# (Machine) Learning the mutation signatures of SARS-CoV-2: a primer for predictive prognosis

**DOI:** 10.1101/2021.08.30.458244

**Authors:** Sunil Nagpal, Nishal Kumar Pinna, Divyanshu Srivastava, Rohan Singh, Sharmila S. Mande

## Abstract

**Motivation:** Continuous emergence of new variants through appearance, accumulation and disappearance of mutations in viruses is a hallmark of many viral diseases. SARS-CoV-2 and its variants have particularly exerted tremendous pressure on global healthcare system owing to their life threatening and debilitating implications. The sheer plurality of the variants and huge scale of genome sequence data available for Covid19 have added to the challenges of traceability of mutations of concern. The latter however provides an opportunity to utilize SARS-CoV-2 genomes and the mutations therein as ‘big data records’ to comprehensively classify the variants through the (machine) learning of mutation patterns. The unprecedented sequencing effort and tracing of disease outcomes provide an excellent ground for identifying important mutations by developing machine learnt models or severity classifiers using mutation profile of SARS-CoV-2. This is expected to provide a significant impetus to the efforts towards not only identifying the mutations of concern but also exploring the potential of mutation driven predictive prognosis of SARS-CoV-2.

**Results:** We describe how a graduated approach of building various severity specific machine learning classifiers, using only the mutation corpus of SARS-CoV-2 genomes, can potentially lead to the identification of important mutations and guide potential prognosis of infection. We demonstrate the applicability of model derived important mutations and use of Shapley values in order to identify the significant mutations of concern as well as for developing sparse models of outcome classification. A total of 77,284 outcome traced SARS-CoV-2 genomes were employed in this study which represented a total corpus of 30346 unique nucleotide mutations and 18647 amino acid mutations. Machine learning models pertaining to graduated classifiers of target outcomes namely ‘Asymptomatic, Mild, Symptomatic/Moderate, Severe and Fatal’ were built considering the TRIPOD guidelines for predictive prognosis. Shapley values for model linked important mutations were employed to select significant mutations leading to identification of less than 20 outcome driving mutations from each classifier. We additionally describe the significance of adopting a ‘temporal modeling approach’ to benchmark the predictive prognosis linked with continuously evolving pathogens. A chronologically distinct sampling is important in evaluating the performance of models trained on ‘past data’ in accurately classifying prognosis linked with genomes of future (observed with new mutations). We conclude that while machine learning approach can play a vital role in identifying relevant mutations, caution should be exercised in using the mutation signatures for predictive prognosis in cases where new mutations have accumulated along with the previously observed mutations of concern.

**Supplementary information:** Supplementary data are enclosed.

## 1 Introduction

Continuous evolution of SARS-CoV-2 and emergence of virulent variants have burdened the global healthcare system at unprecedented levels. With more than 200 million reported cases and over 4 million causalities (worldwide) within the last one year, Covid-19 continues to challenge the adequacy of global healthcare infrastructure (WHO Coronavirus (COVID-19) Dashboard: https://covid19.who.int/, accessed 30 Aug 2021). This has been further complicated by the lack of knowledge pertaining to the factors driving the severity of the SARS-CoV-2 infection. Previously, attempts have been made to predict the infection prognosis using classical machine learning methods based on the symptom profile and co-morbidities of infected individuals (Zoabi et al., 2021). Such efforts are important as they lay the ground for a much-needed thought towards predictive prognosis which may aid in mitigating the potential burden on healthcare system. Mutations in the SARS-CoV-2 genome have a link to the Covid-19 virulence. While the severity of an infection is rightly attributed to host immunity, it is well founded that certain variants of concern (VoCs) are more infectious owing to their mutational peculiarity. Identification of the key mutations and variants of SARS-CoV-2 has therefore become one of the major goals of global genome sequencing efforts. The latter has been exceptional in the entire history of infectious diseases as close to 3 million genome sequences have already been deposited to public repositories like Global initiative on sharing all influenza data (GISAID) (https://www.gisaid.org/, accessed 30 Aug 2021). The traceability of health status of sequencing sample donor is also appreciable, which is reflected in the large cohort of more than 100,000 such samples (and corresponding sequence data) deposited globally with GISAID alone (https://www.epicov.org/epi3/, accessed 30 Aug 2021). Given the large scale of such ‘labelled’ datasets, an ample ground for obtaining clinical intelligence by employing biology agnostic methods is eminent (Zahn, 2021; Nagpal et al., 2020). Supervised machine learning approaches can potentially learn important mutation signatures from these labelled sequences of SARS-CoV-2 genomes and guide prediction of infection severity based on observed mutation signatures (Nagy et al., 2021). Concerted efforts are therefore required to utilize not only the existing methods rooted in biology (e.g., symptom profile, family history, genetic predisposition, sequence analysis, phylogenetics, structural biology, etc.), but also apply unconventional data driven approaches that have conventionally and consistently been proven to yield actionable intelligence in a domain agnostic fashion (Carvalho et al., 2019; Collins et al., 2015; Yadaw et al., 2020). As rightly quoted in a news piece published in Nature lastear, “scientists can spot mutations faster than they can make sense of them” (Callaway, 2020). This situation has only aggravated further with more than 100,000 unique amino acid mutations already identified in 3,134,790 SARS-CoV-2 genome sequences shared by researchers from across the globe through GISAID as on Aug 30, 2021 (Shu and McCauley, 2017).

Like humans, machines or computers can learn from experience. For machines, this experience is derived from the data, which could be labelled or unlabeled. While labelled data refers to the data which is well annotated (e.g., blood biochemistry of diseased and healthy individuals), unlabeled data refers to a data without any ancillary information (e.g., blood biochemistry of unknown samples). These two forms of data availabilities drive the two important types of machine learning approaches, namely, unsupervised and supervised machine learning methods. Unsupervised algorithms, like principal component analysis (PCA) and t-distributed stochastic neighbor embedding (t-SNE), aim to decipher unobserved patterns in the unlabeled data and potentially group the input data points based on patterns of similarity. On the other hand, supervised algorithms, like decision trees and logistic regression, are built on an assumption that there exists a relationship between the input data and their labels, and are therefore aimed at inferring the said relationship. The latter class of machine learning algorithms are therefore cornerstone of predictive analytics and through this article we intend to highlight the possibilities and bottlenecks of predictive prognosis of Covid-19 infection by exploiting the large scale ‘labelled genome sequence’ data. Figure 1 provides a graphical summary of the underlying idea of (machine) learning the labelled genome sequence (and mutation) data of SARS-CoV-2 and developing a severity predictor.

**Figure 1.**
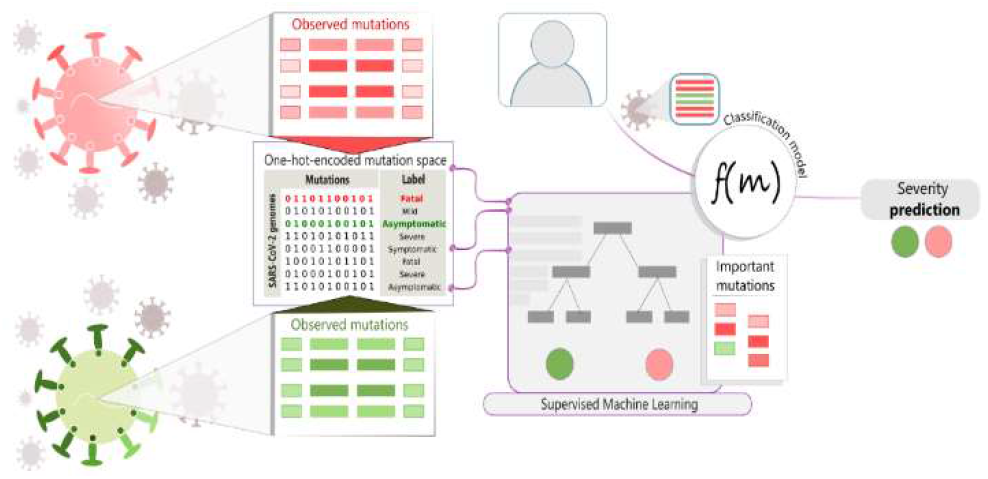
Machine learning the mutation signatures of SARS-CoV-2 for predicting severity outcomes. Mutation profile in a given SARS-CoV-2 genome is an important feature that may drive the course of infection. A one-hot-encoded (presence-absence) matrix of observed mutations across all sequenced genomes can serve as an input data for machine learning (ML) methods. The traced status of patient health associated with each sequence or mutation profile is an important label to train ML algorithms. Supervised machine learning can therefore potentially enable prediction of infection severity by analyzing the patterns of important mutations in the large number of sequenced genomes and in the process enable identification of key mutations that drive the prediction.

However, caution must be exercised in reporting the accuracies and clinical applicability of predictive models, especially where model features (mutations or symptoms) are not expected to exhibit a temporally stable profile (Nagy et al., 2021; Collins et al., 2015; Yadaw et al., 2020). Current approaches, in addition to over speculating the goals of predictive model development, under-utilize the large label space for infection outcomes (Nagy et al., 2021). While the former leads to over-ambitious speculation on clinical applicability of machine learnt models (trained using reported mutations or symptoms in the past) in predicting infection severity; the latter (under-utilized label space) under-estimates the span of significant mutations of concern. It is therefore prudent to acknowledge the limitations of a predictive prognosis exercise while trusting its ability to guide the prediction goals by identifying the mutations of concern from reported variants of SARS-CoV-2.

We adopted a graduated approach of building multiple machine learning classifiers to gauge the predictive power of mutation signatures of SARS-CoV-2 towards prognosis of a Covid-19 infection in terms of all pairs of multiple target labels, namely, (i) Asymptomatic (ii) Mild (iii) Symptomatic (iv) Severe and (v) Fatal. Notably, each binary classifier for different combinations of outcome, was able to classify its target classes with > 85% accuracy and > 0.90 ROC AUC. The mutation signatures identified from the high accuracy models were validated based on the literature evidence for variants of concern (and mutations thereof) and exhibited an encouraging overlap with reported mutations thereof. An additional temporal modeling exercise benchmarked the suitability of non-temporal validation strategies which are currently being adopted to report mutation based predictive prognosis (ML based) methods. We argue that while nontemporal machine learning methods are well adept for identifying the mutation signatures, their applicability for predictive prognosis should be cautiously reported (and adopted).

## 2 Methods

### 2.1. Mutation profiles

Approximately 80,000 SARS-CoV-2 sequences labeled with patient status information were obtained from Global Initiative on Sharing Avian Influenza Data (GISAID)(accessed: July 16, 2021). The complete genome sequence of coronavirus-2 isolate (Wuhan-Hu-1) corresponding to NCBI Genbank accession NC_045512 (GISAID ID EPI_ISL_402125) was employed as the reference (REF.fa) for the purpose of mutation profiling. Fasta files corresponding to each of the downloaded individual genomes (INPUT.fa) were mapped on the reference genome using minimap2 (Li, 2018) with the following flags:

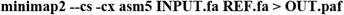

The generated PAF (pairwise alignment format) files were subsequently used for variant calling through the paftools.js module in minimap2 package using the following command in a Linux environment:

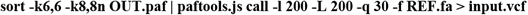

Amino acid changes corresponding to the identified nucleotide variations were predicted using BCFtools/csq program (Danecek and McCarthy, 2017). In total, 30,436 unique nucleotide mutations were identified in 77,284 high quality genome sequences downloaded from GISAID (fulfilling the high coverage, complete sequence and low coverage exclusion criteria of GISAID). The mutation information for all the genomes was one-hot-encoded (binary transformation into 0 and 1) to create a 77284 × 30346 matrix of nucleotide mutation data.

### 2.2. Choice of target outcomes for prediction

Based on the observation of Covid-19 severity, we sought to predict 4 unambiguous target outcomes using machine learning, namely, Asymptomatic, Mild, Moderate and Severe. Additionally, given the good availability of data (2326 genome sequences) corresponding to the ‘fatal’ class of outcome, this class was also included for predictive prognosis of the fatal outcome of Covid-19 infection. The four types of infection outcomes sought to be predicted in this study therefore included - (I) Asymptomatic (II) Mild/Moderate (III) Severe and (IV) Fatal. It may be noted that ambiguous labels (like Symptomatic, Hospitalized, Inpatient, Outpatient, Clinical signs, etc.), that do not provide conclusive indication of health status, were primarily ignored for the purpose of target prediction. ‘Symptomatic’ label was the only apparently ambiguous class of sample that was employed (in later phases of study, post data analysis). This was done after observing a high classification accuracy between Symptomatic class and the unambiguous labels (especially Asymptomatic, Mild and Fatal outcomes), hinting towards a potential employability of this class as Moderate outcome (a label which has not been used often in the patient status data). This led to the total target space of prediction to five labels or disease outcomes: (I) Asymptomatic (II) Mild (III) Symptomatic/Moderate (IV) Severe and (V) Fatal. Caution however must be exercised in conclusive interpretation of ‘Symptomatic’ label and it is recommended that unambiguous labeling be preferred over ambiguous labels.

### 2.3. Choice of machine learning strategy

#### 2.3.1. Unsupervised machine learning exercise

In addition to the primary goal of supervised machine learning based mutation inference and predictive prognosis, a preliminary unsupervised learning of segmentation between genome groups (based on associated disease outcome) was attempted using the non-linear t-Distributed Stochastic Neighbor Embedding (t-SNE)(van der Maaten and Hinton, 2008). The purpose of this exercise was to explore and visualize the large number of genomes and to obtain an initial intuition for the role of mutation signatures in segregating genomes in the space of reduced dimensions.

#### 2.3.2. Supervised machine learning exercise

Given the multi-class nature of patient-status labels, we adopted two approaches for developing unified model(s) to probe predictive power for disease outcomes:

##### a. Using the One-vs-One and One-vs-Rest

We employed the two commonly adopted strategies for arriving at multiclass predictors (Student and Fujarewicz, 2012). The first strategy uses One-vs-One(OVO) approach, wherein discriminant functions are developed for all possible binary combinations of classes (n(n – 1)/2 or 5(5-1)/2 = 10 models in present case). On the other hand, in the second strategy of utilizing One-vs-Rest (OVR) approach, discriminant functions are developed for each individual class by treating rest of the data as opposing single-class of samples (n or 5 models in the present study). Both these approaches aim to develop a single model for predicting one among all the target classes by ensembling the underlying binary predictors. Given the non-linear nature of the one-hot-encoded mutation data, we chose decision tree learning approach for model development and used the well-founded highly efficient gradient boosted tree system of XGboost algorithm (Chen and Guestrin, 2016). The choice of XGboost algorithm, apart from its efficiency, flexibility and portability, is also rooted in the optimized and fast integration of Shapley (Lundberg and Lee, 2017) value assessment for feature importance extraction from the XgBoost trees. The latter, as introduced later, is critical in inferring mutations of concern which is the primary goal of this study.

##### b. Repurposing regression for predicting disease outcomes

The choice of target disease outcomes for prediction has inherently been observed to be incremental in terms of disease severity. For example, the entire range of prediction space falls between Asymptomatic and Fatal classes. This categorical space can potentially be transformed into incremental numeric space (0-4, 0 for asymptomatic, 1 for mild, 2 for symptomatic, 3 for severe and 4 for fatal) and the problem of prediction can be converted into one rooted in regression rather than classification. We therefore transformed the target labels into incremental numerical outcomes and employed XGBoost regressor for developing a regression-based severity estimator. Prediction errors were plotted using a scatter plot between each true level of severity (0-3) on X axis and predicted level of severity on Y axis. The R-squared goodness of fit (coefficient of determination) for the regression model was annotated in the same chart (Supplementary Figure 1).

### 2.4. Training and evaluation

Throughout the study, it was ensured that sample size distribution was equated to the size of minority class population during the model development process. The data with equal proportion of all classes was split into training and testing sets using stratified splitting into 80:20 proportion. In other words, while models were built using 80% of the data, testing of models was performed based on the remaining 20% held out testing set. A stratified 10-fold cross validation was also performed for each model (using the 80% training data) to evaluate the model performance and to ensure that models are not overfitted. Accuracy (average accuracy for cross validation), Precision, Recall, ROC AUC, F1-score and the confusion matrix were assessed to evaluate the models in terms of quantifiable metrics. Classification reports were generated for each of the model consisting of important features (mutations) contributing to model accuracy, confusion matrix, precision-recall-f1 report for each outcome and AUC ROC plot.

### 2.5 Identifying mutations that guide the prediction

Inferring mutations of concern requires identification of mutations (features) that contribute towards the outcome of the model. Manually developed (outside the one-vs-one framework of sci-kit learn library of python) individual binary models/classifiers for all possible pairs of disease outcomes were employed for this purpose. We employed a two-step strategy to identify important mutations for each of the models. The first strategy included creation of a union of model reported important mutations from each iteration of 10-fold cross-validation, in which multiple models were developed across 10 iterations and union of mutations with non-zero model linked importance was performed. This helped in identifying a significantly smaller but important set of features that control the predictive capability of the final model. Once a sparse set of mutations were identified, in the second strategy, Shapley (Lundberg and Lee, 2017) values for each of the features were computed. The concept of Shapley values is originally from coalitional game theory for optimal distribution of game-pay-out to the team players (Lundberg and Lee, 2017). However, this concept has grown popular in the domain of machine learning for assigning outcome contributions to the constituting features (players) of the model towards a given prediction (model payout) (Molnar, 2019; Messalas et al., 2019; Elshawi et al., 2019; Rodríguez-Pérez and Bajorath, 2020; Lundberg and Lee, 2017). Shapley values > 0 were therefore used for identifying the features (mutations) contributing to the positive outcome and values < 0 were used to get important features contributing to the negative outcome. In order to arrive at mutations with consistent but strong contribution, two sets of mutation tables were first created. While the set-1 contained the negative SHAP values for each of the mutation, the set-2 contained positive SHAP values for each of the mutation. Thereafter, for each set, we filtered and retained the mutations that were observed to have an absolute SHAP value of more than or equal to 0.5 in at least 50 genomes (i.e., at least 50 instances of an absolute SHAP value of >=0.5). Subsequently, all SHAP values of the selected mutations were plotted using a Bee-swarm plot for visual inspection of the contribution of each mutation to the disease outcome. Additionally, a compositional assessment of model specific mutations contributing to a given outcome was performed to identify consensus mutations of concern from all the models. The same was visualized using Venn diagram. For example, the mutations selected from the Asymptomatic-Fatal model were compared with the mutations selected using Symptomatic-Fatal model to arrive at reliable set of mutations that deserve attention.

### 2.6. Temporal validation

As discussed previously, a reliable model for predictive prognosis should ideally be benchmarked on unobserved ‘chronologically recent’ data which was not included in the training of the model. This would validate the suitability of proposed models for clinical implementation wherein the viral genome is continuously expected to evolve and accumulate new mutations. Given that it is well founded that SARS-CoV-2 has been evolving with time, an unconditional applicability of models learnt on past data (mutation profiles) must not be assumed.

We therefore devised a chronological data sampling technique with incrementally increasing time windows to test models trained on historical data against a held out unobserved data from a future time-period. For this purpose, entire data (specific to the target outcomes) was first sorted according to the date of collection of samples and multiple held-out test-datasets were created using the chronologically recent subset of data windows. The incremental time window approach was used to create the future test data for observing the effect of time-gap on model performance (where time gap refers to the time-duration between the sample collection day of latest data record used in the training data and the oldest data record of test data). Increase in time gap was approximated by increasing the number of recent samples in the test data without changing the size of training data. It was important not to change the size of the training data to ensure that variations in the performance of model are only time driven (and not training data size driven). Given the best performance observed for Asymptomatic and Fatal outcomes in non-chronological data sampling approach, binary model specific to Asymptomatic-Fatal combination was employed for temporal validation. ROC AUC values for each time-gap based model development exercise were compared for assessing the significance of time as a confounding factor in developing accurate models of predicting the prognosis of SARS-CoV-2 infection using mutation signatures. Five held out test-datasets of 100 samples (genomes) each were created, each with a greater average time gap with respect to the most recent sample in the training dataset.

### 2.8. Databases, tools and implementation

Details of employed software, standard packages, version requirements and cross-validation strategies have been provided in (Supplementary table 1) to facilitate the reproducibility efforts as well as research and discovery of mutation linked prognosis of Covid 19 (and even other diseases).

## 3 Results

### 3.1. Unsupervised learning provides cues to role of mutations in discriminating disease outcome

Figure 2A and 2D represent the outcomes and corresponding sample sizes. Number of genomes considered for each outcome were equal to the size of the minority class/outcome. While Figure 2A-C are representatives of an all-inclusive tSNE, Figure 2D-F represent learning by the exclusion of minority class (Severe: 168 samples). Partially distinct spatial distribution, albeit without conclusive segmentation, of genomes was observed when tSNE was performed using the entire/unfiltered mutation set in the (one-hot-encoded) matrix of genomes pertaining to the target disease outcomes (Figure 2B, 2E). Given that this analysis is based on the mutation (presence-absence) data, the distant points in the space of tSNE plot may be viewed as different variants distanced based on the number of distinct mutations between them. We also overlayed the geographical affiliation of the genomes to the tSNE plots, providing insights about the source of such genome specific data submissions (Figure 2C, F). The said plots do not provide any indication of origin of any variant, which requires complex contact tracing and documentation. The preliminary unsupervised learning task serves to probe initial cues towards utility of mutation signatures alone (or in surrogate to existing methods) in performing a severity risk-assessment of a diagnosed SARS-CoV-2 infection. It was therefore interesting to analyze if mutation-based machine learnt models can be built for guiding the prediction of prognosis or at the least infer mutations contributing to the various outcomes of an infection.

**Figure 2.**
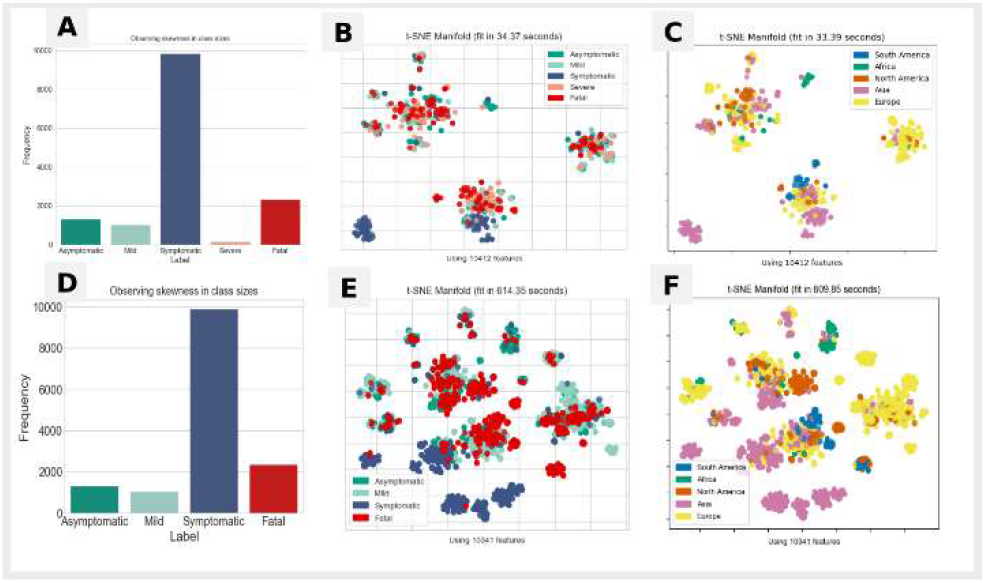
tSNE plot to explore spatial grouping of genomes based on associated infection outcome. Panel (A): a bar plot indicating the class size of various ‘patient status’ specific genome sequences selected for the unfiltered machine learning task Panel (B): the tSNE plot generated using stratified sampling for all chosen outcome labels (randomly picked 168 genomes, i.e. minority class size, from each of category of genome sequences) Panel (c): the plots for genomes presented in panel B, but overlayed with geographical information about location of data submission Panel (D): the class size of samples after excluding the minority class/outcome (Severe). Panel (E) represents the tSNE plot using stratified sampling for labels other than severity group (smallest class of 168 genomes) with geographical information overlayed in Panel (F).

### 3.2. Supervised machine learning for unified model shows limited success, binary models show encouraging signals of discriminative power

#### 3.2.1. Single model for multi-class classification, regression and validation thereof

A single model developed using One-vs-Rest (OvR) approach for all five target classes (Asymptomatic, Symptomatic, Mild, Severe and Fatal) didn’t yield a high accuracy (0.582 ± 0.035, using 10-fold cross validation). Accuracy however cannot be considered as a perfect metric for a multi-class (5 classes in this case) predictor. Specifically, this classifier was trained and tested using 168 samples from each class (as the size of each class was reduced to the minority (Severe) class size to enable an unbiased/stratified learning. Nevertheless, it was encouraging to observe the majority prediction for each of the target outcomes wasn’t significantly skewed (confusion matrix in Figure 3A). An ROC AUC macro-average of 0.833 ± 0.015 through 10-fold cross validation indicated a good degree of separability of individual classes from rest of the data (Supplementary Table 2, ROC plots in Figure 3A). ROC AUC plots in Figure 3 were plotted using the held-out test data (20% of 168 for each class). It may be noted that while Figure 3 represents the performance of model trained on unfiltered mutation set, the comparable results of model performance, generated using only the important mutations (obtained after 10-fold cross validation) are additionally presented in Supplementary Table 2 and 3. Given the expectation that Severe class (168 samples) could be driving the sub-optimal training of otherwise large data size for each of the other target classes (greater than 1000 samples each, Figure 2A), we therefore sought to ask if there is a possibility of improvement in learning of discriminating function by omitting this minority class. The OvR classifier was therefore re-trained using only Asymptomatic, Symptomatic, Mild and Fatal classes (Figure 3C). Omitting Severe class, yielded significantly improved accuracy (0.752 ± 0.017) and macro-average of ROC AUC (0.920 ± 0.010) (Supplementary Table 2, Figure 3C). Model developed using One-vs-One (OvO) approach yielded comparable results (Figure 3B, 3D) on held-out data as well as 10-fold cross validation accuracy (Supplementary Table 3). While these results indicated a need for caution while developing an ambitious single model to predict multiple outcomes of Covid-19 severity, it was encouraging to observe latent signals of mutation peculiarity in the high accuracy (> 0.85) and ROC AUC (> 0.88) for the contributing models of the unified OvR model (Supplementary Table 3, Figure 3C), i.e. Rest Vs Asymptomatic, Rest Vs Mild, Rest vs Symptomatic and Rest vs Fatal. Similarly, the individual models of unified OvO classifier also exhibited high power of discrimination with greater than 0.85 accuracy and 0.90 ROC AUC for most pairs of infection outcomes (Supplementary Table 2). Importantly, the recall value for the more severe outcome was consistently observed to be > 90%, strengthening the assumption that mutational peculiarity not only drives severity but can also be identified using machine learning, thereby guiding the exercise of predictive prognosis, especially in proxy to the other methods (e.g., clinical symptoms and medical history), if not alone. Furthermore, quick identification of the important mutations is crucial to trace the evolution of the virus without missing the hitherto unobserved variants through traditional exercise of variant tracing rooted in epidemiology and phylogenetics. Machine learning can potentially aid this task.

**Figure 3.**
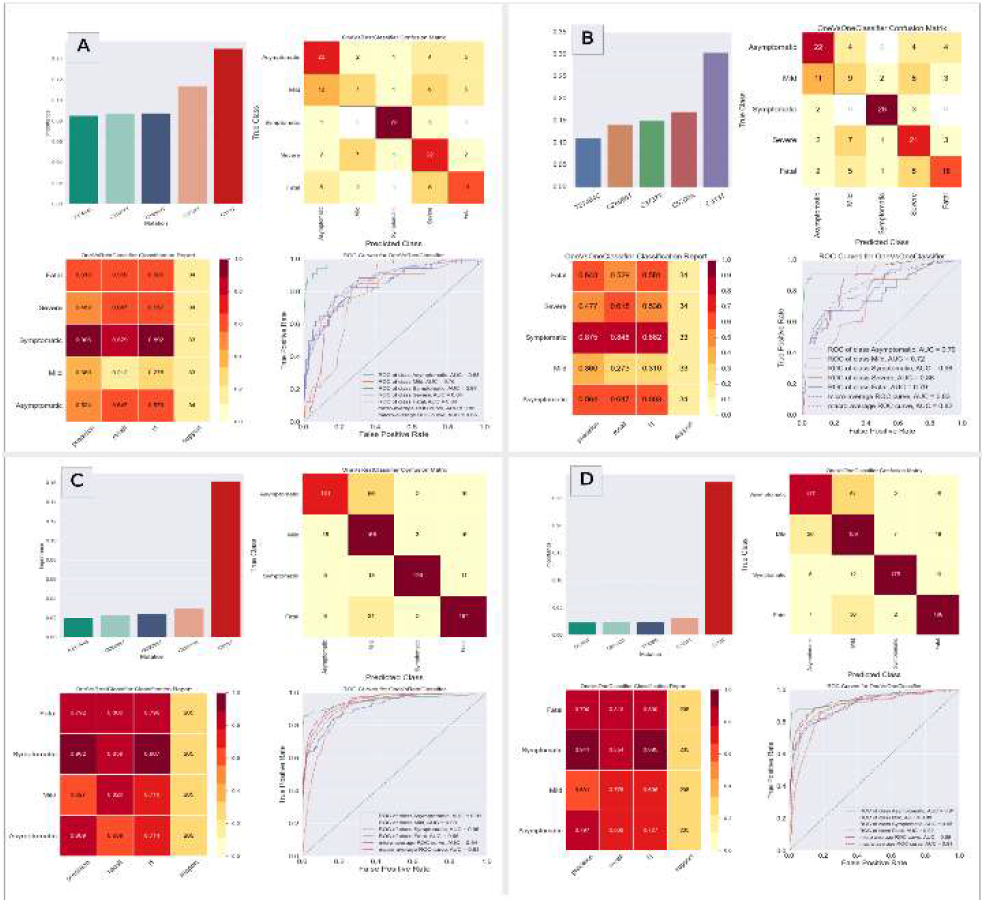
Classification reports generated for the OvR and OvO models consisting of important features (mutations) contributing to model accuracy, confusion matrix, precision-recall-f1 report for each outcome and AUC ROC plot. Panels (A and B) represent the classification report for OvR and OvO models respectively trained for all five target outcomes for prediction. Panels (C-D) represent the classification report of OvR and OvO models respectively trained for all outcomes other than the minority Severe class

The repurposing of regression-based machine learning for predicting severity outcome yielded encouraging results with an R-squared goodness of fit (coefficient of determination) value of 0.625 (∼0.791 coefficient of correlation). These results were obtained by omitting the Severe class outcome for an efficient training using large but stratified sample sizes (∼1000 samples for each class). However, as expected, the results were not encouraging when ‘Severe class’ with 168 samples was included in the training dataset, resulting in the need for stratification of all outcome classes to the minority class size (Supplementary Figure 1). The prediction errors without omission (Supplementary Figure 1A) were 1.25 (RMSE) as compared to 0.68 for training with omission of Severe class (Supplementary Figure 1B). The errors as shown in Supplementary Figure 1 were plotted using a scatter plot between predicted level of severity (Y axis) and original level of severity (X axis).

#### 3.2.2. Identification of mutations of concern

A total of 59 unique genic mutations were selected by using the importance linked SHAP value approach described in methods section. Supplementary Table 4 provides a detailed summary of the selected significant mutations identified for all model outcome pairs. The SHAP value-based contribution inclination of each of the selected mutation is summarized as a Bee-swarm plot in Figure 4 (excluding Severe class due to small sample size), where density of samples with SHAP>0 presents contribution of the presence/absence of the said mutation to the higher severity outcome and SHAP<0 indicates contribution towards less severe outcome. The top five mutations (among the set of total corpora of important mutations) identified from each model which were shortlisted for SHAP value-based mutation filtration are presented in the model summary reports in Supplementary File 2. As summarized in Table 1, the comparison of the outcome linked important mutations indicated a consistent observation of the presence of G25088T (Spike-V1176F) in fatality linked outcome in all the high accuracy models. Similarly, A20268G (NSP15-L216L) and C313T (a synonymous mutation at NSP1-L16) were observed to contribute to the ‘Symptomatic’ outcome across all high accuracy models tested. Mild outcome was observed to be contributed by the presence of G28739T (Nucleocapsid-A156S), while the presence of mutations G11083T (NSP6-L37F), C26456T (Envelope-P71L) and C26885A (Membrane-N121K) were contributing to Asymptomatic outcome, as observed in all tested binary classification models in which Asymptomatic was one of the outcomes (Supplementary Table 4 and Figure 4). Similarly, while absence of C14408T (NSP12-P323L) was consistently observed to contribute to Asymptomatic outcome, there was no mutation whose disappearance was observed to consistently contribute to high levels of severity (Mild, Symptomatic or Fatal). This indicates the rational nature of the process of evolution of the virus wherein variants of concern/interest emerge out of unique or new mutations.

**Figure 4.**
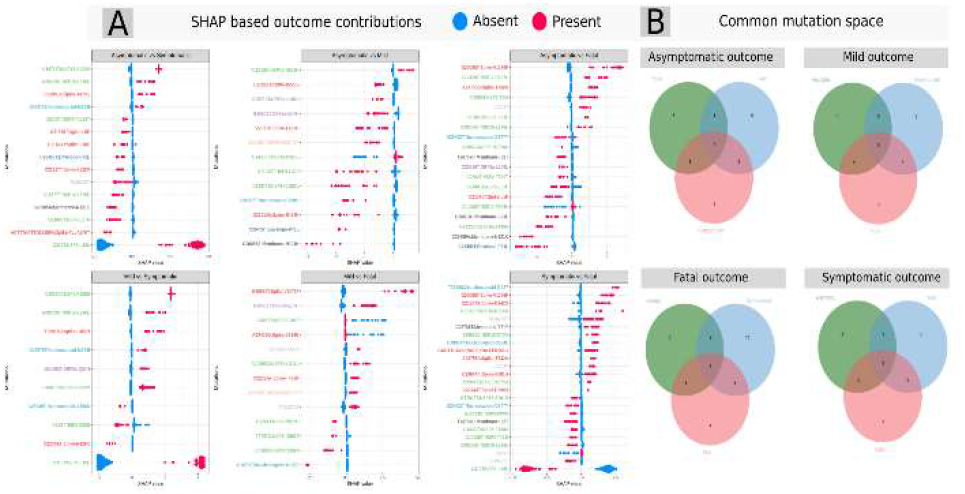
Mutations of concern/interest selected using model importance coupled SHAP value assessment. Bee-swarm plots in panel A indicate the contribution of presence/absence of a mutation towards an outcome across various binary classifiers trained using XgBoost. Values greater than zero indicate contribution towards a more severe outcome, while SHAP values less than zero indicate contribution to less severe outcome of the model. The Venn diagrams in panel B represent the mutational overlap space for each outcome across different models (only the presence linked outcomes are considered in the Venn diagrams). Additionally, the labels of the mutations have been colored according to the protein in which the said mutations appear.

**Table 1.**
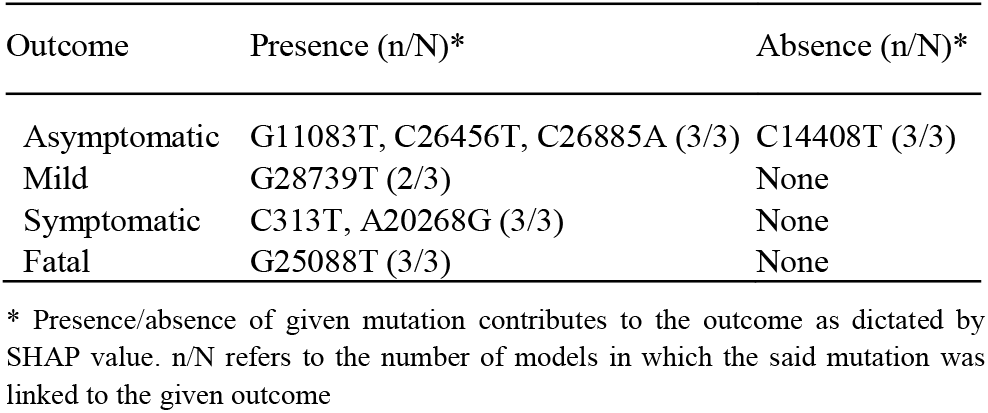
Machine learnt important mutation landscape for Covid-19 outcomes obtained through compositional comparison of individual outcome linked mutations across all trained binary models

| Outcome | Presence (n/N)* | Absence (n/N)* |
| --- | --- | --- |
| Asymptomatic | G11083T, C26456T, C26885A (3/3) | C14408T (3/3) |
| Mild | G28739T (2/3) | None |
| Symptomatic | C313T, A20268G (3/3) | None |
| Fatal | G25088T (3/3) | None |

\* Presence/absence of given mutation contributes to the outcome as dictated by SHAP value. n/N refers to the number of models in which the said mutation was linked to the given outcome

#### 3.2.2. Identification of mutations of concern

A total of 59 unique genic mutations were selected by using the importance linked SHAP value approach described in methods section. Supplementary **Table 4** provides a detailed summary of the selected significant mutations identified for all model outcome pairs. The SHAP value-based contribution inclination of each of the selected mutation is summarized as a Bee-swarm plot in Figure 4 (excluding Severe class due to small sample size), where density of samples with SHAP>0 presents contribution of the presence/absence of the said mutation to the higher severity outcome and SHAP<0 indicates contribution towards less severe outcome. The top five mutations (among the set of total corpora of important mutations) identified from each model which were shortlisted for SHAP value-based mutation filtration are presented in the model summary reports in Supplementary File 2. As summarized in Table 1, the comparison of the outcome linked important mutations indicated a consistent observation of the presence of G25088T (Spike-V1176F) in fatality linked outcome in all the high accuracy models. Similarly, A20268G (NSP15-L216L) and C313T (a synonymous mutation at NSP1-L16) were observed to contribute to the ‘Symptomatic’ outcome across all high accuracy models tested. Mild outcome was observed to be contributed by the presence of G28739T (Nucleocapsid-A156S), while the presence of mutations G11083T (NSP6-L37F), C26456T (Envelope-P71L) and C26885A (Membrane-N121K) were contributing to Asymptomatic outcome, as observed in all tested binary classification models in which Asymptomatic was one of the outcomes (Supplementary Table 4 and Figure 4). Similarly, while absence of C14408T (NSP12-P323L) was consistently observed to contribute to Asymptomatic outcome, there was no mutation whose disappearance was observed to consistently contribute to high levels of severity (Mild, Symptomatic or Fatal). This indicates the rational nature of the process of evolution of the virus wherein variants of concern/interest emerge out of unique or new mutations.

### 3.3. Comparing identified mutation signatures against known variants of SARS-CoV-2

World Health Organization, in global collaboration with researchers, institutions and its partners has been attempting to characterize the virus into certain and variants of concern (VoCs). These characterizations have primarily been rooted in identification of molecular ‘signals’ that indicate a potential to increase the virulence (https://www.who.int/en/activities/tracking-SARS-CoV-2-variants), in addition to tracing the spread, infectivity, hospitalizations linked with given variant. We sought to compare the machine learnt signatures with the mutation profile of the WHO characterized VOCs. A manual survey of the lineages of currently known variants of concern indicated that 24 of these selected 59 mutations have been observed in the said variants with more than 75% prevalence as published on https://www.outbreak.info. The said mapping has been provided under ‘Mapped AA mutation in VoC’, ‘WHO_labcl of VoC’ and ‘Lineages with mutation prevalence >=75 %’ columns of Supplementary Table 4. Interestingly, only 2 out of 8 important mutations obtained through the comparison of outcome linked mutations in all binary models (Table 1) were found with more than 75% prevalence in at least one of the known variants of concern. Among these it was encouraging to observe that the fatality linked G25088T (Spike-V1176F) has already been recognized in gamma variant of concern (Supplementary Table 4). The Asymptomatic outcome linked mutation C26456T (Envelope-P71L) was however traced to be present in an alpha variant (B.1.1.7), which was a counter-intuitive observation, but raises the question regarding the need for probing the effect of co-occurring prevalent mutations. Nevertheless, we opine that the overlap as well as a lack thereof between machine learnt mutations and those found in characterized VoCs provides hints towards the suitability of employing machine learning techniques in not only identifying the mutations of concern but also supporting the existing methods rooted in phylogeny and epidemiology, in the process of variant classification.

### 3.4. Temporal validation

The incremental time window approach of creating five chronologically recent held-out test datasets revealed the rational limitation of mutation based predictive prognosis models. With increase in the time gap between the constant training data and the chronological test datasets, model performance was observed to drop (Figure 5, Supplementary Table 5). While the test data set consisting of 100 samples (50 of Asymptomatic and 50 of Fatal class) chronologically closest to the most recent sample of training data yielded an ROC AUC of 0.95 and accuracy of 0.90 (window 1 on x axis of Figure 5, Supplementary Table 5), the distant window (which pertained to 100 most recent samples and hence chronologically farthest from training data, yielded a held-out ROC AUC of 0.83 and accuracy of 0.70. This leads to the below mentioned three important inferences -

I. as long as the virus is mutating, it may be over-speculative to propose models rooted in mutation signature for prognosis in a clinical setting
II. a judicious use of predictive models can however take place where reliability of the prediction is indexed by the fraction of mutations that are already accounted for in the model (as indicated in the last window wherein model performance is slightly better due to observation of previously learnt mutation signatures)
III. the role of machine learning in identifying the important mutations among the large existing corpus of SARS-CoV-2 mutations should not be ignored as this can significantly aid the ongoing activities of tracing variants of concern.

**Figure 5.**
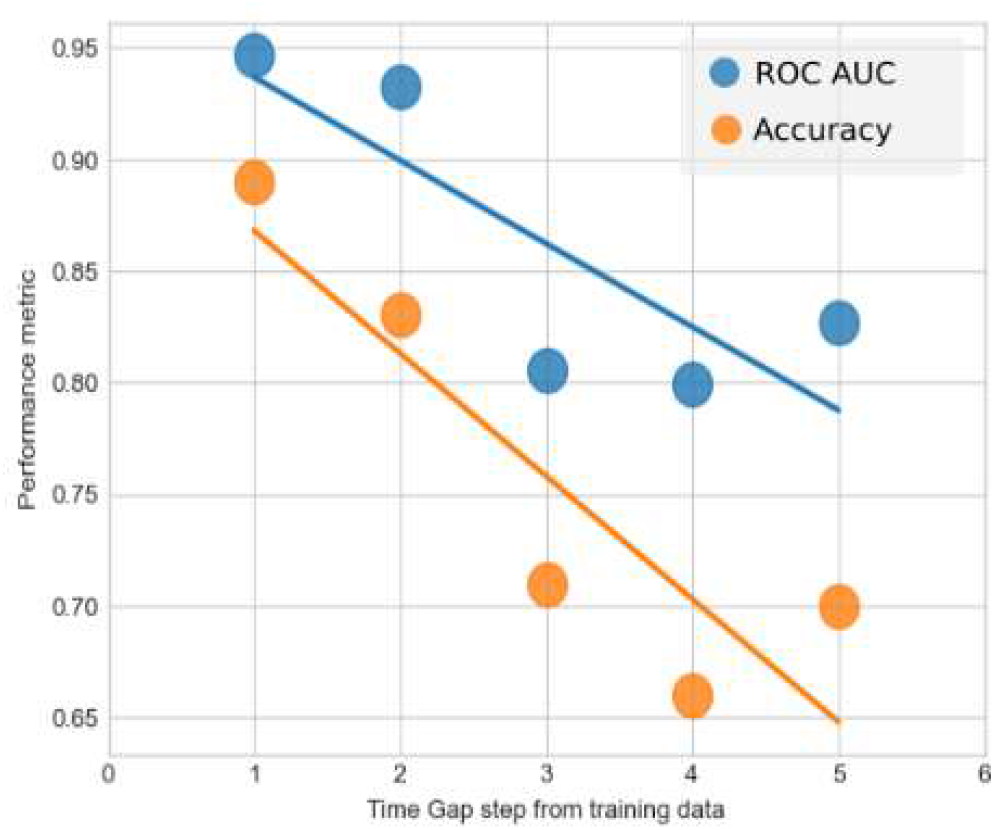
Temporal validation of Asymptomatic-Fatal predictive model. X axis represents the time gap window (wherein window 1 contained 100 test samples which were chronologically closest to the most recent genome in training data, while window 5 contained 100 most distant samples). Y axis represents the value of two performance metrices namely, ROC AUC and accuracy of the model.

## 4 Caveats & Conclusions

Machine learning models are rarely perfect. The imperfection is attributed to fractional representation of information in the chosen datasets (i.e. complete data for any case/event/population is rarely available). Consequently, there is always a scope for improving the learnt models by incorporating new data to the machine learning framework. This limitation is particularly pronounced for viral genomes which are continuously evolving. New data will always be useful in updating the mutation feature profile of the models which will help in improving the accuracy of the prospective predictions. A well streamlined machine learning framework rather simplifies the process of accommodating new data, updating the predictive models and for obtaining quick insights into the new found mutations of concern. Through this study, we not only attempted to provide evidence towards suitability of using SARS-CoV-2 mutation data to develop machine learning methods of severity classification but also tried to develop an approach towards identification of mutations of concern. Importantly, we emphasize on the need for concerted efforts in the direction of building dynamic workflows which can conveniently be reused/improved for generating new models as and when new data is available, which can greatly support the ongoing efforts of variant identification and potentially, predictive prognosis

## Acknowledgements

We gratefully acknowledge all the Authors from the Originating laboratories responsible for obtaining the specimens and the Submitting laboratories where genetic sequence data were generated and shared via the GISAID Initiative, on which this research is based. Genome sequences and meta-data should be downloaded from https://www.gisaid.org. We gratefully acknowledge the original contributors of the virus genome sequences used in this study in Supplementary File 3. SN would like to thank Dr. Bhupesh Taneja (PhD supervisor) for encouraging the work on building a ML framework for important feature recognition.

## Author contribution

SN conceived the idea, designed the study and finalized algorithm. NP generated nucleotide mutation data. SN and NP processed the data. SN coded the workflows which were reviewed by DS, RS, NP and SSM. SN, NP, RS, DS and SSM analysed the results. SN wrote first draft of manuscript. SSM supervised the entire work and finalized manuscript. All authors reviewed and approved the submission.

## References

Callaway, E. (2020) The coronavirus is mutating - does it matter? Nature, 585.

Carvalho, D. v. et al. (2019) Machine learning interpretability: A survey on methods and metrics. Electronics (Switzerland), 8.

Chen, T. and Guestrin, C. (2016) XGBoost.

Collins, G.S. et al. (2015) Transparent reporting of a multivariable prediction model for individual prognosis or diagnosis (TRIPOD): The TRIPOD Statement. European Urology, 67.

Danecek, P. and McCarthy, S.A. (2017) BCFtools/csq: Haplotype-aware variant consequences. Bioinformatics, 33.

Elshawi, R. et al. (2019) On the interpretability of machine learning-based model for predicting hypertension. BMC Medical Informatics and Decision Making, 19.

Li, H. (2018) Minimap2: Pairwise alignment for nucleotide sequences. Bioinformatics, 34.

Lundberg, S.M. and Lee, S.I. (2017) A unified approach to interpreting model predictions. In, Advances in Neural Information Processing Systems.

van der Maaten, L. and Hinton, G. (2008) Visualizing data using t-SNE. Journal of Machine Learning Research, 9.

Messalas, A. et al. (2019) Model-Agnostic Interpretability with Shapley Values. In, 10th International Conference on Information, Intelligence, Systems and Applications, IISA 2019.

Molnar, C. (2019) Interpretable Machine Learning. A Guide for Making Black Box Models Explainable. Book.

Nagpal, S. et al. (2020) What if we perceive SARS-CoV-2 genomes as documents? Topic modelling using Latent Dirichlet Allocation to identify mutation signatures and classify SARS-CoV-2 genomes (preprint). bioRxiv.

Nagy, Á. et al. (2021) COVIDOUTCOME - Estimating COVID severity based on mutation signatures in the SARS-CoV-2 genome. Database, 2021.

Rodríguez-Pérez, R. and Bajorath, J. (2020) Interpretation of machine learning models using shapley values: application to compound potency and multi-target activity predictions. Journal of Computer-Aided Molecular Design, 34.

Shu, Y. and McCauley, J. (2017) GISAID: Global initiative on sharing all influenza data - from vision to reality. Euro surveillance : bulletin Europecn sur les maladies transmissibles = European communicable disease bulletin, 22, 30494.

Student, S. and Fujarewicz, K. (2012) Stable feature selection and classification algorithms for multiclass microarray data. Biology Direct, 7.

Yadaw, A.S. et al. (2020) Clinical features of COVID-19 mortality: development and validation of a clinical prediction model. The Lancet Digital Health, 2.

Zahn, L.M. (2021) Natural language predicts viral escape. Science, 371.

Zoabi, Y. et al. (2021) Machine learning-based prediction of COVID-19 diagnosis based on symptoms. npj Digital Medicine, 4.

