## Supplementary Table 1 for "(Machine) Learning the mutation signatures of SARS-CoV-2: a primer for predictive prognosis"

**Supplementary Table 1:** Details of various key resources including tools and packages employed in the study.

| **Resource** | **Purpose** | **Source link** | **Authors** |
| --- | --- | --- | --- |
| Genome sequences and metadata | Input data and labels | <https://www.gisaid.org/> | (Shu and McCauley, 2017) |
| Reference genome | Variant calling | <https://www.ncbi.nlm.nih.gov/genbank> | (Sayers *et al.*, 2021) |
| ***minimap2, paftools*** | Variant calling | <https://github.com/lh3/minimap2> | (Li, 2018) |
| Pandas library v 1.1.3 (python) | Data handling and structuring | <https://pandas.pydata.org/> | (McKinney, 2011) |
| sklearn library v 0.23.2 (python) | Machine learning framework, conventional algorithms, recursive feature elimination, multiclass prediction strategies (ovr, ovo) and evaluation metrics | <https://scikit-learn.org/stable/> | (Pedregosa *et al.*, 2011) |
| XGBoost library v 1.3.3 (python) | Machine learning algorithm | <https://xgboost.readthedocs.io/> | (Chen and Guestrin, 2016) |
| SHAP library v 0.39.0 (python) | Shapley value computation | <https://shap.readthedocs.io/> | (Lundberg and Lee, 2017) |
| Matplotlib v 3.3.2 and  Yellowbrick 1.3.post1 (python) | Data visualization | <https://matplotlib.org/>  <https://www.scikit-yb.org/> | (Hunter, 2007; Bengfort and Bilbro, 2019) |
