## Supplementary Table 5 for "(Machine) Learning the mutation signatures of SARS-CoV-2: a primer for predictive prognosis"

**Supplementary Table 5.** Temporal validation of Asymptomatic-Fatal predictive model. Column 1 represents the time gap window (wherein T1 gap contained 100 test samples which were chronologically closest to the most recent genome in training data, while window 5 contained 100 most distant samples).

| **Model** | **Accuracy** | **ROC AUC** |
| --- | --- | --- |
| T1 Gap | 0.89 | 0.95 |
| T2 Gap | 0.83 | 0.93 |
| T3 Gap | 0.71 | 0.81 |
| T4 Gap | 0.66 | 0.80 |
| T5 Gap | 0.70 | 0.83 |
