## Supplementary File 3 for "(Machine) Learning the mutation signatures of SARS-CoV-2: a primer for predictive prognosis"

We gratefully acknowledge the following Authors from the Originating laboratories responsible for obtaining the specimens, as well as the Submitting laboratories where the genome data were generated and shared via GISAID, on which this research is based.

All Submitters of data may be contacted directly via www.gisaid.org

Authors are sorted alphabetically.

| Accession ID | Originating Laboratory | Submitting Laboratory | Authors |
|---|---|---|---|
| EPI_ISL_1001044, EPI_ISL_1001052, EPI_ISL_1001059, EPI_ISL_1001062 | Centre Pasteur du Cameroun | Institut Pasteur de Dakar | Njouom Richard |
| EPI_ISL_1007662, EPI_ISL_1007663, EPI_ISL_1007664, EPI_ISL_1007665, EPI_ISL_1007666, EPI_ISL_1007667, EPI_ISL_1007668, EPI_ISL_1007669, EPI_ISL_1007670, EPI_ISL_1007671 | Department of Internal Medicine, College of Medicine, Chosun University | Department of Internal Medicine, College of Medicine, Chosun University | Dong-Min Kim |
| EPI_ISL_1014677, EPI_ISL_1014682 | National Influenza Center, Virology Department | National Influenza Center | NZ Shafiei Jandaghi, V Salimi, A Nejati, K Sadeghi, J Yavarian, N Ghavvami,F Ajaminejad and T Mokhtari Azad |
| EPI_ISL_1014683 | National Influenza Center, Virology Department | National Influenza Center | J Yavarian,K Sadeghi, NZ Shafiei Jandaghi, V Salimi, A Nejati, N Ghavvami,F Ajaminejad and T Mokhtari Azad |
| EPI_ISL_1014684 | National Influenza Center, Virology Department | National Influenza Center | A Nejati, J Yavarian,K Sadeghi, NZ Shafiei Jandaghi, V Salimi, N Ghavvami,F Ajaminejad and T Mokhtari Azad |
| EPI_ISL_1020171, EPI_ISL_1020172, EPI_ISL_1020173, EPI_ISL_1020174 | AZ Klina | AZ Klina | Dr. C. Vael |
| EPI_ISL_1027658, EPI_ISL_1027659, EPI_ISL_1027660, EPI_ISL_1027661, EPI_ISL_1027662, EPI_ISL_1027663, EPI_ISL_1027664, EPI_ISL_1027665, EPI_ISL_1027666 | Mako Medical Laboratories | Mako Medical Laboratories | "Lauren Moon, Matthew Tugwell" |
| EPI_ISL_1029957, EPI_ISL_1029958, EPI_ISL_1029959, EPI_ISL_1029960, EPI_ISL_1029961, EPI_ISL_1029962, EPI_ISL_1029963, EPI_ISL_1029964, EPI_ISL_1029965, EPI_ISL_1029966, EPI_ISL_1029968, EPI_ISL_1029969, EPI_ISL_1029971, EPI_ISL_1029972, EPI_ISL_1029973 | | | |
| see above | Cliniques universitaires Saint-Luc | UCLouvain/IREC/MBLG | Jean Ruelle, Lysa Pinsmaye, Benoit Kabamba Mukadi |
| EPI_ISL_1034305 | Laboratorio de Ecologia de Doencas Transmissiveis na Amazonia, Instituto Leonidas e Maria Deane - Fiocruz Amazonia | Laboratorio de Ecologia de Doencas Transmissiveis na Amazonia, Instituto Leonidas e Maria Deane - Fiocruz Amazonia | Valdinete Nascimento, Victor Souza, André Corado, Fernanda Nascimento, George Silva, Ágatha Costa, Debora Duarte, Karina Pessoa, Matilde Mejía, Luciana Gonçalves, Maria Júlia Brandão, Michele Jesus, Felipe Naveca on behalf of the Fiocruz COVID-19 Genomic Surveillance Network |
| EPI_ISL_1039249 | Virus Research and Diagnostic Laboratory, Medical College Level, Kasturba Hospital for Infectious Diseases | Indian Council of Medical Research-National Institute of Virology, Microbial Containment Complex | Pragya D. Yadav, Jayanti Shastri |
| EPI_ISL_1040031, EPI_ISL_1040033, EPI_ISL_1040034, EPI_ISL_1040036, EPI_ISL_1040037, EPI_ISL_1040040, EPI_ISL_1040041, EPI_ISL_1040042, EPI_ISL_1040043, EPI_ISL_1040045, EPI_ISL_1040046, EPI_ISL_1040047, EPI_ISL_1040048 | | | |
| see above | PCR Laboratory, The First Affiliated Hospital of Zhengzhou University, Zhengzhou, Henan, China | PCR Laboratory, The First Affiliated Hospital of Zhengzhou University, Zhengzhou, Henan, China | Guanglin Cui, Shijie Zhang, Li Sun, Xianchun Meng, Ting Sun, Junhu Wan, Lijuan Rong, Xiangbo Huang, Lei Zheng, Hao Guo, Meijing Long |
| EPI_ISL_1055383 | Virus Research and Diagnostic Laboratory, Medical College Level, Kasturba Hospital for Infectious Diseases | Indian Council of Medical Research-National Institute of Virology, Microbial Containment Complex | Pragya D. Yadav, Jayanti Shastri |
| EPI_ISL_1055384, EPI_ISL_1055768, EPI_ISL_1055770, EPI_ISL_1055771, EPI_ISL_1055785, EPI_ISL_1055788, EPI_ISL_1055789, EPI_ISL_1063400, EPI_ISL_1063401, EPI_ISL_1063494, EPI_ISL_1063526, EPI_ISL_1063527, EPI_ISL_1063529, EPI_ISL_1063531 | | | |
| see above | Virus Research and Diagnostic Laboratory, Medical College Level, Kasturba Hospital for Infectious Diseases | National Institute of Virology-Microbial Containment Complex, Indian Council of Medical Research | Pragya D. Yadav, Jayanti Shastri |
| EPI_ISL_1068593 | Microbiological Laboratory | Microbiological Laboratory,Lu`an Center for Disease Control and Prevention | Yang Wei, Gao Dawei, Chen Zhichao, Chang Hongwei, Zhang Feng, Zhang Limei, Chen Beilei, Zhang Qin, Li Zhaoyang, Zhu Rui |
| EPI_ISL_1068682 | Micrbiological Laboratory | Micrbiological Laboratory,Lu`an Center for Disease Control and Prevention | Yang Wei,Gao Dawei, Chen Zhichao,Chang Hongwei,Zhang Feng,Zhang Limei,Chen Beilei,Zhang Qin,Li Zhaoyang,Zhu Rui |
| EPI_ISL_1069173 | Micrbiological Laboratory,Lu`an Center for Disease Control and Prevention | Micrbiological Laboratory,Lu`an Center for Disease Control and Prevention | Yang Wei,Gao Dawei, Chen Zhichao,Chang Hongwei,Zhang Feng,Zhang Limei,Chen Beilei,Zhang Qin,Li Zhaoyang,Zhu Rui |
| EPI_ISL_1069192 | Micrbiological Laboratory,Lu`an Center for Disease Control and Prevention | No.409,Gaocheng Middle Road,Lu`an City,Anhui Provice | Yang Wei,Gao Dawei, Chen Zhichao,Chang Hongwei,Zhang Feng,Zhang Limei,Chen Beilei,Zhang Qin,Li Zhaoyang,Zhu Rui |
| EPI_ISL_1069193 | Micrbiology Laboratory,Lu`an Center for Disease Control and Prevention | Micrbiology Laboratory,Lu`an Center for Disease Control and Prevention | Yang Wei,Gao Dawei, Chen Zhichao,Chang Hongwei,Zhang Feng,Zhang Limei,Chen Beilei,Zhang Qin,Li Zhaoyang,Zhu Rui |
| EPI_ISL_1069198, EPI_ISL_1069199, EPI_ISL_1069200, EPI_ISL_1069201, EPI_ISL_1069203, EPI_ISL_1069205, EPI_ISL_1069206, EPI_ISL_1069207, EPI_ISL_1069208, EPI_ISL_1069210, EPI_ISL_1069211, EPI_ISL_1069213, EPI_ISL_1069215 | | | |
| see above | Microbiology Laboratory,Lu`an Center for Disease Control and Prevention | Microbiology Laboratory,Lu`an Center for Disease Control and Prevention | Yang Wei,Gao Dawei, Chen Zhichao,Chang Hongwei,Zhang Feng,Zhang Limei,Chen Beilei,Zhang Qin,Li Zhaoyang,Zhu Rui |
| EPI_ISL_1069216 | Microbiology Laboratory,Lu`an Center for Disease Control and Prevention | Microbiology Laboratory,Lu`an Center for Disease Control and Prevention | Microbiology Laboratory,Lu`an Center for Disease Control and Prevention |
| EPI_ISL_1069219, EPI_ISL_1081193, EPI_ISL_1081194, EPI_ISL_1081195, EPI_ISL_1081196, EPI_ISL_1081197, EPI_ISL_1081198, EPI_ISL_1081199, EPI_ISL_1081260, EPI_ISL_1081291, EPI_ISL_1081310, EPI_ISL_1081311, EPI_ISL_1081324, EPI_ISL_1081325, EPI_ISL_1081326, EPI_ISL_1081331, EPI_ISL_1081340 | | | |
| see above | Microbiology Laboratory,Lu`an Center for Disease Control and Prevention | Microbiology Laboratory,Lu`an Center for Disease Control and Prevention | Yang Wei,Gao Dawei, Chen Zhichao,Chang Hongwei,Zhang Feng,Zhang Limei,Chen Beilei,Zhang Qin,Li Zhaoyang,Zhu Rui |
| EPI_ISL_1111075, EPI_ISL_1111076, EPI_ISL_1111077, EPI_ISL_1111078, EPI_ISL_1111079, EPI_ISL_1111080, EPI_ISL_1111081, EPI_ISL_1111086, EPI_ISL_1111087, EPI_ISL_1111088, EPI_ISL_1111089, EPI_ISL_1111090, EPI_ISL_1111091, EPI_ISL_1111092, EPI_ISL_1111093, EPI_ISL_1111094, EPI_ISL_1111095, EPI_ISL_1111096, EPI_ISL_1111097, EPI_ISL_1111098, EPI_ISL_1111100, EPI_ISL_1111101, EPI_ISL_1111102, EPI_ISL_1111103, EPI_ISL_1111104, EPI_ISL_1111105, EPI_ISL_1111106, EPI_ISL_1111107, EPI_ISL_1111108, EPI_ISL_1111109, EPI_ISL_1111110, EPI_ISL_1111111, EPI_ISL_1111112, EPI_ISL_1111113, EPI_ISL_1111114, EPI_ISL_1111115, EPI_ISL_1111116, EPI_ISL_1111117, EPI_ISL_1111118, EPI_ISL_1111119, EPI_ISL_1111120, EPI_ISL_1111122, EPI_ISL_1111123, EPI_ISL_1111124, EPI_ISL_1111125, EPI_ISL_1111126, EPI_ISL_1111162, EPI_ISL_1111163, EPI_ISL_1111164, EPI_ISL_1111169, EPI_ISL_1111170, EPI_ISL_1111171, EPI_ISL_1111172, EPI_ISL_1111174, EPI_ISL_1111175, EPI_ISL_1111176, EPI_ISL_1111177, EPI_ISL_1111178, EPI_ISL_1111179, EPI_ISL_1111180, EPI_ISL_1111181, EPI_ISL_1111182, EPI_ISL_1111183, EPI_ISL_1111184, EPI_ISL_1111185, EPI_ISL_1111186, EPI_ISL_1111187, EPI_ISL_1111188, EPI_ISL_1111189, EPI_ISL_1111190, EPI_ISL_1111192, EPI_ISL_1111193, EPI_ISL_1111194, EPI_ISL_1111195, EPI_ISL_1111196, EPI_ISL_1111197, EPI_ISL_1111198, EPI_ISL_1111199, EPI_ISL_1111200, EPI_ISL_1111201, EPI_ISL_1111202, EPI_ISL_1111203, EPI_ISL_1111204, EPI_ISL_1111205, EPI_ISL_1111206, EPI_ISL_1111207, EPI_ISL_1111208, EPI_ISL_1111209, EPI_ISL_1111210, EPI_ISL_1111211, EPI_ISL_1111212, EPI_ISL_1111213, EPI_ISL_1111214, EPI_ISL_1111215, EPI_ISL_1111216, EPI_ISL_1111217, EPI_ISL_1111218, EPI_ISL_1111219, EPI_ISL_1111220, EPI_ISL_1111221, EPI_ISL_1111222, EPI_ISL_1111223, EPI_ISL_1111224, EPI_ISL_1111225, EPI_ISL_1111226, EPI_ISL_1111227, EPI_ISL_1111228, EPI_ISL_1111229, EPI_ISL_1111230, EPI_ISL_1111231, EPI_ISL_1111232, EPI_ISL_1111233, EPI_ISL_1111234, EPI_ISL_1111235, EPI_ISL_1111236, EPI_ISL_1111237, EPI_ISL_1111238, EPI_ISL_1111239, EPI_ISL_1111240, EPI_ISL_1111267, EPI_ISL_1111268, EPI_ISL_1111269, EPI_ISL_1111270, EPI_ISL_1111271, EPI_ISL_1111272, EPI_ISL_1111273, EPI_ISL_1111274, EPI_ISL_1111275, EPI_ISL_1111276, EPI_ISL_1111277, EPI_ISL_1111278, EPI_ISL_1111279, EPI_ISL_1111280, EPI_ISL_1111313, EPI_ISL_1111333, EPI_ISL_1111342, EPI_ISL_1111352, EPI_ISL_1111356, EPI_ISL_1111358, EPI_ISL_1111366, EPI_ISL_1111367, EPI_ISL_1111368, EPI_ISL_1111369, EPI_ISL_1111370, EPI_ISL_1111371, EPI_ISL_1111372, EPI_ISL_1111373, EPI_ISL_1111374, EPI_ISL_1111375, EPI_ISL_1111376, EPI_ISL_1111377, EPI_ISL_1111378, EPI_ISL_1111379, EPI_ISL_1111380, EPI_ISL_1111381, EPI_ISL_1111382 | | | |
| see above | Laboratorio de Referencia Nacional de Virus Respiratorio. Instituto Nacional de Salud Perú | Laboratorio de Referencia Nacional de Enteropatógenos. Instituto Nacional de Salud del Perú | Ronnie Gavilan Chavez, Junior Caro Castro, Willi Quino Sifuentes, Veronica Hurtado Vela, Iris Silva Molina, Fiorella Orellana Peralta |

| | | | |
|---|---|---|---|
| EPI_ISL_1112947, EPI_ISL_1112948, EPI_ISL_1112949, EPI_ISL_1112950 | Baylor College of Medicine/ GCID | Baylor College of Medicine/ GCID | Vasanthi Avadhanula, Erin Nicholson, David Henke, Pedro Piedra, Harsha Doddapaneni, Donna Muzny, Qingchang Meng, Hsu Chao, Zeineen Momin, Hua Shen, George Weissenberger, Kavya Kottapalli, Yimiti Meiheerguli, Sejal Salvi, Ginger Metcalf, Vipin Menon, Sara J.J. Cregeen, Matthew C. Ross, Tulin Ayvaz, Richard Sucgang, Kristi L. Hoffman, Matthew Wong, Joseph F. Petrosino |
| EPI_ISL_1138989 | Rajendra Institute of Medical Sciences, Ranchi, Jharkhand | Indian Council of Medical Research-National Institute of Virology, Microbial Containment Complex | Pragya D. Yadav, Manoj Kumar |
| EPI_ISL_1139056, EPI_ISL_1139057, EPI_ISL_1139058, EPI_ISL_1139060, EPI_ISL_1139061, EPI_ISL_1139062, EPI_ISL_1139063, EPI_ISL_1139064, EPI_ISL_1139065, EPI_ISL_1139066, EPI_ISL_1139067, EPI_ISL_1139068 | | | |
| see above | LACEN do Mato Grosso do Sul | Instituto Adolfo Lutz, Interdiciplinary Procedures Center, Strategic Laboratory | Claudio Tavares Sacchi, Claudia Regina Gonçalves, Erica Valessa Ramos Gomes, Karoline Rodrigues Campos, Caio Vinicius Dias Lopes |
| EPI_ISL_1139114, EPI_ISL_1139115, EPI_ISL_1139116, EPI_ISL_1139117, EPI_ISL_1139118, EPI_ISL_1139119, EPI_ISL_1139120, EPI_ISL_1139121 | ICMR-RMRC, Dibrugarh | Indian Council of Medical Research-National Institute of Virology, Microbial Containment Complex | Pragya D. Yadav, Biswajyoti Borkakoty |
| EPI_ISL_1163693, EPI_ISL_1163694, EPI_ISL_1163695, EPI_ISL_1163696, EPI_ISL_1163697, EPI_ISL_1163698, EPI_ISL_1163699, EPI_ISL_1163700 | LABCOVID_HCPA | LABRESIS_HCPA | Martins AF, Wink PL, Volpato F, Rosset C, de Paris F, Monteiro F, Zavascki AP, Barth AL |
| EPI_ISL_1163896, EPI_ISL_1163897, EPI_ISL_1163898, EPI_ISL_1163899, EPI_ISL_1163900, EPI_ISL_1163901, EPI_ISL_1163902, EPI_ISL_1163903, EPI_ISL_1163904, EPI_ISL_1163905, EPI_ISL_1163906, EPI_ISL_1163907, EPI_ISL_1163908, EPI_ISL_1163909 | | | |
| see above | IZSM | IZSM-U.O.C. Virologia | Maurizio Viscardi, Lorena Cardillo, Luigi Atripaldi,Giovanna Fusco, Esterina De Carlo, Antonio Limone |
| EPI_ISL_1164932, EPI_ISL_1164933 | B.J. Medical College and Civil hospital | Indian Council of Medical Research-National Institute of Virology, Microbial Containment Complex | Pragya D. Yadav, Disha Patel |
| EPI_ISL_1165082, EPI_ISL_1165083 | Medical Ain Shams Research Institute (MASRI), Ain Shams University | Medical Ain Shams Research Institute (MASRI), Ain Shams University | Hesham Elghazaly, Sara Hassan Agwa, Hala Hafez , Fatma Ebied, Manal Hamdy Elsaid, Mohamed Elhadidi, Aya Mohamed , Shimaa Moustafa, Reham Kassab, Iman Foda , Samia Abdou Girgis, Hoda Ezz Elarab, Hagar Elshora, Osama Mansour. |
| EPI_ISL_1165088, EPI_ISL_1165089, EPI_ISL_1165090, EPI_ISL_1165091, EPI_ISL_1165092 | Government Medical College, Patiala, Punjab | Indian Council of Medical Research-National Institute of Virology, Microbial Containment Complex | Pragya D. Yadav, Rupinder Bakshi |
| EPI_ISL_1180182, EPI_ISL_1180183, EPI_ISL_1180184, EPI_ISL_1180185 | Dept. of Microbiology, All India Institute of Medical Sciences, Jodhpur | Indian Council of Medical Research-National Institute of Virology, Microbial Containment Complex | Pragya D. Yadav, Vijaylakshmi Nag |
| EPI_ISL_1180188, EPI_ISL_1180189, EPI_ISL_1180190, EPI_ISL_1180191, EPI_ISL_1180192, EPI_ISL_1180193, EPI_ISL_1180194 | VRDL, GMC, Haldwani , Nainital | Indian Council of Medical Research-National Institute of Virology, Microbial Containment Complex | Pragya D. Yadav, Vinita Rawat |
| EPI_ISL_1180935 | LHUB-ULB | LHUB-ULB | Charlotte Michel, Olivier Vandenberg, Marie Hallin |
| EPI_ISL_1181614 | Laboratorio Central de Saude Publica do Estado do Parana (LACEN-PR) | Laboratory of Respiratory Viruses and Measles, Oswaldo Cruz Institute, FIOCRUZ | Paola Resende, Luciana Appolinario, Fernando Motta, Ana Carolina Paixao, Ana Carolina Mendonca, Alice Sampaio Rocha, Renata Serrano Lopes, Maria do Carmo Debur, Irina Nastassja Riediger, Marilda Siqueira on behalf of the Fiocruz COVID-19 Genomic Surveillance Network |
| EPI_ISL_1182103, EPI_ISL_1182104 | Adolfo Lutz Institute Santos Regional Center Brazil | Retrovirus Laboratory Adolfo Lutz Institute | Gabriela Bastos Cabral, Giselle I S Lopez-Lopes, Cintia Ahagon, Audrey Cilli, Paula Morena Guimaraes, Igor Mohamed Hussein, Luis Brigido |
| EPI_ISL_1190402 | Institute of Life Sciences | Institute of Life Sciences | Sanchari Chatterjee, Eshna Laha, Ankita Datey, Saikat De, Supriya Suman Keshry, Soumyajit Ghosh, Arup Ghosh, Atimukta Jha, Safal Rajesh Walia, Sunil Raghav, Amol Suryawanshi, Rupesh Dash, Shantibhushan Senapati, Tushar K. Beuria, Rajeeb Swain, Gulam Hussain Syed, Punit Prasad, Soma Chattopadhyay ,Odisha COVID-19 Study Group, ILS COVID-19 Team and Ajay Parida. |
| EPI_ISL_1192100, EPI_ISL_1192114, EPI_ISL_1192115, EPI_ISL_1192116, EPI_ISL_1192117, EPI_ISL_1192118, EPI_ISL_1192119, EPI_ISL_1192120 | Sri Venkateswara Institute of Medical Sciences SVIMS, Tirupati | Indian Council of Medical Research-National Institute of Virology, Microbial Containment Complex | Pragya D. Yadav, Usha Kalawat |
| EPI_ISL_1192149, EPI_ISL_1192150 | Viral Research and Diagnostic Laboratory, Dr. R.P. Govt. Medical College,Kangra | Indian Council of Medical Research-National Institute of Virology, Microbial Containment Complex | Pragya D. Yadav, S.C. Jaryal |
| EPI_ISL_1196305 | Institute of Life Sciences | Institute of Life Sciences | Sanchari Chatterjee, Eshna Laha, Ankita Datey, Saikat De, Supriya Suman Keshry, Soumyajit Ghosh, Arup Ghosh, Atimukta Jha, Safal Rajesh Walia, Sunil Raghav, Amol Suryawanshi, Rupesh Dash, Shantibhushan Senapati, Tushar K. Beuria, Rajeeb Swain, Gulam Hussain Syed, Punit Prasad, Soma Chattopadhyay ,Odisha COVID-19 Study Group, ILS COVID-19 Team and Ajay Parida. |
| EPI_ISL_1196930 | Institute for Medical Research, Infectious Disease Research Centre, National Institutes of Health, Ministry of Health Malaysia | Institute for Medical Research, Infectious Disease Research Centre, National Institutes of Health, Ministry of Health Malaysia | Suppiah J, Kamel K, Mohd Zawawi Z, Ramly N, Robert F, Thayan R |
| EPI_ISL_1201883 | LACEN do Mato Grosso do Sul | Instituto Adolfo Lutz, Interdiciplinary Procedures Center, Strategic Laboratory | Claudio Tavares Sacchi, Claudia Regina Gonçalves, Erica Valessa Ramos Gomes, Karoline Rodrigues Campos, Caio Vinicius Dias Lopes |
| EPI_ISL_1231311, EPI_ISL_1231312, EPI_ISL_1231313 | Depratment of Microbiology, Viral Research and Diagnostic Laboratory | Indian Council of Medical Research-National Institute of Virology, Microbial Containment Complex | Pragya D. Yadav, Tapan Majumdar |
| EPI_ISL_1251267 | Sri Venkateswara Institute of Medical Sciences SVIMS, Tirupati | Indian Council of Medical Research-National Institute of Virology, Microbial Containment Complex | Pragya D. Yadav, Usha Kalawat |
| EPI_ISL_1259400, EPI_ISL_1260834, EPI_ISL_1260836 | BSL-3 Lab, National Institute for Viral Disease Control and Prevention, Chinese Center for Disease Control and Prevention | BSL-3 Lab, National Institute for Viral Disease Control and Prevention, Chinese Center for Disease Control and Prevention | Peipei Liu, Xiang Zhao, Shumei Zou, Jing Zhang, Zhixiao Chen, Yuchao Wu, Dayan Wang, Jun Han, George F. Gao, Guizhen Wu |
| EPI_ISL_1289435, EPI_ISL_1289436, EPI_ISL_1289437, EPI_ISL_1289438, EPI_ISL_1289439, EPI_ISL_1289440, EPI_ISL_1289441, EPI_ISL_1289442, EPI_ISL_1289443, EPI_ISL_1289444, EPI_ISL_1289445, EPI_ISL_1289446 | | | |
| see above | Department of Health Technology and Informatics, The Hong Kong Polytechnic University | Department of Health Technology and Informatics, The Hong Kong Polytechnic University | Gilman Kit-Hang Siu, Lam-Kwong Lee, Kenneth Siu-Sing Leung, Jake Siu-Lun Leung, Timothy Ting-Leung Ng, Chloe Toi-Mei Chan, Kingsley King-Gee Tam, Hiu-Yin Lao, Denise Sze-Hang Wong, Alan Ka-Lun Wu, Miranda Chong-Yee Yau, Yvette Wai-Man Lai, Kitty Sau-Chun Fung, Sandy Ka-Yee Chau, Barry Kin-Chung Wong, Wing-Kin To, Kristine Luk, Alex Yat-Man Ho, Tak-Lun Que, Kam-Tong Yip, Wing Cheong Yam, David Ho-Keung Shum, Shea Ping Yip |
| EPI_ISL_1293351 | Institut National d'Hygiène (INH) | Unité Mixte Internationale TransVIHMI (UMI 233 IRD - U1175 INSERM - Université de Montpellier)IRD (Institut de recherche pour le développement) | Mounerou SALOU, Christelle BUTEL, Wembo A. HALATOKO,Issaka Maman, Abla A. KONOU, Amivi EHLAN, Adodo SADJI, Kokou TEGUENI,Sidonie A.M.KAGNISSODE, Akoélé SILIADIN, Alassane OURO-MEDELI, Messanh DOUFFAN,Déléma MABA,Sika DOSSIM, Améyo DORKENOO, Mireille PRINCE-DAVID,Anoumou DAGNRA,Laetitia SERRANO,Ahidjo AYOUBA,Eric DELAPORTE, Martine PEETERS |
| EPI_ISL_1301550 | Instituto Nacional de Enfermedades Respiratorias (INER) | Instituto de Biotecnología de la UNAM | Authors from IBT, IMSS, InDRE and INER (in alphabetical order): Carlos F. Arias, Santiago Ávila-Ríos, Gisela Barrera-Badillo, Eduardo Becerril-Vargas, Celia Boukadida, Natividad Cruz-Ortiz, Larissa Fernandes-Matano, Ricardo Grande, Lucia Hernandez-Rivas, Alejandra Hernández-Terán, Pavel Isa, Irma Lopez-Martinez, José Arturo Martínez-Orozco, Margarita Matías-Florentino, Fidencio Mejía-Nepomuceno, Edgar Mendieta-Condado, Mario Mújica-Sánchez, José Esteban Muñoz-Medina, Tatiana Nunez-Garcia, Luis Alberto Ochoa-Carrera, Hector Esteban Paz-Juárez, Francisco Pulido, José Ernesto Ramírez-González, Alma Rincón-Rubio, Teresita Rojas-Mendoza, Jorge Salas-Hernández, Alejandro Sanchez-Flores, Clara Esperanza Santacruz-Tinoco, Andrea Santos Coy-Arechavaleta, Blanca Taboada, Gloria Vazquez, Joel Armando Vázquez-Pérez, Jerome Jean Verleyen |
| EPI_ISL_1301552, EPI_ISL_1301553 | Laboratorio Central de Epidemiología IMSS | Instituto de Biotecnología de la UNAM | Authors from IBT, IMSS, InDRE and INER (in alphabetical order): Carlos F. Arias, Santiago Ávila-Ríos, Gisela Barrera-Badillo, Eduardo Becerril-Vargas, Celia |

| | | | |
|---|---|---|---|
| | | | Boukadida, Natividad Cruz-Ortiz, Larissa Fernandes-Matano, Ricardo Grande, Lucia Hernandez-Rivas, Alejandra Hernández-Terán, Pavel Isa, Irma Lopez-Martinez, José Arturo Martínez-Orozco, Margarita Matías-Florentino, Fidencio Mejía-Nepomuceno, Edgar Mendieta-Condado, Mario Mújica-Sánchez, José Esteban Muñoz-Medina, Tatiana Nunez-Garcia, Luis Alberto Ochoa-Carrera, Hector Esteban Paz-Juárez, Francisco Pulido, José Ernesto Ramírez-González, Alma Rincón-Rubio, Teresita Rojas-Mendoza, Jorge Salas-Hernández, Alejandro Sanchez-Flores, Clara Esperanza Santacruz-Tinoco, Andrea Santos Coy-Arechavaleta, Blanca Taboada, Gloria Vazquez, Joel Armando Vázquez-Pérez, Jerome Jean Verleyen |
| EPI_ISL_1301575, EPI_ISL_1301576, EPI_ISL_1301577, EPI_ISL_1301578, EPI_ISL_1301579, EPI_ISL_1301580, EPI_ISL_1301581, EPI_ISL_1301582, EPI_ISL_1301583, EPI_ISL_1301584 | Instituto de Diagnostico y Referencia Epidemiologicos InDRE_RNLSP | Instituto de Biotecnología de la UNAM | Authors from IBT, IMSS, InDRE and INER (in alphabetical order): Carlos F. Arias, Santiago Ávila-Ríos, Gisela Barrera-Badillo, Eduardo Becerril-Vargas, Celia Boukadida, Natividad Cruz-Ortiz, Larissa Fernandes-Matano, Ricardo Grande, Lucia Hernandez-Rivas, Alejandra Hernández-Terán, Pavel Isa, Irma Lopez-Martinez, José Arturo Martínez-Orozco, Margarita Matías-Florentino, Fidencio Mejía-Nepomuceno, Edgar Mendieta-Condado, Mario Mújica-Sánchez, José Esteban Muñoz-Medina, Tatiana Nunez-Garcia, Luis Alberto Ochoa-Carrera, Hector Esteban Paz-Juárez, Francisco Pulido, José Ernesto Ramírez-González, Alma Rincón-Rubio, Teresita Rojas-Mendoza, Jorge Salas-Hernández, Alejandro Sanchez-Flores, Clara Esperanza Santacruz-Tinoco, Andrea Santos Coy-Arechavaleta, Blanca Taboada, Gloria Vazquez, Joel Armando Vázquez-Pérez, Jerome Jean Verleyen |
| EPI_ISL_1301593, EPI_ISL_1301594, EPI_ISL_1301595, EPI_ISL_1301596 | Laboratorio Central de Epidemiología IMSS | Instituto de Biotecnología de la UNAM | Authors from IBT, IMSS, InDRE and INER (in alphabetical order): Carlos F. Arias, Santiago Ávila-Ríos, Gisela Barrera-Badillo, Eduardo Becerril-Vargas, Celia Boukadida, Natividad Cruz-Ortiz, Larissa Fernandes-Matano, Ricardo Grande, Lucia Hernandez-Rivas, Alejandra Hernández-Terán, Pavel Isa, Irma Lopez-Martinez, José Arturo Martínez-Orozco, Margarita Matías-Florentino, Fidencio Mejía-Nepomuceno, Edgar Mendieta-Condado, Mario Mújica-Sánchez, José Esteban Muñoz-Medina, Tatiana Nunez-Garcia, Luis Alberto Ochoa-Carrera, Hector Esteban Paz-Juárez, Francisco Pulido, José Ernesto Ramírez-González, Alma Rincón-Rubio, Teresita Rojas-Mendoza, Jorge Salas-Hernández, Alejandro Sanchez-Flores, Clara Esperanza Santacruz-Tinoco, Andrea Santos Coy-Arechavaleta, Blanca Taboada, Gloria Vazquez, Joel Armando Vázquez-Pérez, Jerome Jean Verleyen |
| EPI_ISL_1301598, EPI_ISL_1301600, EPI_ISL_1301605, EPI_ISL_1301606, EPI_ISL_1301607, EPI_ISL_1301608, EPI_ISL_1301609, EPI_ISL_1301610, EPI_ISL_1301611, EPI_ISL_1301612, EPI_ISL_1301613, EPI_ISL_1301614, EPI_ISL_1301615, EPI_ISL_1301616, EPI_ISL_1301617 | | | |
| see above | Instituto Nacional de Enfermedades Respiratorias (INER) | Instituto de Biotecnología de la UNAM | Authors from IBT, IMSS, InDRE and INER (in alphabetical order): Carlos F. Arias, Santiago Ávila-Ríos, Gisela Barrera-Badillo, Eduardo Becerril-Vargas, Celia Boukadida, Natividad Cruz-Ortiz, Larissa Fernandes-Matano, Ricardo Grande, Lucia Hernandez-Rivas, Alejandra Hernández-Terán, Pavel Isa, Irma Lopez-Martinez, José Arturo Martínez-Orozco, Margarita Matías-Florentino, Fidencio Mejía-Nepomuceno, Edgar Mendieta-Condado, Mario Mújica-Sánchez, José Esteban Muñoz-Medina, Tatiana Nunez-Garcia, Luis Alberto Ochoa-Carrera, Hector Esteban Paz-Juárez, Francisco Pulido, José Ernesto Ramírez-González, Alma Rincón-Rubio, Teresita Rojas-Mendoza, Jorge Salas-Hernández, Alejandro Sanchez-Flores, Clara Esperanza Santacruz-Tinoco, Andrea Santos Coy-Arechavaleta, Blanca Taboada, Gloria Vazquez, Joel Armando Vázquez-Pérez, Jerome Jean Verleyen |
| EPI_ISL_1301620, EPI_ISL_1301621, EPI_ISL_1301622, EPI_ISL_1301623, EPI_ISL_1301624, EPI_ISL_1301625, EPI_ISL_1301626, EPI_ISL_1301627, EPI_ISL_1301628, EPI_ISL_1301629, EPI_ISL_1301630, EPI_ISL_1301631, EPI_ISL_1301632, EPI_ISL_1301633, EPI_ISL_1301634, EPI_ISL_1301635, EPI_ISL_1301636, EPI_ISL_1301637, EPI_ISL_1301638, EPI_ISL_1301639, EPI_ISL_1301640, EPI_ISL_1301641, EPI_ISL_1301642, EPI_ISL_1301643, EPI_ISL_1301644, EPI_ISL_1301645, EPI_ISL_1301646, EPI_ISL_1301647, EPI_ISL_1301648, EPI_ISL_1301649, EPI_ISL_1301650, EPI_ISL_1301651, EPI_ISL_1301652, EPI_ISL_1301653, EPI_ISL_1301654, EPI_ISL_1301655, EPI_ISL_1301656, EPI_ISL_1301657, EPI_ISL_1301658, EPI_ISL_1301659, EPI_ISL_1301660, EPI_ISL_1301661, EPI_ISL_1301662, EPI_ISL_1301663, EPI_ISL_1301664, EPI_ISL_1301665, EPI_ISL_1301666, EPI_ISL_1301667, EPI_ISL_1301668, EPI_ISL_1301669, EPI_ISL_1301670, EPI_ISL_1301671 | | | |
| see above | Laboratorio Central de Epidemiología IMSS | Instituto de Biotecnología de la UNAM | Authors from IBT, IMSS, InDRE and INER (in alphabetical order): Carlos F. Arias, Santiago Ávila-Ríos, Gisela Barrera-Badillo, Eduardo Becerril-Vargas, Celia Boukadida, Natividad Cruz-Ortiz, Larissa Fernandes-Matano, Ricardo Grande, Lucia Hernandez-Rivas, Alejandra Hernández-Terán, Pavel Isa, Irma Lopez-Martinez, José Arturo Martínez-Orozco, Margarita Matías-Florentino, Fidencio Mejía-Nepomuceno, Edgar Mendieta-Condado, Mario Mújica-Sánchez, José Esteban Muñoz-Medina, Tatiana Nunez-Garcia, Luis Alberto Ochoa-Carrera, Hector Esteban Paz-Juárez, Francisco Pulido, José Ernesto Ramírez-González, Alma Rincón-Rubio, Teresita Rojas-Mendoza, Jorge Salas-Hernández, Alejandro Sanchez-Flores, Clara Esperanza Santacruz-Tinoco, Andrea Santos Coy-Arechavaleta, Blanca Taboada, Gloria Vazquez, Joel Armando Vázquez-Pérez, Jerome Jean Verleyen |
| EPI_ISL_1301679, EPI_ISL_1301680, EPI_ISL_1301681, EPI_ISL_1301682, EPI_ISL_1301684, EPI_ISL_1301685, EPI_ISL_1301686, EPI_ISL_1301687, EPI_ISL_1301690, EPI_ISL_1301709 | Instituto de Diagnostico y Referencia Epidemiologicos InDRE_RNLSP | Instituto de Biotecnología de la UNAM | Authors from IBT, IMSS, InDRE and INER (in alphabetical order): Carlos F. Arias, Santiago Ávila-Ríos, Gisela Barrera-Badillo, Eduardo Becerril-Vargas, Celia Boukadida, Natividad Cruz-Ortiz, Larissa Fernandes-Matano, Ricardo Grande, Lucia Hernandez-Rivas, Alejandra Hernández-Terán, Pavel Isa, Irma Lopez-Martinez, José Arturo Martínez-Orozco, Margarita Matías-Florentino, Fidencio Mejía-Nepomuceno, Edgar Mendieta-Condado, Mario Mújica-Sánchez, José Esteban Muñoz-Medina, Tatiana Nunez-Garcia, Luis Alberto Ochoa-Carrera, Hector Esteban Paz-Juárez, Francisco Pulido, José Ernesto Ramírez-González, Alma Rincón-Rubio, Teresita Rojas-Mendoza, Jorge Salas-Hernández, Alejandro Sanchez-Flores, Clara Esperanza Santacruz-Tinoco, Andrea Santos Coy-Arechavaleta, Blanca Taboada, Gloria Vazquez, Joel Armando Vázquez-Pérez, Jerome Jean Verleyen |
| EPI_ISL_1301714, EPI_ISL_1301715, EPI_ISL_1301716 | Laboratorio Central de Epidemiología IMSS | Instituto de Biotecnología de la UNAM | Authors from IBT, IMSS, InDRE and INER (in alphabetical order): Carlos F. Arias, Santiago Ávila-Ríos, Gisela Barrera-Badillo, Eduardo Becerril-Vargas, Celia Boukadida, Natividad Cruz-Ortiz, Larissa Fernandes-Matano, Ricardo Grande, Lucia Hernandez-Rivas, Alejandra Hernández-Terán, Pavel Isa, Irma Lopez-Martinez, José Arturo Martínez-Orozco, Margarita Matías-Florentino, Fidencio Mejía-Nepomuceno, Edgar Mendieta-Condado, Mario Mújica-Sánchez, José Esteban Muñoz-Medina, Tatiana Nunez-Garcia, Luis Alberto Ochoa-Carrera, Hector Esteban Paz-Juárez, Francisco Pulido, José Ernesto Ramírez-González, Alma Rincón-Rubio, Teresita Rojas-Mendoza, Jorge Salas-Hernández, Alejandro Sanchez-Flores, Clara Esperanza Santacruz-Tinoco, Andrea Santos Coy-Arechavaleta, Blanca Taboada, Gloria Vazquez, Joel Armando Vázquez-Pérez, Jerome Jean Verleyen |
| EPI_ISL_1301717, EPI_ISL_1301718, EPI_ISL_1301719, EPI_ISL_1301720, EPI_ISL_1301721, EPI_ISL_1301722, EPI_ISL_1301723, EPI_ISL_1301724, EPI_ISL_1301731 | Instituto Nacional de Enfermedades Respiratorias (INER) | Instituto de Biotecnología de la UNAM | Authors from IBT, IMSS, InDRE and INER (in alphabetical order): Carlos F. Arias, Santiago Ávila-Ríos, Gisela Barrera-Badillo, Eduardo Becerril-Vargas, Celia Boukadida, Natividad Cruz-Ortiz, Larissa Fernandes-Matano, Ricardo Grande, Lucia Hernandez-Rivas, Alejandra Hernández-Terán, Pavel Isa, Irma Lopez-Martinez, José Arturo Martínez-Orozco, Margarita Matías-Florentino, Fidencio Mejía-Nepomuceno, Edgar Mendieta-Condado, Mario Mújica-Sánchez, José Esteban Muñoz-Medina, Tatiana Nunez-Garcia, Luis Alberto Ochoa-Carrera, Hector Esteban Paz-Juárez, Francisco Pulido, José Ernesto Ramírez-González, Alma Rincón-Rubio, Teresita Rojas-Mendoza, Jorge Salas-Hernández, Alejandro Sanchez-Flores, Clara Esperanza Santacruz-Tinoco, Andrea Santos Coy-Arechavaleta, Blanca Taboada, Gloria Vazquez, Joel Armando Vázquez-Pérez, Jerome Jean Verleyen |
| EPI_ISL_1303031 | Institut National d'Hygiène (INH) | Unité Mixte Internationale TransVIHMI (UMI 233 IRD - U1175 INSERM - Université de Montpellier)IRD (Institut de recherche pour le développement) | Mounerou SALOU, Christelle BUTEL, Wembo A. HALATOKO,Issaka Maman, Abla A. KONOU, Amivi EHLAN, Adodo SADJI, Kokou TEGUENI,Sidonie A.M.KAGNISSODE, Akoélé SILIADIN, Alassane OURO-MEDELI, Messanh DOUFFAN,Déléma MABA,Sika DOSSIM, Améyo DORKENOO, Mireille PRINCE-DAVID,Anoumou DAGNRA,Laetitia SERRANO,Ahidjo AYOUBA,Eric DELAPORTE, Martine PEETERS |
| EPI_ISL_1315064 | Armed Forces Medical Research Laboratories and Blood Bank, Egypt | Department of Microbial Biotechnology, Genetic Engineering Division, National Research Centre, | Reham Dawood, Mohamed Khalifa, Ahmed Taha, Ayman Farghaly, Ahmed Elsayed, Mohamed Shemis |
| EPI_ISL_1315065 | Armed Forces Medical Research Laboratories and Blood Bank, Egypt | Department of Microbial Biotechnology, Genetic Engineering Division, National Research Centre | Reham Dawood, Mohamed Khalifa, Ahmed Taha, Ayman Farghaly, Ahmed Elsayed, Mohamed Shemis |
| EPI_ISL_1321604 | Institut National d'Hygiène (INH) | Unité Mixte Internationale TransVIHMI (UMI 233 IRD - U1175 INSERM - Université de Montpellier)IRD (Institut de recherche pour le développement) | Mounerou SALOU, Christelle BUTEL, Wembo A. HALATOKO,Issaka Maman, Abla A. KONOU, Amivi EHLAN, Adodo SADJI, Kokou TEGUENI,Sidonie A.M.KAGNISSODE, Akoélé SILIADIN, Alassane OURO-MEDELI, Messanh DOUFFAN,Déléma MABA,Sika DOSSIM, Améyo DORKENOO, Mireille PRINCE-DAVID,Anoumou DAGNRA,Laetitia SERRANO,Ahidjo AYOUBA,Eric DELAPORTE, Martine PEETERS |
| EPI_ISL_1340764 | Departamento de Virologia, Laboratorio Central de Salud Pública, Avenida Venezuela y Teniente Escurra, Asunción, Paraguay | Laboratory of Respiratory Viruses and Measles, Oswaldo Cruz Institute, FIOCRUZ | Paola Resende, Cynthia Vazquez, Luciana Appolinario, Fernando Motta, Anna Carolina Paixao, Ana Carolina Mendonca, Alice Sampaio Rocha, Renata Serrano Lopes, Marilda Siqueira on behalf of the Fiocruz COVID-19 Genomic Surveillance Network |
| EPI_ISL_1379197 | Department of Molecular and Translational Medicine, Section of Microbiology, University of Brescia, ASST Spedali Civili, Brescia | Department of Molecular and Translational Medicine, Section of Microbiology, University of Brescia, ASST Spedali Civili, Brescia | Francesca Caccuri, Alberto Zani, Serena Messali, Antonella Bugatti, Giovanni Campisi, Federica Filippini, Erika Scaltriti, Simona Fiorentini, Arnaldo Caruso |
| EPI_ISL_1398937 | Emam Ali Hospital | Razi Vaccine and Serum Research Institute | Amir Kaffashi, Jiabin Huang, S Reza Banihashemi, Mohammad Hossein Fallah Mehrabadi, Mohsen Lotfi, Morteza Taghizadeh, Akbar Khorasani, Mohsen Bashashati, Sayed Hamidreza Mozhgan |
| EPI_ISL_1404615 | Institut National d'Hygiène | Unité Mixte Internationale TransVIHMI (UMI 233 IRD - U1175 INSERM - Université de Montpellier) IRD (Institut de recherche pour le développement) | Mounerou SALOU, Christelle BUTEL, Wembo A. HALATOKO,Issaka Maman, Abla A. KONOU, Amivi EHLAN, Adodo SADJI, Kokou TEGUENI,Sidonie A.M.KAGNISSODE, Akoélé SILIADIN, Alassane OURO-MEDELI, Messanh DOUFFAN,Déléma MABA,Sika DOSSIM, Améyo DORKENOO, Mireille PRINCE-DAVID,Anoumou DAGNRA,Laetitia SERRANO,Ahidjo AYOUBA,Eric DELAPORTE, Martine PEETERS |
| EPI_ISL_1406144, EPI_ISL_1406177 | Institut National d'Hygiène (INH) | Unité Mixte Internationale TransVIHMI (UMI 233 IRD - U1175 INSERM - Université de Montpellier)IRD (Institut de recherche pour le développement) | Mounerou SALOU, Christelle BUTEL, Wembo A. HALATOKO,Issaka Maman, Abla A. KONOU, Amivi EHLAN, Adodo SADJI, Kokou TEGUENI,Sidonie A.M.KAGNISSODE, Akoélé SILIADIN, Alassane OURO-MEDELI, Messanh DOUFFAN,Déléma MABA,Sika DOSSIM, Améyo DORKENOO, Mireille PRINCE-DAVID,Anoumou DAGNRA,Laetitia SERRANO,Ahidjo AYOUBA,Eric DELAPORTE, Martine PEETERS |
| EPI_ISL_1415293, EPI_ISL_1415425, EPI_ISL_1415428 | Institut National d'hygiène | "Unité Mixte Internationale TransVIHMI (UMI 233 IRD - U1175 INSERM - Université de Montpellier) IRD (Institut de | Mounerou SALOU, Christelle BUTEL, Wembo A. HALATOKO,Issaka Maman, Abla A. KONOU, Amivi EHLAN, Adodo SADJI, Kokou TEGUENI,Sidonie A.M.KAGNISSODE, Akoélé SILIADIN, Alassane OURO-MEDELI, Messanh DOUFFAN,Déléma MABA,Sika DOSSIM, Améyo DORKENOO, Mireille |

| | | | |
|---|---|---|---|
| | | recherche pour le développement)" | PRINCE-DAVID,Anoumou DAGNRA,Laetitia SERRANO,Ahidjo AYOUBA,Eric DELAPORTE, Martine PEETERS |
| EPI_ISL_1439647, EPI_ISL_1440879 | Institut National d'Hygiène | Unité Mixte Internationale TransVIHMI (UMI 233 IRD - U1175 INSERM - Université de Montpellier) IRD (Institut de recherche pour le développement) | Mounerou SALOU, Christelle BUTEL, Wembo A. HALATOKO,Issaka Maman, Abla A. KONOU, Amivi EHLAN, Adodo SADJI, Kokou TEGUENI,Sidonie A.M.KAGNISSODE, Akoélé SILIADIN, Alassane OURO-MEDELI, Messanh DOUFFAN,Déléma MABA,Sika DOSSIM, Améyo DORKENOO, Mireille PRINCE-DAVID,Anoumou DAGNRA,Laetitia SERRANO,Ahidjo AYOUBA,Eric DELAPORTE, Martine PEETERS |
| EPI_ISL_1442993, EPI_ISL_1443000, EPI_ISL_1443001 | Institut National d'hygiène | "Unité Mixte Internationale TransVIHMI (UMI 233 IRD - U1175 INSERM - Université de Montpellier) IRD (Institut de recherche pour le développement)" | Mounerou SALOU, Christelle BUTEL, Wembo A. HALATOKO,Issaka Maman, Abla A. KONOU, Amivi EHLAN, Adodo SADJI, Kokou TEGUENI,Sidonie A.M.KAGNISSODE, Akoélé SILIADIN, Alassane OURO-MEDELI, Messanh DOUFFAN,Déléma MABA,Sika DOSSIM, Améyo DORKENOO, Mireille PRINCE-DAVID,Anoumou DAGNRA,Laetitia SERRANO,Ahidjo AYOUBA,Eric DELAPORTE, Martine PEETERS |
| EPI_ISL_1447336 | Diagnostic and Research Center of Infectious Diseases, Medical Faculty, Andalas University | Diagnostic and Research Center of Infectious Diseases, Medical Faculty, Andalas University | Andani Eka Putra, Syafrizayanti, Ikhwan R. Sudji, Linosefa, Dede Rahman Agustian, Ayu Novita Trisnawati, Siskalil Fahma, Nia Ayuni Putri, Sekar Asri Tresnaningtyas, SM Rezvi, Fauzul Azhim, Dessy Arisanty, Gestina Aliska, Desmawati, Nita Afriani ,Syandrez Prima Putra, Juane Plantika Menra, Yolani Syaputri, Mutia Lailani |
| EPI_ISL_1469668 | Hospital Unimed Vale do Caí | Epiclin | Fernando Hayashi Sant'Anna, Ana Paula Muterle, Janira Prichula, Juliana Comerlato, Carolina Comerlato, Eliana Márcia Da Ros Wendland |
| EPI_ISL_1469752 | Fundação Hospitalar de Sapucaia do Sul | Epiclin | Fernando Hayashi Sant'Anna, Ana Paula Muterle, Janira Prichula, Juliana Comerlato, Carolina Comerlato, Eliana Márcia Da Ros Wendland |
| EPI_ISL_1469799 | Hospital Regina | Epiclin | Fernando Hayashi Sant'Anna, Ana Paula Muterle, Janira Prichula, Juliana Comerlato, Carolina Comerlato, Eliana Márcia Da Ros Wendland |
| EPI_ISL_1469959 | CUB Hopital Erasme Laboratoire d'Anatomie Pathologique | CUB Hopital Erasme Laboratoire d'Anatomie Pathologique | Prof. Isabelle Salmon, Dr. Nicky D'Haene |
| EPI_ISL_1499504 | Laboratorio di Microbiologia e Virologia, Università Vita-Salute San Raffaele, Milano | Laboratorio di Microbiologia e Virologia, Università Vita-Salute San Raffaele, Milano | Roberto Ferrarese, Roberta Antonia Diotti, Elena Criscuolo, Matteo Castelli, Nicasio Mancini, Massimo Clementi |
| EPI_ISL_1499572 | Laboratorio di Microbiologia e Virologia, Università Vita-Salute San Raffaele, Milano | Laboratorio di Microbiologia e Virologia, Università Vita-Salute San Raffaele, Milano | Matteo Castelli, Elena Criscuolo, Roberta Antonia Diotti, Roberto Ferrarese, Nicasio Mancini, Massimo Clementi, Nicola Clementi on behalf of SARS-CoV-2 ITALIAN RESEARCH ENTERPRISE-(SCIRE) Collaborative Group |
| EPI_ISL_1499593 | Laboratorio di Microbiologia e Virologia, Università Vita-Salute San Raffaele, Milan | Laboratorio di Microbiologia e Virologia, Università Vita-Salute San Raffaele, Milan | Elena Criscuolo, Roberta Antonia Diotti, Roberto Ferrarese, Matteo Castelli, Massimo Clementi, Nicola Clementi, Nicasio Mancini on behalf of SARS-CoV-2 ITALIAN RESEARCH ENTERPRISE-(SCIRE) Collaborative Group |
| EPI_ISL_1502818, EPI_ISL_1502819, EPI_ISL_1502820 | Hospital Regional Anita Moreno Los Santos | Gorgas Memorial Laboratory of Health Studies | Gonzalez Claudia, Best Dayana, Espino Juliee, Leyda Abrego, Moreno Ambar, Oris Chavarria, Jessica Gondola, Marlenne Castillo, Ortiz Alma, Castillo Jorge, Moreno Brechla, Franco Danilo, Lopez-Verges Sandra, Martinez Alexander |
| EPI_ISL_1502821, EPI_ISL_1502822, EPI_ISL_1502823 | Hospital Regional Dr. Luis "Chicho" Fábrega | Gorgas Memorial Laboratory of Health Studies | Gonzalez Claudia, Herrera Melissa, Herrera Genarino, Leyda Abrego, Moreno Ambar, Oris Chavarria, Jessica Gondola, Marlenne Castillo, Ortiz Alma, Castillo Jorge, Moreno Brechla, Franco Danilo, Lopez-Verges Sandra, Martinez Alexander |
| EPI_ISL_1502824 | Hospital Regional Anita Moreno Los Santos | Gorgas Memorial Laboratory of Health Studies | Gonzalez Claudia, Best Dayana, Espino Juliee, Leyda Abrego, Moreno Ambar, Oris Chavarria, Jessica Gondola, Marlenne Castillo, Ortiz Alma, Castillo Jorge, Moreno Brechla, Franco Danilo, Lopez-Verges Sandra, Martinez Alexander |
| EPI_ISL_1502826, EPI_ISL_1502828, EPI_ISL_1502829, EPI_ISL_1502831, EPI_ISL_1502832, EPI_ISL_1502833, EPI_ISL_1502834 | Hospital Regional Dr. Luis "Chicho" Fábrega | Gorgas Memorial Laboratory of Health Studies | Gonzalez Claudia, Herrera Melissa, Herrera Genarino, Leyda Abrego, Moreno Ambar, Oris Chavarria, Jessica Gondola, Marlenne Castillo, Ortiz Alma, Castillo Jorge, Moreno Brechla, Franco Danilo, Lopez-Verges Sandra, Martinez Alexander |
| EPI_ISL_1502836 | Hospital Regional Anita Moreno Los Santos | Gorgas Memorial Laboratory of Health Studies | Gonzalez Claudia, Best Dayana, Espino Juliee, Leyda Abrego, Moreno Ambar, Oris Chavarria, Jessica Gondola, Marlenne Castillo, Ortiz Alma, Castillo Jorge, Moreno Brechla, Franco Danilo, Lopez-Verges Sandra, Martinez Alexander |
| EPI_ISL_1502837, EPI_ISL_1502838, EPI_ISL_1502839, EPI_ISL_1502840, EPI_ISL_1502841 | Hospital Regional Dr. Luis "Chicho" Fábrega | Gorgas Memorial Laboratory of Health Studies | Gonzalez Claudia, Herrera Melissa, Herrera Genarino, Leyda Abrego, Moreno Ambar, Oris Chavarria, Jessica Gondola, Marlenne Castillo, Ortiz Alma, Castillo Jorge, Moreno Brechla, Franco Danilo, Lopez-Verges Sandra, Martinez Alexander |
| EPI_ISL_1502842, EPI_ISL_1502843, EPI_ISL_1502844, EPI_ISL_1502845, EPI_ISL_1502846, EPI_ISL_1502847 | Gorgas Memorial Laboratory of Health Studies | Gorgas Memorial Laboratory of Health Studies | Gonzalez Claudia, Leyda Abrego, Moreno Ambar, Oris Chavarria, Jessica Gondola, Marlenne Castillo, Ortiz Alma, Castillo Jorge, Moreno Brechla, Franco Danilo, Lopez-Verges Sandra, Martinez Alexander |
| EPI_ISL_1510666 | University of Rome Tor Vergata: Departm Experim Medicine Chair of Virology | University of Rome Tor Vergata: Departm Experim Medicine Chair of Virology | Maria Concetta Bellocchi |
| EPI_ISL_1511268 | Medicina Genómica | Medicina Genómica | Ana Laura Sánchez-Sandoval, María Eligia Juárez-Torres, Mariana Villalba-De la Peña, Siddhartha Ríos-Zaragoza, Guadalupe Mercedes Lucía Guerrero-Avendaño, Eira Valeria Barrón-Palma |
| EPI_ISL_1511269, EPI_ISL_1511270 | Hospital General de México, Medicina Genómica | Hospital General de México, Medicina Genómica | Ana Laura Sánchez-Sandoval, María Eligia Juárez-Torres, Mariana Villalba-De la Peña, Siddhartha Ríos-Zaragoza, Guadalupe Mercedes Lucía Guerrero-Avendaño, Eira Valeria Barrón-Palma |
| EPI_ISL_1511641 | Laboratorio de Patologia Clinica - UNICAMP | Laboratorio de Estudos de Virus Emergentes | Mariene R. Amorim, William M. Souza, Antonio C. G. Carlos Jr, Daniel A. Toledo-Teixeira, Karina Bispo-dos-Santos, Camila L. Simeoni, Pierina L. Parise, Aline Vieira, Julia Forato, Ingra M. Claro, Luciana S. Mofatto, Natalia S. Brunetti, Emerson S.S. França, Gisele A. Pedroso, Barbara F. N. Carvalho, Tania R. Zaccariotto, Kamila C. S. Krywacz, André S. Vieira, Marcelo A. Mori, Alessandro S. Farias, Maria H. P. Pavan, Luís Felipe Bachur, Luís G. O. Cardoso, Fernando R. Spilki, Ester C. Sabino, Nuno R. Faria, Magnun N. N. Santos, Rodrigo Angerami, Patricia A. F. Leme, Angelica Schreiber, Maria L. Moretti, Fabiana Granja, José Luiz Proenca-Modena |
| EPI_ISL_1533610 | Shiraz University | Shiraz University | Abozar Ghorbani, Hakimeh Ahmadi, Hamed Gouklani, Ebrahim Eftekhar |
| EPI_ISL_1550878 | Instituto Oswaldo Cruz | Laboratorio de Virologia, Faculdade de Medicina, Universidade Federal de Mato Grosso, campus Cuiabá | Maria de Fatima Ferrerira, Renata Dezengrini Slhessarenko, Janeth Aracely Ramirez Pavon, Luciano Nakazato, Valeria Dutra , Rosane Hahn |
| EPI_ISL_1655937 | State Key Laboratory of Emerging Infectious Diseases, The University of Hong Kong | State Key Laboratory of Emerging Infectious Diseases, The University of Hong Kong | Jia Wang, Lifeng Li, Yiu Man Cheung, Huachen Zhu, Yi Guan |
| EPI_ISL_1678584, EPI_ISL_1690551, EPI_ISL_1694627, EPI_ISL_1694628, EPI_ISL_1694631, EPI_ISL_1694632, EPI_ISL_1694634, EPI_ISL_1695909 | Instituto Estadual do Cérebro Paulo Niemayer (IECPN) | Laboratório de Imunofarmacologia | Thiago Moreno Lopes Souza |
| EPI_ISL_1718321 | SCIENSANO National Influenza Centre | PathoSense BV | Nick Vereecke |
| EPI_ISL_1752476, EPI_ISL_1752480, EPI_ISL_1752484, EPI_ISL_1752495, EPI_ISL_1752496, EPI_ISL_1752497, EPI_ISL_1752498, EPI_ISL_1752499, EPI_ISL_1752500, EPI_ISL_1752501, EPI_ISL_1752502, EPI_ISL_1752503, EPI_ISL_1752504, EPI_ISL_1752505, EPI_ISL_1752506, EPI_ISL_1752507, EPI_ISL_1752508, EPI_ISL_1752509 | | | |
| see above | LHUB-ULB | LHUB-ULB | Charlotte Michel, Olivier Vandenberg, Marie Hallin |
| EPI_ISL_1797550, EPI_ISL_1797552, EPI_ISL_1797553, EPI_ISL_1797554, EPI_ISL_1797556, EPI_ISL_1797558, EPI_ISL_1797560, EPI_ISL_1797561, EPI_ISL_1797563, EPI_ISL_1797564, EPI_ISL_1797565, EPI_ISL_1797566, EPI_ISL_1797567, EPI_ISL_1797568, EPI_ISL_1797570, EPI_ISL_1797571, EPI_ISL_1797572, EPI_ISL_1797573, EPI_ISL_1797575, EPI_ISL_1797576, EPI_ISL_1797578, EPI_ISL_1797579, EPI_ISL_1797582, EPI_ISL_1797583, EPI_ISL_1797584, EPI_ISL_1797585, EPI_ISL_1797586, EPI_ISL_1797588, EPI_ISL_1797589, EPI_ISL_1797590, EPI_ISL_1797591, EPI_ISL_1797592, EPI_ISL_1797595, EPI_ISL_1797597, EPI_ISL_1797598, EPI_ISL_1797599, EPI_ISL_1797600, EPI_ISL_1797601 | | | |
| see above | Medical Center, Aircraft Carrier | IRBA, 2MI | Franck de Laval, Hervé Chaudet, Constance Lacrosse, Joffrey Marchi, Aissata Dia, Gaëtan Texier, Olivier Gorgé, Bakridine Mmadi Mrenda, Vanessa Marbac, Flavie Letois, Anthony Levasseur, Jacques Cobola, Flora Nolent, Fabien Dutasta, Frédéric Janvier, Meynard Jean-Baptiste, Vincent Pommier de Santi |
| EPI_ISL_1805651 | National Center for Communicable Diseases (NCCD) National Influenza Center | National Center for Communicable Diseases (NCCD) National Influenza Center | Naranzul Ts,Bayasgalan N,Khishigmunkh Ch,Seiichiro F,Hideka M,Mina N,Shinji W,Ankhbayar S,Tsogzolmaa G,Darmaa B,Battur L,Nymadawa P |
| EPI_ISL_1859756 | City Clinical Hospital named after S.I. Spasokukotsky of Moscow Healthcare Department, Academic Consortium | FEDERAL RESEARCH AND CLINICAL CENTER OF PHYSICAL-CHEMICAL MEDICINE OF FEDERAL MEDICAL BIOLOGICAL AGENCY, Academic Consortium | Vladislav Babenko, Ramiz Bakhtyev, Vladimir Baklaushev, Larisa Balykova, Pavel Bashkirov, Anna Blagonravova, Daria Boldyreva, Dmitry Fedorov, Ilshat Gafurov, Raushaniya Gaifullina, Yulia Galeeva, Elena Galova, Vadim Govorun, Elena Ilina, Konstantin Ivanov, Daria Kharlampieva, Polina Khromova, Ksenia Klimina, Konstantin Kolontarev, Nadezhda Kolyshkina, Andrey Koritsky, Vyacheslav Kuropatkin, Vasily Lazarev, Alexander Manolov, Valentin Manuvera, Daria |

| | | | |
|---|---|---|---|
| | | | Matyushkina, Ekaterina Moskaleva, Varvara Musarova, Oleg Ogarkov, Elizaveta Orlova, Alexander Pavlenko, Alla Petrova, Natalia Pozhenko, Dmitry Pushkar, Alexander Rumyantsev, Sergey Rumyantsev, Vladimir Rumyantsev, Lyubov Rychkova, Alexander Samoilov, Irina Shirokova, Vyacheslav Sinkov, Svetlana Solovieva, Polina Tikhonova, Galina Trifonova, Alexander Troitsky, Alexander Tulichev, Yuri Udalov, Anna Varizhuk, Alexander Vasiliev, Vladimir Veselovsky, Alexey Volnukhin, Gaukhar Yusubalieva |
| EPI_ISL_1910396 | Centre Hospitalier Universitaire Clermont-Ferrand | CHU Clermont-Ferrand, service de virologie | Bisseux Maxime, Mirand Audrey, Combes Patricia, Henquell Cécile |
| EPI_ISL_1972889, EPI_ISL_1972890, EPI_ISL_1972892, EPI_ISL_1972893, EPI_ISL_1972895 | National Public Health Laboratory Malaysia | National Public Health Laboratory Malaysia | Noorliza MN, JL Tan, Chong CK, Norhayati R, Yukie C, Norazimah T Rehan SAB Selvanesan S, Hannah PYP, NurulAina MCA, W.NurAfiza WMA Zirwatul AA, Hani MH, Nurulsyahida I, Aziyati O, Ushananthiny R, Kamal HKZ, Mohd AY, YF Ngeow |
| EPI_ISL_2000551, EPI_ISL_2000552, EPI_ISL_2000553, EPI_ISL_2000554, EPI_ISL_2000555, EPI_ISL_2000556, EPI_ISL_2000557, EPI_ISL_2000558, EPI_ISL_2000559, EPI_ISL_2000560, EPI_ISL_2000561, EPI_ISL_2000562, EPI_ISL_2000564, EPI_ISL_2000565, EPI_ISL_2000566, EPI_ISL_2000567, EPI_ISL_2000570, EPI_ISL_2000571, EPI_ISL_2000572 | | | |
| see above | Tropical Medicine Institute Pedro Kouri | Respiratory Infections Laboratory | Yahisel Tejero Suarez, Lissette Perez Santos, Yanet Pintos Saavedra, Daylin Mederos, Odalys Valdes, Mirtha Elisa Aguado, Vivian Kouri Cardella |
| EPI_ISL_2001114, EPI_ISL_2001115, EPI_ISL_2001213, EPI_ISL_2001429, EPI_ISL_2001430, EPI_ISL_2001643, EPI_ISL_2001644, EPI_ISL_2001655, EPI_ISL_2001656, EPI_ISL_2001659 | National Public Health Laboratory Malaysia | National Public Health Laboratory Malaysia | Noorliza MN, JL Tan, Chong CK, Norhayati R, Yukie C, Norazimah T Rehan SAB Selvanesan S, Hannah PYP, NurulAina MCA, W.NurAfiza WMA Zirwatul AA, Hani MH, Nurulsyahida I, Aziyati O, Ushananthiny R, Kamal HKZ, Mohd AY, YF Ngeow |
| EPI_ISL_2009116 | National Public Health Laboratory Malaysia | National Public Health Laboratory Malaysia | Noorliza MN, JL Tan, Chong CK, Norhayati R, Yukie C, Norazimah T, Rehan SAB, Selvanesan S, Hannah PYP, NurulAina MCA, W.NurAfiza WMA Zirwatul AA, Hani MH, Nurulsyahida I, Aziyati O, Ushananthiny R, Kamal HKZ, Mohd AY, YF Ngeow |
| EPI_ISL_2009170, EPI_ISL_2009171, EPI_ISL_2009172, EPI_ISL_2009173 | MSK Microbiology Lab | MSK Microbiology Lab | Esther Babady, Tracy McMillen, Krupa Jani |
| EPI_ISL_2013323, EPI_ISL_2015112, EPI_ISL_2015113, EPI_ISL_2015114, EPI_ISL_2015118, EPI_ISL_2015119, EPI_ISL_2015120, EPI_ISL_2015123, EPI_ISL_2015127, EPI_ISL_2015128, EPI_ISL_2015129, EPI_ISL_2015130, EPI_ISL_2015131, EPI_ISL_2015132, EPI_ISL_2015133, EPI_ISL_2015134, EPI_ISL_2015135, EPI_ISL_2015136, EPI_ISL_2015137, EPI_ISL_2015138 | | | |
| see above | IU-Cerrahpasa, Cerrahpasa School of Medicine, COVID-19 Lab | IU-Cerrahpasa, Cerrahpasa School of Medicine, COVID-19 Lab | Mert Kuskucu, Yesim Tuyji Tok, Kenan Midilli |
| EPI_ISL_2134584 | Laboratorio de Virologia Molecular / UFRJ | Bioinformatics Laboratory / LNCC | Alice L. Herlinger, Fábio L. Monteiro, Mirela D'arc, Filipe R.R. Moreira, Harrison J. Westgarth, Rafael M. Galliez, Diana Mariani, Luciana J. da Costa, Luiz Almeida, Carolina M. Voloch, Covid19-UFRJ Workgroup, Adriana S.O. Melo, Renato S. Aguiar, André F.A. dos Santos, Terezinha Marta P. P. Castiñeiras, Ana Tereza R. Vasconcelos, Esaú C. João Filho, Claudia C. Escosteguy, Orlando C. Ferreira Jr, Amilcar Tanuri, Luiza M. Higa |
| EPI_ISL_2138923 | Laboratorio HUB -Azienda Ospedaliero Universitaria - AOU - Cagliari | Laboratorio SPOKE Biologia Molecolare -Azienda Ospedaliero Universitaria - AOU - Cagliari | Germano Orrù, Alessandra Scano, Sara Fais, Valentina Medda, Miriam Loddo, Riccardo Cappai, Ferdinando Coghe |
| EPI_ISL_2138924 | Laboratorio HUB -Azienda Ospedaliero Universitaria - AOU - Cagliari | Laboratorio SPOKE Biologia Molecolare -Azienda Ospedaliero Universitaria - AOU - Cagliari | Alessandra Scano, Germano Orrù, Sara Fais, Valentina Medda, Miriam Loddo, Riccardo Cappai, Ferdinando Coghe |
| EPI_ISL_2140172, EPI_ISL_2140173, EPI_ISL_2140188, EPI_ISL_2140189 | Wits VIDA | Wits VIDA | Vicky Baillie, Shabir Madhi et al. |
| EPI_ISL_2156767 | Valenzuela Hope Molecular Laboratory | Philippine Genome Center | Francis A. Tablizo, Kenneth M. Kim, Carlo M. Lapid, Marc Jerrone R. Castro, Maria Sofia L. Yangzon, Elcid Aaron R. Pangilinan, Benedict A. Maralit, Marc Edsel C. Ayes, Eva Maria Cutiongco-de la Paz, Alethea R. de Guzman, Jan Michael C. Yap, Jo-Hannah S. Llames, Sheila Mae M. Araiza, Kris P. Punayan, Irish Coleen A. Asin, Candice Francheska B. Tambaoan, Asia Louisa U. Chong, Karol Sophia Agape R. Padilla, Rianna Patricia S. Cruz, El King D. Morado, Joshua Gregor A. Dizon, Razel Nikka M. Hao, Arianne A. Zamora, Devon Ray Pacial, Juan Antonio R. Magalang, Marissa Alejandria, Celia Carlos, Anna Ong-Lim, Edsel Maurice Salvaña, John Q. Wong, Jaime C. Montoya, Maria Rosario Singh-Vergeire and Cynthia P. Saloma |
| EPI_ISL_2158598, EPI_ISL_2158600 | Laboratório de Biologia Molecular Jean Piaget | MRCG at LSHTM, Genomics lab | Aladje Balde, Abdul Karim Sesay, Abdoulie Kanteh, Bakary Sanyang, Simão Tchuda Bióté, Rui Inndi, Adul Candé, Faatu Cassama, Milanca Agostinho Cá, Rei José Pereira, Erica Luis Maria Magalhães, Aicha Balde, Bubacar Delgado Pinto Embaló, Edmira Maria da Costa, Paulina Joãozinho da Costa Jarra Manneh, Mariama Kujabi, Dabiri Damilari, Sainabou Laye Ndure |
| EPI_ISL_2254725, EPI_ISL_2254726, EPI_ISL_2255181 | Regional Authority of Public Health Banská Bystrica | Veterinary institute in Zvolen Slovakia | Dirbáková Z., Maarová L., Mancoš M., Strhársky J., Sujová S., Mokryšová S., Tinák M., Mojžiš M. |
| EPI_ISL_2271902, EPI_ISL_2271903, EPI_ISL_2271904 | Hospital General de México, Medicina Genómica | Hospital General de México, Medicina Genómica | Ana Laura Sánchez-Sandoval, María Eligia Juárez-Torres, Mariana Villalba-De la Peña, Siddhartha Ríos-Zaragoza, Guadalupe Mercedes Lucía Guerrero-Avendaño, Erika Anabel Alvarado-Silva, Eira Valeria Barrón-Palma |
| EPI_ISL_2284873 | National Hospital | Institute of Tropical Disease, Universitas Airlangga | Jezzy R Dewantari, Rima R Prasetya, Krisnoadi Rahardjo, Aldise M Nastri, Azwan H Lubis, Gatot Soegiarto, Laksmi Wulandari, Resti Yudhawati, Yasuko Mori, Soetjipto, Kazufumi Shimizu, Maria I Lusida |
| EPI_ISL_2285148, EPI_ISL_2285149, EPI_ISL_2285152 | Microbiology Department. Complexo Hospitalario Universitario de Vigo | Microbiology Department. Complexo Hospitalario Universitario de Vigo | Alfaya N, Alonso I, Alvarez M, Cabrera JJ, Carballo R, Cores O, Cortizo S, del-Campo V, Martinez L, Mediero G, Perez S, Potel C, Regueiro B, Rey S, Vasallo FJ |
| EPI_ISL_2304304, EPI_ISL_2307338, EPI_ISL_2307339, EPI_ISL_2307340, EPI_ISL_2307341 | Genome Analysis Center, Yamanashi Central Hospital | Genome Analysis Center, Yamanashi Central Hospital | Yosuke Hirotsu |
| EPI_ISL_2350785 | CHU Purpan - Laboratoire de Virologie - Institut Fédératif de Biologie | Ecole Veterinaire de Toulouse UMR1225 IHAP Interactions hôtes-agents pathogènes | Thomas Figueroa, Charlotte Foret-Lucas, Pierre Bessière, Romain Volmer, Mathilda Walch, Guillaume Croville, Jean-Luc Guérin, Jacques Izopet |
| EPI_ISL_2350793, EPI_ISL_2350794, EPI_ISL_2350795, EPI_ISL_2350796, EPI_ISL_2350797, EPI_ISL_2350798, EPI_ISL_2350799, EPI_ISL_2350800, EPI_ISL_2350801, EPI_ISL_2350803, EPI_ISL_2350804, EPI_ISL_2350805, EPI_ISL_2350806, EPI_ISL_2350807, EPI_ISL_2350808 | | | |
| see above | Molecular Lab, Evercare hospital Dhaka | International Institute for Zoonosis Control, Hokkaido university | Junya Yamagishi, Mizanur Rahman |
| EPI_ISL_2364506, EPI_ISL_2364507, EPI_ISL_2364508 | Genome Analysis Center, Yamanashi Central Hospital | Genome Analysis Center, Yamanashi Central Hospital | Yosuke Hirotsu |
| EPI_ISL_2382417, EPI_ISL_2382418, EPI_ISL_2382419, EPI_ISL_2382420, EPI_ISL_2398291, EPI_ISL_2398293, EPI_ISL_2398294 | Centre for Human Virology and Genomics, Microbiology Department, Nigerian Institute of Medical Research | Central Research Laboratory, Nigerian Institute of Medical Research | Chika Kingsley Onwuamah, Ayorinde Babatunde James, Joseph Ojonugwa Shaibu, Rahaman A. Ahmed, Azuka Patrick Okwuraiwe, Sola Ajibaye, Sharon Abimbola, Olufemi Samuel Amoo, Muinah Adenike Fowora, Phasha-Muchemenye Mmatshepho, Joy Ayoola, Nyam Itse Yusuf, Grace Oni, Yusuf Jimoh, Josiah Ayoola Isong, Uyi Emokpae, Ngozi Mirabel Otuonye, Rosemary Ajuma Audu, Bamidele Iwalokun, Babatunde Lawal Salako |
| EPI_ISL_2433280, EPI_ISL_2433281, EPI_ISL_2433282, EPI_ISL_2433283, EPI_ISL_2433284 | Fourth affiliated hospital, college of medicine, zhejiang university | Fourth affiliated hospital, college of medicine, zhejiang university | yingping wu,shufa zheng, tian liu, xueke liu, huina tang, yutong he, wei xu, lele li, wenxu yu, ke xing,xiaoping xia |
| EPI_ISL_2443718, EPI_ISL_2443725, EPI_ISL_2443752, EPI_ISL_2443763, EPI_ISL_2443769 | Orange County Public Health Laboratory | University of California, Irvine | Amanda N. Pinski, Ilhem Messaoudi |
| EPI_ISL_2484141 | Mater Misericordiae University Hospital Dublin | University College Dublin, CFS, School of Public Health | Maureen Lynch, Guerrino Macori, Séamus Fanning, Edel O' Regan, Eoin Hunt, Dermot O'Callaghan, Brian McCullagh, Cormac Jennings, Anne Fortune |
| EPI_ISL_2484209, EPI_ISL_2484210, EPI_ISL_2484211, EPI_ISL_2484212, EPI_ISL_2484213, EPI_ISL_2484214 | University of Brescia - Dept. of Molecular and Translational Medicine | Istituto Zooprofilattico Sperimentale della Lombardia e dell'Emilia Romagna (IZSLER), Risk Analysis and Genomic Epidemiology Unit | Caruso Arnaldo, Fiorentini Simona, Caccuri Francesca, Messali Serena, Zani Alberto, Marina Morganti, Erika Scaltriti, Stefano Pongolini |

| | | | |
|---|---|---|---|
| EPI_ISL_2501081, EPI_ISL_2501084, EPI_ISL_2501087 | Centre for Human Virology and Genomics, Microbiology Department, Nigerian Institute of Medical Research | Central Research Laboratory, Nigerian Institute of Medical Research | Chika Kingsley Onwuamah, Ayorinde Babatunde James, Joseph Ojonugwa Shaibu, Rahaman A. Ahmed, Azuka Patrick Okwuraiwe, Sola Ajibaye, Sharon Abimbola, Olufemi Samuel Amoo, Muinah Adenike Fowora, Phasha-Muchemenye Mmatshepho, Joy Ayoola, Nyam Itse Yusuf, Grace Oni, Yusuf Jimoh, Josiah Ayoola Isong, Uyi Emokpae, Ngozi Mirabel Otuonye, Rosemary Ajuma Audu, Bamidele Iwalokun, Babatunde Lawal Salako |
| EPI_ISL_2501394, EPI_ISL_2501395 | Microbiology Department. Complexo Hospitalario Universitario de Vigo | Microbiology Department. Complexo Hospitalario Universitario de Vigo | Alfaya N, Alonso I, Alvarez M, Cabrera JJ, Carballo R, Cores O, Cortizo S, del-Campo V, Martinez L, Mediero G, Perez S, Potel C, Regueiro B, Rey S, Vasallo FJ |
| EPI_ISL_2502015, EPI_ISL_2502016, EPI_ISL_2502017, EPI_ISL_2502023, EPI_ISL_2502024 | Centre for Human Virology and Genomics, Microbiology Department, Nigerian Institute of Medical Research | Central Research Laboratory, Nigerian Institute of Medical Research | Chika Kingsley Onwuamah, Ayorinde Babatunde James, Joseph Ojonugwa Shaibu, Rahaman A. Ahmed, Azuka Patrick Okwuraiwe, Sola Ajibaye, Sharon Abimbola, Olufemi Samuel Amoo, Muinah Adenike Fowora, Phasha-Muchemenye Mmatshepho, Joy Ayoola, Nyam Itse Yusuf, Grace Oni, Yusuf Jimoh, Josiah Ayoola Isong, Uyi Emokpae, Ngozi Mirabel Otuonye, Rosemary Ajuma Audu, Bamidele Iwalokun, Babatunde Lawal Salako |
| EPI_ISL_2502632, EPI_ISL_2502633, EPI_ISL_2502634, EPI_ISL_2502637, EPI_ISL_2502640, EPI_ISL_2502643, EPI_ISL_2502644, EPI_ISL_2502647, EPI_ISL_2502657, EPI_ISL_2502658, EPI_ISL_2502659, EPI_ISL_2502660, EPI_ISL_2502662, EPI_ISL_2502663, EPI_ISL_2502664, EPI_ISL_2502665, EPI_ISL_2502666 | | | |
| see above | All India Institute of Medical Sciences Delhi Hospital | Virology Laboratory, AIIMS Delhi | Aashish Choudhary, Sumedha Bagga, Lata Rani, Dibyabhaba Pradhan, Pooja Pandey, Jyoti Jethani, Shivram Dhakad, Nazneen Arif, Deepankar Srigyan, Megha Brijwal, Lalit Dar, Manish Soneja, Rakesh Lodha, Puneet Kaur, Ritu Gupta, Subrata Sinha, Chitra Sarkar, Randeep Guleria |
| EPI_ISL_2565184 | SK-Roy Romanow Provincial Laboratory | National Microbiology Laboratory (NML) | Anna Majer, Shari Tyson, Grace Seo, Philip Mabon, Elsie Grudeski, Rhiannon Huzarewich, Russell Mandes, Anneliese Landgraff, Jennifer Tanner, Natalie Knox, Morag Graham, Gary Van Domselaar, Ryan McDonald, Amanda Lang, Rachel DePaulo, Jessica Minion, Nathalie Bastien, Yan Li, Timothy Booth, Darian Hole, Madison Chapel, Kirsten Biggar, CanCOGeN's metadata curation team, Public Health Agency of Canada CanCOGeN team |
| EPI_ISL_2603247, EPI_ISL_2603249, EPI_ISL_2603250 | Microbiology Department. Complexo Hospitalario Universitario de Vigo | Microbiology Department. Complexo Hospitalario Universitario de Vigo | Alfaya N, Alonso I, Alvarez M, Cabrera JJ, Carballo R, Cores O, Cortizo S, del-Campo V, Martinez L, Mediero G, Perez S, Potel C, Regueiro B, Rey S, Vasallo FJ |
| EPI_ISL_2614387 | Adolfo Lutz Institute Sao Paulo Virology Center | Retrovirus Laboratory Adolfo Lutz Institute | Gabriela Bastos Cabral, Giselle I S Lopez-Lopes, Cintia Ahagon, Audrey Cilli, Paula Morena Guimaraes, Igor Mohamed Hussein, Luis Brigido |
| EPI_ISL_2663160 | Ochsner Health | Bioinfoexperts, LLC | Judy Crabtree, Amy Feehan, Rebecca Rose, David J. Nolan, Tessa LaFleur, Julia-Garcia-Diaz, Lucio Miele, Susanna L. Lamers |
| EPI_ISL_2671835 | Juntendo University Hospital | Juntendo University Hospital | Koji Tuchiya, Yoshie Hosaka |
| EPI_ISL_2690150 | Ochsner Health | Bioinfoexperts, LLC | Judy Crabtree, Amy Feehan, Rebecca Rose, David J. Nolan, Tessa LaFleur, Julia-Garcia-Diaz, Lucio Miele, Susanna L. Lamers |
| EPI_ISL_2691710 | Department of pathology, school of medicine, Imam Hospital complex, Tehran university of medical sciences, Tehran, Iran | Genetics Research Center, University of Social Welfare and Rehabilitation Sciences | Zohreh Fattahi, Marzieh Mohseni, Khadijeh Jalalvand, Azam Ghaziasadi, Seyedeh elham Mortazavi, Azar Hadadi, Alireza Abdollahi, Ali Jafarpour, Azam Ghaziasad, Seyedeh elham Mortazavi, Saber Soltani, Reza Najafipour, Kimia Kahrizi, Seyed Mohammad Jazayeri, Hossein Najmabadi. |
| EPI_ISL_2692885, EPI_ISL_2692888 | Pars Hospital Laboratory, Rasht, Iran. | Genetics Research Center, University of Social Welfare and Rehabilitation Sciences | Zohreh Fattahi, Marzieh Mohseni, Khadijeh Jalalvand, Issa Jahanzad ,Kimia Kahrizi, Hossein Najmabadi. |
| EPI_ISL_2693672 | Department of Virology, School of Medicine, Ahvaz Jundishapur University of Medical Sciences, Ahvaz, Iran | Genetics Research Center, University of Social Welfare and Rehabilitation Sciences | Zohreh Fattahi, Marzieh Mohseni, Khadijeh Jalalvand, Azam Ghaziasadi, Seyedeh elham Mortazavi, Ali Jafarpour, Azarakhsh Azaran, Farid Yousefi, Gholam Abbas Kaydani, Kimia Kahrizi, Seyed Mohammad Jazayeri, Hossein Najmabadi. |
| EPI_ISL_2693673 | Department of Virology, School of Medicine, Ahvaz Jundishapur University of Medical Sciences, Ahvaz, Iran | Genetics Research Center, University of Social Welfare and Rehabilitation Sciences | Zohreh Fattahi, Marzieh Mohseni, Khadijeh Jalalvand, Azam Ghaziasadi, Seyedeh elham Mortazavi, Ali Jafarpour, Azarakhsh Azaran, Farid Yousefi, Gholam Abbas Kaydani, Kimia Kahrizi, Seyed Mohammad Jazayeri, Hossein Najmabadi. |
| EPI_ISL_2693674 | Department of Virology, School of Medicine, Ahvaz Jundishapur University of Medical Sciences, Ahvaz, Iran | Genetics Research Center, University of Social Welfare and Rehabilitation Sciences | Zohreh Fattahi, Marzieh Mohseni, Khadijeh Jalalvand, Azam Ghaziasadi, Seyedeh elham Mortazavi, Ali Jafarpour, Azarakhsh Azaran, Farid Yousefi, Gholam Abbas Kaydani, Kimia Kahrizi, Seyed Mohammad Jazayeri, Hossein Najmabadi. |
| EPI_ISL_2693675 | Department of Molecular Diagnostic, Emam Ali Educational and Therapeutic Center, Alborz University of Medical Sciences, Karaj, Iran | Genetics Research Center, University of Social Welfare and Rehabilitation Sciences | Zohreh Fattahi, Marzieh Mohseni, Khadijeh Jalalvand, Azam Ghaziasadi, Seyedeh elham Mortazavi, Ali Jafarpour, Zakiye Mokhames Mokhames, Alireza Soleimani, Kimia Kahrizi, Seyed Mohammad Jazayeri, Hossein Najmabadi. |
| EPI_ISL_2693676 | Department of Molecular Diagnostic, Emam Ali Educational and Therapeutic Center, Alborz University of Medical Sciences, Karaj, Iran | Genetics Research Center, University of Social Welfare and Rehabilitation Sciences | Zohreh Fattahi, Marzieh Mohseni, Khadijeh Jalalvand, Azam Ghaziasadi, Seyedeh elham Mortazavi, Ali Jafarpour, Zakiye Mokhames, Alireza Soleimani, Kimia Kahrizi, Seyed Mohammad Jazayeri, Hossein Najmabadi. |
| EPI_ISL_2693905, EPI_ISL_2693906 | Infectious Disease and Tropical Medicine Research Center, Resistant Tuberculosis Institute, Zahedan Univerisry of Medical Sciences, Zahedan, Iran | Genetics Research Center, University of Social Welfare and Rehabilitation Sciences | Zohreh Fattahi, Marzieh Mohseni, Khadijeh Jalalvand, Azam Ghaziasadi, Seyedeh elham Mortazavi, Ali Jafarpour, Ebrahim Kord, Seyed Mohammad Hashemi-Shahri, Kimia Kahrizi, Seyed Mohammad Jazayeri, Hossein Najmabadi. |
| EPI_ISL_2693907 | Infectious Diseases Research Center, Birjand University of Medical Sciences, Birjand, Iran | Genetics Research Center, University of Social Welfare and Rehabilitation Sciences | Zohreh Fattahi, Marzieh Mohseni, Khadijeh Jalalvand, Azam Ghaziasadi, Seyedeh elham Mortazavi, Ali Jafarpour, Shokouh Ghafari, Masood Ziaee, Kimia Kahrizi, Seyed Mohammad Jazayeri, Hossein Najmabadi. |
| EPI_ISL_2693908 | Department of Infectious Disease, School of Medicine, Sina Hospital, Tehran University of Medical Sciences | Genetics Research Center, University of Social Welfare and Rehabilitation Sciences | Zohreh Fattahi, Marzieh Mohseni, Khadijeh Jalalvand, Jila Yavarian, Kimia Kahrizi, Talat Mokhtari-Azad, Hossein Najmabadi. |
| EPI_ISL_2693972, EPI_ISL_2694195, EPI_ISL_2694587 | Virology Centre, School of public health, Tehran University of Medical Sciences, Tehran, Iran | Genetics Research Center, University of Social Welfare and Rehabilitation Sciences | Zohreh Fattahi, Marzieh Mohseni, Khadijeh Jalalvand, Jila Yavarian, Kimia Kahrizi, Talat Mokhtari-Azad, Hossein Najmabadi. |
| EPI_ISL_2710021 | Ochsner Health | Bioinfoexperts, LLC | Judy Crabtree, Amy Feehan, Rebecca Rose, David J. Nolan, Tessa LaFleur, Julia-Garcia-Diaz, Lucio Miele, Susanna L. Lamers |
| EPI_ISL_2710810, EPI_ISL_2710812, EPI_ISL_2710814, EPI_ISL_2710815, EPI_ISL_2710817, EPI_ISL_2710818, EPI_ISL_2710820, EPI_ISL_2710821, EPI_ISL_2710823, EPI_ISL_2710825, EPI_ISL_2710827, EPI_ISL_2710831, EPI_ISL_2710837, EPI_ISL_2710838, EPI_ISL_2710872, EPI_ISL_2710873, EPI_ISL_2710874, EPI_ISL_2710875, EPI_ISL_2710876, EPI_ISL_2710877, EPI_ISL_2710878, EPI_ISL_2710879, EPI_ISL_2710880, EPI_ISL_2710881, EPI_ISL_2710882, EPI_ISL_2713077, EPI_ISL_2713078 | | | |
| see above | Department of Health Technology and Informatics, The Hong Kong Polytechnic University | Department of Health Technology and Informatics, The Hong Kong Polytechnic University | Gilman Kit-Hang Siu, Lam-Kwong Lee, Kenneth Siu-Sing Leung, Jake Siu-Lun Leung, Wing-Hei Lo, Timothy Ting-Leung Ng, Chloe Toi-Mei Chan, Hoi-Ching Jim, Ivan Tak-Fai Wong, Kingsley King-Gee Tam, Hiu-Yin Lao, Denise Sze-Hang Wong, Alan Ka-Lun Wu, Miranda Chong-Yee Yau, Yvette Wai-Man Lai, Kitty Sau-Chun Fung, Sandy Ka-Yee Chau, Barry Kin-Chung Wong, Wing-Kin To, Kristine Luk, Alex Yat-Man Ho, Tak-Lun Que, Kam-Tong Yip, Wing Cheong Yam, David Ho-Keung Shum, Shea Ping Yip |
| EPI_ISL_2727891, EPI_ISL_2727899 | All India Institute of Medical Sciences Delhi Hospital | Virology Laboratory, AIIMS Delhi | Aashish Choudhary, Sumedha Bagga, Lata Rani, Dibyabhaba Pradhan, Pooja Pandey, Jyoti Jethani, Shivram Dhakad, Nazneen Arif, Deepankar Srigyan, Megha Brijwal, Lalit Dar, Manish Soneja, Rakesh Lodha, Puneet Kaur, Ritu Gupta, Subrata Sinha, Chitra Sarkar, Randeep Guleria |
| EPI_ISL_2732373 | Diagnostic Laboratory, Virology Erlangen | Armin Ensser | Alexandra Herrmann, Armin Ensser |
| EPI_ISL_2779638 | State Key Laboratory of Emerging Infectious Diseases, The University of Hong Kong | The University of Hong Kong | Lifeng Li, Xianfan Wu, Yiu Man Cheung, Huachen Zhu, Yi Guan |
| EPI_ISL_2790002, EPI_ISL_2790003 | All India Institute of Medical Sciences Delhi Hospital | Virology Laboratory, AIIMS Delhi | Aashish Choudhary, Sumedha Bagga, Lata Rani, Dibyabhaba Pradhan, Pooja Pandey, Jyoti Jethani, Shivram Dhakad, Nazneen Arif, Deepankar Srigyan, Megha Brijwal, Lalit Dar, Manish Soneja, Rakesh Lodha, Puneet Kaur, Ritu Gupta, Subrata Sinha, Chitra Sarkar, Randeep Guleria |
| EPI_ISL_2790800, EPI_ISL_2790801, EPI_ISL_2790802, EPI_ISL_2790803 | Armies | National Reference Center for Viruses of Respiratory Infections, Institut Pasteur, Paris | Marion Barbet, Sylvie Behillil, Méline Bizard, Angela Brisebarre, Camille Capel, Vincent Enouf, Louise Lefrançois, Frédéric Lemoine, Christophe Malabat, Corinne Maufrais, Emmanuelle Permal, Etienne Simon-Lorière, Maud Vanpeene, Sylvie Van der Werf ,Christine Bigaillon |
| EPI_ISL_2790804, EPI_ISL_2790805, EPI_ISL_2790806, EPI_ISL_2790807, EPI_ISL_2790808 | Hospital | National Reference Center for Viruses of Respiratory Infections, Institut Pasteur, Paris | Marion Barbet, Sylvie Behillil, Méline Bizard, Angela Brisebarre, Camille Capel, Vincent Enouf, Louise Lefrançois, Frédéric Lemoine, Christophe Malabat, Corinne Maufrais, Emmanuelle Permal, Etienne Simon-Lorière, Maud Vanpeene, Sylvie Van der Werf ,Christine Lambert |
| EPI_ISL_2790809 | Hospital | National Reference Center for Viruses of Respiratory Infections, Institut Pasteur, Paris | Marion Barbet, Sylvie Behillil, Méline Bizard, Angela Brisebarre, Camille Capel, Vincent Enouf, Louise Lefrançois, Frédéric Lemoine, Christophe Malabat, Corinne Maufrais, Emmanuelle Permal, Etienne Simon-Lorière, Maud Vanpeene, Sylvie Van der Werf |
| EPI_ISL_2790810, EPI_ISL_2790811, EPI_ISL_2790812, EPI_ISL_2790813, EPI_ISL_2790814 | Hospital | National Reference Center for Viruses of Respiratory Infections, Institut Pasteur, Paris | Marion Barbet, Sylvie Behillil, Méline Bizard, Angela Brisebarre, Camille Capel, Vincent Enouf, Louise Lefrançois, Frédéric Lemoine, Christophe Malabat, Corinne Maufrais, Emmanuelle Permal, Etienne Simon-Lorière, Maud Vanpeene, Sylvie Van der Werf ,Olivia Raulin |
| EPI_ISL_2790815, EPI_ISL_2790816 | Hospital | National Reference Center for Viruses of Respiratory Infections, Institut Pasteur, Paris | Marion Barbet, Sylvie Behillil, Méline Bizard, Angela Brisebarre, Camille Capel, Vincent Enouf, Louise Lefrançois, Frédéric Lemoine, Christophe Malabat, Corinne Maufrais, Emmanuelle Permal, Etienne Simon-Lorière, Maud Vanpeene, Sylvie Van der Werf ,Christine Lambert |

| | | | |
|---|---|---|---|
| EPI_ISL_2790817, EPI_ISL_2790818 | Hospital | National Reference Center for Viruses of Respiratory Infections, Institut Pasteur, Paris | Marion Barbet, Sylvie Behillil, Méline Bizard, Angela Brisebarre, Camille Capel, Vincent Enouf, Louise Lefrançois, Frédéric Lemoine, Christophe Malabat, Corinne Maufrais, Emmanuelle Permal, Etienne Simon-Lorière, Maud Vanpeene, Sylvie Van der Werf |
| EPI_ISL_2790819, EPI_ISL_2790820, EPI_ISL_2790821, EPI_ISL_2790822, EPI_ISL_2790823, EPI_ISL_2790824, EPI_ISL_2790825, EPI_ISL_2790826 | Hospital | National Reference Center for Viruses of Respiratory Infections, Institut Pasteur, Paris | Marion Barbet, Sylvie Behillil, Méline Bizard, Angela Brisebarre, Camille Capel, Vincent Enouf, Louise Lefrançois, Frédéric Lemoine, Christophe Malabat, Corinne Maufrais, Emmanuelle Permal, Etienne Simon-Lorière, Maud Vanpeene, Sylvie Van der Werf ,Olivia Raulin |
| EPI_ISL_2790827, EPI_ISL_2790828, EPI_ISL_2790829, EPI_ISL_2790830, EPI_ISL_2790831, EPI_ISL_2790832, EPI_ISL_2790833 | Hospital | National Reference Center for Viruses of Respiratory Infections, Institut Pasteur, Paris | Marion Barbet, Sylvie Behillil, Méline Bizard, Angela Brisebarre, Camille Capel, Vincent Enouf, Louise Lefrançois, Frédéric Lemoine, Christophe Malabat, Corinne Maufrais, Emmanuelle Permal, Etienne Simon-Lorière, Maud Vanpeene, Sylvie Van der Werf |
| EPI_ISL_2790834, EPI_ISL_2790835, EPI_ISL_2790836, EPI_ISL_2790837 | Hospital | National Reference Center for Viruses of Respiratory Infections, Institut Pasteur, Paris | Marion Barbet, Sylvie Behillil, Méline Bizard, Angela Brisebarre, Camille Capel, Vincent Enouf, Louise Lefrançois, Frédéric Lemoine, Christophe Malabat, Corinne Maufrais, Emmanuelle Permal, Etienne Simon-Lorière, Maud Vanpeene, Sylvie Van der Werf ,Olivia Raulin |
| EPI_ISL_2790838, EPI_ISL_2790839, EPI_ISL_2790840, EPI_ISL_2790841, EPI_ISL_2790842, EPI_ISL_2790843 | Hospital | National Reference Center for Viruses of Respiratory Infections, Institut Pasteur, Paris | Marion Barbet, Sylvie Behillil, Méline Bizard, Angela Brisebarre, Camille Capel, Vincent Enouf, Louise Lefrançois, Frédéric Lemoine, Christophe Malabat, Corinne Maufrais, Emmanuelle Permal, Etienne Simon-Lorière, Maud Vanpeene, Sylvie Van der Werf |
| EPI_ISL_2790844 | Hospital | National Reference Center for Viruses of Respiratory Infections, Institut Pasteur, Paris | Marion Barbet, Sylvie Behillil, Méline Bizard, Angela Brisebarre, Camille Capel, Vincent Enouf, Louise Lefrançois, Frédéric Lemoine, Christophe Malabat, Corinne Maufrais, Emmanuelle Permal, Etienne Simon-Lorière, Maud Vanpeene, Sylvie Van der Werf ,LéA Pilorge |
| EPI_ISL_2790845, EPI_ISL_2790846, EPI_ISL_2790847 | Hospital | National Reference Center for Viruses of Respiratory Infections, Institut Pasteur, Paris | Marion Barbet, Sylvie Behillil, Méline Bizard, Angela Brisebarre, Camille Capel, Vincent Enouf, Louise Lefrançois, Frédéric Lemoine, Christophe Malabat, Corinne Maufrais, Emmanuelle Permal, Etienne Simon-Lorière, Maud Vanpeene, Sylvie Van der Werf ,Patricia Barbut |
| EPI_ISL_2790848 | Hospital | National Reference Center for Viruses of Respiratory Infections, Institut Pasteur, Paris | Marion Barbet, Sylvie Behillil, Méline Bizard, Angela Brisebarre, Camille Capel, Vincent Enouf, Louise Lefrançois, Frédéric Lemoine, Christophe Malabat, Corinne Maufrais, Emmanuelle Permal, Etienne Simon-Lorière, Maud Vanpeene, Sylvie Van der Werf ,Vincent Esteve |
| EPI_ISL_2790849 | Hospital | National Reference Center for Viruses of Respiratory Infections, Institut Pasteur, Paris | Marion Barbet, Sylvie Behillil, Méline Bizard, Angela Brisebarre, Camille Capel, Vincent Enouf, Louise Lefrançois, Frédéric Lemoine, Christophe Malabat, Corinne Maufrais, Emmanuelle Permal, Etienne Simon-Lorière, Maud Vanpeene, Sylvie Van der Werf ,Patricia Barbut |
| EPI_ISL_2790850, EPI_ISL_2790851 | Hospital | National Reference Center for Viruses of Respiratory Infections, Institut Pasteur, Paris | Marion Barbet, Sylvie Behillil, Méline Bizard, Angela Brisebarre, Camille Capel, Vincent Enouf, Louise Lefrançois, Frédéric Lemoine, Christophe Malabat, Corinne Maufrais, Emmanuelle Permal, Etienne Simon-Lorière, Maud Vanpeene, Sylvie Van der Werf ,Olivia Raulin |
| EPI_ISL_2790852, EPI_ISL_2790853, EPI_ISL_2790854, EPI_ISL_2790855, EPI_ISL_2790856 | Hospital | National Reference Center for Viruses of Respiratory Infections, Institut Pasteur, Paris | Marion Barbet, Sylvie Behillil, Méline Bizard, Angela Brisebarre, Camille Capel, Vincent Enouf, Louise Lefrançois, Frédéric Lemoine, Christophe Malabat, Corinne Maufrais, Emmanuelle Permal, Etienne Simon-Lorière, Maud Vanpeene, Sylvie Van der Werf ,Patricia Barbut |
| EPI_ISL_2790857, EPI_ISL_2790858 | EHPAD_IRA | National Reference Center for Viruses of Respiratory Infections, Institut Pasteur, Paris | Marion Barbet, Sylvie Behillil, Méline Bizard, Angela Brisebarre, Camille Capel, Vincent Enouf, Louise Lefrançois, Frédéric Lemoine, Christophe Malabat, Corinne Maufrais, Emmanuelle Permal, Etienne Simon-Lorière, Maud Vanpeene, Sylvie Van der Werf ,La Villa Papyri |
| EPI_ISL_2790859 | EHPAD_IRA | National Reference Center for Viruses of Respiratory Infections, Institut Pasteur, Paris | Marion Barbet, Sylvie Behillil, Méline Bizard, Angela Brisebarre, Camille Capel, Vincent Enouf, Louise Lefrançois, Frédéric Lemoine, Christophe Malabat, Corinne Maufrais, Emmanuelle Permal, Etienne Simon-Lorière, Maud Vanpeene, Sylvie Van der Werf ,Plaisance |
| EPI_ISL_2790904 | Hospital | National Reference Center for Viruses of Respiratory Infections, Institut Pasteur, Paris | Marion Barbet, Sylvie Behillil, Méline Bizard, Angela Brisebarre, Camille Capel, Vincent Enouf, Louise Lefrançois, Frédéric Lemoine, Christophe Malabat, Corinne Maufrais, Emmanuelle Permal, Etienne Simon-Lorière, Maud Vanpeene, Sylvie Van der Werf ,StéPhanie Marque Juillet |
| EPI_ISL_2790992 | Hospital | National Reference Center for Viruses of Respiratory Infections, Institut Pasteur, Paris | Marion Barbet, Sylvie Behillil, Méline Bizard, Angela Brisebarre, Camille Capel, Vincent Enouf, Louise Lefrançois, Frédéric Lemoine, Christophe Malabat, Corinne Maufrais, Emmanuelle Permal, Etienne Simon-Lorière, Maud Vanpeene, Sylvie Van der Werf ,Olivia Raulin |
| EPI_ISL_2790993 | Hospital | National Reference Center for Viruses of Respiratory Infections, Institut Pasteur, Paris | Marion Barbet, Sylvie Behillil, Méline Bizard, Angela Brisebarre, Camille Capel, Vincent Enouf, Louise Lefrançois, Frédéric Lemoine, Christophe Malabat, Corinne Maufrais, Emmanuelle Permal, Etienne Simon-Lorière, Maud Vanpeene, Sylvie Van der Werf ,Laurent Andreoletti |
| EPI_ISL_2790994 | Hospital | National Reference Center for Viruses of Respiratory Infections, Institut Pasteur, Paris | Marion Barbet, Sylvie Behillil, Méline Bizard, Angela Brisebarre, Camille Capel, Vincent Enouf, Louise Lefrançois, Frédéric Lemoine, Christophe Malabat, Corinne Maufrais, Emmanuelle Permal, Etienne Simon-Lorière, Maud Vanpeene, Sylvie Van der Werf ,JéRome Guinard |
| EPI_ISL_2790995, EPI_ISL_2790996, EPI_ISL_2790997 | EHPAD_IRA | National Reference Center for Viruses of Respiratory Infections, Institut Pasteur, Paris | Marion Barbet, Sylvie Behillil, Méline Bizard, Angela Brisebarre, Camille Capel, Vincent Enouf, Louise Lefrançois, Frédéric Lemoine, Christophe Malabat, Corinne Maufrais, Emmanuelle Permal, Etienne Simon-Lorière, Maud Vanpeene, Sylvie Van der Werf ,Vill'Alizé |
| EPI_ISL_2790998, EPI_ISL_2790999, EPI_ISL_2791000, EPI_ISL_2791001, EPI_ISL_2791002 | Hospital | National Reference Center for Viruses of Respiratory Infections, Institut Pasteur, Paris | Marion Barbet, Sylvie Behillil, Méline Bizard, Angela Brisebarre, Camille Capel, Vincent Enouf, Louise Lefrançois, Frédéric Lemoine, Christophe Malabat, Corinne Maufrais, Emmanuelle Permal, Etienne Simon-Lorière, Maud Vanpeene, Sylvie Van der Werf ,Olivia Raulin |
| EPI_ISL_2791003 | Hospital | National Reference Center for Viruses of Respiratory Infections, Institut Pasteur, Paris | Marion Barbet, Sylvie Behillil, Méline Bizard, Angela Brisebarre, Camille Capel, Vincent Enouf, Louise Lefrançois, Frédéric Lemoine, Christophe Malabat, Corinne Maufrais, Emmanuelle Permal, Etienne Simon-Lorière, Maud Vanpeene, Sylvie Van der Werf ,JéRome Guinard |
| EPI_ISL_2791004, EPI_ISL_2791005, EPI_ISL_2791006, EPI_ISL_2791007 | Hospital | National Reference Center for Viruses of Respiratory Infections, Institut Pasteur, Paris | Marion Barbet, Sylvie Behillil, Méline Bizard, Angela Brisebarre, Camille Capel, Vincent Enouf, Louise Lefrançois, Frédéric Lemoine, Christophe Malabat, Corinne Maufrais, Emmanuelle Permal, Etienne Simon-Lorière, Maud Vanpeene, Sylvie Van der Werf ,Olivia Raulin |
| EPI_ISL_2791008, EPI_ISL_2791009 | Hospital | National Reference Center for Viruses of Respiratory Infections, Institut Pasteur, Paris | Marion Barbet, Sylvie Behillil, Méline Bizard, Angela Brisebarre, Camille Capel, Vincent Enouf, Louise Lefrançois, Frédéric Lemoine, Christophe Malabat, Corinne Maufrais, Emmanuelle Permal, Etienne Simon-Lorière, Maud Vanpeene, Sylvie Van der Werf ,Alice Cadennes |
| EPI_ISL_2791010 | Hospital | National Reference Center for Viruses of Respiratory Infections, Institut Pasteur, Paris | Marion Barbet, Sylvie Behillil, Méline Bizard, Angela Brisebarre, Camille Capel, Vincent Enouf, Louise Lefrançois, Frédéric Lemoine, Christophe Malabat, Corinne Maufrais, Emmanuelle Permal, Etienne Simon-Lorière, Maud Vanpeene, Sylvie Van der Werf ,Christine Lambert |
| EPI_ISL_2791011, EPI_ISL_2791012, EPI_ISL_2791013, EPI_ISL_2791014 | Hospital | National Reference Center for Viruses of Respiratory Infections, Institut Pasteur, Paris | Marion Barbet, Sylvie Behillil, Méline Bizard, Angela Brisebarre, Camille Capel, Vincent Enouf, Louise Lefrançois, Frédéric Lemoine, Christophe Malabat, Corinne Maufrais, Emmanuelle Permal, Etienne Simon-Lorière, Maud Vanpeene, Sylvie Van der Werf ,Olivia Raulin |
| EPI_ISL_2791015 | Hospital | National Reference Center for Viruses of Respiratory Infections, Institut Pasteur, Paris | Marion Barbet, Sylvie Behillil, Méline Bizard, Angela Brisebarre, Camille Capel, Vincent Enouf, Louise Lefrançois, Frédéric Lemoine, Christophe Malabat, Corinne Maufrais, Emmanuelle Permal, Etienne Simon-Lorière, Maud Vanpeene, Sylvie Van der Werf ,JéRome Guinard |
| EPI_ISL_2791016, EPI_ISL_2791017, EPI_ISL_2791018, EPI_ISL_2791019, EPI_ISL_2791020 | Hospital | National Reference Center for Viruses of Respiratory Infections, Institut Pasteur, Paris | Marion Barbet, Sylvie Behillil, Méline Bizard, Angela Brisebarre, Camille Capel, Vincent Enouf, Louise Lefrançois, Frédéric Lemoine, Christophe Malabat, Corinne Maufrais, Emmanuelle Permal, Etienne Simon-Lorière, Maud Vanpeene, Sylvie Van der Werf ,Olivia Raulin |
| EPI_ISL_2791021, EPI_ISL_2791022, EPI_ISL_2791023, EPI_ISL_2791024, EPI_ISL_2791025, EPI_ISL_2791026, EPI_ISL_2791027, EPI_ISL_2791028, EPI_ISL_2791029, EPI_ISL_2791030, EPI_ISL_2791031, EPI_ISL_2791032, EPI_ISL_2791033, EPI_ISL_2791034, EPI_ISL_2791035 | | | |
| see above | Armies | National Reference Center for Viruses of Respiratory Infections, Institut Pasteur, Paris | Marion Barbet, Sylvie Behillil, Méline Bizard, Angela Brisebarre, Camille Capel, Vincent Enouf, Louise Lefrançois, Frédéric Lemoine, Christophe Malabat, Corinne Maufrais, Emmanuelle Permal, Etienne Simon-Lorière, Maud Vanpeene, Sylvie Van der Werf ,Christine Bigaillon |
| EPI_ISL_2811805 | Jining Center for Disease Control and Prevention | Jining Center for Disease Control and Prevention | Jiao Bo-yan |
| EPI_ISL_2835273, EPI_ISL_2835274, EPI_ISL_2835275, EPI_ISL_2835276, EPI_ISL_2835277, EPI_ISL_2835278, EPI_ISL_2835279, EPI_ISL_2835280, EPI_ISL_2835281, EPI_ISL_2835282, EPI_ISL_2835283, EPI_ISL_2835284, EPI_ISL_2835285, EPI_ISL_2835286, EPI_ISL_2835287, EPI_ISL_2835288, EPI_ISL_2835289, EPI_ISL_2835290, EPI_ISL_2835291, EPI_ISL_2835292, EPI_ISL_2835293, EPI_ISL_2835294, EPI_ISL_2835295, EPI_ISL_2835296, EPI_ISL_2835297, EPI_ISL_2835298, EPI_ISL_2835299, EPI_ISL_2835300, EPI_ISL_2835301, EPI_ISL_2835302, EPI_ISL_2835303, EPI_ISL_2835304, EPI_ISL_2835305, EPI_ISL_2835306, EPI_ISL_2835307, EPI_ISL_2835308, EPI_ISL_2835309, EPI_ISL_2835310, EPI_ISL_2835311, EPI_ISL_2835312, EPI_ISL_2835313, EPI_ISL_2835314, EPI_ISL_2835315, EPI_ISL_2835316, EPI_ISL_2835317, EPI_ISL_2835318, EPI_ISL_2835319, EPI_ISL_2835320, EPI_ISL_2835321, EPI_ISL_2835322, EPI_ISL_2835323, EPI_ISL_2835324, EPI_ISL_2835325, EPI_ISL_2835326, EPI_ISL_2835327, EPI_ISL_2835328, EPI_ISL_2835329, EPI_ISL_2835330, EPI_ISL_2835331, EPI_ISL_2835332, EPI_ISL_2835333, EPI_ISL_2835334, EPI_ISL_2835335, EPI_ISL_2835336, EPI_ISL_2835337, EPI_ISL_2835338, EPI_ISL_2835339, EPI_ISL_2835340, EPI_ISL_2835341, EPI_ISL_2835342, EPI_ISL_2835343, EPI_ISL_2835344, EPI_ISL_2835345, EPI_ISL_2835346, EPI_ISL_2835347, EPI_ISL_2835348, EPI_ISL_2835349, EPI_ISL_2835351, EPI_ISL_2835352, EPI_ISL_2835353, EPI_ISL_2835354, EPI_ISL_2835355, EPI_ISL_2835356, EPI_ISL_2835357, EPI_ISL_2835358, EPI_ISL_2835359, EPI_ISL_2835360, EPI_ISL_2835361, EPI_ISL_2835362, EPI_ISL_2835363, EPI_ISL_2835365, EPI_ISL_2835366, EPI_ISL_2835371, EPI_ISL_2835372, EPI_ISL_2835373, EPI_ISL_2835375, EPI_ISL_2835376, EPI_ISL_2835377, EPI_ISL_2835378, EPI_ISL_2835379, EPI_ISL_2835406, EPI_ISL_2835418, EPI_ISL_2835419 | | | |
| see above | Instituto de Diagnostico y Referencia Epidemiologicos (InDRE) | Andersen lab at Scripps Research | SEARCH Alliance with Gisela Barrera Badillo, Jose Ernesto Ramirez Gonzalez, Maria Natividad Cruz Ortiz, Nervain Benjamin Contreras Gonzalez, Tatiana Ernestina Nunez Garcia, Dayanira Sarith Arellano Suarez, Irma Lopez Martinez, Rita Flores Leon, Beatriz Olivares Flores |

| | | | |
|---|---|---|---|
| EPI_ISL_2835566 | Sharp HealthCare Laboratory | Andersen lab at Scripps Research | SEARCH Alliance San Diego with Aaron Harding, Jacquelyn Berumen, Cathy Woerle, Liam McGinnis, Art Mendoza, Omid Bakhtar |
| EPI_ISL_2877292, EPI_ISL_2877293 | Hospital | National Reference Center for Viruses of Respiratory Infections, Institut Pasteur, Paris | Marion Barbet, Sylvie Behillil, Méline Bizard, Angela Brisebarre, Camille Capel, Vincent Enouf, Louise Lefrançois, Frédéric Lemoine, Christophe Malabat, Corinne Maufrais, Etienne Simon-Lorière, Maud Vanpeene, Sylvie Van der Werf |
| EPI_ISL_2877294, EPI_ISL_2877295, EPI_ISL_2877296, EPI_ISL_2877297 | Hospital | National Reference Center for Viruses of Respiratory Infections, Institut Pasteur, Paris | Marion Barbet, Sylvie Behillil, Méline Bizard, Angela Brisebarre, Camille Capel, Vincent Enouf, Louise Lefrançois, Frédéric Lemoine, Christophe Malabat, Corinne Maufrais, Etienne Simon-Lorière, Maud Vanpeene, Sylvie Van der Werf ,Olivia Raulin |
| EPI_ISL_2877298 | Hospital | National Reference Center for Viruses of Respiratory Infections, Institut Pasteur, Paris | Marion Barbet, Sylvie Behillil, Méline Bizard, Angela Brisebarre, Camille Capel, Vincent Enouf, Louise Lefrançois, Frédéric Lemoine, Christophe Malabat, Corinne Maufrais, Etienne Simon-Lorière, Maud Vanpeene, Sylvie Van der Werf |
| EPI_ISL_2877299, EPI_ISL_2877300 | EHPAD_IRA | National Reference Center for Viruses of Respiratory Infections, Institut Pasteur, Paris | Marion Barbet, Sylvie Behillil, Méline Bizard, Angela Brisebarre, Camille Capel, Vincent Enouf, Louise Lefrançois, Frédéric Lemoine, Christophe Malabat, Corinne Maufrais, Etienne Simon-Lorière, Maud Vanpeene, Sylvie Van der Werf |
| EPI_ISL_2877301, EPI_ISL_2877302, EPI_ISL_2877303 | Hospital | National Reference Center for Viruses of Respiratory Infections, Institut Pasteur, Paris | Marion Barbet, Sylvie Behillil, Méline Bizard, Angela Brisebarre, Camille Capel, Vincent Enouf, Louise Lefrançois, Frédéric Lemoine, Christophe Malabat, Corinne Maufrais, Etienne Simon-Lorière, Maud Vanpeene, Sylvie Van der Werf |
| EPI_ISL_2877304, EPI_ISL_2877305, EPI_ISL_2877306, EPI_ISL_2877307, EPI_ISL_2877308, EPI_ISL_2877309, EPI_ISL_2877310, EPI_ISL_2877311, EPI_ISL_2877312 | EHPAD_IRA | National Reference Center for Viruses of Respiratory Infections, Institut Pasteur, Paris | Marion Barbet, Sylvie Behillil, Méline Bizard, Angela Brisebarre, Camille Capel, Vincent Enouf, Louise Lefrançois, Frédéric Lemoine, Christophe Malabat, Corinne Maufrais, Etienne Simon-Lorière, Maud Vanpeene, Sylvie Van der Werf |
| EPI_ISL_2877313, EPI_ISL_2877314, EPI_ISL_2877315, EPI_ISL_2877316 | Hospital | National Reference Center for Viruses of Respiratory Infections, Institut Pasteur, Paris | Marion Barbet, Sylvie Behillil, Méline Bizard, Angela Brisebarre, Camille Capel, Vincent Enouf, Louise Lefrançois, Frédéric Lemoine, Christophe Malabat, Corinne Maufrais, Etienne Simon-Lorière, Maud Vanpeene, Sylvie Van der Werf ,Pascale Martres |
| EPI_ISL_2877317, EPI_ISL_2877318, EPI_ISL_2877319 | Hospital | National Reference Center for Viruses of Respiratory Infections, Institut Pasteur, Paris | Marion Barbet, Sylvie Behillil, Méline Bizard, Angela Brisebarre, Camille Capel, Vincent Enouf, Louise Lefrançois, Frédéric Lemoine, Christophe Malabat, Corinne Maufrais, Etienne Simon-Lorière, Maud Vanpeene, Sylvie Van der Werf ,Olivia Raulin |
| EPI_ISL_2877320, EPI_ISL_2877321, EPI_ISL_2877322, EPI_ISL_2877323, EPI_ISL_2877324, EPI_ISL_2877325, EPI_ISL_2877326, EPI_ISL_2877327, EPI_ISL_2877328, EPI_ISL_2877329, EPI_ISL_2877330, EPI_ISL_2877331 | | | |
| see above | Hospital | National Reference Center for Viruses of Respiratory Infections, Institut Pasteur, Paris | Marion Barbet, Sylvie Behillil, Méline Bizard, Angela Brisebarre, Camille Capel, Vincent Enouf, Louise Lefrançois, Frédéric Lemoine, Christophe Malabat, Corinne Maufrais, Etienne Simon-Lorière, Maud Vanpeene, Sylvie Van der Werf |
| EPI_ISL_2877332 | Hospital | National Reference Center for Viruses of Respiratory Infections, Institut Pasteur, Paris | Marion Barbet, Sylvie Behillil, Méline Bizard, Angela Brisebarre, Camille Capel, Vincent Enouf, Louise Lefrançois, Frédéric Lemoine, Christophe Malabat, Corinne Maufrais, Etienne Simon-Lorière, Maud Vanpeene, Sylvie Van der Werf ,S Kouhil |
| EPI_ISL_2877333, EPI_ISL_2877334, EPI_ISL_2877335, EPI_ISL_2877336, EPI_ISL_2877337, EPI_ISL_2877338, EPI_ISL_2877339, EPI_ISL_2877340 | Hospital | National Reference Center for Viruses of Respiratory Infections, Institut Pasteur, Paris | Marion Barbet, Sylvie Behillil, Méline Bizard, Angela Brisebarre, Camille Capel, Vincent Enouf, Louise Lefrançois, Frédéric Lemoine, Christophe Malabat, Corinne Maufrais, Etienne Simon-Lorière, Maud Vanpeene, Sylvie Van der Werf ,Olivia Raulin |
| EPI_ISL_2877341, EPI_ISL_2877342, EPI_ISL_2877343, EPI_ISL_2877344, EPI_ISL_2877345, EPI_ISL_2877346, EPI_ISL_2877347 | Hospital | National Reference Center for Viruses of Respiratory Infections, Institut Pasteur, Paris | Marion Barbet, Sylvie Behillil, Méline Bizard, Angela Brisebarre, Camille Capel, Vincent Enouf, Louise Lefrançois, Frédéric Lemoine, Christophe Malabat, Corinne Maufrais, Etienne Simon-Lorière, Maud Vanpeene, Sylvie Van der Werf |
| EPI_ISL_2877348, EPI_ISL_2877349, EPI_ISL_2877350, EPI_ISL_2877351, EPI_ISL_2877353 | EHPAD_IRA | National Reference Center for Viruses of Respiratory Infections, Institut Pasteur, Paris | Marion Barbet, Sylvie Behillil, Méline Bizard, Angela Brisebarre, Camille Capel, Vincent Enouf, Louise Lefrançois, Frédéric Lemoine, Christophe Malabat, Corinne Maufrais, Etienne Simon-Lorière, Maud Vanpeene, Sylvie Van der Werf |
| EPI_ISL_2877355, EPI_ISL_2877357, EPI_ISL_2877359, EPI_ISL_2877361, EPI_ISL_2877363, EPI_ISL_2877364, EPI_ISL_2877366, EPI_ISL_2877368 | Hospital | National Reference Center for Viruses of Respiratory Infections, Institut Pasteur, Paris | Marion Barbet, Sylvie Behillil, Méline Bizard, Angela Brisebarre, Camille Capel, Vincent Enouf, Louise Lefrançois, Frédéric Lemoine, Christophe Malabat, Corinne Maufrais, Etienne Simon-Lorière, Maud Vanpeene, Sylvie Van der Werf |
| EPI_ISL_2877370 | EHPAD_IRA | National Reference Center for Viruses of Respiratory Infections, Institut Pasteur, Paris | Marion Barbet, Sylvie Behillil, Méline Bizard, Angela Brisebarre, Camille Capel, Vincent Enouf, Louise Lefrançois, Frédéric Lemoine, Christophe Malabat, Corinne Maufrais, Etienne Simon-Lorière, Maud Vanpeene, Sylvie Van der Werf |
| EPI_ISL_2877371, EPI_ISL_2877373, EPI_ISL_2877375, EPI_ISL_2877377, EPI_ISL_2877379, EPI_ISL_2877381, EPI_ISL_2877383 | Hospital | National Reference Center for Viruses of Respiratory Infections, Institut Pasteur, Paris | Marion Barbet, Sylvie Behillil, Méline Bizard, Angela Brisebarre, Camille Capel, Vincent Enouf, Louise Lefrançois, Frédéric Lemoine, Christophe Malabat, Corinne Maufrais, Etienne Simon-Lorière, Maud Vanpeene, Sylvie Van der Werf ,Patricia Barbut |
| EPI_ISL_2877384, EPI_ISL_2877386, EPI_ISL_2877388 | Hospital | National Reference Center for Viruses of Respiratory Infections, Institut Pasteur, Paris | Marion Barbet, Sylvie Behillil, Méline Bizard, Angela Brisebarre, Camille Capel, Vincent Enouf, Louise Lefrançois, Frédéric Lemoine, Christophe Malabat, Corinne Maufrais, Etienne Simon-Lorière, Maud Vanpeene, Sylvie Van der Werf ,Vincent Esteve |
| EPI_ISL_2877390, EPI_ISL_2877392, EPI_ISL_2877394, EPI_ISL_2877395, EPI_ISL_2877397, EPI_ISL_2877399, EPI_ISL_2877401, EPI_ISL_2877403, EPI_ISL_2877405, EPI_ISL_2877407, EPI_ISL_2877408, EPI_ISL_2877410, EPI_ISL_2877412, EPI_ISL_2877414, EPI_ISL_2877416, EPI_ISL_2877418, EPI_ISL_2877420, EPI_ISL_2877422 | | | |
| see above | Hospital | National Reference Center for Viruses of Respiratory Infections, Institut Pasteur, Paris | Marion Barbet, Sylvie Behillil, Méline Bizard, Angela Brisebarre, Camille Capel, Vincent Enouf, Louise Lefrançois, Frédéric Lemoine, Christophe Malabat, Corinne Maufrais, Etienne Simon-Lorière, Maud Vanpeene, Sylvie Van der Werf ,Olivia Raulin |
| EPI_ISL_2877423, EPI_ISL_2877425, EPI_ISL_2877427, EPI_ISL_2877429 | Hospital | National Reference Center for Viruses of Respiratory Infections, Institut Pasteur, Paris | Marion Barbet, Sylvie Behillil, Méline Bizard, Angela Brisebarre, Camille Capel, Vincent Enouf, Louise Lefrançois, Frédéric Lemoine, Christophe Malabat, Corinne Maufrais, Etienne Simon-Lorière, Maud Vanpeene, Sylvie Van der Werf ,Patricia Barbut |
| EPI_ISL_2877431, EPI_ISL_2877433, EPI_ISL_2877435, EPI_ISL_2877437, EPI_ISL_2877438 | EHPAD_IRA | National Reference Center for Viruses of Respiratory Infections, Institut Pasteur, Paris | Marion Barbet, Sylvie Behillil, Méline Bizard, Angela Brisebarre, Camille Capel, Vincent Enouf, Louise Lefrançois, Frédéric Lemoine, Christophe Malabat, Corinne Maufrais, Etienne Simon-Lorière, Maud Vanpeene, Sylvie Van der Werf |
| EPI_ISL_2877440 | Hospital | National Reference Center for Viruses of Respiratory Infections, Institut Pasteur, Paris | Marion Barbet, Sylvie Behillil, Méline Bizard, Angela Brisebarre, Camille Capel, Vincent Enouf, Louise Lefrançois, Frédéric Lemoine, Christophe Malabat, Corinne Maufrais, Etienne Simon-Lorière, Maud Vanpeene, Sylvie Van der Werf ,Patricia Barbut |
| EPI_ISL_2877442 | Hospital | National Reference Center for Viruses of Respiratory Infections, Institut Pasteur, Paris | Marion Barbet, Sylvie Behillil, Méline Bizard, Angela Brisebarre, Camille Capel, Vincent Enouf, Louise Lefrançois, Frédéric Lemoine, Christophe Malabat, Corinne Maufrais, Etienne Simon-Lorière, Maud Vanpeene, Sylvie Van der Werf ,Olivia Raulin |
| EPI_ISL_2877444 | Hospital | National Reference Center for Viruses of Respiratory Infections, Institut Pasteur, Paris | Marion Barbet, Sylvie Behillil, Méline Bizard, Angela Brisebarre, Camille Capel, Vincent Enouf, Louise Lefrançois, Frédéric Lemoine, Christophe Malabat, Corinne Maufrais, Etienne Simon-Lorière, Maud Vanpeene, Sylvie Van der Werf ,Vincent Esteve |
| EPI_ISL_2877446 | Hospital | National Reference Center for Viruses of Respiratory Infections, Institut Pasteur, Paris | Marion Barbet, Sylvie Behillil, Méline Bizard, Angela Brisebarre, Camille Capel, Vincent Enouf, Louise Lefrançois, Frédéric Lemoine, Christophe Malabat, Corinne Maufrais, Etienne Simon-Lorière, Maud Vanpeene, Sylvie Van der Werf ,Olivia Raulin |
| EPI_ISL_2877448, EPI_ISL_2877450, EPI_ISL_2877451, EPI_ISL_2877452, EPI_ISL_2877453 | Hospital | National Reference Center for Viruses of Respiratory Infections, Institut Pasteur, Paris | Marion Barbet, Sylvie Behillil, Méline Bizard, Angela Brisebarre, Camille Capel, Vincent Enouf, Louise Lefrançois, Frédéric Lemoine, Christophe Malabat, Corinne Maufrais, Etienne Simon-Lorière, Maud Vanpeene, Sylvie Van der Werf ,Patricia Barbut |
| EPI_ISL_2877454, EPI_ISL_2877455, EPI_ISL_2877456, EPI_ISL_2877457, EPI_ISL_2877458 | Hospital | National Reference Center for Viruses of Respiratory Infections, Institut Pasteur, Paris | Marion Barbet, Sylvie Behillil, Méline Bizard, Angela Brisebarre, Camille Capel, Vincent Enouf, Louise Lefrançois, Frédéric Lemoine, Christophe Malabat, Corinne Maufrais, Etienne Simon-Lorière, Maud Vanpeene, Sylvie Van der Werf ,Olivia Raulin |
| EPI_ISL_2877459 | EHPAD_IRA | National Reference Center for Viruses of Respiratory Infections, Institut Pasteur, Paris | Marion Barbet, Sylvie Behillil, Méline Bizard, Angela Brisebarre, Camille Capel, Vincent Enouf, Louise Lefrançois, Frédéric Lemoine, Christophe Malabat, Corinne Maufrais, Etienne Simon-Lorière, Maud Vanpeene, Sylvie Van der Werf |

| | | | |
|---|---|---|---|
| EPI_ISL_2877460 | Hospital | National Reference Center for Viruses of Respiratory Infections, Institut Pasteur, Paris | Marion Barbet, Sylvie Behillil, Méline Bizard, Angela Brisebarre, Camille Capel, Vincent Enouf, Louise Lefrançois, Frédéric Lemoine, Christophe Malabat, Corinne Maufrais, Etienne Simon-Lorière, Maud Vanpeene, Sylvie Van der Werf ,S Kouhil |
| EPI_ISL_2877461 | Hospital | National Reference Center for Viruses of Respiratory Infections, Institut Pasteur, Paris | Marion Barbet, Sylvie Behillil, Méline Bizard, Angela Brisebarre, Camille Capel, Vincent Enouf, Louise Lefrançois, Frédéric Lemoine, Christophe Malabat, Corinne Maufrais, Etienne Simon-Lorière, Maud Vanpeene, Sylvie Van der Werf ,Patricia Barbut |
| EPI_ISL_2877462, EPI_ISL_2877463, EPI_ISL_2877464 | Hospital | National Reference Center for Viruses of Respiratory Infections, Institut Pasteur, Paris | Marion Barbet, Sylvie Behillil, Méline Bizard, Angela Brisebarre, Camille Capel, Vincent Enouf, Louise Lefrançois, Frédéric Lemoine, Christophe Malabat, Corinne Maufrais, Etienne Simon-Lorière, Maud Vanpeene, Sylvie Van der Werf ,Olivia Raulin |
| EPI_ISL_2877465, EPI_ISL_2877466 | EHPAD_IRA | National Reference Center for Viruses of Respiratory Infections, Institut Pasteur, Paris | Marion Barbet, Sylvie Behillil, Méline Bizard, Angela Brisebarre, Camille Capel, Vincent Enouf, Louise Lefrançois, Frédéric Lemoine, Christophe Malabat, Corinne Maufrais, Etienne Simon-Lorière, Maud Vanpeene, Sylvie Van der Werf |
| EPI_ISL_2877467 | Sentinelles Province | National Reference Center for Viruses of Respiratory Infections, Institut Pasteur, Paris | Marion Barbet, Sylvie Behillil, Méline Bizard, Angela Brisebarre, Camille Capel, Vincent Enouf, Louise Lefrançois, Frédéric Lemoine, Christophe Malabat, Corinne Maufrais, Etienne Simon-Lorière, Maud Vanpeene, Sylvie Van der Werf ,LaurèNe Millet-Malingrey |
| EPI_ISL_2877468 | Sentinelles Province | National Reference Center for Viruses of Respiratory Infections, Institut Pasteur, Paris | Marion Barbet, Sylvie Behillil, Méline Bizard, Angela Brisebarre, Camille Capel, Vincent Enouf, Louise Lefrançois, Frédéric Lemoine, Christophe Malabat, Corinne Maufrais, Etienne Simon-Lorière, Maud Vanpeene, Sylvie Van der Werf ,Isabelle Husson |
| EPI_ISL_2877469, EPI_ISL_2877470, EPI_ISL_2877471 | Hospital | National Reference Center for Viruses of Respiratory Infections, Institut Pasteur, Paris | Marion Barbet, Sylvie Behillil, Méline Bizard, Angela Brisebarre, Camille Capel, Vincent Enouf, Louise Lefrançois, Frédéric Lemoine, Christophe Malabat, Corinne Maufrais, Etienne Simon-Lorière, Maud Vanpeene, Sylvie Van der Werf ,Patricia Barbut |
| EPI_ISL_2877472 | Hospital | National Reference Center for Viruses of Respiratory Infections, Institut Pasteur, Paris | Marion Barbet, Sylvie Behillil, Méline Bizard, Angela Brisebarre, Camille Capel, Vincent Enouf, Louise Lefrançois, Frédéric Lemoine, Christophe Malabat, Corinne Maufrais, Etienne Simon-Lorière, Maud Vanpeene, Sylvie Van der Werf ,Vincent Esteve |
| EPI_ISL_2877473, EPI_ISL_2877474, EPI_ISL_2877475, EPI_ISL_2877476, EPI_ISL_2877477, EPI_ISL_2877478, EPI_ISL_2877479, EPI_ISL_2877480, EPI_ISL_2877481, EPI_ISL_2877482, EPI_ISL_2877483, EPI_ISL_2877484, EPI_ISL_2877485, EPI_ISL_2877486 | | | |
| see above | Hospital | National Reference Center for Viruses of Respiratory Infections, Institut Pasteur, Paris | Marion Barbet, Sylvie Behillil, Méline Bizard, Angela Brisebarre, Camille Capel, Vincent Enouf, Louise Lefrançois, Frédéric Lemoine, Christophe Malabat, Corinne Maufrais, Etienne Simon-Lorière, Maud Vanpeene, Sylvie Van der Werf ,Olivia Raulin |
| EPI_ISL_2877487 | Sentinelles Idf | National Reference Center for Viruses of Respiratory Infections, Institut Pasteur, Paris | Marion Barbet, Sylvie Behillil, Méline Bizard, Angela Brisebarre, Camille Capel, Vincent Enouf, Louise Lefrançois, Frédéric Lemoine, Christophe Malabat, Corinne Maufrais, Etienne Simon-Lorière, Maud Vanpeene, Sylvie Van der Werf ,Christine Aubry |
| EPI_ISL_2877488, EPI_ISL_2877489, EPI_ISL_2877490, EPI_ISL_2877491, EPI_ISL_2877492, EPI_ISL_2877493, EPI_ISL_2877494, EPI_ISL_2877495, EPI_ISL_2877496, EPI_ISL_2877497 | Hospital | National Reference Center for Viruses of Respiratory Infections, Institut Pasteur, Paris | Marion Barbet, Sylvie Behillil, Méline Bizard, Angela Brisebarre, Camille Capel, Vincent Enouf, Louise Lefrançois, Frédéric Lemoine, Christophe Malabat, Corinne Maufrais, Etienne Simon-Lorière, Maud Vanpeene, Sylvie Van der Werf ,Ilka Engelmann |
| EPI_ISL_402124 | Wuhan Jinyintan Hospital | Wuhan Institute of Virology, Chinese Academy of Sciences | Peng Zhou, Xing-Lou Yang, Ding-Yu Zhang, Lei Zhang, Yan Zhu, Hao-Rui Si, Zhengli Shi |
| EPI_ISL_402126 | Dept. of Virology III, National Institute of Infectious Diseases | Dept. of Virology III, National Institute of Infectious Diseases | Naganori Nao, Kazuya Shirato, Shutoku Matsuyama, Makoto Takeda |
| EPI_ISL_402127, EPI_ISL_402128, EPI_ISL_402129, EPI_ISL_402130 | Wuhan Jinyintan Hospital | Wuhan Institute of Virology, Chinese Academy of Sciences | Peng Zhou, Xing-Lou Yang, Ding-Yu Zhang, Lei Zhang, Yan Zhu, Hao-Rui Si, Zhengli Shi |
| EPI_ISL_402132 | Wuhan Jinyintan Hospital | Hubei Provincial Center for Disease Control and Prevention | Bin Fang, Xiang Li, Xiao Yu, Linlin Liu, Bo Yang, Faxian Zhan, Guojun Ye, Xixiang Huo, Junqiang Xu, Bo Yu, Kun Cai, Jing Li, Yongzhong Jiang. |
| EPI_ISL_403962, EPI_ISL_403963 | Bamrasnaradura Hospital | 1. Department of Medical Sciences, Ministry of Public Health, Thailand 2. Thai Red Cross Emerging Infectious Diseases - Health Science Centre 3. Department of Disease Control, Ministry of Public Health, Thailand | Pilailuk,Okada; Siripaporn,Phuygun; Thanutsapa,Thanadachakul; Supaporn,Wacharapluesadee; Sittiporn,Parnmen; Warawan,Wongboot; Sunthareeya,Waicharoen; Rome,Buathong; Malinee,Chittaganpitch; Nanthawan,Mekha |
| EPI_ISL_404227 | Zhejiang Provincial Center for Disease Control and Prevention | Department of Microbiology, Zhejiang Provincial Center for Disease Control and Prevention | Yin Chen, Yanjun Zhang, Haiyan Mao, Junhang Pan, Xiuyu Lou, Yiyu Lu, Juying Yan, Hanping Zhu, Jian Gao, Yan Feng, Yi Sun, Hao Yan, Zhen Li, Yisheng Sun, Liming Gong, Qiong Ge, Wen Shi, Xinying Wang, Wenwu Yao, Zhangnv Yang, Fang Xu, Chen Chen, Enfu Chen, Zhen Wang, Zhiping Chen, Jianmin Jiang, Chonggao Hu |
| EPI_ISL_404228 | Zhejiang Provincial Center for Disease Control and Prevention | Department of Microbiology, Zhejiang Provincial Center for Disease Control and Prevention | Yanjun Zhang, Yin Chen, Haiyan Mao, Junhang Pan, Xiuyu Lou, Yiyu Lu, Juying Yan, Hanping Zhu, Jian Gao, Yan Feng, Yi Sun, Hao Yan, Zhen Li, Yisheng Sun, Liming Gong, Qiong Ge, Wen Shi, Xinying Wang, Wenwu Yao, Zhangnv Yang, Fang Xu, Chen Chen, Enfu Chen, Zhen Wang, Zhiping Chen, Jianmin Jiang, Chonggao Hu |
| EPI_ISL_406592 | Shenzhen Third People's Hospital | Shenzhen Key Laboratory of Pathogen and Immunity, National Clinical Research Center for Infectious Disease,Shenzhen Third People's Hospital | Yang Yang, Chenguang Shen, Li Xing, Zhixiang Xu, Haixia Zheng, Yingxia Liu |
| EPI_ISL_406593, EPI_ISL_406594, EPI_ISL_406595 | Shenzhen Key Laboratory of Pathogen and Immunity, National Clinical Research Center for Infectious Disease, Shenzhen Third People's Hospital | Shenzhen Key Laboratory of Pathogen and Immunity, National Clinical Research Center for Infectious Disease, Shenzhen Third People's Hospital | Yang Yang, Chenguang Shen, Li Xing, Zhixiang Xu, Haixia Zheng, Yingxia Liu |
| EPI_ISL_406862 | Charité Universitätsmedizin Berlin, Institute of Virology; Institut für Mikrobiologie der Bundeswehr, Munich | Charité Universitätsmedizin Berlin, Institute of Virology | Victor M Corman, Julia Schneider, Talitha Veith, Barbara Mühlemann, Markus Antwerpen, Christian Drosten, Roman Wölfel |
| EPI_ISL_406970 | Hangzhou Center for Disease and Control Microbiology Lab | Hangzhou Center for Disease and Control Microbiology Lab | Yu Hua, Wang Haoqiu, Li Jun, Yu Xinfeng |
| EPI_ISL_407079 | Lapland Central Hospital | Department of Virology, University of Helsinki and Helsinki University Hospital, Helsinki, Finland | Teemu Smura, Suvi Kuivanen, Hannimari Kallio-Kokko, Olli Vapalahti |
| EPI_ISL_407313 | Hangzhou Center for Disease Control and Prevention | Hangzhou Center for Disease Control and Prevention | Jun Li, Haoqiu Wang, Hua Yu, Lingfeng Mao, Xinfen Yu, Zhou Sun, Qingxin Kong, Xin Qian, Shuchang Chen, Xuchu Wang |
| EPI_ISL_407893 | Centre for Infectious Diseases and Microbiology Laboratory Services | NSW Health Pathology - Institute of Clinical Pathology and Medical Research; Westmead Hospital; University of Sydney | Eden J-S, Carter I, Rahman H, Holmes EC, Rockett R, O'Sullivan MV, Sintchenko V, Chen SC, Maddocks S, Kok J and Dwyer DE for the 2019-nCoV Study Group |
| EPI_ISL_407976 | KU Leuven, Clinical and Epidemiological Virology | KU Leuven, Clinical and Epidemiological Virology | Bert Vanmechelen, Elke Wollants, Annabel Rector, Els Keyaerts, Lies Laenen, Marc Van Ranst, and Piet Maes |
| EPI_ISL_408068 | Virology Laboratory National Institute for Infectious Diseases 'Lazzaro Spallanzani' IRCCS | Virology Laboratory National Institute for Infectious Diseases 'Lazzaro Spallanzani' IRCCS | Capobianchi,M.R., Carletti,F., Lalle,E., Bordi,L., Marsella,P.,Colavita,F., Matusali,G., Nicastri,E., Ippolito,G. and Castilletti,C. |
| EPI_ISL_408430 | Department of Infectious and Tropical Diseases, Bichat Claude Bernard Hospital, Paris | National Reference Center for Viruses of Respiratory Infections, Institut Pasteur, Paris | Mélanie Albert, Marion Barbet, Sylvie Behillil, Méline Bizard, Angela Brisebarre, Flora Donati, Vincent Enouf, Maud Vanpeene, Sylvie van der Werf, Yazdan Yazdanpanah, Xavier Lescure |
| EPI_ISL_408431 | Sorbonne Université, Inserm et Assistance Publique-Hôpitaux de Paris (Pitié Salpétrière) | National Reference Center for Viruses of Respiratory Infections, Institut Pasteur, Paris | Mélanie Albert, Marion Barbet, Sylvie Behillil, Méline Bizard, Angela Brisebarre, Flora Donati, Vincent Enouf, Maud Vanpeene, Sylvie van der Werf, Sonia Burrel, Anne-Geneviève Marcelin, Vincent Calvez, David Boutolleau, Elise Klément, Valérie Pourcher, Eric Caumes. |
| EPI_ISL_408480 | National Institute for Viral Disease Control and Prevention, China CDC | National Institute for Viral Disease Control & Prevention, CCDC | Wenjie TanXiaoqing FuXiang ZhaoWenling Wang Peihua NiuRoujian Lu,Yanhong SunBaoying HuangLi ZhaoFei YeWenbo XuGeorge F. GaoGuizhen Wu |
| EPI_ISL_408481 | National Institute for Viral Disease Control and Prevention, China CDC | National Institute for Viral Disease Control & Prevention, CCDC | Wenjie Tan, Hengqin Wang, Xiang Zhao, Wenling Wang, Peihua Niu, Roujian Lu, Sheng Ye, Baoying Huang, Li Zhao, Fei Ye, Wenbo Xu, George F. Gao, Guizhen Wu |
| EPI_ISL_408482 | National Institute for Viral Disease Control and Prevention, China CDC | National Institute for Viral Disease Control & Prevention, CCDC | Wenjie Tan, Zhaoguo Wang, Xiang Zhao, Wenling Wang, Peihua Niu, Roujian Lu, Ti Liu, Baoying Huang, Li Zhao, Fei Ye, Wenbo Xu, George F. Gao, Guizhen Wu |
| EPI_ISL_408484 | National Institute for Viral Disease Control and Prevention, China CDC | National Institute for Viral Disease Control & Prevention, CCDC | Wenjie Tan, Jianan Xu, Wenling Wang, Peihua Niu, Roujian Lu, Huiping Yang, Xiang Zhao, Baoying Huang, Li Zhao, Fei Ye, Wenbo Xu, George F. Gao, Guizhen Wu |

| | | | |
|---|---|---|---|
| EPI_ISL_408485 | National Institute for Viral Disease Control and Prevention, China CDC | National Institute for Viral Disease Control & Prevention, CCDC | Wenjie Tan,Quanyi Wang,Wenling Wang, Peihua Niu,Roujian Lu,Yang Pan,Xiang Zhao,Baoying Huang,Li Zhao,Fei Ye,Wenbo Xu,George F. Gao,Guizhen Wu |
| EPI_ISL_408486 | National Institute for Viral Disease Control and Prevention, China CDC | National Institute for Viral Disease Control & Prevention, CCDC | Wenjie Tan, Yong Shi, Wenling Wang, Peihua Niu, Roujian Lu, Jianxiong Li, Xiang Zhao, Baoying Huang, Li Zhao, Fei Ye, Wenbo Xu, George F. Gao, Guizhen Wu |
| EPI_ISL_408488 | National Institute for Viral Disease Control and Prevention, China CDC | National Institute for Viral Disease Control & Prevention, CCDC | Wenjie Tan, Shenjiao Wang, Wenling Wang, Peihua Niu, Roujian Lu, Kangchen Zhao, Xiang Zhao, Baoying Huang, Li Zhao, Fei Ye, Wenbo Xu, George F. Gao, Guizhen Wu |
| EPI_ISL_408489 | Department of Laboratory Medicine, National Taiwan University Hospital | Microbial Genomics Core Lab, National Taiwan University Centers of Genomic and Precision Medicine | Shiou-Hwei Yeh, You-Yu Lin, Ya-Yun Lai, Chiao-Ling Li, Shan-Chwen Chang, Pei-Jer Chen, Sui-Yuan Chang |
| EPI_ISL_408666, EPI_ISL_408669 | Dept. of Virology III, National Institute of Infectious Diseases | Pathogen Genomics Center, National Institute of Infectious Diseases | Tsuyoshi Sekizuka, Shutoku Matsuyama, Naganori Nao, Kazuya Shirato, Makoto Takeda, Makoto Kuroda |
| EPI_ISL_408977 | Serology, Virology and OTDS Laboratories (SAViD), NSW Health Pathology Randwick | NSW Health Pathology - Institute of Clinical Pathology and Medical Research; Centre for Infectious Diseases and Microbiology Laboratory Services; Westmead Hospital; University of Sydney | Eden J-S, Carter I, Rahman H, Rawlinson W, Holmes EC, Rockett R, O'Sullivan MV, Sintchenko V, Chen SC, Maddocks S, Kok J and Dwyer DE for the 2019-nCoV Study Group* |
| EPI_ISL_410218 | Department of Laboratory Medicine, National Taiwan University Hospital | Microbial Genomics Core Lab, National Taiwan University Centers of Genomic and Precision Medicine | Shiou-Hwei Yeh, You-Yu Lin, Ya-Yun Lai, Chiao-Ling Li, Shan-Chwen Chang, Pei-Jer Chen, Sui-Yuan Chang |
| EPI_ISL_410301 | National Influenza Centre, National Public Health Laboratory, Kathmandu, Nepal | The University of Hong Kong | Ranjit Sah , Runa Jha, Daniel Chu, Haogao Gu, Malik Peiris, Anup Bastola, Alfonso J. Rodriguez-Morales, Bibek Kumar Lal, Basu Dev Pandey, Leo Poon |
| EPI_ISL_410531, EPI_ISL_410532 | Dept. of Pathology, National Institute of Infectious Diseases | Pathogen Genomics Center, National Institute of Infectious Diseases | Tsuyoshi Sekizuka, Harutaka Katano, Shutoku Matsuyama, Naganori Nao, Kazuya Shirato, Motoi Suzuki, Hideki Hasegawa, Takaji Wakita, Makoto Takeda, Tadaki Suzuki, Makoto Kuroda |
| EPI_ISL_410545 | INMI Lazzaro Spallanzani IRCCS | Laboratory of Virology, INMI Lazzaro Spallanzani IRCCS | Maria R. Capobianchi, Cesare E. M. Gruber, Martina Rueca, Barbara Bartolini, Francesco Messina, Emanuela Giombini, Francesca Colavita, Concetta Castilletti, Eleonora Lalle, Fabrizio Carletti, Emanuele Nicastri, Giuseppe Ippolito. |
| EPI_ISL_410546 | INMI Lazzaro Spallanzani IRCCS | Laboratory of Virology, INMI Lazzaro Spallanzani IRCCS | Maria R. Capobianchi, Cesare E. M. Gruber, Martina Rueca, Fabrizio Carletti, Barbara Bartolini, Francesco Messina, Emanuela Giombini, Francesca Colavita, Concetta Castilletti, Eleonora Lalle, Emanuele Nicastri, Giuseppe Ippolito. |
| EPI_ISL_410984 | Department of Infectious and Tropical Diseases, Bichat Claude Bernard Hospital, Paris | National Reference Center for Viruses of Respiratory Infections, Institut Pasteur, Paris | Mélanie Albert, Marion Barbet, Sylvie Behillil, Méline Bizard, Angela Brisebarre, Flora Donati, Vincent Enouf, Maud Vanpeene, Sylvie van der Werf, Yazdan Yazdanpanah, Xavier Lescure |
| EPI_ISL_411218 | Department of Infectious and Tropical Diseases, Bichat Claude Bernard Hospital, Paris | Laboratoire Virpath, CIRI U111, UCBL1, INSERM, CNRS, ENS Lyon | Olivier Terrier, Aurélien Traversier, Julien Fouret, Yazdan Yazdanpanah, Xavier Lescure, Catherine Legras-Lachuer, Alexandre Gaymard, Bruno Lina, Manuel Rosa-Calatrava |
| EPI_ISL_411219, EPI_ISL_411220 | Department of Infectious and Tropical Diseases, Bichat Claude Bernard Hospital, Paris | Laboratoire Virpath, CIRI U111, UCBL1, INSERM, CNRS, ENS Lyon | Olivier Terrier, Aurélien Traversier, Julien Fouret, Yazdan Yazdanpanah, Xavier Lescure, Alexandre Gaymard, Bruno Lina, Manuel Rosa-Calatrava |
| EPI_ISL_411902 | Virology Unit, Institut Pasteur du Cambodge. | Virology Unit, Institut Pasteur du Cambodge (Sequencing done by: Jessica E Manning/Jennifer A Bohl at Malaria and Vector Research Research Laboratory, National Institute of Allergy and Infectious Diseases and Vida Ahyong from Chan-Zuckerberg Biohub) | Erik A Karlsson, Jennifer A Bohl, Vida Ahyong, Veasna Duong, Philippe Dussart, Jessica E Manning. |
| EPI_ISL_412386 | Beijing Ditan Hospital, Capital Medical University | National Institute for Communicable Disease Control and Prevention, Chinese Center for Disease Control and Prevention | Xinmin Xu, Xin Lu, Pan Xiang, Haijian Zhou, Biao Kan, Yajie Wang, Jingyuan Liu, Yanwen Xiong, Huizhu Wang, Ruihong Li, Fangfang Jin, Jie Gong, Xiaoping Chen, Lili Gao, Haofeng Xiong, Lin Pu, Chuansheng Li, Ming Zhang, Jianbo Tan, Yao Sun, Yufeng Liu, Hebing Guo, Jingjing Hao |
| EPI_ISL_412459 | Jingzhou Center for Disease Control and Prevention | Hubei Provincial Center for Disease Control and Prevention | Bin Fang, Xiang Li, Xiao Yu, Linlin Liu, Bo Yang, Faxian Zhan, Guojun Ye, Xixiang Huo, Junqiang Xu, Bo Yu, Kun Cai, Jing Li, Maoyi Chen,Jie Hu, Chunlin Mao, Yongzhong Jiang. |
| EPI_ISL_412964 | Hospital Israelita Albert Einstein | Instituto Adolfo Lutz Interdisciplinary Procedures Center Strategic Laboratory | Jaqueline Goes de Jesus, Claudio Tavares Sacchi, Daniela Bernardes Borges da Silva, Ingra Morales Claro, Flávia Cristina da Silva Sales, Claudia Regina Gonçalves, Joshua Quick, Maria do Carmo, Sampaio Tavares Timenetsky, Nicholas James Loman, Andrew Rambaut, Ester Cerdeira Sabino, Nuno Rodrigues Faria |
| EPI_ISL_412970 | Washington State Department of Health | Seattle Flu Study | Helen Chu, Michael Boeckh, Janet Englund, Michael Famulare, Barry Lutz, Deborah Nickerson, Mark Rieder, Lea Starita, Matthew Thompson, Jay Shendure, and Trevor Bedford |
| EPI_ISL_412972 | Instituto Nacional de Enfermedades Respiratorias | Instituto de Diagnostico y Referencia Epidemiologicos (INDRE) | Ramirez-Gonzalez Ernesto, Garces-Ayala Fabiola, Araiza-Rodriguez Adnan, Mendieta-Condado Edgar, Rodriguez-Maldonado Abril, Wong-Arambula Claudia, Vazquez-Perez Joel, Martinez Arturo, Boukadida Celia, Munoz-Medina Esteban, Sanchez Alejandro, Isa Pavel, Taboada Blanca, Lopez Susana, Arias Carlos, Barrera-Badillo Gisela, Hernandez-Rivas Lucia, Lopez-Martinez Irma |
| EPI_ISL_412975 | Centre for Infectious Diseases and Microbiology Laboratory Services | NSW Health Pathology - Institute of Clinical Pathology and Medical Research; Westmead Hospital; University of Sydney | Eden J-S, Carter I, Rahman H, Holmes EC, Rockett R, O'Sullivan MV, Sintchenko V, Chen SC, Maddocks S, Kok J and Dwyer DE for the 2019-nCoV Study Group |
| EPI_ISL_412978 | The Central Hospital Of Wuhan | Hubei Provincial Center for Disease Control and Prevention | Bin Fang, Xiang Li, Xiao Yu, Linlin Liu, Bo Yang, Faxian Zhan, Guojun Ye, Xixiang Huo, Junqiang Xu, Bo Yu, Kun Cai, Jing Li, Yongzhong Jiang. |
| EPI_ISL_412979, EPI_ISL_412980 | Union Hospital of Tongji Medical College, Huazhong University of Science and Technology | Hubei Provincial Center for Disease Control and Prevention | Bin Fang, Xiang Li, Xiao Yu, Linlin Liu, Bo Yang, Faxian Zhan, Guojun Ye, Xixiang Huo, Junqiang Xu, Bo Yu, Kun Cai, Jing Li, Yongzhong Jiang. |
| EPI_ISL_412981 | CR&WISCO GENERAL HOSPITAL | Hubei Provincial Center for Disease Control and Prevention | Bin Fang, Xiang Li, Xiao Yu, Linlin Liu, Bo Yang, Faxian Zhan, Guojun Ye, Xixiang Huo, Junqiang Xu, Bo Yu, Kun Cai, Jing Li, Yongzhong Jiang. |
| EPI_ISL_412982 | Wuhan Lung Hospital | Hubei Provincial Center for Disease Control and Prevention | Bin Fang, Xiang Li, Xiao Yu, Linlin Liu, Bo Yang, Faxian Zhan, Guojun Ye, Xixiang Huo, Junqiang Xu, Bo Yu, Kun Cai, Jing Li, Yongzhong Jiang. |
| EPI_ISL_412983 | Tianmen Center for Disease Control and Prevention | Hubei Provincial Center for Disease Control and Prevention | Bin Fang, Xiang Li, Xiao Yu, Linlin Liu, Bo Yang, Faxian Zhan, Guojun Ye, Xixiang Huo, Junqiang Xu, Bo Yu, Kun Cai, Jing Li, YiFa Zhu, Yangyang Tao,Xierong Li,Yongzhong Jiang. |
| EPI_ISL_413016 | Hospital Israelita Albert Einstein | Instituto Adolfo Lutz, Interdisciplinary Procedures Center, Strategic Laboratory | Jaqueline Goes de Jesus, Claudio Tavares Sacchi, Fabiana Cristina Pereira dos Santos, Ingra Morales Claro, Flávia Cristina da Silva Sales, Claudia Regina Gonçalves, Joshua Quick, Maria do Carmo Sampaio Tavares Timenetsky, Nicholas James Loman, Andrew Rambaut, Ester Cerdeira Sabino, Nuno Rodrigues Faria |
| EPI_ISL_413213, EPI_ISL_413214 | Centre for Infectious Diseases and Microbiology Laboratory Services | NSW Health Pathology - Institute of Clinical Pathology and Medical Research; Westmead Hospital; University of Sydney | Eden J-S, Carter I, Rahman H, Holmes EC, Rockett R, O'Sullivan MV, Sintchenko V, Chen SC, Maddocks S, Kok J and Dwyer DE for the 2019-nCoV Study Group* |
| EPI_ISL_413219 | National Institute of Health Research and Development | National Institute of Health Research and Development | Setiawaty,V; Subangkit; Pawestri,HA; Puspa,KD; Ikawati,HD; Nugraha,AA; Hariastuti,NI; Ramadhany,R; Susilarini,NK; Pratiwi,E; Agustiningsih; Kurniawati,J; Siswanto |
| EPI_ISL_413459 | Department of Pathology, Toshima Hospital | Pathogen Genomics Center, National Institute of Infectious Diseases | Tsuyoshi Sekizuka, Kentaro Itokawa, Takuya Adachi, Masahiro Sano, Jun Yamazaki, Ippei Miyamoto, Haruka Nishioka, Ja-Mun Chong, Noriko Nakajima, Yuko Sato, Minoru Tobiume, Harutaka Katano, Tadaki Suzuki, Makoto Kuroda |
| EPI_ISL_413490 | Auckland Hospital | Institute of Environmental Science and Research (ESR) | Matt Storey, Xiaoyun Ren, Gary McAuliffe, Sally Roberts, Matthew Blakiston, Erasmus Smit, Lauren Jelly, Joep de Ligt |
| EPI_ISL_413522 | Indian Council of Medical Research - National Institute of Virology | National Influenza Center, Indian Council of Medical Research - National Institute of Virology | Potdar V, Yadav PD, Choudhary ML, Shete-Aich A |

| | | | |
|---|---|---|---|
| EPI_ISL_413523 | Indian Council of Medical Research-National Institute of Virology | National Influenza Center, Indian Council of Medical Research-National Institute of Virology | Potdar V, Yadav PD, Choudhary ML, Shete-Aich A |
| EPI_ISL_413553 | Iran National Influenza Center | Iran National Influenza Center | Jila Yavarian, Nazanin Zahra Shafiei Jandaghi, Kaveh Sadeghi, Fatemeh Ajaminejad, Nastaran Ghavvami and Talat Mokhtari Azad |
| EPI_ISL_413554 | Iran National Influenza Center | Iran National Influenza Center | Jila Yavarian, Nazanin Zahra Shafiei Jandaghi, Kaveh Sadeghi, Nastaran Ghavvami, Fatemeh Ajami Nejad, Fatemeh Saadatmand and Talat Mokhtari Azad |
| EPI_ISL_413594 | Centre for Infectious Diseases and Microbiology Laboratory Services | NSW Health Pathology - Institute of Clinical Pathology and Medical Research; Westmead Hospital; University of Sydney | Rockett R, Eden J-S, Lam C, Gray K, Timms, V, Gall, M, Alicia, A, Carter I, Rahman H, Holmes EC, , O'Sullivan MV, Sintchenko V, Chen SC, Maddocks S, Kok J and Dwyer DE for the 2019-nCoV Study Group* |
| EPI_ISL_413595 | Centre for Infectious Diseases and Microbiology Laboratory Services | NSW Health Pathology - Institute of Clinical Pathology and Medical Research; Westmead Hospital; University of Sydney | Rockett R, Eden J-S, Lam C, Gray K, Timms, V, Gall, M, Carter I, Rahman H, Holmes EC, O'Sullivan MV, Sintchenko V, Chen SC, Maddocks S, Kok J and Dwyer DE for the 2019-nCoV Study Group* |
| EPI_ISL_413596 | Centre for Infectious Diseases and Microbiology - Public Health | NSW Health Pathology - Institute of Clinical Pathology and Medical Research; Westmead Hospital; University of Sydney | Rockett R, Eden J-S, Lam C, Gray K, Timms, V, Gall, M, Carter I, Rahman H, Holmes EC, O'Sullivan MV, Sintchenko V, Chen SC, Maddocks S, Kok J and Dwyer DE for the 2019-nCoV Study Group* |
| EPI_ISL_413597 | Centre for Infectious Diseases and Microbiology- Public Health | NSW Health Pathology - Institute of Clinical Pathology and Medical Research; Westmead Hospital; University of Sydney | Lam C, Eden J-S, Rockett R, Gray K, Timms, V, Gall, M, Carter I, Rahman H, Holmes EC, O'Sullivan MV, Sintchenko V, Chen SC, Maddocks S, Kok J and Dwyer DE for the 2019-nCoV Study Group* |
| EPI_ISL_413598 | Centre for Infectious Diseases and Microbiology - Public Health | NSW Health Pathology - Institute of Clinical Pathology and Medical Research; Westmead Hospital; University of Sydney | Gray K, Eden J-S, Lam C, Rockett R, Timms, V, Gall, M, Carter I, Rahman H, Holmes EC, O'Sullivan MV, Sintchenko V, Chen SC, Maddocks S, Kok J and Dwyer DE for the 2019-nCoV Study Group* |
| EPI_ISL_413599 | Centre for Infectious Diseases and Microbiology - Public Health | NSW Health Pathology - Institute of Clinical Pathology and Medical Research; Westmead Hospital; University of Sydney | Timms, V, Eden J-S, Lam C, Gray K, Rockett R, Gall, M, Carter I, Rahman H, Holmes EC, O'Sullivan MV, Sintchenko V, Chen SC, Maddocks S, Kok J and Dwyer DE for the 2019-nCoV Study Group* |
| EPI_ISL_413600 | Centre for Infectious Diseases and Microbiology - Public Health | NSW Health Pathology - Institute of Clinical Pathology and Medical Research; Westmead Hospital; University of Sydney | Gall, M, Eden J-S, Lam C, Gray K, Timms, V, Rockett R, Carter I, Rahman H, Holmes EC, O'Sullivan MV, Sintchenko V, Chen SC, Maddocks S, Kok J and Dwyer DE for the 2019-nCoV Study Group* |
| EPI_ISL_413647 | Centro Hospital do Porto, E.P.E. - H. Geral de Santo Antonio | Instituto Nacional de Saude (INSA) | Raquel Guiomar, Inês Costa, Pedro Pechirra, Joana Mendonça, Luís Vieira, Helena Ramos, Joana Isidro, Vítor Borges, João Paulo Gomes |
| EPI_ISL_413648 | Centro Hospitalar e Universitário de Sao Joao, Porto | Instituto Nacional de Saude (INSA) | Raquel Guiomar, Inês Costa, Pedro Pechirra, Joana Mendonça, Luís Vieira, João Tiago Guimarães, Joana Isidro, Vítor Borges, João Paulo Gomes |
| EPI_ISL_413904 | Iran National Influenza Center | Iran National Influenza Center | Nazanin Zahra Shafiei Jandaghi,Jila Yavarian,Fatemeh AjamiNejad, Nastaran Ghavvami,Kaveh Sadeghi and Talat Mokhtari Azad |
| EPI_ISL_414014 | Hospital Israelita Albert Einstein | Instituto Adolfo Lutz, Interdiciplinary Procedures Center, Strategic Laboratory | Claudio Tavares Sacchi, Claudia Regina Gonçalves, Katia Correia dos Santos, Carlos Henrique Camargo, Maria do Carmo Sampaito Tavares Timenetsky, Terezinha Maria de Paiva, Ester Cerdeira Sabino |
| EPI_ISL_414015 | Hospital São Joaquim Beneficencia Portuguesa | Instituto Adolfo Lutz, Interdiciplinary Procedures Center, Strategic Laboratory | Claudio Tavares Sacchi, Claudia Regina Gonçalves, SimoneGuadagnucci Morillo, Carlos Henrique Camargo, Maria do Carmo Sampaito Tavares Timenetsky, Fabiana Cristina Pereira dos Santos Terezinha Maria de Paiva, Ester Cerdeira Sabino |
| EPI_ISL_414016 | Hospital São Joaquim Beneficencia Portuguesa | Instituto Adolfo Lutz, Interdiciplinary Procedures Center, Strategic Laboratory | Claudio Tavares Sacchi, Claudia Regina Gonçalves, Audrey Cilli, Carlos Henrique Camargo, Maria do Carmo Sampaito Tavares Timenetsky, Daniela Bernardes Borges da Silva, Terezinha Maria de Paiva, Ester Cerdeira Sabino |
| EPI_ISL_414017 | Hospital São Joaquim Beneficencia Portuguesa | Instituto Adolfo Lutz, Interdiciplinary Procedures Center, Strategic Laboratory | Claudio Tavares Sacchi, Claudia Regina Gonçalves, Fabiana Cristina Pereira dos Santos, Carlos Henrique Camargo, Maria do Carmo Sampaito Tavares Timenetsky, Daniela Bernardes Borges da Silva, Terezinha Maria de Paiva, Ester Cerdeira Sabino |
| EPI_ISL_414372 | Iran National Influenza Center | Iran National Influenza Center | Jila Yavarian, Nazanin Zahra Shafiei Jandaghi, Kaveh Sadeghi, Fatemeh Ajaminejad, Fatemeh Saadatmand and Talat Mokhtari Azad |
| EPI_ISL_414373 | Iran National Influenza Center | Iran National Influenza Center | Kaveh Sadeghi, Jila Yavarian, Nazanin Zahra Shafiei Jandaghi, Ahmad Nejati, Najmeh Parhizghari, Soad Ghabeshi and Talat Mokhtari Azad |
| EPI_ISL_414375, EPI_ISL_414376, EPI_ISL_414377 | National Institute of Health Research and Development | National Institute of Health Research and Development | Setiawaty,V; Subangkit; Puspa,KD; Ikawati,HD; Nugraha, AA; Susilarini, NK; Agustiningsih; Ramadhany,R; Pratiwi,E; Hariastuti,NI; Kurniawati,J; Pawestri,HA; Siswanto |
| EPI_ISL_414477 | The National Institute of Public Health Center for Epidemiology and Microbiology | State Veterinary Institute Prague | Alexander Nagy, Oldrich Bartos, Helena Jirincova, Klara Labska, Ludmila Novakova, Olga Storkanova, Dusan Trnka, Jaromira Vecerova |
| EPI_ISL_414495 | Servicio Microbiología. Hospital Clínico Universitario. Valencia. | Sequencing and Bioinformatics Service. Molecular Epidemiology Laboratory. FISABIO-Public Health | David Navarro, Maria Alma Bracho, Giuseppe D'Auria, Griselda De Marco, Neris Garcia-Gonzalez, Fernando Gonzalez-Candelas |
| EPI_ISL_414496 | Servicio Microbiologia. Hospital Clinico Universitario. Valencia. | Sequencing and Bioinformatics Service. Molecular Epidemiology Laboratory. FISABIO-Public Health | David Navarro, María Alma Bracho, Giuseppe D'Auria, Griselda De Marco, Neris Garcia-Gonzalez, Fernando Gonzalez-Candelas |
| EPI_ISL_414512 | Dipartimento di Medicina e Chirurgia, University of Insubria and Ospedale di Circolo e Fondazione Macchi | Dipartimento di Medicina e Chirurgia, University of Insubria and Ospedale di Circolo e Fondazione Macchi | Genoni,A.P., Baj,A. and Rossi,A. |
| EPI_ISL_414520, EPI_ISL_414521 | Bundeswehr Institute of Microbiology | Bundeswehr Institute of Microbiology | Mathias C Walter, Markus H Antwerpen and Roman Wölfel |
| EPI_ISL_414568 | Pasteur Institute of Iran | Pasteur Institute of Iran | Arash Arashkia, Kayhan Azadmanesh, Mohammad Hassan Pouriayevali, Tahmineh Jalali, Zahra Ahmadi, Mohammad Sadegh Shams Nosrati, Ali Maleki, Zabihollah Shoja, Sanam Azad-Mazjiri, Neda Amin, Mehdi Rohani, Saber Esmaeili, Ahmad Ghasemi, Amir Hesam Nemati, Ahmad Mahmoudi, Zahra Fereydouni, Mahsa Tavakolirad, Tahereh Mohammadi, Sahar Khakifirouz, Mehdi Fazlalipour, Maryam Rostamtabar, Arezoo Parikhani, Hesam Karimi, Kazem Baesi, Seyed Dawood Mousavi Nasab, Mahmood Barati, Mohammad Reza Asadi Karam, Mehri Habibi, Fatemeh Fotouhi-Chahooki, Neda Afzali, Ali Torabi, Azita Eshratkhah mohammadnejad, Seyedeh Sahar Bathaeian, Mina Bahri, Mohamad Mahdi Mortazavipour, Seyedeh Atefe Hosseini, Farideh Niknam, Parastoo Yekta Sanati, Hadiseh Shokouhi, Azam Amirian, Afsaneh zokaei, Hajarossadat Ghaderi, Mahboobeh Rafigh, Elmira Vadaye kheiri, Akram Agharezaei, Akram Abouie Mehrizi, seyedeh Zahra Moravej, Mostafa Salehi-Vaziri |
| EPI_ISL_414577 | Hospital de Talca, Chile | Instituto de Salud Publica de Chile | Andrés E. Castillo, Bárbara Parra, Paz Tapia, Alejandra Acevedo, Jaime Lagos, Winston Andrade, Loredana Arata, Gabriel Leal, Gisselle Barra, Carolina Tambley, Javier Tognarelli, Patricia Bustos, Soledad Ulloa, Rodrigo Fasce, Jorge Fernández. |
| EPI_ISL_414598 | Servicio Microbiologia, Hospital Clinico Universitario, Valencia | Sequencing and Bioinformatics Service and Molecular Epidemiology Research Group. FISABIO-Public Health. | David Navarro, Maria Alma Bracho, Giuseppe D'Auria, Griselda De Marco, Neris Garcia-Gonzalez, Fernando Gonzalez-Candelas |
| EPI_ISL_414623 | Laboratoire de Virologie Institut de Virologie - INSERM U 1109 Hôpitaux Universitaires de Strasbourg | National Reference Center for Viruses of Respiratory Infections, Institut Pasteur, Paris | Mélnie Albert, Marion Barbet, Sylvie Behillil, Méline Bizard, Angela Brisebarre, Flora Donati Vincent Enouf, Maud Vanpeene, Sylvie van der Werf, Samira Fafi-Kremer |
| EPI_ISL_414624 | Centre Hositalier Universitaire de Rouen Laboratoire de Virologie | National Reference Center for Viruses of Respiratory Infections, Institut Pasteur, Paris | Mélnie Albert, Marion Barbet, Sylvie Behillil, Méline Bizard, Angela Brisebarre, Flora Donati Vincent Enouf, Maud Vanpeene, Sylvie van der Werf, Jean-Christophe Plantier |
| EPI_ISL_414625 | Centre Hospitalier Régional Universitaire de Nantes Laboratoire de Virologie | National Reference Center for Viruses of Respiratory Infections, Institut Pasteur, Paris | Mélnie Albert, Marion Barbet, Sylvie Behillil, Méline Bizard, Angela Brisebarre, Flora Donati Vincent Enouf, Maud Vanpeene, Sylvie van der Werf, Marianne Coste-Burel |
| EPI_ISL_414626 | unknown | National Reference Center for Viruses of Respiratory Infections, Institut Pasteur, Paris | Mélnie Albert, Marion Barbet, Sylvie Behillil, Méline Bizard, Angela Brisebarre, Flora Donati Vincent Enouf, Maud Vanpeene, Sylvie van der Werf |
| EPI_ISL_414627, EPI_ISL_414628, EPI_ISL_414629, EPI_ISL_414630 | Centre Hospitalier Compiègne Laboratoire de Biologie | National Reference Center for Viruses of Respiratory Infections, Institut Pasteur, Paris | Mélnie Albert, Marion Barbet, Sylvie Behillil, Méline Bizard, Angela Brisebarre, Flora Donati Vincent Enouf, Maud Vanpeene, Sylvie van der Werf, Raulin Olivia |
| EPI_ISL_414631, EPI_ISL_414632 | Hôpital Robert Debré Laboratoire de Virologie | National Reference Center for Viruses of Respiratory | Mélnie Albert, Marion Barbet, Sylvie Behillil, Méline Bizard, Angela Brisebarre, Flora Donati Vincent Enouf, Maud Vanpeene, Sylvie van der Werf, Laurent |

| | | | |
|---|---|---|---|
| | | Infections, Institut Pasteur, Paris | Andreoletti |
| EPI_ISL_414633 | Centre Hospitalier René Dubois Laboratoire de Microbiologie - Bât A | National Reference Center for Viruses of Respiratory Infections, Institut Pasteur, Paris | Mélnie Albert, Marion Barbet, Sylvie Behillil, Méline Bizard, Angela Brisebarre, Flora Donati Vincent Enouf, Maud Vanpeene, Sylvie van der Werf, Pascale Martres |
| EPI_ISL_414634, EPI_ISL_414635, EPI_ISL_414636, EPI_ISL_414637, EPI_ISL_414638 | Centre Hospitalier Compiègne Laboratoire de Biologie | National Reference Center for Viruses of Respiratory Infections, Institut Pasteur, Paris | Mélnie Albert, Marion Barbet, Sylvie Behillil, Méline Bizard, Angela Brisebarre, Flora Donati Vincent Enouf, Maud Vanpeene, Sylvie van der Werf, Raulin Olivia |
| EPI_ISL_414663, EPI_ISL_414691 | State Key Laboratory of Respiratory Disease, National Clinical Research Center for Respiratory Disease, Guangzhou Institute of Respiratory Health, the First Affiliated Hospital of Guangzhou Medical University | The First Affiliated Hospital of Guangzhou Medical University & BGI-Shenzhen | Zhao et al |
| EPI_ISL_414945 | Iran National Influenza Center | Iran National Influenza Center | Nazanin Zahra Shafiei Jandaghi, Jila Yavarian,Kaveh Sadeghi, Vahid Salimi, Simin Abbasi, Saeedeh Mahfozi and Talat Mokhtari Azad |
| EPI_ISL_415153 | KU Leuven, Clinical and Epidemiological Virology | KU Leuven, Clinical and Epidemiological Virology | Bert Vanmechelen, Joan Marti-Carreras, Tony Wawina, Marc Van Ranst, Piet Maes |
| EPI_ISL_415154 | KU Leuven, Clincal and Epidemiological Virology | KU Leuven, Clincal and Epidemiological Virology | Bert Vanmechelen, Joan Marti-Careras, Tony Wawina, Marc Van Ranst, Piet Maes. |
| EPI_ISL_415155 | KU Leuven, Clinical and Epidemiological Virology | KU Leuven, Clinical and Epidemiological Virology | Bert Vanmechelen, Joan Marti-Carreras, Tony Wawina, Marc Van Ranst, Piet Maes |
| EPI_ISL_415156, EPI_ISL_415157, EPI_ISL_415158, EPI_ISL_415159 | KU Leuven, Clinical and Epidemiological Virology | KU Leuven, Clinical and Epidemiological Virology | Bert Vanmechelen, Joan Marti-Carreras, Tony Wawina, Piet Maes |
| EPI_ISL_415641, EPI_ISL_415642, EPI_ISL_415643, EPI_ISL_415644 | R. G. Lugar Center for Public Health Research, National Center for Disease Control and Public Health (NCDC) of Georgia. | R. G. Lugar Center for Public Health Research, National Center for Disease Control and Public Health (NCDC) of Georgia. | Nato Kotaria, Marine Murtskhvaladze, Ann Machablishvili, Lela Sabadze, Mari Gavashelidze, Ana Papkiauri, Meri Pantsulaia, Gvantsa Brachveli, Tata Imnadze, Tamar Jashiashvili, Tea Tevdoradze, Ketevan Sidamonidze, Ekaterine Khmaladze, Ekaterine Zhghenti, Roena Sukhiashvili, Mariam Zakalashvili, Lela Urushadze, Magda Dgebuadze, Giorgi Tomashvili, Davit Tsaguria, Ekaterine Zangaladze, Nino Berishvili, Gvantsa Chanturia, Adam Kotorashvili, Maia Alkhazashvili, Irma Burjanadze, Anna Kasradze, Khatuna Zakhashvili, Paata Imnadze, Amiran Gamkrelidze. |
| EPI_ISL_415649 | unknown | National Reference Center for Viruses of Respiratory Infections, Institut Pasteur, Paris | Mélnie Albert, Marion Barbet, Sylvie Behillil, Méline Bizard, Angela Brisebarre, Flora Donati Vincent Enouf, Maud Vanpeene, Sylvie van der Werf |
| EPI_ISL_415650 | Hôpital Instruction des Armées - BEGIN | National Reference Center for Viruses of Respiratory Infections, Institut Pasteur, Paris | Mélnie Albert, Marion Barbet, Sylvie Behillil, Méline Bizard, Angela Brisebarre, Flora Donati Vincent Enouf, Maud Vanpeene, Sylvie van der Werf, Christine Bigaillon |
| EPI_ISL_415652 | unknown | National Reference Center for Viruses of Respiratory Infections, Institut Pasteur, Paris | Mélnie Albert, Marion Barbet, Sylvie Behillil, Méline Bizard, Angela Brisebarre, Flora Donati Vincent Enouf, Maud Vanpeene, Sylvie van der Werf |
| EPI_ISL_415654 | Centre Hospitalier Compiègne Laboratoire de Biologie | National Reference Center for Viruses of Respiratory Infections, Institut Pasteur, Paris | Mélnie Albert, Marion Barbet, Sylvie Behillil, Méline Bizard, Angela Brisebarre, Flora Donati Vincent Enouf, Maud Vanpeene, Sylvie van der Werf, Raulin Olivia |
| EPI_ISL_415709 | State Key Laboratory for Diagnosis and Treatment of Infectious Diseases, National Clinical Research Center for Infectious Diseases, First Affiliated Hospital, Zhejiang University School of Medicine, Hangzhou, China. 310003 | State Key Laboratory for Diagnosis and Treatment of Infectious Diseases, National Clinical Research Center for Infectious Diseases, First Affiliated Hospital, Zhejiang University School of Medicine, Hangzhou, China. 310003 | Hangping Yao, Nanping Wu, Chao Jiang, Xiangyun Lu, Linfang Cheng, Fumin Liu, Zhigang Wu, Haibo Wu, Changzhong Jin, Min Zheng, Lanjuan Li |
| EPI_ISL_415710 | WHO National Influenza Centre Russian Federation | WHO National Influenza Centre Russian Federation | Andrey Komissarov, Artem Fadeev, Anna Ivanova, Daria Danilenko |
| EPI_ISL_415711 | State Key Laboratory for Diagnosis and Treatment of Infectious Diseases, National Clinical Research Center for Infectious Diseases, First Affiliated Hospital, Zhejiang University School of Medicine, Hangzhou, China. 310003 | State Key Laboratory for Diagnosis and Treatment of Infectious Diseases, National Clinical Research Center for Infectious Diseases, First Affiliated Hospital, Zhejiang University School of Medicine, Hangzhou, China. 310003 | Hangping Yao, Nanping Wu, Chao Jiang, Xiangyun Lu, Linfang Cheng, Fumin Liu, Zhigang Wu, Haibo Wu, Changzhong Jin, Min Zheng, Lanjuan Li |
| EPI_ISL_416028 | National Influenza Center - Instituto Adolfo Lutz | Instituto Adolfo Lutz, Interdiciplinary Procedures Center, Strategic Laboratory | Claudio Tavares Sacchi, Claudia Regina Gonçalves, Carlos Henrique Camargo, Fabiana Cristina Pereira dos Santos, Daniela Bernardes Borges da Silva, Simone Guadagnucci Morillo, Adriano Abbud, Adriana Bugno, Maria do Carmo Sampaio Tavares Timenetsky, Terezinha Maria de Paiva |
| EPI_ISL_416029 | Laboiratório Fleury | Instituto Adolfo Lutz, Interdiciplinary Procedures Center, Strategic Laboratory | Claudio Tavares Sacchi, Claudia Regina Gonçalves, Carlos Henrique Camargo, Fabiana Cristina Pereira dos Santos, Daniela Bernardes Borges da Silva, Simone Guadagnucci Morillo, Adriano Abbud, Adriana Bugno, Maria do Carmo Sampaio Tavares Timenetsky, Terezinha Maria de Paiva |
| EPI_ISL_416031, EPI_ISL_416032 | National Influenza Center - Instituto Adolfo Lutz | Instituto Adolfo Lutz, Interdiciplinary Procedures Center, Strategic Laboratory | Claudio Tavares Sacchi, Claudia Regina Gonçalves, Carlos Henrique Camargo, Fabiana Cristina Pereira dos Santos, Daniela Bernardes Borges da Silva, Simone Guadagnucci Morillo, Adriano Abbud, Adriana Bugno, Maria do Carmo Sampaio Tavares Timenetsky, Terezinha Maria de Paiva |
| EPI_ISL_416033, EPI_ISL_416034 | Hospital Israelita Albert Einstein | Instituto Adolfo Lutz, Interdiciplinary Procedures Center, Strategic Laboratory | Claudio Tavares Sacchi, Claudia Regina Gonçalves, Carlos Henrique Camargo, Erica Valessa Ramos Gomes, Fabiana Cristina Pereira dos Santos, Daniela Bernardes Borges da Silva, Simone Guadagnucci Morillo, Adriano Abbud, Adriana Bugno, Maria do Carmo Sampaio Tavares Timenetsky, Terezinha Maria de Paiva |
| EPI_ISL_416035, EPI_ISL_416036 | National Influenza Center - Instituto Adolfo Lutz | Instituto Adolfo Lutz, Interdiciplinary Procedures Center, Strategic Laboratory | Claudio Tavares Sacchi, Claudia Regina Gonçalves, Carlos Henrique Camargo, Erica Valessa Ramos Gomes, Fabiana Cristina Pereira dos Santos, Daniela Bernardes Borges da Silva, Simone Guadagnucci Morillo, Adriano Abbud, Adriana Bugno, Maria do Carmo Sampaio Tavares Timenetsky, Terezinha Maria de Paiva |
| EPI_ISL_416042 | State Key Laboratory for Diagnosis and Treatment of Infectious Diseases, National Clinical Research Center for Infectious Diseases, First Affiliated Hospital, Zhejiang University School of Medicine, Hangzhou, China. 310003 | State Key Laboratory for Diagnosis and Treatment of Infectious Diseases, National Clinical Research Center for Infectious Diseases, First Affiliated Hospital, Zhejiang University School of Medicine, Hangzhou, China. 310003 | Hangping Yao, Nanping Wu, Chao Jiang, Xiangyun Lu, Linfang Cheng, Fumin Liu, Zhigang Wu, Haibo Wu, Changzhong Jin, Min Zheng, Lanjuan Li |
| EPI_ISL_416044, EPI_ISL_416046, EPI_ISL_416047 | State Key Laboratory for Diagnosis and Treatment of Infectious Diseases, National Clinical Research Center for Infectious Diseases, First Affiliated Hospital, Zhejiang University School of Medicine, Hangzhou, China 310003 | State Key Laboratory for Diagnosis and Treatment of Infectious Diseases, National Clinical Research Center for Infectious Diseases, First Affiliated Hospital, Zhejiang University School of Medicine, Hangzhou, China 310003 | Hangping Yao, Nanping Wu, Chao Jiang, Xiangyun Lu, Linfang Cheng, Fumin Liu, Zhigang Wu, Haibo Wu, Changzhong Jin, Min Zheng, Lanjuan Li |
| EPI_ISL_416142 | Department of Virus and Microbiological Special diagnostics, Statens Serum Institut, Copenhagen, Denmark. | Statens Serum Institute | Morten Rasmussen, Maiken Worsoe Rosenstierne , Anders Fomsgaard |
| EPI_ISL_416143, EPI_ISL_416144 | Department of Virus and Microbiological Special diagnostics, Statens Serum Institut, Copenhagen, Denmark. | ViFU | Morten Rasmussen, Maiken Worsoe Rosenstierne , Anders Fomsgaard |
| EPI_ISL_416425 | State Key Laboratory for Diagnosis and Treatment of Infectious Diseases, National Clinical Research Center for Infectious Diseases, First Affiliated Hospital, Zhejiang University School of Medicine, Hangzhou, China 310003 | State Key Laboratory for Diagnosis and Treatment of Infectious Diseases, National Clinical Research Center for Infectious Diseases, First Affiliated Hospital, Zhejiang University School of Medicine, Hangzhou, China 310003 | Hangping Yao, Nanping Wu, Chao Jiang, Xiangyun Lu, Linfang Cheng, Fumin Liu, Zhigang Wu, Haibo Wu, Changzhong Jin, Min Zheng, Lanjuan Li |
| EPI_ISL_416426 | Virological Research Group, Szentágothai Research Centre, University of Pécs | Bioinformatics Research Group, Szentágothai Research Centre, University of Pécs | Péter Urbán, Endre Gábor Tóth, Gábor Kemenesi, Róbert Herczeg, Attila Gyenesei, Ferenc Jakab |
| EPI_ISL_416458 | Virology laboratory Ministry of Health Kuwait sequenced at Dasman Diabetes Institute | Dasman Diabetes Institute | Fahd Al-Mulla, Sumi John, Sara Alqabandi, Rasheeba iqbal, Motasem Melhem, Ebaa alOzairi, Qais Al-Duwairi |
| EPI_ISL_416469, EPI_ISL_416470, EPI_ISL_416471, EPI_ISL_416472 | KU Leuven, Clinical and Epidemiological Virology | KU Leuven, Clinical and Epidemiological Virology | Bert Vanmechelen, Tony Wawina, Joan Marti-Carreras, Piet Maes |
| EPI_ISL_416473, EPI_ISL_416474 | State Key Laboratory for Diagnosis and Treatment of Infectious Diseases, National Clinical Research Center for | State Key Laboratory for Diagnosis and Treatment of Infectious Diseases, National Clinical Research Center for | Hangping Yao, Nanping Wu, Chao Jiang, Xiangyun Lu, Linfang Cheng, Fumin Liu, Zhigang Wu, Haibo Wu, Changzhong Jin, Min Zheng, Lanjuan Li |

| | | | |
|---|---|---|---|
| | Infectious Diseases, First Affiliated Hospital, Zhejiang University School of Medicine, Hangzhou, China 310003 | Infectious Diseases, First Affiliated Hospital, Zhejiang University School of Medicine, Hangzhou, China 310003 | |
| EPI_ISL_416475 | KU Leuven, Clinical and Epidemiological Virology | KU Leuven, Clinical and Epidemiological Virology | Bert Vanmechelen, Tony Wawina, Joan Marti-Carreras, Piet Maes |
| EPI_ISL_416485 | Servicio de Microbiologia. Consorcio Hospital General Universitario de Valencia | Sequencing and Bioinformatics Service and Molecular Epidemiology Research Group. FISABIO-Public Health | Griselda De Marco, Neris Garcia-Gonzalez, Maria Alma Bracho, Maria Dolores Ocete, Concepcion Gimeno, Giuseppe D'Auria, Fernando Gonzalez-Candelas |
| EPI_ISL_416486 | Servicio de Microbiología. Consorcio Hospital General Universitario de Valencia | Sequencing and Bioinformatics Service and Molecular Epidemiology Research Group. FISABIO-Public Health | Neris Garcia-Gonzalez, Maria Alma Bracho, Maria Dolores Ocete, Concepcion Gimeno, Giuseppe D'Auria, Griselda De Marco, Fernando Gonzalez-Candelas |
| EPI_ISL_416487 | Servicio de Microbiología. Consorcio Hospital General Universitario de Valencia | Sequencing and Bioinformatics Service and Molecular Epidemiology Research Group. FISABIO-Public Health | Giuseppe D'Auria, Griselda De Marco, Neris Garcia-Gonzalez, Maria Alma Bracho, Maria Dolores Ocete, Concepcion Gimeno, Fernando Gonzalez-Candelas |
| EPI_ISL_416488 | ViroGenetics - BSL3 Laboratory of Virology; Human Genome Variation Research Group & Genomics Centre MCB; Bioinformatics Research Group Department of Virology | ViroGenetics - BSL3 Laboratory of Virology; Human Genome Variation Research Group & Genomics Centre MCB; Bioinformatics Research Group Department of Virology | Aleksandra Milewska, Ewelina Popiech, Agata Jarosz, Adrianna Klajmon, Kamila Marszaek, Katarzyna Pancer, Magdalena Rzeczkowska, Tomasz Wokowicz, Katarzyna Zacharczuk, Agnieszka Koakowska-Kulesza, Natalia Wolaniuk, Ewelina Hallman-Szeliska, Pawe P abaj, Wojciech Branicki, Krzysztof Pyr |
| EPI_ISL_416493 | CH Jean de Navarre Laboratoire de Biologie | National Reference Center for Viruses of Respiratory Infections, Institut Pasteur, Paris | Mélnie Albert, Marion Barbet, Sylvie Behillil, Méline Bizard, Angela Brisebarre, Flora Donati, Etienne Simon-Lorière, Vincent Enouf, Maud Vanpeene, Sylvie van der Werf |
| EPI_ISL_416494 | Centre Hositalier Universitaire de Rouen Laboratoire de Virologie | National Reference Center for Viruses of Respiratory Infections, Institut Pasteur, Paris | Mélnie Albert, Marion Barbet, Sylvie Behillil, Méline Bizard, Angela Brisebarre, Flora Donati, Etienne Simon-Lorière, Vincent Enouf, Maud Vanpeene, Sylvie van der Werf, Jean-Christophe Plantier |
| EPI_ISL_416495, EPI_ISL_416496, EPI_ISL_416497 | Centre Hospitalier Compiègne Laboratoire de Biologie | National Reference Center for Viruses of Respiratory Infections, Institut Pasteur, Paris | Mélnie Albert, Marion Barbet, Sylvie Behillil, Méline Bizard, Angela Brisebarre, Flora Donati, Etienne Simon-Lorière, Vincent Enouf, Maud Vanpeene, Sylvie van der Werf, Raulin Olivia |
| EPI_ISL_416498 | Institut Médico légal- Hop R. Poincaré | National Reference Center for Viruses of Respiratory Infections, Institut Pasteur, Paris | Mélnie Albert, Marion Barbet, Sylvie Behillil, Méline Bizard, Angela Brisebarre, Flora Donati, Etienne Simon-Lorière, Vincent Enouf, Maud Vanpeene, Sylvie van der Werf |
| EPI_ISL_416499, EPI_ISL_416500 | LABM GH nord Essonne | National Reference Center for Viruses of Respiratory Infections, Institut Pasteur, Paris | Mélnie Albert, Marion Barbet, Sylvie Behillil, Méline Bizard, Angela Brisebarre, Flora Donati, Etienne Simon-Lorière, Vincent Enouf, Maud Vanpeene, Sylvie van der Werf |
| EPI_ISL_416501 | Hopital franco britannique - Service des Urgences | National Reference Center for Viruses of Respiratory Infections, Institut Pasteur, Paris | Mélnie Albert, Marion Barbet, Sylvie Behillil, Méline Bizard, Angela Brisebarre, Flora Donati, Etienne Simon-Lorière, Vincent Enouf, Maud Vanpeene, Sylvie van der Werf |
| EPI_ISL_416502, EPI_ISL_416503, EPI_ISL_416504, EPI_ISL_416505, EPI_ISL_416506, EPI_ISL_416507, EPI_ISL_416508, EPI_ISL_416509, EPI_ISL_416510, EPI_ISL_416511, EPI_ISL_416512, EPI_ISL_416513 | | | |
| see above | CHRU Pontchaillou - Laboratoire de Virologie | National Reference Center for Viruses of Respiratory Infections, Institut Pasteur, Paris | Mélnie Albert, Marion Barbet, Sylvie Behillil, Méline Bizard, Angela Brisebarre, Flora Donati, Etienne Simon-Lorière, Vincent Enouf, Maud Vanpeene, Sylvie van der Werf, Gisèle Lagathu |
| EPI_ISL_416565, EPI_ISL_416566, EPI_ISL_416567, EPI_ISL_416569, EPI_ISL_416570, EPI_ISL_416571, EPI_ISL_416572, EPI_ISL_416573, EPI_ISL_416574, EPI_ISL_416575, EPI_ISL_416576, EPI_ISL_416577, EPI_ISL_416578, EPI_ISL_416579, EPI_ISL_416580, EPI_ISL_416581, EPI_ISL_416582, EPI_ISL_416583, EPI_ISL_416585, EPI_ISL_416586, EPI_ISL_416587, EPI_ISL_416588, EPI_ISL_416589, EPI_ISL_416590, EPI_ISL_416591, EPI_ISL_416592, EPI_ISL_416593, EPI_ISL_416594, EPI_ISL_416595, EPI_ISL_416596, EPI_ISL_416597, EPI_ISL_416598, EPI_ISL_416599, EPI_ISL_416600, EPI_ISL_416601, EPI_ISL_416602, EPI_ISL_416603, EPI_ISL_416604, EPI_ISL_416605, EPI_ISL_416606, EPI_ISL_416607, EPI_ISL_416608, EPI_ISL_416609, EPI_ISL_416610, EPI_ISL_416611, EPI_ISL_416612, EPI_ISL_416613, EPI_ISL_416614, EPI_ISL_416615, EPI_ISL_416617, EPI_ISL_416618, EPI_ISL_416619, EPI_ISL_416620, EPI_ISL_416621, EPI_ISL_416622, EPI_ISL_416624, EPI_ISL_416625, EPI_ISL_416626, EPI_ISL_416627, EPI_ISL_416628, EPI_ISL_416629, EPI_ISL_416630, EPI_ISL_416631, EPI_ISL_416632, EPI_ISL_416633, EPI_ISL_416634 | | | |
| see above | Japanese Quarantine Stations | Pathogen Genomics Center, National Institute of Infectious Diseases | Tsuyoshi Sekizuka, Kentaro Itokawa, Rina Tanaka, Masanori Hashino, Tsutomu Kageyama, Shinji Saito, Ikuyo Takayama, Hideki Hasegawa, Takuri Takahashi, Hajime Kamiya, Takuya Yamagishi, Motoi Suzuki, Takaji Wakita, Makoto Kuroda |
| EPI_ISL_416744 | Virological Research Group, Szentágothai Research Centre | Bioinformatics Research Group, Szentágothai Research Centre | Péter Urbán, Endre Gábor Tóth, Gábor Kemenesi, Róbert Herczeg, Attila Gyenesei, Ferenc Jakab |
| EPI_ISL_416829 | National Public Health Laboratory | Malaysia Genome Institute | Mohd Noor Mat Isa, Irni Suhayu Sapian, Yusuf Muhammad Noor, Nurhezreen Md Iqbal, Mohd Faizal Abu Bakar, Enizza Kasim, Shamsidar Sopie, Siti Noraini Othman, Azrin Ahmad, Nor Azfa Johari, Norazimah Tajudin, Noorliza Mohamad Noordin, W Afiza W Mohd Arifin, Rehan Shuhada Abu Bakar, Yu Kie Chem, Selvanesan Sengol, Hani Mat Hussin, Shahrul Hisham Zainal Ariffin |
| EPI_ISL_416866 | National Public Health Laboratory | Malaysia Genome Institute | Mohd Noor Mat Isa, Irni Suhayu Sapian, Yusuf Muhammad Noor, Nurhezreen Md Iqbal, Mohd Faizal Abu Bakar, Enizza Kasim, Shamsidar Sopie, Siti Noraini Othman, Azrin Ahmad, Nor Azfa Johari, Norazimah Tajudin, Noorliza Mohamad Noordin, W Afiza W Mohd Arifin, Rehan Shuhada Abu Bakar, Yu Kie Chem, Selvanesan Sengol, Hani Mat Hussin, Shahrul Hisham Zainal Ariffin |
| EPI_ISL_416885, EPI_ISL_416886, EPI_ISL_416907 | National Public Health Laboratory | Malaysia Genome Institute | Mohd Noor Mat Isa, Irni Suhayu Sapian, Yusuf Muhammad Noor, Nurhezreen Md Iqbal, Mohd Faizal Abu Bakar, Enizza Kasim, Shamsidar Sopie, Siti Noraini Othman, Azrin Ahmad, Nor Azfa Johari, Norazimah Tajudin, Noorliza Mohamad Noordin, W Afiza W Mohd Arifin, Rehan Shuhada Abu Bakar, Yu Kie Chem, Selvanesan Sengol, Hani Mat Hussin, Shahrul Hisham Zainal Ariffin |
| EPI_ISL_416994 | COMPLEJO ASISTENCIAL UNIVERSITARIO DE BURGOS | Instituto de Salud Carlos III | Iglesias-Caballero, M. Molinero Calamita, M. González-Esguevillas, M. Camarero S. Pozo F. Casas I. Jiménez P. Jiménez M. Zaballos A. Monzón, S. Varona, S. Juliá M. Cuesta I. Megias Lobón, G. Hospital: ------ |
| EPI_ISL_417007 | HOSPITAL SANTA MARIA NAI | Instituto de Salud Carlos III | Iglesias-Caballero, M. Molinero Calamita, M. González-Esguevillas, M. Camarero S. Pozo F. Casas I. Jiménez, P. Jiménez, M. Zaballos, A. Monzón, S. Varona, S. Juliá, M. Cuesta, I. García Costa, J. |
| EPI_ISL_417010 | FUNDACION JIMENEZ DIAZ | Instituto de Salud Carlos III | Iglesias-Caballero, M. Molinero Calamita, M. González-Esguevillas, M. Camarero, S. Pozo, F. Casas, I. Jiménez, P. Jiménez, M. Zaballos, A. Monzón, S. Varona, S. Juliá, M. Cuesta, I. Fernández Roblas, R. |
| EPI_ISL_417015, EPI_ISL_417016, EPI_ISL_417018, EPI_ISL_417019 | Department of Clinical Microbiology | GIGA Medical Genomics | Durkin Keith, Artesi Maria, Bontems Sébastien, Boreux Raphaël, Meex Cécile, Melin Pierrette, Hayette Marie-Pierre, Bours Vincent. |
| EPI_ISL_417030 | Centre for Infectious Diseases and Microbiology Laboratory Services | NSW Health Pathology - Institute of Clinical Pathology and Medical Research; Westmead Hospital; University of Sydney | Eden J-S, Rockett R, Carter I, Rahman H, Holmes EC, O'Sullivan MV, Sintchenko V, Chen SC, Maddocks S, Kok J and Dwyer DE for the 2019-nCoV Study Group* |
| EPI_ISL_417034 | Laboratorio de Ecologia de Doencas Transmissiveis na Amazonia, Instituto Leonidas e Maria Deane - Fiocruz Amazonia | Laboratorio de Ecologia de Doencas Transmissiveis na Amazonia, Instituto Leonidas e Maria Deane - Fiocruz Amazonia | Valdinete Nascimento, André Corado, Fernanda Nascimento, Ágatha Costa, Debora Duarte, Luciana Gonçalves, Michele Jesus, Sérgio Luz, Felipe Naveca on behalf of the Fiocruz COVID-19 Genomic Surveillance Network |
| EPI_ISL_417176 | Department of Pathology, Princess Margaret Hospital | Department of Health Technology and Informatics, Faculty of Health and Social Science, The Hong Kong Polytechnic University | Kenneth Siu-Sing LEUNG, Timothy Ting-Leung NG, Alan Ka-Lun WU, Miranda Chong-Yee YAU, Hiu-Yin LAO, Ming-Pan CHOI, Kingsley King-Gee TAM, Lam-Kwong LEE, Barry Kin-Chung WONG, Alex Yat-Man HO, Kam-Tong Yip, Kwok-Cheung LUNG, Raymond Wai-To LIU, Eugene Yuk-Keung TSO, Wai-Shing LEUNG, Man-Chun CHAN, Yuk-Yung NG, Kit-Man SIN, Kitty Sau-Chun FUNG, Sandy Ka-Yee CHAU, Wing-Kin TO, Tak-Lun Que, David Ho-Keung SHUM, Shea Ping YIP, Wing Cheong YAM, Gilman Kit-Hang SIU |
| EPI_ISL_417181, EPI_ISL_417185 | Department of Pathology, United Christian Hospital | Department of Health Technology and Informatics, Faculty of Health and Social Science, The Hong Kong Polytechnic University | Kenneth Siu-Sing LEUNG, Timothy Ting-Leung NG, Alan Ka-Lun WU, Miranda Chong-Yee YAU, Hiu-Yin LAO, Ming-Pan CHOI, Kingsley King-Gee TAM, Lam-Kwong LEE, Barry Kin-Chung WONG, Alex Yat-Man HO, Kam-Tong Yip, Kwok-Cheung LUNG, Raymond Wai-To LIU, Eugene Yuk-Keung TSO, Wai-Shing LEUNG, Man-Chun CHAN, Yuk-Yung NG, Kit-Man SIN, Kitty Sau-Chun FUNG, Sandy Ka-Yee CHAU, Wing-Kin TO, Tak-Lun Que, David Ho-Keung SHUM, Shea Ping YIP, Wing Cheong YAM, Gilman Kit-Hang SIU |
| EPI_ISL_417186 | National Institute for Communicable Diseases of the National Health Laboratory Service | National Institute for Communicable Diseases of the National Health Laboratory Service | Allam M, Kwenda S, van Heusden P, Khumalo Z, Mohale T, Subramoney K, von Gottberg, A, Ismail A, Bhiman JN |
| EPI_ISL_417187, EPI_ISL_417188, EPI_ISL_417193 | Department of Clinical Pathology, Pamela Youde Nethersole Eastern Hospital | Department of Health Technology and Informatics, Faculty of Health and Social Science, The Hong Kong Polytechnic University | Kenneth Siu-Sing LEUNG, Timothy Ting-Leung NG, Alan Ka-Lun WU, Miranda Chong-Yee YAU, Hiu-Yin LAO, Ming-Pan CHOI, Kingsley King-Gee TAM, Lam-Kwong LEE, Barry Kin-Chung WONG, Alex Yat-Man HO, Kam-Tong Yip, Kwok-Cheung LUNG, Raymond Wai-To LIU, Eugene Yuk-Keung TSO, Wai-Shing LEUNG, Man-Chun CHAN, Yuk-Yung NG, Kit-Man SIN, Kitty Sau-Chun FUNG, Sandy Ka-Yee CHAU, Wing-Kin TO, Tak-Lun Que, David Ho-Keung SHUM, Shea Ping YIP, Wing Cheong YAM, Gilman Kit-Hang SIU |

| | | | |
|---|---|---|---|
| EPI_ISL_417333, EPI_ISL_417334, EPI_ISL_417335, EPI_ISL_417336, EPI_ISL_417337, EPI_ISL_417339 | Institut des Agents Infectieux (IAI), Hospices Civils de Lyon | CNR Virus des Infections Respiratoires - France SUD | Antonin Bal, Gregory Destras, Gwendolyne Burfin, Solenne Brun, Carine Moustaud, Raphaelle Lamy, Alexandre Gaymard, Maude Bouscambert-Duchamp, Florence Morfin-Sherpa, Martine Valette, Laurence Josset, Bruno Lina |
| EPI_ISL_417443 | State Key Laboratory for Emerging Infectious Diseases Department of Microbiology Li Ka Shing Faculty of Medicine The University of Hong Kong | State Key Laboratory for Emerging Infectious Diseases Department of Microbiology Li Ka Shing Faculty of Medicine The University of Hong Kong | Pui Wang, Siu-Ying Lau, Shaofeng Deng, Bobo Wing-Yee Mok, Wenjun Song, Kwok-Yung Yuen, Honglin Chen |
| EPI_ISL_417491 | Virology Laboratory, Department of Biomedical Sciences and Public Health, University Politecnica delle Marche | Virology and Legal Medicine Laboratories, Department of Biomedical Sciences and Public Health, University Politecnica delle Marche | Bagnarelli,P., Caucci,S., Di Sante,L., Menzo,S., Alessandrini,F., Onofri,V., Turchi,C., Tagliabracci,A. |
| EPI_ISL_417877, EPI_ISL_417878, EPI_ISL_417879, EPI_ISL_417880 | Institute of Virology, Biomedical Research Center of the Slovak Academy of Sciences, Bratislava; Public Health Authority of the Slovak Republic, Bratislava | Institute of Virology, Biomedical Research Center of the Slovak Academy of Sciences, Bratislava; Comenius University Science Park, Bratislava | Monika Slávikova, Martina Liková, Sabina Fumaová Havlíková, Juraj Koi, Juraj Kopáek, Elena Tichá, Edita Staroová, Jaroslav Budiš, Werner Krampl, Miroslav Böhmer, Diana Rusáková, Tomáš Szemeš, Boris Klempa |
| EPI_ISL_417924 | Secretaría de Salud Medellín | Instituto Nacional de Salud, Universidad Cooperativa de Colombia, Instituto Alexander von Humboldt, Imperial College-London, London School of Hygiene & Tropical Medicine | Marcela Mercado-Reyes, Katherine Laiton-Donato, Diego A. Álvarez-Díaz, Carlos Franco-Muñoz, Jose A. Usme-Ciro, Gloria Puerto, Nicolás D. Franco-Sierra, Mailyn A. Gonzalez, Zulma M. Cucunubá, Christian Julian VillabonaArenas, Liz Villabona-Arenas, Sussy Echeverria-Londoño, Astrid C. Flórez, Sergio Gomez Rangel, Luz Dary Rodriguez, Juliana Barbosa, Erika Ospitia, Diana Marcela Walteros-Acero, Martha Lucia Ospina Martinez |
| EPI_ISL_418183 | Virological Research Group, Szentágothai Research Centre | Bioinformatics Research Group, Szentágothai Research Centre | Péter Urbán, Endre Gábor Tóth, Gábor Kemenesi, Róbert Herczeg, Attila Gyenesei, Ferenc Jakab |
| EPI_ISL_418206, EPI_ISL_418207, EPI_ISL_418208, EPI_ISL_418209, EPI_ISL_418210, EPI_ISL_418211 | Institut Pasteur Dakar | Institut Pasteur de Dakar | Ndongo Dia, Ousmane Faye, Amadou Alpha Sall |
| EPI_ISL_418212 | Institut Pasteur Dakar | Institut Pasteur de Dakar | Ndongo Dia, Ousmane Faye, Amadou Alpha sall |
| EPI_ISL_418213 | Institut Pasteur Dakar | Institut Pasteur de Dakar | Ndongo Dia, Ousmane Faye, Amadou Alpha Sall |
| EPI_ISL_418215 | Instirut Pasteur Dakar | Institut Pasteur de Dakar | Ndongo Dia, Ousmane Faye, Amadou Alpha Sall |
| EPI_ISL_418216, EPI_ISL_418217 | Institut Pasteur Dakar | Institut Pasteur de Dakar | Ndongo Dia, Ousmane Faye, Amadou Alpha Sall |
| EPI_ISL_418218 | Centre Hospitalier Compiègne Laboratoire de Biologie | National Reference Center for Viruses of Respiratory Infections, Institut Pasteur, Paris | Mélanie Albert, Marion Barbet, Sylvie Behillil, Méline Bizard, Angela Brisebarre, Flora Donati, Fabiana Gambaro, Etienne Simon-Lorière, Vincent Enouf, Maud Vanpeene, Sylvie van der Werf, Raulin Olivia |
| EPI_ISL_418219 | CHU - Hôpital Cavale Blanche - Labo. de Virologie | National Reference Center for Viruses of Respiratory Infections, Institut Pasteur, Paris | Mélanie Albert, Marion Barbet, Sylvie Behillil, Méline Bizard, Angela Brisebarre, Flora Donati, Fabiana Gambaro, Etienne Simon-Lorière, Vincent Enouf, Maud Vanpeene, Sylvie van der Werf, Léa Pilorge |
| EPI_ISL_418220, EPI_ISL_418221 | Centre Hospitalier Compiègne Laboratoire de Biologie | National Reference Center for Viruses of Respiratory Infections, Institut Pasteur, Paris | Mélanie Albert, Marion Barbet, Sylvie Behillil, Méline Bizard, Angela Brisebarre, Flora Donati, Fabiana Gambaro, Etienne Simon-Lorière, Vincent Enouf, Maud Vanpeene, Sylvie van der Werf, Raulin Olivia |
| EPI_ISL_418222 | CHRU Bretonneau - Serv. Bacterio-Virol. | National Reference Center for Viruses of Respiratory Infections, Institut Pasteur, Paris | Mélanie Albert, Marion Barbet, Sylvie Behillil, Méline Bizard, Angela Brisebarre, Flora Donati, Fabiana Gambaro, Etienne Simon-Lorière, Vincent Enouf, Maud Vanpeene, Sylvie van der Werf, Julien Marlet |
| EPI_ISL_418223, EPI_ISL_418224, EPI_ISL_418225 | Centre Hospitalier Compiègne Laboratoire de Biologie | National Reference Center for Viruses of Respiratory Infections, Institut Pasteur, Paris | Mélanie Albert, Marion Barbet, Sylvie Behillil, Méline Bizard, Angela Brisebarre, Flora Donati, Fabiana Gambaro, Etienne Simon-Lorière, Vincent Enouf, Maud Vanpeene, Sylvie van der Werf, Raulin Olivia |
| EPI_ISL_418226 | EHPAD - Résidences les Cèdres | National Reference Center for Viruses of Respiratory Infections, Institut Pasteur, Paris | Mélanie Albert, Marion Barbet, Sylvie Behillil, Méline Bizard, Angela Brisebarre, Flora Donati, Etienne Simon-Lorière, Vincent Enouf, Maud Vanpeene, Sylvie van der Werf |
| EPI_ISL_418227, EPI_ISL_418228 | Centre Hospitalier Compiègne Laboratoire de Biologie | National Reference Center for Viruses of Respiratory Infections, Institut Pasteur, Paris | Mélanie Albert, Marion Barbet, Sylvie Behillil, Méline Bizard, Angela Brisebarre, Flora Donati, Etienne Simon-Lorière, Vincent Enouf, Maud Vanpeene, Sylvie van der Werf, Raulin Olivia |
| EPI_ISL_418229 | Hopital franco britannique - Laboratoire | National Reference Center for Viruses of Respiratory Infections, Institut Pasteur, Paris | Mélanie Albert, Marion Barbet, Sylvie Behillil, Méline Bizard, Angela Brisebarre, Flora Donati, Etienne Simon-Lorière, Vincent Enouf, Maud Vanpeene, Sylvie van der Werf, Marianne Asso Bonnet |
| EPI_ISL_418230 | Clinique AVERAY LA BROUSTE, Med. Polyvalente | National Reference Center for Viruses of Respiratory Infections, Institut Pasteur, Paris | Mélanie Albert, Marion Barbet, Sylvie Behillil, Méline Bizard, Angela Brisebarre, Flora Donati, Etienne Simon-Lorière, Vincent Enouf, Maud Vanpeene, Sylvie van der Werf, Elsa Ngwem |
| EPI_ISL_418231 | Centre Hospitalier Compiègne Laboratoire de Biologie | National Reference Center for Viruses of Respiratory Infections, Institut Pasteur, Paris | Mélanie Albert, Marion Barbet, Sylvie Behillil, Méline Bizard, Angela Brisebarre, Flora Donati, Etienne Simon-Lorière, Vincent Enouf, Maud Vanpeene, Sylvie van der Werf, Raulin Olivia |
| EPI_ISL_418232, EPI_ISL_418233 | Service des Urgences | National Reference Center for Viruses of Respiratory Infections, Institut Pasteur, Paris | Mélanie Albert, Marion Barbet, Sylvie Behillil, Méline Bizard, Angela Brisebarre, Flora Donati, Etienne Simon-Lorière, Vincent Enouf, Maud Vanpeene, Sylvie van der Werf, Boubkeur |
| EPI_ISL_418234 | LABM GH nord Essonne | National Reference Center for Viruses of Respiratory Infections, Institut Pasteur, Paris | Mélanie Albert, Marion Barbet, Sylvie Behillil, Méline Bizard, Angela Brisebarre, Flora Donati, Etienne Simon-Lorière, Vincent Enouf, Maud Vanpeene, Sylvie van der Werf, Christine Lambert |
| EPI_ISL_418236, EPI_ISL_418237, EPI_ISL_418238, EPI_ISL_418239 | Centre Hospitalier Compiègne Laboratoire de Biologie | National Reference Center for Viruses of Respiratory Infections, Institut Pasteur, Paris | Mélanie Albert, Marion Barbet, Sylvie Behillil, Méline Bizard, Angela Brisebarre, Flora Donati, Etienne Simon-Lorière, Vincent Enouf, Maud Vanpeene, Sylvie van der Werf, Raulin Olivia |
| EPI_ISL_418240 | LABM GH nord Essonne | National Reference Center for Viruses of Respiratory Infections, Institut Pasteur, Paris | Mélanie Albert, Marion Barbet, Sylvie Behillil, Méline Bizard, Angela Brisebarre, Flora Donati, Etienne Simon-Lorière, Vincent Enouf, Maud Vanpeene, Sylvie van der Werf, Christine Lambert |
| EPI_ISL_418241, EPI_ISL_418242 | NIC Viral Respiratory Unit - Institut Pasteur of Algeria | National Reference Center for Viruses of Respiratory Infections, Institut Pasteur, Paris | Mélanie Albert, Marion Barbet, Sylvie Behillil, Méline Bizard, Angela Brisebarre, Flora Donati, Etienne Simon-Lorière, Vincent Enouf, Maud Vanpeene, Sylvie van der Werf, Fawzi Derrar |
| EPI_ISL_418243, EPI_ISL_418244 | HOSPITAL UNIVERSITARIO VIRGEN DE LAS NIEVES | Instituto de Salud Carlos III | Iglesias-Caballero, M. Molinero Calamita, M. González-Esguevillas, M. Camarero, S. Pozo, F. Casas, I. Jiménez, P. Jiménez, M. Zaballos, A. Monzón, S. Varona, S. Juliá, M. Cuesta, I. Sanbonmatsu S. |
| EPI_ISL_418245, EPI_ISL_418246 | Hospital General y Universitario de Guadalajara | Instituto de Salud Carlos III | Iglesias-Caballero, M. Molinero Calamita, M. González-Esguevillas, M. Camarero, S. Pozo, F. Casas, I. Jiménez, P. Jiménez, M. Zaballos, A. Monzón, S. Varona, S. Juliá, M. Cuesta, I. Gonzalez-Praetorius A. |
| EPI_ISL_418247 | HOSPITAL GENERAL DE SEGOVIA | Instituto de Salud Carlos III | Iglesias-Caballero, M. Molinero Calamita, M. González-Esguevillas, M. Camarero, S. Pozo, F. Casas, I. Jiménez, P. Jiménez, M. Zaballos, A. Monzón, S. Varona, S. Juliá, M. Cuesta, I. Hernando-Real S. |
| EPI_ISL_418248, EPI_ISL_418249 | COMPLEJO ASISTENCIAL UNIVERSITARIO DE BURGOS | Instituto de Salud Carlos III | Iglesias-Caballero, M. Molinero Calamita, M. González-Esguevillas, M. Camarero, S. Pozo, F. Casas, I. Jiménez, P. Jiménez, M. Zaballos, A. Monzón, S. Varona, S. Juliá, M. Cuesta, I. Megias-Lobon G. |
| EPI_ISL_418250 | HOSPITAL CLINIC | Instituto de Salud Carlos III | Iglesias-Caballero, M. Molinero Calamita, M. González-Esguevillas, M. Camarero, S. Pozo, F. Casas, I. Jiménez, P. Jiménez, M. Zaballos, A. Monzón, S. Varona, S. Juliá, M. Cuesta, I. Marcos M.A |
| EPI_ISL_418251 | HOSPITAL UNIVERSITARIO LA PAZ | Instituto de Salud Carlos III | Iglesias-Caballero, M. Molinero Calamita, M. González-Esguevillas, M. Camarero, S. Pozo, F. Casas, I. Jiménez, P. Jiménez, M. Zaballos, A. Monzón, S. Varona, S. Juliá, M. Cuesta, I. Romero P. |
| EPI_ISL_418252 | FUNDACION JIMENEZ DIAZ | Instituto de Salud Carlos III | Iglesias-Caballero, M. Molinero Calamita, M. González-Esguevillas, M. Camarero, S. Pozo, F. Casas, I. Jiménez, P. Jiménez, M. Zaballos, A. Monzón, S. Varona, S. Juliá, M. Cuesta, I. Fernández Roblas, R. |
| EPI_ISL_418253 | HOSPITAL TXAGORRITXU | Instituto de Salud Carlos III | Iglesias-Caballero, M. Molinero Calamita, M. González-Esguevillas, M. Camarero, S. Pozo, F. Casas, I. Jiménez, P. Jiménez, M. Zaballos, A. Monzón, S. Varona, S. Juliá, M. Cuesta, I. Gomez-Gonzalez C. |

| | | | |
|---|---|---|---|
| EPI_ISL_418256 | Ospedale "San Liberatore" di Atri | Istituto Zooprofilattico Sperimentale dell'Abruzzo e Molise "G. Caporale" | Lorusso A, Marcacci M, Di Domenico M, Puglia I, Curini V, Ancora M, Di Pasquale A, Rinaldi A, Mangone I, Cammà C, Savini G. |
| EPI_ISL_418257 | Ospedale Civile Giuseppe Mazzini, Teramo | Istituto Zooprofilattico Sperimentale dell'Abruzzo e Molise "G.Caporale" | Lorusso A, Marcacci M, Di Domenico M, Puglia I, Curini V, Ancora M, Di Pasquale A, Rinaldi A, Mangone I, Cammà C, Savini G. |
| EPI_ISL_418262 | Instituto Nacional de Salud | Instituto Nacional de Salud Universidad Cooperativa de Colombia Instituto Alexander von Humboldt Imperial College-London London School of Hygiene & Tropical Medicine | Marcela Mercado-Reyes, Katherine Laiton-Donato, Diego A. Álvarez-Díaz, Carlos Franco-Muñoz, Jose A. Usme-Ciro, Gloria Puerto, Nicolas D. Franco-Sierra, Mailyn A.Gonzalez, Zulma M. Cucunubá, Christian Julian VillabonaArenas, Liz Villabona-Arenas, Sussy Echeverria, Astrid C. Flórez, Sergio Gomez Rangel, Luz Dary Rodriguez, Juliana Barbosa, Erika Ospitia, Diana Marcela Walteros-Acero, Nuno Rodrigues Faria, Martha Lucia Ospina Martinez |
| EPI_ISL_418624, EPI_ISL_418633, EPI_ISL_418645, EPI_ISL_418646, EPI_ISL_418648, EPI_ISL_418649, EPI_ISL_418650, EPI_ISL_418651, EPI_ISL_418652, EPI_ISL_418659, EPI_ISL_418660, EPI_ISL_418664 | | | |
| see above | Department of Clinical Microbiology | GIGA Medical Genomics | Keith Durkin, Maria Artesi, Sébastien Bontems, Raphaël Boreux, Cécile Meex, Pierrette Melin, Marie-Pierre Hayette, Vincent Bours. |
| EPI_ISL_418800 | KU Leuven, Clinical and Epidemiological Virology | KU Leuven, Clinical and Epidemiological Virology | Bert Vanmechelen, Joan Marti-Carreras, Tony Wawina, Piet Maes |
| EPI_ISL_418815 | Department of Clinical Pathology, Pamela Youde Nethersole Eastern Hospital | Department of Health Technology and Informatics, Faculty of Health and Social Science, The Hong Kong Polytechnic University | Kenneth Siu-Sing LEUNG, Timothy Ting-Leung NG, Alan Ka-Lun WU, Miranda Chong-Yee YAU, Hiu-Yin LAO, Ming-Pan CHOI, Kingsley King-Gee TAM, Lam-Kwong LEE, Barry Kin-Chung WONG, Alex Yat-Man HO, Kam-Tong YIP, Kwok-Cheung LUNG, Raymond Wai-To LIU, Eugene Yuk-Keung TSO, Wai-Shing LEUNG, Man-Chun CHAN, Yuk-Yung NG, Kit-Man SIN, Kitty Sau-Chun FUNG, Sandy Ka-Yee CHAU, Wing-Kin TO, Tak-Lun QUE, David Ho-Keung SHUM, Shea Ping YIP, Wing Cheong YAM, Gilman Kit-Hang SIU |
| EPI_ISL_418990, EPI_ISL_418991 | State Key Laboratory for Diagnosis and Treatment of Infectious Diseases, National Clinical Research Center for Infectious Diseases, First Affiliated Hospital, Zhejiang University School of Medicine, Hangzhou, China 310003 | State Key Laboratory for Diagnosis and Treatment of Infectious Diseases, National Clinical Research Center for Infectious Diseases, First Affiliated Hospital, Zhejiang University School of Medicine, Hangzhou, China 310003 | Hangping Yao, Nanping Wu, Chao Jiang, Xiangyun Lu, Linfang Cheng, Fumin Liu, Zhigang Wu, Haibo Wu, Changzhong Jin, Min Zheng, Lanjuan Li |
| EPI_ISL_419214, EPI_ISL_419215, EPI_ISL_419216 | Department of Clinical Pathology, Pamela Youde Nethersole Eastern Hospital | Department of Health Technology and Informatics, Faculty of Health and Social Science, The Hong Kong Polytechnic University | Kenneth Siu-Sing LEUNG, Timothy Ting-Leung NG, Alan Ka-Lun WU, Miranda Chong-Yee YAU, Hiu-Yin LAO, Ming-Pan CHOI, Kingsley King-Gee TAM, Lam-Kwong LEE, Barry Kin-Chung WONG, Alex Yat-Man HO, Kam-Tong YIP, Kwok-Cheung LUNG, Raymond Wai-To LIU, Eugene Yuk-Keung TSO, Wai-Shing LEUNG, Man-Chun CHAN, Yuk-Yung NG, Kit-Man SIN, Kitty Sau-Chun FUNG, Sandy Ka-Yee CHAU, Wing-Kin TO, Tak-Lun QUE, David Ho-Keung SHUM, Shea Ping YIP, Wing Cheong YAM, Gilman Kit-Hang SIU |
| EPI_ISL_419217 | Department of Pathology, Princess Margaret Hospital | Department of Health Technology and Informatics, Faculty of Health and Social Science, The Hong Kong Polytechnic University | Kenneth Siu-Sing LEUNG, Timothy Ting-Leung NG, Alan Ka-Lun WU, Miranda Chong-Yee YAU, Hiu-Yin LAO, Ming-Pan CHOI, Kingsley King-Gee TAM, Lam-Kwong LEE, Barry Kin-Chung WONG, Alex Yat-Man HO, Kam-Tong YIP, Kwok-Cheung LUNG, Raymond Wai-To LIU, Eugene Yuk-Keung TSO, Wai-Shing LEUNG, Man-Chun CHAN, Yuk-Yung NG, Kit-Man SIN, Kitty Sau-Chun FUNG, Sandy Ka-Yee CHAU, Wing-Kin TO, Tak-Lun QUE, David Ho-Keung SHUM, Shea Ping YIP, Wing Cheong YAM, Gilman Kit-Hang SIU |
| EPI_ISL_419219 | Department of Clinical Pathology, Pamela Youde Nethersole Eastern Hospital | Department of Health Technology and Informatics, Faculty of Health and Social Science, The Hong Kong Polytechnic University | Kenneth Siu-Sing LEUNG, Timothy Ting-Leung NG, Alan Ka-Lun WU, Miranda Chong-Yee YAU, Hiu-Yin LAO, Ming-Pan CHOI, Kingsley King-Gee TAM, Lam-Kwong LEE, Barry Kin-Chung WONG, Alex Yat-Man HO, Kam-Tong YIP, Kwok-Cheung LUNG, Raymond Wai-To LIU, Eugene Yuk-Keung TSO, Wai-Shing LEUNG, Man-Chun CHAN, Yuk-Yung NG, Kit-Man SIN, Kitty Sau-Chun FUNG, Sandy Ka-Yee CHAU, Wing-Kin TO, Tak-Lun QUE, David Ho-Keung SHUM, Shea Ping YIP, Wing Cheong YAM, Gilman Kit-Hang SIU |
| EPI_ISL_419221 | Department of Pathology, United Christian Hospital | Department of Health Technology and Informatics, Faculty of Health and Social Science, The Hong Kong Polytechnic University | Kenneth Siu-Sing LEUNG, Timothy Ting-Leung NG, Alan Ka-Lun WU, Miranda Chong-Yee YAU, Hiu-Yin LAO, Ming-Pan CHOI, Kingsley King-Gee TAM, Lam-Kwong LEE, Barry Kin-Chung WONG, Alex Yat-Man HO, Kam-Tong YIP, Kwok-Cheung LUNG, Raymond Wai-To LIU, Eugene Yuk-Keung TSO, Wai-Shing LEUNG, Man-Chun CHAN, Yuk-Yung NG, Kit-Man SIN, Kitty Sau-Chun FUNG, Sandy Ka-Yee CHAU, Wing-Kin TO, Tak-Lun QUE, David Ho-Keung SHUM, Shea Ping YIP, Wing Cheong YAM, Gilman Kit-Hang SIU |
| EPI_ISL_419222 | Department of Pathology, Princess Margaret Hospital | Department of Health Technology and Informatics, Faculty of Health and Social Science, The Hong Kong Polytechnic University | Kenneth Siu-Sing LEUNG, Timothy Ting-Leung NG, Alan Ka-Lun WU, Miranda Chong-Yee YAU, Hiu-Yin LAO, Ming-Pan CHOI, Kingsley King-Gee TAM, Lam-Kwong LEE, Barry Kin-Chung WONG, Alex Yat-Man HO, Kam-Tong YIP, Kwok-Cheung LUNG, Raymond Wai-To LIU, Eugene Yuk-Keung TSO, Wai-Shing LEUNG, Man-Chun CHAN, Yuk-Yung NG, Kit-Man SIN, Kitty Sau-Chun FUNG, Sandy Ka-Yee CHAU, Wing-Kin TO, Tak-Lun QUE, David Ho-Keung SHUM, Shea Ping YIP, Wing Cheong YAM, Gilman Kit-Hang SIU |
| EPI_ISL_419224, EPI_ISL_419225, EPI_ISL_419226, EPI_ISL_419227, EPI_ISL_419228, EPI_ISL_419229 | Department of Clinical Pathology, Pamela Youde Nethersole Eastern Hospital | Department of Health Technology and Informatics, Faculty of Health and Social Science, The Hong Kong Polytechnic University | Kenneth Siu-Sing LEUNG, Timothy Ting-Leung NG, Alan Ka-Lun WU, Miranda Chong-Yee YAU, Hiu-Yin LAO, Ming-Pan CHOI, Kingsley King-Gee TAM, Lam-Kwong LEE, Barry Kin-Chung WONG, Alex Yat-Man HO, Kam-Tong YIP, Kwok-Cheung LUNG, Raymond Wai-To LIU, Eugene Yuk-Keung TSO, Wai-Shing LEUNG, Man-Chun CHAN, Yuk-Yung NG, Kit-Man SIN, Kitty Sau-Chun FUNG, Sandy Ka-Yee CHAU, Wing-Kin TO, Tak-Lun QUE, David Ho-Keung SHUM, Shea Ping YIP, Wing Cheong YAM, Gilman Kit-Hang SIU |
| EPI_ISL_419230 | Hospital Universitario Virgen de las Nieves | Instituto de Salud Carlos III | Iglesias-Caballero, M.; Molinero Calamita, M.; González-Esguevillas, M.; Camarero, S.; Pozo, F.; Casas, I.; Jiménez, P.; Jiménez, M.; Zaballos, A.; Monzón, S.; Varona, S.; Juliá, M.; Cuesta, I.; Sanbonmatsu, S. |
| EPI_ISL_419231 | Department of Clinical Pathology, Tuen Mun Hospital, 23 Tsing Chung Koon Road, Tuen Mun, N.T. | Department of Health Technology and Informatics, Faculty of Health and Social Science, The Hong Kong Polytechnic University | Kenneth Siu-Sing LEUNG, Timothy Ting-Leung NG, Alan Ka-Lun WU, Miranda Chong-Yee YAU, Hiu-Yin LAO, Ming-Pan CHOI, Kingsley King-Gee TAM, Lam-Kwong LEE, Barry Kin-Chung WONG, Alex Yat-Man HO, Kam-Tong YIP, Kwok-Cheung LUNG, Raymond Wai-To LIU, Eugene Yuk-Keung TSO, Wai-Shing LEUNG, Man-Chun CHAN, Yuk-Yung NG, Kit-Man SIN, Kitty Sau-Chun FUNG, Sandy Ka-Yee CHAU, Wing-Kin TO, Tak-Lun QUE, David Ho-Keung SHUM, Shea Ping YIP, Wing Cheong YAM, Gilman Kit-Hang SIU |
| EPI_ISL_419232 | Department of Clinical Pathology, Pamela Youde Nethersole Eastern Hospital | Department of Health Technology and Informatics, Faculty of Health and Social Science, The Hong Kong Polytechnic University | Kenneth Siu-Sing LEUNG, Timothy Ting-Leung NG, Alan Ka-Lun WU, Miranda Chong-Yee YAU, Hiu-Yin LAO, Ming-Pan CHOI, Kingsley King-Gee TAM, Lam-Kwong LEE, Barry Kin-Chung WONG, Alex Yat-Man HO, Kam-Tong YIP, Kwok-Cheung LUNG, Raymond Wai-To LIU, Eugene Yuk-Keung TSO, Wai-Shing LEUNG, Man-Chun CHAN, Yuk-Yung NG, Kit-Man SIN, Kitty Sau-Chun FUNG, Sandy Ka-Yee CHAU, Wing-Kin TO, Tak-Lun QUE, David Ho-Keung SHUM, Shea Ping YIP, Wing Cheong YAM, Gilman Kit-Hang SIU |
| EPI_ISL_419233 | Hospital Universitario de Canarias | Instituto de Salud Carlos III | Iglesias-Caballero, M.; Molinero Calamita, M.; González-Esguevillas, M.; Camarero, S.; Pozo, F.; Casas, I.; Jiménez, P.; Jiménez, M.; Zaballos, A.; Monzón, S.; Varona, S.; Juliá, M.; Cuesta, I.; Castro, B. |
| EPI_ISL_419234 | Hospital San Pedro | Instituto de Salud Carlos III | Iglesias-Caballero, M.; Molinero Calamita, M.; González-Esguevillas, M.; Camarero, S.; Pozo, F.; Casas, I.; Jiménez, P.; Jiménez, M.; Zaballos, A.; Monzón, S.; Varona, S.; Juliá, M.; Cuesta, I.; Alonso, C. |
| EPI_ISL_419235, EPI_ISL_419236, EPI_ISL_419237 | Fundacion Jimenez Diaz | Instituto de Salud Carlos III | Iglesias-Caballero, M.; Molinero Calamita, M.; González-Esguevillas, M.; Camarero, S.; Pozo, F.; Casas, I.; Jiménez, P.; Jiménez, M.; Zaballos, A.; Monzón, S.; Varona, S.; Juliá, M.; Cuesta, I.; Fernández, R. |
| EPI_ISL_419238 | HOSPITAL DE CRUCES. | Instituto de Salud Carlos III | Iglesias-Caballero, M. Molinero Calamita, M. González-Esguevillas, M. Camarero, S. Pozo, F. Casas, I. Jiménez, P. Jiménez, M. Zaballos, A. Monzón, S. Varona, S. Juliá, M. Cuesta, I. Aranzamendi, M. |
| EPI_ISL_419240 | HOSPITAL TXAGORRITXU | Instituto de Salud Carlos III | Iglesias-Caballero, M. Molinero Calamita, M. González-Esguevillas, M. Camarero, S. Pozo, F. Casas, I. Jiménez, P. Jiménez, M. Zaballos, A. Monzón, S. Varona, S. Juliá, M. Cuesta, I. Gómez, C |
| EPI_ISL_419245, EPI_ISL_419247, EPI_ISL_419250, EPI_ISL_419252 | Department of Clinical Pathology, Pamela Youde Nethersole Eastern Hospital | Department of Health Technology and Informatics, Faculty of Health and Social Science, The Hong Kong Polytechnic University | Kenneth Siu-Sing LEUNG, Timothy Ting-Leung NG, Alan Ka-Lun WU, Miranda Chong-Yee YAU, Hiu-Yin LAO, Ming-Pan CHOI, Kingsley King-Gee TAM, Lam-Kwong LEE, Barry Kin-Chung WONG, Alex Yat-Man HO, Kam-Tong YIP, Kwok-Cheung LUNG, Raymond Wai-To LIU, Eugene Yuk-Keung TSO, Wai-Shing LEUNG, Man-Chun CHAN, Yuk-Yung NG, Kit-Man SIN, Kitty Sau-Chun FUNG, Sandy Ka-Yee CHAU, Wing-Kin TO, Tak-Lun QUE, David Ho-Keung SHUM, Shea Ping YIP, Wing Cheong YAM, Gilman Kit-Hang SIU |
| EPI_ISL_419296 | Kochi Prefectural Institute of Public Health | Pathogen Genomics Center, National Institute of Infectious Diseases | Tsuyoshi Sekizuka, Akihiko Tokaji, Kentaro Itokawa, Rina Tanaka, Masanori Hashino, Hajime Kamiya, Motoi Suzuki, Makoto Kuroda |
| EPI_ISL_419297, EPI_ISL_419298 | Chiba Prefectural Institute of Public Health | Pathogen Genomics Center, National Institute of Infectious Diseases | Tsuyoshi Sekizuka, Masakatsu Taira, Yushi Hachisu, Kentaro Itokawa, Rina Tanaka, Masanori Hashino, Hajime Kamiya, Motoi Suzuki, Makoto Kuroda |
| EPI_ISL_419299, EPI_ISL_419300 | Ishikawa Prefectural Institute of Public Health and Environmental Science | Pathogen Genomics Center, National Institute of Infectious Diseases | Tsuyoshi Sekizuka, Sanae Kuramoto, Eri Nariai, Kentaro Itokawa, Rina Tanaka, Masanori Hashino, Hajime Kamiya, Motoi Suzuki, Makoto Kuroda |
| EPI_ISL_419301, EPI_ISL_419302, | Saitama Prefectural Institute of Public Health | Pathogen Genomics Center, National Institute of Infectious | Tsuyoshi Sekizuka, Michiyo Shinohara, Tsuyoshi Kishimoto, Kentaro Itokawa, Rina Tanaka, Masanori Hashino, Hajime Kamiya, Motoi Suzuki, Makoto Kuroda |

| | | | |
|---|---|---|---|
| EPI_ISL_419303, EPI_ISL_419304, EPI_ISL_419305, EPI_ISL_419306, EPI_ISL_419307, EPI_ISL_419308 | | Diseases | |
| EPI_ISL_419309, EPI_ISL_419310, EPI_ISL_419311 | Chiba Prefectural Institute of Public Health | Pathogen Genomics Center, National Institute of Infectious Diseases | Tsuyoshi Sekizuka, Masakatsu Taira, Yushi Hachisu, Kentaro Itokawa, Rina Tanaka, Masanori Hashino, Hajime Kamiya, Motoi Suzuki, Makoto Kuroda |
| EPI_ISL_419662, EPI_ISL_419664 | Center for Virology, Medical University of Vienna | Bergthaler laboratory, CeMM Research Center for Molecular Medicine of the Austrian Academy of Sciences | Alexandra Popa, Benedikt Agerer, Henrique Colaco, Lukas Endler, Jakob-Wendelin Genger, Alexander Lercher, Mark Smyth, Thomas Penz, Michael Schuster, Judith Aberle, Stephan Aberle, Elisabeth Puchhammer-Stöckl, Christoph Bock, Andreas Bergthaler |
| EPI_ISL_419707 | HOSPITAL CLINIC | Instituto de Salud Carlos III | Iglesias-Caballero, M. Molinero Calamita, M. González-Esguevillas, M. Camarero S. Pozo F. Casas I. Jiménez, P. Jiménez, M. Zaballos, A. Monzón, S. Varona, S. Juliá, M. Cuesta, I. Marcos, M.A |
| EPI_ISL_419709 | HOSPITAL TXAGORRITXU | Instituto de Salud Carlos III | Iglesias-Caballero, M. Molinero Calamita, M. González-Esguevillas, M. Camarero S. Pozo F. Casas I. Jiménez, P. Jiménez, M. Zaballos, A. Monzón, S. Varona, S. Juliá, M. Cuesta, I. Gómez, C. |
| EPI_ISL_420037 | NIC Viral Respiratory Unit - Institut Pasteur of Algeria | National Reference Center for Viruses of Respiratory Infections, Institut Pasteur, Paris | Mélanie Albert, Marion Barbet, Sylvie Behillil, Méline Bizard, Angela Brisebarre, Flora Donati, Etienne Simon-Lorière, Vincent Enouf, Maud Vanpeene, Sylvie van der Werf, Fawzi Derrar |
| EPI_ISL_420039, EPI_ISL_420040 | L'Air du Temps | National Reference Center for Viruses of Respiratory Infections, Institut Pasteur, Paris | Mélanie Albert, Marion Barbet, Sylvie Behillil, Méline Bizard, Angela Brisebarre, Flora Donati, Etienne Simon-Lorière, Vincent Enouf, Maud Vanpeene, Sylvie van der Werf |
| EPI_ISL_420041 | CH Compiègne Laboratoire de Biologie | National Reference Center for Viruses of Respiratory Infections, Institut Pasteur, Paris | Mélanie Albert, Marion Barbet, Sylvie Behillil, Méline Bizard, Angela Brisebarre, Flora Donati, Etienne Simon-Lorière, Vincent Enouf, Maud Vanpeene, Sylvie van der Werf, Raulin Olivia |
| EPI_ISL_420042 | Service de Biologie clinique | National Reference Center for Viruses of Respiratory Infections, Institut Pasteur, Paris | Mélanie Albert, Marion Barbet, Sylvie Behillil, Méline Bizard, Angela Brisebarre, Flora Donati, Etienne Simon-Lorière, Vincent Enouf, Maud Vanpeene, Sylvie van der Werf |
| EPI_ISL_420044 | CH Jean de Navarre Laboratoire de Biologie | National Reference Center for Viruses of Respiratory Infections, Institut Pasteur, Paris | Mélanie Albert, Marion Barbet, Sylvie Behillil, Méline Bizard, Angela Brisebarre, Flora Donati, Etienne Simon-Lorière, Vincent Enouf, Maud Vanpeene, Sylvie van der Werf |
| EPI_ISL_420046, EPI_ISL_420047 | Résidence Villa Caroline | National Reference Center for Viruses of Respiratory Infections, Institut Pasteur, Paris | Mélanie Albert, Marion Barbet, Sylvie Behillil, Méline Bizard, Angela Brisebarre, Flora Donati, Etienne Simon-Lorière, Vincent Enouf, Maud Vanpeene, Sylvie van der Werf |
| EPI_ISL_420048 | Service de Biologie Médicale - BP 125 | National Reference Center for Viruses of Respiratory Infections, Institut Pasteur, Paris | Mélanie Albert, Marion Barbet, Sylvie Behillil, Méline Bizard, Angela Brisebarre, Flora Donati, Etienne Simon-Lorière, Vincent Enouf, Maud Vanpeene, Sylvie van der Werf, Christine Lambert |
| EPI_ISL_420049, EPI_ISL_420050 | CH Compiègne Laboratoire de Biologie | National Reference Center for Viruses of Respiratory Infections, Institut Pasteur, Paris | Mélanie Albert, Marion Barbet, Sylvie Behillil, Méline Bizard, Angela Brisebarre, Flora Donati, Etienne Simon-Lorière, Vincent Enouf, Maud Vanpeene, Sylvie van der Werf, Raulin Olivia |
| EPI_ISL_420051 | Résidence Eleusis | National Reference Center for Viruses of Respiratory Infections, Institut Pasteur, Paris | Mélanie Albert, Marion Barbet, Sylvie Behillil, Méline Bizard, Angela Brisebarre, Flora Donati, Etienne Simon-Lorière, Vincent Enouf, Maud Vanpeene, Sylvie van der Werf |
| EPI_ISL_420052 | Résidence les Marines | National Reference Center for Viruses of Respiratory Infections, Institut Pasteur, Paris | Mélanie Albert, Marion Barbet, Sylvie Behillil, Méline Bizard, Angela Brisebarre, Flora Donati, Etienne Simon-Lorière, Vincent Enouf, Maud Vanpeene, Sylvie van der Werf |
| EPI_ISL_420053 | CH Jean de Navarre Laboratoire de Biologie | National Reference Center for Viruses of Respiratory Infections, Institut Pasteur, Paris | Mélanie Albert, Marion Barbet, Sylvie Behillil, Méline Bizard, Angela Brisebarre, Flora Donati, Etienne Simon-Lorière, Vincent Enouf, Maud Vanpeene, Sylvie van der Werf |
| EPI_ISL_420056, EPI_ISL_420057 | CH Compiègne Laboratoire de Biologie | National Reference Center for Viruses of Respiratory Infections, Institut Pasteur, Paris | Mélanie Albert, Marion Barbet, Sylvie Behillil, Méline Bizard, Angela Brisebarre, Flora Donati, Etienne Simon-Lorière, Vincent Enouf, Maud Vanpeene, Sylvie van der Werf, Raulin Olivia |
| EPI_ISL_420058, EPI_ISL_420059, EPI_ISL_420060 | Service de Biologie Médicale - BP 125 | National Reference Center for Viruses of Respiratory Infections, Institut Pasteur, Paris | Mélanie Albert, Marion Barbet, Sylvie Behillil, Méline Bizard, Angela Brisebarre, Flora Donati, Etienne Simon-Lorière, Vincent Enouf, Maud Vanpeene, Sylvie van der Werf, Christine Lambert |
| EPI_ISL_420063 | Labo BM - Site de Juvisy - Hopital Général | National Reference Center for Viruses of Respiratory Infections, Institut Pasteur, Paris | Mélanie Albert, Marion Barbet, Sylvie Behillil, Méline Bizard, Angela Brisebarre, Flora Donati, Etienne Simon-Lorière, Vincent Enouf, Maud Vanpeene, Sylvie van der Werf |
| EPI_ISL_420064 | Service de Biologie Médicale - BP 125 | National Reference Center for Viruses of Respiratory Infections, Institut Pasteur, Paris | Mélanie Albert, Marion Barbet, Sylvie Behillil, Méline Bizard, Angela Brisebarre, Flora Donati, Etienne Simon-Lorière, Vincent Enouf, Maud Vanpeene, Sylvie van der Werf, Christine Lambert |
| EPI_ISL_420069, EPI_ISL_420070 | Institut Pasteur Dakar | Institut Pasteur de Dakar | Ndongo Dia, Moussa Moise Diagne, Mamadou Diop, Ousmane Faye, Amadou Alpha Sall |
| EPI_ISL_420072, EPI_ISL_420073, EPI_ISL_420074 | Institut Pasteur Dakar | Institut Pasteur de Dakar | Ndongo Dia, Moussa Moise Diagne, Mamadou Diop, Ousmane Faye , Amadou Alpha Sall |
| EPI_ISL_420076 | Institut Pasteur Dakar | Institut Pasteur de Dakar | Ndongo Dia, Moussa Moise Diagne, Mamadou Diop, Ousmane Faye , Ndongo Dia |
| EPI_ISL_420077, EPI_ISL_420078 | Institut Pasteur Dakar | Institut Pasteur de Dakar | Ndongo Dia, Moussa Moise Diagne, Mamadou Diop, Ousmane Faye , Amadou Alpha Sall |
| EPI_ISL_420080 | WHO National Influenza Centre Russian Federation | WHO National Influenza Centre Russian Federation | Andrey Komissarov, Artem Fadeev, Anna Ivanova, Daria Danilenko |
| EPI_ISL_420081 | WHO National Influenza Centre Russian Federation | WHO National Influenza Centre Russian Federation | Andrey Komissarov, Artem Fadeev, Mariia Sergeeva, Anna Ivanova, Daria Danilenko |
| EPI_ISL_421182, EPI_ISL_421183, EPI_ISL_421184, EPI_ISL_421185, EPI_ISL_421186, EPI_ISL_421187, EPI_ISL_421188, EPI_ISL_421189, EPI_ISL_421190, EPI_ISL_421191, EPI_ISL_421192, EPI_ISL_421193, EPI_ISL_421194, EPI_ISL_421195, EPI_ISL_421196 | | | |
| see above | Department of Clinical Microbiology | GIGA Medical Genomics | Keith Durkin, Maria Artesi, Sébastien Bontems, Raphaël Boreux, Cécile Meex, Pierrette Melin, Marie-Pierre Hayette, Vincent Bours. |
| EPI_ISL_421221, EPI_ISL_421222, EPI_ISL_421224, EPI_ISL_421225, EPI_ISL_421226, EPI_ISL_421227, EPI_ISL_421228, EPI_ISL_421229, EPI_ISL_421230, EPI_ISL_421231, EPI_ISL_421232, EPI_ISL_421233, EPI_ISL_421234, EPI_ISL_421235, EPI_ISL_421236 | | | |
| see above | Hangzhou Center for Diseases Control and Prevention | Hangzhou Center for Diseases Control and Prevention | Jun Li, Haoqiu Wang, Lingfeng Mao, Hua Yu, Xinfen Yu, Zhou Sun, Xin Qian, Shuchang Chen, Junfang Chen, Xuchu Wang |
| EPI_ISL_421275 | Russian State Collection of Viruses | Pathogenic Microorganisms Variability Laboratory | Alexey Shchetinin, Maria Nikiforova, Nadezhda Kuznetsova, Ekaterina Aksenova, Marina Kunda, Natalia Ryzhova, Olga Voronina, Inna Dolzhikova, Daria Grousova, Andrey Botikov, Denis Logunov, Alexander Gintsburg, Vladimir Gushchin |
| EPI_ISL_421500 | CH Compiègne Laboratoire de Biologie | National Reference Center for Viruses of Respiratory Infections, Institut Pasteur, Paris | Mélanie Albert, Marion Barbet, Sylvie Behillil, Méline Bizard, Angela Brisebarre, Flora Donati, Etienne Simon-Lorière, Vincent Enouf, Maud Vanpeene, Sylvie van der Werf, Raulin Olivia |
| EPI_ISL_421501 | Service de Biologie Médicale - BP 125 | National Reference Center for Viruses of Respiratory Infections, Institut Pasteur, Paris | Mélanie Albert, Marion Barbet, Sylvie Behillil, Méline Bizard, Angela Brisebarre, Flora Donati, Etienne Simon-Lorière, Vincent Enouf, Maud Vanpeene, Sylvie van der Werf, Christine Lambert |
| EPI_ISL_421502, EPI_ISL_421503 | Parc des Dames | National Reference Center for Viruses of Respiratory Infections, Institut Pasteur, Paris | Mélanie Albert, Marion Barbet, Sylvie Behillil, Méline Bizard, Angela Brisebarre, Flora Donati, Etienne Simon-Lorière, Vincent Enouf, Maud Vanpeene, Sylvie van der Werf |
| EPI_ISL_421504, EPI_ISL_421505, EPI_ISL_421506 | Service de Biologie Médicale - BP 125 | National Reference Center for Viruses of Respiratory Infections, Institut Pasteur, Paris | Mélanie Albert, Marion Barbet, Sylvie Behillil, Méline Bizard, Angela Brisebarre, Flora Donati, Etienne Simon-Lorière, Vincent Enouf, Maud Vanpeene, Sylvie van der Werf, Christine Lambert |
| EPI_ISL_421507, EPI_ISL_421508 | Le Château de Seine-Port | National Reference Center for Viruses of Respiratory Infections, Institut Pasteur, Paris | Mélanie Albert, Marion Barbet, Sylvie Behillil, Méline Bizard, Angela Brisebarre, Flora Donati, Etienne Simon-Lorière, Vincent Enouf, Maud Vanpeene, Sylvie van der Werf |
| EPI_ISL_421509, EPI_ISL_421510, EPI_ISL_421511 | CH Compiègne Laboratoire de Biologie | National Reference Center for Viruses of Respiratory Infections, Institut Pasteur, Paris | Mélanie Albert, Marion Barbet, Sylvie Behillil, Méline Bizard, Angela Brisebarre, Flora Donati, Etienne Simon-Lorière, Vincent Enouf, Maud Vanpeene, Sylvie van der Werf, Raulin Olivia |
| EPI_ISL_421512 | Service de Biologie Médicale - BP 125 | National Reference Center for Viruses of Respiratory Infections, Institut Pasteur, Paris | Mélanie Albert, Marion Barbet, Sylvie Behillil, Méline Bizard, Angela Brisebarre, Flora Donati, Etienne Simon-Lorière, Vincent Enouf, Maud Vanpeene, Sylvie van der Werf, Christine Lambert |
| EPI_ISL_421513 | Service de Biologie clinique | National Reference Center for Viruses of Respiratory | Mélanie Albert, Marion Barbet, Sylvie Behillil, Méline Bizard, Angela Brisebarre, Flora Donati, Etienne Simon-Lorière, Vincent Enouf, Maud Vanpeene, Sylvie van |

| | | | |
|---|---|---|---|
| | | Infections, Institut Pasteur, Paris | der Werf, Christine Lambert |
| EPI_ISL_421574 | Molecular Diagnostic Services | KRISP, KZN Research Innovation and Sequencing Platform | Giandhari J, Pillay S, Ngcapu S, Samsunder N, Lessells R, Chimukangara B, Deforche K, Tegally H, Wilkinson E, de Oliveira T |
| EPI_ISL_421652 | Dasman Diabetes Institute | Dasman Diabetes Institute | Fahd Al-Mulla, Rasheeba Iqbal, Sumi John, Ebaa Al-Ozairi, Qais Al-Duwairi |
| EPI_ISL_422407, EPI_ISL_422408, EPI_ISL_422409, EPI_ISL_422410, EPI_ISL_422411, EPI_ISL_422412, EPI_ISL_422413, EPI_ISL_422414, EPI_ISL_422415, EPI_ISL_422416, EPI_ISL_422417, EPI_ISL_422418, EPI_ISL_422419, EPI_ISL_422420, EPI_ISL_422421, EPI_ISL_422422 | | | |
| see above | Department of Laboratory Medicine, National Taiwan University Hospital | Microbial Genomics Core Lab, National Taiwan University Centers of Genomic and Precision Medicine | Shiou-Hwei Yeh, You-Yu Lin, Ya-Yun Lai, Chiao-Ling Li, Shan-Chwen Chang, Pei-Jer Chen, Sui-Yuan Chang |
| EPI_ISL_422424 | Jaber Al Ahmad Al Sabah Hospital | Dasman diabetes Institute | Fahd Al-Mulla, Rasheeba Iqbal, Sumi John, Ebaa Al-Ozairi, Qais Al-Duwairi |
| EPI_ISL_422427 | JABER AL AHMAD AL SABAH HOSPITAL - KUWAIT CITY | Dasman Diabetes Institute | Fahd Al-Mulla, Rasheeba Iqbal, Sumi John, Ebaa Al-Ozairi, Qais Al-Duwairi |
| EPI_ISL_422636 | The National Institute of Public Health Center for Epidemiology and Microbiology | State Veterinary Institute Prague | Alexander Nagy, Helena Jirincova, Klara Labska, Ludmila Novakova, Olga Storkanova, Dusan Trnka, Jaromira Vecerova |
| EPI_ISL_424366 | Vaccine Research, Development and Application Center, Erciyes University | Gen Era Diagnostics Inc. | Shaikh Terkis Islam Pavel, Hazel Yetiskin, Gunsu Aydin, Can Holyavkin, Muhammet Ali Uygut, Zehra B Dursun, Ihami Celik, Alper Iseri, Aykut Ozdarendeli |
| EPI_ISL_424629, EPI_ISL_424630, EPI_ISL_424631, EPI_ISL_424633, EPI_ISL_424636, EPI_ISL_424637, EPI_ISL_424640, EPI_ISL_424641, EPI_ISL_424642, EPI_ISL_424643, EPI_ISL_424644, EPI_ISL_424647, EPI_ISL_424649, EPI_ISL_424650, EPI_ISL_424652, EPI_ISL_424653, EPI_ISL_424654, EPI_ISL_424655, EPI_ISL_424657, EPI_ISL_424658, EPI_ISL_424660, EPI_ISL_424661, EPI_ISL_424664 | | | |
| see above | Department of Clinical Microbiology | GIGA Medical Genomics | Keith Durkin, Maria Artesi, Sébastien Bontems, Raphaël Boreux, Cécile Meex, Pierrette Melin, Marie-Pierre Hayette, Vincent Bours. |
| EPI_ISL_426364 | Instituto Nacional de Ciencias Medicas y Nutricion Salvador Zubiran | Instituto Nacional de Ciencias Medicas y Nutricion | Guillermo M. Ruiz-Palacios, Pilar Ramos Cervantes, Violeta Ibarra Gonzalez, Fernando Ledesma Barrientos, Luis Alberto García Andrade, Alfredo Ponce de León Garduño, Irma López Martínez, Lucia Hernández Rivas, Gisela Barrera Badillo, Edgar Mendieta Condado, Fabiola Garcés Ayala, Adnan Araiza Rodríguez, José Ernesto Ramírez González, Celia Boukadida, Santiago Avila Ríos, Mario Mújica Sánchez, José Arturo Martínez Orozco, Eduardo Becerril Vargas, Joel Armando Vázquez Pérez, Victor Hugo Borja Aburto, Concepción Grajales Muñiz, Cesar Raúl González Bonilla, Carolina González Torres, Francisco Javier Gaytán Cervantes, José Esteban Muñoz Medina, Blanca Taboada, Alejandro Sánchez, Pavel Isa, Ricardo Grande, Gloria Vázquez, Francisco Pulido, Carlos F. Arias |
| EPI_ISL_426414 | Sir M P Shah Government Medical College | Gujarat Biotechnology Research Centre | Ramesh Pandit, Tejas Shah, Ankit Hinsu, Pritesh Sabara, Apurvasinh Puvar, Janvi Raval, Monika Gandhi, Pinal Trivedi, Maharshi Pandya, Amit Kanani, Akanksha Verma, Nitin Savaliya, Raghawendra Kumar, Dinesh Kumar, Zubair Saiyed, Dipa Kinariwala, Disha Patel, Binita Aring, Geeta Vaghela, Sonia Barve, Bhavesh Modi, Kairavi Joshi, Nidhi Sood, Pranay Shah, R D Dixit, Snehal Bagatharia, Madhvi Joshi, Chaitanya Joshi |
| EPI_ISL_426415 | Sir M P Shah Government Medical College, Jamnagar | Gujarat Biotechnology Research Centre, Gandhinagar | Ramesh Pandit, Tejas Shah, Ankit Hinsu, Pritesh Sabara, Apurvasinh Puvar, Janvi Raval, Monika Gandhi, Pinal Trivedi, Maharshi Pandya, Amit Kanani, Akanksha Verma, Nitin Savaliya, Raghvendra Kumar, Dinesh Kumar, Zuber Saiyed, Dipa Kinariwala, Disha Patel, Binita Aring, Geeta Vaghela, Sonia Barve, Bhavesh Modi, Kairavi Joshi, Nidhi Sood, Pranay Shah, R D Dixit, Snehal Bagatharia, Madhvi Joshi, Chaitanya Joshi |
| EPI_ISL_426580 | Instituto Sabin | Laboratory of Virology | Fernando L Melo,Gustavo Barra, Ticiane H Santa-Rita, Pedro G Mesquita, Ikaro A Andrade, Tatsuya Nagata, Bergmann M Ribeiro |
| EPI_ISL_427307, EPI_ISL_427308, EPI_ISL_427309, EPI_ISL_427310, EPI_ISL_427311, EPI_ISL_427312, EPI_ISL_427314, EPI_ISL_427315, EPI_ISL_427319, EPI_ISL_427320, EPI_ISL_427321, EPI_ISL_427322, EPI_ISL_427323, EPI_ISL_427337, EPI_ISL_427338, EPI_ISL_427339 | | | |
| see above | WHO National Influenza Centre Russian Federation | WHO National Influenza Centre Russian Federation | Andrey Komissarov, Artem Fadeev, Mariia Sergeeva, Anna Ivanova, Daria Danilenko |
| EPI_ISL_427349, EPI_ISL_427350, EPI_ISL_427351, EPI_ISL_427352, EPI_ISL_427354, EPI_ISL_427357, EPI_ISL_427358, EPI_ISL_427359, EPI_ISL_427364, EPI_ISL_427366, EPI_ISL_427370, EPI_ISL_427371, EPI_ISL_427372, EPI_ISL_427379, EPI_ISL_427385, EPI_ISL_427388, EPI_ISL_427390 | | | |
| see above | Department of Clinical Microbiology | GIGA Medical Genomics | Keith Durkin, Maria Artesi, Sébastien Bontems, Raphaël Boreux, Cécile Meex, Pierrette Melin, Marie-Pierre Hayette, Vincent Bours. |
| EPI_ISL_427391 | Genomic Laboratory (GLAB) (Conjoint lab of Health Directorate of Istanbul and Istanbul Technical University) | Genomic Laboratory (GLAB), Istanbul Technical University | Ilker Karacan, Tugba Kizilboga Akgun, Bugra Agaoglu, Gizem Alkurt, Jale Yildiz, Betsi Köse, Elifnaz Çelik, Mehtap Aydn, Levent Doganay, Gizem Dinler Doganay |
| EPI_ISL_428207, EPI_ISL_428208 | National Institute of Health Research and Development | National Institute of Health Research and Development | Setiawaty,V;Subangkit;Puspa,KD;Ikawati,HD;Nugraha,AA;Hariastuti,NI;Ramadhany,R;Susilarini,NK;Pratiwi,E;Agustiningsih;Kurniawati,J;Pawestri,HA;Siswanto |
| EPI_ISL_428209 | Laboratory of Molecular Biology, Diagnostyka sp. z o.o. | Laboratory of Recombinant Vaccines | Lukasz Rabalski, Anna Piotrowska-Mietelska, Boguslaw Szewczyk, Krystyna Bienkowska-Szewczyk |
| EPI_ISL_428229 | TSGH-CP molecular lab | TSGH-CP molecular lab | Cherng-Lih Perng, Ming-Jr Jian, Chih-Kai Chang, Jung-Chung Lin, Kuo-Ming Yeh, Chien-Wen Chen, Sheng-Kang Chiu, Hsing-Yi Chung, Shih-Hung Tsai, Kuo-Sheng Hung, Tien-Yao Chang, Feng-Yee Chang, Hung-Sheng Shang |
| EPI_ISL_428346 | Genomic Laboratory (GLAB) (Conjoint lab of Health Directorate of Istanbul and Istanbul Technical University) | Genomic Laboratory (GLAB), Istanbul Technical University | Ilker Karacan, Tugba Kizilboga Akgun, Bugra Agaoglu, Gizem Alkurt, Jale Yildiz, Betsi Köse, Elifnaz Çelik, Arzu Irvem, Yasemin Kendir Demirkol, Ozlem Akgun Dogan, Mehtap Aydn, Levent Doganay, Gizem Dinler Doganay |
| EPI_ISL_428347, EPI_ISL_428349 | Service de Biologie Médicale - BP 125 | National Reference Center for Viruses of Respiratory Infections, Institut Pasteur, Paris | Mélanie Albert, Marion Barbet, Sylvie Behillil, Méline Bizard, Angela Brisebarre, Flora Donati, Etienne Simon-Lorière, Vincent Enouf, Maud Vanpeene, Sylvie van der Werf |
| EPI_ISL_428350 | CH Jean de Navarre Laboratoire de Biologie | National Reference Center for Viruses of Respiratory Infections, Institut Pasteur, Paris | Mélanie Albert, Marion Barbet, Sylvie Behillil, Méline Bizard, Angela Brisebarre, Flora Donati, Etienne Simon-Lorière, Vincent Enouf, Maud Vanpeene, Sylvie van der Werf |
| EPI_ISL_428351, EPI_ISL_428352 | GH Nord Essonne Service de Biologie clinique | National Reference Center for Viruses of Respiratory Infections, Institut Pasteur, Paris | Mélanie Albert, Marion Barbet, Sylvie Behillil, Méline Bizard, Angela Brisebarre, Flora Donati, Etienne Simon-Lorière, Vincent Enouf, Maud Vanpeene, Sylvie van der Werf |
| EPI_ISL_428353 | CH Compiègne Laboratoire de Biologie | National Reference Center for Viruses of Respiratory Infections, Institut Pasteur, Paris | Mélanie Albert, Marion Barbet, Sylvie Behillil, Méline Bizard, Angela Brisebarre, Flora Donati, Etienne Simon-Lorière, Vincent Enouf, Maud Vanpeene, Sylvie van der Werf |
| EPI_ISL_428354 | LABM GH nord Essonne de Longjumeau - BP 125 | National Reference Center for Viruses of Respiratory Infections, Institut Pasteur, Paris | Mélanie Albert, Marion Barbet, Sylvie Behillil, Méline Bizard, Angela Brisebarre, Flora Donati, Etienne Simon-Lorière, Vincent Enouf, Maud Vanpeene, Sylvie van der Werf |
| EPI_ISL_428355, EPI_ISL_428356, EPI_ISL_428357 | Institut Médico légal- Hop R. Poincaré | National Reference Center for Viruses of Respiratory Infections, Institut Pasteur, Paris | Mélanie Albert, Marion Barbet, Sylvie Behillil, Méline Bizard, Angela Brisebarre, Flora Donati, Etienne Simon-Lorière, Vincent Enouf, Maud Vanpeene, Sylvie van der Werf |
| EPI_ISL_428358 | CH Jeanne de Navarre Laboratoire de Biologie | National Reference Center for Viruses of Respiratory Infections, Institut Pasteur, Paris | Mélanie Albert, Marion Barbet, Sylvie Behillil, Méline Bizard, Angela Brisebarre, Flora Donati, Etienne Simon-Lorière, Vincent Enouf, Maud Vanpeene, Sylvie van der Werf |
| EPI_ISL_428359, EPI_ISL_428360 | CH Compiègne Laboratoire de Biologie | National Reference Center for Viruses of Respiratory Infections, Institut Pasteur, Paris | Mélanie Albert, Marion Barbet, Sylvie Behillil, Méline Bizard, Angela Brisebarre, Flora Donati, Etienne Simon-Lorière, Vincent Enouf, Maud Vanpeene, Sylvie van der Werf |
| EPI_ISL_428361 | LABM GH nord Essonne de Longjumeau - BP 125 | National Reference Center for Viruses of Respiratory Infections, Institut Pasteur, Paris | Mélanie Albert, Marion Barbet, Sylvie Behillil, Méline Bizard, Angela Brisebarre, Flora Donati, Etienne Simon-Lorière, Vincent Enouf, Maud Vanpeene, Sylvie van der Werf |
| EPI_ISL_428363 | GH Nord Essonne Service de Biologie clinique | National Reference Center for Viruses of Respiratory Infections, Institut Pasteur, Paris | Mélanie Albert, Marion Barbet, Sylvie Behillil, Méline Bizard, Angela Brisebarre, Flora Donati, Etienne Simon-Lorière, Vincent Enouf, Maud Vanpeene, Sylvie van der Werf |
| EPI_ISL_428365 | LABM GH nord Essonne de Longjumeau - BP 125 | National Reference Center for Viruses of Respiratory Infections, Institut Pasteur, Paris | Mélanie Albert, Marion Barbet, Sylvie Behillil, Méline Bizard, Angela Brisebarre, Flora Donati, Etienne Simon-Lorière, Vincent Enouf, Maud Vanpeene, Sylvie van der Werf |
| EPI_ISL_428366 | CH Jeanne de Navarre Laboratoire de Biologie | National Reference Center for Viruses of Respiratory Infections, Institut Pasteur, Paris | Mélanie Albert, Marion Barbet, Sylvie Behillil, Méline Bizard, Angela Brisebarre, Flora Donati, Etienne Simon-Lorière, Vincent Enouf, Maud Vanpeene, Sylvie van der Werf |
| EPI_ISL_428368 | Genomic Laboratory (GLAB) (Conjoint lab of Health Directorate of Istanbul and Istanbul Technical University) | Genomic Laboratory (GLAB), Istanbul Technical University | Ilker Karacan, Tugba Kizilboga Akgun, Bugra Agaoglu, Gizem Alkurt, Jale Yildiz, Betsi Köse, Elifnaz Çelik, Arzu Irvem, Yasemin Kendir Demirkol, Ozlem Akgun Dogan, Mehtap Aydn, Levent Doganay, Gizem Dinler Doganay |
| EPI_ISL_428670 | Centre for Dengue Research | Centre for Dengue Research | Chandima Jeewandara, Dinuka Ariyaratne, Laksiri Gomes, Deshni Jayathilaka, Ananda Wijewickrama, Eranga Narangoda, Damayanthi Idampitiya, Neelika Malaige |
| EPI_ISL_428671 | Centre for Dengue Research | Centre for Dengue Research | Chandima Jeewandara, Dinuka Ariyatane, Laksiri Gomes, Deshni Jayathilaka, Diyanath Ranasinghe, Ananda Wijewickrama, Eranga Narangoda, Damayanthi |

| | | | |
|---|---|---|---|
| | | | Tdampitiya, Neelika Malavige |
| EPI_ISL_428672, EPI_ISL_428673 | Centre for Dengue Research | Centre for Dengue Research | Chandima Jeewandara, Dinuka Ariyaratne, Laksiri Gomes, Deshni Jayathilaka, Diyanath Ranasinghe, Ananda Wijewickrama, Eranga Narangoda, Damayanthi Idampitiya, Neelika Malavige |
| EPI_ISL_428851, EPI_ISL_428852 | FSBSI "Chumakov Federal Scientific Center for Research and Development of Immune-and-Biological Products of Russian Academy of Sciences" | FSBSI "Chumakov Federal Scientific Center for Research and Development of Immune-and-Biological Products of Russian Academy of Sciences" & NRC "Kurchatov institute" | Liubov Kozlovskaya, Anastasia Piniaeva, Georgy Ignatyev, Anna Shishova, Aydar Ishmukhametov, Mikhail Rychev, Egor Prokhorchuk, Denis Protsenko, Anastasia Berestovskaya |
| EPI_ISL_428855 | MRCG at LSHTM Geomics lab | MRCG at LSHTM Genomics lab | Sesay et al |
| EPI_ISL_428856 | MRCG at LSHTM Genomics Lab | MRCG at LSHTM Genomics lab | Sesay et al |
| EPI_ISL_429256 | Health Sciences Technology Park, Avicena, 8, 18016 Granada. Spain | Sequencing and Bioinformatics Service FISABIO-Public Health | Joaquín Mendoza, Almudena Rojas, Pablo Mendoza |
| EPI_ISL_429852, EPI_ISL_429853, EPI_ISL_429854 | Centers for Disease Control and Prevention of Lishui | Department of InspectionCenters for Disease Control and Prevention of Lishui | Wang Xiaoguang,Ji Qiaoying,Ji Jiansong,Ye Bifeng,Ye Ling |
| EPI_ISL_429968 | Centre Hospitalier Compiègne Laboratoire de Biologie | National Reference Center for Viruses of Respiratory Infections, Institut Pasteur, Paris | Mélanie Albert, Marion Barbet, Sylvie Behillil, Méline Bizard, Angela Brisebarre, Flora Donati, Fabiana Gambaro, Etienne Simon-Lorière, Vincent Enouf, Maud Vanpeene, Sylvie van der Werf, Raulin Olivia |
| EPI_ISL_430067, EPI_ISL_430068, EPI_ISL_430069, EPI_ISL_430070, EPI_ISL_430071, EPI_ISL_430072, EPI_ISL_430073, EPI_ISL_430074, EPI_ISL_430075, EPI_ISL_430076, EPI_ISL_430077, EPI_ISL_430078, EPI_ISL_430079, EPI_ISL_430080, EPI_ISL_430081, EPI_ISL_430082, EPI_ISL_430083, EPI_ISL_430084, EPI_ISL_430085, EPI_ISL_430086, EPI_ISL_430087, EPI_ISL_430088, EPI_ISL_430089, EPI_ISL_430090, EPI_ISL_430100, EPI_ISL_430101, EPI_ISL_430102, EPI_ISL_430103, EPI_ISL_430104, EPI_ISL_430105, EPI_ISL_430106, EPI_ISL_430107, EPI_ISL_430108, EPI_ISL_430109, EPI_ISL_430110, EPI_ISL_430111, EPI_ISL_430112 | | | |
| see above | WHO National Influenza Centre Russian Federation | WHO National Influenza Centre Russian Federation | Andrey Komissarov, Artem Fadeev, Mariia Sergeeva, Anna Ivanova, Daria Danilenko |
| EPI_ISL_430297 | National Institute for Communicable Diseases of the National Health Laboratory Service | National Institute for Communicable Diseases of the National Health Laboratory Service | Allam M, Kwenda S, van Heusden P, Khumalo Z, Mohale T, Subramoney K, von Gottberg, A, Ismail A, Bhiman JN |
| EPI_ISL_430439 | Institute for Medical Research, Infectious Disease Research Centre, National Institutes of Health, Ministry of Health Malaysia | Institute for Medical Research Infectious Disease Research Centre, National Institutes of Health, Ministry of Health Malaysia | Suppiah.J, Mohd-Zawawi.Z, Kalyanasundram.J, Azizan.M-A, Mat-Sharani.S, Hisham.H-A, Tan.L-P, Abdul-Wahid.M-Z, Mohd-Zain.R, Ahmad.N, Thayan.R |
| EPI_ISL_430440 | Institute for Medical Research, Infectious Disease Research Centre, National Institutes of Health, Ministry of Health Malaysia | Institute for Medical Research, Infectious Disease Research Centre, National Institutes of Health, Ministry of Health Malaysia | Suppiah.J, Mohd-Zawawi.Z, Kalyanasundram.J, Azizan.M-A, Mat-Sharani.S, Hisham.H-A, Tan.L-P, Abdul-Wahid.M-Z, Tengku-Abd-Rashid.T-R, Mohd-Zain.R, Ahmad.N, Thayan.R |
| EPI_ISL_430441, EPI_ISL_430442, EPI_ISL_430443, EPI_ISL_430444 | Institute for Medical Research, Infectious Disease Research Centre, National Institutes of Health, Ministry of Health Malaysia | Institute for Medical Research, Infectious Disease Research Centre, National Institutes of Health, Ministry of Health Malaysia | Suppiah.J, Mohd-Zawawi.Z, Kalyanasundram.J, Azizan.M-A, Mat-Sharani.S, Hisham.H-A, Tan.L-P, Abdul-Wahid.M-Z, Tengku-Rogayah.TAR, Mohd-Zain.R, Ahmad.N, Thayan.R |
| EPI_ISL_430469 | Hellenic Pasteur Institute, Public Health Laboratories | Hellenic Pasteur Institute, Public Health Laboratories, Unit of Bioinformatics and Applied Genomics | Vasiliki Pogka, Timokratis Karamitros, Athanasios Kossyvakis, Antonios Kalliaropoulos, Horefti Elina, Evangelidou Maria, Androniki Voulgari-Kokota, Aspasia Kontou, Andreas Mentis |
| EPI_ISL_430722, EPI_ISL_430724, EPI_ISL_430725, EPI_ISL_430728, EPI_ISL_430729, EPI_ISL_430730, EPI_ISL_430731, EPI_ISL_430733, EPI_ISL_430734, EPI_ISL_430735, EPI_ISL_430736, EPI_ISL_430737, EPI_ISL_430738, EPI_ISL_430740, EPI_ISL_430741, EPI_ISL_430742 | | | |
| see above | Chinese PLA Institute for Disease Control and Prevention | Chinese PLA Institute for Disease Control and Prevention | Peng LiJinhui Li, Lizhong Li |
| EPI_ISL_430819 | Center of Scientific Excellence for Influenza Viruses,National Research Centre (NRC), Egypt. | Center of Scientific Excellence for Influenza Viruses,National Research Centre (NRC), Egypt. | Mohamed Ahmed Ali, Ahmed Kandeil, Ahmed Mostafa, Rabeh El-Shesheny, Mahmoud Shehata, Wael Roshdy, Shymaa Showky Ahmed , Amal Naguib, Nancy M. El Guindy, Mokhtar Gomaa, Ahmed El-Taweel, Ahmed E Kayed, Yassmin Moatasim, Omnia Kutkat, Sara Mahmoud, Mina Kamel, Abo Shama, M Noura, Mohamed El Sayes |
| EPI_ISL_430820 | Center of Scientific Excellence for Influenza Viruses, National Research Centre (NRC), Egypt. | Center of Scientific Excellence for Influenza Viruses, National Research Centre (NRC), Egypt. | Mohamed Ahmed Ali, Ahmed Kandeil, Ahmed Mostafa, Rabeh El-Shesheny, Mahmoud Shehata, Wael Roshdy, Shymaa Showky Ahmed , Amal Naguib, Mokhtar Gomaa, Ahmed El-Taweel, Ahmed E Kayed, Yassmin Moatasim, Omnia Kutkat, Sara Mahmoud, Mina Kamel, Abo Shama, M Noura, Mohamed El Sayes, Nancy M. El Guindy |
| EPI_ISL_431101 | Department of Microbiology,Gandhi Medical College and Hospital | Virus Research Laboratory, Department of Zoology, Osmania University,Hyderabad,India | Muttineni Radhakrishna, Nagamani K, Thrilok Chander B, Raja Rao M, Kalyani Putty, Ravikumar P, Sunitha P, Pankaj Singh D, Anand Kumar K, Amit A. Upadhyay Steven E. Bosinger, Rama Amara |
| EPI_ISL_431102 | Department of MicroBiology,Gandhi Medical College and Hospital,Secendrabad,Hyderabad,India | Department of Microbiology, Gandhi Medical College and Hospital, Secendrabad, Hyderabad | Nagamani K, Muttineni Radhakrishna, Thrilok Chander B, Raja Rao M, Kalyani Putty, Ravikumar P, Sunitha P, Pankaj Singh D, Anand Kumar K, Amit A. Upadhyay, Steven E. Bosinger, Rama Amara |
| EPI_ISL_431103 | Department of Microbiology, Gandhi Medical College and Hospital, Secendrabad, Hyderabad, India | Department of Microbiology, Gandhi Medical College and Hospital, Secendrabad, Hyderabad, India | Nagamani K, Muttineni Radhakrishna, Thrilok Chander B, Raja Rao M, Kalyani Putty, Ravikumar P, Sunitha P, Pankaj Singh D, Anand Kumar K, Amit A. Upadhyay, Steven E. Bosinger, Rama Amara |
| EPI_ISL_431117 | Department of Microbiology, Gandhi Medical College and Hospital, Secendrabad, Hyderabad, India | Department of Microbiology, Gandhi Medical College and Hospital, Secendrabad, Hyderabad, India | Thrilok Chander B, Muttineni Radhakrishna, Nagamani K, Raja Rao M, Kalyani Putty, Ravikumar P, Sunitha P, Pankaj Singh D, Anand Kumar K, Amit A. Upadhyay, Steven E.Bosinger, Rama Amara |
| EPI_ISL_431118, EPI_ISL_431180, EPI_ISL_431240, EPI_ISL_431782, EPI_ISL_431783, EPI_ISL_431784, EPI_ISL_431785 | Fujian Center for Disease Control and Prevention | Fujian Center for Disease Control and Prevention | Lin Qi, Huang Zhimiao, Zhang Yanhua, Weng Yuwei |
| EPI_ISL_434366, EPI_ISL_434369, EPI_ISL_434371 | Hospital AZ Rivierenland | Institute of Tropical Medicine | Philippe Selhorst, Colin Anthony |
| EPI_ISL_434372, EPI_ISL_434373, EPI_ISL_434375, EPI_ISL_434376, EPI_ISL_434377, EPI_ISL_434378, EPI_ISL_434379, EPI_ISL_434380, EPI_ISL_434381, EPI_ISL_434383 | Hospital AZ Rivierenland | Institute of Tropical Medicine | Philippe Selhorst, Colin Anthony, |
| EPI_ISL_434384 | Hospital AZ Rivierenland | Institute of Tropical Medicine | Philippe Selhorst, Colin Anthony |
| EPI_ISL_434533 | Area de Salud Alajuela Sur | Inciensa, Instituto Costarricense de Investigación y Enseñanza en Nutrición y Salud | Francisco Duarte, Hebleen Porras, Claudio Soto-Garita, Estela Cordero, Adriana Godinez & Melany Calderon |
| EPI_ISL_434534 | National Institute for Viral Disease Control and Prevention, China CDC | National Institute for Viral Disease Control and Prevention, China CDC, Yunnan Provincial CDC | Wenjie Tan, Roujian Lu, Wenling Wang, Peihua Niu, Huijuan Wang, Baoying Huang, Li Zhao, Fei Ye, Guizhen Wu |
| EPI_ISL_434535 | Area de Salud Alajuela Sur | Inciensa, Instituto Costarricense de Investigación y Enseñanza en Nutrición y Salud | Francisco Duarte, Hebleen Porras, Claudio Soto-Garita, Estela Cordero, Adriana Godinez & Melany Calderon |
| EPI_ISL_434538 | COOPESAIN | Inciensa, Instituto Costarricense de Investigación y Enseñanza en Nutrición y Salud | Francisco Duarte, Hebleen Porras, Claudio Soto-Garita, Estela Cordero, Adriana Godinez & Melany Calderon |
| EPI_ISL_434539 | Area de Salud Orotina | Inciensa, Instituto Costarricense de Investigación y Enseñanza en Nutrición y Salud | Francisco Duarte, Hebleen Porras, Claudio Soto-Garita, Estela Cordero, Adriana Godinez & Melany Calderon |
| EPI_ISL_434541, EPI_ISL_434543, EPI_ISL_434545, EPI_ISL_434546, EPI_ISL_434547, EPI_ISL_434548, | Puerto Rico Department of Health | Centers for Disease Control and Prevention, Dengue Branch | Gilberto A. Santiago, Glenda Gonzalez, Betzabel Flores, Keyla Charriez, Fabiola Cruz, Chaney Kalinich, Joseph Fauver, Jessica I. Falcon, Nathan Grubaugh, Jorge L. Munoz-Jordan |

| | | | |
|---|---|---|---|
| EPI_ISL_434549, EPI_ISL_434551 | | | |
| EPI_ISL_434572 | The National Institute of Public Health Center for Epidemiology and Microbiology | The National Institute of Public Health Center for Epidemiology and Microbiology | Alexander Nagy, Helena Jirincova, Ludmila Novakova, Dusan Trnka, Jaromira Vecerova |
| EPI_ISL_435045 | Laboratory of Applied Genetics | RSE "National Center for Biotechnology" | Alexandr Shevtsov, Ilyas Akhmetollayev, Viktoriya Lutsay, Asylulan Amirgazin, Ruslan Kalendar, Yerlan Ramanculov |
| EPI_ISL_435049 | B.J. Medical College and Civil hospital | Gujarat Biotechnology Research Centre | Pinal Trivedi, Maharshi Pandya, Amit Kanani, Akanksha Verma, Nitin Savaliya, Raghawendra Kumar, Dinesh Kumar, Zuber Saiyed, Dipa Kinariwala, Disha Patel, Binita Aring, Geeta Vaghela, Sonia Barve, Bhavesh Modi, Kairavi Joshi, Gaurishankar Shrimali, Nidhi Sood, Pranay Shah, R D Dixit, Snehal Bagatharia, Kamlesh J Upadhyay, Ramesh Pandit, Tejas Shah, Ankit Hinsu, Pritesh Sabara, Apurvasinh Puvar, Janvi Raval, Monika Gandhi, Neha Rajpara, Chaitanya Joshi, Madhvi Joshi |
| EPI_ISL_435050 | B.J. Medical College and Civil hospital | Gujarat Biotechnology Research Centre | Ankit Hinsu, Pritesh Sabara, Apurvasinh Puvar, Janvi Raval, Monika Gandhi, Pinal Trivedi, Maharshi Pandya, Amit Kanani, Akanksha Verma, Nitin Savaliya, Raghawendra Kumar, Dinesh Kumar, Zuber Saiyed, Dipa Kinariwala, Disha Patel, Binita Aring, Geeta Vaghela, Sonia Barve, Bhavesh Modi, Kairavi Joshi, Gaurishankar Shrimali, Nidhi Sood, Pranay Shah, R D Dixit, Snehal Bagatharia, Kamlesh J Upadhyay, Ramesh Pandit, Tejas Shah, Dipeshwari Shewale, Chaitanya Joshi, Madhvi Joshi |
| EPI_ISL_435051 | B.J. Medical College and Civil hospital | Gujarat Biotechnology Research Centre | Pritesh Sabara, Apurvasinh Puvar, Janvi Raval, Monika Gandhi, Pinal Trivedi, Maharshi Pandya, Amit Kanani, Akanksha Verma, Nitin Savaliya, Raghawendra Kumar, Dinesh Kumar, Zuber Saiyed, Dipa Kinariwala, Disha Patel, Binita Aring, Geeta Vaghela, Sonia Barve, Bhavesh Modi, Kairavi Joshi, Gaurishankar Shrimali, Nidhi Sood, Pranay Shah, R D Dixit, Snehal Bagatharia, Kamlesh J Upadhyay, Ramesh Pandit, Tejas Shah, Ankit Hinsu, Vasudha Sharma, Chaitanya Joshi, Madhvi Joshi |
| EPI_ISL_435052 | B.J. Medical College and Civil hospital | Gujarat Biotechnology Research Centre | Apurvasinh Puvar, Janvi Raval, Monika Gandhi, Pinal Trivedi, Maharshi Pandya, Amit Kanani, Akanksha Verma, Nitin Savaliya, Raghawendra Kumar, Dinesh Kumar, Zuber Saiyed, Dipa Kinariwala, Disha Patel, Binita Aring, Geeta Vaghela, Sonia Barve, Bhavesh Modi, Kairavi Joshi, Gaurishankar Shrimali, Nidhi Sood, Pranay Shah, R D Dixit, Snehal Bagatharia, Kamlesh J Upadhyay, Ramesh Pandit, Tejas Shah, Ankit Hinsu, Pritesh Sabara, Pooja P Doshi, Chaitanya Joshi, Madhvi Joshi |
| EPI_ISL_435053 | B.J. Medical College and Civil hospital | Gujarat Biotechnology Research Centre | Janvi Raval, Monika Gandhi, Pinal Trivedi, Maharshi Pandya, Amit Kanani, Akanksha Verma, Nitin Savaliya, Raghawendra Kumar, Dinesh Kumar, Zuber Saiyed, Dipa Kinariwala, Disha Patel, Binita Aring, Geeta Vaghela, Sonia Barve, Bhavesh Modi, Kairavi Joshi, Gaurishankar Shrimali, Nidhi Sood, Pranay Shah, R D Dixit, Snehal Bagatharia, Kamlesh J Upadhyay, Ramesh Pandit, Tejas Shah, Ankit Hinsu, Pritesh Sabara, Apurvasinh Puvar, Nidhi Patel, Chaitanya Joshi, Madhvi Joshi |
| EPI_ISL_435055 | Gujarat Biotechnology Research Centre | Gujarat Biotechnology Research Centre | Tejas Shah, Ankit Hinsu, Pritesh Sabara, Apurvasinh Puvar, Janvi Raval, Monika Gandhi, Pinal Trivedi, Maharshi Pandya, Amit Kanani, Akanksha Verma, Nitin Savaliya, Raghawendra Kumar, Dinesh Kumar, Zuber Saiyed, Dipa Kinariwala, Disha Patel, Binita Aring, Geeta Vaghela, Sonia Barve, Bhavesh Modi, Kairavi Joshi, Gaurishankar Shrimali, Nidhi Sood, Pranay Shah, R D Dixit, Snehal Bagatharia, Kamlesh J Upadhyay, Ramesh Pandit, Anjali Rajwal, Chaitanya Joshi, Madhvi Joshi |
| EPI_ISL_435056 | Gujarat Biotechnology Research Centre | Gujarat Biotechnology Research Centre | Maharshi Pandya, Amit Kanani, Akanksha Verma, Nitin Savaliya, Raghawendra Kumar, Dinesh Kumar, Zuber Saiyed, Dipa Kinariwala, Disha Patel, Binita Aring, Geeta Vaghela, Sonia Barve, Bhavesh Modi, Kairavi Joshi, Gaurishankar Shrimali, Nidhi Sood, Pranay Shah, R D Dixit, Snehal Bagatharia, Kamlesh J Upadhyay, Ramesh Pandit, Tejas Shah, Ankit Hinsu, Pritesh Sabara, Apurvasinh Puvar, Janvi Raval, Monika Gandhi, Pinal Trivedi, Afzal Ansari, Chaitanya Joshi, Madhvi Joshi |
| EPI_ISL_435145 | Ospedale Civile Giuseppe Mazzini | Istituto Zooprofilattico Sperimentale dell'Abruzzo e Molise "G.Caporale" | Lorusso A, Marcacci M, Di Domenico M, Ancora M, Curini V, Mangone I, Rinaldi A, Di Pasquale A, Cammà C, Puglia I, Savini G |
| EPI_ISL_435146, EPI_ISL_435147 | Villa Serena del Dr. Leonardo Petruzzi | Istituto Zooprofilattico Sperimentale dell'Abruzzo e Molise "G.Caporale" | Lorusso A, Marcacci M, Di Domenico M, Ancora M, Curini V, Mangone I, Rinaldi A, Di Pasquale A, Cammà C, Puglia I, Savini G |
| EPI_ISL_435148 | Ospedale SS Annunziata | Istituto Zooprofilattico Sperimentale dell'Abruzzo e Molise "G.Caporale" | Lorusso A, Marcacci M, Di Domenico M, Ancora M, Curini V, Mangone I, Rinaldi A, Di Pasquale A, Cammà C, Puglia I, Savini G |
| EPI_ISL_435149 | SERVIZIO DI IGIENE E SANITÀ PUBBLICA ASL Teramo | Istituto Zooprofilattico Sperimentale dell'Abruzzo e Molise "G.Caporale" | Lorusso A, Marcacci M, Di Domenico M, Ancora M, Curini V, Mangone I, Rinaldi A, Di Pasquale A, Cammà C, Puglia I, Savini G |
| EPI_ISL_435150, EPI_ISL_435151 | Ospedale SS Annunziata | Istituto Zooprofilattico Sperimentale dell'Abruzzo e Molise "G.Caporale" | Lorusso A, Marcacci M, Di Domenico M, Ancora M, Curini V, Mangone I, Rinaldi A, Di Pasquale A, Cammà C, Puglia I, Savini G |
| EPI_ISL_435152 | Servizio di Igiene, Epidemiologia e Sanità Pubblica (SIESP) Avezzano | Istituto Zooprofilattico Sperimentale dell'Abruzzo e Molise "G.Caporale" | Lorusso A, Marcacci M, Di Domenico M, Ancora M, Curini V, Mangone I, Rinaldi A, Di Pasquale A, Cammà C, Puglia I, Savini G |
| EPI_ISL_435153, EPI_ISL_435154, EPI_ISL_435155 | SERVIZIO DI IGIENE E SANITÀ PUBBLICA ASL Teramo | Istituto Zooprofilattico Sperimentale dell'Abruzzo e Molise "G.Caporale" | Lorusso A, Marcacci M, Di Domenico M, Ancora M, Curini V, Mangone I, Rinaldi A, Di Pasquale A, Cammà C, Puglia I, Savini G |
| EPI_ISL_435281 | Medistra Hospital Jakarta | Eijkman Institute for Molecular Biology, Ministry of Research and Technology/National Agency for Research and Innovation | Edison Johar, Frilasita A Yudhaputri, Hidayat Trimarsanto, David H Muljono, Safarina G Malik, Khin Saw Myint, Amin Soebandrio |
| EPI_ISL_435282, EPI_ISL_435283 | RS Pondok Indah Hospital - Pondok Indah | Eijkman Institute for Molecular Biology, Ministry of Research and Technology/National Agency for Research and Innovation | Edison Johar, Frilasita A Yudhaputri, Hidayat Trimarsanto, David H Muljono, Safarina G Malik, Khin Saw Myint, Amin Soebandrio |
| EPI_ISL_435284 | Central Virology Laboratory, Israel Ministry of Health | Central Virology Laboratory, Israel Ministry of Health | Neta Zuckerman, Efrat Bucris, Oran Erster, Danit Sofer, Orna Mor, Ella Mendelson, Michal Mandelboim |
| EPI_ISL_435286 | Central Virology Laboratory, Israel Ministry of Health | Central Virology Laboratory, Israel Ministry of Health | eta Zuckerman, Efrat Bucris, Oran Erster, Orna Mor, Ella Mendelson, Michal Mandelboim, Danit Sofer |
| EPI_ISL_435289, EPI_ISL_435291 | Central Virology Laboratory, Israel Ministry of Health | Central Virology Laboratory, Israel Ministry of Health | Neta Zuckerman, Efrat Bucris, Oran Erster, Danit Sofer, Orna Mor, Ella Mendelson, Michal Mandelboim |
| EPI_ISL_435292 | Central Virology Laboratory, Israel Ministry of Health | Central Virology Laboratory, Israel Ministry of Health | Neta Zuckerman, Efrat Bucris, Oran Erster, Danit Sofer, Ella Mendelson, Michal Mandelboim, Orna Mor |
| EPI_ISL_435303 | National Hospital of Tropical Diseases | Oxford University Clinical Research Unit, Hanoi, Vietnam | Nguyen Thi Tam, Van Dinh Trang, Nguyen Thu Trang, Nguyen Thi Ngoc Diep, Le Nguyen Minh Hoa, Pham Ngoc Thach, H.Rogier van Doorn, on behalf of the OUCRU COVID-19 research group |
| EPI_ISL_435305 | National Hospital of Tropical Diseases | Oxford University Clinical Research Unit, Hanoi, Vietnam | Nguyen Thi Tam, Van Dinh Trang, Nguyen Thu Trang, Nguyen Thi Ngoc Diep, Le Nguyen Minh Hoa, Pham Ngoc Thach, H. Rogier van Doorn, on behalf of the OUCRU COVID-19 research group |
| EPI_ISL_435405, EPI_ISL_435407, EPI_ISL_435408, EPI_ISL_435410, EPI_ISL_435412, EPI_ISL_435414, EPI_ISL_435416, EPI_ISL_435419, EPI_ISL_435422, EPI_ISL_435423, EPI_ISL_435424, EPI_ISL_435425, EPI_ISL_435430, EPI_ISL_435431 | | | |
| see above | Virological Research Group, Szentágothai Research Centre | Bioinformatics Research Group, Szentágothai Research Centre | Péter Urbán, Endre Gábor Tóth, Gábor Kemenesi, Róbert Herczeg, Attila Gyenesei, Ferenc Jakab |
| EPI_ISL_436099 | TSGH-CP molecular lab | TSGH-CP molecular lab | Cherng-Lih Perng, Ming-Jr JIAN, Chih-Kai Chang, Jung-Chung Lin, Kuo-Ming Yeh, Chien-Wen Chen, Sheng-Kang Chiu, Hsing-Yi Chung, Shih-Hung Tsai, Kuo-Sheng Hung, Tien-Yao Chang, Feng-Yee Chang, Hung-Sheng Shang |
| EPI_ISL_436100 | TSGH-CP molecular lab | TSGH-CP molecular lab | Cherng-Lih Perng, Ming-Jr Jian, Chih-Kai Chang, Jung-Chung Lin, Kuo-Ming Yeh, Chien-Wen Chen, Sheng-Kang Chiu, Hsing-Yi Chung, Shih-Hung Tsai, Kuo-Sheng Hung, Tien-Yao Chang, Feng-Yee Chang, Hung-Sheng Shang |
| EPI_ISL_436101, EPI_ISL_436102, EPI_ISL_436103, EPI_ISL_436104, EPI_ISL_436106, EPI_ISL_436107, EPI_ISL_436108 | TSGH-CP molecular lab | TSGH-CP molecular lab | Cherng-Lih Perng, Ming-Jr JIAN, Chih-Kai Chang, Jung-Chung Lin, Kuo-Ming Yeh, Chien-Wen Chen, Sheng-Kang Chiu, Hsing-Yi Chung, Shih-Hung Tsai, Kuo-Sheng Hung, Tien-Yao Chang, Feng-Yee Chang, Hung-Sheng Shang |
| EPI_ISL_436157 | District Surveillance Unit | Department of Neurovirology, National Institute of Mental Health and Neuroscience (NIMHANS) | Chitra Pattabiraman, Vijayalakshmi Reddy, Harsha PK, Risha Rasheed, Shafeeq S Hameed, Manjunatha Venkataswamy, Anita Desai, Ravi Vasanthapuram |

| | | | |
|---|---|---|---|
| EPI_ISL_436412 | Viral Respiratory Lab, National Institute for Biomedical Research (INRB) | Pathogen Sequencing Lab, National Institute for Biomedical Research (INRB) | Placide Mbala-Kingebeni, Edith Nkwembe, Eddy Kinganda-Lusamaki, Amuri Aziza, Francisca Muyembe Mawete, Catherine Pratt, Matthias Pauthner, Josh Quick, Allison Black, James Hadfield, Trevor Bedford, Ian Goodfellow, Andrew Rambaut, Nick Loman, Kristian Andersen, Michael Wiley, Steve Ahuka-Mundeke, Jean-Jacques Muyembe Tamfum |
| EPI_ISL_436684, EPI_ISL_436686 | KRISP, KZN Research Innovation and Sequencing Platform | KRISP, KZN Research Innovation and Sequencing Platform | Giandhari J, Pillay S, Lessells R, Chimukangara B, Deforche K, Tegally H, Wilkinson E, de Oliveira T |
| EPI_ISL_436715, EPI_ISL_436716, EPI_ISL_436717 | Genomics and Computational Biology Lab, Scientific Research Institute of Physical-Chemical Medicine, FMBA of Russia | Genomics and Computational Biology Lab, Scientific Research Institute of Physical-Chemical Medicine, FMBA of Russia | A. Pavlenko, O. Guskova, K. Klimina, V. Veselovsky, A. Manolov, D. Fedorov, V. Govorun and E. Ilina |
| EPI_ISL_436718 | Ospedale Regionale San Salvatore | Istituto Zooprofilattico Sperimentale dell'Abruzzo e Molise "G.Caporale" | Lorusso A, Marcacci M, Di Domenico M, Ancora M, Curini V, Mangone I, Rinaldi A, Di Pasquale A, Cammà C, Puglia I, Savini G |
| EPI_ISL_436719, EPI_ISL_436720, EPI_ISL_436721, EPI_ISL_436722 | Ospedale Civile S. Liberatore di Atri | Istituto Zooprofilattico Sperimentale dell'Abruzzo e Molise "G.Caporale" | Lorusso A, Marcacci M, Di Domenico M, Ancora M, Curini V, Mangone I, Rinaldi A, Di Pasquale A, Cammà C, Puglia I, Savini G |
| EPI_ISL_436723 | Ospedale Civile Giuseppe Mazzini | Istituto Zooprofilattico Sperimentale dell'Abruzzo e Molise "G.Caporale" | Lorusso A, Marcacci M, Di Domenico M, Ancora M, Curini V, Mangone I, Rinaldi A, Di Pasquale A, Cammà C, Puglia I, Savini G |
| EPI_ISL_436724 | Ospedale Civile S. Liberatore di Atri | Istituto Zooprofilattico Sperimentale dell'Abruzzo e Molise "G.Caporale" | Lorusso A, Marcacci M, Di Domenico M, Ancora M, Curini V, Mangone I, Rinaldi A, Di Pasquale A, Cammà C, Puglia I, Savini G |
| EPI_ISL_436725 | RSA/RP Villa San Giovanni - Gruppo Edos | Istituto Zooprofilattico Sperimentale dell'Abruzzo e Molise "G.Caporale" | Lorusso A, Marcacci M, Di Domenico M, Ancora M, Curini V, Mangone I, Rinaldi A, Di Pasquale A, Cammà C, Puglia I, Savini G |
| EPI_ISL_436726, EPI_ISL_436727, EPI_ISL_436729 | SERVIZIO DI IGIENE E SANITÀ PUBBLICA ASL Teramo | Istituto Zooprofilattico Sperimentale dell'Abruzzo e Molise "G.Caporale" | Lorusso A, Marcacci M, Di Domenico M, Ancora M, Curini V, Mangone I, Rinaldi A, Di Pasquale A, Cammà C, Puglia I, Savini G |
| EPI_ISL_436730 | Servizio di igiene epidemiologia e sanità pubblica (Siesp) Chieti | Istituto Zooprofilattico Sperimentale dell'Abruzzo e Molise "G.Caporale" | Lorusso A, Marcacci M, Di Domenico M, Ancora M, Curini V, Mangone I, Rinaldi A, Di Pasquale A, Cammà C, Puglia I, Savini G |
| EPI_ISL_436731, EPI_ISL_436732 | Ospedale Civile S. Liberatore di Atri | Istituto Zooprofilattico Sperimentale dell'Abruzzo e Molise "G.Caporale" | Lorusso A, Marcacci M, Di Domenico M, Ancora M, Curini V, Mangone I, Rinaldi A, Di Pasquale A, Cammà C, Puglia I, Savini G |
| EPI_ISL_436939, EPI_ISL_436942, EPI_ISL_436943, EPI_ISL_436944, EPI_ISL_436945, EPI_ISL_436946, EPI_ISL_436947, EPI_ISL_436948, EPI_ISL_436949, EPI_ISL_436950, EPI_ISL_436952, EPI_ISL_436953, EPI_ISL_436954, EPI_ISL_436956, EPI_ISL_436957, EPI_ISL_436958, EPI_ISL_436959, EPI_ISL_436960, EPI_ISL_436961 | | | |
| see above | Ochsner Health | Bioinfoexperts, LLC | Amy Feehan, David J. Nolan, Rebecca Rose, Sissy Cross, David Moraga Amador, Tong Yang, Luke Caruso, Wayra Navia, Lydia Von Borstel, Xiao Hui Zhou, Julia-Garcia-Diaz, Susanna L. Lamers |
| EPI_ISL_437187 | Siloam Hospitals | Institute of Tropical Disease, Universitas Airlangga | Kazufumi Shimizu, Krisnoadi Rahardjo, Aldise M Nastri, Jezzy R Dewantari, Rima R Prasetya, Maria M Padmidewi, Gatot Soegiarto, Laksmi Wulandari, Retno A Setyoningrum, Resti Y Meliana, Yohko K Shimizu, Mitsuhiro Nishimura, Yasuko Mori, Soetjipto, Maria I Lusida |
| EPI_ISL_437188 | RSUD Dr. Soetomo | Institute of Tropical Disease, Universitas Airlangga | Krisnoadi Rahardjo, Aldise M Nastri, Jezzy R Dewantari, Rima R Prasetya, Joni Wahyuhadi, Gatot Soegiarto, Laksmi Wulandari, Retno A Setyoningrum, Resti Y Meliana, Yohko K Shimizu, Mitsuhiro Nishimura, Yasuko Mori, Soetjipto, Kazufumi Shimizu, Maria I Lusida |
| EPI_ISL_437189 | Pusat Pertamina Hospital | Eijkman Institute for Molecular Biology, Ministry of Research and Technology/National Agency for Research and Innovation | Edison Johar, Frilasita A Yudhaputri, Hidayat Trimarsanto, David H Muljono, Safarina G Malik, Khin Saw Myint, Amin Soebandrio |
| EPI_ISL_437190, EPI_ISL_437191 | RS Pondok Indah Hospital - Pondok Indah | Eijkman Institute for Molecular Biology, Ministry of Research and Technology/National Agency for Research and Innovation | Edison Johar, Frilasita A Yudhaputri, Hidayat Trimarsanto, David H Muljono, Safarina G Malik, Khin Saw Myint, Amin Soebandrio |
| EPI_ISL_437192 | Mitra Keluarga Kelapa Gading Hospital | Eijkman Institute for Molecular Biology, Ministry of Research and Technology/National Agency for Research and Innovation | Edison Johar, Frilasita A Yudhaputri, Hidayat Trimarsanto, David H Muljono, Safarina G Malik, Khin Saw Myint, Amin Soebandrio |
| EPI_ISL_437194 | Viral Respiratory Lab, National Institute for Biomedical Research (INRB) | Pathogen Sequencing Lab, National Institute for Biomedical Research (INRB) | Placide Mbala-Kingebeni, Edith Nkwembe, Eddy Kinganda-Lusamaki, Amuri Aziza, Francisca Muyembe Mawete, Catherine Pratt, Matthias Pauthner, Josh Quick, Allison Black, James Hadfield, Trevor Bedford, Ian Goodfellow, Andrew Rambaut, Nick Loman, Kristian Andersen, Michael Wiley, Steve Ahuka-Mundeke, Jean-Jacques Muyembe Tamfum |
| EPI_ISL_437197, EPI_ISL_437198, EPI_ISL_437199, EPI_ISL_437200, EPI_ISL_437201, EPI_ISL_437202, EPI_ISL_437203, EPI_ISL_437298, EPI_ISL_437299, EPI_ISL_437300, EPI_ISL_437301, EPI_ISL_437302, EPI_ISL_437303 | | | |
| see above | Diagnostic- and Research Institute of Pathology, Medical University of Graz | Diagnostic- and Research Institute of Pathology, Medical University of Graz | Karl Kashofer, Peter Regitnig, Martin Zacharias, Gregor Gorkiewicz |
| EPI_ISL_437339 | Viral Respiratory Lab, National Institute for Biomedical Research (INRB) | Pathogen Sequencing Lab, National Institute for Biomedical Research (INRB) | Placide Mbala-Kingebeni, Edith Nkwembe, Eddy Kinganda-Lusamaki, Amuri Aziza, Francisca Muyembe Mawete, Catherine Pratt, Matthias Pauthner, Josh Quick, Allison Black, James Hadfield, Trevor Bedford, Ian Goodfellow, Andrew Rambaut, Nick Loman, Kristian Andersen, Michael Wiley, Steve Ahuka-Mundeke, Jean-Jacques Muyembe Tamfum |
| EPI_ISL_437433 | Bozeman Health Deaconess Hospital | Wiedenheft lab, Montana State University | Artem Nemudryi, Anna Nemudraia, Kevin Surya, Tanner Wiegand, Murat Buyukyoruk, Royce Wilkinson, Blake Wiedenheft |
| EPI_ISL_437440 | Department of MicroBiology, Government Medical College, Surat | Gujarat Biotechnology Research Centre | Nitin Savaliya, Raghawendra Kumar, Dinesh Kumar, Zuber Saiyed, Dipa Kinariwala, Disha Patel, Binita Aring, Neeta Khandelwal, Geeta Vaghela, Sonia Barve, Bhavesh Modi, Kairavi Joshi, Gaurishankar Shrimali, Nidhi Sood, Pranay Shah, R D Dixit, Snehal Bagatharia, Kamlesh J Upadhyay, Ramesh Pandit, Tejas Shah, Ankit Hinsu, Pritesh Sabara, Apurvasinh Puvar, Janvi Raval, Monika Gandhi, Pinal Trivedi, Maharshi Pandya, Amit Kanani, Akanksha Verma, Bhavya Jindal, Chaitanya Joshi, Madhvi Joshi |
| EPI_ISL_437442 | Department of MicroBiology, Government Medical College, Surat | Gujarat Biotechnology Research Centre | Dinesh Kumar, Zuber Saiyed, Dipa Kinariwala, Disha Patel, Binita Aring, Neeta Khandelwal, Geeta Vaghela, Sonia Barve, Bhavesh Modi, Kairavi Joshi, Gaurishankar Shrimali, Nidhi Sood, Pranay Shah, R D Dixit, Snehal Bagatharia, Kamlesh J Upadhyay, Ramesh Pandit, Tejas Shah, Ankit Hinsu, Pritesh Sabara, Apurvasinh Puvar, Janvi Raval, Monika Gandhi, Pinal Trivedi, Maharshi Pandya, Amit Kanani, Akanksha Verma, Nitin Savaliya, Raghawendra Kumar, Dipeshwari Shewale, Chaitanya Joshi, Madhvi Joshi |
| EPI_ISL_437445 | B.J. Medical College and Civil hospital | Gujarat Biotechnology Research Centre | Disha Patel, Binita Aring, Neeta Khandelwal, Geeta Vaghela, Sonia Barve, Bhavesh Modi, Kairavi Joshi, Gaurishankar Shrimali, Nidhi Sood, Pranay Shah, R D Dixit, Snehal Bagatharia, Kamlesh J Upadhyay, Ramesh Pandit, Tejas Shah, Ankit Hinsu, Pritesh Sabara, Apurvasinh Puvar, Janvi Raval, Monika Gandhi, Pinal Trivedi, Maharshi Pandya, Amit Kanani, Akanksha Verma, Nitin Savaliya, Raghawendra Kumar, Dinesh Kumar, Zuber Saiyed, Dipa Kinariwala, Nidhi Patel, Chaitanya Joshi, Madhvi Joshi |
| EPI_ISL_437446 | B.J. Medical College and Civil hospital | Gujarat Biotechnology Research Centre | Binita Aring, Neeta Khandelwal, Geeta Vaghela, Sonia Barve, Bhavesh Modi, Kairavi Joshi, Gaurishankar Shrimali, Nidhi Sood, Pranay Shah, R D Dixit, Snehal Bagatharia, Kamlesh J Upadhyay, Ramesh Pandit, Tejas Shah, Ankit Hinsu, Pritesh Sabara, Apurvasinh Puvar, Janvi Raval, Monika Gandhi, Pinal Trivedi, Maharshi Pandya, Amit Kanani, Akanksha Verma, Nitin Savaliya, Raghawendra Kumar, Dinesh Kumar, Zuber Saiyed, Dipa Kinariwala, Disha Patel, Priti Pandita, Chaitanya Joshi, Madhvi Joshi |
| EPI_ISL_437447 | B.J. Medical College and Civil hospital | Gujarat Biotechnology Research Centre | Neeta Khandelwal, Geeta Vaghela, Sonia Barve, Bhavesh Modi, Kairavi Joshi, Gaurishankar Shrimali, Nidhi Sood, Pranay Shah, R D Dixit, Snehal Bagatharia, Kamlesh J Upadhyay, Ramesh Pandit, Tejas Shah, Ankit Hinsu, Pritesh Sabara, Apurvasinh Puvar, Janvi Raval, Monika Gandhi, Pinal Trivedi, Maharshi Pandya, Amit Kanani, Akanksha Verma, Nitin Savaliya, Raghawendra Kumar, Dinesh Kumar, Zuber Saiyed, Dipa Kinariwala, Disha Patel, Binita Aring, Neha Rajpara, Chaitanya Joshi, Madhvi Joshi |
| EPI_ISL_437448 | B.J. Medical College and Civil hospital | Gujarat Biotechnology Research Centre | Geeta Vaghela, Sonia Barve, Bhavesh Modi, Kairavi Joshi, Gaurishankar Shrimali, Nidhi Sood, Pranay Shah, R D Dixit, Snehal Bagatharia, Kamlesh J Upadhyay, Ramesh Pandit, Tejas Shah, Ankit Hinsu, Pritesh Sabara, Apurvasinh Puvar, Janvi Raval, Monika Gandhi, Pinal Trivedi, Maharshi Pandya, Amit Kanani, Akanksha Verma, Nitin Savaliya, Raghawendra Kumar, Dinesh Kumar, Zuber Saiyed, Dipa Kinariwala, Disha Patel, Binita Aring, Neeta Khandelwal, |

| | | | |
|---|---|---|---|
| | | | Afzal Ansari, Chaitanya Joshi, Madhvi Joshi |
| EPI_ISL_437449 | B.J. Medical College and Civil hospital | Gujarat Biotechnology Research Centre | Sonia Barve, Bhavesh Modi, Kairavi Joshi, Gaurishankar Shrimali, Nidhi Sood, Pranay Shah, R D Dixit, Snehal Bagatharia, Kamlesh J Upadhyay, Ramesh Pandit, Tejas Shah, Ankit Hinsu, Pritesh Sabara, Apurvasinh Puvar, Janvi Raval, Monika Gandhi, Pinal Trivedi, Maharshi Pandya, Amit Kanani, Akanksha Verma, Nitin Savaliya, Raghawendra Kumar, Dinesh Kumar, Zuber Saiyed, Dipa Kinariwala, Disha Patel, Binita Aring, Neeta Khandelwal, Geeta Vaghela, Neelam Nathani, Chaitanya Joshi, Madhvi Joshi |
| EPI_ISL_437450 | B.J. Medical College and Civil hospital | Gujarat Biotechnology Research Centre | Bhavesh Modi, Kairavi Joshi, Gaurishankar Shrimali, Nidhi Sood, Pranay Shah, R D Dixit, Snehal Bagatharia, Kamlesh J Upadhyay, Ramesh Pandit, Tejas Shah, Ankit Hinsu, Pritesh Sabara, Apurvasinh Puvar, Janvi Raval, Monika Gandhi, Pinal Trivedi, Maharshi Pandya, Amit Kanani, Akanksha Verma, Nitin Savaliya, Raghawendra Kumar, Dinesh Kumar, Zuber Saiyed, Dipa Kinariwala, Disha Patel, Binita Aring, Neeta Khandelwal, Geeta Vaghela, Sonia Barve, Armi Chaudhari, Chaitanya Joshi, Madhvi Joshi |
| EPI_ISL_437452 | B.J. Medical College and Civil hospital | Gujarat Biotechnology Research Centre | Gaurishankar Shrimali, Nidhi Sood, Pranay Shah, R D Dixit, Snehal Bagatharia, Kamlesh J Upadhyay, Ramesh Pandit, Tejas Shah, Ankit Hinsu, Pritesh Sabara, Apurvasinh Puvar, Janvi Raval, Monika Gandhi, Pinal Trivedi, Maharshi Pandya, Amit Kanani, Akanksha Verma, Nitin Savaliya, Raghawendra Kumar, Dinesh Kumar, Zuber Saiyed, Dipa Kinariwala, Disha Patel, Binita Aring, Neeta Khandelwal, Geeta Vaghela, Sonia Barve, Bhavesh Modi, Kairavi Joshi, Chaitanya Joshi, Anjali Rajwar, Madhvi Joshi |
| EPI_ISL_437453 | B.J. Medical College and Civil hospital | Gujarat Biotechnology Research Centre | Nidhi Sood, Pranay Shah, R D Dixit, Snehal Bagatharia, Kamlesh J Upadhyay, Ramesh Pandit, Tejas Shah, Ankit Hinsu, Pritesh Sabara, Apurvasinh Puvar, Janvi Raval, Monika Gandhi, Pinal Trivedi, Maharshi Pandya, Amit Kanani, Akanksha Verma, Nitin Savaliya, Raghawendra Kumar, Dinesh Kumar, Zuber Saiyed, Dipa Kinariwala, Disha Patel, Binita Aring, Neeta Khandelwal, Geeta Vaghela, Sonia Barve, Bhavesh Modi, Kairavi Joshi, Gaurishankar Shrimali, Chaitanya Joshi, Dipeshwari Shewale, Madhvi Joshi |
| EPI_ISL_437454 | B.J. Medical College and Civil hospital | Gujarat Biotechnology Research Centre | Pranay Shah, R D Dixit, Snehal Bagatharia, Kamlesh J Upadhyay, Ramesh Pandit, Tejas Shah, Ankit Hinsu, Pritesh Sabara, Apurvasinh Puvar, Janvi Raval, Monika Gandhi, Pinal Trivedi, Maharshi Pandya, Amit Kanani, Akanksha Verma, Nitin Savaliya, Raghawendra Kumar, Dinesh Kumar, Zuber Saiyed, Dipa Kinariwala, Disha Patel, Binita Aring, Neeta Khandelwal, Geeta Vaghela, Sonia Barve, Bhavesh Modi, Kairavi Joshi, Gaurishankar Shrimali, Nidhi Sood, Chaitanya Joshi, Sharmistha Majumdar, Madhvi Joshi |
| EPI_ISL_437459, EPI_ISL_437460, EPI_ISL_437461, EPI_ISL_437462, EPI_ISL_437463, EPI_ISL_437464, EPI_ISL_437465, EPI_ISL_437466, EPI_ISL_437468, EPI_ISL_437469, EPI_ISL_437470, EPI_ISL_437471, EPI_ISL_437472, EPI_ISL_437473, EPI_ISL_437474, EPI_ISL_437475, EPI_ISL_437477, EPI_ISL_437479 | | | |
| see above | Pathogen Genomics Lab King Abdullah University of Science and Technology(KAUST) | Pathogen Genomics Lab King Abdullah University of Science and Technology(KAUST) | Sharif Hala,Raeece Naeem,Sara Mfarrej,Arnab Pain |
| EPI_ISL_437481, EPI_ISL_437482, EPI_ISL_437483, EPI_ISL_437484, EPI_ISL_437485, EPI_ISL_437486, EPI_ISL_437487, EPI_ISL_437488, EPI_ISL_437489, EPI_ISL_437490, EPI_ISL_437491, EPI_ISL_437492, EPI_ISL_437493, EPI_ISL_437494, EPI_ISL_437495, EPI_ISL_437496, EPI_ISL_437497 | | | |
| see above | Pathogen Genomics Lab King Abdullah University of Science and Technology(KAUST) | Pathogen Genomics Lab King Abdullah University of Science and Technology(KAUST) | Sara Mfarrej,Raeece Naeem,Sharif Hala,Amit Subudhi,Fathia Rached,Arnab Pain |
| EPI_ISL_437498, EPI_ISL_437499 | Biotechnology Center for Advanced Technologies | Biotechnology Center for Advanced Technologies | Abdullaev,A., Abdurakhimov,A., Muminov,M., Nuriddinov,S., Dalimova,D., Tsoy,V., Tsay,E., Bozorov,S., Charishnikova,O., Dalimova,D. and Turdikulova,S. |
| EPI_ISL_437626 | Department of Microbiology,Gandhi Medical College and Hospital | Department of Veterinary Biotechnalogy, College of Veterinary Science, Rajendranagar, PV Narsimha Rao Telengana Veterinary University | Kalyani Putty, Muttineni Radhakrishna, Nagamani K, Thrilok Chander B, Raja Rao M, Ravikumar P, Sunitha P, Pankaj Singh D, Anand Kumar K, Amit A. Upadhyay, Steven Bosinger, Rama Amara |
| EPI_ISL_437691, EPI_ISL_437692, EPI_ISL_437693, EPI_ISL_437694, EPI_ISL_437695, EPI_ISL_437696, EPI_ISL_437697, EPI_ISL_437698, EPI_ISL_437699, EPI_ISL_437700, EPI_ISL_437701, EPI_ISL_437702, EPI_ISL_437703, EPI_ISL_437704, EPI_ISL_437705, EPI_ISL_437706, EPI_ISL_437707, EPI_ISL_437708, EPI_ISL_437709, EPI_ISL_437710, EPI_ISL_437711, EPI_ISL_437712, EPI_ISL_437713, EPI_ISL_437714, EPI_ISL_437715, EPI_ISL_437716, EPI_ISL_437717, EPI_ISL_437718, EPI_ISL_437719, EPI_ISL_437720, EPI_ISL_437721, EPI_ISL_437722, EPI_ISL_437723, EPI_ISL_437724, EPI_ISL_437725, EPI_ISL_437726, EPI_ISL_437727, EPI_ISL_437728, EPI_ISL_437729, EPI_ISL_437730, EPI_ISL_437731, EPI_ISL_437732, EPI_ISL_437733, EPI_ISL_437734, EPI_ISL_437735, EPI_ISL_437736, EPI_ISL_437737, EPI_ISL_437738, EPI_ISL_437739, EPI_ISL_437740, EPI_ISL_437741, EPI_ISL_437742, EPI_ISL_437743, EPI_ISL_437744, EPI_ISL_437745, EPI_ISL_437746, EPI_ISL_437747, EPI_ISL_437748, EPI_ISL_437749, EPI_ISL_437750, EPI_ISL_437751, EPI_ISL_437752, EPI_ISL_437753, EPI_ISL_437754, EPI_ISL_437755, EPI_ISL_437756, EPI_ISL_437757, EPI_ISL_437758, EPI_ISL_437759, EPI_ISL_437760, EPI_ISL_437761, EPI_ISL_437762 | | | |
| see above | Pathogen Genomics Lab King Abdullah University of Science and Technology(KAUST) | Pathogen Genomics Lab King Abdullah University of Science and Technology(KAUST) | Sharif Hala,Fadwa Alofi,Afrah Alsomali, Asim Khogeer, Sara Mfarrej, Khaled Algithami,Raeece Naeem, Amit Kumar Subudhi,Fathia Ben-Rached, Rahul Salunke, Anwar Hashem, Naif Almontashiri, Arnab Pain |
| EPI_ISL_438138 | Department of Microbiology,Gandhi Medical College and Hospital | Department of Microbiology, Gandhi Medical College and Hospital Secendrabad, Hyderbad, India | Raja Rao Mesipogu, Muttineni Radhakrishna, Nagamani K, Thrilok Chander B, Kalyani Putty, Ravikumar P, Sunitha P, Pankaj Singh D, Anand Kumar K, Amit A. Upadhyay, Steven Bosinger, Rama Amara |
| EPI_ISL_438139 | Department of Microbiology,Gandhi Medical College and Hospital,Hyderabad | Virus Research Laboratory, Department of Zoology, Osmania University, Hyderabad, India | Muttineni Radhakrishna, Nagamani K, Thrilok Chander B, Raja Rao M, Kalyani Putty, Ravikumar P, Sunitha P, Pankaj Singh D, Anand Kumar K, Amit A. Upadhyay, Steven Bosinger, Rama Amara |
| EPI_ISL_438546, EPI_ISL_438548 | Siloam Hospital Lippo Village | Mochtar Riady Institute for Nanotechnology, Universitas Pelita Harapan | Aksar C Lages, David Rustandi, Febi Andriani, Ivet M Suriapranata, Riska N Taufik, Tri Shinta Kurniasih, Irawan Yusuf |
| EPI_ISL_442044 | Kawsar Human Genetic Research Center | Kawsar Human Genetic Research Center | Mohammad Ali Khosravi, Maryam Abbasalipour Bashash, Sirous Zeinali, Solmaz Sabeghi, Yeganeh Keshvar, Fatemeh Hosseini, Yeganeh Haghdoost |
| EPI_ISL_442523 | Pasteur Institute of Iran | Kawsar Human Genetic Research Company | Sirous Zeinali, Mohammad Ali Khosravi,Maryam Abbasalipour Bashash, Sanaz Mostafavi Jabbari, Maraym Firoozi, Sormeh Pourtavakoli, Elmira Khateri, Razieh Zeinali and Fahimeh Hoseini |
| EPI_ISL_443258, EPI_ISL_443259 | Résidence Ornano | National Reference Center for Viruses of Respiratory Infections, Institut Pasteur, Paris | Mélanie Albert, Marion Barbet, Sylvie Behillil, Méline Bizard, Angela Brisebarre, Flora Donati, Etienne Simon-Lorière, Vincent Enouf, Maud Vanpeene, Sylvie van der Werf |
| EPI_ISL_443260 | LABM GH nord Essonne de Longjumeau - BP 125 | National Reference Center for Viruses of Respiratory Infections, Institut Pasteur, Paris | Mélanie Albert, Marion Barbet, Sylvie Behillil, Méline Bizard, Angela Brisebarre, Flora Donati, Etienne Simon-Lorière, Vincent Enouf, Maud Vanpeene, Sylvie van der Werf |
| EPI_ISL_443261, EPI_ISL_443262, EPI_ISL_443263, EPI_ISL_443264 | CHU de Dijon - Laboratoire de Virologie | National Reference Center for Viruses of Respiratory Infections, Institut Pasteur, Paris | Mélanie Albert, Marion Barbet, Sylvie Behillil, Méline Bizard, Angela Brisebarre, Flora Donati, Etienne Simon-Lorière, Vincent Enouf, Maud Vanpeene, Sylvie van der Werf, Jean-Baptiste Bour |
| EPI_ISL_443265, EPI_ISL_443266, EPI_ISL_443267, EPI_ISL_443268, EPI_ISL_443269, EPI_ISL_443270, EPI_ISL_443271, EPI_ISL_443272, EPI_ISL_443273, EPI_ISL_443274, EPI_ISL_443275, EPI_ISL_443276, EPI_ISL_443277, EPI_ISL_443278, EPI_ISL_443279, EPI_ISL_443280, EPI_ISL_443281, EPI_ISL_443282, EPI_ISL_443283 | | | |
| see above | CHU - Hôpital Cavale Blanche - Labo. de Virologie | National Reference Center for Viruses of Respiratory Infections, Institut Pasteur, Paris | Mélanie Albert, Marion Barbet, Sylvie Behillil, Méline Bizard, Angela Brisebarre, Flora Donati, Etienne Simon-Lorière, Vincent Enouf, Maud Vanpeene, Sylvie van der Werf, Léa Pilorge |
| EPI_ISL_443284, EPI_ISL_443285, EPI_ISL_443286, EPI_ISL_443287, EPI_ISL_443288 | Laboratoire de Microbiologie - Bât A - CH René Dubois | National Reference Center for Viruses of Respiratory Infections, Institut Pasteur, Paris | Mélanie Albert, Marion Barbet, Sylvie Behillil, Méline Bizard, Angela Brisebarre, Flora Donati, Etienne Simon-Lorière, Vincent Enouf, Maud Vanpeene, Sylvie van der Werf, Pascale Martres |
| EPI_ISL_443289, EPI_ISL_443290, EPI_ISL_443291, EPI_ISL_443292, EPI_ISL_443293, EPI_ISL_443294 | CHRU Pontchaillou - Laboratoire de Virologie | National Reference Center for Viruses of Respiratory Infections, Institut Pasteur, Paris | Mélanie Albert, Marion Barbet, Sylvie Behillil, Méline Bizard, Angela Brisebarre, Flora Donati, Etienne Simon-Lorière, Vincent Enouf, Maud Vanpeene, Sylvie van der Werf, Gisèle Lagathu |
| EPI_ISL_443295, EPI_ISL_443296, EPI_ISL_443298, EPI_ISL_443299 | Hôpital Necker - Enfants - Malades Laboratoire de Virologie | National Reference Center for Viruses of Respiratory Infections, Institut Pasteur, Paris | Mélanie Albert, Marion Barbet, Sylvie Behillil, Méline Bizard, Angela Brisebarre, Flora Donati, Etienne Simon-Lorière, Vincent Enouf, Maud Vanpeene, Sylvie van der Werf, Marianne Leruez-Ville |
| EPI_ISL_443303 | Résidence Les Marines | National Reference Center for Viruses of Respiratory Infections, Institut Pasteur, Paris | Mélanie Albert, Marion Barbet, Sylvie Behillil, Méline Bizard, Angela Brisebarre, Flora Donati, Etienne Simon-Lorière, Vincent Enouf, Maud Vanpeene, Sylvie van der Werf |
| EPI_ISL_443305 | LABM GH nord Essonne de Longjumeau - BP 125 | National Reference Center for Viruses of Respiratory Infections, Institut Pasteur, Paris | Mélanie Albert, Marion Barbet, Sylvie Behillil, Méline Bizard, Angela Brisebarre, Flora Donati, Etienne Simon-Lorière, Vincent Enouf, Maud Vanpeene, Sylvie van der Werf |
| EPI_ISL_443307 | La Villa Papyri | National Reference Center for Viruses of Respiratory Infections, Institut Pasteur, Paris | Mélanie Albert, Marion Barbet, Sylvie Behillil, Méline Bizard, Angela Brisebarre, Flora Donati, Etienne Simon-Lorière, Vincent Enouf, Maud Vanpeene, Sylvie van der Werf |
| EPI_ISL_443308 | Plaisance | National Reference Center for Viruses of Respiratory Infections, Institut Pasteur, Paris | Mélanie Albert, Marion Barbet, Sylvie Behillil, Méline Bizard, Angela Brisebarre, Flora Donati, Etienne Simon-Lorière, Vincent Enouf, Maud Vanpeene, Sylvie van der Werf |

| | | | |
|---|---|---|---|
| EPI_ISL_443309 | CH Compiègne Laboratoire de Biologie | National Reference Center for Viruses of Respiratory Infections, Institut Pasteur, Paris | Mélanie Albert, Marion Barbet, Sylvie Behillil, Méline Bizard, Angela Brisebarre, Flora Donati, Etienne Simon-Lorière, Vincent Enouf, Maud Vanpeene, Sylvie van der Werf |
| EPI_ISL_443314 | LABM GH nord Essonne de Longjumeau - BP 125 | National Reference Center for Viruses of Respiratory Infections, Institut Pasteur, Paris | Mélanie Albert, Marion Barbet, Sylvie Behillil, Méline Bizard, Angela Brisebarre, Flora Donati, Etienne Simon-Lorière, Vincent Enouf, Maud Vanpeene, Sylvie van der Werf |
| EPI_ISL_443315 | Château de la Source | National Reference Center for Viruses of Respiratory Infections, Institut Pasteur, Paris | Mélanie Albert, Marion Barbet, Sylvie Behillil, Méline Bizard, Angela Brisebarre, Flora Donati, Etienne Simon-Lorière, Vincent Enouf, Maud Vanpeene, Sylvie van der Werf |
| EPI_ISL_443316 | CH Compiègne Laboratoire de Biologie | National Reference Center for Viruses of Respiratory Infections, Institut Pasteur, Paris | Mélanie Albert, Marion Barbet, Sylvie Behillil, Méline Bizard, Angela Brisebarre, Flora Donati, Etienne Simon-Lorière, Vincent Enouf, Maud Vanpeene, Sylvie van der Werf, Olivia Raulin |
| EPI_ISL_444022 | Baylor College of Medicine | Baylor College of Medicine: HGSC | Vasanthi Avadhanula, Erin Nicholson, David Henke, Pedro Piedra, Harsha Doddapaneni, Donna Muzny, Qingchang Meng, Hsu Chao, Zeineen Momin, Hua Shen, George Weissenberger, Kavya Kottapalli, Yimiti Meiheerguli, Sejal Salvi, Ginger Metcalf, Vipin Menon, Sara J.J. Cregeen, Matthew C. Ross, Tulin Ayvaz, Richard Sucgang, Kristi L. Hoffman, Matthew Wong, Joseph F. Petrosino |
| EPI_ISL_444027 | Pamela Youde Nethersole Eastern Hospital | Hong Kong Department of Health | Mak Gannon C.K., Cheng Peter K.C., Lam Edman T.K., Chan Rickjason C.W., Tsang Dominic N.C. |
| EPI_ISL_444028 | Queen Elizabeth Hospital | Hong Kong Department of Health | Mak Gannon C.K., Cheng Peter K.C., Lam Edman T.K., Chan Rickjason C.W., Tsang Dominic N.C. |
| EPI_ISL_444030 | United Christian Hospital | Hong Kong Department of Health | Mak Gannon C.K., Cheng Peter K.C., Lam Edman T.K., Chan Rickjason C.W., Tsang Dominic N.C. |
| EPI_ISL_444031 | Queen Mary Hospital | Hong Kong Department of Health | Mak Gannon C.K., Cheng Peter K.C., Lam Edman T.K., Chan Rickjason C.W., Tsang Dominic N.C. |
| EPI_ISL_444032 | North Lantau Hospital | Hong Kong Department of Health | Mak Gannon C.K., Cheng Peter K.C., Lam Edman T.K., Chan Rickjason C.W., Tsang Dominic N.C. |
| EPI_ISL_444033 | Queen Mary Hospital | Hong Kong Department of Health | Mak Gannon C.K., Cheng Peter K.C., Lam Edman T.K., Chan Rickjason C.W., Tsang Dominic N.C. |
| EPI_ISL_444034 | Prince of Wales Hospital | Hong Kong Department of Health | Mak Gannon C.K., Cheng Peter K.C., Lam Edman T.K., Chan Rickjason C.W., Tsang Dominic N.C. |
| EPI_ISL_444035 | Princess Margaret Hospital | Hong Kong Department of Health | Mak Gannon C.K., Cheng Peter K.C., Lam Edman T.K., Chan Rickjason C.W., Tsang Dominic N.C. |
| EPI_ISL_444036 | North Lantau Hospital | Hong Kong Department of Health | Mak Gannon C.K., Cheng Peter K.C., Lam Edman T.K., Chan Rickjason C.W., Tsang Dominic N.C. |
| EPI_ISL_444038 | Princess Margaret Hospital | Hong Kong Department of Health | Mak Gannon C.K., Cheng Peter K.C., Lam Edman T.K., Chan Rickjason C.W., Tsang Dominic N.C. |
| EPI_ISL_444040 | Queen Elizabeth Hospital | Hong Kong Department of Health | Mak Gannon C.K., Cheng Peter K.C., Lam Edman T.K., Chan Rickjason C.W., Tsang Dominic N.C. |
| EPI_ISL_444041 | Tuen Mun Hospital | Hong Kong Department of Health | Mak Gannon C.K., Cheng Peter K.C., Lam Edman T.K., Chan Rickjason C.W., Tsang Dominic N.C. |
| EPI_ISL_444042, EPI_ISL_444043 | Pamela Youde Nethersole Eastern Hospital | Hong Kong Department of Health | Mak Gannon C.K., Cheng Peter K.C., Lam Edman T.K., Chan Rickjason C.W., Tsang Dominic N.C. |
| EPI_ISL_444044 | Yan Chai Hospital | Hong Kong Department of Health | Mak Gannon C.K., Cheng Peter K.C., Lam Edman T.K., Chan Rickjason C.W., Tsang Dominic N.C. |
| EPI_ISL_444045, EPI_ISL_444046 | Queen Elizabeth Hospital | Hong Kong Department of Health | Mak Gannon C.K., Cheng Peter K.C., Lam Edman T.K., Chan Rickjason C.W., Tsang Dominic N.C. |
| EPI_ISL_444047 | United Christian Hospital | Hong Kong Department of Health | Mak Gannon C.K., Cheng Peter K.C., Lam Edman T.K., Chan Rickjason C.W., Tsang Dominic N.C. |
| EPI_ISL_444048 | Queen Elizabeth Hospital | Hong Kong Department of Health | Mak Gannon C.K., Cheng Peter K.C., Lam Edman T.K., Chan Rickjason C.W., Tsang Dominic N.C. |
| EPI_ISL_444049 | North Lantau Hospital | Hong Kong Department of Health | Mak Gannon C.K., Cheng Peter K.C., Lam Edman T.K., Chan Rickjason C.W., Tsang Dominic N.C. |
| EPI_ISL_444050 | Shek Wu Hui Jockey Club General Out-patient Clinic | Hong Kong Department of Health | Mak Gannon C.K., Cheng Peter K.C., Lam Edman T.K., Chan Rickjason C.W., Tsang Dominic N.C. |
| EPI_ISL_444456 | B.J. Medical College and Civil hospital | Gujarat Biotechnology Research Centre | R D Dixit, Snehal Bagatharia, Kamlesh J Upadhyay, Ramesh Pandit, Tejas Shah, Ankit Hinsu, Pritesh Sabara, Apurvasinh Puvar, Janvi Raval, Monika Gandhi, Pinal Trivedi, Maharshi Pandya, Amit Kanani, Akanksha Verma, Nitin Savaliya, Raghawendra Kumar, Dinesh Kumar, Zuber Saiyed, Dipa Kinariwala, Disha Patel, Binita Aring, Neeta Khandelwal, Geeta Vaghela, Sonia Barve, Bhavesh Modi, Kairavi Joshi, Gaurishankar Shrimali, Nidhi Sood, Pranay Shah, Pooja P Doshi, Chaitanya Joshi, Madhvi Joshi |
| EPI_ISL_444457 | B.J. Medical College and Civil hospital | Gujarat Biotechnology Research Centre | Snehal Bagatharia, Kamlesh J Upadhyay, Ramesh Pandit, Tejas Shah, Ankit Hinsu, Pritesh Sabara, Apurvasinh Puvar, Janvi Raval, Monika Gandhi, Pinal Trivedi, Maharshi Pandya, Amit Kanani, Akanksha Verma, Nitin Savaliya, Raghawendra Kumar, Dinesh Kumar, Zuber Saiyed, Dipa Kinariwala, Disha Patel, Binita Aring, Neeta Khandelwal, Geeta Vaghela, Sonia Barve, Bhavesh Modi, Kairavi Joshi, Gaurishankar Shrimali, Nidhi Sood, Pranay Shah, R D Dixit, Nidhi Patel, Chaitanya Joshi, Madhvi Joshi |
| EPI_ISL_444458 | B.J. Medical College and Civil hospital | Gujarat Biotechnology Research Centre | Kamlesh J Upadhyay, Ramesh Pandit, Tejas Shah, Ankit Hinsu, Pritesh Sabara, Apurvasinh Puvar, Janvi Raval, Monika Gandhi, Pinal Trivedi, Maharshi Pandya, Amit Kanani, Akanksha Verma, Nitin Savaliya, Raghawendra Kumar, Dinesh Kumar, Zuber Saiyed, Dipa Kinariwala, Disha Patel, Binita Aring, Neeta Khandelwal, Geeta Vaghela, Sonia Barve, Bhavesh Modi, Kairavi Joshi, Gaurishankar Shrimali, Nidhi Sood, Pranay Shah, R D Dixit, Snehal Bagatharia, Priti Pandita, Chaitanya Joshi, Madhvi Joshi |
| EPI_ISL_444459 | B.J. Medical College and Civil hospital | Gujarat Biotechnology Research Centre | Ramesh Pandit, Tejas Shah, Ankit Hinsu, Pritesh Sabara, Apurvasinh Puvar, Janvi Raval, Monika Gandhi, Pinal Trivedi, Maharshi Pandya, Amit Kanani, Akanksha Verma, Nitin Savaliya, Raghawendra Kumar, Dinesh Kumar, Zuber Saiyed, Dipa Kinariwala, Disha Patel, Binita Aring, Neeta Khandelwal, Geeta Vaghela, Sonia Barve, Bhavesh Modi, Kairavi Joshi, Gaurishankar Shrimali, Nidhi Sood, Pranay Shah, R D Dixit, Snehal Bagatharia, Kamlesh J Upadhyay, Neha Rajpara, Chaitanya Joshi, Madhvi Joshi |
| EPI_ISL_444460 | B.J. Medical College and Civil hospital | Gujarat Biotechnology Research Centre | Tejas Shah, Ankit Hinsu, Pritesh Sabara, Apurvasinh Puvar, Janvi Raval, Monika Gandhi, Pinal Trivedi, Maharshi Pandya, Amit Kanani, Akanksha Verma, Nitin Savaliya, Raghawendra Kumar, Dinesh Kumar, Zuber Saiyed, Dipa Kinariwala, Disha Patel, Binita Aring, Neeta Khandelwal, Geeta Vaghela, Sonia Barve, Bhavesh Modi, Kairavi Joshi, Gaurishankar Shrimali, Nidhi Sood, Pranay Shah, R D Dixit, Snehal Bagatharia, Kamlesh J Upadhyay, Ramesh Pandit, Afzal Ansari, Chaitanya Joshi, Madhvi Joshi |
| EPI_ISL_444462 | B.J. Medical College and Civil hospital | Gujarat Biotechnology Research Centre | Pritesh Sabara, Apurvasinh Puvar, Janvi Raval, Monika Gandhi, Pinal Trivedi, Maharshi Pandya, Amit Kanani, Akanksha Verma, Nitin Savaliya, Raghawendra Kumar, Dinesh Kumar, Zuber Saiyed, Dipa Kinariwala, Disha Patel, Binita Aring, Neeta Khandelwal, Geeta Vaghela, Sonia Barve, Bhavesh Modi, Kairavi Joshi, Gaurishankar Shrimali, Nidhi Sood, Pranay Shah, R D Dixit, Snehal Bagatharia, Kamlesh J Upadhyay, Ramesh Pandit, Tejas Shah, Ankit Hinsu, Armi Chaudhari, Chaitanya Joshi, Madhvi Joshi |
| EPI_ISL_444465 | B.J. Medical College and Civil hospital | Gujarat Biotechnology Research Centre | Monika Gandhi, Pinal Trivedi, Maharshi Pandya, Amit Kanani, Akanksha Verma, Nitin Savaliya, Raghawendra Kumar, Dinesh Kumar, Zuber Saiyed, Dipa Kinariwala, Disha Patel, Binita Aring, Neeta Khandelwal, Geeta Vaghela, Sonia Barve, Bhavesh Modi, Kairavi Joshi, Gaurishankar Shrimali, Nidhi Sood, Pranay Shah, R D Dixit, Snehal Bagatharia, Kamlesh J Upadhyay, Ramesh Pandit, Tejas Shah, Ankit Hinsu, Pritesh Sabara, Apurvasinh Puvar, Janvi Raval, Anjali Rajwar, Chaitanya Joshi, Madhvi Joshi |
| EPI_ISL_444466 | B.J. Medical College and Civil hospital | Gujarat Biotechnology Research Centre | Pinal Trivedi, Maharshi Pandya, Amit Kanani, Akanksha Verma, Nitin Savaliya, Raghawendra Kumar, Dinesh Kumar, Zuber Saiyed, Dipa Kinariwala, Disha Patel, Binita Aring, Neeta Khandelwal, Geeta Vaghela, Sonia Barve, Bhavesh Modi, Kairavi Joshi, Gaurishankar Shrimali, Nidhi Sood, Pranay Shah, R D Dixit, Snehal Bagatharia, Kamlesh J Upadhyay, Ramesh Pandit, Tejas Shah, Ankit Hinsu, Pritesh Sabara, Apurvasinh Puvar, Janvi Raval, Monika Gandhi, Sharmistha Majumdar, Chaitanya Joshi, Madhvi Joshi |
| EPI_ISL_444468 | B.J. Medical College and Civil hospital | Gujarat Biotechnology Research Centre | Amit Kanani, Akanksha Verma, Nitin Savaliya, Raghawendra Kumar, Dinesh Kumar, Zuber Saiyed, Dipa Kinariwala, Disha Patel, Binita Aring, Neeta Khandelwal, Geeta Vaghela, Sonia Barve, Bhavesh Modi, Kairavi Joshi, Gaurishankar Shrimali, Nidhi Sood, Pranay Shah, R D Dixit, Snehal Bagatharia, Kamlesh J Upadhyay, Ramesh Pandit, Tejas Shah, Ankit Hinsu, Pritesh Sabara, Apurvasinh Puvar, Janvi Raval, Monika Gandhi, Pinal Trivedi, Maharshi Pandya, Nidhi Patel, Chaitanya Joshi, Madhvi Joshi |
| EPI_ISL_444469 | B.J. Medical College and Civil hospital | Gujarat Biotechnology Research Centre | Akanksha Verma, Nitin Savaliya, Raghawendra Kumar, Dinesh Kumar, Zuber Saiyed, Dipa Kinariwala, Disha Patel, Binita Aring, Neeta Khandelwal, Geeta Vaghela, Sonia Barve, Bhavesh Modi, Kairavi Joshi, Gaurishankar Shrimali, Nidhi Sood, Pranay Shah, R D Dixit, Snehal Bagatharia, Kamlesh J Upadhyay, Ramesh Pandit, Tejas Shah, Ankit Hinsu, Pritesh Sabara, Apurvasinh Puvar, Janvi Raval, Monika Gandhi, Pinal Trivedi, Maharshi Pandya, Amit Kanani, Priti Pandita, Chaitanya Joshi, Madhvi Joshi |
| EPI_ISL_444471 | B.J. Medical College and Civil hospital | Gujarat Biotechnology Research Centre | Raghawendra Kumar, Dinesh Kumar, Zuber Saiyed, Dipa Kinariwala, Disha Patel, Binita Aring, Neeta Khandelwal, Geeta Vaghela, Sonia Barve, Bhavesh Modi, |

| | | | |
|---|---|---|---|
| | | | Kairavi Joshi, Gaurishankar Shrimali, Nidhi Sood, Pranay Shah, R D Dixit, Snehal Bagatharia, Kamlesh J Upadhyay, Ramesh Pandit, Tejas Shah, Ankit Hinsu, Pritesh Sabara, Apurvasinh Puvar, Janvi Raval, Monika Gandhi, Pinal Trivedi, Maharshi Pandya, Amit Kanani, Akanksha Verma, Nitin Savaliya, Afzal Ansari, Chaitanya Joshi, Madhvi Joshi |
| EPI_ISL_444474 | B.J. Medical College and Civil hospital | Gujarat Biotechnology Research Centre | Dipa Kinariwala, Disha Patel, Binita Aring, Neeta Khandelwal, Geeta Vaghela, Sonia Barve, Bhavesh Modi, Kairavi Joshi, Gaurishankar Shrimali, Nidhi Sood, Pranay Shah, R D Dixit, Snehal Bagatharia, Kamlesh J Upadhyay, Ramesh Pandit, Tejas Shah, Ankit Hinsu, Pritesh Sabara, Apurvasinh Puvar, Janvi Raval, Monika Gandhi, Pinal Trivedi, Maharshi Pandya, Amit Kanani, Akanksha Verma, Nitin Savaliya, Raghawendra Kumar, Dinesh Kumar, Zuber Saiyed, Bhavya Jindal, Chaitanya Joshi, Madhvi Joshi |
| EPI_ISL_444475 | B.J. Medical College and Civil hospital | Gujarat Biotechnology Research Centre | Disha Patel, Binita Aring, Neeta Khandelwal, Geeta Vaghela, Sonia Barve, Bhavesh Modi, Kairavi Joshi, Gaurishankar Shrimali, Nidhi Sood, Pranay Shah, R D Dixit, Snehal Bagatharia, Kamlesh J Upadhyay, Ramesh Pandit, Tejas Shah, Ankit Hinsu, Pritesh Sabara, Apurvasinh Puvar, Janvi Raval, Monika Gandhi, Pinal Trivedi, Maharshi Pandya, Amit Kanani, Akanksha Verma, Nitin Savaliya, Raghawendra Kumar, Dinesh Kumar, Zuber Saiyed, Dipa Kinariwala, Dipeshwari Shewale, Chaitanya Joshi, Madhvi Joshi |
| EPI_ISL_444476 | B.J. Medical College and Civil hospital | Gujarat Biotechnology Research Centre | Binita Aring, Neeta Khandelwal, Geeta Vaghela, Sonia Barve, Bhavesh Modi, Kairavi Joshi, Gaurishankar Shrimali, Nidhi Sood, Pranay Shah, R D Dixit, Snehal Bagatharia, Kamlesh J Upadhyay, Ramesh Pandit, Tejas Shah, Ankit Hinsu, Pritesh Sabara, Apurvasinh Puvar, Janvi Raval, Monika Gandhi, Pinal Trivedi, Maharshi Pandya, Amit Kanani, Akanksha Verma, Nitin Savaliya, Raghawendra Kumar, Dinesh Kumar, Zuber Saiyed, Dipa Kinariwala, Disha Patel, Chaitanya Joshi, Madhvi Joshi, Dipeshwari Shewale |
| EPI_ISL_444477 | B.J. Medical College and Civil hospital | Gujarat Biotechnology Research Centre | Neeta Khandelwal, Geeta Vaghela, Sonia Barve, Bhavesh Modi, Kairavi Joshi, Gaurishankar Shrimali, Nidhi Sood, Pranay Shah, R D Dixit, Snehal Bagatharia, Kamlesh J Upadhyay, Ramesh Pandit, Tejas Shah, Ankit Hinsu, Pritesh Sabara, Apurvasinh Puvar, Janvi Raval, Monika Gandhi, Pinal Trivedi, Maharshi Pandya, Amit Kanani, Akanksha Verma, Nitin Savaliya, Raghawendra Kumar, Dinesh Kumar, Zuber Saiyed, Dipa Kinariwala, Disha Patel, Binita Aring, Sharmistha Majumdar, Chaitanya Joshi, Madhvi Joshi |
| EPI_ISL_444480 | B.J. Medical College and Civil hospital | Gujarat Biotechnology Research Centre | Bhavesh Modi, Kairavi Joshi, Gaurishankar Shrimali, Nidhi Sood, Pranay Shah, R D Dixit, Snehal Bagatharia, Kamlesh J Upadhyay, Ramesh Pandit, Tejas Shah, Ankit Hinsu, Pritesh Sabara, Apurvasinh Puvar, Janvi Raval, Monika Gandhi, Pinal Trivedi, Maharshi Pandya, Amit Kanani, Akanksha Verma, Nitin Savaliya, Raghawendra Kumar, Dinesh Kumar, Zuber Saiyed, Dipa Kinariwala, Disha Patel, Binita Aring, Neeta Khandelwal, Geeta Vaghela, Sonia Barve, Priti Pandita, Chaitanya Joshi, Madhvi Joshi |
| EPI_ISL_444481 | B.J. Medical College and Civil hospital | Gujarat Biotechnology Research Centre | Kairavi Joshi, Gaurishankar Shrimali, Nidhi Sood, Pranay Shah, R D Dixit, Snehal Bagatharia, Kamlesh J Upadhyay, Ramesh Pandit, Tejas Shah, Ankit Hinsu, Pritesh Sabara, Apurvasinh Puvar, Janvi Raval, Monika Gandhi, Pinal Trivedi, Maharshi Pandya, Amit Kanani, Akanksha Verma, Nitin Savaliya, Raghawendra Kumar, Dinesh Kumar, Zuber Saiyed, Dipa Kinariwala, Disha Patel, Binita Aring, Neeta Khandelwal, Geeta Vaghela, Sonia Barve, Bhavesh Modi, Neha Rajpara, Chaitanya Joshi, Madhvi Joshi |
| EPI_ISL_444482 | Gujarat Biotechnology Research Centre | Gujarat Biotechnology Research Centre | Gaurishankar Shrimali, Nidhi Sood, Pranay Shah, R D Dixit, Snehal Bagatharia, Kamlesh J Upadhyay, Ramesh Pandit, Tejas Shah, Ankit Hinsu, Pritesh Sabara, Apurvasinh Puvar, Janvi Raval, Monika Gandhi, Pinal Trivedi, Maharshi Pandya, Amit Kanani, Akanksha Verma, Nitin Savaliya, Raghawendra Kumar, Dinesh Kumar, Zuber Saiyed, Dipa Kinariwala, Disha Patel, Binita Aring, Neeta Khandelwal, Geeta Vaghela, Sonia Barve, Bhavesh Modi, Kairavi Joshi, Afzal Ansari, Chaitanya Joshi, Madhvi Joshi |
| EPI_ISL_444483 | Gujarat Biotechnology Research Centre | Gujarat Biotechnology Research Centre | Nidhi Sood, Pranay Shah, R D Dixit, Snehal Bagatharia, Kamlesh J Upadhyay, Ramesh Pandit, Tejas Shah, Ankit Hinsu, Pritesh Sabara, Apurvasinh Puvar, Janvi Raval, Monika Gandhi, Pinal Trivedi, Maharshi Pandya, Amit Kanani, Akanksha Verma, Nitin Savaliya, Raghawendra Kumar, Dinesh Kumar, Zuber Saiyed, Dipa Kinariwala, Disha Patel, Binita Aring, Neeta Khandelwal, Geeta Vaghela, Sonia Barve, Bhavesh Modi, Kairavi Joshi, Gaurishankar Shrimali, Neelam Nathani, Chaitanya Joshi, Madhvi Joshi |
| EPI_ISL_444484 | Gujarat Biotechnology Research Centre | Gujarat Biotechnology Research Centre | Pranay Shah, R D Dixit, Snehal Bagatharia, Kamlesh J Upadhyay, Ramesh Pandit, Tejas Shah, Ankit Hinsu, Pritesh Sabara, Apurvasinh Puvar, Janvi Raval, Monika Gandhi, Pinal Trivedi, Maharshi Pandya, Amit Kanani, Akanksha Verma, Nitin Savaliya, Raghawendra Kumar, Dinesh Kumar, Zuber Saiyed, Dipa Kinariwala, Disha Patel, Binita Aring, Neeta Khandelwal, Geeta Vaghela, Sonia Barve, Bhavesh Modi, Kairavi Joshi, Gaurishankar Shrimali, Nidhi Sood, Armi Chaudhari, Chaitanya Joshi, Madhvi Joshi |
| EPI_ISL_444486 | Gujarat Biotechnology Research Centre | Gujarat Biotechnology Research Centre | Snehal Bagatharia, Kamlesh J Upadhyay, Ramesh Pandit, Tejas Shah, Ankit Hinsu, Pritesh Sabara, Apurvasinh Puvar, Janvi Raval, Monika Gandhi, Pinal Trivedi, Maharshi Pandya, Amit Kanani, Akanksha Verma, Nitin Savaliya, Raghawendra Kumar, Dinesh Kumar, Zuber Saiyed, Dipa Kinariwala, Disha Patel, Binita Aring, Neeta Khandelwal, Geeta Vaghela, Sonia Barve, Bhavesh Modi, Kairavi Joshi, Gaurishankar Shrimali, Nidhi Sood, Pranay Shah, R D Dixit, Dipeshwari Shewale, Chaitanya Joshi, Madhvi Joshi |
| EPI_ISL_444493 | Departamento de Laboratorios de Salud Publica (DLSP, Division Epidemiologia, Ministerio de Salud Publica) | Facultad de Ciencias (Sección Genética Evolutiva, Sección Virología). | Panzera,Y., Delfraro,A., Ramos,N., Frabasile,S., Calleros,L., Techera,C., Grecco,S., Fuques,E., Goni,N., Coppola,L., Ramos,V., Chiparelli,H., Arbiza,J. and Perez,R. |
| EPI_ISL_444969 | Guangzhou Eighth People's Hospital (Jiahe Sector) | Institute of Human Virology, Zhongshan School of Medicine, Sun Yat-sen University | Junsong Zhang, Fei Yu, Jun Liu, Huimin Fan, Ruosu Ying, Feng Huang, Ting Pan, Bingfeng Liu,Yiwen Zhang, Xu Zhang, Mang Shi, Fengyu Hu, Fang Li, Kai Deng, Hui Zhang |
| EPI_ISL_444994, EPI_ISL_444995, EPI_ISL_444996, EPI_ISL_444998, EPI_ISL_445000 | Naval Health Research Center | Naval Medical Research Center Biological Defense Research Directorate | Logan Voegtly, Regina Cer, Dessiree Pena-Gomez, Adrian Paskey,Kyle Long, Roger Pan, Melinda Balansay-Ames, Chris Myers, Ewell Hollis, Nathaniel Christy, Kimberly Bishop-Lilly |
| EPI_ISL_445078, EPI_ISL_445079, EPI_ISL_445080, EPI_ISL_445081, EPI_ISL_445082, EPI_ISL_445083, EPI_ISL_445084 | Baylor College of Medicine | Baylor College of Medicine: HGSC | Vasanthi Avadhanula, Erin Nicholson, David Henke, Pedro Piedra, Harsha Doddapaneni, Donna Muzny, Qingchang Meng, Hsu Chao, Zeineen Momin, Hua Shen, George Weissenberger, Kavya Kottapalli, Yimiti Meiheerguli, Sejal Salvi, Ginger Metcalf, Vipin Menon, Sara J.J. Cregeen, Matthew C. Ross, Tulin Ayvaz, Richard Sucgang, Kristi L. Hoffman, Matthew Wong, Joseph F. Petrosino |
| EPI_ISL_445214, EPI_ISL_445215, EPI_ISL_445216 | DNA Solution Ltd. | DNA Solution Ltd. | Md. Imran Khan, Kazi Nadim Hasan, Abu Sufian, Mohammed Nafiz Imtiaz Polol, Abdul Khaleque, Mizanur Rahman, MSM Chowdhury, Hasan Ul Haider, Mamudul Hasan Razu, Mala Khan, Mohammad Fazle Alam Rabbi |
| EPI_ISL_445219 | Universidad del Valle, Laboratorio de Microbiologia, VIREM | Universidad del Valle, Universidad Nacional de Colombia-Sede Palmira, International Center for Tropical Agriculture | Beatriz Parra, Diana López-Alvarez, Wilmer J. Cuellar |
| EPI_ISL_445220 | Laborarory for Respiratory Viruses, "Cantacuzino" National Military-Medical Institute for Resararch and Development | Cantacuzino Institute | M.Lazar, L.Ustea, A.Cretu |
| EPI_ISL_445243 | Laboratory for Respiratory Viruses, Cantacuzino National Military-Medical Institute for Research and Development | Cantacuzino Institute | M.Lazar, L.Ustea, A.Cretu |
| EPI_ISL_447030 | B.J. Medical College and Civil hospital | Gujarat Biotechnology Research Centre | Kamlesh J Upadhyay, Ramesh Pandit, Tejas Shah, Ankit Hinsu, Pritesh Sabara, Apurvasinh Puvar, Janvi Raval, Monika Gandhi, Pinal Trivedi, Maharshi Pandya, Amit Kanani, Akanksha Verma, Nitin Savaliya, Raghawendra Kumar, Dinesh Kumar, Zuber Saiyed, Dipa Kinariwala, Disha Patel, Binita Aring, Neeta Khandelwal, Geeta Vaghela, Sonia Barve, Bhavesh Modi, Kairavi Joshi, Gaurishankar Shrimali, Nidhi Sood, Pranay Shah, R D Dixit, Snehal Bagatharia, Anjali Rajwar, Chaitanya Joshi, Madhvi Joshi |
| EPI_ISL_447031 | B.J. Medical College and Civil hospital | Gujarat Biotechnology Research Centre | Ramesh Pandit, Tejas Shah, Ankit Hinsu, Pritesh Sabara, Apurvasinh Puvar, Janvi Raval, Monika Gandhi, Pinal Trivedi, Maharshi Pandya, Amit Kanani, Akanksha Verma, Nitin Savaliya, Raghawendra Kumar, Dinesh Kumar, Zuber Saiyed, Dipa Kinariwala, Disha Patel, Binita Aring, Neeta Khandelwal, Geeta Vaghela, Sonia Barve, Bhavesh Modi, Kairavi Joshi, Gaurishankar Shrimali, Nidhi Sood, Pranay Shah, R D Dixit, Snehal Bagatharia, Kamlesh J Upadhyay, Sharmistha Majumdar, Chaitanya Joshi, Madhvi Joshi |
| EPI_ISL_447033 | B.J. Medical College and Civil hospital | Gujarat Biotechnology Research Centre | Ankit Hinsu, Pritesh Sabara, Apurvasinh Puvar, Janvi Raval, Monika Gandhi, Pinal Trivedi, Maharshi Pandya, Amit Kanani, Akanksha Verma, Nitin Savaliya, Raghawendra Kumar, Dinesh Kumar, Zuber Saiyed, Dipa Kinariwala, Disha Patel, Binita Aring, Neeta Khandelwal, Geeta Vaghela, Sonia Barve, Bhavesh Modi, Kairavi Joshi, Gaurishankar Shrimali, Nidhi Sood, Pranay Shah, R D Dixit, Snehal Bagatharia, Kamlesh J Upadhyay, Ramesh Pandit, Tejas Shah, Nidhi Patel, Chaitanya Joshi, Madhvi Joshi |
| EPI_ISL_447034 | B.J. Medical College and Civil hospital | Gujarat Biotechnology Research Centre | Pritesh Sabara, Apurvasinh Puvar, Janvi Raval, Monika Gandhi, Pinal Trivedi, Maharshi Pandya, Amit Kanani, Akanksha Verma, Nitin Savaliya, Raghawendra |

| | | | |
|---|---|---|---|
| | | | Kumar, Dinesh Kumar, Zuber Saiyed, Dipa Kinariwala, Disha Patel, Binita Aring, Neeta Khandelwal, Geeta Vaghela, Sonia Barve, Bhavesh Modi, Kairavi Joshi, Gaurishankar Shrimali, Nidhi Sood, Pranay Shah, R D Dixit, Snehal Bagatharia, Kamlesh J Upadhyay, Ramesh Pandit, Tejas Shah, Ankit Hinsu, Priti Pandita, Chaitanya Joshi, Madhvi Joshi |
| EPI_ISL_447035 | B.J. Medical College and Civil hospital | Gujarat Biotechnology Research Centre | Apurvasinh Puvar, Janvi Raval, Monika Gandhi, Pinal Trivedi, Maharshi Pandya, Amit Kanani, Akanksha Verma, Nitin Savaliya, Raghawendra Kumar, Dinesh Kumar, Zuber Saiyed, Dipa Kinariwala, Disha Patel, Binita Aring, Neeta Khandelwal, Geeta Vaghela, Sonia Barve, Bhavesh Modi, Kairavi Joshi, Gaurishankar Shrimali, Nidhi Sood, Pranay Shah, R D Dixit, Snehal Bagatharia, Kamlesh J Upadhyay, Ramesh Pandit, Tejas Shah, Ankit Hinsu, Pritesh Sabara, Neha Rajpara, Chaitanya Joshi, Madhvi Joshi |
| EPI_ISL_447038 | B.J. Medical College and Civil hospital | Gujarat Biotechnology Research Centre | Pinal Trivedi, Maharshi Pandya, Amit Kanani, Akanksha Verma, Nitin Savaliya, Raghawendra Kumar, Dinesh Kumar, Zuber Saiyed, Dipa Kinariwala, Disha Patel, Binita Aring, Neeta Khandelwal, Geeta Vaghela, Sonia Barve, Bhavesh Modi, Kairavi Joshi, Gaurishankar Shrimali, Nidhi Sood, Pranay Shah, R D Dixit, Snehal Bagatharia, Kamlesh J Upadhyay, Ramesh Pandit, Tejas Shah, Ankit Hinsu, Pritesh Sabara, Apurvasinh Puvar, Janvi Raval, Monika Gandhi, Armi Chaudhari, Chaitanya Joshi, Madhvi Joshi |
| EPI_ISL_447039 | B.J. Medical College and Civil hospital | Gujarat Biotechnology Research Centre | Maharshi Pandya, Amit Kanani, Akanksha Verma, Nitin Savaliya, Raghawendra Kumar, Dinesh Kumar, Zuber Saiyed, Dipa Kinariwala, Disha Patel, Binita Aring, Neeta Khandelwal, Geeta Vaghela, Sonia Barve, Bhavesh Modi, Kairavi Joshi, Gaurishankar Shrimali, Nidhi Sood, Pranay Shah, R D Dixit, Snehal Bagatharia, Kamlesh J Upadhyay, Ramesh Pandit, Tejas Shah, Ankit Hinsu, Pritesh Sabara, Apurvasinh Puvar, Janvi Raval, Monika Gandhi, Pinal Trivedi, Bhavya Jindal, Chaitanya Joshi, Madhvi Joshi |
| EPI_ISL_447042 | B.J. Medical College and Civil hospital | Gujarat Biotechnology Research Centre | Nitin Savaliya, Raghawendra Kumar, Dinesh Kumar, Zuber Saiyed, Dipa Kinariwala, Disha Patel, Binita Aring, Neeta Khandelwal, Geeta Vaghela, Sonia Barve, Bhavesh Modi, Kairavi Joshi, Gaurishankar Shrimali, Nidhi Sood, Pranay Shah, R D Dixit, Snehal Bagatharia, Kamlesh J Upadhyay, Ramesh Pandit, Tejas Shah, Ankit Hinsu, Pritesh Sabara, Apurvasinh Puvar, Janvi Raval, Monika Gandhi, Pinal Trivedi, Maharshi Pandya, Amit Kanani, Akanksha Verma, Pooja P Doshi, Chaitanya Joshi, Madhvi Joshi |
| EPI_ISL_447043 | B.J. Medical College and Civil hospital | Gujarat Biotechnology Research Centre | Raghawendra Kumar, Dinesh Kumar, Zuber Saiyed, Dipa Kinariwala, Disha Patel, Binita Aring, Neeta Khandelwal, Geeta Vaghela, Sonia Barve, Bhavesh Modi, Kairavi Joshi, Gaurishankar Shrimali, Nidhi Sood, Pranay Shah, R D Dixit, Snehal Bagatharia, Kamlesh J Upadhyay, Ramesh Pandit, Tejas Shah, Ankit Hinsu, Pritesh Sabara, Apurvasinh Puvar, Janvi Raval, Monika Gandhi, Pinal Trivedi, Maharshi Pandya, Amit Kanani, Akanksha Verma, Nitin Savaliya, Nidhi Patel, Chaitanya Joshi, Madhvi Joshi |
| EPI_ISL_447044 | B.J. Medical College and Civil hospital | Gujarat Biotechnology Research Centre | Dinesh Kumar, Zuber Saiyed, Dipa Kinariwala, Disha Patel, Binita Aring, Neeta Khandelwal, Geeta Vaghela, Sonia Barve, Bhavesh Modi, Kairavi Joshi, Gaurishankar Shrimali, Nidhi Sood, Pranay Shah, R D Dixit, Snehal Bagatharia, Kamlesh J Upadhyay, Ramesh Pandit, Tejas Shah, Ankit Hinsu, Pritesh Sabara, Apurvasinh Puvar, Janvi Raval, Monika Gandhi, Pinal Trivedi, Maharshi Pandya, Amit Kanani, Akanksha Verma, Nitin Savaliya, Raghawendra Kumar, Priti Pandita, Chaitanya Joshi, Madhvi Joshi |
| EPI_ISL_447045 | B.J. Medical College and Civil hospital | Gujarat Biotechnology Research Centre | Zuber Saiyed, Dipa Kinariwala, Disha Patel, Binita Aring, Neeta Khandelwal, Geeta Vaghela, Sonia Barve, Bhavesh Modi, Kairavi Joshi, Gaurishankar Shrimali, Nidhi Sood, Pranay Shah, R D Dixit, Snehal Bagatharia, Kamlesh J Upadhyay, Ramesh Pandit, Tejas Shah, Ankit Hinsu, Pritesh Sabara, Apurvasinh Puvar, Janvi Raval, Monika Gandhi, Pinal Trivedi, Maharshi Pandya, Amit Kanani, Akanksha Verma, Nitin Savaliya, Raghawendra Kumar, Dinesh Kumar, Neha Rajpara, Chaitanya Joshi, Madhvi Joshi |
| EPI_ISL_447048 | GMERS Medical College and Hospital, Gandhinagar | Gujarat Biotechnology Research Centre | Binita Aring, Neeta Khandelwal, Geeta Vaghela, Sonia Barve, Bhavesh Modi, Kairavi Joshi, Gaurishankar Shrimali, Nidhi Sood, Pranay Shah, R D Dixit, Snehal Bagatharia, Kamlesh J Upadhyay, Ramesh Pandit, Tejas Shah, Ankit Hinsu, Pritesh Sabara, Apurvasinh Puvar, Janvi Raval, Monika Gandhi, Pinal Trivedi, Maharshi Pandya, Amit Kanani, Akanksha Verma, Nitin Savaliya, Raghawendra Kumar, Dinesh Kumar, Zuber Saiyed, Dipa Kinariwala, Disha Patel, Armi Chaudhari, Chaitanya Joshi, Madhvi Joshi |
| EPI_ISL_447053 | GMERS Medical College and Hospital, Gandhinagar | Gujarat Biotechnology Research Centre | Kairavi Joshi, Gaurishankar Shrimali, Nidhi Sood, Pranay Shah, R D Dixit, Snehal Bagatharia, Kamlesh J Upadhyay, Ramesh Pandit, Tejas Shah, Ankit Hinsu, Pritesh Sabara, Apurvasinh Puvar, Janvi Raval, Monika Gandhi, Pinal Trivedi, Maharshi Pandya, Amit Kanani, Akanksha Verma, Nitin Savaliya, Raghawendra Kumar, Dinesh Kumar, Zuber Saiyed, Dipa Kinariwala, Disha Patel, Binita Aring, Neeta Khandelwal, Geeta Vaghela, Sonia Barve, Bhavesh Modi, Pooja P Doshi, Chaitanya Joshi, Madhvi Joshi |
| EPI_ISL_447054 | Cantacuzino National Military-Medical Institute for Research and Development | Cantacuzino Institute | M.Lazar, L.Ustea, A.Cretu |
| EPI_ISL_447251 | Central Virology Laboratory | Central Virology Laboratory | Neta Zuckerman, Efrat Bucris, Oran Erster, Danit Sofer, Orna Mor, Ella Mendelson, Michal Mandelboim |
| EPI_ISL_447252, EPI_ISL_447253, EPI_ISL_447255, EPI_ISL_447257 | TSGH-CP molecular lab | TSGH-CP molecular lab | Cherng-Lih Perng, Ming-Jr JIAN, Chih-Kai Chang, Jung-Chung Lin, Kuo-Ming Yeh, Chien-Wen Chen, Sheng-Kang Chiu, Hsing-Yi Chung, Shih-Hung Tsai, Kuo-Sheng Hung, Tien-Yao Chang, Feng-Yee Chang, Hung-Sheng Shang |
| EPI_ISL_447281, EPI_ISL_447282, EPI_ISL_447283, EPI_ISL_447284, EPI_ISL_447285, EPI_ISL_447286, EPI_ISL_447287, EPI_ISL_447288, EPI_ISL_447289, EPI_ISL_447290, EPI_ISL_447291, EPI_ISL_447292, EPI_ISL_447293, EPI_ISL_447294, EPI_ISL_447295, EPI_ISL_447296, EPI_ISL_447297, EPI_ISL_447299, EPI_ISL_447300, EPI_ISL_447301, EPI_ISL_447302, EPI_ISL_447303, EPI_ISL_447305, EPI_ISL_447306, EPI_ISL_447307, EPI_ISL_447308, EPI_ISL_447309, EPI_ISL_447310 | | | |
| see above | Microbiology Division, Barzilai University Medical Center | Stern Lab | Stern Lab |
| EPI_ISL_447534 | Gujarat Biotechnology Research Centre | Gujarat Biotechnology Research Centre | Gaurishankar Shrimali, Nidhi Sood, Pranay Shah, R D Dixit, Snehal Bagatharia, Kamlesh J Upadhyay, Ramesh Pandit, Tejas Shah, Ankit Hinsu, Pritesh Sabara, Apurvasinh Puvar, Janvi Raval, Monika Gandhi, Pinal Trivedi, Maharshi Pandya, Amit Kanani, Akanksha Verma, Nitin Savaliya, Raghawendra Kumar, Dinesh Kumar, Zuber Saiyed, Dipa Kinariwala, Disha Patel, Binita Aring, Neeta Khandelwal, Geeta Vaghela, Sonia Barve, Bhavesh Modi, Kairavi Joshi, Nidhi Patel, Chaitanya Joshi, Madhvi Joshi |
| EPI_ISL_447535 | Gujarat Biotechnology Research Centre | Gujarat Biotechnology Research Centre | Nidhi Sood, Pranay Shah, R D Dixit, Snehal Bagatharia, Kamlesh J Upadhyay, Ramesh Pandit, Tejas Shah, Ankit Hinsu, Pritesh Sabara, Apurvasinh Puvar, Janvi Raval, Monika Gandhi, Pinal Trivedi, Maharshi Pandya, Amit Kanani, Akanksha Verma, Nitin Savaliya, Raghawendra Kumar, Dinesh Kumar, Zuber Saiyed, Dipa Kinariwala, Disha Patel, Binita Aring, Neeta Khandelwal, Geeta Vaghela, Sonia Barve, Bhavesh Modi, Kairavi Joshi, Gaurishankar Shrimali, Priti Pandita, Chaitanya Joshi, Madhvi Joshi |
| EPI_ISL_447536 | Gujarat Biotechnology Research Centre | Gujarat Biotechnology Research Centre | Pranay Shah, R D Dixit, Snehal Bagatharia, Kamlesh J Upadhyay, Ramesh Pandit, Tejas Shah, Ankit Hinsu, Pritesh Sabara, Apurvasinh Puvar, Janvi Raval, Monika Gandhi, Pinal Trivedi, Maharshi Pandya, Amit Kanani, Akanksha Verma, Nitin Savaliya, Raghawendra Kumar, Dinesh Kumar, Zuber Saiyed, Dipa Kinariwala, Disha Patel, Binita Aring, Neeta Khandelwal, Geeta Vaghela, Sonia Barve, Bhavesh Modi, Kairavi Joshi, Gaurishankar Shrimali, Nidhi Sood, Neha Rajpara, Chaitanya Joshi, Madhvi Joshi |
| EPI_ISL_447537 | Gujarat Biotechnology Research Centre | Gujarat Biotechnology Research Centre | R D Dixit, Snehal Bagatharia, Kamlesh J Upadhyay, Ramesh Pandit, Tejas Shah, Ankit Hinsu, Pritesh Sabara, Apurvasinh Puvar, Janvi Raval, Monika Gandhi, Pinal Trivedi, Maharshi Pandya, Amit Kanani, Akanksha Verma, Nitin Savaliya, Raghawendra Kumar, Dinesh Kumar, Zuber Saiyed, Dipa Kinariwala, Disha Patel, Binita Aring, Neeta Khandelwal, Geeta Vaghela, Sonia Barve, Bhavesh Modi, Kairavi Joshi, Gaurishankar Shrimali, Nidhi Sood, Pranay Shah, Afzal Ansari, Chaitanya Joshi, Madhvi Joshi |
| EPI_ISL_447542 | Gujarat Biotechnology Research Centre | Gujarat Biotechnology Research Centre | Ankit Hinsu, Pritesh Sabara, Apurvasinh Puvar, Janvi Raval, Monika Gandhi, Pinal Trivedi, Maharshi Pandya, Amit Kanani, Akanksha Verma, Nitin Savaliya, Raghawendra Kumar, Dinesh Kumar, Zuber Saiyed, Dipa Kinariwala, Disha Patel, Binita Aring, Neeta Khandelwal, Geeta Vaghela, Sonia Barve, Bhavesh Modi, Kairavi Joshi, Gaurishankar Shrimali, Nidhi Sood, Pranay Shah, R D Dixit, Snehal Bagatharia, Kamlesh J Upadhyay, Ramesh Pandit, Tejas Shah, Dipeshwari Shewale, Chaitanya Joshi, Madhvi Joshi |
| EPI_ISL_447543 | Gujarat Biotechnology Research Centre | Gujarat Biotechnology Research Centre | Pritesh Sabara, Apurvasinh Puvar, Janvi Raval, Monika Gandhi, Pinal Trivedi, Maharshi Pandya, Amit Kanani, Akanksha Verma, Nitin Savaliya, Raghawendra Kumar, Dinesh Kumar, Zuber Saiyed, Dipa Kinariwala, Disha Patel, Binita Aring, Neeta Khandelwal, Geeta Vaghela, Sonia Barve, Bhavesh Modi, Kairavi Joshi, Gaurishankar Shrimali, Nidhi Sood, Pranay Shah, R D Dixit, Snehal Bagatharia, Kamlesh J Upadhyay, Ramesh Pandit, Tejas Shah, Ankit Hinsu, Sharmistha Majumdar, Chaitanya Joshi, Madhvi Joshi |
| EPI_ISL_447544 | Gujarat Biotechnology Research Centre | Gujarat Biotechnology Research Centre | Apurvasinh Puvar, Janvi Raval, Monika Gandhi, Pinal Trivedi, Maharshi Pandya, Amit Kanani, Akanksha Verma, Nitin Savaliya, Raghawendra Kumar, Dinesh Kumar, Zuber Saiyed, Dipa Kinariwala, Disha Patel, Binita Aring, Neeta Khandelwal, Geeta Vaghela, Sonia Barve, Bhavesh Modi, Kairavi Joshi, Gaurishankar Shrimali, Nidhi Sood, Pranay Shah, R D Dixit, Snehal Bagatharia, Kamlesh J Upadhyay, Ramesh Pandit, Tejas Shah, Ankit Hinsu, Pritesh Sabara, Pooja P Doshi, Chaitanya Joshi, Madhvi Joshi |
| EPI_ISL_447545 | Gujarat Biotechnology Research Centre | Gujarat Biotechnology Research Centre | Janvi Raval, Monika Gandhi, Pinal Trivedi, Maharshi Pandya, Amit Kanani, Akanksha Verma, Nitin Savaliya, Raghawendra Kumar, Dinesh Kumar, Zuber |

| | | | |
|---|---|---|---|
| | | | Saiyed, Dipa Kinariwala, Disha Patel, Binita Aring, Neeta Khandelwal, Geeta Vaghela, Sonia Barve, Bhavesh Modi, Kairavi Joshi, Gaurishankar Shrimali, Nidhi Sood, Pranay Shah, R D Dixit, Snehal Bagatharia, Kamlesh J Upadhyay, Ramesh Pandit, Tejas Shah, Ankit Hinsu, Pritesh Sabara, Apurvasinh Puvar, Nidhi Patel, Chaitanya Joshi, Madhvi Joshi |
| EPI_ISL_447546 | Gujarat Biotechnology Research Centre | Gujarat Biotechnology Research Centre | Monika Gandhi, Pinal Trivedi, Maharshi Pandya, Amit Kanani, Akanksha Verma, Nitin Savaliya, Raghawendra Kumar, Dinesh Kumar, Zuber Saiyed, Dipa Kinariwala, Disha Patel, Binita Aring, Neeta Khandelwal, Geeta Vaghela, Sonia Barve, Bhavesh Modi, Kairavi Joshi, Gaurishankar Shrimali, Nidhi Sood, Pranay Shah, R D Dixit, Snehal Bagatharia, Kamlesh J Upadhyay, Ramesh Pandit, Tejas Shah, Ankit Hinsu, Pritesh Sabara, Apurvasinh Puvar, Janvi Raval, Priti Pandita, Chaitanya Joshi, Madhvi Joshi |
| EPI_ISL_447548 | GMERS Medical College and Hospital, Gandhinagar | Gujarat Biotechnology Research Centre | Maharshi Pandya, Amit Kanani, Akanksha Verma, Nitin Savaliya, Raghawendra Kumar, Dinesh Kumar, Zuber Saiyed, Dipa Kinariwala, Disha Patel, Binita Aring, Neeta Khandelwal, Geeta Vaghela, Sonia Barve, Bhavesh Modi, Kairavi Joshi, Gaurishankar Shrimali, Nidhi Sood, Pranay Shah, R D Dixit, Snehal Bagatharia, Kamlesh J Upadhyay, Ramesh Pandit, Tejas Shah, Ankit Hinsu, Pritesh Sabara, Apurvasinh Puvar, Janvi Raval, Monika Gandhi, Pinal Trivedi, Afzal Ansari, Chaitanya Joshi, Madhvi Joshi |
| EPI_ISL_447551 | GMERS Medical College and Hospital, Gandhinagar | Gujarat Biotechnology Research Centre | Nitin Savaliya, Raghawendra Kumar, Dinesh Kumar, Zuber Saiyed, Dipa Kinariwala, Disha Patel, Binita Aring, Neeta Khandelwal, Geeta Vaghela, Sonia Barve, Bhavesh Modi, Kairavi Joshi, Gaurishankar Shrimali, Nidhi Sood, Pranay Shah, R D Dixit, Snehal Bagatharia, Kamlesh J Upadhyay, Ramesh Pandit, Tejas Shah, Ankit Hinsu, Pritesh Sabara, Apurvasinh Puvar, Janvi Raval, Monika Gandhi, Pinal Trivedi, Maharshi Pandya, Amit Kanani, Akanksha Verma, Bhavya Jindal, Chaitanya Joshi, Madhvi Joshi |
| EPI_ISL_447552 | GMERS Medical College and Hospital, Gandhinagar | Gujarat Biotechnology Research Centre | Raghawendra Kumar, Dinesh Kumar, Zuber Saiyed, Dipa Kinariwala, Disha Patel, Binita Aring, Neeta Khandelwal, Geeta Vaghela, Sonia Barve, Bhavesh Modi, Kairavi Joshi, Gaurishankar Shrimali, Nidhi Sood, Pranay Shah, R D Dixit, Snehal Bagatharia, Kamlesh J Upadhyay, Ramesh Pandit, Tejas Shah, Ankit Hinsu, Pritesh Sabara, Apurvasinh Puvar, Janvi Raval, Monika Gandhi, Pinal Trivedi, Maharshi Pandya, Amit Kanani, Akanksha Verma, Nitin Savaliya, Anjali Rajwar, Chaitanya Joshi, Madhvi Joshi |
| EPI_ISL_447553 | GMERS Medical College and Hospital, Gandhinagar | Gujarat Biotechnology Research Centre | Dinesh Kumar, Zuber Saiyed, Dipa Kinariwala, Disha Patel, Binita Aring, Neeta Khandelwal, Geeta Vaghela, Sonia Barve, Bhavesh Modi, Kairavi Joshi, Gaurishankar Shrimali, Nidhi Sood, Pranay Shah, R D Dixit, Snehal Bagatharia, Kamlesh J Upadhyay, Ramesh Pandit, Tejas Shah, Ankit Hinsu, Pritesh Sabara, Apurvasinh Puvar, Janvi Raval, Monika Gandhi, Pinal Trivedi, Maharshi Pandya, Amit Kanani, Akanksha Verma, Nitin Savaliya, Raghawendra Kumar, Dipeshwari Shewale, Chaitanya Joshi, Madhvi Joshi |
| EPI_ISL_447554 | GMERS Medical College and Hospital, Gandhinagar | Gujarat Biotechnology Research Centre | Zuber Saiyed, Dipa Kinariwala, Disha Patel, Binita Aring, Neeta Khandelwal, Geeta Vaghela, Sonia Barve, Bhavesh Modi, Kairavi Joshi, Gaurishankar Shrimali, Nidhi Sood, Pranay Shah, R D Dixit, Snehal Bagatharia, Kamlesh J Upadhyay, Ramesh Pandit, Tejas Shah, Ankit Hinsu, Pritesh Sabara, Apurvasinh Puvar, Janvi Raval, Monika Gandhi, Pinal Trivedi, Maharshi Pandya, Amit Kanani, Akanksha Verma, Nitin Savaliya, Raghawendra Kumar, Dinesh Kumar, Sharmistha Majumdar, Chaitanya Joshi, Madhvi Joshi |
| EPI_ISL_447555 | GMERS Medical College and Hospital, Gandhinagar | Gujarat Biotechnology Research Centre | Dipa Kinariwala, Disha Patel, Binita Aring, Neeta Khandelwal, Geeta Vaghela, Sonia Barve, Bhavesh Modi, Kairavi Joshi, Gaurishankar Shrimali, Nidhi Sood, Pranay Shah, R D Dixit, Snehal Bagatharia, Kamlesh J Upadhyay, Ramesh Pandit, Tejas Shah, Ankit Hinsu, Pritesh Sabara, Apurvasinh Puvar, Janvi Raval, Monika Gandhi, Pinal Trivedi, Maharshi Pandya, Amit Kanani, Akanksha Verma, Nitin Savaliya, Raghawendra Kumar, Dinesh Kumar, Zuber Saiyed, Pooja P Doshi, Chaitanya Joshi, Madhvi Joshi |
| EPI_ISL_447577 | CSIR-Centre for Cellular and Molecular Biology | CSIR-Centre for Cellular and Molecular Biology | Namami Gaur, Sakshi Shambhavi, Lamuk Zaveri, Shagufta Khan, Tulasi Nagabandi, Purushotham Vodnala, Payel Mukherjee, Sofia Banu, Priya Singh, Dhiviya Vedagiri, Divya Gupta, Vishal Sah, Santosh Kumar Kuncha, Krishnan Harinivas Harshan, Archana Bharadwaj Siva, Karthik Bharadwaj Tallapaka, Rakesh K Mishra, Divya Tej Sowpati |
| EPI_ISL_447592, EPI_ISL_447593 | TSGH-CP molecular lab | TSGH-CP molecular lab | Cherng-Lih Perng, Ming-Jr JIAN, Chih-Kai Chang, Jung-Chung Lin, Kuo-Ming Yeh, Chien-Wen Chen, Sheng-Kang Chiu, Hsing-Yi Chung, Shih-Hung Tsai, Kuo-Sheng Hung, Tien-Yao Chang, Feng-Yee Chang, Hung-Sheng Shang |
| EPI_ISL_447614, EPI_ISL_447615, EPI_ISL_447616, EPI_ISL_447617, EPI_ISL_447618, EPI_ISL_447619, EPI_ISL_447620, EPI_ISL_447621, EPI_ISL_447622 | Department of Laboratory Medicine, National Taiwan University Hospital | Microbial Genomics Core Lab, National Taiwan University Centers of Genomic and Precision Medicine | Shiou-Hwei Yeh, You-Yu Lin, Ya-Yun Lai, Chiao-Ling Li, Shan-Chwen Chang, Pei-Jer Chen, Sui-Yuan Chang |
| EPI_ISL_447734, EPI_ISL_447735, EPI_ISL_447736, EPI_ISL_447738, EPI_ISL_447739, EPI_ISL_447740, EPI_ISL_447741, EPI_ISL_447742, EPI_ISL_447743, EPI_ISL_447744, EPI_ISL_447745, EPI_ISL_447746, EPI_ISL_447747, EPI_ISL_447748, EPI_ISL_447750, EPI_ISL_447754 | | | |
| see above | Grupo de Investigaciones Microbiológicas-UR (GIMUR), Departamento de Biología, Facultad de Ciencias Naturales, Universidad del Rosario, Bogotá, Colombia | Grupo de Investigaciones Microbiológicas-UR (GIMUR), Departamento de Biología, Facultad de Ciencias Naturales, Universidad del Rosario, Bogotá, Colombia Instituto Nacional de Salud, Bogotá, Colombia Icahn School of Medicine at Mount Sinai, New York, USA | Juan David Ramírez, Carolina Florez, Marina Muñoz, Carolina Hernandez, Adriana Castillo, Sergio Castañeda, Nathalia Ballesteros, David Martínez, Laura Vega, Jesús E. Jaimes, Sergio Gomez, Angelica Rico, Lisseth Pardo, Esther C. Barros, Martha L. Ospina, Aníbal A. Teherán, Ana S. Gonzalez-Reiche, Matthew M. Hernandez, Emilia Mia Sordillo, Viviana Simon, Harm van Bakel, Alberto Paniz-Mondolfi |
| EPI_ISL_447755, EPI_ISL_447756, EPI_ISL_447757, EPI_ISL_447759, EPI_ISL_447760, EPI_ISL_447761, EPI_ISL_447762, EPI_ISL_447763, EPI_ISL_447765, EPI_ISL_447766, EPI_ISL_447767, EPI_ISL_447768, EPI_ISL_447769, EPI_ISL_447771, EPI_ISL_447774, EPI_ISL_447775, EPI_ISL_447777, EPI_ISL_447778, EPI_ISL_447779, EPI_ISL_447780, EPI_ISL_447781, EPI_ISL_447785, EPI_ISL_447786, EPI_ISL_447789, EPI_ISL_447790, EPI_ISL_447791, EPI_ISL_447792, EPI_ISL_447793, EPI_ISL_447794, EPI_ISL_447795, EPI_ISL_447796, EPI_ISL_447797, EPI_ISL_447798, EPI_ISL_447800, EPI_ISL_447802, EPI_ISL_447803, EPI_ISL_447804, EPI_ISL_447805, EPI_ISL_447806, EPI_ISL_447807, EPI_ISL_447808, EPI_ISL_447809, EPI_ISL_447810, EPI_ISL_447812, EPI_ISL_447813, EPI_ISL_447816, EPI_ISL_447817 | | | |
| see above | Instituto Nacional de Salud, Bogotá, Colombia | Grupo de Investigaciones Microbiológicas-UR (GIMUR), Departamento de Biología, Facultad de Ciencias Naturales, Universidad del Rosario, Bogotá, Colombia Instituto Nacional de Salud, Bogotá, Colombia Icahn School of Medicine at Mount Sinai, New York, USA | Juan David Ramírez, Carolina Florez, Marina Muñoz, Carolina Hernandez, Adriana Castillo, Sergio Castañeda, Nathalia Ballesteros, David Martínez, Laura Vega, Jesús E. Jaimes, Sergio Gomez, Angelica Rico, Lisseth Pardo, Esther C. Barros, Martha L. Ospina, Aníbal A. Teherán, Ana S. Gonzalez-Reiche, Matthew M. Hernandez, Emilia Mia Sordillo, Viviana Simon, Harm van Bakel, Alberto Paniz-Mondolfi |
| EPI_ISL_447854 | CSIR-Centre for Cellular and Molecular Biology | CSIR-Centre for Cellular and Molecular Biology | Payel Mukherjee, Sofia Banu, Priya Singh, Dhiviya Vedagiri, Divya Gupta, Vishal Sah, Santosh Kumar Kuncha, Krishnan Harinivas Harshan, Archana Bharadwaj Siva, Karthik Bharadwaj Tallapaka, Shagufta Khan, Lamuk Zaveri, Namami Gaur, Sakshi Shambhavi, Tulasi Nagabandi, Purushotham Vodnala, Rakesh K Mishra, Divya Tej Sowpati |
| EPI_ISL_447855 | CSIR-Centre for Cellular and Molecular Biology | CSIR-Centre for Cellular and Molecular Biology | Lamuk Zaveri, Shagufta Khan, Namami Gaur, Sakshi Shambhavi, Tulasi Nagabandi, Purushotham Vodnala, Payel Mukherjee, Sofia Banu, Priya Singh, Dhiviya Vedagiri, Divya Gupta, Vishal Sah, Santosh Kumar Kuncha, Krishnan Harinivas Harshan, Archana Bharadwaj Siva, Karthik Bharadwaj Tallapaka, Rakesh K Mishra, Divya Tej Sowpati |
| EPI_ISL_448222 | Pasteur Insitute Ho Chi Minh City | National Key Laboratory of Gene Technology, Institute of Biotechnology, Vietnam Academy of Science and Technology | Le Tung Lam, Nguyen Trung Hieu, Nguyen Hong Trang, Ho Thi Thuong, Nguyen Thi Ngoc Thao, Huynh Thi Kim Loan, Luu Thuy Tien, Tran Huyen Linh, Pham Duy Quang, Luong Chan Quang, Cao Minh Thang, Nguyen Vu Thuong, Hoang Ha, Chu Hoang Ha, Phan Trong Lan, Truong Nam Hai |
| EPI_ISL_450241, EPI_ISL_450242, EPI_ISL_450243, EPI_ISL_450244, EPI_ISL_450245, EPI_ISL_450248, EPI_ISL_450249, EPI_ISL_450250, EPI_ISL_450251, EPI_ISL_450252, EPI_ISL_450258, EPI_ISL_450259, EPI_ISL_450260, EPI_ISL_450261, EPI_ISL_450262, EPI_ISL_450263, EPI_ISL_450264, EPI_ISL_450265, EPI_ISL_450266, EPI_ISL_450267, EPI_ISL_450271, EPI_ISL_450272, EPI_ISL_450273, EPI_ISL_450274, EPI_ISL_450275, EPI_ISL_450276, EPI_ISL_450277, EPI_ISL_450278, EPI_ISL_450279, EPI_ISL_450280, EPI_ISL_450281, EPI_ISL_450282, EPI_ISL_450283, EPI_ISL_450284, EPI_ISL_450285, EPI_ISL_450286, EPI_ISL_450287, EPI_ISL_450288, EPI_ISL_450289, EPI_ISL_450290, EPI_ISL_450292, EPI_ISL_450293 | | | |
| see above | WHO National Influenza Centre Russian Federation | WHO National Influenza Centre Russian Federation | Andrey Komissarov, Artem Fadeev, Mariia Sergeeva, Anna Ivanova, Tamila Musaeva, Ksenia Komissarova, Mariia Timofeeva, Veronica Eder, Mariia Pisareva, Daria Danilenko |
| EPI_ISL_450321 | NIV Pune | CSIR-Centre for Cellular and Molecular Biology | Dr V A Potdar, Dr ML Choudhary,Dr Priya Abraham,V. Vipat, S. Jadhav, U. Saha, H. Kengle, A. Awhale, A. Jagtap, A. Gondhalikar, V Malik, N Srivastava, S. Digraskar, P. Malsane, S. Hundekar, K. Patel, Yogesh Balakartik, M. Kakade, S. Jadhav, R. Gunjikar, V. Awtade, S. Bhorekar, P Shinde, S. Salve, B. Minhas S. Bharadwaj, H Kaushal Y. Gurav, S. Tomar,Payel Mukherjee, Sofia Banu, Priya Singh, Dhiviya Vedagiri, Divya Gupta, Vishal Sah, Santosh Kumar Kuncha, Krishnan Harinivas Harshan, Archana Bharadwaj Siva, Karthik Bharadwaj Tallapaka, Shagufta Khan, Lamuk Zaveri, Namami Gaur, Sakshi Shambhavi, Tulasi Nagabandi, Purushotham Vodnala,G. Aditya Kumar, Koushick Sivakumar, Pooja Ramesh Gupta, Rajan Kumar Jha, Shraddha Vijay Lahoti, Deepak Kumar, Devi Prasad Vijayashankara, Disha Nanda, Divya Das, Jotin Gogoi, Manish |
| EPI_ISL_450322 | NIV Pune | CSIR-Centre for Cellular and Molecular Biology | Dr V A Potdar, Dr ML Choudhary,Dr Priya Abraham,V. Vipat, S. Jadhav, U. Saha, H. Kengle, A. Awhale, A. Jagtap, A. Gondhalikar, V Malik, N Srivastava, S. Digraskar, P. Malsane, S. Hundekar, K. Patel, Yogesh Balakartik, M. Kakade, S. Jadhav, R. Gunjikar, V. Awtade, S. Bhorekar, P Shinde, S. Salve, B. Minhas S. |

| | | | |
|---|---|---|---|
| | | | Bharadwaj, H Kaushal Y. Gurav, S. Tomar,Sofia Banu, Payel Mukherjee, Priya Singh, Dhiviya Vedagiri, Divya Gupta, Vishal Sah, Santosh Kumar Kuncha, Krishnan Harinivas Harshan, Archana Bharadwaj Siva, Karthik Bharadwaj Tallapaka, Shagufta Khan, Lamuk Zaveri, Namami Gaur, Sakshi Shambhavi, Tulasi Nagabandi, Purushotham Vodnala, Disha Nanda, Divya Das, Jotin Gogoi, Manish Bhattacharjee, Ravi Prasad Mukku, Renu Sudhakar, Somesh Gorde, Gangumala Srinivas Reddy, Sujoy Deb, Swati Bayyana, Zeba Rizvi, Rakesh K Mishra |
| EPI_ISL_450323 | NIV Pune | CSIR-Centre for Cellular and Molecular Biology | Dr V A Potdar, Dr ML Choudhary,Dr Priya Abraham,V. Vipat, S. Jadhav, U. Saha, H. Kengle, A. Awhale, A. Jagtap, A. Gondhalikar, V Malik, N Srivastava, S. Digraskar, P. Malsane, S. Hundekar, K. Patel, Yogesh Balakartik, M. Kakade, S. Jadhav, R. Gunjikar, V. Awtade, S. Bhorekar, P Shinde, S. Salve, B. Minhas S. Bharadwaj, H Kaushal Y. Gurav, S. Tomar,Payel Mukherjee, Sofia Banu, Priya Singh, Dhiviya Vedagiri, Divya Gupta, Vishal Sah, Santosh Kumar Kuncha, Krishnan Harinivas Harshan, Archana Bharadwaj Siva, Karthik Bharadwaj Tallapaka, Shagufta Khan, Lamuk Zaveri, Namami Gaur, Sakshi Shambhavi, Tulasi Nagabandi, Purushotham Vodnala,G. Aditya Kumar, Koushick Sivakumar, Pooja Ramesh Gupta, Rajan Kumar Jha, Shraddha Vijay Lahoti, Deepak Kumar, Devi Prasad Vijayashankara, Disha Nanda, Divya Das, Jotin Gogoi, Manish |
| EPI_ISL_450324 | NIV Pune | CSIR-Centre for Cellular and Molecular Biology | Dr V A Potdar, Dr ML Choudhary,Dr Priya Abraham,V. Vipat, S. Jadhav, U. Saha, H. Kengle, A. Awhale, A. Jagtap, A. Gondhalikar, V Malik, N Srivastava, S. Digraskar, P. Malsane, S. Hundekar, K. Patel, Yogesh Balakartik, M. Kakade, S. Jadhav, R. Gunjikar, V. Awtade, S. Bhorekar, P Shinde, S. Salve, B. Minhas S. Bharadwaj, H Kaushal Y. Gurav, S. Tomar,Sofia Banu, Payel Mukherjee, Priya Singh, Dhiviya Vedagiri, Divya Gupta, Vishal Sah, Santosh Kumar Kuncha, Krishnan Harinivas Harshan, Archana Bharadwaj Siva, Karthik Bharadwaj Tallapaka, Shagufta Khan, Lamuk Zaveri, Namami Gaur, Sakshi Shambhavi, Tulasi Nagabandi, Purushotham Vodnala, Disha Nanda, Divya Das, Jotin Gogoi, Manish Bhattacharjee, Ravi Prasad Mukku, Renu Sudhakar, Somesh Gorde, Gangumala Srinivas Reddy, Sujoy Deb, Swati Bayyana, Zeba Rizvi, Rakesh K Mishra |
| EPI_ISL_450325 | NIV Pune | CSIR-Centre for Cellular and Molecular Biology | Dr V A Potdar, Dr ML Choudhary,Dr Priya Abraham,V. Vipat, S. Jadhav, U. Saha, H. Kengle, A. Awhale, A. Jagtap, A. Gondhalikar, V Malik, N Srivastava, S. Digraskar, P. Malsane, S. Hundekar, K. Patel, Yogesh Balakartik, M. Kakade, S. Jadhav, R. Gunjikar, V. Awtade, S. Bhorekar, P Shinde, S. Salve, B. Minhas S. Bharadwaj, H Kaushal Y. Gurav, S. Tomar,Payel Mukherjee, Sofia Banu, Priya Singh, Dhiviya Vedagiri, Divya Gupta, Vishal Sah, Santosh Kumar Kuncha, Krishnan Harinivas Harshan, Archana Bharadwaj Siva, Karthik Bharadwaj Tallapaka, Shagufta Khan, Lamuk Zaveri, Namami Gaur, Sakshi Shambhavi, Tulasi Nagabandi, Purushotham Vodnala,G. Aditya Kumar, Koushick Sivakumar, Pooja Ramesh Gupta, Rajan Kumar Jha, Shraddha Vijay Lahoti, Deepak Kumar, Devi Prasad Vijayashankara, Disha Nanda, Divya Das, Jotin Gogoi, Manish |
| EPI_ISL_450340 | Bangladesh Institute of Tropical & Infectious Diseases, COVID-19 Testing Laboratory | Basic and Applied Research on Jute Project | Rasel Ahmed, Md. Sabbir Hossain, Shah Md Tamim Kabir, Emdadul Mannan Emdad, Md. Nazmul Haq Rony, Eaftekhar Ahmed Rana, Paritous Kumar Biswas, M A Hassan Chowdhury, Md. Shakeel Ahmed, Md. Samiul Haque, Md. Monjurul Alam, Md. Sharifur Rahman, A S M Anwarul Huq, Md. Shahidul Islam, Goutam Buddha Das, AMAM Zonaed Siddiki |
| EPI_ISL_450341 | Bangladesh Institute of Tropical & Infectious Diseases, COVID-19 Testing Laboratory | Basic and Applied Research on Jute Project | Md. Sabbir Hossain, Rasel Ahmed, Shah Md Tamim Kabir, Emdadul Mannan Emdad, Md. Nazmul Haq Rony, Eaftekhar Ahmed Rana, Paritous Kumar Biswas, M A Hassan Chowdhury, Md. Shakeel Ahmed, Md. Samiul Haque, Md. Monjurul Alam, Md. Sharifur Rahman, A S M Anwarul Huq, Md. Shahidul Islam, Goutam Buddha Das, AMAM Zonaed Siddiki |
| EPI_ISL_450342 | Bangladesh Institute of Tropical & Infectious Diseases, COVID-19 Testing Laboratory | Basic and Applied Research on Jute Project | Rasel Ahmed, Md. Sabbir Hossain, Shah Md Tamim Kabir, Emdadul Mannan Emdad, Md. Nazmul Haq Rony, Eaftekhar Ahmed Rana, Paritous Kumar Biswas, M A Hassan Chowdhury, Md. Shakeel Ahmed, Md. Samiul Haque, Md. Monjurul Alam, Md. Sharifur Rahman, A S M Anwarul Huq, Md. Shahidul Islam, Goutam Buddha Das, AMAM Zonaed Siddiki |
| EPI_ISL_450343 | Bangladesh Institute of Tropical & Infectious Diseases, COVID-19 Testing Laboratory | Basic and Applied Research on Jute Project | Md. Sabbir Hossain, Rasel Ahmed, Shah Md Tamim Kabir, Emdadul Mannan Emdad, Md. Nazmul Haq Rony, Eaftekhar Ahmed Rana, Paritous Kumar Biswas, M A Hassan Chowdhury, Md. Shakeel Ahmed, Md. Samiul Haque, Md. Monjurul Alam, Md. Sharifur Rahman, A S M Anwarul Huq, Md. Shahidul Islam, Goutam Buddha Das, AMAM Zonaed Siddiki |
| EPI_ISL_450508, EPI_ISL_450509, EPI_ISL_450511, EPI_ISL_450512, EPI_ISL_450515 | Rafik Hariri University Hospital | Rafik Hariri University Hospital | Rita Feghali |
| EPI_ISL_450724, EPI_ISL_450725, EPI_ISL_450726, EPI_ISL_450727, EPI_ISL_450728, EPI_ISL_450729, EPI_ISL_450730 | Hospital AZ Rivierenland | Institute of Tropical Medicine | Philippe Selhorst, Colin Anthony |
| EPI_ISL_450746 | Laboratory of Molecular Biology, Diagnostyka sp. z o.o. | Laboratory of Recombinant Vaccines | Lukasz Rabalski, Anna Piotrowska-Mietelska, Maciej Kosinski, Boguslaw Szewczyk, Krystyna Bienkowska-Szewczyk |
| EPI_ISL_450781 | Government Medical College-Bhavnagar | Gujarat Biotechnology Research Centre | Kairavi Desai, Saklain Malek, Shirish Patel, Ramesh Pandit, Tejas Shah, Ankit Hinsu, Pritesh Sabara, Apurvasinh Puvar, Janvi Raval, Zarna Patel, Monika Gandhi, Pinal Trivedi, Maharshi Pandya, Amit Kanani, Nidhi Patel, Nitin Savaliya, Raghawendra Kumar, Dinesh Kumar, Zuber Saiyed, Komal Patel, Labdhi Pandya, Snehal Bagatharia, Bhavesh Modi, Gaurishankar Shrimali, R D Dixit, A M Kadri, Akanksha Verma, Chaitanya Joshi, Madhvi Joshi |
| EPI_ISL_450783 | Government Medical College-Bhavnagar | Gujarat Biotechnology Research Centre | Shirish Patel, Kairavi Desai, Saklain Malek, Ankit Hinsu, Pritesh Sabara, Apurvasinh Puvar, Janvi Raval, Zarna Patel, Monika Gandhi, Pinal Trivedi, Maharshi Pandya, Amit Kanani, Nidhi Patel, Nitin Savaliya, Raghawendra Kumar, Dinesh Kumar, Zuber Saiyed, Komal Patel, Labdhi Pandya, Snehal Bagatharia, Ramesh Pandit, Tejas Shah, Bhavesh Modi, Gaurishankar Shrimali, R D Dixit, A M Kadri, Neha Rajpara, Chaitanya Joshi, Madhvi Joshi |
| EPI_ISL_450784 | Government Medical College-Bhavnagar | Gujarat Biotechnology Research Centre | Zarna Patel, Ramesh Pandit, Tejas Shah, Ankit Hinsu, Pritesh Sabara, Apurvasinh Puvar, Janvi Raval, Monika Gandhi, Pinal Trivedi, Maharshi Pandya, Amit Kanani, Nidhi Patel, Nitin Savaliya, Raghawendra Kumar, Dinesh Kumar, Zuber Saiyed, Komal Patel, Labdhi Pandya, Snehal Bagatharia, Kairavi Desai, Saklain Malek, Shirish Patel, Bhavesh Modi, Gaurishankar Shrimali, R D Dixit, A M Kadri, Afzal Ansari, Chaitanya Joshi, Madhvi Joshi |
| EPI_ISL_450785 | Pandit Deendayal Upadhyay Government Medical College, Rajkot | Gujarat Biotechnology Research Centre | Prakash Modi, Sejul Antala, Manish Pattani, Apurvasinh Puvar, Janvi Raval, Zarna Patel, Monika Gandhi, Pinal Trivedi, Maharshi Pandya, Amit Kanani, Nidhi Patel, Nitin Savaliya, Raghawendra Kumar, Dinesh Kumar, Zuber Saiyed, Komal Patel, Labdhi Pandya, Snehal Bagatharia, Ramesh Pandit, Tejas Shah, Ankit Hinsu, Pritesh Sabara, Bhavesh Modi, Gaurishankar Shrimali, R D Dixit, A M Kadri, Neelam Nathani, Chaitanya Joshi, Madhvi Joshi |
| EPI_ISL_450786 | Pandit Deendayal Upadhyay Government Medical College, Rajkot | Gujarat Biotechnology Research Centre | Sejul Antala, Manish Pattani, Prakash Modi, Janvi Raval, Zarna Patel, Monika Gandhi, Pinal Trivedi, Maharshi Pandya, Amit Kanani, Nidhi Patel, Nitin Savaliya, Raghawendra Kumar, Dinesh Kumar, Zuber Saiyed, Komal Patel, Labdhi Pandya, Snehal Bagatharia, Ramesh Pandit, Tejas Shah, Ankit Hinsu, Pritesh Sabara, Apurvasinh Puvar, Bhavesh Modi, Gaurishankar Shrimali, R D Dixit, A M Kadri, Armi Chaudhari, Chaitanya Joshi, Madhvi Joshi |
| EPI_ISL_450788 | Pandit Deendayal Upadhyay Government Medical College, Rajkot | Gujarat Biotechnology Research Centre | Zarna Patel, Tejas Shah, Ankit Hinsu, Pritesh Sabara, Apurvasinh Puvar, Janvi Raval, Monika Gandhi, Pinal Trivedi, Maharshi Pandya, Amit Kanani, Nidhi Patel, Nitin Savaliya, Raghawendra Kumar, Dinesh Kumar, Zuber Saiyed, Komal Patel, Labdhi Pandya, Snehal Bagatharia, Prakash Modi, Sejul Antala, Manish Pattani, Ramesh Pandit, Bhavesh Modi, Gaurishankar Shrimali, R D Dixit, A M Kadri, Camellia Chakraborty, Chaitanya Joshi, Madhvi Joshi |
| EPI_ISL_450789 | Pandit Deendayal Upadhyay Government Medical College, Rajkot | Gujarat Biotechnology Research Centre | Ankit Hinsu, Pritesh Sabara, Apurvasinh Puvar, Janvi Raval, Zarna Patel, Monika Gandhi, Pinal Trivedi, Maharshi Pandya, Amit Kanani, Nidhi Patel, Nitin Savaliya, Raghawendra Kumar, Dinesh Kumar, Zuber Saiyed, Komal Patel, Labdhi Pandya, Snehal Bagatharia, Prakash Modi, Sejul Antala, Manish Pattani, Ramesh Pandit, Tejas Shah, Bhavesh Modi, Gaurishankar Shrimali, R D Dixit, A M Kadri, Siddhant Kumar, Chaitanya Joshi, Madhvi Joshi |
| EPI_ISL_450790 | Pandit Deendayal Upadhyay Government Medical College, Rajkot | Gujarat Biotechnology Research Centre | Zarna Patel, Pritesh Sabara, Apurvasinh Puvar, Janvi Raval, Monika Gandhi, Pinal Trivedi, Maharshi Pandya, Amit Kanani, Nidhi Patel, Nitin Savaliya, Raghawendra Kumar, Dinesh Kumar, Zuber Saiyed, Komal Patel, Labdhi Pandya, Snehal Bagatharia, Prakash Modi, Sejul Antala, Manish Pattani, Ramesh Pandit, Tejas Shah, Ankit Hinsu, Bhavesh Modi, Gaurishankar Shrimali, R D Dixit, A M Kadri, Sharmistha Majumdar, Chaitanya Joshi, Madhvi Joshi |
| EPI_ISL_450791 | Pandit Deendayal Upadhyay Government Medical College, Rajkot | Gujarat Biotechnology Research Centre | Zarna Patel, Apurvasinh Puvar, Janvi Raval, Monika Gandhi, Pinal Trivedi, Maharshi Pandya, Amit Kanani, Nidhi Patel, Nitin Savaliya, Raghawendra Kumar, Dinesh Kumar, Zuber Saiyed, Komal Patel, Labdhi Pandya, Snehal Bagatharia, Prakash Modi, Sejul Antala, Manish Pattani, Ramesh Pandit, Tejas Shah, Ankit Hinsu, Pritesh Sabara, Bhavesh Modi, Gaurishankar Shrimali, R D Dixit, A M Kadri, Pooja P Doshi, Chaitanya Joshi, Madhvi Joshi |
| EPI_ISL_450839 | COVID-19 Laboratory Centre for Advanced Research in Sciences (CARS), University of Dhaka, Dhaka-1000, Bangladesh | DNA Solution Ltd | Sharif Akhteruzzaman, Zeba Islam Seraj, Nazmul Ahsan, Md Imdadul Hoque, MA Malek, Shahryar Nabi, Sabrina Moriom Elius, ABM Khademul Islam, Richard Malo, Imran Khan, Abu Sufian, Sabita Rezwana Rahman, Habibul Bari Shozib, Mamun Ahmed, AHM Nurun Nabi, Mohammad Riazul Islam, Md Mizanur Rahman, Md Ismail Hosen, Latiful Bari, Gazi Nurun Nahar, Haseena Khan, M Anwar Hossain. |
| EPI_ISL_450840 | COVID-19 Laboratory | DNA Solution Ltd. L-5 | Sharif Akhteruzzaman, Zeba Islam Seraj, Nazmul Ahsan, Md Imdadul Hoque, MA Malek, Shahryar Nabi, Sabrina Moriom Elius, ABM Khademul Islam, Richard Malo, Imran Khan, Abu Sufian, Sabita Rezwana Rahman, Habibul Bari Shozib, Mamun Ahmed, AHM Nurun Nabi, Mohammad Riazul Islam, Md Mizanur Rahman, Md Ismail Hosen, Latiful Bari, Gazi Nurun Nahar, Haseena Khan, M Anwar Hossain. |
| EPI_ISL_450841 | COVID-19 Laboratory | DNA Solution Ltd | Sharif Akhteruzzaman, Zeba Islam Seraj, Nazmul Ahsan, Md Imdadul Hoque, MA Malek, Shahryar Nabi, Sabrina Moriom Elius, ABM Khademul Islam, Richard |

| | | | |
|---|---|---|---|
| | | | Malo, Imran Khan, Abu Sufian, Sabita Rezwana Rahman, Habibul Bari Shozib, Mamun Ahmed, AHM Nurun Nabi, Mohammad Riazul Islam, Md Mizanur Rahman, Md Ismail Hosen, Latiful Bari, Gazi Nurun Nahar, Haseena Khan, M Anwar Hossain. |
| EPI_ISL_450843 | COVID-19 Laboratory | DNA Solution Ltd. | Sharif Akhteruzzaman, Zeba Islam Seraj, Nazmul Ahsan, Md Imdadul Hoque, MA Malek, Shahryar Nabi, Sabrina Moriom Elius, ABM Khademul Islam, Richard Malo, Imran Khan, Abu Sufian, Sabita Rezwana Rahman, Habibul Bari Shozib, Mamun Ahmed, AHM Nurun Nabi, Mohammad Riazul Islam, Md Mizanur Rahman, Md Ismail Hosen, Latiful Bari, Gazi Nurun Nahar, Haseena Khan, M Anwar Hossain. |
| EPI_ISL_451076 | West China Hospital of Sichuan University | State Key Laboratory of Biotherapy of Sichuan University | Baowen Du, Minjin Wang, Chao Tanga, Chuan Chena, Yongzhao Zhou, Mingxia Yu, Han-Cheng Wei, Weimin Li, Jing-wen Lin, Jia Geng, Binwu Ying, Lu Chen |
| EPI_ISL_451149 | M.P Shah Government Medocal college Jamnagar | Gujarat Biotechnology Research Centre | Janvi Raval, Zarna Patel, Monika Gandhi, Pinal Trivedi, Maharshi Pandya, Amit Kanani, Nidhi Patel, Nitin Savaliya, Raghawendra Kumar, Dinesh Kumar, Zuber Saiyed, Komal Patel, Labdhi Pandya, Snehal Bagatharia, Ramesh Pandit, Tejas Shah, Ankit Hinsu, Pritesh Sabara, Apurvasinh Puvar, Binita Aring, Bhavesh Modi, Gaurishankar Shrimali, R D Dixit, A M Kadri, Priti Pandita, Chaitanya Joshi, Madhvi Joshi, |
| EPI_ISL_451150 | M.P Shah Government Medocal college Jamnagar | Gujarat Biotechnology Research Centre | Zarna Patel, Monika Gandhi, Pinal Trivedi, Maharshi Pandya, Amit Kanani, Nidhi Patel, Nitin Savaliya, Raghawendra Kumar, Dinesh Kumar, Zuber Saiyed, Komal Patel, Labdhi Pandya, Snehal Bagatharia, Ramesh Pandit, Tejas Shah, Ankit Hinsu, Pritesh Sabara, Apurvasinh Puvar, Binita Aring, Janvi Raval, Bhavesh Modi, Gaurishankar Shrimali, R D Dixit, A M Kadri, Pragya Sharma, Chaitanya Joshi, Madhvi Joshi, |
| EPI_ISL_451151 | M.P Shah Government Medocal college Jamnagar | Gujarat Biotechnology Research Centre | Monika Gandhi, Pinal Trivedi, Maharshi Pandya, Amit Kanani, Nidhi Patel, Nitin Savaliya, Raghawendra Kumar, Dinesh Kumar, Zuber Saiyed, Komal Patel, Labdhi Pandya, Snehal Bagatharia, Ramesh Pandit, Tejas Shah, Ankit Hinsu, Pritesh Sabara, Apurvasinh Puvar, Binita Aring, Janvi Raval, Zarna Patel, Bhavesh Modi, Gaurishankar Shrimali, R D Dixit, A M Kadri, Neha Rajpara, Chaitanya Joshi, Madhvi Joshi, |
| EPI_ISL_451152 | M.P Shah Government Medocal college Jamnagar | Gujarat Biotechnology Research Centre | Pinal Trivedi, Maharshi Pandya, Amit Kanani, Nidhi Patel, Nitin Savaliya, Raghawendra Kumar, Dinesh Kumar, Zuber Saiyed, Komal Patel, Labdhi Pandya, Snehal Bagatharia, Ramesh Pandit, Tejas Shah, Ankit Hinsu, Pritesh Sabara, Apurvasinh Puvar, Binita Aring, Janvi Raval, Zarna Patel, Monika Gandhi, Bhavesh Modi, Gaurishankar Shrimali, R D Dixit, A M Kadri, Afzal Ansari, Chaitanya Joshi, Madhvi Joshi, |
| EPI_ISL_451157 | Government Medical College, Vadodara | Gujarat Biotechnology Research Centre | Nidhi Patel, Nitin Savaliya, Raghawendra Kumar, Dinesh Kumar, Zuber Saiyed, Komal Patel, Labdhi Pandya, Snehal Bagatharia, Ramesh Pandit, Tejas Shah, Ankit Hinsu, Pritesh Sabara, Apurvasinh Puvar, Janvi Raval, Zarna Patel, Monika Gandhi, Pinal Trivedi, Maharshi Pandya, Manish Pattani, Tanuja Javadekar , Amit Kanani, Bhavesh Modi, Gaurishankar Shrimali, R D Dixit, A M Kadri, Camellia Chakraborty, Chaitanya Joshi, Madhvi Joshi |
| EPI_ISL_451158 | Government Medical College, Vadodara | Gujarat Biotechnology Research Centre | Nitin Savaliya, Raghawendra Kumar, Dinesh Kumar, Zuber Saiyed, Komal Patel, Labdhi Pandya, Snehal Bagatharia, Ramesh Pandit, Tejas Shah, Ankit Hinsu, Pritesh Sabara, Apurvasinh Puvar, Janvi Raval, Zarna Patel, Monika Gandhi, Pinal Trivedi, Maharshi Pandya, Manish Pattani, Tanuja Javadekar , Amit Kanani, Nidhi Patel, Bhavesh Modi, Gaurishankar Shrimali, R D Dixit, A M Kadri, Siddhant Kumar, Chaitanya Joshi, Madhvi Joshi |
| EPI_ISL_451161 | Government Medical College, Vadodara | Gujarat Biotechnology Research Centre | Zuber Saiyed, Komal Patel, Labdhi Pandya, Snehal Bagatharia, Ramesh Pandit, Tejas Shah, Ankit Hinsu, Pritesh Sabara, Apurvasinh Puvar, Janvi Raval, Zarna Patel, Monika Gandhi, Pinal Trivedi, Maharshi Pandya, Manish Pattani, Tanuja Javadekar , Amit Kanani, Nidhi Patel, Nitin Savaliya, Raghawendra Kumar, Dinesh Kumar, Bhavesh Modi, Gaurishankar Shrimali, R D Dixit, A M Kadri, Akanksha Verma, Chaitanya Joshi, Madhvi Joshi |
| EPI_ISL_451162 | Government Medical College, Vadodara | Gujarat Biotechnology Research Centre | Komal Patel, Labdhi Pandya, Snehal Bagatharia, Ramesh Pandit, Tejas Shah, Ankit Hinsu, Pritesh Sabara, Apurvasinh Puvar, Janvi Raval, Zarna Patel, Monika Gandhi, Pinal Trivedi, Maharshi Pandya, Manish Pattani, Tanuja Javadekar , Amit Kanani, Nidhi Patel, Nitin Savaliya, Raghawendra Kumar, Dinesh Kumar, Zuber Saiyed, Bhavesh Modi, Gaurishankar Shrimali, R D Dixit, A M Kadri, Priti Pandita, Chaitanya Joshi, Madhvi Joshi |
| EPI_ISL_451298 | Laboratory of Virology, INMI Lazzaro Spallanzani IRCCS | Laboratory of Virology, INMI Lazzaro Spallanzani IRCCS | Cesare E.M. Gruber, Martina Rueca, Barbara Bartolini, Francesco Messina, Antonino Di Caro, Maria R. Capobianchi, Giuseppe Ippolito |
| EPI_ISL_451299 | Laboratory of Virology, INMI Lazzaro Spallanzani IRCCS | Laboratory of Virology, INMI Lazzaro Spallanzani IRCCS | Martina Rueca, Cesare E.M. Gruber, Barbara Bartolini, Francesco Messina, Antonino Di Caro, Maria R. Capobianchi, Giuseppe Ippolito |
| EPI_ISL_451300 | Laboratory of Virology, INMI Lazzaro Spallanzani IRCCS | Laboratory of Virology, INMI Lazzaro Spallanzani IRCCS | Cesare E.M. Gruber, Martina Rueca, Barbara Bartolini, Francesco Messina, Antonino Di Caro, Maria R. Capobianchi, Giuseppe Ippolito |
| EPI_ISL_451301 | Laboratory of Virology, INMI Lazzaro Spallanzani IRCCS | Laboratory of Virology, INMI Lazzaro Spallanzani IRCCS | Martina Rueca, Cesare E.M. Gruber, Barbara Bartolini, Francesco Messina, Antonino Di Caro, Maria R. Capobianchi, Giuseppe Ippolito |
| EPI_ISL_451302 | Laboratory of Virology, INMI Lazzaro Spallanzani IRCCS | Laboratory of Virology, INMI Lazzaro Spallanzani IRCCS | Cesare E.M. Gruber, Martina Rueca, Barbara Bartolini, Francesco Messina, Antonino Di Caro, Maria R. Capobianchi, Giuseppe Ippolito |
| EPI_ISL_451303 | Laboratory of Virology, INMI Lazzaro Spallanzani IRCCS | Laboratory of Virology, INMI Lazzaro Spallanzani IRCCS | Martina Rueca, Cesare E.M. Gruber, Barbara Bartolini, Francesco Messina, Antonino Di Caro, Maria R. Capobianchi, Giuseppe Ippolito |
| EPI_ISL_451304 | Laboratory of Virology, INMI Lazzaro Spallanzani IRCCS | Laboratory of Virology, INMI Lazzaro Spallanzani IRCCS | Cesare E.M. Gruber, Martina Rueca, Barbara Bartolini, Francesco Messina, Antonino Di Caro, Maria R. Capobianchi, Giuseppe Ippolito |
| EPI_ISL_451306 | Molecular Virology Unit, Fondazione IRCCS Policlinico San Matteo , Pavia | Laboratory of Virology, INMI Lazzaro Spallanzani IRCCS | Antonio Piralla, Fausto Baldanti, Martina Rueca, Antonino Di Caro, Maria R. Capobianchi, Cesare E.M. Gruber, Barbara Bartolini |
| EPI_ISL_451307 | Molecular Virology Unit, Fondazione IRCCS Policlinico San Matteo , Pavia | Laboratory of Virology, INMI Lazzaro Spallanzani IRCCS | Fausto Baldanti, Antonio Piralla, Antonino Di Caro, Cesare E.M. Gruber, Martina Rueca, Barbara Bartolini, Maria R. Capobianchi |
| EPI_ISL_451308 | Molecular Virology Unit, Fondazione IRCCS Policlinico San Matteo , Pavia | Laboratory of Virology, INMI Lazzaro Spallanzani IRCCS | Antonio Piralla, Fausto Baldanti, Maria R. Capobianchi, Cesare E.M. Gruber, Martina Rueca, Barbara Bartolini, Antonino Di Caro |
| EPI_ISL_451309 | Molecular Virology Unit, Fondazione IRCCS Policlinico San Matteo , Pavia | Laboratory of Virology, INMI Lazzaro Spallanzani IRCCS | Fausto Baldanti, Antonio Piralla, Cesare E.M. Gruber, Maria R. Capobianchi, Antonino Di Caro, Martina Rueca, Barbara Bartolini |
| EPI_ISL_451310, EPI_ISL_451311 | Hellenic Pasteur Institute, National Influenza Reference laboratory of Southern Greece & Unit of Bioinformatics and Applied Genomics | Hellenic Pasteur Institute, National Influenza Reference laboratory of Southern Greece & Unit of Bioinformatics and Applied Genomics | Vasiliki Pogka, Timokratis Karamitros, Athanasios Kossyvakis, Antonios Kalliaropoulos, Horefti Elina, Evangelidou Maria, Androniki Voulgari-Kokota, Aspasia Kontou, Andreas Mentis |
| EPI_ISL_451313, EPI_ISL_451314, EPI_ISL_451315, EPI_ISL_451316, EPI_ISL_451318, EPI_ISL_451319, EPI_ISL_451320, EPI_ISL_451321, EPI_ISL_451322, EPI_ISL_451325, EPI_ISL_451326, EPI_ISL_451327, EPI_ISL_451328, EPI_ISL_451329, EPI_ISL_451330, EPI_ISL_451331, EPI_ISL_451334, EPI_ISL_451337, EPI_ISL_451338, EPI_ISL_451344, EPI_ISL_451345, EPI_ISL_451346, EPI_ISL_451348, EPI_ISL_451353, EPI_ISL_451354, EPI_ISL_451356, EPI_ISL_451357, EPI_ISL_451359, EPI_ISL_451360, EPI_ISL_451365, EPI_ISL_451369, EPI_ISL_451370, EPI_ISL_451371, EPI_ISL_451374, EPI_ISL_451376, EPI_ISL_451377, EPI_ISL_451378, EPI_ISL_451379, EPI_ISL_451380, EPI_ISL_451381, EPI_ISL_451382, EPI_ISL_451383, EPI_ISL_451384, EPI_ISL_451385, EPI_ISL_451386, EPI_ISL_451387, EPI_ISL_451388, EPI_ISL_451389, EPI_ISL_451390, EPI_ISL_451391, EPI_ISL_451392, EPI_ISL_451393, EPI_ISL_451394, EPI_ISL_451395, EPI_ISL_451398 | | | |
| see above | West China Hospital of Sichuan University | State Key Laboratory of Biotherapy of Sichuan University | Baowen Du, Minjin Wang, Chao Tang, Chuan Chen, Yongzhao Zhou, Mingxia Yu, Hancheng Wei, Weimin Li, Jing-wen Lin, Jia Geng, Binwu Ying, Lu Chen |
| EPI_ISL_451644, EPI_ISL_451645, EPI_ISL_451646, EPI_ISL_451647 | Laboratory of Molecular Biology, Diagnostyka sp. z o.o. | Laboratory of Recombinant Vaccines | Lukasz Rabalski, Anna Piotrowska-Mietelska, Maciej Kosinski, Boguslaw Szewczyk, Krystyna Bienkowska-Szewczyk |
| EPI_ISL_451655, EPI_ISL_451656, EPI_ISL_451657, EPI_ISL_451658, EPI_ISL_451659, EPI_ISL_451660, EPI_ISL_451661, EPI_ISL_451662, EPI_ISL_451663, EPI_ISL_451664 | State Sanitary Inspectorate | Laboratory of Recombinant Vaccines | Lukasz Rabalski, Boguslaw Szewczyk, Krystyna Bienkowska-Szewczyk, Jaroslaw Pinkas |
| EPI_ISL_451666 | M.P Shah Government Medocal college Jamnagar | Gujarat Biotechnology Research Centre | Binita Aring, Janvi Raval, Zarna Patel, Monika Gandhi, Pinal Trivedi, Maharshi Pandya, Amit Kanani, Nidhi Patel, Nitin Savaliya, Raghawendra Kumar, Dinesh Kumar, Zuber Saiyed, Komal Patel, Labdhi Pandya, Snehal Bagatharia, Ramesh Pandit, Tejas Shah, Ankit Hinsu, Pritesh Sabara, Apurvasinh Puvar, Bhavesh Modi, Gaurishankar Shrimali, R D Dixit, A M Kadri, Akanksha Verma, Chaitanya Joshi, Madhvi Joshi, |
| EPI_ISL_451934 | Research Unit, University Hospital for Infectious Diseases "Dr. Fran Mihaljevi" | Cicin Sain lab, Helmholtz Centre for Infection Research | Zeeshan Chaudhry, Kathrin Eschke, Željka Maak Šafranko, Ivan-Christian Kurolt |
| EPI_ISL_451935 | CUB Hopital Erasme Laboratoire d'Anatomie Pathologique | CUB Hopital Erasme Laboratoire d'Anatomie Pathologique | Prof. Isabelle Salmon, Dr. Nicky D'Haene |
| EPI_ISL_451961 | Istituto Zooprofilattico Sperimentale Puglia e Basilicata; Dipartimento di Bioscienze, Biotecnologie e Biofarmaceutica dell'Università degli Studi di Bari "A.Moro"; Istituto di Biomembrane. Bioenergetica e Biotecnologie Molecolari del Consiglio Nazionale delle Ricerche di Bari | Beaconlab (Bioinformatics Evolution and Comparative Genomics lab), Dept of Biosciences, University of Milan | Parisi A.,Pesole G., Manzari C., Chiara M. |
| EPI_ISL_451962 | Istituto Zooprofilattico Sperimentale Puglia e Basilicata; | Beaconlab (Bioinformatics, Evolution and Comparative | Parisi A.,Pesole G., Manzari C., Chiara M. |

| | | | |
|---|---|---|---|
| | Dipartimento di Bioscienze, Biotecnologie e Biofarmaceutica dell'Università degli Studi di Bari "A.Moro"; Istituto di Biomembrane. Bioenergetica e Biotecnologie Molecolari del Consiglio Nazionale delle Ricerche di Bari | Genomics lab), Dept of Biosciences, University on Milan | |
| EPI_ISL_452139 | Instituto de Diagnostico y Referencia Epidemiologicos (INDRE) | Instituto de diagnóstico y Referencia Epidemiologicos (INDRE) | Ramirez-Gonzalez Ernesto, Garces-Ayala Fabiola, Araiza-Rodriguez Adnan, Mendieta-Condado Edgar, Rodriguez-Maldonado Abril, Wong-Arambula Claudia, Barrera-Badillo Gisela, Hernandez-Rivas Lucia, Lopez-Martinez Irma |
| EPI_ISL_452140 | CUB Hopital Erasme Laboratoire d'Anatomie Pathologique | CUB Hopital Erasme Laboratoire d'Anatomie Pathologique | Isabelle Salmon, Nicky D'Haene |
| EPI_ISL_452142 | CUB Hopital Erasme Laboratoire d'Anatomie Pathologique | CUB Hopital Erasme Laboratoire d'Anatomie Pathologique | Isabelle Salmon, Nikcy D'Haene |
| EPI_ISL_452148, EPI_ISL_452149 | CUB Hopital Erasme Laboratoire d'Anatomie Pathologique | CUB Hopital Erasme Laboratoire d'Anatomie Pathologique | Isabelle Salmon, Nicky D'Haene |
| EPI_ISL_452150 | CUB Hopital Erasme Laboratoire d'Anatomie Pathologique | CUB Hopital Erasme Laboratoire d'Anatomie Pathologique | Prof. Isabelle Salmon, Dr Nicky D'Haene |
| EPI_ISL_452151 | CUB Hopital Erasme Laboratoire d'Anatomie Pathologique | CUB Hopital Erasme Laboratoire d'Anatomie Pathologique | Prof. Isabelle Salmon, Dr.Nikcy D'Haene |
| EPI_ISL_452152 | CUB Hopital Erasme Laboratoire d'Anatomie Pathologique | CUB Hopital Erasme Laboratoire d'Anatomie Pathologique | Prof. Isabelle Salmon, Dr Nicky D'Haene |
| EPI_ISL_452192, EPI_ISL_452193, EPI_ISL_452194, EPI_ISL_452195, EPI_ISL_452196, EPI_ISL_452197, EPI_ISL_452198, EPI_ISL_452199, EPI_ISL_452200, EPI_ISL_452201, EPI_ISL_452202, EPI_ISL_452203, EPI_ISL_452204, EPI_ISL_452205, EPI_ISL_452206, EPI_ISL_452207, EPI_ISL_452208, EPI_ISL_452209, EPI_ISL_452210, EPI_ISL_452211, EPI_ISL_452212, EPI_ISL_452213 | | | |
| see above | NIV Influenza | NIV Influenza | Potdar V |
| EPI_ISL_452218, EPI_ISL_452219, EPI_ISL_452220, EPI_ISL_452221, EPI_ISL_452222, EPI_ISL_452223 | Goethe University Hospital Frankfurt | Institute for Medical Virology, Goethe University Hospital Frankfurt | Tuna Toptan, Sebastian Hoehl, Sandra Westhaus, Denisa Bojkova, Annemarie Berger, Björn Rotter, Klaus Hoffmeier, Jindrich Cinatl, Sandra Ciesek, and Marek Widera |
| EPI_ISL_452327, EPI_ISL_452328, EPI_ISL_452329, EPI_ISL_452330, EPI_ISL_452331, EPI_ISL_452332, EPI_ISL_452333, EPI_ISL_452334, EPI_ISL_452335, EPI_ISL_452336, EPI_ISL_452337, EPI_ISL_452338, EPI_ISL_452339, EPI_ISL_452340, EPI_ISL_452341, EPI_ISL_452342, EPI_ISL_452343, EPI_ISL_452344, EPI_ISL_452345, EPI_ISL_452346, EPI_ISL_452347, EPI_ISL_452348, EPI_ISL_452349, EPI_ISL_452350, EPI_ISL_452351, EPI_ISL_452352, EPI_ISL_452353, EPI_ISL_452354, EPI_ISL_452355, EPI_ISL_452356, EPI_ISL_452357, EPI_ISL_452358, EPI_ISL_452359, EPI_ISL_452360, EPI_ISL_452361, EPI_ISL_452362, EPI_ISL_452363, EPI_ISL_452364 | | | |
| see above | Laboratory of Infectious Diseases Center of Beijing Ditan Hospital | Laboratory of Infectious Diseases Center of Beijing Ditan Hospital | Siyuan Yang, Chengjie Jie, Fengting Yu, Yunxia Tang, Liting Yan, Linghang Wang |
| EPI_ISL_452473, EPI_ISL_452480, EPI_ISL_452481, EPI_ISL_452484, EPI_ISL_452487, EPI_ISL_452488, EPI_ISL_452492, EPI_ISL_452496, EPI_ISL_452498, EPI_ISL_452500, EPI_ISL_452503, EPI_ISL_452509, EPI_ISL_452521, EPI_ISL_452534 | | | |
| see above | Clínica Universidad de Navarra. Servicio de Enfermedades Infecciosas y Microbiología clínica | SeqCOVID-SPAIN consortium/IBV(CSIC) | Mirian Fernández-Alonso, Jose Luis del Pozo and SeqCOVID-SPAIN consortium |
| EPI_ISL_454497, EPI_ISL_454498, EPI_ISL_454499, EPI_ISL_454500, EPI_ISL_454501, EPI_ISL_454502, EPI_ISL_454503, EPI_ISL_454504, EPI_ISL_454505, EPI_ISL_454506, EPI_ISL_454507, EPI_ISL_454508, EPI_ISL_454509, EPI_ISL_454510, EPI_ISL_454511, EPI_ISL_454512, EPI_ISL_454513, EPI_ISL_454514, EPI_ISL_454515, EPI_ISL_454516, EPI_ISL_454517, EPI_ISL_454518, EPI_ISL_454519, EPI_ISL_454520 | | | |
| see above | RSE "National Center for Biotechnology" | RSE "National Center for Biotechnology" | Alexandr Shevtsov, Ilyas Akhmetollayev, Viktoriya Lutsay, Asylulan Amirgazin, Askar Abdaliyev, Akbota Rakhmetova, Zabira Aushakhmetova, Ruslan Kalendar, Yerlan Ramankulov |
| EPI_ISL_454521, EPI_ISL_454522, EPI_ISL_454524, EPI_ISL_454525, EPI_ISL_454526, EPI_ISL_454527, EPI_ISL_454528, EPI_ISL_454529, EPI_ISL_454530, EPI_ISL_454531, EPI_ISL_454532, EPI_ISL_454533, EPI_ISL_454534, EPI_ISL_454536, EPI_ISL_454537, EPI_ISL_454540, EPI_ISL_454542, EPI_ISL_454543, EPI_ISL_454544, EPI_ISL_454546, EPI_ISL_454547, EPI_ISL_454549, EPI_ISL_454551, EPI_ISL_454552, EPI_ISL_454556, EPI_ISL_454557, EPI_ISL_454558, EPI_ISL_454560, EPI_ISL_454561, EPI_ISL_454562 | | | |
| see above | NIV Influenza | NIV Influenza | Potdar V |
| EPI_ISL_454574 | nstitute for Public Health | Laboratory for advanced genomics | Filip Roki, Lovro Trgovec-Greif, Neven Sui, Tomislav Rukavina, Igor Jurak, Oliver Vugrek |
| EPI_ISL_454578, EPI_ISL_454581, EPI_ISL_454583, EPI_ISL_454588, EPI_ISL_454592 | University Hospital for Infectious Diseases "Dr. Fran Mihaljevi", Research Unit | University of Zagreb, Centre for research and knowledge transfer in biotechnology | Ivan-Christian Kurolt, Jelena Ivancic Jelecki, Anamarija Slovic |
| EPI_ISL_454602 | Croatian Institute of Public Health | University of Zagreb, Centre for research and knowledge transfer in biotechnology | Irena Tabain, Tatjana Vilibic-Cavlek, Jelena Ivancic Jelecki, Anamarija Slovic |
| EPI_ISL_454605, EPI_ISL_454606 | Institute for Public Health | Laboratory for advanced genomics | Filip Roki, Lovro Trgovec-Greif, Neven Sui, Tomislav Rukavina, Igor Jurak, Oliver Vugrek |
| EPI_ISL_454732 | Russian State Collection of Viruses | Pathogenic Microorganisms Variability Laboratory | Denis Protsenko, Alexey Shchetinin, Maria Nikiforova, Elena Shidlovskaya, Nadezhda Kuznetsova, Vladimir Gushchin, Inna Dolzhikova, Daria Grousova, Andrey Botikov, Denis Logunov, Alexander Gintsburg, Alexey Mazus |
| EPI_ISL_454749 | Japanese Quarantine Stations | Pathogen Genomics Center, National Institute of Infectious Diseases | Tsuyoshi Sekizuka, Kentaro Itokawa, Rina Tanaka, Masanori Hashino, Tsutomu Kageyama, Shinji Saito, Ikuyo Takayama, Hideki Hasegawa, Takuri Takahashi, Hajime Kamiya, Takuya Yamagishi, Motoi Suzuki, Takaji Wakita, Makoto Kuroda |
| EPI_ISL_454830, EPI_ISL_454832, EPI_ISL_454833 | SMS Medical College, Jaipur | CSIR Institute of Genomics and Integrative Biology | Sudhir Bhandari, Rahul Bhoyar, Mohammed Imran, Mohit Divakar, Disha Sharma, Anshul Kumar, Bani Jolly, Rahul Sahlot, Abhinav Jain, Paras Sehgal, Gyan Ranjan, Vinod Scaria, Sridhar Sivasubbu, Sandeep K Mathur |
| EPI_ISL_454858, EPI_ISL_454862, EPI_ISL_454865, EPI_ISL_454866, EPI_ISL_454867 | Translational Health Science and Technology Institute -ESIC medical college and hospital, Faridabad | THSTI Bioassay laboratory | Saurabh Kumar, Jigme Wangchuk, Anil Kumar Pandey, Asim Das, Guruprasad R. Medigeshi |
| EPI_ISL_455015 | Pandit Deendayal Upadhyay Government Medical College, Rajkot | Gujarat Biotechnology Research Centre | Snehal Bagatharia, Prakash Modi, Sejul Antala, Manish Pattani, Ramesh Pandit, Tejas Shah, Ankit Hinsu, Pritesh Sabara, Apurvasinh Puvar, Janvi Raval, Zarna Patel, Monika Gandhi, Pinal Trivedi, Maharshi Pandya, Amit Kanani, Nidhi Patel, Nitin Savaliya, Raghawendra Kumar, Dinesh Kumar, Zuber Saiyed, Komal Patel, Labdhi Pandya, Neha Rajpara, Bhavesh Modi, Gaurishankar Shrimali, R D Dixit, A M Kadri, Umang Mishra, Chaitanya Joshi, Madhvi Joshi |
| EPI_ISL_455016 | Pandit Deendayal Upadhyay Government Medical College, Rajkot | Gujarat Biotechnology Research Centre | Prakash Modi, Sejul Antala, Manish Pattani, Ramesh Pandit, Tejas Shah, Ankit Hinsu, Pritesh Sabara, Apurvasinh Puvar, Janvi Raval, Zarna Patel, Monika Gandhi, Pinal Trivedi, Maharshi Pandya, Amit Kanani, Nidhi Patel, Nitin Savaliya, Raghawendra Kumar, Dinesh Kumar, Zuber Saiyed, Komal Patel, Labdhi Pandya, Snehal Bagatharia, Afzal Ansari, Bhavesh Modi, Gaurishankar Shrimali, R D Dixit, A M Kadri, Umang Mishra, Chaitanya Joshi, Madhvi Joshi |
| EPI_ISL_455017 | Government Medical College, Vadodara | Gujarat Biotechnology Research Centre | Tanuja Javadekar , R N Daveshwar, Ramesh Pandit, Tejas Shah, Ankit Hinsu, Pritesh Sabara, Apurvasinh Puvar, Janvi Raval, Zarna Patel, Monika Gandhi, Pinal Trivedi, Maharshi Pandya, Amit Kanani, Nidhi Patel, Nitin Savaliya, Raghawendra Kumar, Dinesh Kumar, Zuber Saiyed, Komal Patel, Labdhi Pandya, Snehal Bagatharia, Fenil Patel, Bhavesh Modi, Gaurishankar Shrimali, R D Dixit, A M Kadri, Umang Mishra, Chaitanya Joshi, Madhvi Joshi, |
| EPI_ISL_455018 | Government Medical College, Vadodara | Gujarat Biotechnology Research Centre | R N Daveshwar, Ramesh Pandit, Tejas Shah, Ankit Hinsu, Pritesh Sabara, Apurvasinh Puvar, Janvi Raval, Zarna Patel, Monika Gandhi, Pinal Trivedi, Maharshi Pandya, Amit Kanani, Nidhi Patel, Nitin Savaliya, Raghawendra Kumar, Dinesh Kumar, Zuber Saiyed, Komal Patel, Labdhi Pandya, Snehal Bagatharia, Tanuja Javadekar , Neelam Nathani, Bhavesh Modi, Gaurishankar Shrimali, R D Dixit, A M Kadri, Umang Mishra, Chaitanya Joshi, Madhvi Joshi, |
| EPI_ISL_455019 | Government Medical College, Vadodara | Gujarat Biotechnology Research Centre | Ramesh Pandit, Tejas Shah, Ankit Hinsu, Pritesh Sabara, Apurvasinh Puvar, Janvi Raval, Zarna Patel, Monika Gandhi, Pinal Trivedi, Maharshi Pandya, Amit Kanani, Nidhi Patel, Nitin Savaliya, Raghawendra Kumar, Dinesh Kumar, Zuber Saiyed, Komal Patel, Labdhi Pandya, Snehal Bagatharia, Tanuja Javadekar , R N Daveshwar, Armi Chaudhari, Bhavesh Modi, Gaurishankar Shrimali, R D Dixit, A M Kadri, Umang Mishra, Chaitanya Joshi, Madhvi Joshi, |
| EPI_ISL_455020 | Government Medical College, Vadodara | Gujarat Biotechnology Research Centre | Tejas Shah, Ankit Hinsu, Pritesh Sabara, Apurvasinh Puvar, Janvi Raval, Zarna Patel, Monika Gandhi, Pinal Trivedi, Maharshi Pandya, Amit Kanani, Nidhi Patel, Nitin Savaliya, Raghawendra Kumar, Dinesh Kumar, Zuber Saiyed, Komal Patel, Labdhi Pandya, Snehal Bagatharia, Tanuja Javadekar , R N Daveshwar, Ramesh Pandit, Bhavya Jindal, Bhavesh Modi, Gaurishankar Shrimali, R D Dixit, A M Kadri, Umang Mishra, Chaitanya Joshi, Madhvi Joshi, |
| EPI_ISL_455021 | Government Medical College, Vadodara | Gujarat Biotechnology Research Centre | Ankit Hinsu, Pritesh Sabara, Apurvasinh Puvar, Janvi Raval, Zarna Patel, Monika Gandhi, Pinal Trivedi, Maharshi Pandya, Amit Kanani, Nidhi Patel, Nitin Savaliya, Raghawendra Kumar, Dinesh Kumar, Zuber Saiyed, Komal Patel, Labdhi Pandya, Snehal Bagatharia, Tanuja Javadekar , R N Daveshwar, Ramesh Pandit, Tejas Shah, Camellia Chakraborty, Bhavesh Modi, Gaurishankar Shrimali, R D Dixit, A M Kadri, Umang Mishra, Chaitanya Joshi, Madhvi Joshi, |
| EPI_ISL_455022 | Government Medical College, Vadodara | Gujarat Biotechnology Research Centre | Pritesh Sabara, Apurvasinh Puvar, Janvi Raval, Zarna Patel, Monika Gandhi, Pinal Trivedi, Maharshi Pandya, Amit Kanani, Nidhi Patel, Nitin Savaliya, |

| | | | |
|---|---|---|---|
| | | | Raghawendra Kumar, Dinesh Kumar, Zuber Saiyed, Komal Patel, Labdhi Pandya, Snehal Bagatharia, Tanuja Javadekar , R N Daveshwar, Ramesh Pandit, Tejas Shah, Ankit Hinsu, Siddhant Kumar, Bhavesh Modi, Gaurishankar Shrimali, R D Dixit, A M Kadri, Umang Mishra, Chaitanya Joshi, Madhvi Joshi, |
| EPI_ISL_455023 | Government Medical College, Vadodara | Gujarat Biotechnology Research Centre | Apurvasinh Puvar, Janvi Raval, Zarna Patel, Monika Gandhi, Pinal Trivedi, Maharshi Pandya, Amit Kanani, Nidhi Patel, Nitin Savaliya, Raghawendra Kumar, Dinesh Kumar, Zuber Saiyed, Komal Patel, Labdhi Pandya, Snehal Bagatharia, Tanuja Javadekar , R N Daveshwar, Ramesh Pandit, Tejas Shah, Ankit Hinsu, Pritesh Sabara, Priyanka P Vatsa, Bhavesh Modi, Gaurishankar Shrimali, R D Dixit, A M Kadri, Umang Mishra, Chaitanya Joshi, Madhvi Joshi, |
| EPI_ISL_455024 | Government Medical College, Vadodara | Gujarat Biotechnology Research Centre | Janvi Raval, Zarna Patel, Monika Gandhi, Pinal Trivedi, Maharshi Pandya, Amit Kanani, Nidhi Patel, Nitin Savaliya, Raghawendra Kumar, Dinesh Kumar, Zuber Saiyed, Komal Patel, Labdhi Pandya, Snehal Bagatharia, Tanuja Javadekar , R N Daveshwar, Ramesh Pandit, Tejas Shah, Ankit Hinsu, Pritesh Sabara, Apurvasinh Puvar, Pooja P Doshi, Bhavesh Modi, Gaurishankar Shrimali, R D Dixit, A M Kadri, Umang Mishra, Chaitanya Joshi, Madhvi Joshi, |
| EPI_ISL_455025 | Government Medical College, Vadodara | Gujarat Biotechnology Research Centre | Zarna Patel, Monika Gandhi, Pinal Trivedi, Maharshi Pandya, Amit Kanani, Nidhi Patel, Nitin Savaliya, Raghawendra Kumar, Dinesh Kumar, Zuber Saiyed, Komal Patel, Labdhi Pandya, Snehal Bagatharia, Tanuja Javadekar , R N Daveshwar, Ramesh Pandit, Tejas Shah, Ankit Hinsu, Pritesh Sabara, Apurvasinh Puvar, Janvi Raval, Akanksha Verma, Bhavesh Modi, Gaurishankar Shrimali, R D Dixit, A M Kadri, Umang Mishra, Chaitanya Joshi, Madhvi Joshi, |
| EPI_ISL_455026 | Government Medical College, Vadodara | Gujarat Biotechnology Research Centre | Monika Gandhi, Pinal Trivedi, Maharshi Pandya, Amit Kanani, Nidhi Patel, Nitin Savaliya, Raghawendra Kumar, Dinesh Kumar, Zuber Saiyed, Komal Patel, Labdhi Pandya, Snehal Bagatharia, Tanuja Javadekar , R N Daveshwar, Ramesh Pandit, Tejas Shah, Ankit Hinsu, Pritesh Sabara, Apurvasinh Puvar, Janvi Raval, Zarna Patel, Priti Pandita, Bhavesh Modi, Gaurishankar Shrimali, R D Dixit, A M Kadri, Umang Mishra, Chaitanya Joshi, Madhvi Joshi, |
| EPI_ISL_455027 | Government Medical College, Vadodara | Gujarat Biotechnology Research Centre | Pinal Trivedi, Maharshi Pandya, Amit Kanani, Nidhi Patel, Nitin Savaliya, Raghawendra Kumar, Dinesh Kumar, Zuber Saiyed, Komal Patel, Labdhi Pandya, Snehal Bagatharia, Tanuja Javadekar , R N Daveshwar, Ramesh Pandit, Tejas Shah, Ankit Hinsu, Pritesh Sabara, Apurvasinh Puvar, Janvi Raval, Zarna Patel, Monika Gandhi, Pragya Sharma, Bhavesh Modi, Gaurishankar Shrimali, R D Dixit, A M Kadri, Umang Mishra, Chaitanya Joshi, Madhvi Joshi, |
| EPI_ISL_455312 | Microbiology Unit, Department of Pathology & Laboratory Medicine, IIUM Medical Centre | SEA Microbiome Unit, Faculty of Industrial Sciences & Technology, Universiti Malaysia Pahang | Norhidayah Binti Kamarudin, Ahmad Hafiz Bin Zulkifly, Hajar Fauzan Ahmad, Muhammad Adam Lee Abdullah, Mohd Fazli Farida Asras, Ahmad Mahfuz Gazali, Mohd Nazli Bin Kamarulzaman, IIUM Medical Centre Covid19 Taskforce, UMP Covid19 Team |
| EPI_ISL_455313 | Microbiology Unit, Department of Pathology & Laboratory Medicine, IIUM Medical Centre | SEA Microbiome Unit, Faculty of Industrial Sciences & Technology, Universiti Malaysia Pahang | Hajar Fauzan Ahmad, Norhidayah Kamarudin, Ahmad Hafiz Zulkifly, IIUM Medical Centre Covid19 Taskforce, UMP Covid19 Team |
| EPI_ISL_455314 | Hospital Virgen del Rocio | Instituto de Salud Carlos III | Iglesias-Caballero, M. Molinero Calamita, M. González-Esguevillas, M. Camarero, S. Pozo, F. Casas, I. Jiménez, P. Jiménez, M. Zaballos, A. Monzón, S. Varona, S. Juliá, M. Cuesta, I, J. Lepe |
| EPI_ISL_455315, EPI_ISL_455316, EPI_ISL_455317, EPI_ISL_455318, EPI_ISL_455319, EPI_ISL_455320, EPI_ISL_455321, EPI_ISL_455322, EPI_ISL_455324 | Hospital Virgen de las Nieves | Instituto de Salud Carlos III | Iglesias-Caballero, M. Molinero Calamita, M. González-Esguevillas, M. Camarero, S. Pozo, F. Casas, I. Jiménez, P. Jiménez, M. Zaballos, A. Monzón, S. Varona, S. Juliá, M. Cuesta, I, S. Sanbonmatsu |
| EPI_ISL_455325 | Hospital Universitario de Canarias | Instituto de Salud Carlos III | Iglesias-Caballero, M. Molinero Calamita, M. González-Esguevillas, M. Camarero, S. Pozo, F. Casas, I. Jiménez, P. Jiménez, M. Zaballos, A. Monzón, S. Varona, S. Juliá, M. Cuesta, I, B. Castro |
| EPI_ISL_455326 | Hospital Universitario Insular de Gran Canaria | Instituto de Salud Carlos III | Iglesias-Caballero, M. Molinero Calamita, M. González-Esguevillas, M. Camarero, S. Pozo, F. Casas, I. Jiménez, P. Jiménez, M. Zaballos, A. Monzón, S. Varona, S. Juliá, M. Cuesta, I, A. Hernández |
| EPI_ISL_455327 | Consejeria de Sanidad y Asuntos Sociales | Instituto de Salud Carlos III | Iglesias-Caballero, M. Molinero Calamita, M. González-Esguevillas, M. Camarero, S. Pozo, F. Casas, I. Jiménez, P. Jiménez, M. Zaballos, A. Monzón, S. Varona, S. Juliá, M. Cuesta, I, G. Gutiérrez |
| EPI_ISL_455328, EPI_ISL_455329, EPI_ISL_455330, EPI_ISL_455331 | Complejo Hospitalario Universitario La Coruna | Instituto de Salud Carlos III | Iglesias-Caballero, M. Molinero Calamita, M. González-Esguevillas, M. Camarero, S. Pozo, F. Casas, I. Jiménez, P. Jiménez, M. Zaballos, A. Monzón, S. Varona, S. Juliá, M. Cuesta, I, M.A Canizares |
| EPI_ISL_455332 | Xerencia de Xestión Integrada de Pontevedra e o Salnés | Instituto de Salud Carlos III | Iglesias-Caballero, M. Molinero Calamita, M. González-Esguevillas, M. Camarero, S. Pozo, F. Casas, I. Jiménez, P. Jiménez, M. Zaballos, A. Monzón, S. Varona, S. Juliá, M. Cuesta, I, M. Garcia |
| EPI_ISL_455333 | Complejo Hospitalario Universitario de Santiago | Instituto de Salud Carlos III | Iglesias-Caballero, M. Molinero Calamita, M. González-Esguevillas, M. Camarero, S. Pozo, F. Casas, I. Jiménez, P. Jiménez, M. Zaballos, A. Monzón, S. Varona, S. Juliá, M. Cuesta, I, J. Llovo |
| EPI_ISL_455334, EPI_ISL_455335 | Complejo Hospitalario de Orense | Instituto de Salud Carlos III | Iglesias-Caballero, M. Molinero Calamita, M. González-Esguevillas, M. Camarero, S. Pozo, F. Casas, I. Jiménez, P. Jiménez, M. Zaballos, A. Monzón, S. Varona, S. Juliá, M. Cuesta, I, M. Paz |
| EPI_ISL_455336, EPI_ISL_455337, EPI_ISL_455338, EPI_ISL_455339, EPI_ISL_455340, EPI_ISL_455341, EPI_ISL_455342, EPI_ISL_455343 | Hospital San Pedro | Instituto de Salud Carlos III | Iglesias-Caballero, M. Molinero Calamita, M. González-Esguevillas, M. Camarero, S. Pozo, F. Casas, I. Jiménez, P. Jiménez, M. Zaballos, A. Monzón, S. Varona, S. Juliá, M. Cuesta, I, C. Alonso |
| EPI_ISL_455344, EPI_ISL_455345, EPI_ISL_455346, EPI_ISL_455347, EPI_ISL_455348, EPI_ISL_455349 | Hospital Comarcal de Melilla | Instituto de Salud Carlos III | Iglesias-Caballero, M. Molinero Calamita, M. González-Esguevillas, M. Camarero, S. Pozo, F. Casas, I. Jiménez, P. Jiménez, M. Zaballos, A. Monzón, S. Varona, S. Juliá, M. Cuesta, I, I. Pérez |
| EPI_ISL_455350, EPI_ISL_455351 | Hospital Txagorritxu | Instituto de Salud Carlos III | Iglesias-Caballero, M. Molinero Calamita, M. González-Esguevillas, M. Camarero, S. Pozo, F. Casas, I. Jiménez, P. Jiménez, M. Zaballos, A. Monzón, S. Varona, S. Juliá, M. Cuesta, I, C. Gómez |
| EPI_ISL_455352, EPI_ISL_455353, EPI_ISL_455354 | Hospital de Cruces | Instituto de Salud Carlos III | Iglesias-Caballero, M. Molinero Calamita, M. González-Esguevillas, M. Camarero, S. Pozo, F. Casas, I. Jiménez, P. Jiménez, M. Zaballos, A. Monzón, S. Varona, S. Juliá, M. Cuesta, I, M. Aranzamendi |
| EPI_ISL_455362 | Nigeria Centre for Disease Control (NCDC) | African Centre of Excellence for Genomics of Infectious Diseases (ACEGID), Redeemer's University, Ede, Osun State, Nigeria | Oluniyi P.E., Ajogbasile F.V., Kayode A., Olawoye I., Uwanibe J., Oguzie J., Olumade T., Folarin O.A., Ihekweazu C., Happi C.T. |
| EPI_ISL_455412, EPI_ISL_455413, EPI_ISL_455419 | Nigeria Centre for Disease Control (NCDC) | African Centre of Excellence for Genomics of Infectious Diseases (ACEGID), Redeemer's University, Ede, Osun State, Nigeria | Oluniyi P.E., Ajogbasile F.V., Kayode A., Oguzie J., Olawoye I., Uwanibe J., Olumade T., Folarin O.A., Ihekweazu C., Happi C.T. |
| EPI_ISL_455422 | Nigeria Centre for Disease Control | African Centre of Excellence for Genomics of Infectious Diseases (ACEGID), Redeemer's University, Ede, Osun State, Nigeria | Oluniyi P.E., Ajogbasile F.V., Kayode A., Oguzie J., Olawoye I., Uwanibe J., Olumade T., Folarin O.A., Ihekweazu C., Happi C.T. |
| EPI_ISL_455423, EPI_ISL_455424 | Nigeria Centre for Disease Control (NCDC) | African Centre of Excellence for Genomics of Infectious Diseases (ACEGID), Redeemer's University, Ede, Osun State, Nigeria | Oluniyi P.E., Ajogbasile F.V., Kayode A., Oguzie J., Olawoye I., Uwanibe J., Olumade T., Folarin O.A., Ihekweazu C., Happi C.T. |
| EPI_ISL_455426 | Nigeria Centre for Disease Control | African Centre of Excellence for Genomics of Infectious Diseases (ACEGID), Redeemer's University, Ede, Osun State, Nigeria | Oluniyi P.E., Ajogbasile F.V., Kayode A., Oguzie J., Olawoye I., Uwanibe J., Olumade T., Folarin O.A., Ihekweazu C., Happi C.T. |
| EPI_ISL_455429, EPI_ISL_455431 | Nigeria Centre for Disease Control (NCDC) | African Centre of Excellence for Genomics of Infectious Diseases (ACEGID), Redeemer's University, Ede, Osun State, Nigeria | Oluniyi P.E., Ajogbasile F.V., Kayode A., Oguzie J., Olawoye I., Uwanibe J., Olumade T., Folarin O.A., Ihekweazu C., Happi C.T. |
| EPI_ISL_455434 | Instituto de Diagnostico y Referencia Epidemiologicos (INDRE) | Instituto de Diagnostico y Referencia Epidemiologicos (INDRE) | Taboada Ramírez Blanca. Ramirez-Gonzalez Ernesto, Garces-Ayala Fabiola, Araiza-Rodriguez Adnan, Mendieta-Condado Edgar, Rodriguez-Maldonado Abril, Wong-Arambula Claudia, Barrera-Badillo Gisela, Hernandez-Rivas Lucia, Lopez-Martinez Irma. |

| | | | |
|---|---|---|---|
| EPI_ISL_455435 | Instituto de Diagnostico y Referencia Epidemiologicos (INDRE) | Instituto de Diagnostico y Referencia Epidemiologicos (INDRE) | Garces-Ayala Fabiola. Taboada Ramírez Blanca. Ramirez-Gonzalez Ernesto, Araiza-Rodriguez Adnan , Mendieta-Condado Edgar, Rodriguez-Maldonado Abril, Wong-Arambula Claudia, Barrera-Badillo Gisela, Hernandez-Rivas Lucia, Lopez-Martinez Irma |
| EPI_ISL_455436 | Instituto de Diagnostico y Referencia Epidemiologicos (INDRE) | Instituto de Diagnostico y Referencia Epidemiologicos (INDRE) | Garces-Ayala Fabiola. Taboada Ramírez Blanca. Ramirez-Gonzalez Ernesto, Araiza-Rodriguez Adnan, Mendieta-Condado Edgar, Rodriguez-Maldonado Abril, Wong-Arambula Claudia, Barrera-Badillo Gisela, Hernandez-Rivas Lucia, Lopez-Martinez Irma. |
| EPI_ISL_455437, EPI_ISL_455438 | Instituto de Diagnostico y Referencia Epidemiologicos (INDRE) | Instituto de Diagnostico y Referencia Epidemiologicos (INDRE) | Araiza-Rodriguez Adnan, Garces-Ayala Fabiola. Ramirez-Gonzalez Ernesto, Mendieta-Condado Edgar, Rodriguez-Maldonado Abril, Wong-Arambula Claudia, Barrera-Badillo Gisela, Hernandez-Rivas Lucia, Lopez-Martinez Irma, Taboada Ramírez Blanca. |
| EPI_ISL_455439 | Instituto de Diagnostico y Referencia Epidemiologicos (INDRE) | Instituto de Diagnostico y Referencia Epidemiologicos (INDRE) | Mendieta-Condado Edgar, Araiza-Rodriguez Adnan, Garces-Ayala Fabiola, Rodriguez-Maldonado Abril, Wong-Arambula Claudia, Barrera-Badillo Gisela, Hernandez-Rivas Lucia, Lopez-Martinez Irma, Taboada Ramirez Blanca, Ramirez-Gonzalez Ernesto. |
| EPI_ISL_455455 | Instituto de Diagnostico y Referencia Epidemiologicos (INDRE) | Instituto de Diagnostico y Referencia Epidemiologicos (INDRE) | Rodriguez-Maldonado Abril, Mendieta-Condado Edgar, Araiza-Rodriguez Adnan, Garces-Ayala Fabiola. Taboada Ramirez Blanca. Ramirez-Gonzalez Ernesto, Barrera-Badillo Gisela, Hernandez-Rivas Lucia, Lopez-Martinez Irma, Wong-Arambula Claudia. |
| EPI_ISL_455456 | Instituto de Diagnostico y Referencia Epidemiologicos (INDRE) | Instituto de Diagnostico y Referencia Epidemiologicos (INDRE) | Rodriguez-Maldonado Abril, Mendieta-Condado Edgar, Araiza-Rodriguez Adnan, Garces-Ayala Fabiola. Taboada Ramírez Blanca. Ramirez-Gonzalez Ernesto, , Barrera-Badillo Gisela, Hernandez-Rivas Lucia, Lopez-Martinez Irma, Wong-Arambula Claudia. |
| EPI_ISL_455460, EPI_ISL_455461, EPI_ISL_455462, EPI_ISL_455463, EPI_ISL_455464, EPI_ISL_455465, EPI_ISL_455466, EPI_ISL_455467 | Jiangxi Province Center for Disease Control and Prevention | Jiangxi Province Center for Disease Control and Prevention | JianXiong Li,Ying Xiong,Tian Gong,Yong Shi,Jun Zhou,Fang Xiao,ShiWen Liu,XiaoQing Liu,Gang Xu,DaJin Xiao,Xin Ran,YanNi Zhang |
| EPI_ISL_455468, EPI_ISL_455469, EPI_ISL_455470, EPI_ISL_455471, EPI_ISL_455472, EPI_ISL_455473, EPI_ISL_455474 | Laboratory for Respiratory Viruses, Cantacuzino National Military-Medical Institute for Research and Development | Cantacuzino Institute | M.Lazar, L.Ustea, A.Cretu, Tim Durfee |
| EPI_ISL_455475, EPI_ISL_455477 | Laboratory for Respiratory Viruses, Cantacuzino National Military-Medical Institute for Research and Development | Cantacuzino Institute | M.Lazar, L.Ustea, A.Cretu, T.Durfee |
| EPI_ISL_455566, EPI_ISL_455567 | Institute for Public Health | Laboratory for advanced genomics | Filip Roki, Lovro Trgovec-Greif, Neven Sui, Tomislav Rukavina, Igor Jurak, Oliver Vugrek |
| EPI_ISL_455694, EPI_ISL_455695, EPI_ISL_455696, EPI_ISL_455697, EPI_ISL_455698, EPI_ISL_455699, EPI_ISL_455700, EPI_ISL_455701, EPI_ISL_455702, EPI_ISL_455703, EPI_ISL_455704, EPI_ISL_455705, EPI_ISL_455706, EPI_ISL_455707, EPI_ISL_455708, EPI_ISL_455709, EPI_ISL_455710, EPI_ISL_455711, EPI_ISL_455712 | | | |
| see above | National Hospital of Tropical Diseases | Oxford University Clinical Research Unit, Hanoi, Vietnam | Nguyen Thi Tam, Van Dinh Trang, Nguyen Thu Trang, Nguyen Thi Ngoc Diep, Le Nguyen Minh Hoa, Pham Ngoc Thach, H. Rogier van Doorn, on behalf of the OUCRU COVID-19 research group |
| EPI_ISL_455714, EPI_ISL_455716, EPI_ISL_455717, EPI_ISL_455718 | National Hospital of Tropical Diseases | Oxford University Clinical Research Unit, Hanoi, Vietnam | Nguyen Thi Tam, Van Dinh Trang, Nguyen Thi Hong Thuong, Vu Thi Ngoc Bich, Nguyen Thu Trang, Nguyen Thi Ngoc Diep, Le Nguyen Minh Hoa, Pham Ngoc Thach, H. Rogier van Doorn, on behalf of the OUCRU COVID-19 research group |
| EPI_ISL_455790, EPI_ISL_455791 | Institute for Medical Research, Infectious Disease Research Centre, National Institutes of Health, Ministry of Health Malaysia | Malaysia Genome Institute | Mohd Noor Mat Isa, Irni Suhayu Sapian, Yusuf Muhammad Noor, Jeyanthi Suppiah, Nurhezreen Md Iqbal, Enizza Kasim, Zarina Mohd Zawawi, Siti Noraini Othman, Mohd Faizal Abu Bakar, Shamsidar Sopie, Azrin Ahmad, Ravindran Thayan, Norazah Ahmad, Tahir Aris, Shahrul Hisham Zainal Ariffin |
| EPI_ISL_455792, EPI_ISL_455793 | Institute for Medical Research, Infectious Disease Research Centre, National Institutes of Health, Ministry of Health Malaysia | Malaysia Genome Institute | Mohd Noor Mat Isa, Irni Suhayu Sapian, Yusuf Muhammad Noor, Jeyanthi Suppiah, Nurhezreen Md Iqbal, Enizza Kasim, Zarina Mohd Zawawi, Siti Noraini Othman, Mohd Faizal Abu Bakar, Shamsidar Sopie, Azrin Ahmad, Ravindran Thayan, Norazah Ahmad, Tahir Aris, Shahrul Hisham Zainal Ariffin |
| EPI_ISL_456117, EPI_ISL_456119, EPI_ISL_456120, EPI_ISL_456126, EPI_ISL_456127, EPI_ISL_456138, EPI_ISL_456145, EPI_ISL_456146, EPI_ISL_456147, EPI_ISL_456148, EPI_ISL_456149, EPI_ISL_456150, EPI_ISL_456151, EPI_ISL_456152, EPI_ISL_456155 | | | |
| see above | Instituto Nacional de Salud - Unidad de Secuenciación y Análisis Genómico | Instituto Nacional de Salud, Universidad Cooperativa de Colombia, Instituto Alexander von Humboldt, Imperial College-London, London School of Hygiene & Tropical Medicine | Katherine Laiton-Donato, Diego A. Álvarez-Díaz, Carlos Franco-Muñoz, Jose A. Usme-Ciro, Gloria Puerto, Nicolas D. Franco-Sierra, Mailyn A.Gonzalez, Zulma M. Cucunubá, Christian Julian VillabonaArenas, Liz Villabona-Arenas, Sussy Echeverria, Astrid C. Flórez, Sergio Gomez-Rangel, Luz Dary Rodriguez, Juliana Barbosa, Erika Ospitia, Diana Marcela Walteros-Acero, Martha Lucia Ospina Martinez, Marcela Mercado-Reyes. |
| EPI_ISL_457702 | Oman-NIC | Microbiology laboratory- Sultan Qaboos University Hospital | Fahad Zadjali, Samira Al-Maruqi, Amina Al Jardani, Khulood Al-Mammary, Hanan Al-kindi, Fatma BaAlawi, Hamida AL Barwani, Zeyana AL-Dahmani, Intisar Al-Shukri, Aisha Al-Busaidi, Aisha Al-Amri, Ahlam Al-Amri, Mohammed Al-Tobi, Samiha Al Kharusi, Abdulla Balkhair |
| EPI_ISL_457703 | Oman-NIC | Department of Microbiology and Immunology- SQUH | Fahad Zadjali, Samira Al-Maruqi, Amina Al Jardani, Khulood Al-Mammary, Hanan Al-kindi, Fatma BaAlawi, Hamida AL Barwani, Zeyana AL-Dahmani, Intisar Al-Shukri, Aisha Al-Busaidi, Aisha Al-Amri, Ahlam Al-Amri, Mohammed Al-Tobi, Samiha Al Kharusi, Abdulla Balkhair |
| EPI_ISL_457705 | OMAN-NIC | Department of Microbiology and Immunology- SQUH | Fahad Zadjali, Samira Al-Maruqi, Amina Al Jardani, Khulood Al-Mammary, Hanan Al-kindi, Fatma BaAlawi, Hamida AL Barwani, Zeyana AL-Dahmani, Intisar Al-Shukri, Aisha Al-Busaidi, Aisha Al-Amri, Ahlam Al-Amri, Mohammed Al-Tobi, Samiha Al Kharusi, Abdulla Balkhair |
| EPI_ISL_457707 | Oman-NIC | Department of Microbiology and Immunology- SQUH | Fahad Zadjali, Samira Al-Maruqi, Amina Al Jardani, Khulood Al-Mammary, Hanan Al-kindi, Fatma BaAlawi, Hamida AL Barwani, Zeyana AL-Dahmani, Intisar Al-Shukri, Aisha Al-Busaidi, Aisha Al-Amri, Ahlam Al-Amri, Mohammed Al-Tobi, Samiha Al Kharusi, Abdulla Balkhair |
| EPI_ISL_457726, EPI_ISL_457730 | TSGH-CP molecular lab | TSGH-CP molecular lab | Cherng-Lih Perng, Ming-Jr JIAN, Chih-Kai Chang, Jung-Chung Lin, Kuo-Ming Yeh, Chien-Wen Chen, Sheng-Kang Chiu, Hsing-Yi Chung, Shih-Hung Tsai, Kuo-Sheng Hung, Tien-Yao Chang, Feng-Yee Chang, Hung-Sheng Shang |
| EPI_ISL_457750 | Centogene AG | Centogene AG | Prof. Dr. Peter Bauer, Dr. Krishna Kumar Kandaswamy |
| EPI_ISL_457824 | Bezmialem Vakif University, Dept Microbiology, Medical School, Fatih, Istanbul, Turkey | Bezmialem Vakif University, Medical School & Beykoz Institute of Life Sciences & Biotechnology | Mehmet Z. Doymaz, Merve Kalkan, Nesibe Cetin, Elif Karaaslan, Bilge Sumbul, Filiz Guney |
| EPI_ISL_457997 | Oman-NIC | Oman-NIC | Samira Al-Maruqi, Fahad Zadjali, Amina Al Jardani, Khulood Al-Mammary, Hanan Al-kindi, Fatma BaAlawi, Hamida AL Barwani, Zeyana AL-Dahmani, Intisar Al-Shukri, Aisha Al-Busaidi, Aisha Al-Amri, Ahlam Al-Amri, Mohammed Al-Tobi, Samiha Al Kharusi, Abdulla Balkhair |
| EPI_ISL_458029 | TSGH-CP molecular lab | TSGH-CP molecular lab | Cherng-Lih Perng, Ming-Jr JIAN, Chih-Kai Chang, Jung-Chung Lin, Kuo-Ming Yeh, Chien-Wen Chen, Sheng-Kang Chiu, Hsing-Yi Chung, Shih-Hung Tsai, Kuo-Sheng Hung, Tien-Yao Chang, Feng-Yee Chang, Hung-Sheng Shang |
| EPI_ISL_458079 | Mitra Keluarga Hospital Kenjeran | Institute of Tropical Disease, Universitas Airlangga | Aldise M Nastri, Jezzy R Dewantari, Rima R Prasetya, Krisnoadi Rahardjo, Anastasia W Jefuna, Gatot Soegiarto, Laksmi Wulandari, Retno A Setyoningrum, Resti Yudhawati, Yohko K Shimizu, Mitsuhiro Nishimura, Yasuko Mori, Soetjipto, Kazufumi Shimizu, Maria I Lusida |
| EPI_ISL_458081 | RSUD Bangil Pasuruan | Institute of Tropical Disease, Universitas Airlangga | Jezzy R Dewantari, Rima R Prasetya, Krisnoadi Rahardjo, Aldise M Nastri, Arma Roosalina, Gatot Soegiarto, Laksmi Wulandari, Retno A Setyoningrum, Resti Yudhawati, Yohko K Shimizu, Mitsuhiro Nishimura, Yasuko Mori, Soetjipto, Kazufumi Shimizu, Maria I Lusida |
| EPI_ISL_458083 | Adi Husada Undaan Hospital | Institute of Tropical Disease, Universitas Airlangga | Rima R Prasetya, Krisnoadi Rahardjo, Aldise M Nastri, Jezzy R Dewantari, Irawati Marga, Gatot Soegiarto, Laksmi Wulandari, Retno A Setyoningrum, Resti Yudhawati, Yohko K Shimizu, Mitsuhiro Nishimura, Yasuko Mori, Soetjipto, Kazufumi Shimizu, Maria I Lusida |
| EPI_ISL_458103 | Gujarat Biotechnology Research Centre | Gujarat Biotechnology Research Centre | Ramesh Pandit, Tejas Shah, Ankit Hinsu, Pritesh Sabara, Apurvasinh Puvar, Janvi Raval, Zarna Patel, Monika Gandhi, Pinal Trivedi, Maharshi Pandya, Amit Kanani, Nidhi Patel, Nitin Savaliya, Raghawendra Kumar, Dinesh Kumar, Zuber Saiyed, Komal Patel, Labdhi Pandya, Snehal Bagatharia, Armi Chaudhari, Bhavesh Modi, Gaurishankar Shrimali, R D Dixit, A M Kadri, Umang Mishra, Chaitanya Joshi, Madhvi Joshi, , , , , |
| EPI_ISL_458104 | Gujarat Biotechnology Research Centre | Gujarat Biotechnology Research Centre | Tejas Shah, Ankit Hinsu, Pritesh Sabara, Apurvasinh Puvar, Janvi Raval, Zarna Patel, Monika Gandhi, Pinal Trivedi, Maharshi Pandya, Amit Kanani, Nidhi Patel, Nitin Savaliya, Raghawendra Kumar, Dinesh Kumar, Zuber Saiyed, Komal Patel, Labdhi Pandya, Snehal Bagatharia, Ramesh Pandit, Bhavya Jindal, Bhavesh Modi, Gaurishankar Shrimali, R D Dixit, A M Kadri, Umang Mishra, Chaitanya Joshi, Madhvi Joshi, , , , , |
| EPI_ISL_458105 | Gujarat Biotechnology Research Centre | Gujarat Biotechnology Research Centre | Ankit Hinsu, Pritesh Sabara, Apurvasinh Puvar, Janvi Raval, Zarna Patel, Monika Gandhi, Pinal Trivedi, Maharshi Pandya, Amit Kanani, Nidhi Patel, Nitin Savaliya, Raghawendra Kumar, Dinesh Kumar, Zuber Saiyed, Komal Patel, Labdhi Pandya, Snehal Bagatharia, Ramesh Pandit, Tejas Shah, Camellia Chakraborty, Bhavesh Modi, Gaurishankar Shrimali, R D Dixit, A M Kadri, Umang Mishra, Chaitanya Joshi, Madhvi Joshi, , , , , |

| | | | |
|---|---|---|---|
| EPI_ISL_458116, EPI_ISL_458117, EPI_ISL_458118, EPI_ISL_458120 | Oman National Influenza Centre | Department of Microbiology and Immunology-SQUH | Fahad Zadjali, Samira Al-Maruqi, Amina Al Jardani, Khulood Al-Mammary, Hanan Al-kindi, Fatma BaAlawi, Hamida AL Barwani, Zeyana AL-Dahmani, Intisar Al-Shukri, Aisha Al-Busaidi, Aisha Al-Amri, Ahlam Al-Amri, Mohammed Al-Tobi, Samiha Al Kharusi, Abdulla Balkhair |
| EPI_ISL_458133 | National Institute of Biotechnology | Bioinformatics Division, National Institute of Biotechnology | Mohammad Uzzal Hossain, Md. Moniruzzaman, Md. Salim Khan, Md. Nazrul Islam, Md. Hadisur Rahman, Arittra Bhattacharjee, Md. Ruhul Amin, Asif Rashid, Chaman Ara Keya, Keshob Chandra Das, Md. Salimullah |
| EPI_ISL_458139 | Evandro Chagas Institute | Evandro Chagas Institute | Santos, M.C.; Silva, A.M.; Junior, W.D.C.; Barbagelata, L.S.; Ferreira, J.A.; Sousa, E.M.A.; da Silva, P.S.; Resque, H.R; Martins, L.C.; Sousa Junior, E.C.;Viana, G.M.R |
| EPI_ISL_459866, EPI_ISL_459867, EPI_ISL_459868, EPI_ISL_459869, EPI_ISL_459871, EPI_ISL_459872, EPI_ISL_459873, EPI_ISL_459874, EPI_ISL_459875, EPI_ISL_459877, EPI_ISL_459879, EPI_ISL_459880, EPI_ISL_459881, EPI_ISL_459882, EPI_ISL_459883, EPI_ISL_459885, EPI_ISL_459886, EPI_ISL_459887, EPI_ISL_459888, EPI_ISL_459889, EPI_ISL_459890, EPI_ISL_459891, EPI_ISL_459892 | | | |
| see above | Kingston Health Sciences Center | Queen's Genomics Lab at Ongwanada (Q-GLO) | Sjaarda CP, Rustom N, Huang D, Perez-Patrigeon S, Hudson ML, Wong H,Guan H, Ayub M, Soares CN, Colautti R, Evans GA, Sheth P |
| EPI_ISL_459911 | Devki Devi Foundation, a unit of Max Healthcare | CSIR-IGIB/Max | Rajesh Pandey#, Samreen Siddiqui, Pooja Sharma, Bansidhar Tarai, Vivekanand A, Bharathram Uppili, Saruchi Wadhwa, Nishu Tyagi, Mitali Mukerji, Poonam Das, Sujeet Jha, Mohammed Faruq, Vinita Jha, Anurag Agrawal |
| EPI_ISL_459914, EPI_ISL_459915, EPI_ISL_459918, EPI_ISL_459921, EPI_ISL_459922, EPI_ISL_459923, EPI_ISL_459932, EPI_ISL_459933, EPI_ISL_459934, EPI_ISL_459935, EPI_ISL_459936, EPI_ISL_459937, EPI_ISL_459938, EPI_ISL_459939, EPI_ISL_459940, EPI_ISL_459941, EPI_ISL_459942, EPI_ISL_459943 | | | |
| see above | Devki Devi Foundation, a unit of Max Healthcare | CSIR-IGIB/Max | Rajesh Pandey#, Samreen Siddiqui, Pooja Sharma, Bansidhar Tarai, Vivekanand A, Bharathram Uppili, Saruchi Wadhwa, Nishu Tyagi, Mitali Mukerji, Bansidhar Tarai, Poonam Das, Sujeet Jha, Mohammed Faruq, Vinita Jha, Anurag Agrawal |
| EPI_ISL_459953 | Institute for Medical Research, Infectious Disease Research Centre, National Institutes of Health, Ministry of Health Malaysia | Institute for Medical Research Infectious Disease Research Centre, National Institutes of Health, Ministry of Health Malaysia | Suppiah J, Mohd-Zawawi Z, Kamel KA, Ellan K, Kalyanasundram J, Mohd-Zain R, Thayan R |
| EPI_ISL_459954, EPI_ISL_459955, EPI_ISL_459956 | Institute for Medical Research, Infectious Disease Research Centre, National Institutes of Health, Ministry of Health Malaysia | Institute for Medical Research, Infectious Disease Research Centre, National Institutes of Health, Ministry of Health Malaysia | Suppiah J, Mohd-Zawawi Z, Kamel KA, Ellan K, Kalyanasundram J, Mohd-Zain R, Thayan R |
| EPI_ISL_459962, EPI_ISL_459963, EPI_ISL_459964 | Centogene AG | Centogene AG | Prof. Dr. Peter Bauer, Dr. Krishna Kumar Kandaswamy |
| EPI_ISL_460079 | Molecular Virology Unit, Fondazione IRCCS Policlinico San Matteo , Pavia | Laboratory of Virology, INMI Lazzaro Spallanzani IRCCS | Barbara Bartolini, Cesare E.M. Gruber, Maria R. Capobianchi, Martina Rueca, Antonio Piralla, Fausto Baldanti, Antonino Di Caro |
| EPI_ISL_460080 | Molecular Virology Unit, Fondazione IRCCS Policlinico San Matteo , Pavia | Laboratory of Virology, INMI Lazzaro Spallanzani IRCCS | Antonio Piralla, Barbara Bartolini, Fausto Baldanti, Martina Rueca, Antonino Di Caro, Cesare E.M. Gruber, Maria R. Capobianchi |
| EPI_ISL_460081 | Molecular Virology Unit, Fondazione IRCCS Policlinico San Matteo , Pavia | Laboratory of Virology, INMI Lazzaro Spallanzani IRCCS | Fausto Baldanti, Martina Rueca, Antonio Piralla, Antonino Di Caro, Maria R. Capobianchi, Cesare E.M. Gruber, Barbara Bartolini |
| EPI_ISL_460082 | Molecular Virology Unit, Fondazione IRCCS Policlinico San Matteo , Pavia | Laboratory of Virology, INMI Lazzaro Spallanzani IRCCS | Martina Rueca, Cesare E.M. Gruber, Antonio Piralla, Antonino Di Caro, Barbara Bartolini, Maria R. Capobianchi, Fausto Baldanti |
| EPI_ISL_460083 | Molecular Virology Unit, Fondazione IRCCS Policlinico San Matteo , Pavia | Laboratory of Virology, INMI Lazzaro Spallanzani IRCCS | Martina Rueca, Antonino Di Caro, Cesare E.M. Gruber, Barbara Bartolini, Fausto Baldanti, Antonio Piralla, Maria R. Capobianchi |
| EPI_ISL_460084 | Molecular Virology Unit, Fondazione IRCCS Policlinico San Matteo , Pavia | Laboratory of Virology, INMI Lazzaro Spallanzani IRCCS | Fausto Baldanti, Antonio Piralla, Martina Rueca, Barbara Bartolini, Maria R. Capobianchi, Cesare E.M. Gruber, Antonino Di Caro |
| EPI_ISL_460085 | Molecular Virology Unit, Fondazione IRCCS Policlinico San Matteo , Pavia | Laboratory of Virology, INMI Lazzaro Spallanzani IRCCS | Cesare E.M. Gruber, Maria R. Capobianchi, Barbara Bartolini, Fausto Baldanti, Martina Rueca, Antonio Piralla, Antonino Di Caro |
| EPI_ISL_460086 | Molecular Virology Unit, Fondazione IRCCS Policlinico San Matteo , Pavia | Laboratory of Virology, INMI Lazzaro Spallanzani IRCCS | Maria R. Capobianchi, Fausto Baldanti, Antonio Piralla, Antonino Di Caro, Barbara Bartolini, Cesare E.M. Gruber, Martina Rueca |
| EPI_ISL_460087 | Molecular Virology Unit, Fondazione IRCCS Policlinico San Matteo , Pavia | Laboratory of Virology, INMI Lazzaro Spallanzani IRCCS | Cesare E.M. Gruber, Maria R. Capobianchi, Martina Rueca, Barbara Bartolini, Antonino Di Caro, Antonio Piralla, Fausto Baldanti |
| EPI_ISL_460088 | Molecular Virology Unit, Fondazione IRCCS Policlinico San Matteo , Pavia | Laboratory of Virology, INMI Lazzaro Spallanzani IRCCS | Martina Rueca, Barbara Bartolini, Fausto Baldanti, Maria R. Capobianchi, Cesare E.M. Gruber, Antonino Di Caro, Antonio Piralla |
| EPI_ISL_460089 | Molecular Virology Unit, Fondazione IRCCS Policlinico San Matteo , Pavia | Laboratory of Virology, INMI Lazzaro Spallanzani IRCCS | Antonino Di Caro, Barbara Bartolini, Martina Rueca, Cesare E.M. Gruber, Antonio Piralla, Fausto Baldanti, Maria R. Capobianchi |
| EPI_ISL_460091 | Molecular Virology Unit, Fondazione IRCCS Policlinico San Matteo , Pavia | Laboratory of Virology, INMI Lazzaro Spallanzani IRCCS | Antonino Di Caro, Antonio Piralla, Martina Rueca, Fausto Baldanti, Barbara Bartolini, Maria R. Capobianchi, Cesare E.M. Gruber |
| EPI_ISL_460092 | Molecular Virology Unit, Fondazione IRCCS Policlinico San Matteo , Pavia | Laboratory of Virology, INMI Lazzaro Spallanzani IRCCS | Cesare E.M. Gruber, Martina Rueca, Maria R. Capobianchi, Antonino Di Caro, Antonio Piralla, Barbara Bartolini, Fausto Baldanti |
| EPI_ISL_460093 | Molecular Virology Unit, Fondazione IRCCS Policlinico San Matteo , Pavia | Laboratory of Virology, INMI Lazzaro Spallanzani IRCCS | Maria R. Capobianchi, Antonio Piralla, Antonino Di Caro, Fausto Baldanti, Martina Rueca, Cesare E.M. Gruber, Barbara Bartolini |
| EPI_ISL_460094 | Molecular Virology Unit, Fondazione IRCCS Policlinico San Matteo , Pavia | Laboratory of Virology, INMI Lazzaro Spallanzani IRCCS | Barbara Bartolini, Maria R. Capobianchi, Antonino Di Caro, Antonio Piralla, Cesare E.M. Gruber, Martina Rueca, Fausto Baldanti |
| EPI_ISL_461478 | Government Medical College, Vadodara | Gujarat Biotechnology Research Centre | Fenil Patel, Nidhi Patel, Nitin Savaliya, Raghawendra Kumar, Dinesh Kumar, Zuber Saiyed, Komal Patel, Labdhi Pandya, Snehal Bagatharia, Tanuja Javadekar , R N Daveshwar, Tejas Shah, Ankit Hinsu, Pritesh Sabara, Apurvasinh Puvar, Janvi Raval, Zarna Patel, Monika Gandhi, Pinal Trivedi, Maharshi Pandya, R D Dixit, A M Kadri, Harsh Bakshi, Chaitanya Joshi, Madhvi Joshi, |
| EPI_ISL_461479 | Government Medical College, Vadodara | Gujarat Biotechnology Research Centre | Neelam Nathani, Nitin Savaliya, Raghawendra Kumar, Dinesh Kumar, Zuber Saiyed, Komal Patel, Labdhi Pandya, Snehal Bagatharia, Tanuja Javadekar , R N Daveshwar, Tejas Shah, Ankit Hinsu, Pritesh Sabara, Apurvasinh Puvar, Janvi Raval, Zarna Patel, Monika Gandhi, Pinal Trivedi, Maharshi Pandya, Nidhi Patel, R D Dixit, A M Kadri, Harsh Bakshi, Chaitanya Joshi, Madhvi Joshi, |
| EPI_ISL_461480 | Government Medical College, Vadodara | Gujarat Biotechnology Research Centre | Armi Chaudhari, Raghawendra Kumar, Dinesh Kumar, Zuber Saiyed, Komal Patel, Labdhi Pandya, Snehal Bagatharia, Tanuja Javadekar , R N Daveshwar, Tejas Shah, Ankit Hinsu, Pritesh Sabara, Apurvasinh Puvar, Janvi Raval, Zarna Patel, Monika Gandhi, Pinal Trivedi, Maharshi Pandya, Nidhi Patel, Nitin Savaliya, R D Dixit, A M Kadri, Harsh Bakshi, Chaitanya Joshi, Madhvi Joshi, |
| EPI_ISL_461481 | Pandit Deendayal Upadhyay Government Medical College, Rajkot | Gujarat Biotechnology Research Centre | Bhavya Jindal, Dinesh Kumar, Zuber Saiyed, Komal Patel, Labdhi Pandya, Snehal Bagatharia, Prakash Modi, Sejul Antala, Manish Pattani, Tejas Shah, Ankit Hinsu, Pritesh Sabara, Apurvasinh Puvar, Janvi Raval, Zarna Patel, Monika Gandhi, Pinal Trivedi, Maharshi Pandya, Nidhi Patel, Nitin Savaliya, Raghawendra Kumar, R D Dixit, A M Kadri, Harsh Bakshi, Chaitanya Joshi, Madhvi Joshi |
| EPI_ISL_461482 | Pandit Deendayal Upadhyay Government Medical College, Rajkot | Gujarat Biotechnology Research Centre | Anjali Rajwar, Zuber Saiyed, Komal Patel, Labdhi Pandya, Snehal Bagatharia, Prakash Modi, Sejul Antala, Manish Pattani, Tejas Shah, Ankit Hinsu, Pritesh Sabara, Apurvasinh Puvar, Janvi Raval, Zarna Patel, Monika Gandhi, Pinal Trivedi, Maharshi Pandya, Nidhi Patel, Nitin Savaliya, Raghawendra Kumar, Dinesh Kumar, R D Dixit, A M Kadri, Harsh Bakshi, Chaitanya Joshi, Madhvi Joshi |
| EPI_ISL_462447, EPI_ISL_462448 | Fundació Lluita contra la SIDA (FLSida)/Hospital Universitari Germans Trias i Pujol | IrsiCaixa AIDS Research Lab | Marc Noguera-Julian, Mariona Parera, Maria Pilar Armengol, Marc Corbacho, Maria Ubals, Oriol Mitjà, Lidia Ruiz, Nuria Izquierdo, Jorge Carrillo, Roger Paredes, Julia Blanco, Joaquim Segalés, Bonaventura Clotet |
| EPI_ISL_462449 | Fundació Lluita contra la SIDA (FLSida)/Hospital Universitari Germans Trias i Pujol | IrsiCaixa AIDS Research Lab | Marc Noguera-Julian, Mariona Parera, Maria Pilar Armengol, Marc Corbacho, Maria Ubals, Oriol Mitjà, Lidia Ruiz, Nuria Izquierdo, Jorge Carrillo, Roger Paredes, Julia Blanco, Bonaventura Clotet |

| | | | |
|---|---|---|---|
| EPI_ISL_462478 | Fundación Jiménez Díaz | Instituto de Salud Carlos III | Iglesias-Caballero, M. Molinero Calamita, M. González-Esguevillas, M. Camarero, S. Pozo, F. Casas, I. Jiménez, P. Jiménez, M. Zaballos, A. Monzón, S. Varona, S. Juliá, M. Cuesta, I, R. Fernández |
| EPI_ISL_462479 | Hospital Clinic | Instituto de Salud Carlos III | Iglesias-Caballero, M. Molinero Calamita, M. González-Esguevillas, M. Camarero, S. Pozo, F. Casas, I. Jiménez, P. Jiménez, M. Zaballos, A. Monzón, S. Varona, S. Juliá, M. Cuesta, I, M.A Marcos |
| EPI_ISL_463007 | Department of Laboratory Medicine, National Taiwan University Hospital | Microbial Genomics Core Lab, National Taiwan University Centers of Genomic and Precision Medicine | Shiou-Hwei Yeh, You-Yu Lin, Ya-Yun Lai, Chiao-Ling Li, Shan-Chwen Chang, Pei-Jer Chen, Sui-Yuan Chang |
| EPI_ISL_463008 | Institute of Molecular Virology, University Münster | Institute of Molecular Virology, University Münster | Angeles Mecate Zambrano, Linda Brunotte, Stephan Ludwig, Joachim Kühn, Alexander Mellmann |
| EPI_ISL_463277, EPI_ISL_463278, EPI_ISL_463279, EPI_ISL_463280, EPI_ISL_463281, EPI_ISL_463282, EPI_ISL_463283, EPI_ISL_463284, EPI_ISL_463285, EPI_ISL_463286, EPI_ISL_463287, EPI_ISL_463288, EPI_ISL_463289, EPI_ISL_463290, EPI_ISL_463291, EPI_ISL_463292, EPI_ISL_463293, EPI_ISL_463294, EPI_ISL_463295, EPI_ISL_463296, EPI_ISL_463297, EPI_ISL_463298, EPI_ISL_463299, EPI_ISL_463300 | | | |
| see above | Ochsner Health | Bioinfoexperts, LLC | Susanna L. Lamers, David J. Nolan, Rebecca Rose, Sissy Cross, David Moraga Amador, Tong Yang, Luke Caruso, Wayra Navia, Lydia Von Borstel, Xiao Hui Zhou, Amy Feehan, Julia-Garcia-Diaz |
| EPI_ISL_463301 | Tuen Mun Hospital | Hong Kong Department of Health | Mak Gannon C.K., Cheng Peter K.C., Lam Edman T.K., Chan Rickjason C.W., Tsang Dominic N.C. |
| EPI_ISL_463302, EPI_ISL_463303 | United Christian Hospital | Hong Kong Department of Health | Mak Gannon C.K., Cheng Peter K.C., Lam Edman T.K., Chan Rickjason C.W., Tsang Dominic N.C. |
| EPI_ISL_463304 | Mrs Wu York Yu GOPC | Hong Kong Department of Health | Mak Gannon C.K., Cheng Peter K.C., Lam Edman T.K., Chan Rickjason C.W., Tsang Dominic N.C. |
| EPI_ISL_463305 | Queen Elizabeth Hospital | Hong Kong Department of Health | Mak Gannon C.K., Cheng Peter K.C., Lam Edman T.K., Chan Rickjason C.W., Tsang Dominic N.C. |
| EPI_ISL_463306 | Prince of Wales Hospital | Hong Kong Department of Health | Mak Gannon C.K., Cheng Peter K.C., Lam Edman T.K., Chan Rickjason C.W., Tsang Dominic N.C. |
| EPI_ISL_463307 | HK Molecular Pathology Disgnostic Centre | Hong Kong Department of Health | Mak Gannon C.K., Cheng Peter K.C., Lam Edman T.K., Chan Rickjason C.W., Tsang Dominic N.C. |
| EPI_ISL_463309 | Prince of Wales Hospital | Hong Kong Department of Health | Mak Gannon C.K., Cheng Peter K.C., Lam Edman T.K., Chan Rickjason C.W., Tsang Dominic N.C. |
| EPI_ISL_463310 | Tseung Kwan O Hospital | Hong Kong Department of Health | Mak Gannon C.K., Cheng Peter K.C., Lam Edman T.K., Chan Rickjason C.W., Tsang Dominic N.C. |
| EPI_ISL_463311, EPI_ISL_463312 | Queen Elizabeth Hospital | Hong Kong Department of Health | Mak Gannon C.K., Cheng Peter K.C., Lam Edman T.K., Chan Rickjason C.W., Tsang Dominic N.C. |
| EPI_ISL_463313 | Central Health Medical Practice | Hong Kong Department of Health | Mak Gannon C.K., Cheng Peter K.C., Lam Edman T.K., Chan Rickjason C.W., Tsang Dominic N.C. |
| EPI_ISL_463314 | Queen Elizabeth Hospital | Hong Kong Department of Health | Mak Gannon C.K., Cheng Peter K.C., Lam Edman T.K., Chan Rickjason C.W., Tsang Dominic N.C. |
| EPI_ISL_463315 | Princess Margaret Hospital | Hong Kong Department of Health | Mak Gannon C.K., Cheng Peter K.C., Lam Edman T.K., Chan Rickjason C.W., Tsang Dominic N.C. |
| EPI_ISL_463316, EPI_ISL_463317 | Yan Chai Hospital | Hong Kong Department of Health | Mak Gannon C.K., Cheng Peter K.C., Lam Edman T.K., Chan Rickjason C.W., Tsang Dominic N.C. |
| EPI_ISL_463318 | Pamela Youde Nethersole Eastern Hospital | Hong Kong Department of Health | Mak Gannon C.K., Cheng Peter K.C., Lam Edman T.K., Chan Rickjason C.W., Tsang Dominic N.C. |
| EPI_ISL_463319 | Prince of Wales Hospital | Hong Kong Department of Health | Mak Gannon C.K., Cheng Peter K.C., Lam Edman T.K., Chan Rickjason C.W., Tsang Dominic N.C. |
| EPI_ISL_463320 | Tseung Kwan O Hospital | Hong Kong Department of Health | Mak Gannon C.K., Cheng Peter K.C., Lam Edman T.K., Chan Rickjason C.W., Tsang Dominic N.C. |
| EPI_ISL_463321 | Queen Mary Hospital | Hong Kong Department of Health | Mak Gannon C.K., Cheng Peter K.C., Lam Edman T.K., Chan Rickjason C.W., Tsang Dominic N.C. |
| EPI_ISL_463322 | North Lantau Hospital | Hong Kong Department of Health | Mak Gannon C.K., Cheng Peter K.C., Lam Edman T.K., Chan Rickjason C.W., Tsang Dominic N.C. |
| EPI_ISL_463323 | Yuen Long JC GOPC | Hong Kong Department of Health | Mak Gannon C.K., Cheng Peter K.C., Lam Edman T.K., Chan Rickjason C.W., Tsang Dominic N.C. |
| EPI_ISL_463741, EPI_ISL_463742, EPI_ISL_463743, EPI_ISL_463744, EPI_ISL_463745, EPI_ISL_463746, EPI_ISL_463747, EPI_ISL_463748 | Department of Molecular Virology, Cyprus Institute of Neurology and Genetics | Department of Molecular Virology, Cyprus Institute of Neurology and Genetics | Jan Richter, George Krashias, Christina Tryfonos, Stavros Bashiardes, Dana Koptides, Christina Christodoulou |
| EPI_ISL_467300 | General Hospital "Abdulah Nakas" | Alea Genetic Center | Rijad Konjhodzic; Lana Salihefendic; Teufik Goletic; Sead Jazic; Dino Pecar; Nihad Fejzic; Damir Marjanovic; Enis Kandic |
| EPI_ISL_467352, EPI_ISL_467353, EPI_ISL_467367, EPI_ISL_467369 | Laboratory of Respiratory Viruses and Measles, Oswaldo Cruz Institute, FIOCRUZ | Laboratory of Respiratory Viruses and Measles, Oswaldo Cruz Institute, FIOCRUZ | Paola Resende, Luciana Appolinario, Fernando Motta, Anna Carolina Paixão, Ana Carolina Mendonça, Aline Mattos, Milene Miranda, Cristiana Garcia, Braulia Caetano, Maria Ogrzewalska, Jonathan Lopes, Marilda Siqueira on behalf of the Fiocruz COVID-19 Genomic Surveillance Network |
| EPI_ISL_467374 | Dinkes Samarinda | Eijkman Institute for Molecular Biology, Ministry of Research and Technology/National Agency for Research and Innovation | Edison Johar, Frilasita A Yudhaputri, Hidayat Trimarsanto, David H Muljono, Safarina G Malik, Khin Saw Myint, Amin Soebandrio |
| EPI_ISL_467375 | RSUP Prof. Dr. R. Kandou Manado | Eijkman Institute for Molecular Biology, Ministry of Research and Technology/National Agency for Research and Innovation | Edison Johar, Frilasita A Yudhaputri, Hidayat Trimarsanto, David H Muljono, Safarina G Malik, Khin Saw Myint, Amin Soebandrio |
| EPI_ISL_467376 | RSUP Fatmawati | Eijkman Institute for Molecular Biology, Ministry of Research and Technology/National Agency for Research and Innovation | Edison Johar, Frilasita A Yudhaputri, Hidayat Trimarsanto, David H Muljono, Safarina G Malik, Khin Saw Myint, Amin Soebandrio |
| EPI_ISL_467774, EPI_ISL_467775 | Molecular diagnostic laboratory of Federal Budget Institution of Science "Central Research Institute of Epidemiology" of The Federal Service on Customers' Rights Protection and Human Well-being Surveillance | Group of Genomics and Postgenomic Technologies of Central Research Institute of Epidemiology | Speranskaya AS, Kaptelova VV, Samoilov AE, Korneenko EV, Sizova TV, Tivanova EV, Shipulina OY, Akimkin VG |
| EPI_ISL_467809 | Cedars-Sinai Medical Center, Department of Pathology & Laboratory Medicine, Molecular Pathology Laboratory | Cedars-Sinai Medical Center, Molecular Pathology Laboratory of Department of Pathology & Laboratory Medicine and Genomic Core | Wenjuan Zhang, John Paul Govindavari, Brian Davis, Stephanie Chen, Jong Taek Kim, Jianbo Song, Jean Lopategui, Jasmine T Plummer, Eric Vail |
| EPI_ISL_468134, EPI_ISL_468135, EPI_ISL_468136, EPI_ISL_468137, EPI_ISL_468138, EPI_ISL_468139, EPI_ISL_468140, EPI_ISL_468141, EPI_ISL_468142, EPI_ISL_468143, EPI_ISL_468144, EPI_ISL_468145, EPI_ISL_468146, EPI_ISL_468147, EPI_ISL_468148, EPI_ISL_468149, EPI_ISL_468150, EPI_ISL_468151, EPI_ISL_468153, EPI_ISL_468154, EPI_ISL_468155, EPI_ISL_468156 | | | |
| see above | [Romania, Bucharest] National Institute for Infectious Diseases "Prof. Dr. Matei Bal" | [Romania, Bucharest] National Institute for Infectious Diseases "Prof. Dr. Matei Bal" | Leontina Banica, Marius Cotic, Corina Casangiu, Marius Surleac, Simona Paraschiv |
| EPI_ISL_468305, EPI_ISL_468307 | Centro de Vigilancia a Saude de Diadema | Instituto Adolfo Lutz, Interdiciplinary Procedures Center, Strategic Laboratory | Claudio Tavares Sacchi, Claudia Regina Gonçalves, Erica Valessa Ramos Gomes |
| EPI_ISL_468308 | Hospital Municipal do Tatuape Carmino Caricchio | Instituto Adolfo Lutz, Interdiciplinary Procedures Center, Strategic Laboratory | Claudio Tavares Sacchi, Claudia Regina Gonçalves, Erica Valessa Ramos Gomes |
| EPI_ISL_468310 | Hospital Sao Paulo de Ensino da UNIFESP | Instituto Adolfo Lutz, Interdiciplinary Procedures Center, Strategic Laboratory | Claudio Tavares Sacchi, Claudia Regina Gonçalves, Erica Valessa Ramos Gomes |
| EPI_ISL_468311, EPI_ISL_468312 | Hospital Municipal Dr Ignacio Proenca de Gouvea | Instituto Adolfo Lutz, Interdiciplinary Procedures Center, Strategic Laboratory | Claudio Tavares Sacchi, Claudia Regina Gonçalves, Erica Valessa Ramos Gomes |
| EPI_ISL_468313 | Vigilancia Epidemiologica de São Bernardo do Campo | Instituto Adolfo Lutz, Interdiciplinary Procedures Center, Strategic Laboratory | Claudio Tavares Sacchi, Claudia Regina Gonçalves, Erica Valessa Ramos Gomes |
| EPI_ISL_468314 | CTA Centro de Testagem e Aconselhamento | Instituto Adolfo Lutz, Interdiciplinary Procedures Center, | Claudio Tavares Sacchi, Claudia Regina Gonçalves, Erica Valessa Ramos Gomes |

| | | | |
|---|---|---|---|
| | | Strategic Laboratory | |
| EPI_ISL_468315 | Hospital Municipal do Tatuape Carmino Caricchio | Instituto Adolfo Lutz, Interdiciplinary Procedures Center, Strategic Laboratory | Claudio Tavares Sacchi, Claudia Regina Gonçalves, Erica Valessa Ramos Gomes |
| EPI_ISL_468316 | UPA Vila Assis | Instituto Adolfo Lutz, Interdiciplinary Procedures Center, Strategic Laboratory | Claudio Tavares Sacchi, Claudia Regina Gonçalves, Erica Valessa Ramos Gomes |
| EPI_ISL_468318 | Hospital Universitario da USP | Instituto Adolfo Lutz, Interdiciplinary Procedures Center, Strategic Laboratory | Claudio Tavares Sacchi, Claudia Regina Gonçalves, Erica Valessa Ramos Gomes |
| EPI_ISL_468319 | Vigilancia Epidemiologica de São Bernardo do Campo | Instituto Adolfo Lutz, Interdiciplinary Procedures Center, Strategic Laboratory | Claudio Tavares Sacchi, Claudia Regina Gonçalves, Erica Valessa Ramos Gomes |
| EPI_ISL_468320 | Secretaria Municipal de Saude de Hortolandia | Instituto Adolfo Lutz, Interdiciplinary Procedures Center, Strategic Laboratory | Claudio Tavares Sacchi, Claudia Regina Gonçalves, Erica Valessa Ramos Gomes |
| EPI_ISL_468321 | Hospital Universitario da USP | Instituto Adolfo Lutz, Interdiciplinary Procedures Center, Strategic Laboratory | Claudio Tavares Sacchi, Claudia Regina Gonçalves, Erica Valessa Ramos Gomes |
| EPI_ISL_468591, EPI_ISL_468656 | Institute for Public Health | Laboratory for advanced genomics | Filip Roki, Lovro Trgovec-Greif, Neven Sui, Tomislav Rukavina, Igor Jurak, Oliver Vugrek |
| EPI_ISL_468765, EPI_ISL_468766, EPI_ISL_468768, EPI_ISL_468771, EPI_ISL_468772, EPI_ISL_468774, EPI_ISL_468775, EPI_ISL_468776, EPI_ISL_468780, EPI_ISL_468782, EPI_ISL_468788, EPI_ISL_468791, EPI_ISL_468793, EPI_ISL_468795, EPI_ISL_468798, EPI_ISL_468800, EPI_ISL_468801, EPI_ISL_468804, EPI_ISL_468805, EPI_ISL_468808, EPI_ISL_468809, EPI_ISL_468810, EPI_ISL_468812, EPI_ISL_468814, EPI_ISL_468815, EPI_ISL_468817, EPI_ISL_468820, EPI_ISL_468829, EPI_ISL_468830, EPI_ISL_468832, EPI_ISL_468834, EPI_ISL_468835, EPI_ISL_468841, EPI_ISL_468849, EPI_ISL_468851, EPI_ISL_468855, EPI_ISL_468858, EPI_ISL_468859 | | | |
| see above | Servicio de Microbiología, Hospital Miguel Servet, Zaragoza | SeqCOVID-SPAIN consortium/IBV(CSIC) | Antonio Rezusta López, Alexander Tristancho Baró, Ana Milagro, Yolanda Gracia Grataloup, Nieves Martínez Cameo and SeqCOVID-SPAIN consortium |
| EPI_ISL_468914, EPI_ISL_469016, EPI_ISL_469018, EPI_ISL_469019, EPI_ISL_469020, EPI_ISL_469021, EPI_ISL_469022 | Istituto Zooprofilattico Sperimentale Puglia e Basilicata; Dipartimento di Bioscienze, Biotecnologie e Biofarmaceutica dell'Università degli Studi di Bari "A.Moro"; Istituto di Biomembrane. Bioenergetica e Biotecnologie Molecolari del Consiglio Nazionale delle Ricerche di Bari | Beaconlab (Bioinformatics, Evolution and Comparative Genomics lab), Dept of Biosciences, University on Milan | Parisi A.,Pesole G., Manzari C., Chiara M. |
| EPI_ISL_469023 | Istituto Zooprofilattico Sperimentale Puglia e Basilicata; Dipartimento di Bioscienze, Biotecnologie e Biofarmaceutica dell'Università degli Studi di Bari "A.Moro"; Istituto di Biomembrane. Bioenergetica e Biotecnologie Molecolari del Consiglio Nazionale delle Ricerche di Bari | Beaconlab (Bioinformatics, Evolution and Comparative Genomics lab), Dept of Biosciences, University on Milan | Parisi A.,Pesole G., Manzari C., Chiara M |
| EPI_ISL_469275 | Egyptian National Cancer Institute (ENCI) | Human Genome Center | Zekri, Abdel Rahman N, Amer,K.E., Ahmed,O.S., Soliman,H.K., Hafez,M.M., Bahnassy,A.A., Abdelhamid,W., Gad,A., Ali,M., Hassan,W., Samir,M., Raouf,A., Hamdy,M.S., Soliman,M.S., Elsissy,M.H., Elkhateeb,S.M., Ezzelarab,M.H., Abouelhoda, Mohamed |
| EPI_ISL_469282 | Service de Virologie Hôpital Saint-Louis | Laboratory of Cell Biology of viral infection, Unit INSERM-U944 | Laurent Meertens, Lucie Bonnet-Madin, Constance Delaugerre, Ali Amara |
| EPI_ISL_469283, EPI_ISL_469284 | Service de Virologie Hôpital Saint-Louis | Laboratory Cell Biology of Viral Infection-INSERM unit 944 | Laurent Meertens, Lucie Bonnet-Madin, Séverine Mercier-Delarue, Maud SALMONA, Constance Delaugerre, Ali Amara |
| EPI_ISL_470896 | Russian State Collection of Viruses | Pathogenic Microorganisms Variability Laboratory | Alexey Shchetinin, Maria Nikiforova, Elena Shidlovskaya, Nadezhda Kuznetsova, Inna Dolzhikova, Daria Grousova, Andrey Botikov, Denis Logunov, Alexander Gintsburg, Vladimir Gushchin |
| EPI_ISL_470897, EPI_ISL_470898, EPI_ISL_470899 | Pathogenic Microorganisms Variability Laboratory | Pathogenic Microorganisms Variability Laboratory | Alexey Shchetinin, Maria Nikiforova, Elena Shidlovskaya, Nadezhda Kuznetsova, Andrey Botikov, Alexander Gintsburg, Vladimir Gushchin |
| EPI_ISL_470900, EPI_ISL_470901, EPI_ISL_470902 | Influenza etiology and epidemiology laboratory | Pathogenic Microorganisms Variability Laboratory | Alexey Shchetinin, Maria Nikiforova, Elena Shidlovskaya, Nadezhda Kuznetsova, Vladimir Gushchin, Inna Dolzhikova, Daria Grousova, Andrey Botikov, Denis Logunov, Kirill Krasnoslobotsev, Svetlana Trushakova, Elena Burtseva, Ludmila Kolobukhina, Svetlana Smetanina, Alexander Gintsburg |
| EPI_ISL_470903, EPI_ISL_470904 | Influenza etiology and epidemiology laboratory | Pathogenic Microorganisms Variability Laboratory | Alexey Shchetinin, Maria Nikiforova, Elena Shidlovskaya, Nadezhda Kuznetsova, Vladimir Gushchin, Inna Dolzhikova, Daria Grousova, Andrey Botikov, Denis Logunov, Anna Ignatjeva, Evgeniya Mukasheva, Elena Burtseva, Ludmila Kolobukhina, Svetlana Smetanina, Alexander Gintsburg |
| EPI_ISL_471158, EPI_ISL_471163, EPI_ISL_471164, EPI_ISL_471167, EPI_ISL_471171 | MRCG at LSHTM Genomics lab | MRCG at LSHTM Genomics lab | Sesay et al |
| EPI_ISL_471267 | Hospital IESS Babahoyo | Institute of Microbiology, Universidad San Francisco de Quito | Sully Márquez, Belén Prado-Vivar, Juan José Guadalupe, Bernardo Gutiérrez, Francisco Cordova, Ninfa Henriquez, Killen Briones-Zamora, Killen Briones-Claudette, Verónica Barragán, Patricio Rojas-Silva, Gabriel Trueba, Michelle Grunauer, Paúl Cárdenas |
| EPI_ISL_471456, EPI_ISL_471457, EPI_ISL_471458, EPI_ISL_471459, EPI_ISL_471460 | Centre de Virologie des Maladies Tropicales | Functional Genomic Platform/Service Analyses Biologique/UATRS/ Centre National Pour la Recherche Scientifique Et Technique (CNRST) | Hicham EL ANNAZ, Elmostafa EL FAHIME, Marouane MELLOUL, Youssef AKHOUAD, Mly Abdelaziz ELALAOUI, Ahmed REGGAD, Sanaa ALAOUI-Amine, Rachid ABI, Rida TAGAJDID, Zohour KASMY, Safae ELKOCHRI, Nadia TOUIL, Farida HILALI, Abdelkader LAATIRIS, Abdelillah LARAQUI, Tahar BAJJOU , Yassine SEKHSOKH , Idriss-Amine LAHLOU, Mostafa ELOUENNASS, Khalid ENNIBI |
| EPI_ISL_471472 | Hospital Universitari Germans Trias i Pujol(HUGTiP)/Fundació Lluita contra la SIDA (FLSida)/IRTA-CReSA | IrsiCaixa AIDS Research Lab | Marc Noguera-Julian, Pilar Armengol, Jordi Rodón, Julia Vergara, Lidia Ruiz, Nuria Izquierdo, Jorge Carrillo, Roger Paredes, Albert Bensaid, Julia Blanco, Joaquim Segalés, Bonaventura Clotet |
| EPI_ISL_471528, EPI_ISL_471530 | The National Institute of Public Health | State Veterinary Institute Prague and The National Institute of Public Health | Nagy,A;Jirincova,H;Novakova,L;Trnka,D;Vecerova,J |
| EPI_ISL_471539 | Hospital Universitario da USP Sao Paulo | Instituto Adolfo Lutz, Interdiciplinary Procedures Center, Strategic Laboratory | Claudio Tavares Sacchi, Claudia Regina Gonçalves, Erica Valessa Ramos Gomes |
| EPI_ISL_471540 | The National Institute of Public Health | State Veterinary Institute Prague and The National Institute of Public Health | Nagy,A;Jirincova,H;Novakova,L;Trnka,D;Vecerova,J |
| EPI_ISL_471541 | Hospital Geral Santa Marcelina | Instituto Adolfo Lutz, Interdiciplinary Procedures Center, Strategic Laboratory | Claudio Tavares Sacchi, Claudia Regina Gonçalves, Erica Valessa Ramos Gomes |
| EPI_ISL_471542 | Secretaria de Saude de Mogi das Cruzes | Instituto Adolfo Lutz, Interdiciplinary Procedures Center, Strategic Laboratory | Claudio Tavares Sacchi, Claudia Regina Gonçalves, Erica Valessa Ramos Gomes |
| EPI_ISL_471543 | Centro de Saude I Tacito Leite de Carvalho e Silva | Instituto Adolfo Lutz, Interdiciplinary Procedures Center, Strategic Laboratory | Claudio Tavares Sacchi, Claudia Regina Gonçalves, Erica Valessa Ramos Gomes |
| EPI_ISL_471544 | The National Institute of Public Health | State Veterinary Institute Prague and The National Institute of Public Health | Nagy,A;Jirincova,H;Novakova,L;Trnka,D;Vecerova,J |
| EPI_ISL_471545 | Hospital Sao Paulo de Ensino da Unifesp | Instituto Adolfo Lutz, Interdiciplinary Procedures Center, Strategic Laboratory | Claudio Tavares Sacchi, Claudia Regina Gonçalves, Erica Valessa Ramos Gomes |
| EPI_ISL_471546 | AMA DR Jose Soares Hungria | Instituto Adolfo Lutz, Interdiciplinary Procedures Center, Strategic Laboratory | Claudio Tavares Sacchi, Claudia Regina Gonçalves, Erica Valessa Ramos Gomes |
| EPI_ISL_471547 | The National Institute of Public Health | State Veterinary Institute Prague and The National Institute of Public Health | Nagy,A;Jirincova,H;Novakova,L;Trnka,D;Vecerova,J |

| | | | |
|---|---|---|---|
| EPI_ISL_471548 | Hospital do Servidor Público Estadual Francisco Morato de Oliveira | Instituto Adolfo Lutz, Interdiciplinary Procedures Center, Strategic Laboratory | Claudio Tavares Sacchi, Claudia Regina Gonçalves, Erica Valessa Ramos Gomes |
| EPI_ISL_471549 | Hospital Municipal Carmen Prudente | Instituto Adolfo Lutz, Interdiciplinary Procedures Center, Strategic Laboratory | Claudio Tavares Sacchi, Claudia Regina Gonçalves, Erica Valessa Ramos Gomes |
| EPI_ISL_471550 | The National Institute of Public Health | State Veterinary Institute Prague and The National Institute of Public Health | Nagy,A;Jirincova,H;Novakova,L;Trnka,D;Vecerova,J |
| EPI_ISL_471551 | Hospital Sao Paulo de Ensino da Unifesp | Instituto Adolfo Lutz, Interdiciplinary Procedures Center, Strategic Laboratory | Claudio Tavares Sacchi, Claudia Regina Gonçalves, Erica Valessa Ramos Gomes |
| EPI_ISL_471552 | Hospital Sancta Maggiore | Instituto Adolfo Lutz, Interdiciplinary Procedures Center, Strategic Laboratory | Claudio Tavares Sacchi, Claudia Regina Gonçalves, Erica Valessa Ramos Gomes |
| EPI_ISL_471553 | The National Institute of Public Health | State Veterinary Institute Prague and The National Institute of Public Health | Nagy,A;Jirincova,H;Novakova,L;Trnka,D;Vecerova,J |
| EPI_ISL_471554 | Hospital Bosque da Saúde | Instituto Adolfo Lutz, Interdiciplinary Procedures Center, Strategic Laboratory | Claudio Tavares Sacchi, Claudia Regina Gonçalves, Erica Valessa Ramos Gomes |
| EPI_ISL_471555 | The National Institute of Public Health | State Veterinary Institute Prague and The National Institute of Public Health | Nagy,A;Jirincova,H;Novakova,L;Trnka,D;Vecerova,J |
| EPI_ISL_471556 | Pronto Socorro Jose Ibrahin | Instituto Adolfo Lutz, Interdiciplinary Procedures Center, Strategic Laboratory | Claudio Tavares Sacchi, Claudia Regina Gonçalves, Erica Valessa Ramos Gomes |
| EPI_ISL_471562, EPI_ISL_471581, EPI_ISL_471582 | Hosp. Municipal Prof. Dr. Alípio Corrêa Netto | Instituto Adolfo Lutz, Interdiciplinary Procedures Center, Strategic Laboratory | Claudio Tavares Sacchi, Claudia Regina Gonçalves, Erica Valessa Ramos Gomes |
| EPI_ISL_471647 | Hospital Municipal de Barueri Dr. Francisco Moran | Instituto Adolfo Lutz, Interdiciplinary Procedures Center, Strategic Laboratory | Claudio Tavares Sacchi, Claudia Regina Gonçalves, Erica Valessa Ramos Gomes |
| EPI_ISL_471648 | UBS e Pronto Socorro Jd. Jacira | Instituto Adolfo Lutz, Interdiciplinary Procedures Center, Strategic Laboratory | Claudio Tavares Sacchi, Claudia Regina Gonçalves, Erica Valessa Ramos Gomes |
| EPI_ISL_475574, EPI_ISL_475576, EPI_ISL_475577, EPI_ISL_475578, EPI_ISL_475579, EPI_ISL_475580, EPI_ISL_475582, EPI_ISL_475583, EPI_ISL_475585, EPI_ISL_475588, EPI_ISL_475590, EPI_ISL_475593, EPI_ISL_475594, EPI_ISL_475595, EPI_ISL_475596, EPI_ISL_475598, EPI_ISL_475599, EPI_ISL_475600, EPI_ISL_475601, EPI_ISL_475602, EPI_ISL_475603, EPI_ISL_475604, EPI_ISL_475605, EPI_ISL_475606, EPI_ISL_475607, EPI_ISL_475608, EPI_ISL_475609, EPI_ISL_475610, EPI_ISL_475611, EPI_ISL_475612, EPI_ISL_475613, EPI_ISL_475614, EPI_ISL_475615, EPI_ISL_475616, EPI_ISL_475617, EPI_ISL_475618, EPI_ISL_475619, EPI_ISL_475621, EPI_ISL_475622, EPI_ISL_475625, EPI_ISL_475627, EPI_ISL_475628, EPI_ISL_475629, EPI_ISL_475630, EPI_ISL_475631, EPI_ISL_475632, EPI_ISL_475633, EPI_ISL_475634, EPI_ISL_475635, EPI_ISL_475636, EPI_ISL_475637, EPI_ISL_475638, EPI_ISL_475639, EPI_ISL_475640, EPI_ISL_475641, EPI_ISL_475643, EPI_ISL_475644, EPI_ISL_475645, EPI_ISL_475646, EPI_ISL_475647, EPI_ISL_475648, EPI_ISL_475650, EPI_ISL_475651, EPI_ISL_475652, EPI_ISL_475653, EPI_ISL_475654, EPI_ISL_475655, EPI_ISL_475656, EPI_ISL_475657, EPI_ISL_475658, EPI_ISL_475659, EPI_ISL_475660, EPI_ISL_475661, EPI_ISL_475662, EPI_ISL_475663, EPI_ISL_475664, EPI_ISL_475666, EPI_ISL_475667, EPI_ISL_475668, EPI_ISL_475669, EPI_ISL_475670, EPI_ISL_475671, EPI_ISL_475673, EPI_ISL_475674, EPI_ISL_475675, EPI_ISL_475677, EPI_ISL_475678, EPI_ISL_475679, EPI_ISL_475680, EPI_ISL_475681, EPI_ISL_475682, EPI_ISL_475683, EPI_ISL_475684, EPI_ISL_475685, EPI_ISL_475686, EPI_ISL_475687, EPI_ISL_475688, EPI_ISL_475689, EPI_ISL_475690, EPI_ISL_475691, EPI_ISL_475692, EPI_ISL_475694, EPI_ISL_475695, EPI_ISL_475697, EPI_ISL_475698, EPI_ISL_475699, EPI_ISL_475700, EPI_ISL_475701, EPI_ISL_475702, EPI_ISL_475703, EPI_ISL_475705, EPI_ISL_475706, EPI_ISL_475708, EPI_ISL_475709, EPI_ISL_475710, EPI_ISL_475711, EPI_ISL_475712, EPI_ISL_475713, EPI_ISL_475714, EPI_ISL_475715, EPI_ISL_475716 | | | |
| see above | Cedars-Sinai Medical Center, Department of Pathology & Laboratory Medicine, Molecular Pathology Laboratory | Cedars-Sinai Medical Center, Molecular Pathology Laboratory of Department of Pathology & Laboratory Medicine and Genomic Core | Wenjuan Zhang, John Paul Govindavari, Brian Davis, Stephanie Chen, Jong Taek Kim, Jianbo Song, Jean Lopategui, Jasmine T Plummer, Eric Vail |
| EPI_ISL_475745, EPI_ISL_475746, EPI_ISL_475747, EPI_ISL_475748, EPI_ISL_475749, EPI_ISL_475751, EPI_ISL_475752, EPI_ISL_475753 | Medical Ain Shams Research Institute (MASRI), Ain Shams University | Medical Ain Shams Research Institute (MASRI), Ain Shams University | Hesham Elghazaly , Sara Hassan Agwa, Mahmoud Elmeteini , Ahmad Moustafa , Ashraf Omar, Osama Mansour, Samia Abdo, Hala Hafez, Ghada Ismael , Shaimaa Moustafa , Aya Mohamed, Reham Mamdouh , Hoda Abd Elsatar, Manal Hamdy Elsaid, Fatma Ebied |
| EPI_ISL_476022 | Defence Research & Development Establishment | Defence Research & Development Establishment | Shashi Sharma, Paban Kumar Dash, Jyoti S Kumar, Sushil Kumar Sharma, Ambuj Shrivastava |
| EPI_ISL_476023 | Defence Research & Development Establishment (DRDE) | Defence Research & Development Establishment (DRDE) | Shashi Sharma, Paban Kumar Dash, Sushil Kumar Sharma, Ambuj Shrivastava, Jyoti S. Kumar |
| EPI_ISL_476067 | The National Institute of Public Health | State Veterinary Institute Prague and The National Institute of Public Health | Nagy,A;Jirincova,H;Novakova,L;Trnka,D;Vecerova,J |
| EPI_ISL_476702, EPI_ISL_476704 | Incubadora Venezolana de Ciencia, Venezuela | Incubadora Venezolana de Ciencia, Venezuela / Instituto Nacional de Salud, Bogotá, Colombia / Grupo de Investigaciones Microbiológicas-UR (GIMUR), Departamento de Biología, Facultad de Ciencias Naturales, Universidad del Rosario, Bogotá, Colombia / Icahn School of Medicine at Mount Sinai, New York, USA | Alberto Paniz-Mondolfi, Marina Muñoz, Luis Perez-Garcia, Lourdes Delgado, Carolina Florez, Sergio Gomez, Angelica Rico, Lisseth Pardo, Esther C. Barros, Carolina Hernández, Jesús E. Jaimes, Aníbal A. Teherán, Ana S. Gonzalez-Reiche, Matthew M. Hernandez, Emilia Mia Sordillo, Viviana Simon, Harm van Bakel, Juan David Ramírez |
| EPI_ISL_476801, EPI_ISL_476802, EPI_ISL_476803, EPI_ISL_476804 | Hong Kong Department of Health | School of Public Health, The University of Hong Kong | Dominic N.C. Tsang, Daniel K.W. Chu, Leo L.M. Poon, Malik Peiris |
| EPI_ISL_476814, EPI_ISL_476818, EPI_ISL_476819 | Department of Laboratory Medicine, Tan Tock Seng Hospital | Department of Laboratory Medicine, Tan Tock Seng Hospital | Chen YYC, Zair X, Li C, Tang WY, Maurer-Stroh S, Barkham TMS, Nagarajan N, Sessions OM |
| EPI_ISL_476832 | Medical Biology Department, Kocaeli University | Medical Genetics Department, Kocaeli University | Savli H, Cine N, Sunnetci-Akkoyunlu D, Eren-Keskin S, Ilgazli A, Akhan S, Karadenizli A, Kasap M, Sayan M, Akpinar G, Canturk NZ. |
| EPI_ISL_476840, EPI_ISL_476844, EPI_ISL_476846, EPI_ISL_476848, EPI_ISL_476849, EPI_ISL_476850, EPI_ISL_476852, EPI_ISL_476853, EPI_ISL_476854, EPI_ISL_476883 | Defence Research & Development Establishment (DRDE) | Defence Research & Development Establishment (DRDE) | Shashi Sharma, Paban Kumar Dash, Sushil Kumar Sharma, Ambuj Shrivastava, Jyoti S. Kumar |
| EPI_ISL_477163 | Laboratory of Dr. John Lednicky | University of Florida | John A. Lednicky, Maha A. Elbadry, Kuttichantran Subramaniam, Thomas B. Waltzek, John Glenn Morris, Jr. |
| EPI_ISL_477164 | Department of Virology, Public Health Laboratories Division, National Institute of Health | Department of Virology, Public Health Laboratories Division, National Institute of Health | Nazish Badar, Aamer Ikram, Muhammad Salman, Hamza Ahmed Mirza, Abdul Ahad, Yasir Arshad, Massab Umair |
| EPI_ISL_477165, EPI_ISL_477167 | Department of Virology, Public Health Laboratories Division, National Institute of Health | Department of Virology, Public Health Laboratories Division, National Institute of Health | Nazish Badar,Aamer Ikram, Muhammad Salman, Massab Umair, Hamza Ahmed Mirza, Abdul Ahad, Yasir Arshad |
| EPI_ISL_477170 | Department of Laboratory, Medicine Tan Tock Seng Hospital | Department of Laboratory Medicine Tan Tock Seng Hospital | Chen YYC, Zair X, Li C, Tang WY, Maurer-Stroh S, Barkham TMS, Nagarajan N, Sessions OM |
| EPI_ISL_477171 | Department of Laboratory, Medicine Tan Tock Seng Hospital | Department of Laboratory, Medicine Tan Tock Seng Hospital | Chen YYC, Zair X, Li C, Tang WY, Maurer-Stroh S, Barkham TMS, Nagarajan N, Sessions OM |
| EPI_ISL_477172, EPI_ISL_477174, EPI_ISL_477175, EPI_ISL_477177, EPI_ISL_477178, EPI_ISL_477182, EPI_ISL_477184, EPI_ISL_477187, EPI_ISL_477188, EPI_ISL_477189, EPI_ISL_477190, EPI_ISL_477191, EPI_ISL_477192 | | | |
| see above | Department of Laboratory Medicine Tan Tock Seng Hospital | Department of Laboratory Medicine Tan Tock Seng Hospital | Chen YYC, Zair X, Li C, Tang WY, Maurer-Stroh S, Barkham TMS, Nagarajan N, Sessions OM |
| EPI_ISL_477193, EPI_ISL_477194, EPI_ISL_477202 | Istituto Zooprofilattico Sperimentale Puglia e Basilicata; | Beaconlab (Bioinformatics, Evolution and Comparative Genomics lab), Dept of Biosciences, University on Mila | Parisi A.,Pesole G., Manzari C., Chiara M. |
| EPI_ISL_479482, EPI_ISL_479483, EPI_ISL_479484, EPI_ISL_479485, EPI_ISL_479486, EPI_ISL_479487, EPI_ISL_479488, EPI_ISL_479489, EPI_ISL_479490, EPI_ISL_479491, EPI_ISL_479492 | | | |

| | | | |
|---|---|---|---|
| see above | Department of Laboratory Medicine Tan Tock Seng Hospital | Department of Laboratory Medicine Tan Tock Seng Hospital | Chen YYC, Zair X, Li C, Tang WY, Maurer-Stroh S, Barkham TMS, Nagarajan N, Sessions OM |
| EPI_ISL_479493, EPI_ISL_479495, EPI_ISL_479498, EPI_ISL_479499, EPI_ISL_479501, EPI_ISL_479502, EPI_ISL_479505, EPI_ISL_479507, EPI_ISL_479509, EPI_ISL_479510, EPI_ISL_479512, EPI_ISL_479513, EPI_ISL_479514, EPI_ISL_479516, EPI_ISL_479517, EPI_ISL_479519, EPI_ISL_479520, EPI_ISL_479521, EPI_ISL_479522, EPI_ISL_479523, EPI_ISL_479526, EPI_ISL_479527, EPI_ISL_479528, EPI_ISL_479529, EPI_ISL_479530, EPI_ISL_479531, EPI_ISL_479532, EPI_ISL_479533, EPI_ISL_479572 | | | |
| see above | NIV Influenza | NIV Influenza | Potdar V |
| EPI_ISL_479616, EPI_ISL_479617 | Laboratory of Molecular Virology of the International Centre for Genetic Engineering and Biotechnology (ICGEB) | ARGO Open Lab Platform for Genome Sequencing | Licastro, D, Rajasekharan S, Dal Monego S, Segat L, D'Agaro P, Salton F, Confalonieri P, Confalonieri M Marcello A |
| EPI_ISL_479618, EPI_ISL_479619 | Laboratory of Molecular Virology of the International Centre for Genetic Engineering and Biotechnology (ICGEB) | ARGO Open Lab Platform for Genome Sequencing | Licastro, D, Rajasekharan S, Dal Monego S, Segat L, D'Agaro P, Salton F, Confalonieri P, Confalonieri M, Marcello A |
| EPI_ISL_479620, EPI_ISL_479621, EPI_ISL_479622, EPI_ISL_479623, EPI_ISL_479624 | Molecular diagnostic laboratory of Federal Budget Institution of Science "Central Research Institute of Epidemiology" of The Federal Service on Customers' Rights Protection and Human Well-being Surveillance | Group of Genomics and Postgenomic Technologies of Central Research Institute of Epidemiology | Speranskaya AS, Kaptelova VV,Valdokhina AV, Bulanenko VP, Samoilov AE, Korneenko EV, Sizova TV, Tivanova EV, Shipulina OY, Akimkin VG |
| EPI_ISL_479657 | NIV Influenza | NIV Influenza | Potdar V |
| EPI_ISL_479756, EPI_ISL_479757, EPI_ISL_479758 | National Institute of Hygiene and Epidemiology (NIHE) | National Key Laboratory of Gene Technology, Institute of Biotechnology (IBT) | Le Tung Lam, Nguyen Hong Trang, Ho Thi Thuong, Tran Huyen Linh, Ung Thi Hong Trang, Le Thi Thanh, Nguyen Vu Son, Vuong Duc Cuong, Tran Thu Huong, Pham Thi Hien, Nguyen Phuong Anh, Nguyen Le Khanh Hang, Hoang Vu Mai Phuong, Hoang Ha, Taichiro Takemura, Futoshi Hasebe, Chu Hoang Ha, Le Quynh Mai, Dang Duc Anh, Truong Nam Hai |
| EPI_ISL_479790, EPI_ISL_479791 | Laboratory of Molecular Virology of the International Centre for Genetic Engineering and Biotechnology (ICGEB) | ARGO Open Lab Platform for Genome Sequencing | Licastro, D, Rajasekharan S, Dal Monego S, Segat L, D'Agaro P, Salton F, Confalonieri P, Confalonieri M, Marcello A |
| EPI_ISL_479792, EPI_ISL_479793, EPI_ISL_479794, EPI_ISL_479795 | Hokkaido Institute of Public Health | Pathogen Genomics Center, National Institute of Infectious Diseases | Tsuyoshi Sekizuka, Rika Komagome, Kentaro Itokawa, Rina Tanaka, Masanori Hashino, Hajime Kamiya, Motoi Suzuki, Makoto Kuroda |
| EPI_ISL_479796 | Ishikawa Prefectural Institute of Public Health and Environmental Science | Pathogen Genomics Center, National Institute of Infectious Diseases | Tsuyoshi Sekizuka, Sanae Kuramoto, Eri Nariai, Kentaro Itokawa, Rina Tanaka, Masanori Hashino, Hajime Kamiya, Motoi Suzuki, Makoto Kuroda |
| EPI_ISL_479797, EPI_ISL_479798 | Sagamihara City Public Health Research Institute | Pathogen Genomics Center, National Institute of Infectious Diseases | Tsuyoshi Sekizuka, Hiroshi Nakamura, Kentaro Itokawa, Rina Tanaka, Masanori Hashino, Hajime Kamiya, Motoi Suzuki, Makoto Kuroda |
| EPI_ISL_479799, EPI_ISL_479800 | Sapporo City Institute of Public Health | Pathogen Genomics Center, National Institute of Infectious Diseases | Tsuyoshi Sekizuka, Asami Ohnishi, Kentaro Itokawa, Rina Tanaka, Masanori Hashino, Hajime Kamiya, Motoi Suzuki, Makoto Kuroda |
| EPI_ISL_479801 | Hokkaido Institute of Public Health | Pathogen Genomics Center, National Institute of Infectious Diseases | Tsuyoshi Sekizuka, Rika Komagome, Kentaro Itokawa, Rina Tanaka, Masanori Hashino, Hajime Kamiya, Motoi Suzuki, Makoto Kuroda |
| EPI_ISL_479802, EPI_ISL_479803, EPI_ISL_479804 | Sagamihara City Public Health Research Institute | Pathogen Genomics Center, National Institute of Infectious Diseases | Tsuyoshi Sekizuka, Hiroshi Nakamura, Kentaro Itokawa, Rina Tanaka, Masanori Hashino, Hajime Kamiya, Motoi Suzuki, Makoto Kuroda |
| EPI_ISL_479805, EPI_ISL_479806, EPI_ISL_479807, EPI_ISL_479808 | Saitama Prefectural Institute of Public Health | Pathogen Genomics Center, National Institute of Infectious Diseases | Tsuyoshi Sekizuka, Hayato Ehara, Kentaro Itokawa, Rina Tanaka, Masanori Hashino, Hajime Kamiya, Motoi Suzuki, Makoto Kuroda |
| EPI_ISL_479809, EPI_ISL_479810, EPI_ISL_479811 | Chiba Prefectural Institute of Public Health | Pathogen Genomics Center, National Institute of Infectious Diseases | Tsuyoshi Sekizuka, Masakatsu Taira, Kentaro Itokawa, Rina Tanaka, Masanori Hashino, Hajime Kamiya, Motoi Suzuki, Makoto Kuroda |
| EPI_ISL_479812, EPI_ISL_479813, EPI_ISL_479814, EPI_ISL_479815, EPI_ISL_479816, EPI_ISL_479817, EPI_ISL_479818, EPI_ISL_479819, EPI_ISL_479820 | Hokkaido Institute of Public Health | Pathogen Genomics Center, National Institute of Infectious Diseases | Tsuyoshi Sekizuka, Rika Komagome, Kentaro Itokawa, Rina Tanaka, Masanori Hashino, Hajime Kamiya, Motoi Suzuki, Makoto Kuroda |
| EPI_ISL_479821, EPI_ISL_479822 | Department of Infectious Diseases, Kobe Institute of Health | Pathogen Genomics Center, National Institute of Infectious Diseases | Tsuyoshi Sekizuka, Ryohei Nomoto, Kentaro Itokawa, Rina Tanaka, Masanori Hashino, Hajime Kamiya, Motoi Suzuki, Makoto Kuroda |
| EPI_ISL_479823 | Kochi Prefectural Institute of Public Health | Pathogen Genomics Center, National Institute of Infectious Diseases | Tsuyoshi Sekizuka, Akihiko Tokaji, Kentaro Itokawa, Rina Tanaka, Masanori Hashino, Hajime Kamiya, Motoi Suzuki, Makoto Kuroda |
| EPI_ISL_479824 | Kumamoto PrefecturalInstituteof Public-Health and Environmental Science | Pathogen Genomics Center, National Institute of Infectious Diseases | Tsuyoshi Sekizuka, Shunsuke Yahiro, Kentaro Itokawa, Rina Tanaka, Masanori Hashino, Hajime Kamiya, Motoi Suzuki, Makoto Kuroda |
| EPI_ISL_479825 | Tokyo Metropolitan Institute of Public Health | Pathogen Genomics Center, National Institute of Infectious Diseases | Tsuyoshi Sekizuka, Kenji Sadamasu, Takashi Chiba, Mami Nagashima, Kentaro Itokawa, Rina Tanaka, Masanori Hashino, Hajime Kamiya, Motoi Suzuki, Makoto Kuroda |
| EPI_ISL_479826, EPI_ISL_479827, EPI_ISL_479828, EPI_ISL_479829, EPI_ISL_479830, EPI_ISL_479831, EPI_ISL_479832, EPI_ISL_479833, EPI_ISL_479834, EPI_ISL_479835, EPI_ISL_479836, EPI_ISL_479837, EPI_ISL_479838, EPI_ISL_479839, EPI_ISL_479840, EPI_ISL_479841, EPI_ISL_479842, EPI_ISL_479843, EPI_ISL_479844, EPI_ISL_479845, EPI_ISL_479846, EPI_ISL_479847, EPI_ISL_479848, EPI_ISL_479849 | | | |
| see above | Sapporo City Institute of Public Health | Pathogen Genomics Center, National Institute of Infectious Diseases | Tsuyoshi Sekizuka, Asami Ohnishi, Kentaro Itokawa, Rina Tanaka, Masanori Hashino, Hajime Kamiya, Motoi Suzuki, Makoto Kuroda |
| EPI_ISL_479850, EPI_ISL_479851, EPI_ISL_479852, EPI_ISL_479853, EPI_ISL_479854 | Gunma Prefectural Institute of Public Health and Environmental Sciences | Pathogen Genomics Center, National Institute of Infectious Diseases | Tsuyoshi Sekizuka, Hiroyuki Tsukagoshi, Kentaro Itokawa, Rina Tanaka, Masanori Hashino, Hajime Kamiya, Motoi Suzuki, Makoto Kuroda |
| EPI_ISL_479855, EPI_ISL_479856, EPI_ISL_479857, EPI_ISL_479858, EPI_ISL_479859, EPI_ISL_479860, EPI_ISL_479861 | Department of Infectious Diseases, Kobe Institute of Health | Pathogen Genomics Center, National Institute of Infectious Diseases | Tsuyoshi Sekizuka, Ryohei Nomoto, Kentaro Itokawa, Rina Tanaka, Masanori Hashino, Hajime Kamiya, Motoi Suzuki, Makoto Kuroda |
| EPI_ISL_479862, EPI_ISL_479863, EPI_ISL_479864, EPI_ISL_479865, EPI_ISL_479866, EPI_ISL_479867 | Wakayama Prefectural Research Center of Environment and Public Health | Pathogen Genomics Center, National Institute of Infectious Diseases | Tsuyoshi Sekizuka, Fumio Terasoma, Yosuke Hamajima, Kentaro Itokawa, Rina Tanaka, Masanori Hashino, Hajime Kamiya, Motoi Suzuki, Makoto Kuroda |
| EPI_ISL_479868 | Department of Infectious Diseases, Kobe Institute of Health | Pathogen Genomics Center, National Institute of Infectious Diseases | Tsuyoshi Sekizuka, Ryohei Nomoto, Kentaro Itokawa, Rina Tanaka, Masanori Hashino, Hajime Kamiya, Motoi Suzuki, Makoto Kuroda |
| EPI_ISL_479869 | Niigata Prefectural Institute of Public Health and Environmental Sciences | Pathogen Genomics Center, National Institute of Infectious Diseases | Tsuyoshi Sekizuka, Reiko Arai, Kentaro Itokawa, Rina Tanaka, Masanori Hashino, Hajime Kamiya, Motoi Suzuki, Makoto Kuroda |
| EPI_ISL_479870, EPI_ISL_479871 | Sagamihara City Public Health Research Institute | Pathogen Genomics Center, National Institute of Infectious Diseases | Tsuyoshi Sekizuka, Hiroshi Nakamura, Kentaro Itokawa, Rina Tanaka, Masanori Hashino, Hajime Kamiya, Motoi Suzuki, Makoto Kuroda |
| EPI_ISL_479872, EPI_ISL_479873, EPI_ISL_479874, EPI_ISL_479875, EPI_ISL_479876, EPI_ISL_479877, EPI_ISL_479878, EPI_ISL_479879, EPI_ISL_479880, EPI_ISL_479881, EPI_ISL_479882, EPI_ISL_479883, EPI_ISL_479884, EPI_ISL_479885 | | | |
| see above | Sapporo City Institute of Public Health | Pathogen Genomics Center, National Institute of Infectious Diseases | Tsuyoshi Sekizuka, Asami Ohnishi, Kentaro Itokawa, Rina Tanaka, Masanori Hashino, Hajime Kamiya, Motoi Suzuki, Makoto Kuroda |
| EPI_ISL_479886, EPI_ISL_479887, EPI_ISL_479888, EPI_ISL_479889, | Tokyo Metropolitan Institute of Public Health | Pathogen Genomics Center, National Institute of Infectious Diseases | Tsuyoshi Sekizuka, Kenji Sadamasu, Takashi Chiba, Mami Nagashima, Kentaro Itokawa, Rina Tanaka, Masanori Hashino, Hajime Kamiya, Motoi Suzuki, Makoto Kuroda |

| | | | |
|---|---|---|---|
| EPI_ISL_479890, EPI_ISL_479891, EPI_ISL_479892, EPI_ISL_479893, EPI_ISL_479894, EPI_ISL_479895 | | | |
| EPI_ISL_479896, EPI_ISL_479897, EPI_ISL_479898, EPI_ISL_479899, EPI_ISL_479900, EPI_ISL_479901 | Gunma Prefectural Institute of Public Health and Environmental Sciences | Pathogen Genomics Center, National Institute of Infectious Diseases | Tsuyoshi Sekizuka, Hiroyuki Tsukagoshi, Kentaro Itokawa, Rina Tanaka, Masanori Hashino, Hajime Kamiya, Motoi Suzuki, Makoto Kuroda |
| EPI_ISL_479902 | Niigata Prefectural Institute of Public Health and Environmental Sciences | Pathogen Genomics Center, National Institute of Infectious Diseases | Tsuyoshi Sekizuka, Reiko Arai, Kentaro Itokawa, Rina Tanaka, Masanori Hashino, Hajime Kamiya, Motoi Suzuki, Makoto Kuroda |
| EPI_ISL_479903, EPI_ISL_479904, EPI_ISL_479905, EPI_ISL_479906, EPI_ISL_479907, EPI_ISL_479908, EPI_ISL_479909, EPI_ISL_479910, EPI_ISL_479911, EPI_ISL_479912 | Himeji City Institute of Environment and Health | Pathogen Genomics Center, National Institute of Infectious Diseases | Tsuyoshi Sekizuka, Kentaro Itokawa, Rina Tanaka, Masanori Hashino, Hajime Kamiya, Motoi Suzuki, Makoto Kuroda |
| EPI_ISL_479913, EPI_ISL_479914, EPI_ISL_479915, EPI_ISL_479916, EPI_ISL_479917, EPI_ISL_479918, EPI_ISL_479919, EPI_ISL_479920, EPI_ISL_479921, EPI_ISL_479922, EPI_ISL_479923, EPI_ISL_479924 | | | |
| see above | Niigata City Public Health Research Institute | Pathogen Genomics Center, National Institute of Infectious Diseases | Tsuyoshi Sekizuka, Yurie Takahashi, Kentaro Itokawa, Rina Tanaka, Masanori Hashino, Hajime Kamiya, Motoi Suzuki, Makoto Kuroda |
| EPI_ISL_479925, EPI_ISL_479926, EPI_ISL_479927 | Sakai City Institute of Public Health | Pathogen Genomics Center, National Institute of Infectious Diseases | Tsuyoshi Sekizuka, Tatsuya Miyoshi, Kentaro Itokawa, Rina Tanaka, Masanori Hashino, Hajime Kamiya, Motoi Suzuki, Makoto Kuroda |
| EPI_ISL_479928, EPI_ISL_479929, EPI_ISL_479930, EPI_ISL_479931, EPI_ISL_479932, EPI_ISL_479933, EPI_ISL_479934, EPI_ISL_479935 | Saitama Prefectural Institute of Public Health | Pathogen Genomics Center, National Institute of Infectious Diseases | Tsuyoshi Sekizuka, Hayato Ehara, Kentaro Itokawa, Rina Tanaka, Masanori Hashino, Hajime Kamiya, Motoi Suzuki, Makoto Kuroda |
| EPI_ISL_479936, EPI_ISL_479937, EPI_ISL_479938, EPI_ISL_479939, EPI_ISL_479940, EPI_ISL_479941, EPI_ISL_479942, EPI_ISL_479943 | Ibaraki Prefectural Institute of Public Health | Pathogen Genomics Center, National Institute of Infectious Diseases | Tsuyoshi Sekizuka, Keiko Goto, Kentaro Itokawa, Rina Tanaka, Masanori Hashino, Hajime Kamiya, Motoi Suzuki, Makoto Kuroda |
| EPI_ISL_479944, EPI_ISL_479945, EPI_ISL_479946, EPI_ISL_479947, EPI_ISL_479948, EPI_ISL_479949, EPI_ISL_479950, EPI_ISL_479951, EPI_ISL_479952, EPI_ISL_479953, EPI_ISL_479954, EPI_ISL_479955, EPI_ISL_479956, EPI_ISL_479957, EPI_ISL_479958 | | | |
| see above | Osaka Institute of Public Health | Pathogen Genomics Center, National Institute of Infectious Diseases | Tsuyoshi Sekizuka, Satoshi Hiroi, Saeko Morikawa, Kazushi Motomura, Kentaro Itokawa, Rina Tanaka, Masanori Hashino, Hajime Kamiya, Motoi Suzuki, Makoto Kuroda |
| EPI_ISL_479959, EPI_ISL_479960, EPI_ISL_479961, EPI_ISL_479962, EPI_ISL_479963, EPI_ISL_479964, EPI_ISL_479965 | Tokyo Metropolitan Institute of Public Health | Pathogen Genomics Center, National Institute of Infectious Diseases | Tsuyoshi Sekizuka, Kenji Sadamasu, Takashi Chiba, Mami Nagashima, Kentaro Itokawa, Rina Tanaka, Masanori Hashino, Hajime Kamiya, Motoi Suzuki, Makoto Kuroda |
| EPI_ISL_479966 | Osaka Institute of Public Health | Pathogen Genomics Center, National Institute of Infectious Diseases | Tsuyoshi Sekizuka, Satoshi Hiroi, Saeko Morikawa, Kazushi Motomura, Kentaro Itokawa, Rina Tanaka, Masanori Hashino, Hajime Kamiya, Motoi Suzuki, Makoto Kuroda |
| EPI_ISL_479967, EPI_ISL_479968, EPI_ISL_479969, EPI_ISL_479970, EPI_ISL_479971, EPI_ISL_479972, EPI_ISL_479973, EPI_ISL_479974, EPI_ISL_479975, EPI_ISL_479976, EPI_ISL_479977, EPI_ISL_479978 | | | |
| see above | Fukui Prefectural Institute of Public Health and Environmental Science | Pathogen Genomics Center, National Institute of Infectious Diseases | Tsuyoshi Sekizuka, Miho Toho, Kentaro Itokawa, Rina Tanaka, Masanori Hashino, Hajime Kamiya, Motoi Suzuki, Makoto Kuroda |
| EPI_ISL_479979, EPI_ISL_479980, EPI_ISL_479981, EPI_ISL_479982, EPI_ISL_479983, EPI_ISL_479984 | Oita Prefectural Institute of Public Health and Environmental Science | Pathogen Genomics Center, National Institute of Infectious Diseases | Tsuyoshi Sekizuka, Mari Sasaki, Kentaro Itokawa, Rina Tanaka, Masanori Hashino, Hajime Kamiya, Motoi Suzuki, Makoto Kuroda |
| EPI_ISL_479985 | Ibaraki Prefectural Institute of Public Health | Pathogen Genomics Center, National Institute of Infectious Diseases | Tsuyoshi Sekizuka, Keiko Goto, Kentaro Itokawa, Rina Tanaka, Masanori Hashino, Hajime Kamiya, Motoi Suzuki, Makoto Kuroda |
| EPI_ISL_479986, EPI_ISL_479987, EPI_ISL_479988, EPI_ISL_479989 | Department of Infectious Diseases, Kobe Institute of Health | Pathogen Genomics Center, National Institute of Infectious Diseases | Tsuyoshi Sekizuka, Ryohei Nomoto, Kentaro Itokawa, Rina Tanaka, Masanori Hashino, Hajime Kamiya, Motoi Suzuki, Makoto Kuroda |
| EPI_ISL_479990 | Kitakyushu City Institute of Health and Environmental Sciences | Pathogen Genomics Center, National Institute of Infectious Diseases | Tsuyoshi Sekizuka, Katsuya Obata, Asuka Kikuchi Kentaro Itokawa, Rina Tanaka, Masanori Hashino, Hajime Kamiya, Motoi Suzuki, Makoto Kuroda |
| EPI_ISL_479991, EPI_ISL_479992, EPI_ISL_479993, EPI_ISL_479994, EPI_ISL_479995, EPI_ISL_479996 | Kumamoto City Public Health Research Institute | Pathogen Genomics Center, National Institute of Infectious Diseases | Tsuyoshi Sekizuka, Kaori Tashiro, Kentaro Itokawa, Rina Tanaka, Masanori Hashino, Hajime Kamiya, Motoi Suzuki, Makoto Kuroda |
| EPI_ISL_479997, EPI_ISL_479998, EPI_ISL_479999, EPI_ISL_480000, EPI_ISL_480001 | Nagano Environmental Conservation Research Institute | Pathogen Genomics Center, National Institute of Infectious Diseases | Tsuyoshi Sekizuka, Naoko Shimodaira, Kentaro Itokawa, Rina Tanaka, Masanori Hashino, Hajime Kamiya, Motoi Suzuki, Makoto Kuroda |
| EPI_ISL_480002, EPI_ISL_480003 | Nagasaki Prefectural Institute for Environmental Research and Public Health | Pathogen Genomics Center, National Institute of Infectious Diseases | Tsuyoshi Sekizuka, Fumiaki Matsumoto, Kentaro Itokawa, Rina Tanaka, Masanori Hashino, Hajime Kamiya, Motoi Suzuki, Makoto Kuroda |
| EPI_ISL_480004, EPI_ISL_480005, EPI_ISL_480006, EPI_ISL_480007, EPI_ISL_480008, EPI_ISL_480009, EPI_ISL_480010, EPI_ISL_480011, EPI_ISL_480012, EPI_ISL_480013, EPI_ISL_480014 | | | |
| see above | Chiba Prefectural Institute of Public Health | Pathogen Genomics Center, National Institute of Infectious Diseases | Tsuyoshi Sekizuka, Masakatsu Taira, Kentaro Itokawa, Rina Tanaka, Masanori Hashino, Hajime Kamiya, Motoi Suzuki, Makoto Kuroda |
| EPI_ISL_480015, EPI_ISL_480016, EPI_ISL_480017, EPI_ISL_480018, EPI_ISL_480019, EPI_ISL_480020 | Gunma Prefectural Institute of Public Health and Environmental Sciences | Pathogen Genomics Center, National Institute of Infectious Diseases | Tsuyoshi Sekizuka, Hiroyuki Tsukagoshi, Kentaro Itokawa, Rina Tanaka, Masanori Hashino, Hajime Kamiya, Motoi Suzuki, Makoto Kuroda |
| EPI_ISL_480021, EPI_ISL_480022, EPI_ISL_480023, EPI_ISL_480024, EPI_ISL_480025, EPI_ISL_480026, EPI_ISL_480027, EPI_ISL_480028, EPI_ISL_480029 | Ibaraki Prefectural Institute of Public Health | Pathogen Genomics Center, National Institute of Infectious Diseases | Tsuyoshi Sekizuka, Keiko Goto, Kentaro Itokawa, Rina Tanaka, Masanori Hashino, Hajime Kamiya, Motoi Suzuki, Makoto Kuroda |
| EPI_ISL_480030, EPI_ISL_480031, EPI_ISL_480032, EPI_ISL_480033, EPI_ISL_480034, EPI_ISL_480035, EPI_ISL_480036, EPI_ISL_480037, EPI_ISL_480038, EPI_ISL_480039, EPI_ISL_480040, EPI_ISL_480041 | | | |
| see above | Tochigi Prefectural Institute of Public Health and Environmental Science | Pathogen Genomics Center, National Institute of Infectious Diseases | Tsuyoshi Sekizuka, Ako Nakajima, Kentaro Itokawa, Rina Tanaka, Masanori Hashino, Hajime Kamiya, Motoi Suzuki, Makoto Kuroda |
| EPI_ISL_480042, EPI_ISL_480043, EPI_ISL_480044, EPI_ISL_480045, EPI_ISL_480046, EPI_ISL_480047, EPI_ISL_480048, EPI_ISL_480049, EPI_ISL_480050, EPI_ISL_480051, EPI_ISL_480052, EPI_ISL_480053, EPI_ISL_480054, EPI_ISL_480055, EPI_ISL_480056, EPI_ISL_480057, EPI_ISL_480058, EPI_ISL_480059, EPI_ISL_480060, EPI_ISL_480061, EPI_ISL_480062, EPI_ISL_480063, EPI_ISL_480064 | | | |
| see above | Nagoya City Public Health Research Institute | Pathogen Genomics Center, National Institute of Infectious Diseases | Tsuyoshi Sekizuka, Takuya Miki, Shinichiro Shibata, Kentaro Itokawa, Rina Tanaka, Masanori Hashino, Hajime Kamiya, Motoi Suzuki, Makoto Kuroda |

| | | | |
|---|---|---|---|
| EPI_ISL_480065, EPI_ISL_480066, EPI_ISL_480067, EPI_ISL_480068, EPI_ISL_480069, EPI_ISL_480070, EPI_ISL_480071, EPI_ISL_480072 | Sakai City Institute of Public Health | Pathogen Genomics Center, National Institute of Infectious Diseases | Tsuyoshi Sekizuka, Tatsuya Miyoshi, Kentaro Itokawa, Rina Tanaka, Masanori Hashino, Hajime Kamiya, Motoi Suzuki, Makoto Kuroda |
| EPI_ISL_480073 | Tochigi Prefectural Institute of Public Health and Environmental Science | Pathogen Genomics Center, National Institute of Infectious Diseases | Tsuyoshi Sekizuka, Ako Nakajima, Kentaro Itokawa, Rina Tanaka, Masanori Hashino, Hajime Kamiya, Motoi Suzuki, Makoto Kuroda |
| EPI_ISL_480074, EPI_ISL_480075, EPI_ISL_480076, EPI_ISL_480077, EPI_ISL_480078, EPI_ISL_480079, EPI_ISL_480080, EPI_ISL_480081, EPI_ISL_480082 | Shizuoka City Institute of Environmental Sciences and Public Health | Pathogen Genomics Center, National Institute of Infectious Diseases | Tsuyoshi Sekizuka, Takaharu Maehata,Sou Okamura,Yuji Kanazawa,Kenji Yagi, Kentaro Itokawa, Rina Tanaka, Masanori Hashino, Hajime Kamiya, Motoi Suzuki, Makoto Kuroda |
| EPI_ISL_480083, EPI_ISL_480084, EPI_ISL_480085, EPI_ISL_480086, EPI_ISL_480087, EPI_ISL_480088, EPI_ISL_480089 | Gifu Prefectural Institute of Public Health and Environmental Sciences | Pathogen Genomics Center, National Institute of Infectious Diseases | Tsuyoshi Sekizuka, Yoshihiko Kameyama, Kentaro Itokawa, Rina Tanaka, Masanori Hashino, Hajime Kamiya, Motoi Suzuki, Makoto Kuroda |
| EPI_ISL_480090, EPI_ISL_480091, EPI_ISL_480092, EPI_ISL_480093, EPI_ISL_480094, EPI_ISL_480095, EPI_ISL_480097, EPI_ISL_480098, EPI_ISL_480099, EPI_ISL_480100, EPI_ISL_480101, EPI_ISL_480102 | | | |
| see above | Department of Infectious Diseases, Kobe Institute of Health | Pathogen Genomics Center, National Institute of Infectious Diseases | Tsuyoshi Sekizuka, Ryohei Nomoto, Kentaro Itokawa, Rina Tanaka, Masanori Hashino, Hajime Kamiya, Motoi Suzuki, Makoto Kuroda |
| EPI_ISL_480103, EPI_ISL_480104, EPI_ISL_480105, EPI_ISL_480106, EPI_ISL_480107, EPI_ISL_480108 | Koshigaya City Public Health Center | Pathogen Genomics Center, National Institute of Infectious Diseases | Tsuyoshi Sekizuka, Yuka Furui, Aya Tamura, Kyohei Sakata, Takumi Daimon, Yoko Togawa, Yoshiko Hamada, Kentaro Itokawa, Rina Tanaka, Masanori Hashino, Hajime Kamiya, Motoi Suzuki, Makoto Kuroda |
| EPI_ISL_480109, EPI_ISL_480110, EPI_ISL_480111, EPI_ISL_480112, EPI_ISL_480113, EPI_ISL_480114, EPI_ISL_480115, EPI_ISL_480116, EPI_ISL_480117, EPI_ISL_480118, EPI_ISL_480119 | | | |
| see above | Oita Prefectural Institute of Public Health and Environmental Science | Pathogen Genomics Center, National Institute of Infectious Diseases | Tsuyoshi Sekizuka, Mari Sasaki, Kentaro Itokawa, Rina Tanaka, Masanori Hashino, Hajime Kamiya, Motoi Suzuki, Makoto Kuroda |
| EPI_ISL_480120, EPI_ISL_480121, EPI_ISL_480122, EPI_ISL_480123, EPI_ISL_480124, EPI_ISL_480125, EPI_ISL_480126, EPI_ISL_480127, EPI_ISL_480128, EPI_ISL_480129, EPI_ISL_480130, EPI_ISL_480131, EPI_ISL_480132, EPI_ISL_480133, EPI_ISL_480134, EPI_ISL_480135, EPI_ISL_480136, EPI_ISL_480137, EPI_ISL_480138, EPI_ISL_480139, EPI_ISL_480140, EPI_ISL_480141, EPI_ISL_480142, EPI_ISL_480143, EPI_ISL_480144, EPI_ISL_480145, EPI_ISL_480146, EPI_ISL_480147, EPI_ISL_480148, EPI_ISL_480149, EPI_ISL_480150, EPI_ISL_480151, EPI_ISL_480152, EPI_ISL_480153, EPI_ISL_480154, EPI_ISL_480155, EPI_ISL_480156, EPI_ISL_480157, EPI_ISL_480158, EPI_ISL_480159, EPI_ISL_480160, EPI_ISL_480161, EPI_ISL_480162, EPI_ISL_480163, EPI_ISL_480164, EPI_ISL_480165, EPI_ISL_480166, EPI_ISL_480167, EPI_ISL_480168 | | | |
| see above | Fukui Prefectural Institute of Public Health and Environmental Science | Pathogen Genomics Center, National Institute of Infectious Diseases | Tsuyoshi Sekizuka, Miho Toho, Kentaro Itokawa, Rina Tanaka, Masanori Hashino, Hajime Kamiya, Motoi Suzuki, Makoto Kuroda |
| EPI_ISL_480169, EPI_ISL_480170, EPI_ISL_480171, EPI_ISL_480172, EPI_ISL_480173, EPI_ISL_480174, EPI_ISL_480175, EPI_ISL_480176, EPI_ISL_480177 | Gunma Prefectural Institute of Public Health and Environmental Sciences | Pathogen Genomics Center, National Institute of Infectious Diseases | Tsuyoshi Sekizuka, Hiroyuki Tsukagoshi, Kentaro Itokawa, Rina Tanaka, Masanori Hashino, Hajime Kamiya, Motoi Suzuki, Makoto Kuroda |
| EPI_ISL_480178, EPI_ISL_480179 | Hiroshima City Institute of Public Health | Pathogen Genomics Center, National Institute of Infectious Diseases | Tsuyoshi Sekizuka, Kota Noritsune, Kentaro Itokawa, Rina Tanaka, Masanori Hashino, Hajime Kamiya, Motoi Suzuki, Makoto Kuroda |
| EPI_ISL_480180, EPI_ISL_480181, EPI_ISL_480182, EPI_ISL_480183, EPI_ISL_480184, EPI_ISL_480185, EPI_ISL_480186, EPI_ISL_480187, EPI_ISL_480188, EPI_ISL_480189 | Ibaraki Prefectural Institute of Public Health | Pathogen Genomics Center, National Institute of Infectious Diseases | Tsuyoshi Sekizuka, Keiko Goto, Kentaro Itokawa, Rina Tanaka, Masanori Hashino, Hajime Kamiya, Motoi Suzuki, Makoto Kuroda |
| EPI_ISL_480190, EPI_ISL_480191, EPI_ISL_480192, EPI_ISL_480193, EPI_ISL_480194, EPI_ISL_480195 | Ota Health Center Welfare Section | Pathogen Genomics Center, National Institute of Infectious Diseases | Tsuyoshi Sekizuka, Chika Takahashi, Kentaro Itokawa, Rina Tanaka, Masanori Hashino, Hajime Kamiya, Motoi Suzuki, Makoto Kuroda |
| EPI_ISL_480196, EPI_ISL_480197, EPI_ISL_480198, EPI_ISL_480199, EPI_ISL_480200, EPI_ISL_480201, EPI_ISL_480202, EPI_ISL_480203 | Toyama Institute of Health | Pathogen Genomics Center, National Institute of Infectious Diseases | Tsuyoshi Sekizuka, Masae Itamochi, Kazunori Oishi, Kentaro Itokawa, Rina Tanaka, Masanori Hashino, Hajime Kamiya, Motoi Suzuki, Makoto Kuroda |
| EPI_ISL_480204 | Akita City Public Health Center | Pathogen Genomics Center, National Institute of Infectious Diseases | Tsuyoshi Sekizuka, Koichi Ito, Kentaro Itokawa, Rina Tanaka, Masanori Hashino, Hajime Kamiya, Motoi Suzuki, Makoto Kuroda |
| EPI_ISL_480205, EPI_ISL_480206, EPI_ISL_480207 | Department of Infectious Diseases, Kobe Institute of Health | Pathogen Genomics Center, National Institute of Infectious Diseases | Tsuyoshi Sekizuka, Ryohei Nomoto, Kentaro Itokawa, Rina Tanaka, Masanori Hashino, Hajime Kamiya, Motoi Suzuki, Makoto Kuroda |
| EPI_ISL_480208 | Pathogen Genomics Center, National Institute of Infectious Diseases | Pathogen Genomics Center, National Institute of Infectious Diseases | Tsuyoshi Sekizuka, Ryohei Nomoto, Kentaro Itokawa, Rina Tanaka, Masanori Hashino, Hajime Kamiya, Motoi Suzuki, Makoto Kuroda |
| EPI_ISL_480209, EPI_ISL_480210, EPI_ISL_480211, EPI_ISL_480212, EPI_ISL_480213, EPI_ISL_480214, EPI_ISL_480215, EPI_ISL_480216, EPI_ISL_480217, EPI_ISL_480218, EPI_ISL_480219, EPI_ISL_480220 | | | |
| see above | Department of Infectious Diseases, Kobe Institute of Health | Pathogen Genomics Center, National Institute of Infectious Diseases | Tsuyoshi Sekizuka, Ryohei Nomoto, Kentaro Itokawa, Rina Tanaka, Masanori Hashino, Hajime Kamiya, Motoi Suzuki, Makoto Kuroda |
| EPI_ISL_480221, EPI_ISL_480222, EPI_ISL_480223 | Koshigaya City Public Health Center | Pathogen Genomics Center, National Institute of Infectious Diseases | Tsuyoshi Sekizuka, Yuka Furui, Aya Tamura, Kyohei Sakata, Takumi Daimon, Yoko Togawa, Yoshiko Hamada, Kentaro Itokawa, Rina Tanaka, Masanori Hashino, Hajime Kamiya, Motoi Suzuki, Makoto Kuroda |
| EPI_ISL_480228, EPI_ISL_480229, EPI_ISL_480230, EPI_ISL_480231, EPI_ISL_480232, EPI_ISL_480233, EPI_ISL_480234, EPI_ISL_480235, EPI_ISL_480236, EPI_ISL_480237, EPI_ISL_480238, EPI_ISL_480239, EPI_ISL_480240, EPI_ISL_480241, EPI_ISL_480242, EPI_ISL_480243, EPI_ISL_480244, EPI_ISL_480245, EPI_ISL_480246, EPI_ISL_480247, EPI_ISL_480248, EPI_ISL_480249, EPI_ISL_480250, EPI_ISL_480252, EPI_ISL_480253, EPI_ISL_480254, EPI_ISL_480255, EPI_ISL_480256, EPI_ISL_480257, EPI_ISL_480258, EPI_ISL_480259, EPI_ISL_480260, EPI_ISL_480261, EPI_ISL_480262, EPI_ISL_480263, EPI_ISL_480264, EPI_ISL_480265, EPI_ISL_480266, EPI_ISL_480267, EPI_ISL_480268, EPI_ISL_480269, EPI_ISL_480270, EPI_ISL_480271, EPI_ISL_480272, EPI_ISL_480273, EPI_ISL_480274, EPI_ISL_480277, EPI_ISL_480278, EPI_ISL_480281, EPI_ISL_480282, EPI_ISL_480283, EPI_ISL_480284, EPI_ISL_480285, EPI_ISL_480286, EPI_ISL_480289, EPI_ISL_480291, EPI_ISL_480292 | | | |
| see above | Genomic Laboratory (GLAB) (Conjoint lab of Health Directorate of Istanbul and Istanbul Technical University) | Genomic Laboratory (GLAB), Istanbul Technical University | Ilker Karacan, Tugba Kizilboga Akgun, Bugra Agaoglu, Gizem Alkurt, Jale Yildiz, Betsi Köse, Elifnaz Çelik, Arzu Irvem, Yasemin Kendir Demirkol, Ozlem Akgun Dogan, Mehtap Aydn, Levent Doganay, Gizem Dinler Doganay |
| EPI_ISL_480310 | National Reference Laboratory "Influenza and acute respiratory diseases" | NRL-HIV | Ivan Ivanov, Ivailo Alexiev, Ivva Philipova |
| EPI_ISL_480554, EPI_ISL_480556, EPI_ISL_480783, EPI_ISL_480787, EPI_ISL_480789, EPI_ISL_481235, EPI_ISL_481236, EPI_ISL_481237, EPI_ISL_481239, EPI_ISL_481240, EPI_ISL_481243 | | | |
| see above | Institut Pasteur Dakar | Institut Pasteur de Dakar | Ndongo Dia, Moussa Moise Diagne, Mamadou Diop, Marie Henriette Dior Ndione, Mamadou Malado Jallow, Safietou Sanke, Ousmane Faye, Amadou Alpha Sall. |
| EPI_ISL_482575, EPI_ISL_482576, EPI_ISL_482577, EPI_ISL_482578, EPI_ISL_482579, EPI_ISL_482580, EPI_ISL_482581, EPI_ISL_482582, EPI_ISL_482583, EPI_ISL_482584, EPI_ISL_482585, EPI_ISL_482586 | | | |
| see above | Hangzhou Center for Diseases Control and Prevention | Hangzhou Center for Diseases Control and Prevention | Jun Li, Haoqiu Wang, Lingfeng Mao, Hua Yu, Xinfen Yu, Zhou Sun, Xin Qian, Shuchang Chen, Junfang Chen, Xuchu Wang |
| EPI_ISL_482672, EPI_ISL_482673, | Singapore General Hospital | Department of Microbiology | Nurdyana Abdul Rahman, Kun Lee Lim, Chenhao Li, Kian Sing Chan, Lynette Oon, Kern Rei Chng, Niranjan Nagarajan, Karrie Ko |

| | | | |
|---|---|---|---|
| EPI_ISL_482674, EPI_ISL_482675, EPI_ISL_482676, EPI_ISL_482677, EPI_ISL_482680 | | | |
| EPI_ISL_482759, EPI_ISL_482760, EPI_ISL_482761, EPI_ISL_482762, EPI_ISL_482763, EPI_ISL_482764, EPI_ISL_482765, EPI_ISL_482766, EPI_ISL_482767, EPI_ISL_482768, EPI_ISL_482769, EPI_ISL_482770, EPI_ISL_482771, EPI_ISL_482772, EPI_ISL_482773, EPI_ISL_482774 | | | |
| see above | Medical Ain Shams Research Institute (MASRI), Ain Shams University | Medical Ain Shams Research Institute (MASRI), Ain Shams University | Hesham Elghazaly, Sara Hassan Agwa, Ahmad Moustafa, Hala Hafez, Sara Elnakeep, Shaimaa Moustafa, Aya Mohamed, Reham Mamdouh, Ghada Ismael, Ashraf Omar, Osama Mansour, Mahmoud Elmeitini |
| EPI_ISL_482777 | Queen Elizabeth Hospital | Hong Kong Department of Health | Mak Gannon C.K., Cheng Peter K.C., Lam Edman T.K., Chan Rickjason C.W., Tsang Dominic N.C. |
| EPI_ISL_483035, EPI_ISL_483036, EPI_ISL_483038 | Medical Ain Shams Research Institute (MASRI), Ain Shams University | Medical Ain Shams Research Institute (MASRI), Ain Shams University | Hesham Elghazaly, Sara Hassan Agwa, Ahmad Moustafa, Hala Hafez, Sara Elnakeep, Shaimaa Moustafa, Aya Mohamed, Reham Mamdouh, Ghada Ismael, Ashraf Omar, Osama Mansour, Mahmoud Elmeitini |
| EPI_ISL_483059 | Hospital Universitari Germans Trias i Pujol | IrsiCaixa AIDS Research Lab | J. Segalés, M. Puig, J. Rodon, C. Avila-Nieto, J. Carrillo, G. Cantero, M.T. Terrón, S. Cruz, M. Parera ,M. Noguera-Julián, N. Izquierdo-Useros, V. Guallar, E. Vidal, A. Valencia, I. Blanco, J. Blanco, B. Clotet, J. Vergara-Alert |
| EPI_ISL_483637 | National Laboratory of Virology, Szentágothai Research Centre | National Laboratory of Virology, Szentágothai Research Centre | Endre Gábor Tóth, Balázs Somogyi, Ferenc Jakab, Gábor Kemenesi |
| EPI_ISL_486382 | District Surveillence Unit | Department of Neurovirology, National Institute of Mental Health and Neuroscience (NIMHANS) | Chitra Pattabiraman, Vijayalakshmi Reddy, Harsha PK, Risha Rasheed, Shafeeq S Hameed, Manjunatha Venkataswamy, Anita Desai, Ravi Vasanthapuram |
| EPI_ISL_486383 | CV Raman Hospital | Department of Neurovirology, National Institute of Mental Health and Neuroscience (NIMHANS) | Chitra Pattabiraman, Vijayalakshmi Reddy, Harsha PK, Risha Rasheed, Shafeeq S Hameed, Manjunatha Venkataswamy, Anita Desai, Ravi Vasanthapuram |
| EPI_ISL_486388, EPI_ISL_486389 | DH | Department of Neurovirology, National Institute of Mental Health and Neuroscience (NIMHANS) | Chitra Pattabiraman, Vijayalakshmi Reddy, Harsha PK, Risha Rasheed, Shafeeq S Hameed, Manjunatha Venkataswamy, Anita Desai, Ravi Vasanthapuram |
| EPI_ISL_486392 | Victoria Hospital | Department of Neurovirology, National Institute of Mental Health and Neuroscience (NIMHANS) | Chitra Pattabiraman, Vijayalakshmi Reddy, Harsha PK, Risha Rasheed, Shafeeq S Hameed, Manjunatha Venkataswamy, Anita Desai, Ravi Vasanthapuram |
| EPI_ISL_486394 | MIMS | Department of Neurovirology, National Institute of Mental Health and Neuroscience (NIMHANS) | Chitra Pattabiraman, Vijayalakshmi Reddy, Harsha PK, Risha Rasheed, Shafeeq S Hameed, Manjunatha Venkataswamy, Anita Desai, Ravi Vasanthapuram |
| EPI_ISL_486395 | BIMS | Department of Neurovirology, National Institute of Mental Health and Neuroscience (NIMHANS) | Chitra Pattabiraman, Vijayalakshmi Reddy, Harsha PK, Risha Rasheed, Shafeeq S Hameed, Manjunatha Venkataswamy, Anita Desai, Ravi Vasanthapuram |
| EPI_ISL_486397 | KC General Hospital | Department of Neurovirology, National Institute of Mental Health and Neuroscience (NIMHANS) | Chitra Pattabiraman, Vijayalakshmi Reddy, Harsha PK, Risha Rasheed, Shafeeq S Hameed, Manjunatha Venkataswamy, Anita Desai, Ravi Vasanthapuram |
| EPI_ISL_486398, EPI_ISL_486399 | MIMS | Department of Neurovirology, National Institute of Mental Health and Neuroscience (NIMHANS) | Chitra Pattabiraman, Vijayalakshmi Reddy, Harsha PK, Risha Rasheed, Shafeeq S Hameed, Manjunatha Venkataswamy, Anita Desai, Ravi Vasanthapuram |
| EPI_ISL_486400 | Victoria Hospital | Department of Neurovirology, National Institute of Mental Health and Neuroscience (NIMHANS) | Chitra Pattabiraman, Vijayalakshmi Reddy, Harsha PK, Risha Rasheed, Shafeeq S Hameed, Manjunatha Venkataswamy, Anita Desai, Ravi Vasanthapuram |
| EPI_ISL_486405, EPI_ISL_486408, EPI_ISL_486409 | DH | Department of Neurovirology, National Institute of Mental Health and Neuroscience (NIMHANS) | Chitra Pattabiraman, Vijayalakshmi Reddy, Harsha PK, Risha Rasheed, Shafeeq S Hameed, Manjunatha Venkataswamy, Anita Desai, Ravi Vasanthapuram |
| EPI_ISL_486815, EPI_ISL_486816, EPI_ISL_486817, EPI_ISL_486818, EPI_ISL_486819, EPI_ISL_486820, EPI_ISL_486821, EPI_ISL_486822, EPI_ISL_486823, EPI_ISL_486824, EPI_ISL_486825, EPI_ISL_486826, EPI_ISL_486827, EPI_ISL_486828, EPI_ISL_486829 | | | |
| see above | Molecular diagnostic laboratory of Federal Budget Institution of Science "Central Research Institute of Epidemiology" of The Federal Service on Customers' Rights Protection and Human Well-being Surveillance | Group of Genomics and Postgenomic Technologies of Central Research Institute of Epidemilology | Speranskaya AS, Kaptelova VV, Valdokhina AV, Bulanenko VP, Samoilov AE, Korneenko EV, Tivanova EV, Shipulina OY, Akimkin VG |
| EPI_ISL_486856 | Emergency County Hospital | Stefan cel Mare, University Metagenomics lab | Lobiuc Andrei et al. |
| EPI_ISL_486859, EPI_ISL_486860, EPI_ISL_486861, EPI_ISL_486862, EPI_ISL_486863, EPI_ISL_486864, EPI_ISL_486865, EPI_ISL_486866, EPI_ISL_486867, EPI_ISL_486868, EPI_ISL_486870, EPI_ISL_486871, EPI_ISL_486872, EPI_ISL_486873 | | | |
| see above | Institut Pasteur Dakar | Institut Pasteur de Dakar | Ndongo Dia, Moussa Moise Diagne, Mamadou Diop, Marie Henriette Dior Ndione, Mamadou Malado Jallow, Safietou Sanke, Ousmane Faye, Amadou Alpha Sall. |
| EPI_ISL_486881 | CV Raman Hospital | Department of Neurovirology, National Institute of Mental Health and Neuroscience (NIMHANS) | Chitra Pattabiraman, Vijayalakshmi Reddy, Harsha PK, Risha Rasheed, Shafeeq S Hameed, Manjunatha Venkataswamy, Anita Desai, Ravi Vasanthapuram |
| EPI_ISL_487107 | Nigeria Centre for Disease Control (NCDC) | African Centre of Excellence for Genomics of Infectious Diseases (ACEGID), Redeemer's University, Ede, Osun State, Nigeria | Oluniyi P.E., Ajogbasile F.V., Kayode A., Oguzie J., Olawoye I., Uwanibe J., Olumade T., Folarin O.A., Ihekweazu C., Happi C.T. |
| EPI_ISL_487377, EPI_ISL_487381 | Hellenic Pasteur Institute, National Influenza Reference laboratory of Southern Greece & Unit of Bioinformatics and Applied Genomics | Hellenic Pasteur Institute, National Influenza Reference laboratory of Southern Greece & Unit of Bioinformatics and Applied Genomics | Vasiliki Pogka, Timokratis Karamitros, Athanasios Kossyvakis, Antonios Kalliaropoulos, Horefti Elina, Evangelidou Maria, Androniki Voulgari-Kokota, Aspasia Kontou, Andreas Mentis |
| EPI_ISL_487435 | Queen Astrid Military Hospital | Institute of Tropical Medicine | Philippe Selhorst, Colin Anthony |
| EPI_ISL_489708, EPI_ISL_489709 | The National Institute of Public Health | The National Institute of Public Health and State Veterinary Institute Prague | Nagy,A;Jirincova,H;Novakova,L;Trnka,D;Vecerova,J |
| EPI_ISL_489995, EPI_ISL_490106 | CSIR-CDRI/SGPGI, Lucknow | CSIR-CDRI/SGPGI, Lucknow | Saumya Sarkar, Dharam Veer Singh, Rahul Vishvkarma, Ujjala Ghoshal, Uday Ghoshal, Ravishankar Ramachandran, Tapas Kumar Kundu, Rajender Singh |
| EPI_ISL_490112 | The National Institute of Public Health | The National Institute of Public Health and State Veterinary Institute Prague | Nagy,A;Jirincova,H;Novakova,L;Trnka,D;Vecerova,J |
| EPI_ISL_491092 | The National Institute of Public Health | State Veterinary Institute Prague | Nagy,A;Jirincova,H;Novakova,L;Trnka,D;Vecerova,J |
| EPI_ISL_491093, EPI_ISL_491094, EPI_ISL_491095 | The National Institute of Public Health | The National Institute of Public Health and State Veterinary Institute Prague | Nagy,A; Jirincova,H; Novakova,L; Trnka,D; Vecerova,J |
| EPI_ISL_491096 | CSIR-CDRI/SGPGI, Lucknow | CSIR-CDRI/SGPGI, Lucknow | Saumya Sarkar, Dharam Veer Singh, Rahul Vishvkarma, Ujjala Ghoshal, Uday Ghoshal, Ravishankar Ramachandran, Tapas Kumar Kundu, Rajender Singh |
| EPI_ISL_491115 | Cicin-Sain Lab | Cicin-Sain Lab | M. Zeeshan Chaudhry, Kathrin Eschke, Yeonsu Kim, Luka Cicin-Sain |
| EPI_ISL_491117, EPI_ISL_491118 | The National Institute of Public Health | The National Institute of Public Health and State Veterinary Institute Prague | Nagy,A;Jirincova,H;Novakova,L;Trnka,D;Vecerova,J |
| EPI_ISL_491436 | Laboratorio de Referencia Nacional de Virus Respiratorio. Instituto Nacional de Salud Perú | Laboratorio de Referencia Nacional de Biotecnología y Biología Molecular. Instituto Nacional de Salud Perú | Carlos Padilla Rojas, Karolyn Vega Chozo, Priscila Lope Pari, Omar Caceres Rey, Marco Galarza Perez, Maribel Huaringa Nuñez, Johanna Balbuena Torres, Henri Bailon Calderon, Nancy Rojas Serrano |
| EPI_ISL_491437 | Area de Salud Escazu (Coopesana) | Inciensa, Instituto Costarricense de Investigación y Enseñanza en Nutrición y Salud | Francisco Duarte, Hebleen Brenes, Claudio Soto-Garita, Estela Cordero, Adriana Godinez & Melany Calderon |
| EPI_ISL_491438 | Hospital San Rafael de Alajuela | Inciensa, Instituto Costarricense de Investigación y Enseñanza en Nutrición y Salud | Francisco Duarte, Hebleen Brenes, Claudio Soto-Garita, Estela Cordero, Adriana Godinez & Melany Calderon |

| | | | |
|---|---|---|---|
| EPI_ISL_491439 | Hospital Calderon Guardia | Inciensa, Instituto Costarricense de Investigación y Enseñanza en Nutrición y Salud | Francisco Duarte, Hebleen Brenes, Claudio Soto-Garita, Estela Cordero, Adriana Godinez & Melany Calderon |
| EPI_ISL_491440 | Hospital San Rafael de Alajuela | Inciensa, Instituto Costarricense de Investigación y Enseñanza en Nutrición y Salud | Francisco Duarte, Hebleen Brenes, Claudio Soto-Garita, Estela Cordero, Adriana Godinez & Melany Calderon |
| EPI_ISL_491441, EPI_ISL_491442 | Hospital Clinica Biblica | Inciensa, Instituto Costarricense de Investigación y Enseñanza en Nutrición y Salud | Francisco Duarte, Hebleen Brenes, Claudio Soto-Garita, Estela Cordero, Adriana Godinez & Melany Calderon |
| EPI_ISL_491443 | Hospital Fernando Escalante Pradilla | Inciensa, Instituto Costarricense de Investigación y Enseñanza en Nutrición y Salud | Francisco Duarte, Hebleen Brenes, Claudio Soto-Garita, Estela Cordero, Adriana Godinez & Melany Calderon |
| EPI_ISL_491444 | Area de Salud Escazu (Coopesana) | Inciensa, Instituto Costarricense de Investigación y Enseñanza en Nutrición y Salud | Francisco Duarte, Hebleen Brenes, Claudio Soto-Garita, Estela Cordero, Adriana Godinez & Melany Calderon |
| EPI_ISL_491445 | Area de Salud Mata Redonda | Inciensa, Instituto Costarricense de Investigación y Enseñanza en Nutrición y Salud | Francisco Duarte, Hebleen Brenes, Claudio Soto-Garita, Estela Cordero, Adriana Godinez & Melany Calderon |
| EPI_ISL_491446 | Area de Salud Alajuela Central | Inciensa, Instituto Costarricense de Investigación y Enseñanza en Nutrición y Salud | Francisco Duarte, Hebleen Brenes, Claudio Soto-Garita, Estela Cordero, Adriana Godinez & Melany Calderon |
| EPI_ISL_491447 | Hospital Fernando Escalante Pradilla | Inciensa, Instituto Costarricense de Investigación y Enseñanza en Nutrición y Salud | Francisco Duarte, Hebleen Brenes, Claudio Soto-Garita, Estela Cordero, Adriana Godinez & Melany Calderon |
| EPI_ISL_491448 | Hospital San Rafael de Alajuela | Inciensa, Instituto Costarricense de Investigación y Enseñanza en Nutrición y Salud | Francisco Duarte, Hebleen Brenes, Claudio Soto-Garita, Estela Cordero, Adriana Godinez & Melany Calderon |
| EPI_ISL_491449 | Area de Salud Alajuela Sur | Inciensa, Instituto Costarricense de Investigación y Enseñanza en Nutrición y Salud | Francisco Duarte, Hebleen Brenes, Claudio Soto-Garita, Estela Cordero, Adriana Godinez & Melany Calderon |
| EPI_ISL_491450 | Hospital San Juan de Dios | Inciensa, Instituto Costarricense de Investigación y Enseñanza en Nutrición y Salud | Francisco Duarte, Hebleen Brenes, Claudio Soto-Garita, Estela Cordero, Adriana Godinez & Melany Calderon |
| EPI_ISL_491451 | Hospital México | Inciensa, Instituto Costarricense de Investigación y Enseñanza en Nutrición y Salud | Francisco Duarte, Hebleen Brenes, Claudio Soto-Garita, Estela Cordero, Adriana Godinez & Melany Calderon |
| EPI_ISL_491452 | Hospital San Rafael de Alajuela | Inciensa, Instituto Costarricense de Investigación y Enseñanza en Nutrición y Salud | Francisco Duarte, Hebleen Brenes, Claudio Soto-Garita, Estela Cordero, Adriana Godinez & Melany Calderon |
| EPI_ISL_491453 | Hospital México | Inciensa, Instituto Costarricense de Investigación y Enseñanza en Nutrición y Salud | Francisco Duarte, Hebleen Brenes, Claudio Soto-Garita, Estela Cordero, Adriana Godinez & Melany Calderon |
| EPI_ISL_491454 | Hospital San Juan de Dios | Inciensa, Instituto Costarricense de Investigación y Enseñanza en Nutrición y Salud | Francisco Duarte, Hebleen Brenes, Claudio Soto-Garita, Estela Cordero, Adriana Godinez & Melany Calderon |
| EPI_ISL_491455 | Hospital Clinica Biblica | Inciensa, Instituto Costarricense de Investigación y Enseñanza en Nutrición y Salud | Francisco Duarte, Hebleen Brenes, Claudio Soto-Garita, Estela Cordero, Adriana Godinez & Melany Calderon |
| EPI_ISL_491456 | Hospital San Juan de Dios | Inciensa, Instituto Costarricense de Investigación y Enseñanza en Nutrición y Salud | Francisco Duarte, Hebleen Brenes, Claudio Soto-Garita, Estela Cordero, Adriana Godinez & Melany Calderon |
| EPI_ISL_491457 | Area de Salud Los Santos | Inciensa, Instituto Costarricense de Investigación y Enseñanza en Nutrición y Salud | Francisco Duarte, Hebleen Brenes, Claudio Soto-Garita, Estela Cordero, Adriana Godinez & Melany Calderon |
| EPI_ISL_491463 | Laboratorio de Referencia Nacional de Virus Respiratorio. Instituto Nacional de Salud Perú | Laboratorio de Referencia Nacional de Biotecnología y Biología Molecular. Instituto Nacional de Salud Perú | Carlos Padilla Rojas, Karolyn Vega Chozo, Priscila Lope Pari, Omar Caceres Rey, Marco Galarza Perez, Maribel Huaringa Nuñez, Johanna Balbuena Torres, Henri Bailon Calderon, Nancy Rojas Serrano |
| EPI_ISL_491476 | BSL3 Lab, Pendik Veterinary Control Enstitue | Genomic Laboratory (GLAB), Istanbul Technical University | Mustafa HASOKSUZ, Fahriye SARAC, Osman ERGANIS, Serdar UZAR, Hakan ENUL, Cumhur ADIAY, Ahmet SAIT, Orbay SAYI, Kadir YESILBAG, Oguz KARABEY |
| EPI_ISL_491968, EPI_ISL_491969, EPI_ISL_491970, EPI_ISL_491971, EPI_ISL_491972, EPI_ISL_491973, EPI_ISL_491974 | Oman-NIC | Department of Microbiology and Immunology-SQUH | Fahad Zadjali, Samira Al-Maruqi, Amina Al Jardani, Khulood Al-Mammary, Hanan Al-kindi, Fatma BaAlawi, Hamida AL Barwani, Zeyana AL-Dahmani, Intisar Al-Shukri, Aisha Al-Busaidi, Aisha Al-Amri, Ahlam Al-Amri, Mohammed Al-Tobi, Samiha Al Kharusi, Abdulla Balkhair |
| EPI_ISL_492033 | Instituto de Biologia do Exército | Laboratório Metabolismo Macromolecular FirminoTorres de Castro, Instituto de Biofísica Carlos Chagas Filho, Universidade Federal do Rio de Janeiro | Bianca Catarina Azevedo Cabral, Aline Rosa Vianna de Souza, Caleb GM Santos, Marcos Dornelas-Ribeiro, Tatiana LS Nogueira, Nádia Vaez Gonçalves da Cruz, Elizabeth Valentin, Marcio da Costa Cipitelli, Virginia Sara Grancieri do Amaral, Rodrigo Soares de Moura Neto, Clarissa Damaso, Rosane Silva |
| EPI_ISL_492034, EPI_ISL_492038 | Instituto de Biologia do Exército | Laboratório Metabolismo Macromolecular FirminoTorres de Castro, Instituto de Biofísica Carlos Chagas Filho, Universidade Federal do Rio de Janeiro | Bianca Catarina Azevedo Cabral, Aline Rosa Vianna de Souza, Nádia Vaez Gonçalves da Cruz, Caleb GM Santos, Marcos Dornelas-Ribeiro, Tatiana LS Nogueira, Elizabeth Valentin, Marcio da Costa Cipitelli, Virginia Sara Grancieri do Amaral, Rodrigo Soares de Moura Neto, Clarissa Damaso, Rosane Silva |
| EPI_ISL_492039 | Instituto de Biologia do Exército | Laboratório Metabolismo Macromolecular FirminoTorres de Castro, Instituto de Biofísica Carlos Chagas Filho, Universidade Federal do Rio de Janeiro | Bianca Catarina Azevedo Cabral, Aline Rosa Vianna de Souza, Tatiana LS Nogueira, Nádia Vaez Gonçalves da Cruz, Caleb GM Santos, Marcos Dornelas-Ribeiro, Elizabeth Valentin, Marcio da Costa Cipitelli, Virginia Sara Grancieri do Amaral, Rodrigo Soares de Moura Neto, Clarissa Damaso, Rosane Silva |
| EPI_ISL_492041 | Instituto de Biologia do Exército | Laboratório Metabolismo Macromolecular FirminoTorres de Castro, Instituto de Biofísica Carlos Chagas Filho, Universidade Federal do Rio de Janeiro | Bianca Catarina Azevedo Cabral, Aline Rosa Vianna de Souza, Caleb GM Santos, Marcos Dornelas-Ribeiro, Tatiana LS Nogueira, Nádia Vaez Gonçalves da Cruz, Elizabeth Valentin, Marcio da Costa Cipitelli, Virginia Sara Grancieri do Amaral, Rodrigo Soares de Moura Neto, Clarissa Damaso, Rosane Silva |
| EPI_ISL_492042 | Instituto de Biologia do Exército | Laboratório Metabolismo Macromolecular FirminoTorres de Castro, Instituto de Biofísica Carlos Chagas Filho, Universidade Federal do Rio de Janeiro | Bianca Catarina Azevedo Cabral, Aline Rosa Vianna de Souza, Nádia Vaez Gonçalves da Cruz, Caleb GM Santos, Marcos Dornelas-Ribeiro, Tatiana LS Nogueira, Elizabeth Valentin, Marcio da Costa Cipitelli, Virginia Sara Grancieri do Amaral, Rodrigo Soares de Moura Neto, Clarissa Damaso, Rosane Silva |
| EPI_ISL_492184 | INT Fondazione Pascale | INT Fondazione Pascale | Pascale |
| EPI_ISL_492978, EPI_ISL_492979 | Department of Laboratory Medicine Tan Tock Seng Hospital | Department of Laboratory Medicine Tan Tock Seng Hospital | Chen YYC, Zair X, Li C, Tang WY, Maurer-Stroh S, Barkham TMS, Nagarajan N, Sessions OM |
| EPI_ISL_492982, EPI_ISL_492983, EPI_ISL_492984, EPI_ISL_492985, EPI_ISL_492986, EPI_ISL_492987 | IRCCS Sacro Cuore Don Calabria Hospital, Department of Infectious, Tropical Diseases & Microbiology | University of Verona, Department of Biotechnology | Antonio Mori, Michela Deiana, Elena Pomari, Chiara Piubelli; Giulia Lopatriello, Luca Marcolungo, Cristina Beltrami, Chiara Degli Esposti, Emanuela Cosentino, Massimo Delledonne |
| EPI_ISL_493198, EPI_ISL_493199, EPI_ISL_493200, EPI_ISL_493202, EPI_ISL_493205, EPI_ISL_493206, EPI_ISL_493207 | Virology Lab,Department of Pathology, National Cheng Kung University Hospital | Virology Lab,Department of Pathology, National Cheng Kung University Hospital | Huey-Pin Tsai, et al |
| EPI_ISL_493332 | Istituto Zooprofilattico Sperimentale del Mezzogiorno | INMI Lazzaro Spallanzani IRCCS | Cesare E.M. Gruber, Martina Rueca, Barbara Bartolini, Francesco Messina, Antonino Di Caro, Giovanna Fusco, Maurizio Viscardi, Giorgia Borriello, Maria R. Capobianchi |
| EPI_ISL_493333 | Istituto Zooprofilattico Sperimentale del Mezzogiorno | INMI Lazzaro Spallanzani IRCCS | Barbara Bartolini, Martina Rueca, Cesare E.M. Gruber, Francesco Messina, Antonino Di Caro, Giovanna Fusco, Maurizio Viscardi, Giorgia Borriello, Maria R. Capobianchi |

| | | | |
|---|---|---|---|
| EPI_ISL_493334, EPI_ISL_493335, EPI_ISL_493336 | Instituto de Diagnostico y Referencia Epidemiologicos (INDRE) | Instituto de Diagnostico y Referencia Epidemiologicos (INDRE) | Gisela Barrera-Badillo , Abril Rodriguez-Maldonado, Claudia Wong-Arambula , Natividad Cruz-Ortiz, Tatiana Nunez-Garcia, Dayanira Arellano-Suarez, Fabiola Garces-Ayala, Edgar Mendieta-Condado, Lucia Hernandez-Rivas, Irma Lopez-Martinez, Ernesto Ramirez-Gonzalez. |
| EPI_ISL_494757, EPI_ISL_494759, EPI_ISL_494761, EPI_ISL_494771 | INT Fondazione Pascale | INT Fondazione Pascale | INT Fondazione Pascale |
| EPI_ISL_495414, EPI_ISL_495417, EPI_ISL_495423, EPI_ISL_495426, EPI_ISL_495428, EPI_ISL_495432, EPI_ISL_495434, EPI_ISL_495435, EPI_ISL_495437, EPI_ISL_495439, EPI_ISL_495442, EPI_ISL_495443, EPI_ISL_495448, EPI_ISL_495455, EPI_ISL_495457 | | | |
| see above | Kafkas University, Faculty of Medicine, Department of Medical Microbiology | Kafkas University, Faculty of Medicine, Department of Medical Microbiology | Murat Karamese, Didem Ozgur, E. Ediz Tutuncu |
| EPI_ISL_495459 | Centers for Disease Control and Prevention of Lishui | Department of InspectionCenters for Disease Control and Prevention of Lishui | Wang Xiaoguang,Ji Qiaoying,Ji Jiansong,Ye Bifeng,Ye Ling |
| EPI_ISL_496534, EPI_ISL_496548, EPI_ISL_496549, EPI_ISL_496551 | B.J. Govt. Medical College | National Centre For Cell Science | Dhiraj Paul, Kunal Jani, Radha Chauhan, Janesh Kumar, Vasudevan Seshadri, Girdhari Lal, Rajesh Karyakarte, Suvarna Joshi, Murlidhar Tambe, Sourav Sen, Santosh Karade, Kavita Bala Anand, Shelinder Pal Singh Shergill, Rajiv Mohan Gupta, Manoj Kumar Bhat, Arvind Sahu, Maharashtra COVID-19 Study Group, DBT's PAN-INDIA 1000 SARS-CoV2 RNA genome sequencing consortium, Yogesh S Shouche |
| EPI_ISL_496555, EPI_ISL_496556, EPI_ISL_496557, EPI_ISL_496587 | National Centre For Cell Science | National Centre For Cell Science | Dhiraj Paul, Kunal Jani, Radha Chauhan, Janesh Kumar, Vasudevan Seshadri, Girdhari Lal, Rajesh Karyakarte, Suvarna Joshi, Murlidhar Tambe, Sourav Sen, Santosh Karade, Kavita Bala Anand, Shelinder Pal Singh Shergill, Rajiv Mohan Gupta, Manoj Kumar Bhat, Arvind Sahu, Maharashtra COVID-19 Study Group, DBT's PAN-INDIA 1000 SARS-CoV2 RNA genome sequencing consortium, Yogesh S Shouche |
| EPI_ISL_497736, EPI_ISL_497738, EPI_ISL_497744, EPI_ISL_497745 | Instituto Nacional de Salud, Bogotá, Colombia | Instituto Nacional de Salud, Bogotá, Colombia | Katherine Laiton-Donato, Diego A. Álvarez-Díaz, Carlos Franco-Muñoz, Jonathan Reales, Diego Andrés Prada, Jose A. Usme-Ciro, Nicolas D. Franco-Sierra, Zulma M. Cucunubá, Christian Julian VillabonaArenas, Liz Villabona-Arenas, Sussy Echeverria, Astrid C. Flórez, Carolina Ferro, Diana Marcela Walteros-Acero, Franklin Prieto, Carlos Andrés Durán, Martha Lucia Ospina Martinez, Marcela Mercado-Reyes |
| EPI_ISL_497758, EPI_ISL_497760 | CSIR-CDRI/SGPGI, Lucknow | CSIr-CDRI/SGPGI, Lucknow | Saumya Sarkar, Dharam Veer Singh, Rahul Vishvkarma, Ujjala Ghoshal, Uday Ghoshal, Ravishankar Ramachandran, Tapas Kumar Kundu, Rajender Singh |
| EPI_ISL_497763, EPI_ISL_497765 | CSIR-CDRI/SGPGI, Lucknow | CSIR-CDRI/SGPGI, Lucknow | Saumya Sarkar, Dharam Veer Singh, Rahul Vishvkarma, Ujjala Ghoshal, Uday Ghoshal, Ravishankar Ramachandran, Tapas Kumar Kundu, Rajender Singh |
| EPI_ISL_497766 | CSIR-CDRI/SGPGI, Lucknow | CSIR-CDRi/SGPGI, Lucknow | Saumya Sarkar, Dharam Veer Singh, Rahul Vishvkarma, Ujjala Ghoshal, Uday Ghoshal, Ravishankar Ramachandran, Tapas Kumar Kundu, Rajender Singh |
| EPI_ISL_498152, EPI_ISL_498153, EPI_ISL_498154, EPI_ISL_498155, EPI_ISL_498156, EPI_ISL_498157, EPI_ISL_498158, EPI_ISL_498159, EPI_ISL_498160, EPI_ISL_498161, EPI_ISL_498162, EPI_ISL_498163, EPI_ISL_498164, EPI_ISL_498165, EPI_ISL_498166, EPI_ISL_498167, EPI_ISL_498168, EPI_ISL_498169, EPI_ISL_498170 | | | |
| see above | Instituto Nacional de Salud, Bogotá, Colombia | Instituto Nacional de Salud, Bogotá, Colombia | Katherine Laiton-Donato, Diego A. Álvarez-Díaz, Carlos Franco-Muñoz, Jonathan Reales, Diego Andrés Prada, Jose A. Usme-Ciro, Nicolas D. Franco-Sierra, Zulma M. Cucunubá, Christian Julian VillabonaArenas, Liz Villabona-Arenas, Sussy Echeverria, Astrid C. Flórez, Carolina Ferro, Diana Marcela Walteros-Acero, Franklin Prieto, Carlos Andrés Durán, Martha Lucia Ospina Martinez, Marcela Mercado-Reyes |
| EPI_ISL_498171, EPI_ISL_498172, EPI_ISL_498173, EPI_ISL_498174, EPI_ISL_498175, EPI_ISL_498176, EPI_ISL_498177, EPI_ISL_498178, EPI_ISL_498179, EPI_ISL_498180, EPI_ISL_498181, EPI_ISL_498182, EPI_ISL_498183, EPI_ISL_498184, EPI_ISL_498185, EPI_ISL_498186, EPI_ISL_498187, EPI_ISL_498188, EPI_ISL_498189, EPI_ISL_498190, EPI_ISL_498191, EPI_ISL_498192 | | | |
| see above | OUCRU | OUCRU | Nguyen Van Vinh Chau, Nguyen Thi Thu Hong, Nguyen Thi Han Ny, Le Nguyen Truc Nhu, Nghiem My Ngoc, Vo Thanh Lam, Nguyen Thanh Dung, Lam Minh Yen, Ngo Ngoc Quang Minh, Le Manh Hung, Nguyen Tri Dung, Dinh Nguyen Huy Man, Lam Anh Nguyet, Tran Chanh Xuan, Tran Tinh Hien, Nguyen Thanh Phong, Tran Nguyen Hoang Tu, Tran Tan Thanh, Nguyen Thanh Truong, Nguyen Tan Binh, Tang Chi Thuong, Guy Thwaites, and Le Van Tan, for OUCRU COVID-19 research group* |
| EPI_ISL_498229, EPI_ISL_498230, EPI_ISL_498236 | Institut Pasteur de Dakar | Institut Pasteur de Dakar | Ndongo Dia, Moussa Moise Diagne, Mamadou Diop, Marie Henriette Dior Ndione, Mamadou Malado Jallow, Safietou Sankhe Mbengue, Ousmane Faye, Amadou Alpha Sall. |
| EPI_ISL_498551, EPI_ISL_498552, EPI_ISL_498554 | Lebanese American University | Lebanese American University | Abi Habib,W., Abdallah,J., El Shesheny,R., Mokhbat,J., Webby,R.J., Goldstein,J. and Kayali,G. |
| EPI_ISL_499083 | Instituto de Virologia "Dr. J. M. Vanella", Facultad de Ciencias Medicas, Universidad Nacional de Cordoba. Laboratorio Central de la Provincia de Cordoba, Argentina. Ministerio de Salud de la provincia de Cordoba, Argentina. | Laboratorio de Virología, Hospital de Niños Ricardo Gutiérrez, CABA, Argentina. | Sandra Gallego, Brenda Konigheim, Sebastian Blanco, Lorena Spinsanti, Javier Aguilar, Adrian Diaz, Gonzalo Castro, Gabriela Barbas, Mercedes Nabaes, Stephanie Goya, Monica Natale, Silvina Lusso, Mariana Viegas. |
| EPI_ISL_500541 | Singapore General Hospital | Department of Microbiology | Nurdyana Abdul Rahman, Kun Lee Lim, Chenhao Li, Kian Sing Chan, Lynette Oon, Kern Rei Chng, Niranjan Nagarajan, Karrie Ko |
| EPI_ISL_500716 | BSL3 Lab Pendik Veterinary Control Institute | Department of Medicinal Genetics, Bursa Uluda University, Faculty of medicine By Sehime Gülsün Temel, Adem Alemdar, Kadir Yeilba | Mustafa HASOKSUZ, Fahriye SARAC, Osman ERGANIS, Serdar UZAR, Hakan ENUL, Cumhur ADIAY, Ahmet SAIT, Orbay SAYI, Kadir YESILBAG, Oguz KARABEY |
| EPI_ISL_501167, EPI_ISL_501168, EPI_ISL_501169, EPI_ISL_501170, EPI_ISL_501171, EPI_ISL_501172, EPI_ISL_501173, EPI_ISL_501174 | Baylor College of Medicine | Baylor College of Medicine: HGSC | Vasanthi Avadhanula, Erin Nicholson, David Henke, Pedro Piedra, Harsha Doddapaneni, Donna Muzny, Qingchang Meng, Hsu Chao, Zeineen Momin, Hua Shen, George Weissenberger, Kavya Kottapalli, Yimiti Meiheerguli, Sejal Salvi, Ginger Metcalf, Vipin Menon, Sara J.J. Cregeen, Matthew C. Ross, Tulin Ayvaz, Richard Sucgang, Kristi L. Hoffman, Matthew Wong, Joseph F. Petrosino |
| EPI_ISL_501176, EPI_ISL_501177, EPI_ISL_501178, EPI_ISL_501179, EPI_ISL_501180, EPI_ISL_501181, EPI_ISL_501182, EPI_ISL_501183, EPI_ISL_501184, EPI_ISL_501185, EPI_ISL_501186, EPI_ISL_501187, EPI_ISL_501188, EPI_ISL_501189, EPI_ISL_501190, EPI_ISL_501191, EPI_ISL_501192, EPI_ISL_501193, EPI_ISL_501194, EPI_ISL_501195, EPI_ISL_501196, EPI_ISL_501197, EPI_ISL_501198, EPI_ISL_501199, EPI_ISL_501200, EPI_ISL_501201, EPI_ISL_501202, EPI_ISL_501203, EPI_ISL_501204, EPI_ISL_501205, EPI_ISL_501206, EPI_ISL_501207, EPI_ISL_501208, EPI_ISL_501209, EPI_ISL_501210, EPI_ISL_501211, EPI_ISL_501212, EPI_ISL_501213, EPI_ISL_501214, EPI_ISL_501215, EPI_ISL_501216, EPI_ISL_501217, EPI_ISL_501218, EPI_ISL_501219, EPI_ISL_501220, EPI_ISL_501221, EPI_ISL_501222, EPI_ISL_501223, EPI_ISL_501224, EPI_ISL_501225, EPI_ISL_501226, EPI_ISL_501227, EPI_ISL_501228 | | | |
| see above | Department of Medical Microbiology, University Malaya Medical Centre | Department of Medical Microbiology, Faculty of Medicine, University of Malaya | Yoong Min CHONG, Jennifer Chong, I-Ching SAM, Yoke Fun CHAN, University Malaya Medical Centre COVID Team |
| EPI_ISL_501230 | Hellenic Pasteur Institute, Public Health Laboratories | Hellenic Pasteur Institute, National Influenza Reference laboratory of Southern Greece & Unit of Bioinformatics and Applied Genomics | Vasiliki Pogka, Timokratis Karamitros, Athanasios Kossyvakis, Antonios Kalliaropoulos, Horefti Elina, Evangelidou Maria, Androniki Voulgari-Kokota, Aspasia Kontou, Andreas Mentis |
| EPI_ISL_501233, EPI_ISL_501236, EPI_ISL_501248, EPI_ISL_501249, EPI_ISL_501250, EPI_ISL_501251 | Hellenic Pasteur Institute, National Influenza Reference laboratory of Southern Greece & Unit of Bioinformatics and Applied Genomics | Hellenic Pasteur Institute, National Influenza Reference laboratory of Southern Greece & Unit of Bioinformatics and Applied Genomics | Vasiliki Pogka, Timokratis Karamitros, Athanasios Kossyvakis, Antonios Kalliaropoulos, Horefti Elina, Evangelidou Maria, Androniki Voulgari-Kokota, Aspasia Kontou, Andreas Mentis |
| EPI_ISL_505003 | Biology Dpt | Microbiology and Infections Diseases | Emmanuelle Billon-Denis, Audrey Ferrier-Rembert, Annabelle Garnier, Laurence Cheutin, Clarisse Vigne, Emilie Tessier, Jessica Denis, Olivier Gorgé, Flora Nolent, Isabelle Drouet, Noémie Verguet, Olivier Ferraris, Jean-Nicolas Tournier |
| EPI_ISL_506996 | Department of Medical Microbiology, University Malaya Medical Centre | Department of Medical Microbiology, Faculty of Medicine, University of Malaya | Yoong Min CHONG, Jennifer Chong, I-Ching SAM, Yoke Fun CHAN, University Malaya Medical Centre COVID Team |
| EPI_ISL_507002, EPI_ISL_507003, EPI_ISL_507004, EPI_ISL_507005, EPI_ISL_507006 | Department of Laboratory Medicine, Tan Tock Seng Hospital | Department of Laboratory Medicine, Tan Tock Seng Hospital | Chen YYC, Zair X, Li C, Tang WY, Maurer-Stroh S, Barkham TMS, Nagarajan N, Sessions OM |
| EPI_ISL_507008 | National Veterinary Institute | National Veterinary Institute | Siamak Zohari |
| EPI_ISL_507206, EPI_ISL_507207, EPI_ISL_507208, EPI_ISL_507209, EPI_ISL_507210, EPI_ISL_507211, EPI_ISL_507212, EPI_ISL_507213, EPI_ISL_507214, EPI_ISL_507215 | Department of Experimental Modeling and Pathogenesis of Infectious Diseases | WHO National Influenza Centre Russian Federation | Andrey Komissarov, Artem Fadeev, Mariia Sergeeva, Anna Ivanova, Daria Danilenko |
| EPI_ISL_507225, EPI_ISL_507226, | WHO National Influenza Centre Russian Federation | WHO National Influenza Centre Russian Federation | Andrey Komissarov, Artem Fadeev, Mariia Sergeeva, Anna Ivanova, Daria Danilenko |

| | | | |
|---|---|---|---|
| EPI_ISL_507261, EPI_ISL_507262, EPI_ISL_507263, EPI_ISL_507264, EPI_ISL_507291, EPI_ISL_507292, EPI_ISL_507293, EPI_ISL_507294 | | | |
| EPI_ISL_508862, EPI_ISL_508863 | Virology Unit, Institut Pasteur de Madagascar | Virology Unit, Institut Pasteur de Madagascar | Christian Ranaivoson, Cara Brook, Norosoa Razanajatovo, Vida Ahyong, Tsiry Randriambolamanantsoa, Michelle Tan, Vololoniaina Raharinosy, Helisoa Razafimanjato, Cristina M. Tato, Joseph L. DeRisi, Soa Fy Andriamandimby, Jean-Michel Heraud |
| EPI_ISL_509413, EPI_ISL_509416 | Acibadem Labcell Cellular Therapy Laboratory | Acibadem Mehmet Ali Aydinlar University School of Medicine, Medical Genetics Department | Ozden Hatirnaz Ng, Sezer Akyoney, Ilayda Sahin, Gunseli Bayram Akcapinar, Ozkan Ozdemir, Derya Dilek Kancagi, Gozde Sir Karakus, Bulut Yurtsever, Cihan Tastan, Ercument Ovali, Ugur Ozbek |
| EPI_ISL_510689 | Hospital Universitari Germans Trias i Pujol(HUGTiP)/Fundació Lluita contra la SIDA (FLSida)/IRTA-CReSA | IrsiCaixa AIDS Research Lab | Pilar Armengol, Marc Noguera-Julian, Jordi Rodón, Julia Vergara, Lidia Ruiz, Nuria Izquierdo, Jorge Carrillo, Roger Paredes, Albert Bensaid, Julia Blanco, Joaquim Segalés, Bonaventura Clotet |
| EPI_ISL_511891, EPI_ISL_511892, EPI_ISL_511893, EPI_ISL_511894, EPI_ISL_511895, EPI_ISL_511896, EPI_ISL_511897, EPI_ISL_511898 | National Hospital of Tropical Diseases | Oxford University Clinical Research Unit, Hanoi, Vietnam | Nguyen Thi Tam, Van Dinh Trang, Nguyen Thi Hong Thuong, Vu Thi Ngoc Bich, Nguyen Thu Trang, Nguyen Thi Ngoc Diep, Le Nguyen Minh Hoa, Pham Ngoc Thach, H. Rogier van Doorn, on behalf of the OUCRU COVID-19 research group |
| EPI_ISL_512653 | Area De Salud Desamparados 1 - Clinica Dr. Marcial Fallas [Grifo Alto/Desampara | Inciensa, Instituto Costarricense de Investigación y Enseñanza en Nutrición y Salud | Francisco Duarte, Hebleen Porras, Claudio Soto-Garita, Estela Cordero, Adriana Godinez & Melany Calderon |
| EPI_ISL_512844 | Department of Medical Research | DMR_Myanmar | Myat Htut Nyunt, Hnin Ohnmar Soe, Kay Thi Aye, Wah Wah Aung,Yi Yi Kyaw, Aung Kyaw Kyaw, Theingi Win Myat, Phyu Win Ei, Aung Zaw Latt, Nan Aye Thida Oo, Lai Lai San, Su Mon Win, Ni Ni Zaw, Htin Lin, Hlaing Myat Thu, Zaw Than Htun |
| EPI_ISL_512908, EPI_ISL_512909, EPI_ISL_512911, EPI_ISL_512912, EPI_ISL_512913, EPI_ISL_512914, EPI_ISL_512915, EPI_ISL_512916, EPI_ISL_512917 | Pathogen Genomics Lab King Abdullah University of Science and Technology(KAUST) | Pathogen Genomics Lab King Abdullah University of Science and Technology(KAUST) | Sharif Hala, Fadwa Alofi, Sara Mfarrej, Amit Kumar Subudhi, Rahul P Salunke, Fathia Ben Rached, Amanda Ooi, Luke Esau, Afrah Alsomali, Asim Khogeer, Jumana Taha, Abdulaziz Alahmadi, Kahled Algithami, Raeece Naeem, Anwar Hashem, Naif Almontashiri, Arnab Pain |
| EPI_ISL_512918, EPI_ISL_512919, EPI_ISL_512928, EPI_ISL_512929, EPI_ISL_512930, EPI_ISL_512931, EPI_ISL_512932, EPI_ISL_512933, EPI_ISL_512934, EPI_ISL_512935, EPI_ISL_512936, EPI_ISL_512937, EPI_ISL_512938, EPI_ISL_512939, EPI_ISL_512941, EPI_ISL_512942, EPI_ISL_512943, EPI_ISL_512944, EPI_ISL_512945 | | | |
| see above | Pathogen Genomics Lab King Abdullah University of Science and Technology(KAUST) | Pathogen Genomics Lab King Abdullah University of Science and Technology(KAUST) | Fadwa Alofi, Sharif Hala, Rahul P Salunke, Sara Mfarrej, Amit Kumar Subudhi, Fathia Ben Rached, Amanda, Luke, Afrah Alsomali, Asim Khogeer, Jumana Taha, Abdulaziz Alahmadi, Kahled Algithami, Raeece Naeem, Anwar Hashem, Naif Almontashiri, Arnab Pain |
| EPI_ISL_512946, EPI_ISL_512947, EPI_ISL_512948, EPI_ISL_512949, EPI_ISL_512950, EPI_ISL_512951, EPI_ISL_512952, EPI_ISL_512953, EPI_ISL_512954, EPI_ISL_512955, EPI_ISL_512956, EPI_ISL_512957, EPI_ISL_512959, EPI_ISL_512963, EPI_ISL_512964, EPI_ISL_512966, EPI_ISL_512967, EPI_ISL_512969, EPI_ISL_512970, EPI_ISL_512971, EPI_ISL_512973, EPI_ISL_512974, EPI_ISL_512975, EPI_ISL_512976, EPI_ISL_512977, EPI_ISL_512979, EPI_ISL_512980, EPI_ISL_512981, EPI_ISL_512982, EPI_ISL_512983, EPI_ISL_512984, EPI_ISL_512986, EPI_ISL_512987, EPI_ISL_512988 | | | |
| see above | Pathogen Genomics Lab King Abdullah University of Science and Technology(KAUST) | Pathogen Genomics Lab King Abdullah University of Science and Technology(KAUST) | Sara Mfarrej, Raeece Naeem, Rahul P Salunke, Sharif Hala, Fadwa Alofi, Amit Kumar Subudhi, Fathia Ben Rached, Afrah Alsomali, Jumana Taha, Abdulaziz Alahmadi, Asim Khogeer, Nashwa Al-khotani, Anwar Hashem, Naif Almontashiri, Arnab Pain |
| EPI_ISL_512989, EPI_ISL_512990, EPI_ISL_512991, EPI_ISL_512992, EPI_ISL_512993, EPI_ISL_512994, EPI_ISL_512995, EPI_ISL_512996, EPI_ISL_512997, EPI_ISL_512998, EPI_ISL_512999, EPI_ISL_513000, EPI_ISL_513001 | | | |
| see above | Pathogen Genomics Lab King Abdullah University of Science and Technology(KAUST) | Pathogen Genomics Lab King Abdullah University of Science and Technology(KAUST) | Amit Kumar Subudhi, Rahul P Salunke, Sara Mfarrej, Sharif Hala, Fadwa Alofi, Fathia Ben Rached, Afrah Alsomali, Asim Khogeer, Nashwa Al-khotani, Raeece Naeem, Anwar Hashem, Naif Almontashiri, Arnab Pain |
| EPI_ISL_513041, EPI_ISL_513042, EPI_ISL_513043, EPI_ISL_513044, EPI_ISL_513045, EPI_ISL_513046, EPI_ISL_513047, EPI_ISL_513048, EPI_ISL_513049, EPI_ISL_513050, EPI_ISL_513051, EPI_ISL_513052, EPI_ISL_513053, EPI_ISL_513054, EPI_ISL_513055, EPI_ISL_513056, EPI_ISL_513057, EPI_ISL_513058, EPI_ISL_513059, EPI_ISL_513060, EPI_ISL_513061, EPI_ISL_513062 | | | |
| see above | Pathogen Genomics Lab King Abdullah University of Science and Technology(KAUST) | Pathogen Genomics Lab King Abdullah University of Science and Technology(KAUST) | Rahul P Salunke, Sharif Hala, Raeece Naeem, Sara Mfarrej, Amit Kumar Subudhi, Amanda Ooi, Luke Esau, Fadwa Alofi, Fathia Ben Rached, Afrah Alsomali, Asim Khogeer, Ahmad Bakur Mahmoud, Anwar Hashem, Naif Almontashiri, Arnab Pain |
| EPI_ISL_513122, EPI_ISL_513123, EPI_ISL_513124, EPI_ISL_513125, EPI_ISL_513126, EPI_ISL_513127, EPI_ISL_513128, EPI_ISL_513129, EPI_ISL_513131, EPI_ISL_513132, EPI_ISL_513133, EPI_ISL_513134, EPI_ISL_513135, EPI_ISL_513137, EPI_ISL_513138, EPI_ISL_513139, EPI_ISL_513141, EPI_ISL_513142, EPI_ISL_513143, EPI_ISL_513144, EPI_ISL_513145, EPI_ISL_513146 | | | |
| see above | Pathogen Genomics Lab King Abdullah University of Science and Technology(KAUST) | Pathogen Genomics Lab King Abdullah University of Science and Technology(KAUST) | Sharif Hala, Fadwa Alofi, Sara Mfarrej, Amit Kumar Subudhi, Rahul P Salunke, Fathia Ben Rached, Amanda Ooi, Luke Esau, Afrah Alsomali, Asim Khogeer, Jumana Taha, Abdulaziz Alahmadi, Kahled Algithami, Raeece Naeem, Anwar Hashem, Naif Almontashiri, Arnab Pain |
| EPI_ISL_513147, EPI_ISL_513148, EPI_ISL_513149, EPI_ISL_513150, EPI_ISL_513151, EPI_ISL_513152, EPI_ISL_513153, EPI_ISL_513154, EPI_ISL_513156, EPI_ISL_513157, EPI_ISL_513158, EPI_ISL_513159, EPI_ISL_513160, EPI_ISL_513161, EPI_ISL_513162, EPI_ISL_513163, EPI_ISL_513164, EPI_ISL_513165, EPI_ISL_513166, EPI_ISL_513167 | | | |
| see above | Pathogen Genomics Lab King Abdullah University of Science and Technology(KAUST) | Pathogen Genomics Lab King Abdullah University of Science and Technology(KAUST) | Fadwa Alofi, Sharif Hala, Rahul P Salunke, Sara Mfarrej, Amit Kumar Subudhi, Fathia Ben Rached, Amanda, Luke, Afrah Alsomali, Asim Khogeer, Jumana Taha, Abdulaziz Alahmadi, Kahled Algithami, Raeece Naeem, Anwar Hashem, Naif Almontashiri, Arnab Pain |
| EPI_ISL_513168, EPI_ISL_513169, EPI_ISL_513170, EPI_ISL_513171, EPI_ISL_513172, EPI_ISL_513173, EPI_ISL_513174 | Pathogen Genomics Lab King Abdullah University of Science and Technology(KAUST) | Pathogen Genomics Lab King Abdullah University of Science and Technology(KAUST) | Sara Mfarrej, Raeece Naeem, Rahul P Salunke, Sharif Hala, Fadwa Alofi, Amit Kumar Subudhi, Fathia Ben Rached, Afrah Alsomali, Jumana Taha, Abdulaziz Alahmadi, Asim Khogeer, Nashwa Al-khotani, Anwar Hashem, Naif Almontashiri, Arnab Pain |
| EPI_ISL_513175, EPI_ISL_513176, EPI_ISL_513177, EPI_ISL_513178, EPI_ISL_513179, EPI_ISL_513180, EPI_ISL_513181, EPI_ISL_513182, EPI_ISL_513183, EPI_ISL_513184, EPI_ISL_513185, EPI_ISL_513186, EPI_ISL_513187, EPI_ISL_513188, EPI_ISL_513189 | | | |
| see above | Pathogen Genomics Lab King Abdullah University of Science and Technology(KAUST) | Pathogen Genomics Lab King Abdullah University of Science and Technology(KAUST) | Amit Kumar Subudhi, Rahul P Salunke, Sara Mfarrej, Sharif Hala, Fadwa Alofi, Fathia Ben Rached, Afrah Alsomali, Asim Khogeer, Nashwa Al-khotani, Raeece Naeem, Anwar Hashem, Naif Almontashiri, Arnab Pain |
| EPI_ISL_513195 | Pathogen Genomics Lab King Abdullah University of Science and Technology(KAUST) | Pathogen Genomics Lab King Abdullah University of Science and Technology(KAUST) | Afrah Alsomali, Fathia Ben Rached, Raeece Naeem, Sharif Hala,Rahul P Salunke, Amanda Ooi, Luke Esau, Sara Mfarrej, Amit Kumar Subudhi, Fadwa Alofi, Asim Khogeer, Kahled Algithami, Anwar Hashem, Naif Almontashiri, Arnab Pain |
| EPI_ISL_513198, EPI_ISL_513205, EPI_ISL_513206 | Pathogen Genomics Lab King Abdullah University of Science and Technology(KAUST) | Pathogen Genomics Lab King Abdullah University of Science and Technology(KAUST) | Rahul P Salunke, Sharif Hala, Raeece Naeem, Sara Mfarrej, Amit Kumar Subudhi, Amanda Ooi, Luke Esau, Fadwa Alofi, Fathia Ben Rached, Afrah Alsomali, Asim Khogeer, Ahmad Bakur Mahmoud, Anwar Hashem, Naif Almontashiri, Arnab Pain |
| EPI_ISL_513208, EPI_ISL_513209, EPI_ISL_513211 | Pathogen Genomics Lab King Abdullah University of Science and Technology(KAUST) | Pathogen Genomics Lab King Abdullah University of Science and Technology(KAUST) | Raeece Naeem, Rahul P Salunke, Sharif Hala, Sara Mfarrej, Amit Kumar Subudhi, Fadwa Alofi, Fathia Ben Rached, Afrah Alsomali, Asim Khogeer, Ahmad Bakur Mahmoud, Anwar Hashem, Naif Almontashiri, Arnab Pain |
| EPI_ISL_513214, EPI_ISL_513215, EPI_ISL_513216, EPI_ISL_513217, EPI_ISL_513218, EPI_ISL_513219, EPI_ISL_513221, EPI_ISL_513222, EPI_ISL_513224, EPI_ISL_513226 | Pathogen Genomics Lab King Abdullah University of Science and Technology(KAUST) | Pathogen Genomics Lab King Abdullah University of Science and Technology(KAUST) | Sharif Hala, Fadwa Alofi, Sara Mfarrej, Amit Kumar Subudhi, Rahul P Salunke, Fathia Ben Rached, Amanda Ooi, Luke Esau, Afrah Alsomali, Asim Khogeer, Jumana Taha, Abdulaziz Alahmadi, Kahled Algithami, Raeece Naeem, Anwar Hashem, Naif Almontashiri, Arnab Pain |
| EPI_ISL_513228, EPI_ISL_513229, EPI_ISL_513230, EPI_ISL_513231, EPI_ISL_513233, EPI_ISL_513234, EPI_ISL_513235, EPI_ISL_513236, EPI_ISL_513238, EPI_ISL_513240, EPI_ISL_513241, EPI_ISL_513242, EPI_ISL_513243, EPI_ISL_513244, EPI_ISL_513245, EPI_ISL_513246, EPI_ISL_513247, EPI_ISL_513248 | | | |
| see above | Pathogen Genomics Lab King Abdullah University of Science and Technology(KAUST) | Pathogen Genomics Lab King Abdullah University of Science and Technology(KAUST) | Amit Kumar Subudhi, Rahul P Salunke, Sara Mfarrej, Sharif Hala, Fadwa Alofi, Fathia Ben Rached, Afrah Alsomali, Asim Khogeer, Nashwa Al-khotani, Raeece Naeem, Anwar Hashem, Naif Almontashiri, Arnab Pain |
| EPI_ISL_513249, EPI_ISL_513250, EPI_ISL_513251, EPI_ISL_513252, EPI_ISL_513253, EPI_ISL_513254, EPI_ISL_513257, EPI_ISL_513258, EPI_ISL_513259, EPI_ISL_513260, EPI_ISL_513261, EPI_ISL_513262, EPI_ISL_513263, EPI_ISL_513264 | | | |
| see above | Pathogen Genomics Lab King Abdullah University of Science and Technology(KAUST) | Pathogen Genomics Lab King Abdullah University of Science and Technology(KAUST) | Afrah Alsomali, Fathia Ben Rached, Raeece Naeem, Sharif Hala,Rahul P Salunke, Amanda Ooi, Luke Esau, Sara Mfarrej, Amit Kumar Subudhi, Fadwa Alofi, Asim Khogeer, Kahled Algithami, Anwar Hashem, Naif Almontashiri, Arnab Pain |
| EPI_ISL_513314, EPI_ISL_513511, | The National Institute of Public Health | The National Institute of Public Health and State Veterinary | Nagy,A;Jirincova,H;Novakova,L;Trnka,D;Vecerova,J |

| | | | |
|---|---|---|---|
| EPI_ISL_513512 | | Institute Prague | |
| EPI_ISL_513513, EPI_ISL_513514, EPI_ISL_513515, EPI_ISL_513516, EPI_ISL_513518, EPI_ISL_513519, EPI_ISL_513520, EPI_ISL_513521, EPI_ISL_513522, EPI_ISL_513524, EPI_ISL_513525, EPI_ISL_513526, EPI_ISL_513527, EPI_ISL_513529, EPI_ISL_513530, EPI_ISL_513531, EPI_ISL_513532, EPI_ISL_513533, EPI_ISL_513534, EPI_ISL_513535, EPI_ISL_513536, EPI_ISL_513537, EPI_ISL_513538, EPI_ISL_513539, EPI_ISL_513540, EPI_ISL_513541, EPI_ISL_513542, EPI_ISL_513543, EPI_ISL_513544, EPI_ISL_513545, EPI_ISL_513546, EPI_ISL_513547, EPI_ISL_513548, EPI_ISL_513549, EPI_ISL_513550, EPI_ISL_513551, EPI_ISL_513553, EPI_ISL_513556, EPI_ISL_513557, EPI_ISL_513558, EPI_ISL_513559, EPI_ISL_513560, EPI_ISL_513561, EPI_ISL_513562, EPI_ISL_513563, EPI_ISL_513564, EPI_ISL_513567, EPI_ISL_513568, EPI_ISL_513571, EPI_ISL_513572, EPI_ISL_513573, EPI_ISL_513574, EPI_ISL_513575, EPI_ISL_513576, EPI_ISL_513577, EPI_ISL_513578, EPI_ISL_513579, EPI_ISL_513580, EPI_ISL_513581, EPI_ISL_513582, EPI_ISL_513583 | | | |
| see above | Programa de Oncovirologia, Instituto Nacional de Câncer | Programa de Oncovirologia, Instituto Nacional de Câncer | Juliana D. Siqueira, Livia R. Goes, Brunna M. Alves, Claudia Cicala,James Arthos, João P.B. Viola, Andreia C. de Melo, Marcelo A. Soares |
| EPI_ISL_514126 | National Scientific Center for Especially Dangerous Infections (NSCEDI) | Kazakh National Agrarian University (KazNAU) TreeGene LLP Genetic Laboratory | Tabynov Kaissar, Beloussov Vyacheslav, Strochkov Vitaliy, Sandybayev Nurlan, Tabynov Kairat, Turebekov Nurkeldy, Granica Joanna, Solomadin Maksim, Yerubayev Toktassyn, Yespolov Tlektes |
| EPI_ISL_514131 | Rondônia Central Public Health Laboratory (LACEN/RO), vinculated to State Health Secretariat of Rondônia (SESAU/RO) | Molecular Virology Laboratory of Oswaldo Cruz Foundation of Rondônia | Luan Felipo Botelho-Souza, Felipe Souza Nogueira-Lima, Tárcio Peixoto Roca, Alcione de Oliveira dos Santos, Felipe Gomes Naveca, Adriana Cristina Salvador Maia, Cicileia Correia da Silva, Aline Linhares Ferreira de Melo Mendonça, Celina Aparecida Bertoni Lugtenburg, Camila Flávia Gomes Azzi, Juliana Loca Furtado, Suelen Cavalcante, Rita de Cássia Pontello Rampazzo, Caio Henrique Nemeth Santos, Alice Paula Di Sabatino Guimarães, Jansen Fernandes de Medeiros, Fernando Rodrigues Máximo, Juan Miguel Vilallobos-Salcedo and Deusilene Souza Vieira1 |
| EPI_ISL_514132 | Rondônia Central Public Health Laboratory (LACEN/RO), vinculated to State Health Secretariat of Rondônia (SESAU/RO) | Molecular Virology Laboratory of Oswaldo Cruz Foundation of Rondônia | Luan Felipo Botelho-Souza, Felipe Souza Nogueira-Lima, Tárcio Peixoto Roca, Alcione de Oliveira dos Santos, Felipe Gomes Naveca, Adriana Cristina Salvador Maia, Cicileia Correia da Silva, Aline Linhares Ferreira de Melo Mendonça, Celina Aparecida Bertoni Lugtenburg, Camila Flávia Gomes Azzi, Juliana Loca Furtado, Suelen Cavalcante, Rita de Cássia Pontello Rampazzo, Caio Henrique Nemeth Santos, Alice Paula Di Sabatino Guimarães, Jansen Fernandes de Medeiros, Fernando Rodrigues Máximo, Juan Miguel Vilallobos-Salcedo and Deusilene Souza Vieira. |
| EPI_ISL_514133, EPI_ISL_514134, EPI_ISL_514135, EPI_ISL_514136, EPI_ISL_514137 | Rondônia Central Public Health Laboratory (LACEN/RO), vinculated to State Health Secretariat of Rondônia (SESAU/RO) | Molecular Virology Laboratory of Oswaldo Cruz Foundation of Rondônia | Luan Felipo Botelho-Souza, Felipe Souza Nogueira-Lima, Tárcio Peixoto Roca, Alcione de Oliveira dos Santos, Felipe Gomes Naveca, Adriana Cristina Salvador Maia, Cicileia Correia da Silva, Aline Linhares Ferreira de Melo Mendonça, Celina Aparecida Bertoni Lugtenburg, Camila Flávia Gomes Azzi, Juliana Loca Furtado, Suelen Cavalcante, Rita de Cássia Pontello Rampazzo, Caio Henrique Nemeth Santos, Alice Paula Di Sabatino Guimarães, Jansen Fernandes de Medeiros, Fernando Rodrigues Máximo, Juan Miguel Vilallobos-Salcedo and Deusilene Souza Vieira |
| EPI_ISL_515520 | Hospital Municipal do Tatuape Carmino Caricchio | Instituto Adolfo Lutz, Interdiciplinary Procedures Center, Strategic Laboratory | Claudio Tavares Sacchi, Claudia Regina Gonçalves, Erica Valessa Ramos Gomes |
| EPI_ISL_515521 | Hospital Municipal Dr Waldemar Tebaldi | Instituto Adolfo Lutz, Interdiciplinary Procedures Center, Strategic Laboratory | Claudio Tavares Sacchi, Claudia Regina Gonçalves, Erica Valessa Ramos Gomes |
| EPI_ISL_515522 | UPA 24HS de Itatiba | Instituto Adolfo Lutz, Interdiciplinary Procedures Center, Strategic Laboratory | Claudio Tavares Sacchi, Claudia Regina Gonçalves, Erica Valessa Ramos Gomes |
| EPI_ISL_515523, EPI_ISL_515524 | PS Municipal Dr Lauro Ribas Braga | Instituto Adolfo Lutz, Interdiciplinary Procedures Center, Strategic Laboratory | Claudio Tavares Sacchi, Claudia Regina Gonçalves, Erica Valessa Ramos Gomes |
| EPI_ISL_515526 | Hospital Municipal do Tatuape Carmino Caricchio | Instituto Adolfo Lutz, Interdiciplinary Procedures Center, Strategic Laboratory | Claudio Tavares Sacchi, Claudia Regina Gonçalves, Erica Valessa Ramos Gomes |
| EPI_ISL_515527 | Hospital Santa Clara | Instituto Adolfo Lutz, Interdiciplinary Procedures Center, Strategic Laboratory | Claudio Tavares Sacchi, Claudia Regina Gonçalves, Erica Valessa Ramos Gomes |
| EPI_ISL_515528 | Hospital Sao Paulo de Ensino da Unifesp | Instituto Adolfo Lutz, Interdiciplinary Procedures Center, Strategic Laboratory | Claudio Tavares Sacchi, Claudia Regina Gonçalves, Erica Valessa Ramos Gomes |
| EPI_ISL_515529 | Pronto Socorro Municipal Julio Tupy | Instituto Adolfo Lutz, Interdiciplinary Procedures Center, Strategic Laboratory | Claudio Tavares Sacchi, Claudia Regina Gonçalves, Erica Valessa Ramos Gomes |
| EPI_ISL_515541 | Hospital Montemagno | Instituto Adolfo Lutz, Interdiciplinary Procedures Center, Strategic Laboratory | Claudio Tavares Sacchi, Claudia Regina Gonçalves, Erica Valessa Ramos Gomes |
| EPI_ISL_515542 | Vigilância Epidemiológica de Leme | Instituto Adolfo Lutz, Interdiciplinary Procedures Center, Strategic Laboratory | Claudio Tavares Sacchi, Claudia Regina Gonçalves, Erica Valessa Ramos Gomes |
| EPI_ISL_515543 | Serviço de Vigilância Sanitária e Epidemiológica | Instituto Adolfo Lutz, Interdiciplinary Procedures Center, Strategic Laboratory | Claudio Tavares Sacchi, Claudia Regina Gonçalves, Erica Valessa Ramos Gomes |
| EPI_ISL_515544 | Ama Dr Jose Soares Hungria | Instituto Adolfo Lutz, Interdiciplinary Procedures Center, Strategic Laboratory | Claudio Tavares Sacchi, Claudia Regina Gonçalves, Erica Valessa Ramos Gomes |
| EPI_ISL_515545 | Hospital Sao Paulo de Ensino da Unifesp | Instituto Adolfo Lutz, Interdiciplinary Procedures Center, Strategic Laboratory | Claudio Tavares Sacchi, Claudia Regina Gonçalves, Erica Valessa Ramos Gomes |
| EPI_ISL_515546 | Hospital Municipal do Tatuape Carmino Caricchio | Instituto Adolfo Lutz, Interdiciplinary Procedures Center, Strategic Laboratory | Claudio Tavares Sacchi, Claudia Regina Gonçalves, Erica Valessa Ramos Gomes |
| EPI_ISL_515547 | Centro Medico da Policia Militar do Estado de Sao Paulo | Instituto Adolfo Lutz, Interdiciplinary Procedures Center, Strategic Laboratory | Claudio Tavares Sacchi, Claudia Regina Gonçalves, Erica Valessa Ramos Gomes |
| EPI_ISL_515548 | Hospital Municipal Dr. Jose Soares Hungria | Instituto Adolfo Lutz, Interdiciplinary Procedures Center, Strategic Laboratory | Claudio Tavares Sacchi, Claudia Regina Gonçalves, Erica Valessa Ramos Gomes |
| EPI_ISL_515550 | UPA Vila Santa Catarina | Instituto Adolfo Lutz, Interdiciplinary Procedures Center, Strategic Laboratory | Claudio Tavares Sacchi, Claudia Regina Gonçalves, Erica Valessa Ramos Gomes |
| EPI_ISL_515551, EPI_ISL_515552 | Hospital Municipal do Tatuape Carmino Caricchio | Instituto Adolfo Lutz, Interdiciplinary Procedures Center, Strategic Laboratory | Claudio Tavares Sacchi, Claudia Regina Gonçalves, Erica Valessa Ramos Gomes |
| EPI_ISL_515553 | Hospital Municipal Dr. Ignacio Proença de Gouvea | Instituto Adolfo Lutz, Interdiciplinary Procedures Center, Strategic Laboratory | Claudio Tavares Sacchi, Claudia Regina Gonçalves, Erica Valessa Ramos Gomes |
| EPI_ISL_515554 | Pronto Socorro Municipal de Perus | Instituto Adolfo Lutz, Interdiciplinary Procedures Center, Strategic Laboratory | Claudio Tavares Sacchi, Claudia Regina Gonçalves, Erica Valessa Ramos Gomes |
| EPI_ISL_515555 | Hospital Geral de Vila Nova Cachoeirinha | Instituto Adolfo Lutz, Interdiciplinary Procedures Center, Strategic Laboratory | Claudio Tavares Sacchi, Claudia Regina Gonçalves, Erica Valessa Ramos Gomes |
| EPI_ISL_515557 | Hospital Municipal Dr. Moysés Deutsch | Instituto Adolfo Lutz, Interdiciplinary Procedures Center, Strategic Laboratory | Claudio Tavares Sacchi, Claudia Regina Gonçalves, Erica Valessa Ramos Gomes |
| EPI_ISL_515559, EPI_ISL_515560 | Hospital Sao Paulo de Ensino da Unifesp | Instituto Adolfo Lutz, Interdiciplinary Procedures Center, Strategic Laboratory | Claudio Tavares Sacchi, Claudia Regina Gonçalves, Erica Valessa Ramos Gomes |
| EPI_ISL_515561 | Hospital Montemagno | Instituto Adolfo Lutz, Interdiciplinary Procedures Center, Strategic Laboratory | Claudio Tavares Sacchi, Claudia Regina Gonçalves, Erica Valessa Ramos Gomes |
| EPI_ISL_515562 | Hospital Municipal Doutor Alexandre Zaio | Instituto Adolfo Lutz, Interdiciplinary Procedures Center, Strategic Laboratory | Claudio Tavares Sacchi, Claudia Regina Gonçalves, Erica Valessa Ramos Gomes |
| EPI_ISL_515563 | Hospital Municipal Dr. Jose Soares Hungria | Instituto Adolfo Lutz, Interdiciplinary Procedures Center, Strategic Laboratory | Claudio Tavares Sacchi, Claudia Regina Gonçalves, Erica Valessa Ramos Gomes |

| | | | |
|---|---|---|---|
| EPI_ISL_515564 | Hosp. Municipal Prof. Dr. Alípio Corrêa Netto | Instituto Adolfo Lutz, Interdiciplinary Procedures Center, Strategic Laboratory | Claudio Tavares Sacchi, Claudia Regina Gonçalves, Erica Valessa Ramos Gomes |
| EPI_ISL_515565 | Hospital do Servidor Público Estadual Francisco Morato de Oliveira | Instituto Adolfo Lutz, Interdiciplinary Procedures Center, Strategic Laboratory | Claudio Tavares Sacchi, Claudia Regina Gonçalves, Erica Valessa Ramos Gomes |
| EPI_ISL_515566 | PS Municipal Dr Lauro Ribas Braga | Instituto Adolfo Lutz, Interdiciplinary Procedures Center, Strategic Laboratory | Claudio Tavares Sacchi, Claudia Regina Gonçalves, Erica Valessa Ramos Gomes |
| EPI_ISL_515938, EPI_ISL_515941, EPI_ISL_515942 | CV RAMAN HOSPITAL | Department of Neurovirology, National Institute of Mental Health and Neuroscience (NIMHANS) | Chitra Pattabiraman,Vijayalakshmi Reddy, Harsha PK, Risha Rasheed, Pramada Prasad, Shafeeq S Hameed, Manjunatha Venkataswamy, Anita Desai, Ravi Vasanthapuram |
| EPI_ISL_515945, EPI_ISL_515947 | DH | Department of Neurovirology, National Institute of Mental Health and Neuroscience (NIMHANS) | Chitra Pattabiraman,Vijayalakshmi Reddy, Harsha PK, Risha Rasheed, Pramada Prasad, Shafeeq S Hameed, Manjunatha Venkataswamy, Anita Desai, Ravi Vasanthapuram |
| EPI_ISL_515952 | VICTORIA HOSPITAL | Department of Neurovirology, National Institute of Mental Health and Neuroscience (NIMHANS) | Chitra Pattabiraman,Vijayalakshmi Reddy, Harsha PK, Risha Rasheed, Pramada Prasad, Shafeeq S Hameed, Manjunatha Venkataswamy, Anita Desai, Ravi Vasanthapuram |
| EPI_ISL_515954, EPI_ISL_515957, EPI_ISL_515958, EPI_ISL_515959, EPI_ISL_515961 | DH | Department of Neurovirology, National Institute of Mental Health and Neuroscience (NIMHANS) | Chitra Pattabiraman,Vijayalakshmi Reddy, Harsha PK, Risha Rasheed, Pramada Prasad, Shafeeq S Hameed, Manjunatha Venkataswamy, Anita Desai, Ravi Vasanthapuram |
| EPI_ISL_515963, EPI_ISL_515964, EPI_ISL_515966, EPI_ISL_515967 | VICTORIA HOSPITAL | Department of Neurovirology, National Institute of Mental Health and Neuroscience (NIMHANS) | Chitra Pattabiraman,Vijayalakshmi Reddy, Harsha PK, Risha Rasheed, Pramada Prasad, Shafeeq S Hameed, Manjunatha Venkataswamy, Anita Desai, Ravi Vasanthapuram |
| EPI_ISL_516622 | Instituto de Diagnostico y Referencia Epidemiologicos (INDRE) | Instituto de Diagnostico y Referencia Epidemiologicos (INDRE) | Gisela Barrera-Badillo , Abril Rodriguez-Maldonado, Claudia Wong-Arambula , Natividad Cruz-Ortiz, Tatiana Nunez-Garcia, Dayanira Arellano-Suarez, Fabiola Garces-Ayala, Edgar Mendieta-Condado, Lucia Hernandez-Rivas, Irma Lopez-Martinez, Ernesto Ramirez-Gonzalez. |
| EPI_ISL_516801, EPI_ISL_516802, EPI_ISL_516803, EPI_ISL_516804, EPI_ISL_516805 | Department of Laboratory Medicine, Tan Tock Seng Hospital | Department of Laboratory Medicine, Tan Tock Seng Hospital | Chen YYC, Zair X, Li C, Tang WY, Maurer-Stroh S, Barkham TMS, Nagarajan N, Sessions OM |
| EPI_ISL_516940, EPI_ISL_516942, EPI_ISL_516943, EPI_ISL_516946, EPI_ISL_516948, EPI_ISL_516949, EPI_ISL_516969, EPI_ISL_516974 | King Georges Medical University | CSIR-National Botanical Research Institute | Priti Prasad, Shantanu Prakash, Kishan Sahu, Babita Singh, Suruchi Shukla, Hricha Mishra, Danish Nasar Khan , Om Prakash, MLB Bhatt, SK Barik, Mehar H.Asif,Samir V. Sawant,Amita Jain, Sumit Kr. Bag |
| EPI_ISL_522547, EPI_ISL_522548 | Platforme CYROI | UMR PIMIT Université de La Réunion | David Wilkinson, Camille Lebarbenchon, Patrick Mavingui |
| EPI_ISL_523811 | Universidad Iberoamericana, Instituto de Medicina Tropical & Salud Global | International Centre for Genetic Engineering and Biotechnology (ICGEB) and ARGO Open Lab Platform | Robert Paulino-Ramirez, Eileen Riego, Alejandro Vallejo Degaudenzi, Victor Virgilio Calderon, Leandro Tapia, Danilo Licastro, Simeone Dal Monego, Sreejith Rajasekharan and Alessandro Marcello. |
| EPI_ISL_523955 | Hospital Municipal do Tatuape Carmino Caricchio | Instituto Adolfo Lutz, Interdiciplinary Procedures Center, Strategic Laboratory | Claudio Tavares Sacchi, Claudia Regina Gonçalves, Erica Valessa Ramos Gomes |
| EPI_ISL_523956 | Hospital Regional de Assis | Instituto Adolfo Lutz, Interdiciplinary Procedures Center, Strategic Laboratory | Claudio Tavares Sacchi, Claudia Regina Gonçalves, Erica Valessa Ramos Gomes |
| EPI_ISL_523957 | Hospital Itamaraty | Instituto Adolfo Lutz, Interdiciplinary Procedures Center, Strategic Laboratory | Claudio Tavares Sacchi, Claudia Regina Gonçalves, Erica Valessa Ramos Gomes |
| EPI_ISL_523958 | Pronto Socorro Municipal de Perus | Instituto Adolfo Lutz, Interdiciplinary Procedures Center, Strategic Laboratory | Claudio Tavares Sacchi, Claudia Regina Gonçalves, Erica Valessa Ramos Gomes |
| EPI_ISL_523963 | UBS Vila Silvia | Instituto Adolfo Lutz, Interdiciplinary Procedures Center, Strategic Laboratory | Claudio Tavares Sacchi, Claudia Regina Gonçalves, Erica Valessa Ramos Gomes |
| EPI_ISL_523965 | Hospital do Servidor Público Estadual Francisco Morato de Oliveira | Instituto Adolfo Lutz, Interdiciplinary Procedures Center, Strategic Laboratory | Claudio Tavares Sacchi, Claudia Regina Gonçalves, Erica Valessa Ramos Gomes |
| EPI_ISL_523967 | Hospital Sancta Maggiore | Instituto Adolfo Lutz, Interdiciplinary Procedures Center, Strategic Laboratory | Claudio Tavares Sacchi, Claudia Regina Gonçalves, Erica Valessa Ramos Gomes |
| EPI_ISL_523969 | Hospital Sao Paulo de Ensino da Unifesp | Instituto Adolfo Lutz, Interdiciplinary Procedures Center, Strategic Laboratory | Claudio Tavares Sacchi, Claudia Regina Gonçalves, Erica Valessa Ramos Gomes |
| EPI_ISL_523970 | Conjunto Hospitalar do Mandaqui | Instituto Adolfo Lutz, Interdiciplinary Procedures Center, Strategic Laboratory | Claudio Tavares Sacchi, Claudia Regina Gonçalves, Erica Valessa Ramos Gomes |
| EPI_ISL_523971 | Hospital Geral Santa Marcelina | Instituto Adolfo Lutz, Interdiciplinary Procedures Center, Strategic Laboratory | Claudio Tavares Sacchi, Claudia Regina Gonçalves, Erica Valessa Ramos Gomes |
| EPI_ISL_523972 | Hospital do Servidor Público Estadual Francisco Morato de Oliveira | Instituto Adolfo Lutz, Interdiciplinary Procedures Center, Strategic Laboratory | Claudio Tavares Sacchi, Claudia Regina Gonçalves, Erica Valessa Ramos Gomes |
| EPI_ISL_523973 | PS Municipal Dona Maria Antonieta Ferreira de Barros | Instituto Adolfo Lutz, Interdiciplinary Procedures Center, Strategic Laboratory | Claudio Tavares Sacchi, Claudia Regina Gonçalves, Erica Valessa Ramos Gomes |
| EPI_ISL_523974 | Hospital Municipal do Tatuape Carmino Caricchio | Instituto Adolfo Lutz, Interdiciplinary Procedures Center, Strategic Laboratory | Claudio Tavares Sacchi, Claudia Regina Gonçalves, Erica Valessa Ramos Gomes |
| EPI_ISL_523975 | UPA Tito Lopes | Instituto Adolfo Lutz, Interdiciplinary Procedures Center, Strategic Laboratory | Claudio Tavares Sacchi, Claudia Regina Gonçalves, Erica Valessa Ramos Gomes |
| EPI_ISL_523976 | Hospital Municipal do Tatuape Carmino Caricchio | Instituto Adolfo Lutz, Interdiciplinary Procedures Center, Strategic Laboratory | Claudio Tavares Sacchi, Claudia Regina Gonçalves, Erica Valessa Ramos Gomes |
| EPI_ISL_523977 | Hosp. Municipal Prof. Dr. Alípio Corrêa Netto | Instituto Adolfo Lutz, Interdiciplinary Procedures Center, Strategic Laboratory | Claudio Tavares Sacchi, Claudia Regina Gonçalves, Erica Valessa Ramos Gomes |
| EPI_ISL_523978 | Hospital do Servidor Público Estadual Francisco Morato de Oliveira | Instituto Adolfo Lutz, Interdiciplinary Procedures Center, Strategic Laboratory | Claudio Tavares Sacchi, Claudia Regina Gonçalves, Erica Valessa Ramos Gomes |
| EPI_ISL_523980 | UPA Tito Lopes | Instituto Adolfo Lutz, Interdiciplinary Procedures Center, Strategic Laboratory | Claudio Tavares Sacchi, Claudia Regina Gonçalves, Erica Valessa Ramos Gomes |
| EPI_ISL_523981 | Hospital Sao Paulo de Ensino da Unifesp | Instituto Adolfo Lutz, Interdiciplinary Procedures Center, Strategic Laboratory | Claudio Tavares Sacchi, Claudia Regina Gonçalves, Erica Valessa Ramos Gomes |
| EPI_ISL_523982 | Hospital do Servidor Público Estadual Francisco Morato de Oliveira | Instituto Adolfo Lutz, Interdiciplinary Procedures Center, Strategic Laboratory | Claudio Tavares Sacchi, Claudia Regina Gonçalves, Erica Valessa Ramos Gomes |
| EPI_ISL_523983 | UPA Campo Limpo | Instituto Adolfo Lutz, Interdiciplinary Procedures Center, Strategic Laboratory | Claudio Tavares Sacchi, Claudia Regina Gonçalves, Erica Valessa Ramos Gomes |

| | | | |
|---|---|---|---|
| EPI_ISL_523984 | Ama Dr Jose Soares Hungria | Instituto Adolfo Lutz, Interdiciplinary Procedures Center, Strategic Laboratory | Claudio Tavares Sacchi, Claudia Regina Gonçalves, Erica Valessa Ramos Gomes |
| EPI_ISL_523985 | Hospital Municipal Dr. Benedicto Montenegro | Instituto Adolfo Lutz, Interdiciplinary Procedures Center, Strategic Laboratory | Claudio Tavares Sacchi, Claudia Regina Gonçalves, Erica Valessa Ramos Gomes |
| EPI_ISL_523986 | Ama Dr Jose Soares Hungria | Instituto Adolfo Lutz, Interdiciplinary Procedures Center, Strategic Laboratory | Claudio Tavares Sacchi, Claudia Regina Gonçalves, Erica Valessa Ramos Gomes |
| EPI_ISL_523988 | Hospital Sao Paulo de Ensino da Unifesp | Instituto Adolfo Lutz, Interdiciplinary Procedures Center, Strategic Laboratory | Claudio Tavares Sacchi, Claudia Regina Gonçalves, Erica Valessa Ramos Gomes |
| EPI_ISL_523989 | AMA Jardim Joamar | Instituto Adolfo Lutz, Interdiciplinary Procedures Center, Strategic Laboratory | Claudio Tavares Sacchi, Claudia Regina Gonçalves, Erica Valessa Ramos Gomes |
| EPI_ISL_523990 | AMA Jardim Peri | Instituto Adolfo Lutz, Interdiciplinary Procedures Center, Strategic Laboratory | Claudio Tavares Sacchi, Claudia Regina Gonçalves, Erica Valessa Ramos Gomes |
| EPI_ISL_523991, EPI_ISL_523992 | Hospital Municipal Carmen Prudente | Instituto Adolfo Lutz, Interdiciplinary Procedures Center, Strategic Laboratory | Claudio Tavares Sacchi, Claudia Regina Gonçalves, Erica Valessa Ramos Gomes |
| EPI_ISL_524462 | Hospital Metropolitano | Instituto Adolfo Lutz, Interdiciplinary Procedures Center, Strategic Laboratory | Claudio Tavares Sacchi, Claudia Regina Gonçalves, Erica Valessa Ramos Gomes |
| EPI_ISL_524463 | Hospital Regional de Cotia | Instituto Adolfo Lutz, Interdiciplinary Procedures Center, Strategic Laboratory | Claudio Tavares Sacchi, Claudia Regina Gonçalves, Erica Valessa Ramos Gomes |
| EPI_ISL_524464 | Santa Casa de Santa Isabel | Instituto Adolfo Lutz, Interdiciplinary Procedures Center, Strategic Laboratory | Claudio Tavares Sacchi, Claudia Regina Gonçalves, Erica Valessa Ramos Gomes |
| EPI_ISL_524465 | PS Municipal Dr. Caetano Virgílio Neto | Instituto Adolfo Lutz, Interdiciplinary Procedures Center, Strategic Laboratory | Claudio Tavares Sacchi, Claudia Regina Gonçalves, Erica Valessa Ramos Gomes |
| EPI_ISL_524466 | PS Municipal Dr Lauro Ribas Braga | Instituto Adolfo Lutz, Interdiciplinary Procedures Center, Strategic Laboratory | Claudio Tavares Sacchi, Claudia Regina Gonçalves, Erica Valessa Ramos Gomes |
| EPI_ISL_524467 | Hospital Municipal Dr. Moysés Deutsch | Instituto Adolfo Lutz, Interdiciplinary Procedures Center, Strategic Laboratory | Claudio Tavares Sacchi, Claudia Regina Gonçalves, Erica Valessa Ramos Gomes |
| EPI_ISL_524468 | Hospital Municipal Vereador Jose Storopolli | Instituto Adolfo Lutz, Interdiciplinary Procedures Center, Strategic Laboratory | Claudio Tavares Sacchi, Claudia Regina Gonçalves, Erica Valessa Ramos Gomes |
| EPI_ISL_524469 | Santa Casa de Misericordia de Sao Paulo | Instituto Adolfo Lutz, Interdiciplinary Procedures Center, Strategic Laboratory | Claudio Tavares Sacchi, Claudia Regina Gonçalves, Erica Valessa Ramos Gomes |
| EPI_ISL_524470 | Hospital do Servidor Público Estadual Francisco Morato de Oliveira | Instituto Adolfo Lutz, Interdiciplinary Procedures Center, Strategic Laboratory | Claudio Tavares Sacchi, Claudia Regina Gonçalves, Erica Valessa Ramos Gomes |
| EPI_ISL_524800 | Evandro Chagas Institute | Evandro Chagas Institute | Santos, M.C.; Silva, A.M.; Junior, W.D.C.; Barbagelata, L.S.; Ferreira, J.A.; Sousa, E.M.A.; da Silva, P.S.; Resque, H.R; Martins, L.C.; Sousa Junior, E.C.;Viana, G.M.R |
| EPI_ISL_525467 | Universidad Iberoamericana | International Centre for Genetic Engineering and Biotechnology (ICGEB) and ARGO Open Lab Platform | Robert Paulino-Ramirez, Eileen Riego, Alejandro Vallejo Degaudenzi, Victor Virgilio Calderon, Leandro Tapia, Patricia Leon, Danilo Licastro, Simeone Dal Monego, Sreejith Rajasekharan and Alessandro Marcello. |
| EPI_ISL_525468 | Universidad Iberoamericana | International Centre for Genetic Engineering and Biotechnology (ICGEB) and ARGO Open Lab Platform | Robert Paulino-Ramirez, Eileen Riego, Alejandro Vallejo Degaudenzi, Victor Virgilio Calderon, Leandro Tapia,Patricia Leon,Danilo Licastro, Simeone Dal Monego, Sreejith Rajasekharan and Alessandro Marcello. |
| EPI_ISL_525469 | Universidad Iberoamericana | International Centre for Genetic Engineering and Biotechnology (ICGEB) and ARGO Open Lab Platform | Robert Paulino-Ramirez, Eileen Riego, Alejandro Vallejo Degaudenzi, Victor Virgilio Calderon, Leandro Tapia, Patricia Leon, Danilo Licastro, Simeone Dal Monego, Sreejith Rajasekharan and Alessandro Marcello. |
| EPI_ISL_525474 | Centre for Dengue Research | Centre for Dengue Research, USJ, SL | Chandima Jeewandara, Deshni Jayathilaka, Dinuka Ariyaratne, Laksiri Gomes, Diyanath Ranasinghe, Dinuka Guruge, Ruwan Wijayamuni, Gathsaurie Neelika Malavige |
| EPI_ISL_525476 | Centre for Dengue Research | Centre for Dengue Research | Chandima Jeewandara, Deshni Jayathilaka, Dinuka Ariyaratne, Laksiri Gomes, Diyanath Ranasinghe, Dinuka Guruge, Ruwan Wijayamuni, Gathsaurie Neelika Malavige |
| EPI_ISL_525478, EPI_ISL_525479 | Centre for Dengue Research | Centre for Dengue Research | Chandima Jeewandara, Deshni Jayathilaka, Dinuka Ariyaratne, Laksiri Gomes, Diyanath Ranasinghe, Ananda Wijewickrama, Eranga Narangoda, Damayanthi Idampitiya, Gathsaurie Neelika Malavige |
| EPI_ISL_525553, EPI_ISL_525556, EPI_ISL_525557, EPI_ISL_525570, EPI_ISL_525571 | Istituto Zooprofilattico Sperimentale Puglia e Basilicata; Dipartimento di Bioscienze, Biotecnologie e Biofarmaceutica dell'Università degli Studi di Bari "A.Moro"; Istituto di Biomembrane. Bioenergetica e Biotecnologie Molecolari del Consiglio Nazionale delle Ricerche di Bari | Beaconlab (Bioinformatics, Evolution and Comparative Genomics lab), Dept of Biosciences, University on Milan | Parisi A.,Pesole G., Manzari C., Chiara M |
| EPI_ISL_525578, EPI_ISL_525621, EPI_ISL_525627, EPI_ISL_525631, EPI_ISL_525638, EPI_ISL_525639, EPI_ISL_525640, EPI_ISL_525641, EPI_ISL_525642, EPI_ISL_525646, EPI_ISL_525647, EPI_ISL_525648, EPI_ISL_525649, EPI_ISL_525650, EPI_ISL_525654, EPI_ISL_525658, EPI_ISL_525659, EPI_ISL_525676, EPI_ISL_525677, EPI_ISL_525678, EPI_ISL_525679, EPI_ISL_525680, EPI_ISL_525681, EPI_ISL_525682, EPI_ISL_525683, EPI_ISL_525684, EPI_ISL_525685, EPI_ISL_525686 | | | |
| see above | Wadsworth Center, New York State Department of Health | Wadsworth Center, New York State Department of Health | Kirsten St. George, Daryl M. Lamson, Sara Griesemer, Jonathan Plitnick, Navjot Singh, Matthew D. Shudt, Erica Lasek-Nesselquist |
| EPI_ISL_526215, EPI_ISL_526216, EPI_ISL_526217, EPI_ISL_526218, EPI_ISL_526219, EPI_ISL_526220, EPI_ISL_526221, EPI_ISL_526222, EPI_ISL_526226, EPI_ISL_526227, EPI_ISL_526228, EPI_ISL_526229, EPI_ISL_526230, EPI_ISL_526231, EPI_ISL_526232, EPI_ISL_526233, EPI_ISL_526234, EPI_ISL_526235, EPI_ISL_526236, EPI_ISL_526237, EPI_ISL_526238 | | | |
| see above | Hungarian Defence Forces Military Medical Centre | National Laboratory of Virology, Szentágothai Research Centre | Endre Gábor Tóth, Balázs Somogyi, Bálint Eszenyi, Ferenc Jakab, Gábor Kemenesi |
| EPI_ISL_526686, EPI_ISL_526687, EPI_ISL_526688 | Faith Laboratory, Immunology Institute, Icahn School of Medicine at Mount Sinai | van Bakel Laboratory, Genetics and Genomics Sciences, Icahn School of Medicine at Mount Sinai | Graham J. Britton, Alice Chen-Liaw, Francesca Cossarini, Alexandra Livanos, Matthew P. Spindler, Tamar Plitt, Joseph Eggers, Ilaria Mogno, Ana S. Gonzalez-Reiche, Sophia Sui, Michael Tankelevich, Lauren Tal Grinspan, Rebekah E. Dixon, Divya Jha, Gustavo Martinez-Delgado, Fatima Amanat, Daisy Hoagland, Benjamin R. tenOever, Marla C. Dubinsky, Miriam Merad, Harm Van Bakel, Florian Krammer, Gerold Bongers, Saurabh Mehandru and Jeremiah J. Faith |
| EPI_ISL_526933, EPI_ISL_526934 | Instituto Nacional de Salud, Bogotá, Colombia | Instituto Nacional de Salud, Bogotá, Colombia | Katherine Laiton-Donato, Diego A. Álvarez-Díaz, Carlos Franco-Muñoz, Mauricio Pacheco-Montealegre, Jonathan Reales, Diego Andrés Prada, Jose A. Usme-Ciro, Zulma M. Cucunubá, Christian Julian VillabonaArenas, Liz Villabona-Arenas, Sussy Echeverria, Astrid C. Flórez, Carolina Ferro, Diana Marcela Walteros-Acero, Franklin Prieto, Carlos Andrés Durán, Martha Lucia Ospina Martinez, Marcela Mercado-Reyes |
| EPI_ISL_526937, EPI_ISL_526938, EPI_ISL_526939, EPI_ISL_526940, EPI_ISL_526941, EPI_ISL_526943, EPI_ISL_526946 | Faroese National Reference Laboratory for Fish and Animal Diseases | Faroese National Reference Laboratory for Fish and Animal Diseases | Maria Marjunardóttir Dahl, Petra Elisabeth Petersen, Debes Hammershaimb Christiansen |
| EPI_ISL_526949, EPI_ISL_526950, EPI_ISL_526951, EPI_ISL_526955, EPI_ISL_526956, EPI_ISL_526959 | Instituto Nacional de Salud, Bogotá, Colombia | Instituto Nacional de Salud, Bogotá, Colombia | Katherine Laiton-Donato, Diego A. Álvarez-Díaz, Carlos Franco-Muñoz, Mauricio Pacheco-Montealegre, Jonathan Reales, Diego Andrés Prada, Jose A. Usme-Ciro, Zulma M. Cucunubá, Christian Julian VillabonaArenas, Liz Villabona-Arenas, Sussy Echeverria, Astrid C. Flórez, Carolina Ferro, Diana Marcela Walteros-Acero, Franklin Prieto, Carlos Andrés Durán, Martha Lucia Ospina Martinez, Marcela Mercado-Reyes |

| | | | |
|---|---|---|---|
| EPI_ISL_526963, EPI_ISL_526964, EPI_ISL_526965 | Instituto Nacional de Salud, Bogotá, Colombia | Instituto Nacional de Salud, Bogotá, Colombia | Katherine Laiton-Donato, Diego A. Álvarez-Díaz, Carlos Franco-Muñoz, Jonathan Reales, Diego Andrés Prada, Jeadran Malagón-Rojas, Felix Betzler, Wendy K. Jo, Edmilson F. de Oliveira-Filho, Carolina Ferro, Diana Marcela Walteros-Acero, Franklin Prieto, Carlos Andrés Durán, Martha Lucia Ospina Martinez, Marcela Mercado-Reyes |
| EPI_ISL_527819 | Centro de Investigaciones, Universidad de Especialidades Espíritu Santo | Institute of Microbiology, Universidad San Francisco de Quito | Derly Andrade, Juan Carlos Fernandez, Belén Prado-Vivar, Sully Márquez, Juan José Guadalupe, Monica Becerra-Wong, Bernardo Gutiérrez, Gabriel Morey, Ruben Armas, Jose Pedro Barberan, Fernando Espinoza, Edith Lopez, Verónica Barragán, Patricio Rojas-Silva, Gabriel Trueba, Michelle Grunauer, Paúl Cárdenas |
| EPI_ISL_527856 | Hospital Municipal Prof. Waldomiro de Paula | Instituto Adolfo Lutz, Interdiciplinary Procedures Center, Strategic Laboratory | Claudio Tavares Sacchi, Claudia Regina Gonçalves, Erica Valessa Ramos Gomes |
| EPI_ISL_527857 | Hospital Regional Vale do Ribeira | Instituto Adolfo Lutz, Interdiciplinary Procedures Center, Strategic Laboratory | Claudio Tavares Sacchi, Claudia Regina Gonçalves, Erica Valessa Ramos Gomes |
| EPI_ISL_527858 | Pronto Atendimento Sancta Maggiore Jardim Paulista | Instituto Adolfo Lutz, Interdiciplinary Procedures Center, Strategic Laboratory | Claudio Tavares Sacchi, Claudia Regina Gonçalves, Erica Valessa Ramos Gomes |
| EPI_ISL_527859 | Hospital Municipal Vereador Jose Storopolli | Instituto Adolfo Lutz, Interdiciplinary Procedures Center, Strategic Laboratory | Claudio Tavares Sacchi, Claudia Regina Gonçalves, Erica Valessa Ramos Gomes |
| EPI_ISL_527860 | Hospital Municipal de Parelheiros Josanias Castanha Braga | Instituto Adolfo Lutz, Interdiciplinary Procedures Center, Strategic Laboratory | Claudio Tavares Sacchi, Claudia Regina Gonçalves, Erica Valessa Ramos Gomes |
| EPI_ISL_527861 | Hospital e Maternidade Celso Pierro | Instituto Adolfo Lutz, Interdiciplinary Procedures Center, Strategic Laboratory | Av. Dr. Arnaldo, 355 - Brazil, Cerqueira Cesar, São Paulo - SP, 01246-1301 |
| EPI_ISL_527862 | Hospital Municipal de Urgência | Instituto Adolfo Lutz, Interdiciplinary Procedures Center, Strategic Laboratory | Claudio Tavares Sacchi, Claudia Regina Gonçalves, Erica Valessa Ramos Gomes |
| EPI_ISL_527863 | Hospital Municipal do Tatuape Carmino Caricchio | Instituto Adolfo Lutz, Interdiciplinary Procedures Center, Strategic Laboratory | Claudio Tavares Sacchi, Claudia Regina Gonçalves, Erica Valessa Ramos Gomes |
| EPI_ISL_527864 | Hospital e Pronto Socorro Comunitário Vila Iolanda | Instituto Adolfo Lutz, Interdiciplinary Procedures Center, Strategic Laboratory | Claudio Tavares Sacchi, Claudia Regina Gonçalves, Erica Valessa Ramos Gomes |
| EPI_ISL_527865 | Hospital e Maternidade São Cristóvão | Instituto Adolfo Lutz, Interdiciplinary Procedures Center, Strategic Laboratory | Claudio Tavares Sacchi, Claudia Regina Gonçalves, Erica Valessa Ramos Gomes |
| EPI_ISL_527866 | PS Municipal Dr Lauro Ribas Braga | Instituto Adolfo Lutz, Interdiciplinary Procedures Center, Strategic Laboratory | Av. Dr. Arnaldo, 355 - Brazil, Cerqueira Cesar, São Paulo - SP, 01246-1301 |
| EPI_ISL_527867 | Pronto Socorro Municipal - Balneario São José | Instituto Adolfo Lutz, Interdiciplinary Procedures Center, Strategic Laboratory | Claudio Tavares Sacchi, Claudia Regina Gonçalves, Erica Valessa Ramos Gomes |
| EPI_ISL_527868 | Hospital e Maternidade do Braz | Instituto Adolfo Lutz, Interdiciplinary Procedures Center, Strategic Laboratory | Claudio Tavares Sacchi, Claudia Regina Gonçalves, Erica Valessa Ramos Gomes |
| EPI_ISL_527869 | Hospital Municipal Carmen Prudente | Instituto Adolfo Lutz, Interdiciplinary Procedures Center, Strategic Laboratory | Claudio Tavares Sacchi, Claudia Regina Gonçalves, Erica Valessa Ramos Gomes |
| EPI_ISL_527870 | Hospital Municipal Mário Gatti | Instituto Adolfo Lutz, Interdiciplinary Procedures Center, Strategic Laboratory | Claudio Tavares Sacchi, Claudia Regina Gonçalves, Erica Valessa Ramos Gomes |
| EPI_ISL_528382, EPI_ISL_528383, EPI_ISL_528384, EPI_ISL_528385 | Translational Health Science and Technology Institute -ESIC medical college and hospital, Faridabad | THSTI Bioassay laboratory | Saurabh Kumar, Jigme Wangchuk, Anil Kumar Pandey, Asim Das, Guruprasad R. Medigeshi |
| EPI_ISL_528538 | Alsafar | Alsafar | Andreas Henschel, Gihan Elsir Ahmed Daw Elbait, Samuel Feng, Rifat, Ernesto Damiani, Guan Tay, Habiba Alsafar |
| EPI_ISL_528539 | LVM/UFRJ | LNCC | Gustavo M. Romário M. de Souza; Bruno B. Bezerra; Lucio A. Caldas; Fabio Limonte; Elena Cobos; Sharton V. A. Coelho; Luiz Almeida; Luiza Higga; Isadora A. Correa; Diana Marianni; Luciana B. Arruda; Marcelo Bozza; Orlando Ferreira; Wanderley de Souza; Ana Teresa R. Vasconcelos; Terezinha M. Castineiras; Amilcar Tanuri; Luciana J. Costa. |
| EPI_ISL_528637, EPI_ISL_528638 | LVM/UFRJ | Bioinformatics Laboratory / LNCC | Gustavo D. P. Silva; M. Romário M. de Souza; Bruno B. Bezerra; Lucio A. Caldas; Fabio Limonte; Elena Cobos; Sharton V. A. Coelho; Luiz Almeida; Luiza Higga; Isadora A. Correa; Diana Marianni; Luciana B. Arruda; Marcelo Bozza; Orlando Ferreira; Wanderley de Souza; Ana Teresa R. Vasconcelos; Terezinha M. Castineiras; Amilcar Tanuri; Luciana J. Costa |
| EPI_ISL_528687, EPI_ISL_528689, EPI_ISL_528692, EPI_ISL_528693, EPI_ISL_528698, EPI_ISL_528699, EPI_ISL_528701, EPI_ISL_528704, EPI_ISL_528706, EPI_ISL_528709, EPI_ISL_528710, EPI_ISL_528711, EPI_ISL_528712, EPI_ISL_528713, EPI_ISL_528714, EPI_ISL_528715, EPI_ISL_528716, EPI_ISL_528717, EPI_ISL_528718, EPI_ISL_528719, EPI_ISL_528720, EPI_ISL_528721 | | | |
| see above | Alsafar - Khalifa University Abu Dhabi | Alsafar - Khalifa University Abu Dhabi | Andreas Henschel, Gihan Daw Elbait, Samuel Feng, Rifat Hamoudi, Ernesto Damiani, Guan Tay, Habiba Alsafar |
| EPI_ISL_528738, EPI_ISL_528739, EPI_ISL_528740, EPI_ISL_528741 | Malaysia Genome Institute | Malaysia Genome Institute | Mohd Noor Mat Isa, Irni Suhayu Sapian, Yusuf Muhammad Noor, Nurhezreen Md Iqbal, Mohd Faizal Abu Bakar, Enizza Kasim, Shamsidar Sopie, Siti Noraini Othman, Azrin Ahmad, Nor Azfa Johari, Shahrul Hisham Zainal Ariffin |
| EPI_ISL_528742 | Malaysia Genome Institute | Malaysia Genome Institute | Mohd Noor Mat Isa, Irni Suhayu Sapian, Gan Han Ming, Yusuf Muhammad Noor, Tan Ju Lin, Nurhezreen Md Iqbal, Mohd Faizal Abu Bakar, Enizza Kasim, Shamsidar Sopie, Siti Noraini Othman, Azrin Ahmad, Nor Azfa Johari, Shahrul Hisham Zainal Ariffin |
| EPI_ISL_528749 | Santosa Hospital Bandung Central | School of Life Sciences and Technology & School of Pharmacy-Institut Teknologi Bandung; Molecular Genetics Laboratory-Faculty of Medicine-Universitas Padjadjaran; Laboratorium Kesehatan Provinsi Jawa Barat | Husna Nugrahapraja, Azzania Fibriani, Catur Riani, Marselina Irasonia Tan, Yunia Sribudiani, Tarwadi, Ema Rahmawati, Savira Ekawardhani, Hesti Lina Wiraswati, Ryan Bayusantika Ristandi, Rifky Waluyajati Rachman, Cut Nur Cinthia Alamanda, Lia Faridah, Davin H. E. Setiamarga, Rizki Mardian, Hammam Riza, Sony Solistia Wirawan, Agung Eru Wibowo, Irvan Faizal |
| EPI_ISL_528919 | Ospedale Civile S. Liberatore-Atri | Istituto Zooprofilattico Sperimentale dell'Abruzzo e Molise "G.Caporale" | Lorusso A, Marcacci M, Di Domenico M, Curini V, Ancora M, Cammà C, Rinaldi A, Mangone I, Di Pasquale A, Puglia I, Savini G. |
| EPI_ISL_528920, EPI_ISL_528921 | Presidio Ospedaliero "Santo Spirito"-Pescara | Istituto Zooprofilattico Sperimentale dell'Abruzzo e Molise "G.Caporale" | Lorusso A, Marcacci M, Di Domenico M, Curini V, Ancora M, Cammà C, Rinaldi A, Mangone I, Di Pasquale A, Puglia I, Savini G. |
| EPI_ISL_528922, EPI_ISL_528924 | Ospedale "Giuseppe Mazzini"-Teramo | Istituto Zooprofilattico Sperimentale dell'Abruzzo e Molise "G.Caporale" | Lorusso A, Marcacci M, Di Domenico M, Curini V, Ancora M, Cammà C, Rinaldi A, Mangone I, Di Pasquale A, Puglia I, Savini G. |
| EPI_ISL_528925 | Ospedale Regionale San Salvatore-L'Aquila | Istituto Zooprofilattico Sperimentale dell'Abruzzo e Molise "G.Caporale" | Lorusso A, Marcacci M, Di Domenico M, Curini V, Ancora M, Cammà C, Rinaldi A, Mangone I, Di Pasquale A, Puglia I, Savini G. |
| EPI_ISL_528926, EPI_ISL_528927, EPI_ISL_528928 | Ospedale "Giuseppe Mazzini"-Teramo | Istituto Zooprofilattico Sperimentale dell'Abruzzo e Molise "G.Caporale" | Lorusso A, Marcacci M, Di Domenico M, Curini V, Ancora M, Cammà C, Rinaldi A, Mangone I, Di Pasquale A, Puglia I, Savini G. |
| EPI_ISL_528929 | Ospedale Civile S. Liberatore-Atri | Istituto Zooprofilattico Sperimentale dell'Abruzzo e Molise "G.Caporale" | Lorusso A, Marcacci M, Di Domenico M, Curini V, Ancora M, Cammà C, Rinaldi A, Mangone I, Di Pasquale A, Puglia I, Savini G. |
| EPI_ISL_528934, EPI_ISL_528935, EPI_ISL_528936, EPI_ISL_528937, EPI_ISL_528938, EPI_ISL_528939, EPI_ISL_528940, EPI_ISL_528941, EPI_ISL_528943, EPI_ISL_528944, EPI_ISL_528945, EPI_ISL_528946, EPI_ISL_528947, EPI_ISL_528948, EPI_ISL_528949 | | | |
| see above | Agenzia di Tutela della Salute di Bergamo | Istituto Zooprofilattico Sperimentale dell'Abruzzo e Molise "G.Caporale" | Lorusso A, Marcacci M, Di Domenico M, Curini V, Ancora M, Cammà C, Rinaldi A, Mangone I, Di Pasquale A, Puglia I, Savini G. |

| | | | |
|---|---|---|---|
| EPI_ISL_528990 | Ospedale Civile Maria SS. dello Splendore | Istituto Zooprofilattico Sperimentale dell'Abruzzo e Molise "G.Caporale" | Lorusso A, Marcacci M, Di Domenico M, Curini V, Ancora M, Cammà C, Rinaldi A, Mangone I, Di Pasquale A, Puglia I, Savini G. |
| EPI_ISL_528991 | Ospedale SS Annunziata-Sulmona | Istituto Zooprofilattico Sperimentale dell'Abruzzo e Molise "G.Caporale" | Lorusso A, Marcacci M, Di Domenico M, Curini V, Ancora M, Cammà C, Rinaldi A, Mangone I, Di Pasquale A, Puglia I, Savini G. |
| EPI_ISL_528993 | Ospedale Civile S. Liberatore-Atri | Istituto Zooprofilattico Sperimentale dell'Abruzzo e Molise "G.Caporale" | Lorusso A, Marcacci M, Di Domenico M, Curini V, Ancora M, Cammà C, Rinaldi A, Mangone I, Di Pasquale A, Puglia I, Savini G. |
| EPI_ISL_528994, EPI_ISL_528995, EPI_ISL_528996, EPI_ISL_528997, EPI_ISL_528998, EPI_ISL_528999, EPI_ISL_529000, EPI_ISL_529001 | Servizio di igiene epidemiologia e sanità pubblica (SIESP)-Chieti | Istituto Zooprofilattico Sperimentale dell'Abruzzo e Molise "G.Caporale" | Lorusso A, Marcacci M, Di Domenico M, Curini V, Ancora M, Cammà C, Rinaldi A, Mangone I, Di Pasquale A, Puglia I, Savini G. |
| EPI_ISL_529009 | Ospedale Civile S. Liberatore-Atri | Istituto Zooprofilattico Sperimentale dell'Abruzzo e Molise "G.Caporale" | Lorusso A, Marcacci M, Di Domenico M, Curini V, Ancora M, Cammà C, Rinaldi A, Mangone I, Di Pasquale A, Puglia I, Savini G. |
| EPI_ISL_529016 | Ospedale SS Annunziata-Sulmona | Istituto Zooprofilattico Sperimentale dell'Abruzzo e Molise "G.Caporale" | Lorusso A, Marcacci M, Di Domenico M, Curini V, Ancora M, Cammà C, Rinaldi A, Mangone I, Di Pasquale A, Puglia I, Savini G. |
| EPI_ISL_529018 | Ospedale "Giuseppe Mazzini"-Teramo | Istituto Zooprofilattico Sperimentale dell'Abruzzo e Molise "G.Caporale" | Lorusso A, Marcacci M, Di Domenico M, Curini V, Ancora M, Cammà C, Rinaldi A, Mangone I, Di Pasquale A, Puglia I, Savini G. |
| EPI_ISL_529019 | RSA/RP Villa San Giovanni - Gruppo Edos | Istituto Zooprofilattico Sperimentale dell'Abruzzo e Molise "G.Caporale" | Lorusso A, Marcacci M, Di Domenico M, Curini V, Ancora M, Cammà C, Rinaldi A, Mangone I, Di Pasquale A, Puglia I, Savini G. |
| EPI_ISL_529020, EPI_ISL_529021 | Ospedale Civile S. Liberatore-Atri | Istituto Zooprofilattico Sperimentale dell'Abruzzo e Molise "G.Caporale" | Lorusso A, Marcacci M, Di Domenico M, Curini V, Ancora M, Cammà C, Rinaldi A, Mangone I, Di Pasquale A, Puglia I, Savini G. |
| EPI_ISL_529022 | Ospedale "Ss. Annunziata" | Istituto Zooprofilattico Sperimentale dell'Abruzzo e Molise "G.Caporale" | Lorusso A, Marcacci M, Di Domenico M, Curini V, Ancora M, Cammà C, Rinaldi A, Mangone I, Di Pasquale A, Puglia I, Savini G. |
| EPI_ISL_529026 | Ospedale "Giuseppe Mazzini"-Teramo | Istituto Zooprofilattico Sperimentale dell'Abruzzo e Molise "G.Caporale" | Lorusso A, Marcacci M, Di Domenico M, Curini V, Ancora M, Cammà C, Rinaldi A, Mangone I, Di Pasquale A, Puglia I, Savini G. |
| EPI_ISL_529035, EPI_ISL_529036 | Wadsworth Center, New York State Department of Health | Wadsworth Center, New York State Department of Health | Kirsten St. George, Daryl M. Lamson, Sara Griesemer, Jonathan Plitnick, Navjot Singh, Matthew D. Shudt, Erica Lasek-Nesselquist |
| EPI_ISL_529138 | RSAL Dr. Ramelan Surabaya | Institute of Tropical Disease, Universitas Airlangga | Jezzy R Dewantari, Rima R Prasetya, Krisnoadi Rahardjo, Aldise M Nastri, Radito Soesanto, Gatot Soegiarto, Laksmi Wulandari, Retno A Setyoningrum, Resti Yudhawati, Yohko K Shimizu, Mitsuhiro Nishimura, Yasuko Mori, Soetjipto, Kazufumi Shimizu, Maria I Lusida |
| EPI_ISL_529961 | Husada Utama Hospital | Institute of Tropical Disease, Universitas Airlangga | Krisnoadi Rahardjo, Aldise M Nastri, Jezzy R Dewantari, Rima R Prasetya, Didi Dewanto, Gatot Soegiarto, Laksmi Wulandari, Retno A Setyoningrum, Resti Yudhawati, Yohko K Shimizu, Mitsuhiro Nishimura, Yasuko Mori, Soetjipto, Kazufumi Shimizu, Maria I Lusida |
| EPI_ISL_529962 | Universitas Airlangga Hospital | Institute of Tropical Disease, Universitas Airlangga | Jezzy R Dewantari, Rima R Prasetya, Krisnoadi Rahardjo, Aldise M Nastri, Nasronudin, Gatot Soegiarto, Laksmi Wulandari, Retno A Setyoningrum, Resti Yudhawati, Yohko K Shimizu, Mitsuhiro Nishimura, Yasuko Mori, Soetjipto, Kazufumi Shimizu, Maria I Lusida |
| EPI_ISL_529964 | RSUD Bangil Pasuruan | Institute of Tropical Disease, Universitas Airlangga | Aldise M Nastri, Jezzy R Dewantari, Rima R Prasetya, Krisnoadi Rahardjo, Arma Roosalina, Gatot Soegiarto, Laksmi Wulandari, Retno A Setyoningrum, Resti Yudhawati, Yohko K Shimizu, Mitsuhiro Nishimura, Yasuko Mori, Soetjipto, Kazufumi Shimizu, Maria I Lusida |
| EPI_ISL_529965 | RSUD Dr. Soetomo | Institute of Tropical Disease, Universitas Airlangga | Krisnoadi Rahardjo, Aldise M Nastri, Jezzy R Dewantari, Rima R Prasetya, Joni Wahyuhadi, Gatot Soegiarto, Laksmi Wulandari, Retno A Setyoningrum, Resti Yudhawati, Yohko K Shimizu, Mitsuhiro Nishimura, Yasuko Mori, Soetjipto, Kazufumi Shimizu, Maria I Lusida |
| EPI_ISL_529966 | RSUD Sidoarjo | Institute of Tropical Disease, Universitas Airlangga | Kazufumi Shimizu, Krisnoadi Rahardjo, Aldise M Nastri, Jezzy R Dewantari, Rima R Prasetya, Atok Irawan, Gatot Soegiarto, Laksmi Wulandari, Retno A Setyoningrum, Resti Yudhawati, Yohko K Shimizu, Mitsuhiro Nishimura, Yasuko Mori, Soetjipto, Maria I Lusida |
| EPI_ISL_530347, EPI_ISL_530348, EPI_ISL_530349, EPI_ISL_530350 | The National Institute of Public Health | State Veterinary Institute Prague | Nagy,A;Jirincova,H;Novakova,L;Trnka,D;Vecerova,J |
| EPI_ISL_534311 | UPA III 26 de Agosto | Instituto Adolfo Lutz, Interdiciplinary Procedures Center, Strategic Laboratory | Claudio Tavares Sacchi, Claudia Regina Gonçalves, Erica Valessa Ramos Gomes |
| EPI_ISL_534312 | Distrito Sanitario Sul | Instituto Adolfo Lutz, Interdiciplinary Procedures Center, Strategic Laboratory | Claudio Tavares Sacchi, Claudia Regina Gonçalves, Erica Valessa Ramos Gomes |
| EPI_ISL_534313 | Hospital da Sta Casa de Sto Amaro | Instituto Adolfo Lutz, Interdiciplinary Procedures Center, Strategic Laboratory | Claudio Tavares Sacchi, Claudia Regina Gonçalves, Erica Valessa Ramos Gomes |
| EPI_ISL_534314 | Hospital Universitario da USP de SP | Instituto Adolfo Lutz, Interdiciplinary Procedures Center, Strategic Laboratory | Claudio Tavares Sacchi, Claudia Regina Gonçalves, Erica Valessa Ramos Gomes |
| EPI_ISL_534315 | Serviço de Verificação de Óbitos SVO Guaruulhos | Instituto Adolfo Lutz, Interdiciplinary Procedures Center, Strategic Laboratory | Claudio Tavares Sacchi, Claudia Regina Gonçalves, Erica Valessa Ramos Gomes |
| EPI_ISL_534316 | OS Mun Santana Lauro Ribas Braga | Instituto Adolfo Lutz, Interdiciplinary Procedures Center, Strategic Laboratory | Claudio Tavares Sacchi, Claudia Regina Gonçalves, Erica Valessa Ramos Gomes |
| EPI_ISL_534317 | Hospital Geral de Itapevi | Instituto Adolfo Lutz, Interdiciplinary Procedures Center, Strategic Laboratory | Claudio Tavares Sacchi, Claudia Regina Gonçalves, Erica Valessa Ramos Gomes |
| EPI_ISL_534318 | Hospital Municipal Antonio Giglio | Instituto Adolfo Lutz, Interdiciplinary Procedures Center, Strategic Laboratory | Claudio Tavares Sacchi, Claudia Regina Gonçalves, Erica Valessa Ramos Gomes |
| EPI_ISL_534319, EPI_ISL_534320 | Hospital do Serv Pub ESTAFCO Morato de Oliveira | Instituto Adolfo Lutz, Interdiciplinary Procedures Center, Strategic Laboratory | Claudio Tavares Sacchi, Claudia Regina Gonçalves, Erica Valessa Ramos Gomes |
| EPI_ISL_534321 | PS e Maternidade Nair Fonseca Leitao Arantes | Instituto Adolfo Lutz, Interdiciplinary Procedures Center, Strategic Laboratory | Claudio Tavares Sacchi, Claudia Regina Gonçalves, Erica Valessa Ramos Gomes |
| EPI_ISL_534322 | PS Mun Julio Tupy | Instituto Adolfo Lutz, Interdiciplinary Procedures Center, Strategic Laboratory | Claudio Tavares Sacchi, Claudia Regina Gonçalves, Erica Valessa Ramos Gomes |
| EPI_ISL_534323 | Hospital e Pronto Socorro Comunitario Vila Yolanda | Instituto Adolfo Lutz, Interdiciplinary Procedures Center, Strategic Laboratory | Claudio Tavares Sacchi, Claudia Regina Gonçalves, Erica Valessa Ramos Gomes |
| EPI_ISL_534324 | Hospital Mun Ver Jose Storopolli | Instituto Adolfo Lutz, Interdiciplinary Procedures Center, Strategic Laboratory | Claudio Tavares Sacchi, Claudia Regina Gonçalves, Erica Valessa Ramos Gomes |
| EPI_ISL_534325 | Unidade de Vigilancia em Saude de Guarulhos | Instituto Adolfo Lutz, Interdiciplinary Procedures Center, Strategic Laboratory | Claudio Tavares Sacchi, Claudia Regina Gonçalves, Erica Valessa Ramos Gomes |
| EPI_ISL_534326 | Notre Dame Intermedica Saude AS | Instituto Adolfo Lutz, Interdiciplinary Procedures Center, Strategic Laboratory | Claudio Tavares Sacchi, Claudia Regina Gonçalves, Erica Valessa Ramos Gomes |
| EPI_ISL_534346, EPI_ISL_534347, | Molecular diagnostic laboratory of Federal Budget Institution | Group of Genomics and Postgenomic Technologies of | Speranskaya AS, Kaptelova VV, Valdokhina AV, Bulanenko VP, Samoilov AE, Korneenko EV, Tivanova EV, Shipulina OY, Akimkin VG |

| | | | |
|---|---|---|---|
| EPI_ISL_534348, EPI_ISL_534349 | of Science "Central Research Institute of Epidemiology" of The Federal Service on Customers' Rights Protection and Human Well-being Surveillance | Central Research Institute of Epidemilology | |
| EPI_ISL_536399 | Laboratory of Immunovirology. Universidad de Antioquia | Instituto Nacional de Salud - Unidad de Secuenciación y Genómica | Wbeimar Aguilar-Jimenez, Lizdany Flórez, Francisco J. Díaz, Katherine Laiton-Donato, Carlos Franco-Muñoz, Diego Álvarez-Diaz and Marcela Mercado-Reyes |
| EPI_ISL_538435, EPI_ISL_538436, EPI_ISL_538437, EPI_ISL_538438, EPI_ISL_538439, EPI_ISL_538440, EPI_ISL_538441, EPI_ISL_538442, EPI_ISL_538443, EPI_ISL_538444, EPI_ISL_538445, EPI_ISL_538446, EPI_ISL_538447, EPI_ISL_538448, EPI_ISL_538449, EPI_ISL_538450, EPI_ISL_538451, EPI_ISL_538452, EPI_ISL_538453, EPI_ISL_538454, EPI_ISL_538455, EPI_ISL_538456, EPI_ISL_538457, EPI_ISL_538458, EPI_ISL_538459, EPI_ISL_538460, EPI_ISL_538461, EPI_ISL_538462, EPI_ISL_538463, EPI_ISL_538464, EPI_ISL_538465, EPI_ISL_538466, EPI_ISL_538467, EPI_ISL_538468, EPI_ISL_538469, EPI_ISL_538470, EPI_ISL_538471, EPI_ISL_538472, EPI_ISL_538473, EPI_ISL_538474, EPI_ISL_538475, EPI_ISL_538476, EPI_ISL_538477, EPI_ISL_538478, EPI_ISL_538479, EPI_ISL_538480, EPI_ISL_538481, EPI_ISL_538482, EPI_ISL_538483, EPI_ISL_538484, EPI_ISL_538485, EPI_ISL_538486, EPI_ISL_538487, EPI_ISL_538488, EPI_ISL_538489, EPI_ISL_538490, EPI_ISL_538491, EPI_ISL_538492, EPI_ISL_538493, EPI_ISL_538494, EPI_ISL_538495, EPI_ISL_538496, EPI_ISL_538497 | | | |
| see above | Department of Laboratory Medicine, Tan Tock Seng Hospital | Department of Laboratory Medicine, Tan Tock Seng Hospital | Chen YYC, Zair X, Lim JX, Li C, Tang WY, Maurer-Stroh S, Barkham TMS, Nagarajan N, Sessions OM |
| EPI_ISL_539493, EPI_ISL_539494 | The National Institute of Public Health | State Veterinary Institute Prague | Nagy,A;Jirincova,H;Novakova,L;Trnka,D;Vecerova,J |
| EPI_ISL_539496 | Hospital Nostra Senyora de Meritxell | Instituto de Salud Carlos III | Iglesias-Caballero, M. Molinero Calamita, M. González-Esguevillas, M. Camarero, S. Pozo, F. Casas, I. Jiménez, P. Jiménez, M. Zaballos, A. Monzón, S. Varona, S. Juliá, M. Cuesta, I, F. Fernández |
| EPI_ISL_539497, EPI_ISL_539498 | Hospital Virgen de las Nieves | Instituto de Salud Carlos III | Iglesias-Caballero, M. Molinero Calamita, M. González-Esguevillas, M. Camarero, S. Pozo, F. Casas, I. Jiménez, P. Jiménez, M. Zaballos, A. Monzón, S. Varona, S. Juliá, M. Cuesta, I, J. Lepe |
| EPI_ISL_539500, EPI_ISL_539501, EPI_ISL_539502 | Hospital Clínico Universitario Lozano Blesa | Instituto de Salud Carlos III | Iglesias-Caballero, M. Molinero Calamita, M. González-Esguevillas, M. Camarero, S. Pozo, F. Casas, I. Jiménez, P. Jiménez, M. Zaballos, A. Monzón, S. Varona, S. Juliá, M. Cuesta, I, R. Benito |
| EPI_ISL_539503 | Hospital Universitario Miguel Servet | Instituto de Salud Carlos III | Iglesias-Caballero, M. Molinero Calamita, M. González-Esguevillas, M. Camarero, S. Pozo, F. Casas, I. Jiménez, P. Jiménez, M. Zaballos, A. Monzón, S. Varona, S. Juliá, M. Cuesta, I, A. Rezusta |
| EPI_ISL_539504 | Hospital Universitario Miguel Servet | Instituto de Salud Carlos III | Iglesias-Caballero, M. Molinero Calamita, M. González-Esguevillas, M. Camarero, S. Pozo, F. Casas, I. Jiménez, P. Jiménez, M. Zaballos, A. Monzón, S. Varona, S. Juliá, M. Cuesta, I, R. Benito |
| EPI_ISL_539505, EPI_ISL_539506, EPI_ISL_539507, EPI_ISL_539508 | Hospital Universitario Miguel Servet | Instituto de Salud Carlos III | Iglesias-Caballero, M. Molinero Calamita, M. González-Esguevillas, M. Camarero, S. Pozo, F. Casas, I. Jiménez, P. Jiménez, M. Zaballos, A. Monzón, S. Varona, S. Juliá, M. Cuesta, I, A. Rezusta |
| EPI_ISL_539509, EPI_ISL_539510, EPI_ISL_539511, EPI_ISL_539512 | Hospital Clínico Universitario Lozano Blesa | Instituto de Salud Carlos III | Iglesias-Caballero, M. Molinero Calamita, M. González-Esguevillas, M. Camarero, S. Pozo, F. Casas, I. Jiménez, P. Jiménez, M. Zaballos, A. Monzón, S. Varona, S. Juliá, M. Cuesta, I, R. Benito |
| EPI_ISL_539513, EPI_ISL_539514, EPI_ISL_539515, EPI_ISL_539516, EPI_ISL_539517, EPI_ISL_539518, EPI_ISL_539519 | Hospital Universitario Miguel Servet | Instituto de Salud Carlos III | Iglesias-Caballero, M. Molinero Calamita, M. González-Esguevillas, M. Camarero, S. Pozo, F. Casas, I. Jiménez, P. Jiménez, M. Zaballos, A. Monzón, S. Varona, S. Juliá, M. Cuesta, I, A. Rezusta |
| EPI_ISL_539520, EPI_ISL_539521 | Hospital Universitario de Ceuta | Instituto de Salud Carlos III | Iglesias-Caballero, M. Molinero Calamita, M. González-Esguevillas, M. Camarero, S. Pozo, F. Casas, I. Jiménez, P. Jiménez, M. Zaballos, A. Monzón, S. Varona, S. Juliá, M. Cuesta, I, J. López |
| EPI_ISL_539522 | Hospital Universitario de Ceuta | Instituto de Salud Carlos III | Iglesias-Caballero, M. Molinero Calamita, M. González-Esguevillas, M. Camarero, S. Pozo, F. Casas, I. Jiménez, P. Jiménez, M. Zaballos, A. Monzón, S. Varona, S. Juliá, M. Cuesta, I, G. Sánchez |
| EPI_ISL_539523 | Hospital General de Segovia | Instituto de Salud Carlos III | Iglesias-Caballero, M. Molinero Calamita, M. González-Esguevillas, M. Camarero, S. Pozo, F. Casas, I. Jiménez, P. Jiménez, M. Zaballos, A. Monzón, S. Varona, S. Juliá, M. Cuesta, I, S. Hernando |
| EPI_ISL_539524 | Gerencia de Asistencia Sanitaria de Soria | Instituto de Salud Carlos III | Iglesias-Caballero, M. Molinero Calamita, M. González-Esguevillas, M. Camarero, S. Pozo, F. Casas, I. Jiménez, P. Jiménez, M. Zaballos, A. Monzón, S. Varona, S. Juliá, M. Cuesta, I, C. Aldea |
| EPI_ISL_539525 | Hospital Nuestra Señora de Sonsoles | Instituto de Salud Carlos III | Iglesias-Caballero, M. Molinero Calamita, M. González-Esguevillas, M. Camarero, S. Pozo, F. Casas, I. Jiménez, P. Jiménez, M. Zaballos, A. Monzón, S. Varona, S. Juliá, M. Cuesta, I, A. San Pedro |
| EPI_ISL_539526, EPI_ISL_539527, EPI_ISL_539528, EPI_ISL_539529, EPI_ISL_539530 | Consejeria de Sanidad y Asuntos Sociales | Instituto de Salud Carlos III | Iglesias-Caballero, M. Molinero Calamita, M. González-Esguevillas, M. Camarero, S. Pozo, F. Casas, I. Jiménez, P. Jiménez, M. Zaballos, A. Monzón, S. Varona, S. Juliá, M. Cuesta, I, G. Gutiérrez |
| EPI_ISL_539531 | C.H.U Nuestra Señora de Candelaria | Instituto de Salud Carlos III | Iglesias-Caballero, M. Molinero Calamita, M. González-Esguevillas, M. Camarero, S. Pozo, F. Casas, I. Jiménez, P. Jiménez, M. Zaballos, A. Monzón, S. Varona, S. Juliá, M. Cuesta, I, O. Díez |
| EPI_ISL_539532 | Hospital Universitario de Canarias | Instituto de Salud Carlos III | Iglesias-Caballero, M. Molinero Calamita, M. González-Esguevillas, M. Camarero, S. Pozo, F. Casas, I. Jiménez, P. Jiménez, M. Zaballos, A. Monzón, S. Varona, S. Juliá, M. Cuesta, I, B. Castro |
| EPI_ISL_539534, EPI_ISL_539535, EPI_ISL_539536, EPI_ISL_539537, EPI_ISL_539549, EPI_ISL_539550, EPI_ISL_539551, EPI_ISL_539552, EPI_ISL_539553, EPI_ISL_539554, EPI_ISL_539555, EPI_ISL_539556 | | | |
| see above | Hospital Clínic | Instituto de Salud Carlos III | Iglesias-Caballero, M. Molinero Calamita, M. González-Esguevillas, M. Camarero, S. Pozo, F. Casas, I. Jiménez, P. Jiménez, M. Zaballos, A. Monzón, S. Varona, S. Juliá, M. Cuesta, I, M.A Marcos |
| EPI_ISL_539557 | Gerencia del área de salud de Badajoz, Llerena y Zafra | Instituto de Salud Carlos III | Iglesias-Caballero, M. Molinero Calamita, M. González-Esguevillas, M. Camarero, S. Pozo, F. Casas, I. Jiménez, P. Jiménez, M. Zaballos, A. Monzón, S. Varona, S. Juliá, M. Cuesta, I, C. Pazos |
| EPI_ISL_539558 | Hospital San Pedro de Alcántara | Instituto de Salud Carlos III | Iglesias-Caballero, M. Molinero Calamita, M. González-Esguevillas, M. Camarero, S. Pozo, F. Casas, I. Jiménez, P. Jiménez, M. Zaballos, A. Monzón, S. Varona, S. Juliá, M. Cuesta, I, E. Cerro |
| EPI_ISL_539559 | Hospital Campo Arañuelo | Instituto de Salud Carlos III | Iglesias-Caballero, M. Molinero Calamita, M. González-Esguevillas, M. Camarero, S. Pozo, F. Casas, I. Jiménez, P. Jiménez, M. Zaballos, A. Monzón, S. Varona, S. Juliá, M. Cuesta, I, J. López |
| EPI_ISL_539560 | Hospital Campo Arañuelo | Instituto de Salud Carlos III | Iglesias-Caballero, M. Molinero Calamita, M. González-Esguevillas, M. Camarero, S. Pozo, F. Casas, I. Jiménez, P. Jiménez, M. Zaballos, A. Monzón, S. Varona, S. Juliá, M. Cuesta, I, G. Rodríguez |
| EPI_ISL_539561 | Hospital San Pedro de Alcántara | Instituto de Salud Carlos III | Iglesias-Caballero, M. Molinero Calamita, M. González-Esguevillas, M. Camarero, S. Pozo, F. Casas, I. Jiménez, P. Jiménez, M. Zaballos, A. Monzón, S. Varona, S. Juliá, M. Cuesta, I, J. López |
| EPI_ISL_539562 | Hospital San Pedro de Alcántara | Instituto de Salud Carlos III | Iglesias-Caballero, M. Molinero Calamita, M. González-Esguevillas, M. Camarero, S. Pozo, F. Casas, I. Jiménez, P. Jiménez, M. Zaballos, A. Monzón, S. Varona, S. Juliá, M. Cuesta, I, M.A Cañizares |
| EPI_ISL_539569, EPI_ISL_539570, EPI_ISL_539571, EPI_ISL_539572 | Complejo Hospitalario de Navarra | Instituto de Salud Carlos III | Iglesias-Caballero, M. Molinero Calamita, M. González-Esguevillas, M. Camarero, S. Pozo, F. Casas, I. Jiménez, P. Jiménez, M. Zaballos, A. Monzón, S. Varona, S. Juliá, M. Cuesta, I, J. López |
| EPI_ISL_539776 | National Institute of Public Health (Czech Republic) | State Veterinary Institute Prague | Nagy, A; Jirincova, H; Novakova, L; Trnka, D; Vecerova, J. |
| EPI_ISL_539777 | The National Institute of Public Health | State Veterinary Institute Prague | Nagy,A;Jirincova,H;Novakova,L;Trnka,D;Vecerova,J |
| EPI_ISL_539778, EPI_ISL_539779, EPI_ISL_539780, EPI_ISL_539781 | National Institute of Public Health (Czech Republic) | State Veterinary Institute Prague | Nagy, A; Jirincova, H; Novakova, L; Trnka, D; Vecerova, J. |
| EPI_ISL_539820 | Queen Mary Hospital | Hong Kong Department of Health | Alan K.L. Tsang, Peter C.W. Yip, Edman T.K. Lam, Rickjason C.W. Chan, Dominic N.C. Tsang |
| EPI_ISL_539850, EPI_ISL_539851 | Pok Oi Hospital | Hong Kong Department of Health | Alan K.L. Tsang, Peter C.W. Yip, Edman T.K. Lam, Rickjason C.W. Chan, Dominic N.C. Tsang |

| | | | |
|---|---|---|---|
| EPI_ISL_540993, EPI_ISL_540994, EPI_ISL_540995, EPI_ISL_540996, EPI_ISL_540997, EPI_ISL_540998, EPI_ISL_540999, EPI_ISL_541000, EPI_ISL_541001 | Health and Environmental Research Institute of Gwangju Metropolitan city | Health and Environmental Research Institute of Gwangju Metropolitan city | Min Ji Kim, Ji-eun Lee |
| EPI_ISL_541138 | The National Institute of Public Health | State Veterinary Institute Prague | Nagy,A; Jirincova,H; Novakova,L; Trnka,D; Vecerova,J |
| EPI_ISL_541335 | The National Institute of Public Health | Sídlištní 136/24 165 03, Prague Czech Republic | Nagy,A; Jirincova,H; Novakova,L; Trnka,D; Vecerova,J |
| EPI_ISL_541352 | Laboratory of Respiratory Viruses and Measles, Oswaldo Cruz Institute, FIOCRUZ | Laboratory of Respiratory Viruses and Measles, Oswaldo Cruz Institute, FIOCRUZ | Paola Resende, Roxana Loayza, Cinthia Avila, Luciana Appolinario, Fernando Motta, Anna Carolina Paixao, Ana Carolina Mendonca, Marilda Siqueira on behalf of the Fiocruz COVID-19 Genomic Surveillance Network |
| EPI_ISL_541372, EPI_ISL_541374, EPI_ISL_541375, EPI_ISL_541376, EPI_ISL_541379, EPI_ISL_541383, EPI_ISL_541385 | Laboratório Central de Saúde Pública do Estado de Sergipe (LACEN-SE) | Laboratory of Respiratory Viruses and Measles, Oswaldo Cruz Institute, FIOCRUZ | Paola Resende, Luciana Appolinario, Fernando Motta, Anna Carolina Paixão, Ana Carolina Mendonça, Jonathan Lopes, Clioma Santos, Marilda Siqueira on behalf of the Fiocruz COVID-19 Genomic Surveillance Network |
| EPI_ISL_541970 | Influenza Centre, University of Bergen | Norwegian Institute of Public Health, Department of Virology | Fan Zhou, Rebecca J Cox, Karl A Brokstad, Bjørn Blomberg, Kathrine Stene-Johansen, Kamilla Heddeland Instefjord, Hilde Elshaug, Rasmus Riis Kopperud, Hilde Synnøve Vollan, Karoline Bragstad, Olav Hungnes |
| EPI_ISL_542099, EPI_ISL_542101, EPI_ISL_542104, EPI_ISL_542106, EPI_ISL_542107, EPI_ISL_542108, EPI_ISL_542110, EPI_ISL_542112, EPI_ISL_542114, EPI_ISL_542116, EPI_ISL_542118, EPI_ISL_542119, EPI_ISL_542120, EPI_ISL_542121, EPI_ISL_542122, EPI_ISL_542124, EPI_ISL_542128, EPI_ISL_542130, EPI_ISL_542131, EPI_ISL_542133, EPI_ISL_542145, EPI_ISL_542146, EPI_ISL_542152, EPI_ISL_542153, EPI_ISL_542154, EPI_ISL_542155, EPI_ISL_542156, EPI_ISL_542158, EPI_ISL_542159, EPI_ISL_542162, EPI_ISL_542165, EPI_ISL_542166, EPI_ISL_542169, EPI_ISL_542170, EPI_ISL_542172, EPI_ISL_542173, EPI_ISL_542174, EPI_ISL_542175, EPI_ISL_542178, EPI_ISL_542180, EPI_ISL_542181, EPI_ISL_542183, EPI_ISL_542192, EPI_ISL_542193, EPI_ISL_542194, EPI_ISL_542195, EPI_ISL_542196, EPI_ISL_542197, EPI_ISL_542198, EPI_ISL_542199, EPI_ISL_542200, EPI_ISL_542201, EPI_ISL_542202, EPI_ISL_542203, EPI_ISL_542204, EPI_ISL_542205, EPI_ISL_542206, EPI_ISL_542207, EPI_ISL_542208, EPI_ISL_542209, EPI_ISL_542210, EPI_ISL_542211, EPI_ISL_542212, EPI_ISL_542213, EPI_ISL_542214, EPI_ISL_542215, EPI_ISL_542216, EPI_ISL_542217, EPI_ISL_542218, EPI_ISL_542219, EPI_ISL_542220, EPI_ISL_542221, EPI_ISL_542222, EPI_ISL_542223, EPI_ISL_542224, EPI_ISL_542225, EPI_ISL_542226, EPI_ISL_542228, EPI_ISL_542229, EPI_ISL_542230, EPI_ISL_542231, EPI_ISL_542232, EPI_ISL_542233, EPI_ISL_542234, EPI_ISL_542236, EPI_ISL_542237, EPI_ISL_542238, EPI_ISL_542239, EPI_ISL_542240, EPI_ISL_542241, EPI_ISL_542242, EPI_ISL_542243, EPI_ISL_542248, EPI_ISL_542249, EPI_ISL_542252, EPI_ISL_542253, EPI_ISL_542254, EPI_ISL_542255, EPI_ISL_542256, EPI_ISL_542257, EPI_ISL_542258, EPI_ISL_542260, EPI_ISL_542261, EPI_ISL_542262, EPI_ISL_542263, EPI_ISL_542264, EPI_ISL_542274, EPI_ISL_542275, EPI_ISL_542276, EPI_ISL_542277 | | | |
| see above | ASST GOM Niguarda | Dep. Of Oncology and Hemato-Oncology University of Milan | Claudia Alteri, Valeria Cento, Antonio Piralla, Valentino Costabile, Monica Tallarita, Luna Colagrossi, Silvia Renica, Federica Giardina, Federica Novazzi, Stefano Gaiarsa, Elisa Matarazzo, Maria Antonello, Chiara Vismara, Roberto Fumagalli, Oscar Massimiliano Epis, Massimo Puoti, Carlo Federico Perno, Fausto Baldanti |
| EPI_ISL_542278, EPI_ISL_542279, EPI_ISL_542280, EPI_ISL_542281, EPI_ISL_542282, EPI_ISL_542283, EPI_ISL_542284, EPI_ISL_542285, EPI_ISL_542286, EPI_ISL_542287, EPI_ISL_542288, EPI_ISL_542289, EPI_ISL_542290, EPI_ISL_542291, EPI_ISL_542292, EPI_ISL_542294, EPI_ISL_542297, EPI_ISL_542299, EPI_ISL_542302, EPI_ISL_542303, EPI_ISL_542304, EPI_ISL_542305, EPI_ISL_542306, EPI_ISL_542307, EPI_ISL_542308, EPI_ISL_542309, EPI_ISL_542310, EPI_ISL_542311, EPI_ISL_542312, EPI_ISL_542313, EPI_ISL_542314, EPI_ISL_542315, EPI_ISL_542316, EPI_ISL_542317, EPI_ISL_542318, EPI_ISL_542319, EPI_ISL_542320, EPI_ISL_542321, EPI_ISL_542322, EPI_ISL_542323, EPI_ISL_542324, EPI_ISL_542325, EPI_ISL_542326, EPI_ISL_542327, EPI_ISL_542328, EPI_ISL_542329, EPI_ISL_542330, EPI_ISL_542331, EPI_ISL_542332, EPI_ISL_542333, EPI_ISL_542334, EPI_ISL_542335, EPI_ISL_542336, EPI_ISL_542337, EPI_ISL_542338, EPI_ISL_542339, EPI_ISL_542340, EPI_ISL_542341, EPI_ISL_542342, EPI_ISL_542343, EPI_ISL_542344, EPI_ISL_542345, EPI_ISL_542346, EPI_ISL_542347, EPI_ISL_542348, EPI_ISL_542349, EPI_ISL_542350, EPI_ISL_542351, EPI_ISL_542352, EPI_ISL_542353, EPI_ISL_542354, EPI_ISL_542355, EPI_ISL_542357, EPI_ISL_542358, EPI_ISL_542359, EPI_ISL_542360, EPI_ISL_542361, EPI_ISL_542362, EPI_ISL_542363, EPI_ISL_542364, EPI_ISL_542365, EPI_ISL_542366, EPI_ISL_542367, EPI_ISL_542369, EPI_ISL_542370, EPI_ISL_542371, EPI_ISL_542372, EPI_ISL_542373, EPI_ISL_542374, EPI_ISL_542375, EPI_ISL_542376, EPI_ISL_542377, EPI_ISL_542378, EPI_ISL_542379, EPI_ISL_542381, EPI_ISL_542382, EPI_ISL_542383, EPI_ISL_542384, EPI_ISL_542385, EPI_ISL_542386, EPI_ISL_542387, EPI_ISL_542388, EPI_ISL_542389, EPI_ISL_542390, EPI_ISL_542391, EPI_ISL_542392 | | | |
| see above | San Matteo Hospital Pavia | Dep. Of Oncology and Hemato-Oncology University of Milan | Claudia Alteri, Valeria Cento, Antonio Piralla, Valentino Costabile, Monica Tallarita, Luna Colagrossi, Silvia Renica, Federica Giardina, Federica Novazzi, Stefano Gaiarsa, Elisa Matarazzo, Maria Antonello, Chiara Vismara, Roberto Fumagalli, Oscar Massimiliano Epis, Massimo Puoti, Carlo Federico Perno, Fausto Baldanti |
| EPI_ISL_542413, EPI_ISL_542415, EPI_ISL_542419, EPI_ISL_542420, EPI_ISL_542421, EPI_ISL_542422, EPI_ISL_542423, EPI_ISL_542424, EPI_ISL_542425, EPI_ISL_542426, EPI_ISL_542427, EPI_ISL_542428, EPI_ISL_542429, EPI_ISL_542431, EPI_ISL_542432, EPI_ISL_542433, EPI_ISL_542435, EPI_ISL_542440, EPI_ISL_542442, EPI_ISL_542443 | | | |
| see above | ASST GOM Niguarda | Dep. Of Oncology and Hemato-Oncology University of Milan | Claudia Alteri, Valeria Cento, Antonio Piralla, Valentino Costabile, Monica Tallarita, Luna Colagrossi, Silvia Renica, Federica Giardina, Federica Novazzi, Stefano Gaiarsa, Elisa Matarazzo, Maria Antonello, Chiara Vismara, Roberto Fumagalli, Oscar Massimiliano Epis, Massimo Puoti, Carlo Federico Perno, Fausto Baldanti |
| EPI_ISL_547570 | PS Municipal Dr Augusto Gomes de Mattos | Instituto Adolfo Lutz, Interdiciplinary Procedures Center, Strategic Laboratory | Claudio Tavares Sacchi, Claudia Regina Gonçalves, Erica Valessa Ramos Gomes, Karoline Rodrigues Campos |
| EPI_ISL_548208, EPI_ISL_548214 | Laboratoire de Virologie, HUG | Swiss National Reference Centre for Influenza | LAUBSCHER F. |
| EPI_ISL_548243, EPI_ISL_548244 | Faith Laboratory, Immunology Institute, Icahn School of Medicine at Mount Sinai | van Bakel Laboratory, Genetics and Genomics Sciences, Icahn School of Medicine at Mount Sinai | Graham J. Britton, Alice Chen-Liaw, Francesca Cossarini, Alexandra Livanos, Matthew P. Spindler, Tamar Plitt, Joseph Eggers, Ilaria Mogno, Ana S. Gonzalez-Reiche, Sophia Sui, Michael Tankelevich, Lauren Tal Grinspan, Rebekah E. Dixon, Divya Jha, Gustavo Martinez-Delgado, Fatima Amanat, Daisy Hoagland, Benjamin R. tenOever, Marla C. Dubinsky, Miriam Merad, Harm Van Bakel, Florian Krammer, Gerold Bongers, Saurabh Mehandru and Jeremiah J. Faith |
| EPI_ISL_548966, EPI_ISL_548967, EPI_ISL_548968, EPI_ISL_548970, EPI_ISL_548971 | Expo2020 Emergency Center | Agiomix | Walaa Allam, Cherif Ben Hamada, Cengiz Yakicier, Walid Dridi, Rashid Mohammed, Tamer Degheidy |
| EPI_ISL_560407 | Istituto Zooprofilattico Sperimentale del Mezzogiorno | INMI Lazzaro Spallanzani IRCCS | Barbara Bartolini, Cesare E.M. Gruber, Martina Rueca, Francesco Messina, Antonino Di Caro, Giovanna Fusco, Maurizio Viscardi, Giorgia Borriello, Sergio Brandi, Maria R. Capobianchi |
| EPI_ISL_560568, EPI_ISL_560569, EPI_ISL_560570, EPI_ISL_560571, EPI_ISL_560572, EPI_ISL_560573, EPI_ISL_560574, EPI_ISL_560575, EPI_ISL_560576, EPI_ISL_560577, EPI_ISL_560578, EPI_ISL_560579, EPI_ISL_560580 | | | |
| see above | hôpital | National Reference Center for Viruses of Respiratory Infections, Institut Pasteur, Paris | Sylvie Behillil, Fabiana Gambaro, Etienne Simon-Lorière, Vincent Enouf, Maud Vanpeene, Sylvie van der Werf |
| EPI_ISL_560581, EPI_ISL_560582, EPI_ISL_560583, EPI_ISL_560584, EPI_ISL_560585, EPI_ISL_560586 | Hopital | National Reference Center for Viruses of Respiratory Infections, Institut Pasteur, Paris | Sylvie Behillil, Fabiana Gambaro, Etienne Simon-Lorière, Vincent Enouf, Maud Vanpeene, Sylvie van der Werf |
| EPI_ISL_560591, EPI_ISL_560592, EPI_ISL_560593, EPI_ISL_560594, EPI_ISL_560595, EPI_ISL_560596, EPI_ISL_560597 | hopital | National Reference Center for Viruses of Respiratory Infections, Institut Pasteur, Paris | Sylvie Behillil, Fabiana Gambaro, Etienne Simon-Lorière, Vincent Enouf, Maud Vanpeene, Sylvie van der Werf |
| EPI_ISL_560620, EPI_ISL_560621, EPI_ISL_560622, EPI_ISL_560623, EPI_ISL_560624, EPI_ISL_560625, EPI_ISL_560626, EPI_ISL_560627, EPI_ISL_560628, EPI_ISL_560629, EPI_ISL_560630, EPI_ISL_560631, EPI_ISL_560632, EPI_ISL_560633, EPI_ISL_560634, EPI_ISL_560644 | | | |
| see above | Hospital | National Reference Center for Viruses of Respiratory Infections, Institut Pasteur, Paris | Sylvie Behillil, Fabiana Gambaro, Etienne Simon-Lorière, Vincent Enouf, Maud Vanpeene, Sylvie van der Werf |
| EPI_ISL_560991 | RSUD Dr. Soetomo | Institute of Tropical Disease, Universitas Airlangga | Rima R Prasetya, Krisnoadi Rahardjo, Aldise M Nastri, Jezzy R Dewantari, Joni Wahyuhadi, Gatot Soegiarto, Laksmi Wulandari, Retno A Setyoningrum, Resti Yudhawati, Yohko K Shimizu, Mitsuhiro Nishimura, Yasuko Mori, Soetjipto, Kazufumi Shimizu, Maria I Lusida |
| EPI_ISL_568579 | Virus Molecular Laboratory of the Microbiology and Virology Department | INMI Lazzaro Spallanzani IRCCS | Cesare E.M. Gruber, Martina Rueca, Barbara Bartolini, Francesco Messina, Silvia Meschi, Francesca Colavita, Concetta Castilletti, Elena Percivalle, Irene Cassaniti, Edoardo Vecchio Nepita, Fausto Baldanti, Maria R. Capobianchi, Antonino Di Caro |
| EPI_ISL_568686 | RSUD Prof. DR. Margono Soekarjo | Eijkman Institute for Molecular Biology, Ministry of Research and Technology/National Agency for Research and Innovation | Frilasita A Yudhaputri, Edison Johar, Hidayat Trimarsanto, Iskandar A Adnan, Willy Agustine, David H Muljono, Safarina G Malik, Herawati Sudoyo, Khin Saw Myint, Amin Soebandrio |
| EPI_ISL_568687 | RSUP Fatmawati | Eijkman Institute for Molecular Biology, Ministry of Research and Technology/National Agency for Research and Innovation | Frilasita A Yudhaputri, Edison Johar, Hidayat Trimarsanto, Iskandar A Adnan, Willy Agustine, David H Muljono, Safarina G Malik, Herawati Sudoyo, Khin Saw Myint, Amin Soebandrio |

| | | | |
|---|---|---|---|
| EPI_ISL_568688 | RS Pelni | Eijkman Institute for Molecular Biology, Ministry of Research and Technology/National Agency for Research and Innovation | Frilasita A Yudhaputri, Edison Johar, Hidayat Trimarsanto, Iskandar A Adnan, Willy Agustine, David H Muljono, Safarina G Malik, Herawati Sudoyo, Khin Saw Myint, Amin Soebandrio |
| EPI_ISL_568693 | RS Freeport Tembagapura | Eijkman Institute for Molecular Biology, Ministry of Research and Technology/National Agency for Research and Innovation | Frilasita A Yudhaputri, Edison Johar, Hidayat Trimarsanto, Iskandar A Adnan, Willy Agustine, David H Muljono, Safarina G Malik, Herawati Sudoyo, Khin Saw Myint, Amin Soebandrio |
| EPI_ISL_568694 | RSUP Prof. Dr. R. Kandou Manado | Eijkman Institute for Molecular Biology, Ministry of Research and Technology/National Agency for Research and Innovation | Frilasita A Yudhaputri, Edison Johar, Hidayat Trimarsanto, Iskandar A Adnan, Willy Agustine, David H Muljono, Safarina G Malik, Herawati Sudoyo, Khin Saw Myint, Amin Soebandrio |
| EPI_ISL_568873, EPI_ISL_568874, EPI_ISL_568875 | Malaysia Genome Institute | Malaysia Genome Institute | Mohd Noor Mat Isa, Irni Suhayu Sapian, Yusuf Muhammad Noor, Nurhezreen Md Iqbal, Mohd Faizal Abu Bakar, Enizza Kasim, Shamsidar Sopie, Siti Noraini Othman, Azrin Ahmad, Nor Azfa Johari, Shahrul Hisham Zainal Ariffin |
| EPI_ISL_569865, EPI_ISL_569866, EPI_ISL_569867, EPI_ISL_569868, EPI_ISL_569869, EPI_ISL_569870, EPI_ISL_569871, EPI_ISL_569872, EPI_ISL_569873, EPI_ISL_569875, EPI_ISL_569876, EPI_ISL_569877, EPI_ISL_569878, EPI_ISL_569879, EPI_ISL_569880, EPI_ISL_569881, EPI_ISL_569882, EPI_ISL_569883, EPI_ISL_569884, EPI_ISL_569885, EPI_ISL_569886 | | | |
| see above | Amedeo di savoia | Crosetto lab, Karolinska Institutet, SciLifeLab | Michele Simonetti, Maria Grazia Milia, Luuk Harbers, Ning Zhang, Anna Sapino, Valeria Ghisetti, Nicola Crosetto |
| EPI_ISL_572330 | Institute for Virology, University Hospital Duesseldorf, Medical Faculty, Heinrich-Heine-University Duesseldorf | Institute for Virology, University Hospital Duesseldorf, Medical Faculty, Heinrich-Heine-University Duesseldorf | Maximilian Damagnez, Verena Keitel, Björn Jensen, Nadine Lübke, Lisa Müller, Philipp Ostermann, Tina Senff, Ortwin Adams, Philipp Albrecht, Gerald Antoch, Johannes Bode, Edwin Bölke, Saskia Elben, Torsten Feldt, Johannes C. Fischer, , Anselm Kunstein, Caroline Klindt, Alexander Killer, Tom Lüdde, Annemarie Mohring, Jennifer Neubert, Heiner Schaal, Ansgar Schulz, Jörg Timm, Andreas Walker |
| EPI_ISL_572351, EPI_ISL_572386 | Laboratório Central de Saúde Pública do Estado de Pernambuco (LACEN-PE) | WallauLab, Aggeu Magalhaes Institute | Marcelo Henrique Santos Paiva, Duschinka Ribeiro Duarte Guedes, Cássia Docena, Matheus Filgueira Bezerra, Filipe Zimmer Dezordi, Laís Ceschini Machado, Larissa Krokovsky, Elisama Helvecio, Alexandre Freitas da Silva, Luydson Richardson Silva Vasconcelos, Antonio Mauro Rezende, Severino Jefferson Ribeiro da Silva, Kamila Gaudêncio da Silva Sales, Bruna Santos Lima Figueiredo de Sá, Derciliano Lopes da Cruz, Claudio Eduardo Cavalcanti, Armando de Menezes Neto, Caroline Targino Alves da Silva, Renata Pessôa Germano Mendes, Maria Almerice Lopes da Silva, Tiago Gräf, Paola Cristina Resende, Gonzalo Bello, Michelle da Silva Barros, Wheverton Ricardo Correia do Nascimento, Rodrigo Moraes Loyo Arcoverde, Luciane Caroline Albuquerque Bezerra, Sinval Pinto Brandão Filho, Constância Flávia Junqueira Ayres, Gabriel Luz Wallau on behalf of the Fiocruz COVID-19 Genomic Surveillance Network |
| EPI_ISL_574577 | Hospital Municipal Dr. Ignacio Proença de Gouvea | Instituto Adolfo Lutz, Interdiciplinary Procedures Center, Strategic Laboratory | Claudio Tavares Sacchi, Claudia Regina Gonçalves, Erica Valessa Ramos Gomes, Karoline Rodrigues Campos |
| EPI_ISL_574578 | Hospital Municipal Mário Gatti | Instituto Adolfo Lutz, Interdiciplinary Procedures Center, Strategic Laboratory | Claudio Tavares Sacchi, Claudia Regina Gonçalves, Erica Valessa Ramos Gomes, Karoline Rodrigues Campos |
| EPI_ISL_574579 | Hospital Municipal Dr. Ignacio Proença de Gouvea | Instituto Adolfo Lutz, Interdiciplinary Procedures Center, Strategic Laboratory | Claudio Tavares Sacchi, Claudia Regina Gonçalves, Erica Valessa Ramos Gomes, Karoline Rodrigues Campos |
| EPI_ISL_574580 | Hospital Cidade Tiradentes Carmen Prudente | Instituto Adolfo Lutz, Interdiciplinary Procedures Center, Strategic Laboratory | Claudio Tavares Sacchi, Claudia Regina Gonçalves, Erica Valessa Ramos Gomes, Karoline Rodrigues Campos |
| EPI_ISL_574582 | Hospital Municipal Dr. Jose Soares Hungria | Instituto Adolfo Lutz, Interdiciplinary Procedures Center, Strategic Laboratory | Claudio Tavares Sacchi, Claudia Regina Gonçalves, Erica Valessa Ramos Gomes, Karoline Rodrigues Campos |
| EPI_ISL_574583 | Secretaria Municipal de Saude de Jandira | Instituto Adolfo Lutz, Interdiciplinary Procedures Center, Strategic Laboratory | Claudio Tavares Sacchi, Claudia Regina Gonçalves, Erica Valessa Ramos Gomes, Karoline Rodrigues Campos |
| EPI_ISL_574588 | Hospital Estadual Sumare | Instituto Adolfo Lutz, Interdiciplinary Procedures Center, Strategic Laboratory | Claudio Tavares Sacchi, Claudia Regina Gonçalves, Erica Valessa Ramos Gomes, Karoline Rodrigues Campos |
| EPI_ISL_574589 | Hospital Municipal Dr. Jose Soares Hungria | Instituto Adolfo Lutz, Interdiciplinary Procedures Center, Strategic Laboratory | Claudio Tavares Sacchi, Claudia Regina Gonçalves, Erica Valessa Ramos Gomes, Karoline Rodrigues Campos |
| EPI_ISL_574590 | Unidade de Printo Atendimento UPA I Santa Isabel | Instituto Adolfo Lutz, Interdiciplinary Procedures Center, Strategic Laboratory | Claudio Tavares Sacchi, Claudia Regina Gonçalves, Erica Valessa Ramos Gomes, Karoline Rodrigues Campos |
| EPI_ISL_574591, EPI_ISL_574592 | Hospital Domingos Leonardo Ceravolo Presidente Prudente | Instituto Adolfo Lutz, Interdiciplinary Procedures Center, Strategic Laboratory | Claudio Tavares Sacchi, Claudia Regina Gonçalves, Erica Valessa Ramos Gomes, Karoline Rodrigues Campos |
| EPI_ISL_574593 | CS II Dr. Antonio Vicoso Moreira de Rezende Sumare | Instituto Adolfo Lutz, Interdiciplinary Procedures Center, Strategic Laboratory | Claudio Tavares Sacchi, Claudia Regina Gonçalves, Erica Valessa Ramos Gomes, Karoline Rodrigues Campos |
| EPI_ISL_574595 | Hospital Geral de Vila Penteado Dr. Jose Pamgella | Instituto Adolfo Lutz, Interdiciplinary Procedures Center, Strategic Laboratory | Claudio Tavares Sacchi, Claudia Regina Gonçalves, Erica Valessa Ramos Gomes, Karoline Rodrigues Campos |
| EPI_ISL_574603 | RS Freeport Tembagapura | Eijkman Institute for Molecular Biology, Ministry of Research and Technology/National Agency for Research and Innovation | Frilasita A Yudhaputri, Edison Johar, Hidayat Trimarsanto, Iskandar A Adnan, Willy Agustine, David H Muljono, Safarina G Malik, Herawati Sudoyo, Khin Saw Myint, Amin Soebandrio |
| EPI_ISL_574604 | RSUP Prof. Dr. R. Kandou Manado | Eijkman Institute for Molecular Biology, Ministry of Research and Technology/National Agency for Research and Innovation | Frilasita A Yudhaputri, Edison Johar, Hidayat Trimarsanto, Iskandar A Adnan, Willy Agustine, David H Muljono, Safarina G Malik, Herawati Sudoyo, Khin Saw Myint, Amin Soebandrio |
| EPI_ISL_574605 | RSUP dr. Sardjito | Eijkman Institute for Molecular Biology, Ministry of Research and Technology/National Agency for Research and Innovation | Frilasita A Yudhaputri, Edison Johar, Hidayat Trimarsanto, Iskandar A Adnan, Willy Agustine, David H Muljono, Safarina G Malik, Herawati Sudoyo, Khin Saw Myint, Amin Soebandrio |
| EPI_ISL_574606 | RS Siloam Purwakarta | Eijkman Institute for Molecular Biology, Ministry of Research and Technology/National Agency for Research and Innovation | Frilasita A Yudhaputri, Edison Johar, Hidayat Trimarsanto, Iskandar A Adnan, Willy Agustine, David H Muljono, Safarina G Malik, Herawati Sudoyo, Khin Saw Myint, Amin Soebandrio |
| EPI_ISL_574607 | RS Husada | Eijkman Institute for Molecular Biology, Ministry of Research and Technology/National Agency for Research and Innovation | Frilasita A Yudhaputri, Edison Johar, Hidayat Trimarsanto, Iskandar A Adnan, Willy Agustine, David H Muljono, Safarina G Malik, Herawati Sudoyo, Khin Saw Myint, Amin Soebandrio |
| EPI_ISL_574608 | RS Mitra Keluarga Kelapa Gading | Eijkman Institute for Molecular Biology, Ministry of Research and Technology/National Agency for Research and Innovation | Frilasita A Yudhaputri, Edison Johar, Hidayat Trimarsanto, Iskandar A Adnan, Willy Agustine, David H Muljono, Safarina G Malik, Herawati Sudoyo, Khin Saw Myint, Amin Soebandrio |
| EPI_ISL_574609 | RSJPD Harapan Kita | Eijkman Institute for Molecular Biology, Ministry of Research and Technology/National Agency for Research and Innovation | Frilasita A Yudhaputri, Edison Johar, Hidayat Trimarsanto, Iskandar A Adnan, Willy Agustine, David H Muljono, Safarina G Malik, Herawati Sudoyo, Khin Saw Myint, Amin Soebandrio |
| EPI_ISL_574610 | Puskesmas Tambora | Eijkman Institute for Molecular Biology, Ministry of Research and Technology/National Agency for Research and Innovation | Frilasita A Yudhaputri, Edison Johar, Hidayat Trimarsanto, Iskandar A Adnan, Willy Agustine, David H Muljono, Safarina G Malik, Herawati Sudoyo, Khin Saw Myint, Amin Soebandrio |
| EPI_ISL_574611 | RS Premier Bintaro | Eijkman Institute for Molecular Biology, Ministry of Research and Technology/National Agency for Research and | Frilasita A Yudhaputri, Edison Johar, Hidayat Trimarsanto, Iskandar A Adnan, Willy Agustine, David H Muljono, Safarina G Malik, Herawati Sudoyo, Khin Saw Myint, Amin Soebandrio |

| | | | |
|---|---|---|---|
| | | Innovation | |
| EPI_ISL_575330 | National Institute for Viral Disease Control and Prevention, China CDC | National Institute for Viral Disease Control and Prevention, China CDC | Rongbao Gao, Kang Xiao, Qinqin Song, ZhiQiang Xia, Dong Xia, Juan Song, Haijun Du, Yuan He, Shuai Pang, Xuancheng Lu, Guizhen Wu, Geogia Fu Gao, Jun Han |
| EPI_ISL_577629, EPI_ISL_577631 | The National Institute of Public Health | State Veterinary Institute Prague | Nagy,A;Jirincova,H;Novakova,L;Trnka,D;Vecerova,J |
| EPI_ISL_578186 | Hospital General Juan Ramón Jiménez | Instituto de Salud Carlos III | Iglesias-Caballero, M. Molinero Calamita, M. González-Esguevillas, M. Camarero, S. Pozo, F. Casas, I. Jiménez, P. Jiménez, M. Zaballos, A. Monzón, S. Varona, S. Juliá, M. Cuesta, I, J. Saavedra |
| EPI_ISL_578189 | Hospital Universitario Miguel Servet | Instituto de Salud Carlos III | Iglesias-Caballero, M. Molinero Calamita, M. González-Esguevillas, M. Camarero, S. Pozo, F. Casas, I. Jiménez, P. Jiménez, M. Zaballos, A. Monzón, S. Varona, S. Juliá, M. Cuesta, I, A. Rezusta |
| EPI_ISL_578190, EPI_ISL_578191 | Hospital Clínico Universitario Lozano Blesa | Instituto de Salud Carlos III | Iglesias-Caballero, M. Molinero Calamita, M. González-Esguevillas, M. Camarero, S. Pozo, F. Casas, I. Jiménez, P. Jiménez, M. Zaballos, A. Monzón, S. Varona, S. Juliá, M. Cuesta, I, R. Benito |
| EPI_ISL_578192, EPI_ISL_578193 | Hospital Universitario Miguel Servet | Instituto de Salud Carlos III | Iglesias-Caballero, M. Molinero Calamita, M. González-Esguevillas, M. Camarero, S. Pozo, F. Casas, I. Jiménez, P. Jiménez, M. Zaballos, A. Monzón, S. Varona, S. Juliá, M. Cuesta, I, A. Rezusta |
| EPI_ISL_578194 | Complejo Hospitalario Universitario La Coruna | Instituto de Salud Carlos III | Iglesias-Caballero, M. Molinero Calamita, M. González-Esguevillas, M. Camarero, S. Pozo, F. Casas, I. Jiménez, P. Jiménez, M. Zaballos, A. Monzón, S. Varona, S. Juliá, M. Cuesta, I, J. López |
| EPI_ISL_578195 | Complejo Hospitalario de Orense | Instituto de Salud Carlos III | Iglesias-Caballero, M. Molinero Calamita, M. González-Esguevillas, M. Camarero, S. Pozo, F. Casas, I. Jiménez, P. Jiménez, M. Zaballos, A. Monzón, S. Varona, S. Juliá, M. Cuesta, I, M. García |
| EPI_ISL_578196 | Hospital San Pedro | Instituto de Salud Carlos III | Iglesias-Caballero, M. Molinero Calamita, M. González-Esguevillas, M. Camarero, S. Pozo, F. Casas, I. Jiménez, P. Jiménez, M. Zaballos, A. Monzón, S. Varona, S. Juliá, M. Cuesta, I, J.M Azcona |
| EPI_ISL_578197 | Hospital San Pedro | Instituto de Salud Carlos III | Iglesias-Caballero, M. Molinero Calamita, M. González-Esguevillas, M. Camarero, S. Pozo, F. Casas, I. Jiménez, P. Jiménez, M. Zaballos, A. Monzón, S. Varona, S. Juliá, M. Cuesta, I, C. Alonso |
| EPI_ISL_578198, EPI_ISL_578199 | Hospital San Pedro | Instituto de Salud Carlos III | Iglesias-Caballero, M. Molinero Calamita, M. González-Esguevillas, M. Camarero, S. Pozo, F. Casas, I. Jiménez, P. Jiménez, M. Zaballos, A. Monzón, S. Varona, S. Juliá, M. Cuesta, I, M. Blasco |
| EPI_ISL_578200, EPI_ISL_578201 | Hospital San Pedro | Instituto de Salud Carlos III | Iglesias-Caballero, M. Molinero Calamita, M. González-Esguevillas, M. Camarero, S. Pozo, F. Casas, I. Jiménez, P. Jiménez, M. Zaballos, A. Monzón, S. Varona, S. Juliá, M. Cuesta, I, C. Alonso |
| EPI_ISL_582033 | Department of Pathology, School of Medicine, Imam Khomeini Hospital, Tehran University of Medical Sciences | Genetics Research Center. University Of Social Welfare And Rehabilitation Sciences | Zohreh Fattahi, Marzieh Mohseni, Khadijeh Jalalvand, Azam Ghaziasadi, Seyedeh elham Mortazavi, Ali Jafarpour, Azar Hadadi, Alireza Abdollahi, Ali Jafarpour, Azam Ghaziasad, Seyedeh elham Mortazavi, Saber Soltani, Reza Najafipour , Kimia Kahrizi, Seyed Mohammad Jazayeri, Hossein Najmabadi |
| EPI_ISL_582125, EPI_ISL_582126 | Sheikh Khalifa Medical City | Molecular Surveillance lab Sheikh Khalifa Medical City | Amirtharaj Francis, Sajeed Abdul, Hala Imambaccus, Sahar Almarzooqi, Hiba Saud, Stefan Weber |
| EPI_ISL_582127 | Antwerp University Hospital | Institute of Tropical Medicine | Philippe Selhorst, Colin Anthony |
| EPI_ISL_582608, EPI_ISL_582613, EPI_ISL_582620, EPI_ISL_582624, EPI_ISL_582625, EPI_ISL_582632 | Sheikh Khalifa Medical City | Molecular/Surveillance lab Sheikh Khalifa Medical City | Amirtharaj Francis, Sajeed Abdul, Hala Imambaccus, Sahar Almarzooqi, Hiba Saud, Stefan Weber |
| EPI_ISL_583426 | Garcia-Sastre Laboratory, Department of Microbiology, Icahn School of Medicine at Mount Sinai | van Bakel Laboratory, Genetics and Genomics Sciences, Icahn School of Medicine at Mount Sinai | Teresa Aydillo, Ana S. Gonzalez-Reiche, Sadaf Aslam, Adriana van de Guchte, Zenab Khan, Ajay Obla, Jayeeta Dutta, Harm van Bakel, Judith Aberg, Adolfo García-Sastre, Gunjan Shah, Tobias Hohl, Genovefa Papanicolaou, Miguel-Angel Perales, Kent Sepkowitz, Ngolela Esther Babady, and Mini Kamboj |
| EPI_ISL_583427, EPI_ISL_583428 | Memorial Sloan Kettering Cancer Center | van Bakel Laboratory, Genetics and Genomics Sciences, Icahn School of Medicine at Mount Sinai | Teresa Aydillo, Ana S. Gonzalez-Reiche, Sadaf Aslam, Adriana van de Guchte, Zenab Khan, Ajay Obla, Jayeeta Dutta, Harm van Bakel, Judith Aberg, Adolfo García-Sastre, Gunjan Shah, Tobias Hohl, Genovefa Papanicolaou, Miguel-Angel Perales, Kent Sepkowitz, Ngolela Esther Babady, and Mini Kamboj |
| EPI_ISL_583429 | Garcia-Sastre Laboratory, Department of Microbiology, Icahn School of Medicine at Mount Sinai | van Bakel Laboratory, Genetics and Genomics Sciences, Icahn School of Medicine at Mount Sinai | Teresa Aydillo, Ana S. Gonzalez-Reiche, Sadaf Aslam, Adriana van de Guchte, Zenab Khan, Ajay Obla, Jayeeta Dutta, Harm van Bakel, Judith Aberg, Adolfo García-Sastre, Gunjan Shah, Tobias Hohl, Genovefa Papanicolaou, Miguel-Angel Perales, Kent Sepkowitz, Ngolela Esther Babady, and Mini Kamboj |
| EPI_ISL_583430, EPI_ISL_583431, EPI_ISL_583432, EPI_ISL_583433 | Memorial Sloan Kettering Cancer Center | van Bakel Laboratory, Genetics and Genomics Sciences, Icahn School of Medicine at Mount Sinai | Teresa Aydillo, Ana S. Gonzalez-Reiche, Sadaf Aslam, Adriana van de Guchte, Zenab Khan, Ajay Obla, Jayeeta Dutta, Harm van Bakel, Judith Aberg, Adolfo García-Sastre, Gunjan Shah, Tobias Hohl, Genovefa Papanicolaou, Miguel-Angel Perales, Kent Sepkowitz, Ngolela Esther Babady, and Mini Kamboj |
| EPI_ISL_583434, EPI_ISL_583435, EPI_ISL_583436 | Garcia-Sastre Laboratory, Department of Microbiology, Icahn School of Medicine at Mount Sinai | van Bakel Laboratory, Genetics and Genomics Sciences, Icahn School of Medicine at Mount Sinai | Teresa Aydillo, Ana S. Gonzalez-Reiche, Sadaf Aslam, Adriana van de Guchte, Zenab Khan, Ajay Obla, Jayeeta Dutta, Harm van Bakel, Judith Aberg, Adolfo García-Sastre, Gunjan Shah, Tobias Hohl, Genovefa Papanicolaou, Miguel-Angel Perales, Kent Sepkowitz, Ngolela Esther Babady, and Mini Kamboj |
| EPI_ISL_583437, EPI_ISL_583438, EPI_ISL_583439, EPI_ISL_583440 | Memorial Sloan Kettering Cancer Center | van Bakel Laboratory, Genetics and Genomics Sciences, Icahn School of Medicine at Mount Sinai | Teresa Aydillo, Ana S. Gonzalez-Reiche, Sadaf Aslam, Adriana van de Guchte, Zenab Khan, Ajay Obla, Jayeeta Dutta, Harm van Bakel, Judith Aberg, Adolfo García-Sastre, Gunjan Shah, Tobias Hohl, Genovefa Papanicolaou, Miguel-Angel Perales, Kent Sepkowitz, Ngolela Esther Babady, and Mini Kamboj |
| EPI_ISL_583441, EPI_ISL_583442 | Garcia-Sastre Laboratory, Department of Microbiology, Icahn School of Medicine at Mount Sinai | van Bakel Laboratory, Genetics and Genomics Sciences, Icahn School of Medicine at Mount Sinai | Teresa Aydillo, Ana S. Gonzalez-Reiche, Sadaf Aslam, Adriana van de Guchte, Zenab Khan, Ajay Obla, Jayeeta Dutta, Harm van Bakel, Judith Aberg, Adolfo García-Sastre, Gunjan Shah, Tobias Hohl, Genovefa Papanicolaou, Miguel-Angel Perales, Kent Sepkowitz, Ngolela Esther Babady, and Mini Kamboj |
| EPI_ISL_583445, EPI_ISL_583446, EPI_ISL_583447, EPI_ISL_583448, EPI_ISL_583456, EPI_ISL_583457 | Memorial Sloan Kettering Cancer Center | van Bakel Laboratory, Genetics and Genomics Sciences, Icahn School of Medicine at Mount Sinai | Teresa Aydillo, Ana S. Gonzalez-Reiche, Sadaf Aslam, Adriana van de Guchte, Zenab Khan, Ajay Obla, Jayeeta Dutta, Harm van Bakel, Judith Aberg, Adolfo García-Sastre, Gunjan Shah, Tobias Hohl, Genovefa Papanicolaou, Miguel-Angel Perales, Kent Sepkowitz, Ngolela Esther Babady, and Mini Kamboj |
| EPI_ISL_583458, EPI_ISL_583459 | Garcia-Sastre Laboratory, Department of Microbiology, Icahn School of Medicine at Mount Sinai | van Bakel Laboratory, Genetics and Genomics Sciences, Icahn School of Medicine at Mount Sinai | Teresa Aydillo, Ana S. Gonzalez-Reiche, Sadaf Aslam, Adriana van de Guchte, Zenab Khan, Ajay Obla, Jayeeta Dutta, Harm van Bakel, Judith Aberg, Adolfo García-Sastre, Gunjan Shah, Tobias Hohl, Genovefa Papanicolaou, Miguel-Angel Perales, Kent Sepkowitz, Ngolela Esther Babady, and Mini Kamboj |
| EPI_ISL_583460, EPI_ISL_583461, EPI_ISL_583462 | Memorial Sloan Kettering Cancer Center | van Bakel Laboratory, Genetics and Genomics Sciences, Icahn School of Medicine at Mount Sinai | Teresa Aydillo, Ana S. Gonzalez-Reiche, Sadaf Aslam, Adriana van de Guchte, Zenab Khan, Ajay Obla, Jayeeta Dutta, Harm van Bakel, Judith Aberg, Adolfo García-Sastre, Gunjan Shah, Tobias Hohl, Genovefa Papanicolaou, Miguel-Angel Perales, Kent Sepkowitz, Ngolela Esther Babady, and Mini Kamboj |
| EPI_ISL_583463, EPI_ISL_583464 | Garcia-Sastre Laboratory, Department of Microbiology, Icahn School of Medicine at Mount Sinai | van Bakel Laboratory, Genetics and Genomics Sciences, Icahn School of Medicine at Mount Sinai | Teresa Aydillo, Ana S. Gonzalez-Reiche, Sadaf Aslam, Adriana van de Guchte, Zenab Khan, Ajay Obla, Jayeeta Dutta, Harm van Bakel, Judith Aberg, Adolfo García-Sastre, Gunjan Shah, Tobias Hohl, Genovefa Papanicolaou, Miguel-Angel Perales, Kent Sepkowitz, Ngolela Esther Babady, and Mini Kamboj |
| EPI_ISL_583465, EPI_ISL_583466 | Memorial Sloan Kettering Cancer Center | van Bakel Laboratory, Genetics and Genomics Sciences, Icahn School of Medicine at Mount Sinai | Teresa Aydillo, Ana S. Gonzalez-Reiche, Sadaf Aslam, Adriana van de Guchte, Zenab Khan, Ajay Obla, Jayeeta Dutta, Harm van Bakel, Judith Aberg, Adolfo García-Sastre, Gunjan Shah, Tobias Hohl, Genovefa Papanicolaou, Miguel-Angel Perales, Kent Sepkowitz, Ngolela Esther Babady, and Mini Kamboj |
| EPI_ISL_583467 | Garcia-Sastre Laboratory, Department of Microbiology, Icahn School of Medicine at Mount Sinai | van Bakel Laboratory, Genetics and Genomics Sciences, Icahn School of Medicine at Mount Sinai | Teresa Aydillo, Ana S. Gonzalez-Reiche, Sadaf Aslam, Adriana van de Guchte, Zenab Khan, Ajay Obla, Jayeeta Dutta, Harm van Bakel, Judith Aberg, Adolfo García-Sastre, Gunjan Shah, Tobias Hohl, Genovefa Papanicolaou, Miguel-Angel Perales, Kent Sepkowitz, Ngolela Esther Babady, and Mini Kamboj |
| EPI_ISL_583468, EPI_ISL_583469, EPI_ISL_583470, EPI_ISL_583472, EPI_ISL_583475, EPI_ISL_583478, EPI_ISL_583479, EPI_ISL_583480 | Memorial Sloan Kettering Cancer Center | van Bakel Laboratory, Genetics and Genomics Sciences, Icahn School of Medicine at Mount Sinai | Teresa Aydillo, Ana S. Gonzalez-Reiche, Sadaf Aslam, Adriana van de Guchte, Zenab Khan, Ajay Obla, Jayeeta Dutta, Harm van Bakel, Judith Aberg, Adolfo García-Sastre, Gunjan Shah, Tobias Hohl, Genovefa Papanicolaou, Miguel-Angel Perales, Kent Sepkowitz, Ngolela Esther Babady, and Mini Kamboj |
| EPI_ISL_583490 | Hospital Estadual Sumare | Instituto Adolfo Lutz, Interdiciplinary Procedures Center, Strategic Laboratory | Claudio Tavares Sacchi, Claudia Regina Gonçalves, Erica Valessa Ramos Gomes, Karoline Rodrigues Campos |
| EPI_ISL_583491 | Centro de Saude Esf IV Zona Rual Domingos de SJ Rio Pardo | Instituto Adolfo Lutz, Interdiciplinary Procedures Center, Strategic Laboratory | Claudio Tavares Sacchi, Claudia Regina Gonçalves, Erica Valessa Ramos Gomes, Karoline Rodrigues Campos |
| EPI_ISL_583492 | Santa Casa Anna Cintra | Instituto Adolfo Lutz, Interdiciplinary Procedures Center, Strategic Laboratory | Claudio Tavares Sacchi, Claudia Regina Gonçalves, Erica Valessa Ramos Gomes, Karoline Rodrigues Campos |
| EPI_ISL_583493 | Vigilância em Saúde de Cajamar | Instituto Adolfo Lutz, Interdiciplinary Procedures Center, Strategic Laboratory | Claudio Tavares Sacchi, Claudia Regina Gonçalves, Erica Valessa Ramos Gomes, Karoline Rodrigues Campos |

| | | | |
|---|---|---|---|
| EPI_ISL_583494 | CS II Dr. Antonio Vicoso Moreira de Rezende Sumare | Instituto Adolfo Lutz, Interdiciplinary Procedures Center, Strategic Laboratory | Claudio Tavares Sacchi, Claudia Regina Gonçalves, Erica Valessa Ramos Gomes, Karoline Rodrigues Campos |
| EPI_ISL_584048 | Laboratory of Molecular Virology, Department of Biomedical, Surgical and Dental Sciences University of Milano | Laboratory of Molecular Virology, Department of Biomedical, Surgical and Dental Sciences University of Milano | Delbue,S., Modenese,A., Bianchi,M., Fattori,M., D'Alessandro,S., Pariani,E., Basilico,N., Galli,C. and Ferrante,P. |
| EPI_ISL_584049 | Laboratory of Molecular Virology, Department of Biomedical, Surgical and Dental Sciences University of Milano | Laboratory of Molecular Virology, Department of Biomedical, Surgical and Dental Sciences University of Milano | Delbue,S., Modenese,A., Bianchi,M., Fattori,M., D'Alessandro,S.,Pariani,E., Basilico,N., Galli,C. and Ferrante,P. |
| EPI_ISL_584052 | Laboratory of Molecular Virology, Department of Biomedical, Surgical and Dental Sciences University of Milano | Laboratory of Molecular Virology, Department of Biomedical, Surgical and Dental Sciences University of Milano | Delbue,S., D'Alessandro,S., Modenese,A., Signorini,L., Parapini,S.,Dolci,M., Binda,S., Primache,V., Taramelli,D., Incorvaia,B. and Ferrante,P. |
| EPI_ISL_584073, EPI_ISL_584079 | The National Institute of Public Health | State Veterinary Institute Prague | Nagy,A;Jirincova,H;Novakova,L;Trnka,D;Vecerova,J |
| EPI_ISL_591471, EPI_ISL_591472, EPI_ISL_591473, EPI_ISL_591474, EPI_ISL_591475, EPI_ISL_591476, EPI_ISL_591477, EPI_ISL_591478, EPI_ISL_591479, EPI_ISL_591480, EPI_ISL_591481, EPI_ISL_591482, EPI_ISL_591483 | | | |
| see above | Pathogen Genomics Center, National Institute of Infectious Diseases | Pathogen Genomics Center, National Institute of Infectious Diseases | Tsuyoshi Sekizuka, Kentaro Itokawa, Rina Tanaka, Masanori Hashino, Hajime Kamiya, Tomoe Shimada, Makoto Kuroda |
| EPI_ISL_593478, EPI_ISL_593479 | Brigham and Women's Hospital | Jonathan Li Laboratory | Manish C. Choudhary, James Regan, Jonathan Z. Li |
| EPI_ISL_594185, EPI_ISL_594186, EPI_ISL_594187 | Department of Pathology, School of Medicine, Imam Khomeini Hospital, Tehran University of Medical Sciences | Genetics Research Center, University of Social Welfare and Rehabilitation Sciences | Zohreh Fattahi, Marzieh Mohseni, Khadijeh Jalalvand, Azam Ghaziasadi, Seyedeh elham Mortazavi, Ali Jafarpour, Azar Hadadi, Alireza Abdollahi, Ali Jafarpour, Azam Ghaziasad, Seyedeh elham Mortazavi, Saber Soltani, Reza Najafipour, Kimia Kahrizi, Seyed Mohammad Jazayeri, Hossein Najmabadi |
| EPI_ISL_596453 | Booali laboratory, Qom, Iran. Department of Virology, School of Public Health, Tehran University of Medical Sciences, Tehran, Iran. | Genetics Research Center, University of Social Welfare and Rehabilitation Sciences | Zohreh Fattahi, Marzieh Mohseni, Khadijeh Jalalvand, Azam Ghaziasadi, Seyedeh elham Mortazavi, Ali Jafarpour, Mohammad Khazeni, Seyed Amir Momeni, Kimia Kahrizi, Seyed Mohammad Jazayeri, Hossein Najmbadi |
| EPI_ISL_596454 | Infectious Disease and Tropical Medicine Research Center, Resistant Tuberculosis Institute, Zahedan University of Medical Sciences, Zahedan, Iran. | Genetics Research Center, University of Social Welfare and Rehabilitation Sciences | Zohreh Fattahi, Marzieh Mohseni, Khadijeh Jalalvand, Azam Ghaziasadi, Seyedeh elham Mortazavi, Ali Jafarpour, Ebrahim Kord, Seyed Mohammad Hashemi-Shahri, Kimia Kahrizi, Seyed Mohammad Jazayeri, Hossein Najmabadi |
| EPI_ISL_602412 | WHO National Influenza Centre Russian Federation | WHO National Influenza Centre Russian Federation | Andrey Komissarov, Artem Fadeev, Kseniya Komissarova, Anna Ivanova, Dmitry Bazhenov, Daria Danilenko |
| EPI_ISL_605799, EPI_ISL_605800, EPI_ISL_605801, EPI_ISL_605802, EPI_ISL_605803, EPI_ISL_605804, EPI_ISL_605805, EPI_ISL_605806, EPI_ISL_605807, EPI_ISL_605808, EPI_ISL_605809, EPI_ISL_605810, EPI_ISL_605811, EPI_ISL_605812, EPI_ISL_605813, EPI_ISL_605814, EPI_ISL_605815, EPI_ISL_605816 | | | |
| see above | Clinical Virology Laboratory, Institute of Liver and Biliary Sciences | ILBS - IGIB | Ekta Gupta, Sheetalnath Rooge, Abhishek Padhi, Reshu Agarwal, Jaswinder Singh Maras, Shridhar Sivasubbu, Vinod Scaria, Shvetank Sharma |
| EPI_ISL_610237, EPI_ISL_610238 | Molecular diagnostic laboratory of Federal Budget Institution of Science "Central Research Institute of Epidemiology" of The Federal Service on Customers' Rights Protection and Human Well-being Surveillance | Group of Genomics and Postgenomic Technologies of Central Research Institute of Epidemilology | Samoilov AE, Kaptelova VV, Bukharina AY, Speranskaya AS, Tivanova EV, Shipulina OY, Akimkin VG |
| EPI_ISL_610249 | Virologisches Institut, Universitätsklinikum Erlangen | Virologisches Institut, Universitätsklinikum Erlangen | Armin Ensser, Klaus Korn, Klaus Überla |
| EPI_ISL_613419, EPI_ISL_613420, EPI_ISL_613421, EPI_ISL_613422, EPI_ISL_613423, EPI_ISL_613424, EPI_ISL_613425, EPI_ISL_613426, EPI_ISL_613429, EPI_ISL_613431, EPI_ISL_613432, EPI_ISL_613433, EPI_ISL_613434, EPI_ISL_613435, EPI_ISL_613436, EPI_ISL_613437, EPI_ISL_613439, EPI_ISL_613440, EPI_ISL_613445, EPI_ISL_613446 | | | |
| see above | Institut Pasteur de la Guadeloupe | Institut Pasteur de la Guadeloupe | Marion Barbet, Sylvie Behillil, Méline Bizard, Angela Brisebarre, Camille Capel, Etienne Simon-Lorière, Vincent Enouf, Maud Vanpeene, Sylvie van der Werf, Stéphanie Guyomard, Sébastien Breurec, Antoine Talarmin |
| EPI_ISL_613447, EPI_ISL_613448, EPI_ISL_613449, EPI_ISL_613450, EPI_ISL_613451, EPI_ISL_613452, EPI_ISL_613453, EPI_ISL_613454, EPI_ISL_613455, EPI_ISL_613456 | Institut Pasteur de la Guadeloupe | Institut Pasteur de la Guadeloupe | Marion Barbet, Sylvie Behillil, Méline Bizard, Angela Brisebarre, Camille Capel, Etienne Simon-Lorière, Vincent Enouf, Maud Vanpeene, Sylvie van der Werf, Stéphanie Guyomard, Sébastien Breurec, Radjin Steingrover, Cherina Fleming, Antoine Talarmin |
| EPI_ISL_613560 | Laboratorio Biologia Molecolare Sars Cov2 - UOC Laboratorio Analisi - Servizio Medicina di Laboratorio, Ospedale "San Francesco" - ATS-ASSL Nuoro Via Mannironi 1, 08100 Nuoro | Laboratorio specialistico UOC Ematologia - Ospedale "San Francesco" - ATS-ASSL Nuoro Nuoro | Piras Giovanna, Fancello Tatiana, Asproni Rosanna, Fiamma Maura, Monne Maria Itria, Toja Alessandro, Sanna Filomena, Floris Anna Rita, Sulis Vincenzo, Palmas Angelo Domenico, Casu Gavino, Lo Maglio Iana, Mameli Giuseppe |
| EPI_ISL_613563, EPI_ISL_613564, EPI_ISL_613707, EPI_ISL_613708 | Laboratory of Molecular Biology, Blood Center of Ribeirão Preto | Laboratory of Molecular Biology, Blood Center of Ribeirão Preto, Faculty of Medicine of Ribeirão Preto, University of São Paulo | Svetoslav N Slavov, Marta Giovanetti, Vagner Fonseca, Elaine V Santos, Evandra S Rodrigues, Talita Adelino, Joilson Xavier, Glauco de Carvalho Pereira, Aparecida Y Yamamoto, Diego Villa Clé, Rodrigo T Calado; Dimas T Covas, Luiz CJ Alcantara, Simone Kashima |
| EPI_ISL_613709, EPI_ISL_613951 | Laboratory of Molecular Biology, Blood Center of Ribeirão Preto, Faculty of Medicine of Ribeirão Preto, University of São Paulo | Laboratory of Molecular Biology, Blood Center of Ribeirão Preto, Faculty of Medicine of Ribeirão Preto, University of São Paulo | Svetoslav N Slavov, Marta Giovanetti, Vagner Fonseca, Elaine V Santos, Evandra S Rodrigues, Talita Adelino, Joilson Xavier, Glauco de Carvalho Pereira, Aparecida Y Yamamoto, Diego Villa Clé, Rodrigo T Calado; Dimas T Covas, Luiz CJ Alcantara, Simone Kashima |
| EPI_ISL_613955 | Laboratorio Biologia Molecolare Sars Cov2 - UOC Laboratorio Analisi - Servizio Medicina di Laboratorio, Ospedale "San Francesco" - ATS-ASSL Nuoro | Laboratorio specialistico UOC Ematologia - Ospedale "San Francesco" - ATS-ASSL Nuoro | Piras Giovanna, Fancello Tatiana, Asproni Rosanna, Fiamma Maura, Monne Maria Itria, Toja Alessandro, Sanna Filomena, Floris Anna Rita, Sulis Vincenzo, Palmas Angelo Domenico, Casu Gavino, Lo Maglio Iana, Mameli Giuseppe |
| EPI_ISL_614281 | CHRU Pontchaillou - Laboratoire de Virologie 2, rue Henri Le Guilloux | National Reference Center for Viruses of Respiratory Infections, Institut Pasteur, Paris | Marion Barbet, Sylvie Behillil, Méline Bizard, Angela Brisebarre, Camille Capel, Etienne Simon-Lorière, Vincent Enouf, Maud Vanpeene, Sylvie van der Werf, Gisèle Lagathu |
| EPI_ISL_614396, EPI_ISL_614397, EPI_ISL_614398, EPI_ISL_614889 | Laboratorio Biologia Molecolare Sars Cov2 - UOC Laboratorio Analisi - Servizio Medicina di Laboratorio, Ospedale "San Francesco" - ATS-ASSL Nuoro | Laboratorio specialistico UOC Ematologia - Ospedale "San Francesco" - ATS-ASSL Nuoro | Piras Giovanna, Fancello Tatiana, Asproni Rosanna, Fiamma Maura, Monne Maria Itria, Toja Alessandro, Sanna Filomena, Floris Anna Rita, Sulis Vincenzo, Palmas Angelo Domenico, Casu Gavino, Lo Maglio Iana, Mameli Giuseppe |
| EPI_ISL_623108, EPI_ISL_623110, EPI_ISL_623112, EPI_ISL_623114, EPI_ISL_623116, EPI_ISL_623118, EPI_ISL_623119, EPI_ISL_623120, EPI_ISL_623124, EPI_ISL_623126, EPI_ISL_623129, EPI_ISL_623130, EPI_ISL_623132, EPI_ISL_623134, EPI_ISL_623136, EPI_ISL_623138, EPI_ISL_623140, EPI_ISL_623142, EPI_ISL_623143, EPI_ISL_623144, EPI_ISL_623146, EPI_ISL_623148, EPI_ISL_623149, EPI_ISL_623152, EPI_ISL_623154, EPI_ISL_623157, EPI_ISL_623158, EPI_ISL_623160, EPI_ISL_623161, EPI_ISL_623162, EPI_ISL_623164, EPI_ISL_623165, EPI_ISL_623166, EPI_ISL_623168 | | | |
| see above | Laboratorio de Virologia Molecular / UFRJ | Bioinformatics Laboratory / LNCC | Carolina M Voloch, Ronaldo S Francisco Jr, Luiz G P de Almeida, Otavio J. Brustolini, Cynthia C Cardoso, Alexandra L Gerber, Ana Paula de C Guimarães, Diana Mariani, Covid19-UFRJ Workgroup, Luís Cristóvão Pôrto, Renato S Aguiar, Terezinha M P P Castiñeiras, Orlando C. Ferreira, Amilcar Tanuri, Ana Tereza R de Vasconcelos |
| EPI_ISL_625456 | Virology Unit, Institut Pasteur de Madagascar | Virology Unit, Institut Pasteur de Madagascar | Christian Ranaivoson, Cara Brook, Norosoa Razanajatovo, Vida Ahyong, Tsiry Randriambolamanantsoa, Michelle Tan, Vololoniaina Raharinosy, Helisoa Razafimanjato, Cristina M. Tato, Joseph L. DeRisi, Soa Fy Andriamandimby, Jean-Michel Heraud, Philippe Dussart |
| EPI_ISL_626570, EPI_ISL_626593 | The National Institute of Public Health | State Veterinary Institute Prague | Nagy,A;Jirincova,H;Novakova,L;Trnka,D;Vecerova,J |
| EPI_ISL_631504 | Wyckoff Heights Medical Center | New York City Public Health Laboratory | Jade Wang, et al. |
| EPI_ISL_631510, EPI_ISL_631511 | New York Presbyterian/ Weill Cornell Medical Center | New York City Public Health Laboratory | Jade Wang, et al. |
| EPI_ISL_631513 | Montefiore Medical Center | New York City Public Health Laboratory | Jade Wang, et al. |
| EPI_ISL_631538 | Maimonides Medical Center | New York City Public Health Laboratory | Jade Wang, et al. |

| | | | |
|---|---|---|---|
| EPI_ISL_631542 | New York Presbyterian Lower Manhattan | New York City Public Health Laboratory | Jade Wang, et al. |
| EPI_ISL_631546 | Maimonides Medical Center | New York City Public Health Laboratory | Jade Wang, et al. |
| EPI_ISL_631554 | Jamaica Hospital Medical Center | New York City Public Health Laboratory | Jade Wang, et al. |
| EPI_ISL_631556 | NYC HH Jacobi Medical Center | New York City Public Health Laboratory | Jade Wang, et al. |
| EPI_ISL_631558 | NYU Langone Health | New York City Public Health Laboratory | Jade Wang, et al. |
| EPI_ISL_631561 | Montefiore Medical Center | New York City Public Health Laboratory | Jade Wang, et al. |
| EPI_ISL_631562 | Brookdale University Hospital Medical Center | New York City Public Health Laboratory | Jade Wang, et al. |
| EPI_ISL_631568 | New York Presbyterian Queens | New York City Public Health Laboratory | Jade Wang, et al. |
| EPI_ISL_631570 | OCME Office Of Chief Medical Examiner | New York City Public Health Laboratory | Jade Wang, et al. |
| EPI_ISL_631573 | Brookdale University Hospital Medical Center | New York City Public Health Laboratory | Jade Wang, et al. |
| EPI_ISL_631574 | Jamaica Hospital Medical Center | New York City Public Health Laboratory | Jade Wang, et al. |
| EPI_ISL_631577, EPI_ISL_631584, EPI_ISL_631587 | NYC HH Lincoln Medical And Mental Health Center | New York City Public Health Laboratory | Jade Wang, et al. |
| EPI_ISL_631591 | OCME Office Of Chief Medical Examiner | New York City Public Health Laboratory | Jade Wang, et al. |
| EPI_ISL_631593 | Jamaica Hospital Medical Center | New York City Public Health Laboratory | Jade Wang, et al. |
| EPI_ISL_631594 | OCME Office Of Chief Medical Examiner | New York City Public Health Laboratory | Jade Wang, et al. |
| EPI_ISL_631595 | Jamaica Hospital Medical Center | New York City Public Health Laboratory | Jade Wang, et al. |
| EPI_ISL_631596 | James J. Peters VA Medical Center | New York City Public Health Laboratory | Jade Wang, et al. |
| EPI_ISL_631597 | Jamaica Hospital Medical Center | New York City Public Health Laboratory | Jade Wang, et al. |
| EPI_ISL_631598 | OCME Office Of Chief Medical Examiner | New York City Public Health Laboratory | Jade Wang, et al. |
| EPI_ISL_631602 | Flushing Hospital Medical Center | New York City Public Health Laboratory | Jade Wang, et al. |
| EPI_ISL_631603 | NYC HH Lincoln Medical And Mental Health Center | New York City Public Health Laboratory | Jade Wang, et al. |
| EPI_ISL_631605 | OCME Office Of Chief Medical Examiner | New York City Public Health Laboratory | Jade Wang, et al. |
| EPI_ISL_631606 | Flushing Hospital Medical Center | New York City Public Health Laboratory | Jade Wang, et al. |
| EPI_ISL_631607 | NYC HH Lincoln Medical And Mental Health Center | New York City Public Health Laboratory | Jade Wang, et al. |
| EPI_ISL_631610, EPI_ISL_631613, EPI_ISL_631615 | Jamaica Hospital Medical Center | New York City Public Health Laboratory | Jade Wang, et al. |
| EPI_ISL_631616 | Flushing Hospital Medical Center | New York City Public Health Laboratory | Jade Wang, et al. |
| EPI_ISL_631617, EPI_ISL_631618, EPI_ISL_631619 | OCME Office Of Chief Medical Examiner | New York City Public Health Laboratory | Jade Wang, et al. |
| EPI_ISL_631624, EPI_ISL_631625, EPI_ISL_631628, EPI_ISL_631629, EPI_ISL_631633 | Richmond University Medical Center | New York City Public Health Laboratory | Jade Wang, et al. |
| EPI_ISL_631637 | Jamaica Hospital Medical Center | New York City Public Health Laboratory | Jade Wang, et al. |
| EPI_ISL_631686 | NYC HH Lincoln Medical And Mental Health Center | New York City Public Health Laboratory | Jade Wang, et al. |
| EPI_ISL_631687 | OCME Office Of Chief Medical Examiner | New York City Public Health Laboratory | Jade Wang, et al. |
| EPI_ISL_631689, EPI_ISL_631690 | Lincoln Medical Center | New York City Public Health Laboratory | Jade Wang, et al. |
| EPI_ISL_631694, EPI_ISL_631695 | OCME Office Of Chief Medical Examiner | New York City Public Health Laboratory | Jade Wang, et al. |
| EPI_ISL_631697, EPI_ISL_631698 | Jamaica Hospital Medical Center | New York City Public Health Laboratory | Jade Wang, et al. |
| EPI_ISL_631701 | Flushing Hospital Medical Center | New York City Public Health Laboratory | Jade Wang, et al. |
| EPI_ISL_631703, EPI_ISL_631704, EPI_ISL_631705, EPI_ISL_631706 | Jamaica Hospital Medical Center | New York City Public Health Laboratory | Jade Wang, et al. |
| EPI_ISL_631710 | Flushing Hospital Medical Center | New York City Public Health Laboratory | Jade Wang, et al. |
| EPI_ISL_631711, EPI_ISL_631712, EPI_ISL_631716, EPI_ISL_631719, EPI_ISL_631720 | Jamaica Hospital Medical Center | New York City Public Health Laboratory | Jade Wang, et al. |
| EPI_ISL_631724 | Flushing Hospital Medical Center | New York City Public Health Laboratory | Jade Wang, et al. |
| EPI_ISL_631726, EPI_ISL_631728 | Jamaica Hospital Medical Center | New York City Public Health Laboratory | Jade Wang, et al. |
| EPI_ISL_631732 | Flushing Hospital Medical Center | New York City Public Health Laboratory | Jade Wang, et al. |
| EPI_ISL_631733, EPI_ISL_631735, EPI_ISL_631736, EPI_ISL_631740 | Jamaica Hospital Medical Center | New York City Public Health Laboratory | Jade Wang, et al. |
| EPI_ISL_631754 | NYC HH Lincoln Medical And Mental Health Center | New York City Public Health Laboratory | Jade Wang, et al. |
| EPI_ISL_631755, EPI_ISL_631757 | Richmond University Medical Center | New York City Public Health Laboratory | Jade Wang, et al. |
| EPI_ISL_631758, EPI_ISL_631759, EPI_ISL_631760, EPI_ISL_631761, EPI_ISL_631762 | OCME Office Of Chief Medical Examiner | New York City Public Health Laboratory | Jade Wang, et al. |
| EPI_ISL_631767, EPI_ISL_631768 | Jamaica Hospital Medical Center | New York City Public Health Laboratory | Jade Wang, et al. |
| EPI_ISL_631769, EPI_ISL_631770 | Flushing Hospital Medical Center | New York City Public Health Laboratory | Jade Wang, et al. |
| EPI_ISL_631772 | Jamaica Hospital Medical Center | New York City Public Health Laboratory | Jade Wang, et al. |
| EPI_ISL_631773, EPI_ISL_631774 | Flushing Hospital Medical Center | New York City Public Health Laboratory | Jade Wang, et al. |
| EPI_ISL_631775, EPI_ISL_631776, EPI_ISL_631777 | Jamaica Hospital Medical Center | New York City Public Health Laboratory | Jade Wang, et al. |

| | | | |
|---|---|---|---|
| EPI_ISL_631778, EPI_ISL_631781, EPI_ISL_631782 | St Barnabas Hospital | New York City Public Health Laboratory | Jade Wang, et al. |
| EPI_ISL_631797 | OCME Office Of Chief Medical Examiner | New York City Public Health Laboratory | Jade Wang, et al. |
| EPI_ISL_631801 | Richmond University Medical Center | New York City Public Health Laboratory | Jade Wang, et al. |
| EPI_ISL_631803, EPI_ISL_631806 | Brookdale University Hospital Medical Center | New York City Public Health Laboratory | Jade Wang, et al. |
| EPI_ISL_631808, EPI_ISL_631811 | Wyckoff Heights Medical Center | New York City Public Health Laboratory | Jade Wang, et al. |
| EPI_ISL_631812, EPI_ISL_631813 | Jamaica Hospital Medical Center | New York City Public Health Laboratory | Jade Wang, et al. |
| EPI_ISL_631815 | Richmond University Medical Center | New York City Public Health Laboratory | Jade Wang, et al. |
| EPI_ISL_631829 | Brookdale University Hospital Medical Center | New York City Public Health Laboratory | Jade Wang, et al. |
| EPI_ISL_631831 | VA NY Harbor Healthcare System | New York City Public Health Laboratory | Jade Wang, et al. |
| EPI_ISL_631835, EPI_ISL_631836 | Jamaica Hospital Medical Center | New York City Public Health Laboratory | Jade Wang, et al. |
| EPI_ISL_631840, EPI_ISL_631841, EPI_ISL_631845 | OCME Office Of Chief Medical Examiner | New York City Public Health Laboratory | Jade Wang, et al. |
| EPI_ISL_631866, EPI_ISL_631867 | Jamaica Hospital Medical Center | New York City Public Health Laboratory | Jade Wang, et al. |
| EPI_ISL_632313 | NU-sjukvården | Clinical Microbiology, Sahlgrenska University Hospital, | Johan Ringlander, Josefin Olausson, Hedvig Engström Jakobsson, Magnus Lindh |
| EPI_ISL_636476, EPI_ISL_636477, EPI_ISL_636485 | CNR Virus des Infections Respiratoires - France SUD | CNR Virus des Infections Respiratoires - France SUD | Antonin Bal, Gregory Destras, Gwendolyne Burfin, Hadrien Règue, Alexandre Gaymard, Maude Bouscambert-Duchamp, Florence Morfin-Sherpa, Martine Valette, Bruno Lina, Laurence Josset |
| EPI_ISL_636834 | Laboratório de Imunofarmacologia - Instituto Oswaldo Cruz | Laboratório de Imunofarmacologia - Instituto Oswaldo Cruz | Souza,T.M., Fintelman-Rodrigues,N., De Paula,A.D., Saraiva,F.B., Ferreira,M.A. and Sacramento,C.Q. |
| EPI_ISL_636964 | Pathogen Genomics Lab King Abdullah University of Science and Technology(KAUST) | Pathogen Genomics Lab King Abdullah University of Science and Technology(KAUST) | Rahul P Salunke, Sharif Hala, Raeece Naeem, Sara Mfarrej, Amit Kumar Subudhi, Amanda Ooi, Luke Esau, Fadwa Alofi, Fathia Ben Rached, Afrah Alsomali, Asim Khogeer, Ahmad Bakur Mahmoud, Anwar Hashem, Naif Almontashiri, Arnab Pain |
| EPI_ISL_636969, EPI_ISL_636970, EPI_ISL_636971 | Etlik Veterinary Control Central Research Institute | Etlik Veterinary Control Central Research Institute | Sabri Hacioglu, Ahu Pakdemirli, Dilek Dulger, Erdem Danyer, Ummu Sena Sari, Cevdet Yarali, Ozcan Yildirim |
| EPI_ISL_637108, EPI_ISL_637109 | Laboratorio Biologia Molecolare Sars Cov2 - UOC Laboratorio Analisi - Servizio Medicina di Laboratorio, Ospedale "San Francesco" - ATS-ASSL Nuoro | Laboratorio specialistico UOC Ematologia - Ospedale "San Francesco" - ATS-ASSL Nuoro | Piras Giovanna, Fancello Tatiana, Asproni Rosanna, Fiamma Maura, Monne Maria Itria, Toja Alessandro, Sanna Filomena, Floris Anna Rita, Sulis Vincenzo, Palmas Angelo Domenico, Casu Gavino, Lo Maglio Iana, Mameli Giuseppe |
| EPI_ISL_648043, EPI_ISL_648044, EPI_ISL_648045, EPI_ISL_648046, EPI_ISL_648047, EPI_ISL_648048, EPI_ISL_648049, EPI_ISL_648050, EPI_ISL_648051, EPI_ISL_648052, EPI_ISL_648053, EPI_ISL_648054, EPI_ISL_648055, EPI_ISL_648056, EPI_ISL_648057, EPI_ISL_648058, EPI_ISL_648059, EPI_ISL_648060, EPI_ISL_648061, EPI_ISL_648062, EPI_ISL_648063, EPI_ISL_648064, EPI_ISL_648065, EPI_ISL_648066, EPI_ISL_648067, EPI_ISL_648068, EPI_ISL_648069, EPI_ISL_648070, EPI_ISL_648071, EPI_ISL_648072, EPI_ISL_648073, EPI_ISL_648074, EPI_ISL_648075, EPI_ISL_648076, EPI_ISL_648077, EPI_ISL_648078, EPI_ISL_648079, EPI_ISL_648080, EPI_ISL_648081, EPI_ISL_648082, EPI_ISL_648083, EPI_ISL_648084, EPI_ISL_648085, EPI_ISL_648086, EPI_ISL_648087, EPI_ISL_648088, EPI_ISL_648089, EPI_ISL_648090, EPI_ISL_648091, EPI_ISL_648092, EPI_ISL_648093, EPI_ISL_648094, EPI_ISL_648095, EPI_ISL_648096, EPI_ISL_648097, EPI_ISL_648098, EPI_ISL_648099, EPI_ISL_648100, EPI_ISL_648101, EPI_ISL_648102, EPI_ISL_648103, EPI_ISL_648104, EPI_ISL_648105, EPI_ISL_648106, EPI_ISL_648107, EPI_ISL_648108, EPI_ISL_648109, EPI_ISL_648110, EPI_ISL_648111, EPI_ISL_648112, EPI_ISL_648113, EPI_ISL_648114, EPI_ISL_648115, EPI_ISL_648116, EPI_ISL_648117, EPI_ISL_648118, EPI_ISL_648119, EPI_ISL_648120, EPI_ISL_648121 | | | |
| see above | Department of Laboratory Medicine, Tan Tock Seng Hospital | Department of Laboratory Medicine, Tan Tock Seng Hospital | Chen YYC, Zair X, Lim JX, Li C, Tang WY, Maurer-Stroh S, Barkham TMS, Nagarajan N, Sessions OM |
| EPI_ISL_648265, EPI_ISL_648267, EPI_ISL_648270 | Brigham and Women's Hospital | Jonathan Li laboratory | Jonathan Z. Li, Manish C. Choudhary, Upasana D. Adhikari, George Eng, Douglas S Kwon |
| EPI_ISL_648610, EPI_ISL_648611, EPI_ISL_648612, EPI_ISL_648613, EPI_ISL_648614, EPI_ISL_648615, EPI_ISL_648616, EPI_ISL_648617, EPI_ISL_648618, EPI_ISL_648619, EPI_ISL_648620, EPI_ISL_648621, EPI_ISL_648622, EPI_ISL_648623, EPI_ISL_648624, EPI_ISL_648625, EPI_ISL_648626, EPI_ISL_648627, EPI_ISL_648628, EPI_ISL_648629, EPI_ISL_648630, EPI_ISL_648631, EPI_ISL_648632, EPI_ISL_648633, EPI_ISL_648634, EPI_ISL_648635, EPI_ISL_648636, EPI_ISL_648637, EPI_ISL_648638, EPI_ISL_648639, EPI_ISL_648640, EPI_ISL_648641, EPI_ISL_648642, EPI_ISL_648643, EPI_ISL_648644, EPI_ISL_648645, EPI_ISL_648646, EPI_ISL_648647, EPI_ISL_648648, EPI_ISL_648649, EPI_ISL_648650, EPI_ISL_648651, EPI_ISL_648652, EPI_ISL_648653, EPI_ISL_648654, EPI_ISL_648655, EPI_ISL_648656, EPI_ISL_648657, EPI_ISL_648658, EPI_ISL_648659, EPI_ISL_648660, EPI_ISL_648661, EPI_ISL_648662, EPI_ISL_648663, EPI_ISL_648664, EPI_ISL_648665, EPI_ISL_648666, EPI_ISL_648667, EPI_ISL_648668, EPI_ISL_648669, EPI_ISL_648670, EPI_ISL_648671, EPI_ISL_648672, EPI_ISL_648673, EPI_ISL_648674, EPI_ISL_648675, EPI_ISL_648676, EPI_ISL_648742, EPI_ISL_648743, EPI_ISL_648744, EPI_ISL_648745, EPI_ISL_648746, EPI_ISL_648747, EPI_ISL_648748, EPI_ISL_648749, EPI_ISL_648750, EPI_ISL_648751, EPI_ISL_648752, EPI_ISL_648753, EPI_ISL_648754, EPI_ISL_648755, EPI_ISL_648756, EPI_ISL_648757, EPI_ISL_648758, EPI_ISL_648759, EPI_ISL_648760, EPI_ISL_648761, EPI_ISL_648762, EPI_ISL_648763, EPI_ISL_648764, EPI_ISL_648765, EPI_ISL_648766, EPI_ISL_648767, EPI_ISL_648768, EPI_ISL_648769, EPI_ISL_648770, EPI_ISL_648771, EPI_ISL_648772, EPI_ISL_648773, EPI_ISL_648774, EPI_ISL_648775, EPI_ISL_648776, EPI_ISL_648777, EPI_ISL_648778, EPI_ISL_648779, EPI_ISL_648780, EPI_ISL_648781, EPI_ISL_648782, EPI_ISL_648783, EPI_ISL_648784, EPI_ISL_648785, EPI_ISL_648786, EPI_ISL_648787, EPI_ISL_648788, EPI_ISL_648789, EPI_ISL_648790, EPI_ISL_648791 | | | |
| see above | Department of Laboratory Medicine, Tan Tock Seng Hospital | Department of Laboratory Medicine, Tan Tock Seng Hospital | Chen YYC, Zair X, Lim JX, Li C, Tang WY, Maurer-Stroh S, Barkham TMS, Nagarajan N, Sessions OM |
| EPI_ISL_649189 | Istituto Zooprofilattico Sperimentale della Puglia e della Basilicata | Istituto Zooprofilattico Sperimentale della Puglia e della Basilicata | Parisi A., Bianco A., Capozzi L., Del Sambro L., Manzulli V, Rondinone V., Pace L., Galante D., Cipolletta D. |
| EPI_ISL_653813 | Istituto Zooprofilattico Sperimentale della Puglia e della Basilicata | Istituto Zooprofilattico Sperimentale della Puglia e della Basilicata | Parisi A., Bianco A., Capozzi L., Del Sambro L., Manzulli V, Rondinone V., Pace L., Cipolletta D., Galante D. |
| EPI_ISL_653814, EPI_ISL_653815, EPI_ISL_653816, EPI_ISL_653817, EPI_ISL_653818 | I.R.C.C.S. "S. De Bellis" - Ente Ospedaliero | Istituto Zooprofilattico Sperimentale della Puglia e della Basilicata | Parisi A., Bianco A., Capozzi L., Del Sambro L., Lippolis A., Notarnicola M., Cipolletta D., Galante D. |
| EPI_ISL_654016 | Laboratory of Microbiology, National Reference Lab, Charles Nicolle Hospital; 2-University of Tunis ElManar, Faculty of Medicine of Tunis, LR99ES09, Tunis, Tunisia | 1-Clinical and Experimental Pharmacology Lab, LR16SP02, National Center of Pharmacovigilance, University of Tunis El Manar, Tunis, Tunisia. 2-Neurodegenerative diseases and psychiatric troubles, LR18SP03, Razi Hospital, University of Tunis El Manar, Tunis, Tunisia. 3- Ministry of Health, National Observatory of New and Emerging Diseases, 1006, Tunis, Tunisia | Ilhem Boutiba-Ben Boubaker, Sameh Trabelsi, Nissaf Ben Alaya, Maher Kharrat, Alia Ben Kahla, Jalila Ben Khelil, Salma Abid, Sana Ferjani, Mouna Ben Sassi, Mouna Safer, Imen Mkada, Imen Kacem, Gaies Emna, Soumaya Rammeh, Riadh Daghfous, Riadh Gouider. |
| EPI_ISL_654867 | Pasteur Institute in Ho Chi Minh city | Department of Microbiology and Immunology - Pasteur Institute in Ho Chi Minh city | Hoàng Minh, Cao Minh Thng, Phm Th Thu Hng, V Phm Hng Nhung, ào Huy Mnh, Hunh Th Kim Loan, Nguyn Hoàng Quân, Hunh Phng Tho, Hoàng Nh ào, Nguyn Trung Hiu, Nguyn Hoàng Anh, Nguyn Thu Ngc, Phm Th Nhung, La Hoàng Huy, ng Thanh Giang, Nguyn Th Thanh Thng, Trn Th Hng Kim, Nguyn Thanh Long, Phm Duy Quang, Lng Chn Quang, Hoàng Quc Cng, Phan Trng Lân, Nguyn V Thng |
| EPI_ISL_654868 | Pasteur Institute in Ho Chi Minh city | Department of Microbiology and Immunology - Pasteur Institute in Ho Chi Minh city | ào Huy Mnh, Phm Th Thu Hng, V Phm Hng Nhung, Cao Minh Thng, Hunh Th Kim Loan, Nguyn Hoàng Quân, Hunh Phng Tho, Hoàng Nh ào, Nguyn Trung Hiu, Nguyn Hoàng Anh, Nguyn Thu Ngc, Phm Th Nhung, Nguyn Th Ngc Tho, Hà Bích Ngc, Nguyn Th Thanh Thng, Hoàng Minh, Trn Th Hng Kim, Nguyn Thanh Long, Phm Duy Quang, Lng Chn Quang, Hoàng Quc Cng, Nguyn V Thng, Phan Trng Lân |
| EPI_ISL_654869 | Pasteur Institute in Ho Chi Minh city | Department of Microbiology and Immunology - Pasteur Institute in Ho Chi Minh city | Phm Th Nhung, ào Huy Mnh, Phm Th Thu Hng, V Phm Hng Nhung, Cao Minh Thng, Hunh Th Kim Loan, Nguyn Hoàng Quân, Hunh Phng Tho, Hoàng Nh ào, Nguyn Trung Hiu, Nguyn Hoàng Anh, Nguyn Thu Ngc, Nguyn Th Ngc Tho, ng Thanh Giang, Võ Th Trang ài, Hoàng Minh, Trn Th Hng Kim, Nguyn Thanh Long, Phm Duy Quang, Lng Chn Quang, Hoàng Quc Cng, Phan Trng Lân, Nguyn V Thng |
| EPI_ISL_654870 | Pasteur Institute in Ho Chi Minh city | Department of Microbiology and Immunology - Pasteur Institute in Ho Chi Minh city | Trn Th Hng Kim, Phm Th Thu Hng, ào Huy Mnh, V Phm Hng Nhung, Cao Minh Thng, Hunh Th Kim Loan, Nguyn Hoàng Quân, Hunh Phng Tho, Hoàng Nh ào, Nguyn Trung Hiu, Nguyn Hoàng Anh, Nguyn Thu Ngc, Phm Th Nhung, Nguyn Th Ngc Tho, ng Thanh Giang, Nguyn Th Thanh Thng, Hoàng Kim Loan, Nguyn Thanh Long, Phm Duy Quang, Lng Chn Quang, Hoàng Quc Cng, Nguyn V Thng, Phan Trng Lân |
| EPI_ISL_654872 | Pasteur Institute in Ho Chi Minh city | Department of Microbiology and Immunology - Pasteur Institute in Ho Chi Minh city | Phm Th Thu Hng, ào Huy Mnh, V Phm Hng Nhung, Cao Minh Thng, Hunh Th Kim Loan, Nguyn Hoàng Quân, Hunh Phng Tho, Hoàng Nh ào, Nguyn Trung Hiu, Nguyn Hoàng Anh, Nguyn Thu Ngc, Phm Th Nhung, Nguyn Th Ngc Tho, ng Thanh Giang, Nguyn Th Thanh Thng, Hoàng Minh, Nguyn Hoàng V, Nguyn Thanh Long, Phm Duy Quang, Lng Chn Quang, Hoàng Quc Cng, Phan Trng Lân, Nguyn V Thng |

| | | | |
|---|---|---|---|
| EPI_ISL_654873 | Pasteur Institute in Ho Chi Minh city | Department of Microbiology and Immunology - Pasteur Institute in Ho Chi Minh city | Phm Th Thu Hng, ào Huy Mnh, V Phm Hng Nhung, Cao Minh Thng, Hunh Th Kim Loan, Nguyn Hoàng Quân, Hunh Phng Tho, Hoàng Nh ào, Nguyn Trung Hiu, Nguyn Th Thanh Tho, Nguyn Thu Ngc, Phm Th Nhung, Nguyn Th Ngc Tho, ng Thanh Giang, Nguyn Th Thanh Thng, Hoàng Minh, Trn Th Hng Kim, Nguyn Thanh Long, Phm Duy Quang, Lng Chn Quang, Hoàng Quc Cng, Nguyn V Thng, Phan Trng Lân |
| EPI_ISL_654874 | Pasteur Institute in Ho Chi Minh city | Department of Microbiology and Immunology - Pasteur Institute in Ho Chi Minh city | Nguyn Th Ngc Tho, V Phm Hng Nhung, ào Huy Mnh, Phm Th Thu Hng, Cao Minh Thng, Hunh Th Kim Loan, Nguyn Hoàng Quân, Hunh Phng Tho, Hoàng Nh ào, Trn Tôn, Nguyn Hoàng Anh, Nguyn Thu Ngc, Phm Th Nhung, ng Thanh Giang, Nguyn Th Thanh Thng, Hoàng Minh, Trn Th Hng Kim, Nguyn Thanh Long, Phm Duy Quang, Lng Chn Quang, Hoàng Quc Cng, Phan Trng Lân, Nguyn V Thng |
| EPI_ISL_654875 | Pasteur Institute in Ho Chi Minh city | Department of Microbiology and Immunology - Pasteur Institute in Ho Chi Minh city | V Phm Hng Nhung, ào Huy Mnh, Phm Th Thu Hng, Cao Minh Thng, Hunh Th Kim Loan, Nguyn Hoàng Quân, Hunh Phng Tho, Hoàng Nh ào, Nguyn Trung Hiu, Nguyn Hoàng Anh, Nguyn Vit Thinh, Phm Th Nhung, Nguyn Th Ngc Tho, ng Thanh Giang, Nguyn Th Thanh Thng, Hoàng Minh, Trn Th Hng Kim, Nguyn Thanh Long, Phm Duy Quang, Lng Chn Quang, Hoàng Quc Cng, Nguyn V Thng, Phan Trng Lân |
| EPI_ISL_654876 | Pasteur Institute in Ho Chi Minh city | Department of Microbiology and Immunology - Pasteur Institute in Ho Chi Minh city | V Phm Hng Nhung, ào Huy Mnh, Phm Th Thu Hng, Cao Minh Thng, Hunh Th Kim Loan, Nguyn Hoàng Quân, Hunh Phng Tho, Hoàng Nh ào, Nguyn Trung Hiu, Nguyn Hoàng Anh, Nguyn Thu Ngc, Trng Công Hiu, Nguyn Th Ngc Tho, ng Thanh Giang, Nguyn Th Thanh Thng, Hoàng Minh, Trn Th Hng Kim, Nguyn Thanh Long, Phm Duy Quang, Lng Chn Quang, Hoàng Quc Cng, Phan Trng Lân, Nguyn V Thng |
| EPI_ISL_654877 | Pasteur Institute in Ho Chi Minh city | Department of Microbiology and Immunology - Pasteur Institute in Ho Chi Minh city | ng Thanh Giang, Phm Duy Quang, ào Huy Mnh, Phm Th Thu Hng, V Phm Hng Nhung, Cao Minh Thng, Hunh Th Kim Loan, Nguyn Hoàng Quân, Hunh Phng Tho, Hoàng Nh ào, Nguyn Trung Hiu, Nguyn Hoàng Anh, Nguyn Thu Ngc, Phm Th Nhung, Trng Thùy Dung, Nguyn Th Thanh Thng, Hoàng Minh, Trn Th Hng Kim, Nguyn Thanh Long, Lng Chn Quang, Hoàng Quc Cng, Nguyn V Thng, Phan Trng Lân |
| EPI_ISL_654878 | Pasteur Institute in Ho Chi Minh city | Department of Microbiology and Immunology - Pasteur Institute in Ho Chi Minh city | Phm Duy Quang, ào Huy Mnh, Phm Th Thu Hng, V Phm Hng Nhung, Cao Minh Thng, Hunh Th Kim Loan, Nguyn Hoàng Quân, Hunh Phng Tho, Hoàng Nh ào, Nguyn Trung Hiu, Nguyn Hoàng Anh, Nguyn Thu Ngc, Phm Th Nhung, Nguyn Th Ngc Tho, Viên ng Khánh Linh, Nguyn Th Thanh Thng, Hoàng Minh, Trn Th Hng Kim, Nguyn Thanh Long, Lng Chn Quang, Hoàng Quc Cng, Phan Trng Lân, Nguyn V Thng |
| EPI_ISL_654881 | Pasteur Institute in Ho Chi Minh city | Department of Microbiology and Immunology - Pasteur Institute in Ho Chi Minh city | Lng Chn Quang, ào Huy Mnh, Phm Th Thu Hng, V Phm Hng Nhung, Cao Minh Thng, Hunh Th Kim Loan, Nguyn Hoàng Quân, Hunh Phng Tho, Hoàng Nh ào, Nguyn Trung Hiu, Nguyn Hoàng Anh, Nguyn Thu Ngc, Phm Th Nhung, Nguyn Th Ngc Tho, ng Thanh Giang, Nguyn Th Thanh Thng, Hoàng Minh, Nguyn Th Phng Thúy, Nguyn Thanh Long, Phm Duy Quang, Hoàng Quc Cng, Nguyn V Thng, Phan Trng Lân |
| EPI_ISL_654882 | Pasteur Institute in Ho Chi Minh city | Department of Microbiology and Immunology - Pasteur Institute in Ho Chi Minh city | Lng Chn Quang, ào Huy Mnh, Phm Th Thu Hng, V Phm Hng Nhung, Cao Minh Thng, Hunh Th Kim Loan, Nguyn Hoàng Quân, Hunh Phng Tho, Hoàng Nh ào, Nguyn Trung Hiu, Nguyn Hoàng Anh, Nguyn Ngc Anh Tun, Phm Th Nhung, Nguyn Th Ngc Tho, ng Thanh Giang, Nguyn Th Thanh Thng, Hoàng Minh, Trn Th Hng Kim, Nguyn Thanh Long, Phm Duy Quang, Hoàng Quc Cng, Phan Trng Lân, Nguyn V Thng |
| EPI_ISL_658904 | Instituto de Diagnostico y Referencia Epidemiologicos (INDRE) | Instituto de Diagnostico y Referencia Epidemiologicos (INDRE) | Ernesto Ramirez-Gonzalez, Abril Rodriguez-Maldonado, Claudia Wong-Arambula , Natividad Cruz-Ortiz, Tatiana Nunez-Garcia, Dayanira Arellano-Suarez, Fabiola Garces-Ayala, Lucia Hernandez-Rivas, Irma Lopez-Martinez, Gisela Barrera-Badillo, Marisol Galindo-Galindo, Linda Andarade-Sanchez, Octavio Ruiz-Muñiz, Pamela Ramirez-Medina, Roman Canul-Aguilar, Gaudalupe Herrera-Ramirez, Jazmin Galicia-Hernandez. |
| EPI_ISL_666870 | Laboratoire de Virologie, CHU de Caen, Normandie, France. | GRAM2.0, Université de Caen Normandie | Meriadeg LE GOUIL, Januel VOUTEAU, Juliette ECHE, Estelle LEPERCHOIS, Astrid VABRET |
| EPI_ISL_677673 | Wadsworth Center, New York State Department.of Health | Wadsworth Center, New York State Department.of Health | Kirsten St. George, Daryl M. Lamson, Alexis Russel, Jonathan Plitnick, Navjot Singh, John Kelly, Sara Griesemer, Erasmus Schneider, Erica Lasek-Nesselquist |
| EPI_ISL_677678, EPI_ISL_677679, EPI_ISL_677680, EPI_ISL_677681, EPI_ISL_677682, EPI_ISL_677683, EPI_ISL_677684, EPI_ISL_677685, EPI_ISL_677686, EPI_ISL_677687, EPI_ISL_677688, EPI_ISL_677689 | | | |
| see above | Vanda Pharmaceuticals Clinical Site | Vanda Pharmaceuticals | Vanda Pharmaceuticals |
| EPI_ISL_681702, EPI_ISL_681703 | Zurita & Zurita Laboratorios | Zurita & Zurita Laboratorios | Gabriela Sevillano, Camilo Zurita-Salinas, Jeannete Zurita |
| EPI_ISL_684110, EPI_ISL_684111, EPI_ISL_684113, EPI_ISL_684114, EPI_ISL_684115, EPI_ISL_684116, EPI_ISL_684118, EPI_ISL_684119, EPI_ISL_684120, EPI_ISL_684121, EPI_ISL_684122, EPI_ISL_684123, EPI_ISL_684124, EPI_ISL_684125, EPI_ISL_684126, EPI_ISL_684128, EPI_ISL_684129, EPI_ISL_684130, EPI_ISL_684132, EPI_ISL_684133 | | | |
| see above | Pathogen Genomics Center, National Institute of Infectious Diseases | Pathogen Genomics Center, National Institute of Infectious Diseases | Tsuyoshi Sekizuka, Kentaro Itokawa, Rina Tanaka, Masanori Hashino, Makoto Kuroda |
| EPI_ISL_684136 | Tokyo Metropolitan Institute of Public Health | Pathogen Genomics Center, National Institute of Infectious Diseases | Tsuyoshi Sekizuka, Kentaro Itokawa, Rina Tanaka, Masanori Hashino, Makoto Kuroda |
| EPI_ISL_684139, EPI_ISL_684142, EPI_ISL_684143, EPI_ISL_684145, EPI_ISL_684146 | Pathogen Genomics Center, National Institute of Infectious Diseases | Pathogen Genomics Center, National Institute of Infectious Diseases | Tsuyoshi Sekizuka, Kentaro Itokawa, Rina Tanaka, Masanori Hashino, Makoto Kuroda |
| EPI_ISL_684149 | Saitama Prefectural Institute of Public Health | Pathogen Genomics Center, National Institute of Infectious Diseases | Tsuyoshi Sekizuka, Kentaro Itokawa, Rina Tanaka, Masanori Hashino, Makoto Kuroda |
| EPI_ISL_684150, EPI_ISL_684151, EPI_ISL_684152, EPI_ISL_684153 | Pathogen Genomics Center, National Institute of Infectious Diseases | Pathogen Genomics Center, National Institute of Infectious Diseases | Tsuyoshi Sekizuka, Kentaro Itokawa, Rina Tanaka, Masanori Hashino, Makoto Kuroda |
| EPI_ISL_684154, EPI_ISL_684155 | Tokyo Metropolitan Institute of Public Health | Pathogen Genomics Center, National Institute of Infectious Diseases | Tsuyoshi Sekizuka, Kentaro Itokawa, Rina Tanaka, Masanori Hashino, Makoto Kuroda |
| EPI_ISL_684156, EPI_ISL_684157, EPI_ISL_684158 | Pathogen Genomics Center, National Institute of Infectious Diseases | Pathogen Genomics Center, National Institute of Infectious Diseases | Tsuyoshi Sekizuka, Kentaro Itokawa, Rina Tanaka, Masanori Hashino, Makoto Kuroda |
| EPI_ISL_684159, EPI_ISL_684160, EPI_ISL_684161 | Chiba Prefectural Institute of Public Health | Pathogen Genomics Center, National Institute of Infectious Diseases | Tsuyoshi Sekizuka, Kentaro Itokawa, Rina Tanaka, Masanori Hashino, Makoto Kuroda |
| EPI_ISL_684162, EPI_ISL_684163, EPI_ISL_684164, EPI_ISL_684166, EPI_ISL_684167, EPI_ISL_684168, EPI_ISL_684170, EPI_ISL_684171, EPI_ISL_684172, EPI_ISL_684173, EPI_ISL_684174, EPI_ISL_684175, EPI_ISL_684176, EPI_ISL_684177, EPI_ISL_684178, EPI_ISL_684179, EPI_ISL_684180, EPI_ISL_684181, EPI_ISL_684182, EPI_ISL_684183, EPI_ISL_684184, EPI_ISL_684185, EPI_ISL_684186, EPI_ISL_684187, EPI_ISL_684188, EPI_ISL_684189, EPI_ISL_684190, EPI_ISL_684191, EPI_ISL_684192, EPI_ISL_684193, EPI_ISL_684194, EPI_ISL_684195, EPI_ISL_684196, EPI_ISL_684197, EPI_ISL_684198, EPI_ISL_684199, EPI_ISL_684200, EPI_ISL_684201, EPI_ISL_684202, EPI_ISL_684203, EPI_ISL_684204, EPI_ISL_684205, EPI_ISL_684206, EPI_ISL_684207, EPI_ISL_684208, EPI_ISL_684209, EPI_ISL_684210, EPI_ISL_684211, EPI_ISL_684212, EPI_ISL_684213, EPI_ISL_684214, EPI_ISL_684215, EPI_ISL_684216, EPI_ISL_684217, EPI_ISL_684218, EPI_ISL_684219 | | | |
| see above | Pathogen Genomics Center, National Institute of Infectious Diseases | Pathogen Genomics Center, National Institute of Infectious Diseases | Tsuyoshi Sekizuka, Kentaro Itokawa, Rina Tanaka, Masanori Hashino, Makoto Kuroda |
| EPI_ISL_684220, EPI_ISL_684221, EPI_ISL_684222, EPI_ISL_684223, EPI_ISL_684224, EPI_ISL_684225, EPI_ISL_684226, EPI_ISL_684227, EPI_ISL_684228, EPI_ISL_684229, EPI_ISL_684230, EPI_ISL_684231, EPI_ISL_684232, EPI_ISL_684233, EPI_ISL_684234, EPI_ISL_684235, EPI_ISL_684236, EPI_ISL_684237, EPI_ISL_684238, EPI_ISL_684239, EPI_ISL_684240, EPI_ISL_684241, EPI_ISL_684242, EPI_ISL_684243, EPI_ISL_684244, EPI_ISL_684245, EPI_ISL_684246, EPI_ISL_684247, EPI_ISL_684248, EPI_ISL_684249, EPI_ISL_684250, EPI_ISL_684251, EPI_ISL_684252, EPI_ISL_684253, EPI_ISL_684254, EPI_ISL_684255, EPI_ISL_684256, EPI_ISL_684257, EPI_ISL_684258, EPI_ISL_684259, EPI_ISL_684260, EPI_ISL_684261, EPI_ISL_684262, EPI_ISL_684263, EPI_ISL_684264, EPI_ISL_684265, EPI_ISL_684266, EPI_ISL_684267, EPI_ISL_684268, EPI_ISL_684269, EPI_ISL_684270, EPI_ISL_684271, EPI_ISL_684272, EPI_ISL_684273, EPI_ISL_684274, EPI_ISL_684275, EPI_ISL_684276, EPI_ISL_684277, EPI_ISL_684278, EPI_ISL_684279, EPI_ISL_684280, EPI_ISL_684281, EPI_ISL_684282, EPI_ISL_684283, EPI_ISL_684284, EPI_ISL_684285, EPI_ISL_684286, EPI_ISL_684287, EPI_ISL_684288 | | | |
| see above | Gunma Prefectural Institute of Public Health and Environmental Sciences | Pathogen Genomics Center, National Institute of Infectious Diseases | Tsuyoshi Sekizuka, Kentaro Itokawa, Rina Tanaka, Masanori Hashino, Makoto Kuroda |
| EPI_ISL_684289, EPI_ISL_684290, EPI_ISL_684291, EPI_ISL_684292, EPI_ISL_684293, EPI_ISL_684294, EPI_ISL_684295, EPI_ISL_684296, EPI_ISL_684297, EPI_ISL_684298, EPI_ISL_684299, EPI_ISL_684300, EPI_ISL_684301, EPI_ISL_684302, EPI_ISL_684303, EPI_ISL_684304, EPI_ISL_684305, EPI_ISL_684306, EPI_ISL_684307, EPI_ISL_684308, EPI_ISL_684309, EPI_ISL_684310, EPI_ISL_684311, EPI_ISL_684312, EPI_ISL_684315, EPI_ISL_684316, EPI_ISL_684317, EPI_ISL_684319, EPI_ISL_684320 | | | |
| see above | Pathogen Genomics Center, National Institute of Infectious Diseases | Pathogen Genomics Center, National Institute of Infectious Diseases | Tsuyoshi Sekizuka, Kentaro Itokawa, Rina Tanaka, Masanori Hashino, Makoto Kuroda |
| EPI_ISL_684325, EPI_ISL_684326, EPI_ISL_684327, EPI_ISL_684328, EPI_ISL_684329, EPI_ISL_684330, EPI_ISL_684331, EPI_ISL_684332, EPI_ISL_684333, EPI_ISL_684334, EPI_ISL_684335, EPI_ISL_684336, EPI_ISL_684337, EPI_ISL_684338 | | | |
| see above | Gunma Prefectural Institute of Public Health and Environmental Sciences | Pathogen Genomics Center, National Institute of Infectious Diseases | Tsuyoshi Sekizuka, Kentaro Itokawa, Rina Tanaka, Masanori Hashino, Makoto Kuroda |
| EPI_ISL_684339, EPI_ISL_684340, EPI_ISL_684341, EPI_ISL_684342, EPI_ISL_684343, EPI_ISL_684344, EPI_ISL_684345, EPI_ISL_684346, EPI_ISL_684347, EPI_ISL_684348, EPI_ISL_684349, EPI_ISL_684350, EPI_ISL_684351, EPI_ISL_684352, EPI_ISL_684353, EPI_ISL_684354, EPI_ISL_684355, EPI_ISL_684356, EPI_ISL_684357, EPI_ISL_684358, EPI_ISL_684359, EPI_ISL_684360, EPI_ISL_684361, EPI_ISL_684362, EPI_ISL_684363, EPI_ISL_684365, EPI_ISL_684366, EPI_ISL_684367, EPI_ISL_684368, EPI_ISL_684369, EPI_ISL_684370, EPI_ISL_684371, EPI_ISL_684372, EPI_ISL_684373, EPI_ISL_684374, EPI_ISL_684375, | | | |

| | | | |
|---|---|---|---|
| EPI_ISL_684376, EPI_ISL_684377, EPI_ISL_684378, EPI_ISL_684379, EPI_ISL_684381, EPI_ISL_684382, EPI_ISL_684383, EPI_ISL_684384, EPI_ISL_684385, EPI_ISL_684386, EPI_ISL_684387, EPI_ISL_684388, EPI_ISL_684389, EPI_ISL_684390, EPI_ISL_684391, EPI_ISL_684392, EPI_ISL_684393, EPI_ISL_684394, EPI_ISL_684395, EPI_ISL_684396, EPI_ISL_684397, EPI_ISL_684398, EPI_ISL_684399, EPI_ISL_684400, EPI_ISL_684401, EPI_ISL_684402, EPI_ISL_684403, EPI_ISL_684404, EPI_ISL_684405, EPI_ISL_684406, EPI_ISL_684407 | | | |
| see above | Pathogen Genomics Center, National Institute of Infectious Diseases | Pathogen Genomics Center, National Institute of Infectious Diseases | Tsuyoshi Sekizuka, Kentaro Itokawa, Rina Tanaka, Masanori Hashino, Makoto Kuroda |
| EPI_ISL_684408, EPI_ISL_684409, EPI_ISL_684410, EPI_ISL_684411, EPI_ISL_684412, EPI_ISL_684413, EPI_ISL_684414, EPI_ISL_684415, EPI_ISL_684416, EPI_ISL_684417, EPI_ISL_684418, EPI_ISL_684419, EPI_ISL_684420, EPI_ISL_684421, EPI_ISL_684422, EPI_ISL_684423, EPI_ISL_684424, EPI_ISL_684425, EPI_ISL_684426, EPI_ISL_684427, EPI_ISL_684428, EPI_ISL_684429, EPI_ISL_684430, EPI_ISL_684431, EPI_ISL_684432, EPI_ISL_684433, EPI_ISL_684434, EPI_ISL_684435, EPI_ISL_684436, EPI_ISL_684437, EPI_ISL_684438, EPI_ISL_684439, EPI_ISL_684440, EPI_ISL_684441, EPI_ISL_684442, EPI_ISL_684443, EPI_ISL_684444, EPI_ISL_684445, EPI_ISL_684446, EPI_ISL_684447, EPI_ISL_684448, EPI_ISL_684449, EPI_ISL_684450, EPI_ISL_684451, EPI_ISL_684452, EPI_ISL_684453, EPI_ISL_684454, EPI_ISL_684455, EPI_ISL_684456, EPI_ISL_684457, EPI_ISL_684458, EPI_ISL_684459, EPI_ISL_684460, EPI_ISL_684461, EPI_ISL_684462, EPI_ISL_684463, EPI_ISL_684464, EPI_ISL_684465, EPI_ISL_684466, EPI_ISL_684467, EPI_ISL_684468, EPI_ISL_684469, EPI_ISL_684470, EPI_ISL_684471, EPI_ISL_684472, EPI_ISL_684473, EPI_ISL_684474, EPI_ISL_684475, EPI_ISL_684476, EPI_ISL_684477, EPI_ISL_684478, EPI_ISL_684479, EPI_ISL_684480, EPI_ISL_684481, EPI_ISL_684482, EPI_ISL_684483 | | | |
| see above | Fukuoka Institute of Health and Environmental Sciences | Pathogen Genomics Center, National Institute of Infectious Diseases | Tsuyoshi Sekizuka, Kentaro Itokawa, Rina Tanaka, Masanori Hashino, Makoto Kuroda |
| EPI_ISL_684484, EPI_ISL_684485, EPI_ISL_684486, EPI_ISL_684487, EPI_ISL_684488, EPI_ISL_684489, EPI_ISL_684490, EPI_ISL_684491, EPI_ISL_684492, EPI_ISL_684493, EPI_ISL_684494, EPI_ISL_684495, EPI_ISL_684496, EPI_ISL_684497, EPI_ISL_684498, EPI_ISL_684499, EPI_ISL_684500, EPI_ISL_684501, EPI_ISL_684502, EPI_ISL_684503, EPI_ISL_684504, EPI_ISL_684505 | | | |
| see above | Pathogen Genomics Center, National Institute of Infectious Diseases | Pathogen Genomics Center, National Institute of Infectious Diseases | Tsuyoshi Sekizuka, Kentaro Itokawa, Rina Tanaka, Masanori Hashino, Makoto Kuroda |
| EPI_ISL_684506, EPI_ISL_684507, EPI_ISL_684508, EPI_ISL_684509, EPI_ISL_684510, EPI_ISL_684511, EPI_ISL_684512, EPI_ISL_684513, EPI_ISL_684514, EPI_ISL_684515, EPI_ISL_684516, EPI_ISL_684517, EPI_ISL_684518, EPI_ISL_684519, EPI_ISL_684520, EPI_ISL_684521, EPI_ISL_684522 | | | |
| see above | Tokyo Metropolitan Institute of Public Health | Pathogen Genomics Center, National Institute of Infectious Diseases | Tsuyoshi Sekizuka, Kentaro Itokawa, Rina Tanaka, Masanori Hashino, Makoto Kuroda |
| EPI_ISL_684523, EPI_ISL_684524, EPI_ISL_684525, EPI_ISL_684526, EPI_ISL_684527, EPI_ISL_684529, EPI_ISL_684530, EPI_ISL_684531, EPI_ISL_684532, EPI_ISL_684533, EPI_ISL_684534, EPI_ISL_684535, EPI_ISL_684536, EPI_ISL_684537, EPI_ISL_684538, EPI_ISL_684539, EPI_ISL_684540, EPI_ISL_684541, EPI_ISL_684542, EPI_ISL_684543, EPI_ISL_684544, EPI_ISL_684545, EPI_ISL_684546, EPI_ISL_684547, EPI_ISL_684548, EPI_ISL_684549, EPI_ISL_684550, EPI_ISL_684551, EPI_ISL_684552, EPI_ISL_684553, EPI_ISL_684554, EPI_ISL_684555 | | | |
| see above | Pathogen Genomics Center, National Institute of Infectious Diseases | Pathogen Genomics Center, National Institute of Infectious Diseases | Tsuyoshi Sekizuka, Kentaro Itokawa, Rina Tanaka, Masanori Hashino, Makoto Kuroda |
| EPI_ISL_684556, EPI_ISL_684557 | Tokyo Metropolitan Institute of Public Health | Pathogen Genomics Center, National Institute of Infectious Diseases | Tsuyoshi Sekizuka, Kentaro Itokawa, Rina Tanaka, Masanori Hashino, Makoto Kuroda |
| EPI_ISL_684558, EPI_ISL_684559, EPI_ISL_684560, EPI_ISL_684561, EPI_ISL_684562, EPI_ISL_684563, EPI_ISL_684564, EPI_ISL_684565 | Pathogen Genomics Center, National Institute of Infectious Diseases | Pathogen Genomics Center, National Institute of Infectious Diseases | Tsuyoshi Sekizuka, Kentaro Itokawa, Rina Tanaka, Masanori Hashino, Makoto Kuroda |
| EPI_ISL_684566, EPI_ISL_684567, EPI_ISL_684568, EPI_ISL_684569, EPI_ISL_684570, EPI_ISL_684571, EPI_ISL_684572, EPI_ISL_684573, EPI_ISL_684574, EPI_ISL_684575, EPI_ISL_684576 | | | |
| see above | Gunma Prefectural Institute of Public Health and Environmental Sciences | Pathogen Genomics Center, National Institute of Infectious Diseases | Tsuyoshi Sekizuka, Kentaro Itokawa, Rina Tanaka, Masanori Hashino, Makoto Kuroda |
| EPI_ISL_684578, EPI_ISL_684579, EPI_ISL_684580, EPI_ISL_684581, EPI_ISL_684582, EPI_ISL_684586, EPI_ISL_684587, EPI_ISL_684588, EPI_ISL_684589, EPI_ISL_684590, EPI_ISL_684591, EPI_ISL_684592, EPI_ISL_684593, EPI_ISL_684596, EPI_ISL_684597, EPI_ISL_684598, EPI_ISL_684599, EPI_ISL_684600, EPI_ISL_684601, EPI_ISL_684602, EPI_ISL_684603, EPI_ISL_684604, EPI_ISL_684605, EPI_ISL_684606, EPI_ISL_684607, EPI_ISL_684608, EPI_ISL_684609, EPI_ISL_684610, EPI_ISL_684611, EPI_ISL_684614 | | | |
| see above | Pathogen Genomics Center, National Institute of Infectious Diseases | Pathogen Genomics Center, National Institute of Infectious Diseases | Tsuyoshi Sekizuka, Kentaro Itokawa, Rina Tanaka, Masanori Hashino, Makoto Kuroda |
| EPI_ISL_684615, EPI_ISL_684616, EPI_ISL_684617, EPI_ISL_684619, EPI_ISL_684620, EPI_ISL_684622, EPI_ISL_684623, EPI_ISL_684624, EPI_ISL_684625, EPI_ISL_684626, EPI_ISL_684627, EPI_ISL_684628, EPI_ISL_684629, EPI_ISL_684630, EPI_ISL_684631, EPI_ISL_684632, EPI_ISL_684633 | | | |
| see above | Tokyo Metropolitan Institute of Public Health | Pathogen Genomics Center, National Institute of Infectious Diseases | Tsuyoshi Sekizuka, Kentaro Itokawa, Rina Tanaka, Masanori Hashino, Makoto Kuroda |
| EPI_ISL_684634, EPI_ISL_684635, EPI_ISL_684636, EPI_ISL_684637, EPI_ISL_684638, EPI_ISL_684640, EPI_ISL_684641, EPI_ISL_684642, EPI_ISL_684643, EPI_ISL_684644, EPI_ISL_684645, EPI_ISL_684646, EPI_ISL_684647, EPI_ISL_684648, EPI_ISL_684649, EPI_ISL_684650, EPI_ISL_684651, EPI_ISL_684652, EPI_ISL_684653, EPI_ISL_684654, EPI_ISL_684655, EPI_ISL_684656, EPI_ISL_684657, EPI_ISL_684658, EPI_ISL_684659, EPI_ISL_684660, EPI_ISL_684661, EPI_ISL_684662, EPI_ISL_684663, EPI_ISL_684664, EPI_ISL_684665, EPI_ISL_684666, EPI_ISL_684667, EPI_ISL_684668, EPI_ISL_684669, EPI_ISL_684670, EPI_ISL_684671, EPI_ISL_684672, EPI_ISL_684673, EPI_ISL_684674, EPI_ISL_684675, EPI_ISL_684676, EPI_ISL_684677, EPI_ISL_684678, EPI_ISL_684679, EPI_ISL_684680, EPI_ISL_684681, EPI_ISL_684682, EPI_ISL_684683, EPI_ISL_684684, EPI_ISL_684685, EPI_ISL_684686, EPI_ISL_684687, EPI_ISL_684688, EPI_ISL_684689, EPI_ISL_684690, EPI_ISL_684691, EPI_ISL_684692 | | | |
| see above | Fukuoka Institute of Health and Environmental Sciences | Pathogen Genomics Center, National Institute of Infectious Diseases | Tsuyoshi Sekizuka, Kentaro Itokawa, Rina Tanaka, Masanori Hashino, Makoto Kuroda |
| EPI_ISL_684693, EPI_ISL_684694, EPI_ISL_684695, EPI_ISL_684696, EPI_ISL_684697, EPI_ISL_684698, EPI_ISL_684699, EPI_ISL_684700, EPI_ISL_684701, EPI_ISL_684702, EPI_ISL_684703, EPI_ISL_684704, EPI_ISL_684705, EPI_ISL_684706, EPI_ISL_684707, EPI_ISL_684708, EPI_ISL_684709, EPI_ISL_684710, EPI_ISL_684711, EPI_ISL_684712, EPI_ISL_684713, EPI_ISL_684714, EPI_ISL_684715, EPI_ISL_684716, EPI_ISL_684717 | | | |
| see above | Pathogen Genomics Center, National Institute of Infectious Diseases | Pathogen Genomics Center, National Institute of Infectious Diseases | Tsuyoshi Sekizuka, Kentaro Itokawa, Rina Tanaka, Masanori Hashino, Makoto Kuroda |
| EPI_ISL_684723, EPI_ISL_684724, EPI_ISL_684725, EPI_ISL_684726, EPI_ISL_684727 | Fukuoka Institute of Health and Environmental Sciences | Pathogen Genomics Center, National Institute of Infectious Diseases | Tsuyoshi Sekizuka, Kentaro Itokawa, Rina Tanaka, Masanori Hashino, Makoto Kuroda |
| EPI_ISL_684728, EPI_ISL_684729, EPI_ISL_684730, EPI_ISL_684731, EPI_ISL_684732, EPI_ISL_684734, EPI_ISL_684735 | Tokyo Metropolitan Institute of Public Health | Pathogen Genomics Center, National Institute of Infectious Diseases | Tsuyoshi Sekizuka, Kentaro Itokawa, Rina Tanaka, Masanori Hashino, Makoto Kuroda |
| EPI_ISL_684736, EPI_ISL_684737, EPI_ISL_684738, EPI_ISL_684739, EPI_ISL_684740, EPI_ISL_684741, EPI_ISL_684742, EPI_ISL_684743, EPI_ISL_684744, EPI_ISL_684745, EPI_ISL_684746, EPI_ISL_684747, EPI_ISL_684748, EPI_ISL_684749, EPI_ISL_684750, EPI_ISL_684751, EPI_ISL_684752, EPI_ISL_684753, EPI_ISL_684754, EPI_ISL_684755, EPI_ISL_684756, EPI_ISL_684757, EPI_ISL_684758, EPI_ISL_684759, EPI_ISL_684760, EPI_ISL_684761, EPI_ISL_684762, EPI_ISL_684763, EPI_ISL_684764, EPI_ISL_684765, EPI_ISL_684766, EPI_ISL_684767, EPI_ISL_684768, EPI_ISL_684769, EPI_ISL_684770, EPI_ISL_684771, EPI_ISL_684772, EPI_ISL_684773, EPI_ISL_684774, EPI_ISL_684775, EPI_ISL_684776, EPI_ISL_684777, EPI_ISL_684778, EPI_ISL_684779, EPI_ISL_684780, EPI_ISL_684781, EPI_ISL_684782, EPI_ISL_684783, EPI_ISL_684784, EPI_ISL_684785, EPI_ISL_684786, EPI_ISL_684787, EPI_ISL_684788, EPI_ISL_684789, EPI_ISL_684790, EPI_ISL_684791, EPI_ISL_684792, EPI_ISL_684793, EPI_ISL_684794, EPI_ISL_684795, EPI_ISL_684796, EPI_ISL_684797 | | | |
| see above | Pathogen Genomics Center, National Institute of Infectious Diseases | Pathogen Genomics Center, National Institute of Infectious Diseases | Tsuyoshi Sekizuka, Kentaro Itokawa, Rina Tanaka, Masanori Hashino, Makoto Kuroda |
| EPI_ISL_684798, EPI_ISL_684799, EPI_ISL_684800, EPI_ISL_684801, EPI_ISL_684802, EPI_ISL_684803, EPI_ISL_684804, EPI_ISL_684805, EPI_ISL_684806, EPI_ISL_684807, EPI_ISL_684808, EPI_ISL_684809, EPI_ISL_684810, EPI_ISL_684811, EPI_ISL_684812, EPI_ISL_684813, EPI_ISL_684815, EPI_ISL_684816, EPI_ISL_684817, EPI_ISL_684818, EPI_ISL_684819, EPI_ISL_684820, EPI_ISL_684821, EPI_ISL_684822, EPI_ISL_684823, EPI_ISL_684824, EPI_ISL_684825, EPI_ISL_684826, EPI_ISL_684827, EPI_ISL_684828, EPI_ISL_684829, EPI_ISL_684830, EPI_ISL_684831, EPI_ISL_684832, EPI_ISL_684833, EPI_ISL_684834, EPI_ISL_684835, EPI_ISL_684836, EPI_ISL_684837, EPI_ISL_684838, EPI_ISL_684839, EPI_ISL_684840, EPI_ISL_684841, EPI_ISL_684842, EPI_ISL_684843, EPI_ISL_684844, EPI_ISL_684845, EPI_ISL_684846, EPI_ISL_684847, EPI_ISL_684848, EPI_ISL_684849, EPI_ISL_684850, EPI_ISL_684851, EPI_ISL_684852, EPI_ISL_684853, EPI_ISL_684854, EPI_ISL_684855, EPI_ISL_684856 | | | |
| see above | Yamagata Prefectural Institute of Public Health | Pathogen Genomics Center, National Institute of Infectious Diseases | Tsuyoshi Sekizuka, Kentaro Itokawa, Rina Tanaka, Masanori Hashino, Makoto Kuroda |
| EPI_ISL_684857, EPI_ISL_684858, EPI_ISL_684859, EPI_ISL_684860, EPI_ISL_684861, EPI_ISL_684862, EPI_ISL_684863, EPI_ISL_684864, EPI_ISL_684865, EPI_ISL_684866, EPI_ISL_684867, EPI_ISL_684868, EPI_ISL_684869, EPI_ISL_684870, EPI_ISL_684871, EPI_ISL_684872, EPI_ISL_684873, EPI_ISL_684874, EPI_ISL_684875, EPI_ISL_684876, EPI_ISL_684877, EPI_ISL_684878, EPI_ISL_684879, EPI_ISL_684880 | | | |
| see above | Pathogen Genomics Center, National Institute of Infectious Diseases | Pathogen Genomics Center, National Institute of Infectious Diseases | Tsuyoshi Sekizuka, Kentaro Itokawa, Rina Tanaka, Masanori Hashino, Makoto Kuroda |
| EPI_ISL_684881, EPI_ISL_684882, EPI_ISL_684883, EPI_ISL_684884, EPI_ISL_684885, EPI_ISL_684886, EPI_ISL_684887, EPI_ISL_684888, EPI_ISL_684889, EPI_ISL_684890, EPI_ISL_684891, EPI_ISL_684892, EPI_ISL_684893, EPI_ISL_684894, EPI_ISL_684895, EPI_ISL_684896, EPI_ISL_684897, EPI_ISL_684898, EPI_ISL_684899, EPI_ISL_684900, EPI_ISL_684901, EPI_ISL_684902, EPI_ISL_684903, EPI_ISL_684904, EPI_ISL_684905, EPI_ISL_684906, EPI_ISL_684907, EPI_ISL_684908, EPI_ISL_684909, EPI_ISL_684910, EPI_ISL_684911, EPI_ISL_684912, EPI_ISL_684913, EPI_ISL_684914, EPI_ISL_684915, EPI_ISL_684916, | | | |

| | | | |
|---|---|---|---|
| EPI_ISL_684917, EPI_ISL_684918, EPI_ISL_684919, EPI_ISL_684920, EPI_ISL_684921, EPI_ISL_684922, EPI_ISL_684923, EPI_ISL_684924, EPI_ISL_684925, EPI_ISL_684926, EPI_ISL_684927, EPI_ISL_684928, EPI_ISL_684929, EPI_ISL_684930, EPI_ISL_684931, EPI_ISL_684932, EPI_ISL_684933, EPI_ISL_684934, EPI_ISL_684935, EPI_ISL_684936, EPI_ISL_684937, EPI_ISL_684938, EPI_ISL_684939, EPI_ISL_684940, EPI_ISL_684941, EPI_ISL_684942, EPI_ISL_684943, EPI_ISL_684944, EPI_ISL_684945, EPI_ISL_684946, EPI_ISL_684947, EPI_ISL_684948, EPI_ISL_684949, EPI_ISL_684950, EPI_ISL_684951, EPI_ISL_684952, EPI_ISL_684953, EPI_ISL_684954, EPI_ISL_684955, EPI_ISL_684956, EPI_ISL_684957, EPI_ISL_684958, EPI_ISL_684959, EPI_ISL_684960, EPI_ISL_684961, EPI_ISL_684962, EPI_ISL_684963, EPI_ISL_684965, EPI_ISL_684966, EPI_ISL_684967, EPI_ISL_684968, EPI_ISL_684969, EPI_ISL_684970, EPI_ISL_684971, EPI_ISL_684972, EPI_ISL_684973, EPI_ISL_684974, EPI_ISL_684975, EPI_ISL_684976, EPI_ISL_684977, EPI_ISL_684978, EPI_ISL_684979, EPI_ISL_684980, EPI_ISL_684981, EPI_ISL_684982, EPI_ISL_684983, EPI_ISL_684984, EPI_ISL_684985, EPI_ISL_684986, EPI_ISL_684987, EPI_ISL_684988, EPI_ISL_684989, EPI_ISL_684990, EPI_ISL_684991, EPI_ISL_684992, EPI_ISL_684993, EPI_ISL_684994, EPI_ISL_684995, EPI_ISL_684996, EPI_ISL_684997, EPI_ISL_684998, EPI_ISL_684999, EPI_ISL_685000, EPI_ISL_685001, EPI_ISL_685002, EPI_ISL_685003, EPI_ISL_685004, EPI_ISL_685005, EPI_ISL_685006, EPI_ISL_685007, EPI_ISL_685008, EPI_ISL_685009, EPI_ISL_685010, EPI_ISL_685011, EPI_ISL_685012, EPI_ISL_685013, EPI_ISL_685014, EPI_ISL_685015, EPI_ISL_685016, EPI_ISL_685017, EPI_ISL_685018, EPI_ISL_685019, EPI_ISL_685020, EPI_ISL_685021, EPI_ISL_685022, EPI_ISL_685023, EPI_ISL_685024, EPI_ISL_685025, EPI_ISL_685026, EPI_ISL_685027, EPI_ISL_685028, EPI_ISL_685029, EPI_ISL_685030, EPI_ISL_685031, EPI_ISL_685032, EPI_ISL_685033, EPI_ISL_685034, EPI_ISL_685035, EPI_ISL_685036, EPI_ISL_685037, EPI_ISL_685038, EPI_ISL_685039, EPI_ISL_685040, EPI_ISL_685041, EPI_ISL_685042, EPI_ISL_685043, EPI_ISL_685044, EPI_ISL_685045, EPI_ISL_685046, EPI_ISL_685047, EPI_ISL_685048, EPI_ISL_685049, EPI_ISL_685050, EPI_ISL_685051, EPI_ISL_685052, EPI_ISL_685053, EPI_ISL_685054, EPI_ISL_685055, EPI_ISL_685056, EPI_ISL_685057, EPI_ISL_685058, EPI_ISL_685059, EPI_ISL_685060, EPI_ISL_685061, EPI_ISL_685062, EPI_ISL_685063, EPI_ISL_685064, EPI_ISL_685065, EPI_ISL_685066, EPI_ISL_685067, EPI_ISL_685068, EPI_ISL_685069, EPI_ISL_685070, EPI_ISL_685071, EPI_ISL_685072, EPI_ISL_685073, EPI_ISL_685074, EPI_ISL_685075, EPI_ISL_685076, EPI_ISL_685077, EPI_ISL_685078, EPI_ISL_685079, EPI_ISL_685080, EPI_ISL_685081, EPI_ISL_685082, EPI_ISL_685083, EPI_ISL_685084, EPI_ISL_685085, EPI_ISL_685086, EPI_ISL_685087, EPI_ISL_685088, EPI_ISL_685089, EPI_ISL_685090, EPI_ISL_685091, EPI_ISL_685092, EPI_ISL_685093, EPI_ISL_685094, EPI_ISL_685095, EPI_ISL_685096, EPI_ISL_685097, EPI_ISL_685098, EPI_ISL_685099, EPI_ISL_685100, EPI_ISL_685101, EPI_ISL_685102, EPI_ISL_685103, EPI_ISL_685104, EPI_ISL_685105, EPI_ISL_685106, EPI_ISL_685107, EPI_ISL_685108, EPI_ISL_685109, EPI_ISL_685110, EPI_ISL_685111, EPI_ISL_685112, EPI_ISL_685113, EPI_ISL_685114, EPI_ISL_685115, EPI_ISL_685116, EPI_ISL_685117, EPI_ISL_685118, EPI_ISL_685119, EPI_ISL_685120, EPI_ISL_685121, EPI_ISL_685122, EPI_ISL_685123, EPI_ISL_685124, EPI_ISL_685126, EPI_ISL_685127, EPI_ISL_685128, EPI_ISL_685129, EPI_ISL_685130, EPI_ISL_685131, EPI_ISL_685132 | | | |
| see above | Saitama Prefectural Institute of Public Health | Pathogen Genomics Center, National Institute of Infectious Diseases | Tsuyoshi Sekizuka, Kentaro Itokawa, Rina Tanaka, Masanori Hashino, Makoto Kuroda |
| EPI_ISL_685133, EPI_ISL_685134, EPI_ISL_685135, EPI_ISL_685136, EPI_ISL_685137, EPI_ISL_685138, EPI_ISL_685139, EPI_ISL_685140, EPI_ISL_685141, EPI_ISL_685142, EPI_ISL_685143, EPI_ISL_685144, EPI_ISL_685145, EPI_ISL_685146, EPI_ISL_685147, EPI_ISL_685148, EPI_ISL_685149, EPI_ISL_685150, EPI_ISL_685151, EPI_ISL_685152, EPI_ISL_685153, EPI_ISL_685154, EPI_ISL_685155, EPI_ISL_685156, EPI_ISL_685157, EPI_ISL_685158, EPI_ISL_685159, EPI_ISL_685160, EPI_ISL_685161, EPI_ISL_685162, EPI_ISL_685163, EPI_ISL_685164, EPI_ISL_685165, EPI_ISL_685166, EPI_ISL_685167, EPI_ISL_685168, EPI_ISL_685169, EPI_ISL_685170, EPI_ISL_685171, EPI_ISL_685172, EPI_ISL_685173, EPI_ISL_685174, EPI_ISL_685175, EPI_ISL_685176, EPI_ISL_685177, EPI_ISL_685178, EPI_ISL_685179, EPI_ISL_685180, EPI_ISL_685181, EPI_ISL_685182, EPI_ISL_685183, EPI_ISL_685184, EPI_ISL_685185, EPI_ISL_685186, EPI_ISL_685187, EPI_ISL_685188, EPI_ISL_685189, EPI_ISL_685190, EPI_ISL_685191, EPI_ISL_685192, EPI_ISL_685193, EPI_ISL_685194, EPI_ISL_685195, EPI_ISL_685196, EPI_ISL_685197, EPI_ISL_685198, EPI_ISL_685199, EPI_ISL_685200, EPI_ISL_685201, EPI_ISL_685202, EPI_ISL_685203, EPI_ISL_685204, EPI_ISL_685205, EPI_ISL_685206, EPI_ISL_685207, EPI_ISL_685208, EPI_ISL_685209, EPI_ISL_685210, EPI_ISL_685211, EPI_ISL_685212, EPI_ISL_685213, EPI_ISL_685214, EPI_ISL_685215, EPI_ISL_685216, EPI_ISL_685217, EPI_ISL_685218, EPI_ISL_685219 | | | |
| see above | Pathogen Genomics Center, National Institute of Infectious Diseases | Pathogen Genomics Center, National Institute of Infectious Diseases | Tsuyoshi Sekizuka, Kentaro Itokawa, Rina Tanaka, Masanori Hashino, Makoto Kuroda |
| EPI_ISL_685220, EPI_ISL_685221, EPI_ISL_685222, EPI_ISL_685223, EPI_ISL_685224, EPI_ISL_685225, EPI_ISL_685226, EPI_ISL_685227, EPI_ISL_685228, EPI_ISL_685229, EPI_ISL_685230, EPI_ISL_685231, EPI_ISL_685232, EPI_ISL_685233, EPI_ISL_685234, EPI_ISL_685235, EPI_ISL_685236, EPI_ISL_685237 | | | |
| see above | Tokyo Metropolitan Institute of Public Health | Pathogen Genomics Center, National Institute of Infectious Diseases | Tsuyoshi Sekizuka, Kentaro Itokawa, Rina Tanaka, Masanori Hashino, Makoto Kuroda |
| EPI_ISL_685238, EPI_ISL_685240, EPI_ISL_685241, EPI_ISL_685242, EPI_ISL_685243, EPI_ISL_685244, EPI_ISL_685245, EPI_ISL_685246, EPI_ISL_685247, EPI_ISL_685248, EPI_ISL_685249, EPI_ISL_685250, EPI_ISL_685251, EPI_ISL_685252, EPI_ISL_685253, EPI_ISL_685254, EPI_ISL_685255, EPI_ISL_685256, EPI_ISL_685257, EPI_ISL_685258, EPI_ISL_685259, EPI_ISL_685260, EPI_ISL_685261, EPI_ISL_685262, EPI_ISL_685263, EPI_ISL_685264, EPI_ISL_685265, EPI_ISL_685266, EPI_ISL_685267, EPI_ISL_685268, EPI_ISL_685269, EPI_ISL_685270, EPI_ISL_685271, EPI_ISL_685272, EPI_ISL_685273, EPI_ISL_685274, EPI_ISL_685275, EPI_ISL_685276, EPI_ISL_685277, EPI_ISL_685278 | | | |
| see above | Pathogen Genomics Center, National Institute of Infectious Diseases | Pathogen Genomics Center, National Institute of Infectious Diseases | Tsuyoshi Sekizuka, Kentaro Itokawa, Rina Tanaka, Masanori Hashino, Makoto Kuroda |
| EPI_ISL_685279, EPI_ISL_685280, EPI_ISL_685281 | Gunma Prefectural Institute of Public Health and Environmental Sciences | Pathogen Genomics Center, National Institute of Infectious Diseases | Tsuyoshi Sekizuka, Kentaro Itokawa, Rina Tanaka, Masanori Hashino, Makoto Kuroda |
| EPI_ISL_685283, EPI_ISL_685284, EPI_ISL_685285, EPI_ISL_685286, EPI_ISL_685287, EPI_ISL_685288, EPI_ISL_685289, EPI_ISL_685290, EPI_ISL_685291, EPI_ISL_685292, EPI_ISL_685295 | | | |
| see above | Pathogen Genomics Center, National Institute of Infectious Diseases | Pathogen Genomics Center, National Institute of Infectious Diseases | Tsuyoshi Sekizuka, Kentaro Itokawa, Rina Tanaka, Masanori Hashino, Makoto Kuroda |
| EPI_ISL_685297, EPI_ISL_685298, EPI_ISL_685299, EPI_ISL_685300, EPI_ISL_685301, EPI_ISL_685302, EPI_ISL_685303, EPI_ISL_685304, EPI_ISL_685305, EPI_ISL_685306, EPI_ISL_685307, EPI_ISL_685308, EPI_ISL_685309, EPI_ISL_685310 | | | |
| see above | Niigata Prefectural Institute of Public Health and Environmental Sciences | Pathogen Genomics Center, National Institute of Infectious Diseases | Tsuyoshi Sekizuka, Kentaro Itokawa, Rina Tanaka, Masanori Hashino, Makoto Kuroda |
| EPI_ISL_685311, EPI_ISL_685312, EPI_ISL_685313, EPI_ISL_685314, EPI_ISL_685315, EPI_ISL_685316, EPI_ISL_685317, EPI_ISL_685318, EPI_ISL_685319, EPI_ISL_685320, EPI_ISL_685321, EPI_ISL_685322, EPI_ISL_685323, EPI_ISL_685324, EPI_ISL_685325, EPI_ISL_685326, EPI_ISL_685327, EPI_ISL_685328, EPI_ISL_685329, EPI_ISL_685330, EPI_ISL_685331, EPI_ISL_685332, EPI_ISL_685333, EPI_ISL_685334, EPI_ISL_685335, EPI_ISL_685336, EPI_ISL_685337, EPI_ISL_685338, EPI_ISL_685339, EPI_ISL_685340, EPI_ISL_685341, EPI_ISL_685342, EPI_ISL_685343, EPI_ISL_685344, EPI_ISL_685345, EPI_ISL_685346, EPI_ISL_685347, EPI_ISL_685348, EPI_ISL_685349, EPI_ISL_685350, EPI_ISL_685351, EPI_ISL_685352, EPI_ISL_685353, EPI_ISL_685354, EPI_ISL_685355, EPI_ISL_685356, EPI_ISL_685357, EPI_ISL_685358, EPI_ISL_685359, EPI_ISL_685360, EPI_ISL_685361, EPI_ISL_685362, EPI_ISL_685363, EPI_ISL_685364, EPI_ISL_685365, EPI_ISL_685366, EPI_ISL_685367, EPI_ISL_685368, EPI_ISL_685369, EPI_ISL_685370, EPI_ISL_685371, EPI_ISL_685372, EPI_ISL_685373, EPI_ISL_685374, EPI_ISL_685375, EPI_ISL_685376, EPI_ISL_685377, EPI_ISL_685378, EPI_ISL_685379, EPI_ISL_685380, EPI_ISL_685381, EPI_ISL_685382, EPI_ISL_685383, EPI_ISL_685384, EPI_ISL_685385, EPI_ISL_685386, EPI_ISL_685387, EPI_ISL_685388, EPI_ISL_685389, EPI_ISL_685390, EPI_ISL_685391, EPI_ISL_685392, EPI_ISL_685393, EPI_ISL_685394, EPI_ISL_685395, EPI_ISL_685396, EPI_ISL_685397, EPI_ISL_685400, EPI_ISL_685401, EPI_ISL_685402, EPI_ISL_685403, EPI_ISL_685404, EPI_ISL_685405, EPI_ISL_685406 | | | |
| see above | Pathogen Genomics Center, National Institute of Infectious Diseases | Pathogen Genomics Center, National Institute of Infectious Diseases | Tsuyoshi Sekizuka, Kentaro Itokawa, Rina Tanaka, Masanori Hashino, Makoto Kuroda |
| EPI_ISL_685407, EPI_ISL_685408, EPI_ISL_685409, EPI_ISL_685410, EPI_ISL_685411, EPI_ISL_685412, EPI_ISL_685413, EPI_ISL_685414, EPI_ISL_685415, EPI_ISL_685416, EPI_ISL_685417, EPI_ISL_685418, EPI_ISL_685419, EPI_ISL_685420, EPI_ISL_685421, EPI_ISL_685422, EPI_ISL_685423, EPI_ISL_685424, EPI_ISL_685425, EPI_ISL_685426, EPI_ISL_685427, EPI_ISL_685428, EPI_ISL_685429, EPI_ISL_685430, EPI_ISL_685431, EPI_ISL_685432, EPI_ISL_685433, EPI_ISL_685434, EPI_ISL_685435, EPI_ISL_685436, EPI_ISL_685437, EPI_ISL_685438, EPI_ISL_685439, EPI_ISL_685440, EPI_ISL_685441, EPI_ISL_685442, EPI_ISL_685443, EPI_ISL_685444, EPI_ISL_685445, EPI_ISL_685446, EPI_ISL_685447, EPI_ISL_685448, EPI_ISL_685449, EPI_ISL_685450, EPI_ISL_685451, EPI_ISL_685452, EPI_ISL_685453, EPI_ISL_685454, EPI_ISL_685455, EPI_ISL_685456, EPI_ISL_685457, EPI_ISL_685458, EPI_ISL_685459, EPI_ISL_685460, EPI_ISL_685461, EPI_ISL_685462, EPI_ISL_685463, EPI_ISL_685464, EPI_ISL_685465, EPI_ISL_685466, EPI_ISL_685467, EPI_ISL_685468, EPI_ISL_685469, EPI_ISL_685470, EPI_ISL_685471, EPI_ISL_685472, EPI_ISL_685473, EPI_ISL_685474, EPI_ISL_685475, EPI_ISL_685476, EPI_ISL_685477, EPI_ISL_685478, EPI_ISL_685479, EPI_ISL_685480, EPI_ISL_685481, EPI_ISL_685482, EPI_ISL_685483, EPI_ISL_685484, EPI_ISL_685485, EPI_ISL_685486, EPI_ISL_685487, EPI_ISL_685488, EPI_ISL_685489, EPI_ISL_685490, EPI_ISL_685491, EPI_ISL_685492, EPI_ISL_685493, EPI_ISL_685494, EPI_ISL_685495, EPI_ISL_685496, EPI_ISL_685497 | | | |
| see above | Tokyo Metropolitan Institute of Public Health | Pathogen Genomics Center, National Institute of Infectious Diseases | Tsuyoshi Sekizuka, Kentaro Itokawa, Rina Tanaka, Masanori Hashino, Makoto Kuroda |
| EPI_ISL_685498, EPI_ISL_685499, EPI_ISL_685500, EPI_ISL_685503, EPI_ISL_685504 | Yamagata Prefectural Institute of Public Health | Pathogen Genomics Center, National Institute of Infectious Diseases | Tsuyoshi Sekizuka, Kentaro Itokawa, Rina Tanaka, Masanori Hashino, Makoto Kuroda |
| EPI_ISL_685505, EPI_ISL_685506, EPI_ISL_685507, EPI_ISL_685508, EPI_ISL_685509, EPI_ISL_685510, EPI_ISL_685511, EPI_ISL_685512, EPI_ISL_685513, EPI_ISL_685514, EPI_ISL_685515, EPI_ISL_685516, EPI_ISL_685517, EPI_ISL_685518, EPI_ISL_685519, EPI_ISL_685520, EPI_ISL_685521, EPI_ISL_685522, EPI_ISL_685523, EPI_ISL_685524, EPI_ISL_685525, EPI_ISL_685526, EPI_ISL_685527, EPI_ISL_685528, EPI_ISL_685529, EPI_ISL_685530, EPI_ISL_685531, EPI_ISL_685532, EPI_ISL_685533, EPI_ISL_685534, EPI_ISL_685535, EPI_ISL_685536, EPI_ISL_685537, EPI_ISL_685538, EPI_ISL_685539, EPI_ISL_685540, EPI_ISL_685541, EPI_ISL_685542, EPI_ISL_685543, EPI_ISL_685544, EPI_ISL_685545, EPI_ISL_685546, EPI_ISL_685547, EPI_ISL_685548, EPI_ISL_685549, EPI_ISL_685550, EPI_ISL_685551, EPI_ISL_685552, EPI_ISL_685553, EPI_ISL_685554, EPI_ISL_685555, EPI_ISL_685556, EPI_ISL_685557, EPI_ISL_685558, EPI_ISL_685559, EPI_ISL_685560, EPI_ISL_685561, EPI_ISL_685562, EPI_ISL_685563, EPI_ISL_685564, EPI_ISL_685565, EPI_ISL_685566, EPI_ISL_685567, EPI_ISL_685568, EPI_ISL_685569, EPI_ISL_685570, EPI_ISL_685571, EPI_ISL_685572, EPI_ISL_685573, EPI_ISL_685574, EPI_ISL_685575, EPI_ISL_685576, EPI_ISL_685577, EPI_ISL_685578, EPI_ISL_685579, EPI_ISL_685580, EPI_ISL_685581, EPI_ISL_685582, EPI_ISL_685584, EPI_ISL_685585, EPI_ISL_685586, EPI_ISL_685587, EPI_ISL_685588, EPI_ISL_685589, EPI_ISL_685590, EPI_ISL_685591, EPI_ISL_685592, EPI_ISL_685593, EPI_ISL_685594, EPI_ISL_685595, EPI_ISL_685596, EPI_ISL_685597, EPI_ISL_685598, EPI_ISL_685599, EPI_ISL_685600, EPI_ISL_685601, EPI_ISL_685602, EPI_ISL_685603, EPI_ISL_685604, EPI_ISL_685605 | | | |
| see above | Pathogen Genomics Center, National Institute of Infectious Diseases | Pathogen Genomics Center, National Institute of Infectious Diseases | Tsuyoshi Sekizuka, Kentaro Itokawa, Rina Tanaka, Masanori Hashino, Makoto Kuroda |
| EPI_ISL_685606, EPI_ISL_685607, EPI_ISL_685608, EPI_ISL_685609, EPI_ISL_685610, EPI_ISL_685611, EPI_ISL_685612, EPI_ISL_685613, EPI_ISL_685614, EPI_ISL_685615, EPI_ISL_685616 | | | |
| see above | Tokyo Metropolitan Institute of Public Health | Pathogen Genomics Center, National Institute of Infectious Diseases | Tsuyoshi Sekizuka, Kentaro Itokawa, Rina Tanaka, Masanori Hashino, Makoto Kuroda |
| EPI_ISL_685617, EPI_ISL_685618, EPI_ISL_685619, EPI_ISL_685620, EPI_ISL_685621, EPI_ISL_685622, EPI_ISL_685623, EPI_ISL_685624, EPI_ISL_685625, EPI_ISL_685626, EPI_ISL_685627, EPI_ISL_685628, EPI_ISL_685629, EPI_ISL_685630, EPI_ISL_685631, EPI_ISL_685632, EPI_ISL_685633, EPI_ISL_685634, EPI_ISL_685635, EPI_ISL_685636, EPI_ISL_685637, EPI_ISL_685638, EPI_ISL_685639, EPI_ISL_685640, EPI_ISL_685641, EPI_ISL_685642, EPI_ISL_685643, EPI_ISL_685644, EPI_ISL_685645, EPI_ISL_685646, EPI_ISL_685647, EPI_ISL_685648, EPI_ISL_685649, EPI_ISL_685650, EPI_ISL_685651, EPI_ISL_685652, EPI_ISL_685653, EPI_ISL_685654, EPI_ISL_685655, EPI_ISL_685656, EPI_ISL_685657, EPI_ISL_685658, EPI_ISL_685659, EPI_ISL_685660, EPI_ISL_685661, EPI_ISL_685662, EPI_ISL_685663, EPI_ISL_685664, EPI_ISL_685665 | | | |
| see above | Pathogen Genomics Center, National Institute of Infectious Diseases | Pathogen Genomics Center, National Institute of Infectious Diseases | Tsuyoshi Sekizuka, Kentaro Itokawa, Rina Tanaka, Masanori Hashino, Makoto Kuroda |
| EPI_ISL_685666, EPI_ISL_685667, EPI_ISL_685668, EPI_ISL_685669, EPI_ISL_685670, EPI_ISL_685671, EPI_ISL_685672, EPI_ISL_685673, EPI_ISL_685674, EPI_ISL_685675, EPI_ISL_685676, EPI_ISL_685677, EPI_ISL_685678, EPI_ISL_685679, EPI_ISL_685680, EPI_ISL_685681, EPI_ISL_685682, EPI_ISL_685683, EPI_ISL_685684, EPI_ISL_685685, EPI_ISL_685686, EPI_ISL_685687, EPI_ISL_685688, EPI_ISL_685689, EPI_ISL_685690, EPI_ISL_685691, EPI_ISL_685692, EPI_ISL_685693, EPI_ISL_685694 | | | |
| see above | Tokyo Metropolitan Institute of Public Health | Pathogen Genomics Center, National Institute of Infectious | Tsuyoshi Sekizuka, Kentaro Itokawa, Rina Tanaka, Masanori Hashino, Makoto Kuroda |

| | | | |
|---|---|---|---|
| | | Diseases | |
| EPI_ISL_685695, EPI_ISL_685696, EPI_ISL_685697, EPI_ISL_685698, EPI_ISL_685699, EPI_ISL_685700, EPI_ISL_685701, EPI_ISL_685702, EPI_ISL_685703, EPI_ISL_685704, EPI_ISL_685705, EPI_ISL_685706, EPI_ISL_685707, EPI_ISL_685708, EPI_ISL_685709, EPI_ISL_685710, EPI_ISL_685711, EPI_ISL_685712, EPI_ISL_685713, EPI_ISL_685714, EPI_ISL_685715, EPI_ISL_685716, EPI_ISL_685717, EPI_ISL_685718, EPI_ISL_685719, EPI_ISL_685720, EPI_ISL_685721, EPI_ISL_685722, EPI_ISL_685723, EPI_ISL_685724, EPI_ISL_685725, EPI_ISL_685726, EPI_ISL_685727, EPI_ISL_685728, EPI_ISL_685729, EPI_ISL_685730, EPI_ISL_685733, EPI_ISL_685734, EPI_ISL_685735, EPI_ISL_685736, EPI_ISL_685737, EPI_ISL_685738, EPI_ISL_685739, EPI_ISL_685740, EPI_ISL_685741, EPI_ISL_685742, EPI_ISL_685743, EPI_ISL_685744, EPI_ISL_685745, EPI_ISL_685746, EPI_ISL_685747, EPI_ISL_685748, EPI_ISL_685749, EPI_ISL_685750, EPI_ISL_685751, EPI_ISL_685752, EPI_ISL_685753, EPI_ISL_685754, EPI_ISL_685755, EPI_ISL_685756, EPI_ISL_685757, EPI_ISL_685758, EPI_ISL_685759, EPI_ISL_685760, EPI_ISL_685761, EPI_ISL_685762, EPI_ISL_685763, EPI_ISL_685764, EPI_ISL_685765, EPI_ISL_685766, EPI_ISL_685767, EPI_ISL_685768, EPI_ISL_685769, EPI_ISL_685770, EPI_ISL_685771, EPI_ISL_685772, EPI_ISL_685773, EPI_ISL_685774, EPI_ISL_685775, EPI_ISL_685776, EPI_ISL_685777, EPI_ISL_685778, EPI_ISL_685779, EPI_ISL_685780, EPI_ISL_685781, EPI_ISL_685782, EPI_ISL_685783, EPI_ISL_685784, EPI_ISL_685785, EPI_ISL_685786, EPI_ISL_685787, EPI_ISL_685788, EPI_ISL_685789, EPI_ISL_685790, EPI_ISL_685791, EPI_ISL_685792, EPI_ISL_685793, EPI_ISL_685794, EPI_ISL_685795, EPI_ISL_685796, EPI_ISL_685797, EPI_ISL_685798, EPI_ISL_685799, EPI_ISL_685800, EPI_ISL_685801, EPI_ISL_685802, EPI_ISL_685803, EPI_ISL_685804, EPI_ISL_685805, EPI_ISL_685806, EPI_ISL_685807, EPI_ISL_685808, EPI_ISL_685809, EPI_ISL_685810, EPI_ISL_685811, EPI_ISL_685812, EPI_ISL_685813, EPI_ISL_685814, EPI_ISL_685815, EPI_ISL_685816, EPI_ISL_685817, EPI_ISL_685818, EPI_ISL_685819, EPI_ISL_685820, EPI_ISL_685821, EPI_ISL_685822, EPI_ISL_685823, EPI_ISL_685824, EPI_ISL_685825, EPI_ISL_685826, EPI_ISL_685827, EPI_ISL_685828, EPI_ISL_685829, EPI_ISL_685830, EPI_ISL_685831, EPI_ISL_685832, EPI_ISL_685833, EPI_ISL_685834, EPI_ISL_685835, EPI_ISL_685836, EPI_ISL_685837, EPI_ISL_685838, EPI_ISL_685839, EPI_ISL_685840, EPI_ISL_685841, EPI_ISL_685842, EPI_ISL_685843, EPI_ISL_685844, EPI_ISL_685845, EPI_ISL_685846, EPI_ISL_685847, EPI_ISL_685848, EPI_ISL_685849, EPI_ISL_685850, EPI_ISL_685851, EPI_ISL_685852, EPI_ISL_685853, EPI_ISL_685854, EPI_ISL_685855, EPI_ISL_685856, EPI_ISL_685857, EPI_ISL_685858, EPI_ISL_685859, EPI_ISL_685860, EPI_ISL_685861, EPI_ISL_685862, EPI_ISL_685863, EPI_ISL_685864, EPI_ISL_685865, EPI_ISL_685866, EPI_ISL_685867, EPI_ISL_685868, EPI_ISL_685869, EPI_ISL_685870, EPI_ISL_685871, EPI_ISL_685872, EPI_ISL_685873, EPI_ISL_685874, EPI_ISL_685875, EPI_ISL_685876, EPI_ISL_685877, EPI_ISL_685878, EPI_ISL_685879, EPI_ISL_685880, EPI_ISL_685881, EPI_ISL_685882, EPI_ISL_685883, EPI_ISL_685884, EPI_ISL_685885, EPI_ISL_685886, EPI_ISL_685887, EPI_ISL_685888, EPI_ISL_685889, EPI_ISL_685890, EPI_ISL_685891, EPI_ISL_685892, EPI_ISL_685893, EPI_ISL_685894, EPI_ISL_685895, EPI_ISL_685896, EPI_ISL_685897, EPI_ISL_685898, EPI_ISL_685899, EPI_ISL_685900, EPI_ISL_685901, EPI_ISL_685902, EPI_ISL_685903, EPI_ISL_685904, EPI_ISL_685905, EPI_ISL_685906, EPI_ISL_685907, EPI_ISL_685908, EPI_ISL_685909, EPI_ISL_685911, EPI_ISL_685912, EPI_ISL_685913, EPI_ISL_685914, EPI_ISL_685915, EPI_ISL_685916 | | | |
| see above | Pathogen Genomics Center, National Institute of Infectious Diseases | Pathogen Genomics Center, National Institute of Infectious Diseases | Tsuyoshi Sekizuka, Kentaro Itokawa, Rina Tanaka, Masanori Hashino, Makoto Kuroda |
| EPI_ISL_685918, EPI_ISL_685919, EPI_ISL_685920, EPI_ISL_685921, EPI_ISL_685922, EPI_ISL_685924 | Fukuoka Institute of Health and Environmental Sciences | Pathogen Genomics Center, National Institute of Infectious Diseases | Tsuyoshi Sekizuka, Kentaro Itokawa, Rina Tanaka, Masanori Hashino, Makoto Kuroda |
| EPI_ISL_685925, EPI_ISL_685926, EPI_ISL_685927, EPI_ISL_685928, EPI_ISL_685929, EPI_ISL_685930, EPI_ISL_685931, EPI_ISL_685932, EPI_ISL_685933, EPI_ISL_685934, EPI_ISL_685935, EPI_ISL_685936, EPI_ISL_685937, EPI_ISL_685947, EPI_ISL_685949, EPI_ISL_685950, EPI_ISL_685953, EPI_ISL_685954, EPI_ISL_685955, EPI_ISL_685957, EPI_ISL_685958, EPI_ISL_685959, EPI_ISL_685960, EPI_ISL_685961, EPI_ISL_685962 | | | |
| see above | Pathogen Genomics Center, National Institute of Infectious Diseases | Pathogen Genomics Center, National Institute of Infectious Diseases | Tsuyoshi Sekizuka, Kentaro Itokawa, Rina Tanaka, Masanori Hashino, Makoto Kuroda |
| EPI_ISL_685987 | Tokyo Metropolitan Institute of Public Health | Pathogen Genomics Center, National Institute of Infectious Diseases | Tsuyoshi Sekizuka, Kentaro Itokawa, Rina Tanaka, Masanori Hashino, Makoto Kuroda |
| EPI_ISL_685988, EPI_ISL_685989, EPI_ISL_685990, EPI_ISL_685991, EPI_ISL_685992, EPI_ISL_685993, EPI_ISL_685994, EPI_ISL_685995, EPI_ISL_685996, EPI_ISL_685997, EPI_ISL_685998, EPI_ISL_685999, EPI_ISL_686000, EPI_ISL_686001, EPI_ISL_686002, EPI_ISL_686003, EPI_ISL_686004, EPI_ISL_686005, EPI_ISL_686006, EPI_ISL_686007, EPI_ISL_686008, EPI_ISL_686009, EPI_ISL_686010, EPI_ISL_686011, EPI_ISL_686012, EPI_ISL_686013, EPI_ISL_686014, EPI_ISL_686015, EPI_ISL_686016, EPI_ISL_686017, EPI_ISL_686018, EPI_ISL_686019, EPI_ISL_686020, EPI_ISL_686021, EPI_ISL_686022, EPI_ISL_686023, EPI_ISL_686024, EPI_ISL_686025, EPI_ISL_686026, EPI_ISL_686027, EPI_ISL_686028, EPI_ISL_686029, EPI_ISL_686030, EPI_ISL_686031, EPI_ISL_686032, EPI_ISL_686033, EPI_ISL_686034, EPI_ISL_686035, EPI_ISL_686036, EPI_ISL_686037, EPI_ISL_686038, EPI_ISL_686039, EPI_ISL_686040, EPI_ISL_686041, EPI_ISL_686042, EPI_ISL_686043, EPI_ISL_686044, EPI_ISL_686045, EPI_ISL_686046, EPI_ISL_686047, EPI_ISL_686048, EPI_ISL_686049, EPI_ISL_686050, EPI_ISL_686051, EPI_ISL_686052, EPI_ISL_686053, EPI_ISL_686054, EPI_ISL_686055, EPI_ISL_686056, EPI_ISL_686057, EPI_ISL_686058, EPI_ISL_686059, EPI_ISL_686060, EPI_ISL_686061, EPI_ISL_686062, EPI_ISL_686063, EPI_ISL_686064, EPI_ISL_686065, EPI_ISL_686066, EPI_ISL_686067, EPI_ISL_686068, EPI_ISL_686069, EPI_ISL_686070, EPI_ISL_686071, EPI_ISL_686072, EPI_ISL_686073, EPI_ISL_686074, EPI_ISL_686075 | | | |
| see above | Pathogen Genomics Center, National Institute of Infectious Diseases | Pathogen Genomics Center, National Institute of Infectious Diseases | Tsuyoshi Sekizuka, Kentaro Itokawa, Rina Tanaka, Masanori Hashino, Makoto Kuroda |
| EPI_ISL_686083, EPI_ISL_686084, EPI_ISL_686085, EPI_ISL_686086, EPI_ISL_686087, EPI_ISL_686088, EPI_ISL_686089, EPI_ISL_686090, EPI_ISL_686091, EPI_ISL_686092, EPI_ISL_686093, EPI_ISL_686094, EPI_ISL_686095, EPI_ISL_686096, EPI_ISL_686097, EPI_ISL_686098, EPI_ISL_686099, EPI_ISL_686100, EPI_ISL_686101, EPI_ISL_686102, EPI_ISL_686103, EPI_ISL_686104, EPI_ISL_686105, EPI_ISL_686106, EPI_ISL_686107 | | | |
| see above | Saitama Prefectural Institute of Public Health | Pathogen Genomics Center, National Institute of Infectious Diseases | Tsuyoshi Sekizuka, Kentaro Itokawa, Rina Tanaka, Masanori Hashino, Makoto Kuroda |
| EPI_ISL_686845, EPI_ISL_686846, EPI_ISL_686847, EPI_ISL_686848, EPI_ISL_686849, EPI_ISL_686850, EPI_ISL_686851, EPI_ISL_686852, EPI_ISL_686853, EPI_ISL_686854, EPI_ISL_686855, EPI_ISL_686856, EPI_ISL_686857, EPI_ISL_686858, EPI_ISL_686859, EPI_ISL_686860, EPI_ISL_686861, EPI_ISL_686862, EPI_ISL_686863, EPI_ISL_686864, EPI_ISL_686865, EPI_ISL_686866, EPI_ISL_686867, EPI_ISL_686868, EPI_ISL_686869, EPI_ISL_686870, EPI_ISL_686871, EPI_ISL_686872, EPI_ISL_686873 | | | |
| see above | Pathogen Genomics Center, National Institute of Infectious Diseases | Pathogen Genomics Center, National Institute of Infectious Diseases | Tsuyoshi Sekizuka, Kentaro Itokawa, Rina Tanaka, Masanori Hashino, Makoto Kuroda |
| EPI_ISL_687043, EPI_ISL_687044, EPI_ISL_687045, EPI_ISL_687046, EPI_ISL_687047, EPI_ISL_687048, EPI_ISL_687049, EPI_ISL_687050, EPI_ISL_687051, EPI_ISL_687052, EPI_ISL_687053, EPI_ISL_687054, EPI_ISL_687055, EPI_ISL_687056, EPI_ISL_687057, EPI_ISL_687058, EPI_ISL_687059, EPI_ISL_687060, EPI_ISL_687061, EPI_ISL_687062, EPI_ISL_687063, EPI_ISL_687064, EPI_ISL_687065, EPI_ISL_687066, EPI_ISL_687067, EPI_ISL_687068, EPI_ISL_687069, EPI_ISL_687070 | | | |
| see above | Kanagawa Prefectural Institute of Public Health | Pathogen Genomics Center, National Institute of Infectious Diseases | Tsuyoshi Sekizuka, Kentaro Itokawa, Rina Tanaka, Masanori Hashino, Makoto Kuroda |
| EPI_ISL_687618, EPI_ISL_687921, EPI_ISL_687922, EPI_ISL_687923, EPI_ISL_687924, EPI_ISL_687925, EPI_ISL_687926, EPI_ISL_687927, EPI_ISL_687928, EPI_ISL_687929, EPI_ISL_687930, EPI_ISL_687931, EPI_ISL_688084, EPI_ISL_688085, EPI_ISL_688624, EPI_ISL_688757, EPI_ISL_688758, EPI_ISL_688759, EPI_ISL_688760, EPI_ISL_688761, EPI_ISL_688762, EPI_ISL_688763, EPI_ISL_688764, EPI_ISL_688765, EPI_ISL_688766, EPI_ISL_688767, EPI_ISL_688768, EPI_ISL_688769, EPI_ISL_688770, EPI_ISL_688771, EPI_ISL_688772, EPI_ISL_688773, EPI_ISL_688774, EPI_ISL_688775, EPI_ISL_688776, EPI_ISL_688777, EPI_ISL_688778, EPI_ISL_688779, EPI_ISL_688780, EPI_ISL_688781, EPI_ISL_688782, EPI_ISL_688783, EPI_ISL_688784, EPI_ISL_688785, EPI_ISL_688786, EPI_ISL_688787, EPI_ISL_688788, EPI_ISL_688789, EPI_ISL_688790, EPI_ISL_688791, EPI_ISL_688792, EPI_ISL_688793, EPI_ISL_688794, EPI_ISL_688795, EPI_ISL_688796, EPI_ISL_688797, EPI_ISL_688798, EPI_ISL_688799, EPI_ISL_688800, EPI_ISL_688801, EPI_ISL_688802, EPI_ISL_688803, EPI_ISL_688804, EPI_ISL_688805, EPI_ISL_688806, EPI_ISL_688807, EPI_ISL_688808, EPI_ISL_688809, EPI_ISL_688810, EPI_ISL_688811, EPI_ISL_688812, EPI_ISL_688813 | | | |
| see above | Pathogen Genomics Center, National Institute of Infectious Diseases | Pathogen Genomics Center, National Institute of Infectious Diseases | Tsuyoshi Sekizuka, Kentaro Itokawa, Rina Tanaka, Masanori Hashino, Makoto Kuroda |
| EPI_ISL_689047, EPI_ISL_689048, EPI_ISL_689049, EPI_ISL_689050, EPI_ISL_689051, EPI_ISL_689052, EPI_ISL_689053, EPI_ISL_689054, EPI_ISL_689055, EPI_ISL_689056, EPI_ISL_689057, EPI_ISL_689058, EPI_ISL_689059, EPI_ISL_689060, EPI_ISL_689061, EPI_ISL_689062, EPI_ISL_689063, EPI_ISL_689064, EPI_ISL_689065, EPI_ISL_689066, EPI_ISL_689067, EPI_ISL_689068, EPI_ISL_689069, EPI_ISL_689070, EPI_ISL_689071, EPI_ISL_689072, EPI_ISL_689073, EPI_ISL_689074, EPI_ISL_689075, EPI_ISL_689076 | | | |
| see above | Gunma Prefectural Institute of Public Health and Environmental Sciences | Pathogen Genomics Center, National Institute of Infectious Diseases | Tsuyoshi Sekizuka, Kentaro Itokawa, Rina Tanaka, Masanori Hashino, Makoto Kuroda |
| EPI_ISL_689643, EPI_ISL_689680 | Pathogen Genomics Center, National Institute of Infectious Diseases | Pathogen Genomics Center, National Institute of Infectious Diseases | Tsuyoshi Sekizuka, Kentaro Itokawa, Rina Tanaka, Masanori Hashino, Makoto Kuroda |
| EPI_ISL_689686, EPI_ISL_689687, EPI_ISL_689688, EPI_ISL_689689, EPI_ISL_689690, EPI_ISL_689691 | Gunma Prefectural Institute of Public Health and Environmental Sciences | Pathogen Genomics Center, National Institute of Infectious Diseases | Tsuyoshi Sekizuka, Kentaro Itokawa, Rina Tanaka, Masanori Hashino, Makoto Kuroda |
| EPI_ISL_690614, EPI_ISL_690615, EPI_ISL_690617 | Pathogen Genomics Center, National Institute of Infectious Diseases | Pathogen Genomics Center, National Institute of Infectious Diseases | Tsuyoshi Sekizuka, Kentaro Itokawa, Rina Tanaka, Masanori Hashino, Makoto Kuroda |
| EPI_ISL_690619 | Saitama Prefectural Institute of Public Health | Pathogen Genomics Center, National Institute of Infectious Diseases | Tsuyoshi Sekizuka, Kentaro Itokawa, Rina Tanaka, Masanori Hashino, Makoto Kuroda |
| EPI_ISL_690620, EPI_ISL_690621, EPI_ISL_690622, EPI_ISL_690623, EPI_ISL_690625 | Pathogen Genomics Center, National Institute of Infectious Diseases | Pathogen Genomics Center, National Institute of Infectious Diseases | Tsuyoshi Sekizuka, Kentaro Itokawa, Rina Tanaka, Masanori Hashino, Makoto Kuroda |
| EPI_ISL_690631 | Saitama Prefectural Institute of Public Health | Pathogen Genomics Center, National Institute of Infectious Diseases | Tsuyoshi Sekizuka, Kentaro Itokawa, Rina Tanaka, Masanori Hashino, Makoto Kuroda |
| EPI_ISL_690636, EPI_ISL_690637 | Pathogen Genomics Center, National Institute of Infectious Diseases | Pathogen Genomics Center, National Institute of Infectious Diseases | Tsuyoshi Sekizuka, Kentaro Itokawa, Rina Tanaka, Masanori Hashino, Makoto Kuroda |
| EPI_ISL_690641, EPI_ISL_690642, EPI_ISL_690643, EPI_ISL_690644, EPI_ISL_690645, EPI_ISL_690646, EPI_ISL_690647, EPI_ISL_690648, EPI_ISL_690649, EPI_ISL_690650, EPI_ISL_690651, EPI_ISL_690652, EPI_ISL_690653, EPI_ISL_690654, EPI_ISL_690655, EPI_ISL_690656, EPI_ISL_690657, EPI_ISL_690658, EPI_ISL_690659, EPI_ISL_690660, EPI_ISL_690661, EPI_ISL_690662, EPI_ISL_690663, EPI_ISL_690664, EPI_ISL_690665, EPI_ISL_690666, EPI_ISL_690667, EPI_ISL_690668, EPI_ISL_690669, EPI_ISL_690670, EPI_ISL_690671, EPI_ISL_690672, EPI_ISL_690673, EPI_ISL_690674, EPI_ISL_690675, EPI_ISL_690676, EPI_ISL_690677, EPI_ISL_690678, EPI_ISL_690679, EPI_ISL_690680 | | | |
| see above | Saitama Prefectural Institute of Public Health | Pathogen Genomics Center, National Institute of Infectious | Tsuyoshi Sekizuka, Kentaro Itokawa, Rina Tanaka, Masanori Hashino, Makoto Kuroda |

| | | | |
|---|---|---|---|
| | | Diseases | |
| EPI_ISL_690684, EPI_ISL_690685, EPI_ISL_690686, EPI_ISL_690708, EPI_ISL_690709, EPI_ISL_690710, EPI_ISL_690711, EPI_ISL_690712, EPI_ISL_690713, EPI_ISL_690714, EPI_ISL_690715, EPI_ISL_690716, EPI_ISL_690717, EPI_ISL_690718, EPI_ISL_690719, EPI_ISL_690720, EPI_ISL_690721, EPI_ISL_690722, EPI_ISL_690723, EPI_ISL_690724, EPI_ISL_690725, EPI_ISL_690726, EPI_ISL_690727, EPI_ISL_690728, EPI_ISL_690729, EPI_ISL_690730, EPI_ISL_690731, EPI_ISL_690732, EPI_ISL_690733, EPI_ISL_690734, EPI_ISL_690735, EPI_ISL_690736, EPI_ISL_690737, EPI_ISL_690738, EPI_ISL_690739, EPI_ISL_690740, EPI_ISL_690741, EPI_ISL_690742, EPI_ISL_690743, EPI_ISL_690745, EPI_ISL_690746, EPI_ISL_690747, EPI_ISL_690748, EPI_ISL_690749, EPI_ISL_690750, EPI_ISL_690751, EPI_ISL_690752, EPI_ISL_690753, EPI_ISL_690754, EPI_ISL_690755, EPI_ISL_690756, EPI_ISL_690757, EPI_ISL_690758, EPI_ISL_690759, EPI_ISL_690760, EPI_ISL_690761, EPI_ISL_690762, EPI_ISL_690763, EPI_ISL_690764, EPI_ISL_690765, EPI_ISL_690766, EPI_ISL_690767, EPI_ISL_690768, EPI_ISL_690769, EPI_ISL_690770, EPI_ISL_690771, EPI_ISL_690772, EPI_ISL_690773, EPI_ISL_690774, EPI_ISL_690775, EPI_ISL_690776, EPI_ISL_690777, EPI_ISL_690779, EPI_ISL_690780, EPI_ISL_690781, EPI_ISL_690782, EPI_ISL_690783, EPI_ISL_690784, EPI_ISL_690785, EPI_ISL_690786, EPI_ISL_690787, EPI_ISL_690788, EPI_ISL_690789, EPI_ISL_690790, EPI_ISL_690791 | | | |
| see above | Pathogen Genomics Center, National Institute of Infectious Diseases | Pathogen Genomics Center, National Institute of Infectious Diseases | Tsuyoshi Sekizuka, Kentaro Itokawa, Rina Tanaka, Masanori Hashino, Makoto Kuroda |
| EPI_ISL_690792, EPI_ISL_690793 | Tokyo Metropolitan Institute of Public Health | Pathogen Genomics Center, National Institute of Infectious Diseases | Tsuyoshi Sekizuka, Kentaro Itokawa, Rina Tanaka, Masanori Hashino, Makoto Kuroda |
| EPI_ISL_690818, EPI_ISL_690819, EPI_ISL_690820, EPI_ISL_690821, EPI_ISL_690822, EPI_ISL_690823, EPI_ISL_690824, EPI_ISL_690825, EPI_ISL_690826, EPI_ISL_690827, EPI_ISL_690828, EPI_ISL_690829, EPI_ISL_690830, EPI_ISL_690831, EPI_ISL_690832, EPI_ISL_690833, EPI_ISL_690834, EPI_ISL_690835, EPI_ISL_690836 | | | |
| see above | Kanagawa Prefectural Institute of Public Health | Pathogen Genomics Center, National Institute of Infectious Diseases | Tsuyoshi Sekizuka, Kentaro Itokawa, Rina Tanaka, Masanori Hashino, Makoto Kuroda |
| EPI_ISL_690837, EPI_ISL_690838, EPI_ISL_690839, EPI_ISL_690840, EPI_ISL_690841, EPI_ISL_690842, EPI_ISL_690843 | Sakai City Institute of Public Health | Pathogen Genomics Center, National Institute of Infectious Diseases | Tsuyoshi Sekizuka, Kentaro Itokawa, Rina Tanaka, Masanori Hashino, Makoto Kuroda |
| EPI_ISL_690845, EPI_ISL_690846, EPI_ISL_690847, EPI_ISL_690848, EPI_ISL_690849, EPI_ISL_690850, EPI_ISL_690851, EPI_ISL_690852, EPI_ISL_690853, EPI_ISL_690854, EPI_ISL_690855, EPI_ISL_690856, EPI_ISL_690857, EPI_ISL_690858, EPI_ISL_690859, EPI_ISL_690860, EPI_ISL_690861, EPI_ISL_690862, EPI_ISL_690863, EPI_ISL_690864, EPI_ISL_690865, EPI_ISL_690866, EPI_ISL_690867, EPI_ISL_690868, EPI_ISL_690869, EPI_ISL_690870, EPI_ISL_690871, EPI_ISL_690872, EPI_ISL_690873, EPI_ISL_690874, EPI_ISL_690875, EPI_ISL_690876, EPI_ISL_690877, EPI_ISL_690878, EPI_ISL_690879, EPI_ISL_690880, EPI_ISL_690881, EPI_ISL_690882, EPI_ISL_690883, EPI_ISL_690884, EPI_ISL_690885, EPI_ISL_690886, EPI_ISL_690887, EPI_ISL_690888, EPI_ISL_690889, EPI_ISL_690890, EPI_ISL_690891, EPI_ISL_690892, EPI_ISL_690893, EPI_ISL_690894, EPI_ISL_690895, EPI_ISL_690896, EPI_ISL_690897, EPI_ISL_690898, EPI_ISL_690899, EPI_ISL_690900, EPI_ISL_690901, EPI_ISL_690909, EPI_ISL_690910, EPI_ISL_690911, EPI_ISL_690912, EPI_ISL_690913, EPI_ISL_690914, EPI_ISL_690915, EPI_ISL_690916, EPI_ISL_690917, EPI_ISL_690918, EPI_ISL_690919, EPI_ISL_690920, EPI_ISL_690921, EPI_ISL_690922, EPI_ISL_690923, EPI_ISL_690924, EPI_ISL_690925, EPI_ISL_690926, EPI_ISL_690927, EPI_ISL_690928, EPI_ISL_690929, EPI_ISL_690930, EPI_ISL_690931, EPI_ISL_690932, EPI_ISL_690933, EPI_ISL_690934, EPI_ISL_690935, EPI_ISL_690936, EPI_ISL_690937, EPI_ISL_690938, EPI_ISL_690939, EPI_ISL_690940, EPI_ISL_690941, EPI_ISL_690942, EPI_ISL_690943, EPI_ISL_690944, EPI_ISL_690945, EPI_ISL_690946, EPI_ISL_690947, EPI_ISL_690948, EPI_ISL_690949, EPI_ISL_690950, EPI_ISL_690951, EPI_ISL_690952, EPI_ISL_690953, EPI_ISL_690954, EPI_ISL_690955, EPI_ISL_690956, EPI_ISL_690957, EPI_ISL_690958, EPI_ISL_690959, EPI_ISL_690960, EPI_ISL_690961, EPI_ISL_690962, EPI_ISL_690963, EPI_ISL_690964, EPI_ISL_690965, EPI_ISL_690966, EPI_ISL_690967, EPI_ISL_690968, EPI_ISL_690969, EPI_ISL_690970, EPI_ISL_690971, EPI_ISL_690972, EPI_ISL_690973, EPI_ISL_690974, EPI_ISL_690975, EPI_ISL_690976, EPI_ISL_690977, EPI_ISL_690978, EPI_ISL_690979, EPI_ISL_690980 | | | |
| see above | Pathogen Genomics Center, National Institute of Infectious Diseases | Pathogen Genomics Center, National Institute of Infectious Diseases | Tsuyoshi Sekizuka, Kentaro Itokawa, Rina Tanaka, Masanori Hashino, Makoto Kuroda |
| EPI_ISL_690982, EPI_ISL_690983, EPI_ISL_690984, EPI_ISL_690985, EPI_ISL_690986 | Fukuoka Institute of Health and Environmental Sciences | Pathogen Genomics Center, National Institute of Infectious Diseases | Tsuyoshi Sekizuka, Kentaro Itokawa, Rina Tanaka, Masanori Hashino, Makoto Kuroda |
| EPI_ISL_690987, EPI_ISL_690988, EPI_ISL_690989, EPI_ISL_690990, EPI_ISL_690991, EPI_ISL_690992, EPI_ISL_690993, EPI_ISL_690994, EPI_ISL_690995, EPI_ISL_690996, EPI_ISL_690997, EPI_ISL_690998, EPI_ISL_690999, EPI_ISL_691000, EPI_ISL_691001, EPI_ISL_691002, EPI_ISL_691003, EPI_ISL_691004, EPI_ISL_691005, EPI_ISL_691006, EPI_ISL_691007, EPI_ISL_691008, EPI_ISL_691009, EPI_ISL_691010, EPI_ISL_691011, EPI_ISL_691012, EPI_ISL_691013, EPI_ISL_691014, EPI_ISL_691015, EPI_ISL_691016, EPI_ISL_691017, EPI_ISL_691018, EPI_ISL_691019, EPI_ISL_691020, EPI_ISL_691021, EPI_ISL_691022, EPI_ISL_691023, EPI_ISL_691024, EPI_ISL_691025, EPI_ISL_691026, EPI_ISL_691027, EPI_ISL_691028, EPI_ISL_691029, EPI_ISL_691030, EPI_ISL_691031, EPI_ISL_691032, EPI_ISL_691033, EPI_ISL_691034, EPI_ISL_691035, EPI_ISL_691036, EPI_ISL_691037, EPI_ISL_691038, EPI_ISL_691039, EPI_ISL_691040, EPI_ISL_691041, EPI_ISL_691042, EPI_ISL_691043, EPI_ISL_691044, EPI_ISL_691045, EPI_ISL_691046, EPI_ISL_691047, EPI_ISL_691048, EPI_ISL_691049, EPI_ISL_691050, EPI_ISL_691051, EPI_ISL_691052, EPI_ISL_691053, EPI_ISL_691054, EPI_ISL_691055, EPI_ISL_691056, EPI_ISL_691057, EPI_ISL_691058, EPI_ISL_691059, EPI_ISL_691060, EPI_ISL_691061, EPI_ISL_691062, EPI_ISL_691063, EPI_ISL_691064, EPI_ISL_691065, EPI_ISL_691066, EPI_ISL_691067, EPI_ISL_691068, EPI_ISL_691069, EPI_ISL_691070, EPI_ISL_691071, EPI_ISL_691072, EPI_ISL_691073, EPI_ISL_691074, EPI_ISL_691075, EPI_ISL_691076, EPI_ISL_691077, EPI_ISL_691078, EPI_ISL_691079, EPI_ISL_691080, EPI_ISL_691081, EPI_ISL_691082, EPI_ISL_691083, EPI_ISL_691084, EPI_ISL_691085, EPI_ISL_691086, EPI_ISL_691087, EPI_ISL_691088, EPI_ISL_691089, EPI_ISL_691090, EPI_ISL_691091, EPI_ISL_691092, EPI_ISL_691093, EPI_ISL_691094, EPI_ISL_691095, EPI_ISL_691096, EPI_ISL_691097, EPI_ISL_691098, EPI_ISL_691099, EPI_ISL_691100, EPI_ISL_691101, EPI_ISL_691102, EPI_ISL_691103, EPI_ISL_691104, EPI_ISL_691105, EPI_ISL_691106, EPI_ISL_691107, EPI_ISL_691109, EPI_ISL_691110, EPI_ISL_691111, EPI_ISL_691112, EPI_ISL_691113, EPI_ISL_691114, EPI_ISL_691115, EPI_ISL_691116, EPI_ISL_691117, EPI_ISL_691118, EPI_ISL_691119, EPI_ISL_691120, EPI_ISL_691121, EPI_ISL_691122, EPI_ISL_691123, EPI_ISL_691124, EPI_ISL_691125, EPI_ISL_691126, EPI_ISL_691127, EPI_ISL_691128, EPI_ISL_691129, EPI_ISL_691130, EPI_ISL_691131, EPI_ISL_691132, EPI_ISL_691133, EPI_ISL_691134, EPI_ISL_691135, EPI_ISL_691136, EPI_ISL_691137, EPI_ISL_691138, EPI_ISL_691139, EPI_ISL_691140, EPI_ISL_691141, EPI_ISL_691142, EPI_ISL_691143, EPI_ISL_691144, EPI_ISL_691145, EPI_ISL_691146, EPI_ISL_691147, EPI_ISL_691148, EPI_ISL_691149, EPI_ISL_691150, EPI_ISL_691151, EPI_ISL_691152, EPI_ISL_691153, EPI_ISL_691154, EPI_ISL_691155, EPI_ISL_691156, EPI_ISL_691157, EPI_ISL_691158, EPI_ISL_691159, EPI_ISL_691160, EPI_ISL_691161, EPI_ISL_691162, EPI_ISL_691163, EPI_ISL_691164, EPI_ISL_691165, EPI_ISL_691166, EPI_ISL_691167, EPI_ISL_691168, EPI_ISL_691169, EPI_ISL_691170, EPI_ISL_691171, EPI_ISL_691172, EPI_ISL_691173, EPI_ISL_691174, EPI_ISL_691175, EPI_ISL_691176, EPI_ISL_691177, EPI_ISL_691178, EPI_ISL_691179, EPI_ISL_691180, EPI_ISL_691181, EPI_ISL_691182, EPI_ISL_691183, EPI_ISL_691184, EPI_ISL_691185, EPI_ISL_691186, EPI_ISL_691187, EPI_ISL_691188, EPI_ISL_691189, EPI_ISL_691190, EPI_ISL_691191, EPI_ISL_691192, EPI_ISL_691193, EPI_ISL_691194, EPI_ISL_691195, EPI_ISL_691196, EPI_ISL_691197, EPI_ISL_691198, EPI_ISL_691199, EPI_ISL_691200, EPI_ISL_691201, EPI_ISL_691202, EPI_ISL_691203, EPI_ISL_691204, EPI_ISL_691205, EPI_ISL_691206, EPI_ISL_691207, EPI_ISL_691208, EPI_ISL_691209, EPI_ISL_691210, EPI_ISL_691211, EPI_ISL_691212, EPI_ISL_691213, EPI_ISL_691214, EPI_ISL_691215, EPI_ISL_691216, EPI_ISL_691217, EPI_ISL_691218, EPI_ISL_691219, EPI_ISL_691220, EPI_ISL_691221, EPI_ISL_691222, EPI_ISL_691223, EPI_ISL_691224, EPI_ISL_691225, EPI_ISL_691226, EPI_ISL_691227, EPI_ISL_691228, EPI_ISL_691229, EPI_ISL_691230, EPI_ISL_691231, EPI_ISL_691232, EPI_ISL_691233, EPI_ISL_691234, EPI_ISL_691235, EPI_ISL_691236, EPI_ISL_691237, EPI_ISL_691238, EPI_ISL_691240, EPI_ISL_691241, EPI_ISL_691242, EPI_ISL_691243, EPI_ISL_691244, EPI_ISL_691245, EPI_ISL_691246, EPI_ISL_691247, EPI_ISL_691249, EPI_ISL_691250, EPI_ISL_691251, EPI_ISL_691257, EPI_ISL_691258, EPI_ISL_691259, EPI_ISL_691273, EPI_ISL_691274, EPI_ISL_691289, EPI_ISL_691290, EPI_ISL_691291, EPI_ISL_691292, EPI_ISL_691293, EPI_ISL_691294, EPI_ISL_691295, EPI_ISL_691296, EPI_ISL_691297, EPI_ISL_691298, EPI_ISL_691299, EPI_ISL_691300, EPI_ISL_691301, EPI_ISL_691302, EPI_ISL_691303, EPI_ISL_691304, EPI_ISL_691305, EPI_ISL_691306, EPI_ISL_691307, EPI_ISL_691308, EPI_ISL_691309, EPI_ISL_691310, EPI_ISL_691311, EPI_ISL_691312, EPI_ISL_691313, EPI_ISL_691314, EPI_ISL_691315, EPI_ISL_691316, EPI_ISL_691317, EPI_ISL_691318, EPI_ISL_691319, EPI_ISL_691320, EPI_ISL_691321, EPI_ISL_691322, EPI_ISL_691323, EPI_ISL_691324, EPI_ISL_691325, EPI_ISL_691326, EPI_ISL_691327, EPI_ISL_691328, EPI_ISL_691329, EPI_ISL_691330, EPI_ISL_691331, EPI_ISL_691332, EPI_ISL_691333, EPI_ISL_691334, EPI_ISL_691335, EPI_ISL_691336, EPI_ISL_691337, EPI_ISL_691338, EPI_ISL_691339, EPI_ISL_691340, EPI_ISL_691341, EPI_ISL_691342, EPI_ISL_691343, EPI_ISL_691344, EPI_ISL_691345, EPI_ISL_691346, EPI_ISL_691347, EPI_ISL_691348, EPI_ISL_691349, EPI_ISL_691350, EPI_ISL_691351, EPI_ISL_691352, EPI_ISL_691353, EPI_ISL_691354, EPI_ISL_691355, EPI_ISL_691356, EPI_ISL_691357, EPI_ISL_691358, EPI_ISL_691359, EPI_ISL_691360, EPI_ISL_691361, EPI_ISL_691362, EPI_ISL_691363, EPI_ISL_691379, EPI_ISL_691380, EPI_ISL_691381, EPI_ISL_691382, EPI_ISL_691383, EPI_ISL_691384, EPI_ISL_691385, EPI_ISL_691395, EPI_ISL_691396, EPI_ISL_691397, EPI_ISL_691398, EPI_ISL_691399, EPI_ISL_691400, EPI_ISL_691401, EPI_ISL_691402, EPI_ISL_691403, EPI_ISL_691404, EPI_ISL_691405, EPI_ISL_691406, EPI_ISL_691407, EPI_ISL_691408, EPI_ISL_691409, EPI_ISL_691410, EPI_ISL_691411, EPI_ISL_691412, EPI_ISL_691413, EPI_ISL_691414, EPI_ISL_691415, EPI_ISL_691416, EPI_ISL_691417, EPI_ISL_691418, EPI_ISL_691419, EPI_ISL_691420, EPI_ISL_691421, EPI_ISL_691422, EPI_ISL_691423, EPI_ISL_691424, EPI_ISL_691425, EPI_ISL_691426, EPI_ISL_691427, EPI_ISL_691428, EPI_ISL_691429, EPI_ISL_691430, EPI_ISL_691431, EPI_ISL_691432, EPI_ISL_691433, EPI_ISL_691434, EPI_ISL_691435, EPI_ISL_691436, EPI_ISL_691437, EPI_ISL_691438, EPI_ISL_691439, EPI_ISL_691440, EPI_ISL_691441, EPI_ISL_691442, EPI_ISL_691443, EPI_ISL_691444, EPI_ISL_691445, EPI_ISL_691446, EPI_ISL_691447, EPI_ISL_691448, EPI_ISL_691449, EPI_ISL_691450, EPI_ISL_691451, EPI_ISL_691452, EPI_ISL_691453, EPI_ISL_691454, EPI_ISL_691455, EPI_ISL_691456, EPI_ISL_691457, EPI_ISL_691458, EPI_ISL_691459, EPI_ISL_691460, EPI_ISL_691461, EPI_ISL_691462, EPI_ISL_691463, EPI_ISL_691464, EPI_ISL_691465, EPI_ISL_691466, EPI_ISL_691467, EPI_ISL_691468, EPI_ISL_691469, EPI_ISL_691470, EPI_ISL_691471, EPI_ISL_691472, EPI_ISL_691473, EPI_ISL_691474, EPI_ISL_691475, EPI_ISL_691476, EPI_ISL_691477, EPI_ISL_691478, EPI_ISL_691479, EPI_ISL_691480, EPI_ISL_691481, EPI_ISL_691482, EPI_ISL_691483, EPI_ISL_691484, EPI_ISL_691485, EPI_ISL_691486, EPI_ISL_691487, EPI_ISL_691488, EPI_ISL_691489, EPI_ISL_691490, EPI_ISL_691491, EPI_ISL_691492, EPI_ISL_691493, EPI_ISL_691494, EPI_ISL_691495, EPI_ISL_691496, EPI_ISL_691497, EPI_ISL_691498, EPI_ISL_691499, EPI_ISL_691500, EPI_ISL_691501, EPI_ISL_691507, EPI_ISL_691508 | | | |
| see above | Pathogen Genomics Center, National Institute of Infectious Diseases | Pathogen Genomics Center, National Institute of Infectious Diseases | Tsuyoshi Sekizuka, Kentaro Itokawa, Rina Tanaka, Masanori Hashino, Makoto Kuroda |
| EPI_ISL_691509, EPI_ISL_691510, EPI_ISL_691511, EPI_ISL_691512, EPI_ISL_691513, EPI_ISL_691514, EPI_ISL_691515, EPI_ISL_691516, EPI_ISL_691517, EPI_ISL_691518, EPI_ISL_691519, EPI_ISL_691520, EPI_ISL_691521, EPI_ISL_691522, EPI_ISL_691523, EPI_ISL_691524, EPI_ISL_691525, EPI_ISL_691526, EPI_ISL_691527, EPI_ISL_691528, EPI_ISL_691529, EPI_ISL_691530, EPI_ISL_691531, EPI_ISL_691532, EPI_ISL_691533, EPI_ISL_691534, EPI_ISL_691535, EPI_ISL_691536, EPI_ISL_691537, EPI_ISL_691538, EPI_ISL_691539, EPI_ISL_691540, EPI_ISL_691541, EPI_ISL_691542 | | | |
| see above | Sakai City Institute of Public Health | Pathogen Genomics Center, National Institute of Infectious Diseases | Tsuyoshi Sekizuka, Kentaro Itokawa, Rina Tanaka, Masanori Hashino, Makoto Kuroda |
| EPI_ISL_691681 | Consejeria de Sanidad y Asuntos Sociales | Instituto de Salud Carlos III | Iglesias-Caballero, M. Camarero, S. Molinero Calamita, M. González-Esguevillas, M. Pozo, F. Casas, I. Jiménez, P. Jiménez, M. Zaballos, A. Monzón, S. Varona, S. Juliá, M. Cuesta, I. Gutiérrez, G. |
| EPI_ISL_691682 | Hospital General Universitario de Ciudad Real | Instituto de Salud Carlos III | Iglesias-Caballero, M. Camarero, S. Molinero Calamita, M. González-Esguevillas, M. Pozo, F. Casas, I. Jiménez, P. Jiménez, M. Zaballos, A. Monzón, S. Varona, S. Juliá, M. Cuesta, I. Jarilla, M. |
| EPI_ISL_691683 | Hospital General Universitario de Ciudad Real | Instituto de Salud Carlos III | Iglesias-Caballero, M. Camarero, S. Molinero Calamita, M. González-Esguevillas, M. Pozo, F. Casas, I. Jiménez, P. Jiménez, M. Zaballos, A. Monzón, S. Varona, S. Juliá, M. Cuesta, I. Madrigal, M. |
| EPI_ISL_691684 | Complejo Hospitalario Universitario de Albacete | Instituto de Salud Carlos III | Iglesias-Caballero, M. Camarero, S. Molinero Calamita, M. González-Esguevillas, M. Pozo, F. Casas, I. Jiménez, P. Jiménez, M. Zaballos, A. Monzón, S. Varona, S. Juliá, M. Cuesta, I. Martínez, E. |
| EPI_ISL_691757, EPI_ISL_691758, EPI_ISL_691767, EPI_ISL_691861, EPI_ISL_691872, EPI_ISL_691873, EPI_ISL_691874, EPI_ISL_691875, EPI_ISL_691876, EPI_ISL_691877, EPI_ISL_691878, EPI_ISL_691879, EPI_ISL_691880, EPI_ISL_691881, EPI_ISL_691882, EPI_ISL_691883, EPI_ISL_691884, EPI_ISL_691885, EPI_ISL_691886, EPI_ISL_691887, EPI_ISL_691888, EPI_ISL_691889, EPI_ISL_691890, EPI_ISL_691891, EPI_ISL_691892, EPI_ISL_691893, EPI_ISL_691894, EPI_ISL_691895, EPI_ISL_691896, EPI_ISL_691897, EPI_ISL_691898, EPI_ISL_691899, EPI_ISL_691900, EPI_ISL_691901, EPI_ISL_691902, EPI_ISL_691903, EPI_ISL_691904, EPI_ISL_691905, EPI_ISL_691906, EPI_ISL_691907, EPI_ISL_691908, EPI_ISL_691909, EPI_ISL_691910, EPI_ISL_691911, EPI_ISL_691912, EPI_ISL_691913, EPI_ISL_691914, EPI_ISL_691915, EPI_ISL_691916, EPI_ISL_691917, EPI_ISL_691918, EPI_ISL_691919, EPI_ISL_691920, EPI_ISL_691921, EPI_ISL_691922, EPI_ISL_691923, EPI_ISL_691924, EPI_ISL_691925, EPI_ISL_691926, EPI_ISL_691927, EPI_ISL_691928, EPI_ISL_691929, EPI_ISL_691930, EPI_ISL_691931, EPI_ISL_691932, EPI_ISL_691933, EPI_ISL_691934, EPI_ISL_691935, EPI_ISL_691936, EPI_ISL_691937, EPI_ISL_691938, EPI_ISL_691939, EPI_ISL_691940, EPI_ISL_691941, EPI_ISL_691942, EPI_ISL_691943, EPI_ISL_691944, EPI_ISL_691945, EPI_ISL_691946, EPI_ISL_691947, EPI_ISL_691948, EPI_ISL_691949, EPI_ISL_691950, EPI_ISL_691951, EPI_ISL_691952, EPI_ISL_691953, EPI_ISL_691954, EPI_ISL_691955, EPI_ISL_691956, EPI_ISL_691957, EPI_ISL_691958, EPI_ISL_691959, EPI_ISL_691960, EPI_ISL_691961, EPI_ISL_691962, EPI_ISL_691963, EPI_ISL_691964, EPI_ISL_691965, EPI_ISL_691966, EPI_ISL_691967, EPI_ISL_691968, EPI_ISL_691969, EPI_ISL_691970, EPI_ISL_691971, EPI_ISL_691972, EPI_ISL_691973, EPI_ISL_691974, EPI_ISL_691975, EPI_ISL_691976, EPI_ISL_691977, EPI_ISL_691978, EPI_ISL_691979, EPI_ISL_691980, EPI_ISL_691981, EPI_ISL_691982, EPI_ISL_691983, EPI_ISL_691984, EPI_ISL_691985, EPI_ISL_691986, EPI_ISL_691987, EPI_ISL_691988, EPI_ISL_691989, EPI_ISL_691990, EPI_ISL_691991, EPI_ISL_691992, EPI_ISL_691993, | | | |

| | | | |
|---|---|---|---|
| EPI_ISL_691994, EPI_ISL_691995, EPI_ISL_691996, EPI_ISL_691997, EPI_ISL_691998, EPI_ISL_691999, EPI_ISL_692000, EPI_ISL_692001, EPI_ISL_692002, EPI_ISL_692003, EPI_ISL_692004, EPI_ISL_692005, EPI_ISL_692006, EPI_ISL_692007, EPI_ISL_692008, EPI_ISL_692009, EPI_ISL_692010, EPI_ISL_692011, EPI_ISL_692012, EPI_ISL_692013, EPI_ISL_692014, EPI_ISL_692015, EPI_ISL_692016, EPI_ISL_692017, EPI_ISL_692018, EPI_ISL_692019, EPI_ISL_692020, EPI_ISL_692021, EPI_ISL_692022, EPI_ISL_692023, EPI_ISL_692024, EPI_ISL_692025, EPI_ISL_692026, EPI_ISL_692027, EPI_ISL_692028, EPI_ISL_692029, EPI_ISL_692030, EPI_ISL_692031, EPI_ISL_692032, EPI_ISL_692033, EPI_ISL_692034, EPI_ISL_692035, EPI_ISL_692036, EPI_ISL_692037, EPI_ISL_692038, EPI_ISL_692039, EPI_ISL_692040, EPI_ISL_692041, EPI_ISL_692042, EPI_ISL_692043, EPI_ISL_692044, EPI_ISL_692045, EPI_ISL_692046, EPI_ISL_692059, EPI_ISL_692060, EPI_ISL_692197, EPI_ISL_692527, EPI_ISL_692597 | | | |
| see above | Pathogen Genomics Center, National Institute of Infectious Diseases | Pathogen Genomics Center, National Institute of Infectious Diseases | Tsuyoshi Sekizuka, Kentaro Itokawa, Rina Tanaka, Masanori Hashino, Makoto Kuroda |
| EPI_ISL_693195 | Hospital e Pronto Socorro Portinari | Instituto Adolfo Lutz, Interdiciplinary Procedures Center, Strategic Laboratory | Claudio Tavares Sacchi, Claudia Regina Gonçalves, Erica Valessa Ramos Gomes, Karoline Rodrigues Campos |
| EPI_ISL_693196 | Hospital Santa Clara | Instituto Adolfo Lutz, Interdiciplinary Procedures Center, Strategic Laboratory | Claudio Tavares Sacchi, Claudia Regina Gonçalves, Erica Valessa Ramos Gomes, Karoline Rodrigues Campos |
| EPI_ISL_693198 | Santa Casa de Misericordia de Sao Paulo - Hospital Central | Instituto Adolfo Lutz, Interdiciplinary Procedures Center, Strategic Laboratory | Claudio Tavares Sacchi, Claudia Regina Gonçalves, Erica Valessa Ramos Gomes, Karoline Rodrigues Campos |
| EPI_ISL_693200 | Hospital e Maternidade Mairipora | Instituto Adolfo Lutz, Interdiciplinary Procedures Center, Strategic Laboratory | Claudio Tavares Sacchi, Claudia Regina Gonçalves, Erica Valessa Ramos Gomes, Karoline Rodrigues Campos |
| EPI_ISL_693201 | Hospital Sao Paulo de Ensino da Unifesp | Instituto Adolfo Lutz, Interdiciplinary Procedures Center, Strategic Laboratory | Claudio Tavares Sacchi, Claudia Regina Gonçalves, Erica Valessa Ramos Gomes, Karoline Rodrigues Campos |
| EPI_ISL_693202 | Pronto Socorro Municipal Prof. Joao Catarin Mezomo | Instituto Adolfo Lutz, Interdiciplinary Procedures Center, Strategic Laboratory | Claudio Tavares Sacchi, Claudia Regina Gonçalves, Erica Valessa Ramos Gomes, Karoline Rodrigues Campos |
| EPI_ISL_693203 | Hospital Municipal Doutor Arthur Ribeiro de Saboya | Instituto Adolfo Lutz, Interdiciplinary Procedures Center, Strategic Laboratory | Claudio Tavares Sacchi, Claudia Regina Gonçalves, Erica Valessa Ramos Gomes, Karoline Rodrigues Campos |
| EPI_ISL_699656, EPI_ISL_699657 | 1-Laboratory of Microbiology, National Reference Lab, Charles Nicolle Hospital; 2-University of Tunis ElManar, Faculty of Medicine of Tunis, LR99ES09, Tunis, Tunisia | 1-Clinical and Experimental Pharmacology Lab, LR16SP02, National Center of Pharmacovigilance, University of Tunis El Manar, Tunis, Tunisia. 2-Neurodegenerative diseases and psychiatric troubles, LR18SP03, Razi Hospital, University of Tunis El Manar, Tunis, Tunisia. 3- Ministry of Health, National Observatory of New and Emerging Diseases, 1006, Tunis, Tunisia | Ilhem Boutiba-Ben Boubaker, Sameh Trabelsi, Nissaf Ben Alaya, Maher Kharrat, Alia Ben Kahla, Jalila Ben Khelil, Salma Abid, Sana Ferjani, Asma Ferjani, Mouna Ben Sassi, Mouna Safer, Guedi Berrabeh,Salwa Mrabet, Hanen ElJebari, Gaies Emna, Riadh Daghfous, Riadh Gouider. |
| EPI_ISL_699658, EPI_ISL_699659, EPI_ISL_699660, EPI_ISL_699661, EPI_ISL_699662, EPI_ISL_699663, EPI_ISL_699664, EPI_ISL_699665, EPI_ISL_699666, EPI_ISL_699667, EPI_ISL_699668, EPI_ISL_699669, EPI_ISL_699670, EPI_ISL_699671, EPI_ISL_699672, EPI_ISL_699674, EPI_ISL_699675, EPI_ISL_699676, EPI_ISL_699677, EPI_ISL_699678, EPI_ISL_699680, EPI_ISL_699681, EPI_ISL_699682, EPI_ISL_699683, EPI_ISL_699684, EPI_ISL_699685, EPI_ISL_699686, EPI_ISL_699687, EPI_ISL_699688, EPI_ISL_699691, EPI_ISL_699692, EPI_ISL_699693, EPI_ISL_699694, EPI_ISL_699695, EPI_ISL_699696, EPI_ISL_699697, EPI_ISL_699698, EPI_ISL_699699, EPI_ISL_699700, EPI_ISL_699701, EPI_ISL_699702, EPI_ISL_699703, EPI_ISL_699704, EPI_ISL_699705, EPI_ISL_699706, EPI_ISL_699709, EPI_ISL_699711, EPI_ISL_699712, EPI_ISL_699713, EPI_ISL_699714, EPI_ISL_699715, EPI_ISL_699723 | | | |
| see above | Hematopathology Laboratory, ACTREC, TMC | Hematopathology Laboratory, ACTREC, TMC | Hematopathology Laboratory, ACTREC |
| EPI_ISL_700371 | CHU Nantes | CNR Virus des Infections Respiratoires - France SUD | Antonin Bal, Louise Castain, Gregory Destras, Gwendolyne Burfin, Hadrien Règue, Quentin Semanas, Martine Valette, Bruno Lina, Virginie Ferré, Celine Bressollette, Laurence Josset |
| EPI_ISL_707791 | 1-Laboratory of Microbiology, National Reference Lab, Charles Nicolle Hospital; 2-University of Tunis ElManar, Faculty of Medicine of Tunis, LR99ES09, Tunis, Tunisia | 1-Clinical and Experimental Pharmacology Lab, LR16SP02, National Center of Pharmacovigilance, University of Tunis El Manar, Tunis, Tunisia. 2-Neurodegenerative diseases and psychiatric troubles, LR18SP03, Razi Hospital, University of Tunis El Manar, Tunis, Tunisia. 3- Ministry of Health, National Observatory of New and Emerging Diseases, 1006, Tunis, Tunisia | Ilhem Boutiba-Ben Boubaker, Sameh Trabelsi, Nissaf Ben Alaya, Maher Kharrat, Alia Ben Kahla, Jalila Ben Khelil, Salma Abid, Sana Ferjani, Mouna Ben Sassi, Mouna Safer, Awatef El MOussi, Habiba Ben Romdhane, Souissi Amira, Ines Mdini, Hanen El Jebari, Asma Ferjani, Gaies Emna, Riadh Daghfous, Riadh Gouider. |
| EPI_ISL_718132, EPI_ISL_718133, EPI_ISL_718136, EPI_ISL_718137, EPI_ISL_718138, EPI_ISL_718139, EPI_ISL_718140, EPI_ISL_718141, EPI_ISL_718142, EPI_ISL_718144, EPI_ISL_718145, EPI_ISL_718148, EPI_ISL_718149, EPI_ISL_718150, EPI_ISL_718151, EPI_ISL_718152, EPI_ISL_718153, EPI_ISL_718154, EPI_ISL_718155, EPI_ISL_718157, EPI_ISL_718158, EPI_ISL_718159, EPI_ISL_718160, EPI_ISL_718161, EPI_ISL_718162, EPI_ISL_718163, EPI_ISL_718164, EPI_ISL_718165, EPI_ISL_718166, EPI_ISL_718221 | | | |
| see above | Ministry of Health Hospitals | Institute of Health and Community Medicine | David Perera, Ooi Mong How, Chua Hock Hin, Tonnii Sia Loong Loong, Wong Jyn Shan, Wong Kiing Aik, Chan Chia Jui |
| EPI_ISL_718273 | Fujita Health University Okazaki Medical Center | Department of Virology and Parasitology, Fujita Health University School of Medicine | Takayuki Murata, Aki Sakurai, Satoshi Komoto, Tomihiko Ide,Takuma Ishihara, Masahiro suzuki, Yohei Doi |
| EPI_ISL_722899 | Dipartimento di Scienze Biomediche e Oncologia Umana - Azienda Ospedaliero Universitaria Consorziale Policlinico | Istituto Zooprofilattico Sperimentale della Puglia e della Basilicata | Parisi A., Bianco A., Capozzi L., Del Sambro L., Chironna M., Loconsole D. |
| EPI_ISL_727998, EPI_ISL_727999 | Fujita Health University Okazaki Medical Center | Department of Virology and Parasitology, Fujita Health University School of Medicine | Takayuki Murata, Yohei Doi, Takuma Ishihara, Masahiro suzuki, Aki Sakurai, Satoshi Komoto, Tomihiko Ide |
| EPI_ISL_728000 | Fujita Health University Okazaki Medical Center | Department of Virology and Parasitology, Fujita Health University School of Medicin | Takayuki Murata, Yohei Doi, Takuma Ishihara, Masahiro suzuki, Aki Sakurai, Satoshi Komoto, Tomihiko Ide |
| EPI_ISL_728153, EPI_ISL_728154, EPI_ISL_728155 | Fujita Health University Okazaki Medical Center | Department of Virology and Parasitology, Fujita Health University School of Medicine | Takayuki Murata, Yohei Doi, Takuma Ishihara, Masahiro suzuki, Aki Sakurai, Satoshi Komoto, Tomihiko Ide |
| EPI_ISL_728156, EPI_ISL_728159 | Fujita Health University Okazaki Medical Center | Department of Virology and Parasitology, Fujita Health University School of Medicine | Takayuki Murata, Aki Sakurai, Satoshi Komoto, Tomihiko Ide,Takuma Ishihara, Masahiro suzuki, Yohei Doi |
| EPI_ISL_728206, EPI_ISL_728207, EPI_ISL_728208 | Fujita Health University Okazaki Medical Center | Fujita health University, Department of Microbiology | Takayuki Murata, Aki Sakurai, Satoshi Komoto, Tomihiko Ide,Takuma Ishihara, Masahiro suzuki, Yohei Doi |
| EPI_ISL_729797, EPI_ISL_729798 | Laboratório Central de Saúde Pública do Estado do Rio Grande do Sul (LACEN-RS) | Laboratory of Respiratory Viruses and Measles, Oswaldo Cruz Institute, FIOCRUZ | Paola Resende, Luciana Appolinario, Fernando Motta, Anna Carolina Paixão, Ana Carolina Mendonça, Tatiana Schaffer Gregianini, Marilda Tereza Mar da Rosa, Marilda Siqueira on behalf of the Fiocruz COVID-19 Genomic Surveillance Network |
| EPI_ISL_732556, EPI_ISL_732557, EPI_ISL_732558 | Bundeswehr Institute of Microbiology | Bundeswehr Institute of Microbiology | Elham Khatamzas, Markus Antwerpen, Mathias Walter, Alexandra Rehn, Sabine Zange, Enrico Georgi, Michael von Bergwelt-Baildon, Roman Wölfel |
| EPI_ISL_735260, EPI_ISL_735261, EPI_ISL_735262, EPI_ISL_735263, EPI_ISL_735264, EPI_ISL_735265, EPI_ISL_735266, EPI_ISL_735267, EPI_ISL_735268, EPI_ISL_735269, EPI_ISL_735270, EPI_ISL_735271, EPI_ISL_735272, EPI_ISL_735273, EPI_ISL_735274, EPI_ISL_735275, EPI_ISL_735276, EPI_ISL_735277, EPI_ISL_735278, EPI_ISL_735279, EPI_ISL_735280, EPI_ISL_735281, EPI_ISL_735282, EPI_ISL_735283, EPI_ISL_735284, EPI_ISL_735285, EPI_ISL_735286, EPI_ISL_735288, EPI_ISL_735289, EPI_ISL_735290, EPI_ISL_735292, EPI_ISL_735293, EPI_ISL_735294, EPI_ISL_735295, EPI_ISL_735296, EPI_ISL_735297, EPI_ISL_735298, EPI_ISL_735299, EPI_ISL_735300, EPI_ISL_735301, EPI_ISL_735302, EPI_ISL_735303, EPI_ISL_735304, EPI_ISL_735305 | | | |
| see above | Genomic Laboratory (GLAB) (Conjoint lab of Health Directorate of Istanbul and Istanbul Technical University) | Genomic Laboratory (GLAB), Istanbul Technical University | Ilker Karacan, Tugba Kizilboga Akgun, Nihat Bugra Agaoglu, Payam Zolfagharian, Mehtap Aydin, Gizem Alkurt, Jale Yildiz, Betsi Kose, Nisan Denizce Can, Ayse Serra Ozel, Nilsun Altunal, Arzu Irvem, Yasemin Kendir Demirkol, Ozlem Akgun Dogan, Levent Doganay, Gizem Dinler Doganay |
| EPI_ISL_735399 | Hospital Municiplal Dr Ignacio de gouvea | Instituto Adolfo Lutz, Interdiciplinary Procedures Center, Strategic Laboratory | Claudio Tavares Sacchi, Claudia Regina Gonçalves, Erica Valessa Ramos Gomes, Karoline Rodrigues Campos |
| EPI_ISL_738147, EPI_ISL_738194, EPI_ISL_738243 | Microbiology and Virology Unit, Florence Careggi University Hospital | Microbiology and Virology Unit, Florence Careggi University Hospital | Vincenzo Di Pilato, Marco Coppi, Alberto Antonelli, Simona Pollini, Gian Maria Rossolini |
| EPI_ISL_751297, EPI_ISL_751301 | Devki Devi Foundation, a unit of Max Healthcare | CSIR-IGIB/Max | Rajesh Pandey#, Samreen Siddiqui, Janani Srinivasa Vasudevan, Akshay Kanakan, Ranjeet Maurya, Uzma Shamim, Bansidhar Tarai, Akansha Tyagi, Mitali Mukerji, Poonam Das, Sujeet Jha, Mohammed Faruq, Anurag Agrawal |
| EPI_ISL_751320, EPI_ISL_751322, EPI_ISL_751333, EPI_ISL_751334, EPI_ISL_751336, EPI_ISL_751347, EPI_ISL_751348, EPI_ISL_751349, EPI_ISL_751350, EPI_ISL_751351, EPI_ISL_751352, EPI_ISL_751353, EPI_ISL_751355, EPI_ISL_751356, EPI_ISL_751358, EPI_ISL_751359, EPI_ISL_751360, EPI_ISL_751361, | | | |

EPI_ISL_751362, EPI_ISL_751363, EPI_ISL_751364, EPI_ISL_751365, EPI_ISL_751366, EPI_ISL_751367, EPI_ISL_751368, EPI_ISL_751369, EPI_ISL_751370, EPI_ISL_751371, EPI_ISL_751372, EPI_ISL_751373, EPI_ISL_751374, EPI_ISL_751375, EPI_ISL_751376, EPI_ISL_751377, EPI_ISL_751378, EPI_ISL_751380, EPI_ISL_751381, EPI_ISL_751382, EPI_ISL_751383, EPI_ISL_751384, EPI_ISL_751386, EPI_ISL_751387, EPI_ISL_751388, EPI_ISL_751389, EPI_ISL_751390, EPI_ISL_751391, EPI_ISL_751392, EPI_ISL_751393, EPI_ISL_751394, EPI_ISL_751395, EPI_ISL_751397, EPI_ISL_751398, EPI_ISL_751399, EPI_ISL_751400, EPI_ISL_751401, EPI_ISL_751402, EPI_ISL_751403, EPI_ISL_751404, EPI_ISL_751405, EPI_ISL_751407, EPI_ISL_751408, EPI_ISL_751409, EPI_ISL_751410, EPI_ISL_751411, EPI_ISL_751413, EPI_ISL_751414, EPI_ISL_751415, EPI_ISL_751416, EPI_ISL_751417, EPI_ISL_751418, EPI_ISL_751419, EPI_ISL_751420, EPI_ISL_751421, EPI_ISL_751422, EPI_ISL_751423, EPI_ISL_751427, EPI_ISL_751429, EPI_ISL_751430, EPI_ISL_751431, EPI_ISL_751435, EPI_ISL_751439, EPI_ISL_751440, EPI_ISL_751442, EPI_ISL_751444, EPI_ISL_751445, EPI_ISL_751446

| | | | |
|---|---|---|---|
| see above | IRCCS Sacro Cuore Don Calabria Hospital, Department of Infectious, Tropical Diseases & Microbiology | University of Verona, Department of Biotechnology | Antonio Mori, Michela Deiana, Elena Pomari, Chiara Piubelli; Giulia Lopatriello, Luca Marcolungo, Cristina Beltrami, Chiara Degli Esposti, Emanuela Cosentino, Massimo Delledonne |
| EPI_ISL_759955 | Primasatya Husada Citra Hospital | Institute of Tropical Disease, Universitas Airlangga | Aldise M Nastri, Jezzy R Dewantari, Rima R Prasetya, Krisnoadi Rahardjo, Pudji Djanuartono, Gatot Soegiarto, Laksmi Wulandari, Resti Yudhawati, Soetjipto, Yasuko Mori, Maria I Lusida, Kazufumi Shimizu |
| EPI_ISL_759956 | Husada Utama Hospital | Institute of Tropical Disease, Universitas Airlangga | Jezzy R Dewantari, Rima R Prasetya, Krisnoadi Rahardjo, Aldise M Nastri, Didi Dewanto, Gatot Soegiarto, Laksmi Wulandari, Resti Yudhawati, Soetjipto, Yasuko Mori, Maria I Lusida, Kazufumi Shimizu |
| EPI_ISL_759957 | Siti Khodijah Hospital | Institute of Tropical Disease, Universitas Airlangga | Rima R Prasetya, Krisnoadi Rahardjo, Aldise M Nastri, Jezzy R Dewantari, Muhammad Hamdan, Gatot Soegiarto, Laksmi Wulandari, Resti Yudhawati, Soetjipto, Yasuko Mori, Maria I Lusida, Kazufumi Shimizu |
| EPI_ISL_759958 | Universitas Airlangga Hospital | Institute of Tropical Disease, Universitas Airlangga | Maria I Lusida, Krisnoadi Rahardjo, Aldise M Nastri, Jezzy R Dewantari, Rima R Prasetya, Nasronudin, Gatot Soegiarto, Laksmi Wulandari, Resti Yudhawati, Soetjipto, Yasuko Mori, Kazufumi Shimizu |
| EPI_ISL_759959 | Premier Hospital Surabaya | Institute of Tropical Disease, Universitas Airlangga | Krisnoadi Rahardjo, Aldise M Nastri, Jezzy R Dewantari, Rima R Prasetya, Hartono Tanto, Gatot Soegiarto, Laksmi Wulandari, Resti Yudhawati, Soetjipto, Yasuko Mori, Maria I Lusida, Kazufumi Shimizu |
| EPI_ISL_759961 | Universitas Airlangga Hospital | Institute of Tropical Disease, Universitas Airlangga | Jezzy R Dewantari, Rima R Prasetya, Krisnoadi Rahardjo, Aldise M Nastri, Nasronudin, Gatot Soegiarto, Laksmi Wulandari, Resti Yudhawati, Soetjipto, Yasuko Mori, Maria I Lusida, Kazufumi Shimizu |
| EPI_ISL_759962 | RSUD Dr. Soetomo | Institute of Tropical Disease, Universitas Airlangga | Krisnoadi Rahardjo, Aldise M Nastri, Jezzy R Dewantari, Rima R Prasetya, Joni Wahyuhadi, Gatot Soegiarto, Laksmi Wulandari, Resti Yudhawati, Yasuko Mori, Soetjipto, Kazufumi Shimizu ,Maria I Lusida |
| EPI_ISL_759964 | Mitra Keluarga Hospital Waru | Institute of Tropical Disease, Universitas Airlangga | Jezzy R Dewantari, Rima R Prasetya, Krisnoadi Rahardjo, Aldise M Nastri, Christina Dian Anggraeni, Gatot Soegiarto, Laksmi Wulandari, Resti Yudhawati, Yasuko Mori, Soetjipto, Kazufumi Shimizu ,Maria I Lusida |
| EPI_ISL_759965 | RSUD Dr. Soetomo | Institute of Tropical Disease, Universitas Airlangga | Rima R Prasetya, Krisnoadi Rahardjo, Aldise M Nastri, Jezzy R Dewantari, Joni Wahyuhadi, Gatot Soegiarto, Laksmi Wulandari, Resti Yudhawati, Yasuko Mori, Soetjipto, Kazufumi Shimizu ,Maria I Lusida |
| EPI_ISL_759966 | RSUD Sidoarjo | Institute of Tropical Disease, Universitas Airlangga | Krisnoadi Rahardjo, Aldise M Nastri, Jezzy R Dewantari, Rima R Prasetya, Atok Irawan, Gatot Soegiarto, Laksmi Wulandari, Resti Yudhawati, Yasuko Mori, Soetjipto, Kazufumi Shimizu ,Maria I Lusida |
| EPI_ISL_759967 | RSUD Dr. Iskak Tulungagung | Institute of Tropical Disease, Universitas Airlangga | Rima R Prasetya, Krisnoadi Rahardjo, Aldise M Nastri, Jezzy R Dewantari, Supriyanto Dharmoredjo, Gatot Soegiarto, Laksmi Wulandari, Resti Yudhawati, Soetjipto, Yasuko Mori, Maria I Lusida, Kazufumi Shimizu |
| EPI_ISL_766032 | RSUP Dr. Sardjito | Eijkman Institute for Molecular Biology, Ministry of Research and Technology/National Agency for Research and Innovation | Frilasita A Yudhaputri, Edison Johar, Hidayat Trimarsanto, David H Muljono, Safarina G Malik, Khin Saw Myint, Amin Soebandrio |
| EPI_ISL_766034 | RS Mitra Keluarga Bintaro | Eijkman Institute for Molecular Biology, Ministry of Research and Technology/National Agency for Research and Innovation | Frilasita A Yudhaputri, Edison Johar, Hidayat Trimarsanto, David H Muljono, Safarina G Malik, Khin Saw Myint, Amin Soebandrio |
| EPI_ISL_766035 | RS Bhayangkara Brimob Kelapa Dua | Eijkman Institute for Molecular Biology, Ministry of Research and Technology/National Agency for Research and Innovation | Frilasita A Yudhaputri, Edison Johar, Hidayat Trimarsanto, David H Muljono, Safarina G Malik, Khin Saw Myint, Amin Soebandrio |
| EPI_ISL_766036 | RS Mitra Keluarga Bintaro | Eijkman Institute for Molecular Biology, Ministry of Research and Technology/National Agency for Research and Innovation | Frilasita A Yudhaputri, Edison Johar, Hidayat Trimarsanto, David H Muljono, Safarina G Malik, Khin Saw Myint, Amin Soebandrio |
| EPI_ISL_766037 | Puskesmas Palmerah | Eijkman Institute for Molecular Biology, Ministry of Research and Technology/National Agency for Research and Innovation | Frilasita A Yudhaputri, Edison Johar, Hidayat Trimarsanto, David H Muljono, Safarina G Malik, Khin Saw Myint, Amin Soebandrio |
| EPI_ISL_766038 | RSJPD Harapan Kita | Eijkman Institute for Molecular Biology, Ministry of Research and Technology/National Agency for Research and Innovation | Frilasita A Yudhaputri, Edison Johar, Hidayat Trimarsanto, David H Muljono, Safarina G Malik, Khin Saw Myint, Amin Soebandrio |
| EPI_ISL_766039 | RS Bhayangkara Tk. III | Eijkman Institute for Molecular Biology, Ministry of Research and Technology/National Agency for Research and Innovation | Frilasita A Yudhaputri, Edison Johar, Hidayat Trimarsanto, David H Muljono, Safarina G Malik, Khin Saw Myint, Amin Soebandrio |
| EPI_ISL_766040 | RS Tingkat II Moh. Ridwan Meuraksa | Eijkman Institute for Molecular Biology, Ministry of Research and Technology/National Agency for Research and Innovation | Frilasita A Yudhaputri, Edison Johar, Hidayat Trimarsanto, David H Muljono, Safarina G Malik, Khin Saw Myint, Amin Soebandrio |
| EPI_ISL_766041 | RS Polri | Eijkman Institute for Molecular Biology, Ministry of Research and Technology/National Agency for Research and Innovation | Frilasita A Yudhaputri, Edison Johar, Hidayat Trimarsanto, David H Muljono, Safarina G Malik, Khin Saw Myint, Amin Soebandrio |
| EPI_ISL_766042 | RS Sari Asih Ciledug | Eijkman Institute for Molecular Biology, Ministry of Research and Technology/National Agency for Research and Innovation | Frilasita A Yudhaputri, Edison Johar, Hidayat Trimarsanto, David H Muljono, Safarina G Malik, Khin Saw Myint, Amin Soebandrio |
| EPI_ISL_766043 | RS Pelni | Eijkman Institute for Molecular Biology, Ministry of Research and Technology/National Agency for Research and Innovation | Frilasita A Yudhaputri, Edison Johar, Hidayat Trimarsanto, David H Muljono, Safarina G Malik, Khin Saw Myint, Amin Soebandrio |
| EPI_ISL_766044 | RSUP Fatmawati | Eijkman Institute for Molecular Biology, Ministry of Research and Technology/National Agency for Research and Innovation | Frilasita A Yudhaputri, Edison Johar, Hidayat Trimarsanto, David H Muljono, Safarina G Malik, Khin Saw Myint, Amin Soebandrio |
| EPI_ISL_766045 | RS Pelni | Eijkman Institute for Molecular Biology, Ministry of Research and Technology/National Agency for Research and Innovation | Frilasita A Yudhaputri, Edison Johar, Hidayat Trimarsanto, David H Muljono, Safarina G Malik, Khin Saw Myint, Amin Soebandrio |
| EPI_ISL_766048 | RS Bhayangkara Tk. III | Eijkman Institute for Molecular Biology, Ministry of Research and Technology/National Agency for Research and Innovation | Frilasita A Yudhaputri, Edison Johar, Hidayat Trimarsanto, David H Muljono, Safarina G Malik, Khin Saw Myint, Amin Soebandrio |
| EPI_ISL_767043 | University of Bari Biomedical Sciences and Human Oncology | University of Bari Biomedical Sciences and Human Oncology | Maria Chironna, Anna Sallustio, Daniela Loconsole, Marisa Accogli |
| EPI_ISL_767405 | Ochsner Health | BioInfo Experts, LLC | Rebecca Rose, Amy Feehan, David J. Nolan, Sissy Cross, David Moraga Amador, Tong Yang, Luke Caruso, Wayra Navia, Lydia Von Borstel, Xiao Hui Zhou, |

| | | | |
|---|---|---|---|
| | | | Julia-Garcia-Diaz, Susanna L. Lamers |
| EPI_ISL_775259 | INT Fondazione Pascale | INT Fondazione Pascale | INT Fondazione Pascale |
| EPI_ISL_776750, EPI_ISL_776751, EPI_ISL_776752, EPI_ISL_776753, EPI_ISL_776754, EPI_ISL_776755, EPI_ISL_776756 | Instituto Adolfo Lutz - Central | Instituto Adolfo Lutz, Interdiciplinary Procedures Center, Strategic Laboratory | Claudio Tavares Sacchi, Claudia Regina Gonçalves, Erica Valessa Ramos Gomes, Karoline Rodrigues Campos |
| EPI_ISL_776757, EPI_ISL_776758 | Instituto Adolfo Lutz - Regional de Marilia | Instituto Adolfo Lutz, Interdiciplinary Procedures Center, Strategic Laboratory | Claudio Tavares Sacchi, Claudia Regina Gonçalves, Erica Valessa Ramos Gomes, Karoline Rodrigues Campos |
| EPI_ISL_776759, EPI_ISL_776760, EPI_ISL_776761, EPI_ISL_776762, EPI_ISL_776763 | Instituto Adolfo Lutz - Central | Instituto Adolfo Lutz, Interdiciplinary Procedures Center, Strategic Laboratory | Claudio Tavares Sacchi, Claudia Regina Gonçalves, Erica Valessa Ramos Gomes, Karoline Rodrigues Campos |
| EPI_ISL_779704, EPI_ISL_779709, EPI_ISL_779712, EPI_ISL_779713, EPI_ISL_779714, EPI_ISL_780378, EPI_ISL_780381, EPI_ISL_780413 | Laboratory of Infectious Diseases, Department of Biomedical and Clinical Sciences L. Sacco, University of Milan | Laboratory of Infectious Diseases, Department of Biomedical and Clinical Sciences L. Sacco, University of Milan | Alessia Lai, Annalisa Bergna, Carla Della Ventura, Claudia Balotta, Massimo Galli, Gianguglielmo Zehender on behalf of SARS-CoV-2 ITALIAN RESEARCH ENTERPRISE-(SCIRE) Collaborative Group |
| EPI_ISL_788944, EPI_ISL_788945, EPI_ISL_788947, EPI_ISL_788962, EPI_ISL_788963, EPI_ISL_788964, EPI_ISL_788966, EPI_ISL_788967 | Ospedale "Di Venere" | Beaconlab (Bioinformatics, Evolution and Comparative Genomics lab), Dept of Biosciences, University on Milan | Iacobellis M, d'Avenia M, Piluscio R, Parisi A, Chiara M, Manzari C, Pesole G |
| EPI_ISL_792101, EPI_ISL_792102 | Instituto Adolfo Lutz - Central | Instituto Adolfo Lutz, Interdiciplinary Procedures Center, Strategic Laboratory | Claudio Tavares Sacchi, Claudia Regina Gonçalves, Erica Valessa Ramos Gomes, Karoline Rodrigues Campos |
| EPI_ISL_792103 | Instituto Adolfo Lutz - Regional de Santo Andre | Instituto Adolfo Lutz, Interdiciplinary Procedures Center, Strategic Laboratory | Claudio Tavares Sacchi, Claudia Regina Gonçalves, Erica Valessa Ramos Gomes, Karoline Rodrigues Campos |
| EPI_ISL_792104, EPI_ISL_792105, EPI_ISL_792106, EPI_ISL_792107, EPI_ISL_792108, EPI_ISL_792109, EPI_ISL_792110, EPI_ISL_792111, EPI_ISL_792112, EPI_ISL_792113, EPI_ISL_792114 | | | |
| see above | Instituto Adolfo Lutz - Central | Instituto Adolfo Lutz, Interdiciplinary Procedures Center, Strategic Laboratory | Claudio Tavares Sacchi, Claudia Regina Gonçalves, Erica Valessa Ramos Gomes, Karoline Rodrigues Campos |
| EPI_ISL_794745 | Istituto Zooprofilattico Sperimentale della Puglia e della Basilicata | Istituto Zooprofilattico Sperimentale della Puglia e della Basilicata | Parisi A., Bianco A., Capozzi L., Del Sambro L., Manzulli V, Rondinone V., Pace L., Cipolletta D., Galante D. |
| EPI_ISL_807150 | Makmal Kesihatan Awam Kebangsaan, Kementerian Kesihatan Malaysia. | Integrative Pharmacogenomics Institute (iPROMISE) | Mohd Zaki Salleh, Teh Lay Kek, Ariza Adnan, Sazzli Shahlan Kassim, Siti Hamimah Sheikh Abdul Kadir, Fadzilah Mohd Nor, Mohd Nur Fakhruzzaman Noorizhab, Siti Farah Alwani Mohd Nawi, Wang Seok Mui, Hani Mat Hussin, Nor Zahrin Hasran, Lim Wai Feng |
| EPI_ISL_807151, EPI_ISL_807153 | Makmal Kesihatan Awam Kebangsaan, Kementerian Kesihatan Malaysia | Integrative Pharmacogenomics Institute (iPROMISE) | Mohd Zaki Salleh, Teh Lay Kek, Ariza Adnan, Sazzli Shahlan Kassim, Siti Hamimah Sheikh Abdul Kadir, Fadzilah Mohd Nor, Mohd Nur Fakhruzzaman Noorizhab, Siti Farah Alwani Mohd Nawi, Wang Seok Mui, Hani Mat Hussin, Nor Zahrin Hasran, Lim Wai Feng |
| EPI_ISL_811148 | Laboratorio de Ecologia de Doencas Transmissiveis na Amazonia, Instituto Leonidas e Maria Deane - Fiocruz Amazonia | Laboratorio de Ecologia de Doencas Transmissiveis na Amazonia, Instituto Leonidas e Maria Deane - Fiocruz Amazonia | Valdinete Nascimento, Victor Souza, André Corado, Fernanda Nascimento, George Silva, Ágatha Costa, Debora Duarte, Luciana Gonçalves, Matilde Mejía, Karina Pessoa, Maria Júlia Brandão, Michele Jesus, Felipe Naveca on behalf of the Fiocruz COVID-19 Genomic Surveillance Network |
| EPI_ISL_832178 | Hospital | National Reference Center for Viruses of Respiratory Infections, Institut Pasteur, Paris | Marion Barbet, Sylvie Behillil, Méline Bizard, Angela Brisebarre, Camille Capel, Etienne Simon-Lorière, Vincent Enouf, Maud Vanpeene, Sylvie van der Werf, Laurent Roudiere |
| EPI_ISL_832198 | Laboratory Analyses Med | National Reference Center for Viruses of Respiratory Infections, Institut Pasteur, Paris | Marion Barbet, Sylvie Behillil, Méline Bizard, Angela Brisebarre, Camille Capel, Etienne Simon-Lorière, Vincent Enouf, Maud Vanpeene, Sylvie van der Werf, Elise Liscoet |
| EPI_ISL_832202 | Hospital | National Reference Center for Viruses of Respiratory Infections, Institut Pasteur, Paris | Marion Barbet, Sylvie Behillil, Méline Bizard, Angela Brisebarre, Camille Capel, Etienne Simon-Lorière, Vincent Enouf, Maud Vanpeene, Sylvie van der Werf, Patricia Stoessel-Thouvenin |
| EPI_ISL_833152, EPI_ISL_833153, EPI_ISL_833154, EPI_ISL_833155 | Instituto Adolfo Lutz - Central | Instituto Adolfo Lutz, Interdiciplinary Procedures Center, Strategic Laboratory | Claudio Tavares Sacchi, Claudia Regina Gonçalves, Erica Valessa Ramos Gomes, Karoline Rodrigues Campos |
| EPI_ISL_833200 | Hôpital Bichat Claude Bernard, Laboratoire de Virologie | IAME UMR1137 Inserm, Université de Paris, Hôpital Bichat | Antoine Bridier, Amélie Recoing, Quentin Le Hingrat, Lena Daniel, Siham Hamri, Gilles Collin, Alexandre Storto, Mélanie Bertine, Charlotte Charpentier, Nadhira Houhou-Fidouh, Diane Descamps, Benoit Visseaux |
| EPI_ISL_837565, EPI_ISL_837566, EPI_ISL_837568, EPI_ISL_837570, EPI_ISL_837574, EPI_ISL_837575, EPI_ISL_837577 | Centro Nacional de Enfermedades Tropicales (CENETROP) | Laboratory of Respiratory Viruses and Measles, Oswaldo Cruz Institute, FIOCRUZ | Paola Resende, Roxana Loayza, Cinthia Avila, Luciana Appolinario, Fernando Motta, Anna Carolina Paixao, Ana Carolina Mendonca, Marilda Siqueira on behalf of the Fiocruz COVID-19 Genomic Surveillance Network |
| EPI_ISL_837604, EPI_ISL_837605, EPI_ISL_837606, EPI_ISL_837608, EPI_ISL_837609, EPI_ISL_837610, EPI_ISL_837611, EPI_ISL_837612, EPI_ISL_837613, EPI_ISL_837614, EPI_ISL_837615, EPI_ISL_837616, EPI_ISL_837617, EPI_ISL_837618, EPI_ISL_837619, EPI_ISL_837620, EPI_ISL_837621, EPI_ISL_837622, EPI_ISL_837623, EPI_ISL_837624, EPI_ISL_837625, EPI_ISL_837626, EPI_ISL_837627, EPI_ISL_837628, EPI_ISL_837629, EPI_ISL_837630, EPI_ISL_837631, EPI_ISL_837632, EPI_ISL_837633, EPI_ISL_837634, EPI_ISL_837635, EPI_ISL_837636, EPI_ISL_837637, EPI_ISL_837638, EPI_ISL_837639, EPI_ISL_837640, EPI_ISL_837641, EPI_ISL_837642, EPI_ISL_837643, EPI_ISL_837644, EPI_ISL_837645, EPI_ISL_837646, EPI_ISL_837647, EPI_ISL_837648, EPI_ISL_837649, EPI_ISL_837650, EPI_ISL_837651, EPI_ISL_837652, EPI_ISL_837653, EPI_ISL_837654, EPI_ISL_837655, EPI_ISL_837656, EPI_ISL_837657, EPI_ISL_837658, EPI_ISL_837659, EPI_ISL_837660, EPI_ISL_837661, EPI_ISL_837662, EPI_ISL_837663, EPI_ISL_837664, EPI_ISL_837665, EPI_ISL_837666, EPI_ISL_837667, EPI_ISL_837669, EPI_ISL_837670, EPI_ISL_837671, EPI_ISL_837672, EPI_ISL_837673, EPI_ISL_837674, EPI_ISL_837675, EPI_ISL_837676, EPI_ISL_837677, EPI_ISL_837678, EPI_ISL_837679, EPI_ISL_837680, EPI_ISL_837681, EPI_ISL_837682, EPI_ISL_837683, EPI_ISL_837684, EPI_ISL_837685, EPI_ISL_837686, EPI_ISL_837687, EPI_ISL_837688, EPI_ISL_837689, EPI_ISL_837690, EPI_ISL_837691, EPI_ISL_837692, EPI_ISL_837693, EPI_ISL_837694, EPI_ISL_837695, EPI_ISL_837696, EPI_ISL_837697 | | | |
| see above | Instituto Nacional de Enfermedades Respiratorias (INER) | Instituto Nacional de Enfermedades Respiratorias (INER) | Celia Boukadida, Margarita Matías-Florentino, Alma Rincón-Rubio, Hector Esteban Paz-Juárez, Olivia Briceño, Edgar Sevilla-Reyes, Fidencio Mejía-Nepomuceno, Mario Mújica-Sánchez, Eduardo Becerril-Vargas, José Arturo Martínez-Orozco, Alejandra Hernández-Terán, Jorge Salas-Hernández, Santiago Ávila-Ríos, Joel Armando Vázquez-Pérez |
| EPI_ISL_848624 | Evandro Chagas Institute | Evandro Chagas Institute | Santos, M.C.; Silva, A.M.; Junior, W.D.C.; Barbagelata, L.S.; Ferreira, J.A.; Sousa, E.M.A.; da Silva, P.S.; Pinheiro, K.C.; L.C.; Sousa Junior, E.C. |
| EPI_ISL_851048 | Gandhi Medical College and Hospital | CSIR-Centre for Cellular and Molecular Biology | Nagamani Kammili, Winnie Thomas, Madhavi Latha Manolla, Shailaja VV, Sudhamadhuri Devara, Vanisree Rajoli, Archana GJ, Sushma Rajyalakshmi Gudiseva, Sunitha Pakalapati, Manisha Rani, Amrithesh Kumar, Raja Rao Mesepogu, Vinay Shekar Reddy, Thrilok Chander Bingi, Sofia Banu, Divya Tej Sowpati |
| EPI_ISL_851049 | Gandhi Medical College and Hospital | CSIR-Centre for Cellular and Molecular Biology | Nagamani Kammili, Madhavi Latha Manolla, Winnie Thomas, Shailaja VV, Sudhamadhuri Devara, Vanisree Rajoli, Archana GJ,Sushma Rajyalakshmi Gudiseva, Sunitha Pakalapati, Manisha Rani, Amrithesh Kumar, Raja Rao Mesepogu, Vinay Shekar Reddy, Thrilok Chander Bingi, Sofia Banu, Divya Tej Sowpati |
| EPI_ISL_853820, EPI_ISL_853821 | Department of Pathology, Landeskrankenhaus Graz II, Medical University Graz | Bergthaler laboratory, CeMM Research Center for Molecular Medicine of the Austrian Academy of Sciences | Lukas Endler, Alexandra Popa, Benedikt Agerer, Jakob-Wendelin Genger, Alexander Lercher, Anna Schedl, Thomas Penz, Michael Schuster, Jan Laine, Martin Senekowitsch, Christoph Bock, Andreas Bergthaler |
| EPI_ISL_856695 | University of Zambia, School of Veterinary Medicine, Disease Control | University of Zambia, School of Veterinary Medicine, Disease Control | Simulundu,E., Kapata,N., Mupeta,F., Kapata,P.C., Saasa,N., Changula,K., Muleya,W., Chitanga,S., Chambaro,H., Mubemba,B., Masahiro,K., Chanda,D., Mulenga,L., Fwoloshi,S., Shibemba,A.L., Kapaya,F., Zulu,P., Musonda,K., Monze,M., Sinyange,N., Liwewe,M.M., Kapin'a,M., Chipimo,P.J., Ngosa,W., Morales,A.N., Kayeyi,N., Malama,K., Tembo,J., Bates,M., Sawa,H., Takada,A., Nalubamba,K.S., Mukonka,V., Chilufya,C. and Zumla,A. |
| EPI_ISL_856971 | Laboratory of Histology-Embryology, Molecular Carcinogenesis Group Medical School National and Kapodistrian University of Athens | Prof. Gorgoulis Lab | Gorgoulis V.G. |
| EPI_ISL_861419 | Instituto de Diagnostico y Referencia Epidemiologicos | Instituto de Diagnostico y Referencia Epidemiologicos | Fabiola Garces-Ayala, Gisela Barrera-Badillo , Abril Rodriguez-Maldonado, Claudia Wong-Arambula , Natividad Cruz-Ortiz, Tatiana Nunez-Garcia, Dayanira |

| | | | |
|---|---|---|---|
| | (INDRE) | (INDRE) | Arellano-Suarez, Adnan-Araiza Rodríguez, Edgar Mendieta-Condado, Lucia Hernandez-Rivas, Irma Lopez-Martinez, Ernesto Ramirez-Gonzalez. |
| EPI_ISL_861420 | Instituto de Diagnostico y Referencia Epidemiologicos (INDRE) | Instituto de Diagnostico y Referencia Epidemiologicos (INDRE) | Abril Rodriguez-Maldonado, Gisela Barrera-Badillo ,Claudia Wong-Arambula , Natividad Cruz-Ortiz, Tatiana Nunez-Garcia, Dayanira Arellano-Suarez, Fabiola Garces-Ayala, Adnan-Araiza Rodríguez, Edgar Mendieta-Condado, Lucia Hernandez-Rivas, Irma Lopez-Martinez, Ernesto Ramirez-Gonzalez. |
| EPI_ISL_861625, EPI_ISL_861626, EPI_ISL_861627 | Instituto Adolfo Lutz - Central | Instituto Adolfo Lutz, Interdiciplinary Procedures Center, Strategic Laboratory | Claudio Tavares Sacchi, Claudia Regina Gonçalves, Erica Valessa Ramos Gomes, Karoline Rodrigues Campos |
| EPI_ISL_861628 | Laboratorio Municipal de Guarulhos | Instituto Adolfo Lutz, Interdiciplinary Procedures Center, Strategic Laboratory | Claudio Tavares Sacchi, Claudia Regina Gonçalves, Erica Valessa Ramos Gomes, Karoline Rodrigues Campos |
| EPI_ISL_861629, EPI_ISL_861630, EPI_ISL_861631, EPI_ISL_861632, EPI_ISL_861633, EPI_ISL_861634 | Instituto Adolfo Lutz - Central | Instituto Adolfo Lutz, Interdiciplinary Procedures Center, Strategic Laboratory | Claudio Tavares Sacchi, Claudia Regina Gonçalves, Erica Valessa Ramos Gomes, Karoline Rodrigues Campos |
| EPI_ISL_861635 | Hospital e Maternidade Madre Theodora | Instituto Adolfo Lutz, Interdiciplinary Procedures Center, Strategic Laboratory | Claudio Tavares Sacchi, Claudia Regina Gonçalves, Erica Valessa Ramos Gomes, Karoline Rodrigues Campos |
| EPI_ISL_861636 | Hospital Geral de Sao Mateus São Paulo | Instituto Adolfo Lutz, Interdiciplinary Procedures Center, Strategic Laboratory | Claudio Tavares Sacchi, Claudia Regina Gonçalves, Erica Valessa Ramos Gomes, Karoline Rodrigues Campos |
| EPI_ISL_861637 | Hospital Nipo Brasileiro | Instituto Adolfo Lutz, Interdiciplinary Procedures Center, Strategic Laboratory | Claudio Tavares Sacchi, Claudia Regina Gonçalves, Erica Valessa Ramos Gomes, Karoline Rodrigues Campos |
| EPI_ISL_861638 | Hospital Municipal Dr. Moyses Deutsch | Instituto Adolfo Lutz, Interdiciplinary Procedures Center, Strategic Laboratory | Claudio Tavares Sacchi, Claudia Regina Gonçalves, Erica Valessa Ramos Gomes, Karoline Rodrigues Campos |
| EPI_ISL_861639 | Hospital Sao Paulo de Ensino da Unifesp | Instituto Adolfo Lutz, Interdiciplinary Procedures Center, Strategic Laboratory | Claudio Tavares Sacchi, Claudia Regina Gonçalves, Erica Valessa Ramos Gomes, Karoline Rodrigues Campos |
| EPI_ISL_861640, EPI_ISL_861641 | Hospital Municipal Dr. Moyses Deutsch | Instituto Adolfo Lutz, Interdiciplinary Procedures Center, Strategic Laboratory | Claudio Tavares Sacchi, Claudia Regina Gonçalves, Erica Valessa Ramos Gomes, Karoline Rodrigues Campos |
| EPI_ISL_861642, EPI_ISL_861643 | Instituto Adolfo Lutz - Central | Instituto Adolfo Lutz, Interdiciplinary Procedures Center, Strategic Laboratory | Claudio Tavares Sacchi, Claudia Regina Gonçalves, Erica Valessa Ramos Gomes, Karoline Rodrigues Campos |
| EPI_ISL_861644 | Hospital Santa Virginia | Instituto Adolfo Lutz, Interdiciplinary Procedures Center, Strategic Laboratory | Claudio Tavares Sacchi, Claudia Regina Gonçalves, Erica Valessa Ramos Gomes, Karoline Rodrigues Campos |
| EPI_ISL_861645 | Hospital e Pronto Socorro Comunitario Vila Iolanda | Instituto Adolfo Lutz, Interdiciplinary Procedures Center, Strategic Laboratory | Claudio Tavares Sacchi, Claudia Regina Gonçalves, Erica Valessa Ramos Gomes, Karoline Rodrigues Campos |
| EPI_ISL_861650 | Hospital Santa Marcelina Sao Paulo | Instituto Adolfo Lutz, Interdiciplinary Procedures Center, Strategic Laboratory | Claudio Tavares Sacchi, Claudia Regina Gonçalves, Erica Valessa Ramos Gomes, Karoline Rodrigues Campos |
| EPI_ISL_861651 | AMA Jardim Brasil | Instituto Adolfo Lutz, Interdiciplinary Procedures Center, Strategic Laboratory | Claudio Tavares Sacchi, Claudia Regina Gonçalves, Erica Valessa Ramos Gomes, Karoline Rodrigues Campos |
| EPI_ISL_861652 | AMA Wamberto Dias da Costa | Instituto Adolfo Lutz, Interdiciplinary Procedures Center, Strategic Laboratory | Claudio Tavares Sacchi, Claudia Regina Gonçalves, Erica Valessa Ramos Gomes, Karoline Rodrigues Campos |
| EPI_ISL_861653 | Hospital Santa Virgínia | Instituto Adolfo Lutz, Interdiciplinary Procedures Center, Strategic Laboratory | Claudio Tavares Sacchi, Claudia Regina Gonçalves, Erica Valessa Ramos Gomes, Karoline Rodrigues Campos |
| EPI_ISL_861654, EPI_ISL_861655 | Hospital Santa Marcelina Sao Paulo | Instituto Adolfo Lutz, Interdiciplinary Procedures Center, Strategic Laboratory | Claudio Tavares Sacchi, Claudia Regina Gonçalves, Erica Valessa Ramos Gomes, Karoline Rodrigues Campos |
| EPI_ISL_861656 | UPA de Jandira | Instituto Adolfo Lutz, Interdiciplinary Procedures Center, Strategic Laboratory | Claudio Tavares Sacchi, Claudia Regina Gonçalves, Erica Valessa Ramos Gomes, Karoline Rodrigues Campos |
| EPI_ISL_861659, EPI_ISL_861661 | PS e Maternidade Nair Fonseca Leitao Arantes | Instituto Adolfo Lutz, Interdiciplinary Procedures Center, Strategic Laboratory | Claudio Tavares Sacchi, Claudia Regina Gonçalves, Erica Valessa Ramos Gomes, Karoline Rodrigues Campos |
| EPI_ISL_861663 | Instituto Adolfo Lutz - Central | Instituto Adolfo Lutz, Interdiciplinary Procedures Center, Strategic Laboratory | Claudio Tavares Sacchi, Claudia Regina Gonçalves, Erica Valessa Ramos Gomes, Karoline Rodrigues Campos |
| EPI_ISL_862077 | National Influenza Center, Virology Department | National Influenza Center | J Yavarian,K Sadeghi, NZ Shafiei Jandaghi, V Salimi, A Nejati, N Ghavvami,F Ajaminejad and T Mokhtari Azad |
| EPI_ISL_862078 | National Influenza Center, Virology Department | National Influenza Center | A Nejati, J Yavarian,K Sadeghi, NZ Shafiei Jandaghi, V Salimi, N Ghavvami,F Ajaminejad and T Mokhtari Azad |
| EPI_ISL_862567 | Gerencia de Asistencia Sanitaria de Soria | Instituto de Salud Carlos III | Iglesias-Caballero, M. Molinero Calamita, M. González-Esguevillas, M. Camarero, S. Pozo, F. Casas, I. Jiménez, P. Jiménez, M. Zaballos, A. Monzón, S. Varona, S. Juliá, M. Cuesta, I, C. Aldea |
| EPI_ISL_862627, EPI_ISL_862628, EPI_ISL_862629, EPI_ISL_862630, EPI_ISL_862631, EPI_ISL_862632, EPI_ISL_862633, EPI_ISL_862634, EPI_ISL_862635, EPI_ISL_862636 | Hospital Clínic | Instituto de Salud Carlos III | Iglesias-Caballero, M.Camarero, S. Molinero Calamita, M. González-Esguevillas, M. Pozo, F. Casas, I. Jiménez, P. Jiménez, M. Zaballos, A. Monzón, S. Varona, S. Juliá, M. Cuesta, I. Marcos, M.A. |
| EPI_ISL_862637, EPI_ISL_862638, EPI_ISL_862639 | Hospital Nuestra Señora de Sonsoles | Instituto de Salud Carlos III | Iglesias-Caballero, M. Molinero Calamita, M. González-Esguevillas, M. Camarero, S. Pozo, F. Casas, I. Jiménez, P. Jiménez, M. Zaballos, A. Monzón, S. Varona, S. Juliá, M. Cuesta, I, A. San Pedro |
| EPI_ISL_862640 | Gerencia de Asistencia Sanitaria de Soria | Instituto de Salud Carlos III | Iglesias-Caballero, M. Molinero Calamita, M. González-Esguevillas, M. Camarero, S. Pozo, F. Casas, I. Jiménez, P. Jiménez, M. Zaballos, A. Monzón, S. Varona, S. Juliá, M. Cuesta, I, C. Aldea |
| EPI_ISL_862641, EPI_ISL_862642, EPI_ISL_862643, EPI_ISL_862644 | Consejería de Sanidad y Asuntos Sociales | Instituto de Salud Carlos III | Iglesias-Caballero, M. Camarero, S. Molinero Calamita, M. González-Esguevillas, M. Pozo, F. Casas, I. Jiménez, P. Jiménez, M. Zaballos, A. Monzón, S. Varona, S. Juliá, M. Cuesta, I. Gutiérrez, G. |
| EPI_ISL_862645, EPI_ISL_862646, EPI_ISL_862647, EPI_ISL_862648, EPI_ISL_862649, EPI_ISL_862650, EPI_ISL_862651 | Complejo Hospitalario de Navarra | Instituto de Salud Carlos III | Iglesias-Caballero, M.Camarero, S. Molinero Calamita, M. González-Esguevillas, M. Pozo, F. Casas, I. Jiménez, P. Jiménez, M. Zaballos, A. Monzón, S. Varona, S. Juliá, M. Cuesta, I. Ezpeleta, C. |
| EPI_ISL_862652, EPI_ISL_862653 | Clínica Universidad de Navarra | Instituto de Salud Carlos III | Iglesias-Caballero, M.Camarero, S. Molinero Calamita, M. González-Esguevillas, M. Pozo, F. Casas, I. Jiménez, P. Jiménez, M. Zaballos, A. Monzón, S. Varona, S. Juliá, M. Cuesta, I. Fernández, M. |
| EPI_ISL_862654 | Hospital San Pedro de Alcántara | Instituto de Salud Carlos III | Iglesias-Caballero, M.Camarero, S. Molinero Calamita, M. González-Esguevillas, M. Pozo, F. Casas, I. Jiménez, P. Jiménez, M. Zaballos, A. Monzón, S. Varona, S. Juliá, M. Cuesta, I. Rodríguez, G. |
| EPI_ISL_862655, EPI_ISL_862656, EPI_ISL_862657, EPI_ISL_862658 | Hospital Universitario de Ceuta | Instituto de Salud Carlos III | Iglesias-Caballero, M. Camarero, S. Molinero Calamita, M. González-Esguevillas, M. Pozo, F. Casas, I. Jiménez, P. Jiménez, M. Zaballos, A. Monzón, S. Varona, S. Juliá, M. Cuesta, I. Hijano, S. |
| EPI_ISL_862837, EPI_ISL_862838 | Fujita Health University Hospital | Fujita health University, School of Medicine, Department of Microbiology | Masahiro suzuki, Yohei Doi |

| | | | |
|---|---|---|---|
| EPI_ISL_862839 | Fujita Health University Bantane Hospital | Fujita health University, School of Medicine, Department of Microbiology | Masahiro suzuki, Yohei Doi |
| EPI_ISL_862840, EPI_ISL_862841, EPI_ISL_862842 | Fujita Health University Hospital | Fujita health University, School of Medicine, Department of Microbiology | Masahiro suzuki, Yohei Doi |
| EPI_ISL_862843, EPI_ISL_862844, EPI_ISL_862845, EPI_ISL_862846, EPI_ISL_862847 | Nara Medical University Hospital | Fujita health University, School of Medicine, Department of Microbiology | Masahiro suzuki, Yohei Doi |
| EPI_ISL_862848 | Japanese Red Cross Narita Hospital | Fujita health University, School of Medicine, Department of Microbiology | Masahiro suzuki, Yohei Doi |
| EPI_ISL_862849, EPI_ISL_862850, EPI_ISL_862851 | Ehime Prefectural Central Hospital | Fujita health University, School of Medicine, Department of Microbiology | Masahiro suzuki, Yohei Doi |
| EPI_ISL_862852, EPI_ISL_862853, EPI_ISL_862854 | Tosei General Hospital | Fujita health University, School of Medicine, Department of Microbiology | Masahiro suzuki, Yohei Doi |
| EPI_ISL_862855 | National Hospital Organization Kanazawa Medical Center | Fujita health University, School of Medicine, Department of Microbiology | Masahiro suzuki, Yohei Doi |
| EPI_ISL_862856 | Japanese Red Cross Narita Hospital | Fujita health University, School of Medicine, Department of Microbiology | Masahiro suzuki, Yohei Doi |
| EPI_ISL_862857 | Fujita Health University Bantane Hospital | Fujita health University, School of Medicine, Department of Microbiology | Masahiro suzuki, Yohei Doi |
| EPI_ISL_862858 | Isehara Kyodo Hospital | Fujita health University, School of Medicine, Department of Microbiology | Masahiro suzuki, Yohei Doi |
| EPI_ISL_862859, EPI_ISL_862860 | Nara Medical University Hospital | Fujita health University, School of Medicine, Department of Microbiology | Masahiro suzuki, Yohei Doi |
| EPI_ISL_862861, EPI_ISL_862862 | Sagamihara Kyodo Hospital | Fujita health University, School of Medicine, Department of Microbiology | Masahiro suzuki, Yohei Doi |
| EPI_ISL_862863, EPI_ISL_862864, EPI_ISL_862865 | University Hospital Kyoto Prefectural University of Medicine | Fujita health University, School of Medicine, Department of Microbiology | Masahiro suzuki, Yohei Doi |
| EPI_ISL_862866 | Nara Medical University Hospital | Fujita health University, School of Medicine, Department of Microbiology | Masahiro suzuki, Yohei Doi |
| EPI_ISL_862867 | Yokohama City University Hospital | Fujita health University, School of Medicine, Department of Microbiology | Masahiro suzuki, Yohei Doi |
| EPI_ISL_862868 | Fujita Health University Bantane Hospital | Fujita health University, School of Medicine, Department of Microbiology | Masahiro suzuki, Yohei Doi |
| EPI_ISL_862869 | Saiseikai Niigata Hospital | Fujita health University, School of Medicine, Department of Microbiology | Masahiro suzuki, Yohei Doi |
| EPI_ISL_862871 | Nara Medical University Hospital | Fujita health University, School of Medicine, Department of Microbiology | Masahiro suzuki, Yohei Doi |
| EPI_ISL_862872, EPI_ISL_862873 | Japanese Red Cross Narita Hospital | Fujita health University, School of Medicine, Department of Microbiology | Masahiro suzuki, Yohei Doi |
| EPI_ISL_862874 | Japanese Red Cross Nagoya Daiichi Hospital | Fujita health University, School of Medicine, Department of Microbiology | Masahiro suzuki, Yohei Doi |
| EPI_ISL_862875 | Komatsu Municipal Hospital | Fujita health University, School of Medicine, Department of Microbiology | Masahiro suzuki, Yohei Doi |
| EPI_ISL_862876 | NTT East Sapporo Hospital | Fujita health University, School of Medicine, Department of Microbiology | Masahiro suzuki, Yohei Doi |
| EPI_ISL_862877, EPI_ISL_862878 | Nara Medical University Hospital | Fujita health University, School of Medicine, Department of Microbiology | Masahiro suzuki, Yohei Doi |
| EPI_ISL_862879 | Ehime Prefectural Central Hospital | Fujita health University, School of Medicine, Department of Microbiology | Masahiro suzuki, Yohei Doi |
| EPI_ISL_862880 | Nara Medical University Hospital | Fujita health University, School of Medicine, Department of Microbiology | Masahiro suzuki, Yohei Doi |
| EPI_ISL_862881 | Kitakyushu Municipal Medical Center | Fujita health University, School of Medicine, Department of Microbiology | Masahiro suzuki, Yohei Doi |
| EPI_ISL_862882, EPI_ISL_862883, EPI_ISL_862884 | University Hospital Kyoto Prefectural University of Medicine | Fujita health University, School of Medicine, Department of Microbiology | Masahiro suzuki, Yohei Doi |
| EPI_ISL_862885 | Japanese Red Cross Narita Hospital | Fujita health University, School of Medicine, Department of Microbiology | Masahiro suzuki, Yohei Doi |
| EPI_ISL_871973 | Hospital Universitario Severo Ochoa | Instituto de Salud Carlos III | Iglesias-Caballero, M. Camarero, S. Molinero Calamita, M. González-Esguevillas, M. Pozo, F. Casas, I. Jiménez, P. Jiménez, M. Zaballos, A. Monzón, S. Varona, S. Juliá, M. Cuesta, I. García, M.L. |
| EPI_ISL_871974, EPI_ISL_871975 | Hospital Infanta Cristina | Instituto de Salud Carlos III | Iglesias-Caballero, M. Camarero, S. Molinero Calamita, M. González-Esguevillas, M. Pozo, F. Casas, I. Jiménez, P. Jiménez, M. Zaballos, A. Monzón, S. Varona, S. Juliá, M. Cuesta, I. García, P. |
| EPI_ISL_871976, EPI_ISL_871977 | Hospital Universitario Lozano Bielsa | Instituto de Salud Carlos III | Iglesias-Caballero, M. Camarero, S. Molinero Calamita, M. González-Esguevillas, M. Pozo, F. Casas, I. Jiménez, P. Jiménez, M. Zaballos, A. Monzón, S. Varona, S. Juliá, M. Cuesta, I. Benito, R. |
| EPI_ISL_871978 | Hospital Universitario Miguel Servet | Instituto de Salud Carlos III | Iglesias-Caballero, M. Camarero, S. Molinero Calamita, M. González-Esguevillas, M. Pozo, F. Casas, I. Jiménez, P. Jiménez, M. Zaballos, A. Monzón, S. Varona, S. Juliá, M. Cuesta, I. Rezusta, A. |
| EPI_ISL_871979 | Hospital Infanta Cristina | Instituto de Salud Carlos III | Iglesias-Caballero, M. Camarero, S. Molinero Calamita, M. González-Esguevillas, M. Pozo, F. Casas, I. Jiménez, P. Jiménez, M. Zaballos, A. Monzón, S. Varona, S. Juliá, M. Cuesta, I. García, P. |
| EPI_ISL_871980, EPI_ISL_871981, EPI_ISL_871982, EPI_ISL_871983 | Hospital Universitario Miguel Servet | Instituto de Salud Carlos III | Iglesias-Caballero, M. Camarero, S. Molinero Calamita, M. González-Esguevillas, M. Pozo, F. Casas, I. Jiménez, P. Jiménez, M. Zaballos, A. Monzón, S. Varona, S. Juliá, M. Cuesta, I. Rezusta, A. |
| EPI_ISL_872027, EPI_ISL_872028, EPI_ISL_872029, EPI_ISL_872030, | Ospedale "Di Venere" | Beaconlab (Bioinformatics, Evolution and Comparative Genomics lab), Dept of Biosciences, University on Milan | Iacobellis M, d'Avenia M, Piluscio R, Parisi A, Chiara M, Manzari C, Pesole G |

| | | | |
|---|---|---|---|
| EPI_ISL_872031, EPI_ISL_872032, EPI_ISL_872033, EPI_ISL_872034, EPI_ISL_872035, EPI_ISL_872036 | | | |
| EPI_ISL_872088 | Instituto de Diagnostico y Referencia Epidemiologicos (INDRE) | Instituto de Diagnostico y Referencia Epidemiologicos (INDRE) | Fabiola Garces-Ayala, Gisela Barrera-Badillo , Abril Rodriguez-Maldonado, Claudia Wong-Arambula , Natividad Cruz-Ortiz, Tatiana Nunez-Garcia, Dayanira Arellano-Suarez, Adnan-Araiza Rodríguez, Edgar Mendieta-Condado, Lucia Hernandez-Rivas, Irma Lopez-Martinez, Ernesto Ramirez-Gonzalez. |
| EPI_ISL_872089 | Instituto de Diagnostico y Referencia Epidemiologicos (INDRE) | Instituto de Diagnostico y Referencia Epidemiologicos (INDRE) | Abril Rodriguez-Maldonado, Gisela Barrera-Badillo ,Claudia Wong-Arambula , Natividad Cruz-Ortiz, Tatiana Nunez-Garcia, Dayanira Arellano-Suarez, Fabiola Garces-Ayala, Adnan-Araiza Rodríguez, Edgar Mendieta-Condado, Lucia Hernandez-Rivas, Irma Lopez-Martinez, Ernesto Ramirez-Gonzalez. |
| EPI_ISL_872091 | Instituto de Diagnostico y Referencia Epidemiologicos (INDRE) | Instituto de Diagnostico y Referencia Epidemiologicos (INDRE) | Adnan-Araiza Rodríguez,Claudia Wong-Arambula, Gisela Barrera-Badillo , Abril Rodriguez-Maldonado, Natividad Cruz-Ortiz, Tatiana Nunez-Garcia, Dayanira Arellano-Suarez, Fabiola Garces-Ayala, Edgar Mendieta-Condado, Lucia Hernandez-Rivas, Irma Lopez-Martinez, Ernesto Ramirez-Gonzalez. |
| EPI_ISL_872092 | Instituto de Diagnostico y Referencia Epidemiologicos (INDRE) | Instituto de Diagnostico y Referencia Epidemiologicos (INDRE) | Ernesto Ramirez-Gonzalez, Adnan-Araiza Rodríguez,Claudia Wong-Arambula, Gisela Barrera-Badillo , Abril Rodriguez-Maldonado, Natividad Cruz-Ortiz, Tatiana Nunez-Garcia, Dayanira Arellano-Suarez, Fabiola Garces-Ayala, Edgar Mendieta-Condado, Lucia Hernandez-Rivas, Irma Lopez-Martinez. |
| EPI_ISL_872093 | Instituto de Diagnostico y Referencia Epidemiologicos (INDRE) | Instituto de Diagnostico y Referencia Epidemiologicos (INDRE) | Edgar Mendieta-Condado, Ernesto Ramirez-Gonzalez, Adnan-Araiza Rodríguez,Claudia Wong-Arambula, Gisela Barrera-Badillo , Abril Rodriguez-Maldonado, Natividad Cruz-Ortiz, Tatiana Nunez-Garcia, Dayanira Arellano-Suarez, Fabiola Garces-Ayala, Lucia Hernandez-Rivas, Irma Lopez-Martinez. |
| EPI_ISL_872097 | Instituto de Diagnostico y Referencia Epidemiologicos (INDRE) | Instituto de Diagnostico y Referencia Epidemiologicos (INDRE) | Abril Rodriguez-Maldonado, Gisela Barrera-Badillo ,Claudia Wong-Arambula , Natividad Cruz-Ortiz, Tatiana Nunez-Garcia, Dayanira Arellano-Suarez, Fabiola Garces-Ayala, Adnan-Araiza Rodríguez, Edgar Mendieta-Condado, Lucia Hernandez-Rivas, Irma Lopez-Martinez, Ernesto Ramirez-Gonzalez. |
| EPI_ISL_872099 | Instituto de Diagnostico y Referencia Epidemiologicos (INDRE) | Instituto de Diagnostico y Referencia Epidemiologicos (INDRE) | Fabiola Garces-Ayala, Gisela Barrera-Badillo , Abril Rodriguez-Maldonado, Claudia Wong-Arambula , Natividad Cruz-Ortiz, Tatiana Nunez-Garcia, Dayanira Arellano-Suarez, Adnan-Araiza Rodríguez, Edgar Mendieta-Condado, Lucia Hernandez-Rivas, Irma Lopez-Martinez, Ernesto Ramirez-Gonzalez. |
| EPI_ISL_876811, EPI_ISL_876812, EPI_ISL_876813 | Istituto Zooprofilattico Sperimentale della Puglia e della Basilicata | Istituto Zooprofilattico Sperimentale della Puglia e della Basilicata | Parisi A., Bianco A., Capozzi L., Del Sambro L., Manzulli V, Rondinone V., Pace L., Cipolletta D., Galante D. |
| EPI_ISL_877036 | 1.AO Universitaria 'S. Giovanni di Dio e Ruggi D'Aragona, Scuola Medica Salernitana' Hospital / 2.UOC di Virologia e Microbiologia, Università della Campania 'L. Vanvitelli' / 3.AO Universitaria 'Federico II' Napoli Hospital / 4.AORN 'San Giuseppe Moscati' Avellino Hospital / 5.AO 'San Pio - presidio G. Rummo' Benevento Hospital / 6.AO 'Sant'Anna e San Sebastiano' Caserta Hospital / 7.PO 'Maria Santissima Addolorata' Eboli Hospital / 8.Biogem Istituto di Ricerche Genetiche | 1. Genome Research Center for Health (CRGS) / 2. Laboratory of Molecular Medicine and Genomics(LMMGe) / 3. Center for Research in Pure and Applied Mathematics (CRMPA) | Giorgio Giurato (Corresponding Author), Francesca Rizzo (Corresponding Author), Alessandro Weisz (Corresponding Author), Gianluigi Franci, Giovanni Nassa, Pasquale Pagliano, Roberta Tarallo, Elena Alexandrova, Ylenia D'Agostino, Carlo Ferravante, Jessica Lamberti, Viola Melone, Domenico Memoli, Valeria Mirici Cappa, Domenico Palumbo, Giovanni Pecoraro, Assunta Sellitto, Oriana Strianese, Ilaria Terenzi, Giuseppe Fenza, Aniello Gentile, Antonello Saccomanno, Sonia Amabile, Teresa Rocco, Annamaria Salvati, Emilia Vaccaro, Massimiliano Galdiero, Michele Cennamo, Giuseppe Portella, Maria Grazia Foti, Mariarosaria Ingino, Maria Landi, Maurizio Fumi, Vincenzo Rocco, Rita Greco, Vittoria Letizia, Arnolfo Petruzziello, Maddalena Schioppa, Gregorio Goffredi, Francesca Marciano, Michele Caraglia, Alessia Cossu, Marianna Scrima |
| EPI_ISL_888678 | Laboratory for Respiratory Viruses, Cantacuzino National Military-Medical Institute for Research and Development | Cantacuzino Institute Virology | Luiza Ustea, Nicoleta Paraschiv, Mihaela Lazar |
| EPI_ISL_888982 | RS Pondok Indah Hospital - Pondok Indah | Eijkman Institute for Molecular Biology, Ministry of Research and Technology/National Agency for Research and Innovation | Edison Johar, Frilasita A Yudhaputri, Hidayat Trimarsanto, Iskandar Adnan, Lydia V. Panggalo, Sukma Oktavianthi, Willy Agustine, Safarina G Malik, Khin Saw Myint, Amin Soebandrio |
| EPI_ISL_889373, EPI_ISL_889374, EPI_ISL_889375, EPI_ISL_889377, EPI_ISL_889378, EPI_ISL_889379, EPI_ISL_889380, EPI_ISL_889384, EPI_ISL_889385, EPI_ISL_889386, EPI_ISL_889387, EPI_ISL_889388, EPI_ISL_889389, EPI_ISL_889390, EPI_ISL_889391, EPI_ISL_889392, EPI_ISL_889393 | | | |
| see above | Olomouc University Hospital | Institute of Applied Biotechnologies a.s. | Petr Klempt, Ondej Brzo, Martin Kašný, Kateina Kvapilová, Marián Hajdúch, Petr Kvapil |
| EPI_ISL_895747, EPI_ISL_895748, EPI_ISL_895768, EPI_ISL_895769, EPI_ISL_895789, EPI_ISL_895803, EPI_ISL_895805, EPI_ISL_895818, EPI_ISL_895829, EPI_ISL_895833, EPI_ISL_895836, EPI_ISL_895838, EPI_ISL_895839, EPI_ISL_895840, EPI_ISL_895843, EPI_ISL_895844, EPI_ISL_895845, EPI_ISL_895846 | | | |
| see above | Molecular biology division, Institute of Clinical Biochemistry and Diagnostics, Charles University, Faculty of Medicine in Hradec Králové and University Hospital Hradec Králové | Molecular biology division, Institute of Clinical Biochemistry and Diagnostics, Charles University, Faculty of Medicine in Hradec Králové and University Hospital Hradec Králové | Helena Kovaíková, Petr Brož, Ivana Baranová, Kateina Hrochová, Tereza Baková, Jitka Novotná, Kateina Pehlíková, Vladimír Palika. Cooperation project with BioVendor-R&D and bioinformatics company BIOXSYS s r.o. |
| EPI_ISL_900726 | Bozeman Health Deaconess Hospital | Wiedenheft lab, Montana State University | Artem Nemudryi, Anna Nemudraia, Tanner Wiegand, Joseph Nichols, Deann T. Snyder, Jodi F. Hedges, Calvin Cicha, Helen Lee, Karl K. Vanderwood, Diane Bimczok, Mark A. Jutila and Blake Wiedenheft |
| EPI_ISL_906065 | Day Hospital de Ermelino Matarazzo | Instituto Adolfo Lutz, Interdiciplinary Procedures Center, Strategic Laboratory | Claudio Tavares Sacchi, Claudia Regina Gonçalves, Erica Valessa Ramos Gomes, Karoline Rodrigues Campos |
| EPI_ISL_906066 | Hospital Nipo Brasileiro | Instituto Adolfo Lutz, Interdiciplinary Procedures Center, Strategic Laboratory | Claudio Tavares Sacchi, Claudia Regina Gonçalves, Erica Valessa Ramos Gomes, Karoline Rodrigues Campos |
| EPI_ISL_906067 | PS e Maternidade Nair Fonseca Leitao Arantes | Instituto Adolfo Lutz, Interdiciplinary Procedures Center, Strategic Laboratory | Claudio Tavares Sacchi, Claudia Regina Gonçalves, Erica Valessa Ramos Gomes, Karoline Rodrigues Campos |
| EPI_ISL_906769, EPI_ISL_906775 | Dr. Leonard A. Miller Centre for Health Services | National Microbiology Laboratory (NML) | Anna Majer, Shari Tyson, Grace Seo, Philip Mabon, Elsie Grudeski, Rhiannon Huzarewich, Russell Mandes, Anneliese Landgraff, Jennifer Tanner, Natalie Knox, Morag Graham, Gary Van Domselaar, Robert Needle, Yang Yu, Laura Gilbert, George Zahariadis, Nathalie Bastien, Yan Li, Timothy Booth, Darian Hole, Madison Chapel, Kerri Smith, CanCOGeN's metadata curation team, Public Health Agency of Canada CanCOGeN team |
| EPI_ISL_911513 | Laboratoire de Virologie, CHU de Caen, Normandie, France. | GRAM2.0, Université de Caen Normandie Laboratoire de Virologie, CHU de Caen | Meriadeg LE GOUIL, Januel VOUTEAU, Juliette ECHE, Estelle LEPERCHOIS, Astrid VABRET |
| EPI_ISL_940753 | University of Bari Biomedical Sciences and Human Oncology | University of Bari Biomedical Sciences and Human Oncology | Chironna M., Sallustio A., Loconsole D., Accogli M. |
| EPI_ISL_941104, EPI_ISL_941105, EPI_ISL_941106, EPI_ISL_941107, EPI_ISL_941108, EPI_ISL_941109, EPI_ISL_941110, EPI_ISL_941111, EPI_ISL_941112, EPI_ISL_941113, EPI_ISL_941114, EPI_ISL_941115, EPI_ISL_941116, EPI_ISL_941117, EPI_ISL_941118, EPI_ISL_941119, EPI_ISL_941120, EPI_ISL_941121, EPI_ISL_941122, EPI_ISL_941123, EPI_ISL_941125, EPI_ISL_941126, EPI_ISL_941127, EPI_ISL_941129, EPI_ISL_941130, EPI_ISL_941131, EPI_ISL_941132, EPI_ISL_941133, EPI_ISL_941134, EPI_ISL_941135, EPI_ISL_941137, EPI_ISL_941138, EPI_ISL_941139, EPI_ISL_941140, EPI_ISL_941141, EPI_ISL_941142, EPI_ISL_941143, EPI_ISL_941144, EPI_ISL_941145, EPI_ISL_941146, EPI_ISL_941147, EPI_ISL_941148, EPI_ISL_941149, EPI_ISL_941150, EPI_ISL_941151, EPI_ISL_941152, EPI_ISL_941153, EPI_ISL_941154, EPI_ISL_941155, EPI_ISL_941156, EPI_ISL_941157, EPI_ISL_941158, EPI_ISL_941159, EPI_ISL_941160, EPI_ISL_941161, EPI_ISL_941162 | | | |
| see above | Centro de Investigaciones en Microbiología y Biotecnología-UR (CIMBIUR), Facultad de Ciencias Naturales, Universidad del Rosario, Bogotá, Colombia | Centro de Investigaciones en Microbiología y Biotecnología-UR (CIMBIUR), Facultad de Ciencias Naturales, Universidad del Rosario, Bogotá, Colombia Icahn School of Medicine at Mount Sinai, New York, USA | Nathalia Ballesteros, Marina Muñoz, Luz Helena Patiño, Carolina Hernández, Felipe González-Casabianca, Iván Carroll, Mauricio Santos-Vega, Jaime Cascante, Andrés Angel, Alejandro Feged-Rivadeneira, Mónica Palma-Cuero, Carolina Flórez, Sergio Gomez, Adriana van de Guchte, Zenab Khan, Jayeeta Dutta, Hala Alejel Alshammary, Ana S. Gonzalez-Reiche, Matthew M. Hernandez, Emilia Mia Sordillo, Viviana Simon, Harm van Bakel, Alberto Paniz-Mondolfi, Juan David Ramírez |
| EPI_ISL_941952, EPI_ISL_941964, EPI_ISL_941965, EPI_ISL_941966, EPI_ISL_941967, EPI_ISL_941968, EPI_ISL_941969, EPI_ISL_941970, EPI_ISL_941971, EPI_ISL_941972, EPI_ISL_941973, EPI_ISL_941974, EPI_ISL_941975, EPI_ISL_941976, EPI_ISL_941977, EPI_ISL_941978, EPI_ISL_941979, EPI_ISL_941980, EPI_ISL_941981, EPI_ISL_941982, EPI_ISL_941983 | | | |
| see above | Centro de Investigaciones en Microbiología y Biotecnología-UR (CIMBIUR), Facultad de Ciencias Naturales, Universidad del Rosario, Bogotá, Colombia | Centro de Investigaciones en Microbiología y Biotecnología-UR (CIMBIUR), Facultad de Ciencias Naturales, Universidad del Rosario, Bogotá, Colombia Instituto Nacional de Salud, Bogotá, Colombia Icahn School of Medicine at Mount Sinai, New York, USA | Luz Helena Patiño, Marina Muñoz, Nathalia Ballesteros, Carolina Hernández, Carolina Flórez, Sergio Gomez, Adriana van de Guchte, Zenab Khan, Jayeeta Dutta, Hala Alejel Alshammary, Ana S. Gonzalez-Reiche, Matthew M. Hernandez, Emilia Mia Sordillo, Viviana Simon, Harm van Bakel, Alberto Paniz-Mondolfi, Juan David Ramírez |
| EPI_ISL_941984, EPI_ISL_941985 | Instituto Nacional de Salud, Bogotá, Colombia | Centro de Investigaciones en Microbiología y Biotecnología-UR (CIMBIUR), Facultad de Ciencias Naturales, Universidad del Rosario, Bogotá, Colombia Instituto Nacional de Salud, Bogotá, Colombia Icahn School of Medicine at Mount Sinai, New York, USA | Luz Helena Patiño, Marina Muñoz, Nathalia Ballesteros, Carolina Hernández, Carolina Flórez, Sergio Gomez, Adriana van de Guchte, Zenab Khan, Jayeeta Dutta, Hala Alejel Alshammary, Ana S. Gonzalez-Reiche, Matthew M. Hernandez, Emilia Mia Sordillo, Viviana Simon, Harm van Bakel, Alberto Paniz-Mondolfi, Juan David Ramírez |

| | | | |
|---|---|---|---|
| EPI_ISL_941999, EPI_ISL_942000, EPI_ISL_942001, EPI_ISL_942002, EPI_ISL_944635, EPI_ISL_944638 | Centro de Investigaciones en Microbiología y Biotecnología-UR (CIMBIUR), Facultad de Ciencias Naturales, Universidad del Rosario, Bogotá, Colombia | Centro de Investigaciones en Microbiología y Biotecnología-UR (CIMBIUR), Facultad de Ciencias Naturales, Universidad del Rosario, Bogotá, Colombia Instituto Nacional de Salud, Bogotá, Colombia Icahn School of Medicine at Mount Sinai, New York, USA | Luz Helena Patiño, Marina Muñoz, Nathalia Ballesteros, Carolina Hernández, Carolina Flórez, Sergio Gomez, Adriana van de Guchte, Zenab Khan, Jayeeta Dutta, Hala Alejel Alshammary, Ana S. Gonzalez-Reiche, Matthew M. Hernandez, Emilia Mia Sordillo, Viviana Simon, Harm van Bakel, Alberto Paniz-Mondolfi, Juan David Ramírez |
| EPI_ISL_949244, EPI_ISL_949245, EPI_ISL_949246, EPI_ISL_949247 | Cliniques universitaires Saint-Luc | UCLouvain/IREC/MBLG | Jean Ruelle, Lysa Pinsmaye, Benoit Kabamba Mukadi |
| EPI_ISL_949248 | Cliniques Saint-Pierre Ottignies | UCLouvain/IREC/MBLG | Jean Ruelle, Lysa Pinsmaye, Benoit Kabamba Mukadi |
| EPI_ISL_949249, EPI_ISL_949250, EPI_ISL_949252 | Cliniques universitaires Saint-Luc | UCLouvain/IREC/MBLG | Jean Ruelle, Lysa Pinsmaye, Benoit Kabamba Mukadi |
| EPI_ISL_955234 | HE CMN LA RAZA | BIOBANCO / COCTI | Borja-Aburto VH, Grajales-Muñiz C, Santacruz Tinoco CE, Rojas-Mendoza T, Ochoa Carrera LA, Sánchez A, Grande R, Isa P, Taboada B, López S, Arias C, Muñoz-Medina JE |
| EPI_ISL_955235 | UMF 23 UNIDAD MORELOS | BIOBANCO / COCTI | Borja-Aburto VH, Grajales-Muñiz C, Santacruz Tinoco CE, Rojas-Mendoza T, Ochoa Carrera LA, Sánchez A, Grande R, Isa P, Taboada B, López S, Arias C, Muñoz-Medina JE |
| EPI_ISL_955236 | HGR 1 VICENTE GUERRERO | BIOBANCO / COCTI | Borja-Aburto VH, Grajales-Muñiz C, Santacruz Tinoco CE, Rojas-Mendoza T, Ochoa Carrera LA, Sánchez A, Grande R, Isa P, Taboada B, López S, Arias C, Muñoz-Medina JE |
| EPI_ISL_955237 | UMF 75 NETZAHUALCOYOTL | BIOBANCO / COCTI | Borja-Aburto VH, Grajales-Muñiz C, Santacruz Tinoco CE, Rojas-Mendoza T, Ochoa Carrera LA, Sánchez A, Grande R, Isa P, Taboada B, López S, Arias C, Muñoz-Medina JE |
| EPI_ISL_955238 | HI CMN LA RAZA | BIOBANCO / COCTI | Borja-Aburto VH, Grajales-Muñiz C, Santacruz Tinoco CE, Rojas-Mendoza T, Ochoa Carrera LA, Sánchez A, Grande R, Isa P, Taboada B, López S, Arias C, Muñoz-Medina JE |
| EPI_ISL_955239 | HE CMN LA RAZA | BIOBANCO / COCTI | Borja-Aburto VH, Grajales-Muñiz C, Santacruz Tinoco CE, Rojas-Mendoza T, Ochoa Carrera LA, Sánchez A, Grande R, Isa P, Taboada B, López S, Arias C, Muñoz-Medina JE |
| EPI_ISL_955240, EPI_ISL_955241 | HGZ 30 MEXICALI | BIOBANCO / COCTI | Borja-Aburto VH, Grajales-Muñiz C, Santacruz Tinoco CE, Rojas-Mendoza T, Ochoa Carrera LA, Sánchez A, Grande R, Isa P, Taboada B, López S, Arias C, Muñoz-Medina JE |
| EPI_ISL_955242, EPI_ISL_955243 | HGSMF 26 CABO SAN LUCAS | BIOBANCO / COCTI | Borja-Aburto VH, Grajales-Muñiz C, Santacruz Tinoco CE, Rojas-Mendoza T, Ochoa Carrera LA, Sánchez A, Grande R, Isa P, Taboada B, López S, Arias C, Muñoz-Medina JE |
| EPI_ISL_955244 | UMF 26 ACAPULCO | BIOBANCO / COCTI | Borja-Aburto VH, Grajales-Muñiz C, Santacruz Tinoco CE, Rojas-Mendoza T, Ochoa Carrera LA, Sánchez A, Grande R, Isa P, Taboada B, López S, Arias C, Muñoz-Medina JE |
| EPI_ISL_955245 | HGR 1 VICENTE GUERRERO | BIOBANCO / COCTI | Borja-Aburto VH, Grajales-Muñiz C, Santacruz Tinoco CE, Rojas-Mendoza T, Ochoa Carrera LA, Sánchez A, Grande R, Isa P, Taboada B, López S, Arias C, Muñoz-Medina JE |
| EPI_ISL_955246 | HGOMF 60 TLANEPANTLA | BIOBANCO / COCTI | Borja-Aburto VH, Grajales-Muñiz C, Santacruz Tinoco CE, Rojas-Mendoza T, Ochoa Carrera LA, Sánchez A, Grande R, Isa P, Taboada B, López S, Arias C, Muñoz-Medina JE |
| EPI_ISL_955247 | HGR 200 TECAMAC | BIOBANCO / COCTI | Borja-Aburto VH, Grajales-Muñiz C, Santacruz Tinoco CE, Rojas-Mendoza T, Ochoa Carrera LA, Sánchez A, Grande R, Isa P, Taboada B, López S, Arias C, Muñoz-Medina JE |
| EPI_ISL_955248 | BCO SANGRE CM LA RAZA | BIOBANCO / COCTI | Borja-Aburto VH, Grajales-Muñiz C, Santacruz Tinoco CE, Rojas-Mendoza T, Ochoa Carrera LA, Sánchez A, Grande R, Isa P, Taboada B, López S, Arias C, Muñoz-Medina JE |
| EPI_ISL_955249 | HGZ 2A PASO-TRONCOSO | BIOBANCO / COCTI | Borja-Aburto VH, Grajales-Muñiz C, Santacruz Tinoco CE, Rojas-Mendoza T, Ochoa Carrera LA, Sánchez A, Grande R, Isa P, Taboada B, López S, Arias C, Muñoz-Medina JE |
| EPI_ISL_955250, EPI_ISL_955251, EPI_ISL_955252, EPI_ISL_955253 | HGSMF 26 CABO SAN LUCAS | BIOBANCO / COCTI | Borja-Aburto VH, Grajales-Muñiz C, Santacruz Tinoco CE, Rojas-Mendoza T, Ochoa Carrera LA, Sánchez A, Grande R, Isa P, Taboada B, López S, Arias C, Muñoz-Medina JE |
| EPI_ISL_955254 | UMF 28 DEL VALLE | BIOBANCO / COCTI | Borja-Aburto VH, Grajales-Muñiz C, Santacruz Tinoco CE, Rojas-Mendoza T, Ochoa Carrera LA, Sánchez A, Grande R, Isa P, Taboada B, López S, Arias C, Muñoz-Medina JE |
| EPI_ISL_955255 | HGSMF 26 CABO SAN LUCAS | BIOBANCO / COCTI | Borja-Aburto VH, Grajales-Muñiz C, Santacruz Tinoco CE, Rojas-Mendoza T, Ochoa Carrera LA, Sánchez A, Grande R, Isa P, Taboada B, López S, Arias C, Muñoz-Medina JE |
| EPI_ISL_955256 | HGZ 1 CARLOS MC GREGOR | BIOBANCO / COCTI | Borja-Aburto VH, Grajales-Muñiz C, Santacruz Tinoco CE, Rojas-Mendoza T, Ochoa Carrera LA, Sánchez A, Grande R, Isa P, Taboada B, López S, Arias C, Muñoz-Medina JE |
| EPI_ISL_955257 | HGZ 32 VILLA COAPA | BIOBANCO / COCTI | Borja-Aburto VH, Grajales-Muñiz C, Santacruz Tinoco CE, Rojas-Mendoza T, Ochoa Carrera LA, Sánchez A, Grande R, Isa P, Taboada B, López S, Arias C, Muñoz-Medina JE |
| EPI_ISL_956270 | Universitas Airlangga Hospital | Institute of Tropical Disease, Universitas Airlangga | Aldise M Nastri, Jezzy R Dewantari, Rima R Prasetya, Krisnoadi Rahardjo, Nasronudin, Gatot Soegiarto, Laksmi Wulandari, Resti Yudhawati, Soetjipto, Yasuko Mori, Maria I Lusida, Kazufumi Shimizu |
| EPI_ISL_956307 | Primasatya Husada Citra Hospital | Institute of Tropical Disease, Universitas Airlangga | Krisnoadi Rahardjo, Aldise M Nastri, Jezzy R Dewantari, Rima R Prasetya, Pudji Djanuartono, Gatot Soegiarto, Laksmi Wulandari, Resti Yudhawati, Yasuko Mori, Soetjipto, Kazufumi Shimizu, Maria I Lusida |
| EPI_ISL_956308 | RSUD Dr. M. Soewandhie | Institute of Tropical Disease, Universitas Airlangga | Jezzy R Dewantari, Rima R Prasetya, Krisnoadi Rahardjo, Aldise M Nastri, Febria Rachmanita, Gatot Soegiarto, Laksmi Wulandari, Resti Yudhawati, Yasuko Mori, Soetjipto, Kazufumi Shimizu, Maria I Lusida |
| EPI_ISL_956384 | Isolation - Virology Unit, Institut Pasteur du Cambodge; Sequencing - US National Institute of Allergy and Infectious Diseases Cambodia, US Naval Medical Research Unit -2, National Public Health Laboratory, National Institute for Public Health, Cambodia | Virology Unit, Institut Pasteur du Cambodge | Vireak Heang, Jennifer Bohl, Sophana Chea, Sreyngim Lay, Ly Sovann, Kraing Sidonn, Yi Sengdoeurn, Chin Savuth, Chau Darapheak, Veasna Duong, Jose A Garcia-Rivera, Jessica Manning, Erik A Karlsson |
| EPI_ISL_956385, EPI_ISL_956386, EPI_ISL_956387, EPI_ISL_956388, EPI_ISL_956389, EPI_ISL_956390, EPI_ISL_956391, EPI_ISL_956392, EPI_ISL_956393, EPI_ISL_956394, EPI_ISL_956395, EPI_ISL_956396, EPI_ISL_956397, EPI_ISL_956398, EPI_ISL_956399, EPI_ISL_956400, EPI_ISL_956401 | | | |
| see above | Isolation - Virology Unit, Institut Pasteur du Cambodge; Sequencing - US National Institute of Allergy and Infectious Diseases Cambodia, US Naval Medical Research Unit -2, Cambodia National Institute for Public Health | Virology Unit, Institut Pasteur du Cambodge | Vireak Heang, Jennifer Bohl, Sophana Chea, Sreyngim Lay, Ly Sovann, Kraing Sidonn, Yi Sengdoeurn, Chin Savuth, Chau Darapheak, Veasna Duong, Jose A Garcia-Rivera, Jessica Manning, Erik A Karlsson |
| EPI_ISL_959275 | National Influenza Center, Virology Department | National Influenza Center | V Salimi,NZ Shafiei Jandaghi, J Yavarian, A Nejati, K Sadeghi, N Ghavvami,F Ajaminejad and T Mokhtari Azad |
| EPI_ISL_959276 | National Influenza Center, Virology Department | National Influenza Center | NZ Shafiei Jandaghi, V Salimi, A Nejati, K Sadeghi, J Yavarian, N Ghavvami,F Ajaminejad and T Mokhtari Azad |
| EPI_ISL_959280 | National Influenza Center, Virology Department | National Influenza Center | J Yavarian, NZ Shafiei Jandaghi, V Salimi, A Nejati, K Sadeghi, F Ajaminejad,N Ghavvami and T Mokhtari Azad |
| EPI_ISL_959283 | National Influenza Center, Virology Department | National Influenza Center | NZ Shafiei Jandaghi, V Salimi, A Nejati, K Sadeghi, J Yavarian, F Ajaminejad,N Ghavvami and T Mokhtari Azad |
| EPI_ISL_962528, EPI_ISL_962529, EPI_ISL_962530, EPI_ISL_962531, EPI_ISL_962532, EPI_ISL_962533, EPI_ISL_962534, EPI_ISL_962535, EPI_ISL_962536, EPI_ISL_962537, EPI_ISL_962538, EPI_ISL_962539, EPI_ISL_962540, EPI_ISL_962541, EPI_ISL_962542, EPI_ISL_962543, EPI_ISL_962544, EPI_ISL_962545, EPI_ISL_962546, EPI_ISL_962547, EPI_ISL_962548, EPI_ISL_962549, EPI_ISL_962550, EPI_ISL_962551, EPI_ISL_962552, EPI_ISL_962553, EPI_ISL_962554, EPI_ISL_962555, EPI_ISL_962556, EPI_ISL_962557, EPI_ISL_962558, EPI_ISL_962559, EPI_ISL_962560, EPI_ISL_962561, EPI_ISL_962562, EPI_ISL_962563, EPI_ISL_962564, EPI_ISL_962565, EPI_ISL_962566, EPI_ISL_962567, EPI_ISL_962568, EPI_ISL_962569, EPI_ISL_962570, EPI_ISL_962571, EPI_ISL_962572, EPI_ISL_962573, EPI_ISL_962574, EPI_ISL_962575, EPI_ISL_962576, EPI_ISL_962577, EPI_ISL_962578, EPI_ISL_962579, EPI_ISL_962580, EPI_ISL_962581, | | | |

EPI_ISL_962582, EPI_ISL_962583, EPI_ISL_962584, EPI_ISL_962868, EPI_ISL_962869, EPI_ISL_962870, EPI_ISL_962871, EPI_ISL_962873, EPI_ISL_962874, EPI_ISL_962875

| | | | |
|---|---|---|---|
| see above | Infectious Disease Control and Prevention Institute | Infectious Disease Control and Prevention Institute | Xiaolin Jiang, Mingxiao Yao, Yuwei Zhang, Yujie He, Zengqiang Kou, Ti Liu, Yan Li, Jianxing Wang, Bo Pang, Julong Wu |
| EPI_ISL_964878, EPI_ISL_964879 | Institute of Virology Ulm University Medical Center Prof. Dr. Thomas Stamminger | Münch Lab / Ulm University Medical Center Kirchhoff Lab / Ulm University Medical Center Sparrer Lab / Ulm University Medical Center Blum Lab / LMU Munich | Thomas Stamminger, Helmut Blum, Stefan Krebs, Alexander Graf, Rüdiger Gross, Janis Müller, Carina Conzelmann, Jan Münch, Frank Kirchhoff, Konstantin Sparrer |
| EPI_ISL_977357, EPI_ISL_977411 | University of Zambia, School of Veterinary Medicine | UNZAVET and PATH | Mulenga Mwenda-Chimfwembe, Ngonda Saasa, Daniel Bridges |
| EPI_ISL_982182 | Hôpital Henri Mondor | Department of Virology, Henri Mondor University Hospital, Assistance Publique Hôpitaux de Paris, Université Paris-Est Créteil, INSERM U955 | Christophe Rodriguez, Slim Fourati, Vanessa Demontant, Guillaume Gricourt, Melissa N'Debi, Alexandre Soulier, Elisabeth Trawinski, Jean-Michel Pawlotsky |
| EPI_ISL_984277 | National Institute of Health Research and Development | National Institute of Health Research and Development | Kindi Adam, Holy Arif Wibowo, Ririn Ramadhany, Yuni Rukminiati, Agustiningsih, Hana Apsari Pawestri, Subangkit, Kartika Dewi Puspa, Arie Ardiansyah Nugraha, Hartanti Dian Ikawati, Krisna Nur Andriana Pangesti, Ni Ketut Susilarini, Nur Ika Hariastuti, Uly Alfi Nikmah, Mursinah, Asri Febriyani, Reni Herman, Nike Susanti, Herna, Tati Febriyanti, Natalie Laurencia Kipuw, Fauzul Muna, Irene Lorinda Indalao, Aulia Rizki, Nelly Puspandari, Vivi Setiawaty |
| EPI_ISL_985034 | Omics Sciences Lab | Omics Sciences Lab | Derly Andrade Molina, Rubén Armas González, Gabriel Morey León, Darlyn Amaya, Katheryn Sacheri Viteri, Edith Lopez Montanero, Pedro Barberán, Fernando Espinoza Fuentes, Juan Carlos Fernández Cadena |
| EPI_ISL_985394 | Department of Internal Medicine, College of Medicine, Chosun University | Department of Internal Medicine, College of Medicine, Chosun University | Dong-Min Kim |

We gratefully acknowledge the following Authors from the Originating laboratories responsible for obtaining the specimens, as well as the Submitting laboratories where the genome data were generated and shared via GISAID, on which this research is based.

All Submitters of data may be contacted directly via www.gisaid.org

Authors are sorted alphabetically.

| Accession ID | Originating Laboratory | Submitting Laboratory | Authors |
|---|---|---|---|
| EPI_ISL_1007672, EPI_ISL_1007673, EPI_ISL_1007674, EPI_ISL_1007675, EPI_ISL_1007676 | Department of Internal Medicine, College of Medicine, Chosun University | Department of Internal Medicine, College of Medicine, Chosun University | Dong-Min Kim |
| EPI_ISL_1014678 | National Influenza Center, Virology Department | National Influenza Center | J Yavarian,K Sadeghi, NZ Shafiei Jandaghi, V Salimi, A Nejati, N Ghavvami,F Ajaminejad and T Mokhtari Azad |
| EPI_ISL_1014679 | National Influenza Center, Virology Department | National Influenza Center | A Nejati, J Yavarian,K Sadeghi, NZ Shafiei Jandaghi, V Salimi, N Ghavvami,F Ajaminejad and T Mokhtari Azad |
| EPI_ISL_1014680 | National Influenza Center, Virology Department | National Influenza Center | K Sadeghi, A Nejati, J Yavarian, NZ Shafiei Jandaghi, V Salimi, F Ajaminejad,N Ghavvami and T Mokhtari Azad |
| EPI_ISL_1014681 | National Influenza Center, Virology Department | National Influenza Center | V Salimi,NZ Shafiei Jandaghi, J Yavarian, A Nejati, K Sadeghi, N Ghavvami,F Ajaminejad and T Mokhtari Azad |
| EPI_ISL_1014687 | National Influenza Center, Virology Department | National Influenza Center | J Yavarian,K Sadeghi, NZ Shafiei Jandaghi, V Salimi, A Nejati, N Ghavvami,F Ajaminejad and T Mokhtari Azad |
| EPI_ISL_1015362 | Pathogenic Microorganisms Variability Laboratory | Pathogenic Microorganisms Variability Laboratory | Alexey Shchetinin, Olesya Venchakova, Maria Nikiforova, Andrei Siniavin, Nadezhda Kuznetsova, Elena Shidlovskaya, Elizaveta Divisenko, Kirill Krasnoslobotsev, Evgeniya Mukasheva, Anna Ignatieva, Svetlana Trushakova, Andrey Pochtovyy, Valeria Bacalin, Evgeny Usachev, Olga Burgasova, Ludmila Kolobukhina, Svetlana Smetanina, Elena Burtseva, Denis Logunov, Vladimir Gushchin, Alexander Gintsburg |
| EPI_ISL_1018118, EPI_ISL_1018119, EPI_ISL_1018120, EPI_ISL_1018121, EPI_ISL_1018122, EPI_ISL_1018125, EPI_ISL_1018126, EPI_ISL_1018127, EPI_ISL_1018128, EPI_ISL_1018129, EPI_ISL_1018130, EPI_ISL_1018131, EPI_ISL_1018132, EPI_ISL_1018133, EPI_ISL_1018134, EPI_ISL_1018135, EPI_ISL_1018136, EPI_ISL_1018137, EPI_ISL_1018138, EPI_ISL_1018139, EPI_ISL_1018140, EPI_ISL_1018141, EPI_ISL_1018142, EPI_ISL_1018143, EPI_ISL_1018144, EPI_ISL_1018145, EPI_ISL_1018146, EPI_ISL_1018147, EPI_ISL_1018148, EPI_ISL_1018149, EPI_ISL_1018150, EPI_ISL_1018151, EPI_ISL_1018152, EPI_ISL_1018153, EPI_ISL_1018154, EPI_ISL_1018155, EPI_ISL_1018156, EPI_ISL_1018157, EPI_ISL_1018158, EPI_ISL_1018159, EPI_ISL_1018160, EPI_ISL_1018161, EPI_ISL_1018162, EPI_ISL_1018163, EPI_ISL_1018164, EPI_ISL_1018165, EPI_ISL_1018166, EPI_ISL_1018167, EPI_ISL_1018168, EPI_ISL_1018169, EPI_ISL_1018170, EPI_ISL_1018171, EPI_ISL_1018172, EPI_ISL_1018173, EPI_ISL_1018174, EPI_ISL_1018175, EPI_ISL_1018176, EPI_ISL_1018177, EPI_ISL_1018178, EPI_ISL_1018179, EPI_ISL_1018180, EPI_ISL_1018181, EPI_ISL_1018182, EPI_ISL_1018183, EPI_ISL_1018184, EPI_ISL_1018185, EPI_ISL_1018186, EPI_ISL_1018187, EPI_ISL_1018188, EPI_ISL_1018189, EPI_ISL_1018190, EPI_ISL_1018191, EPI_ISL_1018192, EPI_ISL_1018197, EPI_ISL_1018198, EPI_ISL_1018199, EPI_ISL_1018200, EPI_ISL_1018201, EPI_ISL_1018203, EPI_ISL_1018204, EPI_ISL_1018206, EPI_ISL_1018207, EPI_ISL_1018208, EPI_ISL_1018209, EPI_ISL_1018210, EPI_ISL_1018211, EPI_ISL_1018212, EPI_ISL_1018213, EPI_ISL_1018214, EPI_ISL_1018215, EPI_ISL_1018216, EPI_ISL_1018217, EPI_ISL_1018230, EPI_ISL_1018231, EPI_ISL_1018232, EPI_ISL_1018233, EPI_ISL_1018236, EPI_ISL_1018237, EPI_ISL_1018238, EPI_ISL_1018239, EPI_ISL_1018240, EPI_ISL_1018243, EPI_ISL_1018244, EPI_ISL_1018245, EPI_ISL_1018246, EPI_ISL_1018247, EPI_ISL_1018248, EPI_ISL_1018249, EPI_ISL_1018250, EPI_ISL_1018251, EPI_ISL_1018252, EPI_ISL_1018253, EPI_ISL_1018254, EPI_ISL_1018255, EPI_ISL_1018256, EPI_ISL_1018257, EPI_ISL_1018259, EPI_ISL_1018260, EPI_ISL_1018262, EPI_ISL_1018263, EPI_ISL_1018264, EPI_ISL_1018265, EPI_ISL_1018266, EPI_ISL_1018268, EPI_ISL_1018269, EPI_ISL_1018270, EPI_ISL_1018271, EPI_ISL_1018273, EPI_ISL_1018274, EPI_ISL_1018275, EPI_ISL_1018276, EPI_ISL_1018279, EPI_ISL_1018280, EPI_ISL_1018281, EPI_ISL_1018282, EPI_ISL_1018284, EPI_ISL_1018286 | | | |
| see above | Department of Health Technology and Informatics, The Hong Kong Polytechnic University | Department of Health Technology and Informatics, The Hong Kong Polytechnic University | Gilman Kit-Hang Siu, Lam-Kwong Lee, Kenneth Siu-Sing Leung, Jake Siu-Lun Leung, Timothy Ting-Leung Ng, Chloe Toi-Mei Chan, Kingsley King-Gee Tam, Hiu-Yin Lao, Denise Sze-Hang Wong, Alan Ka-Lun Wu, Miranda Chong-Yee Yau, Yvette Wai-Man Lai, Kitty Sau-Chun Fung, Sandy Ka-Yee Chau, Barry Kin-Chung Wong, Wing-Kin To, Kristine Luk, Alex Yat-Man Ho, Tak-Lun Que, Kam-Tong Yip, Wing Cheong Yam, David Ho-Keung Shum, Shea Ping Yip |
| EPI_ISL_1023841 | Centro de Desenvolvimento Tecnológico em Saúde - CDTS | Centro de Desenvolvimento Tecnológico em Saúde - CDTS | Souza,T.M., Fintelman-Rodrigues,N., De Paula,A.D., Saraiva,F.B., Ferreira,M.A. and Sacramento,C.Q. |
| EPI_ISL_1039696 | Instituto Adolfo Lutz - Regional de Presidente Prudente | Instituto Adolfo Lutz, Interdiciplinary Procedures Center, Strategic Laboratory | Claudio Tavares Sacchi, Claudia Regina Gonçalves, Erica Valessa Ramos Gomes, Karoline Rodrigues Campos |
| EPI_ISL_1039702 | Instituto Adolfo Lutz - Regional de Aracatuba | Instituto Adolfo Lutz, Interdiciplinary Procedures Center, Strategic Laboratory | Claudio Tavares Sacchi, Claudia Regina Gonçalves, Erica Valessa Ramos Gomes, Karoline Rodrigues Campos |
| EPI_ISL_1040841, EPI_ISL_1040842, EPI_ISL_1040843, EPI_ISL_1040844, EPI_ISL_1040845, EPI_ISL_1040846, EPI_ISL_1040847, EPI_ISL_1040848, EPI_ISL_1040849, EPI_ISL_1040850, EPI_ISL_1040851 | | | |
| see above | LACEN do Mato Grosso do Sul | Instituto Adolfo Lutz, Interdiciplinary Procedures Center, Strategic Laboratory | Claudio Tavares Sacchi, Claudia Regina Gonçalves, Erica Valessa Ramos Gomes, Karoline Rodrigues Campos |
| EPI_ISL_1040923 | Grupo de Investigación en Enfermedades Tropicales del Ejército (GINETEJ), Laboratorio de Referencia e Investigación, Dirección de Sanidad Ejército, Bogotá, Colombia | Centro de Investigaciones en Microbiología y Biotecnología-UR (CIMBIUR), Facultad de Ciencias Naturales, Universidad del Rosario, Bogotá, Colombia | Juan David Ramírez, Marina Muñoz, Nathalia Ballesteros, Luz H. Patiño, Sergio Castañeda, Carlos A. Rincón, Claudia P. Mendez, Magda C. Oliveros, Julie J. Perez, Elizabeth K. Marquez, Frank de los Santos Ortiz, Camilo Correa-Cardenas, María C. Duque, Alberto Paniz-Mondolfi |
| EPI_ISL_1054940, EPI_ISL_1054944, EPI_ISL_1054947, EPI_ISL_1054960, EPI_ISL_1054964, EPI_ISL_1054968, EPI_ISL_1054978, EPI_ISL_1054980, EPI_ISL_1054983, EPI_ISL_1054988, EPI_ISL_1054995 | | | |
| see above | Instituto de Diagnostico y Referencia Epidemiologicos INDRE_RNLSP | Instituto de Diagnostico y Referencia Epidemiologicos (INDRE) | Claudia Wong-Arambula, Abril Rodriguez-Maldonado, Fabiola Garces-Ayala, Adnan Araiza-Rodriguez, David Fragoso-Fonseca, Sergio Rangel-Guerrero, Mayra Jimenez-Morales, Nancy Munoz-Hernandez, Natividad Cruz-Ortiz, Tatiana Nunez-Garcia, Gisela Barrera-Badillo, Lucia Hernandez-Rivas, Irma Lopez-Martinez, Ernesto Ramirez-Gonzalez. |
| EPI_ISL_1060735, EPI_ISL_1060736, EPI_ISL_1060743, EPI_ISL_1060752, EPI_ISL_1060757, EPI_ISL_1060758, EPI_ISL_1060760, EPI_ISL_1060762 | Instituto de Diagnostico y Referencia Epidemiologicos (INDRE)_RNLSP | Instituto de Diagnostico y Referencia Epidemiologicos (INDRE) | Claudia Wong-Arambula, Abril Rodriguez-Maldonado, Fabiola Garces-Ayala, Adnan Araiza-Rodriguez, David Fragoso-Fonseca, Sergio Rangel-Guerrero, Mayra Jimenez-Morales, Nancy Munoz-Hernandez, Natividad Cruz-Ortiz, Tatiana Nunez-Garcia, Gisela Barrera-Badillo, Lucia Hernandez-Rivas, Irma Lopez-Martinez, Ernesto Ramirez-Gonzalez. |
| EPI_ISL_1081834, EPI_ISL_1081835 | Virus Research and Diagnostic Laboratory, Medical College Level, Kasturba Hospital for Infectious Diseases | National Institute of Virology-Microbial Containment Complex, Indian Council of Medical Research | Pragya D. Yadav, Jayanti Shastri |
| EPI_ISL_1091786 | LDSP | Universidad Nacional de Colombia - Laboratorio Genómico One Health | Andres F. Cardona-Rios, Daniel O. Maldonado-Perez, Laura Silvana Perez, Karl A Ciuoderis, Maria Angélica Maya, Idabely Betancur Ortiz, Sandra Ines Cano, Diego A. Álvarez-Díaz, Carlos Franco-Muñoz, Marcela Mercado-Reyes, Jorge E. Osorio, Juan P. Hernandez-Ortiz |
| EPI_ISL_1098839 | Medical Ain Shams Research Institute (MASRI), Ain Shams University | Medical Ain Shams Research Institute (MASRI), Ain Shams University | Hesham Elghazaly, Sara Hassan Agwa, Hala Hafez , Fatma Ebied, Manal Hamdy Elsaid, Mohamed Elhadidi, Aya Mohamed , Shimaa Moustafa, Reham Kassab, Iman Foda , Samia Abdou Girgis, Hoda Ezz Elarab, Hagar Elshora, Osama Mansour. |
| EPI_ISL_1103556 | Kurnool Medical College (KMC) | CSIR Institute of Genomics and Integrative Biology | Pallavali Roja Rani, Mohamed Imran, Juturu Vijaya Lakshmi, Bani Jolly, Abhinav Jain, Avileli Surekha, Vigneshwar Senthivel, Pulala Chandrasekhar, Mohit Kumar Divakar, Damam Srinivasulu, Rahul C Bhoyar, PR Vanaja, Vinod Scaria, Sridhar Sivasubbu |
| EPI_ISL_1111068, EPI_ISL_1111070, EPI_ISL_1111071, EPI_ISL_1111072, EPI_ISL_1111073 | Laboratorio de Referencia Nacional de Enteropatógenos. Instituto Nacional de Salud del Perú | Laboratorio de Referencia Nacional de Enteropatógenos. Instituto Nacional de Salud del Perú | Ronnie Gavilan Chavez, Junior Caro Castro, Willi Quino Sifuentes, Veronica Hurtado Vela, Iris Silva Molina, Fiorella Orellana Peralta |
| EPI_ISL_1111074, EPI_ISL_1111082, EPI_ISL_1111083, EPI_ISL_1111084, EPI_ISL_1111085, EPI_ISL_1111099, EPI_ISL_1111121, EPI_ISL_1111165, EPI_ISL_1111166, EPI_ISL_1111167, EPI_ISL_1111168, EPI_ISL_1111173, EPI_ISL_1111191, EPI_ISL_1111241, EPI_ISL_1111242, EPI_ISL_1111243, EPI_ISL_1111244, EPI_ISL_1111245, EPI_ISL_1111246, EPI_ISL_1111247, EPI_ISL_1111248, EPI_ISL_1111249, EPI_ISL_1111250, EPI_ISL_1111251, EPI_ISL_1111252, EPI_ISL_1111253, EPI_ISL_1111254, EPI_ISL_1111255, EPI_ISL_1111256, EPI_ISL_1111257, EPI_ISL_1111258, EPI_ISL_1111259, EPI_ISL_1111260, EPI_ISL_1111261, EPI_ISL_1111262, EPI_ISL_1111263, EPI_ISL_1111264, EPI_ISL_1111265, EPI_ISL_1111266, EPI_ISL_1111311, EPI_ISL_1111357, EPI_ISL_1111359, EPI_ISL_1111360, EPI_ISL_1111361, EPI_ISL_1111362, EPI_ISL_1111363, EPI_ISL_1111364, EPI_ISL_1111365, EPI_ISL_1111383, EPI_ISL_1111384 | | | |
| see above | Laboratorio de Referencia Nacional de Virus Respiratorio. Instituto Nacional de Salud Perú | Laboratorio de Referencia Nacional de Enteropatógenos. Instituto Nacional de Salud del Perú | Ronnie Gavilan Chavez, Junior Caro Castro, Willi Quino Sifuentes, Veronica Hurtado Vela, Iris Silva Molina, Fiorella Orellana Peralta |
| EPI_ISL_1111385, EPI_ISL_1111386, EPI_ISL_1111387, EPI_ISL_1111388, EPI_ISL_1111389, EPI_ISL_1111390, EPI_ISL_1111391, EPI_ISL_1111392, EPI_ISL_1111393, EPI_ISL_1111394, EPI_ISL_1111395, EPI_ISL_1111396, EPI_ISL_1111397, EPI_ISL_1111398, EPI_ISL_1111399, EPI_ISL_1111400, EPI_ISL_1111401, EPI_ISL_1111402, EPI_ISL_1111403, EPI_ISL_1111404, EPI_ISL_1111405, EPI_ISL_1111406, EPI_ISL_1111407, EPI_ISL_1111408, EPI_ISL_1111409, EPI_ISL_1111410, EPI_ISL_1111411, EPI_ISL_1111412, EPI_ISL_1111413, EPI_ISL_1111414, EPI_ISL_1111415, EPI_ISL_1111416, EPI_ISL_1111417, EPI_ISL_1111418, EPI_ISL_1111419, EPI_ISL_1111420, EPI_ISL_1111421, EPI_ISL_1111422, EPI_ISL_1111423, EPI_ISL_1111424, EPI_ISL_1111425, EPI_ISL_1111426, EPI_ISL_1111427, EPI_ISL_1111428, EPI_ISL_1111429, EPI_ISL_1111430, EPI_ISL_1111431, EPI_ISL_1111432, EPI_ISL_1111433, EPI_ISL_1111434, EPI_ISL_1111435, EPI_ISL_1111436, EPI_ISL_1111437, EPI_ISL_1111438, EPI_ISL_1111439, EPI_ISL_1111440, EPI_ISL_1111441, EPI_ISL_1111442, EPI_ISL_1111443, EPI_ISL_1111444, EPI_ISL_1111445 | | | |

| | | | |
|---|---|---|---|
| see above | Laboratorio de Referencia Nacional de Enteropatógenos. Instituto Nacional de Salud del Perú | Laboratorio de Referencia Nacional de Enteropatógenos. Instituto Nacional de Salud del Perú | Ronnie Gavilan Chavez, Junior Caro Castro, Willi Quino Sifuentes, Veronica Hurtado Vela, Iris Silva Molina, Fiorella Orellana Peralta |
| EPI_ISL_1112951, EPI_ISL_1112952, EPI_ISL_1112953, EPI_ISL_1112954, EPI_ISL_1112955, EPI_ISL_1112957, EPI_ISL_1112958, EPI_ISL_1112959, EPI_ISL_1112960, EPI_ISL_1112961, EPI_ISL_1112962, EPI_ISL_1112963, EPI_ISL_1112964, EPI_ISL_1112965, EPI_ISL_1112966, EPI_ISL_1112967, EPI_ISL_1112968, EPI_ISL_1112969, EPI_ISL_1112970, EPI_ISL_1112971, EPI_ISL_1112972, EPI_ISL_1112973, EPI_ISL_1112974, EPI_ISL_1112975, EPI_ISL_1112976, EPI_ISL_1112977, EPI_ISL_1112978, EPI_ISL_1112979, EPI_ISL_1112980, EPI_ISL_1112981, EPI_ISL_1112982, EPI_ISL_1112983, EPI_ISL_1112984, EPI_ISL_1112985, EPI_ISL_1112986, EPI_ISL_1112987, EPI_ISL_1112988, EPI_ISL_1112989, EPI_ISL_1112990, EPI_ISL_1112991, EPI_ISL_1112993, EPI_ISL_1112994, EPI_ISL_1112995, EPI_ISL_1112996, EPI_ISL_1112997, EPI_ISL_1112998, EPI_ISL_1112999, EPI_ISL_1113000, EPI_ISL_1113001, EPI_ISL_1113002, EPI_ISL_1113003, EPI_ISL_1113004, EPI_ISL_1113005, EPI_ISL_1113006, EPI_ISL_1113007, EPI_ISL_1113008, EPI_ISL_1113009, EPI_ISL_1113010, EPI_ISL_1113011, EPI_ISL_1113012, EPI_ISL_1113013, EPI_ISL_1113014, EPI_ISL_1113015, EPI_ISL_1113016, EPI_ISL_1113017, EPI_ISL_1113018, EPI_ISL_1113019, EPI_ISL_1113020, EPI_ISL_1113021, EPI_ISL_1113022, EPI_ISL_1113023, EPI_ISL_1113024, EPI_ISL_1113025, EPI_ISL_1113026, EPI_ISL_1113027, EPI_ISL_1113028, EPI_ISL_1113029, EPI_ISL_1113030, EPI_ISL_1113031, EPI_ISL_1113032, EPI_ISL_1113033, EPI_ISL_1113034, EPI_ISL_1113035, EPI_ISL_1113036, EPI_ISL_1113037, EPI_ISL_1113038, EPI_ISL_1113039, EPI_ISL_1113040, EPI_ISL_1113041 | | | |
| see above | Baylor College of Medicine/ GCID | Baylor College of Medicine/ GCID | Vasanthi Avadhanula, Erin Nicholson, David Henke, Pedro Piedra, Harsha Doddapaneni, Donna Muzny, Qingchang Meng, Hsu Chao, Zeineen Momin, Hua Shen, George Weissenberger, Kavya Kottapalli, Yimiti Meiheerguli, Sejal Salvi, Ginger Metcalf, Vipin Menon, Sara J.J. Cregeen, Matthew C. Ross, Tulin Ayvaz, Richard Sucgang, Kristi L. Hoffman, Matthew Wong, Joseph F. Petrosino |
| EPI_ISL_1118929 | Institute of Clinical Hygiene, Medical Microbiology and Infectiology, General Hospital Nürnberg, Paracelsus Medical University, Nuremberg, Germany | Department of Molecular & Medical Virology, Ruhr University Bochum, 44801 Bochum, Germany \| Institute of Virology, Charité - Universitätsmedizin Berlin, corporate member of Freie Universität Berlin, Humboldt-Universität zu Berlin and Berlin Institute of Health (BIH), Berlin, Germany | Daniel Todt, Toni Luise Meister, Baxolele Mhlekude, Christine Goffinet, Jörg Steinmann, Stephanie Pfaender, Yannick Brüggemann, Eike Steinmann |
| EPI_ISL_1121326 | IAL Regional de Bauru | Instituto Adolfo Lutz, Interdiciplinary Procedures Center, Strategic Laboratory | Claudio Tavares Sacchi, Claudia Regina Gonçalves, Erica Valessa Ramos Gomes, Karoline Rodrigues Campos, Caio Vinicius Dias Lopes |
| EPI_ISL_1121327, EPI_ISL_1121328, EPI_ISL_1121329, EPI_ISL_1121330 | LACEN do Mato Grosso do Sul | Instituto Adolfo Lutz, Interdiciplinary Procedures Center, Strategic Laboratory | Claudio Tavares Sacchi, Claudia Regina Gonçalves, Erica Valessa Ramos Gomes, Karoline Rodrigues Campos, Caio Vinicius Dias Lopes |
| EPI_ISL_1129216, EPI_ISL_1129217, EPI_ISL_1129218, EPI_ISL_1129219, EPI_ISL_1129220, EPI_ISL_1129221, EPI_ISL_1129222, EPI_ISL_1129223, EPI_ISL_1129224, EPI_ISL_1129225, EPI_ISL_1129226, EPI_ISL_1129227 | | | |
| see above | Virus Research and Diagnostic Laboratory, Medical College Level, Kasturba Hospital for Infectious Diseases | National Institute of Virology-Microbial Containment Complex, Indian Council of Medical Research | Pragya D. Yadav, Jayanti Shastri |
| EPI_ISL_1138997, EPI_ISL_1138999 | Rajendra Institute of Medical Sciences, Ranchi, Jharkhand | Indian Council of Medical Research-National Institute of Virology, Microbial Containment Complex | Pragya D. Yadav, Manoj Kumar |
| EPI_ISL_1139052, EPI_ISL_1139053, EPI_ISL_1139054, EPI_ISL_1139055 | LACEN do Mato Grosso do Sul | Instituto Adolfo Lutz, Interdiciplinary Procedures Center, Strategic Laboratory | Claudio Tavares Sacchi, Claudia Regina Gonçalves, Erica Valessa Ramos Gomes, Karoline Rodrigues Campos, Caio Vinicius Dias Lopes |
| EPI_ISL_1139122, EPI_ISL_1139123, EPI_ISL_1139124, EPI_ISL_1139125, EPI_ISL_1139126, EPI_ISL_1139127, EPI_ISL_1139128, EPI_ISL_1139129, EPI_ISL_1139130, EPI_ISL_1139131, EPI_ISL_1139132, EPI_ISL_1139133, EPI_ISL_1139134, EPI_ISL_1139135, EPI_ISL_1139136, EPI_ISL_1139137, EPI_ISL_1139138, EPI_ISL_1139139, EPI_ISL_1139140, EPI_ISL_1139141, EPI_ISL_1139142, EPI_ISL_1139143, EPI_ISL_1139144, EPI_ISL_1139145 | | | |
| see above | ICMR-RMRC, Dibrugarh | Indian Council of Medical Research-National Institute of Virology, Microbial Containment Complex | Pragya D. Yadav, Biswajyoti Borkakoty |
| EPI_ISL_1163701, EPI_ISL_1163702, EPI_ISL_1163703, EPI_ISL_1163704, EPI_ISL_1163705, EPI_ISL_1163706, EPI_ISL_1163707, EPI_ISL_1163708 | LABCOVID_HCPA | LABRESIS_HCPA | Martins AF, Wink PL, Volpato F, Rosset C, de Paris F, Monteiro F, Zavascki AP, Barth AL |
| EPI_ISL_1164934, EPI_ISL_1164935 | B.J. Medical College and Civil hospital | Indian Council of Medical Research-National Institute of Virology, Microbial Containment Complex | Pragya D. Yadav, Disha Patel |
| EPI_ISL_1165067 | Siti Khodijah Hospital | Institute of Tropical Disease, Universitas Airlangga | Aldise M Nastri, Jezzy R Dewantari, Rima R Prasetya, Krisnoadi Rahardjo, Muhammad Hamdan, Gatot Soegiarto, Laksmi Wulandari, Resti Yudhawati, Yasuko Mori, Soetjipto, Kazufumi Shimizu, Maria I Lusida |
| EPI_ISL_1165068 | Anwar Medika General Hospital | Institute of Tropical Disease, Universitas Airlangga | Jezzy R Dewantari, Rima R Prasetya, Krisnoadi Rahardjo, Aldise M Nastri, Nungky Taniasari ,Gatot Soegiarto, Laksmi Wulandari, Resti Yudhawati, Yasuko Mori, Soetjipto, Kazufumi Shimizu, Maria I Lusida |
| EPI_ISL_1165069 | Siti Khodijah Hospital | Institute of Tropical Disease, Universitas Airlangga | Rima R Prasetya, Krisnoadi Rahardjo, Aldise M Nastri, Jezzy R Dewantari, Muhammad Hamdan, Gatot Soegiarto, Laksmi Wulandari, Resti Yudhawati, Yasuko Mori, Soetjipto, Kazufumi Shimizu, Maria I Lusida |
| EPI_ISL_1165075 | Paru Hospital | Institute of Tropical Disease, Universitas Airlangga | Jezzy R Dewantari, Rima R Prasetya, Krisnoadi Rahardjo, Aldise M Nastri, Dyah Retno, Gatot Soegiarto, Laksmi Wulandari, Resti Yudhawati, Yasuko Mori, Soetjipto, Kazufumi Shimizu, Maria I Lusida |
| EPI_ISL_1165093, EPI_ISL_1165094, EPI_ISL_1165095, EPI_ISL_1165096, EPI_ISL_1165097, EPI_ISL_1165098, EPI_ISL_1165099, EPI_ISL_1165100, EPI_ISL_1165101, EPI_ISL_1165102, EPI_ISL_1165103, EPI_ISL_1165104, EPI_ISL_1165105, EPI_ISL_1165106, EPI_ISL_1165107, EPI_ISL_1165108, EPI_ISL_1165109, EPI_ISL_1165110, EPI_ISL_1165111, EPI_ISL_1165112, EPI_ISL_1165113, EPI_ISL_1165114, EPI_ISL_1165115, EPI_ISL_1165116, EPI_ISL_1165117, EPI_ISL_1165118, EPI_ISL_1165119, EPI_ISL_1165120, EPI_ISL_1165121, EPI_ISL_1165122, EPI_ISL_1165123, EPI_ISL_1165124, EPI_ISL_1165125, EPI_ISL_1165126, EPI_ISL_1165127, EPI_ISL_1165128, EPI_ISL_1165129, EPI_ISL_1165130, EPI_ISL_1165132, EPI_ISL_1165133, EPI_ISL_1165134, EPI_ISL_1165136, EPI_ISL_1165137 | | | |
| see above | Government Medical College, Patiala, Punjab | Indian Council of Medical Research-National Institute of Virology, Microbial Containment Complex | Pragya D. Yadav, Rupinder Bakshi |
| EPI_ISL_1168693, EPI_ISL_1168694, EPI_ISL_1168695 | Puerto Rico Department of Health | Centers for Disease Control and Prevention, Dengue Branch | Gilberto A. Santiago, Glenda Gonzalez, Betzabel Flores, Keyla Charriez, Gabriela Paz-Bailey, Jorge L. Munoz-Jordan |
| EPI_ISL_1171620 | Instituto Adolfo Lutz Central | Instituto Adolfo Lutz, Interdiciplinary Procedures Center, Strategic Laboratory | Claudio Tavares Sacchi, Claudia Regina Gonçalves, Erica Valessa Ramos Gomes, Karoline Rodrigues Campos, Caio Vinicius Dias Lopes |
| EPI_ISL_1173511 | Mubarak Al-Kabeer Hospital | Virology Unit, Department of Microbiology, Faculty of Medicine, Kuwait | Nada Madi, Mohammed Sadeq, Anfal Al-Adwani |
| EPI_ISL_1180186, EPI_ISL_1180187 | Dept. of Microbiology, All India Institute of Medical Sciences, Jodhpur | Indian Council of Medical Research-National Institute of Virology, Microbial Containment Complex | Pragya D. Yadav, Vijaylakshmi Nag |
| EPI_ISL_1180195, EPI_ISL_1180196, EPI_ISL_1180197, EPI_ISL_1180198, EPI_ISL_1180199, EPI_ISL_1180200, EPI_ISL_1180201, EPI_ISL_1180202, EPI_ISL_1180203, EPI_ISL_1180204, EPI_ISL_1180205, EPI_ISL_1180206, EPI_ISL_1180207 | | | |
| see above | VRDL, GMC, Haldwani , Nainital | Indian Council of Medical Research-National Institute of Virology, Microbial Containment Complex | Pragya D. Yadav, Vinita Rawat |
| EPI_ISL_1181363, EPI_ISL_1181385, EPI_ISL_1181386, EPI_ISL_1181387, EPI_ISL_1181388, EPI_ISL_1181459 | Laboratory of Respiratory Viruses and Measles, Oswaldo Cruz Institute, FIOCRUZ | Laboratory of Respiratory Viruses and Measles, Oswaldo Cruz Institute, FIOCRUZ | Paola Resende, Luciana Appolinario, Fernando Motta, Anna Carolina Paixao, Ana Carolina Mendonca, Alice Sampaio Rocha, Renata Serrano Lopes, Marilda Siqueira on behalf of the Fiocruz COVID-19 Genomic Surveillance Network |
| EPI_ISL_1181474 | Laboratorio Central de Saude Publica do Estado do Parana (LACEN-PR) | Laboratory of Respiratory Viruses and Measles, Oswaldo Cruz Institute, FIOCRUZ | Paola Resende, Luciana Appolinario, Fernando Motta, Anna Carolina Paixao, Ana Carolina Mendonca, Alice Sampaio Rocha, Renata Serrano Lopes, Maria do Carmo Debur, Irina Nastassja Riediger, Marilda Siqueira on behalf of the Fiocruz COVID-19 Genomic Surveillance Network |
| EPI_ISL_1181476 | Laboratory of Respiratory Viruses and Measles, Oswaldo Cruz Institute, FIOCRUZ | Laboratory of Respiratory Viruses and Measles, Oswaldo Cruz Institute, FIOCRUZ | Paola Resende, Luciana Appolinario, Fernando Motta, Anna Carolina Paixao, Ana Carolina Mendonca, Alice Sampaio Rocha, Renata Serrano Lopes, Marilda Siqueira on behalf of the Fiocruz COVID-19 Genomic Surveillance Network |
| EPI_ISL_1181479, EPI_ISL_1181487 | Laboratorio de Imunologia de Transplantes de Goias LTDA (HLAGYN) | Laboratory of Respiratory Viruses and Measles, Oswaldo Cruz Institute, FIOCRUZ | Paola Resende, Luciana Appolinario, Fernando Motta, Anna Carolina Paixao, Ana Carolina Mendonca, Alice Sampaio Rocha, Renata Serrano Lopes , Alessandro Leonardo Alvares Magalhaes, Fernando Antonio Vinhal dos Santos, Erika Lopes Rocha Batista, Marilda Siqueira on behalf of the Fiocruz COVID-19 Genomic Surveillance Network |
| EPI_ISL_1181491, EPI_ISL_1181518 | Laboratory of Respiratory Viruses and Measles, Oswaldo | Laboratory of Respiratory Viruses and Measles, Oswaldo | Paola Resende, Luciana Appolinario, Fernando Motta, Anna Carolina Paixao, Ana Carolina Mendonca, Alice Sampaio Rocha, Renata Serrano Lopes, Marilda |

| | | | |
|---|---|---|---|
| | Cruz Institute, FIOCRUZ | Cruz Institute, FIOCRUZ | Siqueira on behalf of the Fiocruz COVID-19 Genomic Surveillance Network |
| EPI_ISL_1181530 | Laboratorio Central de Saude Publica do Estado do Parana (LACEN-PR) | Laboratory of Respiratory Viruses and Measles, Oswaldo Cruz Institute, FIOCRUZ | Paola Resende, Luciana Appolinario, Fernando Motta, Anna Carolina Paixao, Ana Carolina Mendonca, Alice Sampaio Rocha, Renata Serrano Lopes, Maria do Carmo Debur, Irina Nastassja Riediger, Marilda Siqueira on behalf of the Fiocruz COVID-19 Genomic Surveillance Network |
| EPI_ISL_1181532 | Laboratorio de Imunologia de Transplantes de Goias LTDA (HLAGYN) | Laboratory of Respiratory Viruses and Measles, Oswaldo Cruz Institute, FIOCRUZ | Paola Resende, Luciana Appolinario, Fernando Motta, Anna Carolina Paixao, Ana Carolina Mendonca, Alice Sampaio Rocha, Renata Serrano Lopes , Alessandro Leonardo Alvares Magalhaes, Fernando Antonio Vinhal dos Santos, Erika Lopes Rocha Batista, Marilda Siqueira on behalf of the Fiocruz COVID-19 Genomic Surveillance Network |
| EPI_ISL_1181536, EPI_ISL_1181576 | Laboratorio Central de Saude Publica do Estado do Parana (LACEN-PR) | Laboratory of Respiratory Viruses and Measles, Oswaldo Cruz Institute, FIOCRUZ | Paola Resende, Luciana Appolinario, Fernando Motta, Anna Carolina Paixao, Ana Carolina Mendonca, Alice Sampaio Rocha, Renata Serrano Lopes, Maria do Carmo Debur, Irina Nastassja Riediger, Marilda Siqueira on behalf of the Fiocruz COVID-19 Genomic Surveillance Network |
| EPI_ISL_1191738, EPI_ISL_1191739 | Laboratorio Biologia Molecolare Sars Cov2 - UOC Laboratorio Analisi - Servizio Medicina di Laboratorio, Ospedale "San Francesco" - ATS-ASSL Nuoro | Laboratorio specialistico UOC Ematologia - Ospedale "San Francesco" - ATS-ASSL Nuoro | Piras Giovanna, Asproni Rosanna,,Fiamma Maura, Monne Maria Itria, Palmas Angelo Domenico, Lo Maglio Iana, Mameli Giuseppe |
| EPI_ISL_1191781 | Virology Center Adolfo Lutz Institute Sao Paulo Brazil | Retrovirus Laboratory Adolfo Lutz Institute | Gabriela Bastos Cabral, Giselle I S Lopez-Lopes, Cintia Ahagon, Audrey Cilli, Paula Morena Guimaraes, Igor Mohamed Hussein, Luis Brigido |
| EPI_ISL_1192077, EPI_ISL_1192078 | Sri Venkateswara Institute of Medical Sciences SVIMS, Tirupati | Indian Council of Medical Research-National Institute of Virology, Microbial Containment Complex | Pragya D. Yadav, Usha Kalawat |
| EPI_ISL_1192079 | Viral Research and Diagnostic Laboratory, Dr. R.P. Govt. Medical College,Kangra | Indian Council of Medical Research-National Institute of Virology, Microbial Containment Complex | Pragya D. Yadav, S.C. Jaryal |
| EPI_ISL_1192080, EPI_ISL_1192081, EPI_ISL_1192082, EPI_ISL_1192083, EPI_ISL_1192084 | Sri Venkateswara Institute of Medical Sciences SVIMS, Tirupati | Indian Council of Medical Research-National Institute of Virology, Microbial Containment Complex | Pragya D. Yadav, Usha Kalawat |
| EPI_ISL_1192092 | Indira Gandhi Medical College & Hospital, Shimla | Indian Council of Medical Research-National Institute of Virology, Microbial Containment Complex | Pragya D. Yadav, Santwana Verma |
| EPI_ISL_1192093, EPI_ISL_1192094 | Sri Venkateswara Institute of Medical Sciences SVIMS, Tirupati | Indian Council of Medical Research-National Institute of Virology, Microbial Containment Complex | Pragya D. Yadav, Usha Kalawat |
| EPI_ISL_1192098, EPI_ISL_1192099 | Indira Gandhi Medical College & Hospital, Shimla | Indian Council of Medical Research-National Institute of Virology, Microbial Containment Complex | Pragya D. Yadav, Santwana Verma |
| EPI_ISL_1192101, EPI_ISL_1192102, EPI_ISL_1192103, EPI_ISL_1192104, EPI_ISL_1192105, EPI_ISL_1192106, EPI_ISL_1192107, EPI_ISL_1192108, EPI_ISL_1192109, EPI_ISL_1192110, EPI_ISL_1192111, EPI_ISL_1192112, EPI_ISL_1192113, EPI_ISL_1192121, EPI_ISL_1192122, EPI_ISL_1192123, EPI_ISL_1192124 | | | |
| see above | Sri Venkateswara Institute of Medical Sciences SVIMS, Tirupati | Indian Council of Medical Research-National Institute of Virology, Microbial Containment Complex | Pragya D. Yadav, Usha Kalawat |
| EPI_ISL_1192134, EPI_ISL_1192135, EPI_ISL_1192136, EPI_ISL_1192137, EPI_ISL_1192138, EPI_ISL_1192139, EPI_ISL_1192140, EPI_ISL_1192141, EPI_ISL_1192142, EPI_ISL_1192143, EPI_ISL_1192144, EPI_ISL_1192145, EPI_ISL_1192146, EPI_ISL_1192147, EPI_ISL_1192148 | | | |
| see above | Indira Gandhi Medical College & Hospital, Shimla | Indian Council of Medical Research-National Institute of Virology, Microbial Containment Complex | Pragya D. Yadav, Santwana Verma |
| EPI_ISL_1192151, EPI_ISL_1192152, EPI_ISL_1192153, EPI_ISL_1192154, EPI_ISL_1192155, EPI_ISL_1192156, EPI_ISL_1192157, EPI_ISL_1192158 | Viral Research and Diagnostic Laboratory, Dr. R.P. Govt. Medical College,Kangra | Indian Council of Medical Research-National Institute of Virology, Microbial Containment Complex | Pragya D. Yadav, S.C. Jaryal |
| EPI_ISL_1196283, EPI_ISL_1196284 | LACEN do Mato Grosso do Sul | Instituto Adolfo Lutz, Interdiciplinary Procedures Center, Strategic Laboratory | Claudio Tavares Sacchi, Claudia Regina Gonçalves, Erica Valessa Ramos Gomes, Karoline Rodrigues Campos, Caio Vinicius Dias Lopes |
| EPI_ISL_1231314, EPI_ISL_1231315, EPI_ISL_1231316, EPI_ISL_1231317, EPI_ISL_1231318, EPI_ISL_1231319, EPI_ISL_1231320, EPI_ISL_1231321, EPI_ISL_1231322, EPI_ISL_1231323, EPI_ISL_1231324 | | | |
| see above | Depratment of Microbiology, Viral Research and Diagnostic Laboratory | Indian Council of Medical Research-National Institute of Virology, Microbial Containment Complex | Pragya D. Yadav, Tapan Majumdar |
| EPI_ISL_1238787, EPI_ISL_1238788, EPI_ISL_1238789, EPI_ISL_1238790, EPI_ISL_1238791 | Ecological and Evolutionary Genomics. UGA-CINVESTAV | Unidad Universitaria de Secuenciación Masiva y Bioinformática (UUSMB). IBT-UNAM | Angelica Cibrian |
| EPI_ISL_1241861 | Institute of Life Sciences | Institute of Life Sciences | Sanchari Chatterjee, Ankita Datey, Soumya Sengupta, Sandhya Suranjika, Gargee Bhattacharya, Arup Ghosh, Atimukta Jha, Safal Rajesh Walia, Supriya Suman Keshry, Eshna Laha, Sunil Raghav, Amol Suryawanshi, Rupesh Dash, Shantibhushan Senapati, Tushar K. Beuria, Rajeeb Swain, Gulam Hussain Syed, Punit Prasad, Soma Chattopadhyay ,Odisha COVID-19 Study Group, ILS COVID-19 Team and Ajay Parida. |
| EPI_ISL_1242007 | Institute of Life Sciences | Institute of Life Sciences | Sanchari Chatterjee, Ankita Datey, Soumya Sengupta, Sandhya Suranjika, Gargee Bhattacharya, Arup Ghosh, Atimukta Jha, Safal Rajesh Walia, Supriya Suman Keshry, Eshna Laha, Sunil Raghav, Amol Suryawanshi, Rupesh Dash, Shantibhushan Senapati, Tushar K. Beuria, Rajeeb Swain, Gulam Hussain Syed, Punit Prasad, Soma Chattopadhyay ,Odisha COVID-19 Study Group, ILS COVID-19 Team and Ajay Parida |
| EPI_ISL_1251264 | Government Medical College, Patiala, Punjab | Indian Council of Medical Research-National Institute of Virology, Microbial Containment Complex | Pragya D. Yadav, Rupinder Bakshi |
| EPI_ISL_1251269 | Viral Research and Diagnostic Laboratory, Dr. R.P. Govt. Medical College,Kangra | Indian Council of Medical Research-National Institute of Virology, Microbial Containment Complex | Pragya D. Yadav, S.C. Jaryal |
| EPI_ISL_1251270, EPI_ISL_1251271 | Depratment of Microbiology, Viral Research and Diagnostic Laboratory | Indian Council of Medical Research-National Institute of Virology, Microbial Containment Complex | Pragya D. Yadav, Tapan Majumdar |
| EPI_ISL_1255113, EPI_ISL_1255114, EPI_ISL_1255115, EPI_ISL_1255117, EPI_ISL_1255228, EPI_ISL_1255242, EPI_ISL_1255244, EPI_ISL_1255245, EPI_ISL_1255246, EPI_ISL_1255247, EPI_ISL_1255248, EPI_ISL_1255249 | | | |
| see above | West African Centre for Cell Biology of Infectious Pathogens (WACCBIP), University of Ghana, Accra, Ghana | West African Centre for Cell Biology of Infectious Pathogens (WACCBIP), University of Ghana, Volta Road, Legon-Accra, Ghana | Collins M. Morang'a, Joyce M. Ngoi, Evelyn B. Quansah, Samirah Saiid, Dominic S.Y. Amuzu, Vincent Appiah, Philip M. Soglo, Vanessa Magnussen, Aisha Mohammed, Kesego Tapela, Nelson Kibinge, Abdoulaye B Diallo, Frederick Kumi-Ansah, Theophilus Odoom, Oliver D Boakye5, Emmanuella Amoako4, Abdul-Karim Abass, , Samuel Kaba Akoriyea, Frederick Tei-Maya, Lucas N. Amenga-Etego, Dam Kenneth Mibut, Yaw Bediako, Benjamin Demah Nuertey, Gordon A Awandare, Peter K Quashie, Gordon A Awandare, Yaw Bediako |
| EPI_ISL_1255389, EPI_ISL_1255390 | Institute of Life Sciences | Institute of Life Sciences | Sanchari Chatterjee, Ankita Datey, Soumya Sengupta, Sandhya Suranjika, Gargee Bhattacharya, Arup Ghosh, Atimukta Jha, Safal Rajesh Walia, Supriya Suman Keshry, Eshna Laha, Sunil Raghav, Amol Suryawanshi, Rupesh Dash, Shantibhushan Senapati, Tushar K. Beuria, Rajeeb Swain, Gulam Hussain Syed, Punit Prasad, Soma Chattopadhyay ,Odisha COVID-19 Study Group, ILS COVID-19 Team and Ajay Parida |
| EPI_ISL_1260838, EPI_ISL_1260839 | VRDL, GMC, Haldwani , Nainital | Indian Council of Medical Research-National Institute of Virology, Microbial Containment Complex | Pragya D. Yadav, Vinita Rawat |
| EPI_ISL_1279947 | National Institute of Infectious Diseases-Prof. Dr. Matei Bals Molecular Diagnostics Laboratory | National Institute of Infectious Diseases-Prof. Dr. Matei Bals Molecular Diagnostics Laboratory | Leontina Banica, Marius Surleac, Corina Casangiu, Petre Milu, Andreea Tudor, Simona Paraschiv, Dan Otelea |
| EPI_ISL_1301549 | Instituto de Diagnostico y Referencia Epidemiologicos InDRE_RNLSP | Instituto de Biotecnología de la UNAM | Authors from IBT, IMSS, InDRE and INER (in alphabetical order): Carlos F. Arias, Santiago Ávila-Ríos, Gisela Barrera-Badillo, Eduardo Becerril-Vargas, Celia Boukadida, Natividad Cruz-Ortiz, Larissa Fernandes-Matano, Ricardo Grande, Lucia Hernandez-Rivas, Alejandra Hernández-Terán, Pavel Isa, Irma Lopez-Martinez, José Arturo Martínez-Orozco, Margarita Matías-Florentino, Fidencio Mejía-Nepomuceno, Edgar Mendieta-Condado, Mario Mújica-Sánchez, |

| | | | |
|---|---|---|---|
| | | | José Esteban Muñoz-Medina, Tatiana Nunez-Garcia, Luis Alberto Ochoa-Carrera, Hector Esteban Paz-Juárez, Francisco Pulido, José Ernesto Ramírez-González, Alma Rincón-Rubio, Teresita Rojas-Mendoza, Jorge Salas-Hernández, Alejandro Sanchez-Flores, Clara Esperanza Santacruz-Tinoco, Andrea Santos Coy-Arechavaleta, Blanca Taboada, Gloria Vazquez, Joel Armando Vázquez-Pérez, Jerome Jean Verleyen |
| EPI_ISL_1301551, EPI_ISL_1301554 | Instituto Nacional de Enfermedades Respiratorias (INER) | Instituto de Biotecnología de la UNAM | Authors from IBT, IMSS, InDRE and INER (in alphabetical order): Carlos F. Arias, Santiago Ávila-Ríos, Gisela Barrera-Badillo, Eduardo Becerril-Vargas, Celia Boukadida, Natividad Cruz-Ortiz, Larissa Fernandes-Matano, Ricardo Grande, Lucia Hernandez-Rivas, Alejandra Hernández-Terán, Pavel Isa, Irma Lopez-Martinez, José Arturo Martínez-Orozco, Margarita Matías-Florentino, Fidencio Mejía-Nepomuceno, Edgar Mendieta-Condado, Mario Mújica-Sánchez, José Esteban Muñoz-Medina, Tatiana Nunez-Garcia, Luis Alberto Ochoa-Carrera, Hector Esteban Paz-Juárez, Francisco Pulido, José Ernesto Ramírez-González, Alma Rincón-Rubio, Teresita Rojas-Mendoza, Jorge Salas-Hernández, Alejandro Sanchez-Flores, Clara Esperanza Santacruz-Tinoco, Andrea Santos Coy-Arechavaleta, Blanca Taboada, Gloria Vazquez, Joel Armando Vázquez-Pérez, Jerome Jean Verleyen |
| EPI_ISL_1301555, EPI_ISL_1301556, EPI_ISL_1301557, EPI_ISL_1301558, EPI_ISL_1301559, EPI_ISL_1301560, EPI_ISL_1301561, EPI_ISL_1301562, EPI_ISL_1301563, EPI_ISL_1301565, EPI_ISL_1301566, EPI_ISL_1301567, EPI_ISL_1301568, EPI_ISL_1301569, EPI_ISL_1301570, EPI_ISL_1301571, EPI_ISL_1301572, EPI_ISL_1301573, EPI_ISL_1301574, EPI_ISL_1301585, EPI_ISL_1301586, EPI_ISL_1301587, EPI_ISL_1301588, EPI_ISL_1301589, EPI_ISL_1301590, EPI_ISL_1301592 | | | |
| see above | Instituto de Diagnostico y Referencia Epidemiologicos InDRE_RNLSP | Instituto de Biotecnología de la UNAM | Authors from IBT, IMSS, InDRE and INER (in alphabetical order): Carlos F. Arias, Santiago Ávila-Ríos, Gisela Barrera-Badillo, Eduardo Becerril-Vargas, Celia Boukadida, Natividad Cruz-Ortiz, Larissa Fernandes-Matano, Ricardo Grande, Lucia Hernandez-Rivas, Alejandra Hernández-Terán, Pavel Isa, Irma Lopez-Martinez, José Arturo Martínez-Orozco, Margarita Matías-Florentino, Fidencio Mejía-Nepomuceno, Edgar Mendieta-Condado, Mario Mújica-Sánchez, José Esteban Muñoz-Medina, Tatiana Nunez-Garcia, Luis Alberto Ochoa-Carrera, Hector Esteban Paz-Juárez, Francisco Pulido, José Ernesto Ramírez-González, Alma Rincón-Rubio, Teresita Rojas-Mendoza, Jorge Salas-Hernández, Alejandro Sanchez-Flores, Clara Esperanza Santacruz-Tinoco, Andrea Santos Coy-Arechavaleta, Blanca Taboada, Gloria Vazquez, Joel Armando Vázquez-Pérez, Jerome Jean Verleyen |
| EPI_ISL_1301597 | Laboratorio Central de Epidemiología IMSS | Instituto de Biotecnología de la UNAM | Authors from IBT, IMSS, InDRE and INER (in alphabetical order): Carlos F. Arias, Santiago Ávila-Ríos, Gisela Barrera-Badillo, Eduardo Becerril-Vargas, Celia Boukadida, Natividad Cruz-Ortiz, Larissa Fernandes-Matano, Ricardo Grande, Lucia Hernandez-Rivas, Alejandra Hernández-Terán, Pavel Isa, Irma Lopez-Martinez, José Arturo Martínez-Orozco, Margarita Matías-Florentino, Fidencio Mejía-Nepomuceno, Edgar Mendieta-Condado, Mario Mújica-Sánchez, José Esteban Muñoz-Medina, Tatiana Nunez-Garcia, Luis Alberto Ochoa-Carrera, Hector Esteban Paz-Juárez, Francisco Pulido, José Ernesto Ramírez-González, Alma Rincón-Rubio, Teresita Rojas-Mendoza, Jorge Salas-Hernández, Alejandro Sanchez-Flores, Clara Esperanza Santacruz-Tinoco, Andrea Santos Coy-Arechavaleta, Blanca Taboada, Gloria Vazquez, Joel Armando Vázquez-Pérez, Jerome Jean Verleyen |
| EPI_ISL_1301602, EPI_ISL_1301603, EPI_ISL_1301604, EPI_ISL_1301618, EPI_ISL_1301619 | Instituto Nacional de Enfermedades Respiratorias (INER) | Instituto de Biotecnología de la UNAM | Authors from IBT, IMSS, InDRE and INER (in alphabetical order): Carlos F. Arias, Santiago Ávila-Ríos, Gisela Barrera-Badillo, Eduardo Becerril-Vargas, Celia Boukadida, Natividad Cruz-Ortiz, Larissa Fernandes-Matano, Ricardo Grande, Lucia Hernandez-Rivas, Alejandra Hernández-Terán, Pavel Isa, Irma Lopez-Martinez, José Arturo Martínez-Orozco, Margarita Matías-Florentino, Fidencio Mejía-Nepomuceno, Edgar Mendieta-Condado, Mario Mújica-Sánchez, José Esteban Muñoz-Medina, Tatiana Nunez-Garcia, Luis Alberto Ochoa-Carrera, Hector Esteban Paz-Juárez, Francisco Pulido, José Ernesto Ramírez-González, Alma Rincón-Rubio, Teresita Rojas-Mendoza, Jorge Salas-Hernández, Alejandro Sanchez-Flores, Clara Esperanza Santacruz-Tinoco, Andrea Santos Coy-Arechavaleta, Blanca Taboada, Gloria Vazquez, Joel Armando Vázquez-Pérez, Jerome Jean Verleyen |
| EPI_ISL_1301672, EPI_ISL_1301673, EPI_ISL_1301674, EPI_ISL_1301675, EPI_ISL_1301676, EPI_ISL_1301677, EPI_ISL_1301678 | Laboratorio Central de Epidemiología IMSS | Instituto de Biotecnología de la UNAM | Authors from IBT, IMSS, InDRE and INER (in alphabetical order): Carlos F. Arias, Santiago Ávila-Ríos, Gisela Barrera-Badillo, Eduardo Becerril-Vargas, Celia Boukadida, Natividad Cruz-Ortiz, Larissa Fernandes-Matano, Ricardo Grande, Lucia Hernandez-Rivas, Alejandra Hernández-Terán, Pavel Isa, Irma Lopez-Martinez, José Arturo Martínez-Orozco, Margarita Matías-Florentino, Fidencio Mejía-Nepomuceno, Edgar Mendieta-Condado, Mario Mújica-Sánchez, José Esteban Muñoz-Medina, Tatiana Nunez-Garcia, Luis Alberto Ochoa-Carrera, Hector Esteban Paz-Juárez, Francisco Pulido, José Ernesto Ramírez-González, Alma Rincón-Rubio, Teresita Rojas-Mendoza, Jorge Salas-Hernández, Alejandro Sanchez-Flores, Clara Esperanza Santacruz-Tinoco, Andrea Santos Coy-Arechavaleta, Blanca Taboada, Gloria Vazquez, Joel Armando Vázquez-Pérez, Jerome Jean Verleyen |
| EPI_ISL_1301688, EPI_ISL_1301689, EPI_ISL_1301691, EPI_ISL_1301692, EPI_ISL_1301693, EPI_ISL_1301695, EPI_ISL_1301696, EPI_ISL_1301697, EPI_ISL_1301698, EPI_ISL_1301699, EPI_ISL_1301700, EPI_ISL_1301701, EPI_ISL_1301702, EPI_ISL_1301703, EPI_ISL_1301704, EPI_ISL_1301705, EPI_ISL_1301706, EPI_ISL_1301707, EPI_ISL_1301708, EPI_ISL_1301711, EPI_ISL_1301712, EPI_ISL_1301713 | | | |
| see above | Instituto de Diagnostico y Referencia Epidemiologicos InDRE_RNLSP | Instituto de Biotecnología de la UNAM | Authors from IBT, IMSS, InDRE and INER (in alphabetical order): Carlos F. Arias, Santiago Ávila-Ríos, Gisela Barrera-Badillo, Eduardo Becerril-Vargas, Celia Boukadida, Natividad Cruz-Ortiz, Larissa Fernandes-Matano, Ricardo Grande, Lucia Hernandez-Rivas, Alejandra Hernández-Terán, Pavel Isa, Irma Lopez-Martinez, José Arturo Martínez-Orozco, Margarita Matías-Florentino, Fidencio Mejía-Nepomuceno, Edgar Mendieta-Condado, Mario Mújica-Sánchez, José Esteban Muñoz-Medina, Tatiana Nunez-Garcia, Luis Alberto Ochoa-Carrera, Hector Esteban Paz-Juárez, Francisco Pulido, José Ernesto Ramírez-González, Alma Rincón-Rubio, Teresita Rojas-Mendoza, Jorge Salas-Hernández, Alejandro Sanchez-Flores, Clara Esperanza Santacruz-Tinoco, Andrea Santos Coy-Arechavaleta, Blanca Taboada, Gloria Vazquez, Joel Armando Vázquez-Pérez, Jerome Jean Verleyen |
| EPI_ISL_1301725, EPI_ISL_1301726, EPI_ISL_1301727, EPI_ISL_1301728, EPI_ISL_1301729 | Instituto Nacional de Enfermedades Respiratorias (INER) | Instituto de Biotecnología de la UNAM | Authors from IBT, IMSS, InDRE and INER (in alphabetical order): Carlos F. Arias, Santiago Ávila-Ríos, Gisela Barrera-Badillo, Eduardo Becerril-Vargas, Celia Boukadida, Natividad Cruz-Ortiz, Larissa Fernandes-Matano, Ricardo Grande, Lucia Hernandez-Rivas, Alejandra Hernández-Terán, Pavel Isa, Irma Lopez-Martinez, José Arturo Martínez-Orozco, Margarita Matías-Florentino, Fidencio Mejía-Nepomuceno, Edgar Mendieta-Condado, Mario Mújica-Sánchez, José Esteban Muñoz-Medina, Tatiana Nunez-Garcia, Luis Alberto Ochoa-Carrera, Hector Esteban Paz-Juárez, Francisco Pulido, José Ernesto Ramírez-González, Alma Rincón-Rubio, Teresita Rojas-Mendoza, Jorge Salas-Hernández, Alejandro Sanchez-Flores, Clara Esperanza Santacruz-Tinoco, Andrea Santos Coy-Arechavaleta, Blanca Taboada, Gloria Vazquez, Joel Armando Vázquez-Pérez, Jerome Jean Verleyen |
| EPI_ISL_1301730 | Instituto de Diagnostico y Referencia Epidemiologicos InDRE_RNLSP | Instituto de Biotecnología de la UNAM | Authors from IBT, IMSS, InDRE and INER (in alphabetical order): Carlos F. Arias, Santiago Ávila-Ríos, Gisela Barrera-Badillo, Eduardo Becerril-Vargas, Celia Boukadida, Natividad Cruz-Ortiz, Larissa Fernandes-Matano, Ricardo Grande, Lucia Hernandez-Rivas, Alejandra Hernández-Terán, Pavel Isa, Irma Lopez-Martinez, José Arturo Martínez-Orozco, Margarita Matías-Florentino, Fidencio Mejía-Nepomuceno, Edgar Mendieta-Condado, Mario Mújica-Sánchez, José Esteban Muñoz-Medina, Tatiana Nunez-Garcia, Luis Alberto Ochoa-Carrera, Hector Esteban Paz-Juárez, Francisco Pulido, José Ernesto Ramírez-González, Alma Rincón-Rubio, Teresita Rojas-Mendoza, Jorge Salas-Hernández, Alejandro Sanchez-Flores, Clara Esperanza Santacruz-Tinoco, Andrea Santos Coy-Arechavaleta, Blanca Taboada, Gloria Vazquez, Joel Armando Vázquez-Pérez, Jerome Jean Verleyen |
| EPI_ISL_1301732, EPI_ISL_1301733 | Instituto Nacional de Enfermedades Respiratorias (INER) | Instituto de Biotecnología de la UNAM | Authors from IBT, IMSS, InDRE and INER (in alphabetical order): Carlos F. Arias, Santiago Ávila-Ríos, Gisela Barrera-Badillo, Eduardo Becerril-Vargas, Celia Boukadida, Natividad Cruz-Ortiz, Larissa Fernandes-Matano, Ricardo Grande, Lucia Hernandez-Rivas, Alejandra Hernández-Terán, Pavel Isa, Irma Lopez-Martinez, José Arturo Martínez-Orozco, Margarita Matías-Florentino, Fidencio Mejía-Nepomuceno, Edgar Mendieta-Condado, Mario Mújica-Sánchez, José Esteban Muñoz-Medina, Tatiana Nunez-Garcia, Luis Alberto Ochoa-Carrera, Hector Esteban Paz-Juárez, Francisco Pulido, José Ernesto Ramírez-González, Alma Rincón-Rubio, Teresita Rojas-Mendoza, Jorge Salas-Hernández, Alejandro Sanchez-Flores, Clara Esperanza Santacruz-Tinoco, Andrea Santos Coy-Arechavaleta, Blanca Taboada, Gloria Vazquez, Joel Armando Vázquez-Pérez, Jerome Jean Verleyen |
| EPI_ISL_1335896 | Columbia Asia Hospital Medan | Faculty of Medicine, Universitas Sumatera Utara; Institute of Tropical Disease, Universitas Airlangga | Aldise M Nastri, Jezzy R Dewantari, Rima R Prasetya, Krisnoadi Rahardjo, Yasuko Mori, Soetjipto, Kazufumi Shimizu, Maria I Lusida, Meliani, R Andika D Cahyadi, Mirzan Hasibuan, Irbah R Nainggolan, Ramadhan Bestari, Franciscus Ginting, Muhammad Ichwan, R Lia Kusumawati, Inke N D Lubis |
| EPI_ISL_1340760, EPI_ISL_1340763 | Departamento de Virologia, Laboratorio Central de Salud Pública, Avenida Venezuela y Teniente Escurra, Asunción, Paraguay | Laboratory of Respiratory Viruses and Measles, Oswaldo Cruz Institute, FIOCRUZ | Paola Resende, Cynthia Vazquez, Luciana Appolinario, Fernando Motta, Anna Carolina Paixao, Ana Carolina Mendonca, Alice Sampaio Rocha, Renata Serrano Lopes, Marilda Siqueira on behalf of the Fiocruz COVID-19 Genomic Surveillance Network |
| EPI_ISL_1357605 | Dr. Ann Chahroudi's Lab | Dr. Thomas Vanderford's Lab | Thomas Vanderford, Maud Mavigner, Nils Schoof, Ann Chahroudi |
| EPI_ISL_1372187 | Cliniques universitaires Saint-Luc | UCLouvain/IREC/MBLG | Jean Ruelle, Lysa Pinsmaye, Eleonore Ngyuvula Mantu, Benoit Kabamba Mukadi |
| EPI_ISL_1398364 | Emam Ali Hospital | Razi Vaccine and Serum Research Institute | Amir Kaffashi, Jiabin Huang, S Reza Banihashemi, Mohammad Hossein Fallah Mehrabadi, Mohsen Lotfi, Morteza Taghizadeh, Akbar Khorasani, Mohsen Bashashati, Sayed Hamidreza Mozhgani |
| EPI_ISL_1398925 | Emam Ali Hospital | Razi Vaccine and Serum Research Institute | Amir Kaffashi, Jiabin Huang, S Reza Banihashemi, Mohammad Hossein Fallah Mehrabadi, Mohsen Lotfi, Morteza Taghizadeh, Akbar Khorasani, Mohsen Bashashati, Sayed Hamidreza Mozhgani |
| EPI_ISL_1404131, EPI_ISL_1404132, EPI_ISL_1404133, EPI_ISL_1404134, EPI_ISL_1404135, EPI_ISL_1404136, EPI_ISL_1404137, EPI_ISL_1404138, EPI_ISL_1404139, EPI_ISL_1404140, EPI_ISL_1404141, EPI_ISL_1404142 | | | |
| see above | HCRC | HCRC | Laurens Lambrechts, Jolien Van Cleemput, Nick Vereecke, Sebastiaan Theuns, Janneke Cox, Piet Maes, Linos Vandekerckhove |
| EPI_ISL_1406696 | Laboratory of Respiratory Viruses Teaching and Clinical Center of the Medical University of Lodz | Biobank Lab, Department of Molecular Biophysics, Faculty of Biology and Environmental Protection, University of Lodz | Strapagiel Dominik, Sobalska-Kwapis Marta, Dród Izabela, Tymoniuk Bogusaw, Traczyk-Borszyska Magdalena, Dastych Jaroslaw, Dziadek Jarosaw, Paradowska Edyta, Borowiec Maciej. |
| EPI_ISL_1416165, EPI_ISL_1416192, | Institut National d'hygiène | "Unité Mixte Internationale TransVIHMI (UMI 233 IRD - | Mounerou SALOU, Christelle BUTEL, Wembo A. HALATOKO,Issaka Maman, Abla A. KONOU, Amivi EHLAN, Adodo SADJI, Kokou TEGUENI,Sidonie |

| | | | |
|---|---|---|---|
| EPI_ISL_1416222 | | U1175 INSERM - Université de Montpellier) IRD (Institut de recherche pour le développement)" | A.M.KAGNISSODE, Akoélé SILIADIN, Alassane OURO-MEDELI, Messanh DOUFFAN,Déléma MABA,Sika DOSSIM, Améyo DORKENOO, Mireille PRINCE-DAVID,Anoumou DAGNRA,Laetitia SERRANO,Ahidjo AYOUBA,Eric DELAPORTE, Martine PEETERS |
| EPI_ISL_1424060, EPI_ISL_1424063 | Instituto Nacional de Salud- Dirección de Investigación en Salud Pública | Instituto Nacional de Salud- Dirección de Investigación en Salud Pública | Katherine Laiton-Donato, Carlos Franco-Muñoz, Diego A. Álvarez-Díaz, Mauricio Pacheco, Jhonnatan Reales-González, Diego Andrés Prada, Sheryll Corchuelo, Maria T. Herrera-Sepúlveda, Julian Naizaque, Gerardo Santamaría, Magdalena Wiesner, Martha Lucia Ospina Martínez, Marcela Mercado-Reyes. |
| EPI_ISL_1433132, EPI_ISL_1434443, EPI_ISL_1435763, EPI_ISL_1436817 | Institut National d'Hygiène | Unité Mixte Internationale TransVIHMI (UMI 233 IRD - U1175 INSERM - Université de Montpellier) IRD (Institut de recherche pour le développement) | Mounerou SALOU, Christelle BUTEL, Wembo A. HALATOKO,Issaka Maman, Abla A. KONOU, Amivi EHLAN, Adodo SADJI, Kokou TEGUENI,Sidonie A.M.KAGNISSODE, Akoélé SILIADIN, Alassane OURO-MEDELI, Messanh DOUFFAN,Déléma MABA,Sika DOSSIM, Améyo DORKENOO, Mireille PRINCE-DAVID,Anoumou DAGNRA,Laetitia SERRANO,Ahidjo AYOUBA,Eric DELAPORTE, Martine PEETERS |
| EPI_ISL_1447347 | Diagnostic and Research Center of Infectious Diseases, Medical Faculty, Andalas University | Diagnostic and Research Center of Infectious Diseases, Medical Faculty, Andalas University | Andani Eka Putra, Syafrizayanti, Ikhwan R. Sudji, Linosefa, Dede Rahman Agustian, Ayu Novita Trisnawati, Siskalil Fahma, Nia Ayuni Putri, Sekar Asri Tresnaningtyas, SM Rezvi, Fauzul Azhim, Dessy Arisanty, Gestina Aliska, Desmawati, Nita Afriani ,Syandrez Prima Putra, Juane Plantika Menra, Yolani Syaputri, Mutia Lailani |
| EPI_ISL_1468432, EPI_ISL_1468434 | LACEN do Mato Grosso do Sul | Instituto Adolfo Lutz, Interdiciplinary Procedures Center, Strategic Laboratory | Claudio Tavares Sacchi, Claudia Regina Gonçalves, Erica Valessa Ramos Gomes, Karoline Rodrigues Campos, Caio Vinicius Dias Lopes |
| EPI_ISL_1469587 | Hospital Sapiranga | Epiclin | Fernando Hayashi Sant'Anna, Ana Paula Muterle, Janira Prichula, Juliana Comerlato, Carolina Comerlato, Eliana Márcia Da Ros Wendland |
| EPI_ISL_1469713, EPI_ISL_1469724 | Fundação Hospitalar de Sapucaia do Sul | Epiclin | Fernando Hayashi Sant'Anna, Ana Paula Muterle, Janira Prichula, Juliana Comerlato, Carolina Comerlato, Eliana Márcia Da Ros Wendland |
| EPI_ISL_1469840 | Hospital Sapiranga | Epiclin | Fernando Hayashi Sant'Anna, Ana Paula Muterle, Janira Prichula, Juliana Comerlato, Carolina Comerlato, Eliana Márcia Da Ros Wendland |
| EPI_ISL_1489924, EPI_ISL_1489925, EPI_ISL_1489926 | Laboratory of Respiratory Viruses Teaching and Clinical Center of the Medical University of Lodz | Biobank Lab, Department of Molecular Biophysics, Faculty of Biology and Environmental Protection, University of Lodz | Strapagiel Dominik, Sobalska-Kwapis Marta, Dród Izabela, Tymoniuk Bogusaw, Traczyk-Borszyska Magdalena, Dastych Jaroslaw, Dziadek Jarosaw, Paradowska Edyta, Borowiec Maciej. |
| EPI_ISL_1494717 | Centro de Investigación en Ciencias de la Salud y Biomedicina, U.A.S.L.P | Centro de Investigación en Ciencias de la Salud y Biomedicina, U.A.S.L.P | MD PhD Sofía Bernal Silva |
| EPI_ISL_1494725, EPI_ISL_1494726, EPI_ISL_1494727 | Centro de Investigación en Ciencias de la Salud y Biomedicina, U.A.S.L.P. | Centro de Investigación en Ciencias de la Salud y Biomedicina, U.A.S.L.P. | MD PhD Sofía Bernal Silva |
| EPI_ISL_1495043, EPI_ISL_1495045, EPI_ISL_1495047, EPI_ISL_1495049, EPI_ISL_1495051, EPI_ISL_1495053, EPI_ISL_1495054 | Laboratory of Respiratory Viruses Teaching and Clinical Center of the Medical University of Lodz | Biobank Lab, Department of Molecular Biophysics, Faculty of Biology and Environmental Protection, University of Lodz | Strapagiel Dominik, Sobalska-Kwapis Marta, Dród Izabela, Tymoniuk Bogusaw, Traczyk-Borszyska Magdalena, Dastych Jaroslaw, Dziadek Jarosaw, Paradowska Edyta, Borowiec Maciej. |
| EPI_ISL_1500866 | Centro de Investigación en Ciencias de la Salud y Biomedicina, U.A.S.L.P. | Centro de Investigación en Ciencias de la Salud y Biomedicina, U.A.S.L.P. | MD PhD Sofía Bernal Silva |
| EPI_ISL_1502848, EPI_ISL_1502850, EPI_ISL_1502851, EPI_ISL_1502852, EPI_ISL_1502853, EPI_ISL_1502854, EPI_ISL_1502855, EPI_ISL_1502856, EPI_ISL_1502857, EPI_ISL_1502858, EPI_ISL_1502859, EPI_ISL_1502860 | | | |
| see above | Gorgas Memorial Laboratory of Health Studies | Gorgas Memorial Laboratory of Health Studies | Gonzalez Claudia, Leyda Abrego, Moreno Ambar, Oris Chavarria, Jessica Gondola, Marlenne Castillo, Ortiz Alma, Castillo Jorge, Moreno Brechla, Franco Danilo, Lopez-Verges Sandra, Martinez Alexander |
| EPI_ISL_1502865 | Hospital Regional Dr. Luis "Chicho" Fábrega | Gorgas Memorial Laboratory of Health Studies | Gonzalez Claudia, Herrera Melissa, Herrera Genarino, Leyda Abrego, Moreno Ambar, Oris Chavarria, Jessica Gondola, Marlenne Castillo, Ortiz Alma, Castillo Jorge, Moreno Brechla, Franco Danilo, Lopez-Verges Sandra, Martinez Alexander |
| EPI_ISL_1502866, EPI_ISL_1502871, EPI_ISL_1502874, EPI_ISL_1502876, EPI_ISL_1502877, EPI_ISL_1502879, EPI_ISL_1502881, EPI_ISL_1502884, EPI_ISL_1502885, EPI_ISL_1502886 | Gorgas Memorial Laboratory of Health Studies | Gorgas Memorial Laboratory of Health Studies | Gonzalez Claudia, Leyda Abrego, Moreno Ambar, Oris Chavarria, Jessica Gondola, Marlenne Castillo, Ortiz Alma, Castillo Jorge, Moreno Brechla, Franco Danilo, Lopez-Verges Sandra, Martinez Alexander |
| EPI_ISL_1510667, EPI_ISL_1510668 | University of Rome Tor Vergata: Departm Experim Medicine Chair of Virology | University of Rome Tor Vergata: Departm Experim Medicine Chair of Virology | Maria Concetta Bellocchi |
| EPI_ISL_1517392 | Inciensa, Instituto Costarricense de Investigación y Enseñanza en Nutrición y Salud | Inciensa, Instituto Costarricense de Investigación y Enseñanza en Nutrición y Salud | Cristian Peréz-Corrales, Valeria Peralta-Barquero & Patricia Salas |
| EPI_ISL_1517401, EPI_ISL_1517402, EPI_ISL_1517403 | Inciensa, Instituto Costarricense de Investigación y Enseñanza en Nutrición y Salud | Inciensa, Instituto Costarricense de Investigación y Enseñanza en Nutrición y Salud | Cristian Peréz-Corrales & Valeria Peralta-Barquero |
| EPI_ISL_1517404 | Inciensa, Instituto Costarricense de Investigación y Enseñanza en Nutrición y Salud | Inciensa, Instituto Costarricense de Investigación y Enseñanza en Nutrición y Salud | Cristian Peréz-Corrales, Valeria Peralta-Barquero & Mauricio Bolaños |
| EPI_ISL_1517405, EPI_ISL_1517406, EPI_ISL_1517407 | Inciensa, Instituto Costarricense de Investigación y Enseñanza en Nutrición y Salud | Inciensa, Instituto Costarricense de Investigación y Enseñanza en Nutrición y Salud | Cristian Peréz-Corrales, Valeria Peralta-Barquero & Gallegos-Carrillo B |
| EPI_ISL_1527373, EPI_ISL_1527374, EPI_ISL_1527375, EPI_ISL_1527376, EPI_ISL_1527377, EPI_ISL_1527378, EPI_ISL_1527379, EPI_ISL_1527380, EPI_ISL_1527381 | Molecular diagnostic laboratory of Federal Budget Institution of Science "Central Research Institute of Epidemiology" of The Federal Service on Customers' Rights Protection and Human Well-being Surveillance | Group of Genomics and Postgenomic Technologies of Central Research Institute of Epidemilology | Samoilov AE, Kaptelova VV, Korneenko EV, Saenko SS, Shipulina OY, Speranskaya AS, Tivanova EV, Kondrasheva LY, Akimkin VG |
| EPI_ISL_1532282, EPI_ISL_1532283, EPI_ISL_1532284, EPI_ISL_1532285 | Oman-National Influenza Center | Biotechnology & OMICs Laboratory | Ahmed Al Harrasi, Aisha Al-Amri, Intisar Al-Shukri, Amal Al-Maani, Sajjad Asaf, Bilal Hussain, Samiya Al-Zadjali, Ahmed N Al-Rawahi, Saqib Bilal, Abdul Latif Khan, Samira Al-Mahruqi, Ahmed Al-Rawahi, Hanan Al-Kindi, Amina Al-Jardani. |
| EPI_ISL_1532286, EPI_ISL_1532287, EPI_ISL_1532288, EPI_ISL_1532289 | Oman-National Influenza Center | Biotechnology & OMICs Laboratory | Intisar Al-Shukri, Ahmed Al Harrasi, Aisha Al-Amri, Amal Al-Maani, Sajjad Asaf, Fahad Al-Balushi, Laila Al-Balushi, Ahmed N Al-Rawahi, Saqib Bilal, Abdul Latif Khan, Samira Al-Mahruqi, Ahmed Al-Rawahi, Hanan Al-Kindi, Amina Al-Jardani. |
| EPI_ISL_1532290 | Oman-National Influenza Center | Biotechnology & OMICs Laboratory | Aisha Al-Amri, Ahmed Al Harrasi, Intisar Al-Shukri, Amal Al-Maani, Sajjad Asaf, Bilal Hussain, Samiya Al-Zadjali, Ahmed N Al-Rawahi, Saqib Bilal, Abdul Latif Khan, Samira Al-Mahruqi, Ahmed Al-Rawahi, Hanan Al-Kindi, Amina Al-Jardani. |
| EPI_ISL_1532302 | Oman-National Influenza Center | Biotechnology & OMICs Laboratory | Intisar Al-Shukri, Ahmed Al Harrasi, Aisha Al-Amri, Amal Al-Maani, Sajjad Asaf, Fahad Al-Balushi, Laila Al-Balushi, Ahmed N Al-Rawahi, Saqib Bilal, Abdul Latif Khan, Samira Al-Mahruqi, Ahmed Al-Rawahi, Hanan Al-Kindi, Amina Al-Jardani. |
| EPI_ISL_1532303 | Oman-National Influenza Center | Biotechnology & OMICs Laboratory | Aisha Al-Amri, Ahmed Al Harrasi, Intisar Al-Shukri, Amal Al-Maani, Sajjad Asaf, Bilal Hussain, Samiya Al-Zadjali, Ahmed N Al-Rawahi, Saqib Bilal, Abdul Latif Khan, Samira Al-Mahruqi, Ahmed Al-Rawahi, Hanan Al-Kindi, Amina Al-Jardani. |
| EPI_ISL_1532305 | Oman-National Influenza Center | Biotechnology & OMICs Laboratory | Abdul Latif Khan, Aisha Al-Amri, Intisar Al-Shukri, Amal Al-Maani, Sajjad Asaf, Bilal Hussain, Samiya Al-Zadjali, Ahmed N Al-Rawahi, Saqib Bilal, Samira Al-Mahruqi, Ahmed Al-Rawahi, Hanan Al-Kindi, Amina Al-Jardani, Ahmed Al Harrasi. |
| EPI_ISL_1532794 | Microbiology Department. Complexo Hospitalario Universitario de Vigo | Microbiology Department. Complexo Hospitalario Universitario de Vigo | Alfaya N, Alonso I, Alvarez M, Cabrera JJ, Carballo R, Cores O, Cortizo S, del-Campo V, Martinez L, Mediero G, Perez S, Potel C, Regueiro B, Rey S, Vasallo FJ |
| EPI_ISL_1533066, EPI_ISL_1533077, EPI_ISL_1533094 | Diagnostic and Research Center of Infectious Diseases, Medical Faculty, Andalas University | Diagnostic and Research Center of Infectious Diseases, Medical Faculty, Andalas University | Andani Eka Putra, Syafrizayanti, Ikhwan R. Sudji, Linosefa, Dede Rahman Agustian, Ayu Novita Trisnawati, Siskalil Fahma, Nia Ayuni Putri, Sekar Asri Tresnaningtyas, SM Rezvi, Fauzul Azhim, Dessy Arisanty, Gestina Aliska, Desmawati, Nita Afriani ,Syandrez Prima Putra, Juane Plantika Menra, Yolani Syaputri, Mutia Lailani |
| EPI_ISL_1543934 | Laboratory of Respiratory Viruses Teaching and Clinical Center of the Medical University of Lodz | Laboratory of Respiratory Viruses Teaching and Clinical Center of the Medical University of Lodz | Borowiec Maciej, Strapagiel Dominik, Sobalska-Kwapis Marta, Dród Izabela, Tymoniuk Bogusaw, Traczyk-Borszyska Magdalena, Dastych Jaroslaw, Dziadek Jarosaw, Paradowska Edyta |
| EPI_ISL_1543937, EPI_ISL_1543985, | Laboratory of Respiratory Viruses Teaching and Clinical | Laboratory of Respiratory Viruses Teaching and Clinical | Borowiec Maciej, Strapagiel Dominik, Sobalska-Kwapis Marta, Dród Izabela, Tymoniuk Bogusaw, Traczyk-Borszyska Magdalena, Dastych Jaroslaw, Dziadek |

| | | | |
|---|---|---|---|
| EPI_ISL_1543987 | Center of the Medical University of Lodz | Center of the Medical University of Lodz | Jarosaw, Paradowska Edyta, |
| EPI_ISL_1550388 | Instituto Oswaldo Cruz | Laboratorio de Virologia, Faculdade de Medicina, Universidade Federal de Mato Grosso, campus Cuiabá | María de Fátima Ferreira, Renata Dezengrini Slhessarenko, Janeth Aracely Ramirez Pavon, Luciano Nakazato, Valeria Dutra, Rosane Christiane Hahn, Paola Cristina Resende |
| EPI_ISL_1626483, EPI_ISL_1626485, EPI_ISL_1626486, EPI_ISL_1626487, EPI_ISL_1626491, EPI_ISL_1626492, EPI_ISL_1626494, EPI_ISL_1626496, EPI_ISL_1626497, EPI_ISL_1626498, EPI_ISL_1626503, EPI_ISL_1626504, EPI_ISL_1626505, EPI_ISL_1626506, EPI_ISL_1626507, EPI_ISL_1626508, EPI_ISL_1626509, EPI_ISL_1626510, EPI_ISL_1626511, EPI_ISL_1626513, EPI_ISL_1626518 | | | |
| see above | NSTU COVID-19 Diagnostic Center | NSU Genome Research Institute (NGRI) | Maqsud Hossain, Tahrima Saiha Huq, Aura Rahman, Md. Aminul Islam, Syeda Naushin Tabassum, Kazi Nadim Hasan, Abdul Khaleque, Abdus Sadique, Mohammad Salim Hossain, Newaz Mohammed Bahadur, Firoz Ahmed, Hasan Mahmud Reza |
| EPI_ISL_1672647 | ID and BG Hospital / CSIR - Indian Institute of Chemical Biology (CSIR-IICB) | CSIR Institute of Genomics and Integrative Biology (CSIR-IGIB) | Rajesh Pandey, Janani SV, Ranjeet Maurya, Akshay Kanakan, Neha Jha, Priyanka Mehta, Partha Chattopadhyay, Priti Devi, Shweta Sahni, Azka Khan, Sachin Sharma, Yogiraj Ray, Shekhar Ranjan Paul, Sandip Paul, Debaleena Bhowmik, Dipyaman Ganguly |
| EPI_ISL_1701311 | Laboratório de diagnóstico molecular da COVID-19 - Bahia, Santo Antônio de Jesus, Brazil | Laboratório Baculovírus | Josilene R Pinheiro, Jaime H Amorim, Aline Belmok, Fernando L Melo, Bergmann M Ribeiro |
| EPI_ISL_1708505, EPI_ISL_1708506 | Collection of WHO National Influenza Center, N.F. Gamaleya National Research Center for Epidemiology and Microbiology of the Ministry of Health of the Russian Federation | Laboratory of genome analysis, N.F. Gamaleya National Research Center for Epidemiology and Microbiology of the Ministry of Health of the Russian Federation | Olga Voronina, Ekaterina Aksenova, Natalia Ryzhova, Marina Kunda, Kirill Krasnoslobodtsev, Elena Burtseva, Lyudmila Kolobukhina, Svetlana Smetanina, Denis Logunov |
| EPI_ISL_1708507, EPI_ISL_1708510 | State collection of viruses, N.F. Gamaleya National Research Center for Epidemiology and Microbiology of the Ministry of Health of the Russian Federation | Laboratory of genome analysis, N.F. Gamaleya National Research Center for Epidemiology and Microbiology of the Ministry of Health of the Russian Federation | Olga Voronina, Ekaterina Aksenova, Natalia Ryzhova, Marina Kunda, Inna Dolzhikova, Andrey Botikov, Daria Grousova, Anna Kovyrshina, Denis Logunov |
| EPI_ISL_1716880, EPI_ISL_1752589, EPI_ISL_1752590, EPI_ISL_1752591 | Showa University PCR Center | Showa University PCR Center | Fumihiro Ishikawa, Hideto Oyamada, Yuko Udaka, Keiko Ishino, Yuji Kiuchi |
| EPI_ISL_1761308, EPI_ISL_1761309 | Tokyo Medical and Dental University | Tokyo Medical and Dental University | Kousuke Tanimoto, Yukie Tanaka, Yoko Nukui, Akinori Kimura, Hiroaki Takeuchi |
| EPI_ISL_1795043, EPI_ISL_1795044, EPI_ISL_1795045, EPI_ISL_1795046, EPI_ISL_1795047 | Oman-NIC | Department of Microbiology and Immunology-SQUH | Fahad Zadjali, Samira Al-Maruqi, Amina Al Jardani, Khulood Al-Mammary, Hanan Al-kindi, Fatma BaAlawi, Hamida AL Barwani, Zeyana AL-Dahmani, Intisar Al-Shukri, Aisha Al-Busaidi, Aisha Al-Amri, Ahlam Al-Amri, Mohammed Al-Tobi, Samiha Al Kharusi, Abdulla Balkhair |
| EPI_ISL_1795048 | Oman-NIC | Department of Microbiology and Immunology-SQUH | Samira Al-Maruqi,Fahad Zadjali, Amina Al Jardani, Khulood Al-Mammary, Hanan Al-kindi, Fatma BaAlawi, Hamida AL Barwani, Zeyana AL-Dahmani, Intisar Al-Shukri, Aisha Al-Busaidi, Aisha Al-Amri, Ahlam Al-Amri, Mohammed Al-Tobi, Samiha Al Kharusi, Abdulla Balkhair |
| EPI_ISL_1795049, EPI_ISL_1795050, EPI_ISL_1795051, EPI_ISL_1795052 | Oman-NIC | Oman-National Influenza Center-Department of Microbiology and Immunology-SQUH | Samira Al-Maruqi,Fahad Zadjali, Amina Al Jardani, Khulood Al-Mammary, Hanan Al-kindi, Fatma BaAlawi, Hamida AL Barwani, Zeyana AL-Dahmani, Intisar Al-Shukri, Aisha Al-Busaidi, Aisha Al-Amri, Ahlam Al-Amri, Mohammed Al-Tobi, Samiha Al Kharusi, Abdulla Balkhair |
| EPI_ISL_1799498 | Laboratório de Microbiologia Molecular - Universidade FEEVALE | Molecular Microbiology Laboratory | Alana Witt Hansen, Fágner Henrique Heldt, Fernando Rosado Spilki, Flávio Silveira, Juliana Schons Gularte, Juliane Deise Fleck, Mariana Soares da Silva, Meriane Demoliner, Matheus Nunes Weber, Paula Rodrigues de Almeida, Micheli Filippi. |
| EPI_ISL_1799499, EPI_ISL_1799501, EPI_ISL_1799502, EPI_ISL_1799505, EPI_ISL_1799507 | Laboratório de Microbiologia Molecular - Universidade FEEVALE | Molecular Microbiology Laboratory | Alana Witt Hansen, Fágner Henrique Heldt, Fernando Rosado Spilki, Flávio Silveira, Juliana Schons Gulart, Juliane Deise Fleck, Mariana Soares da Silva, Meriane Demoliner, Matheus Nunes Weber, Paula Rodrigues de Almeida, Michele Filippi. |
| EPI_ISL_1819072, EPI_ISL_1819073, EPI_ISL_1819074, EPI_ISL_1819075, EPI_ISL_1819076, EPI_ISL_1819077 | Oman-NIC | Oman-National Influenza Center-Department of Microbiology and Immunology-SQUH | Samira Al-Maruqi,Fahad Zadjali, Amina Al Jardani, Khulood Al-Mammary, Hanan Al-kindi, Fatma BaAlawi, Hamida AL Barwani, Zeyana AL-Dahmani, Intisar Al-Shukri, Aisha Al-Busaidi, Aisha Al-Amri, Ahlam Al-Amri, Mohammed Al-Tobi, Samiha Al Kharusi, Abdulla Balkhair |
| EPI_ISL_1859361 | Federal State Budgetary Educational Institution of Higher Education «Privolzhsky Research Medical University» of the Ministry of Health of the Russian Federation, Academic Consortium | FEDERAL RESEARCH AND CLINICAL CENTER OF PHYSICAL-CHEMICAL MEDICINE OF FEDERAL MEDICAL BIOLOGICAL AGENCY, Academic Consortium | Vladislav Babenko, Ramiz Bakhtyev, Vladimir Baklaushev, Larisa Balykova, Pavel Bashkirov, Anna Blagonravova, Daria Boldyreva, Dmitry Fedorov, Ilshat Gafurov, Raushaniya Gaifullina, Yulia Galeeva, Elena Galova, Vadim Govorun, Elena Ilina, Konstantin Ivanov, Daria Kharlampieva, Polina Khromova, Ksenia Klimina, Konstantin Kolontarev, Nadezhda Kolyshkina, Andrey Koritsky, Vyacheslav Kuropatkin, Vasily Lazarev, Alexander Manolov, Valentin Manuvera, Daria Matyushkina, Ekaterina Moskaleva, Varvara Musarova, Oleg Ogarkov, Elizaveta Orlova, Alexander Pavlenko, Alla Petrova, Natalia Pozhenko, Dmitry Pushkar, Alexander Rumyantsev, Sergey Rumyantsev, Vladimir Rumyantsev, Lyubov Rychkova, Alexander Samoilov, Irina Shirokova, Vyacheslav Sinkov, Svetlana Solovieva, Polina Tikhonova, Galina Trifonova, Alexander Troitsky, Alexander Tulichev, Yuri Udalov, Anna Varizhuk, Alexander Vasiliev, Vladimir Veselovsky, Alexey Volnukhin, Gaukhar Yusubalieva |
| EPI_ISL_1859397 | FEDERAL STATE BUDGETARY INSTITUTION OF HEALTHCARE HOSPITAL OF THE RUSSIAN ACADEMY OF SCIENCES (TROITSK), Academic Consortium | FEDERAL RESEARCH AND CLINICAL CENTER OF PHYSICAL-CHEMICAL MEDICINE OF FEDERAL MEDICAL BIOLOGICAL AGENCY, Academic Consortium | Vladislav Babenko, Ramiz Bakhtyev, Vladimir Baklaushev, Larisa Balykova, Pavel Bashkirov, Anna Blagonravova, Daria Boldyreva, Dmitry Fedorov, Ilshat Gafurov, Raushaniya Gaifullina, Yulia Galeeva, Elena Galova, Vadim Govorun, Elena Ilina, Konstantin Ivanov, Daria Kharlampieva, Polina Khromova, Ksenia Klimina, Konstantin Kolontarev, Nadezhda Kolyshkina, Andrey Koritsky, Vyacheslav Kuropatkin, Vasily Lazarev, Alexander Manolov, Valentin Manuvera, Daria Matyushkina, Ekaterina Moskaleva, Varvara Musarova, Oleg Ogarkov, Elizaveta Orlova, Alexander Pavlenko, Alla Petrova, Natalia Pozhenko, Dmitry Pushkar, Alexander Rumyantsev, Sergey Rumyantsev, Vladimir Rumyantsev, Lyubov Rychkova, Alexander Samoilov, Irina Shirokova, Vyacheslav Sinkov, Svetlana Solovieva, Polina Tikhonova, Galina Trifonova, Alexander Troitsky, Alexander Tulichev, Yuri Udalov, Anna Varizhuk, Alexander Vasiliev, Vladimir Veselovsky, Alexey Volnukhin, Gaukhar Yusubalieva |
| EPI_ISL_1859412 | Federal State Budgetary Educational Institution of Higher Education «Privolzhsky Research Medical University» of the Ministry of Health of the Russian Federation, Academic Consortium | FEDERAL RESEARCH AND CLINICAL CENTER OF PHYSICAL-CHEMICAL MEDICINE OF FEDERAL MEDICAL BIOLOGICAL AGENCY, Academic Consortium | Vladislav Babenko, Ramiz Bakhtyev, Vladimir Baklaushev, Larisa Balykova, Pavel Bashkirov, Anna Blagonravova, Daria Boldyreva, Dmitry Fedorov, Ilshat Gafurov, Raushaniya Gaifullina, Yulia Galeeva, Elena Galova, Vadim Govorun, Elena Ilina, Konstantin Ivanov, Daria Kharlampieva, Polina Khromova, Ksenia Klimina, Konstantin Kolontarev, Nadezhda Kolyshkina, Andrey Koritsky, Vyacheslav Kuropatkin, Vasily Lazarev, Alexander Manolov, Valentin Manuvera, Daria Matyushkina, Ekaterina Moskaleva, Varvara Musarova, Oleg Ogarkov, Elizaveta Orlova, Alexander Pavlenko, Alla Petrova, Natalia Pozhenko, Dmitry Pushkar, Alexander Rumyantsev, Sergey Rumyantsev, Vladimir Rumyantsev, Lyubov Rychkova, Alexander Samoilov, Irina Shirokova, Vyacheslav Sinkov, Svetlana Solovieva, Polina Tikhonova, Galina Trifonova, Alexander Troitsky, Alexander Tulichev, Yuri Udalov, Anna Varizhuk, Alexander Vasiliev, Vladimir Veselovsky, Alexey Volnukhin, Gaukhar Yusubalieva |
| EPI_ISL_1859425 | Federal State Public Scientific Institution «Scientific entre for Family Health and Human Reproduction Problems», Academic Consortium | FEDERAL RESEARCH AND CLINICAL CENTER OF PHYSICAL-CHEMICAL MEDICINE OF FEDERAL MEDICAL BIOLOGICAL AGENCY, Academic Consortium | Vladislav Babenko, Ramiz Bakhtyev, Vladimir Baklaushev, Larisa Balykova, Pavel Bashkirov, Anna Blagonravova, Daria Boldyreva, Dmitry Fedorov, Ilshat Gafurov, Raushaniya Gaifullina, Yulia Galeeva, Elena Galova, Vadim Govorun, Elena Ilina, Konstantin Ivanov, Daria Kharlampieva, Polina Khromova, Ksenia Klimina, Konstantin Kolontarev, Nadezhda Kolyshkina, Andrey Koritsky, Vyacheslav Kuropatkin, Vasily Lazarev, Alexander Manolov, Valentin Manuvera, Daria Matyushkina, Ekaterina Moskaleva, Varvara Musarova, Oleg Ogarkov, Elizaveta Orlova, Alexander Pavlenko, Alla Petrova, Natalia Pozhenko, Dmitry Pushkar, Alexander Rumyantsev, Sergey Rumyantsev, Vladimir Rumyantsev, Lyubov Rychkova, Alexander Samoilov, Irina Shirokova, Vyacheslav Sinkov, Svetlana Solovieva, Polina Tikhonova, Galina Trifonova, Alexander Troitsky, Alexander Tulichev, Yuri Udalov, Anna Varizhuk, Alexander Vasiliev, Vladimir Veselovsky, Alexey Volnukhin, Gaukhar Yusubalieva |
| EPI_ISL_1859429 | FEDERAL STATE BUDGETARY INSTITUTION OF HEALTHCARE HOSPITAL OF THE RUSSIAN ACADEMY OF SCIENCES (TROITSK), Academic Consortium | FEDERAL RESEARCH AND CLINICAL CENTER OF PHYSICAL-CHEMICAL MEDICINE OF FEDERAL MEDICAL BIOLOGICAL AGENCY, Academic Consortium | Vladislav Babenko, Ramiz Bakhtyev, Vladimir Baklaushev, Larisa Balykova, Pavel Bashkirov, Anna Blagonravova, Daria Boldyreva, Dmitry Fedorov, Ilshat Gafurov, Raushaniya Gaifullina, Yulia Galeeva, Elena Galova, Vadim Govorun, Elena Ilina, Konstantin Ivanov, Daria Kharlampieva, Polina Khromova, Ksenia Klimina, Konstantin Kolontarev, Nadezhda Kolyshkina, Andrey Koritsky, Vyacheslav Kuropatkin, Vasily Lazarev, Alexander Manolov, Valentin Manuvera, Daria Matyushkina, Ekaterina Moskaleva, Varvara Musarova, Oleg Ogarkov, Elizaveta Orlova, Alexander Pavlenko, Alla Petrova, Natalia Pozhenko, Dmitry Pushkar, Alexander Rumyantsev, Sergey Rumyantsev, Vladimir Rumyantsev, Lyubov Rychkova, Alexander Samoilov, Irina Shirokova, Vyacheslav Sinkov, Svetlana Solovieva, Polina Tikhonova, Galina Trifonova, Alexander Troitsky, Alexander Tulichev, Yuri Udalov, Anna Varizhuk, Alexander Vasiliev, Vladimir Veselovsky, Alexey Volnukhin, Gaukhar Yusubalieva |
| EPI_ISL_1859435 | Federal State Public Scientific Institution «Scientific entre for Family Health and Human Reproduction Problems», Academic Consortium | FEDERAL RESEARCH AND CLINICAL CENTER OF PHYSICAL-CHEMICAL MEDICINE OF FEDERAL MEDICAL BIOLOGICAL AGENCY, Academic Consortium | Vladislav Babenko, Ramiz Bakhtyev, Vladimir Baklaushev, Larisa Balykova, Pavel Bashkirov, Anna Blagonravova, Daria Boldyreva, Dmitry Fedorov, Ilshat Gafurov, Raushaniya Gaifullina, Yulia Galeeva, Elena Galova, Vadim Govorun, Elena Ilina, Konstantin Ivanov, Daria Kharlampieva, Polina Khromova, Ksenia Klimina, Konstantin Kolontarev, Nadezhda Kolyshkina, Andrey Koritsky, Vyacheslav Kuropatkin, Vasily Lazarev, Alexander Manolov, Valentin Manuvera, Daria Matyushkina, Ekaterina Moskaleva, Varvara Musarova, Oleg Ogarkov, Elizaveta Orlova, Alexander Pavlenko, Alla Petrova, Natalia Pozhenko, Dmitry Pushkar, Alexander Rumyantsev, Sergey Rumyantsev, Vladimir Rumyantsev, Lyubov Rychkova, Alexander Samoilov, Irina Shirokova, Vyacheslav Sinkov, Svetlana Solovieva, Polina Tikhonova, Galina Trifonova, Alexander Troitsky, Alexander Tulichev, Yuri Udalov, Anna Varizhuk, Alexander Vasiliev, Vladimir Veselovsky, Alexey Volnukhin, Gaukhar Yusubalieva |

| | | | |
|---|---|---|---|
| EPI_ISL_1859447 | City Clinical Hospital named after S.I. Spasokukotsky of Moscow Healthcare Department, Academic Consortium | FEDERAL RESEARCH AND CLINICAL CENTER OF PHYSICAL-CHEMICAL MEDICINE OF FEDERAL MEDICAL BIOLOGICAL AGENCY, Academic Consortium | Vladislav Babenko, Ramiz Bakhtyev, Vladimir Baklaushev, Larisa Balykova, Pavel Bashkirov, Anna Blagonravova, Daria Boldyreva, Dmitry Fedorov, Ilshat Gafurov, Raushaniya Gaifullina, Yulia Galeeva, Elena Galova, Vadim Govorun, Elena Ilina, Konstantin Ivanov, Daria Kharlampieva, Polina Khromova, Ksenia Klimina, Konstantin Kolontarev, Nadezhda Kolyshkina, Andrey Koritsky, Vyacheslav Kuropatkin, Vasily Lazarev, Alexander Manolov, Valentin Manuvera, Daria Matyushkina, Ekaterina Moskaleva, Varvara Musarova, Oleg Ogarkov, Elizaveta Orlova, Alexander Pavlenko, Alla Petrova, Natalia Pozhenko, Dmitry Pushkar, Alexander Rumyantsev, Sergey Rumyantsev, Vladimir Rumyantsev, Lyubov Rychkova, Alexander Samoilov, Irina Shirokova, Vyacheslav Sinkov, Svetlana Solovieva, Polina Tikhonova, Galina Trifonova, Alexander Troitsky, Alexander Tulichev, Yuri Udalov, Anna Varizhuk, Alexander Vasiliev, Vladimir Veselovsky, Alexey Volnukhin, Gaukhar Yusubalieva |
| EPI_ISL_1859449 | Federal State Budgetary Educational Institution of Higher Education «Privolzhsky Research Medical University» of the Ministry of Health of the Russian Federation, Academic Consortium | FEDERAL RESEARCH AND CLINICAL CENTER OF PHYSICAL-CHEMICAL MEDICINE OF FEDERAL MEDICAL BIOLOGICAL AGENCY, Academic Consortium | Vladislav Babenko, Ramiz Bakhtyev, Vladimir Baklaushev, Larisa Balykova, Pavel Bashkirov, Anna Blagonravova, Daria Boldyreva, Dmitry Fedorov, Ilshat Gafurov, Raushaniya Gaifullina, Yulia Galeeva, Elena Galova, Vadim Govorun, Elena Ilina, Konstantin Ivanov, Daria Kharlampieva, Polina Khromova, Ksenia Klimina, Konstantin Kolontarev, Nadezhda Kolyshkina, Andrey Koritsky, Vyacheslav Kuropatkin, Vasily Lazarev, Alexander Manolov, Valentin Manuvera, Daria Matyushkina, Ekaterina Moskaleva, Varvara Musarova, Oleg Ogarkov, Elizaveta Orlova, Alexander Pavlenko, Alla Petrova, Natalia Pozhenko, Dmitry Pushkar, Alexander Rumyantsev, Sergey Rumyantsev, Vladimir Rumyantsev, Lyubov Rychkova, Alexander Samoilov, Irina Shirokova, Vyacheslav Sinkov, Svetlana Solovieva, Polina Tikhonova, Galina Trifonova, Alexander Troitsky, Alexander Tulichev, Yuri Udalov, Anna Varizhuk, Alexander Vasiliev, Vladimir Veselovsky, Alexey Volnukhin, Gaukhar Yusubalieva |
| EPI_ISL_1859456, EPI_ISL_1859488, EPI_ISL_1859492, EPI_ISL_1859500 | Federal State Public Scientific Institution «Scientific entre for Family Health and Human Reproduction Problems», Academic Consortium | FEDERAL RESEARCH AND CLINICAL CENTER OF PHYSICAL-CHEMICAL MEDICINE OF FEDERAL MEDICAL BIOLOGICAL AGENCY, Academic Consortium | Vladislav Babenko, Ramiz Bakhtyev, Vladimir Baklaushev, Larisa Balykova, Pavel Bashkirov, Anna Blagonravova, Daria Boldyreva, Dmitry Fedorov, Ilshat Gafurov, Raushaniya Gaifullina, Yulia Galeeva, Elena Galova, Vadim Govorun, Elena Ilina, Konstantin Ivanov, Daria Kharlampieva, Polina Khromova, Ksenia Klimina, Konstantin Kolontarev, Nadezhda Kolyshkina, Andrey Koritsky, Vyacheslav Kuropatkin, Vasily Lazarev, Alexander Manolov, Valentin Manuvera, Daria Matyushkina, Ekaterina Moskaleva, Varvara Musarova, Oleg Ogarkov, Elizaveta Orlova, Alexander Pavlenko, Alla Petrova, Natalia Pozhenko, Dmitry Pushkar, Alexander Rumyantsev, Sergey Rumyantsev, Vladimir Rumyantsev, Lyubov Rychkova, Alexander Samoilov, Irina Shirokova, Vyacheslav Sinkov, Svetlana Solovieva, Polina Tikhonova, Galina Trifonova, Alexander Troitsky, Alexander Tulichev, Yuri Udalov, Anna Varizhuk, Alexander Vasiliev, Vladimir Veselovsky, Alexey Volnukhin, Gaukhar Yusubalieva |
| EPI_ISL_1859505 | Federal State Budgetary Educational Institution of Higher Education «Privolzhsky Research Medical University» of the Ministry of Health of the Russian Federation, Academic Consortium | FEDERAL RESEARCH AND CLINICAL CENTER OF PHYSICAL-CHEMICAL MEDICINE OF FEDERAL MEDICAL BIOLOGICAL AGENCY, Academic Consortium | Vladislav Babenko, Ramiz Bakhtyev, Vladimir Baklaushev, Larisa Balykova, Pavel Bashkirov, Anna Blagonravova, Daria Boldyreva, Dmitry Fedorov, Ilshat Gafurov, Raushaniya Gaifullina, Yulia Galeeva, Elena Galova, Vadim Govorun, Elena Ilina, Konstantin Ivanov, Daria Kharlampieva, Polina Khromova, Ksenia Klimina, Konstantin Kolontarev, Nadezhda Kolyshkina, Andrey Koritsky, Vyacheslav Kuropatkin, Vasily Lazarev, Alexander Manolov, Valentin Manuvera, Daria Matyushkina, Ekaterina Moskaleva, Varvara Musarova, Oleg Ogarkov, Elizaveta Orlova, Alexander Pavlenko, Alla Petrova, Natalia Pozhenko, Dmitry Pushkar, Alexander Rumyantsev, Sergey Rumyantsev, Vladimir Rumyantsev, Lyubov Rychkova, Alexander Samoilov, Irina Shirokova, Vyacheslav Sinkov, Svetlana Solovieva, Polina Tikhonova, Galina Trifonova, Alexander Troitsky, Alexander Tulichev, Yuri Udalov, Anna Varizhuk, Alexander Vasiliev, Vladimir Veselovsky, Alexey Volnukhin, Gaukhar Yusubalieva |
| EPI_ISL_1859507 | Federal Research and Clinical Center of Specialized Medical Care and Medical Technologies, Federal Biomedical Agency of the Russian Federation, Academic Consortium | FEDERAL RESEARCH AND CLINICAL CENTER OF PHYSICAL-CHEMICAL MEDICINE OF FEDERAL MEDICAL BIOLOGICAL AGENCY, Academic Consortium | Vladislav Babenko, Ramiz Bakhtyev, Vladimir Baklaushev, Larisa Balykova, Pavel Bashkirov, Anna Blagonravova, Daria Boldyreva, Dmitry Fedorov, Ilshat Gafurov, Raushaniya Gaifullina, Yulia Galeeva, Elena Galova, Vadim Govorun, Elena Ilina, Konstantin Ivanov, Daria Kharlampieva, Polina Khromova, Ksenia Klimina, Konstantin Kolontarev, Nadezhda Kolyshkina, Andrey Koritsky, Vyacheslav Kuropatkin, Vasily Lazarev, Alexander Manolov, Valentin Manuvera, Daria Matyushkina, Ekaterina Moskaleva, Varvara Musarova, Oleg Ogarkov, Elizaveta Orlova, Alexander Pavlenko, Alla Petrova, Natalia Pozhenko, Dmitry Pushkar, Alexander Rumyantsev, Sergey Rumyantsev, Vladimir Rumyantsev, Lyubov Rychkova, Alexander Samoilov, Irina Shirokova, Vyacheslav Sinkov, Svetlana Solovieva, Polina Tikhonova, Galina Trifonova, Alexander Troitsky, Alexander Tulichev, Yuri Udalov, Anna Varizhuk, Alexander Vasiliev, Vladimir Veselovsky, Alexey Volnukhin, Gaukhar Yusubalieva |
| EPI_ISL_1859537 | City Clinical Hospital named after S.I. Spasokukotsky of Moscow Healthcare Department, Academic Consortium | FEDERAL RESEARCH AND CLINICAL CENTER OF PHYSICAL-CHEMICAL MEDICINE OF FEDERAL MEDICAL BIOLOGICAL AGENCY, Academic Consortium | Vladislav Babenko, Ramiz Bakhtyev, Vladimir Baklaushev, Larisa Balykova, Pavel Bashkirov, Anna Blagonravova, Daria Boldyreva, Dmitry Fedorov, Ilshat Gafurov, Raushaniya Gaifullina, Yulia Galeeva, Elena Galova, Vadim Govorun, Elena Ilina, Konstantin Ivanov, Daria Kharlampieva, Polina Khromova, Ksenia Klimina, Konstantin Kolontarev, Nadezhda Kolyshkina, Andrey Koritsky, Vyacheslav Kuropatkin, Vasily Lazarev, Alexander Manolov, Valentin Manuvera, Daria Matyushkina, Ekaterina Moskaleva, Varvara Musarova, Oleg Ogarkov, Elizaveta Orlova, Alexander Pavlenko, Alla Petrova, Natalia Pozhenko, Dmitry Pushkar, Alexander Rumyantsev, Sergey Rumyantsev, Vladimir Rumyantsev, Lyubov Rychkova, Alexander Samoilov, Irina Shirokova, Vyacheslav Sinkov, Svetlana Solovieva, Polina Tikhonova, Galina Trifonova, Alexander Troitsky, Alexander Tulichev, Yuri Udalov, Anna Varizhuk, Alexander Vasiliev, Vladimir Veselovsky, Alexey Volnukhin, Gaukhar Yusubalieva |
| EPI_ISL_1859553 | Federal Research and Clinical Center of Specialized Medical Care and Medical Technologies, Federal Biomedical Agency of the Russian Federation, Academic Consortium | FEDERAL RESEARCH AND CLINICAL CENTER OF PHYSICAL-CHEMICAL MEDICINE OF FEDERAL MEDICAL BIOLOGICAL AGENCY, Academic Consortium | Vladislav Babenko, Ramiz Bakhtyev, Vladimir Baklaushev, Larisa Balykova, Pavel Bashkirov, Anna Blagonravova, Daria Boldyreva, Dmitry Fedorov, Ilshat Gafurov, Raushaniya Gaifullina, Yulia Galeeva, Elena Galova, Vadim Govorun, Elena Ilina, Konstantin Ivanov, Daria Kharlampieva, Polina Khromova, Ksenia Klimina, Konstantin Kolontarev, Nadezhda Kolyshkina, Andrey Koritsky, Vyacheslav Kuropatkin, Vasily Lazarev, Alexander Manolov, Valentin Manuvera, Daria Matyushkina, Ekaterina Moskaleva, Varvara Musarova, Oleg Ogarkov, Elizaveta Orlova, Alexander Pavlenko, Alla Petrova, Natalia Pozhenko, Dmitry Pushkar, Alexander Rumyantsev, Sergey Rumyantsev, Vladimir Rumyantsev, Lyubov Rychkova, Alexander Samoilov, Irina Shirokova, Vyacheslav Sinkov, Svetlana Solovieva, Polina Tikhonova, Galina Trifonova, Alexander Troitsky, Alexander Tulichev, Yuri Udalov, Anna Varizhuk, Alexander Vasiliev, Vladimir Veselovsky, Alexey Volnukhin, Gaukhar Yusubalieva |
| EPI_ISL_1859564, EPI_ISL_1859566 | Federal State Public Scientific Institution «Scientific entre for Family Health and Human Reproduction Problems», Academic Consortium | FEDERAL RESEARCH AND CLINICAL CENTER OF PHYSICAL-CHEMICAL MEDICINE OF FEDERAL MEDICAL BIOLOGICAL AGENCY, Academic Consortium | Vladislav Babenko, Ramiz Bakhtyev, Vladimir Baklaushev, Larisa Balykova, Pavel Bashkirov, Anna Blagonravova, Daria Boldyreva, Dmitry Fedorov, Ilshat Gafurov, Raushaniya Gaifullina, Yulia Galeeva, Elena Galova, Vadim Govorun, Elena Ilina, Konstantin Ivanov, Daria Kharlampieva, Polina Khromova, Ksenia Klimina, Konstantin Kolontarev, Nadezhda Kolyshkina, Andrey Koritsky, Vyacheslav Kuropatkin, Vasily Lazarev, Alexander Manolov, Valentin Manuvera, Daria Matyushkina, Ekaterina Moskaleva, Varvara Musarova, Oleg Ogarkov, Elizaveta Orlova, Alexander Pavlenko, Alla Petrova, Natalia Pozhenko, Dmitry Pushkar, Alexander Rumyantsev, Sergey Rumyantsev, Vladimir Rumyantsev, Lyubov Rychkova, Alexander Samoilov, Irina Shirokova, Vyacheslav Sinkov, Svetlana Solovieva, Polina Tikhonova, Galina Trifonova, Alexander Troitsky, Alexander Tulichev, Yuri Udalov, Anna Varizhuk, Alexander Vasiliev, Vladimir Veselovsky, Alexey Volnukhin, Gaukhar Yusubalieva |
| EPI_ISL_1859581 | City Clinical Hospital named after S.I. Spasokukotsky of Moscow Healthcare Department, Academic Consortium | FEDERAL RESEARCH AND CLINICAL CENTER OF PHYSICAL-CHEMICAL MEDICINE OF FEDERAL MEDICAL BIOLOGICAL AGENCY, Academic Consortium | Vladislav Babenko, Ramiz Bakhtyev, Vladimir Baklaushev, Larisa Balykova, Pavel Bashkirov, Anna Blagonravova, Daria Boldyreva, Dmitry Fedorov, Ilshat Gafurov, Raushaniya Gaifullina, Yulia Galeeva, Elena Galova, Vadim Govorun, Elena Ilina, Konstantin Ivanov, Daria Kharlampieva, Polina Khromova, Ksenia Klimina, Konstantin Kolontarev, Nadezhda Kolyshkina, Andrey Koritsky, Vyacheslav Kuropatkin, Vasily Lazarev, Alexander Manolov, Valentin Manuvera, Daria Matyushkina, Ekaterina Moskaleva, Varvara Musarova, Oleg Ogarkov, Elizaveta Orlova, Alexander Pavlenko, Alla Petrova, Natalia Pozhenko, Dmitry Pushkar, Alexander Rumyantsev, Sergey Rumyantsev, Vladimir Rumyantsev, Lyubov Rychkova, Alexander Samoilov, Irina Shirokova, Vyacheslav Sinkov, Svetlana Solovieva, Polina Tikhonova, Galina Trifonova, Alexander Troitsky, Alexander Tulichev, Yuri Udalov, Anna Varizhuk, Alexander Vasiliev, Vladimir Veselovsky, Alexey Volnukhin, Gaukhar Yusubalieva |
| EPI_ISL_1859586, EPI_ISL_1859592 | FEDERAL STATE BUDGETARY INSTITUTION OF HEALTHCARE HOSPITAL OF THE RUSSIAN ACADEMY OF SCIENCES (TROITSK), Academic Consortium | FEDERAL RESEARCH AND CLINICAL CENTER OF PHYSICAL-CHEMICAL MEDICINE OF FEDERAL MEDICAL BIOLOGICAL AGENCY, Academic Consortium | Vladislav Babenko, Ramiz Bakhtyev, Vladimir Baklaushev, Larisa Balykova, Pavel Bashkirov, Anna Blagonravova, Daria Boldyreva, Dmitry Fedorov, Ilshat Gafurov, Raushaniya Gaifullina, Yulia Galeeva, Elena Galova, Vadim Govorun, Elena Ilina, Konstantin Ivanov, Daria Kharlampieva, Polina Khromova, Ksenia Klimina, Konstantin Kolontarev, Nadezhda Kolyshkina, Andrey Koritsky, Vyacheslav Kuropatkin, Vasily Lazarev, Alexander Manolov, Valentin Manuvera, Daria Matyushkina, Ekaterina Moskaleva, Varvara Musarova, Oleg Ogarkov, Elizaveta Orlova, Alexander Pavlenko, Alla Petrova, Natalia Pozhenko, Dmitry Pushkar, Alexander Rumyantsev, Sergey Rumyantsev, Vladimir Rumyantsev, Lyubov Rychkova, Alexander Samoilov, Irina Shirokova, Vyacheslav Sinkov, Svetlana Solovieva, Polina Tikhonova, Galina Trifonova, Alexander Troitsky, Alexander Tulichev, Yuri Udalov, Anna Varizhuk, Alexander Vasiliev, Vladimir Veselovsky, Alexey Volnukhin, Gaukhar Yusubalieva |
| EPI_ISL_1859609 | Federal State Budgetary Educational Institution of Higher Education «Privolzhsky Research Medical University» of the Ministry of Health of the Russian Federation, Academic Consortium | FEDERAL RESEARCH AND CLINICAL CENTER OF PHYSICAL-CHEMICAL MEDICINE OF FEDERAL MEDICAL BIOLOGICAL AGENCY, Academic Consortium | Vladislav Babenko, Ramiz Bakhtyev, Vladimir Baklaushev, Larisa Balykova, Pavel Bashkirov, Anna Blagonravova, Daria Boldyreva, Dmitry Fedorov, Ilshat Gafurov, Raushaniya Gaifullina, Yulia Galeeva, Elena Galova, Vadim Govorun, Elena Ilina, Konstantin Ivanov, Daria Kharlampieva, Polina Khromova, Ksenia Klimina, Konstantin Kolontarev, Nadezhda Kolyshkina, Andrey Koritsky, Vyacheslav Kuropatkin, Vasily Lazarev, Alexander Manolov, Valentin Manuvera, Daria Matyushkina, Ekaterina Moskaleva, Varvara Musarova, Oleg Ogarkov, Elizaveta Orlova, Alexander Pavlenko, Alla Petrova, Natalia Pozhenko, Dmitry Pushkar, Alexander Rumyantsev, Sergey Rumyantsev, Vladimir Rumyantsev, Lyubov Rychkova, Alexander Samoilov, Irina Shirokova, Vyacheslav Sinkov, Svetlana Solovieva, Polina Tikhonova, Galina Trifonova, Alexander Troitsky, Alexander Tulichev, Yuri Udalov, Anna Varizhuk, Alexander Vasiliev, Vladimir Veselovsky, Alexey Volnukhin, Gaukhar Yusubalieva |
| EPI_ISL_1859615, EPI_ISL_1859617 | City Clinical Hospital named after S.I. Spasokukotsky of Moscow Healthcare Department, Academic Consortium | FEDERAL RESEARCH AND CLINICAL CENTER OF PHYSICAL-CHEMICAL MEDICINE OF FEDERAL MEDICAL BIOLOGICAL AGENCY, Academic Consortium | Vladislav Babenko, Ramiz Bakhtyev, Vladimir Baklaushev, Larisa Balykova, Pavel Bashkirov, Anna Blagonravova, Daria Boldyreva, Dmitry Fedorov, Ilshat Gafurov, Raushaniya Gaifullina, Yulia Galeeva, Elena Galova, Vadim Govorun, Elena Ilina, Konstantin Ivanov, Daria Kharlampieva, Polina Khromova, Ksenia Klimina, Konstantin Kolontarev, Nadezhda Kolyshkina, Andrey Koritsky, Vyacheslav Kuropatkin, Vasily Lazarev, Alexander Manolov, Valentin Manuvera, Daria Matyushkina, Ekaterina Moskaleva, Varvara Musarova, Oleg Ogarkov, Elizaveta Orlova, Alexander Pavlenko, Alla Petrova, Natalia Pozhenko, Dmitry Pushkar, Alexander Rumyantsev, Sergey Rumyantsev, Vladimir Rumyantsev, Lyubov Rychkova, Alexander Samoilov, Irina Shirokova, Vyacheslav Sinkov, |

| | | | |
|---|---|---|---|
| | | | Svetlana Solovieva, Polina Tikhonova, Galina Trifonova, Alexander Troitsky, Alexander Tulichev, Yuri Udalov, Anna Varizhuk, Alexander Vasiliev, Vladimir Veselovsky, Alexey Volnukhin, Gaukhar Yusubalieva |
| EPI_ISL_1859619 | Federal State Public Scientific Institution «Scientific entre for Family Health and Human Reproduction Problems», Academic Consortium | FEDERAL RESEARCH AND CLINICAL CENTER OF PHYSICAL-CHEMICAL MEDICINE OF FEDERAL MEDICAL BIOLOGICAL AGENCY, Academic Consortium | Vladislav Babenko, Ramiz Bakhtyev, Vladimir Baklaushev, Larisa Balykova, Pavel Bashkirov, Anna Blagonravova, Daria Boldyreva, Dmitry Fedorov, Ilshat Gafurov, Raushaniya Gaifullina, Yulia Galeeva, Elena Galova, Vadim Govorun, Elena Ilina, Konstantin Ivanov, Daria Kharlampieva, Polina Khromova, Ksenia Klimina, Konstantin Kolontarev, Nadezhda Kolyshkina, Andrey Koritsky, Vyacheslav Kuropatkin, Vasily Lazarev, Alexander Manolov, Valentin Manuvera, Daria Matyushkina, Ekaterina Moskaleva, Varvara Musarova, Oleg Ogarkov, Elizaveta Orlova, Alexander Pavlenko, Alla Petrova, Natalia Pozhenko, Dmitry Pushkar, Alexander Rumyantsev, Sergey Rumyantsev, Vladimir Rumyantsev, Lyubov Rychkova, Alexander Samoilov, Irina Shirokova, Vyacheslav Sinkov, Svetlana Solovieva, Polina Tikhonova, Galina Trifonova, Alexander Troitsky, Alexander Tulichev, Yuri Udalov, Anna Varizhuk, Alexander Vasiliev, Vladimir Veselovsky, Alexey Volnukhin, Gaukhar Yusubalieva |
| EPI_ISL_1859622 | City Clinical Hospital named after S.I. Spasokukotsky of Moscow Healthcare Department, Academic Consortium | FEDERAL RESEARCH AND CLINICAL CENTER OF PHYSICAL-CHEMICAL MEDICINE OF FEDERAL MEDICAL BIOLOGICAL AGENCY, Academic Consortium | Vladislav Babenko, Ramiz Bakhtyev, Vladimir Baklaushev, Larisa Balykova, Pavel Bashkirov, Anna Blagonravova, Daria Boldyreva, Dmitry Fedorov, Ilshat Gafurov, Raushaniya Gaifullina, Yulia Galeeva, Elena Galova, Vadim Govorun, Elena Ilina, Konstantin Ivanov, Daria Kharlampieva, Polina Khromova, Ksenia Klimina, Konstantin Kolontarev, Nadezhda Kolyshkina, Andrey Koritsky, Vyacheslav Kuropatkin, Vasily Lazarev, Alexander Manolov, Valentin Manuvera, Daria Matyushkina, Ekaterina Moskaleva, Varvara Musarova, Oleg Ogarkov, Elizaveta Orlova, Alexander Pavlenko, Alla Petrova, Natalia Pozhenko, Dmitry Pushkar, Alexander Rumyantsev, Sergey Rumyantsev, Vladimir Rumyantsev, Lyubov Rychkova, Alexander Samoilov, Irina Shirokova, Vyacheslav Sinkov, Svetlana Solovieva, Polina Tikhonova, Galina Trifonova, Alexander Troitsky, Alexander Tulichev, Yuri Udalov, Anna Varizhuk, Alexander Vasiliev, Vladimir Veselovsky, Alexey Volnukhin, Gaukhar Yusubalieva |
| EPI_ISL_1859624, EPI_ISL_1859632 | Federal State Public Scientific Institution «Scientific entre for Family Health and Human Reproduction Problems», Academic Consortium | FEDERAL RESEARCH AND CLINICAL CENTER OF PHYSICAL-CHEMICAL MEDICINE OF FEDERAL MEDICAL BIOLOGICAL AGENCY, Academic Consortium | Vladislav Babenko, Ramiz Bakhtyev, Vladimir Baklaushev, Larisa Balykova, Pavel Bashkirov, Anna Blagonravova, Daria Boldyreva, Dmitry Fedorov, Ilshat Gafurov, Raushaniya Gaifullina, Yulia Galeeva, Elena Galova, Vadim Govorun, Elena Ilina, Konstantin Ivanov, Daria Kharlampieva, Polina Khromova, Ksenia Klimina, Konstantin Kolontarev, Nadezhda Kolyshkina, Andrey Koritsky, Vyacheslav Kuropatkin, Vasily Lazarev, Alexander Manolov, Valentin Manuvera, Daria Matyushkina, Ekaterina Moskaleva, Varvara Musarova, Oleg Ogarkov, Elizaveta Orlova, Alexander Pavlenko, Alla Petrova, Natalia Pozhenko, Dmitry Pushkar, Alexander Rumyantsev, Sergey Rumyantsev, Vladimir Rumyantsev, Lyubov Rychkova, Alexander Samoilov, Irina Shirokova, Vyacheslav Sinkov, Svetlana Solovieva, Polina Tikhonova, Galina Trifonova, Alexander Troitsky, Alexander Tulichev, Yuri Udalov, Anna Varizhuk, Alexander Vasiliev, Vladimir Veselovsky, Alexey Volnukhin, Gaukhar Yusubalieva |
| EPI_ISL_1859635 | Federal Research and Clinical Center of Specialized Medical Care and Medical Technologies, Federal Biomedical Agency of the Russian Federation, Academic Consortium | FEDERAL RESEARCH AND CLINICAL CENTER OF PHYSICAL-CHEMICAL MEDICINE OF FEDERAL MEDICAL BIOLOGICAL AGENCY, Academic Consortium | Vladislav Babenko, Ramiz Bakhtyev, Vladimir Baklaushev, Larisa Balykova, Pavel Bashkirov, Anna Blagonravova, Daria Boldyreva, Dmitry Fedorov, Ilshat Gafurov, Raushaniya Gaifullina, Yulia Galeeva, Elena Galova, Vadim Govorun, Elena Ilina, Konstantin Ivanov, Daria Kharlampieva, Polina Khromova, Ksenia Klimina, Konstantin Kolontarev, Nadezhda Kolyshkina, Andrey Koritsky, Vyacheslav Kuropatkin, Vasily Lazarev, Alexander Manolov, Valentin Manuvera, Daria Matyushkina, Ekaterina Moskaleva, Varvara Musarova, Oleg Ogarkov, Elizaveta Orlova, Alexander Pavlenko, Alla Petrova, Natalia Pozhenko, Dmitry Pushkar, Alexander Rumyantsev, Sergey Rumyantsev, Vladimir Rumyantsev, Lyubov Rychkova, Alexander Samoilov, Irina Shirokova, Vyacheslav Sinkov, Svetlana Solovieva, Polina Tikhonova, Galina Trifonova, Alexander Troitsky, Alexander Tulichev, Yuri Udalov, Anna Varizhuk, Alexander Vasiliev, Vladimir Veselovsky, Alexey Volnukhin, Gaukhar Yusubalieva |
| EPI_ISL_1859645 | FEDERAL STATE BUDGETARY INSTITUTION OF HEALTHCARE HOSPITAL OF THE RUSSIAN ACADEMY OF SCIENCES (TROITSK), Academic Consortium | FEDERAL RESEARCH AND CLINICAL CENTER OF PHYSICAL-CHEMICAL MEDICINE OF FEDERAL MEDICAL BIOLOGICAL AGENCY, Academic Consortium | Vladislav Babenko, Ramiz Bakhtyev, Vladimir Baklaushev, Larisa Balykova, Pavel Bashkirov, Anna Blagonravova, Daria Boldyreva, Dmitry Fedorov, Ilshat Gafurov, Raushaniya Gaifullina, Yulia Galeeva, Elena Galova, Vadim Govorun, Elena Ilina, Konstantin Ivanov, Daria Kharlampieva, Polina Khromova, Ksenia Klimina, Konstantin Kolontarev, Nadezhda Kolyshkina, Andrey Koritsky, Vyacheslav Kuropatkin, Vasily Lazarev, Alexander Manolov, Valentin Manuvera, Daria Matyushkina, Ekaterina Moskaleva, Varvara Musarova, Oleg Ogarkov, Elizaveta Orlova, Alexander Pavlenko, Alla Petrova, Natalia Pozhenko, Dmitry Pushkar, Alexander Rumyantsev, Sergey Rumyantsev, Vladimir Rumyantsev, Lyubov Rychkova, Alexander Samoilov, Irina Shirokova, Vyacheslav Sinkov, Svetlana Solovieva, Polina Tikhonova, Galina Trifonova, Alexander Troitsky, Alexander Tulichev, Yuri Udalov, Anna Varizhuk, Alexander Vasiliev, Vladimir Veselovsky, Alexey Volnukhin, Gaukhar Yusubalieva |
| EPI_ISL_1859663 | Federal State Public Scientific Institution «Scientific entre for Family Health and Human Reproduction Problems», Academic Consortium | FEDERAL RESEARCH AND CLINICAL CENTER OF PHYSICAL-CHEMICAL MEDICINE OF FEDERAL MEDICAL BIOLOGICAL AGENCY, Academic Consortium | Vladislav Babenko, Ramiz Bakhtyev, Vladimir Baklaushev, Larisa Balykova, Pavel Bashkirov, Anna Blagonravova, Daria Boldyreva, Dmitry Fedorov, Ilshat Gafurov, Raushaniya Gaifullina, Yulia Galeeva, Elena Galova, Vadim Govorun, Elena Ilina, Konstantin Ivanov, Daria Kharlampieva, Polina Khromova, Ksenia Klimina, Konstantin Kolontarev, Nadezhda Kolyshkina, Andrey Koritsky, Vyacheslav Kuropatkin, Vasily Lazarev, Alexander Manolov, Valentin Manuvera, Daria Matyushkina, Ekaterina Moskaleva, Varvara Musarova, Oleg Ogarkov, Elizaveta Orlova, Alexander Pavlenko, Alla Petrova, Natalia Pozhenko, Dmitry Pushkar, Alexander Rumyantsev, Sergey Rumyantsev, Vladimir Rumyantsev, Lyubov Rychkova, Alexander Samoilov, Irina Shirokova, Vyacheslav Sinkov, Svetlana Solovieva, Polina Tikhonova, Galina Trifonova, Alexander Troitsky, Alexander Tulichev, Yuri Udalov, Anna Varizhuk, Alexander Vasiliev, Vladimir Veselovsky, Alexey Volnukhin, Gaukhar Yusubalieva |
| EPI_ISL_1859669 | Federal State Budgetary Educational Institution of Higher Education «Privolzhsky Research Medical University» of the Ministry of Health of the Russian Federation, Academic Consortium | FEDERAL RESEARCH AND CLINICAL CENTER OF PHYSICAL-CHEMICAL MEDICINE OF FEDERAL MEDICAL BIOLOGICAL AGENCY, Academic Consortium | Vladislav Babenko, Ramiz Bakhtyev, Vladimir Baklaushev, Larisa Balykova, Pavel Bashkirov, Anna Blagonravova, Daria Boldyreva, Dmitry Fedorov, Ilshat Gafurov, Raushaniya Gaifullina, Yulia Galeeva, Elena Galova, Vadim Govorun, Elena Ilina, Konstantin Ivanov, Daria Kharlampieva, Polina Khromova, Ksenia Klimina, Konstantin Kolontarev, Nadezhda Kolyshkina, Andrey Koritsky, Vyacheslav Kuropatkin, Vasily Lazarev, Alexander Manolov, Valentin Manuvera, Daria Matyushkina, Ekaterina Moskaleva, Varvara Musarova, Oleg Ogarkov, Elizaveta Orlova, Alexander Pavlenko, Alla Petrova, Natalia Pozhenko, Dmitry Pushkar, Alexander Rumyantsev, Sergey Rumyantsev, Vladimir Rumyantsev, Lyubov Rychkova, Alexander Samoilov, Irina Shirokova, Vyacheslav Sinkov, Svetlana Solovieva, Polina Tikhonova, Galina Trifonova, Alexander Troitsky, Alexander Tulichev, Yuri Udalov, Anna Varizhuk, Alexander Vasiliev, Vladimir Veselovsky, Alexey Volnukhin, Gaukhar Yusubalieva |
| EPI_ISL_1859688, EPI_ISL_1859720, EPI_ISL_1859742 | Federal State Public Scientific Institution «Scientific entre for Family Health and Human Reproduction Problems», Academic Consortium | FEDERAL RESEARCH AND CLINICAL CENTER OF PHYSICAL-CHEMICAL MEDICINE OF FEDERAL MEDICAL BIOLOGICAL AGENCY, Academic Consortium | Vladislav Babenko, Ramiz Bakhtyev, Vladimir Baklaushev, Larisa Balykova, Pavel Bashkirov, Anna Blagonravova, Daria Boldyreva, Dmitry Fedorov, Ilshat Gafurov, Raushaniya Gaifullina, Yulia Galeeva, Elena Galova, Vadim Govorun, Elena Ilina, Konstantin Ivanov, Daria Kharlampieva, Polina Khromova, Ksenia Klimina, Konstantin Kolontarev, Nadezhda Kolyshkina, Andrey Koritsky, Vyacheslav Kuropatkin, Vasily Lazarev, Alexander Manolov, Valentin Manuvera, Daria Matyushkina, Ekaterina Moskaleva, Varvara Musarova, Oleg Ogarkov, Elizaveta Orlova, Alexander Pavlenko, Alla Petrova, Natalia Pozhenko, Dmitry Pushkar, Alexander Rumyantsev, Sergey Rumyantsev, Vladimir Rumyantsev, Lyubov Rychkova, Alexander Samoilov, Irina Shirokova, Vyacheslav Sinkov, Svetlana Solovieva, Polina Tikhonova, Galina Trifonova, Alexander Troitsky, Alexander Tulichev, Yuri Udalov, Anna Varizhuk, Alexander Vasiliev, Vladimir Veselovsky, Alexey Volnukhin, Gaukhar Yusubalieva |
| EPI_ISL_1859762 | Federal State Budgetary Educational Institution of Higher Education «Privolzhsky Research Medical University» of the Ministry of Health of the Russian Federation, Academic Consortium | FEDERAL RESEARCH AND CLINICAL CENTER OF PHYSICAL-CHEMICAL MEDICINE OF FEDERAL MEDICAL BIOLOGICAL AGENCY, Academic Consortium | Vladislav Babenko, Ramiz Bakhtyev, Vladimir Baklaushev, Larisa Balykova, Pavel Bashkirov, Anna Blagonravova, Daria Boldyreva, Dmitry Fedorov, Ilshat Gafurov, Raushaniya Gaifullina, Yulia Galeeva, Elena Galova, Vadim Govorun, Elena Ilina, Konstantin Ivanov, Daria Kharlampieva, Polina Khromova, Ksenia Klimina, Konstantin Kolontarev, Nadezhda Kolyshkina, Andrey Koritsky, Vyacheslav Kuropatkin, Vasily Lazarev, Alexander Manolov, Valentin Manuvera, Daria Matyushkina, Ekaterina Moskaleva, Varvara Musarova, Oleg Ogarkov, Elizaveta Orlova, Alexander Pavlenko, Alla Petrova, Natalia Pozhenko, Dmitry Pushkar, Alexander Rumyantsev, Sergey Rumyantsev, Vladimir Rumyantsev, Lyubov Rychkova, Alexander Samoilov, Irina Shirokova, Vyacheslav Sinkov, Svetlana Solovieva, Polina Tikhonova, Galina Trifonova, Alexander Troitsky, Alexander Tulichev, Yuri Udalov, Anna Varizhuk, Alexander Vasiliev, Vladimir Veselovsky, Alexey Volnukhin, Gaukhar Yusubalieva |
| EPI_ISL_1910397, EPI_ISL_1910398 | Centre Hospitalier Universitaire Clermont-Ferrand | CHU Clermont-Ferrand, service de virologie | Bisseux Maxime, Mirand Audrey, Combes Patricia, Henquell Cécile |
| EPI_ISL_1972894 | National Public Health Laboratory Malaysia | National Public Health Laboratory Malaysia | Noorliza MN, JL Tan, Chong CK, Norhayati R, Yukie C, Norazimah T Rehan SAB Selvanesan S, Hannah PYP, NurulAina MCA, W.NurAfiza WMA Zirwatul AA, Hani MH, Nurulsyahida I, Aziyati O, Ushananthiny R, Kamal HKZ, Mohd AY, YF Ngeow |
| EPI_ISL_2000568, EPI_ISL_2000569 | Tropical Medicine Institute Pedro Kouri | Respiratory Infections Laboratory | Yahisel Tejero Suarez, Lissette Perez Santos, Yanet Pintos Saavedra, Daylin Mederos, Odalys Valdes, Mirtha Elisa Aguado, Vivian Kouri Cardella |
| EPI_ISL_2001481, EPI_ISL_2001482 | National Public Health Laboratory Malaysia | National Public Health Laboratory Malaysia | Noorliza MN, JL Tan, Chong CK, Norhayati R, Yukie C, Norazimah T Rehan SAB Selvanesan S, Hannah PYP, NurulAina MCA, W.NurAfiza WMA Zirwatul AA, Hani MH, Nurulsyahida I, Aziyati O, Ushananthiny R, Kamal HKZ, Mohd AY, YF Ngeow |
| EPI_ISL_2013325, EPI_ISL_2015101, EPI_ISL_2015102, EPI_ISL_2015105, EPI_ISL_2015106, EPI_ISL_2015107, EPI_ISL_2015108, EPI_ISL_2015109, EPI_ISL_2015110, EPI_ISL_2015111, EPI_ISL_2015115, EPI_ISL_2015124, EPI_ISL_2015126, EPI_ISL_2015141 | | | |
| see above | IU-Cerrahpasa, Cerrahpasa School of Medicine, COVID-19 Lab | IU-Cerrahpasa, Cerrahpasa School of Medicine, COVID-19 Lab | Mert Kuskucu, Yesim Tuyji Tok, Kenan Midilli |
| EPI_ISL_2040230 | Cliniques universitaires Saint-Luc | UCLouvain/IREC/MBLG | Jean Ruelle, Eleonore Ngyuvula, Benoit Kabamba Mukadi |
| EPI_ISL_2081876, EPI_ISL_2081878, EPI_ISL_2081886, EPI_ISL_2081887, EPI_ISL_2081902, EPI_ISL_2081905, EPI_ISL_2081909, EPI_ISL_2081910, EPI_ISL_2081912, EPI_ISL_2081915, EPI_ISL_2081929, EPI_ISL_2081930 | | | |
| see above | National Center of Infectious and Parasitic Diseases | National Center of Infectious and Parasitic Diseases | Alexiev, Ivanov, Korsun, Stoitsova, Philipova, Dimitrova, Grigorova L., Hristova, Donchev, Stoykov, Trifonova, Dobrinov, Grigorova I., Kantardjiev |
| EPI_ISL_2103371, EPI_ISL_2103373 | HOSPITAL SAN JUAN DE DIOS | Inciensa, Instituto Costarricense de Investigación y | Francisco Duarte, Hebleen Brenes, Claudio Soto-Garita, Estela Cordero, Adriana Godinez & Melany Calderon |

| | | | |
|---|---|---|---|
| | | Enseñanza en Nutrición y Salud | |
| EPI_ISL_2148602 | Clínica Meta | Instituto de Virología-Universidad El Bosque | L. Johana Madroñero, Eliana P. Calvo, Maria Fernanda Montañez-Suarez, Adriana Franco, Andres F. Cardona-Rios, Maria Stella López, Jhann Andrés Arturo, Lilia Bernal-Cepeda, Sonia P. Bohórquez, María Angélica Calderón-Peláez, Sigrid J. Camacho-Ortega, Carolina Coronel-Ruiz, Felix G. Delgado, Lady Lopez, Myriam Lucia Velandia-Romero, Laura Silvana Pérez, Karl A Ciuoderis, Jorge E. Osorio, Juan P. Hernandez-Ortiz, Jaime E. Castellanos |
| EPI_ISL_2171040 | Dhulikhel Hospital, Kathmandu University Hospital | Molecular and Genomics Research Lab, Dhulikhel Hospital, Kathmandu University Hospital | Rajeev Shrestha, Nishan Katuwal, Navin Adhikari, Manu Vanaerschot, Dipesh Tamrakar, Meghnath Dhimal, Pradip Gyanwali, Saroj Bhattarai, Surendra Kumar Madhup |
| EPI_ISL_2196362 | Lboratorio Central de Saude Publica do Estado do Parana (LACEN/PR) | Laboratory of Respiratory Viruses and Measles, Oswaldo Cruz Institute, FIOCRUZ | Paola Resende, Luciana Appolinario, Fernando Motta, Anna Carolina Paixao, Ana Carolina Mendonca, Alice Sampaio Rocha, Taina Venas, Elisa Cavalcante Pereira, Renata Serrano Lopes, Irina Riediger, Marilda Siqueira on behalf of the Fiocruz COVID-19 Genomic Surveillance Network |
| EPI_ISL_2226216 | Department of Internal Medicine, College of Medicine, Chosun University | Department of Internal Medicine, College of Medicine, Chosun University | Dong-Min Kim |
| EPI_ISL_2231475, EPI_ISL_2231478, EPI_ISL_2231485 | San Diego County Public Health Laboratory | Andersen lab at Scripps Research | SEARCH Alliance San Diego with Ashleigh Murphy, Jovan Shephard, Brett Austin |
| EPI_ISL_2276072, EPI_ISL_2276073, EPI_ISL_2276074, EPI_ISL_2276076 | Regional Authority of Public Health Trenín | Veterinary institute in Zvolen Slovakia | Dirbáková Z., Buková L., Reizigová L., Beová K., Sujová S., Mokryšová S., Tinák M., Mojžiš M. |
| EPI_ISL_2278896 | Institut National d'Hygiène (INH) | Unité Mixte Internationale TransVIHMI (UMI 233 IRD - U1175 INSERM - Université de Montpellier)IRD (Institut de recherche pour le développement) | Mounerou SALOU, Christelle BUTEL, Wembo A. HALATOKO,Issaka Maman, Abla A. KONOU, Amivi EHLAN, Adodo SADJI, Kokou TEGUENI,Sidonie A.M.KAGNISSODE, Akoélé SILIADIN, Alassane OURO-MEDELI, Messanh DOUFFAN,Déléma MABA,Sika DOSSIM, Améyo DORKENOO, Mireille PRINCE-DAVID,Anoumou DAGNRA,Laetitia SERRANO,Ahidjo AYOUBA,Eric DELAPORTE, Martine PEETERS |
| EPI_ISL_2285105, EPI_ISL_2285126, EPI_ISL_2285155 | Microbiology Department. Complexo Hospitalario Universitario de Vigo | Microbiology Department. Complexo Hospitalario Universitario de Vigo | Alfaya N, Alonso I, Alvarez M, Cabrera JJ, Carballo R, Cores O, Cortizo S, del-Campo V, Martinez L, Mediero G, Perez S, Potel C, Regueiro B, Rey S, Vasallo FJ |
| EPI_ISL_2322456, EPI_ISL_2322457, EPI_ISL_2322458, EPI_ISL_2322459, EPI_ISL_2322460, EPI_ISL_2322461, EPI_ISL_2322462, EPI_ISL_2322463, EPI_ISL_2322464 | Genome Analysis Center, Yamanashi Central Hospital | Genome Analysis Center, Yamanashi Central Hospital | Yosuke Hirotsu |
| EPI_ISL_2339878, EPI_ISL_2339890, EPI_ISL_2339907 | Centro de Estudio de Enfermedades Autoinmunes (CREA), Universidad del Rosario, Bogota, Colombia | Centro de Investigaciones en Microbiología y Biotecnología-UR (CIMBIUR), Facultad de Ciencias Naturales, Universidad del Rosario, Bogotá, Colombia | Juan-Manuel Anaya, Gustavo Salguero, Juan Esteban Gallo, Carolina Ramírez-Santana, Sergio Castañeda, Nathalia Ballesteros, Marina Muñoz, Luz H. Patiño, Juan David Ramírez |
| EPI_ISL_2350809, EPI_ISL_2350810, EPI_ISL_2350811, EPI_ISL_2350812, EPI_ISL_2350813, EPI_ISL_2350814, EPI_ISL_2350815, EPI_ISL_2350816 | Molecular Lab, Evercare hospital Dhaka | International Institute for Zoonosis Control, Hokkaido university | Junya Yamagishi, Mizanur Rahman |
| EPI_ISL_2351932 | Centre for Human Virology and Genomics, Microbiology Department, Nigerian Institute of Medical Research | Central Research Laboratory, Nigerian Institute of Medical Research | Chika Kingsley Onwuamah, Ayorinde Babatunde James, Joseph Ojonugwa Shaibu, Rahaman A. Ahmed, Azuka Patrick Okwuraiwe, Olufemi Samuel Amoo, Sola Ajibaye, Bamidele Iwalokun, Rosemary Ajuma Audu, Babatunde Lawal Salako |
| EPI_ISL_2364491, EPI_ISL_2364492, EPI_ISL_2364493, EPI_ISL_2364494, EPI_ISL_2364495, EPI_ISL_2364496, EPI_ISL_2364497, EPI_ISL_2364498, EPI_ISL_2364499 | Genome Analysis Center, Yamanashi Central Hospital | Genome Analysis Center, Yamanashi Central Hospital | Yosuke Hirotsu |
| EPI_ISL_2365883 | Microbiology Department. Complexo Hospitalario Universitario de Vigo | Microbiology Department. Complexo Hospitalario Universitario de Vigo | Alfaya N, Alonso I, Alvarez M, Cabrera JJ, Carballo R, Cores O, Cortizo S, del-Campo V, Martinez L, Mediero G, Perez S, Potel C, Regueiro B, Rey S, Vasallo FJ |
| EPI_ISL_2382422, EPI_ISL_2382423 | Centre for Human Virology and Genomics, Microbiology Department, Nigerian Institute of Medical Research | Central Research Laboratory, Nigerian Institute of Medical Research | Chika Kingsley Onwuamah, Ayorinde Babatunde James, Joseph Ojonugwa Shaibu, Rahaman A. Ahmed, Azuka Patrick Okwuraiwe, Sola Ajibaye, Sharon Abimbola, Olufemi Samuel Amoo, Muinah Adenike Fowora, Phasha-Muchemenye Mmatshepho, Joy Ayoola, Nyam Itse Yusuf, Grace Oni, Yusuf Jimoh, Josiah Ayoola Isong, Uyi Emokpae, Ngozi Mirabel Otuonye, Rosemary Ajuma Audu, Bamidele Iwalokun, Babatunde Lawal Salako |
| EPI_ISL_2397307 | NHLS-IALCH | KRISP, KZN Research Innovation and Sequencing Platform | Karim F, Moosa MYS, Gosnell BI, Cele S, Giandhari J, Pillay S, Tegally H, Wilkinson E, San JE, Msomi N, Mlisana K, Khan K, Bernstein M, Manickchund N, Singh L, Ramphal U, COMMIT-KZN Team, Hanekom W, Lessells RJ, Sigal A, de Oliveira T |
| EPI_ISL_2398288, EPI_ISL_2398290 | Centre for Human Virology and Genomics, Microbiology Department, Nigerian Institute of Medical Research | Central Research Laboratory, Nigerian Institute of Medical Research | Chika Kingsley Onwuamah, Ayorinde Babatunde James, Joseph Ojonugwa Shaibu, Rahaman A. Ahmed, Azuka Patrick Okwuraiwe, Sola Ajibaye, Sharon Abimbola, Olufemi Samuel Amoo, Muinah Adenike Fowora, Phasha-Muchemenye Mmatshepho, Joy Ayoola, Nyam Itse Yusuf, Grace Oni, Yusuf Jimoh, Josiah Ayoola Isong, Uyi Emokpae, Ngozi Mirabel Otuonye, Rosemary Ajuma Audu, Bamidele Iwalokun, Babatunde Lawal Salako |
| EPI_ISL_2448653 | Molecular diagnostic laboratory of Federal Budget Institution of Science "Central Research Institute of Epidemiology" of The Federal Service on Customers' Rights Protection and Human Well-being Surveillance | Group of Genomics and Postgenomic Technologies of Central Research Institute of Epidemiology | Samoilov A.E., Kaptelova V.V., Korneenko E.V., Valdohina A.V., Bulanenko V.P., Saenko S.S., Sinitsyn S.O., Golubeva A.G., Speranskaya A.S., Tivanova E.V., Shipulina O.Y., Akimkin V.G. |
| EPI_ISL_2458065 | Adolfo Lutz Institute Santos Regional Center | Retrovirus Laboratory Adolfo Lutz Institute | Gabriela Bastos Cabral, Giselle I S Lopez-Lopes, Cintia Ahagon, Audrey Cilli, Paula Morena Guimaraes, Igor Mohamed Hussein, Luis Brigido |
| EPI_ISL_2476401 | Department of Molecular and Translational Medicine, Section of Microbiology, University of Brescia, ASST Spedali Civili, Brescia | Department of Molecular and Translational Medicine, Section of Microbiology, University of Brescia, ASST Spedali Civili, Brescia | Simona Fiorentini, Serena Messali, Alberto Zani, Francesca Caccuri, Arnaldo Caruso |
| EPI_ISL_2484147 | Mater Misericordiae University Hospital Dublin | University College Dublin, CFS, School of Public Health | Maureen Lynch, Guerrino Macori, Séamus Fanning, Edel O' Regan, Eoin Hunt, Dermot O'Callaghan, Brian McCullagh, Cormac Jennings, Anne Fortune |
| EPI_ISL_2484215, EPI_ISL_2484216, EPI_ISL_2484217 | University of Brescia - Dept. of Molecular and Translational Medicine | Istituto Zooprofilattico Sperimentale della Lombardia e dell'Emilia Romagna (IZSLER), Risk Analysis and Genomic Epidemiology Unit | Caruso Arnaldo, Fiorentini Simona, Caccuri Francesca, Messali Serena, Zani Alberto, Marina Morganti, Erika Scaltriti, Stefano Pongolini |
| EPI_ISL_2501079, EPI_ISL_2501080, EPI_ISL_2501082, EPI_ISL_2501083, EPI_ISL_2501085, EPI_ISL_2501086, EPI_ISL_2502018, EPI_ISL_2502020, EPI_ISL_2502021, EPI_ISL_2502022, EPI_ISL_2502025 | | | |
| see above | Centre for Human Virology and Genomics, Microbiology Department, Nigerian Institute of Medical Research | Central Research Laboratory, Nigerian Institute of Medical Research | Chika Kingsley Onwuamah, Ayorinde Babatunde James, Joseph Ojonugwa Shaibu, Rahaman A. Ahmed, Azuka Patrick Okwuraiwe, Sola Ajibaye, Sharon Abimbola, Olufemi Samuel Amoo, Muinah Adenike Fowora, Phasha-Muchemenye Mmatshepho, Joy Ayoola, Nyam Itse Yusuf, Grace Oni, Yusuf Jimoh, Josiah Ayoola Isong, Uyi Emokpae, Ngozi Mirabel Otuonye, Rosemary Ajuma Audu, Bamidele Iwalokun, Babatunde Lawal Salako |
| EPI_ISL_2502630, EPI_ISL_2502635, EPI_ISL_2502636, EPI_ISL_2502638, EPI_ISL_2502649, EPI_ISL_2502656, EPI_ISL_2502661 | All India Institute of Medical Sciences Delhi Hospital | Virology Laboratory, AIIMS Delhi | Aashish Choudhary, Sumedha Bagga, Lata Rani, Dibyabhaba Pradhan, Pooja Pandey, Jyoti Jethani, Shivram Dhakad, Nazneen Arif, Deepankar Srigyan, Megha Brijwal, Lalit Dar, Manish Soneja, Rakesh Lodha, Puneet Kaur, Ritu Gupta, Subrata Sinha, Chitra Sarkar, Randeep Guleria |
| EPI_ISL_2603237, EPI_ISL_2603251, EPI_ISL_2603255 | Microbiology Department. Complexo Hospitalario Universitario de Vigo | Microbiology Department. Complexo Hospitalario Universitario de Vigo | Alfaya N, Alonso I, Alvarez M, Cabrera JJ, Carballo R, Cores O, Cortizo S, del-Campo V, Martinez L, Mediero G, Perez S, Potel C, Regueiro B, Rey S, Vasallo FJ |
| EPI_ISL_2603816 | Oman-National Influenza Center | Biotechnology & OMICs Laboratory | Intisar Al-Shukri, Ahmed Al Harrasi, Aisha Al-Amri, Amal Al-Maani, Sajjad Asaf, Fahad Al-Balushi, Laila Al-Balushi, Ahmed N Al-Rawahi, Saqib Bilal, Abdul Latif Khan, Samira Al-Mahruqi, Ahmed Al-Rawahi, Hanan Al-Kindi, Amina Al-Jardani. |
| EPI_ISL_2614543 | IdenGene Medicina Diagnostica SA | Instituto Adolfo Lutz, Interdiciplinary Procedures Center, | Claudio Tavares Sacchi, Claudia Regina Gonçalves, Erica Valessa Ramos Gomes, Karoline Rodrigues Campos, Caio Vinicius Dias Lopes, Leonardo Jose |

| | | | |
|---|---|---|---|
| | | Strategic Laboratory | Tadeu de Araujo |
| EPI_ISL_2625925 | Microbiology Unit ASST Grande Ospedale Metropolitano Niguarda, Milan, Italy | Microbiology Unit ASST Grande Ospedale Metropolitano Niguarda, Milan, Italy | Merli M. et al. |
| EPI_ISL_2671836, EPI_ISL_2671846, EPI_ISL_2671847, EPI_ISL_2671848 | Juntendo University Hospital | Juntendo University Hospital | Koji Tuchiya, Yoshie Hosaka |
| EPI_ISL_2690146, EPI_ISL_2690147, EPI_ISL_2690148, EPI_ISL_2690149, EPI_ISL_2690180, EPI_ISL_2690181, EPI_ISL_2690182 | Ochsner Health | Bioinfoexperts, LLC | Judy Crabtree, Amy Feehan, Rebecca Rose, David J. Nolan, Tessa LaFleur, Julia-Garcia-Diaz, Lucio Miele, Susanna L. Lamers |
| EPI_ISL_2692890 | Department of Infectious Disease, Ardabil University of Medical Sciences, Ardabil, Iran | Genetics Research Center, University of Social Welfare and Rehabilitation Sciences | Zohreh Fattahi, Marzieh Mohseni, Khadijeh Jalalvand, Azam Ghaziasadi, Seyedeh elham Mortazavi, Ali Jafarpour, Shahram Habibzadeh, Farhad Jeddi, Kimia Kahrizi, Seyed Mohammad Jazayeri, Hossein Najmabadi. |
| EPI_ISL_2692893 | Pars Hospital Laboratory, Rasht, Iran. | Genetics Research Center, University of Social Welfare and Rehabilitation Sciences | Zohreh Fattahi, Marzieh Mohseni, Khadijeh Jalalvand, Issa Jahanzad, Kimia Kahrizi, Hossein Najmabadi. |
| EPI_ISL_2692895 | Department of pathology, school of medicine, Imam Hospital complex, Tehran university of medical sciences, Tehran, Iran | Genetics Research Center, University of Social Welfare and Rehabilitation Sciences | Zohreh Fattahi, Marzieh Mohseni, Khadijeh Jalalvand, Azam Ghaziasadi, Seyedeh elham Mortazavi, Ali Jafarpour, Azar Hadadi, Alireza Abdollahi, Seyedeh elham Mortazavi, Saber Soltani, Reza Najafipour, Kimia Kahrizi, Seyed Mohammad Jazayeri, Hossein Najmabadi. |
| EPI_ISL_2692901 | Department of pathology, school of medicine, Imam Hospital complex, Tehran university of medical sciences, Tehran,Iran | Genetics Research Center, University of Social Welfare and Rehabilitation Sciences | Zohreh Fattahi, Marzieh Mohseni, Khadijeh Jalalvand, Azam Ghaziasadi, Seyedeh elham Mortazavi, Ali Jafarpour, Azar Hadadi, Alireza Abdollahi, Saber Soltani, Reza Najafipour, Kimia Kahrizi, Seyed Mohammad Jazayeri, Hossein Najmabadi. |
| EPI_ISL_2692904 | Pars Hospital Laboratory, Rasht, Iran. | Genetics Research Center, University of Social Welfare and Rehabilitation Sciences | Zohreh Fattahi, Marzieh Mohseni, Khadijeh Jalalvand, Issa Jahanzad, Kimia Kahrizi, Hossein Najmabadi. |
| EPI_ISL_2692906, EPI_ISL_2692907, EPI_ISL_2692908 | Department of pathology, school of medicine, Imam Hospital complex, Tehran university of medical sciences, Tehran, Iran | Genetics Research Center, University of Social Welfare and Rehabilitation Sciences | Zohreh Fattahi, Marzieh Mohseni, Khadijeh Jalalvand, Azam Ghaziasadi, Seyedeh elham Mortazavi, Ali Jafarpour, Azar Hadadi, Alireza Abdollahi, Saber Soltani, Reza Najafipour, Kimia Kahrizi, Seyed Mohammad Jazayeri, Hossein Najmabadi. |
| EPI_ISL_2692910 | Pars Hospital Laboratory, Rasht, Iran. | Genetics Research Center, University of Social Welfare and Rehabilitation Sciences | Zohreh Fattahi, Marzieh Mohseni, Khadijeh Jalalvand, Issa Jahanzad ,Kimia Kahrizi, Hossein Najmabadi. |
| EPI_ISL_2692953 | Pars Hospital Laboratory, Rasht, Iran. | Genetics Research Center, University of Social Welfare and Rehabilitation Sciences | Zohreh Fattahi, Marzieh Mohseni, Khadijeh Jalalvand, Issa Jahanzad, Kimia Kahrizi, Hossein Najmabadi. |
| EPI_ISL_2692982, EPI_ISL_2692984, EPI_ISL_2692985, EPI_ISL_2692988 | Department of pathology, school of medicine, Imam Hospital complex, Tehran university of medical sciences, Tehran, Iran | Genetics Research Center, University of Social Welfare and Rehabilitation Sciences | Zohreh Fattahi, Marzieh Mohseni, Khadijeh Jalalvand, Azam Ghaziasadi, Seyedeh elham Mortazavi, Ali Jafarpour, Azar Hadadi, Alireza Abdollahi, Saber Soltani, Reza Najafipour, Kimia Kahrizi, Seyed Mohammad Jazayeri, Hossein Najmabadi. |
| EPI_ISL_2692996 | Oman-National Influenza Center | Biotechnology & OMICs Laboratory | Ahmed Al Harrasi, Aisha Al-Amri, Intisar Al-Shukri, Amal Al-Maani, Sajjad Asaf, Bilal Hussain, Samiya Al-Zadjali, Ahmed N Al-Rawahi, Saqib Bilal, Abdul Latif Khan, Samira Al-Mahruqi, Ahmed Al-Rawahi, Hanan Al-Kindi, |
| EPI_ISL_2692997 | Oman-National Influenza Center | Biotechnology & OMICs Laboratory | Aisha Al-Amri, Ahmed Al Harrasi, Intisar Al-Shukri, Amal Al-Maani, Sajjad Asaf, Bilal Hussain, Samiya Al-Zadjali, Ahmed N Al-Rawahi, Saqib Bilal, Abdul Latif Khan, Samira Al-Mahruqi, Ahmed Al-Rawahi, Hanan Al-Kindi,Amina Al-Jardani. |
| EPI_ISL_2694588, EPI_ISL_2694589, EPI_ISL_2694590 | Virology Centre, School of public health, Tehran University of Medical Sciences, Tehran, Iran | Genetics Research Center, University of Social Welfare and Rehabilitation Sciences | Zohreh Fattahi, Marzieh Mohseni, Khadijeh Jalalvand, Jila Yavarian, Kimia Kahrizi, Talat Mokhtari-Azad, Hossein Najmabadi. |
| EPI_ISL_2694591 | Department of pathology, school of medicine, Imam Hospital complex, Tehran university of medical sciences, Tehran, Iran | Genetics Research Center, University of Social Welfare and Rehabilitation Sciences | Zohreh Fattahi, Marzieh Mohseni, Khadijeh Jalalvand, Jila Yavarian, Kimia Kahrizi, Talat Mokhtari-Azad, Hossein Najmabadi. |
| EPI_ISL_2694592 | Virology Centre, School of public health, Tehran University of Medical Sciences, Tehran, Iran | Genetics Research Center, University of Social Welfare and Rehabilitation Sciences | Zohreh Fattahi, Marzieh Mohseni, Khadijeh Jalalvand, Jila Yavarian, Kimia Kahrizi, Talat Mokhtari-Azad, Hossein Najmabadi. |
| EPI_ISL_2696940, EPI_ISL_2696941 | All India Institute of Medical Sciences Delhi Hospital | Virology Laboratory, AIIMS Delhi | Aashish Choudhary, Sumedha Bagga, Lata Rani, Dibyabhaba Pradhan, Pooja Pandey, Jyoti Jethani, Shivram Dhakad, Nazneen Arif, Deepankar Srigyan, Megha Brijwal, Lalit Dar, Manish Soneja, Rakesh Lodha, Puneet Kaur, Ritu Gupta, Subrata Sinha, Chitra Sarkar, Randeep Guleria |
| EPI_ISL_2710839, EPI_ISL_2710840, EPI_ISL_2710841, EPI_ISL_2710842, EPI_ISL_2710843, EPI_ISL_2710844, EPI_ISL_2710845, EPI_ISL_2710846, EPI_ISL_2710847, EPI_ISL_2710848, EPI_ISL_2710849, EPI_ISL_2710850, EPI_ISL_2710851, EPI_ISL_2710852, EPI_ISL_2710853, EPI_ISL_2710854, EPI_ISL_2710855, EPI_ISL_2710856, EPI_ISL_2710857, EPI_ISL_2710858, EPI_ISL_2710859, EPI_ISL_2710860, EPI_ISL_2710861, EPI_ISL_2710862, EPI_ISL_2710863, EPI_ISL_2710864, EPI_ISL_2710865, EPI_ISL_2710866, EPI_ISL_2710867, EPI_ISL_2710868, EPI_ISL_2710869, EPI_ISL_2710870, EPI_ISL_2710871, EPI_ISL_2710883, EPI_ISL_2713076, EPI_ISL_2713119 | | | |
| see above | Department of Health Technology and Informatics, The Hong Kong Polytechnic University | Department of Health Technology and Informatics, The Hong Kong Polytechnic University | Gilman Kit-Hang Siu, Lam-Kwong Lee, Kenneth Siu-Sing Leung, Jake Siu-Lun Leung, Wing-Hei Lo, Timothy Ting-Leung Ng, Chloe Toi-Mei Chan, Hoi-Ching Jim, Victor Tak-Fai Wong, Kingsley King-Gee Tam, Hiu-Yin Lao, Denise Sze-Hang Wong, Alan Ka-Lun Wu, Miranda Chong-Yee Yau, Yvette Wai-Man Lai, Kitty Sau-Chun Fung, Sandy Ka-Yee Chau, Barry Kin-Chung Wong, Wing-Kin To, Kristine Luk, Alex Yat-Man Ho, Tak-Lun Que, Kam-Tong Yip, Wing Cheong Yam, David Ho-Keung Shum, Shea Ping Yip |
| EPI_ISL_2727892, EPI_ISL_2727893, EPI_ISL_2727894 | All India Institute of Medical Sciences Delhi Hospital | Virology Laboratory, AIIMS Delhi | Aashish Choudhary, Sumedha Bagga, Lata Rani, Dibyabhaba Pradhan, Pooja Pandey, Jyoti Jethani, Shivram Dhakad, Nazneen Arif, Deepankar Srigyan, Megha Brijwal, Lalit Dar, Manish Soneja, Rakesh Lodha, Puneet Kaur, Ritu Gupta, Subrata Sinha, Chitra Sarkar, Randeep Guleria |
| EPI_ISL_2731461, EPI_ISL_2731466, EPI_ISL_2731467 | Lboratorio Central de Saude Publica do Estado do Parana (LACEN/PR) | Laboratory of Respiratory Viruses and Measles, Oswaldo Cruz Institute, FIOCRUZ | Paola Resende, Luciana Appolinario, Fernando Motta, Anna Carolina Paixao, Ana Carolina Mendonca, Alice Sampaio Rocha, Taina Venas, Elisa Cavalcante Pereira, Renata Serrano Lopes, Irina Riediger, Marilda Siqueira on behalf of the Fiocruz COVID-19 Genomic Surveillance Network |
| EPI_ISL_2779398, EPI_ISL_2779399 | MOH-K | USAMRD-A, Basic Science Laboratory | Gathii Kimita, Josphat Nyataya, Esther Omuseni, Faith Sigei, Alan Lemtudo, Eric Muthanje, Brian Andika, Rehema Liyai, Rachel Githii, Clement Masakwe, Stephen Ochola, George Awinda, Carol Kifude, Beth Mutai, John Waitumbi |
| EPI_ISL_2779418, EPI_ISL_2779421, EPI_ISL_2779424, EPI_ISL_2779425, EPI_ISL_2779426, EPI_ISL_2779427, EPI_ISL_2779428, EPI_ISL_2779429, EPI_ISL_2779430 | Port Health Clinic, Busia border point | USAMRD-A, Basic Science Laboratory | Gathii Kimita, Josphat Nyataya, Esther Omuseni, Faith Sigei, Alan Lemtudo, Eric Muthanje, Brian Andika, Rehema Liyai, Rachel Githii, Clement Masakwe, Stephen Ochola, George Awinda, Carol Kifude, Beth Mutai, John Waitumbi |
| EPI_ISL_2779431, EPI_ISL_2779432, EPI_ISL_2779433 | DFMH | USAMRD-A, Basic Science Laboratory | Gathii Kimita, Josphat Nyataya, Esther Omuseni, Faith Sigei, Alan Lemtudo, Eric Muthanje, Brian Andika, Rehema Liyai, Rachel Githii, Clement Masakwe, Stephen Ochola, George Awinda, Carol Kifude, Beth Mutai, John Waitumbi |
| EPI_ISL_2779435, EPI_ISL_2779436, EPI_ISL_2779438, EPI_ISL_2779439, EPI_ISL_2779441, EPI_ISL_2779442, EPI_ISL_2779443, EPI_ISL_2779444, EPI_ISL_2779445, EPI_ISL_2779446, EPI_ISL_2779448 | | | |
| see above | MOH-K | USAMRD-A, Basic Science Laboratory | Gathii Kimita, Josphat Nyataya, Esther Omuseni, Faith Sigei, Alan Lemtudo, Eric Muthanje, Brian Andika, Rehema Liyai, Rachel Githii, Clement Masakwe, Stephen Ochola, George Awinda, Carol Kifude, Beth Mutai, John Waitumbi |
| EPI_ISL_2779451, EPI_ISL_2779452 | MOH-HQ | USAMRD-A, Basic Science Laboratory | Gathii Kimita, Josphat Nyataya, Esther Omuseni, Faith Sigei, Alan Lemtudo, Eric Muthanje, Brian Andika, Rehema Liyai, Rachel Githii, Clement Masakwe, Stephen Ochola, George Awinda, Carol Kifude, Beth Mutai, John Waitumbi |
| EPI_ISL_2779454 | MOH-K | USAMRD-A, Basic Science Laboratory | Gathii Kimita, Josphat Nyataya, Esther Omuseni, Faith Sigei, Alan Lemtudo, Eric Muthanje, Brian Andika, Rehema Liyai, Rachel Githii, Clement Masakwe, Stephen Ochola, George Awinda, Carol Kifude, Beth Mutai, John Waitumbi |
| EPI_ISL_2779455 | KCRH KEMRI/DEID Surveillance Site | USAMRD-A, Basic Science Laboratory | Gathii Kimita, Josphat Nyataya, Esther Omuseni, Faith Sigei, Alan Lemtudo, Eric Muthanje, Brian Andika, Rehema Liyai, Rachel Githii, Clement Masakwe, Stephen Ochola, George Awinda, Carol Kifude, Beth Mutai, John Waitumbi |
| EPI_ISL_2779467, EPI_ISL_2779469, EPI_ISL_2779470, EPI_ISL_2779471, EPI_ISL_2779472, EPI_ISL_2779473 | MOH-K | USAMRD-A, Basic Science Laboratory | Gathii Kimita, Josphat Nyataya, Esther Omuseni, Faith Sigei, Alan Lemtudo, Eric Muthanje, Brian Andika, Rehema Liyai, Rachel Githii, Clement Masakwe, Stephen Ochola, George Awinda, Carol Kifude, Beth Mutai, John Waitumbi |
| EPI_ISL_2779474 | DFMH | USAMRD-A, Basic Science Laboratory | Gathii Kimita, Josphat Nyataya, Esther Omuseni, Faith Sigei, Alan Lemtudo, Eric Muthanje, Brian Andika, Rehema Liyai, Rachel Githii, Clement Masakwe, |

| | | | |
|---|---|---|---|
| | | | Stephen Ochola, George Awinda, Carol Kifude, Beth Mutai, John Waitumbi |
| EPI_ISL_2779475 | MOH-K | USAMRD-A, Basic Science Laboratory | Gathii Kimita, Josphat Nyataya, Esther Omuseni, Faith Sigei, Alan Lemtudo, Eric Muthanje, Brian Andika, Rehema Liyai, Rachel Githii, Clement Masakwe, Stephen Ochola, George Awinda, Carol Kifude, Beth Mutai, John Waitumbi |
| EPI_ISL_2779482 | MOH-HQ | USAMRD-A, Basic Science Laboratory | Gathii Kimita, Josphat Nyataya, Esther Omuseni, Faith Sigei, Alan Lemtudo, Eric Muthanje, Brian Andika, Rehema Liyai, Rachel Githii, Clement Masakwe, Stephen Ochola, George Awinda, Carol Kifude, Beth Mutai, John Waitumbi |
| EPI_ISL_2779484, EPI_ISL_2779485 | DFMH | USAMRD-A, Basic Science Laboratory | Gathii Kimita, Josphat Nyataya, Esther Omuseni, Faith Sigei, Alan Lemtudo, Eric Muthanje, Brian Andika, Rehema Liyai, Rachel Githii, Clement Masakwe, Stephen Ochola, George Awinda, Carol Kifude, Beth Mutai, John Waitumbi |
| EPI_ISL_2779489, EPI_ISL_2779490 | Lanet hospital | USAMRD-A, Basic Science Laboratory | Gathii Kimita, Josphat Nyataya, Esther Omuseni, Faith Sigei, Alan Lemtudo, Eric Muthanje, Brian Andika, Rehema Liyai, Rachel Githii, Clement Masakwe, Stephen Ochola, George Awinda, Carol Kifude, Beth Mutai, John Waitumbi |
| EPI_ISL_2779495, EPI_ISL_2779496, EPI_ISL_2779497, EPI_ISL_2779498, EPI_ISL_2779499, EPI_ISL_2779500, EPI_ISL_2779501, EPI_ISL_2779502, EPI_ISL_2779503, EPI_ISL_2779504, EPI_ISL_2779505, EPI_ISL_2779506, EPI_ISL_2779507, EPI_ISL_2779508 | | | |
| see above | Port Health Clinic, Malaba border point | USAMRD-A, Basic Science Laboratory | Gathii Kimita, Josphat Nyataya, Esther Omuseni, Faith Sigei, Alan Lemtudo, Eric Muthanje, Brian Andika, Rehema Liyai, Rachel Githii, Clement Masakwe, Stephen Ochola, George Awinda, Carol Kifude, Beth Mutai, John Waitumbi |
| EPI_ISL_2779522, EPI_ISL_2779523 | MOH-HQS | USAMRD-A, Basic Science Laboratory | Gathii Kimita, Josphat Nyataya, Esther Omuseni, Faith Sigei, Alan Lemtudo, Eric Muthanje, Brian Andika, Rehema Liyai, Rachel Githii, Clement Masakwe, Stephen Ochola, George Awinda, Carol Kifude, Beth Mutai, John Waitumbi |
| EPI_ISL_2779524 | DFMH | USAMRD-A, Basic Science Laboratory | Gathii Kimita, Josphat Nyataya, Esther Omuseni, Faith Sigei, Alan Lemtudo, Eric Muthanje, Brian Andika, Rehema Liyai, Rachel Githii, Clement Masakwe, Stephen Ochola, George Awinda, Carol Kifude, Beth Mutai, John Waitumbi |
| EPI_ISL_2779525 | LAB-BMC | USAMRD-A, Basic Science Laboratory | Gathii Kimita, Josphat Nyataya, Esther Omuseni, Faith Sigei, Alan Lemtudo, Eric Muthanje, Brian Andika, Rehema Liyai, Rachel Githii, Clement Masakwe, Stephen Ochola, George Awinda, Carol Kifude, Beth Mutai, John Waitumbi |
| EPI_ISL_2779538, EPI_ISL_2779540, EPI_ISL_2779541 | MOH-HQ | USAMRD-A, Basic Science Laboratory | Gathii Kimita, Josphat Nyataya, Esther Omuseni, Faith Sigei, Alan Lemtudo, Eric Muthanje, Brian Andika, Rehema Liyai, Rachel Githii, Clement Masakwe, Stephen Ochola, George Awinda, Carol Kifude, Beth Mutai, John Waitumbi |
| EPI_ISL_2779542, EPI_ISL_2779543 | DFMH | USAMRD-A, Basic Science Laboratory | Gathii Kimita, Josphat Nyataya, Esther Omuseni, Faith Sigei, Alan Lemtudo, Eric Muthanje, Brian Andika, Rehema Liyai, Rachel Githii, Clement Masakwe, Stephen Ochola, George Awinda, Carol Kifude, Beth Mutai, John Waitumbi |
| EPI_ISL_2779549 | MOH-K | USAMRD-A, Basic Science Laboratory | Gathii Kimita, Josphat Nyataya, Esther Omuseni, Faith Sigei, Alan Lemtudo, Eric Muthanje, Brian Andika, Rehema Liyai, Rachel Githii, Clement Masakwe, Stephen Ochola, George Awinda, Carol Kifude, Beth Mutai, John Waitumbi |
| EPI_ISL_2790004 | All India Institute of Medical Sciences Delhi Hospital | Virology Laboratory, AIIMS Delhi | Aashish Choudhary, Sumedha Bagga, Lata Rani, Dibyabhaba Pradhan, Pooja Pandey, Jyoti Jethani, Shivram Dhakad, Nazneen Arif, Deepankar Srigyan, Megha Brijwal, Lalit Dar, Manish Soneja, Rakesh Lodha, Puneet Kaur, Ritu Gupta, Subrata Sinha, Chitra Sarkar, Randeep Guleria |
| EPI_ISL_2790881, EPI_ISL_2790882 | Labo Analyses Med | National Reference Center for Viruses of Respiratory Infections, Institut Pasteur, Paris | Marion Barbet, Sylvie Behillil, Méline Bizard, Angela Brisebarre, Camille Capel, Vincent Enouf, Louise Lefrançois, Frédéric Lemoine, Christophe Malabat, Corinne Maufrais, Emmanuelle Permal, Etienne Simon-Lorière, Maud Vanpeene, Sylvie Van der Werf ,StéPhane Romand |
| EPI_ISL_2790889, EPI_ISL_2790890, EPI_ISL_2790891, EPI_ISL_2790892, EPI_ISL_2790893, EPI_ISL_2790894, EPI_ISL_2790895, EPI_ISL_2790896, EPI_ISL_2790897, EPI_ISL_2790898, EPI_ISL_2790899, EPI_ISL_2790900 | | | |
| see above | Labo Analyses Med | National Reference Center for Viruses of Respiratory Infections, Institut Pasteur, Paris | Marion Barbet, Sylvie Behillil, Méline Bizard, Angela Brisebarre, Camille Capel, Vincent Enouf, Louise Lefrançois, Frédéric Lemoine, Christophe Malabat, Corinne Maufrais, Emmanuelle Permal, Etienne Simon-Lorière, Maud Vanpeene, Sylvie Van der Werf ,Brieuc Gestin |
| EPI_ISL_2835364, EPI_ISL_2835367, EPI_ISL_2835368, EPI_ISL_2835369, EPI_ISL_2835370, EPI_ISL_2835374, EPI_ISL_2835380, EPI_ISL_2835381, EPI_ISL_2835382, EPI_ISL_2835383, EPI_ISL_2835384, EPI_ISL_2835385, EPI_ISL_2835386, EPI_ISL_2835387, EPI_ISL_2835388, EPI_ISL_2835389, EPI_ISL_2835390, EPI_ISL_2835391, EPI_ISL_2835392, EPI_ISL_2835393, EPI_ISL_2835394, EPI_ISL_2835395, EPI_ISL_2835396, EPI_ISL_2835397, EPI_ISL_2835398, EPI_ISL_2835399, EPI_ISL_2835400, EPI_ISL_2835401, EPI_ISL_2835402, EPI_ISL_2835403, EPI_ISL_2835404, EPI_ISL_2835405, EPI_ISL_2835407, EPI_ISL_2835408, EPI_ISL_2835409, EPI_ISL_2835410, EPI_ISL_2835411, EPI_ISL_2835412, EPI_ISL_2835413, EPI_ISL_2835414, EPI_ISL_2835415, EPI_ISL_2835416, EPI_ISL_2835417, EPI_ISL_2835420, EPI_ISL_2835421, EPI_ISL_2835422, EPI_ISL_2835423, EPI_ISL_2835424, EPI_ISL_2835425, EPI_ISL_2835426, EPI_ISL_2835427, EPI_ISL_2835429, EPI_ISL_2835430, EPI_ISL_2835431, EPI_ISL_2835432, EPI_ISL_2835433, EPI_ISL_2835434, EPI_ISL_2835435, EPI_ISL_2835436, EPI_ISL_2835437, EPI_ISL_2835439, EPI_ISL_2835440, EPI_ISL_2835441, EPI_ISL_2835442, EPI_ISL_2835443, EPI_ISL_2835444, EPI_ISL_2835445, EPI_ISL_2835446, EPI_ISL_2835447, EPI_ISL_2835448, EPI_ISL_2835449, EPI_ISL_2835450, EPI_ISL_2835451, EPI_ISL_2835452, EPI_ISL_2835453, EPI_ISL_2835454, EPI_ISL_2835455, EPI_ISL_2835456, EPI_ISL_2835457, EPI_ISL_2835458, EPI_ISL_2835459, EPI_ISL_2835460, EPI_ISL_2835461, EPI_ISL_2835462, EPI_ISL_2835463, EPI_ISL_2835464, EPI_ISL_2835465, EPI_ISL_2835466, EPI_ISL_2835467, EPI_ISL_2835468, EPI_ISL_2835469, EPI_ISL_2835470, EPI_ISL_2835471, EPI_ISL_2835472, EPI_ISL_2835473, EPI_ISL_2835474, EPI_ISL_2835475, EPI_ISL_2835476, EPI_ISL_2835477, EPI_ISL_2835478, EPI_ISL_2835479, EPI_ISL_2835480, EPI_ISL_2835481, EPI_ISL_2835482, EPI_ISL_2835483, EPI_ISL_2835484, EPI_ISL_2835485, EPI_ISL_2835486, EPI_ISL_2835487, EPI_ISL_2835488, EPI_ISL_2835489 | | | |
| see above | Instituto de Diagnostico y Referencia Epidemiologicos (InDRE) | Andersen lab at Scripps Research | SEARCH Alliance with Gisela Barrera Badillo, Jose Ernesto Ramirez Gonzalez, Maria Natividad Cruz Ortiz, Nervain Benjamin Contreras Gonzalez, Tatiana Ernestina Nunez Garcia, Dayanira Sarith Arellano Suarez, Irma Lopez Martinez, Rita Flores Leon, Beatriz Olivares Flores |
| EPI_ISL_2860774, EPI_ISL_2860775, EPI_ISL_2860776 | Department of Health Technology and Informatics, The Hong Kong Polytechnic University | Department of Health Technology and Informatics, The Hong Kong Polytechnic University | Gilman Kit-Hang Siu, Lam-Kwong Lee, Kenneth Siu-Sing Leung, Jake Siu-Lun Leung, Wing-Hei Lo, Timothy Ting-Leung Ng, Chloe Toi-Mei Chan, Hoi-Ching Jim, Ivan Tak-Fai Wong, Kingsley King-Gee Tam, Hiu-Yin Lao, Denise Sze-Hang Wong, Alan Ka-Lun Wu, Miranda Chong-Yee Yau, Yvette Wai-Man Lai, Kitty Sau-Chun Fung, Sandy Ka-Yee Chau, Barry Kin-Chung Wong, Wing-Kin To, Kristine Luk, Alex Yat-Man Ho, Tak-Lun Que, Kam-Tong Yip, Wing Cheong Yam, David Ho-Keung Shum, Shea Ping Yip |
| EPI_ISL_455476 | Laboratory for Respiratory Viruses, Cantacuzino National Military-Medical Institute for Research and Development | Cantacuzino Institute | M.Lazar, L.Ustea, A.Cretu, T.Durfee |
| EPI_ISL_458086 | B.J. Medical College and Civil hospital | Gujarat Biotechnology Research Centre | Dhaval Vaghela, Ramesh Patel, Pranay Shah, Kamlesh J Upadhyay, Ramesh Pandit, Tejas Shah, Ankit Hinsu, Pritesh Sabara, Apurvasinh Puvar, Janvi Raval, Zarna Patel, Monika Gandhi, Pinal Trivedi, Maharshi Pandya, Amit Kanani, Nidhi Patel, Nitin Savaliya, Raghawendra Kumar, Dinesh Kumar, Zuber Saiyed, Komal Patel, Labdhi Pandya, Snehal Bagatharia, Neha Rajpara, Bhavesh Modi, Gaurishankar Shrimali, R D Dixit, A M Kadri, Umang Mishra, Chaitanya Joshi, Madhvi Joshi |
| EPI_ISL_458087 | B.J. Medical College and Civil hospital | Gujarat Biotechnology Research Centre | Ramesh Patel, Pranay Shah, Kamlesh J Upadhyay, Ramesh Pandit, Tejas Shah, Ankit Hinsu, Pritesh Sabara, Apurvasinh Puvar, Janvi Raval, Zarna Patel, Monika Gandhi, Pinal Trivedi, Maharshi Pandya, Amit Kanani, Nidhi Patel, Nitin Savaliya, Raghawendra Kumar, Dinesh Kumar, Zuber Saiyed, Komal Patel, Labdhi Pandya, Snehal Bagatharia, Dhaval Vaghela, Afzal Ansari, Bhavesh Modi, Gaurishankar Shrimali, R D Dixit, A M Kadri, Umang Mishra, Chaitanya Joshi, Madhvi Joshi |
| EPI_ISL_458088 | B.J. Medical College and Civil hospital | Gujarat Biotechnology Research Centre | Pranay Shah, Kamlesh J Upadhyay, Ramesh Pandit, Tejas Shah, Ankit Hinsu, Pritesh Sabara, Apurvasinh Puvar, Janvi Raval, Zarna Patel, Monika Gandhi, Pinal Trivedi, Maharshi Pandya, Amit Kanani, Nidhi Patel, Nitin Savaliya, Raghawendra Kumar, Dinesh Kumar, Zuber Saiyed, Komal Patel, Labdhi Pandya, Snehal Bagatharia, Dhaval Vaghela, Ramesh Patel, Fenil Patel, Bhavesh Modi, Gaurishankar Shrimali, R D Dixit, A M Kadri, Umang Mishra, Chaitanya Joshi, Madhvi Joshi |
| EPI_ISL_458089 | B.J. Medical College and Civil hospital | Gujarat Biotechnology Research Centre | Pranay Shah, Kamlesh J Upadhyay, Ramesh Pandit, Tejas Shah, Ankit Hinsu, Pritesh Sabara, Apurvasinh Puvar, Janvi Raval, Zarna Patel, Monika Gandhi, Pinal Trivedi, Maharshi Pandya, Amit Kanani, Nidhi Patel, Nitin Savaliya, Raghawendra Kumar, Dinesh Kumar, Zuber Saiyed, Komal Patel, Labdhi Pandya, Snehal Bagatharia, Dhaval Vaghela, Ramesh Patel, Neelam Nathani, Bhavesh Modi, Gaurishankar Shrimali, R D Dixit, A M Kadri, Umang Mishra, Chaitanya Joshi, Madhvi Joshi |
| EPI_ISL_458090 | B.J. Medical College and Civil hospital | Gujarat Biotechnology Research Centre | Kamlesh J Upadhyay, Ramesh Pandit, Tejas Shah, Ankit Hinsu, Pritesh Sabara, Apurvasinh Puvar, Janvi Raval, Zarna Patel, Monika Gandhi, Pinal Trivedi, Maharshi Pandya, Amit Kanani, Nidhi Patel, Nitin Savaliya, Raghawendra Kumar, Dinesh Kumar, Zuber Saiyed, Komal Patel, Labdhi Pandya, Snehal Bagatharia, Dhaval Vaghela, Ramesh Patel, Pranay Shah, Armi Chaudhari, Bhavesh Modi, Gaurishankar Shrimali, R D Dixit, A M Kadri, Umang Mishra, Chaitanya Joshi, Madhvi Joshi |
| EPI_ISL_458091 | B.J. Medical College and Civil hospital | Gujarat Biotechnology Research Centre | Maharshi Pandya, Amit Kanani, Nidhi Patel, Nitin Savaliya, Raghawendra Kumar, Dinesh Kumar, Zuber Saiyed, Komal Patel, Labdhi Pandya, Snehal Bagatharia, Dhaval Vaghela, Ramesh Patel, Pranay Shah, Kamlesh J Upadhyay, Ramesh Pandit, Tejas Shah, Ankit Hinsu, Pritesh Sabara, Apurvasinh Puvar, Janvi Raval, Zarna Patel, Monika Gandhi, Pinal Trivedi, Bhavya Jindal, Bhavesh Modi, Gaurishankar Shrimali, R D Dixit, A M Kadri, Umang Mishra, Chaitanya Joshi, Madhvi Joshi |
| EPI_ISL_458092 | B.J. Medical College and Civil hospital | Gujarat Biotechnology Research Centre | Amit Kanani, Nidhi Patel, Nitin Savaliya, Raghawendra Kumar, Dinesh Kumar, Zuber Saiyed, Komal Patel, Labdhi Pandya, Snehal Bagatharia, Dhaval Vaghela, Ramesh Patel, Pranay Shah, Kamlesh J Upadhyay, Ramesh Pandit, Tejas Shah, Ankit Hinsu, Pritesh Sabara, Apurvasinh Puvar, Janvi Raval, Zarna Patel, Monika Gandhi, Pinal Trivedi, Maharshi Pandya, Camellia Chakraborty, Bhavesh Modi, Gaurishankar Shrimali, R D Dixit, A M Kadri, Umang Mishra, Chaitanya Joshi, Madhvi Joshi |

| | | | |
|---|---|---|---|
| EPI_ISL_458093 | B.J. Medical College and Civil hospital | Gujarat Biotechnology Research Centre | Nidhi Patel, Nitin Savaliya, Raghawendra Kumar, Dinesh Kumar, Zuber Saiyed, Komal Patel, Labdhi Pandya, Snehal Bagatharia, Dhaval Vaghela, Ramesh Patel, Pranay Shah, Kamlesh J Upadhyay, Ramesh Pandit, Tejas Shah, Ankit Hinsu, Pritesh Sabara, Apurvasinh Puvar, Janvi Raval, Zarna Patel, Monika Gandhi, Pinal Trivedi, Maharshi Pandya, Amit Kanani, Siddhant Kumar, Bhavesh Modi, Gaurishankar Shrimali, R D Dixit, A M Kadri, Umang Mishra, Chaitanya Joshi, Madhvi Joshi |
| EPI_ISL_458094 | B.J. Medical College and Civil hospital | Gujarat Biotechnology Research Centre | Nitin Savaliya, Raghawendra Kumar, Dinesh Kumar, Zuber Saiyed, Komal Patel, Labdhi Pandya, Snehal Bagatharia, Dhaval Vaghela, Ramesh Patel, Pranay Shah, Kamlesh J Upadhyay, Ramesh Pandit, Tejas Shah, Ankit Hinsu, Pritesh Sabara, Apurvasinh Puvar, Janvi Raval, Zarna Patel, Monika Gandhi, Pinal Trivedi, Maharshi Pandya, Amit Kanani, Nidhi Patel, Priyanka P Vatsa, Bhavesh Modi, Gaurishankar Shrimali, R D Dixit, A M Kadri, Umang Mishra, Chaitanya Joshi, Madhvi Joshi |
| EPI_ISL_458095 | B.J. Medical College and Civil hospital | Gujarat Biotechnology Research Centre | Raghawendra Kumar, Dinesh Kumar, Zuber Saiyed, Komal Patel, Labdhi Pandya, Snehal Bagatharia, Dhaval Vaghela, Ramesh Patel, Pranay Shah, Kamlesh J Upadhyay, Ramesh Pandit, Tejas Shah, Ankit Hinsu, Pritesh Sabara, Apurvasinh Puvar, Janvi Raval, Zarna Patel, Monika Gandhi, Pinal Trivedi, Maharshi Pandya, Amit Kanani, Nidhi Patel, Nitin Savaliya, Pooja P Doshi, Bhavesh Modi, Gaurishankar Shrimali, R D Dixit, A M Kadri, Umang Mishra, Chaitanya Joshi, Madhvi Joshi |
| EPI_ISL_458096 | B.J. Medical College and Civil hospital | Gujarat Biotechnology Research Centre | Dinesh Kumar, Zuber Saiyed, Komal Patel, Labdhi Pandya, Snehal Bagatharia, Dhaval Vaghela, Ramesh Patel, Pranay Shah, Kamlesh J Upadhyay, Ramesh Pandit, Tejas Shah, Ankit Hinsu, Pritesh Sabara, Apurvasinh Puvar, Janvi Raval, Zarna Patel, Monika Gandhi, Pinal Trivedi, Maharshi Pandya, Amit Kanani, Nidhi Patel, Nitin Savaliya, Raghawendra Kumar, Akanksha Verma, Bhavesh Modi, Gaurishankar Shrimali, R D Dixit, A M Kadri, Umang Mishra, Chaitanya Joshi, Madhvi Joshi |
| EPI_ISL_458097 | B.J. Medical College and Civil hospital | Gujarat Biotechnology Research Centre | Zuber Saiyed, Komal Patel, Labdhi Pandya, Snehal Bagatharia, Dhaval Vaghela, Ramesh Patel, Pranay Shah, Kamlesh J Upadhyay, Ramesh Pandit, Tejas Shah, Ankit Hinsu, Pritesh Sabara, Apurvasinh Puvar, Janvi Raval, Zarna Patel, Monika Gandhi, Pinal Trivedi, Maharshi Pandya, Amit Kanani, Nidhi Patel, Nitin Savaliya, Raghawendra Kumar, Dinesh Kumar, Priti Pandita, Bhavesh Modi, Gaurishankar Shrimali, R D Dixit, A M Kadri, Umang Mishra, Chaitanya Joshi, Madhvi Joshi |
| EPI_ISL_458098 | B.J. Medical College and Civil hospital | Gujarat Biotechnology Research Centre | Komal Patel, Labdhi Pandya, Snehal Bagatharia, Dhaval Vaghela, Ramesh Patel, Pranay Shah, Kamlesh J Upadhyay, Ramesh Pandit, Tejas Shah, Ankit Hinsu, Pritesh Sabara, Apurvasinh Puvar, Janvi Raval, Zarna Patel, Monika Gandhi, Pinal Trivedi, Maharshi Pandya, Amit Kanani, Nidhi Patel, Nitin Savaliya, Raghawendra Kumar, Dinesh Kumar, Zuber Saiyed, Pragya Sharma, Bhavesh Modi, Gaurishankar Shrimali, R D Dixit, A M Kadri, Umang Mishra, Chaitanya Joshi, Madhvi Joshi |
| EPI_ISL_458099 | B.J. Medical College and Civil hospital | Gujarat Biotechnology Research Centre | Labdhi Pandya, Snehal Bagatharia, Dhaval Vaghela, Ramesh Patel, Pranay Shah, Kamlesh J Upadhyay, Ramesh Pandit, Tejas Shah, Ankit Hinsu, Pritesh Sabara, Apurvasinh Puvar, Janvi Raval, Zarna Patel, Monika Gandhi, Pinal Trivedi, Maharshi Pandya, Amit Kanani, Nidhi Patel, Nitin Savaliya, Raghawendra Kumar, Dinesh Kumar, Zuber Saiyed, Komal Patel, Neha Rajpara, Bhavesh Modi, Gaurishankar Shrimali, R D Dixit, A M Kadri, Umang Mishra, Chaitanya Joshi, Madhvi Joshi |
| EPI_ISL_458100 | B.J. Medical College and Civil hospital | Gujarat Biotechnology Research Centre | Snehal Bagatharia, Dhaval Vaghela, Ramesh Patel, Pranay Shah, Kamlesh J Upadhyay, Ramesh Pandit, Tejas Shah, Ankit Hinsu, Pritesh Sabara, Apurvasinh Puvar, Janvi Raval, Zarna Patel, Monika Gandhi, Pinal Trivedi, Maharshi Pandya, Amit Kanani, Nidhi Patel, Nitin Savaliya, Raghawendra Kumar, Dinesh Kumar, Zuber Saiyed, Komal Patel, Labdhi Pandya, Afzal Ansari, Bhavesh Modi, Gaurishankar Shrimali, R D Dixit, A M Kadri, Umang Mishra, Chaitanya Joshi, Madhvi Joshi |
| EPI_ISL_458101 | B.J. Medical College and Civil hospital | Gujarat Biotechnology Research Centre | Dhaval Vaghela, Ramesh Patel, Pranay Shah, Kamlesh J Upadhyay, Ramesh Pandit, Tejas Shah, Ankit Hinsu, Pritesh Sabara, Apurvasinh Puvar, Janvi Raval, Zarna Patel, Monika Gandhi, Pinal Trivedi, Maharshi Pandya, Amit Kanani, Nidhi Patel, Nitin Savaliya, Raghawendra Kumar, Dinesh Kumar, Zuber Saiyed, Komal Patel, Labdhi Pandya, Snehal Bagatharia, Fenil Patel, Bhavesh Modi, Gaurishankar Shrimali, R D Dixit, A M Kadri, Umang Mishra, Chaitanya Joshi, Madhvi Joshi |
| EPI_ISL_458102 | B.J. Medical College and Civil hospital | Gujarat Biotechnology Research Centre | Ramesh Patel, Pranay Shah, Kamlesh J Upadhyay, Ramesh Pandit, Tejas Shah, Ankit Hinsu, Pritesh Sabara, Apurvasinh Puvar, Janvi Raval, Zarna Patel, Monika Gandhi, Pinal Trivedi, Maharshi Pandya, Amit Kanani, Nidhi Patel, Nitin Savaliya, Raghawendra Kumar, Dinesh Kumar, Zuber Saiyed, Komal Patel, Labdhi Pandya, Snehal Bagatharia, Dhaval Vaghela, Neelam Nathani, Bhavesh Modi, Gaurishankar Shrimali, R D Dixit, A M Kadri, Umang Mishra, Chaitanya Joshi, Madhvi Joshi |
| EPI_ISL_458106 | Gujarat Biotechnology Research Centre | Gujarat Biotechnology Research Centre | Pritesh Sabara, Apurvasinh Puvar, Janvi Raval, Zarna Patel, Monika Gandhi, Pinal Trivedi, Maharshi Pandya, Amit Kanani, Nidhi Patel, Nitin Savaliya, Raghawendra Kumar, Dinesh Kumar, Zuber Saiyed, Komal Patel, Labdhi Pandya, Snehal Bagatharia, Ramesh Pandit, Tejas Shah, Ankit Hinsu, Siddhant Kumar, Bhavesh Modi, Gaurishankar Shrimali, R D Dixit, A M Kadri, Umang Mishra, Chaitanya Joshi, Madhvi Joshi, , , , , |
| EPI_ISL_458107 | Gujarat Biotechnology Research Centre | Gujarat Biotechnology Research Centre | Apurvasinh Puvar, Janvi Raval, Zarna Patel, Monika Gandhi, Pinal Trivedi, Maharshi Pandya, Amit Kanani, Nidhi Patel, Nitin Savaliya, Raghawendra Kumar, Dinesh Kumar, Zuber Saiyed, Komal Patel, Labdhi Pandya, Snehal Bagatharia, Ramesh Pandit, Tejas Shah, Ankit Hinsu, Pritesh Sabara, Priyanka P Vatsa, Bhavesh Modi, Gaurishankar Shrimali, R D Dixit, A M Kadri, Umang Mishra, Chaitanya Joshi, Madhvi Joshi, , , , , |
| EPI_ISL_458111 | Gujarat Biotechnology Research Centre | Gujarat Biotechnology Research Centre | Pinal Trivedi, Maharshi Pandya, Amit Kanani, Nidhi Patel, Nitin Savaliya, Raghawendra Kumar, Dinesh Kumar, Zuber Saiyed, Komal Patel, Labdhi Pandya, Snehal Bagatharia, Ramesh Pandit, Tejas Shah, Ankit Hinsu, Pritesh Sabara, Apurvasinh Puvar, Janvi Raval, Zarna Patel, Monika Gandhi, Pragya Sharma, Bhavesh Modi, Gaurishankar Shrimali, R D Dixit, A M Kadri, Umang Mishra, Chaitanya Joshi, Madhvi Joshi, , , , , |
| EPI_ISL_458112 | Gujarat Biotechnology Research Centre | Gujarat Biotechnology Research Centre | Maharshi Pandya, Amit Kanani, Nidhi Patel, Nitin Savaliya, Raghawendra Kumar, Dinesh Kumar, Zuber Saiyed, Komal Patel, Labdhi Pandya, Snehal Bagatharia, Ramesh Pandit, Tejas Shah, Ankit Hinsu, Pritesh Sabara, Apurvasinh Puvar, Janvi Raval, Zarna Patel, Monika Gandhi, Pinal Trivedi, Neha Rajpara, Bhavesh Modi, Gaurishankar Shrimali, R D Dixit, A M Kadri, Umang Mishra, Chaitanya Joshi, Madhvi Joshi, , , , , |
| EPI_ISL_458113 | Gujarat Biotechnology Research Centre | Gujarat Biotechnology Research Centre | Amit Kanani, Nidhi Patel, Nitin Savaliya, Raghawendra Kumar, Dinesh Kumar, Zuber Saiyed, Komal Patel, Labdhi Pandya, Snehal Bagatharia, Ramesh Pandit, Tejas Shah, Ankit Hinsu, Pritesh Sabara, Apurvasinh Puvar, Janvi Raval, Zarna Patel, Monika Gandhi, Pinal Trivedi, Maharshi Pandya, Afzal Ansari, Bhavesh Modi, Gaurishankar Shrimali, R D Dixit, A M Kadri, Umang Mishra, Chaitanya Joshi, Madhvi Joshi, , , , , |
| EPI_ISL_459919, EPI_ISL_459920 | Devki Devi Foundation, a unit of Max Healthcare | CSIR-IGIB/Max | Rajesh Pandey#, Samreen Siddiqui, Pooja Sharma, Bansidhar Tarai, Vivekanand A, Bharathram Uppili, Saruchi Wadhwa, Nishu Tyagi, Mitali Mukerji, Bansidhar Tarai, Poonam Das, Sujeet Jha, Mohammed Faruq, Vinita Jha, Anurag Agrawal |
| EPI_ISL_461483 | B.J. Medical College and Civil hospital | Gujarat Biotechnology Research Centre | Dipeshwari Shewale, Komal Patel, Labdhi Pandya, Snehal Bagatharia, Pranay Shah, Kamlesh J Upadhyay, Tejas Shah, Ankit Hinsu, Pritesh Sabara, Apurvasinh Puvar, Janvi Raval, Zarna Patel, Monika Gandhi, Pinal Trivedi, Maharshi Pandya, Nidhi Patel, Nitin Savaliya, Raghawendra Kumar, Dinesh Kumar, Zuber Saiyed, R D Dixit, A M Kadri, Harsh Bakshi, Chaitanya Joshi, Madhvi Joshi, |
| EPI_ISL_461484 | B.J. Medical College and Civil hospital | Gujarat Biotechnology Research Centre | Priyanka P Vatsa, Labdhi Pandya, Snehal Bagatharia, Pranay Shah, Kamlesh J Upadhyay, Tejas Shah, Ankit Hinsu, Pritesh Sabara, Apurvasinh Puvar, Janvi Raval, Zarna Patel, Monika Gandhi, Pinal Trivedi, Maharshi Pandya, Nidhi Patel, Nitin Savaliya, Raghawendra Kumar, Dinesh Kumar, Zuber Saiyed, Komal Patel, R D Dixit, A M Kadri, Harsh Bakshi, Chaitanya Joshi, Madhvi Joshi, |
| EPI_ISL_461485 | B.J. Medical College and Civil hospital | Gujarat Biotechnology Research Centre | Pooja P Doshi, Snehal Bagatharia, Pranay Shah, Kamlesh J Upadhyay, Tejas Shah, Ankit Hinsu, Pritesh Sabara, Apurvasinh Puvar, Janvi Raval, Zarna Patel, Monika Gandhi, Pinal Trivedi, Maharshi Pandya, Nidhi Patel, Nitin Savaliya, Raghawendra Kumar, Dinesh Kumar, Zuber Saiyed, Komal Patel, Labdhi Pandya, R D Dixit, A M Kadri, Harsh Bakshi, Chaitanya Joshi, Madhvi Joshi, |
| EPI_ISL_461486 | B.J. Medical College and Civil hospital | Gujarat Biotechnology Research Centre | Akanksha Verma, Pranay Shah, Kamlesh J Upadhyay, Tejas Shah, Ankit Hinsu, Pritesh Sabara, Apurvasinh Puvar, Janvi Raval, Zarna Patel, Monika Gandhi, Pinal Trivedi, Maharshi Pandya, Nidhi Patel, Nitin Savaliya, Raghawendra Kumar, Dinesh Kumar, Zuber Saiyed, Komal Patel, Labdhi Pandya, Snehal Bagatharia, R D Dixit, A M Kadri, Harsh Bakshi, Chaitanya Joshi, Madhvi Joshi, |
| EPI_ISL_461487 | B.J. Medical College and Civil hospital | Gujarat Biotechnology Research Centre | Priti Pandita, Kamlesh J Upadhyay, Tejas Shah, Ankit Hinsu, Pritesh Sabara, Apurvasinh Puvar, Janvi Raval, Zarna Patel, Monika Gandhi, Pinal Trivedi, Maharshi Pandya, Nidhi Patel, Nitin Savaliya, Raghawendra Kumar, Dinesh Kumar, Zuber Saiyed, Komal Patel, Labdhi Pandya, Snehal Bagatharia, Pranay Shah, R D Dixit, A M Kadri, Harsh Bakshi, Chaitanya Joshi, Madhvi Joshi, |
| EPI_ISL_461488 | B.J. Medical College and Civil hospital | Gujarat Biotechnology Research Centre | Pragya Sharma, Tejas Shah, Ankit Hinsu, Pritesh Sabara, Apurvasinh Puvar, Janvi Raval, Zarna Patel, Monika Gandhi, Pinal Trivedi, Maharshi Pandya, Nidhi Patel, Nitin Savaliya, Raghawendra Kumar, Dinesh Kumar, Zuber Saiyed, Komal Patel, Labdhi Pandya, Snehal Bagatharia, Pranay Shah, Kamlesh J Upadhyay, R D Dixit, A M Kadri, Harsh Bakshi, Chaitanya Joshi, Madhvi Joshi, |
| EPI_ISL_461489 | B.J. Medical College and Civil hospital | Gujarat Biotechnology Research Centre | Neha Rajpara, Ankit Hinsu, Pritesh Sabara, Apurvasinh Puvar, Janvi Raval, Zarna Patel, Monika Gandhi, Pinal Trivedi, Maharshi Pandya, Nidhi Patel, Nitin |

| | | | |
|---|---|---|---|
| | | | Savaliya, Raghawendra Kumar, Dinesh Kumar, Zuber Saiyed, Komal Patel, Labdhi Pandya, Snehal Bagatharia, Pranay Shah, Kamlesh J Upadhyay, Tejas Shah, R D Dixit, A M Kadri, Harsh Bakshi, Chaitanya Joshi, Madhvi Joshi, |
| EPI_ISL_461490 | B.J. Medical College and Civil hospital | Gujarat Biotechnology Research Centre | Afzal Ansari, Pritesh Sabara, Apurvasinh Puvar, Janvi Raval, Zarna Patel, Monika Gandhi, Pinal Trivedi, Maharshi Pandya, Nidhi Patel, Nitin Savaliya, Raghawendra Kumar, Dinesh Kumar, Zuber Saiyed, Komal Patel, Labdhi Pandya, Snehal Bagatharia, Pranay Shah, Kamlesh J Upadhyay, Tejas Shah, Ankit Hinsu, R D Dixit, A M Kadri, Harsh Bakshi, Chaitanya Joshi, Madhvi Joshi, |
| EPI_ISL_461491 | B.J. Medical College and Civil hospital | Gujarat Biotechnology Research Centre | Fenil Patel, Apurvasinh Puvar, Janvi Raval, Zarna Patel, Monika Gandhi, Pinal Trivedi, Maharshi Pandya, Nidhi Patel, Nitin Savaliya, Raghawendra Kumar, Dinesh Kumar, Zuber Saiyed, Komal Patel, Labdhi Pandya, Snehal Bagatharia, Pranay Shah, Kamlesh J Upadhyay, Tejas Shah, Ankit Hinsu, Pritesh Sabara, R D Dixit, A M Kadri, Harsh Bakshi, Chaitanya Joshi, Madhvi Joshi, |
| EPI_ISL_461492 | B.J. Medical College and Civil hospital | Gujarat Biotechnology Research Centre | Neelam Nathani, Janvi Raval, Zarna Patel, Monika Gandhi, Pinal Trivedi, Maharshi Pandya, Nidhi Patel, Nitin Savaliya, Raghawendra Kumar, Dinesh Kumar, Zuber Saiyed, Komal Patel, Labdhi Pandya, Snehal Bagatharia, Pranay Shah, Kamlesh J Upadhyay, Tejas Shah, Ankit Hinsu, Pritesh Sabara, Apurvasinh Puvar, R D Dixit, A M Kadri, Harsh Bakshi, Chaitanya Joshi, Madhvi Joshi, |
| EPI_ISL_461493 | B.J. Medical College and Civil hospital | Gujarat Biotechnology Research Centre | Armi Chaudhari, Zarna Patel, Monika Gandhi, Pinal Trivedi, Maharshi Pandya, Nidhi Patel, Nitin Savaliya, Raghawendra Kumar, Dinesh Kumar, Zuber Saiyed, Komal Patel, Labdhi Pandya, Snehal Bagatharia, Pranay Shah, Kamlesh J Upadhyay, Tejas Shah, Ankit Hinsu, Pritesh Sabara, Apurvasinh Puvar, Janvi Raval, R D Dixit, A M Kadri, Harsh Bakshi, Chaitanya Joshi, Madhvi Joshi, |
| EPI_ISL_461494 | B.J. Medical College and Civil hospital | Gujarat Biotechnology Research Centre | Bhavya Jindal, Monika Gandhi, Pinal Trivedi, Maharshi Pandya, Nidhi Patel, Nitin Savaliya, Raghawendra Kumar, Dinesh Kumar, Zuber Saiyed, Komal Patel, Labdhi Pandya, Snehal Bagatharia, Pranay Shah, Kamlesh J Upadhyay, Tejas Shah, Ankit Hinsu, Pritesh Sabara, Apurvasinh Puvar, Janvi Raval, Zarna Patel, R D Dixit, A M Kadri, Harsh Bakshi, Chaitanya Joshi, Madhvi Joshi, |
| EPI_ISL_461495 | B.J. Medical College and Civil hospital | Gujarat Biotechnology Research Centre | Anjali Rajwar, Pinal Trivedi, Maharshi Pandya, Nidhi Patel, Nitin Savaliya, Raghawendra Kumar, Dinesh Kumar, Zuber Saiyed, Komal Patel, Labdhi Pandya, Snehal Bagatharia, Pranay Shah, Kamlesh J Upadhyay, Tejas Shah, Ankit Hinsu, Pritesh Sabara, Apurvasinh Puvar, Janvi Raval, Zarna Patel, Monika Gandhi, R D Dixit, A M Kadri, Harsh Bakshi, Chaitanya Joshi, Madhvi Joshi, |
| EPI_ISL_461496 | B.J. Medical College and Civil hospital | Gujarat Biotechnology Research Centre | Dipeshwari Shewale, Maharshi Pandya, Nidhi Patel, Nitin Savaliya, Raghawendra Kumar, Dinesh Kumar, Zuber Saiyed, Komal Patel, Labdhi Pandya, Snehal Bagatharia, Pranay Shah, Kamlesh J Upadhyay, Tejas Shah, Ankit Hinsu, Pritesh Sabara, Apurvasinh Puvar, Janvi Raval, Zarna Patel, Monika Gandhi, Pinal Trivedi, R D Dixit, A M Kadri, Harsh Bakshi, Chaitanya Joshi, Madhvi Joshi, |
| EPI_ISL_461497 | B.J. Medical College and Civil hospital | Gujarat Biotechnology Research Centre | Priyanka P Vatsa, Nidhi Patel, Nitin Savaliya, Raghawendra Kumar, Dinesh Kumar, Zuber Saiyed, Komal Patel, Labdhi Pandya, Snehal Bagatharia, Pranay Shah, Kamlesh J Upadhyay, Tejas Shah, Ankit Hinsu, Pritesh Sabara, Apurvasinh Puvar, Janvi Raval, Zarna Patel, Monika Gandhi, Pinal Trivedi, Maharshi Pandya, R D Dixit, A M Kadri, Harsh Bakshi, Chaitanya Joshi, Madhvi Joshi, |
| EPI_ISL_461498 | B.J. Medical College and Civil hospital | Gujarat Biotechnology Research Centre | Pooja P Doshi, Nitin Savaliya, Raghawendra Kumar, Dinesh Kumar, Zuber Saiyed, Komal Patel, Labdhi Pandya, Snehal Bagatharia, Pranay Shah, Kamlesh J Upadhyay, Tejas Shah, Ankit Hinsu, Pritesh Sabara, Apurvasinh Puvar, Janvi Raval, Zarna Patel, Monika Gandhi, Pinal Trivedi, Maharshi Pandya, Nidhi Patel, R D Dixit, A M Kadri, Harsh Bakshi, Chaitanya Joshi, Madhvi Joshi, |
| EPI_ISL_461499 | B.J. Medical College and Civil hospital | Gujarat Biotechnology Research Centre | Akanksha Verma, Raghawendra Kumar, Dinesh Kumar, Zuber Saiyed, Komal Patel, Labdhi Pandya, Snehal Bagatharia, Pranay Shah, Kamlesh J Upadhyay, Tejas Shah, Ankit Hinsu, Pritesh Sabara, Apurvasinh Puvar, Janvi Raval, Zarna Patel, Monika Gandhi, Pinal Trivedi, Maharshi Pandya, Nidhi Patel, Nitin Savaliya, R D Dixit, A M Kadri, Harsh Bakshi, Chaitanya Joshi, Madhvi Joshi, |
| EPI_ISL_461500 | B.J. Medical College and Civil hospital | Gujarat Biotechnology Research Centre | Priti Pandita, Dinesh Kumar, Zuber Saiyed, Komal Patel, Labdhi Pandya, Snehal Bagatharia, Pranay Shah, Kamlesh J Upadhyay, Tejas Shah, Ankit Hinsu, Pritesh Sabara, Apurvasinh Puvar, Janvi Raval, Zarna Patel, Monika Gandhi, Pinal Trivedi, Maharshi Pandya, Nidhi Patel, Nitin Savaliya, Raghawendra Kumar, R D Dixit, A M Kadri, Harsh Bakshi, Chaitanya Joshi, Madhvi Joshi, |
| EPI_ISL_461501 | B.J. Medical College and Civil hospital | Gujarat Biotechnology Research Centre | Pragya Sharma, Zuber Saiyed, Komal Patel, Labdhi Pandya, Snehal Bagatharia, Pranay Shah, Kamlesh J Upadhyay, Tejas Shah, Ankit Hinsu, Pritesh Sabara, Apurvasinh Puvar, Janvi Raval, Zarna Patel, Monika Gandhi, Pinal Trivedi, Maharshi Pandya, Nidhi Patel, Nitin Savaliya, Raghawendra Kumar, Dinesh Kumar, R D Dixit, A M Kadri, Harsh Bakshi, Chaitanya Joshi, Madhvi Joshi, |
| EPI_ISL_461502 | B.J. Medical College and Civil hospital | Gujarat Biotechnology Research Centre | Neha Rajpara, Komal Patel, Labdhi Pandya, Snehal Bagatharia, Pranay Shah, Kamlesh J Upadhyay, Tejas Shah, Ankit Hinsu, Pritesh Sabara, Apurvasinh Puvar, Janvi Raval, Zarna Patel, Monika Gandhi, Pinal Trivedi, Maharshi Pandya, Nidhi Patel, Nitin Savaliya, Raghawendra Kumar, Dinesh Kumar, Zuber Saiyed, R D Dixit, A M Kadri, Harsh Bakshi, Chaitanya Joshi, Madhvi Joshi, |
| EPI_ISL_461503 | B.J. Medical College and Civil hospital | Gujarat Biotechnology Research Centre | Afzal Ansari, Labdhi Pandya, Snehal Bagatharia, Pranay Shah, Kamlesh J Upadhyay, Tejas Shah, Ankit Hinsu, Pritesh Sabara, Apurvasinh Puvar, Janvi Raval, Zarna Patel, Monika Gandhi, Pinal Trivedi, Maharshi Pandya, Nidhi Patel, Nitin Savaliya, Raghawendra Kumar, Dinesh Kumar, Zuber Saiyed, Komal Patel, R D Dixit, A M Kadri, Harsh Bakshi, Chaitanya Joshi, Madhvi Joshi, |
| EPI_ISL_461504 | B.J. Medical College and Civil hospital | Gujarat Biotechnology Research Centre | Snehal Bagatharia, Pranay Shah, Kamlesh J Upadhyay, Tejas Shah, Ankit Hinsu, Pritesh Sabara, Apurvasinh Puvar, Janvi Raval, Zarna Patel, Monika Gandhi, Pinal Trivedi, Maharshi Pandya, Nidhi Patel, Nitin Savaliya, Raghawendra Kumar, Dinesh Kumar, Zuber Saiyed, Komal Patel, Labdhi Pandya, Fenil Patel, R D Dixit, A M Kadri, Harsh Bakshi, Chaitanya Joshi, Madhvi Joshi, |
| EPI_ISL_461505 | B.J. Medical College and Civil hospital | Gujarat Biotechnology Research Centre | Pranay Shah, Kamlesh J Upadhyay, Tejas Shah, Ankit Hinsu, Pritesh Sabara, Apurvasinh Puvar, Janvi Raval, Zarna Patel, Monika Gandhi, Pinal Trivedi, Maharshi Pandya, Nidhi Patel, Nitin Savaliya, Raghawendra Kumar, Dinesh Kumar, Zuber Saiyed, Komal Patel, Labdhi Pandya, Snehal Bagatharia, Neelam Nathani, R D Dixit, A M Kadri, Harsh Bakshi, Chaitanya Joshi, Madhvi Joshi, |
| EPI_ISL_461506 | B.J. Medical College and Civil hospital | Gujarat Biotechnology Research Centre | Kamlesh J Upadhyay, Tejas Shah, Ankit Hinsu, Pritesh Sabara, Apurvasinh Puvar, Janvi Raval, Zarna Patel, Monika Gandhi, Pinal Trivedi, Maharshi Pandya, Nidhi Patel, Nitin Savaliya, Raghawendra Kumar, Dinesh Kumar, Zuber Saiyed, Komal Patel, Labdhi Pandya, Snehal Bagatharia, Pranay Shah, Armi Chaudhari, R D Dixit, A M Kadri, Harsh Bakshi, Chaitanya Joshi, Madhvi Joshi, |
| EPI_ISL_462753 | University Clinical Hospital of Mostar | University of Sarajevo Veterinary Faculty | Goletic, T., Softic, A., Goletic, S., Ostojic, M., Hukic, M., Eterovic, T., Seho-Alic, A. |
| EPI_ISL_467029 | GMERS Medical College and Hospital, Gandhinagar | Gujarat Biotechnology Research Centre | Seema Bhatt, Gaurishankar Shrimali, Bhavesh Modi, Bharti Rajani, Tejas Shah, Ankit Hinsu, Pritesh Sabara, Apurvasinh Puvar, Janvi Raval, Zarna Patel, Monika Gandhi, Pinal Trivedi, Maharshi Pandya, Nidhi Patel, Nitin Savaliya, Raghawendra Kumar, Dinesh Kumar, Zuber Saiyed, Komal Patel, Labdhi Pandya, Snehal Bagatharia, Bhavya Jindal, R D Dixit, A M Kadri, Harsh Bakshi, Chaitanya Joshi, Madhvi Joshi |
| EPI_ISL_467030 | GMERS Medical College and Hospital, Gandhinagar | Gujarat Biotechnology Research Centre | Gaurishankar Shrimali, Bhavesh Modi, Bharti Rajani, Tejas Shah, Ankit Hinsu, Pritesh Sabara, Apurvasinh Puvar, Janvi Raval, Zarna Patel, Monika Gandhi, Pinal Trivedi, Maharshi Pandya, Nidhi Patel, Nitin Savaliya, Raghawendra Kumar, Dinesh Kumar, Zuber Saiyed, Komal Patel, Labdhi Pandya, Snehal Bagatharia, Seema Bhatt, Priyanka P Vatsa, R D Dixit, A M Kadri, Harsh Bakshi, Chaitanya Joshi, Madhvi Joshi |
| EPI_ISL_467031 | GMERS Medical College and Hospital, Gandhinagar | Gujarat Biotechnology Research Centre | Bhavesh Modi, Bharti Rajani, Tejas Shah, Ankit Hinsu, Pritesh Sabara, Apurvasinh Puvar, Janvi Raval, Zarna Patel, Monika Gandhi, Pinal Trivedi, Maharshi Pandya, Nidhi Patel, Nitin Savaliya, Raghawendra Kumar, Dinesh Kumar, Zuber Saiyed, Komal Patel, Labdhi Pandya, Snehal Bagatharia, Seema Bhatt, Gaurishankar Shrimali, Pooja P Doshi, R D Dixit, A M Kadri, Harsh Bakshi, Chaitanya Joshi, Madhvi Joshi |
| EPI_ISL_467032 | GMERS Medical College and Hospital, Gandhinagar | Gujarat Biotechnology Research Centre | Bharti Rajani, Tejas Shah, Ankit Hinsu, Pritesh Sabara, Apurvasinh Puvar, Janvi Raval, Zarna Patel, Monika Gandhi, Pinal Trivedi, Maharshi Pandya, Nidhi Patel, Nitin Savaliya, Raghawendra Kumar, Dinesh Kumar, Zuber Saiyed, Komal Patel, Labdhi Pandya, Snehal Bagatharia, Seema Bhatt, Gaurishankar Shrimali, Bhavesh Modi, Akanksha Verma, R D Dixit, A M Kadri, Harsh Bakshi, Chaitanya Joshi, Madhvi Joshi |
| EPI_ISL_467033 | GMERS Medical College and Hospital, Gandhinagar | Gujarat Biotechnology Research Centre | Tejas Shah, Ankit Hinsu, Pritesh Sabara, Apurvasinh Puvar, Janvi Raval, Zarna Patel, Monika Gandhi, Pinal Trivedi, Maharshi Pandya, Nidhi Patel, Nitin Savaliya, Raghawendra Kumar, Dinesh Kumar, Zuber Saiyed, Komal Patel, Labdhi Pandya, Snehal Bagatharia, Seema Bhatt, Gaurishankar Shrimali, Bhavesh Modi, Bharti Rajani, Priti Pandita, R D Dixit, A M Kadri, Harsh Bakshi, Chaitanya Joshi, Madhvi Joshi |
| EPI_ISL_467034 | GMERS Medical College and Hospital, Gandhinagar | Gujarat Biotechnology Research Centre | Ankit Hinsu, Pritesh Sabara, Apurvasinh Puvar, Janvi Raval, Zarna Patel, Monika Gandhi, Pinal Trivedi, Maharshi Pandya, Nidhi Patel, Nitin Savaliya, Raghawendra Kumar, Dinesh Kumar, Zuber Saiyed, Komal Patel, Labdhi Pandya, Snehal Bagatharia, Seema Bhatt, Gaurishankar Shrimali, Bhavesh Modi, Bharti Rajani, Tejas Shah, Pragya Sharma, R D Dixit, A M Kadri, Harsh Bakshi, Chaitanya Joshi, Madhvi Joshi |
| EPI_ISL_467035 | GMERS Medical College and Hospital, Gandhinagar | Gujarat Biotechnology Research Centre | Pritesh Sabara, Apurvasinh Puvar, Janvi Raval, Zarna Patel, Monika Gandhi, Pinal Trivedi, Maharshi Pandya, Nidhi Patel, Nitin Savaliya, Raghawendra Kumar, Dinesh Kumar, Zuber Saiyed, Komal Patel, Labdhi Pandya, Snehal Bagatharia, Seema Bhatt, Gaurishankar Shrimali, Bhavesh Modi, Bharti Rajani, Tejas Shah, Ankit Hinsu, Neha Rajpara, R D Dixit, A M Kadri, Harsh Bakshi, Chaitanya Joshi, Madhvi Joshi |

| | | | |
|---|---|---|---|
| EPI_ISL_467036 | GMERS Medical College and Hospital, Gandhinagar | Gujarat Biotechnology Research Centre | Apurvasinh Puvar, Janvi Raval, Zarna Patel, Monika Gandhi, Pinal Trivedi, Maharshi Pandya, Nidhi Patel, Nitin Savaliya, Raghawendra Kumar, Dinesh Kumar, Zuber Saiyed, Komal Patel, Labdhi Pandya, Snehal Bagatharia, Seema Bhatt, Gaurishankar Shrimali, Bhavesh Modi, Bharti Rajani, Tejas Shah, Ankit Hinsu, Pritesh Sabara, Afzal Ansari, R D Dixit, A M Kadri, Harsh Bakshi, Chaitanya Joshi, Madhvi Joshi |
| EPI_ISL_467037 | GMERS Medical College and Hospital, Gandhinagar | Gujarat Biotechnology Research Centre | Janvi Raval, Zarna Patel, Monika Gandhi, Pinal Trivedi, Maharshi Pandya, Nidhi Patel, Nitin Savaliya, Raghawendra Kumar, Dinesh Kumar, Zuber Saiyed, Komal Patel, Labdhi Pandya, Snehal Bagatharia, Seema Bhatt, Gaurishankar Shrimali, Bhavesh Modi, Bharti Rajani, Tejas Shah, Ankit Hinsu, Pritesh Sabara, Apurvasinh Puvar, Fenil Patel, R D Dixit, A M Kadri, Harsh Bakshi, Chaitanya Joshi, Madhvi Joshi |
| EPI_ISL_467038 | GMERS Medical College and Hospital, Gandhinagar | Gujarat Biotechnology Research Centre | Zarna Patel, Monika Gandhi, Pinal Trivedi, Maharshi Pandya, Nidhi Patel, Nitin Savaliya, Raghawendra Kumar, Dinesh Kumar, Zuber Saiyed, Komal Patel, Labdhi Pandya, Snehal Bagatharia, Seema Bhatt, Gaurishankar Shrimali, Bhavesh Modi, Bharti Rajani, Tejas Shah, Ankit Hinsu, Pritesh Sabara, Apurvasinh Puvar, Janvi Raval, Neelam Nathani, R D Dixit, A M Kadri, Harsh Bakshi, Chaitanya Joshi, Madhvi Joshi |
| EPI_ISL_467039 | Government Medical College, Vadodara | Gujarat Biotechnology Research Centre | Meenakshi Shah, Neena Doshi, Varsha Godbole, Tejas Shah, Ankit Hinsu, Pritesh Sabara, Apurvasinh Puvar, Janvi Raval, Zarna Patel, Monika Gandhi, Pinal Trivedi, Maharshi Pandya, Nidhi Patel, Nitin Savaliya, Raghawendra Kumar, Dinesh Kumar, Zuber Saiyed, Komal Patel, Labdhi Pandya, Snehal Bagatharia, Armi Chaudhari, R D Dixit, A M Kadri, Harsh Bakshi, Chaitanya Joshi, Madhvi Joshi, |
| EPI_ISL_467040 | Government Medical College, Vadodara | Gujarat Biotechnology Research Centre | Neena Doshi, Varsha Godbole, Tejas Shah, Ankit Hinsu, Pritesh Sabara, Apurvasinh Puvar, Janvi Raval, Zarna Patel, Monika Gandhi, Pinal Trivedi, Maharshi Pandya, Nidhi Patel, Nitin Savaliya, Raghawendra Kumar, Dinesh Kumar, Zuber Saiyed, Komal Patel, Labdhi Pandya, Snehal Bagatharia, Meenakshi Shah, Bhavya Jindal, R D Dixit, A M Kadri, Harsh Bakshi, Chaitanya Joshi, Madhvi Joshi, |
| EPI_ISL_467041 | B.J. Medical College and Civil hospital | Gujarat Biotechnology Research Centre | Monika Gandhi, Pinal Trivedi, Maharshi Pandya, Nidhi Patel, Nitin Savaliya, Raghawendra Kumar, Dinesh Kumar, Zuber Saiyed, Komal Patel, Labdhi Pandya, Snehal Bagatharia, Pranay Shah, Kamlesh J Upadhyay, Nirav Mungalpara, Tejas Shah, Ankit Hinsu, Pritesh Sabara, Apurvasinh Puvar, Janvi Raval, Zarna Patel, Priyanka P Vatsa, R D Dixit, A M Kadri, Harsh Bakshi, Chaitanya Joshi, Madhvi Joshi, |
| EPI_ISL_467042 | B.J. Medical College and Civil hospital | Gujarat Biotechnology Research Centre | Pinal Trivedi, Maharshi Pandya, Nidhi Patel, Nitin Savaliya, Raghawendra Kumar, Dinesh Kumar, Zuber Saiyed, Komal Patel, Labdhi Pandya, Snehal Bagatharia, Pranay Shah, Kamlesh J Upadhyay, Nirav Mungalpara, Tejas Shah, Ankit Hinsu, Pritesh Sabara, Apurvasinh Puvar, Janvi Raval, Zarna Patel, Monika Gandhi, Pooja P Doshi, R D Dixit, A M Kadri, Harsh Bakshi, Chaitanya Joshi, Madhvi Joshi, |
| EPI_ISL_467043 | B.J. Medical College and Civil hospital | Gujarat Biotechnology Research Centre | Maharshi Pandya, Nidhi Patel, Nitin Savaliya, Raghawendra Kumar, Dinesh Kumar, Zuber Saiyed, Komal Patel, Labdhi Pandya, Snehal Bagatharia, Pranay Shah, Kamlesh J Upadhyay, Nirav Mungalpara, Tejas Shah, Ankit Hinsu, Pritesh Sabara, Apurvasinh Puvar, Janvi Raval, Zarna Patel, Monika Gandhi, Pinal Trivedi, Akanksha Verma, R D Dixit, A M Kadri, Harsh Bakshi, Chaitanya Joshi, Madhvi Joshi, |
| EPI_ISL_467044 | B.J. Medical College and Civil hospital | Gujarat Biotechnology Research Centre | Nidhi Patel, Nitin Savaliya, Raghawendra Kumar, Dinesh Kumar, Zuber Saiyed, Komal Patel, Labdhi Pandya, Snehal Bagatharia, Pranay Shah, Kamlesh J Upadhyay, Nirav Mungalpara, Tejas Shah, Ankit Hinsu, Pritesh Sabara, Apurvasinh Puvar, Janvi Raval, Zarna Patel, Monika Gandhi, Pinal Trivedi, Maharshi Pandya, Priti Pandita, R D Dixit, A M Kadri, Harsh Bakshi, Chaitanya Joshi, Madhvi Joshi, |
| EPI_ISL_467045 | B.J. Medical College and Civil hospital | Gujarat Biotechnology Research Centre | Nitin Savaliya, Raghawendra Kumar, Dinesh Kumar, Zuber Saiyed, Komal Patel, Labdhi Pandya, Snehal Bagatharia, Pranay Shah, Kamlesh J Upadhyay, Nirav Mungalpara, Tejas Shah, Ankit Hinsu, Pritesh Sabara, Apurvasinh Puvar, Janvi Raval, Zarna Patel, Monika Gandhi, Pinal Trivedi, Maharshi Pandya, Nidhi Patel, Pragya Sharma, R D Dixit, A M Kadri, Harsh Bakshi, Chaitanya Joshi, Madhvi Joshi, |
| EPI_ISL_467046 | B.J. Medical College and Civil hospital | Gujarat Biotechnology Research Centre | Raghawendra Kumar, Dinesh Kumar, Zuber Saiyed, Komal Patel, Labdhi Pandya, Snehal Bagatharia, Pranay Shah, Kamlesh J Upadhyay, Nirav Mungalpara, Tejas Shah, Ankit Hinsu, Pritesh Sabara, Apurvasinh Puvar, Janvi Raval, Zarna Patel, Monika Gandhi, Pinal Trivedi, Maharshi Pandya, Nidhi Patel, Nitin Savaliya, Neha Rajpara, R D Dixit, A M Kadri, Harsh Bakshi, Chaitanya Joshi, Madhvi Joshi, |
| EPI_ISL_467047 | B.J. Medical College and Civil hospital | Gujarat Biotechnology Research Centre | Dinesh Kumar, Zuber Saiyed, Komal Patel, Labdhi Pandya, Snehal Bagatharia, Pranay Shah, Kamlesh J Upadhyay, Nirav Mungalpara, Tejas Shah, Ankit Hinsu, Pritesh Sabara, Apurvasinh Puvar, Janvi Raval, Zarna Patel, Monika Gandhi, Pinal Trivedi, Maharshi Pandya, Nidhi Patel, Nitin Savaliya, Raghawendra Kumar, Afzal Ansari, R D Dixit, A M Kadri, Harsh Bakshi, Chaitanya Joshi, Madhvi Joshi, |
| EPI_ISL_467048 | B.J. Medical College and Civil hospital | Gujarat Biotechnology Research Centre | Zuber Saiyed, Komal Patel, Labdhi Pandya, Snehal Bagatharia, Pranay Shah, Kamlesh J Upadhyay, Nirav Mungalpara, Tejas Shah, Ankit Hinsu, Pritesh Sabara, Apurvasinh Puvar, Janvi Raval, Zarna Patel, Monika Gandhi, Pinal Trivedi, Maharshi Pandya, Nidhi Patel, Nitin Savaliya, Raghawendra Kumar, Dinesh Kumar, Fenil Patel, R D Dixit, A M Kadri, Harsh Bakshi, Chaitanya Joshi, Madhvi Joshi, |
| EPI_ISL_467049 | B.J. Medical College and Civil hospital | Gujarat Biotechnology Research Centre | Komal Patel, Labdhi Pandya, Snehal Bagatharia, Pranay Shah, Kamlesh J Upadhyay, Nirav Mungalpara, Tejas Shah, Ankit Hinsu, Pritesh Sabara, Apurvasinh Puvar, Janvi Raval, Zarna Patel, Monika Gandhi, Pinal Trivedi, Maharshi Pandya, Nidhi Patel, Nitin Savaliya, Raghawendra Kumar, Dinesh Kumar, Zuber Saiyed, Neelam Nathani, R D Dixit, A M Kadri, Harsh Bakshi, Chaitanya Joshi, Madhvi Joshi, |
| EPI_ISL_467050 | B.J. Medical College and Civil hospital | Gujarat Biotechnology Research Centre | Labdhi Pandya, Snehal Bagatharia, Pranay Shah, Kamlesh J Upadhyay, Nirav Mungalpara, Tejas Shah, Ankit Hinsu, Pritesh Sabara, Apurvasinh Puvar, Janvi Raval, Zarna Patel, Monika Gandhi, Pinal Trivedi, Maharshi Pandya, Nidhi Patel, Nitin Savaliya, Raghawendra Kumar, Dinesh Kumar, Zuber Saiyed, Komal Patel, Armi Chaudhari, R D Dixit, A M Kadri, Harsh Bakshi, Chaitanya Joshi, Madhvi Joshi, |
| EPI_ISL_467051 | B.J. Medical College and Civil hospital | Gujarat Biotechnology Research Centre | Snehal Bagatharia, Pranay Shah, Kamlesh J Upadhyay, Nirav Mungalpara, Tejas Shah, Ankit Hinsu, Pritesh Sabara, Apurvasinh Puvar, Janvi Raval, Zarna Patel, Monika Gandhi, Pinal Trivedi, Maharshi Pandya, Nidhi Patel, Nitin Savaliya, Raghawendra Kumar, Dinesh Kumar, Zuber Saiyed, Komal Patel, Labdhi Pandya, Bhavya Jindal, R D Dixit, A M Kadri, Harsh Bakshi, Chaitanya Joshi, Madhvi Joshi, |
| EPI_ISL_467052 | B.J. Medical College and Civil hospital | Gujarat Biotechnology Research Centre | Pranay Shah, Kamlesh J Upadhyay, Nirav Mungalpara, Tejas Shah, Ankit Hinsu, Pritesh Sabara, Apurvasinh Puvar, Janvi Raval, Zarna Patel, Monika Gandhi, Pinal Trivedi, Maharshi Pandya, Nidhi Patel, Nitin Savaliya, Raghawendra Kumar, Dinesh Kumar, Zuber Saiyed, Komal Patel, Labdhi Pandya, Snehal Bagatharia, Priyanka P Vatsa, R D Dixit, A M Kadri, Harsh Bakshi, Chaitanya Joshi, Madhvi Joshi, |
| EPI_ISL_467053 | B.J. Medical College and Civil hospital | Gujarat Biotechnology Research Centre | Kamlesh J Upadhyay, Nirav Mungalpara, Tejas Shah, Ankit Hinsu, Pritesh Sabara, Apurvasinh Puvar, Janvi Raval, Zarna Patel, Monika Gandhi, Pinal Trivedi, Maharshi Pandya, Nidhi Patel, Nitin Savaliya, Raghawendra Kumar, Dinesh Kumar, Zuber Saiyed, Komal Patel, Labdhi Pandya, Snehal Bagatharia, Pranay Shah, Pooja P Doshi, R D Dixit, A M Kadri, Harsh Bakshi, Chaitanya Joshi, Madhvi Joshi, |
| EPI_ISL_467054 | B.J. Medical College and Civil hospital | Gujarat Biotechnology Research Centre | Nirav Mungalpara, Tejas Shah, Ankit Hinsu, Pritesh Sabara, Apurvasinh Puvar, Janvi Raval, Zarna Patel, Monika Gandhi, Pinal Trivedi, Maharshi Pandya, Nidhi Patel, Nitin Savaliya, Raghawendra Kumar, Dinesh Kumar, Zuber Saiyed, Komal Patel, Labdhi Pandya, Snehal Bagatharia, Pranay Shah, Kamlesh J Upadhyay, Akanksha Verma, R D Dixit, A M Kadri, Harsh Bakshi, Chaitanya Joshi, Madhvi Joshi, |
| EPI_ISL_468158 | [Romania, Bucharest] National Institute for Infectious Diseases "Prof. Dr. Matei Bal" | [Romania, Bucharest] National Institute for Infectious Diseases "Prof. Dr. Matei Bal" | Leontina Banica, Marius Cotic, Corina Casangiu, Marius Surleac, Simona Paraschiv |
| EPI_ISL_469024 | B.J. Medical College and Civil hospital | Gujarat Biotechnology Research Centre | Tejas Shah, Ankit Hinsu, Pritesh Sabara, Apurvasinh Puvar, Janvi Raval, Zarna Patel, Monika Gandhi, Pinal Trivedi, Maharshi Pandya, Nidhi Patel, Nitin Savaliya, Raghawendra Kumar, Dinesh Kumar, Zuber Saiyed, Komal Patel, Labdhi Pandya, Snehal Bagatharia, Pranay Shah, Kamlesh J Upadhyay, Nirav Mungalpara, Priti Pandita, R D Dixit, A M Kadri, Harsh Bakshi, Chaitanya Joshi, Madhvi Joshi |
| EPI_ISL_469025 | B.J. Medical College and Civil hospital | Gujarat Biotechnology Research Centre | Ankit Hinsu, Pritesh Sabara, Apurvasinh Puvar, Janvi Raval, Zarna Patel, Monika Gandhi, Pinal Trivedi, Maharshi Pandya, Nidhi Patel, Nitin Savaliya, Raghawendra Kumar, Dinesh Kumar, Zuber Saiyed, Komal Patel, Labdhi Pandya, Snehal Bagatharia, Pranay Shah, Kamlesh J Upadhyay, Nirav Mungalpara, Tejas Shah, Pragya Sharma, R D Dixit, A M Kadri, Harsh Bakshi, Chaitanya Joshi, Madhvi Joshi |
| EPI_ISL_469026 | B.J. Medical College and Civil hospital | Gujarat Biotechnology Research Centre | Pritesh Sabara, Apurvasinh Puvar, Janvi Raval, Zarna Patel, Monika Gandhi, Pinal Trivedi, Maharshi Pandya, Nidhi Patel, Nitin Savaliya, Raghawendra Kumar, Dinesh Kumar, Zuber Saiyed, Komal Patel, Labdhi Pandya, Snehal Bagatharia, Pranay Shah, Kamlesh J Upadhyay, Nirav Mungalpara, Tejas Shah, Ankit Hinsu, Neha Rajpara, R D Dixit, A M Kadri, Harsh Bakshi, Chaitanya Joshi, Madhvi Joshi |
| EPI_ISL_469027 | B.J. Medical College and Civil hospital | Gujarat Biotechnology Research Centre | Apurvasinh Puvar, Janvi Raval, Zarna Patel, Monika Gandhi, Pinal Trivedi, Maharshi Pandya, Nidhi Patel, Nitin Savaliya, Raghawendra Kumar, Dinesh Kumar, Zuber Saiyed, Komal Patel, Labdhi Pandya, Snehal Bagatharia, Pranay Shah, Kamlesh J Upadhyay, Nirav Mungalpara, Tejas Shah, Ankit Hinsu, Pritesh Sabara, Afzal Ansari, R D Dixit, A M Kadri, Harsh Bakshi, Chaitanya Joshi, Madhvi Joshi |
| EPI_ISL_469028 | B.J. Medical College and Civil hospital | Gujarat Biotechnology Research Centre | Janvi Raval, Zarna Patel, Monika Gandhi, Pinal Trivedi, Maharshi Pandya, Nidhi Patel, Nitin Savaliya, Raghawendra Kumar, Dinesh Kumar, Zuber Saiyed, Komal Patel, Labdhi Pandya, Snehal Bagatharia, Pranay Shah, Kamlesh J Upadhyay, Nirav Mungalpara, Tejas Shah, Ankit Hinsu, Pritesh Sabara, Apurvasinh Puvar, Fenil Patel, R D Dixit, A M Kadri, Harsh Bakshi, Chaitanya Joshi, Madhvi Joshi |
| EPI_ISL_469029 | Government Medical College, Vadodara | Gujarat Biotechnology Research Centre | Zarna Patel, Monika Gandhi, Pinal Trivedi, Maharshi Pandya, Nidhi Patel, Nitin Savaliya, Raghawendra Kumar, Dinesh Kumar, Zuber Saiyed, Komal Patel, |

| | | | |
|---|---|---|---|
| | | | Labdhi Pandya, Snehal Bagatharia, Meenakshi Shah, Neena Doshi, Varsha Godbole, Tejas Shah, Ankit Hinsu, Pritesh Sabara, Apurvasinh Puvar, Janvi Raval, Neelam Nathani, R D Dixit, A M Kadri, Harsh Bakshi, Chaitanya Joshi, Madhvi Joshi |
| EPI_ISL_469030 | Government Medical College, Vadodara | Gujarat Biotechnology Research Centre | Monika Gandhi, Pinal Trivedi, Maharshi Pandya, Nidhi Patel, Nitin Savaliya, Raghawendra Kumar, Dinesh Kumar, Zuber Saiyed, Komal Patel, Labdhi Pandya, Snehal Bagatharia, Meenakshi Shah, Neena Doshi, Varsha Godbole, Tejas Shah, Ankit Hinsu, Pritesh Sabara, Apurvasinh Puvar, Janvi Raval, Zarna Patel, Armi Chaudhari, R D Dixit, A M Kadri, Harsh Bakshi, Chaitanya Joshi, Madhvi Joshi |
| EPI_ISL_469031 | Government Medical College, Vadodara | Gujarat Biotechnology Research Centre | Pinal Trivedi, Maharshi Pandya, Nidhi Patel, Nitin Savaliya, Raghawendra Kumar, Dinesh Kumar, Zuber Saiyed, Komal Patel, Labdhi Pandya, Snehal Bagatharia, Meenakshi Shah, Neena Doshi, Varsha Godbole, Tejas Shah, Ankit Hinsu, Pritesh Sabara, Apurvasinh Puvar, Janvi Raval, Zarna Patel, Monika Gandhi, Bhavya Jindal, R D Dixit, A M Kadri, Harsh Bakshi, Chaitanya Joshi, Madhvi Joshi |
| EPI_ISL_469032 | Government Medical College, Vadodara | Gujarat Biotechnology Research Centre | Maharshi Pandya, Nidhi Patel, Nitin Savaliya, Raghawendra Kumar, Dinesh Kumar, Zuber Saiyed, Komal Patel, Labdhi Pandya, Snehal Bagatharia, Meenakshi Shah, Neena Doshi, Varsha Godbole, Tejas Shah, Ankit Hinsu, Pritesh Sabara, Apurvasinh Puvar, Janvi Raval, Zarna Patel, Monika Gandhi, Pinal Trivedi, Pragya Sharma, R D Dixit, A M Kadri, Harsh Bakshi, Chaitanya Joshi, Madhvi Joshi |
| EPI_ISL_469033 | Government Medical College, Vadodara | Gujarat Biotechnology Research Centre | Nidhi Patel, Nitin Savaliya, Raghawendra Kumar, Dinesh Kumar, Zuber Saiyed, Komal Patel, Labdhi Pandya, Snehal Bagatharia, Meenakshi Shah, Neena Doshi, Varsha Godbole, Tejas Shah, Ankit Hinsu, Pritesh Sabara, Apurvasinh Puvar, Janvi Raval, Zarna Patel, Monika Gandhi, Pinal Trivedi, Maharshi Pandya, Priyanka P Vatsa, R D Dixit, A M Kadri, Harsh Bakshi, Chaitanya Joshi, Madhvi Joshi |
| EPI_ISL_469034 | Government Medical College, Vadodara | Gujarat Biotechnology Research Centre | Nitin Savaliya, Raghawendra Kumar, Dinesh Kumar, Zuber Saiyed, Komal Patel, Labdhi Pandya, Snehal Bagatharia, Meenakshi Shah, Neena Doshi, Varsha Godbole, Tejas Shah, Ankit Hinsu, Pritesh Sabara, Apurvasinh Puvar, Janvi Raval, Zarna Patel, Monika Gandhi, Pinal Trivedi, Maharshi Pandya, Nidhi Patel, Pooja P Doshi, R D Dixit, A M Kadri, Harsh Bakshi, Chaitanya Joshi, Madhvi Joshi |
| EPI_ISL_469035 | Government Medical College, Vadodara | Gujarat Biotechnology Research Centre | Raghawendra Kumar, Dinesh Kumar, Zuber Saiyed, Komal Patel, Labdhi Pandya, Snehal Bagatharia, Meenakshi Shah, Neena Doshi, Varsha Godbole, Tejas Shah, Ankit Hinsu, Pritesh Sabara, Apurvasinh Puvar, Janvi Raval, Zarna Patel, Monika Gandhi, Pinal Trivedi, Maharshi Pandya, Nidhi Patel, Nitin Savaliya, Akanksha Verma, R D Dixit, A M Kadri, Harsh Bakshi, Chaitanya Joshi, Madhvi Joshi |
| EPI_ISL_469036 | Government Medical College, Vadodara | Gujarat Biotechnology Research Centre | Dinesh Kumar, Zuber Saiyed, Komal Patel, Labdhi Pandya, Snehal Bagatharia, Meenakshi Shah, Neena Doshi, Varsha Godbole, Tejas Shah, Ankit Hinsu, Pritesh Sabara, Apurvasinh Puvar, Janvi Raval, Zarna Patel, Monika Gandhi, Pinal Trivedi, Maharshi Pandya, Nidhi Patel, Nitin Savaliya, Raghawendra Kumar, Priti Pandita, R D Dixit, A M Kadri, Harsh Bakshi, Chaitanya Joshi, Madhvi Joshi |
| EPI_ISL_469037 | GMERS Medical College & Hospital | Gujarat Biotechnology Research Centre | Zuber Saiyed, Komal Patel, Labdhi Pandya, Snehal Bagatharia, Meenakshi Shah, Neena Doshi, Varsha Godbole, Tejas Shah, Ankit Hinsu, Pritesh Sabara, Apurvasinh Puvar, Janvi Raval, Zarna Patel, Monika Gandhi, Pinal Trivedi, Maharshi Pandya, Nidhi Patel, Nitin Savaliya, Raghawendra Kumar, Dinesh Kumar, Pragya Sharma, R D Dixit, A M Kadri, Harsh Bakshi, Chaitanya Joshi, Madhvi Joshi |
| EPI_ISL_469038 | GMERS Medical College & Hospital | Gujarat Biotechnology Research Centre | Komal Patel, Labdhi Pandya, Snehal Bagatharia, Meenakshi Shah, Neena Doshi, Varsha Godbole, Tejas Shah, Ankit Hinsu, Pritesh Sabara, Apurvasinh Puvar, Janvi Raval, Zarna Patel, Monika Gandhi, Pinal Trivedi, Maharshi Pandya, Nidhi Patel, Nitin Savaliya, Raghawendra Kumar, Dinesh Kumar, Zuber Saiyed, Neha Rajpara, R D Dixit, A M Kadri, Harsh Bakshi, Chaitanya Joshi, Madhvi Joshi |
| EPI_ISL_469039 | GMERS Medical College & Hospital | Gujarat Biotechnology Research Centre | Labdhi Pandya, Snehal Bagatharia, Meenakshi Shah, Neena Doshi, Varsha Godbole, Tejas Shah, Ankit Hinsu, Pritesh Sabara, Apurvasinh Puvar, Janvi Raval, Zarna Patel, Monika Gandhi, Pinal Trivedi, Maharshi Pandya, Nidhi Patel, Nitin Savaliya, Raghawendra Kumar, Dinesh Kumar, Zuber Saiyed, Komal Patel, Afzal Ansari, R D Dixit, A M Kadri, Harsh Bakshi, Chaitanya Joshi, Madhvi Joshi |
| EPI_ISL_469040 | GMERS Medical College & Hospital | Gujarat Biotechnology Research Centre | Snehal Bagatharia, Meenakshi Shah, Neena Doshi, Varsha Godbole, Tejas Shah, Ankit Hinsu, Pritesh Sabara, Apurvasinh Puvar, Janvi Raval, Zarna Patel, Monika Gandhi, Pinal Trivedi, Maharshi Pandya, Nidhi Patel, Nitin Savaliya, Raghawendra Kumar, Dinesh Kumar, Zuber Saiyed, Komal Patel, Labdhi Pandya, Fenil Patel, R D Dixit, A M Kadri, Harsh Bakshi, Chaitanya Joshi, Madhvi Joshi |
| EPI_ISL_469041 | GMERS Medical College & Hospital | Gujarat Biotechnology Research Centre | Meenakshi Shah, Neena Doshi, Varsha Godbole, Tejas Shah, Ankit Hinsu, Pritesh Sabara, Apurvasinh Puvar, Janvi Raval, Zarna Patel, Monika Gandhi, Pinal Trivedi, Maharshi Pandya, Nidhi Patel, Nitin Savaliya, Raghawendra Kumar, Dinesh Kumar, Zuber Saiyed, Komal Patel, Labdhi Pandya, Snehal Bagatharia, Neelam Nathani, R D Dixit, A M Kadri, Harsh Bakshi, Chaitanya Joshi, Madhvi Joshi |
| EPI_ISL_469042 | GMERS Medical College & Hospital | Gujarat Biotechnology Research Centre | Neena Doshi, Varsha Godbole, Tejas Shah, Ankit Hinsu, Pritesh Sabara, Apurvasinh Puvar, Janvi Raval, Zarna Patel, Monika Gandhi, Pinal Trivedi, Maharshi Pandya, Nidhi Patel, Nitin Savaliya, Raghawendra Kumar, Dinesh Kumar, Zuber Saiyed, Komal Patel, Labdhi Pandya, Snehal Bagatharia, Meenakshi Shah, Armi Chaudhari, R D Dixit, A M Kadri, Harsh Bakshi, Chaitanya Joshi, Madhvi Joshi |
| EPI_ISL_469043 | Dr. N. D. Desai Medical College & Hospital | Gujarat Biotechnology Research Centre | J G Buch, Jigar Gusani, Supreet Prabhu, Tejas Shah, Ankit Hinsu, Pritesh Sabara, Apurvasinh Puvar, Janvi Raval, Zarna Patel, Monika Gandhi, Pinal Trivedi, Maharshi Pandya, Nidhi Patel, Nitin Savaliya, Raghawendra Kumar, Dinesh Kumar, Zuber Saiyed, Komal Patel, Labdhi Pandya, Snehal Bagatharia, Bhavya Jindal, R D Dixit, A M Kadri, Harsh Bakshi, Chaitanya Joshi, Madhvi Joshi |
| EPI_ISL_469044 | Dr. N. D. Desai Medical College & Hospital | Gujarat Biotechnology Research Centre | Jigar Gusani, Supreet Prabhu, Tejas Shah, Ankit Hinsu, Pritesh Sabara, Apurvasinh Puvar, Janvi Raval, Zarna Patel, Monika Gandhi, Pinal Trivedi, Maharshi Pandya, Nidhi Patel, Nitin Savaliya, Raghawendra Kumar, Dinesh Kumar, Zuber Saiyed, Komal Patel, Labdhi Pandya, Snehal Bagatharia, J G Buch, Neha Rajpara, R D Dixit, A M Kadri, Harsh Bakshi, Chaitanya Joshi, Madhvi Joshi |
| EPI_ISL_469045 | Dr. N. D. Desai Medical College & Hospital | Gujarat Biotechnology Research Centre | Supreet Prabhu, Tejas Shah, Ankit Hinsu, Pritesh Sabara, Apurvasinh Puvar, Janvi Raval, Zarna Patel, Monika Gandhi, Pinal Trivedi, Maharshi Pandya, Nidhi Patel, Nitin Savaliya, Raghawendra Kumar, Dinesh Kumar, Zuber Saiyed, Komal Patel, Labdhi Pandya, Snehal Bagatharia, J G Buch, Jigar Gusani, Priyanka P Vatsa, R D Dixit, A M Kadri, Harsh Bakshi, Chaitanya Joshi, Madhvi Joshi |
| EPI_ISL_469046 | Dr. N. D. Desai Medical College & Hospital | Gujarat Biotechnology Research Centre | Tejas Shah, Ankit Hinsu, Pritesh Sabara, Apurvasinh Puvar, Janvi Raval, Zarna Patel, Monika Gandhi, Pinal Trivedi, Maharshi Pandya, Nidhi Patel, Nitin Savaliya, Raghawendra Kumar, Dinesh Kumar, Zuber Saiyed, Komal Patel, Labdhi Pandya, Snehal Bagatharia, J G Buch, Jigar Gusani, Supreet Prabhu, Pooja P Doshi, R D Dixit, A M Kadri, Harsh Bakshi, Chaitanya Joshi, Madhvi Joshi |
| EPI_ISL_469047 | Dr. N. D. Desai Medical College & Hospital | Gujarat Biotechnology Research Centre | Ankit Hinsu, Pritesh Sabara, Apurvasinh Puvar, Janvi Raval, Zarna Patel, Monika Gandhi, Pinal Trivedi, Maharshi Pandya, Nidhi Patel, Nitin Savaliya, Raghawendra Kumar, Dinesh Kumar, Zuber Saiyed, Komal Patel, Labdhi Pandya, Snehal Bagatharia, J G Buch, Jigar Gusani, Supreet Prabhu, Tejas Shah, Akanksha Verma, R D Dixit, A M Kadri, Harsh Bakshi, Chaitanya Joshi, Madhvi Joshi |
| EPI_ISL_469048 | Banas Medical College and Research Institute | Gujarat Biotechnology Research Centre | Radhika Khara, Sunil R Joshi, Viren s Doshi, Zarna Patel, Monika Gandhi, Pinal Trivedi, Maharshi Pandya, Nidhi Patel, Nitin Savaliya, Raghawendra Kumar, Dinesh Kumar, Zuber Saiyed, Komal Patel, Labdhi Pandya, Snehal Bagatharia, Tejas Shah, Ankit Hinsu, Pritesh Sabara, Apurvasinh Puvar, Janvi Raval, Priti Pandita, R D Dixit, A M Kadri, Harsh Bakshi, Chaitanya Joshi, Madhvi Joshi |
| EPI_ISL_471269, EPI_ISL_471270, EPI_ISL_471271 | Hospital Oncológico Solca Núcleo de Quito | Institute of Microbiology, Universidad San Francisco de Quito | Sully Márquez, Belén Prado-Vivar, Juan José Guadalupe, Bernardo Gutiérrez, Marcos Di Stefano, Grace Salazar, Verónica Barragán, Patricio Rojas-Silva, Gabriel Trueba, Michelle Grunauer, Paúl Cárdenas |
| EPI_ISL_471416, EPI_ISL_471417, EPI_ISL_471419, EPI_ISL_471420, EPI_ISL_471422, EPI_ISL_471423, EPI_ISL_471424 | Laboratory for Respiratory Viruses, National Influenza Centre, Cantacuzino National Military-Medical Institute for Research and Development | Cantacuzino Institute | Luiza Ustea, Nicoleta Paraschiv, Tim Durfee, Mihaela Lazar |
| EPI_ISL_475026 | Banas Medical College and Research Institute | Gujarat Biotechnology Research Centre | Sunil R Joshi, Viren s Doshi, Pritesh Sabara, Apurvasinh Puvar, Janvi Raval, Zarna Patel, Monika Gandhi, Pinal Trivedi, Maharshi Pandya, Nidhi Patel, Nitin Savaliya, Raghawendra Kumar, Dinesh Kumar, Zuber Saiyed, Komal Patel, Labdhi Pandya, Snehal Bagatharia, Radhika Khara, Neha Rajpara, R D Dixit, A M Kadri, Harsh Bakshi, Chaitanya Joshi, Madhvi Joshi |
| EPI_ISL_475027 | Banas Medical College and Research Institute | Gujarat Biotechnology Research Centre | Viren s Doshi, Pritesh Sabara, Apurvasinh Puvar, Janvi Raval, Zarna Patel, Monika Gandhi, Pinal Trivedi, Maharshi Pandya, Nidhi Patel, Nitin Savaliya, Raghawendra Kumar, Dinesh Kumar, Zuber Saiyed, Komal Patel, Labdhi Pandya, Snehal Bagatharia, Radhika Khara, Sunil R Joshi, Afzal Ansari, R D Dixit, A M Kadri, Harsh Bakshi, Chaitanya Joshi, Madhvi Joshi |
| EPI_ISL_475028 | Banas Medical College and Research Institute | Gujarat Biotechnology Research Centre | Pritesh Sabara, Apurvasinh Puvar, Janvi Raval, Zarna Patel, Monika Gandhi, Pinal Trivedi, Maharshi Pandya, Nidhi Patel, Nitin Savaliya, Raghawendra Kumar, Dinesh Kumar, Zuber Saiyed, Komal Patel, Labdhi Pandya, Snehal Bagatharia, Radhika Khara, Sunil R Joshi, Viren s Doshi, Fenil Patel, R D Dixit, A M Kadri, Harsh Bakshi, Chaitanya Joshi, Madhvi Joshi |
| EPI_ISL_475029 | Banas Medical College and Research Institute | Gujarat Biotechnology Research Centre | Apurvasinh Puvar, Janvi Raval, Zarna Patel, Monika Gandhi, Pinal Trivedi, Maharshi Pandya, Nidhi Patel, Nitin Savaliya, Raghawendra Kumar, Dinesh |

| | | | |
|---|---|---|---|
| | | | Kumar, Zuber Saiyed, Komal Patel, Labdhi Pandya, Snehal Bagatharia, Radhika Khara, Sunil R Joshi, Viren s Doshi, Pritesh Sabara, Neelam Nathani, R D Dixit, A M Kadri, Harsh Bakshi, Chaitanya Joshi, Madhvi Joshi |
| EPI_ISL_475030 | Department of MicroBiology, Government Medical College, Surat | Gujarat Biotechnology Research Centre | Janvi Raval, Zarna Patel, Monika Gandhi, Pinal Trivedi, Maharshi Pandya, Nidhi Patel, Nitin Savaliya, Raghawendra Kumar, Dinesh Kumar, Zuber Saiyed, Komal Patel, Labdhi Pandya, Snehal Bagatharia, Naresh Chauhan, Summaiya Mullan, Amit gamit, Pritesh Sabara, Apurvasinh Puvar, Armi Chaudhari, R D Dixit, A M Kadri, Harsh Bakshi, Chaitanya Joshi, Madhvi Joshi |
| EPI_ISL_475031 | Department of MicroBiology, Government Medical College, Surat | Gujarat Biotechnology Research Centre | Zarna Patel, Monika Gandhi, Pinal Trivedi, Maharshi Pandya, Nidhi Patel, Nitin Savaliya, Raghawendra Kumar, Dinesh Kumar, Zuber Saiyed, Komal Patel, Labdhi Pandya, Snehal Bagatharia, Naresh Chauhan, Summaiya Mullan, Amit gamit, Pritesh Sabara, Apurvasinh Puvar, Janvi Raval, Bhavya Jindal, R D Dixit, A M Kadri, Harsh Bakshi, Chaitanya Joshi, Madhvi Joshi |
| EPI_ISL_475032 | Department of MicroBiology, Government Medical College, Surat | Gujarat Biotechnology Research Centre | Monika Gandhi, Pinal Trivedi, Maharshi Pandya, Nidhi Patel, Nitin Savaliya, Raghawendra Kumar, Dinesh Kumar, Zuber Saiyed, Komal Patel, Labdhi Pandya, Snehal Bagatharia, Naresh Chauhan, Summaiya Mullan, Amit gamit, Pritesh Sabara, Apurvasinh Puvar, Janvi Raval, Zarna Patel, Priyanka P Vatsa, R D Dixit, A M Kadri, Harsh Bakshi, Chaitanya Joshi, Madhvi Joshi |
| EPI_ISL_475033 | Department of MicroBiology, Government Medical College, Surat | Gujarat Biotechnology Research Centre | Pinal Trivedi, Maharshi Pandya, Nidhi Patel, Nitin Savaliya, Raghawendra Kumar, Dinesh Kumar, Zuber Saiyed, Komal Patel, Labdhi Pandya, Snehal Bagatharia, Naresh Chauhan, Summaiya Mullan, Amit gamit, Pritesh Sabara, Apurvasinh Puvar, Janvi Raval, Zarna Patel, Monika Gandhi, Pooja P Doshi, R D Dixit, A M Kadri, Harsh Bakshi, Chaitanya Joshi, Madhvi Joshi |
| EPI_ISL_475034 | Department of MicroBiology, Government Medical College, Surat | Gujarat Biotechnology Research Centre | Maharshi Pandya, Nidhi Patel, Nitin Savaliya, Raghawendra Kumar, Dinesh Kumar, Zuber Saiyed, Komal Patel, Labdhi Pandya, Snehal Bagatharia, Naresh Chauhan, Summaiya Mullan, Amit gamit, Pritesh Sabara, Apurvasinh Puvar, Janvi Raval, Zarna Patel, Monika Gandhi, Pinal Trivedi, Akanksha Verma, R D Dixit, A M Kadri, Harsh Bakshi, Chaitanya Joshi, Madhvi Joshi |
| EPI_ISL_475035 | Department of MicroBiology, Government Medical College, Surat | Gujarat Biotechnology Research Centre | Nidhi Patel, Nitin Savaliya, Raghawendra Kumar, Dinesh Kumar, Zuber Saiyed, Komal Patel, Labdhi Pandya, Snehal Bagatharia, Naresh Chauhan, Summaiya Mullan, Amit gamit, Pritesh Sabara, Apurvasinh Puvar, Janvi Raval, Zarna Patel, Monika Gandhi, Pinal Trivedi, Maharshi Pandya, Priti Pandita, R D Dixit, A M Kadri, Harsh Bakshi, Chaitanya Joshi, Madhvi Joshi |
| EPI_ISL_475036 | Department of MicroBiology, Government Medical College, Surat | Gujarat Biotechnology Research Centre | Nitin Savaliya, Raghawendra Kumar, Dinesh Kumar, Zuber Saiyed, Komal Patel, Labdhi Pandya, Snehal Bagatharia, Naresh Chauhan, Summaiya Mullan, Amit gamit, Pritesh Sabara, Apurvasinh Puvar, Janvi Raval, Zarna Patel, Monika Gandhi, Pinal Trivedi, Maharshi Pandya, Nidhi Patel, Pragya Sharma, R D Dixit, A M Kadri, Harsh Bakshi, Chaitanya Joshi, Madhvi Joshi |
| EPI_ISL_475037 | Department of MicroBiology, Government Medical College, Surat | Gujarat Biotechnology Research Centre | Raghawendra Kumar, Dinesh Kumar, Zuber Saiyed, Komal Patel, Labdhi Pandya, Snehal Bagatharia, Naresh Chauhan, Summaiya Mullan, Amit gamit, Pritesh Sabara, Apurvasinh Puvar, Janvi Raval, Zarna Patel, Monika Gandhi, Pinal Trivedi, Maharshi Pandya, Nidhi Patel, Nitin Savaliya, Neha Rajpara, R D Dixit, A M Kadri, Harsh Bakshi, Chaitanya Joshi, Madhvi Joshi |
| EPI_ISL_475038 | Department of MicroBiology, Government Medical College, Surat | Gujarat Biotechnology Research Centre | Dinesh Kumar, Zuber Saiyed, Komal Patel, Labdhi Pandya, Snehal Bagatharia, Naresh Chauhan, Summaiya Mullan, Amit gamit, Pritesh Sabara, Apurvasinh Puvar, Janvi Raval, Zarna Patel, Monika Gandhi, Pinal Trivedi, Maharshi Pandya, Nidhi Patel, Nitin Savaliya, Raghawendra Kumar, Afzal Ansari, R D Dixit, A M Kadri, Harsh Bakshi, Chaitanya Joshi, Madhvi Joshi |
| EPI_ISL_475039 | Department of MicroBiology, Government Medical College, Surat | Gujarat Biotechnology Research Centre | Zuber Saiyed, Komal Patel, Labdhi Pandya, Snehal Bagatharia, Naresh Chauhan, Summaiya Mullan, Amit gamit, Pritesh Sabara, Apurvasinh Puvar, Janvi Raval, Zarna Patel, Monika Gandhi, Pinal Trivedi, Maharshi Pandya, Nidhi Patel, Nitin Savaliya, Raghawendra Kumar, Dinesh Kumar, Fenil Patel, R D Dixit, A M Kadri, Harsh Bakshi, Chaitanya Joshi, Madhvi Joshi |
| EPI_ISL_475040 | Department of MicroBiology, Government Medical College, Surat | Gujarat Biotechnology Research Centre | Komal Patel, Labdhi Pandya, Snehal Bagatharia, Naresh Chauhan, Summaiya Mullan, Amit gamit, Pritesh Sabara, Apurvasinh Puvar, Janvi Raval, Zarna Patel, Monika Gandhi, Pinal Trivedi, Maharshi Pandya, Nidhi Patel, Nitin Savaliya, Raghawendra Kumar, Dinesh Kumar, Zuber Saiyed, Neelam Nathani, R D Dixit, A M Kadri, Harsh Bakshi, Chaitanya Joshi, Madhvi Joshi |
| EPI_ISL_475043 | Department of MicroBiology, Government Medical College, Surat | Gujarat Biotechnology Research Centre | Naresh Chauhan, Summaiya Mullan, Amit gamit, Pritesh Sabara, Apurvasinh Puvar, Janvi Raval, Zarna Patel, Monika Gandhi, Pinal Trivedi, Maharshi Pandya, Nidhi Patel, Nitin Savaliya, Raghawendra Kumar, Dinesh Kumar, Zuber Saiyed, Komal Patel, Labdhi Pandya, Snehal Bagatharia, Priyanka P Vatsa, R D Dixit, A M Kadri, Harsh Bakshi, Chaitanya Joshi, Madhvi Joshi |
| EPI_ISL_475044 | Department of MicroBiology, Government Medical College, Surat | Gujarat Biotechnology Research Centre | Summaiya Mullan, Amit gamit, Pritesh Sabara, Apurvasinh Puvar, Janvi Raval, Zarna Patel, Monika Gandhi, Pinal Trivedi, Maharshi Pandya, Nidhi Patel, Nitin Savaliya, Raghawendra Kumar, Dinesh Kumar, Zuber Saiyed, Komal Patel, Labdhi Pandya, Snehal Bagatharia, Naresh Chauhan, Pooja P Doshi, R D Dixit, A M Kadri, Harsh Bakshi, Chaitanya Joshi, Madhvi Joshi |
| EPI_ISL_475045 | Department of MicroBiology, Government Medical College, Surat | Gujarat Biotechnology Research Centre | Amit gamit, Pritesh Sabara, Apurvasinh Puvar, Janvi Raval, Zarna Patel, Monika Gandhi, Pinal Trivedi, Maharshi Pandya, Nidhi Patel, Nitin Savaliya, Raghawendra Kumar, Dinesh Kumar, Zuber Saiyed, Komal Patel, Labdhi Pandya, Snehal Bagatharia, Naresh Chauhan, Summaiya Mullan, Akanksha Verma, R D Dixit, A M Kadri, Harsh Bakshi, Chaitanya Joshi, Madhvi Joshi |
| EPI_ISL_475046 | Department of MicroBiology, Government Medical College, Surat | Gujarat Biotechnology Research Centre | Pritesh Sabara, Apurvasinh Puvar, Janvi Raval, Zarna Patel, Monika Gandhi, Pinal Trivedi, Maharshi Pandya, Nidhi Patel, Nitin Savaliya, Raghawendra Kumar, Dinesh Kumar, Zuber Saiyed, Komal Patel, Labdhi Pandya, Snehal Bagatharia, Naresh Chauhan, Summaiya Mullan, Amit gamit, Priti Pandita, R D Dixit, A M Kadri, Harsh Bakshi, Chaitanya Joshi, Madhvi Joshi |
| EPI_ISL_475047 | GMERS Medical College & Hospital | Gujarat Biotechnology Research Centre | Apurvasinh Puvar, Janvi Raval, Zarna Patel, Monika Gandhi, Pinal Trivedi, Maharshi Pandya, Nidhi Patel, Nitin Savaliya, Raghawendra Kumar, Dinesh Kumar, Zuber Saiyed, Komal Patel, Labdhi Pandya, Snehal Bagatharia, Meenakshi Shah, Neena Doshi, Varsha Godbole, Pritesh Sabara, Pragya Sharma, R D Dixit, A M Kadri, Harsh Bakshi, Chaitanya Joshi, Madhvi Joshi |
| EPI_ISL_475048 | GMERS Medical College & Hospital | Gujarat Biotechnology Research Centre | Janvi Raval, Zarna Patel, Monika Gandhi, Pinal Trivedi, Maharshi Pandya, Nidhi Patel, Nitin Savaliya, Raghawendra Kumar, Dinesh Kumar, Zuber Saiyed, Komal Patel, Labdhi Pandya, Snehal Bagatharia, Meenakshi Shah, Neena Doshi, Varsha Godbole, Pritesh Sabara, Apurvasinh Puvar, Neha Rajpara, R D Dixit, A M Kadri, Harsh Bakshi, Chaitanya Joshi, Madhvi Joshi |
| EPI_ISL_475050 | GMERS Medical College & Hospital | Gujarat Biotechnology Research Centre | Monika Gandhi, Pinal Trivedi, Maharshi Pandya, Nidhi Patel, Nitin Savaliya, Raghawendra Kumar, Dinesh Kumar, Zuber Saiyed, Komal Patel, Labdhi Pandya, Snehal Bagatharia, Meenakshi Shah, Neena Doshi, Varsha Godbole, Pritesh Sabara, Apurvasinh Puvar, Janvi Raval, Zarna Patel, Fenil Patel, R D Dixit, A M Kadri, Harsh Bakshi, Chaitanya Joshi, Madhvi Joshi |
| EPI_ISL_475051 | GMERS Medical College & Hospital | Gujarat Biotechnology Research Centre | Pinal Trivedi, Maharshi Pandya, Nidhi Patel, Nitin Savaliya, Raghawendra Kumar, Dinesh Kumar, Zuber Saiyed, Komal Patel, Labdhi Pandya, Snehal Bagatharia, Meenakshi Shah, Neena Doshi, Varsha Godbole, Pritesh Sabara, Apurvasinh Puvar, Janvi Raval, Zarna Patel, Monika Gandhi, Neelam Nathani, R D Dixit, A M Kadri, Harsh Bakshi, Chaitanya Joshi, Madhvi Joshi |
| EPI_ISL_475052 | GMERS Medical College & Hospital | Gujarat Biotechnology Research Centre | Maharshi Pandya, Nidhi Patel, Nitin Savaliya, Raghawendra Kumar, Dinesh Kumar, Zuber Saiyed, Komal Patel, Labdhi Pandya, Snehal Bagatharia, Meenakshi Shah, Neena Doshi, Varsha Godbole, Pritesh Sabara, Apurvasinh Puvar, Janvi Raval, Zarna Patel, Monika Gandhi, Pinal Trivedi, Armi Chaudhari, R D Dixit, A M Kadri, Harsh Bakshi, Chaitanya Joshi, Madhvi Joshi |
| EPI_ISL_475053 | GMERS Medical College & Hospital | Gujarat Biotechnology Research Centre | Nidhi Patel, Nitin Savaliya, Raghawendra Kumar, Dinesh Kumar, Zuber Saiyed, Komal Patel, Labdhi Pandya, Snehal Bagatharia, Meenakshi Shah, Neena Doshi, Varsha Godbole, Pritesh Sabara, Apurvasinh Puvar, Janvi Raval, Zarna Patel, Monika Gandhi, Pinal Trivedi, Maharshi Pandya, Bhavya Jindal, R D Dixit, A M Kadri, Harsh Bakshi, Chaitanya Joshi, Madhvi Joshi |
| EPI_ISL_475056 | Dr. N. D. Desai Medical College & Hospital | Gujarat Biotechnology Research Centre | Dinesh Kumar, Zuber Saiyed, Komal Patel, Labdhi Pandya, Snehal Bagatharia, J G Buch, Jigar Gusani, Supreet Prabhu, Pritesh Sabara, Apurvasinh Puvar, Janvi Raval, Zarna Patel, Monika Gandhi, Pinal Trivedi, Maharshi Pandya, Nidhi Patel, Nitin Savaliya, Raghawendra Kumar, Akanksha Verma, R D Dixit, A M Kadri, Harsh Bakshi, Chaitanya Joshi, Madhvi Joshi |
| EPI_ISL_475057 | Dr. N. D. Desai Medical College & Hospital | Gujarat Biotechnology Research Centre | Zuber Saiyed, Komal Patel, Labdhi Pandya, Supreet Prabhu, Snehal Bagatharia, Jigar Gusani, J G Buch, Pritesh Sabara, Apurvasinh Puvar, Janvi Raval, Zarna Patel, Monika Gandhi, Pinal Trivedi, Maharshi Pandya, Nidhi Patel, Nitin Savaliya, Raghawendra Kumar, Dinesh Kumar, Priti Pandita, R D Dixit, A M Kadri, Harsh Bakshi, Chaitanya Joshi, Madhvi Joshi |
| EPI_ISL_475059 | GAIMS & G K General Hospital | Gujarat Biotechnology Research Centre | Hitesh Assudani, Babulal Babhoria, Labdhi Pandya, Snehal Bagatharia, Pritesh Sabara, Apurvasinh Puvar, Janvi Raval, Zarna Patel, Monika Gandhi, Pinal Trivedi, Maharshi Pandya, Nidhi Patel, Nitin Savaliya, Raghawendra Kumar, Dinesh Kumar, Zuber Saiyed, Komal Patel, Neha Rajpara, R D Dixit, A M Kadri, Harsh Bakshi, Chaitanya Joshi, Madhvi Joshi |
| EPI_ISL_476148, EPI_ISL_476149, | Institut Pasteur Dakar | Institut Pasteur de Dakar | Ndongo Dia, Moussa Moise Diagne, Mamadou Diop, Ousmane Faye, Amadou Alpha Sall |

| | | | |
|---|---|---|---|
| EPI_ISL_476492 | | | |
| EPI_ISL_476493 | Institut Pasteur Dakar | Institut Pasteur de Dakar | Ndongo Dia, Moussa Moise Diagne, Mamadou Diop, Ousmane Faye, Amadou alpha Sall |
| EPI_ISL_476494 | Institut Pasteur Dakar | Institut Pasteur de Dakar | Ndongo Dia, Moussa Moise Diagne, Mamadou Diop, Ousmane Faye, Amadou Alpha Sall |
| EPI_ISL_476495 | Institut Pasteur Dakar | Institut Pasteur de Dakar | Ndongo Dia, Moussa Moise Diagne, Mamadou Diop, Ousmane Faye, Amadou alpha Sall |
| EPI_ISL_476514 | Institut Pasteur Dakar | Institut Pasteur de Dakar | Ndongo Dia, Moussa Moise Diagne, Mamadou Diop, Ousmane Faye, Amadou Alpha Sall |
| EPI_ISL_476516 | Institut Pasteur Dakar | Institut Pasteur de Dakar | Ndongo Dia, Moussa Moise Diagne, mamadou Diop, Ousmane Faye, Amadou Alpha Sall |
| EPI_ISL_476558, EPI_ISL_476562 | Institut Pasteur Dakar | Institut Pasteur de Dakar | Ndongo Dia, Moussa Moise Diagne, Mamadou Diop, Ousmane Faye, Amadou Alpha Sall |
| EPI_ISL_476564 | Institut Pasteur Dakar | Institut Pasteur de Dakar | Ndongo Dia, Moussa Moise Diagne, Mamadou diop, Ousmane Faye, Amadou alpha Sall |
| EPI_ISL_476569 | Institut Pasteur Dakar | Institut Pasteur de Dakar | Ndongo Dia, Moussa Moise, Mamadou Diop, Ousmane Faye, Amadou Alpha Sall |
| EPI_ISL_476572, EPI_ISL_476574 | Institut Pasteur Dakar | Institut Pasteur de Dakar | Ndongo Dia, Moussa Moise Diagne, Mamadou Diop, Ousmane Faye, Amadou Alpha Sall |
| EPI_ISL_476855 | GMERS Medical College & Hospital, Gotri, Vadodara | Gujarat Biotechnology Research Centre | Apurvasinh Puvar, Janvi Raval, Zarna Patel, Monika Gandhi, Pinal Trivedi, Maharshi Pandya, Nidhi Patel, Nitin Savaliya, Raghawendra Kumar, Dinesh Kumar, Zuber Saiyed, Komal Patel, Labdhi Pandya, Afzal Ansari, Nikha Trivedi, Meenakshi Shah, Neena Doshi, Varsha Godbole, R D Dixit, A M Kadri, Harsh Bakshi, Chaitanya Joshi, Madhvi Joshi |
| EPI_ISL_476856 | GMERS Medical College & Hospital, Gotri, Vadodara | Gujarat Biotechnology Research Centre | Janvi Raval, Zarna Patel, Monika Gandhi, Pinal Trivedi, Maharshi Pandya, Nidhi Patel, Nitin Savaliya, Raghawendra Kumar, Dinesh Kumar, Zuber Saiyed, Komal Patel, Labdhi Pandya, Afzal Ansari, Nikha Trivedi, Meenakshi Shah, Neena Doshi, Varsha Godbole, Apurvasinh Puvar, R D Dixit, A M Kadri, Harsh Bakshi, Chaitanya Joshi, Madhvi Joshi |
| EPI_ISL_476857 | GMERS Medical College & Hospital, Gotri, Vadodara | Gujarat Biotechnology Research Centre | Zarna Patel, Monika Gandhi, Pinal Trivedi, Maharshi Pandya, Nidhi Patel, Nitin Savaliya, Raghawendra Kumar, Dinesh Kumar, Zuber Saiyed, Komal Patel, Labdhi Pandya, Afzal Ansari, Nikha Trivedi, Meenakshi Shah, Neena Doshi, Varsha Godbole, Apurvasinh Puvar, Janvi Raval, R D Dixit, A M Kadri, Harsh Bakshi, Chaitanya Joshi, Madhvi Joshi |
| EPI_ISL_476858 | GMERS Medical College & Hospital, Gotri, Vadodara | Gujarat Biotechnology Research Centre | Monika Gandhi, Pinal Trivedi, Maharshi Pandya, Nidhi Patel, Nitin Savaliya, Raghawendra Kumar, Dinesh Kumar, Zuber Saiyed, Komal Patel, Labdhi Pandya, Afzal Ansari, Nikha Trivedi, Meenakshi Shah, Neena Doshi, Varsha Godbole, Apurvasinh Puvar, Janvi Raval, Zarna Patel, R D Dixit, A M Kadri, Harsh Bakshi, Chaitanya Joshi, Madhvi Joshi |
| EPI_ISL_476859 | GMERS Medical College & Hospital, Gotri, Vadodara | Gujarat Biotechnology Research Centre | Pinal Trivedi, Maharshi Pandya, Nidhi Patel, Nitin Savaliya, Raghawendra Kumar, Dinesh Kumar, Zuber Saiyed, Komal Patel, Labdhi Pandya, Afzal Ansari, Nikha Trivedi, Meenakshi Shah, Neena Doshi, Varsha Godbole, Apurvasinh Puvar, Janvi Raval, Zarna Patel, Monika Gandhi, R D Dixit, A M Kadri, Harsh Bakshi, Chaitanya Joshi, Madhvi Joshi |
| EPI_ISL_476860 | GMERS Medical College & Hospital, Gotri, Vadodara | Gujarat Biotechnology Research Centre | Maharshi Pandya, Nidhi Patel, Nitin Savaliya, Raghawendra Kumar, Dinesh Kumar, Zuber Saiyed, Komal Patel, Labdhi Pandya, Afzal Ansari, Nikha Trivedi, Meenakshi Shah, Neena Doshi, Varsha Godbole, Apurvasinh Puvar, Janvi Raval, Zarna Patel, Monika Gandhi, Pinal Trivedi, R D Dixit, A M Kadri, Harsh Bakshi, Chaitanya Joshi, Madhvi Joshi |
| EPI_ISL_476861 | GMERS Medical College & Hospital, Gotri, Vadodara | Gujarat Biotechnology Research Centre | Nidhi Patel, Nitin Savaliya, Raghawendra Kumar, Dinesh Kumar, Zuber Saiyed, Komal Patel, Labdhi Pandya, Afzal Ansari, Nikha Trivedi, Meenakshi Shah, Neena Doshi, Varsha Godbole, Apurvasinh Puvar, Janvi Raval, Zarna Patel, Monika Gandhi, Pinal Trivedi, Maharshi Pandya, R D Dixit, A M Kadri, Harsh Bakshi, Chaitanya Joshi, Madhvi Joshi |
| EPI_ISL_476862 | GMERS Medical College & Hospital, Gotri, Vadodara | Gujarat Biotechnology Research Centre | Nitin Savaliya, Raghawendra Kumar, Dinesh Kumar, Zuber Saiyed, Komal Patel, Labdhi Pandya, Afzal Ansari, Nikha Trivedi, Meenakshi Shah, Neena Doshi, Varsha Godbole, Apurvasinh Puvar, Janvi Raval, Zarna Patel, Monika Gandhi, Pinal Trivedi, Maharshi Pandya, Nidhi Patel, R D Dixit, A M Kadri, Harsh Bakshi, Chaitanya Joshi, Madhvi Joshi |
| EPI_ISL_476863 | GMERS Medical College and Hospital, Gandhinagar | Gujarat Biotechnology Research Centre | Raghawendra Kumar, Dinesh Kumar, Zuber Saiyed, Komal Patel, Labdhi Pandya, Afzal Ansari, Nikha Trivedi, Seema Bhatt, Gaurishankar Shrimali, Bhavesh Modi, Bharti Rajani, Apurvasinh Puvar, Janvi Raval, Zarna Patel, Monika Gandhi, Pinal Trivedi, Maharshi Pandya, Nidhi Patel, Nitin Savaliya, R D Dixit, A M Kadri, Harsh Bakshi, Chaitanya Joshi, Madhvi Joshi |
| EPI_ISL_476864 | GMERS Medical College and Hospital, Gandhinagar | Gujarat Biotechnology Research Centre | Dinesh Kumar, Zuber Saiyed, Komal Patel, Labdhi Pandya, Afzal Ansari, Nikha Trivedi, Seema Bhatt, Gaurishankar Shrimali, Bhavesh Modi, Bharti Rajani, Apurvasinh Puvar, Janvi Raval, Zarna Patel, Monika Gandhi, Pinal Trivedi, Maharshi Pandya, Nidhi Patel, Nitin Savaliya, Raghawendra Kumar, R D Dixit, A M Kadri, Harsh Bakshi, Chaitanya Joshi, Madhvi Joshi |
| EPI_ISL_476865 | GMERS Medical College and Hospital, Gandhinagar | Gujarat Biotechnology Research Centre | Zuber Saiyed, Komal Patel, Labdhi Pandya, Afzal Ansari, Nikha Trivedi, Seema Bhatt, Gaurishankar Shrimali, Bhavesh Modi, Bharti Rajani, Apurvasinh Puvar, Janvi Raval, Zarna Patel, Monika Gandhi, Pinal Trivedi, Maharshi Pandya, Nidhi Patel, Nitin Savaliya, Raghawendra Kumar, Dinesh Kumar, R D Dixit, A M Kadri, Harsh Bakshi, Chaitanya Joshi, Madhvi Joshi |
| EPI_ISL_476866 | GMERS Medical College and Hospital, Gandhinagar | Gujarat Biotechnology Research Centre | Komal Patel, Labdhi Pandya, Afzal Ansari, Nikha Trivedi, Seema Bhatt, Gaurishankar Shrimali, Bhavesh Modi, Bharti Rajani, Apurvasinh Puvar, Janvi Raval, Zarna Patel, Monika Gandhi, Pinal Trivedi, Maharshi Pandya, Nidhi Patel, Nitin Savaliya, Raghawendra Kumar, Dinesh Kumar, Zuber Saiyed, R D Dixit, A M Kadri, Harsh Bakshi, Chaitanya Joshi, Madhvi Joshi |
| EPI_ISL_476867 | Banas Medical College and Research Institute | Gujarat Biotechnology Research Centre | Labdhi Pandya, Afzal Ansari, Nikha Trivedi, Radhika Khara, Sunil R Joshi, Viren s Doshi, Apurvasinh Puvar, Janvi Raval, Zarna Patel, Monika Gandhi, Pinal Trivedi, Maharshi Pandya, Nidhi Patel, Nitin Savaliya, Raghawendra Kumar, Dinesh Kumar, Zuber Saiyed, Komal Patel, R D Dixit, A M Kadri, Harsh Bakshi, Chaitanya Joshi, Madhvi Joshi |
| EPI_ISL_476868 | Banas Medical College and Research Institute | Gujarat Biotechnology Research Centre | Afzal Ansari, Nikha Trivedi, Radhika Khara, Sunil R Joshi, Viren s Doshi, Apurvasinh Puvar, Janvi Raval, Zarna Patel, Monika Gandhi, Pinal Trivedi, Maharshi Pandya, Nidhi Patel, Nitin Savaliya, Raghawendra Kumar, Dinesh Kumar, Zuber Saiyed, Komal Patel, Labdhi Pandya, R D Dixit, A M Kadri, Harsh Bakshi, Chaitanya Joshi, Madhvi Joshi |
| EPI_ISL_476869 | Department of MicroBiology, Government Medical College, Surat | Gujarat Biotechnology Research Centre | Nikha Trivedi, Naresh Chauhan, Summaiya Mullan, Amit gamit, Apurvasinh Puvar, Janvi Raval, Zarna Patel, Monika Gandhi, Pinal Trivedi, Maharshi Pandya, Nidhi Patel, Nitin Savaliya, Raghawendra Kumar, Dinesh Kumar, Zuber Saiyed, Komal Patel, Labdhi Pandya, Afzal Ansari, R D Dixit, A M Kadri, Harsh Bakshi, Chaitanya Joshi, Madhvi Joshi |
| EPI_ISL_476870 | Department of MicroBiology, Government Medical College, Surat | Gujarat Biotechnology Research Centre | Naresh Chauhan, Summaiya Mullan, Amit gamit, Apurvasinh Puvar, Janvi Raval, Zarna Patel, Monika Gandhi, Pinal Trivedi, Maharshi Pandya, Nidhi Patel, Nitin Savaliya, Raghawendra Kumar, Dinesh Kumar, Zuber Saiyed, Komal Patel, Labdhi Pandya, Afzal Ansari, Nikha Trivedi, R D Dixit, A M Kadri, Harsh Bakshi, Chaitanya Joshi, Madhvi Joshi |
| EPI_ISL_476871 | Department of MicroBiology, Government Medical College, Surat | Gujarat Biotechnology Research Centre | Summaiya Mullan, Amit gamit, Apurvasinh Puvar, Janvi Raval, Zarna Patel, Monika Gandhi, Pinal Trivedi, Maharshi Pandya, Nidhi Patel, Nitin Savaliya, Raghawendra Kumar, Dinesh Kumar, Zuber Saiyed, Komal Patel, Labdhi Pandya, Afzal Ansari, Nikha Trivedi, Naresh Chauhan, R D Dixit, A M Kadri, Harsh Bakshi, Chaitanya Joshi, Madhvi Joshi |
| EPI_ISL_476872 | Department of MicroBiology, Government Medical College, Surat | Gujarat Biotechnology Research Centre | Amit gamit, Apurvasinh Puvar, Janvi Raval, Zarna Patel, Monika Gandhi, Pinal Trivedi, Maharshi Pandya, Nidhi Patel, Nitin Savaliya, Raghawendra Kumar, Dinesh Kumar, Zuber Saiyed, Komal Patel, Labdhi Pandya, Afzal Ansari, Nikha Trivedi, Naresh Chauhan, Summaiya Mullan, R D Dixit, A M Kadri, Harsh Bakshi, Chaitanya Joshi, Madhvi Joshi |
| EPI_ISL_476873 | Department of MicroBiology, Government Medical College, Surat | Gujarat Biotechnology Research Centre | Apurvasinh Puvar, Janvi Raval, Zarna Patel, Monika Gandhi, Pinal Trivedi, Maharshi Pandya, Nidhi Patel, Nitin Savaliya, Raghawendra Kumar, Dinesh Kumar, Zuber Saiyed, Komal Patel, Labdhi Pandya, Afzal Ansari, Nikha Trivedi, Naresh Chauhan, Summaiya Mullan, Amit gamit, R D Dixit, A M Kadri, Harsh Bakshi, Chaitanya Joshi, Madhvi Joshi |
| EPI_ISL_476874 | Department of MicroBiology, Government Medical College, Surat | Gujarat Biotechnology Research Centre | Janvi Raval, Zarna Patel, Monika Gandhi, Pinal Trivedi, Maharshi Pandya, Nidhi Patel, Nitin Savaliya, Raghawendra Kumar, Dinesh Kumar, Zuber Saiyed, Komal Patel, Labdhi Pandya, Afzal Ansari, Nikha Trivedi, Naresh Chauhan, Summaiya Mullan, Amit gamit, Apurvasinh Puvar, R D Dixit, A M Kadri, Harsh Bakshi, Chaitanya Joshi, Madhvi Joshi |
| EPI_ISL_476875 | Department of MicroBiology, Government Medical College, Surat | Gujarat Biotechnology Research Centre | Zarna Patel, Monika Gandhi, Pinal Trivedi, Maharshi Pandya, Nidhi Patel, Nitin Savaliya, Raghawendra Kumar, Dinesh Kumar, Zuber Saiyed, Komal Patel, Labdhi Pandya, Afzal Ansari, Nikha Trivedi, Naresh Chauhan, Summaiya Mullan, Amit gamit, Apurvasinh Puvar, Janvi Raval, R D Dixit, A M Kadri, Harsh Bakshi, Chaitanya Joshi, Madhvi Joshi |

| | | | |
|---|---|---|---|
| EPI_ISL_476876 | Department of MicroBiology, Government Medical College, Surat | Gujarat Biotechnology Research Centre | Pinal Trivedi, Maharshi Pandya, Nidhi Patel, Nitin Savaliya, Raghawendra Kumar, Dinesh Kumar, Zuber Saiyed, Komal Patel, Labdhi Pandya, Afzal Ansari, Nikha Trivedi, Naresh Chauhan, Summaiya Mullan, Amit gamit, Apurvasinh Puvar, Janvi Raval, Zarna Patel, Monika Gandhi, R D Dixit, A M Kadri, Harsh Bakshi, Chaitanya Joshi, Madhvi Joshi |
| EPI_ISL_476877 | Department of MicroBiology, Government Medical College, Surat | Gujarat Biotechnology Research Centre | Maharshi Pandya, Nidhi Patel, Nitin Savaliya, Raghawendra Kumar, Dinesh Kumar, Zuber Saiyed, Komal Patel, Labdhi Pandya, Afzal Ansari, Nikha Trivedi, Naresh Chauhan, Summaiya Mullan, Amit gamit, Apurvasinh Puvar, Janvi Raval, Zarna Patel, Monika Gandhi, Pinal Trivedi, R D Dixit, A M Kadri, Harsh Bakshi, Chaitanya Joshi, Madhvi Joshi |
| EPI_ISL_476878 | Department of MicroBiology, Government Medical College, Surat | Gujarat Biotechnology Research Centre | Nidhi Patel, Nitin Savaliya, Raghawendra Kumar, Dinesh Kumar, Zuber Saiyed, Komal Patel, Labdhi Pandya, Afzal Ansari, Nikha Trivedi, Naresh Chauhan, Summaiya Mullan, Amit gamit, Apurvasinh Puvar, Janvi Raval, Zarna Patel, Monika Gandhi, Pinal Trivedi, Maharshi Pandya, R D Dixit, A M Kadri, Harsh Bakshi, Chaitanya Joshi, Madhvi Joshi |
| EPI_ISL_476879 | Department of MicroBiology, Government Medical College, Surat | Gujarat Biotechnology Research Centre | Nitin Savaliya, Raghawendra Kumar, Dinesh Kumar, Zuber Saiyed, Komal Patel, Labdhi Pandya, Afzal Ansari, Nikha Trivedi, Naresh Chauhan, Summaiya Mullan, Amit gamit, Apurvasinh Puvar, Janvi Raval, Zarna Patel, Monika Gandhi, Pinal Trivedi, Maharshi Pandya, Nidhi Patel, R D Dixit, A M Kadri, Harsh Bakshi, Chaitanya Joshi, Madhvi Joshi |
| EPI_ISL_476880 | Department of MicroBiology, Government Medical College, Surat | Gujarat Biotechnology Research Centre | Raghawendra Kumar, Dinesh Kumar, Zuber Saiyed, Komal Patel, Labdhi Pandya, Afzal Ansari, Nikha Trivedi, Naresh Chauhan, Summaiya Mullan, Amit gamit, Apurvasinh Puvar, Janvi Raval, Zarna Patel, Monika Gandhi, Pinal Trivedi, Maharshi Pandya, Nidhi Patel, Nitin Savaliya, R D Dixit, A M Kadri, Harsh Bakshi, Chaitanya Joshi, Madhvi Joshi |
| EPI_ISL_476881 | Department of MicroBiology, Government Medical College, Surat | Gujarat Biotechnology Research Centre | Dinesh Kumar, Zuber Saiyed, Komal Patel, Labdhi Pandya, Afzal Ansari, Nikha Trivedi, Naresh Chauhan, Summaiya Mullan, Amit gamit, Apurvasinh Puvar, Janvi Raval, Zarna Patel, Monika Gandhi, Pinal Trivedi, Maharshi Pandya, Nidhi Patel, Nitin Savaliya, Raghawendra Kumar, R D Dixit, A M Kadri, Harsh Bakshi, Chaitanya Joshi, Madhvi Joshi |
| EPI_ISL_476882 | Department of MicroBiology, Government Medical College, Surat | Gujarat Biotechnology Research Centre | Zuber Saiyed, Komal Patel, Labdhi Pandya, Afzal Ansari, Nikha Trivedi, Naresh Chauhan, Summaiya Mullan, Amit gamit, Apurvasinh Puvar, Janvi Raval, Zarna Patel, Monika Gandhi, Pinal Trivedi, Maharshi Pandya, Nidhi Patel, Nitin Savaliya, Raghawendra Kumar, Dinesh Kumar, R D Dixit, A M Kadri, Harsh Bakshi, Chaitanya Joshi, Madhvi Joshi |
| EPI_ISL_476884, EPI_ISL_476885, EPI_ISL_476886, EPI_ISL_476887, EPI_ISL_476888, EPI_ISL_476889, EPI_ISL_476890, EPI_ISL_476891, EPI_ISL_476892, EPI_ISL_476893, EPI_ISL_476894, EPI_ISL_476895, EPI_ISL_476896 | | | |
| see above | Defence Research & Development Establishment (DRDE) | Defence Research & Development Establishment (DRDE) | Shashi Sharma, Paban Kumar Dash, Sushil Kumar Sharma, Ambuj Shrivastava, Jyoti S. Kumar |
| EPI_ISL_477015 | Institute of Microbiology, Universidad San Francisco de Quito | Institute of Microbiology, Universidad San Francisco de Quito | Sully Márquez, Belén Prado-Vivar, Juan José Guadalupe, Monica Becerra-Wong, Carla Torres, Bernardo Gutiérrez, Jorge Luis Velez, Verónica Barragán, Patricio Rojas-Silva, Gabriel Trueba, Michelle Grunauer, Paúl Cárdenas |
| EPI_ISL_477016 | Institute of Microbiology, Universidad San Francisco de Quito | Institute of Microbiology, Universidad San Francisco de Quito | Juan José Guadalupe, Sully Márquez, Belén Prado-Vivar, Monica Becerra-Wong, Carla Torres, Bernardo Gutiérrez, Jorge Luis Velez, Verónica Barragán, Patricio Rojas-Silva, Gabriel Trueba, Michelle Grunauer, Paúl Cárdenas |
| EPI_ISL_477143, EPI_ISL_477150, EPI_ISL_477155, EPI_ISL_477156 | Institut Pasteur Dakar | Institut Pasteur de Dakar | Ndongo Dia, Moussa Moise Diagne, Mamadou Diop, Mamadou Malado Jallow, Marie Henriette Dior Ndione, Safietou Sankhe, Ousmane Faye, Amadou Alpha Sall. |
| EPI_ISL_477180 | Department of Laboratory Medicine Tan Tock Seng Hospital | Department of Laboratory Medicine Tan Tock Seng Hospital | Chen YYC, Zair X, Li C, Tang WY, Maurer-Stroh S, Barkham TMS, Nagarajan N, Sessions OM |
| EPI_ISL_477183 | Department of MicroBiology, Government Medical College, Surat | Gujarat Biotechnology Research Centre | Monika Gandhi, Pinal Trivedi, Maharshi Pandya, Nidhi Patel, Nitin Savaliya, Raghawendra Kumar, Dinesh Kumar, Zuber Saiyed, Komal Patel, Labdhi Pandya, Afzal Ansari, Nikha Trivedi, Naresh Chauhan, Summaiya Mullan, Amit gamit, Apurvasinh Puvar, Janvi Raval, Zarna Patel, R D Dixit, A M Kadri, Harsh Bakshi, Chaitanya Joshi, Madhvi Joshi |
| EPI_ISL_479534, EPI_ISL_479535, EPI_ISL_479536, EPI_ISL_479537, EPI_ISL_479538, EPI_ISL_479539, EPI_ISL_479540, EPI_ISL_479541, EPI_ISL_479543, EPI_ISL_479544, EPI_ISL_479546, EPI_ISL_479547, EPI_ISL_479548, EPI_ISL_479550, EPI_ISL_479551, EPI_ISL_479552, EPI_ISL_479553 | | | |
| see above | NIV Influenza | NIV Influenza | Potdar V |
| EPI_ISL_482682, EPI_ISL_482683, EPI_ISL_482684, EPI_ISL_482685, EPI_ISL_482686, EPI_ISL_482687, EPI_ISL_482688, EPI_ISL_482689, EPI_ISL_482690, EPI_ISL_482691, EPI_ISL_482692, EPI_ISL_482693, EPI_ISL_482694, EPI_ISL_482695, EPI_ISL_482696, EPI_ISL_482697, EPI_ISL_482698, EPI_ISL_482699 | | | |
| see above | Singapore General Hospital | Department of Microbiology | Nurdyana Abdul Rahman, Kun Lee Lim, Chenhao Li, Kian Sing Chan, Lynette Oon, Kern Rei Chng, Niranjan Nagarajan, Karrie Ko |
| EPI_ISL_482702, EPI_ISL_482704, EPI_ISL_482705, EPI_ISL_482708, EPI_ISL_482709 | Molecular Diagnostics Services (MDS) | KRISP, KZN Research Innovation and Sequencing Platform | Giandhari J, Pillay S, Lessells R, Chimukangara B, Mdlalose K, York D, Khan S, Tegally H, Wilkinson E, de Oliveira T |
| EPI_ISL_482710, EPI_ISL_482711, EPI_ISL_482712 | NHLS-IALCH | KRISP, KZN Research Innovation and Sequencing Platform | Giandhari J, Pillay S, Lessells R, Chimukangara B, Mdlalose K, York D, Khan S, Tegally H, Wilkinson E, de Oliveira T |
| EPI_ISL_482716, EPI_ISL_482717, EPI_ISL_482718, EPI_ISL_482719, EPI_ISL_482720, EPI_ISL_482721, EPI_ISL_482722, EPI_ISL_482723 | Molecular Diagnostics Services (MDS) | KRISP, KZN Research Innovation and Sequencing Platform | Giandhari J, Pillay S, Lessells R, Chimukangara B, Mdlalose K, York D, Khan S, Tegally H, Wilkinson E, de Oliveira T |
| EPI_ISL_482726 | NHLS-IALCH | KRISP, KZN Research Innovation and Sequencing Platform | Giandhari J, Pillay S, Lessells R, Chimukangara B, Mdlalose K, York D, Khan S, Tegally H, Wilkinson E, de Oliveira T |
| EPI_ISL_482778, EPI_ISL_482782, EPI_ISL_482783 | Tuen Mun Hospital | Hong Kong Department of Health | Mak Gannon C.K., Cheng Peter K.C., Lam Edman T.K., Chan Rickjason C.W., Tsang Dominic N.C. |
| EPI_ISL_482852, EPI_ISL_482853, EPI_ISL_482862, EPI_ISL_482867 | Molecular Diagnostics Services (MDS) | KRISP, KZN Research Innovation and Sequencing Platform | Giandhari J, Pillay S, Lessells R, Chimukangara B, Mdlalose K, York D, Khan S, Tegally H, Wilkinson E, de Oliveira T |
| EPI_ISL_482874, EPI_ISL_482875, EPI_ISL_482876, EPI_ISL_482877, EPI_ISL_482878 | Institut Pasteur Dakar | Institut Pasteur de Dakar | Ndongo Dia, Moussa Moise Diagne, Mamadou Diop, Marie Henriette Dior Ndione, Mamadou malado Jallow, Safietou Sankhe, Ousmane Faye, Amadou Alpha Sall. |
| EPI_ISL_483566 | Clinical Microbiology Laboratory- Basurto University Hospital | Biocruces-Bizkaia | Mikel J. Urrutikoetxea-Gutierrez, Ana Belén Belén de la Hoz, Matxalen Vidal-García, Mº Carmen Nieto Toboso, Estibaliz Ugalde-Zarraga, José Luis Díaz de Tuesta del Arco |
| EPI_ISL_483570 | Clinical Microbiology Laboratory- Basurto University Hospita | Biocruces-Bizkaia | Mikel J. Urrutikoetxea-Gutierrez, Ana Belén Belén de la Hoz, Matxalen Vidal-García, Mº Carmen Nieto Toboso, Estibaliz Ugalde-Zarraga, José Luis Díaz de Tuesta del Arco |
| EPI_ISL_483571, EPI_ISL_483572, EPI_ISL_483573 | Clinical Microbiology Laboratory- Basurto University Hospital | Biocruces-Bizkaia | Mikel J. Urrutikoetxea-Gutierrez, Ana Belén Belén de la Hoz, Matxalen Vidal-García, Mº Carmen Nieto Toboso, Estibaliz Ugalde-Zarraga, José Luis Díaz de Tuesta del Arco |
| EPI_ISL_483820 | GMERS Medical College and Hospital, Gandhinagar | Gujarat Biotechnology Research Centre | Komal Patel, Labdhi Pandya, Afzal Ansari, Nikha Trivedi, Seema Bhatt, Gaurishankar Shrimali, Bhavesh Modi, Bharti Rajani, Apurvasinh Puvar, Janvi Raval, Zarna Patel, Monika Gandhi, Pinal Trivedi, Maharshi Pandya, Nidhi Patel, Nitin Savaliya, Raghawendra Kumar, Dinesh Kumar, Zuber Saiyed, R D Dixit, A M Kadri, Harsh Bakshi, Chaitanya Joshi, Madhvi Joshi |
| EPI_ISL_483821 | Government Medical College, Vadodara | Gujarat Biotechnology Research Centre | Labdhi Pandya, Afzal Ansari, Nikha Trivedi, Meenakshi Shah, Neena Doshi, Varsha Godbole, Apurvasinh Puvar, Janvi Raval, Zarna Patel, Monika Gandhi, Pinal Trivedi, Maharshi Pandya, Nidhi Patel, Nitin Savaliya, Raghawendra Kumar, Dinesh Kumar, Zuber Saiyed, Komal Patel, R D Dixit, A M Kadri, Harsh Bakshi, Chaitanya Joshi, Madhvi Joshi |
| EPI_ISL_483822 | Government Medical College, Vadodara | Gujarat Biotechnology Research Centre | Afzal Ansari, Nikha Trivedi, Meenakshi Shah, Neena Doshi, Varsha Godbole, Apurvasinh Puvar, Janvi Raval, Zarna Patel, Monika Gandhi, Pinal Trivedi, Maharshi Pandya, Nidhi Patel, Nitin Savaliya, Raghawendra Kumar, Dinesh Kumar, Zuber Saiyed, Komal Patel, Labdhi Pandya, R D Dixit, A M Kadri, Harsh Bakshi, Chaitanya Joshi, Madhvi Joshi |
| EPI_ISL_483823 | GMERS Medical College Himmatnagar | Gujarat Biotechnology Research Centre | Nikha Trivedi, Himanshu Khatri, Mayur Gandhi, Apurvasinh Puvar, Janvi Raval, Zarna Patel, Monika Gandhi, Pinal Trivedi, Maharshi Pandya, Nidhi Patel, Nitin Savaliya, Raghawendra Kumar, Dinesh Kumar, Zuber Saiyed, Komal Patel, Labdhi Pandya, Afzal Ansari, R D Dixit, A M Kadri, Harsh Bakshi, Chaitanya Joshi, Madhvi Joshi |

| | | | |
|---|---|---|---|
| EPI_ISL_483824 | GMERS Medical College Himmatnagar | Gujarat Biotechnology Research Centre | Himanshu Khatri, Mayur Gandhi, Apurvasinh Puvar, Janvi Raval, Zarna Patel, Monika Gandhi, Pinal Trivedi, Maharshi Pandya, Nidhi Patel, Nitin Savaliya, Raghawendra Kumar, Dinesh Kumar, Zuber Saiyed, Komal Patel, Labdhi Pandya, Afzal Ansari, Nikha Trivedi, R D Dixit, A M Kadri, Harsh Bakshi, Chaitanya Joshi, Madhvi Joshi |
| EPI_ISL_483825 | GMERS Medical College Himmatnagar | Gujarat Biotechnology Research Centre | Mayur Gandhi, Apurvasinh Puvar, Janvi Raval, Zarna Patel, Monika Gandhi, Pinal Trivedi, Maharshi Pandya, Nidhi Patel, Nitin Savaliya, Raghawendra Kumar, Dinesh Kumar, Zuber Saiyed, Komal Patel, Labdhi Pandya, Afzal Ansari, Nikha Trivedi, Himanshu Khatri, R D Dixit, A M Kadri, Harsh Bakshi, Chaitanya Joshi, Madhvi Joshi |
| EPI_ISL_483826 | GMERS Medical College Himmatnagar | Gujarat Biotechnology Research Centre | Apurvasinh Puvar, Janvi Raval, Zarna Patel, Monika Gandhi, Pinal Trivedi, Maharshi Pandya, Nidhi Patel, Nitin Savaliya, Raghawendra Kumar, Dinesh Kumar, Zuber Saiyed, Komal Patel, Labdhi Pandya, Afzal Ansari, Nikha Trivedi, Himanshu Khatri, Mayur Gandhi, R D Dixit, A M Kadri, Harsh Bakshi, Chaitanya Joshi, Madhvi Joshi |
| EPI_ISL_483827 | GMERS Medical College Himmatnagar | Gujarat Biotechnology Research Centre | Janvi Raval, Zarna Patel, Monika Gandhi, Pinal Trivedi, Maharshi Pandya, Nidhi Patel, Nitin Savaliya, Raghawendra Kumar, Dinesh Kumar, Zuber Saiyed, Komal Patel, Labdhi Pandya, Afzal Ansari, Nikha Trivedi, Himanshu Khatri, Mayur Gandhi, Apurvasinh Puvar, R D Dixit, A M Kadri, Harsh Bakshi, Chaitanya Joshi, Madhvi Joshi |
| EPI_ISL_483828 | GMERS Medical College Himmatnagar | Gujarat Biotechnology Research Centre | Zarna Patel, Monika Gandhi, Pinal Trivedi, Maharshi Pandya, Nidhi Patel, Nitin Savaliya, Raghawendra Kumar, Dinesh Kumar, Zuber Saiyed, Komal Patel, Labdhi Pandya, Afzal Ansari, Nikha Trivedi, Himanshu Khatri, Mayur Gandhi, Apurvasinh Puvar, Janvi Raval, R D Dixit, A M Kadri, Harsh Bakshi, Chaitanya Joshi, Madhvi Joshi |
| EPI_ISL_483829 | Pandit Deendayal Upadhyay Government Medical College, Rajkot | Gujarat Biotechnology Research Centre | Monika Gandhi, Pinal Trivedi, Maharshi Pandya, Nidhi Patel, Nitin Savaliya, Raghawendra Kumar, Dinesh Kumar, Zuber Saiyed, Komal Patel, Labdhi Pandya, Afzal Ansari, Nikha Trivedi, Gauravi Dhruv, Arti Trivedi, Apurvasinh Puvar, Janvi Raval, Zarna Patel, R D Dixit, A M Kadri, Harsh Bakshi, Chaitanya Joshi, Madhvi Joshi |
| EPI_ISL_483830 | Pandit Deendayal Upadhyay Government Medical College, Rajkot | Gujarat Biotechnology Research Centre | Pinal Trivedi, Maharshi Pandya, Nidhi Patel, Nitin Savaliya, Raghawendra Kumar, Dinesh Kumar, Zuber Saiyed, Komal Patel, Labdhi Pandya, Afzal Ansari, Nikha Trivedi, Gauravi Dhruv, Arti Trivedi, Apurvasinh Puvar, Janvi Raval, Zarna Patel, Monika Gandhi, R D Dixit, A M Kadri, Harsh Bakshi, Chaitanya Joshi, Madhvi Joshi |
| EPI_ISL_483831 | Pandit Deendayal Upadhyay Government Medical College, Rajkot | Gujarat Biotechnology Research Centre | Maharshi Pandya, Nidhi Patel, Nitin Savaliya, Raghawendra Kumar, Dinesh Kumar, Zuber Saiyed, Komal Patel, Labdhi Pandya, Afzal Ansari, Nikha Trivedi, Gauravi Dhruv, Arti Trivedi, Apurvasinh Puvar, Janvi Raval, Zarna Patel, Monika Gandhi, Pinal Trivedi, R D Dixit, A M Kadri, Harsh Bakshi, Chaitanya Joshi, Madhvi Joshi |
| EPI_ISL_483832 | Pandit Deendayal Upadhyay Government Medical College, Rajkot | Gujarat Biotechnology Research Centre | Nidhi Patel, Nitin Savaliya, Raghawendra Kumar, Dinesh Kumar, Zuber Saiyed, Komal Patel, Labdhi Pandya, Afzal Ansari, Nikha Trivedi, Gauravi Dhruv, Arti Trivedi, Apurvasinh Puvar, Janvi Raval, Zarna Patel, Monika Gandhi, Pinal Trivedi, Maharshi Pandya, R D Dixit, A M Kadri, Harsh Bakshi, Chaitanya Joshi, Madhvi Joshi |
| EPI_ISL_483833 | Pandit Deendayal Upadhyay Government Medical College, Rajkot | Gujarat Biotechnology Research Centre | Nitin Savaliya, Raghawendra Kumar, Dinesh Kumar, Zuber Saiyed, Komal Patel, Labdhi Pandya, Afzal Ansari, Nikha Trivedi, Gauravi Dhruv, Arti Trivedi, Apurvasinh Puvar, Janvi Raval, Zarna Patel, Monika Gandhi, Pinal Trivedi, Maharshi Pandya, Nidhi Patel, R D Dixit, A M Kadri, Harsh Bakshi, Chaitanya Joshi, Madhvi Joshi |
| EPI_ISL_483834 | Pandit Deendayal Upadhyay Government Medical College, Rajkot | Gujarat Biotechnology Research Centre | Raghawendra Kumar, Dinesh Kumar, Zuber Saiyed, Komal Patel, Labdhi Pandya, Afzal Ansari, Nikha Trivedi, Gauravi Dhruv, Arti Trivedi, Apurvasinh Puvar, Janvi Raval, Zarna Patel, Monika Gandhi, Pinal Trivedi, Maharshi Pandya, Nidhi Patel, Nitin Savaliya, R D Dixit, A M Kadri, Harsh Bakshi, Chaitanya Joshi, Madhvi Joshi |
| EPI_ISL_483835 | Pandit Deendayal Upadhyay Government Medical College, Rajkot | Gujarat Biotechnology Research Centre | Dinesh Kumar, Zuber Saiyed, Komal Patel, Labdhi Pandya, Afzal Ansari, Nikha Trivedi, Gauravi Dhruv, Arti Trivedi, Apurvasinh Puvar, Janvi Raval, Zarna Patel, Monika Gandhi, Pinal Trivedi, Maharshi Pandya, Nidhi Patel, Nitin Savaliya, Raghawendra Kumar, R D Dixit, A M Kadri, Harsh Bakshi, Chaitanya Joshi, Madhvi Joshi |
| EPI_ISL_483836 | Pandit Deendayal Upadhyay Government Medical College, Rajkot | Gujarat Biotechnology Research Centre | Zuber Saiyed, Komal Patel, Labdhi Pandya, Afzal Ansari, Nikha Trivedi, Gauravi Dhruv, Arti Trivedi, Apurvasinh Puvar, Janvi Raval, Zarna Patel, Monika Gandhi, Pinal Trivedi, Maharshi Pandya, Nidhi Patel, Nitin Savaliya, Raghawendra Kumar, Dinesh Kumar, R D Dixit, A M Kadri, Harsh Bakshi, Chaitanya Joshi, Madhvi Joshi |
| EPI_ISL_483837 | Pandit Deendayal Upadhyay Government Medical College, Rajkot | Gujarat Biotechnology Research Centre | Komal Patel, Labdhi Pandya, Afzal Ansari, Nikha Trivedi, Gauravi Dhruv, Arti Trivedi, Apurvasinh Puvar, Janvi Raval, Zarna Patel, Monika Gandhi, Pinal Trivedi, Maharshi Pandya, Nidhi Patel, Nitin Savaliya, Raghawendra Kumar, Dinesh Kumar, Zuber Saiyed, R D Dixit, A M Kadri, Harsh Bakshi, Chaitanya Joshi, Madhvi Joshi |
| EPI_ISL_483838 | Pandit Deendayal Upadhyay Government Medical College, Rajkot | Gujarat Biotechnology Research Centre | Labdhi Pandya, Afzal Ansari, Nikha Trivedi, Gauravi Dhruv, Arti Trivedi, Apurvasinh Puvar, Janvi Raval, Zarna Patel, Monika Gandhi, Pinal Trivedi, Maharshi Pandya, Nidhi Patel, Nitin Savaliya, Raghawendra Kumar, Dinesh Kumar, Zuber Saiyed, Komal Patel, R D Dixit, A M Kadri, Harsh Bakshi, Chaitanya Joshi, Madhvi Joshi |
| EPI_ISL_483839 | Pandit Deendayal Upadhyay Government Medical College, Rajkot | Gujarat Biotechnology Research Centre | Afzal Ansari, Nikha Trivedi, Gauravi Dhruv, Arti Trivedi, Apurvasinh Puvar, Janvi Raval, Zarna Patel, Monika Gandhi, Pinal Trivedi, Maharshi Pandya, Nidhi Patel, Nitin Savaliya, Raghawendra Kumar, Dinesh Kumar, Zuber Saiyed, Komal Patel, Labdhi Pandya, R D Dixit, A M Kadri, Harsh Bakshi, Chaitanya Joshi, Madhvi Joshi |
| EPI_ISL_483840 | Pandit Deendayal Upadhyay Government Medical College, Rajkot | Gujarat Biotechnology Research Centre | Nikha Trivedi, Gauravi Dhruv, Arti Trivedi, Apurvasinh Puvar, Janvi Raval, Zarna Patel, Monika Gandhi, Pinal Trivedi, Maharshi Pandya, Nidhi Patel, Nitin Savaliya, Raghawendra Kumar, Dinesh Kumar, Zuber Saiyed, Komal Patel, Labdhi Pandya, Afzal Ansari, R D Dixit, A M Kadri, Harsh Bakshi, Chaitanya Joshi, Madhvi Joshi |
| EPI_ISL_483841 | Pandit Deendayal Upadhyay Government Medical College, Rajkot | Gujarat Biotechnology Research Centre | Gauravi Dhruv, Arti Trivedi, Apurvasinh Puvar, Janvi Raval, Zarna Patel, Monika Gandhi, Pinal Trivedi, Maharshi Pandya, Nidhi Patel, Nitin Savaliya, Raghawendra Kumar, Dinesh Kumar, Zuber Saiyed, Komal Patel, Labdhi Pandya, Afzal Ansari, Nikha Trivedi, R D Dixit, A M Kadri, Harsh Bakshi, Chaitanya Joshi, Madhvi Joshi |
| EPI_ISL_483842 | Pandit Deendayal Upadhyay Government Medical College, Rajkot | Gujarat Biotechnology Research Centre | Arti Trivedi, Apurvasinh Puvar, Janvi Raval, Zarna Patel, Monika Gandhi, Pinal Trivedi, Maharshi Pandya, Nidhi Patel, Nitin Savaliya, Raghawendra Kumar, Dinesh Kumar, Zuber Saiyed, Komal Patel, Labdhi Pandya, Afzal Ansari, Nikha Trivedi, Gauravi Dhruv, R D Dixit, A M Kadri, Harsh Bakshi, Chaitanya Joshi, Madhvi Joshi |
| EPI_ISL_483843 | Pandit Deendayal Upadhyay Government Medical College, Rajkot | Gujarat Biotechnology Research Centre | Apurvasinh Puvar, Janvi Raval, Zarna Patel, Monika Gandhi, Pinal Trivedi, Maharshi Pandya, Nidhi Patel, Nitin Savaliya, Raghawendra Kumar, Dinesh Kumar, Zuber Saiyed, Komal Patel, Labdhi Pandya, Afzal Ansari, Nikha Trivedi, Gauravi Dhruv, Arti Trivedi, R D Dixit, A M Kadri, Harsh Bakshi, Chaitanya Joshi, Madhvi Joshi |
| EPI_ISL_483844 | Pandit Deendayal Upadhyay Government Medical College, Rajkot | Gujarat Biotechnology Research Centre | Janvi Raval, Zarna Patel, Monika Gandhi, Pinal Trivedi, Maharshi Pandya, Nidhi Patel, Nitin Savaliya, Raghawendra Kumar, Dinesh Kumar, Zuber Saiyed, Komal Patel, Labdhi Pandya, Afzal Ansari, Nikha Trivedi, Gauravi Dhruv, Arti Trivedi, Apurvasinh Puvar, R D Dixit, A M Kadri, Harsh Bakshi, Chaitanya Joshi, Madhvi Joshi |
| EPI_ISL_483845 | Pandit Deendayal Upadhyay Government Medical College, Rajkot | Gujarat Biotechnology Research Centre | Zarna Patel, Monika Gandhi, Pinal Trivedi, Maharshi Pandya, Nidhi Patel, Nitin Savaliya, Raghawendra Kumar, Dinesh Kumar, Zuber Saiyed, Komal Patel, Labdhi Pandya, Afzal Ansari, Nikha Trivedi, Gauravi Dhruv, Arti Trivedi, Apurvasinh Puvar, Janvi Raval, R D Dixit, A M Kadri, Harsh Bakshi, Chaitanya Joshi, Madhvi Joshi |
| EPI_ISL_483846 | Pandit Deendayal Upadhyay Government Medical College, Rajkot | Gujarat Biotechnology Research Centre | Monika Gandhi, Pinal Trivedi, Maharshi Pandya, Nidhi Patel, Nitin Savaliya, Raghawendra Kumar, Dinesh Kumar, Zuber Saiyed, Komal Patel, Labdhi Pandya, Afzal Ansari, Nikha Trivedi, Gauravi Dhruv, Arti Trivedi, Apurvasinh Puvar, Janvi Raval, Zarna Patel, R D Dixit, A M Kadri, Harsh Bakshi, Chaitanya Joshi, Madhvi Joshi |
| EPI_ISL_483847 | Pandit Deendayal Upadhyay Government Medical College, Rajkot | Gujarat Biotechnology Research Centre | Pinal Trivedi, Maharshi Pandya, Nidhi Patel, Nitin Savaliya, Raghawendra Kumar, Dinesh Kumar, Zuber Saiyed, Komal Patel, Labdhi Pandya, Afzal Ansari, Nikha Trivedi, Gauravi Dhruv, Arti Trivedi, Apurvasinh Puvar, Janvi Raval, Zarna Patel, Monika Gandhi, R D Dixit, A M Kadri, Harsh Bakshi, Chaitanya Joshi, Madhvi Joshi |
| EPI_ISL_483848 | Pandit Deendayal Upadhyay Government Medical College, Rajkot | Gujarat Biotechnology Research Centre | Maharshi Pandya, Nidhi Patel, Nitin Savaliya, Raghawendra Kumar, Dinesh Kumar, Zuber Saiyed, Komal Patel, Labdhi Pandya, Afzal Ansari, Nikha Trivedi, Gauravi Dhruv, Arti Trivedi, Apurvasinh Puvar, Janvi Raval, Zarna Patel, Monika Gandhi, Pinal Trivedi, R D Dixit, A M Kadri, Harsh Bakshi, Chaitanya Joshi, Madhvi Joshi |

| | | | |
|---|---|---|---|
| EPI_ISL_483849 | Pandit Deendayal Upadhyay Government Medical College, Rajkot | Gujarat Biotechnology Research Centre | Nidhi Patel, Nitin Savaliya, Raghawendra Kumar, Dinesh Kumar, Zuber Saiyed, Komal Patel, Labdhi Pandya, Afzal Ansari, Nikha Trivedi, Gauravi Dhruv, Arti Trivedi, Apurvasinh Puvar, Janvi Raval, Zarna Patel, Monika Gandhi, Pinal Trivedi, Maharshi Pandya, R D Dixit, A M Kadri, Harsh Bakshi, Chaitanya Joshi, Madhvi Joshi |
| EPI_ISL_483850 | Department of Microbiology, Government Medical College, Surat | Gujarat Biotechnology Research Centre | Nitin Savaliya, Raghawendra Kumar, Dinesh Kumar, Zuber Saiyed, Komal Patel, Labdhi Pandya, Afzal Ansari, Nikha Trivedi, Naresh Chauhan, Summaiya Mullan, Amit gamit, Apurvasinh Puvar, Janvi Raval, Zarna Patel, Monika Gandhi, Pinal Trivedi, Maharshi Pandya, Nidhi Patel, R D Dixit, A M Kadri, Harsh Bakshi, Chaitanya Joshi, Madhvi Joshi |
| EPI_ISL_483851 | Department of Microbiology, Government Medical College, Surat | Gujarat Biotechnology Research Centre | Raghawendra Kumar, Dinesh Kumar, Zuber Saiyed, Komal Patel, Labdhi Pandya, Afzal Ansari, Nikha Trivedi, Naresh Chauhan, Summaiya Mullan, Amit gamit, Apurvasinh Puvar, Janvi Raval, Zarna Patel, Monika Gandhi, Pinal Trivedi, Maharshi Pandya, Nidhi Patel, Nitin Savaliya, R D Dixit, A M Kadri, Harsh Bakshi, Chaitanya Joshi, Madhvi Joshi |
| EPI_ISL_483852 | Department of Microbiology, Government Medical College, Surat | Gujarat Biotechnology Research Centre | Dinesh Kumar, Zuber Saiyed, Komal Patel, Labdhi Pandya, Afzal Ansari, Nikha Trivedi, Naresh Chauhan, Summaiya Mullan, Amit gamit, Apurvasinh Puvar, Janvi Raval, Zarna Patel, Monika Gandhi, Pinal Trivedi, Maharshi Pandya, Nidhi Patel, Nitin Savaliya, Raghawendra Kumar, R D Dixit, A M Kadri, Harsh Bakshi, Chaitanya Joshi, Madhvi Joshi |
| EPI_ISL_483853 | Department of Microbiology, Government Medical College, Surat | Gujarat Biotechnology Research Centre | Zuber Saiyed, Komal Patel, Labdhi Pandya, Afzal Ansari, Nikha Trivedi, Naresh Chauhan, Summaiya Mullan, Amit gamit, Apurvasinh Puvar, Janvi Raval, Zarna Patel, Monika Gandhi, Pinal Trivedi, Maharshi Pandya, Nidhi Patel, Nitin Savaliya, Raghawendra Kumar, Dinesh Kumar, R D Dixit, A M Kadri, Harsh Bakshi, Chaitanya Joshi, Madhvi Joshi |
| EPI_ISL_483854 | Department of Microbiology, Government Medical College, Surat | Gujarat Biotechnology Research Centre | Komal Patel, Labdhi Pandya, Afzal Ansari, Nikha Trivedi, Naresh Chauhan, Summaiya Mullan, Amit gamit, Apurvasinh Puvar, Janvi Raval, Zarna Patel, Monika Gandhi, Pinal Trivedi, Maharshi Pandya, Nidhi Patel, Nitin Savaliya, Raghawendra Kumar, Dinesh Kumar, Zuber Saiyed, R D Dixit, A M Kadri, Harsh Bakshi, Chaitanya Joshi, Madhvi Joshi |
| EPI_ISL_483855 | Department of Microbiology, Government Medical College, Surat | Gujarat Biotechnology Research Centre | Labdhi Pandya, Afzal Ansari, Nikha Trivedi, Naresh Chauhan, Summaiya Mullan, Amit gamit, Apurvasinh Puvar, Janvi Raval, Zarna Patel, Monika Gandhi, Pinal Trivedi, Maharshi Pandya, Nidhi Patel, Nitin Savaliya, Raghawendra Kumar, Dinesh Kumar, Zuber Saiyed, Komal Patel, R D Dixit, A M Kadri, Harsh Bakshi, Chaitanya Joshi, Madhvi Joshi |
| EPI_ISL_483856 | Department of Microbiology, Government Medical College, Surat | Gujarat Biotechnology Research Centre | Afzal Ansari, Nikha Trivedi, Naresh Chauhan, Summaiya Mullan, Amit gamit, Apurvasinh Puvar, Janvi Raval, Zarna Patel, Monika Gandhi, Pinal Trivedi, Maharshi Pandya, Nidhi Patel, Nitin Savaliya, Raghawendra Kumar, Dinesh Kumar, Zuber Saiyed, Komal Patel, Labdhi Pandya, R D Dixit, A M Kadri, Harsh Bakshi, Chaitanya Joshi, Madhvi Joshi |
| EPI_ISL_483857 | Department of Microbiology, Government Medical College, Surat | Gujarat Biotechnology Research Centre | Nikha Trivedi, Naresh Chauhan, Summaiya Mullan, Amit gamit, Apurvasinh Puvar, Janvi Raval, Zarna Patel, Monika Gandhi, Pinal Trivedi, Maharshi Pandya, Nidhi Patel, Nitin Savaliya, Raghawendra Kumar, Dinesh Kumar, Zuber Saiyed, Komal Patel, Labdhi Pandya, Afzal Ansari, R D Dixit, A M Kadri, Harsh Bakshi, Chaitanya Joshi, Madhvi Joshi |
| EPI_ISL_483858 | Department of Microbiology, Government Medical College, Surat | Gujarat Biotechnology Research Centre | Naresh Chauhan, Summaiya Mullan, Amit gamit, Apurvasinh Puvar, Janvi Raval, Zarna Patel, Monika Gandhi, Pinal Trivedi, Maharshi Pandya, Nidhi Patel, Nitin Savaliya, Raghawendra Kumar, Dinesh Kumar, Zuber Saiyed, Komal Patel, Labdhi Pandya, Afzal Ansari, Nikha Trivedi, R D Dixit, A M Kadri, Harsh Bakshi, Chaitanya Joshi, Madhvi Joshi |
| EPI_ISL_483859 | Department of Microbiology, Government Medical College, Surat | Gujarat Biotechnology Research Centre | Summaiya Mullan, Amit gamit, Apurvasinh Puvar, Janvi Raval, Zarna Patel, Monika Gandhi, Pinal Trivedi, Maharshi Pandya, Nidhi Patel, Nitin Savaliya, Raghawendra Kumar, Dinesh Kumar, Zuber Saiyed, Komal Patel, Labdhi Pandya, Afzal Ansari, Nikha Trivedi, Naresh Chauhan, R D Dixit, A M Kadri, Harsh Bakshi, Chaitanya Joshi, Madhvi Joshi |
| EPI_ISL_483860 | Department of Microbiology, Government Medical College, Surat | Gujarat Biotechnology Research Centre | Amit gamit, Apurvasinh Puvar, Janvi Raval, Zarna Patel, Monika Gandhi, Pinal Trivedi, Maharshi Pandya, Nidhi Patel, Nitin Savaliya, Raghawendra Kumar, Dinesh Kumar, Zuber Saiyed, Komal Patel, Labdhi Pandya, Afzal Ansari, Nikha Trivedi, Naresh Chauhan, Summaiya Mullan, R D Dixit, A M Kadri, Harsh Bakshi, Chaitanya Joshi, Madhvi Joshi |
| EPI_ISL_483861 | Department of Microbiology, Government Medical College, Surat | Gujarat Biotechnology Research Centre | Apurvasinh Puvar, Janvi Raval, Zarna Patel, Monika Gandhi, Pinal Trivedi, Maharshi Pandya, Nidhi Patel, Nitin Savaliya, Raghawendra Kumar, Dinesh Kumar, Zuber Saiyed, Komal Patel, Labdhi Pandya, Afzal Ansari, Nikha Trivedi, Naresh Chauhan, Summaiya Mullan, Amit gamit, R D Dixit, A M Kadri, Harsh Bakshi, Chaitanya Joshi, Madhvi Joshi |
| EPI_ISL_483862 | Department of Microbiology, Government Medical College, Surat | Gujarat Biotechnology Research Centre | Janvi Raval, Zarna Patel, Monika Gandhi, Pinal Trivedi, Maharshi Pandya, Nidhi Patel, Nitin Savaliya, Raghawendra Kumar, Dinesh Kumar, Zuber Saiyed, Komal Patel, Labdhi Pandya, Afzal Ansari, Nikha Trivedi, Naresh Chauhan, Summaiya Mullan, Amit gamit, Apurvasinh Puvar, R D Dixit, A M Kadri, Harsh Bakshi, Chaitanya Joshi, Madhvi Joshi |
| EPI_ISL_483863 | Department of Microbiology, Government Medical College, Surat | Gujarat Biotechnology Research Centre | Zarna Patel, Monika Gandhi, Pinal Trivedi, Maharshi Pandya, Nidhi Patel, Nitin Savaliya, Raghawendra Kumar, Dinesh Kumar, Zuber Saiyed, Komal Patel, Labdhi Pandya, Afzal Ansari, Nikha Trivedi, Naresh Chauhan, Summaiya Mullan, Amit gamit, Apurvasinh Puvar, Janvi Raval, R D Dixit, A M Kadri, Harsh Bakshi, Chaitanya Joshi, Madhvi Joshi |
| EPI_ISL_483864 | Department of Microbiology, Government Medical College, Surat | Gujarat Biotechnology Research Centre | Monika Gandhi, Pinal Trivedi, Maharshi Pandya, Nidhi Patel, Nitin Savaliya, Raghawendra Kumar, Dinesh Kumar, Zuber Saiyed, Komal Patel, Labdhi Pandya, Afzal Ansari, Nikha Trivedi, Naresh Chauhan, Summaiya Mullan, Amit gamit, Apurvasinh Puvar, Janvi Raval, Zarna Patel, R D Dixit, A M Kadri, Harsh Bakshi, Chaitanya Joshi, Madhvi Joshi |
| EPI_ISL_483865 | Department of Microbiology, Government Medical College, Surat | Gujarat Biotechnology Research Centre | Pinal Trivedi, Maharshi Pandya, Nidhi Patel, Nitin Savaliya, Raghawendra Kumar, Dinesh Kumar, Zuber Saiyed, Komal Patel, Labdhi Pandya, Afzal Ansari, Nikha Trivedi, Naresh Chauhan, Summaiya Mullan, Amit gamit, Apurvasinh Puvar, Janvi Raval, Zarna Patel, Monika Gandhi, R D Dixit, A M Kadri, Harsh Bakshi, Chaitanya Joshi, Madhvi Joshi |
| EPI_ISL_483866 | Department of Microbiology, Government Medical College, Surat | Gujarat Biotechnology Research Centre | Maharshi Pandya, Nidhi Patel, Nitin Savaliya, Raghawendra Kumar, Dinesh Kumar, Zuber Saiyed, Komal Patel, Labdhi Pandya, Afzal Ansari, Nikha Trivedi, Naresh Chauhan, Summaiya Mullan, Amit gamit, Apurvasinh Puvar, Janvi Raval, Zarna Patel, Monika Gandhi, Pinal Trivedi, R D Dixit, A M Kadri, Harsh Bakshi, Chaitanya Joshi, Madhvi Joshi |
| EPI_ISL_483867 | Department of Microbiology, Government Medical College, Surat | Gujarat Biotechnology Research Centre | Nidhi Patel, Nitin Savaliya, Raghawendra Kumar, Dinesh Kumar, Zuber Saiyed, Komal Patel, Labdhi Pandya, Afzal Ansari, Nikha Trivedi, Naresh Chauhan, Summaiya Mullan, Amit gamit, Apurvasinh Puvar, Janvi Raval, Zarna Patel, Monika Gandhi, Pinal Trivedi, Maharshi Pandya, R D Dixit, A M Kadri, Harsh Bakshi, Chaitanya Joshi, Madhvi Joshi |
| EPI_ISL_483868 | Department of Microbiology, Government Medical College, Surat | Gujarat Biotechnology Research Centre | Nitin Savaliya, Raghawendra Kumar, Dinesh Kumar, Zuber Saiyed, Komal Patel, Labdhi Pandya, Afzal Ansari, Nikha Trivedi, Naresh Chauhan, Summaiya Mullan, Amit gamit, Apurvasinh Puvar, Janvi Raval, Zarna Patel, Monika Gandhi, Pinal Trivedi, Maharshi Pandya, Nidhi Patel, R D Dixit, A M Kadri, Harsh Bakshi, Chaitanya Joshi, Madhvi Joshi |
| EPI_ISL_483869 | Department of Microbiology, Government Medical College, Surat | Gujarat Biotechnology Research Centre | Raghawendra Kumar, Dinesh Kumar, Zuber Saiyed, Komal Patel, Labdhi Pandya, Afzal Ansari, Nikha Trivedi, Naresh Chauhan, Summaiya Mullan, Amit gamit, Apurvasinh Puvar, Janvi Raval, Zarna Patel, Monika Gandhi, Pinal Trivedi, Maharshi Pandya, Nidhi Patel, Nitin Savaliya, R D Dixit, A M Kadri, Harsh Bakshi, Chaitanya Joshi, Madhvi Joshi |
| EPI_ISL_483870 | Department of Microbiology, Government Medical College, Surat | Gujarat Biotechnology Research Centre | Dinesh Kumar, Zuber Saiyed, Komal Patel, Labdhi Pandya, Afzal Ansari, Nikha Trivedi, Naresh Chauhan, Summaiya Mullan, Amit gamit, Apurvasinh Puvar, Janvi Raval, Zarna Patel, Monika Gandhi, Pinal Trivedi, Maharshi Pandya, Nidhi Patel, Nitin Savaliya, Raghawendra Kumar, R D Dixit, A M Kadri, Harsh Bakshi, Chaitanya Joshi, Madhvi Joshi |
| EPI_ISL_483871 | Department of Microbiology, Government Medical College, Surat | Gujarat Biotechnology Research Centre | Zuber Saiyed, Komal Patel, Labdhi Pandya, Afzal Ansari, Nikha Trivedi, Naresh Chauhan, Summaiya Mullan, Amit gamit, Apurvasinh Puvar, Janvi Raval, Zarna Patel, Monika Gandhi, Pinal Trivedi, Maharshi Pandya, Nidhi Patel, Nitin Savaliya, Raghawendra Kumar, Dinesh Kumar, R D Dixit, A M Kadri, Harsh Bakshi, Chaitanya Joshi, Madhvi Joshi |
| EPI_ISL_483872 | Department of Microbiology, Government Medical College, Surat | Gujarat Biotechnology Research Centre | Komal Patel, Labdhi Pandya, Afzal Ansari, Nikha Trivedi, Naresh Chauhan, Summaiya Mullan, Amit gamit, Apurvasinh Puvar, Janvi Raval, Zarna Patel, Monika Gandhi, Pinal Trivedi, Maharshi Pandya, Nidhi Patel, Nitin Savaliya, Raghawendra Kumar, Dinesh Kumar, Zuber Saiyed, R D Dixit, A M Kadri, Harsh Bakshi, Chaitanya Joshi, Madhvi Joshi |
| EPI_ISL_483873 | Department of Microbiology, Government Medical College, Surat | Gujarat Biotechnology Research Centre | Labdhi Pandya, Afzal Ansari, Nikha Trivedi, Naresh Chauhan, Summaiya Mullan, Amit gamit, Apurvasinh Puvar, Janvi Raval, Zarna Patel, Monika Gandhi, Pinal Trivedi, Maharshi Pandya, Nidhi Patel, Nitin Savaliya, Raghawendra Kumar, Dinesh Kumar, Zuber Saiyed, Komal Patel, R D Dixit, A M Kadri, Harsh Bakshi, Chaitanya Joshi, Madhvi Joshi |

| | | | |
|---|---|---|---|
| EPI_ISL_483874 | Department of Microbiology, Government Medical College, Surat | Gujarat Biotechnology Research Centre | Afzal Ansari, Nikha Trivedi, Naresh Chauhan, Summaiya Mullan, Amit gamit, Apurvasinh Puvar, Janvi Raval, Zarna Patel, Monika Gandhi, Pinal Trivedi, Maharshi Pandya, Nidhi Patel, Nitin Savaliya, Raghawendra Kumar, Dinesh Kumar, Zuber Saiyed, Komal Patel, Labdhi Pandya, R D Dixit, A M Kadri, Harsh Bakshi, Chaitanya Joshi, Madhvi Joshi |
| EPI_ISL_483875 | Department of Microbiology, Government Medical College, Surat | Gujarat Biotechnology Research Centre | Nikha Trivedi, Naresh Chauhan, Summaiya Mullan, Amit gamit, Apurvasinh Puvar, Janvi Raval, Zarna Patel, Monika Gandhi, Pinal Trivedi, Maharshi Pandya, Nidhi Patel, Nitin Savaliya, Raghawendra Kumar, Dinesh Kumar, Zuber Saiyed, Komal Patel, Labdhi Pandya, Afzal Ansari, R D Dixit, A M Kadri, Harsh Bakshi, Chaitanya Joshi, Madhvi Joshi |
| EPI_ISL_483876 | Department of Microbiology, Government Medical College, Surat | Gujarat Biotechnology Research Centre | Naresh Chauhan, Summaiya Mullan, Amit gamit, Apurvasinh Puvar, Janvi Raval, Zarna Patel, Monika Gandhi, Pinal Trivedi, Maharshi Pandya, Nidhi Patel, Nitin Savaliya, Raghawendra Kumar, Dinesh Kumar, Zuber Saiyed, Komal Patel, Labdhi Pandya, Afzal Ansari, Nikha Trivedi, R D Dixit, A M Kadri, Harsh Bakshi, Chaitanya Joshi, Madhvi Joshi |
| EPI_ISL_483877 | Department of Microbiology, Government Medical College, Surat | Gujarat Biotechnology Research Centre | Summaiya Mullan, Amit gamit, Apurvasinh Puvar, Janvi Raval, Zarna Patel, Monika Gandhi, Pinal Trivedi, Maharshi Pandya, Nidhi Patel, Nitin Savaliya, Raghawendra Kumar, Dinesh Kumar, Zuber Saiyed, Komal Patel, Labdhi Pandya, Afzal Ansari, Nikha Trivedi, Naresh Chauhan, R D Dixit, A M Kadri, Harsh Bakshi, Chaitanya Joshi, Madhvi Joshi |
| EPI_ISL_483878 | Department of Microbiology, Government Medical College, Surat | Gujarat Biotechnology Research Centre | Amit gamit, Apurvasinh Puvar, Janvi Raval, Zarna Patel, Monika Gandhi, Pinal Trivedi, Maharshi Pandya, Nidhi Patel, Nitin Savaliya, Raghawendra Kumar, Dinesh Kumar, Zuber Saiyed, Komal Patel, Labdhi Pandya, Afzal Ansari, Nikha Trivedi, Naresh Chauhan, Summaiya Mullan, R D Dixit, A M Kadri, Harsh Bakshi, Chaitanya Joshi, Madhvi Joshi |
| EPI_ISL_483879 | Department of Microbiology, Government Medical College, Surat | Gujarat Biotechnology Research Centre | Apurvasinh Puvar, Janvi Raval, Zarna Patel, Monika Gandhi, Pinal Trivedi, Maharshi Pandya, Nidhi Patel, Nitin Savaliya, Raghawendra Kumar, Dinesh Kumar, Zuber Saiyed, Komal Patel, Labdhi Pandya, Afzal Ansari, Nikha Trivedi, Naresh Chauhan, Summaiya Mullan, Amit gamit, R D Dixit, A M Kadri, Harsh Bakshi, Chaitanya Joshi, Madhvi Joshi |
| EPI_ISL_485635, EPI_ISL_485708, EPI_ISL_485710 | Institut Pasteur Dakar | Institut Pasteur de Dakar | Ndongo Dia, Moussa Moise Diagne, Mamadou diop, Marie Henriette Dior Ndione, Mamadou Malado Jallow, Safietou Sanke, Ousmane Faye, Amadou Alpha Sall. |
| EPI_ISL_485712 | Institut Pasteur | Institut Pasteur de Dakar | Ndongo Dia, Moussa Moise Diagne, Mamadou diop, Marie Henriette Dior Ndione, Mamadou Malado Jallow, Safietou Sanke, Ousmane Faye, Amadou Alpha Sall. |
| EPI_ISL_485713, EPI_ISL_485715, EPI_ISL_485716, EPI_ISL_485717 | Institut Pasteur Dakar | Institut Pasteur de Dakar | Ndongo Dia, Moussa Moise Diagne, Mamadou diop, Marie Henriette Dior Ndione, Mamadou Malado Jallow, Safietou Sanke, Ousmane Faye, Amadou Alpha Sall. |
| EPI_ISL_486834 | Suceava County Energency Hospital "Sf. Ioan cel Nou" | SMU Metagenomics lab | Lobiuc Andrei, Antoniadis Panagiotis |
| EPI_ISL_486842, EPI_ISL_486843, EPI_ISL_486844 | Institute of Microbiology, Universidad San Francisco de Quito | Institute of Microbiology, Universidad San Francisco de Quito | Belén Prado-Vivar, Sully Márquez, Juan José Guadalupe, Monica Becerra-Wong, Carla Torres, Bernardo Gutiérrez, Fausto Maldonado, Geovanny Carzola, Verónica Barragán, Patricio Rojas-Silva, Gabriel Trueba, Michelle Grunauer, Paúl Cárdenas |
| EPI_ISL_486845, EPI_ISL_486846, EPI_ISL_486847, EPI_ISL_486848, EPI_ISL_486849, EPI_ISL_486850, EPI_ISL_486851 | Institute of Microbiology, Universidad San Francisco de Quito | Institute of Microbiology, Universidad San Francisco de Quito | Belén Prado-Vivar, Sully Márquez, Juan José Guadalupe, Monica Becerra-Wong, Carla Torres, Bernardo Gutiérrez, Jonathan Araujo, Verónica Barragán, Patricio Rojas-Silva, Gabriel Trueba, Michelle Grunauer, Paúl Cárdenas |
| EPI_ISL_486852 | CDRI/SGPGI | CSIR-CDRI/SGPGI | Saumya Sarkar, Dharam Veer Singh, Rahul Vishvkarma, Ujjala Ghoshal, Uday Ghoshal, Ravishankar Ramachandran, Tapas Kumar Kundu, Rajender Singh |
| EPI_ISL_486853 | CSIR-CDRI/SGPGI | CSIR-CDRI/SGPGI | Saumya Sarkar, Dharam Veer Singh, Rahul Vishvkarma, Ujjala Ghoshal, Uday Ghoshal, Ravishankar Ramachandran, Tapas Kumar Kundu, Rajender Singh |
| EPI_ISL_486854 | Emergency County Hospital Suceava | Stefan cel Mare, University Metagenomics lab | Lobiuc Andrei et al. |
| EPI_ISL_486876 | Clinical Microbiology Laboratory- Basurto University Hospital | Biocruces-Bizkaia | Mikel J. Urrutikoetxea-Gutierrez, Ana Belén Belén de la Hoz, Matxalen Vidal-García, Mº Carmen Nieto Toboso, Estibaliz Ugalde-Zarraga, José Luis Díaz de Tuesta del Arco |
| EPI_ISL_487277, EPI_ISL_487280, EPI_ISL_487281, EPI_ISL_487288, EPI_ISL_487295, EPI_ISL_487297, EPI_ISL_487304, EPI_ISL_487308, EPI_ISL_487311, EPI_ISL_487312, EPI_ISL_487313, EPI_ISL_487314, EPI_ISL_487316, EPI_ISL_487318, EPI_ISL_487319, EPI_ISL_487320, EPI_ISL_487321, EPI_ISL_487322, EPI_ISL_487324, EPI_ISL_487325, EPI_ISL_487328 | | | |
| see above | NHLS-IALCH | KRISP, KZN Research Innovation and Sequencing Platform | Giandhari J, Pillay S, Lessells R, Chimukangara B, Mdlalose K, York D, Khan S, Tegally H, Wilkinson E, de Oliveira T |
| EPI_ISL_487329, EPI_ISL_487330, EPI_ISL_487332, EPI_ISL_487334, EPI_ISL_487336, EPI_ISL_487337, EPI_ISL_487338, EPI_ISL_487339, EPI_ISL_487340 | Molecular Diagnostics Services (MDS) | KRISP, KZN Research Innovation and Sequencing Platform | Giandhari J, Pillay S, Lessells R, Chimukangara B, Mdlalose K, York D, Khan S, Tegally H, Wilkinson E, de Oliveira T |
| EPI_ISL_487432, EPI_ISL_487433, EPI_ISL_487434, EPI_ISL_487436 | Queen Astrid Military Hospital | Institute of Tropical Medicine | Philippe Selhorst, Colin Anthony |
| EPI_ISL_489833, EPI_ISL_489834, EPI_ISL_489835 | Clinical Microbiology Laboratory- Basurto University Hospital | Biocruces-Bizkaia | Mikel J. Urrutikoetxea-Gutierrez, Ana Belén Belén de la Hoz, Matxalen Vidal-García, Mº Carmen Nieto Toboso, Estibaliz Ugalde-Zarraga, José Luis Díaz de Tuesta del Arco |
| EPI_ISL_490101, EPI_ISL_490103 | Institute for Medical Research, Infectious Disease Research Centre, National Institutes of Health, Ministry of Health Malaysia | Institute for Medical Research, Infectious Disease Research Centre, National Institutes of Health, Ministry of Health Malaysia | Suppiah J, Mohd-Zawawi Z, Kamel K, Kalyanasundram J, Thayan R |
| EPI_ISL_490202, EPI_ISL_490203, EPI_ISL_490204, EPI_ISL_490977 | Clinical Microbiology Laboratory- Basurto University Hospital | Biocruces-Bizkaia | Mikel J. Urrutikoetxea-Gutierrez, Ana Belén Belén de la Hoz, Matxalen Vidal-García, Mº Carmen Nieto Toboso, Estibaliz Ugalde-Zarraga, José Luis Díaz de Tuesta del Arco |
| EPI_ISL_491976, EPI_ISL_491977, EPI_ISL_491978, EPI_ISL_491979, EPI_ISL_491980, EPI_ISL_491981, EPI_ISL_491982, EPI_ISL_491983, EPI_ISL_491984, EPI_ISL_491985, EPI_ISL_491986, EPI_ISL_491988, EPI_ISL_491989, EPI_ISL_491990, EPI_ISL_491991, EPI_ISL_491992, EPI_ISL_491994, EPI_ISL_491995, EPI_ISL_491996, EPI_ISL_491997, EPI_ISL_491998, EPI_ISL_491999, EPI_ISL_492000, EPI_ISL_492001, EPI_ISL_492002, EPI_ISL_492003, EPI_ISL_492005, EPI_ISL_492006, EPI_ISL_492007, EPI_ISL_492008, EPI_ISL_492009, EPI_ISL_492010, EPI_ISL_492011, EPI_ISL_492012, EPI_ISL_492014, EPI_ISL_492016, EPI_ISL_492017, EPI_ISL_492019, EPI_ISL_492020, EPI_ISL_492021, EPI_ISL_492022, EPI_ISL_492023, EPI_ISL_492024, EPI_ISL_492025, EPI_ISL_492026 | | | |
| see above | Oman-NIC | Department of Microbiology and Immunology-SQUH | Fahad Zadjali, Samira Al-Maruqi, Amina Al Jardani, Khulood Al-Mammary, Hanan Al-kindi, Fatma BaAlawi, Hamida AL Barwani, Zeyana AL-Dahmani, Intisar Al-Shukri, Aisha Al-Busaidi, Aisha Al-Amri, Ahlam Al-Amri, Mohammed Al-Tobi, Samiha Al Kharusi, Abdulla Balkhair |
| EPI_ISL_492035 | Instituto de Biologia do Exército | Laboratório Metabolismo Macromolecular FirminoTorres de Castro, Instituto de Biofísica Carlos Chagas Filho, Universidade Federal do Rio de Janeiro | Bianca Catarina Azevedo Cabral, Aline Rosa Vianna de Souza, Tatiana LS Nogueira, Nádia Vaez Gonçalves da Cruz, Caleb GM Santos, Marcos Dornelas-Ribeiro, Elizabeth Valentin, Marcio da Costa Cipitelli, Virginia Sara Grancieri do Amaral, Rodrigo Soares de Moura Neto, Clarissa Damaso, Rosane Silva |
| EPI_ISL_492036 | Instituto de Biologia do Exército | Laboratório Metabolismo Macromolecular FirminoTorres de Castro, Instituto de Biofísica Carlos Chagas Filho, Universidade Federal do Rio de Janeiro | Bianca Catarina Azevedo Cabral, Aline Rosa Vianna de Souza , Marcos Dornelas-Ribeiro, Tatiana LS Nogueira, Nádia Vaez Gonçalves da Cruz, Caleb GM Santos, Elizabeth Valentin, Marcio da Costa Cipitelli, Virginia Sara Grancieri do Amaral, Rodrigo Soares de Moura Neto, Clarissa Damaso, Rosane Silva |
| EPI_ISL_492043 | Instituto de Biologia do Exército | Laboratório Metabolismo Macromolecular FirminoTorres de Castro, Instituto de Biofísica Carlos Chagas Filho, Universidade Federal do Rio de Janeiro | Bianca Catarina Azevedo Cabral, Aline Rosa Vianna de Souza, Tatiana LS Nogueira, Nádia Vaez Gonçalves da Cruz, Caleb GM Santos, Marcos Dornelas-Ribeiro, Elizabeth Valentin, Marcio da Costa Cipitelli, Virginia Sara Grancieri do Amaral, Rodrigo Soares de Moura Neto, Clarissa Damaso, Rosane Silva |
| EPI_ISL_492044 | Instituto de Biologia do Exército | Laboratório Metabolismo Macromolecular FirminoTorres de Castro, Instituto de Biofísica Carlos Chagas Filho, Universidade Federal do Rio de Janeiro | Bianca Catarina Azevedo Cabral, Aline Rosa Vianna de Souza , Marcos Dornelas-Ribeiro, Tatiana LS Nogueira, Nádia Vaez Gonçalves da Cruz, Caleb GM Santos, Elizabeth Valentin, Marcio da Costa Cipitelli, Virginia Sara Grancieri do Amaral, Rodrigo Soares de Moura Neto, Clarissa Damaso, Rosane Silva |
| EPI_ISL_492045 | Instituto de Biologia do Exército | Laboratório Metabolismo Macromolecular FirminoTorres de Castro, Instituto de Biofísica Carlos Chagas Filho, Universidade Federal do Rio de Janeiro | Bianca Catarina Azevedo Cabral, Aline Rosa Vianna de Souza, Caleb GM Santos, Marcos Dornelas-Ribeiro, Tatiana LS Nogueira, Nádia Vaez Gonçalves da Cruz, Elizabeth Valentin, Marcio da Costa Cipitelli, Virginia Sara Grancieri do Amaral, Rodrigo Soares de Moura Neto, Clarissa Damaso, Rosane Silva |

| | | | |
|---|---|---|---|
| EPI_ISL_492048 | Instituto de Biologia do Exército | Laboratório Metabolismo Macromolecular FirminoTorres de Castro, Instituto de Biofísica Carlos Chagas Filho, Universidade Federal do Rio de Janeiro | Bianca Catarina Azevedo Cabral, Aline Rosa Vianna de Souza , Marcos Dornelas-Ribeiro, Tatiana LS Nogueira, Nádia Vaez Gonçalves da Cruz, Caleb GM Santos, Elizabeth Valentin, Marcio da Costa Cipitelli, Virginia Sara Grancieri do Amaral, Rodrigo Soares de Moura Neto, Clarissa Damaso, Rosane Silva |
| EPI_ISL_493346, EPI_ISL_493348 | Instituto de Diagnostico y Referencia Epidemiologicos (INDRE) | Instituto de Diagnostico y Referencia Epidemiologicos (INDRE) | Ernesto Ramirez-Gonzalez, Abril Rodriguez-Maldonado, Claudia Wong-Arambula , Natividad Cruz-Ortiz, Tatiana Nunez-Garcia, Dayanira Arellano-Suarez, Adnan Araiza-Rodriguez, Fabiola Garces-Ayala, Lucia Hernandez-Rivas, Irma Lopez-Martinez, Gisela Barrera-Badillo. |
| EPI_ISL_495014 | B.J. Medical College and Civil hospital | Gujarat Biotechnology Research Centre | Janvi Raval, Zarna Patel, Monika Gandhi, Pinal Trivedi, Maharshi Pandya, Nidhi Patel, Nitin Savaliya, Raghawendra Kumar, Dinesh Kumar, Zuber Saiyed, Komal Patel, Labdhi Pandya, Afzal Ansari, Nikha Trivedi, Pranay Shah, Kamlesh J Upadhyay, Sanjay Kapadia, Apurvasinh Puvar, R D Dixit, A M Kadri, Harsh Bakshi, Chaitanya Joshi, Madhvi Joshi |
| EPI_ISL_495015 | B.J. Medical College and Civil hospital | Gujarat Biotechnology Research Centre | Zarna Patel, Monika Gandhi, Pinal Trivedi, Maharshi Pandya, Nidhi Patel, Nitin Savaliya, Raghawendra Kumar, Dinesh Kumar, Zuber Saiyed, Komal Patel, Labdhi Pandya, Afzal Ansari, Nikha Trivedi, Pranay Shah, Kamlesh J Upadhyay, Sanjay Kapadia, Apurvasinh Puvar, Janvi Raval, R D Dixit, A M Kadri, Harsh Bakshi, Chaitanya Joshi, Madhvi Joshi |
| EPI_ISL_495016 | B.J. Medical College and Civil hospital | Gujarat Biotechnology Research Centre | Monika Gandhi, Pinal Trivedi, Maharshi Pandya, Nidhi Patel, Nitin Savaliya, Raghawendra Kumar, Dinesh Kumar, Zuber Saiyed, Komal Patel, Labdhi Pandya, Afzal Ansari, Nikha Trivedi, Pranay Shah, Kamlesh J Upadhyay, Sanjay Kapadia, Apurvasinh Puvar, Janvi Raval, Zarna Patel, R D Dixit, A M Kadri, Harsh Bakshi, Chaitanya Joshi, Madhvi Joshi |
| EPI_ISL_495017 | B.J. Medical College and Civil hospital | Gujarat Biotechnology Research Centre | Pinal Trivedi, Maharshi Pandya, Nidhi Patel, Nitin Savaliya, Raghawendra Kumar, Dinesh Kumar, Zuber Saiyed, Komal Patel, Labdhi Pandya, Afzal Ansari, Nikha Trivedi, Pranay Shah, Kamlesh J Upadhyay, Sanjay Kapadia, Apurvasinh Puvar, Janvi Raval, Zarna Patel, Monika Gandhi, R D Dixit, A M Kadri, Harsh Bakshi, Chaitanya Joshi, Madhvi Joshi |
| EPI_ISL_495018 | B.J. Medical College and Civil hospital | Gujarat Biotechnology Research Centre | Maharshi Pandya, Nidhi Patel, Nitin Savaliya, Raghawendra Kumar, Dinesh Kumar, Zuber Saiyed, Komal Patel, Labdhi Pandya, Afzal Ansari, Nikha Trivedi, Pranay Shah, Kamlesh J Upadhyay, Sanjay Kapadia, Apurvasinh Puvar, Janvi Raval, Zarna Patel, Monika Gandhi, Pinal Trivedi, R D Dixit, A M Kadri, Harsh Bakshi, Chaitanya Joshi, Madhvi Joshi |
| EPI_ISL_495019 | B.J. Medical College and Civil hospital | Gujarat Biotechnology Research Centre | Nidhi Patel, Nitin Savaliya, Raghawendra Kumar, Dinesh Kumar, Zuber Saiyed, Komal Patel, Labdhi Pandya, Afzal Ansari, Nikha Trivedi, Pranay Shah, Kamlesh J Upadhyay, Sanjay Kapadia, Apurvasinh Puvar, Janvi Raval, Zarna Patel, Monika Gandhi, Pinal Trivedi, Maharshi Pandya, R D Dixit, A M Kadri, Harsh Bakshi, Chaitanya Joshi, Madhvi Joshi |
| EPI_ISL_495020 | Government Medical College, Bhavnagar | Gujarat Biotechnology Research Centre | Kairavi Desai, Saklin Malek, Shirish Patel, Nitin Savaliya, Raghawendra Kumar, Dinesh Kumar, Zuber Saiyed, Komal Patel, Labdhi Pandya, Afzal Ansari, Nikha Trivedi, Apurvasinh Puvar, Janvi Raval, Zarna Patel, Monika Gandhi, Pinal Trivedi, Maharshi Pandya, Nidhi Patel, R D Dixit, A M Kadri, Harsh Bakshi, Chaitanya Joshi, Madhvi Joshi |
| EPI_ISL_495021 | Government Medical College, Bhavnagar | Gujarat Biotechnology Research Centre | Saklin Malek, Shirish Patel, Kairavi Desai, Raghawendra Kumar, Dinesh Kumar, Zuber Saiyed, Komal Patel, Labdhi Pandya, Afzal Ansari, Nikha Trivedi, Apurvasinh Puvar, Janvi Raval, Zarna Patel, Monika Gandhi, Pinal Trivedi, Maharshi Pandya, Nidhi Patel, Nitin Savaliya, R D Dixit, A M Kadri, Harsh Bakshi, Chaitanya Joshi, Madhvi Joshi |
| EPI_ISL_495022 | Government Medical College, Bhavnagar | Gujarat Biotechnology Research Centre | Shirish Patel, Kairavi Desai, Saklin Malek, Dinesh Kumar, Zuber Saiyed, Komal Patel, Labdhi Pandya, Afzal Ansari, Nikha Trivedi, Apurvasinh Puvar, Janvi Raval, Zarna Patel, Monika Gandhi, Pinal Trivedi, Maharshi Pandya, Nidhi Patel, Nitin Savaliya, Raghawendra Kumar, R D Dixit, A M Kadri, Harsh Bakshi, Chaitanya Joshi, Madhvi Joshi |
| EPI_ISL_495023 | Government Medical College, Bhavnagar | Gujarat Biotechnology Research Centre | Nitin Savaliya, Raghawendra Kumar, Dinesh Kumar, Zuber Saiyed, Komal Patel, Labdhi Pandya, Afzal Ansari, Nikha Trivedi, Kairavi Desai, Saklin Malek, Shirish Patel, Apurvasinh Puvar, Janvi Raval, Zarna Patel, Monika Gandhi, Pinal Trivedi, Maharshi Pandya, Nidhi Patel, R D Dixit, A M Kadri, Harsh Bakshi, Chaitanya Joshi, Madhvi Joshi |
| EPI_ISL_495024 | Government Medical College, Bhavnagar | Gujarat Biotechnology Research Centre | Raghawendra Kumar, Dinesh Kumar, Zuber Saiyed, Komal Patel, Labdhi Pandya, Afzal Ansari, Nikha Trivedi, Kairavi Desai, Saklin Malek, Shirish Patel, Apurvasinh Puvar, Janvi Raval, Zarna Patel, Monika Gandhi, Pinal Trivedi, Maharshi Pandya, Nidhi Patel, Nitin Savaliya, R D Dixit, A M Kadri, Harsh Bakshi, Chaitanya Joshi, Madhvi Joshi |
| EPI_ISL_495025 | Government Medical College, Bhavnagar | Gujarat Biotechnology Research Centre | Dinesh Kumar, Zuber Saiyed, Komal Patel, Labdhi Pandya, Afzal Ansari, Nikha Trivedi, Kairavi Desai, Saklin Malek, Shirish Patel, Apurvasinh Puvar, Janvi Raval, Zarna Patel, Monika Gandhi, Pinal Trivedi, Maharshi Pandya, Nidhi Patel, Nitin Savaliya, Raghawendra Kumar, R D Dixit, A M Kadri, Harsh Bakshi, Chaitanya Joshi, Madhvi Joshi |
| EPI_ISL_495026 | GMERS Medical College & Hospital, Gotri, Vadodara | Gujarat Biotechnology Research Centre | Komal Patel, Labdhi Pandya, Afzal Ansari, Nikha Trivedi, Meenakshi Shah, Neena Doshi, Varsha Godbole, Apurvasinh Puvar, Janvi Raval, Zarna Patel, Monika Gandhi, Pinal Trivedi, Maharshi Pandya, Nidhi Patel, Nitin Savaliya, Raghawendra Kumar, Dinesh Kumar, Zuber Saiyed, R D Dixit, A M Kadri, Harsh Bakshi, Chaitanya Joshi, Madhvi Joshi |
| EPI_ISL_495027 | GMERS Medical College & Hospital, Gotri, Vadodara | Gujarat Biotechnology Research Centre | Labdhi Pandya, Afzal Ansari, Nikha Trivedi, Meenakshi Shah, Neena Doshi, Varsha Godbole, Apurvasinh Puvar, Janvi Raval, Zarna Patel, Monika Gandhi, Pinal Trivedi, Maharshi Pandya, Nidhi Patel, Nitin Savaliya, Raghawendra Kumar, Dinesh Kumar, Zuber Saiyed, Komal Patel, R D Dixit, A M Kadri, Harsh Bakshi, Chaitanya Joshi, Madhvi Joshi |
| EPI_ISL_495028 | GMERS Medical College & Hospital, Gotri, Vadodara | Gujarat Biotechnology Research Centre | Afzal Ansari, Nikha Trivedi, Meenakshi Shah, Neena Doshi, Varsha Godbole, Apurvasinh Puvar, Janvi Raval, Zarna Patel, Monika Gandhi, Pinal Trivedi, Maharshi Pandya, Nidhi Patel, Nitin Savaliya, Raghawendra Kumar, Dinesh Kumar, Zuber Saiyed, Komal Patel, Labdhi Pandya, R D Dixit, A M Kadri, Harsh Bakshi, Chaitanya Joshi, Madhvi Joshi |
| EPI_ISL_495029 | GMERS Medical College & Hospital, Gotri, Vadodara | Gujarat Biotechnology Research Centre | Nikha Trivedi, Meenakshi Shah, Neena Doshi, Varsha Godbole, Apurvasinh Puvar, Janvi Raval, Zarna Patel, Monika Gandhi, Pinal Trivedi, Maharshi Pandya, Nidhi Patel, Nitin Savaliya, Raghawendra Kumar, Dinesh Kumar, Zuber Saiyed, Komal Patel, Labdhi Pandya, Afzal Ansari, R D Dixit, A M Kadri, Harsh Bakshi, Chaitanya Joshi, Madhvi Joshi |
| EPI_ISL_495030 | GMERS Medical College & Hospital, Gotri, Vadodara | Gujarat Biotechnology Research Centre | Meenakshi Shah, Neena Doshi, Varsha Godbole, Apurvasinh Puvar, Janvi Raval, Zarna Patel, Monika Gandhi, Pinal Trivedi, Maharshi Pandya, Nidhi Patel, Nitin Savaliya, Raghawendra Kumar, Dinesh Kumar, Zuber Saiyed, Komal Patel, Labdhi Pandya, Afzal Ansari, Nikha Trivedi, R D Dixit, A M Kadri, Harsh Bakshi, Chaitanya Joshi, Madhvi Joshi |
| EPI_ISL_495031 | GMERS Medical College & Hospital, Gotri, Vadodara | Gujarat Biotechnology Research Centre | Neena Doshi, Varsha Godbole, Apurvasinh Puvar, Janvi Raval, Zarna Patel, Monika Gandhi, Pinal Trivedi, Maharshi Pandya, Nidhi Patel, Nitin Savaliya, Raghawendra Kumar, Dinesh Kumar, Zuber Saiyed, Komal Patel, Labdhi Pandya, Afzal Ansari, Nikha Trivedi, Meenakshi Shah, R D Dixit, A M Kadri, Harsh Bakshi, Chaitanya Joshi, Madhvi Joshi |
| EPI_ISL_495032 | GMERS Medical College & Hospital, Gotri, Vadodara | Gujarat Biotechnology Research Centre | Varsha Godbole, Apurvasinh Puvar, Janvi Raval, Zarna Patel, Monika Gandhi, Pinal Trivedi, Maharshi Pandya, Nidhi Patel, Nitin Savaliya, Raghawendra Kumar, Dinesh Kumar, Zuber Saiyed, Komal Patel, Labdhi Pandya, Afzal Ansari, Nikha Trivedi, Meenakshi Shah, Neena Doshi, R D Dixit, A M Kadri, Harsh Bakshi, Chaitanya Joshi, Madhvi Joshi |
| EPI_ISL_495033 | GMERS Medical College & Hospital, Gotri, Vadodara | Gujarat Biotechnology Research Centre | Apurvasinh Puvar, Janvi Raval, Zarna Patel, Monika Gandhi, Pinal Trivedi, Maharshi Pandya, Nidhi Patel, Nitin Savaliya, Raghawendra Kumar, Dinesh Kumar, Zuber Saiyed, Komal Patel, Labdhi Pandya, Afzal Ansari, Nikha Trivedi, Meenakshi Shah, Neena Doshi, Varsha Godbole, R D Dixit, A M Kadri, Harsh Bakshi, Chaitanya Joshi, Madhvi Joshi |
| EPI_ISL_495034 | GMERS Medical College & Hospital, Gotri, Vadodara | Gujarat Biotechnology Research Centre | Janvi Raval, Zarna Patel, Monika Gandhi, Pinal Trivedi, Maharshi Pandya, Nidhi Patel, Nitin Savaliya, Raghawendra Kumar, Dinesh Kumar, Zuber Saiyed, Komal Patel, Labdhi Pandya, Afzal Ansari, Nikha Trivedi, Meenakshi Shah, Neena Doshi, Varsha Godbole, Apurvasinh Puvar, R D Dixit, A M Kadri, Harsh Bakshi, Chaitanya Joshi, Madhvi Joshi |
| EPI_ISL_495035 | GMERS Medical College & Hospital, Gotri, Vadodara | Gujarat Biotechnology Research Centre | Zarna Patel, Monika Gandhi, Pinal Trivedi, Maharshi Pandya, Nidhi Patel, Nitin Savaliya, Raghawendra Kumar, Dinesh Kumar, Zuber Saiyed, Komal Patel, Labdhi Pandya, Afzal Ansari, Nikha Trivedi, Meenakshi Shah, Neena Doshi, Varsha Godbole, Apurvasinh Puvar, Janvi Raval, R D Dixit, A M Kadri, Harsh Bakshi, Chaitanya Joshi, Madhvi Joshi |
| EPI_ISL_495038 | GMERS Medical College & Hospital, Gotri, Vadodara | Gujarat Biotechnology Research Centre | Maharshi Pandya, Nidhi Patel, Nitin Savaliya, Raghawendra Kumar, Dinesh Kumar, Zuber Saiyed, Komal Patel, Labdhi Pandya, Afzal Ansari, Nikha Trivedi, Meenakshi Shah, Neena Doshi, Varsha Godbole, Apurvasinh Puvar, Janvi Raval, Zarna Patel, Monika Gandhi, Pinal Trivedi, R D Dixit, A M Kadri, Harsh Bakshi, Chaitanya Joshi, Madhvi Joshi |
| EPI_ISL_495039 | GMERS Medical College & Hospital, Gotri, Vadodara | Gujarat Biotechnology Research Centre | Nidhi Patel, Nitin Savaliya, Raghawendra Kumar, Dinesh Kumar, Zuber Saiyed, Komal Patel, Labdhi Pandya, Afzal Ansari, Nikha Trivedi, Meenakshi Shah, |

| | | | |
|---|---|---|---|
| | | | Neena Doshi, Varsha Godbole, Apurvasinh Puvar, Janvi Raval, Zarna Patel, Monika Gandhi, Pinal Trivedi, Maharshi Pandya, R D Dixit, A M Kadri, Harsh Bakshi, Chaitanya Joshi, Madhvi Joshi |
| EPI_ISL_495042 | GMERS Medical College & Hospital, Gotri, Vadodara | Gujarat Biotechnology Research Centre | Dinesh Kumar, Zuber Saiyed, Komal Patel, Labdhi Pandya, Afzal Ansari, Nikha Trivedi, Meenakshi Shah, Neena Doshi, Varsha Godbole, Apurvasinh Puvar, Janvi Raval, Zarna Patel, Monika Gandhi, Pinal Trivedi, Maharshi Pandya, Nidhi Patel, Nitin Savaliya, Raghawendra Kumar, R D Dixit, A M Kadri, Harsh Bakshi, Chaitanya Joshi, Madhvi Joshi |
| EPI_ISL_495043 | GMERS Medical College & Hospital, Gotri, Vadodara | Gujarat Biotechnology Research Centre | Zuber Saiyed, Komal Patel, Labdhi Pandya, Afzal Ansari, Nikha Trivedi, Meenakshi Shah, Neena Doshi, Varsha Godbole, Apurvasinh Puvar, Janvi Raval, Zarna Patel, Monika Gandhi, Pinal Trivedi, Maharshi Pandya, Nidhi Patel, Nitin Savaliya, Raghawendra Kumar, Dinesh Kumar, R D Dixit, A M Kadri, Harsh Bakshi, Chaitanya Joshi, Madhvi Joshi |
| EPI_ISL_495044 | GMERS Medical College & Hospital, Gotri, Vadodara | Gujarat Biotechnology Research Centre | Komal Patel, Labdhi Pandya, Afzal Ansari, Nikha Trivedi, Meenakshi Shah, Neena Doshi, Varsha Godbole, Apurvasinh Puvar, Janvi Raval, Zarna Patel, Monika Gandhi, Pinal Trivedi, Maharshi Pandya, Nidhi Patel, Nitin Savaliya, Raghawendra Kumar, Dinesh Kumar, Zuber Saiyed, R D Dixit, A M Kadri, Harsh Bakshi, Chaitanya Joshi, Madhvi Joshi |
| EPI_ISL_495045 | GMERS Medical College & Hospital, Gotri, Vadodara | Gujarat Biotechnology Research Centre | Labdhi Pandya, Afzal Ansari, Nikha Trivedi, Meenakshi Shah, Neena Doshi, Varsha Godbole, Apurvasinh Puvar, Janvi Raval, Zarna Patel, Monika Gandhi, Pinal Trivedi, Maharshi Pandya, Nidhi Patel, Nitin Savaliya, Raghawendra Kumar, Dinesh Kumar, Zuber Saiyed, Komal Patel, R D Dixit, A M Kadri, Harsh Bakshi, Chaitanya Joshi, Madhvi Joshi |
| EPI_ISL_495046 | GMERS Medical College & Hospital, Gotri, Vadodara | Gujarat Biotechnology Research Centre | Afzal Ansari, Nikha Trivedi, Meenakshi Shah, Neena Doshi, Varsha Godbole, Apurvasinh Puvar, Janvi Raval, Zarna Patel, Monika Gandhi, Pinal Trivedi, Maharshi Pandya, Nidhi Patel, Nitin Savaliya, Raghawendra Kumar, Dinesh Kumar, Zuber Saiyed, Komal Patel, Labdhi Pandya, R D Dixit, A M Kadri, Harsh Bakshi, Chaitanya Joshi, Madhvi Joshi |
| EPI_ISL_495047 | GMERS Medical College & Hospital, Gotri, Vadodara | Gujarat Biotechnology Research Centre | Nikha Trivedi, Meenakshi Shah, Neena Doshi, Varsha Godbole, Apurvasinh Puvar, Janvi Raval, Zarna Patel, Monika Gandhi, Pinal Trivedi, Maharshi Pandya, Nidhi Patel, Nitin Savaliya, Raghawendra Kumar, Dinesh Kumar, Zuber Saiyed, Komal Patel, Labdhi Pandya, Afzal Ansari, R D Dixit, A M Kadri, Harsh Bakshi, Chaitanya Joshi, Madhvi Joshi |
| EPI_ISL_495048 | GMERS Medical College & Hospital, Gotri, Vadodara | Gujarat Biotechnology Research Centre | Meenakshi Shah, Neena Doshi, Varsha Godbole, Apurvasinh Puvar, Janvi Raval, Zarna Patel, Monika Gandhi, Pinal Trivedi, Maharshi Pandya, Nidhi Patel, Nitin Savaliya, Raghawendra Kumar, Dinesh Kumar, Zuber Saiyed, Komal Patel, Labdhi Pandya, Afzal Ansari, Nikha Trivedi, R D Dixit, A M Kadri, Harsh Bakshi, Chaitanya Joshi, Madhvi Joshi |
| EPI_ISL_495049 | GMERS Medical College & Hospital, Gotri, Vadodara | Gujarat Biotechnology Research Centre | Neena Doshi, Varsha Godbole, Apurvasinh Puvar, Janvi Raval, Zarna Patel, Monika Gandhi, Pinal Trivedi, Maharshi Pandya, Nidhi Patel, Nitin Savaliya, Raghawendra Kumar, Dinesh Kumar, Zuber Saiyed, Komal Patel, Labdhi Pandya, Afzal Ansari, Nikha Trivedi, Meenakshi Shah, R D Dixit, A M Kadri, Harsh Bakshi, Chaitanya Joshi, Madhvi Joshi |
| EPI_ISL_495050 | GMERS Medical College & Hospital, Gotri, Vadodara | Gujarat Biotechnology Research Centre | Varsha Godbole, Apurvasinh Puvar, Janvi Raval, Zarna Patel, Monika Gandhi, Pinal Trivedi, Maharshi Pandya, Nidhi Patel, Nitin Savaliya, Raghawendra Kumar, Dinesh Kumar, Zuber Saiyed, Komal Patel, Labdhi Pandya, Afzal Ansari, Nikha Trivedi, Meenakshi Shah, Neena Doshi, R D Dixit, A M Kadri, Harsh Bakshi, Chaitanya Joshi, Madhvi Joshi |
| EPI_ISL_495051 | GMERS Medical College & Hospital, Gotri, Vadodara | Gujarat Biotechnology Research Centre | Apurvasinh Puvar, Janvi Raval, Zarna Patel, Monika Gandhi, Pinal Trivedi, Maharshi Pandya, Nidhi Patel, Nitin Savaliya, Raghawendra Kumar, Dinesh Kumar, Zuber Saiyed, Komal Patel, Labdhi Pandya, Afzal Ansari, Nikha Trivedi, Meenakshi Shah, Neena Doshi, Varsha Godbole, R D Dixit, A M Kadri, Harsh Bakshi, Chaitanya Joshi, Madhvi Joshi |
| EPI_ISL_495052 | GMERS Medical College & Hospital, Gotri, Vadodara | Gujarat Biotechnology Research Centre | Janvi Raval, Zarna Patel, Monika Gandhi, Pinal Trivedi, Maharshi Pandya, Nidhi Patel, Nitin Savaliya, Raghawendra Kumar, Dinesh Kumar, Zuber Saiyed, Komal Patel, Labdhi Pandya, Afzal Ansari, Nikha Trivedi, Meenakshi Shah, Neena Doshi, Varsha Godbole, Apurvasinh Puvar, R D Dixit, A M Kadri, Harsh Bakshi, Chaitanya Joshi, Madhvi Joshi |
| EPI_ISL_495053 | GMERS Medical College & Hospital, Gotri, Vadodara | Gujarat Biotechnology Research Centre | Zarna Patel, Monika Gandhi, Pinal Trivedi, Maharshi Pandya, Nidhi Patel, Nitin Savaliya, Raghawendra Kumar, Dinesh Kumar, Zuber Saiyed, Komal Patel, Labdhi Pandya, Afzal Ansari, Nikha Trivedi, Meenakshi Shah, Neena Doshi, Varsha Godbole, Apurvasinh Puvar, Janvi Raval, R D Dixit, A M Kadri, Harsh Bakshi, Chaitanya Joshi, Madhvi Joshi |
| EPI_ISL_495054 | GMERS Medical College & Hospital, Gotri, Vadodara | Gujarat Biotechnology Research Centre | Monika Gandhi, Pinal Trivedi, Maharshi Pandya, Nidhi Patel, Nitin Savaliya, Raghawendra Kumar, Dinesh Kumar, Zuber Saiyed, Komal Patel, Labdhi Pandya, Afzal Ansari, Nikha Trivedi, Meenakshi Shah, Neena Doshi, Varsha Godbole, Apurvasinh Puvar, Janvi Raval, Zarna Patel, R D Dixit, A M Kadri, Harsh Bakshi, Chaitanya Joshi, Madhvi Joshi |
| EPI_ISL_495055 | GMERS Medical College & Hospital, Gotri, Vadodara | Gujarat Biotechnology Research Centre | Pinal Trivedi, Maharshi Pandya, Nidhi Patel, Nitin Savaliya, Raghawendra Kumar, Dinesh Kumar, Zuber Saiyed, Komal Patel, Labdhi Pandya, Afzal Ansari, Nikha Trivedi, Meenakshi Shah, Neena Doshi, Varsha Godbole, Apurvasinh Puvar, Janvi Raval, Zarna Patel, Monika Gandhi, R D Dixit, A M Kadri, Harsh Bakshi, Chaitanya Joshi, Madhvi Joshi |
| EPI_ISL_495056 | GMERS Medical College & Hospital, Gotri, Vadodara | Gujarat Biotechnology Research Centre | Maharshi Pandya, Nidhi Patel, Nitin Savaliya, Raghawendra Kumar, Dinesh Kumar, Zuber Saiyed, Komal Patel, Labdhi Pandya, Afzal Ansari, Nikha Trivedi, Meenakshi Shah, Neena Doshi, Varsha Godbole, Apurvasinh Puvar, Janvi Raval, Zarna Patel, Monika Gandhi, Pinal Trivedi, R D Dixit, A M Kadri, Harsh Bakshi, Chaitanya Joshi, Madhvi Joshi |
| EPI_ISL_495057 | GMERS Medical College & Hospital, Gotri, Vadodara | Gujarat Biotechnology Research Centre | Nidhi Patel, Nitin Savaliya, Raghawendra Kumar, Dinesh Kumar, Zuber Saiyed, Komal Patel, Labdhi Pandya, Afzal Ansari, Nikha Trivedi, Meenakshi Shah, Neena Doshi, Varsha Godbole, Apurvasinh Puvar, Janvi Raval, Zarna Patel, Monika Gandhi, Pinal Trivedi, Maharshi Pandya, R D Dixit, A M Kadri, Harsh Bakshi, Chaitanya Joshi, Madhvi Joshi |
| EPI_ISL_495058 | GMERS Medical College & Hospital, Gotri, Vadodara | Gujarat Biotechnology Research Centre | Nitin Savaliya, Raghawendra Kumar, Dinesh Kumar, Zuber Saiyed, Komal Patel, Labdhi Pandya, Afzal Ansari, Nikha Trivedi, Meenakshi Shah, Neena Doshi, Varsha Godbole, Apurvasinh Puvar, Janvi Raval, Zarna Patel, Monika Gandhi, Pinal Trivedi, Maharshi Pandya, Nidhi Patel, R D Dixit, A M Kadri, Harsh Bakshi, Chaitanya Joshi, Madhvi Joshi |
| EPI_ISL_495059 | GMERS Medical College & Hospital, Gotri, Vadodara | Gujarat Biotechnology Research Centre | Raghawendra Kumar, Dinesh Kumar, Zuber Saiyed, Komal Patel, Labdhi Pandya, Afzal Ansari, Nikha Trivedi, Meenakshi Shah, Neena Doshi, Varsha Godbole, Apurvasinh Puvar, Janvi Raval, Zarna Patel, Monika Gandhi, Pinal Trivedi, Maharshi Pandya, Nidhi Patel, Nitin Savaliya, R D Dixit, A M Kadri, Harsh Bakshi, Chaitanya Joshi, Madhvi Joshi |
| EPI_ISL_495060 | GMERS Medical College & Hospital, Gotri, Vadodara | Gujarat Biotechnology Research Centre | Dinesh Kumar, Zuber Saiyed, Komal Patel, Labdhi Pandya, Afzal Ansari, Nikha Trivedi, Meenakshi Shah, Neena Doshi, Varsha Godbole, Apurvasinh Puvar, Janvi Raval, Zarna Patel, Monika Gandhi, Pinal Trivedi, Maharshi Pandya, Nidhi Patel, Nitin Savaliya, Raghawendra Kumar, R D Dixit, A M Kadri, Harsh Bakshi, Chaitanya Joshi, Madhvi Joshi |
| EPI_ISL_495061 | GMERS Medical College & Hospital, Gotri, Vadodara | Gujarat Biotechnology Research Centre | Zuber Saiyed, Komal Patel, Labdhi Pandya, Afzal Ansari, Nikha Trivedi, Meenakshi Shah, Neena Doshi, Varsha Godbole, Apurvasinh Puvar, Janvi Raval, Zarna Patel, Monika Gandhi, Pinal Trivedi, Maharshi Pandya, Nidhi Patel, Nitin Savaliya, Raghawendra Kumar, Dinesh Kumar, R D Dixit, A M Kadri, Harsh Bakshi, Chaitanya Joshi, Madhvi Joshi |
| EPI_ISL_495062 | GMERS Medical College & Hospital, Gotri, Vadodara | Gujarat Biotechnology Research Centre | Komal Patel, Labdhi Pandya, Afzal Ansari, Nikha Trivedi, Meenakshi Shah, Neena Doshi, Varsha Godbole, Apurvasinh Puvar, Janvi Raval, Zarna Patel, Monika Gandhi, Pinal Trivedi, Maharshi Pandya, Nidhi Patel, Nitin Savaliya, Raghawendra Kumar, Dinesh Kumar, Zuber Saiyed, R D Dixit, A M Kadri, Harsh Bakshi, Chaitanya Joshi, Madhvi Joshi |
| EPI_ISL_495063 | GMERS Medical College & Hospital, Himmatnagar | Gujarat Biotechnology Research Centre | Labdhi Pandya, Afzal Ansari, Nikha Trivedi, Himanshu Khatri, Mayur Gandhi, Apurvasinh Puvar, Janvi Raval, Zarna Patel, Monika Gandhi, Pinal Trivedi, Maharshi Pandya, Nidhi Patel, Nitin Savaliya, Raghawendra Kumar, Dinesh Kumar, Zuber Saiyed, Komal Patel, R D Dixit, A M Kadri, Harsh Bakshi, Chaitanya Joshi, Madhvi Joshi, |
| EPI_ISL_495064 | GMERS Medical College & Hospital, Himmatnagar | Gujarat Biotechnology Research Centre | Afzal Ansari, Nikha Trivedi, Himanshu Khatri, Mayur Gandhi, Apurvasinh Puvar, Janvi Raval, Zarna Patel, Monika Gandhi, Pinal Trivedi, Maharshi Pandya, Nidhi Patel, Nitin Savaliya, Raghawendra Kumar, Dinesh Kumar, Zuber Saiyed, Komal Patel, Labdhi Pandya, R D Dixit, A M Kadri, Harsh Bakshi, Chaitanya Joshi, Madhvi Joshi, |
| EPI_ISL_495065 | GAIMS & G K General Hospital | Gujarat Biotechnology Research Centre | Nikha Trivedi, Babulal Babhoria, Hitesh Assudani, Apurvasinh Puvar, Janvi Raval, Zarna Patel, Monika Gandhi, Pinal Trivedi, Maharshi Pandya, Nidhi Patel, Nitin Savaliya, Raghawendra Kumar, Dinesh Kumar, Zuber Saiyed, Komal Patel, Labdhi Pandya, Afzal Ansari, R D Dixit, A M Kadri, Harsh Bakshi, Chaitanya Joshi, Madhvi Joshi, |
| EPI_ISL_495066 | GAIMS & G K General Hospital | Gujarat Biotechnology Research Centre | Babulal Babhoria, Hitesh Assudani, Apurvasinh Puvar, Janvi Raval, Zarna Patel, Monika Gandhi, Pinal Trivedi, Maharshi Pandya, Nidhi Patel, Nitin Savaliya, |

| | | | |
|---|---|---|---|
| | | | Raghawendra Kumar, Dinesh Kumar, Zuber Saiyed, Komal Patel, Labdhi Pandya, Afzal Ansari, Nikha Trivedi, R D Dixit, A M Kadri, Harsh Bakshi, Chaitanya Joshi, Madhvi Joshi, |
| EPI_ISL_495067 | Dr. N. D. Desai Medical College & Hospital | Gujarat Biotechnology Research Centre | J G Buch, Jigar Gusani, Supreet Prabhu, Apurvasinh Puvar, Janvi Raval, Zarna Patel, Monika Gandhi, Pinal Trivedi, Maharshi Pandya, Nidhi Patel, Nitin Savaliya, Raghawendra Kumar, Dinesh Kumar, Zuber Saiyed, Komal Patel, Labdhi Pandya, Afzal Ansari, Nikha Trivedi, R D Dixit, A M Kadri, Harsh Bakshi, Chaitanya Joshi, Madhvi Joshi |
| EPI_ISL_495068 | Dr. N. D. Desai Medical College & Hospital | Gujarat Biotechnology Research Centre | Jigar Gusani, Supreet Prabhu, Apurvasinh Puvar, Janvi Raval, Zarna Patel, Monika Gandhi, Pinal Trivedi, Maharshi Pandya, Nidhi Patel, Nitin Savaliya, Raghawendra Kumar, Dinesh Kumar, Zuber Saiyed, Komal Patel, Labdhi Pandya, Afzal Ansari, Nikha Trivedi, J G Buch, R D Dixit, A M Kadri, Harsh Bakshi, Chaitanya Joshi, Madhvi Joshi |
| EPI_ISL_495069 | Dr. N. D. Desai Medical College & Hospital | Gujarat Biotechnology Research Centre | Supreet Prabhu, Apurvasinh Puvar, Janvi Raval, Zarna Patel, Monika Gandhi, Pinal Trivedi, Maharshi Pandya, Nidhi Patel, Nitin Savaliya, Raghawendra Kumar, Dinesh Kumar, Zuber Saiyed, Komal Patel, Labdhi Pandya, Afzal Ansari, Nikha Trivedi, J G Buch, Jigar Gusani, R D Dixit, A M Kadri, Harsh Bakshi, Chaitanya Joshi, Madhvi Joshi |
| EPI_ISL_495070 | Department of MicroBiology, Government Medical College, Surat | Gujarat Biotechnology Research Centre | Apurvasinh Puvar, Janvi Raval, Zarna Patel, Monika Gandhi, Pinal Trivedi, Maharshi Pandya, Nidhi Patel, Nitin Savaliya, Raghawendra Kumar, Dinesh Kumar, Zuber Saiyed, Komal Patel, Labdhi Pandya, Afzal Ansari, Nikha Trivedi, Naresh Chauhan, Summaiya Mullan, Amit gamit, R D Dixit, A M Kadri, Harsh Bakshi, Chaitanya Joshi, Madhvi Joshi |
| EPI_ISL_495071 | Department of MicroBiology, Government Medical College, Surat | Gujarat Biotechnology Research Centre | Janvi Raval, Zarna Patel, Monika Gandhi, Pinal Trivedi, Maharshi Pandya, Nidhi Patel, Nitin Savaliya, Raghawendra Kumar, Dinesh Kumar, Zuber Saiyed, Komal Patel, Labdhi Pandya, Afzal Ansari, Nikha Trivedi, Naresh Chauhan, Summaiya Mullan, Amit gamit, Apurvasinh Puvar, R D Dixit, A M Kadri, Harsh Bakshi, Chaitanya Joshi, Madhvi Joshi |
| EPI_ISL_495072 | Department of MicroBiology, Government Medical College, Surat | Gujarat Biotechnology Research Centre | Zarna Patel, Monika Gandhi, Pinal Trivedi, Maharshi Pandya, Nidhi Patel, Nitin Savaliya, Raghawendra Kumar, Dinesh Kumar, Zuber Saiyed, Komal Patel, Labdhi Pandya, Afzal Ansari, Nikha Trivedi, Naresh Chauhan, Summaiya Mullan, Amit gamit, Apurvasinh Puvar, Janvi Raval, R D Dixit, A M Kadri, Harsh Bakshi, Chaitanya Joshi, Madhvi Joshi |
| EPI_ISL_495073 | Department of MicroBiology, Government Medical College, Surat | Gujarat Biotechnology Research Centre | Monika Gandhi, Pinal Trivedi, Maharshi Pandya, Nidhi Patel, Nitin Savaliya, Raghawendra Kumar, Dinesh Kumar, Zuber Saiyed, Komal Patel, Labdhi Pandya, Afzal Ansari, Nikha Trivedi, Naresh Chauhan, Summaiya Mullan, Amit gamit, Apurvasinh Puvar, Janvi Raval, Zarna Patel, R D Dixit, A M Kadri, Harsh Bakshi, Chaitanya Joshi, Madhvi Joshi |
| EPI_ISL_495074 | Department of MicroBiology, Government Medical College, Surat | Gujarat Biotechnology Research Centre | Pinal Trivedi, Maharshi Pandya, Nidhi Patel, Nitin Savaliya, Raghawendra Kumar, Dinesh Kumar, Zuber Saiyed, Komal Patel, Labdhi Pandya, Afzal Ansari, Nikha Trivedi, Naresh Chauhan, Summaiya Mullan, Amit gamit, Apurvasinh Puvar, Janvi Raval, Zarna Patel, Monika Gandhi, R D Dixit, A M Kadri, Harsh Bakshi, Chaitanya Joshi, Madhvi Joshi |
| EPI_ISL_495075 | Department of MicroBiology, Government Medical College, Surat | Gujarat Biotechnology Research Centre | Nidhi Patel, Nitin Savaliya, Raghawendra Kumar, Dinesh Kumar, Zuber Saiyed, Komal Patel, Labdhi Pandya, Afzal Ansari, Nikha Trivedi, Naresh Chauhan, Summaiya Mullan, Amit gamit, Apurvasinh Puvar, Janvi Raval, Zarna Patel, Monika Gandhi, Pinal Trivedi, Maharshi Pandya, R D Dixit, A M Kadri, Harsh Bakshi, Chaitanya Joshi, Madhvi Joshi |
| EPI_ISL_495076 | Department of MicroBiology, Government Medical College, Surat | Gujarat Biotechnology Research Centre | Nitin Savaliya, Raghawendra Kumar, Dinesh Kumar, Zuber Saiyed, Komal Patel, Labdhi Pandya, Afzal Ansari, Nikha Trivedi, Naresh Chauhan, Summaiya Mullan, Amit gamit, Apurvasinh Puvar, Janvi Raval, Zarna Patel, Monika Gandhi, Pinal Trivedi, Maharshi Pandya, Nidhi Patel, R D Dixit, A M Kadri, Harsh Bakshi, Chaitanya Joshi, Madhvi Joshi |
| EPI_ISL_495077 | Department of MicroBiology, Government Medical College, Surat | Gujarat Biotechnology Research Centre | Raghawendra Kumar, Dinesh Kumar, Zuber Saiyed, Komal Patel, Labdhi Pandya, Afzal Ansari, Nikha Trivedi, Naresh Chauhan, Summaiya Mullan, Amit gamit, Apurvasinh Puvar, Janvi Raval, Zarna Patel, Monika Gandhi, Pinal Trivedi, Maharshi Pandya, Nidhi Patel, Nitin Savaliya, R D Dixit, A M Kadri, Harsh Bakshi, Chaitanya Joshi, Madhvi Joshi |
| EPI_ISL_495078 | Department of MicroBiology, Government Medical College, Surat | Gujarat Biotechnology Research Centre | Dinesh Kumar, Zuber Saiyed, Komal Patel, Labdhi Pandya, Afzal Ansari, Nikha Trivedi, Naresh Chauhan, Summaiya Mullan, Amit gamit, Apurvasinh Puvar, Janvi Raval, Zarna Patel, Monika Gandhi, Pinal Trivedi, Maharshi Pandya, Nidhi Patel, Nitin Savaliya, Raghawendra Kumar, R D Dixit, A M Kadri, Harsh Bakshi, Chaitanya Joshi, Madhvi Joshi |
| EPI_ISL_495079 | Department of MicroBiology, Government Medical College, Surat | Gujarat Biotechnology Research Centre | Afzal Ansari, Nikha Trivedi, Naresh Chauhan, Summaiya Mullan, Amit gamit, Apurvasinh Puvar, Janvi Raval, Zarna Patel, Monika Gandhi, Pinal Trivedi, Maharshi Pandya, Nidhi Patel, Nitin Savaliya, Raghawendra Kumar, Dinesh Kumar, Zuber Saiyed, Komal Patel, Labdhi Pandya, R D Dixit, A M Kadri, Harsh Bakshi, Chaitanya Joshi, Madhvi Joshi |
| EPI_ISL_495080 | Department of MicroBiology, Government Medical College, Surat | Gujarat Biotechnology Research Centre | Nikha Trivedi, Naresh Chauhan, Summaiya Mullan, Amit gamit, Apurvasinh Puvar, Janvi Raval, Zarna Patel, Monika Gandhi, Pinal Trivedi, Maharshi Pandya, Nidhi Patel, Nitin Savaliya, Raghawendra Kumar, Dinesh Kumar, Zuber Saiyed, Komal Patel, Labdhi Pandya, Afzal Ansari, R D Dixit, A M Kadri, Harsh Bakshi, Chaitanya Joshi, Madhvi Joshi |
| EPI_ISL_495411, EPI_ISL_495412, EPI_ISL_495416, EPI_ISL_495418, EPI_ISL_495420, EPI_ISL_495421, EPI_ISL_495422, EPI_ISL_495425, EPI_ISL_495429, EPI_ISL_495431, EPI_ISL_495433, EPI_ISL_495436, EPI_ISL_495438, EPI_ISL_495444, EPI_ISL_495445, EPI_ISL_495447, EPI_ISL_495449, EPI_ISL_495450, EPI_ISL_495451, EPI_ISL_495452, EPI_ISL_495453, EPI_ISL_495454, EPI_ISL_495456 | | | |
| see above | Kafkas University, Faculty of Medicine, Department of Medical Microbiology | Kafkas University, Faculty of Medicine, Department of Medical Microbiology | Murat Karamese, Didem Ozgur, E. Ediz Tutuncu |
| EPI_ISL_495520, EPI_ISL_495527, EPI_ISL_495528, EPI_ISL_495529, EPI_ISL_495530, EPI_ISL_495534 | NHLS-IALCH | KRISP, KZN Research Innovation and Sequencing Platform | Giandhari J, Pillay S, Lessells R, Chimukangara B, Mdlalose K, York D, Khan S, Tegally H, Wilkinson E, de Oliveira T |
| EPI_ISL_495535, EPI_ISL_495536, EPI_ISL_495537, EPI_ISL_495538, EPI_ISL_495539, EPI_ISL_495540, EPI_ISL_495541, EPI_ISL_495542 | Medical Disagnostics Services (MDS) | KRISP, KZN Research Innovation and Sequencing Platform | Giandhari J, Pillay S, Lessells R, Chimukangara B, Mdlalose K, York D, Khan S, Tegally H, Wilkinson E, de Oliveira T |
| EPI_ISL_495543, EPI_ISL_495544, EPI_ISL_495546, EPI_ISL_495549, EPI_ISL_495550, EPI_ISL_495557, EPI_ISL_495558, EPI_ISL_495559, EPI_ISL_495561, EPI_ISL_495562 | NHLS-IALCH | KRISP, KZN Research Innovation and Sequencing Platform | Giandhari J, Pillay S, Lessells R, Chimukangara B, Mdlalose K, York D, Khan S, Tegally H, Wilkinson E, de Oliveira T |
| EPI_ISL_496520, EPI_ISL_496529, EPI_ISL_496533, EPI_ISL_496537, EPI_ISL_496540, EPI_ISL_496541, EPI_ISL_496542, EPI_ISL_496543, EPI_ISL_496544, EPI_ISL_496545 | Armed Forces Medical College | National Centre For Cell Science | Dhiraj Paul, Kunal Jani, Radha Chauhan, Janesh Kumar, Vasudevan Seshadri, Girdhari Lal, Rajesh Karyakarte, Suvarna Joshi, Murlidhar Tambe, Sourav Sen, Santosh Karade, Kavita Bala Anand, Shelinder Pal Singh Shergill, Rajiv Mohan Gupta, Manoj Kumar Bhat, Arvind Sahu, Maharashtra COVID-19 Study Group, DBT's PAN-INDIA 1000 SARS-CoV2 RNA genome sequencing consortium, Yogesh S Shouche |
| EPI_ISL_496559, EPI_ISL_496560, EPI_ISL_496571, EPI_ISL_496581 | National Centre For Cell Science | National Centre For Cell Science | Dhiraj Paul, Kunal Jani, Radha Chauhan, Janesh Kumar, Vasudevan Seshadri, Girdhari Lal, Rajesh Karyakarte, Suvarna Joshi, Murlidhar Tambe, Sourav Sen, Santosh Karade, Kavita Bala Anand, Shelinder Pal Singh Shergill, Rajiv Mohan Gupta, Manoj Kumar Bhat, Arvind Sahu, Maharashtra COVID-19 Study Group, DBT's PAN-INDIA 1000 SARS-CoV2 RNA genome sequencing consortium, Yogesh S Shouche |
| EPI_ISL_497950 | Shaoxing CDC | Zhejiang Provincial Center for Disease Control and Prevention | Yin Chen, Yanjun Zhang, Haiyan Mao, Junhang Pan, Xiuyu Lou, Yi Sun, Hao Yan, Zhen Li, Wen Shi |
| EPI_ISL_498055, EPI_ISL_498060, EPI_ISL_498061, EPI_ISL_498062, EPI_ISL_498064, EPI_ISL_498065, EPI_ISL_498066, EPI_ISL_498067, EPI_ISL_498069, EPI_ISL_498070, EPI_ISL_498071, EPI_ISL_498072, EPI_ISL_498073, EPI_ISL_498074, EPI_ISL_498075, EPI_ISL_498077, EPI_ISL_498078, EPI_ISL_498079, EPI_ISL_498081, EPI_ISL_498082, EPI_ISL_498084, EPI_ISL_498085, EPI_ISL_498086, EPI_ISL_498087, EPI_ISL_498088, EPI_ISL_498089, EPI_ISL_498090, EPI_ISL_498091, EPI_ISL_498092, EPI_ISL_498093, EPI_ISL_498094, EPI_ISL_498095, EPI_ISL_498096, EPI_ISL_498097, EPI_ISL_498098, EPI_ISL_498099, EPI_ISL_498100, EPI_ISL_498101, EPI_ISL_498102, EPI_ISL_498103, EPI_ISL_498104, EPI_ISL_498105, EPI_ISL_498106, EPI_ISL_498107, EPI_ISL_498108, EPI_ISL_498109, EPI_ISL_498110, EPI_ISL_498111, EPI_ISL_498112, EPI_ISL_498113, EPI_ISL_498114, EPI_ISL_498115, EPI_ISL_498116, EPI_ISL_498117, EPI_ISL_498118, EPI_ISL_498119, EPI_ISL_498120, EPI_ISL_498121, EPI_ISL_498122, EPI_ISL_498123, EPI_ISL_498124, EPI_ISL_498125, EPI_ISL_498126 | | | |
| see above | NHLS-IALCH | KRISP, KZN Research Innovation and Sequencing Platform | Giandhari J, Pillay S, Lessells R, Chimukangara B, Mdlalose K, York D, Khan S, Tegally H, Wilkinson E, de Oliveira T |

| | | | |
|---|---|---|---|
| EPI_ISL_498238, EPI_ISL_498242, EPI_ISL_498243, EPI_ISL_498249, EPI_ISL_498250 | Institut Pasteur de Dakar | Institut Pasteur de Dakar | Ndongo Dia, Moussa Moise Diagne, Mamadou Diop, Marie Henriette Dior Ndione, Mamadou Malado Jallow, Safietou Sankhe Mbengue, Ousmane Faye, Amadou Alpha Sall. |
| EPI_ISL_498558, EPI_ISL_498559, EPI_ISL_498560, EPI_ISL_498561, EPI_ISL_498562, EPI_ISL_498563 | Laboratory of Molecular Virology International Center for Genetic Engineering and Biotechnology (ICGEB) | ARGO Open Lab Platform for Genome Sequencing | Licastro D, Rajasekharan S, Dal Monego S, Segat L, D'Agaro P, Marcello A |
| EPI_ISL_500539, EPI_ISL_500540, EPI_ISL_500542, EPI_ISL_500543, EPI_ISL_500544, EPI_ISL_500545, EPI_ISL_500546, EPI_ISL_500547, EPI_ISL_500548, EPI_ISL_500550, EPI_ISL_500551, EPI_ISL_500552, EPI_ISL_500553, EPI_ISL_500554, EPI_ISL_500555, EPI_ISL_500556, EPI_ISL_500557, EPI_ISL_500558, EPI_ISL_500559, EPI_ISL_500560, EPI_ISL_500561, EPI_ISL_500562, EPI_ISL_500563, EPI_ISL_500564, EPI_ISL_500565, EPI_ISL_500566, EPI_ISL_500567, EPI_ISL_500568, EPI_ISL_500569, EPI_ISL_500570, EPI_ISL_500571, EPI_ISL_500572 | | | |
| see above | Singapore General Hospital | Department of Microbiology | Nurdyana Abdul Rahman, Kun Lee Lim, Chenhao Li, Kian Sing Chan, Lynette Oon, Kern Rei Chng, Niranjan Nagarajan, Karrie Ko |
| EPI_ISL_500946 | GMERS Medical College & Hospital, Gotri, Vadodara | Gujarat Biotechnology Research Centre | Zuber Saiyed, Komal Patel, Labdhi Pandya, Afzal Ansari, Nikha Trivedi, Meenakshi Shah, Neena Doshi, Varsha Godbole, Apurvasinh Puvar, Janvi Raval, Zarna Patel, Monika Gandhi, Pinal Trivedi, Maharshi Pandya, Nidhi Patel, Nitin Savaliya, Raghawendra Kumar, Dinesh Kumar, R D Dixit, A M Kadri, Harsh Bakshi, Chaitanya Joshi, Madhvi Joshi |
| EPI_ISL_500947 | Department of MicroBiology, Government Medical College, Surat | Gujarat Biotechnology Research Centre | Maharshi Pandya, Nidhi Patel, Nitin Savaliya, Raghawendra Kumar, Dinesh Kumar, Zuber Saiyed, Komal Patel, Labdhi Pandya, Afzal Ansari, Nikha Trivedi, Naresh Chauhan, Summaiya Mullan, Amit gamit, Apurvasinh Puvar, Janvi Raval, Zarna Patel, Monika Gandhi, Pinal Trivedi, R D Dixit, A M Kadri, Harsh Bakshi, Chaitanya Joshi, Madhvi Joshi |
| EPI_ISL_500948 | Department of MicroBiology, Government Medical College, Surat | Gujarat Biotechnology Research Centre | Zuber Saiyed, Komal Patel, Labdhi Pandya, Afzal Ansari, Nikha Trivedi, Naresh Chauhan, Summaiya Mullan, Amit gamit, Apurvasinh Puvar, Janvi Raval, Zarna Patel, Monika Gandhi, Pinal Trivedi, Maharshi Pandya, Nidhi Patel, Nitin Savaliya, Raghawendra Kumar, Dinesh Kumar, R D Dixit, A M Kadri, Harsh Bakshi, Chaitanya Joshi, Madhvi Joshi |
| EPI_ISL_500949 | Department of MicroBiology, Government Medical College, Surat | Gujarat Biotechnology Research Centre | Komal Patel, Labdhi Pandya, Afzal Ansari, Nikha Trivedi, Naresh Chauhan, Summaiya Mullan, Amit gamit, Apurvasinh Puvar, Janvi Raval, Zarna Patel, Monika Gandhi, Pinal Trivedi, Maharshi Pandya, Nidhi Patel, Nitin Savaliya, Raghawendra Kumar, Dinesh Kumar, Zuber Saiyed, R D Dixit, A M Kadri, Harsh Bakshi, Chaitanya Joshi, Madhvi Joshi |
| EPI_ISL_500950 | Department of MicroBiology, Government Medical College, Surat | Gujarat Biotechnology Research Centre | Labdhi Pandya, Afzal Ansari, Nikha Trivedi, Naresh Chauhan, Summaiya Mullan, Amit gamit, Apurvasinh Puvar, Janvi Raval, Zarna Patel, Monika Gandhi, Pinal Trivedi, Maharshi Pandya, Nidhi Patel, Nitin Savaliya, Raghawendra Kumar, Dinesh Kumar, Zuber Saiyed, Komal Patel, R D Dixit, A M Kadri, Harsh Bakshi, Chaitanya Joshi, Madhvi Joshi |
| EPI_ISL_507274, EPI_ISL_507279 | WHO National Influenza Centre Russian Federation | WHO National Influenza Centre Russian Federation | Andrey Komissarov, Artem Fadeev, Mariia Sergeeva, Anna Ivanova, Daria Danilenko |
| EPI_ISL_509223, EPI_ISL_509224, EPI_ISL_509225, EPI_ISL_509226, EPI_ISL_509228, EPI_ISL_509229, EPI_ISL_509230, EPI_ISL_509231, EPI_ISL_509232, EPI_ISL_509233, EPI_ISL_509234, EPI_ISL_509235, EPI_ISL_509236, EPI_ISL_509237, EPI_ISL_509238, EPI_ISL_509239, EPI_ISL_509240, EPI_ISL_509241, EPI_ISL_509242, EPI_ISL_509243, EPI_ISL_509244, EPI_ISL_509245, EPI_ISL_509246, EPI_ISL_509247, EPI_ISL_509248, EPI_ISL_509249, EPI_ISL_509250, EPI_ISL_509251, EPI_ISL_509252, EPI_ISL_509253, EPI_ISL_509254, EPI_ISL_509255, EPI_ISL_509256, EPI_ISL_509257, EPI_ISL_509258, EPI_ISL_509260, EPI_ISL_509261, EPI_ISL_509262, EPI_ISL_509263, EPI_ISL_509264, EPI_ISL_509265, EPI_ISL_509266, EPI_ISL_509267, EPI_ISL_509269, EPI_ISL_509270, EPI_ISL_509271, EPI_ISL_509274, EPI_ISL_509275, EPI_ISL_509276, EPI_ISL_509279, EPI_ISL_509281, EPI_ISL_509283, EPI_ISL_509284, EPI_ISL_509285, EPI_ISL_509286, EPI_ISL_509287, EPI_ISL_509288, EPI_ISL_509289, EPI_ISL_509290, EPI_ISL_509292, EPI_ISL_509293, EPI_ISL_509294, EPI_ISL_509295, EPI_ISL_509296, EPI_ISL_509297, EPI_ISL_509299, EPI_ISL_509300, EPI_ISL_509301, EPI_ISL_509302, EPI_ISL_509303, EPI_ISL_509304, EPI_ISL_509305, EPI_ISL_509306, EPI_ISL_509307, EPI_ISL_509308, EPI_ISL_509309, EPI_ISL_509310, EPI_ISL_509311, EPI_ISL_509312, EPI_ISL_509313, EPI_ISL_509314, EPI_ISL_509315, EPI_ISL_509316, EPI_ISL_509317, EPI_ISL_509318, EPI_ISL_509319, EPI_ISL_509320, EPI_ISL_509321, EPI_ISL_509322, EPI_ISL_509323, EPI_ISL_509324, EPI_ISL_509325, EPI_ISL_509326, EPI_ISL_509327, EPI_ISL_509328, EPI_ISL_509329, EPI_ISL_509330, EPI_ISL_509331, EPI_ISL_509332, EPI_ISL_509333, EPI_ISL_509334, EPI_ISL_509335, EPI_ISL_509336, EPI_ISL_509337, EPI_ISL_509338, EPI_ISL_509339, EPI_ISL_509340, EPI_ISL_509341, EPI_ISL_509342, EPI_ISL_509343, EPI_ISL_509344, EPI_ISL_509345, EPI_ISL_509346, EPI_ISL_509347, EPI_ISL_509348, EPI_ISL_509349, EPI_ISL_509350, EPI_ISL_509351, EPI_ISL_509352, EPI_ISL_509353, EPI_ISL_509354, EPI_ISL_509355, EPI_ISL_509356, EPI_ISL_509357, EPI_ISL_509358, EPI_ISL_509360, EPI_ISL_509361, EPI_ISL_509362, EPI_ISL_509363, EPI_ISL_509364, EPI_ISL_509365, EPI_ISL_509366, EPI_ISL_509367, EPI_ISL_509368, EPI_ISL_509369 | | | |
| see above | NHLS-IALCH | KRISP, KZN Research Innovation and Sequencing Platform | Giandhari J, Pillay S, Lessells R, Mdlalose K, York D, Tegally H, Wilkinson E, de Oliveira T |
| EPI_ISL_509373, EPI_ISL_509374, EPI_ISL_509375, EPI_ISL_509377, EPI_ISL_509378, EPI_ISL_509379, EPI_ISL_509380, EPI_ISL_509381, EPI_ISL_509382, EPI_ISL_509383, EPI_ISL_509384, EPI_ISL_509385, EPI_ISL_509386, EPI_ISL_509387, EPI_ISL_509388, EPI_ISL_509389, EPI_ISL_509390, EPI_ISL_509391, EPI_ISL_509392, EPI_ISL_509393, EPI_ISL_509394 | | | |
| see above | Singapore General Hospital | Department of Microbiology | Nurdyana Abdul Rahman, Kun Lee Lim, Chenhao Li, Kian Sing Chan, Lynette Oon, Kern Rei Chng, Niranjan Nagarajan, Karrie Ko |
| EPI_ISL_510081, EPI_ISL_510082 | Communicable Disease Branch | Hong Kong Department of Health | Mak Gannon C.K., Lam Edman T.K., Chan Rickjason C.W., Tsang Dominic N.C. |
| EPI_ISL_510083 | Princess Margaret Hospital | Hong Kong Department of Health | Mak Gannon C.K., Lam Edman T.K., Chan Rickjason C.W., Tsang Dominic N.C. |
| EPI_ISL_510084 | Prince of Wales Hospital | Hong Kong Department of Health | Mak Gannon C.K., Lam Edman T.K., Chan Rickjason C.W., Tsang Dominic N.C. |
| EPI_ISL_510085 | Yan Chai Hospital | Hong Kong Department of Health | Mak Gannon C.K., Lam Edman T.K., Chan Rickjason C.W., Tsang Dominic N.C. |
| EPI_ISL_510086 | Kwong Wah Hospital | Hong Kong Department of Health | Mak Gannon C.K., Lam Edman T.K., Chan Rickjason C.W., Tsang Dominic N.C. |
| EPI_ISL_510087 | Private medical practitioner | Hong Kong Department of Health | Mak Gannon C.K., Lam Edman T.K., Chan Rickjason C.W., Tsang Dominic N.C. |
| EPI_ISL_510088 | United Christian Hospital | Hong Kong Department of Health | Mak Gannon C.K., Lam Edman T.K., Chan Rickjason C.W., Tsang Dominic N.C. |
| EPI_ISL_510089 | Queen Elizabeth Hospital | Hong Kong Department of Health | Mak Gannon C.K., Lam Edman T.K., Chan Rickjason C.W., Tsang Dominic N.C. |
| EPI_ISL_510090 | Hong Kong Baptist Hospital | Hong Kong Department of Health | Mak Gannon C.K., Lam Edman T.K., Chan Rickjason C.W., Tsang Dominic N.C. |
| EPI_ISL_510091 | Kwong Wah Hospital | Hong Kong Department of Health | Mak Gannon C.K., Lam Edman T.K., Chan Rickjason C.W., Tsang Dominic N.C. |
| EPI_ISL_510092, EPI_ISL_510093, EPI_ISL_510094, EPI_ISL_510095, EPI_ISL_510096 | Queen Elizabeth Hospital | Hong Kong Department of Health | Mak Gannon C.K., Lam Edman T.K., Chan Rickjason C.W., Tsang Dominic N.C. |
| EPI_ISL_510097, EPI_ISL_510098 | United Christian Hospital | Hong Kong Department of Health | Mak Gannon C.K., Lam Edman T.K., Chan Rickjason C.W., Tsang Dominic N.C. |
| EPI_ISL_510099 | Queen Mary Hospital | Hong Kong Department of Health | Mak Gannon C.K., Lam Edman T.K., Chan Rickjason C.W., Tsang Dominic N.C. |
| EPI_ISL_510100 | Tuen Mun Hospital | Hong Kong Department of Health | Mak Gannon C.K., Lam Edman T.K., Chan Rickjason C.W., Tsang Dominic N.C. |
| EPI_ISL_510101 | Pamela Youde Nethersole Eastern Hospital | Hong Kong Department of Health | Mak Gannon C.K., Lam Edman T.K., Chan Rickjason C.W., Tsang Dominic N.C. |
| EPI_ISL_511879 | Laboratorium Kesehatan Provinsi Jawa Barat | Molecular Genetics Laboratory-Faculty of Medicine-Universitas Padjadjaran; School of Life Sciences and Technology & School of Pharmacy-Institut Teknologi Bandung; Laboratorium Kesehatan Provinsi Jawa Barat | Marselina Irasonia Tan, Yunia Sribudiani, Catur Riani, Azzania Fibriani, Husna Nugrahapraja, Tarwadi, Ema Rahmawati, Hesti Lina Wiraswati, Lia Faridah, Savira Ekawardhani, Ryan Bayusantika Ristandi, Rifky Waluyajati Rachman, Cut Nur Cinthia Alamanda, Hammam Riza, Soni Solistia Wirawan, Agung Eru Wibowo |
| EPI_ISL_512058 | B.J. Medical College and Civil hospital, Ahmedabad | Gujarat Biotechnology Research Centre | Monika Gandhi, Pinal Trivedi, Maharshi Pandya, Nidhi Patel, Nitin Savaliya, Raghawendra Kumar, Dinesh Kumar, Zuber Saiyed, Komal Patel, Labdhi Pandya, Afzal Ansari, Nikha Trivedi, Pranay Shah, Kamlesh J Upadhyay, Sanjay Kapadia, Apurvasinh Puvar, Janvi Raval, Zarna Patel, R D Dixit, A M Kadri, Harsh Bakshi, Chaitanya Joshi, Madhvi Joshi |
| EPI_ISL_512059 | B.J. Medical College and Civil hospital, Ahmedabad | Gujarat Biotechnology Research Centre | Pinal Trivedi, Maharshi Pandya, Nidhi Patel, Nitin Savaliya, Raghawendra Kumar, Dinesh Kumar, Zuber Saiyed, Komal Patel, Labdhi Pandya, Afzal Ansari, Nikha Trivedi, Pranay Shah, Kamlesh J Upadhyay, Sanjay Kapadia, Apurvasinh Puvar, Janvi Raval, Zarna Patel, Monika Gandhi, R D Dixit, A M Kadri, Harsh Bakshi, Chaitanya Joshi, Madhvi Joshi |
| EPI_ISL_512060 | B.J. Medical College and Civil hospital, Ahmedabad | Gujarat Biotechnology Research Centre | Maharshi Pandya, Nidhi Patel, Nitin Savaliya, Raghawendra Kumar, Dinesh Kumar, Zuber Saiyed, Komal Patel, Labdhi Pandya, Afzal Ansari, Nikha Trivedi, Pranay Shah, Kamlesh J Upadhyay, Sanjay Kapadia, Apurvasinh Puvar, Janvi Raval, Zarna Patel, Monika Gandhi, Pinal Trivedi, R D Dixit, A M Kadri, Harsh Bakshi, Chaitanya Joshi, Madhvi Joshi |
| EPI_ISL_512063 | B.J. Medical College and Civil hospital, Ahmedabad | Gujarat Biotechnology Research Centre | Raghawendra Kumar, Dinesh Kumar, Zuber Saiyed, Komal Patel, Labdhi Pandya, Afzal Ansari, Nikha Trivedi, Pranay Shah, Kamlesh J Upadhyay, Sanjay Kapadia, Apurvasinh Puvar, Janvi Raval, Zarna Patel, Monika Gandhi, Pinal Trivedi, Maharshi Pandya, Nidhi Patel, Nitin Savaliya, R D Dixit, A M Kadri, Harsh Bakshi, Chaitanya Joshi, Madhvi Joshi |
| EPI_ISL_512064 | B.J. Medical College and Civil hospital, Ahmedabad | Gujarat Biotechnology Research Centre | Dinesh Kumar, Zuber Saiyed, Komal Patel, Labdhi Pandya, Afzal Ansari, Nikha Trivedi, Pranay Shah, Kamlesh J Upadhyay, Sanjay Kapadia, Apurvasinh |

| | | | |
|---|---|---|---|
| | | | Puvar, Janvi Raval, Zarna Patel, Monika Gandhi, Pinal Trivedi, Maharshi Pandya, Nidhi Patel, Nitin Savaliya, Raghawendra Kumar, R D Dixit, A M Kadri, Harsh Bakshi, Chaitanya Joshi, Madhvi Joshi |
| EPI_ISL_512065 | B.J. Medical College and Civil hospital, Ahmedabad | Gujarat Biotechnology Research Centre | Zuber Saiyed, Komal Patel, Labdhi Pandya, Afzal Ansari, Nikha Trivedi, Pranay Shah, Kamlesh J Upadhyay, Sanjay Kapadia, Apurvasinh Puvar, Janvi Raval, Zarna Patel, Monika Gandhi, Pinal Trivedi, Maharshi Pandya, Nidhi Patel, Nitin Savaliya, Raghawendra Kumar, Dinesh Kumar, R D Dixit, A M Kadri, Harsh Bakshi, Chaitanya Joshi, Madhvi Joshi |
| EPI_ISL_512066 | Sardar Vallabhbhai Patel Institute of Medical Sciences & Research | Gujarat Biotechnology Research Centre | Komal Patel, Labdhi Pandya, Afzal Ansari, Nikha Trivedi, Pranay Shah, Kamlesh J Upadhyay, Sanjay Kapadia, Apurvasinh Puvar, Janvi Raval, Zarna Patel, Monika Gandhi, Pinal Trivedi, Maharshi Pandya, Nidhi Patel, Nitin Savaliya, Raghawendra Kumar, Dinesh Kumar, Zuber Saiyed, R D Dixit, A M Kadri, Harsh Bakshi, Chaitanya Joshi, Madhvi Joshi |
| EPI_ISL_512067 | Sardar Vallabhbhai Patel Institute of Medical Sciences & Research | Gujarat Biotechnology Research Centre | Labdhi Pandya, Afzal Ansari, Nikha Trivedi, Pranay Shah, Kamlesh J Upadhyay, Sanjay Kapadia, Apurvasinh Puvar, Janvi Raval, Zarna Patel, Monika Gandhi, Pinal Trivedi, Maharshi Pandya, Nidhi Patel, Nitin Savaliya, Raghawendra Kumar, Dinesh Kumar, Zuber Saiyed, Komal Patel, R D Dixit, A M Kadri, Harsh Bakshi, Chaitanya Joshi, Madhvi Joshi |
| EPI_ISL_512068 | Sardar Vallabhbhai Patel Institute of Medical Sciences & Research | Gujarat Biotechnology Research Centre | Afzal Ansari, Nikha Trivedi, Pranay Shah, Kamlesh J Upadhyay, Sanjay Kapadia, Apurvasinh Puvar, Janvi Raval, Zarna Patel, Monika Gandhi, Pinal Trivedi, Maharshi Pandya, Nidhi Patel, Nitin Savaliya, Raghawendra Kumar, Dinesh Kumar, Zuber Saiyed, Komal Patel, Labdhi Pandya, R D Dixit, A M Kadri, Harsh Bakshi, Chaitanya Joshi, Madhvi Joshi |
| EPI_ISL_512069 | Sardar Vallabhbhai Patel Institute of Medical Sciences & Research | Gujarat Biotechnology Research Centre | Nikha Trivedi, Pranay Shah, Kamlesh J Upadhyay, Sanjay Kapadia, Apurvasinh Puvar, Janvi Raval, Zarna Patel, Monika Gandhi, Pinal Trivedi, Maharshi Pandya, Nidhi Patel, Nitin Savaliya, Raghawendra Kumar, Dinesh Kumar, Zuber Saiyed, Komal Patel, Labdhi Pandya, Afzal Ansari, R D Dixit, A M Kadri, Harsh Bakshi, Chaitanya Joshi, Madhvi Joshi |
| EPI_ISL_512070 | Department of MicroBiology, Government Medical College, Surat | Gujarat Biotechnology Research Centre | Naresh Chauhan, Summaiya Mullan, Amit gamit, Apurvasinh Puvar, Janvi Raval, Zarna Patel, Monika Gandhi, Pinal Trivedi, Maharshi Pandya, Nidhi Patel, Nitin Savaliya, Raghawendra Kumar, Dinesh Kumar, Zuber Saiyed, Komal Patel, Labdhi Pandya, Afzal Ansari, Nikha Trivedi, R D Dixit, A M Kadri, Harsh Bakshi, Chaitanya Joshi, Madhvi Joshi |
| EPI_ISL_512071 | Department of MicroBiology, Government Medical College, Surat | Gujarat Biotechnology Research Centre | Summaiya Mullan, Amit gamit, Apurvasinh Puvar, Janvi Raval, Zarna Patel, Monika Gandhi, Pinal Trivedi, Maharshi Pandya, Nidhi Patel, Nitin Savaliya, Raghawendra Kumar, Dinesh Kumar, Zuber Saiyed, Komal Patel, Labdhi Pandya, Afzal Ansari, Nikha Trivedi, Naresh Chauhan, R D Dixit, A M Kadri, Harsh Bakshi, Chaitanya Joshi, Madhvi Joshi |
| EPI_ISL_512072 | Saikrishna Hospital, Mehsana | Gujarat Biotechnology Research Centre | Harshadbhai Parmar, Apurvasinh Puvar, Janvi Raval, Zarna Patel, Monika Gandhi, Pinal Trivedi, Maharshi Pandya, Nidhi Patel, Nitin Savaliya, Raghawendra Kumar, Dinesh Kumar, Zuber Saiyed, Komal Patel, Labdhi Pandya, Afzal Ansari, Nikha Trivedi, R D Dixit, A M Kadri, Harsh Bakshi, Chaitanya Joshi, Madhvi Joshi |
| EPI_ISL_512073 | Saikrishna Hospital, Mehsana | Gujarat Biotechnology Research Centre | Apurvasinh Puvar, Janvi Raval, Zarna Patel, Monika Gandhi, Pinal Trivedi, Maharshi Pandya, Nidhi Patel, Nitin Savaliya, Raghawendra Kumar, Dinesh Kumar, Zuber Saiyed, Komal Patel, Labdhi Pandya, Afzal Ansari, Nikha Trivedi, Harshadbhai Parmar, R D Dixit, A M Kadri, Harsh Bakshi, Chaitanya Joshi, Madhvi Joshi |
| EPI_ISL_512074 | Saikrishna Hospital, Mehsana | Gujarat Biotechnology Research Centre | Janvi Raval, Zarna Patel, Monika Gandhi, Pinal Trivedi, Maharshi Pandya, Nidhi Patel, Nitin Savaliya, Raghawendra Kumar, Dinesh Kumar, Zuber Saiyed, Komal Patel, Labdhi Pandya, Afzal Ansari, Nikha Trivedi, Harshadbhai Parmar, Apurvasinh Puvar, R D Dixit, A M Kadri, Harsh Bakshi, Chaitanya Joshi, Madhvi Joshi |
| EPI_ISL_512075 | Dr. RSS Hospital, Modasa | Gujarat Biotechnology Research Centre | Monika Gandhi, Pinal Trivedi, Maharshi Pandya, Nidhi Patel, Nitin Savaliya, Raghawendra Kumar, Dinesh Kumar, Zuber Saiyed, Komal Patel, Labdhi Pandya, Afzal Ansari, Nikha Trivedi, Harsh Chaudhari, Apurvasinh Puvar, Janvi Raval, Zarna Patel, R D Dixit, A M Kadri, Harsh Bakshi, Chaitanya Joshi, Madhvi Joshi |
| EPI_ISL_512076 | Dr. RSS Hospital, Modasa | Gujarat Biotechnology Research Centre | Pinal Trivedi, Maharshi Pandya, Nidhi Patel, Nitin Savaliya, Raghawendra Kumar, Dinesh Kumar, Zuber Saiyed, Komal Patel, Labdhi Pandya, Afzal Ansari, Nikha Trivedi, Harsh Chaudhari, Apurvasinh Puvar, Janvi Raval, Zarna Patel, Monika Gandhi, R D Dixit, A M Kadri, Harsh Bakshi, Chaitanya Joshi, Madhvi Joshi |
| EPI_ISL_512077 | Dr. RSS Hospital, Modasa | Gujarat Biotechnology Research Centre | Maharshi Pandya, Nidhi Patel, Nitin Savaliya, Raghawendra Kumar, Dinesh Kumar, Zuber Saiyed, Komal Patel, Labdhi Pandya, Afzal Ansari, Nikha Trivedi, Harsh Chaudhari, Apurvasinh Puvar, Janvi Raval, Zarna Patel, Monika Gandhi, Pinal Trivedi, R D Dixit, A M Kadri, Harsh Bakshi, Chaitanya Joshi, Madhvi Joshi |
| EPI_ISL_512670 | Centro Nacional De Rehabilitacion Humberto Araya Rojas (Cenare) | Inciensa, Instituto Costarricense de Investigación y Enseñanza en Nutrición y Salud | Francisco Duarte, Hebleen Porras, Claudio Soto-Garita, Estela Cordero, Adriana Godinez & Melany Calderon |
| EPI_ISL_512846 | O.I.J. MORGUE JUDICIAL | Inciensa, Instituto Costarricense de Investigación y Enseñanza en Nutrición y Salud | Francisco Duarte, Hebleen Porras, Claudio Soto-Garita, Estela Cordero, Adriana Godinez & Melany Calderon |
| EPI_ISL_512874, EPI_ISL_512875, EPI_ISL_512876, EPI_ISL_512877, EPI_ISL_512878, EPI_ISL_512879, EPI_ISL_512880, EPI_ISL_512881, EPI_ISL_512882, EPI_ISL_512883, EPI_ISL_512884, EPI_ISL_512885, EPI_ISL_512886, EPI_ISL_512887, EPI_ISL_512888, EPI_ISL_512889, EPI_ISL_512890, EPI_ISL_512891, EPI_ISL_512892, EPI_ISL_512893, EPI_ISL_512894, EPI_ISL_512895, EPI_ISL_512896, EPI_ISL_512897, EPI_ISL_512898, EPI_ISL_512899, EPI_ISL_512900, EPI_ISL_512901, EPI_ISL_512902, EPI_ISL_512903 | | | |
| see above | Pathogen Genomics Lab King Abdullah University of Science and Technology(KAUST) | Pathogen Genomics Lab King Abdullah University of Science and Technology(KAUST) | Raeece Naeem, Rahul P Salunke, Sharif Hala, Sara Mfarrej, Amit Kumar Subudhi, Fadwa Alofi, Fathia Ben Rached, Afrah Alsomali, Asim Khogeer, Ahmad Bakur Mahmoud, Anwar Hashem, Naif Almontashiri, Arnab Pain |
| EPI_ISL_512904, EPI_ISL_512905, EPI_ISL_512906, EPI_ISL_512907 | Pathogen Genomics Lab King Abdullah University of Science and Technology(KAUST) | Pathogen Genomics Lab King Abdullah University of Science and Technology(KAUST) | Fathia Ben Rached, Raeece Naeem, Sharif Hala, Fadwa Alofi, Rahul P Salunke, Sara Mfarrej, Amit Kumar Subudhi, Afrah Alsomali, Asim Khogeer, Ahmad Bakur Mahmoud, Anwar Hashem, Naif Almontashiri, Arnab Pain |
| EPI_ISL_512920, EPI_ISL_512921, EPI_ISL_512922, EPI_ISL_512923, EPI_ISL_512924, EPI_ISL_512926, EPI_ISL_512927 | Pathogen Genomics Lab King Abdullah University of Science and Technology(KAUST) | Pathogen Genomics Lab King Abdullah University of Science and Technology(KAUST) | Fadwa Alofi, Sharif Hala, Rahul P Salunke, Sara Mfarrej, Amit Kumar Subudhi, Fathia Ben Rached, Amanda, Luke, Afrah Alsomali, Asim Khogeer, Jumana Taha, Abdulaziz Alahmadi, Kahled Algithami, Raeece Naeem, Anwar Hashem, Naif Almontashiri, Arnab Pain |
| EPI_ISL_513002, EPI_ISL_513003, EPI_ISL_513004, EPI_ISL_513005, EPI_ISL_513006, EPI_ISL_513008, EPI_ISL_513009, EPI_ISL_513010, EPI_ISL_513012, EPI_ISL_513013, EPI_ISL_513014, EPI_ISL_513015, EPI_ISL_513016, EPI_ISL_513017, EPI_ISL_513018, EPI_ISL_513019, EPI_ISL_513020, EPI_ISL_513021, EPI_ISL_513022, EPI_ISL_513023, EPI_ISL_513024, EPI_ISL_513025, EPI_ISL_513026 | | | |
| see above | Pathogen Genomics Lab King Abdullah University of Science and Technology(KAUST) | Pathogen Genomics Lab King Abdullah University of Science and Technology(KAUST) | Afrah Alsomali, Fathia Ben Rached, Raeece Naeem, Sharif Hala,Rahul P Salunke, Amanda Ooi, Luke Esau, Sara Mfarrej, Amit Kumar Subudhi, Fadwa Alofi, Asim Khogeer, Kahled Algithami, Anwar Hashem, Naif Almontashiri, Arnab Pain |
| EPI_ISL_513027, EPI_ISL_513028, EPI_ISL_513029, EPI_ISL_513032, EPI_ISL_513033, EPI_ISL_513034, EPI_ISL_513035, EPI_ISL_513036, EPI_ISL_513037, EPI_ISL_513038, EPI_ISL_513039, EPI_ISL_513040, EPI_ISL_513063 | | | |
| see above | Pathogen Genomics Lab King Abdullah University of Science and Technology(KAUST) | Pathogen Genomics Lab King Abdullah University of Science and Technology(KAUST) | Rahul P Salunke, Sharif Hala, Raeece Naeem, Sara Mfarrej, Amit Kumar Subudhi, Amanda Ooi, Luke Esau, Fadwa Alofi, Fathia Ben Rached, Afrah Alsomali, Asim Khogeer, Ahmad Bakur Mahmoud, Anwar Hashem, Naif Almontashiri, Arnab Pain |
| EPI_ISL_513064, EPI_ISL_513065, EPI_ISL_513066, EPI_ISL_513067, EPI_ISL_513068, EPI_ISL_513069, EPI_ISL_513070, EPI_ISL_513071, EPI_ISL_513072, EPI_ISL_513073, EPI_ISL_513074, EPI_ISL_513075, EPI_ISL_513076 | | | |
| see above | Pathogen Genomics Lab King Abdullah University of Science and Technology(KAUST) | Pathogen Genomics Lab King Abdullah University of Science and Technology(KAUST) | Raeece Naeem, Rahul P Salunke, Sharif Hala, Sara Mfarrej, Amit Kumar Subudhi, Fadwa Alofi, Fathia Ben Rached, Afrah Alsomali, Asim Khogeer, Ahmad Bakur Mahmoud, Anwar Hashem, Naif Almontashiri, Arnab Pain |
| EPI_ISL_513078, EPI_ISL_513079, EPI_ISL_513081, EPI_ISL_513082, EPI_ISL_513083, EPI_ISL_513086, EPI_ISL_513087, EPI_ISL_513089, EPI_ISL_513090, EPI_ISL_513092, EPI_ISL_513093, EPI_ISL_513094, EPI_ISL_513095, EPI_ISL_513096, EPI_ISL_513097, EPI_ISL_513098, EPI_ISL_513099, EPI_ISL_513100, EPI_ISL_513101, EPI_ISL_513102, EPI_ISL_513103, EPI_ISL_513104, EPI_ISL_513105, EPI_ISL_513106, EPI_ISL_513107, EPI_ISL_513108, EPI_ISL_513109, EPI_ISL_513111, EPI_ISL_513112, EPI_ISL_513113, EPI_ISL_513114, EPI_ISL_513115, EPI_ISL_513116, EPI_ISL_513117, EPI_ISL_513118, EPI_ISL_513119, EPI_ISL_513120 | | | |
| see above | Pathogen Genomics Lab King Abdullah University of Science and Technology(KAUST) | Pathogen Genomics Lab King Abdullah University of Science and Technology(KAUST) | Fathia Ben Rached, Raeece Naeem, Sharif Hala, Fadwa Alofi, Rahul P Salunke, Sara Mfarrej, Amit Kumar Subudhi, Afrah Alsomali, Asim Khogeer, Ahmad Bakur Mahmoud, Anwar Hashem, Naif Almontashiri, Arnab Pain |
| EPI_ISL_513190, EPI_ISL_513191, EPI_ISL_513192, EPI_ISL_513193, EPI_ISL_513194, EPI_ISL_513196, EPI_ISL_513197 | Pathogen Genomics Lab King Abdullah University of Science and Technology(KAUST) | Pathogen Genomics Lab King Abdullah University of Science and Technology(KAUST) | Afrah Alsomali, Fathia Ben Rached, Raeece Naeem, Sharif Hala,Rahul P Salunke, Amanda Ooi, Luke Esau, Sara Mfarrej, Amit Kumar Subudhi, Fadwa Alofi, Asim Khogeer, Kahled Algithami, Anwar Hashem, Naif Almontashiri, Arnab Pain |

| | | | |
|---|---|---|---|
| EPI_ISL_513199, EPI_ISL_513200, EPI_ISL_513201, EPI_ISL_513202, EPI_ISL_513203, EPI_ISL_513204, EPI_ISL_513207 | Pathogen Genomics Lab King Abdullah University of Science and Technology(KAUST) | Pathogen Genomics Lab King Abdullah University of Science and Technology(KAUST) | Rahul P Salunke, Sharif Hala, Raeece Naeem, Sara Mfarrej, Amit Kumar Subudhi, Amanda Ooi, Luke Esau, Fadwa Alofi, Fathia Ben Rached, Afrah Alsomali, Asim Khogeer, Ahmad Bakur Mahmoud, Anwar Hashem, Naif Almontashiri, Arnab Pain |
| EPI_ISL_513210, EPI_ISL_513212, EPI_ISL_513213 | Pathogen Genomics Lab King Abdullah University of Science and Technology(KAUST) | Pathogen Genomics Lab King Abdullah University of Science and Technology(KAUST) | Raeece Naeem, Rahul P Salunke, Sharif Hala, Sara Mfarrej, Amit Kumar Subudhi, Fadwa Alofi, Fathia Ben Rached, Afrah Alsomali, Asim Khogeer, Ahmad Bakur Mahmoud, Anwar Hashem, Naif Almontashiri, Arnab Pain |
| EPI_ISL_514093 | National Scientific Center for Especially Dangerous Infections (NSCEDI) | Kazakh National Agrarian University (KazNAU) TreeGene LLP Genetic Laboratory | Beloussov Vyacheslav, Strochkov Vitaliy, Tabynov Kaissar, Sandybayev Nurlan, Tabynov Kairat, Turebekov Nurkeldy, Granica Joanna, Solomadin Maksim, Yerubayev Toktassyn, Yespolov Tlektes |
| EPI_ISL_514127 | National Scientific Center for Especially Dangerous Infections (NSCEDI) | Kazakh National Agrarian University (KazNAU) TreeGene LLP Genetic Laboratory | Tabynov Kairat, Beloussov Vyacheslav, Strochkov Vitaliy, Sandybayev Nurlan, Tabynov Kaissar, Turebekov Nurkeldy, Granica Joanna, Solomadin Maksim, Yerubayev Toktassyn, Yespolov Tlektes |
| EPI_ISL_514138 | Rondônia Central Public Health Laboratory (LACEN/RO), vinculated to State Health Secretariat of Rondônia (SESAU/RO) | Molecular Virology Laboratory of Oswaldo Cruz Foundation of Rondônia | Luan Felipo Botelho-Souza, Felipe Souza Nogueira-Lima, Tárcio Peixoto Roca, Alcione de Oliveira dos Santos, Felipe Gomes Naveca, Adriana Cristina Salvador Maia, Cicileia Correia da Silva, Aline Linhares Ferreira de Melo Mendonça, Celina Aparecida Bertoni Lugtenburg, Camila Flávia Gomes Azzi, Juliana Loca Furtado, Suelen Cavalcante, Rita de Cássia Pontello Rampazzo, Caio Henrique Nemeth Santos, Alice Paula Di Sabatino Guimarães, Jansen Fernandes de Medeiros, Fernando Rodrigues Máximo, Juan Miguel Vilallobos-Salcedo and Deusilene Souza Vieira |
| EPI_ISL_514434 | NSTU COVID-19 Diagnostic Center, | NSU Genome Research Institute (NGRI), North South University | Dr. Muhammad Maqsud Hossain, Aura Rahman, Prof. Firoz Ahmed, Tahrima Huq, Abdus Sadique, Jahidul Alam, Md Aminul Islam, Prof. Md. Didar-Ul-Alam, Prof. Kazi Nadim Hasan, Prof. Abdul Khaleque, Prof, Hasan Mahmud Reza |
| EPI_ISL_514435 | Dr. RSS Hospital, Modasa | Gujarat Biotechnology Research Centre | Harsh Chaudhari, Zarna Patel, Monika Gandhi, Pinal Trivedi, Maharshi Pandya, Nidhi Patel, Nitin Savaliya, Raghawendra Kumar, Dinesh Kumar, Zuber Saiyed, Komal Patel, Labdhi Pandya, Afzal Ansari, Nikha Trivedi, Apurvasinh Puvar, Janvi Raval, R D Dixit, A M Kadri, Harsh Bakshi, Chaitanya Joshi, Madhvi Joshi |
| EPI_ISL_514436 | Dr. RSS Hospital, Modasa | Gujarat Biotechnology Research Centre | Zarna Patel, Monika Gandhi, Pinal Trivedi, Maharshi Pandya, Nidhi Patel, Nitin Savaliya, Raghawendra Kumar, Dinesh Kumar, Zuber Saiyed, Komal Patel, Labdhi Pandya, Afzal Ansari, Nikha Trivedi, Harsh Chaudhari, Apurvasinh Puvar, Janvi Raval, R D Dixit, A M Kadri, Harsh Bakshi, Chaitanya Joshi, Madhvi Joshi |
| EPI_ISL_514437 | Dr. RSS Hospital, Modasa | Gujarat Biotechnology Research Centre | Nidhi Patel, Nitin Savaliya, Raghawendra Kumar, Dinesh Kumar, Zuber Saiyed, Komal Patel, Labdhi Pandya, Afzal Ansari, Nikha Trivedi, Harsh Chaudhari, Apurvasinh Puvar, Janvi Raval, Zarna Patel, Monika Gandhi, Pinal Trivedi, Maharshi Pandya, R D Dixit, A M Kadri, Harsh Bakshi, Chaitanya Joshi, Madhvi Joshi |
| EPI_ISL_514440 | NSTU COVID-19 Diagnostic Center | NSU Genome Research Institute (NGRI), North South University | Dr. Muhammad Maqsud Hossain, Aura Rahman, Prof. Firoz Ahmed, Tahrima Huq, Abdus Sadique, Tamanna Afroze, Jahidul Alam, Md Aminul Islam, Prof. Md. Didar-Ul-Alam, Prof. Kazi Nadim Hasan, Prof. Abdul Khaleque, Prof, Hasan Mahmud Reza |
| EPI_ISL_514441 | NSTU COVID-19 Diagnostic Center | NSU Genome Research Institute (NGRI), North South University | Dr. Muhammad Maqsud Hossain, Aura Rahman, Prof. Firoz Ahmed, Tahrima Huq, Abdus Sadique, Jahidul Alam, Md Aminul Islam, Prof. Md. Didar-Ul-Alam, Prof. Kazi Nadim Hasan, Prof. Abdul Khaleque, Prof, Hasan Mahmud Reza |
| EPI_ISL_514442, EPI_ISL_514580 | NSTU COVID-19 Diagnostic Center | NSU Genome Research Institute (NGRI), North South University | Dr. Muhammad Maqsud Hossain, Aura Rahman, Prof. Firoz Ahmed, Tahrima Huq, Abdus Sadique, Jahidul Alam, Tamanna Afroze, Md Aminul Islam, Prof. Md. Didar-Ul-Alam, Prof. Kazi Nadim Hasan, Prof. Abdul Khaleque, Prof, Hasan Mahmud Reza |
| EPI_ISL_514581 | B.J. Medical College and Civil hospital, Ahmedabad | Gujarat Biotechnology Research Centre | Nitin Savaliya, Raghawendra Kumar, Dinesh Kumar, Zuber Saiyed, Komal Patel, Labdhi Pandya, Afzal Ansari, Nikha Trivedi, Pranay Shah, Kamlesh J Upadhyay, Sanjay Kapadia, Apurvasinh Puvar, Janvi Raval, Zarna Patel, Monika Gandhi, Pinal Trivedi, Maharshi Pandya, Nidhi Patel, R D Dixit, A M Kadri, Harsh Bakshi, Chaitanya Joshi, Madhvi Joshi |
| EPI_ISL_514582 | B.J. Medical College and Civil hospital, Ahmedabad | Gujarat Biotechnology Research Centre | Raghawendra Kumar, Dinesh Kumar, Zuber Saiyed, Komal Patel, Labdhi Pandya, Afzal Ansari, Nikha Trivedi, Pranay Shah, Kamlesh J Upadhyay, Sanjay Kapadia, Apurvasinh Puvar, Janvi Raval, Zarna Patel, Monika Gandhi, Pinal Trivedi, Maharshi Pandya, Nidhi Patel, Nitin Savaliya, R D Dixit, A M Kadri, Harsh Bakshi, Chaitanya Joshi, Madhvi Joshi |
| EPI_ISL_514583 | B.J. Medical College and Civil hospital, Ahmedabad | Gujarat Biotechnology Research Centre | Dinesh Kumar, Zuber Saiyed, Komal Patel, Labdhi Pandya, Afzal Ansari, Nikha Trivedi, Pranay Shah, Kamlesh J Upadhyay, Sanjay Kapadia, Apurvasinh Puvar, Janvi Raval, Zarna Patel, Monika Gandhi, Pinal Trivedi, Maharshi Pandya, Nidhi Patel, Nitin Savaliya, Raghawendra Kumar, R D Dixit, A M Kadri, Harsh Bakshi, Chaitanya Joshi, Madhvi Joshi |
| EPI_ISL_514584 | B.J. Medical College and Civil hospital, Ahmedabad | Gujarat Biotechnology Research Centre | Zuber Saiyed, Komal Patel, Labdhi Pandya, Afzal Ansari, Nikha Trivedi, Pranay Shah, Kamlesh J Upadhyay, Sanjay Kapadia, Apurvasinh Puvar, Janvi Raval, Zarna Patel, Monika Gandhi, Pinal Trivedi, Maharshi Pandya, Nidhi Patel, Nitin Savaliya, Raghawendra Kumar, Dinesh Kumar, R D Dixit, A M Kadri, Harsh Bakshi, Chaitanya Joshi, Madhvi Joshi |
| EPI_ISL_514585 | B.J. Medical College and Civil hospital, Ahmedabad | Gujarat Biotechnology Research Centre | Komal Patel, Labdhi Pandya, Afzal Ansari, Nikha Trivedi, Pranay Shah, Kamlesh J Upadhyay, Sanjay Kapadia, Apurvasinh Puvar, Janvi Raval, Zarna Patel, Monika Gandhi, Pinal Trivedi, Maharshi Pandya, Nidhi Patel, Nitin Savaliya, Raghawendra Kumar, Dinesh Kumar, Zuber Saiyed, R D Dixit, A M Kadri, Harsh Bakshi, Chaitanya Joshi, Madhvi Joshi |
| EPI_ISL_514586 | B.J. Medical College and Civil hospital, Ahmedabad | Gujarat Biotechnology Research Centre | Labdhi Pandya, Afzal Ansari, Nikha Trivedi, Pranay Shah, Kamlesh J Upadhyay, Sanjay Kapadia, Apurvasinh Puvar, Janvi Raval, Zarna Patel, Monika Gandhi, Pinal Trivedi, Maharshi Pandya, Nidhi Patel, Nitin Savaliya, Raghawendra Kumar, Dinesh Kumar, Zuber Saiyed, Komal Patel, R D Dixit, A M Kadri, Harsh Bakshi, Chaitanya Joshi, Madhvi Joshi |
| EPI_ISL_514587 | B.J. Medical College and Civil hospital, Ahmedabad | Gujarat Biotechnology Research Centre | Afzal Ansari, Nikha Trivedi, Pranay Shah, Kamlesh J Upadhyay, Sanjay Kapadia, Apurvasinh Puvar, Janvi Raval, Zarna Patel, Monika Gandhi, Pinal Trivedi, Maharshi Pandya, Nidhi Patel, Nitin Savaliya, Raghawendra Kumar, Dinesh Kumar, Zuber Saiyed, Komal Patel, Labdhi Pandya, R D Dixit, A M Kadri, Harsh Bakshi, Chaitanya Joshi, Madhvi Joshi |
| EPI_ISL_514588 | B.J. Medical College and Civil hospital, Ahmedabad | Gujarat Biotechnology Research Centre | Nikha Trivedi, Pranay Shah, Kamlesh J Upadhyay, Sanjay Kapadia, Apurvasinh Puvar, Janvi Raval, Zarna Patel, Monika Gandhi, Pinal Trivedi, Maharshi Pandya, Nidhi Patel, Nitin Savaliya, Raghawendra Kumar, Dinesh Kumar, Zuber Saiyed, Komal Patel, Labdhi Pandya, Afzal Ansari, R D Dixit, A M Kadri, Harsh Bakshi, Chaitanya Joshi, Madhvi Joshi |
| EPI_ISL_514589 | B.J. Medical College and Civil hospital, Ahmedabad | Gujarat Biotechnology Research Centre | Pranay Shah, Kamlesh J Upadhyay, Sanjay Kapadia, Apurvasinh Puvar, Janvi Raval, Zarna Patel, Monika Gandhi, Pinal Trivedi, Maharshi Pandya, Nidhi Patel, Nitin Savaliya, Raghawendra Kumar, Dinesh Kumar, Zuber Saiyed, Komal Patel, Labdhi Pandya, Afzal Ansari, Nikha Trivedi, R D Dixit, A M Kadri, Harsh Bakshi, Chaitanya Joshi, Madhvi Joshi |
| EPI_ISL_514590 | B.J. Medical College and Civil hospital, Ahmedabad | Gujarat Biotechnology Research Centre | Kamlesh J Upadhyay, Sanjay Kapadia, Apurvasinh Puvar, Janvi Raval, Zarna Patel, Monika Gandhi, Pinal Trivedi, Maharshi Pandya, Nidhi Patel, Nitin Savaliya, Raghawendra Kumar, Dinesh Kumar, Zuber Saiyed, Komal Patel, Labdhi Pandya, Afzal Ansari, Nikha Trivedi, Pranay Shah, R D Dixit, A M Kadri, Harsh Bakshi, Chaitanya Joshi, Madhvi Joshi |
| EPI_ISL_514591 | B.J. Medical College and Civil hospital, Ahmedabad | Gujarat Biotechnology Research Centre | Sanjay Kapadia,Apurvasinh Puvar,Janvi Raval,Zarna Patel, Monika Gandhi, Pinal Trivedi, Maharshi Pandya, Nidhi Patel, Nitin Savaliya, Raghawendra Kumar, Dinesh Kumar, Zuber Saiyed, Komal Patel, Labdhi Pandya, Afzal Ansari, Nikha Trivedi, Pranay Shah, Kamlesh J Upadhyay, R D Dixit, A M Kadri, Harsh Bakshi, Chaitanya Joshi, Madhvi Joshi |
| EPI_ISL_514592 | B.J. Medical College and Civil hospital, Ahmedabad | Gujarat Biotechnology Research Centre | Apurvasinh Puvar, Janvi Raval, Zarna Patel, Monika Gandhi, Pinal Trivedi, Maharshi Pandya, Nidhi Patel, Nitin Savaliya, Raghawendra Kumar, Dinesh Kumar, Zuber Saiyed, Komal Patel, Labdhi Pandya, Afzal Ansari, Nikha Trivedi, Pranay Shah, Kamlesh J Upadhyay, Sanjay Kapadia, R D Dixit, A M Kadri, Harsh Bakshi, Chaitanya Joshi, Madhvi Joshi |
| EPI_ISL_514593 | B.J. Medical College and Civil hospital, Ahmedabad | Gujarat Biotechnology Research Centre | Janvi Raval, Zarna Patel, Monika Gandhi, Pinal Trivedi, Maharshi Pandya, Nidhi Patel, Nitin Savaliya, Raghawendra Kumar, Dinesh Kumar, Zuber Saiyed, Komal Patel, Labdhi Pandya, Afzal Ansari, Nikha Trivedi, Pranay Shah, Kamlesh J Upadhyay, Sanjay Kapadia, Apurvasinh Puvar, R D Dixit, A M Kadri, Harsh Bakshi, Chaitanya Joshi, Madhvi Joshi |
| EPI_ISL_514594 | B.J. Medical College and Civil hospital, Ahmedabad | Gujarat Biotechnology Research Centre | Zarna Patel, Monika Gandhi, Pinal Trivedi, Maharshi Pandya, Nidhi Patel, Nitin Savaliya, Raghawendra Kumar, Dinesh Kumar, Zuber Saiyed, Komal Patel, Labdhi Pandya, Afzal Ansari, Nikha Trivedi, Pranay Shah, Kamlesh J Upadhyay, Sanjay Kapadia, Apurvasinh Puvar, Janvi Raval, R D Dixit, A M Kadri, Harsh Bakshi, Chaitanya Joshi, Madhvi Joshi |
| EPI_ISL_514595 | B.J. Medical College and Civil hospital, Ahmedabad | Gujarat Biotechnology Research Centre | Monika Gandhi, Pinal Trivedi, Maharshi Pandya, Nidhi Patel, Nitin Savaliya, Raghawendra Kumar, Dinesh Kumar, Zuber Saiyed, Komal Patel, Labdhi Pandya, |

| | | | |
|---|---|---|---|
| | | | Afzal Ansari, Nikha Trivedi, Pranay Shah, Kamlesh J Upadhyay, Sanjay Kapadia, Apurvasinh Puvar, Janvi Raval, Zarna Patel, R D Dixit, A M Kadri, Harsh Bakshi, Chaitanya Joshi, Madhvi Joshi |
| EPI_ISL_514596 | B.J. Medical College and Civil hospital, Ahmedabad | Gujarat Biotechnology Research Centre | Pinal Trivedi, Maharshi Pandya, Nidhi Patel, Nitin Savaliya, Raghawendra Kumar, Dinesh Kumar, Zuber Saiyed, Komal Patel, Labdhi Pandya, Afzal Ansari, Nikha Trivedi, Pranay Shah, Kamlesh J Upadhyay, Sanjay Kapadia, Apurvasinh Puvar, Janvi Raval, Zarna Patel, Monika Gandhi, R D Dixit, A M Kadri, Harsh Bakshi, Chaitanya Joshi, Madhvi Joshi |
| EPI_ISL_514597 | B.J. Medical College and Civil hospital, Ahmedabad | Gujarat Biotechnology Research Centre | Maharshi Pandya, Nidhi Patel, Nitin Savaliya, Raghawendra Kumar, Dinesh Kumar, Zuber Saiyed, Komal Patel, Labdhi Pandya, Afzal Ansari, Nikha Trivedi, Pranay Shah, Kamlesh J Upadhyay, Sanjay Kapadia, Apurvasinh Puvar, Janvi Raval, Zarna Patel, Monika Gandhi, Pinal Trivedi, R D Dixit, A M Kadri, Harsh Bakshi, Chaitanya Joshi, Madhvi Joshi |
| EPI_ISL_514598 | B.J. Medical College and Civil hospital, Ahmedabad | Gujarat Biotechnology Research Centre | Nidhi Patel, Nitin Savaliya, Raghawendra Kumar, Dinesh Kumar, Zuber Saiyed, Komal Patel, Labdhi Pandya, Afzal Ansari, Nikha Trivedi, Pranay Shah, Kamlesh J Upadhyay, Sanjay Kapadia, Apurvasinh Puvar, Janvi Raval, Zarna Patel, Monika Gandhi, Pinal Trivedi, Maharshi Pandya, R D Dixit, A M Kadri, Harsh Bakshi, Chaitanya Joshi, Madhvi Joshi |
| EPI_ISL_514599 | B.J. Medical College and Civil hospital, Ahmedabad | Gujarat Biotechnology Research Centre | Nitin Savaliya, Raghawendra Kumar, Dinesh Kumar, Zuber Saiyed, Komal Patel, Labdhi Pandya, Afzal Ansari, Nikha Trivedi, Pranay Shah, Kamlesh J Upadhyay, Sanjay Kapadia, Apurvasinh Puvar, Janvi Raval, Zarna Patel, Monika Gandhi, Pinal Trivedi, Maharshi Pandya, Nidhi Patel, R D Dixit, A M Kadri, Harsh Bakshi, Chaitanya Joshi, Madhvi Joshi |
| EPI_ISL_514600 | B.J. Medical College and Civil hospital, Ahmedabad | Gujarat Biotechnology Research Centre | Raghawendra Kumar, Dinesh Kumar, Zuber Saiyed, Komal Patel, Labdhi Pandya, Afzal Ansari, Nikha Trivedi, Pranay Shah, Kamlesh J Upadhyay, Sanjay Kapadia, Apurvasinh Puvar, Janvi Raval, Zarna Patel, Monika Gandhi, Pinal Trivedi, Maharshi Pandya, Nidhi Patel, Nitin Savaliya, R D Dixit, A M Kadri, Harsh Bakshi, Chaitanya Joshi, Madhvi Joshi |
| EPI_ISL_514601 | B.J. Medical College and Civil hospital, Ahmedabad | Gujarat Biotechnology Research Centre | Dinesh Kumar, Zuber Saiyed, Komal Patel, Labdhi Pandya, Afzal Ansari, Nikha Trivedi, Pranay Shah, Kamlesh J Upadhyay, Sanjay Kapadia, Apurvasinh Puvar, Janvi Raval, Zarna Patel, Monika Gandhi, Pinal Trivedi, Maharshi Pandya, Nidhi Patel, Nitin Savaliya, Raghawendra Kumar, R D Dixit, A M Kadri, Harsh Bakshi, Chaitanya Joshi, Madhvi Joshi |
| EPI_ISL_514602 | B.J. Medical College and Civil hospital, Ahmedabad | Gujarat Biotechnology Research Centre | Zuber Saiyed, Komal Patel, Labdhi Pandya, Afzal Ansari, Nikha Trivedi, Pranay Shah, Kamlesh J Upadhyay, Sanjay Kapadia, Apurvasinh Puvar, Janvi Raval, Zarna Patel, Monika Gandhi, Pinal Trivedi, Maharshi Pandya, Nidhi Patel, Nitin Savaliya, Raghawendra Kumar, Dinesh Kumar, R D Dixit, A M Kadri, Harsh Bakshi, Chaitanya Joshi, Madhvi Joshi |
| EPI_ISL_514603 | B.J. Medical College and Civil hospital, Ahmedabad | Gujarat Biotechnology Research Centre | Komal Patel, Labdhi Pandya, Afzal Ansari, Nikha Trivedi, Pranay Shah, Kamlesh J Upadhyay, Sanjay Kapadia, Apurvasinh Puvar, Janvi Raval, Zarna Patel, Monika Gandhi, Pinal Trivedi, Maharshi Pandya, Nidhi Patel, Nitin Savaliya, Raghawendra Kumar, Dinesh Kumar, Zuber Saiyed, R D Dixit, A M Kadri, Harsh Bakshi, Chaitanya Joshi, Madhvi Joshi |
| EPI_ISL_514604 | B.J. Medical College and Civil hospital, Ahmedabad | Gujarat Biotechnology Research Centre | Labdhi Pandya, Afzal Ansari, Nikha Trivedi, Pranay Shah, Kamlesh J Upadhyay, Sanjay Kapadia, Apurvasinh Puvar, Janvi Raval, Zarna Patel, Monika Gandhi, Pinal Trivedi, Maharshi Pandya, Nidhi Patel, Nitin Savaliya, Raghawendra Kumar, Dinesh Kumar, Zuber Saiyed, Komal Patel, R D Dixit, A M Kadri, Harsh Bakshi, Chaitanya Joshi, Madhvi Joshi |
| EPI_ISL_514605 | B.J. Medical College and Civil hospital, Ahmedabad | Gujarat Biotechnology Research Centre | Afzal Ansari, Nikha Trivedi, Pranay Shah, Kamlesh J Upadhyay, Sanjay Kapadia, Apurvasinh Puvar, Janvi Raval, Zarna Patel, Monika Gandhi, Pinal Trivedi, Maharshi Pandya, Nidhi Patel, Nitin Savaliya, Raghawendra Kumar, Dinesh Kumar, Zuber Saiyed, Komal Patel, Labdhi Pandya, R D Dixit, A M Kadri, Harsh Bakshi, Chaitanya Joshi, Madhvi Joshi |
| EPI_ISL_514606 | B.J. Medical College and Civil hospital, Ahmedabad | Gujarat Biotechnology Research Centre | Nikha Trivedi, Pranay Shah, Kamlesh J Upadhyay, Sanjay Kapadia, Apurvasinh Puvar, Janvi Raval, Zarna Patel, Monika Gandhi, Pinal Trivedi, Maharshi Pandya, Nidhi Patel, Nitin Savaliya, Raghawendra Kumar, Dinesh Kumar, Zuber Saiyed, Komal Patel, Labdhi Pandya, Afzal Ansari, R D Dixit, A M Kadri, Harsh Bakshi, Chaitanya Joshi, Madhvi Joshi |
| EPI_ISL_514607 | B.J. Medical College and Civil hospital, Ahmedabad | Gujarat Biotechnology Research Centre | Pranay Shah, Kamlesh J Upadhyay, Sanjay Kapadia, Apurvasinh Puvar, Janvi Raval, Zarna Patel, Monika Gandhi, Pinal Trivedi, Maharshi Pandya, Nidhi Patel, Nitin Savaliya, Raghawendra Kumar, Dinesh Kumar, Zuber Saiyed, Komal Patel, Labdhi Pandya, Afzal Ansari, Nikha Trivedi, R D Dixit, A M Kadri, Harsh Bakshi, Chaitanya Joshi, Madhvi Joshi |
| EPI_ISL_514608 | B.J. Medical College and Civil hospital, Ahmedabad | Gujarat Biotechnology Research Centre | Kamlesh J Upadhyay, Sanjay Kapadia, Apurvasinh Puvar, Janvi Raval, Zarna Patel, Monika Gandhi, Pinal Trivedi, Maharshi Pandya, Nidhi Patel, Nitin Savaliya, Raghawendra Kumar, Dinesh Kumar, Zuber Saiyed, Komal Patel, Labdhi Pandya, Afzal Ansari, Nikha Trivedi, Pranay Shah, R D Dixit, A M Kadri, Harsh Bakshi, Chaitanya Joshi, Madhvi Joshi |
| EPI_ISL_514609 | B.J. Medical College and Civil hospital, Ahmedabad | Gujarat Biotechnology Research Centre | Sanjay Kapadia, Apurvasinh Puvar, Janvi Raval, Zarna Patel, Monika Gandhi, Pinal Trivedi, Maharshi Pandya, Nidhi Patel, Nitin Savaliya, Raghawendra Kumar, Dinesh Kumar, Zuber Saiyed, Komal Patel, Labdhi Pandya, Afzal Ansari, Nikha Trivedi, Pranay Shah, Kamlesh J Upadhyay, R D Dixit, A M Kadri, Harsh Bakshi, Chaitanya Joshi, Madhvi Joshi |
| EPI_ISL_514610 | GMERS Medical College & Hospital, Gotri, Vadodara | Gujarat Biotechnology Research Centre | Meenakshi Shah, Neena Doshi, Varsha Godbole, Apurvasinh Puvar, Janvi Raval, Zarna Patel, Monika Gandhi, Pinal Trivedi, Maharshi Pandya, Nidhi Patel, Nitin Savaliya, Raghawendra Kumar, Dinesh Kumar, Zuber Saiyed, Komal Patel, Labdhi Pandya, Afzal Ansari, Nikha Trivedi, Pranay Shah, Kamlesh J Upadhyay, Sanjay Kapadia, R D Dixit, A M Kadri, Harsh Bakshi, Chaitanya Joshi, Madhvi Joshi, |
| EPI_ISL_514611 | GMERS Medical College and Hospital, Gandhinagar | Gujarat Biotechnology Research Centre | Seema Bhatt, Gaurishankar Shrimali, Bhavesh Modi, Bharti Rajani, Apurvasinh Puvar, Janvi Raval, Zarna Patel, Monika Gandhi, Pinal Trivedi, Maharshi Pandya, Nidhi Patel, Nitin Savaliya, Raghawendra Kumar, Dinesh Kumar, Zuber Saiyed, Komal Patel, Labdhi Pandya, Afzal Ansari, Nikha Trivedi, Pranay Shah, Kamlesh J Upadhyay, Sanjay Kapadia, R D Dixit, A M Kadri, Harsh Bakshi, Chaitanya Joshi, Madhvi Joshi |
| EPI_ISL_514612 | GMERS Medical College and Hospital, Gandhinagar | Gujarat Biotechnology Research Centre | Gaurishankar Shrimali, Bhavesh Modi, Bharti Rajani, Apurvasinh Puvar, Janvi Raval, Zarna Patel, Monika Gandhi, Pinal Trivedi, Maharshi Pandya, Nidhi Patel, Nitin Savaliya, Raghawendra Kumar, Dinesh Kumar, Zuber Saiyed, Komal Patel, Labdhi Pandya, Afzal Ansari, Nikha Trivedi, Pranay Shah, Kamlesh J Upadhyay, Sanjay Kapadia, Seema Bhatt, R D Dixit, A M Kadri, Harsh Bakshi, Chaitanya Joshi, Madhvi Joshi |
| EPI_ISL_514613, EPI_ISL_514615 | NSTU COVID-19 Diagnostic Center | NSU Genome Research Institute (NGRI), North South University | Dr. Muhammad Maqsud Hossain, Aura Rahman, Prof. Firoz Ahmed, Tahrima Huq, Abdus Sadique, Jahidul Alam, Tamanna Afroze, Md Aminul Islam, Prof. Md. Didar-Ul-Alam, Prof. Kazi Nadim Hasan, Prof. Abdul Khaleque, Prof, Hasan Mahmud Reza |
| EPI_ISL_515466 | Discovery DNA | Discovery DNA | Dustin Hittel, Marina Kerr, Leo Dimnik, Desmond Koo, Alice Li, Aneal Khan |
| EPI_ISL_515525 | National Influenza Center - Instituto Adolfo Lutz | Instituto Adolfo Lutz, Interdiciplinary Procedures Center, Strategic Laboratory | Claudio Tavares Sacchi, Claudia Regina Gonçalves, Erica Valessa Ramos Gomes |
| EPI_ISL_515568, EPI_ISL_515569, EPI_ISL_515570, EPI_ISL_515573, EPI_ISL_515574, EPI_ISL_515575, EPI_ISL_515579, EPI_ISL_515580, EPI_ISL_515582, EPI_ISL_515584, EPI_ISL_515585, EPI_ISL_515586, EPI_ISL_515589, EPI_ISL_515592, EPI_ISL_515593, EPI_ISL_515594, EPI_ISL_515596, EPI_ISL_515597, EPI_ISL_515600, EPI_ISL_515601, EPI_ISL_515602, EPI_ISL_515603, EPI_ISL_515604, EPI_ISL_515606, EPI_ISL_515607, EPI_ISL_515610, EPI_ISL_515612, EPI_ISL_515613, EPI_ISL_515615, EPI_ISL_515616, EPI_ISL_515619, EPI_ISL_515622, EPI_ISL_515623, EPI_ISL_515627, EPI_ISL_515628, EPI_ISL_515631, EPI_ISL_515633, EPI_ISL_515634, EPI_ISL_515635, EPI_ISL_515639, EPI_ISL_515640, EPI_ISL_515641, EPI_ISL_515642, EPI_ISL_515643, EPI_ISL_515644, EPI_ISL_515645, EPI_ISL_515647, EPI_ISL_515649, EPI_ISL_515650, EPI_ISL_515651, EPI_ISL_515652, EPI_ISL_515653, EPI_ISL_515654, EPI_ISL_515655, EPI_ISL_515656, EPI_ISL_515657, EPI_ISL_515658, EPI_ISL_515659, EPI_ISL_515660, EPI_ISL_515661, EPI_ISL_515662, EPI_ISL_515665, EPI_ISL_515666, EPI_ISL_515668, EPI_ISL_515669, EPI_ISL_515671, EPI_ISL_515672, EPI_ISL_515674, EPI_ISL_515675, EPI_ISL_515676, EPI_ISL_515678, EPI_ISL_515679, EPI_ISL_515680, EPI_ISL_515681, EPI_ISL_515682, EPI_ISL_515683, EPI_ISL_515684, EPI_ISL_515685, EPI_ISL_515686, EPI_ISL_515687, EPI_ISL_515688, EPI_ISL_515689, EPI_ISL_515690, EPI_ISL_515691, EPI_ISL_515692, EPI_ISL_515693, EPI_ISL_515694, EPI_ISL_515696, EPI_ISL_515697, EPI_ISL_515698, EPI_ISL_515699, EPI_ISL_515700, EPI_ISL_515701, EPI_ISL_515702, EPI_ISL_515703, EPI_ISL_515704, EPI_ISL_515705, EPI_ISL_515706, EPI_ISL_515707, EPI_ISL_515708, EPI_ISL_515709, EPI_ISL_515710, EPI_ISL_515711, EPI_ISL_515712, EPI_ISL_515714, EPI_ISL_515715, EPI_ISL_515716, EPI_ISL_515718, EPI_ISL_515719, EPI_ISL_515720, EPI_ISL_515721, EPI_ISL_515722, EPI_ISL_515723, EPI_ISL_515724, EPI_ISL_515725, EPI_ISL_515728, EPI_ISL_515729, EPI_ISL_515730, EPI_ISL_515731, EPI_ISL_515732, EPI_ISL_515733, EPI_ISL_515734, EPI_ISL_515736, EPI_ISL_515737, EPI_ISL_515738, EPI_ISL_515739, EPI_ISL_515740, EPI_ISL_515741, EPI_ISL_515743, EPI_ISL_515744, EPI_ISL_515745, EPI_ISL_515747, EPI_ISL_515748, EPI_ISL_515749, EPI_ISL_515751, EPI_ISL_515752, EPI_ISL_515753, EPI_ISL_515754, EPI_ISL_515756, EPI_ISL_515757, EPI_ISL_515758, EPI_ISL_515759, EPI_ISL_515760, EPI_ISL_515762, EPI_ISL_515763, EPI_ISL_515764, EPI_ISL_515767, EPI_ISL_515769, EPI_ISL_515771, EPI_ISL_515773, EPI_ISL_515775, EPI_ISL_515776, EPI_ISL_515777, EPI_ISL_515778, EPI_ISL_515780, EPI_ISL_515781, EPI_ISL_515782, EPI_ISL_515783, EPI_ISL_515784, EPI_ISL_515787, EPI_ISL_515789, EPI_ISL_515790, EPI_ISL_515791, EPI_ISL_515792, EPI_ISL_515794, EPI_ISL_515796, EPI_ISL_515797, EPI_ISL_515799 | | | |
| see above | NHLS-IALCH | KRISP, KZN Research Innovation and Sequencing Platform | Giandhari J, Pillay S, Lessells R, Mdlalose K, York D, Khan S, Tegally H, Wilkinson E, de Oliveira T |
| EPI_ISL_515801, EPI_ISL_515802, EPI_ISL_515803, EPI_ISL_515804, EPI_ISL_515805, EPI_ISL_515806, EPI_ISL_515807, EPI_ISL_515808, EPI_ISL_515809, EPI_ISL_515810, EPI_ISL_515811, EPI_ISL_515812, EPI_ISL_515813, EPI_ISL_515814, EPI_ISL_515815, EPI_ISL_515816, EPI_ISL_515817, EPI_ISL_515818, EPI_ISL_515819, EPI_ISL_515820, EPI_ISL_515821, EPI_ISL_515822, EPI_ISL_515823, EPI_ISL_515824, EPI_ISL_515825, EPI_ISL_515826, EPI_ISL_515827, EPI_ISL_515828, EPI_ISL_515829, EPI_ISL_515830, EPI_ISL_515831, EPI_ISL_515832, EPI_ISL_515833, EPI_ISL_515835, EPI_ISL_515836, EPI_ISL_515837, EPI_ISL_515838, EPI_ISL_515839, EPI_ISL_515840, EPI_ISL_515841, EPI_ISL_515842, EPI_ISL_515843, EPI_ISL_515844, EPI_ISL_515845, EPI_ISL_515846, EPI_ISL_515847, EPI_ISL_515848, EPI_ISL_515849, EPI_ISL_515850, EPI_ISL_515851, EPI_ISL_515852, EPI_ISL_515853, EPI_ISL_515854, EPI_ISL_515855, EPI_ISL_515856, EPI_ISL_515857, EPI_ISL_515858, EPI_ISL_515859, EPI_ISL_515860, EPI_ISL_515861, EPI_ISL_515862, EPI_ISL_515863, EPI_ISL_515864, EPI_ISL_515865, EPI_ISL_515866, EPI_ISL_515867, EPI_ISL_515868, EPI_ISL_515869, EPI_ISL_515870, EPI_ISL_515871, EPI_ISL_515872, EPI_ISL_515873, EPI_ISL_515874, EPI_ISL_515875, EPI_ISL_515876, EPI_ISL_515877, EPI_ISL_515878, EPI_ISL_515880, EPI_ISL_515881, EPI_ISL_515882, EPI_ISL_515883, EPI_ISL_515884, EPI_ISL_515885, EPI_ISL_515886, EPI_ISL_515887, EPI_ISL_515888, EPI_ISL_515889, EPI_ISL_515890, EPI_ISL_515891, EPI_ISL_515892, EPI_ISL_515893 | | | |

| | | | |
|---|---|---|---|
| see above | Medical Disagnostics Services (MDS) | KRISP, KZN Research Innovation and Sequencing Platform | Giandhari J, Pillay S, Lessells R, ChimukangaraB, Mdlalose K, York D, Khan S, Tegally H, Wilkinson E, de Oliveira T |
| EPI_ISL_515968, EPI_ISL_515969, EPI_ISL_515970, EPI_ISL_515971, EPI_ISL_515972, EPI_ISL_515973 | MIMS | Department of Neurovirology, National Institute of Mental Health and Neuroscience (NIMHANS) | Chitra Pattabiraman,Vijayalakshmi Reddy, Harsha PK, Risha Rasheed, Pramada Prasad, Shafeeq S Hameed, Manjunatha Venkataswamy, Anita Desai, Ravi Vasanthapuram |
| EPI_ISL_516609 | Instituto de Diagnostico y Referencia Epidemiologicos (INDRE) | Instituto de Diagnostico y Referencia Epidemiologicos (INDRE) | Ernesto Ramirez-Gonzalez, Abril Rodriguez-Maldonado, Claudia Wong-Arambula , Natividad Cruz-Ortiz, Tatiana Nunez-Garcia, Dayanira Arellano-Suarez, Adnan Araiza-Rodriguez, Fabiola Garces-Ayala, Lucia Hernandez-Rivas, Irma Lopez-Martinez, Gisela Barrera-Badillo. |
| EPI_ISL_516611 | Instituto de Diagnostico y Referencia Epidemiologicos (INDRE) | Instituto de Diagnostico y Referencia Epidemiologicos (INDRE) | Gisela Barrera-Badillo , Abril Rodriguez-Maldonado, Claudia Wong-Arambula , Natividad Cruz-Ortiz, Tatiana Nunez-Garcia, Dayanira Arellano-Suarez, Fabiola Garces-Ayala, Edgar Mendieta-Condado, Lucia Hernandez-Rivas, Irma Lopez-Martinez, Ernesto Ramirez-Gonzalez. |
| EPI_ISL_516613 | Instituto de Diagnostico y Referencia Epidemiologicos (INDRE) | Instituto de Diagnostico y Referencia Epidemiologicos (INDRE) | Ernesto Ramirez-Gonzalez, Abril Rodriguez-Maldonado, Claudia Wong-Arambula , Natividad Cruz-Ortiz, Tatiana Nunez-Garcia, Dayanira Arellano-Suarez, Adnan Araiza-Rodriguez, Edgar Mendieta-Condado, Lucia Hernandez-Rivas, Irma Lopez-Martinez, Gisela Barrera-Badillo. |
| EPI_ISL_516618 | Instituto de Diagnostico y Referencia Epidemiologicos (INDRE) | Instituto de Diagnostico y Referencia Epidemiologicos (INDRE) | Gisela Barrera-Badillo , Abril Rodriguez-Maldonado, Claudia Wong-Arambula , Natividad Cruz-Ortiz, Tatiana Nunez-Garcia, Dayanira Arellano-Suarez, Fabiola Garces-Ayala, Edgar Mendieta-Condado, Lucia Hernandez-Rivas, Irma Lopez-Martinez, Ernesto Ramirez-Gonzalez. |
| EPI_ISL_516620 | Instituto de Diagnostico y Referencia Epidemiologicos (INDRE) | Instituto de Diagnostico y Referencia Epidemiologicos (INDRE) | Gisela Barrera-Badillo , Abril Rodriguez-Maldonado, Claudia Wong-Arambula , Natividad Cruz-Ortiz, Tatiana Nunez-Garcia, Dayanira Arellano-Suarez, Adnan Araiza-Rodriguez, Edgar Mendieta-Condado, Lucia Hernandez-Rivas, Irma Lopez-Martinez, Ernesto Ramirez-Gonzalez. |
| EPI_ISL_516624, EPI_ISL_516625 | Instituto de Diagnostico y Referencia Epidemiologicos (INDRE) | Instituto de Diagnostico y Referencia Epidemiologicos (INDRE) | Gisela Barrera-Badillo , Abril Rodriguez-Maldonado, Claudia Wong-Arambula , Natividad Cruz-Ortiz, Tatiana Nunez-Garcia, Dayanira Arellano-Suarez, Fabiola Garces-Ayala, Edgar Mendieta-Condado, Lucia Hernandez-Rivas, Irma Lopez-Martinez, Ernesto Ramirez-Gonzalez. |
| EPI_ISL_516748, EPI_ISL_516749, EPI_ISL_516750, EPI_ISL_516752, EPI_ISL_516753, EPI_ISL_516755, EPI_ISL_516756, EPI_ISL_516757, EPI_ISL_516758, EPI_ISL_516762, EPI_ISL_516763, EPI_ISL_516764, EPI_ISL_516765, EPI_ISL_516766, EPI_ISL_516767, EPI_ISL_516768, EPI_ISL_516770, EPI_ISL_516771, EPI_ISL_516772, EPI_ISL_516773, EPI_ISL_516774, EPI_ISL_516775, EPI_ISL_516776, EPI_ISL_516777, EPI_ISL_516778, EPI_ISL_516779 | | | |
| see above | van Bakel Laboratory, Genetics and Genomics Sciences, Icahn School of Medicine at Mount Sinai | van Bakel Laboratory, Genetics and Genomics Sciences, Icahn School of Medicine at Mount Sinai | Andrew G. Letizia, Irene Ramos, Ajay Obla, Carl Goforth, Dawn Weir, Yongchao Ge, Marcas M. Bamman, Jayeeta Dutta, Ethan Ellis, Luis Estrella, Mary-Catherine George, Ana S. Gonzalez-Reiche, Darnell Graham, Adriana van de Guchte, Ramiro Gutierrez, Franca Jones, Aspasia Kalomoiri, Rhonda Lizewski, Stephen Lizewski, Jan Marayag, Nada Marjanovic, Eugene V. Millar, Venugopalan Nair, German Nudelman, Edgar Nunez, Brian Pike, James Regeimbal, Stas Rirak , Ernesto Santa Ana, Rachel S. Gelernter Sealfon, Robert Sebra, Mark Simons, Alessandra Soares-Schanoski, Michael Termini, Sindhu Vangeti, Carlos Williams, Harm van Bakel, Stuart C. Sealfon |
| EPI_ISL_516800 | Rumah Sakit Akademik Universitas Gadjah Mada | Genetics Working Group (Pokja Genetik) Faculty of Medicine, Public Health and Nursing Universitas Gadjah Mada (FK-KMK UGM); Disease Investigation Center Wates Ministry of Agriculture Indonesia; Department of Microbiology FK-KMK UGM; Laboratorium Diagnostik Yayasan Tahija World Mosquito Program (WMP) Yogyakarta Center for Tropical Medicine FK-KMK UGM; Integrated Research center FK-KMK UGM; Department of Computer Science and Electronics FMIPA UGM | Gunadi, Hendra Wibawa, . Marcellus, Mohamad S. Hakim, Edwin W. Daniwijaya, Ludhang P. Rizki, Endah Supriyati, Eggi Arguni, Titik Nuryastuti, Tri Wibawa, Dwi AA Nugrahaningsih, Afiahayati, Siswanto, Alvin S. Kalim, Desyifa Mursalin |
| EPI_ISL_516806 | Rumah Sakit PKU Gamping | Genetics Working Group (Pokja Genetik) Faculty of Medicine, Public Health and Nursing Universitas Gadjah Mada (FK-KMK UGM); Disease Investigation Center Wates Ministry of Agriculture Indonesia; Department of Microbiology FK-KMK UGM; Laboratorium Diagnostik Yayasan Tahija World Mosquito Program (WMP) Yogyakarta Center for Tropical Medicine FK-KMK UGM; Integrated Research center FK-KMK UGM; Department of Computer Science and Electronics FMIPA UGM | Gunadi, Hendra Wibawa, . Marcellus, Mohamad S. Hakim, Edwin W. Daniwijaya, Ludhang P. Rizki, Endah Supriyati, Eggi Arguni, Titik Nuryastuti, Tri Wibawa, Dwi AA Nugrahaningsih, . Afiahayati , . Siswanto, Ardorisye Saptaty Fornia, Kemala Athollah |
| EPI_ISL_516829 | RSUD Nyi Ageng Serang | Genetics Working Group (Pokja Genetik) Faculty of Medicine, Public Health and Nursing Universitas Gadjah Mada (FK-KMK UGM), Disease Investigation Center Wates Ministry of Agriculture Indonesia, Department of Microbiology FK-KMK UGM, Laboratorium Diagnostik Yayasan Tahija World Mosquito Program (WMP) Yogyakarta Center for Tropical Medicine FK-KMK UGM, Integrated Research Center FK-KMK UGM, Department of Computer Science and Electronics FMIPA UGM | Gunadi, Hendra Wibawa, . Marcellus, Mohamad S. Hakim, Edwin W. Daniwijaya, Ludhang P. Rizki, Endah Supriyati, Eggi Arguni, Titik Nuryastuti, Tri Wibawa, Dwi AA Nugrahaningsih, Afiahayati , . Siswanto, Beby Dewi Sartika, Dyah Ayu Puspitarani |
| EPI_ISL_516934, EPI_ISL_516935, EPI_ISL_516938 | Nicolae Testemitanu State University of Medicine and Pharmacy | International Centre for Genetic Engineering and Biotechnology (ICGEB) and ARGO Open Lab Platform for Genome Sequencing | Ulinici M, Licastro D, Dal Monego S, Rajasekharan S, Marcello A |
| EPI_ISL_516976, EPI_ISL_516977, EPI_ISL_516980, EPI_ISL_516981, EPI_ISL_516982, EPI_ISL_516983, EPI_ISL_516984, EPI_ISL_516986 | King Georges Medical University | CSIR-National Botanical Research Institute | Priti Prasad, Shantanu Prakash, Kishan Sahu, Babita Singh, Suruchi Shukla, Hricha Mishra, Danish Nasar Khan , Om Prakash, MLB Bhatt, SK Barik, Mehar H.Asif,Samir V. Sawant,Amita Jain, Sumit Kr. Bag |
| EPI_ISL_517996, EPI_ISL_517997, EPI_ISL_517998, EPI_ISL_517999, EPI_ISL_518001, EPI_ISL_518002, EPI_ISL_518003, EPI_ISL_518004, EPI_ISL_518005, EPI_ISL_518006, EPI_ISL_518007, EPI_ISL_518008, EPI_ISL_518009, EPI_ISL_518010, EPI_ISL_518011, EPI_ISL_518013, EPI_ISL_518014, EPI_ISL_518015, EPI_ISL_518016 | | | |
| see above | Singapore General Hospital | Department of Microbiology | Nurdyana Abdul Rahman, Kun Lee Lim, Chenhao Li, Kian Sing Chan, Lynette Oon, Kern Rei Chng, Niranjan Nagarajan, Karrie Ko |
| EPI_ISL_518034, EPI_ISL_518035, EPI_ISL_518037, EPI_ISL_518038, EPI_ISL_518039, EPI_ISL_518040, EPI_ISL_518041, EPI_ISL_518042, EPI_ISL_518043, EPI_ISL_518044, EPI_ISL_518045, EPI_ISL_518046, EPI_ISL_518047, EPI_ISL_518048, EPI_ISL_518049, EPI_ISL_518050, EPI_ISL_518051, EPI_ISL_518052 | | | |
| see above | NHLS-IALCH | KRISP, KZN Research Innovation and Sequencing Platform | Giandhari J, Pillay S, Lessells R, Mdlalose K, York D, Khan S, Tegally H, Wilkinson E, de Oliveira T |
| EPI_ISL_518819 | Qadr Hospital, Tangerang, Banten | Biosafety Level-3 Laboratory, Indonesian Institute of Sciences (LIPI) | Anik Budhi Dharmayanthi, Syam Budi Iryanto, Andri Wardiana, Anggia Prasetyoputri, Isa Nuryana, Ade Andriani, Ario Betha Juanssilfero, Asep M Ridwanuloh, Ahmad Fathoni, Rifki Sadikin, Ratih Asmana Ningrum, Wien Kusharyoto, Puspita Lisdiyanti |
| EPI_ISL_522350 | KU Leuven, Rega Institute, Clinical and Epidemiological Virology | KU Leuven, Rega Institute, Clinical and Epidemiological Virology | Tony Wawina-Bokalanga, Joan Marti-Carerras, Bert Vanmechelen, Piet Maes |
| EPI_ISL_522874, EPI_ISL_522877 | Instituto Nacional de Medicina Genomica | Instituto Nacional de Medicina Genomica | Hidalgo-Miranda A, Mendoza-Vargas A, Reyes-Grajeda JP, Cisneros-Villanueva M, Cedro-Tanda A, Hurtado-Cordova E, Peñaloza-Figueroa F, Herrera-Montalvo LA |
| EPI_ISL_522878 | Instituto Nacional de Medicina Genomica | Instituto Nacional de Medicina Genomica | Hidalgo-Miranda A, Mendoza-Vargas A, Reyes-Grajeda JP, Cisneros-Villanueva M, Hurtado-Cordova E, Cedro-Tanda A, Peñaloza-Figueroa F, Herrera-Montalvo LA |
| EPI_ISL_522880, EPI_ISL_522940, EPI_ISL_522941, EPI_ISL_522942, EPI_ISL_522978, EPI_ISL_522980, EPI_ISL_522981, EPI_ISL_522982, EPI_ISL_522984, EPI_ISL_522986, EPI_ISL_523500 | | | |
| see above | Instituto Nacional de Medicina Genómica | Instituto Nacional de Medicina Genómica | Hidalgo-Miranda A, Mendoza-Vargas A, Reyes-Grajeda JP, Cisneros-Villanueva M, Cedro-Tanda A, Hurtado-Cordova E, Peñaloza-Figueroa F, Herrera-Montalvo LA |
| EPI_ISL_523824 | Communicable Disease Branch | Hong Kong Department of Health | Mak Gannon C.K., Lam Edman T.K., Chan Rickjason C.W., Tsang Dominic N.C. |

| | | | |
|---|---|---|---|
| EPI_ISL_523825 | United Christian Hospital | Hong Kong Department of Health | Mak Gannon C.K., Lam Edman T.K., Chan Rickjason C.W., Tsang Dominic N.C. |
| EPI_ISL_523826, EPI_ISL_523827 | Queen Mary Hospital | Hong Kong Department of Health | Mak Gannon C.K., Lam Edman T.K., Chan Rickjason C.W., Tsang Dominic N.C. |
| EPI_ISL_523828, EPI_ISL_523829 | Tuen Mun Hospital | Hong Kong Department of Health | Mak Gannon C.K., Lam Edman T.K., Chan Rickjason C.W., Tsang Dominic N.C. |
| EPI_ISL_523830, EPI_ISL_523831 | Our Lady of Maryknoll Hospital | Hong Kong Department of Health | Mak Gannon C.K., Lam Edman T.K., Chan Rickjason C.W., Tsang Dominic N.C. |
| EPI_ISL_523832, EPI_ISL_523834 | Tuen Mun Hospital | Hong Kong Department of Health | Mak Gannon C.K., Lam Edman T.K., Chan Rickjason C.W., Tsang Dominic N.C. |
| EPI_ISL_523835, EPI_ISL_523836 | Queen Elizabeth Hospital | Hong Kong Department of Health | Mak Gannon C.K., Lam Edman T.K., Chan Rickjason C.W., Tsang Dominic N.C. |
| EPI_ISL_523837 | Yan Chai Hospital | Hong Kong Department of Health | Mak Gannon C.K., Lam Edman T.K., Chan Rickjason C.W., Tsang Dominic N.C. |
| EPI_ISL_523838 | Princess Margaret Hospital | Hong Kong Department of Health | Mak Gannon C.K., Lam Edman T.K., Chan Rickjason C.W., Tsang Dominic N.C. |
| EPI_ISL_523839 | Tseung Kwan O Hospital | Hong Kong Department of Health | Mak Gannon C.K., Lam Edman T.K., Chan Rickjason C.W., Tsang Dominic N.C. |
| EPI_ISL_523840 | Our Lady of Maryknoll Hospital | Hong Kong Department of Health | Mak Gannon C.K., Lam Edman T.K., Chan Rickjason C.W., Tsang Dominic N.C. |
| EPI_ISL_523841 | Queen Elizabeth Hospital | Hong Kong Department of Health | Mak Gannon C.K., Lam Edman T.K., Chan Rickjason C.W., Tsang Dominic N.C. |
| EPI_ISL_523842 | Kwong Wah Hospital | Hong Kong Department of Health | Mak Gannon C.K., Lam Edman T.K., Chan Rickjason C.W., Tsang Dominic N.C. |
| EPI_ISL_523843, EPI_ISL_523844, EPI_ISL_523845, EPI_ISL_523846, EPI_ISL_523847 | Queen Elizabeth Hospital | Hong Kong Department of Health | Mak Gannon C.K., Lam Edman T.K., Chan Rickjason C.W., Tsang Dominic N.C. |
| EPI_ISL_523848, EPI_ISL_523849 | United Christian Hospital | Hong Kong Department of Health | Mak Gannon C.K., Lam Edman T.K., Chan Rickjason C.W., Tsang Dominic N.C. |
| EPI_ISL_523850, EPI_ISL_523851, EPI_ISL_523852 | Tseung Kwan O Hospital | Hong Kong Department of Health | Mak Gannon C.K., Lam Edman T.K., Chan Rickjason C.W., Tsang Dominic N.C. |
| EPI_ISL_523853 | Hong Kong Baptist Hospital | Hong Kong Department of Health | Mak Gannon C.K., Lam Edman T.K., Chan Rickjason C.W., Tsang Dominic N.C. |
| EPI_ISL_523854 | Alice Ho Miu Ling Nethersole Hospital | Hong Kong Department of Health | Mak Gannon C.K., Lam Edman T.K., Chan Rickjason C.W., Tsang Dominic N.C. |
| EPI_ISL_523855 | Kwong Wah Hospital | Hong Kong Department of Health | Mak Gannon C.K., Lam Edman T.K., Chan Rickjason C.W., Tsang Dominic N.C. |
| EPI_ISL_523959 | Pronto Socorro Municipal de Perus | Instituto Adolfo Lutz, Interdiciplinary Procedures Center, Strategic Laboratory | Claudio Tavares Sacchi, Claudia Regina Gonçalves, Erica Valessa Ramos Gomes |
| EPI_ISL_524445, EPI_ISL_524447, EPI_ISL_524448, EPI_ISL_524449 | Singapore General Hospital | Department of Microbiology | Nurdyana Abdul Rahman, Kun Lee Lim, Chenhao Li, Kian Sing Chan, Lynette Oon, Kern Rei Chng, Niranjan Nagarajan, Karrie Ko |
| EPI_ISL_524713 | B.J. Medical College and Civil hospital, Ahmedabad | Gujarat Biotechnology Research Centre | Apurvasinh Puvar, Janvi Raval, Zarna Patel, Monika Gandhi, Pinal Trivedi, Maharshi Pandya, Nidhi Patel, Nitin Savaliya, Raghawendra Kumar, Dinesh Kumar, Zuber Saiyed, Komal Patel, Labdhi Pandya, Afzal Ansari, Nikha Trivedi, Pranay Shah, Kamlesh J Upadhyay, Sanjay Kapadia, R D Dixit, A M Kadri, Harsh Bakshi, Chaitanya Joshi, Madhvi Joshi |
| EPI_ISL_524714 | B.J. Medical College and Civil hospital, Ahmedabad | Gujarat Biotechnology Research Centre | Janvi Raval, Zarna Patel, Monika Gandhi, Pinal Trivedi, Maharshi Pandya, Nidhi Patel, Nitin Savaliya, Raghawendra Kumar, Dinesh Kumar, Zuber Saiyed, Komal Patel, Labdhi Pandya, Afzal Ansari, Nikha Trivedi, Pranay Shah, Kamlesh J Upadhyay, Sanjay Kapadia, Apurvasinh Puvar, R D Dixit, A M Kadri, Harsh Bakshi, Chaitanya Joshi, Madhvi Joshi |
| EPI_ISL_524715 | B.J. Medical College and Civil hospital, Ahmedabad | Gujarat Biotechnology Research Centre | Zarna Patel, Monika Gandhi, Pinal Trivedi, Maharshi Pandya, Nidhi Patel, Nitin Savaliya, Raghawendra Kumar, Dinesh Kumar, Zuber Saiyed, Komal Patel, Labdhi Pandya, Afzal Ansari, Nikha Trivedi, Pranay Shah, Kamlesh J Upadhyay, Sanjay Kapadia, Apurvasinh Puvar, Janvi Raval, R D Dixit, A M Kadri, Harsh Bakshi, Chaitanya Joshi, Madhvi Joshi |
| EPI_ISL_524716 | B.J. Medical College and Civil hospital, Ahmedabad | Gujarat Biotechnology Research Centre | Monika Gandhi, Pinal Trivedi, Maharshi Pandya, Nidhi Patel, Nitin Savaliya, Raghawendra Kumar, Dinesh Kumar, Zuber Saiyed, Komal Patel, Labdhi Pandya, Afzal Ansari, Nikha Trivedi, Pranay Shah, Kamlesh J Upadhyay, Sanjay Kapadia, Apurvasinh Puvar, Janvi Raval, Zarna Patel, R D Dixit, A M Kadri, Harsh Bakshi, Chaitanya Joshi, Madhvi Joshi |
| EPI_ISL_524717 | B.J. Medical College and Civil hospital, Ahmedabad | Gujarat Biotechnology Research Centre | Pinal Trivedi, Maharshi Pandya, Nidhi Patel, Nitin Savaliya, Raghawendra Kumar, Dinesh Kumar, Zuber Saiyed, Komal Patel, Labdhi Pandya, Afzal Ansari, Nikha Trivedi, Pranay Shah, Kamlesh J Upadhyay, Sanjay Kapadia, Apurvasinh Puvar, Janvi Raval, Zarna Patel, Monika Gandhi, R D Dixit, A M Kadri, Harsh Bakshi, Chaitanya Joshi, Madhvi Joshi |
| EPI_ISL_524718 | B.J. Medical College and Civil hospital, Ahmedabad | Gujarat Biotechnology Research Centre | Maharshi Pandya, Nidhi Patel, Nitin Savaliya, Raghawendra Kumar, Dinesh Kumar, Zuber Saiyed, Komal Patel, Labdhi Pandya, Afzal Ansari, Nikha Trivedi, Pranay Shah, Kamlesh J Upadhyay, Sanjay Kapadia, Apurvasinh Puvar, Janvi Raval, Zarna Patel, Monika Gandhi, Pinal Trivedi, R D Dixit, A M Kadri, Harsh Bakshi, Chaitanya Joshi, Madhvi Joshi |
| EPI_ISL_524719 | B.J. Medical College and Civil hospital, Ahmedabad | Gujarat Biotechnology Research Centre | Nidhi Patel, Nitin Savaliya, Raghawendra Kumar, Dinesh Kumar, Zuber Saiyed, Komal Patel, Labdhi Pandya, Afzal Ansari, Nikha Trivedi, Pranay Shah, Kamlesh J Upadhyay, Sanjay Kapadia, Apurvasinh Puvar, Janvi Raval, Zarna Patel, Monika Gandhi, Pinal Trivedi, Maharshi Pandya, R D Dixit, A M Kadri, Harsh Bakshi, Chaitanya Joshi, Madhvi Joshi |
| EPI_ISL_524720 | B.J. Medical College and Civil hospital, Ahmedabad | Gujarat Biotechnology Research Centre | Nitin Savaliya, Raghawendra Kumar, Dinesh Kumar, Zuber Saiyed, Komal Patel, Labdhi Pandya, Afzal Ansari, Nikha Trivedi, Pranay Shah, Kamlesh J Upadhyay, Sanjay Kapadia, Apurvasinh Puvar, Janvi Raval, Zarna Patel, Monika Gandhi, Pinal Trivedi, Maharshi Pandya, Nidhi Patel, R D Dixit, A M Kadri, Harsh Bakshi, Chaitanya Joshi, Madhvi Joshi |
| EPI_ISL_524721 | B.J. Medical College and Civil hospital, Ahmedabad | Gujarat Biotechnology Research Centre | Raghawendra Kumar, Dinesh Kumar, Zuber Saiyed, Komal Patel, Labdhi Pandya, Afzal Ansari, Nikha Trivedi, Pranay Shah, Kamlesh J Upadhyay, Sanjay Kapadia, Apurvasinh Puvar, Janvi Raval, Zarna Patel, Monika Gandhi, Pinal Trivedi, Maharshi Pandya, Nidhi Patel, Nitin Savaliya, R D Dixit, A M Kadri, Harsh Bakshi, Chaitanya Joshi, Madhvi Joshi |
| EPI_ISL_524722 | B.J. Medical College and Civil hospital, Ahmedabad | Gujarat Biotechnology Research Centre | Dinesh Kumar, Zuber Saiyed, Komal Patel, Labdhi Pandya, Afzal Ansari, Nikha Trivedi, Pranay Shah, Kamlesh J Upadhyay, Sanjay Kapadia, Apurvasinh Puvar, Janvi Raval, Zarna Patel, Monika Gandhi, Pinal Trivedi, Maharshi Pandya, Nidhi Patel, Nitin Savaliya, Raghawendra Kumar, R D Dixit, A M Kadri, Harsh Bakshi, Chaitanya Joshi, Madhvi Joshi |
| EPI_ISL_524723 | B.J. Medical College and Civil hospital, Ahmedabad | Gujarat Biotechnology Research Centre | Zuber Saiyed, Komal Patel, Labdhi Pandya, Afzal Ansari, Nikha Trivedi, Pranay Shah, Kamlesh J Upadhyay, Sanjay Kapadia, Apurvasinh Puvar, Janvi Raval, Zarna Patel, Monika Gandhi, Pinal Trivedi, Maharshi Pandya, Nidhi Patel, Nitin Savaliya, Raghawendra Kumar, Dinesh Kumar, R D Dixit, A M Kadri, Harsh Bakshi, Chaitanya Joshi, Madhvi Joshi |
| EPI_ISL_524724 | B.J. Medical College and Civil hospital, Ahmedabad | Gujarat Biotechnology Research Centre | Komal Patel, Labdhi Pandya, Afzal Ansari, Nikha Trivedi, Pranay Shah, Kamlesh J Upadhyay, Sanjay Kapadia, Apurvasinh Puvar, Janvi Raval, Zarna Patel, Monika Gandhi, Pinal Trivedi, Maharshi Pandya, Nidhi Patel, Nitin Savaliya, Raghawendra Kumar, Dinesh Kumar, Zuber Saiyed, R D Dixit, A M Kadri, Harsh Bakshi, Chaitanya Joshi, Madhvi Joshi |
| EPI_ISL_524725 | B.J. Medical College and Civil hospital, Ahmedabad | Gujarat Biotechnology Research Centre | Labdhi Pandya, Afzal Ansari, Nikha Trivedi, Pranay Shah, Kamlesh J Upadhyay, Sanjay Kapadia, Apurvasinh Puvar, Janvi Raval, Zarna Patel, Monika Gandhi, Pinal Trivedi, Maharshi Pandya, Nidhi Patel, Nitin Savaliya, Raghawendra Kumar, Dinesh Kumar, Zuber Saiyed, Komal Patel, R D Dixit, A M Kadri, Harsh Bakshi, Chaitanya Joshi, Madhvi Joshi |
| EPI_ISL_524726 | B.J. Medical College and Civil hospital, Ahmedabad | Gujarat Biotechnology Research Centre | Afzal Ansari, Nikha Trivedi, Pranay Shah, Kamlesh J Upadhyay, Sanjay Kapadia, Apurvasinh Puvar, Janvi Raval, Zarna Patel, Monika Gandhi, Pinal Trivedi, Maharshi Pandya, Nidhi Patel, Nitin Savaliya, Raghawendra Kumar, Dinesh Kumar, Zuber Saiyed, Komal Patel, Labdhi Pandya, R D Dixit, A M Kadri, Harsh Bakshi, Chaitanya Joshi, Madhvi Joshi |
| EPI_ISL_524727 | B.J. Medical College and Civil hospital, Ahmedabad | Gujarat Biotechnology Research Centre | Nikha Trivedi, Pranay Shah, Kamlesh J Upadhyay, Sanjay Kapadia, Apurvasinh Puvar, Janvi Raval, Zarna Patel, Monika Gandhi, Pinal Trivedi, Maharshi Pandya, Nidhi Patel, Nitin Savaliya, Raghawendra Kumar, Dinesh Kumar, Zuber Saiyed, Komal Patel, Labdhi Pandya, Afzal Ansari, R D Dixit, A M Kadri, Harsh Bakshi, Chaitanya Joshi, Madhvi Joshi |

| | | | |
|---|---|---|---|
| EPI_ISL_524728 | B.J. Medical College and Civil hospital, Ahmedabad | Gujarat Biotechnology Research Centre | Pranay Shah, Kamlesh J Upadhyay, Sanjay Kapadia, Apurvasinh Puvar, Janvi Raval, Zarna Patel, Monika Gandhi, Pinal Trivedi, Maharshi Pandya, Nidhi Patel, Nitin Savaliya, Raghawendra Kumar, Dinesh Kumar, Zuber Saiyed, Komal Patel, Labdhi Pandya, Afzal Ansari, Nikha Trivedi, R D Dixit, A M Kadri, Harsh Bakshi, Chaitanya Joshi, Madhvi Joshi |
| EPI_ISL_524729 | B.J. Medical College and Civil hospital, Ahmedabad | Gujarat Biotechnology Research Centre | Kamlesh J Upadhyay, Sanjay Kapadia, Apurvasinh Puvar, Janvi Raval, Zarna Patel, Monika Gandhi, Pinal Trivedi, Maharshi Pandya, Nidhi Patel, Nitin Savaliya, Raghawendra Kumar, Dinesh Kumar, Zuber Saiyed, Komal Patel, Labdhi Pandya, Afzal Ansari, Nikha Trivedi, Pranay Shah, R D Dixit, A M Kadri, Harsh Bakshi, Chaitanya Joshi, Madhvi Joshi |
| EPI_ISL_524730 | GMERS Medical College & Hospital, Gotri, Vadodara | Gujarat Biotechnology Research Centre | Meenakshi Shah, Neena Doshi, Varsha Godbole, Apurvasinh Puvar, Janvi Raval, Zarna Patel, Monika Gandhi, Pinal Trivedi, Maharshi Pandya, Nidhi Patel, Nitin Savaliya, Raghawendra Kumar, Dinesh Kumar, Zuber Saiyed, Komal Patel, Labdhi Pandya, Afzal Ansari, Nikha Trivedi, R D Dixit, A M Kadri, Harsh Bakshi, Chaitanya Joshi, Madhvi Joshi |
| EPI_ISL_524731 | GMERS Medical College & Hospital, Gotri, Vadodara | Gujarat Biotechnology Research Centre | Neena Doshi, Varsha Godbole, Apurvasinh Puvar, Janvi Raval, Zarna Patel, Monika Gandhi, Pinal Trivedi, Maharshi Pandya, Nidhi Patel, Nitin Savaliya, Raghawendra Kumar, Dinesh Kumar, Zuber Saiyed, Komal Patel, Labdhi Pandya, Afzal Ansari, Nikha Trivedi, Meenakshi Shah, R D Dixit, A M Kadri, Harsh Bakshi, Chaitanya Joshi, Madhvi Joshi |
| EPI_ISL_524732 | Dr. RSS Hospital, Modasa | Gujarat Biotechnology Research Centre | Harsh Chaudhari, Apurvasinh Puvar, Janvi Raval, Zarna Patel, Monika Gandhi, Pinal Trivedi, Maharshi Pandya, Nidhi Patel, Nitin Savaliya, Raghawendra Kumar, Dinesh Kumar, Zuber Saiyed, Komal Patel, Labdhi Pandya, Afzal Ansari, Nikha Trivedi, R D Dixit, A M Kadri, Harsh Bakshi, Chaitanya Joshi, Madhvi Joshi, Neena Doshi, Varsha Godbole |
| EPI_ISL_524733 | GMERS Medical College and Hospital, Dharpur, Patan | Gujarat Biotechnology Research Centre | A N Parmar, Apurvasinh Puvar, Janvi Raval, Zarna Patel, Monika Gandhi, Pinal Trivedi, Maharshi Pandya, Nidhi Patel, Nitin Savaliya, Raghawendra Kumar, Dinesh Kumar, Zuber Saiyed, Komal Patel, Labdhi Pandya, Afzal Ansari, Nikha Trivedi, R D Dixit, A M Kadri, Harsh Bakshi, Chaitanya Joshi, Madhvi Joshi |
| EPI_ISL_524734 | GMERS Medical College and Hospital, Dharpur, Patan | Gujarat Biotechnology Research Centre | Apurvasinh Puvar, Janvi Raval, Zarna Patel, Monika Gandhi, Pinal Trivedi, Maharshi Pandya, Nidhi Patel, Nitin Savaliya, Raghawendra Kumar, Dinesh Kumar, Zuber Saiyed, Komal Patel, Labdhi Pandya, Afzal Ansari, Nikha Trivedi, A N Parmar, R D Dixit, A M Kadri, Harsh Bakshi, Chaitanya Joshi, Madhvi Joshi |
| EPI_ISL_524735 | Government General Hospital, Jam Khambhaliya, Devbhoomi Dwarka | Gujarat Biotechnology Research Centre | Harish Matani, Janvi Raval, Zarna Patel, Monika Gandhi, Pinal Trivedi, Maharshi Pandya, Nidhi Patel, Nitin Savaliya, Raghawendra Kumar, Dinesh Kumar, Zuber Saiyed, Komal Patel, Labdhi Pandya, Afzal Ansari, Nikha Trivedi, Apurvasinh Puvar, R D Dixit, A M Kadri, Harsh Bakshi, Chaitanya Joshi, Madhvi Joshi |
| EPI_ISL_524736 | Government General Hospital, Jam Khambhaliya, Devbhoomi Dwarka | Gujarat Biotechnology Research Centre | Janvi Raval, Zarna Patel, Monika Gandhi, Pinal Trivedi, Maharshi Pandya, Nidhi Patel, Nitin Savaliya, Raghawendra Kumar, Dinesh Kumar, Zuber Saiyed, Komal Patel, Labdhi Pandya, Afzal Ansari, Nikha Trivedi, Harish Matani, Apurvasinh Puvar, R D Dixit, A M Kadri, Harsh Bakshi, Chaitanya Joshi, Madhvi Joshi |
| EPI_ISL_524737 | GMERS Medical College and Hospital, Gandhinagar | Gujarat Biotechnology Research Centre | Zarna Patel, Monika Gandhi, Pinal Trivedi, Maharshi Pandya, Nidhi Patel, Nitin Savaliya, Raghawendra Kumar, Dinesh Kumar, Zuber Saiyed, Komal Patel, Labdhi Pandya, Afzal Ansari, Nikha Trivedi, Seema Bhatt, Gaurishankar Shrimali, Bhavesh Modi, Bharti Rajani, Apurvasinh Puvar, Janvi Raval, R D Dixit, A M Kadri, Harsh Bakshi, Chaitanya Joshi, Madhvi Joshi |
| EPI_ISL_524738 | GMERS Medical College and Hospital, Gandhinagar | Gujarat Biotechnology Research Centre | Monika Gandhi, Pinal Trivedi, Maharshi Pandya, Nidhi Patel, Nitin Savaliya, Raghawendra Kumar, Dinesh Kumar, Zuber Saiyed, Komal Patel, Labdhi Pandya, Afzal Ansari, Nikha Trivedi, Seema Bhatt, Gaurishankar Shrimali, Bhavesh Modi, Bharti Rajani, Apurvasinh Puvar, Janvi Raval, Zarna Patel, R D Dixit, A M Kadri, Harsh Bakshi, Chaitanya Joshi, Madhvi Joshi |
| EPI_ISL_524739 | GMERS Medical College and Hospital, Gandhinagar | Gujarat Biotechnology Research Centre | Pinal Trivedi, Maharshi Pandya, Nidhi Patel, Nitin Savaliya, Raghawendra Kumar, Dinesh Kumar, Zuber Saiyed, Komal Patel, Labdhi Pandya, Afzal Ansari, Nikha Trivedi, Seema Bhatt, Gaurishankar Shrimali, Bhavesh Modi, Bharti Rajani, Apurvasinh Puvar, Janvi Raval, Zarna Patel, Monika Gandhi, R D Dixit, A M Kadri, Harsh Bakshi, Chaitanya Joshi, Madhvi Joshi |
| EPI_ISL_524740 | GMERS Medical College and Hospital, Gandhinagar | Gujarat Biotechnology Research Centre | Maharshi Pandya, Nidhi Patel, Nitin Savaliya, Raghawendra Kumar, Dinesh Kumar, Zuber Saiyed, Komal Patel, Labdhi Pandya, Afzal Ansari, Nikha Trivedi, Seema Bhatt, Gaurishankar Shrimali, Bhavesh Modi, Bharti Rajani, Apurvasinh Puvar, Janvi Raval, Zarna Patel, Monika Gandhi, Pinal Trivedi, R D Dixit, A M Kadri, Harsh Bakshi, Chaitanya Joshi, Madhvi Joshi |
| EPI_ISL_524741 | GMERS Medical College and Hospital, Gandhinagar | Gujarat Biotechnology Research Centre | Nidhi Patel, Nitin Savaliya, Raghawendra Kumar, Dinesh Kumar, Zuber Saiyed, Komal Patel, Labdhi Pandya, Afzal Ansari, Nikha Trivedi, Seema Bhatt, Gaurishankar Shrimali, Bhavesh Modi, Bharti Rajani, Apurvasinh Puvar, Janvi Raval, Zarna Patel, Monika Gandhi, Pinal Trivedi, Maharshi Pandya, R D Dixit, A M Kadri, Harsh Bakshi, Chaitanya Joshi, Madhvi Joshi |
| EPI_ISL_524742 | GMERS Medical College and Hospital, Gandhinagar | Gujarat Biotechnology Research Centre | Dinesh Kumar, Zuber Saiyed, Komal Patel, Labdhi Pandya, Afzal Ansari, Nikha Trivedi, Seema Bhatt, Gaurishankar Shrimali, Bhavesh Modi, Bharti Rajani, Apurvasinh Puvar, Janvi Raval, Zarna Patel, Monika Gandhi, Pinal Trivedi, Maharshi Pandya, Nidhi Patel, Nitin Savaliya, Raghawendra Kumar, R D Dixit, A M Kadri, Harsh Bakshi, Chaitanya Joshi, Madhvi Joshi |
| EPI_ISL_524743 | GMERS Medical College and Hospital, Gandhinagar | Gujarat Biotechnology Research Centre | Zuber Saiyed, Komal Patel, Labdhi Pandya, Afzal Ansari, Nikha Trivedi, Seema Bhatt, Gaurishankar Shrimali, Bhavesh Modi, Bharti Rajani, Apurvasinh Puvar, Janvi Raval, Zarna Patel, Monika Gandhi, Pinal Trivedi, Maharshi Pandya, Nidhi Patel, Nitin Savaliya, Raghawendra Kumar, Dinesh Kumar, R D Dixit, A M Kadri, Harsh Bakshi, Chaitanya Joshi, Madhvi Joshi |
| EPI_ISL_524744 | GMERS Medical College and Hospital, Gandhinagar | Gujarat Biotechnology Research Centre | Komal Patel, Labdhi Pandya, Afzal Ansari, Nikha Trivedi, Seema Bhatt, Gaurishankar Shrimali, Bhavesh Modi, Bharti Rajani, Apurvasinh Puvar, Janvi Raval, Zarna Patel, Monika Gandhi, Pinal Trivedi, Maharshi Pandya, Nidhi Patel, Nitin Savaliya, Raghawendra Kumar, Dinesh Kumar, Zuber Saiyed, R D Dixit, A M Kadri, Harsh Bakshi, Chaitanya Joshi, Madhvi Joshi |
| EPI_ISL_524745 | GMERS Medical College and Hospital, Gandhinagar | Gujarat Biotechnology Research Centre | Labdhi Pandya, Afzal Ansari, Nikha Trivedi, Seema Bhatt, Gaurishankar Shrimali, Bhavesh Modi, Bharti Rajani, Apurvasinh Puvar, Janvi Raval, Zarna Patel, Monika Gandhi, Pinal Trivedi, Maharshi Pandya, Nidhi Patel, Nitin Savaliya, Raghawendra Kumar, Dinesh Kumar, Zuber Saiyed, Komal Patel, R D Dixit, A M Kadri, Harsh Bakshi, Chaitanya Joshi, Madhvi Joshi |
| EPI_ISL_524746 | Sir M P Shah Government Medical College, Jamnagar | Gujarat Biotechnology Research Centre | Binita Aring, Afzal Ansari, Nikha Trivedi, Apurvasinh Puvar, Janvi Raval, Zarna Patel, Monika Gandhi, Pinal Trivedi, Maharshi Pandya, Nidhi Patel, Nitin Savaliya, Raghawendra Kumar, Dinesh Kumar, Zuber Saiyed, Komal Patel, Labdhi Pandya, R D Dixit, A M Kadri, Harsh Bakshi, Chaitanya Joshi, Madhvi Joshi |
| EPI_ISL_524747 | Sir M P Shah Government Medical College, Jamnagar | Gujarat Biotechnology Research Centre | Afzal Ansari, Nikha Trivedi, Binita Aring, Apurvasinh Puvar, Janvi Raval, Zarna Patel, Monika Gandhi, Pinal Trivedi, Maharshi Pandya, Nidhi Patel, Nitin Savaliya, Raghawendra Kumar, Dinesh Kumar, Zuber Saiyed, Komal Patel, Labdhi Pandya, R D Dixit, A M Kadri, Harsh Bakshi, Chaitanya Joshi, Madhvi Joshi |
| EPI_ISL_524748 | Sir M P Shah Government Medical College, Jamnagar | Gujarat Biotechnology Research Centre | Nikha Trivedi, Apurvasinh Puvar, Janvi Raval, Binita Aring, Zarna Patel, Monika Gandhi, Pinal Trivedi, Maharshi Pandya, Nidhi Patel, Nitin Savaliya, Raghawendra Kumar, Dinesh Kumar, Zuber Saiyed, Komal Patel, Labdhi Pandya, Afzal Ansari, R D Dixit, A M Kadri, Harsh Bakshi, Chaitanya Joshi, Madhvi Joshi |
| EPI_ISL_524749 | Sir M P Shah Government Medical College, Jamnagar | Gujarat Biotechnology Research Centre | Apurvasinh Puvar, Janvi Raval, Zarna Patel, Monika Gandhi, Pinal Trivedi, Maharshi Pandya, Nidhi Patel, Nitin Savaliya, Raghawendra Kumar, Dinesh Kumar, Zuber Saiyed, Komal Patel, Labdhi Pandya, Afzal Ansari, Nikha Trivedi, Binita Aring, R D Dixit, A M Kadri, Harsh Bakshi, Chaitanya Joshi, Madhvi Joshi |
| EPI_ISL_524750 | Sir M P Shah Government Medical College, Jamnagar | Gujarat Biotechnology Research Centre | Janvi Raval, Zarna Patel, Monika Gandhi, Pinal Trivedi, Maharshi Pandya, Nidhi Patel, Nitin Savaliya, Raghawendra Kumar, Dinesh Kumar, Zuber Saiyed, Komal Patel, Labdhi Pandya, Afzal Ansari, Nikha Trivedi, Binita Aring, Apurvasinh Puvar, R D Dixit, A M Kadri, Harsh Bakshi, Chaitanya Joshi, Madhvi Joshi |
| EPI_ISL_524751 | Sir M P Shah Government Medical College, Jamnagar | Gujarat Biotechnology Research Centre | Zarna Patel, Monika Gandhi, Pinal Trivedi, Maharshi Pandya, Nidhi Patel, Nitin Savaliya, Raghawendra Kumar, Dinesh Kumar, Zuber Saiyed, Komal Patel, Labdhi Pandya, Afzal Ansari, Nikha Trivedi, Binita Aring, Apurvasinh Puvar, Janvi Raval, R D Dixit, A M Kadri, Harsh Bakshi, Chaitanya Joshi, Madhvi Joshi |
| EPI_ISL_524752 | GMERS Medical College and Hospital, Dharpur, Patan | Gujarat Biotechnology Research Centre | Monika Gandhi, Pinal Trivedi, Maharshi Pandya, Nidhi Patel, Nitin Savaliya, Raghawendra Kumar, Dinesh Kumar, Zuber Saiyed, Komal Patel, Labdhi Pandya, Afzal Ansari, Nikha Trivedi, A N Parmar, Apurvasinh Puvar, Janvi Raval, Zarna Patel, R D Dixit, A M Kadri, Harsh Bakshi, Chaitanya Joshi, Madhvi Joshi |
| EPI_ISL_524753 | GMERS Medical College and Hospital, Dharpur, Patan | Gujarat Biotechnology Research Centre | Pinal Trivedi, Maharshi Pandya, Nidhi Patel, Nitin Savaliya, Raghawendra Kumar, Dinesh Kumar, Zuber Saiyed, Komal Patel, Labdhi Pandya, Afzal Ansari, Nikha Trivedi, A N Parmar, Apurvasinh Puvar, Janvi Raval, Zarna Patel, Monika Gandhi, R D Dixit, A M Kadri, Harsh Bakshi, Chaitanya Joshi, Madhvi Joshi |
| EPI_ISL_524754 | GMERS Medical College and Hospital, Dharpur, Patan | Gujarat Biotechnology Research Centre | Maharshi Pandya, Nidhi Patel, Nitin Savaliya, Raghawendra Kumar, Dinesh Kumar, Zuber Saiyed, Komal Patel, Labdhi Pandya, Afzal Ansari, Nikha Trivedi, A N Parmar, Apurvasinh Puvar, Janvi Raval, Zarna Patel, Monika Gandhi, Pinal Trivedi, R D Dixit, A M Kadri, Harsh Bakshi, Chaitanya Joshi, Madhvi Joshi |
| EPI_ISL_524755 | GMERS Medical College and Hospital, Dharpur, Patan | Gujarat Biotechnology Research Centre | Nidhi Patel, Nitin Savaliya, Raghawendra Kumar, Dinesh Kumar, Zuber Saiyed, Komal Patel, Labdhi Pandya, Afzal Ansari, Nikha Trivedi, A N Parmar, |

| | | | |
|---|---|---|---|
| | | | Apurvasinh Puvar, Janvi Raval, Zarna Patel, Monika Gandhi, Pinal Trivedi, Maharshi Pandya, R D Dixit, A M Kadri, Harsh Bakshi, Chaitanya Joshi, Madhvi Joshi |
| EPI_ISL_524756 | GMERS Medical College and Hospital, Dharpur, Patan | Gujarat Biotechnology Research Centre | Nitin Savaliya, Raghawendra Kumar, Dinesh Kumar, Zuber Saiyed, Komal Patel, Labdhi Pandya, Afzal Ansari, Nikha Trivedi, A N Parmar, Apurvasinh Puvar, Janvi Raval, Zarna Patel, Monika Gandhi, Pinal Trivedi, Maharshi Pandya, Nidhi Patel, R D Dixit, A M Kadri, Harsh Bakshi, Chaitanya Joshi, Madhvi Joshi |
| EPI_ISL_524757 | GMERS Medical College and Hospital, Dharpur, Patan | Gujarat Biotechnology Research Centre | Raghawendra Kumar, Dinesh Kumar, Zuber Saiyed, Komal Patel, Labdhi Pandya, Afzal Ansari, Nikha Trivedi, A N Parmar, Apurvasinh Puvar, Janvi Raval, Zarna Patel, Monika Gandhi, Pinal Trivedi, Maharshi Pandya, Nidhi Patel, Nitin Savaliya, R D Dixit, A M Kadri, Harsh Bakshi, Chaitanya Joshi, Madhvi Joshi |
| EPI_ISL_524758 | GMERS Medical College and Hospital, Dharpur, Patan | Gujarat Biotechnology Research Centre | Dinesh Kumar, Zuber Saiyed, Komal Patel, Labdhi Pandya, Afzal Ansari, Nikha Trivedi, A N Parmar, Apurvasinh Puvar, Janvi Raval, Zarna Patel, Monika Gandhi, Pinal Trivedi, Maharshi Pandya, Nidhi Patel, Nitin Savaliya, Raghawendra Kumar, R D Dixit, A M Kadri, Harsh Bakshi, Chaitanya Joshi, Madhvi Joshi |
| EPI_ISL_524759 | GMERS Medical College and Hospital, Dharpur, Patan | Gujarat Biotechnology Research Centre | Zuber Saiyed, Komal Patel, Labdhi Pandya, Afzal Ansari, Nikha Trivedi, A N Parmar, Apurvasinh Puvar, Janvi Raval, Zarna Patel, Monika Gandhi, Pinal Trivedi, Maharshi Pandya, Nidhi Patel, Nitin Savaliya, Raghawendra Kumar, Dinesh Kumar, R D Dixit, A M Kadri, Harsh Bakshi, Chaitanya Joshi, Madhvi Joshi |
| EPI_ISL_524760 | GMERS Medical College and Hospital, Dharpur, Patan | Gujarat Biotechnology Research Centre | Komal Patel, Labdhi Pandya, Afzal Ansari, Nikha Trivedi, A N Parmar, Apurvasinh Puvar, Janvi Raval, Zarna Patel, Monika Gandhi, Pinal Trivedi, Maharshi Pandya, Nidhi Patel, Nitin Savaliya, Raghawendra Kumar, Dinesh Kumar, Zuber Saiyed, R D Dixit, A M Kadri, Harsh Bakshi, Chaitanya Joshi, Madhvi Joshi |
| EPI_ISL_524761 | GMERS Medical College and Hospital, Dharpur, Patan | Gujarat Biotechnology Research Centre | Labdhi Pandya, Afzal Ansari, Nikha Trivedi, A N Parmar, Apurvasinh Puvar, Janvi Raval, Zarna Patel, Monika Gandhi, Pinal Trivedi, Maharshi Pandya, Nidhi Patel, Nitin Savaliya, Raghawendra Kumar, Dinesh Kumar, Zuber Saiyed, Komal Patel, R D Dixit, A M Kadri, Harsh Bakshi, Chaitanya Joshi, Madhvi Joshi |
| EPI_ISL_524762 | GMERS Medical College and Hospital, Dharpur, Patan | Gujarat Biotechnology Research Centre | Afzal Ansari, Nikha Trivedi, A N Parmar, Apurvasinh Puvar, Janvi Raval, Zarna Patel, Monika Gandhi, Pinal Trivedi, Maharshi Pandya, Nidhi Patel, Nitin Savaliya, Raghawendra Kumar, Dinesh Kumar, Zuber Saiyed, Komal Patel, Labdhi Pandya, R D Dixit, A M Kadri, Harsh Bakshi, Chaitanya Joshi, Madhvi Joshi |
| EPI_ISL_524763 | GMERS Medical College and Hospital, Dharpur, Patan | Gujarat Biotechnology Research Centre | Nikha Trivedi, A N Parmar, Apurvasinh Puvar, Janvi Raval, Zarna Patel, Monika Gandhi, Pinal Trivedi, Maharshi Pandya, Nidhi Patel, Nitin Savaliya, Raghawendra Kumar, Dinesh Kumar, Zuber Saiyed, Komal Patel, Labdhi Pandya, Afzal Ansari, R D Dixit, A M Kadri, Harsh Bakshi, Chaitanya Joshi, Madhvi Joshi |
| EPI_ISL_524764 | GMERS Medical College and Hospital, Dharpur, Patan | Gujarat Biotechnology Research Centre | A N Parmar, Apurvasinh Puvar, Janvi Raval, Zarna Patel, Monika Gandhi, Pinal Trivedi, Maharshi Pandya, Nidhi Patel, Nitin Savaliya, Raghawendra Kumar, Dinesh Kumar, Zuber Saiyed, Komal Patel, Labdhi Pandya, Afzal Ansari, Nikha Trivedi, R D Dixit, A M Kadri, Harsh Bakshi, Chaitanya Joshi, Madhvi Joshi |
| EPI_ISL_524765 | GMERS Medical College and Hospital, Dharpur, Patan | Gujarat Biotechnology Research Centre | Apurvasinh Puvar, Janvi Raval, Zarna Patel, Monika Gandhi, Pinal Trivedi, Maharshi Pandya, Nidhi Patel, Nitin Savaliya, Raghawendra Kumar, Dinesh Kumar, Zuber Saiyed, Komal Patel, Labdhi Pandya, Afzal Ansari, Nikha Trivedi, A N Parmar, R D Dixit, A M Kadri, Harsh Bakshi, Chaitanya Joshi, Madhvi Joshi |
| EPI_ISL_524766 | GMERS Medical College and Hospital, Dharpur, Patan | Gujarat Biotechnology Research Centre | Janvi Raval, Zarna Patel, Monika Gandhi, Pinal Trivedi, Maharshi Pandya, Nidhi Patel, Nitin Savaliya, Raghawendra Kumar, Dinesh Kumar, Zuber Saiyed, Komal Patel, Labdhi Pandya, Afzal Ansari, Nikha Trivedi, Apurvasinh Puvar, A N Parmar, R D Dixit, A M Kadri, Harsh Bakshi, Chaitanya Joshi, Madhvi Joshi |
| EPI_ISL_525419 | GMERS Medical College and Hospital, Gandhinagar | Gujarat Biotechnology Research Centre | Nitin Savaliya, Raghawendra Kumar, Dinesh Kumar, Zuber Saiyed, Komal Patel, Labdhi Pandya, Afzal Ansari, Nikha Trivedi, Seema Bhatt, Gaurishankar Shrimali, Bhavesh Modi, Bharti Rajani, Apurvasinh Puvar, Janvi Raval, Zarna Patel, Monika Gandhi, Pinal Trivedi, Maharshi Pandya, Nidhi Patel, R D Dixit, A M Kadri, Harsh Bakshi, Chaitanya Joshi, Madhvi Joshi |
| EPI_ISL_525420 | GMERS Medical College and Hospital, Gandhinagar | Gujarat Biotechnology Research Centre | Raghawendra Kumar, Dinesh Kumar, Zuber Saiyed, Komal Patel, Labdhi Pandya, Afzal Ansari, Nikha Trivedi, Seema Bhatt, Gaurishankar Shrimali, Bhavesh Modi, Bharti Rajani, Apurvasinh Puvar, Janvi Raval, Zarna Patel, Monika Gandhi, Pinal Trivedi, Maharshi Pandya, Nidhi Patel, Nitin Savaliya, R D Dixit, A M Kadri, Harsh Bakshi, Chaitanya Joshi, Madhvi Joshi |
| EPI_ISL_525421 | B.J. Medical College and Civil hospital, Ahmedabad | Gujarat Biotechnology Research Centre | Zarna Patel, Monika Gandhi, Pinal Trivedi, Maharshi Pandya, Nidhi Patel, Nitin Savaliya, Raghawendra Kumar, Dinesh Kumar, Zuber Saiyed, Komal Patel, Labdhi Pandya, Afzal Ansari, Nikha Trivedi, Pranay Shah, Kamlesh J Upadhyay, Sanjay Kapadia, Apurvasinh Puvar, Janvi Raval, R D Dixit, A M Kadri, Harsh Bakshi, Chaitanya Joshi, Madhvi Joshi |
| EPI_ISL_525422 | B.J. Medical College and Civil hospital, Ahmedabad | Gujarat Biotechnology Research Centre | Monika Gandhi, Pinal Trivedi, Maharshi Pandya, Nidhi Patel, Nitin Savaliya, Raghawendra Kumar, Dinesh Kumar, Zuber Saiyed, Komal Patel, Labdhi Pandya, Afzal Ansari, Nikha Trivedi, Pranay Shah, Kamlesh J Upadhyay, Sanjay Kapadia, Apurvasinh Puvar, Janvi Raval, Zarna Patel, R D Dixit, A M Kadri, Harsh Bakshi, Chaitanya Joshi, Madhvi Joshi |
| EPI_ISL_525430 | Institute of Microbiology, Universidad San Francisco de Quito | Institute of Microbiology, Universidad San Francisco de Quito | Juan José Guadalupe, Monica Becerra-Wong, Belén Prado-Vivar, Sully Márquez, Bernardo Gutiérrez, Verónica Barragán, Patricio Rojas-Silva, Gabriel Trueba, Michelle Grunauer, Paúl Cárdenas |
| EPI_ISL_525481 | Centre for Dengue Research | Centre for Dengue Research | Chandima Jeewandara, Deshni Jayathilaka, Dinuka Ariyaratne, Laksiri Gomes, Diyanath Ranasinghe, Ananda Wijewickrama, Eranga Narangoda, Damayanthi Idampitiya, Gathsaurie Neelika Malavige |
| EPI_ISL_525486, EPI_ISL_525489 | Centre for Dengue Research | Centre for Dengue Research | Chandima Jeewandara, Deshni Jayathilaka, Dinuka Ariyaratne, Laksiri Gomes, Diyanath Ranasinghe, Ananda Wijewickrama, Malika Karunaratne, Eranga Narangoda, Damayanthi Idampitiya, Gathsaurie Neelika Malavige |
| EPI_ISL_525492 | RSUP dr. SOERADJI TIRTONEGORO | Genetics Working Group (Pokja Genetik) Faculty of Medicine, Public Health and Nursing Universitas Gadjah Mada (FK-KMK UGM), Disease Investigation Center Wates Ministry of Agriculture Indonesia, Department of Microbiology FK-KMK UGM, Laboratorium Diagnostik Yayasan Tahija World Mosquito Program (WMP) Yogyakarta Center for Tropical Medicine FK-KMK UGM, Integrated Research Center FK-KMK UGM, Department of Computer Science and Electronics FMIPA UGM | Gunadi, Hendra Wibawa, . Marcellus, Mohamad S. Hakim, Edwin W. Daniwijaya, Ludhang P. Rizki, Endah Supriyati, Eggi Arguni, Titik Nuryastuti, Tri Wibawa, Dwi AA Nugrahaningsih, Afiahayati , . Siswanto, Kurniyanto, Indah Juliana, Alvin S. Kalim, Dwiki Afandy |
| EPI_ISL_525495, EPI_ISL_525496 | Laboratory of Molecular Virology of the International Centre for Genetic Engineering and Biotechnology (ICGEB) | ARGO Open Lab Platform for Genome Sequencing | Licastro D, Rajasekharan S, Dal Monego S, Segat L, D'Agaro P, Marcello A |
| EPI_ISL_525572 | Istituto Zooprofilattico Sperimentale Puglia e Basilicata; Dipartimento di Bioscienze, Biotecnologie e Biofarmaceutica dell'Università degli Studi di Bari "A.Moro"; Istituto di Biomembrane. Bioenergetica e Biotecnologie Molecolari del Consiglio Nazionale delle Ricerche di Bari | Beaconlab (Bioinformatics, Evolution and Comparative Genomics lab), Dept of Biosciences, University on Milan | Parisi A.,Pesole G., Manzari C., Chiara M |
| EPI_ISL_526223, EPI_ISL_526224, EPI_ISL_526225 | Hungarian Defence Forces Military Medical Centre | National Laboratory of Virology, Szentágothai Research Centre | Endre Gábor Tóth, Balázs Somogyi, Bálint Eszenyi, Ferenc Jakab, Gábor Kemenesi |
| EPI_ISL_526958, EPI_ISL_526967, EPI_ISL_526971 | Instituto Nacional de Salud, Bogotá, Colombia | Instituto Nacional de Salud, Bogotá, Colombia | Katherine Laiton-Donato, Diego A. Álvarez-Díaz, Carlos Franco-Muñoz, Mauricio Pacheco-Montealegre, Jonathan Reales, Diego Andrés Prada, Jose A. Usme-Ciro, Zulma M. Cucunubá, Christian Julian VillabonaArenas, Liz Villabona-Arenas, Sussy Echeverria, Astrid C. Flórez, Carolina Ferro, Diana Marcela Walteros-Acero, Franklin Prieto, Carlos Andrés Durán, Martha Lucia Ospina Martinez, Marcela Mercado-Reyes |
| EPI_ISL_527742 | Centro Nacional De Rehabilitacion Humberto Araya Rojas (Cenare) | Inciensa, Instituto Costarricense de Investigación y Enseñanza en Nutrición y Salud | Francisco Duarte, Hebleen Porras, Claudio Soto-Garita, Estela Cordero, Adriana Godinez & Melany Calderon |
| EPI_ISL_527753 | Hospital San Vicente De Paul | Inciensa, Instituto Costarricense de Investigación y Enseñanza en Nutrición y Salud | Francisco Duarte, Hebleen Porras, Claudio Soto-Garita, Estela Cordero, Adriana Godinez & Melany Calderon |
| EPI_ISL_527818 | Centro de Investigaciones, Universidad de Especialidades Espíritu Santo | Institute of Microbiology, Universidad San Francisco de Quito | Derly Andrade, Juan Carlos Fernandez, Belén Prado-Vivar, Sully Márquez, Juan José Guadalupe, Monica Becerra-Wong, Bernardo Gutiérrez, Gabriel Morey, Ruben Armas, Jose Pedro Barberan, Fernando Espinoza, Edith Lopez, Verónica Barragán, Patricio Rojas-Silva, Gabriel Trueba, Michelle Grunauer, Paúl Cárdenas |

| | | | |
|---|---|---|---|
| EPI_ISL_528691, EPI_ISL_528694, EPI_ISL_528695, EPI_ISL_528696, EPI_ISL_528700, EPI_ISL_528702, EPI_ISL_528703, EPI_ISL_528705, EPI_ISL_528707, EPI_ISL_528708 | Alsafar - Khalifa University Abu Dhabi | Alsafar - Khalifa University Abu Dhabi | Andreas Henschel, Gihan Daw Elbait, Samuel Feng, Rifat Hamoudi, Ernesto Damiani, Guan Tay, Habiba Alsafar |
| EPI_ISL_528743, EPI_ISL_528744 | Malaysia Genome Institute | Malaysia Genome Institute | Mohd Noor Mat Isa, Irni Suhayu Sapian, Yusuf Muhammad Noor, Nurhezreen Md Iqbal, Mohd Faizal Abu Bakar, Enizza Kasim, Shamsidar Sopie, Siti Noraini Othman, Azrin Ahmad, Nor Azfa Johari, Shahrul Hisham Zainal Ariffin |
| EPI_ISL_528745 | Laboratorium Kesehatan Provinsi Jawa Barat | School of Life Sciences and Technology & School of Pharmacy-Institut Teknologi Bandung; Molecular Genetics Laboratory-Faculty of Medicine-Universitas Padjadjaran; Laboratorium Kesehatan Provinsi Jawa Barat | Marselina Irasonia Tan, Yunia Sribudiani, Catur Riani, Azzania Fibriani, Husna Nugrahapraja, Tarwadi, Ema Rahmawati, Savira Ekawardhani, Hesti Lina Wiraswati, Ryan Bayusantika Ristandi, Rifky Waluyajati Rachman, Cut Nur Cinthia Alamanda, Lia Faridah, Tri Hanggono Achmad, Mas Rizky A.A. Syamsunarno, Fensi Amalina, Hammam Riza, Sony Solistia Wirawan, Agung Eru Wibowo, Irvan Faizal |
| EPI_ISL_528746 | Immanuel Hospital | Molecular Genetics Laboratory-Faculty of Medicine-Universitas Padjadjaran; School of Life Sciences and Technology & School of Pharmacy-Institut Teknologi Bandung; Laboratorium Kesehatan Provinsi Jawa Barat | Yunia Sribudiani,Tri Hanggono Achmad, Mas Rizky A.A. Syamsunarno, Fensi Amalina, Catur Riani, Azzania Fibriani, Husna Nugrahapraja, Marselina Irasonia Tan, Tarwadi, Ema Rahmawati, Savira Ekawardhani, Hesti Lina Wiraswati, Ryan Bayusantika Ristandi, Rifky Waluyajati Rachman, Cut Nur Cinthia Alamanda, Lia Faridah, Hammam Riza, Sony Solistia Wirawan, Agung Eru Wibowo, Irvan Faizal |
| EPI_ISL_528747 | Santo Borromeus Hospital | School of Pharmacy & School of Life Sciences and Technology - Institut Teknologi Bandung; Molecular Genetics Laboratory-Faculty of Medicine-Universitas Padjadjaran; Laboratorium Kesehatan Provinsi Jawa Barat | Catur Riani, Marselina Irasonia Tan, Yunia Sribudiani, Azzania Fibriani, Husna Nugrahapraja, Tarwadi, Ema Rahmawati, Savira Ekawardhani, Hesti Lina Wiraswati, Ryan Bayusantika Ristandi, Rifky Waluyajati Rachman, Cut Nur Cinthia Alamanda, Lia Faridah, Miftahul Faridl, Karimatu Khoirunnisa, Hammam Riza, Sony Solistia Wirawan, Agung Eru Wibowo, Irvan Faizal |
| EPI_ISL_528748 | Dinkes Provinsi Jawa Barat | School of Life Sciences and Technology & School of Pharmacy-Institut Teknologi Bandung; Molecular Genetics Laboratory-Faculty of Medicine-Universitas Padjadjaran; Laboratorium Kesehatan Provinsi Jawa Barat | Azzania Fibriani, Catur Riani, Marselina Irasonia Tan, Yunia Sribudiani, Husna Nugrahapraja, Tarwadi, Ema Rahmawati, Savira Ekawardhani, Hesti Lina Wiraswati, Ryan Bayusantika Ristandi, Rifky Waluyajati Rachman, Cut Nur Cinthia Alamanda, Lia Faridah, Gusti Ayu Prani Pradani, Adelina Khristiani Rahayu, Hammam Riza, Sony Solistia Wirawan, Agung Eru Wibowo, Irvan Faizal |
| EPI_ISL_528750 | Santo Borromeus Hospital | School of Life Sciences and Technology & School of Pharmacy-Institut Teknologi Bandung; Molecular Genetics Laboratory-Faculty of Medicine-Universitas Padjadjaran; Laboratorium Kesehatan Provinsi Jawa Barat | Marselina Irasonia Tan, Yunia Sribudiani, Catur Riani, Azzania Fibriani, Husna Nugrahapraja, Tarwadi, Ema Rahmawati, Savira Ekawardhani, Hesti Lina Wiraswati, Ryan Bayusantika Ristandi, Rifky Waluyajati Rachman, Cut Nur Cinthia Alamanda, Lia Faridah, Miftahul Faridl, Karimatu Khoirunnisa, Hammam Riza, Sony Solistia Wirawan, Agung Eru Wibowo, Irvan Faizal |
| EPI_ISL_528751 | Santo Borromeus Hospital | Molecular Genetics Laboratory-Faculty of Medicine-Universitas Padjadjaran; School of Life Sciences and Technology & School of Pharmacy-Institut Teknologi Bandung; Laboratorium Kesehatan Provinsi Jawa Barat | Yunia Sribudiani, Tri Hanggono Achmad, Mas Rizky A.A. Syamsunarno, Fensi Amalina, Catur Riani, Azzania Fibriani, Husna Nugrahapraja, Marselina Irasonia Tan, Tarwadi, Ema Rahmawati, Savira Ekawardhani, Hesti Lina Wiraswati, Ryan Bayusantika Ristandi, Rifky Waluyajati Rachman, Cut Nur Cinthia Alamanda, Lia Faridah, Hammam Riza, Sony Solistia Wirawan, Agung Eru Wibowo, Irvan Faizal |
| EPI_ISL_528752 | Dr. H. A. Rotinsulu Lung Hospital | School of Pharmacy & School of Life Sciences and Technology - Institut Teknologi Bandung; Molecular Genetics Laboratory-Faculty of Medicine-Universitas Padjadjaran; Laboratorium Kesehatan Provinsi Jawa Barat | Catur Riani, Marselina Irasonia Tan, Yunia Sribudiani, Azzania Fibriani, Husna Nugrahapraja, Tarwadi, Ema Rahmawati, Savira Ekawardhani, Hesti Lina Wiraswati, Ryan Bayusantika Ristandi, Rifky Waluyajati Rachman, Cut Nur Cinthia Alamanda, Lia Faridah, Gusti Ayu Prani Pradani, Adelina Khristiani Rahayu, Hammam Riza, Sony Solistia Wirawan, Agung Eru Wibowo, Irvan Faizal |
| EPI_ISL_528753 | Dinkes Kota Bogor | School of Life Sciences and Technology & School of Pharmacy-Institut Teknologi Bandung; Molecular Genetics Laboratory-Faculty of Medicine-Universitas Padjadjaran; Laboratorium Kesehatan Provinsi Jawa Barat | Azzania Fibriani, Catur Riani, Marselina Irasonia Tan, Yunia Sribudiani, Husna Nugrahapraja, Tarwadi, Ema Rahmawati, Savira Ekawardhani, Hesti Lina Wiraswati, Ryan Bayusantika Ristandi, Rifky Waluyajati Rachman, Cut Nur Cinthia Alamanda, Lia Faridah, Davin H. E. Setiamarga, Rizki Mardian , Hammam Riza, Sony Solistia Wirawan, Agung Eru Wibowo, Irvan Faizal |
| EPI_ISL_528759 | Santo Borromeus Hospital | School of Life Sciences and Technology & School of Pharmacy-Institut Teknologi Bandung; Molecular Genetics Laboratory-Faculty of Medicine-Universitas Padjadjaran; Laboratorium Kesehatan Provinsi Jawa Barat | Husna Nugrahapraja, Azzania Fibriani, Catur Riani, Marselina Irasonia Tan, Yunia Sribudiani, Tarwadi, Ema Rahmawati, Savira Ekawardhani, Hesti Lina Wiraswati, Ryan Bayusantika Ristandi, Rifky Waluyajati Rachman, Cut Nur Cinthia Alamanda, Lia Faridah, Tri Hanggono Achmad, Mas Rizky A.A. Syamsunarno, Fensi Amalina, Hammam Riza, Sony Solistia Wirawan, Agung Eru Wibowo, Irvan Faizal |
| EPI_ISL_528809 | Department of Medicine, Gandhi hospital, Hyderabad | CSIR-Centre for Cellular and Molecular Biology | Rajarao Mesipogu , Thrilok Chander Bingi ,Vinayasekhar Aedula,Tulasi Nagabandi, Namami Gaur, Sakshi Shambhavi, Lamuk Zaveri, Shagufta Khan, Nikhil Hajirnis, M Soujanya Reddy, Pratheusa Maccha, Purushotham Vodnala, Payel Mukherjee, Sofia Banu, Priya Singh, Onkar Kulkarni, Dhiviya Vedagiri, Divya Gupta, Vishal Sah, Santosh Kumar Kuncha, Krishnan Harinivas Harshan, Archana Bharadwaj Siva, Karthik Bharadwaj Tallapaka,G. Aditya Kumar, Koushick Sivakumar, Pooja Ramesh Gupta, Rajan Kumar Jha, Shraddha Vijay Lahoti, Rakesh K Mishra, Divya Tej Sowpati |
| EPI_ISL_528810, EPI_ISL_528811, EPI_ISL_528812 | Department of Medicine, Gandhi hospital, Hyderabad | CSIR-Centre for Cellular and Molecular Biology | Thrilok Chander Bingi,Rajarao Mesipogu ,Vinayasekhar Aedula,Lamuk Zaveri, Shagufta Khan, Namami Gaur, Sakshi Shambhavi, Nikhil Hajirnis, M Soujanya Reddy, Pratheusa Maccha, Tulasi Nagabandi, Purushotham Vodnala, Payel Mukherjee, Sofia Banu, Priya Singh, Onkar Kulkarni, Dhiviya Vedagiri, Divya Gupta, Vishal Sah, Santosh Kumar Kuncha, Krishnan Harinivas Harshan, Archana Bharadwaj Siva, Karthik Bharadwaj Tallapaka, Renu Sudhakar, Somesh Gorde, Gangumala Srinivas Reddy, Sujoy Deb, Swati Bayyana, Rakesh K Mishra, Divya Tej Sowpati |
| EPI_ISL_528813 | Department of Medicine, Gandhi hospital, Hyderabad | CSIR-Centre for Cellular and Molecular Biology | Vinayasekhar Aedula,Thrilok Chander Bingi, Rajarao Mesipogu, Shagufta Khan, Lamuk Zaveri, Namami Gaur, Sakshi Shambhavi, Nikhil Hajirnis, M Soujanya Reddy, Pratheusa Maccha,Tulasi Nagabandi, Purushotham Vodnala, Payel Mukherjee, Sofia Banu, Priya Singh, Onkar Kulkarni, Dhiviya Vedagiri, Divya Gupta, Vishal Sah, Santosh Kumar Kuncha, Krishnan Harinivas Harshan, Archana Bharadwaj Siva, Karthik Bharadwaj Tallapaka,Umesh Kumar, Unis Ahmad Bhat, Ajay Sarawagi, Priyanka Pant, Rajkanwar Nathawat, Rakesh K Mishra, Divya Tej Sowpati |
| EPI_ISL_528814 | Department of Medicine, Gandhi hospital, Hyderabad | CSIR-Centre for Cellular and Molecular Biology | Rajarao Mesipogu , Thrilok Chander Bingi ,Vinayasekhar Aedula,Tulasi Nagabandi, Namami Gaur, Sakshi Shambhavi, Lamuk Zaveri, Shagufta Khan, Nikhil Hajirnis, M Soujanya Reddy, Pratheusa Maccha, Purushotham Vodnala, Payel Mukherjee, Sofia Banu, Priya Singh, Onkar Kulkarni, Dhiviya Vedagiri, Divya Gupta, Vishal Sah, Santosh Kumar Kuncha, Krishnan Harinivas Harshan, Archana Bharadwaj Siva, Karthik Bharadwaj Tallapaka,G. Aditya Kumar, Koushick Sivakumar, Pooja Ramesh Gupta, Rajan Kumar Jha, Shraddha Vijay Lahoti, Rakesh K Mishra, Divya Tej Sowpati |
| EPI_ISL_528815 | Department of Medicine, Gandhi hospital, Hyderabad | CSIR-Centre for Cellular and Molecular Biology | Vinayasekhar Aedula,Thrilok Chander Bingi, Rajarao Mesipogu, Shagufta Khan, Lamuk Zaveri, Namami Gaur, Sakshi Shambhavi, Nikhil Hajirnis, M Soujanya Reddy, Pratheusa Maccha,Tulasi Nagabandi, Purushotham Vodnala, Payel Mukherjee, Sofia Banu, Priya Singh, Onkar Kulkarni, Dhiviya Vedagiri, Divya Gupta, Vishal Sah, Santosh Kumar Kuncha, Krishnan Harinivas Harshan, Archana Bharadwaj Siva, Karthik Bharadwaj Tallapaka,Umesh Kumar, Unis Ahmad Bhat, Ajay Sarawagi, Priyanka Pant, Rajkanwar Nathawat, Rakesh K Mishra, Divya Tej Sowpati |
| EPI_ISL_528816 | Department of Medicine, Gandhi hospital, Hyderabad | CSIR-Centre for Cellular and Molecular Biology | Rajarao Mesipogu , Thrilok Chander Bingi ,Vinayasekhar Aedula,Tulasi Nagabandi, Namami Gaur, Sakshi Shambhavi, Lamuk Zaveri, Shagufta Khan, Nikhil Hajirnis, M Soujanya Reddy, Pratheusa Maccha, Purushotham Vodnala, Payel Mukherjee, Sofia Banu, Priya Singh, Onkar Kulkarni, Dhiviya Vedagiri, Divya Gupta, Vishal Sah, Santosh Kumar Kuncha, Krishnan Harinivas Harshan, Archana Bharadwaj Siva, Karthik Bharadwaj Tallapaka,G. Aditya Kumar, Koushick Sivakumar, Pooja Ramesh Gupta, Rajan Kumar Jha, Shraddha Vijay Lahoti, Rakesh K Mishra, Divya Tej Sowpati |
| EPI_ISL_528817 | Department of Medicine, Gandhi hospital, Hyderabad | CSIR-Centre for Cellular and Molecular Biology | Thrilok Chander Bingi,Rajarao Mesipogu ,Vinayasekhar Aedula,Lamuk Zaveri, Shagufta Khan, Namami Gaur, Sakshi Shambhavi, Nikhil Hajirnis, M Soujanya Reddy, Pratheusa Maccha, Tulasi Nagabandi, Purushotham Vodnala, Payel Mukherjee, Sofia Banu, Priya Singh, Onkar Kulkarni, Dhiviya Vedagiri, Divya Gupta, Vishal Sah, Santosh Kumar Kuncha, Krishnan Harinivas Harshan, Archana Bharadwaj Siva, Karthik Bharadwaj Tallapaka, Renu Sudhakar, Somesh Gorde, Gangumala Srinivas Reddy, Sujoy Deb, Swati Bayyana, Rakesh K Mishra, Divya Tej Sowpati |
| EPI_ISL_528818 | Department of Medicine, Gandhi hospital, Hyderabad | CSIR-Centre for Cellular and Molecular Biology | Vinayasekhar Aedula,Thrilok Chander Bingi, Rajarao Mesipogu, Shagufta Khan, Lamuk Zaveri, Namami Gaur, Sakshi Shambhavi, Nikhil Hajirnis, M Soujanya Reddy, Pratheusa Maccha,Tulasi Nagabandi, Purushotham Vodnala, Payel Mukherjee, Sofia Banu, Priya Singh, Onkar Kulkarni, Dhiviya Vedagiri, Divya Gupta, Vishal Sah, Santosh Kumar Kuncha, Krishnan Harinivas Harshan, Archana Bharadwaj Siva, Karthik Bharadwaj Tallapaka,Umesh Kumar, Unis Ahmad Bhat, Ajay Sarawagi, Priyanka Pant, Rajkanwar Nathawat, Rakesh K Mishra, Divya Tej Sowpati |
| EPI_ISL_528819, EPI_ISL_528820 | Department of Medicine, Gandhi hospital, Hyderabad | CSIR-Centre for Cellular and Molecular Biology | Rajarao Mesipogu , Thrilok Chander Bingi ,Vinayasekhar Aedula,Tulasi Nagabandi, Namami Gaur, Sakshi Shambhavi, Lamuk Zaveri, Shagufta Khan, Nikhil Hajirnis, M Soujanya Reddy, Pratheusa Maccha, Purushotham Vodnala, Payel Mukherjee, Sofia Banu, Priya Singh, Onkar Kulkarni, Dhiviya Vedagiri, Divya Gupta, Vishal Sah, Santosh Kumar Kuncha, Krishnan Harinivas Harshan, Archana Bharadwaj Siva, Karthik Bharadwaj Tallapaka,G. Aditya Kumar, Koushick |

| | | | |
|---|---|---|---|
| | | | Sivakumar, Pooja Ramesh Gupta, Rajan Kumar Jha, Shraddha Vijay Lahoti, Rakesh K Mishra, Divya Tej Sowpati |
| EPI_ISL_528821 | Department of Medicine, Gandhi hospital, Hyderabad | CSIR-Centre for Cellular and Molecular Biology | Thrilok Chander Bingi,Rajarao Mesipogu ,Vinayasekhar Aedula,Lamuk Zaveri, Shagufta Khan, Namami Gaur, Sakshi Shambhavi, Nikhil Hajirnis, M Soujanya Reddy, Pratheusa Maccha, Tulasi Nagabandi, Purushotham Vodnala, Payel Mukherjee, Sofia Banu, Priya Singh, Onkar Kulkarni, Dhiviya Vedagiri, Divya Gupta, Vishal Sah, Santosh Kumar Kuncha, Krishnan Harinivas Harshan, Archana Bharadwaj Siva, Karthik Bharadwaj Tallapaka, Renu Sudhakar, Somesh Gorde, Gangumala Srinivas Reddy, Sujoy Deb, Swati Bayyana, Rakesh K Mishra, Divya Tej Sowpati |
| EPI_ISL_528822 | Department of Medicine, Gandhi hospital, Hyderabad | CSIR-Centre for Cellular and Molecular Biology | Vinayasekhar Aedula,Thrilok Chander Bingi, Rajarao Mesipogu, Shagufta Khan, Lamuk Zaveri, Namami Gaur, Sakshi Shambhavi, Nikhil Hajirnis, M Soujanya Reddy, Pratheusa Maccha,Tulasi Nagabandi, Purushotham Vodnala, Payel Mukherjee, Sofia Banu, Priya Singh, Onkar Kulkarni, Dhiviya Vedagiri, Divya Gupta, Vishal Sah, Santosh Kumar Kuncha, Krishnan Harinivas Harshan, Archana Bharadwaj Siva, Karthik Bharadwaj Tallapaka,Umesh Kumar, Unis Ahmad Bhat, Ajay Sarawagi, Priyanka Pant, Rajkanwar Nathawat, Rakesh K Mishra, Divya Tej Sowpati |
| EPI_ISL_529003, EPI_ISL_529004, EPI_ISL_529005 | Servizio di igiene epidemiologia e sanità pubblica (SIESP)-Chieti | Istituto Zooprofilattico Sperimentale dell'Abruzzo e Molise "G.Caporale" | Lorusso A, Marcacci M, Di Domenico M, Curini V, Ancora M, Cammà C, Rinaldi A, Mangone I, Di Pasquale A, Puglia I, Savini G. |
| EPI_ISL_529006 | Servizio di igiene e sanità pubblica (SIESP)-Teramo | Istituto Zooprofilattico Sperimentale dell'Abruzzo e Molise "G.Caporale" | Lorusso A, Marcacci M, Di Domenico M, Curini V, Ancora M, Cammà C, Rinaldi A, Mangone I, Di Pasquale A, Puglia I, Savini G. |
| EPI_ISL_529007 | Ospedale Civile S. Liberatore-Atri | Istituto Zooprofilattico Sperimentale dell'Abruzzo e Molise "G.Caporale" | Lorusso A, Marcacci M, Di Domenico M, Curini V, Ancora M, Cammà C, Rinaldi A, Mangone I, Di Pasquale A, Puglia I, Savini G. |
| EPI_ISL_529010, EPI_ISL_529011, EPI_ISL_529012 | Servizio Igiene Epidemiologia e Sanità Pubblica (SIESP)-L'Aquila | Istituto Zooprofilattico Sperimentale dell'Abruzzo e Molise "G.Caporale" | Lorusso A, Marcacci M, Di Domenico M, Curini V, Ancora M, Cammà C, Rinaldi A, Mangone I, Di Pasquale A, Puglia I, Savini G. |
| EPI_ISL_529013 | Presidio Ospedaliero "S.Filippo e Nicola"-Avezzano | Istituto Zooprofilattico Sperimentale dell'Abruzzo e Molise "G.Caporale" | Lorusso A, Marcacci M, Di Domenico M, Curini V, Ancora M, Cammà C, Rinaldi A, Mangone I, Di Pasquale A, Puglia I, Savini G. |
| EPI_ISL_529014, EPI_ISL_529015 | Ospedale "Ss. Annunziata" | Istituto Zooprofilattico Sperimentale dell'Abruzzo e Molise "G.Caporale" | Lorusso A, Marcacci M, Di Domenico M, Curini V, Ancora M, Cammà C, Rinaldi A, Mangone I, Di Pasquale A, Puglia I, Savini G. |
| EPI_ISL_529031 | Central Molecular Microbiology Laboratory, Clinical and Chemical Pathology Department, Faculty of Medicine, CAIRO UNIVERSITY | Next Generation Sequencing Reference Laboratory, Faculty of Medicine, Cairo University and The Center for Genome and Microbiome Research, Faculty of Pharmacy, CAIRO UNIVERSITY | May Sherif Soliman, May Abdelfattah, Ramy Karam Aziz |
| EPI_ISL_529032 | Central Molecular Microbiology Laboratory and Next Generation Sequencing Reference Laboratory, Clinical and Chemical Pathology Department, Faculty of Medicine, CAIRO UNIVERSITY | Next Generation Sequencing Reference Laboratory, Faculty of Medicine, CAIRO UNIVERSITY and The Center for Genome and Microbiome Research, Faculty of Pharmacy, CAIRO UNIVERSITY | May Sherif Soliman, May Abdelfattah, Ramy Karam Aziz |
| EPI_ISL_529720, EPI_ISL_529721, EPI_ISL_529723, EPI_ISL_529724, EPI_ISL_529726, EPI_ISL_529728, EPI_ISL_529729, EPI_ISL_529730, EPI_ISL_529731, EPI_ISL_529732, EPI_ISL_529733, EPI_ISL_529734, EPI_ISL_529735, EPI_ISL_529736, EPI_ISL_529737, EPI_ISL_529738, EPI_ISL_529739, EPI_ISL_529740, EPI_ISL_529741, EPI_ISL_529742, EPI_ISL_529743, EPI_ISL_529744, EPI_ISL_529745, EPI_ISL_529747, EPI_ISL_529748, EPI_ISL_529749, EPI_ISL_529750, EPI_ISL_529751, EPI_ISL_529752, EPI_ISL_529753, EPI_ISL_529755, EPI_ISL_529756, EPI_ISL_529757, EPI_ISL_529758, EPI_ISL_529759, EPI_ISL_529760, EPI_ISL_529761, EPI_ISL_529762, EPI_ISL_529763, EPI_ISL_529764, EPI_ISL_529765, EPI_ISL_529766, EPI_ISL_529767, EPI_ISL_529769, EPI_ISL_529770, EPI_ISL_529771, EPI_ISL_529772, EPI_ISL_529773, EPI_ISL_529774, EPI_ISL_529775, EPI_ISL_529776, EPI_ISL_529777, EPI_ISL_529778, EPI_ISL_529779, EPI_ISL_529780, EPI_ISL_529782, EPI_ISL_529783, EPI_ISL_529784, EPI_ISL_529785, EPI_ISL_529786, EPI_ISL_529787, EPI_ISL_529788, EPI_ISL_529789, EPI_ISL_529790, EPI_ISL_529791, EPI_ISL_529792, EPI_ISL_529793, EPI_ISL_529794, EPI_ISL_529796, EPI_ISL_529797, EPI_ISL_529798, EPI_ISL_529799, EPI_ISL_529800, EPI_ISL_529801, EPI_ISL_529802 | | | |
| see above | NHLS-IALCH | KRISP, KZN Research Innovation and Sequencing Platform | Giandhari J, Pillay S, Lessells R, Mdlalose K, York D, Khan S, Tegally H, Wilkinson E, de Oliveira T |
| EPI_ISL_529963 | Universitas Airlangga Hospital | Institute of Tropical Disease, Universitas Airlangga | Rima R Prasetya, Krisnoadi Rahardjo, Aldise M Nastri, Jezzy R Dewantari, Nasronudin, Gatot Soegiarto, Laksmi Wulandari, Retno A Setyoningrum, Resti Yudhawati, Yohko K Shimizu, Mitsuhiro Nishimura, Yasuko Mori, Soetjipto, Kazufumi Shimizu, Maria I Lusida |
| EPI_ISL_529967 | RSUD Sidoarjo | Institute of Tropical Disease, Universitas Airlangga | Aldise M Nastri, Jezzy R Dewantari, Rima R Prasetya, Krisnoadi Rahardjo, Atok Irawan, Gatot Soegiarto, Laksmi Wulandari, Retno A Setyoningrum, Resti Yudhawati, Yohko K Shimizu, Mitsuhiro Nishimura, Yasuko Mori, Soetjipto, Kazufumi Shimizu, Maria I Lusida |
| EPI_ISL_534336 | Department of Laboratory Medicine, National Taiwan University Hospital | Microbial Genomics Core Lab, National Taiwan University Centers of Genomic and Precision Medicine | Shiou-Hwei Yeh, You-Yu Lin, Ya-Yun Lai, Chiao-Ling Li, Shan-Chwen Chang, Pei-Jer Chen, Sui-Yuan Chang |
| EPI_ISL_534337, EPI_ISL_534338, EPI_ISL_534339, EPI_ISL_534340, EPI_ISL_534341, EPI_ISL_534342, EPI_ISL_534343, EPI_ISL_534344, EPI_ISL_534345 | Molecular diagnostic laboratory of Federal Budget Institution of Science "Central Research Institute of Epidemiology" of The Federal Service on Customers' Rights Protection and Human Well-being Surveillance | Group of Genomics and Postgenomic Technologies of Central Research Institute of Epidemilology | Speranskaya AS, Kaptelova VV, Samoilov AE, Valdokhina AV, Bulanenko VP, Bukharina A.Y., Tivanova EV, Shipulina OY, Akimkin VG |
| EPI_ISL_535390, EPI_ISL_535392, EPI_ISL_535393, EPI_ISL_535394, EPI_ISL_535395, EPI_ISL_535396, EPI_ISL_535397, EPI_ISL_535398, EPI_ISL_535399, EPI_ISL_535400, EPI_ISL_535403, EPI_ISL_535404, EPI_ISL_535405, EPI_ISL_535406, EPI_ISL_535408, EPI_ISL_535410, EPI_ISL_535412, EPI_ISL_535413, EPI_ISL_535414, EPI_ISL_535415, EPI_ISL_535416, EPI_ISL_535417, EPI_ISL_535418, EPI_ISL_535420, EPI_ISL_535421, EPI_ISL_535422, EPI_ISL_535424, EPI_ISL_535425, EPI_ISL_535426, EPI_ISL_535427, EPI_ISL_535428, EPI_ISL_535430, EPI_ISL_535431, EPI_ISL_535432, EPI_ISL_535433, EPI_ISL_535434, EPI_ISL_535435, EPI_ISL_535436, EPI_ISL_535438, EPI_ISL_535439, EPI_ISL_535440, EPI_ISL_535441, EPI_ISL_535442, EPI_ISL_535443, EPI_ISL_535444, EPI_ISL_535445, EPI_ISL_535446, EPI_ISL_535447, EPI_ISL_535448, EPI_ISL_535449, EPI_ISL_535450, EPI_ISL_535451, EPI_ISL_535452, EPI_ISL_535454, EPI_ISL_535455, EPI_ISL_535456, EPI_ISL_535457, EPI_ISL_535458, EPI_ISL_535459, EPI_ISL_535460, EPI_ISL_535461, EPI_ISL_535462, EPI_ISL_535463, EPI_ISL_535464, EPI_ISL_535465, EPI_ISL_535466, EPI_ISL_535467, EPI_ISL_535468, EPI_ISL_535469, EPI_ISL_535470, EPI_ISL_535471, EPI_ISL_535472, EPI_ISL_535473, EPI_ISL_535475, EPI_ISL_535476, EPI_ISL_535477, EPI_ISL_535478, EPI_ISL_535479, EPI_ISL_535480, EPI_ISL_535481, EPI_ISL_535483, EPI_ISL_535484, EPI_ISL_535485, EPI_ISL_535487, EPI_ISL_535488, EPI_ISL_535496, EPI_ISL_535497, EPI_ISL_535498, EPI_ISL_535499, EPI_ISL_535500, EPI_ISL_535501, EPI_ISL_535502, EPI_ISL_535503, EPI_ISL_535504, EPI_ISL_535505, EPI_ISL_535506, EPI_ISL_535507, EPI_ISL_535508, EPI_ISL_535509, EPI_ISL_535510, EPI_ISL_535511, EPI_ISL_535512, EPI_ISL_535513, EPI_ISL_535514, EPI_ISL_535515, EPI_ISL_535516, EPI_ISL_535518, EPI_ISL_535519, EPI_ISL_535520, EPI_ISL_535522, EPI_ISL_535523, EPI_ISL_535526, EPI_ISL_535527, EPI_ISL_535529, EPI_ISL_535530, EPI_ISL_535531, EPI_ISL_535532, EPI_ISL_535533, EPI_ISL_535534, EPI_ISL_535536, EPI_ISL_535537, EPI_ISL_535539, EPI_ISL_535540, EPI_ISL_535541, EPI_ISL_535542, EPI_ISL_535543, EPI_ISL_535544, EPI_ISL_535545, EPI_ISL_535546, EPI_ISL_535547, EPI_ISL_535551, EPI_ISL_535552, EPI_ISL_535553, EPI_ISL_535554, EPI_ISL_535555, EPI_ISL_535556, EPI_ISL_535557, EPI_ISL_535558, EPI_ISL_535559, EPI_ISL_535560, EPI_ISL_535561, EPI_ISL_535562, EPI_ISL_535563, EPI_ISL_535564, EPI_ISL_535565, EPI_ISL_535566, EPI_ISL_535567, EPI_ISL_535568, EPI_ISL_535569, EPI_ISL_535570, EPI_ISL_535571, EPI_ISL_535572 | | | |
| see above | NHLS-IALCH | KRISP, KZN Research Innovation and Sequencing Platform | Giandhari J, Pillay S, Lessells R, Mdlalose K, York D, Khan S, Tegally H, Wilkinson E, de Oliveira T |
| EPI_ISL_535575 | Hospital Universitari Germans Trias i Pujol(HUGTiP)/Fundació Lluita contra la SIDA (FLSida) | IrsiCaixa AIDS Research Lab | Marc Noguera-Julian, Mariona Parera, Maria Pilar Armengol, Marta Massanella, Ester Ballana, Lidia Ruiz, Nuria Izquierdo, Jorge Carrillo, Roger Paredes, Julia Blanco, Joaquim Segalés, Bonaventura Clotet |
| EPI_ISL_536411, EPI_ISL_536793 | Medtimes Molecular Laboratory | Medtimes Molecular Laboratory | Eric Chan, Winsome Wong, Jacqueline Tam, Isaac Chow |
| EPI_ISL_538500, EPI_ISL_538501 | RS Prima Husada Cipta Medan North Sumatra | National Institute of Health Research and Development | Pawestri, HA; Subangkit; Puspa, KD; Nugraha, AA; Ikawati, HD; Pangesti, KNA; Soekarso, T; Paisal; Setiawaty,V. |
| EPI_ISL_538502 | RSUD Sultan Imanudin Pangkalan Bun Center Kalimantan | National Institute of Health Research and Development | Pawestri, HA; Subangkit; Puspa, KD; Nugraha, AA; Ikawati, HD; Pangesti, KNA; Soekarso, T; Paisal; Setiawaty,V. |
| EPI_ISL_538552 | Hospital Universitari Germans Trias i Pujol(HUGTiP)/Fundació Lluita contra la SIDA (FLSida) | IrsiCaixa AIDS Research Lab | Marc Noguera-Julian, Mariona Parera, Maria Pilar Armengol, Marta Massanella, Ester Ballana, Lidia Ruiz, Nuria Izquierdo, Jorge Carrillo, Roger Paredes, Julia Blanco, Joaquim Segalés, Bonaventura Clotet |
| EPI_ISL_539495 | Centers for Disease Control and Prevention, Dengue Branch | Centers for Disease Control and Prevention, Dengue Branch | Gilberto A. Santiago, Glenda Gonzalez, Betzabel Flores, Keyla Charriez, Jorge L. Munoz-Jordan, Gabriela Paz-Bailey, Janice Perez, Vanessa Rivera-Amill, Diego Sainz de la Peña, Jorge Bertran |
| EPI_ISL_539538, EPI_ISL_539539, EPI_ISL_539540, EPI_ISL_539541, EPI_ISL_539542, EPI_ISL_539543, EPI_ISL_539544, EPI_ISL_539547, EPI_ISL_539548 | Hospital Clínic | Instituto de Salud Carlos III | Iglesias-Caballero, M. Molinero Calamita, M. González-Esguevillas, M. Camarero, S. Pozo, F. Casas, I. Jiménez, P. Jiménez, M. Zaballos, A. Monzón, S. Varona, S. Juliá, M. Cuesta, I, M.A Marcos |
| EPI_ISL_539567 | Hospital Comarcal de Melilla | Instituto de Salud Carlos III | Iglesias-Caballero, M. Molinero Calamita, M. González-Esguevillas, M. Camarero, S. Pozo, F. Casas, I. Jiménez, P. Jiménez, M. Zaballos, A. Monzón, S. Varona, S. Juliá, M. Cuesta, I, J. López |
| EPI_ISL_539568 | Hospital Comarcal de Melilla | Instituto de Salud Carlos III | Iglesias-Caballero, M. Molinero Calamita, M. González-Esguevillas, M. Camarero, S. Pozo, F. Casas, I. Jiménez, P. Jiménez, M. Zaballos, A. Monzón, S. Varona, S. Juliá, M. Cuesta, I, C. Ezpeleta |

| | | | |
|---|---|---|---|
| EPI_ISL_539782 | National Institute of Public Health (Czech Republic) | State Veterinary Institute Prague | Nagy, A; Jirincova, H; Novakova, L; Trnka, D; Vecerova, J. |
| EPI_ISL_539803 | Queen Mary Hospital | Hong Kong Department of Health | Alan K.L. Tsang, Peter C.W. Yip, Edman T.K. Lam, Rickjason C.W. Chan, Dominic N.C. Tsang |
| EPI_ISL_539804 | Yan Chai Hospital | Hong Kong Department of Health | Alan K.L. Tsang, Peter C.W. Yip, Edman T.K. Lam, Rickjason C.W. Chan, Dominic N.C. Tsang |
| EPI_ISL_539805 | Kwong Wah Hospital | Hong Kong Department of Health | Alan K.L. Tsang, Peter C.W. Yip, Edman T.K. Lam, Rickjason C.W. Chan, Dominic N.C. Tsang |
| EPI_ISL_539806, EPI_ISL_539807, EPI_ISL_539808 | Princess Margaret Hospital | Hong Kong Department of Health | Alan K.L. Tsang, Peter C.W. Yip, Edman T.K. Lam, Rickjason C.W. Chan, Dominic N.C. Tsang |
| EPI_ISL_539809, EPI_ISL_539811, EPI_ISL_539812 | Asiaworld Expo Command Post | Hong Kong Department of Health | Alan K.L. Tsang, Peter C.W. Yip, Edman T.K. Lam, Rickjason C.W. Chan, Dominic N.C. Tsang |
| EPI_ISL_539813 | Prince of Wales Hospital | Hong Kong Department of Health | Alan K.L. Tsang, Peter C.W. Yip, Edman T.K. Lam, Rickjason C.W. Chan, Dominic N.C. Tsang |
| EPI_ISL_539814 | Tuen Mun Hospital | Hong Kong Department of Health | Alan K.L. Tsang, Peter C.W. Yip, Edman T.K. Lam, Rickjason C.W. Chan, Dominic N.C. Tsang |
| EPI_ISL_539815 | Queen Elizabeth Hospital | Hong Kong Department of Health | Alan K.L. Tsang, Peter C.W. Yip, Edman T.K. Lam, Rickjason C.W. Chan, Dominic N.C. Tsang |
| EPI_ISL_539816, EPI_ISL_539817 | Queen Mary Hospital | Hong Kong Department of Health | Alan K.L. Tsang, Peter C.W. Yip, Edman T.K. Lam, Rickjason C.W. Chan, Dominic N.C. Tsang |
| EPI_ISL_539818, EPI_ISL_539819 | Tuen Mun Hospital | Hong Kong Department of Health | Alan K.L. Tsang, Peter C.W. Yip, Edman T.K. Lam, Rickjason C.W. Chan, Dominic N.C. Tsang |
| EPI_ISL_539821 | Asiaworld Expo Command Post | Hong Kong Department of Health | Alan K.L. Tsang, Peter C.W. Yip, Edman T.K. Lam, Rickjason C.W. Chan, Dominic N.C. Tsang |
| EPI_ISL_539822, EPI_ISL_539823 | Communicable Disease Branch | Hong Kong Department of Health | Alan K.L. Tsang, Peter C.W. Yip, Edman T.K. Lam, Rickjason C.W. Chan, Dominic N.C. Tsang |
| EPI_ISL_541002, EPI_ISL_541003, EPI_ISL_541004, EPI_ISL_541005, EPI_ISL_541006, EPI_ISL_541007 | Health and Environmental Research Institute of Gwangju Metropolitan city | Health and Environmental Research Institute of Gwangju Metropolitan city | Min Ji Kim, Ji-eun Lee |
| EPI_ISL_541082 | The National Institute of Public Health | State Veterinary Institute Prague | Nagy,A;Jirincova,H;Novakova,L;Trnka,D;Vecerova,J |
| EPI_ISL_541332, EPI_ISL_541333, EPI_ISL_541334, EPI_ISL_541336, EPI_ISL_541337 | The National Institute of Public Health | State Veterinary Institute Prague | Nagy,A; Jirincova,H; Novakova,L; Trnka,D; Vecerova,J |
| EPI_ISL_545580, EPI_ISL_545954, EPI_ISL_545956 | The National Institute of Public Health | State Veterinary Institute Prague | Nagy,A;Jirincova,H;Novakova,L;Trnka,D;Vecerova,J |
| EPI_ISL_545957, EPI_ISL_546436, EPI_ISL_546935, EPI_ISL_546936 | The National Institute of Public Health | State Veterinary Institute Prague | Nagy,A; Jirincova,H; Novakova,L; Trnka,D; Vecerova,J |
| EPI_ISL_547571 | Hospital Municipal Antônio Giglio | Instituto Adolfo Lutz, Interdiciplinary Procedures Center, Strategic Laboratory | Claudio Tavares Sacchi, Claudia Regina Gonçalves, Erica Valessa Ramos Gomes, Karoline Rodrigues Campos |
| EPI_ISL_547573 | Vigilância em Saúde de Cajamar | Instituto Adolfo Lutz, Interdiciplinary Procedures Center, Strategic Laboratory | Claudio Tavares Sacchi, Claudia Regina Gonçalves, Erica Valessa Ramos Gomes, Karoline Rodrigues Campos |
| EPI_ISL_547574 | Hospital Universitario da USP | Instituto Adolfo Lutz, Interdiciplinary Procedures Center, Strategic Laboratory | Claudio Tavares Sacchi, Claudia Regina Gonçalves, Erica Valessa Ramos Gomes, Karoline Rodrigues Campos |
| EPI_ISL_547575 | SVO Jundiaí | Instituto Adolfo Lutz, Interdiciplinary Procedures Center, Strategic Laboratory | Claudio Tavares Sacchi, Claudia Regina Gonçalves, Erica Valessa Ramos Gomes, Karoline Rodrigues Campos |
| EPI_ISL_547576 | Secretaria Municipal de Saúde | Instituto Adolfo Lutz, Interdiciplinary Procedures Center, Strategic Laboratory | Claudio Tavares Sacchi, Claudia Regina Gonçalves, Erica Valessa Ramos Gomes, Karoline Rodrigues Campos |
| EPI_ISL_547577 | Hospital e Maternidade Nossa Senhora das Graças | Instituto Adolfo Lutz, Interdiciplinary Procedures Center, Strategic Laboratory | Claudio Tavares Sacchi, Claudia Regina Gonçalves, Erica Valessa Ramos Gomes, Karoline Rodrigues Campos |
| EPI_ISL_547578 | Hospital Doutor Domingos Leonardo Cerávolo | Instituto Adolfo Lutz, Interdiciplinary Procedures Center, Strategic Laboratory | Claudio Tavares Sacchi, Claudia Regina Gonçalves, Erica Valessa Ramos Gomes, Karoline Rodrigues Campos |
| EPI_ISL_547579 | Santa Casa de Misericórdia de Araçatuba | Instituto Adolfo Lutz, Interdiciplinary Procedures Center, Strategic Laboratory | Claudio Tavares Sacchi, Claudia Regina Gonçalves, Erica Valessa Ramos Gomes, Karoline Rodrigues Campos |
| EPI_ISL_547580 | Santa Casa da Misericórdia de Presidente Prudente | Instituto Adolfo Lutz, Interdiciplinary Procedures Center, Strategic Laboratory | Claudio Tavares Sacchi, Claudia Regina Gonçalves, Erica Valessa Ramos Gomes, Karoline Rodrigues Campos |
| EPI_ISL_547966, EPI_ISL_547967, EPI_ISL_547968 | The National Institute of Public Health | State Veterinary Institute Prague | Nagy,A;Jirincova,H;Novakova,L;Trnka,D;Vecerova,J |
| EPI_ISL_560408, EPI_ISL_560409, EPI_ISL_560410, EPI_ISL_560411, EPI_ISL_560412 | The National Institute of Public Health | State Veterinary Institute Prague | Nagy, A; Jirincova, H; Novakova, L; Trnka, D; Vecerova, J |
| EPI_ISL_560587, EPI_ISL_560588, EPI_ISL_560589, EPI_ISL_560590 | Hopital | National Reference Center for Viruses of Respiratory Infections, Institut Pasteur, Paris | Sylvie Behillil, Fabiana Gambaro, Etienne Simon-Lorière, Vincent Enouf, Maud Vanpeene, Sylvie van der Werf |
| EPI_ISL_560598, EPI_ISL_560599, EPI_ISL_560600, EPI_ISL_560601, EPI_ISL_560602, EPI_ISL_560603, EPI_ISL_560604, EPI_ISL_560605, EPI_ISL_560606, EPI_ISL_560607, EPI_ISL_560608, EPI_ISL_560609, EPI_ISL_560610, EPI_ISL_560611, EPI_ISL_560612, EPI_ISL_560613, EPI_ISL_560614, EPI_ISL_560615, EPI_ISL_560616, EPI_ISL_560617, EPI_ISL_560618, EPI_ISL_560636 | | | |
| see above | Hospital | National Reference Center for Viruses of Respiratory Infections, Institut Pasteur, Paris | Sylvie Behillil, Fabiana Gambaro, Etienne Simon-Lorière, Vincent Enouf, Maud Vanpeene, Sylvie van der Werf |
| EPI_ISL_560637, EPI_ISL_560638, EPI_ISL_560639, EPI_ISL_560640, EPI_ISL_560641, EPI_ISL_560642 | Labo Analyses Med | National Reference Center for Viruses of Respiratory Infections, Institut Pasteur, Paris | Sylvie Behillil, Fabiana Gambaro, Etienne Simon-Lorière, Vincent Enouf, Maud Vanpeene, Sylvie van der Werf |
| EPI_ISL_560643, EPI_ISL_560645, EPI_ISL_560646 | Hospital | National Reference Center for Viruses of Respiratory Infections, Institut Pasteur, Paris | Sylvie Behillil, Fabiana Gambaro, Etienne Simon-Lorière, Vincent Enouf, Maud Vanpeene, Sylvie van der Werf |
| EPI_ISL_566077, EPI_ISL_566078, EPI_ISL_566079, EPI_ISL_566080, EPI_ISL_566081, EPI_ISL_566082, EPI_ISL_566083 | Pathogenic Microorganisms Variability Laboratory | WHO National Influenza Centre Russian Federation | Andrey Komissarov, Artem Fadeev, Anna Ivanova, Kseniya Komissarova, Daria Danilenko, Dmitry Lioznov, Nadezhda Kuznetsova, Elena Shidlovskaya, Elizaveta Divisenko, Ekaterina Milashenko, Kirill Krasnoslobotsev, Evgeniya Mukasheva, Anna Ignatieva, Svetlana Trushakova, Alexey Shchetinin, Maria Nikiforova, Andrey Pochtovyy, Valeria Bacalin, Evgeny Usachev, Olga Burgasova, Ludmila Kolobukhina, Svetlana Smetanina, Elena Burtseva, Artem Tkachuk, Vladimir Gushchin, Alexander Gintsburg |
| EPI_ISL_568689 | RS Pondok Indah Puri Indah | Eijkman Institute for Molecular Biology, Ministry of Research and Technology/National Agency for Research and Innovation | Frilasita A Yudhaputri, Edison Johar, Hidayat Trimarsanto, Iskandar A Adnan, Willy Agustine, David H Muljono, Safarina G Malik, Herawati Sudoyo, Khin Saw Myint, Amin Soebandrio |
| EPI_ISL_568691 | RSUP Fatmawati | Eijkman Institute for Molecular Biology, Ministry of Research and Technology/National Agency for Research and Innovation | Frilasita A Yudhaputri, Edison Johar, Hidayat Trimarsanto, Iskandar A Adnan, Willy Agustine, David H Muljono, Safarina G Malik, Herawati Sudoyo, Khin Saw Myint, Amin Soebandrio |

| | | | |
|---|---|---|---|
| EPI_ISL_568692 | RS Kramat 128 | Eijkman Institute for Molecular Biology, Ministry of Research and Technology/National Agency for Research and Innovation | Frilasita A Yudhaputri, Edison Johar, Hidayat Trimarsanto, Iskandar A Adnan, Willy Agustine, David H Muljono, Safarina G Malik, Herawati Sudoyo, Khin Saw Myint, Amin Soebandrio |
| EPI_ISL_572331, EPI_ISL_572397 | Institute for Virology, University Hospital Duesseldorf, Medical Faculty, Heinrich-Heine-University Duesseldorf | Institute for Virology, University Hospital Duesseldorf, Medical Faculty, Heinrich-Heine-University Duesseldorf | Maximilian Damagnez, Verena Keitel, Björn Jensen, Nadine Lübke, Lisa Müller, Philipp Ostermann, Tina Senff, Ortwin Adams, Philipp Albrecht, Gerald Antoch, Johannes Bode, Edwin Bölke, Saskia Elben, Torsten Feldt, Johannes C. Fischer, , Anselm Kunstein, Caroline Klindt, Alexander Killer, Tom Lüdde, Annemarie Mohring, Jennifer Neubert, Heiner Schaal, Ansgar Schulz, Jörg Timm, Andreas Walker |
| EPI_ISL_572398 | Laboratory of Biology and Identification of Arboviruses | Pathogenic Microorganisms Variability Laboratory | Alexey Shchetinin, Maria Nikiforova, Andrei Siniavin, Victor Larichev, Alina Kozlova, Muhammad Saifullin, Alexey Prilipov, Vladimir Gushchin, Alexander Gintsburg |
| EPI_ISL_573152, EPI_ISL_574259 | Institute for Virology, University Hospital Duesseldorf, Medical Faculty, Heinrich-Heine-University Duesseldorf | Institute for Virology, University Hospital Duesseldorf, Medical Faculty, Heinrich-Heine-University Duesseldorf | Maximilian Damagnez, Verena Keitel, Björn Jensen, Nadine Lübke, Lisa Müller, Philipp Ostermann, Tina Senff, Ortwin Adams, Philipp Albrecht, Gerald Antoch, Johannes Bode, Edwin Bölke, Saskia Elben, Torsten Feldt, Johannes C. Fischer, , Anselm Kunstein, Caroline Klindt, Alexander Killer, Tom Lüdde, Annemarie Mohring, Jennifer Neubert, Heiner Schaal, Ansgar Schulz, Jörg Timm, Andreas Walker |
| EPI_ISL_574594 | Hospital Escola da Universidade de Taubate | Instituto Adolfo Lutz, Interdiciplinary Procedures Center, Strategic Laboratory | Claudio Tavares Sacchi, Claudia Regina Gonçalves, Erica Valessa Ramos Gomes, Karoline Rodrigues Campos |
| EPI_ISL_574596 | CS II Dr. Antonio Vicoso Moreira de Rezende Sumare | Instituto Adolfo Lutz, Interdiciplinary Procedures Center, Strategic Laboratory | Claudio Tavares Sacchi, Claudia Regina Gonçalves, Erica Valessa Ramos Gomes, Karoline Rodrigues Campos |
| EPI_ISL_574597 | Secretaria Municipal de Saude de Jarinu | Instituto Adolfo Lutz, Interdiciplinary Procedures Center, Strategic Laboratory | Claudio Tavares Sacchi, Claudia Regina Gonçalves, Erica Valessa Ramos Gomes, Karoline Rodrigues Campos |
| EPI_ISL_574598 | Servico de Verificacao de Obito SVO | Instituto Adolfo Lutz, Interdiciplinary Procedures Center, Strategic Laboratory | Claudio Tavares Sacchi, Claudia Regina Gonçalves, Erica Valessa Ramos Gomes, Karoline Rodrigues Campos |
| EPI_ISL_574612 | RS Hermina Mekarsari | Eijkman Institute for Molecular Biology, Ministry of Research and Technology/National Agency for Research and Innovation | Frilasita A Yudhaputri, Edison Johar, Hidayat Trimarsanto, Iskandar A Adnan, Willy Agustine, David H Muljono, Safarina G Malik, Herawati Sudoyo, Khin Saw Myint, Amin Soebandrio |
| EPI_ISL_574614 | RS Kramat 128 | Eijkman Institute for Molecular Biology, Ministry of Research and Technology/National Agency for Research and Innovation | Frilasita A Yudhaputri, Edison Johar, Hidayat Trimarsanto, Iskandar A Adnan, Willy Agustine, David H Muljono, Safarina G Malik, Herawati Sudoyo, Khin Saw Myint, Amin Soebandrio |
| EPI_ISL_574615 | RS Harapan Bunda | Eijkman Institute for Molecular Biology, Ministry of Research and Technology/National Agency for Research and Innovation | Frilasita A Yudhaputri, Edison Johar, Hidayat Trimarsanto, Iskandar A Adnan, Willy Agustine, David H Muljono, Safarina G Malik, Herawati Sudoyo, Khin Saw Myint, Amin Soebandrio |
| EPI_ISL_574616, EPI_ISL_574617 | RS Kramat 128 | Eijkman Institute for Molecular Biology, Ministry of Research and Technology/National Agency for Research and Innovation | Frilasita A Yudhaputri, Edison Johar, Hidayat Trimarsanto, Iskandar A Adnan, Willy Agustine, David H Muljono, Safarina G Malik, Herawati Sudoyo, Khin Saw Myint, Amin Soebandrio |
| EPI_ISL_574618 | RS Kramat 128 | Eijkman Institute for Molecular Biology, Ministry of Research and Technology/National Agency for Research and Innovation | Hidayat Trimarsanto, Frilasita A Yudhaputri, Edison Johar, Iskandar A Adnan, Willy Agustine, David H Muljono, Safarina G Malik, Herawati Sudoyo, Khin Saw Myint, Amin Soebandrio |
| EPI_ISL_574619, EPI_ISL_574620, EPI_ISL_574621, EPI_ISL_574622, EPI_ISL_574623 | BTKLPP Kelas I Manado | Eijkman Institute for Molecular Biology, Ministry of Research and Technology/National Agency for Research and Innovation | Hidayat Trimarsanto, Frilasita A Yudhaputri, Edison Johar, Iskandar A Adnan, Willy Agustine, David H Muljono, Safarina G Malik, Herawati Sudoyo, Khin Saw Myint, Amin Soebandrio |
| EPI_ISL_575331 | RSUD Wates | Genetics Working Group (Pokja Genetik) Faculty of Medicine, Public Health and Nursing Universitas Gadjah Mada (FK-KMK UGM); Disease Investigation Center Wates Ministry of Agriculture Indonesia; Department of Microbiology FK-KMK UGM; Laboratorium Diagnostik Yayasan Tahija World Mosquito Program (WMP) Yogyakarta Center for Tropical Medicine FK-KMK UGM; Integrated Research Center FK-KMK UGM; Department of Computer Science and Electronics FMIPA UGM | Gunadi, Hendra Wibawa, . Marcellus, Mohamad S. Hakim, Edwin W. Daniwijaya, Ludhang P. Rizki, Endah Supriyati, Eggi Arguni, Titik Nuryastuti, Tri Wibawa, Dwi AA Nugrahaningsih, Afiahayati, Siswanto, Kristy Iskandar, Nungki Anggorowati, Bastiano Kusumajaya, Zumrati Ahmad, Alvin S. Kalim, Susan Simanjaya |
| EPI_ISL_576113 | RSUP Dr. Sardjito | Genetics Working Group (Pokja Genetik) Faculty of Medicine, Public Health and Nursing Universitas Gadjah Mada (FK-KMK UGM); Disease Investigation Center Wates Ministry of Agriculture Indonesia; Department of Microbiology FK-KMK UGM; Laboratorium Diagnostik Yayasan Tahija World Mosquito Program (WMP) Yogyakarta Center for Tropical Medicine FK-KMK UGM; Integrated Research Center FK-KMK UGM; Department of Computer Science and Electronics FMIPA UGM | Gunadi, Hendra Wibawa, . Marcellus, Mohamad S. Hakim, Edwin W. Daniwijaya, Ludhang P. Rizki, Endah Supriyati, Eggi Arguni, Titik Nuryastuti, Tri Wibawa, Dwi AA Nugrahaningsih, Afiahayati, Siswanto, Kristy Iskandar, Nungki Anggorowati, Ika Trisnawati, Riat El Khair, William Widitjiarso, Fadil Fahri |
| EPI_ISL_576114 | RSUP Dr. Sardjito | Genetics Working Group (Pokja Genetik) Faculty of Medicine, Public Health and Nursing Universitas Gadjah Mada (FK-KMK UGM); Disease Investigation Center Wates Ministry of Agriculture Indonesia; Department of Microbiology FK-KMK UGM; Laboratorium Diagnostik Yayasan Tahija World Mosquito Program (WMP) Yogyakarta Center for Tropical Medicine FK-KMK UGM; Integrated Research Center FK-KMK UGM; Department of Computer Science and Electronics FMIPA UGM | Gunadi, Hendra Wibawa, . Marcellus, Mohamad S. Hakim, Edwin W. Daniwijaya, Ludhang P. Rizki, Endah Supriyati, Eggi Arguni, Titik Nuryastuti, Tri Wibawa, Dwi AA Nugrahaningsih, Afiahayati, Siswanto, Kristy Iskandar, Nungki Anggorowati, Ika Trisnawati, Riat El Khair, Audric Kenny Tedja, Aditya Rifqi Fauzi |
| EPI_ISL_576115 | RSUP Dr Sardjito | Genetics Working Group (Pokja Genetik) Faculty of Medicine, Public Health and Nursing Universitas Gadjah Mada (FK-KMK UGM); Disease Investigation Center Wates Ministry of Agriculture Indonesia; Department of Microbiology FK-KMK UGM; Laboratorium Diagnostik Yayasan Tahija World Mosquito Program (WMP) Yogyakarta Center for Tropical Medicine FK-KMK UGM; Integrated Research Center FK-KMK UGM; Department of Computer Science and Electronics FMIPA UGM | Gunadi, Hendra Wibawa, . Marcellus, Mohamad S. Hakim, Edwin W. Daniwijaya, Ludhang P. Rizki, Endah Supriyati, Eggi Arguni, Titik Nuryastuti, Tri Wibawa, Dwi AA Nugrahaningsih, Afiahayati, Siswanto, Kristy Iskandar, Nungki Anggorowati, Ika Trisnawati, Riat El Khair, Yunika Puspadewi, Osman Sianipar, Umi Solekhah Intansari, Elizabeth Henny Herningtiyas, Ira Puspitawati, Nur Imma Fatimah Harahap, Untung Wirawan, Maria Patricia Inggriani |
| EPI_ISL_576116 | RSUP Dr. Sardjito | Genetics Working Group (Pokja Genetik) Faculty of Medicine, Public Health and Nursing Universitas Gadjah Mada (FK-KMK UGM); Disease Investigation Center Wates Ministry of Agriculture Indonesia; Department of Microbiology FK-KMK UGM; Laboratorium Diagnostik Yayasan Tahija | Gunadi, Hendra Wibawa, . Marcellus, Mohamad S. Hakim, Edwin W. Daniwijaya, Ludhang P. Rizki, Endah Supriyati, Eggi Arguni, Titik Nuryastuti, Tri Wibawa, Dwi AA Nugrahaningsih, Afiahayati, Siswanto, Kristy Iskandar, Nungki Anggorowati, Ika Trisnawati, Sumardi, Eko Budiono, Bambang Sigit Riyanto, Heni Retnowulan, Munawar Gani, Satria Maulana, Nur Rahmi Ananda, Riat El Khair, Yunika Puspadewi, Osman Sianipar, Umi Solekhah Intansari, Elizabeth Henny Herningtiyas, Ira Puspitawati, Nur Imma Fatimah Harahap, Ririn Enggy Yuliyanti, Alvin Santoso Kalim |

| | | | |
|---|---|---|---|
| | | World Mosquito Program (WMP) Yogyakarta Center for Tropical Medicine FK-KMK UGM; Integrated Research Center FK-KMK UGM; Department of Computer Science and Electronics FMIPA UGM | |
| EPI_ISL_576128 | RSUP Dr. Sardjito | Genetics Working Group (Pokja Genetik) Faculty of Medicine, Public Health and Nursing Universitas Gadjah Mada (FK-KMK UGM); Disease Investigation Center Wates Ministry of Agriculture Indonesia; Department of Microbiology FK-KMK UGM; Laboratorium Diagnostik Yayasan Tahija World Mosquito Program (WMP) Yogyakarta Center for Tropical Medicine FK-KMK UGM; Integrated Research Center FK-KMK UGM; Department of Computer Science and Electronics FMIPA UGM | Gunadi, Hendra Wibawa, Marcellus, Mohamad S. Hakim, Edwin W. Daniwijaya, Ludhang P. Rizki, Endah Supriyati, Eggi Arguni, Titik Nuryastuti, Tri Wibawa, Dwi AA Nugrahaningsih, Afiahayati, Siswanto, Kristy Iskandar, Nungki Anggorowati, Ika Trisnawati, Sumardi, Eko Budiono, Bambang Sigit Riyanto, Heni Retnowulan, Munawar Gani, Satria Maulana, Nur Rahmi Ananda, Riat El Khair, Yunika Puspadewi, Osman Sianipar, Umi Solekhah Intansari, Elizabeth Henny Herningtiyas, Ira Puspitawati, Nur Imma Fatimah Harahap, Dyah Ayu Puspitarani, Kemala Athollah |
| EPI_ISL_576130 | RSUP Dr. Sardjito | Genetics Working Group (Pokja Genetik) Faculty of Medicine, Public Health and Nursing Universitas Gadjah Mada (FK-KMK UGM); Disease Investigation Center Wates Ministry of Agriculture Indonesia; Department of Microbiology FK-KMK UGM; Laboratorium Diagnostik Yayasan Tahija World Mosquito Program (WMP) Yogyakarta Center for Tropical Medicine FK-KMK UGM; Integrated Research Center FK-KMK UGM; Department of Computer Science and Electronics FMIPA UGM | Gunadi, Hendra Wibawa, Marcellus, Mohamad S. Hakim, Edwin W. Daniwijaya, Ludhang P. Rizki, Endah Supriyati, Eggi Arguni, Titik Nuryastuti, Tri Wibawa, Dwi AA Nugrahaningsih, Afiahayati, Siswanto, Kristy Iskandar, Nungki Anggorowati, Rina Triasih, Amalia Setyati, Dwikisworo Setyowireni, Ida Safitri Laksanawati, Cahya Dewi Satria, Titis Widowati, Elisabeth Siti Herini, Riat El Khair, Yunika Puspadewi, Osman Sianipar, Umi Solekhah Intansari, Elizabeth Henny Herningtiyas, Ira Puspitawati, Nur Imma Fatimah Harahap, Alvin S. Kalim, Susan Simanjaya, Desyifa Mursalin |
| EPI_ISL_576145 | RSA Universitas Gadjah Mada | Genetics Working Group (Pokja Genetik) Faculty of Medicine, Public Health and Nursing Universitas Gadjah Mada (FK-KMK UGM); Disease Investigation Center Wates Ministry of Agriculture Indonesia; Department of Microbiology FK-KMK UGM; Laboratorium Diagnostik Yayasan Tahija World Mosquito Program (WMP) Yogyakarta Center for Tropical Medicine FK-KMK UGM; Integrated Research Center FK-KMK UGM; Department of Computer Science and Electronics FMIPA UGM | Gunadi, Hendra Wibawa, Marcellus, Mohamad S. Hakim, Edwin W. Daniwijaya, Ludhang P. Rizki, Endah Supriyati, Eggi Arguni, Titik Nuryastuti, Tri Wibawa, Dwi AA Nugrahaningsih, Afiahayati, Siswanto, Kristy Iskandar, Nungki Anggorowati, William Widitjiarso, Fadil Fahri |
| EPI_ISL_576146, EPI_ISL_576147, EPI_ISL_576148 | Department of Respiratory & Other Viral Infections of L.V. Gromashevsky Institute of Epidemiology & Infectious Diseases NAMS of Ukraine | Department of Respiratory & Other Viral Infections of L.V. Gromashevsky Institute of Epidemiology & Infectious Diseases NAMS of Ukraine, JSC "Farmak" | Alla Mironenko, Ihor Kravchuk, Liudmyla Bolotova, Larysa Radchenko, Nataliia Teteriuk |
| EPI_ISL_576149 | Department of Respiratory & Other Viral Infections of L.V. Gromashevsky Institute of Epidemiology & Infectious Diseases NAMS of Ukraine | Department of Respiratory & Other Viral Infections of L.V. Gromashevsky Institute of Epidemiology & Infectious Diseases NAMS of Ukraine, JSC "Farmak" | Alla Mironenko, Andriy Goy, Ihor Kravchuk, Liudmyla Bolotova, Larysa Radchenko, Nataliia Teteriuk |
| EPI_ISL_576258, EPI_ISL_576259, EPI_ISL_576260 | Instituto de Diagnostico y Referencia Epidemiologicos (INDRE) | Instituto de Diagnostico y Referencia Epidemiologicos (INDRE) | Gisela Barrera-Badillo , Abril Rodriguez-Maldonado, Claudia Wong-Arambula , Natividad Cruz-Ortiz, Tatiana Nunez-Garcia, Dayanira Arellano-Suarez, Fabiola Garces-Ayala, Edgar Mendieta-Condado, Lucia Hernandez-Rivas, Irma Lopez-Martinez, Ernesto Ramirez-Gonzalez. |
| EPI_ISL_576261, EPI_ISL_576262, EPI_ISL_576263 | Instituto de Diagnostico y Referencia Epidemiologicos (INDRE) | Instituto de Diagnostico y Referencia Epidemiologicos (INDRE) | Gisela Barrera-Badillo , Abril Rodriguez-Maldonado, Claudia Wong-Arambula , Natividad Cruz-Ortiz, Tatiana Nunez-Garcia, Dayanira Arellano-Suarez, Adnan Araiza-Rodriguez, Edgar Mendieta-Condado, Lucia Hernandez-Rivas, Irma Lopez-Martinez, Ernesto Ramirez-Gonzalez. |
| EPI_ISL_576264, EPI_ISL_576265 | Instituto de Diagnostico y Referencia Epidemiologicos (INDRE) | Instituto de Diagnostico y Referencia Epidemiologicos (INDRE) | Ernesto Ramirez-Gonzalez, Abril Rodriguez-Maldonado, Claudia Wong-Arambula , Natividad Cruz-Ortiz, Tatiana Nunez-Garcia, Dayanira Arellano-Suarez, Adnan Araiza-Rodriguez, Edgar Mendieta-Condado, Lucia Hernandez-Rivas, Irma Lopez-Martinez, Gisela Barrera-Badillo. |
| EPI_ISL_576266, EPI_ISL_576267, EPI_ISL_576268, EPI_ISL_576269, EPI_ISL_576270, EPI_ISL_576271 | Instituto de Diagnostico y Referencia Epidemiologicos (INDRE) | Instituto de Diagnostico y Referencia Epidemiologicos (INDRE) | Ernesto Ramirez-Gonzalez, Abril Rodriguez-Maldonado, Claudia Wong-Arambula , Natividad Cruz-Ortiz, Tatiana Nunez-Garcia, Dayanira Arellano-Suarez, Adnan Araiza-Rodriguez, Fabiola Garces-Ayala, Lucia Hernandez-Rivas, Irma Lopez-Martinez, Gisela Barrera-Badillo. |
| EPI_ISL_576272 | Instituto de Diagnostico y Referencia Epidemiologicos (INDRE) | Instituto de Diagnostico y Referencia Epidemiologicos (INDRE) | Gisela Barrera-Badillo , Abril Rodriguez-Maldonado, Claudia Wong-Arambula , Natividad Cruz-Ortiz, Tatiana Nunez-Garcia, Dayanira Arellano-Suarez, Fabiola Garces-Ayala, Edgar Mendieta-Condado, Lucia Hernandez-Rivas, Irma Lopez-Martinez, Ernesto Ramirez-Gonzalez. |
| EPI_ISL_576273 | Instituto de Diagnostico y Referencia Epidemiologicos (INDRE) | Instituto de Diagnostico y Referencia Epidemiologicos (INDRE) | Ernesto Ramirez-Gonzalez, Abril Rodriguez-Maldonado, Claudia Wong-Arambula , Natividad Cruz-Ortiz, Tatiana Nunez-Garcia, Dayanira Arellano-Suarez, Adnan Araiza-Rodriguez, Fabiola Garces-Ayala, Lucia Hernandez-Rivas, Irma Lopez-Martinez, Gisela Barrera-Badillo. |
| EPI_ISL_576274, EPI_ISL_576275, EPI_ISL_576276 | Instituto de Diagnostico y Referencia Epidemiologicos (INDRE) | Instituto de Diagnostico y Referencia Epidemiologicos (INDRE) | Gisela Barrera-Badillo , Abril Rodriguez-Maldonado, Claudia Wong-Arambula , Natividad Cruz-Ortiz, Tatiana Nunez-Garcia, Dayanira Arellano-Suarez, Adnan Araiza-Rodriguez, Edgar Mendieta-Condado, Lucia Hernandez-Rivas, Irma Lopez-Martinez, Ernesto Ramirez-Gonzalez. |
| EPI_ISL_576277, EPI_ISL_576278 | Instituto de Diagnostico y Referencia Epidemiologicos (INDRE) | Instituto de Diagnostico y Referencia Epidemiologicos (INDRE) | Ernesto Ramirez-Gonzalez, Abril Rodriguez-Maldonado, Claudia Wong-Arambula , Natividad Cruz-Ortiz, Tatiana Nunez-Garcia, Dayanira Arellano-Suarez, Adnan Araiza-Rodriguez, Edgar Mendieta-Condado, Lucia Hernandez-Rivas, Irma Lopez-Martinez, Gisela Barrera-Badillo. |
| EPI_ISL_576279 | Instituto de Diagnostico y Referencia Epidemiologicos (INDRE) | Instituto de Diagnostico y Referencia Epidemiologicos (INDRE) | Ernesto Ramirez-Gonzalez, Abril Rodriguez-Maldonado, Claudia Wong-Arambula , Natividad Cruz-Ortiz, Tatiana Nunez-Garcia, Dayanira Arellano-Suarez, Adnan Araiza-Rodriguez, Fabiola Garces-Ayala, Lucia Hernandez-Rivas, Irma Lopez-Martinez, Gisela Barrera-Badillo. |
| EPI_ISL_577624, EPI_ISL_577625, EPI_ISL_577626, EPI_ISL_577627, EPI_ISL_577628, EPI_ISL_577632, EPI_ISL_577633, EPI_ISL_577634, EPI_ISL_577635, EPI_ISL_577636, EPI_ISL_577637, EPI_ISL_577638 | | | |
| see above | The National Institute of Public Health | State Veterinary Institute Prague | Nagy,A;Jirincova,H;Novakova,L;Trnka,D;Vecerova,J |
| EPI_ISL_578187, EPI_ISL_578188 | Hospital Virgen de las Nieves | Instituto de Salud Carlos III | Iglesias-Caballero, M. Molinero Calamita, M. González-Esguevillas, M. Camarero, S. Pozo, F. Casas, I. Jiménez, P. Jiménez, M. Zaballos, A. Monzón, S. Varona, S. Juliá, M. Cuesta, I, J.M Navarro |
| EPI_ISL_582030 | Biology Department, College of Science, Al-Muthanna University | International Centre for Genetic Engineering and Biotechnology (ICGEB) and ARGO Open Lab Platform | Nihad Al-Rashedi, Danilo Licastro, Sreejith Rajasekharan, Simeone Dal Monego, Alessandro Marcello |
| EPI_ISL_582124 | Malaysia Genome Institute | Malaysia Genome Institute | Mohd Noor Mat Isa, Irni Suhayu Sapian, Yusuf Muhammad Noor, Nurhezreen Md Iqbal, Mohd Faizal Abu Bakar, Enizza Kasim, Shamsidar Sopie, Siti Noraini Othman, Azrin Ahmad, Nor Azfa Johari, Shahrul Hisham Zainal Ariffin |
| EPI_ISL_582509 | Department of Respiratory and other Viral Infections of L.V.Gromashevsky Institute of Epidemiology & Infectious Diseases NAMS of Ukraine | Department of Respiratory and other Viral Infections of L.V.Gromashevsky Institute of Epidemiology & Infectious Diseases NAMS of Ukraine, JSC "Farmak" | Alla Mironenko, Andriy Goy, Ihor Kravchuk, Ludmyla Bolotova, Larysa Radchenko, Nataliia Teteriuk |
| EPI_ISL_582510 | Department of Respiratory and other Viral Infections of L.V.Gromashevsky Institute of Epidemiology & Infectious Diseases NAMS of Ukraine | Department of Respiratory and other Viral Infections of L.V.Gromashevsky Institute of Epidemiology & Infectious Diseases NAMS of Ukrain, JSC "Farmak" | Alla Mironenko, Andriy Goy, Ihor Kravchuk, Ludmyla Bolotova, Larysa Radchenko, Nataliia Teteriuk |
| EPI_ISL_582511, EPI_ISL_582512, EPI_ISL_582513 | Department of Respiratory and other Viral Infections of L.V.Gromashevsky Institute of Epidemiology & Infectious Diseases NAMS of Ukrain | Department of Respiratory and other Viral Infections of L.V.Gromashevsky Institute of Epidemiology & Infectious Diseases NAMS of Ukrain, JSC "Farmak" | Alla Mironenko, Andriy Goy, Ihor Kravchuk, Ludmyla Bolotova, Larysa Radchenko, Nataliia Teteriuk |
| EPI_ISL_582642, EPI_ISL_582643, EPI_ISL_582644, EPI_ISL_582645, EPI_ISL_582646, EPI_ISL_582647, EPI_ISL_582648, EPI_ISL_582649, EPI_ISL_582650, EPI_ISL_582651, EPI_ISL_582652, EPI_ISL_582653, EPI_ISL_582657 | | | |
| see above | Sheikh Khalifa Medical City | Molecular/Surveillance lab Sheikh Khalifa Medical City | Amirtharaj Francis, Sajeed Abdul, Hala Imambaccus, Sahar Almarzooqi, Hiba Saud, Stefan Weber |

| | | | |
|---|---|---|---|
| EPI_ISL_583443, EPI_ISL_583449, EPI_ISL_583450, EPI_ISL_583451 | Memorial Sloan Kettering Cancer Center | van Bakel Laboratory, Genetics and Genomics Sciences, Icahn School of Medicine at Mount Sinai | Teresa Aydillo, Ana S. Gonzalez-Reiche, Sadaf Aslam, Adriana van de Guchte, Zenab Khan, Ajay Obla, Jayeeta Dutta, Harm van Bakel, Judith Aberg, Adolfo García-Sastre, Gunjan Shah, Tobias Hohl, Genovefa Papanicolaou, Miguel-Angel Perales, Kent Sepkowitz, Ngolela Esther Babady, and Mini Kamboj |
| EPI_ISL_583452, EPI_ISL_583453, EPI_ISL_583454, EPI_ISL_583473 | Garcia-Sastre Laboratory, Department of Microbiology, Icahn School of Medicine at Mount Sinai | van Bakel Laboratory, Genetics and Genomics Sciences, Icahn School of Medicine at Mount Sinai | Teresa Aydillo, Ana S. Gonzalez-Reiche, Sadaf Aslam, Adriana van de Guchte, Zenab Khan, Ajay Obla, Jayeeta Dutta, Harm van Bakel, Judith Aberg, Adolfo García-Sastre, Gunjan Shah, Tobias Hohl, Genovefa Papanicolaou, Miguel-Angel Perales, Kent Sepkowitz, Ngolela Esther Babady, and Mini Kamboj |
| EPI_ISL_583476 | Memorial Sloan Kettering Cancer Center | van Bakel Laboratory, Genetics and Genomics Sciences, Icahn School of Medicine at Mount Sinai | Teresa Aydillo, Ana S. Gonzalez-Reiche, Sadaf Aslam, Adriana van de Guchte, Zenab Khan, Ajay Obla, Jayeeta Dutta, Harm van Bakel, Judith Aberg, Adolfo García-Sastre, Gunjan Shah, Tobias Hohl, Genovefa Papanicolaou, Miguel-Angel Perales, Kent Sepkowitz, Ngolela Esther Babady, and Mini Kamboj |
| EPI_ISL_583477 | Garcia-Sastre Laboratory, Department of Microbiology, Icahn School of Medicine at Mount Sinai | van Bakel Laboratory, Genetics and Genomics Sciences, Icahn School of Medicine at Mount Sinai | Teresa Aydillo, Ana S. Gonzalez-Reiche, Sadaf Aslam, Adriana van de Guchte, Zenab Khan, Ajay Obla, Jayeeta Dutta, Harm van Bakel, Judith Aberg, Adolfo García-Sastre, Gunjan Shah, Tobias Hohl, Genovefa Papanicolaou, Miguel-Angel Perales, Kent Sepkowitz, Ngolela Esther Babady, and Mini Kamboj |
| EPI_ISL_583495 | Serviço de Verificação de Óbitos SVO Guaruulhos | Instituto Adolfo Lutz, Interdiciplinary Procedures Center, Strategic Laboratory | Claudio Tavares Sacchi, Claudia Regina Gonçalves, Erica Valessa Ramos Gomes, Karoline Rodrigues Campos |
| EPI_ISL_583496 | UPA Jandira | Instituto Adolfo Lutz, Interdiciplinary Procedures Center, Strategic Laboratory | Claudio Tavares Sacchi, Claudia Regina Gonçalves, Erica Valessa Ramos Gomes, Karoline Rodrigues Campos |
| EPI_ISL_583497 | Complexo Hospitalar Ouro Verde de Campinas | Instituto Adolfo Lutz, Interdiciplinary Procedures Center, Strategic Laboratory | Claudio Tavares Sacchi, Claudia Regina Gonçalves, Erica Valessa Ramos Gomes, Karoline Rodrigues Campos |
| EPI_ISL_583498 | Hospital Municipal Dr. Waldemar Tebaldi | Instituto Adolfo Lutz, Interdiciplinary Procedures Center, Strategic Laboratory | Claudio Tavares Sacchi, Claudia Regina Gonçalves, Erica Valessa Ramos Gomes, Karoline Rodrigues Campos |
| EPI_ISL_583499 | Distrito Sanitario Sul Campinas | Instituto Adolfo Lutz, Interdiciplinary Procedures Center, Strategic Laboratory | Claudio Tavares Sacchi, Claudia Regina Gonçalves, Erica Valessa Ramos Gomes, Karoline Rodrigues Campos |
| EPI_ISL_583500 | Centro de Saude I Tacito Leite de Carvalho e Silva | Instituto Adolfo Lutz, Interdiciplinary Procedures Center, Strategic Laboratory | Claudio Tavares Sacchi, Claudia Regina Gonçalves, Erica Valessa Ramos Gomes, Karoline Rodrigues Campos |
| EPI_ISL_583501 | Hospital Estadual de CampanhaCOVID 19 Barradas | Instituto Adolfo Lutz, Interdiciplinary Procedures Center, Strategic Laboratory | Claudio Tavares Sacchi, Claudia Regina Gonçalves, Erica Valessa Ramos Gomes, Karoline Rodrigues Campos |
| EPI_ISL_583502 | Serv de Vig Sanitaria Epidemio e CTRL de Zoonoses Guaruja | Instituto Adolfo Lutz, Interdiciplinary Procedures Center, Strategic Laboratory | Claudio Tavares Sacchi, Claudia Regina Gonçalves, Erica Valessa Ramos Gomes, Karoline Rodrigues Campos |
| EPI_ISL_583503 | CTA Centro de Testagem e Aconselhamento | Instituto Adolfo Lutz, Interdiciplinary Procedures Center, Strategic Laboratory | Claudio Tavares Sacchi, Claudia Regina Gonçalves, Erica Valessa Ramos Gomes, Karoline Rodrigues Campos |
| EPI_ISL_583504, EPI_ISL_583505 | Casa de Saude Stella Maris | Instituto Adolfo Lutz, Interdiciplinary Procedures Center, Strategic Laboratory | Claudio Tavares Sacchi, Claudia Regina Gonçalves, Erica Valessa Ramos Gomes, Karoline Rodrigues Campos |
| EPI_ISL_583894, EPI_ISL_583895, EPI_ISL_583896, EPI_ISL_583897, EPI_ISL_583898, EPI_ISL_583899, EPI_ISL_583900 | Singapore General Hospital | Department of Microbiology | Nurdyana Abdul Rahman, Kun Lee Lim, Chenhao Li, Sui Sin Goh, Kenneth Xin Long Chan, Kian Sing Chan, Lynette Oon, Kern Rei Chng, Niranjan Nagarajan, Karrie Ko |
| EPI_ISL_584074, EPI_ISL_584075, EPI_ISL_584076, EPI_ISL_584077, EPI_ISL_584078, EPI_ISL_584080 | The National Institute of Public Health | State Veterinary Institute Prague | Nagy,A;Jirincova,H;Novakova,L;Trnka,D;Vecerova,J |
| EPI_ISL_586513 | B.J. Medical College and Civil hospital, Ahmedabad | Gujarat Biotechnology Research Centre | Pinal Trivedi, Maharshi Pandya, Nidhi Patel, Nitin Savaliya, Raghawendra Kumar, Dinesh Kumar, Zuber Saiyed, Komal Patel, Labdhi Pandya, Afzal Ansari, Nikha Trivedi, Pranay Shah, Kamlesh J Upadhyay, Sanjay Kapadia, Apurvasinh Puvar, Janvi Raval, Zarna Patel, Monika Gandhi, R D Dixit, A M Kadri, Harsh Bakshi, Chaitanya Joshi, Madhvi Joshi |
| EPI_ISL_586514 | B.J. Medical College and Civil hospital, Ahmedabad | Gujarat Biotechnology Research Centre | Maharshi Pandya, Nidhi Patel, Nitin Savaliya, Raghawendra Kumar, Dinesh Kumar, Zuber Saiyed, Komal Patel, Labdhi Pandya, Afzal Ansari, Nikha Trivedi, Pranay Shah, Kamlesh J Upadhyay, Sanjay Kapadia, Apurvasinh Puvar, Janvi Raval, Zarna Patel, Monika Gandhi, Pinal Trivedi, R D Dixit, A M Kadri, Harsh Bakshi, Chaitanya Joshi, Madhvi Joshi |
| EPI_ISL_586515 | B.J. Medical College and Civil hospital, Ahmedabad | Gujarat Biotechnology Research Centre | Nidhi Patel, Nitin Savaliya, Raghawendra Kumar, Dinesh Kumar, Zuber Saiyed, Komal Patel, Labdhi Pandya, Afzal Ansari, Nikha Trivedi, Pranay Shah, Kamlesh J Upadhyay, Sanjay Kapadia, Apurvasinh Puvar, Janvi Raval, Zarna Patel, Monika Gandhi, Pinal Trivedi, Maharshi Pandya, R D Dixit, A M Kadri, Harsh Bakshi, Chaitanya Joshi, Madhvi Joshi |
| EPI_ISL_586516 | B.J. Medical College and Civil hospital, Ahmedabad | Gujarat Biotechnology Research Centre | Nitin Savaliya, Raghawendra Kumar, Dinesh Kumar, Zuber Saiyed, Komal Patel, Labdhi Pandya, Afzal Ansari, Nikha Trivedi, Pranay Shah, Kamlesh J Upadhyay, Sanjay Kapadia, Apurvasinh Puvar, Janvi Raval, Zarna Patel, Monika Gandhi, Pinal Trivedi, Maharshi Pandya, Nidhi Patel, R D Dixit, A M Kadri, Harsh Bakshi, Chaitanya Joshi, Madhvi Joshi |
| EPI_ISL_586517 | B.J. Medical College and Civil hospital, Ahmedabad | Gujarat Biotechnology Research Centre | Raghawendra Kumar, Dinesh Kumar, Zuber Saiyed, Komal Patel, Labdhi Pandya, Afzal Ansari, Nikha Trivedi, Pranay Shah, Kamlesh J Upadhyay, Sanjay Kapadia, Apurvasinh Puvar, Janvi Raval, Zarna Patel, Monika Gandhi, Pinal Trivedi, Maharshi Pandya, Nidhi Patel, Nitin Savaliya, R D Dixit, A M Kadri, Harsh Bakshi, Chaitanya Joshi, Madhvi Joshi |
| EPI_ISL_586518 | B.J. Medical College and Civil hospital, Ahmedabad | Gujarat Biotechnology Research Centre | Dinesh Kumar, Zuber Saiyed, Komal Patel, Labdhi Pandya, Afzal Ansari, Nikha Trivedi, Pranay Shah, Kamlesh J Upadhyay, Sanjay Kapadia, Apurvasinh Puvar, Janvi Raval, Zarna Patel, Monika Gandhi, Pinal Trivedi, Maharshi Pandya, Nidhi Patel, Nitin Savaliya, Raghawendra Kumar, R D Dixit, A M Kadri, Harsh Bakshi, Chaitanya Joshi, Madhvi Joshi |
| EPI_ISL_586519 | B.J. Medical College and Civil hospital, Ahmedabad | Gujarat Biotechnology Research Centre | Zuber Saiyed, Komal Patel, Labdhi Pandya, Afzal Ansari, Nikha Trivedi, Pranay Shah, Kamlesh J Upadhyay, Sanjay Kapadia, Apurvasinh Puvar, Janvi Raval, Zarna Patel, Monika Gandhi, Pinal Trivedi, Maharshi Pandya, Nidhi Patel, Nitin Savaliya, Raghawendra Kumar, Dinesh Kumar, R D Dixit, A M Kadri, Harsh Bakshi, Chaitanya Joshi, Madhvi Joshi |
| EPI_ISL_586520 | B.J. Medical College and Civil hospital, Ahmedabad | Gujarat Biotechnology Research Centre | Komal Patel, Labdhi Pandya, Afzal Ansari, Nikha Trivedi, Pranay Shah, Kamlesh J Upadhyay, Sanjay Kapadia, Apurvasinh Puvar, Janvi Raval, Zarna Patel, Monika Gandhi, Pinal Trivedi, Maharshi Pandya, Nidhi Patel, Nitin Savaliya, Raghawendra Kumar, Dinesh Kumar, Zuber Saiyed, R D Dixit, A M Kadri, Harsh Bakshi, Chaitanya Joshi, Madhvi Joshi |
| EPI_ISL_586521 | B.J. Medical College and Civil hospital, Ahmedabad | Gujarat Biotechnology Research Centre | Labdhi Pandya, Afzal Ansari, Nikha Trivedi, Pranay Shah, Kamlesh J Upadhyay, Sanjay Kapadia, Apurvasinh Puvar, Janvi Raval, Zarna Patel, Monika Gandhi, Pinal Trivedi, Maharshi Pandya, Nidhi Patel, Nitin Savaliya, Raghawendra Kumar, Dinesh Kumar, Zuber Saiyed, Komal Patel, R D Dixit, A M Kadri, Harsh Bakshi, Chaitanya Joshi, Madhvi Joshi |
| EPI_ISL_586522 | B.J. Medical College and Civil hospital, Ahmedabad | Gujarat Biotechnology Research Centre | Afzal Ansari, Nikha Trivedi, Pranay Shah, Kamlesh J Upadhyay, Sanjay Kapadia, Apurvasinh Puvar, Janvi Raval, Zarna Patel, Monika Gandhi, Pinal Trivedi, Maharshi Pandya, Nidhi Patel, Nitin Savaliya, Raghawendra Kumar, Dinesh Kumar, Zuber Saiyed, Komal Patel, Labdhi Pandya, R D Dixit, A M Kadri, Harsh Bakshi, Chaitanya Joshi, Madhvi Joshi |
| EPI_ISL_586523 | B.J. Medical College and Civil hospital, Ahmedabad | Gujarat Biotechnology Research Centre | Nikha Trivedi, Pranay Shah, Kamlesh J Upadhyay, Sanjay Kapadia, Apurvasinh Puvar, Janvi Raval, Zarna Patel, Monika Gandhi, Pinal Trivedi, Maharshi Pandya, Nidhi Patel, Nitin Savaliya, Raghawendra Kumar, Dinesh Kumar, Zuber Saiyed, Komal Patel, Labdhi Pandya, Afzal Ansari, R D Dixit, A M Kadri, Harsh Bakshi, Chaitanya Joshi, Madhvi Joshi |
| EPI_ISL_586524 | B.J. Medical College and Civil hospital, Ahmedabad | Gujarat Biotechnology Research Centre | Pranay Shah, Kamlesh J Upadhyay, Sanjay Kapadia, Apurvasinh Puvar, Janvi Raval, Zarna Patel, Monika Gandhi, Pinal Trivedi, Maharshi Pandya, Nidhi Patel, Nitin Savaliya, Raghawendra Kumar, Dinesh Kumar, Zuber Saiyed, Komal Patel, Labdhi Pandya, Afzal Ansari, Nikha Trivedi, R D Dixit, A M Kadri, Harsh Bakshi, Chaitanya Joshi, Madhvi Joshi |
| EPI_ISL_586525 | B.J. Medical College and Civil hospital, Ahmedabad | Gujarat Biotechnology Research Centre | Kamlesh J Upadhyay, Sanjay Kapadia, Apurvasinh Puvar, Janvi Raval, Zarna Patel, Monika Gandhi, Pinal Trivedi, Maharshi Pandya, Nidhi Patel, Nitin Savaliya, Raghawendra Kumar, Dinesh Kumar, Zuber Saiyed, Komal Patel, Labdhi Pandya, Afzal Ansari, Nikha Trivedi, Pranay Shah, R D Dixit, A M Kadri, |

| | | | |
|---|---|---|---|
| | | | Harsh Bakshi, Chaitanya Joshi, Madhvi Joshi |
| EPI_ISL_586526 | B.J. Medical College and Civil hospital, Ahmedabad | Gujarat Biotechnology Research Centre | Sanjay Kapadia, Apurvasinh Puvar, Janvi Raval, Zarna Patel, Monika Gandhi, Pinal Trivedi, Maharshi Pandya, Nidhi Patel, Nitin Savaliya, Raghawendra Kumar, Dinesh Kumar, Zuber Saiyed, Komal Patel, Labdhi Pandya, Afzal Ansari, Nikha Trivedi, Pranay Shah, Kamlesh J Upadhyay, R D Dixit, A M Kadri, Harsh Bakshi, Chaitanya Joshi, Madhvi Joshi |
| EPI_ISL_586527 | B.J. Medical College and Civil hospital, Ahmedabad | Gujarat Biotechnology Research Centre | Apurvasinh Puvar, Janvi Raval, Zarna Patel, Monika Gandhi, Pinal Trivedi, Maharshi Pandya, Nidhi Patel, Nitin Savaliya, Raghawendra Kumar, Dinesh Kumar, Zuber Saiyed, Komal Patel, Labdhi Pandya, Afzal Ansari, Nikha Trivedi, Pranay Shah, Kamlesh J Upadhyay, Sanjay Kapadia, R D Dixit, A M Kadri, Harsh Bakshi, Chaitanya Joshi, Madhvi Joshi |
| EPI_ISL_586528 | B.J. Medical College and Civil hospital, Ahmedabad | Gujarat Biotechnology Research Centre | Zarna Patel, Monika Gandhi, Pinal Trivedi, Maharshi Pandya, Nidhi Patel, Nitin Savaliya, Raghawendra Kumar, Dinesh Kumar, Zuber Saiyed, Komal Patel, Labdhi Pandya, Afzal Ansari, Nikha Trivedi, Pranay Shah, Kamlesh J Upadhyay, Sanjay Kapadia, Apurvasinh Puvar, Janvi Raval, R D Dixit, A M Kadri, Harsh Bakshi, Chaitanya Joshi, Madhvi Joshi |
| EPI_ISL_586529 | B.J. Medical College and Civil hospital, Ahmedabad | Gujarat Biotechnology Research Centre | Monika Gandhi, Pinal Trivedi, Maharshi Pandya, Nidhi Patel, Nitin Savaliya, Raghawendra Kumar, Dinesh Kumar, Zuber Saiyed, Komal Patel, Labdhi Pandya, Afzal Ansari, Nikha Trivedi, Pranay Shah, Kamlesh J Upadhyay, Sanjay Kapadia, Apurvasinh Puvar, Janvi Raval, Zarna Patel, R D Dixit, A M Kadri, Harsh Bakshi, Chaitanya Joshi, Madhvi Joshi |
| EPI_ISL_586530 | B.J. Medical College and Civil hospital, Ahmedabad | Gujarat Biotechnology Research Centre | Pinal Trivedi, Maharshi Pandya, Nidhi Patel, Nitin Savaliya, Raghawendra Kumar, Dinesh Kumar, Zuber Saiyed, Komal Patel, Labdhi Pandya, Afzal Ansari, Nikha Trivedi, Pranay Shah, Kamlesh J Upadhyay, Sanjay Kapadia, Apurvasinh Puvar, Janvi Raval, Zarna Patel, Monika Gandhi, R D Dixit, A M Kadri, Harsh Bakshi, Chaitanya Joshi, Madhvi Joshi |
| EPI_ISL_586531 | General Hospital,Chhotaudepur | Gujarat Biotechnology Research Centre | Yogesh Parmar, Arti Thakur, Dharak Pandya, Maharshi Pandya, Nidhi Patel, Nitin Savaliya, Raghawendra Kumar, Dinesh Kumar, Zuber Saiyed, Komal Patel, Labdhi Pandya, Afzal Ansari, Nikha Trivedi, Apurvasinh Puvar, Janvi Raval, Zarna Patel, Monika Gandhi, Pinal Trivedi, R D Dixit, A M Kadri, Harsh Bakshi, Chaitanya Joshi, Madhvi Joshi |
| EPI_ISL_586532 | General Hospital,Chhotaudepur | Gujarat Biotechnology Research Centre | Arti Thakur, Dharak Pandya, Maharshi Pandya, Nidhi Patel, Nitin Savaliya, Raghawendra Kumar, Dinesh Kumar, Zuber Saiyed, Komal Patel, Labdhi Pandya, Afzal Ansari, Nikha Trivedi, Apurvasinh Puvar, Janvi Raval, Zarna Patel, Monika Gandhi, Pinal Trivedi, Yogesh Parmar, R D Dixit, A M Kadri, Harsh Bakshi, Chaitanya Joshi, Madhvi Joshi |
| EPI_ISL_586533 | General Hospital,Chhotaudepur | Gujarat Biotechnology Research Centre | Dharak Pandya, Maharshi Pandya, Nidhi Patel, Nitin Savaliya, Raghawendra Kumar, Dinesh Kumar, Zuber Saiyed, Komal Patel, Labdhi Pandya, Afzal Ansari, Nikha Trivedi, Apurvasinh Puvar, Janvi Raval, Zarna Patel, Monika Gandhi, Pinal Trivedi, Yogesh Parmar, Arti Thakur, R D Dixit, A M Kadri, Harsh Bakshi, Chaitanya Joshi, Madhvi Joshi |
| EPI_ISL_586534 | General Hospital,Chhotaudepur | Gujarat Biotechnology Research Centre | Maharshi Pandya, Nidhi Patel, Nitin Savaliya, Raghawendra Kumar, Dinesh Kumar, Zuber Saiyed, Komal Patel, Labdhi Pandya, Afzal Ansari, Nikha Trivedi, Apurvasinh Puvar, Janvi Raval, Zarna Patel, Monika Gandhi, Pinal Trivedi, Yogesh Parmar, Arti Thakur, Dharak Pandya, R D Dixit, A M Kadri, Harsh Bakshi, Chaitanya Joshi, Madhvi Joshi |
| EPI_ISL_586535 | General Hospital,Chhotaudepur | Gujarat Biotechnology Research Centre | Nidhi Patel, Nitin Savaliya, Raghawendra Kumar, Dinesh Kumar, Zuber Saiyed, Komal Patel, Labdhi Pandya, Afzal Ansari, Nikha Trivedi, Apurvasinh Puvar, Janvi Raval, Zarna Patel, Monika Gandhi, Pinal Trivedi, Yogesh Parmar, Arti Thakur, Dharak Pandya, Maharshi Pandya, R D Dixit, A M Kadri, Harsh Bakshi, Chaitanya Joshi, Madhvi Joshi |
| EPI_ISL_586536 | Zydus Medical Hospital,Dahod | Gujarat Biotechnology Research Centre | Kalpesh Labana, Mohit Desai, Nitin Savaliya, Raghawendra Kumar, Dinesh Kumar, Zuber Saiyed, Komal Patel, Labdhi Pandya, Afzal Ansari, Nikha Trivedi, Apurvasinh Puvar, Janvi Raval, Zarna Patel, Monika Gandhi, Pinal Trivedi, Maharshi Pandya, Nidhi Patel, R D Dixit, A M Kadri, Harsh Bakshi, Chaitanya Joshi, Madhvi Joshi |
| EPI_ISL_586537 | Zydus Medical Hospital,Dahod | Gujarat Biotechnology Research Centre | Mohit Desai, Nitin Savaliya, Raghawendra Kumar, Dinesh Kumar, Zuber Saiyed, Komal Patel, Labdhi Pandya, Afzal Ansari, Nikha Trivedi, Apurvasinh Puvar, Janvi Raval, Zarna Patel, Monika Gandhi, Pinal Trivedi, Maharshi Pandya, Nidhi Patel, Kalpesh Labana, R D Dixit, A M Kadri, Harsh Bakshi, Chaitanya Joshi, Madhvi Joshi |
| EPI_ISL_586538 | Zydus Medical Hospital,Dahod | Gujarat Biotechnology Research Centre | Nitin Savaliya, Raghawendra Kumar, Dinesh Kumar, Zuber Saiyed, Komal Patel, Labdhi Pandya, Afzal Ansari, Nikha Trivedi, Apurvasinh Puvar, Janvi Raval, Zarna Patel, Monika Gandhi, Pinal Trivedi, Maharshi Pandya, Nidhi Patel, Kalpesh Labana, Mohit Desai, R D Dixit, A M Kadri, Harsh Bakshi, Chaitanya Joshi, Madhvi Joshi |
| EPI_ISL_586539 | Zydus Medical Hospital,Dahod | Gujarat Biotechnology Research Centre | Raghawendra Kumar, Dinesh Kumar, Zuber Saiyed, Komal Patel, Labdhi Pandya, Afzal Ansari, Nikha Trivedi, Apurvasinh Puvar, Janvi Raval, Zarna Patel, Monika Gandhi, Pinal Trivedi, Maharshi Pandya, Nidhi Patel, Kalpesh Labana, Mohit Desai, Nitin Savaliya, R D Dixit, A M Kadri, Harsh Bakshi, Chaitanya Joshi, Madhvi Joshi |
| EPI_ISL_586540 | Zydus Medical Hospital,Dahod | Gujarat Biotechnology Research Centre | Dinesh Kumar, Zuber Saiyed, Komal Patel, Labdhi Pandya, Afzal Ansari, Nikha Trivedi, Apurvasinh Puvar, Janvi Raval, Zarna Patel, Monika Gandhi, Pinal Trivedi, Maharshi Pandya, Nidhi Patel, Kalpesh Labana, Mohit Desai, Nitin Savaliya, Raghawendra Kumar, R D Dixit, A M Kadri, Harsh Bakshi, Chaitanya Joshi, Madhvi Joshi |
| EPI_ISL_586541 | Zydus Medical Hospital,Dahod | Gujarat Biotechnology Research Centre | Zuber Saiyed, Komal Patel, Labdhi Pandya, Afzal Ansari, Nikha Trivedi, Apurvasinh Puvar, Janvi Raval, Zarna Patel, Monika Gandhi, Pinal Trivedi, Maharshi Pandya, Nidhi Patel, Kalpesh Labana, Mohit Desai, Nitin Savaliya, Raghawendra Kumar, Dinesh Kumar, R D Dixit, A M Kadri, Harsh Bakshi, Chaitanya Joshi, Madhvi Joshi |
| EPI_ISL_586542 | Government General Hospital, Jam Khambhaliya, Devbhoomi Dwarka | Gujarat Biotechnology Research Centre | Komal Patel, Labdhi Pandya, Afzal Ansari, Nikha Trivedi, Harish Matani, Apurvasinh Puvar, Janvi Raval, Zarna Patel, Monika Gandhi, Pinal Trivedi, Maharshi Pandya, Nidhi Patel, Nitin Savaliya, Raghawendra Kumar, Dinesh Kumar, Zuber Saiyed, R D Dixit, A M Kadri, Harsh Bakshi, Chaitanya Joshi, Madhvi Joshi |
| EPI_ISL_586543 | Government General Hospital, Jam Khambhaliya, Devbhoomi Dwarka | Gujarat Biotechnology Research Centre | Labdhi Pandya, Afzal Ansari, Nikha Trivedi, Harish Matani, Apurvasinh Puvar, Janvi Raval, Zarna Patel, Monika Gandhi, Pinal Trivedi, Maharshi Pandya, Nidhi Patel, Nitin Savaliya, Raghawendra Kumar, Dinesh Kumar, Zuber Saiyed, Komal Patel, R D Dixit, A M Kadri, Harsh Bakshi, Chaitanya Joshi, Madhvi Joshi |
| EPI_ISL_586544 | Government General Hospital, Jam Khambhaliya, Devbhoomi Dwarka | Gujarat Biotechnology Research Centre | Afzal Ansari, Nikha Trivedi, Harish Matani, Apurvasinh Puvar, Janvi Raval, Zarna Patel, Monika Gandhi, Pinal Trivedi, Maharshi Pandya, Nidhi Patel, Nitin Savaliya, Raghawendra Kumar, Dinesh Kumar, Zuber Saiyed, Komal Patel, Labdhi Pandya, R D Dixit, A M Kadri, Harsh Bakshi, Chaitanya Joshi, Madhvi Joshi |
| EPI_ISL_586545 | Government General Hospital, Jam Khambhaliya, Devbhoomi Dwarka | Gujarat Biotechnology Research Centre | Nikha Trivedi, Harish Matani, Apurvasinh Puvar, Janvi Raval, Zarna Patel, Monika Gandhi, Pinal Trivedi, Maharshi Pandya, Nidhi Patel, Nitin Savaliya, Raghawendra Kumar, Dinesh Kumar, Zuber Saiyed, Komal Patel, Labdhi Pandya, Afzal Ansari, R D Dixit, A M Kadri, Harsh Bakshi, Chaitanya Joshi, Madhvi Joshi |
| EPI_ISL_586546 | General Hospital,Godhra | Gujarat Biotechnology Research Centre | Neela Katara, Pinal Gandhi, Sangeeta Kumar, Apurvasinh Puvar, Janvi Raval, Zarna Patel, Monika Gandhi, Pinal Trivedi, Maharshi Pandya, Nidhi Patel, Nitin Savaliya, Raghawendra Kumar, Dinesh Kumar, Zuber Saiyed, Komal Patel, Labdhi Pandya, Afzal Ansari, Nikha Trivedi, R D Dixit, A M Kadri, Harsh Bakshi, Chaitanya Joshi, Madhvi Joshi |
| EPI_ISL_586547 | General Hospital,Godhra | Gujarat Biotechnology Research Centre | Pinal Gandhi, Sangeeta Kumar, Apurvasinh Puvar, Janvi Raval, Zarna Patel, Monika Gandhi, Pinal Trivedi, Maharshi Pandya, Nidhi Patel, Nitin Savaliya, Raghawendra Kumar, Dinesh Kumar, Zuber Saiyed, Komal Patel, Labdhi Pandya, Afzal Ansari, Nikha Trivedi, Neela Katara, R D Dixit, A M Kadri, Harsh Bakshi, Chaitanya Joshi, Madhvi Joshi |
| EPI_ISL_586548 | General Hospital,Godhra | Gujarat Biotechnology Research Centre | Sangeeta Kumar, Apurvasinh Puvar, Janvi Raval, Zarna Patel, Monika Gandhi, Pinal Trivedi, Maharshi Pandya, Nidhi Patel, Nitin Savaliya, Raghawendra Kumar, Dinesh Kumar, Zuber Saiyed, Komal Patel, Labdhi Pandya, Afzal Ansari, Nikha Trivedi, Neela Katara, Pinal Gandhi, R D Dixit, A M Kadri, Harsh Bakshi, Chaitanya Joshi, Madhvi Joshi |
| EPI_ISL_586549 | CCC,Veraval | Gujarat Biotechnology Research Centre | Jignesh Parmar, Jitendra Bamrotia, Apurvasinh Puvar, Janvi Raval, Zarna Patel, Monika Gandhi, Pinal Trivedi, Maharshi Pandya, Nidhi Patel, Nitin Savaliya, Raghawendra Kumar, Dinesh Kumar, Zuber Saiyed, Komal Patel, Labdhi Pandya, Afzal Ansari, Nikha Trivedi, R D Dixit, A M Kadri, Harsh Bakshi, Chaitanya Joshi, Madhvi Joshi |
| EPI_ISL_586550 | CCC,Veraval | Gujarat Biotechnology Research Centre | Jitendra Bamrotia, Apurvasinh Puvar, Janvi Raval, Zarna Patel, Monika Gandhi, Pinal Trivedi, Maharshi Pandya, Nidhi Patel, Nitin Savaliya, Raghawendra Kumar, Dinesh Kumar, Zuber Saiyed, Komal Patel, Labdhi Pandya, Afzal Ansari, Nikha Trivedi, Jignesh Parmar, R D Dixit, A M Kadri, Harsh Bakshi, Chaitanya Joshi, Madhvi Joshi |

| | | | |
|---|---|---|---|
| EPI_ISL_586551 | Government Hospital,Veraval | Gujarat Biotechnology Research Centre | Monika Gandhi, Pinal Trivedi, Maharshi Pandya, Nidhi Patel, Nitin Savaliya, Raghawendra Kumar, Dinesh Kumar, Zuber Saiyed, Komal Patel, Labdhi Pandya, Afzal Ansari, Nikha Trivedi, Jignesh Parmar, Jitendra Bamrotia, Apurvasinh Puvar, Janvi Raval, Zarna Patel, R D Dixit, A M Kadri, Harsh Bakshi, Chaitanya Joshi, Madhvi Joshi |
| EPI_ISL_586552 | Government Hospital,Veraval | Gujarat Biotechnology Research Centre | Pinal Trivedi, Maharshi Pandya, Nidhi Patel, Nitin Savaliya, Raghawendra Kumar, Dinesh Kumar, Zuber Saiyed, Komal Patel, Labdhi Pandya, Afzal Ansari, Nikha Trivedi, Jignesh Parmar, Jitendra Bamrotia, Apurvasinh Puvar, Janvi Raval, Zarna Patel, Monika Gandhi, R D Dixit, A M Kadri, Harsh Bakshi, Chaitanya Joshi, Madhvi Joshi |
| EPI_ISL_586553 | Government Hospital,Veraval | Gujarat Biotechnology Research Centre | Maharshi Pandya, Nidhi Patel, Nitin Savaliya, Raghawendra Kumar, Dinesh Kumar, Zuber Saiyed, Komal Patel, Labdhi Pandya, Afzal Ansari, Nikha Trivedi, Jignesh Parmar, Jitendra Bamrotia, Apurvasinh Puvar, Janvi Raval, Zarna Patel, Monika Gandhi, Pinal Trivedi, R D Dixit, A M Kadri, Harsh Bakshi, Chaitanya Joshi, Madhvi Joshi |
| EPI_ISL_586554 | Government Hospital,Veraval | Gujarat Biotechnology Research Centre | Nidhi Patel, Nitin Savaliya, Raghawendra Kumar, Dinesh Kumar, Zuber Saiyed, Komal Patel, Labdhi Pandya, Afzal Ansari, Nikha Trivedi, Jignesh Parmar, Jitendra Bamrotia, Apurvasinh Puvar, Janvi Raval, Zarna Patel, Monika Gandhi, Pinal Trivedi, Maharshi Pandya, R D Dixit, A M Kadri, Harsh Bakshi, Chaitanya Joshi, Madhvi Joshi |
| EPI_ISL_586555 | CCC,Veraval | Gujarat Biotechnology Research Centre | Nitin Savaliya, Raghawendra Kumar, Dinesh Kumar, Zuber Saiyed, Komal Patel, Labdhi Pandya, Afzal Ansari, Nikha Trivedi, Jignesh Parmar, Jitendra Bamrotia, Apurvasinh Puvar, Janvi Raval, Zarna Patel, Monika Gandhi, Pinal Trivedi, Maharshi Pandya, Nidhi Patel, R D Dixit, A M Kadri, Harsh Bakshi, Chaitanya Joshi, Madhvi Joshi |
| EPI_ISL_586556 | CCC,Veraval | Gujarat Biotechnology Research Centre | Raghawendra Kumar, Dinesh Kumar, Zuber Saiyed, Komal Patel, Labdhi Pandya, Afzal Ansari, Nikha Trivedi, Jignesh Parmar, Jitendra Bamrotia, Apurvasinh Puvar, Janvi Raval, Zarna Patel, Monika Gandhi, Pinal Trivedi, Maharshi Pandya, Nidhi Patel, Nitin Savaliya, R D Dixit, A M Kadri, Harsh Bakshi, Chaitanya Joshi, Madhvi Joshi |
| EPI_ISL_586557 | CCC,Veraval | Gujarat Biotechnology Research Centre | Dinesh Kumar, Zuber Saiyed, Komal Patel, Labdhi Pandya, Afzal Ansari, Nikha Trivedi, Jignesh Parmar, Jitendra Bamrotia, Apurvasinh Puvar, Janvi Raval, Zarna Patel, Monika Gandhi, Pinal Trivedi, Maharshi Pandya, Nidhi Patel, Nitin Savaliya, Raghawendra Kumar, R D Dixit, A M Kadri, Harsh Bakshi, Chaitanya Joshi, Madhvi Joshi |
| EPI_ISL_586558 | CCC,Veraval | Gujarat Biotechnology Research Centre | Zuber Saiyed, Komal Patel, Labdhi Pandya, Afzal Ansari, Nikha Trivedi, Jignesh Parmar, Jitendra Bamrotia, Apurvasinh Puvar, Janvi Raval, Zarna Patel, Monika Gandhi, Pinal Trivedi, Maharshi Pandya, Nidhi Patel, Nitin Savaliya, Raghawendra Kumar, Dinesh Kumar, R D Dixit, A M Kadri, Harsh Bakshi, Chaitanya Joshi, Madhvi Joshi |
| EPI_ISL_586559 | CCC,Veraval | Gujarat Biotechnology Research Centre | Komal Patel, Labdhi Pandya, Afzal Ansari, Nikha Trivedi, Jignesh Parmar, Jitendra Bamrotia, Apurvasinh Puvar, Janvi Raval, Zarna Patel, Monika Gandhi, Pinal Trivedi, Maharshi Pandya, Nidhi Patel, Nitin Savaliya, Raghawendra Kumar, Dinesh Kumar, Zuber Saiyed, R D Dixit, A M Kadri, Harsh Bakshi, Chaitanya Joshi, Madhvi Joshi |
| EPI_ISL_586560 | CCC,Veraval | Gujarat Biotechnology Research Centre | Labdhi Pandya, Afzal Ansari, Nikha Trivedi, Jignesh Parmar, Jitendra Bamrotia, Apurvasinh Puvar, Janvi Raval, Zarna Patel, Monika Gandhi, Pinal Trivedi, Maharshi Pandya, Nidhi Patel, Nitin Savaliya, Raghawendra Kumar, Dinesh Kumar, Zuber Saiyed, Komal Patel, R D Dixit, A M Kadri, Harsh Bakshi, Chaitanya Joshi, Madhvi Joshi |
| EPI_ISL_586561 | CCC,Veraval | Gujarat Biotechnology Research Centre | Afzal Ansari, Nikha Trivedi, Jignesh Parmar, Jitendra Bamrotia, Apurvasinh Puvar, Janvi Raval, Zarna Patel, Monika Gandhi, Pinal Trivedi, Maharshi Pandya, Nidhi Patel, Nitin Savaliya, Raghawendra Kumar, Dinesh Kumar, Zuber Saiyed, Komal Patel, Labdhi Pandya, R D Dixit, A M Kadri, Harsh Bakshi, Chaitanya Joshi, Madhvi Joshi |
| EPI_ISL_586562 | GMERS GMERS Medical College and Hospital, Junagadh | Gujarat Biotechnology Research Centre | Nikha Trivedi, Bhavesh Bagda , Ravi Dedaniya, Apurvasinh Puvar, Janvi Raval, Zarna Patel, Monika Gandhi, Pinal Trivedi, Maharshi Pandya, Nidhi Patel, Nitin Savaliya, Raghawendra Kumar, Dinesh Kumar, Zuber Saiyed, Komal Patel, Labdhi Pandya, Afzal Ansari, R D Dixit, A M Kadri, Harsh Bakshi, Chaitanya Joshi, Madhvi Joshi |
| EPI_ISL_586563 | GMERS GMERS Medical College and Hospital, Junagadh | Gujarat Biotechnology Research Centre | Bhavesh Bagda , Ravi Dedaniya, Apurvasinh Puvar, Janvi Raval, Zarna Patel, Monika Gandhi, Pinal Trivedi, Maharshi Pandya, Nidhi Patel, Nitin Savaliya, Raghawendra Kumar, Dinesh Kumar, Zuber Saiyed, Komal Patel, Labdhi Pandya, Afzal Ansari, Nikha Trivedi, R D Dixit, A M Kadri, Harsh Bakshi, Chaitanya Joshi, Madhvi Joshi |
| EPI_ISL_586564 | GMERS GMERS Medical College and Hospital, Junagadh | Gujarat Biotechnology Research Centre | Ravi Dedaniya, Apurvasinh Puvar, Janvi Raval, Zarna Patel, Monika Gandhi, Pinal Trivedi, Maharshi Pandya, Nidhi Patel, Nitin Savaliya, Raghawendra Kumar, Dinesh Kumar, Zuber Saiyed, Komal Patel, Labdhi Pandya, Afzal Ansari, Nikha Trivedi, Bhavesh Bagda , R D Dixit, A M Kadri, Harsh Bakshi, Chaitanya Joshi, Madhvi Joshi |
| EPI_ISL_586565 | GMERS GMERS Medical College and Hospital, Junagadh | Gujarat Biotechnology Research Centre | Apurvasinh Puvar, Janvi Raval, Zarna Patel, Monika Gandhi, Pinal Trivedi, Maharshi Pandya, Nidhi Patel, Nitin Savaliya, Raghawendra Kumar, Dinesh Kumar, Zuber Saiyed, Komal Patel, Labdhi Pandya, Afzal Ansari, Nikha Trivedi, Bhavesh Bagda , Ravi Dedaniya, R D Dixit, A M Kadri, Harsh Bakshi, Chaitanya Joshi, Madhvi Joshi |
| EPI_ISL_586566 | GMERS GMERS Medical College and Hospital, Junagadh | Gujarat Biotechnology Research Centre | Janvi Raval, Zarna Patel, Monika Gandhi, Pinal Trivedi, Maharshi Pandya, Nidhi Patel, Nitin Savaliya, Raghawendra Kumar, Dinesh Kumar, Zuber Saiyed, Komal Patel, Labdhi Pandya, Afzal Ansari, Nikha Trivedi, Bhavesh Bagda , Ravi Dedaniya, Apurvasinh Puvar, R D Dixit, A M Kadri, Harsh Bakshi, Chaitanya Joshi, Madhvi Joshi |
| EPI_ISL_586567, EPI_ISL_586568 | Saikrishna Hospital,Mehsana | Gujarat Biotechnology Research Centre | Zarna Patel, Monika Gandhi, Pinal Trivedi, Maharshi Pandya, Nidhi Patel, Nitin Savaliya, Raghawendra Kumar, Dinesh Kumar, Zuber Saiyed, Komal Patel, Labdhi Pandya, Afzal Ansari, Nikha Trivedi, Harshadbhai Parmar, Apurvasinh Puvar, Janvi Raval, R D Dixit, A M Kadri, Harsh Bakshi, Chaitanya Joshi, Madhvi Joshi |
| EPI_ISL_590693 | INMI Lazzaro Spallanzani IRCCS | INMI Lazzaro Spallanzani IRCCS | Martina Rueca, Barbara Bartolini, Cesare E.M. Gruber, Francesco Messina, Emanuela Giombini, Beatrice Valli, Eleonora Lalle, Simone Lanini, Francesco Vairo, Maria R. Capobianchi, Antonino Di Caro |
| EPI_ISL_590694 | INMI Lazzaro Spallanzani IRCCS | INMI Lazzaro Spallanzani IRCCS | Barbara Bartolini, Martina Rueca, Francesco Messina, Cesare E.M. Gruber, Emanuela Giombini, Beatrice Valli, Eleonora Lalle, Simone Lanini, Francesco Vairo, Maria R. Capobianchi, Antonino Di Caro |
| EPI_ISL_590695 | INMI Lazzaro Spallanzani IRCCS | INMI Lazzaro Spallanzani IRCCS | Cesare E.M. Gruber, Francesco Messina, Barbara Bartolini, Martina Rueca, Emanuela Giombini, Beatrice Valli, Eleonora Lalle, Simone Lanini, Francesco Vairo, Antonino Di Caro, Maria R. Capobianchi |
| EPI_ISL_590696 | INMI Lazzaro Spallanzani IRCCS | INMI Lazzaro Spallanzani IRCCS | Cesare E.M. Gruber, Barbara Bartolini, Francesco Messina, Martina Rueca, Emanuela Giombini, Beatrice Valli, Eleonora Lalle, Simone Lanini, Francesco Vairo, Antonino Di Caro, Maria R. Capobianchi |
| EPI_ISL_590697 | INMI Lazzaro Spallanzani IRCCS | INMI Lazzaro Spallanzani IRCCS | Martina Rueca, Cesare E.M. Gruber, Barbara Bartolini, Francesco Messina, Emanuela Giombini, Beatrice Valli, Eleonora Lalle, Simone Lanini, Francesco Vairo, Antonino Di Caro, Maria R. Capobianchi |
| EPI_ISL_590698 | INMI Lazzaro Spallanzani IRCCS | INMI Lazzaro Spallanzani IRCCS | Barbara Bartolini, Francesco Messina, Cesare E.M. Gruber, Martina Rueca, Emanuela Giombini, Beatrice Valli, Eleonora Lalle, Simone Lanini, Francesco Vairo, Maria R. Capobianchi, Antonino Di Caro |
| EPI_ISL_593480, EPI_ISL_593553, EPI_ISL_593554, EPI_ISL_593555, EPI_ISL_593556 | Brigham and Women's Hospital | Jonathan Li Laboratory | Manish C. Choudhary, James Regan, Jonathan Z. Li |
| EPI_ISL_593596 | Swedish Medical Center, Seattle, WA, USA | Institute for Systems Biology | Jason D. Goldman, Kai Wang, Katharina Röltgen, Sandra C. A. Nielsen, Jared C. Roach, Samia N. Naccache, Fan Yang, Oliver F. Wirz, Kathryn E. Yost, Ji-Yeun Lee, Kelly Chun, Terri Wrin,, Christos J. Petropoulos, Inyoul Lee, Shannon Fallen, Paula M. Manner, Julie A. Wallick, Heather A. Algren, RN, Kim M. Murray, Yapeng Su, Jennifer Hadlock, Joshua Jeharajah, William R. Berrington, George P. Pappas, Sonam T. Nyatsatsang, Alexander L. Greninger, Ansuman T. Satpathy, John S. Pauk, Scott D. Boyd, James R. Heath |
| EPI_ISL_593912 | Hospital, Marange Silvange | National Reference Center for Viruses of Respiratory Infections, Institut Pasteur, Paris | Sylvie Behillil, Fabiana Gambaro, Etienne Simon-Lorière, Vincent Enouf, Maud Vanpeene, Sylvie van der Werf |
| EPI_ISL_593917 | Hospital, Montigny les Metz | National Reference Center for Viruses of Respiratory Infections, Institut Pasteur, Paris | Sylvie Behillil, Fabiana Gambaro, Etienne Simon-Lorière, Vincent Enouf, Maud Vanpeene, Sylvie van der Werf |

| | | | |
|---|---|---|---|
| EPI_ISL_593932 | Labo Analyses Med, Puteaux | National Reference Center for Viruses of Respiratory Infections, Institut Pasteur, Paris | Sylvie Behillil, Fabiana Gambaro, Etienne Simon-Lorière, Vincent Enouf, Maud Vanpeene, Sylvie van der Werf |
| EPI_ISL_594118 | Centro de Investigaciones, Universidad de Especialidades Espíritu Santo | Institute of Microbiology, Universidad San Francisco de Quito | Derly Andrade, Juan Carlos Fernandez, Belén Prado-Vivar, Sully Márquez, Juan José Guadalupe, Monica Becerra-Wong, Bernardo Gutiérrez, Gabriel Morey, Ruben Armas, Jose Pedro Barberan, Fernando Espinoza, Edith Lopez, Verónica Barragán, Patricio Rojas-Silva, Gabriel Trueba, Michelle Grunauer, Paúl Cárdenas |
| EPI_ISL_594188 | Department of Pathology, School of Medicine, Imam Khomeini Hospital, Tehran University of Medical Sciences | Genetics Research Center, University of Social Welfare and Rehabilitation Sciences | Zohreh Fattahi, Marzieh Mohseni, Khadijeh Jalalvand, Azam Ghaziasadi, Seyedeh elham Mortazavi, Ali Jafarpour, Azar Hadadi, Alireza Abdollahi, Ali Jafarpour, Azam Ghaziasad, Seyedeh elham Mortazavi, Saber Soltani, Reza Najafipour, Kimia Kahrizi, Seyed Mohammad Jazayeri, Hossein Najmabadi |
| EPI_ISL_596228 | WHO National Influenza Centre Russian Federation | WHO National Influenza Centre Russian Federation | Andrey Komissarov, Artem Fadeev, Anna Ivanova, Kseniya Komissarova, Dmitry Bazhenov, Daria Danilenko, Ksenia Safina, Elena Nabieva, Georgii Bazykin, Dmitry Lioznov |
| EPI_ISL_596264, EPI_ISL_596265 | WHO National Influenza Centre Russian Federation | WHO National Influenza Centre Russian Federation | Andrey Komissarov, Artem Fadeev, Anna Ivanova, Mariia Sergeeva, Kseniya Komissarova, Dmitry Bazhenov, Daria Danilenko |
| EPI_ISL_596266, EPI_ISL_596267 | WHO National Influenza Centre Russian Federation | WHO National Influenza Centre Russian Federation | Andrey Komissarov, Artem Fadeev, Anna Ivanova, Kseniya Komissarova, Dmitry Bazhenov, Daria Danilenko |
| EPI_ISL_596449 | Institute for Medical Research, Infectious Disease Research Centre, National Institutes of Health, Ministry of Health Malaysia | Institute for Medical Research, Infectious Disease Research Centre, National Institutes of Health, Ministry of Health Malaysia | Suppiah J, Kamel K, Mohd-Zawawi Z, Thayan R |
| EPI_ISL_596451, EPI_ISL_596455 | Department of Pathology, School of Medicine, Imam Khomeini Hospital, Tehran University of Medical Sciences | Genetics Research Center, University of Social Welfare and Rehabilitation Sciences | Zohreh Fattahi, Marzieh Mohseni, Khadijeh Jalalvand, Azam Ghaziasadi, Seyedeh elham Mortazavi, Ali Jafarpour, Azar Hadadi, Alireza Abdollahi, Ali Jafarpour, Azam Ghaziasad, Seyedeh elham Mortazavi, Saber Soltani, Reza Najafipour, Kimia Kahrizi, Seyed Mohammad Jazayeri, Hossein Najmabadi |
| EPI_ISL_602582, EPI_ISL_602583 | Private medical practitioner | Hong Kong Department of Health | Mak Gannon C.K., Lam Edman T.K., Chan Rickjason C.W., Tsang Dominic N.C. |
| EPI_ISL_602584 | Queen Mary Hospital | Hong Kong Department of Health | Mak Gannon C.K., Lam Edman T.K., Chan Rickjason C.W., Tsang Dominic N.C. |
| EPI_ISL_602585 | Private medical practitioner | Hong Kong Department of Health | Mak Gannon C.K., Lam Edman T.K., Chan Rickjason C.W., Tsang Dominic N.C. |
| EPI_ISL_602586 | Queen Elizabeth Hospital | Hong Kong Department of Health | Mak Gannon C.K., Lam Edman T.K., Chan Rickjason C.W., Tsang Dominic N.C. |
| EPI_ISL_602587 | Private medical practitioner | Hong Kong Department of Health | Mak Gannon C.K., Lam Edman T.K., Chan Rickjason C.W., Tsang Dominic N.C. |
| EPI_ISL_602588 | United Christian Hospital | Hong Kong Department of Health | Mak Gannon C.K., Lam Edman T.K., Chan Rickjason C.W., Tsang Dominic N.C. |
| EPI_ISL_602589 | Queen Elizabeth Hospital | Hong Kong Department of Health | Mak Gannon C.K., Lam Edman T.K., Chan Rickjason C.W., Tsang Dominic N.C. |
| EPI_ISL_602590 | Caritas Medical Centre | Hong Kong Department of Health | Mak Gannon C.K., Lam Edman T.K., Chan Rickjason C.W., Tsang Dominic N.C. |
| EPI_ISL_602591 | Queen Elizabeth Hospital | Hong Kong Department of Health | Mak Gannon C.K., Lam Edman T.K., Chan Rickjason C.W., Tsang Dominic N.C. |
| EPI_ISL_602592 | United Christian Hospital | Hong Kong Department of Health | Mak Gannon C.K., Lam Edman T.K., Chan Rickjason C.W., Tsang Dominic N.C. |
| EPI_ISL_602593 | Queen Mary Hospital | Hong Kong Department of Health | Mak Gannon C.K., Lam Edman T.K., Chan Rickjason C.W., Tsang Dominic N.C. |
| EPI_ISL_602594 | Prince of Wales Hospital | Hong Kong Department of Health | Mak Gannon C.K., Lam Edman T.K., Chan Rickjason C.W., Tsang Dominic N.C. |
| EPI_ISL_602595, EPI_ISL_602596 | Pamela Youde Nethersole Eastern Hospital | Hong Kong Department of Health | Mak Gannon C.K., Lam Edman T.K., Chan Rickjason C.W., Tsang Dominic N.C. |
| EPI_ISL_602597 | Private medical practitioner | Hong Kong Department of Health | Mak Gannon C.K., Lam Edman T.K., Chan Rickjason C.W., Tsang Dominic N.C. |
| EPI_ISL_602598 | Our Lady of Maryknoll Hospital | Hong Kong Department of Health | Mak Gannon C.K., Lam Edman T.K., Chan Rickjason C.W., Tsang Dominic N.C. |
| EPI_ISL_602599 | Tseung Kwan O Hospital | Hong Kong Department of Health | Mak Gannon C.K., Lam Edman T.K., Chan Rickjason C.W., Tsang Dominic N.C. |
| EPI_ISL_602600 | United Christian Hospital | Hong Kong Department of Health | Mak Gannon C.K., Lam Edman T.K., Chan Rickjason C.W., Tsang Dominic N.C. |
| EPI_ISL_602601 | Tseung Kwan O Hospital | Hong Kong Department of Health | Mak Gannon C.K., Lam Edman T.K., Chan Rickjason C.W., Tsang Dominic N.C. |
| EPI_ISL_602602 | Ruttonjee Hospital | Hong Kong Department of Health | Mak Gannon C.K., Lam Edman T.K., Chan Rickjason C.W., Tsang Dominic N.C. |
| EPI_ISL_602603, EPI_ISL_602604 | Queen Mary Hospital | Hong Kong Department of Health | Mak Gannon C.K., Lam Edman T.K., Chan Rickjason C.W., Tsang Dominic N.C. |
| EPI_ISL_602605 | Pamela Youde Nethersole Eastern Hospital | Hong Kong Department of Health | Mak Gannon C.K., Lam Edman T.K., Chan Rickjason C.W., Tsang Dominic N.C. |
| EPI_ISL_602606 | Queen Elizabeth Hospital | Hong Kong Department of Health | Mak Gannon C.K., Lam Edman T.K., Chan Rickjason C.W., Tsang Dominic N.C. |
| EPI_ISL_602607 | Tuen Mun Hospital | Hong Kong Department of Health | Mak Gannon C.K., Lam Edman T.K., Chan Rickjason C.W., Tsang Dominic N.C. |
| EPI_ISL_602608 | United Christian Hospital | Hong Kong Department of Health | Mak Gannon C.K., Lam Edman T.K., Chan Rickjason C.W., Tsang Dominic N.C. |
| EPI_ISL_602609 | Pamela Youde Nethersole Eastern Hospital | Hong Kong Department of Health | Mak Gannon C.K., Lam Edman T.K., Chan Rickjason C.W., Tsang Dominic N.C. |
| EPI_ISL_602610 | Queen Mary Hospital | Hong Kong Department of Health | Mak Gannon C.K., Lam Edman T.K., Chan Rickjason C.W., Tsang Dominic N.C. |
| EPI_ISL_602611 | Queen Elizabeth Hospital | Hong Kong Department of Health | Mak Gannon C.K., Lam Edman T.K., Chan Rickjason C.W., Tsang Dominic N.C. |
| EPI_ISL_602612 | Tseung Kwan O Hospital | Hong Kong Department of Health | Mak Gannon C.K., Lam Edman T.K., Chan Rickjason C.W., Tsang Dominic N.C. |
| EPI_ISL_602613, EPI_ISL_602614, EPI_ISL_602615 | Pamela Youde Nethersole Eastern Hospital | Hong Kong Department of Health | Mak Gannon C.K., Lam Edman T.K., Chan Rickjason C.W., Tsang Dominic N.C. |
| EPI_ISL_602616 | Kwong Wah Hospital | Hong Kong Department of Health | Mak Gannon C.K., Lam Edman T.K., Chan Rickjason C.W., Tsang Dominic N.C. |
| EPI_ISL_602617, EPI_ISL_602618 | Queen Mary Hospital | Hong Kong Department of Health | Mak Gannon C.K., Lam Edman T.K., Chan Rickjason C.W., Tsang Dominic N.C. |
| EPI_ISL_602619 | United Christian Hospital | Hong Kong Department of Health | Mak Gannon C.K., Lam Edman T.K., Chan Rickjason C.W., Tsang Dominic N.C. |
| EPI_ISL_602620 | Queen Elizabeth Hospital | Hong Kong Department of Health | Mak Gannon C.K., Lam Edman T.K., Chan Rickjason C.W., Tsang Dominic N.C. |
| EPI_ISL_602621 | GLENEAGLES HONG KONG (CLINICAL LAB) | Hong Kong Department of Health | Mak Gannon C.K., Lam Edman T.K., Chan Rickjason C.W., Tsang Dominic N.C. |
| EPI_ISL_602622 | AHRI-Sigal | KRISP, KZN Research Innovation and Sequencing Platform | Gazy I, Sigla, Karim F, Cele S, Giandhari J, Pillay S, Tegally H, Wilkinson E, de Oliveira T |
| EPI_ISL_602624, EPI_ISL_602625, EPI_ISL_602626, EPI_ISL_602627, EPI_ISL_602628, EPI_ISL_602629, EPI_ISL_602630 | AHRI-Sigal | KRISP, KZN Research Innovation and Sequencing Platform | Gazy I, Sigl A, Karim F, Cele S, Giandhari J, Pillay S, Tegally H, Wilkinson E, de Oliveira T |
| EPI_ISL_602632, EPI_ISL_602633, EPI_ISL_602634, EPI_ISL_602635, EPI_ISL_602636, EPI_ISL_602637, EPI_ISL_602638, EPI_ISL_602640, EPI_ISL_602641, EPI_ISL_602644, EPI_ISL_602645, EPI_ISL_602650, EPI_ISL_602654, EPI_ISL_602659, EPI_ISL_602660, EPI_ISL_602661, EPI_ISL_602666, EPI_ISL_602667, EPI_ISL_602668, EPI_ISL_602669, EPI_ISL_602670, EPI_ISL_602671, EPI_ISL_602672, EPI_ISL_602673, EPI_ISL_602674, EPI_ISL_602675, EPI_ISL_602676, EPI_ISL_602677, EPI_ISL_602678, EPI_ISL_602679, EPI_ISL_602680, EPI_ISL_602681 | | | |
| see above | NHLS-IALCH | KRISP, KZN Research Innovation and Sequencing Platform | Giandhari J, Pillay S, Lessells R, Mdlalose K, York D, Khan S, Tegally H, Wilkinson E, de Oliveira T |
| EPI_ISL_603021 | Pronto Socorro Dr. Conrado Cesarino Nuvolini | Instituto Adolfo Lutz, Interdiciplinary Procedures Center, Strategic Laboratory | Claudio Tavares Sacchi, Claudia Regina Gonçalves, Erica Valessa Ramos Gomes, Karoline Rodrigues Campos |
| EPI_ISL_603022 | Departamento de Vigilância à Saúde | Instituto Adolfo Lutz, Interdiciplinary Procedures Center, Strategic Laboratory | Claudio Tavares Sacchi, Claudia Regina Gonçalves, Erica Valessa Ramos Gomes, Karoline Rodrigues Campos |
| EPI_ISL_603023 | Vigilância em Saúde Visa Sul | Instituto Adolfo Lutz, Interdiciplinary Procedures Center, | Claudio Tavares Sacchi, Claudia Regina Gonçalves, Erica Valessa Ramos Gomes, Karoline Rodrigues Campos |

| | | | |
|---|---|---|---|
| | | Strategic Laboratory | |
| EPI_ISL_603024 | Santa Casa de Misericordia de Araçatuba | Instituto Adolfo Lutz, Interdiciplinary Procedures Center, Strategic Laboratory | Claudio Tavares Sacchi, Claudia Regina Gonçalves, Erica Valessa Ramos Gomes, Karoline Rodrigues Campos |
| EPI_ISL_603025 | UPA Central de Caraguatatuba | Instituto Adolfo Lutz, Interdiciplinary Procedures Center, Strategic Laboratory | Claudio Tavares Sacchi, Claudia Regina Gonçalves, Erica Valessa Ramos Gomes, Karoline Rodrigues Campos |
| EPI_ISL_603026 | Santa Casa da Misericórdia de Presidente Prudente | Instituto Adolfo Lutz, Interdiciplinary Procedures Center, Strategic Laboratory | Claudio Tavares Sacchi, Claudia Regina Gonçalves, Erica Valessa Ramos Gomes, Karoline Rodrigues Campos |
| EPI_ISL_603027 | Santa Casa de Misericordia de Araçatuba | Instituto Adolfo Lutz, Interdiciplinary Procedures Center, Strategic Laboratory | Claudio Tavares Sacchi, Claudia Regina Gonçalves, Erica Valessa Ramos Gomes, Karoline Rodrigues Campos |
| EPI_ISL_603028 | Hospital Municipal Santa Ana | Instituto Adolfo Lutz, Interdiciplinary Procedures Center, Strategic Laboratory | Claudio Tavares Sacchi, Claudia Regina Gonçalves, Erica Valessa Ramos Gomes, Karoline Rodrigues Campos |
| EPI_ISL_603029 | Hospital Municipal Mário Gatti | Instituto Adolfo Lutz, Interdiciplinary Procedures Center, Strategic Laboratory | Claudio Tavares Sacchi, Claudia Regina Gonçalves, Erica Valessa Ramos Gomes, Karoline Rodrigues Campos |
| EPI_ISL_603030 | Hospital Domingos Leonardo Ceravolo Presidente Prudente | Instituto Adolfo Lutz, Interdiciplinary Procedures Center, Strategic Laboratory | Claudio Tavares Sacchi, Claudia Regina Gonçalves, Erica Valessa Ramos Gomes, Karoline Rodrigues Campos |
| EPI_ISL_603031 | Santa Casa de Presidente Epitácio | Instituto Adolfo Lutz, Interdiciplinary Procedures Center, Strategic Laboratory | Claudio Tavares Sacchi, Claudia Regina Gonçalves, Erica Valessa Ramos Gomes, Karoline Rodrigues Campos |
| EPI_ISL_603032 | Santa Casa da Misericórdia de Presidente Prudente | Instituto Adolfo Lutz, Interdiciplinary Procedures Center, Strategic Laboratory | Claudio Tavares Sacchi, Claudia Regina Gonçalves, Erica Valessa Ramos Gomes, Karoline Rodrigues Campos |
| EPI_ISL_603033 | Vigilancia Epidemiologica de São Bernardo do Campo | Instituto Adolfo Lutz, Interdiciplinary Procedures Center, Strategic Laboratory | Claudio Tavares Sacchi, Claudia Regina Gonçalves, Erica Valessa Ramos Gomes, Karoline Rodrigues Campos |
| EPI_ISL_603034 | Departamento de Vigilância à Saúde | Instituto Adolfo Lutz, Interdiciplinary Procedures Center, Strategic Laboratory | Claudio Tavares Sacchi, Claudia Regina Gonçalves, Erica Valessa Ramos Gomes, Karoline Rodrigues Campos |
| EPI_ISL_603035 | Secretaria Municipal de Saúde | Instituto Adolfo Lutz, Interdiciplinary Procedures Center, Strategic Laboratory | Claudio Tavares Sacchi, Claudia Regina Gonçalves, Erica Valessa Ramos Gomes, Karoline Rodrigues Campos |
| EPI_ISL_603036 | Hospital Santa Ana | Instituto Adolfo Lutz, Interdiciplinary Procedures Center, Strategic Laboratory | Claudio Tavares Sacchi, Claudia Regina Gonçalves, Erica Valessa Ramos Gomes, Karoline Rodrigues Campos |
| EPI_ISL_603037 | Hospital Geral de Pedreira | Instituto Adolfo Lutz, Interdiciplinary Procedures Center, Strategic Laboratory | Claudio Tavares Sacchi, Claudia Regina Gonçalves, Erica Valessa Ramos Gomes, Karoline Rodrigues Campos |
| EPI_ISL_603038 | Santa Casa de Misericordia de Araçatuba | Instituto Adolfo Lutz, Interdiciplinary Procedures Center, Strategic Laboratory | Claudio Tavares Sacchi, Claudia Regina Gonçalves, Erica Valessa Ramos Gomes, Karoline Rodrigues Campos |
| EPI_ISL_603039 | Hospital Municipal Mário Gatti | Instituto Adolfo Lutz, Interdiciplinary Procedures Center, Strategic Laboratory | Claudio Tavares Sacchi, Claudia Regina Gonçalves, Erica Valessa Ramos Gomes, Karoline Rodrigues Campos |
| EPI_ISL_603137 | INMI Lazzaro Spallanzani IRCCS | INMI Lazzaro Spallanzani IRCCS | Cesare E.M. Gruber, Martina Rueca, Barbara Bartolini, Francesco Messina, Emanuela Giombini, Simone Lanini, Antonino Di Caro, Maria R. Capobianchi |
| EPI_ISL_603138 | INMI Lazzaro Spallanzani IRCCS | INMI Lazzaro Spallanzani IRCCS | Martina Rueca, Francesco Messina, Barbara Bartolini, Cesare E.M. Gruber, Emanuela Giombini, Simone Lanini, Antonino Di Caro, Maria R. Capobianchi |
| EPI_ISL_603139 | INMI Lazzaro Spallanzani IRCCS | INMI Lazzaro Spallanzani IRCCS | Martina Rueca, Cesare E.M. Gruber, Barbara Bartolini, Francesco Messina, Emanuela Giombini, Simone Lanini, Antonino Di Caro, Maria R. Capobianchi |
| EPI_ISL_603140 | INMI Lazzaro Spallanzani IRCCS | INMI Lazzaro Spallanzani IRCCS | Martina Rueca, Cesare E.M. Gruber, Francesco Messina, Barbara Bartolini, Emanuela Giombini, Simone Lanini, Antonino Di Caro, Maria R. Capobianchi |
| EPI_ISL_603141 | INMI Lazzaro Spallanzani IRCCS | INMI Lazzaro Spallanzani IRCCS | Francesco Messina, Cesare E.M. Gruber, Martina Rueca, Barbara Bartolini, Emanuela Giombini, Simone Lanini, Maria R. Capobianchi, Antonino Di Caro |
| EPI_ISL_603142 | INMI Lazzaro Spallanzani IRCCS | INMI Lazzaro Spallanzani IRCCS | Barbara Bartolini, Cesare E.M. Gruber, Francesco Messina, Martina Rueca, Simone Lanini, Emanuela Giombini, Maria R. Capobianchi, Antonino Di Caro |
| EPI_ISL_603143 | INMI Lazzaro Spallanzani IRCCS | INMI Lazzaro Spallanzani IRCCS | Martina Rueca, Francesco Messina, Cesare E.M. Gruber, Barbara Bartolini, Emanuela Giombini, Simone Lanini, Antonino Di Caro, Maria R. Capobianchi |
| EPI_ISL_603144 | INMI Lazzaro Spallanzani IRCCS | INMI Lazzaro Spallanzani IRCCS | Cesare E.M. Gruber, Martina Rueca, Francesco Messina, Barbara Bartolini, Emanuela Giombini, Simone Lanini, Maria R. Capobianchi, Antonino Di Caro |
| EPI_ISL_603145 | INMI Lazzaro Spallanzani IRCCS | INMI Lazzaro Spallanzani IRCCS | Francesco Messina, Cesare E.M. Gruber, Martina Rueca, Barbara Bartolini, Emanuela Giombini, Simone Lanini, Antonino Di Caro, Maria R. Capobianchi |
| EPI_ISL_603146 | INMI Lazzaro Spallanzani IRCCS | INMI Lazzaro Spallanzani IRCCS | Cesare E.M. Gruber, Martina Rueca, Barbara Bartolini, Emanuela Giombini, Francesco Messina, Simone Lanini, Maria R. Capobianchi, Antonino Di Caro |
| EPI_ISL_603147 | INMI Lazzaro Spallanzani IRCCS | INMI Lazzaro Spallanzani IRCCS | Cesare E.M. Gruber, Barbara Bartolini, Francesco Messina, Martina Rueca, Simone Lanini, Emanuela Giombini, Antonino Di Caro, Maria R. Capobianchi |
| EPI_ISL_603148 | INMI Lazzaro Spallanzani IRCCS | INMI Lazzaro Spallanzani IRCCS | Cesare E.M. Gruber, Francesco Messina, Martina Rueca, Simone Lanini, Emanuela Giombini, Barbara Bartolini, Maria R. Capobianchi, Antonino Di Caro |
| EPI_ISL_603149 | INMI Lazzaro Spallanzani IRCCS | INMI Lazzaro Spallanzani IRCCS | Francesco Messina, Barbara Bartolini, Cesare E.M. Gruber, Martina Rueca, Simone Lanini, Emanuela Giombini, Patrizia Massarelli, Antonino Di Caro, Maria R. Capobianchi |
| EPI_ISL_603150 | INMI Lazzaro Spallanzani IRCCS | INMI Lazzaro Spallanzani IRCCS | Barbara Bartolini, Francesco Messina, Martina Rueca, Cesare E.M. Gruber, Simone Lanini, Emanuela Giombini, Patrizia Massarelli, Maria R. Capobianchi, Antonino Di Caro |
| EPI_ISL_603151 | INMI Lazzaro Spallanzani IRCCS | INMI Lazzaro Spallanzani IRCCS | Francesco Messina, Barbara Bartolini, Cesare E.M. Gruber, Martina Rueca, Emanuela Giombini, Simone Lanini, Patrizia Massarelli, Maria R. Capobianchi, Antonino Di Caro |
| EPI_ISL_603152 | INMI Lazzaro Spallanzani IRCCS | INMI Lazzaro Spallanzani IRCCS | Barbara Bartolini, Cesare E.M. Gruber, Francesco Messina, Martina Rueca, Emanuela Giombini, Simone Lanini, Patrizia Massarelli, Antonino Di Caro, Maria R. Capobianchi |
| EPI_ISL_603153 | INMI Lazzaro Spallanzani IRCCS | INMI Lazzaro Spallanzani IRCCS | Francesco Messina, Cesare E.M. Gruber, Barbara Bartolini, Simone Lanini, Emanuela Giombini, Martina Rueca, Marcello Meledandri, Maria Letizia Schiavone, Annunziata Tamburro, Antonino Di Caro, Maria R. Capobianchi |
| EPI_ISL_603154 | INMI Lazzaro Spallanzani IRCCS | INMI Lazzaro Spallanzani IRCCS | Martina Rueca, Francesco Messina, Cesare E.M. Gruber, Barbara Bartolini, Emanuela Giombini, Simone Lanini, Antonino Di Caro, Maria R. Capobianchi |
| EPI_ISL_603155 | INMI Lazzaro Spallanzani IRCCS | INMI Lazzaro Spallanzani IRCCS | Martina Rueca, Francesco Messina, Barbara Bartolini, Cesare E.M. Gruber, Simone Lanini, Emanuela Giombini, Maria R. Capobianchi, Antonino Di Caro |
| EPI_ISL_603156 | INMI Lazzaro Spallanzani IRCCS | INMI Lazzaro Spallanzani IRCCS | Barbara Bartolini, Francesco Messina, Cesare E.M. Gruber, Martina Rueca, Emanuela Giombini, Simone Lanini, Maria R. Capobianchi, Antonino Di Caro |
| EPI_ISL_603157 | INMI Lazzaro Spallanzani IRCCS | INMI Lazzaro Spallanzani IRCCS | Francesco Messina, Martina Rueca, Barbara Bartolini, Cesare E.M. Gruber, Simone Lanini, Emanuela Giombini, Fulvia Pimpinelli, Antonino Di Caro, Maria R. Capobianchi |
| EPI_ISL_603158 | INMI Lazzaro Spallanzani IRCCS | INMI Lazzaro Spallanzani IRCCS | Barbara Bartolini, Cesare E.M. Gruber, Francesco Messina, Martina Rueca, Emanuela Giombini, Simone Lanini, Fulvia Pimpinelli, Maria R. Capobianchi, Antonino Di Caro |
| EPI_ISL_603159 | INMI Lazzaro Spallanzani IRCCS | INMI Lazzaro Spallanzani IRCCS | Barbara Bartolini, Francesco Messina, Cesare E.M. Gruber, Martina Rueca, Simone Lanini, Emanuela Giombini, Fulvia Pimpinelli, Maria R. Capobianchi, Antonino Di Caro |
| EPI_ISL_603160 | INMI Lazzaro Spallanzani IRCCS | INMI Lazzaro Spallanzani IRCCS | Barbara Bartolini, Simone Lanini, Emanuela Giombini, Francesco Messina, Cesare E.M. Gruber, Martina Rueca, Fulvia Pimpinelli, Maria R. Capobianchi, Antonino Di Caro |
| EPI_ISL_603161 | INMI Lazzaro Spallanzani IRCCS | INMI Lazzaro Spallanzani IRCCS | Cesare E.M. Gruber, Barbara Bartolini, Francesco Messina, Martina Rueca, Emanuela Giombini, Simone Lanini, Fulvia Pimpinelli, Antonino Di Caro, Maria R. |

| | | | |
|---|---|---|---|
| | | | Capobianchi |
| EPI_ISL_603162 | INMI Lazzaro Spallanzani IRCCS | INMI Lazzaro Spallanzani IRCCS | Cesare E.M. Gruber, Francesco Messina, Martina Rueca, Barbara Bartolini, Emanuela Giombini, Simone Lanini, Fulvia Pimpinelli, Antonino Di Caro, Maria R. Capobianchi |
| EPI_ISL_603163 | INMI Lazzaro Spallanzani IRCCS | INMI Lazzaro Spallanzani IRCCS | Cesare E.M. Gruber, Martina Rueca, Emanuela Giombini, Simone Lanini, Francesco Messina, Barbara Bartolini, Fulvia Pimpinelli, Maria R. Capobianchi, Antonino Di Caro |
| EPI_ISL_603164 | INMI Lazzaro Spallanzani IRCCS | INMI Lazzaro Spallanzani IRCCS | Cesare E.M. Gruber, Francesco Messina, Martina Rueca, Barbara Bartolini, Emanuela Giombini, Simone Lanini, Fulvia Pimpinelli, Antonino Di Caro, Maria R. Capobianchi |
| EPI_ISL_603165 | INMI Lazzaro Spallanzani IRCCS | INMI Lazzaro Spallanzani IRCCS | Martina Rueca, Francesco Messina, Barbara Bartolini, Cesare E.M. Gruber, Emanuela Giombini, Simone Lanini, Fulvia Pimpinelli, Maria R. Capobianchi, Antonino Di Caro |
| EPI_ISL_603166 | INMI Lazzaro Spallanzani IRCCS | INMI Lazzaro Spallanzani IRCCS | Martina Rueca, Cesare E.M. Gruber, Barbara Bartolini, Francesco Messina, Simone Lanini, Emanuela Giombini, Fulvia Pimpinelli, Maria R. Capobianchi, Antonino Di Caro |
| EPI_ISL_603167 | INMI Lazzaro Spallanzani IRCCS | INMI Lazzaro Spallanzani IRCCS | Martina Rueca, Cesare E.M. Gruber, Francesco Messina, Barbara Bartolini, Simone Lanini, Emanuela Giombini, Fulvia Pimpinelli, Maria R. Capobianchi, Antonino Di Caro |
| EPI_ISL_603168 | INMI Lazzaro Spallanzani IRCCS | INMI Lazzaro Spallanzani IRCCS | Martina Rueca, Barbara Bartolini, Cesare E.M. Gruber, Simone Lanini, Emanuela Giombini, Francesco Messina, Fulvia Pimpinelli, Antonino Di Caro, Maria R. Capobianchi |
| EPI_ISL_603169 | INMI Lazzaro Spallanzani IRCCS | INMI Lazzaro Spallanzani IRCCS | Francesco Messina, Barbara Bartolini, Cesare E.M. Gruber, Martina Rueca, Simone Lanini, Emanuela Giombini, Fulvia Pimpinelli, Maria R. Capobianchi, Antonino Di Caro |
| EPI_ISL_603170 | INMI Lazzaro Spallanzani IRCCS | INMI Lazzaro Spallanzani IRCCS | Martina Rueca, Barbara Bartolini, Cesare E.M. Gruber, Francesco Messina, Simone Lanini, Emanuela Giombini, Maria R. Capobianchi, Antonino Di Caro |
| EPI_ISL_603171 | INMI Lazzaro Spallanzani IRCCS | INMI Lazzaro Spallanzani IRCCS | Barbara Bartolini, Cesare E.M. Gruber, Francesco Messina, Martina Rueca, Emanuela Giombini, Simone Lanini, Mariarosa Gaudio, Antonino Di Caro, Maria R. Capobianchi |
| EPI_ISL_603172 | INMI Lazzaro Spallanzani IRCCS | INMI Lazzaro Spallanzani IRCCS | Cesare E.M. Gruber, Martina Rueca, Barbara Bartolini, Simone Lanini, Francesco Messina, Emanuela Giombini, Mariarosa Gaudio, Antonino Di Caro, Maria R. Capobianchi |
| EPI_ISL_603173 | INMI Lazzaro Spallanzani IRCCS | INMI Lazzaro Spallanzani IRCCS | Barbara Bartolini, Cesare E.M. Gruber, Martina Rueca, Francesco Messina, Simone Lanini, Emanuela Giombini, Giuseppina Cappiello, Maria R. Capobianchi, Antonino Di Caro |
| EPI_ISL_603174 | INMI Lazzaro Spallanzani IRCCS | INMI Lazzaro Spallanzani IRCCS | Francesco Messina, Barbara Bartolini, Martina Rueca, Emanuela Giombini, Simone Lanini, Cesare E.M. Gruber, Mariarosa Gaudio, Maria R. Capobianchi, Antonino Di Caro |
| EPI_ISL_603175 | INMI Lazzaro Spallanzani IRCCS | INMI Lazzaro Spallanzani IRCCS | Barbara Bartolini, Cesare E.M. Gruber, Simone Lanini, Emanuela Giombini, Francesco Messina, Martina Rueca, Mariarosa Gaudio, Antonino Di Caro, Maria R. Capobianchi |
| EPI_ISL_603176 | INMI Lazzaro Spallanzani IRCCS | INMI Lazzaro Spallanzani IRCCS | Cesare E.M. Gruber, Barbara Bartolini, Francesco Messina, Martina Rueca, Simone Lanini, Emanuela Giombini, Antonino Di Caro, Maria R. Capobianchi |
| EPI_ISL_603223 | National Institute of Laboratory Medicine and Referral Center | Genomic Research Lab, BCSIR | Md. Murshed Hasan Sarkar, Abu Sayeed Mohammad Mahmud, Mohammad Samir Uzzaman, Eshrar Osman, Md. Ahashan Habib, Shahina Akter, Tanjina Akhter Banu, Barna Goswami, Iffat Jahan, Md. Saddam Hossain, Tasnim Nafisa, Md. Maruf Ahmed Molla, Mahmuda Yeasmin, Asish Kumar Ghosh, A. K. M. Shamsuzzaman, Monira Parveen, Md. Masum Hossain Arif, Md. Salim Khan |
| EPI_ISL_610162 | RSUD Dr. Tjitrowardojo | Genetics Working Group (Pokja Genetik) Faculty of Medicine, Public Health and Nursing Universitas Gadjah Mada (FK-KMK UGM); Disease Investigation Center Wates Ministry of Agriculture Indonesia; Department of Microbiology FK-KMK UGM; Laboratorium Diagnostik Yayasan Tahija World Mosquito Program (WMP) Yogyakarta Center for Tropical Medicine FK-KMK UGM; Integrated Research Center FK-KMK UGM; Department of Computer Science and Electronics FMIPA UGM; Balai Besar Teknik Kesehatan Lingkungan dan Pengendalian Penyakit (BBTKLPP) Yogyakarta | Gunadi, Hendra Wibawa, Marcellus, Mohamad S. Hakim, Edwin W. Daniwijaya, Ludhang P. Rizki, Endah Supriyati, Eggi Arguni, Titik Nuryastuti, Tri Wibawa, Dwi AA Nugrahaningsih, Afiahayati, Siswanto, Kristy Iskandar, Nungki Anggorowati, Irene, Indaryati, Havid Setyawan, Wuryanto, Susan Simanjaya, Alvin Santoso Kalim |
| EPI_ISL_610227, EPI_ISL_610228, EPI_ISL_610229, EPI_ISL_610230, EPI_ISL_610231, EPI_ISL_610232, EPI_ISL_610233, EPI_ISL_610234, EPI_ISL_610235, EPI_ISL_610236, EPI_ISL_610239, EPI_ISL_610240, EPI_ISL_610241, EPI_ISL_610242, EPI_ISL_610243, EPI_ISL_610244 | | | |
| see above | Molecular diagnostic laboratory of Federal Budget Institution of Science "Central Research Institute of Epidemiology" of The Federal Service on Customers' Rights Protection and Human Well-being Surveillance | Group of Genomics and Postgenomic Technologies of Central Research Institute of Epidemilology | Samoilov AE, Kaptelova VV, Valdokhina AV, Bulanenko VP, Speranskaya AS, Tivanova EV, Shipulina OY, Akimkin VG |
| EPI_ISL_613457 | QUARANTINE CAMP | Hong Kong Department of Health | Alan K.L. Tsang, Peter C.W. Yip, Edman T.K. Lam, Rickjason C.W. Chan, Dominic N.C. Tsang |
| EPI_ISL_613545, EPI_ISL_613546, EPI_ISL_613547, EPI_ISL_613548, EPI_ISL_613550 | CHRU Pontchaillou - Laboratoire de Virologie 2, rue Henri Le Guilloux | National Reference Center for Viruses of Respiratory Infections, Institut Pasteur, Paris | Marion Barbet, Sylvie Behillil, Méline Bizard, Angela Brisebarre, Camille Capel, Etienne Simon-Lorière, Vincent Enouf, Maud Vanpeene, Sylvie van der Werf, Gisèle Lagathu |
| EPI_ISL_614265, EPI_ISL_614266, EPI_ISL_614267, EPI_ISL_614268, EPI_ISL_614269, EPI_ISL_614270, EPI_ISL_614271, EPI_ISL_614272, EPI_ISL_614273, EPI_ISL_614274, EPI_ISL_614275, EPI_ISL_614276, EPI_ISL_614277, EPI_ISL_614278, EPI_ISL_614279, EPI_ISL_614280 | | | |
| see above | Eurofins | National Reference Center for Viruses of Respiratory Infections, Institut Pasteur, Paris | Marion Barbet, Sylvie Behillil, Méline Bizard, Angela Brisebarre, Camille Capel, Etienne Simon-Lorière, Vincent Enouf, Maud Vanpeene, Sylvie van der Werf |
| EPI_ISL_614307, EPI_ISL_614308, EPI_ISL_614310, EPI_ISL_614311, EPI_ISL_614312 | Faroese National Reference Laboratory for Fish and Animal Diseases | Faroese National Reference Laboratory for Fish and Animal Diseases | Maria Marjunardóttir Dahl, Petra Elisabeth Petersen, Debes Hammershaimb Christiansen |
| EPI_ISL_623145, EPI_ISL_623147 | Laboratorio de Virologia Molecular / UFRJ | Bioinformatics Laboratory / LNCC | Carolina M Voloch, Ronaldo S Francisco Jr, Luiz G P de Almeida, Otavio J. Brustolini, Cynthia C Cardoso, Alexandra L Gerber, Ana Paula de C Guimarães, Diana Mariani, Covid19-UFRJ Workgroup, Luís Cristóvão Pôrto, Renato S Aguiar, Terezinha M P P Castiñeiras, Orlando C. Ferreira, Amilcar Tanuri, Ana Tereza R de Vasconcelos |
| EPI_ISL_632312 | NU-sjukvården | Clinical microbiology, Sahlgrenska University Hospital | Johan Ringlander, Josefin Olausson, Hedvig Engström Jakobsson, Magnus Lindh |
| EPI_ISL_632936 | Puskesmas Mlati 1 Sleman | Genetics Working Group (Pokja Genetik) Faculty of Medicine, Public Health and Nursing Universitas Gadjah Mada (FK-KMK UGM); Disease Investigation Center Wates Ministry of Agriculture Indonesia; Department of Microbiology FK-KMK UGM; Laboratorium Diagnostik Yayasan Tahija World Mosquito Program (WMP) Yogyakarta Center for Tropical Medicine FK-KMK UGM; Integrated Research Center FK-KMK UGM; Department of Computer Science and Electronics FMIPA UGM; RSUP Dr. Sardjito | Gunadi, Hendra Wibawa, Marcellus, Mohamad S. Hakim, Edwin W. Daniwijaya, Ludhang P. Rizki, Endah Supriyati, Eggi Arguni, Titik Nuryastuti, Tri Wibawa, Dwi AA Nugrahaningsih, Afiahayati, Siswanto, Kristy Iskandar, Nungki Anggorowati, Ika Trisnawati, Sumardi, Eko Budiono, Bambang Sigit Riyanto, Heni Retnowulan, Munawar Gani, Satria Maulana, Nur Rahmi Ananda, Riat El Khair, Yunika Puspadewi, Osman Sianipar, Umi Solekhah Intansari, Elizabeth Henny Herningtiyas, Ira Puspitawati, Nur Imma Fatimah Harahap, Ernawati, Untung Wirawan , William Widitjiarso |
| EPI_ISL_636737, EPI_ISL_636835, EPI_ISL_636837 | Laboratório de Imunofarmacologia - Instituto Oswaldo Cruz | Laboratório de Imunofarmacologia - Instituto Oswaldo Cruz | Souza,T.M., Fintelman-Rodrigues,N., De Paula,A.D., Saraiva,F.B., Ferreira,M.A. and Sacramento,C.Q. |

| | | | |
|---|---|---|---|
| EPI_ISL_636965 | Pathogen Genomics Lab King Abdullah University of Science and Technology(KAUST) | Pathogen Genomics Lab King Abdullah University of Science and Technology(KAUST) | Raeece Naeem, Rahul P Salunke, Sharif Hala, Sara Mfarrej, Amit Kumar Subudhi, Fadwa Alofi, Fathia Ben Rached, Afrah Alsomali, Asim Khogeer, Ahmad Bakur Mahmoud, Anwar Hashem, Naif Almontashiri, Arnab Pain |
| EPI_ISL_636966, EPI_ISL_636967 | Pathogen Genomics Lab King Abdullah University of Science and Technology(KAUST) | Pathogen Genomics Lab King Abdullah University of Science and Technology(KAUST) | Fathia Ben Rached, Raeece Naeem, Sharif Hala, Fadwa Alofi, Rahul P Salunke, Sara Mfarrej, Amit Kumar Subudhi, Afrah Alsomali, Asim Khogeer, Ahmad Bakur Mahmoud, Anwar Hashem, Naif Almontashiri, Arnab Pain |
| EPI_ISL_636972 | Pathogen Genomics Lab King Abdullah University of Science and Technology (KAUST) | Pathogen Genomics Lab King Abdullah University of Science and Technology (KAUST) | Afrah Alsomali, Fathia Ben Rached, Raeece Naeem, Sharif Hala,Rahul P Salunke, Amanda Ooi, Luke Esau, Sara Mfarrej, Amit Kumar Subudhi, Fadwa Alofi, Asim Khogeer, Kahled Algithami, Anwar Hashem, Naif Almontashiri, Arnab Pain |
| EPI_ISL_636973 | Public Health Lab | Public Health Lab | Alwasti, H |
| EPI_ISL_636977 | HP Pemba | KRISP, KZN Research Innovation and Sequencing Platform | Ismael N, Giandhari J, Pillay S, Tegally H, Wilkinson E, de Oliveira T, Nadia Sitoe, Paulo Arnaldo, Nedio Mabunda |
| EPI_ISL_636980 | CS Xai Xai | KRISP, KZN Research Innovation and Sequencing Platform | Ismael N, Giandhari J, Pillay S, Tegally H, Wilkinson E, de Oliveira T, Nadia Sitoe, Paulo Arnaldo, Nedio Mabunda |
| EPI_ISL_637022 | Tin Shui Wai Hospital | Hong Kong Department of Health | Mak Gannon C.K., Lam Edman T.K., Chan Rickjason C.W., Tsang Dominic N.C. |
| EPI_ISL_637023 | Queen Elizabeth Hospital | Hong Kong Department of Health | Mak Gannon C.K., Lam Edman T.K., Chan Rickjason C.W., Tsang Dominic N.C. |
| EPI_ISL_637024 | Pamela Youde Nethersole Eastern Hospital | Hong Kong Department of Health | Mak Gannon C.K., Lam Edman T.K., Chan Rickjason C.W., Tsang Dominic N.C. |
| EPI_ISL_637025 | Queen Elizabeth Hospital | Hong Kong Department of Health | Mak Gannon C.K., Lam Edman T.K., Chan Rickjason C.W., Tsang Dominic N.C. |
| EPI_ISL_637026 | Kwong Wah Hospital | Hong Kong Department of Health | Mak Gannon C.K., Lam Edman T.K., Chan Rickjason C.W., Tsang Dominic N.C. |
| EPI_ISL_637027 | Hong Kong Baptist Hospital | Hong Kong Department of Health | Mak Gannon C.K., Lam Edman T.K., Chan Rickjason C.W., Tsang Dominic N.C. |
| EPI_ISL_637028 | Queen Mary Hospital | Hong Kong Department of Health | Mak Gannon C.K., Lam Edman T.K., Chan Rickjason C.W., Tsang Dominic N.C. |
| EPI_ISL_637029 | Caritas Medical Centre | Hong Kong Department of Health | Mak Gannon C.K., Lam Edman T.K., Chan Rickjason C.W., Tsang Dominic N.C. |
| EPI_ISL_637030, EPI_ISL_637031 | Prince of Wales Hospital | Hong Kong Department of Health | Mak Gannon C.K., Lam Edman T.K., Chan Rickjason C.W., Tsang Dominic N.C. |
| EPI_ISL_637032 | Private medical practitioner | Hong Kong Department of Health | Mak Gannon C.K., Lam Edman T.K., Chan Rickjason C.W., Tsang Dominic N.C. |
| EPI_ISL_637033 | Chun Yeung Estate Quarantine Center | Hong Kong Department of Health | Mak Gannon C.K., Lam Edman T.K., Chan Rickjason C.W., Tsang Dominic N.C. |
| EPI_ISL_637034 | Queen Elizabeth Hospital | Hong Kong Department of Health | Mak Gannon C.K., Lam Edman T.K., Chan Rickjason C.W., Tsang Dominic N.C. |
| EPI_ISL_637035 | Princess Margaret Hospital | Hong Kong Department of Health | Mak Gannon C.K., Lam Edman T.K., Chan Rickjason C.W., Tsang Dominic N.C. |
| EPI_ISL_637036 | Tseung Kwan O Hospital | Hong Kong Department of Health | Mak Gannon C.K., Lam Edman T.K., Chan Rickjason C.W., Tsang Dominic N.C. |
| EPI_ISL_637037, EPI_ISL_637038, EPI_ISL_637039 | Queen Elizabeth Hospital | Hong Kong Department of Health | Mak Gannon C.K., Lam Edman T.K., Chan Rickjason C.W., Tsang Dominic N.C. |
| EPI_ISL_637040 | Prince of Wales Hospital | Hong Kong Department of Health | Mak Gannon C.K., Lam Edman T.K., Chan Rickjason C.W., Tsang Dominic N.C. |
| EPI_ISL_637041 | Private medical practitioner | Hong Kong Department of Health | Mak Gannon C.K., Lam Edman T.K., Chan Rickjason C.W., Tsang Dominic N.C. |
| EPI_ISL_637042 | Prince of Wales Hospital | Hong Kong Department of Health | Mak Gannon C.K., Lam Edman T.K., Chan Rickjason C.W., Tsang Dominic N.C. |
| EPI_ISL_637043 | Temporary Specimen Collection Centre at the AsiaWorld-Expo | Hong Kong Department of Health | Mak Gannon C.K., Lam Edman T.K., Chan Rickjason C.W., Tsang Dominic N.C. |
| EPI_ISL_648217 | SILAB | Laboratorio Mixto de Biotecnología Acuática (LMBA) | Joaquín Ezpeleta, Ignacio García Labari, Victoria Posner, Vanina Villanova, Pablo Casal, Sofía Lavista Llanos, Federico Remes Lenicov, Ana Paletta, Flavio Spetale, Agustina Cerri, Silvana Spinelli, Elisa Bolatti, Diego Chouhy, María Re, Gastón Viarengo, Ana Cavatorta, Julian Acosta, Javier Murillo, Laura Angelone, Leandro Ciappina, Pilar Bulacio, Adriana Giri, Silvia Arranz, Elizabeth Tapia |
| EPI_ISL_648678, EPI_ISL_648679, EPI_ISL_648680, EPI_ISL_648681, EPI_ISL_648682, EPI_ISL_648683, EPI_ISL_648684, EPI_ISL_648685, EPI_ISL_648686, EPI_ISL_648687, EPI_ISL_648688, EPI_ISL_648689, EPI_ISL_648690, EPI_ISL_648691, EPI_ISL_648692, EPI_ISL_648693, EPI_ISL_648694, EPI_ISL_648695, EPI_ISL_648696, EPI_ISL_648697, EPI_ISL_648698, EPI_ISL_648699, EPI_ISL_648700, EPI_ISL_648701, EPI_ISL_648702, EPI_ISL_648703, EPI_ISL_648704, EPI_ISL_648705, EPI_ISL_648706, EPI_ISL_648707, EPI_ISL_648708, EPI_ISL_648709, EPI_ISL_648710, EPI_ISL_648711, EPI_ISL_648712, EPI_ISL_648713, EPI_ISL_648714, EPI_ISL_648715, EPI_ISL_648716, EPI_ISL_648717, EPI_ISL_648718, EPI_ISL_648719, EPI_ISL_648720, EPI_ISL_648721, EPI_ISL_648722, EPI_ISL_648723, EPI_ISL_648724, EPI_ISL_648725, EPI_ISL_648726, EPI_ISL_648727, EPI_ISL_648728, EPI_ISL_648729, EPI_ISL_648730, EPI_ISL_648731, EPI_ISL_648732, EPI_ISL_648733, EPI_ISL_648734, EPI_ISL_648735, EPI_ISL_648736, EPI_ISL_648737, EPI_ISL_648738, EPI_ISL_648739, EPI_ISL_648740, EPI_ISL_648741, EPI_ISL_648792, EPI_ISL_648793, EPI_ISL_648794, EPI_ISL_648795, EPI_ISL_648796, EPI_ISL_648797, EPI_ISL_648798, EPI_ISL_648799, EPI_ISL_648800, EPI_ISL_648801, EPI_ISL_648802, EPI_ISL_648803, EPI_ISL_648804, EPI_ISL_648805, EPI_ISL_648806, EPI_ISL_648807, EPI_ISL_648808, EPI_ISL_648809, EPI_ISL_648810, EPI_ISL_648811, EPI_ISL_648812, EPI_ISL_648813, EPI_ISL_648814, EPI_ISL_648815, EPI_ISL_648816, EPI_ISL_648817, EPI_ISL_648818, EPI_ISL_648819, EPI_ISL_648820 | | | |
| see above | Department of Laboratory Medicine, Tan Tock Seng Hospital | Department of Laboratory Medicine, Tan Tock Seng Hospital | Chen YYC, Zair X, Lim JX, Li C, Tang WY, Maurer-Stroh S, Barkham TMS, Nagarajan N, Sessions OM |
| EPI_ISL_649154 | Queen Astrid Military Hospital | Institute of Tropical Medicine | Philippe Selhorst, Colin Anthony |
| EPI_ISL_649190 | Istituto Zooprofilattico Sperimentale della Puglia e della Basilicata | Istituto Zooprofilattico Sperimentale della Puglia e della Basilicata | Parisi A., Bianco A., Capozzi L., Del Sambro L., Manzulli V, Rondinone V., Pace L., Galante D., Cipolletta D. |
| EPI_ISL_649191 | Istituto Zooprofilattico Sperimentale della Puglia e della Basilicata | Istituto Zooprofilattico Sperimentale della Puglia e della Basilicata | Parisi A., Bianco A., Capozzi L., Del Sambro L., Manzulli V, Rondinone V., Pace L., Cipolletta D., Galante D. |
| EPI_ISL_649938 | I.R.C.C.S. "S. De Bellis" - Ente Ospedaliero | Istituto Zooprofilattico Sperimentale della Puglia e della Basilicata | Parisi A., Bianco A., Capozzi L., Del Sambro L., Lippolis A., Notarnicola M., Cipolletta D., Galante D. |
| EPI_ISL_649939 | I.R.C.C.S. "S. De Bellis" - Ente Ospedaliero | Istituto Zooprofilattico Sperimentale della Puglia e della Basilicata | Parisi A., Bianco A., Capozzi L., Del Sambro L., Lippolis A., Notarnicola M., Manzulli V, Rondinone V., Pace L. |
| EPI_ISL_649940 | Istituto Zooprofilattico Sperimentale della Puglia e della Basilicata | Istituto Zooprofilattico Sperimentale della Puglia e della Basilicata | Parisi A., Bianco A., Capozzi L., Del Sambro L., Manzulli V, Rondinone V., Pace L., Cipolletta D., Galante D. |
| EPI_ISL_653746, EPI_ISL_653747, EPI_ISL_653755, EPI_ISL_653756, EPI_ISL_653757 | Instituto Nacional de Salud, Bogotá, Colombia | Instituto Nacional de Salud, Bogotá, Colombia | Katherine Laiton-Donato, Diego A. Álvarez-Díaz, Carlos Franco-Muñoz, Mauricio Pacheco-Montealegre, Jonathan Reales, Diego Andrés Prada, Jose A. Usme-Ciro, Zulma M. Cucunubá, Christian Julian VillabonaArenas, Liz Villabona-Arenas, Sussy Echeverria, Astrid C. Flórez, Carolina Ferro, Diana Marcela Walteros-Acero, Franklin Prieto, Carlos Andrés Durán, Martha Lucia Ospina Martinez, Marcela Mercado-Reyes |
| EPI_ISL_653763, EPI_ISL_653764 | I.R.C.C.S. "S. De Bellis" - Ente Ospedaliero | Istituto Zooprofilattico Sperimentale della Puglia e della Basilicata | Parisi A., Bianco A., Capozzi L., Del Sambro L., Lippolis A., Notarnicola M., Manzulli V, Rondinone V., Pace L. |
| EPI_ISL_653823 | I.R.C.C.S. "S. De Bellis" - Ente Ospedaliero | Istituto Zooprofilattico Sperimentale della Puglia e della Basilicata | Parisi A., Bianco A., Capozzi L., Del Sambro L., Lippolis A., Notarnicola M., Cipolletta D., Galante D. |
| EPI_ISL_653922, EPI_ISL_653923, EPI_ISL_653924, EPI_ISL_653925, EPI_ISL_653926, EPI_ISL_653927, EPI_ISL_653928, EPI_ISL_653929, EPI_ISL_653930, EPI_ISL_653931, EPI_ISL_653932 | | | |
| see above | Molecular diagnostic laboratory of Federal Budget Institution of Science "Central Research Institute of Epidemiology" of The Federal Service on Customers' Rights Protection and Human Well-being Surveillance | Group of Genomics and Postgenomic Technologies of Central Research Institute of Epidemilology | Samoilov AE, Kaptelova VV, Dudorova A.V., Speranskaya AS, Tivanova EV, Shipulina OY, Akimkin VG |
| EPI_ISL_654818 | Department of Respiratory and other Viral Infections of L.V.Gromashevsky Institute of Epidemiology & Infectious Diseases NAMS of Ukrain | Department of Respiratory and other Viral Infections of L.V.Gromashevsky Institute of Epidemiology & Infectious Diseases NAMS of Ukrain, JSC "Farmak" | Alla Mironenko, Andriy Goy, Ihor Kravchuk, Ludmyla Bolotova, Larysa Radchenko, Nataliia Teteriuk |
| EPI_ISL_654819, EPI_ISL_654820 | Department of Respiratory and other Viral Infections of L.V.Gromashevsky Institute of Epidemiology & Infectious | Department of Respiratory and other Viral Infections of L.V.Gromashevsky Institute of Epidemiology & Infectious | Alla Mironenko, Andriy Goy, Ihor Kravchuk, Ludmyla Bolotova, Larysa Radchenko, Nataliia Teteriuk |

| | | | |
|---|---|---|---|
| | Diseases NAMS of Ukrain | Diseases NAMS of Ukrain, JSC "Farmak" | |
| EPI_ISL_658896 | Instituto de Diagnostico y Referencia Epidemiologicos (INDRE) | Instituto de Diagnostico y Referencia Epidemiologicos (INDRE) | Ernesto Ramirez-Gonzalez, Abril Rodriguez-Maldonado, Claudia Wong-Arambula , Natividad Cruz-Ortiz, Tatiana Nunez-Garcia, Dayanira Arellano-Suarez, Fabiola Garces-Ayala, Lucia Hernandez-Rivas, Irma Lopez-Martinez, Gisela Barrera-Badillo. |
| EPI_ISL_660069 | Zurita & Zurita Laboratorios | Zurita & Zurita Laboratorios | Gabriela Sevillano Camilo Zurita-Salinas Karen Loaiza David Ortega-Paredes Jeannete Zurita |
| EPI_ISL_660109, EPI_ISL_660110 | Division of Infectious Diseases, Department of Internal Medicine, Yonsei University College of Medicine | Department of biotechnology, Yonsei University | Minwoo Kim, Youn-Jung Lee, Jae sun Yoon, Jin Young Ahn, Jung Ho Kim, Jun Yong Choi*, and Jong-Won Oh* |
| EPI_ISL_660165, EPI_ISL_660166, EPI_ISL_660168, EPI_ISL_660169, EPI_ISL_660171, EPI_ISL_660172, EPI_ISL_660173, EPI_ISL_660174, EPI_ISL_660175, EPI_ISL_660176, EPI_ISL_660181 | | | |
| see above | NHLS-IALCH | KRISP, KZN Research Innovation and Sequencing Platform | Gazy I, Sigal A, Karim F, Cele S, Giandhari J, Pillay S, Tegally H, Wilkinson E, de Oliveira T |
| EPI_ISL_660435, EPI_ISL_660436, EPI_ISL_660437, EPI_ISL_660438 | Molecular diagnostic laboratory of Federal Budget Institution of Science "Central Research Institute of Epidemiology" of The Federal Service on Customers' Rights Protection and Human Well-being Surveillance | Group of Genomics and Postgenomic Technologies of Central Research Institute of Epidemilology | Samoilov AE, Kaptelova VV, Valdokhina AV, Bulanenko VP, Speranskaya AS, Tivanova EV, Shipulina OY, Akimkin VG |
| EPI_ISL_671419, EPI_ISL_671420, EPI_ISL_671421, EPI_ISL_671423, EPI_ISL_671424, EPI_ISL_671425, EPI_ISL_671427, EPI_ISL_671429, EPI_ISL_671437 | University of Debrecen, Department of Medical Microbiology | National Laboratory of Virology, Szentágothai Research Centre | Endre Gábor Tóth, Balázs Somogyi, Brigitta Zana, Eszter Csoma, Ferenc Jakab, Gábor Kemenesi |
| EPI_ISL_677728, EPI_ISL_677729, EPI_ISL_677730, EPI_ISL_677731, EPI_ISL_677732, EPI_ISL_677733, EPI_ISL_677734, EPI_ISL_677735, EPI_ISL_677736, EPI_ISL_677737, EPI_ISL_677738 | | | |
| see above | University of Szeged, Institute of Clinical Microbiology | National Laboratory of Virology, Szentágothai Research Centre | Endre Gábor Tóth, Balázs Somogyi, Brigitta, Gabriella Terhes, Ferenc Jakab, Gábor Kemenesi |
| EPI_ISL_682242 | HOSPITAL SAN VICENTE DE PAUL | Inciensa, Instituto Costarricense de Investigación y Enseñanza en Nutrición y Salud | Francisco Duarte, Hebleen Porras, Claudio Soto-Garita, Estela Cordero, Adriana Godinez, Melany Calderon & Mariel López |
| EPI_ISL_682250, EPI_ISL_682252 | AREA DE SALUD ESCAZU (COOPESANA) | Inciensa, Instituto Costarricense de Investigación y Enseñanza en Nutrición y Salud | Francisco Duarte, Hebleen Porras, Claudio Soto-Garita, Estela Cordero, Adriana Godinez & Melany Calderon |
| EPI_ISL_682255 | HOSPITAL SAN VICENTE DE PAUL | Inciensa, Instituto Costarricense de Investigación y Enseñanza en Nutrición y Salud | Francisco Duarte, Hebleen Porras, Claudio Soto-Garita, Estela Cordero, Adriana Godinez & Melany Calderon |
| EPI_ISL_682257 | HOSPITAL SAN JUAN DE DIOS | Inciensa, Instituto Costarricense de Investigación y Enseñanza en Nutrición y Salud | Francisco Duarte, Hebleen Porras, Claudio Soto-Garita, Estela Cordero, Adriana Godinez & Melany Calderon |
| EPI_ISL_682261 | TAMIZAJE COMUNITARIO- PASO CANOAS | Inciensa, Instituto Costarricense de Investigación y Enseñanza en Nutrición y Salud | Francisco Duarte, Hebleen Porras, Claudio Soto-Garita, Estela Cordero, Adriana Godinez & Melany Calderon |
| EPI_ISL_682262, EPI_ISL_682264, EPI_ISL_682266, EPI_ISL_682273 | HOSPITAL SAN JUAN DE DIOS | Inciensa, Instituto Costarricense de Investigación y Enseñanza en Nutrición y Salud | Francisco Duarte, Hebleen Porras, Claudio Soto-Garita, Estela Cordero, Adriana Godinez & Melany Calderon |
| EPI_ISL_685731, EPI_ISL_685732, EPI_ISL_685910, EPI_ISL_685917, EPI_ISL_685938, EPI_ISL_685939, EPI_ISL_685940, EPI_ISL_685941, EPI_ISL_685942, EPI_ISL_685943, EPI_ISL_685944, EPI_ISL_685945, EPI_ISL_685946, EPI_ISL_685948, EPI_ISL_685951, EPI_ISL_685952 | | | |
| see above | Pathogen Genomics Center, National Institute of Infectious Diseases | Pathogen Genomics Center, National Institute of Infectious Diseases | Tsuyoshi Sekizuka, Kentaro Itokawa, Rina Tanaka, Masanori Hashino, Makoto Kuroda |
| EPI_ISL_685963, EPI_ISL_685964, EPI_ISL_685965, EPI_ISL_685966, EPI_ISL_685967, EPI_ISL_685968, EPI_ISL_685969, EPI_ISL_685970, EPI_ISL_685971, EPI_ISL_685972, EPI_ISL_685973, EPI_ISL_685974, EPI_ISL_685975, EPI_ISL_685976, EPI_ISL_685977, EPI_ISL_685978, EPI_ISL_685979, EPI_ISL_685980, EPI_ISL_685981, EPI_ISL_685982, EPI_ISL_685983, EPI_ISL_685984, EPI_ISL_685985, EPI_ISL_685986 | | | |
| see above | Tokyo Metropolitan Institute of Public Health | Pathogen Genomics Center, National Institute of Infectious Diseases | Tsuyoshi Sekizuka, Kentaro Itokawa, Rina Tanaka, Masanori Hashino, Makoto Kuroda |
| EPI_ISL_686076, EPI_ISL_686077, EPI_ISL_686078, EPI_ISL_686079, EPI_ISL_686080, EPI_ISL_686081, EPI_ISL_686082, EPI_ISL_686109, EPI_ISL_686110, EPI_ISL_686111, EPI_ISL_686112, EPI_ISL_686113, EPI_ISL_686114, EPI_ISL_686115, EPI_ISL_686116, EPI_ISL_686117, EPI_ISL_686118, EPI_ISL_686119, EPI_ISL_686120, EPI_ISL_686121, EPI_ISL_686122, EPI_ISL_686123, EPI_ISL_686124, EPI_ISL_686125, EPI_ISL_686126, EPI_ISL_686127, EPI_ISL_686128, EPI_ISL_686129, EPI_ISL_686130, EPI_ISL_686131, EPI_ISL_686132, EPI_ISL_686133, EPI_ISL_686134, EPI_ISL_686135, EPI_ISL_686136, EPI_ISL_686137, EPI_ISL_686138, EPI_ISL_686139, EPI_ISL_686140, EPI_ISL_686141, EPI_ISL_686142, EPI_ISL_686143, EPI_ISL_686144, EPI_ISL_686145, EPI_ISL_686146, EPI_ISL_686147, EPI_ISL_686148, EPI_ISL_686149, EPI_ISL_686150, EPI_ISL_686151, EPI_ISL_686152, EPI_ISL_686153, EPI_ISL_686154, EPI_ISL_686155, EPI_ISL_686156, EPI_ISL_686157, EPI_ISL_686158, EPI_ISL_686159, EPI_ISL_686160, EPI_ISL_686161, EPI_ISL_686162, EPI_ISL_686163, EPI_ISL_686164, EPI_ISL_686165, EPI_ISL_686166, EPI_ISL_686167, EPI_ISL_686168, EPI_ISL_686169, EPI_ISL_686170, EPI_ISL_686171, EPI_ISL_686172, EPI_ISL_686173, EPI_ISL_686174, EPI_ISL_686175, EPI_ISL_686176, EPI_ISL_686177, EPI_ISL_686178, EPI_ISL_686179, EPI_ISL_686180, EPI_ISL_686181, EPI_ISL_686182, EPI_ISL_686183, EPI_ISL_686184, EPI_ISL_686185, EPI_ISL_686186, EPI_ISL_686187, EPI_ISL_686188, EPI_ISL_686189, EPI_ISL_686191, EPI_ISL_686192, EPI_ISL_686193, EPI_ISL_686194, EPI_ISL_686195, EPI_ISL_686196, EPI_ISL_686197, EPI_ISL_686198, EPI_ISL_686199, EPI_ISL_686200, EPI_ISL_686201, EPI_ISL_686202, EPI_ISL_686203, EPI_ISL_686204, EPI_ISL_686205, EPI_ISL_686206, EPI_ISL_686207, EPI_ISL_686208, EPI_ISL_686209, EPI_ISL_686210, EPI_ISL_686211, EPI_ISL_686212, EPI_ISL_686213, EPI_ISL_686215, EPI_ISL_686216, EPI_ISL_686223, EPI_ISL_686224, EPI_ISL_686225, EPI_ISL_686226, EPI_ISL_686227, EPI_ISL_686228, EPI_ISL_686283, EPI_ISL_686284, EPI_ISL_686285, EPI_ISL_686286, EPI_ISL_686287, EPI_ISL_686288, EPI_ISL_686289, EPI_ISL_686290, EPI_ISL_686291, EPI_ISL_686292, EPI_ISL_686293, EPI_ISL_686294, EPI_ISL_686295, EPI_ISL_686296, EPI_ISL_686297, EPI_ISL_686298, EPI_ISL_686299, EPI_ISL_686301, EPI_ISL_686302, EPI_ISL_686303, EPI_ISL_686304, EPI_ISL_686305, EPI_ISL_686306, EPI_ISL_686307, EPI_ISL_686308, EPI_ISL_686309, EPI_ISL_686310, EPI_ISL_686311, EPI_ISL_686312, EPI_ISL_686313, EPI_ISL_686314, EPI_ISL_686315, EPI_ISL_686316, EPI_ISL_686317, EPI_ISL_686318, EPI_ISL_686319, EPI_ISL_686320, EPI_ISL_686321, EPI_ISL_686322, EPI_ISL_686323, EPI_ISL_686324, EPI_ISL_686325, EPI_ISL_686326, EPI_ISL_686327, EPI_ISL_686328, EPI_ISL_686329, EPI_ISL_686330, EPI_ISL_686331, EPI_ISL_686332, EPI_ISL_686333, EPI_ISL_686334, EPI_ISL_686335, EPI_ISL_686367, EPI_ISL_686368, EPI_ISL_686369, EPI_ISL_686370, EPI_ISL_686371, EPI_ISL_686372, EPI_ISL_686373, EPI_ISL_686374, EPI_ISL_686375, EPI_ISL_686376, EPI_ISL_686377, EPI_ISL_686378, EPI_ISL_686379, EPI_ISL_686380, EPI_ISL_686381, EPI_ISL_686382, EPI_ISL_686383, EPI_ISL_686384, EPI_ISL_686385, EPI_ISL_686386, EPI_ISL_686387, EPI_ISL_686388, EPI_ISL_686389, EPI_ISL_686390, EPI_ISL_686391, EPI_ISL_686392, EPI_ISL_686393, EPI_ISL_686394, EPI_ISL_686395, EPI_ISL_686396, EPI_ISL_686397, EPI_ISL_686398, EPI_ISL_686399, EPI_ISL_686400, EPI_ISL_686401, EPI_ISL_686402, EPI_ISL_686403, EPI_ISL_686404, EPI_ISL_686405, EPI_ISL_686406, EPI_ISL_686407, EPI_ISL_686408, EPI_ISL_686410, EPI_ISL_686411, EPI_ISL_686412, EPI_ISL_686413, EPI_ISL_686414, EPI_ISL_686415, EPI_ISL_686418, EPI_ISL_686419, EPI_ISL_686420, EPI_ISL_686421, EPI_ISL_686422, EPI_ISL_686423, EPI_ISL_686424, EPI_ISL_686425, EPI_ISL_686426, EPI_ISL_686427, EPI_ISL_686429, EPI_ISL_686430, EPI_ISL_686431, EPI_ISL_686432, EPI_ISL_686433, EPI_ISL_686434, EPI_ISL_686435, EPI_ISL_686436, EPI_ISL_686437, EPI_ISL_686438, EPI_ISL_686439, EPI_ISL_686440, EPI_ISL_686441, EPI_ISL_686442, EPI_ISL_686443, EPI_ISL_686444, EPI_ISL_686445, EPI_ISL_686446, EPI_ISL_686447, EPI_ISL_686469, EPI_ISL_686470, EPI_ISL_686471, EPI_ISL_686472, EPI_ISL_686473, EPI_ISL_686474, EPI_ISL_686475, EPI_ISL_686476, EPI_ISL_686477, EPI_ISL_686478, EPI_ISL_686479, EPI_ISL_686480, EPI_ISL_686481, EPI_ISL_686482, EPI_ISL_686483, EPI_ISL_686484, EPI_ISL_686485, EPI_ISL_686486, EPI_ISL_686487, EPI_ISL_686488, EPI_ISL_686489, EPI_ISL_686490, EPI_ISL_686491, EPI_ISL_686492, EPI_ISL_686493 | | | |
| see above | Pathogen Genomics Center, National Institute of Infectious Diseases | Pathogen Genomics Center, National Institute of Infectious Diseases | Tsuyoshi Sekizuka, Kentaro Itokawa, Rina Tanaka, Masanori Hashino, Makoto Kuroda |
| EPI_ISL_686521 | Sakai City Institute of Public Health | Pathogen Genomics Center, National Institute of Infectious Diseases | Tsuyoshi Sekizuka, Kentaro Itokawa, Rina Tanaka, Masanori Hashino, Makoto Kuroda |
| EPI_ISL_686612, EPI_ISL_686613, EPI_ISL_686614, EPI_ISL_686615, EPI_ISL_686616, EPI_ISL_686617, EPI_ISL_686618, EPI_ISL_686619, EPI_ISL_686620, EPI_ISL_686621, EPI_ISL_686622, EPI_ISL_686623, EPI_ISL_686624, EPI_ISL_686625, EPI_ISL_686626, EPI_ISL_686627, EPI_ISL_686628, EPI_ISL_686629, EPI_ISL_686630, EPI_ISL_686631, EPI_ISL_686632, EPI_ISL_686633, EPI_ISL_686634, EPI_ISL_686635, EPI_ISL_686636, EPI_ISL_686637, EPI_ISL_686638, EPI_ISL_686639, EPI_ISL_686640, EPI_ISL_686641, EPI_ISL_686642, EPI_ISL_686643, EPI_ISL_686644, EPI_ISL_686645, EPI_ISL_686646, EPI_ISL_686647, EPI_ISL_686648, EPI_ISL_686649, EPI_ISL_686650, EPI_ISL_686651, EPI_ISL_686652, EPI_ISL_686653 | | | |
| see above | Pathogen Genomics Center, National Institute of Infectious Diseases | Pathogen Genomics Center, National Institute of Infectious Diseases | Tsuyoshi Sekizuka, Kentaro Itokawa, Rina Tanaka, Masanori Hashino, Makoto Kuroda |
| EPI_ISL_686659, EPI_ISL_686660, EPI_ISL_686661, EPI_ISL_686662, EPI_ISL_686663, EPI_ISL_686664, EPI_ISL_686665, EPI_ISL_686666, EPI_ISL_686667, EPI_ISL_686668, EPI_ISL_686669, EPI_ISL_686670, EPI_ISL_686671, EPI_ISL_686672, EPI_ISL_686673, EPI_ISL_686674, EPI_ISL_686675, EPI_ISL_686676, EPI_ISL_686677, EPI_ISL_686678, EPI_ISL_686679, EPI_ISL_686680, EPI_ISL_686681, EPI_ISL_686682, EPI_ISL_686683, EPI_ISL_686684, EPI_ISL_686685, EPI_ISL_686686, EPI_ISL_686687, EPI_ISL_686688, EPI_ISL_686689, EPI_ISL_686690, EPI_ISL_686691, EPI_ISL_686692, EPI_ISL_686693, EPI_ISL_686694, EPI_ISL_686695, EPI_ISL_686696, EPI_ISL_686697, EPI_ISL_686698, EPI_ISL_686699, EPI_ISL_686700, EPI_ISL_686701, EPI_ISL_686702, EPI_ISL_686703, EPI_ISL_686704, EPI_ISL_686705, EPI_ISL_686706, EPI_ISL_686707, EPI_ISL_686708, EPI_ISL_686709, EPI_ISL_686710, EPI_ISL_686711, EPI_ISL_686712, EPI_ISL_686713, EPI_ISL_686714, EPI_ISL_686715, EPI_ISL_686716, EPI_ISL_686717, EPI_ISL_686718, EPI_ISL_686719, EPI_ISL_686720, EPI_ISL_686721, EPI_ISL_686722, EPI_ISL_686723, EPI_ISL_686724, EPI_ISL_686725, EPI_ISL_686726, EPI_ISL_686727, EPI_ISL_686728, EPI_ISL_686729, EPI_ISL_686730, EPI_ISL_686731, EPI_ISL_686732, EPI_ISL_686733, EPI_ISL_686734, EPI_ISL_686735, EPI_ISL_686736, EPI_ISL_686737, EPI_ISL_686738, EPI_ISL_686739, EPI_ISL_686740, EPI_ISL_686741, EPI_ISL_686742, EPI_ISL_686743, EPI_ISL_686744, EPI_ISL_686745, EPI_ISL_686746, EPI_ISL_686747, EPI_ISL_686748, EPI_ISL_686749, EPI_ISL_686750, EPI_ISL_686751, EPI_ISL_686752, EPI_ISL_686753, EPI_ISL_686754, EPI_ISL_686755, EPI_ISL_686756, EPI_ISL_686757, EPI_ISL_686758, EPI_ISL_686759, EPI_ISL_686760, EPI_ISL_686761, EPI_ISL_686762, EPI_ISL_686763, EPI_ISL_686764, EPI_ISL_686765, EPI_ISL_686766, EPI_ISL_686767, EPI_ISL_686768, EPI_ISL_686769, EPI_ISL_686770, EPI_ISL_686771, EPI_ISL_686772, EPI_ISL_686773, EPI_ISL_686774, EPI_ISL_686775, EPI_ISL_686776, EPI_ISL_686777, EPI_ISL_686778, EPI_ISL_686779, EPI_ISL_686780, EPI_ISL_686781, EPI_ISL_686782, EPI_ISL_686783, EPI_ISL_686784, EPI_ISL_686785, EPI_ISL_686786, EPI_ISL_686787, EPI_ISL_686788, EPI_ISL_686789, EPI_ISL_686790, EPI_ISL_686791, EPI_ISL_686792, EPI_ISL_686793, EPI_ISL_686794, EPI_ISL_686795, EPI_ISL_686796, EPI_ISL_686797, EPI_ISL_686798, EPI_ISL_686799, EPI_ISL_686800, EPI_ISL_686801, EPI_ISL_686802, EPI_ISL_686803, EPI_ISL_686804, EPI_ISL_686805, EPI_ISL_686806, EPI_ISL_686807, EPI_ISL_686808, EPI_ISL_686809, EPI_ISL_686810, EPI_ISL_686811, EPI_ISL_686812, EPI_ISL_686813, EPI_ISL_686814, EPI_ISL_686815, EPI_ISL_686816, EPI_ISL_686817, EPI_ISL_686818, EPI_ISL_686819, EPI_ISL_686820, EPI_ISL_686821, EPI_ISL_686822, EPI_ISL_686823, EPI_ISL_686824, EPI_ISL_686825, EPI_ISL_686826, EPI_ISL_686827, EPI_ISL_686828, EPI_ISL_686829, EPI_ISL_686830 | | | |

| | | | |
|---|---|---|---|
| see above | Chiba Prefectural Institute of Public Health | Pathogen Genomics Center, National Institute of Infectious Diseases | Tsuyoshi Sekizuka, Kentaro Itokawa, Rina Tanaka, Masanori Hashino, Makoto Kuroda |
| EPI_ISL_686874, EPI_ISL_686875, EPI_ISL_686876, EPI_ISL_686877, EPI_ISL_686878, EPI_ISL_686879, EPI_ISL_686880, EPI_ISL_686881, EPI_ISL_686882, EPI_ISL_686883, EPI_ISL_686884, EPI_ISL_686885, EPI_ISL_686886, EPI_ISL_686887, EPI_ISL_686888, EPI_ISL_686889, EPI_ISL_686890, EPI_ISL_686891, EPI_ISL_686892, EPI_ISL_686893, EPI_ISL_686894, EPI_ISL_686895, EPI_ISL_686896, EPI_ISL_686897, EPI_ISL_686898, EPI_ISL_686899, EPI_ISL_686900, EPI_ISL_686901, EPI_ISL_686902, EPI_ISL_686903, EPI_ISL_686904, EPI_ISL_686905, EPI_ISL_686906, EPI_ISL_686907, EPI_ISL_686908, EPI_ISL_686909, EPI_ISL_686910, EPI_ISL_686911, EPI_ISL_686912, EPI_ISL_686913, EPI_ISL_686914, EPI_ISL_686915, EPI_ISL_686916, EPI_ISL_686917, EPI_ISL_686918, EPI_ISL_686919, EPI_ISL_686920, EPI_ISL_686921, EPI_ISL_686922, EPI_ISL_686923, EPI_ISL_686924, EPI_ISL_686925, EPI_ISL_686926, EPI_ISL_686927, EPI_ISL_686928, EPI_ISL_686929, EPI_ISL_686930, EPI_ISL_686931, EPI_ISL_686932, EPI_ISL_686933, EPI_ISL_686934, EPI_ISL_686935, EPI_ISL_686936, EPI_ISL_686937, EPI_ISL_686938, EPI_ISL_686939, EPI_ISL_686940, EPI_ISL_686941, EPI_ISL_686942, EPI_ISL_686943, EPI_ISL_686944, EPI_ISL_686945, EPI_ISL_686946, EPI_ISL_686947, EPI_ISL_686948, EPI_ISL_686949, EPI_ISL_686950, EPI_ISL_686951, EPI_ISL_686952, EPI_ISL_686953, EPI_ISL_686954, EPI_ISL_686955, EPI_ISL_686956, EPI_ISL_686957, EPI_ISL_686958, EPI_ISL_686959, EPI_ISL_686960, EPI_ISL_686961, EPI_ISL_686962, EPI_ISL_686963, EPI_ISL_686964, EPI_ISL_686965, EPI_ISL_686966, EPI_ISL_686967, EPI_ISL_686968, EPI_ISL_686969, EPI_ISL_686970, EPI_ISL_686971, EPI_ISL_686972, EPI_ISL_686973, EPI_ISL_686974, EPI_ISL_686975, EPI_ISL_686976, EPI_ISL_686977, EPI_ISL_686978, EPI_ISL_686979, EPI_ISL_686980, EPI_ISL_686981, EPI_ISL_686982, EPI_ISL_686983, EPI_ISL_686984, EPI_ISL_686985, EPI_ISL_686986, EPI_ISL_686987, EPI_ISL_686988, EPI_ISL_686989, EPI_ISL_686990, EPI_ISL_686991, EPI_ISL_686992, EPI_ISL_686993, EPI_ISL_686994, EPI_ISL_686995, EPI_ISL_686996, EPI_ISL_686997, EPI_ISL_686998, EPI_ISL_686999, EPI_ISL_687000, EPI_ISL_687001, EPI_ISL_687002, EPI_ISL_687003, EPI_ISL_687004, EPI_ISL_687005, EPI_ISL_687006, EPI_ISL_687007, EPI_ISL_687008, EPI_ISL_687009, EPI_ISL_687010, EPI_ISL_687011, EPI_ISL_687012, EPI_ISL_687013, EPI_ISL_687014, EPI_ISL_687015, EPI_ISL_687016, EPI_ISL_687017, EPI_ISL_687018, EPI_ISL_687019, EPI_ISL_687020, EPI_ISL_687021, EPI_ISL_687022, EPI_ISL_687023, EPI_ISL_687024, EPI_ISL_687025, EPI_ISL_687026, EPI_ISL_687027, EPI_ISL_687028, EPI_ISL_687029, EPI_ISL_687030, EPI_ISL_687031, EPI_ISL_687032, EPI_ISL_687033, EPI_ISL_687034, EPI_ISL_687035, EPI_ISL_687036, EPI_ISL_687037, EPI_ISL_687038, EPI_ISL_687039, EPI_ISL_687040, EPI_ISL_687041, EPI_ISL_687042 | | | |
| see above | Pathogen Genomics Center, National Institute of Infectious Diseases | Pathogen Genomics Center, National Institute of Infectious Diseases | Tsuyoshi Sekizuka, Kentaro Itokawa, Rina Tanaka, Masanori Hashino, Makoto Kuroda |
| EPI_ISL_687071, EPI_ISL_687072, EPI_ISL_687073, EPI_ISL_687074, EPI_ISL_687075, EPI_ISL_687076, EPI_ISL_687077, EPI_ISL_687078 | Kanagawa Prefectural Institute of Public Health | Pathogen Genomics Center, National Institute of Infectious Diseases | Tsuyoshi Sekizuka, Kentaro Itokawa, Rina Tanaka, Masanori Hashino, Makoto Kuroda |
| EPI_ISL_687079, EPI_ISL_687080, EPI_ISL_687081, EPI_ISL_687082, EPI_ISL_687083, EPI_ISL_687084, EPI_ISL_687085, EPI_ISL_687086, EPI_ISL_687087, EPI_ISL_687088, EPI_ISL_687089, EPI_ISL_687090, EPI_ISL_687091, EPI_ISL_687092, EPI_ISL_687093, EPI_ISL_687094, EPI_ISL_687095, EPI_ISL_687096, EPI_ISL_687097, EPI_ISL_687098, EPI_ISL_687100, EPI_ISL_687101, EPI_ISL_687102, EPI_ISL_687103, EPI_ISL_687104, EPI_ISL_687105, EPI_ISL_687106, EPI_ISL_687107, EPI_ISL_687108, EPI_ISL_687109, EPI_ISL_687110, EPI_ISL_687111, EPI_ISL_687112, EPI_ISL_687113, EPI_ISL_687114, EPI_ISL_687115, EPI_ISL_687116, EPI_ISL_687117, EPI_ISL_687118, EPI_ISL_687119, EPI_ISL_687120, EPI_ISL_687121, EPI_ISL_687122, EPI_ISL_687123, EPI_ISL_687124, EPI_ISL_687125, EPI_ISL_687126, EPI_ISL_687127, EPI_ISL_687128, EPI_ISL_687129, EPI_ISL_687130, EPI_ISL_687131, EPI_ISL_687132, EPI_ISL_687133, EPI_ISL_687134, EPI_ISL_687135, EPI_ISL_687136, EPI_ISL_687137, EPI_ISL_687139, EPI_ISL_687140, EPI_ISL_687141, EPI_ISL_687142, EPI_ISL_687143, EPI_ISL_687145, EPI_ISL_687146, EPI_ISL_687147, EPI_ISL_687152, EPI_ISL_687153, EPI_ISL_687154, EPI_ISL_687155, EPI_ISL_687156, EPI_ISL_687157, EPI_ISL_687163, EPI_ISL_687164, EPI_ISL_687165, EPI_ISL_687166, EPI_ISL_687172, EPI_ISL_687173, EPI_ISL_687174, EPI_ISL_687177, EPI_ISL_687179, EPI_ISL_687181, EPI_ISL_687182, EPI_ISL_687184, EPI_ISL_687185, EPI_ISL_687188, EPI_ISL_687189, EPI_ISL_687190, EPI_ISL_687191, EPI_ISL_687192, EPI_ISL_687193, EPI_ISL_687194, EPI_ISL_687195, EPI_ISL_687196, EPI_ISL_687197, EPI_ISL_687198, EPI_ISL_687199, EPI_ISL_687200, EPI_ISL_687201, EPI_ISL_687202, EPI_ISL_687203, EPI_ISL_687204, EPI_ISL_687205, EPI_ISL_687206, EPI_ISL_687207, EPI_ISL_687208, EPI_ISL_687209, EPI_ISL_687210, EPI_ISL_687211, EPI_ISL_687212, EPI_ISL_687213, EPI_ISL_687214, EPI_ISL_687215, EPI_ISL_687216, EPI_ISL_687217, EPI_ISL_687218, EPI_ISL_687219, EPI_ISL_687220, EPI_ISL_687221, EPI_ISL_687222, EPI_ISL_687223, EPI_ISL_687224, EPI_ISL_687225, EPI_ISL_687226, EPI_ISL_687227, EPI_ISL_687228, EPI_ISL_687229, EPI_ISL_687230, EPI_ISL_687231, EPI_ISL_687232, EPI_ISL_687233, EPI_ISL_687234, EPI_ISL_687235, EPI_ISL_687236, EPI_ISL_687237, EPI_ISL_687238, EPI_ISL_687239, EPI_ISL_687240, EPI_ISL_687241, EPI_ISL_687242, EPI_ISL_687243, EPI_ISL_687244, EPI_ISL_687245, EPI_ISL_687255, EPI_ISL_687256, EPI_ISL_687257, EPI_ISL_687258, EPI_ISL_687259, EPI_ISL_687260, EPI_ISL_687261, EPI_ISL_687262, EPI_ISL_687263, EPI_ISL_687264, EPI_ISL_687265, EPI_ISL_687266, EPI_ISL_687267, EPI_ISL_687268, EPI_ISL_687269, EPI_ISL_687270, EPI_ISL_687271, EPI_ISL_687272, EPI_ISL_687273, EPI_ISL_687274, EPI_ISL_687275, EPI_ISL_687276, EPI_ISL_687277, EPI_ISL_687278, EPI_ISL_687279, EPI_ISL_687280, EPI_ISL_687281, EPI_ISL_687282, EPI_ISL_687283, EPI_ISL_687284, EPI_ISL_687285, EPI_ISL_687286, EPI_ISL_687287, EPI_ISL_687288, EPI_ISL_687289, EPI_ISL_687290, EPI_ISL_687291, EPI_ISL_687292, EPI_ISL_687293, EPI_ISL_687294, EPI_ISL_687295, EPI_ISL_687296, EPI_ISL_687297, EPI_ISL_687298, EPI_ISL_687299, EPI_ISL_687300, EPI_ISL_687301, EPI_ISL_687302, EPI_ISL_687303, EPI_ISL_687304, EPI_ISL_687305, EPI_ISL_687306, EPI_ISL_687307, EPI_ISL_687308, EPI_ISL_687309, EPI_ISL_687310, EPI_ISL_687311, EPI_ISL_687312, EPI_ISL_687313, EPI_ISL_687314, EPI_ISL_687315, EPI_ISL_687316, EPI_ISL_687317, EPI_ISL_687318, EPI_ISL_687319, EPI_ISL_687320, EPI_ISL_687321, EPI_ISL_687322, EPI_ISL_687323, EPI_ISL_687324, EPI_ISL_687325, EPI_ISL_687326, EPI_ISL_687327, EPI_ISL_687328, EPI_ISL_687329, EPI_ISL_687330, EPI_ISL_687331, EPI_ISL_687332, EPI_ISL_687333, EPI_ISL_687334, EPI_ISL_687335, EPI_ISL_687336, EPI_ISL_687337, EPI_ISL_687338, EPI_ISL_687339, EPI_ISL_687340, EPI_ISL_687341, EPI_ISL_687342, EPI_ISL_687343, EPI_ISL_687344, EPI_ISL_687345, EPI_ISL_687346, EPI_ISL_687347, EPI_ISL_687348, EPI_ISL_687349, EPI_ISL_687350, EPI_ISL_687351, EPI_ISL_687352, EPI_ISL_687353, EPI_ISL_687354, EPI_ISL_687355, EPI_ISL_687356, EPI_ISL_687357, EPI_ISL_687358, EPI_ISL_687359, EPI_ISL_687361, EPI_ISL_687362, EPI_ISL_687363, EPI_ISL_687364, EPI_ISL_687365, EPI_ISL_687366, EPI_ISL_687367, EPI_ISL_687368, EPI_ISL_687369, EPI_ISL_687370, EPI_ISL_687371, EPI_ISL_687372, EPI_ISL_687373, EPI_ISL_687374, EPI_ISL_687375, EPI_ISL_687376, EPI_ISL_687377, EPI_ISL_687378, EPI_ISL_687379, EPI_ISL_687380, EPI_ISL_687381, EPI_ISL_687382, EPI_ISL_687383, EPI_ISL_687384, EPI_ISL_687385, EPI_ISL_687386, EPI_ISL_687387, EPI_ISL_687388, EPI_ISL_687389, EPI_ISL_687390, EPI_ISL_687391, EPI_ISL_687392, EPI_ISL_687393, EPI_ISL_687394, EPI_ISL_687395, EPI_ISL_687396, EPI_ISL_687397, EPI_ISL_687398, EPI_ISL_687399, EPI_ISL_687400, EPI_ISL_687401, EPI_ISL_687402, EPI_ISL_687403, EPI_ISL_687404, EPI_ISL_687405, EPI_ISL_687406, EPI_ISL_687407, EPI_ISL_687408, EPI_ISL_687409, EPI_ISL_687410, EPI_ISL_687411, EPI_ISL_687412, EPI_ISL_687413, EPI_ISL_687414, EPI_ISL_687415, EPI_ISL_687416, EPI_ISL_687417, EPI_ISL_687418, EPI_ISL_687419, EPI_ISL_687420, EPI_ISL_687421, EPI_ISL_687422, EPI_ISL_687423, EPI_ISL_687424, EPI_ISL_687425, EPI_ISL_687426, EPI_ISL_687427, EPI_ISL_687428, EPI_ISL_687429, EPI_ISL_687430, EPI_ISL_687431, EPI_ISL_687432, EPI_ISL_687433, EPI_ISL_687434, EPI_ISL_687435, EPI_ISL_687436, EPI_ISL_687437, EPI_ISL_687438, EPI_ISL_687439, EPI_ISL_687440, EPI_ISL_687441, EPI_ISL_687442, EPI_ISL_687443, EPI_ISL_687444, EPI_ISL_687445, EPI_ISL_687446, EPI_ISL_687447, EPI_ISL_687448, EPI_ISL_687449, EPI_ISL_687450, EPI_ISL_687451, EPI_ISL_687452, EPI_ISL_687453, EPI_ISL_687454, EPI_ISL_687455, EPI_ISL_687456, EPI_ISL_687459, EPI_ISL_687460, EPI_ISL_687461, EPI_ISL_687462, EPI_ISL_687463, EPI_ISL_687464, EPI_ISL_687465, EPI_ISL_687466, EPI_ISL_687467, EPI_ISL_687468, EPI_ISL_687469, EPI_ISL_687470, EPI_ISL_687471, EPI_ISL_687472, EPI_ISL_687473, EPI_ISL_687474, EPI_ISL_687475, EPI_ISL_687476, EPI_ISL_687477, EPI_ISL_687478, EPI_ISL_687479, EPI_ISL_687480, EPI_ISL_687481, EPI_ISL_687482, EPI_ISL_687483, EPI_ISL_687484, EPI_ISL_687485, EPI_ISL_687486, EPI_ISL_687487, EPI_ISL_687488, EPI_ISL_687489, EPI_ISL_687490, EPI_ISL_687491, EPI_ISL_687492, EPI_ISL_687493, EPI_ISL_687494, EPI_ISL_687495, EPI_ISL_687496, EPI_ISL_687497, EPI_ISL_687498, EPI_ISL_687499, EPI_ISL_687500, EPI_ISL_687501, EPI_ISL_687502, EPI_ISL_687503, EPI_ISL_687504, EPI_ISL_687505, EPI_ISL_687506, EPI_ISL_687507, EPI_ISL_687508, EPI_ISL_687509, EPI_ISL_687510, EPI_ISL_687511, EPI_ISL_687512, EPI_ISL_687513, EPI_ISL_687514, EPI_ISL_687515, EPI_ISL_687516, EPI_ISL_687517, EPI_ISL_687518, EPI_ISL_687519, EPI_ISL_687520, EPI_ISL_687521, EPI_ISL_687522, EPI_ISL_687561, EPI_ISL_687562, EPI_ISL_687563, EPI_ISL_687564, EPI_ISL_687565, EPI_ISL_687566, EPI_ISL_687567, EPI_ISL_687568, EPI_ISL_687569, EPI_ISL_687570, EPI_ISL_687572, EPI_ISL_687573, EPI_ISL_687574, EPI_ISL_687575, EPI_ISL_687576, EPI_ISL_687577, EPI_ISL_687578, EPI_ISL_687579, EPI_ISL_687580, EPI_ISL_687581, EPI_ISL_687582, EPI_ISL_687583, EPI_ISL_687584, EPI_ISL_687585, EPI_ISL_687586, EPI_ISL_687587, EPI_ISL_687588, EPI_ISL_687589, EPI_ISL_687590, EPI_ISL_687591, EPI_ISL_687592, EPI_ISL_687593, EPI_ISL_687594, EPI_ISL_687595, EPI_ISL_687596, EPI_ISL_687597, EPI_ISL_687598, EPI_ISL_687599, EPI_ISL_687600, EPI_ISL_687609, EPI_ISL_687610, EPI_ISL_687611, EPI_ISL_687614, EPI_ISL_687616, EPI_ISL_687617 | | | |
| see above | Pathogen Genomics Center, National Institute of Infectious Diseases | Pathogen Genomics Center, National Institute of Infectious Diseases | Tsuyoshi Sekizuka, Kentaro Itokawa, Rina Tanaka, Masanori Hashino, Makoto Kuroda |
| EPI_ISL_687619, EPI_ISL_687620 | Saitama Prefectural Institute of Public Health | Pathogen Genomics Center, National Institute of Infectious Diseases | Tsuyoshi Sekizuka, Kentaro Itokawa, Rina Tanaka, Masanori Hashino, Makoto Kuroda |
| EPI_ISL_687624, EPI_ISL_687625, EPI_ISL_687626, EPI_ISL_687627, EPI_ISL_687628, EPI_ISL_687629, EPI_ISL_687630, EPI_ISL_687631, EPI_ISL_687632, EPI_ISL_687633, EPI_ISL_687634, EPI_ISL_687635, EPI_ISL_687636, EPI_ISL_687637, EPI_ISL_687638, EPI_ISL_687639, EPI_ISL_687640, EPI_ISL_687641, EPI_ISL_687642, EPI_ISL_687643, EPI_ISL_687644, EPI_ISL_687645, EPI_ISL_687646, EPI_ISL_687647, EPI_ISL_687648, EPI_ISL_687649, EPI_ISL_687650, EPI_ISL_687651, EPI_ISL_687652, EPI_ISL_687653, EPI_ISL_687654, EPI_ISL_687655, EPI_ISL_687659, EPI_ISL_687660, EPI_ISL_687661, EPI_ISL_687662, EPI_ISL_687663, EPI_ISL_687664, EPI_ISL_687665, EPI_ISL_687666, EPI_ISL_687667, EPI_ISL_687668, EPI_ISL_687673, EPI_ISL_687674, EPI_ISL_687675 | | | |
| see above | Pathogen Genomics Center, National Institute of Infectious Diseases | Pathogen Genomics Center, National Institute of Infectious Diseases | Tsuyoshi Sekizuka, Kentaro Itokawa, Rina Tanaka, Masanori Hashino, Makoto Kuroda |
| EPI_ISL_687699, EPI_ISL_687700, EPI_ISL_687701, EPI_ISL_687702, EPI_ISL_687703, EPI_ISL_687704, EPI_ISL_687705, EPI_ISL_687706, EPI_ISL_687707, EPI_ISL_687708, EPI_ISL_687709, EPI_ISL_687710, EPI_ISL_687712, EPI_ISL_687714 | | | |
| see above | Niigata Prefectural Institute of Public Health and Environmental Sciences | Pathogen Genomics Center, National Institute of Infectious Diseases | Tsuyoshi Sekizuka, Kentaro Itokawa, Rina Tanaka, Masanori Hashino, Makoto Kuroda |
| EPI_ISL_687718, EPI_ISL_687719, EPI_ISL_687720, EPI_ISL_687721, EPI_ISL_687722, EPI_ISL_687723, EPI_ISL_687724, EPI_ISL_687725, EPI_ISL_687726, EPI_ISL_687727, EPI_ISL_687728, EPI_ISL_687729, EPI_ISL_687730, EPI_ISL_687731, EPI_ISL_687732, EPI_ISL_687733, EPI_ISL_687734, EPI_ISL_687735, EPI_ISL_687736, EPI_ISL_687737, EPI_ISL_687738, EPI_ISL_687739, EPI_ISL_687740, EPI_ISL_687741, EPI_ISL_687742, EPI_ISL_687746, EPI_ISL_687747, EPI_ISL_687748, EPI_ISL_687749, EPI_ISL_687750, EPI_ISL_687751, EPI_ISL_687752, EPI_ISL_687753, EPI_ISL_687754, EPI_ISL_687755, EPI_ISL_687756, EPI_ISL_687757, EPI_ISL_687758, EPI_ISL_687759, EPI_ISL_687760, EPI_ISL_687761, EPI_ISL_687762, EPI_ISL_687763, EPI_ISL_687764, EPI_ISL_687765, EPI_ISL_687766, EPI_ISL_687767, EPI_ISL_687768, EPI_ISL_687769, EPI_ISL_687770, EPI_ISL_687771, EPI_ISL_687772, EPI_ISL_687773, EPI_ISL_687774, EPI_ISL_687775, EPI_ISL_687776, EPI_ISL_687777, EPI_ISL_687778, EPI_ISL_687779, EPI_ISL_687780, EPI_ISL_687781, EPI_ISL_687782, EPI_ISL_687783, EPI_ISL_687784, EPI_ISL_687785, EPI_ISL_687786, EPI_ISL_687787, EPI_ISL_687788, EPI_ISL_687789, EPI_ISL_687790, EPI_ISL_687791, EPI_ISL_687792, EPI_ISL_687793, EPI_ISL_687794, EPI_ISL_687795, EPI_ISL_687796, EPI_ISL_687797, EPI_ISL_687798, EPI_ISL_687799, EPI_ISL_687800, EPI_ISL_687801, EPI_ISL_687802, EPI_ISL_687803, EPI_ISL_687804, EPI_ISL_687805, EPI_ISL_687806, EPI_ISL_687807, EPI_ISL_687808, EPI_ISL_687809, EPI_ISL_687810, EPI_ISL_687811, EPI_ISL_687812, EPI_ISL_687813, EPI_ISL_687814, EPI_ISL_687815, EPI_ISL_687816, EPI_ISL_687817, EPI_ISL_687818, EPI_ISL_687819, EPI_ISL_687820, EPI_ISL_687821, EPI_ISL_687822, EPI_ISL_687823, EPI_ISL_687824, EPI_ISL_687825, EPI_ISL_687826, EPI_ISL_687827, EPI_ISL_687828, EPI_ISL_687829, EPI_ISL_687830, EPI_ISL_687831, EPI_ISL_687832, EPI_ISL_687833, EPI_ISL_687834, EPI_ISL_687835, EPI_ISL_687836, EPI_ISL_687837, EPI_ISL_687838, EPI_ISL_687839, EPI_ISL_687840, EPI_ISL_687841, EPI_ISL_687842, EPI_ISL_687843, EPI_ISL_687844, EPI_ISL_687845, EPI_ISL_687846, EPI_ISL_687847, EPI_ISL_687848, EPI_ISL_687849, EPI_ISL_687850, EPI_ISL_687851, EPI_ISL_687852, EPI_ISL_687853, EPI_ISL_687854, EPI_ISL_687855, EPI_ISL_687856, EPI_ISL_687857, EPI_ISL_687858, EPI_ISL_687859, EPI_ISL_687860, EPI_ISL_687861, EPI_ISL_687862, EPI_ISL_687863, EPI_ISL_687864, EPI_ISL_687865, EPI_ISL_687866, EPI_ISL_687867, EPI_ISL_687868, EPI_ISL_687869, EPI_ISL_687870, EPI_ISL_687871, EPI_ISL_687872, EPI_ISL_687873, EPI_ISL_687874, EPI_ISL_687875, EPI_ISL_687876, EPI_ISL_687877, EPI_ISL_687878, EPI_ISL_687879, EPI_ISL_687880, EPI_ISL_687881, EPI_ISL_687882, EPI_ISL_687883, EPI_ISL_687884, EPI_ISL_687885, EPI_ISL_687886, EPI_ISL_687887, EPI_ISL_687888, EPI_ISL_687889, EPI_ISL_687890, EPI_ISL_687891, EPI_ISL_687892, EPI_ISL_687893, EPI_ISL_687894, EPI_ISL_687895, EPI_ISL_687896, EPI_ISL_687897, EPI_ISL_687898, EPI_ISL_687899, EPI_ISL_687900, EPI_ISL_687901, EPI_ISL_687902, EPI_ISL_687903, EPI_ISL_687904, EPI_ISL_687905, EPI_ISL_687906, EPI_ISL_687907, EPI_ISL_687908, EPI_ISL_687909, EPI_ISL_687910, EPI_ISL_687911, EPI_ISL_687912, EPI_ISL_687913, EPI_ISL_687914, EPI_ISL_687915, EPI_ISL_687916, EPI_ISL_687917, EPI_ISL_687918, EPI_ISL_687919, EPI_ISL_687932, EPI_ISL_687933, EPI_ISL_687934, EPI_ISL_687935, EPI_ISL_687936, EPI_ISL_687937, EPI_ISL_687938, EPI_ISL_687939, EPI_ISL_687940, EPI_ISL_687941, EPI_ISL_687942, EPI_ISL_687943, EPI_ISL_687973, EPI_ISL_687974, EPI_ISL_687975, EPI_ISL_687976, EPI_ISL_687977, EPI_ISL_687978, EPI_ISL_687979, EPI_ISL_687980, EPI_ISL_687981, EPI_ISL_687982, EPI_ISL_687983, EPI_ISL_687984, EPI_ISL_687985, EPI_ISL_687986, EPI_ISL_687987, EPI_ISL_687988, EPI_ISL_687989, EPI_ISL_687990, EPI_ISL_687991, EPI_ISL_687992, EPI_ISL_687993, EPI_ISL_687994, EPI_ISL_687995, EPI_ISL_687996, EPI_ISL_687997, EPI_ISL_687998, EPI_ISL_687999, EPI_ISL_688000, EPI_ISL_688001, EPI_ISL_688002, EPI_ISL_688003, EPI_ISL_688004, EPI_ISL_688005, EPI_ISL_688006, EPI_ISL_688056, EPI_ISL_688057, EPI_ISL_688058, EPI_ISL_688059, EPI_ISL_688060, EPI_ISL_688061, EPI_ISL_688062, EPI_ISL_688063, EPI_ISL_688064, EPI_ISL_688065, EPI_ISL_688066, EPI_ISL_688067, EPI_ISL_688068, EPI_ISL_688069, EPI_ISL_688070, EPI_ISL_688071, EPI_ISL_688072, EPI_ISL_688073, EPI_ISL_688074, EPI_ISL_688075, EPI_ISL_688076, EPI_ISL_688077, EPI_ISL_688078, EPI_ISL_688079, EPI_ISL_688080, EPI_ISL_688081, EPI_ISL_688082, EPI_ISL_688083, EPI_ISL_688086, EPI_ISL_688087, EPI_ISL_688088, EPI_ISL_688089, EPI_ISL_688090, EPI_ISL_688091, EPI_ISL_688092, EPI_ISL_688093, EPI_ISL_688094, EPI_ISL_688095, EPI_ISL_688096, EPI_ISL_688097, EPI_ISL_688098, EPI_ISL_688099, EPI_ISL_688100, EPI_ISL_688109, EPI_ISL_688110, EPI_ISL_688111, EPI_ISL_688112, EPI_ISL_688113, EPI_ISL_688114, EPI_ISL_688115, EPI_ISL_688116, EPI_ISL_688117, EPI_ISL_688118, EPI_ISL_688119, EPI_ISL_688120, EPI_ISL_688121, EPI_ISL_688122, EPI_ISL_688123, EPI_ISL_688124, EPI_ISL_688125, EPI_ISL_688126, EPI_ISL_688127, EPI_ISL_688128, EPI_ISL_688129, EPI_ISL_688130, EPI_ISL_688131, EPI_ISL_688132, EPI_ISL_688133, EPI_ISL_688134, EPI_ISL_688135, EPI_ISL_688136, EPI_ISL_688137, EPI_ISL_688138, EPI_ISL_688139, EPI_ISL_688140, EPI_ISL_688141, EPI_ISL_688142, EPI_ISL_688143, EPI_ISL_688144, EPI_ISL_688145, EPI_ISL_688146, EPI_ISL_688147, EPI_ISL_688148, EPI_ISL_688149, EPI_ISL_688150, EPI_ISL_688151, EPI_ISL_688152, EPI_ISL_688153, EPI_ISL_688154, EPI_ISL_688155, EPI_ISL_688156, EPI_ISL_688157, EPI_ISL_688158, EPI_ISL_688159, EPI_ISL_688160, EPI_ISL_688161, EPI_ISL_688162, EPI_ISL_688163, EPI_ISL_688164, EPI_ISL_688165, EPI_ISL_688166, EPI_ISL_688167, EPI_ISL_688168, EPI_ISL_688169, EPI_ISL_688170, EPI_ISL_688171, EPI_ISL_688172, EPI_ISL_688173, EPI_ISL_688174, EPI_ISL_688175, EPI_ISL_688176, EPI_ISL_688177, EPI_ISL_688178, EPI_ISL_688179, EPI_ISL_688180, EPI_ISL_688181, EPI_ISL_688182, EPI_ISL_688183, EPI_ISL_688184, EPI_ISL_688185, EPI_ISL_688186, EPI_ISL_688187, EPI_ISL_688188, EPI_ISL_688189, EPI_ISL_688190, EPI_ISL_688191, EPI_ISL_688192, EPI_ISL_688193, EPI_ISL_688194, EPI_ISL_688195, EPI_ISL_688196, EPI_ISL_688197, EPI_ISL_688198, EPI_ISL_688199, EPI_ISL_688200, EPI_ISL_688201, EPI_ISL_688202, EPI_ISL_688203, EPI_ISL_688204, EPI_ISL_688205, EPI_ISL_688206, EPI_ISL_688207, EPI_ISL_688208, EPI_ISL_688209, EPI_ISL_688210, EPI_ISL_688211, EPI_ISL_688212, EPI_ISL_688213, EPI_ISL_688214, EPI_ISL_688215, EPI_ISL_688216, EPI_ISL_688217, EPI_ISL_688218, EPI_ISL_688219, EPI_ISL_688220, EPI_ISL_688221, EPI_ISL_688222, EPI_ISL_688223, EPI_ISL_688224, EPI_ISL_688225, EPI_ISL_688226, EPI_ISL_688227, EPI_ISL_688228, EPI_ISL_688229, EPI_ISL_688230, EPI_ISL_688231, EPI_ISL_688232, EPI_ISL_688233, EPI_ISL_688234, EPI_ISL_688235, EPI_ISL_688236, EPI_ISL_688237, EPI_ISL_688238, EPI_ISL_688239, EPI_ISL_688240, EPI_ISL_688241, EPI_ISL_688242, EPI_ISL_688243, EPI_ISL_688244, EPI_ISL_688245, EPI_ISL_688246, EPI_ISL_688247, EPI_ISL_688248, EPI_ISL_688249, EPI_ISL_688250, EPI_ISL_688269, EPI_ISL_688270, EPI_ISL_688271, EPI_ISL_688272, EPI_ISL_688273, EPI_ISL_688274, EPI_ISL_688275, EPI_ISL_688276, EPI_ISL_688277, EPI_ISL_688278, EPI_ISL_688279, EPI_ISL_688280, EPI_ISL_688281, EPI_ISL_688282, EPI_ISL_688283 | | | |

| | | | |
|---|---|---|---|
| see above | Pathogen Genomics Center, National Institute of Infectious Diseases | Pathogen Genomics Center, National Institute of Infectious Diseases | Tsuyoshi Sekizuka, Kentaro Itokawa, Rina Tanaka, Masanori Hashino, Makoto Kuroda |
| EPI_ISL_688284, EPI_ISL_688285, EPI_ISL_688286, EPI_ISL_688287, EPI_ISL_688288, EPI_ISL_688289, EPI_ISL_688290, EPI_ISL_688291, EPI_ISL_688292, EPI_ISL_688293, EPI_ISL_688294, EPI_ISL_688295, EPI_ISL_688296, EPI_ISL_688297, EPI_ISL_688298, EPI_ISL_688299, EPI_ISL_688300, EPI_ISL_688301, EPI_ISL_688302, EPI_ISL_688303, EPI_ISL_688304, EPI_ISL_688305, EPI_ISL_688306, EPI_ISL_688307, EPI_ISL_688308, EPI_ISL_688309, EPI_ISL_688310, EPI_ISL_688311, EPI_ISL_688312, EPI_ISL_688313, EPI_ISL_688314, EPI_ISL_688315, EPI_ISL_688316, EPI_ISL_688317, EPI_ISL_688318, EPI_ISL_688319, EPI_ISL_688320, EPI_ISL_688321, EPI_ISL_688322, EPI_ISL_688323, EPI_ISL_688324, EPI_ISL_688325, EPI_ISL_688326, EPI_ISL_688327, EPI_ISL_688328, EPI_ISL_688329, EPI_ISL_688330, EPI_ISL_688331, EPI_ISL_688332, EPI_ISL_688333, EPI_ISL_688334, EPI_ISL_688335, EPI_ISL_688336, EPI_ISL_688337, EPI_ISL_688338, EPI_ISL_688339, EPI_ISL_688340, EPI_ISL_688341, EPI_ISL_688342, EPI_ISL_688343, EPI_ISL_688344, EPI_ISL_688345, EPI_ISL_688346, EPI_ISL_688347, EPI_ISL_688348, EPI_ISL_688349, EPI_ISL_688350, EPI_ISL_688351, EPI_ISL_688352, EPI_ISL_688353, EPI_ISL_688354, EPI_ISL_688355, EPI_ISL_688356, EPI_ISL_688357, EPI_ISL_688358, EPI_ISL_688359, EPI_ISL_688360, EPI_ISL_688361, EPI_ISL_688362, EPI_ISL_688363, EPI_ISL_688364, EPI_ISL_688365, EPI_ISL_688366, EPI_ISL_688367, EPI_ISL_688368, EPI_ISL_688369, EPI_ISL_688370, EPI_ISL_688371, EPI_ISL_688372, EPI_ISL_688373, EPI_ISL_688374, EPI_ISL_688375, EPI_ISL_688376, EPI_ISL_688377, EPI_ISL_688378, EPI_ISL_688379, EPI_ISL_688380, EPI_ISL_688381, EPI_ISL_688382, EPI_ISL_688383, EPI_ISL_688384, EPI_ISL_688385, EPI_ISL_688386, EPI_ISL_688387, EPI_ISL_688388, EPI_ISL_688389, EPI_ISL_688390, EPI_ISL_688391, EPI_ISL_688392, EPI_ISL_688393, EPI_ISL_688394, EPI_ISL_688395, EPI_ISL_688396, EPI_ISL_688397, EPI_ISL_688398, EPI_ISL_688399, EPI_ISL_688400, EPI_ISL_688401 | | | |
| see above | Saitama Prefectural Institute of Public Health | Pathogen Genomics Center, National Institute of Infectious Diseases | Tsuyoshi Sekizuka, Kentaro Itokawa, Rina Tanaka, Masanori Hashino, Makoto Kuroda |
| EPI_ISL_688402, EPI_ISL_688403, EPI_ISL_688404, EPI_ISL_688405, EPI_ISL_688406, EPI_ISL_688407, EPI_ISL_688408, EPI_ISL_688409, EPI_ISL_688410, EPI_ISL_688411, EPI_ISL_688412, EPI_ISL_688413, EPI_ISL_688414, EPI_ISL_688415, EPI_ISL_688416, EPI_ISL_688417, EPI_ISL_688418, EPI_ISL_688419, EPI_ISL_688420, EPI_ISL_688421, EPI_ISL_688422, EPI_ISL_688423, EPI_ISL_688424, EPI_ISL_688425, EPI_ISL_688426, EPI_ISL_688427, EPI_ISL_688428, EPI_ISL_688429, EPI_ISL_688430, EPI_ISL_688431, EPI_ISL_688432, EPI_ISL_688433, EPI_ISL_688434, EPI_ISL_688435, EPI_ISL_688436, EPI_ISL_688437, EPI_ISL_688438, EPI_ISL_688439, EPI_ISL_688440, EPI_ISL_688441, EPI_ISL_688442, EPI_ISL_688443, EPI_ISL_688444, EPI_ISL_688445, EPI_ISL_688446, EPI_ISL_688447, EPI_ISL_688448, EPI_ISL_688449, EPI_ISL_688450, EPI_ISL_688451, EPI_ISL_688452, EPI_ISL_688453, EPI_ISL_688454, EPI_ISL_688455, EPI_ISL_688456, EPI_ISL_688457, EPI_ISL_688458, EPI_ISL_688459, EPI_ISL_688460, EPI_ISL_688461, EPI_ISL_688462, EPI_ISL_688463, EPI_ISL_688464, EPI_ISL_688465, EPI_ISL_688466, EPI_ISL_688467, EPI_ISL_688468, EPI_ISL_688469, EPI_ISL_688470, EPI_ISL_688471, EPI_ISL_688472, EPI_ISL_688473, EPI_ISL_688474, EPI_ISL_688475, EPI_ISL_688476, EPI_ISL_688477, EPI_ISL_688478, EPI_ISL_688479, EPI_ISL_688480, EPI_ISL_688481, EPI_ISL_688482, EPI_ISL_688483, EPI_ISL_688484, EPI_ISL_688485, EPI_ISL_688486, EPI_ISL_688487, EPI_ISL_688488, EPI_ISL_688489, EPI_ISL_688490, EPI_ISL_688491, EPI_ISL_688492, EPI_ISL_688493, EPI_ISL_688494, EPI_ISL_688495, EPI_ISL_688496, EPI_ISL_688497, EPI_ISL_688498, EPI_ISL_688499, EPI_ISL_688500, EPI_ISL_688501, EPI_ISL_688502, EPI_ISL_688503, EPI_ISL_688504, EPI_ISL_688505, EPI_ISL_688506, EPI_ISL_688507, EPI_ISL_688508, EPI_ISL_688509, EPI_ISL_688510, EPI_ISL_688511, EPI_ISL_688512, EPI_ISL_688513, EPI_ISL_688514, EPI_ISL_688515, EPI_ISL_688516, EPI_ISL_688517, EPI_ISL_688518, EPI_ISL_688519, EPI_ISL_688520, EPI_ISL_688521, EPI_ISL_688522, EPI_ISL_688523, EPI_ISL_688524, EPI_ISL_688525, EPI_ISL_688526, EPI_ISL_688527, EPI_ISL_688528, EPI_ISL_688529, EPI_ISL_688530, EPI_ISL_688531, EPI_ISL_688532, EPI_ISL_688533, EPI_ISL_688534, EPI_ISL_688535, EPI_ISL_688536, EPI_ISL_688537, EPI_ISL_688538, EPI_ISL_688539, EPI_ISL_688540, EPI_ISL_688541, EPI_ISL_688542, EPI_ISL_688543, EPI_ISL_688544, EPI_ISL_688545, EPI_ISL_688546, EPI_ISL_688547, EPI_ISL_688548, EPI_ISL_688549, EPI_ISL_688550, EPI_ISL_688551, EPI_ISL_688552, EPI_ISL_688553, EPI_ISL_688554, EPI_ISL_688555, EPI_ISL_688556, EPI_ISL_688557, EPI_ISL_688558, EPI_ISL_688559, EPI_ISL_688560, EPI_ISL_688561, EPI_ISL_688562, EPI_ISL_688563, EPI_ISL_688564, EPI_ISL_688565, EPI_ISL_688566, EPI_ISL_688567, EPI_ISL_688568, EPI_ISL_688569, EPI_ISL_688570, EPI_ISL_688571, EPI_ISL_688572, EPI_ISL_688573, EPI_ISL_688574, EPI_ISL_688575, EPI_ISL_688576, EPI_ISL_688577, EPI_ISL_688578, EPI_ISL_688579, EPI_ISL_688580, EPI_ISL_688581, EPI_ISL_688582 | | | |
| see above | Pathogen Genomics Center, National Institute of Infectious Diseases | Pathogen Genomics Center, National Institute of Infectious Diseases | Tsuyoshi Sekizuka, Kentaro Itokawa, Rina Tanaka, Masanori Hashino, Makoto Kuroda |
| EPI_ISL_688583, EPI_ISL_688584, EPI_ISL_688585, EPI_ISL_688586, EPI_ISL_688587, EPI_ISL_688588, EPI_ISL_688589, EPI_ISL_688590, EPI_ISL_688591, EPI_ISL_688592, EPI_ISL_688593, EPI_ISL_688594, EPI_ISL_688595, EPI_ISL_688596, EPI_ISL_688597, EPI_ISL_688598, EPI_ISL_688599, EPI_ISL_688600, EPI_ISL_688601, EPI_ISL_688602, EPI_ISL_688603, EPI_ISL_688604 | | | |
| see above | Tokyo Metropolitan Institute of Public Health | Pathogen Genomics Center, National Institute of Infectious Diseases | Tsuyoshi Sekizuka, Kentaro Itokawa, Rina Tanaka, Masanori Hashino, Makoto Kuroda |
| EPI_ISL_688605, EPI_ISL_688606, EPI_ISL_688607, EPI_ISL_688608, EPI_ISL_688609, EPI_ISL_688610, EPI_ISL_688611 | Pathogen Genomics Center, National Institute of Infectious Diseases | Pathogen Genomics Center, National Institute of Infectious Diseases | Tsuyoshi Sekizuka, Kentaro Itokawa, Rina Tanaka, Masanori Hashino, Makoto Kuroda |
| EPI_ISL_688612 | Fukuoka Institute of Health and Environmental Sciences | Pathogen Genomics Center, National Institute of Infectious Diseases | Tsuyoshi Sekizuka, Kentaro Itokawa, Rina Tanaka, Masanori Hashino, Makoto Kuroda |
| EPI_ISL_688613, EPI_ISL_688616, EPI_ISL_688617, EPI_ISL_688618 | Pathogen Genomics Center, National Institute of Infectious Diseases | Pathogen Genomics Center, National Institute of Infectious Diseases | Tsuyoshi Sekizuka, Kentaro Itokawa, Rina Tanaka, Masanori Hashino, Makoto Kuroda |
| EPI_ISL_688619, EPI_ISL_688620, EPI_ISL_688621, EPI_ISL_688622, EPI_ISL_688623 | Fukuoka Institute of Health and Environmental Sciences | Pathogen Genomics Center, National Institute of Infectious Diseases | Tsuyoshi Sekizuka, Kentaro Itokawa, Rina Tanaka, Masanori Hashino, Makoto Kuroda |
| EPI_ISL_688625, EPI_ISL_688632, EPI_ISL_688633 | Pathogen Genomics Center, National Institute of Infectious Diseases | Pathogen Genomics Center, National Institute of Infectious Diseases | Tsuyoshi Sekizuka, Kentaro Itokawa, Rina Tanaka, Masanori Hashino, Makoto Kuroda |
| EPI_ISL_688634 | Fukuoka Institute of Health and Environmental Sciences | Pathogen Genomics Center, National Institute of Infectious Diseases | Tsuyoshi Sekizuka, Kentaro Itokawa, Rina Tanaka, Masanori Hashino, Makoto Kuroda |
| EPI_ISL_688635, EPI_ISL_688636, EPI_ISL_688637, EPI_ISL_688638, EPI_ISL_688639, EPI_ISL_688640, EPI_ISL_688641, EPI_ISL_688642, EPI_ISL_688643, EPI_ISL_688644, EPI_ISL_688645, EPI_ISL_688646, EPI_ISL_688647, EPI_ISL_688648, EPI_ISL_688649, EPI_ISL_688650, EPI_ISL_688651, EPI_ISL_688652, EPI_ISL_688653, EPI_ISL_688654, EPI_ISL_688655, EPI_ISL_688656, EPI_ISL_688657, EPI_ISL_688658, EPI_ISL_688659, EPI_ISL_688660, EPI_ISL_688661, EPI_ISL_688662, EPI_ISL_688663, EPI_ISL_688664, EPI_ISL_688665, EPI_ISL_688666, EPI_ISL_688667, EPI_ISL_688668, EPI_ISL_688669, EPI_ISL_688670, EPI_ISL_688671, EPI_ISL_688672, EPI_ISL_688673, EPI_ISL_688674, EPI_ISL_688675, EPI_ISL_688676, EPI_ISL_688677, EPI_ISL_688678, EPI_ISL_688679, EPI_ISL_688680, EPI_ISL_688681, EPI_ISL_688682, EPI_ISL_688683, EPI_ISL_688684, EPI_ISL_688685, EPI_ISL_688686, EPI_ISL_688687, EPI_ISL_688688, EPI_ISL_688689, EPI_ISL_688690, EPI_ISL_688691, EPI_ISL_688692, EPI_ISL_688693, EPI_ISL_688694, EPI_ISL_688695, EPI_ISL_688696, EPI_ISL_688697, EPI_ISL_688698, EPI_ISL_688699 | | | |
| see above | Pathogen Genomics Center, National Institute of Infectious Diseases | Pathogen Genomics Center, National Institute of Infectious Diseases | Tsuyoshi Sekizuka, Kentaro Itokawa, Rina Tanaka, Masanori Hashino, Makoto Kuroda |
| EPI_ISL_688700, EPI_ISL_688701, EPI_ISL_688702, EPI_ISL_688703, EPI_ISL_688704, EPI_ISL_688705, EPI_ISL_688706, EPI_ISL_688707, EPI_ISL_688708, EPI_ISL_688709 | Sakai City Institute of Public Health | Pathogen Genomics Center, National Institute of Infectious Diseases | Tsuyoshi Sekizuka, Kentaro Itokawa, Rina Tanaka, Masanori Hashino, Makoto Kuroda |
| EPI_ISL_688710, EPI_ISL_688711, EPI_ISL_688712, EPI_ISL_688713, EPI_ISL_688714, EPI_ISL_688715, EPI_ISL_688716, EPI_ISL_688717, EPI_ISL_688718, EPI_ISL_688719, EPI_ISL_688720, EPI_ISL_688721, EPI_ISL_688722, EPI_ISL_688723, EPI_ISL_688724, EPI_ISL_688725, EPI_ISL_688726, EPI_ISL_688727, EPI_ISL_688728, EPI_ISL_688729, EPI_ISL_688730, EPI_ISL_688731, EPI_ISL_688732, EPI_ISL_688733, EPI_ISL_688734, EPI_ISL_688735, EPI_ISL_688736, EPI_ISL_688737, EPI_ISL_688738, EPI_ISL_688739, EPI_ISL_688740, EPI_ISL_688741, EPI_ISL_688742, EPI_ISL_688743, EPI_ISL_688744, EPI_ISL_688745, EPI_ISL_688747, EPI_ISL_688748, EPI_ISL_688749, EPI_ISL_688750, EPI_ISL_688751, EPI_ISL_688752, EPI_ISL_688753, EPI_ISL_688754, EPI_ISL_688755, EPI_ISL_688756, EPI_ISL_688814, EPI_ISL_688815, EPI_ISL_688816, EPI_ISL_688817, EPI_ISL_688818, EPI_ISL_688819, EPI_ISL_688820, EPI_ISL_688821, EPI_ISL_688822, EPI_ISL_688823, EPI_ISL_688824, EPI_ISL_688825, EPI_ISL_688826, EPI_ISL_688827, EPI_ISL_688828, EPI_ISL_688829, EPI_ISL_688830, EPI_ISL_688831, EPI_ISL_688832, EPI_ISL_688833, EPI_ISL_688834, EPI_ISL_688835, EPI_ISL_688836, EPI_ISL_688837, EPI_ISL_688838, EPI_ISL_688839, EPI_ISL_688840, EPI_ISL_688841, EPI_ISL_688842, EPI_ISL_688843, EPI_ISL_688844, EPI_ISL_688845, EPI_ISL_688846, EPI_ISL_688847, EPI_ISL_688848, EPI_ISL_688849, EPI_ISL_688850, EPI_ISL_688851, EPI_ISL_688852, EPI_ISL_688853, EPI_ISL_688854, EPI_ISL_688855, EPI_ISL_688856, EPI_ISL_688857, EPI_ISL_688858, EPI_ISL_688859, EPI_ISL_688860, EPI_ISL_688861, EPI_ISL_688862, EPI_ISL_688863, EPI_ISL_688864, EPI_ISL_688865, EPI_ISL_688866, EPI_ISL_688867, EPI_ISL_688868, EPI_ISL_688869, EPI_ISL_688870, EPI_ISL_688871, EPI_ISL_688872, EPI_ISL_688873, EPI_ISL_688874, EPI_ISL_688875, EPI_ISL_688876, EPI_ISL_688877, EPI_ISL_688878, EPI_ISL_688879, EPI_ISL_688880, EPI_ISL_688881, EPI_ISL_688882, EPI_ISL_688883, EPI_ISL_688884, EPI_ISL_688885, EPI_ISL_688886, EPI_ISL_688887, EPI_ISL_688888, EPI_ISL_688889, EPI_ISL_688890, EPI_ISL_688891, EPI_ISL_688892, EPI_ISL_688893, EPI_ISL_688894, EPI_ISL_688895, EPI_ISL_688896 | | | |
| see above | Pathogen Genomics Center, National Institute of Infectious Diseases | Pathogen Genomics Center, National Institute of Infectious Diseases | Tsuyoshi Sekizuka, Kentaro Itokawa, Rina Tanaka, Masanori Hashino, Makoto Kuroda |
| EPI_ISL_688897, EPI_ISL_688898, EPI_ISL_688899, EPI_ISL_688900, EPI_ISL_688901, EPI_ISL_688902, EPI_ISL_688903, EPI_ISL_688904, EPI_ISL_688905, EPI_ISL_688906, EPI_ISL_688907, EPI_ISL_688908, EPI_ISL_688909, EPI_ISL_688910, EPI_ISL_688911, EPI_ISL_688912, EPI_ISL_688913, EPI_ISL_688914, EPI_ISL_688915, EPI_ISL_688916, EPI_ISL_688917, EPI_ISL_688918, EPI_ISL_688919, EPI_ISL_688920, EPI_ISL_688921, EPI_ISL_688922, EPI_ISL_688923, EPI_ISL_688924, EPI_ISL_688925, EPI_ISL_688926, EPI_ISL_688927, EPI_ISL_688928, EPI_ISL_688929, EPI_ISL_688930, EPI_ISL_688931, EPI_ISL_688932, EPI_ISL_688933, EPI_ISL_688934, EPI_ISL_688935, EPI_ISL_688936, EPI_ISL_688937, EPI_ISL_688938, EPI_ISL_688939, EPI_ISL_688940, EPI_ISL_688941, EPI_ISL_688942, EPI_ISL_688943, EPI_ISL_688944, EPI_ISL_688945, EPI_ISL_688946, EPI_ISL_688947, EPI_ISL_688948, EPI_ISL_688949, EPI_ISL_688950, EPI_ISL_688951, EPI_ISL_688952, EPI_ISL_688953, EPI_ISL_688954, EPI_ISL_688955, EPI_ISL_688956, EPI_ISL_688957, EPI_ISL_688958, EPI_ISL_688959, EPI_ISL_688960, EPI_ISL_688961, EPI_ISL_688962, EPI_ISL_688963, EPI_ISL_688964, EPI_ISL_688965, EPI_ISL_688966, EPI_ISL_688967, EPI_ISL_688968 | | | |
| see above | Fukuoka Institute of Health and Environmental Sciences | Pathogen Genomics Center, National Institute of Infectious Diseases | Tsuyoshi Sekizuka, Kentaro Itokawa, Rina Tanaka, Masanori Hashino, Makoto Kuroda |
| EPI_ISL_688969, EPI_ISL_688970, EPI_ISL_688971, EPI_ISL_688972, EPI_ISL_688973, EPI_ISL_688974, EPI_ISL_688975, EPI_ISL_688976, EPI_ISL_688977, EPI_ISL_688978, EPI_ISL_688979, EPI_ISL_688980, EPI_ISL_688981, EPI_ISL_688982, EPI_ISL_688983, EPI_ISL_688984, EPI_ISL_688985, EPI_ISL_688986, EPI_ISL_688987, EPI_ISL_688988, EPI_ISL_688989, EPI_ISL_688990, EPI_ISL_688991, EPI_ISL_688992, EPI_ISL_688993, EPI_ISL_688994, EPI_ISL_688995, EPI_ISL_688996, EPI_ISL_688997, EPI_ISL_688998, EPI_ISL_688999, EPI_ISL_689000, EPI_ISL_689001, EPI_ISL_689002, EPI_ISL_689003, EPI_ISL_689004, EPI_ISL_689005, EPI_ISL_689006, EPI_ISL_689007, EPI_ISL_689008, EPI_ISL_689009, EPI_ISL_689010, EPI_ISL_689011, EPI_ISL_689012, EPI_ISL_689013, EPI_ISL_689014, EPI_ISL_689015, EPI_ISL_689016, EPI_ISL_689017, EPI_ISL_689018, EPI_ISL_689019, EPI_ISL_689020, EPI_ISL_689021, EPI_ISL_689022, EPI_ISL_689023, EPI_ISL_689024, EPI_ISL_689025, EPI_ISL_689026, EPI_ISL_689027, EPI_ISL_689028, EPI_ISL_689029, EPI_ISL_689030, EPI_ISL_689031, EPI_ISL_689032, EPI_ISL_689033, EPI_ISL_689034, EPI_ISL_689035, EPI_ISL_689036, EPI_ISL_689037, EPI_ISL_689038, EPI_ISL_689039, EPI_ISL_689040, EPI_ISL_689041, EPI_ISL_689042, EPI_ISL_689043, EPI_ISL_689044, EPI_ISL_689045, EPI_ISL_689046 | | | |
| see above | Pathogen Genomics Center, National Institute of Infectious Diseases | Pathogen Genomics Center, National Institute of Infectious Diseases | Tsuyoshi Sekizuka, Kentaro Itokawa, Rina Tanaka, Masanori Hashino, Makoto Kuroda |

EPI_ISL_689077, EPI_ISL_689078, EPI_ISL_689079, EPI_ISL_689080, EPI_ISL_689081, EPI_ISL_689082, EPI_ISL_689083, EPI_ISL_689084, EPI_ISL_689085, EPI_ISL_689086, EPI_ISL_689087, EPI_ISL_689088

| | | | |
|---|---|---|---|
| see above | Tokyo Metropolitan Institute of Public Health | Pathogen Genomics Center, National Institute of Infectious Diseases | Tsuyoshi Sekizuka, Kentaro Itokawa, Rina Tanaka, Masanori Hashino, Makoto Kuroda |

EPI_ISL_689089, EPI_ISL_689090, EPI_ISL_689091, EPI_ISL_689092, EPI_ISL_689093, EPI_ISL_689094, EPI_ISL_689095, EPI_ISL_689096, EPI_ISL_689097, EPI_ISL_689098, EPI_ISL_689099, EPI_ISL_689100, EPI_ISL_689101, EPI_ISL_689102, EPI_ISL_689103, EPI_ISL_689104, EPI_ISL_689105, EPI_ISL_689106, EPI_ISL_689107, EPI_ISL_689108, EPI_ISL_689109, EPI_ISL_689110, EPI_ISL_689111, EPI_ISL_689112, EPI_ISL_689113, EPI_ISL_689114, EPI_ISL_689115, EPI_ISL_689116, EPI_ISL_689117, EPI_ISL_689118, EPI_ISL_689119, EPI_ISL_689120, EPI_ISL_689121, EPI_ISL_689122, EPI_ISL_689123, EPI_ISL_689124, EPI_ISL_689125, EPI_ISL_689126, EPI_ISL_689127, EPI_ISL_689128, EPI_ISL_689129, EPI_ISL_689130, EPI_ISL_689131, EPI_ISL_689132, EPI_ISL_689133, EPI_ISL_689134, EPI_ISL_689135, EPI_ISL_689136, EPI_ISL_689137, EPI_ISL_689138, EPI_ISL_689139, EPI_ISL_689140, EPI_ISL_689141, EPI_ISL_689142, EPI_ISL_689143, EPI_ISL_689144, EPI_ISL_689145, EPI_ISL_689146, EPI_ISL_689147, EPI_ISL_689148, EPI_ISL_689149, EPI_ISL_689150, EPI_ISL_689151, EPI_ISL_689152, EPI_ISL_689153, EPI_ISL_689154, EPI_ISL_689155, EPI_ISL_689156, EPI_ISL_689157, EPI_ISL_689158, EPI_ISL_689159, EPI_ISL_689160, EPI_ISL_689161, EPI_ISL_689162, EPI_ISL_689163, EPI_ISL_689164, EPI_ISL_689165, EPI_ISL_689166, EPI_ISL_689167, EPI_ISL_689168, EPI_ISL_689169, EPI_ISL_689170, EPI_ISL_689171, EPI_ISL_689172, EPI_ISL_689173, EPI_ISL_689174, EPI_ISL_689175, EPI_ISL_689176, EPI_ISL_689177, EPI_ISL_689178, EPI_ISL_689179, EPI_ISL_689180, EPI_ISL_689181, EPI_ISL_689182, EPI_ISL_689183, EPI_ISL_689184, EPI_ISL_689185, EPI_ISL_689186, EPI_ISL_689187, EPI_ISL_689188, EPI_ISL_689189, EPI_ISL_689190, EPI_ISL_689191, EPI_ISL_689192, EPI_ISL_689193, EPI_ISL_689194, EPI_ISL_689195, EPI_ISL_689196, EPI_ISL_689197, EPI_ISL_689198, EPI_ISL_689199, EPI_ISL_689200, EPI_ISL_689201, EPI_ISL_689202, EPI_ISL_689203, EPI_ISL_689204, EPI_ISL_689205, EPI_ISL_689206, EPI_ISL_689207, EPI_ISL_689208, EPI_ISL_689209, EPI_ISL_689210, EPI_ISL_689211, EPI_ISL_689212, EPI_ISL_689213, EPI_ISL_689214, EPI_ISL_689215, EPI_ISL_689216, EPI_ISL_689217, EPI_ISL_689218, EPI_ISL_689219, EPI_ISL_689220, EPI_ISL_689221, EPI_ISL_689222, EPI_ISL_689223, EPI_ISL_689224, EPI_ISL_689225, EPI_ISL_689226, EPI_ISL_689227, EPI_ISL_689228, EPI_ISL_689229, EPI_ISL_689230, EPI_ISL_689231, EPI_ISL_689232, EPI_ISL_689233, EPI_ISL_689234, EPI_ISL_689235, EPI_ISL_689236, EPI_ISL_689237, EPI_ISL_689238, EPI_ISL_689239, EPI_ISL_689240, EPI_ISL_689241, EPI_ISL_689242, EPI_ISL_689243, EPI_ISL_689244, EPI_ISL_689245, EPI_ISL_689246, EPI_ISL_689247, EPI_ISL_689248, EPI_ISL_689249, EPI_ISL_689250, EPI_ISL_689251, EPI_ISL_689252, EPI_ISL_689253, EPI_ISL_689254, EPI_ISL_689255, EPI_ISL_689256, EPI_ISL_689257, EPI_ISL_689258, EPI_ISL_689259, EPI_ISL_689260, EPI_ISL_689261, EPI_ISL_689262, EPI_ISL_689263, EPI_ISL_689264, EPI_ISL_689265, EPI_ISL_689266, EPI_ISL_689267, EPI_ISL_689268, EPI_ISL_689269, EPI_ISL_689270, EPI_ISL_689271, EPI_ISL_689272, EPI_ISL_689273, EPI_ISL_689274, EPI_ISL_689275, EPI_ISL_689276, EPI_ISL_689277, EPI_ISL_689278, EPI_ISL_689279, EPI_ISL_689280, EPI_ISL_689281, EPI_ISL_689282, EPI_ISL_689283, EPI_ISL_689284, EPI_ISL_689285, EPI_ISL_689286, EPI_ISL_689287, EPI_ISL_689288, EPI_ISL_689289, EPI_ISL_689290, EPI_ISL_689291, EPI_ISL_689292, EPI_ISL_689293, EPI_ISL_689294, EPI_ISL_689295, EPI_ISL_689296, EPI_ISL_689297, EPI_ISL_689298, EPI_ISL_689299, EPI_ISL_689300, EPI_ISL_689301, EPI_ISL_689302, EPI_ISL_689303, EPI_ISL_689304, EPI_ISL_689305, EPI_ISL_689306, EPI_ISL_689307, EPI_ISL_689308, EPI_ISL_689310, EPI_ISL_689311, EPI_ISL_689312, EPI_ISL_689313, EPI_ISL_689314, EPI_ISL_689315, EPI_ISL_689316, EPI_ISL_689317, EPI_ISL_689318, EPI_ISL_689319, EPI_ISL_689320, EPI_ISL_689321, EPI_ISL_689322, EPI_ISL_689323, EPI_ISL_689324, EPI_ISL_689325, EPI_ISL_689326, EPI_ISL_689327, EPI_ISL_689328, EPI_ISL_689329, EPI_ISL_689330, EPI_ISL_689331, EPI_ISL_689332, EPI_ISL_689333, EPI_ISL_689334, EPI_ISL_689335, EPI_ISL_689336, EPI_ISL_689337, EPI_ISL_689338, EPI_ISL_689339, EPI_ISL_689340, EPI_ISL_689341, EPI_ISL_689342, EPI_ISL_689343, EPI_ISL_689344, EPI_ISL_689345, EPI_ISL_689346, EPI_ISL_689347, EPI_ISL_689348, EPI_ISL_689349, EPI_ISL_689350, EPI_ISL_689351, EPI_ISL_689352, EPI_ISL_689353, EPI_ISL_689354, EPI_ISL_689355, EPI_ISL_689356, EPI_ISL_689357, EPI_ISL_689358, EPI_ISL_689359, EPI_ISL_689360, EPI_ISL_689361, EPI_ISL_689362, EPI_ISL_689363, EPI_ISL_689364, EPI_ISL_689365, EPI_ISL_689366, EPI_ISL_689367, EPI_ISL_689368, EPI_ISL_689369, EPI_ISL_689370, EPI_ISL_689371, EPI_ISL_689372, EPI_ISL_689373, EPI_ISL_689374, EPI_ISL_689375, EPI_ISL_689376, EPI_ISL_689377, EPI_ISL_689378, EPI_ISL_689379, EPI_ISL_689380, EPI_ISL_689381, EPI_ISL_689382, EPI_ISL_689383, EPI_ISL_689384, EPI_ISL_689385, EPI_ISL_689386, EPI_ISL_689387, EPI_ISL_689388, EPI_ISL_689389, EPI_ISL_689390, EPI_ISL_689391, EPI_ISL_689392, EPI_ISL_689393, EPI_ISL_689394, EPI_ISL_689395, EPI_ISL_689396, EPI_ISL_689397, EPI_ISL_689398, EPI_ISL_689399, EPI_ISL_689400, EPI_ISL_689401, EPI_ISL_689402, EPI_ISL_689403, EPI_ISL_689404, EPI_ISL_689405, EPI_ISL_689406, EPI_ISL_689407, EPI_ISL_689408, EPI_ISL_689409, EPI_ISL_689410, EPI_ISL_689411, EPI_ISL_689412, EPI_ISL_689413, EPI_ISL_689414, EPI_ISL_689415, EPI_ISL_689416, EPI_ISL_689417, EPI_ISL_689418, EPI_ISL_689419, EPI_ISL_689420, EPI_ISL_689421, EPI_ISL_689422, EPI_ISL_689423, EPI_ISL_689424, EPI_ISL_689425, EPI_ISL_689426, EPI_ISL_689427, EPI_ISL_689428, EPI_ISL_689429, EPI_ISL_689430, EPI_ISL_689431, EPI_ISL_689432, EPI_ISL_689433, EPI_ISL_689434, EPI_ISL_689435, EPI_ISL_689436, EPI_ISL_689437, EPI_ISL_689438, EPI_ISL_689439, EPI_ISL_689440, EPI_ISL_689441, EPI_ISL_689442, EPI_ISL_689443, EPI_ISL_689444, EPI_ISL_689445, EPI_ISL_689446, EPI_ISL_689447, EPI_ISL_689448, EPI_ISL_689449, EPI_ISL_689450, EPI_ISL_689451, EPI_ISL_689452, EPI_ISL_689453, EPI_ISL_689454, EPI_ISL_689455, EPI_ISL_689456, EPI_ISL_689457, EPI_ISL_689458, EPI_ISL_689459

| | | | |
|---|---|---|---|
| see above | Pathogen Genomics Center, National Institute of Infectious Diseases | Pathogen Genomics Center, National Institute of Infectious Diseases | Tsuyoshi Sekizuka, Kentaro Itokawa, Rina Tanaka, Masanori Hashino, Makoto Kuroda |

EPI_ISL_689460, EPI_ISL_689461, EPI_ISL_689462, EPI_ISL_689463, EPI_ISL_689464, EPI_ISL_689465, EPI_ISL_689466, EPI_ISL_689467, EPI_ISL_689468, EPI_ISL_689469, EPI_ISL_689470, EPI_ISL_689471, EPI_ISL_689472, EPI_ISL_689473, EPI_ISL_689474, EPI_ISL_689475, EPI_ISL_689476, EPI_ISL_689477, EPI_ISL_689478, EPI_ISL_689479, EPI_ISL_689480, EPI_ISL_689481, EPI_ISL_689482, EPI_ISL_689483, EPI_ISL_689484, EPI_ISL_689485, EPI_ISL_689486, EPI_ISL_689487, EPI_ISL_689488, EPI_ISL_689489, EPI_ISL_689490, EPI_ISL_689491, EPI_ISL_689492, EPI_ISL_689493, EPI_ISL_689494, EPI_ISL_689495, EPI_ISL_689496

| | | | |
|---|---|---|---|
| see above | Fukuoka Institute of Health and Environmental Sciences | Pathogen Genomics Center, National Institute of Infectious Diseases | Tsuyoshi Sekizuka, Kentaro Itokawa, Rina Tanaka, Masanori Hashino, Makoto Kuroda |

EPI_ISL_689497, EPI_ISL_689498, EPI_ISL_689499, EPI_ISL_689500, EPI_ISL_689501, EPI_ISL_689502, EPI_ISL_689503, EPI_ISL_689504, EPI_ISL_689505, EPI_ISL_689506, EPI_ISL_689507, EPI_ISL_689508, EPI_ISL_689509, EPI_ISL_689510, EPI_ISL_689511, EPI_ISL_689512, EPI_ISL_689513, EPI_ISL_689514, EPI_ISL_689515, EPI_ISL_689516, EPI_ISL_689517, EPI_ISL_689518, EPI_ISL_689519, EPI_ISL_689520, EPI_ISL_689521, EPI_ISL_689522, EPI_ISL_689523, EPI_ISL_689524, EPI_ISL_689525, EPI_ISL_689526, EPI_ISL_689527, EPI_ISL_689528, EPI_ISL_689529, EPI_ISL_689530, EPI_ISL_689531, EPI_ISL_689532, EPI_ISL_689533, EPI_ISL_689534, EPI_ISL_689535, EPI_ISL_689536, EPI_ISL_689537, EPI_ISL_689538, EPI_ISL_689539, EPI_ISL_689540, EPI_ISL_689541, EPI_ISL_689542, EPI_ISL_689543, EPI_ISL_689544, EPI_ISL_689545, EPI_ISL_689546, EPI_ISL_689547, EPI_ISL_689548, EPI_ISL_689551, EPI_ISL_689552, EPI_ISL_689553, EPI_ISL_689554, EPI_ISL_689555, EPI_ISL_689556, EPI_ISL_689557, EPI_ISL_689558, EPI_ISL_689559, EPI_ISL_689560, EPI_ISL_689561, EPI_ISL_689562, EPI_ISL_689563, EPI_ISL_689564, EPI_ISL_689565, EPI_ISL_689566, EPI_ISL_689567, EPI_ISL_689568, EPI_ISL_689569, EPI_ISL_689570, EPI_ISL_689571, EPI_ISL_689572, EPI_ISL_689573, EPI_ISL_689574, EPI_ISL_689575, EPI_ISL_689576, EPI_ISL_689577, EPI_ISL_689578, EPI_ISL_689579, EPI_ISL_689580, EPI_ISL_689581, EPI_ISL_689582, EPI_ISL_689583, EPI_ISL_689584, EPI_ISL_689585, EPI_ISL_689586, EPI_ISL_689587, EPI_ISL_689588, EPI_ISL_689589, EPI_ISL_689590, EPI_ISL_689591, EPI_ISL_689601, EPI_ISL_689602, EPI_ISL_689603, EPI_ISL_689604, EPI_ISL_689605, EPI_ISL_689606, EPI_ISL_689607, EPI_ISL_689608, EPI_ISL_689609, EPI_ISL_689610, EPI_ISL_689611, EPI_ISL_689624, EPI_ISL_689625, EPI_ISL_689626, EPI_ISL_689627, EPI_ISL_689628, EPI_ISL_689629, EPI_ISL_689630, EPI_ISL_689631, EPI_ISL_689632, EPI_ISL_689633, EPI_ISL_689634, EPI_ISL_689635, EPI_ISL_689636, EPI_ISL_689637, EPI_ISL_689638, EPI_ISL_689639, EPI_ISL_689641, EPI_ISL_689642, EPI_ISL_689644, EPI_ISL_689645, EPI_ISL_689647, EPI_ISL_689648, EPI_ISL_689649, EPI_ISL_689650, EPI_ISL_689651, EPI_ISL_689652, EPI_ISL_689653, EPI_ISL_689654, EPI_ISL_689655, EPI_ISL_689656, EPI_ISL_689657, EPI_ISL_689658, EPI_ISL_689659, EPI_ISL_689660, EPI_ISL_689661, EPI_ISL_689662, EPI_ISL_689663, EPI_ISL_689664, EPI_ISL_689665, EPI_ISL_689666, EPI_ISL_689667, EPI_ISL_689668, EPI_ISL_689669, EPI_ISL_689670, EPI_ISL_689671, EPI_ISL_689672, EPI_ISL_689673, EPI_ISL_689674, EPI_ISL_689675, EPI_ISL_689676, EPI_ISL_689677, EPI_ISL_689678, EPI_ISL_689679, EPI_ISL_689681, EPI_ISL_689682, EPI_ISL_689683, EPI_ISL_689684, EPI_ISL_689685

| | | | |
|---|---|---|---|
| see above | Pathogen Genomics Center, National Institute of Infectious Diseases | Pathogen Genomics Center, National Institute of Infectious Diseases | Tsuyoshi Sekizuka, Kentaro Itokawa, Rina Tanaka, Masanori Hashino, Makoto Kuroda |
| EPI_ISL_689692, EPI_ISL_689693, EPI_ISL_689694, EPI_ISL_689695, EPI_ISL_689696, EPI_ISL_689697 | Fukuoka Institute of Health and Environmental Sciences | Pathogen Genomics Center, National Institute of Infectious Diseases | Tsuyoshi Sekizuka, Kentaro Itokawa, Rina Tanaka, Masanori Hashino, Makoto Kuroda |

EPI_ISL_689698, EPI_ISL_689699, EPI_ISL_689700, EPI_ISL_689701, EPI_ISL_689702, EPI_ISL_689703, EPI_ISL_689704, EPI_ISL_689705, EPI_ISL_689706, EPI_ISL_689707, EPI_ISL_689708, EPI_ISL_689709, EPI_ISL_689710, EPI_ISL_689711, EPI_ISL_689712, EPI_ISL_689713, EPI_ISL_689714, EPI_ISL_689715, EPI_ISL_689716, EPI_ISL_689717, EPI_ISL_689718, EPI_ISL_689719, EPI_ISL_689720, EPI_ISL_689721, EPI_ISL_689722, EPI_ISL_689723, EPI_ISL_689724, EPI_ISL_689725, EPI_ISL_689726, EPI_ISL_689727, EPI_ISL_689728, EPI_ISL_689729, EPI_ISL_689730, EPI_ISL_689731, EPI_ISL_689732, EPI_ISL_689733, EPI_ISL_689734, EPI_ISL_689735, EPI_ISL_689736, EPI_ISL_689737, EPI_ISL_689738, EPI_ISL_689739, EPI_ISL_689740, EPI_ISL_689741, EPI_ISL_689742, EPI_ISL_689743, EPI_ISL_689744, EPI_ISL_689745, EPI_ISL_689746, EPI_ISL_689747, EPI_ISL_689748, EPI_ISL_689749, EPI_ISL_689750, EPI_ISL_689751, EPI_ISL_689752, EPI_ISL_689753, EPI_ISL_689754, EPI_ISL_689755, EPI_ISL_689756, EPI_ISL_689757, EPI_ISL_689758

| | | | |
|---|---|---|---|
| see above | Pathogen Genomics Center, National Institute of Infectious Diseases | Pathogen Genomics Center, National Institute of Infectious Diseases | Tsuyoshi Sekizuka, Kentaro Itokawa, Rina Tanaka, Masanori Hashino, Makoto Kuroda |

EPI_ISL_689759, EPI_ISL_689760, EPI_ISL_689761, EPI_ISL_689762, EPI_ISL_689763, EPI_ISL_689764, EPI_ISL_689765, EPI_ISL_689766, EPI_ISL_689767, EPI_ISL_689768, EPI_ISL_689769, EPI_ISL_689770, EPI_ISL_689771, EPI_ISL_689772, EPI_ISL_689773, EPI_ISL_689774, EPI_ISL_689775, EPI_ISL_689776, EPI_ISL_689777, EPI_ISL_689778, EPI_ISL_689779, EPI_ISL_689780, EPI_ISL_689781, EPI_ISL_689782, EPI_ISL_689783, EPI_ISL_689784, EPI_ISL_689785, EPI_ISL_689786, EPI_ISL_689787, EPI_ISL_689788, EPI_ISL_689789, EPI_ISL_689790, EPI_ISL_689791, EPI_ISL_689792, EPI_ISL_689793, EPI_ISL_689794, EPI_ISL_689795, EPI_ISL_689796, EPI_ISL_689797, EPI_ISL_689798, EPI_ISL_689799, EPI_ISL_689800, EPI_ISL_689801, EPI_ISL_689802, EPI_ISL_689803, EPI_ISL_689804, EPI_ISL_689805, EPI_ISL_689806, EPI_ISL_689807, EPI_ISL_689808, EPI_ISL_689809, EPI_ISL_689810, EPI_ISL_689811, EPI_ISL_689812, EPI_ISL_689813, EPI_ISL_689814, EPI_ISL_689815, EPI_ISL_689816, EPI_ISL_689817, EPI_ISL_689818, EPI_ISL_689819, EPI_ISL_689820, EPI_ISL_689821, EPI_ISL_689822, EPI_ISL_689823, EPI_ISL_689824, EPI_ISL_689825

| | | | |
|---|---|---|---|
| see above | Fukuoka Institute of Health and Environmental Sciences | Pathogen Genomics Center, National Institute of Infectious Diseases | Tsuyoshi Sekizuka, Kentaro Itokawa, Rina Tanaka, Masanori Hashino, Makoto Kuroda |

EPI_ISL_689826, EPI_ISL_689827, EPI_ISL_689828, EPI_ISL_689829, EPI_ISL_689830, EPI_ISL_689831, EPI_ISL_689832, EPI_ISL_689833, EPI_ISL_689834, EPI_ISL_689835, EPI_ISL_689836, EPI_ISL_689837, EPI_ISL_689838, EPI_ISL_689839, EPI_ISL_689840, EPI_ISL_689841, EPI_ISL_689842, EPI_ISL_689843, EPI_ISL_689844, EPI_ISL_689845, EPI_ISL_689846, EPI_ISL_689847, EPI_ISL_689848, EPI_ISL_689849, EPI_ISL_689850, EPI_ISL_689851, EPI_ISL_689852, EPI_ISL_689853, EPI_ISL_689854, EPI_ISL_689855, EPI_ISL_689856, EPI_ISL_689857, EPI_ISL_689858, EPI_ISL_689859, EPI_ISL_689860, EPI_ISL_689861, EPI_ISL_689862, EPI_ISL_689863, EPI_ISL_689864, EPI_ISL_689865, EPI_ISL_689866, EPI_ISL_689867, EPI_ISL_689868, EPI_ISL_689869, EPI_ISL_689870, EPI_ISL_689871, EPI_ISL_689872, EPI_ISL_689873, EPI_ISL_689874, EPI_ISL_689875, EPI_ISL_689876, EPI_ISL_689877, EPI_ISL_689878, EPI_ISL_689879, EPI_ISL_689880, EPI_ISL_689881, EPI_ISL_689882, EPI_ISL_689883, EPI_ISL_689884, EPI_ISL_689885, EPI_ISL_689886, EPI_ISL_689887, EPI_ISL_689888, EPI_ISL_689889, EPI_ISL_689890, EPI_ISL_689891, EPI_ISL_689892, EPI_ISL_689893, EPI_ISL_689894, EPI_ISL_689895, EPI_ISL_689896, EPI_ISL_689897, EPI_ISL_689898, EPI_ISL_689899, EPI_ISL_689900, EPI_ISL_689901, EPI_ISL_689902, EPI_ISL_689903, EPI_ISL_689904, EPI_ISL_689905, EPI_ISL_689906, EPI_ISL_689907, EPI_ISL_689908, EPI_ISL_689909, EPI_ISL_689910, EPI_ISL_689911, EPI_ISL_689912, EPI_ISL_689913, EPI_ISL_689914, EPI_ISL_689915, EPI_ISL_689916, EPI_ISL_689917, EPI_ISL_689918, EPI_ISL_689919, EPI_ISL_689920, EPI_ISL_689921, EPI_ISL_689922, EPI_ISL_689923, EPI_ISL_689924, EPI_ISL_689925, EPI_ISL_689926, EPI_ISL_689927, EPI_ISL_689928, EPI_ISL_689929, EPI_ISL_689930, EPI_ISL_689931, EPI_ISL_689932, EPI_ISL_689933, EPI_ISL_689934, EPI_ISL_689935, EPI_ISL_689936, EPI_ISL_689937, EPI_ISL_689938, EPI_ISL_689939, EPI_ISL_689940, EPI_ISL_689941, EPI_ISL_689942, EPI_ISL_689943, EPI_ISL_689944, EPI_ISL_689945, EPI_ISL_689946, EPI_ISL_689947, EPI_ISL_689948, EPI_ISL_689949, EPI_ISL_689950, EPI_ISL_689951, EPI_ISL_689952, EPI_ISL_689953, EPI_ISL_689954, EPI_ISL_689955, EPI_ISL_689956, EPI_ISL_689957, EPI_ISL_689958, EPI_ISL_689959, EPI_ISL_689960, EPI_ISL_689961, EPI_ISL_689962, EPI_ISL_689963, EPI_ISL_689964, EPI_ISL_689965, EPI_ISL_689966, EPI_ISL_689967, EPI_ISL_689968, EPI_ISL_689969, EPI_ISL_689970, EPI_ISL_689971, EPI_ISL_689972, EPI_ISL_689973, EPI_ISL_689974, EPI_ISL_689975, EPI_ISL_689976, EPI_ISL_689977, EPI_ISL_689978, EPI_ISL_689979, EPI_ISL_689980, EPI_ISL_689981, EPI_ISL_689982, EPI_ISL_689983, EPI_ISL_689984, EPI_ISL_689985, EPI_ISL_689986, EPI_ISL_689987, EPI_ISL_689988, EPI_ISL_689989, EPI_ISL_689990, EPI_ISL_689991, EPI_ISL_689992, EPI_ISL_689993, EPI_ISL_689994, EPI_ISL_689995, EPI_ISL_689996, EPI_ISL_689997, EPI_ISL_689998, EPI_ISL_689999, EPI_ISL_690000, EPI_ISL_690001, EPI_ISL_690002, EPI_ISL_690003, EPI_ISL_690004, EPI_ISL_690005, EPI_ISL_690006, EPI_ISL_690007, EPI_ISL_690008, EPI_ISL_690009, EPI_ISL_690010

| | | | |
|---|---|---|---|
| see above | Pathogen Genomics Center, National Institute of Infectious Diseases | Pathogen Genomics Center, National Institute of Infectious Diseases | Tsuyoshi Sekizuka, Kentaro Itokawa, Rina Tanaka, Masanori Hashino, Makoto Kuroda |

EPI_ISL_690011, EPI_ISL_690012, EPI_ISL_690013, EPI_ISL_690014, EPI_ISL_690015, EPI_ISL_690016, EPI_ISL_690017, EPI_ISL_690018, EPI_ISL_690019, EPI_ISL_690020, EPI_ISL_690021, EPI_ISL_690022, EPI_ISL_690023, EPI_ISL_690024, EPI_ISL_690025, EPI_ISL_690026, EPI_ISL_690027, EPI_ISL_690028

| | | | |
|---|---|---|---|
| see above | Sakai City Institute of Public Health | Pathogen Genomics Center, National Institute of Infectious Diseases | Tsuyoshi Sekizuka, Kentaro Itokawa, Rina Tanaka, Masanori Hashino, Makoto Kuroda |

EPI_ISL_690029, EPI_ISL_690030, EPI_ISL_690031, EPI_ISL_690032, EPI_ISL_690033, EPI_ISL_690034, EPI_ISL_690035, EPI_ISL_690036, EPI_ISL_690037, EPI_ISL_690038, EPI_ISL_690039, EPI_ISL_690040, EPI_ISL_690041, EPI_ISL_690042, EPI_ISL_690043, EPI_ISL_690044, EPI_ISL_690045, EPI_ISL_690046, EPI_ISL_690047, EPI_ISL_690048, EPI_ISL_690049, EPI_ISL_690050, EPI_ISL_690051, EPI_ISL_690052, EPI_ISL_690053, EPI_ISL_690054, EPI_ISL_690055, EPI_ISL_690056, EPI_ISL_690057, EPI_ISL_690058, EPI_ISL_690059, EPI_ISL_690060, EPI_ISL_690061, EPI_ISL_690062, EPI_ISL_690063, EPI_ISL_690064, EPI_ISL_690065, EPI_ISL_690066, EPI_ISL_690067, EPI_ISL_690068, EPI_ISL_690069, EPI_ISL_690070, EPI_ISL_690071, EPI_ISL_690072, EPI_ISL_690073, EPI_ISL_690074, EPI_ISL_690075, EPI_ISL_690076, EPI_ISL_690077, EPI_ISL_690078, EPI_ISL_690079, EPI_ISL_690080, EPI_ISL_690081, EPI_ISL_690082, EPI_ISL_690083, EPI_ISL_690084, EPI_ISL_690085, EPI_ISL_690086, EPI_ISL_690087, EPI_ISL_690088, EPI_ISL_690089, EPI_ISL_690090, EPI_ISL_690091, EPI_ISL_690092, EPI_ISL_690093, EPI_ISL_690094, EPI_ISL_690095, EPI_ISL_690096, EPI_ISL_690097, EPI_ISL_690098, EPI_ISL_690099, EPI_ISL_690100, EPI_ISL_690101, EPI_ISL_690102, EPI_ISL_690103, EPI_ISL_690104, EPI_ISL_690105, EPI_ISL_690106, EPI_ISL_690107, EPI_ISL_690108, EPI_ISL_690109, EPI_ISL_690110, EPI_ISL_690111, EPI_ISL_690112, EPI_ISL_690113, EPI_ISL_690114, EPI_ISL_690115, EPI_ISL_690116, EPI_ISL_690117, EPI_ISL_690118, EPI_ISL_690119, EPI_ISL_690120, EPI_ISL_690121, EPI_ISL_690122, EPI_ISL_690123, EPI_ISL_690124, EPI_ISL_690125, EPI_ISL_690126, EPI_ISL_690127, EPI_ISL_690128, EPI_ISL_690129, EPI_ISL_690130, EPI_ISL_690131, EPI_ISL_690132, EPI_ISL_690133, EPI_ISL_690134, EPI_ISL_690135, EPI_ISL_690136,

| | | | |
|---|---|---|---|
| EPI_ISL_690137, EPI_ISL_690138, EPI_ISL_690139, EPI_ISL_690140, EPI_ISL_690141, EPI_ISL_690142, EPI_ISL_690143, EPI_ISL_690144, EPI_ISL_690145, EPI_ISL_690146, EPI_ISL_690147, EPI_ISL_690148, EPI_ISL_690149, EPI_ISL_690150, EPI_ISL_690151, EPI_ISL_690152, EPI_ISL_690153, EPI_ISL_690154, EPI_ISL_690155, EPI_ISL_690156, EPI_ISL_690157, EPI_ISL_690158, EPI_ISL_690159, EPI_ISL_690160, EPI_ISL_690161, EPI_ISL_690162, EPI_ISL_690163, EPI_ISL_690164, EPI_ISL_690165, EPI_ISL_690166, EPI_ISL_690167, EPI_ISL_690168, EPI_ISL_690169, EPI_ISL_690170, EPI_ISL_690171, EPI_ISL_690172, EPI_ISL_690173, EPI_ISL_690174, EPI_ISL_690175, EPI_ISL_690176, EPI_ISL_690177, EPI_ISL_690178, EPI_ISL_690179, EPI_ISL_690180, EPI_ISL_690181, EPI_ISL_690182, EPI_ISL_690183, EPI_ISL_690184, EPI_ISL_690185, EPI_ISL_690186, EPI_ISL_690187, EPI_ISL_690188, EPI_ISL_690189, EPI_ISL_690190, EPI_ISL_690191, EPI_ISL_690192, EPI_ISL_690193, EPI_ISL_690194, EPI_ISL_690195, EPI_ISL_690196, EPI_ISL_690197, EPI_ISL_690198, EPI_ISL_690199, EPI_ISL_690200, EPI_ISL_690201, EPI_ISL_690202, EPI_ISL_690203, EPI_ISL_690204, EPI_ISL_690205, EPI_ISL_690206, EPI_ISL_690207, EPI_ISL_690208, EPI_ISL_690209 | | | |
| see above | Pathogen Genomics Center, National Institute of Infectious Diseases | Pathogen Genomics Center, National Institute of Infectious Diseases | Tsuyoshi Sekizuka, Kentaro Itokawa, Rina Tanaka, Masanori Hashino, Makoto Kuroda |
| EPI_ISL_690210, EPI_ISL_690211, EPI_ISL_690212, EPI_ISL_690213, EPI_ISL_690214, EPI_ISL_690215, EPI_ISL_690216, EPI_ISL_690217, EPI_ISL_690218, EPI_ISL_690219, EPI_ISL_690220, EPI_ISL_690221, EPI_ISL_690222, EPI_ISL_690223, EPI_ISL_690224, EPI_ISL_690225, EPI_ISL_690226, EPI_ISL_690227, EPI_ISL_690228, EPI_ISL_690229, EPI_ISL_690230, EPI_ISL_690231, EPI_ISL_690232, EPI_ISL_690233, EPI_ISL_690234, EPI_ISL_690235, EPI_ISL_690236, EPI_ISL_690237, EPI_ISL_690238, EPI_ISL_690239, EPI_ISL_690240, EPI_ISL_690241, EPI_ISL_690242, EPI_ISL_690243, EPI_ISL_690244, EPI_ISL_690245, EPI_ISL_690246, EPI_ISL_690247, EPI_ISL_690248, EPI_ISL_690249, EPI_ISL_690250, EPI_ISL_690251, EPI_ISL_690252, EPI_ISL_690253 | | | |
| see above | Gunma Prefectural Institute of Public Health and Environmental Sciences | Pathogen Genomics Center, National Institute of Infectious Diseases | Tsuyoshi Sekizuka, Kentaro Itokawa, Rina Tanaka, Masanori Hashino, Makoto Kuroda |
| EPI_ISL_690254, EPI_ISL_690255, EPI_ISL_690256, EPI_ISL_690257, EPI_ISL_690258, EPI_ISL_690259, EPI_ISL_690260, EPI_ISL_690261, EPI_ISL_690262, EPI_ISL_690263, EPI_ISL_690264, EPI_ISL_690265, EPI_ISL_690266, EPI_ISL_690267, EPI_ISL_690268, EPI_ISL_690269, EPI_ISL_690270, EPI_ISL_690271, EPI_ISL_690272, EPI_ISL_690273, EPI_ISL_690274, EPI_ISL_690275, EPI_ISL_690276, EPI_ISL_690277, EPI_ISL_690278, EPI_ISL_690279, EPI_ISL_690280, EPI_ISL_690281, EPI_ISL_690282, EPI_ISL_690283, EPI_ISL_690284, EPI_ISL_690285, EPI_ISL_690286, EPI_ISL_690287, EPI_ISL_690288, EPI_ISL_690289, EPI_ISL_690290, EPI_ISL_690291, EPI_ISL_690292, EPI_ISL_690293, EPI_ISL_690294, EPI_ISL_690295, EPI_ISL_690296, EPI_ISL_690297, EPI_ISL_690298, EPI_ISL_690299, EPI_ISL_690300, EPI_ISL_690301, EPI_ISL_690302, EPI_ISL_690303, EPI_ISL_690304, EPI_ISL_690305, EPI_ISL_690306, EPI_ISL_690307, EPI_ISL_690308, EPI_ISL_690309, EPI_ISL_690310, EPI_ISL_690311, EPI_ISL_690312, EPI_ISL_690313, EPI_ISL_690314, EPI_ISL_690315, EPI_ISL_690316, EPI_ISL_690317, EPI_ISL_690318, EPI_ISL_690319, EPI_ISL_690320, EPI_ISL_690321, EPI_ISL_690322, EPI_ISL_690323, EPI_ISL_690324, EPI_ISL_690325, EPI_ISL_690326, EPI_ISL_690327, EPI_ISL_690328, EPI_ISL_690329, EPI_ISL_690330, EPI_ISL_690331, EPI_ISL_690332, EPI_ISL_690333, EPI_ISL_690334, EPI_ISL_690335, EPI_ISL_690336, EPI_ISL_690337, EPI_ISL_690338, EPI_ISL_690339, EPI_ISL_690340, EPI_ISL_690341, EPI_ISL_690342, EPI_ISL_690343, EPI_ISL_690344, EPI_ISL_690345, EPI_ISL_690346, EPI_ISL_690347, EPI_ISL_690348, EPI_ISL_690349, EPI_ISL_690350, EPI_ISL_690351, EPI_ISL_690352, EPI_ISL_690353, EPI_ISL_690354, EPI_ISL_690355, EPI_ISL_690356, EPI_ISL_690357, EPI_ISL_690358, EPI_ISL_690359, EPI_ISL_690360, EPI_ISL_690361, EPI_ISL_690362, EPI_ISL_690363, EPI_ISL_690364, EPI_ISL_690365, EPI_ISL_690366, EPI_ISL_690367, EPI_ISL_690368, EPI_ISL_690369, EPI_ISL_690370, EPI_ISL_690371, EPI_ISL_690372, EPI_ISL_690373, EPI_ISL_690374, EPI_ISL_690375, EPI_ISL_690376, EPI_ISL_690377, EPI_ISL_690378, EPI_ISL_690379, EPI_ISL_690380, EPI_ISL_690381, EPI_ISL_690382, EPI_ISL_690383, EPI_ISL_690384, EPI_ISL_690385, EPI_ISL_690386, EPI_ISL_690387, EPI_ISL_690388, EPI_ISL_690389, EPI_ISL_690390, EPI_ISL_690391, EPI_ISL_690392, EPI_ISL_690393, EPI_ISL_690394, EPI_ISL_690395, EPI_ISL_690396, EPI_ISL_690397, EPI_ISL_690398, EPI_ISL_690399, EPI_ISL_690400, EPI_ISL_690401, EPI_ISL_690402, EPI_ISL_690403, EPI_ISL_690404, EPI_ISL_690405, EPI_ISL_690406, EPI_ISL_690407, EPI_ISL_690408, EPI_ISL_690409, EPI_ISL_690410, EPI_ISL_690411, EPI_ISL_690412, EPI_ISL_690413, EPI_ISL_690414, EPI_ISL_690415, EPI_ISL_690416, EPI_ISL_690417, EPI_ISL_690418, EPI_ISL_690419, EPI_ISL_690420, EPI_ISL_690421, EPI_ISL_690422, EPI_ISL_690423, EPI_ISL_690424, EPI_ISL_690425, EPI_ISL_690426, EPI_ISL_690427, EPI_ISL_690428, EPI_ISL_690429, EPI_ISL_690430, EPI_ISL_690431 | | | |
| see above | Pathogen Genomics Center, National Institute of Infectious Diseases | Pathogen Genomics Center, National Institute of Infectious Diseases | Tsuyoshi Sekizuka, Kentaro Itokawa, Rina Tanaka, Masanori Hashino, Makoto Kuroda |
| EPI_ISL_690432, EPI_ISL_690433, EPI_ISL_690434, EPI_ISL_690435, EPI_ISL_690436, EPI_ISL_690437, EPI_ISL_690438, EPI_ISL_690439, EPI_ISL_690440, EPI_ISL_690441, EPI_ISL_690442, EPI_ISL_690443, EPI_ISL_690444, EPI_ISL_690445, EPI_ISL_690446, EPI_ISL_690447, EPI_ISL_690448, EPI_ISL_690449, EPI_ISL_690450, EPI_ISL_690451, EPI_ISL_690452, EPI_ISL_690453, EPI_ISL_690454, EPI_ISL_690455, EPI_ISL_690456, EPI_ISL_690457, EPI_ISL_690458, EPI_ISL_690459, EPI_ISL_690460, EPI_ISL_690461, EPI_ISL_690462, EPI_ISL_690463, EPI_ISL_690464, EPI_ISL_690465, EPI_ISL_690466, EPI_ISL_690467, EPI_ISL_690468, EPI_ISL_690469, EPI_ISL_690470, EPI_ISL_690471, EPI_ISL_690472, EPI_ISL_690473, EPI_ISL_690474, EPI_ISL_690475, EPI_ISL_690476, EPI_ISL_690477, EPI_ISL_690478, EPI_ISL_690479, EPI_ISL_690480, EPI_ISL_690481, EPI_ISL_690482, EPI_ISL_690483, EPI_ISL_690484, EPI_ISL_690485, EPI_ISL_690486, EPI_ISL_690487, EPI_ISL_690488, EPI_ISL_690489, EPI_ISL_690490, EPI_ISL_690491, EPI_ISL_690492, EPI_ISL_690493, EPI_ISL_690494, EPI_ISL_690495, EPI_ISL_690496, EPI_ISL_690497, EPI_ISL_690498, EPI_ISL_690499, EPI_ISL_690500, EPI_ISL_690501, EPI_ISL_690502, EPI_ISL_690503, EPI_ISL_690504, EPI_ISL_690505, EPI_ISL_690506, EPI_ISL_690507, EPI_ISL_690508, EPI_ISL_690509, EPI_ISL_690510, EPI_ISL_690511, EPI_ISL_690512, EPI_ISL_690513, EPI_ISL_690514, EPI_ISL_690515, EPI_ISL_690516, EPI_ISL_690517, EPI_ISL_690518, EPI_ISL_690519, EPI_ISL_690520, EPI_ISL_690521, EPI_ISL_690522, EPI_ISL_690523, EPI_ISL_690524, EPI_ISL_690525, EPI_ISL_690526 | | | |
| see above | Fukuoka Institute of Health and Environmental Sciences | Pathogen Genomics Center, National Institute of Infectious Diseases | Tsuyoshi Sekizuka, Kentaro Itokawa, Rina Tanaka, Masanori Hashino, Makoto Kuroda |
| EPI_ISL_690527, EPI_ISL_690528, EPI_ISL_690529, EPI_ISL_690530, EPI_ISL_690531, EPI_ISL_690532, EPI_ISL_690533, EPI_ISL_690534, EPI_ISL_690535, EPI_ISL_690536, EPI_ISL_690537, EPI_ISL_690538, EPI_ISL_690539, EPI_ISL_690540, EPI_ISL_690541, EPI_ISL_690542, EPI_ISL_690543, EPI_ISL_690544, EPI_ISL_690545, EPI_ISL_690546, EPI_ISL_690547, EPI_ISL_690548, EPI_ISL_690549, EPI_ISL_690550, EPI_ISL_690551, EPI_ISL_690552, EPI_ISL_690553, EPI_ISL_690554, EPI_ISL_690555, EPI_ISL_690556, EPI_ISL_690557, EPI_ISL_690558, EPI_ISL_690559, EPI_ISL_690560, EPI_ISL_690561, EPI_ISL_690562, EPI_ISL_690563, EPI_ISL_690564, EPI_ISL_690565, EPI_ISL_690566, EPI_ISL_690567, EPI_ISL_690568, EPI_ISL_690569, EPI_ISL_690570, EPI_ISL_690571, EPI_ISL_690572, EPI_ISL_690573, EPI_ISL_690574, EPI_ISL_690575, EPI_ISL_690576, EPI_ISL_690577, EPI_ISL_690578, EPI_ISL_690579, EPI_ISL_690580, EPI_ISL_690581, EPI_ISL_690582, EPI_ISL_690583, EPI_ISL_690584, EPI_ISL_690585, EPI_ISL_690586, EPI_ISL_690587, EPI_ISL_690588, EPI_ISL_690589, EPI_ISL_690590, EPI_ISL_690591, EPI_ISL_690592, EPI_ISL_690593, EPI_ISL_690594, EPI_ISL_690595, EPI_ISL_690596, EPI_ISL_690597, EPI_ISL_690598, EPI_ISL_690599, EPI_ISL_690600, EPI_ISL_690601, EPI_ISL_690602, EPI_ISL_690603, EPI_ISL_690604, EPI_ISL_690605, EPI_ISL_690606, EPI_ISL_690607, EPI_ISL_690608, EPI_ISL_690609, EPI_ISL_690610, EPI_ISL_690611, EPI_ISL_690616, EPI_ISL_690624, EPI_ISL_690632, EPI_ISL_690633, EPI_ISL_690634, EPI_ISL_690635 | | | |
| see above | Pathogen Genomics Center, National Institute of Infectious Diseases | Pathogen Genomics Center, National Institute of Infectious Diseases | Tsuyoshi Sekizuka, Kentaro Itokawa, Rina Tanaka, Masanori Hashino, Makoto Kuroda |
| EPI_ISL_690638, EPI_ISL_690639, EPI_ISL_690640 | Tokyo Metropolitan Institute of Public Health | Pathogen Genomics Center, National Institute of Infectious Diseases | Tsuyoshi Sekizuka, Kentaro Itokawa, Rina Tanaka, Masanori Hashino, Makoto Kuroda |
| EPI_ISL_690681, EPI_ISL_690682, EPI_ISL_690683 | Saitama Prefectural Institute of Public Health | Pathogen Genomics Center, National Institute of Infectious Diseases | Tsuyoshi Sekizuka, Kentaro Itokawa, Rina Tanaka, Masanori Hashino, Makoto Kuroda |
| EPI_ISL_690687, EPI_ISL_690688, EPI_ISL_690689, EPI_ISL_690690, EPI_ISL_690691, EPI_ISL_690692, EPI_ISL_690693, EPI_ISL_690694, EPI_ISL_690695, EPI_ISL_690696, EPI_ISL_690697, EPI_ISL_690698, EPI_ISL_690699, EPI_ISL_690700, EPI_ISL_690701, EPI_ISL_690702, EPI_ISL_690703, EPI_ISL_690704, EPI_ISL_690705, EPI_ISL_690706, EPI_ISL_690707, EPI_ISL_690744, EPI_ISL_690778 | | | |
| see above | Pathogen Genomics Center, National Institute of Infectious Diseases | Pathogen Genomics Center, National Institute of Infectious Diseases | Tsuyoshi Sekizuka, Kentaro Itokawa, Rina Tanaka, Masanori Hashino, Makoto Kuroda |
| EPI_ISL_690794, EPI_ISL_690795, EPI_ISL_690796, EPI_ISL_690797, EPI_ISL_690798, EPI_ISL_690799, EPI_ISL_690800, EPI_ISL_690801, EPI_ISL_690802, EPI_ISL_690803, EPI_ISL_690804, EPI_ISL_690805, EPI_ISL_690806, EPI_ISL_690807, EPI_ISL_690808, EPI_ISL_690809, EPI_ISL_690810, EPI_ISL_690811, EPI_ISL_690812, EPI_ISL_690813, EPI_ISL_690814, EPI_ISL_690815, EPI_ISL_690816, EPI_ISL_690817 | | | |
| see above | Tokyo Metropolitan Institute of Public Health | Pathogen Genomics Center, National Institute of Infectious Diseases | Tsuyoshi Sekizuka, Kentaro Itokawa, Rina Tanaka, Masanori Hashino, Makoto Kuroda |
| EPI_ISL_690844 | Sakai City Institute of Public Health | Pathogen Genomics Center, National Institute of Infectious Diseases | Tsuyoshi Sekizuka, Kentaro Itokawa, Rina Tanaka, Masanori Hashino, Makoto Kuroda |
| EPI_ISL_690902, EPI_ISL_690903, EPI_ISL_690904, EPI_ISL_690905, EPI_ISL_690906, EPI_ISL_690907, EPI_ISL_690908 | Tokyo Metropolitan Institute of Public Health | Pathogen Genomics Center, National Institute of Infectious Diseases | Tsuyoshi Sekizuka, Kentaro Itokawa, Rina Tanaka, Masanori Hashino, Makoto Kuroda |
| EPI_ISL_690981, EPI_ISL_691108, EPI_ISL_691239, EPI_ISL_691248, EPI_ISL_691252, EPI_ISL_691253, EPI_ISL_691254, EPI_ISL_691255, EPI_ISL_691256, EPI_ISL_691260, EPI_ISL_691261, EPI_ISL_691262, EPI_ISL_691263, EPI_ISL_691264, EPI_ISL_691265, EPI_ISL_691266, EPI_ISL_691267, EPI_ISL_691268, EPI_ISL_691269, EPI_ISL_691270, EPI_ISL_691271, EPI_ISL_691272, EPI_ISL_691275, EPI_ISL_691276, EPI_ISL_691277, EPI_ISL_691278, EPI_ISL_691279, EPI_ISL_691280, EPI_ISL_691281, EPI_ISL_691282, EPI_ISL_691283 | | | |
| see above | Pathogen Genomics Center, National Institute of Infectious Diseases | Pathogen Genomics Center, National Institute of Infectious Diseases | Tsuyoshi Sekizuka, Kentaro Itokawa, Rina Tanaka, Masanori Hashino, Makoto Kuroda |
| EPI_ISL_691284, EPI_ISL_691285, EPI_ISL_691286, EPI_ISL_691287, EPI_ISL_691288 | Chiba Prefectural Institute of Public Health | Pathogen Genomics Center, National Institute of Infectious Diseases | Tsuyoshi Sekizuka, Kentaro Itokawa, Rina Tanaka, Masanori Hashino, Makoto Kuroda |
| EPI_ISL_691364, EPI_ISL_691365, EPI_ISL_691366, EPI_ISL_691367, EPI_ISL_691368, EPI_ISL_691369, EPI_ISL_691370, EPI_ISL_691371, EPI_ISL_691372, EPI_ISL_691373, EPI_ISL_691374, EPI_ISL_691375, EPI_ISL_691376, EPI_ISL_691377, EPI_ISL_691378 | | | |
| see above | Saitama Prefectural Institute of Public Health | Pathogen Genomics Center, National Institute of Infectious Diseases | Tsuyoshi Sekizuka, Kentaro Itokawa, Rina Tanaka, Masanori Hashino, Makoto Kuroda |
| EPI_ISL_691386, EPI_ISL_691387, EPI_ISL_691388, EPI_ISL_691389, EPI_ISL_691390, EPI_ISL_691391, EPI_ISL_691392, EPI_ISL_691393, EPI_ISL_691394, EPI_ISL_691502, EPI_ISL_691503, EPI_ISL_691504, EPI_ISL_691505, EPI_ISL_691506 | | | |
| see above | Pathogen Genomics Center, National Institute of Infectious Diseases | Pathogen Genomics Center, National Institute of Infectious Diseases | Tsuyoshi Sekizuka, Kentaro Itokawa, Rina Tanaka, Masanori Hashino, Makoto Kuroda |
| EPI_ISL_691543, EPI_ISL_691544, EPI_ISL_691545 | Sakai City Institute of Public Health | Pathogen Genomics Center, National Institute of Infectious Diseases | Tsuyoshi Sekizuka, Kentaro Itokawa, Rina Tanaka, Masanori Hashino, Makoto Kuroda |
| EPI_ISL_691546, EPI_ISL_691547, | Pathogen Genomics Center, National Institute of Infectious | Pathogen Genomics Center, National Institute of Infectious | Tsuyoshi Sekizuka, Kentaro Itokawa, Rina Tanaka, Masanori Hashino, Makoto Kuroda |

| | | | |
|---|---|---|---|
| EPI_ISL_691548, EPI_ISL_691549 | Diseases | Diseases | |
| EPI_ISL_691550, EPI_ISL_691551 | Fukuoka Institute of Health and Environmental Sciences | Pathogen Genomics Center, National Institute of Infectious Diseases | Tsuyoshi Sekizuka, Kentaro Itokawa, Rina Tanaka, Masanori Hashino, Makoto Kuroda |
| EPI_ISL_691552, EPI_ISL_691553, EPI_ISL_691554, EPI_ISL_691555, EPI_ISL_691556, EPI_ISL_691557, EPI_ISL_691558, EPI_ISL_691559, EPI_ISL_691560, EPI_ISL_691561, EPI_ISL_691562, EPI_ISL_691563, EPI_ISL_691564, EPI_ISL_691565, EPI_ISL_691566, EPI_ISL_691567, EPI_ISL_691568, EPI_ISL_691569, EPI_ISL_691570, EPI_ISL_691571, EPI_ISL_691572, EPI_ISL_691573, EPI_ISL_691574, EPI_ISL_691575, EPI_ISL_691576, EPI_ISL_691577, EPI_ISL_691578, EPI_ISL_691579, EPI_ISL_691580, EPI_ISL_691581, EPI_ISL_691582, EPI_ISL_691583, EPI_ISL_691584, EPI_ISL_691585, EPI_ISL_691586, EPI_ISL_691587, EPI_ISL_691588, EPI_ISL_691589, EPI_ISL_691590, EPI_ISL_691591, EPI_ISL_691592, EPI_ISL_691593, EPI_ISL_691594, EPI_ISL_691595, EPI_ISL_691596, EPI_ISL_691597, EPI_ISL_691598, EPI_ISL_691599, EPI_ISL_691600, EPI_ISL_691601, EPI_ISL_691602, EPI_ISL_691603, EPI_ISL_691604, EPI_ISL_691605, EPI_ISL_691606, EPI_ISL_691607, EPI_ISL_691608, EPI_ISL_691609, EPI_ISL_691610, EPI_ISL_691611 | | | |
| see above | Pathogen Genomics Center, National Institute of Infectious Diseases | Pathogen Genomics Center, National Institute of Infectious Diseases | Tsuyoshi Sekizuka, Kentaro Itokawa, Rina Tanaka, Masanori Hashino, Makoto Kuroda |
| EPI_ISL_691700, EPI_ISL_691701, EPI_ISL_691702, EPI_ISL_691703, EPI_ISL_691704, EPI_ISL_691705, EPI_ISL_691706, EPI_ISL_691707, EPI_ISL_691708, EPI_ISL_691709, EPI_ISL_691710, EPI_ISL_691711, EPI_ISL_691712, EPI_ISL_691713, EPI_ISL_691714, EPI_ISL_691715, EPI_ISL_691716, EPI_ISL_691717, EPI_ISL_691718, EPI_ISL_691719, EPI_ISL_691720, EPI_ISL_691721, EPI_ISL_691722, EPI_ISL_691723, EPI_ISL_691724, EPI_ISL_691725, EPI_ISL_691726, EPI_ISL_691727, EPI_ISL_691728 | | | |
| see above | Hospital Clínico San Carlos | Instituto de Salud Carlos III | Iglesias-Caballero, M. Camarero, S. Molinero Calamita, M. González-Esguevillas, M. Pozo, F. Casas, I. Jiménez, P. Jiménez, M. Zaballos, A. Monzón, S. Varona, S. Juliá, M. Cuesta, I. Rodríguez, I. |
| EPI_ISL_691744, EPI_ISL_691748, EPI_ISL_691752 | Pathogen Genomics Center, National Institute of Infectious Diseases | Pathogen Genomics Center, National Institute of Infectious Diseases | Tsuyoshi Sekizuka, Kentaro Itokawa, Rina Tanaka, Masanori Hashino, Makoto Kuroda |
| EPI_ISL_691753 | Sakai City Institute of Public Health | Pathogen Genomics Center, National Institute of Infectious Diseases | Tsuyoshi Sekizuka, Kentaro Itokawa, Rina Tanaka, Masanori Hashino, Makoto Kuroda |
| EPI_ISL_691754, EPI_ISL_691759, EPI_ISL_691760, EPI_ISL_691761, EPI_ISL_691762, EPI_ISL_691763 | Pathogen Genomics Center, National Institute of Infectious Diseases | Pathogen Genomics Center, National Institute of Infectious Diseases | Tsuyoshi Sekizuka, Kentaro Itokawa, Rina Tanaka, Masanori Hashino, Makoto Kuroda |
| EPI_ISL_691764, EPI_ISL_691765 | Fukuoka Institute of Health and Environmental Sciences | Pathogen Genomics Center, National Institute of Infectious Diseases | Tsuyoshi Sekizuka, Kentaro Itokawa, Rina Tanaka, Masanori Hashino, Makoto Kuroda |
| EPI_ISL_691766 | Saitama Prefectural Institute of Public Health | Pathogen Genomics Center, National Institute of Infectious Diseases | Tsuyoshi Sekizuka, Kentaro Itokawa, Rina Tanaka, Masanori Hashino, Makoto Kuroda |
| EPI_ISL_691768, EPI_ISL_691769, EPI_ISL_691770, EPI_ISL_691771, EPI_ISL_691772, EPI_ISL_691773, EPI_ISL_691774, EPI_ISL_691775, EPI_ISL_691776, EPI_ISL_691777, EPI_ISL_691778, EPI_ISL_691779, EPI_ISL_691780, EPI_ISL_691781, EPI_ISL_691782, EPI_ISL_691783, EPI_ISL_691784, EPI_ISL_691785, EPI_ISL_691786, EPI_ISL_691787, EPI_ISL_691788, EPI_ISL_691789, EPI_ISL_691790, EPI_ISL_691791, EPI_ISL_691792, EPI_ISL_691793, EPI_ISL_691794, EPI_ISL_691795, EPI_ISL_691796, EPI_ISL_691797, EPI_ISL_691798, EPI_ISL_691799, EPI_ISL_691800, EPI_ISL_691801, EPI_ISL_691802, EPI_ISL_691803, EPI_ISL_691804, EPI_ISL_691805, EPI_ISL_691806, EPI_ISL_691807, EPI_ISL_691808, EPI_ISL_691809, EPI_ISL_691810, EPI_ISL_691811, EPI_ISL_691812, EPI_ISL_691813, EPI_ISL_691814, EPI_ISL_691815, EPI_ISL_691816, EPI_ISL_691817, EPI_ISL_691818, EPI_ISL_691819, EPI_ISL_691820, EPI_ISL_691821, EPI_ISL_691822, EPI_ISL_691823, EPI_ISL_691824, EPI_ISL_691825, EPI_ISL_691826, EPI_ISL_691827, EPI_ISL_691828, EPI_ISL_691829, EPI_ISL_691830, EPI_ISL_691831, EPI_ISL_691832, EPI_ISL_691833, EPI_ISL_691834, EPI_ISL_691835, EPI_ISL_691836, EPI_ISL_691837, EPI_ISL_691838, EPI_ISL_691839, EPI_ISL_691840, EPI_ISL_691841, EPI_ISL_691842, EPI_ISL_691843, EPI_ISL_691844, EPI_ISL_691845, EPI_ISL_691846, EPI_ISL_691847, EPI_ISL_691848, EPI_ISL_691849, EPI_ISL_691850, EPI_ISL_691851, EPI_ISL_691852, EPI_ISL_691853, EPI_ISL_691854, EPI_ISL_691855, EPI_ISL_691856, EPI_ISL_691857, EPI_ISL_691858, EPI_ISL_691859, EPI_ISL_691860, EPI_ISL_691862, EPI_ISL_691863, EPI_ISL_691864, EPI_ISL_691865, EPI_ISL_691866, EPI_ISL_691867, EPI_ISL_691868, EPI_ISL_691869, EPI_ISL_691870, EPI_ISL_691871 | | | |
| see above | Pathogen Genomics Center, National Institute of Infectious Diseases | Pathogen Genomics Center, National Institute of Infectious Diseases | Tsuyoshi Sekizuka, Kentaro Itokawa, Rina Tanaka, Masanori Hashino, Makoto Kuroda |
| EPI_ISL_692047 | Gunma Prefectural Institute of Public Health and Environmental Sciences | Pathogen Genomics Center, National Institute of Infectious Diseases | Tsuyoshi Sekizuka, Kentaro Itokawa, Rina Tanaka, Masanori Hashino, Makoto Kuroda |
| EPI_ISL_692048, EPI_ISL_692049, EPI_ISL_692050 | Fukuoka Institute of Health and Environmental Sciences | Pathogen Genomics Center, National Institute of Infectious Diseases | Tsuyoshi Sekizuka, Kentaro Itokawa, Rina Tanaka, Masanori Hashino, Makoto Kuroda |
| EPI_ISL_692051, EPI_ISL_692052 | Niigata Prefectural Institute of Public Health and Environmental Sciences | Pathogen Genomics Center, National Institute of Infectious Diseases | Tsuyoshi Sekizuka, Kentaro Itokawa, Rina Tanaka, Masanori Hashino, Makoto Kuroda |
| EPI_ISL_692053, EPI_ISL_692054, EPI_ISL_692055, EPI_ISL_692056, EPI_ISL_692057, EPI_ISL_692058 | Sakai City Institute of Public Health | Pathogen Genomics Center, National Institute of Infectious Diseases | Tsuyoshi Sekizuka, Kentaro Itokawa, Rina Tanaka, Masanori Hashino, Makoto Kuroda |
| EPI_ISL_692061, EPI_ISL_692062, EPI_ISL_692063, EPI_ISL_692064, EPI_ISL_692065, EPI_ISL_692066, EPI_ISL_692067, EPI_ISL_692068, EPI_ISL_692069, EPI_ISL_692070, EPI_ISL_692071, EPI_ISL_692072, EPI_ISL_692073, EPI_ISL_692074, EPI_ISL_692075, EPI_ISL_692076, EPI_ISL_692077, EPI_ISL_692078, EPI_ISL_692079, EPI_ISL_692080, EPI_ISL_692081, EPI_ISL_692082, EPI_ISL_692083, EPI_ISL_692084, EPI_ISL_692085, EPI_ISL_692086, EPI_ISL_692087, EPI_ISL_692088, EPI_ISL_692089, EPI_ISL_692090, EPI_ISL_692091, EPI_ISL_692092, EPI_ISL_692093 | | | |
| see above | Pathogen Genomics Center, National Institute of Infectious Diseases | Pathogen Genomics Center, National Institute of Infectious Diseases | Tsuyoshi Sekizuka, Kentaro Itokawa, Rina Tanaka, Masanori Hashino, Makoto Kuroda |
| EPI_ISL_692094, EPI_ISL_692095, EPI_ISL_692096, EPI_ISL_692097 | Yamagata Prefectural Institute of Public Health | Pathogen Genomics Center, National Institute of Infectious Diseases | Tsuyoshi Sekizuka, Kentaro Itokawa, Rina Tanaka, Masanori Hashino, Makoto Kuroda |
| EPI_ISL_692098, EPI_ISL_692099, EPI_ISL_692100, EPI_ISL_692101, EPI_ISL_692102, EPI_ISL_692103, EPI_ISL_692104, EPI_ISL_692105, EPI_ISL_692106, EPI_ISL_692107, EPI_ISL_692108, EPI_ISL_692109, EPI_ISL_692110, EPI_ISL_692111, EPI_ISL_692112, EPI_ISL_692113, EPI_ISL_692114, EPI_ISL_692115, EPI_ISL_692116, EPI_ISL_692117, EPI_ISL_692118 | | | |
| see above | Pathogen Genomics Center, National Institute of Infectious Diseases | Pathogen Genomics Center, National Institute of Infectious Diseases | Tsuyoshi Sekizuka, Kentaro Itokawa, Rina Tanaka, Masanori Hashino, Makoto Kuroda |
| EPI_ISL_692119, EPI_ISL_692120, EPI_ISL_692121, EPI_ISL_692122, EPI_ISL_692123, EPI_ISL_692124, EPI_ISL_692125, EPI_ISL_692126, EPI_ISL_692127, EPI_ISL_692128 | Tokyo Metropolitan Institute of Public Health | Pathogen Genomics Center, National Institute of Infectious Diseases | Tsuyoshi Sekizuka, Kentaro Itokawa, Rina Tanaka, Masanori Hashino, Makoto Kuroda |
| EPI_ISL_692129, EPI_ISL_692130, EPI_ISL_692131, EPI_ISL_692132, EPI_ISL_692133, EPI_ISL_692134, EPI_ISL_692135, EPI_ISL_692136, EPI_ISL_692137, EPI_ISL_692138, EPI_ISL_692139, EPI_ISL_692140, EPI_ISL_692141, EPI_ISL_692142, EPI_ISL_692143, EPI_ISL_692144, EPI_ISL_692145, EPI_ISL_692146, EPI_ISL_692147, EPI_ISL_692148, EPI_ISL_692149, EPI_ISL_692150, EPI_ISL_692151, EPI_ISL_692152, EPI_ISL_692153, EPI_ISL_692154, EPI_ISL_692155, EPI_ISL_692156, EPI_ISL_692157, EPI_ISL_692158, EPI_ISL_692159, EPI_ISL_692160, EPI_ISL_692161 | | | |
| see above | Pathogen Genomics Center, National Institute of Infectious Diseases | Pathogen Genomics Center, National Institute of Infectious Diseases | Tsuyoshi Sekizuka, Kentaro Itokawa, Rina Tanaka, Masanori Hashino, Makoto Kuroda |
| EPI_ISL_692162, EPI_ISL_692163 | Tokyo Metropolitan Institute of Public Health | Pathogen Genomics Center, National Institute of Infectious Diseases | Tsuyoshi Sekizuka, Kentaro Itokawa, Rina Tanaka, Masanori Hashino, Makoto Kuroda |
| EPI_ISL_692164, EPI_ISL_692165, EPI_ISL_692166, EPI_ISL_692167, EPI_ISL_692168, EPI_ISL_692169, EPI_ISL_692170, EPI_ISL_692171, EPI_ISL_692172, EPI_ISL_692173, EPI_ISL_692174, EPI_ISL_692175, EPI_ISL_692176, EPI_ISL_692177, EPI_ISL_692178, EPI_ISL_692179, EPI_ISL_692180, EPI_ISL_692181, EPI_ISL_692182, EPI_ISL_692183, EPI_ISL_692184, EPI_ISL_692185, EPI_ISL_692186, EPI_ISL_692187, EPI_ISL_692188, EPI_ISL_692189, EPI_ISL_692190, EPI_ISL_692191, EPI_ISL_692192, EPI_ISL_692193, EPI_ISL_692194, EPI_ISL_692195, EPI_ISL_692196, EPI_ISL_692198, EPI_ISL_692199, EPI_ISL_692200, EPI_ISL_692201, EPI_ISL_692202, EPI_ISL_692203, EPI_ISL_692204, EPI_ISL_692205, EPI_ISL_692206, EPI_ISL_692207, EPI_ISL_692208, EPI_ISL_692209, EPI_ISL_692210, EPI_ISL_692211, EPI_ISL_692212, EPI_ISL_692213, EPI_ISL_692214, EPI_ISL_692215, EPI_ISL_692216, EPI_ISL_692217, EPI_ISL_692218 | | | |
| see above | Pathogen Genomics Center, National Institute of Infectious Diseases | Pathogen Genomics Center, National Institute of Infectious Diseases | Tsuyoshi Sekizuka, Kentaro Itokawa, Rina Tanaka, Masanori Hashino, Makoto Kuroda |
| EPI_ISL_692219, EPI_ISL_692220, EPI_ISL_692221, EPI_ISL_692222, EPI_ISL_692223, EPI_ISL_692224, EPI_ISL_692225, EPI_ISL_692226, EPI_ISL_692227, EPI_ISL_692228, EPI_ISL_692229 | | | |
| see above | Fukuoka Institute of Health and Environmental Sciences | Pathogen Genomics Center, National Institute of Infectious Diseases | Tsuyoshi Sekizuka, Kentaro Itokawa, Rina Tanaka, Masanori Hashino, Makoto Kuroda |
| EPI_ISL_692230, EPI_ISL_692231, EPI_ISL_692232, EPI_ISL_692233, EPI_ISL_692234, EPI_ISL_692235, EPI_ISL_692236, EPI_ISL_692237, EPI_ISL_692238, EPI_ISL_692239, EPI_ISL_692240, EPI_ISL_692241, EPI_ISL_692242, EPI_ISL_692243, EPI_ISL_692244, EPI_ISL_692245, EPI_ISL_692246, EPI_ISL_692247, EPI_ISL_692248, EPI_ISL_692249, EPI_ISL_692250, EPI_ISL_692251, EPI_ISL_692252, EPI_ISL_692253, EPI_ISL_692254, EPI_ISL_692255, EPI_ISL_692256, EPI_ISL_692257, EPI_ISL_692258, EPI_ISL_692259, EPI_ISL_692260, EPI_ISL_692261, EPI_ISL_692262, EPI_ISL_692263, EPI_ISL_692264, EPI_ISL_692265, EPI_ISL_692266, EPI_ISL_692267, EPI_ISL_692268, EPI_ISL_692269, EPI_ISL_692270, EPI_ISL_692271, EPI_ISL_692272, EPI_ISL_692273, EPI_ISL_692274, EPI_ISL_692275, EPI_ISL_692276, EPI_ISL_692277, EPI_ISL_692278, EPI_ISL_692279, EPI_ISL_692280, EPI_ISL_692281, EPI_ISL_692282, EPI_ISL_692283 | | | |

| | | | |
|---|---|---|---|
| see above | Pathogen Genomics Center, National Institute of Infectious Diseases | Pathogen Genomics Center, National Institute of Infectious Diseases | Tsuyoshi Sekizuka, Kentaro Itokawa, Rina Tanaka, Masanori Hashino, Makoto Kuroda |
| EPI_ISL_692284, EPI_ISL_692285, EPI_ISL_692286, EPI_ISL_692287, EPI_ISL_692288, EPI_ISL_692289, EPI_ISL_692290, EPI_ISL_692291, EPI_ISL_692292, EPI_ISL_692293, EPI_ISL_692294, EPI_ISL_692295, EPI_ISL_692296, EPI_ISL_692297, EPI_ISL_692298, EPI_ISL_692299, EPI_ISL_692300, EPI_ISL_692301, EPI_ISL_692302, EPI_ISL_692303, EPI_ISL_692304, EPI_ISL_692305, EPI_ISL_692306, EPI_ISL_692307, EPI_ISL_692308, EPI_ISL_692309, EPI_ISL_692310, EPI_ISL_692311, EPI_ISL_692312, EPI_ISL_692313, EPI_ISL_692314, EPI_ISL_692315 | | | |
| see above | Sakai City Institute of Public Health | Pathogen Genomics Center, National Institute of Infectious Diseases | Tsuyoshi Sekizuka, Kentaro Itokawa, Rina Tanaka, Masanori Hashino, Makoto Kuroda |
| EPI_ISL_692316, EPI_ISL_692317, EPI_ISL_692318, EPI_ISL_692319, EPI_ISL_692320, EPI_ISL_692321, EPI_ISL_692322, EPI_ISL_692323, EPI_ISL_692324, EPI_ISL_692325, EPI_ISL_692326, EPI_ISL_692327 | | | |
| see above | Gunma Prefectural Institute of Public Health and Environmental Sciences | Pathogen Genomics Center, National Institute of Infectious Diseases | Tsuyoshi Sekizuka, Kentaro Itokawa, Rina Tanaka, Masanori Hashino, Makoto Kuroda |
| EPI_ISL_692328, EPI_ISL_692329, EPI_ISL_692330, EPI_ISL_692331, EPI_ISL_692332, EPI_ISL_692333, EPI_ISL_692334, EPI_ISL_692335, EPI_ISL_692336, EPI_ISL_692337, EPI_ISL_692338, EPI_ISL_692339, EPI_ISL_692340, EPI_ISL_692341, EPI_ISL_692342, EPI_ISL_692343, EPI_ISL_692344, EPI_ISL_692345, EPI_ISL_692346, EPI_ISL_692347, EPI_ISL_692348, EPI_ISL_692349, EPI_ISL_692350, EPI_ISL_692351, EPI_ISL_692352, EPI_ISL_692353, EPI_ISL_692354, EPI_ISL_692355, EPI_ISL_692356, EPI_ISL_692357, EPI_ISL_692358, EPI_ISL_692359, EPI_ISL_692360, EPI_ISL_692361, EPI_ISL_692362, EPI_ISL_692363, EPI_ISL_692364, EPI_ISL_692365, EPI_ISL_692366, EPI_ISL_692367, EPI_ISL_692368, EPI_ISL_692369, EPI_ISL_692370, EPI_ISL_692371, EPI_ISL_692372, EPI_ISL_692373, EPI_ISL_692374, EPI_ISL_692375, EPI_ISL_692376, EPI_ISL_692377, EPI_ISL_692378, EPI_ISL_692379, EPI_ISL_692380, EPI_ISL_692381, EPI_ISL_692382, EPI_ISL_692383, EPI_ISL_692384, EPI_ISL_692385, EPI_ISL_692386, EPI_ISL_692387, EPI_ISL_692388, EPI_ISL_692389, EPI_ISL_692390, EPI_ISL_692391, EPI_ISL_692392, EPI_ISL_692393, EPI_ISL_692394, EPI_ISL_692395, EPI_ISL_692396, EPI_ISL_692397, EPI_ISL_692398, EPI_ISL_692399, EPI_ISL_692400, EPI_ISL_692401, EPI_ISL_692402, EPI_ISL_692403, EPI_ISL_692404, EPI_ISL_692405, EPI_ISL_692406, EPI_ISL_692407, EPI_ISL_692408, EPI_ISL_692409, EPI_ISL_692410, EPI_ISL_692411, EPI_ISL_692412, EPI_ISL_692413, EPI_ISL_692414, EPI_ISL_692415, EPI_ISL_692416, EPI_ISL_692417, EPI_ISL_692418, EPI_ISL_692419, EPI_ISL_692420, EPI_ISL_692421, EPI_ISL_692422, EPI_ISL_692423, EPI_ISL_692424, EPI_ISL_692425, EPI_ISL_692426, EPI_ISL_692427, EPI_ISL_692428, EPI_ISL_692429, EPI_ISL_692430, EPI_ISL_692431, EPI_ISL_692432, EPI_ISL_692433, EPI_ISL_692434, EPI_ISL_692435, EPI_ISL_692436, EPI_ISL_692437, EPI_ISL_692438, EPI_ISL_692439, EPI_ISL_692440, EPI_ISL_692441 | | | |
| see above | Pathogen Genomics Center, National Institute of Infectious Diseases | Pathogen Genomics Center, National Institute of Infectious Diseases | Tsuyoshi Sekizuka, Kentaro Itokawa, Rina Tanaka, Masanori Hashino, Makoto Kuroda |
| EPI_ISL_692442, EPI_ISL_692443, EPI_ISL_692444, EPI_ISL_692445, EPI_ISL_692446, EPI_ISL_692447, EPI_ISL_692448, EPI_ISL_692449, EPI_ISL_692450, EPI_ISL_692451, EPI_ISL_692452, EPI_ISL_692453, EPI_ISL_692454, EPI_ISL_692455, EPI_ISL_692456, EPI_ISL_692457, EPI_ISL_692458, EPI_ISL_692459, EPI_ISL_692460, EPI_ISL_692461, EPI_ISL_692462, EPI_ISL_692463, EPI_ISL_692464, EPI_ISL_692465, EPI_ISL_692466, EPI_ISL_692467, EPI_ISL_692468, EPI_ISL_692469, EPI_ISL_692470, EPI_ISL_692471, EPI_ISL_692472, EPI_ISL_692473, EPI_ISL_692474, EPI_ISL_692475, EPI_ISL_692476, EPI_ISL_692477, EPI_ISL_692478, EPI_ISL_692479, EPI_ISL_692480, EPI_ISL_692481, EPI_ISL_692482, EPI_ISL_692483, EPI_ISL_692484, EPI_ISL_692485, EPI_ISL_692486, EPI_ISL_692487, EPI_ISL_692488, EPI_ISL_692489, EPI_ISL_692490, EPI_ISL_692491, EPI_ISL_692492, EPI_ISL_692493, EPI_ISL_692494, EPI_ISL_692495, EPI_ISL_692496, EPI_ISL_692497 | | | |
| see above | Fukuoka Institute of Health and Environmental Sciences | Pathogen Genomics Center, National Institute of Infectious Diseases | Tsuyoshi Sekizuka, Kentaro Itokawa, Rina Tanaka, Masanori Hashino, Makoto Kuroda |
| EPI_ISL_692498, EPI_ISL_692499, EPI_ISL_692500, EPI_ISL_692501, EPI_ISL_692502, EPI_ISL_692503, EPI_ISL_692504, EPI_ISL_692505, EPI_ISL_692506, EPI_ISL_692507, EPI_ISL_692508, EPI_ISL_692509, EPI_ISL_692510, EPI_ISL_692511, EPI_ISL_692512, EPI_ISL_692513, EPI_ISL_692514, EPI_ISL_692515, EPI_ISL_692516, EPI_ISL_692517, EPI_ISL_692518, EPI_ISL_692519, EPI_ISL_692520, EPI_ISL_692521, EPI_ISL_692522, EPI_ISL_692523, EPI_ISL_692524, EPI_ISL_692525, EPI_ISL_692526, EPI_ISL_692528, EPI_ISL_692529, EPI_ISL_692530, EPI_ISL_692531, EPI_ISL_692532, EPI_ISL_692533, EPI_ISL_692534, EPI_ISL_692535, EPI_ISL_692536, EPI_ISL_692537, EPI_ISL_692538, EPI_ISL_692539, EPI_ISL_692540, EPI_ISL_692541, EPI_ISL_692542, EPI_ISL_692543, EPI_ISL_692544, EPI_ISL_692545, EPI_ISL_692546, EPI_ISL_692547, EPI_ISL_692548, EPI_ISL_692549, EPI_ISL_692550, EPI_ISL_692551, EPI_ISL_692552, EPI_ISL_692553, EPI_ISL_692554, EPI_ISL_692555, EPI_ISL_692556, EPI_ISL_692557, EPI_ISL_692558, EPI_ISL_692559, EPI_ISL_692560, EPI_ISL_692561, EPI_ISL_692562, EPI_ISL_692563, EPI_ISL_692564, EPI_ISL_692565, EPI_ISL_692566, EPI_ISL_692567, EPI_ISL_692568, EPI_ISL_692569, EPI_ISL_692570, EPI_ISL_692571, EPI_ISL_692572, EPI_ISL_692573, EPI_ISL_692574, EPI_ISL_692575, EPI_ISL_692576, EPI_ISL_692577, EPI_ISL_692578, EPI_ISL_692579, EPI_ISL_692580, EPI_ISL_692581, EPI_ISL_692582, EPI_ISL_692583, EPI_ISL_692584, EPI_ISL_692585, EPI_ISL_692586, EPI_ISL_692587, EPI_ISL_692588, EPI_ISL_692589, EPI_ISL_692590, EPI_ISL_692591, EPI_ISL_692592, EPI_ISL_692593, EPI_ISL_692594, EPI_ISL_692595, EPI_ISL_692596, EPI_ISL_692598, EPI_ISL_692599, EPI_ISL_692600, EPI_ISL_692601, EPI_ISL_692602, EPI_ISL_692603, EPI_ISL_692604, EPI_ISL_692605, EPI_ISL_692606, EPI_ISL_692607, EPI_ISL_692608, EPI_ISL_692609, EPI_ISL_692610, EPI_ISL_692611, EPI_ISL_692612, EPI_ISL_692613, EPI_ISL_692614, EPI_ISL_692615, EPI_ISL_692616, EPI_ISL_692617, EPI_ISL_692618, EPI_ISL_692619, EPI_ISL_692620, EPI_ISL_692621, EPI_ISL_692622, EPI_ISL_692623, EPI_ISL_692624, EPI_ISL_692625, EPI_ISL_692626, EPI_ISL_692627, EPI_ISL_692628, EPI_ISL_692629, EPI_ISL_692630, EPI_ISL_692631, EPI_ISL_692632, EPI_ISL_692633, EPI_ISL_692634, EPI_ISL_692635, EPI_ISL_692636, EPI_ISL_692637, EPI_ISL_692638, EPI_ISL_692639, EPI_ISL_692640, EPI_ISL_692641, EPI_ISL_692642, EPI_ISL_692643, EPI_ISL_692644, EPI_ISL_692645, EPI_ISL_692646, EPI_ISL_692647, EPI_ISL_692648, EPI_ISL_692649, EPI_ISL_692650, EPI_ISL_692651, EPI_ISL_692652 | | | |
| see above | Pathogen Genomics Center, National Institute of Infectious Diseases | Pathogen Genomics Center, National Institute of Infectious Diseases | Tsuyoshi Sekizuka, Kentaro Itokawa, Rina Tanaka, Masanori Hashino, Makoto Kuroda |
| EPI_ISL_692653, EPI_ISL_692654, EPI_ISL_692655, EPI_ISL_692656, EPI_ISL_692657, EPI_ISL_692658, EPI_ISL_692659, EPI_ISL_692660, EPI_ISL_692661, EPI_ISL_692662, EPI_ISL_692663, EPI_ISL_692664, EPI_ISL_692665, EPI_ISL_692666, EPI_ISL_692667, EPI_ISL_692668, EPI_ISL_692669, EPI_ISL_692670, EPI_ISL_692671, EPI_ISL_692672, EPI_ISL_692673, EPI_ISL_692674, EPI_ISL_692675, EPI_ISL_692676, EPI_ISL_692677, EPI_ISL_692678, EPI_ISL_692679 | | | |
| see above | Sakai City Institute of Public Health | Pathogen Genomics Center, National Institute of Infectious Diseases | Tsuyoshi Sekizuka, Kentaro Itokawa, Rina Tanaka, Masanori Hashino, Makoto Kuroda |
| EPI_ISL_692680, EPI_ISL_692681, EPI_ISL_692682, EPI_ISL_692683, EPI_ISL_692684, EPI_ISL_692685, EPI_ISL_692686, EPI_ISL_692687, EPI_ISL_692688, EPI_ISL_692689, EPI_ISL_692690, EPI_ISL_692691, EPI_ISL_692692, EPI_ISL_692693, EPI_ISL_692694, EPI_ISL_692695, EPI_ISL_692696, EPI_ISL_692697, EPI_ISL_692698, EPI_ISL_692699, EPI_ISL_692700, EPI_ISL_692701, EPI_ISL_692702, EPI_ISL_692703, EPI_ISL_692704, EPI_ISL_692705, EPI_ISL_692706, EPI_ISL_692707, EPI_ISL_692708, EPI_ISL_692709, EPI_ISL_692710, EPI_ISL_692711, EPI_ISL_692712, EPI_ISL_692713, EPI_ISL_692714, EPI_ISL_692715, EPI_ISL_692716, EPI_ISL_692717, EPI_ISL_692718, EPI_ISL_692719, EPI_ISL_692720, EPI_ISL_692721, EPI_ISL_692722, EPI_ISL_692723, EPI_ISL_692724, EPI_ISL_692725, EPI_ISL_692726, EPI_ISL_692727, EPI_ISL_692728, EPI_ISL_692729, EPI_ISL_692730, EPI_ISL_692731, EPI_ISL_692732 | | | |
| see above | Pathogen Genomics Center, National Institute of Infectious Diseases | Pathogen Genomics Center, National Institute of Infectious Diseases | Tsuyoshi Sekizuka, Kentaro Itokawa, Rina Tanaka, Masanori Hashino, Makoto Kuroda |
| EPI_ISL_693199 | Hospital do Servidor Publico Estadual Francisco Morato de Oliveira | Instituto Adolfo Lutz, Interdiciplinary Procedures Center, Strategic Laboratory | Claudio Tavares Sacchi, Claudia Regina Gonçalves, Erica Valessa Ramos Gomes, Karoline Rodrigues Campos |
| EPI_ISL_693204 | Pronto Socorro Dr. Conrado Cesarino Nuvolini | Instituto Adolfo Lutz, Interdiciplinary Procedures Center, Strategic Laboratory | Claudio Tavares Sacchi, Claudia Regina Gonçalves, Erica Valessa Ramos Gomes, Karoline Rodrigues Campos |
| EPI_ISL_693205 | Hospital de Campanha Covid-19 Assis | Instituto Adolfo Lutz, Interdiciplinary Procedures Center, Strategic Laboratory | Claudio Tavares Sacchi, Claudia Regina Gonçalves, Erica Valessa Ramos Gomes, Karoline Rodrigues Campos |
| EPI_ISL_693206 | Hospital Municipal Mario Gatti | Instituto Adolfo Lutz, Interdiciplinary Procedures Center, Strategic Laboratory | Claudio Tavares Sacchi, Claudia Regina Gonçalves, Erica Valessa Ramos Gomes, Karoline Rodrigues Campos |
| EPI_ISL_693207 | Cs II Doutor Antonio Vicoso Moreira de Rezende | Instituto Adolfo Lutz, Interdiciplinary Procedures Center, Strategic Laboratory | Claudio Tavares Sacchi, Claudia Regina Gonçalves, Erica Valessa Ramos Gomes, Karoline Rodrigues Campos |
| EPI_ISL_693208, EPI_ISL_693209 | Hospital Municipal Antonio Giglio | Instituto Adolfo Lutz, Interdiciplinary Procedures Center, Strategic Laboratory | Claudio Tavares Sacchi, Claudia Regina Gonçalves, Erica Valessa Ramos Gomes, Karoline Rodrigues Campos |
| EPI_ISL_693210 | Pronto-Socorro Dr. Osmar Mesquita | Instituto Adolfo Lutz, Interdiciplinary Procedures Center, Strategic Laboratory | Claudio Tavares Sacchi, Claudia Regina Gonçalves, Erica Valessa Ramos Gomes, Karoline Rodrigues Campos |
| EPI_ISL_693211 | Santa Casa de Misericordia e Maternidade | Instituto Adolfo Lutz, Interdiciplinary Procedures Center, Strategic Laboratory | Claudio Tavares Sacchi, Claudia Regina Gonçalves, Erica Valessa Ramos Gomes, Karoline Rodrigues Campos |
| EPI_ISL_693212 | Santa Casa de Misericordia de Braganca Paulista | Instituto Adolfo Lutz, Interdiciplinary Procedures Center, Strategic Laboratory | Claudio Tavares Sacchi, Claudia Regina Gonçalves, Erica Valessa Ramos Gomes, Karoline Rodrigues Campos |
| EPI_ISL_693213 | Hospital E Maternidade Municipal Governador Mario Covas | Instituto Adolfo Lutz, Interdiciplinary Procedures Center, Strategic Laboratory | Claudio Tavares Sacchi, Claudia Regina Gonçalves, Erica Valessa Ramos Gomes, Karoline Rodrigues Campos |
| EPI_ISL_693214 | Unidade de Pronto Atendimento Central de Caraguatatuba | Instituto Adolfo Lutz, Interdiciplinary Procedures Center, Strategic Laboratory | Claudio Tavares Sacchi, Claudia Regina Gonçalves, Erica Valessa Ramos Gomes, Karoline Rodrigues Campos |
| EPI_ISL_693215 | Secretaria Municipal de Saúde de Iracemapolis | Instituto Adolfo Lutz, Interdiciplinary Procedures Center, Strategic Laboratory | Claudio Tavares Sacchi, Claudia Regina Gonçalves, Erica Valessa Ramos Gomes, Karoline Rodrigues Campos |
| EPI_ISL_693216, EPI_ISL_693217 | Unidade de Vigilância Epiudemiológica de Araras | Instituto Adolfo Lutz, Interdiciplinary Procedures Center, Strategic Laboratory | Claudio Tavares Sacchi, Claudia Regina Gonçalves, Erica Valessa Ramos Gomes, Karoline Rodrigues Campos |
| EPI_ISL_693218 | Hospital Domingos Leonardo Ceravolo Presidente Prudente | Instituto Adolfo Lutz, Interdiciplinary Procedures Center, Strategic Laboratory | Claudio Tavares Sacchi, Claudia Regina Gonçalves, Erica Valessa Ramos Gomes, Karoline Rodrigues Campos |
| EPI_ISL_693219 | Santa Casa da Misericórdia de Presidente Prudente | Instituto Adolfo Lutz, Interdiciplinary Procedures Center, | Claudio Tavares Sacchi, Claudia Regina Gonçalves, Erica Valessa Ramos Gomes, Karoline Rodrigues Campos |

| | | | |
|---|---|---|---|
| | | Strategic Laboratory | |
| EPI_ISL_693220 | Laboratório Municipal de Piracicaba | Instituto Adolfo Lutz, Interdiciplinary Procedures Center, Strategic Laboratory | Claudio Tavares Sacchi, Claudia Regina Gonçalves, Erica Valessa Ramos Gomes, Karoline Rodrigues Campos |
| EPI_ISL_693221, EPI_ISL_693222 | Secretaria Municipal de Saúde de Birigui | Instituto Adolfo Lutz, Interdiciplinary Procedures Center, Strategic Laboratory | Claudio Tavares Sacchi, Claudia Regina Gonçalves, Erica Valessa Ramos Gomes, Karoline Rodrigues Campos |
| EPI_ISL_693223, EPI_ISL_693224 | Laboratório Municipal de Piracicaba | Instituto Adolfo Lutz, Interdiciplinary Procedures Center, Strategic Laboratory | Claudio Tavares Sacchi, Claudia Regina Gonçalves, Erica Valessa Ramos Gomes, Karoline Rodrigues Campos |
| EPI_ISL_693225 | Ubs Vila Rosa - Olímpia Gomes De Almeida | Instituto Adolfo Lutz, Interdiciplinary Procedures Center, Strategic Laboratory | Claudio Tavares Sacchi, Claudia Regina Gonçalves, Erica Valessa Ramos Gomes, Karoline Rodrigues Campos |
| EPI_ISL_693226 | Unidade de Pronto Atendimento Sao José | Instituto Adolfo Lutz, Interdiciplinary Procedures Center, Strategic Laboratory | Claudio Tavares Sacchi, Claudia Regina Gonçalves, Erica Valessa Ramos Gomes, Karoline Rodrigues Campos |
| EPI_ISL_693227 | UBS Vila Marchi | Instituto Adolfo Lutz, Interdiciplinary Procedures Center, Strategic Laboratory | Claudio Tavares Sacchi, Claudia Regina Gonçalves, Erica Valessa Ramos Gomes, Karoline Rodrigues Campos |
| EPI_ISL_693228 | Secretaria Municipal de Sorocaba | Instituto Adolfo Lutz, Interdiciplinary Procedures Center, Strategic Laboratory | Claudio Tavares Sacchi, Claudia Regina Gonçalves, Erica Valessa Ramos Gomes, Karoline Rodrigues Campos |
| EPI_ISL_693229 | Hospital 8 de Maio | Instituto Adolfo Lutz, Interdiciplinary Procedures Center, Strategic Laboratory | Claudio Tavares Sacchi, Claudia Regina Gonçalves, Erica Valessa Ramos Gomes, Karoline Rodrigues Campos |
| EPI_ISL_693230 | Hospital e Pronto Socorro Portinari | Instituto Adolfo Lutz, Interdiciplinary Procedures Center, Strategic Laboratory | Claudio Tavares Sacchi, Claudia Regina Gonçalves, Erica Valessa Ramos Gomes, Karoline Rodrigues Campos |
| EPI_ISL_693231 | Pronto Socorro Municipal de Santa Branca | Instituto Adolfo Lutz, Interdiciplinary Procedures Center, Strategic Laboratory | Claudio Tavares Sacchi, Claudia Regina Gonçalves, Erica Valessa Ramos Gomes, Karoline Rodrigues Campos |
| EPI_ISL_693232 | Hospital e Pronto Socorro Portinari | Instituto Adolfo Lutz, Interdiciplinary Procedures Center, Strategic Laboratory | Claudio Tavares Sacchi, Claudia Regina Gonçalves, Erica Valessa Ramos Gomes, Karoline Rodrigues Campos |
| EPI_ISL_693233 | Hospital Santa Cruz | Instituto Adolfo Lutz, Interdiciplinary Procedures Center, Strategic Laboratory | Claudio Tavares Sacchi, Claudia Regina Gonçalves, Erica Valessa Ramos Gomes, Karoline Rodrigues Campos |
| EPI_ISL_693234 | Upa Vereador Jose Da Rocha Goncalves | Instituto Adolfo Lutz, Interdiciplinary Procedures Center, Strategic Laboratory | Claudio Tavares Sacchi, Claudia Regina Gonçalves, Erica Valessa Ramos Gomes, Karoline Rodrigues Campos |
| EPI_ISL_693235 | Casmi Centro Atendimento Saude da Mulher e Infancia | Instituto Adolfo Lutz, Interdiciplinary Procedures Center, Strategic Laboratory | Claudio Tavares Sacchi, Claudia Regina Gonçalves, Erica Valessa Ramos Gomes, Karoline Rodrigues Campos |
| EPI_ISL_693236 | Hospital Santa Marcelina Sao Paulo | Instituto Adolfo Lutz, Interdiciplinary Procedures Center, Strategic Laboratory | Claudio Tavares Sacchi, Claudia Regina Gonçalves, Erica Valessa Ramos Gomes, Karoline Rodrigues Campos |
| EPI_ISL_693237 | UPA Santa Isabel | Instituto Adolfo Lutz, Interdiciplinary Procedures Center, Strategic Laboratory | Claudio Tavares Sacchi, Claudia Regina Gonçalves, Erica Valessa Ramos Gomes, Karoline Rodrigues Campos |
| EPI_ISL_693238, EPI_ISL_693239 | Secao Centro de Diagnostico Secedi | Instituto Adolfo Lutz, Interdiciplinary Procedures Center, Strategic Laboratory | Claudio Tavares Sacchi, Claudia Regina Gonçalves, Erica Valessa Ramos Gomes, Karoline Rodrigues Campos |
| EPI_ISL_693240 | Centro de Vigilância a Saude de Diadema | Instituto Adolfo Lutz, Interdiciplinary Procedures Center, Strategic Laboratory | Claudio Tavares Sacchi, Claudia Regina Gonçalves, Erica Valessa Ramos Gomes, Karoline Rodrigues Campos |
| EPI_ISL_693241 | Hospital e Maternidade Sao Lucas | Instituto Adolfo Lutz, Interdiciplinary Procedures Center, Strategic Laboratory | Claudio Tavares Sacchi, Claudia Regina Gonçalves, Erica Valessa Ramos Gomes, Karoline Rodrigues Campos |
| EPI_ISL_693242 | Centro de Vigilância a Saude de Diadema | Instituto Adolfo Lutz, Interdiciplinary Procedures Center, Strategic Laboratory | Claudio Tavares Sacchi, Claudia Regina Gonçalves, Erica Valessa Ramos Gomes, Karoline Rodrigues Campos |
| EPI_ISL_693243 | Laboratório Municipal de Piracicaba | Instituto Adolfo Lutz, Interdiciplinary Procedures Center, Strategic Laboratory | Claudio Tavares Sacchi, Claudia Regina Gonçalves, Erica Valessa Ramos Gomes, Karoline Rodrigues Campos |
| EPI_ISL_693244 | Centro Médico da Polícia Militar do Estado de Sao Paulo | Instituto Adolfo Lutz, Interdiciplinary Procedures Center, Strategic Laboratory | Claudio Tavares Sacchi, Claudia Regina Gonçalves, Erica Valessa Ramos Gomes, Karoline Rodrigues Campos |
| EPI_ISL_693245 | UPA Santa Isabel | Instituto Adolfo Lutz, Interdiciplinary Procedures Center, Strategic Laboratory | Claudio Tavares Sacchi, Claudia Regina Gonçalves, Erica Valessa Ramos Gomes, Karoline Rodrigues Campos |
| EPI_ISL_693246 | Laboratorio Municipal de Rio Grande da Serra | Instituto Adolfo Lutz, Interdiciplinary Procedures Center, Strategic Laboratory | Claudio Tavares Sacchi, Claudia Regina Gonçalves, Erica Valessa Ramos Gomes, Karoline Rodrigues Campos |
| EPI_ISL_693247 | Secao Centro de Diagnostico Secedi | Instituto Adolfo Lutz, Interdiciplinary Procedures Center, Strategic Laboratory | Claudio Tavares Sacchi, Claudia Regina Gonçalves, Erica Valessa Ramos Gomes, Karoline Rodrigues Campos |
| EPI_ISL_693248 | Centro Municipal de Epidemiologia e Imunizações | Instituto Adolfo Lutz, Interdiciplinary Procedures Center, Strategic Laboratory | Claudio Tavares Sacchi, Claudia Regina Gonçalves, Erica Valessa Ramos Gomes, Karoline Rodrigues Campos |
| EPI_ISL_699716, EPI_ISL_699717, EPI_ISL_699718, EPI_ISL_699719, EPI_ISL_699720, EPI_ISL_699721, EPI_ISL_699724, EPI_ISL_699725, EPI_ISL_699726, EPI_ISL_699727, EPI_ISL_699728, EPI_ISL_699729, EPI_ISL_699730, EPI_ISL_699731, EPI_ISL_699732, EPI_ISL_699733, EPI_ISL_699735, EPI_ISL_699736, EPI_ISL_699737, EPI_ISL_699738, EPI_ISL_699739, EPI_ISL_699740, EPI_ISL_699741, EPI_ISL_699742, EPI_ISL_699743, EPI_ISL_699744, EPI_ISL_699745, EPI_ISL_699746, EPI_ISL_699747, EPI_ISL_699748, EPI_ISL_699749, EPI_ISL_699750, EPI_ISL_699751, EPI_ISL_699754, EPI_ISL_699759, EPI_ISL_699760, EPI_ISL_699761, EPI_ISL_699763, EPI_ISL_699764, EPI_ISL_699765, EPI_ISL_699766, EPI_ISL_699769, EPI_ISL_699770, EPI_ISL_699771, EPI_ISL_699772, EPI_ISL_699773, EPI_ISL_699774, EPI_ISL_699775, EPI_ISL_699776, EPI_ISL_699777, EPI_ISL_699778, EPI_ISL_699779, EPI_ISL_699780, EPI_ISL_699782, EPI_ISL_699783, EPI_ISL_699785, EPI_ISL_699786, EPI_ISL_699787, EPI_ISL_699788, EPI_ISL_699790, EPI_ISL_699791, EPI_ISL_699792, EPI_ISL_699793, EPI_ISL_699794, EPI_ISL_699795, EPI_ISL_699796, EPI_ISL_699797, EPI_ISL_699798, EPI_ISL_699799, EPI_ISL_699802, EPI_ISL_699803, EPI_ISL_699804, EPI_ISL_699805, EPI_ISL_699806, EPI_ISL_699807, EPI_ISL_699808, EPI_ISL_699809, EPI_ISL_699811, EPI_ISL_699813, EPI_ISL_699815, EPI_ISL_699816, EPI_ISL_699818, EPI_ISL_699820, EPI_ISL_699821, EPI_ISL_699822, EPI_ISL_699823, EPI_ISL_699826, EPI_ISL_699828, EPI_ISL_699829, EPI_ISL_699830, EPI_ISL_699832, EPI_ISL_699833, EPI_ISL_699834, EPI_ISL_699835, EPI_ISL_699838, EPI_ISL_699839, EPI_ISL_699840, EPI_ISL_699841, EPI_ISL_699843, EPI_ISL_699844, EPI_ISL_699845, EPI_ISL_699846, EPI_ISL_699847, EPI_ISL_699851, EPI_ISL_699852, EPI_ISL_699853, EPI_ISL_699854, EPI_ISL_699855, EPI_ISL_699856, EPI_ISL_699857, EPI_ISL_699858, EPI_ISL_699859, EPI_ISL_699860, EPI_ISL_699861, EPI_ISL_699862, EPI_ISL_699863, EPI_ISL_699864, EPI_ISL_699865, EPI_ISL_699866, EPI_ISL_699867, EPI_ISL_699868, EPI_ISL_699869, EPI_ISL_699870, EPI_ISL_699871, EPI_ISL_699872, EPI_ISL_699873, EPI_ISL_699874, EPI_ISL_699878, EPI_ISL_699879, EPI_ISL_699880, EPI_ISL_699882, EPI_ISL_699883, EPI_ISL_699884, EPI_ISL_699886, EPI_ISL_699887, EPI_ISL_699888, EPI_ISL_699889, EPI_ISL_699890, EPI_ISL_699891, EPI_ISL_699893, EPI_ISL_699894, EPI_ISL_699897, EPI_ISL_699898, EPI_ISL_699899, EPI_ISL_699902, EPI_ISL_699904, EPI_ISL_699906, EPI_ISL_699907, EPI_ISL_699909, EPI_ISL_699910, EPI_ISL_699911, EPI_ISL_699912, EPI_ISL_699914, EPI_ISL_699915, EPI_ISL_699916, EPI_ISL_699917, EPI_ISL_699918, EPI_ISL_699919, EPI_ISL_699920, EPI_ISL_699921, EPI_ISL_699922, EPI_ISL_699923, EPI_ISL_699924, EPI_ISL_699925, EPI_ISL_699926, EPI_ISL_699927, EPI_ISL_699928, EPI_ISL_699930, EPI_ISL_699931, EPI_ISL_699932, EPI_ISL_699933, EPI_ISL_699934, EPI_ISL_699935, EPI_ISL_699936, EPI_ISL_699937, EPI_ISL_699938, EPI_ISL_699939, EPI_ISL_699940, EPI_ISL_699941, EPI_ISL_699942, EPI_ISL_699943, EPI_ISL_699944, EPI_ISL_699945, EPI_ISL_699946, EPI_ISL_699947, EPI_ISL_699948, EPI_ISL_699949, EPI_ISL_699950, EPI_ISL_699951, EPI_ISL_699952, EPI_ISL_699953, EPI_ISL_699954, EPI_ISL_699955, EPI_ISL_699956, EPI_ISL_699957, EPI_ISL_699958, EPI_ISL_699961, EPI_ISL_699962, EPI_ISL_699963, EPI_ISL_699964, EPI_ISL_699966, EPI_ISL_699967, EPI_ISL_699968, EPI_ISL_699970, EPI_ISL_699971, EPI_ISL_699972, EPI_ISL_699973, EPI_ISL_699974, EPI_ISL_699975, EPI_ISL_699979, EPI_ISL_699980, EPI_ISL_699981, EPI_ISL_699983, EPI_ISL_699984, EPI_ISL_699985, EPI_ISL_699986, EPI_ISL_699987, EPI_ISL_699988, EPI_ISL_699989, EPI_ISL_699990, EPI_ISL_699991, EPI_ISL_699993, EPI_ISL_699994, EPI_ISL_699996, EPI_ISL_699997, EPI_ISL_699998, EPI_ISL_699999, EPI_ISL_700000, EPI_ISL_700002, EPI_ISL_700003, EPI_ISL_700004, EPI_ISL_700005, EPI_ISL_700006, EPI_ISL_700009, EPI_ISL_700010, EPI_ISL_700011, EPI_ISL_700012, EPI_ISL_700013, EPI_ISL_700016, EPI_ISL_700017, EPI_ISL_700018, EPI_ISL_700020, EPI_ISL_700021, EPI_ISL_700024, EPI_ISL_700025, EPI_ISL_700027, EPI_ISL_700028, EPI_ISL_700030, EPI_ISL_700031, EPI_ISL_700033, EPI_ISL_700034, EPI_ISL_700036, EPI_ISL_700037, EPI_ISL_700038, EPI_ISL_700039, EPI_ISL_700040, EPI_ISL_700041, EPI_ISL_700043, EPI_ISL_700044, EPI_ISL_700046, EPI_ISL_700047, EPI_ISL_700048, EPI_ISL_700049, EPI_ISL_700050, EPI_ISL_700051, EPI_ISL_700052, EPI_ISL_700053, EPI_ISL_700054, EPI_ISL_700055, EPI_ISL_700056, EPI_ISL_700057, EPI_ISL_700058, EPI_ISL_700059, EPI_ISL_700061, EPI_ISL_700064, EPI_ISL_700065, EPI_ISL_700066, EPI_ISL_700067, EPI_ISL_700068, EPI_ISL_700069, EPI_ISL_700070, EPI_ISL_700071, EPI_ISL_700072, EPI_ISL_700073, EPI_ISL_700074, EPI_ISL_700076, EPI_ISL_700077, EPI_ISL_700078, EPI_ISL_700079, EPI_ISL_700080, EPI_ISL_700081, EPI_ISL_700082, EPI_ISL_700083, EPI_ISL_700084, EPI_ISL_700085, EPI_ISL_700086, EPI_ISL_700088, EPI_ISL_700089, EPI_ISL_700091, EPI_ISL_700092, EPI_ISL_700094, EPI_ISL_700096, EPI_ISL_700097, EPI_ISL_700098, EPI_ISL_700099, EPI_ISL_700100, EPI_ISL_700101, EPI_ISL_700102, EPI_ISL_700103, EPI_ISL_700104, EPI_ISL_700107, EPI_ISL_700108, EPI_ISL_700109, EPI_ISL_700110, EPI_ISL_700111, EPI_ISL_700112, EPI_ISL_700113, EPI_ISL_700114, EPI_ISL_700115, EPI_ISL_700116, EPI_ISL_700118, EPI_ISL_700120, EPI_ISL_700123, EPI_ISL_700124, EPI_ISL_700125, EPI_ISL_700127, EPI_ISL_700128, EPI_ISL_700129, EPI_ISL_700131, EPI_ISL_700132, EPI_ISL_700136, EPI_ISL_700138, EPI_ISL_700140, EPI_ISL_700143, EPI_ISL_700144, EPI_ISL_700145, EPI_ISL_700146, EPI_ISL_700147, EPI_ISL_700149, EPI_ISL_700150, EPI_ISL_700151, EPI_ISL_700154, | | | |

| | | | |
|---|---|---|---|
| EPI_ISL_700155, EPI_ISL_700156, EPI_ISL_700157, EPI_ISL_700158, EPI_ISL_700160, EPI_ISL_700161, EPI_ISL_700162, EPI_ISL_700164, EPI_ISL_700166, EPI_ISL_700167, EPI_ISL_700168, EPI_ISL_700169, EPI_ISL_700172, EPI_ISL_700176, EPI_ISL_700177, EPI_ISL_700179, EPI_ISL_700180, EPI_ISL_700181, EPI_ISL_700183, EPI_ISL_700184, EPI_ISL_700185, EPI_ISL_700187, EPI_ISL_700188, EPI_ISL_700189, EPI_ISL_700190, EPI_ISL_700191, EPI_ISL_700192, EPI_ISL_700193, EPI_ISL_700194, EPI_ISL_700195, EPI_ISL_700196, EPI_ISL_700197, EPI_ISL_700198, EPI_ISL_700199, EPI_ISL_700200, EPI_ISL_700201, EPI_ISL_700202, EPI_ISL_700203, EPI_ISL_700204, EPI_ISL_700205, EPI_ISL_700206, EPI_ISL_700208, EPI_ISL_700209, EPI_ISL_700210, EPI_ISL_700211, EPI_ISL_700212, EPI_ISL_700214, EPI_ISL_700215, EPI_ISL_700216, EPI_ISL_700217 | | | |
| see above | Hematopathology Laboratory, ACTREC, TMC | Hematopathology Laboratory, ACTREC, TMC | Hematopathology Laboratory, ACTREC |
| EPI_ISL_707928 | Fujita Health University Hospital | Fujita Health University School of Medicine, Department of Microbiology | Masahiro Suzuki, Aki Sakurai, Yohei Doi |
| EPI_ISL_707936, EPI_ISL_707937, EPI_ISL_707938, EPI_ISL_707939, EPI_ISL_708184, EPI_ISL_708185, EPI_ISL_708186, EPI_ISL_708188, EPI_ISL_708189, EPI_ISL_708190, EPI_ISL_708191, EPI_ISL_708192, EPI_ISL_708193, EPI_ISL_708194, EPI_ISL_708195 | | | |
| see above | Pamukkale University Hospital | Pamukkale University Department of Medical Genetics | Onur TOKGUN et al. |
| EPI_ISL_708529 | Secretária Municipal de Saude de Fernandópolis | Instituto Adolfo Lutz, Interdiciplinary Procedures Center, Strategic Laboratory | Claudio Tavares Sacchi, Claudia Regina Gonçalves, Erica Valessa Ramos Gomes, Carlos Henrique Camargo, Karoline Rodrigues Campos, Fernanda Modesto Tolentino Binhardi, Maricelia Navarro Pinheiro Flores, Marcia Maria Costa Nunes Soares, Janaina Other Martins Montanha |
| EPI_ISL_708839, EPI_ISL_708840 | National Institute of Blood Diseases (NIBD), Molecular Biology Lab | Genomics Lab NIBD | Samina Naz Mukry, Sayed Ali Raza, Shariq Ahmed, Aneeta Shahni, Gul Sufaida, Arshi Naz , Tahir Sultan Shamsi |
| EPI_ISL_718134, EPI_ISL_718135, EPI_ISL_718143, EPI_ISL_718146, EPI_ISL_718156 | Ministry of Health Hospitals | Institute of Health and Community Medicine | David Perera, Ooi Mong How, Chua Hock Hin, Tonnii Sia Loong Loong, Wong Jyn Shan, Wong Kiing Aik, Chan Chia Jui |
| EPI_ISL_718267, EPI_ISL_718268, EPI_ISL_718271, EPI_ISL_718272, EPI_ISL_718274, EPI_ISL_718275, EPI_ISL_718277, EPI_ISL_718279 | Institute for Medical Research, Infectious Disease Research Centre, National Institutes of Health, Ministry of Health Malaysia | Institute for Medical Research, Infectious Disease Research Centre, National Institutes of Health, Ministry of Health Malaysia | Suppiah J, Kamel K, Mohd-Zawawi Z, Thayan R |
| EPI_ISL_722851 | I.R.C.C.S. "S. De Bellis" - Ente Ospedaliero | Istituto Zooprofilattico Sperimentale della Puglia e della Basilicata | Parisi A., Bianco A., Capozzi L., Del Sambro L., Lippolis A., Notarnicola M., Manzulli V, Rondinone V., Pace L. |
| EPI_ISL_722852 | I.R.C.C.S. "S. De Bellis" - Ente Ospedaliero | Istituto Zooprofilattico Sperimentale della Puglia e della Basilicata | Parisi A., Bianco A., Capozzi L., Del Sambro L., Lippolis A., Notarnicola M., Cipolletta D., Galante D. |
| EPI_ISL_722853 | I.R.C.C.S. "S. De Bellis" - Ente Ospedaliero | Istituto Zooprofilattico Sperimentale della Puglia e della Basilicata | Parisi A., Bianco A., Capozzi L., Del Sambro L., Lippolis A., Notarnicola M., Manzulli V, Rondinone V., Pace L. |
| EPI_ISL_722854 | I.R.C.C.S. "S. De Bellis" - Ente Ospedaliero | Istituto Zooprofilattico Sperimentale della Puglia e della Basilicata | Parisi A., Bianco A., Capozzi L., Del Sambro L., Lippolis A., Notarnicola M., Cipolletta D., Galante D. |
| EPI_ISL_722855, EPI_ISL_722856, EPI_ISL_722857, EPI_ISL_722858 | Dipartimento di Scienze Biomediche e Oncologia Umana - Azienda Ospedaliero Universitaria Consorziale Policlinico | Istituto Zooprofilattico Sperimentale della Puglia e della Basilicata | Parisi A., Bianco A., Capozzi L., Del Sambro L., Chironna M., Loconsole D. |
| EPI_ISL_722859, EPI_ISL_722860 | I.R.C.C.S. "S. De Bellis" - Ente Ospedaliero | Istituto Zooprofilattico Sperimentale della Puglia e della Basilicata | Parisi A., Bianco A., Capozzi L., Del Sambro L., Lippolis A., Notarnicola M., Manzulli V, Rondinone V., Pace L. |
| EPI_ISL_722861 | I.R.C.C.S. "S. De Bellis" - Ente Ospedaliero | Istituto Zooprofilattico Sperimentale della Puglia e della Basilicata | Parisi A., Bianco A., Capozzi L., Del Sambro L., Lippolis A., Notarnicola M., Cipolletta D., Galante D. |
| EPI_ISL_722862 | I.R.C.C.S. "S. De Bellis" - Ente Ospedaliero | Istituto Zooprofilattico Sperimentale della Puglia e della Basilicata | Parisi A., Bianco A., Capozzi L., Del Sambro L., Lippolis A., Notarnicola M., Manzulli V, Rondinone V., Pace L. |
| EPI_ISL_722863 | I.R.C.C.S. "S. De Bellis" - Ente Ospedaliero | Istituto Zooprofilattico Sperimentale della Puglia e della Basilicata | Parisi A., Bianco A., Capozzi L., Del Sambro L., Lippolis A., Notarnicola M., Cipolletta D., Galante D. |
| EPI_ISL_722864, EPI_ISL_722865 | I.R.C.C.S. "S. De Bellis" - Ente Ospedaliero | Istituto Zooprofilattico Sperimentale della Puglia e della Basilicata | Parisi A., Bianco A., Capozzi L., Del Sambro L., Lippolis A., Notarnicola M., Manzulli V, Rondinone V., Pace L. |
| EPI_ISL_722866 | I.R.C.C.S. "S. De Bellis" - Ente Ospedaliero | Istituto Zooprofilattico Sperimentale della Puglia e della Basilicata | Parisi A., Bianco A., Capozzi L., Del Sambro L., Lippolis A., Notarnicola M., Cipolletta D., Galante D. |
| EPI_ISL_722867, EPI_ISL_722868 | I.R.C.C.S. "S. De Bellis" - Ente Ospedaliero | Istituto Zooprofilattico Sperimentale della Puglia e della Basilicata | Parisi A., Bianco A., Capozzi L., Del Sambro L., Lippolis A., Notarnicola M., Manzulli V, Rondinone V., Pace L. |
| EPI_ISL_722869 | I.R.C.C.S. "S. De Bellis" - Ente Ospedaliero | Istituto Zooprofilattico Sperimentale della Puglia e della Basilicata | Parisi A., Bianco A., Capozzi L., Del Sambro L., Lippolis A., Notarnicola M., Cipolletta D., Galante D. |
| EPI_ISL_722870 | I.R.C.C.S. "S. De Bellis" - Ente Ospedaliero | Istituto Zooprofilattico Sperimentale della Puglia e della Basilicata | Parisi A., Bianco A., Capozzi L., Del Sambro L., Lippolis A., Notarnicola M., Manzulli V, Rondinone V., Pace L. |
| EPI_ISL_722871 | I.R.C.C.S. "S. De Bellis" - Ente Ospedaliero | Istituto Zooprofilattico Sperimentale della Puglia e della Basilicata | Parisi A., Bianco A., Capozzi L., Del Sambro L., Lippolis A., Notarnicola M., Cipolletta D., Galante D. |
| EPI_ISL_722872 | Dipartimento di Scienze Biomediche e Oncologia Umana - Azienda Ospedaliero Universitaria Consorziale Policlinico | Istituto Zooprofilattico Sperimentale della Puglia e della Basilicata | Parisi A., Bianco A., Capozzi L., Del Sambro L., Chironna M., Loconsole D. |
| EPI_ISL_722892 | Istituto Zooprofilattico Sperimentale della Puglia e della Basilicata | Istituto Zooprofilattico Sperimentale della Puglia e della Basilicata | Parisi A., Bianco A., Capozzi L., Del Sambro L., Manzulli V, Rondinone V., Pace L., Cipolletta D., Galante D. |
| EPI_ISL_722895 | I.R.C.C.S. "S. De Bellis" - Ente Ospedaliero | Istituto Zooprofilattico Sperimentale della Puglia e della Basilicata | Parisi A., Bianco A., Capozzi L., Del Sambro L., Lippolis A., Notarnicola M., Cipolletta D., Galante D. |
| EPI_ISL_722897 | Dipartimento di Scienze Biomediche e Oncologia Umana - Azienda Ospedaliero Universitaria Consorziale Policlinico | Istituto Zooprofilattico Sperimentale della Puglia e della Basilicata | Parisi A., Bianco A., Capozzi L., Del Sambro L., Chironna M., Loconsole D. |
| EPI_ISL_722898, EPI_ISL_722900 | I.R.C.C.S. "S. De Bellis" - Ente Ospedaliero | Istituto Zooprofilattico Sperimentale della Puglia e della Basilicata | Parisi A., Bianco A., Capozzi L., Del Sambro L., Lippolis A., Notarnicola M., Cipolletta D., Galante D. |
| EPI_ISL_729800, EPI_ISL_729848, EPI_ISL_729849 | Laboratório Central de Saúde Pública do Estado do Rio Grande do Sul (LACEN-RS) | Laboratory of Respiratory Viruses and Measles, Oswaldo Cruz Institute, FIOCRUZ | Paola Resende, Luciana Appolinario, Fernando Motta, Anna Carolina Paixão, Ana Carolina Mendonça, Tatiana Schaffer Gregianini, Marilda Tereza Mar da Rosa, Marilda Siqueira on behalf of the Fiocruz COVID-19 Genomic Surveillance Network |
| EPI_ISL_730578 | Home Quarantine Taskforce | Hong Kong Department of Health | Mak Gannon C.K., Lam Edman T.K., Chan Rickjason C.W., Tsang Dominic N.C. |
| EPI_ISL_730601 | Temporary Specimen Collection Centre at the AsiaWorld-Expo | Hong Kong Department of Health | Mak Gannon C.K., Lam Edman T.K., Chan Rickjason C.W., Tsang Dominic N.C. |
| EPI_ISL_732531, EPI_ISL_732532, EPI_ISL_732533, EPI_ISL_732534, EPI_ISL_732535, EPI_ISL_732536, EPI_ISL_732537, EPI_ISL_732538, EPI_ISL_732539, EPI_ISL_732540, EPI_ISL_732559, EPI_ISL_732560 | | | |
| see above | Bundeswehr Institute of Microbiology | Bundeswehr Institute of Microbiology | Elham Khatamzas, Markus Antwerpen, Mathias Walter, Alexandra Rehn, Sabine Zange, Enrico Georgi, Michael von Bergwelt-Baildon, Roman Wölfel |
| EPI_ISL_735306, EPI_ISL_735307, EPI_ISL_735308, EPI_ISL_735309, EPI_ISL_735310, EPI_ISL_735311, EPI_ISL_735312, EPI_ISL_735313, EPI_ISL_735314, EPI_ISL_735315, EPI_ISL_735316, EPI_ISL_735317, EPI_ISL_735318, EPI_ISL_735319, EPI_ISL_735320, EPI_ISL_735321, EPI_ISL_735322, EPI_ISL_735323, EPI_ISL_735324, EPI_ISL_735325, EPI_ISL_735326, EPI_ISL_735327, EPI_ISL_735328, EPI_ISL_735329, EPI_ISL_735330, EPI_ISL_735331, EPI_ISL_735332, EPI_ISL_735333, EPI_ISL_735334, EPI_ISL_735335, EPI_ISL_735336, EPI_ISL_735337, EPI_ISL_735338, EPI_ISL_735339, EPI_ISL_735340, EPI_ISL_735342, EPI_ISL_735343, EPI_ISL_735344, EPI_ISL_735345, EPI_ISL_735346, EPI_ISL_735347, EPI_ISL_735348, EPI_ISL_735349, EPI_ISL_735350, EPI_ISL_735351, EPI_ISL_735352, EPI_ISL_735353, EPI_ISL_735354, EPI_ISL_735355, EPI_ISL_735356, EPI_ISL_735357, EPI_ISL_735358, EPI_ISL_735359, EPI_ISL_735360, EPI_ISL_735361, EPI_ISL_735362, EPI_ISL_735363, EPI_ISL_735364, EPI_ISL_735365 | | | |

| | | | |
|---|---|---|---|
| see above | Genomic Laboratory (GLAB) (Conjoint lab of Health Directorate of Istanbul and Istanbul Technical University) | Genomic Laboratory (GLAB), Istanbul Technical University | Ilker Karacan, Tugba Kizilboga Akgun, Nihat Bugra Agaoglu, Payam Zolfagharian, Mehtap Aydin, Gizem Alkurt, Jale Yildiz, Betsi Kose, Nisan Denizce Can, Ayse Serra Ozel, Nilsun Altunal, Arzu Irvem, Yasemin Kendir Demirkol, Ozlem Akgun Dogan, Levent Doganay, Gizem Dinler Doganay |
| EPI_ISL_735396 | Hospital de Camplanha COVID 19 SER | Instituto Adolfo Lutz, Interdiciplinary Procedures Center, Strategic Laboratory | Claudio Tavares Sacchi, Claudia Regina Gonçalves, Erica Valessa Ramos Gomes, Karoline Rodrigues Campos |
| EPI_ISL_735397 | Unidade Respiratória Nova Hortolandia | Instituto Adolfo Lutz, Interdiciplinary Procedures Center, Strategic Laboratory | Claudio Tavares Sacchi, Claudia Regina Gonçalves, Erica Valessa Ramos Gomes, Karoline Rodrigues Campos |
| EPI_ISL_735398 | Laboratorio Fleury | Instituto Adolfo Lutz, Interdiciplinary Procedures Center, Strategic Laboratory | Claudio Tavares Sacchi, Claudia Regina Gonçalves, Erica Valessa Ramos Gomes, Karoline Rodrigues Campos |
| EPI_ISL_735400 | Instituto Adolfo Lutz - Regional de Santos | Instituto Adolfo Lutz, Interdiciplinary Procedures Center, Strategic Laboratory | Claudio Tavares Sacchi, Claudia Regina Gonçalves, Erica Valessa Ramos Gomes, Karoline Rodrigues Campos |
| EPI_ISL_735401, EPI_ISL_735402, EPI_ISL_735403, EPI_ISL_735404 | Instituto Adolfo Lutz - Regional de Rio Claro | Instituto Adolfo Lutz, Interdiciplinary Procedures Center, Strategic Laboratory | Claudio Tavares Sacchi, Claudia Regina Gonçalves, Erica Valessa Ramos Gomes, Karoline Rodrigues Campos |
| EPI_ISL_735405 | Secretaria Minucipal de Saude de Birigui | Instituto Adolfo Lutz, Interdiciplinary Procedures Center, Strategic Laboratory | Claudio Tavares Sacchi, Claudia Regina Gonçalves, Erica Valessa Ramos Gomes, Karoline Rodrigues Campos |
| EPI_ISL_735406 | Unidade de Pronto Atendimento UPA I Sta Isabel | Instituto Adolfo Lutz, Interdiciplinary Procedures Center, Strategic Laboratory | Claudio Tavares Sacchi, Claudia Regina Gonçalves, Erica Valessa Ramos Gomes, Karoline Rodrigues Campos |
| EPI_ISL_735407 | Santa Casa de Marilia | Instituto Adolfo Lutz, Interdiciplinary Procedures Center, Strategic Laboratory | Claudio Tavares Sacchi, Claudia Regina Gonçalves, Erica Valessa Ramos Gomes, Karoline Rodrigues Campos |
| EPI_ISL_735408 | COVID 19 Centro de Combate ao Coronavirus CCC Jandira | Instituto Adolfo Lutz, Interdiciplinary Procedures Center, Strategic Laboratory | Claudio Tavares Sacchi, Claudia Regina Gonçalves, Erica Valessa Ramos Gomes, Karoline Rodrigues Campos |
| EPI_ISL_735409 | Unidade de Pronto Atendimento Carlos Lourenco | Instituto Adolfo Lutz, Interdiciplinary Procedures Center, Strategic Laboratory | Claudio Tavares Sacchi, Claudia Regina Gonçalves, Erica Valessa Ramos Gomes, Karoline Rodrigues Campos |
| EPI_ISL_735410 | Instituto Adolfo Lutz - Regional de Rio Claro | Instituto Adolfo Lutz, Interdiciplinary Procedures Center, Strategic Laboratory | Claudio Tavares Sacchi, Claudia Regina Gonçalves, Erica Valessa Ramos Gomes, Karoline Rodrigues Campos |
| EPI_ISL_735411 | Centro de Vigilancia a Saude de Diadema | Instituto Adolfo Lutz, Interdiciplinary Procedures Center, Strategic Laboratory | Claudio Tavares Sacchi, Claudia Regina Gonçalves, Erica Valessa Ramos Gomes, Karoline Rodrigues Campos |
| EPI_ISL_735412 | Hospital e Pronto Socorro Portinari | Instituto Adolfo Lutz, Interdiciplinary Procedures Center, Strategic Laboratory | Claudio Tavares Sacchi, Claudia Regina Gonçalves, Erica Valessa Ramos Gomes, Karoline Rodrigues Campos |
| EPI_ISL_735413 | Militello Centro de Diagnosticos e Biopesquisa Clinica | Instituto Adolfo Lutz, Interdiciplinary Procedures Center, Strategic Laboratory | Claudio Tavares Sacchi, Claudia Regina Gonçalves, Erica Valessa Ramos Gomes, Karoline Rodrigues Campos |
| EPI_ISL_735414, EPI_ISL_735415 | Unidade de Pronto Atendimento de Agenor de Campos | Instituto Adolfo Lutz, Interdiciplinary Procedures Center, Strategic Laboratory | Claudio Tavares Sacchi, Claudia Regina Gonçalves, Erica Valessa Ramos Gomes, Karoline Rodrigues Campos |
| EPI_ISL_735416 | Centro de Saude II Dr Jose Paione Mococa | Instituto Adolfo Lutz, Interdiciplinary Procedures Center, Strategic Laboratory | Claudio Tavares Sacchi, Claudia Regina Gonçalves, Erica Valessa Ramos Gomes, Karoline Rodrigues Campos |
| EPI_ISL_735417 | Unidade de Pronto Atendimento de Agenor de Campos | Instituto Adolfo Lutz, Interdiciplinary Procedures Center, Strategic Laboratory | Claudio Tavares Sacchi, Claudia Regina Gonçalves, Erica Valessa Ramos Gomes, Karoline Rodrigues Campos |
| EPI_ISL_735418 | Hospital Regional do Vale do Paraiba | Instituto Adolfo Lutz, Interdiciplinary Procedures Center, Strategic Laboratory | Claudio Tavares Sacchi, Claudia Regina Gonçalves, Erica Valessa Ramos Gomes, Karoline Rodrigues Campos |
| EPI_ISL_735419 | UBS Alvarenga | Instituto Adolfo Lutz, Interdiciplinary Procedures Center, Strategic Laboratory | Claudio Tavares Sacchi, Claudia Regina Gonçalves, Erica Valessa Ramos Gomes, Karoline Rodrigues Campos |
| EPI_ISL_735420 | UBS Riacho Grande | Instituto Adolfo Lutz, Interdiciplinary Procedures Center, Strategic Laboratory | Claudio Tavares Sacchi, Claudia Regina Gonçalves, Erica Valessa Ramos Gomes, Karoline Rodrigues Campos |
| EPI_ISL_735421 | UBS Sta Terezinha | Instituto Adolfo Lutz, Interdiciplinary Procedures Center, Strategic Laboratory | Claudio Tavares Sacchi, Claudia Regina Gonçalves, Erica Valessa Ramos Gomes, Karoline Rodrigues Campos |
| EPI_ISL_735422 | UBS Dematchi | Instituto Adolfo Lutz, Interdiciplinary Procedures Center, Strategic Laboratory | Claudio Tavares Sacchi, Claudia Regina Gonçalves, Erica Valessa Ramos Gomes, Karoline Rodrigues Campos |
| EPI_ISL_735423, EPI_ISL_735424 | Centro de Vigilancia a Saude de Diadema | Instituto Adolfo Lutz, Interdiciplinary Procedures Center, Strategic Laboratory | Claudio Tavares Sacchi, Claudia Regina Gonçalves, Erica Valessa Ramos Gomes, Karoline Rodrigues Campos |
| EPI_ISL_735425 | Hospital e Maternidade Sao Lucas | Instituto Adolfo Lutz, Interdiciplinary Procedures Center, Strategic Laboratory | Claudio Tavares Sacchi, Claudia Regina Gonçalves, Erica Valessa Ramos Gomes, Karoline Rodrigues Campos |
| EPI_ISL_735426 | Centro de Vigilancia a Saude de Diadema | Instituto Adolfo Lutz, Interdiciplinary Procedures Center, Strategic Laboratory | Claudio Tavares Sacchi, Claudia Regina Gonçalves, Erica Valessa Ramos Gomes, Karoline Rodrigues Campos |
| EPI_ISL_735427 | Instituto Adolfo Lutz - Regional de Santos | Instituto Adolfo Lutz, Interdiciplinary Procedures Center, Strategic Laboratory | Claudio Tavares Sacchi, Claudia Regina Gonçalves, Erica Valessa Ramos Gomes, Karoline Rodrigues Campos |
| EPI_ISL_735428, EPI_ISL_735429 | Hospital Nipo Brasileiro | Instituto Adolfo Lutz, Interdiciplinary Procedures Center, Strategic Laboratory | Claudio Tavares Sacchi, Claudia Regina Gonçalves, Erica Valessa Ramos Gomes, Karoline Rodrigues Campos |
| EPI_ISL_735430 | Instituto Adolfo Lutz - Regional de Santos | Instituto Adolfo Lutz, Interdiciplinary Procedures Center, Strategic Laboratory | Claudio Tavares Sacchi, Claudia Regina Gonçalves, Erica Valessa Ramos Gomes, Karoline Rodrigues Campos |
| EPI_ISL_735431, EPI_ISL_735432 | Hospital Nipo Brasileiro | Instituto Adolfo Lutz, Interdiciplinary Procedures Center, Strategic Laboratory | Claudio Tavares Sacchi, Claudia Regina Gonçalves, Erica Valessa Ramos Gomes, Karoline Rodrigues Campos |
| EPI_ISL_735433 | Posto de Atendimento Saude Cidade Pasc Cajati | Instituto Adolfo Lutz, Interdiciplinary Procedures Center, Strategic Laboratory | Claudio Tavares Sacchi, Claudia Regina Gonçalves, Erica Valessa Ramos Gomes, Karoline Rodrigues Campos |
| EPI_ISL_735436 | Nucleic Acid Testing - Rwanda National Reference Laboratory | GIGA Medical Genomics | Yvan Butera,Keith Durkin, Maria Artesi, Bouchra Boujemla, Robert Rutayisire, Patrick Tuyisenge, Esperence Umumararungu, Sébastien Bontems, Marie-Pierre Hayette, Swaibu Gatare, Jacob Souopgui, Sabin Nsanzimana, Vincent Bours, Léon Mutesa |
| EPI_ISL_735437 | Nucleic Acid Testing - Rwanda National Reference Laboratory | GIGA Medical Genomics | Yvan Butera, Keith Durkin, Maria Artesi, Bouchra Boujemla, Robert Rutayisire, Patrick Tuyisenge, Esperence Umumararungu, Sébastien Bontems, Marie-Pierre Hayette, Swaibu Gatare, Jacob Souopgui, Sabin Nsanzimana, Vincent Bours, Léon Mutesa |
| EPI_ISL_735438 | Nuclei Acid Testing - Rwanda National Reference Laboratory | GIGA Medical Genomics | Yvan Butera, Keith Durkin, Maria Artesi, Bouchra Boujemla, Robert Rutayisire, Patrick Tuyisenge, Esperence Umumararungu, Sébastien Bontems, Marie-Pierre Hayette, Swaibu Gatare, Jacob Souopgui, Sabin Nsanzimana, Vincent Bours, Léon Mutesa |
| EPI_ISL_737201 | National Reference Laboratory, Nigeria Centre for Disease Control. | National Reference Laboratory, Nigeria Centre for Disease Control, Gaduwa,Abuja, Nigeria | Dr Ndodo Nnaemeka, Olusola Akanbi, Chimaobi Chukwu, Dr Adesuyi Omoare, oluwaseyi Ajegbe, Grace Esebanmen, Nwando Mba, Shirlee Wohl, Anthony Ahumibe, Celestina Obiekea, Catherine Okoi, Akinpelu Afolabi, Kingsley Njoku, Dr Sikiru Badaru, Dr Chinwe Ochu, Dr Chikwe Ihekweazu |

| | | | |
|---|---|---|---|
| EPI_ISL_738122, EPI_ISL_738123, EPI_ISL_738124, EPI_ISL_738125, EPI_ISL_738126, EPI_ISL_738127, EPI_ISL_738128, EPI_ISL_738130 | IZSM-U.O.C. Virologia | Istituto Zooprofilattico Sperimentale del Mezzogiorno | Maurizio Viscardi, Lorena Cardillo, Giovanna Fusco |
| EPI_ISL_745191 | Mubarak Al-Kabeer Hospital | Virology Unit, Microbiology Department, Faculty of Medicine, Kuwait University | Nada Madi |
| EPI_ISL_745223, EPI_ISL_745259 | Molecular diagnostic laboratory of Federal Budget Institution of Science "Central Research Institute of Epidemiology" of The Federal Service on Customers' Rights Protection and Human Well-being Surveillance | Group of Genomics and Postgenomic Technologies of Central Research Institute of Epidemilology | Samoilov AE, Kaptelova VV, Korneenko EV, Dudorova AV, Saenko SS, Speranskaya AS, Tivanova EV, Shipulina OY, Akimkin VG |
| EPI_ISL_747234 | National Institute of Health Research and Development | National Institute of Health Research and Development | Subangkit; Indrasari,ND; Wulandari,D; Monika,M; Pawestri,HA; Puspa,KD; Nugraha,AA; Ikawati,HD; Pangesti,KNA; Soekarso,T; Paisal; Setiawaty,Vivi |
| EPI_ISL_747242 | Pathogen Lab (BSL3), Biomedical Innovation Department, Applied and Experimental Biology Division, Scientific Research Center and High Education from Ensenada (CICESE) | Pathogen Laboratory (BSL3), Biomedical Innovation Department, Experimental and Applied Biology Division, Scientific Research Center and High Education from Ensenada (CICESE) | Cervantes-Luevano K, Martinez M, Saavedra A, Galindo C and Licea-Navarro A |
| EPI_ISL_751245, EPI_ISL_751246, EPI_ISL_751277, EPI_ISL_751278, EPI_ISL_751279, EPI_ISL_751280, EPI_ISL_751281, EPI_ISL_751282, EPI_ISL_751283, EPI_ISL_751284, EPI_ISL_751285, EPI_ISL_751286, EPI_ISL_751287, EPI_ISL_751288, EPI_ISL_751289, EPI_ISL_751290, EPI_ISL_751291, EPI_ISL_751292, EPI_ISL_751293, EPI_ISL_751294, EPI_ISL_751295, EPI_ISL_751296, EPI_ISL_751298, EPI_ISL_751299, EPI_ISL_751300, EPI_ISL_751302, EPI_ISL_751303, EPI_ISL_751304, EPI_ISL_751305, EPI_ISL_751306, EPI_ISL_751307, EPI_ISL_751308, EPI_ISL_751309, EPI_ISL_751310, EPI_ISL_751311, EPI_ISL_751312, EPI_ISL_751313, EPI_ISL_751315, EPI_ISL_751317 | | | |
| see above | Devki Devi Foundation, a unit of Max Healthcare | CSIR-IGIB/Max | Rajesh Pandey#, Samreen Siddiqui, Janani Srinivasa Vasudevan, Akshay Kanakan, Ranjeet Maurya, Uzma Shamim, Bansidhar Tarai, Akansha Tyagi, Mitali Mukerji, Poonam Das, Sujeet Jha, Mohammed Faruq, Anurag Agrawal |
| EPI_ISL_751337 | IRCCS Sacro Cuore Don Calabria Hospital, Department of Infectious, Tropical Diseases & Microbiology | University of Verona, Department of Biotechnology | Antonio Mori, Michela Deiana, Elena Pomari, Chiara Piubelli; Giulia Lopatriello, Luca Marcolungo, Cristina Beltrami, Chiara Degli Esposti, Emanuela Cosentino, Massimo Delledonne |
| EPI_ISL_753699 | RSUD Ulin Banjarmasin Kalimantan Selatan | National Institute of Health Research and Development | Subangkit;Pawestri,HA;Puspa,KD;Nugraha,AA;Ikawati,HD;Pangesti,KNA;Soekarso,T;Puspandari,N;Pasaribu,M;Setiawaty,V |
| EPI_ISL_754239 | Santosa Hospital Bandung Central | "School of Life Sciences and Technology & School of Pharmacy-Institut Teknologi Bandung; Molecular Genetics Laboratory-Faculty of Medicine-Universitas Padjadjaran; Laboratorium Kesehatan Provinsi Jawa Barat" | Marselina Irasonia Tan, Yunia Sribudiani, Catur Riani, Azzania Fibriani, Husna Nugrahapraja, Tarwadi, Ema Rahmawati, Savira Ekawardhani, Hesti Lina Wiraswati, Ryan Bayusantika Ristandi, Rifky Waluyajati Rachman, Cut Nur Cinthia Alamanda, Lia Faridah, Miftahul Faridl, Karimatu Khoirunnisa, Hammam Riza, Soni Solistia Wirawan, Agung Eru Wibowo, Irvan Faizal |
| EPI_ISL_754241 | Dinkes Tasikmalaya | "School of Life Sciences and Technology & School of Pharmacy-Institut Teknologi Bandung; Molecular Genetics Laboratory-Faculty of Medicine-Universitas Padjadjaran; Laboratorium Kesehatan Provinsi Jawa Barat" | Husna Nugrahapraja, Marselina Irasonia Tan, Yunia Sribudiani, Catur Riani, Azzania Fibriani, Tarwadi, Ema Rahmawati, Savira Ekawardhani, Hesti Lina Wiraswati, Ryan Bayusantika Ristandi, Rifky Waluyajati Rachman, Cut Nur Cinthia Alamanda, Lia Faridah, Miftahul Faridl, Karimatu Khoirunnisa, Hammam Riza, Soni Solistia Wirawan, Agung Eru Wibowo, Irvan Faizal |
| EPI_ISL_756293 | Center for Biotechnology and Cell Therapy, São Rafael Hospital, Salvador, Brazil | Center for Biotechnology and Cell Therapy, São Rafael Hospital, Salvador, Brazil | Carolina Kymie Vasques Nonaka, Marília Miranda Franco, Tiago Gräf, Ana Verena Almeida Mendes, Renato Santana de Aguiar, Marta Giovanetti, Bruno Solano de Freitas Souza |
| EPI_ISL_756401 | National Institute of Health Research and Development | National Institute of Health Research and Development | Adam,K; Wibowo,HA; Ramadhany,R; Rukminiati,Y; Agustiningsih; Pawestri,HA; Subangkit; Puspa,KD; Nugraha,AA; Ikawati,HD; Pangesti,KNA; Soekarso,T; Susilarini,NK; Hariastuti,NI; Nikmah,UA; Mursinah; Febriyani,A; Herman,R; Susanti,N; Herna; Febriyanti,T; Nurhadi,M; Kurniawati,J; Kipuw,NL; Muna,F; Indalao,IL; Rizki,A; Puspandari,N; Setiawaty,Vivi |
| EPI_ISL_759960 | Surabaya Public Health Office | Institute of Tropical Disease, Universitas Airlangga | Aldise M Nastri, Jezzy R Dewantari, Rima R Prasetya, Krisnoadi Rahardjo, Febria Rachmanita, Gatot Soegiarto, Laksmi Wulandari, Resti Yudhawati, Soetjipto, Yasuko Mori, Maria I Lusida, Kazufumi Shimizu |
| EPI_ISL_759963 | RSUD Sidoarjo | Institute of Tropical Disease, Universitas Airlangga | Aldise M Nastri, Jezzy R Dewantari, Rima R Prasetya, Krisnoadi Rahardjo, Atok Irawan, Gatot Soegiarto, Laksmi Wulandari, Resti Yudhawati, Yasuko Mori, Soetjipto, Kazufumi Shimizu ,Maria I Lusida |
| EPI_ISL_760031 | Hong Kong Department of Health | School of Public Health, The University of Hong Kong | Daniel Chu, Haogao Gu, Pavithra Krishnan, Daisy Ng, Gigi Liu, Carrie Wan, Malik Peiris, Leo Poon |
| EPI_ISL_770009 | Area De Salud Escazu (Coopesana) | Inciensa, Instituto Costarricense de Investigación y Enseñanza en Nutrición y Salud | Francisco Duarte, Hebleen Porras, Claudio Soto-Garita, Estela Cordero, Adriana Godínez & Melany Calderón |
| EPI_ISL_776764, EPI_ISL_776765 | Instituto Adolfo Lutz - Regional de Santo Andre | Instituto Adolfo Lutz, Interdiciplinary Procedures Center, Strategic Laboratory | Claudio Tavares Sacchi, Claudia Regina Gonçalves, Erica Valessa Ramos Gomes, Karoline Rodrigues Campos |
| EPI_ISL_779383, EPI_ISL_779391, EPI_ISL_779395 | Siloam Hospital Lippo Village | Mochtar Riady Institute for Nanotechnology-Universitas Pelita Harapan | Aksar C Lages, David Rustandi, Ivet M Suriapranata, Rinaldy Kusuma, Febi Andriani, Tri Shinta Kurniasih, Riska N Taufik, Young O Larasati, Rury M Wahyuni, Irawan Yusuf |
| EPI_ISL_780418 | CERBA LAB | CERBA LAB | Roquebert B; Costa JM; Hedbaut E; Trombert S; Lecorche E; Verdurme L; Malek Ramdane, Olivi M; Haïm-Boukobza S |
| EPI_ISL_788933, EPI_ISL_788936 | Centre de Recherche et de Formation en Infectiologie Guinée | TransVIHMI, IRD/INSERM/Monpellier University | Alpha Kabinet KEITA, Abdoul Karim SOUMAH, Abdoulaye TOURE, Moriba POVOGUI, Joel KOIVOGUI, Jean-louis MONEMOU, Mamadou Saliou SOW, Penda Malhado DIALLO, Mamadou Bhoye KEITA, Alimou CAMARA, Kaba KOUROUMA, Mandiou DIAKITE, Mamadou Saliou BAH, Sakoba KEITA, Bouna Yatassaye, Christelle BUTEL, Laetitia SERRANO, Ahidjo AYOUBA, Eric DELAPORTE, Martine PEETERS |
| EPI_ISL_788937 | Centre de Recherche et de Formation en Infectiologie Guinée | TransVIHMI, IRD/INSERM/Monpellier University | Alpha Kabinet KEITA, Abdoul Karim SOUMAH, Abdoulaye TOURE, Moriba POVOGUI, Joel KOIVOGUI, Jean-louis MONEMOU, Mamadou Saliou SOW, Penda Malhado DIALLO, Mamadou Bhoye KEITA, Alimou CAMARA, Kaba KOUROUMA, Mandiou DIAKITE, Mamadou Saliou BAH, Sakoba KEITA, Bouna |
| EPI_ISL_788938 | Centre de Recherche et de Formation en Infectiologie Guinée | TransVIHMI, IRD/INSERM/Monpellier University | Alpha Kabinet KEITA, Abdoul Karim SOUMAH, Abdoulaye TOURE, Moriba POVOGUI, Joel KOIVOGUI, Jean-louis MONEMOU, Mamadou Saliou SOW, Penda Malhado DIALLO, Mamadou Bhoye KEITA, Alimou CAMARA, Kaba KOUROUMA, Mandiou DIAKITE, Mamadou Saliou BAH, Sakoba KEITA, Bouna Yatassaye, Christelle BUTEL, Laetitia SERRANO, Ahidjo AYOUBA, Eric DELAPORTE, Martine PEETERS |
| EPI_ISL_788953, EPI_ISL_788954, EPI_ISL_788955, EPI_ISL_788956, EPI_ISL_788957, EPI_ISL_788958, EPI_ISL_788959, EPI_ISL_788960, EPI_ISL_788961, EPI_ISL_788970, EPI_ISL_788971, EPI_ISL_788972, EPI_ISL_788973, EPI_ISL_788974, EPI_ISL_788975, EPI_ISL_788976, EPI_ISL_788977, EPI_ISL_788978 | | | |
| see above | Ospedale "Di Venere" | Beaconlab (Bioinformatics, Evolution and Comparative Genomics lab), Dept of Biosciences, University on Milan | Iacobellis M, d'Avenia M, Piluscio R, Parisi A, Chiara M, Manzari C, Pesole G |
| EPI_ISL_792115, EPI_ISL_792116 | Instituto Adolfo Lutz - Regional de Taubate | Instituto Adolfo Lutz, Interdiciplinary Procedures Center, Strategic Laboratory | Claudio Tavares Sacchi, Claudia Regina Gonçalves, Erica Valessa Ramos Gomes, Karoline Rodrigues Campos |
| EPI_ISL_804949, EPI_ISL_804950, EPI_ISL_804951, EPI_ISL_804952, EPI_ISL_804953, EPI_ISL_804954, EPI_ISL_804955, EPI_ISL_804956, EPI_ISL_804957, EPI_ISL_804958, EPI_ISL_804959, EPI_ISL_804960, EPI_ISL_804961, EPI_ISL_804962, EPI_ISL_804965, EPI_ISL_804966, EPI_ISL_804967, EPI_ISL_804970, EPI_ISL_804977 | | | |
| see above | Hospital Comarcal de Melilla | Instituto de Salud Carlos III | Iglesias-Caballero, M. Molinero Calamita, M. González-Esguevillas, M. Camarero, S. Pozo, F. Casas, I. Jiménez, P. Jiménez, M. Zaballos, A. Monzón, S. Varona, S. Juliá, M. Cuesta, I, J. López |
| EPI_ISL_807157 | Mubarak Al-Kabeer Hospital | Virology Unit, Department of Microbiology, Faculty of Medicine | Dr. Nada Madi, Dr. Mohammed Sadeq, Mrs. Anfal Al-Adwani |
| EPI_ISL_811161, EPI_ISL_811162, EPI_ISL_811163, EPI_ISL_811164, EPI_ISL_811165, EPI_ISL_811166, EPI_ISL_811167, EPI_ISL_811168, EPI_ISL_811169, EPI_ISL_811170, EPI_ISL_811171, EPI_ISL_811172, EPI_ISL_811173, EPI_ISL_811174, EPI_ISL_811175, EPI_ISL_811176, EPI_ISL_811177, EPI_ISL_811178, EPI_ISL_811179, EPI_ISL_811180, EPI_ISL_811181, EPI_ISL_811182, EPI_ISL_811183, EPI_ISL_811184, EPI_ISL_811185, EPI_ISL_811186, EPI_ISL_811187, EPI_ISL_811188, EPI_ISL_811189, EPI_ISL_811190, EPI_ISL_811191, EPI_ISL_811192, EPI_ISL_811193 | | | |
| see above | Dharwad | CSIR Institute of Genomics and Integrative Biology | Dr. Shivarudrapp B Bhairappanavar, Rahul Bhoyar, Mohammed Imran, Mohit Divakar, Disha Sharma, Dr. Vijay A Yenagi, Dr. Suresh B Arakera, Dr. Amit Ugargol, Dr. Rgavendra B Nayak, Bani Jolly, Abhinav Jain, Paras Sehgal, Gyan Ranjan, Vinod Scaria, Sridhar Sivasubbu |
| EPI_ISL_826520 | University of Bari Biomedical Sciences and Human Oncology | University of Bari Biomedical Sciences and Human Oncology | Chironna M, Sallustio A., Loconsole D., Accogli M. |

| | | | |
|---|---|---|---|
| EPI_ISL_831474, EPI_ISL_831645, EPI_ISL_831646, EPI_ISL_831660, EPI_ISL_831678, EPI_ISL_831681, EPI_ISL_831683, EPI_ISL_831685, EPI_ISL_831688, EPI_ISL_831689, EPI_ISL_831892, EPI_ISL_831898, EPI_ISL_831913, EPI_ISL_831938 | | | |
| see above | Laboratório de Microbiologia Molecular - Universidade FEEVALE | Universidade Federal de Ciências da Saúde de Porto Alegre | Vinícius Bonetti Franceschi, Amanda de Menezes Mayer, Gabriel Dickin Caldana, Carla Andretta Moreira Neves, Patrícia Aline Gröhs Ferrareze, Gabriela Bettella Cybis, Ricardo Ariel Zimerman, Lívia Kmetzsch, Fernando Rosado Spilki, Claudia Elizabeth Thompson |
| EPI_ISL_833156 | Instituto Adolfo Lutz - Regional de Sorocaba | Instituto Adolfo Lutz, Interdiciplinary Procedures Center, Strategic Laboratory | Claudio Tavares Sacchi, Claudia Regina Gonçalves, Erica Valessa Ramos Gomes, Karoline Rodrigues Campos |
| EPI_ISL_833191, EPI_ISL_833192, EPI_ISL_833193, EPI_ISL_833194, EPI_ISL_833195, EPI_ISL_833197, EPI_ISL_833198, EPI_ISL_833199 | Hôpital Bichat Claude Bernard, Laboratoire de Virologie | IAME UMR1137 Inserm, Université de Paris, Hôpital Bichat | Antoine Bridier, Amélie Recoing, Quentin Le Hingrat, Lena Daniel, Siham Hamri, Gilles Collin, Alexandre Storto, Mélanie Bertine, Charlotte Charpentier, Nadhira Houhou-Fidouh, Diane Descamps, Benoit Visseaux |
| EPI_ISL_833391 | Klinik Apotek Dein, Jakarta, Indonesia | Biosafety Level-3 Laboratory, Indonesian Institute of Sciences (LIPI) | Andri Wardiana, Anggia Prasetyoputri, Isa Nuryana, Ade Andriani, Anik Budhi Dharmayanthi, Syam Budi Iryanto, Ahmad Fathoni, Herman Irawan, Dadang Supriatna, Ratih Asmana Ningrum |
| EPI_ISL_833392 | RSU Sayang Bunda, Makasar,Indonesia | Biosafety Level-3 Laboratory, Indonesian Institute of Sciences (LIPI) | Anggia Prasetyoputri, Isa Nuryana, Ade Andriani, Anik Budhi Dharmayanthi, Syam Budi Iryanto, Andri Wardiana, Ahmad Fathoni, Dini Nurdiani, Wahyuni, Ratih Asmana Ningrum |
| EPI_ISL_833493 | RS PMI Bogor, Indonesia | Biosafety Level-3 Laboratory, Indonesian Institute of Sciences (LIPI) | Isa Nuryana, Ade Andriani, Anik Budhi Dharmayanthi, Syam Budi Iryanto, Andri Wardiana, Anggia Prasetyoputri, Ahmad Fathoni, Indriawati, Nurulliawaty, Ratih Asmana Ningrum |
| EPI_ISL_833494 | RS PMI Bogor, Indonesia | Biosafety Level-3 Laboratory, Indonesian Institute of Sciences (LIPI) | Ade Andriani, Anik Budhi Dharmayanthi, Syam Budi Iryanto, Andri Wardiana, Anggia Prasetyoputri, Isa Nuryana, Ahmad Fathoni, Maritsa Nurfatwa, Anky Zannati, Ratih Asmana Ningrum |
| EPI_ISL_833495 | Klinik Apotek Dein, Jakarta, Indonesia | Biosafety Level-3 Laboratory, Indonesian Institute of Sciences (LIPI) | Ade Andriani, Anik Budhi Dharmayanthi, Syam Budi Iryanto, Andri Wardiana, Anggia Prasetyoputri, Isa Nuryana, Ahmad Fathoni, Hariyatun, Akhirta Atikana, Ratih Asmana Ningrum |
| EPI_ISL_833497 | Klinik Apotek Dein, Jakarta, Indonesia | Biosafety Level-3 Laboratory, Indonesian Institute of Sciences (LIPI) | Syam Budi Iryanto, Andri Wardiana, Anggia Prasetyoputri, Anik Budhi Dharmayanthi, Isa Nuryana, Ade Andriani, Ahmad Fathoni, Saiful anwar, Pamella Apriliana, Ratih Asmana Ningrum |
| EPI_ISL_833498 | Kartika Pulomas Hospital, Jakarta, Indonesia | Biosafety Level-3 Laboratory, Indonesian Institute of Sciences (LIPI) | Anggia Prasetyoputri, Isa Nuryana, Ade Andriani, Anik Budhi Dharmayanthi, Syam Budi Iryanto, Andri Wardiana, Ahmad Fathoni, Winda Tasia, Pinka Alisa Diena Shafa, Ratih Asmana Ningrum |
| EPI_ISL_833499 | RS Qadr, Tangerang, Indonesia | Biosafety Level-3 Laboratory, Indonesian Institute of Sciences (LIPI) | Isa Nuryana, Ade Andriani, Ahmad Fathoni, Anik Budhi Dharmayanthi, Syam Budi Iryanto, Andri Wardiana, Anggia Prasetyoputri, Ratna Dwi Ramadani, Sugiyono Saputra, Ratih Asmana Ningrum |
| EPI_ISL_833500 | Klinik Apotek Dein, Jakarta, Indonesia | Biosafety Level-3 Laboratory, Indonesian Institute of Sciences (LIPI) | Anik Budhi Dharmayanthi, Syam Budi Iryanto, Andri Wardiana, Anggia Prasetyoputri, Isa Nuryana, Ade Andriani, Ahmad Fathoni, Ratih Asmana Ningrum |
| EPI_ISL_833503 | RS MMC, Jakarta, Indonesia | Biosafety Level-3 Laboratory, Indonesian Institute of Sciences (LIPI) | Anggia Prasetyoputri, Isa Nuryana, Anik Budhi Dharmayanthi, Syam Budi Iryanto, Andri Wardiana, Ade Andriani, Ahmad Fathoni, Idris, Ruby Setiawan, Ratih Asmana Ningrum |
| EPI_ISL_833504 | RS MMC, Jakarta, Indonesia | Biosafety Level-3 Laboratory, Indonesian Institute of Sciences (LIPI) | Isa Nuryana, Anik Budhi Dharmayanthi, Syam Budi Iryanto, Andri Wardiana, Anggia Prasetyoputri, Ade Andriani, Ahmad Fathoni, Ahmad Randy, Listiana Oktavia, Ratih Asmana Ningrum |
| EPI_ISL_833506 | RS PMI, Bogor, Indonesia | Biosafety Level-3 Laboratory, Indonesian Institute of Sciences (LIPI) | Isa Nuryana, Ade Andriani, Ahmad Fathoni, Anik Budhi Dharmayanthi, Syam Budi Iryanto, Andri Wardiana, Anggia Prasetyoputri, Satrio Haryo Benowo, Gilang Akbariani, Ratih Asmana Ningrum |
| EPI_ISL_833571 | RS Qadr, Tangerang, Indonesia | Biosafety Level-3 Laboratory, Indonesian Institute of Sciences (LIPI) | Ade Andriani, Ahmad Fathoni, Anik Budhi Dharmayanthi, Syam Budi Iryanto, Andri Wardiana, Anggia Prasetyoputri, Isa Nuryana, Erwin Fajar Hasrianda, Ryan Haryo Setiawan, Ratih Asmana Ningrum |
| EPI_ISL_837564, EPI_ISL_837569 | Centro Nacional de Enfermedades Tropicales (CENETROP) | Laboratory of Respiratory Viruses and Measles, Oswaldo Cruz Institute, FIOCRUZ | Paola Resende, Roxana Loayza, Cinthia Avila, Luciana Appolinario, Fernando Motta, Anna Carolina Paixao, Ana Carolina Mendonca, Marilda Siqueira on behalf of the Fiocruz COVID-19 Genomic Surveillance Network |
| EPI_ISL_837602, EPI_ISL_837603, EPI_ISL_837607, EPI_ISL_837699, EPI_ISL_837700, EPI_ISL_837701, EPI_ISL_837702, EPI_ISL_837703, EPI_ISL_837704, EPI_ISL_837705, EPI_ISL_837706, EPI_ISL_837707, EPI_ISL_837708, EPI_ISL_837709, EPI_ISL_837710, EPI_ISL_837711, EPI_ISL_837712, EPI_ISL_837713, EPI_ISL_837715, EPI_ISL_837716, EPI_ISL_837717, EPI_ISL_837718, EPI_ISL_837719, EPI_ISL_837720, EPI_ISL_837721, EPI_ISL_837722, EPI_ISL_837723, EPI_ISL_837724, EPI_ISL_837725, EPI_ISL_837726, EPI_ISL_837727, EPI_ISL_837728, EPI_ISL_837729, EPI_ISL_837730, EPI_ISL_837731, EPI_ISL_837732, EPI_ISL_837733, EPI_ISL_837734, EPI_ISL_837735, EPI_ISL_837736, EPI_ISL_837737, EPI_ISL_837738, EPI_ISL_837739, EPI_ISL_837740, EPI_ISL_837741, EPI_ISL_837742, EPI_ISL_837743, EPI_ISL_837745, EPI_ISL_837746, EPI_ISL_837747, EPI_ISL_837748, EPI_ISL_837749, EPI_ISL_837750, EPI_ISL_837751, EPI_ISL_837753, EPI_ISL_837754, EPI_ISL_837755, EPI_ISL_837756, EPI_ISL_837757, EPI_ISL_837758, EPI_ISL_837761, EPI_ISL_837762, EPI_ISL_837763, EPI_ISL_837764, EPI_ISL_837765, EPI_ISL_837766, EPI_ISL_837767, EPI_ISL_837768, EPI_ISL_837769, EPI_ISL_837770, EPI_ISL_837771, EPI_ISL_837772, EPI_ISL_837773, EPI_ISL_837774, EPI_ISL_837775, EPI_ISL_837776, EPI_ISL_837777, EPI_ISL_837778, EPI_ISL_837780, EPI_ISL_837781, EPI_ISL_837782, EPI_ISL_837783, EPI_ISL_837784, EPI_ISL_837785, EPI_ISL_837786, EPI_ISL_837787, EPI_ISL_837789, EPI_ISL_837790, EPI_ISL_837791, EPI_ISL_837793, EPI_ISL_837794, EPI_ISL_837795, EPI_ISL_837796, EPI_ISL_837797, EPI_ISL_837798, EPI_ISL_837799, EPI_ISL_837800, EPI_ISL_837802, EPI_ISL_837803, EPI_ISL_837804, EPI_ISL_837805, EPI_ISL_837806, EPI_ISL_837807, EPI_ISL_837808, EPI_ISL_837809, EPI_ISL_843191, EPI_ISL_843192 | | | |
| see above | Instituto Nacional de Enfermedades Respiratorias (INER) | Instituto Nacional de Enfermedades Respiratorias (INER) | Celia Boukadida, Margarita Matías-Florentino, Alma Rincón-Rubio, Hector Esteban Paz-Juárez, Olivia Briceño, Edgar Sevilla-Reyes, Fidencio Mejía-Nepomuceno, Mario Мújica-Sánchez, Eduardo Becerril-Vargas, José Arturo Martínez-Orozco, Alejandra Hernández-Terán, Jorge Salas-Hernández, Santiago Ávila-Ríos, Joel Armando Vázquez-Pérez |
| EPI_ISL_848574 | Evandro Chagas Institute | Evandro Chagas Institute | Santos, M.C.; Silva, A.M.; Junior, W.D.C.; Barbagelata, L.S.; Ferreira, J.A.; Sousa, E.M.A.; da Silva, P.S.; Pinheiro, K.C.; L.C.; Sousa Junior, E.C. |
| EPI_ISL_849650, EPI_ISL_849652 | Servizio di igiene epidemiologia e sanità pubblica (SIESP)-Chieti | Istituto Zooprofilattico Sperimentale dell'Abruzzo e Molise "G.Caporale" | Lorusso A, Marcacci M, Di Domenico M, Curini V, Ancora M, Cammà C, Rinaldi A, Mangone I, Di Pasquale A, Puglia I, Savini G. |
| EPI_ISL_849653 | Servizio di igiene e sanità pubblica (SIESP)-Teramo | Istituto Zooprofilattico Sperimentale dell'Abruzzo e Molise "G.Caporale" | Lorusso A, Marcacci M, Di Domenico M, Curini V, Ancora M, Cammà C, Rinaldi A, Mangone I, Di Pasquale A, Puglia I, Savini G. |
| EPI_ISL_849654, EPI_ISL_849656, EPI_ISL_849657, EPI_ISL_849659, EPI_ISL_849661 | Servizio Igiene Epidemiologia e Sanità Pubblica (SIESP)-L'Aquila | Istituto Zooprofilattico Sperimentale dell'Abruzzo e Molise "G.Caporale" | Lorusso A, Marcacci M, Di Domenico M, Curini V, Ancora M, Cammà C, Rinaldi A, Mangone I, Di Pasquale A, Puglia I, Savini G. |
| EPI_ISL_849662, EPI_ISL_849663 | Servizio di igiene epidemiologia e sanità pubblica (SIESP)-Chieti | Istituto Zooprofilattico Sperimentale dell'Abruzzo e Molise "G.Caporale" | Lorusso A, Marcacci M, Di Domenico M, Curini V, Ancora M, Cammà C, Rinaldi A, Mangone I, Di Pasquale A, Puglia I, Savini G. |
| EPI_ISL_850507 | Gandhi Medical College and Hospital | VRDL-Gandhi Medical College | Nagamani Kammili, Rajeshwar Rao, Madhavi Latha Manolla, Winnie Thomas, Shailaja VV, Sudhamadhuri Devara, Vanisree Rajoli, Archana GJ,Sushma Rajyalakshmi Gudiseva, Sunitha Pakalapati, Manisha Rani, Amrithesh Kumar, Raja Rao Mesepogu, Vinay Shekar Reddy, Thrilok Chander Bingi, Sofia Banu, Divya Tej Sowpati |
| EPI_ISL_850509 | Gandhi Medical College and Hospital | VRDL-Gandhi Medical College | Nagamani Kammili, Rajeshwar Rao, Winnie Thomas, Madhavi Latha Manolla, Shailaja VV, Sudhamadhuri Devara, Vanisree Rajoli, Archana GJ, Sushma Rajyalakshmi Gudiseva, Sunitha Pakalapati, Manisha Rani, Amrithesh Kumar, Raja Rao Mesepogu, Vinay Shekar Reddy, Thrilok Chander Bingi, Sofia Banu, Divya Tej Sowpati |
| EPI_ISL_850510 | Gandhi Medical College and Hospital | MDRU-DHR,Gandhi Medical College | Nagamani Kammili, Winnie Thomas, Madhavi Latha Manolla, Shailaja VV, Sudhamadhuri Devara, Vanisree Rajoli, Archana GJ, Sushma Rajyalakshmi Gudiseva, Sunitha Pakalapati, Manisha Rani, Amrithesh Kumar, Raja Rao Mesepogu, Vinay Shekar Reddy, Thrilok Chander Bingi, Sofia Banu, Divya Tej Sowpati |
| EPI_ISL_850514 | Gandhi Medical College and Hospital | MDRU-DHR,Gandhi Medical College | Nagamani Kammili, Madhavi Latha Manolla, Winnie Thomas, Shailaja VV, Sudhamadhuri Devara, Vanisree Rajoli, Archana GJ,Sushma Rajyalakshmi Gudiseva, Sunitha Pakalapati, Manisha Rani, Amrithesh Kumar, Raja Rao Mesepogu, Vinay Shekar Reddy, Thrilok Chander Bingi, Sofia Banu, Divya Tej Sowpati |
| EPI_ISL_851050 | Gandhi Medical College and Hospital | CSIR-Centre for Cellular and Molecular Biology | Nagamani Kammili, Winnie Thomas, Madhavi Latha Manolla, Shailaja VV, Sudhamadhuri Devara, Vanisree Rajoli, Archana GJ, Sushma Rajyalakshmi Gudiseva, Sunitha Pakalapati, Manisha Rani, Amrithesh Kumar, Raja Rao Mesepogu, Vinay Shekar Reddy, Thrilok Chander Bingi, Sofia Banu, Divya Tej Sowpati |
| EPI_ISL_851051 | Gandhi Medical College and Hospital | CSIR-Centre for Cellular and Molecular Biology | Nagamani Kammili, Madhavi Latha Manolla, Winnie Thomas, Shailaja VV, Sudhamadhuri Devara, Vanisree Rajoli, Archana GJ,Sushma Rajyalakshmi Gudiseva, Sunitha Pakalapati, Manisha Rani, Amrithesh Kumar, Raja Rao Mesepogu, Vinay Shekar Reddy, Thrilok Chander Bingi, Sofia Banu, Divya Tej |

| | | | |
|---|---|---|---|
| | | | Sowpati |
| EPI_ISL_852580, EPI_ISL_852581, EPI_ISL_852582, EPI_ISL_852586, EPI_ISL_852588, EPI_ISL_852591, EPI_ISL_852592 | Max von Pettenkofer Institute, Virology, National Reference Center for Retroviruses, LMU München | Laboratory for Functional Genome Analysis, Dept. Genomics, Gene Center of the LMU Munich | Max Muenchhoff, Stefan Krebs, Alexander Graf, Oliver Keppler, Helmut Blum |
| EPI_ISL_853819 | Department of Pathology, Landeskrankenhaus Graz II, Medical University Graz | Bergthaler laboratory, CeMM Research Center for Molecular Medicine of the Austrian Academy of Sciences | Lukas Endler, Alexandra Popa, Benedikt Agerer, Jakob-Wendelin Genger, Alexander Lercher, Anna Schedl, Thomas Penz, Michael Schuster, Jan Laine, Martin Senekowitsch, Christoph Bock, Andreas Bergthaler |
| EPI_ISL_861646, EPI_ISL_861647 | Hospital Santa Marcelina Sao Paulo | Instituto Adolfo Lutz, Interdiciplinary Procedures Center, Strategic Laboratory | Claudio Tavares Sacchi, Claudia Regina Gonçalves, Erica Valessa Ramos Gomes, Karoline Rodrigues Campos |
| EPI_ISL_861648 | Hospital e Pronto Socorro Portinari | Instituto Adolfo Lutz, Interdiciplinary Procedures Center, Strategic Laboratory | Claudio Tavares Sacchi, Claudia Regina Gonçalves, Erica Valessa Ramos Gomes, Karoline Rodrigues Campos |
| EPI_ISL_861649 | Hospital Renascença Campinas | Instituto Adolfo Lutz, Interdiciplinary Procedures Center, Strategic Laboratory | Claudio Tavares Sacchi, Claudia Regina Gonçalves, Erica Valessa Ramos Gomes, Karoline Rodrigues Campos |
| EPI_ISL_861657 | Hospital e Maternidade Sino Brasileiro | Instituto Adolfo Lutz, Interdiciplinary Procedures Center, Strategic Laboratory | Claudio Tavares Sacchi, Claudia Regina Gonçalves, Erica Valessa Ramos Gomes, Karoline Rodrigues Campos |
| EPI_ISL_861658 | Hospital Municipal Antônio Giglio | Instituto Adolfo Lutz, Interdiciplinary Procedures Center, Strategic Laboratory | Claudio Tavares Sacchi, Claudia Regina Gonçalves, Erica Valessa Ramos Gomes, Karoline Rodrigues Campos |
| EPI_ISL_861660 | PS e Maternidade Nair Fonseca Leitao Arantes | Instituto Adolfo Lutz, Interdiciplinary Procedures Center, Strategic Laboratory | Claudio Tavares Sacchi, Claudia Regina Gonçalves, Erica Valessa Ramos Gomes, Karoline Rodrigues Campos |
| EPI_ISL_861662 | CS I Tacito Leite de Carvalho e Silva | Instituto Adolfo Lutz, Interdiciplinary Procedures Center, Strategic Laboratory | Claudio Tavares Sacchi, Claudia Regina Gonçalves, Erica Valessa Ramos Gomes, Karoline Rodrigues Campos |
| EPI_ISL_861664 | Instituto Adolfo Lutz - Regional de Campinas | Instituto Adolfo Lutz, Interdiciplinary Procedures Center, Strategic Laboratory | Claudio Tavares Sacchi, Claudia Regina Gonçalves, Erica Valessa Ramos Gomes, Karoline Rodrigues Campos |
| EPI_ISL_861665 | Instituto Adolfo Lutz - Regional de Taubate | Instituto Adolfo Lutz, Interdiciplinary Procedures Center, Strategic Laboratory | Claudio Tavares Sacchi, Claudia Regina Gonçalves, Erica Valessa Ramos Gomes, Karoline Rodrigues Campos |
| EPI_ISL_861666 | PSF Dr. Antonio Pires de Almeida | Instituto Adolfo Lutz, Interdiciplinary Procedures Center, Strategic Laboratory | Claudio Tavares Sacchi, Claudia Regina Gonçalves, Erica Valessa Ramos Gomes, Karoline Rodrigues Campos |
| EPI_ISL_861667 | Instituto Adolfo Lutz - Regional de Rio Claro | Instituto Adolfo Lutz, Interdiciplinary Procedures Center, Strategic Laboratory | Claudio Tavares Sacchi, Claudia Regina Gonçalves, Erica Valessa Ramos Gomes, Karoline Rodrigues Campos |
| EPI_ISL_862594 | Hospital Comarcal de Melilla | Instituto de Salud Carlos III | Iglesias-Caballero, M. Camarero, S. Molinero Calamita, M. González-Esguevillas, M. Pozo, F. Casas, I. Jiménez, P. Jiménez, M. Zaballos, A. Monzón, S. Varona, S. Juliá, M. Cuesta, I. López, J. |
| EPI_ISL_871790 | Multidisciplinary Research Unit, DHR-ICMR, Institute of Medical Sciences, Banaras Hindu University | Multidisciplinary Research Unit, DHR-ICMR, Institute of Medical Sciences, Banaras Hindu University | Royana Singh, Priyoneel Basu, Ashish, Nitish Kumar Singh, Abhay Kumar Yadav, Manpreet Kaur, Arup Acharjee, Deepa Devadas, Chetan Sahni, Gulshan Kumar, Tribhuwan Mohan Mohapatra, Richa Arya, Prashant Singh, Jay Prakash Maurya, Surendra Pratap Mishra |
| EPI_ISL_871984 | Hospital Universitario Severo Ochoa | Instituto de Salud Carlos III | Iglesias-Caballero, M. Camarero, S. Molinero Calamita, M. González-Esguevillas, M. Pozo, F. Casas, I. Jiménez, P. Jiménez, M. Zaballos, A. Monzón, S. Varona, S. Juliá, M. Cuesta, I. García, M.L. |
| EPI_ISL_871985, EPI_ISL_871986, EPI_ISL_871987, EPI_ISL_871988, EPI_ISL_871989, EPI_ISL_871990 | Hospital Clínico San Carlos | Instituto de Salud Carlos III | Iglesias-Caballero, M. Camarero, S. Molinero Calamita, M. González-Esguevillas, M. Pozo, F. Casas, I. Jiménez, P. Jiménez, M. Zaballos, A. Monzón, S. Varona, S. Juliá, M. Cuesta, I. Rodríguez, I. |
| EPI_ISL_871991 | Hospital Universitario Severo Ochoa | Instituto de Salud Carlos III | Iglesias-Caballero, M. Camarero, S. Molinero Calamita, M. González-Esguevillas, M. Pozo, F. Casas, I. Jiménez, P. Jiménez, M. Zaballos, A. Monzón, S. Varona, S. Juliá, M. Cuesta, I. García, M.L. |
| EPI_ISL_871992, EPI_ISL_871993 | Hospital Clínico San Carlos | Instituto de Salud Carlos III | Iglesias-Caballero, M. Camarero, S. Molinero Calamita, M. González-Esguevillas, M. Pozo, F. Casas, I. Jiménez, P. Jiménez, M. Zaballos, A. Monzón, S. Varona, S. Juliá, M. Cuesta, I. Rodríguez, I. |
| EPI_ISL_872009, EPI_ISL_872010, EPI_ISL_872011, EPI_ISL_872012, EPI_ISL_872014, EPI_ISL_872016, EPI_ISL_872017, EPI_ISL_872018, EPI_ISL_872019, EPI_ISL_872020, EPI_ISL_872021, EPI_ISL_872022, EPI_ISL_872023, EPI_ISL_872024, EPI_ISL_872025, EPI_ISL_872026 | | | |
| see above | Ospedale "Di Venere" | Beaconlab (Bioinformatics, Evolution and Comparative Genomics lab), Dept of Biosciences, University on Milan | Iacobellis M, d'Avenia M, Piluscio R, Parisi A, Chiara M, Manzari C, Pesole G |
| EPI_ISL_872628, EPI_ISL_872629, EPI_ISL_872630, EPI_ISL_872631, EPI_ISL_872632, EPI_ISL_872633, EPI_ISL_872634, EPI_ISL_872635, EPI_ISL_872636, EPI_ISL_872637, EPI_ISL_872638, EPI_ISL_872639, EPI_ISL_872640 | | | |
| see above | Pathogenic Microorganisms Variability Laboratory | Pathogenic Microorganisms Variability Laboratory | Alexey Shchetinin, Olesya Venchakova, Maria Nikiforova, Andrei Siniavin, Nadezhda Kuznetsova, Elena Shidlovskaya, Elizaveta Divisenko, Kirill Krasnoslobotsev, Evgeniya Mukasheva, Anna Ignatieva, Svetlana Trushakova, Andrey Pochtovyy, Valeria Bacalin, Evgeny Usachev, Olga Burgasova, Ludmila Kolobukhina, Svetlana Smetanina, Elena Burtseva, Denis Logunov, Vladimir Gushchin, Alexander Gintsburg |
| EPI_ISL_872641, EPI_ISL_872642, EPI_ISL_875515 | Influenza etiology and epidemiology laboratory | Pathogenic Microorganisms Variability Laboratory | Alexey Shchetinin, Olesya Venchakova, Maria Nikiforova, Andrei Siniavin, Nadezhda Kuznetsova, Elena Shidlovskaya, Elizaveta Divisenko, Kirill Krasnoslobotsev, Evgeniya Mukasheva, Anna Ignatieva, Svetlana Trushakova, Andrey Pochtovyy, Valeria Bacalin, Evgeny Usachev, Olga Burgasova, Ludmila Kolobukhina, Svetlana Smetanina, Elena Burtseva, Denis Logunov, Vladimir Gushchin, Alexander Gintsburg |
| EPI_ISL_882657 | LACEN do Estado do Piaui, Dr. Costa Alvarenga | Instituto Adolfo Lutz, Interdiciplinary Procedures Center, Strategic Laboratory | Claudio Tavares Sacchi, Claudia Regina Gonçalves, Erica Valessa Ramos Gomes, Karoline Rodrigues Campos |
| EPI_ISL_889357, EPI_ISL_889358, EPI_ISL_889359, EPI_ISL_889360, EPI_ISL_889362, EPI_ISL_889363, EPI_ISL_889364, EPI_ISL_889365 | Motol University Hospital | Institute of Applied Biotechnologies a.s. | Petr Klempt, Ondej Brzo, Martin Kašný, Kateina Kvapilová, Pavel Devínek, Petr Kvapil |
| EPI_ISL_895834 | Molecular biology division, Institute of Clinical Biochemistry and Diagnostics, Charles University, Faculty of Medicine in Hradec Králové and University Hospital Hradec Králové | Molecular biology division, Institute of Clinical Biochemistry and Diagnostics, Charles University, Faculty of Medicine in Hradec Králové and University Hospital Hradec Králové | Helena Kovaíková, Petr Brož, Ivana Baranová, Kateina Hrochová, Tereza Baková, Jitka Novotná, Kateina Pehlíková, Vladimír Palika. Cooperation project with BioVendor-R&D and bioinformatics company BIOXSYS s r.o. |
| EPI_ISL_900691, EPI_ISL_900692, EPI_ISL_900693, EPI_ISL_900694, EPI_ISL_900695, EPI_ISL_900696, EPI_ISL_900697, EPI_ISL_900698, EPI_ISL_900699, EPI_ISL_900700, EPI_ISL_900701, EPI_ISL_900702, EPI_ISL_900704, EPI_ISL_900705, EPI_ISL_900706, EPI_ISL_900707, EPI_ISL_900708, EPI_ISL_900709, EPI_ISL_900710, EPI_ISL_900711, EPI_ISL_900712, EPI_ISL_900713, EPI_ISL_900714, EPI_ISL_900715, EPI_ISL_900716, EPI_ISL_900717, EPI_ISL_900718, EPI_ISL_900719, EPI_ISL_900720, EPI_ISL_900721, EPI_ISL_900722, EPI_ISL_900723, EPI_ISL_900724, EPI_ISL_900725, EPI_ISL_900727, EPI_ISL_900728, EPI_ISL_900729, EPI_ISL_900730, EPI_ISL_900731, EPI_ISL_900732, EPI_ISL_900733, EPI_ISL_900734, EPI_ISL_900735, EPI_ISL_900736, EPI_ISL_900737, EPI_ISL_900738, EPI_ISL_900740, EPI_ISL_900741 | | | |
| see above | Bozeman Health Deaconess Hospital | Wiedenheft lab, Montana State University | Artem Nemudryi, Anna Nemudraia, Tanner Wiegand, Joseph Nichols, Deann T. Snyder, Jodi F. Hedges, Calvin Cicha, Helen Lee, Karl K. Vanderwood, Diane Bimczok, Mark A. Jutila and Blake Wiedenheft |
| EPI_ISL_902917 | Tanjungpura University Hospital | Tanjungpura University Hospital | Andriani; Mahyarudin; Novianry V; Liana DF; Shofiyah SS Astuti P; Kahtan MI; Rialita A; Putri EA; Windarti W; Sastriawan W |
| EPI_ISL_903339, EPI_ISL_903340, EPI_ISL_903341, EPI_ISL_903342, EPI_ISL_903343, EPI_ISL_903344, EPI_ISL_903345, EPI_ISL_903346 | Bozeman Health Deaconess Hospital | Wiedenheft lab, Montana State University | Artem Nemudryi, Anna Nemudraia, Tanner Wiegand, Joseph Nichols, Deann T. Snyder, Jodi F. Hedges, Calvin Cicha, Helen Lee, Karl K. Vanderwood, Diane Bimczok, Mark A. Jutila and Blake Wiedenheft |
| EPI_ISL_906060 | Tilia Laboratories s.r.o. | Tilia Laboratories s.r.o. | Sona Pekova, MD, PhD. |

| | | | |
|---|---|---|---|
| EPI_ISL_913914 | Instituto de Diagnostico y Referencia Epidemiologicos INDRE_RNLSP | Instituto de Diagnostico y Referencia Epidemiologicos (INDRE) | Claudia Wong-Arambula, Abril Rodriguez-Maldonado, Fabiola Garces-Ayala, Adnan Araiza-Rodriguez, David Fragoso-Fonseca, Sergio Rangel-Guerrero, Mayra Jimenez-Morales, Nancy Munoz-Hernandez, Natividad Cruz-Ortiz, Tatiana Nunez-Garcia, Gisela Barrera-Badillo, Lucia Hernandez-Rivas, Irma Lopez-Martinez, Ernesto Ramirez-Gonzalez. |
| EPI_ISL_930855, EPI_ISL_930857, EPI_ISL_930858 | Central Laboratory of Public Health of Rio Grande do Sul (Lacen-RS) | State Center for Health Surveillance of the Health Department of the State of Rio Grande do Sul (CEVS/SES-RS) | Barcellos R, Campos A, Dornelles C, Godinho F, Gonzalez A, Gregianini T, Molina C, Salvato R, Schaurich A, |
| EPI_ISL_940608 | Laboratório Sao Lucas | Instituto Adolfo Lutz, Interdicíplinary Procedures Center, Strategic Laboratory | Claudio Tavares Sacchi, Claudia Regina Gonçalves, Erica Valessa Ramos Gomes, Karoline Rodrigues Campos |
| EPI_ISL_940756, EPI_ISL_940895, EPI_ISL_940896, EPI_ISL_940897, EPI_ISL_940898, EPI_ISL_940899, EPI_ISL_940900, EPI_ISL_940902, EPI_ISL_940903, EPI_ISL_940904, EPI_ISL_940906, EPI_ISL_940907, EPI_ISL_940908, EPI_ISL_940909, EPI_ISL_940910, EPI_ISL_940911, EPI_ISL_940912, EPI_ISL_940913, EPI_ISL_940914, EPI_ISL_940916, EPI_ISL_940918, EPI_ISL_940919, EPI_ISL_940921, EPI_ISL_940922, EPI_ISL_940923, EPI_ISL_940924, EPI_ISL_940925, EPI_ISL_940927, EPI_ISL_940928, EPI_ISL_940929, EPI_ISL_940930, EPI_ISL_940931, EPI_ISL_940932, EPI_ISL_940933, EPI_ISL_940934, EPI_ISL_940935, EPI_ISL_940950, EPI_ISL_940951, EPI_ISL_940952, EPI_ISL_940953, EPI_ISL_940955, EPI_ISL_940956, EPI_ISL_940957, EPI_ISL_940958, EPI_ISL_940959 | | | |
| see above | Centers for Disease Control and Prevention, Dengue Branch | Centers for Disease Control and Prevention, Dengue Branch | Gilberto A. Santiago, Glenda Gonzalez, Betzabel Flores, Keyla Charriez, Gabriela Paz-Bailey, Jorge L. Munoz-Jordan |
| EPI_ISL_941986, EPI_ISL_941987, EPI_ISL_941988, EPI_ISL_941989, EPI_ISL_941990, EPI_ISL_941991, EPI_ISL_941992, EPI_ISL_941993, EPI_ISL_941994 | Instituto Nacional de Salud, Bogotá, Colombia | Centro de Investigaciones en Microbiología y Biotecnología-UR (CIMBIUR), Facultad de Ciencias Naturales, Universidad del Rosario, Bogotá, Colombia Instituto Nacional de Salud, Bogotá, Colombia Icahn School of Medicine at Mount Sinai, New York, USA | Luz Helena Patiño, Marina Muñoz, Nathalia Ballesteros, Carolina Hernández, Carolina Flórez, Sergio Gomez, Adriana van de Guchte, Zenab Khan, Jayeeta Dutta, Hala Alejel Alshammary, Ana S. Gonzalez-Reiche, Matthew M. Hernandez, Emilia Mia Sordillo, Viviana Simon, Harm van Bakel, Alberto Paniz-Mondolfi, Juan David Ramírez |
| EPI_ISL_941995, EPI_ISL_941996, EPI_ISL_941997, EPI_ISL_941998 | Centro de Investigaciones en Microbiología y Biotecnología-UR (CIMBIUR), Facultad de Ciencias Naturales, Universidad del Rosario, Bogotá, Colombia | Centro de Investigaciones en Microbiología y Biotecnología-UR (CIMBIUR), Facultad de Ciencias Naturales, Universidad del Rosario, Bogotá, Colombia Instituto Nacional de Salud, Bogotá, Colombia Icahn School of Medicine at Mount Sinai, New York, USA | Luz Helena Patiño, Marina Muñoz, Nathalia Ballesteros, Carolina Hernández, Carolina Flórez, Sergio Gomez, Adriana van de Guchte, Zenab Khan, Jayeeta Dutta, Hala Alejel Alshammary, Ana S. Gonzalez-Reiche, Matthew M. Hernandez, Emilia Mia Sordillo, Viviana Simon, Harm van Bakel, Alberto Paniz-Mondolfi, Juan David Ramírez |
| EPI_ISL_942003 | Instituto Nacional de Salud, Bogotá, Colombia | Centro de Investigaciones en Microbiología y Biotecnología-UR (CIMBIUR), Facultad de Ciencias Naturales, Universidad del Rosario, Bogotá, Colombia Instituto Nacional de Salud, Bogotá, Colombia Icahn School of Medicine at Mount Sinai, New York, USA | Luz Helena Patiño, Marina Muñoz, Nathalia Ballesteros, Carolina Hernández, Carolina Flórez, Sergio Gomez, Adriana van de Guchte, Zenab Khan, Jayeeta Dutta, Hala Alejel Alshammary, Ana S. Gonzalez-Reiche, Matthew M. Hernandez, Emilia Mia Sordillo, Viviana Simon, Harm van Bakel, Alberto Paniz-Mondolfi, Juan David Ramírez |
| EPI_ISL_942004 | Centro de Investigaciones en Microbiología y Biotecnología-UR (CIMBIUR), Facultad de Ciencias Naturales, Universidad del Rosario, Bogotá, Colombia | Centro de Investigaciones en Microbiología y Biotecnología-UR (CIMBIUR), Facultad de Ciencias Naturales, Universidad del Rosario, Bogotá, Colombia Instituto Nacional de Salud, Bogotá, Colombia Icahn School of Medicine at Mount Sinai, New York, USA | Luz Helena Patiño, Marina Muñoz, Nathalia Ballesteros, Carolina Hernández, Carolina Flórez, Sergio Gomez, Adriana van de Guchte, Zenab Khan, Jayeeta Dutta, Hala Alejel Alshammary, Ana S. Gonzalez-Reiche, Matthew M. Hernandez, Emilia Mia Sordillo, Viviana Simon, Harm van Bakel, Alberto Paniz-Mondolfi, Juan David Ramírez |
| EPI_ISL_942375, EPI_ISL_942897, EPI_ISL_942898 | Central Laboratory of Public Health of Rio Grande do Sul (Lacen-RS) | State Center for Health Surveillance of the Health Department of the State of Rio Grande do Sul (CEVS/SES-RS) | Barcellos R, Campos A, Crescente L, Da Silva A, Dornelles C, Fonseca V, Garay L, Godinho F, Gonzalez A, Gregianini T, Molina C, Salvato R, Schaurich A |
| EPI_ISL_942929 | Instituto de Diagnostico y Referencia Epidemiologicos INDRE_RNLSP | Instituto de Diagnostico y Referencia Epidemiologicos (INDRE) | Claudia Wong-Arambula, Abril Rodriguez-Maldonado, Fabiola Garces-Ayala, Natividad Cruz-Ortiz, Tatiana Nunez-Garcia, Gisela Barrera-Badillo, Lucia Hernandez-Rivas, Irma Lopez-Martinez, Ernesto Ramirez-Gonzalez. |
| EPI_ISL_942930, EPI_ISL_942931 | Central Laboratory of Public Health of Rio Grande do Sul (Lacen-RS) | State Center for Health Surveillance of the Health Department of the State of Rio Grande do Sul (CEVS/SES-RS) | Barcellos R, Campos A, Crescente L, Da Silva A, Dornelles C, Fonseca V, Garay L, Godinho F, Gonzalez A, Gregianini T, Molina C, Salvato R, Schaurich A |
| EPI_ISL_943574, EPI_ISL_943575, EPI_ISL_943576, EPI_ISL_943577, EPI_ISL_943581, EPI_ISL_943582, EPI_ISL_943584, EPI_ISL_943586, EPI_ISL_943587, EPI_ISL_943588, EPI_ISL_943590 | | | |
| see above | Central Laboratory of Public Health of Rio Grande do Sul (Lacen-RS) | State Center for Health Surveillance of the Health Department of the State of Rio Grande do Sul (CEVS/SES-RS) | Aline Campos, Amanda da Silva, Anelise Schaurich, Claudia Dornelles, Cynthia Molina, Fernanda Godinho, Lara Crescente, Leticia Garay, Regina Barcellos, Richard Salvato, Tatiana Gregianini, Vagner Fonseca |
| EPI_ISL_943988 | LACEN do Estado de Goias | Instituto Adolfo Lutz, Interdicíplinary Procedures Center, Strategic Laboratory | Claudio Tavares Sacchi, Claudia Regina Gonçalves, Erica Valessa Ramos Gomes, Karoline Rodrigues Campos |
| EPI_ISL_954810 | Hospital General Universitario de Ciudad Real | Instituto de Salud Carlos III | Iglesias-Caballero, M. Camarero, S. Sandonís,V. Vázquez, S. Pozo, F. Casas, I. Jiménez, P. Zaballos, A. Monzón, S. Varona, S. Cuesta, I. Illescas, S. |
| EPI_ISL_955142 | University of Sarajevo, Veterinary Faculty, Laboratory for Molecular Diagnostic and Research Laboratory | University of Sarajevo, Veterinary Faculty, Laboratory for Molecular Diagnostic and Research Laboratory | Goleti T., Goleti Š., Softi A., Ali-Šeho A., Šabi E., Jaži A., Nicevi M., Hodži A., Terzi I. |
| EPI_ISL_955144 | University of Sarajevo, Veterinary Faculty, Laboratory for Molecular Diagnostic and Research Laboratory | University of Sarajevo, Veterinary Faculty, Laboratory for Molecular Diagnostic and Research Laboratory | Goleti T., Goleti Š., Softi A., Ali-Šeho A., Hodži A., Šabi E., Terzi I. Jaži A., Nicevi M. |
| EPI_ISL_956271 | Adi Husada Undaan Hospital | Institute of Tropical Disease, Universitas Airlangga | Aldise M Nastri, Jezzy R Dewantari, Rima R Prasetya, Krisnoadi Rahardjo, Irawati Marga, Gatot Soegiarto, Laksmi Wulandari, Retno A Setyoningrum, Resti Yudhawati, Yohko K Shimizu, Mitsuhiro Nishimura, Soetjipto, Yasuko Mori, Maria I Lusida, Kazufumi Shimizu |
| EPI_ISL_956273 | Universitas Airlangga Hospital | Institute of Tropical Disease, Universitas Airlangga | Jezzy R Dewantari, Rima R Prasetya, Krisnoadi Rahardjo, Aldise M Nastri, Nasronudin, Gatot Soegiarto, Laksmi Wulandari, Resti Yudhawati, Soetjipto, Yasuko Mori, Maria I Lusida, Kazufumi Shimizu |
| EPI_ISL_956274 | RSUD Sidoarjo | Institute of Tropical Disease, Universitas Airlangga | Rima R Prasetya, Krisnoadi Rahardjo, Aldise M Nastri, Jezzy R Dewantari, Atok Irawan, Gatot Soegiarto, Laksmi Wulandari, Resti Yudhawati, Soetjipto, Yasuko Mori, Maria I Lusida, Kazufumi Shimizu |
| EPI_ISL_956276 | Mitra Keluarga Hospital Waru | Institute of Tropical Disease, Universitas Airlangga | Maria I Lusida, Krisnoadi Rahardjo, Aldise M Nastri, Jezzy R Dewantari, Rima R Prasetya, Christina Dian Anggraeni, Gatot Soegiarto, Laksmi Wulandari, Resti Yudhawati, Soetjipto, Yasuko Mori, Kazufumi Shimizu |
| EPI_ISL_956277 | Paru Hospital | Institute of Tropical Disease, Universitas Airlangga | Krisnoadi Rahardjo, Aldise M Nastri, Jezzy R Dewantari, Rima R Prasetya, Dyah Retno, Gatot Soegiarto, Laksmi Wulandari, Resti Yudhawati, Soetjipto, Yasuko Mori, Maria I Lusida, Kazufumi Shimizu |
| EPI_ISL_956278 | Siti Khodijah Hospital | Institute of Tropical Disease, Universitas Airlangga | Aldise M Nastri, Jezzy R Dewantari, Rima R Prasetya, Krisnoadi Rahardjo, Muhammad Hamdan, Gatot Soegiarto, Laksmi Wulandari, Resti Yudhawati, Soetjipto, Yasuko Mori, Maria I Lusida, Kazufumi Shimizu |
| EPI_ISL_956309 | RSUD Sidoarjo | Institute of Tropical Disease, Universitas Airlangga | Rima R Prasetya, Krisnoadi Rahardjo, Aldise M Nastri, Jezzy R Dewantari, Atok Irawan, Gatot Soegiarto, Laksmi Wulandari, Resti Yudhawati, Yasuko Mori, Soetjipto, Kazufumi Shimizu, Maria I Lusida |
| EPI_ISL_956310 | RSUD Dr. M. Soewandhie | Institute of Tropical Disease, Universitas Airlangga | Aldise M Nastri, Jezzy R Dewantari, Rima R Prasetya, Krisnoadi Rahardjo, Febria Rachmanita, Gatot Soegiarto, Laksmi Wulandari, Resti Yudhawati, Yasuko Mori, Soetjipto, Kazufumi Shimizu, Maria I Lusida |
| EPI_ISL_956311 | Islamic Hospital | Institute of Tropical Disease, Universitas Airlangga | Kazufumi Shimizu, Krisnoadi Rahardjo, Aldise M Nastri, Jezzy R Dewantari, Rima R Prasetya, Dodo Anondo, Gatot Soegiarto, Laksmi Wulandari, Resti Yudhawati, Yasuko Mori, Soetjipto, Maria I Lusida |
| EPI_ISL_959277 | National Influenza Center, Virology Department | National Influenza Center | J Yavarian,K Sadeghi, NZ Shafiei Jandaghi, V Salimi, A Nejati, N Ghavvami,F Ajaminejad and T Mokhtari Azad |
| EPI_ISL_959278 | National Influenza Center, Virology Department | National Influenza Center | A Nejati, J Yavarian,K Sadeghi, NZ Shafiei Jandaghi, V Salimi, N Ghavvami,F Ajaminejad and T Mokhtari Azad |
| EPI_ISL_959281 | National Influenza Center, Virology Department | National Influenza Center | NZ Shafiei Jandaghi, V Salimi, A Nejati, K Sadeghi, J Yavarian, F Ajaminejad,N Ghavvami and T Mokhtari Azad |

| | | | |
|---|---|---|---|
| EPI_ISL_959282 | National Influenza Center, Virology Department | National Influenza Center | J Yavarian, NZ Shafiei Jandaghi, V Salimi, A Nejati, K Sadeghi, F Ajaminejad,N Ghavvami and T Mokhtari Azad |
| EPI_ISL_961761, EPI_ISL_961771, EPI_ISL_961780 | Laboratorio de Infectología, Servicio de Infectología, Hospital Universitario Dr. José Eleuterio González - Universidad Autónoma de Nuevo León | Laboratorio de Infectología Molecular, Departamento de Bioquímica y Medicina Molecular,Facultad de Medicina - Universidad Autónoma de Nuevo León | Kame A. Galán-Huerta, María F. Herrera-Saldivar, Natalia Martínez-Acuña, Sonia A. Lozano-Sepúlveda, Daniel Arellanos-Soto, Ana M. Rivas-Estilla, Paola Bocanegra-Ibarias, Samantha M. Flores-Treviño, Elvira Garza-González, Eduardo Perez-Alba, Laura Nuzzolo-Shihadeh, Adrian Camacho-Ortiz |
| EPI_ISL_969509, EPI_ISL_969740 | Indian Council of Medical Research-National Institute of Virology, Pune | Indian Council of Medical Research-National Institute of Virology, Maximum Containment Laboratory | Pragya D. Yadav, Anita Shete-Aich, Dimpal A. Nyayanit, Abhinendra Kumar, Savita Patil, Triparna D. Majumdar |
| EPI_ISL_971451 | Cell culture Unit at CV-MIT belonging to HIMMV | Functional Genomic Platform UATRS-biology, CNRST | Nadia Touil, Abderrazzak Rfak, Mouhssine Hemlali, Sanaa ALAOUI-Amine, Marouane MELLOUL, Mly Abdelaziz ELALAOUI, Hicham Elannaz, Amine Idriss Lahlou, Mostafa ELOUENNASS, Khalid ENNIBI, Elmostafa EL FAHIME |
| EPI_ISL_977471 | Instituto Adolfo Lutz - Regional de Presidente Prudente | Instituto Adolfo Lutz, Interdiciplinary Procedures Center, Strategic Laboratory | Claudio Tavares Sacchi, Claudia Regina Gonçalves, Erica Valessa Ramos Gomes, Karoline Rodrigues Campos |
| EPI_ISL_977472, EPI_ISL_977473, EPI_ISL_977474 | Instituto Adolfo Lutz Central | Instituto Adolfo Lutz, Interdiciplinary Procedures Center, Strategic Laboratory | Claudio Tavares Sacchi, Claudia Regina Gonçalves, Erica Valessa Ramos Gomes, Karoline Rodrigues Campos |
| EPI_ISL_977475, EPI_ISL_977478 | Instituto Adolfo Lutz - Regional de Presidente Prudente | Instituto Adolfo Lutz, Interdiciplinary Procedures Center, Strategic Laboratory | Claudio Tavares Sacchi, Claudia Regina Gonçalves, Erica Valessa Ramos Gomes, Karoline Rodrigues Campos |
| EPI_ISL_977479 | Lab Loc - Itapecerica da Serra | Instituto Adolfo Lutz, Interdiciplinary Procedures Center, Strategic Laboratory | Claudio Tavares Sacchi, Claudia Regina Gonçalves, Erica Valessa Ramos Gomes, Karoline Rodrigues Campos |
| EPI_ISL_977480, EPI_ISL_977481 | Instituto Adolfo Lutz - Regional de Presidente Prudente | Instituto Adolfo Lutz, Interdiciplinary Procedures Center, Strategic Laboratory | Claudio Tavares Sacchi, Claudia Regina Gonçalves, Erica Valessa Ramos Gomes, Karoline Rodrigues Campos |
| EPI_ISL_977483, EPI_ISL_977484 | Instituto Adolfo Lutz Central | Instituto Adolfo Lutz, Interdiciplinary Procedures Center, Strategic Laboratory | Claudio Tavares Sacchi, Claudia Regina Gonçalves, Erica Valessa Ramos Gomes, Karoline Rodrigues Campos |
| EPI_ISL_977485 | Instituto Adolfo Lutz - Regional de Presidente Prudente | Instituto Adolfo Lutz, Interdiciplinary Procedures Center, Strategic Laboratory | Claudio Tavares Sacchi, Claudia Regina Gonçalves, Erica Valessa Ramos Gomes, Karoline Rodrigues Campos |
| EPI_ISL_977486 | Instituto Adolfo Lutz - Regional de Santo Andre | Instituto Adolfo Lutz, Interdiciplinary Procedures Center, Strategic Laboratory | Claudio Tavares Sacchi, Claudia Regina Gonçalves, Erica Valessa Ramos Gomes, Karoline Rodrigues Campos |
| EPI_ISL_977488 | Instituto Adolfo Lutz - Regional de Presidente Prudente | Instituto Adolfo Lutz, Interdiciplinary Procedures Center, Strategic Laboratory | Claudio Tavares Sacchi, Claudia Regina Gonçalves, Erica Valessa Ramos Gomes, Karoline Rodrigues Campos |
| EPI_ISL_979970, EPI_ISL_979971, EPI_ISL_979972, EPI_ISL_979973 | Department of Respiratory and other Viral Infections of L.V.Gromashevsky Institute of Epidemiology & Infectious Diseases NAMS of Ukrain | Department of Respiratory and other Viral Infections of L.V.Gromashevsky Institute of Epidemiology & Infectious Diseases NAMS of Ukrain, JSC "Farmak" | Alla Mironenko, Andriy Goy, Ihor Kravchuk, Ludmyla Bolotova, Larysa Radchenko, Nataliia Teteriuk |
| EPI_ISL_984242 | Instituto Adolfo Lutz Central | Instituto Adolfo Lutz, Interdiciplinary Procedures Center, Strategic Laboratory | Claudio Tavares Sacchi, Claudia Regina Gonçalves, Erica Valessa Ramos Gomes, Karoline Rodrigues Campos |
| EPI_ISL_984243, EPI_ISL_984244 | Instituto Adolfo Lutz - Regional de Marilia | Instituto Adolfo Lutz, Interdiciplinary Procedures Center, Strategic Laboratory | Claudio Tavares Sacchi, Claudia Regina Gonçalves, Erica Valessa Ramos Gomes, Karoline Rodrigues Campos |
| EPI_ISL_984246 | Instituto Adolfo Lutz Central | Instituto Adolfo Lutz, Interdiciplinary Procedures Center, Strategic Laboratory | Claudio Tavares Sacchi, Claudia Regina Gonçalves, Erica Valessa Ramos Gomes, Karoline Rodrigues Campos |
| EPI_ISL_985170 | Instituto Adolfo Lutz - Regional de Presidente Prudente | Instituto Adolfo Lutz, Interdiciplinary Procedures Center, Strategic Laboratory | Claudio Tavares Sacchi, Claudia Regina Gonçalves, Erica Valessa Ramos Gomes, Karoline Rodrigues Campos |
| EPI_ISL_985171, EPI_ISL_985172, EPI_ISL_985173, EPI_ISL_985174 | Instituto Adolfo Lutz - Regional de Taubate | Instituto Adolfo Lutz, Interdiciplinary Procedures Center, Strategic Laboratory | Claudio Tavares Sacchi, Claudia Regina Gonçalves, Erica Valessa Ramos Gomes, Karoline Rodrigues Campos |
| EPI_ISL_985175, EPI_ISL_985176, EPI_ISL_985177 | Instituto Adolfo Lutz Central | Instituto Adolfo Lutz, Interdiciplinary Procedures Center, Strategic Laboratory | Claudio Tavares Sacchi, Claudia Regina Gonçalves, Erica Valessa Ramos Gomes, Karoline Rodrigues Campos |
| EPI_ISL_985178 | Lab Loc - Itapecerica da Serra | Instituto Adolfo Lutz, Interdiciplinary Procedures Center, Strategic Laboratory | Claudio Tavares Sacchi, Claudia Regina Gonçalves, Erica Valessa Ramos Gomes, Karoline Rodrigues Campos |

We gratefully acknowledge the following Authors from the Originating laboratories responsible for obtaining the specimens, as well as the Submitting laboratories where the genome data were generated and shared via GISAID, on which this research is based.

All Submitters of data may be contacted directly via www.gisaid.org

Authors are sorted alphabetically.

| Accession ID | Originating Laboratory | Submitting Laboratory | Authors |
|---|---|---|---|
| EPI_ISL_1001050 | Centre Pasteur du Cameroun | Institut Pasteur de Dakar | Njouom Richard |
| EPI_ISL_1001353 | hopital | National Reference Center for Viruses of Respiratory Infections, Institut Pasteur, Paris | Marion Barbet, Sylvie Behillil, Méline Bizard, Angela Brisebarre, Camille Capel, Etienne Simon-Lorière, Vincent Enouf, Maud Vanpeene, Sylvie van der Werf,Stoessel-Thouvenin Patricia |
| EPI_ISL_1001376, EPI_ISL_1001377, EPI_ISL_1001378, EPI_ISL_1001379, EPI_ISL_1001380 | Labo Analyses Med | National Reference Center for Viruses of Respiratory Infections, Institut Pasteur, Paris | Marion Barbet, Sylvie Behillil, Méline Bizard, Angela Brisebarre, Camille Capel, Etienne Simon-Lorière, Vincent Enouf, Maud Vanpeene, Sylvie van der Werf,Besson J |
| EPI_ISL_1001456 | Hopital | National Reference Center for Viruses of Respiratory Infections, Institut Pasteur, Paris | Marion Barbet, Sylvie Behillil, Méline Bizard, Angela Brisebarre, Camille Capel, Etienne Simon-Lorière, Vincent Enouf, Maud Vanpeene, Sylvie van der Werf,Ducancelle Alexandra |
| EPI_ISL_1001458 | Gorgas memorial Institute for Health Studies | Gorgas memorial Institute for Health Studies | Díaz Y, Castillo D, Moreno B, Castillo M,González C, Gondola J, Moreno A, Pitti Y, Chavarria O, Franco D, Saenz L, Gaitan M, Arauz D, Martinez AA, Lopez-Verges S. |
| EPI_ISL_1005696 | RSUD Dr. Tjitrowardojo | Genetics Working Group (Pokja Genetik) Faculty of Medicine, Public Health and Nursing Universitas Gadjah Mada (FK-KMK UGM); Disease Investigation Center Wates Ministry of Agriculture Indonesia; Department of Microbiology FK-KMK UGM; Laboratorium Diagnostik Yayasan Tahija World Mosquito Program (WMP) Yogyakarta Center for Tropical Medicine FK-KMK UGM; Integrated Research Center FK-KMK UGM; Department of Computer Science and Electronics FMIPA UGM; Balai Besar Teknik Kesehatan Lingkungan dan Pengendalian Penyakit (BBTKLPP) Yogyakarta | Gunadi, Hendra Wibawa, Marcellus, Mohamad S. Hakim, Edwin W. Daniwijaya, Ludhang P. Rizki, Endah Supriyati, Eggi Arguni, Titik Nuryastuti, Tri Wibawa, Dwi AA Nugrahaningsih, Afiahayati, Siswanto, Kristy Iskandar, Nungki Anggorowati, Irene, Indaryati, Havid Setyawan, Arry Setyowati, Chusni Mubarakh, Try Nirmala Sari, Susan Simanjaya, Alvin Santoso Kalim |
| EPI_ISL_1005697 | RSUD Dr. Tjitrowardojo | Genetics Working Group (Pokja Genetik) Faculty of Medicine, Public Health and Nursing Universitas Gadjah Mada (FK-KMK UGM); Disease Investigation Center Wates Ministry of Agriculture Indonesia; Department of Microbiology FK-KMK UGM; Laboratorium Diagnostik Yayasan Tahija World Mosquito Program (WMP) Yogyakarta Center for Tropical Medicine FK-KMK UGM; Integrated Research Center FK-KMK UGM; Department of Computer Science and Electronics FMIPA UGM; Balai Besar Teknik Kesehatan Lingkungan dan Pengendalian Penyakit (BBTKLPP) Yogyakarta | Gunadi, Hendra Wibawa, Marcellus, Mohamad S. Hakim, Edwin W. Daniwijaya, Ludhang P. Rizki, Endah Supriyati, Eggi Arguni, Titik Nuryastuti, Tri Wibawa, Dwi AA Nugrahaningsih, Afiahayati, Siswanto, Kristy Iskandar, Nungki Anggorowati, Irene, Indaryati, Havid Setyawan, Arry Setyowati, Chusni Mubarakh, Try Nirmala Sari, Kemala Athollah, Dyah Ayu Puspitarani |
| EPI_ISL_1005698 | RSUD Dr. Tjitrowardojo | Genetics Working Group (Pokja Genetik) Faculty of Medicine, Public Health and Nursing Universitas Gadjah Mada (FK-KMK UGM); Disease Investigation Center Wates Ministry of Agriculture Indonesia; Department of Microbiology FK-KMK UGM; Laboratorium Diagnostik Yayasan Tahija World Mosquito Program (WMP) Yogyakarta Center for Tropical Medicine FK-KMK UGM; Integrated Research Center FK-KMK UGM; Department of Computer Science and Electronics FMIPA UGM; Balai Besar Teknik Kesehatan Lingkungan dan Pengendalian Penyakit (BBTKLPP) Yogyakarta | Gunadi, Hendra Wibawa, Marcellus, Mohamad S. Hakim, Edwin W. Daniwijaya, Ludhang P. Rizki, Endah Supriyati, Eggi Arguni, Titik Nuryastuti, Tri Wibawa, Dwi AA Nugrahaningsih, Afiahayati, Siswanto, Kristy Iskandar, Nungki Anggorowati, Irene, Indaryati, Havid Setyawan, Arry Setyowati, Chusni Mubarakh, Try Nirmala Sari, Aditya Rifqi Fauzi, Fadil Fahri |
| EPI_ISL_1007677, EPI_ISL_1007678, EPI_ISL_1007679 | Department of Internal Medicine, College of Medicine, Chosun University | Department of Internal Medicine, College of Medicine, Chosun University | Dong-Min Kim |
| EPI_ISL_1008703 | Presidio di Brindisi Di Summa - Perrino | Istituto Zooprofilattico Sperimentale della Puglia e della Basilicata | Parisi A., Bianco A., Capozzi L., Del Sambro L., Simone D., Chironna M., Loconsole D., Sallustio A., Ridolfi D. |
| EPI_ISL_1010728, EPI_ISL_1010729, EPI_ISL_1010730, EPI_ISL_1010731 | Department of Laboratory Medicine, National Taiwan University Hospital | Microbial Genomics Core Lab, National Taiwan University Centers of Genomic and Precision Medicine | Shiou-Hwei Yeh, You-Yu Lin, Ya-Yun Lai, Chiao-Ling Li, Shan-Chwen Chang, Pei-Jer Chen, Sui-Yuan Chang |
| EPI_ISL_1013094 | Imelda Hospital | Imelda Hospital | Johan Frans, Hanne Valgaeren, Dagmar Obbels |
| EPI_ISL_1013099, EPI_ISL_1013100, EPI_ISL_1013101, EPI_ISL_1013102, EPI_ISL_1013103, EPI_ISL_1013104, EPI_ISL_1013105, EPI_ISL_1013106, EPI_ISL_1013107, EPI_ISL_1013108, EPI_ISL_1013109, EPI_ISL_1013110, EPI_ISL_1013111, EPI_ISL_1013112, EPI_ISL_1013113, EPI_ISL_1013114, EPI_ISL_1013115, EPI_ISL_1013116, EPI_ISL_1013117, EPI_ISL_1013118, EPI_ISL_1013119, EPI_ISL_1013120, EPI_ISL_1013121, EPI_ISL_1013122, EPI_ISL_1013123, EPI_ISL_1013124, EPI_ISL_1013125, EPI_ISL_1013126, EPI_ISL_1013127, EPI_ISL_1013128, EPI_ISL_1013129, EPI_ISL_1013130, EPI_ISL_1013131, EPI_ISL_1013132, EPI_ISL_1013133, EPI_ISL_1013134, EPI_ISL_1013135, EPI_ISL_1013136, EPI_ISL_1013137, EPI_ISL_1013138, EPI_ISL_1013139, EPI_ISL_1013140, EPI_ISL_1013141, EPI_ISL_1013142, EPI_ISL_1013143, EPI_ISL_1013144, EPI_ISL_1013145, EPI_ISL_1013146, EPI_ISL_1013147, EPI_ISL_1013148, EPI_ISL_1013149, EPI_ISL_1013150, EPI_ISL_1013151, EPI_ISL_1013152, EPI_ISL_1013153, EPI_ISL_1013154, EPI_ISL_1013155, EPI_ISL_1013156, EPI_ISL_1013157, EPI_ISL_1013158, EPI_ISL_1013159, EPI_ISL_1013160, EPI_ISL_1013161, EPI_ISL_1013162, EPI_ISL_1013163, EPI_ISL_1013164, EPI_ISL_1013165, EPI_ISL_1013166, EPI_ISL_1013167, EPI_ISL_1013168, EPI_ISL_1013169, EPI_ISL_1013170, EPI_ISL_1013171, EPI_ISL_1013172, EPI_ISL_1013173, EPI_ISL_1013174, EPI_ISL_1013175, EPI_ISL_1013176, EPI_ISL_1013177, EPI_ISL_1013178, EPI_ISL_1013179, EPI_ISL_1013180, EPI_ISL_1013181, EPI_ISL_1013182, EPI_ISL_1013183, EPI_ISL_1013184, EPI_ISL_1013185, EPI_ISL_1013186, EPI_ISL_1013187, EPI_ISL_1013188, EPI_ISL_1013189, EPI_ISL_1013190, EPI_ISL_1013191, EPI_ISL_1013192, EPI_ISL_1013193, EPI_ISL_1013194, EPI_ISL_1013195, EPI_ISL_1013196, EPI_ISL_1013197, EPI_ISL_1013198, EPI_ISL_1013199, EPI_ISL_1013200, EPI_ISL_1013201, EPI_ISL_1013202, EPI_ISL_1013203, EPI_ISL_1013204, EPI_ISL_1013205, EPI_ISL_1013206, EPI_ISL_1013207, EPI_ISL_1013208 | | | |
| see above | CH de Mayotte | National Reference Center for Viruses of Respiratory Infections, Institut Pasteur, Paris | Marion Barbet, Sylvie Behillil, Méline Bizard, Angela Brisebarre, Camille Capel, Etienne Simon-Lorière, Vincent Enouf, Maud Vanpeene, Sylvie van der Werf,Combe Patrice |
| EPI_ISL_1013437 | Institut Pasteur de Guadeloupe | National Reference Center for Viruses of Respiratory Infections, Institut Pasteur, Paris | Marion Barbet, Sylvie Behillil, Méline Bizard, Angela Brisebarre, Camille Capel, Etienne Simon-Lorière, Vincent Enouf, Maud Vanpeene, Sylvie van der Werf,Talarmin Antoine |
| EPI_ISL_1013444 | Laboratoire associé au CNR des Virus Influenzae | National Reference Center for Viruses of Respiratory Infections, Institut Pasteur, Paris | Marion Barbet, Sylvie Behillil, Méline Bizard, Angela Brisebarre, Camille Capel, Etienne Simon-Lorière, Vincent Enouf, Maud Vanpeene, Sylvie van der Werf,Rousset (Guy) Dominique |
| EPI_ISL_1013779, EPI_ISL_1014102, EPI_ISL_1014238 | Zurita & Zurita Laboratorios | Zurita & Zurita Laboratorios | Jeannete Zurita, Camilo Zurita-Salinas, Gabriela Sevillano |
| EPI_ISL_1014676 | National Influenza Center, Virology Department | National Influenza Center | V Salimi,NZ Shafiei Jandaghi, J Yavarian, A Nejati, K Sadeghi, N Ghavvami,F Ajaminejad and T Mokhtari Azad |

| | | | |
|---|---|---|---|
| EPI_ISL_1014685 | National Influenza Center, Virology Department | National Influenza Center | K Sadeghi, A Nejati, J Yavarian, NZ Shafiei Jandaghi, V Salimi, F Ajaminejad,N Ghavvami and T Mokhtari Azad |
| EPI_ISL_1014686 | National Influenza Center, Virology Department | National Influenza Center | NZ Shafiei Jandaghi, V Salimi, A Nejati, K Sadeghi, J Yavarian, N Ghavvami,F Ajaminejad and T Mokhtari Azad |
| EPI_ISL_1016690 | University of Sarajevo, Veterinary Faculty, Laboratory for Molecular Diagnostic and Research Laboratory | University of Sarajevo, Veterinary Faculty, Laboratory for Molecular Diagnostic and Research Laboratory | Goletic T., Goletic S., Softic A., Alic-Seho A., Sabic E., Terzic I., Jazic A., Nicevic M., Hodzic A. |
| EPI_ISL_1016822 | University of Sarajevo, Veterinary Faculty, Laboratory for Molecular Diagnostic and Research Laboratory | University of Sarajevo, Veterinary Faculty, Laboratory for Molecular Diagnostic and Research Laboratory | Goletic T., Goletic S., Softic A., Alic-Seho A., Hodzic A., Sabic E., Nicevic M., Terzic I., Jazic A. |
| EPI_ISL_1018287, EPI_ISL_1018289, EPI_ISL_1018290, EPI_ISL_1018291, EPI_ISL_1018292, EPI_ISL_1018294, EPI_ISL_1018295, EPI_ISL_1018296, EPI_ISL_1018297, EPI_ISL_1018299, EPI_ISL_1018300, EPI_ISL_1018303, EPI_ISL_1018305, EPI_ISL_1018306, EPI_ISL_1018307, EPI_ISL_1018308, EPI_ISL_1018310, EPI_ISL_1018312, EPI_ISL_1018313, EPI_ISL_1018315, EPI_ISL_1018316, EPI_ISL_1018318, EPI_ISL_1018319, EPI_ISL_1018321, EPI_ISL_1018322, EPI_ISL_1018323, EPI_ISL_1018324, EPI_ISL_1018325, EPI_ISL_1018326, EPI_ISL_1018327, EPI_ISL_1018328, EPI_ISL_1019637, EPI_ISL_1019642, EPI_ISL_1019645, EPI_ISL_1019647, EPI_ISL_1019650, EPI_ISL_1019653, EPI_ISL_1019655, EPI_ISL_1019658, EPI_ISL_1019660, EPI_ISL_1019663, EPI_ISL_1019665, EPI_ISL_1019668, EPI_ISL_1019671, EPI_ISL_1019674, EPI_ISL_1019678, EPI_ISL_1019681, EPI_ISL_1019683, EPI_ISL_1019688, EPI_ISL_1019693, EPI_ISL_1019699, EPI_ISL_1019702, EPI_ISL_1019705, EPI_ISL_1019720, EPI_ISL_1019724, EPI_ISL_1019726, EPI_ISL_1019737, EPI_ISL_1019739, EPI_ISL_1019742, EPI_ISL_1019748, EPI_ISL_1019750, EPI_ISL_1019753, EPI_ISL_1019756, EPI_ISL_1019762, EPI_ISL_1019767, EPI_ISL_1019775, EPI_ISL_1019778, EPI_ISL_1019783, EPI_ISL_1019785, EPI_ISL_1019788, EPI_ISL_1019790, EPI_ISL_1019793, EPI_ISL_1019795, EPI_ISL_1019798, EPI_ISL_1019800, EPI_ISL_1019803, EPI_ISL_1019805, EPI_ISL_1019810, EPI_ISL_1019813, EPI_ISL_1019815, EPI_ISL_1019828, EPI_ISL_1019838, EPI_ISL_1019846, EPI_ISL_1019849, EPI_ISL_1019851, EPI_ISL_1019853, EPI_ISL_1019855, EPI_ISL_1019858, EPI_ISL_1019861, EPI_ISL_1019863, EPI_ISL_1019866, EPI_ISL_1019869, EPI_ISL_1019872, EPI_ISL_1019875, EPI_ISL_1019877, EPI_ISL_1019880, EPI_ISL_1019882, EPI_ISL_1019885, EPI_ISL_1019888, EPI_ISL_1019890, EPI_ISL_1019893, EPI_ISL_1020022, EPI_ISL_1020023, EPI_ISL_1020024, EPI_ISL_1020025, EPI_ISL_1020026, EPI_ISL_1020027, EPI_ISL_1020028, EPI_ISL_1020029, EPI_ISL_1020030, EPI_ISL_1020031, EPI_ISL_1020032, EPI_ISL_1020033, EPI_ISL_1020034, EPI_ISL_1020035, EPI_ISL_1020036, EPI_ISL_1020037, EPI_ISL_1020038, EPI_ISL_1020039, EPI_ISL_1020040, EPI_ISL_1020041, EPI_ISL_1020042, EPI_ISL_1020043, EPI_ISL_1020044, EPI_ISL_1020045, EPI_ISL_1020046, EPI_ISL_1020047, EPI_ISL_1020048, EPI_ISL_1020049, EPI_ISL_1020050, EPI_ISL_1020051, EPI_ISL_1020052, EPI_ISL_1020053, EPI_ISL_1020054, EPI_ISL_1020055, EPI_ISL_1020056, EPI_ISL_1020058, EPI_ISL_1020059, EPI_ISL_1020060, EPI_ISL_1020061, EPI_ISL_1020062, EPI_ISL_1020063, EPI_ISL_1020064, EPI_ISL_1020065, EPI_ISL_1020066, EPI_ISL_1020067, EPI_ISL_1020068, EPI_ISL_1020069, EPI_ISL_1020070, EPI_ISL_1020071, EPI_ISL_1020072, EPI_ISL_1020073, EPI_ISL_1020074, EPI_ISL_1020075, EPI_ISL_1020076, EPI_ISL_1020077, EPI_ISL_1020078, EPI_ISL_1020079, EPI_ISL_1020080, EPI_ISL_1020081, EPI_ISL_1020083, EPI_ISL_1020084, EPI_ISL_1020086, EPI_ISL_1020087, EPI_ISL_1020088, EPI_ISL_1020089, EPI_ISL_1020091, EPI_ISL_1020092, EPI_ISL_1020093, EPI_ISL_1020094, EPI_ISL_1020095, EPI_ISL_1020096, EPI_ISL_1020098, EPI_ISL_1020099, EPI_ISL_1020101, EPI_ISL_1020102, EPI_ISL_1020104, EPI_ISL_1020105, EPI_ISL_1020106, EPI_ISL_1020107, EPI_ISL_1020108, EPI_ISL_1020109, EPI_ISL_1020110, EPI_ISL_1020111, EPI_ISL_1020112, EPI_ISL_1020113, EPI_ISL_1020114, EPI_ISL_1020115, EPI_ISL_1020116, EPI_ISL_1020117, EPI_ISL_1020118, EPI_ISL_1020119, EPI_ISL_1020120 | | | |
| see above | Department of Health Technology and Informatics, The Hong Kong Polytechnic University | Department of Health Technology and Informatics, The Hong Kong Polytechnic University | Gilman Kit-Hang Siu, Lam-Kwong Lee, Kenneth Siu-Sing Leung, Jake Siu-Lun Leung, Timothy Ting-Leung Ng, Chloe Toi-Mei Chan, Kingsley King-Gee Tam, Hiu-Yin Lao, Denise Sze-Hang Wong, Alan Ka-Lun Wu, Miranda Chong-Yee Yau, Yvette Wai-Man Lai, Kitty Sau-Chun Fung, Sandy Ka-Yee Chau, Barry Kin-Chung Wong, Wing-Kin To, Kristine Luk, Alex Yat-Man Ho, Tak-Lun Que, Kam-Tong Yip, Wing Cheong Yam, David Ho-Keung Shum, Shea Ping Yip |
| EPI_ISL_1020192 | Melania Hospital, Bagor , west Java | Biosafety Level-3 Laboratory, Indonesian Institute of Sciences (LIPI) | Anik Budhi Dharmayanthi, Ade Andriani, Hariyatun, Asep M. Ridwanuloh, Syam Budi Iryanto, Eko Wahyu Putro, Gita Syahputra, Susanti, Inswasti Cahyani, Matthew W. Loose, Mohammad Ilyas, Wien Kusharyoto |
| EPI_ISL_1020194 | UMMI Hospital, Bogor, West Java | Biosafety Level-3 Laboratory, Indonesian Institute of Sciences (LIPI) | Anggia Prasetyoputri, Isa Nuryana, Sri Swasthikawati, Sugiyono Saputra, Syam Budi Iryanto, Gita Syahputra, Ratih Asmana Ningrum |
| EPI_ISL_1020195 | Melania Hospital, Bogor , West Java | Biosafety Level-3 Laboratory, Indonesian Institute of Sciences (LIPI) | Anggia Prasetyoputri, Erwin Hasrianda, Herjuno A. Nugroho, Nova Dhilla Yanthi, Fahrurrozi, Syam Budi Iryanto, Gita Syahputra, Ratih Asmana Ningrum |
| EPI_ISL_1020197 | Dinkes, Bogor, West Java | Biosafety Level-3 Laboratory, Indonesian Institute of Sciences (LIPI) | Anik Budhi Dharmayanthi, Ade Andriani, Hariyatun, Asep M. Ridwanuloh, Syam Budi Iryanto, Eko Wahyu Putro, Gita Syahputra, Susanti, Inswasti Cahyani, Matthew W. Loose, Mohammad Ilyas, Wien Kusharyoto |
| EPI_ISL_1020198 | RS Trimitra, Bogor, West java | Biosafety Level-3 Laboratory, Indonesian Institute of Sciences (LIPI) | Sri Swasthikawati, Linda Sukmarini, Idris, Masrukhin, Alfi T. Fathurahman, Eko Wahyu Putro, Ratih Asmana Ningrum, Sugiyono Saputra |
| EPI_ISL_1020203 | Klinik Apotik Dien, Jakarta | Biosafety Level-3 Laboratory, Indonesian Institute of Sciences (LIPI) | Anik Budhi Dharmayanthi, Ade Andriani, Hariyatun, Asep M. Ridwanuloh, Syam Budi Iryanto, Eko Wahyu Putro, Gita Syahputra, Susanti, Inswasti Cahyani, Matthew W. Loose, Mohammad Ilyas, Wien Kusharyoto |
| EPI_ISL_1020206 | Qadr Hospital, Tangerang, Banten | Biosafety Level-3 Laboratory, Indonesian Institute of Sciences (LIPI) | Anggia Prasetyoputri, Isa Nuryana, Sri Swasthikawati, Sugiyono Saputra, Syam Budi Iryanto, Eko Wahyu Putro, Gita Syahputra, Ari Sulistyo Wulandari, Ratih Asmana Ningrum |
| EPI_ISL_1020207 | AZ Klina | AZ Klina | Dr. C. Vael |
| EPI_ISL_1020214, EPI_ISL_1020215, EPI_ISL_1020216, EPI_ISL_1020217, EPI_ISL_1020218, EPI_ISL_1020219, EPI_ISL_1020220, EPI_ISL_1020221, EPI_ISL_1020222, EPI_ISL_1020223, EPI_ISL_1020224, EPI_ISL_1020225, EPI_ISL_1020226, EPI_ISL_1020227, EPI_ISL_1020228, EPI_ISL_1020229, EPI_ISL_1020230, EPI_ISL_1020231, EPI_ISL_1020232, EPI_ISL_1020233, EPI_ISL_1020234, EPI_ISL_1020235, EPI_ISL_1020236, EPI_ISL_1020237, EPI_ISL_1020238, EPI_ISL_1020239, EPI_ISL_1020240, EPI_ISL_1020241, EPI_ISL_1020242, EPI_ISL_1020243, EPI_ISL_1020244, EPI_ISL_1020245, EPI_ISL_1020246, EPI_ISL_1020247, EPI_ISL_1020248, EPI_ISL_1020249, EPI_ISL_1020250, EPI_ISL_1020251, EPI_ISL_1020252, EPI_ISL_1020253, EPI_ISL_1020254, EPI_ISL_1020255, EPI_ISL_1020256, EPI_ISL_1020257, EPI_ISL_1020258, EPI_ISL_1020259, EPI_ISL_1020260, EPI_ISL_1020261, EPI_ISL_1020262, EPI_ISL_1020263, EPI_ISL_1020264, EPI_ISL_1020265, EPI_ISL_1020266, EPI_ISL_1020267, EPI_ISL_1020268, EPI_ISL_1020269, EPI_ISL_1020270, EPI_ISL_1020271, EPI_ISL_1020272, EPI_ISL_1020273, EPI_ISL_1020274, EPI_ISL_1020275, EPI_ISL_1020276, EPI_ISL_1020277, EPI_ISL_1020278, EPI_ISL_1020279, EPI_ISL_1020280, EPI_ISL_1020281, EPI_ISL_1020282, EPI_ISL_1020283, EPI_ISL_1020284 | | | |
| see above | Department of Health Technology and Informatics, The Hong Kong Polytechnic University | Department of Health Technology and Informatics, The Hong Kong Polytechnic University | Gilman Kit-Hang Siu, Lam-Kwong Lee, Kenneth Siu-Sing Leung, Jake Siu-Lun Leung, Timothy Ting-Leung Ng, Chloe Toi-Mei Chan, Kingsley King-Gee Tam, Hiu-Yin Lao, Denise Sze-Hang Wong, Alan Ka-Lun Wu, Miranda Chong-Yee Yau, Yvette Wai-Man Lai, Kitty Sau-Chun Fung, Sandy Ka-Yee Chau, Barry Kin-Chung Wong, Wing-Kin To, Kristine Luk, Alex Yat-Man Ho, Tak-Lun Que, Kam-Tong Yip, Wing Cheong Yam, David Ho-Keung Shum, Shea Ping Yip |
| EPI_ISL_1020285, EPI_ISL_1020286, EPI_ISL_1020287, EPI_ISL_1020289, EPI_ISL_1020290, EPI_ISL_1020291, EPI_ISL_1020292, EPI_ISL_1020293 | AZ Klina | AZ Klina | Dr. C. Vael |
| EPI_ISL_1027629, EPI_ISL_1027630, EPI_ISL_1027631, EPI_ISL_1027632, EPI_ISL_1027633, EPI_ISL_1027634, EPI_ISL_1027635, EPI_ISL_1027636 | Mako Medical Laboratories | Mako Medical Laboratories | "Lauren Moon, Matthew Tugwell" |
| EPI_ISL_1029967 | Cliniques universitaires Saint-Luc | UCLouvain/IREC/MBLG | Jean Ruelle, Lysa Pinsmaye, Benoit Kabamba Mukadi |
| EPI_ISL_1035889, EPI_ISL_1035890, EPI_ISL_1035891 | Istituto Zooprofilattico Sperimentale del Mezzogiorno | IZSM-U.O.C. Virologia | Maurizio Viscardi, Lorena Cardillo, Sergio Brandi, Pellegrino Cerino, Massimo Zollo,Giovanna Fusco, Esterina De Carlo, Antonio Limone |
| EPI_ISL_1035892 | Istituto Zooprofilattico Sperimentale del Mezzogiorno | IZSM-U.O.C. Virologia | Maurizio Viscardi, Lorena Cardillo, Claudio de Martinis, Giovanna Fusco, Esterina De Carlo, Antonio Limone |
| EPI_ISL_1035903, EPI_ISL_1035949, EPI_ISL_1035950, EPI_ISL_1035951 | Outre Mer CH de Mayotte | National Reference Center for Viruses of Respiratory Infections, Institut Pasteur, Paris | Marion Barbet, Sylvie Behillil, Méline Bizard, Angela Brisebarre, Camille Capel, Etienne Simon-Lorière, Vincent Enouf, Maud Vanpeene, Sylvie van der Werf,Combe Patrice |
| EPI_ISL_1036213, EPI_ISL_1036214, EPI_ISL_1036218, EPI_ISL_1036219, EPI_ISL_1036220 | SIESP TERAMO | Istituto Zooprofilattico Sperimentale dell'Abruzzo e Molise "G. Caporale" | Lorusso A, Marcacci M, Di Domenico M, Ancora M, Curini V, Mangone I, Rinaldi A, Scialabba S, Di Pasquale A, Cammà C, Puglia I, Calistri P, Savini G |
| EPI_ISL_1039008, EPI_ISL_1039009, EPI_ISL_1039010, EPI_ISL_1039011, EPI_ISL_1039157, EPI_ISL_1039158 | WHO National Influenza Centre Russian Federation | WHO National Influenza Centre Russian Federation | Andrey Komissarov, Artem Fadeev, Anna Ivanova, Kseniya Komissarova, Dmitry Bazhenov, Tamila Musaeva, Maria Timofeeva, Veronika Eder, Maria Pisareva, Daria Danilenko, Ksenia Safina, Elena Nabieva, Georgii Bazykin, Dmitry Lioznov |
| EPI_ISL_1039160 | Department of Laboratory Medicine, National Taiwan University Hospital | Microbial Genomics Core Lab, National Taiwan University Centers of Genomic and Precision Medicine | Shiou-Hwei Yeh, You-Yu Lin, Ya-Yun Lai, Chiao-Ling Li, Shan-Chwen Chang, Pei-Jer Chen, Sui-Yuan Chang |
| EPI_ISL_1039697 | Instituto Adolfo Lutz Central | Instituto Adolfo Lutz, Interdiciplinary Procedures Center, Strategic Laboratory | Claudio Tavares Sacchi, Claudia Regina Gonçalves, Erica Valessa Ramos Gomes, Karoline Rodrigues Campos |
| EPI_ISL_1039698 | Lab Loc - Itapecerica da Serra | Instituto Adolfo Lutz, Interdiciplinary Procedures Center, Strategic Laboratory | Claudio Tavares Sacchi, Claudia Regina Gonçalves, Erica Valessa Ramos Gomes, Karoline Rodrigues Campos |
| EPI_ISL_1039699 | Instituto Adolfo Lutz - Regional de Taubate | Instituto Adolfo Lutz, Interdiciplinary Procedures Center, Strategic Laboratory | Claudio Tavares Sacchi, Claudia Regina Gonçalves, Erica Valessa Ramos Gomes, Karoline Rodrigues Campos |
| EPI_ISL_1039700, EPI_ISL_1039701 | Instituto Adolfo Lutz Central | Instituto Adolfo Lutz, Interdiciplinary Procedures Center, | Claudio Tavares Sacchi, Claudia Regina Gonçalves, Erica Valessa Ramos Gomes, Karoline Rodrigues Campos |

| | | | |
|---|---|---|---|
| | | Strategic Laboratory | |
| EPI_ISL_1039703 | Instituto Adolfo Lutz - Regional de Taubate | Instituto Adolfo Lutz, Interdiciplinary Procedures Center, Strategic Laboratory | Claudio Tavares Sacchi, Claudia Regina Gonçalves, Erica Valessa Ramos Gomes, Karoline Rodrigues Campos |
| EPI_ISL_1039704 | Lab Loc - Itapecerica da Serra | Instituto Adolfo Lutz, Interdiciplinary Procedures Center, Strategic Laboratory | Claudio Tavares Sacchi, Claudia Regina Gonçalves, Erica Valessa Ramos Gomes, Karoline Rodrigues Campos |
| EPI_ISL_1039705, EPI_ISL_1039706, EPI_ISL_1039707, EPI_ISL_1039708, EPI_ISL_1039709, EPI_ISL_1039710 | Instituto Adolfo Lutz Central | Instituto Adolfo Lutz, Interdiciplinary Procedures Center, Strategic Laboratory | Claudio Tavares Sacchi, Claudia Regina Gonçalves, Erica Valessa Ramos Gomes, Karoline Rodrigues Campos |
| EPI_ISL_1040028, EPI_ISL_1040029 | Original detection - Virology Unit, Institut Pasteur du Cambodge; Sequencing - US National Institute of Allergy and Infectious Diseases Cambodia | Virology Unit, Institut Pasteur du Cambodge | Jennifer Bohl, Sophana Chea, Sreyngim Lay, Ly Sovann, Kraing Sidonn, Yi Sengdoeurn, Chin Savuth, Chau Darapheak, Veasna Duong, Jessica Manning, Erik A Karlsson |
| EPI_ISL_1040825, EPI_ISL_1040826, EPI_ISL_1040827, EPI_ISL_1040828, EPI_ISL_1040829, EPI_ISL_1040830, EPI_ISL_1040831, EPI_ISL_1040832, EPI_ISL_1040833, EPI_ISL_1040834, EPI_ISL_1040835, EPI_ISL_1040836, EPI_ISL_1040837, EPI_ISL_1040838, EPI_ISL_1040839, EPI_ISL_1040840 | | | |
| see above | LACEN do Mato Grosso do Sul | Instituto Adolfo Lutz, Interdiciplinary Procedures Center, Strategic Laboratory | Claudio Tavares Sacchi, Claudia Regina Gonçalves, Erica Valessa Ramos Gomes, Karoline Rodrigues Campos |
| EPI_ISL_1041030 | Dr. Cipto Mangunkusumo (RSCM) Hospital | Faculty of Medicine, Universitas Indonesia | Rina La Distia Nora, Mei Riasanti, Rani Dwi Ningtias, Ari Fahrial Syam |
| EPI_ISL_1041195, EPI_ISL_1041196, EPI_ISL_1041197, EPI_ISL_1041198 | Hungarian Defence Forces Military Medical Centre | National Laboratory of Virology, Szentágothai Research Centre | Endre Gábor Tóth, Balázs Somogyi, Ágnes Balázs-Nagy, Csaba Pereszlényi,Ferenc Jakab, Gábor Kemenesi |
| EPI_ISL_1041957, EPI_ISL_1041958 | Department of Laboratory Medicine, National Taiwan University Hospital | Microbial Genomics Core Lab, National Taiwan University Centers of Genomic and Precision Medicine | Shiou-Hwei Yeh, You-Yu Lin, Ya-Yun Lai, Chiao-Ling Li, Shan-Chwen Chang, Pei-Jer Chen, Sui-Yuan Chang |
| EPI_ISL_1048372 | Farmalab Indoutama Jakarta | National Institute of Health Research and Development | Hana Apsari Pawestri, Subangkit, Kartika Dewi Puspa, Arie Ardiansyah Nugraha, Hartanti Dian Ikawati, Krisna Nur Andriana Pangesti, Nelly Puspandari, Vivi Setiawaty. |
| EPI_ISL_1048406 | Balai Penelitian dan Pengembangan Biomedis Papua | National Institute of Health Research and Development | Hana Apsari Pawestri, Subangkit, Kartika Dewi Puspa, Arie Ardiansyah Nugraha, Hartanti Dian Ikawati, Krisna Nur Andriana Pangesti, Nelly Puspandari, Anton Oktavianus, Hotma Sitompul, Vivi Setiawaty. |
| EPI_ISL_1048806, EPI_ISL_1048807, EPI_ISL_1048808, EPI_ISL_1048809, EPI_ISL_1048810, EPI_ISL_1048811, EPI_ISL_1048812, EPI_ISL_1048813 | Laboratory of Clinical Microbiology, Virology and Bioemergencies - Fatebenefratelli Sacco Hospital | Virology Laboratory, Scientific Department, Army Medical Center | Valeria Micheli, Alessandro Mancon, Alberto Rizzo, Maria Rita Gismondo, Silvia Fillo, Anella Monte, Anna Anselmo, Vanessa Vera Fain, Giovanni Faggioni, Riccardo De Sanctis, Florigio Lista |
| EPI_ISL_1055191 | Temporary Specimen Collection Centre at the AsiaWorld-Expo | Hong Kong Department of Health | Mak Gannon C.K., Lam Edman T.K., Chan Rickjason C.W., Tsang Dominic N.C. |
| EPI_ISL_1055192 | Queen Mary Hospital | Hong Kong Department of Health | Mak Gannon C.K., Lam Edman T.K., Chan Rickjason C.W., Tsang Dominic N.C. |
| EPI_ISL_1055193 | Prince of Wales Hospital | Hong Kong Department of Health | Mak Gannon C.K., Lam Edman T.K., Chan Rickjason C.W., Tsang Dominic N.C. |
| EPI_ISL_1055194, EPI_ISL_1055195 | United Christian Hospital | Hong Kong Department of Health | Mak Gannon C.K., Lam Edman T.K., Chan Rickjason C.W., Tsang Dominic N.C. |
| EPI_ISL_1055196 | Queen Mary Hospital | Hong Kong Department of Health | Mak Gannon C.K., Lam Edman T.K., Chan Rickjason C.W., Tsang Dominic N.C. |
| EPI_ISL_1055197, EPI_ISL_1055198 | Prince of Wales Hospital | Hong Kong Department of Health | Mak Gannon C.K., Lam Edman T.K., Chan Rickjason C.W., Tsang Dominic N.C. |
| EPI_ISL_1055199 | Princess Margaret Hospital | Hong Kong Department of Health | Mak Gannon C.K., Lam Edman T.K., Chan Rickjason C.W., Tsang Dominic N.C. |
| EPI_ISL_1055200 | Queen Mary Hospital | Hong Kong Department of Health | Mak Gannon C.K., Lam Edman T.K., Chan Rickjason C.W., Tsang Dominic N.C. |
| EPI_ISL_1055201 | Prince of Wales Hospital | Hong Kong Department of Health | Mak Gannon C.K., Lam Edman T.K., Chan Rickjason C.W., Tsang Dominic N.C. |
| EPI_ISL_1055202 | Hong Kong Adventist Hospital - Tsuen Wan | Hong Kong Department of Health | Mak Gannon C.K., Lam Edman T.K., Chan Rickjason C.W., Tsang Dominic N.C. |
| EPI_ISL_1055203 | Private medical practitioner | Hong Kong Department of Health | Mak Gannon C.K., Lam Edman T.K., Chan Rickjason C.W., Tsang Dominic N.C. |
| EPI_ISL_1055204 | Queen Elizabeth Hospital | Hong Kong Department of Health | Mak Gannon C.K., Lam Edman T.K., Chan Rickjason C.W., Tsang Dominic N.C. |
| EPI_ISL_1055205 | Queen Mary Hospital | Hong Kong Department of Health | Mak Gannon C.K., Lam Edman T.K., Chan Rickjason C.W., Tsang Dominic N.C. |
| EPI_ISL_1055206 | Princess Margaret Hospital | Hong Kong Department of Health | Mak Gannon C.K., Lam Edman T.K., Chan Rickjason C.W., Tsang Dominic N.C. |
| EPI_ISL_1055207 | Pamela Youde Nethersole Eastern Hospital | Hong Kong Department of Health | Mak Gannon C.K., Lam Edman T.K., Chan Rickjason C.W., Tsang Dominic N.C. |
| EPI_ISL_1055208 | Private medical practitioner | Hong Kong Department of Health | Mak Gannon C.K., Lam Edman T.K., Chan Rickjason C.W., Tsang Dominic N.C. |
| EPI_ISL_1055209 | Tuen Mun Hospital | Hong Kong Department of Health | Mak Gannon C.K., Lam Edman T.K., Chan Rickjason C.W., Tsang Dominic N.C. |
| EPI_ISL_1055210 | Hong Kong Adventist Hospital - Tsuen Wan | Hong Kong Department of Health | Mak Gannon C.K., Lam Edman T.K., Chan Rickjason C.W., Tsang Dominic N.C. |
| EPI_ISL_1055211, EPI_ISL_1055212 | Community Testing Centre | Hong Kong Department of Health | Mak Gannon C.K., Lam Edman T.K., Chan Rickjason C.W., Tsang Dominic N.C. |
| EPI_ISL_1055213 | Queen Mary Hospital | Hong Kong Department of Health | Mak Gannon C.K., Lam Edman T.K., Chan Rickjason C.W., Tsang Dominic N.C. |
| EPI_ISL_1055214 | Prince of Wales Hospital | Hong Kong Department of Health | Mak Gannon C.K., Lam Edman T.K., Chan Rickjason C.W., Tsang Dominic N.C. |
| EPI_ISL_1055215 | Private medical practitioner | Hong Kong Department of Health | Mak Gannon C.K., Lam Edman T.K., Chan Rickjason C.W., Tsang Dominic N.C. |
| EPI_ISL_1055216 | Community Testing Centre | Hong Kong Department of Health | Mak Gannon C.K., Lam Edman T.K., Chan Rickjason C.W., Tsang Dominic N.C. |
| EPI_ISL_1055217 | Private medical practitioner | Hong Kong Department of Health | Mak Gannon C.K., Lam Edman T.K., Chan Rickjason C.W., Tsang Dominic N.C. |
| EPI_ISL_1055218 | Tseung Kwan O Hospital | Hong Kong Department of Health | Mak Gannon C.K., Lam Edman T.K., Chan Rickjason C.W., Tsang Dominic N.C. |
| EPI_ISL_1055219 | Princess Margaret Hospital | Hong Kong Department of Health | Mak Gannon C.K., Lam Edman T.K., Chan Rickjason C.W., Tsang Dominic N.C. |
| EPI_ISL_1055220 | Community Testing Centre | Hong Kong Department of Health | Mak Gannon C.K., Lam Edman T.K., Chan Rickjason C.W., Tsang Dominic N.C. |
| EPI_ISL_1055221 | Queen Mary Hospital | Hong Kong Department of Health | Mak Gannon C.K., Lam Edman T.K., Chan Rickjason C.W., Tsang Dominic N.C. |
| EPI_ISL_1055222, EPI_ISL_1055223 | Queen Elizabeth Hospital | Hong Kong Department of Health | Mak Gannon C.K., Lam Edman T.K., Chan Rickjason C.W., Tsang Dominic N.C. |
| EPI_ISL_1055224 | United Christian Hospital | Hong Kong Department of Health | Mak Gannon C.K., Lam Edman T.K., Chan Rickjason C.W., Tsang Dominic N.C. |
| EPI_ISL_1055225 | Community Testing Centre | Hong Kong Department of Health | Mak Gannon C.K., Lam Edman T.K., Chan Rickjason C.W., Tsang Dominic N.C. |
| EPI_ISL_1055226 | Queen Mary Hospital | Hong Kong Department of Health | Mak Gannon C.K., Lam Edman T.K., Chan Rickjason C.W., Tsang Dominic N.C. |
| EPI_ISL_1055227 | Hong Kong Police Force | Hong Kong Department of Health | Mak Gannon C.K., Lam Edman T.K., Chan Rickjason C.W., Tsang Dominic N.C. |
| EPI_ISL_1055228 | Queen Mary Hospital | Hong Kong Department of Health | Mak Gannon C.K., Lam Edman T.K., Chan Rickjason C.W., Tsang Dominic N.C. |
| EPI_ISL_1055229 | Queen Elizabeth Hospital | Hong Kong Department of Health | Mak Gannon C.K., Lam Edman T.K., Chan Rickjason C.W., Tsang Dominic N.C. |
| EPI_ISL_1055230 | Prince of Wales Hospital | Hong Kong Department of Health | Mak Gannon C.K., Lam Edman T.K., Chan Rickjason C.W., Tsang Dominic N.C. |

| | | | |
|---|---|---|---|
| EPI_ISL_1055231 | Private medical practitioner | Hong Kong Department of Health | Mak Gannon C.K., Lam Edman T.K., Chan Rickjason C.W., Tsang Dominic N.C. |
| EPI_ISL_1055232 | Hong Kong Baptist Hospital | Hong Kong Department of Health | Mak Gannon C.K., Lam Edman T.K., Chan Rickjason C.W., Tsang Dominic N.C. |
| EPI_ISL_1055233 | Pamela Youde Nethersole Eastern Hospital | Hong Kong Department of Health | Mak Gannon C.K., Lam Edman T.K., Chan Rickjason C.W., Tsang Dominic N.C. |
| EPI_ISL_1055234 | Queen Elizabeth Hospital | Hong Kong Department of Health | Mak Gannon C.K., Lam Edman T.K., Chan Rickjason C.W., Tsang Dominic N.C. |
| EPI_ISL_1055235 | Pamela Youde Nethersole Eastern Hospital | Hong Kong Department of Health | Mak Gannon C.K., Lam Edman T.K., Chan Rickjason C.W., Tsang Dominic N.C. |
| EPI_ISL_1055236 | Prince of Wales Hospital | Hong Kong Department of Health | Mak Gannon C.K., Lam Edman T.K., Chan Rickjason C.W., Tsang Dominic N.C. |
| EPI_ISL_1055237 | Queen Mary Hospital | Hong Kong Department of Health | Mak Gannon C.K., Lam Edman T.K., Chan Rickjason C.W., Tsang Dominic N.C. |
| EPI_ISL_1055238 | United Christian Hospital | Hong Kong Department of Health | Mak Gannon C.K., Lam Edman T.K., Chan Rickjason C.W., Tsang Dominic N.C. |
| EPI_ISL_1055239 | Community Testing Centre | Hong Kong Department of Health | Mak Gannon C.K., Lam Edman T.K., Chan Rickjason C.W., Tsang Dominic N.C. |
| EPI_ISL_1055240 | Pamela Youde Nethersole Eastern Hospital | Hong Kong Department of Health | Mak Gannon C.K., Lam Edman T.K., Chan Rickjason C.W., Tsang Dominic N.C. |
| EPI_ISL_1055241 | United Christian Hospital | Hong Kong Department of Health | Mak Gannon C.K., Lam Edman T.K., Chan Rickjason C.W., Tsang Dominic N.C. |
| EPI_ISL_1055242 | Queen Mary Hospital | Hong Kong Department of Health | Mak Gannon C.K., Lam Edman T.K., Chan Rickjason C.W., Tsang Dominic N.C. |
| EPI_ISL_1055243 | Prince of Wales Hospital | Hong Kong Department of Health | Mak Gannon C.K., Lam Edman T.K., Chan Rickjason C.W., Tsang Dominic N.C. |
| EPI_ISL_1055244 | Precious Blood Hospital | Hong Kong Department of Health | Mak Gannon C.K., Lam Edman T.K., Chan Rickjason C.W., Tsang Dominic N.C. |
| EPI_ISL_1055245 | Queen Mary Hospital | Hong Kong Department of Health | Mak Gannon C.K., Lam Edman T.K., Chan Rickjason C.W., Tsang Dominic N.C. |
| EPI_ISL_1055246 | Pok Oi Hospital | Hong Kong Department of Health | Mak Gannon C.K., Lam Edman T.K., Chan Rickjason C.W., Tsang Dominic N.C. |
| EPI_ISL_1055247 | Queen Elizabeth Hospital | Hong Kong Department of Health | Mak Gannon C.K., Lam Edman T.K., Chan Rickjason C.W., Tsang Dominic N.C. |
| EPI_ISL_1055248 | Prince of Wales Hospital | Hong Kong Department of Health | Mak Gannon C.K., Lam Edman T.K., Chan Rickjason C.W., Tsang Dominic N.C. |
| EPI_ISL_1055249 | Caritas Medical Centre | Hong Kong Department of Health | Mak Gannon C.K., Lam Edman T.K., Chan Rickjason C.W., Tsang Dominic N.C. |
| EPI_ISL_1055250 | Community Testing Centre | Hong Kong Department of Health | Mak Gannon C.K., Lam Edman T.K., Chan Rickjason C.W., Tsang Dominic N.C. |
| EPI_ISL_1055251 | Princess Margaret Hospital | Hong Kong Department of Health | Mak Gannon C.K., Lam Edman T.K., Chan Rickjason C.W., Tsang Dominic N.C. |
| EPI_ISL_1055252, EPI_ISL_1055253 | Community Testing Centre | Hong Kong Department of Health | Mak Gannon C.K., Lam Edman T.K., Chan Rickjason C.W., Tsang Dominic N.C. |
| EPI_ISL_1055254 | Princess Margaret Hospital | Hong Kong Department of Health | Mak Gannon C.K., Lam Edman T.K., Chan Rickjason C.W., Tsang Dominic N.C. |
| EPI_ISL_1055255 | Tuen Mun Hospital | Hong Kong Department of Health | Mak Gannon C.K., Lam Edman T.K., Chan Rickjason C.W., Tsang Dominic N.C. |
| EPI_ISL_1055256 | Home Quarantine Taskforce | Hong Kong Department of Health | Mak Gannon C.K., Lam Edman T.K., Chan Rickjason C.W., Tsang Dominic N.C. |
| EPI_ISL_1055257 | Pamela Youde Nethersole Eastern Hospital | Hong Kong Department of Health | Mak Gannon C.K., Lam Edman T.K., Chan Rickjason C.W., Tsang Dominic N.C. |
| EPI_ISL_1055258 | Tuen Mun Hospital | Hong Kong Department of Health | Mak Gannon C.K., Lam Edman T.K., Chan Rickjason C.W., Tsang Dominic N.C. |
| EPI_ISL_1055259 | United Christian Hospital | Hong Kong Department of Health | Mak Gannon C.K., Lam Edman T.K., Chan Rickjason C.W., Tsang Dominic N.C. |
| EPI_ISL_1055260 | Caritas Medical Centre | Hong Kong Department of Health | Mak Gannon C.K., Lam Edman T.K., Chan Rickjason C.W., Tsang Dominic N.C. |
| EPI_ISL_1055263 | Institute for Medical Research, Infectious Disease Research Centre, National Institutes of Health, Ministry of Health Malaysia | Institute for Medical Research, Infectious Disease Research Centre, National Institutes of Health, Ministry of Health Malaysia | Suppiah J, Kamel K,Moh Zawawi Z, Ramly N, Robert F, Thayan R |
| EPI_ISL_1055755, EPI_ISL_1055756, EPI_ISL_1055757, EPI_ISL_1055758 | Microbiology Department. Complexo Hospitalario Universitario de Santiago | Microbiology Department. Complexo Hospitalario Universitario de Vigo | Microbiology Department, Complexo Hospitalario Universitario de Vigo (CHUVI). Complexo Hospitalario Universitario de Santiago (CHUS). |
| EPI_ISL_1055759, EPI_ISL_1055760 | Vithas Lab. Hospital Vithas Vigo | Microbiology Department. Complexo Hospitalario Universitario de Vigo | Microbiology Department, Complexo Hospitalario Universitario de Vigo (CHUVI). Vithas Lab, Hospital Vithas Vigo |
| EPI_ISL_1055816 | Ospedale SS Annunziata Chieti | Istituto Zooprofilattico Sperimentale dell'Abruzzo e Molise "G. Caporale" | Lorusso A, Marcacci M, Di Domenico M, Ancora M, Curini V, Mangone I, Rinaldi A, Delli Compagni E, Scialabba S, Di Pasquale A, Cammà C, Puglia I, Calistri P, Savini G |
| EPI_ISL_1055817 | SIESP DIPARTIMENTO DI PREVENZIONE CHIETI | Istituto Zooprofilattico Sperimentale dell'Abruzzo e Molise "G. Caporale" | Lorusso A, Marcacci M, Di Domenico M, Ancora M, Curini V, Mangone I, Rinaldi A, Delli Compagni E, Scialabba S, Di Pasquale A, Cammà C, Puglia I, Calistri P, Savini G |
| EPI_ISL_1055818 | DIP. PREV. AVEZZANO SERVIZIO DI IGIENE EPIDEMIOLOGIAE SANITA' PUBBLICA | Istituto Zooprofilattico Sperimentale dell'Abruzzo e Molise "G. Caporale" | Lorusso A, Marcacci M, Di Domenico M, Ancora M, Curini V, Mangone I, Rinaldi A, Delli Compagni E, Scialabba S, Di Pasquale A, Cammà C, Puglia I, Calistri P, Savini G |
| EPI_ISL_1055819 | SIESP DIPARTIMENTO DI PREVENZIONE SULMONA | Istituto Zooprofilattico Sperimentale dell'Abruzzo e Molise "G. Caporale" | Lorusso A, Marcacci M, Di Domenico M, Ancora M, Curini V, Mangone I, Rinaldi A, Delli Compagni E, Scialabba S, Di Pasquale A, Cammà C, Puglia I, Calistri P, Savini G |
| EPI_ISL_1058285, EPI_ISL_1058286, EPI_ISL_1058287, EPI_ISL_1058288, EPI_ISL_1058289, EPI_ISL_1058290, EPI_ISL_1058291, EPI_ISL_1058292, EPI_ISL_1058293, EPI_ISL_1058294, EPI_ISL_1058295, EPI_ISL_1058296, EPI_ISL_1058297, EPI_ISL_1058298, EPI_ISL_1058299, EPI_ISL_1058300, EPI_ISL_1058301, EPI_ISL_1058302, EPI_ISL_1058303, EPI_ISL_1058304, EPI_ISL_1058305, EPI_ISL_1058306, EPI_ISL_1058307, EPI_ISL_1058308, EPI_ISL_1058309, EPI_ISL_1058310, EPI_ISL_1058311, EPI_ISL_1058312, EPI_ISL_1058313, EPI_ISL_1058314, EPI_ISL_1058315, EPI_ISL_1058316, EPI_ISL_1058317, EPI_ISL_1058318, EPI_ISL_1058319, EPI_ISL_1058320, EPI_ISL_1058321, EPI_ISL_1058322, EPI_ISL_1058323, EPI_ISL_1058324, EPI_ISL_1058325, EPI_ISL_1058326, EPI_ISL_1058327, EPI_ISL_1058328, EPI_ISL_1058329, EPI_ISL_1058330, EPI_ISL_1058331, EPI_ISL_1058332, EPI_ISL_1058333, EPI_ISL_1058334, EPI_ISL_1058335, EPI_ISL_1058336, EPI_ISL_1058337, EPI_ISL_1058338, EPI_ISL_1058339, EPI_ISL_1058340, EPI_ISL_1058341, EPI_ISL_1058342, EPI_ISL_1058343, EPI_ISL_1058344, EPI_ISL_1058345, EPI_ISL_1058346, EPI_ISL_1058347, EPI_ISL_1058348, EPI_ISL_1058349, EPI_ISL_1058350, EPI_ISL_1058351, EPI_ISL_1058352, EPI_ISL_1058353, EPI_ISL_1058354, EPI_ISL_1058355, EPI_ISL_1058356, EPI_ISL_1058357, EPI_ISL_1058358, EPI_ISL_1058359, EPI_ISL_1058360, EPI_ISL_1058361, EPI_ISL_1058362, EPI_ISL_1058363, EPI_ISL_1058364, EPI_ISL_1058365, EPI_ISL_1058366, EPI_ISL_1058367, EPI_ISL_1058368, EPI_ISL_1058369, EPI_ISL_1058370, EPI_ISL_1058371, EPI_ISL_1058372, EPI_ISL_1058373, EPI_ISL_1058374, EPI_ISL_1058375, EPI_ISL_1058376, EPI_ISL_1058377, EPI_ISL_1058378, EPI_ISL_1058379, EPI_ISL_1058380, EPI_ISL_1058381, EPI_ISL_1058382, EPI_ISL_1058383, EPI_ISL_1058384, EPI_ISL_1058385, EPI_ISL_1058386, EPI_ISL_1058387, EPI_ISL_1058388, EPI_ISL_1058389, EPI_ISL_1058390, EPI_ISL_1058391, EPI_ISL_1058392, EPI_ISL_1058393, EPI_ISL_1058394, EPI_ISL_1058395, EPI_ISL_1058396, EPI_ISL_1058397, EPI_ISL_1058398, EPI_ISL_1058399, EPI_ISL_1058400, EPI_ISL_1058401, EPI_ISL_1058402, EPI_ISL_1058403, EPI_ISL_1058404, EPI_ISL_1058405, EPI_ISL_1058406, EPI_ISL_1058407, EPI_ISL_1058408, EPI_ISL_1058409, EPI_ISL_1058410, EPI_ISL_1058411, EPI_ISL_1058412, EPI_ISL_1058413, EPI_ISL_1058414, EPI_ISL_1058415, EPI_ISL_1058416, EPI_ISL_1058417, EPI_ISL_1058418, EPI_ISL_1058419, EPI_ISL_1058420, EPI_ISL_1058421, EPI_ISL_1058422, EPI_ISL_1058423, EPI_ISL_1058424, EPI_ISL_1058425, EPI_ISL_1058426, EPI_ISL_1058427, EPI_ISL_1058428, EPI_ISL_1058429, EPI_ISL_1058430, EPI_ISL_1058431, EPI_ISL_1058432, EPI_ISL_1058433, EPI_ISL_1058434, EPI_ISL_1058435, EPI_ISL_1058436 | | | |
| see above | Microbiology Department. Complexo Hospitalario Universitario de Vigo | Microbiology Department. Complexo Hospitalario Universitario de Vigo | Microbiology Department, Complexo Hospitalario Universitario de Vigo (CHUVI). |
| EPI_ISL_1059900, EPI_ISL_1059901 | Malaysia Genome Institute | Malaysia Genome Institute | Mohd Noor Mat Isa, Irni Suhayu Sapian, Yusuf Muhammad Noor, Nurhezreen Md Iqbal, Mohd Faizal Abu Bakar, Enizza Kasim, Shamsidar Sopie, Siti Noraini Othman, Azrin Ahmad, Nor Azfa Johari. |
| EPI_ISL_1059902 | DNA Laboratories Sdn. Bhd. | Malaysia Genome Institute | Mohd Noor Mat Isa, Irni Suhayu Sapian, Yusuf Muhammad Noor, Nurhezreen Md Iqbal, Mohd Faizal Abu Bakar, Enizza Kasim, Shamsidar Sopie, Siti Noraini Othman, Azrin Ahmad, Wong Yong Wee, Nor Azfa Johari. |
| EPI_ISL_1060359, EPI_ISL_1060360, EPI_ISL_1060361, EPI_ISL_1060362, EPI_ISL_1060363, EPI_ISL_1060364, EPI_ISL_1060365, EPI_ISL_1060366, EPI_ISL_1060367 | Hopital | National Reference Center for Viruses of Respiratory Infections, Institut Pasteur, Paris | Marion Barbet, Sylvie Behillil, Méline Bizard, Angela Brisebarre, Camille Capel, Etienne Simon-Lorière, Vincent Enouf, Maud Vanpeene, Sylvie van der Werf,Fourgeaud Jacques |
| EPI_ISL_1060395 | Labo Analyses Med | National Reference Center for Viruses of Respiratory Infections, Institut Pasteur, Paris | Marion Barbet, Sylvie Behillil, Méline Bizard, Angela Brisebarre, Camille Capel, Etienne Simon-Lorière, Vincent Enouf, Maud Vanpeene, Sylvie van der Werf,Chauvet |

| | | | |
|---|---|---|---|
| EPI_ISL_1060398, EPI_ISL_1060433, EPI_ISL_1060461, EPI_ISL_1060462 | Hopital | National Reference Center for Viruses of Respiratory Infections, Institut Pasteur, Paris | Marion Barbet, Sylvie Behillil, Méline Bizard, Angela Brisebarre, Camille Capel, Etienne Simon-Lorière, Vincent Enouf, Maud Vanpeene, Sylvie van der Werf,Fourgeaud Jacques |
| EPI_ISL_1060475 | Labo Analyses Med | National Reference Center for Viruses of Respiratory Infections, Institut Pasteur, Paris | Marion Barbet, Sylvie Behillil, Méline Bizard, Angela Brisebarre, Camille Capel, Etienne Simon-Lorière, Vincent Enouf, Maud Vanpeene, Sylvie van der Werf,Fiorina Jc |
| EPI_ISL_1060561, EPI_ISL_1060563, EPI_ISL_1060564, EPI_ISL_1060565, EPI_ISL_1060566 | Hopital | National Reference Center for Viruses of Respiratory Infections, Institut Pasteur, Paris | Marion Barbet, Sylvie Behillil, Méline Bizard, Angela Brisebarre, Camille Capel, Etienne Simon-Lorière, Vincent Enouf, Maud Vanpeene, Sylvie van der Werf,Fourgeaud Jacques |
| EPI_ISL_1060727 | Instituto de Diagnostico y Referencia Epidemiologicos (INDRE)_RNLSP | Instituto de Diagnostico y Referencia Epidemiologicos (INDRE) | Claudia Wong-Arambula, Abril Rodriguez-Maldonado, Fabiola Garces-Ayala, Adnan Araiza-Rodriguez, David Fragoso-Fonseca, Sergio Rangel-Guerrero, Mayra Jimenez-Morales, Nancy Munoz-Hernandez, Natividad Cruz-Ortiz, Tatiana Nunez-Garcia, Gisela Barrera-Badillo, Lucia Hernandez-Rivas, Irma Lopez-Martinez, Ernesto Ramirez-Gonzalez. |
| EPI_ISL_1061404, EPI_ISL_1061405 | Queen Elizabeth Hospital | Hong Kong Children's Hospital | Herman Tse, Garnet Kwan-Yue Choi, Sally Cheuk-Ying Wong, David Christopher Lung |
| EPI_ISL_1061406 | Kwong Wah Hospital | Hong Kong Children's Hospital | Herman Tse, Garnet Kwan-Yue Choi, Sally Cheuk-Ying Wong, David Christopher Lung |
| EPI_ISL_1061407 | Queen Elizabeth Hospital | Hong Kong Children's Hospital | Herman Tse, Garnet Kwan-Yue Choi, Sally Cheuk-Ying Wong, David Christopher Lung |
| EPI_ISL_1061408 | Kwong Wah Hospital | Hong Kong Children's Hospital | Herman Tse, Garnet Kwan-Yue Choi, Sally Cheuk-Ying Wong, David Christopher Lung |
| EPI_ISL_1061409, EPI_ISL_1061410, EPI_ISL_1061411 | Queen Elizabeth Hospital | Hong Kong Children's Hospital | Herman Tse, Garnet Kwan-Yue Choi, Sally Cheuk-Ying Wong, David Christopher Lung |
| EPI_ISL_1063784 | OSP SS ANNIUNZIATA CHIETI, PRONTO SOCCORSO | Istituto Zooprofilattico Sperimentale dell'Abruzzo e Molise "G. Caporale" | Lorusso A, Marcacci M, Di Domenico M, Ancora M, Curini V, Mangone I, Rinaldi A, Scialabba S, Di Pasquale A, Cammà C, Puglia I, Calistri P, Savini G |
| EPI_ISL_1078981, EPI_ISL_1078983, EPI_ISL_1078984, EPI_ISL_1078986, EPI_ISL_1078987, EPI_ISL_1078988, EPI_ISL_1078989, EPI_ISL_1078992 | IAL Regional de Bauru | Instituto Adolfo Lutz, Interdiciplinary Procedures Center, Strategic Laboratory | Claudio Tavares Sacchi, Claudia Regina Gonçalves, Erica Valessa Ramos Gomes, Karoline Rodrigues Campos |
| EPI_ISL_1080375, EPI_ISL_1080376, EPI_ISL_1080378, EPI_ISL_1080379, EPI_ISL_1080380, EPI_ISL_1080381, EPI_ISL_1080382, EPI_ISL_1080384, EPI_ISL_1080385, EPI_ISL_1080386, EPI_ISL_1080387, EPI_ISL_1080388, EPI_ISL_1080389, EPI_ISL_1080390, EPI_ISL_1080391, EPI_ISL_1080491, EPI_ISL_1080492 | | | |
| see above | Istituto Zooprofilattico Sperimentale del Mezzogiorno | Istituto Zooprofilattico Sperimentale del Mezzogiorno-U.O.C. Virologia | Maurizio Viscardi, Lorena Cardillo, Loredana Cozzolino, Pellegrino Cerino, Massimo Zollo,Giovanna Fusco, Esterina De Carlo, Antonio Limone |
| EPI_ISL_1080580, EPI_ISL_1080581, EPI_ISL_1080582, EPI_ISL_1080583, EPI_ISL_1080584, EPI_ISL_1080585, EPI_ISL_1080586, EPI_ISL_1080587, EPI_ISL_1080588, EPI_ISL_1080589, EPI_ISL_1080590, EPI_ISL_1080591, EPI_ISL_1080592, EPI_ISL_1080593, EPI_ISL_1080594, EPI_ISL_1080595, EPI_ISL_1080596, EPI_ISL_1080597, EPI_ISL_1080598, EPI_ISL_1080599, EPI_ISL_1080600, EPI_ISL_1080601, EPI_ISL_1080602 | | | |
| see above | Istituto Zooprofilattico Sperimentale del Mezzogiorno | Istituto Zooprofilattico Sperimentale del Mezzogiorno-U.O.C. Virologia | Maurizio Viscardi, Lorena Cardillo, Loredana Cozzolino Pellegrino Cerino, Massimo Zollo,Giovanna Fusco, Esterina De Carlo, Antonio Limone |
| EPI_ISL_1080857, EPI_ISL_1081150, EPI_ISL_1081151, EPI_ISL_1081171, EPI_ISL_1081172, EPI_ISL_1081174, EPI_ISL_1081175, EPI_ISL_1081180, EPI_ISL_1081185, EPI_ISL_1081186, EPI_ISL_1081187, EPI_ISL_1081188, EPI_ISL_1081190, EPI_ISL_1081191 | | | |
| see above | Baylor Scott & White-Temple | Baylor Scott & White-Temple | Ari Rao, Linden Morales, Kimberly Walker, Marcus Volz, Shelby Hendrickson |
| EPI_ISL_1081333 | Institute for Medical Research, Infectious Disease Research Centre, National Institutes of Health, Ministry of Health Malaysia | Institute for Medical Research, Infectious Disease Research Centre, National Institutes of Health, Ministry of Health Malaysia | Suppiah J, Kamel K, Azizan MA, Thayan R |
| EPI_ISL_1081954 | "Stefan S. Nicolau"Institute of Virology | "Stefan S. Nicolau"Institute of Virology | Anca Botezatu, Iulia Virginia Iancu, Coralia Bleotu, Mihaela Economescu, Laura Necula, Denisa Dragu, Lilia Matei, Ioana Pitica, Marinela Bostan, Mirela Mihaila, Laura Grecu, Camelia Sultana, Simona Ruta, Ana Iulia Neagu, Adriana Plesa, Saviana Nedeianu, Alina Nastasie, Cristina Mambet, Gabriela Anton, Carmen Cristina Diaconu. |
| EPI_ISL_1081958, EPI_ISL_1081959, EPI_ISL_1081961, EPI_ISL_1082252, EPI_ISL_1082290, EPI_ISL_1082291, EPI_ISL_1082294 | "Stefan S. Nicolau"Institute of Virology | "Stefan S. Nicolau"Institute of Virology | Anca Botezatu, Iulia Virginia Iancu, Coralia Bleotu, Mihaela Economescu, Laura Necula, Denisa Dragu, Lilia Matei, Ioana Pitica, Marinela Bostan, Mirela Mihaila, Laura Grecu, Camelia Sultana, Simona Ruta, Ana Iulia Neagu, Adriana Plesa, Saviana Nedeianu, Alina Nastasie, Cristina Mambet,Gabriela Anton, Carmen Cristina Diaconu |
| EPI_ISL_1082296 | "Stefan S. Nicolau"Institute of Virology | 285 Mihai Bravu Ave, Bucharest, Romania | Anca Botezatu, Iulia Virginia Iancu, Coralia Bleotu, Mihaela Economescu, Laura Necula, Denisa Dragu, Lilia Matei, Ioana Pitica, Marinela Bostan, Mirela Mihaila, Laura Grecu, Camelia Sultana, Simona Ruta, Ana Iulia Neagu, Adriana Plesa, Saviana Nedeianu, Alina Nastasie, Cristina Mambet,Gabriela Anton, Carmen Cristina Diaconu |
| EPI_ISL_1085013 | Laboratorio Biologia Molecolare Sars Cov2 - UOC Laboratorio Analisi - Servizio Medicina di Laboratorio, Ospedale "San Francesco" - ATS-ASSL Nuoro | Laboratorio Biologia Molecolare Sars Cov2 - UOC Laboratorio Analisi - Servizio Medicina di Laboratorio, Ospedale "San Francesco" - ATS-ASSL Nuoro | Piras Giovanna, Asproni Rosanna, Monne Maria Itria, Fiamma Maura, , Palmas Angelo Domenico, Lo Maglio Iana, Mameli Giuseppe |
| EPI_ISL_1085534, EPI_ISL_1085674, EPI_ISL_1085675 | Hôpital Henri Mondor | Department of Virology, Henri Mondor University Hospital, Assistance Publique Hôpitaux de Paris, Université Paris-Est Créteil, INSERM U955 | Christophe Rodriguez, Slim Fourati, Vanessa Demontant, Guillaume Gricourt, Melissa N'Debi, Alexandre Soulier, Elisabeth Trawinski, Jean-Michel Pawlotsky |
| EPI_ISL_1085676 | University of Oregon COVID-19 MAP Laboratory | University of Oregon Genomics and Cell Characterization Core Facility (GC3F) | Douglas Turnbull, Ariana White, Jeff Bishop, Jason Carriere |
| EPI_ISL_1085677 | University of Oregon COVID-19 MAP Laboratory | University of Oregon Genomics and Cell Characterization Core Facility (GC3F) | Douglas Turnbull, Ariana White, Jason Carriere, Jeff Bishop |
| EPI_ISL_1086048, EPI_ISL_1086049 | IAL Regional de Bauru | Instituto Adolfo Lutz, Interdiciplinary Procedures Center, Strategic Laboratory | Claudio Tavares Sacchi, Claudia Regina Gonçalves, Erica Valessa Ramos Gomes, Karoline Rodrigues Campos, Caio Vinicius Dias Lopes |
| EPI_ISL_1091781 | ICMT | Universidad Nacional de Colombia - Laboratorio Genómico One Health | Andres F. Cardona-Rios, Daniel O. Maldonado-Perez, Laura Silvana Perez, Karl A Ciuoderis, Idabely Betancur Ortiz, Sandra Ines Cano, Diego A. Álvarez-Díaz, Carlos Franco-Muñoz, Marcela Mercado-Reyes, Jorge E. Osorio, Juan P. Hernandez-Ortiz |
| EPI_ISL_1091784, EPI_ISL_1091785 | LDSP | Universidad Nacional de Colombia - Laboratorio Genómico One Health | Andres F. Cardona-Rios, Daniel O. Maldonado-Perez, Laura Silvana Perez, Karl A Ciuoderis, Maria Angélica Maya, Idabely Betancur Ortiz, Sandra Ines Cano, Diego A. Álvarez-Díaz, Carlos Franco-Muñoz, Marcela Mercado-Reyes, Jorge E. Osorio, Juan P. Hernandez-Ortiz |
| EPI_ISL_1092005 | Laboratorio de Salud Publica de Cesar | Instituto Nacional de Salud- Dirección de Investigación en Salud Pública | Katherine Laiton-Donato, Diego A. Álvarez-Díaz, Carlos Franco-Muñoz, Hector Alejandro Ruiz-Moreno, Maria T. Herrera-Sepúlveda, Diego Andrés Prada, Jhonnatan Reales-González, Sheryll Corchuelo, Julian Naizaque, Gerardo Santamaría, Magdalena Wiesner, Martha Lucia Ospina Martinez, Marcela Mercado-Reyes |
| EPI_ISL_1092006 | Laboratorio IMAT | Instituto Nacional de Salud- Dirección de Investigación en Salud Pública | Katherine Laiton-Donato, Diego A. Álvarez-Díaz, Carlos Franco-Muñoz, Hector Alejandro Ruiz-Moreno, Maria T. Herrera-Sepúlveda, Diego Andrés Prada, Jhonnatan Reales-González, Sheryll Corchuelo, Julian Naizaque, Gerardo Santamaría, Magdalena Wiesner, Martha Lucia Ospina Martinez, Marcela Mercado-Reyes |
| EPI_ISL_1092007 | Laboratorio Continental | Instituto Nacional de Salud- Dirección de Investigación en Salud Pública | Katherine Laiton-Donato, Diego A. Álvarez-Díaz, Carlos Franco-Muñoz, Hector Alejandro Ruiz-Moreno, Maria T. Herrera-Sepúlveda, Diego Andrés Prada, Jhonnatan Reales-González, Sheryll Corchuelo, Julian Naizaque, Gerardo Santamaría, Magdalena Wiesner, Martha Lucia Ospina Martinez, Marcela Mercado-Reyes |
| EPI_ISL_1096120 | IAL Regional de Bauru | Instituto Adolfo Lutz, Interdiciplinary Procedures Center, Strategic Laboratory | Claudio Tavares Sacchi, Claudia Regina Gonçalves, Erica Valessa Ramos Gomes, Karoline Rodrigues Campos, Caio Vinicius Dias Lopes |
| EPI_ISL_1096249, EPI_ISL_1096261, EPI_ISL_1096278, EPI_ISL_1096280, EPI_ISL_1096283, EPI_ISL_1096284, EPI_ISL_1096286, EPI_ISL_1096289, EPI_ISL_1096292, EPI_ISL_1096294, EPI_ISL_1096297, EPI_ISL_1096299 | | | |

| | | | |
|---|---|---|---|
| see above | Hopital | National Reference Center for Viruses of Respiratory Infections, Institut Pasteur, Paris | Marion Barbet, Sylvie Behillil, Méline Bizard, Angela Brisebarre, Camille Capel, Etienne Simon-Lorière, Vincent Enouf, Maud Vanpeene, Sylvie van der Werf,Guinard JéRome |
| EPI_ISL_1096331 | Research | National Reference Center for Viruses of Respiratory Infections, Institut Pasteur, Paris | Marion Barbet, Sylvie Behillil, Méline Bizard, Angela Brisebarre, Camille Capel, Etienne Simon-Lorière, Vincent Enouf, Maud Vanpeene, Sylvie van der Werf,Leruez-Ville Marianne |
| EPI_ISL_1097313, EPI_ISL_1097314, EPI_ISL_1097315, EPI_ISL_1097316, EPI_ISL_1097317, EPI_ISL_1097321, EPI_ISL_1097322, EPI_ISL_1097323, EPI_ISL_1097324, EPI_ISL_1097325, EPI_ISL_1097326, EPI_ISL_1097327, EPI_ISL_1097328, EPI_ISL_1097329, EPI_ISL_1097330, EPI_ISL_1097331, EPI_ISL_1097332, EPI_ISL_1097333, EPI_ISL_1097334, EPI_ISL_1097335, EPI_ISL_1097336, EPI_ISL_1097337, EPI_ISL_1097338, EPI_ISL_1097339, EPI_ISL_1097340, EPI_ISL_1097341, EPI_ISL_1097342, EPI_ISL_1097343, EPI_ISL_1097344, EPI_ISL_1097345, EPI_ISL_1097346 | | | |
| see above | University of Oregon COVID-19 MAP Laboratory | University of Oregon Genomics and Cell Characterization Core Facility (GC3F) | Douglas Turnbull, Ariana White, Jeff Bishop, Jason Carriere |
| EPI_ISL_1103557 | Kurnool Medical College (KMC) | CSIR Institute of Genomics and Integrative Biology | Pallavali Roja Rani, Mohamed Imran, Juturu Vijaya Lakshmi, Bani Jolly, Abhinav Jain, Avileli Surekha, Vigneshwar Senthivel, Pulala Chandrasekhar, Mohit Kumar Divakar, Damam Srinivasulu, Rahul C Bhoyar, PR Vanaja, Vinod Scaria, Sridhar Sivasubbu |
| EPI_ISL_1109637 | Althaia. Xarxa Assistencial Universitària de Manresa | IrsiCaixa - Can Ruti CovidSeq | Marc Noguera-Julian, Mariona Parera, Maria Casadellà, Pilar Armengol, Francesc Catala-Moll, Roger Paredes, Bonaventura Clotet Gloria Trujillo, Rafel Perez Vidal, Jaume Trape Pujol, Carolina Gonzalez Fernandez, Roger Paredes, Eulalia Grau, Bonaventura Clotet |
| EPI_ISL_1110291 | Baylor Scott & White-Temple | Baylor Scott & White-Temple | Ari Rao, Linden Morales, Kimberly Walker, Marcus Volz, Shelby Hendrickson |
| EPI_ISL_1110410, EPI_ISL_1110411, EPI_ISL_1110412, EPI_ISL_1110413 | Hopital | National Reference Center for Viruses of Respiratory Infections, Institut Pasteur, Paris | Marion Barbet, Sylvie Behillil, Méline Bizard, Angela Brisebarre, Camille Capel, Etienne Simon-Lorière, Vincent Enouf, Maud Vanpeene, Sylvie van der Werf,Pilorge LéA |
| EPI_ISL_1110429 | Labo Analyses Med | National Reference Center for Viruses of Respiratory Infections, Institut Pasteur, Paris | Marion Barbet, Sylvie Behillil, Méline Bizard, Angela Brisebarre, Camille Capel, Etienne Simon-Lorière, Vincent Enouf, Maud Vanpeene, Sylvie van der Werf,Etienne Elodie |
| EPI_ISL_1110435, EPI_ISL_1110437, EPI_ISL_1110438, EPI_ISL_1110440 | Hopital | National Reference Center for Viruses of Respiratory Infections, Institut Pasteur, Paris | Marion Barbet, Sylvie Behillil, Méline Bizard, Angela Brisebarre, Camille Capel, Etienne Simon-Lorière, Vincent Enouf, Maud Vanpeene, Sylvie van der Werf,Pilorge LéA |
| EPI_ISL_1110463 | Labo Analyses Med | National Reference Center for Viruses of Respiratory Infections, Institut Pasteur, Paris | Marion Barbet, Sylvie Behillil, Méline Bizard, Angela Brisebarre, Camille Capel, Etienne Simon-Lorière, Vincent Enouf, Maud Vanpeene, Sylvie van der Werf,Morvan Odile |
| EPI_ISL_1110472 | Labo Analyses Med | National Reference Center for Viruses of Respiratory Infections, Institut Pasteur, Paris | Marion Barbet, Sylvie Behillil, Méline Bizard, Angela Brisebarre, Camille Capel, Etienne Simon-Lorière, Vincent Enouf, Maud Vanpeene, Sylvie van der Werf,Etienne Elodie |
| EPI_ISL_1110492, EPI_ISL_1110493, EPI_ISL_1110494, EPI_ISL_1110495, EPI_ISL_1110496, EPI_ISL_1110497, EPI_ISL_1110498, EPI_ISL_1110499, EPI_ISL_1110500, EPI_ISL_1110501, EPI_ISL_1110502, EPI_ISL_1110503, EPI_ISL_1110504, EPI_ISL_1110505, EPI_ISL_1110506, EPI_ISL_1110507, EPI_ISL_1110508, EPI_ISL_1110509, EPI_ISL_1110510 | | | |
| see above | Hopital | National Reference Center for Viruses of Respiratory Infections, Institut Pasteur, Paris | Marion Barbet, Sylvie Behillil, Méline Bizard, Angela Brisebarre, Camille Capel, Etienne Simon-Lorière, Vincent Enouf, Maud Vanpeene, Sylvie van der Werf,Pilorge LéA |
| EPI_ISL_1110952 | Research | National Reference Center for Viruses of Respiratory Infections, Institut Pasteur, Paris | Marion Barbet, Sylvie Behillil, Méline Bizard, Angela Brisebarre, Camille Capel, Etienne Simon-Lorière, Vincent Enouf, Maud Vanpeene, Sylvie van der Werf,Leruez-Ville Marianne |
| EPI_ISL_1110967 | Hopital | National Reference Center for Viruses of Respiratory Infections, Institut Pasteur, Paris | Marion Barbet, Sylvie Behillil, Méline Bizard, Angela Brisebarre, Camille Capel, Etienne Simon-Lorière, Vincent Enouf, Maud Vanpeene, Sylvie van der Werf,Pilorge LéA |
| EPI_ISL_1111069, EPI_ISL_1111132, EPI_ISL_1111133, EPI_ISL_1111134, EPI_ISL_1111135, EPI_ISL_1111136, EPI_ISL_1111137, EPI_ISL_1111138, EPI_ISL_1111139, EPI_ISL_1111140, EPI_ISL_1111141, EPI_ISL_1111142, EPI_ISL_1111143, EPI_ISL_1111144, EPI_ISL_1111145, EPI_ISL_1111146, EPI_ISL_1111147, EPI_ISL_1111148, EPI_ISL_1111149, EPI_ISL_1111150, EPI_ISL_1111151, EPI_ISL_1111152, EPI_ISL_1111153, EPI_ISL_1111154, EPI_ISL_1111155, EPI_ISL_1111156, EPI_ISL_1111157, EPI_ISL_1111158, EPI_ISL_1111159, EPI_ISL_1111160, EPI_ISL_1111161, EPI_ISL_1111308, EPI_ISL_1111309, EPI_ISL_1111310, EPI_ISL_1111312, EPI_ISL_1111314, EPI_ISL_1111315, EPI_ISL_1111446, EPI_ISL_1111457, EPI_ISL_1111458, EPI_ISL_1111459, EPI_ISL_1111460, EPI_ISL_1111461, EPI_ISL_1111462, EPI_ISL_1111463, EPI_ISL_1111464, EPI_ISL_1111465, EPI_ISL_1111466, EPI_ISL_1111467, EPI_ISL_1111469, EPI_ISL_1111470, EPI_ISL_1111471, EPI_ISL_1111472, EPI_ISL_1111473, EPI_ISL_1111474, EPI_ISL_1111475, EPI_ISL_1111476, EPI_ISL_1111477, EPI_ISL_1111478, EPI_ISL_1111479, EPI_ISL_1111480, EPI_ISL_1111481, EPI_ISL_1111482, EPI_ISL_1111483, EPI_ISL_1111484, EPI_ISL_1111485, EPI_ISL_1111486, EPI_ISL_1111487, EPI_ISL_1111488, EPI_ISL_1111489, EPI_ISL_1111490, EPI_ISL_1111491, EPI_ISL_1111492, EPI_ISL_1111493, EPI_ISL_1111494, EPI_ISL_1111495, EPI_ISL_1111496, EPI_ISL_1111497, EPI_ISL_1111498, EPI_ISL_1111499, EPI_ISL_1111500, EPI_ISL_1111501, EPI_ISL_1111502, EPI_ISL_1111503, EPI_ISL_1111504, EPI_ISL_1111505 | | | |
| see above | Laboratorio de Referencia Nacional de Virus Respiratorio. Instituto Nacional de Salud Perú | Laboratorio de Referencia Nacional de Enteropatógenos. Instituto Nacional de Salud del Perú | Ronnie Gavilan Chavez, Junior Caro Castro, Willi Quino Sifuentes, Veronica Hurtado Vela, Iris Silva Molina, Fiorella Orellana Peralta |
| EPI_ISL_1112227, EPI_ISL_1112231 | Istituto Zooprofilattico Sperimentale della Puglia e della Basilicata | Istituto Zooprofilattico Sperimentale della Puglia e della Basilicata | Parisi A., Bianco A., Capozzi L., Del Sambro L., Simone D., Giannico A., Ridolfi D. |
| EPI_ISL_1114726 | Institute for Medical Research, Infectious Disease Research Centre, National Institutes of Health, Ministry of Health Malaysia | Institute for Medical Research, Infectious Disease Research Centre, National Institutes of Health, Ministry of Health Malaysia | Suppiah J, Kamel K, Azizan MA, Thayan R |
| EPI_ISL_1117376 | Beijing Center for Disease Prevention and Control | Beijing Center for Disease Prevention and Control | Yang Pan, Shujuan Cui, Fu Li, Zhaomin Feng, Bing Lyu, Zhichao Liang, Lijuan Chen, Daitao Zhang, Quanyi Wang |
| EPI_ISL_1117377 | Beijing Center for Disease Prevention and Control | Beijing Center for Disease Prevention and Control | Yang Pan, Fu Li, Zhaomin Feng, Shujuan Cui, Bing Lyu, Zhichao Liang, Lijuan Chen, Daitao Zhang, Quanyi Wang |
| EPI_ISL_1117378 | Beijing Center for Disease Prevention and Control | Beijing Center for Disease Prevention and Control | Yang Pan, Zhaomin Feng, Fu Li, Shujuan Cui, Bing Lyu, Zhichao Liang, Lijuan Chen, Daitao Zhang, Quanyi Wang |
| EPI_ISL_1117450 | Melania Hospital, Bagor , west Java | Biosafety Level-3 Laboratory, Indonesian Institute of Sciences (LIPI) | Anggia Prasetyoputri, Isa Nuryana, Sri Swasthikawati, Sugiyono Saputra, Syam Budi Iryanto, Eko Wahyu Putro, Gita Syahputra, Ari Sulistyo Wulandari, Ratih Asmana Ningrum |
| EPI_ISL_1117457 | Klinik Apotik Dien, Jakarta | Biosafety Level-3 Laboratory, Indonesian Institute of Sciences (LIPI) | Anik Budhi Dharmayanthi, Ade Andriani, Hariyatun, Asep M. Ridwanuloh, Syam Budi Iryanto, Eko Wahyu Putro, Gita Syahputra, Susanti, Inswasti Cahyani, Matthew W. Loose, Mohammad Ilyas, Wien Kusharyoto |
| EPI_ISL_1117590 | RS Trimitra, Bogor, West java | Biosafety Level-3 Laboratory, Indonesian Institute of Sciences (LIPI) | Anik Budhi Dharmayanthi, Ade Andriani, Hariyatun, Asep M. Ridwanuloh, Syam Budi Iryanto, Eko Wahyu Putro, Gita Syahputra, Susanti, Inswasti Cahyani, Matthew W. Loose, Mohammad Ilyas, Wien Kusharyoto |
| EPI_ISL_1117592 | Melania Hospital, Bagor , West Java | Biosafety Level-3 Laboratory, Indonesian Institute of Sciences (LIPI) | Anik Budhi Dharmayanthi, Ade Andriani, Hariyatun, Asep M. Ridwanuloh, Syam Budi Iryanto, Eko Wahyu Putro, Gita Syahputra, Susanti, Inswasti Cahyani, Matthew W. Loose, Mohammad Ilyas, Wien Kusharyoto |
| EPI_ISL_1117937, EPI_ISL_1117938, EPI_ISL_1117939, EPI_ISL_1117940, EPI_ISL_1117941, EPI_ISL_1117942, EPI_ISL_1117943, EPI_ISL_1117944, EPI_ISL_1117945, EPI_ISL_1117946, EPI_ISL_1117947, EPI_ISL_1117948 | | | |
| see above | Austrian Agency for Health and Food Safety (AGES) | Bergthaler laboratory, CeMM Research Center for Molecular Medicine of the Austrian Academy of Sciences | Lukas Endler, Anna Schedl, Thomas Penz, Benedikt Agerer, Maelle Le Moing, Michael Schuster, Bekir Erguner, Jan Laine, Martin Senekowitsch, Christoph Bock, Andreas Bergthaler |
| EPI_ISL_1117953, EPI_ISL_1117954, EPI_ISL_1117961, EPI_ISL_1117969, EPI_ISL_1117981, EPI_ISL_1117982, EPI_ISL_1117983, EPI_ISL_1117986, EPI_ISL_1117987, EPI_ISL_1117989, EPI_ISL_1117990, EPI_ISL_1118020 | | | |
| see above | HG Pharma GmbH | Bergthaler laboratory, CeMM Research Center for Molecular Medicine of the Austrian Academy of Sciences | Lukas Endler, Anna Schedl, Thomas Penz, Benedikt Agerer, Maelle Le Moing, Michael Schuster, Bekir Erguner, Jan Laine, Martin Senekowitsch, Christoph Bock, Andreas Bergthaler |
| EPI_ISL_1118206 | National Institute of Health Research and Development | National Institute of Health Research and Development | Hana Apsari Pawestri, Subangkit, Kartika Dewi Puspa, Arie Ardiansyah Nugraha, Hartanti Dian Ikawati, Krisna Nur Andriana Pangesti, Yuni Rukminiati, Ririn Ramadhany, Agustiningsih, Kindi Adam, Holy Arif Wibowo, Triyani Soekarso, Ni Ketut Susilarini, Nurika Hariastuti, Ully Alfi Nikmah, Reni Herman, Nike Susanti, Herna, Tati Febriyanti, Natalie Laurencia Kipuw, Fauzul Muna, Irene Lorinda Indalao, Nelly Puspandari, Vivi Setiawaty. |
| EPI_ISL_1118280 | Merial Health Hospital, Jakarta | Biosafety Level-3 Laboratory, Indonesian Institute of Sciences (LIPI) | Anik Budhi Dharmayanthi, Ade Andriani, Hariyatun, Asep M. Ridwanuloh, Syam Budi Iryanto, Eko Wahyu Putro, Gita Syahputra, Susanti, Inswasti Cahyani, Matthew W. Loose, Mohammad Ilyas, Wien Kusharyoto |
| EPI_ISL_1118879, EPI_ISL_1118880, EPI_ISL_1118881, EPI_ISL_1118882 | Labo Analyses Med | National Reference Center for Viruses of Respiratory Infections, Institut Pasteur, Paris | Marion Barbet, Sylvie Behillil, Méline Bizard, Angela Brisebarre, Camille Capel, Etienne Simon-Lorière, Vincent Enouf, Maud Vanpeene, Sylvie van der Werf,King Lisa |
| EPI_ISL_1121316 | LACEN do Rio Grande do Sul | Instituto Adolfo Lutz, Interdiciplinary Procedures Center, Strategic Laboratory | Claudio Tavares Sacchi, Claudia Regina Gonçalves, Erica Valessa Ramos Gomes, Karoline Rodrigues Campos, Caio Vinicius Dias Lopes |

| | | | |
|---|---|---|---|
| EPI_ISL_1123375 | IAL Regional de Santos | Instituto Adolfo Lutz, Interdiciplinary Procedures Center, Strategic Laboratory | Claudio Tavares Sacchi, Claudia Regina Gonçalves, Erica Valessa Ramos Gomes, Karoline Rodrigues Campos, Caio Vinicius Dias Lopes |
| EPI_ISL_1133133 | LABCOVID_HCPA | LABRESIS_HCPA | Martins AF, Wink PL, Volpato F, Rosset C, de Paris F, Monteiro F, Barth AL |
| EPI_ISL_1138411 | Hospital HIA Percy, Medical Biology | National Reference Center for Viruses of Respiratory Infections, Institut Pasteur, Paris | Marion Barbet, Sylvie Behillil, Méline Bizard, Angela Brisebarre, Camille Capel, Etienne Simon-Lorière, Vincent Enouf, Maud Vanpeene, Sylvie van der Werf,Vincent Foissaud |
| EPI_ISL_1138512, EPI_ISL_1138513, EPI_ISL_1138514, EPI_ISL_1138515, EPI_ISL_1138516, EPI_ISL_1138517 | SA Pärnu Hospital Laboratory | 1. Laboratory of Communicable Diseases (Estonia); 2. Eurofins Genomics Europe Sequencing GmbH | Liidia Dotsenko |
| EPI_ISL_1139059 | LACEN do Mato Grosso do Sul | Instituto Adolfo Lutz, Interdiciplinary Procedures Center, Strategic Laboratory | Claudio Tavares Sacchi, Claudia Regina Gonçalves, Erica Valessa Ramos Gomes, Karoline Rodrigues Campos, Caio Vinicius Dias Lopes |
| EPI_ISL_1159070, EPI_ISL_1159071, EPI_ISL_1159072, EPI_ISL_1159073, EPI_ISL_1159074, EPI_ISL_1159075, EPI_ISL_1159076, EPI_ISL_1159077, EPI_ISL_1159078, EPI_ISL_1159079, EPI_ISL_1159080, EPI_ISL_1159081, EPI_ISL_1159082, EPI_ISL_1159083, EPI_ISL_1159084, EPI_ISL_1159085, EPI_ISL_1159086, EPI_ISL_1159087, EPI_ISL_1159088, EPI_ISL_1159089, EPI_ISL_1159090, EPI_ISL_1159091, EPI_ISL_1159092, EPI_ISL_1159093, EPI_ISL_1159094, EPI_ISL_1159095, EPI_ISL_1159096, EPI_ISL_1159097, EPI_ISL_1159098, EPI_ISL_1159099, EPI_ISL_1159100, EPI_ISL_1159101, EPI_ISL_1159102, EPI_ISL_1159103, EPI_ISL_1159104, EPI_ISL_1159105, EPI_ISL_1159106 | | | |
| see above | Department of Health Technology and Informatics, The Hong Kong Polytechnic University | Department of Health Technology and Informatics, The Hong Kong Polytechnic University | Gilman Kit-Hang Siu, Lam-Kwong Lee, Kenneth Siu-Sing Leung, Jake Siu-Lun Leung, Timothy Ting-Leung Ng, Chloe Toi-Mei Chan, Kingsley King-Gee Tam, Hiu-Yin Lao, Denise Sze-Hang Wong, Alan Ka-Lun Wu, Miranda Chong-Yee Yau, Yvette Wai-Man Lai, Kitty Sau-Chun Fung, Sandy Ka-Yee Chau, Barry Kin-Chung Wong, Wing-Kin To, Kristine Luk, Alex Yat-Man Ho, Tak-Lun Que, Kam-Tong Yip, Wing Cheong Yam, David Ho-Keung Shum, Shea Ping Yip |
| EPI_ISL_1159382 | Adi Husada Kapasari Hospital | Institute of Tropical Disease, Universitas Airlangga | Maria I Lusida, Krisnoadi Rahardjo, Aldise M Nastri, Jezzy R Dewantari, Rima R Prasetya, Hermanto Wijaya, Gatot Soegiarto, Laksmi Wulandari, Resti Yudhawati, Soetjipto, Yasuko Mori, Kazufumi Shimizu |
| EPI_ISL_1159383 | Siti Khodijah Hospital | Institute of Tropical Disease, Universitas Airlangga | Aldise M Nastri, Jezzy R Dewantari, Rima R Prasetya, Krisnoadi Rahardjo, Muhammad Hamdan, Gatot Soegiarto, Laksmi Wulandari, Resti Yudhawati, Soetjipto, Yasuko Mori, Maria I Lusida, Kazufumi Shimizu |
| EPI_ISL_1159385 | Graha Medika Maternity Hospital | Institute of Tropical Disease, Universitas Airlangga | Rima R Prasetya, Krisnoadi Rahardjo, Aldise M Nastri, Jezzy R Dewantari, Diah R Kusumawati, Gatot Soegiarto, Laksmi Wulandari, Resti Yudhawati, Soetjipto, Yasuko Mori, Maria I Lusida, Kazufumi Shimizu |
| EPI_ISL_1159386 | Siti Khodijah Hospital | Institute of Tropical Disease, Universitas Airlangga | Krisnoadi Rahardjo, Aldise M Nastri, Jezzy R Dewantari, Rima R Prasetya, Muhammad Hamdan, Gatot Soegiarto, Laksmi Wulandari, Resti Yudhawati, Yasuko Mori, Soetjipto, Kazufumi Shimizu, Maria I Lusida |
| EPI_ISL_1159387 | Adi Husada Kapasari Hospital | Institute of Tropical Disease, Universitas Airlangga | Maria I Lusida, Krisnoadi Rahardjo, Aldise M Nastri, Jezzy R Dewantari, Rima R Prasetya, Hermanto Wijaya, Gatot Soegiarto, Laksmi Wulandari, Resti Yudhawati, Soetjipto, Yasuko Mori, Kazufumi Shimizu |
| EPI_ISL_1159388 | Paru Hospital | Institute of Tropical Disease, Universitas Airlangga | Aldise M Nastri, Jezzy R Dewantari, Rima R Prasetya, Krisnoadi Rahardjo, Dyah Retno, Gatot Soegiarto, Laksmi Wulandari, Resti Yudhawati, Soetjipto, Yasuko Mori, Maria I Lusida, Kazufumi Shimizu |
| EPI_ISL_1159389 | Darmo Hospital | Institute of Tropical Disease, Universitas Airlangga | Jezzy R Dewantari, Rima R Prasetya, Krisnoadi Rahardjo, Aldise M Nastri, Sulung Budianto, Gatot Soegiarto, Laksmi Wulandari, Resti Yudhawati, Soetjipto, Yasuko Mori, Maria I Lusida, Kazufumi Shimizu |
| EPI_ISL_1161401, EPI_ISL_1161402, EPI_ISL_1161403, EPI_ISL_1161404, EPI_ISL_1161406, EPI_ISL_1161407, EPI_ISL_1161408, EPI_ISL_1161409, EPI_ISL_1163712, EPI_ISL_1163713, EPI_ISL_1163714, EPI_ISL_1163715, EPI_ISL_1163716 | | | |
| see above | LABCOVID_HCPA | LABRESIS_HCPA | Martins AF, Wink PL, Volpato F, Rosset C, de Paris F, Monteiro F, Zavascki AP, Barth AL |
| EPI_ISL_1163736 | LABCOVID_HCPA | LABCOVID_HCPA | Martins AF, Wink PL, Volpato F, Rosset C, de Paris F, Monteiro F, Zavascki AP, Barth AL |
| EPI_ISL_1163737 | LABCOVID_HCPA | LABRESIS_HCPA | Martins AF, Wink PL, Volpato F, Rosset C, de Paris F, Monteiro F, Zavascki AP, Barth AL |
| EPI_ISL_1163827 | Università degli Studi di Perugia | Istituto Zooprofilattico Sperimentale dell'Abruzzo e Molise "G. Caporale" | Mencacci A., Camilloni B., Lorusso A, Marcacci M, Di Domenico M, Ancora M, Curini V, Mangone I, Rinaldi A, Scialabba S, Di Pasquale A, Cammà C, Puglia I, Calistri P, Savini G |
| EPI_ISL_1164927, EPI_ISL_1164928, EPI_ISL_1164929 | Genome Analysis Center, Yamanashi Central Hospital | Genome Analysis Center, Yamanashi Central Hospital | Yosuke Hirotsu |
| EPI_ISL_1165070 | Siti Khodijah Hospital | Institute of Tropical Disease, Universitas Airlangga | Rima R Prasetya, Krisnoadi Rahardjo, Aldise M Nastri, Jezzy R Dewantari, Muhammad Hamdan, Gatot Soegiarto, Laksmi Wulandari, Resti Yudhawati, Yasuko Mori, Soetjipto, Kazufumi Shimizu, Maria I Lusida |
| EPI_ISL_1165071 | Anwar Medika General Hospital | Institute of Tropical Disease, Universitas Airlangga | Kazufumi Shimizu, Krisnoadi Rahardjo, Aldise M Nastri, Jezzy R Dewantari, Rima R Prasetya, Nungky Taniasari, Gatot Soegiarto, Laksmi Wulandari, Resti Yudhawati, Yasuko Mori, Soetjipto, Maria I Lusida |
| EPI_ISL_1165073 | Darus Syifa' Islamic Hospital | Institute of Tropical Disease, Universitas Airlangga | Kazufumi Shimizu, Krisnoadi Rahardjo, Aldise M Nastri, Jezzy R Dewantari, Rima R Prasetya, HM Faiz, Gatot Soegiarto, Laksmi Wulandari, Resti Yudhawati, Yasuko Mori, Soetjipto, Maria I Lusida |
| EPI_ISL_1165076 | Rahman Rahim Hospital | Institute of Tropical Disease, Universitas Airlangga | Jezzy R Dewantari, Rima R Prasetya, Krisnoadi Rahardjo, Aldise M Nastri, Rendra E Febrianto, Gatot Soegiarto, Laksmi Wulandari, Resti Yudhawati, Yasuko Mori, Soetjipto, Kazufumi Shimizu, Maria I Lusida |
| EPI_ISL_1166891, EPI_ISL_1166892, EPI_ISL_1166893, EPI_ISL_1166894, EPI_ISL_1166895, EPI_ISL_1166896, EPI_ISL_1166897, EPI_ISL_1166898, EPI_ISL_1166899, EPI_ISL_1166900, EPI_ISL_1166901, EPI_ISL_1166902, EPI_ISL_1166903, EPI_ISL_1166904, EPI_ISL_1166905, EPI_ISL_1166906, EPI_ISL_1166907, EPI_ISL_1166908, EPI_ISL_1166909, EPI_ISL_1166910, EPI_ISL_1166911, EPI_ISL_1166912, EPI_ISL_1166913, EPI_ISL_1166914, EPI_ISL_1166915, EPI_ISL_1166916, EPI_ISL_1166917, EPI_ISL_1166919, EPI_ISL_1166920 | | | |
| see above | Hopital | National Reference Center for Viruses of Respiratory Infections, Institut Pasteur, Paris | Marion Barbet, Sylvie Behillil, Méline Bizard, Angela Brisebarre, Camille Capel, Etienne Simon-Lorière, Vincent Enouf, Maud Vanpeene, Sylvie van der Werf,Combe Patrice |
| EPI_ISL_1167135, EPI_ISL_1167138, EPI_ISL_1167139, EPI_ISL_1167140, EPI_ISL_1167141, EPI_ISL_1167145, EPI_ISL_1167146, EPI_ISL_1167147, EPI_ISL_1167148, EPI_ISL_1167149, EPI_ISL_1167150, EPI_ISL_1167173, EPI_ISL_1167175, EPI_ISL_1167176, EPI_ISL_1167177 | | | |
| see above | Iressef Genomics lab | L'institut de Recherche en Santé, de Surveillance Épidémiologique et de Formation (IRESSEF) | Souleymane MBOUP, Abdou PADANE, Abdoulie KANTEH ,Abdul Karim SESAY , Khadim GUEYE, Papa Alassane DIAW, Birahim Piere NDIAYE, Barada CISSE, Aminata MBOUP, Moustapha MBOW, Ndeye Coumba Toure KANE, Nafisatou LEYE, Gora LO, Ambroise AHOUIDI , Astou Gaye GAYE, Aminata DIA, Yacine DIA |
| EPI_ISL_1168481, EPI_ISL_1168494, EPI_ISL_1168532, EPI_ISL_1168533, EPI_ISL_1168534, EPI_ISL_1168535, EPI_ISL_1168552, EPI_ISL_1168577, EPI_ISL_1168578, EPI_ISL_1168580, EPI_ISL_1168581, EPI_ISL_1168582, EPI_ISL_1168615, EPI_ISL_1168619, EPI_ISL_1168622, EPI_ISL_1168623 | | | |
| see above | Instituto de Diagnostico y Referencia Epidemiologicos INDRE_RNLSP | Instituto de Diagnostico y Referencia Epidemiologicos (INDRE) | Claudia Wong-Arambula, Abril Rodriguez-Maldonado, Vanessa Rivero-Arredondo, Ariadna Medina-Benitez, Joaquin Quiroz-Mercado,David Fragoso-Fonseca, Sergio Rangel-Guerrero, Natividad Cruz-Ortiz, Tatiana Nunez-Garcia, Gisela Barrera-Badillo, Lucia Hernandez-Rivas, Irma Lopez-Martinez, Ernesto Ramirez-Gonzalez. |
| EPI_ISL_1168653, EPI_ISL_1168654, EPI_ISL_1168656, EPI_ISL_1168658, EPI_ISL_1168659, EPI_ISL_1168660, EPI_ISL_1168661, EPI_ISL_1168662, EPI_ISL_1168663, EPI_ISL_1168664, EPI_ISL_1168665, EPI_ISL_1168666, EPI_ISL_1168667, EPI_ISL_1168668, EPI_ISL_1168669, EPI_ISL_1168670, EPI_ISL_1168671, EPI_ISL_1168672, EPI_ISL_1168673, EPI_ISL_1168674, EPI_ISL_1168675, EPI_ISL_1168677, EPI_ISL_1168678, EPI_ISL_1168679, EPI_ISL_1168680, EPI_ISL_1168681 | | | |
| see above | Centers for Disease Control and Prevention, Dengue Branch | Centers for Disease Control and Prevention, Dengue Branch | Gilberto A. Santiago, Glenda Gonzalez, Betzabel Flores, Keyla Charriez, Gabriela Paz-Bailey, Jorge L. Munoz-Jordan |
| EPI_ISL_1168686 | Puerto Rico Department of Health | Centers for Disease Control and Prevention, Dengue Branch | Gilberto A. Santiago, Glenda Gonzalez, Betzabel Flores, Keyla Charriez, Gabriela Paz-Bailey, Jorge L. Munoz-Jordan |
| EPI_ISL_1169047 | RSUP Dr. Mohammad Hoesin Palembang Sumatera Selatan | National Institute of Health Research and Development | Vivi Setiawaty, Hana Apsari Pawestri, Subangkit, Kartika Dewi Puspa, Arie Ardiansyah Nugraha, Hartanti Dian Ikawati, Nelly Puspandari |
| EPI_ISL_1169048 | BBTKLPP Banjarbaru, Kalimantan Selatan | National Institute of Health Research and Development | Vivi Setiawaty, Hana Apsari Pawestri, Subangkit, Kartika Dewi Puspa, Arie Ardiansyah Nugraha, Hartanti Dian Ikawati, Nelly Puspandari |
| EPI_ISL_1169919 | HIV Molecular Laboratory, Ethiopian Public Health Institute | HIV Molecular Laboratory, Ethiopian Public Health Institute | Weldemariam,A.G., Vanaerschot,M., Abichu,G., Kidane,E., Damena,D., Tessema,S.K., Gutema,G., Ahyong,V., Detweiler,A., Tan,M., Neff,N., Fahsbender,E., Kalantar,K., Leta,D. and Tato,C. |
| EPI_ISL_1170955 | National Institute of Health Research and Development | National Institute of Health Research and Development | Yuni Rukminiati, Agustiningsih, Kindi Adam, Holy Arif Wibowo, Ririn Ramadhany, Hana Apsari Pawestri, Subangkit, Kartika Dewi Puspa, Arie Ardiansyah Nugraha, Hartanti Dian Ikawati, Krisna Nur Andriana Pangesti, Ni Ketut Susilarini, Nur Ika Hariastuti, Uly Alfi Nikmah, Mursinah, Asri Febriyani, Reni Herman, Nike Susanti, Herna, Tati Febriyanti, Natalie Laurencia Kipuw, Fauzul Muna, Irene Lorinda Indalao, Aulia Rizki, Nelly Puspandari, Vivi Setiawaty |

| | | | |
|---|---|---|---|
| EPI_ISL_1170956 | HIV Molecular Lab, Ethiopian Public Health Institute | HIV Molecular Lab | Weldemariam,A.G., Vanaerschot,M., Abichu,G., Kidane,E., Damena,D., Tessema,S.K., Gutema,G., Ahyong,V., Detweiler,A., Tan,M., Neff,N., Fahsbender,E., Kalantar,K., Leta,D. and Tato,C. |
| EPI_ISL_1171621 | LACEN do Mato Grosso do Sul | Instituto Adolfo Lutz, Interdiciplinary Procedures Center, Strategic Laboratory | Claudio Tavares Sacchi, Claudia Regina Gonçalves, Erica Valessa Ramos Gomes, Karoline Rodrigues Campos, Caio Vinicius Dias Lopes |
| EPI_ISL_1171622, EPI_ISL_1171623, EPI_ISL_1171624, EPI_ISL_1171625, EPI_ISL_1171627, EPI_ISL_1171629, EPI_ISL_1171630, EPI_ISL_1171631, EPI_ISL_1171632, EPI_ISL_1171633, EPI_ISL_1171635, EPI_ISL_1171636, EPI_ISL_1171637, EPI_ISL_1171639, EPI_ISL_1171640 | | | |
| see above | IAL Regional de Santos | Instituto Adolfo Lutz, Interdiciplinary Procedures Center, Strategic Laboratory | Claudio Tavares Sacchi, Claudia Regina Gonçalves, Erica Valessa Ramos Gomes, Karoline Rodrigues Campos, Caio Vinicius Dias Lopes |
| EPI_ISL_1171648, EPI_ISL_1171649, EPI_ISL_1171650 | Centro de Saude Dr. Jose Paione em Mococa | Instituto Adolfo Lutz, Interdiciplinary Procedures Center, Strategic Laboratory | Claudio Tavares Sacchi, Claudia Regina Gonçalves, Erica Valessa Ramos Gomes, Karoline Rodrigues Campos, Caio Vinicius Dias Lopes |
| EPI_ISL_1171668, EPI_ISL_1171669, EPI_ISL_1171670 | IAL Regional de Presidente Prudente | Instituto Adolfo Lutz, Interdiciplinary Procedures Center, Strategic Laboratory | Claudio Tavares Sacchi, Claudia Regina Gonçalves, Erica Valessa Ramos Gomes, Karoline Rodrigues Campos, Caio Vinicius Dias Lopes |
| EPI_ISL_1173202 | Laboratorio Biologia Molecolare Sars Cov2 - UOC Laboratorio Analisi - Servizio Medicina di Laboratorio, Ospedale "San Francesco" - ATS-ASSL Nuoro | Laboratorio Specialistico Ematologia, UOC Ematologia, Ospedale "San Francesco" - ATS-ASSL Nuoro | Pira Giovanna, Rosu Valentina, Fancello Patrizia, Asproni Rosanna, Carta Franco, Serra Silvia, Piras Giovanna, Monne Maria Itria, Malune Paolo, Fiamma Maura, Palmas Angelo Domenico, Lo Maglio Iana, Mameli Giuseppe |
| EPI_ISL_1180250, EPI_ISL_1180252, EPI_ISL_1180253 | Laboratorio Biologia Molecolare Sars Cov2 - UOC Laboratorio Analisi - Servizio Medicina di Laboratorio, Ospedale "San Francesco" - ATS-ASSL Nuoro | Laboratorio specialistico UOC Ematologia - Ospedale "San Francesco" - ATS-ASSL Nuoro | Piras Giovanna, Asproni Rosanna,,Fiamma Maura, Monne Maria Itria, Palmas Angelo Domenico, Lo Maglio Iana, Mameli Giuseppe |
| EPI_ISL_1180672, EPI_ISL_1180673 | Institut für Pathologie, Salzkammergut Klinikum Vöcklabruck | Bergthaler laboratory, CeMM Research Center for Molecular Medicine of the Austrian Academy of Sciences | Lukas Endler, Anna Schedl, Fabian Amman, Thomas Penz, Benedikt Agerer, Maelle Le Moing, Michael Schuster, Bekir Erguner, Jan Laine, Martin Senekowitsch, Christoph Bock, Andreas Bergthaler |
| EPI_ISL_1180675, EPI_ISL_1180677, EPI_ISL_1180678, EPI_ISL_1180679, EPI_ISL_1180680, EPI_ISL_1180681, EPI_ISL_1180682, EPI_ISL_1180683, EPI_ISL_1180684, EPI_ISL_1180685, EPI_ISL_1180686, EPI_ISL_1180687, EPI_ISL_1180688, EPI_ISL_1180689, EPI_ISL_1180690, EPI_ISL_1180692, EPI_ISL_1180693, EPI_ISL_1180694, EPI_ISL_1180696, EPI_ISL_1180698, EPI_ISL_1180699, EPI_ISL_1180700, EPI_ISL_1180701, EPI_ISL_1180702, EPI_ISL_1180703, EPI_ISL_1180704, EPI_ISL_1180705, EPI_ISL_1180706, EPI_ISL_1180707, EPI_ISL_1180708, EPI_ISL_1180709, EPI_ISL_1180710, EPI_ISL_1180711, EPI_ISL_1180712, EPI_ISL_1180713, EPI_ISL_1180714 | | | |
| see above | Public Health Authority of the Slovak Republic | Bergthaler laboratory, CeMM Research Center for Molecular Medicine of the Austrian Academy of Sciences | Lukas Endler, Anna Schedl, Fabian Amman, Thomas Penz, Benedikt Agerer, Maelle Le Moing, Michael Schuster, Bekir Erguner, Jan Laine, Martin Senekowitsch, Christoph Bock, Andreas Bergthaler |
| EPI_ISL_1180881, EPI_ISL_1180882, EPI_ISL_1180883, EPI_ISL_1180884 | Department of Laboratory Medicine, Division of Clinical Virology, University of Medicine, Vienna | Bergthaler laboratory, CeMM Research Center for Molecular Medicine of the Austrian Academy of Sciences | Lukas Endler, Anna Schedl, Fabian Amman, Thomas Penz, Benedikt Agerer, Maelle Le Moing, Michael Schuster, Bekir Erguner, Jan Laine, Martin Senekowitsch, Christoph Bock, Andreas Bergthaler |
| EPI_ISL_1181360, EPI_ISL_1181381, EPI_ISL_1181382, EPI_ISL_1181383 | Laboratory of Respiratory Viruses and Measles, Oswaldo Cruz Institute, FIOCRUZ | Laboratory of Respiratory Viruses and Measles, Oswaldo Cruz Institute, FIOCRUZ | Paola Resende, Luciana Appolinario, Fernando Motta, Anna Carolina Paixao, Ana Carolina Mendonca, Alice Sampaio Rocha, Renata Serrano Lopes, Marilda Siqueira on behalf of the Fiocruz COVID-19 Genomic Surveillance Network |
| EPI_ISL_1181434 | Laboratorio Central de Saude Publica do Estado do Rio Grande do Sul (LACEN-RS) | Laboratory of Respiratory Viruses and Measles, Oswaldo Cruz Institute, FIOCRUZ | Paola Resende, Luciana Appolinario, Fernando Motta, Anna Carolina Paixao, Ana Carolina Mendonca, Alice Sampaio Rocha, Renata Serrano Lopes, Tatiana Schaffer Gregianini, Marilda Siqueira on behalf of the Fiocruz COVID-19 Genomic Surveillance Network |
| EPI_ISL_1181446 | Laboratory of Respiratory Viruses and Measles, Oswaldo Cruz Institute, FIOCRUZ | Laboratory of Respiratory Viruses and Measles, Oswaldo Cruz Institute, FIOCRUZ | Paola Resende, Luciana Appolinario, Fernando Motta, Anna Carolina Paixao, Ana Carolina Mendonca, Alice Sampaio Rocha, Renata Serrano Lopes, Marilda Siqueira on behalf of the Fiocruz COVID-19 Genomic Surveillance Network |
| EPI_ISL_1181451 | Laboratorio Central de Saude Publica do Estado do Rio Grande do Sul (LACEN-RS) | Laboratory of Respiratory Viruses and Measles, Oswaldo Cruz Institute, FIOCRUZ | Paola Resende, Luciana Appolinario, Fernando Motta, Anna Carolina Paixao, Ana Carolina Mendonca, Alice Sampaio Rocha, Renata Serrano Lopes, Tatiana Schaffer Gregianini, Marilda Siqueira on behalf of the Fiocruz COVID-19 Genomic Surveillance Network |
| EPI_ISL_1186028 | Institute for Medical Research, Infectious Disease Research Centre, National Institutes of Health, Ministry of Health Malaysia | Institute for Medical Research, Infectious Disease Research Centre, National Institutes of Health, Ministry of Health Malaysia | Suppiah J, Kamel K, Mohd Zawawi Z, Ramly N, Robert F, Thayan R |
| EPI_ISL_1190761, EPI_ISL_1190762, EPI_ISL_1190765, EPI_ISL_1190766, EPI_ISL_1190767, EPI_ISL_1190768 | CREMER(Centre de Rechercherches sur les Maladies Emergentes et Ré-émergentes) | TransVIHMI(Recherches Translationnelles sur le VIH et les Maladies Infectieuses) | Celestin Godwe, Christelle Butel, Martin Maidadi Foudi, Laetitia Serrano, Dowbiss Meta Djomsi, Nicole Vidal, Esemu Livo, Rodrigue Kamga, Marie Amougou , Eitel Mpoudi Ngole,Ahidjo Ayouba, Marcel Tongo, Martine Peeters, Eric Delaporte |
| EPI_ISL_1191141, EPI_ISL_1191736, EPI_ISL_1191737 | Laboratorio Biologia Molecolare Sars Cov2 - UOC Laboratorio Analisi - Servizio Medicina di Laboratorio, Ospedale "San Francesco" - ATS-ASSL Nuoro | Laboratorio specialistico UOC Ematologia - Ospedale "San Francesco" - ATS-ASSL Nuoro | Piras Giovanna, Asproni Rosanna,,Fiamma Maura, Monne Maria Itria, Palmas Angelo Domenico, Lo Maglio Iana, Mameli Giuseppe |
| EPI_ISL_1196282 | LACEN do Mato Grosso do Sul | Instituto Adolfo Lutz, Interdiciplinary Procedures Center, Strategic Laboratory | Claudio Tavares Sacchi, Claudia Regina Gonçalves, Erica Valessa Ramos Gomes, Karoline Rodrigues Campos, Caio Vinicius Dias Lopes |
| EPI_ISL_1196285, EPI_ISL_1196286, EPI_ISL_1196287, EPI_ISL_1196288, EPI_ISL_1196289, EPI_ISL_1196290, EPI_ISL_1196291, EPI_ISL_1196292, EPI_ISL_1196293, EPI_ISL_1196294 | LACEN do Distrito Federal | Instituto Adolfo Lutz, Interdiciplinary Procedures Center, Strategic Laboratory | Claudio Tavares Sacchi, Claudia Regina Gonçalves, Erica Valessa Ramos Gomes, Karoline Rodrigues Campos, Caio Vinicius Dias Lopes |
| EPI_ISL_1201427, EPI_ISL_1201428, EPI_ISL_1201429, EPI_ISL_1201430 | Laboratory of Genetics and Personalized Medicine, Zan Mitrev Clinic | Laboratory of Genetics and Personalized Medicine, Zan Mitrev Clinic | Kungulovski G. et al. |
| EPI_ISL_1201480 | AZ St.Jan Brugge-Oostende | AZ SINT-JAN BRUGGE | Marijke Reynders, Patrick Descheemaeker, Laurien Hoornaert, Thomas Van Landschoot, Jorn Hellemans |
| EPI_ISL_1201488, EPI_ISL_1201489, EPI_ISL_1201490, EPI_ISL_1201491, EPI_ISL_1201492, EPI_ISL_1201493, EPI_ISL_1201494, EPI_ISL_1201495, EPI_ISL_1201496, EPI_ISL_1201497, EPI_ISL_1201498, EPI_ISL_1201499, EPI_ISL_1201500, EPI_ISL_1201501, EPI_ISL_1201502, EPI_ISL_1201503, EPI_ISL_1201504, EPI_ISL_1201505, EPI_ISL_1201506 | | | |
| see above | Hopital | National Reference Center for Viruses of Respiratory Infections, Institut Pasteur, Paris | Marion Barbet, Sylvie Behillil, Méline Bizard, Angela Brisebarre, Camille Capel, Etienne Simon-Lorière, Vincent Enouf, Maud Vanpeene, Sylvie van der Werf,Gastli Nabil |
| EPI_ISL_1201535, EPI_ISL_1201536 | Outre Mer | National Reference Center for Viruses of Respiratory Infections, Institut Pasteur, Paris | Marion Barbet, Sylvie Behillil, Méline Bizard, Angela Brisebarre, Camille Capel, Etienne Simon-Lorière, Vincent Enouf, Maud Vanpeene, Sylvie van der Werf,Rousset Dominique |
| EPI_ISL_1201894 | Diagnosticos da America - DASA | Instituto Adolfo Lutz, Interdiciplinary Procedures Center, Strategic Laboratory | Claudio Tavares Sacchi, Claudia Regina Gonçalves, Erica Valessa Ramos Gomes, Karoline Rodrigues Campos, Caio Vinicius Dias Lopes |
| EPI_ISL_1204503 | Institute for Medical Research, Infectious Disease Research Centre, National Institutes of Health, Ministry of Health Malaysia | Institute for Medical Research, Infectious Disease Research Centre, National Institutes of Health, Ministry of Health Malaysia | Suppiah J, Kamel K, Mohd Zawawi Z, Ramly N, Robert F, Thayan R |
| EPI_ISL_1205468 | Institut de Virologie du CHU de Strasbourg | Institut de Virologie du CHU de Strasbourg | Fafi-Kremer Samira, Gallais Floriane, Gantner Pierre, Laugel Elodie, Solis Morgane, Velay Aurélie, Wendling Marie-Josée |
| EPI_ISL_1208949 | "Scientific and Practical Center for Sanitary and Epidemiological Expertise and Monitoring" branch of the RSE on the REU "National Center for Public Health" of the Ministry of Health RK | Kazakh National Agrarian University (KazNAU); TreeGene LLP Genetic Laboratory; National Scientific Center for Especially Dangerous Infections (NSCEDI) | Beloussov Vyacheslav, Strochkov Vitaliy, Tabynov Kaissar, Sandybayev Nurlan, Tabynov Kairat, Turebekov Nurkeldy, Granica Joanna, Yerubayev Toktassyn, Yespolov Tlektes |
| EPI_ISL_1208951 | "Scientific and Practical Center for Sanitary and Epidemiological Expertise and Monitoring" branch of the RSE on the REU "National Center for Public Health" of the Ministry of Health RK | Kazakh National Agrarian University (KazNAU); TreeGene LLP Genetic Laboratory; National Scientific Center for Especially Dangerous Infections (NSCEDI) | Tabynov Kaissar, Sandybayev Nurlan, Tabynov Kairat, Turebekov Nurkeldy, Granica Joanna, Yerubayev Toktassyn, Beloussov Vyacheslav, Strochkov Vitaliy, Yespolov Tlektes |

| | | | |
|---|---|---|---|
| EPI_ISL_1209243, EPI_ISL_1209244, EPI_ISL_1209245, EPI_ISL_1209246, EPI_ISL_1209247, EPI_ISL_1209248, EPI_ISL_1209249, EPI_ISL_1209250, EPI_ISL_1209251, EPI_ISL_1209252, EPI_ISL_1209253, EPI_ISL_1209254, EPI_ISL_1209255, EPI_ISL_1209256, EPI_ISL_1209258, EPI_ISL_1209259, EPI_ISL_1209260, EPI_ISL_1209261, EPI_ISL_1209262 | | | |
| see above | Public Health Authority of the Slovak Republic | Bergthaler laboratory, CeMM Research Center for Molecular Medicine of the Austrian Academy of Sciences | Lukas Endler, Anna Schedl, Fabian Amman, Thomas Penz, Benedikt Agerer, Maelle Le Moing, Michael Schuster, Bekir Erguner, Jan Laine, Martin Senekowitsch, Christoph Bock, Andreas Bergthaler |
| EPI_ISL_1209299, EPI_ISL_1209300, EPI_ISL_1209301, EPI_ISL_1209302, EPI_ISL_1209303, EPI_ISL_1209304, EPI_ISL_1209305, EPI_ISL_1209306, EPI_ISL_1209307, EPI_ISL_1209308, EPI_ISL_1209309, EPI_ISL_1209310, EPI_ISL_1209311, EPI_ISL_1209312, EPI_ISL_1209313, EPI_ISL_1209314, EPI_ISL_1209315, EPI_ISL_1209316, EPI_ISL_1209317, EPI_ISL_1209318, EPI_ISL_1209319, EPI_ISL_1209320, EPI_ISL_1209321, EPI_ISL_1209322, EPI_ISL_1209323, EPI_ISL_1209324, EPI_ISL_1209325, EPI_ISL_1209326, EPI_ISL_1209327, EPI_ISL_1209328, EPI_ISL_1209329, EPI_ISL_1209330, EPI_ISL_1209331, EPI_ISL_1209332, EPI_ISL_1209333, EPI_ISL_1209334 | | | |
| see above | Department of Laboratory Medicine, Division of Clinical Virology, University of Medicine, Vienna | Bergthaler laboratory, CeMM Research Center for Molecular Medicine of the Austrian Academy of Sciences | Lukas Endler, Anna Schedl, Fabian Amman, Thomas Penz, Benedikt Agerer, Maelle Le Moing, Michael Schuster, Bekir Erguner, Jan Laine, Martin Senekowitsch, Christoph Bock, Andreas Bergthaler |
| EPI_ISL_1212673, EPI_ISL_1212675, EPI_ISL_1212677, EPI_ISL_1212679, EPI_ISL_1212681 | Istituto Zooprofilattico Sperimentale Umbria e Marche "Togo Rosati" | Istituto Zooprofilattico Sperimentale dell'Abruzzo e Molise "G. Caporale" | Biagetti M., Giammarioli M., Lorusso A, Marcacci M, Di Domenico M, Ancora M, Curini V, Mangone I, Rinaldi A, Scialabba S, Di Pasquale A, Cammà C, Puglia I, Calistri P, Savini G |
| EPI_ISL_1219707, EPI_ISL_1219710 | Laboratorio Biologia Molecolare Sars Cov2 - UOC Laboratorio Analisi - Servizio Medicina di Laboratorio, Ospedale "San Francesco" - ATS-ASSL Nuoro | Laboratorio specialistico UOC Ematologia - Ospedale "San Francesco" - ATS-ASSL Nuoro | Piras Giovanna, Asproni Rosanna,,Fiamma Maura, Monne Maria Itria, Palmas Angelo Domenico, Lo Maglio Iana, Mameli Giuseppe |
| EPI_ISL_1219714 | CICSaB/UASLP/LESP SLP | Instituto de Diagnostico y Referencia Epidemiologicos (INDRE) | Claudia Wong-Arambula, Abril Rodriguez-Maldonado, Vanessa Rivero-Arredondo, Ariadna Medina-Benitez, Joaquin Quiroz-Mercado,David Fragoso-Fonseca, Sergio Rangel-Guerrero, Andreu Comas-Garcia, Natividad Cruz-Ortiz, Tatiana Nunez-Garcia, Gisela Barrera-Badillo, Lucia Hernandez-Rivas, Irma Lopez-Martinez, Ernesto Ramirez-Gonzalez. |
| EPI_ISL_1219930, EPI_ISL_1219931 | Hopital | National Reference Center for Viruses of Respiratory Infections, Institut Pasteur, Paris | Marion Barbet, Sylvie Behillil, Méline Bizard, Angela Brisebarre, Camille Capel, Etienne Simon-Lorière, Vincent Enouf, Maud Vanpeene, Sylvie van der Werf,Guiard Carine |
| EPI_ISL_1219942 | Hopital | National Reference Center for Viruses of Respiratory Infections, Institut Pasteur, Paris | Marion Barbet, Sylvie Behillil, Méline Bizard, Angela Brisebarre, Camille Capel, Etienne Simon-Lorière, Vincent Enouf, Maud Vanpeene, Sylvie van der Werf,Pilorge LéA |
| EPI_ISL_1219961 | Labo Analyses Med | National Reference Center for Viruses of Respiratory Infections, Institut Pasteur, Paris | Marion Barbet, Sylvie Behillil, Méline Bizard, Angela Brisebarre, Camille Capel, Etienne Simon-Lorière, Vincent Enouf, Maud Vanpeene, Sylvie van der Werf,Oliel Jonathan |
| EPI_ISL_1220057 | LABORATORIO BIOLOGIA MOLECULAR IMAT SAS | Instituto Nacional de Salud- Dirección de Investigación en Salud Pública | Katherine Laiton-Donato, Carlos Franco-Muñoz, Diego A. Álvarez-Díaz, Hector Alejandro Ruiz-Moreno, Jhonnatan Reales-González, Diego Andrés Prada, Sheryll Corchuelo, Maria T. Herrera-Sepúlveda, Julian Naizaque, Gerardo Santamaría, Magdalena Wiesner, Martha Lucia Ospina Martínez, Marcela Mercado-Reyes. |
| EPI_ISL_1220060 | FUNDACION VALLE DEL LILI | Instituto Nacional de Salud- Dirección de Investigación en Salud Pública | Katherine Laiton-Donato, Carlos Franco-Muñoz, Diego A. Álvarez-Díaz, Hector Alejandro Ruiz-Moreno, Jhonnatan Reales-González, Diego Andrés Prada, Sheryll Corchuelo, Maria T. Herrera-Sepúlveda, Julian Naizaque, Gerardo Santamaría, Magdalena Wiesner, Martha Lucia Ospina Martínez, Marcela Mercado-Reyes. |
| EPI_ISL_1222779 | Institut de Virologie du CHU de Strasbourg | Institut de Virologie du CHU de Strasbourg | Fafi-Kremer Samira, Gallais Floriane, Gantner Pierre, Laugel Elodie, Solis Morgane, Velay Aurélie, Wendling Marie-Josée |
| EPI_ISL_1225156, EPI_ISL_1225157, EPI_ISL_1225158, EPI_ISL_1225160, EPI_ISL_1225161 | Turkish Society of Internal Medicine | Gen Era Diagnostics Corporation Life Science Research and Molecular Diagnostics | Serhat Ünal |
| EPI_ISL_1229144, EPI_ISL_1229145, EPI_ISL_1229146, EPI_ISL_1229147, EPI_ISL_1229148, EPI_ISL_1229149 | Laboratorio Biologia Molecolare Sars Cov2 - UOC Laboratorio Analisi - Servizio Medicina di Laboratorio, Ospedale "San Francesco" - ATS-ASSL Nuoro | Laboratorio specialistico UOC Ematologia - Ospedale "San Francesco" - ATS-ASSL Nuoro | Giovanna Piras, Rosanna Asproni, Maura Fiamma, Maria Itria Monne, Angelo Domenico Palmas, Iana Lo Maglio, Giuseppe Mameli |
| EPI_ISL_1233440, EPI_ISL_1233441, EPI_ISL_1233443, EPI_ISL_1233444, EPI_ISL_1233445, EPI_ISL_1233446, EPI_ISL_1233447, EPI_ISL_1233448, EPI_ISL_1233449, EPI_ISL_1233450, EPI_ISL_1233451, EPI_ISL_1233452, EPI_ISL_1233453, EPI_ISL_1233454, EPI_ISL_1233455, EPI_ISL_1233456, EPI_ISL_1233457, EPI_ISL_1233459, EPI_ISL_1233460, EPI_ISL_1233461, EPI_ISL_1233462, EPI_ISL_1233463, EPI_ISL_1233464, EPI_ISL_1233465, EPI_ISL_1233466, EPI_ISL_1233467, EPI_ISL_1233469, EPI_ISL_1233470, EPI_ISL_1233471, EPI_ISL_1233472, EPI_ISL_1233473, EPI_ISL_1233474, EPI_ISL_1233475, EPI_ISL_1233476, EPI_ISL_1233477, EPI_ISL_1233509, EPI_ISL_1233510, EPI_ISL_1233511, EPI_ISL_1233512, EPI_ISL_1233513, EPI_ISL_1233514, EPI_ISL_1233515, EPI_ISL_1233516, EPI_ISL_1233517, EPI_ISL_1233518, EPI_ISL_1233519, EPI_ISL_1233520, EPI_ISL_1233521, EPI_ISL_1233522, EPI_ISL_1233523, EPI_ISL_1233524, EPI_ISL_1233525, EPI_ISL_1233526, EPI_ISL_1233527, EPI_ISL_1233528, EPI_ISL_1233529, EPI_ISL_1233530, EPI_ISL_1233531, EPI_ISL_1233532, EPI_ISL_1233533, EPI_ISL_1233534, EPI_ISL_1233535, EPI_ISL_1233536, EPI_ISL_1233537, EPI_ISL_1233538, EPI_ISL_1233539, EPI_ISL_1233540, EPI_ISL_1233541, EPI_ISL_1233542, EPI_ISL_1233543, EPI_ISL_1233544, EPI_ISL_1233623, EPI_ISL_1233624, EPI_ISL_1233625, EPI_ISL_1233626, EPI_ISL_1233627 | | | |
| see above | National Center for Infectious and Parasitic Diseases (NCIPD) | National Center for Infectious and Parasitic Diseases (NCIPD) | Alexiev et al |
| EPI_ISL_1235678, EPI_ISL_1235682, EPI_ISL_1235683 | Baylor Scott & White-Temple | Baylor Scott & White-Temple | Ari Rao, Linden Morales, Kimberly Walker, Marcus Volz, Shelby Hendrickson, Caitlin Maloney |
| EPI_ISL_1238792, EPI_ISL_1238793 | Ecological and Evolutionary Genomics. UGA-CINVESTAV | Unidad Universitaria de Secuenciación Masiva y Bioinformática (UUSMB). IBT-UNAM | Angelica Cibrian |
| EPI_ISL_1239145, EPI_ISL_1239216, EPI_ISL_1239217, EPI_ISL_1239219 | National Institute of Health Research and Development | National Institute of Health Research and Development | Hana Apsari Pawestri, Subangkit, Kartika Dewi Puspa, Arie Ardiansyah Nugraha, Hartanti Dian Ikawati, Krisna Nur Andriana Pangesti, Yuni Rukminiati, Ririn Ramadhany, Agustiningsih, Kindi Adam, Holy Arif Wibowo, Triyani Soekarso, Ni Ketut Susilarini, Nurika Hariastuti, Ully Alfi Nikmah, Reni Herman, Nike Susanti, Herna, Tati Febriyanti, Natalie Laurencia Kipuw, Fauzul Muna, Irene Lorinda Indalao, Nelly Puspandari, Vivi Setiawaty. |
| EPI_ISL_1239220 | RSUPN Dr. Cipto Mangunkusumo | National Institute of Health Research and Development | Hana Apsari Pawestri, Subangkit, Kartika Dewi Puspa, Arie Ardiansyah Nugraha, Hartanti Dian Ikawati, Krisna Nur Andriana Pangesti, Yuni Rukminiati, Ririn Ramadhany, Agustiningsih, Kindi Adam, Holy Arif Wibowo, Triyani Soekarso, Ni Ketut Susilarini, Nurika Hariastuti, Ully Alfi Nikmah, Reni Herman, Nike Susanti, Herna, Tati Febriyanti, Natalie Laurencia Kipuw, Fauzul Muna, Irene Lorinda Indalao, Nelly Puspandari, Vivi Setiawaty. |
| EPI_ISL_1239221, EPI_ISL_1239222 | National Institute of Health Research and Development | National Institute of Health Research and Development | Hana Apsari Pawestri, Subangkit, Kartika Dewi Puspa, Arie Ardiansyah Nugraha, Hartanti Dian Ikawati, Krisna Nur Andriana Pangesti, Yuni Rukminiati, Ririn Ramadhany, Agustiningsih, Kindi Adam, Holy Arif Wibowo, Triyani Soekarso, Ni Ketut Susilarini, Nurika Hariastuti, Ully Alfi Nikmah, Reni Herman, Nike Susanti, Herna, Tati Febriyanti, Natalie Laurencia Kipuw, Fauzul Muna, Irene Lorinda Indalao, Nelly Puspandari, Vivi Setiawaty. |
| EPI_ISL_1250222 | Clinical Microbiology Lab RSPTN Universitas Hasanuddin | Clinical Microbiology Lab RSPTN Universitas Hasanuddin | Muhammad Nasrum Massi, Rizalinda Sjahril, Gita Vita Soraya, Triwibawa, Handayani Halik, Muhammad Yusuf Usman, Rufika Shari |
| EPI_ISL_1251274, EPI_ISL_1251275 | "Stefan S. Nicolau"Institute of Virology | "Stefan S. Nicolau"Institute of Virology | Anca Botezatu, Iulia Virginia Iancu, Coralia Bleotu, Mihaela Economescu, Laura Necula, Denisa Dragu, Lilia Matei, Ioana Pitica, Marinela Bostan, Mirela Mihaila, Laura Grecu, Camelia Sultana, Simona Ruta, Ana Iulia Neagu, Adriana Plesa, Saviana Nedeianu, Alina Nastasie, Cristina Mambet,Gabriela Anton, Carmen Cristina Diaconu |
| EPI_ISL_1254648 | Ospedale di Venere | Istituto Zooprofilattico Sperimentale della Puglia e della Basilicata | Parisi A., Bianco A., Capozzi L., Del Sambro L., Simone D., Iacobellis M., Ridolfi D., Giannico A. |
| EPI_ISL_1255105 | West African Centre for Cell Biology of Infectious Pathogens (WACCBIP), University of Ghana, Volta Road, Legon-Accra, Ghana | West African Centre for Cell Biology of Infectious Pathogens (WACCBIP), University of Ghana, Volta Road, Legon-Accra, Ghana | Collins M. Morang'a, Joyce M. Ngoi, Evelyn B. Quansah, Samirah Saiid, Dominic S.Y. Amuzu, Vincent Appiah, Philip M. Soglo, Vanessa Magnussen, Aisha Mohammed, Kesego Tapela, Nelson Kibinge, Abdoulaye B Diallo, Frederick Kumi-Ansah, Theophilus Odoom, Oliver D Boakye5, Emmanuella Amoako4, Abdul-Karim Abass, , Samuel Kaba Akoriyea, Frederick Tei-Maya, Lucas N. Amenga-Etego, Dam Kenneth Mibut, Yaw Bediako, Benjamin Demah Nuertey, Gordon A Awandare, Peter K Quashie, Gordon A Awandare, Yaw Bediako |
| EPI_ISL_1255123, EPI_ISL_1255124, EPI_ISL_1255125, EPI_ISL_1255126, EPI_ISL_1255127, EPI_ISL_1255128, EPI_ISL_1255129, EPI_ISL_1255130, EPI_ISL_1255131, EPI_ISL_1255132, EPI_ISL_1255133, EPI_ISL_1255134, EPI_ISL_1255135, EPI_ISL_1255136, EPI_ISL_1255137, EPI_ISL_1255138, EPI_ISL_1255139, EPI_ISL_1255184, EPI_ISL_1255185 | | | |
| see above | West African Centre for Cell Biology of Infectious Pathogens (WACCBIP), University of Ghana, Accra, Ghana | West African Centre for Cell Biology of Infectious Pathogens (WACCBIP), University of Ghana, Volta Road, Legon-Accra, Ghana | Collins M. Morang'a, Joyce M. Ngoi, Evelyn B. Quansah, Samirah Saiid, Dominic S.Y. Amuzu, Vincent Appiah, Philip M. Soglo, Vanessa Magnussen, Aisha Mohammed, Kesego Tapela, Nelson Kibinge, Abdoulaye B Diallo, Frederick Kumi-Ansah, Theophilus Odoom, Oliver D Boakye5, Emmanuella Amoako4, Abdul-Karim Abass, , Samuel Kaba Akoriyea, Frederick Tei-Maya, Lucas N. Amenga-Etego, Dam Kenneth Mibut, Yaw Bediako, Benjamin Demah Nuertey, Gordon A Awandare, Peter K Quashie, Gordon A Awandare, Yaw Bediako |

| | | | |
|---|---|---|---|
| EPI_ISL_1257818, EPI_ISL_1257819, EPI_ISL_1257820, EPI_ISL_1257822 | Balai Litbangkes Papua | National Institute of Health Research and Development | Hana Apsari Pawestri, Subangkit, Kartika Dewi Puspa, Arie Ardiansyah Nugraha, Hartanti Dian Ikawati, Krisna Pangesti, Nelly Puspandari, Hotma Sitompul, Antonius Octavian, Vivi Setiawaty |
| EPI_ISL_1257823, EPI_ISL_1257824, EPI_ISL_1257825, EPI_ISL_1257826 | RSUP DR Mohammad Hoesin | National Institute of Health Research and Development | Hana Apsari Pawestri, Subangkit, Kartika Dewi Puspa, Arie Ardiansyah Nugraha, Hartanti Dian Ikawati, Krisna Pangesti, Nelly Puspandari, Eny Rahmawati, Rogaya, Vivi Setiawaty |
| EPI_ISL_1257832 | BTKL-PP Ambon | National Institute of Health Research and Development | Vivi Setiawaty, Hana Apsari Pawestri, Subangkit, Kartika Dewi Puspa, Arie Ardiansyah Nugraha, Hartanti Dian Ikawati, Nelly Puspandari, Krisna Pangesti |
| EPI_ISL_1257851, EPI_ISL_1257852, EPI_ISL_1257853 | BBTKLPP Banjarbaru | National Institute of Health Research and Development | Hartanti Dian Ikawati, Hana Apsari Pawestri, Subangkit, Kartika Dewi Puspa, Arie Ardiansyah Nugraha, Krisna Pangesti, Nelly Puspandari, Dewi Hermawati, Mutia Raudah, Vivi Setiawaty |
| EPI_ISL_1257860 | Farmalab Indoutama | National Institute of Health Research and Development | Hana Apsari Pawestri, Subangkit, Kartika Dewi Puspa, Arie Ardiansyah Nugraha, Hartanti Dian Ikawati, Nelly Puspandari, Krisna Pangesti, Vivi Setiawaty |
| EPI_ISL_1257863 | RSUP Dr. Sardjito | National Institute of Health Research and Development | Kartika Dewi Puspa, Hana Apsari Pawestri, Subangkit, Arie Ardiansyah Nugraha, Hartanti Dian Ikawati, Nelly Puspandari, Krisna Pangesti, Vivi Setiawaty |
| EPI_ISL_1257864 | BTKL-PP Ambon | National Institute of Health Research and Development | Kartika Dewi Puspa, Hana Apsari Pawestri, Subangkit, Arie Ardiansyah Nugraha, Hartanti Dian Ikawati, Nelly Puspandari, Krisna Pangesti, Vivi Setiawaty |
| EPI_ISL_1257875, EPI_ISL_1257878, EPI_ISL_1257879, EPI_ISL_1257880 | Centre Pasteur du Cameroun | Institut Pasteur de Dakar | Njouom Richard, Diagne Moussa Moïse, Dia Ndongo, Diallo Amadou, Sankhe Safietou, Diop Mamadou, Loucoubar Cheikh, Elisabeth Carniel, Faye Ousmane, Sall Amadou Alpha |
| EPI_ISL_1258423, EPI_ISL_1258424 | Clinical Microbiology Lab RSPTN Universitas Hasanuddin | Clinical Microbiology Lab RSPTN Universitas Hasanuddin | Muhammad Nasrum Massi, Rizalinda Sjahril, Gita Vita Soraya, Triwibawa, Handayani, Muhammad Yusuf Usman, Rufika Shari |
| EPI_ISL_1258453 | National Institute of Infectious Diseases-Prof. Dr. Matei Bals Molecular Diagnostics Laboratory | National Institute of Infectious Diseases-Prof. Dr. Matei Bals Molecular Diagnostics Laboratory | Corina Casangiu, Leontina Banica, Marius Surleac, Ovidiu Vlaicu, Ionelia Nicolae, Simona Paraschiv, Dan Otelea |
| EPI_ISL_1258463, EPI_ISL_1258464, EPI_ISL_1258465 | Clinical Microbiology Lab RSPTN Universitas Hasanuddin | Clinical Microbiology Lab RSPTN Universitas Hasanuddin | Muhammad Nasrum Massi, Rizalinda Sjahril, Gita Vita Soraya, Triwibawa, Handayani Halik, Muhammad Yusuf Usman, Rufika Shari |
| EPI_ISL_1259297 | Hopital | National Reference Center for Viruses of Respiratory Infections, Institut Pasteur, Paris | Marion Barbet, Sylvie Behillil, Méline Bizard, Angela Brisebarre, Camille Capel, Etienne Simon-Lorière, Vincent Enouf, Maud Vanpeene, Sylvie van der Werf,Dufour Mj |
| EPI_ISL_1259304, EPI_ISL_1259305, EPI_ISL_1259306 | Hopital | National Reference Center for Viruses of Respiratory Infections, Institut Pasteur, Paris | Marion Barbet, Sylvie Behillil, Méline Bizard, Angela Brisebarre, Camille Capel, Etienne Simon-Lorière, Vincent Enouf, Maud Vanpeene, Sylvie van der Werf,Gastli Nabil |
| EPI_ISL_1259309 | Labo Analyses Med | National Reference Center for Viruses of Respiratory Infections, Institut Pasteur, Paris | Marion Barbet, Sylvie Behillil, Méline Bizard, Angela Brisebarre, Camille Capel, Etienne Simon-Lorière, Vincent Enouf, Maud Vanpeene, Sylvie van der Werf |
| EPI_ISL_1260862, EPI_ISL_1260871 | "Stefan S. Nicolau"Institute of Virology | "Stefan S. Nicolau"Institute of Virology | Anca Botezatu, Iulia Virginia Iancu, Coralia Bleotu, Mihaela Economescu, Laura Necula, Denisa Dragu, Lilia Matei, Ioana Pitica, Marinela Bostan, Mirela Mihaila, Laura Grecu, Camelia Sultana, Simona Ruta, Ana Iulia Neagu, Adriana Plesa, Saviana Nedeianu, Alina Nastasie, Cristina Mambet,Gabriela Anton, Carmen Cristina Diaconu |
| EPI_ISL_1262760, EPI_ISL_1262761, EPI_ISL_1262762 | Hopital | National Reference Center for Viruses of Respiratory Infections, Institut Pasteur, Paris | Marion Barbet, Sylvie Behillil, Méline Bizard, Angela Brisebarre, Camille Capel, Etienne Simon-Lorière, Vincent Enouf, Maud Vanpeene, Sylvie van der Werf,Pilorge LéA |
| EPI_ISL_1262814 | Labo Analyses Med | National Reference Center for Viruses of Respiratory Infections, Institut Pasteur, Paris | Marion Barbet, Sylvie Behillil, Méline Bizard, Angela Brisebarre, Camille Capel, Etienne Simon-Lorière, Vincent Enouf, Maud Vanpeene, Sylvie van der Werf,Lebaupain Aude |
| EPI_ISL_1262816, EPI_ISL_1262817, EPI_ISL_1262818 | Hopital | National Reference Center for Viruses of Respiratory Infections, Institut Pasteur, Paris | Marion Barbet, Sylvie Behillil, Méline Bizard, Angela Brisebarre, Camille Capel, Etienne Simon-Lorière, Vincent Enouf, Maud Vanpeene, Sylvie van der Werf,Louis Mathilde |
| EPI_ISL_1265373 | "Stefan S. Nicolau"Institute of Virology | "Stefan S. Nicolau"Institute of Virology | Anca Botezatu, Iulia Virginia Iancu, Coralia Bleotu, Mihaela Economescu, Laura Necula, Denisa Dragu, Lilia Matei, Ioana Pitica, Marinela Bostan, Mirela Mihaila, Laura Grecu, Camelia Sultana, Simona Ruta, Ana Iulia Neagu, Adriana Plesa, Saviana Nedeianu, Alina Nastasie, Cristina Mambet,Gabriela Anton, Carmen Cristina Diaconu |
| EPI_ISL_1273443 | Department of Virology, Bangabandhu Sheikh Mujib Medical University | Bioinformatics Division, National Institute of Biotechnology (NIB) | Munira Jahan, Arittra Bhattacharjee, Raad Rahmat, S M Rashedul Islam, Tahmina Akhter, Ishtiaque Ahammad, Mohammad Uzzal Hossain, Md. Salimullah, Saif Ullah Munshi |
| EPI_ISL_1279948, EPI_ISL_1279950, EPI_ISL_1279951, EPI_ISL_1279952, EPI_ISL_1279953, EPI_ISL_1279954, EPI_ISL_1279956, EPI_ISL_1279957, EPI_ISL_1279958, EPI_ISL_1279959, EPI_ISL_1279960, EPI_ISL_1279962, EPI_ISL_1279963, EPI_ISL_1279964, EPI_ISL_1279966, EPI_ISL_1279967, EPI_ISL_1279968, EPI_ISL_1279970, EPI_ISL_1279973, EPI_ISL_1279974, EPI_ISL_1279975, EPI_ISL_1279976, EPI_ISL_1279977 | | | |
| see above | National Institute of Infectious Diseases-Prof. Dr. Matei Bals Molecular Diagnostics Laboratory | National Institute of Infectious Diseases-Prof. Dr. Matei Bals Molecular Diagnostics Laboratory | Leontina Banica, Marius Surleac, Corina Casangiu, Petre Milu, Andreea Tudor, Simona Paraschiv, Dan Otelea |
| EPI_ISL_1281568, EPI_ISL_1281569, EPI_ISL_1281571, EPI_ISL_1281573 | Laboratory of Extremely Dangerous Infections of Federal Budget Health Care Instute "Center of Hygiene and Epidemiology in Sakhalin region" | Group of Genomics and Postgenomic Technologies of Central Research Institute of Epidemiology | Samoilov AE, Kaptelova VV, Korneenko EV, Valdokhina AV, Bulanenko VP, Saenko SS, Arutiunian AE, Byankina MA, Golubeva AG, Speranskaya AS, Tivanova EV, Kondrasheva LV, Akimkin VG |
| EPI_ISL_1281680 | Imelda hospital Bonheiden | Imelda hospital Bonheiden | Johan Frans, Dagmar Obbels, Hanne Valgaeren |
| EPI_ISL_1283923, EPI_ISL_1283924 | RSIA Sayyidah | Eijkman Institute for Molecular Biology, Ministry of Research and Technology/National Agency for Research and Innovation | Edison Johar, Frilasita A Yudhaputri, Hidayat Trimarsanto, Iskandar Adnan, Lydia V. Panggalo, Sukma Oktavianthi, Willy Agustine, Safarina G Malik, Khin Saw Myint, Amin Soebandrio |
| EPI_ISL_1283929 | RS Mitra Keluarga Kelapa Gading | Eijkman Institute for Molecular Biology, Ministry of Research and Technology/National Agency for Research and Innovation | Edison Johar, Frilasita A Yudhaputri, Hidayat Trimarsanto, Iskandar Adnan, Lydia V. Panggalo, Sukma Oktavianthi, Willy Agustine, Safarina G Malik, Khin Saw Myint, Amin Soebandrio |
| EPI_ISL_1283932 | RS Mulya Tangerang | Eijkman Institute for Molecular Biology, Ministry of Research and Technology/National Agency for Research and Innovation | Edison Johar, Frilasita A Yudhaputri, Hidayat Trimarsanto, Iskandar Adnan, Lydia V. Panggalo, Sukma Oktavianthi, Willy Agustine, Safarina G Malik, Khin Saw Myint, Amin Soebandrio |
| EPI_ISL_1283933 | RS Mitra Keluarga Gading Serpong | Eijkman Institute for Molecular Biology, Ministry of Research and Technology/National Agency for Research and Innovation | Edison Johar, Frilasita A Yudhaputri, Hidayat Trimarsanto, Iskandar Adnan, Lydia V. Panggalo, Sukma Oktavianthi, Willy Agustine, Safarina G Malik, Khin Saw Myint, Amin Soebandrio |
| EPI_ISL_1283936 | RS Sentra Medika Cikarang | Eijkman Institute for Molecular Biology, Ministry of Research and Technology/National Agency for Research and Innovation | Edison Johar, Frilasita A Yudhaputri, Hidayat Trimarsanto, Iskandar Adnan, Lydia V. Panggalo, Sukma Oktavianthi, Willy Agustine, Safarina G Malik, Khin Saw Myint, Amin Soebandrio |
| EPI_ISL_1283938 | RSUD Cileungsi | Eijkman Institute for Molecular Biology, Ministry of Research and Technology/National Agency for Research and Innovation | Frilasita A Yudhaputri, Hidayat Trimarsanto, Iskandar Adnan, Lydia V. Panggalo, Sukma Oktavianthi, Willy Agustine, Edison Johar, Safarina G Malik, Khin Saw Myint, Amin Soebandrio |
| EPI_ISL_1283957 | RSIA Selasih Medika | Eijkman Institute for Molecular Biology, Ministry of Research and Technology/National Agency for Research and Innovation | Frilasita A Yudhaputri, Hidayat Trimarsanto, Iskandar Adnan, Lydia V. Panggalo, Sukma Oktavianthi, Willy Agustine, Edison Johar, Safarina G Malik, Khin Saw Myint, Amin Soebandrio |
| EPI_ISL_1283960 | RS Asy-Syifaa Leuwiliang | Eijkman Institute for Molecular Biology, Ministry of Research and Technology/National Agency for Research and Innovation | Hidayat Trimarsanto, Iskandar Adnan, Lydia V. Panggalo, Sukma Oktavianthi, Willy Agustine, Edison Johar, Frilasita A Yudhaputri, Safarina G Malik, Khin Saw Myint, Amin Soebandrio |
| EPI_ISL_1283967 | RS Bina Husada Bogor | Eijkman Institute for Molecular Biology, Ministry of Research and Technology/National Agency for Research and Innovation | Hidayat Trimarsanto, Iskandar Adnan, Lydia V. Panggalo, Sukma Oktavianthi, Willy Agustine, Edison Johar, Frilasita A Yudhaputri, Safarina G Malik, Khin Saw Myint, Amin Soebandrio |

| | | | |
|---|---|---|---|
| EPI_ISL_1283970 | RSIA Selasih Medika | Eijkman Institute for Molecular Biology, Ministry of Research and Technology/National Agency for Research and Innovation | Hidayat Trimarsanto, Iskandar Adnan, Lydia V. Panggalo, Sukma Oktavianthi, Willy Agustine, Edison Johar, Frilasita A Yudhaputri, Safarina G Malik, Khin Saw Myint, Amin Soebandrio |
| EPI_ISL_1283973 | RSU Bunda Mulia Cikarang | Eijkman Institute for Molecular Biology, Ministry of Research and Technology/National Agency for Research and Innovation | Hidayat Trimarsanto, Iskandar Adnan, Lydia V. Panggalo, Sukma Oktavianthi, Willy Agustine, Edison Johar, Frilasita A Yudhaputri, Safarina G Malik, Khin Saw Myint, Amin Soebandrio |
| EPI_ISL_1283979 | RS Cendana | Eijkman Institute for Molecular Biology, Ministry of Research and Technology/National Agency for Research and Innovation | Iskandar Adnan, Lydia V. Panggalo, Sukma Oktavianthi, Willy Agustine, Edison Johar, Hidayat Trimarsanto, Frilasita A Yudhaputri, Safarina G Malik, Khin Saw Myint, Amin Soebandrio |
| EPI_ISL_1283987 | RS Islam Sari Asih Ar-Rahmah | Eijkman Institute for Molecular Biology, Ministry of Research and Technology/National Agency for Research and Innovation | Iskandar Adnan, Lydia V. Panggalo, Sukma Oktavianthi, Willy Agustine, Edison Johar, Hidayat Trimarsanto, Frilasita A Yudhaputri, Safarina G Malik, Khin Saw Myint, Amin Soebandrio |
| EPI_ISL_1283988 | RSUD Cileungsi | Eijkman Institute for Molecular Biology, Ministry of Research and Technology/National Agency for Research and Innovation | Iskandar Adnan, Lydia V. Panggalo, Sukma Oktavianthi, Willy Agustine, Edison Johar, Hidayat Trimarsanto, Frilasita A Yudhaputri, Safarina G Malik, Khin Saw Myint, Amin Soebandrio |
| EPI_ISL_1283996 | RSIA Bunda Sejahtera | Eijkman Institute for Molecular Biology, Ministry of Research and Technology/National Agency for Research and Innovation | Lydia V. Panggalo, Sukma Oktavianthi, Willy Agustine, Edison Johar, Hidayat Trimarsanto, Iskandar Adnan, Frilasita A Yudhaputri, Safarina G Malik, Khin Saw Myint, Amin Soebandrio |
| EPI_ISL_1284000 | RSU HGA Depok | Eijkman Institute for Molecular Biology, Ministry of Research and Technology/National Agency for Research and Innovation | Lydia V. Panggalo, Sukma Oktavianthi, Willy Agustine, Edison Johar, Hidayat Trimarsanto, Iskandar Adnan, Frilasita A Yudhaputri, Safarina G Malik, Khin Saw Myint, Amin Soebandrio |
| EPI_ISL_1284007, EPI_ISL_1284008 | RS Mitra Keluarga Gading Serpong | Eijkman Institute for Molecular Biology, Ministry of Research and Technology/National Agency for Research and Innovation | Sukma Oktavianthi, Willy Agustine, Edison Johar, Hidayat Trimarsanto, Iskandar Adnan, Lydia V. Panggalo, Frilasita A Yudhaputri, Safarina G Malik, Khin Saw Myint, Amin Soebandrio |
| EPI_ISL_1284014 | RSUD Cileungsi | Eijkman Institute for Molecular Biology, Ministry of Research and Technology/National Agency for Research and Innovation | Sukma Oktavianthi, Willy Agustine, Edison Johar, Hidayat Trimarsanto, Iskandar Adnan, Lydia V. Panggalo, Frilasita A Yudhaputri, Safarina G Malik, Khin Saw Myint, Amin Soebandrio |
| EPI_ISL_1284017 | RS Mitra Keluarga Kalideres | Eijkman Institute for Molecular Biology, Ministry of Research and Technology/National Agency for Research and Innovation | Sukma Oktavianthi, Willy Agustine, Edison Johar, Hidayat Trimarsanto, Iskandar Adnan, Lydia V. Panggalo, Frilasita A Yudhaputri, Safarina G Malik, Khin Saw Myint, Amin Soebandrio |
| EPI_ISL_1284021, EPI_ISL_1284025 | RS Mitra Keluarga Gading Serpong | Eijkman Institute for Molecular Biology, Ministry of Research and Technology/National Agency for Research and Innovation | Sukma Oktavianthi, Willy Agustine, Edison Johar, Hidayat Trimarsanto, Iskandar Adnan, Lydia V. Panggalo, Frilasita A Yudhaputri, Safarina G Malik, Khin Saw Myint, Amin Soebandrio |
| EPI_ISL_1284027 | RS Sentra Medika Cikarang | Eijkman Institute for Molecular Biology, Ministry of Research and Technology/National Agency for Research and Innovation | Willy Agustine, Edison Johar, Hidayat Trimarsanto, Iskandar Adnan, Lydia V. Panggalo, Sukma Oktavianthi, Frilasita A Yudhaputri, Safarina G Malik, Khin Saw Myint, Amin Soebandrio |
| EPI_ISL_1284031 | RS OMNI Alam Sutera | Eijkman Institute for Molecular Biology, Ministry of Research and Technology/National Agency for Research and Innovation | Willy Agustine, Edison Johar, Hidayat Trimarsanto, Iskandar Adnan, Lydia V. Panggalo, Sukma Oktavianthi, Frilasita A Yudhaputri, Safarina G Malik, Khin Saw Myint, Amin Soebandrio |
| EPI_ISL_1284035 | RS Taman Harapan Baru | Eijkman Institute for Molecular Biology, Ministry of Research and Technology/National Agency for Research and Innovation | Willy Agustine, Edison Johar, Hidayat Trimarsanto, Iskandar Adnan, Lydia V. Panggalo, Sukma Oktavianthi, Frilasita A Yudhaputri, Safarina G Malik, Khin Saw Myint, Amin Soebandrio |
| EPI_ISL_1284040 | RSU Azra | Eijkman Institute for Molecular Biology, Ministry of Research and Technology/National Agency for Research and Innovation | Willy Agustine, Edison Johar, Hidayat Trimarsanto, Iskandar Adnan, Lydia V. Panggalo, Sukma Oktavianthi, Frilasita A Yudhaputri, Safarina G Malik, Khin Saw Myint, Amin Soebandrio |
| EPI_ISL_1284044, EPI_ISL_1284048 | RS Bhakti Husada Cikarang | Eijkman Institute for Molecular Biology, Ministry of Research and Technology/National Agency for Research and Innovation | Edison Johar, Frilasita A Yudhaputri, Hidayat Trimarsanto, Iskandar Adnan, Lydia V. Panggalo, Sukma Oktavianthi, Willy Agustine, Safarina G Malik, Khin Saw Myint, Amin Soebandrio |
| EPI_ISL_1284049 | RSU Harapan Bunda | Eijkman Institute for Molecular Biology, Ministry of Research and Technology/National Agency for Research and Innovation | Edison Johar, Frilasita A Yudhaputri, Hidayat Trimarsanto, Iskandar Adnan, Lydia V. Panggalo, Sukma Oktavianthi, Willy Agustine, Safarina G Malik, Khin Saw Myint, Amin Soebandrio |
| EPI_ISL_1284052 | RS Rumah Indonesia Sehat | Eijkman Institute for Molecular Biology, Ministry of Research and Technology/National Agency for Research and Innovation | Edison Johar, Frilasita A Yudhaputri, Hidayat Trimarsanto, Iskandar Adnan, Lydia V. Panggalo, Sukma Oktavianthi, Willy Agustine, Safarina G Malik, Khin Saw Myint, Amin Soebandrio |
| EPI_ISL_1284054 | Metro Hospitals Cikarang | Eijkman Institute for Molecular Biology, Ministry of Research and Technology/National Agency for Research and Innovation | Edison Johar, Frilasita A Yudhaputri, Hidayat Trimarsanto, Iskandar Adnan, Lydia V. Panggalo, Sukma Oktavianthi, Willy Agustine, Safarina G Malik, Khin Saw Myint, Amin Soebandrio |
| EPI_ISL_1284057 | RS Primaya Tangerang | Eijkman Institute for Molecular Biology, Ministry of Research and Technology/National Agency for Research and Innovation | Edison Johar, Frilasita A Yudhaputri, Hidayat Trimarsanto, Iskandar Adnan, Lydia V. Panggalo, Sukma Oktavianthi, Willy Agustine, Safarina G Malik, Khin Saw Myint, Amin Soebandrio |
| EPI_ISL_1284063 | RS Mitra Keluarga Gading Serpong | Eijkman Institute for Molecular Biology, Ministry of Research and Technology/National Agency for Research and Innovation | Frilasita A Yudhaputri, Hidayat Trimarsanto, Iskandar Adnan, Lydia V. Panggalo, Sukma Oktavianthi, Willy Agustine, Edison Johar, Safarina G Malik, Khin Saw Myint, Amin Soebandrio |
| EPI_ISL_1284064 | RSU HGA Depok | Eijkman Institute for Molecular Biology, Ministry of Research and Technology/National Agency for Research and Innovation | Frilasita A Yudhaputri, Hidayat Trimarsanto, Iskandar Adnan, Lydia V. Panggalo, Sukma Oktavianthi, Willy Agustine, Edison Johar, Safarina G Malik, Khin Saw Myint, Amin Soebandrio |
| EPI_ISL_1289960 | Laboratory of Virology, Ribeirão Preto General Hospital, Ribeirão Preto Medical School, University of São Paulo | Laboratory of Oncology, Blood Center of Ribeirão Preto, Ribeirão Preto School of Medicine, University of São Paulo | CAMPOS, MR; SANTOS, A.L.P.; YAMAMOTO, A.Y.; COLLI, L.M.; FONSECA, B.A.L.; BELLISSIMO-RODRIGUES, F. |
| EPI_ISL_1293051 | LACEN do Distrito Federal | Instituto Adolfo Lutz, Interdiciplinary Procedures Center, Strategic Laboratory | Claudio Tavares Sacchi, Claudia Regina Gonçalves, Erica Valessa Ramos Gomes, Karoline Rodrigues Campos, Caio Vinicius Dias Lopes |
| EPI_ISL_1293353 | OLVZ Aalst | OLVZ Aalst | Astrid Holderbeke |
| EPI_ISL_1297194 | RSUD Pasarminggu | Stem Cell Lab. Universitas Pembangunan Nasional Veteran Jakarta (UPNVJ) | Andri Pramesyanti Pramono, Yudhi Nugraha, Ria Maria Theresa, Yudi Amiarno, Megadianty Mokoginta |

EPI_ISL_1299195, EPI_ISL_1299196, EPI_ISL_1299197, EPI_ISL_1299198, EPI_ISL_1299199, EPI_ISL_1299200, EPI_ISL_1299201, EPI_ISL_1299202, EPI_ISL_1299203, EPI_ISL_1299204, EPI_ISL_1299205, EPI_ISL_1299206, EPI_ISL_1299207, EPI_ISL_1299208, EPI_ISL_1299209, EPI_ISL_1299210, EPI_ISL_1299211,

| | | | |
|---|---|---|---|
| EPI_ISL_1299212, EPI_ISL_1299213, EPI_ISL_1299214, EPI_ISL_1299215, EPI_ISL_1299216 | | | |
| see above | Department of Virology, Bangabandhu Sheikh Mujib Medical University | Department of Virology, Bangabandhu Sheikh Mujib Medical University | Munira Jahan, Arittra Bhattacharjee, Raad Rahmat, S M Rashedul Islam, Tahmina Akhter, Ishtiaque Ahammad, Mohammad Uzzal Hossain, Md. Salimullah, Saif Ullah Munshi |
| EPI_ISL_1300496 | RSUD Pasarminggu | Stem Cell Lab, Universitas Pembangunan Nasional Veteran Jakarta (UPNVJ) | Andri Pramesyanti Pramono, Yudhi Nugraha, Ria Maria Theresa, Yudi Amiarno, Megadianty Mokoginta |
| EPI_ISL_1300521 | RSUD Pasarminggu | Stem Cell Lab, Universitas Pembangunan Nasional Veteran Jakarta | Andri Pramesyanti Pramono, Yudhi Nugraha, Ria Maria Theresa, Yudi Amiarno, Megadianty Mokoginta |
| EPI_ISL_1300535, EPI_ISL_1300540, EPI_ISL_1300644, EPI_ISL_1300660 | RSUD Pasarminggu | Stem Cell Lab, Universitas Pembangunan Nasional Veteran Jakarta (UPNVJ) | Andri Pramesyanti Pramono, Yudhi Nugraha, Ria Maria Theresa, Yudi Amiarno, Megadianty Mokoginta |
| EPI_ISL_1301830, EPI_ISL_1301832 | Outre Mer | National Reference Center for Viruses of Respiratory Infections, Institut Pasteur, Paris | Marion Barbet, Sylvie Behillil, Méline Bizard, Angela Brisebarre, Camille Capel, Louise Lefrançois, Etienne Simon-Lorière, Vincent Enouf, Maud Vanpeene, Sylvie van der Werf,Combe Patrice |
| EPI_ISL_1301834, EPI_ISL_1301835, EPI_ISL_1301836, EPI_ISL_1301837 | Hopital | National Reference Center for Viruses of Respiratory Infections, Institut Pasteur, Paris | Marion Barbet, Sylvie Behillil, Méline Bizard, Angela Brisebarre, Camille Capel, Louise Lefrançois, Etienne Simon-Lorière, Vincent Enouf, Maud Vanpeene, Sylvie van der Werf,Leruez-Ville Marianne |
| EPI_ISL_1301889 | Outre Mer | National Reference Center for Viruses of Respiratory Infections, Institut Pasteur, Paris | Marion Barbet, Sylvie Behillil, Méline Bizard, Angela Brisebarre, Camille Capel, Louise Lefrançois, Etienne Simon-Lorière, Vincent Enouf, Maud Vanpeene, Sylvie van der Werf,Combe Patrice |
| EPI_ISL_1301895 | Hopital | National Reference Center for Viruses of Respiratory Infections, Institut Pasteur, Paris | Marion Barbet, Sylvie Behillil, Méline Bizard, Angela Brisebarre, Camille Capel, Louise Lefrançois, Etienne Simon-Lorière, Vincent Enouf, Maud Vanpeene, Sylvie van der Werf,Combe Patrice |
| EPI_ISL_1301903, EPI_ISL_1301904, EPI_ISL_1301905, EPI_ISL_1301906, EPI_ISL_1301907, EPI_ISL_1301908, EPI_ISL_1301909, EPI_ISL_1301910, EPI_ISL_1301911, EPI_ISL_1301912, EPI_ISL_1301913, EPI_ISL_1301914 | | | |
| see above | Outre Mer | National Reference Center for Viruses of Respiratory Infections, Institut Pasteur, Paris | Marion Barbet, Sylvie Behillil, Méline Bizard, Angela Brisebarre, Camille Capel, Louise Lefrançois, Etienne Simon-Lorière, Vincent Enouf, Maud Vanpeene, Sylvie van der Werf,Combe Patrice |
| EPI_ISL_1301915 | Labo Analyses Med | National Reference Center for Viruses of Respiratory Infections, Institut Pasteur, Paris | Marion Barbet, Sylvie Behillil, Méline Bizard, Angela Brisebarre, Camille Capel, Louise Lefrançois, Etienne Simon-Lorière, Vincent Enouf, Maud Vanpeene, Sylvie van der Werf,Delaval José |
| EPI_ISL_1301920 | Hopital | National Reference Center for Viruses of Respiratory Infections, Institut Pasteur, Paris | Marion Barbet, Sylvie Behillil, Méline Bizard, Angela Brisebarre, Camille Capel, Louise Lefrançois, Etienne Simon-Lorière, Vincent Enouf, Maud Vanpeene, Sylvie van der Werf,Leruez-Ville Marianne |
| EPI_ISL_1301921 | Hopital | National Reference Center for Viruses of Respiratory Infections, Institut Pasteur, Paris | Marion Barbet, Sylvie Behillil, Méline Bizard, Angela Brisebarre, Camille Capel, Louise Lefrançois, Etienne Simon-Lorière, Vincent Enouf, Maud Vanpeene, Sylvie van der Werf,Ducancelle Alexandra |
| EPI_ISL_1301924, EPI_ISL_1301925, EPI_ISL_1301926, EPI_ISL_1301927, EPI_ISL_1301928, EPI_ISL_1301929 | Hopital | National Reference Center for Viruses of Respiratory Infections, Institut Pasteur, Paris | Marion Barbet, Sylvie Behillil, Méline Bizard, Angela Brisebarre, Camille Capel, Louise Lefrançois, Etienne Simon-Lorière, Vincent Enouf, Maud Vanpeene, Sylvie van der Werf,Leruez-Ville Marianne |
| EPI_ISL_1301981, EPI_ISL_1301982, EPI_ISL_1301983, EPI_ISL_1301984, EPI_ISL_1302090, EPI_ISL_1302094, EPI_ISL_1302098, EPI_ISL_1302099, EPI_ISL_1302100, EPI_ISL_1302101, EPI_ISL_1302102, EPI_ISL_1302103, EPI_ISL_1302104, EPI_ISL_1302105, EPI_ISL_1302106, EPI_ISL_1302107, EPI_ISL_1302108, EPI_ISL_1302109, EPI_ISL_1302122, EPI_ISL_1302123, EPI_ISL_1302124, EPI_ISL_1302125, EPI_ISL_1302126, EPI_ISL_1302127, EPI_ISL_1302128, EPI_ISL_1302129, EPI_ISL_1302130, EPI_ISL_1302131, EPI_ISL_1302132, EPI_ISL_1302133, EPI_ISL_1302134, EPI_ISL_1302135, EPI_ISL_1302136, EPI_ISL_1302137, EPI_ISL_1302138, EPI_ISL_1302139, EPI_ISL_1302140, EPI_ISL_1302141, EPI_ISL_1302142, EPI_ISL_1302143, EPI_ISL_1302144, EPI_ISL_1302145, EPI_ISL_1302146, EPI_ISL_1302147, EPI_ISL_1302148 | | | |
| see above | National Center of Infectious and Parasitic Diseases | National Center of Infectious and Parasitic Diseases | Alexiev et al |
| EPI_ISL_1302395 | Instituto Nacional de Enfermedades Respiratorias (INER) | Instituto de Biotecnología de la UNAM | Authors from IBT, IMSS, InDRE and INER (in alphabetical order): Carlos F. Arias, Santiago Ávila-Ríos, Gisela Barrera-Badillo, Eduardo Becerril-Vargas, Celia Boukadida, Natividad Cruz-Ortiz, Larissa Fernandes-Matano, Ricardo Grande, Lucia Hernandez-Rivas, Alejandra Hernández-Terán, Pavel Isa, Irma Lopez-Martinez, José Arturo Martínez-Orozco, Margarita Matías-Florentino, Fidencio Mejía-Nepomuceno, Edgar Mendieta-Condado, Mario Mújica-Sánchez, José Esteban Muñoz-Medina, Tatiana Nunez-Garcia, Luis Alberto Ochoa-Carrera, Hector Esteban Paz-Juárez, Francisco Pulido, José Ernesto Ramírez-González, Alma Rincón-Rubio, Teresita Rojas-Mendoza, Jorge Salas-Hernández, Alejandro Sanchez-Flores, Clara Esperanza Santacruz-Tinoco, Andrea Santos Coy-Arechavaleta, Blanca Taboada, Gloria Vazquez, Joel Armando Vázquez-Pérez, Jerome Jean Verleyen |
| EPI_ISL_1302396, EPI_ISL_1302397, EPI_ISL_1302398 | Instituto de Diagnostico y Referencia Epidemiologicos InDRE_RNLSP | Instituto de Biotecnología de la UNAM | Authors from IBT, IMSS, InDRE and INER (in alphabetical order): Carlos F. Arias, Santiago Ávila-Ríos, Gisela Barrera-Badillo, Eduardo Becerril-Vargas, Celia Boukadida, Natividad Cruz-Ortiz, Larissa Fernandes-Matano, Ricardo Grande, Lucia Hernandez-Rivas, Alejandra Hernández-Terán, Pavel Isa, Irma Lopez-Martinez, José Arturo Martínez-Orozco, Margarita Matías-Florentino, Fidencio Mejía-Nepomuceno, Edgar Mendieta-Condado, Mario Mújica-Sánchez, José Esteban Muñoz-Medina, Tatiana Nunez-Garcia, Luis Alberto Ochoa-Carrera, Hector Esteban Paz-Juárez, Francisco Pulido, José Ernesto Ramírez-González, Alma Rincón-Rubio, Teresita Rojas-Mendoza, Jorge Salas-Hernández, Alejandro Sanchez-Flores, Clara Esperanza Santacruz-Tinoco, Andrea Santos Coy-Arechavaleta, Blanca Taboada, Gloria Vazquez, Joel Armando Vázquez-Pérez, Jerome Jean Verleyen |
| EPI_ISL_1302399, EPI_ISL_1302400 | Instituto Nacional de Enfermedades Respiratorias (INER) | Instituto de Biotecnología de la UNAM | Authors from IBT, IMSS, InDRE and INER (in alphabetical order): Carlos F. Arias, Santiago Ávila-Ríos, Gisela Barrera-Badillo, Eduardo Becerril-Vargas, Celia Boukadida, Natividad Cruz-Ortiz, Larissa Fernandes-Matano, Ricardo Grande, Lucia Hernandez-Rivas, Alejandra Hernández-Terán, Pavel Isa, Irma Lopez-Martinez, José Arturo Martínez-Orozco, Margarita Matías-Florentino, Fidencio Mejía-Nepomuceno, Edgar Mendieta-Condado, Mario Mújica-Sánchez, José Esteban Muñoz-Medina, Tatiana Nunez-Garcia, Luis Alberto Ochoa-Carrera, Hector Esteban Paz-Juárez, Francisco Pulido, José Ernesto Ramírez-González, Alma Rincón-Rubio, Teresita Rojas-Mendoza, Jorge Salas-Hernández, Alejandro Sanchez-Flores, Clara Esperanza Santacruz-Tinoco, Andrea Santos Coy-Arechavaleta, Blanca Taboada, Gloria Vazquez, Joel Armando Vázquez-Pérez, Jerome Jean Verleyen |
| EPI_ISL_1303368 | Clinica de la Costa | Instituto Nacional de Salud- Dirección de Investigación en Salud Pública | Katherine Laiton-Donato, Diego A. Álvarez-Díaz, Carlos Franco-Muñoz, Hector Alejandro Ruiz-Moreno, Maria T. Herrera-Sepúlveda, Diego Andrés Prada, Jhonnatan Reales-González, Sheryll Corchuelo, Julian Naizaque, Gerardo Santamaría, Sergio Gomez, Lisseth Pardo, Juan Camilo Martinez, Marta Lopez Blanco, Ángela Alarcon Cruz, Diana Malo, Carmen Osorio, Magdalena Wiesner, Martha Lucia Ospina Martinez, Marcela Mercado-Reyes |
| EPI_ISL_1303504 | LACEN de Rondonia | Instituto Adolfo Lutz, Interdiciplinary Procedures Center, Strategic Laboratory | Claudio Tavares Sacchi, Claudia Regina Gonçalves, Erica Valessa Ramos Gomes, Karoline Rodrigues Campos, Caio Vinicius Dias Lopes |
| EPI_ISL_1303513, EPI_ISL_1303515, EPI_ISL_1303516 | LACEN do Estado de Goias | Instituto Adolfo Lutz, Interdiciplinary Procedures Center, Strategic Laboratory | Claudio Tavares Sacchi, Claudia Regina Gonçalves, Erica Valessa Ramos Gomes, Karoline Rodrigues Campos, Caio Vinicius Dias Lopes |
| EPI_ISL_1311863 | Laboratorio Biologia Molecolare Sars Cov2 - UOC Laboratorio Analisi - Servizio Medicina di Laboratorio, Ospedale "San Francesco" - ATS-ASSL Nuoro | Laboratorio specialistico UOC Ematologia - Ospedale "San Francesco" - ATS-ASSL Nuoro | Piras Giovanna, Asproni Rosanna,Fiamma Maura, Monne Maria Itria, Malune Paolo, Palmas Angelo Domenico, Lo Maglio Iana, Mameli Giuseppe |
| EPI_ISL_1318203, EPI_ISL_1318204, EPI_ISL_1318205, EPI_ISL_1318208, EPI_ISL_1318209, EPI_ISL_1318211, EPI_ISL_1318212 | Ministry of Health Hospitals | Institute of Health and Community Medicine | Chan Chia Jui, David Perera, Ooi Mong How, Chua Hock Hin, Tonnii Sia Loong Loong, Wong Jyn Shan, Wong Kiing Aik |
| EPI_ISL_1324766 | Laboratorio de Diagnostico de Enfermedades Emergentes y Reemergentes | Instituto de Diagnostico y Referencia Epidemiologicos (INDRE) | Claudia Wong-Arambula, Abril Rodriguez-Maldonado, Vanessa Rivero-Arredondo, Ariadna Medina-Benitez, Joaquin Quiroz-Mercado, Sergio Rangel-Guerrero,Natali Vega-Magana, Natividad Cruz-Ortiz, Tatiana Nunez-Garcia, Gisela Barrera-Badillo, Lucia Hernandez-Rivas, Irma Lopez-Martinez, Ernesto Ramirez-Gonzalez. |
| EPI_ISL_1335884 | Department of Microbiology, Faculty of Medicine, Universitas Sumatera Utara | Faculty of Medicine, Universitas Sumatera Utara; Institute of Tropical Disease, Universitas Airlangga | Jezzy R Dewantari, Rima R Prasetya, Krisnoadi Rahardjo, Aldise M Nastri, Yasuko Mori, Soetjipto, Kazufumi Shimizu, Maria I Lusida, Meliani, R Andika D Cahyadi, Mirzan Hasibuan, Irbah R Nainggolan, Ramadhan Bestari, Franciscus Ginting, Muhammad Ichwan, R Lia Kusumawati |
| EPI_ISL_1335886 | Sundari Hospital Medan | Faculty of Medicine, Universitas Sumatera Utara; Institute of Tropical Disease, Universitas Airlangga | Kazufumi Shimizu, Krisnoadi Rahardjo, Aldise M Nastri, Jezzy R Dewantari, Rima R Prasetya, Yasuko Mori, Soetjipto, Maria I Lusida, Meliani, R Andika D Cahyadi, Mirzan Hasibuan, Irbah R Nainggolan, Ramadhan Bestari, Franciscus Ginting, Muhammad Ichwan, R Lia Kusumawati, Inke N D Lubis |
| EPI_ISL_1335888 | Martha Friska Multatuli Hospital Medan | Faculty of Medicine, Universitas Sumatera Utara; Institute of | Krisnoadi Rahardjo, Aldise M Nastri, Jezzy R Dewantari, Rima R Prasetya, Yasuko Mori, Soetjipto, Kazufumi Shimizu, Maria I Lusida, Meliani, R Andika D |

| | | | |
|---|---|---|---|
| | | Tropical Disease, Universitas Airlangga | Cahyadi, Mirzan Hasibuan, Irbah R Nainggolan, Ramadhan Bestari, Franciscus Ginting, Muhammad Ichwan, R Lia Kusumawati, Inke N D Lubis |
| EPI_ISL_1335889 | Advent Hospital Medan | Faculty of Medicine, Universitas Sumatera Utara; Institute of Tropical Disease, Universitas Airlangga | Krisnoadi Rahardjo, Aldise M Nastri, Jezzy R Dewantari, Rima R Prasetya, Yasuko Mori, Soetjipto, Kazufumi Shimizu, Maria I Lusida, Meliani, R Andika D Cahyadi, Mirzan Hasibuan, Irbah R Nainggolan, Ramadhan Bestari, Franciscus Ginting, Muhammad Ichwan, R Lia Kusumawati, Inke N D Lubis |
| EPI_ISL_1335890 | Elisabeth Hospital Medan | Faculty of Medicine, Universitas Sumatera Utara; Institute of Tropical Disease, Universitas Airlangga | Aldise M Nastri, Jezzy R Dewantari, Rima R Prasetya, Krisnoadi Rahardjo, Yasuko Mori, Soetjipto, Kazufumi Shimizu, Maria I Lusida, Meliani, R Andika D Cahyadi, Mirzan Hasibuan, Irbah R Nainggolan, Ramadhan Bestari, Franciscus Ginting, Muhammad Ichwan, R Lia Kusumawati, Inke N D Lubis |
| EPI_ISL_1335891 | Advent Hospital Medan | Faculty of Medicine, Universitas Sumatera Utara; Institute of Tropical Disease, Universitas Airlangga | Jezzy R Dewantari, Rima R Prasetya, Krisnoadi Rahardjo, Aldise M Nastri, Yasuko Mori, Soetjipto, Kazufumi Shimizu, Maria I Lusida, Meliani, R Andika D Cahyadi, Mirzan Hasibuan, Irbah R Nainggolan, Ramadhan Bestari, Franciscus Ginting, Muhammad Ichwan, R Lia Kusumawati, Inke N D Lubis |
| EPI_ISL_1335892 | Department of Microbiology, Faculty of Medicine, Universitas Sumatera Utara | Faculty of Medicine, Universitas Sumatera Utara; Institute of Tropical Disease, Universitas Airlangga | Rima R Prasetya, Krisnoadi Rahardjo, Aldise M Nastri, Jezzy R Dewantari, Yasuko Mori, Soetjipto, Kazufumi Shimizu, Maria I Lusida, Meliani, R Andika D Cahyadi, Mirzan Hasibuan, Irbah R Nainggolan, Ramadhan Bestari, Franciscus Ginting, Muhammad Ichwan, R Lia Kusumawati, Inke N D Lubis |
| EPI_ISL_1335893 | Department of Microbiology, Faculty of Medicine, Universitas Sumatera Utara | Faculty of Medicine, Universitas Sumatera Utara; Institute of Tropical Disease, Universitas Airlangga | Kazufumi Shimizu, Krisnoadi Rahardjo, Aldise M Nastri, Jezzy R Dewantari, Rima R Prasetya, Yasuko Mori, Soetjipto, Maria I Lusida, Meliani, R Andika D Cahyadi, Mirzan Hasibuan, Irbah R Nainggolan, Ramadhan Bestari, Franciscus Ginting, Muhammad Ichwan, R Lia Kusumawati, Inke N D Lubis |
| EPI_ISL_1335894 | Department of Microbiology, Faculty of Medicine, Universitas Sumatera Utara | Faculty of Medicine, Universitas Sumatera Utara; Institute of Tropical Disease, Universitas Airlangga | Maria I Lusida, Krisnoadi Rahardjo, Aldise M Nastri, Jezzy R Dewantari, Rima R Prasetya, Yasuko Mori, Soetjipto, Kazufumi Shimizu, Meliani, R Andika D Cahyadi, Mirzan Hasibuan, Irbah R Nainggolan, Ramadhan Bestari, Franciscus Ginting, Muhammad Ichwan, R Lia Kusumawati, Inke N D Lubis |
| EPI_ISL_1335895 | Martha Friska Multatuli Hospital Medan | Faculty of Medicine, Universitas Sumatera Utara; Institute of Tropical Disease, Universitas Airlangga | Krisnoadi Rahardjo, Aldise M Nastri, Jezzy R Dewantari, Rima R Prasetya, Yasuko Mori, Soetjipto, Kazufumi Shimizu, Maria I Lusida, Meliani, R Andika D Cahyadi, Mirzan Hasibuan, Irbah R Nainggolan, Ramadhan Bestari, Franciscus Ginting, Muhammad Ichwan, R Lia Kusumawati, Inke N D Lubis |
| EPI_ISL_1340754, EPI_ISL_1340755, EPI_ISL_1340756 | Departamento de Virologia, Laboratorio Central de Salud Pública, Avenida Venezuela y Teniente Escurra, Asunción, Paraguay | Laboratory of Respiratory Viruses and Measles, Oswaldo Cruz Institute, FIOCRUZ | Paola Resende, Cynthia Vazquez, Luciana Appolinario, Fernando Motta, Anna Carolina Paixao, Ana Carolina Mendonca, Alice Sampaio Rocha, Renata Serrano Lopes, Marilda Siqueira on behalf of the Fiocruz COVID-19 Genomic Surveillance Network |
| EPI_ISL_1340782, EPI_ISL_1340783, EPI_ISL_1340784, EPI_ISL_1340785 | Northwestern Memorial Hospital | Northwestern University - Ozer Lab | Ramon Lorenzo-Redondo, Lacy M. Simons, Taylor J. Dean, Chad J. Achenbach, Lawrence J. Jennings, Chao Qi, Michael G. Ison, Judd F. Hultquist, Egon A. Ozer |
| EPI_ISL_1346626, EPI_ISL_1346628, EPI_ISL_1346630, EPI_ISL_1346631 | Center of Excellence in Clinical Virology | Research Unit of Systems Microbiology | Jiratchaya Puenpa, Pattaraporn Nimsamer, Oraphan Mayuramart, Vorthon Sawaswong, Sunchai Payungporn, Anek Mungaomklang, Ritthideach Yorsaeng, Jira Chansaenroj, Kamolthip Atsawawaranunt, Vichan Pawan, Suwanna Petto, Amornmas Kongklieng, Pakkaporn Panwijitkul, Yong Poovowan |
| EPI_ISL_1358316, EPI_ISL_1358317 | LACEN do Mato Grosso do Sul | Instituto Adolfo Lutz, Interdiciplinary Procedures Center, Strategic Laboratory | Claudio Tavares Sacchi, Claudia Regina Gonçalves, Erica Valessa Ramos Gomes, Karoline Rodrigues Campos, Caio Vinicius Dias Lopes |
| EPI_ISL_1360296 | CSIR-National Environmental Engineering Research Institute | CSIR-Centre for Cellular and Molecular Biology - INSACOG | Onkar Kulkarni, Lamuk Zaveri, Ara Sreenivas, Sofia Banu, Shreekant Verma, Amareshwar Vodapalli , Viswagithe S L, B Himasri, Sharath Chandra Thota, Karthik Bharadwaj Tallapaka, Krishna Khairnar, Rakesh K Mishra,Divya Tej Sowpati |
| EPI_ISL_1360298 | CSIR-National Environmental Engineering Research Institute | CSIR-Centre for Cellular and Molecular Biology - INSACOG | Lamuk Zaveri, Ara Sreenivas, Onkar Kulkarni, Sofia Banu, Shreekant Verma,Amareshwar Vodapalli, Viswagithe S L, B Himasri, Sharath Chandra Thota, Karthik Bharadwaj Tallapaka, Krishna Khairnar, Rakesh K Mishra,Divya Tej Sowpati |
| EPI_ISL_1360299, EPI_ISL_1360300 | CSIR-National Environmental Engineering Research Institute | CSIR-Centre for Cellular and Molecular Biology - INSACOG | Ara Sreenivas, Onkar Kulkarni, Lamuk Zaveri, Sofia Banu, Shreekant Verma, Amareshwar Vodapalli , Viswagithe S L, B Himasri, Sharath Chandra Thota, Karthik Bharadwaj Tallapaka, Krishna Khairnar, Rakesh K Mishra,Divya Tej Sowpati |
| EPI_ISL_1360301, EPI_ISL_1360302 | CSIR-National Environmental Engineering Research Institute | CSIR-Centre for Cellular and Molecular Biology - INSACOG | Sofia Banu, Lamuk Zaveri, Ara Sreenivas, Shreekant Verma, Onkar Kulkarni, Amareshwar Vodapalli ,Viswagithe S L, B Himasri, Sharath Chandra Thota, Karthik Bharadwaj Tallapaka, Krishna Khairnar, Rakesh K Mishra,Divya Tej Sowpati |
| EPI_ISL_1360303, EPI_ISL_1360304 | CSIR-National Environmental Engineering Research Institute | CSIR-Centre for Cellular and Molecular Biology - INSACOG | Onkar Kulkarni, Lamuk Zaveri, Ara Sreenivas, Sofia Banu, Shreekant Verma, Amareshwar Vodapalli , Viswagithe S L, B Himasri, Sharath Chandra Thota, Karthik Bharadwaj Tallapaka, Krishna Khairnar, Rakesh K Mishra,Divya Tej Sowpati |
| EPI_ISL_1360305 | CSIR-National Environmental Engineering Research Institute | CSIR-Centre for Cellular and Molecular Biology - INSACOG | Lamuk Zaveri, Ara Sreenivas, Sofia Banu, Onkar Kulkarni, Shreekant Verma, Amareshwar Vodapalli , Viswagithe S L, B Himasri, Sharath Chandra Thota, Karthik Bharadwaj Tallapaka, Krishna Khairnar, Rakesh K Mishra,Divya Tej Sowpati |
| EPI_ISL_1360306 | CSIR-National Environmental Engineering Research Institute | CSIR-Centre for Cellular and Molecular Biology - INSACOG | Lamuk Zaveri, Ara Sreenivas, Onkar Kulkarni, Sofia Banu, Shreekant Verma,Amareshwar Vodapalli, Viswagithe S L, B Himasri, Sharath Chandra Thota, Karthik Bharadwaj Tallapaka, Krishna Khairnar, Rakesh K Mishra,Divya Tej Sowpati |
| EPI_ISL_1360307, EPI_ISL_1360308 | CSIR-National Environmental Engineering Research Institute | CSIR-Centre for Cellular and Molecular Biology - INSACOG | Ara Sreenivas, Onkar Kulkarni, Lamuk Zaveri, Sofia Banu, Shreekant Verma, Amareshwar Vodapalli , Viswagithe S L, B Himasri, Sharath Chandra Thota, Karthik Bharadwaj Tallapaka, Krishna Khairnar, Rakesh K Mishra,Divya Tej Sowpati |
| EPI_ISL_1360309, EPI_ISL_1360310 | CSIR-National Environmental Engineering Research Institute | CSIR-Centre for Cellular and Molecular Biology - INSACOG | Sofia Banu, Lamuk Zaveri, Ara Sreenivas, Shreekant Verma, Onkar Kulkarni, Amareshwar Vodapalli ,Viswagithe S L, B Himasri, Sharath Chandra Thota, Karthik Bharadwaj Tallapaka, Krishna Khairnar, Rakesh K Mishra,Divya Tej Sowpati |
| EPI_ISL_1360311, EPI_ISL_1360312 | CSIR-National Environmental Engineering Research Institute | CSIR-Centre for Cellular and Molecular Biology - INSACOG | Onkar Kulkarni, Lamuk Zaveri, Ara Sreenivas, Sofia Banu, Shreekant Verma, Amareshwar Vodapalli , Viswagithe S L, B Himasri, Sharath Chandra Thota, Karthik Bharadwaj Tallapaka, Krishna Khairnar, Rakesh K Mishra,Divya Tej Sowpati |
| EPI_ISL_1360313 | CSIR-National Environmental Engineering Research Institute | CSIR-Centre for Cellular and Molecular Biology - INSACOG | Lamuk Zaveri, Ara Sreenivas, Sofia Banu, Onkar Kulkarni, Shreekant Verma, Amareshwar Vodapalli , Viswagithe S L, B Himasri, Sharath Chandra Thota, Karthik Bharadwaj Tallapaka, Krishna Khairnar, Rakesh K Mishra,Divya Tej Sowpati |
| EPI_ISL_1360314 | CSIR-National Environmental Engineering Research Institute | CSIR-Centre for Cellular and Molecular Biology - INSACOG | Lamuk Zaveri, Ara Sreenivas, Onkar Kulkarni, Sofia Banu, Shreekant Verma,Amareshwar Vodapalli, Viswagithe S L, B Himasri, Sharath Chandra Thota, Karthik Bharadwaj Tallapaka, Krishna Khairnar, Rakesh K Mishra,Divya Tej Sowpati |
| EPI_ISL_1360315, EPI_ISL_1360316 | CSIR-National Environmental Engineering Research Institute | CSIR-Centre for Cellular and Molecular Biology - INSACOG | Ara Sreenivas, Onkar Kulkarni, Lamuk Zaveri, Sofia Banu, Shreekant Verma, Amareshwar Vodapalli , Viswagithe S L, B Himasri, Sharath Chandra Thota, Karthik Bharadwaj Tallapaka, Krishna Khairnar, Rakesh K Mishra,Divya Tej Sowpati |
| EPI_ISL_1360317, EPI_ISL_1360318 | CSIR-National Environmental Engineering Research Institute | CSIR-Centre for Cellular and Molecular Biology - INSACOG | Sofia Banu, Lamuk Zaveri, Ara Sreenivas, Shreekant Verma, Onkar Kulkarni, Amareshwar Vodapalli ,Viswagithe S L, B Himasri, Sharath Chandra Thota, Karthik Bharadwaj Tallapaka, Krishna Khairnar, Rakesh K Mishra,Divya Tej Sowpati |
| EPI_ISL_1360319, EPI_ISL_1360320 | CSIR-National Environmental Engineering Research Institute | CSIR-Centre for Cellular and Molecular Biology - INSACOG | Onkar Kulkarni, Lamuk Zaveri, Ara Sreenivas, Sofia Banu, Shreekant Verma, Amareshwar Vodapalli , Viswagithe S L, B Himasri, Sharath Chandra Thota, Karthik Bharadwaj Tallapaka, Krishna Khairnar, Rakesh K Mishra,Divya Tej Sowpati |
| EPI_ISL_1360321 | CSIR-National Environmental Engineering Research Institute | CSIR-Centre for Cellular and Molecular Biology - INSACOG | Lamuk Zaveri, Ara Sreenivas, Sofia Banu, Onkar Kulkarni, Shreekant Verma, Amareshwar Vodapalli , Viswagithe S L, B Himasri, Sharath Chandra Thota, Karthik Bharadwaj Tallapaka, Krishna Khairnar, Rakesh K Mishra,Divya Tej Sowpati |
| EPI_ISL_1360322 | CSIR-National Environmental Engineering Research Institute | CSIR-Centre for Cellular and Molecular Biology - INSACOG | Lamuk Zaveri, Ara Sreenivas, Onkar Kulkarni, Sofia Banu, Shreekant Verma,Amareshwar Vodapalli, Viswagithe S L, B Himasri, Sharath Chandra Thota, Karthik Bharadwaj Tallapaka, Krishna Khairnar, Rakesh K Mishra,Divya Tej Sowpati |
| EPI_ISL_1360323, EPI_ISL_1360324 | CSIR-National Environmental Engineering Research Institute | CSIR-Centre for Cellular and Molecular Biology - INSACOG | Ara Sreenivas, Onkar Kulkarni, Lamuk Zaveri, Sofia Banu, Shreekant Verma, Amareshwar Vodapalli , Viswagithe S L, B Himasri, Sharath Chandra Thota, Karthik Bharadwaj Tallapaka, Krishna Khairnar, Rakesh K Mishra,Divya Tej Sowpati |
| EPI_ISL_1360325, EPI_ISL_1360326 | CSIR-National Environmental Engineering Research Institute | CSIR-Centre for Cellular and Molecular Biology - INSACOG | Sofia Banu, Lamuk Zaveri, Ara Sreenivas, Shreekant Verma, Onkar Kulkarni, Amareshwar Vodapalli ,Viswagithe S L, B Himasri, Sharath Chandra Thota, Karthik Bharadwaj Tallapaka, Krishna Khairnar, Rakesh K Mishra,Divya Tej Sowpati |
| EPI_ISL_1360327, EPI_ISL_1360328 | CSIR-National Environmental Engineering Research Institute | CSIR-Centre for Cellular and Molecular Biology - INSACOG | Onkar Kulkarni, Lamuk Zaveri, Ara Sreenivas, Sofia Banu, Shreekant Verma, Amareshwar Vodapalli , Viswagithe S L, B Himasri, Sharath Chandra Thota, Karthik Bharadwaj Tallapaka, Krishna Khairnar, Rakesh K Mishra,Divya Tej Sowpati |
| EPI_ISL_1360329 | CSIR-National Environmental Engineering Research Institute | CSIR-Centre for Cellular and Molecular Biology - INSACOG | Lamuk Zaveri, Ara Sreenivas, Sofia Banu, Onkar Kulkarni, Shreekant Verma, Amareshwar Vodapalli , Viswagithe S L, B Himasri, Sharath Chandra Thota, Karthik Bharadwaj Tallapaka, Krishna Khairnar, Rakesh K Mishra,Divya Tej Sowpati |
| EPI_ISL_1360330 | CSIR-National Environmental Engineering Research Institute | CSIR-Centre for Cellular and Molecular Biology - INSACOG | Lamuk Zaveri, Ara Sreenivas, Onkar Kulkarni, Sofia Banu, Shreekant Verma,Amareshwar Vodapalli, Viswagithe S L, B Himasri, Sharath Chandra Thota, Karthik Bharadwaj Tallapaka, Krishna Khairnar, Rakesh K Mishra,Divya Tej Sowpati |
| EPI_ISL_1360331, EPI_ISL_1360332 | CSIR-National Environmental Engineering Research Institute | CSIR-Centre for Cellular and Molecular Biology - INSACOG | Ara Sreenivas, Onkar Kulkarni, Lamuk Zaveri, Sofia Banu, Shreekant Verma, Amareshwar Vodapalli , Viswagithe S L, B Himasri, Sharath Chandra Thota, Karthik Bharadwaj Tallapaka, Krishna Khairnar, Rakesh K Mishra,Divya Tej Sowpati |

| | | | |
|---|---|---|---|
| EPI_ISL_1360333 | CSIR-National Environmental Engineering Research Institute | CSIR-Centre for Cellular and Molecular Biology - INSACOG | Sofia Banu, Lamuk Zaveri, Ara Sreenivas, Shreekant Verma, Onkar Kulkarni, Amareshwar Vodapalli ,Viswagithe S L, B Himasri, Sharath Chandra Thota, Karthik Bharadwaj Tallapaka, Krishna Khairnar, Rakesh K Mishra,Divya Tej Sowpati |
| EPI_ISL_1360335, EPI_ISL_1360336 | CSIR-National Environmental Engineering Research Institute | CSIR-Centre for Cellular and Molecular Biology - INSACOG | Onkar Kulkarni, Lamuk Zaveri, Ara Sreenivas, Sofia Banu, Shreekant Verma, Amareshwar Vodapalli , Viswagithe S L, B Himasri, Sharath Chandra Thota, Karthik Bharadwaj Tallapaka, Krishna Khairnar, Rakesh K Mishra,Divya Tej Sowpati |
| EPI_ISL_1360338 | CSIR-National Environmental Engineering Research Institute | CSIR-Centre for Cellular and Molecular Biology - INSACOG | Lamuk Zaveri, Ara Sreenivas, Onkar Kulkarni, Sofia Banu, Shreekant Verma,Amareshwar Vodapalli, Viswagithe S L, B Himasri, Sharath Chandra Thota, Karthik Bharadwaj Tallapaka, Krishna Khairnar, Rakesh K Mishra,Divya Tej Sowpati |
| EPI_ISL_1360407, EPI_ISL_1360412 | Laboratorio de Enfermedades Emergentes y Reemergentes | Laboratorio de Enfermedades Emergentes y Reemergentes | Vega-Magaña Natali, Peña-Rodriguez Marcela, Viera-Segura Oliver, García-Chagollan Mariel, Hernández-Bello Jorge, Muñoz-Valle José Francisco |
| EPI_ISL_1363082, EPI_ISL_1363084, EPI_ISL_1363085, EPI_ISL_1363086, EPI_ISL_1363087, EPI_ISL_1363088, EPI_ISL_1363090, EPI_ISL_1363091, EPI_ISL_1363092, EPI_ISL_1363093, EPI_ISL_1363094, EPI_ISL_1363095, EPI_ISL_1363096, EPI_ISL_1363097, EPI_ISL_1363098, EPI_ISL_1363099, EPI_ISL_1363101, EPI_ISL_1363102, EPI_ISL_1363103, EPI_ISL_1363104, EPI_ISL_1363105, EPI_ISL_1363106, EPI_ISL_1363107, EPI_ISL_1363108, EPI_ISL_1363109, EPI_ISL_1363111, EPI_ISL_1363113 | | | |
| see above | Virological laboratory FBIH "The Center for Hygiene and Epidemiology in the Khabarovsk Region" | Group of Genomics and Postgenomic Technologies of Central Research Institute of Epidemiology | Samoilov AE, Kaptelova VV, Korneenko EV, Valdokhina AV, Bulanenko VP, Saenko SS, Lebedeva LA, Shubina YA, Savosina LV, Zhaleyko ZP, Lukyanov AV, Golubeva AG, Speranskaya AS, Tivanova EV, Kondrasheva LY, Akimkin VG |
| EPI_ISL_1363119, EPI_ISL_1363120, EPI_ISL_1363121, EPI_ISL_1363122, EPI_ISL_1363123 | Borneo Medical Centre | Institute of Health and Community Medicine | Chan Chia Jui, David Perera, Ooi Mong How, Chua Hock Hin, Tonnii Sia Loong Loong, Wong Jyn Shan, Wong Kiing Aik |
| EPI_ISL_1363758 | Departments of Intensive Care Medicine, and Clinical lab, Osaka Women's and Children's Hospital | Department of Developmental Medicine, Research Institute, | Itaru Yanagihara, Yukiko Nakura, Muneyuki Takeuchi, Shinobu Ida |
| EPI_ISL_1363759 | Departments of Intensive Care Medicine, and Clinical lab, Osaka Women's and Children's Hospital | Department of Developmental Medicine, Research Institute, Osaka Women's and Children's Hospital | Itaru Yanagihara, Yukiko Nakura, Muneyuki Takeuchi, Shinobu Ida |
| EPI_ISL_1372283, EPI_ISL_1372286, EPI_ISL_1372301, EPI_ISL_1372304, EPI_ISL_1372307, EPI_ISL_1372309, EPI_ISL_1372310, EPI_ISL_1372312 | WHO National Influenza Centre Russian Federation | WHO National Influenza Centre Russian Federation | Andrey Komissarov, Artem Fadeev, Anna Ivanova, Kseniya Komissarova, Dmitry Bazhenov, Tamila Musaeva, Maria Timofeeva, Veronika Eder, Maria Pisareva, Daria Danilenko, Ksenia Safina, Elena Nabieva, Georgii Bazykin, Dmitry Lioznov |
| EPI_ISL_1372400, EPI_ISL_1372401, EPI_ISL_1372402, EPI_ISL_1372403, EPI_ISL_1372404, EPI_ISL_1372405, EPI_ISL_1372406, EPI_ISL_1372407, EPI_ISL_1372408 | Pathogenic Microorganisms Variability Laboratory | WHO National Influenza Centre Russian Federation | Andrey Komissarov, Artem Fadeev, Anna Ivanova, Kseniya Komissarova, Dmitry Bazhenov, Daria Danilenko, Ksenia Safina, Elena Nabieva, Georgii Bazykin, Nadezhda Kuznetsova, Elena Shidlovskaya, Sergey Alkhovsky, Tatyana Vishnevskaya, Elizaveta Divisenko, Alexey Shchetinin, Maria Nikiforova, Andrey Pochtovyy, Evgeny Usachev, Elena Vokalova, Maxim Rubalsky, Oleg Rubalsky, Artem Tkachuk, Vladimir Gushchin, Alexander Gintsburg, Dmitry Lioznov |
| EPI_ISL_1379391 | Department of Molecular and Translational Medicine, Section of Microbiology, University of Brescia, ASST Spedali Civili, Brescia | Department of Molecular and Translational Medicine, Section of Microbiology, University of Brescia, ASST Spedali Civili, Brescia | Simona Fiorentini, Serena Messali, Alberto Zani, Francesca Caccuri, Arnaldo Caruso |
| EPI_ISL_1379435 | Department of Molecular and Translational Medicine, Section of Microbiology, University of Brescia, ASST Spedali Civili, Brescia | Department of Molecular and Translational Medicine, Section of Microbiology, University of Brescia, ASST Spedali Civili, Brescia | Simona Fiorentini, Serena Messali, Alberto Zani, Erika Scaltriti, Francesca Caccuri, Arnaldo Caruso |
| EPI_ISL_1379733 | Stanford Health Care | Stanford University School of Medicine, Clinical Virology Laboratory | Fumiko Yamamoto, Malaya K. Sahoo, Mamdouh Sibai, Daniel Solis, Michelle Vergese, ChunHong Huang, James Zehnder, and Benjamin A. Pinsky |
| EPI_ISL_1380060, EPI_ISL_1380061, EPI_ISL_1380062, EPI_ISL_1380063, EPI_ISL_1380064, EPI_ISL_1380065 | Laboratorio Biologia Molecolare Sars Cov2 - UOC Laboratorio Analisi - Servizio Medicina di Laboratorio, Ospedale "San Francesco" - ATS-ASSL Nuoro | Laboratorio specialistico UOC Ematologia - Ospedale "San Francesco" - ATS-ASSL Nuoro | Piras Giovanna, Aspronì Rosanna,,Fiamma Maura, Monne Maria Itria, Palmas Angelo Domenico, Lo Maglio Iana, Mameli Giuseppe |
| EPI_ISL_1381066, EPI_ISL_1381067 | LACEN do Mato Grosso do Sul | Instituto Adolfo Lutz, Interdiciplinary Procedures Center, Strategic Laboratory | Claudio Tavares Sacchi, Claudia Regina Gonçalves, Erica Valessa Ramos Gomes, Karoline Rodrigues Campos, Caio Vinicius Dias Lopes |
| EPI_ISL_1381146 | Hospital | National Reference Center for Viruses of Respiratory Infections, Institut Pasteur, Paris | Marion Barbet, Sylvie Behillil, Méline Bizard, Angela Brisebarre, Camille Capel, Louise Lefrançois, Etienne Simon-Lorière, Vincent Enouf, Maud Vanpeene, Sylvie van der Werf,Hecquet Denise |
| EPI_ISL_1381161 | Labo Analyses Med | National Reference Center for Viruses of Respiratory Infections, Institut Pasteur, Paris | Marion Barbet, Sylvie Behillil, Méline Bizard, Angela Brisebarre, Camille Capel, Louise Lefrançois, Etienne Simon-Lorière, Vincent Enouf, Maud Vanpeene, Sylvie van der Werf,Besson J |
| EPI_ISL_1381165, EPI_ISL_1381166, EPI_ISL_1381167, EPI_ISL_1381168, EPI_ISL_1381169, EPI_ISL_1381170, EPI_ISL_1381171, EPI_ISL_1381172, EPI_ISL_1381173, EPI_ISL_1381174, EPI_ISL_1381175, EPI_ISL_1381176, EPI_ISL_1381177, EPI_ISL_1381178, EPI_ISL_1381179, EPI_ISL_1381180, EPI_ISL_1381181, EPI_ISL_1381182, EPI_ISL_1381183, EPI_ISL_1381184, EPI_ISL_1381185, EPI_ISL_1381186, EPI_ISL_1381187, EPI_ISL_1381188, EPI_ISL_1381189, EPI_ISL_1381190, EPI_ISL_1381191, EPI_ISL_1381192, EPI_ISL_1381193, EPI_ISL_1381194, EPI_ISL_1381195, EPI_ISL_1381196, EPI_ISL_1381197, EPI_ISL_1381198, EPI_ISL_1381199, EPI_ISL_1381201, EPI_ISL_1381202 | | | |
| see above | Hospital | National Reference Center for Viruses of Respiratory Infections, Institut Pasteur, Paris | Marion Barbet, Sylvie Behillil, Méline Bizard, Angela Brisebarre, Camille Capel, Louise Lefrançois, Etienne Simon-Lorière, Vincent Enouf, Maud Vanpeene, Sylvie van der Werf,Bressollette CéLine |
| EPI_ISL_1381231, EPI_ISL_1381232 | Hospital | National Reference Center for Viruses of Respiratory Infections, Institut Pasteur, Paris | Marion Barbet, Sylvie Behillil, Méline Bizard, Angela Brisebarre, Camille Capel, Louise Lefrançois, Etienne Simon-Lorière, Vincent Enouf, Maud Vanpeene, Sylvie van der Werf,Pilorge LéA |
| EPI_ISL_1381386 | Centers for Disease Control, R.O.C. (Taiwan) | Centers for Disease Control, R.O.C. (Taiwan) | Ji-Rong Yang, Yu-Chi Lin, Jung-Jung Mu, Ming-Tsan Liu |
| EPI_ISL_1381822, EPI_ISL_1381823, EPI_ISL_1381824, EPI_ISL_1381825, EPI_ISL_1381826, EPI_ISL_1381827 | Hospital | National Reference Center for Viruses of Respiratory Infections, Institut Pasteur, Paris | Marion Barbet, Sylvie Behillil, Méline Bizard, Angela Brisebarre, Camille Capel, Louise Lefrançois, Etienne Simon-Lorière, Vincent Enouf, Maud Vanpeene, Sylvie van der Werf,Bressollette CéLine |
| EPI_ISL_1381831 | Hospital | National Reference Center for Viruses of Respiratory Infections, Institut Pasteur, Paris | Marion Barbet, Sylvie Behillil, Méline Bizard, Angela Brisebarre, Camille Capel, Louise Lefrançois, Etienne Simon-Lorière, Vincent Enouf, Maud Vanpeene, Sylvie van der Werf,Pilorge LéA |
| EPI_ISL_1392889, EPI_ISL_1392890, EPI_ISL_1392891 | Centrum Badawczo-Rozwojowe Invicta (Invicta sp. z o.o.) | Diagtron Laboratoria | Maciej Kosinski, Natalia Derewonko, Lukasz Rabalski, Karolina Gackowska, Celina Cybulska, Krzysztof Lukaszczuk |
| EPI_ISL_1396347 | Laboratorio del Hospital Regional Ushuaia Gdor. Ernesto Campos | Nodo de Secuenciación Tierra del Fuego - Hospital Regional Ushuaia - Centro Austral De Investigaciones Científicas - Universidad Nacional De Tierra Del Fuego on behalf of 'Proyecto Argentino Interinstitucional de genomica de SARS-CoV-2' (PAIS Consortium) | Carina Andrea De Roccis, Gabriel Alejandro Castro, Silvana Beatriz Cáceres, Carolina Beatriz Yulan, Manuel Fabian Boutureira, Alejandro Ezequiel Rojas, Fernando Gallego, Santiago Guillermo Ceballos, Cristina Fernanda Nardi, Ivan Dario Gramundi |
| EPI_ISL_1400529 | WHO National Influenza Centre Russian Federation | WHO National Influenza Centre Russian Federation | Andrey Komissarov, Artem Fadeev, Anna Ivanova, Kseniya Komissarova, Dmitry Bazhenov, Tamila Musaeva, Maria Timofeeva, Veronika Eder, Maria Pisareva, Daria Danilenko, Ksenia Safina, Elena Nabieva, Georgii Bazykin, Dmitry Lioznov |
| EPI_ISL_1400532, EPI_ISL_1400533, EPI_ISL_1400534, EPI_ISL_1400535, EPI_ISL_1400536, EPI_ISL_1400537, EPI_ISL_1400538, EPI_ISL_1400539, EPI_ISL_1400540, EPI_ISL_1400560 | WHO National Influenza Centre Russian Federation | WHO National Influenza Centre Russian Federation | Andrey Komissarov, Artem Fadeev, Anna Ivanova, Kseniya Komissarova, Dmitry Bazhenov, Mikhail Bakaev, Daria Danilenko, Ksenia Safina, Elena Nabieva, Georgii Bazykin, Dmitry Lioznov |
| EPI_ISL_1402423, EPI_ISL_1402425 | Private clinic of Biogen Med, Tashkent, Uzbekistan | Center of Genomics and bioinformatics, Bioinformatics laboratory | Mirzakamol S Ayubov, Zabardast T Buriev, Mukhammadjon H Mirzakhmedov, Abdurakhmon N Yusupov, Shukhrat E Shermatov, Ibrokhim Y Abdurakhmonov. |

| | | | |
|---|---|---|---|
| EPI_ISL_1404375 | Klinisch Laboratorium ZNA | Klinisch Laboratorium ZNA | Verstrepen et al. |
| EPI_ISL_1406651 | HOPITAL DE VICHY | CHU Clermont-Ferrand, service de virologie | Bisseux Maxime, Mirand Audrey, Combes Patricia, Henquell Cécile |
| EPI_ISL_1406681 | Centre Jean Perrin | CHU Clermont-Ferrand, service de virologie | Bisseux Maxime, Mirand Audrey, Combes Patricia, Henquell Cécile |
| EPI_ISL_1406694 | National Center of Infectious and Parasitic Diseases | National Center of Infectious and Parasitic Diseases | Alexiev et al |
| EPI_ISL_1416992 | Outre mer | National Reference Center for Viruses of Respiratory Infections, Institut Pasteur, Paris | Marion Barbet, Sylvie Behillil, Méline Bizard,Frédéric Lemoine,Corinne Maufrais,Christophe Malabat, Angela Brisebarre, Camille Capel, Louise Lefrançois, Etienne Simon-Lorière, Vincent Enouf, Maud Vanpeene, Sylvie van der Werf,Anne Holstein |
| EPI_ISL_1416993, EPI_ISL_1416994 | Outre mer | National Reference Center for Viruses of Respiratory Infections, Institut Pasteur, Paris | Marion Barbet, Sylvie Behillil, Méline Bizard,Frédéric Lemoine,Corinne Maufrais,Christophe Malabat, Angela Brisebarre, Camille Capel, Louise Lefrançois, Etienne Simon-Lorière, Vincent Enouf, Maud Vanpeene, Sylvie van der Werf,GréGoire Potiron |
| EPI_ISL_1416995 | Outre mer | National Reference Center for Viruses of Respiratory Infections, Institut Pasteur, Paris | Marion Barbet, Sylvie Behillil, Méline Bizard,Frédéric Lemoine,Corinne Maufrais,Christophe Malabat, Angela Brisebarre, Camille Capel, Louise Lefrançois, Etienne Simon-Lorière, Vincent Enouf, Maud Vanpeene, Sylvie van der Werf,StéPhanie Guyomard-Rabenirina |
| EPI_ISL_1416996, EPI_ISL_1416997 | Outre mer | National Reference Center for Viruses of Respiratory Infections, Institut Pasteur, Paris | Marion Barbet, Sylvie Behillil, Méline Bizard,Frédéric Lemoine,Corinne Maufrais,Christophe Malabat, Angela Brisebarre, Camille Capel, Louise Lefrançois, Etienne Simon-Lorière, Vincent Enouf, Maud Vanpeene, Sylvie van der Werf,Anne Holstein |
| EPI_ISL_1417067 | Hospital | National Reference Center for Viruses of Respiratory Infections, Institut Pasteur, Paris | Marion Barbet, Sylvie Behillil, Méline Bizard,Frédéric Lemoine,Corinne Maufrais,Christophe Malabat, Angela Brisebarre, Camille Capel, Louise Lefrançois, Etienne Simon-Lorière, Vincent Enouf, Maud Vanpeene, Sylvie van der Werf,OphéLie Said-Delattre |
| EPI_ISL_1417069 | Hospital | National Reference Center for Viruses of Respiratory Infections, Institut Pasteur, Paris | Marion Barbet, Sylvie Behillil, Méline Bizard,Frédéric Lemoine,Corinne Maufrais,Christophe Malabat, Angela Brisebarre, Camille Capel, Louise Lefrançois, Etienne Simon-Lorière, Vincent Enouf, Maud Vanpeene, Sylvie van der Werf,StéPhanie Guyomard-Rabenirina |
| EPI_ISL_1417070 | Hospital | National Reference Center for Viruses of Respiratory Infections, Institut Pasteur, Paris | Marion Barbet, Sylvie Behillil, Méline Bizard,Frédéric Lemoine,Corinne Maufrais,Christophe Malabat, Angela Brisebarre, Camille Capel, Louise Lefrançois, Etienne Simon-Lorière, Vincent Enouf, Maud Vanpeene, Sylvie van der Werf,OphéLie Said-Delattre |
| EPI_ISL_1418308 | National Institute of Infectious Diseases-Prof. Dr. Matei Bals Molecular Diagnostics Laboratory | National Institute of Infectious Diseases-Prof. Dr. Matei Bals Molecular Diagnostics Laboratory | Corina Casangiu, Leontina Banica, Marius Surleac, Ovidiu Vlaicu, Ionelia Nicolae, Simona Paraschiv, Dan Otelea |
| EPI_ISL_1424065 | Institute for Medical Research, Infectious Disease Research Centre, National Institutes of Health, Ministry of Health Malaysia | Institute for Medical Research, Infectious Disease Research Centre, National Institutes of Health, Ministry of Health Malaysia | Suppiah J, Kamel K, Mohd Zawawi Z, Ramly N, Robert F, Thayan R |
| EPI_ISL_1435783, EPI_ISL_1435785, EPI_ISL_1435787, EPI_ISL_1435789, EPI_ISL_1435791, EPI_ISL_1435792, EPI_ISL_1435794, EPI_ISL_1435796, EPI_ISL_1435798, EPI_ISL_1435800 | DNALAB Sdn. Bhd. | Malaysia Genome Institute | Mohd Noor Mat Isa, Irni Suhayu Sapian, Yusuf Muhammad Noor, Nurhezreen Md Iqbal, Mohd Faizal Abu Bakar, Enizza Kasim, Shamsidar Sopie, Siti Noraini Othman, Azrin Ahmad, Wong Yong Wee, Nor Azfa Johari. |
| EPI_ISL_1435802, EPI_ISL_1435804, EPI_ISL_1435806, EPI_ISL_1435808, EPI_ISL_1435810, EPI_ISL_1435812, EPI_ISL_1435814, EPI_ISL_1435816, EPI_ISL_1435817, EPI_ISL_1435820 | Malaysia Genome Institute | Malaysia Genome Institute | Mohd Noor Mat Isa, Irni Suhayu Sapian, Yusuf Muhammad Noor, Nurhezreen Md Iqbal, Mohd Faizal Abu Bakar, Enizza Kasim, Shamsidar Sopie, Siti Noraini Othman, Azrin Ahmad, Nor Azfa Johari. |
| EPI_ISL_1437749 | Institut National d'Hygiène | Unité Mixte Internationale TransVIHMI (UMI 233 IRD - U1175 INSERM - Université de Montpellier) IRD (Institut de recherche pour le développement) | Mounerou SALOU, Christelle BUTEL, Wembo A. HALATOKO,Issaka Maman, Abla A. KONOU, Amivi EHLAN, Adodo SADJI, Kokou TEGUENI,Sidonie A.M.KAGNISSODE, Akoélé SILIADIN, Alassane OURO-MEDELI, Messanh DOUFFAN,Déléma MABA,Sika DOSSIM, Améyo DORKENOO, Mireille PRINCE-DAVID,Anoumou DAGNRA,Laetitia SERRANO,Ahidjo AYOUBA,Eric DELAPORTE, Martine PEETERS |
| EPI_ISL_1443002 | Institut National d'hygiène | "Unité Mixte Internationale TransVIHMI (UMI 233 IRD - U1175 INSERM - Université de Montpellier) IRD (Institut de recherche pour le développement)" | Mounerou SALOU, Christelle BUTEL, Wembo A. HALATOKO,Issaka Maman, Abla A. KONOU, Amivi EHLAN, Adodo SADJI, Kokou TEGUENI,Sidonie A.M.KAGNISSODE, Akoélé SILIADIN, Alassane OURO-MEDELI, Messanh DOUFFAN,Déléma MABA,Sika DOSSIM, Améyo DORKENOO, Mireille PRINCE-DAVID,Anoumou DAGNRA,Laetitia SERRANO,Ahidjo AYOUBA,Eric DELAPORTE, Martine PEETERS |
| EPI_ISL_1443196, EPI_ISL_1443197, EPI_ISL_1443198 | Hospital Aliança | Hospital São Rafael - IDOR | Isadora Cristina de Siqueira, Aquiles Assunção Camelier, Elves A.P. Maciel, Margarida Celia L. C. Neves, Carolina Kymie Vasques Nonaka, Karoline Almeida Félix de Sousa, Victor Costa Araujo, Yasmin Santos Freitas Macêdo, Aurea Angelica Paste, Bruno Solano de Freitas Souza, Tiago Gräf |
| EPI_ISL_1443901, EPI_ISL_1443902 | Hospital | National Reference Center for Viruses of Respiratory Infections, Institut Pasteur, Paris | Marion Barbet, Sylvie Behillil, Méline Bizard,Frédéric Lemoine,Corinne Maufrais,Christophe Malabat, Angela Brisebarre, Camille Capel, Louise Lefrançois, Etienne Simon-Lorière, Vincent Enouf, Maud Vanpeene, Sylvie van der Werf,CéLine Bressollette |
| EPI_ISL_1445275 | Institut National d'Hygiène (INH) | Unité Mixte Internationale TransVIHMI (UMI 233 IRD - U1175 INSERM - Université de Montpellier)IRD (Institut de recherche pour le développement) | Mounerou SALOU, Christelle BUTEL, Wembo A. HALATOKO,Issaka Maman, Abla A. KONOU, Amivi EHLAN, Adodo SADJI, Kokou TEGUENI,Sidonie A.M.KAGNISSODE, Akoélé SILIADIN, Alassane OURO-MEDELI, Messanh DOUFFAN,Déléma MABA,Sika DOSSIM, Améyo DORKENOO, Mireille PRINCE-DAVID,Anoumou DAGNRA,Laetitia SERRANO,Ahidjo AYOUBA,Eric DELAPORTE, Martine PEETERS |
| EPI_ISL_1447339, EPI_ISL_1447341, EPI_ISL_1447343, EPI_ISL_1447348, EPI_ISL_1447349, EPI_ISL_1447350, EPI_ISL_1447353, EPI_ISL_1447354, EPI_ISL_1447355, EPI_ISL_1447356, EPI_ISL_1447357, EPI_ISL_1447358, EPI_ISL_1447359, EPI_ISL_1447360, EPI_ISL_1447361 | | | |
| see above | Diagnostic and Research Center of Infectious Diseases, Medical Faculty, Andalas University | Diagnostic and Research Center of Infectious Diseases, Medical Faculty, Andalas University | Andani Eka Putra, Syafrizayanti, Ikhwan R. Sudji, Linosefa, Dede Rahman Agustian, Ayu Novita Trisnawati, Siskalil Fahma, Nia Ayuni Putri, Sekar Asri Tresnaningtyas, SM Rezvi, Fauzul Azhim, Dessy Arisanty, Gestina Aliska, Desmawati, Nita Afriani ,Syandrez Prima Putra, Juane Plantika Menra, Yolani Syaputri, Mutia Lailani |
| EPI_ISL_1468418 | Secretaria Municipal de Saude de Birigui | Instituto Adolfo Lutz, Interdiciplinary Procedures Center, Strategic Laboratory | Claudio Tavares Sacchi, Claudia Regina Gonçalves, Erica Valessa Ramos Gomes, Karoline Rodrigues Campos, Caio Vinicius Dias Lopes |
| EPI_ISL_1468433, EPI_ISL_1468435, EPI_ISL_1468436, EPI_ISL_1468437, EPI_ISL_1468438, EPI_ISL_1468439, EPI_ISL_1468440, EPI_ISL_1468441 | LACEN do Mato Grosso do Sul | Instituto Adolfo Lutz, Interdiciplinary Procedures Center, Strategic Laboratory | Claudio Tavares Sacchi, Claudia Regina Gonçalves, Erica Valessa Ramos Gomes, Karoline Rodrigues Campos, Caio Vinicius Dias Lopes |
| EPI_ISL_1469236, EPI_ISL_1469237 | Diagnostic and Research Center of Infectious Diseases, Medical Faculty, Andalas University | Diagnostic and Research Center of Infectious Diseases, Medical Faculty, Andalas University | Andani Eka Putra, Syafrizayanti, Ikhwan R. Sudji, Linosefa, Dede Rahman Agustian, Ayu Novita Trisnawati, Siskalil Fahma, Nia Ayuni Putri, Sekar Asri Tresnaningtyas, SM Rezvi, Fauzul Azhim, Dessy Arisanty, Gestina Aliska, Desmawati, Nita Afriani ,Syandrez Prima Putra, Juane Plantika Menra, Yolani Syaputri, Mutia Lailani |
| EPI_ISL_1469279 | RSUD Surakarta | National Institute of Health Research and Development | Kartika Dewi Puspa, Hana Apsari Pawestri, Subangkit, Arie Ardiansyah Nugraha, Hartanti Dian Ikawati, Nelly Puspandari, Krisna Pangesti, Vivi Setiawaty |
| EPI_ISL_1469290, EPI_ISL_1469291 | National Institute Health Research and Development | National Institute of Health Research and Development | Nur Ika Hariastuti, Nike Susanti, Hana Apsari Pawestri, Subangkit, Kartika Dewi Puspa, Arie Ardiansyah Nugraha, Hartanti Dian Ikawati, Nelly Puspandari, Vivi Setiawaty |
| EPI_ISL_1469450, EPI_ISL_1469451, EPI_ISL_1469452, EPI_ISL_1469453, EPI_ISL_1469454, EPI_ISL_1469455, EPI_ISL_1469456, EPI_ISL_1469457, EPI_ISL_1469458, EPI_ISL_1469459, EPI_ISL_1469460, EPI_ISL_1469461, EPI_ISL_1469462, EPI_ISL_1469463, EPI_ISL_1469465, EPI_ISL_1469466, EPI_ISL_1469467, EPI_ISL_1469468, EPI_ISL_1469469, EPI_ISL_1469470, EPI_ISL_1469471, EPI_ISL_1469472, EPI_ISL_1469474, EPI_ISL_1469475, EPI_ISL_1469476, EPI_ISL_1469477, EPI_ISL_1469478, EPI_ISL_1469479, EPI_ISL_1469480, EPI_ISL_1469481, EPI_ISL_1469482, EPI_ISL_1469483, EPI_ISL_1469484, EPI_ISL_1469485, EPI_ISL_1469486, EPI_ISL_1469488, EPI_ISL_1469489, EPI_ISL_1469490, EPI_ISL_1469491, EPI_ISL_1469492 | | | |
| see above | Center for Laboratory Medicine St. Gallen | Center for Laboratory Medicine St. Gallen | Yannick Gerth |
| EPI_ISL_1469549 | FUNDACAO HOSPITALAR DE SAPUCAIA DO SUL | Epiclin | Fernando Hayashi Sant'Anna, Ana Paula Muterle, Janira Prichula, Juliana Comerlato, Carolina Comerlato, Eliana Márcia Da Ros Wendland |
| EPI_ISL_1472367, EPI_ISL_1472368, EPI_ISL_1472369, EPI_ISL_1472370, EPI_ISL_1472371 | Department of Pathology, National Institute of Infectious Diseases | Department of Pathology, National Institute of Infectious Diseases | Tsuyoshi Sekizuka, Kentaro Itokawa, Rina Tanaka, Masanori Hashino, Makoto Kuroda, Harutaka Katano, Tadaki Suzuki |
| EPI_ISL_1477044 | Private clinic of Biogen Med, Tashkent, Uzbekistan | Center of Genomics and bioinformatics, Bioinformatics laboratory | Mirzakamol S Ayubov, Zabardast T Buriev, Mukhammadjon H Mirzakhmedov, Abdurakhmon N Yusupov, Shukhrat E Shermatov, Ibrokhim Y Abdurakhmonov. |

| | | | |
|---|---|---|---|
| EPI_ISL_1478754 | Microbiology Department. Complexo Hospitalario Universitario de Vigo | Microbiology Department. Complexo Hospitalario Universitario de Vigo | Alfaia N, Alonso I, Alvarez M, Cabrera JJ, Carballo R, Cores O, Cortizo S, del-Campo V, Martinez L, Mediero G, Perez S, Potel C, Regueiro B, Rey S, Vasallo FJ |
| EPI_ISL_1483319, EPI_ISL_1483320 | Microbiology Department. Complexo Hospitalario Universitario de Vigo | Microbiology Department. Complexo Hospitalario Universitario de Vigo | Alfaya N, Alonso I, Alvarez M, Cabrera JJ, Carballo R, Cores O, Cortizo S, del-Campo V, Martinez L, Mediero G, Perez S, Potel C, Regueiro B, Rey S, Vasallo FJ |
| EPI_ISL_1491614, EPI_ISL_1491617, EPI_ISL_1491628, EPI_ISL_1491629, EPI_ISL_1491630, EPI_ISL_1491631, EPI_ISL_1491632, EPI_ISL_1491633, EPI_ISL_1491634, EPI_ISL_1491635, EPI_ISL_1491636, EPI_ISL_1491637, EPI_ISL_1491638, EPI_ISL_1491639, EPI_ISL_1491640, EPI_ISL_1491641, EPI_ISL_1491642, EPI_ISL_1491643, EPI_ISL_1491644, EPI_ISL_1491645, EPI_ISL_1491646, EPI_ISL_1491647, EPI_ISL_1491648, EPI_ISL_1491649, EPI_ISL_1491650, EPI_ISL_1491651, EPI_ISL_1491652, EPI_ISL_1491653, EPI_ISL_1491654, EPI_ISL_1491655, EPI_ISL_1491656, EPI_ISL_1491657, EPI_ISL_1491658, EPI_ISL_1491659, EPI_ISL_1491660, EPI_ISL_1491662, EPI_ISL_1491663, EPI_ISL_1491664, EPI_ISL_1491665, EPI_ISL_1491666, EPI_ISL_1491667, EPI_ISL_1491668, EPI_ISL_1491669, EPI_ISL_1491670 | | | |
| see above | Pathogenic Microorganisms Variability Laboratory | WHO National Influenza Centre Russian Federation | Andrey Komissarov, Artem Fadeev, Anna Ivanova, Kseniya Komissarova, Alexey Masharsky, Maria Baturova, Dmitry Bazhenov, Daria Danilenko, Ksenia Safina, Elena Nabieva, Georgii Bazykin, Nadezhda Kuznetsova, Elena Shidlovskaya, Sergey Alkhovsky, Tatyana Vishnevskaya, Elizaveta Divisenko, Alexey Shchetinin, Maria Nikiforova, Andrey Pochtovyy, Evgeny Usachev, Elena Vokalova, Maxim Rubalsky, Oleg Rubalsky, Artem Tkachuk, Vladimir Gushchin, Alexander Gintsburg, Dmitry Lioznov |
| EPI_ISL_1493597, EPI_ISL_1493598, EPI_ISL_1493599, EPI_ISL_1493600 | LACEN do Estado de Rondonia | Instituto Adolfo Lutz, Interdiciplinary Procedures Center, Strategic Laboratory | Claudio Tavares Sacchi, Claudia Regina Gonçalves, Erica Valessa Ramos Gomes, Karoline Rodrigues Campos, Caio Vinicius Dias Lopes |
| EPI_ISL_1494728, EPI_ISL_1494729, EPI_ISL_1494730 | Centro de Investigación en Ciencias de la Salud y Biomedicina, U.A.S.L.P. | Centro de Investigación en Ciencias de la Salud y Biomedicina, U.A.S.L.P. | MD PhD Sofía Bernal Silva |
| EPI_ISL_1500960, EPI_ISL_1500961, EPI_ISL_1500962, EPI_ISL_1500963, EPI_ISL_1500964, EPI_ISL_1500965, EPI_ISL_1500966, EPI_ISL_1500967, EPI_ISL_1500968, EPI_ISL_1500969, EPI_ISL_1500970, EPI_ISL_1500971, EPI_ISL_1500994, EPI_ISL_1500995, EPI_ISL_1500996, EPI_ISL_1500997, EPI_ISL_1500998, EPI_ISL_1500999, EPI_ISL_1501000, EPI_ISL_1501001, EPI_ISL_1501002, EPI_ISL_1501003, EPI_ISL_1501218 | | | |
| see above | HEGP - Laboratoire de Virologie | HEGP - Laboratoire de Virologie | David Veyer |
| EPI_ISL_1502814, EPI_ISL_1502816, EPI_ISL_1502817 | Infectious Diseases Department , Instituto Nacional de Ciencias Medicas y Nutrición | Instituto Nacional de Ciencias Medicas y Nutricion Infectious Diseases | Guillermo M. Ruiz-Palacios, Pilar Ramos Cervantes, Violeta Ibarra Gonzalez, Fernando Ledesma Barrientos, Miriam Arciniega Fuentes , Fernando Arteaga Cabello, Diana Paola Davalos Martinez, Luis Alberto Garcia Andrade, Luz Elena Cervantes Villar |
| EPI_ISL_1502887, EPI_ISL_1502888, EPI_ISL_1502889, EPI_ISL_1502890, EPI_ISL_1502891, EPI_ISL_1502892, EPI_ISL_1502893, EPI_ISL_1502894, EPI_ISL_1502895, EPI_ISL_1502896, EPI_ISL_1502897, EPI_ISL_1502898, EPI_ISL_1502899, EPI_ISL_1502900, EPI_ISL_1502901, EPI_ISL_1502903, EPI_ISL_1502905, EPI_ISL_1502907, EPI_ISL_1502908, EPI_ISL_1502911, EPI_ISL_1502912, EPI_ISL_1502913, EPI_ISL_1502915, EPI_ISL_1502917, EPI_ISL_1502919, EPI_ISL_1502921, EPI_ISL_1502923, EPI_ISL_1502925, EPI_ISL_1502928, EPI_ISL_1502929, EPI_ISL_1502930, EPI_ISL_1502931, EPI_ISL_1502933, EPI_ISL_1502934, EPI_ISL_1502935, EPI_ISL_1502936, EPI_ISL_1502937, EPI_ISL_1502939, EPI_ISL_1502941, EPI_ISL_1502943, EPI_ISL_1502944, EPI_ISL_1502946, EPI_ISL_1502947, EPI_ISL_1502949, EPI_ISL_1502951, EPI_ISL_1502954, EPI_ISL_1502955, EPI_ISL_1502956, EPI_ISL_1502957, EPI_ISL_1502959, EPI_ISL_1502961, EPI_ISL_1502963, EPI_ISL_1502966, EPI_ISL_1502967, EPI_ISL_1502968, EPI_ISL_1502970, EPI_ISL_1502972, EPI_ISL_1502978, EPI_ISL_1502983, EPI_ISL_1502984, EPI_ISL_1502985, EPI_ISL_1502986, EPI_ISL_1502987, EPI_ISL_1502988, EPI_ISL_1502989, EPI_ISL_1502990, EPI_ISL_1502991, EPI_ISL_1502992, EPI_ISL_1502993, EPI_ISL_1502994, EPI_ISL_1502995, EPI_ISL_1502996, EPI_ISL_1502998, EPI_ISL_1502999, EPI_ISL_1503003 | | | |
| see above | Gorgas Memorial Laboratory of Health Studies | Gorgas Memorial Laboratory of Health Studies | Gonzalez Claudia, Leyda Abrego, Moreno Ambar, Oris Chavarria, Jessica Gondola, Marlenne Castillo, Ortiz Alma, Castillo Jorge, Moreno Brechla, Franco Danilo, Lopez-Verges Sandra, Martinez Alexander |
| EPI_ISL_1508848 | Medical Genetics Laboratory, Regional Centre of Medical Genetics, Emergency County Hospital Craiova | Medical Genetics Laboratory, Regional Centre of Medical Genetics, Emergency County Hospital Craiova | Anca-Lelia (Riza) Costache, Ioana Streata, Stefania Dorobantu, Alina Dragos, Mihai Cucu, Ana-Maria Buga, Razvan Plesea, Andrei Pirvu, Elena Plesea, Monica Cara, Mihai Ioana |
| EPI_ISL_1508953 | Institut National d'hygiène | Unité Mixte Internationale TransVIHMI (UMI 233 IRD - U1175 INSERM - Université de Montpellier) IRD (Institut de recherche pour le développement) | Mounerou SALOU, Christelle BUTEL, Wembo A. HALATOKO, Amivi EHLAN, Abla A. KONOU, Issaka Maman,Syntyche DEVATCHAGNI, Adodo SADJI, Kokou TEGUENI,Koku AGBODEKA, Sidonie A.M.KAGNISSODE, Akoélé SILIADIN, Alassane OURO-MEDELI, Messanh DOUFFAN,Déléma MABA,Sika DOSSIM, Améyo DORKENOO, Mireille PRINCE-DAVID,Anoumou DAGNRA, Laetitia SERRANO, Ahidjo AYOUBA, Eric DELAPORTE, Martine PEETERS |
| EPI_ISL_1508956 | Laboratoire Biolim/FSS/UL | Unité Mixte Internationale TransVIHMI (UMI 233 IRD - U1175 INSERM - Université de Montpellier) IRD (Institut de recherche pour le développement) | Mounerou SALOU, Christelle BUTEL, Wembo A. HALATOKO, Amivi EHLAN, Abla A. KONOU, Issaka Maman, Syntyche DEVATCHAGNI, Adodo SADJI, Kokou TEGUENI,Koku AGBODEKA, Sidonie A.M.KAGNISSODE, Akoélé SILIADIN, Alassane OURO-MEDELI, Messanh DOUFFAN,Déléma MABA,Sika DOSSIM, Améyo DORKENOO, Mireille PRINCE-DAVID,Anoumou DAGNRA, Laetitia SERRANO,Ahidjo AYOUBA, Eric DELAPORTE, Martine PEETERS |
| EPI_ISL_1508958 | Institut National d'hygiène | Unité Mixte Internationale TransVIHMI (UMI 233 IRD - U1175 INSERM - Université de Montpellier) IRD (Institut de recherche pour le développement) | Mounerou SALOU, Christelle BUTEL, Wembo A. HALATOKO, Amivi EHLAN, Abla A. KONOU, Issaka Maman,Syntyche DEVATCHAGNI, Adodo SADJI, Kokou TEGUENI,Koku AGBODEKA, Sidonie A.M.KAGNISSODE, Akoélé SILIADIN, Alassane OURO-MEDELI, Messanh DOUFFAN,Déléma MABA,Sika DOSSIM, Améyo DORKENOO, Mireille PRINCE-DAVID,Anoumou DAGNRA, Laetitia SERRANO,Ahidjo AYOUBA, Eric DELAPORTE, Martine PEETERS |
| EPI_ISL_1508960 | Institut National d'hygiène | Unité Mixte Internationale TransVIHMI (UMI 233 IRD - U1175 INSERM - Université de Montpellier) IRD (Institut de recherche pour le développement) | Mounerou SALOU, Christelle BUTEL, Wembo A. HALATOKO, Amivi EHLAN, Abla A. KONOU, Issaka Maman, Syntyche DEVATCHAGNI, Adodo SADJI, Kokou TEGUENI,Koku AGBODEKA, Sidonie A.M.KAGNISSODE, Akoélé SILIADIN, Alassane OURO-MEDELI, Messanh DOUFFAN,Déléma MABA,Sika DOSSIM, Améyo DORKENOO, Mireille PRINCE-DAVID,Anoumou DAGNRA, Laetitia SERRANO,Ahidjo AYOUBA, Eric DELAPORTE, Martine PEETERS |
| EPI_ISL_1510669, EPI_ISL_1510670, EPI_ISL_1510671, EPI_ISL_1510672, EPI_ISL_1510673, EPI_ISL_1510674, EPI_ISL_1510675, EPI_ISL_1510676, EPI_ISL_1510677, EPI_ISL_1510678, EPI_ISL_1510679, EPI_ISL_1510680, EPI_ISL_1510681, EPI_ISL_1510682, EPI_ISL_1510683, EPI_ISL_1510684, EPI_ISL_1510685, EPI_ISL_1510686 | | | |
| see above | University of Rome Tor Vergata: Departm Experim Medicine Chair of Virology | University of Rome Tor Vergata: Departm Experim Medicine Chair of Virology | Maria Concetta Bellocchi |
| EPI_ISL_1511400, EPI_ISL_1511401, EPI_ISL_1511402 | Hospital General de México, Medicina Genómica | Hospital General de México, Medicina Genómica | Ana Laura Sánchez-Sandoval, María Eligia Juárez-Torres, Mariana Villalba-De la Peña, Siddhartha Ríos-Zaragoza, Guadalupe Mercedes Lucía Guerrero-Avendaño, Eira Valeria Barrón-Palma |
| EPI_ISL_1516898 | Hospital | National Reference Center for Viruses of Respiratory Infections, Institut Pasteur, Paris | Marion Barbet, Sylvie Behillil, Méline Bizard, Angela Brisebarre, Camille Capel,Frédéric Lemoine,Corinne Maufrais,Christophe Malabat,Pierre Lechat, Louise Lefrançois, Etienne Simon-Lorière, Vincent Enouf, Maud Vanpeene, Sylvie van der Werf,FrançOise Jouble |
| EPI_ISL_1517037, EPI_ISL_1517038, EPI_ISL_1517040, EPI_ISL_1517042, EPI_ISL_1517044, EPI_ISL_1517048, EPI_ISL_1517050, EPI_ISL_1517053 | Hospital | National Reference Center for Viruses of Respiratory Infections, Institut Pasteur, Paris | Marion Barbet, Sylvie Behillil, Méline Bizard, Angela Brisebarre, Camille Capel,Frédéric Lemoine,Corinne Maufrais,Christophe Malabat, Louise Lefrançois, Etienne Simon-Lorière, Vincent Enouf, Maud Vanpeene, Sylvie van der Werf,LéA Pilorge |
| EPI_ISL_1517085, EPI_ISL_1517088 | Labo Analyses Med | National Reference Center for Viruses of Respiratory Infections, Institut Pasteur, Paris | Marion Barbet, Sylvie Behillil, Méline Bizard, Angela Brisebarre, Camille Capel,Frédéric Lemoine,Corinne Maufrais,Christophe Malabat, Louise Lefrançois, Etienne Simon-Lorière, Vincent Enouf, Maud Vanpeene, Sylvie van der Werf,J Besson |
| EPI_ISL_1517198 | Labo Analyses Med | National Reference Center for Viruses of Respiratory Infections, Institut Pasteur, Paris | Marion Barbet, Sylvie Behillil, Méline Bizard, Angela Brisebarre, Camille Capel,Frédéric Lemoine,Corinne Maufrais,Christophe Malabat, Louise Lefrançois, Etienne Simon-Lorière, Vincent Enouf, Maud Vanpeene, Sylvie van der Werf,Nabil Gastli |
| EPI_ISL_1517387 | Inciensa, Instituto Costarricense de Investigación y Enseñanza en Nutrición y Salud | Inciensa, Instituto Costarricense de Investigación y Enseñanza en Nutrición y Salud | Cristian Peréz-Corrales, Valeria Peralta-Barquero & Patricia Salas |
| EPI_ISL_1517388, EPI_ISL_1517389 | Inciensa, Instituto Costarricense de Investigación y Enseñanza en Nutrición y Salud | Inciensa, Instituto Costarricense de Investigación y Enseñanza en Nutrición y Salud | Cristian Peréz-Corrales & Valeria Peralta-Barquero |
| EPI_ISL_1517390 | Inciensa, Instituto Costarricense de Investigación y Enseñanza en Nutrición y Salud | Inciensa, Instituto Costarricense de Investigación y Enseñanza en Nutrición y Salud | Cristian Peréz-Corrales, Valeria Peralta-Barquero & Blanco-Delgado C |
| EPI_ISL_1517391, EPI_ISL_1517393, EPI_ISL_1517394, EPI_ISL_1517395, EPI_ISL_1517396 | Inciensa, Instituto Costarricense de Investigación y Enseñanza en Nutrición y Salud | Inciensa, Instituto Costarricense de Investigación y Enseñanza en Nutrición y Salud | Cristian Peréz-Corrales & Valeria Peralta-Barquero |
| EPI_ISL_1517397 | Inciensa, Instituto Costarricense de Investigación y Enseñanza en Nutrición y Salud | Inciensa, Instituto Costarricense de Investigación y Enseñanza en Nutrición y Salud | Cristian Peréz-Corrales, Valeria Peralta-Barquero & Patricia Salas |
| EPI_ISL_1517398, EPI_ISL_1517399, EPI_ISL_1517400 | Inciensa, Instituto Costarricense de Investigación y Enseñanza en Nutrición y Salud | Inciensa, Instituto Costarricense de Investigación y Enseñanza en Nutrición y Salud | Cristian Peréz-Corrales & Valeria Peralta-Barquero |
| EPI_ISL_1517411 | Inciensa, Instituto Costarricense de Investigación y Enseñanza en Nutrición y Salud | Inciensa, Instituto Costarricense de Investigación y Enseñanza en Nutrición y Salud | Cristian Peréz-Corrales, Valeria Peralta-Barquero & Kellerman-Montero K |
| EPI_ISL_1517412, EPI_ISL_1517413, | Inciensa, Instituto Costarricense de Investigación y | Inciensa, Instituto Costarricense de Investigación y | Cristian Peréz-Corrales, Valeria Peralta-Barquero & Salas-Abarca P |

| | | | |
|---|---|---|---|
| EPI_ISL_1517414 | Enseñanza en Nutrición y Salud | Enseñanza en Nutrición y Salud | |
| EPI_ISL_1517415, EPI_ISL_1517416, EPI_ISL_1517417, EPI_ISL_1517418, EPI_ISL_1517419 | Inciensa, Instituto Costarricense de Investigación y Enseñanza en Nutrición y Salud | Inciensa, Instituto Costarricense de Investigación y Enseñanza en Nutrición y Salud | Cristian Peréz-Corrales, Valeria Peralta-Barquero & César Cerdas Quesada |
| EPI_ISL_1517420, EPI_ISL_1517421, EPI_ISL_1517422, EPI_ISL_1517423, EPI_ISL_1517424 | Inciensa, Instituto Costarricense de Investigación y Enseñanza en Nutrición y Salud | Inciensa, Instituto Costarricense de Investigación y Enseñanza en Nutrición y Salud | Cristian Peréz-Corrales, Valeria Peralta-Barquero & Gallegos-Carrillo B |
| EPI_ISL_1517425, EPI_ISL_1517426 | Inciensa, Instituto Costarricense de Investigación y Enseñanza en Nutrición y Salud | Inciensa, Instituto Costarricense de Investigación y Enseñanza en Nutrición y Salud | Cristian Peréz-Corrales, Valeria Peralta-Barquero & Salas-Abarca P |
| EPI_ISL_1517427, EPI_ISL_1517428 | Inciensa, Instituto Costarricense de Investigación y Enseñanza en Nutrición y Salud | Inciensa, Instituto Costarricense de Investigación y Enseñanza en Nutrición y Salud | Cristian Peréz-Corrales, Valeria Peralta-Barquero & Kellerman-Montero K |
| EPI_ISL_1520107 | LACEN do Estado de Rondonia | Instituto Adolfo Lutz, Interdiciplinary Procedures Center, Strategic Laboratory | Claudio Tavares Sacchi, Claudia Regina Gonçalves, Erica Valessa Ramos Gomes, Karoline Rodrigues Campos, Caio Vinicius Dias Lopes |
| EPI_ISL_1524332, EPI_ISL_1524344 | Biology Department, College of Science, Al Muthanna University and Public Health Laboratory, Al-Muthanna Health Directorate | Department of Virology, Faculty of Medicine, University of Helsinki, Helsinki, Finland | Nihad Al-Rashedi, Hussein Alburkat, Murad Munahi, Alaa Hameed, Ali Jasim, Olli Vapalahti, Tarja Sironen,Teemu Smura |
| EPI_ISL_1526784 | Hospital | National Reference Center for Viruses of Respiratory Infections, Institut Pasteur, Paris | Marion Barbet, Sylvie Behillil, Méline Bizard, Angela Brisebarre, Camille Capel,Frédéric Lemoine,Corinne Maufrais,Christophe Malabat, Louise Lefrançois, Etienne Simon-Lorière, Vincent Enouf, Maud Vanpeene, Sylvie van der Werf,LéA Pilorge |
| EPI_ISL_1527364, EPI_ISL_1527365, EPI_ISL_1527366, EPI_ISL_1527367, EPI_ISL_1527368, EPI_ISL_1527370, EPI_ISL_1527371, EPI_ISL_1527372 | Molecular diagnostic laboratory of Federal Budget Institution of Science "Central Research Institute of Epidemiology" of The Federal Service on Customers' Rights Protection and Human Well-being Surveillance | Group of Genomics and Postgenomic Technologies of Central Research Institute of Epidemilology | Samoilov AE, Kaptelova VV, Korneenko EV, Saenko SS, Shipulina OY, Speranskaya AS, Tivanova EV, Kondrasheva LY, Akimkin VG |
| EPI_ISL_1527390, EPI_ISL_1527391, EPI_ISL_1527392, EPI_ISL_1527394, EPI_ISL_1527395, EPI_ISL_1527396, EPI_ISL_1527397, EPI_ISL_1527398 | Laboratory of Extremely Dangerous Infections of Federal Budget Health Care Instute "Center of Hygiene and Epidemiology in Sakhalin region" | Group of Genomics and Postgenomic Technologies of Central Research Institute of Epidemiology | Samoilov AE, Kaptelova VV, Korneenko EV, Valdokhina AV, Bulanenko VP, Saenko SS, Arutiunian AE, Byankina MA, Zagorodnov YL, Golubeva AG, Speranskaya AS, Tivanova EV, Kondrasheva LY, Akimkin VG |
| EPI_ISL_1528224, EPI_ISL_1528225 | Labo Analyses Med | National Reference Center for Viruses of Respiratory Infections, Institut Pasteur, Paris | Marion Barbet, Sylvie Behillil, Frédéric Lemoine, Corinne Maufrais, Christophe Malabat, Méline Bizard, Angela Brisebarre, Camille Capel, Louise Lefrançois, Etienne Simon-Lorière, Vincent Enouf, Maud Vanpeene, Sylvie van der Werf,Sarah Schillinger |
| EPI_ISL_1528240 | Hospital | National Reference Center for Viruses of Respiratory Infections, Institut Pasteur, Paris | Marion Barbet, Sylvie Behillil, Frédéric Lemoine, Corinne Maufrais, Christophe Malabat, Méline Bizard, Angela Brisebarre, Camille Capel, Louise Lefrançois, Etienne Simon-Lorière, Vincent Enouf, Maud Vanpeene, Sylvie van der Werf,Eric Farfour |
| EPI_ISL_1532291, EPI_ISL_1532292, EPI_ISL_1532293, EPI_ISL_1532294 | Oman-National Influenza Center | Biotechnology & OMICs Laboratory | Aisha Al-Amri, Ahmed Al Harrasi, Intisar Al-Shukri, Amal Al-Maani, Sajjad Asaf, Bilal Hussain, Samiya Al-Zadjali, Ahmed N Al-Rawahi, Saqib Bilal, Abdul Latif Khan, Samira Al-Mahruqi, Ahmed Al-Rawahi, Hanan Al-Kindi, Amina Al-Jardani. |
| EPI_ISL_1532295, EPI_ISL_1532296, EPI_ISL_1532297, EPI_ISL_1532298, EPI_ISL_1532299, EPI_ISL_1532300 | Oman-National Influenza Center | Biotechnology & OMICs Laboratory | Abdul Latif Khan, Aisha Al-Amri, Intisar Al-Shukri, Amal Al-Maani, Sajjad Asaf, Bilal Hussain, Samiya Al-Zadjali, Ahmed N Al-Rawahi, Saqib Bilal, Samira Al-Mahruqi, Ahmed Al-Rawahi, Hanan Al-Kindi, Amina Al-Jardani, Ahmed Al Harrasi. |
| EPI_ISL_1532301 | Oman-National Influenza Center | Biotechnology & OMICs Laboratory | Ahmed Al Harrasi, Aisha Al-Amri, Intisar Al-Shukri, Amal Al-Maani, Sajjad Asaf, Bilal Hussain, Samiya Al-Zadjali, Ahmed N Al-Rawahi, Saqib Bilal, Abdul Latif Khan, Samira Al-Mahruqi, Ahmed Al-Rawahi, Hanan Al-Kindi, Amina Al-Jardani. |
| EPI_ISL_1532304 | Oman-National Influenza Center | Biotechnology & OMICs Laboratory | Abdul Latif Khan, Aisha Al-Amri, Intisar Al-Shukri, Amal Al-Maani, Sajjad Asaf, Bilal Hussain, Samiya Al-Zadjali, Ahmed N Al-Rawahi, Saqib Bilal, Samira Al-Mahruqi, Ahmed Al-Rawahi, Hanan Al-Kindi, Amina Al-Jardani, Ahmed Al Harrasi. |
| EPI_ISL_1532793, EPI_ISL_1532795 | Microbiology Department. Complexo Hospitalario Universitario de Vigo | Microbiology Department. Complexo Hospitalario Universitario de Vigo | Alfaya N, Alonso I, Alvarez M, Cabrera JJ, Carballo R, Cores O, Cortizo S, del-Campo V, Martinez L, Mediero G, Perez S, Potel C, Regueiro B, Rey S, Vasallo FJ |
| EPI_ISL_1533034, EPI_ISL_1533035, EPI_ISL_1533038, EPI_ISL_1533039, EPI_ISL_1533041, EPI_ISL_1533042, EPI_ISL_1533043, EPI_ISL_1533045, EPI_ISL_1533047, EPI_ISL_1533048, EPI_ISL_1533049, EPI_ISL_1533050, EPI_ISL_1533051, EPI_ISL_1533052, EPI_ISL_1533053, EPI_ISL_1533054, EPI_ISL_1533055, EPI_ISL_1533056, EPI_ISL_1533057, EPI_ISL_1533059, EPI_ISL_1533060, EPI_ISL_1533062, EPI_ISL_1533063, EPI_ISL_1533064, EPI_ISL_1533067, EPI_ISL_1533068, EPI_ISL_1533070, EPI_ISL_1533071, EPI_ISL_1533072, EPI_ISL_1533073, EPI_ISL_1533074, EPI_ISL_1533075, EPI_ISL_1533076, EPI_ISL_1533078, EPI_ISL_1533079, EPI_ISL_1533080, EPI_ISL_1533081, EPI_ISL_1533083, EPI_ISL_1533084, EPI_ISL_1533085, EPI_ISL_1533086, EPI_ISL_1533087, EPI_ISL_1533088, EPI_ISL_1533090, EPI_ISL_1533091, EPI_ISL_1533092, EPI_ISL_1533093, EPI_ISL_1533095, EPI_ISL_1533096, EPI_ISL_1533097, EPI_ISL_1533098, EPI_ISL_1533100, EPI_ISL_1533101 | | | |
| see above | Diagnostic and Research Center of Infectious Diseases, Medical Faculty, Andalas University | Diagnostic and Research Center of Infectious Diseases, Medical Faculty, Andalas University | Andani Eka Putra, Syafrizayanti, Ikhwan R. Sudji, Linosefa, Dede Rahman Agustian, Ayu Novita Trisnawati, Siskalil Fahma, Nia Ayuni Putri, Sekar Asri Tresnaningtyas, SM Rezvi, Fauzul Azhim, Dessy Arisanty, Gestina Aliska, Desmawati, Nita Afriani ,Syandrez Prima Putra, Juane Plantika Menra, Yolani Syaputri, Mutia Lailani |
| EPI_ISL_1540448 | Labo Analyses Med | National Reference Center for Viruses of Respiratory Infections, Institut Pasteur, Paris | Marion Barbet, Sylvie Behillil, Frédéric Lemoine, Corinne Maufrais, Christophe Malabat,Adrien Pain, Méline Bizard, Angela Brisebarre, Camille Capel, Louise Lefrançois, Etienne Simon-Lorière, Vincent Enouf, Maud Vanpeene, Sylvie van der Werf,Sarah Schillinger |
| EPI_ISL_1544118 | Supratech Micropath Laboratory Research Institute Pvt Ltd, Ahmedabad | Gujarat Biotechnology Research Centre | Mahendra Parikh,Zuber Saiyed,Ramesh Pandit,Janvi Raval,Zarna Patel,Nitin Savaliya,Dinesh Kumar,Twinkle Soni,Sonal Sharma ,Umang Mishra,Nitesh Shah,Chaitanya Joshi,Madhvi Joshi |
| EPI_ISL_1544119 | Supratech Micropath Laboratory Research Institute Pvt Ltd, Ahmedabad | Gujarat Biotechnology Research Centre | Mahendra Parikh,Ramesh Pandit,Janvi Raval,Zarna Patel,Nitin Savaliya,Dinesh Kumar,Twinkle Soni,Sonal Sharma ,Zuber Saiyed,Umang Mishra,Nitesh Shah,Chaitanya Joshi,Madhvi Joshi |
| EPI_ISL_1544120 | Supratech Micropath Laboratory Research Institute Pvt Ltd, Ahmedabad | Gujarat Biotechnology Research Centre | Mahendra Parikh,Janvi Raval,Zarna Patel,Nitin Savaliya,Dinesh Kumar,Twinkle Soni,Sonal Sharma ,Zuber Saiyed,Ramesh Pandit,Umang Mishra,Nitesh Shah,Chaitanya Joshi,Madhvi Joshi |
| EPI_ISL_1544121 | Supratech Micropath Laboratory Research Institute Pvt Ltd, Ahmedabad | Gujarat Biotechnology Research Centre | Zarna Patel,Nitin Savaliya,Dinesh Kumar,Twinkle Soni,Sonal Sharma ,Zuber Saiyed,Ramesh Pandit,Janvi Raval,Mahendra Parikh,Umang Mishra,Nitesh Shah,Chaitanya Joshi,Madhvi Joshi |
| EPI_ISL_1544122 | Supratech Micropath Laboratory Research Institute Pvt Ltd, Ahmedabad | Gujarat Biotechnology Research Centre | Nitin Savaliya,Dinesh Kumar,Twinkle Soni,Sonal Sharma ,Zuber Saiyed,Ramesh Pandit,Janvi Raval,Zarna Patel,Mahendra Parikh,Umang Mishra,Nitesh Shah,Chaitanya Joshi,Madhvi Joshi |
| EPI_ISL_1544123 | Supratech Micropath Laboratory Research Institute Pvt Ltd, Ahmedabad | Gujarat Biotechnology Research Centre | Dinesh Kumar,Twinkle Soni,Sonal Sharma ,Zuber Saiyed,Ramesh Pandit,Janvi Raval,Zarna Patel,Nitin Savaliya,Mahendra Parikh,Umang Mishra,Nitesh Shah,Chaitanya Joshi,Madhvi Joshi |
| EPI_ISL_1544124 | Supratech Micropath Laboratory Research Institute Pvt Ltd, Ahmedabad | Gujarat Biotechnology Research Centre | Twinkle Soni,Sonal Sharma ,Zuber Saiyed,Ramesh Pandit,Janvi Raval,Zarna Patel,Nitin Savaliya,Dinesh Kumar,Mahendra Parikh,Umang Mishra,Nitesh Shah,Chaitanya Joshi,Madhvi Joshi |
| EPI_ISL_1544125 | Supratech Micropath Laboratory Research Institute Pvt Ltd, Ahmedabad | Gujarat Biotechnology Research Centre | Sonal Sharma ,Zuber Saiyed,Ramesh Pandit,Janvi Raval,Zarna Patel,Nitin Savaliya,Dinesh Kumar,Twinkle Soni,Mahendra Parikh,Umang Mishra,Nitesh Shah,Chaitanya Joshi,Madhvi Joshi |
| EPI_ISL_1544126 | Supratech Micropath Laboratory Research Institute Pvt Ltd, Ahmedabad | Gujarat Biotechnology Research Centre | Zuber Saiyed,Ramesh Pandit,Janvi Raval,Zarna Patel,Nitin Savaliya,Dinesh Kumar,Twinkle Soni,Sonal Sharma ,Mahendra Parikh,Umang Mishra,Nitesh Shah,Chaitanya Joshi,Madhvi Joshi |
| EPI_ISL_1544127 | Supratech Micropath Laboratory Research Institute Pvt Ltd, Ahmedabad | Gujarat Biotechnology Research Centre | Ramesh Pandit,Janvi Raval,Zarna Patel,Nitin Savaliya,Dinesh Kumar,Twinkle Soni,Sonal Sharma ,Zuber Saiyed,Mahendra Parikh,Umang Mishra,Nitesh Shah,Chaitanya Joshi,Madhvi Joshi |
| EPI_ISL_1544128 | Supratech Micropath Laboratory Research Institute Pvt Ltd, Ahmedabad | Gujarat Biotechnology Research Centre | Janvi Raval,Zarna Patel,Nitin Savaliya,Dinesh Kumar,Twinkle Soni,Sonal Sharma ,Zuber Saiyed,Ramesh Pandit,Mahendra Parikh,Umang Mishra,Nitesh Shah,Chaitanya Joshi,Madhvi Joshi |

| | | | |
|---|---|---|---|
| EPI_ISL_1544129 | Supratech Micropath Laboratory Research Institute Pvt Ltd, Ahmedabad | Gujarat Biotechnology Research Centre | Zarna Patel,Nitin Savaliya,Dinesh Kumar,Twinkle Soni,Sonal Sharma ,Zuber Saiyed,Ramesh Pandit,Janvi Raval,Mahendra Parikh,Umang Mishra,Nitesh Shah,Chaitanya Joshi,Madhvi Joshi |
| EPI_ISL_1544130 | Supratech Micropath Laboratory Research Institute Pvt Ltd, Ahmedabad | Gujarat Biotechnology Research Centre | Nitin Savaliya,Dinesh Kumar,Twinkle Soni,Sonal Sharma ,Zuber Saiyed,Ramesh Pandit,Janvi Raval,Zarna Patel,Mahendra Parikh,Umang Mishra,Nitesh Shah,Chaitanya Joshi,Madhvi Joshi |
| EPI_ISL_1544131 | Supratech Micropath Laboratory Research Institute Pvt Ltd, Ahmedabad | Gujarat Biotechnology Research Centre | Dinesh Kumar,Twinkle Soni,Sonal Sharma ,Zuber Saiyed,Ramesh Pandit,Janvi Raval,Zarna Patel,Nitin Savaliya,Mahendra Parikh,Umang Mishra,Nitesh Shah,Chaitanya Joshi,Madhvi Joshi |
| EPI_ISL_1544132 | Supratech Micropath Laboratory Research Institute Pvt Ltd, Ahmedabad | Gujarat Biotechnology Research Centre | Twinkle Soni,Sonal Sharma ,Zuber Saiyed,Ramesh Pandit,Janvi Raval,Zarna Patel,Nitin Savaliya,Dinesh Kumar,Mahendra Parikh,Umang Mishra,Nitesh Shah,Chaitanya Joshi,Madhvi Joshi |
| EPI_ISL_1544135 | Supratech Micropath Laboratory Research Institute Pvt Ltd, Ahmedabad | Gujarat Biotechnology Research Centre | Ramesh Pandit,Janvi Raval,Zarna Patel,Nitin Savaliya,Dinesh Kumar,Twinkle Soni,Sonal Sharma ,Zuber Saiyed,Mahendra Parikh,Umang Mishra,Nitesh Shah,Chaitanya Joshi,Madhvi Joshi |
| EPI_ISL_1544136 | Supratech Micropath Laboratory Research Institute Pvt Ltd, Ahmedabad | Gujarat Biotechnology Research Centre | Janvi Raval,Zarna Patel,Nitin Savaliya,Dinesh Kumar,Twinkle Soni,Sonal Sharma ,Zuber Saiyed,Ramesh Pandit,Mahendra Parikh,Umang Mishra,Nitesh Shah,Chaitanya Joshi,Madhvi Joshi |
| EPI_ISL_1544137 | Supratech Micropath Laboratory Research Institute Pvt Ltd, Ahmedabad | Gujarat Biotechnology Research Centre | Zarna Patel,Nitin Savaliya,Dinesh Kumar,Twinkle Soni,Sonal Sharma ,Zuber Saiyed,Ramesh Pandit,Janvi Raval,Mahendra Parikh,Umang Mishra,Nitesh Shah,Chaitanya Joshi,Madhvi Joshi |
| EPI_ISL_1544138 | Supratech Micropath Laboratory Research Institute Pvt Ltd, Ahmedabad | Gujarat Biotechnology Research Centre | Nitin Savaliya,Dinesh Kumar,Twinkle Soni,Sonal Sharma ,Zuber Saiyed,Ramesh Pandit,Janvi Raval,Zarna Patel,Mahendra Parikh,Umang Mishra,Nitesh Shah,Chaitanya Joshi,Madhvi Joshi |
| EPI_ISL_1544139 | Supratech Micropath Laboratory Research Institute Pvt Ltd, Ahmedabad | Gujarat Biotechnology Research Centre | Dinesh Kumar,Twinkle Soni,Sonal Sharma ,Zuber Saiyed,Ramesh Pandit,Janvi Raval,Zarna Patel,Nitin Savaliya,Mahendra Parikh,Umang Mishra,Nitesh Shah,Chaitanya Joshi,Madhvi Joshi |
| EPI_ISL_1544140 | Supratech Micropath Laboratory Research Institute Pvt Ltd, Ahmedabad | Gujarat Biotechnology Research Centre | Twinkle Soni,Sonal Sharma ,Zuber Saiyed,Ramesh Pandit,Janvi Raval,Zarna Patel,Nitin Savaliya,Dinesh Kumar,Mahendra Parikh,Umang Mishra,Nitesh Shah,Chaitanya Joshi,Madhvi Joshi |
| EPI_ISL_1544141 | Supratech Micropath Laboratory Research Institute Pvt Ltd, Ahmedabad | Gujarat Biotechnology Research Centre | Sonal Sharma ,Zuber Saiyed,Ramesh Pandit,Janvi Raval,Zarna Patel,Nitin Savaliya,Dinesh Kumar,Twinkle Soni,Mahendra Parikh,Umang Mishra,Nitesh Shah,Chaitanya Joshi,Madhvi Joshi |
| EPI_ISL_1544142 | SSG Medical College, Baroda | Gujarat Biotechnology Research Centre | Tanuja B. Javadekar,Zuber Saiyed,Ramesh Pandit,Janvi Raval,Zarna Patel,Nitin Savaliya,Dinesh Kumar,Twinkle Soni,Sonal Sharma ,Umang Mishra,Nitesh Shah,Chaitanya Joshi,Madhvi Joshi |
| EPI_ISL_1544143 | SSG Medical College, Baroda | Gujarat Biotechnology Research Centre | Tanuja B. Javadekar,Ramesh Pandit,Janvi Raval,Zarna Patel,Nitin Savaliya,Dinesh Kumar,Twinkle Soni,Sonal Sharma ,Zuber Saiyed,Umang Mishra,Nitesh Shah,Chaitanya Joshi,Madhvi Joshi |
| EPI_ISL_1544144 | SSG Medical College, Baroda | Gujarat Biotechnology Research Centre | Tanuja B. Javadekar,Janvi Raval,Zarna Patel,Nitin Savaliya,Dinesh Kumar,Twinkle Soni,Sonal Sharma ,Zuber Saiyed,Ramesh Pandit,Umang Mishra,Nitesh Shah,Chaitanya Joshi,Madhvi Joshi |
| EPI_ISL_1544145 | SSG Medical College, Baroda | Gujarat Biotechnology Research Centre | Zarna Patel,Nitin Savaliya,Dinesh Kumar,Twinkle Soni,Sonal Sharma ,Zuber Saiyed,Ramesh Pandit,Janvi Raval,Tanuja B. Javadekar,Umang Mishra,Nitesh Shah,Chaitanya Joshi,Madhvi Joshi |
| EPI_ISL_1544146 | SSG Medical College, Baroda | Gujarat Biotechnology Research Centre | Nitin Savaliya,Dinesh Kumar,Twinkle Soni,Sonal Sharma ,Zuber Saiyed,Ramesh Pandit,Janvi Raval,Zarna Patel,Tanuja B. Javadekar,Umang Mishra,Nitesh Shah,Chaitanya Joshi,Madhvi Joshi |
| EPI_ISL_1544147 | SSG Medical College, Baroda | Gujarat Biotechnology Research Centre | Dinesh Kumar,Twinkle Soni,Sonal Sharma ,Zuber Saiyed,Ramesh Pandit,Janvi Raval,Zarna Patel,Nitin Savaliya,Tanuja B. Javadekar,Umang Mishra,Nitesh Shah,Chaitanya Joshi,Madhvi Joshi |
| EPI_ISL_1544148 | SSG Medical College, Baroda | Gujarat Biotechnology Research Centre | Twinkle Soni,Sonal Sharma ,Zuber Saiyed,Ramesh Pandit,Janvi Raval,Zarna Patel,Nitin Savaliya,Dinesh Kumar,Tanuja B. Javadekar,Umang Mishra,Nitesh Shah,Chaitanya Joshi,Madhvi Joshi |
| EPI_ISL_1544149 | SSG Medical College, Baroda | Gujarat Biotechnology Research Centre | Sonal Sharma ,Zuber Saiyed,Ramesh Pandit,Janvi Raval,Zarna Patel,Nitin Savaliya,Dinesh Kumar,Twinkle Soni,Tanuja B. Javadekar,Umang Mishra,Nitesh Shah,Chaitanya Joshi,Madhvi Joshi |
| EPI_ISL_1544150 | SSG Medical College, Baroda | Gujarat Biotechnology Research Centre | Zuber Saiyed,Ramesh Pandit,Janvi Raval,Zarna Patel,Nitin Savaliya,Dinesh Kumar,Twinkle Soni,Sonal Sharma ,Tanuja B. Javadekar,Umang Mishra,Nitesh Shah,Chaitanya Joshi,Madhvi Joshi |
| EPI_ISL_1544151 | SSG Medical College, Baroda | Gujarat Biotechnology Research Centre | Ramesh Pandit,Janvi Raval,Zarna Patel,Nitin Savaliya,Dinesh Kumar,Twinkle Soni,Sonal Sharma ,Zuber Saiyed,Tanuja B. Javadekar,Umang Mishra,Nitesh Shah,Chaitanya Joshi,Madhvi Joshi |
| EPI_ISL_1544152 | SSG Medical College, Baroda | Gujarat Biotechnology Research Centre | Janvi Raval,Zarna Patel,Nitin Savaliya,Dinesh Kumar,Twinkle Soni,Sonal Sharma ,Zuber Saiyed,Ramesh Pandit,Tanuja B. Javadekar,Umang Mishra,Nitesh Shah,Chaitanya Joshi,Madhvi Joshi |
| EPI_ISL_1544153 | SSG Medical College, Baroda | Gujarat Biotechnology Research Centre | Zarna Patel,Nitin Savaliya,Dinesh Kumar,Twinkle Soni,Sonal Sharma ,Zuber Saiyed,Ramesh Pandit,Janvi Raval,Tanuja B. Javadekar,Umang Mishra,Nitesh Shah,Chaitanya Joshi,Madhvi Joshi |
| EPI_ISL_1544154 | SSG Medical College, Baroda | Gujarat Biotechnology Research Centre | Nitin Savaliya,Dinesh Kumar,Twinkle Soni,Sonal Sharma ,Zuber Saiyed,Ramesh Pandit,Janvi Raval,Zarna Patel,Tanuja B. Javadekar,Umang Mishra,Nitesh Shah,Chaitanya Joshi,Madhvi Joshi |
| EPI_ISL_1544155 | SSG Medical College, Baroda | Gujarat Biotechnology Research Centre | Dinesh Kumar,Twinkle Soni,Sonal Sharma ,Zuber Saiyed,Ramesh Pandit,Janvi Raval,Zarna Patel,Nitin Savaliya,Tanuja B. Javadekar,Umang Mishra,Nitesh Shah,Chaitanya Joshi,Madhvi Joshi |
| EPI_ISL_1544156 | SSG Medical College, Baroda | Gujarat Biotechnology Research Centre | Twinkle Soni,Sonal Sharma ,Zuber Saiyed,Ramesh Pandit,Janvi Raval,Zarna Patel,Nitin Savaliya,Dinesh Kumar,Tanuja B. Javadekar,Umang Mishra,Nitesh Shah,Chaitanya Joshi,Madhvi Joshi |
| EPI_ISL_1544157 | SSG Medical College, Baroda | Gujarat Biotechnology Research Centre | Sonal Sharma ,Zuber Saiyed,Ramesh Pandit,Janvi Raval,Zarna Patel,Nitin Savaliya,Dinesh Kumar,Twinkle Soni,Tanuja B. Javadekar,Umang Mishra,Nitesh Shah,Chaitanya Joshi,Madhvi Joshi |
| EPI_ISL_1544158 | SSG Medical College, Baroda | Gujarat Biotechnology Research Centre | Zuber Saiyed,Ramesh Pandit,Janvi Raval,Zarna Patel,Nitin Savaliya,Dinesh Kumar,Twinkle Soni,Sonal Sharma ,Tanuja B. Javadekar,Umang Mishra,Nitesh Shah,Chaitanya Joshi,Madhvi Joshi |
| EPI_ISL_1544159 | SSG Medical College, Baroda | Gujarat Biotechnology Research Centre | Ramesh Pandit,Janvi Raval,Zarna Patel,Nitin Savaliya,Dinesh Kumar,Twinkle Soni,Sonal Sharma ,Zuber Saiyed,Tanuja B. Javadekar,Umang Mishra,Nitesh Shah,Chaitanya Joshi,Madhvi Joshi |
| EPI_ISL_1544160 | SSG Medical College, Baroda | Gujarat Biotechnology Research Centre | Janvi Raval,Zarna Patel,Nitin Savaliya,Dinesh Kumar,Twinkle Soni,Sonal Sharma ,Zuber Saiyed,Ramesh Pandit,Tanuja B. Javadekar,Umang Mishra,Nitesh Shah,Chaitanya Joshi,Madhvi Joshi |
| EPI_ISL_1544161 | SSG Medical College, Baroda | Gujarat Biotechnology Research Centre | Zarna Patel,Nitin Savaliya,Dinesh Kumar,Twinkle Soni,Sonal Sharma ,Zuber Saiyed,Ramesh Pandit,Janvi Raval,Tanuja B. Javadekar,Umang Mishra,Nitesh Shah,Chaitanya Joshi,Madhvi Joshi |
| EPI_ISL_1544162 | SSG Medical College, Baroda | Gujarat Biotechnology Research Centre | Nitin Savaliya,Dinesh Kumar,Twinkle Soni,Sonal Sharma ,Zuber Saiyed,Ramesh Pandit,Janvi Raval,Zarna Patel,Tanuja B. Javadekar,Umang Mishra,Nitesh Shah,Chaitanya Joshi,Madhvi Joshi |
| EPI_ISL_1544163 | SSG Medical College, Baroda | Gujarat Biotechnology Research Centre | Dinesh Kumar,Twinkle Soni,Sonal Sharma ,Zuber Saiyed,Ramesh Pandit,Janvi Raval,Zarna Patel,Nitin Savaliya,Tanuja B. Javadekar,Umang Mishra,Nitesh Shah,Chaitanya Joshi,Madhvi Joshi |
| EPI_ISL_1544164 | SSG Medical College, Baroda | Gujarat Biotechnology Research Centre | Twinkle Soni,Sonal Sharma ,Zuber Saiyed,Ramesh Pandit,Janvi Raval,Zarna Patel,Nitin Savaliya,Dinesh Kumar,Tanuja B. Javadekar,Umang Mishra,Nitesh Shah,Chaitanya Joshi,Madhvi Joshi |
| EPI_ISL_1544165 | SSG Medical College, Baroda | Gujarat Biotechnology Research Centre | Sonal Sharma ,Zuber Saiyed,Ramesh Pandit,Janvi Raval,Zarna Patel,Nitin Savaliya,Dinesh Kumar,Twinkle Soni,Tanuja B. Javadekar,Umang Mishra,Nitesh Shah,Chaitanya Joshi,Madhvi Joshi |

| | | | |
|---|---|---|---|
| EPI_ISL_1544166 | Supratech Micropath Laboratory Research Institute Pvt Ltd, Ahmedabad | Gujarat Biotechnology Research Centre | Zuber Saiyed,Ramesh Pandit,Janvi Raval,Zarna Patel,Nitin Savaliya,Dinesh Kumar,Twinkle Soni,Sonal Sharma ,Mahendra Parikh,Umang Mishra,Nitesh Shah,Chaitanya Joshi,Madhvi Joshi |
| EPI_ISL_1544167 | Supratech Micropath Laboratory Research Institute Pvt Ltd, Ahmedabad | Gujarat Biotechnology Research Centre | Ramesh Pandit,Janvi Raval,Zarna Patel,Nitin Savaliya,Dinesh Kumar,Twinkle Soni,Sonal Sharma ,Zuber Saiyed,Mahendra Parikh,Umang Mishra,Nitesh Shah,Chaitanya Joshi,Madhvi Joshi |
| EPI_ISL_1544168 | Supratech Micropath Laboratory Research Institute Pvt Ltd, Ahmedabad | Gujarat Biotechnology Research Centre | Janvi Raval,Zarna Patel,Nitin Savaliya,Dinesh Kumar,Twinkle Soni,Sonal Sharma ,Zuber Saiyed,Ramesh Pandit,Mahendra Parikh,Umang Mishra,Nitesh Shah,Chaitanya Joshi,Madhvi Joshi |
| EPI_ISL_1544169 | Supratech Micropath Laboratory Research Institute Pvt Ltd, Ahmedabad | Gujarat Biotechnology Research Centre | Zarna Patel,Nitin Savaliya,Dinesh Kumar,Twinkle Soni,Sonal Sharma ,Zuber Saiyed,Ramesh Pandit,Janvi Raval,Mahendra Parikh,Umang Mishra,Nitesh Shah,Chaitanya Joshi,Madhvi Joshi |
| EPI_ISL_1544170 | SSG Medical College, Baroda | Gujarat Biotechnology Research Centre | Nitin Savaliya,Dinesh Kumar,Twinkle Soni,Sonal Sharma ,Zuber Saiyed,Ramesh Pandit,Janvi Raval,Zarna Patel,Tanuja B. Javadekar,Umang Mishra,Nitesh Shah,Chaitanya Joshi,Madhvi Joshi |
| EPI_ISL_1544171 | SSG Medical College, Baroda | Gujarat Biotechnology Research Centre | Dinesh Kumar,Twinkle Soni,Sonal Sharma ,Zuber Saiyed,Ramesh Pandit,Janvi Raval,Zarna Patel,Nitin Savaliya,Tanuja B. Javadekar,Umang Mishra,Nitesh Shah,Chaitanya Joshi,Madhvi Joshi |
| EPI_ISL_1544172 | SSG Medical College, Baroda | Gujarat Biotechnology Research Centre | Twinkle Soni,Sonal Sharma ,Zuber Saiyed,Ramesh Pandit,Janvi Raval,Zarna Patel,Nitin Savaliya,Dinesh Kumar,Tanuja B. Javadekar,Umang Mishra,Nitesh Shah,Chaitanya Joshi,Madhvi Joshi |
| EPI_ISL_1544173 | SSG Medical College, Baroda | Gujarat Biotechnology Research Centre | Sonal Sharma ,Zuber Saiyed,Ramesh Pandit,Janvi Raval,Zarna Patel,Nitin Savaliya,Dinesh Kumar,Twinkle Soni,Tanuja B. Javadekar,Umang Mishra,Nitesh Shah,Chaitanya Joshi,Madhvi Joshi |
| EPI_ISL_1544174 | SSG Medical College, Baroda | Gujarat Biotechnology Research Centre | Zuber Saiyed,Ramesh Pandit,Janvi Raval,Zarna Patel,Nitin Savaliya,Dinesh Kumar,Twinkle Soni,Sonal Sharma ,Tanuja B. Javadekar,Umang Mishra,Nitesh Shah,Chaitanya Joshi,Madhvi Joshi |
| EPI_ISL_1544175 | SSG Medical College, Baroda | Gujarat Biotechnology Research Centre | Ramesh Pandit,Janvi Raval,Zarna Patel,Nitin Savaliya,Dinesh Kumar,Twinkle Soni,Sonal Sharma ,Zuber Saiyed,Tanuja B. Javadekar,Umang Mishra,Nitesh Shah,Chaitanya Joshi,Madhvi Joshi |
| EPI_ISL_1544176 | SSG Medical College, Baroda | Gujarat Biotechnology Research Centre | Janvi Raval,Zarna Patel,Nitin Savaliya,Dinesh Kumar,Twinkle Soni,Sonal Sharma ,Zuber Saiyed,Ramesh Pandit,Tanuja B. Javadekar,Umang Mishra,Nitesh Shah,Chaitanya Joshi,Madhvi Joshi |
| EPI_ISL_1544177 | SSG Medical College, Baroda | Gujarat Biotechnology Research Centre | Zarna Patel,Nitin Savaliya,Dinesh Kumar,Twinkle Soni,Sonal Sharma ,Zuber Saiyed,Ramesh Pandit,Janvi Raval,Tanuja B. Javadekar,Umang Mishra,Nitesh Shah,Chaitanya Joshi,Madhvi Joshi |
| EPI_ISL_1544178 | SSG Medical College, Baroda | Gujarat Biotechnology Research Centre | Nitin Savaliya,Dinesh Kumar,Twinkle Soni,Sonal Sharma ,Zuber Saiyed,Ramesh Pandit,Janvi Raval,Zarna Patel,Tanuja B. Javadekar,Umang Mishra,Nitesh Shah,Chaitanya Joshi,Madhvi Joshi |
| EPI_ISL_1544179 | Government Medical College (GMC), Surat | Gujarat Biotechnology Research Centre | Neeta Khandelwal,Dinesh Kumar,Twinkle Soni,Sonal Sharma ,Zuber Saiyed,Ramesh Pandit,Janvi Raval,Zarna Patel,Nitin Savaliya,Umang Mishra,Nitesh Shah,Chaitanya Joshi,Madhvi Joshi |
| EPI_ISL_1544180 | Government Medical College (GMC), Surat | Gujarat Biotechnology Research Centre | Neeta Khandelwal,Twinkle Soni,Sonal Sharma ,Zuber Saiyed,Ramesh Pandit,Janvi Raval,Zarna Patel,Nitin Savaliya,Dinesh Kumar,Umang Mishra,Nitesh Shah,Chaitanya Joshi,Madhvi Joshi |
| EPI_ISL_1544181 | Government Medical College (GMC), Surat | Gujarat Biotechnology Research Centre | Neeta Khandelwal,Sonal Sharma ,Zuber Saiyed,Ramesh Pandit,Janvi Raval,Zarna Patel,Nitin Savaliya,Dinesh Kumar,Twinkle Soni,Umang Mishra,Nitesh Shah,Chaitanya Joshi,Madhvi Joshi |
| EPI_ISL_1544182 | Government Medical College (GMC), Surat | Gujarat Biotechnology Research Centre | Zuber Saiyed,Ramesh Pandit,Janvi Raval,Zarna Patel,Nitin Savaliya,Dinesh Kumar,Twinkle Soni,Sonal Sharma ,Neeta Khandelwal,Umang Mishra,Nitesh Shah,Chaitanya Joshi,Madhvi Joshi |
| EPI_ISL_1544183 | Government Medical College (GMC), Surat | Gujarat Biotechnology Research Centre | Ramesh Pandit,Janvi Raval,Zarna Patel,Nitin Savaliya,Dinesh Kumar,Twinkle Soni,Sonal Sharma ,Zuber Saiyed,Neeta Khandelwal,Umang Mishra,Nitesh Shah,Chaitanya Joshi,Madhvi Joshi |
| EPI_ISL_1544184 | Government Medical College (GMC), Surat | Gujarat Biotechnology Research Centre | Janvi Raval,Zarna Patel,Nitin Savaliya,Dinesh Kumar,Twinkle Soni,Sonal Sharma ,Zuber Saiyed,Ramesh Pandit,Neeta Khandelwal,Umang Mishra,Nitesh Shah,Chaitanya Joshi,Madhvi Joshi |
| EPI_ISL_1544185 | Government Medical College (GMC), Surat | Gujarat Biotechnology Research Centre | Zarna Patel,Nitin Savaliya,Dinesh Kumar,Twinkle Soni,Sonal Sharma ,Zuber Saiyed,Ramesh Pandit,Janvi Raval,Neeta Khandelwal,Umang Mishra,Nitesh Shah,Chaitanya Joshi,Madhvi Joshi |
| EPI_ISL_1544186 | Government Medical College (GMC), Surat | Gujarat Biotechnology Research Centre | Nitin Savaliya,Dinesh Kumar,Twinkle Soni,Sonal Sharma ,Zuber Saiyed,Ramesh Pandit,Janvi Raval,Zarna Patel,Neeta Khandelwal,Umang Mishra,Nitesh Shah,Chaitanya Joshi,Madhvi Joshi |
| EPI_ISL_1544187 | Government Medical College (GMC), Surat | Gujarat Biotechnology Research Centre | Dinesh Kumar,Twinkle Soni,Sonal Sharma ,Zuber Saiyed,Ramesh Pandit,Janvi Raval,Zarna Patel,Nitin Savaliya,Neeta Khandelwal,Umang Mishra,Nitesh Shah,Chaitanya Joshi,Madhvi Joshi |
| EPI_ISL_1544188 | Government Medical College (GMC), Surat | Gujarat Biotechnology Research Centre | Twinkle Soni,Sonal Sharma ,Zuber Saiyed,Ramesh Pandit,Janvi Raval,Zarna Patel,Nitin Savaliya,Dinesh Kumar,Neeta Khandelwal,Umang Mishra,Nitesh Shah,Chaitanya Joshi,Madhvi Joshi |
| EPI_ISL_1544189 | Government Medical College (GMC), Surat | Gujarat Biotechnology Research Centre | Sonal Sharma ,Zuber Saiyed,Ramesh Pandit,Janvi Raval,Zarna Patel,Nitin Savaliya,Dinesh Kumar,Twinkle Soni,Neeta Khandelwal,Umang Mishra,Nitesh Shah,Chaitanya Joshi,Madhvi Joshi |
| EPI_ISL_1544190 | Government Medical College (GMC), Surat | Gujarat Biotechnology Research Centre | Zuber Saiyed,Ramesh Pandit,Janvi Raval,Zarna Patel,Nitin Savaliya,Dinesh Kumar,Twinkle Soni,Sonal Sharma ,Neeta Khandelwal,Umang Mishra,Nitesh Shah,Chaitanya Joshi,Madhvi Joshi |
| EPI_ISL_1544191 | Government Medical College (GMC), Surat | Gujarat Biotechnology Research Centre | Ramesh Pandit,Janvi Raval,Zarna Patel,Nitin Savaliya,Dinesh Kumar,Twinkle Soni,Sonal Sharma ,Zuber Saiyed,Neeta Khandelwal,Umang Mishra,Nitesh Shah,Chaitanya Joshi,Madhvi Joshi |
| EPI_ISL_1544192 | Government Medical College (GMC), Surat | Gujarat Biotechnology Research Centre | Janvi Raval,Zarna Patel,Nitin Savaliya,Dinesh Kumar,Twinkle Soni,Sonal Sharma ,Zuber Saiyed,Ramesh Pandit,Neeta Khandelwal,Umang Mishra,Nitesh Shah,Chaitanya Joshi,Madhvi Joshi |
| EPI_ISL_1544193 | Government Medical College (GMC), Surat | Gujarat Biotechnology Research Centre | Zarna Patel,Nitin Savaliya,Dinesh Kumar,Twinkle Soni,Sonal Sharma ,Zuber Saiyed,Ramesh Pandit,Janvi Raval,Neeta Khandelwal,Umang Mishra,Nitesh Shah,Chaitanya Joshi,Madhvi Joshi |
| EPI_ISL_1544194 | Government Medical College (GMC), Surat | Gujarat Biotechnology Research Centre | Nitin Savaliya,Dinesh Kumar,Twinkle Soni,Sonal Sharma ,Zuber Saiyed,Ramesh Pandit,Janvi Raval,Zarna Patel,Neeta Khandelwal,Umang Mishra,Nitesh Shah,Chaitanya Joshi,Madhvi Joshi |
| EPI_ISL_1544195 | Government Medical College (GMC), Surat | Gujarat Biotechnology Research Centre | Dinesh Kumar,Twinkle Soni,Sonal Sharma ,Zuber Saiyed,Ramesh Pandit,Janvi Raval,Zarna Patel,Nitin Savaliya,Neeta Khandelwal,Umang Mishra,Nitesh Shah,Chaitanya Joshi,Madhvi Joshi |
| EPI_ISL_1548045, EPI_ISL_1548046, EPI_ISL_1548047, EPI_ISL_1548048 | University of Rome Tor Vergata: Departm Experim Medicine Chair of Virology | University of Rome Tor Vergata: Departm Experim Medicine Chair of Virology | Maria Concetta Bellocchi, Luca Carioti, Maria Botticelli, Lorenzo Piermatteo, Francesca Ceccherini-Silberstein |
| EPI_ISL_1575297, EPI_ISL_1575436 | Laboratoire Biolim/FSS/UL | Unité Mixte Internationale TransVIHMI (UMI 233 IRD - U1175 INSERM - Université de Montpellier) IRD (Institut de recherche pour le développement) | Mounerou SALOU, Christelle BUTEL, Wembo A. HALATOKO, Amivi EHLAN, Abla A. KONOU, Issaka Maman, Syntyche DEVATCHAGNI, Adodo SADJI, Kokou TEGUENI,Koku AGBODEKA, Sidonie A.M.KAGNISSODE, Akoélé SILIADIN, Alassane OURO-MEDELI, Messanh DOUFFAN,Déléma MABA,Sika DOSSIM, Améyo DORKENOO, Mireille PRINCE-DAVID,Anoumou DAGNRA, Laetitia SERRANO,Ahidjo AYOUBA, Eric DELAPORTE, Martine PEETERS |
| EPI_ISL_1576835, EPI_ISL_1577026, EPI_ISL_1577182, EPI_ISL_1577390, EPI_ISL_1582589, EPI_ISL_1582590 | Universidad Industrial de Santander (Laboratorio Central de Investigaciones - Clínica Chicamocha). | Universidad Industrial de Santander. | Francisco Martinez-Perez, Cristian E. Cadena-Caballero, Diego Rueda-Plata, Carolina S. Torres-Jiménez, Lizeth J. Forero-Buitrago, Erika Lizarazo-Gutiérrez, Carlos Barrios-Hernández, Lina M. Vera-Cala. |
| EPI_ISL_1582639, EPI_ISL_1582640, EPI_ISL_1582641 | Research | National Reference Center for Viruses of Respiratory Infections, Institut Pasteur, Paris | Marion Barbet, Sylvie Behillil, Frédéric Lemoine, Corinne Maufrais, Christophe Malabat,Gael Millot, Méline Bizard, Angela Brisebarre, Camille Capel, Louise Lefrançois, Etienne Simon-Lorière, Vincent Enouf, Maud Vanpeene, Sylvie van der Werf,Marianne Leruez-Ville |
| EPI_ISL_1582643, EPI_ISL_1582644, | Research | National Reference Center for Viruses of Respiratory | Marion Barbet, Sylvie Behillil, Frédéric Lemoine, Corinne Maufrais, Christophe Malabat, Méline Bizard, Angela Brisebarre, Camille Capel, Louise Lefrançois, |

| | | | |
|---|---|---|---|
| EPI_ISL_1582645, EPI_ISL_1582646 | | Infections, Institut Pasteur, Paris | Etienne Simon-Lorière, Vincent Enouf, Maud Vanpeene, Sylvie van der Werf,Marianne Leruez-Ville |
| EPI_ISL_1582647 | Research | National Reference Center for Viruses of Respiratory Infections, Institut Pasteur, Paris | Marion Barbet, Sylvie Behillil, Frédéric Lemoine, Corinne Maufrais, Christophe Malabat,Gael Millot, Méline Bizard, Angela Brisebarre, Camille Capel, Louise Lefrançois, Etienne Simon-Lorière, Vincent Enouf, Maud Vanpeene, Sylvie van der Werf,Marianne Leruez-Ville |
| EPI_ISL_1582648, EPI_ISL_1582649 | Research | National Reference Center for Viruses of Respiratory Infections, Institut Pasteur, Paris | Marion Barbet, Sylvie Behillil, Frédéric Lemoine, Corinne Maufrais, Christophe Malabat, Méline Bizard, Angela Brisebarre, Camille Capel, Louise Lefrançois, Etienne Simon-Lorière, Vincent Enouf, Maud Vanpeene, Sylvie van der Werf,Marianne Leruez-Ville |
| EPI_ISL_1582650 | Research | National Reference Center for Viruses of Respiratory Infections, Institut Pasteur, Paris | Marion Barbet, Sylvie Behillil, Frédéric Lemoine, Corinne Maufrais, Christophe Malabat,Gael Millot, Méline Bizard, Angela Brisebarre, Camille Capel, Louise Lefrançois, Etienne Simon-Lorière, Vincent Enouf, Maud Vanpeene, Sylvie van der Werf,Marianne Leruez-Ville |
| EPI_ISL_1582651, EPI_ISL_1582652 | Research | National Reference Center for Viruses of Respiratory Infections, Institut Pasteur, Paris | Marion Barbet, Sylvie Behillil, Frédéric Lemoine, Corinne Maufrais, Christophe Malabat, Méline Bizard, Angela Brisebarre, Camille Capel, Louise Lefrançois, Etienne Simon-Lorière, Vincent Enouf, Maud Vanpeene, Sylvie van der Werf,Marianne Leruez-Ville |
| EPI_ISL_1582653 | Research | National Reference Center for Viruses of Respiratory Infections, Institut Pasteur, Paris | Marion Barbet, Sylvie Behillil, Frédéric Lemoine, Corinne Maufrais, Christophe Malabat,Pierre Lechat, Méline Bizard, Angela Brisebarre, Camille Capel, Louise Lefrançois, Etienne Simon-Lorière, Vincent Enouf, Maud Vanpeene, Sylvie van der Werf,Marianne Leruez-Ville |
| EPI_ISL_1582654, EPI_ISL_1582655, EPI_ISL_1582656 | Research | National Reference Center for Viruses of Respiratory Infections, Institut Pasteur, Paris | Marion Barbet, Sylvie Behillil, Frédéric Lemoine, Corinne Maufrais, Christophe Malabat, Méline Bizard, Angela Brisebarre, Camille Capel, Louise Lefrançois, Etienne Simon-Lorière, Vincent Enouf, Maud Vanpeene, Sylvie van der Werf,Marianne Leruez-Ville |
| EPI_ISL_1582756 | Hospital | National Reference Center for Viruses of Respiratory Infections, Institut Pasteur, Paris | Marion Barbet, Sylvie Behillil, Frédéric Lemoine, Corinne Maufrais, Christophe Malabat,Pierre Lechat, Méline Bizard, Angela Brisebarre, Camille Capel, Louise Lefrançois, Etienne Simon-Lorière, Vincent Enouf, Maud Vanpeene, Sylvie van der Werf,Franck Griscelli |
| EPI_ISL_1583053 | Labo Analyses Med | National Reference Center for Viruses of Respiratory Infections, Institut Pasteur, Paris | Marion Barbet, Sylvie Behillil, Frédéric Lemoine, Corinne Maufrais, Christophe Malabat, Méline Bizard, Angela Brisebarre, Camille Capel, Louise Lefrançois, Etienne Simon-Lorière, Vincent Enouf, Maud Vanpeene, Sylvie van der Werf,GréGoire Potiron |
| EPI_ISL_1583074, EPI_ISL_1583078 | Labo analyses Med | National Reference Center for Viruses of Respiratory Infections, Institut Pasteur, Paris | Marion Barbet, Sylvie Behillil, Frédéric Lemoine, Corinne Maufrais, Christophe Malabat, Méline Bizard, Angela Brisebarre, Camille Capel, Louise Lefrançois, Etienne Simon-Lorière, Vincent Enouf, Maud Vanpeene, Sylvie van der Werf,Anne Holstein |
| EPI_ISL_1589116, EPI_ISL_1589117, EPI_ISL_1589118, EPI_ISL_1589119, EPI_ISL_1589127, EPI_ISL_1589128, EPI_ISL_1589129, EPI_ISL_1589130, EPI_ISL_1589131, EPI_ISL_1589132, EPI_ISL_1589133, EPI_ISL_1589134, EPI_ISL_1589135, EPI_ISL_1589136, EPI_ISL_1589137, EPI_ISL_1589138 | | | |
| see above | Department of Medical, Surgical, and Health Sciences, University of Trieste and Institute for Maternal and Child Health "IRCCS Burlo Garofolo" | Department of Biomedical, Surgical and Dental Sciences; University of Milan | Ferrante Pasquale, Delbue Serena, Signorini Lucia, Dolci Maria, Comar Manola, Campisciano Giuseppina, Cason Carolina, Ticozzi Rosalia |
| EPI_ISL_1591101, EPI_ISL_1591102, EPI_ISL_1591103, EPI_ISL_1591104 | Queen Astrid Military Hospital | Institute of Tropical Medicine | Philippe Selhorst |
| EPI_ISL_1594431 | Labo Analyses Med | National Reference Center for Viruses of Respiratory Infections, Institut Pasteur, Paris | Marion Barbet, Sylvie Behillil, Frédéric Lemoine, Corinne Maufrais, Christophe Malabat, Adrien Pain, Méline Bizard, Angela Brisebarre, Camille Capel, Louise Lefrançois, Etienne Simon-Lorière, Vincent Enouf, Maud Vanpeene, Sylvie van der Werf,J Besson |
| EPI_ISL_1595731 | Microbiology Department. Complexo Hospitalario Universitario de Vigo | Microbiology Department. Complexo Hospitalario Universitario de Vigo | Alfaya N, Alonso I, Alvarez M, Cabrera JJ, Carballo R, Cores O, Cortizo S, del-Campo V, Martinez L, Mediero G, Perez S, Potel C, Regueiro B, Rey S, Vasallo FJ |
| EPI_ISL_1620172, EPI_ISL_1620173, EPI_ISL_1620174 | MRCG at LSHTM Genomics lab | MRCG at LSHTM Genomics lab | Abdul Karim sesay, Abdoulie Kanteh, Jarra Manneh, Mariama Kujabi, Bakary Sanyang |
| EPI_ISL_1622360, EPI_ISL_1622361 | RS Mitra Keluarga Gading Serpong | Eijkman Institute for Molecular Biology, Ministry of Research and Technology/National Agency for Research and Innovation | Willy Agustine, Hidayat Trimarsanto, Lydia V. Panggalo, Iskandar Adnan, Sukma Oktavianthi, Edison Johar, Frilasita A Yudhaputri, Muhammad Rezki Rasyak, Safarina G Malik, Khin Saw Myint, Amin Soebandrio |
| EPI_ISL_1622362 | RS Sentra Medika Cikarang | Eijkman Institute for Molecular Biology, Ministry of Research and Technology/National Agency for Research and Innovation | Willy Agustine, Hidayat Trimarsanto, Lydia V. Panggalo, Iskandar Adnan, Sukma Oktavianthi, Edison Johar, Frilasita A Yudhaputri, Muhammad Rezki Rasyak, Safarina G Malik, Khin Saw Myint, Amin Soebandrio |
| EPI_ISL_1622363, EPI_ISL_1622364 | RS Sari Asih Ciledug | Eijkman Institute for Molecular Biology, Ministry of Research and Technology/National Agency for Research and Innovation | Willy Agustine, Hidayat Trimarsanto, Lydia V. Panggalo, Iskandar Adnan, Sukma Oktavianthi, Edison Johar, Frilasita A Yudhaputri, Muhammad Rezki Rasyak, Safarina G Malik, Khin Saw Myint, Amin Soebandrio |
| EPI_ISL_1622365 | RS Bhakti Husada Cikarang | Eijkman Institute for Molecular Biology, Ministry of Research and Technology/National Agency for Research and Innovation | Willy Agustine, Hidayat Trimarsanto, Lydia V. Panggalo, Iskandar Adnan, Sukma Oktavianthi, Edison Johar, Frilasita A Yudhaputri, Muhammad Rezki Rasyak, Safarina G Malik, Khin Saw Myint, Amin Soebandrio |
| EPI_ISL_1622366 | RSUD Cileungsi | Eijkman Institute for Molecular Biology, Ministry of Research and Technology/National Agency for Research and Innovation | Willy Agustine, Hidayat Trimarsanto, Lydia V. Panggalo, Iskandar Adnan, Sukma Oktavianthi, Edison Johar, Frilasita A Yudhaputri, Muhammad Rezki Rasyak, Safarina G Malik, Khin Saw Myint, Amin Soebandrio |
| EPI_ISL_1622367 | RS Harapan Bunda | Eijkman Institute for Molecular Biology, Ministry of Research and Technology/National Agency for Research and Innovation | Willy Agustine, Hidayat Trimarsanto, Lydia V. Panggalo, Iskandar Adnan, Sukma Oktavianthi, Edison Johar, Frilasita A Yudhaputri, Muhammad Rezki Rasyak, Safarina G Malik, Khin Saw Myint, Amin Soebandrio |
| EPI_ISL_1622368 | RSJ Dr. Soeharto Heerdjan | Eijkman Institute for Molecular Biology, Ministry of Research and Technology/National Agency for Research and Innovation | Willy Agustine, Hidayat Trimarsanto, Lydia V. Panggalo, Iskandar Adnan, Sukma Oktavianthi, Edison Johar, Frilasita A Yudhaputri, Muhammad Rezki Rasyak, Safarina G Malik, Khin Saw Myint, Amin Soebandrio |
| EPI_ISL_1622369 | RS Sumber Waras | Eijkman Institute for Molecular Biology, Ministry of Research and Technology/National Agency for Research and Innovation | Willy Agustine, Hidayat Trimarsanto, Lydia V. Panggalo, Iskandar Adnan, Sukma Oktavianthi, Edison Johar, Frilasita A Yudhaputri, Muhammad Rezki Rasyak, Safarina G Malik, Khin Saw Myint, Amin Soebandrio |
| EPI_ISL_1622370 | RSIA Selasih Medika | Eijkman Institute for Molecular Biology, Ministry of Research and Technology/National Agency for Research and Innovation | Willy Agustine, Hidayat Trimarsanto, Lydia V. Panggalo, Iskandar Adnan, Sukma Oktavianthi, Edison Johar, Frilasita A Yudhaputri, Muhammad Rezki Rasyak, Safarina G Malik, Khin Saw Myint, Amin Soebandrio |
| EPI_ISL_1622371 | RS Mitra Keluarga Gading Serpong | Eijkman Institute for Molecular Biology, Ministry of Research and Technology/National Agency for Research and Innovation | Willy Agustine, Hidayat Trimarsanto, Lydia V. Panggalo, Iskandar Adnan, Sukma Oktavianthi, Edison Johar, Frilasita A Yudhaputri, Muhammad Rezki Rasyak, Safarina G Malik, Khin Saw Myint, Amin Soebandrio |
| EPI_ISL_1622372 | RS Mitra Keluarga Gading Serpong | Eijkman Institute for Molecular Biology, Ministry of Research and Technology/National Agency for Research and Innovation | Hidayat Trimarsanto, Lydia V. Panggalo, Iskandar Adnan, Sukma Oktavianthi, Edison Johar, Frilasita A Yudhaputri, Muhammad Rezki Rasyak, Willy Agustine, Safarina G Malik, Khin Saw Myint, Amin Soebandrio |
| EPI_ISL_1622373 | RS Mary Cileungsi Hijau | Eijkman Institute for Molecular Biology, Ministry of Research and Technology/National Agency for Research and Innovation | Hidayat Trimarsanto, Lydia V. Panggalo, Iskandar Adnan, Sukma Oktavianthi, Edison Johar, Frilasita A Yudhaputri, Muhammad Rezki Rasyak, Willy Agustine, Safarina G Malik, Khin Saw Myint, Amin Soebandrio |
| EPI_ISL_1622374 | RS Sentra Medika Cikarang | Eijkman Institute for Molecular Biology, Ministry of Research and Technology/National Agency for Research and Innovation | Hidayat Trimarsanto, Lydia V. Panggalo, Iskandar Adnan, Sukma Oktavianthi, Edison Johar, Frilasita A Yudhaputri, Muhammad Rezki Rasyak, Willy Agustine, Safarina G Malik, Khin Saw Myint, Amin Soebandrio |
| EPI_ISL_1622375 | RS Taman Harapan Baru | Eijkman Institute for Molecular Biology, Ministry of Research and Technology/National Agency for Research and Innovation | Hidayat Trimarsanto, Lydia V. Panggalo, Iskandar Adnan, Sukma Oktavianthi, Edison Johar, Frilasita A Yudhaputri, Muhammad Rezki Rasyak, Willy Agustine, Safarina G Malik, Khin Saw Myint, Amin Soebandrio |

| | | | |
|---|---|---|---|
| EPI_ISL_1622376, EPI_ISL_1622377 | RSAB Harapan Kita | Eijkman Institute for Molecular Biology, Ministry of Research and Technology/National Agency for Research and Innovation | Hidayat Trimarsanto, Lydia V. Panggalo, Iskandar Adnan, Sukma Oktavianthi, Edison Johar, Frilasita A Yudhaputri, Muhammad Rezki Rasyak, Willy Agustine, Safarina G Malik, Khin Saw Myint, Amin Soebandrio |
| EPI_ISL_1622378 | RSIA Bun | Eijkman Institute for Molecular Biology, Ministry of Research and Technology/National Agency for Research and Innovation | Hidayat Trimarsanto, Lydia V. Panggalo, Iskandar Adnan, Sukma Oktavianthi, Edison Johar, Frilasita A Yudhaputri, Muhammad Rezki Rasyak, Willy Agustine, Safarina G Malik, Khin Saw Myint, Amin Soebandrio |
| EPI_ISL_1622379, EPI_ISL_1622380 | RS Mitra Keluarga Gading Serpong | Eijkman Institute for Molecular Biology, Ministry of Research and Technology/National Agency for Research and Innovation | Hidayat Trimarsanto, Lydia V. Panggalo, Iskandar Adnan, Sukma Oktavianthi, Edison Johar, Frilasita A Yudhaputri, Muhammad Rezki Rasyak, Willy Agustine, Safarina G Malik, Khin Saw Myint, Amin Soebandrio |
| EPI_ISL_1622388 | RS Mitra Keluarga Gading Serpong | Eijkman Institute for Molecular Biology, Ministry of Research and Technology/National Agency for Research and Innovation | Lydia V. Panggalo, Iskandar Adnan, Sukma Oktavianthi, Edison Johar, Frilasita A Yudhaputri, Muhammad Rezki Rasyak, Willy Agustine, Hidayat Trimarsanto, Safarina G Malik, Khin Saw Myint, Amin Soebandrio |
| EPI_ISL_1623687 | National Platform bis UMONS/Jolimont | National Platform bis UMONS/Jolimont | François Dufrasne, Guillaume Bayon-Vicente, Gautier Detry, Ruddy Wattiez |
| EPI_ISL_1626009 | Instituto Adolfo Lutz - Regional de Rio Claro | Instituto Adolfo Lutz, Interdiciplinary Procedures Center, Strategic Laboratory | Claudio Tavares Sacchi, Claudia Regina Gonçalves, Erica Valessa Ramos Gomes, Karoline Rodrigues Campos, Caio Vinicius Dias Lopes, Leonardo Jose Tadeu de Araujo, Katia Correa de Oliveira Santos |
| EPI_ISL_1626490, EPI_ISL_1626493, EPI_ISL_1626499, EPI_ISL_1626500, EPI_ISL_1626501, EPI_ISL_1626502, EPI_ISL_1626514, EPI_ISL_1626516, EPI_ISL_1626519, EPI_ISL_1626521, EPI_ISL_1626522, EPI_ISL_1626523, EPI_ISL_1626524, EPI_ISL_1626527 | | | |
| see above | NSTU COVID-19 Diagnostic Center | NSU Genome Research Institute (NGRI) | Maqsud Hossain, Tahrima Saiha Huq, Aura Rahman, Md. Aminul Islam, Syeda Naushin Tabassum, Kazi Nadim Hasan, Abdul Khaleque, Abdus Sadique, Mohammad Salim Hossain, Newaz Mohammed Bahadur, Firoz Ahmed, Hasan Mahmud Reza |
| EPI_ISL_1628345, EPI_ISL_1628346 | Instituto Adolfo Lutz - Regional de Rio Claro | Instituto Adolfo Lutz, Interdiciplinary Procedures Center, Strategic Laboratory | Claudio Tavares Sacchi, Claudia Regina Gonçalves, Erica Valessa Ramos Gomes, Karoline Rodrigues Campos, Caio Vinicius Dias Lopes, Leonardo Jose Tadeu de Araujo, Katia Correa de Oliveira Santos |
| EPI_ISL_1628474 | Universidad Industrial de Santander (Laboratorio Central de Investigaciones - Clínica Chicamocha). | Universidad Industrial de Santander. | Francisco Martinez-Perez, Cristian E. Cadena-Caballero, Diego Rueda-Plata, Carolina S. Torres-Jiménez, Lizeth J. Forero-Buitrago, Erika Lizarazo-Gutiérrez, Carlos Barrios-Hernández, Lina M. Vera-Cala. |
| EPI_ISL_1628499 | Universidad Industrial de Santander (Laboratorio Central de Investigaciones - Clínica Chicamocha). | Universidad Industrial de Santander | Francisco Martinez-Perez, Cristian E. Cadena-Caballero, Diego Rueda-Plata, Carolina S. Torres-Jiménez, Lizeth J. Forero-Buitrago, Erika Lizarazo-Gutiérrez, Carlos Barrios-Hernández, Lina M. Vera-Cala. |
| EPI_ISL_1629382, EPI_ISL_1629387, EPI_ISL_1629388, EPI_ISL_1629397 | CHU | UMR PIMIT | Dr David A Wilkinson, Dr Patrick Mavingui, Dr Camille Lebarbenchon, Magali Turpin |
| EPI_ISL_1629710 | Universidad Industrial de Santander (Laboratorio Central de Investigaciones - Clínica Chicamocha). | Universidad Industrial de Santander. | Francisco Martinez-Perez, Cristian E. Cadena-Caballero, Diego Rueda-Plata, Carolina S. Torres-Jiménez, Lizeth J. Forero-Buitrago, Erika Lizarazo-Gutiérrez, Carlos Barrios-Hernández, Lina M. Vera-Cala. |
| EPI_ISL_1629720 | ICMT | Universidad Nacional de Colombia - Laboratorio Genómico One Health | Maria Stella López, Andres F. Cardona-Rios, Daniel O. Maldonado-Perez, Simón Villegas Velásquez, Laura Silvana Perez, Karl A Ciuoderis, Idabely Betancur Ortiz, Sandra Ines Cano, Eliana Patricia Calvo Tapiero, Diego A. Álvarez-Díaz, Carlos Franco-Muñoz, Hector Alejandro Ruiz-Moreno, Marcela Mercado-Reyes, Jorge E. Osorio, Juan P. Hernandez-Ortiz |
| EPI_ISL_1629724, EPI_ISL_1629725 | SOMER | Universidad Nacional de Colombia - Laboratorio Genómico One Health | Simón Villegas Velásquez, Andres F. Cardona-Rios, Daniel O. Maldonado-Perez, Maria Stella López, Laura Silvana Perez, Karl A Ciuoderis, Idabely Betancur Ortiz, Sandra Ines Cano, Eliana Patricia Calvo Tapiero, Diego A. Álvarez-Díaz, Carlos Franco-Muñoz, Hector Alejandro Ruiz-Moreno, Marcela Mercado-Reyes, Jorge E. Osorio, Juan P. Hernandez-Ortiz |
| EPI_ISL_1629729 | UDEA | Universidad Nacional de Colombia - Laboratorio Genómico One Health | Daniel O. Maldonado-Perez, Andres F. Cardona-Rios, Maria Stella López, Simón Villegas Velásquez, Laura Silvana Perez, Karl A Ciuoderis, Idabely Betancur Ortiz, Sandra Ines Cano, Eliana Patricia Calvo Tapiero, Diego A. Álvarez-Díaz, Carlos Franco-Muñoz, Hector Alejandro Ruiz-Moreno, Marcela Mercado-Reyes, Jorge E. Osorio, Juan P. Hernandez-Ortiz |
| EPI_ISL_1629734 | SURA | Universidad Nacional de Colombia - Laboratorio Genómico One Health | Laura Silvana Perez, Andres F. Cardona-Rios, Daniel O. Maldonado-Perez, Maria Stella López, Simón Villegas Velásquez, Karl A Ciuoderis, Idabely Betancur Ortiz, Sandra Ines Cano, Eliana Patricia Calvo Tapiero, Diego A. Álvarez-Díaz, Carlos Franco-Muñoz, Hector Alejandro Ruiz-Moreno, Marcela Mercado-Reyes, Jorge E. Osorio, Juan P. Hernandez-Ortiz |
| EPI_ISL_1629744, EPI_ISL_1629745 | LABORATORIO ECHAVARRIA | Universidad Nacional de Colombia - Laboratorio Genómico One Health | Andres F. Cardona-Rios, Daniel O. Maldonado-Perez, Maria Stella López, Simón Villegas Velásquez, Laura Silvana Perez, Karl A Ciuoderis, Idabely Betancur Ortiz, Sandra Ines Cano, Eliana Patricia Calvo Tapiero, Diego A. Álvarez-Díaz, Carlos Franco-Muñoz, Hector Alejandro Ruiz-Moreno, Marcela Mercado-Reyes, Jorge E. Osorio, Juan P. Hernandez-Ortiz |
| EPI_ISL_1629752 | SYNLAB | Universidad Nacional de Colombia - Laboratorio Genómico One Health | Andres F. Cardona-Rios, Daniel O. Maldonado-Perez, Maria Stella López, Simón Villegas Velásquez, Laura Silvana Perez, Karl A Ciuoderis, Idabely Betancur Ortiz, Sandra Ines Cano, Eliana Patricia Calvo Tapiero, Diego A. Álvarez-Díaz, Carlos Franco-Muñoz, Hector Alejandro Ruiz-Moreno, Marcela Mercado-Reyes, Jorge E. Osorio, Juan P. Hernandez-Ortiz |
| EPI_ISL_1629753, EPI_ISL_1629754, EPI_ISL_1629755, EPI_ISL_1629757, EPI_ISL_1629759 | H.U SAN VICENTE FUNDACION | Universidad Nacional de Colombia - Laboratorio Genómico One Health | Andres F. Cardona-Rios, Daniel O. Maldonado-Perez, Maria Stella López, Simón Villegas Velásquez, Laura Silvana Perez, Karl A Ciuoderis, Idabely Betancur Ortiz, Sandra Ines Cano, Eliana Patricia Calvo Tapiero, Diego A. Álvarez-Díaz, Carlos Franco-Muñoz, Hector Alejandro Ruiz-Moreno, Marcela Mercado-Reyes, Jorge E. Osorio, Juan P. Hernandez-Ortiz |
| EPI_ISL_1629760, EPI_ISL_1629761 | LABMÉDICO | Universidad Nacional de Colombia - Laboratorio Genómico One Health | Andres F. Cardona-Rios, Daniel O. Maldonado-Perez, Maria Stella López, Simón Villegas Velásquez, Laura Silvana Perez, Karl A Ciuoderis, Idabely Betancur Ortiz, Sandra Ines Cano, Eliana Patricia Calvo Tapiero, Diego A. Álvarez-Díaz, Carlos Franco-Muñoz, Hector Alejandro Ruiz-Moreno, Marcela Mercado-Reyes, Jorge E. Osorio, Juan P. Hernandez-Ortiz |
| EPI_ISL_1629769, EPI_ISL_1629770, EPI_ISL_1629772, EPI_ISL_1629776, EPI_ISL_1629778 | Hospital Carlos Alberto Seguín Escobedo - EsSalud | Laboratorio de Genómica Microbiana, Universidad Peruana Cayetano Heredia | Lenin Maturrano, Pablo Tsukayama, Alejandra Dávila-Barclay, Guillermo Salvatierra, Luis González, Pedro E. Romero, Janet Huancachoque, Pool Marcos, Fernando Sánchez Fragoso, Lin Zevallos Cuarite |
| EPI_ISL_1630158 | Laboratório de Microbiologia Molecular - Universidade FEEVALE | Laboratório de Microbiologia Molecular - Universidade FEEVALE | Alana Witt Hansen, Fágner Henrique Heldt, Fernando Rosado Spilki, Flávio Silveira, Juliana Schons Gularte, Juliane Deise Fleck, Mariana Soares da Silva, Meriane Demoliner, Matheus Nunes Weber, Paula Rodrigues de Almeida, Micheli Filippi. |
| EPI_ISL_1647425, EPI_ISL_1647426, EPI_ISL_1647446, EPI_ISL_1647447, EPI_ISL_1647448, EPI_ISL_1647449, EPI_ISL_1647450, EPI_ISL_1647451, EPI_ISL_1647452 | Indian Council of Medical Research-National Institute of Virology, Microbial Containment Complex | Indian Council of Medical Research-National Institute of Virology, Microbial Containment Complex | Pragya D Yadav |
| EPI_ISL_1652049, EPI_ISL_1652050, EPI_ISL_1652051, EPI_ISL_1652052, EPI_ISL_1652053, EPI_ISL_1652054, EPI_ISL_1652057, EPI_ISL_1652058, EPI_ISL_1652061, EPI_ISL_1652062, EPI_ISL_1652064, EPI_ISL_1652066, EPI_ISL_1652067, EPI_ISL_1652068, EPI_ISL_1652071, EPI_ISL_1652072, EPI_ISL_1652073 | | | |
| see above | Integrated Biorepository of H3Africa Uganda - IBRH3AU | Molecular Biology Laboratory | Savannah Mwesigwa, Eric Katagirya, Gerald Mboowa, David Patrick Kateete, Misaki Wayengera, Emmanuel Nasinghe, Ashaba Fred Katabazi, Edgar Kigozi, Samuel Kirimunda, Rogers Kamulegeya, Moses Luutu, Lwanga Newton, Nsubuga Gideon, Bernard Ssentalo Bagaya, Sarah Stanley, Moses Joloba |
| EPI_ISL_1653804, EPI_ISL_1653805, EPI_ISL_1653807, EPI_ISL_1653809, EPI_ISL_1653811, EPI_ISL_1653812 | Southern Nevada Public Health Laboratory | Southern Nevada Public Health Laboratory | Michael Picker |
| EPI_ISL_1654202, EPI_ISL_1654204, EPI_ISL_1654205 | Research | National Reference Center for Viruses of Respiratory Infections, Institut Pasteur, Paris | Marion Barbet, Sylvie Behillil, Frédéric Lemoine, Corinne Maufrais, Christophe Malabat, Damien Mornico, Méline Bizard, Angela Brisebarre, Camille Capel, Louise Lefrançois, Etienne Simon-Lorière, Vincent Enouf, Maud Vanpeene, Sylvie van der Werf,Marianne Leruez-Ville |
| EPI_ISL_1655075 | Fujian Center for Disease Control and Prevention | Fujian Center for Disease Control and Prevention | Lin Qi, Huang Zhimiao, Weng Yuwei |
| EPI_ISL_1661250, EPI_ISL_1661251, EPI_ISL_1661252 | Laboratorio de Ecologia de Doencas Transmissiveis na Amazonia, Instituto Leonidas e Maria Deane - Fiocruz | Laboratorio de Ecologia de Doencas Transmissiveis na Amazonia, Instituto Leonidas e Maria Deane - Fiocruz | Valdinete Nascimento, Victor Souza, André Corado, Fernanda Nascimento, George Silva, Ágatha Costa, Debora Duarte, Karina Pessoa, Matilde Mejía, Luciana Gonçalves, Maria Júlia Brandão, Michele Jesus, Felipe Naveca |

| | | | |
|---|---|---|---|
| | Amazonia | Amazonia | |
| EPI_ISL_1663017 | Fujian Center for Disease Control and Prevention | Fujian Center for Disease Control and Prevention | Lin Qi, Huang Zhimiao, Weng Yuwei |
| EPI_ISL_1673261, EPI_ISL_1673262 | LABORATORIO LAS AMERICAS | Universidad Nacional de Colombia - Laboratorio Genómico One Health | Maria Stella López, Andres F. Cardona-Rios, Daniel O. Maldonado-Perez, Simón Villegas Velásquez, Lina M. Hurtado, Laura Silvana Perez, Karl A Ciuoderis, Idabely Betancur Ortiz, Sandra Ines Cano, Eliana Patricia Calvo Tapiero, Diego A. Álvarez-Díaz, Carlos Franco-Muñoz, Hector Alejandro Ruiz-Moreno, Marcela Mercado-Reyes, Jorge E. Osorio, Juan P. Hernandez-Ortiz |
| EPI_ISL_1675009 | Labo Analyses Med | National Reference Center for Viruses of Respiratory Infections, Institut Pasteur, Paris | Marion Barbet, Sylvie Behillil, Méline Bizard, Angela Brisebarre, Camille Capel, Vincent Enouf, Louise Lefrançois, Frédéric Lemoine, Christophe Malabat, Corinne Maufrais, Etienne Simon-Lorière, Maud Vanpeene, Sylvie Van der Werf ,Brieuc Gestin |
| EPI_ISL_1675033 | Hospital | National Reference Center for Viruses of Respiratory Infections, Institut Pasteur, Paris | Marion Barbet, Sylvie Behillil, Méline Bizard, Angela Brisebarre, Camille Capel, Vincent Enouf, Louise Lefrançois, Frédéric Lemoine, Christophe Malabat, Corinne Maufrais, Etienne Simon-Lorière, Maud Vanpeene, Sylvie Van der Werf ,Alexandra Ducancelle |
| EPI_ISL_1708512 | Collection of WHO National Influenza Center, N.F. Gamaleya National Research Center for Epidemiology and Microbiology of the Ministry of Health of the Russian Federation | Laboratory of genome analysis, N.F. Gamaleya National Research Center for Epidemiology and Microbiology of the Ministry of Health of the Russian Federation | Olga Voronina, Ekaterina Aksenova, Natalia Ryzhova, Marina Kunda, Kirill Krasnoslobodtsev, Elena Burtseva, Denis Logunov |
| EPI_ISL_1708513, EPI_ISL_1708514, EPI_ISL_1708515, EPI_ISL_1708516, EPI_ISL_1708519 | State collection of viruses, N.F. Gamaleya National Research Center for Epidemiology and Microbiology of the Ministry of Health of the Russian Federation | Laboratory of genome analysis, N.F. Gamaleya National Research Center for Epidemiology and Microbiology of the Ministry of Health of the Russian Federation | Olga Voronina, Ekaterina Aksenova, Natalia Ryzhova, Marina Kunda, Inna Dolzhikova, Andrey Botikov, Daria Grousova, Anna Kovyrshina, Denis Logunov |
| EPI_ISL_1710849, EPI_ISL_1710850, EPI_ISL_1710851, EPI_ISL_1710852, EPI_ISL_1710853, EPI_ISL_1710854 | Pathogenic Microorganisms Variability Laboratory | Pathogenic Microorganisms Variability Laboratory | Alexey Shchetinin, Maria Nikiforova, Andrei Siniavin, Andrey Pochtovyy, Evgeny Usachev, Elena Shidlovskaya, Elizaveta Divisenko, Nadezhda Kuznetsova, Valeria Bacalin, Olga Burgasova, Ludmila Kolobukhina, Svetlana Smetanina, Denis Logunov, Vladimir Gushchin, Alexander Gintsburg |
| EPI_ISL_1714798 | Institute for Medical Research, Infectious Disease Research Centre, National Institutes of Health, Ministry of Health Malaysia | Institute for Medical Research, Infectious Disease Research Centre, National Institutes of Health, Ministry of Health Malaysia | Suppiah J, Kamel K, Azizan MA, Thayan R |
| EPI_ISL_1731043 | Collection of WHO National Influenza Center, N.F. Gamaleya National Research Center for Epidemiology and Microbiology of the Ministry of Health of the Russian Federation | N.F. Gamaleya Research Center for Epidemiology and Microbiology | Olga Voronina, Ekaterina Aksenova, Natalia Ryzhova, Marina Kunda, Kirill Krasnoslobodtsev, Elena Burtseva, Denis Logunov |
| EPI_ISL_1731044, EPI_ISL_1731045 | State collection of viruses, N.F. Gamaleya National Research Center for Epidemiology and Microbiology of the Ministry of Health of the Russian Federation | N.F. Gamaleya Research Center for Epidemiology and Microbiology | Olga Voronina, Ekaterina Aksenova, Natalia Ryzhova, Marina Kunda, Inna Dolzhikova, Andrey Botikov, Daria Grousova, Anna Kovyrshina, Denis Logunov |
| EPI_ISL_1731067, EPI_ISL_1731068 | Molecular diagnostic laboratory of Federal Budget Institution of Science "Central Research Institute of Epidemiology" of The Federal Service on Customers' Rights Protection and Human Well-being Surveillance | Group of Genomics and Postgenomic Technologies of Central Research Institute of Epidemilology | Samoilov AE, Kaptelova VV, Korneenko EV, Saenko SS, Shipulina OY, Speranskaya AS, Tivanova EV, Kondrasheva LY, Akimkin VG |
| EPI_ISL_1731069, EPI_ISL_1731070, EPI_ISL_1731071, EPI_ISL_1731072, EPI_ISL_1731073, EPI_ISL_1731074, EPI_ISL_1731075, EPI_ISL_1731076, EPI_ISL_1731078, EPI_ISL_1731079 | Laboratory of Extremely Dangerous Infections of Federal Budget Health Care Instute "Center of Hygiene and Epidemiology in Sakhalin region" | Group of Genomics and Postgenomic Technologies of Central Research Institute of Epidemiology | Samoilov AE, Kaptelova VV, Korneenko EV, Valdokhina AV, Bulanenko VP, Saenko SS, Arutiunian AE, Byankina MA, Zagorodnov YL, Golubeva AG, Speranskaya AS, Tivanova EV, Kondrasheva LY, Akimkin VG |
| EPI_ISL_1752475, EPI_ISL_1752510 | LHUB-ULB | LHUB-ULB | Charlotte Michel, Olivier Vandenberg, Marie Hallin |
| EPI_ISL_1752592, EPI_ISL_1752593, EPI_ISL_1752594, EPI_ISL_1752595, EPI_ISL_1752596, EPI_ISL_1752597, EPI_ISL_1752598, EPI_ISL_1752599, EPI_ISL_1752600, EPI_ISL_1752601, EPI_ISL_1752602, EPI_ISL_1752603, EPI_ISL_1752604, EPI_ISL_1752605, EPI_ISL_1752606, EPI_ISL_1752607, EPI_ISL_1752608, EPI_ISL_1752609, EPI_ISL_1752610, EPI_ISL_1752611, EPI_ISL_1752612, EPI_ISL_1752613, EPI_ISL_1752614, EPI_ISL_1752615, EPI_ISL_1752616, EPI_ISL_1752617, EPI_ISL_1752618, EPI_ISL_1752619, EPI_ISL_1752620, EPI_ISL_1752632, EPI_ISL_1752633, EPI_ISL_1752634, EPI_ISL_1752635 | | | |
| see above | Showa University PCR Center | Showa University PCR Center | Fumihiro Ishikawa, Hideto Oyamada, Yuko Udaka, Keiko Ishino, Yuji Kiuchi |
| EPI_ISL_1753847 | Servicio Microbiología Hospital La Paz | Servicio Microbiología Hospital La Paz | Fernando Lázaro, Rubén Cáceres, Jesús Mingorance Cruz, Elie Dahdouh |
| EPI_ISL_1760210, EPI_ISL_1760211, EPI_ISL_1760212, EPI_ISL_1760213, EPI_ISL_1760214, EPI_ISL_1760215, EPI_ISL_1760216, EPI_ISL_1760217, EPI_ISL_1760218, EPI_ISL_1760219, EPI_ISL_1760220, EPI_ISL_1760221, EPI_ISL_1760222, EPI_ISL_1760223, EPI_ISL_1760224, EPI_ISL_1760225, EPI_ISL_1760226, EPI_ISL_1760227, EPI_ISL_1760228, EPI_ISL_1760229, EPI_ISL_1760230, EPI_ISL_1760231, EPI_ISL_1760232, EPI_ISL_1760233, EPI_ISL_1760234, EPI_ISL_1760235, EPI_ISL_1760241, EPI_ISL_1760242 | | | |
| see above | Centers for Disease Control and Prevention, Dengue Branch | Centers for Disease Control and Prevention, Dengue Branch | Gilberto A. Santiago, Glenda Gonzalez, Betzabel Flores, Keyla Charriez, Gabriela Paz-Bailey, Jorge L. Munoz-Jordan |
| EPI_ISL_1760615, EPI_ISL_1760616, EPI_ISL_1760617, EPI_ISL_1760618, EPI_ISL_1760619, EPI_ISL_1760621, EPI_ISL_1760623, EPI_ISL_1760624, EPI_ISL_1760625, EPI_ISL_1760626, EPI_ISL_1760627, EPI_ISL_1760628, EPI_ISL_1760629, EPI_ISL_1760631, EPI_ISL_1760632, EPI_ISL_1760633, EPI_ISL_1760634, EPI_ISL_1760635, EPI_ISL_1760636, EPI_ISL_1760637, EPI_ISL_1760638, EPI_ISL_1760639, EPI_ISL_1760640, EPI_ISL_1760641, EPI_ISL_1760642, EPI_ISL_1760643, EPI_ISL_1760644, EPI_ISL_1760645, EPI_ISL_1760646 | | | |
| see above | Clinical specimen | Centre de Recherche et de Formation en Infectiologie Guinée | Alpha Kabinet KEITA, Haby DIALLO, Abdoul Karim SOUMAH, Abdoulaye TOURE, Thibaut Armel Cherif GNIMADI, Joel KOIVOGUI, Jean-louis MONEMOU, Moriba POVOGUI, Mamadou Saliou SOW, Mamadou Bhoye KEITA, Penda Malhado DIALLO, Alimou CAMARA, Kaba KOUROUMA, Mandiou DIAKITE, Mamadou Saliou BAH, Sakoba KEITA, Bouna Yatassaye, Christelle BUTEL, Laetitia SERRANO, Ahidjo AYOUBA, Eric DELAPORTE, Martine PEETERS |
| EPI_ISL_1761310, EPI_ISL_1761311, EPI_ISL_1761312, EPI_ISL_1761313, EPI_ISL_1761314, EPI_ISL_1761315, EPI_ISL_1761316, EPI_ISL_1761317, EPI_ISL_1761318, EPI_ISL_1761319, EPI_ISL_1761320, EPI_ISL_1761321, EPI_ISL_1761323, EPI_ISL_1761324, EPI_ISL_1761325, EPI_ISL_1761326, EPI_ISL_1761327, EPI_ISL_1761328, EPI_ISL_1761329, EPI_ISL_1761330, EPI_ISL_1761331, EPI_ISL_1761332, EPI_ISL_1761333, EPI_ISL_1761334, EPI_ISL_1761335, EPI_ISL_1761336, EPI_ISL_1761337, EPI_ISL_1761338, EPI_ISL_1761339, EPI_ISL_1761340, EPI_ISL_1761363 | | | |
| see above | Tokyo Medical and Dental University | Tokyo Medical and Dental University | Kousuke Tanimoto, Yukie Tanaka, Yoko Nukui, Akinori Kimura, Hiroaki Takeuchi |
| EPI_ISL_1761587 | Hospital de Palamós. Serveis de Salut Integral Baix Empordà (SSIBE) | IrsiCaixa | Marc Noguera-Julian, Mariona Parera, Maria Casadellà, Pilar Armengol, Francesc Catala-Moll, Roger Paredes, Bonaventura Clotet Martin-Urda Diez-Canseco, Anabel; Toda Savall, Mº de la Roca; Marchena Romero, Jose Andres; Rodriguez Jareño, Mari Cruz; Erick Fernando Mayer Arispe; Claver Belver, Natalia |
| EPI_ISL_1785071, EPI_ISL_1785072, EPI_ISL_1785074, EPI_ISL_1785075, EPI_ISL_1785076, EPI_ISL_1785077, EPI_ISL_1785078, EPI_ISL_1785079, EPI_ISL_1785080 | Infectious Agents & Hygiene | GIMAP and Virpath teams-CIRI | Sylvie Pillet, Thomas Bourlet, Julien Fouret, Thomas Julien, Victoria Dulière, Olivier Terrier, Andrés Pizzorno, Manuel Rosa-Calatrava, Bruno Pozzetto, Stéphane Paul |
| EPI_ISL_1789114 | Siti Khodijah Hospital | Institute of Tropical Disease, Universitas Airlangga | Jezzy R Dewantari, Rima R Prasetya, Krisnoadi Rahardjo, Aldise M Nastri, Muhammad Hamdan, Gatot Soegiarto, Laksmi Wulandari, Resti Yudhawati, Soetjipto, Yasuko Mori, Maria I Lusida, Kazufumi Shimizu |
| EPI_ISL_1789116 | Anwar Medika General Hospital | Institute of Tropical Disease, Universitas Airlangga | Maria I Lusida, Krisnoadi Rahardjo, Aldise M Nastri, Jezzy R Dewantari, Rima R Prasetya, Nungky Taniasari, Gatot Soegiarto, Laksmi Wulandari, Resti Yudhawati, Soetjipto, Yasuko Mori, Kazufumi Shimizu |
| EPI_ISL_1789117 | Anwar Medika General Hospital | Institute of Tropical Disease, Universitas Airlangga | Krisnoadi Rahardjo, Aldise M Nastri, Jezzy R Dewantari, Rima R Prasetya, Muhammad Hamdan, Gatot Soegiarto, Laksmi Wulandari, Resti Yudhawati, Yasuko Mori, Soetjipto, Kazufumi Shimizu, Maria I Lusida |
| EPI_ISL_1790080, EPI_ISL_1790082, EPI_ISL_1790084, EPI_ISL_1790086 | Centre Pasteur Annexe de Garoua | Institut Pasteur de Dakar | Njouom Richard, Diagne Moussa Moïse, Dia Ndongo, Diallo Amadou, Sankhe Safietou, Diop Mamadou, Ndiaye Ndack, Loucoubar Cheikh, Carniel Elisabeth, Faye Ousmane, Sall Amadou Alpha |
| EPI_ISL_1795053 | Anwar Medika General Hospital | Institute of Tropical Disease, Universitas Airlangga | Aldise M Nastri, Jezzy R Dewantari, Rima R Prasetya, Krisnoadi Rahardjo, Nungky Taniasari, Gatot Soegiarto, Laksmi Wulandari, Resti Yudhawati, Yasuko Mori, Soetjipto, Kazufumi Shimizu, Maria I Lusida |
| EPI_ISL_1795054 | Darus Syifa' Islamic Hospital | Institute of Tropical Disease, Universitas Airlangga | Krisnoadi Rahardjo, Aldise M Nastri, Jezzy R Dewantari, Rima R Prasetya, HM Faiz, Gatot Soegiarto, Laksmi Wulandari, Resti Yudhawati, Yasuko Mori, Soetjipto, Kazufumi Shimizu, Maria I Lusida |

| | | | |
|---|---|---|---|
| EPI_ISL_1795055 | Muhammadiyah Hospital | Institute of Tropical Disease, Universitas Airlangga | Rima R Prasetya, Krisnoadi Rahardjo, Aldise M Nastri, Jezzy R Dewantari, Imam Suyuthi, Gatot Soegiarto, Laksmi Wulandari, Resti Yudhawati, Yasuko Mori, Soetjipto, Kazufumi Shimizu, Maria I Lusida |
| EPI_ISL_1795056 | Siloam Hospital | Institute of Tropical Disease, Universitas Airlangga | Kazufumi Shimizu, Krisnoadi Rahardjo, Aldise M Nastri, Jezzy R Dewantari, Rima R Prasetya, Maria M Padmidewi, Gatot Soegiarto, Laksmi Wulandari, Retno A Setyoningrum, Resti Yudhawati, Yohko K Shimizu, Mitsuhiro Nishimura, Yasuko Mori, Soetjipto, Maria I Lusida |
| EPI_ISL_1795057 | Muhammadiyah Hospital | Institute of Tropical Disease, Universitas Airlangga | Jezzy R Dewantari, Rima R Prasetya, Krisnoadi Rahardjo, Aldise M Nastri, Imam Suyuthi, Gatot Soegiarto, Laksmi Wulandari, Resti Yudhawati, Yasuko Mori, Soetjipto, Kazufumi Shimizu, Maria I Lusida |
| EPI_ISL_1795059 | Citra Medika Hospital | Institute of Tropical Disease, Universitas Airlangga | Aldise M Nastri, Jezzy R Dewantari, Rima R Prasetya, Krisnoadi Rahardjo, Wahidah Rachmaniyah, Gatot Soegiarto, Laksmi Wulandari, Resti Yudhawati, Yasuko Mori, Soetjipto, Kazufumi Shimizu, Maria I Lusida |
| EPI_ISL_1805697 | National Center for Communicable Diseases (NCCD) National Influenza Center | National Centre for Disease Control (NCDC) National Influenza Center | Naranzul Ts,Bayasgalan N,Khishigmunkh Ch,Seiichiro F,Hideka M,Mina N,Shinji W,Ankhbayar S,Tsogzolmaa G,Darmaa B,Battur L,Nymadawa P |
| EPI_ISL_1805716, EPI_ISL_1805717, EPI_ISL_1805718, EPI_ISL_1805740, EPI_ISL_1805741, EPI_ISL_1805744, EPI_ISL_1805745, EPI_ISL_1805787, EPI_ISL_1805789, EPI_ISL_1805790 | National Center for Communicable Diseases (NCCD) National Influenza Center | National Center for Communicable Diseases (NCCD) National Influenza Center | Naranzul Ts,Bayasgalan N,Khishigmunkh Ch,Seiichiro F,Hideka M,Mina N,Shinji W,Ankhbayar S,Tsogzolmaa G,Darmaa B,Battur L,Nymadawa P |
| EPI_ISL_1821208 | Hospital de Base de Bauru Hospital de Base Setima Regiao | Instituto Adolfo Lutz, Interdiciplinary Procedures Center, Strategic Laboratory | Claudio Tavares Sacchi, Claudia Regina Gonçalves, Erica Valessa Ramos Gomes, Karoline Rodrigues Campos, Caio Vinicius Dias Lopes, Leonardo Jose Tadeu de Araujo |
| EPI_ISL_1823412 | Indiana Animal Disease Diagnostic Laboratory | Carpi Laboratory - Purdue University | Jack Dorman, Ilinca I Ciubotariu, Nicole M Perry, Jobin J Kattoor, Lev Gorenstein, Abebe A Fola, G Kenitra Hendrix, Rebecca P Wilkes, Giovanna Carpi |
| EPI_ISL_1824616 | RSUD Bahteramas Kendari | National Institute of Health Research and Development | Hana Apsari Pawestri, Subangkit, Kartika Dewi Puspa, Arie Ardiansyah Nugraha, Hartanti Dian Ikawati, Krisna Pangesti, Nelly Puspandari, Vivi Setiawaty |
| EPI_ISL_1859356, EPI_ISL_1859367, EPI_ISL_1859368, EPI_ISL_1859376, EPI_ISL_1859378, EPI_ISL_1859382, EPI_ISL_1859384, EPI_ISL_1859385, EPI_ISL_1859401, EPI_ISL_1859406 | Kazan Federal University, Academic Consortium | FEDERAL RESEARCH AND CLINICAL CENTER OF PHYSICAL-CHEMICAL MEDICINE OF FEDERAL MEDICAL BIOLOGICAL AGENCY, Academic Consortium | Vladislav Babenko, Ramiz Bakhtyev, Vladimir Baklaushev, Larisa Balykova, Pavel Bashkirov, Anna Blagonravova, Daria Boldyreva, Dmitry Fedorov, Ilshat Gafurov, Raushaniya Gaifullina, Yulia Galeeva, Elena Galova, Vadim Govorun, Elena Ilina, Konstantin Ivanov, Daria Kharlampieva, Polina Khromova, Ksenia Klimina, Konstantin Kolontarev, Nadezhda Kolyshkina, Andrey Koritsky, Vyacheslav Kuropatkin, Vasily Lazarev, Alexander Manolov, Valentin Manuvera, Daria Matyushkina, Ekaterina Moskaleva, Varvara Musarova, Oleg Ogarkov, Elizaveta Orlova, Alexander Pavlenko, Alla Petrova, Natalia Pozhenko, Dmitry Pushkar, Alexander Rumyantsev, Sergey Rumyantsev, Vladimir Rumyantsev, Lyubov Rychkova, Alexander Samoilov, Irina Shirokova, Vyacheslav Sinkov, Svetlana Solovieva, Polina Tikhonova, Galina Trifonova, Alexander Troitsky, Alexander Tulichev, Yuri Udalov, Anna Varizhuk, Alexander Vasiliev, Vladimir Veselovsky, Alexey Volnukhin, Gaukhar Yusubalieva |
| EPI_ISL_1859410 | Federal State Budgetary Educational Institution of Higher Education «Privolzhsky Research Medical University» of the Ministry of Health of the Russian Federation, Academic Consortium | FEDERAL RESEARCH AND CLINICAL CENTER OF PHYSICAL-CHEMICAL MEDICINE OF FEDERAL MEDICAL BIOLOGICAL AGENCY, Academic Consortium | Vladislav Babenko, Ramiz Bakhtyev, Vladimir Baklaushev, Larisa Balykova, Pavel Bashkirov, Anna Blagonravova, Daria Boldyreva, Dmitry Fedorov, Ilshat Gafurov, Raushaniya Gaifullina, Yulia Galeeva, Elena Galova, Vadim Govorun, Elena Ilina, Konstantin Ivanov, Daria Kharlampieva, Polina Khromova, Ksenia Klimina, Konstantin Kolontarev, Nadezhda Kolyshkina, Andrey Koritsky, Vyacheslav Kuropatkin, Vasily Lazarev, Alexander Manolov, Valentin Manuvera, Daria Matyushkina, Ekaterina Moskaleva, Varvara Musarova, Oleg Ogarkov, Elizaveta Orlova, Alexander Pavlenko, Alla Petrova, Natalia Pozhenko, Dmitry Pushkar, Alexander Rumyantsev, Sergey Rumyantsev, Vladimir Rumyantsev, Lyubov Rychkova, Alexander Samoilov, Irina Shirokova, Vyacheslav Sinkov, Svetlana Solovieva, Polina Tikhonova, Galina Trifonova, Alexander Troitsky, Alexander Tulichev, Yuri Udalov, Anna Varizhuk, Alexander Vasiliev, Vladimir Veselovsky, Alexey Volnukhin, Gaukhar Yusubalieva |
| EPI_ISL_1859424, EPI_ISL_1859427, EPI_ISL_1859433, EPI_ISL_1859475, EPI_ISL_1859481 | Kazan Federal University, Academic Consortium | FEDERAL RESEARCH AND CLINICAL CENTER OF PHYSICAL-CHEMICAL MEDICINE OF FEDERAL MEDICAL BIOLOGICAL AGENCY, Academic Consortium | Vladislav Babenko, Ramiz Bakhtyev, Vladimir Baklaushev, Larisa Balykova, Pavel Bashkirov, Anna Blagonravova, Daria Boldyreva, Dmitry Fedorov, Ilshat Gafurov, Raushaniya Gaifullina, Yulia Galeeva, Elena Galova, Vadim Govorun, Elena Ilina, Konstantin Ivanov, Daria Kharlampieva, Polina Khromova, Ksenia Klimina, Konstantin Kolontarev, Nadezhda Kolyshkina, Andrey Koritsky, Vyacheslav Kuropatkin, Vasily Lazarev, Alexander Manolov, Valentin Manuvera, Daria Matyushkina, Ekaterina Moskaleva, Varvara Musarova, Oleg Ogarkov, Elizaveta Orlova, Alexander Pavlenko, Alla Petrova, Natalia Pozhenko, Dmitry Pushkar, Alexander Rumyantsev, Sergey Rumyantsev, Vladimir Rumyantsev, Lyubov Rychkova, Alexander Samoilov, Irina Shirokova, Vyacheslav Sinkov, Svetlana Solovieva, Polina Tikhonova, Galina Trifonova, Alexander Troitsky, Alexander Tulichev, Yuri Udalov, Anna Varizhuk, Alexander Vasiliev, Vladimir Veselovsky, Alexey Volnukhin, Gaukhar Yusubalieva |
| EPI_ISL_1859498 | City Clinical Hospital named after S.I. Spasokukotsky of Moscow Healthcare Department, Academic Consortium | FEDERAL RESEARCH AND CLINICAL CENTER OF PHYSICAL-CHEMICAL MEDICINE OF FEDERAL MEDICAL BIOLOGICAL AGENCY, Academic Consortium | Vladislav Babenko, Ramiz Bakhtyev, Vladimir Baklaushev, Larisa Balykova, Pavel Bashkirov, Anna Blagonravova, Daria Boldyreva, Dmitry Fedorov, Ilshat Gafurov, Raushaniya Gaifullina, Yulia Galeeva, Elena Galova, Vadim Govorun, Elena Ilina, Konstantin Ivanov, Daria Kharlampieva, Polina Khromova, Ksenia Klimina, Konstantin Kolontarev, Nadezhda Kolyshkina, Andrey Koritsky, Vyacheslav Kuropatkin, Vasily Lazarev, Alexander Manolov, Valentin Manuvera, Daria Matyushkina, Ekaterina Moskaleva, Varvara Musarova, Oleg Ogarkov, Elizaveta Orlova, Alexander Pavlenko, Alla Petrova, Natalia Pozhenko, Dmitry Pushkar, Alexander Rumyantsev, Sergey Rumyantsev, Vladimir Rumyantsev, Lyubov Rychkova, Alexander Samoilov, Irina Shirokova, Vyacheslav Sinkov, Svetlana Solovieva, Polina Tikhonova, Galina Trifonova, Alexander Troitsky, Alexander Tulichev, Yuri Udalov, Anna Varizhuk, Alexander Vasiliev, Vladimir Veselovsky, Alexey Volnukhin, Gaukhar Yusubalieva |
| EPI_ISL_1859513, EPI_ISL_1859530, EPI_ISL_1859547, EPI_ISL_1859562 | Kazan Federal University, Academic Consortium | FEDERAL RESEARCH AND CLINICAL CENTER OF PHYSICAL-CHEMICAL MEDICINE OF FEDERAL MEDICAL BIOLOGICAL AGENCY, Academic Consortium | Vladislav Babenko, Ramiz Bakhtyev, Vladimir Baklaushev, Larisa Balykova, Pavel Bashkirov, Anna Blagonravova, Daria Boldyreva, Dmitry Fedorov, Ilshat Gafurov, Raushaniya Gaifullina, Yulia Galeeva, Elena Galova, Vadim Govorun, Elena Ilina, Konstantin Ivanov, Daria Kharlampieva, Polina Khromova, Ksenia Klimina, Konstantin Kolontarev, Nadezhda Kolyshkina, Andrey Koritsky, Vyacheslav Kuropatkin, Vasily Lazarev, Alexander Manolov, Valentin Manuvera, Daria Matyushkina, Ekaterina Moskaleva, Varvara Musarova, Oleg Ogarkov, Elizaveta Orlova, Alexander Pavlenko, Alla Petrova, Natalia Pozhenko, Dmitry Pushkar, Alexander Rumyantsev, Sergey Rumyantsev, Vladimir Rumyantsev, Lyubov Rychkova, Alexander Samoilov, Irina Shirokova, Vyacheslav Sinkov, Svetlana Solovieva, Polina Tikhonova, Galina Trifonova, Alexander Troitsky, Alexander Tulichev, Yuri Udalov, Anna Varizhuk, Alexander Vasiliev, Vladimir Veselovsky, Alexey Volnukhin, Gaukhar Yusubalieva |
| EPI_ISL_1859572 | City Clinical Hospital named after S.I. Spasokukotsky of Moscow Healthcare Department, Academic Consortium | FEDERAL RESEARCH AND CLINICAL CENTER OF PHYSICAL-CHEMICAL MEDICINE OF FEDERAL MEDICAL BIOLOGICAL AGENCY, Academic Consortium | Vladislav Babenko, Ramiz Bakhtyev, Vladimir Baklaushev, Larisa Balykova, Pavel Bashkirov, Anna Blagonravova, Daria Boldyreva, Dmitry Fedorov, Ilshat Gafurov, Raushaniya Gaifullina, Yulia Galeeva, Elena Galova, Vadim Govorun, Elena Ilina, Konstantin Ivanov, Daria Kharlampieva, Polina Khromova, Ksenia Klimina, Konstantin Kolontarev, Nadezhda Kolyshkina, Andrey Koritsky, Vyacheslav Kuropatkin, Vasily Lazarev, Alexander Manolov, Valentin Manuvera, Daria Matyushkina, Ekaterina Moskaleva, Varvara Musarova, Oleg Ogarkov, Elizaveta Orlova, Alexander Pavlenko, Alla Petrova, Natalia Pozhenko, Dmitry Pushkar, Alexander Rumyantsev, Sergey Rumyantsev, Vladimir Rumyantsev, Lyubov Rychkova, Alexander Samoilov, Irina Shirokova, Vyacheslav Sinkov, Svetlana Solovieva, Polina Tikhonova, Galina Trifonova, Alexander Troitsky, Alexander Tulichev, Yuri Udalov, Anna Varizhuk, Alexander Vasiliev, Vladimir Veselovsky, Alexey Volnukhin, Gaukhar Yusubalieva |
| EPI_ISL_1859600, EPI_ISL_1859602, EPI_ISL_1859626 | Kazan Federal University, Academic Consortium | FEDERAL RESEARCH AND CLINICAL CENTER OF PHYSICAL-CHEMICAL MEDICINE OF FEDERAL MEDICAL BIOLOGICAL AGENCY, Academic Consortium | Vladislav Babenko, Ramiz Bakhtyev, Vladimir Baklaushev, Larisa Balykova, Pavel Bashkirov, Anna Blagonravova, Daria Boldyreva, Dmitry Fedorov, Ilshat Gafurov, Raushaniya Gaifullina, Yulia Galeeva, Elena Galova, Vadim Govorun, Elena Ilina, Konstantin Ivanov, Daria Kharlampieva, Polina Khromova, Ksenia Klimina, Konstantin Kolontarev, Nadezhda Kolyshkina, Andrey Koritsky, Vyacheslav Kuropatkin, Vasily Lazarev, Alexander Manolov, Valentin Manuvera, Daria Matyushkina, Ekaterina Moskaleva, Varvara Musarova, Oleg Ogarkov, Elizaveta Orlova, Alexander Pavlenko, Alla Petrova, Natalia Pozhenko, Dmitry Pushkar, Alexander Rumyantsev, Sergey Rumyantsev, Vladimir Rumyantsev, Lyubov Rychkova, Alexander Samoilov, Irina Shirokova, Vyacheslav Sinkov, Svetlana Solovieva, Polina Tikhonova, Galina Trifonova, Alexander Troitsky, Alexander Tulichev, Yuri Udalov, Anna Varizhuk, Alexander Vasiliev, Vladimir Veselovsky, Alexey Volnukhin, Gaukhar Yusubalieva |
| EPI_ISL_1859654 | FEDERAL RESEARCH AND CLINICAL CENTER OF PHYSICAL-CHEMICAL MEDICINE OF FEDERAL MEDICAL BIOLOGICAL AGENCY, Academic Consortium | FEDERAL RESEARCH AND CLINICAL CENTER OF PHYSICAL-CHEMICAL MEDICINE OF FEDERAL MEDICAL BIOLOGICAL AGENCY, Academic Consortium | Vladislav Babenko, Ramiz Bakhtyev, Vladimir Baklaushev, Larisa Balykova, Pavel Bashkirov, Anna Blagonravova, Daria Boldyreva, Dmitry Fedorov, Ilshat Gafurov, Raushaniya Gaifullina, Yulia Galeeva, Elena Galova, Vadim Govorun, Elena Ilina, Konstantin Ivanov, Daria Kharlampieva, Polina Khromova, Ksenia Klimina, Konstantin Kolontarev, Nadezhda Kolyshkina, Andrey Koritsky, Vyacheslav Kuropatkin, Vasily Lazarev, Alexander Manolov, Valentin Manuvera, Daria Matyushkina, Ekaterina Moskaleva, Varvara Musarova, Oleg Ogarkov, Elizaveta Orlova, Alexander Pavlenko, Alla Petrova, Natalia Pozhenko, Dmitry Pushkar, Alexander Rumyantsev, Sergey Rumyantsev, Vladimir Rumyantsev, Lyubov Rychkova, Alexander Samoilov, Irina Shirokova, Vyacheslav Sinkov, Svetlana Solovieva, Polina Tikhonova, Galina Trifonova, Alexander Troitsky, Alexander Tulichev, Yuri Udalov, Anna Varizhuk, Alexander Vasiliev, Vladimir Veselovsky, Alexey Volnukhin, Gaukhar Yusubalieva |
| EPI_ISL_1859673, EPI_ISL_1859675, EPI_ISL_1859677, EPI_ISL_1859681, EPI_ISL_1859683, EPI_ISL_1859687, | Kazan Federal University, Academic Consortium | FEDERAL RESEARCH AND CLINICAL CENTER OF PHYSICAL-CHEMICAL MEDICINE OF FEDERAL MEDICAL BIOLOGICAL AGENCY, Academic Consortium | Vladislav Babenko, Ramiz Bakhtyev, Vladimir Baklaushev, Larisa Balykova, Pavel Bashkirov, Anna Blagonravova, Daria Boldyreva, Dmitry Fedorov, Ilshat Gafurov, Raushaniya Gaifullina, Yulia Galeeva, Elena Galova, Vadim Govorun, Elena Ilina, Konstantin Ivanov, Daria Kharlampieva, Polina Khromova, Ksenia Klimina, Konstantin Kolontarev, Nadezhda Kolyshkina, Andrey Koritsky, Vyacheslav Kuropatkin, Vasily Lazarev, Alexander Manolov, Valentin Manuvera, Daria |

| | | | |
|---|---|---|---|
| EPI_ISL_1859689, EPI_ISL_1859696, EPI_ISL_1859721 | | | Matyushkina, Ekaterina Moskaleva, Varvara Musarova, Oleg Ogarkov, Elizaveta Orlova, Alexander Pavlenko, Alla Petrova, Natalia Pozhenko, Dmitry Pushkar, Alexander Rumyantsev, Sergey Rumyantsev, Vladimir Rumyantsev, Lyubov Rychkova, Alexander Samoilov, Irina Shirokova, Vyacheslav Sinkov, Svetlana Solovieva, Polina Tikhonova, Galina Trifonova, Alexander Troitsky, Alexander Tulichev, Yuri Udalov, Anna Varizhuk, Alexander Vasiliev, Vladimir Veselovsky, Alexey Volnukhin, Gaukhar Yusubalieva |
| EPI_ISL_1859727 | City Clinical Hospital named after S.I. Spasokukotsky of Moscow Healthcare Department, Academic Consortium | FEDERAL RESEARCH AND CLINICAL CENTER OF PHYSICAL-CHEMICAL MEDICINE OF FEDERAL MEDICAL BIOLOGICAL AGENCY, Academic Consortium | Vladislav Babenko, Ramiz Bakhtyev, Vladimir Baklaushev, Larisa Balykova, Pavel Bashkirov, Anna Blagonravova, Daria Boldyreva, Dmitry Fedorov, Ilshat Gafurov, Raushaniya Gaifullina, Yulia Galeeva, Elena Galova, Vadim Govorun, Elena Ilina, Konstantin Ivanov, Daria Kharlampieva, Polina Khromova, Ksenia Klimina, Konstantin Kolontarev, Nadezhda Kolyshkina, Andrey Koritsky, Vyacheslav Kuropatkin, Vasily Lazarev, Alexander Manolov, Valentin Manuvera, Daria Matyushkina, Ekaterina Moskaleva, Varvara Musarova, Oleg Ogarkov, Elizaveta Orlova, Alexander Pavlenko, Alla Petrova, Natalia Pozhenko, Dmitry Pushkar, Alexander Rumyantsev, Sergey Rumyantsev, Vladimir Rumyantsev, Lyubov Rychkova, Alexander Samoilov, Irina Shirokova, Vyacheslav Sinkov, Svetlana Solovieva, Polina Tikhonova, Galina Trifonova, Alexander Troitsky, Alexander Tulichev, Yuri Udalov, Anna Varizhuk, Alexander Vasiliev, Vladimir Veselovsky, Alexey Volnukhin, Gaukhar Yusubalieva |
| EPI_ISL_1859729, EPI_ISL_1859733, EPI_ISL_1859738, EPI_ISL_1859746, EPI_ISL_1859764 | Kazan Federal University, Academic Consortium | FEDERAL RESEARCH AND CLINICAL CENTER OF PHYSICAL-CHEMICAL MEDICINE OF FEDERAL MEDICAL BIOLOGICAL AGENCY, Academic Consortium | Vladislav Babenko, Ramiz Bakhtyev, Vladimir Baklaushev, Larisa Balykova, Pavel Bashkirov, Anna Blagonravova, Daria Boldyreva, Dmitry Fedorov, Ilshat Gafurov, Raushaniya Gaifullina, Yulia Galeeva, Elena Galova, Vadim Govorun, Elena Ilina, Konstantin Ivanov, Daria Kharlampieva, Polina Khromova, Ksenia Klimina, Konstantin Kolontarev, Nadezhda Kolyshkina, Andrey Koritsky, Vyacheslav Kuropatkin, Vasily Lazarev, Alexander Manolov, Valentin Manuvera, Daria Matyushkina, Ekaterina Moskaleva, Varvara Musarova, Oleg Ogarkov, Elizaveta Orlova, Alexander Pavlenko, Alla Petrova, Natalia Pozhenko, Dmitry Pushkar, Alexander Rumyantsev, Sergey Rumyantsev, Vladimir Rumyantsev, Lyubov Rychkova, Alexander Samoilov, Irina Shirokova, Vyacheslav Sinkov, Svetlana Solovieva, Polina Tikhonova, Galina Trifonova, Alexander Troitsky, Alexander Tulichev, Yuri Udalov, Anna Varizhuk, Alexander Vasiliev, Vladimir Veselovsky, Alexey Volnukhin, Gaukhar Yusubalieva |
| EPI_ISL_1893362, EPI_ISL_1893366, EPI_ISL_1893368, EPI_ISL_1893370, EPI_ISL_1893372, EPI_ISL_1893374, EPI_ISL_1893375, EPI_ISL_1893376, EPI_ISL_1893377, EPI_ISL_1893382, EPI_ISL_1893437, EPI_ISL_1893439, EPI_ISL_1893442, EPI_ISL_1893449, EPI_ISL_1893451, EPI_ISL_1893453, EPI_ISL_1893455, EPI_ISL_1893460, EPI_ISL_1893462, EPI_ISL_1893469, EPI_ISL_1893473, EPI_ISL_1893475, EPI_ISL_1893478, EPI_ISL_1893480, EPI_ISL_1893487, EPI_ISL_1893489, EPI_ISL_1893490, EPI_ISL_1893493, EPI_ISL_1893495, EPI_ISL_1893497, EPI_ISL_1893499, EPI_ISL_1893507, EPI_ISL_1893508, EPI_ISL_1893531, EPI_ISL_1893533, EPI_ISL_1893534, EPI_ISL_1893537, EPI_ISL_1893539, EPI_ISL_1893541, EPI_ISL_1893543, EPI_ISL_1893545, EPI_ISL_1893547, EPI_ISL_1893549, EPI_ISL_1893552, EPI_ISL_1893553, EPI_ISL_1893555, EPI_ISL_1893557, EPI_ISL_1893592, EPI_ISL_1893599, EPI_ISL_1893602, EPI_ISL_1893603, EPI_ISL_1893606, EPI_ISL_1893608, EPI_ISL_1893610 | | | |
| see above | IPK | Respiratory Infections Laboratory | Lissette Perez, Yahisel Tejero, Yanet Pintos, Daylin Mederos, Odalys Valdes, Mirtha Elisa Aguado, Liena Ponce |
| EPI_ISL_1910399 | Centre Hospitalier Universitaire Clermont-Ferrand | CHU Clermont-Ferrand, service de virologie | Bisseux Maxime, Mirand Audrey, Combes Patricia, Henquell Cécile |
| EPI_ISL_1910400 | Centre Jean Perrin | CHU Clermont-Ferrand, service de virologie | Bisseux Maxime, Mirand Audrey, Combes Patricia, Henquell Cécile |
| EPI_ISL_1910401, EPI_ISL_1910402 | Centre Hospitalier Universitaire Clermont-Ferrand | CHU Clermont-Ferrand, service de virologie | Bisseux Maxime, Mirand Audrey, Combes Patricia, Henquell Cécile |
| EPI_ISL_1910403, EPI_ISL_1910404, EPI_ISL_1910405, EPI_ISL_1910406, EPI_ISL_1910407, EPI_ISL_1910408, EPI_ISL_1910409, EPI_ISL_1910410, EPI_ISL_1910411, EPI_ISL_1910412, EPI_ISL_1910413, EPI_ISL_1910414, EPI_ISL_1910415 | | | |
| see above | Centre Hospitalier Billom | CHU Clermont-Ferrand, service de virologie | Bisseux Maxime, Mirand Audrey, Combes Patricia, Henquell Cécile |
| EPI_ISL_1910416 | Centre Hospitalier Paul Ardier | CHU Clermont-Ferrand, service de virologie | Bisseux Maxime, Mirand Audrey, Combes Patricia, Henquell Cécile |
| EPI_ISL_1910417 | HOPITAL DE VICHY | CHU Clermont-Ferrand, service de virologie | Bisseux Maxime, Mirand Audrey, Combes Patricia, Henquell Cécile |
| EPI_ISL_1910418 | Centre Hospitalier Universitaire Clermont-Ferrand | CHU Clermont-Ferrand, service de virologie | Bisseux Maxime, Mirand Audrey, Combes Patricia, Henquell Cécile |
| EPI_ISL_1910419, EPI_ISL_1910420 | Centre Hospitalier Billom | CHU Clermont-Ferrand, service de virologie | Bisseux Maxime, Mirand Audrey, Combes Patricia, Henquell Cécile |
| EPI_ISL_1910421 | Centre Hospitalier Universitaire Clermont-Ferrand | CHU Clermont-Ferrand, service de virologie | Bisseux Maxime, Mirand Audrey, Combes Patricia, Henquell Cécile |
| EPI_ISL_1910422 | Centre Hospitalier Billom | CHU Clermont-Ferrand, service de virologie | Bisseux Maxime, Mirand Audrey, Combes Patricia, Henquell Cécile |
| EPI_ISL_1910423 | Centre Hospitalier Universitaire Clermont-Ferrand | CHU Clermont-Ferrand, service de virologie | Bisseux Maxime, Mirand Audrey, Combes Patricia, Henquell Cécile |
| EPI_ISL_1910424, EPI_ISL_1910425 | Centre Hospitalier Billom | CHU Clermont-Ferrand, service de virologie | Bisseux Maxime, Mirand Audrey, Combes Patricia, Henquell Cécile |
| EPI_ISL_1910426, EPI_ISL_1910427, EPI_ISL_1910428, EPI_ISL_1910429, EPI_ISL_1910430, EPI_ISL_1910431, EPI_ISL_1910432, EPI_ISL_1910433, EPI_ISL_1910434, EPI_ISL_1910435 | Centre Hospitalier Universitaire Clermont-Ferrand | CHU Clermont-Ferrand, service de virologie | Bisseux Maxime, Mirand Audrey, Combes Patricia, Henquell Cécile |
| EPI_ISL_1910436 | Centre Hospitalier Paul Ardier | CHU Clermont-Ferrand, service de virologie | Bisseux Maxime, Mirand Audrey, Combes Patricia, Henquell Cécile |
| EPI_ISL_1910437, EPI_ISL_1910438, EPI_ISL_1910439, EPI_ISL_1910440, EPI_ISL_1910441, EPI_ISL_1910442, EPI_ISL_1910443, EPI_ISL_1910446, EPI_ISL_1910447, EPI_ISL_1910448, EPI_ISL_1910449, EPI_ISL_1910450 | | | |
| see above | Centre Hospitalier Universitaire Clermont-Ferrand | CHU Clermont-Ferrand, service de virologie | Bisseux Maxime, Mirand Audrey, Combes Patricia, Henquell Cécile |
| EPI_ISL_1911013, EPI_ISL_1911014, EPI_ISL_1911015, EPI_ISL_1911016, EPI_ISL_1911017, EPI_ISL_1911018, EPI_ISL_1911019, EPI_ISL_1911020, EPI_ISL_1911021, EPI_ISL_1911022, EPI_ISL_1911023, EPI_ISL_1911024, EPI_ISL_1911025, EPI_ISL_1911026, EPI_ISL_1911027, EPI_ISL_1911028, EPI_ISL_1911029, EPI_ISL_1911030, EPI_ISL_1911031, EPI_ISL_1911032, EPI_ISL_1911034, EPI_ISL_1911035, EPI_ISL_1911036, EPI_ISL_1911037, EPI_ISL_1911038, EPI_ISL_1911039, EPI_ISL_1911040, EPI_ISL_1911041, EPI_ISL_1911042, EPI_ISL_1911043, EPI_ISL_1911044, EPI_ISL_1911045, EPI_ISL_1911047, EPI_ISL_1911048, EPI_ISL_1911049, EPI_ISL_1911050, EPI_ISL_1911051, EPI_ISL_1911052, EPI_ISL_1911053, EPI_ISL_1911054, EPI_ISL_1911055, EPI_ISL_1911056, EPI_ISL_1911057, EPI_ISL_1911058, EPI_ISL_1911059, EPI_ISL_1911060, EPI_ISL_1911061, EPI_ISL_1911062, EPI_ISL_1911063, EPI_ISL_1911064, EPI_ISL_1911065, EPI_ISL_1911066, EPI_ISL_1911067, EPI_ISL_1911068, EPI_ISL_1911069, EPI_ISL_1911070, EPI_ISL_1911071, EPI_ISL_1911073, EPI_ISL_1911074, EPI_ISL_1911075, EPI_ISL_1911076, EPI_ISL_1911077, EPI_ISL_1911078, EPI_ISL_1911079, EPI_ISL_1911080, EPI_ISL_1911081 | | | |
| see above | The Ohio State University Applied Microbiology Services Laboratory | The Ohio State University Applied Microbiology Services Laboratory | Seth A. Faith PhD |
| EPI_ISL_1915141 | Hospital | National Reference Center for Viruses of Respiratory Infections, Institut Pasteur, Paris | Marion Barbet, Sylvie Behillil, Méline Bizard, Angela Brisebarre, Camille Capel, Vincent Enouf, Louise Lefrançois, Frédéric Lemoine, Christophe Malabat, Corinne Maufrais, Emmanuelle Permal, Etienne Simon-Lorière, Maud Vanpeene, Sylvie Van der Werf ,Farida Moreau Benaoudia |
| EPI_ISL_1915189 | Labo Analyses Med | National Reference Center for Viruses of Respiratory Infections, Institut Pasteur, Paris | Marion Barbet, Sylvie Behillil, Méline Bizard, Angela Brisebarre, Camille Capel, Vincent Enouf, Louise Lefrançois, Frédéric Lemoine, Christophe Malabat, Corinne Maufrais, Emmanuelle Permal, Etienne Simon-Lorière, Maud Vanpeene, Sylvie Van der Werf ,Anne Grattard |
| EPI_ISL_1915604 | The Ohio State University Applied Microbiology Services Laboratory | The Ohio State University Applied Microbiology Services Laboratory | Seth A. Faith PhD |
| EPI_ISL_1916277, EPI_ISL_1916278, EPI_ISL_1916280, EPI_ISL_1916281, EPI_ISL_1916290, EPI_ISL_1916293, EPI_ISL_1916296, EPI_ISL_1916305, EPI_ISL_1916306 | CSIR-National Environmental Engineering Research Institute | CSIR-Centre for Cellular and Molecular Biology - INSACOG | Sofia Banu, Onkar Kulkarni, Lamuk Zaveri, Ara Sreenivas, Shreekant Verma, Amareshwar Vodapalli , Viswagithe S L, B Himasri, Sharath Chandra Thota, Karthik Bharadwaj Tallapaka, Krishna Khairnar, Rakesh K Mishra,Divya Tej Sowpati |
| EPI_ISL_1916312, EPI_ISL_1916313, EPI_ISL_1916370, EPI_ISL_1916371, EPI_ISL_1916374, EPI_ISL_1916382, EPI_ISL_1916383, EPI_ISL_1916387, EPI_ISL_1916395 | CSIR-National Environmental Engineering Research Institute | CSIR-Centre for Cellular and Molecular Biology - INSACOG | Lamuk Zaveri, Sofia Banu, Onkar Kulkarni, Ara Sreenivas, Shreekant Verma, Amareshwar Vodapalli , Viswagithe S L, B Himasri, Sharath Chandra Thota, Karthik Bharadwaj Tallapaka, Krishna Khairnar, Rakesh K Mishra,Divya Tej Sowpati |
| EPI_ISL_1916396, EPI_ISL_1916399 | CSIR-National Environmental Engineering Research Institute | CSIR-Centre for Cellular and Molecular Biology - INSACOG | Onkar Kulkarni, Lamuk Zaveri, Ara Sreenivas, Sofia Banu, Shreekant Verma, Amareshwar Vodapalli , Viswagithe S L, B Himasri, Sharath Chandra Thota, Karthik Bharadwaj Tallapaka, Krishna Khairnar, Rakesh K Mishra,Divya Tej Sowpati |
| EPI_ISL_1916491, EPI_ISL_1916492, EPI_ISL_1916493, EPI_ISL_1916494, EPI_ISL_1916495, EPI_ISL_1916496, EPI_ISL_1916497, EPI_ISL_1916498 | CSIR-National Environmental Engineering Research Institute | CSIR-Centre for Cellular and Molecular Biology - INSACOG | Ara Sreenivas, Onkar Kulkarni, Lamuk Zaveri, Sofia Banu, Shreekant Verma, Amareshwar Vodapalli , Viswagithe S L, B Himasri, Sharath Chandra Thota, Karthik Bharadwaj Tallapaka, Krishna Khairnar, Rakesh K Mishra,Divya Tej Sowpati |

| | | | |
|---|---|---|---|
| EPI_ISL_1965722 | 3. Medizinische Abteilung, Hanusch Krankenhaus | Bergthaler laboratory, CeMM Research Center for Molecular Medicine of the Austrian Academy of Sciences | Lukas Endler, Anna Schedl, Fabian Amman, Petr Triska, Thomas Penz, Benedikt Agerer, Maelle Le Moing, Michael Schuster, Bekir Erguner, Jan Laine, Martin Senekowitsch, Christoph Bock, Andreas Bergthaler |
| EPI_ISL_1970625, EPI_ISL_1970626 | CSIR-National Environmental Engineering Research Institute | CSIR-Centre for Cellular and Molecular Biology - INSACOG | Sofia Banu, Onkar Kulkarni, Lamuk Zaveri, Ara Sreenivas, Shreekant Verma, Amareshwar Vodapalli , Viswagithe S L, B Himasri, Sharath Chandra Thota, Karthik Bharadwaj Tallapaka, Krishna Khairnar, Rakesh K Mishra,Divya Tej Sowpati |
| EPI_ISL_2001116 | VRDL, Goverment Medical College (GMC), Bhavnagar | Gujarat Biotechnology Research Centre | Dinesh Kumar,Twinkle Soni,Sonal Sharma ,Zuber Saiyed,Ramesh Pandit,Janvi Raval,Zarna Patel,Nitin Savaliya,Kairavi Jwalantbhai Desai,Umang Mishra,Nitesh Shah,Chaitanya Joshi,Madhvi Joshi |
| EPI_ISL_2001119 | VRDL, Goverment Medical College (GMC), Bhavnagar | Gujarat Biotechnology Research Centre | Zuber Saiyed,Ramesh Pandit,Janvi Raval,Zarna Patel,Nitin Savaliya,Dinesh Kumar,Twinkle Soni,Sonal Sharma ,Kairavi Jwalantbhai Desai,Umang Mishra,Nitesh Shah,Chaitanya Joshi,Madhvi Joshi |
| EPI_ISL_2001120 | VRDL, Goverment Medical College (GMC), Bhavnagar | Gujarat Biotechnology Research Centre | Ramesh Pandit,Janvi Raval,Zarna Patel,Nitin Savaliya,Dinesh Kumar,Twinkle Soni,Sonal Sharma ,Zuber Saiyed,Kairavi Jwalantbhai Desai,Umang Mishra,Nitesh Shah,Chaitanya Joshi,Madhvi Joshi |
| EPI_ISL_2001149 | VRDL, Goverment Medical College (GMC), Bhavnagar | Gujarat Biotechnology Research Centre | Zuber Saiyed,Ramesh Pandit,Janvi Raval,Zarna Patel,Nitin Savaliya,Dinesh Kumar,Twinkle Soni,Sonal Sharma ,Kairavi Jwalantbhai Desai,Umang Mishra,Nitesh Shah,Chaitanya Joshi,Madhvi Joshi |
| EPI_ISL_2001150 | VRDL, Goverment Medical College (GMC), Bhavnagar | Gujarat Biotechnology Research Centre | Ramesh Pandit,Janvi Raval,Zarna Patel,Nitin Savaliya,Dinesh Kumar,Twinkle Soni,Sonal Sharma ,Zuber Saiyed,Kairavi Jwalantbhai Desai,Umang Mishra,Nitesh Shah,Chaitanya Joshi,Madhvi Joshi |
| EPI_ISL_2003113 | Hospital Municipal Dr Mario Gatti Campinas | Instituto Adolfo Lutz, Interdiciplinary Procedures Center, Strategic Laboratory | Claudio Tavares Sacchi, Claudia Regina Gonçalves, Erica Valessa Ramos Gomes, Karoline Rodrigues Campos, Caio Vinicius Dias Lopes, Leonardo Jose Tadeu de Araujo |
| EPI_ISL_2003153, EPI_ISL_2003154, EPI_ISL_2003155, EPI_ISL_2003156, EPI_ISL_2003157, EPI_ISL_2003158, EPI_ISL_2003159, EPI_ISL_2003160, EPI_ISL_2003161 | Instituto Adolfo Lutz Central | Instituto Adolfo Lutz, Interdiciplinary Procedures Center, Strategic Laboratory | Claudio Tavares Sacchi, Claudia Regina Gonçalves, Erica Valessa Ramos Gomes, Karoline Rodrigues Campos, Caio Vinicius Dias Lopes, Leonardo Jose Tadeu de Araujo |
| EPI_ISL_2009174, EPI_ISL_2009175, EPI_ISL_2009178, EPI_ISL_2009179, EPI_ISL_2009180, EPI_ISL_2009181, EPI_ISL_2009182, EPI_ISL_2009183, EPI_ISL_2009184, EPI_ISL_2009185 | MSK Microbiology Lab | MSK Microbiology Lab | Esther Babady, Tracy McMillen, Krupa Jani |
| EPI_ISL_2009232, EPI_ISL_2009233, EPI_ISL_2009234, EPI_ISL_2009305, EPI_ISL_2009530, EPI_ISL_2009595 | National Public Health Laboratory Malaysia | National Public Health Laboratory Malaysia | Noorliza MN, JL Tan, Chong CK, Norhayati R, Yukie C, Norazimah T, Rehan SAB, Selvanesan S, Hannah PYP, NurulAina MCA, W.NurAfiza WMA Zirwatul AA, Hani MH, Nurulsyahida I, Aziyati O, Ushananthiny R, Kamal HKZ, Mohd AY, YF Ngeow |
| EPI_ISL_2013043 | RS Sentra Medika Cikarang | Eijkman Institute for Molecular Biology, National Agency for Research and Innovation | Edison Johar, Frilasita A Yudhaputri, Muhammad Rezki Rasyak, Willy Agustine, Hidayat Trimarsanto, Lydia V. Panggalo, Iskandar Adnan, Sukma Oktavianthi, Safarina G Malik, Khin Saw Myint, Amin Soebandrio |
| EPI_ISL_2013044 | RS Sentra Medika Cikarang | Eijkman Institute for Molecular Biology, National Agency for Research and Innovation | Willy Agustine, Hidayat Trimarsanto, Lydia V. Panggalo, Iskandar Adnan, Sukma Oktavianthi, Edison Johar, Frilasita A Yudhaputri, Muhammad Rezki Rasyak, Safarina G Malik, Khin Saw Myint, Amin Soebandrio |
| EPI_ISL_2013045 | RS Mitra Keluarga Cikarang | Eijkman Institute for Molecular Biology, National Agency for Research and Innovation | Edison Johar, Frilasita A Yudhaputri, Muhammad Rezki Rasyak, Willy Agustine, Hidayat Trimarsanto, Lydia V. Panggalo, Iskandar Adnan, Sukma Oktavianthi, Safarina G Malik, Khin Saw Myint, Amin Soebandrio |
| EPI_ISL_2013046 | RSUD Cileungsi | Eijkman Institute for Molecular Biology, National Agency for Research and Innovation | Lydia V. Panggalo, Iskandar Adnan, Sukma Oktavianthi, Edison Johar, Frilasita A Yudhaputri, Muhammad Rezki Rasyak, Willy Agustine, Hidayat Trimarsanto, Safarina G Malik, Khin Saw Myint, Amin Soebandrio |
| EPI_ISL_2013047 | RS Mitra Keluarga Kelapa Gading | Eijkman Institute for Molecular Biology, National Agency for Research and Innovation | Hidayat Trimarsanto, Lydia V. Panggalo, Iskandar Adnan, Sukma Oktavianthi, Edison Johar, Frilasita A Yudhaputri, Muhammad Rezki Rasyak, Willy Agustine, Safarina G Malik, Khin Saw Myint, Amin Soebandrio |
| EPI_ISL_2013326 | IU-Cerrahpasa, Cerrahpasa School of Medicine, COVID-19 Lab | IU-Cerrahpasa, Cerrahpasa School of Medicine, COVID-19 Lab | Mert Kuskucu, Yesim Tuyji Tok, Kenan Midilli |
| EPI_ISL_2013509 | RS Harapan Bunda | Eijkman Institute for Molecular Biology, National Agency for Research and Innovation | Lydia V. Panggalo, Iskandar Adnan, Sukma Oktavianthi, Edison Johar, Frilasita A Yudhaputri, Muhammad Rezki Rasyak, Willy Agustine, Hidayat Trimarsanto, Safarina G Malik, Khin Saw Myint, Amin Soebandrio |
| EPI_ISL_2015096, EPI_ISL_2015097, EPI_ISL_2015098, EPI_ISL_2015099, EPI_ISL_2015100, EPI_ISL_2015103, EPI_ISL_2015104, EPI_ISL_2015116, EPI_ISL_2015117, EPI_ISL_2015121, EPI_ISL_2015122, EPI_ISL_2015125, EPI_ISL_2015139, EPI_ISL_2015140, EPI_ISL_2015142, EPI_ISL_2015143 | | | |
| see above | IU-Cerrahpasa, Cerrahpasa School of Medicine, COVID-19 Lab | IU-Cerrahpasa, Cerrahpasa School of Medicine, COVID-19 Lab | Mert Kuskucu, Yesim Tuyji Tok, Kenan Midilli |
| EPI_ISL_2020241 | Middlemore Hospital | Institute of Environmental Science and Research (ESR) WHO National Influenza Centre | Nikki Freed, Paula scholes, Susan Lin, Xiaoyun Ren, Matt Storey, Muhammad Faisal, Jing Wang, Hermes Perez, Anja Werno, Antje van der Linden, Arlo Upton, Chris Mansell, David Hammer, Dragana Drinkovic, Gary McAuliffe, Hana Sofia Andersson, James Ussher, Jill Sherwood, Josh Freeman, Julia Howard, Juliet Elvy, Mary DeAlmeida, Matt Blakiston, Matthew Rogers, Max Bloomfield, Michael Addidle, Michelle Balm, Sally Roberts, Sarah Jefferies, Sharmini Muttaiyah, Susan Morpeth, Susan Taylor, Timothy Blackmore, Vani Sathyendran, Veronica Playle, Virginia Hope, Erasmus Smit, Lauren Jelly, Joep de Ligt, Olin Silander |
| EPI_ISL_2029607 | Hospital | National Reference Center for Viruses of Respiratory Infections, Institut Pasteur, Paris | Marion Barbet, Sylvie Behillil, Méline Bizard, Angela Brisebarre, Camille Capel, Vincent Enouf, Louise Lefrançois, Frédéric Lemoine, Christophe Malabat, Corinne Maufrais, Amaury Vaysse, Etienne Simon-Lorière, Maud Vanpeene, Sylvie Van der Werf ,CéLine Bressollette |
| EPI_ISL_2029618, EPI_ISL_2029624 | Labo Analyses Med | National Reference Center for Viruses of Respiratory Infections, Institut Pasteur, Paris | Marion Barbet, Sylvie Behillil, Méline Bizard, Angela Brisebarre, Camille Capel, Vincent Enouf, Louise Lefrançois, Frédéric Lemoine, Christophe Malabat, Corinne Maufrais, Etienne Simon-Lorière, Maud Vanpeene, Sylvie Van der Werf ,Jean-Christophe Denis |
| EPI_ISL_2029723 | Labo Analyses Med | National Reference Center for Viruses of Respiratory Infections, Institut Pasteur, Paris | Marion Barbet, Sylvie Behillil, Méline Bizard, Angela Brisebarre, Camille Capel, Vincent Enouf, Louise Lefrançois, Frédéric Lemoine, Christophe Malabat, Corinne Maufrais, Etienne Simon-Lorière, Maud Vanpeene, Sylvie Van der Werf |
| EPI_ISL_2029793 | Labo Analyses Med | National Reference Center for Viruses of Respiratory Infections, Institut Pasteur, Paris | Marion Barbet, Sylvie Behillil, Méline Bizard, Angela Brisebarre, Camille Capel, Vincent Enouf, Louise Lefrançois, Frédéric Lemoine, Christophe Malabat, Corinne Maufrais, Etienne Simon-Lorière, Maud Vanpeene, Sylvie Van der Werf ,Jean-Christophe Denis |
| EPI_ISL_2029796 | Labo Analyses Med | National Reference Center for Viruses of Respiratory Infections, Institut Pasteur, Paris | Marion Barbet, Sylvie Behillil, Méline Bizard, Angela Brisebarre, Camille Capel, Vincent Enouf, Louise Lefrançois, Frédéric Lemoine, Christophe Malabat, Corinne Maufrais, Emmanuelle Permal, Etienne Simon-Lorière, Maud Vanpeene, Sylvie Van der Werf ,StéPhane Romand |
| EPI_ISL_2029862 | Labo Analyses Med | National Reference Center for Viruses of Respiratory Infections, Institut Pasteur, Paris | Marion Barbet, Sylvie Behillil, Méline Bizard, Angela Brisebarre, Camille Capel, Vincent Enouf, Louise Lefrançois, Frédéric Lemoine, Christophe Malabat, Corinne Maufrais, Etienne Simon-Lorière, Maud Vanpeene, Sylvie Van der Werf ,Jean-Christophe Denis |
| EPI_ISL_2029903 | Labo Analyses Med | National Reference Center for Viruses of Respiratory Infections, Institut Pasteur, Paris | Marion Barbet, Sylvie Behillil, Méline Bizard, Angela Brisebarre, Camille Capel, Vincent Enouf, Louise Lefrançois, Frédéric Lemoine, Christophe Malabat, Corinne Maufrais, Emmanuelle Permal, Etienne Simon-Lorière, Maud Vanpeene, Sylvie Van der Werf ,StéPhane Romand |
| EPI_ISL_2029974 | Labo Analyses Med | National Reference Center for Viruses of Respiratory Infections, Institut Pasteur, Paris | Marion Barbet, Sylvie Behillil, Méline Bizard, Angela Brisebarre, Camille Capel, Vincent Enouf, Louise Lefrançois, Frédéric Lemoine, Christophe Malabat, Corinne Maufrais, Etienne Simon-Lorière, Maud Vanpeene, Sylvie Van der Werf ,Jean-Christophe Denis |
| EPI_ISL_2030109, EPI_ISL_2030171 | Hospital | National Reference Center for Viruses of Respiratory Infections, Institut Pasteur, Paris | Marion Barbet, Sylvie Behillil, Méline Bizard, Angela Brisebarre, Camille Capel, Vincent Enouf, Louise Lefrançois, Frédéric Lemoine, Christophe Malabat, Corinne Maufrais, Etienne Simon-Lorière, Maud Vanpeene, Sylvie Van der Werf ,Ali Si-Mohammed |
| EPI_ISL_2081877, EPI_ISL_2081879, EPI_ISL_2081880, EPI_ISL_2081881, EPI_ISL_2081882, EPI_ISL_2081883, EPI_ISL_2081884, EPI_ISL_2081885, EPI_ISL_2081888, EPI_ISL_2081889, EPI_ISL_2081890, EPI_ISL_2081891, EPI_ISL_2081892, EPI_ISL_2081893, EPI_ISL_2081894, EPI_ISL_2081895, EPI_ISL_2081896, EPI_ISL_2081897, EPI_ISL_2081898, EPI_ISL_2081899, EPI_ISL_2081900, EPI_ISL_2081901, EPI_ISL_2081903, EPI_ISL_2081904, EPI_ISL_2081906, EPI_ISL_2081907, EPI_ISL_2081908, EPI_ISL_2081911, EPI_ISL_2081913, EPI_ISL_2081914, EPI_ISL_2081916, EPI_ISL_2081921, EPI_ISL_2081931, EPI_ISL_2081932, | | | |

| | | | |
|---|---|---|---|
| EPI_ISL_2081933, EPI_ISL_2081934, EPI_ISL_2081935, EPI_ISL_2081936, EPI_ISL_2081938, EPI_ISL_2081939, EPI_ISL_2081940, EPI_ISL_2081941, EPI_ISL_2081942, EPI_ISL_2081943, EPI_ISL_2081944, EPI_ISL_2081945, EPI_ISL_2081946, EPI_ISL_2081947, EPI_ISL_2081948, EPI_ISL_2081958, EPI_ISL_2081959, EPI_ISL_2081960, EPI_ISL_2081961, EPI_ISL_2081962, EPI_ISL_2081963, EPI_ISL_2082013, EPI_ISL_2082014, EPI_ISL_2082015, EPI_ISL_2082016, EPI_ISL_2082017, EPI_ISL_2082018, EPI_ISL_2082019, EPI_ISL_2082020, EPI_ISL_2082021, EPI_ISL_2082022, EPI_ISL_2082023, EPI_ISL_2082024, EPI_ISL_2082030, EPI_ISL_2082031 | | | |
| see above | National Center of Infectious and Parasitic Diseases | National Center of Infectious and Parasitic Diseases | Alexiev, Ivanov, Korsun, Stoitsova, Philipova, Dimitrova, Grigorova L., Hristova, Donchev, Stoykov, Trifonova, Dobrinov, Grigorova I., Kantardjiev |
| EPI_ISL_2086187, EPI_ISL_2086192 | CQRC_QUALITY CONTROL CHEMICAL BIOLOGICAL RISK_AOOR Villa Sofia Cervello Palermo | CQRC_QUALITY CONTROL CHEMICAL BIOLOGICAL RISK_AOOR Villa Sofia Cervello Palermo | Di Gaudio, F., Contino, F., Brunacci, G.. |
| EPI_ISL_2086202, EPI_ISL_2086207, EPI_ISL_2086210 | CQRC_QUALITY CONTROL CHEMICAL BIOLOGICAL RISK_AOOR Villa Sofia Cervello Palermo | CQRC_QUALITY CONTROL CHEMICAL BIOLOGICAL RISK_AOOR Villa Sofia Cervello Palermo | Di Gaudio, F., Brunacci, G., Contino, F. |
| EPI_ISL_2086268 | Laboratory of Molecular Biology of Flavivirus, Oswaldo Cruz Institute, FIOCRUZ | Laboratory of Molecular Biology of Flavivirus, Oswaldo Cruz Institute, FIOCRUZ | Fernando Lopez Tort, Ieda Ribeiro, Lidiane Menezes, Alexandre dos Santos, Marta Pereira, Myrna Bonaldo |
| EPI_ISL_2098849, EPI_ISL_2098851, EPI_ISL_2098852, EPI_ISL_2098853, EPI_ISL_2098854, EPI_ISL_2098857, EPI_ISL_2098859 | CQRC_QUALITY CONTROL CHEMICAL BIOLOGICAL RISK_AOOR Villa Sofia Cervello Palermo | CQRC_QUALITY CONTROL CHEMICAL BIOLOGICAL RISK_AOOR Villa Sofia Cervello Palermo | Di Gaudio, F., Contino, F., Brunacci, G. |
| EPI_ISL_2098860, EPI_ISL_2098862, EPI_ISL_2098863, EPI_ISL_2098864, EPI_ISL_2098865, EPI_ISL_2098866, EPI_ISL_2098867, EPI_ISL_2098868, EPI_ISL_2098869, EPI_ISL_2098871 | CQRC_QUALITY CONTROL CHEMICAL BIOLOGICAL RISK_AOOR Villa Sofia Cervello Palermo | CQRC_QUALITY CONTROL CHEMICAL BIOLOGICAL RISK_AOOR Villa Sofia Cervello Palermo | Di Gaudio, F., Brunacci G., Contino, F. |
| EPI_ISL_2098948 | Laboratorio Biologia Molecolare Sars Cov2 - UOC Laboratorio Analisi - Servizio Medicina di Laboratorio, Ospedale "San Francesco" - ATS-ASSL Nuoro | Laboratorio specialistico UOC Ematologia - Ospedale "San Francesco" - ATS-ASSL Nuoro | Piras Giovanna, Asproni Rosanna,Paolo Malune,Monne Maria Itria, Palmas Angelo Domenico, Fiamma Maura, Garau Maria Cristina, Mameli Giuseppe |
| EPI_ISL_2099949, EPI_ISL_2099950, EPI_ISL_2099951, EPI_ISL_2099952, EPI_ISL_2099953 | CQRC_QUALITY CONTROL CHEMICAL BIOLOGICAL RISK_AOOR Villa Sofia Cervello Palermo | CQRC_QUALITY CONTROL CHEMICAL BIOLOGICAL RISK_AOOR Villa Sofia Cervello Palermo | Di Gaudio, F., Contino, F., Brunacci, G. |
| EPI_ISL_2099955 | CQRC_QUALITY CONTROL CHEMICAL BIOLOGICAL RISK_AOOR Villa Sofia Cervello Palermo | CQRC_QUALITY CONTROL CHEMICAL BIOLOGICAL RISK_AOOR Villa Sofia Cervello Palermo | Di Gaudio, F., Contino, F., Gasparro G., Rinaudo G., Brunacci, G. |
| EPI_ISL_2099956, EPI_ISL_2099958, EPI_ISL_2099959, EPI_ISL_2099960, EPI_ISL_2099961, EPI_ISL_2099962, EPI_ISL_2099963, EPI_ISL_2099964 | CQRC_QUALITY CONTROL CHEMICAL BIOLOGICAL RISK_AOOR Villa Sofia Cervello Palermo | CQRC_QUALITY CONTROL CHEMICAL BIOLOGICAL RISK_AOOR Villa Sofia Cervello Palermo | Di Gaudio, Brunacci, G., F. Contino, F. |
| EPI_ISL_2099965 | CQRC_QUALITY CONTROL CHEMICAL BIOLOGICAL RISK_AOOR Villa Sofia Cervello Palermo | CQRC_QUALITY CONTROL CHEMICAL BIOLOGICAL RISK_AOOR Villa Sofia Cervello Palermo | Di Gaudio, F., Contino, F., Gasparro G., Rinaudo G., Brunacci, G. |
| EPI_ISL_2100701, EPI_ISL_2100702, EPI_ISL_2100703, EPI_ISL_2100704, EPI_ISL_2100705, EPI_ISL_2100706, EPI_ISL_2100707, EPI_ISL_2100708, EPI_ISL_2100710, EPI_ISL_2100711, EPI_ISL_2100712, EPI_ISL_2100713, EPI_ISL_2100714, EPI_ISL_2100715, EPI_ISL_2100716, EPI_ISL_2100717, EPI_ISL_2100718, EPI_ISL_2100719, EPI_ISL_2100720, EPI_ISL_2100721, EPI_ISL_2100722, EPI_ISL_2100723, EPI_ISL_2100724, EPI_ISL_2100725, EPI_ISL_2100726, EPI_ISL_2100727, EPI_ISL_2100728, EPI_ISL_2100729, EPI_ISL_2100730, EPI_ISL_2100731, EPI_ISL_2100732, EPI_ISL_2100733, EPI_ISL_2100734, EPI_ISL_2100735, EPI_ISL_2100736, EPI_ISL_2100737, EPI_ISL_2100738, EPI_ISL_2100739, EPI_ISL_2100740, EPI_ISL_2100741, EPI_ISL_2100742, EPI_ISL_2100744, EPI_ISL_2100745, EPI_ISL_2100746, EPI_ISL_2100747, EPI_ISL_2100748, EPI_ISL_2100749, EPI_ISL_2100750, EPI_ISL_2100751, EPI_ISL_2100752, EPI_ISL_2100753, EPI_ISL_2100754, EPI_ISL_2100755, EPI_ISL_2100756, EPI_ISL_2100757, EPI_ISL_2100758, EPI_ISL_2100759, EPI_ISL_2100760, EPI_ISL_2100761, EPI_ISL_2100762, EPI_ISL_2100763, EPI_ISL_2100764, EPI_ISL_2100765, EPI_ISL_2100766, EPI_ISL_2100767, EPI_ISL_2100768, EPI_ISL_2100769, EPI_ISL_2100770, EPI_ISL_2100771, EPI_ISL_2100772, EPI_ISL_2100773, EPI_ISL_2100775, EPI_ISL_2100776, EPI_ISL_2100777, EPI_ISL_2100778, EPI_ISL_2100779, EPI_ISL_2100780, EPI_ISL_2100781, EPI_ISL_2100782, EPI_ISL_2100783, EPI_ISL_2100784, EPI_ISL_2100785, EPI_ISL_2100786, EPI_ISL_2100789, EPI_ISL_2100791, EPI_ISL_2100792, EPI_ISL_2100794, EPI_ISL_2100796, EPI_ISL_2100797, EPI_ISL_2100798, EPI_ISL_2100799, EPI_ISL_2100800, EPI_ISL_2100801, EPI_ISL_2100802, EPI_ISL_2100804, EPI_ISL_2100805, EPI_ISL_2100806, EPI_ISL_2100807, EPI_ISL_2100808, EPI_ISL_2100810, EPI_ISL_2100811, EPI_ISL_2100812, EPI_ISL_2100813, EPI_ISL_2100814, EPI_ISL_2100816, EPI_ISL_2100818, EPI_ISL_2100819, EPI_ISL_2100820, EPI_ISL_2100822, EPI_ISL_2100824, EPI_ISL_2100825, EPI_ISL_2100826, EPI_ISL_2100827, EPI_ISL_2100828, EPI_ISL_2100830, EPI_ISL_2100831, EPI_ISL_2100832, EPI_ISL_2100833, EPI_ISL_2100834, EPI_ISL_2100835, EPI_ISL_2100836, EPI_ISL_2100839, EPI_ISL_2100840, EPI_ISL_2100841, EPI_ISL_2100842, EPI_ISL_2100843, EPI_ISL_2100845, EPI_ISL_2100846, EPI_ISL_2100847, EPI_ISL_2100848, EPI_ISL_2100849, EPI_ISL_2100850, EPI_ISL_2100851, EPI_ISL_2100852, EPI_ISL_2100854, EPI_ISL_2100855, EPI_ISL_2100857, EPI_ISL_2100858, EPI_ISL_2100859, EPI_ISL_2100860, EPI_ISL_2100861, EPI_ISL_2100862, EPI_ISL_2100865, EPI_ISL_2100867, EPI_ISL_2100868, EPI_ISL_2100869, EPI_ISL_2100883, EPI_ISL_2100884, EPI_ISL_2100885, EPI_ISL_2100886, EPI_ISL_2100887, EPI_ISL_2100888, EPI_ISL_2100889, EPI_ISL_2100890 | | | |
| see above | The Ohio State University Applied Microbiology Services Laboratory | The Ohio State University Applied Microbiology Services Laboratory | Seth A. Faith PhD |
| EPI_ISL_2101022 | Molecular Biology Laboratory, Faculty Medicine and Health Sciences, Warmadewa University | Eijkman Institute for Molecular Biology, National Agency for Research and Innovation; Molecular Biology Laboratory, Faculty Medicine and Health Sciences, Warmadewa University | Frilasita A Yudhaputri, Muhammad Rezki Rasyak, Willy Agustine, Hidayat Trimarsanto, Lydia V. Panggalo, Iskandar Adnan, Sukma Oktavianthi, Lidwina Priliani, Edison Johar, Sri Masyeni, Erly Sintya, Safarina G Malik, Khin Saw Myint, Amin Soebandrio |
| EPI_ISL_2101023 | Molecular Biology Laboratory, Faculty Medicine and Health Sciences, Warmadewa University | Eijkman Institute for Molecular Biology, National Agency for Research and Innovation; Molecular Biology Laboratory, Faculty Medicine and Health Sciences, Warmadewa University | Muhammad Rezki Rasyak, Willy Agustine, Hidayat Trimarsanto, Lydia V. Panggalo, Iskandar Adnan, Sukma Oktavianthi, Lidwina Priliani, Edison Johar, Frilasita A Yudhaputri, Sri Masyeni, Erly Sintya, Safarina G Malik, Khin Saw Myint, Amin Soebandrio |
| EPI_ISL_2101024 | Molecular Biology Laboratory, Faculty Medicine and Health Sciences, Warmadewa University | Eijkman Institute for Molecular Biology, National Agency for Research and Innovation; Molecular Biology Laboratory, Faculty Medicine and Health Sciences, Warmadewa University | Willy Agustine, Hidayat Trimarsanto, Lydia V. Panggalo, Iskandar Adnan, Sukma Oktavianthi, Lidwina Priliani, Edison Johar, Frilasita A Yudhaputri, Muhammad Rezki Rasyak, Sri Masyeni, Erly Sintya, Safarina G Malik, Khin Saw Myint, Amin Soebandrio |
| EPI_ISL_2101025 | Molecular Biology Laboratory, Faculty Medicine and Health Sciences, Warmadewa University | Eijkman Institute for Molecular Biology, National Agency for Research and Innovation; Molecular Biology Laboratory, Faculty Medicine and Health Sciences, Warmadewa University | Hidayat Trimarsanto, Lydia V. Panggalo, Iskandar Adnan, Sukma Oktavianthi, Lidwina Priliani, Edison Johar, Frilasita A Yudhaputri, Muhammad Rezki Rasyak, Willy Agustine, Sri Masyeni, Erly Sintya, Safarina G Malik, Khin Saw Myint, Amin Soebandrio |
| EPI_ISL_2101026 | Molecular Biology Laboratory, Faculty Medicine and Health Sciences, Warmadewa University | Eijkman Institute for Molecular Biology, National Agency for Research and Innovation; Molecular Biology Laboratory, Faculty Medicine and Health Sciences, Warmadewa University | Willy Agustine, Hidayat Trimarsanto, Lydia V. Panggalo, Iskandar Adnan, Sukma Oktavianthi, Lidwina Priliani, Edison Johar, Frilasita A Yudhaputri, Muhammad Rezki Rasyak, Sri Masyeni, Erly Sintya, Safarina G Malik, Khin Saw Myint, Amin Soebandrio |
| EPI_ISL_2101027 | Molecular Biology Laboratory, Faculty Medicine and Health Sciences, Warmadewa University | Eijkman Institute for Molecular Biology, National Agency for Research and Innovation; Molecular Biology Laboratory, Faculty Medicine and Health Sciences, Warmadewa University | Iskandar Adnan, Sukma Oktavianthi, Lidwina Priliani, Edison Johar, Frilasita A Yudhaputri, Muhammad Rezki Rasyak, Willy Agustine, Hidayat Trimarsanto, Lydia V. Panggalo, Sri Masyeni, Erly Sintya, Safarina G Malik, Khin Saw Myint, Amin Soebandrio |
| EPI_ISL_2101028 | Molecular Biology Laboratory, Faculty Medicine and Health Sciences, Warmadewa University | Eijkman Institute for Molecular Biology, National Agency for Research and Innovation; Molecular Biology Laboratory, Faculty Medicine and Health Sciences, Warmadewa University | Edison Johar, Frilasita A Yudhaputri, Muhammad Rezki Rasyak, Willy Agustine, Hidayat Trimarsanto, Lydia V. Panggalo, Iskandar Adnan, Sukma Oktavianthi, Lidwina Priliani, Sri Masyeni, Erly Sintya, Safarina G Malik, Khin Saw Myint, Amin Soebandrio |
| EPI_ISL_2101029 | Molecular Biology Laboratory, Faculty Medicine and Health Sciences, Warmadewa University | Eijkman Institute for Molecular Biology, National Agency for Research and Innovation; Molecular Biology Laboratory, Faculty Medicine and Health Sciences, Warmadewa University | Hidayat Trimarsanto, Lydia V. Panggalo, Iskandar Adnan, Sukma Oktavianthi, Lidwina Priliani, Edison Johar, Frilasita A Yudhaputri, Muhammad Rezki Rasyak, Willy Agustine, Sri Masyeni, Erly Sintya, Safarina G Malik, Khin Saw Myint, Amin Soebandrio |

| | | | |
|---|---|---|---|
| EPI_ISL_2101030 | Molecular Biology Laboratory, Faculty Medicine and Health Sciences, Warmadewa University | Eijkman Institute for Molecular Biology, National Agency for Research and Innovation; Molecular Biology Laboratory, Faculty Medicine and Health Sciences, Warmadewa University | Sukma Oktavianthi, Lidwina Priliani, Edison Johar, Frilasita A Yudhaputri, Muhammad Rezki Rasyak, Willy Agustine, Hidayat Trimarsanto, Lydia V. Panggalo, Iskandar Adnan, Sri Masyeni, Erly Sintya, Safarina G Malik, Khin Saw Myint, Amin Soebandrio |
| EPI_ISL_2101031 | Molecular Biology Laboratory, Faculty Medicine and Health Sciences, Warmadewa University | Eijkman Institute for Molecular Biology, National Agency for Research and Innovation; Molecular Biology Laboratory, Faculty Medicine and Health Sciences, Warmadewa University | Frilasita A Yudhaputri, Muhammad Rezki Rasyak, Willy Agustine, Hidayat Trimarsanto, Lydia V. Panggalo, Iskandar Adnan, Sukma Oktavianthi, Lidwina Priliani, Edison Johar, Sri Masyeni, Erly Sintya, Safarina G Malik, Khin Saw Myint, Amin Soebandrio |
| EPI_ISL_2101032 | Molecular Biology Laboratory, Faculty Medicine and Health Sciences, Warmadewa University | Eijkman Institute for Molecular Biology, National Agency for Research and Innovation; Molecular Biology Laboratory, Faculty Medicine and Health Sciences, Warmadewa University | Lydia V. Panggalo, Iskandar Adnan, Sukma Oktavianthi, Lidwina Priliani, Edison Johar, Frilasita A Yudhaputri, Muhammad Rezki Rasyak, Willy Agustine, Hidayat Trimarsanto, Sri Masyeni, Erly Sintya, Safarina G Malik, Khin Saw Myint, Amin Soebandrio |
| EPI_ISL_2101034 | Molecular Biology Laboratory, Faculty Medicine and Health Sciences, Warmadewa University | Eijkman Institute for Molecular Biology, National Agency for Research and Innovation; Molecular Biology Laboratory, Faculty Medicine and Health Sciences, Warmadewa University | Willy Agustine, Hidayat Trimarsanto, Lydia V. Panggalo, Iskandar Adnan, Sukma Oktavianthi, Lidwina Priliani, Edison Johar, Frilasita A Yudhaputri, Muhammad Rezki Rasyak, Sri Masyeni, Erly Sintya, Safarina G Malik, Khin Saw Myint, Amin Soebandrio |
| EPI_ISL_2101035, EPI_ISL_2101036, EPI_ISL_2101037 | Molecular Biology Laboratory, Faculty Medicine and Health Sciences, Warmadewa University | Eijkman Institute for Molecular Biology, National Agency for Research and Innovation; Molecular Biology Laboratory, Faculty Medicine and Health Sciences, Warmadewa University | Lydia V. Panggalo, Iskandar Adnan, Sukma Oktavianthi, Lidwina Priliani, Edison Johar, Frilasita A Yudhaputri, Muhammad Rezki Rasyak, Willy Agustine, Hidayat Trimarsanto, Sri Masyeni, Erly Sintya, Safarina G Malik, Khin Saw Myint, Amin Soebandrio |
| EPI_ISL_2101038 | Molecular Biology Laboratory, Faculty Medicine and Health Sciences, Warmadewa University | Eijkman Institute for Molecular Biology, National Agency for Research and Innovation; Molecular Biology Laboratory, Faculty Medicine and Health Sciences, Warmadewa University | Iskandar Adnan, Sukma Oktavianthi, Lidwina Priliani, Edison Johar, Frilasita A Yudhaputri, Muhammad Rezki Rasyak, Willy Agustine, Hidayat Trimarsanto, Lydia V. Panggalo, Sri Masyeni, Erly Sintya, Safarina G Malik, Khin Saw Myint, Amin Soebandrio |
| EPI_ISL_2101039 | Molecular Biology Laboratory, Faculty Medicine and Health Sciences, Warmadewa University | Eijkman Institute for Molecular Biology, National Agency for Research and Innovation; Molecular Biology Laboratory, Faculty Medicine and Health Sciences, Warmadewa University | Edison Johar, Frilasita A Yudhaputri, Muhammad Rezki Rasyak, Willy Agustine, Hidayat Trimarsanto, Lydia V. Panggalo, Iskandar Adnan, Sukma Oktavianthi, Lidwina Priliani, Sri Masyeni, Erly Sintya, Safarina G Malik, Khin Saw Myint, Amin Soebandrio |
| EPI_ISL_2101040 | Molecular Biology Laboratory, Faculty Medicine and Health Sciences, Warmadewa University | Eijkman Institute for Molecular Biology, National Agency for Research and Innovation; Molecular Biology Laboratory, Faculty Medicine and Health Sciences, Warmadewa University | Hidayat Trimarsanto, Lydia V. Panggalo, Iskandar Adnan, Sukma Oktavianthi, Lidwina Priliani, Edison Johar, Frilasita A Yudhaputri, Muhammad Rezki Rasyak, Willy Agustine, Sri Masyeni, Erly Sintya, Safarina G Malik, Khin Saw Myint, Amin Soebandrio |
| EPI_ISL_2101041 | Molecular Biology Laboratory, Faculty Medicine and Health Sciences, Warmadewa University | Eijkman Institute for Molecular Biology, National Agency for Research and Innovation; Molecular Biology Laboratory, Faculty Medicine and Health Sciences, Warmadewa University | Sukma Oktavianthi, Lidwina Priliani, Edison Johar, Frilasita A Yudhaputri, Muhammad Rezki Rasyak, Willy Agustine, Hidayat Trimarsanto, Lydia V. Panggalo, Iskandar Adnan, Sri Masyeni, Erly Sintya, Safarina G Malik, Khin Saw Myint, Amin Soebandrio |
| EPI_ISL_2101042 | Molecular Biology Laboratory, Faculty Medicine and Health Sciences, Warmadewa University | Eijkman Institute for Molecular Biology, National Agency for Research and Innovation; Molecular Biology Laboratory, Faculty Medicine and Health Sciences, Warmadewa University | Willy Agustine, Hidayat Trimarsanto, Lydia V. Panggalo, Iskandar Adnan, Sukma Oktavianthi, Lidwina Priliani, Edison Johar, Frilasita A Yudhaputri, Muhammad Rezki Rasyak, Sri Masyeni, Erly Sintya, Safarina G Malik, Khin Saw Myint, Amin Soebandrio |
| EPI_ISL_2101043 | Molecular Biology Laboratory, Faculty Medicine and Health Sciences, Warmadewa University | Eijkman Institute for Molecular Biology, National Agency for Research and Innovation; Molecular Biology Laboratory, Faculty Medicine and Health Sciences, Warmadewa University | Hidayat Trimarsanto, Lydia V. Panggalo, Iskandar Adnan, Sukma Oktavianthi, Lidwina Priliani, Edison Johar, Frilasita A Yudhaputri, Muhammad Rezki Rasyak, Willy Agustine, Sri Masyeni, Erly Sintya, Safarina G Malik, Khin Saw Myint, Amin Soebandrio |
| EPI_ISL_2101044 | Molecular Biology Laboratory, Faculty Medicine and Health Sciences, Warmadewa University | Eijkman Institute for Molecular Biology, National Agency for Research and Innovation; Molecular Biology Laboratory, Faculty Medicine and Health Sciences, Warmadewa University | Lydia V. Panggalo, Iskandar Adnan, Sukma Oktavianthi, Lidwina Priliani, Edison Johar, Frilasita A Yudhaputri, Muhammad Rezki Rasyak, Willy Agustine, Hidayat Trimarsanto, Sri Masyeni, Erly Sintya, Safarina G Malik, Khin Saw Myint, Amin Soebandrio |
| EPI_ISL_2101045 | Molecular Biology Laboratory, Faculty Medicine and Health Sciences, Warmadewa University | Eijkman Institute for Molecular Biology, National Agency for Research and Innovation; Molecular Biology Laboratory, Faculty Medicine and Health Sciences, Warmadewa University | Iskandar Adnan, Sukma Oktavianthi, Lidwina Priliani, Edison Johar, Frilasita A Yudhaputri, Muhammad Rezki Rasyak, Willy Agustine, Hidayat Trimarsanto, Lydia V. Panggalo, Sri Masyeni, Erly Sintya, Safarina G Malik, Khin Saw Myint, Amin Soebandrio |
| EPI_ISL_2101046 | Molecular Biology Laboratory, Faculty Medicine and Health Sciences, Warmadewa University | Eijkman Institute for Molecular Biology, National Agency for Research and Innovation; Molecular Biology Laboratory, Faculty Medicine and Health Sciences, Warmadewa University | Sukma Oktavianthi, Lidwina Priliani, Edison Johar, Frilasita A Yudhaputri, Muhammad Rezki Rasyak, Willy Agustine, Hidayat Trimarsanto, Lydia V. Panggalo, Iskandar Adnan, Sri Masyeni, Erly Sintya, Safarina G Malik, Khin Saw Myint, Amin Soebandrio |
| EPI_ISL_2101047 | Molecular Biology Laboratory, Faculty Medicine and Health Sciences, Warmadewa University | Eijkman Institute for Molecular Biology, National Agency for Research and Innovation; Molecular Biology Laboratory, Faculty Medicine and Health Sciences, Warmadewa University | Lidwina Priliani, Edison Johar, Frilasita A Yudhaputri, Muhammad Rezki Rasyak, Willy Agustine, Hidayat Trimarsanto, Lydia V. Panggalo, Iskandar Adnan, Sukma Oktavianthi, Sri Masyeni, Erly Sintya, Safarina G Malik, Khin Saw Myint, Amin Soebandrio |
| EPI_ISL_2101048 | Molecular Biology Laboratory, Faculty Medicine and Health Sciences, Warmadewa University | Eijkman Institute for Molecular Biology, National Agency for Research and Innovation; Molecular Biology Laboratory, Faculty Medicine and Health Sciences, Warmadewa University | Edison Johar, Frilasita A Yudhaputri, Muhammad Rezki Rasyak, Willy Agustine, Hidayat Trimarsanto, Lydia V. Panggalo, Iskandar Adnan, Sukma Oktavianthi, Lidwina Priliani, Sri Masyeni, Erly Sintya, Safarina G Malik, Khin Saw Myint, Amin Soebandrio |
| EPI_ISL_2101049 | Molecular Biology Laboratory, Faculty Medicine and Health Sciences, Warmadewa University | Eijkman Institute for Molecular Biology, National Agency for Research and Innovation; Molecular Biology Laboratory, Faculty Medicine and Health Sciences, Warmadewa University | Frilasita A Yudhaputri, Muhammad Rezki Rasyak, Willy Agustine, Hidayat Trimarsanto, Lydia V. Panggalo, Iskandar Adnan, Sukma Oktavianthi, Lidwina Priliani, Edison Johar, Sri Masyeni, Erly Sintya, Safarina G Malik, Khin Saw Myint, Amin Soebandrio |
| EPI_ISL_2101050 | Molecular Biology Laboratory, Faculty Medicine and Health Sciences, Warmadewa University | Eijkman Institute for Molecular Biology, National Agency for Research and Innovation; Molecular Biology Laboratory, Faculty Medicine and Health Sciences, Warmadewa University | Willy Agustine, Hidayat Trimarsanto, Lydia V. Panggalo, Iskandar Adnan, Sukma Oktavianthi, Lidwina Priliani, Edison Johar, Frilasita A Yudhaputri, Muhammad Rezki Rasyak, Sri Masyeni, Erly Sintya, Safarina G Malik, Khin Saw Myint, Amin Soebandrio |
| EPI_ISL_2101051 | Molecular Biology Laboratory, Faculty Medicine and Health Sciences, Warmadewa University | Eijkman Institute for Molecular Biology, National Agency for Research and Innovation; Molecular Biology Laboratory, Faculty Medicine and Health Sciences, Warmadewa University | Hidayat Trimarsanto, Lydia V. Panggalo, Iskandar Adnan, Sukma Oktavianthi, Lidwina Priliani, Edison Johar, Frilasita A Yudhaputri, Muhammad Rezki Rasyak, Willy Agustine, Sri Masyeni, Erly Sintya, Safarina G Malik, Khin Saw Myint, Amin Soebandrio |
| EPI_ISL_2101052 | Molecular Biology Laboratory, Faculty Medicine and Health Sciences, Warmadewa University | Eijkman Institute for Molecular Biology, National Agency for Research and Innovation; Molecular Biology Laboratory, | Lydia V. Panggalo, Iskandar Adnan, Sukma Oktavianthi, Lidwina Priliani, Edison Johar, Frilasita A Yudhaputri, Muhammad Rezki Rasyak, Willy Agustine, Hidayat Trimarsanto, Sri Masyeni, Erly Sintya, Safarina G Malik, Khin Saw Myint, Amin Soebandrio |

| | | | |
|---|---|---|---|
| | | Faculty Medicine and Health Sciences, Warmadewa University | |
| EPI_ISL_2101053 | Molecular Biology Laboratory, Faculty Medicine and Health Sciences, Warmadewa University | Eijkman Institute for Molecular Biology, National Agency for Research and Innovation; Molecular Biology Laboratory, Faculty Medicine and Health Sciences, Warmadewa University | Iskandar Adnan, Sukma Oktavianthi, Lidwina Priliani, Edison Johar, Frilasita A Yudhaputri, Muhammad Rezki Rasyak, Willy Agustine, Hidayat Trimarsanto, Lydia V. Panggalo, Sri Masyeni, Erly Sintya, Safarina G Malik, Khin Saw Myint, Amin Soebandrio |
| EPI_ISL_2101054 | Molecular Biology Laboratory, Faculty Medicine and Health Sciences, Warmadewa University | Eijkman Institute for Molecular Biology, National Agency for Research and Innovation; Molecular Biology Laboratory, Faculty Medicine and Health Sciences, Warmadewa University | Sukma Oktavianthi, Lidwina Priliani, Edison Johar, Frilasita A Yudhaputri, Muhammad Rezki Rasyak, Willy Agustine, Hidayat Trimarsanto, Lydia V. Panggalo, Iskandar Adnan, Sri Masyeni, Erly Sintya, Safarina G Malik, Khin Saw Myint, Amin Soebandrio |
| EPI_ISL_2101055 | Molecular Biology Laboratory, Faculty Medicine and Health Sciences, Warmadewa University | Eijkman Institute for Molecular Biology, National Agency for Research and Innovation; Molecular Biology Laboratory, Faculty Medicine and Health Sciences, Warmadewa University | Lidwina Priliani, Edison Johar, Frilasita A Yudhaputri, Muhammad Rezki Rasyak, Willy Agustine, Hidayat Trimarsanto, Lydia V. Panggalo, Iskandar Adnan, Sukma Oktavianthi, Sri Masyeni, Erly Sintya, Safarina G Malik, Khin Saw Myint, Amin Soebandrio |
| EPI_ISL_2101056 | Molecular Biology Laboratory, Faculty Medicine and Health Sciences, Warmadewa University | Eijkman Institute for Molecular Biology, National Agency for Research and Innovation; Molecular Biology Laboratory, Faculty Medicine and Health Sciences, Warmadewa University | Edison Johar, Frilasita A Yudhaputri, Muhammad Rezki Rasyak, Willy Agustine, Hidayat Trimarsanto, Lydia V. Panggalo, Iskandar Adnan, Sukma Oktavianthi, Lidwina Priliani, Sri Masyeni, Erly Sintya, Safarina G Malik, Khin Saw Myint, Amin Soebandrio |
| EPI_ISL_2101057 | Molecular Biology Laboratory, Faculty Medicine and Health Sciences, Warmadewa University | Eijkman Institute for Molecular Biology, National Agency for Research and Innovation; Molecular Biology Laboratory, Faculty Medicine and Health Sciences, Warmadewa University | Frilasita A Yudhaputri, Muhammad Rezki Rasyak, Willy Agustine, Hidayat Trimarsanto, Lydia V. Panggalo, Iskandar Adnan, Sukma Oktavianthi, Lidwina Priliani, Edison Johar, Sri Masyeni, Erly Sintya, Safarina G Malik, Khin Saw Myint, Amin Soebandrio |
| EPI_ISL_2101058 | Molecular Biology Laboratory, Faculty Medicine and Health Sciences, Warmadewa University | Eijkman Institute for Molecular Biology, National Agency for Research and Innovation; Molecular Biology Laboratory, Faculty Medicine and Health Sciences, Warmadewa University | Willy Agustine, Hidayat Trimarsanto, Lydia V. Panggalo, Iskandar Adnan, Sukma Oktavianthi, Lidwina Priliani, Edison Johar, Frilasita A Yudhaputri, Muhammad Rezki Rasyak, Sri Masyeni, Erly Sintya, Safarina G Malik, Khin Saw Myint, Amin Soebandrio |
| EPI_ISL_2101059 | Molecular Biology Laboratory, Faculty Medicine and Health Sciences, Warmadewa University | Eijkman Institute for Molecular Biology, National Agency for Research and Innovation; Molecular Biology Laboratory, Faculty Medicine and Health Sciences, Warmadewa University | Hidayat Trimarsanto, Lydia V. Panggalo, Iskandar Adnan, Sukma Oktavianthi, Lidwina Priliani, Edison Johar, Frilasita A Yudhaputri, Muhammad Rezki Rasyak, Willy Agustine, Sri Masyeni, Erly Sintya, Safarina G Malik, Khin Saw Myint, Amin Soebandrio |
| EPI_ISL_2101060 | Molecular Biology Laboratory, Faculty Medicine and Health Sciences, Warmadewa University | Eijkman Institute for Molecular Biology, National Agency for Research and Innovation; Molecular Biology Laboratory, Faculty Medicine and Health Sciences, Warmadewa University | Lydia V. Panggalo, Iskandar Adnan, Sukma Oktavianthi, Lidwina Priliani, Edison Johar, Frilasita A Yudhaputri, Muhammad Rezki Rasyak, Willy Agustine, Hidayat Trimarsanto, Sri Masyeni, Erly Sintya, Safarina G Malik, Khin Saw Myint, Amin Soebandrio |
| EPI_ISL_2101061 | Molecular Biology Laboratory, Faculty Medicine and Health Sciences, Warmadewa University | Eijkman Institute for Molecular Biology, National Agency for Research and Innovation; Molecular Biology Laboratory, Faculty Medicine and Health Sciences, Warmadewa University | Iskandar Adnan, Sukma Oktavianthi, Lidwina Priliani, Edison Johar, Frilasita A Yudhaputri, Muhammad Rezki Rasyak, Willy Agustine, Hidayat Trimarsanto, Lydia V. Panggalo, Sri Masyeni, Erly Sintya, Safarina G Malik, Khin Saw Myint, Amin Soebandrio |
| EPI_ISL_2101170 | Molecular Biology Laboratory, Faculty Medicine and Health Sciences, Warmadewa University | Eijkman Institute for Molecular Biology, National Agency for Research and Innovation; Molecular Biology Laboratory, Faculty Medicine and Health Sciences, Warmadewa University | Lidwina Priliani, Edison Johar, Frilasita A Yudhaputri, Muhammad Rezki Rasyak, Willy Agustine, Hidayat Trimarsanto, Lydia V. Panggalo, Iskandar Adnan, Sukma Oktavianthi, Sri Masyeni, Erly Sintya, Safarina G Malik, Khin Saw Myint, Amin Soebandrio |
| EPI_ISL_2101171 | Molecular Biology Laboratory, Faculty Medicine and Health Sciences, Warmadewa University | Eijkman Institute for Molecular Biology, National Agency for Research and Innovation; Molecular Biology Laboratory, Faculty Medicine and Health Sciences, Warmadewa University | Edison Johar, Frilasita A Yudhaputri, Muhammad Rezki Rasyak, Willy Agustine, Hidayat Trimarsanto, Lydia V. Panggalo, Iskandar Adnan, Sukma Oktavianthi, Lidwina Priliani, Sri Masyeni, Erly Sintya, Safarina G Malik, Khin Saw Myint, Amin Soebandrio |
| EPI_ISL_2101172 | Molecular Biology Laboratory, Faculty Medicine and Health Sciences, Warmadewa University | Eijkman Institute for Molecular Biology, National Agency for Research and Innovation; Molecular Biology Laboratory, Faculty Medicine and Health Sciences, Warmadewa University | Iskandar Adnan, Sukma Oktavianthi, Lidwina Priliani, Edison Johar, Frilasita A Yudhaputri, Muhammad Rezki Rasyak, Willy Agustine, Hidayat Trimarsanto, Lydia V. Panggalo, Sri Masyeni, Erly Sintya, Safarina G Malik, Khin Saw Myint, Amin Soebandrio |
| EPI_ISL_2101173 | Molecular Biology Laboratory, Faculty Medicine and Health Sciences, Warmadewa University | Eijkman Institute for Molecular Biology, National Agency for Research and Innovation; Molecular Biology Laboratory, Faculty Medicine and Health Sciences, Warmadewa University | Sukma Oktavianthi, Lidwina Priliani, Edison Johar, Frilasita A Yudhaputri, Muhammad Rezki Rasyak, Willy Agustine, Hidayat Trimarsanto, Lydia V. Panggalo, Iskandar Adnan, Sri Masyeni, Erly Sintya, Safarina G Malik, Khin Saw Myint, Amin Soebandrio |
| EPI_ISL_2101174 | Molecular Biology Laboratory, Faculty Medicine and Health Sciences, Warmadewa University | Eijkman Institute for Molecular Biology, National Agency for Research and Innovation; Molecular Biology Laboratory, Faculty Medicine and Health Sciences, Warmadewa University | Lidwina Priliani, Edison Johar, Frilasita A Yudhaputri, Muhammad Rezki Rasyak, Willy Agustine, Hidayat Trimarsanto, Lydia V. Panggalo, Iskandar Adnan, Sukma Oktavianthi, Sri Masyeni, Erly Sintya, Safarina G Malik, Khin Saw Myint, Amin Soebandrio |
| EPI_ISL_2101175 | Molecular Biology Laboratory, Faculty Medicine and Health Sciences, Warmadewa University | Eijkman Institute for Molecular Biology, National Agency for Research and Innovation; Molecular Biology Laboratory, Faculty Medicine and Health Sciences, Warmadewa University | Edison Johar, Frilasita A Yudhaputri, Muhammad Rezki Rasyak, Willy Agustine, Hidayat Trimarsanto, Lydia V. Panggalo, Iskandar Adnan, Sukma Oktavianthi, Lidwina Priliani, Sri Masyeni, Erly Sintya, Safarina G Malik, Khin Saw Myint, Amin Soebandrio |
| EPI_ISL_2101176 | Molecular Biology Laboratory, Faculty Medicine and Health Sciences, Warmadewa University | Eijkman Institute for Molecular Biology, National Agency for Research and Innovation; Molecular Biology Laboratory, Faculty Medicine and Health Sciences, Warmadewa University | Frilasita A Yudhaputri, Muhammad Rezki Rasyak, Willy Agustine, Hidayat Trimarsanto, Lydia V. Panggalo, Iskandar Adnan, Sukma Oktavianthi, Lidwina Priliani, Edison Johar, Sri Masyeni, Erly Sintya, Safarina G Malik, Khin Saw Myint, Amin Soebandrio |
| EPI_ISL_2101177 | Molecular Biology Laboratory, Faculty Medicine and Health Sciences, Warmadewa University | Eijkman Institute for Molecular Biology, National Agency for Research and Innovation; Molecular Biology Laboratory, Faculty Medicine and Health Sciences, Warmadewa University | Willy Agustine, Hidayat Trimarsanto, Lydia V. Panggalo, Iskandar Adnan, Sukma Oktavianthi, Lidwina Priliani, Edison Johar, Frilasita A Yudhaputri, Muhammad Rezki Rasyak, Sri Masyeni, Erly Sintya, Safarina G Malik, Khin Saw Myint, Amin Soebandrio |
| EPI_ISL_2101178 | Molecular Biology Laboratory, Faculty Medicine and Health Sciences, Warmadewa University | Eijkman Institute for Molecular Biology, National Agency for Research and Innovation; Molecular Biology Laboratory, Faculty Medicine and Health Sciences, Warmadewa University | Hidayat Trimarsanto, Lydia V. Panggalo, Iskandar Adnan, Sukma Oktavianthi, Lidwina Priliani, Edison Johar, Frilasita A Yudhaputri, Muhammad Rezki Rasyak, Willy Agustine, Sri Masyeni, Erly Sintya, Safarina G Malik, Khin Saw Myint, Amin Soebandrio |
| EPI_ISL_2101179 | Molecular Biology Laboratory, Faculty Medicine and Health Sciences, Warmadewa University | Eijkman Institute for Molecular Biology, National Agency for Research and Innovation; Molecular Biology Laboratory, Faculty Medicine and Health Sciences, Warmadewa University | Lydia V. Panggalo, Iskandar Adnan, Sukma Oktavianthi, Lidwina Priliani, Edison Johar, Frilasita A Yudhaputri, Muhammad Rezki Rasyak, Willy Agustine, Hidayat Trimarsanto, Sri Masyeni, Erly Sintya, Safarina G Malik, Khin Saw Myint, Amin Soebandrio |

| | | | |
|---|---|---|---|
| EPI_ISL_2101180 | Molecular Biology Laboratory, Faculty Medicine and Health Sciences, Warmadewa University | Eijkman Institute for Molecular Biology, National Agency for Research and Innovation; Molecular Biology Laboratory, Faculty Medicine and Health Sciences, Warmadewa University | Sukma Oktavianthi, Lidwina Priliani, Edison Johar, Frilasita A Yudhaputri, Muhammad Rezki Rasyak, Willy Agustine, Hidayat Trimarsanto, Lydia V. Panggalo, Iskandar Adnan, Sri Masyeni, Erly Sintya, Safarina G Malik, Khin Saw Myint, Amin Soebandrio |
| EPI_ISL_2101181 | Molecular Biology Laboratory, Faculty Medicine and Health Sciences, Warmadewa University | Eijkman Institute for Molecular Biology, National Agency for Research and Innovation; Molecular Biology Laboratory, Faculty Medicine and Health Sciences, Warmadewa University | Lidwina Priliani, Edison Johar, Frilasita A Yudhaputri, Muhammad Rezki Rasyak, Willy Agustine, Hidayat Trimarsanto, Lydia V. Panggalo, Iskandar Adnan, Sukma Oktavianthi, Sri Masyeni, Erly Sintya, Safarina G Malik, Khin Saw Myint, Amin Soebandrio |
| EPI_ISL_2101182 | Molecular Biology Laboratory, Faculty Medicine and Health Sciences, Warmadewa University | Eijkman Institute for Molecular Biology, National Agency for Research and Innovation; Molecular Biology Laboratory, Faculty Medicine and Health Sciences, Warmadewa University | Edison Johar, Frilasita A Yudhaputri, Muhammad Rezki Rasyak, Willy Agustine, Hidayat Trimarsanto, Lydia V. Panggalo, Iskandar Adnan, Sukma Oktavianthi, Lidwina Priliani, Sri Masyeni, Erly Sintya, Safarina G Malik, Khin Saw Myint, Amin Soebandrio |
| EPI_ISL_2101183 | Molecular Biology Laboratory, Faculty Medicine and Health Sciences, Warmadewa University | Eijkman Institute for Molecular Biology, National Agency for Research and Innovation; Molecular Biology Laboratory, Faculty Medicine and Health Sciences, Warmadewa University | Frilasita A Yudhaputri, Muhammad Rezki Rasyak, Willy Agustine, Hidayat Trimarsanto, Lydia V. Panggalo, Iskandar Adnan, Sukma Oktavianthi, Lidwina Priliani, Edison Johar, Sri Masyeni, Erly Sintya, Safarina G Malik, Khin Saw Myint, Amin Soebandrio |
| EPI_ISL_2101184 | Molecular Biology Laboratory, Faculty Medicine and Health Sciences, Warmadewa University | Eijkman Institute for Molecular Biology, National Agency for Research and Innovation; Molecular Biology Laboratory, Faculty Medicine and Health Sciences, Warmadewa University | Willy Agustine, Hidayat Trimarsanto, Lydia V. Panggalo, Iskandar Adnan, Sukma Oktavianthi, Lidwina Priliani, Edison Johar, Frilasita A Yudhaputri, Muhammad Rezki Rasyak, Sri Masyeni, Erly Sintya, Safarina G Malik, Khin Saw Myint, Amin Soebandrio |
| EPI_ISL_2101185 | Molecular Biology Laboratory, Faculty Medicine and Health Sciences, Warmadewa University | Eijkman Institute for Molecular Biology, National Agency for Research and Innovation; Molecular Biology Laboratory, Faculty Medicine and Health Sciences, Warmadewa University | Hidayat Trimarsanto, Lydia V. Panggalo, Iskandar Adnan, Sukma Oktavianthi, Lidwina Priliani, Edison Johar, Frilasita A Yudhaputri, Muhammad Rezki Rasyak, Willy Agustine, Sri Masyeni, Erly Sintya, Safarina G Malik, Khin Saw Myint, Amin Soebandrio |
| EPI_ISL_2101186 | Molecular Biology Laboratory, Faculty Medicine and Health Sciences, Warmadewa University | Eijkman Institute for Molecular Biology, National Agency for Research and Innovation; Molecular Biology Laboratory, Faculty Medicine and Health Sciences, Warmadewa University | Lydia V. Panggalo, Iskandar Adnan, Sukma Oktavianthi, Lidwina Priliani, Edison Johar, Frilasita A Yudhaputri, Muhammad Rezki Rasyak, Willy Agustine, Hidayat Trimarsanto, Sri Masyeni, Erly Sintya, Safarina G Malik, Khin Saw Myint, Amin Soebandrio |
| EPI_ISL_2101187 | Molecular Biology Laboratory, Faculty Medicine and Health Sciences, Warmadewa University | Eijkman Institute for Molecular Biology, National Agency for Research and Innovation; Molecular Biology Laboratory, Faculty Medicine and Health Sciences, Warmadewa University | Iskandar Adnan, Sukma Oktavianthi, Lidwina Priliani, Edison Johar, Frilasita A Yudhaputri, Muhammad Rezki Rasyak, Willy Agustine, Hidayat Trimarsanto, Lydia V. Panggalo, Sri Masyeni, Erly Sintya, Safarina G Malik, Khin Saw Myint, Amin Soebandrio |
| EPI_ISL_2101188 | Molecular Biology Laboratory, Faculty Medicine and Health Sciences, Warmadewa University | Eijkman Institute for Molecular Biology, National Agency for Research and Innovation; Molecular Biology Laboratory, Faculty Medicine and Health Sciences, Warmadewa University | Sukma Oktavianthi, Lidwina Priliani, Edison Johar, Frilasita A Yudhaputri, Muhammad Rezki Rasyak, Willy Agustine, Hidayat Trimarsanto, Lydia V. Panggalo, Iskandar Adnan, Sri Masyeni, Erly Sintya, Safarina G Malik, Khin Saw Myint, Amin Soebandrio |
| EPI_ISL_2101189 | Molecular Biology Laboratory, Faculty Medicine and Health Sciences, Warmadewa University | Eijkman Institute for Molecular Biology, National Agency for Research and Innovation; Molecular Biology Laboratory, Faculty Medicine and Health Sciences, Warmadewa University | Lidwina Priliani, Edison Johar, Frilasita A Yudhaputri, Muhammad Rezki Rasyak, Willy Agustine, Hidayat Trimarsanto, Lydia V. Panggalo, Iskandar Adnan, Sukma Oktavianthi, Sri Masyeni, Erly Sintya, Safarina G Malik, Khin Saw Myint, Amin Soebandrio |
| EPI_ISL_2101190 | Molecular Biology Laboratory, Faculty Medicine and Health Sciences, Warmadewa University | Eijkman Institute for Molecular Biology, National Agency for Research and Innovation; Molecular Biology Laboratory, Faculty Medicine and Health Sciences, Warmadewa University | Frilasita A Yudhaputri, Muhammad Rezki Rasyak, Willy Agustine, Hidayat Trimarsanto, Lydia V. Panggalo, Iskandar Adnan, Sukma Oktavianthi, Lidwina Priliani, Edison Johar, Sri Masyeni, Erly Sintya, Safarina G Malik, Khin Saw Myint, Amin Soebandrio |
| EPI_ISL_2101191 | Molecular Biology Laboratory, Faculty Medicine and Health Sciences, Warmadewa University | Eijkman Institute for Molecular Biology, National Agency for Research and Innovation; Molecular Biology Laboratory, Faculty Medicine and Health Sciences, Warmadewa University | Willy Agustine, Hidayat Trimarsanto, Lydia V. Panggalo, Iskandar Adnan, Sukma Oktavianthi, Lidwina Priliani, Edison Johar, Frilasita A Yudhaputri, Muhammad Rezki Rasyak, Sri Masyeni, Erly Sintya, Safarina G Malik, Khin Saw Myint, Amin Soebandrio |
| EPI_ISL_2101192 | Molecular Biology Laboratory, Faculty Medicine and Health Sciences, Warmadewa University | Eijkman Institute for Molecular Biology, National Agency for Research and Innovation; Molecular Biology Laboratory, Faculty Medicine and Health Sciences, Warmadewa University | Hidayat Trimarsanto, Lydia V. Panggalo, Iskandar Adnan, Sukma Oktavianthi, Lidwina Priliani, Edison Johar, Frilasita A Yudhaputri, Muhammad Rezki Rasyak, Willy Agustine, Sri Masyeni, Erly Sintya, Safarina G Malik, Khin Saw Myint, Amin Soebandrio |
| EPI_ISL_2101193 | Molecular Biology Laboratory, Faculty Medicine and Health Sciences, Warmadewa University | Eijkman Institute for Molecular Biology, National Agency for Research and Innovation; Molecular Biology Laboratory, Faculty Medicine and Health Sciences, Warmadewa University | Lydia V. Panggalo, Iskandar Adnan, Sukma Oktavianthi, Lidwina Priliani, Edison Johar, Frilasita A Yudhaputri, Muhammad Rezki Rasyak, Willy Agustine, Hidayat Trimarsanto, Sri Masyeni, Erly Sintya, Safarina G Malik, Khin Saw Myint, Amin Soebandrio |
| EPI_ISL_2101194 | Molecular Biology Laboratory, Faculty Medicine and Health Sciences, Warmadewa University | Eijkman Institute for Molecular Biology, National Agency for Research and Innovation; Molecular Biology Laboratory, Faculty Medicine and Health Sciences, Warmadewa University | Iskandar Adnan, Sukma Oktavianthi, Lidwina Priliani, Edison Johar, Frilasita A Yudhaputri, Muhammad Rezki Rasyak, Willy Agustine, Hidayat Trimarsanto, Lydia V. Panggalo, Sri Masyeni, Erly Sintya, Safarina G Malik, Khin Saw Myint, Amin Soebandrio |
| EPI_ISL_2101195 | Molecular Biology Laboratory, Faculty Medicine and Health Sciences, Warmadewa University | Eijkman Institute for Molecular Biology, National Agency for Research and Innovation; Molecular Biology Laboratory, Faculty Medicine and Health Sciences, Warmadewa University | Sukma Oktavianthi, Lidwina Priliani, Edison Johar, Frilasita A Yudhaputri, Muhammad Rezki Rasyak, Willy Agustine, Hidayat Trimarsanto, Lydia V. Panggalo, Iskandar Adnan, Sri Masyeni, Erly Sintya, Safarina G Malik, Khin Saw Myint, Amin Soebandrio |
| EPI_ISL_2101196 | Molecular Biology Laboratory, Faculty Medicine and Health Sciences, Warmadewa University | Eijkman Institute for Molecular Biology, National Agency for Research and Innovation; Molecular Biology Laboratory, Faculty Medicine and Health Sciences, Warmadewa University | Lidwina Priliani, Edison Johar, Frilasita A Yudhaputri, Muhammad Rezki Rasyak, Willy Agustine, Hidayat Trimarsanto, Lydia V. Panggalo, Iskandar Adnan, Sukma Oktavianthi, Sri Masyeni, Erly Sintya, Safarina G Malik, Khin Saw Myint, Amin Soebandrio |
| EPI_ISL_2101197 | Molecular Biology Laboratory, Faculty Medicine and Health Sciences, Warmadewa University | Eijkman Institute for Molecular Biology, National Agency for Research and Innovation; Molecular Biology Laboratory, Faculty Medicine and Health Sciences, Warmadewa University | Edison Johar, Frilasita A Yudhaputri, Muhammad Rezki Rasyak, Willy Agustine, Hidayat Trimarsanto, Lydia V. Panggalo, Iskandar Adnan, Sukma Oktavianthi, Lidwina Priliani, Sri Masyeni, Erly Sintya, Safarina G Malik, Khin Saw Myint, Amin Soebandrio |
| EPI_ISL_2101198 | Molecular Biology Laboratory, Faculty Medicine and Health Sciences, Warmadewa University | Eijkman Institute for Molecular Biology, National Agency for Research and Innovation; Molecular Biology Laboratory, Faculty Medicine and Health Sciences, Warmadewa University | Frilasita A Yudhaputri, Muhammad Rezki Rasyak, Willy Agustine, Hidayat Trimarsanto, Lydia V. Panggalo, Iskandar Adnan, Sukma Oktavianthi, Lidwina Priliani, Edison Johar, Sri Masyeni, Erly Sintya, Safarina G Malik, Khin Saw Myint, Amin Soebandrio |
| EPI_ISL_2101199 | Molecular Biology Laboratory, Faculty Medicine and Health Sciences, Warmadewa University | Eijkman Institute for Molecular Biology, National Agency for Research and Innovation; Molecular Biology Laboratory, | Willy Agustine, Hidayat Trimarsanto, Lydia V. Panggalo, Iskandar Adnan, Sukma Oktavianthi, Lidwina Priliani, Edison Johar, Frilasita A Yudhaputri, Muhammad Rezki Rasyak, Sri Masyeni, Erly Sintya, Safarina G Malik, Khin Saw Myint, Amin Soebandrio |

| | | | |
|---|---|---|---|
| | | Faculty Medicine and Health Sciences, Warmadewa University | |
| EPI_ISL_2101200 | Molecular Biology Laboratory, Faculty Medicine and Health Sciences, Warmadewa University | Eijkman Institute for Molecular Biology, National Agency for Research and Innovation; Molecular Biology Laboratory, Faculty Medicine and Health Sciences, Warmadewa University | Lydia V. Panggalo, Iskandar Adnan, Sukma Oktavianthi, Lidwina Priliani, Edison Johar, Frilasita A Yudhaputri, Muhammad Rezki Rasyak, Willy Agustine, Hidayat Trimarsanto, Sri Masyeni, Erly Sintya, Safarina G Malik, Khin Saw Myint, Amin Soebandrio |
| EPI_ISL_2101202 | Molecular Biology Laboratory, Faculty Medicine and Health Sciences, Warmadewa University | Eijkman Institute for Molecular Biology, National Agency for Research and Innovation; Molecular Biology Laboratory, Faculty Medicine and Health Sciences, Warmadewa University | Iskandar Adnan, Sukma Oktavianthi, Lidwina Priliani, Edison Johar, Frilasita A Yudhaputri, Muhammad Rezki Rasyak, Willy Agustine, Hidayat Trimarsanto, Lydia V. Panggalo, Sri Masyeni, Erly Sintya, Safarina G Malik, Khin Saw Myint, Amin Soebandrio |
| EPI_ISL_2101203 | Molecular Biology Laboratory, Faculty Medicine and Health Sciences, Warmadewa University | Eijkman Institute for Molecular Biology, National Agency for Research and Innovation; Molecular Biology Laboratory, Faculty Medicine and Health Sciences, Warmadewa University | Sukma Oktavianthi, Lidwina Priliani, Edison Johar, Frilasita A Yudhaputri, Muhammad Rezki Rasyak, Willy Agustine, Hidayat Trimarsanto, Lydia V. Panggalo, Iskandar Adnan, Sri Masyeni, Erly Sintya, Safarina G Malik, Khin Saw Myint, Amin Soebandrio |
| EPI_ISL_2101204 | Molecular Biology Laboratory, Faculty Medicine and Health Sciences, Warmadewa University | Eijkman Institute for Molecular Biology, National Agency for Research and Innovation; Molecular Biology Laboratory, Faculty Medicine and Health Sciences, Warmadewa University | Lidwina Priliani, Edison Johar, Frilasita A Yudhaputri, Muhammad Rezki Rasyak, Willy Agustine, Hidayat Trimarsanto, Lydia V. Panggalo, Iskandar Adnan, Sukma Oktavianthi, Sri Masyeni, Erly Sintya, Safarina G Malik, Khin Saw Myint, Amin Soebandrio |
| EPI_ISL_2101205 | Molecular Biology Laboratory, Faculty Medicine and Health Sciences, Warmadewa University | Eijkman Institute for Molecular Biology, National Agency for Research and Innovation; Molecular Biology Laboratory, Faculty Medicine and Health Sciences, Warmadewa University | Edison Johar, Frilasita A Yudhaputri, Muhammad Rezki Rasyak, Willy Agustine, Hidayat Trimarsanto, Lydia V. Panggalo, Iskandar Adnan, Sukma Oktavianthi, Lidwina Priliani, Sri Masyeni, Erly Sintya, Safarina G Malik, Khin Saw Myint, Amin Soebandrio |
| EPI_ISL_2101206 | Molecular Biology Laboratory, Faculty Medicine and Health Sciences, Warmadewa University | Eijkman Institute for Molecular Biology, National Agency for Research and Innovation; Molecular Biology Laboratory, Faculty Medicine and Health Sciences, Warmadewa University | Frilasita A Yudhaputri, Muhammad Rezki Rasyak, Willy Agustine, Hidayat Trimarsanto, Lydia V. Panggalo, Iskandar Adnan, Sukma Oktavianthi, Lidwina Priliani, Edison Johar, Sri Masyeni, Erly Sintya, Safarina G Malik, Khin Saw Myint, Amin Soebandrio |
| EPI_ISL_2101207 | Molecular Biology Laboratory, Faculty Medicine and Health Sciences, Warmadewa University | Eijkman Institute for Molecular Biology, National Agency for Research and Innovation; Molecular Biology Laboratory, Faculty Medicine and Health Sciences, Warmadewa University | Willy Agustine, Hidayat Trimarsanto, Lydia V. Panggalo, Iskandar Adnan, Sukma Oktavianthi, Lidwina Priliani, Edison Johar, Frilasita A Yudhaputri, Muhammad Rezki Rasyak, Sri Masyeni, Erly Sintya, Safarina G Malik, Khin Saw Myint, Amin Soebandrio |
| EPI_ISL_2101208 | Molecular Biology Laboratory, Faculty Medicine and Health Sciences, Warmadewa University | Eijkman Institute for Molecular Biology, National Agency for Research and Innovation; Molecular Biology Laboratory, Faculty Medicine and Health Sciences, Warmadewa University | Hidayat Trimarsanto, Lydia V. Panggalo, Iskandar Adnan, Sukma Oktavianthi, Lidwina Priliani, Edison Johar, Frilasita A Yudhaputri, Muhammad Rezki Rasyak, Willy Agustine, Sri Masyeni, Erly Sintya, Safarina G Malik, Khin Saw Myint, Amin Soebandrio |
| EPI_ISL_2101209 | Molecular Biology Laboratory, Faculty Medicine and Health Sciences, Warmadewa University | Eijkman Institute for Molecular Biology, National Agency for Research and Innovation; Molecular Biology Laboratory, Faculty Medicine and Health Sciences, Warmadewa University | Lydia V. Panggalo, Iskandar Adnan, Sukma Oktavianthi, Lidwina Priliani, Edison Johar, Frilasita A Yudhaputri, Muhammad Rezki Rasyak, Willy Agustine, Hidayat Trimarsanto, Sri Masyeni, Erly Sintya, Safarina G Malik, Khin Saw Myint, Amin Soebandrio |
| EPI_ISL_2101210 | Molecular Biology Laboratory, Faculty Medicine and Health Sciences, Warmadewa University | Eijkman Institute for Molecular Biology, National Agency for Research and Innovation; Molecular Biology Laboratory, Faculty Medicine and Health Sciences, Warmadewa University | Iskandar Adnan, Sukma Oktavianthi, Lidwina Priliani, Edison Johar, Frilasita A Yudhaputri, Muhammad Rezki Rasyak, Willy Agustine, Hidayat Trimarsanto, Lydia V. Panggalo, Sri Masyeni, Erly Sintya, Safarina G Malik, Khin Saw Myint, Amin Soebandrio |
| EPI_ISL_2101211 | Molecular Biology Laboratory, Faculty Medicine and Health Sciences, Warmadewa University | Eijkman Institute for Molecular Biology, National Agency for Research and Innovation; Molecular Biology Laboratory, Faculty Medicine and Health Sciences, Warmadewa University | Lidwina Priliani, Edison Johar, Frilasita A Yudhaputri, Muhammad Rezki Rasyak, Willy Agustine, Hidayat Trimarsanto, Lydia V. Panggalo, Iskandar Adnan, Sukma Oktavianthi, Sri Masyeni, Erly Sintya, Safarina G Malik, Khin Saw Myint, Amin Soebandrio |
| EPI_ISL_2101212 | Molecular Biology Laboratory, Faculty Medicine and Health Sciences, Warmadewa University | Eijkman Institute for Molecular Biology, National Agency for Research and Innovation; Molecular Biology Laboratory, Faculty Medicine and Health Sciences, Warmadewa University | Edison Johar, Frilasita A Yudhaputri, Muhammad Rezki Rasyak, Willy Agustine, Hidayat Trimarsanto, Lydia V. Panggalo, Iskandar Adnan, Sukma Oktavianthi, Lidwina Priliani, Sri Masyeni, Erly Sintya, Safarina G Malik, Khin Saw Myint, Amin Soebandrio |
| EPI_ISL_2101213 | Molecular Biology Laboratory, Faculty Medicine and Health Sciences, Warmadewa University | Eijkman Institute for Molecular Biology, National Agency for Research and Innovation; Molecular Biology Laboratory, Faculty Medicine and Health Sciences, Warmadewa University | Frilasita A Yudhaputri, Muhammad Rezki Rasyak, Willy Agustine, Hidayat Trimarsanto, Lydia V. Panggalo, Iskandar Adnan, Sukma Oktavianthi, Lidwina Priliani, Edison Johar, Sri Masyeni, Erly Sintya, Safarina G Malik, Khin Saw Myint, Amin Soebandrio |
| EPI_ISL_2101214 | Molecular Biology Laboratory, Faculty Medicine and Health Sciences, Warmadewa University | Eijkman Institute for Molecular Biology, National Agency for Research and Innovation; Molecular Biology Laboratory, Faculty Medicine and Health Sciences, Warmadewa University | Willy Agustine, Hidayat Trimarsanto, Lydia V. Panggalo, Iskandar Adnan, Sukma Oktavianthi, Lidwina Priliani, Edison Johar, Frilasita A Yudhaputri, Muhammad Rezki Rasyak, Sri Masyeni, Erly Sintya, Safarina G Malik, Khin Saw Myint, Amin Soebandrio |
| EPI_ISL_2101215 | Molecular Biology Laboratory, Faculty Medicine and Health Sciences, Warmadewa University | Eijkman Institute for Molecular Biology, National Agency for Research and Innovation; Molecular Biology Laboratory, Faculty Medicine and Health Sciences, Warmadewa University | Hidayat Trimarsanto, Lydia V. Panggalo, Iskandar Adnan, Sukma Oktavianthi, Lidwina Priliani, Edison Johar, Frilasita A Yudhaputri, Muhammad Rezki Rasyak, Willy Agustine, Sri Masyeni, Erly Sintya, Safarina G Malik, Khin Saw Myint, Amin Soebandrio |
| EPI_ISL_2101236 | RSUD Klungkung | Eijkman Institute for Molecular Biology, National Agency for Research and Innovation; Molecular Biology Laboratory, Faculty Medicine and Health Sciences, Warmadewa University | Frilasita A Yudhaputri, Muhammad Rezki Rasyak, Willy Agustine, Hidayat Trimarsanto, Lydia V. Panggalo, Iskandar Adnan, Sukma Oktavianthi, Lidwina Priliani, Edison Johar, Sri Masyeni, Erly Sintya, I Komang Parwata, Safarina G Malik, Khin Saw Myint, Amin Soebandrio |
| EPI_ISL_2101237 | RSUD Klungkung | Eijkman Institute for Molecular Biology, National Agency for Research and Innovation; Molecular Biology Laboratory, Faculty Medicine and Health Sciences, Warmadewa University | Willy Agustine, Hidayat Trimarsanto, Lydia V. Panggalo, Iskandar Adnan, Sukma Oktavianthi, Lidwina Priliani, Edison Johar, Frilasita A Yudhaputri, Muhammad Rezki Rasyak, Sri Masyeni, Erly Sintya, I Komang Parwata, Safarina G Malik, Khin Saw Myint, Amin Soebandrio |
| EPI_ISL_2101238 | RSUD Klungkung | Eijkman Institute for Molecular Biology, National Agency for Research and Innovation; Molecular Biology Laboratory, Faculty Medicine and Health Sciences, Warmadewa University | Iskandar Adnan, Sukma Oktavianthi, Lidwina Priliani, Edison Johar, Frilasita A Yudhaputri, Muhammad Rezki Rasyak, Willy Agustine, Hidayat Trimarsanto, Lydia V. Panggalo, Sri Masyeni, Erly Sintya, I Komang Parwata, Safarina G Malik, Khin Saw Myint, Amin Soebandrio |
| EPI_ISL_2101239 | RSUD Klungkung | Eijkman Institute for Molecular Biology, National Agency for Research and Innovation; Molecular Biology Laboratory, Faculty Medicine and Health Sciences, Warmadewa University | Lidwina Priliani, Edison Johar, Frilasita A Yudhaputri, Muhammad Rezki Rasyak, Willy Agustine, Hidayat Trimarsanto, Lydia V. Panggalo, Iskandar Adnan, Sukma Oktavianthi, Sri Masyeni, Erly Sintya, I Komang Parwata, Safarina G Malik, Khin Saw Myint, Amin Soebandrio |

| | | | |
|---|---|---|---|
| EPI_ISL_2101240 | RSUD Klungkung | Eijkman Institute for Molecular Biology, National Agency for Research and Innovation; Molecular Biology Laboratory, Faculty Medicine and Health Sciences, Warmadewa University | Edison Johar, Frilasita A Yudhaputri, Muhammad Rezki Rasyak, Willy Agustine, Hidayat Trimarsanto, Lydia V. Panggalo, Iskandar Adnan, Sukma Oktavianthi, Lidwina Priliani, Sri Masyeni, Erly Sintya, I Komang Parwata, Safarina G Malik, Khin Saw Myint, Amin Soebandrio |
| EPI_ISL_2101241 | RSUD Klungkung | Eijkman Institute for Molecular Biology, National Agency for Research and Innovation; Molecular Biology Laboratory, Faculty Medicine and Health Sciences, Warmadewa University | Frilasita A Yudhaputri, Muhammad Rezki Rasyak, Willy Agustine, Hidayat Trimarsanto, Lydia V. Panggalo, Iskandar Adnan, Sukma Oktavianthi, Lidwina Priliani, Edison Johar, Sri Masyeni, Erly Sintya, I Komang Parwata, Safarina G Malik, Khin Saw Myint, Amin Soebandrio |
| EPI_ISL_2101242 | RSUD Klungkung | Eijkman Institute for Molecular Biology, National Agency for Research and Innovation; Molecular Biology Laboratory, Faculty Medicine and Health Sciences, Warmadewa University | Muhammad Rezki Rasyak, Willy Agustine, Hidayat Trimarsanto, Lydia V. Panggalo, Iskandar Adnan, Sukma Oktavianthi, Lidwina Priliani, Edison Johar, Frilasita A Yudhaputri, Sri Masyeni, Erly Sintya, I Komang Parwata, Safarina G Malik, Khin Saw Myint, Amin Soebandrio |
| EPI_ISL_2101243 | RSUD Klungkung | Eijkman Institute for Molecular Biology, National Agency for Research and Innovation; Molecular Biology Laboratory, Faculty Medicine and Health Sciences, Warmadewa University | Willy Agustine, Hidayat Trimarsanto, Lydia V. Panggalo, Iskandar Adnan, Sukma Oktavianthi, Lidwina Priliani, Edison Johar, Frilasita A Yudhaputri, Muhammad Rezki Rasyak, Sri Masyeni, Erly Sintya, I Komang Parwata, Safarina G Malik, Khin Saw Myint, Amin Soebandrio |
| EPI_ISL_2101244 | RSUD Klungkung | Eijkman Institute for Molecular Biology, National Agency for Research and Innovation; Molecular Biology Laboratory, Faculty Medicine and Health Sciences, Warmadewa University | Lydia V. Panggalo, Iskandar Adnan, Sukma Oktavianthi, Lidwina Priliani, Edison Johar, Frilasita A Yudhaputri, Muhammad Rezki Rasyak, Willy Agustine, Hidayat Trimarsanto, Sri Masyeni, Erly Sintya, I Komang Parwata, Safarina G Malik, Khin Saw Myint, Amin Soebandrio |
| EPI_ISL_2101245 | RSUD Klungkung | Eijkman Institute for Molecular Biology, National Agency for Research and Innovation; Molecular Biology Laboratory, Faculty Medicine and Health Sciences, Warmadewa University | Iskandar Adnan, Sukma Oktavianthi, Lidwina Priliani, Edison Johar, Frilasita A Yudhaputri, Muhammad Rezki Rasyak, Willy Agustine, Hidayat Trimarsanto, Lydia V. Panggalo, Sri Masyeni, Erly Sintya, I Komang Parwata, Safarina G Malik, Khin Saw Myint, Amin Soebandrio |
| EPI_ISL_2101256 | RSUD Mangusada | Eijkman Institute for Molecular Biology, National Agency for Research and Innovation; Molecular Biology Laboratory, Faculty Medicine and Health Sciences, Warmadewa University | Edison Johar, Frilasita A Yudhaputri, Muhammad Rezki Rasyak, Willy Agustine, Hidayat Trimarsanto, Lydia V. Panggalo, Iskandar Adnan, Sukma Oktavianthi, Lidwina Priliani, Sri Masyeni, Erly Sintya, Ida Ayu Wayan Mahayani, Safarina G Malik, Khin Saw Myint, Amin Soebandrio |
| EPI_ISL_2101257 | RSUD Mangusada | Eijkman Institute for Molecular Biology, National Agency for Research and Innovation; Molecular Biology Laboratory, Faculty Medicine and Health Sciences, Warmadewa University | Frilasita A Yudhaputri, Muhammad Rezki Rasyak, Willy Agustine, Hidayat Trimarsanto, Lydia V. Panggalo, Iskandar Adnan, Sukma Oktavianthi, Lidwina Priliani, Edison Johar, Sri Masyeni, Erly Sintya, Ida Ayu Wayan Mahayani, Safarina G Malik, Khin Saw Myint, Amin Soebandrio |
| EPI_ISL_2101258 | RSUD Mangusada | Eijkman Institute for Molecular Biology, National Agency for Research and Innovation; Molecular Biology Laboratory, Faculty Medicine and Health Sciences, Warmadewa University | Muhammad Rezki Rasyak, Willy Agustine, Hidayat Trimarsanto, Lydia V. Panggalo, Iskandar Adnan, Sukma Oktavianthi, Lidwina Priliani, Edison Johar, Frilasita A Yudhaputri, Sri Masyeni, Erly Sintya, Ida Ayu Wayan Mahayani, Safarina G Malik, Khin Saw Myint, Amin Soebandrio |
| EPI_ISL_2101259 | RSUD Mangusada | Eijkman Institute for Molecular Biology, National Agency for Research and Innovation; Molecular Biology Laboratory, Faculty Medicine and Health Sciences, Warmadewa University | Willy Agustine, Hidayat Trimarsanto, Lydia V. Panggalo, Iskandar Adnan, Sukma Oktavianthi, Lidwina Priliani, Edison Johar, Frilasita A Yudhaputri, Muhammad Rezki Rasyak, Sri Masyeni, Erly Sintya, Ida Ayu Wayan Mahayani, Safarina G Malik, Khin Saw Myint, Amin Soebandrio |
| EPI_ISL_2101260 | RSUD Mangusada | Eijkman Institute for Molecular Biology, National Agency for Research and Innovation; Molecular Biology Laboratory, Faculty Medicine and Health Sciences, Warmadewa University | Hidayat Trimarsanto, Lydia V. Panggalo, Iskandar Adnan, Sukma Oktavianthi, Lidwina Priliani, Edison Johar, Frilasita A Yudhaputri, Muhammad Rezki Rasyak, Willy Agustine, Sri Masyeni, Erly Sintya, Ida Ayu Wayan Mahayani, Safarina G Malik, Khin Saw Myint, Amin Soebandrio |
| EPI_ISL_2101261 | RSUD Mangusada | Eijkman Institute for Molecular Biology, National Agency for Research and Innovation; Molecular Biology Laboratory, Faculty Medicine and Health Sciences, Warmadewa University | Lydia V. Panggalo, Iskandar Adnan, Sukma Oktavianthi, Lidwina Priliani, Edison Johar, Frilasita A Yudhaputri, Muhammad Rezki Rasyak, Willy Agustine, Hidayat Trimarsanto, Sri Masyeni, Erly Sintya, Ida Ayu Wayan Mahayani, Safarina G Malik, Khin Saw Myint, Amin Soebandrio |
| EPI_ISL_2101262 | RSUD Mangusada | Eijkman Institute for Molecular Biology, National Agency for Research and Innovation; Molecular Biology Laboratory, Faculty Medicine and Health Sciences, Warmadewa University | Iskandar Adnan, Sukma Oktavianthi, Lidwina Priliani, Edison Johar, Frilasita A Yudhaputri, Muhammad Rezki Rasyak, Willy Agustine, Hidayat Trimarsanto, Lydia V. Panggalo, Sri Masyeni, Erly Sintya, Ida Ayu Wayan Mahayani, Safarina G Malik, Khin Saw Myint, Amin Soebandrio |
| EPI_ISL_2101263 | RSUD Mangusada | Eijkman Institute for Molecular Biology, National Agency for Research and Innovation; Molecular Biology Laboratory, Faculty Medicine and Health Sciences, Warmadewa University | Sukma Oktavianthi, Lidwina Priliani, Edison Johar, Frilasita A Yudhaputri, Muhammad Rezki Rasyak, Willy Agustine, Hidayat Trimarsanto, Lydia V. Panggalo, Iskandar Adnan, Sri Masyeni, Erly Sintya, Ida Ayu Wayan Mahayani, Safarina G Malik, Khin Saw Myint, Amin Soebandrio |
| EPI_ISL_2101264 | RSUD Mangusada | Eijkman Institute for Molecular Biology, National Agency for Research and Innovation; Molecular Biology Laboratory, Faculty Medicine and Health Sciences, Warmadewa University | Lidwina Priliani, Edison Johar, Frilasita A Yudhaputri, Muhammad Rezki Rasyak, Willy Agustine, Hidayat Trimarsanto, Lydia V. Panggalo, Iskandar Adnan, Sukma Oktavianthi, Sri Masyeni, Erly Sintya, Ida Ayu Wayan Mahayani, Safarina G Malik, Khin Saw Myint, Amin Soebandrio |
| EPI_ISL_2101265 | RSUD Mangusada | Eijkman Institute for Molecular Biology, National Agency for Research and Innovation; Molecular Biology Laboratory, Faculty Medicine and Health Sciences, Warmadewa University | Edison Johar, Frilasita A Yudhaputri, Muhammad Rezki Rasyak, Willy Agustine, Hidayat Trimarsanto, Lydia V. Panggalo, Iskandar Adnan, Sukma Oktavianthi, Lidwina Priliani, Sri Masyeni, Erly Sintya, Ida Ayu Wayan Mahayani, Safarina G Malik, Khin Saw Myint, Amin Soebandrio |
| EPI_ISL_2101266 | RSUD Mangusada | Eijkman Institute for Molecular Biology, National Agency for Research and Innovation; Molecular Biology Laboratory, Faculty Medicine and Health Sciences, Warmadewa University | Frilasita A Yudhaputri, Muhammad Rezki Rasyak, Willy Agustine, Hidayat Trimarsanto, Lydia V. Panggalo, Iskandar Adnan, Sukma Oktavianthi, Lidwina Priliani, Edison Johar, Sri Masyeni, Erly Sintya, Ida Ayu Wayan Mahayani, Safarina G Malik, Khin Saw Myint, Amin Soebandrio |
| EPI_ISL_2101267 | RSUD Mangusada | Eijkman Institute for Molecular Biology, National Agency for Research and Innovation; Molecular Biology Laboratory, Faculty Medicine and Health Sciences, Warmadewa University | Muhammad Rezki Rasyak, Willy Agustine, Hidayat Trimarsanto, Lydia V. Panggalo, Iskandar Adnan, Sukma Oktavianthi, Lidwina Priliani, Edison Johar, Frilasita A Yudhaputri, Sri Masyeni, Erly Sintya, Ida Ayu Wayan Mahayani, Safarina G Malik, Khin Saw Myint, Amin Soebandrio |
| EPI_ISL_2101268 | RSUD Mangusada | Eijkman Institute for Molecular Biology, National Agency for Research and Innovation; Molecular Biology Laboratory, Faculty Medicine and Health Sciences, Warmadewa University | Willy Agustine, Hidayat Trimarsanto, Lydia V. Panggalo, Iskandar Adnan, Sukma Oktavianthi, Lidwina Priliani, Edison Johar, Frilasita A Yudhaputri, Muhammad Rezki Rasyak, Sri Masyeni, Erly Sintya, Ida Ayu Wayan Mahayani, Safarina G Malik, Khin Saw Myint, Amin Soebandrio |
| EPI_ISL_2101269 | RSUD Mangusada | Eijkman Institute for Molecular Biology, National Agency for Research and Innovation; Molecular Biology Laboratory, | Hidayat Trimarsanto, Lydia V. Panggalo, Iskandar Adnan, Sukma Oktavianthi, Lidwina Priliani, Edison Johar, Frilasita A Yudhaputri, Muhammad Rezki Rasyak, Willy Agustine, Sri Masyeni, Erly Sintya, Ida Ayu Wayan Mahayani, Safarina G Malik, Khin Saw Myint, Amin Soebandrio |

| | | | |
|---|---|---|---|
| | | Faculty Medicine and Health Sciences, Warmadewa University | |
| EPI_ISL_2101270 | RSUD Mangusada | Eijkman Institute for Molecular Biology, National Agency for Research and Innovation; Molecular Biology Laboratory, Faculty Medicine and Health Sciences, Warmadewa University | Lydia V. Panggalo, Iskandar Adnan, Sukma Oktavianthi, Lidwina Priliani, Edison Johar, Frilasita A Yudhaputri, Muhammad Rezki Rasyak, Willy Agustine, Hidayat Trimarsanto, Sri Masyeni, Erly Sintya, Ida Ayu Wayan Mahayani, Safarina G Malik, Khin Saw Myint, Amin Soebandrio |
| EPI_ISL_2101271 | RSUD Mangusada | Eijkman Institute for Molecular Biology, National Agency for Research and Innovation; Molecular Biology Laboratory, Faculty Medicine and Health Sciences, Warmadewa University | Iskandar Adnan, Sukma Oktavianthi, Lidwina Priliani, Edison Johar, Frilasita A Yudhaputri, Muhammad Rezki Rasyak, Willy Agustine, Hidayat Trimarsanto, Lydia V. Panggalo, Sri Masyeni, Erly Sintya, Ida Ayu Wayan Mahayani, Safarina G Malik, Khin Saw Myint, Amin Soebandrio |
| EPI_ISL_2101272 | RSUD Mangusada | Eijkman Institute for Molecular Biology, National Agency for Research and Innovation; Molecular Biology Laboratory, Faculty Medicine and Health Sciences, Warmadewa University | Sukma Oktavianthi, Lidwina Priliani, Edison Johar, Frilasita A Yudhaputri, Muhammad Rezki Rasyak, Willy Agustine, Hidayat Trimarsanto, Lydia V. Panggalo, Iskandar Adnan, Sri Masyeni, Erly Sintya, Ida Ayu Wayan Mahayani, Safarina G Malik, Khin Saw Myint, Amin Soebandrio |
| EPI_ISL_2101273 | RSUD Mangusada | Eijkman Institute for Molecular Biology, National Agency for Research and Innovation; Molecular Biology Laboratory, Faculty Medicine and Health Sciences, Warmadewa University | Lidwina Priliani, Edison Johar, Frilasita A Yudhaputri, Muhammad Rezki Rasyak, Willy Agustine, Hidayat Trimarsanto, Lydia V. Panggalo, Iskandar Adnan, Sukma Oktavianthi, Sri Masyeni, Erly Sintya, Ida Ayu Wayan Mahayani, Safarina G Malik, Khin Saw Myint, Amin Soebandrio |
| EPI_ISL_2101274 | RSUD Mangusada | Eijkman Institute for Molecular Biology, National Agency for Research and Innovation; Molecular Biology Laboratory, Faculty Medicine and Health Sciences, Warmadewa University | Edison Johar, Frilasita A Yudhaputri, Muhammad Rezki Rasyak, Willy Agustine, Hidayat Trimarsanto, Lydia V. Panggalo, Iskandar Adnan, Sukma Oktavianthi, Lidwina Priliani, Sri Masyeni, Erly Sintya, Ida Ayu Wayan Mahayani, Safarina G Malik, Khin Saw Myint, Amin Soebandrio |
| EPI_ISL_2101275 | RSUD Mangusada | Eijkman Institute for Molecular Biology, National Agency for Research and Innovation; Molecular Biology Laboratory, Faculty Medicine and Health Sciences, Warmadewa University | Frilasita A Yudhaputri, Muhammad Rezki Rasyak, Willy Agustine, Hidayat Trimarsanto, Lydia V. Panggalo, Iskandar Adnan, Sukma Oktavianthi, Lidwina Priliani, Edison Johar, Sri Masyeni, Erly Sintya, Ida Ayu Wayan Mahayani, Safarina G Malik, Khin Saw Myint, Amin Soebandrio |
| EPI_ISL_2101276 | RSUD Mangusada | Eijkman Institute for Molecular Biology, National Agency for Research and Innovation; Molecular Biology Laboratory, Faculty Medicine and Health Sciences, Warmadewa University | Muhammad Rezki Rasyak, Willy Agustine, Hidayat Trimarsanto, Lydia V. Panggalo, Iskandar Adnan, Sukma Oktavianthi, Lidwina Priliani, Edison Johar, Frilasita A Yudhaputri, Sri Masyeni, Erly Sintya, Ida Ayu Wayan Mahayani, Safarina G Malik, Khin Saw Myint, Amin Soebandrio |
| EPI_ISL_2101277 | RSUD Mangusada | Eijkman Institute for Molecular Biology, National Agency for Research and Innovation; Molecular Biology Laboratory, Faculty Medicine and Health Sciences, Warmadewa University | Willy Agustine, Hidayat Trimarsanto, Lydia V. Panggalo, Iskandar Adnan, Sukma Oktavianthi, Lidwina Priliani, Edison Johar, Frilasita A Yudhaputri, Muhammad Rezki Rasyak, Sri Masyeni, Erly Sintya, Ida Ayu Wayan Mahayani, Safarina G Malik, Khin Saw Myint, Amin Soebandrio |
| EPI_ISL_2101278 | RSUD Mangusada | Eijkman Institute for Molecular Biology, National Agency for Research and Innovation; Molecular Biology Laboratory, Faculty Medicine and Health Sciences, Warmadewa University | Hidayat Trimarsanto, Lydia V. Panggalo, Iskandar Adnan, Sukma Oktavianthi, Lidwina Priliani, Edison Johar, Frilasita A Yudhaputri, Muhammad Rezki Rasyak, Willy Agustine, Sri Masyeni, Erly Sintya, Ida Ayu Wayan Mahayani, Safarina G Malik, Khin Saw Myint, Amin Soebandrio |
| EPI_ISL_2101279 | RSUD Mangusada | Eijkman Institute for Molecular Biology, National Agency for Research and Innovation; Molecular Biology Laboratory, Faculty Medicine and Health Sciences, Warmadewa University | Lydia V. Panggalo, Iskandar Adnan, Sukma Oktavianthi, Lidwina Priliani, Edison Johar, Frilasita A Yudhaputri, Muhammad Rezki Rasyak, Willy Agustine, Hidayat Trimarsanto, Sri Masyeni, Erly Sintya, Ida Ayu Wayan Mahayani, Safarina G Malik, Khin Saw Myint, Amin Soebandrio |
| EPI_ISL_2101280 | RSUD Mangusada | Eijkman Institute for Molecular Biology, National Agency for Research and Innovation; Molecular Biology Laboratory, Faculty Medicine and Health Sciences, Warmadewa University | Iskandar Adnan, Sukma Oktavianthi, Lidwina Priliani, Edison Johar, Frilasita A Yudhaputri, Muhammad Rezki Rasyak, Willy Agustine, Hidayat Trimarsanto, Lydia V. Panggalo, Sri Masyeni, Erly Sintya, Ida Ayu Wayan Mahayani, Safarina G Malik, Khin Saw Myint, Amin Soebandrio |
| EPI_ISL_2101281 | RSUD Mangusada | Eijkman Institute for Molecular Biology, National Agency for Research and Innovation; Molecular Biology Laboratory, Faculty Medicine and Health Sciences, Warmadewa University | Sukma Oktavianthi, Lidwina Priliani, Edison Johar, Frilasita A Yudhaputri, Muhammad Rezki Rasyak, Willy Agustine, Hidayat Trimarsanto, Lydia V. Panggalo, Iskandar Adnan, Sri Masyeni, Erly Sintya, Ida Ayu Wayan Mahayani, Safarina G Malik, Khin Saw Myint, Amin Soebandrio |
| EPI_ISL_2101282 | RSUD Mangusada | Eijkman Institute for Molecular Biology, National Agency for Research and Innovation; Molecular Biology Laboratory, Faculty Medicine and Health Sciences, Warmadewa University | Lidwina Priliani, Edison Johar, Frilasita A Yudhaputri, Muhammad Rezki Rasyak, Willy Agustine, Hidayat Trimarsanto, Lydia V. Panggalo, Iskandar Adnan, Sukma Oktavianthi, Sri Masyeni, Erly Sintya, Ida Ayu Wayan Mahayani, Safarina G Malik, Khin Saw Myint, Amin Soebandrio |
| EPI_ISL_2101283 | RSUD Mangusada | Eijkman Institute for Molecular Biology, National Agency for Research and Innovation; Molecular Biology Laboratory, Faculty Medicine and Health Sciences, Warmadewa University | Edison Johar, Frilasita A Yudhaputri, Muhammad Rezki Rasyak, Willy Agustine, Hidayat Trimarsanto, Lydia V. Panggalo, Iskandar Adnan, Sukma Oktavianthi, Lidwina Priliani, Sri Masyeni, Erly Sintya, Ida Ayu Wayan Mahayani, Safarina G Malik, Khin Saw Myint, Amin Soebandrio |
| EPI_ISL_2101284 | RSUD Mangusada | Eijkman Institute for Molecular Biology, National Agency for Research and Innovation; Molecular Biology Laboratory, Faculty Medicine and Health Sciences, Warmadewa University | Frilasita A Yudhaputri, Muhammad Rezki Rasyak, Willy Agustine, Hidayat Trimarsanto, Lydia V. Panggalo, Iskandar Adnan, Sukma Oktavianthi, Lidwina Priliani, Edison Johar, Sri Masyeni, Erly Sintya, Ida Ayu Wayan Mahayani, Safarina G Malik, Khin Saw Myint, Amin Soebandrio |
| EPI_ISL_2101285 | RSUD Mangusada | Eijkman Institute for Molecular Biology, National Agency for Research and Innovation; Molecular Biology Laboratory, Faculty Medicine and Health Sciences, Warmadewa University | Muhammad Rezki Rasyak, Willy Agustine, Hidayat Trimarsanto, Lydia V. Panggalo, Iskandar Adnan, Sukma Oktavianthi, Lidwina Priliani, Edison Johar, Frilasita A Yudhaputri, Sri Masyeni, Erly Sintya, Ida Ayu Wayan Mahayani, Safarina G Malik, Khin Saw Myint, Amin Soebandrio |
| EPI_ISL_2101286 | RSUD Mangusada | Eijkman Institute for Molecular Biology, National Agency for Research and Innovation; Molecular Biology Laboratory, Faculty Medicine and Health Sciences, Warmadewa University | Willy Agustine, Hidayat Trimarsanto, Lydia V. Panggalo, Iskandar Adnan, Sukma Oktavianthi, Lidwina Priliani, Edison Johar, Frilasita A Yudhaputri, Muhammad Rezki Rasyak, Sri Masyeni, Erly Sintya, Ida Ayu Wayan Mahayani, Safarina G Malik, Khin Saw Myint, Amin Soebandrio |
| EPI_ISL_2101287 | RSUD Mangusada | Eijkman Institute for Molecular Biology, National Agency for Research and Innovation; Molecular Biology Laboratory, Faculty Medicine and Health Sciences, Warmadewa University | Hidayat Trimarsanto, Lydia V. Panggalo, Iskandar Adnan, Sukma Oktavianthi, Lidwina Priliani, Edison Johar, Frilasita A Yudhaputri, Muhammad Rezki Rasyak, Willy Agustine, Sri Masyeni, Erly Sintya, Ida Ayu Wayan Mahayani, Safarina G Malik, Khin Saw Myint, Amin Soebandrio |
| EPI_ISL_2101288 | RSUD Mangusada | Eijkman Institute for Molecular Biology, National Agency for Research and Innovation; Molecular Biology Laboratory, Faculty Medicine and Health Sciences, Warmadewa University | Lydia V. Panggalo, Iskandar Adnan, Sukma Oktavianthi, Lidwina Priliani, Edison Johar, Frilasita A Yudhaputri, Muhammad Rezki Rasyak, Willy Agustine, Hidayat Trimarsanto, Sri Masyeni, Erly Sintya, Ida Ayu Wayan Mahayani, Safarina G Malik, Khin Saw Myint, Amin Soebandrio |

| | | | |
|---|---|---|---|
| EPI_ISL_2101290 | RSUD Mangusada | Eijkman Institute for Molecular Biology, National Agency for Research and Innovation; Molecular Biology Laboratory, Faculty Medicine and Health Sciences, Warmadewa University | Iskandar Adnan, Sukma Oktavianthi, Lidwina Priliani, Edison Johar, Frilasita A Yudhaputri, Muhammad Rezki Rasyak, Willy Agustine, Hidayat Trimarsanto, Lydia V. Panggalo, Sri Masyeni, Erly Sintya, Ida Ayu Wayan Mahayani, Safarina G Malik, Khin Saw Myint, Amin Soebandrio |
| EPI_ISL_2101291 | RSUD Mangusada | Eijkman Institute for Molecular Biology, National Agency for Research and Innovation; Molecular Biology Laboratory, Faculty Medicine and Health Sciences, Warmadewa University | Sukma Oktavianthi, Lidwina Priliani, Edison Johar, Frilasita A Yudhaputri, Muhammad Rezki Rasyak, Willy Agustine, Hidayat Trimarsanto, Lydia V. Panggalo, Iskandar Adnan, Sri Masyeni, Erly Sintya, Ida Ayu Wayan Mahayani, Safarina G Malik, Khin Saw Myint, Amin Soebandrio |
| EPI_ISL_2105218, EPI_ISL_2105225, EPI_ISL_2105226 | Cagayan Valley Medical Center Molecular Laboratory | Philippine Genome Center | Francis A. Tablizo, Kenneth M. Kim, Carlo M. Lapid, Marc Jerrone R. Castro, Maria Sofia L. Yangzon, Benedict A. Maralit, Marc Edsel C. Ayes, Eva Maria Cutiongco-de la Paz, Alethea R. de Guzman, Jan Michael C. Yap, Jo-Hannah S. Llames, Sheila Mae M. Araiza, Kris P. Punayan, Irish Coleen A. Asin, Candice Francheska B. Tambaoan, Asia Louisa U. Chong, Karol Sophia Agape R. Padilla, Rianna Patricia S. Cruz, El King D. Morado, Joshua Gregor A. Dizon, Razel Nikka M. Hao, Arianne A. Zamora, Devon Ray Pacial, Juan Antonio R. Magalang, Marissa Alejandria, Celia Carlos, Anna Ong-Lim, Edsel Maurice Salvaña, John Q. Wong, Jaime C. Montoya, Maria Rosario Singh-Vergeire and Cynthia P. Saloma |
| EPI_ISL_2105232 | South Super Highway Molecular Diagnostic Laboratory | Philippine Genome Center | Francis A. Tablizo, Kenneth M. Kim, Carlo M. Lapid, Marc Jerrone R. Castro, Maria Sofia L. Yangzon, Benedict A. Maralit, Marc Edsel C. Ayes, Eva Maria Cutiongco-de la Paz, Alethea R. de Guzman, Jan Michael C. Yap, Jo-Hannah S. Llames, Sheila Mae M. Araiza, Kris P. Punayan, Irish Coleen A. Asin, Candice Francheska B. Tambaoan, Asia Louisa U. Chong, Karol Sophia Agape R. Padilla, Rianna Patricia S. Cruz, El King D. Morado, Joshua Gregor A. Dizon, Razel Nikka M. Hao, Arianne A. Zamora, Devon Ray Pacial, Juan Antonio R. Magalang, Marissa Alejandria, Celia Carlos, Anna Ong-Lim, Edsel Maurice Salvaña, John Q. Wong, Jaime C. Montoya, Maria Rosario Singh-Vergeire and Cynthia P. Saloma |
| EPI_ISL_2105242, EPI_ISL_2105245, EPI_ISL_2105253 | Cagayan Valley Medical Center Molecular Laboratory | Philippine Genome Center | Francis A. Tablizo, Kenneth M. Kim, Carlo M. Lapid, Marc Jerrone R. Castro, Maria Sofia L. Yangzon, Benedict A. Maralit, Marc Edsel C. Ayes, Eva Maria Cutiongco-de la Paz, Alethea R. de Guzman, Jan Michael C. Yap, Jo-Hannah S. Llames, Sheila Mae M. Araiza, Kris P. Punayan, Irish Coleen A. Asin, Candice Francheska B. Tambaoan, Asia Louisa U. Chong, Karol Sophia Agape R. Padilla, Rianna Patricia S. Cruz, El King D. Morado, Joshua Gregor A. Dizon, Razel Nikka M. Hao, Arianne A. Zamora, Devon Ray Pacial, Juan Antonio R. Magalang, Marissa Alejandria, Celia Carlos, Anna Ong-Lim, Edsel Maurice Salvaña, John Q. Wong, Jaime C. Montoya, Maria Rosario Singh-Vergeire and Cynthia P. Saloma |
| EPI_ISL_2105264, EPI_ISL_2105266 | Southern Philippines Medical Center (SPMC) | Philippine Genome Center | Francis A. Tablizo, Kenneth M. Kim, Carlo M. Lapid, Marc Jerrone R. Castro, Maria Sofia L. Yangzon, Benedict A. Maralit, Marc Edsel C. Ayes, Eva Maria Cutiongco-de la Paz, Alethea R. de Guzman, Jan Michael C. Yap, Jo-Hannah S. Llames, Sheila Mae M. Araiza, Kris P. Punayan, Irish Coleen A. Asin, Candice Francheska B. Tambaoan, Asia Louisa U. Chong, Karol Sophia Agape R. Padilla, Rianna Patricia S. Cruz, El King D. Morado, Joshua Gregor A. Dizon, Razel Nikka M. Hao, Arianne A. Zamora, Devon Ray Pacial, Juan Antonio R. Magalang, Marissa Alejandria, Celia Carlos, Anna Ong-Lim, Edsel Maurice Salvaña, John Q. Wong, Jaime C. Montoya, Maria Rosario Singh-Vergeire and Cynthia P. Saloma |
| EPI_ISL_2105276 | Cagayan Valley Medical Center Molecular Laboratory | Philippine Genome Center | Francis A. Tablizo, Kenneth M. Kim, Carlo M. Lapid, Marc Jerrone R. Castro, Maria Sofia L. Yangzon, Benedict A. Maralit, Marc Edsel C. Ayes, Eva Maria Cutiongco-de la Paz, Alethea R. de Guzman, Jan Michael C. Yap, Jo-Hannah S. Llames, Sheila Mae M. Araiza, Kris P. Punayan, Irish Coleen A. Asin, Candice Francheska B. Tambaoan, Asia Louisa U. Chong, Karol Sophia Agape R. Padilla, Rianna Patricia S. Cruz, El King D. Morado, Joshua Gregor A. Dizon, Razel Nikka M. Hao, Arianne A. Zamora, Devon Ray Pacial, Juan Antonio R. Magalang, Marissa Alejandria, Celia Carlos, Anna Ong-Lim, Edsel Maurice Salvaña, John Q. Wong, Jaime C. Montoya, Maria Rosario Singh-Vergeire and Cynthia P. Saloma |
| EPI_ISL_2105280 | The Lord's Grace Medical and Industrial Clinic | Philippine Genome Center | Francis A. Tablizo, Kenneth M. Kim, Carlo M. Lapid, Marc Jerrone R. Castro, Maria Sofia L. Yangzon, Benedict A. Maralit, Marc Edsel C. Ayes, Eva Maria Cutiongco-de la Paz, Alethea R. de Guzman, Jan Michael C. Yap, Jo-Hannah S. Llames, Sheila Mae M. Araiza, Kris P. Punayan, Irish Coleen A. Asin, Candice Francheska B. Tambaoan, Asia Louisa U. Chong, Karol Sophia Agape R. Padilla, Rianna Patricia S. Cruz, El King D. Morado, Joshua Gregor A. Dizon, Razel Nikka M. Hao, Arianne A. Zamora, Devon Ray Pacial, Juan Antonio R. Magalang, Marissa Alejandria, Celia Carlos, Anna Ong-Lim, Edsel Maurice Salvaña, John Q. Wong, Jaime C. Montoya, Maria Rosario Singh-Vergeire and Cynthia P. Saloma |
| EPI_ISL_2105282 | Cagayan Valley Medical Center Molecular Laboratory | Philippine Genome Center | Francis A. Tablizo, Kenneth M. Kim, Carlo M. Lapid, Marc Jerrone R. Castro, Maria Sofia L. Yangzon, Benedict A. Maralit, Marc Edsel C. Ayes, Eva Maria Cutiongco-de la Paz, Alethea R. de Guzman, Jan Michael C. Yap, Jo-Hannah S. Llames, Sheila Mae M. Araiza, Kris P. Punayan, Irish Coleen A. Asin, Candice Francheska B. Tambaoan, Asia Louisa U. Chong, Karol Sophia Agape R. Padilla, Rianna Patricia S. Cruz, El King D. Morado, Joshua Gregor A. Dizon, Razel Nikka M. Hao, Arianne A. Zamora, Devon Ray Pacial, Juan Antonio R. Magalang, Marissa Alejandria, Celia Carlos, Anna Ong-Lim, Edsel Maurice Salvaña, John Q. Wong, Jaime C. Montoya, Maria Rosario Singh-Vergeire and Cynthia P. Saloma |
| EPI_ISL_2105289 | Research Institute for Tropical Medicine, Inc. (RITM) | Philippine Genome Center | Francis A. Tablizo, Kenneth M. Kim, Carlo M. Lapid, Marc Jerrone R. Castro, Maria Sofia L. Yangzon, Benedict A. Maralit, Marc Edsel C. Ayes, Eva Maria Cutiongco-de la Paz, Alethea R. de Guzman, Jan Michael C. Yap, Jo-Hannah S. Llames, Sheila Mae M. Araiza, Kris P. Punayan, Irish Coleen A. Asin, Candice Francheska B. Tambaoan, Asia Louisa U. Chong, Karol Sophia Agape R. Padilla, Rianna Patricia S. Cruz, El King D. Morado, Joshua Gregor A. Dizon, Razel Nikka M. Hao, Arianne A. Zamora, Devon Ray Pacial, Juan Antonio R. Magalang, Marissa Alejandria, Celia Carlos, Anna Ong-Lim, Edsel Maurice Salvaña, John Q. Wong, Jaime C. Montoya, Maria Rosario Singh-Vergeire and Cynthia P. Saloma |
| EPI_ISL_2105291 | Cagayan Valley Medical Center Molecular Laboratory | Philippine Genome Center | Francis A. Tablizo, Kenneth M. Kim, Carlo M. Lapid, Marc Jerrone R. Castro, Maria Sofia L. Yangzon, Benedict A. Maralit, Marc Edsel C. Ayes, Eva Maria Cutiongco-de la Paz, Alethea R. de Guzman, Jan Michael C. Yap, Jo-Hannah S. Llames, Sheila Mae M. Araiza, Kris P. Punayan, Irish Coleen A. Asin, Candice Francheska B. Tambaoan, Asia Louisa U. Chong, Karol Sophia Agape R. Padilla, Rianna Patricia S. Cruz, El King D. Morado, Joshua Gregor A. Dizon, Razel Nikka M. Hao, Arianne A. Zamora, Devon Ray Pacial, Juan Antonio R. Magalang, Marissa Alejandria, Celia Carlos, Anna Ong-Lim, Edsel Maurice Salvaña, John Q. Wong, Jaime C. Montoya, Maria Rosario Singh-Vergeire and Cynthia P. Saloma |
| EPI_ISL_2105294, EPI_ISL_2105300, EPI_ISL_2105302, EPI_ISL_2105303 | Southern Philippines Medical Center (SPMC-SNL-MBL) | Philippine Genome Center | Francis A. Tablizo, Kenneth M. Kim, Carlo M. Lapid, Marc Jerrone R. Castro, Maria Sofia L. Yangzon, Benedict A. Maralit, Marc Edsel C. Ayes, Eva Maria Cutiongco-de la Paz, Alethea R. de Guzman, Jan Michael C. Yap, Jo-Hannah S. Llames, Sheila Mae M. Araiza, Kris P. Punayan, Irish Coleen A. Asin, Candice Francheska B. Tambaoan, Asia Louisa U. Chong, Karol Sophia Agape R. Padilla, Rianna Patricia S. Cruz, El King D. Morado, Joshua Gregor A. Dizon, Razel Nikka M. Hao, Arianne A. Zamora, Devon Ray Pacial, Juan Antonio R. Magalang, Marissa Alejandria, Celia Carlos, Anna Ong-Lim, Edsel Maurice Salvaña, John Q. Wong, Jaime C. Montoya, Maria Rosario Singh-Vergeire and Cynthia P. Saloma |
| EPI_ISL_2105306 | Dr. Jorge P. Royeca Hospital | Philippine Genome Center | Francis A. Tablizo, Kenneth M. Kim, Carlo M. Lapid, Marc Jerrone R. Castro, Maria Sofia L. Yangzon, Benedict A. Maralit, Marc Edsel C. Ayes, Eva Maria Cutiongco-de la Paz, Alethea R. de Guzman, Jan Michael C. Yap, Jo-Hannah S. Llames, Sheila Mae M. Araiza, Kris P. Punayan, Irish Coleen A. Asin, Candice Francheska B. Tambaoan, Asia Louisa U. Chong, Karol Sophia Agape R. Padilla, Rianna Patricia S. Cruz, El King D. Morado, Joshua Gregor A. Dizon, Razel Nikka M. Hao, Arianne A. Zamora, Devon Ray Pacial, Juan Antonio R. Magalang, Marissa Alejandria, Celia Carlos, Anna Ong-Lim, Edsel Maurice Salvaña, John Q. Wong, Jaime C. Montoya, Maria Rosario Singh-Vergeire and Cynthia P. Saloma |
| EPI_ISL_2105309, EPI_ISL_2105316, EPI_ISL_2105321, EPI_ISL_2105324, EPI_ISL_2105336 | Southern Philippines Medical Center (SPMC-SNL-MBL) | Philippine Genome Center | Francis A. Tablizo, Kenneth M. Kim, Carlo M. Lapid, Marc Jerrone R. Castro, Maria Sofia L. Yangzon, Benedict A. Maralit, Marc Edsel C. Ayes, Eva Maria Cutiongco-de la Paz, Alethea R. de Guzman, Jan Michael C. Yap, Jo-Hannah S. Llames, Sheila Mae M. Araiza, Kris P. Punayan, Irish Coleen A. Asin, Candice Francheska B. Tambaoan, Asia Louisa U. Chong, Karol Sophia Agape R. Padilla, Rianna Patricia S. Cruz, El King D. Morado, Joshua Gregor A. Dizon, Razel Nikka M. Hao, Arianne A. Zamora, Devon Ray Pacial, Juan Antonio R. Magalang, Marissa Alejandria, Celia Carlos, Anna Ong-Lim, Edsel Maurice Salvaña, John Q. Wong, Jaime C. Montoya, Maria Rosario Singh-Vergeire and Cynthia P. Saloma |
| EPI_ISL_2105337 | Research Institute for Tropical Medicine, Inc. (RITM) | Philippine Genome Center | Francis A. Tablizo, Kenneth M. Kim, Carlo M. Lapid, Marc Jerrone R. Castro, Maria Sofia L. Yangzon, Benedict A. Maralit, Marc Edsel C. Ayes, Eva Maria Cutiongco-de la Paz, Alethea R. de Guzman, Jan Michael C. Yap, Jo-Hannah S. Llames, Sheila Mae M. Araiza, Kris P. Punayan, Irish Coleen A. Asin, Candice Francheska B. Tambaoan, Asia Louisa U. Chong, Karol Sophia Agape R. Padilla, Rianna Patricia S. Cruz, El King D. Morado, Joshua Gregor A. Dizon, Razel Nikka M. Hao, Arianne A. Zamora, Devon Ray Pacial, Juan Antonio R. Magalang, Marissa Alejandria, Celia Carlos, Anna Ong-Lim, Edsel Maurice Salvaña, John Q. Wong, Jaime C. Montoya, Maria Rosario Singh-Vergeire and Cynthia P. Saloma |
| EPI_ISL_2105344, EPI_ISL_2105346, EPI_ISL_2105347, EPI_ISL_2105348, EPI_ISL_2105349, EPI_ISL_2105351, EPI_ISL_2105358 | Southern Philippines Medical Center (SPMC-SNL-MBL) | Philippine Genome Center | Francis A. Tablizo, Kenneth M. Kim, Carlo M. Lapid, Marc Jerrone R. Castro, Maria Sofia L. Yangzon, Benedict A. Maralit, Marc Edsel C. Ayes, Eva Maria Cutiongco-de la Paz, Alethea R. de Guzman, Jan Michael C. Yap, Jo-Hannah S. Llames, Sheila Mae M. Araiza, Kris P. Punayan, Irish Coleen A. Asin, Candice Francheska B. Tambaoan, Asia Louisa U. Chong, Karol Sophia Agape R. Padilla, Rianna Patricia S. Cruz, El King D. Morado, Joshua Gregor A. Dizon, Razel Nikka M. Hao, Arianne A. Zamora, Devon Ray Pacial, Juan Antonio R. Magalang, Marissa Alejandria, Celia Carlos, Anna Ong-Lim, Edsel Maurice Salvaña, John Q. Wong, Jaime C. Montoya, Maria Rosario Singh-Vergeire and Cynthia P. Saloma |
| EPI_ISL_2105360 | Dr. Jorge P. Royeca Hospital | Philippine Genome Center | Francis A. Tablizo, Kenneth M. Kim, Carlo M. Lapid, Marc Jerrone R. Castro, Maria Sofia L. Yangzon, Benedict A. Maralit, Marc Edsel C. Ayes, Eva Maria |

| | | | |
|---|---|---|---|
| | | | Cutiongco-de la Paz, Alethea R. de Guzman, Jan Michael C. Yap, Jo-Hannah S. Llames, Sheila Mae M. Araiza, Kris P. Punayan, Irish Coleen A. Asin, Candice Francheska B. Tambaoan, Asia Louisa U. Chong, Karol Sophia Agape R. Padilla, Rianna Patricia S. Cruz, El King D. Morado, Joshua Gregor A. Dizon, Razel Nikka M. Hao, Arianne A. Zamora, Devon Ray Pacial, Juan Antonio R. Magalang, Marissa Alejandria, Celia Carlos, Anna Ong-Lim, Edsel Maurice Salvaña, John Q. Wong, Jaime C. Montoya, Maria Rosario Singh-Vergeire and Cynthia P. Saloma |
| EPI_ISL_2105362 | Southern Philippines Medical Center (SPMC-SNL-MBL) | Philippine Genome Center | Francis A. Tablizo, Kenneth M. Kim, Carlo M. Lapid, Marc Jerrone R. Castro, Maria Sofia L. Yangzon, Benedict A. Maralit, Marc Edsel C. Ayes, Eva Maria Cutiongco-de la Paz, Alethea R. de Guzman, Jan Michael C. Yap, Jo-Hannah S. Llames, Sheila Mae M. Araiza, Kris P. Punayan, Irish Coleen A. Asin, Candice Francheska B. Tambaoan, Asia Louisa U. Chong, Karol Sophia Agape R. Padilla, Rianna Patricia S. Cruz, El King D. Morado, Joshua Gregor A. Dizon, Razel Nikka M. Hao, Arianne A. Zamora, Devon Ray Pacial, Juan Antonio R. Magalang, Marissa Alejandria, Celia Carlos, Anna Ong-Lim, Edsel Maurice Salvaña, John Q. Wong, Jaime C. Montoya, Maria Rosario Singh-Vergeire and Cynthia P. Saloma |
| EPI_ISL_2105365 | Dr. Jorge P. Royeca Hospital | Philippine Genome Center | Francis A. Tablizo, Kenneth M. Kim, Carlo M. Lapid, Marc Jerrone R. Castro, Maria Sofia L. Yangzon, Benedict A. Maralit, Marc Edsel C. Ayes, Eva Maria Cutiongco-de la Paz, Alethea R. de Guzman, Jan Michael C. Yap, Jo-Hannah S. Llames, Sheila Mae M. Araiza, Kris P. Punayan, Irish Coleen A. Asin, Candice Francheska B. Tambaoan, Asia Louisa U. Chong, Karol Sophia Agape R. Padilla, Rianna Patricia S. Cruz, El King D. Morado, Joshua Gregor A. Dizon, Razel Nikka M. Hao, Arianne A. Zamora, Devon Ray Pacial, Juan Antonio R. Magalang, Marissa Alejandria, Celia Carlos, Anna Ong-Lim, Edsel Maurice Salvaña, John Q. Wong, Jaime C. Montoya, Maria Rosario Singh-Vergeire and Cynthia P. Saloma |
| EPI_ISL_2105366, EPI_ISL_2105374 | Southern Philippines Medical Center (SPMC-SNL-MBL) | Philippine Genome Center | Francis A. Tablizo, Kenneth M. Kim, Carlo M. Lapid, Marc Jerrone R. Castro, Maria Sofia L. Yangzon, Benedict A. Maralit, Marc Edsel C. Ayes, Eva Maria Cutiongco-de la Paz, Alethea R. de Guzman, Jan Michael C. Yap, Jo-Hannah S. Llames, Sheila Mae M. Araiza, Kris P. Punayan, Irish Coleen A. Asin, Candice Francheska B. Tambaoan, Asia Louisa U. Chong, Karol Sophia Agape R. Padilla, Rianna Patricia S. Cruz, El King D. Morado, Joshua Gregor A. Dizon, Razel Nikka M. Hao, Arianne A. Zamora, Devon Ray Pacial, Juan Antonio R. Magalang, Marissa Alejandria, Celia Carlos, Anna Ong-Lim, Edsel Maurice Salvaña, John Q. Wong, Jaime C. Montoya, Maria Rosario Singh-Vergeire and Cynthia P. Saloma |
| EPI_ISL_2105377 | Safeguard DNA Diagnostic, Inc. | Philippine Genome Center | Francis A. Tablizo, Kenneth M. Kim, Carlo M. Lapid, Marc Jerrone R. Castro, Maria Sofia L. Yangzon, Benedict A. Maralit, Marc Edsel C. Ayes, Eva Maria Cutiongco-de la Paz, Alethea R. de Guzman, Jan Michael C. Yap, Jo-Hannah S. Llames, Sheila Mae M. Araiza, Kris P. Punayan, Irish Coleen A. Asin, Candice Francheska B. Tambaoan, Asia Louisa U. Chong, Karol Sophia Agape R. Padilla, Rianna Patricia S. Cruz, El King D. Morado, Joshua Gregor A. Dizon, Razel Nikka M. Hao, Arianne A. Zamora, Devon Ray Pacial, Juan Antonio R. Magalang, Marissa Alejandria, Celia Carlos, Anna Ong-Lim, Edsel Maurice Salvaña, John Q. Wong, Jaime C. Montoya, Maria Rosario Singh-Vergeire and Cynthia P. Saloma |
| EPI_ISL_2105384, EPI_ISL_2105386, EPI_ISL_2105387, EPI_ISL_2105389, EPI_ISL_2105390, EPI_ISL_2105402 | Southern Philippines Medical Center (SPMC-SNL-MBL) | Philippine Genome Center | Francis A. Tablizo, Kenneth M. Kim, Carlo M. Lapid, Marc Jerrone R. Castro, Maria Sofia L. Yangzon, Benedict A. Maralit, Marc Edsel C. Ayes, Eva Maria Cutiongco-de la Paz, Alethea R. de Guzman, Jan Michael C. Yap, Jo-Hannah S. Llames, Sheila Mae M. Araiza, Kris P. Punayan, Irish Coleen A. Asin, Candice Francheska B. Tambaoan, Asia Louisa U. Chong, Karol Sophia Agape R. Padilla, Rianna Patricia S. Cruz, El King D. Morado, Joshua Gregor A. Dizon, Razel Nikka M. Hao, Arianne A. Zamora, Devon Ray Pacial, Juan Antonio R. Magalang, Marissa Alejandria, Celia Carlos, Anna Ong-Lim, Edsel Maurice Salvaña, John Q. Wong, Jaime C. Montoya, Maria Rosario Singh-Vergeire and Cynthia P. Saloma |
| EPI_ISL_2105404 | Negros Oriental Provincial Hospital | Philippine Genome Center | Francis A. Tablizo, Kenneth M. Kim, Carlo M. Lapid, Marc Jerrone R. Castro, Maria Sofia L. Yangzon, Benedict A. Maralit, Marc Edsel C. Ayes, Eva Maria Cutiongco-de la Paz, Alethea R. de Guzman, Jan Michael C. Yap, Jo-Hannah S. Llames, Sheila Mae M. Araiza, Kris P. Punayan, Irish Coleen A. Asin, Candice Francheska B. Tambaoan, Asia Louisa U. Chong, Karol Sophia Agape R. Padilla, Rianna Patricia S. Cruz, El King D. Morado, Joshua Gregor A. Dizon, Razel Nikka M. Hao, Arianne A. Zamora, Devon Ray Pacial, Juan Antonio R. Magalang, Marissa Alejandria, Celia Carlos, Anna Ong-Lim, Edsel Maurice Salvaña, John Q. Wong, Jaime C. Montoya, Maria Rosario Singh-Vergeire and Cynthia P. Saloma |
| EPI_ISL_2105409 | Cagayan Valley Medical Center Molecular Laboratory | Philippine Genome Center | Francis A. Tablizo, Kenneth M. Kim, Carlo M. Lapid, Marc Jerrone R. Castro, Maria Sofia L. Yangzon, Benedict A. Maralit, Marc Edsel C. Ayes, Eva Maria Cutiongco-de la Paz, Alethea R. de Guzman, Jan Michael C. Yap, Jo-Hannah S. Llames, Sheila Mae M. Araiza, Kris P. Punayan, Irish Coleen A. Asin, Candice Francheska B. Tambaoan, Asia Louisa U. Chong, Karol Sophia Agape R. Padilla, Rianna Patricia S. Cruz, El King D. Morado, Joshua Gregor A. Dizon, Razel Nikka M. Hao, Arianne A. Zamora, Devon Ray Pacial, Juan Antonio R. Magalang, Marissa Alejandria, Celia Carlos, Anna Ong-Lim, Edsel Maurice Salvaña, John Q. Wong, Jaime C. Montoya, Maria Rosario Singh-Vergeire and Cynthia P. Saloma |
| EPI_ISL_2105412, EPI_ISL_2105416 | Negros Oriental Provincial Hospital | Philippine Genome Center | Francis A. Tablizo, Kenneth M. Kim, Carlo M. Lapid, Marc Jerrone R. Castro, Maria Sofia L. Yangzon, Benedict A. Maralit, Marc Edsel C. Ayes, Eva Maria Cutiongco-de la Paz, Alethea R. de Guzman, Jan Michael C. Yap, Jo-Hannah S. Llames, Sheila Mae M. Araiza, Kris P. Punayan, Irish Coleen A. Asin, Candice Francheska B. Tambaoan, Asia Louisa U. Chong, Karol Sophia Agape R. Padilla, Rianna Patricia S. Cruz, El King D. Morado, Joshua Gregor A. Dizon, Razel Nikka M. Hao, Arianne A. Zamora, Devon Ray Pacial, Juan Antonio R. Magalang, Marissa Alejandria, Celia Carlos, Anna Ong-Lim, Edsel Maurice Salvaña, John Q. Wong, Jaime C. Montoya, Maria Rosario Singh-Vergeire and Cynthia P. Saloma |
| EPI_ISL_2105418 | Research Institute for Tropical Medicine, Inc. (RITM) | Philippine Genome Center | Francis A. Tablizo, Kenneth M. Kim, Carlo M. Lapid, Marc Jerrone R. Castro, Maria Sofia L. Yangzon, Benedict A. Maralit, Marc Edsel C. Ayes, Eva Maria Cutiongco-de la Paz, Alethea R. de Guzman, Jan Michael C. Yap, Jo-Hannah S. Llames, Sheila Mae M. Araiza, Kris P. Punayan, Irish Coleen A. Asin, Candice Francheska B. Tambaoan, Asia Louisa U. Chong, Karol Sophia Agape R. Padilla, Rianna Patricia S. Cruz, El King D. Morado, Joshua Gregor A. Dizon, Razel Nikka M. Hao, Arianne A. Zamora, Devon Ray Pacial, Juan Antonio R. Magalang, Marissa Alejandria, Celia Carlos, Anna Ong-Lim, Edsel Maurice Salvaña, John Q. Wong, Jaime C. Montoya, Maria Rosario Singh-Vergeire and Cynthia P. Saloma |
| EPI_ISL_2105419, EPI_ISL_2105422, EPI_ISL_2105423 | Negros Oriental Provincial Hospital | Philippine Genome Center | Francis A. Tablizo, Kenneth M. Kim, Carlo M. Lapid, Marc Jerrone R. Castro, Maria Sofia L. Yangzon, Benedict A. Maralit, Marc Edsel C. Ayes, Eva Maria Cutiongco-de la Paz, Alethea R. de Guzman, Jan Michael C. Yap, Jo-Hannah S. Llames, Sheila Mae M. Araiza, Kris P. Punayan, Irish Coleen A. Asin, Candice Francheska B. Tambaoan, Asia Louisa U. Chong, Karol Sophia Agape R. Padilla, Rianna Patricia S. Cruz, El King D. Morado, Joshua Gregor A. Dizon, Razel Nikka M. Hao, Arianne A. Zamora, Devon Ray Pacial, Juan Antonio R. Magalang, Marissa Alejandria, Celia Carlos, Anna Ong-Lim, Edsel Maurice Salvaña, John Q. Wong, Jaime C. Montoya, Maria Rosario Singh-Vergeire and Cynthia P. Saloma |
| EPI_ISL_2105425 | Southern Philippines Medical Center (SPMC-SNL-MBL) | Philippine Genome Center | Francis A. Tablizo, Kenneth M. Kim, Carlo M. Lapid, Marc Jerrone R. Castro, Maria Sofia L. Yangzon, Benedict A. Maralit, Marc Edsel C. Ayes, Eva Maria Cutiongco-de la Paz, Alethea R. de Guzman, Jan Michael C. Yap, Jo-Hannah S. Llames, Sheila Mae M. Araiza, Kris P. Punayan, Irish Coleen A. Asin, Candice Francheska B. Tambaoan, Asia Louisa U. Chong, Karol Sophia Agape R. Padilla, Rianna Patricia S. Cruz, El King D. Morado, Joshua Gregor A. Dizon, Razel Nikka M. Hao, Arianne A. Zamora, Devon Ray Pacial, Juan Antonio R. Magalang, Marissa Alejandria, Celia Carlos, Anna Ong-Lim, Edsel Maurice Salvaña, John Q. Wong, Jaime C. Montoya, Maria Rosario Singh-Vergeire and Cynthia P. Saloma |
| EPI_ISL_2105427, EPI_ISL_2105431 | Negros Oriental Provincial Hospital | Philippine Genome Center | Francis A. Tablizo, Kenneth M. Kim, Carlo M. Lapid, Marc Jerrone R. Castro, Maria Sofia L. Yangzon, Benedict A. Maralit, Marc Edsel C. Ayes, Eva Maria Cutiongco-de la Paz, Alethea R. de Guzman, Jan Michael C. Yap, Jo-Hannah S. Llames, Sheila Mae M. Araiza, Kris P. Punayan, Irish Coleen A. Asin, Candice Francheska B. Tambaoan, Asia Louisa U. Chong, Karol Sophia Agape R. Padilla, Rianna Patricia S. Cruz, El King D. Morado, Joshua Gregor A. Dizon, Razel Nikka M. Hao, Arianne A. Zamora, Devon Ray Pacial, Juan Antonio R. Magalang, Marissa Alejandria, Celia Carlos, Anna Ong-Lim, Edsel Maurice Salvaña, John Q. Wong, Jaime C. Montoya, Maria Rosario Singh-Vergeire and Cynthia P. Saloma |
| EPI_ISL_2105433 | Dr. Jorge P. Royeca Hospital | Philippine Genome Center | Francis A. Tablizo, Kenneth M. Kim, Carlo M. Lapid, Marc Jerrone R. Castro, Maria Sofia L. Yangzon, Benedict A. Maralit, Marc Edsel C. Ayes, Eva Maria Cutiongco-de la Paz, Alethea R. de Guzman, Jan Michael C. Yap, Jo-Hannah S. Llames, Sheila Mae M. Araiza, Kris P. Punayan, Irish Coleen A. Asin, Candice Francheska B. Tambaoan, Asia Louisa U. Chong, Karol Sophia Agape R. Padilla, Rianna Patricia S. Cruz, El King D. Morado, Joshua Gregor A. Dizon, Razel Nikka M. Hao, Arianne A. Zamora, Devon Ray Pacial, Juan Antonio R. Magalang, Marissa Alejandria, Celia Carlos, Anna Ong-Lim, Edsel Maurice Salvaña, John Q. Wong, Jaime C. Montoya, Maria Rosario Singh-Vergeire and Cynthia P. Saloma |
| EPI_ISL_2105435 | Negros Oriental Provincial Hospital | Philippine Genome Center | Francis A. Tablizo, Kenneth M. Kim, Carlo M. Lapid, Marc Jerrone R. Castro, Maria Sofia L. Yangzon, Benedict A. Maralit, Marc Edsel C. Ayes, Eva Maria Cutiongco-de la Paz, Alethea R. de Guzman, Jan Michael C. Yap, Jo-Hannah S. Llames, Sheila Mae M. Araiza, Kris P. Punayan, Irish Coleen A. Asin, Candice Francheska B. Tambaoan, Asia Louisa U. Chong, Karol Sophia Agape R. Padilla, Rianna Patricia S. Cruz, El King D. Morado, Joshua Gregor A. Dizon, Razel Nikka M. Hao, Arianne A. Zamora, Devon Ray Pacial, Juan Antonio R. Magalang, Marissa Alejandria, Celia Carlos, Anna Ong-Lim, Edsel Maurice Salvaña, John Q. Wong, Jaime C. Montoya, Maria Rosario Singh-Vergeire and Cynthia P. Saloma |
| EPI_ISL_2105448 | Cagayan Valley Medical Center Molecular Laboratory | Philippine Genome Center | Francis A. Tablizo, Kenneth M. Kim, Carlo M. Lapid, Marc Jerrone R. Castro, Maria Sofia L. Yangzon, Benedict A. Maralit, Marc Edsel C. Ayes, Eva Maria Cutiongco-de la Paz, Alethea R. de Guzman, Jan Michael C. Yap, Jo-Hannah S. Llames, Sheila Mae M. Araiza, Kris P. Punayan, Irish Coleen A. Asin, Candice Francheska B. Tambaoan, Asia Louisa U. Chong, Karol Sophia Agape R. Padilla, Rianna Patricia S. Cruz, El King D. Morado, Joshua Gregor A. Dizon, Razel Nikka M. Hao, Arianne A. Zamora, Devon Ray Pacial, Juan Antonio R. Magalang, Marissa Alejandria, Celia Carlos, Anna Ong-Lim, Edsel Maurice Salvaña, John Q. Wong, Jaime C. Montoya, Maria Rosario Singh-Vergeire and Cynthia P. Saloma |

| | | | |
|---|---|---|---|
| EPI_ISL_2105450, EPI_ISL_2105451, EPI_ISL_2105452 | Negros Oriental Provincial Hospital | Philippine Genome Center | Francis A. Tablizo, Kenneth M. Kim, Carlo M. Lapid, Marc Jerrone R. Castro, Maria Sofia L. Yangzon, Benedict A. Maralit, Marc Edsel C. Ayes, Eva Maria Cutiongco-de la Paz, Alethea R. de Guzman, Jan Michael C. Yap, Jo-Hannah S. Llames, Sheila Mae M. Araiza, Kris P. Punayan, Irish Coleen A. Asin, Candice Francheska B. Tambaoan, Asia Louisa U. Chong, Karol Sophia Agape R. Padilla, Rianna Patricia S. Cruz, El King D. Morado, Joshua Gregor A. Dizon, Razel Nikka M. Hao, Arianne A. Zamora, Devon Ray Pacial, Juan Antonio R. Magalang, Marissa Alejandria, Celia Carlos, Anna Ong-Lim, Edsel Maurice Salvaña, John Q. Wong, Jaime C. Montoya, Maria Rosario Singh-Vergeire and Cynthia P. Saloma |
| EPI_ISL_2105453 | Cagayan Valley Medical Center Molecular Laboratory | Philippine Genome Center | Francis A. Tablizo, Kenneth M. Kim, Carlo M. Lapid, Marc Jerrone R. Castro, Maria Sofia L. Yangzon, Benedict A. Maralit, Marc Edsel C. Ayes, Eva Maria Cutiongco-de la Paz, Alethea R. de Guzman, Jan Michael C. Yap, Jo-Hannah S. Llames, Sheila Mae M. Araiza, Kris P. Punayan, Irish Coleen A. Asin, Candice Francheska B. Tambaoan, Asia Louisa U. Chong, Karol Sophia Agape R. Padilla, Rianna Patricia S. Cruz, El King D. Morado, Joshua Gregor A. Dizon, Razel Nikka M. Hao, Arianne A. Zamora, Devon Ray Pacial, Juan Antonio R. Magalang, Marissa Alejandria, Celia Carlos, Anna Ong-Lim, Edsel Maurice Salvaña, John Q. Wong, Jaime C. Montoya, Maria Rosario Singh-Vergeire and Cynthia P. Saloma |
| EPI_ISL_2105460 | Negros Oriental Provincial Hospital | Philippine Genome Center | Francis A. Tablizo, Kenneth M. Kim, Carlo M. Lapid, Marc Jerrone R. Castro, Maria Sofia L. Yangzon, Benedict A. Maralit, Marc Edsel C. Ayes, Eva Maria Cutiongco-de la Paz, Alethea R. de Guzman, Jan Michael C. Yap, Jo-Hannah S. Llames, Sheila Mae M. Araiza, Kris P. Punayan, Irish Coleen A. Asin, Candice Francheska B. Tambaoan, Asia Louisa U. Chong, Karol Sophia Agape R. Padilla, Rianna Patricia S. Cruz, El King D. Morado, Joshua Gregor A. Dizon, Razel Nikka M. Hao, Arianne A. Zamora, Devon Ray Pacial, Juan Antonio R. Magalang, Marissa Alejandria, Celia Carlos, Anna Ong-Lim, Edsel Maurice Salvaña, John Q. Wong, Jaime C. Montoya, Maria Rosario Singh-Vergeire and Cynthia P. Saloma |
| EPI_ISL_2105471, EPI_ISL_2105483, EPI_ISL_2105502 | Governor Celestino Gallares Memorial Hospital | Philippine Genome Center | Francis A. Tablizo, Kenneth M. Kim, Carlo M. Lapid, Marc Jerrone R. Castro, Maria Sofia L. Yangzon, Benedict A. Maralit, Marc Edsel C. Ayes, Eva Maria Cutiongco-de la Paz, Alethea R. de Guzman, Jan Michael C. Yap, Jo-Hannah S. Llames, Sheila Mae M. Araiza, Kris P. Punayan, Irish Coleen A. Asin, Candice Francheska B. Tambaoan, Asia Louisa U. Chong, Karol Sophia Agape R. Padilla, Rianna Patricia S. Cruz, El King D. Morado, Joshua Gregor A. Dizon, Razel Nikka M. Hao, Arianne A. Zamora, Devon Ray Pacial, Juan Antonio R. Magalang, Marissa Alejandria, Celia Carlos, Anna Ong-Lim, Edsel Maurice Salvaña, John Q. Wong, Jaime C. Montoya, Maria Rosario Singh-Vergeire and Cynthia P. Saloma |
| EPI_ISL_2105513, EPI_ISL_2105518, EPI_ISL_2105520 | Army General Hospital Molecular Laboratory | Philippine Genome Center | Francis A. Tablizo, Kenneth M. Kim, Carlo M. Lapid, Marc Jerrone R. Castro, Maria Sofia L. Yangzon, Benedict A. Maralit, Marc Edsel C. Ayes, Eva Maria Cutiongco-de la Paz, Alethea R. de Guzman, Jan Michael C. Yap, Jo-Hannah S. Llames, Sheila Mae M. Araiza, Kris P. Punayan, Irish Coleen A. Asin, Candice Francheska B. Tambaoan, Asia Louisa U. Chong, Karol Sophia Agape R. Padilla, Rianna Patricia S. Cruz, El King D. Morado, Joshua Gregor A. Dizon, Razel Nikka M. Hao, Arianne A. Zamora, Devon Ray Pacial, Juan Antonio R. Magalang, Marissa Alejandria, Celia Carlos, Anna Ong-Lim, Edsel Maurice Salvaña, John Q. Wong, Jaime C. Montoya, Maria Rosario Singh-Vergeire and Cynthia P. Saloma |
| EPI_ISL_2105555 | Governor Celestino Gallares Memorial Hospital | Philippine Genome Center | Francis A. Tablizo, Kenneth M. Kim, Carlo M. Lapid, Marc Jerrone R. Castro, Maria Sofia L. Yangzon, Benedict A. Maralit, Marc Edsel C. Ayes, Eva Maria Cutiongco-de la Paz, Alethea R. de Guzman, Jan Michael C. Yap, Jo-Hannah S. Llames, Sheila Mae M. Araiza, Kris P. Punayan, Irish Coleen A. Asin, Candice Francheska B. Tambaoan, Asia Louisa U. Chong, Karol Sophia Agape R. Padilla, Rianna Patricia S. Cruz, El King D. Morado, Joshua Gregor A. Dizon, Razel Nikka M. Hao, Arianne A. Zamora, Devon Ray Pacial, Juan Antonio R. Magalang, Marissa Alejandria, Celia Carlos, Anna Ong-Lim, Edsel Maurice Salvaña, John Q. Wong, Jaime C. Montoya, Maria Rosario Singh-Vergeire and Cynthia P. Saloma |
| EPI_ISL_2107800, EPI_ISL_2107802 | National Center of Infectious and Parasitic Diseases | National Center of Infectious and Parasitic Diseases | Alexiev, Ivanov, Korsun, Stoitsova, Philipova, Dimitrova, Grigorova L., Hristova, Donchev, Stoykov, Trifonova, Dobrinov, Grigorova I., Kantardjiev |
| EPI_ISL_2131619, EPI_ISL_2131803, EPI_ISL_2131835, EPI_ISL_2131836, EPI_ISL_2131837, EPI_ISL_2131840 | Laboratoire de Virologie - CHU François Mitterrand - Dijon - France | INSERM U1231 - Equipe GAD | Ali SI-MOHAMMED, Yannis DUFFOURD, Hélène GIRAUDON, Lionel PIROTH |
| EPI_ISL_2150999 | Laboratório de Biologia Molecular Jean Piaget | MRCG at LSHTM, Genomics lab | Aladje Balde, Abdul Karim Sesay, Abdoulie Kanteh, Bakary Sanyang, Simão Tchuda Bióté, Rui Inndi, Adul Candé, Faatu Cassama, Milanca Agostinho Cá, Rei José Pereira, Erica Luis Maria Magalhães, Aicha Balde, Bubacar Delgado Pinto Embaló, Edmira Maria da Costa, Paulina Joãozinho da Costa Jarra Manneh, Mariama Kujabi, Dabiri Damilari, Sainabou Laye Ndure |
| EPI_ISL_2154027 | Dr. Jorge P. Royeca Hospital | Philippine Genome Center | Francis A. Tablizo, Kenneth M. Kim, Carlo M. Lapid, Marc Jerrone R. Castro, Maria Sofia L. Yangzon, Benedict A. Maralit, Marc Edsel C. Ayes, Eva Maria Cutiongco-de la Paz, Alethea R. de Guzman, Jan Michael C. Yap, Jo-Hannah S. Llames, Sheila Mae M. Araiza, Kris P. Punayan, Irish Coleen A. Asin, Candice Francheska B. Tambaoan, Asia Louisa U. Chong, Karol Sophia Agape R. Padilla, Rianna Patricia S. Cruz, El King D. Morado, Joshua Gregor A. Dizon, Razel Nikka M. Hao, Arianne A. Zamora, Devon Ray Pacial, Juan Antonio R. Magalang, Marissa Alejandria, Celia Carlos, Anna Ong-Lim, Edsel Maurice Salvaña, John Q. Wong, Jaime C. Montoya, Maria Rosario Singh-Vergeire and Cynthia P. Saloma |
| EPI_ISL_2154028 | Philippine Red Cross Molecular Laboratory | Philippine Genome Center | Francis A. Tablizo, Kenneth M. Kim, Carlo M. Lapid, Marc Jerrone R. Castro, Maria Sofia L. Yangzon, Benedict A. Maralit, Marc Edsel C. Ayes, Eva Maria Cutiongco-de la Paz, Alethea R. de Guzman, Jan Michael C. Yap, Jo-Hannah S. Llames, Sheila Mae M. Araiza, Kris P. Punayan, Irish Coleen A. Asin, Candice Francheska B. Tambaoan, Asia Louisa U. Chong, Karol Sophia Agape R. Padilla, Rianna Patricia S. Cruz, El King D. Morado, Joshua Gregor A. Dizon, Razel Nikka M. Hao, Arianne A. Zamora, Devon Ray Pacial, Juan Antonio R. Magalang, Marissa Alejandria, Celia Carlos, Anna Ong-Lim, Edsel Maurice Salvaña, John Q. Wong, Jaime C. Montoya, Maria Rosario Singh-Vergeire and Cynthia P. Saloma |
| EPI_ISL_2154029, EPI_ISL_2154030 | Baguio General Hospital Medical Center (BGHMC) | Philippine Genome Center | Francis A. Tablizo, Kenneth M. Kim, Carlo M. Lapid, Marc Jerrone R. Castro, Maria Sofia L. Yangzon, Benedict A. Maralit, Marc Edsel C. Ayes, Eva Maria Cutiongco-de la Paz, Alethea R. de Guzman, Jan Michael C. Yap, Jo-Hannah S. Llames, Sheila Mae M. Araiza, Kris P. Punayan, Irish Coleen A. Asin, Candice Francheska B. Tambaoan, Asia Louisa U. Chong, Karol Sophia Agape R. Padilla, Rianna Patricia S. Cruz, El King D. Morado, Joshua Gregor A. Dizon, Razel Nikka M. Hao, Arianne A. Zamora, Devon Ray Pacial, Juan Antonio R. Magalang, Marissa Alejandria, Celia Carlos, Anna Ong-Lim, Edsel Maurice Salvaña, John Q. Wong, Jaime C. Montoya, Maria Rosario Singh-Vergeire and Cynthia P. Saloma |
| EPI_ISL_2154031 | Army General Hospital Molecular Laboratory | Philippine Genome Center | Francis A. Tablizo, Kenneth M. Kim, Carlo M. Lapid, Marc Jerrone R. Castro, Maria Sofia L. Yangzon, Benedict A. Maralit, Marc Edsel C. Ayes, Eva Maria Cutiongco-de la Paz, Alethea R. de Guzman, Jan Michael C. Yap, Jo-Hannah S. Llames, Sheila Mae M. Araiza, Kris P. Punayan, Irish Coleen A. Asin, Candice Francheska B. Tambaoan, Asia Louisa U. Chong, Karol Sophia Agape R. Padilla, Rianna Patricia S. Cruz, El King D. Morado, Joshua Gregor A. Dizon, Razel Nikka M. Hao, Arianne A. Zamora, Devon Ray Pacial, Juan Antonio R. Magalang, Marissa Alejandria, Celia Carlos, Anna Ong-Lim, Edsel Maurice Salvaña, John Q. Wong, Jaime C. Montoya, Maria Rosario Singh-Vergeire and Cynthia P. Saloma |
| EPI_ISL_2154032, EPI_ISL_2154033 | Baguio General Hospital Medical Center (BGHMC) | Philippine Genome Center | Francis A. Tablizo, Kenneth M. Kim, Carlo M. Lapid, Marc Jerrone R. Castro, Maria Sofia L. Yangzon, Benedict A. Maralit, Marc Edsel C. Ayes, Eva Maria Cutiongco-de la Paz, Alethea R. de Guzman, Jan Michael C. Yap, Jo-Hannah S. Llames, Sheila Mae M. Araiza, Kris P. Punayan, Irish Coleen A. Asin, Candice Francheska B. Tambaoan, Asia Louisa U. Chong, Karol Sophia Agape R. Padilla, Rianna Patricia S. Cruz, El King D. Morado, Joshua Gregor A. Dizon, Razel Nikka M. Hao, Arianne A. Zamora, Devon Ray Pacial, Juan Antonio R. Magalang, Marissa Alejandria, Celia Carlos, Anna Ong-Lim, Edsel Maurice Salvaña, John Q. Wong, Jaime C. Montoya, Maria Rosario Singh-Vergeire and Cynthia P. Saloma |
| EPI_ISL_2154034 | Army General Hospital Molecular Laboratory | Philippine Genome Center | Francis A. Tablizo, Kenneth M. Kim, Carlo M. Lapid, Marc Jerrone R. Castro, Maria Sofia L. Yangzon, Benedict A. Maralit, Marc Edsel C. Ayes, Eva Maria Cutiongco-de la Paz, Alethea R. de Guzman, Jan Michael C. Yap, Jo-Hannah S. Llames, Sheila Mae M. Araiza, Kris P. Punayan, Irish Coleen A. Asin, Candice Francheska B. Tambaoan, Asia Louisa U. Chong, Karol Sophia Agape R. Padilla, Rianna Patricia S. Cruz, El King D. Morado, Joshua Gregor A. Dizon, Razel Nikka M. Hao, Arianne A. Zamora, Devon Ray Pacial, Juan Antonio R. Magalang, Marissa Alejandria, Celia Carlos, Anna Ong-Lim, Edsel Maurice Salvaña, John Q. Wong, Jaime C. Montoya, Maria Rosario Singh-Vergeire and Cynthia P. Saloma |
| EPI_ISL_2154035, EPI_ISL_2154036, EPI_ISL_2154037 | Baguio General Hospital Medical Center (BGHMC) | Philippine Genome Center | Francis A. Tablizo, Kenneth M. Kim, Carlo M. Lapid, Marc Jerrone R. Castro, Maria Sofia L. Yangzon, Benedict A. Maralit, Marc Edsel C. Ayes, Eva Maria Cutiongco-de la Paz, Alethea R. de Guzman, Jan Michael C. Yap, Jo-Hannah S. Llames, Sheila Mae M. Araiza, Kris P. Punayan, Irish Coleen A. Asin, Candice Francheska B. Tambaoan, Asia Louisa U. Chong, Karol Sophia Agape R. Padilla, Rianna Patricia S. Cruz, El King D. Morado, Joshua Gregor A. Dizon, Razel Nikka M. Hao, Arianne A. Zamora, Devon Ray Pacial, Juan Antonio R. Magalang, Marissa Alejandria, Celia Carlos, Anna Ong-Lim, Edsel Maurice Salvaña, John Q. Wong, Jaime C. Montoya, Maria Rosario Singh-Vergeire and Cynthia P. Saloma |
| EPI_ISL_2154038 | St. Luke's Medical Center- QC | Philippine Genome Center | Francis A. Tablizo, Kenneth M. Kim, Carlo M. Lapid, Marc Jerrone R. Castro, Maria Sofia L. Yangzon, Benedict A. Maralit, Marc Edsel C. Ayes, Eva Maria Cutiongco-de la Paz, Alethea R. de Guzman, Jan Michael C. Yap, Jo-Hannah S. Llames, Sheila Mae M. Araiza, Kris P. Punayan, Irish Coleen A. Asin, Candice Francheska B. Tambaoan, Asia Louisa U. Chong, Karol Sophia Agape R. Padilla, Rianna Patricia S. Cruz, El King D. Morado, Joshua Gregor A. Dizon, Razel Nikka M. Hao, Arianne A. Zamora, Devon Ray Pacial, Juan Antonio R. Magalang, Marissa Alejandria, Celia Carlos, Anna Ong-Lim, Edsel Maurice Salvaña, John Q. Wong, Jaime C. Montoya, Maria Rosario Singh-Vergeire and Cynthia P. Saloma |
| EPI_ISL_2154039, EPI_ISL_2154040 | Baguio General Hospital Medical Center (BGHMC) | Philippine Genome Center | Francis A. Tablizo, Kenneth M. Kim, Carlo M. Lapid, Marc Jerrone R. Castro, Maria Sofia L. Yangzon, Benedict A. Maralit, Marc Edsel C. Ayes, Eva Maria Cutiongco-de la Paz, Alethea R. de Guzman, Jan Michael C. Yap, Jo-Hannah S. Llames, Sheila Mae M. Araiza, Kris P. Punayan, Irish Coleen A. Asin, Candice |

| | | | |
|---|---|---|---|
| | | | Francheska B. Tambaoan, Asia Louisa U. Chong, Karol Sophia Agape R. Padilla, Rianna Patricia S. Cruz, El King D. Morado, Joshua Gregor A. Dizon, Razel Nikka M. Hao, Arianne A. Zamora, Devon Ray Pacial, Juan Antonio R. Magalang, Marissa Alejandria, Celia Carlos, Anna Ong-Lim, Edsel Maurice Salvaña, John Q. Wong, Jaime C. Montoya, Maria Rosario Singh-Vergeire and Cynthia P. Saloma |
| EPI_ISL_2154041 | Army General Hospital Molecular Laboratory | Philippine Genome Center | Francis A. Tablizo, Kenneth M. Kim, Carlo M. Lapid, Marc Jerrone R. Castro, Maria Sofia L. Yangzon, Benedict A. Maralit, Marc Edsel C. Ayes, Eva Maria Cutiongco-de la Paz, Alethea R. de Guzman, Jan Michael C. Yap, Jo-Hannah S. Llames, Sheila Mae M. Araiza, Kris P. Punayan, Irish Coleen A. Asin, Candice Francheska B. Tambaoan, Asia Louisa U. Chong, Karol Sophia Agape R. Padilla, Rianna Patricia S. Cruz, El King D. Morado, Joshua Gregor A. Dizon, Razel Nikka M. Hao, Arianne A. Zamora, Devon Ray Pacial, Juan Antonio R. Magalang, Marissa Alejandria, Celia Carlos, Anna Ong-Lim, Edsel Maurice Salvaña, John Q. Wong, Jaime C. Montoya, Maria Rosario Singh-Vergeire and Cynthia P. Saloma |
| EPI_ISL_2154042, EPI_ISL_2154043 | Baguio General Hospital Medical Center (BGHMC) | Philippine Genome Center | Francis A. Tablizo, Kenneth M. Kim, Carlo M. Lapid, Marc Jerrone R. Castro, Maria Sofia L. Yangzon, Benedict A. Maralit, Marc Edsel C. Ayes, Eva Maria Cutiongco-de la Paz, Alethea R. de Guzman, Jan Michael C. Yap, Jo-Hannah S. Llames, Sheila Mae M. Araiza, Kris P. Punayan, Irish Coleen A. Asin, Candice Francheska B. Tambaoan, Asia Louisa U. Chong, Karol Sophia Agape R. Padilla, Rianna Patricia S. Cruz, El King D. Morado, Joshua Gregor A. Dizon, Razel Nikka M. Hao, Arianne A. Zamora, Devon Ray Pacial, Juan Antonio R. Magalang, Marissa Alejandria, Celia Carlos, Anna Ong-Lim, Edsel Maurice Salvaña, John Q. Wong, Jaime C. Montoya, Maria Rosario Singh-Vergeire and Cynthia P. Saloma |
| EPI_ISL_2154395 | Research Institute for Tropical Medicine, Inc. (RITM) | Philippine Genome Center | Francis A. Tablizo, Kenneth M. Kim, Carlo M. Lapid, Marc Jerrone R. Castro, Maria Sofia L. Yangzon, Benedict A. Maralit, Marc Edsel C. Ayes, Eva Maria Cutiongco-de la Paz, Alethea R. de Guzman, Jan Michael C. Yap, Jo-Hannah S. Llames, Sheila Mae M. Araiza, Kris P. Punayan, Irish Coleen A. Asin, Candice Francheska B. Tambaoan, Asia Louisa U. Chong, Karol Sophia Agape R. Padilla, Rianna Patricia S. Cruz, El King D. Morado, Joshua Gregor A. Dizon, Razel Nikka M. Hao, Arianne A. Zamora, Devon Ray Pacial, Juan Antonio R. Magalang, Marissa Alejandria, Celia Carlos, Anna Ong-Lim, Edsel Maurice Salvaña, John Q. Wong, Jaime C. Montoya, Maria Rosario Singh-Vergeire and Cynthia P. Saloma |
| EPI_ISL_2154415, EPI_ISL_2154419, EPI_ISL_2154422, EPI_ISL_2154424, EPI_ISL_2154427, EPI_ISL_2154431, EPI_ISL_2154459, EPI_ISL_2154472, EPI_ISL_2154478 | Cagayan Valley Medical Center Molecular Laboratory | Philippine Genome Center | Francis A. Tablizo, Kenneth M. Kim, Carlo M. Lapid, Marc Jerrone R. Castro, Maria Sofia L. Yangzon, Benedict A. Maralit, Marc Edsel C. Ayes, Eva Maria Cutiongco-de la Paz, Alethea R. de Guzman, Jan Michael C. Yap, Jo-Hannah S. Llames, Sheila Mae M. Araiza, Kris P. Punayan, Irish Coleen A. Asin, Candice Francheska B. Tambaoan, Asia Louisa U. Chong, Karol Sophia Agape R. Padilla, Rianna Patricia S. Cruz, El King D. Morado, Joshua Gregor A. Dizon, Razel Nikka M. Hao, Arianne A. Zamora, Devon Ray Pacial, Juan Antonio R. Magalang, Marissa Alejandria, Celia Carlos, Anna Ong-Lim, Edsel Maurice Salvaña, John Q. Wong, Jaime C. Montoya, Maria Rosario Singh-Vergeire and Cynthia P. Saloma |
| EPI_ISL_2154490 | Cardinal Santos Medical Center | Philippine Genome Center | Francis A. Tablizo, Kenneth M. Kim, Carlo M. Lapid, Marc Jerrone R. Castro, Maria Sofia L. Yangzon, Benedict A. Maralit, Marc Edsel C. Ayes, Eva Maria Cutiongco-de la Paz, Alethea R. de Guzman, Jan Michael C. Yap, Jo-Hannah S. Llames, Sheila Mae M. Araiza, Kris P. Punayan, Irish Coleen A. Asin, Candice Francheska B. Tambaoan, Asia Louisa U. Chong, Karol Sophia Agape R. Padilla, Rianna Patricia S. Cruz, El King D. Morado, Joshua Gregor A. Dizon, Razel Nikka M. Hao, Arianne A. Zamora, Devon Ray Pacial, Juan Antonio R. Magalang, Marissa Alejandria, Celia Carlos, Anna Ong-Lim, Edsel Maurice Salvaña, John Q. Wong, Jaime C. Montoya, Maria Rosario Singh-Vergeire and Cynthia P. Saloma |
| EPI_ISL_2154497, EPI_ISL_2154499, EPI_ISL_2154501, EPI_ISL_2154505 | Cagayan Valley Medical Center Molecular Laboratory | Philippine Genome Center | Francis A. Tablizo, Kenneth M. Kim, Carlo M. Lapid, Marc Jerrone R. Castro, Maria Sofia L. Yangzon, Benedict A. Maralit, Marc Edsel C. Ayes, Eva Maria Cutiongco-de la Paz, Alethea R. de Guzman, Jan Michael C. Yap, Jo-Hannah S. Llames, Sheila Mae M. Araiza, Kris P. Punayan, Irish Coleen A. Asin, Candice Francheska B. Tambaoan, Asia Louisa U. Chong, Karol Sophia Agape R. Padilla, Rianna Patricia S. Cruz, El King D. Morado, Joshua Gregor A. Dizon, Razel Nikka M. Hao, Arianne A. Zamora, Devon Ray Pacial, Juan Antonio R. Magalang, Marissa Alejandria, Celia Carlos, Anna Ong-Lim, Edsel Maurice Salvaña, John Q. Wong, Jaime C. Montoya, Maria Rosario Singh-Vergeire and Cynthia P. Saloma |
| EPI_ISL_2154547 | Southern Isabela Medical Center Molecular Diagnostic & Research Laboratory (SIMC - MDRL) | Philippine Genome Center | Francis A. Tablizo, Kenneth M. Kim, Carlo M. Lapid, Marc Jerrone R. Castro, Maria Sofia L. Yangzon, Benedict A. Maralit, Marc Edsel C. Ayes, Eva Maria Cutiongco-de la Paz, Alethea R. de Guzman, Jan Michael C. Yap, Jo-Hannah S. Llames, Sheila Mae M. Araiza, Kris P. Punayan, Irish Coleen A. Asin, Candice Francheska B. Tambaoan, Asia Louisa U. Chong, Karol Sophia Agape R. Padilla, Rianna Patricia S. Cruz, El King D. Morado, Joshua Gregor A. Dizon, Razel Nikka M. Hao, Arianne A. Zamora, Devon Ray Pacial, Juan Antonio R. Magalang, Marissa Alejandria, Celia Carlos, Anna Ong-Lim, Edsel Maurice Salvaña, John Q. Wong, Jaime C. Montoya, Maria Rosario Singh-Vergeire and Cynthia P. Saloma |
| EPI_ISL_2154559 | Philippine Red Cross Molecular Laboratory | Philippine Genome Center | Francis A. Tablizo, Kenneth M. Kim, Carlo M. Lapid, Marc Jerrone R. Castro, Maria Sofia L. Yangzon, Benedict A. Maralit, Marc Edsel C. Ayes, Eva Maria Cutiongco-de la Paz, Alethea R. de Guzman, Jan Michael C. Yap, Jo-Hannah S. Llames, Sheila Mae M. Araiza, Kris P. Punayan, Irish Coleen A. Asin, Candice Francheska B. Tambaoan, Asia Louisa U. Chong, Karol Sophia Agape R. Padilla, Rianna Patricia S. Cruz, El King D. Morado, Joshua Gregor A. Dizon, Razel Nikka M. Hao, Arianne A. Zamora, Devon Ray Pacial, Juan Antonio R. Magalang, Marissa Alejandria, Celia Carlos, Anna Ong-Lim, Edsel Maurice Salvaña, John Q. Wong, Jaime C. Montoya, Maria Rosario Singh-Vergeire and Cynthia P. Saloma |
| EPI_ISL_2154584 | Cagayan Valley Medical Center Molecular Laboratory | Philippine Genome Center | Francis A. Tablizo, Kenneth M. Kim, Carlo M. Lapid, Marc Jerrone R. Castro, Maria Sofia L. Yangzon, Benedict A. Maralit, Marc Edsel C. Ayes, Eva Maria Cutiongco-de la Paz, Alethea R. de Guzman, Jan Michael C. Yap, Jo-Hannah S. Llames, Sheila Mae M. Araiza, Kris P. Punayan, Irish Coleen A. Asin, Candice Francheska B. Tambaoan, Asia Louisa U. Chong, Karol Sophia Agape R. Padilla, Rianna Patricia S. Cruz, El King D. Morado, Joshua Gregor A. Dizon, Razel Nikka M. Hao, Arianne A. Zamora, Devon Ray Pacial, Juan Antonio R. Magalang, Marissa Alejandria, Celia Carlos, Anna Ong-Lim, Edsel Maurice Salvaña, John Q. Wong, Jaime C. Montoya, Maria Rosario Singh-Vergeire and Cynthia P. Saloma |
| EPI_ISL_2154597 | Research Institute for Tropical Medicine, Inc. (RITM) | Philippine Genome Center | Francis A. Tablizo, Kenneth M. Kim, Carlo M. Lapid, Marc Jerrone R. Castro, Maria Sofia L. Yangzon, Benedict A. Maralit, Marc Edsel C. Ayes, Eva Maria Cutiongco-de la Paz, Alethea R. de Guzman, Jan Michael C. Yap, Jo-Hannah S. Llames, Sheila Mae M. Araiza, Kris P. Punayan, Irish Coleen A. Asin, Candice Francheska B. Tambaoan, Asia Louisa U. Chong, Karol Sophia Agape R. Padilla, Rianna Patricia S. Cruz, El King D. Morado, Joshua Gregor A. Dizon, Razel Nikka M. Hao, Arianne A. Zamora, Devon Ray Pacial, Juan Antonio R. Magalang, Marissa Alejandria, Celia Carlos, Anna Ong-Lim, Edsel Maurice Salvaña, John Q. Wong, Jaime C. Montoya, Maria Rosario Singh-Vergeire and Cynthia P. Saloma |
| EPI_ISL_2154612 | Detoxicare Molecular Diagnostics Laboratory | Philippine Genome Center | Francis A. Tablizo, Kenneth M. Kim, Carlo M. Lapid, Marc Jerrone R. Castro, Maria Sofia L. Yangzon, Benedict A. Maralit, Marc Edsel C. Ayes, Eva Maria Cutiongco-de la Paz, Alethea R. de Guzman, Jan Michael C. Yap, Jo-Hannah S. Llames, Sheila Mae M. Araiza, Kris P. Punayan, Irish Coleen A. Asin, Candice Francheska B. Tambaoan, Asia Louisa U. Chong, Karol Sophia Agape R. Padilla, Rianna Patricia S. Cruz, El King D. Morado, Joshua Gregor A. Dizon, Razel Nikka M. Hao, Arianne A. Zamora, Devon Ray Pacial, Juan Antonio R. Magalang, Marissa Alejandria, Celia Carlos, Anna Ong-Lim, Edsel Maurice Salvaña, John Q. Wong, Jaime C. Montoya, Maria Rosario Singh-Vergeire and Cynthia P. Saloma |
| EPI_ISL_2154613 | Cagayan Valley Medical Center Molecular Laboratory | Philippine Genome Center | Francis A. Tablizo, Kenneth M. Kim, Carlo M. Lapid, Marc Jerrone R. Castro, Maria Sofia L. Yangzon, Benedict A. Maralit, Marc Edsel C. Ayes, Eva Maria Cutiongco-de la Paz, Alethea R. de Guzman, Jan Michael C. Yap, Jo-Hannah S. Llames, Sheila Mae M. Araiza, Kris P. Punayan, Irish Coleen A. Asin, Candice Francheska B. Tambaoan, Asia Louisa U. Chong, Karol Sophia Agape R. Padilla, Rianna Patricia S. Cruz, El King D. Morado, Joshua Gregor A. Dizon, Razel Nikka M. Hao, Arianne A. Zamora, Devon Ray Pacial, Juan Antonio R. Magalang, Marissa Alejandria, Celia Carlos, Anna Ong-Lim, Edsel Maurice Salvaña, John Q. Wong, Jaime C. Montoya, Maria Rosario Singh-Vergeire and Cynthia P. Saloma |
| EPI_ISL_2154632 | South Super Highway Molecular Diagnostic Laboratory | Philippine Genome Center | Francis A. Tablizo, Kenneth M. Kim, Carlo M. Lapid, Marc Jerrone R. Castro, Maria Sofia L. Yangzon, Benedict A. Maralit, Marc Edsel C. Ayes, Eva Maria Cutiongco-de la Paz, Alethea R. de Guzman, Jan Michael C. Yap, Jo-Hannah S. Llames, Sheila Mae M. Araiza, Kris P. Punayan, Irish Coleen A. Asin, Candice Francheska B. Tambaoan, Asia Louisa U. Chong, Karol Sophia Agape R. Padilla, Rianna Patricia S. Cruz, El King D. Morado, Joshua Gregor A. Dizon, Razel Nikka M. Hao, Arianne A. Zamora, Devon Ray Pacial, Juan Antonio R. Magalang, Marissa Alejandria, Celia Carlos, Anna Ong-Lim, Edsel Maurice Salvaña, John Q. Wong, Jaime C. Montoya, Maria Rosario Singh-Vergeire and Cynthia P. Saloma |
| EPI_ISL_2154650 | Research Institute for Tropical Medicine, Inc. (RITM) | Philippine Genome Center | Francis A. Tablizo, Kenneth M. Kim, Carlo M. Lapid, Marc Jerrone R. Castro, Maria Sofia L. Yangzon, Benedict A. Maralit, Marc Edsel C. Ayes, Eva Maria Cutiongco-de la Paz, Alethea R. de Guzman, Jan Michael C. Yap, Jo-Hannah S. Llames, Sheila Mae M. Araiza, Kris P. Punayan, Irish Coleen A. Asin, Candice Francheska B. Tambaoan, Asia Louisa U. Chong, Karol Sophia Agape R. Padilla, Rianna Patricia S. Cruz, El King D. Morado, Joshua Gregor A. Dizon, Razel Nikka M. Hao, Arianne A. Zamora, Devon Ray Pacial, Juan Antonio R. Magalang, Marissa Alejandria, Celia Carlos, Anna Ong-Lim, Edsel Maurice Salvaña, John Q. Wong, Jaime C. Montoya, Maria Rosario Singh-Vergeire and Cynthia P. Saloma |
| EPI_ISL_2154662 | Philippine Red Cross Molecular Laboratory | Philippine Genome Center | Francis A. Tablizo, Kenneth M. Kim, Carlo M. Lapid, Marc Jerrone R. Castro, Maria Sofia L. Yangzon, Benedict A. Maralit, Marc Edsel C. Ayes, Eva Maria Cutiongco-de la Paz, Alethea R. de Guzman, Jan Michael C. Yap, Jo-Hannah S. Llames, Sheila Mae M. Araiza, Kris P. Punayan, Irish Coleen A. Asin, Candice Francheska B. Tambaoan, Asia Louisa U. Chong, Karol Sophia Agape R. Padilla, Rianna Patricia S. Cruz, El King D. Morado, Joshua Gregor A. Dizon, Razel Nikka M. Hao, Arianne A. Zamora, Devon Ray Pacial, Juan Antonio R. Magalang, Marissa Alejandria, Celia Carlos, Anna Ong-Lim, Edsel Maurice Salvaña, John Q. Wong, Jaime C. Montoya, Maria Rosario Singh-Vergeire and Cynthia P. Saloma |
| EPI_ISL_2154684, EPI_ISL_2154687, | Cagayan Valley Medical Center Molecular Laboratory | Philippine Genome Center | Francis A. Tablizo, Kenneth M. Kim, Carlo M. Lapid, Marc Jerrone R. Castro, Maria Sofia L. Yangzon, Benedict A. Maralit, Marc Edsel C. Ayes, Eva Maria |

| | | | |
|---|---|---|---|
| EPI_ISL_2154688 | | | Cutiongco-de la Paz, Alethea R. de Guzman, Jan Michael C. Yap, Jo-Hannah S. Llames, Sheila Mae M. Araiza, Kris P. Punayan, Irish Coleen A. Asin, Candice Francheska B. Tambaoan, Asia Louisa U. Chong, Karol Sophia Agape R. Padilla, Rianna Patricia S. Cruz, El King D. Morado, Joshua Gregor A. Dizon, Razel Nikka M. Hao, Arianne A. Zamora, Devon Ray Pacial, Juan Antonio R. Magalang, Marissa Alejandria, Celia Carlos, Anna Ong-Lim, Edsel Maurice Salvaña, John Q. Wong, Jaime C. Montoya, Maria Rosario Singh-Vergeire and Cynthia P. Saloma |
| EPI_ISL_2154702 | Health Delivery Systems, Inc. | Philippine Genome Center | Francis A. Tablizo, Kenneth M. Kim, Carlo M. Lapid, Marc Jerrone R. Castro, Maria Sofia L. Yangzon, Benedict A. Maralit, Marc Edsel C. Ayes, Eva Maria Cutiongco-de la Paz, Alethea R. de Guzman, Jan Michael C. Yap, Jo-Hannah S. Llames, Sheila Mae M. Araiza, Kris P. Punayan, Irish Coleen A. Asin, Candice Francheska B. Tambaoan, Asia Louisa U. Chong, Karol Sophia Agape R. Padilla, Rianna Patricia S. Cruz, El King D. Morado, Joshua Gregor A. Dizon, Razel Nikka M. Hao, Arianne A. Zamora, Devon Ray Pacial, Juan Antonio R. Magalang, Marissa Alejandria, Celia Carlos, Anna Ong-Lim, Edsel Maurice Salvaña, John Q. Wong, Jaime C. Montoya, Maria Rosario Singh-Vergeire and Cynthia P. Saloma |
| EPI_ISL_2154704 | Cagayan Valley Medical Center Molecular Laboratory | Philippine Genome Center | Francis A. Tablizo, Kenneth M. Kim, Carlo M. Lapid, Marc Jerrone R. Castro, Maria Sofia L. Yangzon, Benedict A. Maralit, Marc Edsel C. Ayes, Eva Maria Cutiongco-de la Paz, Alethea R. de Guzman, Jan Michael C. Yap, Jo-Hannah S. Llames, Sheila Mae M. Araiza, Kris P. Punayan, Irish Coleen A. Asin, Candice Francheska B. Tambaoan, Asia Louisa U. Chong, Karol Sophia Agape R. Padilla, Rianna Patricia S. Cruz, El King D. Morado, Joshua Gregor A. Dizon, Razel Nikka M. Hao, Arianne A. Zamora, Devon Ray Pacial, Juan Antonio R. Magalang, Marissa Alejandria, Celia Carlos, Anna Ong-Lim, Edsel Maurice Salvaña, John Q. Wong, Jaime C. Montoya, Maria Rosario Singh-Vergeire and Cynthia P. Saloma |
| EPI_ISL_2154712 | South Super Highway Molecular Diagnostic Laboratory | Philippine Genome Center | Francis A. Tablizo, Kenneth M. Kim, Carlo M. Lapid, Marc Jerrone R. Castro, Maria Sofia L. Yangzon, Benedict A. Maralit, Marc Edsel C. Ayes, Eva Maria Cutiongco-de la Paz, Alethea R. de Guzman, Jan Michael C. Yap, Jo-Hannah S. Llames, Sheila Mae M. Araiza, Kris P. Punayan, Irish Coleen A. Asin, Candice Francheska B. Tambaoan, Asia Louisa U. Chong, Karol Sophia Agape R. Padilla, Rianna Patricia S. Cruz, El King D. Morado, Joshua Gregor A. Dizon, Razel Nikka M. Hao, Arianne A. Zamora, Devon Ray Pacial, Juan Antonio R. Magalang, Marissa Alejandria, Celia Carlos, Anna Ong-Lim, Edsel Maurice Salvaña, John Q. Wong, Jaime C. Montoya, Maria Rosario Singh-Vergeire and Cynthia P. Saloma |
| EPI_ISL_2154730, EPI_ISL_2154733, EPI_ISL_2154771, EPI_ISL_2154783, EPI_ISL_2154787 | Cagayan Valley Medical Center Molecular Laboratory | Philippine Genome Center | Francis A. Tablizo, Kenneth M. Kim, Carlo M. Lapid, Marc Jerrone R. Castro, Maria Sofia L. Yangzon, Benedict A. Maralit, Marc Edsel C. Ayes, Eva Maria Cutiongco-de la Paz, Alethea R. de Guzman, Jan Michael C. Yap, Jo-Hannah S. Llames, Sheila Mae M. Araiza, Kris P. Punayan, Irish Coleen A. Asin, Candice Francheska B. Tambaoan, Asia Louisa U. Chong, Karol Sophia Agape R. Padilla, Rianna Patricia S. Cruz, El King D. Morado, Joshua Gregor A. Dizon, Razel Nikka M. Hao, Arianne A. Zamora, Devon Ray Pacial, Juan Antonio R. Magalang, Marissa Alejandria, Celia Carlos, Anna Ong-Lim, Edsel Maurice Salvaña, John Q. Wong, Jaime C. Montoya, Maria Rosario Singh-Vergeire and Cynthia P. Saloma |
| EPI_ISL_2154794 | Research Institute for Tropical Medicine, Inc. (RITM) | Philippine Genome Center | Francis A. Tablizo, Kenneth M. Kim, Carlo M. Lapid, Marc Jerrone R. Castro, Maria Sofia L. Yangzon, Benedict A. Maralit, Marc Edsel C. Ayes, Eva Maria Cutiongco-de la Paz, Alethea R. de Guzman, Jan Michael C. Yap, Jo-Hannah S. Llames, Sheila Mae M. Araiza, Kris P. Punayan, Irish Coleen A. Asin, Candice Francheska B. Tambaoan, Asia Louisa U. Chong, Karol Sophia Agape R. Padilla, Rianna Patricia S. Cruz, El King D. Morado, Joshua Gregor A. Dizon, Razel Nikka M. Hao, Arianne A. Zamora, Devon Ray Pacial, Juan Antonio R. Magalang, Marissa Alejandria, Celia Carlos, Anna Ong-Lim, Edsel Maurice Salvaña, John Q. Wong, Jaime C. Montoya, Maria Rosario Singh-Vergeire and Cynthia P. Saloma |
| EPI_ISL_2154808 | South Super Highway Molecular Diagnostic Laboratory | Philippine Genome Center | Francis A. Tablizo, Kenneth M. Kim, Carlo M. Lapid, Marc Jerrone R. Castro, Maria Sofia L. Yangzon, Benedict A. Maralit, Marc Edsel C. Ayes, Eva Maria Cutiongco-de la Paz, Alethea R. de Guzman, Jan Michael C. Yap, Jo-Hannah S. Llames, Sheila Mae M. Araiza, Kris P. Punayan, Irish Coleen A. Asin, Candice Francheska B. Tambaoan, Asia Louisa U. Chong, Karol Sophia Agape R. Padilla, Rianna Patricia S. Cruz, El King D. Morado, Joshua Gregor A. Dizon, Razel Nikka M. Hao, Arianne A. Zamora, Devon Ray Pacial, Juan Antonio R. Magalang, Marissa Alejandria, Celia Carlos, Anna Ong-Lim, Edsel Maurice Salvaña, John Q. Wong, Jaime C. Montoya, Maria Rosario Singh-Vergeire and Cynthia P. Saloma |
| EPI_ISL_2154814, EPI_ISL_2154823, EPI_ISL_2154825 | Cagayan Valley Medical Center Molecular Laboratory | Philippine Genome Center | Francis A. Tablizo, Kenneth M. Kim, Carlo M. Lapid, Marc Jerrone R. Castro, Maria Sofia L. Yangzon, Benedict A. Maralit, Marc Edsel C. Ayes, Eva Maria Cutiongco-de la Paz, Alethea R. de Guzman, Jan Michael C. Yap, Jo-Hannah S. Llames, Sheila Mae M. Araiza, Kris P. Punayan, Irish Coleen A. Asin, Candice Francheska B. Tambaoan, Asia Louisa U. Chong, Karol Sophia Agape R. Padilla, Rianna Patricia S. Cruz, El King D. Morado, Joshua Gregor A. Dizon, Razel Nikka M. Hao, Arianne A. Zamora, Devon Ray Pacial, Juan Antonio R. Magalang, Marissa Alejandria, Celia Carlos, Anna Ong-Lim, Edsel Maurice Salvaña, John Q. Wong, Jaime C. Montoya, Maria Rosario Singh-Vergeire and Cynthia P. Saloma |
| EPI_ISL_2154835 | Research Institute for Tropical Medicine, Inc. (RITM) | Philippine Genome Center | Francis A. Tablizo, Kenneth M. Kim, Carlo M. Lapid, Marc Jerrone R. Castro, Maria Sofia L. Yangzon, Benedict A. Maralit, Marc Edsel C. Ayes, Eva Maria Cutiongco-de la Paz, Alethea R. de Guzman, Jan Michael C. Yap, Jo-Hannah S. Llames, Sheila Mae M. Araiza, Kris P. Punayan, Irish Coleen A. Asin, Candice Francheska B. Tambaoan, Asia Louisa U. Chong, Karol Sophia Agape R. Padilla, Rianna Patricia S. Cruz, El King D. Morado, Joshua Gregor A. Dizon, Razel Nikka M. Hao, Arianne A. Zamora, Devon Ray Pacial, Juan Antonio R. Magalang, Marissa Alejandria, Celia Carlos, Anna Ong-Lim, Edsel Maurice Salvaña, John Q. Wong, Jaime C. Montoya, Maria Rosario Singh-Vergeire and Cynthia P. Saloma |
| EPI_ISL_2154838, EPI_ISL_2154844, EPI_ISL_2154861, EPI_ISL_2154866, EPI_ISL_2154874 | Cagayan Valley Medical Center Molecular Laboratory | Philippine Genome Center | Francis A. Tablizo, Kenneth M. Kim, Carlo M. Lapid, Marc Jerrone R. Castro, Maria Sofia L. Yangzon, Benedict A. Maralit, Marc Edsel C. Ayes, Eva Maria Cutiongco-de la Paz, Alethea R. de Guzman, Jan Michael C. Yap, Jo-Hannah S. Llames, Sheila Mae M. Araiza, Kris P. Punayan, Irish Coleen A. Asin, Candice Francheska B. Tambaoan, Asia Louisa U. Chong, Karol Sophia Agape R. Padilla, Rianna Patricia S. Cruz, El King D. Morado, Joshua Gregor A. Dizon, Razel Nikka M. Hao, Arianne A. Zamora, Devon Ray Pacial, Juan Antonio R. Magalang, Marissa Alejandria, Celia Carlos, Anna Ong-Lim, Edsel Maurice Salvaña, John Q. Wong, Jaime C. Montoya, Maria Rosario Singh-Vergeire and Cynthia P. Saloma |
| EPI_ISL_2154879 | Baguio General Hospital Medical Center (BGHMC) | Philippine Genome Center | Francis A. Tablizo, Kenneth M. Kim, Carlo M. Lapid, Marc Jerrone R. Castro, Maria Sofia L. Yangzon, Benedict A. Maralit, Marc Edsel C. Ayes, Eva Maria Cutiongco-de la Paz, Alethea R. de Guzman, Jan Michael C. Yap, Jo-Hannah S. Llames, Sheila Mae M. Araiza, Kris P. Punayan, Irish Coleen A. Asin, Candice Francheska B. Tambaoan, Asia Louisa U. Chong, Karol Sophia Agape R. Padilla, Rianna Patricia S. Cruz, El King D. Morado, Joshua Gregor A. Dizon, Razel Nikka M. Hao, Arianne A. Zamora, Devon Ray Pacial, Juan Antonio R. Magalang, Marissa Alejandria, Celia Carlos, Anna Ong-Lim, Edsel Maurice Salvaña, John Q. Wong, Jaime C. Montoya, Maria Rosario Singh-Vergeire and Cynthia P. Saloma |
| EPI_ISL_2154880 | Cagayan Valley Medical Center Molecular Laboratory | Philippine Genome Center | Francis A. Tablizo, Kenneth M. Kim, Carlo M. Lapid, Marc Jerrone R. Castro, Maria Sofia L. Yangzon, Benedict A. Maralit, Marc Edsel C. Ayes, Eva Maria Cutiongco-de la Paz, Alethea R. de Guzman, Jan Michael C. Yap, Jo-Hannah S. Llames, Sheila Mae M. Araiza, Kris P. Punayan, Irish Coleen A. Asin, Candice Francheska B. Tambaoan, Asia Louisa U. Chong, Karol Sophia Agape R. Padilla, Rianna Patricia S. Cruz, El King D. Morado, Joshua Gregor A. Dizon, Razel Nikka M. Hao, Arianne A. Zamora, Devon Ray Pacial, Juan Antonio R. Magalang, Marissa Alejandria, Celia Carlos, Anna Ong-Lim, Edsel Maurice Salvaña, John Q. Wong, Jaime C. Montoya, Maria Rosario Singh-Vergeire and Cynthia P. Saloma |
| EPI_ISL_2154883 | Baguio General Hospital Medical Center (BGHMC) | Philippine Genome Center | Francis A. Tablizo, Kenneth M. Kim, Carlo M. Lapid, Marc Jerrone R. Castro, Maria Sofia L. Yangzon, Benedict A. Maralit, Marc Edsel C. Ayes, Eva Maria Cutiongco-de la Paz, Alethea R. de Guzman, Jan Michael C. Yap, Jo-Hannah S. Llames, Sheila Mae M. Araiza, Kris P. Punayan, Irish Coleen A. Asin, Candice Francheska B. Tambaoan, Asia Louisa U. Chong, Karol Sophia Agape R. Padilla, Rianna Patricia S. Cruz, El King D. Morado, Joshua Gregor A. Dizon, Razel Nikka M. Hao, Arianne A. Zamora, Devon Ray Pacial, Juan Antonio R. Magalang, Marissa Alejandria, Celia Carlos, Anna Ong-Lim, Edsel Maurice Salvaña, John Q. Wong, Jaime C. Montoya, Maria Rosario Singh-Vergeire and Cynthia P. Saloma |
| EPI_ISL_2154894 | Detoxicare Molecular Diagnostics Laboratory | Philippine Genome Center | Francis A. Tablizo, Kenneth M. Kim, Carlo M. Lapid, Marc Jerrone R. Castro, Maria Sofia L. Yangzon, Benedict A. Maralit, Marc Edsel C. Ayes, Eva Maria Cutiongco-de la Paz, Alethea R. de Guzman, Jan Michael C. Yap, Jo-Hannah S. Llames, Sheila Mae M. Araiza, Kris P. Punayan, Irish Coleen A. Asin, Candice Francheska B. Tambaoan, Asia Louisa U. Chong, Karol Sophia Agape R. Padilla, Rianna Patricia S. Cruz, El King D. Morado, Joshua Gregor A. Dizon, Razel Nikka M. Hao, Arianne A. Zamora, Devon Ray Pacial, Juan Antonio R. Magalang, Marissa Alejandria, Celia Carlos, Anna Ong-Lim, Edsel Maurice Salvaña, John Q. Wong, Jaime C. Montoya, Maria Rosario Singh-Vergeire and Cynthia P. Saloma |
| EPI_ISL_2154896 | Cagayan Valley Medical Center Molecular Laboratory | Philippine Genome Center | Francis A. Tablizo, Kenneth M. Kim, Carlo M. Lapid, Marc Jerrone R. Castro, Maria Sofia L. Yangzon, Benedict A. Maralit, Marc Edsel C. Ayes, Eva Maria Cutiongco-de la Paz, Alethea R. de Guzman, Jan Michael C. Yap, Jo-Hannah S. Llames, Sheila Mae M. Araiza, Kris P. Punayan, Irish Coleen A. Asin, Candice Francheska B. Tambaoan, Asia Louisa U. Chong, Karol Sophia Agape R. Padilla, Rianna Patricia S. Cruz, El King D. Morado, Joshua Gregor A. Dizon, Razel Nikka M. Hao, Arianne A. Zamora, Devon Ray Pacial, Juan Antonio R. Magalang, Marissa Alejandria, Celia Carlos, Anna Ong-Lim, Edsel Maurice Salvaña, John Q. Wong, Jaime C. Montoya, Maria Rosario Singh-Vergeire and Cynthia P. Saloma |
| EPI_ISL_2154903, EPI_ISL_2154908, EPI_ISL_2155013 | Baguio General Hospital Medical Center (BGHMC) | Philippine Genome Center | Francis A. Tablizo, Kenneth M. Kim, Carlo M. Lapid, Marc Jerrone R. Castro, Maria Sofia L. Yangzon, Benedict A. Maralit, Marc Edsel C. Ayes, Eva Maria Cutiongco-de la Paz, Alethea R. de Guzman, Jan Michael C. Yap, Jo-Hannah S. Llames, Sheila Mae M. Araiza, Kris P. Punayan, Irish Coleen A. Asin, Candice Francheska B. Tambaoan, Asia Louisa U. Chong, Karol Sophia Agape R. Padilla, Rianna Patricia S. Cruz, El King D. Morado, Joshua Gregor A. Dizon, Razel Nikka M. Hao, Arianne A. Zamora, Devon Ray Pacial, Juan Antonio R. Magalang, Marissa Alejandria, Celia Carlos, Anna Ong-Lim, Edsel Maurice Salvaña, John Q. Wong, Jaime C. Montoya, Maria Rosario Singh-Vergeire and Cynthia P. Saloma |

| | | | |
|---|---|---|---|
| EPI_ISL_2155055, EPI_ISL_2155056 | Research Institute for Tropical Medicine, Inc. (RITM) | Philippine Genome Center | Francis A. Tablizo, Kenneth M. Kim, Carlo M. Lapid, Marc Jerrone R. Castro, Maria Sofia L. Yangzon, Benedict A. Maralit, Marc Edsel C. Ayes, Eva Maria Cutiongco-de la Paz, Alethea R. de Guzman, Jan Michael C. Yap, Jo-Hannah S. Llames, Sheila Mae M. Araiza, Kris P. Punayan, Irish Coleen A. Asin, Candice Francheska B. Tambaoan, Asia Louisa U. Chong, Karol Sophia Agape R. Padilla, Rianna Patricia S. Cruz, El King D. Morado, Joshua Gregor A. Dizon, Razel Nikka M. Hao, Arianne A. Zamora, Devon Ray Pacial, Juan Antonio R. Magalang, Marissa Alejandria, Celia Carlos, Anna Ong-Lim, Edsel Maurice Salvaña, John Q. Wong, Jaime C. Montoya, Maria Rosario Singh-Vergeire and Cynthia P. Saloma |
| EPI_ISL_2155059 | Philippine Red Cross - Port Area | Philippine Genome Center | Francis A. Tablizo, Kenneth M. Kim, Carlo M. Lapid, Marc Jerrone R. Castro, Maria Sofia L. Yangzon, Benedict A. Maralit, Marc Edsel C. Ayes, Eva Maria Cutiongco-de la Paz, Alethea R. de Guzman, Jan Michael C. Yap, Jo-Hannah S. Llames, Sheila Mae M. Araiza, Kris P. Punayan, Irish Coleen A. Asin, Candice Francheska B. Tambaoan, Asia Louisa U. Chong, Karol Sophia Agape R. Padilla, Rianna Patricia S. Cruz, El King D. Morado, Joshua Gregor A. Dizon, Razel Nikka M. Hao, Arianne A. Zamora, Devon Ray Pacial, Juan Antonio R. Magalang, Marissa Alejandria, Celia Carlos, Anna Ong-Lim, Edsel Maurice Salvaña, John Q. Wong, Jaime C. Montoya, Maria Rosario Singh-Vergeire and Cynthia P. Saloma |
| EPI_ISL_2155060 | Research Institute for Tropical Medicine, Inc. (RITM) | Philippine Genome Center | Francis A. Tablizo, Kenneth M. Kim, Carlo M. Lapid, Marc Jerrone R. Castro, Maria Sofia L. Yangzon, Benedict A. Maralit, Marc Edsel C. Ayes, Eva Maria Cutiongco-de la Paz, Alethea R. de Guzman, Jan Michael C. Yap, Jo-Hannah S. Llames, Sheila Mae M. Araiza, Kris P. Punayan, Irish Coleen A. Asin, Candice Francheska B. Tambaoan, Asia Louisa U. Chong, Karol Sophia Agape R. Padilla, Rianna Patricia S. Cruz, El King D. Morado, Joshua Gregor A. Dizon, Razel Nikka M. Hao, Arianne A. Zamora, Devon Ray Pacial, Juan Antonio R. Magalang, Marissa Alejandria, Celia Carlos, Anna Ong-Lim, Edsel Maurice Salvaña, John Q. Wong, Jaime C. Montoya, Maria Rosario Singh-Vergeire and Cynthia P. Saloma |
| EPI_ISL_2155069 | Laguna Provincial Hospital - San Pablo District Hospital | Philippine Genome Center | Francis A. Tablizo, Kenneth M. Kim, Carlo M. Lapid, Marc Jerrone R. Castro, Maria Sofia L. Yangzon, Benedict A. Maralit, Marc Edsel C. Ayes, Eva Maria Cutiongco-de la Paz, Alethea R. de Guzman, Jan Michael C. Yap, Jo-Hannah S. Llames, Sheila Mae M. Araiza, Kris P. Punayan, Irish Coleen A. Asin, Candice Francheska B. Tambaoan, Asia Louisa U. Chong, Karol Sophia Agape R. Padilla, Rianna Patricia S. Cruz, El King D. Morado, Joshua Gregor A. Dizon, Razel Nikka M. Hao, Arianne A. Zamora, Devon Ray Pacial, Juan Antonio R. Magalang, Marissa Alejandria, Celia Carlos, Anna Ong-Lim, Edsel Maurice Salvaña, John Q. Wong, Jaime C. Montoya, Maria Rosario Singh-Vergeire and Cynthia P. Saloma |
| EPI_ISL_2155072 | Research Institute for Tropical Medicine, Inc. (RITM) | Philippine Genome Center | Francis A. Tablizo, Kenneth M. Kim, Carlo M. Lapid, Marc Jerrone R. Castro, Maria Sofia L. Yangzon, Benedict A. Maralit, Marc Edsel C. Ayes, Eva Maria Cutiongco-de la Paz, Alethea R. de Guzman, Jan Michael C. Yap, Jo-Hannah S. Llames, Sheila Mae M. Araiza, Kris P. Punayan, Irish Coleen A. Asin, Candice Francheska B. Tambaoan, Asia Louisa U. Chong, Karol Sophia Agape R. Padilla, Rianna Patricia S. Cruz, El King D. Morado, Joshua Gregor A. Dizon, Razel Nikka M. Hao, Arianne A. Zamora, Devon Ray Pacial, Juan Antonio R. Magalang, Marissa Alejandria, Celia Carlos, Anna Ong-Lim, Edsel Maurice Salvaña, John Q. Wong, Jaime C. Montoya, Maria Rosario Singh-Vergeire and Cynthia P. Saloma |
| EPI_ISL_2155073 | Qualimed Hospital Sta. Rosa | Philippine Genome Center | Francis A. Tablizo, Kenneth M. Kim, Carlo M. Lapid, Marc Jerrone R. Castro, Maria Sofia L. Yangzon, Benedict A. Maralit, Marc Edsel C. Ayes, Eva Maria Cutiongco-de la Paz, Alethea R. de Guzman, Jan Michael C. Yap, Jo-Hannah S. Llames, Sheila Mae M. Araiza, Kris P. Punayan, Irish Coleen A. Asin, Candice Francheska B. Tambaoan, Asia Louisa U. Chong, Karol Sophia Agape R. Padilla, Rianna Patricia S. Cruz, El King D. Morado, Joshua Gregor A. Dizon, Razel Nikka M. Hao, Arianne A. Zamora, Devon Ray Pacial, Juan Antonio R. Magalang, Marissa Alejandria, Celia Carlos, Anna Ong-Lim, Edsel Maurice Salvaña, John Q. Wong, Jaime C. Montoya, Maria Rosario Singh-Vergeire and Cynthia P. Saloma |
| EPI_ISL_2155079 | Research Institute for Tropical Medicine, Inc. (RITM) | Philippine Genome Center | Francis A. Tablizo, Kenneth M. Kim, Carlo M. Lapid, Marc Jerrone R. Castro, Maria Sofia L. Yangzon, Benedict A. Maralit, Marc Edsel C. Ayes, Eva Maria Cutiongco-de la Paz, Alethea R. de Guzman, Jan Michael C. Yap, Jo-Hannah S. Llames, Sheila Mae M. Araiza, Kris P. Punayan, Irish Coleen A. Asin, Candice Francheska B. Tambaoan, Asia Louisa U. Chong, Karol Sophia Agape R. Padilla, Rianna Patricia S. Cruz, El King D. Morado, Joshua Gregor A. Dizon, Razel Nikka M. Hao, Arianne A. Zamora, Devon Ray Pacial, Juan Antonio R. Magalang, Marissa Alejandria, Celia Carlos, Anna Ong-Lim, Edsel Maurice Salvaña, John Q. Wong, Jaime C. Montoya, Maria Rosario Singh-Vergeire and Cynthia P. Saloma |
| EPI_ISL_2155080 | Southern Isabela Medical Center Molecular Diagnostic & Research Laboratory (SIMC - MDRL) | Philippine Genome Center | Francis A. Tablizo, Kenneth M. Kim, Carlo M. Lapid, Marc Jerrone R. Castro, Maria Sofia L. Yangzon, Benedict A. Maralit, Marc Edsel C. Ayes, Eva Maria Cutiongco-de la Paz, Alethea R. de Guzman, Jan Michael C. Yap, Jo-Hannah S. Llames, Sheila Mae M. Araiza, Kris P. Punayan, Irish Coleen A. Asin, Candice Francheska B. Tambaoan, Asia Louisa U. Chong, Karol Sophia Agape R. Padilla, Rianna Patricia S. Cruz, El King D. Morado, Joshua Gregor A. Dizon, Razel Nikka M. Hao, Arianne A. Zamora, Devon Ray Pacial, Juan Antonio R. Magalang, Marissa Alejandria, Celia Carlos, Anna Ong-Lim, Edsel Maurice Salvaña, John Q. Wong, Jaime C. Montoya, Maria Rosario Singh-Vergeire and Cynthia P. Saloma |
| EPI_ISL_2155084 | Qualimed Hospital Sta. Rosa | Philippine Genome Center | Francis A. Tablizo, Kenneth M. Kim, Carlo M. Lapid, Marc Jerrone R. Castro, Maria Sofia L. Yangzon, Benedict A. Maralit, Marc Edsel C. Ayes, Eva Maria Cutiongco-de la Paz, Alethea R. de Guzman, Jan Michael C. Yap, Jo-Hannah S. Llames, Sheila Mae M. Araiza, Kris P. Punayan, Irish Coleen A. Asin, Candice Francheska B. Tambaoan, Asia Louisa U. Chong, Karol Sophia Agape R. Padilla, Rianna Patricia S. Cruz, El King D. Morado, Joshua Gregor A. Dizon, Razel Nikka M. Hao, Arianne A. Zamora, Devon Ray Pacial, Juan Antonio R. Magalang, Marissa Alejandria, Celia Carlos, Anna Ong-Lim, Edsel Maurice Salvaña, John Q. Wong, Jaime C. Montoya, Maria Rosario Singh-Vergeire and Cynthia P. Saloma |
| EPI_ISL_2155095, EPI_ISL_2155099, EPI_ISL_2155102 | Research Institute for Tropical Medicine, Inc. (RITM) | Philippine Genome Center | Francis A. Tablizo, Kenneth M. Kim, Carlo M. Lapid, Marc Jerrone R. Castro, Maria Sofia L. Yangzon, Benedict A. Maralit, Marc Edsel C. Ayes, Eva Maria Cutiongco-de la Paz, Alethea R. de Guzman, Jan Michael C. Yap, Jo-Hannah S. Llames, Sheila Mae M. Araiza, Kris P. Punayan, Irish Coleen A. Asin, Candice Francheska B. Tambaoan, Asia Louisa U. Chong, Karol Sophia Agape R. Padilla, Rianna Patricia S. Cruz, El King D. Morado, Joshua Gregor A. Dizon, Razel Nikka M. Hao, Arianne A. Zamora, Devon Ray Pacial, Juan Antonio R. Magalang, Marissa Alejandria, Celia Carlos, Anna Ong-Lim, Edsel Maurice Salvaña, John Q. Wong, Jaime C. Montoya, Maria Rosario Singh-Vergeire and Cynthia P. Saloma |
| EPI_ISL_2155110 | Philippine Red Cross - Port Area | Philippine Genome Center | Francis A. Tablizo, Kenneth M. Kim, Carlo M. Lapid, Marc Jerrone R. Castro, Maria Sofia L. Yangzon, Benedict A. Maralit, Marc Edsel C. Ayes, Eva Maria Cutiongco-de la Paz, Alethea R. de Guzman, Jan Michael C. Yap, Jo-Hannah S. Llames, Sheila Mae M. Araiza, Kris P. Punayan, Irish Coleen A. Asin, Candice Francheska B. Tambaoan, Asia Louisa U. Chong, Karol Sophia Agape R. Padilla, Rianna Patricia S. Cruz, El King D. Morado, Joshua Gregor A. Dizon, Razel Nikka M. Hao, Arianne A. Zamora, Devon Ray Pacial, Juan Antonio R. Magalang, Marissa Alejandria, Celia Carlos, Anna Ong-Lim, Edsel Maurice Salvaña, John Q. Wong, Jaime C. Montoya, Maria Rosario Singh-Vergeire and Cynthia P. Saloma |
| EPI_ISL_2155111, EPI_ISL_2155116, EPI_ISL_2155120, EPI_ISL_2155122 | Research Institute for Tropical Medicine, Inc. (RITM) | Philippine Genome Center | Francis A. Tablizo, Kenneth M. Kim, Carlo M. Lapid, Marc Jerrone R. Castro, Maria Sofia L. Yangzon, Benedict A. Maralit, Marc Edsel C. Ayes, Eva Maria Cutiongco-de la Paz, Alethea R. de Guzman, Jan Michael C. Yap, Jo-Hannah S. Llames, Sheila Mae M. Araiza, Kris P. Punayan, Irish Coleen A. Asin, Candice Francheska B. Tambaoan, Asia Louisa U. Chong, Karol Sophia Agape R. Padilla, Rianna Patricia S. Cruz, El King D. Morado, Joshua Gregor A. Dizon, Razel Nikka M. Hao, Arianne A. Zamora, Devon Ray Pacial, Juan Antonio R. Magalang, Marissa Alejandria, Celia Carlos, Anna Ong-Lim, Edsel Maurice Salvaña, John Q. Wong, Jaime C. Montoya, Maria Rosario Singh-Vergeire and Cynthia P. Saloma |
| EPI_ISL_2155123 | Cagayan Valley Medical Center Molecular Laboratory | Philippine Genome Center | Francis A. Tablizo, Kenneth M. Kim, Carlo M. Lapid, Marc Jerrone R. Castro, Maria Sofia L. Yangzon, Benedict A. Maralit, Marc Edsel C. Ayes, Eva Maria Cutiongco-de la Paz, Alethea R. de Guzman, Jan Michael C. Yap, Jo-Hannah S. Llames, Sheila Mae M. Araiza, Kris P. Punayan, Irish Coleen A. Asin, Candice Francheska B. Tambaoan, Asia Louisa U. Chong, Karol Sophia Agape R. Padilla, Rianna Patricia S. Cruz, El King D. Morado, Joshua Gregor A. Dizon, Razel Nikka M. Hao, Arianne A. Zamora, Devon Ray Pacial, Juan Antonio R. Magalang, Marissa Alejandria, Celia Carlos, Anna Ong-Lim, Edsel Maurice Salvaña, John Q. Wong, Jaime C. Montoya, Maria Rosario Singh-Vergeire and Cynthia P. Saloma |
| EPI_ISL_2155124 | Research Institute for Tropical Medicine, Inc. (RITM) | Philippine Genome Center | Francis A. Tablizo, Kenneth M. Kim, Carlo M. Lapid, Marc Jerrone R. Castro, Maria Sofia L. Yangzon, Benedict A. Maralit, Marc Edsel C. Ayes, Eva Maria Cutiongco-de la Paz, Alethea R. de Guzman, Jan Michael C. Yap, Jo-Hannah S. Llames, Sheila Mae M. Araiza, Kris P. Punayan, Irish Coleen A. Asin, Candice Francheska B. Tambaoan, Asia Louisa U. Chong, Karol Sophia Agape R. Padilla, Rianna Patricia S. Cruz, El King D. Morado, Joshua Gregor A. Dizon, Razel Nikka M. Hao, Arianne A. Zamora, Devon Ray Pacial, Juan Antonio R. Magalang, Marissa Alejandria, Celia Carlos, Anna Ong-Lim, Edsel Maurice Salvaña, John Q. Wong, Jaime C. Montoya, Maria Rosario Singh-Vergeire and Cynthia P. Saloma |
| EPI_ISL_2155137 | Cagayan Valley Medical Center Molecular Laboratory | Philippine Genome Center | Francis A. Tablizo, Kenneth M. Kim, Carlo M. Lapid, Marc Jerrone R. Castro, Maria Sofia L. Yangzon, Benedict A. Maralit, Marc Edsel C. Ayes, Eva Maria Cutiongco-de la Paz, Alethea R. de Guzman, Jan Michael C. Yap, Jo-Hannah S. Llames, Sheila Mae M. Araiza, Kris P. Punayan, Irish Coleen A. Asin, Candice Francheska B. Tambaoan, Asia Louisa U. Chong, Karol Sophia Agape R. Padilla, Rianna Patricia S. Cruz, El King D. Morado, Joshua Gregor A. Dizon, Razel Nikka M. Hao, Arianne A. Zamora, Devon Ray Pacial, Juan Antonio R. Magalang, Marissa Alejandria, Celia Carlos, Anna Ong-Lim, Edsel Maurice Salvaña, John Q. Wong, Jaime C. Montoya, Maria Rosario Singh-Vergeire and Cynthia P. Saloma |
| EPI_ISL_2155141 | SAN PABLO CITY HEALTH OFFICE | Philippine Genome Center | Francis A. Tablizo, Kenneth M. Kim, Carlo M. Lapid, Marc Jerrone R. Castro, Maria Sofia L. Yangzon, Benedict A. Maralit, Marc Edsel C. Ayes, Eva Maria Cutiongco-de la Paz, Alethea R. de Guzman, Jan Michael C. Yap, Jo-Hannah S. Llames, Sheila Mae M. Araiza, Kris P. Punayan, Irish Coleen A. Asin, Candice Francheska B. Tambaoan, Asia Louisa U. Chong, Karol Sophia Agape R. Padilla, Rianna Patricia S. Cruz, El King D. Morado, Joshua Gregor A. Dizon, Razel Nikka M. Hao, Arianne A. Zamora, Devon Ray Pacial, Juan Antonio R. Magalang, Marissa Alejandria, Celia Carlos, Anna Ong-Lim, Edsel Maurice Salvaña, John Q. Wong, Jaime C. Montoya, Maria Rosario Singh-Vergeire and Cynthia P. Saloma |

| | | | |
|---|---|---|---|
| EPI_ISL_2155143 | Research Institute for Tropical Medicine, Inc. (RITM) | Philippine Genome Center | Francis A. Tablizo, Kenneth M. Kim, Carlo M. Lapid, Marc Jerrone R. Castro, Maria Sofia L. Yangzon, Benedict A. Maralit, Marc Edsel C. Ayes, Eva Maria Cutiongco-de la Paz, Alethea R. de Guzman, Jan Michael C. Yap, Jo-Hannah S. Llames, Sheila Mae M. Araiza, Kris P. Punayan, Irish Coleen A. Asin, Candice Francheska B. Tambaoan, Asia Louisa U. Chong, Karol Sophia Agape R. Padilla, Rianna Patricia S. Cruz, El King D. Morado, Joshua Gregor A. Dizon, Razel Nikka M. Hao, Arianne A. Zamora, Devon Ray Pacial, Juan Antonio R. Magalang, Marissa Alejandria, Celia Carlos, Anna Ong-Lim, Edsel Maurice Salvaña, John Q. Wong, Jaime C. Montoya, Maria Rosario Singh-Vergeire and Cynthia P. Saloma |
| EPI_ISL_2155144 | Cagayan Valley Medical Center Molecular Laboratory | Philippine Genome Center | Francis A. Tablizo, Kenneth M. Kim, Carlo M. Lapid, Marc Jerrone R. Castro, Maria Sofia L. Yangzon, Benedict A. Maralit, Marc Edsel C. Ayes, Eva Maria Cutiongco-de la Paz, Alethea R. de Guzman, Jan Michael C. Yap, Jo-Hannah S. Llames, Sheila Mae M. Araiza, Kris P. Punayan, Irish Coleen A. Asin, Candice Francheska B. Tambaoan, Asia Louisa U. Chong, Karol Sophia Agape R. Padilla, Rianna Patricia S. Cruz, El King D. Morado, Joshua Gregor A. Dizon, Razel Nikka M. Hao, Arianne A. Zamora, Devon Ray Pacial, Juan Antonio R. Magalang, Marissa Alejandria, Celia Carlos, Anna Ong-Lim, Edsel Maurice Salvaña, John Q. Wong, Jaime C. Montoya, Maria Rosario Singh-Vergeire and Cynthia P. Saloma |
| EPI_ISL_2155147 | Philippine Red Cross - Port Area | Philippine Genome Center | Francis A. Tablizo, Kenneth M. Kim, Carlo M. Lapid, Marc Jerrone R. Castro, Maria Sofia L. Yangzon, Benedict A. Maralit, Marc Edsel C. Ayes, Eva Maria Cutiongco-de la Paz, Alethea R. de Guzman, Jan Michael C. Yap, Jo-Hannah S. Llames, Sheila Mae M. Araiza, Kris P. Punayan, Irish Coleen A. Asin, Candice Francheska B. Tambaoan, Asia Louisa U. Chong, Karol Sophia Agape R. Padilla, Rianna Patricia S. Cruz, El King D. Morado, Joshua Gregor A. Dizon, Razel Nikka M. Hao, Arianne A. Zamora, Devon Ray Pacial, Juan Antonio R. Magalang, Marissa Alejandria, Celia Carlos, Anna Ong-Lim, Edsel Maurice Salvaña, John Q. Wong, Jaime C. Montoya, Maria Rosario Singh-Vergeire and Cynthia P. Saloma |
| EPI_ISL_2155149, EPI_ISL_2155152 | Research Institute for Tropical Medicine, Inc. (RITM) | Philippine Genome Center | Francis A. Tablizo, Kenneth M. Kim, Carlo M. Lapid, Marc Jerrone R. Castro, Maria Sofia L. Yangzon, Benedict A. Maralit, Marc Edsel C. Ayes, Eva Maria Cutiongco-de la Paz, Alethea R. de Guzman, Jan Michael C. Yap, Jo-Hannah S. Llames, Sheila Mae M. Araiza, Kris P. Punayan, Irish Coleen A. Asin, Candice Francheska B. Tambaoan, Asia Louisa U. Chong, Karol Sophia Agape R. Padilla, Rianna Patricia S. Cruz, El King D. Morado, Joshua Gregor A. Dizon, Razel Nikka M. Hao, Arianne A. Zamora, Devon Ray Pacial, Juan Antonio R. Magalang, Marissa Alejandria, Celia Carlos, Anna Ong-Lim, Edsel Maurice Salvaña, John Q. Wong, Jaime C. Montoya, Maria Rosario Singh-Vergeire and Cynthia P. Saloma |
| EPI_ISL_2155155 | Philippine Children's Medical Center | Philippine Genome Center | Francis A. Tablizo, Kenneth M. Kim, Carlo M. Lapid, Marc Jerrone R. Castro, Maria Sofia L. Yangzon, Benedict A. Maralit, Marc Edsel C. Ayes, Eva Maria Cutiongco-de la Paz, Alethea R. de Guzman, Jan Michael C. Yap, Jo-Hannah S. Llames, Sheila Mae M. Araiza, Kris P. Punayan, Irish Coleen A. Asin, Candice Francheska B. Tambaoan, Asia Louisa U. Chong, Karol Sophia Agape R. Padilla, Rianna Patricia S. Cruz, El King D. Morado, Joshua Gregor A. Dizon, Razel Nikka M. Hao, Arianne A. Zamora, Devon Ray Pacial, Juan Antonio R. Magalang, Marissa Alejandria, Celia Carlos, Anna Ong-Lim, Edsel Maurice Salvaña, John Q. Wong, Jaime C. Montoya, Maria Rosario Singh-Vergeire and Cynthia P. Saloma |
| EPI_ISL_2155156 | Philippine Red Cross - Port Area | Philippine Genome Center | Francis A. Tablizo, Kenneth M. Kim, Carlo M. Lapid, Marc Jerrone R. Castro, Maria Sofia L. Yangzon, Benedict A. Maralit, Marc Edsel C. Ayes, Eva Maria Cutiongco-de la Paz, Alethea R. de Guzman, Jan Michael C. Yap, Jo-Hannah S. Llames, Sheila Mae M. Araiza, Kris P. Punayan, Irish Coleen A. Asin, Candice Francheska B. Tambaoan, Asia Louisa U. Chong, Karol Sophia Agape R. Padilla, Rianna Patricia S. Cruz, El King D. Morado, Joshua Gregor A. Dizon, Razel Nikka M. Hao, Arianne A. Zamora, Devon Ray Pacial, Juan Antonio R. Magalang, Marissa Alejandria, Celia Carlos, Anna Ong-Lim, Edsel Maurice Salvaña, John Q. Wong, Jaime C. Montoya, Maria Rosario Singh-Vergeire and Cynthia P. Saloma |
| EPI_ISL_2155160 | Laguna Provincial Hospital - San Pablo District Hospital | Philippine Genome Center | Francis A. Tablizo, Kenneth M. Kim, Carlo M. Lapid, Marc Jerrone R. Castro, Maria Sofia L. Yangzon, Benedict A. Maralit, Marc Edsel C. Ayes, Eva Maria Cutiongco-de la Paz, Alethea R. de Guzman, Jan Michael C. Yap, Jo-Hannah S. Llames, Sheila Mae M. Araiza, Kris P. Punayan, Irish Coleen A. Asin, Candice Francheska B. Tambaoan, Asia Louisa U. Chong, Karol Sophia Agape R. Padilla, Rianna Patricia S. Cruz, El King D. Morado, Joshua Gregor A. Dizon, Razel Nikka M. Hao, Arianne A. Zamora, Devon Ray Pacial, Juan Antonio R. Magalang, Marissa Alejandria, Celia Carlos, Anna Ong-Lim, Edsel Maurice Salvaña, John Q. Wong, Jaime C. Montoya, Maria Rosario Singh-Vergeire and Cynthia P. Saloma |
| EPI_ISL_2155162, EPI_ISL_2155164, EPI_ISL_2155167 | Research Institute for Tropical Medicine, Inc. (RITM) | Philippine Genome Center | Francis A. Tablizo, Kenneth M. Kim, Carlo M. Lapid, Marc Jerrone R. Castro, Maria Sofia L. Yangzon, Benedict A. Maralit, Marc Edsel C. Ayes, Eva Maria Cutiongco-de la Paz, Alethea R. de Guzman, Jan Michael C. Yap, Jo-Hannah S. Llames, Sheila Mae M. Araiza, Kris P. Punayan, Irish Coleen A. Asin, Candice Francheska B. Tambaoan, Asia Louisa U. Chong, Karol Sophia Agape R. Padilla, Rianna Patricia S. Cruz, El King D. Morado, Joshua Gregor A. Dizon, Razel Nikka M. Hao, Arianne A. Zamora, Devon Ray Pacial, Juan Antonio R. Magalang, Marissa Alejandria, Celia Carlos, Anna Ong-Lim, Edsel Maurice Salvaña, John Q. Wong, Jaime C. Montoya, Maria Rosario Singh-Vergeire and Cynthia P. Saloma |
| EPI_ISL_2155168 | Qualimed Hospital Sta. Rosa | Philippine Genome Center | Francis A. Tablizo, Kenneth M. Kim, Carlo M. Lapid, Marc Jerrone R. Castro, Maria Sofia L. Yangzon, Benedict A. Maralit, Marc Edsel C. Ayes, Eva Maria Cutiongco-de la Paz, Alethea R. de Guzman, Jan Michael C. Yap, Jo-Hannah S. Llames, Sheila Mae M. Araiza, Kris P. Punayan, Irish Coleen A. Asin, Candice Francheska B. Tambaoan, Asia Louisa U. Chong, Karol Sophia Agape R. Padilla, Rianna Patricia S. Cruz, El King D. Morado, Joshua Gregor A. Dizon, Razel Nikka M. Hao, Arianne A. Zamora, Devon Ray Pacial, Juan Antonio R. Magalang, Marissa Alejandria, Celia Carlos, Anna Ong-Lim, Edsel Maurice Salvaña, John Q. Wong, Jaime C. Montoya, Maria Rosario Singh-Vergeire and Cynthia P. Saloma |
| EPI_ISL_2155177, EPI_ISL_2155178, EPI_ISL_2155180 | Research Institute for Tropical Medicine, Inc. (RITM) | Philippine Genome Center | Francis A. Tablizo, Kenneth M. Kim, Carlo M. Lapid, Marc Jerrone R. Castro, Maria Sofia L. Yangzon, Benedict A. Maralit, Marc Edsel C. Ayes, Eva Maria Cutiongco-de la Paz, Alethea R. de Guzman, Jan Michael C. Yap, Jo-Hannah S. Llames, Sheila Mae M. Araiza, Kris P. Punayan, Irish Coleen A. Asin, Candice Francheska B. Tambaoan, Asia Louisa U. Chong, Karol Sophia Agape R. Padilla, Rianna Patricia S. Cruz, El King D. Morado, Joshua Gregor A. Dizon, Razel Nikka M. Hao, Arianne A. Zamora, Devon Ray Pacial, Juan Antonio R. Magalang, Marissa Alejandria, Celia Carlos, Anna Ong-Lim, Edsel Maurice Salvaña, John Q. Wong, Jaime C. Montoya, Maria Rosario Singh-Vergeire and Cynthia P. Saloma |
| EPI_ISL_2155186 | Qualimed Hospital Sta. Rosa | Philippine Genome Center | Francis A. Tablizo, Kenneth M. Kim, Carlo M. Lapid, Marc Jerrone R. Castro, Maria Sofia L. Yangzon, Benedict A. Maralit, Marc Edsel C. Ayes, Eva Maria Cutiongco-de la Paz, Alethea R. de Guzman, Jan Michael C. Yap, Jo-Hannah S. Llames, Sheila Mae M. Araiza, Kris P. Punayan, Irish Coleen A. Asin, Candice Francheska B. Tambaoan, Asia Louisa U. Chong, Karol Sophia Agape R. Padilla, Rianna Patricia S. Cruz, El King D. Morado, Joshua Gregor A. Dizon, Razel Nikka M. Hao, Arianne A. Zamora, Devon Ray Pacial, Juan Antonio R. Magalang, Marissa Alejandria, Celia Carlos, Anna Ong-Lim, Edsel Maurice Salvaña, John Q. Wong, Jaime C. Montoya, Maria Rosario Singh-Vergeire and Cynthia P. Saloma |
| EPI_ISL_2155188 | Laguna Provincial Hospital - San Pablo District Hospital | Philippine Genome Center | Francis A. Tablizo, Kenneth M. Kim, Carlo M. Lapid, Marc Jerrone R. Castro, Maria Sofia L. Yangzon, Benedict A. Maralit, Marc Edsel C. Ayes, Eva Maria Cutiongco-de la Paz, Alethea R. de Guzman, Jan Michael C. Yap, Jo-Hannah S. Llames, Sheila Mae M. Araiza, Kris P. Punayan, Irish Coleen A. Asin, Candice Francheska B. Tambaoan, Asia Louisa U. Chong, Karol Sophia Agape R. Padilla, Rianna Patricia S. Cruz, El King D. Morado, Joshua Gregor A. Dizon, Razel Nikka M. Hao, Arianne A. Zamora, Devon Ray Pacial, Juan Antonio R. Magalang, Marissa Alejandria, Celia Carlos, Anna Ong-Lim, Edsel Maurice Salvaña, John Q. Wong, Jaime C. Montoya, Maria Rosario Singh-Vergeire and Cynthia P. Saloma |
| EPI_ISL_2155191 | Cagayan Valley Medical Center Molecular Laboratory | Philippine Genome Center | Francis A. Tablizo, Kenneth M. Kim, Carlo M. Lapid, Marc Jerrone R. Castro, Maria Sofia L. Yangzon, Benedict A. Maralit, Marc Edsel C. Ayes, Eva Maria Cutiongco-de la Paz, Alethea R. de Guzman, Jan Michael C. Yap, Jo-Hannah S. Llames, Sheila Mae M. Araiza, Kris P. Punayan, Irish Coleen A. Asin, Candice Francheska B. Tambaoan, Asia Louisa U. Chong, Karol Sophia Agape R. Padilla, Rianna Patricia S. Cruz, El King D. Morado, Joshua Gregor A. Dizon, Razel Nikka M. Hao, Arianne A. Zamora, Devon Ray Pacial, Juan Antonio R. Magalang, Marissa Alejandria, Celia Carlos, Anna Ong-Lim, Edsel Maurice Salvaña, John Q. Wong, Jaime C. Montoya, Maria Rosario Singh-Vergeire and Cynthia P. Saloma |
| EPI_ISL_2155192, EPI_ISL_2155197 | Research Institute for Tropical Medicine, Inc. (RITM) | Philippine Genome Center | Francis A. Tablizo, Kenneth M. Kim, Carlo M. Lapid, Marc Jerrone R. Castro, Maria Sofia L. Yangzon, Benedict A. Maralit, Marc Edsel C. Ayes, Eva Maria Cutiongco-de la Paz, Alethea R. de Guzman, Jan Michael C. Yap, Jo-Hannah S. Llames, Sheila Mae M. Araiza, Kris P. Punayan, Irish Coleen A. Asin, Candice Francheska B. Tambaoan, Asia Louisa U. Chong, Karol Sophia Agape R. Padilla, Rianna Patricia S. Cruz, El King D. Morado, Joshua Gregor A. Dizon, Razel Nikka M. Hao, Arianne A. Zamora, Devon Ray Pacial, Juan Antonio R. Magalang, Marissa Alejandria, Celia Carlos, Anna Ong-Lim, Edsel Maurice Salvaña, John Q. Wong, Jaime C. Montoya, Maria Rosario Singh-Vergeire and Cynthia P. Saloma |
| EPI_ISL_2155199 | Laguna Provincial Hospital - San Pablo District Hospital | Philippine Genome Center | Francis A. Tablizo, Kenneth M. Kim, Carlo M. Lapid, Marc Jerrone R. Castro, Maria Sofia L. Yangzon, Benedict A. Maralit, Marc Edsel C. Ayes, Eva Maria Cutiongco-de la Paz, Alethea R. de Guzman, Jan Michael C. Yap, Jo-Hannah S. Llames, Sheila Mae M. Araiza, Kris P. Punayan, Irish Coleen A. Asin, Candice Francheska B. Tambaoan, Asia Louisa U. Chong, Karol Sophia Agape R. Padilla, Rianna Patricia S. Cruz, El King D. Morado, Joshua Gregor A. Dizon, Razel Nikka M. Hao, Arianne A. Zamora, Devon Ray Pacial, Juan Antonio R. Magalang, Marissa Alejandria, Celia Carlos, Anna Ong-Lim, Edsel Maurice Salvaña, John Q. Wong, Jaime C. Montoya, Maria Rosario Singh-Vergeire and Cynthia P. Saloma |
| EPI_ISL_2155203 | Philippine Red Cross - Port Area | Philippine Genome Center | Francis A. Tablizo, Kenneth M. Kim, Carlo M. Lapid, Marc Jerrone R. Castro, Maria Sofia L. Yangzon, Benedict A. Maralit, Marc Edsel C. Ayes, Eva Maria Cutiongco-de la Paz, Alethea R. de Guzman, Jan Michael C. Yap, Jo-Hannah S. Llames, Sheila Mae M. Araiza, Kris P. Punayan, Irish Coleen A. Asin, Candice Francheska B. Tambaoan, Asia Louisa U. Chong, Karol Sophia Agape R. Padilla, Rianna Patricia S. Cruz, El King D. Morado, Joshua Gregor A. Dizon, Razel Nikka M. Hao, Arianne A. Zamora, Devon Ray Pacial, Juan Antonio R. Magalang, Marissa Alejandria, Celia Carlos, Anna Ong-Lim, Edsel Maurice Salvaña, John Q. Wong, Jaime C. Montoya, Maria Rosario Singh-Vergeire and Cynthia P. Saloma |

| | | | |
|---|---|---|---|
| EPI_ISL_2155206 | Research Institute for Tropical Medicine, Inc. (RITM) | Philippine Genome Center | Francis A. Tablizo, Kenneth M. Kim, Carlo M. Lapid, Marc Jerrone R. Castro, Maria Sofia L. Yangzon, Benedict A. Maralit, Marc Edsel C. Ayes, Eva Maria Cutiongco-de la Paz, Alethea R. de Guzman, Jan Michael C. Yap, Jo-Hannah S. Llames, Sheila Mae M. Araiza, Kris P. Punayan, Irish Coleen A. Asin, Candice Francheska B. Tambaoan, Asia Louisa U. Chong, Karol Sophia Agape R. Padilla, Rianna Patricia S. Cruz, El King D. Morado, Joshua Gregor A. Dizon, Razel Nikka M. Hao, Arianne A. Zamora, Devon Ray Pacial, Juan Antonio R. Magalang, Marissa Alejandria, Celia Carlos, Anna Ong-Lim, Edsel Maurice Salvaña, John Q. Wong, Jaime C. Montoya, Maria Rosario Singh-Vergeire and Cynthia P. Saloma |
| EPI_ISL_2155207 | Qualimed Hospital Sta. Rosa | Philippine Genome Center | Francis A. Tablizo, Kenneth M. Kim, Carlo M. Lapid, Marc Jerrone R. Castro, Maria Sofia L. Yangzon, Benedict A. Maralit, Marc Edsel C. Ayes, Eva Maria Cutiongco-de la Paz, Alethea R. de Guzman, Jan Michael C. Yap, Jo-Hannah S. Llames, Sheila Mae M. Araiza, Kris P. Punayan, Irish Coleen A. Asin, Candice Francheska B. Tambaoan, Asia Louisa U. Chong, Karol Sophia Agape R. Padilla, Rianna Patricia S. Cruz, El King D. Morado, Joshua Gregor A. Dizon, Razel Nikka M. Hao, Arianne A. Zamora, Devon Ray Pacial, Juan Antonio R. Magalang, Marissa Alejandria, Celia Carlos, Anna Ong-Lim, Edsel Maurice Salvaña, John Q. Wong, Jaime C. Montoya, Maria Rosario Singh-Vergeire and Cynthia P. Saloma |
| EPI_ISL_2155210, EPI_ISL_2155221, EPI_ISL_2155223 | Research Institute for Tropical Medicine, Inc. (RITM) | Philippine Genome Center | Francis A. Tablizo, Kenneth M. Kim, Carlo M. Lapid, Marc Jerrone R. Castro, Maria Sofia L. Yangzon, Benedict A. Maralit, Marc Edsel C. Ayes, Eva Maria Cutiongco-de la Paz, Alethea R. de Guzman, Jan Michael C. Yap, Jo-Hannah S. Llames, Sheila Mae M. Araiza, Kris P. Punayan, Irish Coleen A. Asin, Candice Francheska B. Tambaoan, Asia Louisa U. Chong, Karol Sophia Agape R. Padilla, Rianna Patricia S. Cruz, El King D. Morado, Joshua Gregor A. Dizon, Razel Nikka M. Hao, Arianne A. Zamora, Devon Ray Pacial, Juan Antonio R. Magalang, Marissa Alejandria, Celia Carlos, Anna Ong-Lim, Edsel Maurice Salvaña, John Q. Wong, Jaime C. Montoya, Maria Rosario Singh-Vergeire and Cynthia P. Saloma |
| EPI_ISL_2155225 | Philippine Red Cross - Port Area | Philippine Genome Center | Francis A. Tablizo, Kenneth M. Kim, Carlo M. Lapid, Marc Jerrone R. Castro, Maria Sofia L. Yangzon, Benedict A. Maralit, Marc Edsel C. Ayes, Eva Maria Cutiongco-de la Paz, Alethea R. de Guzman, Jan Michael C. Yap, Jo-Hannah S. Llames, Sheila Mae M. Araiza, Kris P. Punayan, Irish Coleen A. Asin, Candice Francheska B. Tambaoan, Asia Louisa U. Chong, Karol Sophia Agape R. Padilla, Rianna Patricia S. Cruz, El King D. Morado, Joshua Gregor A. Dizon, Razel Nikka M. Hao, Arianne A. Zamora, Devon Ray Pacial, Juan Antonio R. Magalang, Marissa Alejandria, Celia Carlos, Anna Ong-Lim, Edsel Maurice Salvaña, John Q. Wong, Jaime C. Montoya, Maria Rosario Singh-Vergeire and Cynthia P. Saloma |
| EPI_ISL_2155227 | Research Institute for Tropical Medicine, Inc. (RITM) | Philippine Genome Center | Francis A. Tablizo, Kenneth M. Kim, Carlo M. Lapid, Marc Jerrone R. Castro, Maria Sofia L. Yangzon, Benedict A. Maralit, Marc Edsel C. Ayes, Eva Maria Cutiongco-de la Paz, Alethea R. de Guzman, Jan Michael C. Yap, Jo-Hannah S. Llames, Sheila Mae M. Araiza, Kris P. Punayan, Irish Coleen A. Asin, Candice Francheska B. Tambaoan, Asia Louisa U. Chong, Karol Sophia Agape R. Padilla, Rianna Patricia S. Cruz, El King D. Morado, Joshua Gregor A. Dizon, Razel Nikka M. Hao, Arianne A. Zamora, Devon Ray Pacial, Juan Antonio R. Magalang, Marissa Alejandria, Celia Carlos, Anna Ong-Lim, Edsel Maurice Salvaña, John Q. Wong, Jaime C. Montoya, Maria Rosario Singh-Vergeire and Cynthia P. Saloma |
| EPI_ISL_2155237 | Philippine Red Cross - Port Area | Philippine Genome Center | Francis A. Tablizo, Kenneth M. Kim, Carlo M. Lapid, Marc Jerrone R. Castro, Maria Sofia L. Yangzon, Benedict A. Maralit, Marc Edsel C. Ayes, Eva Maria Cutiongco-de la Paz, Alethea R. de Guzman, Jan Michael C. Yap, Jo-Hannah S. Llames, Sheila Mae M. Araiza, Kris P. Punayan, Irish Coleen A. Asin, Candice Francheska B. Tambaoan, Asia Louisa U. Chong, Karol Sophia Agape R. Padilla, Rianna Patricia S. Cruz, El King D. Morado, Joshua Gregor A. Dizon, Razel Nikka M. Hao, Arianne A. Zamora, Devon Ray Pacial, Juan Antonio R. Magalang, Marissa Alejandria, Celia Carlos, Anna Ong-Lim, Edsel Maurice Salvaña, John Q. Wong, Jaime C. Montoya, Maria Rosario Singh-Vergeire and Cynthia P. Saloma |
| EPI_ISL_2155238 | Cagayan Valley Medical Center Molecular Laboratory | Philippine Genome Center | Francis A. Tablizo, Kenneth M. Kim, Carlo M. Lapid, Marc Jerrone R. Castro, Maria Sofia L. Yangzon, Benedict A. Maralit, Marc Edsel C. Ayes, Eva Maria Cutiongco-de la Paz, Alethea R. de Guzman, Jan Michael C. Yap, Jo-Hannah S. Llames, Sheila Mae M. Araiza, Kris P. Punayan, Irish Coleen A. Asin, Candice Francheska B. Tambaoan, Asia Louisa U. Chong, Karol Sophia Agape R. Padilla, Rianna Patricia S. Cruz, El King D. Morado, Joshua Gregor A. Dizon, Razel Nikka M. Hao, Arianne A. Zamora, Devon Ray Pacial, Juan Antonio R. Magalang, Marissa Alejandria, Celia Carlos, Anna Ong-Lim, Edsel Maurice Salvaña, John Q. Wong, Jaime C. Montoya, Maria Rosario Singh-Vergeire and Cynthia P. Saloma |
| EPI_ISL_2155241 | Research Institute for Tropical Medicine, Inc. (RITM) | Philippine Genome Center | Francis A. Tablizo, Kenneth M. Kim, Carlo M. Lapid, Marc Jerrone R. Castro, Maria Sofia L. Yangzon, Benedict A. Maralit, Marc Edsel C. Ayes, Eva Maria Cutiongco-de la Paz, Alethea R. de Guzman, Jan Michael C. Yap, Jo-Hannah S. Llames, Sheila Mae M. Araiza, Kris P. Punayan, Irish Coleen A. Asin, Candice Francheska B. Tambaoan, Asia Louisa U. Chong, Karol Sophia Agape R. Padilla, Rianna Patricia S. Cruz, El King D. Morado, Joshua Gregor A. Dizon, Razel Nikka M. Hao, Arianne A. Zamora, Devon Ray Pacial, Juan Antonio R. Magalang, Marissa Alejandria, Celia Carlos, Anna Ong-Lim, Edsel Maurice Salvaña, John Q. Wong, Jaime C. Montoya, Maria Rosario Singh-Vergeire and Cynthia P. Saloma |
| EPI_ISL_2155243 | Qualimed Hospital Sta. Rosa | Philippine Genome Center | Francis A. Tablizo, Kenneth M. Kim, Carlo M. Lapid, Marc Jerrone R. Castro, Maria Sofia L. Yangzon, Benedict A. Maralit, Marc Edsel C. Ayes, Eva Maria Cutiongco-de la Paz, Alethea R. de Guzman, Jan Michael C. Yap, Jo-Hannah S. Llames, Sheila Mae M. Araiza, Kris P. Punayan, Irish Coleen A. Asin, Candice Francheska B. Tambaoan, Asia Louisa U. Chong, Karol Sophia Agape R. Padilla, Rianna Patricia S. Cruz, El King D. Morado, Joshua Gregor A. Dizon, Razel Nikka M. Hao, Arianne A. Zamora, Devon Ray Pacial, Juan Antonio R. Magalang, Marissa Alejandria, Celia Carlos, Anna Ong-Lim, Edsel Maurice Salvaña, John Q. Wong, Jaime C. Montoya, Maria Rosario Singh-Vergeire and Cynthia P. Saloma |
| EPI_ISL_2155244, EPI_ISL_2155248 | Research Institute for Tropical Medicine, Inc. (RITM) | Philippine Genome Center | Francis A. Tablizo, Kenneth M. Kim, Carlo M. Lapid, Marc Jerrone R. Castro, Maria Sofia L. Yangzon, Benedict A. Maralit, Marc Edsel C. Ayes, Eva Maria Cutiongco-de la Paz, Alethea R. de Guzman, Jan Michael C. Yap, Jo-Hannah S. Llames, Sheila Mae M. Araiza, Kris P. Punayan, Irish Coleen A. Asin, Candice Francheska B. Tambaoan, Asia Louisa U. Chong, Karol Sophia Agape R. Padilla, Rianna Patricia S. Cruz, El King D. Morado, Joshua Gregor A. Dizon, Razel Nikka M. Hao, Arianne A. Zamora, Devon Ray Pacial, Juan Antonio R. Magalang, Marissa Alejandria, Celia Carlos, Anna Ong-Lim, Edsel Maurice Salvaña, John Q. Wong, Jaime C. Montoya, Maria Rosario Singh-Vergeire and Cynthia P. Saloma |
| EPI_ISL_2155254, EPI_ISL_2155260 | Philippine Red Cross - Port Area | Philippine Genome Center | Francis A. Tablizo, Kenneth M. Kim, Carlo M. Lapid, Marc Jerrone R. Castro, Maria Sofia L. Yangzon, Benedict A. Maralit, Marc Edsel C. Ayes, Eva Maria Cutiongco-de la Paz, Alethea R. de Guzman, Jan Michael C. Yap, Jo-Hannah S. Llames, Sheila Mae M. Araiza, Kris P. Punayan, Irish Coleen A. Asin, Candice Francheska B. Tambaoan, Asia Louisa U. Chong, Karol Sophia Agape R. Padilla, Rianna Patricia S. Cruz, El King D. Morado, Joshua Gregor A. Dizon, Razel Nikka M. Hao, Arianne A. Zamora, Devon Ray Pacial, Juan Antonio R. Magalang, Marissa Alejandria, Celia Carlos, Anna Ong-Lim, Edsel Maurice Salvaña, John Q. Wong, Jaime C. Montoya, Maria Rosario Singh-Vergeire and Cynthia P. Saloma |
| EPI_ISL_2155261, EPI_ISL_2155266 | Research Institute for Tropical Medicine, Inc. (RITM) | Philippine Genome Center | Francis A. Tablizo, Kenneth M. Kim, Carlo M. Lapid, Marc Jerrone R. Castro, Maria Sofia L. Yangzon, Benedict A. Maralit, Marc Edsel C. Ayes, Eva Maria Cutiongco-de la Paz, Alethea R. de Guzman, Jan Michael C. Yap, Jo-Hannah S. Llames, Sheila Mae M. Araiza, Kris P. Punayan, Irish Coleen A. Asin, Candice Francheska B. Tambaoan, Asia Louisa U. Chong, Karol Sophia Agape R. Padilla, Rianna Patricia S. Cruz, El King D. Morado, Joshua Gregor A. Dizon, Razel Nikka M. Hao, Arianne A. Zamora, Devon Ray Pacial, Juan Antonio R. Magalang, Marissa Alejandria, Celia Carlos, Anna Ong-Lim, Edsel Maurice Salvaña, John Q. Wong, Jaime C. Montoya, Maria Rosario Singh-Vergeire and Cynthia P. Saloma |
| EPI_ISL_2155271 | Laguna Provincial Hospital - San Pablo District Hospital | Philippine Genome Center | Francis A. Tablizo, Kenneth M. Kim, Carlo M. Lapid, Marc Jerrone R. Castro, Maria Sofia L. Yangzon, Benedict A. Maralit, Marc Edsel C. Ayes, Eva Maria Cutiongco-de la Paz, Alethea R. de Guzman, Jan Michael C. Yap, Jo-Hannah S. Llames, Sheila Mae M. Araiza, Kris P. Punayan, Irish Coleen A. Asin, Candice Francheska B. Tambaoan, Asia Louisa U. Chong, Karol Sophia Agape R. Padilla, Rianna Patricia S. Cruz, El King D. Morado, Joshua Gregor A. Dizon, Razel Nikka M. Hao, Arianne A. Zamora, Devon Ray Pacial, Juan Antonio R. Magalang, Marissa Alejandria, Celia Carlos, Anna Ong-Lim, Edsel Maurice Salvaña, John Q. Wong, Jaime C. Montoya, Maria Rosario Singh-Vergeire and Cynthia P. Saloma |
| EPI_ISL_2155273, EPI_ISL_2155281, EPI_ISL_2155287 | Research Institute for Tropical Medicine, Inc. (RITM) | Philippine Genome Center | Francis A. Tablizo, Kenneth M. Kim, Carlo M. Lapid, Marc Jerrone R. Castro, Maria Sofia L. Yangzon, Benedict A. Maralit, Marc Edsel C. Ayes, Eva Maria Cutiongco-de la Paz, Alethea R. de Guzman, Jan Michael C. Yap, Jo-Hannah S. Llames, Sheila Mae M. Araiza, Kris P. Punayan, Irish Coleen A. Asin, Candice Francheska B. Tambaoan, Asia Louisa U. Chong, Karol Sophia Agape R. Padilla, Rianna Patricia S. Cruz, El King D. Morado, Joshua Gregor A. Dizon, Razel Nikka M. Hao, Arianne A. Zamora, Devon Ray Pacial, Juan Antonio R. Magalang, Marissa Alejandria, Celia Carlos, Anna Ong-Lim, Edsel Maurice Salvaña, John Q. Wong, Jaime C. Montoya, Maria Rosario Singh-Vergeire and Cynthia P. Saloma |
| EPI_ISL_2155294 | Philippine Red Cross - Port Area | Philippine Genome Center | Francis A. Tablizo, Kenneth M. Kim, Carlo M. Lapid, Marc Jerrone R. Castro, Maria Sofia L. Yangzon, Benedict A. Maralit, Marc Edsel C. Ayes, Eva Maria Cutiongco-de la Paz, Alethea R. de Guzman, Jan Michael C. Yap, Jo-Hannah S. Llames, Sheila Mae M. Araiza, Kris P. Punayan, Irish Coleen A. Asin, Candice Francheska B. Tambaoan, Asia Louisa U. Chong, Karol Sophia Agape R. Padilla, Rianna Patricia S. Cruz, El King D. Morado, Joshua Gregor A. Dizon, Razel Nikka M. Hao, Arianne A. Zamora, Devon Ray Pacial, Juan Antonio R. Magalang, Marissa Alejandria, Celia Carlos, Anna Ong-Lim, Edsel Maurice Salvaña, John Q. Wong, Jaime C. Montoya, Maria Rosario Singh-Vergeire and Cynthia P. Saloma |
| EPI_ISL_2155296 | Research Institute for Tropical Medicine, Inc. (RITM) | Philippine Genome Center | Francis A. Tablizo, Kenneth M. Kim, Carlo M. Lapid, Marc Jerrone R. Castro, Maria Sofia L. Yangzon, Benedict A. Maralit, Marc Edsel C. Ayes, Eva Maria Cutiongco-de la Paz, Alethea R. de Guzman, Jan Michael C. Yap, Jo-Hannah S. Llames, Sheila Mae M. Araiza, Kris P. Punayan, Irish Coleen A. Asin, Candice Francheska B. Tambaoan, Asia Louisa U. Chong, Karol Sophia Agape R. Padilla, Rianna Patricia S. Cruz, El King D. Morado, Joshua Gregor A. Dizon, Razel Nikka M. Hao, Arianne A. Zamora, Devon Ray Pacial, Juan Antonio R. Magalang, Marissa Alejandria, Celia Carlos, Anna Ong-Lim, Edsel Maurice Salvaña, John Q. Wong, Jaime C. Montoya, Maria Rosario Singh-Vergeire and Cynthia P. Saloma |

| | | | |
|---|---|---|---|
| EPI_ISL_2155302 | Philippine Children's Medical Center | Philippine Genome Center | Francis A. Tablizo, Kenneth M. Kim, Carlo M. Lapid, Marc Jerrone R. Castro, Maria Sofia L. Yangzon, Benedict A. Maralit, Marc Edsel C. Ayes, Eva Maria Cutiongco-de la Paz, Alethea R. de Guzman, Jan Michael C. Yap, Jo-Hannah S. Llames, Sheila Mae M. Araiza, Kris P. Punayan, Irish Coleen A. Asin, Candice Francheska B. Tambaoan, Asia Louisa U. Chong, Karol Sophia Agape R. Padilla, Rianna Patricia S. Cruz, El King D. Morado, Joshua Gregor A. Dizon, Razel Nikka M. Hao, Arianne A. Zamora, Devon Ray Pacial, Juan Antonio R. Magalang, Marissa Alejandria, Celia Carlos, Anna Ong-Lim, Edsel Maurice Salvaña, John Q. Wong, Jaime C. Montoya, Maria Rosario Singh-Vergeire and Cynthia P. Saloma |
| EPI_ISL_2155303 | Philippine Red Cross - National Blood Center | Philippine Genome Center | Francis A. Tablizo, Kenneth M. Kim, Carlo M. Lapid, Marc Jerrone R. Castro, Maria Sofia L. Yangzon, Benedict A. Maralit, Marc Edsel C. Ayes, Eva Maria Cutiongco-de la Paz, Alethea R. de Guzman, Jan Michael C. Yap, Jo-Hannah S. Llames, Sheila Mae M. Araiza, Kris P. Punayan, Irish Coleen A. Asin, Candice Francheska B. Tambaoan, Asia Louisa U. Chong, Karol Sophia Agape R. Padilla, Rianna Patricia S. Cruz, El King D. Morado, Joshua Gregor A. Dizon, Razel Nikka M. Hao, Arianne A. Zamora, Devon Ray Pacial, Juan Antonio R. Magalang, Marissa Alejandria, Celia Carlos, Anna Ong-Lim, Edsel Maurice Salvaña, John Q. Wong, Jaime C. Montoya, Maria Rosario Singh-Vergeire and Cynthia P. Saloma |
| EPI_ISL_2155307, EPI_ISL_2155312, EPI_ISL_2155314, EPI_ISL_2155322 | Research Institute for Tropical Medicine, Inc. (RITM) | Philippine Genome Center | Francis A. Tablizo, Kenneth M. Kim, Carlo M. Lapid, Marc Jerrone R. Castro, Maria Sofia L. Yangzon, Benedict A. Maralit, Marc Edsel C. Ayes, Eva Maria Cutiongco-de la Paz, Alethea R. de Guzman, Jan Michael C. Yap, Jo-Hannah S. Llames, Sheila Mae M. Araiza, Kris P. Punayan, Irish Coleen A. Asin, Candice Francheska B. Tambaoan, Asia Louisa U. Chong, Karol Sophia Agape R. Padilla, Rianna Patricia S. Cruz, El King D. Morado, Joshua Gregor A. Dizon, Razel Nikka M. Hao, Arianne A. Zamora, Devon Ray Pacial, Juan Antonio R. Magalang, Marissa Alejandria, Celia Carlos, Anna Ong-Lim, Edsel Maurice Salvaña, John Q. Wong, Jaime C. Montoya, Maria Rosario Singh-Vergeire and Cynthia P. Saloma |
| EPI_ISL_2155325 | Philippine Children's Medical Center | Philippine Genome Center | Francis A. Tablizo, Kenneth M. Kim, Carlo M. Lapid, Marc Jerrone R. Castro, Maria Sofia L. Yangzon, Benedict A. Maralit, Marc Edsel C. Ayes, Eva Maria Cutiongco-de la Paz, Alethea R. de Guzman, Jan Michael C. Yap, Jo-Hannah S. Llames, Sheila Mae M. Araiza, Kris P. Punayan, Irish Coleen A. Asin, Candice Francheska B. Tambaoan, Asia Louisa U. Chong, Karol Sophia Agape R. Padilla, Rianna Patricia S. Cruz, El King D. Morado, Joshua Gregor A. Dizon, Razel Nikka M. Hao, Arianne A. Zamora, Devon Ray Pacial, Juan Antonio R. Magalang, Marissa Alejandria, Celia Carlos, Anna Ong-Lim, Edsel Maurice Salvaña, John Q. Wong, Jaime C. Montoya, Maria Rosario Singh-Vergeire and Cynthia P. Saloma |
| EPI_ISL_2155326, EPI_ISL_2155332, EPI_ISL_2155334, EPI_ISL_2155335 | Research Institute for Tropical Medicine, Inc. (RITM) | Philippine Genome Center | Francis A. Tablizo, Kenneth M. Kim, Carlo M. Lapid, Marc Jerrone R. Castro, Maria Sofia L. Yangzon, Benedict A. Maralit, Marc Edsel C. Ayes, Eva Maria Cutiongco-de la Paz, Alethea R. de Guzman, Jan Michael C. Yap, Jo-Hannah S. Llames, Sheila Mae M. Araiza, Kris P. Punayan, Irish Coleen A. Asin, Candice Francheska B. Tambaoan, Asia Louisa U. Chong, Karol Sophia Agape R. Padilla, Rianna Patricia S. Cruz, El King D. Morado, Joshua Gregor A. Dizon, Razel Nikka M. Hao, Arianne A. Zamora, Devon Ray Pacial, Juan Antonio R. Magalang, Marissa Alejandria, Celia Carlos, Anna Ong-Lim, Edsel Maurice Salvaña, John Q. Wong, Jaime C. Montoya, Maria Rosario Singh-Vergeire and Cynthia P. Saloma |
| EPI_ISL_2155337 | Cagayan Valley Medical Center Molecular Laboratory | Philippine Genome Center | Francis A. Tablizo, Kenneth M. Kim, Carlo M. Lapid, Marc Jerrone R. Castro, Maria Sofia L. Yangzon, Benedict A. Maralit, Marc Edsel C. Ayes, Eva Maria Cutiongco-de la Paz, Alethea R. de Guzman, Jan Michael C. Yap, Jo-Hannah S. Llames, Sheila Mae M. Araiza, Kris P. Punayan, Irish Coleen A. Asin, Candice Francheska B. Tambaoan, Asia Louisa U. Chong, Karol Sophia Agape R. Padilla, Rianna Patricia S. Cruz, El King D. Morado, Joshua Gregor A. Dizon, Razel Nikka M. Hao, Arianne A. Zamora, Devon Ray Pacial, Juan Antonio R. Magalang, Marissa Alejandria, Celia Carlos, Anna Ong-Lim, Edsel Maurice Salvaña, John Q. Wong, Jaime C. Montoya, Maria Rosario Singh-Vergeire and Cynthia P. Saloma |
| EPI_ISL_2155340 | Research Institute for Tropical Medicine, Inc. (RITM) | Philippine Genome Center | Francis A. Tablizo, Kenneth M. Kim, Carlo M. Lapid, Marc Jerrone R. Castro, Maria Sofia L. Yangzon, Benedict A. Maralit, Marc Edsel C. Ayes, Eva Maria Cutiongco-de la Paz, Alethea R. de Guzman, Jan Michael C. Yap, Jo-Hannah S. Llames, Sheila Mae M. Araiza, Kris P. Punayan, Irish Coleen A. Asin, Candice Francheska B. Tambaoan, Asia Louisa U. Chong, Karol Sophia Agape R. Padilla, Rianna Patricia S. Cruz, El King D. Morado, Joshua Gregor A. Dizon, Razel Nikka M. Hao, Arianne A. Zamora, Devon Ray Pacial, Juan Antonio R. Magalang, Marissa Alejandria, Celia Carlos, Anna Ong-Lim, Edsel Maurice Salvaña, John Q. Wong, Jaime C. Montoya, Maria Rosario Singh-Vergeire and Cynthia P. Saloma |
| EPI_ISL_2155344 | Qualimed Hospital Sta. Rosa | Philippine Genome Center | Francis A. Tablizo, Kenneth M. Kim, Carlo M. Lapid, Marc Jerrone R. Castro, Maria Sofia L. Yangzon, Benedict A. Maralit, Marc Edsel C. Ayes, Eva Maria Cutiongco-de la Paz, Alethea R. de Guzman, Jan Michael C. Yap, Jo-Hannah S. Llames, Sheila Mae M. Araiza, Kris P. Punayan, Irish Coleen A. Asin, Candice Francheska B. Tambaoan, Asia Louisa U. Chong, Karol Sophia Agape R. Padilla, Rianna Patricia S. Cruz, El King D. Morado, Joshua Gregor A. Dizon, Razel Nikka M. Hao, Arianne A. Zamora, Devon Ray Pacial, Juan Antonio R. Magalang, Marissa Alejandria, Celia Carlos, Anna Ong-Lim, Edsel Maurice Salvaña, John Q. Wong, Jaime C. Montoya, Maria Rosario Singh-Vergeire and Cynthia P. Saloma |
| EPI_ISL_2155347 | Research Institute for Tropical Medicine, Inc. (RITM) | Philippine Genome Center | Francis A. Tablizo, Kenneth M. Kim, Carlo M. Lapid, Marc Jerrone R. Castro, Maria Sofia L. Yangzon, Benedict A. Maralit, Marc Edsel C. Ayes, Eva Maria Cutiongco-de la Paz, Alethea R. de Guzman, Jan Michael C. Yap, Jo-Hannah S. Llames, Sheila Mae M. Araiza, Kris P. Punayan, Irish Coleen A. Asin, Candice Francheska B. Tambaoan, Asia Louisa U. Chong, Karol Sophia Agape R. Padilla, Rianna Patricia S. Cruz, El King D. Morado, Joshua Gregor A. Dizon, Razel Nikka M. Hao, Arianne A. Zamora, Devon Ray Pacial, Juan Antonio R. Magalang, Marissa Alejandria, Celia Carlos, Anna Ong-Lim, Edsel Maurice Salvaña, John Q. Wong, Jaime C. Montoya, Maria Rosario Singh-Vergeire and Cynthia P. Saloma |
| EPI_ISL_2155348 | Philippine Children's Medical Center | Philippine Genome Center | Francis A. Tablizo, Kenneth M. Kim, Carlo M. Lapid, Marc Jerrone R. Castro, Maria Sofia L. Yangzon, Benedict A. Maralit, Marc Edsel C. Ayes, Eva Maria Cutiongco-de la Paz, Alethea R. de Guzman, Jan Michael C. Yap, Jo-Hannah S. Llames, Sheila Mae M. Araiza, Kris P. Punayan, Irish Coleen A. Asin, Candice Francheska B. Tambaoan, Asia Louisa U. Chong, Karol Sophia Agape R. Padilla, Rianna Patricia S. Cruz, El King D. Morado, Joshua Gregor A. Dizon, Razel Nikka M. Hao, Arianne A. Zamora, Devon Ray Pacial, Juan Antonio R. Magalang, Marissa Alejandria, Celia Carlos, Anna Ong-Lim, Edsel Maurice Salvaña, John Q. Wong, Jaime C. Montoya, Maria Rosario Singh-Vergeire and Cynthia P. Saloma |
| EPI_ISL_2155349 | Philippine Red Cross - Port Area | Philippine Genome Center | Francis A. Tablizo, Kenneth M. Kim, Carlo M. Lapid, Marc Jerrone R. Castro, Maria Sofia L. Yangzon, Benedict A. Maralit, Marc Edsel C. Ayes, Eva Maria Cutiongco-de la Paz, Alethea R. de Guzman, Jan Michael C. Yap, Jo-Hannah S. Llames, Sheila Mae M. Araiza, Kris P. Punayan, Irish Coleen A. Asin, Candice Francheska B. Tambaoan, Asia Louisa U. Chong, Karol Sophia Agape R. Padilla, Rianna Patricia S. Cruz, El King D. Morado, Joshua Gregor A. Dizon, Razel Nikka M. Hao, Arianne A. Zamora, Devon Ray Pacial, Juan Antonio R. Magalang, Marissa Alejandria, Celia Carlos, Anna Ong-Lim, Edsel Maurice Salvaña, John Q. Wong, Jaime C. Montoya, Maria Rosario Singh-Vergeire and Cynthia P. Saloma |
| EPI_ISL_2155350 | Research Institute for Tropical Medicine, Inc. (RITM) | Philippine Genome Center | Francis A. Tablizo, Kenneth M. Kim, Carlo M. Lapid, Marc Jerrone R. Castro, Maria Sofia L. Yangzon, Benedict A. Maralit, Marc Edsel C. Ayes, Eva Maria Cutiongco-de la Paz, Alethea R. de Guzman, Jan Michael C. Yap, Jo-Hannah S. Llames, Sheila Mae M. Araiza, Kris P. Punayan, Irish Coleen A. Asin, Candice Francheska B. Tambaoan, Asia Louisa U. Chong, Karol Sophia Agape R. Padilla, Rianna Patricia S. Cruz, El King D. Morado, Joshua Gregor A. Dizon, Razel Nikka M. Hao, Arianne A. Zamora, Devon Ray Pacial, Juan Antonio R. Magalang, Marissa Alejandria, Celia Carlos, Anna Ong-Lim, Edsel Maurice Salvaña, John Q. Wong, Jaime C. Montoya, Maria Rosario Singh-Vergeire and Cynthia P. Saloma |
| EPI_ISL_2155357 | Cagayan Valley Medical Center Molecular Laboratory | Philippine Genome Center | Francis A. Tablizo, Kenneth M. Kim, Carlo M. Lapid, Marc Jerrone R. Castro, Maria Sofia L. Yangzon, Benedict A. Maralit, Marc Edsel C. Ayes, Eva Maria Cutiongco-de la Paz, Alethea R. de Guzman, Jan Michael C. Yap, Jo-Hannah S. Llames, Sheila Mae M. Araiza, Kris P. Punayan, Irish Coleen A. Asin, Candice Francheska B. Tambaoan, Asia Louisa U. Chong, Karol Sophia Agape R. Padilla, Rianna Patricia S. Cruz, El King D. Morado, Joshua Gregor A. Dizon, Razel Nikka M. Hao, Arianne A. Zamora, Devon Ray Pacial, Juan Antonio R. Magalang, Marissa Alejandria, Celia Carlos, Anna Ong-Lim, Edsel Maurice Salvaña, John Q. Wong, Jaime C. Montoya, Maria Rosario Singh-Vergeire and Cynthia P. Saloma |
| EPI_ISL_2155412, EPI_ISL_2155414 | Fatima University Medical Center Antipolo Corp. | Philippine Genome Center | Francis A. Tablizo, Kenneth M. Kim, Carlo M. Lapid, Marc Jerrone R. Castro, Maria Sofia L. Yangzon, Benedict A. Maralit, Marc Edsel C. Ayes, Eva Maria Cutiongco-de la Paz, Alethea R. de Guzman, Jan Michael C. Yap, Jo-Hannah S. Llames, Sheila Mae M. Araiza, Kris P. Punayan, Irish Coleen A. Asin, Candice Francheska B. Tambaoan, Asia Louisa U. Chong, Karol Sophia Agape R. Padilla, Rianna Patricia S. Cruz, El King D. Morado, Joshua Gregor A. Dizon, Razel Nikka M. Hao, Arianne A. Zamora, Devon Ray Pacial, Juan Antonio R. Magalang, Marissa Alejandria, Celia Carlos, Anna Ong-Lim, Edsel Maurice Salvaña, John Q. Wong, Jaime C. Montoya, Maria Rosario Singh-Vergeire and Cynthia P. Saloma |
| EPI_ISL_2155438, EPI_ISL_2155446, EPI_ISL_2155447, EPI_ISL_2155450, EPI_ISL_2155452, EPI_ISL_2155458 | Philippine Red Cross - National Blood Center | Philippine Genome Center | Francis A. Tablizo, Kenneth M. Kim, Carlo M. Lapid, Marc Jerrone R. Castro, Maria Sofia L. Yangzon, Benedict A. Maralit, Marc Edsel C. Ayes, Eva Maria Cutiongco-de la Paz, Alethea R. de Guzman, Jan Michael C. Yap, Jo-Hannah S. Llames, Sheila Mae M. Araiza, Kris P. Punayan, Irish Coleen A. Asin, Candice Francheska B. Tambaoan, Asia Louisa U. Chong, Karol Sophia Agape R. Padilla, Rianna Patricia S. Cruz, El King D. Morado, Joshua Gregor A. Dizon, Razel Nikka M. Hao, Arianne A. Zamora, Devon Ray Pacial, Juan Antonio R. Magalang, Marissa Alejandria, Celia Carlos, Anna Ong-Lim, Edsel Maurice Salvaña, John Q. Wong, Jaime C. Montoya, Maria Rosario Singh-Vergeire and Cynthia P. Saloma |
| EPI_ISL_2155525, EPI_ISL_2155527, EPI_ISL_2155531, EPI_ISL_2155532, EPI_ISL_2155554, EPI_ISL_2155555, EPI_ISL_2155561, EPI_ISL_2155569, EPI_ISL_2155570, EPI_ISL_2155577, EPI_ISL_2155579, EPI_ISL_2155580, EPI_ISL_2155581 | | | |
| see above | UPLB Covid-19 Molecular Diagnostic Laboratory | Philippine Genome Center | Francis A. Tablizo, Kenneth M. Kim, Carlo M. Lapid, Marc Jerrone R. Castro, Maria Sofia L. Yangzon, Benedict A. Maralit, Marc Edsel C. Ayes, Eva Maria Cutiongco-de la Paz, Alethea R. de Guzman, Jan Michael C. Yap, Jo-Hannah S. Llames, Sheila Mae M. Araiza, Kris P. Punayan, Irish Coleen A. Asin, Candice Francheska B. Tambaoan, Asia Louisa U. Chong, Karol Sophia Agape R. Padilla, Rianna Patricia S. Cruz, El King D. Morado, Joshua Gregor A. Dizon, Razel |

| | | | |
|---|---|---|---|
| | | | Nikka M. Hao, Arianne A. Zamora, Devon Ray Pacial, Juan Antonio R. Magalang, Marissa Alejandria, Celia Carlos, Anna Ong-Lim, Edsel Maurice Salvaña, John Q. Wong, Jaime C. Montoya, Maria Rosario Singh-Vergeire and Cynthia P. Saloma |
| EPI_ISL_2155611, EPI_ISL_2155614, EPI_ISL_2155616, EPI_ISL_2155618, EPI_ISL_2155619, EPI_ISL_2155620, EPI_ISL_2155626, EPI_ISL_2155628, EPI_ISL_2155631, EPI_ISL_2155632, EPI_ISL_2155634, EPI_ISL_2155637 | | | |
| see above | Research Institute for Tropical Medicine, Inc. (RITM) | Philippine Genome Center | Francis A. Tablizo, Kenneth M. Kim, Carlo M. Lapid, Marc Jerrone R. Castro, Maria Sofia L. Yangzon, Benedict A. Maralit, Marc Edsel C. Ayes, Eva Maria Cutiongco-de la Paz, Alethea R. de Guzman, Jan Michael C. Yap, Jo-Hannah S. Llames, Sheila Mae M. Araiza, Kris P. Punayan, Irish Coleen A. Asin, Candice Francheska B. Tambaoan, Asia Louisa U. Chong, Karol Sophia Agape R. Padilla, Rianna Patricia S. Cruz, El King D. Morado, Joshua Gregor A. Dizon, Razel Nikka M. Hao, Arianne A. Zamora, Devon Ray Pacial, Juan Antonio R. Magalang, Marissa Alejandria, Celia Carlos, Anna Ong-Lim, Edsel Maurice Salvaña, John Q. Wong, Jaime C. Montoya, Maria Rosario Singh-Vergeire and Cynthia P. Saloma |
| EPI_ISL_2155657 | Ilocos Training and Regional Medical Center | Philippine Genome Center | Francis A. Tablizo, Kenneth M. Kim, Carlo M. Lapid, Marc Jerrone R. Castro, Maria Sofia L. Yangzon, Benedict A. Maralit, Marc Edsel C. Ayes, Eva Maria Cutiongco-de la Paz, Alethea R. de Guzman, Jan Michael C. Yap, Jo-Hannah S. Llames, Sheila Mae M. Araiza, Kris P. Punayan, Irish Coleen A. Asin, Candice Francheska B. Tambaoan, Asia Louisa U. Chong, Karol Sophia Agape R. Padilla, Rianna Patricia S. Cruz, El King D. Morado, Joshua Gregor A. Dizon, Razel Nikka M. Hao, Arianne A. Zamora, Devon Ray Pacial, Juan Antonio R. Magalang, Marissa Alejandria, Celia Carlos, Anna Ong-Lim, Edsel Maurice Salvaña, John Q. Wong, Jaime C. Montoya, Maria Rosario Singh-Vergeire and Cynthia P. Saloma |
| EPI_ISL_2155705 | Cagayan Valley Medical Center Molecular Laboratory | Philippine Genome Center | Francis A. Tablizo, Kenneth M. Kim, Carlo M. Lapid, Marc Jerrone R. Castro, Maria Sofia L. Yangzon, Benedict A. Maralit, Marc Edsel C. Ayes, Eva Maria Cutiongco-de la Paz, Alethea R. de Guzman, Jan Michael C. Yap, Jo-Hannah S. Llames, Sheila Mae M. Araiza, Kris P. Punayan, Irish Coleen A. Asin, Candice Francheska B. Tambaoan, Asia Louisa U. Chong, Karol Sophia Agape R. Padilla, Rianna Patricia S. Cruz, El King D. Morado, Joshua Gregor A. Dizon, Razel Nikka M. Hao, Arianne A. Zamora, Devon Ray Pacial, Juan Antonio R. Magalang, Marissa Alejandria, Celia Carlos, Anna Ong-Lim, Edsel Maurice Salvaña, John Q. Wong, Jaime C. Montoya, Maria Rosario Singh-Vergeire and Cynthia P. Saloma |
| EPI_ISL_2155733, EPI_ISL_2155753 | Philippine Red Cross Logistics and Multipurpose Center | Philippine Genome Center | Francis A. Tablizo, Kenneth M. Kim, Carlo M. Lapid, Marc Jerrone R. Castro, Maria Sofia L. Yangzon, Benedict A. Maralit, Marc Edsel C. Ayes, Eva Maria Cutiongco-de la Paz, Alethea R. de Guzman, Jan Michael C. Yap, Jo-Hannah S. Llames, Sheila Mae M. Araiza, Kris P. Punayan, Irish Coleen A. Asin, Candice Francheska B. Tambaoan, Asia Louisa U. Chong, Karol Sophia Agape R. Padilla, Rianna Patricia S. Cruz, El King D. Morado, Joshua Gregor A. Dizon, Razel Nikka M. Hao, Arianne A. Zamora, Devon Ray Pacial, Juan Antonio R. Magalang, Marissa Alejandria, Celia Carlos, Anna Ong-Lim, Edsel Maurice Salvaña, John Q. Wong, Jaime C. Montoya, Maria Rosario Singh-Vergeire and Cynthia P. Saloma |
| EPI_ISL_2155756, EPI_ISL_2155758, EPI_ISL_2155759 | First Aide Diagnostic Center | Philippine Genome Center | Francis A. Tablizo, Kenneth M. Kim, Carlo M. Lapid, Marc Jerrone R. Castro, Maria Sofia L. Yangzon, Benedict A. Maralit, Marc Edsel C. Ayes, Eva Maria Cutiongco-de la Paz, Alethea R. de Guzman, Jan Michael C. Yap, Jo-Hannah S. Llames, Sheila Mae M. Araiza, Kris P. Punayan, Irish Coleen A. Asin, Candice Francheska B. Tambaoan, Asia Louisa U. Chong, Karol Sophia Agape R. Padilla, Rianna Patricia S. Cruz, El King D. Morado, Joshua Gregor A. Dizon, Razel Nikka M. Hao, Arianne A. Zamora, Devon Ray Pacial, Juan Antonio R. Magalang, Marissa Alejandria, Celia Carlos, Anna Ong-Lim, Edsel Maurice Salvaña, John Q. Wong, Jaime C. Montoya, Maria Rosario Singh-Vergeire and Cynthia P. Saloma |
| EPI_ISL_2155760, EPI_ISL_2155761 | Qualimed Hospital Sta. Rosa | Philippine Genome Center | Francis A. Tablizo, Kenneth M. Kim, Carlo M. Lapid, Marc Jerrone R. Castro, Maria Sofia L. Yangzon, Benedict A. Maralit, Marc Edsel C. Ayes, Eva Maria Cutiongco-de la Paz, Alethea R. de Guzman, Jan Michael C. Yap, Jo-Hannah S. Llames, Sheila Mae M. Araiza, Kris P. Punayan, Irish Coleen A. Asin, Candice Francheska B. Tambaoan, Asia Louisa U. Chong, Karol Sophia Agape R. Padilla, Rianna Patricia S. Cruz, El King D. Morado, Joshua Gregor A. Dizon, Razel Nikka M. Hao, Arianne A. Zamora, Devon Ray Pacial, Juan Antonio R. Magalang, Marissa Alejandria, Celia Carlos, Anna Ong-Lim, Edsel Maurice Salvaña, John Q. Wong, Jaime C. Montoya, Maria Rosario Singh-Vergeire and Cynthia P. Saloma |
| EPI_ISL_2155767, EPI_ISL_2155768, EPI_ISL_2155770, EPI_ISL_2155771, EPI_ISL_2155774, EPI_ISL_2155775, EPI_ISL_2155779, EPI_ISL_2155780, EPI_ISL_2155782 | Philippine Red Cross - Port Area | Philippine Genome Center | Francis A. Tablizo, Kenneth M. Kim, Carlo M. Lapid, Marc Jerrone R. Castro, Maria Sofia L. Yangzon, Benedict A. Maralit, Marc Edsel C. Ayes, Eva Maria Cutiongco-de la Paz, Alethea R. de Guzman, Jan Michael C. Yap, Jo-Hannah S. Llames, Sheila Mae M. Araiza, Kris P. Punayan, Irish Coleen A. Asin, Candice Francheska B. Tambaoan, Asia Louisa U. Chong, Karol Sophia Agape R. Padilla, Rianna Patricia S. Cruz, El King D. Morado, Joshua Gregor A. Dizon, Razel Nikka M. Hao, Arianne A. Zamora, Devon Ray Pacial, Juan Antonio R. Magalang, Marissa Alejandria, Celia Carlos, Anna Ong-Lim, Edsel Maurice Salvaña, John Q. Wong, Jaime C. Montoya, Maria Rosario Singh-Vergeire and Cynthia P. Saloma |
| EPI_ISL_2155803, EPI_ISL_2155805, EPI_ISL_2155808 | Safeguard DNA Diagnostic, Inc. | Philippine Genome Center | Francis A. Tablizo, Kenneth M. Kim, Carlo M. Lapid, Marc Jerrone R. Castro, Maria Sofia L. Yangzon, Benedict A. Maralit, Marc Edsel C. Ayes, Eva Maria Cutiongco-de la Paz, Alethea R. de Guzman, Jan Michael C. Yap, Jo-Hannah S. Llames, Sheila Mae M. Araiza, Kris P. Punayan, Irish Coleen A. Asin, Candice Francheska B. Tambaoan, Asia Louisa U. Chong, Karol Sophia Agape R. Padilla, Rianna Patricia S. Cruz, El King D. Morado, Joshua Gregor A. Dizon, Razel Nikka M. Hao, Arianne A. Zamora, Devon Ray Pacial, Juan Antonio R. Magalang, Marissa Alejandria, Celia Carlos, Anna Ong-Lim, Edsel Maurice Salvaña, John Q. Wong, Jaime C. Montoya, Maria Rosario Singh-Vergeire and Cynthia P. Saloma |
| EPI_ISL_2155816 | Southern Isabela Medical Center Molecular Diagnostic & Research Laboratory (SIMC - MDRL) | Philippine Genome Center | Francis A. Tablizo, Kenneth M. Kim, Carlo M. Lapid, Marc Jerrone R. Castro, Maria Sofia L. Yangzon, Benedict A. Maralit, Marc Edsel C. Ayes, Eva Maria Cutiongco-de la Paz, Alethea R. de Guzman, Jan Michael C. Yap, Jo-Hannah S. Llames, Sheila Mae M. Araiza, Kris P. Punayan, Irish Coleen A. Asin, Candice Francheska B. Tambaoan, Asia Louisa U. Chong, Karol Sophia Agape R. Padilla, Rianna Patricia S. Cruz, El King D. Morado, Joshua Gregor A. Dizon, Razel Nikka M. Hao, Arianne A. Zamora, Devon Ray Pacial, Juan Antonio R. Magalang, Marissa Alejandria, Celia Carlos, Anna Ong-Lim, Edsel Maurice Salvaña, John Q. Wong, Jaime C. Montoya, Maria Rosario Singh-Vergeire and Cynthia P. Saloma |
| EPI_ISL_2156120 | Manila Doctors Hospital | Philippine Genome Center | Francis A. Tablizo, Kenneth M. Kim, Carlo M. Lapid, Marc Jerrone R. Castro, Maria Sofia L. Yangzon, Benedict A. Maralit, Marc Edsel C. Ayes, Eva Maria Cutiongco-de la Paz, Alethea R. de Guzman, Jan Michael C. Yap, Jo-Hannah S. Llames, Sheila Mae M. Araiza, Kris P. Punayan, Irish Coleen A. Asin, Candice Francheska B. Tambaoan, Asia Louisa U. Chong, Karol Sophia Agape R. Padilla, Rianna Patricia S. Cruz, El King D. Morado, Joshua Gregor A. Dizon, Razel Nikka M. Hao, Arianne A. Zamora, Devon Ray Pacial, Juan Antonio R. Magalang, Marissa Alejandria, Celia Carlos, Anna Ong-Lim, Edsel Maurice Salvaña, John Q. Wong, Jaime C. Montoya, Maria Rosario Singh-Vergeire and Cynthia P. Saloma |
| EPI_ISL_2156131 | Baguio General Hospital Medical Center (BGHMC) | Philippine Genome Center | Francis A. Tablizo, Kenneth M. Kim, Carlo M. Lapid, Marc Jerrone R. Castro, Maria Sofia L. Yangzon, Benedict A. Maralit, Marc Edsel C. Ayes, Eva Maria Cutiongco-de la Paz, Alethea R. de Guzman, Jan Michael C. Yap, Jo-Hannah S. Llames, Sheila Mae M. Araiza, Kris P. Punayan, Irish Coleen A. Asin, Candice Francheska B. Tambaoan, Asia Louisa U. Chong, Karol Sophia Agape R. Padilla, Rianna Patricia S. Cruz, El King D. Morado, Joshua Gregor A. Dizon, Razel Nikka M. Hao, Arianne A. Zamora, Devon Ray Pacial, Juan Antonio R. Magalang, Marissa Alejandria, Celia Carlos, Anna Ong-Lim, Edsel Maurice Salvaña, John Q. Wong, Jaime C. Montoya, Maria Rosario Singh-Vergeire and Cynthia P. Saloma |
| EPI_ISL_2156170, EPI_ISL_2156171, EPI_ISL_2156175, EPI_ISL_2156180, EPI_ISL_2156182, EPI_ISL_2156183 | Research Institute for Tropical Medicine, Inc. (RITM) | Philippine Genome Center | Francis A. Tablizo, Kenneth M. Kim, Carlo M. Lapid, Marc Jerrone R. Castro, Maria Sofia L. Yangzon, Benedict A. Maralit, Marc Edsel C. Ayes, Eva Maria Cutiongco-de la Paz, Alethea R. de Guzman, Jan Michael C. Yap, Jo-Hannah S. Llames, Sheila Mae M. Araiza, Kris P. Punayan, Irish Coleen A. Asin, Candice Francheska B. Tambaoan, Asia Louisa U. Chong, Karol Sophia Agape R. Padilla, Rianna Patricia S. Cruz, El King D. Morado, Joshua Gregor A. Dizon, Razel Nikka M. Hao, Arianne A. Zamora, Devon Ray Pacial, Juan Antonio R. Magalang, Marissa Alejandria, Celia Carlos, Anna Ong-Lim, Edsel Maurice Salvaña, John Q. Wong, Jaime C. Montoya, Maria Rosario Singh-Vergeire and Cynthia P. Saloma |
| EPI_ISL_2156207, EPI_ISL_2156209, EPI_ISL_2156212, EPI_ISL_2156213, EPI_ISL_2156215, EPI_ISL_2156216 | Cagayan Valley Medical Center Molecular Laboratory | Philippine Genome Center | Francis A. Tablizo, Kenneth M. Kim, Carlo M. Lapid, Marc Jerrone R. Castro, Maria Sofia L. Yangzon, Benedict A. Maralit, Marc Edsel C. Ayes, Eva Maria Cutiongco-de la Paz, Alethea R. de Guzman, Jan Michael C. Yap, Jo-Hannah S. Llames, Sheila Mae M. Araiza, Kris P. Punayan, Irish Coleen A. Asin, Candice Francheska B. Tambaoan, Asia Louisa U. Chong, Karol Sophia Agape R. Padilla, Rianna Patricia S. Cruz, El King D. Morado, Joshua Gregor A. Dizon, Razel Nikka M. Hao, Arianne A. Zamora, Devon Ray Pacial, Juan Antonio R. Magalang, Marissa Alejandria, Celia Carlos, Anna Ong-Lim, Edsel Maurice Salvaña, John Q. Wong, Jaime C. Montoya, Maria Rosario Singh-Vergeire and Cynthia P. Saloma |
| EPI_ISL_2156617 | Research Institute for Tropical Medicine, Inc. (RITM) | Philippine Genome Center | Francis A. Tablizo, Kenneth M. Kim, Carlo M. Lapid, Marc Jerrone R. Castro, Maria Sofia L. Yangzon, Elcid Aaron R. Pangilinan, Benedict A. Maralit, Marc Edsel C. Ayes, Eva Maria Cutiongco-de la Paz, Alethea R. de Guzman, Jan Michael C. Yap, Jo-Hannah S. Llames, Sheila Mae M. Araiza, Kris P. Punayan, Irish Coleen A. Asin, Candice Francheska B. Tambaoan, Asia Louisa U. Chong, Karol Sophia Agape R. Padilla, Rianna Patricia S. Cruz, El King D. Morado, Joshua Gregor A. Dizon, Razel Nikka M. Hao, Arianne A. Zamora, Devon Ray Pacial, Juan Antonio R. Magalang, Marissa Alejandria, Celia Carlos, Anna Ong-Lim, Edsel Maurice Salvaña, John Q. Wong, Jaime C. Montoya, Maria Rosario Singh-Vergeire and Cynthia P. Saloma |
| EPI_ISL_2156623 | Baguio General Hospital Medical Center (BGHMC) | Philippine Genome Center | Francis A. Tablizo, Kenneth M. Kim, Carlo M. Lapid, Marc Jerrone R. Castro, Maria Sofia L. Yangzon, Elcid Aaron R. Pangilinan, Benedict A. Maralit, Marc Edsel C. Ayes, Eva Maria Cutiongco-de la Paz, Alethea R. de Guzman, Jan Michael C. Yap, Jo-Hannah S. Llames, Sheila Mae M. Araiza, Kris P. Punayan, Irish Coleen A. Asin, Candice Francheska B. Tambaoan, Asia Louisa U. Chong, Karol Sophia Agape R. Padilla, Rianna Patricia S. Cruz, El King D. Morado, Joshua Gregor A. Dizon, Razel Nikka M. Hao, Arianne A. Zamora, Devon Ray Pacial, Juan Antonio R. Magalang, Marissa Alejandria, Celia Carlos, Anna Ong-Lim, Edsel Maurice Salvaña, John Q. Wong, Jaime C. Montoya, Maria Rosario Singh-Vergeire and Cynthia P. Saloma |
| EPI_ISL_2156624 | University of the East Ramon Magsaysay Memorial Medical | Philippine Genome Center | Francis A. Tablizo, Kenneth M. Kim, Carlo M. Lapid, Marc Jerrone R. Castro, Maria Sofia L. Yangzon, Elcid Aaron R. Pangilinan, Benedict A. Maralit, Marc |

| | | | |
|---|---|---|---|
| | Center (UERM) | | Edsel C. Ayes, Eva Maria Cutiongco-de la Paz, Alethea R. de Guzman, Jan Michael C. Yap, Jo-Hannah S. Llames, Sheila Mae M. Araiza, Kris P. Punayan, Irish Coleen A. Asin, Candice Francheska B. Tambaoan, Asia Louisa U. Chong, Karol Sophia Agape R. Padilla, Rianna Patricia S. Cruz, El King D. Morado, Joshua Gregor A. Dizon, Razel Nikka M. Hao, Arianne A. Zamora, Devon Ray Pacial, Juan Antonio R. Magalang, Marissa Alejandria, Celia Carlos, Anna Ong-Lim, Edsel Maurice Salvaña, John Q. Wong, Jaime C. Montoya, Maria Rosario Singh-Vergeire and Cynthia P. Saloma |
| EPI_ISL_2156625 | Cagayan Valley Medical Center Molecular Laboratory | Philippine Genome Center | Francis A. Tablizo, Kenneth M. Kim, Carlo M. Lapid, Marc Jerrone R. Castro, Maria Sofia L. Yangzon, Elcid Aaron R. Pangilinan, Benedict A. Maralit, Marc Edsel C. Ayes, Eva Maria Cutiongco-de la Paz, Alethea R. de Guzman, Jan Michael C. Yap, Jo-Hannah S. Llames, Sheila Mae M. Araiza, Kris P. Punayan, Irish Coleen A. Asin, Candice Francheska B. Tambaoan, Asia Louisa U. Chong, Karol Sophia Agape R. Padilla, Rianna Patricia S. Cruz, El King D. Morado, Joshua Gregor A. Dizon, Razel Nikka M. Hao, Arianne A. Zamora, Devon Ray Pacial, Juan Antonio R. Magalang, Marissa Alejandria, Celia Carlos, Anna Ong-Lim, Edsel Maurice Salvaña, John Q. Wong, Jaime C. Montoya, Maria Rosario Singh-Vergeire and Cynthia P. Saloma |
| EPI_ISL_2156629 | Baguio General Hospital Medical Center (BGHMC) | Philippine Genome Center | Francis A. Tablizo, Kenneth M. Kim, Carlo M. Lapid, Marc Jerrone R. Castro, Maria Sofia L. Yangzon, Elcid Aaron R. Pangilinan, Benedict A. Maralit, Marc Edsel C. Ayes, Eva Maria Cutiongco-de la Paz, Alethea R. de Guzman, Jan Michael C. Yap, Jo-Hannah S. Llames, Sheila Mae M. Araiza, Kris P. Punayan, Irish Coleen A. Asin, Candice Francheska B. Tambaoan, Asia Louisa U. Chong, Karol Sophia Agape R. Padilla, Rianna Patricia S. Cruz, El King D. Morado, Joshua Gregor A. Dizon, Razel Nikka M. Hao, Arianne A. Zamora, Devon Ray Pacial, Juan Antonio R. Magalang, Marissa Alejandria, Celia Carlos, Anna Ong-Lim, Edsel Maurice Salvaña, John Q. Wong, Jaime C. Montoya, Maria Rosario Singh-Vergeire and Cynthia P. Saloma |
| EPI_ISL_2156631, EPI_ISL_2156634, EPI_ISL_2156635 | Research Institute for Tropical Medicine, Inc. (RITM) | Philippine Genome Center | Francis A. Tablizo, Kenneth M. Kim, Carlo M. Lapid, Marc Jerrone R. Castro, Maria Sofia L. Yangzon, Elcid Aaron R. Pangilinan, Benedict A. Maralit, Marc Edsel C. Ayes, Eva Maria Cutiongco-de la Paz, Alethea R. de Guzman, Jan Michael C. Yap, Jo-Hannah S. Llames, Sheila Mae M. Araiza, Kris P. Punayan, Irish Coleen A. Asin, Candice Francheska B. Tambaoan, Asia Louisa U. Chong, Karol Sophia Agape R. Padilla, Rianna Patricia S. Cruz, El King D. Morado, Joshua Gregor A. Dizon, Razel Nikka M. Hao, Arianne A. Zamora, Devon Ray Pacial, Juan Antonio R. Magalang, Marissa Alejandria, Celia Carlos, Anna Ong-Lim, Edsel Maurice Salvaña, John Q. Wong, Jaime C. Montoya, Maria Rosario Singh-Vergeire and Cynthia P. Saloma |
| EPI_ISL_2156637 | Laguna Provincial Hospital - San Pablo District Hospital | Philippine Genome Center | Francis A. Tablizo, Kenneth M. Kim, Carlo M. Lapid, Marc Jerrone R. Castro, Maria Sofia L. Yangzon, Elcid Aaron R. Pangilinan, Benedict A. Maralit, Marc Edsel C. Ayes, Eva Maria Cutiongco-de la Paz, Alethea R. de Guzman, Jan Michael C. Yap, Jo-Hannah S. Llames, Sheila Mae M. Araiza, Kris P. Punayan, Irish Coleen A. Asin, Candice Francheska B. Tambaoan, Asia Louisa U. Chong, Karol Sophia Agape R. Padilla, Rianna Patricia S. Cruz, El King D. Morado, Joshua Gregor A. Dizon, Razel Nikka M. Hao, Arianne A. Zamora, Devon Ray Pacial, Juan Antonio R. Magalang, Marissa Alejandria, Celia Carlos, Anna Ong-Lim, Edsel Maurice Salvaña, John Q. Wong, Jaime C. Montoya, Maria Rosario Singh-Vergeire and Cynthia P. Saloma |
| EPI_ISL_2156638, EPI_ISL_2156639, EPI_ISL_2156643, EPI_ISL_2156645, EPI_ISL_2156647, EPI_ISL_2156648, EPI_ISL_2156649 | Research Institute for Tropical Medicine, Inc. (RITM) | Philippine Genome Center | Francis A. Tablizo, Kenneth M. Kim, Carlo M. Lapid, Marc Jerrone R. Castro, Maria Sofia L. Yangzon, Elcid Aaron R. Pangilinan, Benedict A. Maralit, Marc Edsel C. Ayes, Eva Maria Cutiongco-de la Paz, Alethea R. de Guzman, Jan Michael C. Yap, Jo-Hannah S. Llames, Sheila Mae M. Araiza, Kris P. Punayan, Irish Coleen A. Asin, Candice Francheska B. Tambaoan, Asia Louisa U. Chong, Karol Sophia Agape R. Padilla, Rianna Patricia S. Cruz, El King D. Morado, Joshua Gregor A. Dizon, Razel Nikka M. Hao, Arianne A. Zamora, Devon Ray Pacial, Juan Antonio R. Magalang, Marissa Alejandria, Celia Carlos, Anna Ong-Lim, Edsel Maurice Salvaña, John Q. Wong, Jaime C. Montoya, Maria Rosario Singh-Vergeire and Cynthia P. Saloma |
| EPI_ISL_2156650, EPI_ISL_2156652 | Cagayan Valley Medical Center Molecular Laboratory | Philippine Genome Center | Francis A. Tablizo, Kenneth M. Kim, Carlo M. Lapid, Marc Jerrone R. Castro, Maria Sofia L. Yangzon, Elcid Aaron R. Pangilinan, Benedict A. Maralit, Marc Edsel C. Ayes, Eva Maria Cutiongco-de la Paz, Alethea R. de Guzman, Jan Michael C. Yap, Jo-Hannah S. Llames, Sheila Mae M. Araiza, Kris P. Punayan, Irish Coleen A. Asin, Candice Francheska B. Tambaoan, Asia Louisa U. Chong, Karol Sophia Agape R. Padilla, Rianna Patricia S. Cruz, El King D. Morado, Joshua Gregor A. Dizon, Razel Nikka M. Hao, Arianne A. Zamora, Devon Ray Pacial, Juan Antonio R. Magalang, Marissa Alejandria, Celia Carlos, Anna Ong-Lim, Edsel Maurice Salvaña, John Q. Wong, Jaime C. Montoya, Maria Rosario Singh-Vergeire and Cynthia P. Saloma |
| EPI_ISL_2156657, EPI_ISL_2156664, EPI_ISL_2156665 | Research Institute for Tropical Medicine, Inc. (RITM) | Philippine Genome Center | Francis A. Tablizo, Kenneth M. Kim, Carlo M. Lapid, Marc Jerrone R. Castro, Maria Sofia L. Yangzon, Elcid Aaron R. Pangilinan, Benedict A. Maralit, Marc Edsel C. Ayes, Eva Maria Cutiongco-de la Paz, Alethea R. de Guzman, Jan Michael C. Yap, Jo-Hannah S. Llames, Sheila Mae M. Araiza, Kris P. Punayan, Irish Coleen A. Asin, Candice Francheska B. Tambaoan, Asia Louisa U. Chong, Karol Sophia Agape R. Padilla, Rianna Patricia S. Cruz, El King D. Morado, Joshua Gregor A. Dizon, Razel Nikka M. Hao, Arianne A. Zamora, Devon Ray Pacial, Juan Antonio R. Magalang, Marissa Alejandria, Celia Carlos, Anna Ong-Lim, Edsel Maurice Salvaña, John Q. Wong, Jaime C. Montoya, Maria Rosario Singh-Vergeire and Cynthia P. Saloma |
| EPI_ISL_2156666, EPI_ISL_2156668 | Southern Isabela Medical Center Molecular Diagnostic & Research Laboratory (SIMC - MDRL) | Philippine Genome Center | Francis A. Tablizo, Kenneth M. Kim, Carlo M. Lapid, Marc Jerrone R. Castro, Maria Sofia L. Yangzon, Elcid Aaron R. Pangilinan, Benedict A. Maralit, Marc Edsel C. Ayes, Eva Maria Cutiongco-de la Paz, Alethea R. de Guzman, Jan Michael C. Yap, Jo-Hannah S. Llames, Sheila Mae M. Araiza, Kris P. Punayan, Irish Coleen A. Asin, Candice Francheska B. Tambaoan, Asia Louisa U. Chong, Karol Sophia Agape R. Padilla, Rianna Patricia S. Cruz, El King D. Morado, Joshua Gregor A. Dizon, Razel Nikka M. Hao, Arianne A. Zamora, Devon Ray Pacial, Juan Antonio R. Magalang, Marissa Alejandria, Celia Carlos, Anna Ong-Lim, Edsel Maurice Salvaña, John Q. Wong, Jaime C. Montoya, Maria Rosario Singh-Vergeire and Cynthia P. Saloma |
| EPI_ISL_2156670, EPI_ISL_2156674, EPI_ISL_2156675 | Research Institute for Tropical Medicine, Inc. (RITM) | Philippine Genome Center | Francis A. Tablizo, Kenneth M. Kim, Carlo M. Lapid, Marc Jerrone R. Castro, Maria Sofia L. Yangzon, Elcid Aaron R. Pangilinan, Benedict A. Maralit, Marc Edsel C. Ayes, Eva Maria Cutiongco-de la Paz, Alethea R. de Guzman, Jan Michael C. Yap, Jo-Hannah S. Llames, Sheila Mae M. Araiza, Kris P. Punayan, Irish Coleen A. Asin, Candice Francheska B. Tambaoan, Asia Louisa U. Chong, Karol Sophia Agape R. Padilla, Rianna Patricia S. Cruz, El King D. Morado, Joshua Gregor A. Dizon, Razel Nikka M. Hao, Arianne A. Zamora, Devon Ray Pacial, Juan Antonio R. Magalang, Marissa Alejandria, Celia Carlos, Anna Ong-Lim, Edsel Maurice Salvaña, John Q. Wong, Jaime C. Montoya, Maria Rosario Singh-Vergeire and Cynthia P. Saloma |
| EPI_ISL_2156677, EPI_ISL_2156678 | Southern Isabela Medical Center Molecular Diagnostic & Research Laboratory (SIMC - MDRL) | Philippine Genome Center | Francis A. Tablizo, Kenneth M. Kim, Carlo M. Lapid, Marc Jerrone R. Castro, Maria Sofia L. Yangzon, Elcid Aaron R. Pangilinan, Benedict A. Maralit, Marc Edsel C. Ayes, Eva Maria Cutiongco-de la Paz, Alethea R. de Guzman, Jan Michael C. Yap, Jo-Hannah S. Llames, Sheila Mae M. Araiza, Kris P. Punayan, Irish Coleen A. Asin, Candice Francheska B. Tambaoan, Asia Louisa U. Chong, Karol Sophia Agape R. Padilla, Rianna Patricia S. Cruz, El King D. Morado, Joshua Gregor A. Dizon, Razel Nikka M. Hao, Arianne A. Zamora, Devon Ray Pacial, Juan Antonio R. Magalang, Marissa Alejandria, Celia Carlos, Anna Ong-Lim, Edsel Maurice Salvaña, John Q. Wong, Jaime C. Montoya, Maria Rosario Singh-Vergeire and Cynthia P. Saloma |
| EPI_ISL_2156679 | Cagayan Valley Medical Center Molecular Laboratory | Philippine Genome Center | Francis A. Tablizo, Kenneth M. Kim, Carlo M. Lapid, Marc Jerrone R. Castro, Maria Sofia L. Yangzon, Elcid Aaron R. Pangilinan, Benedict A. Maralit, Marc Edsel C. Ayes, Eva Maria Cutiongco-de la Paz, Alethea R. de Guzman, Jan Michael C. Yap, Jo-Hannah S. Llames, Sheila Mae M. Araiza, Kris P. Punayan, Irish Coleen A. Asin, Candice Francheska B. Tambaoan, Asia Louisa U. Chong, Karol Sophia Agape R. Padilla, Rianna Patricia S. Cruz, El King D. Morado, Joshua Gregor A. Dizon, Razel Nikka M. Hao, Arianne A. Zamora, Devon Ray Pacial, Juan Antonio R. Magalang, Marissa Alejandria, Celia Carlos, Anna Ong-Lim, Edsel Maurice Salvaña, John Q. Wong, Jaime C. Montoya, Maria Rosario Singh-Vergeire and Cynthia P. Saloma |
| EPI_ISL_2156681 | Research Institute for Tropical Medicine, Inc. (RITM) | Philippine Genome Center | Francis A. Tablizo, Kenneth M. Kim, Carlo M. Lapid, Marc Jerrone R. Castro, Maria Sofia L. Yangzon, Elcid Aaron R. Pangilinan, Benedict A. Maralit, Marc Edsel C. Ayes, Eva Maria Cutiongco-de la Paz, Alethea R. de Guzman, Jan Michael C. Yap, Jo-Hannah S. Llames, Sheila Mae M. Araiza, Kris P. Punayan, Irish Coleen A. Asin, Candice Francheska B. Tambaoan, Asia Louisa U. Chong, Karol Sophia Agape R. Padilla, Rianna Patricia S. Cruz, El King D. Morado, Joshua Gregor A. Dizon, Razel Nikka M. Hao, Arianne A. Zamora, Devon Ray Pacial, Juan Antonio R. Magalang, Marissa Alejandria, Celia Carlos, Anna Ong-Lim, Edsel Maurice Salvaña, John Q. Wong, Jaime C. Montoya, Maria Rosario Singh-Vergeire and Cynthia P. Saloma |
| EPI_ISL_2156682 | Cagayan Valley Medical Center Molecular Laboratory | Philippine Genome Center | Francis A. Tablizo, Kenneth M. Kim, Carlo M. Lapid, Marc Jerrone R. Castro, Maria Sofia L. Yangzon, Elcid Aaron R. Pangilinan, Benedict A. Maralit, Marc Edsel C. Ayes, Eva Maria Cutiongco-de la Paz, Alethea R. de Guzman, Jan Michael C. Yap, Jo-Hannah S. Llames, Sheila Mae M. Araiza, Kris P. Punayan, Irish Coleen A. Asin, Candice Francheska B. Tambaoan, Asia Louisa U. Chong, Karol Sophia Agape R. Padilla, Rianna Patricia S. Cruz, El King D. Morado, Joshua Gregor A. Dizon, Razel Nikka M. Hao, Arianne A. Zamora, Devon Ray Pacial, Juan Antonio R. Magalang, Marissa Alejandria, Celia Carlos, Anna Ong-Lim, Edsel Maurice Salvaña, John Q. Wong, Jaime C. Montoya, Maria Rosario Singh-Vergeire and Cynthia P. Saloma |
| EPI_ISL_2156687, EPI_ISL_2156688, EPI_ISL_2156693 | Southern Isabela Medical Center Molecular Diagnostic & Research Laboratory (SIMC - MDRL) | Philippine Genome Center | Francis A. Tablizo, Kenneth M. Kim, Carlo M. Lapid, Marc Jerrone R. Castro, Maria Sofia L. Yangzon, Elcid Aaron R. Pangilinan, Benedict A. Maralit, Marc Edsel C. Ayes, Eva Maria Cutiongco-de la Paz, Alethea R. de Guzman, Jan Michael C. Yap, Jo-Hannah S. Llames, Sheila Mae M. Araiza, Kris P. Punayan, Irish Coleen A. Asin, Candice Francheska B. Tambaoan, Asia Louisa U. Chong, Karol Sophia Agape R. Padilla, Rianna Patricia S. Cruz, El King D. Morado, Joshua Gregor A. Dizon, Razel Nikka M. Hao, Arianne A. Zamora, Devon Ray Pacial, Juan Antonio R. Magalang, Marissa Alejandria, Celia Carlos, Anna Ong-Lim, Edsel Maurice Salvaña, John Q. Wong, Jaime C. Montoya, Maria Rosario Singh-Vergeire and Cynthia P. Saloma |
| EPI_ISL_2156697 | Manila Doctors Hospital | Philippine Genome Center | Francis A. Tablizo, Kenneth M. Kim, Carlo M. Lapid, Marc Jerrone R. Castro, Maria Sofia L. Yangzon, Elcid Aaron R. Pangilinan, Benedict A. Maralit, Marc Edsel C. Ayes, Eva Maria Cutiongco-de la Paz, Alethea R. de Guzman, Jan Michael C. Yap, Jo-Hannah S. Llames, Sheila Mae M. Araiza, Kris P. Punayan, Irish Coleen A. Asin, Candice Francheska B. Tambaoan, Asia Louisa U. Chong, Karol Sophia Agape R. Padilla, Rianna Patricia S. Cruz, El King D. Morado, Joshua Gregor A. Dizon, Razel Nikka M. Hao, Arianne A. Zamora, Devon Ray Pacial, Juan Antonio R. Magalang, Marissa Alejandria, Celia Carlos, Anna Ong-Lim, Edsel Maurice Salvaña, John Q. Wong, Jaime C. Montoya, Maria Rosario Singh-Vergeire and Cynthia P. Saloma |

| | | | |
|---|---|---|---|
| EPI_ISL_2156698, EPI_ISL_2156699 | Research Institute for Tropical Medicine, Inc. (RITM) | Philippine Genome Center | Francis A. Tablizo, Kenneth M. Kim, Carlo M. Lapid, Marc Jerrone R. Castro, Maria Sofia L. Yangzon, Elcid Aaron R. Pangilinan, Benedict A. Maralit, Marc Edsel C. Ayes, Eva Maria Cutiongco-de la Paz, Alethea R. de Guzman, Jan Michael C. Yap, Jo-Hannah S. Llames, Sheila Mae M. Araiza, Kris P. Punayan, Irish Coleen A. Asin, Candice Francheska B. Tambaoan, Asia Louisa U. Chong, Karol Sophia Agape R. Padilla, Rianna Patricia S. Cruz, El King D. Morado, Joshua Gregor A. Dizon, Razel Nikka M. Hao, Arianne A. Zamora, Devon Ray Pacial, Juan Antonio R. Magalang, Marissa Alejandria, Celia Carlos, Anna Ong-Lim, Edsel Maurice Salvaña, John Q. Wong, Jaime C. Montoya, Maria Rosario Singh-Vergeire and Cynthia P. Saloma |
| EPI_ISL_2156700, EPI_ISL_2156705 | Cagayan Valley Medical Center Molecular Laboratory | Philippine Genome Center | Francis A. Tablizo, Kenneth M. Kim, Carlo M. Lapid, Marc Jerrone R. Castro, Maria Sofia L. Yangzon, Elcid Aaron R. Pangilinan, Benedict A. Maralit, Marc Edsel C. Ayes, Eva Maria Cutiongco-de la Paz, Alethea R. de Guzman, Jan Michael C. Yap, Jo-Hannah S. Llames, Sheila Mae M. Araiza, Kris P. Punayan, Irish Coleen A. Asin, Candice Francheska B. Tambaoan, Asia Louisa U. Chong, Karol Sophia Agape R. Padilla, Rianna Patricia S. Cruz, El King D. Morado, Joshua Gregor A. Dizon, Razel Nikka M. Hao, Arianne A. Zamora, Devon Ray Pacial, Juan Antonio R. Magalang, Marissa Alejandria, Celia Carlos, Anna Ong-Lim, Edsel Maurice Salvaña, John Q. Wong, Jaime C. Montoya, Maria Rosario Singh-Vergeire and Cynthia P. Saloma |
| EPI_ISL_2156707, EPI_ISL_2156709 | Research Institute for Tropical Medicine, Inc. (RITM) | Philippine Genome Center | Francis A. Tablizo, Kenneth M. Kim, Carlo M. Lapid, Marc Jerrone R. Castro, Maria Sofia L. Yangzon, Elcid Aaron R. Pangilinan, Benedict A. Maralit, Marc Edsel C. Ayes, Eva Maria Cutiongco-de la Paz, Alethea R. de Guzman, Jan Michael C. Yap, Jo-Hannah S. Llames, Sheila Mae M. Araiza, Kris P. Punayan, Irish Coleen A. Asin, Candice Francheska B. Tambaoan, Asia Louisa U. Chong, Karol Sophia Agape R. Padilla, Rianna Patricia S. Cruz, El King D. Morado, Joshua Gregor A. Dizon, Razel Nikka M. Hao, Arianne A. Zamora, Devon Ray Pacial, Juan Antonio R. Magalang, Marissa Alejandria, Celia Carlos, Anna Ong-Lim, Edsel Maurice Salvaña, John Q. Wong, Jaime C. Montoya, Maria Rosario Singh-Vergeire and Cynthia P. Saloma |
| EPI_ISL_2156711 | UPLB Covid-19 Molecular Diagnostic Laboratory | Philippine Genome Center | Francis A. Tablizo, Kenneth M. Kim, Carlo M. Lapid, Marc Jerrone R. Castro, Maria Sofia L. Yangzon, Elcid Aaron R. Pangilinan, Benedict A. Maralit, Marc Edsel C. Ayes, Eva Maria Cutiongco-de la Paz, Alethea R. de Guzman, Jan Michael C. Yap, Jo-Hannah S. Llames, Sheila Mae M. Araiza, Kris P. Punayan, Irish Coleen A. Asin, Candice Francheska B. Tambaoan, Asia Louisa U. Chong, Karol Sophia Agape R. Padilla, Rianna Patricia S. Cruz, El King D. Morado, Joshua Gregor A. Dizon, Razel Nikka M. Hao, Arianne A. Zamora, Devon Ray Pacial, Juan Antonio R. Magalang, Marissa Alejandria, Celia Carlos, Anna Ong-Lim, Edsel Maurice Salvaña, John Q. Wong, Jaime C. Montoya, Maria Rosario Singh-Vergeire and Cynthia P. Saloma |
| EPI_ISL_2156712 | Cagayan Valley Medical Center Molecular Laboratory | Philippine Genome Center | Francis A. Tablizo, Kenneth M. Kim, Carlo M. Lapid, Marc Jerrone R. Castro, Maria Sofia L. Yangzon, Elcid Aaron R. Pangilinan, Benedict A. Maralit, Marc Edsel C. Ayes, Eva Maria Cutiongco-de la Paz, Alethea R. de Guzman, Jan Michael C. Yap, Jo-Hannah S. Llames, Sheila Mae M. Araiza, Kris P. Punayan, Irish Coleen A. Asin, Candice Francheska B. Tambaoan, Asia Louisa U. Chong, Karol Sophia Agape R. Padilla, Rianna Patricia S. Cruz, El King D. Morado, Joshua Gregor A. Dizon, Razel Nikka M. Hao, Arianne A. Zamora, Devon Ray Pacial, Juan Antonio R. Magalang, Marissa Alejandria, Celia Carlos, Anna Ong-Lim, Edsel Maurice Salvaña, John Q. Wong, Jaime C. Montoya, Maria Rosario Singh-Vergeire and Cynthia P. Saloma |
| EPI_ISL_2156716, EPI_ISL_2156721, EPI_ISL_2156726, EPI_ISL_2156728 | Research Institute for Tropical Medicine, Inc. (RITM) | Philippine Genome Center | Francis A. Tablizo, Kenneth M. Kim, Carlo M. Lapid, Marc Jerrone R. Castro, Maria Sofia L. Yangzon, Elcid Aaron R. Pangilinan, Benedict A. Maralit, Marc Edsel C. Ayes, Eva Maria Cutiongco-de la Paz, Alethea R. de Guzman, Jan Michael C. Yap, Jo-Hannah S. Llames, Sheila Mae M. Araiza, Kris P. Punayan, Irish Coleen A. Asin, Candice Francheska B. Tambaoan, Asia Louisa U. Chong, Karol Sophia Agape R. Padilla, Rianna Patricia S. Cruz, El King D. Morado, Joshua Gregor A. Dizon, Razel Nikka M. Hao, Arianne A. Zamora, Devon Ray Pacial, Juan Antonio R. Magalang, Marissa Alejandria, Celia Carlos, Anna Ong-Lim, Edsel Maurice Salvaña, John Q. Wong, Jaime C. Montoya, Maria Rosario Singh-Vergeire and Cynthia P. Saloma |
| EPI_ISL_2156729, EPI_ISL_2156732, EPI_ISL_2156733, EPI_ISL_2156734 | Cagayan Valley Medical Center Molecular Laboratory | Philippine Genome Center | Francis A. Tablizo, Kenneth M. Kim, Carlo M. Lapid, Marc Jerrone R. Castro, Maria Sofia L. Yangzon, Elcid Aaron R. Pangilinan, Benedict A. Maralit, Marc Edsel C. Ayes, Eva Maria Cutiongco-de la Paz, Alethea R. de Guzman, Jan Michael C. Yap, Jo-Hannah S. Llames, Sheila Mae M. Araiza, Kris P. Punayan, Irish Coleen A. Asin, Candice Francheska B. Tambaoan, Asia Louisa U. Chong, Karol Sophia Agape R. Padilla, Rianna Patricia S. Cruz, El King D. Morado, Joshua Gregor A. Dizon, Razel Nikka M. Hao, Arianne A. Zamora, Devon Ray Pacial, Juan Antonio R. Magalang, Marissa Alejandria, Celia Carlos, Anna Ong-Lim, Edsel Maurice Salvaña, John Q. Wong, Jaime C. Montoya, Maria Rosario Singh-Vergeire and Cynthia P. Saloma |
| EPI_ISL_2156736, EPI_ISL_2156740, EPI_ISL_2156753 | Research Institute for Tropical Medicine, Inc. (RITM) | Philippine Genome Center | Francis A. Tablizo, Kenneth M. Kim, Carlo M. Lapid, Marc Jerrone R. Castro, Maria Sofia L. Yangzon, Elcid Aaron R. Pangilinan, Benedict A. Maralit, Marc Edsel C. Ayes, Eva Maria Cutiongco-de la Paz, Alethea R. de Guzman, Jan Michael C. Yap, Jo-Hannah S. Llames, Sheila Mae M. Araiza, Kris P. Punayan, Irish Coleen A. Asin, Candice Francheska B. Tambaoan, Asia Louisa U. Chong, Karol Sophia Agape R. Padilla, Rianna Patricia S. Cruz, El King D. Morado, Joshua Gregor A. Dizon, Razel Nikka M. Hao, Arianne A. Zamora, Devon Ray Pacial, Juan Antonio R. Magalang, Marissa Alejandria, Celia Carlos, Anna Ong-Lim, Edsel Maurice Salvaña, John Q. Wong, Jaime C. Montoya, Maria Rosario Singh-Vergeire and Cynthia P. Saloma |
| EPI_ISL_2156754, EPI_ISL_2156757 | Cagayan Valley Medical Center Molecular Laboratory | Philippine Genome Center | Francis A. Tablizo, Kenneth M. Kim, Carlo M. Lapid, Marc Jerrone R. Castro, Maria Sofia L. Yangzon, Elcid Aaron R. Pangilinan, Benedict A. Maralit, Marc Edsel C. Ayes, Eva Maria Cutiongco-de la Paz, Alethea R. de Guzman, Jan Michael C. Yap, Jo-Hannah S. Llames, Sheila Mae M. Araiza, Kris P. Punayan, Irish Coleen A. Asin, Candice Francheska B. Tambaoan, Asia Louisa U. Chong, Karol Sophia Agape R. Padilla, Rianna Patricia S. Cruz, El King D. Morado, Joshua Gregor A. Dizon, Razel Nikka M. Hao, Arianne A. Zamora, Devon Ray Pacial, Juan Antonio R. Magalang, Marissa Alejandria, Celia Carlos, Anna Ong-Lim, Edsel Maurice Salvaña, John Q. Wong, Jaime C. Montoya, Maria Rosario Singh-Vergeire and Cynthia P. Saloma |
| EPI_ISL_2156760 | University of the Philippines National Institutes of Health (UP NIH) | Philippine Genome Center | Francis A. Tablizo, Kenneth M. Kim, Carlo M. Lapid, Marc Jerrone R. Castro, Maria Sofia L. Yangzon, Elcid Aaron R. Pangilinan, Benedict A. Maralit, Marc Edsel C. Ayes, Eva Maria Cutiongco-de la Paz, Alethea R. de Guzman, Jan Michael C. Yap, Jo-Hannah S. Llames, Sheila Mae M. Araiza, Kris P. Punayan, Irish Coleen A. Asin, Candice Francheska B. Tambaoan, Asia Louisa U. Chong, Karol Sophia Agape R. Padilla, Rianna Patricia S. Cruz, El King D. Morado, Joshua Gregor A. Dizon, Razel Nikka M. Hao, Arianne A. Zamora, Devon Ray Pacial, Juan Antonio R. Magalang, Marissa Alejandria, Celia Carlos, Anna Ong-Lim, Edsel Maurice Salvaña, John Q. Wong, Jaime C. Montoya, Maria Rosario Singh-Vergeire and Cynthia P. Saloma |
| EPI_ISL_2156763 | Cagayan Valley Medical Center Molecular Laboratory | Philippine Genome Center | Francis A. Tablizo, Kenneth M. Kim, Carlo M. Lapid, Marc Jerrone R. Castro, Maria Sofia L. Yangzon, Elcid Aaron R. Pangilinan, Benedict A. Maralit, Marc Edsel C. Ayes, Eva Maria Cutiongco-de la Paz, Alethea R. de Guzman, Jan Michael C. Yap, Jo-Hannah S. Llames, Sheila Mae M. Araiza, Kris P. Punayan, Irish Coleen A. Asin, Candice Francheska B. Tambaoan, Asia Louisa U. Chong, Karol Sophia Agape R. Padilla, Rianna Patricia S. Cruz, El King D. Morado, Joshua Gregor A. Dizon, Razel Nikka M. Hao, Arianne A. Zamora, Devon Ray Pacial, Juan Antonio R. Magalang, Marissa Alejandria, Celia Carlos, Anna Ong-Lim, Edsel Maurice Salvaña, John Q. Wong, Jaime C. Montoya, Maria Rosario Singh-Vergeire and Cynthia P. Saloma |
| EPI_ISL_2156766 | University of the Philippines National Institutes of Health (UP NIH) | Philippine Genome Center | Francis A. Tablizo, Kenneth M. Kim, Carlo M. Lapid, Marc Jerrone R. Castro, Maria Sofia L. Yangzon, Elcid Aaron R. Pangilinan, Benedict A. Maralit, Marc Edsel C. Ayes, Eva Maria Cutiongco-de la Paz, Alethea R. de Guzman, Jan Michael C. Yap, Jo-Hannah S. Llames, Sheila Mae M. Araiza, Kris P. Punayan, Irish Coleen A. Asin, Candice Francheska B. Tambaoan, Asia Louisa U. Chong, Karol Sophia Agape R. Padilla, Rianna Patricia S. Cruz, El King D. Morado, Joshua Gregor A. Dizon, Razel Nikka M. Hao, Arianne A. Zamora, Devon Ray Pacial, Juan Antonio R. Magalang, Marissa Alejandria, Celia Carlos, Anna Ong-Lim, Edsel Maurice Salvaña, John Q. Wong, Jaime C. Montoya, Maria Rosario Singh-Vergeire and Cynthia P. Saloma |
| EPI_ISL_2157345, EPI_ISL_2157346, EPI_ISL_2157577 | Red - Regional de Vigilancia Genómica del COVID-19 | Laboratory of Respiratory Viruses and Measles, Oswaldo Cruz Institute, FIOCRUZ | Paola Resende, Mitzi Castro, Claudia Díaz, Sandra Paola Paz, Luciana Appolinario, Fernando Motta, Anna Carolina Paixao, Ana Carolina Mendonca, Alice Sampaio Rocha, Taina Venas, Eilisa Cavalcante Pereira, Renata Serrano Lopes, Marilda Siqueira on behalf of the Fiocruz COVID-19 Genomic Surveillance Network |
| EPI_ISL_2171041 | Dhulikhel Hospital, Kathmandu University Hospital | Molecular and Genomics Research Lab, Dhulikhel Hospital, Kathmandu University Hospital | Rajeev Shrestha, Nishan Katuwal, Navin Adhikari, Manu Vanaerschot, Dipesh Tamrakar, Meghnath Dhimal, Pradip Gyanwali, Saroj Bhattarai, Surendra Kumar Madhup |
| EPI_ISL_2171266 | Janamaitri Hospital | Molecular and Genomics Research Lab, Dhulikhel Hospital, Kathmandu University Hospital | Rajeev Shrestha, Nishan Katuwal, Navin Adhikari, Dipesh Tamrakar, Meghnath Dhimal, Pradip Gyanwali, Saroj Bhattarai, Surendra Kumar Madhup |
| EPI_ISL_2178470 | Hospital | National Reference Center for Viruses of Respiratory Infections, Institut Pasteur, Paris | Marion Barbet, Sylvie Behillil, Méline Bizard, Angela Brisebarre, Camille Capel, Vincent Enouf, Louise Lefrançois, Frédéric Lemoine, Christophe Malabat, Corinne Maufrais, Pierre Lechat, Etienne Simon-Lorière, Maud Vanpeene, Sylvie Van der Werf ,Laura Djamdjian |
| EPI_ISL_2188323 | Hospital | National Reference Center for Viruses of Respiratory Infections, Institut Pasteur, Paris | Marion Barbet, Sylvie Behillil, Méline Bizard, Angela Brisebarre, Camille Capel, Vincent Enouf, Louise Lefrançois, Frédéric Lemoine, Christophe Malabat, Corinne Maufrais, Etienne Simon-Lorière, Maud Vanpeene, Sylvie Van der Werf ,Alexandra Ducancelle |
| EPI_ISL_2188437, EPI_ISL_2188438 | Labo Analyses Med | National Reference Center for Viruses of Respiratory Infections, Institut Pasteur, Paris | Marion Barbet, Sylvie Behillil, Méline Bizard, Angela Brisebarre, Camille Capel, Vincent Enouf, Louise Lefrançois, Frédéric Lemoine, Christophe Malabat, Corinne Maufrais, Etienne Simon-Lorière, Maud Vanpeene, Sylvie Van der Werf ,M.-J. Blanc-Galiby |
| EPI_ISL_2202673, EPI_ISL_2202675, EPI_ISL_2202678 | The Ohio State University Applied Microbiology Services Laboratory | The Ohio State University Applied Microbiology Services Laboratory | Seth A. Faith PhD |
| EPI_ISL_2214905 | Centre Hospitalier Universitaire Clermont-Ferrand | CHU Clermont-Ferrand, service de virologie | Bisseux Maxime, Mirand Audrey, Combes Patricia, Henquell Cécile |
| EPI_ISL_2214914 | EHPAD | CHU Clermont-Ferrand, service de virologie | Bisseux Maxime, Mirand Audrey, Combes Patricia, Henquell Cécile |

| | | | |
|---|---|---|---|
| EPI_ISL_2226217, EPI_ISL_2226218, EPI_ISL_2226219, EPI_ISL_2226220, EPI_ISL_2226221, EPI_ISL_2226222, EPI_ISL_2226223, EPI_ISL_2226225, EPI_ISL_2226226, EPI_ISL_2226227, EPI_ISL_2226228 | | | |
| see above | Department of Internal Medicine, College of Medicine, Chosun University | Department of Internal Medicine, College of Medicine, Chosun University | Dong-Min Kim |
| EPI_ISL_2226264, EPI_ISL_2226325, EPI_ISL_2226329, EPI_ISL_2226336, EPI_ISL_2226339, EPI_ISL_2226340 | BIO86 CHAUMONT | CHU Poitiers | Caroline MICHAUD, Maxime PICHON, Manon PRAT, Valentin BON-BARET, Birama N'DIAYE, Magali GARCIA, Agnes BEBY-DEFAUX, Nicolas LEVEQUE |
| EPI_ISL_2227286 | Veterinary institute in Zvolen, Slovakia | Veterinary institute in Zvolen | Dirbáková Z., Sujová S., Mokryšová S., Tinák M., Mojžiš M. |
| EPI_ISL_2227287 | Veterinary institute in Zvolen, Pod Drahami 918, 960 86 Zvolen Slovakia | Veterinary institute in Zvolen, Pod Drahami 918, 960 86 Zvolen Slovakia | Dirbáková Z., Sujová S., Mokryšová S., Tinák M., Mojžiš M. |
| EPI_ISL_2227288 | Veterinary institute in Zvolen Slovakia | Veterinary institute in Zvolen Slovakia | Dirbáková Z., Sujová S., Mokryšová S., Tinák M., Mojžiš M. |
| EPI_ISL_2231152, EPI_ISL_2231153 | Centro de Diagnostico COVID-19 UABC Tijuana | Andersen lab at Scripps Research | SEARCH Alliance San Diego with Idanya Rubi Serafin Higuera, Manuel Sanchez Alavez, Jorge Luis Jimenez Niebla, German Ibarra, Jonathan Vincent Baena, Oscar Efren Zazueta Fierro |
| EPI_ISL_2231154 | Sharp HealthCare Laboratory | Andersen lab at Scripps Research | SEARCH Alliance San Diego with Aaron Harding, Jacquelyn Berumen, Cathy Woerle, Liam McGinnis, Art Mendoza, Omid Bakhtar |
| EPI_ISL_2231171, EPI_ISL_2231172 | Centro de Diagnostico COVID-19 UABC Tijuana | Andersen lab at Scripps Research | SEARCH Alliance San Diego with Idanya Rubi Serafin Higuera, Manuel Sanchez Alavez, Jorge Luis Jimenez Niebla, German Ibarra, Jonathan Vincent Baena, Oscar Efren Zazueta Fierro |
| EPI_ISL_2231173 | San Diego County Public Health Laboratory | Andersen lab at Scripps Research | SEARCH Alliance San Diego with Ashleigh Murphy, Jovan Shephard, Brett Austin |
| EPI_ISL_2231174 | Centro de Diagnostico COVID-19 UABC Tijuana | Andersen lab at Scripps Research | SEARCH Alliance San Diego with Idanya Rubi Serafin Higuera, Manuel Sanchez Alavez, Jorge Luis Jimenez Niebla, German Ibarra, Jonathan Vincent Baena, Oscar Efren Zazueta Fierro |
| EPI_ISL_2231175, EPI_ISL_2231186, EPI_ISL_2231188, EPI_ISL_2231189 | Sharp HealthCare Laboratory | Andersen lab at Scripps Research | SEARCH Alliance San Diego with Aaron Harding, Jacquelyn Berumen, Cathy Woerle, Liam McGinnis, Art Mendoza, Omid Bakhtar |
| EPI_ISL_2231190, EPI_ISL_2231191, EPI_ISL_2231192 | Centro de Diagnostico COVID-19 UABC Tijuana | Andersen lab at Scripps Research | SEARCH Alliance San Diego with Idanya Rubi Serafin Higuera, Manuel Sanchez Alavez, Jorge Luis Jimenez Niebla, German Ibarra, Jonathan Vincent Baena, Oscar Efren Zazueta Fierro |
| EPI_ISL_2231193 | San Diego County Public Health Laboratory | Andersen lab at Scripps Research | SEARCH Alliance San Diego with Ashleigh Murphy, Jovan Shephard, Brett Austin |
| EPI_ISL_2231194, EPI_ISL_2231196, EPI_ISL_2231197 | Sharp HealthCare Laboratory | Andersen lab at Scripps Research | SEARCH Alliance San Diego with Aaron Harding, Jacquelyn Berumen, Cathy Woerle, Liam McGinnis, Art Mendoza, Omid Bakhtar |
| EPI_ISL_2231199, EPI_ISL_2231200 | Centro de Diagnostico COVID-19 UABC Tijuana | Andersen lab at Scripps Research | SEARCH Alliance San Diego with Idanya Rubi Serafin Higuera, Manuel Sanchez Alavez, Jorge Luis Jimenez Niebla, German Ibarra, Jonathan Vincent Baena, Oscar Efren Zazueta Fierro |
| EPI_ISL_2231201, EPI_ISL_2231202, EPI_ISL_2231203, EPI_ISL_2231204, EPI_ISL_2231207, EPI_ISL_2231208, EPI_ISL_2231209, EPI_ISL_2231210, EPI_ISL_2231211, EPI_ISL_2231212, EPI_ISL_2231213, EPI_ISL_2231214, EPI_ISL_2231215, EPI_ISL_2231216, EPI_ISL_2231217, EPI_ISL_2231218, EPI_ISL_2231219, EPI_ISL_2231220, EPI_ISL_2231221, EPI_ISL_2231222, EPI_ISL_2231223, EPI_ISL_2231224, EPI_ISL_2231225, EPI_ISL_2231226, EPI_ISL_2231227, EPI_ISL_2231228, EPI_ISL_2231229, EPI_ISL_2231230, EPI_ISL_2231231, EPI_ISL_2231232, EPI_ISL_2231233, EPI_ISL_2231234, EPI_ISL_2231235, EPI_ISL_2231236, EPI_ISL_2231237, EPI_ISL_2231238, EPI_ISL_2231239, EPI_ISL_2231240, EPI_ISL_2231241, EPI_ISL_2231242, EPI_ISL_2231243, EPI_ISL_2231244, EPI_ISL_2231245, EPI_ISL_2231246, EPI_ISL_2231247, EPI_ISL_2231248, EPI_ISL_2231249, EPI_ISL_2231250, EPI_ISL_2231251, EPI_ISL_2231252, EPI_ISL_2231253, EPI_ISL_2231254, EPI_ISL_2231255, EPI_ISL_2231256, EPI_ISL_2231257, EPI_ISL_2231258, EPI_ISL_2231259, EPI_ISL_2231260, EPI_ISL_2231261, EPI_ISL_2231262, EPI_ISL_2231263, EPI_ISL_2231264, EPI_ISL_2231265, EPI_ISL_2231266, EPI_ISL_2231267, EPI_ISL_2231268, EPI_ISL_2231269 | | | |
| see above | San Diego County Public Health Laboratory | Andersen lab at Scripps Research | SEARCH Alliance San Diego with Ashleigh Murphy, Jovan Shephard, Brett Austin |
| EPI_ISL_2231270, EPI_ISL_2231271, EPI_ISL_2231272, EPI_ISL_2231273, EPI_ISL_2231274, EPI_ISL_2231275, EPI_ISL_2231276, EPI_ISL_2231277, EPI_ISL_2231278, EPI_ISL_2231279, EPI_ISL_2231280, EPI_ISL_2231281, EPI_ISL_2231282, EPI_ISL_2231283, EPI_ISL_2231284, EPI_ISL_2231285, EPI_ISL_2231286, EPI_ISL_2231287, EPI_ISL_2231288, EPI_ISL_2231289, EPI_ISL_2231290, EPI_ISL_2231291, EPI_ISL_2231292, EPI_ISL_2231293, EPI_ISL_2231294, EPI_ISL_2231295, EPI_ISL_2231296, EPI_ISL_2231297, EPI_ISL_2231298, EPI_ISL_2231299, EPI_ISL_2231300, EPI_ISL_2231301, EPI_ISL_2231302, EPI_ISL_2231303, EPI_ISL_2231304, EPI_ISL_2231305, EPI_ISL_2231306, EPI_ISL_2231307, EPI_ISL_2231308, EPI_ISL_2231309, EPI_ISL_2231310, EPI_ISL_2231311, EPI_ISL_2231312, EPI_ISL_2231313, EPI_ISL_2231314, EPI_ISL_2231315, EPI_ISL_2231316, EPI_ISL_2231317 | | | |
| see above | Sharp HealthCare Laboratory | Andersen lab at Scripps Research | SEARCH Alliance San Diego with Aaron Harding, Jacquelyn Berumen, Cathy Woerle, Liam McGinnis, Art Mendoza, Omid Bakhtar |
| EPI_ISL_2231321, EPI_ISL_2231322, EPI_ISL_2231323, EPI_ISL_2231324, EPI_ISL_2231325, EPI_ISL_2231326, EPI_ISL_2231327, EPI_ISL_2231328, EPI_ISL_2231329, EPI_ISL_2231330, EPI_ISL_2231331, EPI_ISL_2231332, EPI_ISL_2231333, EPI_ISL_2231334, EPI_ISL_2231335, EPI_ISL_2231336, EPI_ISL_2231337, EPI_ISL_2231338, EPI_ISL_2231339, EPI_ISL_2231340, EPI_ISL_2231341, EPI_ISL_2231342, EPI_ISL_2231343, EPI_ISL_2231344, EPI_ISL_2231345, EPI_ISL_2231346, EPI_ISL_2231347, EPI_ISL_2231348, EPI_ISL_2231349, EPI_ISL_2231350, EPI_ISL_2231351, EPI_ISL_2231352, EPI_ISL_2231353, EPI_ISL_2231354, EPI_ISL_2231355, EPI_ISL_2231356, EPI_ISL_2231357, EPI_ISL_2231358, EPI_ISL_2231359, EPI_ISL_2231360, EPI_ISL_2231361, EPI_ISL_2231362, EPI_ISL_2231363, EPI_ISL_2231364, EPI_ISL_2231365, EPI_ISL_2231366, EPI_ISL_2231367, EPI_ISL_2231368, EPI_ISL_2231369, EPI_ISL_2231370, EPI_ISL_2231371, EPI_ISL_2231372, EPI_ISL_2231373, EPI_ISL_2231374, EPI_ISL_2231375, EPI_ISL_2231376, EPI_ISL_2231377, EPI_ISL_2231378, EPI_ISL_2231379, EPI_ISL_2231380, EPI_ISL_2231381, EPI_ISL_2231382, EPI_ISL_2231383, EPI_ISL_2231384, EPI_ISL_2231385, EPI_ISL_2231386, EPI_ISL_2231387, EPI_ISL_2231388, EPI_ISL_2231389, EPI_ISL_2231390, EPI_ISL_2231391, EPI_ISL_2231392, EPI_ISL_2231393, EPI_ISL_2231394, EPI_ISL_2231395, EPI_ISL_2231396, EPI_ISL_2231397, EPI_ISL_2231398, EPI_ISL_2231399, EPI_ISL_2231400, EPI_ISL_2231401, EPI_ISL_2231402, EPI_ISL_2231403, EPI_ISL_2231404, EPI_ISL_2231405, EPI_ISL_2231406, EPI_ISL_2231407, EPI_ISL_2231408, EPI_ISL_2231409, EPI_ISL_2231410, EPI_ISL_2231411, EPI_ISL_2231412, EPI_ISL_2231413, EPI_ISL_2231414, EPI_ISL_2231415, EPI_ISL_2231416, EPI_ISL_2231417, EPI_ISL_2231418, EPI_ISL_2231419, EPI_ISL_2231420, EPI_ISL_2231421, EPI_ISL_2231422, EPI_ISL_2231423, EPI_ISL_2231424, EPI_ISL_2231425, EPI_ISL_2231426, EPI_ISL_2231427, EPI_ISL_2231428, EPI_ISL_2231429, EPI_ISL_2231430, EPI_ISL_2231431, EPI_ISL_2231432, EPI_ISL_2231433, EPI_ISL_2231434 | | | |
| see above | Centro de Diagnostico COVID-19 UABC Tijuana | Andersen lab at Scripps Research | SEARCH Alliance San Diego with Idanya Rubi Serafin Higuera, Manuel Sanchez Alavez, Jorge Luis Jimenez Niebla, German Ibarra, Jonathan Vincent Baena, Oscar Efren Zazueta Fierro |
| EPI_ISL_2231436, EPI_ISL_2231437, EPI_ISL_2231438, EPI_ISL_2231439, EPI_ISL_2231440, EPI_ISL_2231441, EPI_ISL_2231443, EPI_ISL_2231444, EPI_ISL_2231445, EPI_ISL_2231446, EPI_ISL_2231447, EPI_ISL_2231448, EPI_ISL_2231449, EPI_ISL_2231450, EPI_ISL_2231451, EPI_ISL_2231452, EPI_ISL_2231453, EPI_ISL_2231454, EPI_ISL_2231455, EPI_ISL_2231456, EPI_ISL_2231457, EPI_ISL_2231458, EPI_ISL_2231459, EPI_ISL_2231460, EPI_ISL_2231461, EPI_ISL_2231462, EPI_ISL_2231463, EPI_ISL_2231464, EPI_ISL_2231465 | | | |
| see above | San Diego County Public Health Laboratory | Andersen lab at Scripps Research | SEARCH Alliance San Diego with Ashleigh Murphy, Jovan Shephard, Brett Austin |
| EPI_ISL_2231467 | Centro de Diagnostico COVID-19 UABC Tijuana | Andersen lab at Scripps Research | SEARCH Alliance San Diego with Idanya Rubi Serafin Higuera, Manuel Sanchez Alavez, Jorge Luis Jimenez Niebla, German Ibarra, Jonathan Vincent Baena, Oscar Efren Zazueta Fierro |
| EPI_ISL_2231468, EPI_ISL_2231469, EPI_ISL_2231470, EPI_ISL_2231471, EPI_ISL_2231473, EPI_ISL_2231474, EPI_ISL_2231477, EPI_ISL_2231480, EPI_ISL_2231481, EPI_ISL_2231482, EPI_ISL_2231483, EPI_ISL_2231484, EPI_ISL_2231487, EPI_ISL_2231488, EPI_ISL_2231489, EPI_ISL_2231490, EPI_ISL_2231491, EPI_ISL_2231492, EPI_ISL_2231493, EPI_ISL_2231495, EPI_ISL_2231496, EPI_ISL_2231497, EPI_ISL_2231500, EPI_ISL_2231501, EPI_ISL_2231504, EPI_ISL_2231505, EPI_ISL_2231506, EPI_ISL_2231507 | | | |
| see above | San Diego County Public Health Laboratory | Andersen lab at Scripps Research | SEARCH Alliance San Diego with Ashleigh Murphy, Jovan Shephard, Brett Austin |
| EPI_ISL_2231508 | Centro de Diagnostico COVID-19 UABC Tijuana | Andersen lab at Scripps Research | SEARCH Alliance San Diego with Idanya Rubi Serafin Higuera, Manuel Sanchez Alavez, Jorge Luis Jimenez Niebla, German Ibarra, Jonathan Vincent Baena, Oscar Efren Zazueta Fierro |
| EPI_ISL_2231510, EPI_ISL_2231511, EPI_ISL_2231512, EPI_ISL_2231513, EPI_ISL_2231514, EPI_ISL_2231518, EPI_ISL_2231519, EPI_ISL_2231520, EPI_ISL_2231521, EPI_ISL_2231522 | San Diego County Public Health Laboratory | Andersen lab at Scripps Research | SEARCH Alliance San Diego with Ashleigh Murphy, Jovan Shephard, Brett Austin |
| EPI_ISL_2231528, EPI_ISL_2231529, EPI_ISL_2231531, EPI_ISL_2231532, EPI_ISL_2231533, EPI_ISL_2231534, EPI_ISL_2231535, EPI_ISL_2231536, EPI_ISL_2231537, EPI_ISL_2231538, EPI_ISL_2231539, EPI_ISL_2231540, EPI_ISL_2231541, EPI_ISL_2231542, EPI_ISL_2231543, EPI_ISL_2231544, EPI_ISL_2231545, EPI_ISL_2231546, EPI_ISL_2231547, EPI_ISL_2231548, EPI_ISL_2231549, EPI_ISL_2231550, EPI_ISL_2231551, EPI_ISL_2231552, EPI_ISL_2231553, EPI_ISL_2231554, EPI_ISL_2231555, EPI_ISL_2231556, EPI_ISL_2231557, EPI_ISL_2231558, EPI_ISL_2231559, EPI_ISL_2231560, EPI_ISL_2231561, EPI_ISL_2231562, EPI_ISL_2231563, EPI_ISL_2231564 | | | |
| see above | Sharp HealthCare Laboratory | Andersen lab at Scripps Research | SEARCH Alliance San Diego with Aaron Harding, Jacquelyn Berumen, Cathy Woerle, Liam McGinnis, Art Mendoza, Omid Bakhtar |
| EPI_ISL_2231883, EPI_ISL_2231884, EPI_ISL_2231885, EPI_ISL_2231886, EPI_ISL_2231887, EPI_ISL_2231888, EPI_ISL_2231889, EPI_ISL_2231890 | CQRC_QUALITY CONTROL CHEMICAL BIOLOGICAL RISK_AOOR Villa Sofia Cervello Palermo | CQRC_QUALITY CONTROL CHEMICAL BIOLOGICAL RISK_AOOR Villa Sofia Cervello Palermo | Di Gaudio, F., Contino, F., Brunacci, G. |
| EPI_ISL_2231891, EPI_ISL_2231892, EPI_ISL_2231893 | CQRC_QUALITY CONTROL CHEMICAL BIOLOGICAL RISK_AOOR Villa Sofia Cervello Palermo | CQRC_QUALITY CONTROL CHEMICAL BIOLOGICAL RISK_AOOR Villa Sofia Cervello Palermo | Di Gaudio, F., Brunacci, G., Contino, F. |
| EPI_ISL_2231894 | CQRC_QUALITY CONTROL CHEMICAL BIOLOGICAL | CQRC_QUALITY CONTROL CHEMICAL BIOLOGICAL | Di Gaudio, F., Contino, F., Brunacci, G. |

| | | | |
|---|---|---|---|
| | RISK_AOOR Villa Sofia Cervello Palermo | RISK_AOOR Villa Sofia Cervello Palermo | |
| EPI_ISL_2231899 | CQRC_QUALITY CONTROL CHEMICAL BIOLOGICAL RISK_AOOR Villa Sofia Cervello Palermo | CQRC_QUALITY CONTROL CHEMICAL BIOLOGICAL RISK_AOOR Villa Sofia Cervello Palermo | Di Gaudio, F., Contino, F., Gasparro O., Rinaudo G., Brunacci, G. |
| EPI_ISL_2231900, EPI_ISL_2231901, EPI_ISL_2231902, EPI_ISL_2231903, EPI_ISL_2231904 | CQRC_QUALITY CONTROL CHEMICAL BIOLOGICAL RISK_AOOR Villa Sofia Cervello Palermo | CQRC_QUALITY CONTROL CHEMICAL BIOLOGICAL RISK_AOOR Villa Sofia Cervello Palermo | Di Gaudio, F., Brunacci, G., Contino, F. |
| EPI_ISL_2240801 | Laboratorio Biologia Molecolare Sars Cov2 - UOC Laboratorio Analisi - Servizio Medicina di Laboratorio, Ospedale "San Francesco" - ATS-ASSL Nuoro | Laboratorio specialistico UOC Ematologia - Ospedale "San Francesco" - ATS-ASSL Nuoro | Piras Giovanna, Asproni Rosanna,,Fiamma Maura, Monne Maria Itria, Palmas Angelo Domenico, Lo Maglio Iana, Mameli Giuseppe |
| EPI_ISL_2253214, EPI_ISL_2254130, EPI_ISL_2254496 | Veterinary institute in Zvolen Slovakia | Veterinary institute in Zvolen Slovakia | Dirbáková Z., Sujová S., Mokryšová S., Tinák M., Mojžiš M |
| EPI_ISL_2258200 | Laboratorio Biologia Molecolare Sars Cov2 - UOC Laboratorio Analisi - Servizio Medicina di Laboratorio, Ospedale "San Francesco" - ATS-ASSL Nuoro | Laboratorio specialistico UOC Ematologia - Ospedale "San Francesco" - ATS-ASSL Nuoro | Piras Giovanna, Asproni Rosanna,,Fiamma Maura, Monne Maria Itria, Palmas Angelo Domenico, Lo Maglio Iana, Mameli Giuseppe |
| EPI_ISL_2271905, EPI_ISL_2271910, EPI_ISL_2271911 | Hospital General de México, Medicina Genómica | Hospital General de México, Medicina Genómica | Ana Laura Sánchez-Sandoval, María Eligia Juárez-Torres, Mariana Villalba-De la Peña, Siddhartha Ríos-Zaragoza, Guadalupe Mercedes Lucía Guerrero-Avendaño, Erika Anabel Alvarado-Silva, Eira Valeria Barrón-Palma |
| EPI_ISL_2272477, EPI_ISL_2272479, EPI_ISL_2272480, EPI_ISL_2272481, EPI_ISL_2272482, EPI_ISL_2272483, EPI_ISL_2272484, EPI_ISL_2272485, EPI_ISL_2272486, EPI_ISL_2272487, EPI_ISL_2272488, EPI_ISL_2272489, EPI_ISL_2272490, EPI_ISL_2272491, EPI_ISL_2272492, EPI_ISL_2272493, EPI_ISL_2272494, EPI_ISL_2272495, EPI_ISL_2272496 | | | |
| see above | Indian Council of Medical Research-National Institute of Virology, Microbial Containment Complex | Indian Council of Medical Research-National Institute of Virology, Microbial Containment Complex | Pragya D Yadav |
| EPI_ISL_2279498 | Institut National d'Hygiène (INH) | Unité Mixte Internationale TransVIHMI (UMI 233 IRD - U1175 INSERM - Université de Montpellier)IRD (Institut de recherche pour le développement) | Mounerou SALOU, Christelle BUTEL, Wembo A. HALATOKO,Issaka Maman, Abla A. KONOU, Amivi EHLAN, Adodo SADJI, Kokou TEGUENI,Sidonie A.M.KAGNISSODE, Akoélé SILIADIN, Alassane OURO-MEDELI, Messanh DOUFFAN,Déléma MABA,Sika DOSSIM, Améyo DORKENOO, Mireille PRINCE-DAVID,Anoumou DAGNRA,Laetitia SERRANO,Ahidjo AYOUBA,Eric DELAPORTE, Martine PEETERS |
| EPI_ISL_2283768, EPI_ISL_2283769, EPI_ISL_2283770, EPI_ISL_2283771, EPI_ISL_2283772, EPI_ISL_2283773, EPI_ISL_2283774, EPI_ISL_2283775, EPI_ISL_2283776, EPI_ISL_2283777, EPI_ISL_2283778, EPI_ISL_2283779, EPI_ISL_2283780, EPI_ISL_2283781, EPI_ISL_2283782, EPI_ISL_2283783, EPI_ISL_2283784, EPI_ISL_2283785, EPI_ISL_2283786, EPI_ISL_2283787, EPI_ISL_2283788, EPI_ISL_2283789, EPI_ISL_2283790, EPI_ISL_2283791, EPI_ISL_2283792, EPI_ISL_2283793, EPI_ISL_2283794, EPI_ISL_2283795, EPI_ISL_2283796, EPI_ISL_2283797, EPI_ISL_2283798, EPI_ISL_2283799, EPI_ISL_2283800, EPI_ISL_2283801, EPI_ISL_2283802, EPI_ISL_2283803, EPI_ISL_2283804, EPI_ISL_2283807, EPI_ISL_2283808, EPI_ISL_2283809, EPI_ISL_2283810 | | | |
| see above | Genome Analysis Center, Yamanashi Central Hospital | Genome Analysis Center, Yamanashi Central Hospital | Yosuke Hirotsu |
| EPI_ISL_2284874 | Siti Khodijah Hospital | Institute of Tropical Disease, Universitas Airlangga | Kazufumi Shimizu, Krisnoadi Rahardjo, Aldise M Nastri, Jezzy R Dewantari, Rima R Prasetya, Muhammad Hamdan, Gatot Soegiarto, Laksmi Wulandari, Resti Yudhawati, Yasuko Mori, Soetjipto, Maria I Lusida |
| EPI_ISL_2285106, EPI_ISL_2285150 | Microbiology Department. Complexo Hospitalario Universitario de Vigo | Microbiology Department. Complexo Hospitalario Universitario de Vigo | Alfaya N, Alonso I, Alvarez M, Cabrera JJ, Carballo R, Cores O, Cortizo S, del-Campo V, Martinez L, Mediero G, Perez S, Potel C, Regueiro B, Rey S, Vasallo FJ |
| EPI_ISL_2285723, EPI_ISL_2285732 | Department of Pathology, National Institute of Infectious Diseases | Department of Pathology, National Institute of Infectious Diseases | Harutaka Katano, Takayuki Kanno, Tadaki Suzuki |
| EPI_ISL_2293282 | Labo Analyses Med | National Reference Center for Viruses of Respiratory Infections, Institut Pasteur, Paris | Marion Barbet, Sylvie Behillil, Méline Bizard, Angela Brisebarre, Camille Capel, Vincent Enouf, Louise Lefrançois, Frédéric Lemoine, Christophe Malabat, Corinne Maufrais, Etienne Simon-Lorière, Maud Vanpeene, Sylvie Van der Werf ,Caroline Malderet |
| EPI_ISL_2293421, EPI_ISL_2293422 | Research | National Reference Center for Viruses of Respiratory Infections, Institut Pasteur, Paris | Marion Barbet, Sylvie Behillil, Méline Bizard, Angela Brisebarre, Camille Capel, Vincent Enouf, Louise Lefrançois, Frédéric Lemoine, Christophe Malabat, Corinne Maufrais, Etienne Simon-Lorière, Maud Vanpeene, Sylvie Van der Werf ,Marianne Leruez-Ville |
| EPI_ISL_2296063 | LESP Hidalgo | Instituto de Diagnostico y Referencia Epidemiologicos (INDRE) | Claudia Wong-Arambula, Abril Rodriguez-Maldonado, Vanessa Rivero-Arredondo, Ariadna Medina-Benitez, Joaquin Quiroz-Mercado, Sergio Rangel-Guerrero, Natividad Cruz-Ortiz, Tatiana Nunez-Garcia, Gisela Barrera-Badillo, Lucia Hernandez-Rivas, Irma Lopez-Martinez, Ernesto Ramirez-Gonzalez. |
| EPI_ISL_2304298, EPI_ISL_2304299, EPI_ISL_2304300, EPI_ISL_2304301, EPI_ISL_2304302, EPI_ISL_2304303, EPI_ISL_2307325, EPI_ISL_2307327, EPI_ISL_2307328, EPI_ISL_2307331, EPI_ISL_2307332, EPI_ISL_2307333, EPI_ISL_2307334, EPI_ISL_2307335, EPI_ISL_2307336, EPI_ISL_2307337, EPI_ISL_2307349 | | | |
| see above | Genome Analysis Center, Yamanashi Central Hospital | Genome Analysis Center, Yamanashi Central Hospital | Yosuke Hirotsu |
| EPI_ISL_2308271, EPI_ISL_2308272, EPI_ISL_2308273, EPI_ISL_2308274, EPI_ISL_2308275, EPI_ISL_2308276 | Indian Council of Medical Research-National Institute of Virology, Microbial Containment Complex | Indian Council of Medical Research-National Institute of Virology, Microbial Containment Complex | Pragya D Yadav |
| EPI_ISL_2321226, EPI_ISL_2321230, EPI_ISL_2322453, EPI_ISL_2322454 | Genome Analysis Center, Yamanashi Central Hospital | Genome Analysis Center, Yamanashi Central Hospital | Yosuke Hirotsu |
| EPI_ISL_2322758, EPI_ISL_2322761 | Collection of WHO National Influenza Center, N.F. Gamaleya National Research Center for Epidemiology and Microbiology of the Ministry of Health of the Russian Federation | Laboratory of genome analysis, N.F. Gamaleya National Research Center for Epidemiology and Microbiology of the Ministry of Health of the Russian Federation | Olga Voronina, Ekaterina Aksenova, Natalia Ryzhova, Marina Kunda, Kirill Krasnoslobodtsev, Elena Burtseva, Denis Logunov |
| EPI_ISL_2324194 | Austrian Agency for Health and Food Safety (AGES) | Bergthaler laboratory, CeMM Research Center for Molecular Medicine of the Austrian Academy of Sciences | Lukas Endler, Anna Schedl, Fabian Amman, Petr Triska, Thomas Penz, Benedikt Agerer, Maelle Le Moing, Michael Schuster, Bekir Erguner, Jan Laine, Martin Senekowitsch, Christoph Bock, Andreas Bergthaler |
| EPI_ISL_2324258, EPI_ISL_2324259, EPI_ISL_2324260, EPI_ISL_2324261 | Institut für Virologie am Department für Hygiene, Mikrobiologie und Public Health | Bergthaler laboratory, CeMM Research Center for Molecular Medicine of the Austrian Academy of Sciences | Lukas Endler, Anna Schedl, Fabian Amman, Petr Triska, Thomas Penz, Benedikt Agerer, Maelle Le Moing, Michael Schuster, Bekir Erguner, Jan Laine, Martin Senekowitsch, Christoph Bock, Andreas Bergthaler |
| EPI_ISL_2339875, EPI_ISL_2339903, EPI_ISL_2339904 | Centro de Estudio de Enfermedades Autoinmunes (CREA), Universidad del Rosario, Bogota, Colombia | Centro de Investigaciones en Microbiología y Biotecnología-UR (CIMBIUR), Facultad de Ciencias Naturales, Universidad del Rosario, Bogotá, Colombia | Juan-Manuel Anaya, Gustavo Salguero, Juan Esteban Gallo, Carolina Ramírez-Santana, Sergio Castañeda, Nathalia Ballesteros, Marina Muñoz, Luz H. Patiño, Juan David Ramírez |
| EPI_ISL_2342585, EPI_ISL_2342586, EPI_ISL_2342587, EPI_ISL_2342588, EPI_ISL_2342589, EPI_ISL_2342590, EPI_ISL_2342591, EPI_ISL_2342592, EPI_ISL_2342593, EPI_ISL_2342594, EPI_ISL_2342595, EPI_ISL_2342596, EPI_ISL_2342597, EPI_ISL_2342598, EPI_ISL_2342599, EPI_ISL_2342600 | | | |
| see above | Malaysia Genome Institute | Malaysia Genome Institute | Mohd Noor Mat Isa, Irni Suhayu Sapian, Yusuf Muhammad Noor, Nurhezreen Md Iqbal, Mohd Faizal Abu Bakar, Enizza Kasim, Shamsidar Sopie, Siti Noraini Othman, Azrin Ahmad, Nor Azfa Johari. |
| EPI_ISL_2343010 | LESP Hidalgo | Instituto de Diagnostico y Referencia Epidemiologicos (INDRE) | Claudia Wong-Arambula, Abril Rodriguez-Maldonado, Vanessa Rivero-Arredondo, Ariadna Medina-Benitez, Joaquin Quiroz-Mercado, Sergio Rangel-Guerrero, Natividad Cruz-Ortiz, Tatiana Nunez-Garcia, Gisela Barrera-Badillo, Lucia Hernandez-Rivas, Irma Lopez-Martinez, Ernesto Ramirez-Gonzalez. |
| EPI_ISL_2350786 | Translational Health Science and Technology Institute -ESIC medical college and hospital, Faridabad | THSTI Bioassay laboratory | Naseem Ahmed Khan, Saurabh Kumar, Neha Jha, Anish Maitra, Partha Chattopadhyay, Anil Kumar Pandey, Guruprasad R Medigeshi, Rajesh Pandey |
| EPI_ISL_2364489, EPI_ISL_2364490, EPI_ISL_2364500, EPI_ISL_2364501, EPI_ISL_2364502, EPI_ISL_2364503, EPI_ISL_2364509 | Genome Analysis Center, Yamanashi Central Hospital | Genome Analysis Center, Yamanashi Central Hospital | Yosuke Hirotsu |
| EPI_ISL_2364568 | RS Medirossa Cikarang | Eijkman Institute for Molecular Biology, National Agency for Research and Innovation | Willy Agustine, Hidayat Trimarsanto, Lydia V. Panggalo, Iskandar Adnan, Sukma Oktavianthi, Lidwina Priliani, Edison Johar, Frilasita A Yudhaputri, Muhammad Rezki Rasyak, Safarina G Malik, Khin Saw Myint, Amin Soebandrio |
| EPI_ISL_2364569, EPI_ISL_2364570, EPI_ISL_2364571 | RS Sentra Medika Cikarang | Eijkman Institute for Molecular Biology, National Agency for Research and Innovation | Lydia V. Panggalo, Iskandar Adnan, Sukma Oktavianthi, Lidwina Priliani, Edison Johar, Frilasita A Yudhaputri, Muhammad Rezki Rasyak, Willy Agustine, Hidayat Trimarsanto, Safarina G Malik, Khin Saw Myint, Amin Soebandrio |

| | | | |
|---|---|---|---|
| EPI_ISL_2364572 | RS Sentra Medika Cikarang | Eijkman Institute for Molecular Biology, National Agency for Research and Innovation | Iskandar Adnan, Sukma Oktavianthi, Lidwina Priliani, Edison Johar, Frilasita A Yudhaputri, Muhammad Rezki Rasyak, Willy Agustine, Hidayat Trimarsanto, Lydia V. Panggalo, Safarina G Malik, Khin Saw Myint, Amin Soebandrio |
| EPI_ISL_2364573 | RS Bhakti Husada Cikarang | Eijkman Institute for Molecular Biology, National Agency for Research and Innovation | Iskandar Adnan, Sukma Oktavianthi, Lidwina Priliani, Edison Johar, Frilasita A Yudhaputri, Muhammad Rezki Rasyak, Willy Agustine, Hidayat Trimarsanto, Lydia V. Panggalo, Safarina G Malik, Khin Saw Myint, Amin Soebandrio |
| EPI_ISL_2364574 | RS Bhakti Husada Cikarang | Eijkman Institute for Molecular Biology, National Agency for Research and Innovation | Sukma Oktavianthi, Lidwina Priliani, Edison Johar, Frilasita A Yudhaputri, Muhammad Rezki Rasyak, Willy Agustine, Hidayat Trimarsanto, Lydia V. Panggalo, Iskandar Adnan, Safarina G Malik, Khin Saw Myint, Amin Soebandrio |
| EPI_ISL_2364575 | RS Masmitra | Eijkman Institute for Molecular Biology, National Agency for Research and Innovation | Iskandar Adnan, Sukma Oktavianthi, Lidwina Priliani, Edison Johar, Frilasita A Yudhaputri, Muhammad Rezki Rasyak, Willy Agustine, Hidayat Trimarsanto, Lydia V. Panggalo, Safarina G Malik, Khin Saw Myint, Amin Soebandrio |
| EPI_ISL_2364576 | RS Harapan Bunda | Eijkman Institute for Molecular Biology, National Agency for Research and Innovation | Hidayat Trimarsanto, Lydia V. Panggalo, Iskandar Adnan, Sukma Oktavianthi, Lidwina Priliani, Edison Johar, Frilasita A Yudhaputri, Muhammad Rezki Rasyak, Willy Agustine, Safarina G Malik, Khin Saw Myint, Amin Soebandrio |
| EPI_ISL_2364577 | RS Permata Bekasi | Eijkman Institute for Molecular Biology, National Agency for Research and Innovation | Sukma Oktavianthi, Lidwina Priliani, Edison Johar, Frilasita A Yudhaputri, Muhammad Rezki Rasyak, Willy Agustine, Hidayat Trimarsanto, Lydia V. Panggalo, Iskandar Adnan, Safarina G Malik, Khin Saw Myint, Amin Soebandrio |
| EPI_ISL_2364578 | RS Asy-Syifaa Leuwiliang | Eijkman Institute for Molecular Biology, National Agency for Research and Innovation | Sukma Oktavianthi, Lidwina Priliani, Edison Johar, Frilasita A Yudhaputri, Muhammad Rezki Rasyak, Willy Agustine, Hidayat Trimarsanto, Lydia V. Panggalo, Iskandar Adnan, Safarina G Malik, Khin Saw Myint, Amin Soebandrio |
| EPI_ISL_2364579 | RS Mitra Keluarga Cibubur | Eijkman Institute for Molecular Biology, National Agency for Research and Innovation | Lidwina Priliani, Edison Johar, Frilasita A Yudhaputri, Muhammad Rezki Rasyak, Willy Agustine, Hidayat Trimarsanto, Lydia V. Panggalo, Iskandar Adnan, Sukma Oktavianthi, Safarina G Malik, Khin Saw Myint, Amin Soebandrio |
| EPI_ISL_2364580 | RSUD Cileungsi | Eijkman Institute for Molecular Biology, National Agency for Research and Innovation | Edison Johar, Frilasita A Yudhaputri, Muhammad Rezki Rasyak, Willy Agustine, Hidayat Trimarsanto, Lydia V. Panggalo, Iskandar Adnan, Sukma Oktavianthi, Lidwina Priliani, Safarina G Malik, Khin Saw Myint, Amin Soebandrio |
| EPI_ISL_2364581 | RS Sentra Medika Cikarang | Eijkman Institute for Molecular Biology, National Agency for Research and Innovation | Iskandar Adnan, Sukma Oktavianthi, Lidwina Priliani, Edison Johar, Frilasita A Yudhaputri, Muhammad Rezki Rasyak, Willy Agustine, Hidayat Trimarsanto, Lydia V. Panggalo, Safarina G Malik, Khin Saw Myint, Amin Soebandrio |
| EPI_ISL_2364582 | RS Sentra Medika Cikarang | Eijkman Institute for Molecular Biology, National Agency for Research and Innovation | Frilasita A Yudhaputri, Muhammad Rezki Rasyak, Willy Agustine, Hidayat Trimarsanto, Lydia V. Panggalo, Iskandar Adnan, Sukma Oktavianthi, Lidwina Priliani, Edison Johar, Safarina G Malik, Khin Saw Myint, Amin Soebandrio |
| EPI_ISL_2364583 | RSJ Dr. Soeharto Heerdjan | Eijkman Institute for Molecular Biology, National Agency for Research and Innovation | Muhammad Rezki Rasyak, Willy Agustine, Hidayat Trimarsanto, Lydia V. Panggalo, Iskandar Adnan, Sukma Oktavianthi, Lidwina Priliani, Edison Johar, Frilasita A Yudhaputri, Safarina G Malik, Khin Saw Myint, Amin Soebandrio |
| EPI_ISL_2364584 | Ciputra Hospital Citra Raya Tangerang | Eijkman Institute for Molecular Biology, National Agency for Research and Innovation | Edison Johar, Frilasita A Yudhaputri, Muhammad Rezki Rasyak, Willy Agustine, Hidayat Trimarsanto, Lydia V. Panggalo, Iskandar Adnan, Sukma Oktavianthi, Lidwina Priliani, Safarina G Malik, Khin Saw Myint, Amin Soebandrio |
| EPI_ISL_2364585 | RSUD Cileungsi | Eijkman Institute for Molecular Biology, National Agency for Research and Innovation | Frilasita A Yudhaputri, Muhammad Rezki Rasyak, Willy Agustine, Hidayat Trimarsanto, Lydia V. Panggalo, Iskandar Adnan, Sukma Oktavianthi, Lidwina Priliani, Edison Johar, Safarina G Malik, Khin Saw Myint, Amin Soebandrio |
| EPI_ISL_2364586 | RS Graha Juanda | Eijkman Institute for Molecular Biology, National Agency for Research and Innovation | Muhammad Rezki Rasyak, Willy Agustine, Hidayat Trimarsanto, Lydia V. Panggalo, Iskandar Adnan, Sukma Oktavianthi, Lidwina Priliani, Edison Johar, Frilasita A Yudhaputri, Safarina G Malik, Khin Saw Myint, Amin Soebandrio |
| EPI_ISL_2364588 | RS Mitra Keluarga Gading Serpong | Eijkman Institute for Molecular Biology, National Agency for Research and Innovation | Hidayat Trimarsanto, Lydia V. Panggalo, Iskandar Adnan, Sukma Oktavianthi, Lidwina Priliani, Edison Johar, Frilasita A Yudhaputri, Muhammad Rezki Rasyak, Willy Agustine, Safarina G Malik, Khin Saw Myint, Amin Soebandrio |
| EPI_ISL_2364590 | RS Bhakti Husada Cikarang | Eijkman Institute for Molecular Biology, National Agency for Research and Innovation | Lidwina Priliani, Edison Johar, Frilasita A Yudhaputri, Muhammad Rezki Rasyak, Willy Agustine, Hidayat Trimarsanto, Lydia V. Panggalo, Iskandar Adnan, Sukma Oktavianthi, Safarina G Malik, Khin Saw Myint, Amin Soebandrio |
| EPI_ISL_2364591 | RS Graha Juanda | Eijkman Institute for Molecular Biology, National Agency for Research and Innovation | Edison Johar, Frilasita A Yudhaputri, Muhammad Rezki Rasyak, Willy Agustine, Hidayat Trimarsanto, Lydia V. Panggalo, Iskandar Adnan, Sukma Oktavianthi, Lidwina Priliani, Safarina G Malik, Khin Saw Myint, Amin Soebandrio |
| EPI_ISL_2364592 | RS Mitra Keluarga Gading Serpong | Eijkman Institute for Molecular Biology, National Agency for Research and Innovation | Frilasita A Yudhaputri, Muhammad Rezki Rasyak, Willy Agustine, Hidayat Trimarsanto, Lydia V. Panggalo, Iskandar Adnan, Sukma Oktavianthi, Lidwina Priliani, Edison Johar, Safarina G Malik, Khin Saw Myint, Amin Soebandrio |
| EPI_ISL_2364594 | RSUD Cileungsi | Eijkman Institute for Molecular Biology, National Agency for Research and Innovation | Muhammad Rezki Rasyak, Willy Agustine, Hidayat Trimarsanto, Lydia V. Panggalo, Iskandar Adnan, Sukma Oktavianthi, Lidwina Priliani, Edison Johar, Frilasita A Yudhaputri, Safarina G Malik, Khin Saw Myint, Amin Soebandrio |
| EPI_ISL_2364596 | RS Mitra Keluarga Kelapa Gading | Eijkman Institute for Molecular Biology, National Agency for Research and Innovation | Hidayat Trimarsanto, Lydia V. Panggalo, Iskandar Adnan, Sukma Oktavianthi, Lidwina Priliani, Edison Johar, Frilasita A Yudhaputri, Muhammad Rezki Rasyak, Willy Agustine, Safarina G Malik, Khin Saw Myint, Amin Soebandrio |
| EPI_ISL_2364597 | RS Permata Cibubur | Eijkman Institute for Molecular Biology, National Agency for Research and Innovation | Edison Johar, Frilasita A Yudhaputri, Muhammad Rezki Rasyak, Willy Agustine, Hidayat Trimarsanto, Lydia V. Panggalo, Iskandar Adnan, Sukma Oktavianthi, Lidwina Priliani, Safarina G Malik, Khin Saw Myint, Amin Soebandrio |
| EPI_ISL_2364598 | RSIA Bunda Sejahtera | Eijkman Institute for Molecular Biology, National Agency for Research and Innovation | Muhammad Rezki Rasyak, Willy Agustine, Hidayat Trimarsanto, Lydia V. Panggalo, Iskandar Adnan, Sukma Oktavianthi, Lidwina Priliani, Edison Johar, Frilasita A Yudhaputri, Safarina G Malik, Khin Saw Myint, Amin Soebandrio |
| EPI_ISL_2364600 | RS Mitra Keluarga Cibubur | Eijkman Institute for Molecular Biology, National Agency for Research and Innovation | Willy Agustine, Hidayat Trimarsanto, Lydia V. Panggalo, Iskandar Adnan, Sukma Oktavianthi, Lidwina Priliani, Edison Johar, Frilasita A Yudhaputri, Muhammad Rezki Rasyak, Safarina G Malik, Khin Saw Myint, Amin Soebandrio |
| EPI_ISL_2364601 | RS Mary Cileungsi Hijau | Eijkman Institute for Molecular Biology, National Agency for Research and Innovation | Hidayat Trimarsanto, Lydia V. Panggalo, Iskandar Adnan, Sukma Oktavianthi, Lidwina Priliani, Edison Johar, Frilasita A Yudhaputri, Muhammad Rezki Rasyak, Willy Agustine, Safarina G Malik, Khin Saw Myint, Amin Soebandrio |
| EPI_ISL_2364602 | RS Sumber Waras | Eijkman Institute for Molecular Biology, National Agency for Research and Innovation | Iskandar Adnan, Sukma Oktavianthi, Lidwina Priliani, Edison Johar, Frilasita A Yudhaputri, Muhammad Rezki Rasyak, Willy Agustine, Hidayat Trimarsanto, Lydia V. Panggalo, Safarina G Malik, Khin Saw Myint, Amin Soebandrio |
| EPI_ISL_2364603 | RS Mitra Keluarga Gading Serpong | Eijkman Institute for Molecular Biology, National Agency for Research and Innovation | Sukma Oktavianthi, Lidwina Priliani, Edison Johar, Frilasita A Yudhaputri, Muhammad Rezki Rasyak, Willy Agustine, Hidayat Trimarsanto, Lydia V. Panggalo, Iskandar Adnan, Safarina G Malik, Khin Saw Myint, Amin Soebandrio |
| EPI_ISL_2364605 | RS Islam Sari Asih Ar-Rahmah | Eijkman Institute for Molecular Biology, National Agency for Research and Innovation | Willy Agustine, Hidayat Trimarsanto, Lydia V. Panggalo, Iskandar Adnan, Sukma Oktavianthi, Lidwina Priliani, Edison Johar, Frilasita A Yudhaputri, Muhammad Rezki Rasyak, Safarina G Malik, Khin Saw Myint, Amin Soebandrio |
| EPI_ISL_2364607 | RS Medika Dramaga | Eijkman Institute for Molecular Biology, National Agency for Research and Innovation | Lydia V. Panggalo, Iskandar Adnan, Sukma Oktavianthi, Lidwina Priliani, Edison Johar, Frilasita A Yudhaputri, Muhammad Rezki Rasyak, Willy Agustine, Hidayat Trimarsanto, Safarina G Malik, Khin Saw Myint, Amin Soebandrio |
| EPI_ISL_2364608 | RS Harapan Bunda | Eijkman Institute for Molecular Biology, National Agency for Research and Innovation | Iskandar Adnan, Sukma Oktavianthi, Lidwina Priliani, Edison Johar, Frilasita A Yudhaputri, Muhammad Rezki Rasyak, Willy Agustine, Hidayat Trimarsanto, Lydia V. Panggalo, Safarina G Malik, Khin Saw Myint, Amin Soebandrio |
| EPI_ISL_2364609 | RS Harapan Bunda | Eijkman Institute for Molecular Biology, National Agency for Research and Innovation | Frilasita A Yudhaputri, Muhammad Rezki Rasyak, Willy Agustine, Hidayat Trimarsanto, Lydia V. Panggalo, Iskandar Adnan, Sukma Oktavianthi, Lidwina Priliani, Edison Johar, Safarina G Malik, Khin Saw Myint, Amin Soebandrio |
| EPI_ISL_2364610 | RS Mitra Keluarga Kalideres | Eijkman Institute for Molecular Biology, National Agency for Research and Innovation | Muhammad Rezki Rasyak, Willy Agustine, Hidayat Trimarsanto, Lydia V. Panggalo, Iskandar Adnan, Sukma Oktavianthi, Lidwina Priliani, Edison Johar, Frilasita A Yudhaputri, Safarina G Malik, Khin Saw Myint, Amin Soebandrio |
| EPI_ISL_2364611 | RSUD Cileungsi | Eijkman Institute for Molecular Biology, National Agency for Research and Innovation | Hidayat Trimarsanto, Lydia V. Panggalo, Iskandar Adnan, Sukma Oktavianthi, Lidwina Priliani, Edison Johar, Frilasita A Yudhaputri, Muhammad Rezki Rasyak, Willy Agustine, Safarina G Malik, Khin Saw Myint, Amin Soebandrio |
| EPI_ISL_2364612 | RSUD Cileungsi | Eijkman Institute for Molecular Biology, National Agency for Research and Innovation | Sukma Oktavianthi, Lidwina Priliani, Edison Johar, Frilasita A Yudhaputri, Muhammad Rezki Rasyak, Willy Agustine, Hidayat Trimarsanto, Lydia V. Panggalo, Iskandar Adnan, Safarina G Malik, Khin Saw Myint, Amin Soebandrio |
| EPI_ISL_2364613 | RSUD Cileungsi | Eijkman Institute for Molecular Biology, National Agency for Research and Innovation | Frilasita A Yudhaputri, Muhammad Rezki Rasyak, Willy Agustine, Hidayat Trimarsanto, Lydia V. Panggalo, Iskandar Adnan, Sukma Oktavianthi, Lidwina Priliani, Edison Johar, Safarina G Malik, Khin Saw Myint, Amin Soebandrio |

| | | | |
|---|---|---|---|
| EPI_ISL_2364615 | RS Sentra Medika Cibinong | Eijkman Institute for Molecular Biology, National Agency for Research and Innovation | Lidwina Priliani, Edison Johar, Frilasita A Yudhaputri, Muhammad Rezki Rasyak, Willy Agustine, Hidayat Trimarsanto, Lydia V. Panggalo, Iskandar Adnan, Sukma Oktavianthi, Safarina G Malik, Khin Saw Myint, Amin Soebandrio |
| EPI_ISL_2364616 | RS Mitra Keluarga Gading Serpong | Eijkman Institute for Molecular Biology, National Agency for Research and Innovation | Muhammad Rezki Rasyak, Willy Agustine, Hidayat Trimarsanto, Lydia V. Panggalo, Iskandar Adnan, Sukma Oktavianthi, Lidwina Priliani, Edison Johar, Frilasita A Yudhaputri, Safarina G Malik, Khin Saw Myint, Amin Soebandrio |
| EPI_ISL_2364617 | RS Mitra Keluarga Gading Serpong | Eijkman Institute for Molecular Biology, National Agency for Research and Innovation | Hidayat Trimarsanto, Lydia V. Panggalo, Iskandar Adnan, Sukma Oktavianthi, Lidwina Priliani, Edison Johar, Frilasita A Yudhaputri, Muhammad Rezki Rasyak, Willy Agustine, Safarina G Malik, Khin Saw Myint, Amin Soebandrio |
| EPI_ISL_2364618 | RS Mitra Keluarga Kelapa Gading | Eijkman Institute for Molecular Biology, National Agency for Research and Innovation | Lydia V. Panggalo, Iskandar Adnan, Sukma Oktavianthi, Lidwina Priliani, Edison Johar, Frilasita A Yudhaputri, Muhammad Rezki Rasyak, Willy Agustine, Hidayat Trimarsanto, Safarina G Malik, Khin Saw Myint, Amin Soebandrio |
| EPI_ISL_2364619 | RS Mitra Keluarga Kelapa Gading | Eijkman Institute for Molecular Biology, National Agency for Research and Innovation | Iskandar Adnan, Sukma Oktavianthi, Lidwina Priliani, Edison Johar, Frilasita A Yudhaputri, Muhammad Rezki Rasyak, Willy Agustine, Hidayat Trimarsanto, Lydia V. Panggalo, Safarina G Malik, Khin Saw Myint, Amin Soebandrio |
| EPI_ISL_2364621 | RS Sentra Medika Cikarang | Eijkman Institute for Molecular Biology, National Agency for Research and Innovation | Sukma Oktavianthi, Lidwina Priliani, Edison Johar, Frilasita A Yudhaputri, Muhammad Rezki Rasyak, Willy Agustine, Hidayat Trimarsanto, Lydia V. Panggalo, Iskandar Adnan, Safarina G Malik, Khin Saw Myint, Amin Soebandrio |
| EPI_ISL_2364624 | RS Sentra Medika Cikarang | Eijkman Institute for Molecular Biology, National Agency for Research and Innovation | Lidwina Priliani, Edison Johar, Frilasita A Yudhaputri, Muhammad Rezki Rasyak, Willy Agustine, Hidayat Trimarsanto, Lydia V. Panggalo, Iskandar Adnan, Sukma Oktavianthi, Safarina G Malik, Khin Saw Myint, Amin Soebandrio |
| EPI_ISL_2364626 | RS Sentra Medika Cikarang | Eijkman Institute for Molecular Biology, National Agency for Research and Innovation | Edison Johar, Frilasita A Yudhaputri, Muhammad Rezki Rasyak, Willy Agustine, Hidayat Trimarsanto, Lydia V. Panggalo, Iskandar Adnan, Sukma Oktavianthi, Lidwina Priliani, Safarina G Malik, Khin Saw Myint, Amin Soebandrio |
| EPI_ISL_2364629 | RS Bhakti Husada Cikarang | Eijkman Institute for Molecular Biology, National Agency for Research and Innovation | Frilasita A Yudhaputri, Muhammad Rezki Rasyak, Willy Agustine, Hidayat Trimarsanto, Lydia V. Panggalo, Iskandar Adnan, Sukma Oktavianthi, Lidwina Priliani, Edison Johar, Safarina G Malik, Khin Saw Myint, Amin Soebandrio |
| EPI_ISL_2364632 | RS Mitra Keluarga Bekasi Barat | Eijkman Institute for Molecular Biology, National Agency for Research and Innovation | Edison Johar, Frilasita A Yudhaputri, Muhammad Rezki Rasyak, Willy Agustine, Hidayat Trimarsanto, Lydia V. Panggalo, Iskandar Adnan, Sukma Oktavianthi, Lidwina Priliani, Safarina G Malik, Khin Saw Myint, Amin Soebandrio |
| EPI_ISL_2364635 | RS Harapan Bunda | Eijkman Institute for Molecular Biology, National Agency for Research and Innovation | Iskandar Adnan, Sukma Oktavianthi, Lidwina Priliani, Edison Johar, Frilasita A Yudhaputri, Muhammad Rezki Rasyak, Willy Agustine, Hidayat Trimarsanto, Lydia V. Panggalo, Safarina G Malik, Khin Saw Myint, Amin Soebandrio |
| EPI_ISL_2364638 | RS Mary Cileungsi Hijau | Eijkman Institute for Molecular Biology, National Agency for Research and Innovation | Willy Agustine, Hidayat Trimarsanto, Lydia V. Panggalo, Iskandar Adnan, Sukma Oktavianthi, Lidwina Priliani, Edison Johar, Frilasita A Yudhaputri, Muhammad Rezki Rasyak, Safarina G Malik, Khin Saw Myint, Amin Soebandrio |
| EPI_ISL_2364641 | RS Mary Cileungsi Hijau | Eijkman Institute for Molecular Biology, National Agency for Research and Innovation | Frilasita A Yudhaputri, Muhammad Rezki Rasyak, Willy Agustine, Hidayat Trimarsanto, Lydia V. Panggalo, Iskandar Adnan, Sukma Oktavianthi, Lidwina Priliani, Edison Johar, Safarina G Malik, Khin Saw Myint, Amin Soebandrio |
| EPI_ISL_2364644 | RSUD Cileungsi | Eijkman Institute for Molecular Biology, National Agency for Research and Innovation | Sukma Oktavianthi, Lidwina Priliani, Edison Johar, Frilasita A Yudhaputri, Muhammad Rezki Rasyak, Willy Agustine, Hidayat Trimarsanto, Lydia V. Panggalo, Iskandar Adnan, Safarina G Malik, Khin Saw Myint, Amin Soebandrio |
| EPI_ISL_2364646 | RS Permata Cibubur | Eijkman Institute for Molecular Biology, National Agency for Research and Innovation | Lidwina Priliani, Edison Johar, Frilasita A Yudhaputri, Muhammad Rezki Rasyak, Willy Agustine, Hidayat Trimarsanto, Lydia V. Panggalo, Iskandar Adnan, Sukma Oktavianthi, Safarina G Malik, Khin Saw Myint, Amin Soebandrio |
| EPI_ISL_2364650 | RS Permata Cibubur | Eijkman Institute for Molecular Biology, National Agency for Research and Innovation | Edison Johar, Frilasita A Yudhaputri, Muhammad Rezki Rasyak, Willy Agustine, Hidayat Trimarsanto, Lydia V. Panggalo, Iskandar Adnan, Sukma Oktavianthi, Lidwina Priliani, Safarina G Malik, Khin Saw Myint, Amin Soebandrio |
| EPI_ISL_2364653 | RSU HGA Depok | Eijkman Institute for Molecular Biology, National Agency for Research and Innovation | Willy Agustine, Hidayat Trimarsanto, Lydia V. Panggalo, Iskandar Adnan, Sukma Oktavianthi, Lidwina Priliani, Edison Johar, Frilasita A Yudhaputri, Muhammad Rezki Rasyak, Safarina G Malik, Khin Saw Myint, Amin Soebandrio |
| EPI_ISL_2364656 | RSIA Assyifa | Eijkman Institute for Molecular Biology, National Agency for Research and Innovation | Hidayat Trimarsanto, Lydia V. Panggalo, Iskandar Adnan, Sukma Oktavianthi, Lidwina Priliani, Edison Johar, Frilasita A Yudhaputri, Muhammad Rezki Rasyak, Willy Agustine, Safarina G Malik, Khin Saw Myint, Amin Soebandrio |
| EPI_ISL_2364659 | RS Medirossa Cikarang | Eijkman Institute for Molecular Biology, National Agency for Research and Innovation | Hidayat Trimarsanto, Lydia V. Panggalo, Iskandar Adnan, Sukma Oktavianthi, Lidwina Priliani, Edison Johar, Frilasita A Yudhaputri, Muhammad Rezki Rasyak, Willy Agustine, Safarina G Malik, Khin Saw Myint, Amin Soebandrio |
| EPI_ISL_2364663 | RS Medirossa Cikarang | Eijkman Institute for Molecular Biology, National Agency for Research and Innovation | Frilasita A Yudhaputri, Muhammad Rezki Rasyak, Willy Agustine, Hidayat Trimarsanto, Lydia V. Panggalo, Iskandar Adnan, Sukma Oktavianthi, Lidwina Priliani, Edison Johar, Safarina G Malik, Khin Saw Myint, Amin Soebandrio |
| EPI_ISL_2364666 | RSIA PKU Muhammadiyah | Eijkman Institute for Molecular Biology, National Agency for Research and Innovation | Willy Agustine, Hidayat Trimarsanto, Lydia V. Panggalo, Iskandar Adnan, Sukma Oktavianthi, Lidwina Priliani, Edison Johar, Frilasita A Yudhaputri, Muhammad Rezki Rasyak, Safarina G Malik, Khin Saw Myint, Amin Soebandrio |
| EPI_ISL_2364669 | RSIA PKU Muhammadiyah | Eijkman Institute for Molecular Biology, National Agency for Research and Innovation | Muhammad Rezki Rasyak, Willy Agustine, Hidayat Trimarsanto, Lydia V. Panggalo, Iskandar Adnan, Sukma Oktavianthi, Lidwina Priliani, Edison Johar, Frilasita A Yudhaputri, Safarina G Malik, Khin Saw Myint, Amin Soebandrio |
| EPI_ISL_2365884, EPI_ISL_2365896 | Microbiology Department. Complexo Hospitalario Universitario de Vigo | Microbiology Department. Complexo Hospitalario Universitario de Vigo | Alfaya N, Alonso I, Alvarez M, Cabrera JJ, Carballo R, Cores O, Cortizo S, del-Campo V, Martinez L, Mediero G, Perez S, Potel C, Regueiro B, Rey S, Vasallo FJ |
| EPI_ISL_2374159 | Research | National Reference Center for Viruses of Respiratory Infections, Institut Pasteur, Paris | Marion Barbet, Sylvie Behillil, Méline Bizard, Angela Brisebarre, Camille Capel, Vincent Enouf, Louise Lefrançois, Frédéric Lemoine, Christophe Malabat, Corinne Maufrais, Etienne Simon-Lorière, Maud Vanpeene, Sylvie Van der Werf ,Marianne Leruez-Ville |
| EPI_ISL_2374160 | Research | National Reference Center for Viruses of Respiratory Infections, Institut Pasteur, Paris | Marion Barbet, Sylvie Behillil, Méline Bizard, Angela Brisebarre, Camille Capel, Vincent Enouf, Louise Lefrançois, Frédéric Lemoine, Christophe Malabat, Corinne Maufrais, Emmanuelle Permal, Etienne Simon-Lorière, Maud Vanpeene, Sylvie Van der Werf ,Marianne Leruez-Ville |
| EPI_ISL_2374161 | Research | National Reference Center for Viruses of Respiratory Infections, Institut Pasteur, Paris | Marion Barbet, Sylvie Behillil, Méline Bizard, Angela Brisebarre, Camille Capel, Vincent Enouf, Louise Lefrançois, Frédéric Lemoine, Christophe Malabat, Corinne Maufrais, Etienne Simon-Lorière, Maud Vanpeene, Sylvie Van der Werf ,Marianne Leruez-Ville |
| EPI_ISL_2375848, EPI_ISL_2375849 | Programa de Oncovirologia, Instituto Nacional de Câncer | Programa de Oncovirologia, Instituto Nacional de Câncer | Livia R. Goes, Juliana D. Siqueira, Marianne M. Garrido, Brunna M. Alves, Ana Cristina P. M. Pereira, Claudia Cicala, James Arthos, João P.B. Viola, Marcelo A. Soares |
| EPI_ISL_2383317 | The teaching Baghdad hospital | PCR lab | Abdulhussein Thaer, Fadhil Hula |
| EPI_ISL_2384369 | Cayman Islands Forensic Science Lab | Cayman Islands Forensic Science Lab | Jonathan Smellie, Karla Montes, Brittany Balcewich, Allen Grolla, Sarah Medina |
| EPI_ISL_2398296, EPI_ISL_2399459, EPI_ISL_2399460, EPI_ISL_2399461, EPI_ISL_2399462 | Centre for Human Virology and Genomics, Microbiology Department, Nigerian Institute of Medical Research | Central Research Laboratory, Nigerian Institute of Medical Research | Chika Kingsley Onwuamah, Ayorinde Babatunde James, Joseph Ojonugwa Shaibu, Rahaman A. Ahmed, Azuka Patrick Okwuraiwe, Sola Ajibaye, Sharon Abimbola, Olufemi Samuel Amoo, Muinah Adenike Fowora, Phasha-Muchemenye Mmatshepho, Joy Ayoola, Nyam Itse Yusuf, Grace Oni, Yusuf Jimoh, Josiah Ayoola Isong, Uyi Emokpae, Ngozi Mirabel Otuonye, Rosemary Ajuma Audu, Bamidele Iwalokun, Babatunde Lawal Salako |
| EPI_ISL_2402534 | OLVZ Aalst | OLVZ Aalst | Astrid Holderbeke |
| EPI_ISL_2422576, EPI_ISL_2422580, EPI_ISL_2422584, EPI_ISL_2422587, EPI_ISL_2422590, EPI_ISL_2422592, EPI_ISL_2422596, EPI_ISL_2422599, EPI_ISL_2422606, EPI_ISL_2422613, EPI_ISL_2422616, EPI_ISL_2422620, EPI_ISL_2422623, EPI_ISL_2422626, EPI_ISL_2422629, EPI_ISL_2422632, EPI_ISL_2422633, EPI_ISL_2422634, EPI_ISL_2422635, EPI_ISL_2422638, EPI_ISL_2422640, EPI_ISL_2422641, EPI_ISL_2422642, EPI_ISL_2422643, EPI_ISL_2422644, EPI_ISL_2422645, EPI_ISL_2422647, EPI_ISL_2422648, EPI_ISL_2422649, EPI_ISL_2422651, EPI_ISL_2422656, EPI_ISL_2422657, EPI_ISL_2422661, EPI_ISL_2422662, EPI_ISL_2422664, EPI_ISL_2422665, EPI_ISL_2422667 | | | |
| see above | San Diego County Public Health Laboratory | Andersen lab at Scripps Research | SEARCH Alliance San Diego with Ashleigh Murphy, Jovan Shephard, Brett Austin |
| EPI_ISL_2422669, EPI_ISL_2422670, EPI_ISL_2422671, EPI_ISL_2422672, EPI_ISL_2422674, EPI_ISL_2422675, EPI_ISL_2422676, EPI_ISL_2422677, EPI_ISL_2422678, EPI_ISL_2422679, EPI_ISL_2422680, EPI_ISL_2422681, EPI_ISL_2422682, EPI_ISL_2422683, EPI_ISL_2422684, EPI_ISL_2422685, EPI_ISL_2422686, EPI_ISL_2422687, EPI_ISL_2422688, EPI_ISL_2422689, EPI_ISL_2422690, EPI_ISL_2422691, EPI_ISL_2422692, EPI_ISL_2422693, EPI_ISL_2422694, EPI_ISL_2422695, EPI_ISL_2422696, EPI_ISL_2422697, EPI_ISL_2422698, EPI_ISL_2422699, EPI_ISL_2422700, EPI_ISL_2422701, EPI_ISL_2422702, EPI_ISL_2422703, EPI_ISL_2422704, EPI_ISL_2422705, EPI_ISL_2422706, EPI_ISL_2422707, EPI_ISL_2422708, EPI_ISL_2422709, EPI_ISL_2422710, EPI_ISL_2422711, EPI_ISL_2422712, EPI_ISL_2422713, EPI_ISL_2422714, EPI_ISL_2422715, EPI_ISL_2422716, EPI_ISL_2422717, EPI_ISL_2422718, EPI_ISL_2422719 | | | |
| see above | Sharp HealthCare Laboratory | Andersen lab at Scripps Research | SEARCH Alliance San Diego with Aaron Harding, Jacquelyn Berumen, Cathy Woerle, Liam McGinnis, Art Mendoza, Omid Bakhtar |
| EPI_ISL_2422720, EPI_ISL_2422722, EPI_ISL_2422723, EPI_ISL_2422724, EPI_ISL_2422726, EPI_ISL_2422727, EPI_ISL_2422728, EPI_ISL_2422729, EPI_ISL_2422730, EPI_ISL_2422731, EPI_ISL_2422733, EPI_ISL_2422734, EPI_ISL_2422737, EPI_ISL_2422738, EPI_ISL_2422739, EPI_ISL_2422740, EPI_ISL_2422742, EPI_ISL_2422743, EPI_ISL_2422744, EPI_ISL_2422745, EPI_ISL_2422747, EPI_ISL_2422748, EPI_ISL_2422749, EPI_ISL_2422750, EPI_ISL_2422751, EPI_ISL_2422752, EPI_ISL_2422753, EPI_ISL_2422754, EPI_ISL_2422755, EPI_ISL_2422757, EPI_ISL_2422758, EPI_ISL_2422759, EPI_ISL_2422760, EPI_ISL_2422763, | | | |

| | | | |
|---|---|---|---|
| EPI_ISL_2422764, EPI_ISL_2422765, EPI_ISL_2422766, EPI_ISL_2422767, EPI_ISL_2422768, EPI_ISL_2422769, EPI_ISL_2422770, EPI_ISL_2422771, EPI_ISL_2422772, EPI_ISL_2422774, EPI_ISL_2422776, EPI_ISL_2422778, EPI_ISL_2422779, EPI_ISL_2422780, EPI_ISL_2422781, EPI_ISL_2422782 | | | |
| see above | San Diego County Public Health Laboratory | Andersen lab at Scripps Research | SEARCH Alliance San Diego with Ashleigh Murphy, Jovan Shephard, Brett Austin |
| EPI_ISL_2422783, EPI_ISL_2422784, EPI_ISL_2422785, EPI_ISL_2422786, EPI_ISL_2422787, EPI_ISL_2422788, EPI_ISL_2422789, EPI_ISL_2422790, EPI_ISL_2422791, EPI_ISL_2422792, EPI_ISL_2422793, EPI_ISL_2422794, EPI_ISL_2422795, EPI_ISL_2422796, EPI_ISL_2422797, EPI_ISL_2422798, EPI_ISL_2422799, EPI_ISL_2422800, EPI_ISL_2422801, EPI_ISL_2422802, EPI_ISL_2422803, EPI_ISL_2422804, EPI_ISL_2422805, EPI_ISL_2422806, EPI_ISL_2422807, EPI_ISL_2422808, EPI_ISL_2422809, EPI_ISL_2422810, EPI_ISL_2422811, EPI_ISL_2422812, EPI_ISL_2422813, EPI_ISL_2422814, EPI_ISL_2422828, EPI_ISL_2422829, EPI_ISL_2422830, EPI_ISL_2422834, EPI_ISL_2422835, EPI_ISL_2422836, EPI_ISL_2422837, EPI_ISL_2422838, EPI_ISL_2422839, EPI_ISL_2422840, EPI_ISL_2422841, EPI_ISL_2422842, EPI_ISL_2422843, EPI_ISL_2422844, EPI_ISL_2422845, EPI_ISL_2422846, EPI_ISL_2422847, EPI_ISL_2422848, EPI_ISL_2422849, EPI_ISL_2422850, EPI_ISL_2422851, EPI_ISL_2422852, EPI_ISL_2422853, EPI_ISL_2422854, EPI_ISL_2422855, EPI_ISL_2422856, EPI_ISL_2422857, EPI_ISL_2422858, EPI_ISL_2422859, EPI_ISL_2422860, EPI_ISL_2422861, EPI_ISL_2422862, EPI_ISL_2422863, EPI_ISL_2422864, EPI_ISL_2422865, EPI_ISL_2422866, EPI_ISL_2422867, EPI_ISL_2422868, EPI_ISL_2422869, EPI_ISL_2422870, EPI_ISL_2422871, EPI_ISL_2422872, EPI_ISL_2422873, EPI_ISL_2422874, EPI_ISL_2422875, EPI_ISL_2422876, EPI_ISL_2422877, EPI_ISL_2422878, EPI_ISL_2422879, EPI_ISL_2422880, EPI_ISL_2422881, EPI_ISL_2422882, EPI_ISL_2422883, EPI_ISL_2422884, EPI_ISL_2422885, EPI_ISL_2422886, EPI_ISL_2422887, EPI_ISL_2422888, EPI_ISL_2422889, EPI_ISL_2422890, EPI_ISL_2422891, EPI_ISL_2422892, EPI_ISL_2422893, EPI_ISL_2422894, EPI_ISL_2422895, EPI_ISL_2422896 | | | |
| see above | Sharp HealthCare Laboratory | Andersen lab at Scripps Research | SEARCH Alliance San Diego with Aaron Harding, Jacquelyn Berumen, Cathy Woerle, Liam McGinnis, Art Mendoza, Omid Bakhtar |
| EPI_ISL_2422897, EPI_ISL_2422898, EPI_ISL_2422899, EPI_ISL_2422900, EPI_ISL_2422901, EPI_ISL_2422902, EPI_ISL_2422903, EPI_ISL_2422905, EPI_ISL_2422906, EPI_ISL_2422907, EPI_ISL_2422908, EPI_ISL_2422909, EPI_ISL_2422910, EPI_ISL_2422911, EPI_ISL_2422912, EPI_ISL_2422913, EPI_ISL_2422914, EPI_ISL_2422915, EPI_ISL_2422916, EPI_ISL_2422917, EPI_ISL_2422918, EPI_ISL_2422919, EPI_ISL_2422920, EPI_ISL_2422921, EPI_ISL_2422922, EPI_ISL_2422923, EPI_ISL_2422924, EPI_ISL_2422925, EPI_ISL_2422926, EPI_ISL_2422927, EPI_ISL_2422928, EPI_ISL_2422929, EPI_ISL_2422930, EPI_ISL_2422931, EPI_ISL_2422932, EPI_ISL_2422933, EPI_ISL_2422934, EPI_ISL_2422935, EPI_ISL_2422936, EPI_ISL_2422937, EPI_ISL_2422938, EPI_ISL_2422939, EPI_ISL_2422940, EPI_ISL_2422941, EPI_ISL_2422942, EPI_ISL_2422943, EPI_ISL_2422944, EPI_ISL_2422945, EPI_ISL_2422946, EPI_ISL_2422947, EPI_ISL_2422948, EPI_ISL_2422949, EPI_ISL_2422950, EPI_ISL_2422951, EPI_ISL_2422952, EPI_ISL_2422953, EPI_ISL_2422954, EPI_ISL_2422955, EPI_ISL_2422956, EPI_ISL_2422957, EPI_ISL_2422958, EPI_ISL_2422959, EPI_ISL_2422960, EPI_ISL_2422961, EPI_ISL_2422962, EPI_ISL_2422963, EPI_ISL_2422964, EPI_ISL_2422965, EPI_ISL_2422966, EPI_ISL_2422967, EPI_ISL_2422970, EPI_ISL_2422971, EPI_ISL_2422972, EPI_ISL_2422973, EPI_ISL_2422974, EPI_ISL_2422975 | | | |
| see above | San Diego County Public Health Laboratory | Andersen lab at Scripps Research | SEARCH Alliance San Diego with Ashleigh Murphy, Jovan Shephard, Brett Austin |
| EPI_ISL_2426898, EPI_ISL_2426899, EPI_ISL_2426900, EPI_ISL_2426901, EPI_ISL_2426902, EPI_ISL_2426903, EPI_ISL_2426904, EPI_ISL_2426905, EPI_ISL_2426906, EPI_ISL_2426907, EPI_ISL_2426908, EPI_ISL_2426909, EPI_ISL_2426910, EPI_ISL_2426911, EPI_ISL_2426912, EPI_ISL_2426913, EPI_ISL_2426914, EPI_ISL_2426915, EPI_ISL_2426916, EPI_ISL_2426917, EPI_ISL_2426918, EPI_ISL_2426919, EPI_ISL_2426920, EPI_ISL_2426921, EPI_ISL_2426922, EPI_ISL_2426923, EPI_ISL_2426924, EPI_ISL_2426925, EPI_ISL_2426926, EPI_ISL_2426927, EPI_ISL_2426928, EPI_ISL_2426929, EPI_ISL_2426930, EPI_ISL_2426931, EPI_ISL_2426932, EPI_ISL_2426933, EPI_ISL_2426934, EPI_ISL_2426935, EPI_ISL_2426936, EPI_ISL_2426937, EPI_ISL_2426938, EPI_ISL_2426939, EPI_ISL_2426940, EPI_ISL_2426942, EPI_ISL_2426943 | | | |
| see above | Centers for Disease Control and Prevention, Dengue Branch | Centers for Disease Control and Prevention, Dengue Branch | Gilberto A. Santiago, Glenda Gonzalez, Betzabel Flores, Keyla Charriez, Gabriela Paz-Bailey, Jorge L. Munoz-Jordan |
| EPI_ISL_2438892 | Motol University Hospital | Charles University, Faculty of Science, BIOCEV, OMICS Genomics | Vladimír Hampl, Štpánka Hrdá, Ruth Tachezy, Blanka Hamplová, Sebastian Cristian Treitli, Zoltán Füssy, Magdalena Janáová, Ingrid Poláková, Jana Šmahelová, Jíí Novák |
| EPI_ISL_2443546 | Lboratorio Central de Saude Publica do Estado do Parana (LACEN/PR) | Laboratory of Respiratory Viruses and Measles, Oswaldo Cruz Institute, FIOCRUZ | Paola Resende, Luciana Appolinario, Fernando Motta, Anna Carolina Paixao, Ana Carolina Mendonca, Alice Sampaio Rocha, Taina Venas, Elisa Cavalcante Pereira, Renata Serrano Lopes, Irina Riediger, Marilda Siqueira on behalf of the Fiocruz COVID-19 Genomic Surveillance Network |
| EPI_ISL_2448646, EPI_ISL_2448647, EPI_ISL_2448648, EPI_ISL_2448649, EPI_ISL_2448650, EPI_ISL_2448651, EPI_ISL_2448652, EPI_ISL_2448654, EPI_ISL_2448679, EPI_ISL_2448680, EPI_ISL_2448681 | | | |
| see above | Molecular diagnostic laboratory of Federal Budget Institution of Science "Central Research Institute of Epidemiology" of The Federal Service on Customers' Rights Protection and Human Well-being Surveillance | Group of Genomics and Postgenomic Technologies of Central Research Institute of Epidemiology | Samoilov A.E., Kaptelova V.V., Korneenko E.V., Valdohina A.V., Bulanenko V.P., Saenko S.S., Sinitsyn S.O., Golubeva A.G., Speranskaya A.S., Tivanova E.V., Shipulina O.Y., Akimkin V.G. |
| EPI_ISL_2455237 | CICESE-BSL3 | CICESE-BSL3 | Cervantes Karla, Gonzalez-Sanchez Ricardo, Martinez Miguel, Galindo Clara and Licea Alexei |
| EPI_ISL_2455244 | CICESE-BSL3 | CICESE-BSL3 | Karla Cervantes, Ricardo Gonzalez, Miguel Martinez, Clara Galindo and Alexei Licea |
| EPI_ISL_2458066 | Retrovirus Laboratory Adolfo Lutz Institute_Sao Paulo | Retrovirus Laboratory Adolfo Lutz Institute | Gabriela Bastos Cabral, Giselle I S Lopez-Lopes, Cintia Ahagon, Audrey Cilli, Paula Morena Guimaraes, Igor Mohamed Hussein, Luís Brigido |
| EPI_ISL_2462895 | Hospital | National Reference Center for Viruses of Respiratory Infections, Institut Pasteur, Paris | Marion Barbet, Sylvie Behillil, Méline Bizard, Angela Brisebarre, Camille Capel, Vincent Enouf, Louise Lefrançois, Frédéric Lemoine, Christophe Malabat, Corinne Maufrais, Etienne Simon-Lorière, Maud Vanpeene, Sylvie Van der Werf ,Olivia Raulin |
| EPI_ISL_2467872, EPI_ISL_2467873, EPI_ISL_2467874, EPI_ISL_2467875, EPI_ISL_2467876, EPI_ISL_2467877, EPI_ISL_2467878, EPI_ISL_2467879 | Rockford Ambulatory Surgery Center | BioInfoExperts LLC | Sarah E. Hagarty, David J Nolan, Rebecca Rose, Susanna L Lamers, Ed Yavitz |
| EPI_ISL_2484218, EPI_ISL_2484219 | University of Brescia - Dept. of Molecular and Translational Medicine | Istituto Zooprofilattico Sperimentale della Lombardia e dell'Emilia Romagna (IZSLER), Risk Analysis and Genomic Epidemiology Unit | Caruso Arnaldo, Fiorentini Simona, Caccuri Francesca, Messali Serena, Zani Alberto, Marina Morganti, Erika Scaltriti, Stefano Pongolini |
| EPI_ISL_2485171, EPI_ISL_2485172, EPI_ISL_2485173, EPI_ISL_2485174, EPI_ISL_2485176, EPI_ISL_2485180, EPI_ISL_2485181, EPI_ISL_2485188, EPI_ISL_2485189, EPI_ISL_2485190, EPI_ISL_2485191, EPI_ISL_2485194, EPI_ISL_2485195, EPI_ISL_2485197, EPI_ISL_2485202, EPI_ISL_2485203, EPI_ISL_2485204, EPI_ISL_2485207, EPI_ISL_2485208, EPI_ISL_2485210, EPI_ISL_2485214, EPI_ISL_2485215, EPI_ISL_2485220, EPI_ISL_2485222, EPI_ISL_2485223, EPI_ISL_2485224, EPI_ISL_2485225 | | | |
| see above | ILV Kärnten | Bergthaler laboratory, CeMM Research Center for Molecular Medicine of the Austrian Academy of Sciences | Lukas Endler, Anna Schedl, Fabian Amman, Petr Triska, Thomas Penz, Benedikt Agerer, Maelle Le Moing, Michael Schuster, Bekir Erguner, Jan Laine, Martin Senekowitsch, Christoph Bock, Andreas Bergthaler |
| EPI_ISL_2491665, EPI_ISL_2491666, EPI_ISL_2491668 | Centre Hospitalier Universitaire Clermont-Ferrand | CHU Clermont-Ferrand, service de virologie | Bisseux Maxime, Mirand Audrey, Combes Patricia, Henquell Cécile |
| EPI_ISL_2491669 | Centre De Prelevement COVID RIOM | CHU Clermont-Ferrand, service de virologie | Bisseux Maxime, Mirand Audrey, Combes Patricia, Henquell Cécile |
| EPI_ISL_2491670 | Centre Hospitalier Universitaire Clermont-Ferrand | CHU Clermont-Ferrand, service de virologie | Bisseux Maxime, Mirand Audrey, Combes Patricia, Henquell Cécile |
| EPI_ISL_2495717, EPI_ISL_2495718, EPI_ISL_2495719, EPI_ISL_2495720 | MB-Cadham Provincial laboratory | National Microbiology Laboratory (NML) | Anna Majer, Shari Tyson, Grace Seo, Philip Mabon, Elsie Grudeski, Rhiannon Huzarewich, Russell Mandes, Anneliese Landgraff, Jennifer Tanner, Natalie Knox, Morag Graham, Gary Van Domselaar, Paul Van Caeseele, Jared Bullard, David Alexander, Kerry Dust, Nathalie Bastien, Yan Li, Timothy Booth, Darian Hole, Madison Chapel, Kirsten Biggar, CanCOGeN's metadata curation team, Public Health Agency of Canada CanCOGeN team |
| EPI_ISL_2501088, EPI_ISL_2501089 | Centre for Human Virology and Genomics, Microbiology Department, Nigerian Institute of Medical Research | Central Research Laboratory, Nigerian Institute of Medical Research | Chika Kingsley Onwuamah, Ayorinde Babatunde James, Joseph Ojonugwa Shaibu, Rahaman A. Ahmed, Azuka Patrick Okwuraiwe, Sola Ajibaye, Sharon Abimbola, Olufemi Samuel Amoo, Muinah Adenike Fowora, Phasha-Muchemenye Mmatshepho, Joy Ayoola, Nyam Itse Yusuf, Grace Oni, Yusuf Jimoh, Josiah Ayoola Isong, Uyi Emokpae, Ngozi Mirabel Otuonye, Rosemary Ajuma Audu, Bamidele Iwalokun, Babatunde Lawal Salako |
| EPI_ISL_2501312, EPI_ISL_2501348, EPI_ISL_2501349, EPI_ISL_2501350, EPI_ISL_2501351, EPI_ISL_2501352, EPI_ISL_2501353, EPI_ISL_2501354, EPI_ISL_2501355, EPI_ISL_2501356, EPI_ISL_2501357, EPI_ISL_2501358, EPI_ISL_2501359, EPI_ISL_2501360, EPI_ISL_2501361, EPI_ISL_2501362, EPI_ISL_2501379, EPI_ISL_2501380, EPI_ISL_2501381, EPI_ISL_2501382, EPI_ISL_2501383, EPI_ISL_2501384, EPI_ISL_2501385, EPI_ISL_2501386, EPI_ISL_2501387, EPI_ISL_2501388, EPI_ISL_2501389, EPI_ISL_2501390, EPI_ISL_2501391, EPI_ISL_2501392, EPI_ISL_2501393 | | | |
| see above | Microbiology Department. Complexo Hospitalario Universitario de Vigo | Microbiology Department. Complexo Hospitalario Universitario de Vigo | Alfaya N, Alonso I, Alvarez M, Cabrera JJ, Carballo R, Cores O, Cortizo S, del-Campo V, Martinez L, Mediero G, Perez S, Potel C, Regueiro B, Rey S, Vasallo FJ |
| EPI_ISL_2502019, EPI_ISL_2502026 | Centre for Human Virology and Genomics, Microbiology Department, Nigerian Institute of Medical Research | Central Research Laboratory, Nigerian Institute of Medical Research | Chika Kingsley Onwuamah, Ayorinde Babatunde James, Joseph Ojonugwa Shaibu, Rahaman A. Ahmed, Azuka Patrick Okwuraiwe, Sola Ajibaye, Sharon Abimbola, Olufemi Samuel Amoo, Muinah Adenike Fowora, Phasha-Muchemenye Mmatshepho, Joy Ayoola, Nyam Itse Yusuf, Grace Oni, Yusuf Jimoh, Josiah Ayoola Isong, Uyi Emokpae, Ngozi Mirabel Otuonye, Rosemary Ajuma Audu, Bamidele Iwalokun, Babatunde Lawal Salako |
| EPI_ISL_2502639, EPI_ISL_2502641, EPI_ISL_2502642, EPI_ISL_2502645, EPI_ISL_2502648, EPI_ISL_2502650, EPI_ISL_2502651, EPI_ISL_2502652, EPI_ISL_2502653, EPI_ISL_2502654, EPI_ISL_2502655 | | | |
| see above | All India Institute of Medical Sciences Delhi Hospital | Virology Laboratory, AIIMS Delhi | Aashish Choudhary, Sumedha Bagga, Lata Rani, Dibyabhaba Pradhan, Pooja Pandey, Jyoti Jethani, Shivram Dhakad, Nazneen Arif, Deepankar Srigyan, Megha Brijwal, Lalit Dar, Manish Soneja, Rakesh Lodha, Puneet Kaur, Ritu Gupta, Subrata Sinha, Chitra Sarkar, Randeep Guleria |
| EPI_ISL_2532369 | Labo Analyses Med | National Reference Center for Viruses of Respiratory Infections, Institut Pasteur, Paris | Marion Barbet, Sylvie Behillil, Méline Bizard, Angela Brisebarre, Camille Capel, Vincent Enouf, Louise Lefrançois, Frédéric Lemoine, Christophe Malabat, Corinne Maufrais, Etienne Simon-Lorière, Maud Vanpeene, Sylvie Van der Werf ,Hervé Dupont |
| EPI_ISL_2532467 | Labo Analyses Med | National Reference Center for Viruses of Respiratory Infections, Institut Pasteur, Paris | Marion Barbet, Sylvie Behillil, Méline Bizard, Angela Brisebarre, Camille Capel, Vincent Enouf, Louise Lefrançois, Frédéric Lemoine, Christophe Malabat, Corinne Maufrais, Adrien Pain, Etienne Simon-Lorière, Maud Vanpeene, Sylvie Van der Werf |
| EPI_ISL_2532470 | Labo Analyses Med | National Reference Center for Viruses of Respiratory Infections, Institut Pasteur, Paris | Marion Barbet, Sylvie Behillil, Méline Bizard, Angela Brisebarre, Camille Capel, Vincent Enouf, Louise Lefrançois, Frédéric Lemoine, Christophe Malabat, Corinne Maufrais, Etienne Simon-Lorière, Maud Vanpeene, Sylvie Van der Werf |
| EPI_ISL_2532676 | IZSM | IZSM-U.O.C. Virologia | Maurizio Viscardi, Lorena Cardillo, Claudio de Martinis, Giovanna Fusco, Esterina De Carlo, Antonio Limone |

| | | | |
|---|---|---|---|
| EPI_ISL_2535801 | National Influenza Centre | National Influenza Centre | William K. Ampofo, Michael Marks, Ivy A. Asante, Sharon Hsu, Benjamiin B. Lindsey, Benjamin H. Foulkes, Mildred Adusei-Poku, Linda Boatemaa, Lorreta Kwasah, Joseph Oliver-Commey, Ernest Asiedu, Franklin Asiedu-Bekoe, Gordon Awandare, Joyce Ngoi, Dennis Laryea, Mathew D. Parker, Thushan I de Silva, |
| EPI_ISL_2544854, EPI_ISL_2544855, EPI_ISL_2544879, EPI_ISL_2544880, EPI_ISL_2544883, EPI_ISL_2544884, EPI_ISL_2544885, EPI_ISL_2544886, EPI_ISL_2544888, EPI_ISL_2544890, EPI_ISL_2544892, EPI_ISL_2544893, EPI_ISL_2544894, EPI_ISL_2544895, EPI_ISL_2544898 | | | |
| see above | Laboratorio de Pesquisa em Virologia, FAMERP, SJRP | Laboratorio de Pesquisa em Virologia, FAMERP, SJRP | Cecília Artico Banho; Lívia Sacchetto; Guilherme Campos; Fábio Sossai Possebon; Leila Sabrina Ullmann; Cíntia Bittar; Helena Lage Ferreira; Jorge A. Petroli Marchesi; Maisa C. Pereira Parra; Marília Moraes; Paula Rahal; Paulo Inacio da Costa; João Pessoa Araújo Jr.; Maurício L. Nogueira. |
| EPI_ISL_2566185, EPI_ISL_2566187, EPI_ISL_2566190, EPI_ISL_2603239, EPI_ISL_2603240, EPI_ISL_2603241, EPI_ISL_2603245, EPI_ISL_2603246 | Microbiology Department. Complexo Hospitalario Universitario de Vigo | Microbiology Department. Complexo Hospitalario Universitario de Vigo | Alfaya N, Alonso I, Alvarez M, Carballo R, Cabrera JJ, Cores O, Cortizo S, del-Campo V, Martinez L, Mediero G, Perez S, Potel C, Regueiro B, Rey S, Vasallo FJ |
| EPI_ISL_2612601, EPI_ISL_2612612 | Labo Analyses Med | National Reference Center for Viruses of Respiratory Infections, Institut Pasteur, Paris | Marion Barbet, Sylvie Behillil, Méline Bizard, Angela Brisebarre, Camille Capel, Vincent Enouf, Louise Lefrançois, Frédéric Lemoine, Christophe Malabat, Corinne Maufrais, Etienne Simon-Lorière, Maud Vanpeene, Sylvie Van der Werf ,Pierre-Yves Leonard |
| EPI_ISL_2614351 | Labortorio Central de Saude Publica do Estado do Rio de Janeiro (LACEN/RJ) | Laboratory of Respiratory Viruses and Measles, Oswaldo Cruz Institute, FIOCRUZ | Paola Resende, Luciana Appolinario, Fernando Motta, Anna Carolina Paixao, Ana Carolina Mendonca, Alice Sampaio Rocha, Taina Venas, Elisa Cavalcante Pereira, Renata Serrano Lopes, Andrea Cony Cavalcanti, Marilda Siqueira on behalf of the Fiocruz COVID-19 Genomic Surveillance Network |
| EPI_ISL_2614513, EPI_ISL_2614514, EPI_ISL_2614515, EPI_ISL_2614516, EPI_ISL_2614517 | Instituto Adolfo Lutz Central | Instituto Adolfo Lutz, Interdiciplinary Procedures Center, Strategic Laboratory | Claudio Tavares Sacchi, Claudia Regina Gonçalves, Erica Valessa Ramos Gomes, Karoline Rodrigues Campos, Caio Vinicius Dias Lopes, Leonardo Jose Tadeu de Araujo |
| EPI_ISL_2614540 | IdenGene Medicina Diagnostica SA | Instituto Adolfo Lutz, Interdiciplinary Procedures Center, Strategic Laboratory | Claudio Tavares Sacchi, Claudia Regina Gonçalves, Erica Valessa Ramos Gomes, Karoline Rodrigues Campos, Caio Vinicius Dias Lopes, Leonardo Jose Tadeu de Araujo |
| EPI_ISL_2614541 | LACEN do Estado de Goias | Instituto Adolfo Lutz, Interdiciplinary Procedures Center, Strategic Laboratory | Claudio Tavares Sacchi, Claudia Regina Gonçalves, Erica Valessa Ramos Gomes, Karoline Rodrigues Campos, Caio Vinicius Dias Lopes, Leonardo Jose Tadeu de Araujo |
| EPI_ISL_2617086 | Africa_CDC - Malawi P4 S2 | CERI, Centre for Epidemic Response and Innvoation, Stellenbosch University and KRISP, KZN Research Innovation and Sequencing Platform, UKZN. | Mvula B, Chilima B, Chiwaula M, Mwangomba W, Panja L, Kasambara W, Auld A, Kim L, Kampira E, Kaba M, Wadonda N, Maida A, Giandhari J, Pillay S, Naidoo Y, Lessells R, Emmanuel SJ, Tegally H, Wilkinson E, de Oliveira T |
| EPI_ISL_2617876, EPI_ISL_2617877, EPI_ISL_2617878, EPI_ISL_2617879 | LABCOVID_HCPA | LABRESIS_HCPA | Wink PL, Martins AF, Volpato F, Monteiro F, Zavascki AP, Barth AL |
| EPI_ISL_2627738 | Belo Horizonte center-south emergency care unit - UPA-BH | Laboratório de Virologia Clínica e Molecular | Erick Gustavo Dorlass, Karine Lima Lourenço, Rubens Daniel Miserani Magalhães, Hugo Sato, Alex Fiorini, Renata Peixoto, Helena Perez Coelho, Ana Paula Salles Fernandes, Bruna Larotonda Telezynski, Guilherme Pereira Scagion, Tatiana Ometto, Luciano Matsumiya Thomazelli, Danielle Bruna Leal Oliveira, Edison Luiz Durigon, Flavio Fonseca e Santuza Teixeira |
| EPI_ISL_2627985, EPI_ISL_2627986, EPI_ISL_2627987 | Hospital | National Reference Center for Viruses of Respiratory Infections, Institut Pasteur, Paris | Marion Barbet, Sylvie Behillil, Méline Bizard, Angela Brisebarre, Camille Capel, Vincent Enouf, Louise Lefrançois, Frédéric Lemoine, Christophe Malabat, Corinne Maufrais, Etienne Simon-Lorière, Maud Vanpeene, Sylvie Van der Werf ,CéCile Farrugia |
| EPI_ISL_2629070 | HEGP - Laboratoire de Virologie | HEGP - Laboratoire de Virologie | David Veyer, Sandrine Imbeaud, Julien Puech, Hélène Péré, Maxime Wack, Nicolas Robillard |
| EPI_ISL_2637801 | U.O. Igiene, Ospedale Policlinico San Martino | U.O. Igiene, Ospedale Policlinico San Martino | Bruzzone Bianca, Caligiuri Patrizia, De Pace Vanessa, Domnich Alexander, Orsi Andrea, Ricucci Valentina, Lai Alessia, Bergna Annalisa, Ventura Francesco, Della Ventura Carla, Zehender Gianguglielmo, Icardi Giancarlo on behalf of SARS-CoV-2 ITALIAN RESEARCH ENTERPRISE-(SCIRE) Collaborative Group |
| EPI_ISL_2645531 | Laboratorio Central de Saude Publica do Estado do Espirito Santo (LACEN/ES) | Laboratory of Respiratory Viruses and Measles, Oswaldo Cruz Institute, FIOCRUZ | Paola Resende, Luciana Appolinario, Fernando Motta, Anna Carolina Paixao, Ana Carolina Mendonca, Alice Sampaio Rocha, Taina Venas, Eilisa Cavalcante Pereira, Renata Serrano Lopes, Rodrigo Ribeiro Rodrigues, Marilda Siqueira on behalf of the Fiocruz COVID-19 Genomic Surveillance Network |
| EPI_ISL_2645639 | Laboratorio Central de Saude Publica do Estado de Alagoas (LACEN/AL) | Laboratory of Respiratory Viruses and Measles, Oswaldo Cruz Institute, FIOCRUZ | Paola Resende, Luciana Appolinario, Fernando Motta, Anna Carolina Paixao, Ana Carolina Mendonca, Alice Sampaio Rocha, Taina Venas, Elisa Cavalcante Pereira, Renata Serrano Lopes, Anderson Brandao Leite, Marilda Siqueira on behalf of the Fiocruz COVID-19 Genomic Surveillance Network |
| EPI_ISL_2652117 | AZ Klina | AZ Klina | Carl Vael - Lynsey Berckmans |
| EPI_ISL_2652123 | Molecular diagnostic laboratory of Federal Budget Institution of Science "Central Research Institute of Epidemiology" of The Federal Service on Customers' Rights Protection and Human Well-being Surveillance | Group of Genomics and Postgenomic Technologies of Central Research Institute of Epidemiology | Samoilov A.E., Kaptelova V.V., Korneenko E.V., Valdohina A.V., Bulanenko V.P., Saenko S.S., Golubeva A.G., Lebedeva L.A., Shubina Y.A., Savosina L.V., Zhaleiko Z.P., Lukyanov A.V., Speranskaya A.S., Tivanova E.V., Kondrasheva L.V., Akimkin V.G. |
| EPI_ISL_2652124 | Molecular diagnostic laboratory of Federal Budget Institution of Science "Central Research Institute of Epidemiology" of The Federal Service on Customers' Rights Protection and Human Well-being Surveillance | Group of Genomics and Postgenomic Technologies of Central Research Institute of Epidemiology | Samoilov A.E., Kaptelova V.V., Korneenko E.V., Valdohina A.V., Bulanenko V.P., Saenko S.S., Sinitsyn S.O., Golubeva A.G., Speranskaya A.S., Tivanova E.V., Shipulina O.Y., Akimkin V.G. |
| EPI_ISL_2652133, EPI_ISL_2652134 | Molecular diagnostic laboratory of Federal Budget Institution of Science "Central Research Institute of Epidemiology" of The Federal Service on Customers' Rights Protection and Human Well-being Surveillance | Group of Genomics and Postgenomic Technologies of Central Research Institute of Epidemiology | Samoilov A.E., Kaptelova V.V., Korneenko E.V., Valdohina A.V., Bulanenko V.P., Saenko S.S., Golubeva A.G., Lebedeva L.A., Shubina Y.A., Savosina L.V., Zhaleiko Z.P., Lukyanov A.V., Speranskaya A.S., Tivanova E.V., Kondrasheva L.V., Akimkin V.G. |
| EPI_ISL_2652135, EPI_ISL_2652136 | Molecular diagnostic laboratory of Federal Budget Institution of Science "Central Research Institute of Epidemiology" of The Federal Service on Customers' Rights Protection and Human Well-being Surveillance | Group of Genomics and Postgenomic Technologies of Central Research Institute of Epidemiology | Samoilov A.E., Kaptelova V.V., Korneenko E.V., Valdohina A.V., Bulanenko V.P., Saenko S.S., Sinitsyn S.O., Golubeva A.G., Speranskaya A.S., Tivanova E.V., Shipulina O.Y., Akimkin V.G. |
| EPI_ISL_2660452 | Laboratory of Respiratory Viruses and Measles, Oswaldo Cruz Institute, FIOCRUZ | Laboratory of Respiratory Viruses and Measles, Oswaldo Cruz Institute, FIOCRUZ | Paola Resende, Luciana Appolinario, Fernando Motta, Anna Carolina Paixao, Ana Carolina Mendonca, Alice Sampaio Rocha, Taina Venas, Elisa Cavalcante Pereira, Renata Serrano Lopes, Marilda Siqueira on behalf of the Fiocruz COVID-19 Genomic Surveillance Network |
| EPI_ISL_2661782, EPI_ISL_2661783, EPI_ISL_2661784, EPI_ISL_2661786, EPI_ISL_2661788, EPI_ISL_2661794, EPI_ISL_2661801, EPI_ISL_2661803, EPI_ISL_2661804, EPI_ISL_2661811, EPI_ISL_2661813, EPI_ISL_2661823, EPI_ISL_2661824, EPI_ISL_2661825, EPI_ISL_2661826, EPI_ISL_2661827 | | | |
| see above | Laboratorio Central de Saude Publica do Estado do Rio Grande do Sul (LACEN-RS) | Laboratory of Respiratory Viruses and Measles, Oswaldo Cruz Institute, FIOCRUZ | Paola Resende, Luciana Appolinario, Fernando Motta, Anna Carolina Paixao, Ana Carolina Mendonca, Alice Sampaio Rocha, Taina Venas, Elisa Cavalcante Pereira, Renata Serrano Lopes, Tatiana Schaffer Gregianini, Richard Salvato, Marilda Siqueira on behalf of the Fiocruz COVID-19 Genomic Surveillance Network |
| EPI_ISL_2663115, EPI_ISL_2663117, EPI_ISL_2663118, EPI_ISL_2663120, EPI_ISL_2663121, EPI_ISL_2663123, EPI_ISL_2663162, EPI_ISL_2663163, EPI_ISL_2663165, EPI_ISL_2663166, EPI_ISL_2663168, EPI_ISL_2663171, EPI_ISL_2663173, EPI_ISL_2663174, EPI_ISL_2663179, EPI_ISL_2663181, EPI_ISL_2663182, EPI_ISL_2663184, EPI_ISL_2663185, EPI_ISL_2663187, EPI_ISL_2663188, EPI_ISL_2663190 | | | |
| see above | Ochsner Health | Bioinfoexperts, LLC | Judy Crabtree, Amy Feehan, Rebecca Rose, David J. Nolan, Tessa LaFleur, Julia-Garcia-Diaz, Lucio Miele, Susanna L. Lamers |
| EPI_ISL_2677079 | Labortorio Central de Saude Publica do Estado de Santa Catarina (LACEN/SC) | Laboratory of Respiratory Viruses and Measles, Oswaldo Cruz Institute, FIOCRUZ | Paola Resende, Luciana Appolinario, Fernando Motta, Anna Carolina Paixao, Ana Carolina Mendonca, Alice Sampaio Rocha, Taina Venas, Elisa Cavalcante Pereira, Renata Serrano Lopes, Darcita Buerger Rovaris, Sandra Bianchini Fernandes, Marilda Siqueira on behalf of the Fiocruz COVID-19 Genomic Surveillance Network |
| EPI_ISL_2688267, EPI_ISL_2688277 | Centers for Disease Control and Prevention, Dengue Branch | Centers for Disease Control and Prevention, Dengue Branch | Gilberto A. Santiago, Glenda Gonzalez, Betzabel Flores, Keyla Charriez, Gabriela Paz-Bailey, Jorge L. Munoz-Jordan |
| EPI_ISL_2689147, EPI_ISL_2689148 | MSK Microbiology Lab | MSK Microbiology Lab | Esther Babady, Tracy McMillen, Krupa Jani |
| EPI_ISL_2690129, EPI_ISL_2690130, EPI_ISL_2690131, EPI_ISL_2690132, EPI_ISL_2690151, EPI_ISL_2690152, EPI_ISL_2690153, EPI_ISL_2690154, EPI_ISL_2690155, EPI_ISL_2690156, EPI_ISL_2690158, EPI_ISL_2690159, EPI_ISL_2690160, EPI_ISL_2690162, EPI_ISL_2690163, EPI_ISL_2690164, EPI_ISL_2690165, EPI_ISL_2690166, EPI_ISL_2690167, EPI_ISL_2690173, EPI_ISL_2690176, EPI_ISL_2690177, EPI_ISL_2690185, EPI_ISL_2690186, EPI_ISL_2690187, EPI_ISL_2690188, EPI_ISL_2690189, EPI_ISL_2690190 | | | |
| see above | Ochsner Health | Bioinfoexperts, LLC | Judy Crabtree, Amy Feehan, Rebecca Rose, David J. Nolan, Tessa LaFleur, Julia-Garcia-Diaz, Lucio Miele, Susanna L. Lamers |

| | | | |
|---|---|---|---|
| EPI_ISL_2690433, EPI_ISL_2690434, EPI_ISL_2690444, EPI_ISL_2690572, EPI_ISL_2690576, EPI_ISL_2690580, EPI_ISL_2690595, EPI_ISL_2690596, EPI_ISL_2690602, EPI_ISL_2690608, EPI_ISL_2690609, EPI_ISL_2690611, EPI_ISL_2690612, EPI_ISL_2690614, EPI_ISL_2690621, EPI_ISL_2690624, EPI_ISL_2691003 | | | |
| see above | MSK Microbiology Lab | MSK Microbiology Lab | Esther Babady, Tracy McMillen, Krupa Jani |
| EPI_ISL_2692207 | Clinical Microbiology Laboratory, Faculty of Medicine, Universitas Indonesia | Clinical Microbiology Laboratory, Faculty of Medicine, Universitas Indonesia | Ibrahim F, Natasha A, Yasmon A, Fitrhiyah, Saharman YR, Karuniawati A, Sudarmono P. |
| EPI_ISL_2692886, EPI_ISL_2692887 | Clinical Microbiology Lab RSPTN Universitas Hasanuddin | Clinical Microbiology Lab RSPTN Universitas Hasanuddin | Muhammad Nasrum Massi, Handayani Halik, Rizalinda Sjahril, Gita Vita Soraya, Triwibawa, Muhammad Yusuf Usman, Rufika Shari |
| EPI_ISL_2696943, EPI_ISL_2696944 | All India Institute of Medical Sciences Delhi Hospital | Virology Laboratory, AIIMS Delhi | Aashish Choudhary, Sumedha Bagga, Lata Rani, Dibyabhaba Pradhan, Pooja Pandey, Jyoti Jethani, Shivram Dhakad, Nazneen Arif, Deepankar Srigyan, Megha Brijwal, Lalit Dar, Manish Soneja, Rakesh Lodha, Puneet Kaur, Ritu Gupta, Subrata Sinha, Chitra Sarkar, Randeep Guleria |
| EPI_ISL_2710012, EPI_ISL_2710013, EPI_ISL_2710014, EPI_ISL_2710015, EPI_ISL_2710024, EPI_ISL_2710026 | Ochsner Health | Bioinfoexperts, LLC | Judy Crabtree, Amy Feehan, Rebecca Rose, David J. Nolan, Tessa LaFleur, Julia-Garcia-Diaz, Lucio Miele, Susanna L. Lamers |
| EPI_ISL_2713080, EPI_ISL_2713081, EPI_ISL_2713082, EPI_ISL_2713084, EPI_ISL_2713085, EPI_ISL_2713086, EPI_ISL_2713087, EPI_ISL_2713089, EPI_ISL_2713090, EPI_ISL_2713092, EPI_ISL_2713094, EPI_ISL_2713098, EPI_ISL_2713099, EPI_ISL_2713114, EPI_ISL_2713115, EPI_ISL_2713120 | | | |
| see above | Department of Health Technology and Informatics, The Hong Kong Polytechnic University | Department of Health Technology and Informatics, The Hong Kong Polytechnic University | Gilman Kit-Hang Siu, Lam-Kwong Lee, Kenneth Siu-Sing Leung, Jake Siu-Lun Leung, Wing-Hei Lo, Timothy Ting-Leung Ng, Chloe Toi-Mei Chan, Hoi-Ching Jim, Ivan Tak-Fai Wong, Kingsley King-Gee Tam, Hiu-Yin Lao, Denise Sze-Hang Wong, Alan Ka-Lun Wu, Miranda Chong-Yee Yau, Yvette Wai-Man Lai, Kitty Sau-Chun Fung, Sandy Ka-Yee Chau, Barry Kin-Chung Wong, Wing-Kin To, Kristine Luk, Alex Yat-Man Ho, Tak-Lun Que, Kam-Tong Yip, Wing Cheong Yam, David Ho-Keung Shum, Shea Ping Yip |
| EPI_ISL_2727895, EPI_ISL_2727896, EPI_ISL_2727897, EPI_ISL_2727898 | All India Institute of Medical Sciences Delhi Hospital | Virology Laboratory, AIIMS Delhi | Aashish Choudhary, Sumedha Bagga, Lata Rani, Dibyabhaba Pradhan, Pooja Pandey, Jyoti Jethani, Shivram Dhakad, Nazneen Arif, Deepankar Srigyan, Megha Brijwal, Lalit Dar, Manish Soneja, Rakesh Lodha, Puneet Kaur, Ritu Gupta, Subrata Sinha, Chitra Sarkar, Randeep Guleria |
| EPI_ISL_2728580 | Laboratorio de Infectologia y Virologia Molecular | Laboratory of Molecular Virology, School of Medicine, Pontificia Universidad Catolica de Chile | Catalina Pardo-Roa, Constanza Martinez-Valdevenito, Jennifer Angulo, Tamara Garcia-Salum, Estefany Poblete, Maria Jose Avendano, Leonardo I. Almonacid, Ana Maria Contreras, Carlos Palma, Jorge Levican, Eileen Serrano, Constanza Maldonado, M. Belen Leyton, Erick Salinas, Andres E. Munoz-Marcos, Francisco Melo, Marcela Ferres, Rafael A. Medina |
| EPI_ISL_2756453 | Instituto Adolfo Lutz Central | Instituto Adolfo Lutz, Interdiciplinary Procedures Center, Strategic Laboratory | Claudio Tavares Sacchi, Claudia Regina Gonçalves, Erica Valessa Ramos Gomes, Karoline Rodrigues Campos, Caio Vinicius Dias Lopes, Leonardo Jose Tadeu de Araujo |
| EPI_ISL_2779355, EPI_ISL_2779356, EPI_ISL_2779357, EPI_ISL_2779358, EPI_ISL_2779359, EPI_ISL_2779360, EPI_ISL_2779361, EPI_ISL_2779362, EPI_ISL_2779363, EPI_ISL_2779364 | KEMRI/DEID Surveillance Site (Mtongwe Navy Base Hospital) | USAMRD-A, Basic Science Laboratory | Gathii Kimita, Josphat Nyataya, Esther Omuseni, Faith Sigei, Alan Lemtudo, Eric Muthanje, Brian Andika, Rehema Liyai, Rachel Githii, Clement Masakwe, Stephen Ochola, George Awinda, Carol Kifude, Beth Mutai, John Waitumbi |
| EPI_ISL_2779365, EPI_ISL_2779366, EPI_ISL_2779367, EPI_ISL_2779368, EPI_ISL_2779369, EPI_ISL_2779370 | Jaramogi Oginga Odinga Teaching and Referral Hospital | USAMRD-A, Basic Science Laboratory | Gathii Kimita, Josphat Nyataya, Esther Omuseni, Faith Sigei, Alan Lemtudo, Eric Muthanje, Brian Andika, Rehema Liyai, Rachel Githii, Clement Masakwe, Stephen Ochola, George Awinda, Carol Kifude, Beth Mutai, John Waitumbi |
| EPI_ISL_2779374, EPI_ISL_2779375 | KEMRI/DEID Surveillance Site (Lamu County Hospital) | USAMRD-A, Basic Science Laboratory | Gathii Kimita, Josphat Nyataya, Esther Omuseni, Faith Sigei, Alan Lemtudo, Eric Muthanje, Brian Andika, Rehema Liyai, Rachel Githii, Clement Masakwe, Stephen Ochola, George Awinda, Carol Kifude, Beth Mutai, John Waitumbi |
| EPI_ISL_2779382, EPI_ISL_2779386 | Jaramogi Oginga Odinga Teaching and Referral Hospital | USAMRD-A, Basic Science Laboratory | Gathii Kimita, Josphat Nyataya, Esther Omuseni, Faith Sigei, Alan Lemtudo, Eric Muthanje, Brian Andika, Rehema Liyai, Rachel Githii, Clement Masakwe, Stephen Ochola, George Awinda, Carol Kifude, Beth Mutai, John Waitumbi |
| EPI_ISL_2779387, EPI_ISL_2779396, EPI_ISL_2779397 | Busia County Referral Hospital | USAMRD-A, Basic Science Laboratory | Gathii Kimita, Josphat Nyataya, Esther Omuseni, Faith Sigei, Alan Lemtudo, Eric Muthanje, Brian Andika, Rehema Liyai, Rachel Githii, Clement Masakwe, Stephen Ochola, George Awinda, Carol Kifude, Beth Mutai, John Waitumbi |
| EPI_ISL_2779419 | Port Health Clinic, Busia border point | USAMRD-A, Basic Science Laboratory | Gathii Kimita, Josphat Nyataya, Esther Omuseni, Faith Sigei, Alan Lemtudo, Eric Muthanje, Brian Andika, Rehema Liyai, Rachel Githii, Clement Masakwe, Stephen Ochola, George Awinda, Carol Kifude, Beth Mutai, John Waitumbi |
| EPI_ISL_2779422, EPI_ISL_2779423 | GK Prison Busia (Matayo)Matayos Prison | USAMRD-A, Basic Science Laboratory | Gathii Kimita, Josphat Nyataya, Esther Omuseni, Faith Sigei, Alan Lemtudo, Eric Muthanje, Brian Andika, Rehema Liyai, Rachel Githii, Clement Masakwe, Stephen Ochola, George Awinda, Carol Kifude, Beth Mutai, John Waitumbi |
| EPI_ISL_2779456, EPI_ISL_2779457, EPI_ISL_2779458, EPI_ISL_2779459, EPI_ISL_2779460, EPI_ISL_2779461, EPI_ISL_2779463, EPI_ISL_2779464, EPI_ISL_2779465 | KCRH KEMRI/DEID Surveillance Site | USAMRD-A, Basic Science Laboratory | Gathii Kimita, Josphat Nyataya, Esther Omuseni, Faith Sigei, Alan Lemtudo, Eric Muthanje, Brian Andika, Rehema Liyai, Rachel Githii, Clement Masakwe, Stephen Ochola, George Awinda, Carol Kifude, Beth Mutai, John Waitumbi |
| EPI_ISL_2779477, EPI_ISL_2779478, EPI_ISL_2779479, EPI_ISL_2779480, EPI_ISL_2779491 | MOH-K | USAMRD-A, Basic Science Laboratory | Gathii Kimita, Josphat Nyataya, Esther Omuseni, Faith Sigei, Alan Lemtudo, Eric Muthanje, Brian Andika, Rehema Liyai, Rachel Githii, Clement Masakwe, Stephen Ochola, George Awinda, Carol Kifude, Beth Mutai, John Waitumbi |
| EPI_ISL_2779493 | Port Health Clinic, Malaba border point | USAMRD-A, Basic Science Laboratory | Gathii Kimita, Josphat Nyataya, Esther Omuseni, Faith Sigei, Alan Lemtudo, Eric Muthanje, Brian Andika, Rehema Liyai, Rachel Githii, Clement Masakwe, Stephen Ochola, George Awinda, Carol Kifude, Beth Mutai, John Waitumbi |
| EPI_ISL_2779526, EPI_ISL_2779527, EPI_ISL_2779528, EPI_ISL_2779529, EPI_ISL_2779530, EPI_ISL_2779531, EPI_ISL_2779532, EPI_ISL_2779533, EPI_ISL_2779534 | MOH-Q | USAMRD-A, Basic Science Laboratory | Gathii Kimita, Josphat Nyataya, Esther Omuseni, Faith Sigei, Alan Lemtudo, Eric Muthanje, Brian Andika, Rehema Liyai, Rachel Githii, Clement Masakwe, Stephen Ochola, George Awinda, Carol Kifude, Beth Mutai, John Waitumbi |
| EPI_ISL_2779535 | PSTC | USAMRD-A, Basic Science Laboratory | Gathii Kimita, Josphat Nyataya, Esther Omuseni, Faith Sigei, Alan Lemtudo, Eric Muthanje, Brian Andika, Rehema Liyai, Rachel Githii, Clement Masakwe, Stephen Ochola, George Awinda, Carol Kifude, Beth Mutai, John Waitumbi |
| EPI_ISL_2779544 | KCRH KEMRI/DEID Surveillance Site | USAMRD-A, Basic Science Laboratory | Gathii Kimita, Josphat Nyataya, Esther Omuseni, Faith Sigei, Alan Lemtudo, Eric Muthanje, Brian Andika, Rehema Liyai, Rachel Githii, Clement Masakwe, Stephen Ochola, George Awinda, Carol Kifude, Beth Mutai, John Waitumbi |
| EPI_ISL_2779545 | Port Health Clinic, Malaba border point | USAMRD-A, Basic Science Laboratory | Gathii Kimita, Josphat Nyataya, Esther Omuseni, Faith Sigei, Alan Lemtudo, Eric Muthanje, Brian Andika, Rehema Liyai, Rachel Githii, Clement Masakwe, Stephen Ochola, George Awinda, Carol Kifude, Beth Mutai, John Waitumbi |
| EPI_ISL_2779546, EPI_ISL_2779547, EPI_ISL_2779548 | KCRH KEMRI/DEID Surveillance Site | USAMRD-A, Basic Science Laboratory | Gathii Kimita, Josphat Nyataya, Esther Omuseni, Faith Sigei, Alan Lemtudo, Eric Muthanje, Brian Andika, Rehema Liyai, Rachel Githii, Clement Masakwe, Stephen Ochola, George Awinda, Carol Kifude, Beth Mutai, John Waitumbi |
| EPI_ISL_2789820, EPI_ISL_2789821, EPI_ISL_2789822 | Centre Hospitalier Universitaire Clermont-Ferrand | CHU Clermont-Ferrand, service de virologie | Bisseux Maxime, Mirand Audrey, Combes Patricia, Henquell Cécile |
| EPI_ISL_2789838 | Centre Hospitalier RIOM | CHU Clermont-Ferrand, service de virologie | Bisseux Maxime, Mirand Audrey, Combes Patricia, Henquell Cécile |
| EPI_ISL_2789839, EPI_ISL_2789840, EPI_ISL_2789841, EPI_ISL_2789842, EPI_ISL_2789843, EPI_ISL_2789844, EPI_ISL_2789845, EPI_ISL_2789846, EPI_ISL_2789847, EPI_ISL_2789848, EPI_ISL_2789849, EPI_ISL_2789850, EPI_ISL_2789851, EPI_ISL_2789852, EPI_ISL_2789853, EPI_ISL_2789854, EPI_ISL_2789855, EPI_ISL_2789856, EPI_ISL_2789857, EPI_ISL_2789858, EPI_ISL_2789859, EPI_ISL_2789860, EPI_ISL_2789861, EPI_ISL_2789862, EPI_ISL_2789863, EPI_ISL_2789864, EPI_ISL_2789865, EPI_ISL_2789866, EPI_ISL_2789867, EPI_ISL_2789868, EPI_ISL_2789869, EPI_ISL_2789870, EPI_ISL_2789871, EPI_ISL_2789872, EPI_ISL_2789873, EPI_ISL_2789874, EPI_ISL_2789875 | | | |
| see above | Centre Hospitalier Universitaire Clermont-Ferrand | CHU Clermont-Ferrand, service de virologie | Bisseux Maxime, Mirand Audrey, Combes Patricia, Henquell Cécile |
| EPI_ISL_2790860, EPI_ISL_2790861 | Hospital | National Reference Center for Viruses of Respiratory Infections, Institut Pasteur, Paris | Marion Barbet, Sylvie Behillil, Méline Bizard, Angela Brisebarre, Camille Capel, Vincent Enouf, Louise Lefrançois, Frédéric Lemoine, Christophe Malabat, Corinne Maufrais, Emmanuelle Permal, Etienne Simon-Lorière, Maud Vanpeene, Sylvie Van der Werf ,Pascale Martres |

| | | | |
|---|---|---|---|
| EPI_ISL_2790873, EPI_ISL_2790874, EPI_ISL_2790875, EPI_ISL_2790876, EPI_ISL_2790877, EPI_ISL_2790878, EPI_ISL_2790879, EPI_ISL_2790880, EPI_ISL_2790883, EPI_ISL_2790884, EPI_ISL_2790885, EPI_ISL_2790886, EPI_ISL_2790887 | | | |
| see above | Labo Analyses Med | National Reference Center for Viruses of Respiratory Infections, Institut Pasteur, Paris | Marion Barbet, Sylvie Behillil, Méline Bizard, Angela Brisebarre, Camille Capel, Vincent Enouf, Louise Lefrançois, Frédéric Lemoine, Christophe Malabat, Corinne Maufrais, Emmanuelle Permal, Etienne Simon-Lorière, Maud Vanpeene, Sylvie Van der Werf ,StéPhane Romand |
| EPI_ISL_2790888 | Medecine sentinel network | National Reference Center for Viruses of Respiratory Infections, Institut Pasteur, Paris | Marion Barbet, Sylvie Behillil, Méline Bizard, Angela Brisebarre, Camille Capel, Vincent Enouf, Louise Lefrançois, Frédéric Lemoine, Christophe Malabat, Corinne Maufrais, Emmanuelle Permal, Etienne Simon-Lorière, Maud Vanpeene, Sylvie Van der Werf ,Anne Garrigues |
| EPI_ISL_2790902, EPI_ISL_2790903 | Hospital | National Reference Center for Viruses of Respiratory Infections, Institut Pasteur, Paris | Marion Barbet, Sylvie Behillil, Méline Bizard, Angela Brisebarre, Camille Capel, Vincent Enouf, Louise Lefrançois, Frédéric Lemoine, Christophe Malabat, Corinne Maufrais, Emmanuelle Permal, Etienne Simon-Lorière, Maud Vanpeene, Sylvie Van der Werf ,GisèLe Lagathu |
| EPI_ISL_2790905 | Medecine sentinel network | National Reference Center for Viruses of Respiratory Infections, Institut Pasteur, Paris | Marion Barbet, Sylvie Behillil, Méline Bizard, Angela Brisebarre, Camille Capel, Vincent Enouf, Louise Lefrançois, Frédéric Lemoine, Christophe Malabat, Corinne Maufrais, Emmanuelle Permal, Etienne Simon-Lorière, Maud Vanpeene, Sylvie Van der Werf ,Sophie Langlais |
| EPI_ISL_2795658 | LESP Hidalgo | LESP Hidalgo GENE2LIFE | HERRERA-GOMEZ CONNIE ALEJANDRA; CASTILLO-ORTEGA LAURA SOFIA; GARCIA-REYES MELITO; VIGUERAS-MORALES YAJAIRA SULIM; GARCIA-GONZALEZ OCTAVIO; VILLAFUENTES-TELLEZ HECTOR ALBERTO |
| EPI_ISL_2820471, EPI_ISL_2820473 | The Hong Kong Polytechnic University - Molecular Diagnostic Laboratory | The Hong Kong Polytechnic University - Molecular Diagnostic Laboratory | Hin Fung Tsang, Allen Chi Shing Yu, Aldrin Kay Yuen Yim, Wai Ming Stanley Leung, Jiachi Chiou, Yin Kwan Evelyn Wong, Dominic Ngai Chong Tsang, Peter and SO, Samuel Chun Lap Lo,Wing Tak Wong, Sze Chuen Cesar Wong |
| EPI_ISL_2835003, EPI_ISL_2835004 | Laboratoire Central Analyse Médicale mohammedia | Functional Genomic Platform_UATRS_CNRST_Rabat | Hajjaji Mohammed, Oumaima Ait Si Mohammed, Mouhcine Gardoul, Marouane Melloul, Sanaa Alaoui-Amine, Mouhssine Hemlali, Taha Chouati Omar Askander, Belayachi Lamiae, Hassan Aguenaou, Hasnae Benkirane, Jaafar Heikel, Amina Barkat, Nadia Touil, Elmostafa El fahime |
| EPI_ISL_2835490, EPI_ISL_2835491, EPI_ISL_2835492, EPI_ISL_2835493, EPI_ISL_2835494, EPI_ISL_2835495, EPI_ISL_2835496, EPI_ISL_2835497, EPI_ISL_2835499, EPI_ISL_2835500, EPI_ISL_2835502, EPI_ISL_2835503, EPI_ISL_2835504, EPI_ISL_2835505, EPI_ISL_2835506, EPI_ISL_2835507, EPI_ISL_2835508, EPI_ISL_2835509, EPI_ISL_2835510 | | | |
| see above | Instituto de Diagnostico y Referencia Epidemiologicos (InDRE) | Andersen lab at Scripps Research | SEARCH Alliance with Gisela Barrera Badillo, Jose Ernesto Ramirez Gonzalez, Maria Natividad Cruz Ortiz, Nervain Benjamin Contreras Gonzalez, Tatiana Ernestina Nunez Garcia, Dayanira Sarith Arellano Suarez, Irma Lopez Martinez, Rita Flores Leon, Beatriz Olivares Flores |
| EPI_ISL_2835846, EPI_ISL_2835847, EPI_ISL_2835848 | Hospital | National Reference Center for Viruses of Respiratory Infections, Institut Pasteur, Paris | Marion Barbet, Sylvie Behillil, Méline Bizard, Angela Brisebarre, Camille Capel, Vincent Enouf, Louise Lefrançois, Frédéric Lemoine, Christophe Malabat, Corinne Maufrais, Emmanuelle Permal, Etienne Simon-Lorière, Maud Vanpeene, Sylvie Van der Werf ,Evelyne Schvoerer |
| EPI_ISL_2836970, EPI_ISL_2836971, EPI_ISL_2837247, EPI_ISL_2838053, EPI_ISL_2838054, EPI_ISL_2838055, EPI_ISL_2838353 | Department of Microbiology and Virology, School of Medicine, Mazandaran University of Medical Sciences, Sari, Iran | Genetics Research Center, University of Social Welfare and Rehabilitation Sciences | Zohreh Fattahi, Marzieh Mohseni, Kimia Kahrizi, Mohammad Reza Haghshenas, Hossein Najmabadi. |
| EPI_ISL_2839238, EPI_ISL_2839240, EPI_ISL_2839243, EPI_ISL_2839245, EPI_ISL_2839247, EPI_ISL_2839249, EPI_ISL_2839251, EPI_ISL_2839255, EPI_ISL_2839259, EPI_ISL_2839261, EPI_ISL_2839268, EPI_ISL_2839270, EPI_ISL_2839272, EPI_ISL_2839274, EPI_ISL_2839276, EPI_ISL_2839278, EPI_ISL_2839280, EPI_ISL_2839282, EPI_ISL_2839284, EPI_ISL_2839286, EPI_ISL_2839287, EPI_ISL_2839289, EPI_ISL_2839291, EPI_ISL_2839299, EPI_ISL_2839301, EPI_ISL_2839303, EPI_ISL_2839305, EPI_ISL_2839309, EPI_ISL_2839311, EPI_ISL_2839315, EPI_ISL_2839319, EPI_ISL_2839321, EPI_ISL_2839323, EPI_ISL_2839325, EPI_ISL_2839327, EPI_ISL_2839329, EPI_ISL_2839331, EPI_ISL_2839335, EPI_ISL_2839336, EPI_ISL_2839340, EPI_ISL_2839342, EPI_ISL_2839344, EPI_ISL_2839346, EPI_ISL_2839348, EPI_ISL_2839352, EPI_ISL_2839353, EPI_ISL_2839359, EPI_ISL_2839361, EPI_ISL_2839367, EPI_ISL_2839371, EPI_ISL_2839373, EPI_ISL_2839377, EPI_ISL_2839380, EPI_ISL_2839382 | | | |
| see above | Instituto Nacional de Enfermedades Respiratorias (INER) | Centro de Investigación en Enfermedades Infecciosas (CIENI), Instituto Nacional de Enfermedades Respiratorias (INER) | Consorcio Mexicano de Vigilancia Genómica (CoViGen-Mex). Authors (in alphabetical order): Julio Elias Alvarado-Yaah, Carlos F. Arias, Santiago Ávila-Ríos, Eduardo Becerril-Vargas, Víctor Hugo Borja-Aburto, Celia Boukadida, Cristóbal Cháidez-Quiróz, Juan Bautista Chale-Dzul, Ricardo Ciria Merce, Andreu Comas-García, Célida Duque Molina, Julissa Enciso-Ibarra, José Antonio Enciso-Moreno, Gloria Elena Espinosa-Ayala, Fernando Fontove-Herrera, Daniel Fregoso-Rueda, Victor Eduardo Garcia-Arias, Alejandra Garcia-Gasca, Bruno Gomez-Gil, Jean Pierre González, Irvin González-López, Concepción Grajales-Muñiz, Ricardo Grande, Rosa María Gutierrez Rios, Alejandra Hernández-Terán, Alfredo Herrera-Estrella, Carla Ivón Herrera-Najera, Pavel Isa, Daniel Lira Morales, Susana Lopez, Antonio Loza Román, Brenda Irasema Maldonado-Meza, Bernardo Martínez-Miguel, José Arturo Martínez-Orozco, Célida Martínez- Rodríguez, Margarita Matías-Florentino, Fidencio Mejía-Nepomuceno, María Guadalupe de Jesús Mireles-Rivera, Gloria María Molina-Salinas, Hector Montoya-Fuentes, Mario Mújica-Sánchez, José Esteban Muñoz-Medina, José de Jesús Nuñez-Contreras, Alicia Ocaña-Mondragón, Luis Alberto Ochoa-Carrera, Hector Esteban Paz-Juárez, Marissa Pérez-García, Francisco Pulido, Helen Haydee Fernanda Ramírez-Plascencia, Jorge Salas-Hernández, Angel Gustavo Salas-Lais, Alejandro Sanchez-Flores, Clara Esperanza Santacruz-Tinoco, María Guadalupe Santiago-Mauricio, Selene Zárate, Nelly Sélem-Mojica, Blanca Taboada, Kathia Elizabeth Tapia-Diaz, Gloria Vazquez, Joel Armando Vázquez-Pérez. |
| EPI_ISL_2860777, EPI_ISL_2860778, EPI_ISL_2860779, EPI_ISL_2860780, EPI_ISL_2860782, EPI_ISL_2860783, EPI_ISL_2860787, EPI_ISL_2860788 | Department of Health Technology and Informatics, The Hong Kong Polytechnic University | Department of Health Technology and Informatics, The Hong Kong Polytechnic University | Gilman Kit-Hang Siu, Lam-Kwong Lee, Kenneth Siu-Sing Leung, Jake Siu-Lun Leung, Wing-Hei Lo, Timothy Ting-Leung Ng, Chloe Toi-Mei Chan, Hoi-Ching Jim, Ivan Tak-Fai Wong, Kingsley King-Gee Tam, Hiu-Yin Lao, Denise Sze-Hang Wong, Alan Ka-Lun Wu, Miranda Chong-Yee Yau, Yvette Wai-Man Lai, Kitty Sau-Chun Fung, Sandy Ka-Yee Chau, Barry Kin-Chung Wong, Wing-Kin To, Kristine Luk, Alex Yat-Man Ho, Tak-Lun Que, Kam-Tong Yip, Wing Cheong Yam, David Ho-Keung Shum, Shea Ping Yip |
| EPI_ISL_2864604, EPI_ISL_2864606 | Ga East Hospital | UHAS COVID-19 Lab | Kwabena O. Duedu, Jones Gyamfi, Reuben Ayivor-Djanie, John O. Gyapong and the UHAS COVID-19 Lab Team |
| EPI_ISL_568577 | The National Institute of Public Health | State Veterinary Institute Prague | Nagy, A; Jirincova, H; Novakova, L; Trnka,D; Vecerova, J |
| EPI_ISL_574431 | Hospital IESS Babahoyo | Institute of Microbiology, Universidad San Francisco de Quito | Belén Prado-Vivar, Sully Márquez, Juan José Guadalupe, Monica Becerra-Wong, Fernanda Zurita, Bernardo Gutiérrez, Francisco Cordova, Ninfa Henriquez, Killen Briones-Zamora, Killen Briones-Claudette, Verónica Barragán, Patricio Rojas-Silva, Gabriel Trueba, Michelle Grunauer, Paúl Cárdenas |
| EPI_ISL_576383 | RSUD Budi Rahayu Kota Magelang | Genetics Working Group (Pokja Genetik) Faculty of Medicine, Public Health and Nursing Universitas Gadjah Mada (FK-KMK UGM); Disease Investigation Center Wates Ministry of Agriculture Indonesia; Department of Microbiology FK-KMK UGM; Laboratorium Diagnostik Yayasan Tahija World Mosquito Program (WMP) Yogyakarta Center for Tropical Medicine FK-KMK UGM; Integrated Research Center FK-KMK UGM; Department of Computer Science and Electronics FMIPA UGM; Balai Besar Teknik Kesehatan Lingkungan dan Pengendalian Penyakit (BBTKLPP) Yogyakarta | Gunadi, Hendra Wibawa, Marcellus, Mohamad S. Hakim, Edwin W. Daniwijaya, Ludhang P. Rizki, Endah Supriyati, Eggi Arguni, Titik Nuryastuti, Tri Wibawa, Dwi AA Nugrahaningsih, Afiahayati, Siswanto, Kristy Iskandar, Nungki Anggorowati, Irene, Indaryati, Havid Setyawan, Ari Meliyanti, Merliana Sari Situmeang, Audric Kenny Tedja, Aditya Rifqi Fauzi |
| EPI_ISL_577630 | The National Institute of Public Health | State Veterinary Institute Prague | Nagy,A;Jirincova,H;Novakova,L;Trnka,D;Vecerova,J |
| EPI_ISL_582128, EPI_ISL_582129, EPI_ISL_582130, EPI_ISL_582131, EPI_ISL_582132 | Antwerp University Hospital | Institute of Tropical Medicine | Philippe Selhorst, Colin Anthony |
| EPI_ISL_582659, EPI_ISL_582662, EPI_ISL_582663, EPI_ISL_582674, EPI_ISL_582675, EPI_ISL_582679, EPI_ISL_582680, EPI_ISL_582681, EPI_ISL_582683, EPI_ISL_582688 | Sheikh Khalifa Medical City | Molecular/Surveillance lab Sheikh Khalifa Medical City | Amirtharaj Francis, Sajeed Abdul, Hala Imambaccus, Sahar Almarzooqi, Hiba Saud, Stefan Weber |
| EPI_ISL_583893 | Singapore General Hospital | Department of Microbiology | Nurdyana Abdul Rahman, Kun Lee Lim, Chenhao Li, Sui Sin Goh, Kenneth Xin Long Chan, Kian Sing Chan, Lynette Oon, Kern Rei Chng, Niranjan Nagarajan, Karrie Ko |
| EPI_ISL_584082 | The National Institute of Public Health | State Veterinary Institute Prague | Nagy,A;Jirincova,H;Novakova,L;Trnka,D;Vecerova,J |
| EPI_ISL_586569 | GCRI,Ahmedabad | Gujarat Biotechnology Research Centre | Shashank Pandya, Harsha Panchal, Apurvasinh Puvar, Janvi Raval, Zarna Patel, Monika Gandhi, Pinal Trivedi, Maharshi Pandya, Nidhi Patel, Nitin Savaliya, Raghawendra Kumar, Dinesh Kumar, Zuber Saiyed, Komal Patel, Labdhi Pandya, Afzal Ansari, Nikha Trivedi, R D Dixit, A M Kadri, Harsh Bakshi, Chaitanya Joshi, Madhvi Joshi |
| EPI_ISL_586570 | GCRI,Ahmedabad | Gujarat Biotechnology Research Centre | Harsha Panchal, Apurvasinh Puvar, Janvi Raval, Zarna Patel, Monika Gandhi, Pinal Trivedi, Maharshi Pandya, Nidhi Patel, Nitin Savaliya, Raghawendra |

| | | | |
|---|---|---|---|
| | | | Kumar, Dinesh Kumar, Zuber Saiyed, Komal Patel, Labdhi Pandya, Afzal Ansari, Nikha Trivedi, Shashank Pandya, R D Dixit, A M Kadri, Harsh Bakshi, Chaitanya Joshi, Madhvi Joshi |
| EPI_ISL_593558 | Brigham and Women's Hospital | Jonathan Li Laboratory | Manish C. Choudhary, James Regan, Jonathan Z. Li |
| EPI_ISL_593902 | Sentinelles, Limay | National Reference Center for Viruses of Respiratory Infections, Institut Pasteur, Paris | Sylvie Behillil, Fabiana Gambaro, Etienne Simon-Lorière, Vincent Enouf, Maud Vanpeene, Sylvie van der Werf |
| EPI_ISL_593903 | Hospital, Argenteuil | National Reference Center for Viruses of Respiratory Infections, Institut Pasteur, Paris | Sylvie Behillil, Fabiana Gambaro, Etienne Simon-Lorière, Vincent Enouf, Maud Vanpeene, Sylvie van der Werf |
| EPI_ISL_593904, EPI_ISL_593905, EPI_ISL_593906, EPI_ISL_593907, EPI_ISL_593908, EPI_ISL_593909 | Labo Analyses Med, Sarcelles | National Reference Center for Viruses of Respiratory Infections, Institut Pasteur, Paris | Sylvie Behillil, Fabiana Gambaro, Etienne Simon-Lorière, Vincent Enouf, Maud Vanpeene, Sylvie van der Werf |
| EPI_ISL_593910 | Hospital, Boulogne-Billancourt | National Reference Center for Viruses of Respiratory Infections, Institut Pasteur, Paris | Sylvie Behillil, Fabiana Gambaro, Etienne Simon-Lorière, Vincent Enouf, Maud Vanpeene, Sylvie van der Werf |
| EPI_ISL_593911 | Hospital, Le Chesnay Cédex | National Reference Center for Viruses of Respiratory Infections, Institut Pasteur, Paris | Sylvie Behillil, Fabiana Gambaro, Etienne Simon-Lorière, Vincent Enouf, Maud Vanpeene, Sylvie van der Werf |
| EPI_ISL_593913 | Hospital, Talange | National Reference Center for Viruses of Respiratory Infections, Institut Pasteur, Paris | Sylvie Behillil, Fabiana Gambaro, Etienne Simon-Lorière, Vincent Enouf, Maud Vanpeene, Sylvie van der Werf |
| EPI_ISL_593914 | Hospital, Metz | National Reference Center for Viruses of Respiratory Infections, Institut Pasteur, Paris | Sylvie Behillil, Fabiana Gambaro, Etienne Simon-Lorière, Vincent Enouf, Maud Vanpeene, Sylvie van der Werf |
| EPI_ISL_593915 | Hospital, Amneville | National Reference Center for Viruses of Respiratory Infections, Institut Pasteur, Paris | Sylvie Behillil, Fabiana Gambaro, Etienne Simon-Lorière, Vincent Enouf, Maud Vanpeene, Sylvie van der Werf |
| EPI_ISL_593916 | Hospital, Sarreguemines | National Reference Center for Viruses of Respiratory Infections, Institut Pasteur, Paris | Sylvie Behillil, Fabiana Gambaro, Etienne Simon-Lorière, Vincent Enouf, Maud Vanpeene, Sylvie van der Werf |
| EPI_ISL_593918 | Hospital, Joeuf | National Reference Center for Viruses of Respiratory Infections, Institut Pasteur, Paris | Sylvie Behillil, Fabiana Gambaro, Etienne Simon-Lorière, Vincent Enouf, Maud Vanpeene, Sylvie van der Werf |
| EPI_ISL_593919 | Hospital, Ottonville | National Reference Center for Viruses of Respiratory Infections, Institut Pasteur, Paris | Sylvie Behillil, Fabiana Gambaro, Etienne Simon-Lorière, Vincent Enouf, Maud Vanpeene, Sylvie van der Werf |
| EPI_ISL_593920 | Hospital, Metz | National Reference Center for Viruses of Respiratory Infections, Institut Pasteur, Paris | Sylvie Behillil, Fabiana Gambaro, Etienne Simon-Lorière, Vincent Enouf, Maud Vanpeene, Sylvie van der Werf |
| EPI_ISL_593921 | Hospital, Fameck | National Reference Center for Viruses of Respiratory Infections, Institut Pasteur, Paris | Sylvie Behillil, Fabiana Gambaro, Etienne Simon-Lorière, Vincent Enouf, Maud Vanpeene, Sylvie van der Werf |
| EPI_ISL_593922, EPI_ISL_593923 | Hospital, Maizières les Metz | National Reference Center for Viruses of Respiratory Infections, Institut Pasteur, Paris | Sylvie Behillil, Fabiana Gambaro, Etienne Simon-Lorière, Vincent Enouf, Maud Vanpeene, Sylvie van der Werf |
| EPI_ISL_593924, EPI_ISL_593925, EPI_ISL_593926, EPI_ISL_593927, EPI_ISL_593928, EPI_ISL_593929, EPI_ISL_593930, EPI_ISL_593933 | Labo Analyses Med, Puteaux | National Reference Center for Viruses of Respiratory Infections, Institut Pasteur, Paris | Sylvie Behillil, Fabiana Gambaro, Etienne Simon-Lorière, Vincent Enouf, Maud Vanpeene, Sylvie van der Werf |
| EPI_ISL_593934 | Sentinelles, Chanteloup-En-Brie | National Reference Center for Viruses of Respiratory Infections, Institut Pasteur, Paris | Sylvie Behillil, Fabiana Gambaro, Etienne Simon-Lorière, Vincent Enouf, Maud Vanpeene, Sylvie van der Werf |
| EPI_ISL_593935 | Sentinelles, Plessis-Trevise | National Reference Center for Viruses of Respiratory Infections, Institut Pasteur, Paris | Sylvie Behillil, Fabiana Gambaro, Etienne Simon-Lorière, Vincent Enouf, Maud Vanpeene, Sylvie van der Werf |
| EPI_ISL_593936 | Sentinelles, Fondettes | National Reference Center for Viruses of Respiratory Infections, Institut Pasteur, Paris | Sylvie Behillil, Fabiana Gambaro, Etienne Simon-Lorière, Vincent Enouf, Maud Vanpeene, Sylvie van der Werf |
| EPI_ISL_602413, EPI_ISL_602414, EPI_ISL_602415, EPI_ISL_602416 | WHO National Influenza Centre Russian Federation | WHO National Influenza Centre Russian Federation | Andrey Komissarov, Artem Fadeev, Kseniya Komissarova, Anna Ivanova, Dmitry Bazhenov, Daria Danilenko |
| EPI_ISL_602564 | Centre for Dengue Research, Department of Immunology and Molecular Medicine | Centre for Dengue Research | Chandima Jeewandara, Deshni Jayathilaka, Dinuka Ariyaratne, Diyanath Ranasinghe, Laksiri Gomes, Ananda Wijewickrama, Malika Karunaratne, Gathsaurie Neelika Malavige |

EPI_ISL_602639, EPI_ISL_602642, EPI_ISL_602646, EPI_ISL_602647, EPI_ISL_602648, EPI_ISL_602649, EPI_ISL_602651, EPI_ISL_602652, EPI_ISL_602653, EPI_ISL_602656, EPI_ISL_602657, EPI_ISL_602658, EPI_ISL_602662, EPI_ISL_602663, EPI_ISL_602664, EPI_ISL_602665, EPI_ISL_602682, EPI_ISL_602683, EPI_ISL_602684, EPI_ISL_602685, EPI_ISL_602686, EPI_ISL_602687, EPI_ISL_602688, EPI_ISL_602689, EPI_ISL_602690, EPI_ISL_602691, EPI_ISL_602692, EPI_ISL_602693, EPI_ISL_602694, EPI_ISL_602695, EPI_ISL_602696, EPI_ISL_602697, EPI_ISL_602698, EPI_ISL_602700, EPI_ISL_602701, EPI_ISL_602703, EPI_ISL_602704, EPI_ISL_602705, EPI_ISL_602706, EPI_ISL_602707, EPI_ISL_602708, EPI_ISL_602709, EPI_ISL_602711, EPI_ISL_602712, EPI_ISL_602713, EPI_ISL_602714, EPI_ISL_602715, EPI_ISL_602716, EPI_ISL_602717, EPI_ISL_602718, EPI_ISL_602719, EPI_ISL_602720, EPI_ISL_602721, EPI_ISL_602722, EPI_ISL_602723, EPI_ISL_602724, EPI_ISL_602725, EPI_ISL_602726, EPI_ISL_602727, EPI_ISL_602728, EPI_ISL_602729, EPI_ISL_602730, EPI_ISL_602731, EPI_ISL_602732, EPI_ISL_602733, EPI_ISL_602735, EPI_ISL_602736, EPI_ISL_602737, EPI_ISL_602738, EPI_ISL_602739, EPI_ISL_602740, EPI_ISL_602741, EPI_ISL_602742, EPI_ISL_602743, EPI_ISL_602744, EPI_ISL_602745, EPI_ISL_602746, EPI_ISL_602747, EPI_ISL_602748, EPI_ISL_602749, EPI_ISL_602750, EPI_ISL_602751, EPI_ISL_602752, EPI_ISL_602753, EPI_ISL_602754, EPI_ISL_602755, EPI_ISL_602756, EPI_ISL_602757, EPI_ISL_602758, EPI_ISL_602759, EPI_ISL_602760, EPI_ISL_602761, EPI_ISL_602762, EPI_ISL_602763, EPI_ISL_602764, EPI_ISL_602765, EPI_ISL_602766, EPI_ISL_602767, EPI_ISL_602768, EPI_ISL_602769, EPI_ISL_602770, EPI_ISL_602771, EPI_ISL_602772, EPI_ISL_602775, EPI_ISL_602776, EPI_ISL_602777, EPI_ISL_602778, EPI_ISL_602779, EPI_ISL_602780, EPI_ISL_602781, EPI_ISL_602782, EPI_ISL_602783, EPI_ISL_602784, EPI_ISL_602785, EPI_ISL_602786, EPI_ISL_602787, EPI_ISL_602788, EPI_ISL_602789, EPI_ISL_602790, EPI_ISL_602791, EPI_ISL_602793, EPI_ISL_602795, EPI_ISL_602796, EPI_ISL_602797, EPI_ISL_602798, EPI_ISL_602799, EPI_ISL_602800, EPI_ISL_602801, EPI_ISL_602802, EPI_ISL_602803, EPI_ISL_602804, EPI_ISL_602806, EPI_ISL_602807, EPI_ISL_602808, EPI_ISL_602809, EPI_ISL_602810, EPI_ISL_602811, EPI_ISL_602812, EPI_ISL_602813, EPI_ISL_602814, EPI_ISL_602815, EPI_ISL_602817, EPI_ISL_602818, EPI_ISL_602820, EPI_ISL_602821, EPI_ISL_602822, EPI_ISL_602824, EPI_ISL_602825, EPI_ISL_602826, EPI_ISL_602828, EPI_ISL_602831, EPI_ISL_602832, EPI_ISL_602833, EPI_ISL_602834, EPI_ISL_602835, EPI_ISL_602836, EPI_ISL_602837, EPI_ISL_602838, EPI_ISL_602839, EPI_ISL_602840, EPI_ISL_602841, EPI_ISL_602842, EPI_ISL_602843, EPI_ISL_602844, EPI_ISL_602845, EPI_ISL_602848, EPI_ISL_602849, EPI_ISL_602850, EPI_ISL_602851, EPI_ISL_602853, EPI_ISL_602854, EPI_ISL_602855, EPI_ISL_602858, EPI_ISL_602859, EPI_ISL_602860, EPI_ISL_602861, EPI_ISL_602862, EPI_ISL_602863, EPI_ISL_602864, EPI_ISL_602865, EPI_ISL_602866, EPI_ISL_602867, EPI_ISL_602869, EPI_ISL_602870, EPI_ISL_602871, EPI_ISL_602872, EPI_ISL_602873, EPI_ISL_602875, EPI_ISL_602876, EPI_ISL_602877, EPI_ISL_602878, EPI_ISL_602879, EPI_ISL_602880, EPI_ISL_602881, EPI_ISL_602883, EPI_ISL_602884, EPI_ISL_602887, EPI_ISL_602890, EPI_ISL_602892, EPI_ISL_602894, EPI_ISL_602895, EPI_ISL_602896, EPI_ISL_602899, EPI_ISL_602900, EPI_ISL_602902, EPI_ISL_602904, EPI_ISL_602906, EPI_ISL_602907, EPI_ISL_602908, EPI_ISL_602911, EPI_ISL_602912, EPI_ISL_602913, EPI_ISL_602916, EPI_ISL_602917, EPI_ISL_602919, EPI_ISL_602920, EPI_ISL_602922, EPI_ISL_602923, EPI_ISL_602924, EPI_ISL_602926, EPI_ISL_602928, EPI_ISL_602929, EPI_ISL_602930

| | | | |
|---|---|---|---|
| see above | NHLS-IALCH | KRISP, KZN Research Innovation and Sequencing Platform | Giandhari J, Pillay S, Lessells R, Mdlalose K, York D, Khan S, Tegally H, Wilkinson E, de Oliveira T |
| EPI_ISL_603221, EPI_ISL_603222 | National Institute of Laboratory Medicine and Referral Center | Genomic Research Lab, BCSIR | Abu Sayeed Mohammad Mahmud, Mohammad Samir Uzzaman, Eshrar Osman, Md. Ahashan Habib, Shahina Akter, Tanjina Akhter Banu, Md. Murshed Hasan Sarkar, Barna Goswami, Iffat Jahan, Md. Saddam Hossain, Tasnim Nafisa, Md. Maruf Ahmed Molla, Mahmuda Yeasmin, Asish Kumar Ghosh, A. K. M. Shamsuzzaman, Monira Parveen, Md. Masum Hossain Arif, Md. Salim Khan |
| EPI_ISL_603238, EPI_ISL_603239 | National Institute of Laboratory Medicine and Referral Center | Genomic Research Lab, BCSIR | Shahina Akter, Abu Sayeed Mohammad Mahmud, Mohammad Samir Uzzaman, Eshrar Osman, Md. Ahashan Habib, Tanjina Akhter Banu, Md. Murshed Hasan Sarkar, Barna Goswami, Iffat Jahan, Md. Saddam Hossain, Tasnim Nafisa, Md. Maruf Ahmed Molla, Mahmuda Yeasmin, Asish Kumar Ghosh, A. K. M. Shamsuzzaman, Monira Parveen, Md. Masum Hossain Arif, Md. Salim Khan |
| EPI_ISL_603240, EPI_ISL_603241 | National Institute of Laboratory Medicine and Referral Center | Genomic Research Lab, BCSIR | Tanjina Akhter Banu, Abu Sayeed Mohammad Mahmud, Mohammad Samir Uzzaman, Eshrar Osman, Md. Ahashan Habib, Shahina Akter, Md. Murshed Hasan Sarkar, Barna Goswami, Iffat Jahan, Md. Saddam Hossain, Tasnim Nafisa, Md. Maruf Ahmed Molla, Mahmuda Yeasmin, Asish Kumar Ghosh, A. K. M. Shamsuzzaman, Monira Parveen, Md. Masum Hossain Arif, Md. Salim Khan |
| EPI_ISL_605785, EPI_ISL_605786, EPI_ISL_605787, EPI_ISL_605788, EPI_ISL_605790 | NHLS-IALCH | KRISP, KZN Research Innovation and Sequencing Platform | Giandhari J, Pillay S, Lessells R, Mdlalose K, York D, Khan S, Tegally H, Wilkinson E, de Oliveira T |
| EPI_ISL_609994 | INMI Lazzaro Spallanzani IRCCS | INMI Lazzaro Spallanzani IRCCS | C.E.M Gruber, F Messina, M Rueca, B Bartolini, E Giombini, MR Capobianchi, A Di Caro |

| | | | |
|---|---|---|---|
| EPI_ISL_609995 | INMI Lazzaro Spallanzani IRCCS | INMI Lazzaro Spallanzani IRCCS | E Giombini, C.E.M Gruber, M Rueca, B Bartolini, F Messina, A Di Caro, MR Capobianchi |
| EPI_ISL_609996 | INMI Lazzaro Spallanzani IRCCS | INMI Lazzaro Spallanzani IRCCS | F Messina, M Rueca, B Bartolini, C.E.M Gruber, E Giombini, MR Capobianchi, A Di Caro |
| EPI_ISL_609997 | INMI Lazzaro Spallanzani IRCCS | INMI Lazzaro Spallanzani IRCCS | M Rueca, B Bartolini, C.E.M Gruber, F Messina, E Giombini, A Di Caro, MR Capobianchi |
| EPI_ISL_610152, EPI_ISL_610153, EPI_ISL_610154 | Singapore General Hospital | Department of Microbiology | Nurdyana Abdul Rahman, Kun Lee Lim, Chenhao Li, Sui Sin Goh, Kenneth Xin Long Chan, Kian Sing Chan, Lynette Oon, Kern Rei Chng, Niranjan Nagarajan, Karrie Ko |
| EPI_ISL_610161 | RSUD Dr. Tjitrowardojo | Genetics Working Group (Pokja Genetik) Faculty of Medicine, Public Health and Nursing Universitas Gadjah Mada (FK-KMK UGM); Disease Investigation Center Wates Ministry of Agriculture Indonesia; Department of Microbiology FK-KMK UGM; Laboratorium Diagnostik Yayasan Tahija World Mosquito Program (WMP) Yogyakarta Center for Tropical Medicine FK-KMK UGM; Integrated Research Center FK-KMK UGM; Department of Computer Science and Electronics FMIPA UGM; Balai Besar Teknik Kesehatan Lingkungan dan Pengendalian Penyakit (BBTKLPP) Yogyakarta | Gunadi, Hendra Wibawa, Marcellus, Mohamad S. Hakim, Edwin W. Daniwijaya, Ludhang P. Rizki, Endah Supriyati, Eggi Arguni, Titik Nuryastuti, Tri Wibawa, Dwi AA Nugrahaningsih, Afiahayati, Siswanto, Kristy Iskandar, Nungki Anggorowati, Irene, Indaryati, Havid Setyawan, Wuryanto, Kemala Athollah, Desyifa Mursalin |
| EPI_ISL_613549, EPI_ISL_613551, EPI_ISL_613552, EPI_ISL_613553, EPI_ISL_613554, EPI_ISL_613555, EPI_ISL_613556, EPI_ISL_613557, EPI_ISL_613558, EPI_ISL_613559 | CHRU Pontchaillou - Laboratoire de Virologie 2, rue Henri Le Guilloux | National Reference Center for Viruses of Respiratory Infections, Institut Pasteur, Paris | Marion Barbet, Sylvie Behillil, Méline Bizard, Angela Brisebarre, Camille Capel, Etienne Simon-Lorière, Vincent Enouf, Maud Vanpeene, Sylvie van der Werf, Gisèle Lagathu |
| EPI_ISL_614282, EPI_ISL_614283, EPI_ISL_614284, EPI_ISL_614285, EPI_ISL_614286, EPI_ISL_614287, EPI_ISL_614288, EPI_ISL_614289, EPI_ISL_614290, EPI_ISL_614291, EPI_ISL_614292, EPI_ISL_614293 | | | |
| see above | General practitioner | National Reference Center for Viruses of Respiratory Infections, Institut Pasteur, Paris | Marion Barbet, Sylvie Behillil, Méline Bizard, Angela Brisebarre, Camille Capel, Etienne Simon-Lorière, Vincent Enouf, Maud Vanpeene, Sylvie van der Werf |
| EPI_ISL_614298, EPI_ISL_614304 | Faroese National Reference Laboratory for Fish and Animal Diseases | Faroese National Reference Laboratory for Fish and Animal Diseases | Maria Marjunardóttir Dahl, Petra Elisabeth Petersen, Debes Hammershaimb Christiansen |
| EPI_ISL_623096, EPI_ISL_623097 | General practitioner | National Reference Center for Viruses of Respiratory Infections, Institut Pasteur, Paris | Marion Barbet, Sylvie Behillil, Méline Bizard, Angela Brisebarre, Camille Capel, Etienne Simon-Lorière, Vincent Enouf, Maud Vanpeene, Sylvie van der Werf |
| EPI_ISL_626571, EPI_ISL_626572, EPI_ISL_626573, EPI_ISL_626575, EPI_ISL_626577, EPI_ISL_626578, EPI_ISL_626579, EPI_ISL_626580, EPI_ISL_626581, EPI_ISL_626582, EPI_ISL_626583, EPI_ISL_626585, EPI_ISL_626586, EPI_ISL_626587, EPI_ISL_626588, EPI_ISL_626590, EPI_ISL_626591, EPI_ISL_626592, EPI_ISL_626594, EPI_ISL_626595, EPI_ISL_626596, EPI_ISL_626597, EPI_ISL_626598, EPI_ISL_626599, EPI_ISL_626600, EPI_ISL_626601, EPI_ISL_626602, EPI_ISL_626603, EPI_ISL_626604, EPI_ISL_626605, EPI_ISL_626606, EPI_ISL_626608, EPI_ISL_626609, EPI_ISL_626612, EPI_ISL_626613, EPI_ISL_626614, EPI_ISL_626615, EPI_ISL_626616, EPI_ISL_626617, EPI_ISL_626618, EPI_ISL_626619, EPI_ISL_626620, EPI_ISL_626621, EPI_ISL_626623, EPI_ISL_626624, EPI_ISL_626625 | | | |
| see above | The National Institute of Public Health | State Veterinary Institute Prague | Nagy,A;Jirincova,H;Novakova,L;Trnka,D;Vecerova,J |
| EPI_ISL_632310 | 1-Laboratory of Microbiology, National Reference Lab, Charles Nicolle Hospital; 2-University of Tunis ElManar, Faculty of Medicine of Tunis, LR99ES09, Tunis, Tunisia | 1-Clinical and Experimental Pharmacology Lab, LR16SP02, National Center of Pharmacovigilance, University of Tunis El Manar, Tunis, Tunisia. 2-Neurodegenerative diseases and psychiatric troubles, LR18SP03, Razi Hospital, University of Tunis El Manar, Tunis, Tunisia. 3- Ministry of Health, National Observatory of New and Emerging Diseases, 1006, Tunis, Tunisia | Ilhem Boutiba-Ben Boubaker, Sameh Trabelsi, Nissaf Ben Alaya, Maher Kharrat, Alia Ben Kahla, Jalila Ben Khelil, Salma Abid, Sana Ferjani, Mouna Ben Sassi, Mouna Safer, Imen Mkada, Imen Kacem, Gaies Emna, Soumaya Rammeh, Riadh Daghfous, Riadh Gouider. |
| EPI_ISL_632934 | Department of Acute Infectious Diseases Control and Prevention,Yunnan Provincial Center for Disease Control and Prevention | Department of Acute Infectious Diseases Control and Prevention,Yunnan Provincial Center for Disease Control and Prevention | Meiling Zhang,Jienan Zhou,Senquan Jia,Xiaonan Zhao,Xiaoqing Fu |
| EPI_ISL_632937 | RSUD Saptosari Gunung Kidul | Genetics Working Group (Pokja Genetik) Faculty of Medicine, Public Health and Nursing Universitas Gadjah Mada (FK-KMK UGM); Disease Investigation Center Wates Ministry of Agriculture Indonesia; Department of Microbiology FK-KMK UGM; Laboratorium Diagnostik Yayasan Tahija World Mosquito Program (WMP) Yogyakarta Center for Tropical Medicine FK-KMK UGM; Integrated Research Center FK-KMK UGM; Department of Computer Science and Electronics FMIPA UGM; Balai Besar Teknik Kesehatan Lingkungan dan Pengendalian Penyakit (BBTKLPP) Yogyakarta | Gunadi, Hendra Wibawa, Marcellus, Mohamad S. Hakim, Edwin W. Daniwijaya, Ludhang P. Rizki, Endah Supriyati, Eggi Arguni, Titik Nuryastuti, Tri Wibawa, Dwi AA Nugrahaningsih, Afiahayati, Siswanto, Kristy Iskandar, Nungki Anggorowati, Irene, Indaryati, Havid Setyawan, Eko Darmawan, Maria Patricia Inggriani, Audric Kenny Tedja |
| EPI_ISL_634977 | 1-Laboratory of Microbiology, National Reference Lab, Charles Nicolle Hospital; 2-University of Tunis ElManar, Faculty of Medicine of Tunis, LR99ES09, Tunis, Tunisia | 1-Clinical and Experimental Pharmacology Lab, LR16SP02, National Center of Pharmacovigilance, University of Tunis El Manar, Tunis, Tunisia. 2-Neurodegenerative diseases and psychiatric troubles, LR18SP03, Razi Hospital, University of Tunis El Manar, Tunis, Tunisia. 3- Ministry of Health, National Observatory of New and Emerging Diseases, 1006, Tunis, Tunisia | Ilhem Boutiba-Ben Boubaker, Sameh Trabelsi, Nissaf Ben Alaya, Maher Kharrat, Alia Ben Kahla, Jalila Ben Khelil, Salma Abid, Sana Ferjani, Mouna Ben Sassi, Mouna Safer, Imen Mkada, Imen Kacem, Gaies Emna, Soumaya Rammeh, Riadh Daghfous, Riadh Gouider. |
| EPI_ISL_634978, EPI_ISL_634979, EPI_ISL_634981, EPI_ISL_634983, EPI_ISL_634984, EPI_ISL_634985, EPI_ISL_634986, EPI_ISL_634987, EPI_ISL_634989, EPI_ISL_634990, EPI_ISL_634993, EPI_ISL_634994, EPI_ISL_634995, EPI_ISL_634996, EPI_ISL_634997, EPI_ISL_634998, EPI_ISL_634999, EPI_ISL_635000, EPI_ISL_635001, EPI_ISL_635002, EPI_ISL_635003, EPI_ISL_635004, EPI_ISL_635005, EPI_ISL_635006, EPI_ISL_635007, EPI_ISL_635008, EPI_ISL_635009, EPI_ISL_635010, EPI_ISL_635011, EPI_ISL_635012, EPI_ISL_635013, EPI_ISL_635014, EPI_ISL_635015, EPI_ISL_635016, EPI_ISL_635017, EPI_ISL_635018, EPI_ISL_635019, EPI_ISL_635021, EPI_ISL_635022, EPI_ISL_635024, EPI_ISL_635025, EPI_ISL_635026, EPI_ISL_635027, EPI_ISL_635028, EPI_ISL_635029, EPI_ISL_635030, EPI_ISL_635031, EPI_ISL_635032, EPI_ISL_635033, EPI_ISL_635034, EPI_ISL_635035, EPI_ISL_635036, EPI_ISL_635037, EPI_ISL_635038, EPI_ISL_635039, EPI_ISL_635040, EPI_ISL_635041, EPI_ISL_635043, EPI_ISL_635045, EPI_ISL_635046, EPI_ISL_635047, EPI_ISL_635048, EPI_ISL_635049, EPI_ISL_635050, EPI_ISL_635051, EPI_ISL_635052, EPI_ISL_635053, EPI_ISL_635054, EPI_ISL_635055, EPI_ISL_635056, EPI_ISL_635057, EPI_ISL_635058 | | | |
| see above | National Health Laboratory Service - Inkosi Albert Luthuli Central Hospital (NHLS-IALCH) | KRISP, KZN Research Innovation and Sequencing Platform | Giandhari J, Pillay S, Lessells R, Mdlalose K, York D, Khan S, Tegally H, Wilkinson E, de Oliveira T |
| EPI_ISL_635061, EPI_ISL_635062 | 1-Laboratory of Microbiology, National Reference Lab, Charles Nicolle Hospital; 2-University of Tunis ElManar, Faculty of Medicine of Tunis, LR99ES09, Tunis, Tunisia | 1-Clinical and Experimental Pharmacology Lab, LR16SP02, National Center of Pharmacovigilance, University of Tunis El Manar, Tunis, Tunisia. 2-Neurodegenerative diseases and psychiatric troubles, LR18SP03, Razi Hospital, University of Tunis El Manar, Tunis, Tunisia. 3- Ministry of Health, National Observatory of New and Emerging Diseases, 1006, Tunis, Tunisia | Ilhem Boutiba-Ben Boubaker, Sameh Trabelsi, Nissaf Ben Alaya, Maher Kharrat, Alia Ben Kahla, Jalila Ben Khelil, Salma Abid, Sana Ferjani, Mouna Ben Sassi, Mouna Safer, Imen Mkada, Imen Kacem, Gaies Emna, Soumaya Rammeh, Riadh Daghfous, Riadh Gouider. |
| EPI_ISL_644672 | Institute for Medical Research, Infectious Disease Research Centre, National Institutes of Health, Ministry of Health Malaysia | Institute for Medical Research, Infectious Disease Research Centre, National Institutes of Health, Ministry of Health Malaysia | Suppiah J, Kamel K, Mohd-Zawawi Z, Thayan R |
| EPI_ISL_653754 | Instituto Nacional de Salud, Bogotá, Colombia | Instituto Nacional de Salud, Bogotá, Colombia | Katherine Laiton-Donato, Diego A. Álvarez-Díaz, Carlos Franco-Muñoz, Mauricio Pacheco-Montealegre, Jonathan Reales, Diego Andrés Prada, Jose A. |

| | | | |
|---|---|---|---|
| | | | Usme-Ciro, Zulma M. Cucunubá, Christian Julian VillabonaArenas, Liz Villabona-Arenas, Sussy Echeverria, Astrid C. Flórez, Carolina Ferro, Diana Marcela Walteros-Acero, Franklin Prieto, Carlos Andrés Durán, Martha Lucia Ospina Martinez, Marcela Mercado-Reyes |
| EPI_ISL_653765, EPI_ISL_653766, EPI_ISL_653767, EPI_ISL_653768, EPI_ISL_653769, EPI_ISL_653770, EPI_ISL_653771, EPI_ISL_653772, EPI_ISL_653773, EPI_ISL_653774, EPI_ISL_653775, EPI_ISL_653776, EPI_ISL_653777, EPI_ISL_653778, EPI_ISL_653779, EPI_ISL_653780, EPI_ISL_653781, EPI_ISL_653782, EPI_ISL_653783 | | | |
| see above | I.R.C.C.S. "S. De Bellis" - Ente Ospedaliero | Istituto Zooprofilattico Sperimentale della Puglia e della Basilicata | Parisi A., Bianco A., Capozzi L., Del Sambro L., Lippolis A., Notarnicola M., Manzulli V, Rondinone V., Pace L. |
| EPI_ISL_653784, EPI_ISL_653785, EPI_ISL_653786 | Istituto Zooprofilattico Sperimentale della Puglia e della Basilicata | Istituto Zooprofilattico Sperimentale della Puglia e della Basilicata | Parisi A., Bianco A., Capozzi L., Del Sambro L., Manzulli V, Rondinone V., Pace L., Cipolletta D., Galante D. |
| EPI_ISL_653787, EPI_ISL_653788, EPI_ISL_653789, EPI_ISL_653790, EPI_ISL_653791, EPI_ISL_653792, EPI_ISL_653793 | I.R.C.C.S. "S. De Bellis" - Ente Ospedaliero | Istituto Zooprofilattico Sperimentale della Puglia e della Basilicata | Parisi A., Bianco A., Capozzi L., Del Sambro L., Lippolis A., Notarnicola M., Manzulli V, Rondinone V., Pace L. |
| EPI_ISL_653794, EPI_ISL_653795, EPI_ISL_653796, EPI_ISL_653797, EPI_ISL_653798, EPI_ISL_653799, EPI_ISL_653800, EPI_ISL_653801, EPI_ISL_653802, EPI_ISL_653803, EPI_ISL_653804, EPI_ISL_653805, EPI_ISL_653806, EPI_ISL_653807, EPI_ISL_653808, EPI_ISL_653809, EPI_ISL_653810, EPI_ISL_653811, EPI_ISL_653812, EPI_ISL_653819, EPI_ISL_653820, EPI_ISL_653821, EPI_ISL_653822 | | | |
| see above | I.R.C.C.S. "S. De Bellis" - Ente Ospedaliero | Istituto Zooprofilattico Sperimentale della Puglia e della Basilicata | Parisi A., Bianco A., Capozzi L., Del Sambro L., Lippolis A., Notarnicola M., Cipolletta D., Galante D. |
| EPI_ISL_653916 | Diagnostic- and Research Institute of Pathology, Medical University of Graz | Diagnostic- and Research Institute of Pathology, Medical University of Graz | Karl Kashofer, Peter Regitnig, Martin Zacharias, Gregor Gorkiewicz |
| EPI_ISL_654017, EPI_ISL_654018, EPI_ISL_654019, EPI_ISL_654020 | Laboratory of Microbiology, National Reference Lab, Charles Nicolle Hospital; 2-University of Tunis ElManar, Faculty of Medicine of Tunis, LR99ES09, Tunis, Tunisia | 1-Clinical and Experimental Pharmacology Lab, LR16SP02, National Center of Pharmacovigilance, University of Tunis El Manar, Tunis, Tunisia. 2-Neurodegenerative diseases and psychiatric troubles, LR18SP03, Razi Hospital, University of Tunis El Manar, Tunis, Tunisia. 3- Ministry of Health, National Observatory of New and Emerging Diseases, 1006, Tunis, Tunisia | Ilhem Boutiba-Ben Boubaker, Sameh Trabelsi, Nissaf Ben Alaya, Maher Kharrat, Alia Ben Kahla, Jalila Ben Khelil, Salma Abid, Sana Ferjani, Mouna Ben Sassi, Mouna Safer, Imen Kacem, Gaies Emna, Soumaya Rammeh, Riadh Daghfous, Riadh Gouider. |
| EPI_ISL_658863, EPI_ISL_658865, EPI_ISL_658868, EPI_ISL_658870 | Instituto de Diagnostico y Referencia Epidemiologicos (INDRE) | Instituto de Diagnostico y Referencia Epidemiologicos (INDRE) | Gisela Barrera-Badillo , Abril Rodriguez-Maldonado, Claudia Wong-Arambula , Natividad Cruz-Ortiz, Tatiana Nunez-Garcia, Dayanira Arellano-Suarez, Fabiola Garces-Ayala, Lucia Hernandez-Rivas, Irma Lopez-Martinez, Ernesto Ramirez-Gonzalez. |
| EPI_ISL_658873, EPI_ISL_658875 | Instituto de Diagnostico y Referencia Epidemiologicos (INDRE) | Instituto de Diagnostico y Referencia Epidemiologicos (INDRE) | Gisela Barrera-Badillo , Abril Rodriguez-Maldonado, Claudia Wong-Arambula , Natividad Cruz-Ortiz, Tatiana Nunez-Garcia, Dayanira Arellano-Suarez, Fabiola Garces-Ayala, Lucia Hernandez-Rivas, Irma Lopez-Martinez, Ernesto Ramirez-Gonzalez. |
| EPI_ISL_658878, EPI_ISL_658880, EPI_ISL_658883, EPI_ISL_658886, EPI_ISL_658888, EPI_ISL_658891, EPI_ISL_658893, EPI_ISL_658901 | Instituto de Diagnostico y Referencia Epidemiologicos (INDRE) | Instituto de Diagnostico y Referencia Epidemiologicos (INDRE) | Ernesto Ramirez-Gonzalez, Abril Rodriguez-Maldonado, Claudia Wong-Arambula , Natividad Cruz-Ortiz, Tatiana Nunez-Garcia, Dayanira Arellano-Suarez, Fabiola Garces-Ayala, Lucia Hernandez-Rivas, Irma Lopez-Martinez, Gisela Barrera-Badillo. |
| EPI_ISL_660068 | Instituto de Diagnostico y Referencia Epidemiologicos (INDRE) | Instituto de diagnóstico y Referencia Epidemiologicos (INDRE) Departamento de Virología | Gisela Barrera-Badillo , Abril Rodriguez-Maldonado, Claudia Wong-Arambula , Natividad Cruz-Ortiz, Tatiana Nunez-Garcia, Dayanira Arellano-Suarez, Fabiola Garces-Ayala, Lucia Hernandez-Rivas, Irma Lopez-Martinez, Ernesto Ramirez-Gonzalez. |
| EPI_ISL_660070 | Zurita & Zurita Laboratorios | Zurita & Zurita Laboratorios | Gabriela Sevillano Camilo Zurita-Salinas Karen Loaiza David Ortega-Paredes Jeannete Zurita |
| EPI_ISL_660121, EPI_ISL_660122, EPI_ISL_660124, EPI_ISL_660127 | Ampath | National Health Laboratory Service (NHLS), Tygerberg | Susan Engelbrecht, Draper C, Davis M-A, Siegfried N, Williamson C, Hsiao M, Kayla Delaney, Bronwyn Kleinhans, Houriiyah Tegally, Eduan Wilkindon, Gert van Zyl, Wolfgang Preiser, Tulio de Oliveira |
| EPI_ISL_660139, EPI_ISL_660141 | Hamadi | National Health Laboratory Service (NHLS), Tygerberg | Susan Engelbrecht, Draper C, Davis M-A, Siegfried N, Williamson C, Hsiao M, Kayla Delaney, Bronwyn Kleinhans, Houriiyah Tegally, Eduan Wilkindon, Gert van Zyl, Wolfgang Preiser, Tulio de Oliveira |
| EPI_ISL_660143, EPI_ISL_660144, EPI_ISL_660145, EPI_ISL_660146, EPI_ISL_660148, EPI_ISL_660150, EPI_ISL_660152, EPI_ISL_660154, EPI_ISL_660157, EPI_ISL_660158 | PathCare | National Health Laboratory Service (NHLS), Tygerberg | Susan Engelbrecht, Draper C, Davis M-A, Siegfried N, Williamson C, Hsiao M, Kayla Delaney, Bronwyn Kleinhans, Houriiyah Tegally, Eduan Wilkindon, Gert van Zyl, Wolfgang Preiser, Tulio de Oliveira |
| EPI_ISL_660159, EPI_ISL_660160, EPI_ISL_660161, EPI_ISL_660162, EPI_ISL_660163, EPI_ISL_660164 | NHLS-IALCH | KRISP, KZN Research Innovation and Sequencing Platform | Giandhari J, Pillay S, Lessells R, Mdlalose K, York D, Khan S, Tegally H, Wilkinson E, de Oliveira T |
| EPI_ISL_660185 | NHLS-IALCH | KRISP, KZN Research Innovation and Sequencing Platform | Gazy I, Sigal A, Karim F, Cele S, Giandhari J, Pillay S, Tegally H, Wilkinson E, de Oliveira T |
| EPI_ISL_660189, EPI_ISL_660190, EPI_ISL_660191, EPI_ISL_660192, EPI_ISL_660193, EPI_ISL_660194, EPI_ISL_660195, EPI_ISL_660196, EPI_ISL_660197, EPI_ISL_660198, EPI_ISL_660199, EPI_ISL_660200, EPI_ISL_660201, EPI_ISL_660202, EPI_ISL_660203, EPI_ISL_660204, EPI_ISL_660205, EPI_ISL_660206, EPI_ISL_660207, EPI_ISL_660208, EPI_ISL_660209, EPI_ISL_660210, EPI_ISL_660211, EPI_ISL_660212, EPI_ISL_660213, EPI_ISL_660214, EPI_ISL_660215, EPI_ISL_660216, EPI_ISL_660217, EPI_ISL_660218, EPI_ISL_660219, EPI_ISL_660220 | | | |
| see above | NHLS-IALCH | KRISP, KZN Research Innovation and Sequencing Platform | Giandhari J, Pillay S, Lessells R, Mdlalose K, York D, Khan S, Tegally H, Wilkinson E, de Oliveira T |
| EPI_ISL_660221, EPI_ISL_660222 | KRISP, KZN Research Innovation and Sequencing Platform | KRISP, KZN Research Innovation and Sequencing Platform | Giandhari J, Pillay S, Lessells R, Mdlalose K, York D, Khan S, Tegally H, Wilkinson E, de Oliveira T |
| EPI_ISL_660223, EPI_ISL_660224, EPI_ISL_660225, EPI_ISL_660227 | NHLS-IALCH | KRISP, KZN Research Innovation and Sequencing Platform | Giandhari J, Pillay S, Lessells R, Mdlalose K, York D, Khan S, Tegally H, Wilkinson E, de Oliveira T |
| EPI_ISL_660228, EPI_ISL_660229, EPI_ISL_660230, EPI_ISL_660231, EPI_ISL_660232, EPI_ISL_660233 | KRISP, KZN Research Innovation and Sequencing Platform | KRISP, KZN Research Innovation and Sequencing Platform | Giandhari J, Pillay S, Lessells R, Mdlalose K, York D, Khan S, Tegally H, Wilkinson E, de Oliveira T |
| EPI_ISL_660234, EPI_ISL_660235, EPI_ISL_660236, EPI_ISL_660237, EPI_ISL_660238, EPI_ISL_660239, EPI_ISL_660240, EPI_ISL_660241, EPI_ISL_660242, EPI_ISL_660243, EPI_ISL_660244, EPI_ISL_660245, EPI_ISL_660246, EPI_ISL_660248, EPI_ISL_660249, EPI_ISL_660250, EPI_ISL_660251, EPI_ISL_660252, EPI_ISL_660253, EPI_ISL_660254, EPI_ISL_660255 | | | |
| see above | NHLS-IALCH | KRISP, KZN Research Innovation and Sequencing Platform | Giandhari J, Pillay S, Lessells R, Mdlalose K, York D, Khan S, Tegally H, Wilkinson E, de Oliveira T |
| EPI_ISL_660256, EPI_ISL_660257, EPI_ISL_660258 | KRISP, KZN Research Innovation and Sequencing Platform | KRISP, KZN Research Innovation and Sequencing Platform | Giandhari J, Pillay S, Lessells R, Mdlalose K, York D, Khan S, Tegally H, Wilkinson E, de Oliveira T |
| EPI_ISL_660259, EPI_ISL_660261, EPI_ISL_660262, EPI_ISL_660263 | Molecualr Diagnostic Services (MDS) | KRISP, KZN Research Innovation and Sequencing Platform | Giandhari J, Pillay S, Lessells R, Mdlalose K, York D, Khan S, Tegally H, Wilkinson E, de Oliveira T |
| EPI_ISL_660323 | Hospital | National Reference Center for Viruses of Respiratory Infections, Institut Pasteur, Paris | Marion Barbet, Sylvie Behillil, Méline Bizard, Angela Brisebarre, Camille Capel, Etienne Simon-Lorière, Vincent Enouf, Maud Vanpeene, Sylvie van der Werf, Alexandra Ducancelle |
| EPI_ISL_660324, EPI_ISL_660325 | General practitioner | National Reference Center for Viruses of Respiratory Infections, Institut Pasteur, Paris | Marion Barbet, Sylvie Behillil, Méline Bizard, Angela Brisebarre, Camille Capel, Etienne Simon-Lorière, Vincent Enouf, Maud Vanpeene, Sylvie van der Werf |
| EPI_ISL_660553, EPI_ISL_660554, EPI_ISL_660555, EPI_ISL_660556, EPI_ISL_660557, EPI_ISL_660558, EPI_ISL_660559, EPI_ISL_660560, EPI_ISL_660561, EPI_ISL_660562, EPI_ISL_660563, EPI_ISL_660564, EPI_ISL_660565, EPI_ISL_660566, EPI_ISL_660567, EPI_ISL_660568, EPI_ISL_660569, EPI_ISL_660570, EPI_ISL_660571, EPI_ISL_660572, EPI_ISL_660573 | | | |
| see above | The National Institute of Public Health | State Veterinary Institute Prague | Nagy,A;Jirincova,H;Novakova,L;Trnka,D;Vecerova,J |

| | | | |
|---|---|---|---|
| EPI_ISL_660605, EPI_ISL_660606, EPI_ISL_660609, EPI_ISL_660610, EPI_ISL_660611, EPI_ISL_660613, EPI_ISL_660614, EPI_ISL_660615, EPI_ISL_660618, EPI_ISL_660619, EPI_ISL_660620, EPI_ISL_660621, EPI_ISL_660622, EPI_ISL_660623, EPI_ISL_660624, EPI_ISL_660625, EPI_ISL_660626, EPI_ISL_660627, EPI_ISL_660628, EPI_ISL_660629, EPI_ISL_660630, EPI_ISL_660631, EPI_ISL_660632, EPI_ISL_660633, EPI_ISL_660634, EPI_ISL_660635, EPI_ISL_660636, EPI_ISL_660637, EPI_ISL_660638, EPI_ISL_660639, EPI_ISL_660640, EPI_ISL_660641, EPI_ISL_660642, EPI_ISL_660643, EPI_ISL_660644, EPI_ISL_660645, EPI_ISL_660646, EPI_ISL_660649, EPI_ISL_660650, EPI_ISL_660651, EPI_ISL_660652, EPI_ISL_660654, EPI_ISL_660655, EPI_ISL_660657, EPI_ISL_660659, EPI_ISL_660661, EPI_ISL_660662, EPI_ISL_660663 | | | |
| see above | NHLS-IALCH | KRISP, KZN Research Innovation and Sequencing Platform | Giandhari J, Pillay S, Lessells R, Mdlalose K, York D, Khan S, Tegally H, Wilkinson E, de Oliveira T |
| EPI_ISL_668454 | Centre for Dengue Research, Department of Immunology and Molecular Medicine | Centre for Dengue Research, Department of Immunology and Molecular Medicine | Chandima Jeewandara, Deshni Jayathilaka, Dinuka Ariyaratne, Diyanath Ranasinghe, Laksiri Gomes, Gathsaurie Neelika Malavige |
| EPI_ISL_671414, EPI_ISL_671415, EPI_ISL_671416, EPI_ISL_671417, EPI_ISL_671418 | National Laboratory of Virology, Szentágothai Research Centre | National Laboratory of Virology, Szentágothai Research Centre | Endre Gábor Tóth, Balázs Somogyi, Brigitta, Ferenc Jakab, Gábor Kemenesi |
| EPI_ISL_671428, EPI_ISL_671430, EPI_ISL_671431, EPI_ISL_671432, EPI_ISL_671433, EPI_ISL_671434, EPI_ISL_671435, EPI_ISL_671436, EPI_ISL_671438, EPI_ISL_671440, EPI_ISL_671441, EPI_ISL_671443, EPI_ISL_671444, EPI_ISL_671445, EPI_ISL_671447, EPI_ISL_671448, EPI_ISL_671449, EPI_ISL_671451, EPI_ISL_671452, EPI_ISL_671453, EPI_ISL_671454, EPI_ISL_671455, EPI_ISL_671456, EPI_ISL_671457, EPI_ISL_671458, EPI_ISL_671459, EPI_ISL_671460, EPI_ISL_671461, EPI_ISL_671462, EPI_ISL_671463, EPI_ISL_671464, EPI_ISL_671465, EPI_ISL_671466, EPI_ISL_671467, EPI_ISL_671468, EPI_ISL_671469, EPI_ISL_671470, EPI_ISL_671471, EPI_ISL_671472, EPI_ISL_671473, EPI_ISL_671474, EPI_ISL_671475, EPI_ISL_671476, EPI_ISL_671477, EPI_ISL_671478, EPI_ISL_671479, EPI_ISL_671480, EPI_ISL_671481, EPI_ISL_671482, EPI_ISL_671483, EPI_ISL_671484, EPI_ISL_671485, EPI_ISL_671486, EPI_ISL_671487, EPI_ISL_671488, EPI_ISL_671489, EPI_ISL_671490, EPI_ISL_671491, EPI_ISL_671492, EPI_ISL_671493, EPI_ISL_671494, EPI_ISL_671495 | | | |
| see above | University of Debrecen, Department of Medical Microbiology | National Laboratory of Virology, Szentágothai Research Centre | Endre Gábor Tóth, Balázs Somogyi, Brigitta Zana, Eszter Csoma, Ferenc Jakab, Gábor Kemenesi |
| EPI_ISL_677699 | Vanda Pharmaceuticals Clinical Site | Vanda Pharmaceuticals | Vanda Pharmaceuticals |
| EPI_ISL_677727, EPI_ISL_677739, EPI_ISL_677740, EPI_ISL_677741, EPI_ISL_677742, EPI_ISL_677743, EPI_ISL_677744, EPI_ISL_677745, EPI_ISL_677746, EPI_ISL_677748, EPI_ISL_677749, EPI_ISL_677750, EPI_ISL_677751, EPI_ISL_677752, EPI_ISL_677753, EPI_ISL_677754, EPI_ISL_677755, EPI_ISL_677756, EPI_ISL_677757, EPI_ISL_677758, EPI_ISL_677759, EPI_ISL_677760, EPI_ISL_677761, EPI_ISL_677762, EPI_ISL_677763, EPI_ISL_677764, EPI_ISL_677765, EPI_ISL_677766, EPI_ISL_677767, EPI_ISL_677768, EPI_ISL_677770, EPI_ISL_677771, EPI_ISL_677773, EPI_ISL_677774, EPI_ISL_677775, EPI_ISL_677776, EPI_ISL_677777, EPI_ISL_677780, EPI_ISL_677781, EPI_ISL_677782, EPI_ISL_677783, EPI_ISL_677784, EPI_ISL_677785, EPI_ISL_677786, EPI_ISL_677787, EPI_ISL_677788, EPI_ISL_677789, EPI_ISL_677790, EPI_ISL_677791, EPI_ISL_677792, EPI_ISL_677793, EPI_ISL_677794, EPI_ISL_677795, EPI_ISL_677796, EPI_ISL_677797, EPI_ISL_677798, EPI_ISL_677799, EPI_ISL_677800, EPI_ISL_677801, EPI_ISL_677802, EPI_ISL_677805, EPI_ISL_677806, EPI_ISL_677808, EPI_ISL_677809, EPI_ISL_677810, EPI_ISL_677811, EPI_ISL_677812, EPI_ISL_677813, EPI_ISL_677814, EPI_ISL_677815, EPI_ISL_677816, EPI_ISL_677817, EPI_ISL_677818, EPI_ISL_677819, EPI_ISL_678290, EPI_ISL_678291 | | | |
| see above | University of Szeged, Institute of Clinical Microbiology | National Laboratory of Virology, Szentágothai Research Centre | Endre Gábor Tóth, Balázs Somogyi, Brigitta, Gabriella Terhes, Ferenc Jakab, Gábor Kemenesi |
| EPI_ISL_678387, EPI_ISL_678388, EPI_ISL_678389, EPI_ISL_678390, EPI_ISL_678391, EPI_ISL_678392, EPI_ISL_678393, EPI_ISL_678394, EPI_ISL_678395, EPI_ISL_678396 | University of Debrecen, Department of Medical Microbiology | National Laboratory of Virology, Szentágothai Research Centre | Endre Gábor Tóth, Balázs Somogyi, Brigitta Zana, Eszter Csoma, Ferenc Jakab, Gábor Kemenesi |
| EPI_ISL_686214, EPI_ISL_686217, EPI_ISL_686218, EPI_ISL_686219, EPI_ISL_686220, EPI_ISL_686221, EPI_ISL_686222, EPI_ISL_686229, EPI_ISL_686230, EPI_ISL_686231, EPI_ISL_686232, EPI_ISL_686233, EPI_ISL_686234, EPI_ISL_686235, EPI_ISL_686236, EPI_ISL_686237, EPI_ISL_686238, EPI_ISL_686239, EPI_ISL_686240, EPI_ISL_686241, EPI_ISL_686242, EPI_ISL_686243, EPI_ISL_686244, EPI_ISL_686245, EPI_ISL_686246, EPI_ISL_686247, EPI_ISL_686248, EPI_ISL_686249, EPI_ISL_686250, EPI_ISL_686251, EPI_ISL_686252, EPI_ISL_686253, EPI_ISL_686254, EPI_ISL_686255, EPI_ISL_686256, EPI_ISL_686257, EPI_ISL_686258, EPI_ISL_686259, EPI_ISL_686260, EPI_ISL_686261, EPI_ISL_686262, EPI_ISL_686263, EPI_ISL_686264, EPI_ISL_686265, EPI_ISL_686267, EPI_ISL_686268, EPI_ISL_686269, EPI_ISL_686270, EPI_ISL_686271, EPI_ISL_686272, EPI_ISL_686273, EPI_ISL_686274, EPI_ISL_686275, EPI_ISL_686276, EPI_ISL_686277, EPI_ISL_686278, EPI_ISL_686279, EPI_ISL_686280, EPI_ISL_686281, EPI_ISL_686282, EPI_ISL_686336, EPI_ISL_686337, EPI_ISL_686338, EPI_ISL_686339, EPI_ISL_686340, EPI_ISL_686341, EPI_ISL_686342, EPI_ISL_686343, EPI_ISL_686344, EPI_ISL_686345, EPI_ISL_686346, EPI_ISL_686347, EPI_ISL_686348, EPI_ISL_686349, EPI_ISL_686350, EPI_ISL_686351, EPI_ISL_686353, EPI_ISL_686354, EPI_ISL_686355, EPI_ISL_686356, EPI_ISL_686357, EPI_ISL_686358, EPI_ISL_686359, EPI_ISL_686360, EPI_ISL_686361, EPI_ISL_686362, EPI_ISL_686363, EPI_ISL_686364, EPI_ISL_686365, EPI_ISL_686366, EPI_ISL_686409, EPI_ISL_686416, EPI_ISL_686417, EPI_ISL_686448, EPI_ISL_686449, EPI_ISL_686450, EPI_ISL_686451, EPI_ISL_686452, EPI_ISL_686453, EPI_ISL_686454, EPI_ISL_686455, EPI_ISL_686456, EPI_ISL_686457, EPI_ISL_686458, EPI_ISL_686459, EPI_ISL_686460, EPI_ISL_686461, EPI_ISL_686462, EPI_ISL_686463, EPI_ISL_686464, EPI_ISL_686465, EPI_ISL_686466, EPI_ISL_686467, EPI_ISL_686468, EPI_ISL_686494, EPI_ISL_686495, EPI_ISL_686496, EPI_ISL_686497, EPI_ISL_686498, EPI_ISL_686499, EPI_ISL_686500, EPI_ISL_686501, EPI_ISL_686502, EPI_ISL_686503, EPI_ISL_686504, EPI_ISL_686505, EPI_ISL_686506, EPI_ISL_686507, EPI_ISL_686508, EPI_ISL_686509, EPI_ISL_686510, EPI_ISL_686511, EPI_ISL_686512, EPI_ISL_686513, EPI_ISL_686514, EPI_ISL_686515, EPI_ISL_686516, EPI_ISL_686517, EPI_ISL_686518, EPI_ISL_686519, EPI_ISL_686520 | | | |
| see above | Pathogen Genomics Center, National Institute of Infectious Diseases | Pathogen Genomics Center, National Institute of Infectious Diseases | Tsuyoshi Sekizuka, Kentaro Itokawa, Rina Tanaka, Masanori Hashino, Makoto Kuroda |
| EPI_ISL_686522, EPI_ISL_686523, EPI_ISL_686524, EPI_ISL_686525, EPI_ISL_686526, EPI_ISL_686527, EPI_ISL_686528, EPI_ISL_686529, EPI_ISL_686530, EPI_ISL_686531, EPI_ISL_686532, EPI_ISL_686533, EPI_ISL_686534 | | | |
| see above | Sakai City Institute of Public Health | Pathogen Genomics Center, National Institute of Infectious Diseases | Tsuyoshi Sekizuka, Kentaro Itokawa, Rina Tanaka, Masanori Hashino, Makoto Kuroda |
| EPI_ISL_686654, EPI_ISL_686655, EPI_ISL_686656, EPI_ISL_686657, EPI_ISL_686658 | Pathogen Genomics Center, National Institute of Infectious Diseases | Pathogen Genomics Center, National Institute of Infectious Diseases | Tsuyoshi Sekizuka, Kentaro Itokawa, Rina Tanaka, Masanori Hashino, Makoto Kuroda |
| EPI_ISL_686831, EPI_ISL_686832, EPI_ISL_686833, EPI_ISL_686834, EPI_ISL_686835, EPI_ISL_686836, EPI_ISL_686837, EPI_ISL_686838, EPI_ISL_686839, EPI_ISL_686840, EPI_ISL_686841, EPI_ISL_686842, EPI_ISL_686843, EPI_ISL_686844 | | | |
| see above | Chiba Prefectural Institute of Public Health | Pathogen Genomics Center, National Institute of Infectious Diseases | Tsuyoshi Sekizuka, Kentaro Itokawa, Rina Tanaka, Masanori Hashino, Makoto Kuroda |
| EPI_ISL_687099, EPI_ISL_687138, EPI_ISL_687148, EPI_ISL_687149, EPI_ISL_687150, EPI_ISL_687151, EPI_ISL_687158, EPI_ISL_687159, EPI_ISL_687160, EPI_ISL_687161, EPI_ISL_687162, EPI_ISL_687246, EPI_ISL_687247, EPI_ISL_687248, EPI_ISL_687249, EPI_ISL_687250, EPI_ISL_687251, EPI_ISL_687252, EPI_ISL_687253, EPI_ISL_687254, EPI_ISL_687360, EPI_ISL_687457, EPI_ISL_687458, EPI_ISL_687523, EPI_ISL_687524, EPI_ISL_687525, EPI_ISL_687526, EPI_ISL_687527, EPI_ISL_687528, EPI_ISL_687529, EPI_ISL_687530, EPI_ISL_687531, EPI_ISL_687532 | | | |
| see above | Pathogen Genomics Center, National Institute of Infectious Diseases | Pathogen Genomics Center, National Institute of Infectious Diseases | Tsuyoshi Sekizuka, Kentaro Itokawa, Rina Tanaka, Masanori Hashino, Makoto Kuroda |
| EPI_ISL_687533, EPI_ISL_687534, EPI_ISL_687535, EPI_ISL_687536, EPI_ISL_687537, EPI_ISL_687538, EPI_ISL_687539, EPI_ISL_687540, EPI_ISL_687541, EPI_ISL_687542, EPI_ISL_687543, EPI_ISL_687544, EPI_ISL_687545, EPI_ISL_687546, EPI_ISL_687547, EPI_ISL_687548, EPI_ISL_687549, EPI_ISL_687550, EPI_ISL_687551, EPI_ISL_687552, EPI_ISL_687553, EPI_ISL_687554, EPI_ISL_687555, EPI_ISL_687556, EPI_ISL_687557, EPI_ISL_687558, EPI_ISL_687559, EPI_ISL_687560 | | | |
| see above | Fukuoka Institute of Health and Environmental Sciences | Pathogen Genomics Center, National Institute of Infectious Diseases | Tsuyoshi Sekizuka, Kentaro Itokawa, Rina Tanaka, Masanori Hashino, Makoto Kuroda |
| EPI_ISL_687601, EPI_ISL_687602, EPI_ISL_687603, EPI_ISL_687604, EPI_ISL_687605, EPI_ISL_687606, EPI_ISL_687607, EPI_ISL_687608, EPI_ISL_687621, EPI_ISL_687622, EPI_ISL_687623, EPI_ISL_687669, EPI_ISL_687670, EPI_ISL_687671, EPI_ISL_687672, EPI_ISL_687676, EPI_ISL_687677, EPI_ISL_687678, EPI_ISL_687679, EPI_ISL_687680, EPI_ISL_687681, EPI_ISL_687682, EPI_ISL_687683, EPI_ISL_687684, EPI_ISL_687685, EPI_ISL_687686, EPI_ISL_687687, EPI_ISL_687688, EPI_ISL_687689, EPI_ISL_687690, EPI_ISL_687691, EPI_ISL_687692, EPI_ISL_687693, EPI_ISL_687694, EPI_ISL_687695, EPI_ISL_687696, EPI_ISL_687697, EPI_ISL_687698, EPI_ISL_687743, EPI_ISL_687744, EPI_ISL_687745, EPI_ISL_687920, EPI_ISL_687944, EPI_ISL_687945, EPI_ISL_687946, EPI_ISL_687947, EPI_ISL_687948, EPI_ISL_687949, EPI_ISL_687950, EPI_ISL_687951, EPI_ISL_687952, EPI_ISL_687953, EPI_ISL_687954, EPI_ISL_687955, EPI_ISL_687956, EPI_ISL_687957, EPI_ISL_687958, EPI_ISL_687959, EPI_ISL_687960, EPI_ISL_687961, EPI_ISL_687962, EPI_ISL_687963, EPI_ISL_687964, EPI_ISL_687965, EPI_ISL_687966, EPI_ISL_687967, EPI_ISL_687968, EPI_ISL_687969, EPI_ISL_687970, EPI_ISL_687971, EPI_ISL_687972, EPI_ISL_688007, EPI_ISL_688008, EPI_ISL_688009, EPI_ISL_688010, EPI_ISL_688011, EPI_ISL_688012, EPI_ISL_688013, EPI_ISL_688014, EPI_ISL_688015, EPI_ISL_688016, EPI_ISL_688017, EPI_ISL_688018, EPI_ISL_688019, EPI_ISL_688020, EPI_ISL_688021, EPI_ISL_688022, EPI_ISL_688023, EPI_ISL_688024, EPI_ISL_688025, EPI_ISL_688026, EPI_ISL_688027, EPI_ISL_688028, EPI_ISL_688029, EPI_ISL_688030, EPI_ISL_688031, EPI_ISL_688032, EPI_ISL_688033, EPI_ISL_688034, EPI_ISL_688035, EPI_ISL_688036, EPI_ISL_688037, EPI_ISL_688038, EPI_ISL_688039, EPI_ISL_688040, EPI_ISL_688041, EPI_ISL_688042, EPI_ISL_688043, EPI_ISL_688044, EPI_ISL_688045, EPI_ISL_688046, EPI_ISL_688047, EPI_ISL_688048, EPI_ISL_688049, EPI_ISL_688050, EPI_ISL_688051, EPI_ISL_688052, EPI_ISL_688053, EPI_ISL_688054, EPI_ISL_688055, EPI_ISL_688101, EPI_ISL_688102, EPI_ISL_688103, EPI_ISL_688104, EPI_ISL_688105, EPI_ISL_688106, EPI_ISL_688107, EPI_ISL_688108, EPI_ISL_688251, EPI_ISL_688252, EPI_ISL_688253, EPI_ISL_688254, EPI_ISL_688255, EPI_ISL_688256, EPI_ISL_688257, EPI_ISL_688258, EPI_ISL_688259, EPI_ISL_688260, EPI_ISL_688261, EPI_ISL_688262, EPI_ISL_688263, EPI_ISL_688264, EPI_ISL_688265, EPI_ISL_688266, EPI_ISL_688267, EPI_ISL_688268, EPI_ISL_688614, EPI_ISL_688746, EPI_ISL_689549, EPI_ISL_689550, EPI_ISL_689592, EPI_ISL_689593, EPI_ISL_689594, EPI_ISL_689595, EPI_ISL_689596, EPI_ISL_689597, EPI_ISL_689598, EPI_ISL_689599, EPI_ISL_689600 | | | |
| see above | Pathogen Genomics Center, National Institute of Infectious Diseases | Pathogen Genomics Center, National Institute of Infectious Diseases | Tsuyoshi Sekizuka, Kentaro Itokawa, Rina Tanaka, Masanori Hashino, Makoto Kuroda |
| EPI_ISL_691685 | Hospital de Leon | Instituto de Salud Carlos III | Iglesias-Caballero, M. Camarero, S. Molinero Calamita, M. González-Esguevillas, M. Pozo, F. Casas, I. Jiménez, P. Jiménez, M. Zaballos, A. Monzón, S. Varona, S. Juliá, M. Cuesta, I. Vidan, J. |
| EPI_ISL_691686, EPI_ISL_691687, EPI_ISL_691688, EPI_ISL_691689, EPI_ISL_691690, EPI_ISL_691691, EPI_ISL_691692, EPI_ISL_691693, EPI_ISL_691694, EPI_ISL_691695, EPI_ISL_691696, EPI_ISL_691697, EPI_ISL_691698, EPI_ISL_691699 | | | |
| see above | Hospital Universitario de Ceuta | Instituto de Salud Carlos III | Iglesias-Caballero, M. Camarero, S. Molinero Calamita, M. González-Esguevillas, M. Pozo, F. Casas, I. Jiménez, P. Jiménez, M. Zaballos, A. Monzón, S. Varona, S. Juliá, M. Cuesta, I. Hijano, S. |
| EPI_ISL_691729, EPI_ISL_691730 | Hospital Universitario Severo Ochoa | Instituto de Salud Carlos III | Iglesias-Caballero, M. Camarero, S. Molinero Calamita, M. González-Esguevillas, M. Pozo, F. Casas, I. Jiménez, P. Jiménez, M. Zaballos, A. Monzón, S. Varona, S. Juliá, M. Cuesta, I. García, M. |
| EPI_ISL_691731 | Hospital Universitario Puerta de Hierro | Instituto de Salud Carlos III | Iglesias-Caballero, M. Camarero, S. Molinero Calamita, M. González-Esguevillas, M. Pozo, F. Casas, I. Jiménez, P. Jiménez, M. Zaballos, A. Monzón, S. |

| | | | |
|---|---|---|---|
| | | | Varona, S. Juliá, M. Cuesta, I. Velasco, A. |
| EPI_ISL_691732 | Hospital de Madrid | Instituto de Salud Carlos III | Iglesias-Caballero, M. Camarero, S. Molinero Calamita, M. González-Esguevillas, M. Pozo, F. Casas, I. Jiménez, P. Jiménez, M. Zaballos, A. Monzón, S. Varona, S. Juliá, M. Cuesta, I. Alhambra, A. |
| EPI_ISL_693302, EPI_ISL_693303, EPI_ISL_693304, EPI_ISL_693305, EPI_ISL_693306 | Department of Laboratory Medicine, National Taiwan University Hospital | Microbial Genomics Core Lab, National Taiwan University Centers of Genomic and Precision Medicine | Shiou-Hwei Yeh, You-Yu Lin, Ya-Yun Lai, Chiao-Ling Li, Shan-Chwen Chang, Pei-Jer Chen, Sui-Yuan Chang |
| EPI_ISL_693660, EPI_ISL_693661, EPI_ISL_693663, EPI_ISL_693665, EPI_ISL_693666, EPI_ISL_693667, EPI_ISL_693669, EPI_ISL_693672, EPI_ISL_693674, EPI_ISL_693675, EPI_ISL_693677, EPI_ISL_693678, EPI_ISL_693679, EPI_ISL_693681, EPI_ISL_693682, EPI_ISL_693683, EPI_ISL_693684, EPI_ISL_693685, EPI_ISL_693687 | | | |
| see above | The National Institute of Public Health | State Veterinary Institute Prague | Nagy,A;Jirincova,H;Trnka,D;Vecerova,J |
| EPI_ISL_693758, EPI_ISL_693759, EPI_ISL_693760, EPI_ISL_693761, EPI_ISL_693762, EPI_ISL_693764 | Hospital | National Reference Center for Viruses of Respiratory Infections, Institut Pasteur, Paris | Marion Barbet, Sylvie Behillil, Méline Bizard, Angela Brisebarre, Camille Capel, Etienne Simon-Lorière, Vincent Enouf, Maud Vanpeene, Sylvie van der Werf, Gisèle Lagathu |
| EPI_ISL_693765 | hospital | National Reference Center for Viruses of Respiratory Infections, Institut Pasteur, Paris | Marion Barbet, Sylvie Behillil, Méline Bizard, Angela Brisebarre, Camille Capel, Etienne Simon-Lorière, Vincent Enouf, Maud Vanpeene, Sylvie van der Werf, Gisèle Lagathu |
| EPI_ISL_693766, EPI_ISL_693767, EPI_ISL_693768, EPI_ISL_693769, EPI_ISL_693770, EPI_ISL_693771, EPI_ISL_693772, EPI_ISL_693773, EPI_ISL_693774 | General practitioner | National Reference Center for Viruses of Respiratory Infections, Institut Pasteur, Paris | Marion Barbet, Sylvie Behillil, Méline Bizard, Angela Brisebarre, Camille Capel, Etienne Simon-Lorière, Vincent Enouf, Maud Vanpeene, Sylvie van der Werf |
| EPI_ISL_699655 | 1-Laboratory of Microbiology, National Reference Lab, Charles Nicolle Hospital; 2-University of Tunis ElManar, Faculty of Medicine of Tunis, LR99ES09, Tunis, Tunisia | 1-Clinical and Experimental Pharmacology Lab, LR16SP02, National Center of Pharmacovigilance, University of Tunis El Manar, Tunis, Tunisia. 2-Neurodegenerative diseases and psychiatric troubles, LR18SP03, Razi Hospital, University of Tunis El Manar, Tunis, Tunisia. 3- Ministry of Health, National Observatory of New and Emerging Diseases, 1006, Tunis, Tunisia | Ilhem Boutiba-Ben Boubaker, Sameh Trabelsi, Nissaf Ben Alaya, Maher Kharrat, Alia Ben Kahla, Jalila Ben Khelil, Salma Abid, Sana Ferjani, Asma Ferjani, Mouna Ben Sassi, Mouna Safer, Guedi Berrabeh,Salwa Mrabet, Hanen ElJebari, Gaies Emna, Riadh Daghfous, Riadh Gouider. |
| EPI_ISL_700219, EPI_ISL_700221, EPI_ISL_700224, EPI_ISL_700227, EPI_ISL_700229, EPI_ISL_700230, EPI_ISL_700231, EPI_ISL_700232, EPI_ISL_700234, EPI_ISL_700237, EPI_ISL_700238, EPI_ISL_700240, EPI_ISL_700241, EPI_ISL_700242, EPI_ISL_700245, EPI_ISL_700246, EPI_ISL_700247, EPI_ISL_700248, EPI_ISL_700249, EPI_ISL_700250, EPI_ISL_700251, EPI_ISL_700252, EPI_ISL_700253, EPI_ISL_700254, EPI_ISL_700255, EPI_ISL_700256, EPI_ISL_700258, EPI_ISL_700260, EPI_ISL_700262, EPI_ISL_700263, EPI_ISL_700265, EPI_ISL_700266, EPI_ISL_700267, EPI_ISL_700268, EPI_ISL_700269, EPI_ISL_700270, EPI_ISL_700271, EPI_ISL_700273, EPI_ISL_700274, EPI_ISL_700275, EPI_ISL_700276, EPI_ISL_700277, EPI_ISL_700278, EPI_ISL_700279, EPI_ISL_700280, EPI_ISL_700281, EPI_ISL_700282, EPI_ISL_700283, EPI_ISL_700284, EPI_ISL_700285, EPI_ISL_700286, EPI_ISL_700287, EPI_ISL_700288, EPI_ISL_700289, EPI_ISL_700290, EPI_ISL_700291, EPI_ISL_700293, EPI_ISL_700294, EPI_ISL_700295, EPI_ISL_700296, EPI_ISL_700297, EPI_ISL_700298, EPI_ISL_700299, EPI_ISL_700302, EPI_ISL_700303, EPI_ISL_700304, EPI_ISL_700305, EPI_ISL_700306, EPI_ISL_700307, EPI_ISL_700308, EPI_ISL_700309, EPI_ISL_700310, EPI_ISL_700311, EPI_ISL_700312, EPI_ISL_700313, EPI_ISL_700314, EPI_ISL_700315, EPI_ISL_700316, EPI_ISL_700318, EPI_ISL_700320, EPI_ISL_700321, EPI_ISL_700322, EPI_ISL_700323, EPI_ISL_700324, EPI_ISL_700325 | | | |
| see above | Hematopathology Laboratory, ACTREC, TMC | Hematopathology Laboratory, ACTREC, TMC | Hematopathology Laboratory, ACTREC |
| EPI_ISL_707696 | Medical Research Center, Faculty of Medicine, Syarif Hidayatullah State Islamic University Jakarta | Medical Research Center, Faculty of Medicine, Syarif Hidayatullah State Islamic University Jakarta | Laifa Hendarmin, Chris Adhiyanto, Erike Suwarsono, Zeti Harriyati, Rini Puspitaningrum, Ferania Mela, Dennis Nurjadi |
| EPI_ISL_707697, EPI_ISL_707698, EPI_ISL_707699, EPI_ISL_707700 | 1-Laboratory of Microbiology, National Reference Lab, Charles Nicolle Hospital; 2-University of Tunis ElManar, Faculty of Medicine of Tunis, LR99ES09, Tunis, Tunisia | 1-Clinical and Experimental Pharmacology Lab, LR16SP02, National Center of Pharmacovigilance, University of Tunis El Manar, Tunis, Tunisia. 2-Neurodegenerative diseases and psychiatric troubles, LR18SP03, Razi Hospital, University of Tunis El Manar, Tunis, Tunisia. 3- Ministry of Health, National Observatory of New and Emerging Diseases, 1006, Tunis, Tunisia | Ilhem Boutiba-Ben Boubaker, Sameh Trabelsi, Nissaf Ben Alaya, Maher Kharrat, Alia Ben Kahla, Jalila Ben Khelil, Salma Abid, Sana Ferjani, Mouna Ben Sassi, Mouna Safer, Zaineb Hamzaoui, Habiba Ben Romdhane, Souissi Amira, Rouaa Ben Othman, Hanen El Jebari, Asma Ferjani, Gaies Emna, Riadh Daghfous, Riadh Gouider. |
| EPI_ISL_708530 | Secretaria Municipal de Saude de Fernandópolis | Instituto Adolfo Lutz, Interdiciplinary Procedures Center, Strategic Laboratory | Claudio Tavares Sacchi, Claudia Regina Gonçalves, Erica Valessa Ramos Gomes, Carlos Henrique Camargo, Karoline Rodrigues Campos, Fernanda Modesto Tolentino Binhardi, Maricelia Navarro Pinheiro Flores, Marcia Maria Costa Nunes Soares, Janaina Other Martins Montanha |
| EPI_ISL_708652 | Medical Research Center, Faculty of Medicine, Syarif Hidayatullah State Islamic University Jakarta | Medical Research Center, Faculty of Medicine, Syarif Hidayatullah State Islamic University Jakarta | Chris Adhiyanto, Laifa Hendarmin, Erike A Suwarsono, Zeti Harriyati, Endah Wulandari, Flori R Sari, Hari Hendarto |
| EPI_ISL_710483, EPI_ISL_710484 | Department of Medical Laboratory Sciences, Arab American University | Department of Medical Laboratory Sciences, Arab American University | Al-Jawabreh,A., Nasereddin,A., Dumaidi,K., Al-Jawabreh,H., Ereqat,S. |
| EPI_ISL_710503 | Laboratorio specialistico UOC Ematologia - Ospedale "San Francesco" - ATS-ASSL Nuoro | Laboratorio specialistico UOC Ematologia - Ospedale "San Francesco" - ATS-ASSL Nuoro | Giovanna Piras |
| EPI_ISL_710534, EPI_ISL_710537, EPI_ISL_710540, EPI_ISL_710541, EPI_ISL_710575, EPI_ISL_711057, EPI_ISL_712060 | Hôpital Fattouma-Bourguiba de Monastir | Laboratoire des Procédés de Criblage Moléculaire et Cellulaire-Centre de Biotechnologie de Sfax | Souissi,A., Abid,N., Ben Ayed,I., Gargouri,S., Abdelmoulah,F.,Elargoubi,A., Smeti,I., Bensaid,M., Stambouli,N., Kharat,N., Ajili,F., Fki-berrajah,L., Mhalla,S., Chtourou,A., Gaaloul,I., Nabli,A., Turki,M., Aouni,M., Hammami,A., Mastouri,M., Karray Hakim,H., Kamoun,S., Rebai,A. and Masmoudi,S. |
| EPI_ISL_717965, EPI_ISL_717966, EPI_ISL_717967, EPI_ISL_717968, EPI_ISL_717969, EPI_ISL_717970, EPI_ISL_717971, EPI_ISL_717972, EPI_ISL_717973, EPI_ISL_717974 | Molecular diagnostic laboratory of Federal Budget Institution of Science "Central Research Institute of Epidemiology" of The Federal Service on Customers' Rights Protection and Human Well-being Surveillance | Group of Genomics and Postgenomic Technologies of Central Research Institute of Epidemilology | Samoilov AE, Kaptelova VV, Dudorova AV, Korneenko EV, Speranskaya AS, Tivanova EV, Shipulina OY, Akimkin VG |
| EPI_ISL_718167 | Borneo Medical Centre | Institute of Health and Community Medicine | David Perera, Ooi Mong How, Chua Hock Hin, Tonnii Sia Loong Loong, Wong Jyn Shan, Wong Kiing Aik, Chan Chia Jui |
| EPI_ISL_718168, EPI_ISL_718169, EPI_ISL_718170, EPI_ISL_718171, EPI_ISL_718172, EPI_ISL_718173 | Ministry of Health Hospitals | Institute of Health and Community Medicine | David Perera, Ooi Mong How, Chua Hock Hin, Tonnii Sia Loong Loong, Wong Jyn Shan, Wong Kiing Aik, Chan Chia Jui |
| EPI_ISL_718174 | Borneo Medical Centre | Institute of Health and Community Medicine | David Perera, Ooi Mong How, Chua Hock Hin, Tonnii Sia Loong Loong, Wong Jyn Shan, Wong Kiing Aik, Chan Chia Jui |
| EPI_ISL_718175 | Ministry of Health Hospitals | Institute of Health and Community Medicine | David Perera, Ooi Mong How, Chua Hock Hin, Tonnii Sia Loong Loong, Wong Jyn Shan, Wong Kiing Aik, Chan Chia Jui |
| EPI_ISL_718176 | Borneo Medical Centre | Institute of Health and Community Medicine | David Perera, Ooi Mong How, Chua Hock Hin, Tonnii Sia Loong Loong, Wong Jyn Shan, Wong Kiing Aik, Chan Chia Jui |
| EPI_ISL_718177 | Ministry of Health Hospitals | Institute of Health and Community Medicine | David Perera, Ooi Mong How, Chua Hock Hin, Tonnii Sia Loong Loong, Wong Jyn Shan, Wong Kiing Aik, Chan Chia Jui |
| EPI_ISL_718178 | Borneo Medical Centre | Institute of Health and Community Medicine | David Perera, Ooi Mong How, Chua Hock Hin, Tonnii Sia Loong Loong, Wong Jyn Shan, Wong Kiing Aik, Chan Chia Jui |
| EPI_ISL_718179, EPI_ISL_718180, EPI_ISL_718181 | Ministry of Health Hospitals | Institute of Health and Community Medicine | David Perera, Ooi Mong How, Chua Hock Hin, Tonnii Sia Loong Loong, Wong Jyn Shan, Wong Kiing Aik, Chan Chia Jui |
| EPI_ISL_718182 | Borneo Medical Centre | Institute of Health and Community Medicine | David Perera, Ooi Mong How, Chua Hock Hin, Tonnii Sia Loong Loong, Wong Jyn Shan, Wong Kiing Aik, Chan Chia Jui |
| EPI_ISL_718183, EPI_ISL_718184 | Ministry of Health Hospitals | Institute of Health and Community Medicine | David Perera, Ooi Mong How, Chua Hock Hin, Tonnii Sia Loong Loong, Wong Jyn Shan, Wong Kiing Aik, Chan Chia Jui |
| EPI_ISL_718185 | Borneo Medical Centre | Institute of Health and Community Medicine | David Perera, Ooi Mong How, Chua Hock Hin, Tonnii Sia Loong Loong, Wong Jyn Shan, Wong Kiing Aik, Chan Chia Jui |
| EPI_ISL_718186, EPI_ISL_718187, | Ministry of Health Hospitals | Institute of Health and Community Medicine | David Perera, Ooi Mong How, Chua Hock Hin, Tonnii Sia Loong Loong, Wong Jyn Shan, Wong Kiing Aik, Chan Chia Jui |

| | | | |
|---|---|---|---|
| EPI_ISL_718188, EPI_ISL_718189, EPI_ISL_718190 | | | |
| EPI_ISL_718191, EPI_ISL_718192 | Borneo Medical Centre | Institute of Health and Community Medicine | David Perera, Ooi Mong How, Chua Hock Hin, Tonnii Sia Loong Loong, Wong Jyn Shan, Wong Kiing Aik, Chan Chia Jui |
| EPI_ISL_718193, EPI_ISL_718194, EPI_ISL_718195, EPI_ISL_718196 | Ministry of Health Hospitals | Institute of Health and Community Medicine | David Perera, Ooi Mong How, Chua Hock Hin, Tonnii Sia Loong Loong, Wong Jyn Shan, Wong Kiing Aik, Chan Chia Jui |
| EPI_ISL_718197, EPI_ISL_718198 | Borneo Medical Centre | Institute of Health and Community Medicine | David Perera, Ooi Mong How, Chua Hock Hin, Tonnii Sia Loong Loong, Wong Jyn Shan, Wong Kiing Aik, Chan Chia Jui |
| EPI_ISL_718199, EPI_ISL_718200, EPI_ISL_718201, EPI_ISL_718202 | Ministry of Health Hospitals | Institute of Health and Community Medicine | David Perera, Ooi Mong How, Chua Hock Hin, Tonnii Sia Loong Loong, Wong Jyn Shan, Wong Kiing Aik, Chan Chia Jui |
| EPI_ISL_718203, EPI_ISL_718204, EPI_ISL_718205 | Borneo Medical Centre | Institute of Health and Community Medicine | David Perera, Ooi Mong How, Chua Hock Hin, Tonnii Sia Loong Loong, Wong Jyn Shan, Wong Kiing Aik, Chan Chia Jui |
| EPI_ISL_718206, EPI_ISL_718207, EPI_ISL_718208, EPI_ISL_718209, EPI_ISL_718210, EPI_ISL_718211, EPI_ISL_718212, EPI_ISL_718213, EPI_ISL_718214, EPI_ISL_718215 | Ministry of Health Hospitals | Institute of Health and Community Medicine | David Perera, Ooi Mong How, Chua Hock Hin, Tonnii Sia Loong Loong, Wong Jyn Shan, Wong Kiing Aik, Chan Chia Jui |
| EPI_ISL_718216 | Borneo Medical Centre | Institute of Health and Community Medicine | David Perera, Ooi Mong How, Chua Hock Hin, Tonnii Sia Loong Loong, Wong Jyn Shan, Wong Kiing Aik, Chan Chia Jui |
| EPI_ISL_718217 | Ministry of Health Hospitals | Institute of Health and Community Medicine | David Perera, Ooi Mong How, Chua Hock Hin, Tonnii Sia Loong Loong, Wong Jyn Shan, Wong Kiing Aik, Chan Chia Jui |
| EPI_ISL_718218 | Borneo Medical Centre | Institute of Health and Community Medicine | David Perera, Ooi Mong How, Chua Hock Hin, Tonnii Sia Loong Loong, Wong Jyn Shan, Wong Kiing Aik, Chan Chia Jui |
| EPI_ISL_718220 | Ministry of Health Hospitals | Institute of Health and Community Medicine | David Perera, Ooi Mong How, Chua Hock Hin, Tonnii Sia Loong Loong, Wong Jyn Shan, Wong Kiing Aik, Chan Chia Jui |
| EPI_ISL_718222 | Borneo Medical Centre | Institute of Health and Community Medicine | David Perera, Ooi Mong How, Chua Hock Hin, Tonnii Sia Loong Loong, Wong Jyn Shan, Wong Kiing Aik, Chan Chia Jui |
| EPI_ISL_718226, EPI_ISL_718227, EPI_ISL_718228, EPI_ISL_718229, EPI_ISL_718230, EPI_ISL_718231, EPI_ISL_718232, EPI_ISL_718233, EPI_ISL_718234, EPI_ISL_718235, EPI_ISL_718236, EPI_ISL_718237, EPI_ISL_718238, EPI_ISL_718239, EPI_ISL_718240, EPI_ISL_718241, EPI_ISL_718242, EPI_ISL_718243, EPI_ISL_718244, EPI_ISL_718245, EPI_ISL_718246, EPI_ISL_718247, EPI_ISL_718248, EPI_ISL_718249 | | | |
| see above | Hospital | National Reference Center for Viruses of Respiratory Infections, Institut Pasteur, Paris | Marion Barbet, Sylvie Behillil, Méline Bizard, Angela Brisebarre, Camille Capel, Etienne Simon-Lorière, Vincent Enouf, Maud Vanpeene, Sylvie van der Werf, Gisèle Lagathu |
| EPI_ISL_722873, EPI_ISL_722874, EPI_ISL_722875, EPI_ISL_722876, EPI_ISL_722877, EPI_ISL_722878, EPI_ISL_722879, EPI_ISL_722880, EPI_ISL_722881, EPI_ISL_722882, EPI_ISL_722883, EPI_ISL_722884, EPI_ISL_722885, EPI_ISL_722886, EPI_ISL_722887, EPI_ISL_722888, EPI_ISL_722889, EPI_ISL_722890, EPI_ISL_722891, EPI_ISL_722893, EPI_ISL_722894, EPI_ISL_722901, EPI_ISL_722902, EPI_ISL_722903, EPI_ISL_722904, EPI_ISL_722905, EPI_ISL_722906, EPI_ISL_722907, EPI_ISL_722908, EPI_ISL_722909, EPI_ISL_722910, EPI_ISL_722911 | | | |
| see above | Istituto Zooprofilattico Sperimentale della Puglia e della Basilicata | Istituto Zooprofilattico Sperimentale della Puglia e della Basilicata | Parisi A., Bianco A., Capozzi L., Del Sambro L., Manzulli V, Rondinone V., Pace L., Cipolletta D., Galante D. |
| EPI_ISL_730579, EPI_ISL_730580, EPI_ISL_730581 | Home Quarantine Taskforce | Hong Kong Department of Health | Mak Gannon C.K., Lam Edman T.K., Chan Rickjason C.W., Tsang Dominic N.C. |
| EPI_ISL_730582 | Hong Kong Adventist Hospital - Tsuen Wan | Hong Kong Department of Health | Mak Gannon C.K., Lam Edman T.K., Chan Rickjason C.W., Tsang Dominic N.C. |
| EPI_ISL_730583, EPI_ISL_730584 | Hong Kong Sanatorium & Hospital | Hong Kong Department of Health | Mak Gannon C.K., Lam Edman T.K., Chan Rickjason C.W., Tsang Dominic N.C. |
| EPI_ISL_730585, EPI_ISL_730586, EPI_ISL_730587 | Pamela Youde Nethersole Eastern Hospital | Hong Kong Department of Health | Mak Gannon C.K., Lam Edman T.K., Chan Rickjason C.W., Tsang Dominic N.C. |
| EPI_ISL_730588, EPI_ISL_730589, EPI_ISL_730590 | Prince of Wales Hospital | Hong Kong Department of Health | Mak Gannon C.K., Lam Edman T.K., Chan Rickjason C.W., Tsang Dominic N.C. |
| EPI_ISL_730591, EPI_ISL_730592 | Princess Margaret Hospital | Hong Kong Department of Health | Mak Gannon C.K., Lam Edman T.K., Chan Rickjason C.W., Tsang Dominic N.C. |
| EPI_ISL_730593 | Private medical practitioner | Hong Kong Department of Health | Mak Gannon C.K., Lam Edman T.K., Chan Rickjason C.W., Tsang Dominic N.C. |
| EPI_ISL_730594, EPI_ISL_730595 | Queen Elizabeth Hospital | Hong Kong Department of Health | Mak Gannon C.K., Lam Edman T.K., Chan Rickjason C.W., Tsang Dominic N.C. |
| EPI_ISL_730596, EPI_ISL_730597, EPI_ISL_730598, EPI_ISL_730599, EPI_ISL_730600 | Queen Mary Hospital | Hong Kong Department of Health | Mak Gannon C.K., Lam Edman T.K., Chan Rickjason C.W., Tsang Dominic N.C. |
| EPI_ISL_730602, EPI_ISL_730603 | Tseung Kwan O Hospital | Hong Kong Department of Health | Mak Gannon C.K., Lam Edman T.K., Chan Rickjason C.W., Tsang Dominic N.C. |
| EPI_ISL_730604, EPI_ISL_730605, EPI_ISL_730606, EPI_ISL_730607, EPI_ISL_730608, EPI_ISL_730609, EPI_ISL_730610 | Tuen Mun Hospital | Hong Kong Department of Health | Mak Gannon C.K., Lam Edman T.K., Chan Rickjason C.W., Tsang Dominic N.C. |
| EPI_ISL_730611 | Union Hospital | Hong Kong Department of Health | Mak Gannon C.K., Lam Edman T.K., Chan Rickjason C.W., Tsang Dominic N.C. |
| EPI_ISL_730612, EPI_ISL_730613, EPI_ISL_730614, EPI_ISL_730615, EPI_ISL_730616 | United Christian Hospital | Hong Kong Department of Health | Mak Gannon C.K., Lam Edman T.K., Chan Rickjason C.W., Tsang Dominic N.C. |
| EPI_ISL_730617 | Universal Community Testing Programme | Hong Kong Department of Health | Mak Gannon C.K., Lam Edman T.K., Chan Rickjason C.W., Tsang Dominic N.C. |
| EPI_ISL_730618, EPI_ISL_730619 | Yan Chai Hospital | Hong Kong Department of Health | Mak Gannon C.K., Lam Edman T.K., Chan Rickjason C.W., Tsang Dominic N.C. |
| EPI_ISL_730653 | University of Bari, Valenzano, Italy | Istituto Zooprofilattico Sperimentale dell'Abruzzo e del Molise "G. Caporale". | N. Decaro, E. Lorusso, G. Elia, C. Desario, D. Buonavoglia, V., Martella, C. Buonavoglia, A. Lorusso, C. Cammà, V. Curini |
| EPI_ISL_732563 | Bundeswehr Institute of Microbiology | Bundeswehr Institute of Microbiology | Markus Antwerpen, Alexandra Rehn, Mathias Walter, Malena Bestehorn-Willmann, Sabine Zange, Enrico Georgi, Roman Wölfel |
| EPI_ISL_732658 | Bundeswehr Institute of Microbiology | Bundeswehr Institute of Microbiology | Elham Khatamzas, Markus Antwerpen, Mathias Walter, Alexandra Rehn, Sabine Zange, Enrico Georgi, Michael von Bergwelt-Baildon, Roman Wölfel |
| EPI_ISL_732659, EPI_ISL_732660, EPI_ISL_732661, EPI_ISL_732662, EPI_ISL_732663, EPI_ISL_732664, EPI_ISL_732665, EPI_ISL_732666, EPI_ISL_732667, EPI_ISL_732668, EPI_ISL_732669, EPI_ISL_732670, EPI_ISL_732671, EPI_ISL_732672, EPI_ISL_732673, EPI_ISL_732674, EPI_ISL_732675, EPI_ISL_732676 | | | |
| see above | Molecular diagnostic laboratory of Federal Budget Institution of Science "Central Research Institute of Epidemiology" of The Federal Service on Customers' Rights Protection and Human Well-being Surveillance | Group of Genomics and Postgenomic Technologies of Central Research Institute of Epidemilology | Samoilov AE, Kaptelova VV, Dudorova AV, Korneenko EV, Saenko SS, Speranskaya AS, Tivanova EV, Shipulina OY, Akimkin VG |
| EPI_ISL_732975, EPI_ISL_732976, EPI_ISL_732979, EPI_ISL_732980, EPI_ISL_732982, EPI_ISL_732983, EPI_ISL_732985 | WHO National Influenza Centre Russian Federation | WHO National Influenza Centre Russian Federation | Andrey Komissarov, Artem Fadeev, Anna Ivanova, Kseniya Komissarova, Dmitry Bazhenov, Daria Danilenko, Ksenia Safina, Elena Nabieva, Georgii Bazykin, Dmitry Lioznov |
| EPI_ISL_733569 | CHAI WAN FAMILIES CLINIC | Hong Kong Department of Health | Alan K.L. Tsang, Peter C.W. Yip, Edman T.K. Lam, Rickjason C.W. Chan, Dominic N.C. Tsang |
| EPI_ISL_733570 | POK OI HOSPITAL | Hong Kong Department of Health | Alan K.L. Tsang, Peter C.W. Yip, Edman T.K. Lam, Rickjason C.W. Chan, Dominic N.C. Tsang |

| | | | |
|---|---|---|---|
| EPI_ISL_733571 | Queen Mary Hospital | Hong Kong Department of Health | Alan K.L. Tsang, Peter C.W. Yip, Edman T.K. Lam, Rickjason C.W. Chan, Dominic N.C. Tsang |
| EPI_ISL_733572, EPI_ISL_733573, EPI_ISL_733574, EPI_ISL_733575, EPI_ISL_733576, EPI_ISL_733577, EPI_ISL_733579 | Temporary Specimen Collection Centre | Hong Kong Department of Health | Alan K.L. Tsang, Peter C.W. Yip, Edman T.K. Lam, Rickjason C.W. Chan, Dominic N.C. Tsang |
| EPI_ISL_734168, EPI_ISL_734169 | CHRU Pontchaillou - Laboratoire de Virologie | National Reference Center for Viruses of Respiratory Infections, Institut Pasteur, Paris | Marion Barbet, Sylvie Behillil, Méline Bizard, Angela Brisebarre, Camille Capel, Etienne Simon-Lorière, Vincent Enouf, Maud Vanpeene, Sylvie van der Werf, Gisèle Lagathu |
| EPI_ISL_735444 | Nucleic Acid Testing - Rwanda National Reference Laboratory | GIGA Medical Genomics | Yvan Butera, Keith Durkin, Maria Artesi, Bouchra Boujemla, Robert Rutayisire, Patrick Tuyisenge, Esperence Umumararungu, Sébastien Bontems, Marie-Pierre Hayette, Swaibu Gatare, Jacob Souopgui, Sabin Nsanzimana, Vincent Bours, Léon Mutesa |
| EPI_ISL_735445 | Rwanda National Reference Laboratory | GIGA Medical Genomics | Yvan Butera, Keith Durkin, Maria Artesi, Bouchra Boujemla, Robert Rutayisire, Patrick Tuyisenge, Esperence Umumararungu, Sébastien Bontems, Marie-Pierre Hayette, Swaibu Gatare, Jacob Souopgui, Sabin Nsanzimana, Vincent Bours, Léon Mutesa |
| EPI_ISL_735446, EPI_ISL_735447, EPI_ISL_735448 | Nucleic Acid Testing - Rwanda National Reference Laboratory | GIGA Medical Genomics | Yvan Butera, Keith Durkin, Maria Artesi, Bouchra Boujemla, Robert Rutayisire, Patrick Tuyisenge, Esperence Umumararungu, Sébastien Bontems, Marie-Pierre Hayette, Swaibu Gatare, Jacob Souopgui, Sabin Nsanzimana, Vincent Bours, Léon Mutesa |
| EPI_ISL_735491 | Virology Laboratory, Ospedali Riuniti, Ancona | Virology Laboratory, Ospedali Riuniti, Ancona | Stefano Menzo, Roberta Longo, Laura Di Sante, Sara Caucci, Patrizia Bagnarelli |
| EPI_ISL_736996 | Istituto Zooprofilattico Sperimentale del Mezzogiorno | National Institute for Infectious Diseases, INMI, "L. Spallanzani" IRCCS | E. Giombini, C.E.M Gruber, F. Messina, M. Rueca, B. Bartolini, F. Carletti, P. Cerino, B. Pierri, C. Buonerba, D. Di Concilio, M.C. Cuomo, A. Di Caro, M.R Capobianchi |
| EPI_ISL_736997 | Istituto Zooprofilattico Sperimentale del Mezzogiorno | National Institute for Infectious Diseases, INMI, "L. Spallanzani" IRCCS | F. Messina, C.E.M Gruber, M. Rueca, B. Bartolini, E. Giombini, F. Carletti, P. Cerino, B. Pierri, C. Buonerba, D. Di Concilio, M.C. Cuomo, M.R. Capobianchi, A. Di Caro |
| EPI_ISL_736998, EPI_ISL_737002, EPI_ISL_737003, EPI_ISL_737008, EPI_ISL_737009, EPI_ISL_737010, EPI_ISL_737011, EPI_ISL_737012, EPI_ISL_737013, EPI_ISL_737014, EPI_ISL_737015, EPI_ISL_737016, EPI_ISL_737017, EPI_ISL_737018, EPI_ISL_737019, EPI_ISL_737021, EPI_ISL_737022, EPI_ISL_737023, EPI_ISL_737024, EPI_ISL_737025, EPI_ISL_737026, EPI_ISL_737027, EPI_ISL_737028, EPI_ISL_737029, EPI_ISL_737031, EPI_ISL_737032 | | | |
| see above | The National Institute of Public Health | State Veterinary Institute Prague | Nagy,A;Jirincova,H;Trnka,D;Vecerova,J |
| EPI_ISL_738044 | SIESP CHIETI - DRIVE IN LANCIANO | Istituto Zooprofilattico Sperimentale dell'Abruzzo e Molise "G. Caporale" | Lorusso A, Marcacci M, Di Domenico M, Ancora M, Curini V, Mangone I, Rinaldi A, Di Pasquale A, Cammà C, Puglia I, Savini G |
| EPI_ISL_738045 | SIESP DIPARTIMENTO DI PREVENZIONE CHIETI | Istituto Zooprofilattico Sperimentale dell'Abruzzo e Molise "G. Caporale" | Lorusso A, Marcacci M, Di Domenico M, Ancora M, Curini V, Mangone I, Rinaldi A, Di Pasquale A, Cammà C, Puglia I, Savini G |
| EPI_ISL_738046, EPI_ISL_738047, EPI_ISL_738048 | SIESP CHIETI - DRIVE IN ORTONA | Istituto Zooprofilattico Sperimentale dell'Abruzzo e Molise "G. Caporale" | Lorusso A, Marcacci M, Di Domenico M, Ancora M, Curini V, Mangone I, Rinaldi A, Di Pasquale A, Cammà C, Puglia I, Savini G |
| EPI_ISL_738064, EPI_ISL_738065 | Department of Laboratory Medicine, National Taiwan University Hospital | Microbial Genomics Core Lab, National Taiwan University Centers of Genomic and Precision Medicine | Shiou-Hwei Yeh, You-Yu Lin, Ya-Yun Lai, Chiao-Ling Li, Shan-Chwen Chang, Pei-Jer Chen, Sui-Yuan Chang |
| EPI_ISL_738121, EPI_ISL_738129 | IZSM-U.O.C. Virologia | Istituto Zooprofilattico Sperimentale del Mezzogiorno | Maurizio Viscardi, Lorena Cardillo, Giovanna Fusco |
| EPI_ISL_738144 | University of Bari Biomedical Sciences and Human Oncology | University of Bari Biomedical Sciences and Human Oncology | Maria Chironna, Anna Sallustio, Daniela Loconsole, Marisa Accogli |
| EPI_ISL_739660, EPI_ISL_739661 | Al-Quds Nutrition and Health Research Institute, Al-Quds University | Al-Quds Nutrition and Health Research Institute, Al-Quds University | Nasereddin,A., Ereqat,S., Al-Jawabreh,A., Rishmawi,C. |
| EPI_ISL_739671, EPI_ISL_739672, EPI_ISL_739673 | Instituto Nacional de Salud, Bogotá, Colombia | Instituto Nacional de Salud, Bogotá, Colombia | Katherine Laiton-Donato, Diego A. Álvarez-Díaz, Carlos Franco-Muñoz, Mauricio Pacheco-Montealegre, Jonathan Reales, Diego Andrés Prada, Sheryl Corchuelo, Magdalena Weisner, Martha Lucia Ospina Martinez, Marcela Mercado-Reyes |
| EPI_ISL_740547 | Department of Laboratory Medicine, National Taiwan University Hospital | Microbial Genomics Core Lab, National Taiwan University Centers of Genomic and Precision Medicine | Shiou-Hwei Yeh, You-Yu Lin, Ya-Yun Lai, Chiao-Ling Li, Shan-Chwen Chang, Pei-Jer Chen, Sui-Yuan Chang |
| EPI_ISL_745033 | West Java Health Laboratory | West Java Health Laboratory | Azzania Fibriani, Ema Rahmawati, Ryan Bayusantika Ristandi, Rifky Waluyajati Rachman, Cut Nur Cinthia Alamanda, Isak Solihin, Rini Robiani, Miftahul Faridl, Karimatu Khoirunnisa |
| EPI_ISL_745193 | Ospedale Di Venere | Istituto Zooprofilattico Sperimentale della Puglia e della Basilicata | Parisi A., Bianco A., Capozzi L., Del Sambro L., Chiara M., De Sabato L., Pesole G., Iacobellis M |
| EPI_ISL_745224, EPI_ISL_745230, EPI_ISL_745231, EPI_ISL_745232, EPI_ISL_745233, EPI_ISL_745234, EPI_ISL_745235, EPI_ISL_745236, EPI_ISL_745237, EPI_ISL_745238, EPI_ISL_745239, EPI_ISL_745240, EPI_ISL_745241, EPI_ISL_745242, EPI_ISL_745243, EPI_ISL_745244, EPI_ISL_745245, EPI_ISL_745246, EPI_ISL_745247, EPI_ISL_745248, EPI_ISL_745249, EPI_ISL_745250, EPI_ISL_745251, EPI_ISL_745252, EPI_ISL_745253, EPI_ISL_745254, EPI_ISL_745255, EPI_ISL_745256, EPI_ISL_745257, EPI_ISL_745258 | | | |
| see above | Molecular diagnostic laboratory of Federal Budget Institution of Science "Central Research Institute of Epidemiology" of The Federal Service on Customers' Rights Protection and Human Well-being Surveillance | Group of Genomics and Postgenomic Technologies of Central Research Institute of Epidemilology | Samoilov AE, Kaptelova VV, Korneenko EV, Dudorova AV, Saenko SS, Speranskaya AS, Tivanova EV, Shipulina OY, Akimkin VG |
| EPI_ISL_747235 | Sukabumi Health Laboratory | West Java Health Laboratory; School of Life Sciences and Technology, Institut Teknologi Bandung | Azzania Fibriani, Ema Rahmawati, Ryan Bayusantika Ristandi, Rifky Waluyajati Rachman, Cut Nur Cinthia Alamanda, Isak Solihin, Rini Robiani, Miftahul Faridl, Karimatu Khoirunnisa |
| EPI_ISL_747236 | Sukabumi Public Health | West Java Health Laboratory; School of Life Sciences and Technology, Institut Teknologi Bandung | Azzania Fibriani, Ema Rahmawati, Ryan Bayusantika Ristandi, Rifky Waluyajati Rachman, Cut Nur Cinthia Alamanda, Isak Solihin, Rini Robiani, Miftahul Faridl, Karimatu Khoirunnisa |
| EPI_ISL_747237 | National Institute of Health Research and Development | National Institute of Health Research and Development | Subangkit,Nafiandi,Estaria,V;Pawestri,HA;Puspa,KD;Nugraha,AA;Ikawati,HD;Pangesti,KNA;Soekarso,T;Paisal;Puspandari,N;Setiawaty,V |
| EPI_ISL_747238 | Bogor Public Health | West Java Health Laboratory; School of Life Sciences and Technology, Institut Teknologi Bandung | Azzania Fibriani, Ema Rahmawati, Ryan Bayusantika Ristandi, Rifky Waluyajati Rachman, Cut Nur Cinthia Alamanda, Isak Solihin, Rini Robiani, Miftahul Faridl, Karimatu Khoirunnisa |
| EPI_ISL_747239 | Kuningan Publlic Health Office | West Java Health Laboratory; School of Life Sciences and Technology, Institut Teknologi Bandung | Azzania Fibriani, Ema Rahmawati, Ryan Bayusantika Ristandi, Rifky Waluyajati Rachman, Cut Nur Cinthia Alamanda, Isak Solihin, Rini Robiani, Miftahul Faridl, Karimatu Khoirunnisa |
| EPI_ISL_747240 | Al Islam Hospital | West Java Health Laboratory; School of Life Sciences and Technology, Institut Teknologi Bandung | Azzania Fibriani, Ema Rahmawati, Ryan Bayusantika Ristandi, Rifky Waluyajati Rachman, Cut Nur Cinthia Alamanda, Isak Solihin, Rini Robiani, Miftahul Faridl, Karimatu Khoirunnisa |
| EPI_ISL_747241 | Subang Public Health Office | West Java Health Laboratory; School of Life Sciences and Technology, Institut Teknologi Bandung | Azzania Fibriani, Ema Rahmawati, Ryan Bayusantika Ristandi, Rifky Waluyajati Rachman, Cut Nur Cinthia Alamanda, Isak Solihin, Rini Robiani, Miftahul Faridl, Karimatu Khoirunnisa |
| EPI_ISL_747243 | General Hospital Laboratory | Pathogen Laboratory (BSL3), Biomedical Innovation Department, Experimental and Applied Biology Division, Scientific Research Center and High Education from Ensenada (CICESE) | Cervantes-Luevano K., Martinez M.A., Saavedra-Flores A., Galindo C. and Licea-Navarro A. |
| EPI_ISL_747283 | DNA SOLUTION LTD. | DNA SOLUTION LTD. | Md. Imran Khan, Kazi Nadim Hasan, Abu Sufian, Jannatun Naima, Abdul Khaleque, Mizanur Rahman, Firoz Kabir, Mohammad Fazle Alam Rabbi, Sharif Akhteruzzaman |
| EPI_ISL_754141, EPI_ISL_754142, EPI_ISL_754143, EPI_ISL_754144, | Laboratoire de Microbiologie | National Reference Center for Viruses of Respiratory Infections, Institut Pasteur, Paris | Marion Barbet, Sylvie Behillil, Méline Bizard, Angela Brisebarre, Camille Capel, Etienne Simon-Lorière, Vincent Enouf, Maud Vanpeene, Sylvie van der Werf, Marie-Sarah Fangous |

| | | | |
|---|---|---|---|
| EPI_ISL_754145, EPI_ISL_754147, EPI_ISL_754148, EPI_ISL_754149, EPI_ISL_754150, EPI_ISL_754151 | | | |
| EPI_ISL_754183 | Wyoming Public Health Laboratory | Wyoming Public Health Laborator | Noah Hull, Taylor Fearing, Channing Weber, Ashley Norberg, Bailey Bowcutt, and Wanda Manley |
| EPI_ISL_754196, EPI_ISL_754197, EPI_ISL_754198, EPI_ISL_754199, EPI_ISL_754200, EPI_ISL_754201, EPI_ISL_754202, EPI_ISL_754203, EPI_ISL_754204, EPI_ISL_754205, EPI_ISL_754206, EPI_ISL_754207, EPI_ISL_754208, EPI_ISL_754209, EPI_ISL_754210, EPI_ISL_754211, EPI_ISL_754212, EPI_ISL_754213, EPI_ISL_754214 | | | |
| see above | Molecular diagnostic laboratory of Federal Budget Institution of Science "Central Research Institute of Epidemiology" of The Federal Service on Customers' Rights Protection and Human Well-being Surveillance | Group of Genomics and Postgenomic Technologies of Central Research Institute of Epidemilology | Samoilov AE, Kaptelova VV, Korneenko EV, Saenko SS, Dudorova AV, Speranskaya AS, Tivanova EV, Shipulina OY, Akimkin VG |
| EPI_ISL_754238 | Laboratory for Respiratory Viruses, Cantacuzino National Military-Medical Institute for Research and Development | Cantacuzino Institute Virology | Luiza Ustea, Mühlemann Barbara, Mihaela Lazar |
| EPI_ISL_754843, EPI_ISL_754844, EPI_ISL_754845, EPI_ISL_754846 | Laboratoire de Virologie Hôpital Robert Debré | National Reference Center for Viruses of Respiratory Infections, Institut Pasteur, Paris | Marion Barbet, Sylvie Behillil, Méline Bizard, Angela Brisebarre, Camille Capel, Etienne Simon-Lorière, Vincent Enouf, Maud Vanpeene, Sylvie van der Werf, Andreoletti Laurent |
| EPI_ISL_754847 | Korian Villa d'Azon | National Reference Center for Viruses of Respiratory Infections, Institut Pasteur, Paris | Marion Barbet, Sylvie Behillil, Méline Bizard, Angela Brisebarre, Camille Capel, Etienne Simon-Lorière, Vincent Enouf, Maud Vanpeene, Sylvie van der Werf |
| EPI_ISL_754849, EPI_ISL_754850, EPI_ISL_754851, EPI_ISL_754852 | Laboratoire de Virologie Hôpital Robert Debré | National Reference Center for Viruses of Respiratory Infections, Institut Pasteur, Paris | Marion Barbet, Sylvie Behillil, Méline Bizard, Angela Brisebarre, Camille Capel, Etienne Simon-Lorière, Vincent Enouf, Maud Vanpeene, Sylvie van der Werf, Andreoletti Laurent |
| EPI_ISL_754853, EPI_ISL_754854, EPI_ISL_754855, EPI_ISL_754856 | CHU - Hôpital Cavale Blanche - Labo. de Virologie | National Reference Center for Viruses of Respiratory Infections, Institut Pasteur, Paris | Marion Barbet, Sylvie Behillil, Méline Bizard, Angela Brisebarre, Camille Capel, Etienne Simon-Lorière, Vincent Enouf, Maud Vanpeene, Sylvie van der Werf, Pilorge léa |
| EPI_ISL_755639 | Bundeswehr Institute of Microbiology | Bundeswehr Institute of Microbiology | Malena Bestehorn-Willmann, Mathias Walter, Christina Bugert, Joachim Bugert, Roman Wölfel, Markus Antwerpen |
| EPI_ISL_755640 | Instituto Adolfo Lutz - Central | Instituto Adolfo Lutz, Interdiciplinary Procedures Center, Strategic Laboratory | Claudio Tavares Sacchi, Claudia Regina Gonçalves, Erica Valessa Ramos Gomes, Karoline Rodrigues Campos |
| EPI_ISL_755641 | Instituto Adolfo Lutz - Regional de Santo Andre | Instituto Adolfo Lutz, Interdiciplinary Procedures Center, Strategic Laboratory | Claudio Tavares Sacchi, Claudia Regina Gonçalves, Erica Valessa Ramos Gomes, Karoline Rodrigues Campos |
| EPI_ISL_755642, EPI_ISL_755643 | Instituto Adolfo Lutz - Central | Instituto Adolfo Lutz, Interdiciplinary Procedures Center, Strategic Laboratory | Claudio Tavares Sacchi, Claudia Regina Gonçalves, Erica Valessa Ramos Gomes, Karoline Rodrigues Campos |
| EPI_ISL_755644, EPI_ISL_755645 | Lab LOC - Itapecerica da Serra | Instituto Adolfo Lutz, Interdiciplinary Procedures Center, Strategic Laboratory | Claudio Tavares Sacchi, Claudia Regina Gonçalves, Erica Valessa Ramos Gomes, Karoline Rodrigues Campos |
| EPI_ISL_755646, EPI_ISL_755647 | Instituto Adolfo Lutz - Regional de Santo Andre | Instituto Adolfo Lutz, Interdiciplinary Procedures Center, Strategic Laboratory | Claudio Tavares Sacchi, Claudia Regina Gonçalves, Erica Valessa Ramos Gomes, Karoline Rodrigues Campos |
| EPI_ISL_755648 | Instituto Adolfo Lutz - Regional de Taubate | Instituto Adolfo Lutz, Interdiciplinary Procedures Center, Strategic Laboratory | Claudio Tavares Sacchi, Claudia Regina Gonçalves, Erica Valessa Ramos Gomes, Karoline Rodrigues Campos |
| EPI_ISL_755649 | Instituto Adolfo Lutz - Regional de Santo Andre | Instituto Adolfo Lutz, Interdiciplinary Procedures Center, Strategic Laboratory | Claudio Tavares Sacchi, Claudia Regina Gonçalves, Erica Valessa Ramos Gomes, Karoline Rodrigues Campos |
| EPI_ISL_755650 | Instituto Adolfo Lutz - Regional de Taubate | Instituto Adolfo Lutz, Interdiciplinary Procedures Center, Strategic Laboratory | Claudio Tavares Sacchi, Claudia Regina Gonçalves, Erica Valessa Ramos Gomes, Karoline Rodrigues Campos |
| EPI_ISL_755651 | Instituto Adolfo Lutz - Central | Instituto Adolfo Lutz, Interdiciplinary Procedures Center, Strategic Laboratory | Claudio Tavares Sacchi, Claudia Regina Gonçalves, Erica Valessa Ramos Gomes, Karoline Rodrigues Campos |
| EPI_ISL_755652 | Lab LOC - Itapecerica da Serra | Instituto Adolfo Lutz, Interdiciplinary Procedures Center, Strategic Laboratory | Claudio Tavares Sacchi, Claudia Regina Gonçalves, Erica Valessa Ramos Gomes, Karoline Rodrigues Campos |
| EPI_ISL_755653, EPI_ISL_755654 | Instituto Adolfo Lutz - Central | Instituto Adolfo Lutz, Interdiciplinary Procedures Center, Strategic Laboratory | Claudio Tavares Sacchi, Claudia Regina Gonçalves, Erica Valessa Ramos Gomes, Karoline Rodrigues Campos |
| EPI_ISL_755655 | Instituto Adolfo Lutz - Regional de Campinas | Instituto Adolfo Lutz, Interdiciplinary Procedures Center, Strategic Laboratory | Claudio Tavares Sacchi, Claudia Regina Gonçalves, Erica Valessa Ramos Gomes, Karoline Rodrigues Campos |
| EPI_ISL_756294 | Center for Biotechnology and Cell Therapy, São Rafael Hospital, Salvador, Brazil | Center for Biotechnology and Cell Therapy, São Rafael Hospital, Salvador, Brazil | Carolina Kymie Vasques Nonaka, Marília Miranda Franco, Tiago Gräf, Ana Verena Almeida Mendes, Renato Santana de Aguiar, Marta Giovanetti, Bruno Solano de Freitas Souza |
| EPI_ISL_757285 | Department of Virology, Public Health Laboratories Division | Department of Virology, Public Health Laboratories Division | Massab Umair, Aamer Ikram, Muhammad Salman |
| EPI_ISL_759701 | RS UNS, Surakarta | UNS/RS-UNS, Faculty of Medicine | Betty Suryawati, Yulia Sari, Hartono, Maryani, Afif Avicenna Gufron, HanaHana Apsari Pawestri, Kartika Dewi Puspa, Hartanti Dian Ika, Arie Ardiansyah Nugraha, Vivi Setiawaty. |
| EPI_ISL_759892, EPI_ISL_759893, EPI_ISL_759894 | Temporary Specimen Collection Centre | Hong Kong Department of Health | Alan K.L. Tsang, Peter C.W. Yip, Edman T.K. Lam, Rickjason C.W. Chan, Dominic N.C. Tsang |
| EPI_ISL_759895 | United Christian Hospital | Hong Kong Department of Health | Alan K.L. Tsang, Peter C.W. Yip, Edman T.K. Lam, Rickjason C.W. Chan, Dominic N.C. Tsang |
| EPI_ISL_759896 | Queen Elizabeth Hospital | Hong Kong Department of Health | Alan K.L. Tsang, Peter C.W. Yip, Edman T.K. Lam, Rickjason C.W. Chan, Dominic N.C. Tsang |
| EPI_ISL_759897, EPI_ISL_759898 | United Christian Hospital | Hong Kong Department of Health | Alan K.L. Tsang, Peter C.W. Yip, Edman T.K. Lam, Rickjason C.W. Chan, Dominic N.C. Tsang |
| EPI_ISL_759968 | RSI Darus Syifa' | Institute of Tropical Disease, Universitas Airlangga | Maria I Lusida, Krisnoadi Rahardjo, Aldise M Nastri, Jezzy R Dewantari, Rima R Prasetya, HM Faiz, Gatot Soegiarto, Laksmi Wulandari, Resti Yudhawati, Soetjipto, Yasuko Mori, Kazufumi Shimizu |
| EPI_ISL_760032, EPI_ISL_760033, EPI_ISL_760034, EPI_ISL_760035, EPI_ISL_760036, EPI_ISL_760037, EPI_ISL_760038, EPI_ISL_760039, EPI_ISL_760040, EPI_ISL_760041, EPI_ISL_760042, EPI_ISL_760043, EPI_ISL_760044, EPI_ISL_760045, EPI_ISL_760046, EPI_ISL_760047, EPI_ISL_760048, EPI_ISL_760049, EPI_ISL_760050, EPI_ISL_760051, EPI_ISL_760052, EPI_ISL_760053, EPI_ISL_760054, EPI_ISL_760055, EPI_ISL_760056, EPI_ISL_760057, EPI_ISL_760058 | | | |
| see above | Hong Kong Department of Health | School of Public Health, The University of Hong Kong | Daniel Chu, Haogao Gu, Pavithra Krishnan, Daisy Ng, Gigi Liu, Carrie Wan, Malik Peiris, Leo Poon |
| EPI_ISL_763074, EPI_ISL_763075 | Diagnosticos da America - DASA | Instituto Adolfo Lutz, Interdiciplinary Procedures Center, Strategic Laboratory | Claudio Tavares Sacchi, Claudia Regina Gonçalves, Erica Valessa Ramos Gomes, Karoline Rodrigues Campos |
| EPI_ISL_765478 | EHPAD | National Reference Center for Viruses of Respiratory Infections, Institut Pasteur, Paris | Marion Barbet, Sylvie Behillil, Méline Bizard, Angela Brisebarre, Camille Capel, Etienne Simon-Lorière, Vincent Enouf, Maud Vanpeene, Sylvie van der Werf |
| EPI_ISL_765479 | Laboratoire de Virologie, Hôpital Robert Debré | National Reference Center for Viruses of Respiratory Infections, Institut Pasteur, Paris | Marion Barbet, Sylvie Behillil, Méline Bizard, Angela Brisebarre, Camille Capel, Etienne Simon-Lorière, Vincent Enouf, Maud Vanpeene, Sylvie van der Werf, Andreoletti Laurent |
| EPI_ISL_767013 | University of Bari Biomedical Sciences and Human Oncology | University of Bari Biomedical Sciences and Human Oncology | Maria Chironna, Anna Sallustio, Daniela Loconsole, Marisa Accogli |
| EPI_ISL_768324 | Singapore General Hospital | Department of Microbiology | Nurdyana Abdul Rahman, Kun Lee Lim, Chenhao Li, Sui Sin Goh, Kenneth Xin Long Chan, Kian Sing Chan, Lynette Oon, Kern Rei Chng, Niranjan Nagarajan, Karrie Ko |

| | | | |
|---|---|---|---|
| EPI_ISL_774976 | Gonoshasthya-RNA Molecular Diagnostic and Research Center | Gonoshasthya-RNA Molecular Diagnostic and Research Center | Nihad Adnan, Mohd. Raeed Jamiruddin, Md. Ahsanul Haq, Mohib Ullah Khondoker, Nafisa Azmuda, Firoz Ahmed, Shahana Sharmin, Salma Akter, Taslin Jahan Mou, Mahfuza Marzan, Sayeda Moriam Liza, Nowshin Jahan, Tamanna Ali, Maha Jamiruddin, Mousumi Chaity, Shahad Saif Khandker, Mumtarin Jannat Oishee |
| EPI_ISL_775019 | Gonoshasthya-RNA Molecular Research Center | Gonoshasthya-RNA Molecular Research Center | Nihad Adnan, Mohd. Raeed Jamiruddin, Md. Ahsanul Haq, Mohib Ullah Khondoker, Nafisa Azmuda, Firoz Ahmed, Shahana Sharmin, Salma Akter, Taslin Jahan Mou, Mahfuza Marzan, Sayeda Moriam Liza, Nowshin Jahan, Tamanna Ali, Maha Jamiruddin, Mousumi Chaity, Shahad Saif Khandker, Mumtarin Jannat Oishee |
| EPI_ISL_775020 | Gonoshasthya-RNA Molecular Research Center | Gonoshasthya-RNA Molecular Research Center | Mohd. Raeed Jamiruddin, Nihad Adnan, Md. Ahsanul Haq, Mohib Ullah Khondoker, Nafisa Azmuda, Firoz Ahmed, Shahana Sharmin, Salma Akter, Taslin Jahan Mou, Mahfuza Marzan, Sayeda Moriam Liza, Nowshin Jahan, Tamanna Ali, Shahad Saif Khandker, Maha Jamiruddin, Mousumi Chaity, Mumtarin Jannat Oishee |
| EPI_ISL_775213, EPI_ISL_775215 | Gonoshasthya-RNA Molecular Research Center | Gonoshasthya-RNA Molecular Research Center | Nihad Adnan, Mohd. Raeed Jamiruddin, Md. Ahsanul Haq, Mohib Ullah Khondoker, Nafisa Azmuda, Firoz Ahmed, Shahana Sharmin, Salma Akter, Taslin Jahan Mou, Mahfuza Marzan, Sayeda Moriam Liza, Nowshin Jahan, Tamanna Ali, Maha Jamiruddin, Mousumi Chaity, Shahad Saif Khandker, Mumtarin Jannat Oishee |
| EPI_ISL_775216 | Gonoshasthya-RNA Molecular Research Center | Gonoshasthya-RNA Molecular Research Center | Mohd. Raeed Jamiruddin, Nihad Adnan, Md. Ahsanul Haq, Mohib Ullah Khondoker, Nafisa Azmuda, Firoz Ahmed, Shahana Sharmin, Salma Akter, Taslin Jahan Mou, Mahfuza Marzan, Sayeda Moriam Liza, Nowshin Jahan, Tamanna Ali, Shahad Saif Khandker, Maha Jamiruddin, Mousumi Chaity, Mumtarin Jannat Oishee |
| EPI_ISL_775217 | Gonoshasthya-RNA Molecular Research Center | Gonoshasthya-RNA Molecular Research Center | Nihad Adnan, Mohd. Raeed Jamiruddin, Md. Ahsanul Haq, Mohib Ullah Khondoker, Nafisa Azmuda, Firoz Ahmed, Shahana Sharmin, Salma Akter, Taslin Jahan Mou, Mahfuza Marzan, Sayeda Moriam Liza, Nowshin Jahan, Tamanna Ali, Maha Jamiruddin, Mousumi Chaity, Shahad Saif Khandker, Mumtarin Jannat Oishee |
| EPI_ISL_775218 | Gonoshasthya-RNA Molecular Research Center | Gonoshasthya-RNA Molecular Research Center | Mohd. Raeed Jamiruddin, Nihad Adnan, Md. Ahsanul Haq, Mohib Ullah Khondoker, Nafisa Azmuda, Firoz Ahmed, Shahana Sharmin, Salma Akter, Taslin Jahan Mou, Mahfuza Marzan, Sayeda Moriam Liza, Nowshin Jahan, Tamanna Ali, Shahad Saif Khandker, Maha Jamiruddin, Mousumi Chaity, Mumtarin Jannat Oishee |
| EPI_ISL_776766 | Instituto Adolfo Lutz - Regional de Santo Andre | Instituto Adolfo Lutz, Interdiciplinary Procedures Center, Strategic Laboratory | Claudio Tavares Sacchi, Claudia Regina Gonçalves, Erica Valessa Ramos Gomes, Karoline Rodrigues Campos |
| EPI_ISL_776767 | Instituto Adolfo Lutz - Regional de Marilia | Instituto Adolfo Lutz, Interdiciplinary Procedures Center, Strategic Laboratory | Claudio Tavares Sacchi, Claudia Regina Gonçalves, Erica Valessa Ramos Gomes, Karoline Rodrigues Campos |
| EPI_ISL_776768 | Instituto Adolfo Lutz - Regional de Aracatuba | Instituto Adolfo Lutz, Interdiciplinary Procedures Center, Strategic Laboratory | Claudio Tavares Sacchi, Claudia Regina Gonçalves, Erica Valessa Ramos Gomes, Karoline Rodrigues Campos |
| EPI_ISL_776769 | Instituto Adolfo Lutz - Regional de Santo Andre | Instituto Adolfo Lutz, Interdiciplinary Procedures Center, Strategic Laboratory | Claudio Tavares Sacchi, Claudia Regina Gonçalves, Erica Valessa Ramos Gomes, Karoline Rodrigues Campos |
| EPI_ISL_788943, EPI_ISL_788948, EPI_ISL_788949, EPI_ISL_788950, EPI_ISL_788951, EPI_ISL_788952, EPI_ISL_788968, EPI_ISL_788969 | Ospedale "Di Venere" | Beaconlab (Bioinformatics, Evolution and Comparative Genomics lab), Dept of Biosciences, University on Milan | Iacobellis M, d'Avenia M, Piluscio R, Parisi A, Chiara M, Manzari C, Pesole G |
| EPI_ISL_791087, EPI_ISL_791093, EPI_ISL_791096 | Instituto Nacional de Salud - Unidad de Secuenciación y Análisis Genómico | Instituto Nacional de Salud - Dirección de Investigación en Salud Pública | Katherine Laiton-Donato, Diego A. Álvarez-Díaz, Carlos Franco-Muñoz, Mauricio Pacheco-Montealegre, Jonathan Reales, Sheryl Corchuelo, Maria T. Herrera, Julian Naizaque, Gerardo Santamaría, Paola Muñoz-Laiton, Diego Andrés Prada, Magdalena Wiesner, Martha Lucia Ospina Martinez, Marcela Mercado-Reyes |
| EPI_ISL_791992, EPI_ISL_791994, EPI_ISL_791995, EPI_ISL_791996, EPI_ISL_791997, EPI_ISL_791998 | CHU - Hôpital Cavale Blanche | National Reference Center for Viruses of Respiratory Infections, Institut Pasteur, Paris | Marion Barbet, Sylvie Behillil, Méline Bizard, Angela Brisebarre, Camille Capel, Etienne Simon-Lorière, Vincent Enouf, Maud Vanpeene, Sylvie van der Werf, Léa Pilorgé |
| EPI_ISL_792000, EPI_ISL_792001, EPI_ISL_792002, EPI_ISL_792004, EPI_ISL_792005, EPI_ISL_792006, EPI_ISL_792007 | Laboratoire de Virologie Hôpital Robert Debré | National Reference Center for Viruses of Respiratory Infections, Institut Pasteur, Paris | Marion Barbet, Sylvie Behillil, Méline Bizard, Angela Brisebarre, Camille Capel, Etienne Simon-Lorière, Vincent Enouf, Maud Vanpeene, Sylvie van der Werf, Andreoletti Laurent |
| EPI_ISL_792008, EPI_ISL_792009, EPI_ISL_792010, EPI_ISL_792011, EPI_ISL_792012, EPI_ISL_792013, EPI_ISL_792014, EPI_ISL_792015 | CH de Bethune - Laboratoire de Biologie Médicale | National Reference Center for Viruses of Respiratory Infections, Institut Pasteur, Paris | Marion Barbet, Sylvie Behillil, Méline Bizard, Angela Brisebarre, Camille Capel, Etienne Simon-Lorière, Vincent Enouf, Maud Vanpeene, Sylvie van der Werf, Léa Pilorgé |
| EPI_ISL_792016, EPI_ISL_792017, EPI_ISL_792018, EPI_ISL_792019, EPI_ISL_792020, EPI_ISL_792021 | Plateforme COVID IDF | National Reference Center for Viruses of Respiratory Infections, Institut Pasteur, Paris | Marion Barbet, Sylvie Behillil, Méline Bizard, Angela Brisebarre, Camille Capel, Etienne Simon-Lorière, Vincent Enouf, Maud Vanpeene, Sylvie van der Werf,Jacques Fourgeaud |
| EPI_ISL_792042 | General practitioner | National Reference Center for Viruses of Respiratory Infections, Institut Pasteur, Paris | Marion Barbet, Sylvie Behillil, Méline Bizard, Angela Brisebarre, Camille Capel, Etienne Simon-Lorière, Vincent Enouf, Maud Vanpeene, Sylvie van der Werf, |
| EPI_ISL_792045, EPI_ISL_792046, EPI_ISL_792047, EPI_ISL_792048, EPI_ISL_792050 | General practitioner | National Reference Center for Viruses of Respiratory Infections, Institut Pasteur, Paris | Marion Barbet, Sylvie Behillil, Méline Bizard, Angela Brisebarre, Camille Capel, Etienne Simon-Lorière, Vincent Enouf, Maud Vanpeene, Sylvie van der Werf,Brieuc Lefaure |
| EPI_ISL_792053 | General practitioner | National Reference Center for Viruses of Respiratory Infections, Institut Pasteur, Paris | Marion Barbet, Sylvie Behillil, Méline Bizard, Angela Brisebarre, Camille Capel, Etienne Simon-Lorière, Vincent Enouf, Maud Vanpeene, Sylvie van der Werf |
| EPI_ISL_792055 | hospital | National Reference Center for Viruses of Respiratory Infections, Institut Pasteur, Paris | Marion Barbet, Sylvie Behillil, Méline Bizard, Angela Brisebarre, Camille Capel, Etienne Simon-Lorière, Vincent Enouf, Maud Vanpeene, Sylvie van der Werf,Patricia Stoessel |
| EPI_ISL_792056 | Hopital | National Reference Center for Viruses of Respiratory Infections, Institut Pasteur, Paris | Marion Barbet, Sylvie Behillil, Méline Bizard, Angela Brisebarre, Camille Capel, Etienne Simon-Lorière, Vincent Enouf, Maud Vanpeene, Sylvie van der Werf,Patricia Stoessel |
| EPI_ISL_792560 | Laboratorio de Ecologia de Doencas Transmissiveis na Amazonia, Instituto Leonidas e Maria Deane - Fiocruz Amazonia | Laboratorio de Ecologia de Doencas Transmissiveis na Amazonia, Instituto Leonidas e Maria Deane - Fiocruz Amazonia | Valdinete Nascimento, Victor Souza, André Corado, Fernanda Nascimento, George Silva, Ágatha Costa, Karina Pessoa, Debora Duarte, Luciana Gonçalves, Maria Júlia Brandão, Michele Jesus, Felipe Naveca on behalf of the Fiocruz COVID-19 Genomic Surveillance Network |
| EPI_ISL_792684, EPI_ISL_792685, EPI_ISL_792686, EPI_ISL_792687, EPI_ISL_792688, EPI_ISL_792689, EPI_ISL_792690, EPI_ISL_792691, EPI_ISL_792692, EPI_ISL_792693, EPI_ISL_792694, EPI_ISL_792695, EPI_ISL_792696, EPI_ISL_792697, EPI_ISL_792698, EPI_ISL_792699, EPI_ISL_792700, EPI_ISL_792701, EPI_ISL_792702, EPI_ISL_792703, EPI_ISL_792704 | | | |
| see above | The National Institute of Public Health | State Veterinary Institute Prague | Nagy,A;Jirincova,H;Trnka,D;Vecerova,J |
| EPI_ISL_794592 | Laboratorio Estatal de Salud Pública de Tamaulipas | Instituto de diagnóstico y Referencia Epidemiologicos (INDRE) | Abril Rodriguez-Maldonado, Claudia Wong-Arambula, Fabiola Garces-Ayala, Gisela Barrera-Badillo, Ana Maria Cortez-Calderon, Bernardita Reyes-Berrones, Hilda del Carmen-Selvera, Gloria Molina-Gamboa, Lucia Hernandez-Rivas, Irma Lopez-Martinez, Celia Alpuche-Aranda, Jose Luis Alomia-Zegarra, Hugo Lopez Gatell-Ramirez, Ernesto Ramirez-Gonzalez. |
| EPI_ISL_794649 | Fundación Cardio Infantil | Instituto Nacional de Salud - Dirección de Investigación en Salud Pública | Katherine Laiton-Donato, Diego A. Álvarez-Díaz, Carlos Franco-Muñoz, Mauricio Pacheco-Montealegre, Jonathan Reales, Sheryl Corchuelo, Maria T. Herrera, Julian Naizaque, Gerardo Santamaría, Paola Muñoz-Laiton, Diego Andrés Prada, Magdalena Wiesner, Martha Lucia Ospina Martinez, Marcela Mercado-Reyes |
| EPI_ISL_794654 | Fundación Valle del Lili | Instituto Nacional de Salud - Dirección de Investigación en | Katherine Laiton-Donato, Diego A. Álvarez-Díaz, Carlos Franco-Muñoz, Mauricio Pacheco-Montealegre, Jonathan Reales, Sheryl Corchuelo, Maria T. Herrera, |

| | | | |
|---|---|---|---|
| | | Salud Pública | Julian Naizaque, Gerardo Santamaría, Paola Muñoz-Laiton, Diego Andrés Prada, Magdalena Wiesner, Martha Lucia Ospina Martinez, Marcela Mercado-Reyes |
| EPI_ISL_794659 | HOSPITAL UNIVERSITARIO SAN IGNACIO | Instituto Nacional de Salud - Dirección de Investigación en Salud Pública | Katherine Laiton-Donato, Diego A. Álvarez-Díaz, Carlos Franco-Muñoz, Mauricio Pacheco-Montealegre, Jonathan Reales, Sheryl Corchuelo, Maria T. Herrera, Julian Naizaque, Gerardo Santamaría, Paola Muñoz-Laiton, Diego Andrés Prada, Magdalena Wiesner, Martha Lucia Ospina Martinez, Marcela Mercado-Reyes |
| EPI_ISL_794735, EPI_ISL_794736, EPI_ISL_794738 | 1-Laboratory of Microbiology, National Reference Lab, Charles Nicolle Hospital; 2-University of Tunis ElManar, Faculty of Medicine of Tunis, LR99ES09, Tunis, Tunisia | 1-Clinical and Experimental Pharmacology Lab, LR16SP02, National Center of Pharmacovigilance, University of Tunis El Manar, Tunis, Tunisia. 2-Neurodegenerative diseases and psychiatric troubles, LR18SP03, Razi Hospital, University of Tunis El Manar, Tunis, Tunisia. 3- Ministry of Health, National Observatory of New and Emerging Diseases, 1006, Tunis, Tunisia | Ilhem Boutiba-Ben Boubaker, Sameh Trabelsi, Nissaf Ben Alaya, Maher Kharrat, Alia BenKahla, Jalila Ben Khelil, Salma Abid, Sana Ferjani, Mouna Ben Sassi, Mouna Safer, Zaineb Hamzaoui, Guedi Ali Barreh, Habiba Ben Romdhane, Souissi Amira, Sarra Chamman, Hanen El Jebari, Asma Ferjani, Gaies Emna, Riadh Daghfous, Riadh Gouider. |
| EPI_ISL_794744 | National Institute of Infectious Diseases-Prof. Dr. Matei Bals Molecular Diagnostics Laboratory | National Institute of Infectious Diseases-Prof. Dr. Matei Bals Molecular Diagnostics Laboratory | Leontina Banica, Marius Surleac, Corina Casangiu, Petre Milu, Andreea Tudor, Simona Paraschiv, Dan Otelea |
| EPI_ISL_794746 | SISP ASL BA AREA SUD | Istituto Zooprofilattico Sperimentale della Puglia e della Basilicata | Parisi A., Bianco A., Capozzi L., Del Sambro L., Manzulli V, Rondinone V., Pace L., Cipolletta D., Galante D. |
| EPI_ISL_794747 | Ospedale Vito Fazzi | Istituto Zooprofilattico Sperimentale della Puglia e della Basilicata | Parisi A., Bianco A., Capozzi L., Del Sambro L., Manzulli V, Rondinone V., Pace L., Cipolletta D., Galante D. |
| EPI_ISL_794748 | SISP ASL BA AREA SUD | Istituto Zooprofilattico Sperimentale della Puglia e della Basilicata | Parisi A., Bianco A., Capozzi L., Del Sambro L., Manzulli V, Rondinone V., Pace L., Cipolletta D., Galante D. |
| EPI_ISL_794749 | Istituto Zooprofilattico Sperimentale della Puglia e della Basilicata | Istituto Zooprofilattico Sperimentale della Puglia e della Basilicata | Parisi A., Bianco A., Capozzi L., Del Sambro L., Manzulli V, Rondinone V., Pace L., Cipolletta D., Galante D. |
| EPI_ISL_794750 | Ospedale Santa Caterina Novella | Istituto Zooprofilattico Sperimentale della Puglia e della Basilicata | Parisi A., Bianco A., Capozzi L., Del Sambro L., Manzulli V, Rondinone V., Pace L., Cipolletta D., Galante D. |
| EPI_ISL_794751 | Ospedale Vito Fazzi | Istituto Zooprofilattico Sperimentale della Puglia e della Basilicata | Parisi A., Bianco A., Capozzi L., Del Sambro L., Manzulli V, Rondinone V., Pace L., Cipolletta D., Galante D. |
| EPI_ISL_794752 | Istituto Zooprofilattico Sperimentale della Puglia e della Basilicata | Istituto Zooprofilattico Sperimentale della Puglia e della Basilicata | Parisi A., Bianco A., Capozzi L., Del Sambro L., Manzulli V, Rondinone V., Pace L., Cipolletta D., Galante D. |
| EPI_ISL_794753 | Ospedale Cardinale G. Panico | Istituto Zooprofilattico Sperimentale della Puglia e della Basilicata | Parisi A., Bianco A., Capozzi L., Del Sambro L., Manzulli V, Rondinone V., Pace L., Cipolletta D., Galante D. |
| EPI_ISL_794754, EPI_ISL_794755 | Istituto Zooprofilattico Sperimentale della Puglia e della Basilicata | Istituto Zooprofilattico Sperimentale della Puglia e della Basilicata | Parisi A., Bianco A., Capozzi L., Del Sambro L., Manzulli V, Rondinone V., Pace L., Cipolletta D., Galante D. |
| EPI_ISL_794756, EPI_ISL_794757, EPI_ISL_794758 | Presidio di Brindisi Di Summa - Perrino | Istituto Zooprofilattico Sperimentale della Puglia e della Basilicata | Parisi A., Bianco A., Capozzi L., Del Sambro L., Manzulli V, Rondinone V., Pace L., Cipolletta D., Galante D. |
| EPI_ISL_794759, EPI_ISL_794760, EPI_ISL_794761, EPI_ISL_794762, EPI_ISL_794763, EPI_ISL_794764, EPI_ISL_794765, EPI_ISL_794766, EPI_ISL_794767, EPI_ISL_794768, EPI_ISL_794769, EPI_ISL_794770, EPI_ISL_794771, EPI_ISL_794772, EPI_ISL_794773, EPI_ISL_794774, EPI_ISL_794775, EPI_ISL_794776, EPI_ISL_794777, EPI_ISL_794778, EPI_ISL_794779, EPI_ISL_794780, EPI_ISL_794781, EPI_ISL_794782, EPI_ISL_794783, EPI_ISL_794784, EPI_ISL_794785, EPI_ISL_794786, EPI_ISL_794787, EPI_ISL_794788, EPI_ISL_794789, EPI_ISL_794790, EPI_ISL_794791, EPI_ISL_794792, EPI_ISL_794793, EPI_ISL_794794, EPI_ISL_794795, EPI_ISL_794796, EPI_ISL_794797, EPI_ISL_794798, EPI_ISL_794799, EPI_ISL_794800, EPI_ISL_794801, EPI_ISL_794802, EPI_ISL_794803, EPI_ISL_794804, EPI_ISL_794805, EPI_ISL_794806, EPI_ISL_794807, EPI_ISL_794808, EPI_ISL_794809, EPI_ISL_794810, EPI_ISL_794811, EPI_ISL_794812, EPI_ISL_794813, EPI_ISL_794814, EPI_ISL_794815, EPI_ISL_794816, EPI_ISL_794817 | | | |
| see above | Istituto Zooprofilattico Sperimentale della Puglia e della Basilicata | Istituto Zooprofilattico Sperimentale della Puglia e della Basilicata | Parisi A., Bianco A., Capozzi L., Del Sambro L., Manzulli V, Rondinone V., Pace L., Cipolletta D., Galante D. |
| EPI_ISL_794818, EPI_ISL_794820 | Greek Genome Center, Biomedical Research Foundation of the Academy of Athens (BRFAA) | Greek Genome Center, Biomedical Research Foundation of the Academy of Athens (BRFAA) | Emmanouil Athanasiadis, Ioannis Vatsellas, Thodoris Loupis, Christina Maria Kravvari, Katerina Zoi, Dimitrios Thanos |
| EPI_ISL_796027, EPI_ISL_796028, EPI_ISL_796029, EPI_ISL_796030, EPI_ISL_796031, EPI_ISL_796032, EPI_ISL_796033, EPI_ISL_796034, EPI_ISL_796035, EPI_ISL_796036, EPI_ISL_796037, EPI_ISL_796038, EPI_ISL_796040, EPI_ISL_796054, EPI_ISL_796055, EPI_ISL_796056, EPI_ISL_796057, EPI_ISL_796058, EPI_ISL_796059, EPI_ISL_796060, EPI_ISL_796061 | | | |
| see above | Institute of Virology, University of Cologne | Institute of Virology, University of Cologne | Saleta Sierra, Gibran Rubio, Zevanya Tessalonica, Dominik Aschenmeier, Eva Heger, Elena Knops, Rolf Kaiser, Maximilian Damagnez, Andreas Walker, Jörg Timm, Alexander Dilthey, Martin Däumer, Alex Thielen |
| EPI_ISL_802726, EPI_ISL_802727, EPI_ISL_802728, EPI_ISL_802729, EPI_ISL_802730, EPI_ISL_802731, EPI_ISL_802732, EPI_ISL_802733, EPI_ISL_802734, EPI_ISL_802735, EPI_ISL_802736, EPI_ISL_802737, EPI_ISL_802738, EPI_ISL_802739, EPI_ISL_802740, EPI_ISL_802741, EPI_ISL_802742, EPI_ISL_802743, EPI_ISL_802744, EPI_ISL_802745, EPI_ISL_802746, EPI_ISL_802747, EPI_ISL_802748, EPI_ISL_802749 | | | |
| see above | Hospital Clínic de Barcelona | Instituto de Salud Carlos III | Iglesias-Caballero, M. Molinero Calamita, M. González-Esguevillas, M. Camarero, S. Pozo, F. Casas, I. Jiménez, P. Jiménez, M. Zaballos, A. Monzón, S. Varona, S. Juliá, M. Cuesta, I, M.A Marcos. |
| EPI_ISL_802750, EPI_ISL_802751, EPI_ISL_802752, EPI_ISL_802753, EPI_ISL_802754, EPI_ISL_802755, EPI_ISL_802756, EPI_ISL_802757, EPI_ISL_802758, EPI_ISL_802759, EPI_ISL_802760, EPI_ISL_802761, EPI_ISL_802762, EPI_ISL_802763, EPI_ISL_802764, EPI_ISL_802765, EPI_ISL_802766, EPI_ISL_802767, EPI_ISL_802768, EPI_ISL_802769, EPI_ISL_802770, EPI_ISL_802771, EPI_ISL_802772, EPI_ISL_802773, EPI_ISL_802774, EPI_ISL_802775, EPI_ISL_802776 | | | |
| see above | Hospital Santa María Nai | Instituto de Salud Carlos III | Iglesias-Caballero, M. Molinero Calamita, M. González-Esguevillas, M. Camarero, S. Pozo, F. Casas, I. Jiménez, P. Jiménez, M. Zaballos, A. Monzón, S. Varona, S. Juliá, M. Cuesta, I, J. García. |
| EPI_ISL_803896 | Virology Unit, Pisa University Hospital and Retrovirus Center, University of Pisa | National Institute for Infectious Diseases, INMI, "L. Spallanzani" IRCCS | M Rueca, E Giombini, C.E.M Gruber, B Bartolini, O Butera, F Messina, A Rosellini, P Mazzetti, M Pistello, A Di Caro, MR Capobianchi |
| EPI_ISL_804038, EPI_ISL_804040, EPI_ISL_804041, EPI_ISL_804042, EPI_ISL_804044, EPI_ISL_804048, EPI_ISL_804052 | SC (UCO) Igiene e Sanità Pubblica (funzione integrata con SC Microbiologia e Virologia) e Laboratory of Molecular Virology of the International Centre for Genetic Engineering and Biotechnology (ICGEB) | ARGO Laboratorio Genomica ed Epigenomica | Licastro D, Dal Monego S, Degasperi M, Marcello A, D'Agaro P |
| EPI_ISL_804963, EPI_ISL_804964, EPI_ISL_804968, EPI_ISL_804969, EPI_ISL_804971, EPI_ISL_804972, EPI_ISL_804973, EPI_ISL_804974, EPI_ISL_804975, EPI_ISL_804976, EPI_ISL_804978, EPI_ISL_804979 | | | |
| see above | Hospital Comarcal de Melilla | Instituto de Salud Carlos III | Iglesias-Caballero, M. Molinero Calamita, M. González-Esguevillas, M. Camarero, S. Pozo, F. Casas, I. Jiménez, P. Jiménez, M. Zaballos, A. Monzón, S. Varona, S. Juliá, M. Cuesta, I, J. López |
| EPI_ISL_806544 | INSPI Instituto Nacional de Investigación en Salud Pública | Av. Julián Coronel 905 entre Esmeraldas y José Mascote Av. Juan Tanca Marengo No. 100 y Av. de las Américas | Leandro Patiño, Doménica de Mora, Maritza Olmedo, Andrés Carrazco, Orson Mestanza, Mary Regato, Melissa Zambrano, Manuel González, Alfredo Bruno, Alberto Orlando. |
| EPI_ISL_806730 | Presidio Ospedaliero S.Liberatore Atri | Istituto Zooprofilattico Sperimentale dell'Abruzzo e Molise "G. Caporale" | Lorusso A, Marcacci M, Di Domenico M, Ancora M, Curini V, Mangone I, Rinaldi A, Di Pasquale A, Cammà C, Puglia I, Calistri P, Savini G |
| EPI_ISL_806732 | SIESP DIPARTIMENTO DI PREVENZIONE CHIETI | Istituto Zooprofilattico Sperimentale dell'Abruzzo e Molise "G. Caporale" | Lorusso A, Marcacci M, Di Domenico M, Ancora M, Curini V, Mangone I, Rinaldi A, Di Pasquale A, Cammà C, Puglia I, Calistri P, Savini G |
| EPI_ISL_806733 | SIESP CHIETI - DRIVE IN CHIETI | Istituto Zooprofilattico Sperimentale dell'Abruzzo e Molise "G. Caporale" | Lorusso A, Marcacci M, Di Domenico M, Ancora M, Curini V, Mangone I, Rinaldi A, Di Pasquale A, Cammà C, Puglia I, Calistri P, Savini G |
| EPI_ISL_806734 | SIESP CHIETI - DRIVE IN LANCIANO | Istituto Zooprofilattico Sperimentale dell'Abruzzo e Molise "G. | Lorusso A, Marcacci M, Di Domenico M, Ancora M, Curini V, Mangone I, Rinaldi A, Di Pasquale A, Cammà C, Puglia I, Calistri P, Savini G |

| | | | |
|---|---|---|---|
| | | Caporale" | |
| EPI_ISL_806735, EPI_ISL_806736, EPI_ISL_806737, EPI_ISL_806738, EPI_ISL_806739, EPI_ISL_806740, EPI_ISL_806741, EPI_ISL_806742, EPI_ISL_806743, EPI_ISL_806744 | SIESP DIPARTIMENTO DI PREVENZIONE TERAMO | Istituto Zooprofilattico Sperimentale dell'Abruzzo e Molise "G. Caporale" | Lorusso A, Marcacci M, Di Domenico M, Ancora M, Curini V, Mangone I, Rinaldi A, Di Pasquale A, Cammà C, Puglia I, Calistri P, Savini G |
| EPI_ISL_806745 | RSA Giulianova | Istituto Zooprofilattico Sperimentale dell'Abruzzo e Molise "G. Caporale" | Lorusso A, Marcacci M, Di Domenico M, Ancora M, Curini V, Mangone I, Rinaldi A, Di Pasquale A, Cammà C, Puglia I, Calistri P, Savini G |
| EPI_ISL_806746, EPI_ISL_806747, EPI_ISL_806748 | SIESP DIPARTIMENTO DI PREVENZIONE TERAMO | Istituto Zooprofilattico Sperimentale dell'Abruzzo e Molise "G. Caporale" | Lorusso A, Marcacci M, Di Domenico M, Ancora M, Curini V, Mangone I, Rinaldi A, Di Pasquale A, Cammà C, Puglia I, Calistri P, Savini G |
| EPI_ISL_806749, EPI_ISL_806750, EPI_ISL_806751, EPI_ISL_806752, EPI_ISL_806753, EPI_ISL_806754, EPI_ISL_806755, EPI_ISL_806757 | SIESP DIPARTIMENTO DI PREVENZIONE SULMONA | Istituto Zooprofilattico Sperimentale dell'Abruzzo e Molise "G. Caporale" | Lorusso A, Marcacci M, Di Domenico M, Ancora M, Curini V, Mangone I, Rinaldi A, Di Pasquale A, Cammà C, Puglia I, Calistri P, Savini G |
| EPI_ISL_806758, EPI_ISL_806759, EPI_ISL_806760, EPI_ISL_806761 | DIPARTIMENTO PREVENZIONE AVEZZANO-SERVIZIO DI IGIENE EPIDEMIOLOGIA E SANITA' PUBBLICA | Istituto Zooprofilattico Sperimentale dell'Abruzzo e Molise "G. Caporale" | Lorusso A, Marcacci M, Di Domenico M, Ancora M, Curini V, Mangone I, Rinaldi A, Di Pasquale A, Cammà C, Puglia I, Calistri P, Savini G |
| EPI_ISL_806762 | SIESP CHIETI - DRIVE IN ORTONA | Istituto Zooprofilattico Sperimentale dell'Abruzzo e Molise "G. Caporale" | Lorusso A, Marcacci M, Di Domenico M, Ancora M, Curini V, Mangone I, Rinaldi A, Di Pasquale A, Cammà C, Puglia I, Calistri P, Savini G |
| EPI_ISL_806763, EPI_ISL_806764, EPI_ISL_806765 | DIPARTIMENTO PREVENZIONE AVEZZANO-SERVIZIO DI IGIENE EPIDEMIOLOGIA E SANITA' PUBBLICA | Istituto Zooprofilattico Sperimentale dell'Abruzzo e Molise "G. Caporale" | Lorusso A, Marcacci M, Di Domenico M, Ancora M, Curini V, Mangone I, Rinaldi A, Di Pasquale A, Cammà C, Puglia I, Calistri P, Savini G |
| EPI_ISL_806766, EPI_ISL_806767, EPI_ISL_806768, EPI_ISL_806770, EPI_ISL_806771 | SIESP DIPARTIMENTO DI PREVENZIONE TERAMO | Istituto Zooprofilattico Sperimentale dell'Abruzzo e Molise "G. Caporale" | Lorusso A, Marcacci M, Di Domenico M, Ancora M, Curini V, Mangone I, Rinaldi A, Di Pasquale A, Cammà C, Puglia I, Calistri P, Savini G |
| EPI_ISL_806773, EPI_ISL_806774 | Ospedale S.Salvatore-Medicina Interna L'Aquila | Istituto Zooprofilattico Sperimentale dell'Abruzzo e Molise "G. Caporale" | Lorusso A, Marcacci M, Di Domenico M, Ancora M, Curini V, Mangone I, Rinaldi A, Di Pasquale A, Cammà C, Puglia I, Calistri P, Savini G |
| EPI_ISL_806775, EPI_ISL_806776 | SIESP CHIETI - DRIVE IN LANCIANO | Istituto Zooprofilattico Sperimentale dell'Abruzzo e Molise "G. Caporale" | Lorusso A, Marcacci M, Di Domenico M, Ancora M, Curini V, Mangone I, Rinaldi A, Di Pasquale A, Cammà C, Puglia I, Calistri P, Savini G |
| EPI_ISL_806777, EPI_ISL_806778, EPI_ISL_806779, EPI_ISL_806780 | SIESP DIPARTIMENTO DI PREVENZIONE TERAMO | Istituto Zooprofilattico Sperimentale dell'Abruzzo e Molise "G. Caporale" | Lorusso A, Marcacci M, Di Domenico M, Ancora M, Curini V, Mangone I, Rinaldi A, Di Pasquale A, Cammà C, Puglia I, Calistri P, Savini G |
| EPI_ISL_806781, EPI_ISL_806782 | DIPARTIMENTO PREVENZIONE AVEZZANO-SERVIZIO DI IGIENE EPIDEMIOLOGIA E SANITA' PUBBLICA | Istituto Zooprofilattico Sperimentale dell'Abruzzo e Molise "G. Caporale" | Lorusso A, Marcacci M, Di Domenico M, Ancora M, Curini V, Mangone I, Rinaldi A, Di Pasquale A, Cammà C, Puglia I, Calistri P, Savini G |
| EPI_ISL_806783 | Casa di cura Di Lorenzo- Avezzano | Istituto Zooprofilattico Sperimentale dell'Abruzzo e Molise "G. Caporale" | Lorusso A, Marcacci M, Di Domenico M, Ancora M, Curini V, Mangone I, Rinaldi A, Di Pasquale A, Cammà C, Puglia I, Calistri P, Savini G |
| EPI_ISL_806784, EPI_ISL_806785 | Ospedale Civile G.Mazzini-Teramo | Istituto Zooprofilattico Sperimentale dell'Abruzzo e Molise "G. Caporale" | Lorusso A, Marcacci M, Di Domenico M, Ancora M, Curini V, Mangone I, Rinaldi A, Di Pasquale A, Cammà C, Puglia I, Calistri P, Savini G |
| EPI_ISL_806786, EPI_ISL_806787, EPI_ISL_806788 | SIESP DIPARTIMENTO DI PREVENZIONE TERAMO | Istituto Zooprofilattico Sperimentale dell'Abruzzo e Molise "G. Caporale" | Lorusso A, Marcacci M, Di Domenico M, Ancora M, Curini V, Mangone I, Rinaldi A, Di Pasquale A, Cammà C, Puglia I, Calistri P, Savini G |
| EPI_ISL_806789 | Ospedale Civile Giulianova | Istituto Zooprofilattico Sperimentale dell'Abruzzo e Molise "G. Caporale" | Lorusso A, Marcacci M, Di Domenico M, Ancora M, Curini V, Mangone I, Rinaldi A, Di Pasquale A, Cammà C, Puglia I, Calistri P, Savini G |
| EPI_ISL_806791, EPI_ISL_806792, EPI_ISL_806793, EPI_ISL_806795, EPI_ISL_806796, EPI_ISL_806797, EPI_ISL_806798 | SIESP CHIETI - DRIVE IN ORTONA | Istituto Zooprofilattico Sperimentale dell'Abruzzo e Molise "G. Caporale" | Lorusso A, Marcacci M, Di Domenico M, Ancora M, Curini V, Mangone I, Rinaldi A, Di Pasquale A, Cammà C, Puglia I, Calistri P, Savini G |
| EPI_ISL_806799 | SIESP CHIETI - DRIVE IN LANCIANO | Istituto Zooprofilattico Sperimentale dell'Abruzzo e Molise "G. Caporale" | Lorusso A, Marcacci M, Di Domenico M, Ancora M, Curini V, Mangone I, Rinaldi A, Di Pasquale A, Cammà C, Puglia I, Calistri P, Savini G |
| EPI_ISL_806800, EPI_ISL_806801, EPI_ISL_806802, EPI_ISL_806803 | SIESP CHIETI - DRIVE IN GISSI | Istituto Zooprofilattico Sperimentale dell'Abruzzo e Molise "G. Caporale" | Lorusso A, Marcacci M, Di Domenico M, Ancora M, Curini V, Mangone I, Rinaldi A, Di Pasquale A, Cammà C, Puglia I, Calistri P, Savini G |
| EPI_ISL_806804, EPI_ISL_806805 | SIESP CHIETI - DISTRETTO SANITARIO CHIETI | Istituto Zooprofilattico Sperimentale dell'Abruzzo e Molise "G. Caporale" | Lorusso A, Marcacci M, Di Domenico M, Ancora M, Curini V, Mangone I, Rinaldi A, Di Pasquale A, Cammà C, Puglia I, Calistri P, Savini G |
| EPI_ISL_811139, EPI_ISL_811140, EPI_ISL_811142, EPI_ISL_811143 | Ministry of Health Turkey | Ministry of Health Turkey | Fatma Bayrakdar, Yasemin Cogun, Süleyman Yalcin, Aye Baak Alta, Gülay Korukluolu |
| EPI_ISL_811149 | Laboratorio de Ecologia de Doencas Transmissiveis na Amazonia, Instituto Leonidas e Maria Deane - Fiocruz Amazonia | Laboratorio de Ecologia de Doencas Transmissiveis na Amazonia, Instituto Leonidas e Maria Deane - Fiocruz Amazonia | Valdinete Nascimento, Victor Souza, André Corado, Fernanda Nascimento, George Silva, Ágatha Costa, Debora Duarte, Luciana Gonçalves, Matilde Mejía, Karina Pessoa, Maria Júlia Brandão, Michele Jesus, Felipe Naveca on behalf of the Fiocruz COVID-19 Genomic Surveillance Network |
| EPI_ISL_812359 | Area of Virology, Serology and Virology Division (SAViD), New South Wales Health Pathology Randwick | Virology Research Laboratory; Area of Virology, Serology and Virology Division (SAViD), New South Wales Health Pathology Randwick | Foster, C.; Au, J.; Ruiz Silva, M.; Deveson, I.; Bull, R.; Van Hal, S.; Rawlinson, W. |
| EPI_ISL_812723, EPI_ISL_812724 | DOHMH PHL | New York City Public Health Laboratory | Jade Wang, et al. |
| EPI_ISL_812726 | DOHMH Corona | New York City Public Health Laboratory | Jade Wang, et al. |
| EPI_ISL_812728 | DOHMH Riverside | New York City Public Health Laboratory | Jade Wang, et al. |
| EPI_ISL_812730 | DOHMH Crown Heights | New York City Public Health Laboratory | Jade Wang, et al. |
| EPI_ISL_812731 | DOHMH Jamaica | New York City Public Health Laboratory | Jade Wang, et al. |
| EPI_ISL_812732 | DOHMH Corona | New York City Public Health Laboratory | Jade Wang, et al. |
| EPI_ISL_812734 | DOHMH Crown Heights | New York City Public Health Laboratory | Jade Wang, et al. |
| EPI_ISL_812735, EPI_ISL_812736, EPI_ISL_812737 | DOHMH Morrisania | New York City Public Health Laboratory | Jade Wang, et al. |
| EPI_ISL_812738 | DOHMH Crown Heights | New York City Public Health Laboratory | Jade Wang, et al. |
| EPI_ISL_812739 | DOHMH Central Harlem | New York City Public Health Laboratory | Jade Wang, et al. |
| EPI_ISL_812740, EPI_ISL_812742, EPI_ISL_812743, EPI_ISL_812744 | DOHMH Corona | New York City Public Health Laboratory | Jade Wang, et al. |

| | | | |
|---|---|---|---|
| EPI_ISL_812745 | DOHMH Riverside | New York City Public Health Laboratory | Jade Wang, et al. |
| EPI_ISL_812748 | DOHMH Corona | New York City Public Health Laboratory | Jade Wang, et al. |
| EPI_ISL_812749 | DOHMH Central Harlem | New York City Public Health Laboratory | Jade Wang, et al. |
| EPI_ISL_812750, EPI_ISL_812751 | DOHMH Corona | New York City Public Health Laboratory | Jade Wang, et al. |
| EPI_ISL_812753, EPI_ISL_812754, EPI_ISL_812755, EPI_ISL_812756, EPI_ISL_812757 | DOHMH Morrisania | New York City Public Health Laboratory | Jade Wang, et al. |
| EPI_ISL_812759 | DOHMH Riverside | New York City Public Health Laboratory | Jade Wang, et al. |
| EPI_ISL_812760 | DOHMH Corona | New York City Public Health Laboratory | Jade Wang, et al. |
| EPI_ISL_813974 | Molecular Diagnostics Mater Dei Hospital | Molecular Diagnostics Mater Dei Hospital | Graziella Zahra |
| EPI_ISL_824555, EPI_ISL_824556, EPI_ISL_824557, EPI_ISL_824558, EPI_ISL_824559, EPI_ISL_824560, EPI_ISL_824561, EPI_ISL_824562, EPI_ISL_824563, EPI_ISL_824564, EPI_ISL_824565, EPI_ISL_824566, EPI_ISL_824567, EPI_ISL_824568, EPI_ISL_824569, EPI_ISL_824570, EPI_ISL_824571, EPI_ISL_824572, EPI_ISL_824573, EPI_ISL_824574, EPI_ISL_824575, EPI_ISL_824576, EPI_ISL_824577, EPI_ISL_824578, EPI_ISL_824579, EPI_ISL_824580, EPI_ISL_824581, EPI_ISL_824582, EPI_ISL_824583, EPI_ISL_824584, EPI_ISL_824585, EPI_ISL_824586, EPI_ISL_824587, EPI_ISL_824588, EPI_ISL_824589, EPI_ISL_824590, EPI_ISL_824591, EPI_ISL_824592, EPI_ISL_824593, EPI_ISL_824594, EPI_ISL_824595, EPI_ISL_824596, EPI_ISL_824597, EPI_ISL_824598, EPI_ISL_824599, EPI_ISL_824600, EPI_ISL_824601, EPI_ISL_824602, EPI_ISL_824603, EPI_ISL_824604, EPI_ISL_824605, EPI_ISL_824606, EPI_ISL_824607, EPI_ISL_824608, EPI_ISL_824609, EPI_ISL_824610, EPI_ISL_824611, EPI_ISL_824612, EPI_ISL_824613, EPI_ISL_824614, EPI_ISL_824615, EPI_ISL_824616, EPI_ISL_824618, EPI_ISL_824619, EPI_ISL_824620, EPI_ISL_824621, EPI_ISL_824622, EPI_ISL_824623, EPI_ISL_824624, EPI_ISL_824625, EPI_ISL_824626, EPI_ISL_824627, EPI_ISL_824628, EPI_ISL_824629, EPI_ISL_824630, EPI_ISL_824631, EPI_ISL_824632, EPI_ISL_824633, EPI_ISL_824634, EPI_ISL_824635, EPI_ISL_824636, EPI_ISL_824637, EPI_ISL_824638, EPI_ISL_824639, EPI_ISL_824640, EPI_ISL_824641, EPI_ISL_824642, EPI_ISL_824643, EPI_ISL_824644, EPI_ISL_824645, EPI_ISL_824646, EPI_ISL_824647, EPI_ISL_824648, EPI_ISL_824649, EPI_ISL_824650, EPI_ISL_824651, EPI_ISL_824652, EPI_ISL_824653, EPI_ISL_824654, EPI_ISL_824655, EPI_ISL_824656, EPI_ISL_824658, EPI_ISL_824659, EPI_ISL_824660, EPI_ISL_824661, EPI_ISL_824662, EPI_ISL_824663, EPI_ISL_824664, EPI_ISL_824665, EPI_ISL_824666, EPI_ISL_824667, EPI_ISL_824668, EPI_ISL_824669, EPI_ISL_824670, EPI_ISL_824671, EPI_ISL_824672, EPI_ISL_824673, EPI_ISL_824674, EPI_ISL_824675, EPI_ISL_824676, EPI_ISL_824677, EPI_ISL_824678, EPI_ISL_824679, EPI_ISL_824680, EPI_ISL_824681, EPI_ISL_824682, EPI_ISL_824683, EPI_ISL_824684, EPI_ISL_824685, EPI_ISL_824686, EPI_ISL_824687, EPI_ISL_824688, EPI_ISL_824689, EPI_ISL_824690, EPI_ISL_824691, EPI_ISL_824692, EPI_ISL_824693, EPI_ISL_824694, EPI_ISL_824695, EPI_ISL_824696, EPI_ISL_824697, EPI_ISL_824698, EPI_ISL_824699, EPI_ISL_824700, EPI_ISL_824701, EPI_ISL_824702, EPI_ISL_824703, EPI_ISL_824704, EPI_ISL_824705, EPI_ISL_824706, EPI_ISL_824707, EPI_ISL_824708, EPI_ISL_824709, EPI_ISL_824710, EPI_ISL_824711, EPI_ISL_824712, EPI_ISL_824713, EPI_ISL_824714, EPI_ISL_824715, EPI_ISL_824716, EPI_ISL_824717, EPI_ISL_824718, EPI_ISL_824719, EPI_ISL_824720, EPI_ISL_824721, EPI_ISL_824722, EPI_ISL_824723, EPI_ISL_824724, EPI_ISL_824725, EPI_ISL_824726, EPI_ISL_824727, EPI_ISL_824728, EPI_ISL_824729, EPI_ISL_824730, EPI_ISL_824732, EPI_ISL_824733, EPI_ISL_824734, EPI_ISL_824735, EPI_ISL_824736, EPI_ISL_824737, EPI_ISL_824738, EPI_ISL_824739, EPI_ISL_824740, EPI_ISL_824741 | | | |
| see above | Cedars-Sinai Medical Center, Department of Pathology & Laboratory Medicine, Molecular Pathology Laboratory | Cedars-Sinai Medical Center, Molecular Pathology Laboratory of Department of Pathology & Laboratory Medicine and Genomic Core | Wenjuan Zhang, Brian Davis, Stephanie Chen, Jorge Mario Sincuir Martinez, Jasmine T Plummer, Eric Vail |
| EPI_ISL_825034 | B.J. Medical College and Civil hospital, Ahmedabad | Gujarat Biotechnology Research Centre | Kamlesh J Upadhyay, Zarna Patel, Nitin Savaliya, Dinesh Kumar, Zuber Saiyed, Labdhi Pandya, Afzal Ansari, Nikha Trivedi, Apurvasinh Puvar, Ramesh Pandit, Janvi Raval, Pranay Shah, Nilima Shah, Sanjay Kapadia, Dipa Kinariwala, Chaitanya Joshi, Madhvi Joshi |
| EPI_ISL_825035 | B.J. Medical College and Civil hospital, Ahmedabad | Gujarat Biotechnology Research Centre | Nilima Shah, Sanjay Kapadia, Nitin Savaliya, Dinesh Kumar, Zuber Saiyed, Labdhi Pandya, Afzal Ansari, Nikha Trivedi, Apurvasinh Puvar, Ramesh Pandit, Janvi Raval, Zarna Patel, Kamlesh J Upadhyay, Pranay Shah, Dipa Kinariwala, Chaitanya Joshi, Madhvi Joshi |
| EPI_ISL_825036 | B.J. Medical College and Civil hospital, Ahmedabad | Gujarat Biotechnology Research Centre | Dipa Kinariwala, Dinesh Kumar, Zuber Saiyed, Labdhi Pandya, Afzal Ansari, Nikha Trivedi, Apurvasinh Puvar, Ramesh Pandit, Janvi Raval, Zarna Patel, Nitin Savaliya, Kamlesh J Upadhyay, Nilima Shah, Sanjay Kapadia, Pranay Shah, Chaitanya Joshi, Madhvi Joshi |
| EPI_ISL_825037 | B.J. Medical College and Civil hospital, Ahmedabad | Gujarat Biotechnology Research Centre | Pranay Shah, Janvi Raval, Zarna Patel, Nitin Savaliya, Dinesh Kumar, Zuber Saiyed, Labdhi Pandya, Afzal Ansari, Nikha Trivedi, Apurvasinh Puvar, Ramesh Pandit, Kamlesh J Upadhyay, Nilima Shah, Sanjay Kapadia, Dipa Kinariwala, Chaitanya Joshi, Madhvi Joshi |
| EPI_ISL_825038 | B.J. Medical College and Civil hospital, Ahmedabad | Gujarat Biotechnology Research Centre | Zuber Saiyed, Labdhi Pandya, Afzal Ansari, Nikha Trivedi, Apurvasinh Puvar, Ramesh Pandit, Janvi Raval, Zarna Patel, Nitin Savaliya, Dinesh Kumar, Pranay Shah, Kamlesh J Upadhyay, Sanjay Kapadia, Dipa Kinariwala, Chaitanya Joshi, Madhvi Joshi |
| EPI_ISL_825039 | B.J. Medical College and Civil hospital, Ahmedabad | Gujarat Biotechnology Research Centre | Labdhi Pandya, Afzal Ansari, Nikha Trivedi, Apurvasinh Puvar, Ramesh Pandit, Janvi Raval, Zarna Patel, Nitin Savaliya, Dinesh Kumar, Zuber Saiyed, Pranay Shah, Kamlesh J Upadhyay, Sanjay Kapadia, Dipa Kinariwala, Chaitanya Joshi, Madhvi Joshi |
| EPI_ISL_825040 | B.J. Medical College and Civil hospital, Ahmedabad | Gujarat Biotechnology Research Centre | Afzal Ansari, Nikha Trivedi, Apurvasinh Puvar, Ramesh Pandit, Janvi Raval, Zarna Patel, Nitin Savaliya, Dinesh Kumar, Zuber Saiyed, Labdhi Pandya, Pranay Shah, Kamlesh J Upadhyay, Sanjay Kapadia, Dipa Kinariwala, Chaitanya Joshi, Madhvi Joshi |
| EPI_ISL_825041 | B.J. Medical College and Civil hospital, Ahmedabad | Gujarat Biotechnology Research Centre | Nikha Trivedi, Apurvasinh Puvar, Ramesh Pandit, Janvi Raval, Zarna Patel, Nitin Savaliya, Dinesh Kumar, Zuber Saiyed, Labdhi Pandya, Afzal Ansari, Pranay Shah, Kamlesh J Upadhyay, Sanjay Kapadia, Dipa Kinariwala, Chaitanya Joshi, Madhvi Joshi |
| EPI_ISL_825042 | B.J. Medical College and Civil hospital, Ahmedabad | Gujarat Biotechnology Research Centre | Apurvasinh Puvar, Ramesh Pandit, Janvi Raval, Zarna Patel, Nitin Savaliya, Dinesh Kumar, Zuber Saiyed, Labdhi Pandya, Afzal Ansari, Nikha Trivedi, Pranay Shah, Kamlesh J Upadhyay, Sanjay Kapadia, Dipa Kinariwala, Chaitanya Joshi, Madhvi Joshi |
| EPI_ISL_825044 | NHL Municipal Medical College, Ahmedbad | Gujarat Biotechnology Research Centre | Monila Patel, Janvi Raval, Zarna Patel, Nitin Savaliya, Dinesh Kumar, Zuber Saiyed, Labdhi Pandya, Afzal Ansari, Nikha Trivedi, Apurvasinh Puvar, Ramesh Pandit, Jayshri Pethani, Atit Shah, NM Shaikh, Bimal Chauhan, Tanmay Mehta, Bhavin Prajapati, Chaitanya Joshi, Madhvi Joshi |
| EPI_ISL_825045 | NHL Municipal Medical College, Ahmedbad | Gujarat Biotechnology Research Centre | Atit Shah, Zarna Patel, Nitin Savaliya, Dinesh Kumar, Zuber Saiyed, Labdhi Pandya, Afzal Ansari, Nikha Trivedi, Apurvasinh Puvar, Ramesh Pandit, Janvi Raval, Jayshri Pethani, Monila Patel, NM Shaikh, Bimal Chauhan, Tanmay Mehta, Bhavin Prajapati, Chaitanya Joshi, Madhvi Joshi |
| EPI_ISL_825046 | NHL Municipal Medical College, Ahmedbad | Gujarat Biotechnology Research Centre | NM Shaikh, Nitin Savaliya, Dinesh Kumar, Zuber Saiyed, Labdhi Pandya, Afzal Ansari, Nikha Trivedi, Apurvasinh Puvar, Ramesh Pandit, Janvi Raval, Zarna Patel, Jayshri Pethani, Monila Patel, Atit Shah, Bimal Chauhan, Tanmay Mehta, Bhavin Prajapati, Chaitanya Joshi, Madhvi Joshi |
| EPI_ISL_825047 | NHL Municipal Medical College, Ahmedbad | Gujarat Biotechnology Research Centre | Bimal Chauhan, Dinesh Kumar, Zuber Saiyed, Labdhi Pandya, Afzal Ansari, Nikha Trivedi, Apurvasinh Puvar, Ramesh Pandit, Janvi Raval, Zarna Patel, Nitin Savaliya, Jayshri Pethani, Monila Patel, Atit Shah, NM Shaikh, Tanmay Mehta, Bhavin Prajapati, Chaitanya Joshi, Madhvi Joshi |
| EPI_ISL_825048 | GMERS Medical College and Hospital, Gotri, Vadodara | Gujarat Biotechnology Research Centre | Mitesh Kamothi, Zuber Saiyed, Labdhi Pandya, Afzal Ansari, Nikha Trivedi, Apurvasinh Puvar, Ramesh Pandit, Janvi Raval, Zarna Patel, Nitin Savaliya, Dinesh Kumar, Bithika Duttaroy, Kalpesh Mistry, Chaitanya Joshi, Madhvi Joshi |
| EPI_ISL_825049 | GMERS Medical College and Hospital, Gotri, Vadodara | Gujarat Biotechnology Research Centre | Bithika Duttaroy, Labdhi Pandya, Afzal Ansari, Nikha Trivedi, Apurvasinh Puvar, Ramesh Pandit, Janvi Raval, Zarna Patel, Nitin Savaliya, Dinesh Kumar, Zuber Saiyed, Mitesh Kamothi, Kalpesh Mistry, Chaitanya Joshi, Madhvi Joshi |
| EPI_ISL_825050 | NHL Municipal Medical College, Ahmedbad | Gujarat Biotechnology Research Centre | Tanmay Mehta, Afzal Ansari, Nikha Trivedi, Apurvasinh Puvar, Ramesh Pandit, Janvi Raval, Zarna Patel, Nitin Savaliya, Dinesh Kumar, Zuber Saiyed, Labdhi Pandya, Jayshri Pethani, Monila Patel, Atit Shah, NM Shaikh, Bimal Chauhan, Bhavin Prajapati, Chaitanya Joshi, Madhvi Joshi |
| EPI_ISL_825051 | NHL Municipal Medical College, Ahmedbad | Gujarat Biotechnology Research Centre | Bhavin Prajapati, Nikha Trivedi, Apurvasinh Puvar, Ramesh Pandit, Janvi Raval, Zarna Patel, Nitin Savaliya, Dinesh Kumar, Zuber Saiyed, Labdhi Pandya, Afzal Ansari, Jayshri Pethani, Monila Patel, Atit Shah, NM Shaikh, Bimal Chauhan, Tanmay Mehta, Chaitanya Joshi, Madhvi Joshi |
| EPI_ISL_825052 | NHL Municipal Medical College, Ahmedbad | Gujarat Biotechnology Research Centre | Apurvasinh Puvar, Ramesh Pandit, Janvi Raval, Zarna Patel, Nitin Savaliya, Dinesh Kumar, Zuber Saiyed, Labdhi Pandya, Afzal Ansari, Nikha Trivedi, Jayshri Pethani, Monila Patel, Atit Shah, NM Shaikh, Bimal Chauhan, Tanmay Mehta, Bhavin Prajapati, Chaitanya Joshi, Madhvi Joshi |
| EPI_ISL_825053 | NHL Municipal Medical College, Ahmedbad | Gujarat Biotechnology Research Centre | Ramesh Pandit, Janvi Raval, Zarna Patel, Nitin Savaliya, Dinesh Kumar, Zuber Saiyed, Labdhi Pandya, Afzal Ansari, Nikha Trivedi, Apurvasinh Puvar, Jayshri Pethani, Monila Patel, Atit Shah, NM Shaikh, Bimal Chauhan, Tanmay Mehta, Bhavin Prajapati, Chaitanya Joshi, Madhvi Joshi |
| EPI_ISL_825054 | GMERS Medical College and Hospital, Gotri, Vadodara | Gujarat Biotechnology Research Centre | Kalpesh Mistry, Janvi Raval, Zarna Patel, Nitin Savaliya, Dinesh Kumar, Zuber Saiyed, Afzal Ansari, Nikha Trivedi, Apurvasinh Puvar, Ramesh Pandit, Mitesh Kamothi, Bithika Duttaroy, Chaitanya Joshi, Madhvi Joshi |
| EPI_ISL_825055 | GMERS Medical College and Hospital, Gotri, Vadodara | Gujarat Biotechnology Research Centre | Zarna Patel, Nitin Savaliya, Dinesh Kumar, Zuber Saiyed, Afzal Ansari, Nikha Trivedi, Apurvasinh Puvar, Ramesh Pandit, Janvi Raval, Mitesh Kamothi, Bithika Duttaroy, Kalpesh Mistry, Chaitanya Joshi, Madhvi Joshi |
| EPI_ISL_825056 | GMERS Medical College and Hospital, Gotri, Vadodara | Gujarat Biotechnology Research Centre | Nitin Savaliya, Dinesh Kumar, Zuber Saiyed, Afzal Ansari, Nikha Trivedi, Apurvasinh Puvar, Ramesh Pandit, Janvi Raval, Zarna Patel, Mitesh Kamothi, Bithika Duttaroy, Kalpesh Mistry, Chaitanya Joshi, Madhvi Joshi |
| EPI_ISL_825057 | GMERS Medical College and Hospital, Gotri, Vadodara | Gujarat Biotechnology Research Centre | Dinesh Kumar, Zuber Saiyed, Afzal Ansari, Nikha Trivedi, Apurvasinh Puvar, Ramesh Pandit, Janvi Raval, Zarna Patel, Nitin Savaliya, Mitesh Kamothi, Bithika Duttaroy, Kalpesh Mistry, Chaitanya Joshi, Madhvi Joshi |
| EPI_ISL_825058 | GMERS Medical College and Hospital, Gotri, Vadodara | Gujarat Biotechnology Research Centre | Zuber Saiyed, Afzal Ansari, Nikha Trivedi, Apurvasinh Puvar, Ramesh Pandit, Janvi Raval, Zarna Patel, Nitin Savaliya, Dinesh Kumar, Mitesh Kamothi, Bithika |

| | | | |
|---|---|---|---|
| | | | Duttaroy, Kalpesh Mistry, Chaitanya Joshi, Madhvi Joshi |
| EPI_ISL_825059 | NHL Municipal Medical College, Ahmedbad | Gujarat Biotechnology Research Centre | Afzal Ansari, Nikha Trivedi, Apurvasinh Puvar, Ramesh Pandit, Janvi Raval, Zarna Patel, Nitin Savaliya, Dinesh Kumar, Zuber Saiyed, Jayshri Pethani, Monila Patel, Atit Shah, NM Shaikh, Bimal Chauhan, Tanmay Mehta, Bhavin Prajapati, Chaitanya Joshi, Madhvi Joshi |
| EPI_ISL_825060 | NHL Municipal Medical College, Ahmedbad | Gujarat Biotechnology Research Centre | Nikha Trivedi, Apurvasinh Puvar, Ramesh Pandit, Janvi Raval, Zarna Patel, Nitin Savaliya, Dinesh Kumar, Zuber Saiyed, Afzal Ansari, Jayshri Pethani, Monila Patel, Atit Shah, NM Shaikh, Bimal Chauhan, Tanmay Mehta, Bhavin Prajapati, Chaitanya Joshi, Madhvi Joshi |
| EPI_ISL_825061 | GMERS Medical College and Hospital, Gotri, Vadodara | Gujarat Biotechnology Research Centre | Apurvasinh Puvar, Ramesh Pandit, Janvi Raval, Zarna Patel, Nitin Savaliya, Dinesh Kumar, Zuber Saiyed, Afzal Ansari, Nikha Trivedi, Mitesh Kamothi, Bithika Duttaroy, Kalpesh Mistry, Chaitanya Joshi, Madhvi Joshi |
| EPI_ISL_825062 | GMERS Medical College and Hospital, Gotri, Vadodara | Gujarat Biotechnology Research Centre | Ramesh Pandit, Janvi Raval, Zarna Patel, Nitin Savaliya, Dinesh Kumar, Zuber Saiyed, Afzal Ansari, Nikha Trivedi, Apurvasinh Puvar, Mitesh Kamothi, Bithika Duttaroy, Kalpesh Mistry, Chaitanya Joshi, Madhvi Joshi |
| EPI_ISL_825063 | NHL Municipal Medical College, Ahmedbad | Gujarat Biotechnology Research Centre | Janvi Raval, Zarna Patel, Nitin Savaliya, Dinesh Kumar, Zuber Saiyed, Afzal Ansari, Nikha Trivedi, Apurvasinh Puvar, Ramesh Pandit, Jayshri Pethani, Monila Patel, Atit Shah, NM Shaikh, Bimal Chauhan, Tanmay Mehta, Bhavin Prajapati, Chaitanya Joshi, Madhvi Joshi |
| EPI_ISL_825155, EPI_ISL_825156 | Microbiology and Virology Unit, Florence Careggi University Hospital | Microbiology and Virology Unit, Florence Careggi University Hospital | Vincenzo Di Pilato, Marco Coppi, Fabio Morecchiato, Alberto Antonelli, Emanuele Gori, Gian Maria Rossolini |
| EPI_ISL_826269, EPI_ISL_826270 | University of Debrecen, Department of Medical Microbiology | National Laboratory of Virology, Szentágothai Research Centre | Endre Gábor Tóth, Balázs Somogyi, Brigitta Zana, Eszter Csoma, Ferenc Jakab, Gábor Kemenesi |
| EPI_ISL_826274 | Sterling Hospital, Memnagar, Ahmedabad | Gujarat Biotechnology Research Centre | Naman Shashtri, Zarna Patel, Nitin Savaliya, Dinesh Kumar, Zuber Saiyed, Labdhi Pandya, Afzal Ansari, Nikha Trivedi, Apurvasinh Puvar, Ramesh Pandit, Janvi Raval, Atul K Patel, Chaitanya Joshi, Madhvi Joshi |
| EPI_ISL_826275 | Sterling Hospital, Memnagar, Ahmedabad | Gujarat Biotechnology Research Centre | Atul K Patel, Nitin Savaliya, Dinesh Kumar, Zuber Saiyed, Labdhi Pandya, Afzal Ansari, Nikha Trivedi, Apurvasinh Puvar, Ramesh Pandit, Janvi Raval, Zarna Patel, Naman Shashtri, Chaitanya Joshi, Madhvi Joshi |
| EPI_ISL_826276 | Sterling Hospital, Memnagar, Ahmedabad | Gujarat Biotechnology Research Centre | Dinesh Kumar, Zuber Saiyed, Labdhi Pandya, Afzal Ansari, Nikha Trivedi, Apurvasinh Puvar, Ramesh Pandit, Janvi Raval, Zarna Patel, Nitin Savaliya, Atul K Patel, Naman Shashtri, Chaitanya Joshi, Madhvi Joshi |
| EPI_ISL_826284, EPI_ISL_826458 | University of Bari Biomedical Sciences and Human Oncology | University of Bari Biomedical Sciences and Human Oncology | Chironna Maria, Sallustio Anna, Loconsole Daniela, Accogli Marisa |
| EPI_ISL_827042 | Institute of Virology, University of Cologne | Institute of Virology, University of Cologne | Saleta Sierra, Gibran Rubio, Zevanya Tessalonica, Dominik Aschenmeier, Eva Heger, Elena Knops, Rolf Kaiser, Martin Däumer, Alex Thielen |
| EPI_ISL_831939, EPI_ISL_831940, EPI_ISL_832009, EPI_ISL_832010, EPI_ISL_832011, EPI_ISL_832012, EPI_ISL_832013 | Laboratório de Microbiologia Molecular - Universidade FEEVALE | Universidade Federal de Ciências da Saúde de Porto Alegre | Vinícius Bonetti Franceschi, Amanda de Menezes Mayer, Gabriel Dickin Caldana, Carla Andretta Moreira Neves, Patrícia Aline Gröhs Ferrareze, Gabriela Bettella Cybis, Ricardo Ariel Zimerman, Lívia Kmetzsch, Fernando Rosado Spilki, Claudia Elizabeth Thompson |
| EPI_ISL_832128 | Sentinelles IDF | National Reference Center for Viruses of Respiratory Infections, Institut Pasteur, Paris | Marion Barbet, Sylvie Behillil, Méline Bizard, Angela Brisebarre, Camille Capel, Etienne Simon-Lorière, Vincent Enouf, Maud Vanpeene, Sylvie van der Werf, Patricia Lefebure |
| EPI_ISL_832129 | Sentinelles IDF | National Reference Center for Viruses of Respiratory Infections, Institut Pasteur, Paris | Marion Barbet, Sylvie Behillil, Méline Bizard, Angela Brisebarre, Camille Capel, Etienne Simon-Lorière, Vincent Enouf, Maud Vanpeene, Sylvie van der Werf, Claire Watermez |
| EPI_ISL_832130, EPI_ISL_832131, EPI_ISL_832132, EPI_ISL_832133, EPI_ISL_832134, EPI_ISL_832135, EPI_ISL_832136, EPI_ISL_832137, EPI_ISL_832138, EPI_ISL_832139, EPI_ISL_832140, EPI_ISL_832141, EPI_ISL_832142, EPI_ISL_832143 | | | |
| see above | Hospital | National Reference Center for Viruses of Respiratory Infections, Institut Pasteur, Paris | Marion Barbet, Sylvie Behillil, Méline Bizard, Angela Brisebarre, Camille Capel, Etienne Simon-Lorière, Vincent Enouf, Maud Vanpeene, Sylvie van der Werf, Laurent Andreoletti |
| EPI_ISL_832144, EPI_ISL_832145, EPI_ISL_832146, EPI_ISL_832147, EPI_ISL_832148, EPI_ISL_832149, EPI_ISL_832150, EPI_ISL_832151, EPI_ISL_832152, EPI_ISL_832153, EPI_ISL_832154, EPI_ISL_832155, EPI_ISL_832156, EPI_ISL_832157, EPI_ISL_832158, EPI_ISL_832159, EPI_ISL_832160, EPI_ISL_832161, EPI_ISL_832162, EPI_ISL_832163, EPI_ISL_832164, EPI_ISL_832165, EPI_ISL_832166 | | | |
| see above | Hospital | National Reference Center for Viruses of Respiratory Infections, Institut Pasteur, Paris | Marion Barbet, Sylvie Behillil, Méline Bizard, Angela Brisebarre, Camille Capel, Etienne Simon-Lorière, Vincent Enouf, Maud Vanpeene, Sylvie van der Werf, Léa Pilorge |
| EPI_ISL_832167 | Sentinelles Province | National Reference Center for Viruses of Respiratory Infections, Institut Pasteur, Paris | Marion Barbet, Sylvie Behillil, Méline Bizard, Angela Brisebarre, Camille Capel, Etienne Simon-Lorière, Vincent Enouf, Maud Vanpeene, Sylvie van der Werf, Jean-Michel Monnier |
| EPI_ISL_832168 | Sentinelles Province | National Reference Center for Viruses of Respiratory Infections, Institut Pasteur, Paris | Marion Barbet, Sylvie Behillil, Méline Bizard, Angela Brisebarre, Camille Capel, Etienne Simon-Lorière, Vincent Enouf, Maud Vanpeene, Sylvie van der Werf, Denis Calvet |
| EPI_ISL_832169, EPI_ISL_832170 | Hospital | National Reference Center for Viruses of Respiratory Infections, Institut Pasteur, Paris | Marion Barbet, Sylvie Behillil, Méline Bizard, Angela Brisebarre, Camille Capel, Etienne Simon-Lorière, Vincent Enouf, Maud Vanpeene, Sylvie van der Werf, Clémence Guillaume |
| EPI_ISL_832171, EPI_ISL_832172, EPI_ISL_832173, EPI_ISL_832174, EPI_ISL_832175 | Hospital | National Reference Center for Viruses of Respiratory Infections, Institut Pasteur, Paris | Marion Barbet, Sylvie Behillil, Méline Bizard, Angela Brisebarre, Camille Capel, Etienne Simon-Lorière, Vincent Enouf, Maud Vanpeene, Sylvie van der Werf, Patricia Stoessel-Thouvenin |
| EPI_ISL_832176, EPI_ISL_832177 | Hospital | National Reference Center for Viruses of Respiratory Infections, Institut Pasteur, Paris | Marion Barbet, Sylvie Behillil, Méline Bizard, Angela Brisebarre, Camille Capel, Etienne Simon-Lorière, Vincent Enouf, Maud Vanpeene, Sylvie van der Werf, Aurélie Guigon |
| EPI_ISL_832179, EPI_ISL_832180 | Hospital | National Reference Center for Viruses of Respiratory Infections, Institut Pasteur, Paris | Marion Barbet, Sylvie Behillil, Méline Bizard, Angela Brisebarre, Camille Capel, Etienne Simon-Lorière, Vincent Enouf, Maud Vanpeene, Sylvie van der Werf, Laurent Roudiere |
| EPI_ISL_832182, EPI_ISL_832183, EPI_ISL_832188, EPI_ISL_832190, EPI_ISL_832191, EPI_ISL_832192, EPI_ISL_832193 | Laboratory Analyses Med | National Reference Center for Viruses of Respiratory Infections, Institut Pasteur, Paris | Marion Barbet, Sylvie Behillil, Méline Bizard, Angela Brisebarre, Camille Capel, Etienne Simon-Lorière, Vincent Enouf, Maud Vanpeene, Sylvie van der Werf, Brieuc Lefaure |
| EPI_ISL_832194, EPI_ISL_832195 | Laboratory Analyses Med | National Reference Center for Viruses of Respiratory Infections, Institut Pasteur, Paris | Marion Barbet, Sylvie Behillil, Méline Bizard, Angela Brisebarre, Camille Capel, Etienne Simon-Lorière, Vincent Enouf, Maud Vanpeene, Sylvie van der Werf, Brieuc Gestin |
| EPI_ISL_832197 | Hospital | National Reference Center for Viruses of Respiratory Infections, Institut Pasteur, Paris | Marion Barbet, Sylvie Behillil, Méline Bizard, Angela Brisebarre, Camille Capel, Etienne Simon-Lorière, Vincent Enouf, Maud Vanpeene, Sylvie van der Werf, Esther Gyde |
| EPI_ISL_832201 | Hospital | National Reference Center for Viruses of Respiratory Infections, Institut Pasteur, Paris | Marion Barbet, Sylvie Behillil, Méline Bizard, Angela Brisebarre, Camille Capel, Etienne Simon-Lorière, Vincent Enouf, Maud Vanpeene, Sylvie van der Werf, Patricia Stoessel-Thouvenin |
| EPI_ISL_832378 | Ohio State | James Molecular Lab | Huolin Tu, Matthew R Avenarius, Laura Kubatko, Matthew Hunt, Xiaokang Pan, Peng Ru, Jason Garee, Keelie Thomas, Peter Mohler, Preeti Pancholi, Dan Jones |
| EPI_ISL_832815, EPI_ISL_832816 | SIESP CHIETI - DRIVE IN LANCIANO | Istituto Zooprofilattico Sperimentale dell'Abruzzo e Molise "G.Caporale" | Lorusso A, Marcacci M, Di Domenico M, Curini V, Ancora M, Cammà C, Rinaldi A, Mangone I, Di Pasquale A, Puglia I, Calistri P, Savini G. |
| EPI_ISL_832817 | SIESP DIPARTIMENTO DI PREVENZIONE TERAMO | Istituto Zooprofilattico Sperimentale dell'Abruzzo e Molise "G.Caporale" | Lorusso A, Marcacci M, Di Domenico M, Curini V, Ancora M, Cammà C, Rinaldi A, Mangone I, Di Pasquale A, Puglia I, Calistri P, Savini G. |
| EPI_ISL_832818 | Ospedale Civile Atri | Istituto Zooprofilattico Sperimentale dell'Abruzzo e Molise "G.Caporale" | Lorusso A, Marcacci M, Di Domenico M, Curini V, Ancora M, Cammà C, Rinaldi A, Mangone I, Di Pasquale A, Puglia I, Calistri P, Savini G. |
| EPI_ISL_832819 | SIESP DIPARTIMENTO DI PREVENZIONE SULMONA | Istituto Zooprofilattico Sperimentale dell'Abruzzo e Molise | Lorusso A, Marcacci M, Di Domenico M, Curini V, Ancora M, Cammà C, Rinaldi A, Mangone I, Di Pasquale A, Puglia I, Calistri P, Savini G. |

| | | | |
|---|---|---|---|
| | | "G.Caporale" | |
| EPI_ISL_832820 | Medico Competente P.O. L'Aquila | Istituto Zooprofilattico Sperimentale dell'Abruzzo e Molise "G.Caporale" | Lorusso A, Marcacci M, Di Domenico M, Curini V, Ancora M, Cammà C, Rinaldi A, Mangone I, Di Pasquale A, Puglia I, Calistri P, Savini G. |
| EPI_ISL_832821 | SIESP CHIETI - DRIVE IN CHIETI | Istituto Zooprofilattico Sperimentale dell'Abruzzo e Molise "G.Caporale" | Lorusso A, Marcacci M, Di Domenico M, Curini V, Ancora M, Cammà C, Rinaldi A, Mangone I, Di Pasquale A, Puglia I, Calistri P, Savini G. |
| EPI_ISL_832822 | SIESP CHIETI - DRIVE IN LANCIANO | Istituto Zooprofilattico Sperimentale dell'Abruzzo e Molise "G.Caporale" | Lorusso A, Marcacci M, Di Domenico M, Curini V, Ancora M, Cammà C, Rinaldi A, Mangone I, Di Pasquale A, Puglia I, Calistri P, Savini G. |
| EPI_ISL_833040 | RSUD Dr. Abdul Aziz Singkawang | National Institute of Health Research and Development | Subangkit;Pawestri,HA;Ikawati,HD;Nugraha,AA;Puspa,KD;Pangesti,KNA;Soekarso,T;Junus,HN;Puspandari,N;Setiawaty,V |
| EPI_ISL_833041 | Defence Services Medical Research Center, Biological Research Laboratory | Defence Services Medical Research Center, Biological Research Laboratory | Khine Zaw Oo, Nay Myo Aung, Ko Ko Win, Phyo Kyaw Aung, Zaw Win Htun, Sat Paing Htoo, Kyaw Wanna, Thein Zaw, Kyee Myint, Ko Ko Lwin |
| EPI_ISL_833043 | SIESP DIPARTIMENTO DI PREVENZIONE CHIETI | Istituto Zooprofilattico Sperimentale dell'Abruzzo e Molise "G. Caporale" | Lorusso A, Marcacci M, Di Domenico M, Ancora M, Curini V, Mangone I, Rinaldi A, Delli Compagni E, Di Pasquale A, Cammà C, Puglia I, Calistri P, Savini G |
| EPI_ISL_833044 | SIESP CHIETI - DRIVE IN CHIETI | Istituto Zooprofilattico Sperimentale dell'Abruzzo e Molise "G. Caporale" | Lorusso A, Marcacci M, Di Domenico M, Ancora M, Curini V, Mangone I, Rinaldi A, Delli Compagni E, Di Pasquale A, Cammà C, Puglia I, Calistri P, Savini G |
| EPI_ISL_833045 | SIESP CHIETI - DRIVE IN ORTONA | Istituto Zooprofilattico Sperimentale dell'Abruzzo e Molise "G. Caporale" | Lorusso A, Marcacci M, Di Domenico M, Ancora M, Curini V, Mangone I, Rinaldi A, Delli Compagni E, Di Pasquale A, Cammà C, Puglia I, Calistri P, Savini G |
| EPI_ISL_833046 | FPAM | Istituto Zooprofilattico Sperimentale dell'Abruzzo e Molise "G. Caporale" | Lorusso A, Marcacci M, Di Domenico M, Ancora M, Curini V, Mangone I, Rinaldi A, Delli Compagni E, Di Pasquale A, Cammà C, Puglia I, Calistri P, Savini G |
| EPI_ISL_833047, EPI_ISL_833048 | OSPEDALE CIVILE ATRI | Istituto Zooprofilattico Sperimentale dell'Abruzzo e Molise "G. Caporale" | Lorusso A, Marcacci M, Di Domenico M, Ancora M, Curini V, Mangone I, Rinaldi A, Delli Compagni E, Di Pasquale A, Cammà C, Puglia I, Calistri P, Savini G |
| EPI_ISL_833049 | SIESP CHIETI - DRIVE IN LANCIANO | Istituto Zooprofilattico Sperimentale dell'Abruzzo e Molise "G. Caporale" | Lorusso A, Marcacci M, Di Domenico M, Ancora M, Curini V, Mangone I, Rinaldi A, Delli Compagni E, Di Pasquale A, Cammà C, Puglia I, Calistri P, Savini G |
| EPI_ISL_833050 | DIP. PREV. AVEZZANO SERVIZIO DI IGIENE EPIDEMIOLOGIAE SANITA' PUBBLICA | Istituto Zooprofilattico Sperimentale dell'Abruzzo e Molise "G. Caporale" | Lorusso A, Marcacci M, Di Domenico M, Ancora M, Curini V, Mangone I, Rinaldi A, Delli Compagni E, Di Pasquale A, Cammà C, Puglia I, Calistri P, Savini G |
| EPI_ISL_833052, EPI_ISL_833053, EPI_ISL_833054 | "Presidio Ospedaliero "San Liberatore" Atri | Istituto Zooprofilattico Sperimentale dell'Abruzzo e Molise "G. Caporale" | Lorusso A, Marcacci M, Di Domenico M, Ancora M, Curini V, Mangone I, Rinaldi A, Delli Compagni E, Di Pasquale A, Cammà C, Puglia I, Calistri P, Savini G |
| EPI_ISL_833055, EPI_ISL_833056 | "SIESP DIPARTIMENTO DI PREVENZIONE CHIETi | Istituto Zooprofilattico Sperimentale dell'Abruzzo e Molise "G. Caporale" | Lorusso A, Marcacci M, Di Domenico M, Ancora M, Curini V, Mangone I, Rinaldi A, Delli Compagni E, Di Pasquale A, Cammà C, Puglia I, Calistri P, Savini G |
| EPI_ISL_833057, EPI_ISL_833058, EPI_ISL_833059, EPI_ISL_833060, EPI_ISL_833061, EPI_ISL_833062, EPI_ISL_833063, EPI_ISL_833064, EPI_ISL_833065, EPI_ISL_833066, EPI_ISL_833067 | | | |
| see above | DIP. PREV. AVEZZANO SERVIZIO DI IGIENE EPIDEMIOLOGIAE SANITA' PUBBLICA | Istituto Zooprofilattico Sperimentale dell'Abruzzo e Molise "G. Caporale" | Lorusso A, Marcacci M, Di Domenico M, Ancora M, Curini V, Mangone I, Rinaldi A, Delli Compagni E, Di Pasquale A, Cammà C, Puglia I, Calistri P, Savini G |
| EPI_ISL_833068, EPI_ISL_833069, EPI_ISL_833070, EPI_ISL_833071, EPI_ISL_833072, EPI_ISL_833073, EPI_ISL_833074, EPI_ISL_833075 | FPAM | Istituto Zooprofilattico Sperimentale dell'Abruzzo e Molise "G. Caporale" | Lorusso A, Marcacci M, Di Domenico M, Ancora M, Curini V, Mangone I, Rinaldi A, Delli Compagni E, Di Pasquale A, Cammà C, Puglia I, Calistri P, Savini G |
| EPI_ISL_833076 | IZSAM | Istituto Zooprofilattico Sperimentale dell'Abruzzo e Molise "G. Caporale" | Lorusso A, Marcacci M, Di Domenico M, Ancora M, Curini V, Mangone I, Rinaldi A, Delli Compagni E, Di Pasquale A, Cammà C, Puglia I, Calistri P, Savini G |
| EPI_ISL_833077, EPI_ISL_833078 | OSPEDALE CIVILE ATRI | Istituto Zooprofilattico Sperimentale dell'Abruzzo e Molise "G. Caporale" | Lorusso A, Marcacci M, Di Domenico M, Ancora M, Curini V, Mangone I, Rinaldi A, Delli Compagni E, Di Pasquale A, Cammà C, Puglia I, Calistri P, Savini G |
| EPI_ISL_833079, EPI_ISL_833080 | OSPEDALE CIVILE TERAMO | Istituto Zooprofilattico Sperimentale dell'Abruzzo e Molise "G. Caporale" | Lorusso A, Marcacci M, Di Domenico M, Ancora M, Curini V, Mangone I, Rinaldi A, Delli Compagni E, Di Pasquale A, Cammà C, Puglia I, Calistri P, Savini G |
| EPI_ISL_833081 | OSPEDALE S.S. ANNUNZIATA CHIETI | Istituto Zooprofilattico Sperimentale dell'Abruzzo e Molise "G. Caporale" | Lorusso A, Marcacci M, Di Domenico M, Ancora M, Curini V, Mangone I, Rinaldi A, Delli Compagni E, Di Pasquale A, Cammà C, Puglia I, Calistri P, Savini G |
| EPI_ISL_833082 | OSPEDALE SAN SALVATORE | Istituto Zooprofilattico Sperimentale dell'Abruzzo e Molise "G. Caporale" | Lorusso A, Marcacci M, Di Domenico M, Ancora M, Curini V, Mangone I, Rinaldi A, Delli Compagni E, Di Pasquale A, Cammà C, Puglia I, Calistri P, Savini G |
| EPI_ISL_833083, EPI_ISL_833084 | RPS | Istituto Zooprofilattico Sperimentale dell'Abruzzo e Molise "G. Caporale" | Lorusso A, Marcacci M, Di Domenico M, Ancora M, Curini V, Mangone I, Rinaldi A, Delli Compagni E, Di Pasquale A, Cammà C, Puglia I, Calistri P, Savini G |
| EPI_ISL_833085, EPI_ISL_833086 | Servizio di Igiene Epidemiologia e Sanità Pubblica (SIESP) CHIETI - DRIVE IN GISSI | Istituto Zooprofilattico Sperimentale dell'Abruzzo e Molise "G. Caporale" | Lorusso A, Marcacci M, Di Domenico M, Ancora M, Curini V, Mangone I, Rinaldi A, Delli Compagni E, Di Pasquale A, Cammà C, Puglia I, Calistri P, Savini G |
| EPI_ISL_833087 | Servizio di Igiene Epidemiologia e Sanità Pubblica (SIESP) L'AQUILA | Istituto Zooprofilattico Sperimentale dell'Abruzzo e Molise "G. Caporale" | Lorusso A, Marcacci M, Di Domenico M, Ancora M, Curini V, Mangone I, Rinaldi A, Delli Compagni E, Di Pasquale A, Cammà C, Puglia I, Calistri P, Savini G |
| EPI_ISL_833088, EPI_ISL_833089 | Servizio di Igiene Epidemiologia e Sanità Pubblica (SIESP) SULMONA | Istituto Zooprofilattico Sperimentale dell'Abruzzo e Molise "G. Caporale" | Lorusso A, Marcacci M, Di Domenico M, Ancora M, Curini V, Mangone I, Rinaldi A, Delli Compagni E, Di Pasquale A, Cammà C, Puglia I, Calistri P, Savini G |
| EPI_ISL_833090, EPI_ISL_833091, EPI_ISL_833092, EPI_ISL_833093, EPI_ISL_833094, EPI_ISL_833095, EPI_ISL_833096 | SIESP CHIETI - DRIVE IN CHIETI | Istituto Zooprofilattico Sperimentale dell'Abruzzo e Molise "G. Caporale" | Lorusso A, Marcacci M, Di Domenico M, Ancora M, Curini V, Mangone I, Rinaldi A, Delli Compagni E, Di Pasquale A, Cammà C, Puglia I, Calistri P, Savini G |
| EPI_ISL_833097, EPI_ISL_833098 | SIESP CHIETI - DRIVE IN LANCIANO | Istituto Zooprofilattico Sperimentale dell'Abruzzo e Molise "G. Caporale" | Lorusso A, Marcacci M, Di Domenico M, Ancora M, Curini V, Mangone I, Rinaldi A, Delli Compagni E, Di Pasquale A, Cammà C, Puglia I, Calistri P, Savini G |
| EPI_ISL_833099, EPI_ISL_833100 | SIESP CHIETI - DRIVE IN ORTONA | Istituto Zooprofilattico Sperimentale dell'Abruzzo e Molise "G. Caporale" | Lorusso A, Marcacci M, Di Domenico M, Ancora M, Curini V, Mangone I, Rinaldi A, Delli Compagni E, Di Pasquale A, Cammà C, Puglia I, Calistri P, Savini G |
| EPI_ISL_833101, EPI_ISL_833102 | SIESP DIPARTIMENTO DI PREVENZIONE CHIETI | Istituto Zooprofilattico Sperimentale dell'Abruzzo e Molise "G. Caporale" | Lorusso A, Marcacci M, Di Domenico M, Ancora M, Curini V, Mangone I, Rinaldi A, Delli Compagni E, Di Pasquale A, Cammà C, Puglia I, Calistri P, Savini G |
| EPI_ISL_833103, EPI_ISL_833104, EPI_ISL_833105, EPI_ISL_833106, EPI_ISL_833107, EPI_ISL_833108, EPI_ISL_833109, EPI_ISL_833110, EPI_ISL_833111, EPI_ISL_833112, EPI_ISL_833113, EPI_ISL_833114, EPI_ISL_833115, EPI_ISL_833116, EPI_ISL_833117, EPI_ISL_833118 | | | |
| see above | SIESP DIPARTIMENTO DI PREVENZIONE TERAMO C.DA CASALENA | Istituto Zooprofilattico Sperimentale dell'Abruzzo e Molise "G. Caporale" | Lorusso A, Marcacci M, Di Domenico M, Ancora M, Curini V, Mangone I, Rinaldi A, Delli Compagni E, Di Pasquale A, Cammà C, Puglia I, Calistri P, Savini G |
| EPI_ISL_833119 | SIESP L'AQUILA | Istituto Zooprofilattico Sperimentale dell'Abruzzo e Molise "G. Caporale" | Lorusso A, Marcacci M, Di Domenico M, Ancora M, Curini V, Mangone I, Rinaldi A, Delli Compagni E, Di Pasquale A, Cammà C, Puglia I, Calistri P, Savini G |
| EPI_ISL_833120, EPI_ISL_833121, EPI_ISL_833122 | SIESP SULMONA | Istituto Zooprofilattico Sperimentale dell'Abruzzo e Molise "G. Caporale" | Lorusso A, Marcacci M, Di Domenico M, Ancora M, Curini V, Mangone I, Rinaldi A, Delli Compagni E, Di Pasquale A, Cammà C, Puglia I, Calistri P, Savini G |

| | | | |
|---|---|---|---|
| EPI_ISL_833123 | Usca Pizzoli | Istituto Zooprofilattico Sperimentale dell'Abruzzo e Molise "G. Caporale" | Lorusso A, Marcacci M, Di Domenico M, Ancora M, Curini V, Mangone I, Rinaldi A, Delli Compagni E, Di Pasquale A, Cammà C, Puglia I, Calistri P, Savini G |
| EPI_ISL_833124, EPI_ISL_833125, EPI_ISL_833126, EPI_ISL_833127, EPI_ISL_833128 | USCA Sulmona | Istituto Zooprofilattico Sperimentale dell'Abruzzo e Molise "G. Caporale" | Lorusso A, Marcacci M, Di Domenico M, Ancora M, Curini V, Mangone I, Rinaldi A, Delli Compagni E, Di Pasquale A, Cammà C, Puglia I, Calistri P, Savini G |
| EPI_ISL_833129 | USCA Tagliacozzo | Istituto Zooprofilattico Sperimentale dell'Abruzzo e Molise "G. Caporale" | Lorusso A, Marcacci M, Di Domenico M, Ancora M, Curini V, Mangone I, Rinaldi A, Delli Compagni E, Di Pasquale A, Cammà C, Puglia I, Calistri P, Savini G |
| EPI_ISL_833141, EPI_ISL_833142, EPI_ISL_833143, EPI_ISL_833144, EPI_ISL_833145, EPI_ISL_833147 | Genomic Laboratory (GLAB) (Conjoint lab of Health Directorate of Istanbul and Istanbul Technical University) | Genomic Laboratory (GLAB), Istanbul Technical University | Ilker Karacan, Tugba Kizilboga Akgun, Payam Zolfagharian, Nisan Denizce Can, Pari Sharifli, Levent Doganay, Gizem Dinler Doganay |
| EPI_ISL_833157, EPI_ISL_833158 | Instituto Adolfo Lutz - Regional de Santo Andre | Instituto Adolfo Lutz, Interdiciplinary Procedures Center, Strategic Laboratory | Claudio Tavares Sacchi, Claudia Regina Gonçalves, Erica Valessa Ramos Gomes, Karoline Rodrigues Campos |
| EPI_ISL_833159 | Instituto Adolfo Lutz - Central | Instituto Adolfo Lutz, Interdiciplinary Procedures Center, Strategic Laboratory | Claudio Tavares Sacchi, Claudia Regina Gonçalves, Erica Valessa Ramos Gomes, Karoline Rodrigues Campos |
| EPI_ISL_833160 | Instituto Adolfo Lutz - Regional de Santo Andre | Instituto Adolfo Lutz, Interdiciplinary Procedures Center, Strategic Laboratory | Claudio Tavares Sacchi, Claudia Regina Gonçalves, Erica Valessa Ramos Gomes, Karoline Rodrigues Campos |
| EPI_ISL_833161 | Instituto Adolfo Lutz - Central | Instituto Adolfo Lutz, Interdiciplinary Procedures Center, Strategic Laboratory | Claudio Tavares Sacchi, Claudia Regina Gonçalves, Erica Valessa Ramos Gomes, Karoline Rodrigues Campos |
| EPI_ISL_833162 | Lab LOC - Itapecerica da Serra | Instituto Adolfo Lutz, Interdiciplinary Procedures Center, Strategic Laboratory | Claudio Tavares Sacchi, Claudia Regina Gonçalves, Erica Valessa Ramos Gomes, Karoline Rodrigues Campos |
| EPI_ISL_833163 | Instituto Adolfo Lutz - Regional de Marilia | Instituto Adolfo Lutz, Interdiciplinary Procedures Center, Strategic Laboratory | Claudio Tavares Sacchi, Claudia Regina Gonçalves, Erica Valessa Ramos Gomes, Karoline Rodrigues Campos |
| EPI_ISL_833164 | Secretaria Municipal de Saude de Santa Barbara d'oeste | Instituto Adolfo Lutz, Interdiciplinary Procedures Center, Strategic Laboratory | Claudio Tavares Sacchi, Claudia Regina Gonçalves, Erica Valessa Ramos Gomes, Karoline Rodrigues Campos |
| EPI_ISL_833165, EPI_ISL_833166 | Hospital Samaritano | Instituto Adolfo Lutz, Interdiciplinary Procedures Center, Strategic Laboratory | Claudio Tavares Sacchi, Claudia Regina Gonçalves, Erica Valessa Ramos Gomes, Karoline Rodrigues Campos |
| EPI_ISL_833183 | Microbiology Department. Complexo Hospitalario Universitario de Vigo | Microbiology Department. Complexo Hospitalario Universitario de Vigo | Microbiology Department, Complexo Hospitalario Universitario de Vigo. EPICOVIGAL. |
| EPI_ISL_833184 | Microbiology Department. Hospital Clinico Universitario de Santiago | Microbiology Department. Complexo Hospitalario Universitario de Vigo | Microbiology Departments from Complexo Hospitalario Universitario de Vigo and Hospital Clinico Universitario de Santiago. EPICOVIGAL. |
| EPI_ISL_833228, EPI_ISL_833229 | R.P. GUARDIAGRELE Ospedale di Comunità | Istitituto Zooprofilattico Sperimentale dell'Abruzzo e Molise "G.Caporale" | Lorusso A, Marcacci M, Di Domenico M, Curini V, Ancora M, Cammà C, Rinaldi A, Mangone I, Di Pasquale A, Puglia I, Calistri P, Savini G. |
| EPI_ISL_833230 | OSPEDALE S.S. ANNUNZIATA CHIETI - CLINICA MEDICA (MEDICINA GENERALE 1) | Istitituto Zooprofilattico Sperimentale dell'Abruzzo e Molise "G.Caporale" | Lorusso A, Marcacci M, Di Domenico M, Curini V, Ancora M, Cammà C, Rinaldi A, Mangone I, Di Pasquale A, Puglia I, Calistri P, Savini G. |
| EPI_ISL_833231 | SIESP CHIETI - DRIVE IN LANCIANO | Istitituto Zooprofilattico Sperimentale dell'Abruzzo e Molise "G.Caporale" | Lorusso A, Marcacci M, Di Domenico M, Curini V, Ancora M, Cammà C, Rinaldi A, Mangone I, Di Pasquale A, Puglia I, Calistri P, Savini G. |
| EPI_ISL_833232 | SIESP CHIETI - DRIVE IN CHIETI | Istitituto Zooprofilattico Sperimentale dell'Abruzzo e Molise "G.Caporale" | Lorusso A, Marcacci M, Di Domenico M, Curini V, Ancora M, Cammà C, Rinaldi A, Mangone I, Di Pasquale A, Puglia I, Calistri P, Savini G. |
| EPI_ISL_833233, EPI_ISL_833234, EPI_ISL_833235, EPI_ISL_833236, EPI_ISL_833237, EPI_ISL_833238 | SIESP DIPARTIMENTO DI PREVENZIONE CHIETI | Istitituto Zooprofilattico Sperimentale dell'Abruzzo e Molise "G.Caporale" | Lorusso A, Marcacci M, Di Domenico M, Curini V, Ancora M, Cammà C, Rinaldi A, Mangone I, Di Pasquale A, Puglia I, Calistri P, Savini G. |
| EPI_ISL_833239, EPI_ISL_833240 | SIESP DIPARTIMENTO DI PREVENZIONE TERAMO | Istitituto Zooprofilattico Sperimentale dell'Abruzzo e Molise "G.Caporale" | Lorusso A, Marcacci M, Di Domenico M, Curini V, Ancora M, Cammà C, Rinaldi A, Mangone I, Di Pasquale A, Puglia I, Calistri P, Savini G. |
| EPI_ISL_833241 | SIESP DIPARTIMENTO DI PREVENZIONE CHIETI | Istitituto Zooprofilattico Sperimentale dell'Abruzzo e Molise "G.Caporale" | Lorusso A, Marcacci M, Di Domenico M, Curini V, Ancora M, Cammà C, Rinaldi A, Mangone I, Di Pasquale A, Puglia I, Calistri P, Savini G. |
| EPI_ISL_833242 | SIESP CHIETI - DRIVE IN LANCIANO | Istitituto Zooprofilattico Sperimentale dell'Abruzzo e Molise "G.Caporale" | Lorusso A, Marcacci M, Di Domenico M, Curini V, Ancora M, Cammà C, Rinaldi A, Mangone I, Di Pasquale A, Puglia I, Calistri P, Savini G. |
| EPI_ISL_833243, EPI_ISL_833246 | SIESP DIPARTIMENTO DI PREVENZIONE CHIETI | Istitituto Zooprofilattico Sperimentale dell'Abruzzo e Molise "G.Caporale" | Lorusso A, Marcacci M, Di Domenico M, Curini V, Ancora M, Cammà C, Rinaldi A, Mangone I, Di Pasquale A, Puglia I, Calistri P, Savini G. |
| EPI_ISL_833250 | SIESP CHIETI - DRIVE IN ORTONA | Istituto Zooprofilattico Sperimentale dell'Abruzzo e Molise "G. Caporale" | Lorusso A, Marcacci M, Di Domenico M, Ancora M, Curini V, Mangone I, Rinaldi A, Di Pasquale A, Cammà C, Puglia I, Calistri P, Savini G |
| EPI_ISL_833251 | Ospedale Civile Atri-Medicina Interna | Istituto Zooprofilattico Sperimentale dell'Abruzzo e Molise "G. Caporale" | Lorusso A, Marcacci M, Di Domenico M, Ancora M, Curini V, Mangone I, Rinaldi A, Di Pasquale A, Cammà C, Puglia I, Calistri P, Savini G |
| EPI_ISL_833252 | Ospedale Civile Giulianova-Pronto Soccorso | Istituto Zooprofilattico Sperimentale dell'Abruzzo e Molise "G. Caporale" | Lorusso A, Marcacci M, Di Domenico M, Ancora M, Curini V, Mangone I, Rinaldi A, Di Pasquale A, Cammà C, Puglia I, Calistri P, Savini G |
| EPI_ISL_833253 | USCA-Avezzano | Istituto Zooprofilattico Sperimentale dell'Abruzzo e Molise "G. Caporale" | Lorusso A, Marcacci M, Di Domenico M, Ancora M, Curini V, Mangone I, Rinaldi A, Di Pasquale A, Cammà C, Puglia I, Calistri P, Savini G |
| EPI_ISL_833254 | USCA-Pescina | Istituto Zooprofilattico Sperimentale dell'Abruzzo e Molise "G. Caporale" | Lorusso A, Marcacci M, Di Domenico M, Ancora M, Curini V, Mangone I, Rinaldi A, Di Pasquale A, Cammà C, Puglia I, Calistri P, Savini G |
| EPI_ISL_833255, EPI_ISL_833256, EPI_ISL_833257 | Ospedale Civile Atri | Istituto Zooprofilattico Sperimentale dell'Abruzzo e Molise "G. Caporale" | Lorusso A, Marcacci M, Di Domenico M, Ancora M, Curini V, Mangone I, Rinaldi A, Di Pasquale A, Cammà C, Puglia I, Calistri P, Savini G |
| EPI_ISL_833258, EPI_ISL_833259 | SIESP DIPARTIMENTO DI PREVENZIONE TERAMO | Istituto Zooprofilattico Sperimentale dell'Abruzzo e Molise "G. Caporale" | Lorusso A, Marcacci M, Di Domenico M, Ancora M, Curini V, Mangone I, Rinaldi A, Di Pasquale A, Cammà C, Puglia I, Calistri P, Savini G |
| EPI_ISL_833260, EPI_ISL_833261 | Giulianova | Istituto Zooprofilattico Sperimentale dell'Abruzzo e Molise "G. Caporale" | Lorusso A, Marcacci M, Di Domenico M, Ancora M, Curini V, Mangone I, Rinaldi A, Di Pasquale A, Cammà C, Puglia I, Calistri P, Savini G |
| EPI_ISL_833262, EPI_ISL_833263, EPI_ISL_833264 | SIESP DIPARTIMENTO DI PREVENZIONE CHIETI | Istituto Zooprofilattico Sperimentale dell'Abruzzo e Molise "G. Caporale" | Lorusso A, Marcacci M, Di Domenico M, Ancora M, Curini V, Mangone I, Rinaldi A, Di Pasquale A, Cammà C, Puglia I, Calistri P, Savini G |
| EPI_ISL_833265, EPI_ISL_833266, EPI_ISL_833267, EPI_ISL_833268, EPI_ISL_833269 | Ospedale SS Annunziata Chieti | Istituto Zooprofilattico Sperimentale dell'Abruzzo e Molise "G. Caporale" | Lorusso A, Marcacci M, Di Domenico M, Ancora M, Curini V, Mangone I, Rinaldi A, Di Pasquale A, Cammà C, Puglia I, Calistri P, Savini G |
| EPI_ISL_833270 | SIESP CHIETI - DRIVE IN ORTONA | Istituto Zooprofilattico Sperimentale dell'Abruzzo e Molise "G. | Lorusso A, Marcacci M, Di Domenico M, Ancora M, Curini V, Mangone I, Rinaldi A, Di Pasquale A, Cammà C, Puglia I, Calistri P, Savini G |

| | | | |
|---|---|---|---|
| | | Caporale" | |
| EPI_ISL_833271 | SIESP CHIETI - DRIVE IN GISSI | Istituto Zooprofilattico Sperimentale dell'Abruzzo e Molise "G. Caporale" | Lorusso A, Marcacci M, Di Domenico M, Ancora M, Curini V, Mangone I, Rinaldi A, Di Pasquale A, Cammà C, Puglia I, Calistri P, Savini G |
| EPI_ISL_833272, EPI_ISL_833273, EPI_ISL_833274, EPI_ISL_833275 | SIESP CHIETI - DRIVE IN ORTONA | Istituto Zooprofilattico Sperimentale dell'Abruzzo e Molise "G. Caporale" | Lorusso A, Marcacci M, Di Domenico M, Ancora M, Curini V, Mangone I, Rinaldi A, Di Pasquale A, Cammà C, Puglia I, Calistri P, Savini G |
| EPI_ISL_833276 | Ospedale Civile G.Mazzini-Teramo | Istituto Zooprofilattico Sperimentale dell'Abruzzo e Molise "G. Caporale" | Lorusso A, Marcacci M, Di Domenico M, Ancora M, Curini V, Mangone I, Rinaldi A, Di Pasquale A, Cammà C, Puglia I, Calistri P, Savini G |
| EPI_ISL_833277 | USCA-Sulmona | Istituto Zooprofilattico Sperimentale dell'Abruzzo e Molise "G. Caporale" | Lorusso A, Marcacci M, Di Domenico M, Ancora M, Curini V, Mangone I, Rinaldi A, Di Pasquale A, Cammà C, Puglia I, Calistri P, Savini G |
| EPI_ISL_833278, EPI_ISL_833279, EPI_ISL_833280 | SIESP CHIETI- DRIVE IN CHIETI | Istituto Zooprofilattico Sperimentale dell'Abruzzo e Molise "G. Caporale" | Lorusso A, Marcacci M, Di Domenico M, Ancora M, Curini V, Mangone I, Rinaldi A, Di Pasquale A, Cammà C, Puglia I, Calistri P, Savini G |
| EPI_ISL_833281, EPI_ISL_833282 | SIESP DIPARTIMENTO DI PREVENZIONE TERAMO | Istituto Zooprofilattico Sperimentale dell'Abruzzo e Molise "G. Caporale" | Lorusso A, Marcacci M, Di Domenico M, Ancora M, Curini V, Mangone I, Rinaldi A, Di Pasquale A, Cammà C, Puglia I, Calistri P, Savini G |
| EPI_ISL_833283 | DIPARTIMENTO PREVENZIONE AVEZZANO-SERVIZIO DI IGIENE EPIDEMIOLOGIA E SANITA' PUBBLICA | Istituto Zooprofilattico Sperimentale dell'Abruzzo e Molise "G. Caporale" | Lorusso A, Marcacci M, Di Domenico M, Ancora M, Curini V, Mangone I, Rinaldi A, Di Pasquale A, Cammà C, Puglia I, Calistri P, Savini G |
| EPI_ISL_833284, EPI_ISL_833285, EPI_ISL_833286, EPI_ISL_833287 | SIESP CHIETI - DRIVE IN ORTONA | Istituto Zooprofilattico Sperimentale dell'Abruzzo e Molise "G. Caporale" | Lorusso A, Marcacci M, Di Domenico M, Ancora M, Curini V, Mangone I, Rinaldi A, Di Pasquale A, Cammà C, Puglia I, Calistri P, Savini G |
| EPI_ISL_833288 | PRESIDIO TERRITORIALE CASOLI CH | Istituto Zooprofilattico Sperimentale dell'Abruzzo e Molise "G. Caporale" | Lorusso A, Marcacci M, Di Domenico M, Ancora M, Curini V, Mangone I, Rinaldi A, Di Pasquale A, Cammà C, Puglia I, Calistri P, Savini G |
| EPI_ISL_833289 | Ospedale SS Annunziata Chieti | Istituto Zooprofilattico Sperimentale dell'Abruzzo e Molise "G. Caporale" | Lorusso A, Marcacci M, Di Domenico M, Ancora M, Curini V, Mangone I, Rinaldi A, Di Pasquale A, Cammà C, Puglia I, Calistri P, Savini G |
| EPI_ISL_833290 | PRESIDIO TERRITORIALE CASOLI CH | Istituto Zooprofilattico Sperimentale dell'Abruzzo e Molise "G. Caporale" | Lorusso A, Marcacci M, Di Domenico M, Ancora M, Curini V, Mangone I, Rinaldi A, Di Pasquale A, Cammà C, Puglia I, Calistri P, Savini G |
| EPI_ISL_833291 | SIESP DIPARTIMENTO DI PREVENZIONE DELL'AQUILA | Istituto Zooprofilattico Sperimentale dell'Abruzzo e Molise "G. Caporale" | Lorusso A, Marcacci M, Di Domenico M, Ancora M, Curini V, Mangone I, Rinaldi A, Di Pasquale A, Cammà C, Puglia I, Calistri P, Savini G |
| EPI_ISL_833292, EPI_ISL_833293, EPI_ISL_833294 | SIESP CHIETI - DRIVE IN LANCIANO | Istituto Zooprofilattico Sperimentale dell'Abruzzo e Molise "G. Caporale" | Lorusso A, Marcacci M, Di Domenico M, Ancora M, Curini V, Mangone I, Rinaldi A, Di Pasquale A, Cammà C, Puglia I, Calistri P, Savini G |
| EPI_ISL_833295, EPI_ISL_833296 | SIESP DIPARTIMENTO DI PREVENZIONE TERAMO | Istituto Zooprofilattico Sperimentale dell'Abruzzo e Molise "G. Caporale" | Lorusso A, Marcacci M, Di Domenico M, Ancora M, Curini V, Mangone I, Rinaldi A, Di Pasquale A, Cammà C, Puglia I, Calistri P, Savini G |
| EPI_ISL_833297 | Ospedale Civile Giulianova | Istituto Zooprofilattico Sperimentale dell'Abruzzo e Molise "G. Caporale" | Lorusso A, Marcacci M, Di Domenico M, Ancora M, Curini V, Mangone I, Rinaldi A, Di Pasquale A, Cammà C, Puglia I, Calistri P, Savini G |
| EPI_ISL_833298 | SIESP CHIETI - DRIVE IN LANCIANO | Istituto Zooprofilattico Sperimentale dell'Abruzzo e Molise "G. Caporale" | Lorusso A, Marcacci M, Di Domenico M, Ancora M, Curini V, Mangone I, Rinaldi A, Di Pasquale A, Cammà C, Puglia I, Calistri P, Savini G |
| EPI_ISL_833299 | SIESP DIPARTIMENTO DI PREVENZIONE CHIETI | Istituto Zooprofilattico Sperimentale dell'Abruzzo e Molise "G. Caporale" | Lorusso A, Marcacci M, Di Domenico M, Ancora M, Curini V, Mangone I, Rinaldi A, Di Pasquale A, Cammà C, Puglia I, Calistri P, Savini G |
| EPI_ISL_833300 | SIESP CHIETI - DRIVE IN LANCIANO | Istituto Zooprofilattico Sperimentale dell'Abruzzo e Molise "G. Caporale" | Lorusso A, Marcacci M, Di Domenico M, Ancora M, Curini V, Mangone I, Rinaldi A, Di Pasquale A, Cammà C, Puglia I, Calistri P, Savini G |
| EPI_ISL_833301 | DIPARTIMENTO PREVENZIONE AVEZZANO-SERVIZIO DI IGIENE EPIDEMIOLOGIA E SANITA' PUBBLICA | Istituto Zooprofilattico Sperimentale dell'Abruzzo e Molise "G. Caporale" | Lorusso A, Marcacci M, Di Domenico M, Ancora M, Curini V, Mangone I, Rinaldi A, Di Pasquale A, Cammà C, Puglia I, Calistri P, Savini G |
| EPI_ISL_833302, EPI_ISL_833303, EPI_ISL_833304, EPI_ISL_833305 | SIESP DIPARTIMENTO DI PREVENZIONE TERAMO | Istituto Zooprofilattico Sperimentale dell'Abruzzo e Molise "G. Caporale" | Lorusso A, Marcacci M, Di Domenico M, Ancora M, Curini V, Mangone I, Rinaldi A, Di Pasquale A, Cammà C, Puglia I, Calistri P, Savini G |
| EPI_ISL_833306, EPI_ISL_833307, EPI_ISL_833308, EPI_ISL_833309, EPI_ISL_833310 | SIESP CHIETI - DRIVE IN ORTONA | Istituto Zooprofilattico Sperimentale dell'Abruzzo e Molise "G. Caporale" | Lorusso A, Marcacci M, Di Domenico M, Ancora M, Curini V, Mangone I, Rinaldi A, Di Pasquale A, Cammà C, Puglia I, Calistri P, Savini G |
| EPI_ISL_833311, EPI_ISL_833312 | SIESP CHIETI- DRIVE IN CHIETI | Istituto Zooprofilattico Sperimentale dell'Abruzzo e Molise "G. Caporale" | Lorusso A, Marcacci M, Di Domenico M, Ancora M, Curini V, Mangone I, Rinaldi A, Di Pasquale A, Cammà C, Puglia I, Calistri P, Savini G |
| EPI_ISL_833313, EPI_ISL_833314, EPI_ISL_833315 | SIESP CHIETI - DRIVE IN LANCIANO | Istituto Zooprofilattico Sperimentale dell'Abruzzo e Molise "G. Caporale" | Lorusso A, Marcacci M, Di Domenico M, Ancora M, Curini V, Mangone I, Rinaldi A, Di Pasquale A, Cammà C, Puglia I, Calistri P, Savini G |
| EPI_ISL_833316 | SIESP CHIETI - DRIVE IN GISSI | Istituto Zooprofilattico Sperimentale dell'Abruzzo e Molise "G. Caporale" | Lorusso A, Marcacci M, Di Domenico M, Ancora M, Curini V, Mangone I, Rinaldi A, Di Pasquale A, Cammà C, Puglia I, Calistri P, Savini G |
| EPI_ISL_833319, EPI_ISL_833320 | SIESP DIPARTIMENTO DI PREVENZIONE SULMONA | Istituto Zooprofilattico Sperimentale dell'Abruzzo e Molise "G. Caporale" | Lorusso A, Marcacci M, Di Domenico M, Ancora M, Curini V, Mangone I, Rinaldi A, Di Pasquale A, Cammà C, Puglia I, Calistri P, Savini G |
| EPI_ISL_833321 | SIESP DIPARTIMENTO DI PREVENZIONE CHIETI | Istituto Zooprofilattico Sperimentale dell'Abruzzo e Molise "G. Caporale" | Lorusso A, Marcacci M, Di Domenico M, Ancora M, Curini V, Mangone I, Rinaldi A, Di Pasquale A, Cammà C, Puglia I, Calistri P, Savini G |
| EPI_ISL_833322 | SIESP DIPARTIMENTO DI PREVENZIONE SULMONA | Istituto Zooprofilattico Sperimentale dell'Abruzzo e Molise "G. Caporale" | Lorusso A, Marcacci M, Di Domenico M, Ancora M, Curini V, Mangone I, Rinaldi A, Di Pasquale A, Cammà C, Puglia I, Calistri P, Savini G |
| EPI_ISL_833323 | SIESP CHIETI - DRIVE IN CHIETI | Istituto Zooprofilattico Sperimentale dell'Abruzzo e Molise "G. Caporale" | Lorusso A, Marcacci M, Di Domenico M, Ancora M, Curini V, Mangone I, Rinaldi A, Di Pasquale A, Cammà C, Puglia I, Calistri P, Savini G |
| EPI_ISL_833324 | SIESP DIPARTIMENTO DI PREVENZIONE CHIETI | Istituto Zooprofilattico Sperimentale dell'Abruzzo e Molise "G. Caporale" | Lorusso A, Marcacci M, Di Domenico M, Ancora M, Curini V, Mangone I, Rinaldi A, Di Pasquale A, Cammà C, Puglia I, Calistri P, Savini G |
| EPI_ISL_833325 | Ospedale SS Annunziata Chieti | Istituto Zooprofilattico Sperimentale dell'Abruzzo e Molise "G. Caporale" | Lorusso A, Marcacci M, Di Domenico M, Ancora M, Curini V, Mangone I, Rinaldi A, Di Pasquale A, Cammà C, Puglia I, Calistri P, Savini G |
| EPI_ISL_833326 | SIESP DIPARTIMENTO DI PREVENZIONE CHIETI | Istituto Zooprofilattico Sperimentale dell'Abruzzo e Molise "G. Caporale" | Lorusso A, Marcacci M, Di Domenico M, Ancora M, Curini V, Mangone I, Rinaldi A, Di Pasquale A, Cammà C, Puglia I, Calistri P, Savini G |
| EPI_ISL_833328, EPI_ISL_833329, EPI_ISL_833330 | SIESP CHIETI - DRIVE IN ORTONA | Istituto Zooprofilattico Sperimentale dell'Abruzzo e Molise "G. Caporale" | Lorusso A, Marcacci M, Di Domenico M, Ancora M, Curini V, Mangone I, Rinaldi A, Di Pasquale A, Cammà C, Puglia I, Calistri P, Savini G |
| EPI_ISL_833331 | Ospedale Civile Atri | Istituto Zooprofilattico Sperimentale dell'Abruzzo e Molise "G. Caporale" | Lorusso A, Marcacci M, Di Domenico M, Ancora M, Curini V, Mangone I, Rinaldi A, Di Pasquale A, Cammà C, Puglia I, Calistri P, Savini G |

EPI_ISL_833435, EPI_ISL_833436, EPI_ISL_833437, EPI_ISL_833438, EPI_ISL_833439, EPI_ISL_833440, EPI_ISL_833441, EPI_ISL_833442, EPI_ISL_833443, EPI_ISL_833444, EPI_ISL_833445, EPI_ISL_833446, EPI_ISL_833447, EPI_ISL_833448, EPI_ISL_833449, EPI_ISL_833450, EPI_ISL_833451, EPI_ISL_833453, EPI_ISL_833454, EPI_ISL_833455

| | | | |
|---|---|---|---|
| see above | SC (UCO) Igiene e Sanità Pubblica (funzione integrata con SC Microbiologia e Virologia) e Laboratory of Molecular Virology of the International Centre for Genetic Engineering and Biotechnology (ICGEB) | ARGO Laboratorio Genomica ed Epigenomica | Licastro D, Dal Monego S, Degasperi M, Marcello A, D'Agaro P |
| EPI_ISL_833578 | National Reference Laboratory for COVID-19, Pasteur Institute of Iran | National Reference Laboratory for COVID-19, Pasteur Institute of Iran | Zahra Ahmadi, Marzieh Sadjadi, Tahmineh Jalali, Mohammad Hassan Pouriayevali, Mahsa Tavakoli, Zahra Fereydouni, Setareh Kashanian, Sanam Azad-Manjiri, Tahereh Mohammadi, Kayhan Azadmanesh, Zabihollah Shoja, Parastoo Yekta, Farideh Niknam, Hessam Nemati, Ahmad Ghasemi, Sahar Khakifirouz, Sepideh Gerdooei,Maryam Rostamtabar,Sana Eybpoosh,Mohammad Mehdi Mortazavipour, Mohamad Sadegh Shams Nosrati, Zeynab VeisiZadeh, Mostafa Salehi-Vaziri |
| EPI_ISL_836143 | Hospital de Campanha COVID-19 de Mairipora | Instituto Adolfo Lutz, Interdiciplinary Procedures Center, Strategic Laboratory | Claudio Tavares Sacchi, Claudia Regina Gonçalves, Erica Valessa Ramos Gomes, Karoline Rodrigues Campos |
| EPI_ISL_836977 | Hospital Municiplal Dr. Jose de Carvalho Florence | Instituto Adolfo Lutz, Interdiciplinary Procedures Center, Strategic Laboratory | Claudio Tavares Sacchi, Claudia Regina Gonçalves, Erica Valessa Ramos Gomes, Karoline Rodrigues Campos |
| EPI_ISL_836978 | Irmandade da Santa Casa de Misericordia de Lorena | Instituto Adolfo Lutz, Interdiciplinary Procedures Center, Strategic Laboratory | Claudio Tavares Sacchi, Claudia Regina Gonçalves, Erica Valessa Ramos Gomes, Karoline Rodrigues Campos |
| EPI_ISL_837053 | UBS Darcy Alves e Robalinho | Instituto Adolfo Lutz, Interdiciplinary Procedures Center, Strategic Laboratory | Claudio Tavares Sacchi, Claudia Regina Gonçalves, Erica Valessa Ramos Gomes, Karoline Rodrigues Campos |
| EPI_ISL_837054 | UBS Jose Sabino Ferreira | Instituto Adolfo Lutz, Interdiciplinary Procedures Center, Strategic Laboratory | Claudio Tavares Sacchi, Claudia Regina Gonçalves, Erica Valessa Ramos Gomes, Karoline Rodrigues Campos |
| EPI_ISL_837810, EPI_ISL_837811, EPI_ISL_837812, EPI_ISL_837814 | Instituto Nacional de Enfermedades Respiratorias (INER) | Instituto Nacional de Enfermedades Respiratorias (INER) | Celia Boukadida, Margarita Matías-Florentino, Alma Rincón-Rubio, Hector Esteban Paz-Juárez, Olivia Briceño, Edgar Sevilla-Reyes, Fidencio Mejía-Nepomuceno, Mario Mújica-Sánchez, Eduardo Becerril-Vargas, José Arturo Martínez-Orozco, Alejandra Hernández-Terán, Jorge Salas-Hernández, Santiago Ávila-Ríos, Joel Armando Vázquez-Pérez |
| EPI_ISL_842652 | Laboratorio de Biología Molecular Hospital Pedro de Elizalde | Grupo de Genómica y Bioinformática del Instituto de Investigación de la Cadena Láctea CONICET-INTA on behalf of 'Proyecto Argentino Interinstitucional de genomica de SARS-CoV-2' (PAIS Consortium) | Amadio, AF, Eberhardt, MF; Irazoqui, M; Indart, J; Rocovich, J; Montoto Piazza, L; Wenk, G; Martín, ME; Sanchez, MF; Marchetti, P; Morandi, F; Sueiro, ML; Claps, A; Bressan, L; Torres, FJ; Chamorro, J; Gondolessi, J; Gómez, ML; Diaz, B; Rosales, D; Alegre, F; Zamora, N; Osaba, E; Paez, E; Lorenzo, F; Torres, C; Aulicino, P; König, G; Alexay, S; Natale, M; Valinotto, L; Lusso, S; Goya, S; Nabaes Jodar, MS; Viegas, M. |
| EPI_ISL_846631, EPI_ISL_846632, EPI_ISL_846633, EPI_ISL_846634, EPI_ISL_846635, EPI_ISL_846636 | Hopital | National Reference Center for Viruses of Respiratory Infections, Institut Pasteur, Paris | Marion Barbet, Sylvie Behillil, Méline Bizard, Angela Brisebarre, Camille Capel, Etienne Simon-Lorière, Vincent Enouf, Maud Vanpeene, Sylvie van der Werf,Farfour Eric |
| EPI_ISL_846637, EPI_ISL_846638, EPI_ISL_846639, EPI_ISL_846640 | Hopital | National Reference Center for Viruses of Respiratory Infections, Institut Pasteur, Paris | Marion Barbet, Sylvie Behillil, Méline Bizard, Angela Brisebarre, Camille Capel, Etienne Simon-Lorière, Vincent Enouf, Maud Vanpeene, Sylvie van der Werf,Lagathu Gisele |
| EPI_ISL_846641, EPI_ISL_846642, EPI_ISL_846643 | Labo Analyses Med | National Reference Center for Viruses of Respiratory Infections, Institut Pasteur, Paris | Marion Barbet, Sylvie Behillil, Méline Bizard, Angela Brisebarre, Camille Capel, Etienne Simon-Lorière, Vincent Enouf, Maud Vanpeene, Sylvie van der Werf |
| EPI_ISL_846644 | Hopital | National Reference Center for Viruses of Respiratory Infections, Institut Pasteur, Paris | Marion Barbet, Sylvie Behillil, Méline Bizard, Angela Brisebarre, Camille Capel, Etienne Simon-Lorière, Vincent Enouf, Maud Vanpeene, Sylvie van der Werf |
| EPI_ISL_846645 | Labo Analyses Med | National Reference Center for Viruses of Respiratory Infections, Institut Pasteur, Paris | Marion Barbet, Sylvie Behillil, Méline Bizard, Angela Brisebarre, Camille Capel, Etienne Simon-Lorière, Vincent Enouf, Maud Vanpeene, Sylvie van der Werf |
| EPI_ISL_846646, EPI_ISL_846647 | Labo Analyses Med | National Reference Center for Viruses of Respiratory Infections, Institut Pasteur, Paris | Marion Barbet, Sylvie Behillil, Méline Bizard, Angela Brisebarre, Camille Capel, Etienne Simon-Lorière, Vincent Enouf, Maud Vanpeene, Sylvie van der Werf,Coignard Catherine |
| EPI_ISL_846648 | Labo Analyses Med | National Reference Center for Viruses of Respiratory Infections, Institut Pasteur, Paris | Marion Barbet, Sylvie Behillil, Méline Bizard, Angela Brisebarre, Camille Capel, Etienne Simon-Lorière, Vincent Enouf, Maud Vanpeene, Sylvie van der Werf |
| EPI_ISL_846649, EPI_ISL_846650 | Hopital | National Reference Center for Viruses of Respiratory Infections, Institut Pasteur, Paris | Marion Barbet, Sylvie Behillil, Méline Bizard, Angela Brisebarre, Camille Capel, Etienne Simon-Lorière, Vincent Enouf, Maud Vanpeene, Sylvie van der Werf,Farfour Eric |
| EPI_ISL_847825 | Lab. Microbiologia e Virologia, Cotugno, A.O. dei Colli | Lab. Microbiologia e Virologia, Cotugno, A.O. dei Colli | Luigi Atripaldi, Claudia Tiberio, Anna Perfetti |
| EPI_ISL_848198 | Microbiology Lab | NBCC Sequencing Facility | Dr. Jeff Wrana |
| EPI_ISL_848199 | Microbiology Department. Complexo Hospitalario Universitario de Vigo | Microbiology Department. Complexo Hospitalario Universitario de Vigo | Microbiology Department, Complexo Hospitalario Universitario de Vigo (CHUVI). EPICOVIGAL. |
| EPI_ISL_848201, EPI_ISL_848202 | Charité Universitätsmedizin Berlin, Institute of Virology, Charitéplatz 1, 10117 Berlin, Germany | Charité Universitätsmedizin Berlin, Institute of Virology, Charitéplatz 1, 10117 Berlin, Germany | Victor M Corman, Julia Schneider, Jörn Beheim-Schwarzbach, Tobias Bleicker, Julia Tesch, Barbara Mühlemann, Talitha Veith, Terry Jones, Christian Drosten |
| EPI_ISL_849634 | SIESP DIPARTIMENTO DI PREVENZIONE CHIETI | Istitituto Zooprofilattico Sperimentale dell'Abruzzo e Molise "G.Caporale" | Lorusso A, Marcacci M, Di Domenico M, Curini V, Ancora M, Cammà C, Rinaldi A, Mangone I, Di Pasquale A, Puglia I, Savini G. |
| EPI_ISL_849635, EPI_ISL_849636 | DIPARTIMENTO PREVENZIONE AVEZZANO-SERVIZIO DI IGIENE EPIDEMIOLOGIA E SANITA' PUBBLICA | Istitituto Zooprofilattico Sperimentale dell'Abruzzo e Molise "G.Caporale" | Lorusso A, Marcacci M, Di Domenico M, Curini V, Ancora M, Cammà C, Rinaldi A, Mangone I, Di Pasquale A, Puglia I, Savini G. |
| EPI_ISL_849637, EPI_ISL_849638, EPI_ISL_849639, EPI_ISL_849640, EPI_ISL_849641, EPI_ISL_849642, EPI_ISL_849643, EPI_ISL_849644, EPI_ISL_849645, EPI_ISL_849646 | SIESP DIPARTIMENTO DI PREVENZIONE CHIETI | Istitituto Zooprofilattico Sperimentale dell'Abruzzo e Molise "G.Caporale" | Lorusso A, Marcacci M, Di Domenico M, Curini V, Ancora M, Cammà C, Rinaldi A, Mangone I, Di Pasquale A, Puglia I, Savini G. |
| EPI_ISL_849647, EPI_ISL_849648, EPI_ISL_849649 | SIESP- DRIVE IN CHIETI | Istitituto Zooprofilattico Sperimentale dell'Abruzzo e Molise "G.Caporale" | Lorusso A, Marcacci M, Di Domenico M, Curini V, Ancora M, Cammà C, Rinaldi A, Mangone I, Di Pasquale A, Puglia I, Savini G. |
| EPI_ISL_850663, EPI_ISL_850665, EPI_ISL_850666, EPI_ISL_850667, EPI_ISL_850668, EPI_ISL_850669, EPI_ISL_850670, EPI_ISL_850671, EPI_ISL_850672, EPI_ISL_850673, EPI_ISL_850674 | | | |
| see above | The National Institute of Public Health | State Veterinary Institute Prague | Nagy,A;Jirincova,H;Trnka,D;Vecerova,J;Trinklova,M |
| EPI_ISL_850679, EPI_ISL_850680, EPI_ISL_850681, EPI_ISL_850682, EPI_ISL_850684, EPI_ISL_850685, EPI_ISL_850686 | The National Institute of Public Health | State Veterinary Institute Prague | Nagy,A;Jirincova,H;Trnka,D;Vecerova,J |
| EPI_ISL_852583, EPI_ISL_852584, EPI_ISL_852585, EPI_ISL_852587, EPI_ISL_852589, EPI_ISL_852590, EPI_ISL_852593, EPI_ISL_852594, EPI_ISL_852595, EPI_ISL_852596, EPI_ISL_852597, EPI_ISL_852598, EPI_ISL_852599, EPI_ISL_852600, EPI_ISL_852601, EPI_ISL_852602, EPI_ISL_852603 | | | |
| see above | Max von Pettenkofer Institute, Virology, National Reference Center for Retroviruses, LMU München | Laboratory for Functional Genome Analysis, Dept. Genomics, Gene Center of the LMU Munich | Max Muenchhoff, Stefan Krebs, Alexander Graf, Oliver Keppler, Helmut Blum |
| EPI_ISL_852619, EPI_ISL_852629, EPI_ISL_852630, EPI_ISL_852631, EPI_ISL_852632, EPI_ISL_852813, EPI_ISL_852819, EPI_ISL_852823, EPI_ISL_852825, EPI_ISL_852828 | Lab. Microbiologia e Virologia, Cotugno, A.O. dei Colli | Lab. Microbiologia e Virologia, Cotugno, A.O. dei Colli | Luigi Atripaldi, Claudia Tiberio, Anna Perfetti |

| | | | |
|---|---|---|---|
| EPI_ISL_854594 | Faroese National Reference Laboratory for Fish and Animal Diseases | Faroese National Reference Laboratory for Fish and Animal Diseases | Maria Marjunardóttir Dahl, Petra Elisabeth Petersen, Arnfinnur Kallsberg Junior, Debes Hammershaimb Christiansen |
| EPI_ISL_855353 | Hospital | National Reference Center for Viruses of Respiratory Infections, Institut Pasteur, Paris | Marion Barbet, Sylvie Behillil, Méline Bizard, Angela Brisebarre, Camille Capel, Etienne Simon-Lorière, Vincent Enouf, Maud Vanpeene, Sylvie van der Werf,Pilorge Léa |
| EPI_ISL_855354, EPI_ISL_855355, EPI_ISL_855356 | Hospital | National Reference Center for Viruses of Respiratory Infections, Institut Pasteur, Paris | Marion Barbet, Sylvie Behillil, Méline Bizard, Angela Brisebarre, Camille Capel, Etienne Simon-Lorière, Vincent Enouf, Maud Vanpeene, Sylvie van der Werf |
| EPI_ISL_855357 | Labo Analyses Med | National Reference Center for Viruses of Respiratory Infections, Institut Pasteur, Paris | Marion Barbet, Sylvie Behillil, Méline Bizard, Angela Brisebarre, Camille Capel, Etienne Simon-Lorière, Vincent Enouf, Maud Vanpeene, Sylvie van der Werf,Harich |
| EPI_ISL_855358 | Hospital | National Reference Center for Viruses of Respiratory Infections, Institut Pasteur, Paris | Marion Barbet, Sylvie Behillil, Méline Bizard, Angela Brisebarre, Camille Capel, Etienne Simon-Lorière, Vincent Enouf, Maud Vanpeene, Sylvie van der Werf,Griscelli Franck |
| EPI_ISL_855359 | Hospital | National Reference Center for Viruses of Respiratory Infections, Institut Pasteur, Paris | Marion Barbet, Sylvie Behillil, Méline Bizard, Angela Brisebarre, Camille Capel, Etienne Simon-Lorière, Vincent Enouf, Maud Vanpeene, Sylvie van der Werf,Fourgeaud Jacques |
| EPI_ISL_855360, EPI_ISL_855361, EPI_ISL_855362, EPI_ISL_855363, EPI_ISL_855364, EPI_ISL_855365, EPI_ISL_855366, EPI_ISL_855367, EPI_ISL_855368, EPI_ISL_855369, EPI_ISL_855370, EPI_ISL_855371, EPI_ISL_855372, EPI_ISL_855373, EPI_ISL_855374, EPI_ISL_855375, EPI_ISL_855376, EPI_ISL_855377, EPI_ISL_855378, EPI_ISL_855379, EPI_ISL_855380 | | | |
| see above | Hospital | National Reference Center for Viruses of Respiratory Infections, Institut Pasteur, Paris | Marion Barbet, Sylvie Behillil, Méline Bizard, Angela Brisebarre, Camille Capel, Etienne Simon-Lorière, Vincent Enouf, Maud Vanpeene, Sylvie van der Werf,Combe Patrice |
| EPI_ISL_855381 | Labo Analyses Med | National Reference Center for Viruses of Respiratory Infections, Institut Pasteur, Paris | Marion Barbet, Sylvie Behillil, Méline Bizard, Angela Brisebarre, Camille Capel, Etienne Simon-Lorière, Vincent Enouf, Maud Vanpeene, Sylvie van der Werf |
| EPI_ISL_855388 | Hospital | National Reference Center for Viruses of Respiratory Infections, Institut Pasteur, Paris | Marion Barbet, Sylvie Behillil, Méline Bizard, Angela Brisebarre, Camille Capel, Etienne Simon-Lorière, Vincent Enouf, Maud Vanpeene, Sylvie van der Werf,Ducancelle Alexandra |
| EPI_ISL_855573, EPI_ISL_855611, EPI_ISL_855612, EPI_ISL_855613 | Lab. Microbiologia e Virologia, Cotugno, A.O. dei Colli | Lab. Microbiologia e Virologia, Cotugno, A.O. dei Colli | Luigi Atripaldi, Claudia Tiberio, Anna Perfetti |
| EPI_ISL_857056, EPI_ISL_857057, EPI_ISL_857058, EPI_ISL_857059, EPI_ISL_857061, EPI_ISL_857066, EPI_ISL_857289, EPI_ISL_857291, EPI_ISL_857292, EPI_ISL_857294, EPI_ISL_857295, EPI_ISL_857296, EPI_ISL_857297, EPI_ISL_857299, EPI_ISL_857300, EPI_ISL_857302, EPI_ISL_857303, EPI_ISL_857304, EPI_ISL_857305, EPI_ISL_857307 | | | |
| see above | OCME Office Of Chief Medical Examiner | New York City Public Health Laboratory | Jade Wang, et al. |
| EPI_ISL_857314 | Centers for Disease Control, R.O.C. (Taiwan) | Centers for Disease Control, R.O.C. (Taiwan) | Ji-Rong Yang, Yu-Chi Lin, Jung-Jung-Mu, Ming-Tsan Liu |
| EPI_ISL_857395 | S.M.S.Medical College, Jaipur, Rajasthan | S.M.S.Medical College, Jaipur, Rajasthan | Dr. Bharti Malhotra, Dr. Swati Gautam |
| EPI_ISL_860095, EPI_ISL_860203, EPI_ISL_860204, EPI_ISL_860257, EPI_ISL_860290, EPI_ISL_860297 | Lab. Microbiologia e Virologia, Cotugno, A.O. dei Colli | Lab. Microbiologia e Virologia, Cotugno, A.O. dei Colli | Luigi Atripaldi, Claudia Tiberio, Anna Perfetti |
| EPI_ISL_860317, EPI_ISL_860633 | Instituto Adolfo Lutz - Regional de Sorocaba | Instituto Adolfo Lutz, Interdiciplinary Procedures Center, Strategic Laboratory | Claudio Tavares Sacchi, Claudia Regina Gonçalves, Erica Valessa Ramos Gomes, Karoline Rodrigues Campos |
| EPI_ISL_860716 | Cerballiance Côte d'Azur | CERBA LAB | Roquebert B; Prots.L; Barrieu-Moussat S;Haïm-Boukobza S. |
| EPI_ISL_860833, EPI_ISL_860834, EPI_ISL_860835, EPI_ISL_860836, EPI_ISL_860837, EPI_ISL_860838, EPI_ISL_860839, EPI_ISL_860840 | Labo Analyses Med | National Reference Center for Viruses of Respiratory Infections, Institut Pasteur, Paris | Marion Barbet, Sylvie Behillil, Méline Bizard, Angela Brisebarre, Camille Capel, Etienne Simon-Lorière, Vincent Enouf, Maud Vanpeene, Sylvie van der Werf,Amzalag Jonas |
| EPI_ISL_860841 | Labo Analyses Med | National Reference Center for Viruses of Respiratory Infections, Institut Pasteur, Paris | Marion Barbet, Sylvie Behillil, Méline Bizard, Angela Brisebarre, Camille Capel, Etienne Simon-Lorière, Vincent Enouf, Maud Vanpeene, Sylvie van der Werf, |
| EPI_ISL_860842, EPI_ISL_860843, EPI_ISL_860844, EPI_ISL_860845 | Labo Analyses med | National Reference Center for Viruses of Respiratory Infections, Institut Pasteur, Paris | Marion Barbet, Sylvie Behillil, Méline Bizard, Angela Brisebarre, Camille Capel, Etienne Simon-Lorière, Vincent Enouf, Maud Vanpeene, Sylvie van der Werf,Amzalag Jonas |
| EPI_ISL_860846 | Labo Analyses Med | National Reference Center for Viruses of Respiratory Infections, Institut Pasteur, Paris | Marion Barbet, Sylvie Behillil, Méline Bizard, Angela Brisebarre, Camille Capel, Etienne Simon-Lorière, Vincent Enouf, Maud Vanpeene, Sylvie van der Werf,Amzalag Jonas |
| EPI_ISL_860847 | Centre Hospitalier Compiègne-Noyon - Laboratoire | National Reference Center for Viruses of Respiratory Infections, Institut Pasteur, Paris | Marion Barbet, Sylvie Behillil, Méline Bizard, Angela Brisebarre, Camille Capel, Etienne Simon-Lorière, Vincent Enouf, Maud Vanpeene, Sylvie van der Werf,Foissaud Vincent |
| EPI_ISL_860848 | CH Dunkerque Laboratoire de Biologie | National Reference Center for Viruses of Respiratory Infections, Institut Pasteur, Paris | Marion Barbet, Sylvie Behillil, Méline Bizard, Angela Brisebarre, Camille Capel, Etienne Simon-Lorière, Vincent Enouf, Maud Vanpeene, Sylvie van der Werf,Joly Isabelle |
| EPI_ISL_860849 | HIA BÉGIN Service de Biologie | National Reference Center for Viruses of Respiratory Infections, Institut Pasteur, Paris | Marion Barbet, Sylvie Behillil, Méline Bizard, Angela Brisebarre, Camille Capel, Etienne Simon-Lorière, Vincent Enouf, Maud Vanpeene, Sylvie van der Werf,Merens Audrey |
| EPI_ISL_860850, EPI_ISL_860851 | HIA PERCY, Biologie médicale | National Reference Center for Viruses of Respiratory Infections, Institut Pasteur, Paris | Marion Barbet, Sylvie Behillil, Méline Bizard, Angela Brisebarre, Camille Capel, Etienne Simon-Lorière, Vincent Enouf, Maud Vanpeene, Sylvie van der Werf,Foissaud Vincent |
| EPI_ISL_860852, EPI_ISL_860853, EPI_ISL_860854, EPI_ISL_860855, EPI_ISL_860856, EPI_ISL_860857, EPI_ISL_860858, EPI_ISL_860859 | Labo Analyses Med | National Reference Center for Viruses of Respiratory Infections, Institut Pasteur, Paris | Marion Barbet, Sylvie Behillil, Méline Bizard, Angela Brisebarre, Camille Capel, Etienne Simon-Lorière, Vincent Enouf, Maud Vanpeene, Sylvie van der Werf,Amzalag Jonas |
| EPI_ISL_860860, EPI_ISL_860861, EPI_ISL_860862 | Labo Analyses med | National Reference Center for Viruses of Respiratory Infections, Institut Pasteur, Paris | Marion Barbet, Sylvie Behillil, Méline Bizard, Angela Brisebarre, Camille Capel, Etienne Simon-Lorière, Vincent Enouf, Maud Vanpeene, Sylvie van der Werf,Amzalag Jonas |
| EPI_ISL_860863, EPI_ISL_860865 | CHU de Nantes - Hôtel Dieu - Labo. Virologie | National Reference Center for Viruses of Respiratory Infections, Institut Pasteur, Paris | Marion Barbet, Sylvie Behillil, Méline Bizard, Angela Brisebarre, Camille Capel, Etienne Simon-Lorière, Vincent Enouf, Maud Vanpeene, Sylvie van der Werf,Andre-Garnier Elisabeth |
| EPI_ISL_860867 | CHU Tours - Virologie | National Reference Center for Viruses of Respiratory Infections, Institut Pasteur, Paris | Marion Barbet, Sylvie Behillil, Méline Bizard, Angela Brisebarre, Camille Capel, Etienne Simon-Lorière, Vincent Enouf, Maud Vanpeene, Sylvie van der Werf,Gaudy Graffin Catherine |
| EPI_ISL_860868 | CHU de Nantes - Hôtel Dieu - Labo. Virologie | National Reference Center for Viruses of Respiratory Infections, Institut Pasteur, Paris | Marion Barbet, Sylvie Behillil, Méline Bizard, Angela Brisebarre, Camille Capel, Etienne Simon-Lorière, Vincent Enouf, Maud Vanpeene, Sylvie van der Werf,Andre-Garnier Elisabeth |
| EPI_ISL_860869, EPI_ISL_860870, EPI_ISL_860871, EPI_ISL_860872, EPI_ISL_860873, EPI_ISL_860874, EPI_ISL_860875, EPI_ISL_860876, EPI_ISL_860877, EPI_ISL_860878, EPI_ISL_860879, EPI_ISL_860880, EPI_ISL_860881, EPI_ISL_860882, EPI_ISL_860883, EPI_ISL_860884, EPI_ISL_860885, EPI_ISL_860886, EPI_ISL_860887, EPI_ISL_860888 | | | |
| see above | Labo Analyses Med | National Reference Center for Viruses of Respiratory Infections, Institut Pasteur, Paris | Marion Barbet, Sylvie Behillil, Méline Bizard, Angela Brisebarre, Camille Capel, Etienne Simon-Lorière, Vincent Enouf, Maud Vanpeene, Sylvie van der Werf,Lefaure Brieuc |
| EPI_ISL_860889 | Labo Analyses Med | National Reference Center for Viruses of Respiratory Infections, Institut Pasteur, Paris | Marion Barbet, Sylvie Behillil, Méline Bizard, Angela Brisebarre, Camille Capel, Etienne Simon-Lorière, Vincent Enouf, Maud Vanpeene, Sylvie van der Werf,Amzalag Jonas |
| EPI_ISL_860890 | CHU de Nantes - Hôtel Dieu - Labo. Virologie | National Reference Center for Viruses of Respiratory | Marion Barbet, Sylvie Behillil, Méline Bizard, Angela Brisebarre, Camille Capel, Etienne Simon-Lorière, Vincent Enouf, Maud Vanpeene, Sylvie van der |

| | | | |
|---|---|---|---|
| | | Infections, Institut Pasteur, Paris | Werf,Andre-Garnier Elisabeth |
| EPI_ISL_860891, EPI_ISL_860892, EPI_ISL_860893, EPI_ISL_860894, EPI_ISL_860895, EPI_ISL_860896, EPI_ISL_860897 | CHU Tours - Virologie | National Reference Center for Viruses of Respiratory Infections, Institut Pasteur, Paris | Marion Barbet, Sylvie Behillil, Méline Bizard, Angela Brisebarre, Camille Capel, Etienne Simon-Lorière, Vincent Enouf, Maud Vanpeene, Sylvie van der Werf,Gaudy Graffin Catherine |
| EPI_ISL_860898 | Centre Hospitalier Compiègne-Noyon - Laboratoire | National Reference Center for Viruses of Respiratory Infections, Institut Pasteur, Paris | Marion Barbet, Sylvie Behillil, Méline Bizard, Angela Brisebarre, Camille Capel, Etienne Simon-Lorière, Vincent Enouf, Maud Vanpeene, Sylvie van der Werf,Emond Jean-Philippe |
| EPI_ISL_860899 | HOPITAL PAUL BROUSSE - Service Microbiologie | National Reference Center for Viruses of Respiratory Infections, Institut Pasteur, Paris | Marion Barbet, Sylvie Behillil, Méline Bizard, Angela Brisebarre, Camille Capel, Etienne Simon-Lorière, Vincent Enouf, Maud Vanpeene, Sylvie van der Werf,Roque-Afonso Anne-Marie |
| EPI_ISL_860900, EPI_ISL_860901, EPI_ISL_860902, EPI_ISL_860903, EPI_ISL_860904 | CH du MANS - Lab. Bio. Moléculaire | National Reference Center for Viruses of Respiratory Infections, Institut Pasteur, Paris | Marion Barbet, Sylvie Behillil, Méline Bizard, Angela Brisebarre, Camille Capel, Etienne Simon-Lorière, Vincent Enouf, Maud Vanpeene, Sylvie van der Werf,Ramanantsoa CéLine |
| EPI_ISL_860905, EPI_ISL_860906 | CHU Angers - Dpt des Agents Infectieux | National Reference Center for Viruses of Respiratory Infections, Institut Pasteur, Paris | Marion Barbet, Sylvie Behillil, Méline Bizard, Angela Brisebarre, Camille Capel, Etienne Simon-Lorière, Vincent Enouf, Maud Vanpeene, Sylvie van der Werf,Ducancelle Alexandra |
| EPI_ISL_860907, EPI_ISL_860908, EPI_ISL_860909 | CHU Tours - Virologie | National Reference Center for Viruses of Respiratory Infections, Institut Pasteur, Paris | Marion Barbet, Sylvie Behillil, Méline Bizard, Angela Brisebarre, Camille Capel, Etienne Simon-Lorière, Vincent Enouf, Maud Vanpeene, Sylvie van der Werf,Gaudy Graffin Catherine |
| EPI_ISL_860910 | CH du MANS - Lab. Bio. Moléculaire | National Reference Center for Viruses of Respiratory Infections, Institut Pasteur, Paris | Marion Barbet, Sylvie Behillil, Méline Bizard, Angela Brisebarre, Camille Capel, Etienne Simon-Lorière, Vincent Enouf, Maud Vanpeene, Sylvie van der Werf,Ramanantsoa CéLine |
| EPI_ISL_860911, EPI_ISL_860912 | CHU Angers - Dpt des Agents Infectieux | National Reference Center for Viruses of Respiratory Infections, Institut Pasteur, Paris | Marion Barbet, Sylvie Behillil, Méline Bizard, Angela Brisebarre, Camille Capel, Etienne Simon-Lorière, Vincent Enouf, Maud Vanpeene, Sylvie van der Werf,Ducancelle Alexandra |
| EPI_ISL_860913, EPI_ISL_860914, EPI_ISL_860915, EPI_ISL_860916, EPI_ISL_860917, EPI_ISL_860918, EPI_ISL_860919, EPI_ISL_860920, EPI_ISL_860921, EPI_ISL_860922, EPI_ISL_860923, EPI_ISL_860924, EPI_ISL_860925, EPI_ISL_860926, EPI_ISL_860927, EPI_ISL_860928, EPI_ISL_860929, EPI_ISL_860930 | | | |
| see above | CHU de Nantes - Hôtel Dieu - Labo. Virologie | National Reference Center for Viruses of Respiratory Infections, Institut Pasteur, Paris | Marion Barbet, Sylvie Behillil, Méline Bizard, Angela Brisebarre, Camille Capel, Etienne Simon-Lorière, Vincent Enouf, Maud Vanpeene, Sylvie van der Werf,Andre-Garnier Elisabeth |
| EPI_ISL_860931 | Labo Analyses Med | National Reference Center for Viruses of Respiratory Infections, Institut Pasteur, Paris | Marion Barbet, Sylvie Behillil, Méline Bizard, Angela Brisebarre, Camille Capel, Etienne Simon-Lorière, Vincent Enouf, Maud Vanpeene, Sylvie van der Werf |
| EPI_ISL_861236 | CHU Angers - Dpt des Agents Infectieux | National Reference Center for Viruses of Respiratory Infections, Institut Pasteur, Paris | Marion Barbet, Sylvie Behillil, Méline Bizard, Angela Brisebarre, Camille Capel, Etienne Simon-Lorière, Vincent Enouf, Maud Vanpeene, Sylvie van der Werf,Ducancelle Alexandra |
| EPI_ISL_861239 | Labo Analyses med | National Reference Center for Viruses of Respiratory Infections, Institut Pasteur, Paris | Marion Barbet, Sylvie Behillil, Méline Bizard, Angela Brisebarre, Camille Capel, Etienne Simon-Lorière, Vincent Enouf, Maud Vanpeene, Sylvie van der Werf,Amzalag Jonas |
| EPI_ISL_861503 | Cerballiance Alpes Durance | CERBA LAB | Dyen P; Prots L. |
| EPI_ISL_861504, EPI_ISL_861505 | Cerballiance Provence | CERBA LAB | Delaunay E; Roig JC; Prots L. |
| EPI_ISL_861506 | Cerballiance Provence | CERBA LAB | Roig JC; Delaunay E; Prots L. |
| EPI_ISL_861508 | Cerballiance Haut de France | CERBA LAB | Felloni C; Mainardi A; Lahmidi S. |
| EPI_ISL_861668 | Centro de Triagem Covid19 | Instituto Adolfo Lutz, Interdiciplinary Procedures Center, Strategic Laboratory | Claudio Tavares Sacchi, Claudia Regina Gonçalves, Erica Valessa Ramos Gomes, Karoline Rodrigues Campos |
| EPI_ISL_861669 | Lab LOC - Itapecerica da Serra | Instituto Adolfo Lutz, Interdiciplinary Procedures Center, Strategic Laboratory | Claudio Tavares Sacchi, Claudia Regina Gonçalves, Erica Valessa Ramos Gomes, Karoline Rodrigues Campos |
| EPI_ISL_861670 | Instituto Adolfo Lutz - Regional de Taubate | Instituto Adolfo Lutz, Interdiciplinary Procedures Center, Strategic Laboratory | Claudio Tavares Sacchi, Claudia Regina Gonçalves, Erica Valessa Ramos Gomes, Karoline Rodrigues Campos |
| EPI_ISL_861671 | Hospital Municipal Prefeito Waldemar Costa Filho | Instituto Adolfo Lutz, Interdiciplinary Procedures Center, Strategic Laboratory | Claudio Tavares Sacchi, Claudia Regina Gonçalves, Erica Valessa Ramos Gomes, Karoline Rodrigues Campos |
| EPI_ISL_861672 | Day Hospital de Ermelino Matarazzo | Instituto Adolfo Lutz, Interdiciplinary Procedures Center, Strategic Laboratory | Claudio Tavares Sacchi, Claudia Regina Gonçalves, Erica Valessa Ramos Gomes, Karoline Rodrigues Campos |
| EPI_ISL_861673 | PA Novo Osasco | Instituto Adolfo Lutz, Interdiciplinary Procedures Center, Strategic Laboratory | Claudio Tavares Sacchi, Claudia Regina Gonçalves, Erica Valessa Ramos Gomes, Karoline Rodrigues Campos |
| EPI_ISL_861674, EPI_ISL_861675 | UPA Central de Caraguatatuba | Instituto Adolfo Lutz, Interdiciplinary Procedures Center, Strategic Laboratory | Claudio Tavares Sacchi, Claudia Regina Gonçalves, Erica Valessa Ramos Gomes, Karoline Rodrigues Campos |
| EPI_ISL_861676 | UPA Vila Santa Catarina | Instituto Adolfo Lutz, Interdiciplinary Procedures Center, Strategic Laboratory | Claudio Tavares Sacchi, Claudia Regina Gonçalves, Erica Valessa Ramos Gomes, Karoline Rodrigues Campos |
| EPI_ISL_861677 | Instituto Adolfo Lutz - Central | Instituto Adolfo Lutz, Interdiciplinary Procedures Center, Strategic Laboratory | Claudio Tavares Sacchi, Claudia Regina Gonçalves, Erica Valessa Ramos Gomes, Karoline Rodrigues Campos |
| EPI_ISL_861678 | Laboratorio Municipal de Guarulhos | Instituto Adolfo Lutz, Interdiciplinary Procedures Center, Strategic Laboratory | Claudio Tavares Sacchi, Claudia Regina Gonçalves, Erica Valessa Ramos Gomes, Karoline Rodrigues Campos |
| EPI_ISL_861679 | Instituto Adolfo Lutz - Regional de Taubate | Instituto Adolfo Lutz, Interdiciplinary Procedures Center, Strategic Laboratory | Claudio Tavares Sacchi, Claudia Regina Gonçalves, Erica Valessa Ramos Gomes, Karoline Rodrigues Campos |
| EPI_ISL_861680 | Hospital e Pronto Socorro Portinari | Instituto Adolfo Lutz, Interdiciplinary Procedures Center, Strategic Laboratory | Claudio Tavares Sacchi, Claudia Regina Gonçalves, Erica Valessa Ramos Gomes, Karoline Rodrigues Campos |
| EPI_ISL_861682 | UPA Vila Santa Catarina | Instituto Adolfo Lutz, Interdiciplinary Procedures Center, Strategic Laboratory | Claudio Tavares Sacchi, Claudia Regina Gonçalves, Erica Valessa Ramos Gomes, Karoline Rodrigues Campos |
| EPI_ISL_861683 | Complexo Hospitalar Padre Bento de Guarulhos | Instituto Adolfo Lutz, Interdiciplinary Procedures Center, Strategic Laboratory | Claudio Tavares Sacchi, Claudia Regina Gonçalves, Erica Valessa Ramos Gomes, Karoline Rodrigues Campos |
| EPI_ISL_861684, EPI_ISL_861685 | Hospital Pronto Socorro Itaquera | Instituto Adolfo Lutz, Interdiciplinary Procedures Center, Strategic Laboratory | Claudio Tavares Sacchi, Claudia Regina Gonçalves, Erica Valessa Ramos Gomes, Karoline Rodrigues Campos |
| EPI_ISL_861710 | Laboratory for Respiratory Viruses, Cantacuzino National Military-Medical Institute for Research and Development | Cantacuzino Institute Virology | Mihaela Lazar, Luiza Ustea, Nicoleta Parashiv, Tim Durfee |
| EPI_ISL_861711, EPI_ISL_861712, EPI_ISL_861713 | Laboratory for Respiratory Viruses, Cantacuzino National Military-Medical Institute for Research and Development | Cantacuzino Institute Virology | Mihaela Lazar, Luiza Ustea, Nicoleta Paraschiv, Tim Durfee |
| EPI_ISL_861714 | Cerba | CERBA LAB | Roquebert B; Costa JM; Hedbaut E; Trombert S; Lecorche E; Verdurme L; Malek Ramdane, Olivi M; Haïm-Boukobza S. |
| EPI_ISL_861715 | CERBA | CERBA LAB | Roquebert B; Costa JM; Hedbaut E; Trombert S; Lecorche E; Verdurme L; Malek Ramdane, Olivi M; Haïm-Boukobza S. |

| | | | |
|---|---|---|---|
| EPI_ISL_861716, EPI_ISL_861727, EPI_ISL_861728 | CERBA | CERBA LAB | Roquebert B; Costa JM; Hedbaut E; Trombert S; Lecorche E; Verdurme L; Malek Ramdane; Olivi M; Haïm-Boukobza S. |
| EPI_ISL_862039 | RSUP Dr. Sardjito | Genetics Working Group (Pokja Genetik) Faculty of Medicine, Public Health and Nursing Universitas Gadjah Mada (FK-KMK UGM); Disease Investigation Center Wates Ministry of Agriculture Indonesia; Department of Microbiology FK-KMK UGM; Laboratorium Diagnostik Yayasan Tahija World Mosquito Program (WMP) Yogyakarta Center for Tropical Medicine FK-KMK UGM; Integrated Research Center FK-KMK UGM; Department of Computer Science and Electronics FMIPA UGM; RSUP Dr. Sardjito | Gunadi, Hendra Wibawa, . Marcellus, Mohamad S. Hakim, Edwin W. Daniwijaya, Ludhang P. Rizki, Endah Supriyati, Eggi Arguni, Titik Nuryastuti, Tri Wibawa, Dwi AA Nugrahaningsih, Afiahayati, Siswanto, Kristy Iskandar, Nungki Anggorowati, Ika Trisnawati, Sumardi, Eko Budiono, Bambang Sigit Riyanto, Heni Retnowulan, Munawar Gani, Satria Maulana, Nur Rahmi Ananda, Riat El Khair, Yunika Puspadewi, Osman Sianipar, Umi Solekhah Intansari, Elizabeth Henny Herningtiyas, Ira Puspitawati, Nur Imma Fatimah Harahap,William Widitjiarso, Maria Patricia Inggriani |
| EPI_ISL_862040 | RSUP Dr. Sardjito | Genetics Working Group (Pokja Genetik) Faculty of Medicine, Public Health and Nursing Universitas Gadjah Mada (FK-KMK UGM); Disease Investigation Center Wates Ministry of Agriculture Indonesia; Department of Microbiology FK-KMK UGM; Laboratorium Diagnostik Yayasan Tahija World Mosquito Program (WMP) Yogyakarta Center for Tropical Medicine FK-KMK UGM; Integrated Research Center FK-KMK UGM; Department of Computer Science and Electronics FMIPA UGM; RSUP Dr. Sardjito | Gunadi, Hendra Wibawa, . Marcellus, Mohamad S. Hakim, Edwin W. Daniwijaya, Ludhang P. Rizki, Endah Supriyati, Eggi Arguni, Titik Nuryastuti, Tri Wibawa, Dwi AA Nugrahaningsih, Afiahayati, Siswanto, Kristy Iskandar, Nungki Anggorowati, Ika Trisnawati, Sumardi, Eko Budiono, Bambang Sigit Riyanto, Heni Retnowulan, Munawar Gani, Satria Maulana, Nur Rahmi Ananda, Riat El Khair, Yunika Puspadewi, Osman Sianipar, Umi Solekhah Intansari, Elizabeth Henny Herningtiyas, Ira Puspitawati, Nur Imma Fatimah Harahap,Alvin Santoso Kalim, Untung Riawan |
| EPI_ISL_862041 | RSUP Dr. Sardjito | Genetics Working Group (Pokja Genetik) Faculty of Medicine, Public Health and Nursing Universitas Gadjah Mada (FK-KMK UGM); Disease Investigation Center Wates Ministry of Agriculture Indonesia; Department of Microbiology FK-KMK UGM; Laboratorium Diagnostik Yayasan Tahija World Mosquito Program (WMP) Yogyakarta Center for Tropical Medicine FK-KMK UGM; Integrated Research Center FK-KMK UGM; Department of Computer Science and Electronics FMIPA UGM; RSUP Dr. Sardjito | Gunadi, Hendra Wibawa, . Marcellus, Mohamad S. Hakim, Edwin W. Daniwijaya, Ludhang P. Rizki, Endah Supriyati, Eggi Arguni, Titik Nuryastuti, Tri Wibawa, Dwi AA Nugrahaningsih, Afiahayati, Siswanto, Kristy Iskandar, Nungki Anggorowati, Ika Trisnawati, Sumardi, Eko Budiono, Bambang Sigit Riyanto, Heni Retnowulan, Munawar Gani, Satria Maulana, Nur Rahmi Ananda, Riat El Khair, Yunika Puspadewi, Osman Sianipar, Umi Solekhah Intansari, Elizabeth Henny Herningtiyas, Ira Puspitawati, Nur Imma Fatimah Harahap, Dyah Ayu Puspitarani, Audric Kenny Tedja |
| EPI_ISL_862042, EPI_ISL_862043, EPI_ISL_862045 | CERBA | CERBA LAB | Roquebert B; Costa JM; Hedbaut E; Trombert S; Lecorche E; Verdurme L; Malek Ramdane, Olivi M; Haïm-Boukobza S. |
| EPI_ISL_862046, EPI_ISL_862047 | Cerballiance Val de Loire | CERBA LAB | Abs G. |
| EPI_ISL_862051 | Multidisciplinary Research Unit, DHR-ICMR, Institute of Medical Sciences, Banaras Hindu University | Multidisciplinary Research Unit, DHR-ICMR, Institute of Medical Sciences, Banaras Hindu University | Priyoneel Basu, Ashish, Nitish Kumar Singh, Abhay Kumar Yadav, Manpreet Kaur, Arup Acharjee, Deepa Devadas, Chetan Sahni, Sanjay Kumar, Tribhuwan Mohan Mahapatra, Richa Arya, Prashant Singh, Jay Prakash Maurya, Surendra Pratap Mishra, Royana Singh |
| EPI_ISL_862075 | National Influenza Center, Virology Department | National Influenza Center | V Salimi,NZ Shafiei Jandaghi, J Yavarian, A Nejati, K Sadeghi, N Ghavvami,F Ajaminejad and T Mokhtari Azad |
| EPI_ISL_862076 | National Influenza Center, Virology Department | National Influenza Center | NZ Shafiei Jandaghi, V Salimi, A Nejati, K Sadeghi, J Yavarian, N Ghavvami,F Ajaminejad and T Mokhtari Azad |
| EPI_ISL_862079 | National Influenza Center, Virology Department | National Influenza Center | K Sadeghi, A Nejati, J Yavarian, NZ Shafiei Jandaghi, V Salimi, F Ajaminejad,N Ghavvami and T Mokhtari Azad |
| EPI_ISL_862080 | National Influenza Center, Virology Department | National Influenza Center | J Yavarian, NZ Shafiei Jandaghi, V Salimi, A Nejati, K Sadeghi, F Ajaminejad,N Ghavvami and T Mokhtari Azad |
| EPI_ISL_862081 | National Influenza Center, Virology Department | National Influenza Center | NZ Shafiei Jandaghi, V Salimi, A Nejati, K Sadeghi, J Yavarian, F Ajaminejad,N Ghavvami and T Mokhtari Azad |
| EPI_ISL_862544 | Compejo Hospitalario Xeral-Calde | Instituto de Salud Carlos III | Iglesias-Caballero, M.Camarero, S. Molinero Calamita, M. González-Esguevillas, M. Pozo, F. Casas, I. Jiménez, P. Jiménez, M. Zaballos, A. Monzón, S. Varona, S. Juliá, M. Cuesta, I. Alonso, P. |
| EPI_ISL_862545 | Hospital Universitario Virgen de la Arrixaca | Instituto de Salud Carlos III | Iglesias-Caballero, M. Camarero, S. Molinero Calamita, M. González-Esguevillas, M. Pozo, F. Casas, I. Jiménez, P. Jiménez, M. Zaballos, A. Monzón, S. Varona, S. Juliá, M. Cuesta, I, Moreno, A. |
| EPI_ISL_862546, EPI_ISL_862547 | Consejería de Sanidad y Asuntos Sociales | Instituto de Salud Carlos III | Iglesias-Caballero, M. Camarero, S. Molinero Calamita, M. González-Esguevillas, M. Pozo, F. Casas, I. Jiménez, P. Jiménez, M. Zaballos, A. Monzón, S. Varona, S. Juliá, M. Cuesta, I. Gutiérrez, G. |
| EPI_ISL_862548 | Hospital Clínico San Carlos | Instituto de Salud Carlos III | Iglesias-Caballero, M. Camarero, S. Molinero Calamita, M. González-Esguevillas, M. Pozo, F. Casas, I. Jiménez, P. Jiménez, M. Zaballos, A. Monzón, S. Varona, S. Juliá, M. Cuesta, I. Rodríguez, I. |
| EPI_ISL_862549 | Hospital Universitario Virgen de la Arrixaca | Instituto de Salud Carlos III | Iglesias-Caballero, M. Camarero, S. Molinero Calamita, M. González-Esguevillas, M. Pozo, F. Casas, I. Jiménez, P. Jiménez, M. Zaballos, A. Monzón, S. Varona, S. Juliá, M. Cuesta, I, Moreno, A. |
| EPI_ISL_862550 | Complejo Hospitalario de Navarra | Instituto de Salud Carlos III | Iglesias-Caballero, M.Camarero, S. Molinero Calamita, M. González-Esguevillas, M. Pozo, F. Casas, I. Jiménez, P. Jiménez, M. Zaballos, A. Monzón, S. Varona, S. Juliá, M. Cuesta, I. Ezpeleta, C. |
| EPI_ISL_862551 | Hospital J.M. Morales Meseguer | Instituto de Salud Carlos III | Iglesias-Caballero, M. Camarero, S. Molinero Calamita, M. González-Esguevillas, M. . Pozo, F. Casas, I. Jiménez, P. Jiménez, M. Zaballos, A. Monzón, S. Varona, S. Juliá, M. Cuesta, I. Guerrero, C. |
| EPI_ISL_862552 | Complejo Hospitalario de Navarra | Instituto de Salud Carlos III | Iglesias-Caballero, M.Camarero, S. Molinero Calamita, M. González-Esguevillas, M. Pozo, F. Casas, I. Jiménez, P. Jiménez, M. Zaballos, A. Monzón, S. Varona, S. Juliá, M. Cuesta, I. Ezpeleta, C. |
| EPI_ISL_862553, EPI_ISL_862554, EPI_ISL_862555 | Compejo Hospitalario Xeral-Calde | Instituto de Salud Carlos III | Iglesias-Caballero, M.Camarero, S. Molinero Calamita, M. González-Esguevillas, M. Pozo, F. Casas, I. Jiménez, P. Jiménez, M. Zaballos, A. Monzón, S. Varona, S. Juliá, M. Cuesta, I. Alonso, P. |
| EPI_ISL_862556 | Gerencia de Asistencia Sanitaria de Soria | Instituto de Salud Carlos III | Iglesias-Caballero, M.Camarero, S. Molinero Calamita, M. González-Esguevillas, M. Pozo, F. Casas, I. Jiménez, P. Jiménez, M. Zaballos, A. Monzón, S. Varona, S. Juliá, M. Cuesta, I. Aldea, C. |
| EPI_ISL_862557, EPI_ISL_862558, EPI_ISL_862559, EPI_ISL_862560, EPI_ISL_862561, EPI_ISL_862562, EPI_ISL_862563 | Consejería de Sanidad y Asuntos Sociales | Instituto de Salud Carlos III | Iglesias-Caballero, M. Camarero, S. Molinero Calamita, M. González-Esguevillas, M. Pozo, F. Casas, I. Jiménez, P. Jiménez, M. Zaballos, A. Monzón, S. Varona, S. Juliá, M. Cuesta, I. Gutiérrez, G. |
| EPI_ISL_862564, EPI_ISL_862565 | Gerencia de Asistencia Sanitaria de Soria | Instituto de Salud Carlos III | Iglesias-Caballero, M.Camarero, S. Molinero Calamita, M. González-Esguevillas, M. Pozo, F. Casas, I. Jiménez, P. Jiménez, M. Zaballos, A. Monzón, S. Varona, S. Juliá, M. Cuesta, I. Aldea, C. |
| EPI_ISL_862566 | Consejería de Sanidad y Asuntos Sociales | Instituto de Salud Carlos III | Iglesias-Caballero, M. Camarero, S. Molinero Calamita, M. González-Esguevillas, M. Pozo, F. Casas, I. Jiménez, P. Jiménez, M. Zaballos, A. Monzón, S. Varona, S. Juliá, M. Cuesta, I. Gutiérrez, G. |
| EPI_ISL_862568 | Hospital Universitario Virgen de la Arrixaca | Instituto de Salud Carlos III | Iglesias-Caballero, M. Camarero, S. Molinero Calamita, M. González-Esguevillas, M. Pozo, F. Casas, I. Jiménez, P. Jiménez, M. Zaballos, A. Monzón, S. Varona, S. Juliá, M. Cuesta, I, Moreno, A. |
| EPI_ISL_862569 | Hospital Clínico San Carlos | Instituto de Salud Carlos III | Iglesias-Caballero, M. Camarero, S. Molinero Calamita, M. González-Esguevillas, M. Pozo, F. Casas, I. Jiménez, P. Jiménez, M. Zaballos, A. Monzón, S. Varona, S. Juliá, M. Cuesta, I. Rodríguez, I. |
| EPI_ISL_862570, EPI_ISL_862571, EPI_ISL_862572 | Complejo Hospitalario de Navarra | Instituto de Salud Carlos III | Iglesias-Caballero, M.Camarero, S. Molinero Calamita, M. González-Esguevillas, M. Pozo, F. Casas, I. Jiménez, P. Jiménez, M. Zaballos, A. Monzón, S. Varona, S. Juliá, M. Cuesta, I. Ezpeleta, C. |

| | | | |
|---|---|---|---|
| EPI_ISL_862573, EPI_ISL_862574, EPI_ISL_862575, EPI_ISL_862576, EPI_ISL_862577, EPI_ISL_862578, EPI_ISL_862579, EPI_ISL_862580, EPI_ISL_862581, EPI_ISL_862582, EPI_ISL_862583, EPI_ISL_862584, EPI_ISL_862585, EPI_ISL_862586, EPI_ISL_862587, EPI_ISL_862588 | | | |
| see above | Hospital Clínic | Instituto de Salud Carlos III | Iglesias-Caballero, M.Camarero, S. Molinero Calamita, M. González-Esguevillas, M. Pozo, F. Casas, I. Jiménez, P. Jiménez, M. Zaballos, A. Monzón, S. Varona, S. Juliá, M. Cuesta, I. Marcos, M.A. |
| EPI_ISL_862589, EPI_ISL_862590, EPI_ISL_862591, EPI_ISL_862592, EPI_ISL_862593, EPI_ISL_862595, EPI_ISL_862596, EPI_ISL_862597, EPI_ISL_862598, EPI_ISL_862599, EPI_ISL_862600, EPI_ISL_862601, EPI_ISL_862602, EPI_ISL_862603, EPI_ISL_862604, EPI_ISL_862605, EPI_ISL_862606, EPI_ISL_862607, EPI_ISL_862608, EPI_ISL_862609, EPI_ISL_862610, EPI_ISL_862611, EPI_ISL_862612, EPI_ISL_862613, EPI_ISL_862614, EPI_ISL_862615, EPI_ISL_862616, EPI_ISL_862617, EPI_ISL_862618, EPI_ISL_862619, EPI_ISL_862620, EPI_ISL_862621, EPI_ISL_862622, EPI_ISL_862623, EPI_ISL_862624, EPI_ISL_862625, EPI_ISL_862626 | | | |
| see above | Hospital Comarcal de Melilla | Instituto de Salud Carlos III | Iglesias-Caballero, M. Camarero, S. Molinero Calamita, M. González-Esguevillas, M. Pozo, F. Casas, I. Jiménez, P. Jiménez, M. Zaballos, A. Monzón, S. Varona, S. Juliá, M. Cuesta, I. López, J. |
| EPI_ISL_862815 | Multidisciplinary Research Unit, DHR-ICMR, Institute of Medical Sciences, Banaras Hindu University, Varanasi | Multidisciplinary Research Unit, DHR-ICMR, Institute of Medical Sciences, Banaras Hindu University, Varanasi | Royana Singh, Priyoneel Basu, Ashish, Nitish Kumar Singh, Abhay Kumar Yadav, Manpreet Kaur, Deepa Devadas, Chetan Sahni, Gulshan Kumar, Tribhuwan Mohan Mohapatra, Richa Arya, Prashant Singh, Jay Prakash Maurya, Surendra Pratap Mishra |
| EPI_ISL_862831, EPI_ISL_862832, EPI_ISL_862833, EPI_ISL_862834 | National Institute of Infectious Diseases-Prof. Dr. Matei Bals Molecular Diagnostics Laboratory | National Institute of Infectious Diseases-Prof. Dr. Matei Bals Molecular Diagnostics Laboratory | Leontina Banica, Marius Surleac, Corina Casangiu, Petre Milu, Andreea Tudor, Simona Paraschiv, Dan Otelea |
| EPI_ISL_869241 | Suceava County Emergency Hospital | "Stefan cel Mare" University Metagenomics Laboratory | Lobiuc Andrei, Puscaselu Roxana |
| EPI_ISL_869261 | UMMC-Health | WHO National Influenza Centre Russian Federation | Andrey Komissarov, Artem Fadeev, Anna Ivanova, Kseniya Komissarova, Dmitry Bazhenov, Mikhail Bakaev,Tatiana Platonova, Daria Danilenko, Ksenia Safina, Elena Nabieva, Georgii Bazykin, Dmitry Lioznov |
| EPI_ISL_869262, EPI_ISL_869263, EPI_ISL_869264 | Krasnoyarsk Regional Clinical Hospital | WHO National Influenza Centre Russian Federation | Andrey Komissarov, Artem Fadeev, Anna Ivanova, Kseniya Komissarova, Dmitry Bazhenov, Mikhail Bakaev, Daria Danilenko, Ksenia Safina, Elena Nabieva, Georgii Bazykin, Dmitry Lioznov |
| EPI_ISL_871789 | Cerballiance Normandie | CERBA LAB | LAFOREST D, BRIAND G, LEPIGEON, technicien V; COUDRAY J; MARTINS-AMARAL F; COULON C, THOMAS Johan, VITEL R, HAMEL A. |
| EPI_ISL_871876 | Cerballiance Côte d'Azur | CERBA LAB | Prots L; Barrieu-Moussat S; Roquebert B; Costa JM; Hedbaut E; Trombert S; Lecorche E; Verdurme L; Malek Ramdane, Olivi M; Haïm-Boukobza S. |
| EPI_ISL_871972 | Instituto de Salud Carlos III | Instituto de Salud Carlos III | Iglesias-Caballero, M. Camarero, S. Molinero Calamita, M. González-Esguevillas, M. Pozo, F. Casas, I. Jiménez, P. Jiménez, M. Zaballos, A. Monzón, S. Varona, S. Juliá, M. Cuesta, I. |
| EPI_ISL_871994 | Hospital de Madrid | Instituto de Salud Carlos III | Iglesias-Caballero, M. Camarero, S. Molinero Calamita, M. González-Esguevillas, M. Pozo, F. Casas, I. Jiménez, P. Jiménez, M. Zaballos, A. Monzón, S. Varona, S. Juliá, M. Cuesta, I. Alhambra, A. |
| EPI_ISL_871995 | Hospital Universitario de Ceuta | Instituto de Salud Carlos III | Iglesias-Caballero, M. Camarero, S. Molinero Calamita, M. González-Esguevillas, M. Pozo, F. Casas, I. Jiménez, P. Jiménez, M. Zaballos, A. Monzón, S. Varona, S. Juliá, M. Cuesta, I. Hijano, S. |
| EPI_ISL_871996, EPI_ISL_871997, EPI_ISL_871998 | Instituto de Salud Carlos III | Instituto de Salud Carlos III | Iglesias-Caballero, M. Camarero, S. Molinero Calamita, M. González-Esguevillas, M. Pozo, F. Casas, I. Jiménez, P. Jiménez, M. Zaballos, A. Monzón, S. Varona, S. Juliá, M. Cuesta, I. |
| EPI_ISL_872003, EPI_ISL_872004, EPI_ISL_872005, EPI_ISL_872006, EPI_ISL_872007, EPI_ISL_872008 | Ospedale "Di Venere" | Beaconlab (Bioinformatics, Evolution and Comparative Genomics lab), Dept of Biosciences, University on Milan | Iacobellis M, d'Avenia M, Piluscio R, Parisi A, Chiara M, Manzari C, Pesole G |
| EPI_ISL_872188 | RSUP Dr. Sardjito | Genetics Working Group (Pokja Genetik) Faculty of Medicine, Public Health and Nursing Universitas Gadjah Mada (FK-KMK UGM); Disease Investigation Center Wates Ministry of Agriculture Indonesia; Department of Microbiology FK-KMK UGM; Laboratorium Diagnostik Yayasan Tahija World Mosquito Program (WMP) Yogyakarta Center for Tropical Medicine FK-KMK UGM; Integrated Research Center FK-KMK UGM; Department of Computer Science and Electronics FMIPA UGM; RSUP Dr. Sardjito | Gunadi, Hendra Wibawa, . Marcellus, Mohamad S. Hakim, Edwin W. Daniwijaya, Ludhang P. Rizki, Endah Supriyati, Eggi Arguni, Titik Nuryastuti, Tri Wibawa, Dwi AA Nugrahaningsih, Afiahayati, Siswanto, Kristy Iskandar, Nungki Anggorowati, Ika Trisnawati, Sumardi, Eko Budiono, Bambang Sigit Riyanto, Heni Retnowulan, Munawar Gani, Satria Maulana, Nur Rahmi Ananda, Riat El Khair, Yunika Puspadewi, Osman Sianipar, Umi Solekhah Intansari, Elizabeth Henny Herningtiyas, Ira Puspitawati, Nur Imma Fatimah Harahap, Aditya Rifqi Fauzi, Fadil Fahri |
| EPI_ISL_872189 | RSUP Dr. Sardjito | Genetics Working Group (Pokja Genetik) Faculty of Medicine, Public Health and Nursing Universitas Gadjah Mada (FK-KMK UGM); Disease Investigation Center Wates Ministry of Agriculture Indonesia; Department of Microbiology FK-KMK UGM; Laboratorium Diagnostik Yayasan Tahija World Mosquito Program (WMP) Yogyakarta Center for Tropical Medicine FK-KMK UGM; Integrated Research Center FK-KMK UGM; Department of Computer Science and Electronics FMIPA UGM; RSUP Dr. Sardjito | Gunadi, Hendra Wibawa, . Marcellus, Mohamad S. Hakim, Edwin W. Daniwijaya, Ludhang P. Rizki, Endah Supriyati, Eggi Arguni, Titik Nuryastuti, Tri Wibawa, Dwi AA Nugrahaningsih, Afiahayati, Siswanto, Kristy Iskandar, Nungki Anggorowati, Ika Trisnawati, Sumardi, Eko Budiono, Bambang Sigit Riyanto, Heni Retnowulan, Munawar Gani, Satria Maulana, Nur Rahmi Ananda, Riat El Khair, Yunika Puspadewi, Osman Sianipar, Umi Solekhah Intansari, Elizabeth Henny Herningtiyas, Ira Puspitawati, Nur Imma Fatimah Harahap, Alvin S. Kalim, Dwiki afandy |
| EPI_ISL_872190 | RSUP Dr. Sardjito | Genetics Working Group (Pokja Genetik) Faculty of Medicine, Public Health and Nursing Universitas Gadjah Mada (FK-KMK UGM); Disease Investigation Center Wates Ministry of Agriculture Indonesia; Department of Microbiology FK-KMK UGM; Laboratorium Diagnostik Yayasan Tahija World Mosquito Program (WMP) Yogyakarta Center for Tropical Medicine FK-KMK UGM; Integrated Research Center FK-KMK UGM; Department of Computer Science and Electronics FMIPA UGM; RSUP Dr. Sardjito | Gunadi, Hendra Wibawa, . Marcellus, Mohamad S. Hakim, Edwin W. Daniwijaya, Ludhang P. Rizki, Endah Supriyati, Eggi Arguni, Titik Nuryastuti, Tri Wibawa, Dwi AA Nugrahaningsih, Afiahayati, Siswanto, Kristy Iskandar, Nungki Anggorowati, Ika Trisnawati, Sumardi, Eko Budiono, Bambang Sigit Riyanto, Heni Retnowulan, Munawar Gani, Satria Maulana, Nur Rahmi Ananda, Riat El Khair, Yunika Puspadewi, Osman Sianipar, Umi Solekhah Intansari, Elizabeth Henny Herningtiyas, Ira Puspitawati, Nur Imma Fatimah Harahap, Dyah Ayu Puspitarani, Kemala Athollah |
| EPI_ISL_872191 | Conjunto Hospitalar do Mandaqui de Sao Paulo | Instituto Adolfo Lutz, Interdiciplinary Procedures Center, Strategic Laboratory | Claudio Tavares Sacchi, Claudia Regina Gonçalves, Erica Valessa Ramos Gomes, Karoline Rodrigues Campos, Katia Correa de Oliveira Santos, Ana Lucia de Carvalho Avelino, Fabiana Cristina Pereira dos Santos |
| EPI_ISL_872197 | hopital | National Reference Center for Viruses of Respiratory Infections, Institut Pasteur, Paris | Marion Barbet, Sylvie Behillil, Méline Bizard, Angela Brisebarre, Camille Capel, Etienne Simon-Lorière, Vincent Enouf, Maud Vanpeene, Sylvie van der Werf,Combe Patrice |
| EPI_ISL_872198 | Hopital | National Reference Center for Viruses of Respiratory Infections, Institut Pasteur, Paris | Marion Barbet, Sylvie Behillil, Méline Bizard, Angela Brisebarre, Camille Capel, Etienne Simon-Lorière, Vincent Enouf, Maud Vanpeene, Sylvie van der Werf |
| EPI_ISL_872199 | Hopital | National Reference Center for Viruses of Respiratory Infections, Institut Pasteur, Paris | Marion Barbet, Sylvie Behillil, Méline Bizard, Angela Brisebarre, Camille Capel, Etienne Simon-Lorière, Vincent Enouf, Maud Vanpeene, Sylvie van der Werf,Ramanantsoa CéLine |
| EPI_ISL_872200 | hopital | National Reference Center for Viruses of Respiratory Infections, Institut Pasteur, Paris | Marion Barbet, Sylvie Behillil, Méline Bizard, Angela Brisebarre, Camille Capel, Etienne Simon-Lorière, Vincent Enouf, Maud Vanpeene, Sylvie van der Werf,Combe Patrice |
| EPI_ISL_872201 | Labo Analyses Med | National Reference Center for Viruses of Respiratory Infections, Institut Pasteur, Paris | Marion Barbet, Sylvie Behillil, Méline Bizard, Angela Brisebarre, Camille Capel, Etienne Simon-Lorière, Vincent Enouf, Maud Vanpeene, Sylvie van der Werf,Girard Sophie |
| EPI_ISL_872202 | hopital | National Reference Center for Viruses of Respiratory Infections, Institut Pasteur, Paris | Marion Barbet, Sylvie Behillil, Méline Bizard, Angela Brisebarre, Camille Capel, Etienne Simon-Lorière, Vincent Enouf, Maud Vanpeene, Sylvie van der Werf,Combe Patrice |
| EPI_ISL_872203 | Labo Analyses Med | National Reference Center for Viruses of Respiratory Infections, Institut Pasteur, Paris | Marion Barbet, Sylvie Behillil, Méline Bizard, Angela Brisebarre, Camille Capel, Etienne Simon-Lorière, Vincent Enouf, Maud Vanpeene, Sylvie van der Werf |
| EPI_ISL_872204, EPI_ISL_872205 | Labo Analyses Med | National Reference Center for Viruses of Respiratory | Marion Barbet, Sylvie Behillil, Méline Bizard, Angela Brisebarre, Camille Capel, Etienne Simon-Lorière, Vincent Enouf, Maud Vanpeene, Sylvie van der |

| | | | |
|---|---|---|---|
| | | Infections, Institut Pasteur, Paris | Werf,Girard Sophie |
| EPI_ISL_872206 | Hopital | National Reference Center for Viruses of Respiratory Infections, Institut Pasteur, Paris | Marion Barbet, Sylvie Behillil, Méline Bizard, Angela Brisebarre, Camille Capel, Etienne Simon-Lorière, Vincent Enouf, Maud Vanpeene, Sylvie van der Werf,Morvan Odile |
| EPI_ISL_872207, EPI_ISL_872208, EPI_ISL_872209 | Labo Analyses Med | National Reference Center for Viruses of Respiratory Infections, Institut Pasteur, Paris | Marion Barbet, Sylvie Behillil, Méline Bizard, Angela Brisebarre, Camille Capel, Etienne Simon-Lorière, Vincent Enouf, Maud Vanpeene, Sylvie van der Werf |
| EPI_ISL_872210, EPI_ISL_872211 | Hopital | National Reference Center for Viruses of Respiratory Infections, Institut Pasteur, Paris | Marion Barbet, Sylvie Behillil, Méline Bizard, Angela Brisebarre, Camille Capel, Etienne Simon-Lorière, Vincent Enouf, Maud Vanpeene, Sylvie van der Werf,Guigon AuréLie |
| EPI_ISL_872212 | Labo analyses med | National Reference Center for Viruses of Respiratory Infections, Institut Pasteur, Paris | Marion Barbet, Sylvie Behillil, Méline Bizard, Angela Brisebarre, Camille Capel, Etienne Simon-Lorière, Vincent Enouf, Maud Vanpeene, Sylvie van der Werf,Amzalag Jonas |
| EPI_ISL_872213, EPI_ISL_872214, EPI_ISL_872215, EPI_ISL_872216, EPI_ISL_872217, EPI_ISL_872218, EPI_ISL_872219 | Hopital | National Reference Center for Viruses of Respiratory Infections, Institut Pasteur, Paris | Marion Barbet, Sylvie Behillil, Méline Bizard, Angela Brisebarre, Camille Capel, Etienne Simon-Lorière, Vincent Enouf, Maud Vanpeene, Sylvie van der Werf,Fourgeaud Jacques |
| EPI_ISL_872220 | Hopital | National Reference Center for Viruses of Respiratory Infections, Institut Pasteur, Paris | Marion Barbet, Sylvie Behillil, Méline Bizard, Angela Brisebarre, Camille Capel, Etienne Simon-Lorière, Vincent Enouf, Maud Vanpeene, Sylvie van der Werf,Brichler Ségolène |
| EPI_ISL_872222, EPI_ISL_872224, EPI_ISL_872225, EPI_ISL_872226 | hopital | National Reference Center for Viruses of Respiratory Infections, Institut Pasteur, Paris | Marion Barbet, Sylvie Behillil, Méline Bizard, Angela Brisebarre, Camille Capel, Etienne Simon-Lorière, Vincent Enouf, Maud Vanpeene, Sylvie van der Werf,Combe Patrice |
| EPI_ISL_872227 | Hopital | National Reference Center for Viruses of Respiratory Infections, Institut Pasteur, Paris | Marion Barbet, Sylvie Behillil, Méline Bizard, Angela Brisebarre, Camille Capel, Etienne Simon-Lorière, Vincent Enouf, Maud Vanpeene, Sylvie van der Werf |
| EPI_ISL_872228 | Hopital | National Reference Center for Viruses of Respiratory Infections, Institut Pasteur, Paris | Marion Barbet, Sylvie Behillil, Méline Bizard, Angela Brisebarre, Camille Capel, Etienne Simon-Lorière, Vincent Enouf, Maud Vanpeene, Sylvie van der Werf,Irimia Alix |
| EPI_ISL_872229, EPI_ISL_872230 | Hopital | National Reference Center for Viruses of Respiratory Infections, Institut Pasteur, Paris | Marion Barbet, Sylvie Behillil, Méline Bizard, Angela Brisebarre, Camille Capel, Etienne Simon-Lorière, Vincent Enouf, Maud Vanpeene, Sylvie van der Werf,Ramanantsoa CéLine |
| EPI_ISL_872231 | Hopital | National Reference Center for Viruses of Respiratory Infections, Institut Pasteur, Paris | Marion Barbet, Sylvie Behillil, Méline Bizard, Angela Brisebarre, Camille Capel, Etienne Simon-Lorière, Vincent Enouf, Maud Vanpeene, Sylvie van der Werf,Florin CéCile |
| EPI_ISL_872232 | hopital | National Reference Center for Viruses of Respiratory Infections, Institut Pasteur, Paris | Marion Barbet, Sylvie Behillil, Méline Bizard, Angela Brisebarre, Camille Capel, Etienne Simon-Lorière, Vincent Enouf, Maud Vanpeene, Sylvie van der Werf,Combe Patrice |
| EPI_ISL_872233 | Hopital | National Reference Center for Viruses of Respiratory Infections, Institut Pasteur, Paris | Marion Barbet, Sylvie Behillil, Méline Bizard, Angela Brisebarre, Camille Capel, Etienne Simon-Lorière, Vincent Enouf, Maud Vanpeene, Sylvie van der Werf,Fourgeaud Jacques |
| EPI_ISL_872234 | Labo Analyses Med | National Reference Center for Viruses of Respiratory Infections, Institut Pasteur, Paris | Marion Barbet, Sylvie Behillil, Méline Bizard, Angela Brisebarre, Camille Capel, Etienne Simon-Lorière, Vincent Enouf, Maud Vanpeene, Sylvie van der Werf,Girard Sophie |
| EPI_ISL_872235 | Labo Analyses Med | National Reference Center for Viruses of Respiratory Infections, Institut Pasteur, Paris | Marion Barbet, Sylvie Behillil, Méline Bizard, Angela Brisebarre, Camille Capel, Etienne Simon-Lorière, Vincent Enouf, Maud Vanpeene, Sylvie van der Werf |
| EPI_ISL_872236 | Hopital | National Reference Center for Viruses of Respiratory Infections, Institut Pasteur, Paris | Marion Barbet, Sylvie Behillil, Méline Bizard, Angela Brisebarre, Camille Capel, Etienne Simon-Lorière, Vincent Enouf, Maud Vanpeene, Sylvie van der Werf,Guigon AuréLie |
| EPI_ISL_872237 | Hopital | National Reference Center for Viruses of Respiratory Infections, Institut Pasteur, Paris | Marion Barbet, Sylvie Behillil, Méline Bizard, Angela Brisebarre, Camille Capel, Etienne Simon-Lorière, Vincent Enouf, Maud Vanpeene, Sylvie van der Werf,Florin CéCile |
| EPI_ISL_872238 | Labo analyses med | National Reference Center for Viruses of Respiratory Infections, Institut Pasteur, Paris | Marion Barbet, Sylvie Behillil, Méline Bizard, Angela Brisebarre, Camille Capel, Etienne Simon-Lorière, Vincent Enouf, Maud Vanpeene, Sylvie van der Werf,Amzalag Jonas |
| EPI_ISL_872239 | Labo Analyses Med | National Reference Center for Viruses of Respiratory Infections, Institut Pasteur, Paris | Marion Barbet, Sylvie Behillil, Méline Bizard, Angela Brisebarre, Camille Capel, Etienne Simon-Lorière, Vincent Enouf, Maud Vanpeene, Sylvie van der Werf,Rouah Raquel |
| EPI_ISL_872240, EPI_ISL_872241, EPI_ISL_872242, EPI_ISL_872243, EPI_ISL_872244, EPI_ISL_872245, EPI_ISL_872246, EPI_ISL_872247, EPI_ISL_872248 | hopital | National Reference Center for Viruses of Respiratory Infections, Institut Pasteur, Paris | Marion Barbet, Sylvie Behillil, Méline Bizard, Angela Brisebarre, Camille Capel, Etienne Simon-Lorière, Vincent Enouf, Maud Vanpeene, Sylvie van der Werf,Combe Patrice |
| EPI_ISL_872249, EPI_ISL_872250 | Hopital | National Reference Center for Viruses of Respiratory Infections, Institut Pasteur, Paris | Marion Barbet, Sylvie Behillil, Méline Bizard, Angela Brisebarre, Camille Capel, Etienne Simon-Lorière, Vincent Enouf, Maud Vanpeene, Sylvie van der Werf,Ducancelle Alexandra |
| EPI_ISL_872251 | Hopital | National Reference Center for Viruses of Respiratory Infections, Institut Pasteur, Paris | Marion Barbet, Sylvie Behillil, Méline Bizard, Angela Brisebarre, Camille Capel, Etienne Simon-Lorière, Vincent Enouf, Maud Vanpeene, Sylvie van der Werf,Irimia Alix |
| EPI_ISL_872252 | Hopital | National Reference Center for Viruses of Respiratory Infections, Institut Pasteur, Paris | Marion Barbet, Sylvie Behillil, Méline Bizard, Angela Brisebarre, Camille Capel, Etienne Simon-Lorière, Vincent Enouf, Maud Vanpeene, Sylvie van der Werf,Ramanantsoa CéLine |
| EPI_ISL_872253 | Hopital | National Reference Center for Viruses of Respiratory Infections, Institut Pasteur, Paris | Marion Barbet, Sylvie Behillil, Méline Bizard, Angela Brisebarre, Camille Capel, Etienne Simon-Lorière, Vincent Enouf, Maud Vanpeene, Sylvie van der Werf,Irimia Alix |
| EPI_ISL_872254 | hopital | National Reference Center for Viruses of Respiratory Infections, Institut Pasteur, Paris | Marion Barbet, Sylvie Behillil, Méline Bizard, Angela Brisebarre, Camille Capel, Etienne Simon-Lorière, Vincent Enouf, Maud Vanpeene, Sylvie van der Werf,Ducancelle Alexandra |
| EPI_ISL_872255 | Hopital | National Reference Center for Viruses of Respiratory Infections, Institut Pasteur, Paris | Marion Barbet, Sylvie Behillil, Méline Bizard, Angela Brisebarre, Camille Capel, Etienne Simon-Lorière, Vincent Enouf, Maud Vanpeene, Sylvie van der Werf,Philippi Jean-FrançOis |
| EPI_ISL_872256 | Labo Analyses Med | National Reference Center for Viruses of Respiratory Infections, Institut Pasteur, Paris | Marion Barbet, Sylvie Behillil, Méline Bizard, Angela Brisebarre, Camille Capel, Etienne Simon-Lorière, Vincent Enouf, Maud Vanpeene, Sylvie van der Werf |
| EPI_ISL_872257 | Labo Analyses Med | National Reference Center for Viruses of Respiratory Infections, Institut Pasteur, Paris | Marion Barbet, Sylvie Behillil, Méline Bizard, Angela Brisebarre, Camille Capel, Etienne Simon-Lorière, Vincent Enouf, Maud Vanpeene, Sylvie van der Werf,Bour Jean Baptiste |
| EPI_ISL_872258 | Labo Analyses Med | National Reference Center for Viruses of Respiratory Infections, Institut Pasteur, Paris | Marion Barbet, Sylvie Behillil, Méline Bizard, Angela Brisebarre, Camille Capel, Etienne Simon-Lorière, Vincent Enouf, Maud Vanpeene, Sylvie van der Werf,Girard Sophie |
| EPI_ISL_872259, EPI_ISL_872260 | Labo Analyses Med | National Reference Center for Viruses of Respiratory Infections, Institut Pasteur, Paris | Marion Barbet, Sylvie Behillil, Méline Bizard, Angela Brisebarre, Camille Capel, Etienne Simon-Lorière, Vincent Enouf, Maud Vanpeene, Sylvie van der Werf |
| EPI_ISL_872261 | Hopital | National Reference Center for Viruses of Respiratory Infections, Institut Pasteur, Paris | Marion Barbet, Sylvie Behillil, Méline Bizard, Angela Brisebarre, Camille Capel, Etienne Simon-Lorière, Vincent Enouf, Maud Vanpeene, Sylvie van der Werf,Guigon AuréLie |
| EPI_ISL_872262, EPI_ISL_872263 | Hopital | National Reference Center for Viruses of Respiratory Infections, Institut Pasteur, Paris | Marion Barbet, Sylvie Behillil, Méline Bizard, Angela Brisebarre, Camille Capel, Etienne Simon-Lorière, Vincent Enouf, Maud Vanpeene, Sylvie van der Werf,Florin CéCile |
| EPI_ISL_872264, EPI_ISL_872265 | Labo analyses med | National Reference Center for Viruses of Respiratory | Marion Barbet, Sylvie Behillil, Méline Bizard, Angela Brisebarre, Camille Capel, Etienne Simon-Lorière, Vincent Enouf, Maud Vanpeene, Sylvie van der |

| | | | |
|---|---|---|---|
| | | Infections, Institut Pasteur, Paris | Werf,Amzalag Jonas |
| EPI_ISL_872266, EPI_ISL_872267, EPI_ISL_872268 | Hopital | National Reference Center for Viruses of Respiratory Infections, Institut Pasteur, Paris | Marion Barbet, Sylvie Behillil, Méline Bizard, Angela Brisebarre, Camille Capel, Etienne Simon-Lorière, Vincent Enouf, Maud Vanpeene, Sylvie van der Werf,Fourgeaud Jacques |
| EPI_ISL_872269 | Hopital | National Reference Center for Viruses of Respiratory Infections, Institut Pasteur, Paris | Marion Barbet, Sylvie Behillil, Méline Bizard, Angela Brisebarre, Camille Capel, Etienne Simon-Lorière, Vincent Enouf, Maud Vanpeene, Sylvie van der Werf,Foissaud Vincent |
| EPI_ISL_872270 | Hopital | National Reference Center for Viruses of Respiratory Infections, Institut Pasteur, Paris | Marion Barbet, Sylvie Behillil, Méline Bizard, Angela Brisebarre, Camille Capel, Etienne Simon-Lorière, Vincent Enouf, Maud Vanpeene, Sylvie van der Werf,Fourgeaud Jacques |
| EPI_ISL_872271, EPI_ISL_872272 | hopital | National Reference Center for Viruses of Respiratory Infections, Institut Pasteur, Paris | Marion Barbet, Sylvie Behillil, Méline Bizard, Angela Brisebarre, Camille Capel, Etienne Simon-Lorière, Vincent Enouf, Maud Vanpeene, Sylvie van der Werf,Combe Patrice |
| EPI_ISL_872274 | Hopital | National Reference Center for Viruses of Respiratory Infections, Institut Pasteur, Paris | Marion Barbet, Sylvie Behillil, Méline Bizard, Angela Brisebarre, Camille Capel, Etienne Simon-Lorière, Vincent Enouf, Maud Vanpeene, Sylvie van der Werf,Irimia Alix |
| EPI_ISL_872275, EPI_ISL_872276, EPI_ISL_872278 | Hopital | National Reference Center for Viruses of Respiratory Infections, Institut Pasteur, Paris | Marion Barbet, Sylvie Behillil, Méline Bizard, Angela Brisebarre, Camille Capel, Etienne Simon-Lorière, Vincent Enouf, Maud Vanpeene, Sylvie van der Werf,Ramanantsoa CéLine |
| EPI_ISL_872279, EPI_ISL_872280, EPI_ISL_872281 | Labo Analyses Med | National Reference Center for Viruses of Respiratory Infections, Institut Pasteur, Paris | Marion Barbet, Sylvie Behillil, Méline Bizard, Angela Brisebarre, Camille Capel, Etienne Simon-Lorière, Vincent Enouf, Maud Vanpeene, Sylvie van der Werf,Bour Jean Baptiste |
| EPI_ISL_872282, EPI_ISL_872283 | Labo Analyses Med | National Reference Center for Viruses of Respiratory Infections, Institut Pasteur, Paris | Marion Barbet, Sylvie Behillil, Méline Bizard, Angela Brisebarre, Camille Capel, Etienne Simon-Lorière, Vincent Enouf, Maud Vanpeene, Sylvie van der Werf,Girard Sophie |
| EPI_ISL_872284, EPI_ISL_872285, EPI_ISL_872286, EPI_ISL_872287, EPI_ISL_872288, EPI_ISL_872289, EPI_ISL_872290 | Labo Analyses Med | National Reference Center for Viruses of Respiratory Infections, Institut Pasteur, Paris | Marion Barbet, Sylvie Behillil, Méline Bizard, Angela Brisebarre, Camille Capel, Etienne Simon-Lorière, Vincent Enouf, Maud Vanpeene, Sylvie van der Werf |
| EPI_ISL_872291, EPI_ISL_872292, EPI_ISL_872293, EPI_ISL_872294, EPI_ISL_872295, EPI_ISL_872296, EPI_ISL_872297 | Hopital | National Reference Center for Viruses of Respiratory Infections, Institut Pasteur, Paris | Marion Barbet, Sylvie Behillil, Méline Bizard, Angela Brisebarre, Camille Capel, Etienne Simon-Lorière, Vincent Enouf, Maud Vanpeene, Sylvie van der Werf,Guigon AuréLie |
| EPI_ISL_872298, EPI_ISL_872299, EPI_ISL_872300, EPI_ISL_872301, EPI_ISL_872302, EPI_ISL_872303, EPI_ISL_872304, EPI_ISL_872305 | Hopital | National Reference Center for Viruses of Respiratory Infections, Institut Pasteur, Paris | Marion Barbet, Sylvie Behillil, Méline Bizard, Angela Brisebarre, Camille Capel, Etienne Simon-Lorière, Vincent Enouf, Maud Vanpeene, Sylvie van der Werf,Florin CéCile |
| EPI_ISL_872307 | Hopital | National Reference Center for Viruses of Respiratory Infections, Institut Pasteur, Paris | Marion Barbet, Sylvie Behillil, Méline Bizard, Angela Brisebarre, Camille Capel, Etienne Simon-Lorière, Vincent Enouf, Maud Vanpeene, Sylvie van der Werf,Castelain Sandrine |
| EPI_ISL_872308 | Hopital | National Reference Center for Viruses of Respiratory Infections, Institut Pasteur, Paris | Marion Barbet, Sylvie Behillil, Méline Bizard, Angela Brisebarre, Camille Capel, Etienne Simon-Lorière, Vincent Enouf, Maud Vanpeene, Sylvie van der Werf,Brichler Ségolène |
| EPI_ISL_872309, EPI_ISL_872310, EPI_ISL_872311, EPI_ISL_872312, EPI_ISL_872313, EPI_ISL_872314 | Labo analyses med | National Reference Center for Viruses of Respiratory Infections, Institut Pasteur, Paris | Marion Barbet, Sylvie Behillil, Méline Bizard, Angela Brisebarre, Camille Capel, Etienne Simon-Lorière, Vincent Enouf, Maud Vanpeene, Sylvie van der Werf,Amzalag Jonas |
| EPI_ISL_872315, EPI_ISL_872316, EPI_ISL_872318, EPI_ISL_872319 | Hopital | National Reference Center for Viruses of Respiratory Infections, Institut Pasteur, Paris | Marion Barbet, Sylvie Behillil, Méline Bizard, Angela Brisebarre, Camille Capel, Etienne Simon-Lorière, Vincent Enouf, Maud Vanpeene, Sylvie van der Werf,Fourgeaud Jacques |
| EPI_ISL_872320, EPI_ISL_872321, EPI_ISL_872322 | Labo Analyses Med | National Reference Center for Viruses of Respiratory Infections, Institut Pasteur, Paris | Marion Barbet, Sylvie Behillil, Méline Bizard, Angela Brisebarre, Camille Capel, Etienne Simon-Lorière, Vincent Enouf, Maud Vanpeene, Sylvie van der Werf |
| EPI_ISL_872323 | Labo Analyses Med | National Reference Center for Viruses of Respiratory Infections, Institut Pasteur, Paris | Marion Barbet, Sylvie Behillil, Méline Bizard, Angela Brisebarre, Camille Capel, Etienne Simon-Lorière, Vincent Enouf, Maud Vanpeene, Sylvie van der Werf,Rouah Raquel |
| EPI_ISL_872324 | Labo Analyses Med | National Reference Center for Viruses of Respiratory Infections, Institut Pasteur, Paris | Marion Barbet, Sylvie Behillil, Méline Bizard, Angela Brisebarre, Camille Capel, Etienne Simon-Lorière, Vincent Enouf, Maud Vanpeene, Sylvie van der Werf |
| EPI_ISL_872325 | Hopital | National Reference Center for Viruses of Respiratory Infections, Institut Pasteur, Paris | Marion Barbet, Sylvie Behillil, Méline Bizard, Angela Brisebarre, Camille Capel, Etienne Simon-Lorière, Vincent Enouf, Maud Vanpeene, Sylvie van der Werf,Brichler Ségolène |
| EPI_ISL_872326, EPI_ISL_872327, EPI_ISL_872328, EPI_ISL_872329, EPI_ISL_872330, EPI_ISL_872331, EPI_ISL_872332, EPI_ISL_872333, EPI_ISL_872335, EPI_ISL_872336, EPI_ISL_872337, EPI_ISL_872338 | | | |
| see above | hopital | National Reference Center for Viruses of Respiratory Infections, Institut Pasteur, Paris | Marion Barbet, Sylvie Behillil, Méline Bizard, Angela Brisebarre, Camille Capel, Etienne Simon-Lorière, Vincent Enouf, Maud Vanpeene, Sylvie van der Werf,Combe Patrice |
| EPI_ISL_872339, EPI_ISL_872340, EPI_ISL_872342 | Hopital | National Reference Center for Viruses of Respiratory Infections, Institut Pasteur, Paris | Marion Barbet, Sylvie Behillil, Méline Bizard, Angela Brisebarre, Camille Capel, Etienne Simon-Lorière, Vincent Enouf, Maud Vanpeene, Sylvie van der Werf,Ducancelle Alexandra |
| EPI_ISL_872343 | Hopital | National Reference Center for Viruses of Respiratory Infections, Institut Pasteur, Paris | Marion Barbet, Sylvie Behillil, Méline Bizard, Angela Brisebarre, Camille Capel, Etienne Simon-Lorière, Vincent Enouf, Maud Vanpeene, Sylvie van der Werf,Irimia Alix |
| EPI_ISL_872344, EPI_ISL_872345, EPI_ISL_872346 | Hopital | National Reference Center for Viruses of Respiratory Infections, Institut Pasteur, Paris | Marion Barbet, Sylvie Behillil, Méline Bizard, Angela Brisebarre, Camille Capel, Etienne Simon-Lorière, Vincent Enouf, Maud Vanpeene, Sylvie van der Werf,Ramanantsoa CéLine |
| EPI_ISL_872347, EPI_ISL_872348 | Hopital | National Reference Center for Viruses of Respiratory Infections, Institut Pasteur, Paris | Marion Barbet, Sylvie Behillil, Méline Bizard, Angela Brisebarre, Camille Capel, Etienne Simon-Lorière, Vincent Enouf, Maud Vanpeene, Sylvie van der Werf,Irimia Alix |
| EPI_ISL_872569 | UMMC-Health | WHO National Influenza Centre Russian Federation | Andrey Komissarov, Artem Fadeev, Anna Ivanova, Kseniya Komissarova, Dmitry Bazhenov, Mikhail Bakaev,Tatiana Platonova, Daria Danilenko, Ksenia Safina, Elena Nabieva, Georgii Bazykin, Dmitry Lioznov |
| EPI_ISL_872570 | Krasnoyarsk Regional Clinical Hospital | WHO National Influenza Centre Russian Federation | Andrey Komissarov, Artem Fadeev, Anna Ivanova, Kseniya Komissarova, Dmitry Bazhenov, Mikhail Bakaev, Daria Danilenko, Ksenia Safina, Elena Nabieva, Georgii Bazykin, Dmitry Lioznov |
| EPI_ISL_872591, EPI_ISL_872592, EPI_ISL_872593, EPI_ISL_872594, EPI_ISL_872595, EPI_ISL_872596, EPI_ISL_872597, EPI_ISL_872598, EPI_ISL_872599 | Department of Laboratory Medicine, National Taiwan University Hospital | Microbial Genomics Core Lab, National Taiwan University Centers of Genomic and Precision Medicine | Shiou-Hwei Yeh, You-Yu Lin, Ya-Yun Lai, Chiao-Ling Li, Shan-Chwen Chang, Pei-Jer Chen, Sui-Yuan Chang |
| EPI_ISL_872643 | Pathogenic Microorganisms Variability Laboratory | Pathogenic Microorganisms Variability Laboratory | Alexey Shchetinin, Olesya Venchakova, Maria Nikiforova, Andrei Siniavin, Nadezhda Kuznetsova, Elena Shidlovskaya, Elizaveta Divisenko, Kirill Krasnoslobotsev, Evgeniya Mukasheva, Anna Ignatieva, Svetlana Trushakova, Andrey Pochtovyy, Valeria Bacalin, Evgeny Usachev, Olga Burgasova, Ludmila Kolobukhina, Svetlana Smetanina, Elena Burtseva, Denis Logunov, Vladimir Gushchin, Alexander Gintsburg |
| EPI_ISL_872898, EPI_ISL_872899, EPI_ISL_872901, EPI_ISL_872903, EPI_ISL_872904, EPI_ISL_872905, EPI_ISL_872910, EPI_ISL_872912, EPI_ISL_872914, EPI_ISL_872915, EPI_ISL_872916, EPI_ISL_872917, EPI_ISL_872918, EPI_ISL_872919, EPI_ISL_872921, EPI_ISL_872925, EPI_ISL_872930, EPI_ISL_872931, EPI_ISL_872933, EPI_ISL_872934, EPI_ISL_872935, EPI_ISL_872936, EPI_ISL_872937, EPI_ISL_872939, EPI_ISL_872940, EPI_ISL_872941, EPI_ISL_872943, EPI_ISL_872944, EPI_ISL_872945, EPI_ISL_872946, EPI_ISL_872947, EPI_ISL_872949, EPI_ISL_872950, EPI_ISL_872952, EPI_ISL_872953, EPI_ISL_872954, EPI_ISL_872955, EPI_ISL_872958, EPI_ISL_872961, EPI_ISL_872962, EPI_ISL_872963, EPI_ISL_872965, EPI_ISL_872967, EPI_ISL_872968, EPI_ISL_872969, EPI_ISL_872970, EPI_ISL_872971 | | | |

| | | | |
|---|---|---|---|
| see above | WHO National Influenza Centre Russian Federation | WHO National Influenza Centre Russian Federation | Andrey Komissarov, Artem Fadeev, Anna Ivanova, Kseniya Komissarova, Dmitry Bazhenov, Mikhail Bakaev, Daria Danilenko, Ksenia Safina, Elena Nabieva, Georgii Bazykin, Dmitry Lioznov |
| EPI_ISL_875346 | National Institute of Infectious Diseases-Prof. Dr. Matei Bals Molecular Diagnostics Laboratory | National Institute of Infectious Diseases-Prof. Dr. Matei Bals Molecular Diagnostics Laboratory | Leontina Banica, Marius Surleac, Corina Casangiu, Petre Milu, Andreea Tudor, Simona Paraschiv, Dan Otelea |
| EPI_ISL_875688 | National Influenza Center - Instituto Adolfo Lutz | Instituto Adolfo Lutz, Interdiciplinary Procedures Center, Strategic Laboratory | Claudio Tavares Sacchi, Claudia Regina Gonçalves, Erica Valessa Ramos Gomes, Karoline Rodrigues Campos, Katia Correa de Oliveira Santos, Ana Lucia de Carvalho Avelino, Clovis Roberto Abe Constantino |
| EPI_ISL_875689 | Hospital do Servidor Publico | Instituto Adolfo Lutz, Interdiciplinary Procedures Center, Strategic Laboratory | Claudio Tavares Sacchi, Claudia Regina Gonçalves, Erica Valessa Ramos Gomes, Karoline Rodrigues Campos |
| EPI_ISL_876738, EPI_ISL_876740, EPI_ISL_876741 | Istituto Zooprofilattico Sperimentale della Puglia e della Basilicata | Istituto Zooprofilattico Sperimentale della Puglia e della Basilicata | Parisi A., Bianco A., Capozzi L., Del Sambro L., Manzulli V, Rondinone V., Pace L., Cipolletta D., Galante D. |
| EPI_ISL_876742 | Ospedale Santa Caterina Novella | Istituto Zooprofilattico Sperimentale della Puglia e della Basilicata | Parisi A., Bianco A., Capozzi L., Del Sambro L., Manzulli V, Rondinone V., Pace L., Cipolletta D., Galante D. |
| EPI_ISL_876743 | Presidio di Brindisi Di Summa - Perrino | Istituto Zooprofilattico Sperimentale della Puglia e della Basilicata | Parisi A., Bianco A., Capozzi L., Del Sambro L., Manzulli V, Rondinone V., Pace L., Cipolletta D., Galante D. |
| EPI_ISL_876744, EPI_ISL_876745, EPI_ISL_876746, EPI_ISL_876747, EPI_ISL_876748, EPI_ISL_876749, EPI_ISL_876750, EPI_ISL_876751, EPI_ISL_876752, EPI_ISL_876753, EPI_ISL_876754, EPI_ISL_876755, EPI_ISL_876756, EPI_ISL_876757, EPI_ISL_876758, EPI_ISL_876759, EPI_ISL_876760, EPI_ISL_876761, EPI_ISL_876762, EPI_ISL_876763, EPI_ISL_876764, EPI_ISL_876765, EPI_ISL_876766, EPI_ISL_876767, EPI_ISL_876768, EPI_ISL_876769, EPI_ISL_876770, EPI_ISL_876771, EPI_ISL_876772, EPI_ISL_876773, EPI_ISL_876774, EPI_ISL_876775, EPI_ISL_876776, EPI_ISL_876777, EPI_ISL_876778, EPI_ISL_876779, EPI_ISL_876780, EPI_ISL_876781, EPI_ISL_876782, EPI_ISL_876783, EPI_ISL_876784, EPI_ISL_876785, EPI_ISL_876786, EPI_ISL_876787, EPI_ISL_876788, EPI_ISL_876789, EPI_ISL_876790, EPI_ISL_876791, EPI_ISL_876792, EPI_ISL_876793, EPI_ISL_876794, EPI_ISL_876795, EPI_ISL_876796, EPI_ISL_876797, EPI_ISL_876798, EPI_ISL_876799, EPI_ISL_876800, EPI_ISL_876801, EPI_ISL_876802, EPI_ISL_876803, EPI_ISL_876804, EPI_ISL_876805, EPI_ISL_876806, EPI_ISL_876807, EPI_ISL_876808, EPI_ISL_876809 | | | |
| see above | Istituto Zooprofilattico Sperimentale della Puglia e della Basilicata | Istituto Zooprofilattico Sperimentale della Puglia e della Basilicata | Parisi A., Bianco A., Capozzi L., Del Sambro L., Manzulli V, Rondinone V., Pace L., Cipolletta D., Galante D. |
| EPI_ISL_877126 | Academic Hospital of Gajdah Mada University (RSA UGM) | Genetics Working Group (Pokja Genetik) Faculty of Medicine, Public Health and Nursing Universitas Gadjah Mada (FK-KMK UGM); Disease Investigation Center Wates Ministry of Agriculture Indonesia; Department of Microbiology FK-KMK UGM; Laboratorium Diagnostik Yayasan Tahija World Mosquito Program (WMP) Yogyakarta Center for Tropical Medicine FK-KMK UGM; Integrated Research center FK-KMK UGM; Department of Computer Science and Electronics FMIPA UGM | Gunadi, Hendra Wibawa, Marcellus, Mohamad S. Hakim, Edwin W. Daniwijaya, Ludhang P. Rizki, Endah Supriyati, Eggi Arguni, Titik Nuryastuti, Tri Wibawa, Dwi AA Nugrahaningsih, Afiahayati, Siswanto,Untung Riawan, William Widitjiarso |
| EPI_ISL_877127 | Academic Hospital of Gajdah Mada University (RSA UGM) | Genetics Working Group (Pokja Genetik) Faculty of Medicine, Public Health and Nursing Universitas Gadjah Mada (FK-KMK UGM); Disease Investigation Center Wates Ministry of Agriculture Indonesia; Department of Microbiology FK-KMK UGM; Laboratorium Diagnostik Yayasan Tahija World Mosquito Program (WMP) Yogyakarta Center for Tropical Medicine FK-KMK UGM; Integrated Research center FK-KMK UGM; Department of Computer Science and Electronics FMIPA UGM | Gunadi, Hendra Wibawa, Marcellus, Mohamad S. Hakim, Edwin W. Daniwijaya, Ludhang P. Rizki, Endah Supriyati, Eggi Arguni, Titik Nuryastuti, Tri Wibawa, Dwi AA Nugrahaningsih, Afiahayati, Siswanto,Audric Kenny Tedja, Maria Patricia Inggriani |
| EPI_ISL_877128 | Academic Hospital of Gajdah Mada University (RSA UGM) | Genetics Working Group (Pokja Genetik) Faculty of Medicine, Public Health and Nursing Universitas Gadjah Mada (FK-KMK UGM); Disease Investigation Center Wates Ministry of Agriculture Indonesia; Department of Microbiology FK-KMK UGM; Laboratorium Diagnostik Yayasan Tahija World Mosquito Program (WMP) Yogyakarta Center for Tropical Medicine FK-KMK UGM; Integrated Research center FK-KMK UGM; Department of Computer Science and Electronics FMIPA UGM | Gunadi, Hendra Wibawa, Marcellus, Mohamad S. Hakim, Edwin W. Daniwijaya, Ludhang P. Rizki, Endah Supriyati, Eggi Arguni, Titik Nuryastuti, Tri Wibawa, Dwi AA Nugrahaningsih, Afiahayati, Siswanto, Kristy Iskandar, Nungki Anggorowati, Aditya Rifqi Fauzi, Fadil Fahri |
| EPI_ISL_877129 | Academic Hospital of Gajdah Mada University (RSA UGM) | Genetics Working Group (Pokja Genetik) Faculty of Medicine, Public Health and Nursing Universitas Gadjah Mada (FK-KMK UGM); Disease Investigation Center Wates Ministry of Agriculture Indonesia; Department of Microbiology FK-KMK UGM; Laboratorium Diagnostik Yayasan Tahija World Mosquito Program (WMP) Yogyakarta Center for Tropical Medicine FK-KMK UGM; Integrated Research Center FK-KMK UGM; Department of Computer Science and Electronics FMIPA UGM | Gunadi, Hendra Wibawa, Marcellus, Mohamad S. Hakim, Edwin W. Daniwijaya, Ludhang P. Rizki, Endah Supriyati, Eggi Arguni, Titik Nuryastuti, Tri Wibawa, Dwi AA Nugrahaningsih, Afiahayati, Siswanto, Kristy Iskandar, Nungki Anggorowati, Dwiki afandy, Dyah Ayu Puspitarani |
| EPI_ISL_877130 | Academic Hospital of Gajdah Mada University (RSA UGM) | Genetics Working Group (Pokja Genetik) Faculty of Medicine, Public Health and Nursing Universitas Gadjah Mada (FK-KMK UGM); Disease Investigation Center Wates Ministry of Agriculture Indonesia; Department of Microbiology FK-KMK UGM; Laboratorium Diagnostik Yayasan Tahija World Mosquito Program (WMP) Yogyakarta Center for Tropical Medicine FK-KMK UGM; Integrated Research center FK-KMK UGM; Department of Computer Science and Electronics FMIPA UGM | Gunadi, Hendra Wibawa, Marcellus, Mohamad S. Hakim, Edwin W. Daniwijaya, Ludhang P. Rizki, Endah Supriyati, Eggi Arguni, Titik Nuryastuti, Tri Wibawa, Dwi AA Nugrahaningsih, Afiahayati, Siswanto, Kristy Iskandar, Nungki Anggorowati, Kemala Athollah, Desyifa Mursalin |
| EPI_ISL_877131 | Academic Hospital of Gajdah Mada University (RSA UGM) | Genetics Working Group (Pokja Genetik) Faculty of Medicine, Public Health and Nursing Universitas Gadjah Mada (FK-KMK UGM); Disease Investigation Center Wates Ministry of Agriculture Indonesia; Department of Microbiology FK-KMK UGM; Laboratorium Diagnostik Yayasan Tahija World Mosquito Program (WMP) Yogyakarta Center for Tropical Medicine FK-KMK UGM; Integrated Research center FK-KMK UGM; Department of Computer Science and Electronics FMIPA UGM | Gunadi, Hendra Wibawa, Marcellus, Mohamad S. Hakim, Edwin W. Daniwijaya, Ludhang P. Rizki, Endah Supriyati, Eggi Arguni, Titik Nuryastuti, Tri Wibawa, Dwi AA Nugrahaningsih, Afiahayati, Siswanto, Kristy Iskandar, Nungki Anggorowati, Susan Simanjaya, Alvin Santoso Kalim |
| EPI_ISL_877228 | Institute for Medical Research, Infectious Disease Research Centre, National Institutes of Health, Ministry of Health Malaysia | Institute for Medical Research, Infectious Disease Research Centre, National Institutes of Health, Ministry of Health Malaysia | Suppiah J, Kamel K, Azizan MA, Thayan R |
| EPI_ISL_877230, EPI_ISL_877231, EPI_ISL_877232, EPI_ISL_877233, | University College Sedaya International (UCSI University) | Institute for Medical Research, Infectious Disease Research Centre, National Institutes of Health, Ministry of Health | Suppiah J, Kamel K, Azizan MA, Sekaran SD, Thayan R |

| | | | |
|---|---|---|---|
| EPI_ISL_877234, EPI_ISL_877235, EPI_ISL_877236, EPI_ISL_877329, EPI_ISL_877330 | | Malaysia | |
| EPI_ISL_877419 | Siti Khodijah Hospital | Institute of Tropical Disease, Universitas Airlangga | Krisnoadi Rahardjo, Aldise M Nastri, Jezzy R Dewantari, Rima R Prasetya, Muhammad Hamdan, Gatot Soegiarto, Laksmi Wulandari, Resti Yudhawati, Yasuko Mori, Soetjipto, Kazufumi Shimizu, Maria I Lusida |
| EPI_ISL_877420 | Darus Syifa' Islamic Hospital | Institute of Tropical Disease, Universitas Airlangga | Aldise M Nastri, Jezzy R Dewantari, Rima R Prasetya, Krisnoadi Rahardjo, HM Faiz, Gatot Soegiarto, Laksmi Wulandari, Resti Yudhawati, Soetjipto, Yasuko Mori, Maria I Lusida, Kazufumi Shimizu |
| EPI_ISL_877422 | Citra Medika Hospital | Institute of Tropical Disease, Universitas Airlangga | Rima R Prasetya, Krisnoadi Rahardjo, Aldise M Nastri, Jezzy R Dewantari, Wahidah Rachmaniyah, Gatot Soegiarto, Laksmi Wulandari, Resti Yudhawati, Soetjipto, Yasuko Mori, Maria I Lusida, Kazufumi Shimizu |
| EPI_ISL_877423 | Primasatya Husada Citra Hospital | Institute of Tropical Disease, Universitas Airlangga | Maria I Lusida, Krisnoadi Rahardjo, Aldise M Nastri, Jezzy R Dewantari, Rima R Prasetya, Pudji Djanuartono, Gatot Soegiarto, Laksmi Wulandari, Resti Yudhawati, Soetjipto, Yasuko Mori, Kazufumi Shimizu |
| EPI_ISL_877456 | Primasatya Husada Citra Hospital | Institute of Tropical Disease, Universitas Airlangga | Krisnoadi Rahardjo, Aldise M Nastri, Jezzy R Dewantari, Rima R Prasetya, Pudji Djanuartono, Gatot Soegiarto, Laksmi Wulandari, Resti Yudhawati, Soetjipto, Yasuko Mori, Maria I Lusida, Kazufumi Shimizu |
| EPI_ISL_877457 | Primasatya Husada Citra Hospital | Institute of Tropical Disease, Universitas Airlangga | Aldise M Nastri, Jezzy R Dewantari, Rima R Prasetya, Krisnoadi Rahardjo, Pudji Djanuartono, Gatot Soegiarto, Laksmi Wulandari, Resti Yudhawati, Soetjipto, Yasuko Mori, Maria I Lusida, Kazufumi Shimizu |
| EPI_ISL_877458 | Anwar Medika General Hospital | Institute of Tropical Disease, Universitas Airlangga | Jezzy R Dewantari, Rima R Prasetya, Krisnoadi Rahardjo, Aldise M Nastri, Nungky Taniasari, Gatot Soegiarto, Laksmi Wulandari, Resti Yudhawati, Soetjipto, Yasuko Mori, Maria I Lusida, Kazufumi Shimizu |
| EPI_ISL_877459 | Darmo Hospital | Institute of Tropical Disease, Universitas Airlangga | Rima R Prasetya, Krisnoadi Rahardjo, Aldise M Nastri, Jezzy R Dewantari, Sulung Budianto, Gatot Soegiarto, Laksmi Wulandari, Resti Yudhawati, Soetjipto, Yasuko Mori, Maria I Lusida, Kazufumi Shimizu |
| EPI_ISL_877460 | Siti Khodijah Hospital | Institute of Tropical Disease, Universitas Airlangga | Maria I Lusida, Krisnoadi Rahardjo, Aldise M Nastri, Jezzy R Dewantari, Rima R Prasetya, Muhammad Hamdan, Gatot Soegiarto, Laksmi Wulandari, Resti Yudhawati, Soetjipto, Yasuko Mori, Kazufumi Shimizu |
| EPI_ISL_877538 | Siti Khodijah Hospital | Institute of Tropical Disease, Universitas Airlangga | Krisnoadi Rahardjo, Aldise M Nastri, Jezzy R Dewantari, Rima R Prasetya, Muhammad Hamdan, Gatot Soegiarto, Laksmi Wulandari, Resti Yudhawati, Soetjipto, Yasuko Mori, Maria I Lusida, Kazufumi Shimizu |
| EPI_ISL_882635 | SC (UCO) Igiene e Sanità Pubblica (funzione integrata con SC Microbiologia e Virologia) e Laboratory of Molecular Virology of the International Centre for Genetic Engineering and Biotechnology (ICGEB) | ARGO Laboratorio Genomica ed EpigenomicA | Licastro D, Dal Monego S, Degasperi M, Marcello A, D'Agaro P |
| EPI_ISL_882636 | SC (UCO) Igiene e Sanità Pubblica (funzione integrata con SC Microbiologia e Virologia) e Laboratory of Molecular Virology of the International Centre for Genetic Engineering and Biotechnology (ICGEB) | ARGO Laboratorio Genomica ed Epigenomica | Licastro D, Dal Monego S, Degasperi M, Marcello A, D'Agaro P |
| EPI_ISL_882645 | Medical Research Center, Faculty of Medicine, Syarif Hidayatullah State Islamic University Jakarta | Medical Research Center, Faculty of Medicine, Syarif Hidayatullah State Islamic University Jakarta | Chris Adhiyanto, Erike A Suwarsono, Laifa Hendarmin, Zeti Harriyati, Endah Wulandari, Flori Ratna Sari, Fika Ekayanti, Hari Hendarto |
| EPI_ISL_882646 | Medical Research Center, Faculty of Medicine, Syarif Hidayatullah State Islamic University Jakarta | Medical Research Center, Faculty of Medicine, Syarif Hidayatullah State Islamic University Jakarta | Chris Adhiyanto, Laifa Hendarmin, Zeti Harriyati, Endah Wulandari, Flori Ratna Sari, Fika Ekayanti, Hari Hendarto |
| EPI_ISL_882656 | Medical Research Center, Faculty of Medicine, Syarif Hidayatullah State Islamic University Jakarta | Medical Research Center, Faculty of Medicine, Syarif Hidayatullah State Islamic University Jakarta | Erike A Suwarsono, Chris Adhiyanto, Laifa Hendarmin, Zeti Harriyati, Flori Ratna Sari, Endah Wulandari, Fika Ekayanti |
| EPI_ISL_882658 | Secretaria Municipal de Saude | Instituto Adolfo Lutz, Interdiciplinary Procedures Center, Strategic Laboratory | Claudio Tavares Sacchi, Claudia Regina Gonçalves, Erica Valessa Ramos Gomes, Karoline Rodrigues Campos |
| EPI_ISL_882659 | Centro de Triagem Covid19 | Instituto Adolfo Lutz, Interdiciplinary Procedures Center, Strategic Laboratory | Claudio Tavares Sacchi, Claudia Regina Gonçalves, Erica Valessa Ramos Gomes, Karoline Rodrigues Campos |
| EPI_ISL_882660 | Hospital Municipal Prefeito Waldemar Costa Filho | Instituto Adolfo Lutz, Interdiciplinary Procedures Center, Strategic Laboratory | Claudio Tavares Sacchi, Claudia Regina Gonçalves, Erica Valessa Ramos Gomes, Karoline Rodrigues Campos |
| EPI_ISL_882661, EPI_ISL_882662 | Hospital de Santa Barbara de Goias | Instituto Adolfo Lutz, Interdiciplinary Procedures Center, Strategic Laboratory | Claudio Tavares Sacchi, Claudia Regina Gonçalves, Erica Valessa Ramos Gomes, Karoline Rodrigues Campos |
| EPI_ISL_882663, EPI_ISL_882664 | LACEN do Distrito Federal | Instituto Adolfo Lutz, Interdiciplinary Procedures Center, Strategic Laboratory | Claudio Tavares Sacchi, Claudia Regina Gonçalves, Erica Valessa Ramos Gomes, Karoline Rodrigues Campos |
| EPI_ISL_882665 | Unidade de Pronto Atendimento Dra Zilda Arns | Instituto Adolfo Lutz, Interdiciplinary Procedures Center, Strategic Laboratory | Claudio Tavares Sacchi, Claudia Regina Gonçalves, Erica Valessa Ramos Gomes, Karoline Rodrigues Campos |
| EPI_ISL_882666 | UMS de Juquitiba | Instituto Adolfo Lutz, Interdiciplinary Procedures Center, Strategic Laboratory | Claudio Tavares Sacchi, Claudia Regina Gonçalves, Erica Valessa Ramos Gomes, Karoline Rodrigues Campos |
| EPI_ISL_882667 | Hospital de Clinicas Caieiras | Instituto Adolfo Lutz, Interdiciplinary Procedures Center, Strategic Laboratory | Claudio Tavares Sacchi, Claudia Regina Gonçalves, Erica Valessa Ramos Gomes, Karoline Rodrigues Campos |
| EPI_ISL_882668 | UMS de Juquitiba | Instituto Adolfo Lutz, Interdiciplinary Procedures Center, Strategic Laboratory | Claudio Tavares Sacchi, Claudia Regina Gonçalves, Erica Valessa Ramos Gomes, Karoline Rodrigues Campos |
| EPI_ISL_882669 | UPA Vila Santa Catarina | Instituto Adolfo Lutz, Interdiciplinary Procedures Center, Strategic Laboratory | Claudio Tavares Sacchi, Claudia Regina Gonçalves, Erica Valessa Ramos Gomes, Karoline Rodrigues Campos |
| EPI_ISL_882670 | Hospital Samaritano Paulista | Instituto Adolfo Lutz, Interdiciplinary Procedures Center, Strategic Laboratory | Claudio Tavares Sacchi, Claudia Regina Gonçalves, Erica Valessa Ramos Gomes, Karoline Rodrigues Campos |
| EPI_ISL_882671, EPI_ISL_882672, EPI_ISL_882673 | Hospital Municipal Dr. Guido Guida | Instituto Adolfo Lutz, Interdiciplinary Procedures Center, Strategic Laboratory | Claudio Tavares Sacchi, Claudia Regina Gonçalves, Erica Valessa Ramos Gomes, Karoline Rodrigues Campos |
| EPI_ISL_882770 | Darus Syifa' Islamic Hospital | Institute of Tropical Disease, Universitas Airlangga | Aldise M Nastri, Jezzy R Dewantari, Rima R Prasetya, Krisnoadi Rahardjo, HM Faiz, Gatot Soegiarto, Laksmi Wulandari, Resti Yudhawati, Soetjipto, Yasuko Mori, Kazufumi Shimizu, Maria I Lusida |
| EPI_ISL_882771 | Siti Khodijah Hospital | Institute of Tropical Disease, Universitas Airlangga | Kazufumi Shimizu, Krisnoadi Rahardjo, Aldise M Nastri, Jezzy R Dewantari, Rima R Prasetya, Muhammad Hamdan, Gatot Soegiarto, Laksmi Wulandari, Resti Yudhawati, Yasuko Mori, Soetjipto, Maria I Lusida |
| EPI_ISL_882943, EPI_ISL_882944, EPI_ISL_882945, EPI_ISL_882946, EPI_ISL_882947, EPI_ISL_882948, EPI_ISL_882949, EPI_ISL_882950, EPI_ISL_882952 | The National Institute of Public Health | State Veterinary Institute Prague | Nagy,A;Jirincova,H;Vecerova,JTrnka,D |
| EPI_ISL_882954, EPI_ISL_882955 | State Veterinary Institute Prague | State Veterinary Institute Prague | Nagy,A;Cernikova,L;Stara,M |
| EPI_ISL_883286, EPI_ISL_883287 | SIESP DIPARTIMENTO DI PREVENZIONE CHIE | Istituto Zooprofilattico Sperimentale dell'Abruzzo e Molise "G. | Lorusso A, Marcacci M, Di Domenico M, Ancora M, Curini V, Mangone I, Rinaldi A, Scialabba S, Di Pasquale A, Cammà C, Puglia I, Calistri P, Savini G |

| | | | |
|---|---|---|---|
| | | Caporale" | |
| EPI_ISL_883288, EPI_ISL_883289 | SIESP CHIETI-DRIVE IN ORTONA | Istituto Zooprofilattico Sperimentale dell'Abruzzo e Molise "G. Caporale" | Lorusso A, Marcacci M, Di Domenico M, Ancora M, Curini V, Mangone I, Rinaldi A, Scialabba S, Di Pasquale A, Cammà C, Puglia I, Calistri P, Savini G |
| EPI_ISL_883290 | SIESP CHIETI-DRIVE IN CHIETI | Istituto Zooprofilattico Sperimentale dell'Abruzzo e Molise "G. Caporale" | Lorusso A, Marcacci M, Di Domenico M, Ancora M, Curini V, Mangone I, Rinaldi A, Scialabba S, Di Pasquale A, Cammà C, Puglia I, Calistri P, Savini G |
| EPI_ISL_883291 | RP Guardiagrele-Ospedale di Comunità | Istituto Zooprofilattico Sperimentale dell'Abruzzo e Molise "G. Caporale" | Lorusso A, Marcacci M, Di Domenico M, Ancora M, Curini V, Mangone I, Rinaldi A, Scialabba S, Di Pasquale A, Cammà C, Puglia I, Calistri P, Savini G |
| EPI_ISL_883292 | SIESP CHIETI-DRIVE IN ORTONA | Istituto Zooprofilattico Sperimentale dell'Abruzzo e Molise "G. Caporale" | Lorusso A, Marcacci M, Di Domenico M, Ancora M, Curini V, Mangone I, Rinaldi A, Scialabba S, Di Pasquale A, Cammà C, Puglia I, Calistri P, Savini G |
| EPI_ISL_883293 | SIESP CHIETI-DRIVE IN LANCIANO | Istituto Zooprofilattico Sperimentale dell'Abruzzo e Molise "G. Caporale" | Lorusso A, Marcacci M, Di Domenico M, Ancora M, Curini V, Mangone I, Rinaldi A, Scialabba S, Di Pasquale A, Cammà C, Puglia I, Calistri P, Savini G |
| EPI_ISL_883294, EPI_ISL_883295 | RP Guardiagrele-Ospedale di Comunità | Istituto Zooprofilattico Sperimentale dell'Abruzzo e Molise "G. Caporale" | Lorusso A, Marcacci M, Di Domenico M, Ancora M, Curini V, Mangone I, Rinaldi A, Scialabba S, Di Pasquale A, Cammà C, Puglia I, Calistri P, Savini G |
| EPI_ISL_883296 | SIESP CHIETI-DRIVE IN ORTONA | Istituto Zooprofilattico Sperimentale dell'Abruzzo e Molise "G. Caporale" | Lorusso A, Marcacci M, Di Domenico M, Ancora M, Curini V, Mangone I, Rinaldi A, Scialabba S, Di Pasquale A, Cammà C, Puglia I, Calistri P, Savini G |
| EPI_ISL_883297 | RP Guardiagrele-Ospedale di Comunità | Istituto Zooprofilattico Sperimentale dell'Abruzzo e Molise "G. Caporale" | Lorusso A, Marcacci M, Di Domenico M, Ancora M, Curini V, Mangone I, Rinaldi A, Scialabba S, Di Pasquale A, Cammà C, Puglia I, Calistri P, Savini G |
| EPI_ISL_883298, EPI_ISL_883299 | SIESP DIPARTIMENTO DI PREVENZIONE CHIE | Istituto Zooprofilattico Sperimentale dell'Abruzzo e Molise "G. Caporale" | Lorusso A, Marcacci M, Di Domenico M, Ancora M, Curini V, Mangone I, Rinaldi A, Scialabba S, Di Pasquale A, Cammà C, Puglia I, Calistri P, Savini G |
| EPI_ISL_883301, EPI_ISL_883302 | RP Guardiagrele-Ospedale di Comunità | Istituto Zooprofilattico Sperimentale dell'Abruzzo e Molise "G. Caporale" | Lorusso A, Marcacci M, Di Domenico M, Ancora M, Curini V, Mangone I, Rinaldi A, Scialabba S, Di Pasquale A, Cammà C, Puglia I, Calistri P, Savini G |
| EPI_ISL_883303, EPI_ISL_883304, EPI_ISL_883305 | SIESP DIPARTIMENTO DI PREVENZIONE CHIE | Istituto Zooprofilattico Sperimentale dell'Abruzzo e Molise "G. Caporale" | Lorusso A, Marcacci M, Di Domenico M, Ancora M, Curini V, Mangone I, Rinaldi A, Scialabba S, Di Pasquale A, Cammà C, Puglia I, Calistri P, Savini G |
| EPI_ISL_883328, EPI_ISL_883336, EPI_ISL_883350, EPI_ISL_883351, EPI_ISL_883352, EPI_ISL_883353, EPI_ISL_883390, EPI_ISL_883391, EPI_ISL_883392, EPI_ISL_883393, EPI_ISL_883394, EPI_ISL_883397, EPI_ISL_883398, EPI_ISL_883399, EPI_ISL_883400, EPI_ISL_883401, EPI_ISL_883402, EPI_ISL_883404, EPI_ISL_883405 | | | |
| see above | OCME Office Of Chief Medical Examiner | New York City Public Health Laboratory | Jade Wang, et al. |
| EPI_ISL_883957 | Medical Research Center, Faculty of Medicine, Syarif Hidayatullah State Islamic University Jakarta | Medical Research Center, Faculty of Medicine, Syarif Hidayatullah State Islamic University Jakarta | Erike A Suwarsono, Chris Adhiyanto, Laifa Hendarmin, Zeti Harriyati, Endah Wulandari, Flori Ratna Sari, Fika Ekayanti, Hari Hendarto |
| EPI_ISL_883958 | Hopital | National Reference Center for Viruses of Respiratory Infections, Institut Pasteur, Paris | Marion Barbet, Sylvie Behillil, Méline Bizard, Angela Brisebarre, Camille Capel, Etienne Simon-Lorière, Vincent Enouf, Maud Vanpeene, Sylvie van der Werf,Lagathu GisèLe |
| EPI_ISL_883959 | Labo Analyses Med | National Reference Center for Viruses of Respiratory Infections, Institut Pasteur, Paris | Marion Barbet, Sylvie Behillil, Méline Bizard, Angela Brisebarre, Camille Capel, Etienne Simon-Lorière, Vincent Enouf, Maud Vanpeene, Sylvie van der Werf |
| EPI_ISL_883960, EPI_ISL_883961 | Labo Analyses Med | National Reference Center for Viruses of Respiratory Infections, Institut Pasteur, Paris | Marion Barbet, Sylvie Behillil, Méline Bizard, Angela Brisebarre, Camille Capel, Etienne Simon-Lorière, Vincent Enouf, Maud Vanpeene, Sylvie van der Werf,Jacques Alexandra |
| EPI_ISL_883962, EPI_ISL_883963 | Labo Analyses Med | National Reference Center for Viruses of Respiratory Infections, Institut Pasteur, Paris | Marion Barbet, Sylvie Behillil, Méline Bizard, Angela Brisebarre, Camille Capel, Etienne Simon-Lorière, Vincent Enouf, Maud Vanpeene, Sylvie van der Werf,Amzalag Jonas |
| EPI_ISL_883964, EPI_ISL_883965, EPI_ISL_883966, EPI_ISL_883967, EPI_ISL_883968, EPI_ISL_883969, EPI_ISL_883970, EPI_ISL_883971 | Labo Analyses med | National Reference Center for Viruses of Respiratory Infections, Institut Pasteur, Paris | Marion Barbet, Sylvie Behillil, Méline Bizard, Angela Brisebarre, Camille Capel, Etienne Simon-Lorière, Vincent Enouf, Maud Vanpeene, Sylvie van der Werf,Amzalag Jonas |
| EPI_ISL_883972, EPI_ISL_883973, EPI_ISL_883974, EPI_ISL_883975, EPI_ISL_883976, EPI_ISL_883977, EPI_ISL_883978, EPI_ISL_883979, EPI_ISL_883980, EPI_ISL_883981, EPI_ISL_883982, EPI_ISL_883983, EPI_ISL_883984, EPI_ISL_883985 | | | |
| see above | Hopital | National Reference Center for Viruses of Respiratory Infections, Institut Pasteur, Paris | Marion Barbet, Sylvie Behillil, Méline Bizard, Angela Brisebarre, Camille Capel, Etienne Simon-Lorière, Vincent Enouf, Maud Vanpeene, Sylvie van der Werf,Fourgeaud Jacques |
| EPI_ISL_883986, EPI_ISL_883987, EPI_ISL_883988, EPI_ISL_883989 | Hopital | National Reference Center for Viruses of Respiratory Infections, Institut Pasteur, Paris | Marion Barbet, Sylvie Behillil, Méline Bizard, Angela Brisebarre, Camille Capel, Etienne Simon-Lorière, Vincent Enouf, Maud Vanpeene, Sylvie van der Werf |
| EPI_ISL_883993, EPI_ISL_883994, EPI_ISL_883995, EPI_ISL_883996 | hopital | National Reference Center for Viruses of Respiratory Infections, Institut Pasteur, Paris | Marion Barbet, Sylvie Behillil, Méline Bizard, Angela Brisebarre, Camille Capel, Etienne Simon-Lorière, Vincent Enouf, Maud Vanpeene, Sylvie van der Werf,Andre-Garnier Elisabeth |
| EPI_ISL_883997 | Labo Analyses Med | National Reference Center for Viruses of Respiratory Infections, Institut Pasteur, Paris | Marion Barbet, Sylvie Behillil, Méline Bizard, Angela Brisebarre, Camille Capel, Etienne Simon-Lorière, Vincent Enouf, Maud Vanpeene, Sylvie van der Werf,Selas Labomaine |
| EPI_ISL_883998, EPI_ISL_883999, EPI_ISL_884000, EPI_ISL_884001, EPI_ISL_884002, EPI_ISL_884003, EPI_ISL_884004, EPI_ISL_884005 | Labo Analyses Med | National Reference Center for Viruses of Respiratory Infections, Institut Pasteur, Paris | Marion Barbet, Sylvie Behillil, Méline Bizard, Angela Brisebarre, Camille Capel, Etienne Simon-Lorière, Vincent Enouf, Maud Vanpeene, Sylvie van der Werf,Rousset Dominique |
| EPI_ISL_884006, EPI_ISL_884007 | hopital | National Reference Center for Viruses of Respiratory Infections, Institut Pasteur, Paris | Marion Barbet, Sylvie Behillil, Méline Bizard, Angela Brisebarre, Camille Capel, Etienne Simon-Lorière, Vincent Enouf, Maud Vanpeene, Sylvie van der Werf,Combe Patrice |
| EPI_ISL_884087, EPI_ISL_884088 | Wazed Mia Science Research Center, Jahangirnagar University in collaboration with GRMDRC, Dhaka and Invent Technologies Ltd, Dhaka, Bangladesh | COVID Research Cell (CRC), Wazed Miah Science Research Center | Md. Abdul Matin, Bishajit Sarkar, Md. Asad Ullah, Yusha Araf, Nihad Adnan, Abdullah Al Nahid, Rashedul Islam, Mohammad Shahedur Rahman |
| EPI_ISL_885142 | RSUP Dr. Sardjito | Genetics Working Group (Pokja Genetik) Faculty of Medicine, Public Health and Nursing Universitas Gadjah Mada (FK-KMK UGM); Disease Investigation Center Wates Ministry of Agriculture Indonesia; Department of Microbiology FK-KMK UGM; Laboratorium Diagnostik Yayasan Tahija World Mosquito Program (WMP) Yogyakarta Center for Tropical Medicine FK-KMK UGM; Integrated Research Center FK-KMK UGM; Department of Computer Science and Electronics FMIPA UGM; RSUP Dr. Sardjito | Gunadi, Hendra Wibawa, . Marcellus, Mohamad S. Hakim, Edwin W. Daniwijaya, Ludhang P. Rizki, Endah Supriyati, Eggi Arguni, Titik Nuryastuti, Tri Wibawa, Dwi AA Nugrahaningsih, Afiahayati, Siswanto, Kristy Iskandar, Nungki Anggorowati, Ika Trisnawati, Sumardi, Eko Budiono, Bambang Sigit Riyanto, Heni Retnowulan, Munawar Gani, Satria Maulana, Nur Rahmi Ananda, Riat El Khair, Yunika Puspadewi, Osman Sianipar, Umi Solekhah Intansari, Elizabeth Henny Herningtiyas, Ira Puspitawati, Nur Imma Fatimah Harahap, Alvin Santoso Kalim, Susan Simanjaya |
| EPI_ISL_885146 | Genomic Laboratory (GLAB) (Conjoint lab of Health Directorate of Istanbul and Istanbul Technical University) | Genomic Laboratory (GLAB), Istanbul Technical University | Ilker Karacan, Tugba Kizilboga Akgun, Payam Zolfagharian, Nisan Denizce Can, Pari Sharifli, Levent Doganay, Gizem Dinler Doganay |
| EPI_ISL_888591 | Siti Khodijah Hospital | Institute of Tropical Disease, Universitas Airlangga | Krisnoadi Rahardjo, Aldise M Nastri, Jezzy R Dewantari, Rima R Prasetya, Muhammad Hamdan, Gatot Soegiarto, Laksmi Wulandari, Resti Yudhawati, Yasuko Mori, Soetjipto, Kazufumi Shimizu, Maria I Lusida |
| EPI_ISL_888593 | Siti Khodijah Hospital | Institute of Tropical Disease, Universitas Airlangga | Rima R Prasetya, Krisnoadi Rahardjo, Aldise M Nastri, Jezzy R Dewantari, Muhammad Hamdan, Gatot Soegiarto, Laksmi Wulandari, Resti Yudhawati, |

| | | | |
|---|---|---|---|
| | | | Soetjipto, Yasuko Mori, Kazufumi Shimizu, Maria I Lusida |
| EPI_ISL_888677 | Laboratory for Respiratory Viruses, Cantacuzino National Military-Medical Institute for Research and Development | Cantacuzino Institute Virology | Mihaela Lazar, Luiza Ustea, Nicoleta Paraschiv, Tim Durfee |
| EPI_ISL_888826 | Hospital | National Reference Center for Viruses of Respiratory Infections, Institut Pasteur, Paris | Marion Barbet, Sylvie Behillil, Méline Bizard, Angela Brisebarre, Camille Capel, Etienne Simon-Lorière, Vincent Enouf, Maud Vanpeene, Sylvie van der Werf, Gisèle Lagathu |
| EPI_ISL_888827, EPI_ISL_888828, EPI_ISL_888829 | General practitioner office | National Reference Center for Viruses of Respiratory Infections, Institut Pasteur, Paris | Marion Barbet, Sylvie Behillil, Méline Bizard, Angela Brisebarre, Camille Capel, Etienne Simon-Lorière, Vincent Enouf, Maud Vanpeene, Sylvie van der Werf |
| EPI_ISL_888979 | RS Bhakti Mulia | Eijkman Institute for Molecular Biology, Ministry of Research and Technology/National Agency for Research and Innovation | Willy Agustine, Edison Johar, Hidayat Trimarsanto, Iskandar Adnan, Lydia V. Panggalo, Sukma Oktavianthi, Frilasita A Yudhaputri, Safarina G Malik, Khin Saw Myint, Amin Soebandrio |
| EPI_ISL_888980 | RS Anna | Eijkman Institute for Molecular Biology, Ministry of Research and Technology/National Agency for Research and Innovation | Hidayat Trimarsanto, Iskandar Adnan, Lydia V. Panggalo, Sukma Oktavianthi, Willy Agustine, Edison Johar, Frilasita A Yudhaputri, Safarina G Malik, Khin Saw Myint, Amin Soebandrio |
| EPI_ISL_888981 | RS Omni Pulomas | Eijkman Institute for Molecular Biology, Ministry of Research and Technology/National Agency for Research and Innovation | Iskandar Adnan, Lydia V. Panggalo, Sukma Oktavianthi, Willy Agustine, Edison Johar, Hidayat Trimarsanto, Frilasita A Yudhaputri, Safarina G Malik, Khin Saw Myint, Amin Soebandrio |
| EPI_ISL_888983 | RS Hermina Daan Mogot | Eijkman Institute for Molecular Biology, Ministry of Research and Technology/National Agency for Research and Innovation | Edison Johar, Frilasita A Yudhaputri, Hidayat Trimarsanto, Iskandar Adnan, Lydia V. Panggalo, Sukma Oktavianthi, Willy Agustine, Safarina G Malik, Khin Saw Myint, Amin Soebandrio |
| EPI_ISL_888985 | RS Hermina Serpong | Eijkman Institute for Molecular Biology, Ministry of Research and Technology/National Agency for Research and Innovation | Edison Johar, Frilasita A Yudhaputri, Hidayat Trimarsanto, Iskandar Adnan, Lydia V. Panggalo, Sukma Oktavianthi, Willy Agustine, Safarina G Malik, Khin Saw Myint, Amin Soebandrio |
| EPI_ISL_888986 | RSU Bhakti Kartini | Eijkman Institute for Molecular Biology, Ministry of Research and Technology/National Agency for Research and Innovation | Edison Johar, Frilasita A Yudhaputri, Hidayat Trimarsanto, Iskandar Adnan, Lydia V. Panggalo, Sukma Oktavianthi, Willy Agustine, Safarina G Malik, Khin Saw Myint, Amin Soebandrio |
| EPI_ISL_888987 | RS Citra Medika Depok | Eijkman Institute for Molecular Biology, Ministry of Research and Technology/National Agency for Research and Innovation | Frilasita A Yudhaputri, Hidayat Trimarsanto, Iskandar Adnan, Lydia V. Panggalo, Sukma Oktavianthi, Willy Agustine, Edison Johar, Safarina G Malik, Khin Saw Myint, Amin Soebandrio |
| EPI_ISL_888988 | RS Qadr | Eijkman Institute for Molecular Biology, Ministry of Research and Technology/National Agency for Research and Innovation | Frilasita A Yudhaputri, Hidayat Trimarsanto, Iskandar Adnan, Lydia V. Panggalo, Sukma Oktavianthi, Willy Agustine, Edison Johar, Safarina G Malik, Khin Saw Myint, Amin Soebandrio |
| EPI_ISL_888989, EPI_ISL_888990 | RS Pondok Indah Bintaro Jaya | Eijkman Institute for Molecular Biology, Ministry of Research and Technology/National Agency for Research and Innovation | Frilasita A Yudhaputri, Hidayat Trimarsanto, Iskandar Adnan, Lydia V. Panggalo, Sukma Oktavianthi, Willy Agustine, Edison Johar, Safarina G Malik, Khin Saw Myint, Amin Soebandrio |
| EPI_ISL_888991 | RS Mitra Keluarga Gading Serpong | Eijkman Institute for Molecular Biology, Ministry of Research and Technology/National Agency for Research and Innovation | Frilasita A Yudhaputri, Hidayat Trimarsanto, Iskandar Adnan, Lydia V. Panggalo, Sukma Oktavianthi, Willy Agustine, Edison Johar, Safarina G Malik, Khin Saw Myint, Amin Soebandrio |
| EPI_ISL_888992 | RS Mitra Keluarga Gading Serpong | Eijkman Institute for Molecular Biology, Ministry of Research and Technology/National Agency for Research and Innovation | Hidayat Trimarsanto, Iskandar Adnan, Lydia V. Panggalo, Sukma Oktavianthi, Willy Agustine, Edison Johar, Frilasita A Yudhaputri, Safarina G Malik, Khin Saw Myint, Amin Soebandrio |
| EPI_ISL_888993 | RSU Medirossa | Eijkman Institute for Molecular Biology, Ministry of Research and Technology/National Agency for Research and Innovation | Iskandar Adnan, Lydia V. Panggalo, Sukma Oktavianthi, Willy Agustine, Edison Johar, Hidayat Trimarsanto, Frilasita A Yudhaputri, Safarina G Malik, Khin Saw Myint, Amin Soebandrio |
| EPI_ISL_888994 | RSU MMC | Eijkman Institute for Molecular Biology, Ministry of Research and Technology/National Agency for Research and Innovation | Lydia V. Panggalo, Sukma Oktavianthi, Willy Agustine, Edison Johar, Hidayat Trimarsanto, Iskandar Adnan, Frilasita A Yudhaputri, Safarina G Malik, Khin Saw Myint, Amin Soebandrio |
| EPI_ISL_888996 | RSU Graha Juanda | Eijkman Institute for Molecular Biology, Ministry of Research and Technology/National Agency for Research and Innovation | Lydia V. Panggalo, Sukma Oktavianthi, Willy Agustine, Edison Johar, Hidayat Trimarsanto, Iskandar Adnan, Frilasita A Yudhaputri, Safarina G Malik, Khin Saw Myint, Amin Soebandrio |
| EPI_ISL_888997 | RSU Asysyifaa | Eijkman Institute for Molecular Biology, Ministry of Research and Technology/National Agency for Research and Innovation | Sukma Oktavianthi, Willy Agustine, Edison Johar, Hidayat Trimarsanto, Iskandar Adnan, Lydia V. Panggalo, Frilasita A Yudhaputri, Safarina G Malik, Khin Saw Myint, Amin Soebandrio |
| EPI_ISL_888998 | RSUD Cileungsi | Eijkman Institute for Molecular Biology, Ministry of Research and Technology/National Agency for Research and Innovation | Sukma Oktavianthi, Willy Agustine, Edison Johar, Hidayat Trimarsanto, Iskandar Adnan, Lydia V. Panggalo, Frilasita A Yudhaputri, Safarina G Malik, Khin Saw Myint, Amin Soebandrio |
| EPI_ISL_888999 | RS Mitra Keluarga Cibubur | Eijkman Institute for Molecular Biology, Ministry of Research and Technology/National Agency for Research and Innovation | Sukma Oktavianthi, Willy Agustine, Edison Johar, Hidayat Trimarsanto, Iskandar Adnan, Lydia V. Panggalo, Frilasita A Yudhaputri, Safarina G Malik, Khin Saw Myint, Amin Soebandrio |
| EPI_ISL_889000 | RSUD Cileungsi | Eijkman Institute for Molecular Biology, Ministry of Research and Technology/National Agency for Research and Innovation | Sukma Oktavianthi, Willy Agustine, Edison Johar, Hidayat Trimarsanto, Iskandar Adnan, Lydia V. Panggalo, Frilasita A Yudhaputri, Safarina G Malik, Khin Saw Myint, Amin Soebandrio |
| EPI_ISL_889001 | RSU Azra | Eijkman Institute for Molecular Biology, Ministry of Research and Technology/National Agency for Research and Innovation | Sukma Oktavianthi, Willy Agustine, Edison Johar, Hidayat Trimarsanto, Iskandar Adnan, Lydia V. Panggalo, Frilasita A Yudhaputri, Safarina G Malik, Khin Saw Myint, Amin Soebandrio |
| EPI_ISL_889003, EPI_ISL_889004 | RS Mitra Keluarga Gading Serpong | Eijkman Institute for Molecular Biology, Ministry of Research and Technology/National Agency for Research and Innovation | Willy Agustine, Edison Johar, Hidayat Trimarsanto, Iskandar Adnan, Lydia V. Panggalo, Sukma Oktavianthi, Frilasita A Yudhaputri, Safarina G Malik, Khin Saw Myint, Amin Soebandrio |
| EPI_ISL_889005 | Eijkman Institute for Molecular Biology, Ministry of Research and Technology/National Agency for Research and Innovation | Eijkman Institute for Molecular Biology, Ministry of Research and Technology/National Agency for Research and Innovation | Willy Agustine, Edison Johar, Hidayat Trimarsanto, Iskandar Adnan, Lydia V. Panggalo, Sukma Oktavianthi, Frilasita A Yudhaputri, Safarina G Malik, Khin Saw Myint, Amin Soebandrio |
| EPI_ISL_889006 | RSU Harapan Bunda | Eijkman Institute for Molecular Biology, Ministry of Research and Technology/National Agency for Research and Innovation | Willy Agustine, Edison Johar, Hidayat Trimarsanto, Iskandar Adnan, Lydia V. Panggalo, Sukma Oktavianthi, Frilasita A Yudhaputri, Safarina G Malik, Khin Saw Myint, Amin Soebandrio |
| EPI_ISL_889007 | RSUD Cileungsi | Eijkman Institute for Molecular Biology, Ministry of Research | Willy Agustine, Edison Johar, Hidayat Trimarsanto, Iskandar Adnan, Lydia V. Panggalo, Sukma Oktavianthi, Frilasita A Yudhaputri, Safarina G Malik, Khin Saw |

| | | | |
|---|---|---|---|
| | | and Technology/National Agency for Research and Innovation | Myint, Amin Soebandrio |
| EPI_ISL_889009 | RS Mitra Keluarga Bintaro | Eijkman Institute for Molecular Biology, Ministry of Research and Technology/National Agency for Research and Innovation | Edison Johar, Frilasita A Yudhaputri, Hidayat Trimarsanto, Iskandar Adnan, Lydia V. Panggalo, Sukma Oktavianthi, Willy Agustine, Safarina G Malik, Khin Saw Myint, Amin Soebandrio |
| EPI_ISL_889011 | RSU Permata Cibubur | Eijkman Institute for Molecular Biology, Ministry of Research and Technology/National Agency for Research and Innovation | Hidayat Trimarsanto, Iskandar Adnan, Lydia V. Panggalo, Sukma Oktavianthi, Willy Agustine, Edison Johar, Frilasita A Yudhaputri, Safarina G Malik, Khin Saw Myint, Amin Soebandrio |
| EPI_ISL_889012 | RS Mitra Keluarga Pratama Jatiasih | Eijkman Institute for Molecular Biology, Ministry of Research and Technology/National Agency for Research and Innovation | Hidayat Trimarsanto, Iskandar Adnan, Lydia V. Panggalo, Sukma Oktavianthi, Willy Agustine, Edison Johar, Frilasita A Yudhaputri, Safarina G Malik, Khin Saw Myint, Amin Soebandrio |
| EPI_ISL_889013 | RS Rawa Lumbu | Eijkman Institute for Molecular Biology, Ministry of Research and Technology/National Agency for Research and Innovation | Iskandar Adnan, Lydia V. Panggalo, Sukma Oktavianthi, Willy Agustine, Edison Johar, Hidayat Trimarsanto, Frilasita A Yudhaputri, Safarina G Malik, Khin Saw Myint, Amin Soebandrio |
| EPI_ISL_889014 | RSU Medika Dramaga | Eijkman Institute for Molecular Biology, Ministry of Research and Technology/National Agency for Research and Innovation | Iskandar Adnan, Lydia V. Panggalo, Sukma Oktavianthi, Willy Agustine, Edison Johar, Hidayat Trimarsanto, Frilasita A Yudhaputri, Safarina G Malik, Khin Saw Myint, Amin Soebandrio |
| EPI_ISL_889016 | RS Mitra Husada | Eijkman Institute for Molecular Biology, Ministry of Research and Technology/National Agency for Research and Innovation | Iskandar Adnan, Lydia V. Panggalo, Sukma Oktavianthi, Willy Agustine, Edison Johar, Hidayat Trimarsanto, Frilasita A Yudhaputri, Safarina G Malik, Khin Saw Myint, Amin Soebandrio |
| EPI_ISL_889017 | RS Hermina Tangerang | Eijkman Institute for Molecular Biology, Ministry of Research and Technology/National Agency for Research and Innovation | Lydia V. Panggalo, Sukma Oktavianthi, Willy Agustine, Edison Johar, Hidayat Trimarsanto, Iskandar Adnan, Frilasita A Yudhaputri, Safarina G Malik, Khin Saw Myint, Amin Soebandrio |
| EPI_ISL_889018 | RSU Medika Dramaga | Eijkman Institute for Molecular Biology, Ministry of Research and Technology/National Agency for Research and Innovation | Lydia V. Panggalo, Sukma Oktavianthi, Willy Agustine, Edison Johar, Hidayat Trimarsanto, Iskandar Adnan, Frilasita A Yudhaputri, Safarina G Malik, Khin Saw Myint, Amin Soebandrio |
| EPI_ISL_889019 | RSIA Sayyidah | Eijkman Institute for Molecular Biology, Ministry of Research and Technology/National Agency for Research and Innovation | Lydia V. Panggalo, Sukma Oktavianthi, Willy Agustine, Edison Johar, Hidayat Trimarsanto, Iskandar Adnan, Frilasita A Yudhaputri, Safarina G Malik, Khin Saw Myint, Amin Soebandrio |
| EPI_ISL_889020 | RS Omni Cikarang | Eijkman Institute for Molecular Biology, Ministry of Research and Technology/National Agency for Research and Innovation | Edison Johar, Frilasita A Yudhaputri, Hidayat Trimarsanto, Iskandar Adnan, Lydia V. Panggalo, Sukma Oktavianthi, Willy Agustine, Safarina G Malik, Khin Saw Myint, Amin Soebandrio |
| EPI_ISL_889021 | RSIA Kenari Graha Medika | Eijkman Institute for Molecular Biology, Ministry of Research and Technology/National Agency for Research and Innovation | Hidayat Trimarsanto, Iskandar Adnan, Lydia V. Panggalo, Sukma Oktavianthi, Willy Agustine, Edison Johar, Frilasita A Yudhaputri, Safarina G Malik, Khin Saw Myint, Amin Soebandrio |
| EPI_ISL_889022 | RS Anna | Eijkman Institute for Molecular Biology, Ministry of Research and Technology/National Agency for Research and Innovation | Hidayat Trimarsanto, Iskandar Adnan, Lydia V. Panggalo, Sukma Oktavianthi, Willy Agustine, Edison Johar, Frilasita A Yudhaputri, Safarina G Malik, Khin Saw Myint, Amin Soebandrio |
| EPI_ISL_889398, EPI_ISL_889399, EPI_ISL_889400, EPI_ISL_889401, EPI_ISL_889402, EPI_ISL_889403, EPI_ISL_889404, EPI_ISL_889418, EPI_ISL_889424, EPI_ISL_889432, EPI_ISL_889436 | | | |
| see above | Olomouc University Hospital | Institute of Applied Biotechnologies a.s. | Petr Klempt, Ondej Brzo, Martin Kašný, Kateina Kvapilová, Marián Hajdúch, Petr Kvapil |
| EPI_ISL_890185 | Academic Hospital of Gadjah Mada University (RSA UGM) | Genetics Working Group (Pokja Genetik) Faculty of Medicine, Public Health and Nursing Universitas Gadjah Mada (FK-KMK UGM); Disease Investigation Center Wates Ministry of Agriculture Indonesia; Department of Microbiology FK-KMK UGM; Laboratorium Diagnostik Yayasan Tahija World Mosquito Program (WMP) Yogyakarta Center for Tropical Medicine FK-KMK UGM; Integrated Research center FK-KMK UGM; Department of Computer Science and Electronics FMIPA UGM | Gunadi, Hendra Wibawa, Marcellus, Mohamad S. Hakim, Edwin W. Daniwijaya, Ludhang P. Rizki, Endah Supriyati, Eggi Arguni, Titik Nuryastuti, Tri Wibawa, Dwi AA Nugrahaningsih, Afiahayati, Siswanto, Kristy Iskandar, Nungki Anggorowati, Aditya Rifqi Fauzi, Fadil Fahri |
| EPI_ISL_890186 | Academic Hospital of Gadjah Mada University (RSA UGM) | Genetics Working Group (Pokja Genetik) Faculty of Medicine, Public Health and Nursing Universitas Gadjah Mada (FK-KMK UGM); Disease Investigation Center Wates Ministry of Agriculture Indonesia; Department of Microbiology FK-KMK UGM; Laboratorium Diagnostik Yayasan Tahija World Mosquito Program (WMP) Yogyakarta Center for Tropical Medicine FK-KMK UGM; Integrated Research center FK-KMK UGM; Department of Computer Science and Electronics FMIPA UGM | Gunadi, Hendra Wibawa, Marcellus, Mohamad S. Hakim, Edwin W. Daniwijaya, Ludhang P. Rizki, Endah Supriyati, Eggi Arguni, Titik Nuryastuti, Tri Wibawa, Dwi AA Nugrahaningsih, Afiahayati, Siswanto, Kristy Iskandar, Nungki Anggorowati, Audric Kenny Tedja, Maria Patricia Inggriani |
| EPI_ISL_890187 | Academic Hospital of Gadjah Mada University (RSA UGM) | Genetics Working Group (Pokja Genetik) Faculty of Medicine, Public Health and Nursing Universitas Gadjah Mada (FK-KMK UGM); Disease Investigation Center Wates Ministry of Agriculture Indonesia; Department of Microbiology FK-KMK UGM; Laboratorium Diagnostik Yayasan Tahija World Mosquito Program (WMP) Yogyakarta Center for Tropical Medicine FK-KMK UGM; Integrated Research center FK-KMK UGM; Department of Computer Science and Electronics FMIPA UGM | Gunadi, Hendra Wibawa, Marcellus, Mohamad S. Hakim, Edwin W. Daniwijaya, Ludhang P. Rizki, Endah Supriyati, Eggi Arguni, Titik Nuryastuti, Tri Wibawa, Dwi AA Nugrahaningsih, Afiahayati, Siswanto, Kristy Iskandar, Nungki Anggorowati, Untung Riawan, William Widitjiarso |
| EPI_ISL_890352, EPI_ISL_890353 | Labo Analyses Med | National Reference Center for Viruses of Respiratory Infections, Institut Pasteur, Paris | Marion Barbet, Sylvie Behillil, Méline Bizard, Angela Brisebarre, Camille Capel, Etienne Simon-Lorière, Vincent Enouf, Maud Vanpeene, Sylvie van der Werf,Rousset Dominique |
| EPI_ISL_890354 | Labo Analyses Med | National Reference Center for Viruses of Respiratory Infections, Institut Pasteur, Paris | Marion Barbet, Sylvie Behillil, Méline Bizard, Angela Brisebarre, Camille Capel, Etienne Simon-Lorière, Vincent Enouf, Maud Vanpeene, Sylvie van der Werf,Coignard Catherine |
| EPI_ISL_890355, EPI_ISL_890356, EPI_ISL_890357, EPI_ISL_890358 | Hospital | National Reference Center for Viruses of Respiratory Infections, Institut Pasteur, Paris | Marion Barbet, Sylvie Behillil, Méline Bizard, Angela Brisebarre, Camille Capel, Etienne Simon-Lorière, Vincent Enouf, Maud Vanpeene, Sylvie van der Werf,Combe Patrice |
| EPI_ISL_890359 | Hospital | National Reference Center for Viruses of Respiratory Infections, Institut Pasteur, Paris | Marion Barbet, Sylvie Behillil, Méline Bizard, Angela Brisebarre, Camille Capel, Etienne Simon-Lorière, Vincent Enouf, Maud Vanpeene, Sylvie van der Werf,Guigon AuréLie |
| EPI_ISL_890891 | Hospital | National Reference Center for Viruses of Respiratory | Marion Barbet, Sylvie Behillil, Méline Bizard, Angela Brisebarre, Camille Capel, Etienne Simon-Lorière, Vincent Enouf, Maud Vanpeene, Sylvie van der Werf, |

| | | | |
|---|---|---|---|
| | | Infections, Institut Pasteur, Paris | Patricia Stoessel-Thouvenin |
| EPI_ISL_890892 | Hospital | National Reference Center for Viruses of Respiratory Infections, Institut Pasteur, Paris | Marion Barbet, Sylvie Behillil, Méline Bizard, Angela Brisebarre, Camille Capel, Etienne Simon-Lorière, Vincent Enouf, Maud Vanpeene, Sylvie van der Werf, Léa Pilorge |
| EPI_ISL_890893 | Hospital | National Reference Center for Viruses of Respiratory Infections, Institut Pasteur, Paris | Marion Barbet, Sylvie Behillil, Méline Bizard, Angela Brisebarre, Camille Capel, Etienne Simon-Lorière, Vincent Enouf, Maud Vanpeene, Sylvie van der Werf, Patricia Stoessel-Thouvenin |
| EPI_ISL_890895 | Laboratory Analyses Med | National Reference Center for Viruses of Respiratory Infections, Institut Pasteur, Paris | Marion Barbet, Sylvie Behillil, Méline Bizard, Angela Brisebarre, Camille Capel, Etienne Simon-Lorière, Vincent Enouf, Maud Vanpeene, Sylvie van der Werf, Céline Menaut |
| EPI_ISL_891151 | Academic Hospital of Gadjah Mada University (RSA UGM) | Genetics Working Group (Pokja Genetik) Faculty of Medicine, Public Health and Nursing Universitas Gadjah Mada (FK-KMK UGM); Disease Investigation Center Wates Ministry of Agriculture Indonesia; Department of Microbiology FK-KMK UGM; Laboratorium Diagnostik Yayasan Tahija World Mosquito Program (WMP) Yogyakarta Center for Tropical Medicine FK-KMK UGM; Integrated Research center FK-KMK UGM; Department of Computer Science and Electronics FMIPA UGM | Gunadi, Hendra Wibawa, Marcellus, Mohamad S. Hakim, Edwin W. Daniwijaya, Ludhang P. Rizki, Endah Supriyati, Eggi Arguni, Titik Nuryastuti, Tri Wibawa, Dwi AA Nugrahaningsih, Afiahayati, Siswanto, Kristy Iskandar, Nungki Anggorowati, Alvin Santoso Kalim, Susan Simanjaya |
| EPI_ISL_891152 | SMS Medical College Jaipur | SMS Medical College Jaipur | Bharti Malhotra, Swati Gautam,Himanshu Sharma, Pratibha Sharma, Neha Bhomia, Nivedita Gupta, Pragya Yadav, Varsha Potdar |
| EPI_ISL_891207 | SMS Medical College Jaipur | SMS Medical College jaipur | Bharti Malhotra, Swati Gautam, Himanshu Sharma,Pratibha Sharma, Neha Bhomia, Nivedita Gupta, Pragya Yadav, Varsha Potdar |
| EPI_ISL_891210 | SMS Medical College jaipur | SMS Medical College Jaipur | Bharti Malhotra, Swati Gautam, Himanshu Sharma, Pratibha Sharma, Neha Bhomia, Nivedita Gupta , Pragya Yadav,Varsha Potdar |
| EPI_ISL_891211 | SMS Medical College Jaipur | SMS Medical College jaipur | Bharti Malhotra, Swati Gautam, Himanshu Sharma, Pratibha Sharma, Neha Bhomia, Nivedita Gupta, Pragya Yadav, Varsha Potdar |
| EPI_ISL_891213 | SMS Medical College Jaipur | SMS Medical College Jaipur | Bharti Malhotra, Swati Gautam, Himanshu Sharma, Pratibha Sharma, Neha Bhomia, Nivedita Gupta, Pragya Yadav, Varsha Potdar |
| EPI_ISL_891214 | SMS Medical College Jaipur | SMS Medical College Jaipur | Bharti Malhotra, Swati Gautam, Himanshu Sharma, Pratibha Sharma, Neha Bhomia, Nivedita Gupta, Pragya Yadav,Varsha Potdar |
| EPI_ISL_891219 | SMS Medical College Jaipur | SMS Medical College Jaipur | Bharti Malhotra, Swati Gautam, Himanshu Sharma, Pratibha Sharma, Neha Bhomia, Nivedita Gupta, Pragya Yadav, Varsha Potdar |
| EPI_ISL_894219 | Hopital | National Reference Center for Viruses of Respiratory Infections, Institut Pasteur, Paris | Marion Barbet, Sylvie Behillil, Méline Bizard, Angela Brisebarre, Camille Capel, Etienne Simon-Lorière, Vincent Enouf, Maud Vanpeene, Sylvie van der Werf,Guigon AuréLie |
| EPI_ISL_894220, EPI_ISL_894221, EPI_ISL_894222 | CH de Mayotte - Laboratoire de Biologie | National Reference Center for Viruses of Respiratory Infections, Institut Pasteur, Paris | Marion Barbet, Sylvie Behillil, Méline Bizard, Angela Brisebarre, Camille Capel, Etienne Simon-Lorière, Vincent Enouf, Maud Vanpeene, Sylvie van der Werf,Combe Patrice |
| EPI_ISL_894223 | Labo Analyses med | National Reference Center for Viruses of Respiratory Infections, Institut Pasteur, Paris | Marion Barbet, Sylvie Behillil, Méline Bizard, Angela Brisebarre, Camille Capel, Etienne Simon-Lorière, Vincent Enouf, Maud Vanpeene, Sylvie van der Werf,Amzalag Jonas |
| EPI_ISL_894224, EPI_ISL_894225, EPI_ISL_894226, EPI_ISL_894227, EPI_ISL_894228, EPI_ISL_894229, EPI_ISL_894230, EPI_ISL_894231 | CH de Mayotte - Laboratoire de Biologie | National Reference Center for Viruses of Respiratory Infections, Institut Pasteur, Paris | Marion Barbet, Sylvie Behillil, Méline Bizard, Angela Brisebarre, Camille Capel, Etienne Simon-Lorière, Vincent Enouf, Maud Vanpeene, Sylvie van der Werf,Combe Patrice |
| EPI_ISL_894232 | Labo Analyses med | National Reference Center for Viruses of Respiratory Infections, Institut Pasteur, Paris | Marion Barbet, Sylvie Behillil, Méline Bizard, Angela Brisebarre, Camille Capel, Etienne Simon-Lorière, Vincent Enouf, Maud Vanpeene, Sylvie van der Werf,Amzalag Jonas |
| EPI_ISL_900492, EPI_ISL_900493, EPI_ISL_900494 | Althaia. Xarxa Assistencial Universitària de Manresa | IrsiCaixa - Can Ruti CovidSeq | Fundació irsiCaixa. Hospital Universitari Germans Trias i Pujol(HUGTiP), 2a planta, maternal Ctra Canyet s/n, Badalona Gloria Trujillo, Rafel Perez Vidal, Jaume Trape Pujol, Carolina Gonzalez Fernandez, Roger Paredes, Eulalia Grau, Bonaventura Clotet |
| EPI_ISL_902737 | RSA UGM | Genetics Working Group (Pokja Genetik) Faculty of Medicine, Public Health and Nursing Universitas Gadjah Mada (FK-KMK UGM); Disease Investigation Center Wates Ministry of Agriculture Indonesia; Department of Microbiology FK-KMK UGM; Laboratorium Diagnostik Yayasan Tahija World Mosquito Program (WMP) Yogyakarta Center for Tropical Medicine FK-KMK UGM; Integrated Research center FK-KMK UGM; Department of Computer Science and Electronics FMIPA UGM | Gunadi, Hendra Wibawa, Marcellus, Mohamad S. Hakim, Edwin W. Daniwijaya, Ludhang P. Rizki, Endah Supriyati, Eggi Arguni, Titik Nuryastuti, Tri Wibawa, Dwi AA Nugrahaningsih, Afiahayati, Siswanto, Kristy Iskandar, Nungki Anggorowati, Aditya Rifqi Fauzi, Fadil Fahri |
| EPI_ISL_902749 | RSA UGM | Genetics Working Group (Pokja Genetik) Faculty of Medicine, Public Health and Nursing Universitas Gadjah Mada (FK-KMK UGM); Disease Investigation Center Wates Ministry of Agriculture Indonesia; Department of Microbiology FK-KMK UGM; Laboratorium Diagnostik Yayasan Tahija World Mosquito Program (WMP) Yogyakarta Center for Tropical Medicine FK-KMK UGM; Integrated Research center FK-KMK UGM; Department of Computer Science and Electronics FMIPA UGM | Gunadi, Hendra Wibawa, Marcellus, Mohamad S. Hakim, Edwin W. Daniwijaya, Ludhang P. Rizki, Endah Supriyati, Eggi Arguni, Titik Nuryastuti, Tri Wibawa, Dwi AA Nugrahaningsih, Afiahayati, Siswanto, Kristy Iskandar, Nungki Anggorowati, Dyah Ayu Puspitarani, Dwiki afandy |
| EPI_ISL_902924 | Tanjungpura University Hospital | Tanjungpura University Hospital | Andriani; Mahyarudin; Virhan Novianry; Delima Fajar Liana; Sofi Siti Shofiyah; Puji Astuti ; Muhammad Ibnu Kahtan; Ambar Rialita; Eka Ardiani Putri; Wiwik Windarti; Helmi Sastriawan |
| EPI_ISL_902926 | SC (UCO) Igiene e Sanità Pubblica (funzione integrata con SC Microbiologia e Virologia) e Laboratory of Molecular Virology of the International Centre for Genetic Engineering and Biotechnology (ICGEB) | ARGO Laboratorio Genomica ed Epigenomica | Licastro D, Dal Monego S, Degasperi M, Marcello A, D'Agaro P |
| EPI_ISL_903383, EPI_ISL_903388, EPI_ISL_903389 | MOH - Jaber Al-Ahmad Hospital (Innovation Research Laboratory) | MOH - Jaber Al-Ahmad Hospital (Innovation Research Laboratory) | Salman Al-Sabah , Mohammad Alghounaim |
| EPI_ISL_905731 | Academic Hospital of Gadjah Mada University (RSA UGM) | Genetics Working Group (Pokja Genetik) Faculty of Medicine, Public Health and Nursing Universitas Gadjah Mada (FK-KMK UGM); Disease Investigation Center Wates Ministry of Agriculture Indonesia; Department of Microbiology FK-KMK UGM; Laboratorium Diagnostik Yayasan Tahija World Mosquito Program (WMP) Yogyakarta Center for Tropical Medicine FK-KMK UGM; Integrated Research center FK-KMK UGM; Department of Computer Science and Electronics FMIPA UGM | Gunadi, Hendra Wibawa, Marcellus, Mohamad S. Hakim, Edwin W. Daniwijaya, Ludhang P. Rizki, Endah Supriyati, Eggi Arguni, Titik Nuryastuti, Tri Wibawa, Dwi AA Nugrahaningsih, Afiahayati, Siswanto, Kristy Iskandar,Nungki Anggorowati, Dyah Ayu Puspitarani, Dwiki afandy |
| EPI_ISL_906050 | RSA UGM | Genetics Working Group (Pokja Genetik) Faculty of Medicine, Public Health and Nursing Universitas Gadjah Mada (FK-KMK UGM); Disease Investigation Center Wates Ministry | Gunadi, Hendra Wibawa, Marcellus, Mohamad S. Hakim, Edwin W. Daniwijaya, Ludhang P. Rizki, Endah Supriyati, Eggi Arguni, Titik Nuryastuti, Tri Wibawa, Dwi AA Nugrahaningsih, Afiahayati, Siswanto,Kristy Iskandar,Nungki Anggorowati, Susan Simanjaya, Kemala Athollah |

| | | | |
|---|---|---|---|
| | | of Agriculture Indonesia; Department of Microbiology FK-KMK UGM; Laboratorium Diagnostik Yayasan Tahija World Mosquito Program (WMP) Yogyakarta Center for Tropical Medicine FK-KMK UGM; Integrated Research center FK-KMK UGM; Department of Computer Science and Electronics FMIPA UGM | |
| EPI_ISL_906051 | RSA UGM | Genetics Working Group (Pokja Genetik) Faculty of Medicine, Public Health and Nursing Universitas Gadjah Mada (FK-KMK UGM); Disease Investigation Center Wates Ministry of Agriculture Indonesia; Department of Microbiology FK-KMK UGM; Laboratorium Diagnostik Yayasan Tahija World Mosquito Program (WMP) Yogyakarta Center for Tropical Medicine FK-KMK UGM; Integrated Research center FK-KMK UGM; Department of Computer Science and Electronics FMIPA UGM | Gunadi, Hendra Wibawa, Marcellus, Mohamad S. Hakim, Edwin W. Daniwijaya, Ludhang P. Rizki, Endah Supriyati, Eggi Arguni, Titik Nuryastuti, Tri Wibawa, Dwi AA Nugrahaningsih, Afiahayati, Siswanto, Kristy Iskandar, Nungki Anggorowati, Alvin Santoso Kalim, Dwiki Afandy |
| EPI_ISL_906052 | RSA UGM | Genetics Working Group (Pokja Genetik) Faculty of Medicine, Public Health and Nursing Universitas Gadjah Mada (FK-KMK UGM); Disease Investigation Center Wates Ministry of Agriculture Indonesia; Department of Microbiology FK-KMK UGM; Laboratorium Diagnostik Yayasan Tahija World Mosquito Program (WMP) Yogyakarta Center for Tropical Medicine FK-KMK UGM; Integrated Research center FK-KMK UGM; Department of Computer Science and Electronics FMIPA UGM | Gunadi, Hendra Wibawa, Marcellus, Mohamad S. Hakim, Edwin W. Daniwijaya, Ludhang P. Rizki, Endah Supriyati, Eggi Arguni, Titik Nuryastuti, Tri Wibawa, Dwi AA Nugrahaningsih, Afiahayati, Siswanto, Kristy Iskandar, Nungki Anggorowati, William Widitjiarso, Untung Riawan |
| EPI_ISL_906054 | Tilia Laboratories s.r.o. | Tilia Laboratories s.r.o. | Sona Pekova, MD, PhD. |
| EPI_ISL_906055 | Tilia Laboratories s.r.o. | Tilia Laboratories s.r.o. | Sona Pekova MD, PhD. |
| EPI_ISL_906056, EPI_ISL_906057, EPI_ISL_906058, EPI_ISL_906059, EPI_ISL_906061, EPI_ISL_906062, EPI_ISL_906063, EPI_ISL_906064 | Tilia Laboratories s.r.o. | Tilia Laboratories s.r.o. | Sona Pekova, MD, PhD. |
| EPI_ISL_906068, EPI_ISL_906069 | Instituto Adolfo Lutz - Regional de Campinas | Instituto Adolfo Lutz, Interdiciplinary Procedures Center, Strategic Laboratory | Claudio Tavares Sacchi, Claudia Regina Gonçalves, Erica Valessa Ramos Gomes, Karoline Rodrigues Campos |
| EPI_ISL_909656, EPI_ISL_909657, EPI_ISL_909658, EPI_ISL_909659, EPI_ISL_909660, EPI_ISL_909661, EPI_ISL_909662, EPI_ISL_909663, EPI_ISL_909664 | Labo Analyses med | National Reference Center for Viruses of Respiratory Infections, Institut Pasteur, Paris | Marion Barbet, Sylvie Behillil, Méline Bizard, Angela Brisebarre, Camille Capel, Etienne Simon-Lorière, Vincent Enouf, Maud Vanpeene, Sylvie van der Werf,Amzalag Jonas |
| EPI_ISL_909665 | Labo Analyses Med | National Reference Center for Viruses of Respiratory Infections, Institut Pasteur, Paris | Marion Barbet, Sylvie Behillil, Méline Bizard, Angela Brisebarre, Camille Capel, Etienne Simon-Lorière, Vincent Enouf, Maud Vanpeene, Sylvie van der Werf,Le Vicky |
| EPI_ISL_909666 | Hopital Avicenne. Laboratoire central de bactériologie virologie hygiène | National Reference Center for Viruses of Respiratory Infections, Institut Pasteur, Paris | Marion Barbet, Sylvie Behillil, Méline Bizard, Angela Brisebarre, Camille Capel, Etienne Simon-Lorière, Vincent Enouf, Maud Vanpeene, Sylvie van der Werf,Brichler Ségolène |
| EPI_ISL_909667, EPI_ISL_909668 | Labo Analyses Med | National Reference Center for Viruses of Respiratory Infections, Institut Pasteur, Paris | Marion Barbet, Sylvie Behillil, Méline Bizard, Angela Brisebarre, Camille Capel, Etienne Simon-Lorière, Vincent Enouf, Maud Vanpeene, Sylvie van der Werf,Le Vicky |
| EPI_ISL_909669, EPI_ISL_909670, EPI_ISL_909671, EPI_ISL_909672, EPI_ISL_909673 | Labo Analyses med | National Reference Center for Viruses of Respiratory Infections, Institut Pasteur, Paris | Marion Barbet, Sylvie Behillil, Méline Bizard, Angela Brisebarre, Camille Capel, Etienne Simon-Lorière, Vincent Enouf, Maud Vanpeene, Sylvie van der Werf,Amzalag Jonas |
| EPI_ISL_909674 | CH de Mayotte - Laboratoire de Biologie | National Reference Center for Viruses of Respiratory Infections, Institut Pasteur, Paris | Marion Barbet, Sylvie Behillil, Méline Bizard, Angela Brisebarre, Camille Capel, Etienne Simon-Lorière, Vincent Enouf, Maud Vanpeene, Sylvie van der Werf,Combe Patrice |
| EPI_ISL_909675, EPI_ISL_909676, EPI_ISL_909677, EPI_ISL_909678, EPI_ISL_909679, EPI_ISL_909680, EPI_ISL_909681 | Labo Analyses Med | National Reference Center for Viruses of Respiratory Infections, Institut Pasteur, Paris | Marion Barbet, Sylvie Behillil, Méline Bizard, Angela Brisebarre, Camille Capel, Etienne Simon-Lorière, Vincent Enouf, Maud Vanpeene, Sylvie van der Werf,Le Vicky |
| EPI_ISL_909682, EPI_ISL_909683 | Hopital Avicenne. Laboratoire central de bactériologie virologie hygiène | National Reference Center for Viruses of Respiratory Infections, Institut Pasteur, Paris | Marion Barbet, Sylvie Behillil, Méline Bizard, Angela Brisebarre, Camille Capel, Etienne Simon-Lorière, Vincent Enouf, Maud Vanpeene, Sylvie van der Werf,Brichler Ségolène |
| EPI_ISL_909684, EPI_ISL_909685 | Virologie - Laboratoire de Microbiologie | National Reference Center for Viruses of Respiratory Infections, Institut Pasteur, Paris | Marion Barbet, Sylvie Behillil, Méline Bizard, Angela Brisebarre, Camille Capel, Etienne Simon-Lorière, Vincent Enouf, Maud Vanpeene, Sylvie van der Werf,Schvoerer Evelyne |
| EPI_ISL_909686 | GH Cochin-Broca Hôtel Dieu/Lab Virologie | National Reference Center for Viruses of Respiratory Infections, Institut Pasteur, Paris | Marion Barbet, Sylvie Behillil, Méline Bizard, Angela Brisebarre, Camille Capel, Etienne Simon-Lorière, Vincent Enouf, Maud Vanpeene, Sylvie van der Werf,L'Honneur Anne-Sophie |
| EPI_ISL_909687, EPI_ISL_909688, EPI_ISL_909689, EPI_ISL_909690, EPI_ISL_909691, EPI_ISL_909692, EPI_ISL_909693, EPI_ISL_909694, EPI_ISL_909695, EPI_ISL_909696, EPI_ISL_909697, EPI_ISL_909698, EPI_ISL_909699, EPI_ISL_909700, EPI_ISL_909701 | | | |
| see above | CH de Mayotte - Laboratoire de Biologie | National Reference Center for Viruses of Respiratory Infections, Institut Pasteur, Paris | Marion Barbet, Sylvie Behillil, Méline Bizard, Angela Brisebarre, Camille Capel, Etienne Simon-Lorière, Vincent Enouf, Maud Vanpeene, Sylvie van der Werf,Combe Patrice |
| EPI_ISL_909702 | CHU de Nantes - Hôtel Dieu. Laboratoire de Virologie | National Reference Center for Viruses of Respiratory Infections, Institut Pasteur, Paris | Marion Barbet, Sylvie Behillil, Méline Bizard, Angela Brisebarre, Camille Capel, Etienne Simon-Lorière, Vincent Enouf, Maud Vanpeene, Sylvie van der Werf,Bressollette CéLine |
| EPI_ISL_909703, EPI_ISL_909704, EPI_ISL_909705, EPI_ISL_909706 | Labo Analyses med | National Reference Center for Viruses of Respiratory Infections, Institut Pasteur, Paris | Marion Barbet, Sylvie Behillil, Méline Bizard, Angela Brisebarre, Camille Capel, Etienne Simon-Lorière, Vincent Enouf, Maud Vanpeene, Sylvie van der Werf,Besson J |
| EPI_ISL_909707 | Labo Analyses Med | National Reference Center for Viruses of Respiratory Infections, Institut Pasteur, Paris | Marion Barbet, Sylvie Behillil, Méline Bizard, Angela Brisebarre, Camille Capel, Etienne Simon-Lorière, Vincent Enouf, Maud Vanpeene, Sylvie van der Werf |
| EPI_ISL_909708, EPI_ISL_909709, EPI_ISL_909710, EPI_ISL_909711, EPI_ISL_909712, EPI_ISL_909713, EPI_ISL_909714, EPI_ISL_909715, EPI_ISL_909716, EPI_ISL_909717, EPI_ISL_909718, EPI_ISL_909719, EPI_ISL_909720, EPI_ISL_909721, EPI_ISL_909722, EPI_ISL_909723 | | | |
| see above | CH de Mayotte - Laboratoire de Biologie | National Reference Center for Viruses of Respiratory Infections, Institut Pasteur, Paris | Marion Barbet, Sylvie Behillil, Méline Bizard, Angela Brisebarre, Camille Capel, Etienne Simon-Lorière, Vincent Enouf, Maud Vanpeene, Sylvie van der Werf,Combe Patrice |
| EPI_ISL_909724, EPI_ISL_909725, EPI_ISL_909726, EPI_ISL_909727, EPI_ISL_909728, EPI_ISL_909729, EPI_ISL_909730, EPI_ISL_909731 | Labo Analyses Med | National Reference Center for Viruses of Respiratory Infections, Institut Pasteur, Paris | Marion Barbet, Sylvie Behillil, Méline Bizard, Angela Brisebarre, Camille Capel, Etienne Simon-Lorière, Vincent Enouf, Maud Vanpeene, Sylvie van der Werf,Nizard Patrice |
| EPI_ISL_909732 | Pôle médico-technique Laboratoire | National Reference Center for Viruses of Respiratory | Marion Barbet, Sylvie Behillil, Méline Bizard, Angela Brisebarre, Camille Capel, Etienne Simon-Lorière, Vincent Enouf, Maud Vanpeene, Sylvie van der Werf |

| | | | |
|---|---|---|---|
| | | Infections, Institut Pasteur, Paris | |
| EPI_ISL_909733, EPI_ISL_909734, EPI_ISL_909735 | Labo Analyses Med | National Reference Center for Viruses of Respiratory Infections, Institut Pasteur, Paris | Marion Barbet, Sylvie Behillil, Méline Bizard, Angela Brisebarre, Camille Capel, Etienne Simon-Lorière, Vincent Enouf, Maud Vanpeene, Sylvie van der Werf |
| EPI_ISL_909772 | National Institute of Health Research and Development | National Institute of Health Research and Development | Holy Arif Wibowo; Ririn Ramadhany; Yuni Rukminiati; Agustiningsih; Kindi Adam; Hana Apsari Pawestri; Subangkit; Kartika Dewi Puspa; Arie Ardiansyah Nugraha; Hartanti Dian Ikawati; Krisna Nur Andriana Pangesti; Triyani Soekarso; Ni Ketut Susilarini; Nur Ika Hariastuti; Uli Alfi Nikmah; Mursinah; Asri Febriyani; Reni Herman; Nike Susanti; Herna; Tati Febriyanti; Juwita Kurniawati; Natalie Laurencia Kipuw; Fauzul Muna; Irene Lorinda Indalao; Aulia Rizki; Nelly Puspandari; vivi Setiawaty |
| EPI_ISL_909948, EPI_ISL_909949, EPI_ISL_909950, EPI_ISL_909951, EPI_ISL_909952 | CUSL/UCLouvain COVID testing federal platform | UCLouvain/IREC/MBLG | Jean Ruelle, Lysa Pinsmaye, Benoit Kabamba Mukadi |
| EPI_ISL_910016, EPI_ISL_910029 | Laboratory for Respiratory Viruses, Cantacuzino National Military-Medical Institute for Research and Development | Cantacuzino Institute Virology | Luiza Ustea, Nicoleta Paraschiv, Mihaela Lazar |
| EPI_ISL_910222, EPI_ISL_910324 | Viollier AG | University Hospital Basel, Clinical Bacteriology | Tim Roloff, Madlen Stange, Helena MB Seth-Smith, Alfredo Mari, Karoline Leuzinger, Julia Bielicki, Christiane Beckmann, Manuel Battegay, Hans Hirsch, Adrian Egli |
| EPI_ISL_910335, EPI_ISL_910336, EPI_ISL_910337, EPI_ISL_910338, EPI_ISL_910339 | Laboratory for Respiratory Viruses, Cantacuzino National Military-Medical Institute for Research and Development | Cantacuzino Institute Virology | Luiza Ustea, Nicoleta Paraschiv, Mihaela Lazar |
| EPI_ISL_911248, EPI_ISL_911249, EPI_ISL_911250, EPI_ISL_911251, EPI_ISL_911252, EPI_ISL_911253, EPI_ISL_911254, EPI_ISL_911255, EPI_ISL_911256, EPI_ISL_911257, EPI_ISL_911258, EPI_ISL_911259, EPI_ISL_911260, EPI_ISL_911261, EPI_ISL_911262, EPI_ISL_911263, EPI_ISL_911264, EPI_ISL_911265, EPI_ISL_911266, EPI_ISL_911267, EPI_ISL_911268, EPI_ISL_911269, EPI_ISL_911270, EPI_ISL_911271, EPI_ISL_911272, EPI_ISL_911273, EPI_ISL_911274, EPI_ISL_911275, EPI_ISL_911276, EPI_ISL_911277, EPI_ISL_911278, EPI_ISL_911279, EPI_ISL_911280, EPI_ISL_911281, EPI_ISL_911282, EPI_ISL_911283 | | | |
| see above | AZDelta | AZDelta | Geert Martens; Dieter De Smet |
| EPI_ISL_911524, EPI_ISL_911526, EPI_ISL_911527 | Microbiology and Virology Unit, Florence Careggi University Hospital | Microbiology and Virology Unit, Florence Careggi University Hospital | Vincenzo Di Pilato, Marco Coppi, Fabio Morecchiato, Noemi Aiezza, Ilaria Baccani, Alberto Antonelli, Emanuele Gori, Gian Maria Rossolini |
| EPI_ISL_911707 | Academic Hospital of Gadjah Mada University (RSA UGM) | Genetics Working Group (Pokja Genetik) Faculty of Medicine, Public Health and Nursing Universitas Gadjah Mada (FK-KMK UGM); Disease Investigation Center Wates Ministry of Agriculture Indonesia; Department of Microbiology FK-KMK UGM; Laboratorium Diagnostik Yayasan Tahija World Mosquito Program (WMP) Yogyakarta Center for Tropical Medicine FK-KMK UGM; Integrated Research center FK-KMK UGM; Department of Computer Science and Electronics FMIPA UGM | Gunadi, Hendra Wibawa, Marcellus, Mohamad S. Hakim, Edwin W. Daniwijaya, Ludhang P. Rizki, Endah Supriyati, Eggi Arguni, Titik Nuryastuti, Tri Wibawa, Dwi AA Nugrahaningsih, Afiahayati, Siswanto, Kristy Iskandar, Nungki Anggorowati, Untung Riawan, Kemala Athollah |
| EPI_ISL_911708 | Tanjungpura University Hospital | Tanjungpura University Hospital | Puji Astuti; Andriani; Mahyarudin; Delima Fajar Liana; Virhan Novianry ; Sofi Siti Shofiyah |
| EPI_ISL_911709 | Academic Hospital of Gadjah Mada University (RSA UGM) | Genetics Working Group (Pokja Genetik) Faculty of Medicine, Public Health and Nursing Universitas Gadjah Mada (FK-KMK UGM); Disease Investigation Center Wates Ministry of Agriculture Indonesia; Department of Microbiology FK-KMK UGM; Laboratorium Diagnostik Yayasan Tahija World Mosquito Program (WMP) Yogyakarta Center for Tropical Medicine FK-KMK UGM; Integrated Research center FK-KMK UGM; Department of Computer Science and Electronics FMIPA UGM | Gunadi, Hendra Wibawa, Marcellus, Mohamad S. Hakim, Edwin W. Daniwijaya, Ludhang P. Rizki, Endah Supriyati, Eggi Arguni, Titik Nuryastuti, Tri Wibawa, Dwi AA Nugrahaningsih, Afiahayati, Siswanto, Kristy Iskandar, Nungki Anggorowati, Dyah Ayu Puspitarani, Dwiki afandy |
| EPI_ISL_911750 | Tanjungpura University Hospital | Tanjungpura University Hospital | Virhan Novianry; Delima Fajar Liana; Mahyarudin; Andriani; Sofi Siti Shofiyah; Puji Astuti; Muhammad Kahtan; Ambar Rialita; Eka Ardiani Putri; Wiwik Windarti; Helmi Sastriawan, Rini Andriani |
| EPI_ISL_912397, EPI_ISL_912399, EPI_ISL_912400 | Institute for Medical Research, Infectious Disease Research Centre, National Institutes of Health, Ministry of Health Malaysia | Institute for Medical Research, Infectious Disease Research Centre, National Institutes of Health, Ministry of Health Malaysia | Suppiah J, Kamel K, Azizan MA, Thayan R |
| EPI_ISL_912402, EPI_ISL_912403 | CH de Mayotte - Laboratoire de Biologie | National Reference Center for Viruses of Respiratory Infections, Institut Pasteur, Paris | Marion Barbet, Sylvie Behillil, Méline Bizard, Angela Brisebarre, Camille Capel, Etienne Simon-Lorière, Vincent Enouf, Maud Vanpeene, Sylvie van der Werf,Combe Patrice |
| EPI_ISL_912549 | Labo Analyses Med | National Reference Center for Viruses of Respiratory Infections, Institut Pasteur, Paris | Marion Barbet, Sylvie Behillil, Méline Bizard, Angela Brisebarre, Camille Capel, Etienne Simon-Lorière, Vincent Enouf, Maud Vanpeene, Sylvie van der Werf |
| EPI_ISL_912550, EPI_ISL_912551, EPI_ISL_912552, EPI_ISL_912553 | Labo Analyses Med | National Reference Center for Viruses of Respiratory Infections, Institut Pasteur, Paris | Marion Barbet, Sylvie Behillil, Méline Bizard, Angela Brisebarre, Camille Capel, Etienne Simon-Lorière, Vincent Enouf, Maud Vanpeene, Sylvie van der Werf,Amzalag Jonas |
| EPI_ISL_912554, EPI_ISL_912555, EPI_ISL_912556 | C.H.R. d'Orléans - Hôpital de la Source | National Reference Center for Viruses of Respiratory Infections, Institut Pasteur, Paris | Marion Barbet, Sylvie Behillil, Méline Bizard, Angela Brisebarre, Camille Capel, Etienne Simon-Lorière, Vincent Enouf, Maud Vanpeene, Sylvie van der Werf,Guinard Jérome |
| EPI_ISL_912557, EPI_ISL_912558 | Centre Hospitalier de Gisors Route de Rouen | National Reference Center for Viruses of Respiratory Infections, Institut Pasteur, Paris | Marion Barbet, Sylvie Behillil, Méline Bizard, Angela Brisebarre, Camille Capel, Etienne Simon-Lorière, Vincent Enouf, Maud Vanpeene, Sylvie van der Werf |
| EPI_ISL_912559 | Hopital Avicenne Laboratoire central de bactériologie virologie hygiène | National Reference Center for Viruses of Respiratory Infections, Institut Pasteur, Paris | Marion Barbet, Sylvie Behillil, Méline Bizard, Angela Brisebarre, Camille Capel, Etienne Simon-Lorière, Vincent Enouf, Maud Vanpeene, Sylvie van der Werf,Brichler Ségolène |
| EPI_ISL_912560, EPI_ISL_912561, EPI_ISL_912562 | HOPITAL SIMONE VEIL | National Reference Center for Viruses of Respiratory Infections, Institut Pasteur, Paris | Marion Barbet, Sylvie Behillil, Méline Bizard, Angela Brisebarre, Camille Capel, Etienne Simon-Lorière, Vincent Enouf, Maud Vanpeene, Sylvie van der Werf,Moreau Farida |
| EPI_ISL_912563, EPI_ISL_912564 | Laboratoire Central - American Hospital of Paris | National Reference Center for Viruses of Respiratory Infections, Institut Pasteur, Paris | Marion Barbet, Sylvie Behillil, Méline Bizard, Angela Brisebarre, Camille Capel, Etienne Simon-Lorière, Vincent Enouf, Maud Vanpeene, Sylvie van der Werf,Ramirez José-Manuel |
| EPI_ISL_912565 | Laboratoire de Virologie du CHU de Lille Bâtiment Paul Boulanger | National Reference Center for Viruses of Respiratory Infections, Institut Pasteur, Paris | Marion Barbet, Sylvie Behillil, Méline Bizard, Angela Brisebarre, Camille Capel, Etienne Simon-Lorière, Vincent Enouf, Maud Vanpeene, Sylvie van der Werf,Guigon Aurélie |
| EPI_ISL_912566, EPI_ISL_912567 | Hopital | National Reference Center for Viruses of Respiratory Infections, Institut Pasteur, Paris | Marion Barbet, Sylvie Behillil, Méline Bizard, Angela Brisebarre, Camille Capel, Etienne Simon-Lorière, Vincent Enouf, Maud Vanpeene, Sylvie van der Werf |
| EPI_ISL_912568 | Laboratoire de Virologie du CHU de Lille Bâtiment Paul Boulanger | National Reference Center for Viruses of Respiratory Infections, Institut Pasteur, Paris | Marion Barbet, Sylvie Behillil, Méline Bizard, Angela Brisebarre, Camille Capel, Etienne Simon-Lorière, Vincent Enouf, Maud Vanpeene, Sylvie van der Werf,Guigon Aurélie |
| EPI_ISL_912569 | Labo Analyses Med | National Reference Center for Viruses of Respiratory Infections, Institut Pasteur, Paris | Marion Barbet, Sylvie Behillil, Méline Bizard, Angela Brisebarre, Camille Capel, Etienne Simon-Lorière, Vincent Enouf, Maud Vanpeene, Sylvie van der Werf |
| EPI_ISL_912570, EPI_ISL_912571, EPI_ISL_912572 | Labo Analyses Med | National Reference Center for Viruses of Respiratory Infections, Institut Pasteur, Paris | Marion Barbet, Sylvie Behillil, Méline Bizard, Angela Brisebarre, Camille Capel, Etienne Simon-Lorière, Vincent Enouf, Maud Vanpeene, Sylvie van der Werf,Amzalag Jonas |
| EPI_ISL_912573 | C.H.R. d'Orléans - Hôpital de la Source | National Reference Center for Viruses of Respiratory | Marion Barbet, Sylvie Behillil, Méline Bizard, Angela Brisebarre, Camille Capel, Etienne Simon-Lorière, Vincent Enouf, Maud Vanpeene, Sylvie van der |

| | | | |
|---|---|---|---|
| | | Infections, Institut Pasteur, Paris | Werf,Guinard Jérome |
| EPI_ISL_912574 | Laboratoire Central - American Hospital of Paris | National Reference Center for Viruses of Respiratory Infections, Institut Pasteur, Paris | Marion Barbet, Sylvie Behillil, Méline Bizard, Angela Brisebarre, Camille Capel, Etienne Simon-Lorière, Vincent Enouf, Maud Vanpeene, Sylvie van der Werf,Ramirez José-Manuel |
| EPI_ISL_912575, EPI_ISL_912576, EPI_ISL_912577, EPI_ISL_912578 | Laboratoire de Virologie du CHU de Lille Bâtiment Paul Boulanger | National Reference Center for Viruses of Respiratory Infections, Institut Pasteur, Paris | Marion Barbet, Sylvie Behillil, Méline Bizard, Angela Brisebarre, Camille Capel, Etienne Simon-Lorière, Vincent Enouf, Maud Vanpeene, Sylvie van der Werf,Guigon Aurélie |
| EPI_ISL_912580, EPI_ISL_912581, EPI_ISL_912582, EPI_ISL_912583 | CENTRE HOSPITALIER LEON BINET | National Reference Center for Viruses of Respiratory Infections, Institut Pasteur, Paris | Marion Barbet, Sylvie Behillil, Méline Bizard, Angela Brisebarre, Camille Capel, Etienne Simon-Lorière, Vincent Enouf, Maud Vanpeene, Sylvie van der Werf,Meziane Ilham |
| EPI_ISL_912584, EPI_ISL_912585, EPI_ISL_912586, EPI_ISL_912587, EPI_ISL_912588, EPI_ISL_912589, EPI_ISL_912590, EPI_ISL_912591, EPI_ISL_912592, EPI_ISL_912593, EPI_ISL_912594, EPI_ISL_912595, EPI_ISL_912596, EPI_ISL_912597 | | | |
| see above | Labo Analyses Med | National Reference Center for Viruses of Respiratory Infections, Institut Pasteur, Paris | Marion Barbet, Sylvie Behillil, Méline Bizard, Angela Brisebarre, Camille Capel, Etienne Simon-Lorière, Vincent Enouf, Maud Vanpeene, Sylvie van der Werf,Le Vicky |
| EPI_ISL_912599 | HOPITAL SIMONE VEIL | National Reference Center for Viruses of Respiratory Infections, Institut Pasteur, Paris | Marion Barbet, Sylvie Behillil, Méline Bizard, Angela Brisebarre, Camille Capel, Etienne Simon-Lorière, Vincent Enouf, Maud Vanpeene, Sylvie van der Werf,Moreau Farida |
| EPI_ISL_912600 | Labo Analyses Med | National Reference Center for Viruses of Respiratory Infections, Institut Pasteur, Paris | Marion Barbet, Sylvie Behillil, Méline Bizard, Angela Brisebarre, Camille Capel, Etienne Simon-Lorière, Vincent Enouf, Maud Vanpeene, Sylvie van der Werf,Gaumery Bertrand |
| EPI_ISL_912602 | Laboratoire de Virologie du CHU de Lille Bâtiment Paul Boulanger | National Reference Center for Viruses of Respiratory Infections, Institut Pasteur, Paris | Marion Barbet, Sylvie Behillil, Méline Bizard, Angela Brisebarre, Camille Capel, Etienne Simon-Lorière, Vincent Enouf, Maud Vanpeene, Sylvie van der Werf,Guigon Aurélie |
| EPI_ISL_912603, EPI_ISL_912604, EPI_ISL_912605 | Labo Analyses Med | National Reference Center for Viruses of Respiratory Infections, Institut Pasteur, Paris | Marion Barbet, Sylvie Behillil, Méline Bizard, Angela Brisebarre, Camille Capel, Etienne Simon-Lorière, Vincent Enouf, Maud Vanpeene, Sylvie van der Werf |
| EPI_ISL_912606 | Outre mer | National Reference Center for Viruses of Respiratory Infections, Institut Pasteur, Paris | Marion Barbet, Sylvie Behillil, Méline Bizard, Angela Brisebarre, Camille Capel, Etienne Simon-Lorière, Vincent Enouf, Maud Vanpeene, Sylvie van der Werf,Combe Patrice |
| EPI_ISL_912607, EPI_ISL_912608, EPI_ISL_912609, EPI_ISL_912610, EPI_ISL_912611 | Labo Analyses Med | National Reference Center for Viruses of Respiratory Infections, Institut Pasteur, Paris | Marion Barbet, Sylvie Behillil, Méline Bizard, Angela Brisebarre, Camille Capel, Etienne Simon-Lorière, Vincent Enouf, Maud Vanpeene, Sylvie van der Werf,Holstein Anne |
| EPI_ISL_912612, EPI_ISL_912613 | Centre Hospitalier de Compiègne Laboratoire Compiègne | National Reference Center for Viruses of Respiratory Infections, Institut Pasteur, Paris | Marion Barbet, Sylvie Behillil, Méline Bizard, Angela Brisebarre, Camille Capel, Etienne Simon-Lorière, Vincent Enouf, Maud Vanpeene, Sylvie van der Werf,Romand Sylvie |
| EPI_ISL_912615, EPI_ISL_912616 | Outre mer | National Reference Center for Viruses of Respiratory Infections, Institut Pasteur, Paris | Marion Barbet, Sylvie Behillil, Méline Bizard, Angela Brisebarre, Camille Capel, Etienne Simon-Lorière, Vincent Enouf, Maud Vanpeene, Sylvie van der Werf,Combe Patrice |
| EPI_ISL_912617, EPI_ISL_912618, EPI_ISL_912619, EPI_ISL_912620 | Labo Analyses Med | National Reference Center for Viruses of Respiratory Infections, Institut Pasteur, Paris | Marion Barbet, Sylvie Behillil, Méline Bizard, Angela Brisebarre, Camille Capel, Etienne Simon-Lorière, Vincent Enouf, Maud Vanpeene, Sylvie van der Werf,Le Vicky |
| EPI_ISL_912621 | HOPITAL YVES LE FOLL Laboratoire de Biologie Médicale | National Reference Center for Viruses of Respiratory Infections, Institut Pasteur, Paris | Marion Barbet, Sylvie Behillil, Méline Bizard, Angela Brisebarre, Camille Capel, Etienne Simon-Lorière, Vincent Enouf, Maud Vanpeene, Sylvie van der Werf,Chrevet Laurent |
| EPI_ISL_912622, EPI_ISL_912624 | Labo Analyses Med | National Reference Center for Viruses of Respiratory Infections, Institut Pasteur, Paris | Marion Barbet, Sylvie Behillil, Méline Bizard, Angela Brisebarre, Camille Capel, Etienne Simon-Lorière, Vincent Enouf, Maud Vanpeene, Sylvie van der Werf |
| EPI_ISL_912669, EPI_ISL_912722, EPI_ISL_912724, EPI_ISL_912725, EPI_ISL_912726, EPI_ISL_912727, EPI_ISL_912730, EPI_ISL_912734, EPI_ISL_912735, EPI_ISL_912737, EPI_ISL_912738, EPI_ISL_912739, EPI_ISL_912742, EPI_ISL_912745, EPI_ISL_912746, EPI_ISL_912747, EPI_ISL_912749, EPI_ISL_912810, EPI_ISL_912853, EPI_ISL_912855, EPI_ISL_912856, EPI_ISL_912857, EPI_ISL_912858, EPI_ISL_912863 | | | |
| see above | Hôpital Henri Mondor | Department of Virology, Henri Mondor University Hospital, Assistance Publique Hôpitaux de Paris, Université Paris-Est Créteil, INSERM U955 | Christophe Rodriguez, Slim Fourati, Vanessa Demontant, Guillaume Gricourt, Melissa N'Debi, Alexandre Soulier, Elisabeth Trawinski, Jean-Michel Pawlotsky |
| EPI_ISL_912913 | Hôpital Pitié-Salpêtrière | Department of Virology, Henri Mondor University Hospital, Assistance Publique Hôpitaux de Paris, Université Paris-Est Créteil, INSERM U955 | Christophe Rodriguez, Slim Fourati, Vanessa Demontant, Guillaume Gricourt, Melissa N'Debi, Alexandre Soulier, Elisabeth Trawinski, Jean-Michel Pawlotsky |
| EPI_ISL_912921, EPI_ISL_912922, EPI_ISL_912944, EPI_ISL_912946, EPI_ISL_912947, EPI_ISL_912948, EPI_ISL_912949, EPI_ISL_912950, EPI_ISL_912951, EPI_ISL_912952, EPI_ISL_912953, EPI_ISL_912954, EPI_ISL_912955, EPI_ISL_912956, EPI_ISL_912957, EPI_ISL_912958, EPI_ISL_912964, EPI_ISL_912965, EPI_ISL_912968, EPI_ISL_912970, EPI_ISL_912971, EPI_ISL_912972, EPI_ISL_912974, EPI_ISL_912985, EPI_ISL_912989, EPI_ISL_912990, EPI_ISL_912999, EPI_ISL_913002, EPI_ISL_913003, EPI_ISL_913006, EPI_ISL_913009, EPI_ISL_913010, EPI_ISL_913013, EPI_ISL_913014, EPI_ISL_913015, EPI_ISL_913017 | | | |
| see above | Hôpital Henri Mondor | Department of Virology, Henri Mondor University Hospital, Assistance Publique Hôpitaux de Paris, Université Paris-Est Créteil, INSERM U955 | Christophe Rodriguez, Slim Fourati, Vanessa Demontant, Guillaume Gricourt, Melissa N'Debi, Alexandre Soulier, Elisabeth Trawinski, Jean-Michel Pawlotsky |
| EPI_ISL_913020 | Hospital Universitario de Ceuta | Instituto de Salud Carlos III | Iglesias-Caballero, M. Camarero, S. Sandonís,V. Vázquez, S. Pozo, F. Casas, I. Jiménez, P. Zaballos, A. Monzón, S. Varona, S. Cuesta, I. López, J. |
| EPI_ISL_913021, EPI_ISL_913022, EPI_ISL_913023 | Hospital San Pedro de Alcántara | Instituto de Salud Carlos III | Iglesias-Caballero, M. Camarero, S. Sandonís,V. Vázquez, S. Pozo, F. Casas, I. Jiménez, P. Zaballos, A. Monzón, S. Varona, S. Cuesta, I. Rodríguez, G. |
| EPI_ISL_913024 | Complejo Hospitalario Universitario de Santiago | Instituto de Salud Carlos III | Iglesias-Caballero, M. Camarero, S. Sandonís,V. Vázquez, S. Pozo, F. Casas, I. Jiménez, P. Zaballos, A. Monzón, S. Varona, S. Cuesta, I. Peña,F. |
| EPI_ISL_913026, EPI_ISL_913027, EPI_ISL_913028, EPI_ISL_913030, EPI_ISL_913031 | Hospital los Arcos | Instituto de Salud Carlos III | Iglesias-Caballero, M. Camarero, S. Sandonís,V. Vázquez, S. Pozo, F. Casas, I. Jiménez, P. Zaballos, A. Monzón, S. Varona, S. Cuesta, I. Cámara, M. |
| EPI_ISL_913032, EPI_ISL_913033, EPI_ISL_913034, EPI_ISL_913035 | Hospital San Pedro de Alcántara | Instituto de Salud Carlos III | Iglesias-Caballero, M. Camarero, S. Sandonís,V. Vázquez, S. Pozo, F. Casas, I. Jiménez, P. Zaballos, A. Monzón, S. Varona, S. Cuesta, I. Rodríguez, G. |
| EPI_ISL_913036, EPI_ISL_913037 | Hospital los Arcos | Instituto de Salud Carlos III | Iglesias-Caballero, M. Camarero, S. Sandonís,V. Vázquez, S. Pozo, F. Casas, I. Jiménez, P. Zaballos, A. Monzón, S. Varona, S. Cuesta, I. Cámara, M. |
| EPI_ISL_913038 | Complejo Asistencial Universitario de Burgos | Instituto de Salud Carlos III | Iglesias-Caballero, M. Camarero, S. Sandonís,V. Vázquez, S. Pozo, F. Casas, I. Jiménez, P. Zaballos, A. Monzón, S. Varona, S. Cuesta, I. Megías, G. |
| EPI_ISL_913039 | Complejo Hospitalario Xeral-Calde | Instituto de Salud Carlos III | Iglesias-Caballero, M. Camarero, S. Sandonís,V. Vázquez, S. Pozo, F. Casas, I. Jiménez, P. Zaballos, A. Monzón, S. Varona, S. Cuesta, I. Alonso, P. |
| EPI_ISL_913040 | Hospital Universitario de Guadalajara | Instituto de Salud Carlos III | Iglesias-Caballero, M. Camarero, S. Sandonís,V. Vázquez, S. Pozo, F. Casas, I. Jiménez, P. Zaballos, A. Monzón, S. Varona, S. Cuesta, I. González, A. |
| EPI_ISL_913041 | Consejería de Sanidad y Asuntos Sociales | Instituto de Salud Carlos III | Iglesias-Caballero, M. Camarero, S. Sandonís,V. Vázquez, S. Pozo, F. Casas, I. Jiménez, P. Zaballos, A. Monzón, S. Varona, S. Cuesta, I. Gutiérrez, G. |
| EPI_ISL_913042, EPI_ISL_913043, EPI_ISL_913044, EPI_ISL_913045, EPI_ISL_913046, EPI_ISL_913047, EPI_ISL_913048, EPI_ISL_913049, EPI_ISL_913050, EPI_ISL_913051 | Hospital Comarcal Sierrallana | Instituto de Salud Carlos III | Iglesias-Caballero, M. Camarero, S. Sandonís,V. Vázquez, S. Pozo, F. Casas, I. Jiménez, P. Zaballos, A. Monzón, S. Varona, S. Cuesta, I. De Benito, I. |
| EPI_ISL_913052, EPI_ISL_913053 | Complejo Asistencial Universitario de Salamanca | Instituto de Salud Carlos III | Iglesias-Caballero, M. Camarero, S. Sandonís,V. Vázquez, S. Pozo, F. Casas, I. Jiménez, P. Zaballos, A. Monzón, S. Varona, S. Cuesta, I. Ávila, A. |
| EPI_ISL_913054, EPI_ISL_913055, EPI_ISL_913056 | Hospital Universitario de Ceuta | Instituto de Salud Carlos III | Iglesias-Caballero, M. Camarero, S. Sandonís,V. Vázquez, S. Pozo, F. Casas, I. Jiménez, P. Zaballos, A. Monzón, S. Varona, S. Cuesta, I. López, J. |
| EPI_ISL_913057 | Hospital San Pedro de Alcántara | Instituto de Salud Carlos III | Iglesias-Caballero, M. Camarero, S. Sandonís,V. Vázquez, S. Pozo, F. Casas, I. Jiménez, P. Zaballos, A. Monzón, S. Varona, S. Cuesta, I. Rodríguez, G. |
| EPI_ISL_913058 | Hospital Universitario de Ceuta | Instituto de Salud Carlos III | Iglesias-Caballero, M. Camarero, S. Sandonís,V. Vázquez, S. Pozo, F. Casas, I. Jiménez, P. Zaballos, A. Monzón, S. Varona, S. Cuesta, I. López, J. |

| | | | |
|---|---|---|---|
| EPI_ISL_913059 | Consejería de Sanidad y Asuntos Sociales | Instituto de Salud Carlos III | Iglesias-Caballero, M. Camarero, S. Sandonís,V. Vázquez, S. Pozo, F. Casas, I. Jiménez, P. Zaballos, A. Monzón, S. Varona, S. Cuesta, I. Gutiérrez, G. |
| EPI_ISL_913060 | Hospital Universitario Virgen de la Arrixaca | Instituto de Salud Carlos III | Iglesias-Caballero, M. Camarero, S. Sandonís,V. Vázquez, S. Pozo, F. Casas, I. Jiménez, P. Zaballos, A. Monzón, S. Varona, S. Cuesta, I. Moreno, A. |
| EPI_ISL_913061, EPI_ISL_913062 | Hospital Nuestra Señora de Sonsoles | Instituto de Salud Carlos III | Iglesias-Caballero, M. Camarero, S. Sandonís,V. Vázquez, S. Pozo, F. Casas, I. Jiménez, P. Zaballos, A. Monzón, S. Varona, S. Cuesta, I. Sánchez, R. |
| EPI_ISL_913449 | Istituto Zooprofilattico Sperimentale del Mezzogiorno | INMI Lazzaro Spallanzani IRCCS | Barbara Bartolini, Ornella Butera, Cesare EM Gruber, Martina Rueca, Francesco Messina, Emanuela Giombini, Silvia Meschi, Francesca Colavita, Concetta Castilletti, Antonino Di Caro, Maria R Capobianchi |
| EPI_ISL_913607 | Hopital Avicenne Laboratoire central de bactériologie virologie hygiène | National Reference Center for Viruses of Respiratory Infections, Institut Pasteur, Paris | Marion Barbet, Sylvie Behillil, Méline Bizard, Angela Brisebarre, Camille Capel, Etienne Simon-Lorière, Vincent Enouf, Maud Vanpeene, Sylvie van der Werf,Brichler Ségolène |
| EPI_ISL_913608, EPI_ISL_913609, EPI_ISL_913610 | Laboratoire de Virologie du CHU de Lille Bâtiment Paul Boulanger | National Reference Center for Viruses of Respiratory Infections, Institut Pasteur, Paris | Marion Barbet, Sylvie Behillil, Méline Bizard, Angela Brisebarre, Camille Capel, Etienne Simon-Lorière, Vincent Enouf, Maud Vanpeene, Sylvie van der Werf,Guigon Aurélie |
| EPI_ISL_913926, EPI_ISL_913927, EPI_ISL_913928, EPI_ISL_913929, EPI_ISL_913930, EPI_ISL_913931, EPI_ISL_913932, EPI_ISL_913957, EPI_ISL_913962, EPI_ISL_913967, EPI_ISL_913970, EPI_ISL_913979, EPI_ISL_913980, EPI_ISL_913981, EPI_ISL_913983 | | | |
| see above | Instituto de Diagnostico y Referencia Epidemiologicos INDRE_RNLSP | Instituto de Diagnostico y Referencia Epidemiologicos (INDRE) | Claudia Wong-Arambula, Abril Rodriguez-Maldonado, Fabiola Garces-Ayala, Adnan Araiza-Rodriguez, David Fragoso-Fonseca, Sergio Rangel-Guerrero, Mayra Jimenez-Morales, Nancy Munoz-Hernandez, Natividad Cruz-Ortiz, Tatiana Nunez-Garcia, Gisela Barrera-Badillo, Lucia Hernandez-Rivas, Irma Lopez-Martinez, Ernesto Ramirez-Gonzalez. |
| EPI_ISL_914802 | AREA DE SALUD CATEDRAL NORESTE | Inciensa, Instituto Costarricense de Investigación y Enseñanza en Nutrición y Salud | Francisco Duarte, Hebleen Porras, Claudio Soto-Garita, Estela Cordero, Adriana Godínez, Melany Calderón & Mariel López |
| EPI_ISL_915421 | MRCG at LSHTM Genomics lab | MRCG at LSHTM Genomics lab | Abdul Karim sesay, Abdoulie Kanteh, Jarra Manneh, Mariama Kujabi, Bakary Sanyang |
| EPI_ISL_918165 | UMR190-Unité des virus emergents | UMR190-Unité des virus emergents | Cecile Baronti |
| EPI_ISL_918263 | SIESP DIPARTIMENTO DI PREVENZIONE CHIE | Istituto Zooprofilattico Sperimentale dell'Abruzzo e Molise "G. Caporale" | Lorusso A, Marcacci M, Di Domenico M, Ancora M, Curini V, Mangone I, Rinaldi A, Scialabba S, Di Pasquale A, Cammà C, Puglia I, Calistri P, Savini G |
| EPI_ISL_918264 | SIESP CHIETI-DRIVE IN CHIETI | Istituto Zooprofilattico Sperimentale dell'Abruzzo e Molise "G. Caporale" | Lorusso A, Marcacci M, Di Domenico M, Ancora M, Curini V, Mangone I, Rinaldi A, Scialabba S, Di Pasquale A, Cammà C, Puglia I, Calistri P, Savini G |
| EPI_ISL_918266 | SIESP CHIETI-Drive in Lanciano | Istituto Zooprofilattico Sperimentale dell'Abruzzo e Molise "G. Caporale" | Lorusso A, Marcacci M, Di Domenico M, Ancora M, Curini V, Mangone I, Rinaldi A, Scialabba S, Di Pasquale A, Cammà C, Puglia I, Calistri P, Savini G |
| EPI_ISL_918267 | SIESP CHIETI - DRIVE IN CHIETI | Istituto Zooprofilattico Sperimentale dell'Abruzzo e Molise "G. Caporale" | Lorusso A, Marcacci M, Di Domenico M, Ancora M, Curini V, Mangone I, Rinaldi A, Scialabba S, Di Pasquale A, Cammà C, Puglia I, Calistri P, Savini G |
| EPI_ISL_918268 | SIESP CHIETI - DRIVE IN ORTONA | Istituto Zooprofilattico Sperimentale dell'Abruzzo e Molise "G. Caporale" | Lorusso A, Marcacci M, Di Domenico M, Ancora M, Curini V, Mangone I, Rinaldi A, Scialabba S, Di Pasquale A, Cammà C, Puglia I, Calistri P, Savini G |
| EPI_ISL_918270 | SIESP CHIETI - DRIVE IN LANCIANO | Istituto Zooprofilattico Sperimentale dell'Abruzzo e Molise "G. Caporale" | Lorusso A, Marcacci M, Di Domenico M, Ancora M, Curini V, Mangone I, Rinaldi A, Scialabba S, Di Pasquale A, Cammà C, Puglia I, Calistri P, Savini G |
| EPI_ISL_918271 | SIESP DIPARTIMENTO DI PREVENZIONE CHIETI | Istituto Zooprofilattico Sperimentale dell'Abruzzo e Molise "G. Caporale" | Lorusso A, Marcacci M, Di Domenico M, Ancora M, Curini V, Mangone I, Rinaldi A, Scialabba S, Di Pasquale A, Cammà C, Puglia I, Calistri P, Savini G |
| EPI_ISL_918359, EPI_ISL_918360, EPI_ISL_918361, EPI_ISL_918362, EPI_ISL_918363, EPI_ISL_918364, EPI_ISL_918365, EPI_ISL_918366, EPI_ISL_918367, EPI_ISL_918368, EPI_ISL_918369, EPI_ISL_918370, EPI_ISL_918371, EPI_ISL_918372, EPI_ISL_918373, EPI_ISL_918374, EPI_ISL_918375, EPI_ISL_918376 | | | |
| see above | Virology Unit, Institut Pasteur du Cambodge | Virology Unit, Institut Pasteur du Cambodge | Sokhoun Yann, Ly Sovann, Kraing Sidonn, Yi Sengdoeurn, Chin Savuth, Chau Darapheak, Etienne Simon-Loriere, Veasna Duong, Erik A Karlsson |
| EPI_ISL_918408, EPI_ISL_918409 | Ospedale Santa Caterina Novella | Istituto Zooprofilattico Sperimentale della Puglia e della Basilicata | Parisi A., Bianco A., Capozzi L., Del Sambro L., Simone D., Manzulli V, Rondinone V., Pace L., Cipolletta D., Galante D. |
| EPI_ISL_918489 | CUSL/UCLouvain COVID testing federal platform | UCLouvain/IREC/MBLG | Jean Ruelle, Lysa Pinsmaye, Benoit Kabamba Mukadi |
| EPI_ISL_918516, EPI_ISL_918546, EPI_ISL_918549 | LACEN - Laboratório Central de Saúde Pública do Para | Evandro Chagas Institute | Santos, M.C.; Silva, A.M.; Junior, W.D.C.; Barbagelata, L.S.; Ferreira, J.A.; Sousa, E.M.A.; da Silva, P.S.; Pinheiro, K.C.; L.C.; Sousa Junior, E.C. |
| EPI_ISL_922076 | General Hospital #Abdulah Nakas" Sarajevo | Alea Genetic Centre | Rijad Konjhodzic, Dino Pecar, Lana Salihefendic, Sead Jazic |
| EPI_ISL_933533 | Laboratory for HIV and opportunistic infections diagnosis The Republican Research and Practical Center for Epidemiology and Microbiology (RRPCEM) | Laboratory for HIV and opportunistic infections diagnosis The Republican Research and Practical Center for Epidemiology and Microbiology (RRPCEM) | Elena Gasich, Kirill Bulda, Artur Akhremchuk, Leonid Valentovich, Vladimir Gorbunov |
| EPI_ISL_933780, EPI_ISL_933781, EPI_ISL_933782, EPI_ISL_933783, EPI_ISL_933784, EPI_ISL_933785 | Virology Unit, Institut Pasteur du Cambodge | Virology Unit, Institut Pasteur du Cambodge | Sokhoun Yann, Ly Sovann, Kraing Sidonn, Yi Sengdoeurn, Chin Savuth, Chau Darapheak, Etienne Simon-Loriere, Veasna Duong, Erik A Karlsson |
| EPI_ISL_934989, EPI_ISL_934990, EPI_ISL_934991, EPI_ISL_935000, EPI_ISL_935001, EPI_ISL_935002, EPI_ISL_935003, EPI_ISL_935004, EPI_ISL_935005, EPI_ISL_935006, EPI_ISL_935007, EPI_ISL_935008, EPI_ISL_935009, EPI_ISL_935010, EPI_ISL_935011 | | | |
| see above | ADMED Microbiologie | Genomics and Transcriptomics, Philip Morris International | Reto Lienhard, Marie-Lise Tritten, Emmanuel Guedj, Nicolas Sierro, Rémi Dulize, David Bornand, Mehdi Auberson, Maxime Berthouzoz, Nikolai Ivanov, Manuel Peitsch |
| EPI_ISL_935156, EPI_ISL_935158, EPI_ISL_935159 | SIESP CHIETI | Istituto Zooprofilattico Sperimentale dell'Abruzzo e Molise "G. Caporale" | Lorusso A, Marcacci M, Di Domenico M, Ancora M, Curini V, Mangone I, Rinaldi A, Scialabba S, Di Pasquale A, Cammà C, Puglia I, Calistri P, Savini G |
| EPI_ISL_935160 | SIESP CHIETI-DRIVE IN ORTONA | Istituto Zooprofilattico Sperimentale dell'Abruzzo e Molise "G. Caporale" | Lorusso A, Marcacci M, Di Domenico M, Ancora M, Curini V, Mangone I, Rinaldi A, Scialabba S, Di Pasquale A, Cammà C, Puglia I, Calistri P, Savini G |
| EPI_ISL_935161, EPI_ISL_935162 | SIESP CHIETI | Istituto Zooprofilattico Sperimentale dell'Abruzzo e Molise "G. Caporale" | Lorusso A, Marcacci M, Di Domenico M, Ancora M, Curini V, Mangone I, Rinaldi A, Scialabba S, Di Pasquale A, Cammà C, Puglia I, Calistri P, Savini G |
| EPI_ISL_935163, EPI_ISL_935164 | SIESP CHIETI-DRIVE IN ORTONA | Istituto Zooprofilattico Sperimentale dell'Abruzzo e Molise "G. Caporale" | Lorusso A, Marcacci M, Di Domenico M, Ancora M, Curini V, Mangone I, Rinaldi A, Scialabba S, Di Pasquale A, Cammà C, Puglia I, Calistri P, Savini G |
| EPI_ISL_935165, EPI_ISL_935166, EPI_ISL_935167 | SIESP CHIETI | Istituto Zooprofilattico Sperimentale dell'Abruzzo e Molise "G. Caporale" | Lorusso A, Marcacci M, Di Domenico M, Ancora M, Curini V, Mangone I, Rinaldi A, Scialabba S, Di Pasquale A, Cammà C, Puglia I, Calistri P, Savini G |
| EPI_ISL_935527, EPI_ISL_935528, EPI_ISL_935529, EPI_ISL_935530, EPI_ISL_935531, EPI_ISL_935532 | Labo Analyses Med | National Reference Center for Viruses of Respiratory Infections, Institut Pasteur, Paris | Marion Barbet, Sylvie Behillil, Méline Bizard, Angela Brisebarre, Camille Capel, Etienne Simon-Lorière, Vincent Enouf, Maud Vanpeene, Sylvie van der Werf,Merah Kader |
| EPI_ISL_935537, EPI_ISL_935538 | Labo Analyses Med | National Reference Center for Viruses of Respiratory Infections, Institut Pasteur, Paris | Marion Barbet, Sylvie Behillil, Méline Bizard, Angela Brisebarre, Camille Capel, Etienne Simon-Lorière, Vincent Enouf, Maud Vanpeene, Sylvie van der Werf,Bonnaudet GéRaldine |
| EPI_ISL_935539 | Labo Analyses Med | National Reference Center for Viruses of Respiratory Infections, Institut Pasteur, Paris | Marion Barbet, Sylvie Behillil, Méline Bizard, Angela Brisebarre, Camille Capel, Etienne Simon-Lorière, Vincent Enouf, Maud Vanpeene, Sylvie van der Werf,Tomasi Florent |
| EPI_ISL_935540, EPI_ISL_935541, EPI_ISL_935542, EPI_ISL_935543, EPI_ISL_935544, EPI_ISL_935545 | Hopital | National Reference Center for Viruses of Respiratory Infections, Institut Pasteur, Paris | Marion Barbet, Sylvie Behillil, Méline Bizard, Angela Brisebarre, Camille Capel, Etienne Simon-Lorière, Vincent Enouf, Maud Vanpeene, Sylvie van der Werf,Pilorge LéA |
| EPI_ISL_935547 | Labo Analyses Med | National Reference Center for Viruses of Respiratory | Marion Barbet, Sylvie Behillil, Méline Bizard, Angela Brisebarre, Camille Capel, Etienne Simon-Lorière, Vincent Enouf, Maud Vanpeene, Sylvie van der |

| | | | |
|---|---|---|---|
| | | Infections, Institut Pasteur, Paris | Werf,Besson J |
| EPI_ISL_935548 | Labo Analyses Med | National Reference Center for Viruses of Respiratory Infections, Institut Pasteur, Paris | Marion Barbet, Sylvie Behillil, Méline Bizard, Angela Brisebarre, Camille Capel, Etienne Simon-Lorière, Vincent Enouf, Maud Vanpeene, Sylvie van der Werf,Pasquier |
| EPI_ISL_935552, EPI_ISL_935553, EPI_ISL_935554, EPI_ISL_935555, EPI_ISL_935556, EPI_ISL_935557 | Hopital | National Reference Center for Viruses of Respiratory Infections, Institut Pasteur, Paris | Marion Barbet, Sylvie Behillil, Méline Bizard, Angela Brisebarre, Camille Capel, Etienne Simon-Lorière, Vincent Enouf, Maud Vanpeene, Sylvie van der Werf,Goudeau Alain |
| EPI_ISL_935560 | Labo Analyses Med | National Reference Center for Viruses of Respiratory Infections, Institut Pasteur, Paris | Marion Barbet, Sylvie Behillil, Méline Bizard, Angela Brisebarre, Camille Capel, Etienne Simon-Lorière, Vincent Enouf, Maud Vanpeene, Sylvie van der Werf |
| EPI_ISL_935571, EPI_ISL_935572, EPI_ISL_935573, EPI_ISL_935574, EPI_ISL_935575 | Hopital | National Reference Center for Viruses of Respiratory Infections, Institut Pasteur, Paris | Marion Barbet, Sylvie Behillil, Méline Bizard, Angela Brisebarre, Camille Capel, Etienne Simon-Lorière, Vincent Enouf, Maud Vanpeene, Sylvie van der Werf,Castelain Sandrine |
| EPI_ISL_935581 | Hopital | National Reference Center for Viruses of Respiratory Infections, Institut Pasteur, Paris | Marion Barbet, Sylvie Behillil, Méline Bizard, Angela Brisebarre, Camille Capel, Etienne Simon-Lorière, Vincent Enouf, Maud Vanpeene, Sylvie van der Werf,Brichler SéGolèNe |
| EPI_ISL_935587, EPI_ISL_935589, EPI_ISL_935597, EPI_ISL_935598 | Labo Analyses med | National Reference Center for Viruses of Respiratory Infections, Institut Pasteur, Paris | Marion Barbet, Sylvie Behillil, Méline Bizard, Angela Brisebarre, Camille Capel, Etienne Simon-Lorière, Vincent Enouf, Maud Vanpeene, Sylvie van der Werf,Amzalag Jonas |
| EPI_ISL_935640 | Labo Analyses Med | National Reference Center for Viruses of Respiratory Infections, Institut Pasteur, Paris | Marion Barbet, Sylvie Behillil, Méline Bizard, Angela Brisebarre, Camille Capel, Etienne Simon-Lorière, Vincent Enouf, Maud Vanpeene, Sylvie van der Werf,Le Guillou-Guillemette HéLèNe |
| EPI_ISL_935641 | Labo Analyses Med | National Reference Center for Viruses of Respiratory Infections, Institut Pasteur, Paris | Marion Barbet, Sylvie Behillil, Méline Bizard, Angela Brisebarre, Camille Capel, Etienne Simon-Lorière, Vincent Enouf, Maud Vanpeene, Sylvie van der Werf,Caron Melanie |
| EPI_ISL_935644, EPI_ISL_935650, EPI_ISL_935651, EPI_ISL_935652 | Labo Analyses Med | National Reference Center for Viruses of Respiratory Infections, Institut Pasteur, Paris | Marion Barbet, Sylvie Behillil, Méline Bizard, Angela Brisebarre, Camille Capel, Etienne Simon-Lorière, Vincent Enouf, Maud Vanpeene, Sylvie van der Werf |
| EPI_ISL_936488 | Hospital Queen Elizabeth | Institute for Medical Research, Infectious Disease Research Centre, National Institutes of Health, Ministry of Health Malaysia | Suppiah J, Kamel K, Azizan MA, Thayan R |
| EPI_ISL_936489 | Hospital Raja Perempuan Zainab II | Institute for Medical Research, Infectious Disease Research Centre, National Institutes of Health, Ministry of Health Malaysia | Suppiah J, Kamel K, Azizan MA, Thayan R |
| EPI_ISL_936491 | Institute for Medical Research, Infectious Disease Research Centre, National Institutes of Health, Ministry of Health Malaysia | Institute for Medical Research, Infectious Disease Research Centre, National Institutes of Health, Ministry of Health Malaysia | Suppiah J, Kamel K, Azizan MA, Thayan R |
| EPI_ISL_936494 | Hospital Enche Besar Hajjah Khalsom Kluang | Institute for Medical Research, Infectious Disease Research Centre, National Institutes of Health, Ministry of Health Malaysia | Suppiah J, Kamel K, Azizan MA, Thayan R |
| EPI_ISL_936495 | Sarawak General Hospital | Institute for Medical Research, Infectious Disease Research Centre, National Institutes of Health, Ministry of Health Malaysia | Suppiah J, Kamel K, Azizan MA, Thayan R |
| EPI_ISL_936496 | Institute for Medical Research, Infectious Disease Research Centre, National Institutes of Health, Ministry of Health Malaysia | Institute for Medical Research, Infectious Disease Research Centre, National Institutes of Health, Ministry of Health Malaysia | Suppiah J, Kamel K, Azizan MA, Thayan R |
| EPI_ISL_939628 | Laboratory of Virology and Molecular Diagnostics | Institute of Public Health of Republic of North Macedonia Laboratory of Virology and Molecular Diagnostics | Maja Kuzmanovska. Golubinka Boshevska, Elizabeta Janchevska |
| EPI_ISL_940146 | Laboratory of Virology and Molecular Diagnostics | Institute of Public Health of Republic of North Macedonia Laboratory of Virology and Molecular Diagnostics | Maja Kuzmanovska, Golubinka Boshevska, Elizabeta Janchevska |
| EPI_ISL_940157 | Laboratory of Virology and Molecular Diagnostics | Institute of Public Health of Republic of North Macedonia Laboratory of Virology and Molecular Diagnostics | Maja Kuzmanovska, Golubinka Bosehvska, Elizabeta Janchevska |
| EPI_ISL_940607 | U.O. Genomics, S.S. Genetics and Advanced Omics Techniques, Istituto Zooprofilattico Sperimentale del Piemonte Liguria e Valle d'Aosta | U.O. Genomics, S.S. Genetics and Advanced Omics Techniques, Istituto Zooprofilattico Sperimentale del Piemonte Liguria e Valle d'Aosta | Chiara Beltramo, Francesco Cerutti, Valeria Ghisetti, Elena Bozzetta, Pier Luigi Acutis, Simone Peletto |
| EPI_ISL_940612 | Hospital de Campanha para Enfrentamento do Coronavirus - Goiania | Instituto Adolfo Lutz, Interdiciplinary Procedures Center, Strategic Laboratory | Claudio Tavares Sacchi, Claudia Regina Gonçalves, Erica Valessa Ramos Gomes, Karoline Rodrigues Campos |
| EPI_ISL_940619, EPI_ISL_940620, EPI_ISL_940621, EPI_ISL_940622, EPI_ISL_940623, EPI_ISL_940624 | Hospital Sao Joaquim - Beneficiencia Portuguesa | Instituto Adolfo Lutz, Interdiciplinary Procedures Center, Strategic Laboratory | Claudio Tavares Sacchi, Claudia Regina Gonçalves, Erica Valessa Ramos Gomes, Karoline Rodrigues Campos |
| EPI_ISL_940749 | University of Bari Biomedical Sciences and Human Oncology | University of Bari Biomedical Sciences and Human Oncology | Chironna M., Sallustio A., Loconsole D., Accogli M. |
| EPI_ISL_940766 | Laboratory of Virology and Molecular Diagnostics | Institute of Public Health of Republic of North Macedonia Laboratory of Virology and Molecular Diagnostics | Maja Kuzmanovska, Golubinka Boshevska, Elizabeta Janchevska |
| EPI_ISL_940767 | Heilig Hart Lier | UAntwerp, Laboratory of Medical Microbiology, Campus Drie Eiken S6.26, Universiteitsplein 1, 2610, Wilrijk, Belgium | Basil Britto Xavier, Jasmine Coppens, Marie Le Mercier, Christine Lammens, Veerle Matheeussen, Herman Goossens |
| EPI_ISL_940769 | INSPI-CRN de Influenza y otros virus respiratorios | INSPI-Centro de Investigación Multidisciplinaria de la DTIDI | Leandro Patiño, Doménica de Mora, Maritza Olmedo, Andrés Carrazco-Montalvo, Orson Mestanza, Mary Regato-Arrata, Melissa Zambrano, Manuel González, Alfredo Bruno, Alberto Orlando. |
| EPI_ISL_940858, EPI_ISL_940859, EPI_ISL_940861, EPI_ISL_940862, EPI_ISL_940863, EPI_ISL_940864, EPI_ISL_940877, EPI_ISL_940878, EPI_ISL_940879, EPI_ISL_940880, EPI_ISL_940881, EPI_ISL_940882, EPI_ISL_940883, EPI_ISL_940884, EPI_ISL_940885, EPI_ISL_940886, EPI_ISL_940887, EPI_ISL_940888, EPI_ISL_940889 | | | |
| see above | Vaccines and Infectious Diseases Analytics Research Unit (VIDA) | KRISP, KZN Research Innovation and Sequencing Platform | Baillie Vicky, du Plessis Jeanine, Giandhari Jennifer, Pillay Sureshnee, Naidoo Yeshnee, Tegally Houriiyah, de Oliveira Tulio, Madhi Shabir |
| EPI_ISL_940936, EPI_ISL_940937, EPI_ISL_940938, EPI_ISL_940939, EPI_ISL_940940, EPI_ISL_940942, EPI_ISL_940943, EPI_ISL_940945, EPI_ISL_940946, EPI_ISL_940947, EPI_ISL_940948, EPI_ISL_940949, EPI_ISL_940954, EPI_ISL_940960, EPI_ISL_940961, EPI_ISL_940962, EPI_ISL_940963, EPI_ISL_940964, EPI_ISL_940965, EPI_ISL_940966, EPI_ISL_940967, EPI_ISL_940968, EPI_ISL_940969, EPI_ISL_940970, EPI_ISL_940971, EPI_ISL_940972, EPI_ISL_940973, EPI_ISL_940974, EPI_ISL_940975, EPI_ISL_940976, EPI_ISL_940977, EPI_ISL_940978, EPI_ISL_940979, EPI_ISL_940980, EPI_ISL_940981, EPI_ISL_940982, EPI_ISL_940983, EPI_ISL_940984, EPI_ISL_940985, EPI_ISL_940986, EPI_ISL_940987, EPI_ISL_940988, EPI_ISL_940989, EPI_ISL_940990, EPI_ISL_940991, EPI_ISL_940992, EPI_ISL_940993, EPI_ISL_940994 | | | |
| see above | Centers for Disease Control and Prevention, Dengue Branch | Centers for Disease Control and Prevention, Dengue Branch | Gilberto A. Santiago, Glenda Gonzalez, Betzabel Flores, Keyla Charriez, Gabriela Paz-Bailey, Jorge L. Munoz-Jordan |
| EPI_ISL_940995 | Hopital | National Reference Center for Viruses of Respiratory Infections, Institut Pasteur, Paris | Marion Barbet, Sylvie Behillil, Méline Bizard, Angela Brisebarre, Camille Capel, Etienne Simon-Lorière, Vincent Enouf, Maud Vanpeene, Sylvie van der Werf |
| EPI_ISL_940997, EPI_ISL_940998, | Labo Analyses Med | National Reference Center for Viruses of Respiratory | Marion Barbet, Sylvie Behillil, Méline Bizard, Angela Brisebarre, Camille Capel, Etienne Simon-Lorière, Vincent Enouf, Maud Vanpeene, Sylvie van der Werf, |

| | | | |
|---|---|---|---|
| EPI_ISL_940999 | | Infections, Institut Pasteur, Paris | Gestin (B) Brieuc |
| EPI_ISL_941000, EPI_ISL_941001, EPI_ISL_941002, EPI_ISL_941003 | Hopital | National Reference Center for Viruses of Respiratory Infections, Institut Pasteur, Paris | Marion Barbet, Sylvie Behillil, Méline Bizard, Angela Brisebarre, Camille Capel, Etienne Simon-Lorière, Vincent Enouf, Maud Vanpeene, Sylvie van der Werf, Cron Sophie |
| EPI_ISL_941004, EPI_ISL_941005, EPI_ISL_941006, EPI_ISL_941007, EPI_ISL_941008, EPI_ISL_941009, EPI_ISL_941010, EPI_ISL_941011, EPI_ISL_941012, EPI_ISL_941013, EPI_ISL_941014 | | | |
| see above | Labo Analyses Med | National Reference Center for Viruses of Respiratory Infections, Institut Pasteur, Paris | Marion Barbet, Sylvie Behillil, Méline Bizard, Angela Brisebarre, Camille Capel, Etienne Simon-Lorière, Vincent Enouf, Maud Vanpeene, Sylvie van der Werf, Amzalag Jonas |
| EPI_ISL_941015, EPI_ISL_941016, EPI_ISL_941017, EPI_ISL_941018, EPI_ISL_941019, EPI_ISL_941020, EPI_ISL_941021, EPI_ISL_941022, EPI_ISL_941023, EPI_ISL_941024, EPI_ISL_941025 | | | |
| see above | Hopital | National Reference Center for Viruses of Respiratory Infections, Institut Pasteur, Paris | Marion Barbet, Sylvie Behillil, Méline Bizard, Angela Brisebarre, Camille Capel, Etienne Simon-Lorière, Vincent Enouf, Maud Vanpeene, Sylvie van der Werf, Bour Jean Baptiste |
| EPI_ISL_941028, EPI_ISL_941029, EPI_ISL_941039, EPI_ISL_941042, EPI_ISL_941043, EPI_ISL_941047, EPI_ISL_941052, EPI_ISL_941054, EPI_ISL_941055, EPI_ISL_941059, EPI_ISL_941062, EPI_ISL_941064, EPI_ISL_941067, EPI_ISL_941068, EPI_ISL_941070, EPI_ISL_941072, EPI_ISL_941077, EPI_ISL_941078, EPI_ISL_941079, EPI_ISL_941080, EPI_ISL_941081, EPI_ISL_941082, EPI_ISL_941083, EPI_ISL_941084, EPI_ISL_941085, EPI_ISL_941086, EPI_ISL_941087, EPI_ISL_941088, EPI_ISL_941089, EPI_ISL_941090, EPI_ISL_941091, EPI_ISL_941093, EPI_ISL_941094, EPI_ISL_941095, EPI_ISL_941096, EPI_ISL_941097, EPI_ISL_941098, EPI_ISL_941099, EPI_ISL_941100, EPI_ISL_941101, EPI_ISL_941102, EPI_ISL_941103 | | | |
| see above | Labo Analyses Med | National Reference Center for Viruses of Respiratory Infections, Institut Pasteur, Paris | Marion Barbet, Sylvie Behillil, Méline Bizard, Angela Brisebarre, Camille Capel, Etienne Simon-Lorière, Vincent Enouf, Maud Vanpeene, Sylvie van der Werf, Merah Kader |
| EPI_ISL_941270 | HIA PERCY - lab biologie | National Reference Center for Viruses of Respiratory Infections, Institut Pasteur, Paris | Marion Barbet, Sylvie Behillil, Méline Bizard, Angela Brisebarre, Camille Capel, Etienne Simon-Lorière, Vincent Enouf, Maud Vanpeene, Sylvie van der Werf, Vincent Foissaud |
| EPI_ISL_942007, EPI_ISL_942008, EPI_ISL_942009, EPI_ISL_942010, EPI_ISL_942011, EPI_ISL_942012 | Centers for Disease Control and Prevention, Dengue Branch | Centers for Disease Control and Prevention, Dengue Branch | Gilberto A. Santiago, Glenda Gonzalez, Betzabel Flores, Keyla Charriez, Gabriela Paz-Bailey, Jorge L. Munoz-Jordan |
| EPI_ISL_942928 | Instituto de Diagnostico y Referencia Epidemiologicos INDRE_RNLSP | Instituto de Diagnostico y Referencia Epidemiologicos (INDRE) | Claudia Wong-Arambula, Abril Rodriguez-Maldonado, Fabiola Garces-Ayala, Natividad Cruz-Ortiz, Tatiana Nunez-Garcia, Gisela Barrera-Badillo, Lucia Hernandez-Rivas, Irma Lopez-Martinez, Ernesto Ramirez-Gonzalez. |
| EPI_ISL_942960, EPI_ISL_942971 | National Institute of Health Research and Development | National Institute of Health Research and Development | Subangkit, Hana Apsari Pawestri, Kartika Dewi Puspa, Arie Ardiansyah Nugraha, Hartanti Dian Ikawati, Krisna Nur Andriana Pangesti, Yuni Rukminiati, Ririn Ramadhany, Agustiningsih, Kindi Adam, Holy Arif Wibowo, Triyani Soekarso, Ni Ketut Susilarini, Nurika Hariastuti, Ully Alfi Nikmah, Reni Herman, Nike Susanti, Herna, Tati Febriyanti, Natalie Laurencia Kipuw, Fauzul Muna, Irene Lorinda Indalao, Nelly Puspandari, Vivi Setiawaty. |
| EPI_ISL_943548 | National Institute of Health Research and Development | National Institute of Health Research and Development | Subangkit, Hana Apsari Pawestri, Kartika Dewi Puspa, Arie Ardiansyah Nugraha, Hartanti Dian Ikawati, Krisna Nur Andriana Pangesti, Yuni Rukminiati, Ririn Ramadhany, Agustiningsih, Kindi Adam, Holy Arif Wibowo, Triyani Soekarso, Ni Ketut Susilarini, Nurika Hariastuti, Ully Alfi Nikmah, Reni Herman, Nike Susanti, Herna, Tati Febriyanti, Natalie Laurencia Kipuw, Fauzul Muna, Irene Lorinda Indalao, Nelly Puspandari, Vivi Setiawaty |
| EPI_ISL_943564, EPI_ISL_943565, EPI_ISL_943567, EPI_ISL_943568, EPI_ISL_943573 | Servizo de Microbioloxía. Complexo Hospitalario Universitario de Santiago de Compostela | Servizo de Microbioloxía. Complexo Hospitalario Universitario de Santiago de Compostela | Antonio Aguilera, Gema Barbeito, Amparo Coira, José Costa, Rocio Trastoy, María Luisa Pérez del Molino. |
| EPI_ISL_943608 | Central Laboratory of Public Health of Rio Grande do Sul (Lacen-RS) | State Center for Health Surveillance of the Health Department of the State of Rio Grande do Sul (CEVS/SES-RS) | Aline Campos, Amanda da Silva, Anelise Schaurich, Claudia Dornelles, Cynthia Molina, Fernanda Godinho, Lara Crescente, Leticia Garay, Regina Barcellos, Richard Salvato, Tatiana Gregianini, Vagner Fonseca |
| EPI_ISL_943805, EPI_ISL_943809 | Servizo de Microbioloxía. Complexo Hospitalario Universitario de Santiago de Compostela | Servizo de Microbioloxía. Complexo Hospitalario Universitario de Santiago de Compostela | Antonio Aguilera, Gema Barbeito, Amparo Coira, José Costa, Rocio Trastoy, María Luisa Pérez del Molino. |
| EPI_ISL_943973, EPI_ISL_943974, EPI_ISL_943975, EPI_ISL_943976, EPI_ISL_943977, EPI_ISL_943978, EPI_ISL_943979, EPI_ISL_943980, EPI_ISL_943981, EPI_ISL_943982, EPI_ISL_943983, EPI_ISL_943985, EPI_ISL_943986 | | | |
| see above | LACEN do Estado de Tocantins | Instituto Adolfo Lutz, Interdiciplinary Procedures Center, Strategic Laboratory | Claudio Tavares Sacchi, Claudia Regina Gonçalves, Erica Valessa Ramos Gomes, Karoline Rodrigues Campos |
| EPI_ISL_943989 | LACEN do Estado de Goias | Instituto Adolfo Lutz, Interdiciplinary Procedures Center, Strategic Laboratory | Claudio Tavares Sacchi, Claudia Regina Gonçalves, Erica Valessa Ramos Gomes, Karoline Rodrigues Campos |
| EPI_ISL_943991 | LACEN do Estado de Tocantins | Instituto Adolfo Lutz, Interdiciplinary Procedures Center, Strategic Laboratory | Claudio Tavares Sacchi, Claudia Regina Gonçalves, Erica Valessa Ramos Gomes, Karoline Rodrigues Campos |
| EPI_ISL_944094, EPI_ISL_944096, EPI_ISL_944099, EPI_ISL_944100, EPI_ISL_944102, EPI_ISL_944103, EPI_ISL_944104 | Institute for Medical Research, Infectious Disease Research Centre, National Institutes of Health, Ministry of Health Malaysia | Institute for Medical Research, Infectious Disease Research Centre, National Institutes of Health, Ministry of Health Malaysia | Suppiah J, Kamel K, Azizan MA, Thayan R |
| EPI_ISL_944753 | State Virus Research and Diagnostic Laboratory (VRDL), AIIMS Raipur | State Virus Research and Diagnostic Laboratory (VRDL), AIIMS Raipur | Pushpendra Singh, Kuldeep Sharma, Pragya Agarwala, Priyanka Singh, Somya Sharma, Sanjay Singh Negi, Anudita Bhargava |
| EPI_ISL_947247 | RSU Bunda Mulia | Eijkman Institute for Molecular Biology, Ministry of Research and Technology/National Agency for Research and Innovation | Edison Johar, Frilasita A Yudhaputri, Hidayat Trimarsanto, Iskandar Adnan, Lydia V. Panggalo, Sukma Oktavianthi, Willy Agustine, Safarina G Malik, Khin Saw Myint, Amin Soebandrio |
| EPI_ISL_947249, EPI_ISL_947252 | RS Bina Husada Cibinong | Eijkman Institute for Molecular Biology, Ministry of Research and Technology/National Agency for Research and Innovation | Edison Johar, Frilasita A Yudhaputri, Hidayat Trimarsanto, Iskandar Adnan, Lydia V. Panggalo, Sukma Oktavianthi, Willy Agustine, Safarina G Malik, Khin Saw Myint, Amin Soebandrio |
| EPI_ISL_947255 | RSU Meilia | Eijkman Institute for Molecular Biology, Ministry of Research and Technology/National Agency for Research and Innovation | Edison Johar, Frilasita A Yudhaputri, Hidayat Trimarsanto, Iskandar Adnan, Lydia V. Panggalo, Sukma Oktavianthi, Willy Agustine, Safarina G Malik, Khin Saw Myint, Amin Soebandrio |
| EPI_ISL_947257 | RS Bina Husada Cibinong | Eijkman Institute for Molecular Biology, Ministry of Research and Technology/National Agency for Research and Innovation | Edison Johar, Frilasita A Yudhaputri, Hidayat Trimarsanto, Iskandar Adnan, Lydia V. Panggalo, Sukma Oktavianthi, Willy Agustine, Safarina G Malik, Khin Saw Myint, Amin Soebandrio |
| EPI_ISL_947262 | RS Qadr | Eijkman Institute for Molecular Biology, Ministry of Research and Technology/National Agency for Research and Innovation | Frilasita A Yudhaputri, Hidayat Trimarsanto, Iskandar Adnan, Lydia V. Panggalo, Sukma Oktavianthi, Willy Agustine, Edison Johar, Safarina G Malik, Khin Saw Myint, Amin Soebandrio |
| EPI_ISL_947265 | RS Sari Asih Ciledug | Eijkman Institute for Molecular Biology, Ministry of Research and Technology/National Agency for Research and Innovation | Frilasita A Yudhaputri, Hidayat Trimarsanto, Iskandar Adnan, Lydia V. Panggalo, Sukma Oktavianthi, Willy Agustine, Edison Johar, Safarina G Malik, Khin Saw Myint, Amin Soebandrio |
| EPI_ISL_947268 | RSU Anggrek Mas | Eijkman Institute for Molecular Biology, Ministry of Research and Technology/National Agency for Research and Innovation | Frilasita A Yudhaputri, Hidayat Trimarsanto, Iskandar Adnan, Lydia V. Panggalo, Sukma Oktavianthi, Willy Agustine, Edison Johar, Safarina G Malik, Khin Saw Myint, Amin Soebandrio |
| EPI_ISL_947275 | RS Umum Mulia Pajajaran | Eijkman Institute for Molecular Biology, Ministry of Research and Technology/National Agency for Research and Innovation | Frilasita A Yudhaputri, Hidayat Trimarsanto, Iskandar Adnan, Lydia V. Panggalo, Sukma Oktavianthi, Willy Agustine, Edison Johar, Safarina G Malik, Khin Saw Myint, Amin Soebandrio |
| EPI_ISL_947278 | RS Umum Ciputra Hospital Citragarden City | Eijkman Institute for Molecular Biology, Ministry of Research | Hidayat Trimarsanto, Iskandar Adnan, Lydia V. Panggalo, Sukma Oktavianthi, Willy Agustine, Edison Johar, Frilasita A Yudhaputri, Safarina G Malik, Khin Saw |

| | | | |
|---|---|---|---|
| | | and Technology/National Agency for Research and Innovation | Myint, Amin Soebandrio |
| EPI_ISL_947281 | RSU Medika Dramaga | Eijkman Institute for Molecular Biology, Ministry of Research and Technology/National Agency for Research and Innovation | Hidayat Trimarsanto, Iskandar Adnan, Lydia V. Panggalo, Sukma Oktavianthi, Willy Agustine, Edison Johar, Frilasita A Yudhaputri, Safarina G Malik, Khin Saw Myint, Amin Soebandrio |
| EPI_ISL_947284 | RSIA Bunda Sejati | Eijkman Institute for Molecular Biology, Ministry of Research and Technology/National Agency for Research and Innovation | Hidayat Trimarsanto, Iskandar Adnan, Lydia V. Panggalo, Sukma Oktavianthi, Willy Agustine, Edison Johar, Frilasita A Yudhaputri, Safarina G Malik, Khin Saw Myint, Amin Soebandrio |
| EPI_ISL_947287 | RSIA PKU Muhammadiyah | Eijkman Institute for Molecular Biology, Ministry of Research and Technology/National Agency for Research and Innovation | Hidayat Trimarsanto, Iskandar Adnan, Lydia V. Panggalo, Sukma Oktavianthi, Willy Agustine, Edison Johar, Frilasita A Yudhaputri, Safarina G Malik, Khin Saw Myint, Amin Soebandrio |
| EPI_ISL_947290 | RS Hermina Bitung | Eijkman Institute for Molecular Biology, Ministry of Research and Technology/National Agency for Research and Innovation | Hidayat Trimarsanto, Iskandar Adnan, Lydia V. Panggalo, Sukma Oktavianthi, Willy Agustine, Edison Johar, Frilasita A Yudhaputri, Safarina G Malik, Khin Saw Myint, Amin Soebandrio |
| EPI_ISL_947298 | RSU Medirossa | Eijkman Institute for Molecular Biology, Ministry of Research and Technology/National Agency for Research and Innovation | Iskandar Adnan, Lydia V. Panggalo, Sukma Oktavianthi, Willy Agustine, Edison Johar, Hidayat Trimarsanto, Frilasita A Yudhaputri, Safarina G Malik, Khin Saw Myint, Amin Soebandrio |
| EPI_ISL_947301 | RS Mitra Keluarga Cibubur | Eijkman Institute for Molecular Biology, Ministry of Research and Technology/National Agency for Research and Innovation | Iskandar Adnan, Lydia V. Panggalo, Sukma Oktavianthi, Willy Agustine, Edison Johar, Hidayat Trimarsanto, Frilasita A Yudhaputri, Safarina G Malik, Khin Saw Myint, Amin Soebandrio |
| EPI_ISL_947307 | RS Sentra Medika Cikarang | Eijkman Institute for Molecular Biology, Ministry of Research and Technology/National Agency for Research and Innovation | Lydia V. Panggalo, Sukma Oktavianthi, Willy Agustine, Edison Johar, Hidayat Trimarsanto, Iskandar Adnan, Frilasita A Yudhaputri, Safarina G Malik, Khin Saw Myint, Amin Soebandrio |
| EPI_ISL_947312 | RSU Bunda Mulia | Eijkman Institute for Molecular Biology, Ministry of Research and Technology/National Agency for Research and Innovation | Lydia V. Panggalo, Sukma Oktavianthi, Willy Agustine, Edison Johar, Hidayat Trimarsanto, Iskandar Adnan, Frilasita A Yudhaputri, Safarina G Malik, Khin Saw Myint, Amin Soebandrio |
| EPI_ISL_947315 | RSU Harapan Bunda | Eijkman Institute for Molecular Biology, Ministry of Research and Technology/National Agency for Research and Innovation | Lydia V. Panggalo, Sukma Oktavianthi, Willy Agustine, Edison Johar, Hidayat Trimarsanto, Iskandar Adnan, Frilasita A Yudhaputri, Safarina G Malik, Khin Saw Myint, Amin Soebandrio |
| EPI_ISL_947317 | RSU Medirossa | Eijkman Institute for Molecular Biology, Ministry of Research and Technology/National Agency for Research and Innovation | Lydia V. Panggalo, Sukma Oktavianthi, Willy Agustine, Edison Johar, Hidayat Trimarsanto, Iskandar Adnan, Frilasita A Yudhaputri, Safarina G Malik, Khin Saw Myint, Amin Soebandrio |
| EPI_ISL_947320 | RS Sari Asih Ciledug | Eijkman Institute for Molecular Biology, Ministry of Research and Technology/National Agency for Research and Innovation | Sukma Oktavianthi, Willy Agustine, Edison Johar, Hidayat Trimarsanto, Iskandar Adnan, Lydia V. Panggalo, Frilasita A Yudhaputri, Safarina G Malik, Khin Saw Myint, Amin Soebandrio |
| EPI_ISL_947322 | RS Mitra Keluarga Kelapa Gading | Eijkman Institute for Molecular Biology, Ministry of Research and Technology/National Agency for Research and Innovation | Sukma Oktavianthi, Willy Agustine, Edison Johar, Hidayat Trimarsanto, Iskandar Adnan, Lydia V. Panggalo, Frilasita A Yudhaputri, Safarina G Malik, Khin Saw Myint, Amin Soebandrio |
| EPI_ISL_947324 | RSU Medirossa | Eijkman Institute for Molecular Biology, Ministry of Research and Technology/National Agency for Research and Innovation | Sukma Oktavianthi, Willy Agustine, Edison Johar, Hidayat Trimarsanto, Iskandar Adnan, Lydia V. Panggalo, Frilasita A Yudhaputri, Safarina G Malik, Khin Saw Myint, Amin Soebandrio |
| EPI_ISL_947325 | RSIA PKU Muhammadiyah | Eijkman Institute for Molecular Biology, Ministry of Research and Technology/National Agency for Research and Innovation | Sukma Oktavianthi, Willy Agustine, Edison Johar, Hidayat Trimarsanto, Iskandar Adnan, Lydia V. Panggalo, Frilasita A Yudhaputri, Safarina G Malik, Khin Saw Myint, Amin Soebandrio |
| EPI_ISL_947326, EPI_ISL_947327 | RS Mitra Keluarga Gading Serpong | Eijkman Institute for Molecular Biology, Ministry of Research and Technology/National Agency for Research and Innovation | Willy Agustine, Edison Johar, Hidayat Trimarsanto, Iskandar Adnan, Lydia V. Panggalo, Sukma Oktavianthi, Frilasita A Yudhaputri, Safarina G Malik, Khin Saw Myint, Amin Soebandrio |
| EPI_ISL_947329, EPI_ISL_947330 | RSU Bunda Mulia | Eijkman Institute for Molecular Biology, Ministry of Research and Technology/National Agency for Research and Innovation | Willy Agustine, Edison Johar, Hidayat Trimarsanto, Iskandar Adnan, Lydia V. Panggalo, Sukma Oktavianthi, Frilasita A Yudhaputri, Safarina G Malik, Khin Saw Myint, Amin Soebandrio |
| EPI_ISL_949183 | National Institute of Health Research and Development | National Institute of Health Research and Development | Kindi Adam; Holy Arif Wibowo; Ririn Ramadhany; Yuni Rukminiati; Agustiningsih; Hana Apsari Pawestri; Subangkit; Kartika Dewi Puspa; Arie Ardiansyah Nugraha; Hartanti Dian Ikawati; Krisna Nur Andriana Pangesti; Triyani Soekarso; Ni Ketut Susilarini; Nur Ika Hariastuti; Uly Alfi Nikmah; Mursinah; Asri Febriyani; Reni Herman; Nike Susanti; Herna; Tati Febriyanti; Natalie Laurencia Kipuw; Fauzul Muna; Irene Lorinda Indalao; Aulia Rizki; Nelly Puspandari; Vivi Setiawaty |
| EPI_ISL_949192 | National Institute of Health Research and Development | National Institute of Health Research and Development | Ririn Ramadhany; Yuni Rukminiati; Agustiningsih; Kindi Adam; Holy Arif Wibowo; Hana Apsari Pawestri; Subangkit; Kartika Dewi Puspa; Arie Ardiansyah Nugraha; Hartanti Dian Ikawati; Krisna Nur Andriana Pangesti; Ni Ketut Susilarini; Nur Ika Hariastuti; Uly Alfi Nikmah; Mursinah; Asri Febriyani; Reni Herman; Nike Susanti; Herna; Tati Febriyanti; Natalie Laurentia Kipuw; Fauzul Muna; Irene Lorinda Indalao; Aulia Rizki; Nelly Puspandari; Vivi Setiawaty |
| EPI_ISL_953427 | National Institute of Health Research and Development | National Institute of Health Research and Development | Subangkit, Hana Apsari Pawestri, Kartika Dewi Puspa, Arie Ardiansyah Nugraha, Hartanti Dian Ikawati, Krisna Nur Andriana Pangesti, Yuni Rukminiati, Ririn Ramadhany, Agustiningsih, Kindi Adam, Holy Arif Wibowo, Triyani Soekarso, Ni Ketut Susilarini, Nurika Hariastuti, Ully Alfi Nikmah, Reni Herman, Nike Susanti, Herna, Tati Febriyanti, Natalie Laurentia Kipuw, Fauzul Muna, Irene Lorinda Indalao, Nelly Puspandari, Vivi Setiawaty |
| EPI_ISL_953941 | Hopital | National Reference Center for Viruses of Respiratory Infections, Institut Pasteur, Paris | Marion Barbet, Sylvie Behillil, Méline Bizard, Angela Brisebarre, Camille Capel, Etienne Simon-Lorière, Vincent Enouf, Maud Vanpeene, Sylvie van der Werf,Lagathu GisèLe |
| EPI_ISL_953994 | Labo Analyses Med | National Reference Center for Viruses of Respiratory Infections, Institut Pasteur, Paris | Marion Barbet, Sylvie Behillil, Méline Bizard, Angela Brisebarre, Camille Capel, Etienne Simon-Lorière, Vincent Enouf, Maud Vanpeene, Sylvie van der Werf,Amzalag Jonas |
| EPI_ISL_953995, EPI_ISL_953996, EPI_ISL_953997 | Hopital | National Reference Center for Viruses of Respiratory Infections, Institut Pasteur, Paris | Marion Barbet, Sylvie Behillil, Méline Bizard, Angela Brisebarre, Camille Capel, Etienne Simon-Lorière, Vincent Enouf, Maud Vanpeene, Sylvie van der Werf,Lagathu GisèLe |
| EPI_ISL_953998, EPI_ISL_953999 | Labo Analyses Med | National Reference Center for Viruses of Respiratory Infections, Institut Pasteur, Paris | Marion Barbet, Sylvie Behillil, Méline Bizard, Angela Brisebarre, Camille Capel, Etienne Simon-Lorière, Vincent Enouf, Maud Vanpeene, Sylvie van der Werf,Amzalag Jonas |
| EPI_ISL_954000 | Hopital | National Reference Center for Viruses of Respiratory Infections, Institut Pasteur, Paris | Marion Barbet, Sylvie Behillil, Méline Bizard, Angela Brisebarre, Camille Capel, Etienne Simon-Lorière, Vincent Enouf, Maud Vanpeene, Sylvie van der Werf,Schvoerer Evelyne |
| EPI_ISL_954001 | Hopital | National Reference Center for Viruses of Respiratory Infections, Institut Pasteur, Paris | Marion Barbet, Sylvie Behillil, Méline Bizard, Angela Brisebarre, Camille Capel, Etienne Simon-Lorière, Vincent Enouf, Maud Vanpeene, Sylvie van der Werf,Eloy Clarence |
| EPI_ISL_954002, EPI_ISL_954003 | Hopital | National Reference Center for Viruses of Respiratory Infections, Institut Pasteur, Paris | Marion Barbet, Sylvie Behillil, Méline Bizard, Angela Brisebarre, Camille Capel, Etienne Simon-Lorière, Vincent Enouf, Maud Vanpeene, Sylvie van der Werf,Ducancelle Alexandra |

| | | | |
|---|---|---|---|
| EPI_ISL_954004, EPI_ISL_954005, EPI_ISL_954006 | Labo Analyses Med | National Reference Center for Viruses of Respiratory Infections, Institut Pasteur, Paris | Marion Barbet, Sylvie Behillil, Méline Bizard, Angela Brisebarre, Camille Capel, Etienne Simon-Lorière, Vincent Enouf, Maud Vanpeene, Sylvie van der Werf,Laboratoire Palissy |
| EPI_ISL_954007, EPI_ISL_954008, EPI_ISL_954022, EPI_ISL_954023, EPI_ISL_954024, EPI_ISL_954025, EPI_ISL_954026, EPI_ISL_954027, EPI_ISL_954028, EPI_ISL_954029, EPI_ISL_954031 | | | |
| see above | Labo Analyses Med | National Reference Center for Viruses of Respiratory Infections, Institut Pasteur, Paris | Marion Barbet, Sylvie Behillil, Méline Bizard, Angela Brisebarre, Camille Capel, Etienne Simon-Lorière, Vincent Enouf, Maud Vanpeene, Sylvie van der Werf,Amzalag Jonas |
| EPI_ISL_954035 | Outre Mer | National Reference Center for Viruses of Respiratory Infections, Institut Pasteur, Paris | Marion Barbet, Sylvie Behillil, Méline Bizard, Angela Brisebarre, Camille Capel, Etienne Simon-Lorière, Vincent Enouf, Maud Vanpeene, Sylvie van der Werf,Rousset Dominique |
| EPI_ISL_954114 | Labo Analyses Med | National Reference Center for Viruses of Respiratory Infections, Institut Pasteur, Paris | Marion Barbet, Sylvie Behillil, Méline Bizard, Angela Brisebarre, Camille Capel, Etienne Simon-Lorière, Vincent Enouf, Maud Vanpeene, Sylvie van der Werf,Selas Labomaine |
| EPI_ISL_954115 | Hopital | National Reference Center for Viruses of Respiratory Infections, Institut Pasteur, Paris | Marion Barbet, Sylvie Behillil, Méline Bizard, Angela Brisebarre, Camille Capel, Etienne Simon-Lorière, Vincent Enouf, Maud Vanpeene, Sylvie van der Werf,Lagathu GisèLe |
| EPI_ISL_954116, EPI_ISL_954117, EPI_ISL_954118 | Outre Mer | National Reference Center for Viruses of Respiratory Infections, Institut Pasteur, Paris | Marion Barbet, Sylvie Behillil, Méline Bizard, Angela Brisebarre, Camille Capel, Etienne Simon-Lorière, Vincent Enouf, Maud Vanpeene, Sylvie van der Werf,Rousset Dominique |
| EPI_ISL_954119 | Labo Analyses Med | National Reference Center for Viruses of Respiratory Infections, Institut Pasteur, Paris | Marion Barbet, Sylvie Behillil, Méline Bizard, Angela Brisebarre, Camille Capel, Etienne Simon-Lorière, Vincent Enouf, Maud Vanpeene, Sylvie van der Werf,Takoudju Eve-Marie |
| EPI_ISL_954120 | Hopital | National Reference Center for Viruses of Respiratory Infections, Institut Pasteur, Paris | Marion Barbet, Sylvie Behillil, Méline Bizard, Angela Brisebarre, Camille Capel, Etienne Simon-Lorière, Vincent Enouf, Maud Vanpeene, Sylvie van der Werf,Lagathu GisèLe |
| EPI_ISL_954121 | Labo Analyses Med | National Reference Center for Viruses of Respiratory Infections, Institut Pasteur, Paris | Marion Barbet, Sylvie Behillil, Méline Bizard, Angela Brisebarre, Camille Capel, Etienne Simon-Lorière, Vincent Enouf, Maud Vanpeene, Sylvie van der Werf,Takoudju Eve-Marie |
| EPI_ISL_954122 | Hopital | National Reference Center for Viruses of Respiratory Infections, Institut Pasteur, Paris | Marion Barbet, Sylvie Behillil, Méline Bizard, Angela Brisebarre, Camille Capel, Etienne Simon-Lorière, Vincent Enouf, Maud Vanpeene, Sylvie van der Werf,Lagathu GisèLe |
| EPI_ISL_954123, EPI_ISL_954124 | Labo Analyses Med | National Reference Center for Viruses of Respiratory Infections, Institut Pasteur, Paris | Marion Barbet, Sylvie Behillil, Méline Bizard, Angela Brisebarre, Camille Capel, Etienne Simon-Lorière, Vincent Enouf, Maud Vanpeene, Sylvie van der Werf,Takoudju Eve-Marie |
| EPI_ISL_954125 | Outre Mer | National Reference Center for Viruses of Respiratory Infections, Institut Pasteur, Paris | Marion Barbet, Sylvie Behillil, Méline Bizard, Angela Brisebarre, Camille Capel, Etienne Simon-Lorière, Vincent Enouf, Maud Vanpeene, Sylvie van der Werf,Rousset Dominique |
| EPI_ISL_954126 | Hopital | National Reference Center for Viruses of Respiratory Infections, Institut Pasteur, Paris | Marion Barbet, Sylvie Behillil, Méline Bizard, Angela Brisebarre, Camille Capel, Etienne Simon-Lorière, Vincent Enouf, Maud Vanpeene, Sylvie van der Werf,Lesimple BéAtrice |
| EPI_ISL_954127, EPI_ISL_954128 | Labo Analyses Med | National Reference Center for Viruses of Respiratory Infections, Institut Pasteur, Paris | Marion Barbet, Sylvie Behillil, Méline Bizard, Angela Brisebarre, Camille Capel, Etienne Simon-Lorière, Vincent Enouf, Maud Vanpeene, Sylvie van der Werf,Takoudju Eve-Marie |
| EPI_ISL_954129 | Labo Analyses Med | National Reference Center for Viruses of Respiratory Infections, Institut Pasteur, Paris | Marion Barbet, Sylvie Behillil, Méline Bizard, Angela Brisebarre, Camille Capel, Etienne Simon-Lorière, Vincent Enouf, Maud Vanpeene, Sylvie van der Werf |
| EPI_ISL_954130, EPI_ISL_954131, EPI_ISL_954132, EPI_ISL_954133, EPI_ISL_954134, EPI_ISL_954135 | Labo Analyses Med | National Reference Center for Viruses of Respiratory Infections, Institut Pasteur, Paris | Marion Barbet, Sylvie Behillil, Méline Bizard, Angela Brisebarre, Camille Capel, Etienne Simon-Lorière, Vincent Enouf, Maud Vanpeene, Sylvie van der Werf,Desjardins Nicolas |
| EPI_ISL_954136, EPI_ISL_954137, EPI_ISL_954138 | Labo Analyses Med | National Reference Center for Viruses of Respiratory Infections, Institut Pasteur, Paris | Marion Barbet, Sylvie Behillil, Méline Bizard, Angela Brisebarre, Camille Capel, Etienne Simon-Lorière, Vincent Enouf, Maud Vanpeene, Sylvie van der Werf,Breton-Cazaux JéRôMe |
| EPI_ISL_954780 | COMPLEJO HOSPITALARIO XERAL-CALDE | Instituto de Salud Carlos III | Iglesias-Caballero, M. Camarero, S. Sandonís,V. Vázquez, S. Pozo, F. Casas, I. Jiménez, P. Zaballos, A. Monzón, S. Varona, S. Cuesta, I. Alonso, P. |
| EPI_ISL_954781 | Hospital General Universitario de Ciudad Real | Instituto de Salud Carlos III | Iglesias-Caballero, M. Camarero, S. Sandonís,V. Vázquez, S. Pozo, F. Casas, I. Jiménez, P. Zaballos, A. Monzón, S. Varona, S. Cuesta, I. Illescas, S. |
| EPI_ISL_954784, EPI_ISL_954786 | Complejo Hospitalario de Navarra | Instituto de Salud Carlos III | Iglesias-Caballero, M. Camarero, S. Sandonís,V. Vázquez, S. Pozo, F. Casas, I. Jiménez, P. Zaballos, A. Monzón, S. Varona, S. Cuesta, I. Ezpeleta,C. |
| EPI_ISL_954788 | Servicio Murciano de Salud | Instituto de Salud Carlos III | Iglesias-Caballero, M. Camarero, S. Sandonís,V. Vázquez, S. Pozo, F. Casas, I. Jiménez, P. Zaballos, A. Monzón, S. Varona, S. Cuesta, I. Blázquez, A. |
| EPI_ISL_954790 | Hospital J.M. Morales Meseguer | Instituto de Salud Carlos III | Iglesias-Caballero, M. Camarero, S. Sandonís,V. Vázquez, S. Pozo, F. Casas, I. Jiménez, P. Zaballos, A. Monzón, S. Varona, S. Cuesta, I. Navarro, D. |
| EPI_ISL_954792 | HOSPITAL COMARCAL DE MELILLA | Instituto de Salud Carlos III | Iglesias-Caballero, M. Camarero, S. Sandonís,V. Vázquez, S. Pozo, F. Casas, I. Jiménez, P. Zaballos, A. Monzón, S. Varona, S. Cuesta, I. Roman, S. |
| EPI_ISL_954793 | Hospital Universitario Virgen de la Arrixaca | Instituto de Salud Carlos III | Iglesias-Caballero, M. Camarero, S. Sandonís,V. Vázquez, S. Pozo, F. Casas, I. Jiménez, P. Zaballos, A. Monzón, S. Varona, S. Cuesta, I. Moreno, L. |
| EPI_ISL_954796 | HOSPITAL COMARCAL DE MELILLA | Instituto de Salud Carlos III | Iglesias-Caballero, M. Camarero, S. Sandonís,V. Vázquez, S. Pozo, F. Casas, I. Jiménez, P. Zaballos, A. Monzón, S. Varona, S. Cuesta, I. Illescas, S. |
| EPI_ISL_954798 | Hospital Comarcal de Melilla | Instituto de Salud Carlos III | Iglesias-Caballero, M. Camarero, S. Sandonís,V. Vázquez, S. Pozo, F. Casas, I. Jiménez, P. Zaballos, A. Monzón, S. Varona, S. Cuesta, I. Roman, S. |
| EPI_ISL_954804 | Complejo Hospitalario de Navarra | Instituto de Salud Carlos III | Iglesias-Caballero, M. Camarero, S. Sandonís,V. Vázquez, S. Pozo, F. Casas, I. Jiménez, P. Zaballos, A. Monzón, S. Varona, S. Cuesta, I. Ezpeleta,C. |
| EPI_ISL_954805 | Hospital General Universitario de Ciudad Real | Instituto de Salud Carlos III | Iglesias-Caballero, M. Camarero, S. Sandonís,V. Vázquez, S. Pozo, F. Casas, I. Jiménez, P. Zaballos, A. Monzón, S. Varona, S. Cuesta, I. Illescas, S. |
| EPI_ISL_954806 | Gerencia de Asistencia Sanitaria de Soria | Instituto de Salud Carlos III | Iglesias-Caballero, M. Camarero, S. Sandonís,V. Vázquez, S. Pozo, F. Casas, I. Jiménez, P. Zaballos, A. Monzón, S. Varona, S. Cuesta, I. Aldea, C. |
| EPI_ISL_954807, EPI_ISL_954808 | Complejo Hospitalario Universitario La Coruña | Instituto de Salud Carlos III | Iglesias-Caballero, M. Camarero, S. Sandonís,V. Vázquez, S. Pozo, F. Casas, I. Jiménez, P. Zaballos, A. Monzón, S. Varona, S. Cuesta, I. Peña, F. |
| EPI_ISL_954809 | Hospital General de Lanzarote | Instituto de Salud Carlos III | Iglesias-Caballero, M. Camarero, S. Sandonís,V. Vázquez, S. Pozo, F. Casas, I. Jiménez, P. Zaballos, A. Monzón, S. Varona, S. Cuesta, I. Copado, R. |
| EPI_ISL_954811, EPI_ISL_954813 | HOSPITAL UNIVERSITARIO DE GUADALAJARA | Instituto de Salud Carlos III | Iglesias-Caballero, M. Camarero, S. Sandonís,V. Vázquez, S. Pozo, F. Casas, I. Jiménez, P. Zaballos, A. Monzón, S. Varona, S. Cuesta, I. González, A. |
| EPI_ISL_955146 | University of Sarajevo, Veterinary Faculty, Laboratory for Molecular Diagnostic and Research Laboratory | University of Sarajevo, Veterinary Faculty, Laboratory for Molecular Diagnostic and Research Laboratory | Goleti T., Goleti Š., Softi A., Ali-Šeho A., Nicevi M., Jaži A., Šabi E., Terzi I., Hodži A. |
| EPI_ISL_955148 | University of Sarajevo, Veterinary Faculty, Laboratory for Molecular Diagnostic and Research Laboratory | University of Sarajevo, Veterinary Faculty, Laboratory for Molecular Diagnostic and Research Laboratory | Goleti T., Goleti Š., Softi A., Ali-Šeho A., Jaži A., Šabi E., Terzi I., Nicevi M., Hodži A. |
| EPI_ISL_955150 | University of Sarajevo, Veterinary Faculty, Laboratory for Molecular Diagnostic and Research Laboratory | University of Sarajevo, Veterinary Faculty, Laboratory for Molecular Diagnostic and Research Laboratory | Goleti T., Goleti Š., Softi A., Ali-Šeho A., Hodži A., Šabi E., Jaži A., Nicevi M., Terzi I. |
| EPI_ISL_955151 | University of Sarajevo, Veterinary Faculty, Laboratory for Molecular Diagnostic and Research Laboratory | University of Sarajevo, Veterinary Faculty, Laboratory for Molecular Diagnostic and Research Laboratory | Goleti T., Goleti Š., Softi A., Ali-Šeho A., Terzi I., Šabi E., Jaži A., Nicevi M., Hodži A. |
| EPI_ISL_955155 | University of Sarajevo, Veterinary Faculty, Laboratory for Molecular Diagnostic and Research Laboratory | University of Sarajevo, Veterinary Faculty, Laboratory for Molecular Diagnostic and Research Laboratory | Goleti T., Goleti Š., Softi A., Ali-Šeho A., Hodži A., Šabi E., Jaži A., Nicevi M., Terzi I. |
| EPI_ISL_955156 | University of Sarajevo, Veterinary Faculty, Laboratory for Molecular Diagnostic and Research Laboratory | University of Sarajevo, Veterinary Faculty, Laboratory for Molecular Diagnostic and Research Laboratory | Goleti T., Goleti Š., Softi A., Ali-Šeho A., Jaži A., Šabi E., Nicevi M., Hodži A., Terzi I. |
| EPI_ISL_955169 | University of Sarajevo, Veterinary Faculty, Laboratory for Molecular Diagnostic and Research Laboratory | University of Sarajevo, Veterinary Faculty, Laboratory for Molecular Diagnostic and Research Laboratory | Goleti Š., Goleti T., Softi A., Ali-Šeho A., Šabi E., Jaži A., Nicevi M., Hodži A., Terzi I. |
| EPI_ISL_955170 | University of Sarajevo, Veterinary Faculty, Laboratory for Molecular Diagnostic and Research Laboratory | University of Sarajevo, Veterinary Faculty, Laboratory for Molecular Diagnostic and Research Laboratory | Goleti T., Goleti Š., Ali-Šeho A., Softi A., Šabi E., Jaži A., Nicevi M., Hodži A., Terzi I. |

| | | | |
|---|---|---|---|
| EPI_ISL_955171 | University of Sarajevo, Veterinary Faculty, Laboratory for Molecular Diagnostic and Research Laboratory | University of Sarajevo, Veterinary Faculty, Laboratory for Molecular Diagnostic and Research Laboratory | Goleti Š., Goleti T., Ali-Šeho A., Softi A., Nicevi M., Šabi E., Jaži A., Hodži A., Terzi I. |
| EPI_ISL_955173 | University of Sarajevo, Veterinary Faculty, Laboratory for Molecular Diagnostic and Research Laboratory | University of Sarajevo, Veterinary Faculty, Laboratory for Molecular Diagnostic and Research Laboratory | Goleti Š., Goleti T., Softi A., Ali-Šeho A., Jaži A., Šabi E., Hodži A., Terzi I., Nicevi M. |
| EPI_ISL_955184 | University of Sarajevo, Veterinary Faculty, Laboratory for Molecular Diagnostic and Research Laboratory | University of Sarajevo, Veterinary Faculty, Laboratory for Molecular Diagnostic and Research Laboratory | Goleti Š., Goleti T., Softi A., Ali-Šeho A., Nicevi M., Hodži A., Terzi I., Šabi E., Jaži A. |
| EPI_ISL_955185 | University of Sarajevo, Veterinary Faculty, Laboratory for Molecular Diagnostic and Research Laboratory | University of Sarajevo, Veterinary Faculty, Laboratory for Molecular Diagnostic and Research Laboratory | Goleti Š., Goleti T., Ali-Šeho A., Softi A., Terzi I., Šabi E., Jaži A., Nicevi M., Hodži A. |
| EPI_ISL_955186 | University of Sarajevo, Veterinary Faculty, Laboratory for Molecular Diagnostic and Research Laboratory | University of Sarajevo, Veterinary Faculty, Laboratory for Molecular Diagnostic and Research Laboratory | Goleti Š., Goleti T., Ali-Šeho A., Softi A., Terzi I., Jaži A., Nicevi M., Hodži A., Šabi E. |
| EPI_ISL_955211 | University of Sarajevo, Veterinary Faculty, Laboratory for Molecular Diagnostic and Research Laboratory | University of Sarajevo, Veterinary Faculty, Laboratory for Molecular Diagnostic and Research Laboratory | Goleti Š., Goleti T., Ali-Šeho A., Softi A., Jaži A., Terzi I., Šabi E., Nicevi M., Hodži A. |
| EPI_ISL_955905 | Pamela Youde Nethersole Eastern Hospital | Hong Kong Department of Health | Alan K.L. Tsang, Peter C.W. Yip, Edman T.K. Lam, Rickjason C.W. Chan, Dominic N.C. Tsang |
| EPI_ISL_956279 | Siti Khodijah Hospital | Institute of Tropical Disease, Universitas Airlangga | Jezzy R Dewantari, Rima R Prasetya, Krisnoadi Rahardjo, Aldise M Nastri, Muhammad Hamdan, Gatot Soegiarto, Laksmi Wulandari, Resti Yudhawati, Soetjipto, Yasuko Mori, Maria I Lusida, Kazufumi Shimizu |
| EPI_ISL_956281 | Siti Khodijah Hospital | Institute of Tropical Disease, Universitas Airlangga | Maria I Lusida, Krisnoadi Rahardjo, Aldise M Nastri, Jezzy R Dewantari, Rima R Prasetya, Muhammad Hamdan, Gatot Soegiarto, Laksmi Wulandari, Resti Yudhawati, Soetjipto, Yasuko Mori, Kazufumi Shimizu |
| EPI_ISL_956289, EPI_ISL_956297 | Instituto Nacional de Salud- Dirección de Redes de Laboratorios de Salud Pública | Instituto Nacional de Salud- Dirección de Investigación en Salud Pública | Katherine Laiton-Donato, Diego A. Álvarez-Díaz, Carlos Franco-Muñoz, Mauricio Pacheco-Montealegre, Hector Alejandro Ruiz-Moreno, Maria T. Herrera-Sepúlveda, Diego Andrés Prada, Jhonnatan Reales-González, Sheryll Corchuelo, Julian Naizaque, Gerardo Santamaría, Magdalena Wiesner, Martha Lucia Ospina Martinez, Marcela Mercado-Reyes |
| EPI_ISL_956313 | Siti Khodijah Hospital | Institute of Tropical Disease, Universitas Airlangga | Jezzy R Dewantari, Rima R Prasetya, Krisnoadi Rahardjo, Aldise M Nastri, Muhammad Hamdan, Gatot Soegiarto, Laksmi Wulandari, Resti Yudhawati, Yasuko Mori, Soetjipto, Kazufumi Shimizu, Maria I Lusida |
| EPI_ISL_956314 | Paru Hospital | Institute of Tropical Disease, Universitas Airlangga | Rima R Prasetya, Krisnoadi Rahardjo, Aldise M Nastri, Jezzy R Dewantari, Dyah Retno, Gatot Soegiarto, Laksmi Wulandari, Resti Yudhawati, Yasuko Mori, Soetjipto, Kazufumi Shimizu, Maria I Lusida |
| EPI_ISL_959279 | National Influenza Center, Virology Department | National Influenza Center | K Sadeghi, A Nejati, J Yavarian, NZ Shafiei Jandaghi, V Salimi, F Ajaminejad,N Ghavvami and T Mokhtari Azad |
| EPI_ISL_959284 | National Influenza Center, Virology Department | National Influenza Center | V Salimi,NZ Shafiei Jandaghi, J Yavarian, A Nejati, K Sadeghi, N Ghavvami,F Ajaminejad and T Mokhtari Azad |
| EPI_ISL_960404, EPI_ISL_960405, EPI_ISL_960408, EPI_ISL_960409, EPI_ISL_960410, EPI_ISL_960411, EPI_ISL_960412, EPI_ISL_960413, EPI_ISL_960414, EPI_ISL_960415, EPI_ISL_960416, EPI_ISL_960417, EPI_ISL_960418, EPI_ISL_960419, EPI_ISL_960423, EPI_ISL_960424, EPI_ISL_960425, EPI_ISL_960426, EPI_ISL_960436, EPI_ISL_960437, EPI_ISL_960438, EPI_ISL_960439 | | | |
| see above | The National Institute of Public Health | State Veterinary Institute Prague | Nagy,A;Vecerova,J;Cernikova,L;Stara,M;Jirincova,H;Trnka,D |
| EPI_ISL_961466 | Laboratorios Lister | Instituto de Diagnostico y Referencia Epidemiologicos (INDRE) | Claudia Wong-Arambula, Abril Rodriguez-Maldonado, Fabiola Garces-Ayala, Natividad Cruz-Ortiz, Tatiana Nunez-Garcia, Gisela Barrera-Badillo, Lucia Hernandez-Rivas, Irma Lopez-Martinez, Ernesto Ramirez-Gonzalez. |
| EPI_ISL_962203, EPI_ISL_962204, EPI_ISL_962206 | Institute for Medical Research, Infectious Disease Research Centre, National Institutes of Health, Ministry of Health Malaysia | Institute for Medical Research, Infectious Disease Research Centre, National Institutes of Health, Ministry of Health Malaysia | Suppiah J, Kamel K, Azizan MA, Thayan R |
| EPI_ISL_964916 | Hopital | National Reference Center for Viruses of Respiratory Infections, Institut Pasteur, Paris | Marion Barbet, Sylvie Behillil, Méline Bizard, Angela Brisebarre, Camille Capel, Etienne Simon-Lorière, Vincent Enouf, Maud Vanpeene, Sylvie van der Werf,Smati-Lafarge Mounira |
| EPI_ISL_964949 | Laboratorio de Virologia del HUCA | Laboratorio de Virologia del HUCA | Castelló C, Gómez de Oña J, Boga JA, Rojo S, Alvarez-Arguelles ME, Abreu F, Costales I, Sandoval M, Perez-Martinez Z, Martín-Rodriguez G, Coto E, Melon S |
| EPI_ISL_965029 | Laboratorio de Virologia HUCA | Laboratorio de Virologia HUCA | Castelló C, Gómez de Oña J, Boga JA, Rojo S, Alvarez-Arguelles ME, Abreu F, Costales I, Sandoval M, Perez-Martínez Z, Martín-Rodríguez G, Coto E, Melón S |
| EPI_ISL_965033 | Laboratorio de Virología HUCA | Laboratorio de Virología HUCA | Castelló C, Gómez de Oña J, Boga JA, Rojo S, Alvarez-Arguelles ME, Abreu F, Costales I, Sandoval M, Perez-Martínez Z, Martín-Rodríguez G, Coto E, Melón S |
| EPI_ISL_965179 | Laboratorio Biologia Molecolare SARS-COV2- UOC Laboratorio Analisi- Servizio Medicina di Laboratorio, Ospedale San Francesco - ATS-ASSLNUoro | LABORATORIO SPECIALISTICO UOC EMATOLOGIA-Ospedale San Francesco - ATS-ASSLNUoro | SULIS VINCENZO, PIRAS GIOVANNA, ASPRONI ROSANNA, FANCELLO TATIANA, MONNE MARIA ITRIA, FIAMMA MAURA, TOJA ALESSANDRO, SANNA FILOMENA, FLORE ANNA RITA. PALMAS ANGELO DOMENICO, IANA LO MAGLIO, MAMELI GIUSEPPE |
| EPI_ISL_965912 | Servicio Murciano de Salud | Instituto de Salud Carlos III | Vázquez, S. Iglesias-Caballero, M. Sandonís,V. Camarero, S. Pozo, F. Casas, I. Jiménez, P. Zaballos, A. Monzón, S. Varona, S. Cuesta, I. Blázquez, A. |
| EPI_ISL_965934 | Hospital General Universitario de Ciudad Real | Instituto de Salud Carlos III | Sandonís,V. Vázquez, S. Iglesias-Caballero, M. Camarero, S. Pozo, F. Casas, I. Jiménez, P. Zaballos, A. Monzón, S. Varona, S. Cuesta, I. Illescas, S. |
| EPI_ISL_965938 | Consejería de Sanidad y Asuntos Sociales de Castilla La Mancha | Instituto de Salud Carlos III | Sandonís,V. Vázquez, S. Iglesias-Caballero, M. Camarero, S. Pozo, F. Casas, I. Jiménez, P. Zaballos, A. Monzón, S. Varona, S. Cuesta, I. Gutiérrez, G. |
| EPI_ISL_965952 | Consejería de Sanidad y Asuntos Sociales de Castilla La Mancha | Instituto de Salud Carlos III | Iglesias-Caballero, M. Sandonís,V. Vázquez, S. Camarero, S. Pozo, F. Casas, I. Jiménez, P. Zaballos, A. Monzón, S. Varona, S. Cuesta, I. Gutiérrez, G. |
| EPI_ISL_965959 | Hospital General Universitario de Ciudad Real | Instituto de Salud Carlos III | Iglesias-Caballero, M. Sandonís,V. Vázquez, S. Camarero, S. Pozo, F. Casas, I. Jiménez, P. Zaballos, A. Monzón, S. Varona, S. Cuesta, I. Illescas, S. |
| EPI_ISL_969397 | Indian Council of Medical Research-National Institute of Virology, Pune | Indian Council of Medical Research-National Institute of Virology, Maximum Containment Laboratory | Pragya D. Yadav, Anita Shete-Aich, Dimpal A. Nyayanit, Abhinendra Kumar, Savita Patil, Triparna D. Majumdar |
| EPI_ISL_974150 | Laboratorio de Virología / Hospital Universitario Central de Asturias (HUCA) | Laboratorio de Virología / Hospital Universitario Central de Asturias (HUCA) | Castelló C, Gómez de Oña J, Boga JA, Rojo S, Alvarez-Arguelles ME, Abreu F, Costales I, Sandoval M, Perez-Martínez Z, Martín-Rodríguez G, Coto E, Melón S |
| EPI_ISL_977476, EPI_ISL_977477 | Instituto Adolfo Lutz Central | Instituto Adolfo Lutz, Interdiciplinary Procedures Center, Strategic Laboratory | Claudio Tavares Sacchi, Claudia Regina Gonçalves, Erica Valessa Ramos Gomes, Karoline Rodrigues Campos |
| EPI_ISL_977482 | Instituto Adolfo Lutz - Regional de Aracatuba | Instituto Adolfo Lutz, Interdiciplinary Procedures Center, Strategic Laboratory | Claudio Tavares Sacchi, Claudia Regina Gonçalves, Erica Valessa Ramos Gomes, Karoline Rodrigues Campos |
| EPI_ISL_977487 | Instituto Adolfo Lutz Central | Instituto Adolfo Lutz, Interdiciplinary Procedures Center, Strategic Laboratory | Claudio Tavares Sacchi, Claudia Regina Gonçalves, Erica Valessa Ramos Gomes, Karoline Rodrigues Campos |
| EPI_ISL_977591, EPI_ISL_977592, EPI_ISL_977593 | Malaysia Genome Institute | Malaysia Genome Institute | Mohd Noor Mat Isa, Irni Suhayu Sapian, Yusuf Muhammad Noor, Nurhezreen Md Iqbal, Mohd Faizal Abu Bakar, Enizza Kasim, Shamsidar Sopie, Siti Noraini Othman, Azrin Ahmad, Nor Azfa Johari. |
| EPI_ISL_977661, EPI_ISL_977662, EPI_ISL_977663 | Labo analyses med | National Reference Center for Viruses of Respiratory Infections, Institut Pasteur, Paris | Marion Barbet, Sylvie Behillil, Méline Bizard, Angela Brisebarre, Camille Capel, Etienne Simon-Lorière, Vincent Enouf, Maud Vanpeene, Sylvie van der Werf,Amzalag Jonas |
| EPI_ISL_977664 | Hopital | National Reference Center for Viruses of Respiratory Infections, Institut Pasteur, Paris | Marion Barbet, Sylvie Behillil, Méline Bizard, Angela Brisebarre, Camille Capel, Etienne Simon-Lorière, Vincent Enouf, Maud Vanpeene, Sylvie van der Werf,Combe Patrice |
| EPI_ISL_981006, EPI_ISL_981370 | AZ Klina | AZ Klina | Dr. C. Vael |

| | | | |
|---|---|---|---|
| EPI_ISL_982184, EPI_ISL_982192, EPI_ISL_982193, EPI_ISL_982196, EPI_ISL_982197, EPI_ISL_982198, EPI_ISL_982322, EPI_ISL_982323, EPI_ISL_982328, EPI_ISL_982332, EPI_ISL_982336, EPI_ISL_982342 | | | |
| see above | Hôpital Henri Mondor | Department of Virology, Henri Mondor University Hospital, Assistance Publique Hôpitaux de Paris, Université Paris-Est Créteil, INSERM U955 | Christophe Rodriguez, Slim Fourati, Vanessa Demontant, Guillaume Gricourt, Melissa N'Debi, Alexandre Soulier, Elisabeth Trawinski, Jean-Michel Pawlotsky |
| EPI_ISL_984247, EPI_ISL_984248, EPI_ISL_984249, EPI_ISL_984250, EPI_ISL_984251, EPI_ISL_984252, EPI_ISL_984253, EPI_ISL_984254, EPI_ISL_984255, EPI_ISL_984256, EPI_ISL_984257, EPI_ISL_984258, EPI_ISL_984259, EPI_ISL_984260, EPI_ISL_984261, EPI_ISL_984262 | | | |
| see above | IAL Regional de Marilia | Instituto Adolfo Lutz, Interdiciplinary Procedures Center, Strategic Laboratory | Claudio Tavares Sacchi, Claudia Regina Gonçalves, Erica Valessa Ramos Gomes, Karoline Rodrigues Campos |
| EPI_ISL_984263 | IAL Regional de Bauru | Instituto Adolfo Lutz, Interdiciplinary Procedures Center, Strategic Laboratory | Claudio Tavares Sacchi, Claudia Regina Gonçalves, Erica Valessa Ramos Gomes, Karoline Rodrigues Campos |
| EPI_ISL_984264 | RSUPN Dr. Cipto Mangunkusumo | National Institute of Health Research and Development | Subangkit, Hana Apsari Pawestri, Kartika Dewi Puspa, Arie Ardiansyah Nugraha, Hartanti Dian Ikawati, Krisna Nur Andriana Pangesti, Nuri Dyah Indrasari, Dewi Wulandari, Nelly Puspandari, Vivi Setiawaty |
| EPI_ISL_984276 | National Institute of Health Research and Development | National Institute of Health Research and Development | Holy Arif Wibowo, Ririn Ramadhany, Yuni Rukminiati, Agustiningsih, Kindi Adam, Hana Apsari Pawestri, Subangkit, Kartika Dewi Puspa, Arie Ardiansyah Nugraha, Hartanti Dian Ikawati, Krisna Nur Andriana Pangesti, Ni Ketut Susilarini, Nur Ika Hariastuti, Uly Alfi Nikmah, Mursinah, Asri Febriyani, Reni Herman, Nike Susanti, Herna, Tati Febriyanti, Natalie Laurencia Kipuw, Fauzul Muna, Irene Lorinda Indalao, Aulia Rizki, Nelly Puspandari, Vivi Setiawaty |
| EPI_ISL_984309 | National Institute of Health Research and Development | National Institute of Health Research and Development | Agustiningsih, Kindi Adam, Holy Arif Wibowo, Ririn Ramadhany, Yuni Rukminiati, Hana Apsari Pawestri, Subangkit, Kartika Dewi Puspa, Arie Ardiansyah Nugraha, Hartanti Dian Ikawati, Krisna Nur Andriana Pangesti, Ni Ketut Susilarini, Nur Ika Hariastuti, Uly Alfi Nikmah, Mursinah, Asri Febriyani, Reni Herman, Nike Susanti, Herna, Tati Febriyanti, Natalie Laurencia Kipuw, Fauzul Muna, Irene Lorinda Indalao, Aulia Rizki, Nelly Puspandari, Vivi Setiawaty |
| EPI_ISL_984313 | National Institute of Health Research and Development | National Institute of Health Research and Development | Yuni Rukminiati, Agustiningsih, Kindi Adam, Holy Arif Wibowo, Ririn Ramadhany, Hana Apsari Pawestri, Subangkit, Kartika Dewi Puspa, Arie Ardiansyah Nugraha, Hartanti Dian Ikawati, Krisna Nur Andriana Pangesti, Ni Ketut Susilarini, Nur Ika Hariastuti, Uly Alfi Nikmah, Mursinah, Asri Febriyani, Reni Herman, Nike Susanti, Herna, Tati Febriyanti, Natalie Laurencia Kipuw, Fauzul Muna, Irene Lorinda Indalao, Aulia Rizki, Nelly Puspandari, Vivi Setiawaty |
| EPI_ISL_985058, EPI_ISL_985059, EPI_ISL_985060, EPI_ISL_985061, EPI_ISL_985062, EPI_ISL_985063, EPI_ISL_985064, EPI_ISL_985065, EPI_ISL_985066, EPI_ISL_985067, EPI_ISL_985068, EPI_ISL_985069, EPI_ISL_985070, EPI_ISL_985071, EPI_ISL_985072, EPI_ISL_985073, EPI_ISL_985074, EPI_ISL_985075, EPI_ISL_985076, EPI_ISL_985077, EPI_ISL_985078, EPI_ISL_985079, EPI_ISL_985080, EPI_ISL_985081, EPI_ISL_985082, EPI_ISL_985083, EPI_ISL_985084, EPI_ISL_985085, EPI_ISL_985086, EPI_ISL_985087, EPI_ISL_985088, EPI_ISL_985089, EPI_ISL_985090, EPI_ISL_985091, EPI_ISL_985092, EPI_ISL_985093, EPI_ISL_985094, EPI_ISL_985095, EPI_ISL_985096, EPI_ISL_985097, EPI_ISL_985098, EPI_ISL_985099, EPI_ISL_985100, EPI_ISL_985101, EPI_ISL_985102, EPI_ISL_985103, EPI_ISL_985104, EPI_ISL_985125, EPI_ISL_985126, EPI_ISL_985127, EPI_ISL_985128, EPI_ISL_985129 | | | |
| see above | Biorepository and Clinical Virology Laboratory | Ozer Lab | Ramon Lorenzo-Redondo, Adeola A. Fowotade, Ewean C. Omoruyi, Johnson A. Adeniji, Lacy M. Simons, Judd F. Hultquist, Babafemi O. Taiwo, Olubusuyi M. Adewumi, Egon A. Ozer |
| EPI_ISL_985318, EPI_ISL_985319 | LACEN de Santa Catarina | Instituto Adolfo Lutz, Interdiciplinary Procedures Center, Strategic Laboratory | Claudio Tavares Sacchi, Claudia Regina Gonçalves, Erica Valessa Ramos Gomes, Karoline Rodrigues Campos |
| EPI_ISL_985396 | RSUD Dr. Tjitrowardojo | Genetics Working Group (Pokja Genetik) Faculty of Medicine, Public Health and Nursing Universitas Gadjah Mada (FK-KMK UGM); Disease Investigation Center Wates Ministry of Agriculture Indonesia; Department of Microbiology FK-KMK UGM; Laboratorium Diagnostik Yayasan Tahija World Mosquito Program (WMP) Yogyakarta Center for Tropical Medicine FK-KMK UGM; Integrated Research Center FK-KMK UGM; Department of Computer Science and Electronics FMIPA UGM; Balai Besar Teknik Kesehatan Lingkungan dan Pengendalian Penyakit (BBTKLPP) Yogyakarta | Gunadi, Hendra Wibawa, Marcellus, Mohamad S. Hakim, Edwin W. Daniwijaya, Ludhang P. Rizki, Endah Supriyati, Eggi Arguni, Titik Nuryastuti, Tri Wibawa, Dwi AA Nugrahaningsih, Afiahayati, Siswanto, Kristy Iskandar, Nungki Anggorowati, Irene, Indaryati, Havid Setyawan, Arry Setyowati, Chusni Mubarakh, Try Nirmala Sari, Dwiki afandy, Dyah Ayu Puspitarani |
| EPI_ISL_985397 | RSUD Dr. Tjitrowardojo | Genetics Working Group (Pokja Genetik) Faculty of Medicine, Public Health and Nursing Universitas Gadjah Mada (FK-KMK UGM); Disease Investigation Center Wates Ministry of Agriculture Indonesia; Department of Microbiology FK-KMK UGM; Laboratorium Diagnostik Yayasan Tahija World Mosquito Program (WMP) Yogyakarta Center for Tropical Medicine FK-KMK UGM; Integrated Research Center FK-KMK UGM; Department of Computer Science and Electronics FMIPA UGM; Balai Besar Teknik Kesehatan Lingkungan dan Pengendalian Penyakit (BBTKLPP) Yogyakarta | Gunadi, Hendra Wibawa, Marcellus, Mohamad S. Hakim, Edwin W. Daniwijaya, Ludhang P. Rizki, Endah Supriyati, Eggi Arguni, Titik Nuryastuti, Tri Wibawa, Dwi AA Nugrahaningsih, Afiahayati, Siswanto, Kristy Iskandar, Nungki Anggorowati, Irene, Indaryati, Havid Setyawan, Arry Setyowati, Chusni Mubarakh, Try Nirmala Sari, Untung Riawan, William Widitjiarso |
| EPI_ISL_985398 | RSA UGM | Genetics Working Group (Pokja Genetik) Faculty of Medicine, Public Health and Nursing Universitas Gadjah Mada (FK-KMK UGM); Disease Investigation Center Wates Ministry of Agriculture Indonesia; Department of Microbiology FK-KMK UGM; Laboratorium Diagnostik Yayasan Tahija World Mosquito Program (WMP) Yogyakarta Center for Tropical Medicine FK-KMK UGM; Integrated Research center FK-KMK UGM; Department of Computer Science and Electronics FMIPA UGM | Gunadi, Hendra Wibawa, Marcellus, Mohamad S. Hakim, Edwin W. Daniwijaya, Ludhang P. Rizki, Endah Supriyati, Eggi Arguni, Titik Nuryastuti, Tri Wibawa, Dwi AA Nugrahaningsih, Afiahayati, Siswanto, Kristy Iskandar, Nungki Anggorowati, Susan Simanjaya, Maria Patricia Inggriani |
| EPI_ISL_992956 | Laboratorio de Virología HUCA | Laboratorio de Virología Hospital Universitario Central de Asturias (HUCA) | Sandoval M, Castelló C, Gómez de Oña J, Boga JA, Rojo S, Alvarez-Arguelles ME, Abreu F, Costales I, Perez-Martínez Z, Martín-Rodríguez G, Coto E, Melón S |
| EPI_ISL_993630 | Hospital San Agustín | Laboratorio de Virología Hospital Universitario Central de Asturias (HUCA) | Sandoval M, Castelló C, Gómez de Oña J, Boga JA, Rojo S, Alvarez-Arguelles ME, Abreu F, Costales I, Perez-Martínez Z, Martín-Rodríguez G, Coto E, Melón S |
| EPI_ISL_995446, EPI_ISL_995448 | Outre mer | National Reference Center for Viruses of Respiratory Infections, Institut Pasteur, Paris | Marion Barbet, Sylvie Behillil, Méline Bizard, Angela Brisebarre, Camille Capel, Etienne Simon-Lorière, Vincent Enouf, Maud Vanpeene, Sylvie van der Werf,Combe Patrice |

We gratefully acknowledge the following Authors from the Originating laboratories responsible for obtaining the specimens, as well as the Submitting laboratories where the genome data were generated and shared via GISAID, on which this research is based.

All Submitters of data may be contacted directly via www.gisaid.org

Authors are sorted alphabetically.

| Accession ID | Originating Laboratory | Submitting Laboratory | Authors |
|---|---|---|---|
| EPI_ISL_1000968 | Cerballiance Provence | Cerba | Roig JC, Haim-Boukobza S, Meyer S, Trombert-Paolantoni S, Lecorche E, Roquebert B |
| EPI_ISL_1001301 | Hopital | National Reference Center for Viruses of Respiratory Infections, Institut Pasteur, Paris | Marion Barbet, Sylvie Behillil, Méline Bizard, Angela Brisebarre, Camille Capel, Etienne Simon-Lorière, Vincent Enouf, Maud Vanpeene, Sylvie van der Werf,Lagathu GisèLe |
| EPI_ISL_1001302 | Hopital | National Reference Center for Viruses of Respiratory Infections, Institut Pasteur, Paris | Marion Barbet, Sylvie Behillil, Méline Bizard, Angela Brisebarre, Camille Capel, Etienne Simon-Lorière, Vincent Enouf, Maud Vanpeene, Sylvie van der Werf,Guinard JéRome |
| EPI_ISL_1001303, EPI_ISL_1001304, EPI_ISL_1001305, EPI_ISL_1001306, EPI_ISL_1001307, EPI_ISL_1001308 | Hopital | National Reference Center for Viruses of Respiratory Infections, Institut Pasteur, Paris | Marion Barbet, Sylvie Behillil, Méline Bizard, Angela Brisebarre, Camille Capel, Etienne Simon-Lorière, Vincent Enouf, Maud Vanpeene, Sylvie van der Werf,Fourgeaud Jacques |
| EPI_ISL_1001309 | Hopital | National Reference Center for Viruses of Respiratory Infections, Institut Pasteur, Paris | Marion Barbet, Sylvie Behillil, Méline Bizard, Angela Brisebarre, Camille Capel, Etienne Simon-Lorière, Vincent Enouf, Maud Vanpeene, Sylvie van der Werf,Laplanche Sophie |
| EPI_ISL_1001310 | Labo Analyses Med | National Reference Center for Viruses of Respiratory Infections, Institut Pasteur, Paris | Marion Barbet, Sylvie Behillil, Méline Bizard, Angela Brisebarre, Camille Capel, Etienne Simon-Lorière, Vincent Enouf, Maud Vanpeene, Sylvie van der Werf,Cady Anne |
| EPI_ISL_1001311 | hopital | National Reference Center for Viruses of Respiratory Infections, Institut Pasteur, Paris | Marion Barbet, Sylvie Behillil, Méline Bizard, Angela Brisebarre, Camille Capel, Etienne Simon-Lorière, Vincent Enouf, Maud Vanpeene, Sylvie van der Werf,Stoessel-Thouvenin Patricia |
| EPI_ISL_1001312, EPI_ISL_1001313, EPI_ISL_1001314, EPI_ISL_1001315, EPI_ISL_1001316, EPI_ISL_1001317 | Hopital | National Reference Center for Viruses of Respiratory Infections, Institut Pasteur, Paris | Marion Barbet, Sylvie Behillil, Méline Bizard, Angela Brisebarre, Camille Capel, Etienne Simon-Lorière, Vincent Enouf, Maud Vanpeene, Sylvie van der Werf,Martres Pascale |
| EPI_ISL_1001318 | Labo Analyses Med | National Reference Center for Viruses of Respiratory Infections, Institut Pasteur, Paris | Marion Barbet, Sylvie Behillil, Méline Bizard, Angela Brisebarre, Camille Capel, Etienne Simon-Lorière, Vincent Enouf, Maud Vanpeene, Sylvie van der Werf,Cady Anne |
| EPI_ISL_1001319, EPI_ISL_1001320 | Hopital | National Reference Center for Viruses of Respiratory Infections, Institut Pasteur, Paris | Marion Barbet, Sylvie Behillil, Méline Bizard, Angela Brisebarre, Camille Capel, Etienne Simon-Lorière, Vincent Enouf, Maud Vanpeene, Sylvie van der Werf,Bressollette (C) CéLine |
| EPI_ISL_1001321, EPI_ISL_1001322, EPI_ISL_1001323, EPI_ISL_1001324 | Hopital | National Reference Center for Viruses of Respiratory Infections, Institut Pasteur, Paris | Marion Barbet, Sylvie Behillil, Méline Bizard, Angela Brisebarre, Camille Capel, Etienne Simon-Lorière, Vincent Enouf, Maud Vanpeene, Sylvie van der Werf,Lagathu GisèLe |
| EPI_ISL_1001325 | Hopital | National Reference Center for Viruses of Respiratory Infections, Institut Pasteur, Paris | Marion Barbet, Sylvie Behillil, Méline Bizard, Angela Brisebarre, Camille Capel, Etienne Simon-Lorière, Vincent Enouf, Maud Vanpeene, Sylvie van der Werf,Fourgeaud Jacques |
| EPI_ISL_1001326, EPI_ISL_1001327 | hopital | National Reference Center for Viruses of Respiratory Infections, Institut Pasteur, Paris | Marion Barbet, Sylvie Behillil, Méline Bizard, Angela Brisebarre, Camille Capel, Etienne Simon-Lorière, Vincent Enouf, Maud Vanpeene, Sylvie van der Werf,Stoessel-Thouvenin Patricia |
| EPI_ISL_1001328 | Hopital | National Reference Center for Viruses of Respiratory Infections, Institut Pasteur, Paris | Marion Barbet, Sylvie Behillil, Méline Bizard, Angela Brisebarre, Camille Capel, Etienne Simon-Lorière, Vincent Enouf, Maud Vanpeene, Sylvie van der Werf,Martres Pascale |
| EPI_ISL_1001329 | Labo Analyses Med | National Reference Center for Viruses of Respiratory Infections, Institut Pasteur, Paris | Marion Barbet, Sylvie Behillil, Méline Bizard, Angela Brisebarre, Camille Capel, Etienne Simon-Lorière, Vincent Enouf, Maud Vanpeene, Sylvie van der Werf,Dynalab |
| EPI_ISL_1001330 | Hopital | National Reference Center for Viruses of Respiratory Infections, Institut Pasteur, Paris | Marion Barbet, Sylvie Behillil, Méline Bizard, Angela Brisebarre, Camille Capel, Etienne Simon-Lorière, Vincent Enouf, Maud Vanpeene, Sylvie van der Werf,Bressollette (C) CéLine |
| EPI_ISL_1001331, EPI_ISL_1001332, EPI_ISL_1001333 | hopital | National Reference Center for Viruses of Respiratory Infections, Institut Pasteur, Paris | Marion Barbet, Sylvie Behillil, Méline Bizard, Angela Brisebarre, Camille Capel, Etienne Simon-Lorière, Vincent Enouf, Maud Vanpeene, Sylvie van der Werf,Stoessel-Thouvenin Patricia |
| EPI_ISL_1001334, EPI_ISL_1001335 | hopital | National Reference Center for Viruses of Respiratory Infections, Institut Pasteur, Paris | Marion Barbet, Sylvie Behillil, Méline Bizard, Angela Brisebarre, Camille Capel, Etienne Simon-Lorière, Vincent Enouf, Maud Vanpeene, Sylvie van der Werf,Guigon AuréLie |
| EPI_ISL_1001336, EPI_ISL_1001337 | Hopital | National Reference Center for Viruses of Respiratory Infections, Institut Pasteur, Paris | Marion Barbet, Sylvie Behillil, Méline Bizard, Angela Brisebarre, Camille Capel, Etienne Simon-Lorière, Vincent Enouf, Maud Vanpeene, Sylvie van der Werf,Laplanche Sophie |
| EPI_ISL_1001338, EPI_ISL_1001339 | hopital | National Reference Center for Viruses of Respiratory Infections, Institut Pasteur, Paris | Marion Barbet, Sylvie Behillil, Méline Bizard, Angela Brisebarre, Camille Capel, Etienne Simon-Lorière, Vincent Enouf, Maud Vanpeene, Sylvie van der Werf,Stoessel-Thouvenin Patricia |
| EPI_ISL_1001340, EPI_ISL_1001341, EPI_ISL_1001342, EPI_ISL_1001343 | hopital | National Reference Center for Viruses of Respiratory Infections, Institut Pasteur, Paris | Marion Barbet, Sylvie Behillil, Méline Bizard, Angela Brisebarre, Camille Capel, Etienne Simon-Lorière, Vincent Enouf, Maud Vanpeene, Sylvie van der Werf,Guigon AuréLie |
| EPI_ISL_1001344, EPI_ISL_1001345, EPI_ISL_1001346 | hopital | National Reference Center for Viruses of Respiratory Infections, Institut Pasteur, Paris | Marion Barbet, Sylvie Behillil, Méline Bizard, Angela Brisebarre, Camille Capel, Etienne Simon-Lorière, Vincent Enouf, Maud Vanpeene, Sylvie van der Werf,Stoessel-Thouvenin Patricia |
| EPI_ISL_1001347, EPI_ISL_1001348, EPI_ISL_1001349, EPI_ISL_1001350 | hopital | National Reference Center for Viruses of Respiratory Infections, Institut Pasteur, Paris | Marion Barbet, Sylvie Behillil, Méline Bizard, Angela Brisebarre, Camille Capel, Etienne Simon-Lorière, Vincent Enouf, Maud Vanpeene, Sylvie van der Werf,Guigon AuréLie |
| EPI_ISL_1001351 | hopital | National Reference Center for Viruses of Respiratory Infections, Institut Pasteur, Paris | Marion Barbet, Sylvie Behillil, Méline Bizard, Angela Brisebarre, Camille Capel, Etienne Simon-Lorière, Vincent Enouf, Maud Vanpeene, Sylvie van der Werf,Stoessel-Thouvenin Patricia |
| EPI_ISL_1001352 | Hopital | National Reference Center for Viruses of Respiratory Infections, Institut Pasteur, Paris | Marion Barbet, Sylvie Behillil, Méline Bizard, Angela Brisebarre, Camille Capel, Etienne Simon-Lorière, Vincent Enouf, Maud Vanpeene, Sylvie van der Werf,Laplanche Sophie |
| EPI_ISL_1001354 | Hopital | National Reference Center for Viruses of Respiratory Infections, Institut Pasteur, Paris | Marion Barbet, Sylvie Behillil, Méline Bizard, Angela Brisebarre, Camille Capel, Etienne Simon-Lorière, Vincent Enouf, Maud Vanpeene, Sylvie van der Werf,Griscelli Franck |
| EPI_ISL_1001355 | Hopital | National Reference Center for Viruses of Respiratory Infections, Institut Pasteur, Paris | Marion Barbet, Sylvie Behillil, Méline Bizard, Angela Brisebarre, Camille Capel, Etienne Simon-Lorière, Vincent Enouf, Maud Vanpeene, Sylvie van der Werf,Marque Juillet StéPhanie |
| EPI_ISL_1001356 | Hopital | National Reference Center for Viruses of Respiratory Infections, Institut Pasteur, Paris | Marion Barbet, Sylvie Behillil, Méline Bizard, Angela Brisebarre, Camille Capel, Etienne Simon-Lorière, Vincent Enouf, Maud Vanpeene, Sylvie van der Werf,Guinard JéRome |
| EPI_ISL_1001357, EPI_ISL_1001358, EPI_ISL_1001359, EPI_ISL_1001360, EPI_ISL_1001361, EPI_ISL_1001362, EPI_ISL_1001363, EPI_ISL_1001364 | Hopital | National Reference Center for Viruses of Respiratory Infections, Institut Pasteur, Paris | Marion Barbet, Sylvie Behillil, Méline Bizard, Angela Brisebarre, Camille Capel, Etienne Simon-Lorière, Vincent Enouf, Maud Vanpeene, Sylvie van der Werf,Fourgeaud Jacques |

| | | | |
|---|---|---|---|
| EPI_ISL_1001365, EPI_ISL_1001366 | Hopital | National Reference Center for Viruses of Respiratory Infections, Institut Pasteur, Paris | Marion Barbet, Sylvie Behillil, Méline Bizard, Angela Brisebarre, Camille Capel, Etienne Simon-Lorière, Vincent Enouf, Maud Vanpeene, Sylvie van der Werf,Lagathu GisèLe |
| EPI_ISL_1001367 | Hopital | National Reference Center for Viruses of Respiratory Infections, Institut Pasteur, Paris | Marion Barbet, Sylvie Behillil, Méline Bizard, Angela Brisebarre, Camille Capel, Etienne Simon-Lorière, Vincent Enouf, Maud Vanpeene, Sylvie van der Werf,Laplanche Sophie |
| EPI_ISL_1001368, EPI_ISL_1001369, EPI_ISL_1001370, EPI_ISL_1001371 | hopital | National Reference Center for Viruses of Respiratory Infections, Institut Pasteur, Paris | Marion Barbet, Sylvie Behillil, Méline Bizard, Angela Brisebarre, Camille Capel, Etienne Simon-Lorière, Vincent Enouf, Maud Vanpeene, Sylvie van der Werf,Guigon AuréLie |
| EPI_ISL_1001372, EPI_ISL_1001373 | Hopital | National Reference Center for Viruses of Respiratory Infections, Institut Pasteur, Paris | Marion Barbet, Sylvie Behillil, Méline Bizard, Angela Brisebarre, Camille Capel, Etienne Simon-Lorière, Vincent Enouf, Maud Vanpeene, Sylvie van der Werf,Laplanche Sophie |
| EPI_ISL_1001374, EPI_ISL_1001375, EPI_ISL_1001381, EPI_ISL_1001382, EPI_ISL_1001383 | hopital | National Reference Center for Viruses of Respiratory Infections, Institut Pasteur, Paris | Marion Barbet, Sylvie Behillil, Méline Bizard, Angela Brisebarre, Camille Capel, Etienne Simon-Lorière, Vincent Enouf, Maud Vanpeene, Sylvie van der Werf,Guigon AuréLie |
| EPI_ISL_1001384, EPI_ISL_1001385 | Outre mer | National Reference Center for Viruses of Respiratory Infections, Institut Pasteur, Paris | Marion Barbet, Sylvie Behillil, Méline Bizard, Angela Brisebarre, Camille Capel, Etienne Simon-Lorière, Vincent Enouf, Maud Vanpeene, Sylvie van der Werf,Rousset (Guy) Dominique |
| EPI_ISL_1001386 | Hopital | National Reference Center for Viruses of Respiratory Infections, Institut Pasteur, Paris | Marion Barbet, Sylvie Behillil, Méline Bizard, Angela Brisebarre, Camille Capel, Etienne Simon-Lorière, Vincent Enouf, Maud Vanpeene, Sylvie van der Werf,Griscelli Franck |
| EPI_ISL_1001387 | Hopital | National Reference Center for Viruses of Respiratory Infections, Institut Pasteur, Paris | Marion Barbet, Sylvie Behillil, Méline Bizard, Angela Brisebarre, Camille Capel, Etienne Simon-Lorière, Vincent Enouf, Maud Vanpeene, Sylvie van der Werf,Marque Juillet StéPhanie |
| EPI_ISL_1001388 | Hopital | National Reference Center for Viruses of Respiratory Infections, Institut Pasteur, Paris | Marion Barbet, Sylvie Behillil, Méline Bizard, Angela Brisebarre, Camille Capel, Etienne Simon-Lorière, Vincent Enouf, Maud Vanpeene, Sylvie van der Werf,Fourgeaud Jacques |
| EPI_ISL_1001389 | hopital | National Reference Center for Viruses of Respiratory Infections, Institut Pasteur, Paris | Marion Barbet, Sylvie Behillil, Méline Bizard, Angela Brisebarre, Camille Capel, Etienne Simon-Lorière, Vincent Enouf, Maud Vanpeene, Sylvie van der Werf,Stoessel-Thouvenin Patricia |
| EPI_ISL_1001390 | Hopital | National Reference Center for Viruses of Respiratory Infections, Institut Pasteur, Paris | Marion Barbet, Sylvie Behillil, Méline Bizard, Angela Brisebarre, Camille Capel, Etienne Simon-Lorière, Vincent Enouf, Maud Vanpeene, Sylvie van der Werf,Bigaillon Christine |
| EPI_ISL_1001391 | Hopital | National Reference Center for Viruses of Respiratory Infections, Institut Pasteur, Paris | Marion Barbet, Sylvie Behillil, Méline Bizard, Angela Brisebarre, Camille Capel, Etienne Simon-Lorière, Vincent Enouf, Maud Vanpeene, Sylvie van der Werf,Thellier |
| EPI_ISL_1001392 | Hopital | National Reference Center for Viruses of Respiratory Infections, Institut Pasteur, Paris | Marion Barbet, Sylvie Behillil, Méline Bizard, Angela Brisebarre, Camille Capel, Etienne Simon-Lorière, Vincent Enouf, Maud Vanpeene, Sylvie van der Werf,Lagathu GisèLe |
| EPI_ISL_1001393 | Hopital | National Reference Center for Viruses of Respiratory Infections, Institut Pasteur, Paris | Marion Barbet, Sylvie Behillil, Méline Bizard, Angela Brisebarre, Camille Capel, Etienne Simon-Lorière, Vincent Enouf, Maud Vanpeene, Sylvie van der Werf,Thellier |
| EPI_ISL_1001394 | Labo Analyses Med | National Reference Center for Viruses of Respiratory Infections, Institut Pasteur, Paris | Marion Barbet, Sylvie Behillil, Méline Bizard, Angela Brisebarre, Camille Capel, Etienne Simon-Lorière, Vincent Enouf, Maud Vanpeene, Sylvie van der Werf |
| EPI_ISL_1001395, EPI_ISL_1001396, EPI_ISL_1001397, EPI_ISL_1001398, EPI_ISL_1001399, EPI_ISL_1001400, EPI_ISL_1001401, EPI_ISL_1001402 | Outre mer | National Reference Center for Viruses of Respiratory Infections, Institut Pasteur, Paris | Marion Barbet, Sylvie Behillil, Méline Bizard, Angela Brisebarre, Camille Capel, Etienne Simon-Lorière, Vincent Enouf, Maud Vanpeene, Sylvie van der Werf,Rousset (Guy) Dominique |
| EPI_ISL_1001403, EPI_ISL_1001404, EPI_ISL_1001405, EPI_ISL_1001406, EPI_ISL_1001407 | Hopital | National Reference Center for Viruses of Respiratory Infections, Institut Pasteur, Paris | Marion Barbet, Sylvie Behillil, Méline Bizard, Angela Brisebarre, Camille Capel, Etienne Simon-Lorière, Vincent Enouf, Maud Vanpeene, Sylvie van der Werf,Guinoiseau Thibault |
| EPI_ISL_1001408, EPI_ISL_1001409, EPI_ISL_1001410, EPI_ISL_1001411, EPI_ISL_1001412, EPI_ISL_1001413, EPI_ISL_1001414 | Outre mer | National Reference Center for Viruses of Respiratory Infections, Institut Pasteur, Paris | Marion Barbet, Sylvie Behillil, Méline Bizard, Angela Brisebarre, Camille Capel, Etienne Simon-Lorière, Vincent Enouf, Maud Vanpeene, Sylvie van der Werf,Rousset (Guy) Dominique |
| EPI_ISL_1001415 | Hopital | National Reference Center for Viruses of Respiratory Infections, Institut Pasteur, Paris | Marion Barbet, Sylvie Behillil, Méline Bizard, Angela Brisebarre, Camille Capel, Etienne Simon-Lorière, Vincent Enouf, Maud Vanpeene, Sylvie van der Werf,Lagathu GisèLe |
| EPI_ISL_1001416, EPI_ISL_1001417, EPI_ISL_1001418, EPI_ISL_1001419, EPI_ISL_1001420, EPI_ISL_1001421, EPI_ISL_1001422, EPI_ISL_1001423, EPI_ISL_1001424, EPI_ISL_1001425, EPI_ISL_1001426, EPI_ISL_1001427, EPI_ISL_1001428, EPI_ISL_1001429, EPI_ISL_1001430, EPI_ISL_1001431, EPI_ISL_1001432, EPI_ISL_1001433, EPI_ISL_1001434, EPI_ISL_1001435, EPI_ISL_1001436, EPI_ISL_1001437, EPI_ISL_1001438, EPI_ISL_1001439, EPI_ISL_1001440, EPI_ISL_1001441, EPI_ISL_1001442, EPI_ISL_1001443, EPI_ISL_1001444, EPI_ISL_1001445, EPI_ISL_1001446, EPI_ISL_1001447, EPI_ISL_1001448, EPI_ISL_1001449 | | | |
| see above | Outre mer | National Reference Center for Viruses of Respiratory Infections, Institut Pasteur, Paris | Marion Barbet, Sylvie Behillil, Méline Bizard, Angela Brisebarre, Camille Capel, Etienne Simon-Lorière, Vincent Enouf, Maud Vanpeene, Sylvie van der Werf,Rousset (Guy) Dominique |
| EPI_ISL_1001450, EPI_ISL_1001451 | Hopital | National Reference Center for Viruses of Respiratory Infections, Institut Pasteur, Paris | Marion Barbet, Sylvie Behillil, Méline Bizard, Angela Brisebarre, Camille Capel, Etienne Simon-Lorière, Vincent Enouf, Maud Vanpeene, Sylvie van der Werf,Ducancelle Alexandra |
| EPI_ISL_1001452, EPI_ISL_1001453, EPI_ISL_1001454, EPI_ISL_1001455 | Outre mer | National Reference Center for Viruses of Respiratory Infections, Institut Pasteur, Paris | Marion Barbet, Sylvie Behillil, Méline Bizard, Angela Brisebarre, Camille Capel, Etienne Simon-Lorière, Vincent Enouf, Maud Vanpeene, Sylvie van der Werf,Rousset (Guy) Dominique |
| EPI_ISL_1001459 | Laboratoire Cerballiance Port de Bouc | Cerba | Roig JC, Haim-Boukobza S, Meyer S, Trombert-Paolantoni S, Lecorche E, Roquebert B |
| EPI_ISL_1001461 | Cerballiance | Cerba | Roig JC, Haim-Boukobza S, Meyer S, Trombert-Paolantoni S, Lecorche E, Roquebert B |
| EPI_ISL_1001462 | Cerballiance Provence | Cerba | Roig JC, Haim-Boukobza S, Meyer S, Trombert-Paolantoni S, Lecorche E, Roquebert B |
| EPI_ISL_1001463 | Cerballiance Port de bouc | Cerba | Roig JC, Haim-Boukobza S, Meyer S, Trombert-Paolantoni S, Lecorche E, Roquebert B |
| EPI_ISL_1001464 | Laboratoire Cerballiance du pole santé | Cerba | Roig JC, Haim-Boukobza S, Meyer S, Trombert-Paolantoni S, Lecorche E, Roquebert B |
| EPI_ISL_1001465 | Cerballiance Carpentras CV | Cerba | Roig JC, Haim-Boukobza S, Meyer S, Trombert-Paolantoni S, Lecorche E, Roquebert B |
| EPI_ISL_1001466 | Cerballiance Haifa | Cerba | Roig JC, Haim-Boukobza S, Meyer S, Trombert-Paolantoni S, Lecorche E, Roquebert B |
| EPI_ISL_1001467 | Cerballiance Carpentras CV | Cerba | Roig JC, Haim-Boukobza S, Meyer S, Trombert-Paolantoni S, Lecorche E, Roquebert B |
| EPI_ISL_1001573 | Cerballiance Luynes | Cerba | Roig JC, Haim-Boukobza S, Meyer S, Trombert-Paolantoni S, Lecorche E, Roquebert B |
| EPI_ISL_1002710 | Cerballiance Rouviere | Cerba | Roig JC, Haim-Boukobza S, Meyer S, Trombert-Paolantoni S, Lecorche E, Roquebert B |
| EPI_ISL_1003769 | Cerballiance Endoume | Cerba | Roig JC, Haim-Boukobza S, Meyer S, Trombert-Paolantoni S, Lecorche E, Roquebert B |
| EPI_ISL_1004174 | Cerballiance Port de Bouc | Cerba | Roig JC, Haim-Boukobza S, Meyer S, Trombert-Paolantoni S, Lecorche E, Roquebert B |
| EPI_ISL_1005682 | Martigues hospital | Cerba | Negre V, Vidal B, Olivi M, Benazra M, Haim-Boukobza S, Trombert-Paolantoni S, Lecorche E, Roquebert B |
| EPI_ISL_1005683 | Martigues Hospital | Cerba | Negre V, Vidal B, Olivi M, Benazra M, Haim-Boukobza S, Trombert-Paolantoni S, Lecorche E, Roquebert B |
| EPI_ISL_1005684 | Cerballiance Monthléry | Cerba | Collin T, Lesenne A, Olivi M, Ricard B, Benazra M, Delaplace E, Gresset N, Haim-Boukobza S, Trombert-Paolantoni S, Lecorche E, Roquebert B |
| EPI_ISL_1005685 | Cerballiance Montlhéry | Cerba | Collin T, Lesenne A, Olivi M, Ricard B, Benazra M, Delaplace E, Gresset N, Trombert-Paolantoni S, Lecorche E, Haim-Boukobza S, Roquebert B |

| | | | |
|---|---|---|---|
| EPI_ISL_1005687 | Cerballiance Lardy | Cerba | Collin T, Lesenne A, Olivi M, Ricard B, Benazra M, Delaplace E, Gresset N, Haim-Boukobza S, Trombert-Paolantoni S, Lecorche E, Roquebert B |
| EPI_ISL_1005688 | Cerballiance Antony Velpeau | Cerba | Collin T, Lesenne A, Olivi M, Ricard B, Benazra M, Delaplace E, Gresset N, Haim-Boukobza S, Trombert-Paolantoni S, Lecorche E, Roquebert B |
| EPI_ISL_1007659 | Thai Red Cross Emerging Infectious Diseases Health Science Centre, Chulalongkorn Hospital, Faculty of Medicine, Chulalongkorn University | Thai Red Cross Emerging Infectious Diseases Center and Faculty of Medicine, Chulalongkorn University | Rome Buathong, Pornvimol Leethong, Non Chindavech, Sopon Iamsirithaworn, Sininat Petcharat, Yutthana Joyjinda, Weenassarin Ampoot, Apaporn Rodpan, Thiravat Hemachudha, Supaporn Wacharapluesadee |
| EPI_ISL_1008664 | Ospedale Santa Caterina Novella | Istituto Zooprofilattico Sperimentale della Puglia e della Basilicata | Parisi A., Bianco A., Capozzi L., Del Sambro L., Simone D., Chironna M., Loconsole D., Sallustio A., Ridolfi D. |
| EPI_ISL_1008665, EPI_ISL_1008666, EPI_ISL_1008667, EPI_ISL_1008668 | Istituto Zooprofilattico Sperimentale della Puglia e della Basilicata | Istituto Zooprofilattico Sperimentale della Puglia e della Basilicata | Parisi A., Bianco A., Capozzi L., Del Sambro L., Simone D., Chironna M., Loconsole D., Sallustio A., Ridolfi D. |
| EPI_ISL_1008669, EPI_ISL_1008670, EPI_ISL_1008671, EPI_ISL_1008672, EPI_ISL_1008673, EPI_ISL_1008674, EPI_ISL_1008675, EPI_ISL_1008676, EPI_ISL_1008677 | Istituto Zooprofilattico Sperimentale della Puglia e della Basilicata | Istituto Zooprofilattico Sperimentale della Puglia e della Basilicata | Parisi A., Bianco A., Capozzi L., Del Sambro L., Simone D., Chironna M., Loconsole D., Sallustio A., Giannico A. |
| EPI_ISL_1008678 | Presidio di Brindisi Di Summa - Perrino | Istituto Zooprofilattico Sperimentale della Puglia e della Basilicata | Parisi A., Bianco A., Capozzi L., Del Sambro L., Simone D., Chironna M., Loconsole D., Sallustio A., Ridolfi D. |
| EPI_ISL_1008679, EPI_ISL_1008680, EPI_ISL_1008681, EPI_ISL_1008682, EPI_ISL_1008683, EPI_ISL_1008684, EPI_ISL_1008685, EPI_ISL_1008686, EPI_ISL_1008687 | Istituto Zooprofilattico Sperimentale della Puglia e della Basilicata | Istituto Zooprofilattico Sperimentale della Puglia e della Basilicata | Parisi A., Bianco A., Capozzi L., Del Sambro L., Simone D., Chironna M., Loconsole D., Sallustio A., Ridolfi D. |
| EPI_ISL_1008688, EPI_ISL_1008689, EPI_ISL_1008690, EPI_ISL_1008691, EPI_ISL_1008692, EPI_ISL_1008693, EPI_ISL_1008694 | Istituto Zooprofilattico Sperimentale della Puglia e della Basilicata | Istituto Zooprofilattico Sperimentale della Puglia e della Basilicata | Parisi A., Bianco A., Capozzi L., Del Sambro L., Simone D., Chironna M., Loconsole D., Sallustio A., Giannico A. |
| EPI_ISL_1008700 | Ospedale Santa Caterina Novella | Istituto Zooprofilattico Sperimentale della Puglia e della Basilicata | Parisi A., Bianco A., Capozzi L., Del Sambro L., Simone D., Chironna M., Loconsole D., Sallustio A., Ridolfi D. |
| EPI_ISL_1008701, EPI_ISL_1008702 | Istituto Zooprofilattico Sperimentale della Puglia e della Basilicata | Istituto Zooprofilattico Sperimentale della Puglia e della Basilicata | Parisi A., Bianco A., Capozzi L., Del Sambro L., Simone D., Chironna M., Loconsole D., Sallustio A., Giannico A. |
| EPI_ISL_1008704, EPI_ISL_1008705 | Presidio di Brindisi Di Summa - Perrino | Istituto Zooprofilattico Sperimentale della Puglia e della Basilicata | Parisi A., Bianco A., Capozzi L., Del Sambro L., Simone D., Chironna M., Loconsole D., Sallustio A., Ridolfi D. |
| EPI_ISL_1008706 | Ospedale Santissima Annunziata | Istituto Zooprofilattico Sperimentale della Puglia e della Basilicata | Parisi A., Bianco A., Capozzi L., Del Sambro L., Simone D., Chironna M., Loconsole D., Sallustio A., Ridolfi D. |
| EPI_ISL_1008707, EPI_ISL_1008708, EPI_ISL_1008709, EPI_ISL_1008710, EPI_ISL_1008711, EPI_ISL_1008712 | Presidio di Brindisi Di Summa - Perrino | Istituto Zooprofilattico Sperimentale della Puglia e della Basilicata | Parisi A., Bianco A., Capozzi L., Del Sambro L., Simone D., Chironna M., Loconsole D., Sallustio A., Ridolfi D. |
| EPI_ISL_1008713 | Lister Laboratorio de Referencia S.A. de C.V. | Instituto de diagnóstico y Referencia Epidemiologicos (INDRE) Departamento de Virología | Claudia Wong-Arambula, Abril Rodriguez-Maldonado, Fabiola Garces-Ayala, Natividad Cruz-Ortiz, Tatiana Nunez-Garcia, Gisela Barrera-Badillo, Lucia Hernandez-Rivas, Irma Lopez-Martinez, Ernesto Ramirez-Gonzalez. |
| EPI_ISL_1008714 | Unidad de Patología Clínica | Instituto de diagnóstico y Referencia Epidemiologicos (INDRE) Departamento de Virología | Claudia Wong-Arambula, Abril Rodriguez-Maldonado, Fabiola Garces-Ayala, Natividad Cruz-Ortiz, Tatiana Nunez-Garcia, Gisela Barrera-Badillo, Lucia Hernandez-Rivas, Irma Lopez-Martinez, Ernesto Ramirez-Gonzalez. |
| EPI_ISL_1009618 | Cerballiance Cerrieres-le-Buisson | Cerba | Collin T, Lesenne A, Olivi M, Ricard B, Benazra M, Delaplace E, Gresset N, Haim-Boukobza S, Trombert-Paolantoni S, Lecorche E, Roquebert B |
| EPI_ISL_1009627 | Laboratoire Cerballiance Bretigny | Cerba | Collin T, Lesenne A, Olivi M, Ricard B, Benazra M, Delaplace E, Gresset N, Haim-Boukobza S, Trombert-Paolantoni S, Lecorche E, Roquebert B |
| EPI_ISL_1009667 | Cerballiance Bourg-la-Reine | Cerba | Collin T, Lesenne A, Olivi M, Ricard B, Benazra M, Delaplace E, Gresset N, Haim-Boukobza S, Trombert-Paolantoni S, Lecorche E, Roquebert B |
| EPI_ISL_1009673 | CHI Andre Gregoire | Cerba | Fremont-Goudot G, Penot P, Haïm-Boukobza S, Trombert-Paolantoni S, Olivi M, Benadza M, Lecorche E, Roquebert B |
| EPI_ISL_1009674 | Cerballiance Wilson | Cerba | Haïm-Boukobza S, Merah K, Olivi M, Benazra M, Trombert-Paolantoni S, Lecorche E, Roquebert B |
| EPI_ISL_1011752 | Cerballiance Grigny | Cerba | Collin T, Lesenne A, Olivi M, Ricard B, Benazra M, Delaplace E, Gresset N, Haim-Boukobza S, Trombert-Paolantoni S, Lecorche E, Roquebert B |
| EPI_ISL_1012922 | Somedi | Imelda Hospital | Johan Frans, Hanne Valgaeren, Dagmar Obbels |
| EPI_ISL_1012925 | Hospital Universitari Germans Trias i Pujol(HUGTiP)/Fundació Lluita contra la SIDA (FLSida) | IrsiCaixa - Can Ruti CovidSeq | Marc Noguera-Julian, Mariona Parera, Maria Casadellà, Pilar Armengol, Francesc Catala-Moll, Roger Paredes, Bonaventura Clotet Marta Massanella, Ester Ballana, Lidia Ruiz, Nuria Izquierdo, Jorge Carrillo, Roger Paredes, Julia Blanco, Joaquim Segalés, Bonaventura Clotet |
| EPI_ISL_1013210, EPI_ISL_1013211, EPI_ISL_1013212, EPI_ISL_1013213 | Hôpital Necker | National Reference Center for Viruses of Respiratory Infections, Institut Pasteur, Paris | Marion Barbet, Sylvie Behillil, Méline Bizard, Angela Brisebarre, Camille Capel, Etienne Simon-Lorière, Vincent Enouf, Maud Vanpeene, Sylvie van der Werf,Fourgeaud Jacques |
| EPI_ISL_1013214, EPI_ISL_1013215, EPI_ISL_1013216 | Institut Pasteur de Guadeloupe | National Reference Center for Viruses of Respiratory Infections, Institut Pasteur, Paris | Marion Barbet, Sylvie Behillil, Méline Bizard, Angela Brisebarre, Camille Capel, Etienne Simon-Lorière, Vincent Enouf, Maud Vanpeene, Sylvie van der Werf,Talarmin Antoine |
| EPI_ISL_1013217, EPI_ISL_1013218, EPI_ISL_1013219, EPI_ISL_1013220, EPI_ISL_1013221 | Laboratoire de Microbiologie | National Reference Center for Viruses of Respiratory Infections, Institut Pasteur, Paris | Marion Barbet, Sylvie Behillil, Méline Bizard, Angela Brisebarre, Camille Capel, Etienne Simon-Lorière, Vincent Enouf, Maud Vanpeene, Sylvie van der Werf,Martres Pascale |
| EPI_ISL_1013222, EPI_ISL_1013223 | Service Biologie | National Reference Center for Viruses of Respiratory Infections, Institut Pasteur, Paris | Marion Barbet, Sylvie Behillil, Méline Bizard, Angela Brisebarre, Camille Capel, Etienne Simon-Lorière, Vincent Enouf, Maud Vanpeene, Sylvie van der Werf,Marque Juillet StéPhanie |
| EPI_ISL_1013224, EPI_ISL_1013225, EPI_ISL_1013226, EPI_ISL_1013227, EPI_ISL_1013228, EPI_ISL_1013229, EPI_ISL_1013230, EPI_ISL_1013231, EPI_ISL_1013232, EPI_ISL_1013233, EPI_ISL_1013234 | | | |
| see above | CH de Mayotte | National Reference Center for Viruses of Respiratory Infections, Institut Pasteur, Paris | Marion Barbet, Sylvie Behillil, Méline Bizard, Angela Brisebarre, Camille Capel, Etienne Simon-Lorière, Vincent Enouf, Maud Vanpeene, Sylvie van der Werf,Combe Patrice |
| EPI_ISL_1013235 | Hôpital Necker | National Reference Center for Viruses of Respiratory Infections, Institut Pasteur, Paris | Marion Barbet, Sylvie Behillil, Méline Bizard, Angela Brisebarre, Camille Capel, Etienne Simon-Lorière, Vincent Enouf, Maud Vanpeene, Sylvie van der Werf,Fourgeaud Jacques |
| EPI_ISL_1013236 | Institut Pasteur de Guadeloupe | National Reference Center for Viruses of Respiratory Infections, Institut Pasteur, Paris | Marion Barbet, Sylvie Behillil, Méline Bizard, Angela Brisebarre, Camille Capel, Etienne Simon-Lorière, Vincent Enouf, Maud Vanpeene, Sylvie van der Werf,Talarmin Antoine |
| EPI_ISL_1013237, EPI_ISL_1013238, EPI_ISL_1013239, EPI_ISL_1013244, EPI_ISL_1013245, EPI_ISL_1013246, EPI_ISL_1013247, EPI_ISL_1013248, EPI_ISL_1013249, EPI_ISL_1013250, EPI_ISL_1013251, EPI_ISL_1013252, EPI_ISL_1013253, EPI_ISL_1013254, EPI_ISL_1013255, EPI_ISL_1013256, EPI_ISL_1013257, EPI_ISL_1013258, EPI_ISL_1013259, EPI_ISL_1013260, EPI_ISL_1013261, EPI_ISL_1013262, EPI_ISL_1013263, EPI_ISL_1013264, EPI_ISL_1013265, EPI_ISL_1013266, EPI_ISL_1013267, EPI_ISL_1013268, EPI_ISL_1013269, EPI_ISL_1013270, EPI_ISL_1013271, EPI_ISL_1013272, EPI_ISL_1013273, EPI_ISL_1013274, EPI_ISL_1013275, EPI_ISL_1013276, EPI_ISL_1013277, EPI_ISL_1013278, EPI_ISL_1013279, EPI_ISL_1013280, EPI_ISL_1013281, EPI_ISL_1013282, EPI_ISL_1013283, EPI_ISL_1013284, EPI_ISL_1013285, EPI_ISL_1013286, EPI_ISL_1013287, EPI_ISL_1013288, EPI_ISL_1013289, EPI_ISL_1013290, EPI_ISL_1013291, EPI_ISL_1013292, EPI_ISL_1013293, EPI_ISL_1013294, EPI_ISL_1013295, EPI_ISL_1013296, EPI_ISL_1013297, EPI_ISL_1013298, EPI_ISL_1013299, EPI_ISL_1013300, EPI_ISL_1013301, EPI_ISL_1013302, EPI_ISL_1013303, EPI_ISL_1013304, EPI_ISL_1013305, EPI_ISL_1013306, EPI_ISL_1013307, EPI_ISL_1013308, EPI_ISL_1013309, EPI_ISL_1013310, EPI_ISL_1013311, EPI_ISL_1013312, EPI_ISL_1013313, EPI_ISL_1013314, EPI_ISL_1013315, EPI_ISL_1013316, EPI_ISL_1013317, EPI_ISL_1013318, EPI_ISL_1013319, EPI_ISL_1013320, EPI_ISL_1013321, EPI_ISL_1013322, EPI_ISL_1013323, EPI_ISL_1013324, EPI_ISL_1013325, EPI_ISL_1013326, EPI_ISL_1013327, EPI_ISL_1013328, EPI_ISL_1013329, EPI_ISL_1013330, EPI_ISL_1013331, EPI_ISL_1013332, EPI_ISL_1013333, EPI_ISL_1013334, EPI_ISL_1013335, EPI_ISL_1013336, EPI_ISL_1013337, EPI_ISL_1013338, EPI_ISL_1013339, EPI_ISL_1013340, EPI_ISL_1013341, EPI_ISL_1013342, EPI_ISL_1013343, EPI_ISL_1013344, EPI_ISL_1013345, EPI_ISL_1013346, EPI_ISL_1013347, EPI_ISL_1013348, EPI_ISL_1013349, EPI_ISL_1013350, EPI_ISL_1013351, EPI_ISL_1013352, EPI_ISL_1013353, EPI_ISL_1013354, EPI_ISL_1013355, EPI_ISL_1013356, EPI_ISL_1013357, EPI_ISL_1013358, EPI_ISL_1013359, | | | |

| | | | |
|---|---|---|---|
| EPI_ISL_1013360, EPI_ISL_1013361, EPI_ISL_1013362, EPI_ISL_1013363, EPI_ISL_1013364, EPI_ISL_1013365, EPI_ISL_1013366, EPI_ISL_1013367, EPI_ISL_1013368, EPI_ISL_1013369, EPI_ISL_1013370, EPI_ISL_1013371, EPI_ISL_1013372, EPI_ISL_1013373, EPI_ISL_1013374, EPI_ISL_1013375, EPI_ISL_1013376, EPI_ISL_1013377, EPI_ISL_1013378, EPI_ISL_1013379, EPI_ISL_1013380, EPI_ISL_1013381, EPI_ISL_1013382, EPI_ISL_1013383, EPI_ISL_1013384, EPI_ISL_1013385, EPI_ISL_1013386, EPI_ISL_1013387, EPI_ISL_1013388, EPI_ISL_1013389, EPI_ISL_1013390, EPI_ISL_1013391, EPI_ISL_1013392, EPI_ISL_1013393, EPI_ISL_1013394, EPI_ISL_1013395, EPI_ISL_1013396, EPI_ISL_1013397, EPI_ISL_1013398, EPI_ISL_1013399, EPI_ISL_1013400, EPI_ISL_1013401, EPI_ISL_1013402, EPI_ISL_1013403, EPI_ISL_1013404, EPI_ISL_1013405, EPI_ISL_1013406, EPI_ISL_1013407 | | | |
| see above | CH de Mayotte | National Reference Center for Viruses of Respiratory Infections, Institut Pasteur, Paris | Marion Barbet, Sylvie Behillil, Méline Bizard, Angela Brisebarre, Camille Capel, Etienne Simon-Lorière, Vincent Enouf, Maud Vanpeene, Sylvie van der Werf,Combe Patrice |
| EPI_ISL_1013408, EPI_ISL_1013409, EPI_ISL_1013410, EPI_ISL_1013411, EPI_ISL_1013412, EPI_ISL_1013413, EPI_ISL_1013414 | CHU - Hôpital Cavale Blanche | National Reference Center for Viruses of Respiratory Infections, Institut Pasteur, Paris | Marion Barbet, Sylvie Behillil, Méline Bizard, Angela Brisebarre, Camille Capel, Etienne Simon-Lorière, Vincent Enouf, Maud Vanpeene, Sylvie van der Werf,Pilorge LéA |
| EPI_ISL_1013415 | Hôpital Necker | National Reference Center for Viruses of Respiratory Infections, Institut Pasteur, Paris | Marion Barbet, Sylvie Behillil, Méline Bizard, Angela Brisebarre, Camille Capel, Etienne Simon-Lorière, Vincent Enouf, Maud Vanpeene, Sylvie van der Werf,Fourgeaud Jacques |
| EPI_ISL_1013416 | Hôpital Necker | National Reference Center for Viruses of Respiratory Infections, Institut Pasteur, Paris | Marion Barbet, Sylvie Behillil, Méline Bizard, Angela Brisebarre, Camille Capel, Etienne Simon-Lorière, Vincent Enouf, Maud Vanpeene, Sylvie van der Werf,Leruez-Ville Marianne |
| EPI_ISL_1013417, EPI_ISL_1013418, EPI_ISL_1013419, EPI_ISL_1013420, EPI_ISL_1013421, EPI_ISL_1013422, EPI_ISL_1013423, EPI_ISL_1013424, EPI_ISL_1013425, EPI_ISL_1013426, EPI_ISL_1013427, EPI_ISL_1013428, EPI_ISL_1013429, EPI_ISL_1013430, EPI_ISL_1013431, EPI_ISL_1013432, EPI_ISL_1013433, EPI_ISL_1013434, EPI_ISL_1013435, EPI_ISL_1013436 | | | |
| see above | Institut Pasteur de Guadeloupe | National Reference Center for Viruses of Respiratory Infections, Institut Pasteur, Paris | Marion Barbet, Sylvie Behillil, Méline Bizard, Angela Brisebarre, Camille Capel, Etienne Simon-Lorière, Vincent Enouf, Maud Vanpeene, Sylvie van der Werf,Talarmin Antoine |
| EPI_ISL_1013438, EPI_ISL_1013439, EPI_ISL_1013440, EPI_ISL_1013441, EPI_ISL_1013442, EPI_ISL_1013443, EPI_ISL_1013445, EPI_ISL_1013446, EPI_ISL_1013447, EPI_ISL_1013448, EPI_ISL_1013449 | | | |
| see above | Laboratoire associé au CNR des Virus Influenzae | National Reference Center for Viruses of Respiratory Infections, Institut Pasteur, Paris | Marion Barbet, Sylvie Behillil, Méline Bizard, Angela Brisebarre, Camille Capel, Etienne Simon-Lorière, Vincent Enouf, Maud Vanpeene, Sylvie van der Werf,Rousset (Guy) Dominique |
| EPI_ISL_1013594, EPI_ISL_1013771 | Imelda Hospital | Imelda Hospital | Johan Frans, Hanne Valgaeren, Dagmar Obbels |
| EPI_ISL_1014237, EPI_ISL_1014690 | University of Bari Biomedical Sciences and Human Oncology | University of Bari Biomedical Sciences and Human Oncology | Chironna M., Sallustio A., Loconsole D., Accogli M. |
| EPI_ISL_1016508 | University of Sarajevo, Veterinary Faculty, Laboratory for Molecular Diagnostic and Research Laboratory | University of Sarajevo, Veterinary Faculty, Laboratory for Molecular Diagnostic and Research Laboratory | Goletic T., Goletic Š., Softic A., Alic-Šeho A., Šabic E., Jažic A., Nicevic M., Hodžic A., Terzic I. |
| EPI_ISL_1016842 | University of Sarajevo, Veterinary Faculty, Laboratory for Molecular Diagnostic and Research Laboratory | University of Sarajevo, Veterinary Faculty, Laboratory for Molecular Diagnostic and Research Laboratory | Goletic T., Goletic S., Softic A., Alic-Seho A., Nicevic M., Jazic A., Hodzic A., Terzic I., Sabic E. |
| EPI_ISL_1016843 | University of Sarajevo, Veterinary Faculty, Laboratory for Molecular Diagnostic and Research Laboratory | University of Sarajevo, Veterinary Faculty, Laboratory for Molecular Diagnostic and Research Laboratory | Goletic T., Goletic S., Softic A., Alic-Seho A., Hodzic A., Terzic I, Jazic A., Sabic E., Nicevic M. |
| EPI_ISL_1016884 | University of Sarajevo, Veterinary Faculty, Laboratory for Molecular Diagnostic and Research Laboratory | University of Sarajevo, Veterinary Faculty, Laboratory for Molecular Diagnostic and Research Laboratory | Goletic T., Goletic S., Softic A., Alic-Seho A., Nicevic M., Sabic E., Terzic I., Jazic A., Hodzic A. |
| EPI_ISL_1016885 | University of Sarajevo, Veterinary Faculty, Laboratory for Molecular Diagnostic and Research Laboratory | University of Sarajevo, Veterinary Faculty, Laboratory for Molecular Diagnostic and Research Laboratory | Goletic S., Goletic T., Softic A., Alic-Seho A., Terzic I., Jazic A., Nicevic M., Hodzic A., Sabic E. |
| EPI_ISL_1016968 | University of Sarajevo, Veterinary Faculty, Laboratory for Molecular Diagnostic and Research Laboratory | University of Sarajevo, Veterinary Faculty, Laboratory for Molecular Diagnostic and Research Laboratory | Goletic S., Goletic T., Softic A., Alic-Seho A., Hodzic A., Terzic I., Sabic E., Jazic A., Nicevic M. |
| EPI_ISL_1016969 | University of Sarajevo, Veterinary Faculty, Laboratory for Molecular Diagnostic and Research Laboratory | University of Sarajevo, Veterinary Faculty, Laboratory for Molecular Diagnostic and Research Laboratory | Goletic S., Goletic T., Softic A., Alic-Seho A., Nicevic M., Terzic I., Jazic A., Hodzic A., Sabic E. |
| EPI_ISL_1016970 | University of Sarajevo, Veterinary Faculty, Laboratory for Molecular Diagnostic and Research Laboratory | University of Sarajevo, Veterinary Faculty, Laboratory for Molecular Diagnostic and Research Laboratory | Goletic S., Goletic T., Softic A., Alic-Seho A., Jazic A., Sabic E., Nicevic M., Terzic I., Hodzic A. |
| EPI_ISL_1018110, EPI_ISL_1018112, EPI_ISL_1018114, EPI_ISL_1018116 | AZ Klina | AZ Klina | Dr. C. Vael |
| EPI_ISL_1018220, EPI_ISL_1018222, EPI_ISL_1018223, EPI_ISL_1018224, EPI_ISL_1018225, EPI_ISL_1018226, EPI_ISL_1018227 | Althaia. Xarxa Assistencial Universitària de Manresa | IrsiCaixa - Can Ruti CovidSeq | Marc Noguera-Julian, Mariona Parera, Maria Casadellà, Pilar Armengol, Francesc Catala-Moll, Roger Paredes, Bonaventura Clotet Gloria Trujillo, Rafel Perez Vidal, Jaume Trape Pujol, Carolina Gonzalez Fernandez, Roger Paredes, Eulalia Grau, Bonaventura Clotet |
| EPI_ISL_1020018, EPI_ISL_1020019, EPI_ISL_1020020 | AZ Klina | AZ Klina | Dr. C. Vael |
| EPI_ISL_1020121, EPI_ISL_1020122, EPI_ISL_1020123, EPI_ISL_1020126 | Laboratorio de Virología HUCA | Laboratorio de Virología HUCA | Sandoval M, Castelló C, Gómez de Oña J, Boga JA, Rojo S, Alvarez-Arguelles ME, Abreu F, Costales I, Perez-Martínez Z, Martín-Rodríguez G, Coto E, Melón S |
| EPI_ISL_1020127 | AZ Klina | AZ Klina | Dr. C. Vael |
| EPI_ISL_1020128 | Laboratorio de Virología HUCA | Laboratorio de Virología HUCA | Sandoval M, Castelló C, Gómez de Oña J, Boga JA, Rojo S, Alvarez-Arguelles ME, Abreu F, Costales I, Perez-Martínez Z, Martín-Rodríguez G, Coto E, Melón S |
| EPI_ISL_1020129, EPI_ISL_1020130 | AZ Klina | AZ Klina | Dr. C. Vael |
| EPI_ISL_1020132 | Laboratorio de Virología HUCA | Laboratorio de Virología HUCA | Sandoval M, Castelló C, Gómez de Oña J, Boga JA, Rojo S, Alvarez-Arguelles ME, Abreu F, Costales I, Perez-Martínez Z, Martín-Rodríguez G, Coto E, Melón S |
| EPI_ISL_1020133 | AZ Klina | AZ Klina | Dr. C. Vael |
| EPI_ISL_1020135 | Laboratorio de Virología HUCA | Laboratorio de Virología HUCA | Sandoval M, Castelló C, Gómez de Oña J, Boga JA, Rojo S, Alvarez-Arguelles ME, Abreu F, Costales I, Perez-Martínez Z, Martín-Rodríguez G, Coto E, Melón S |
| EPI_ISL_1020136, EPI_ISL_1020137, EPI_ISL_1020139, EPI_ISL_1020142, EPI_ISL_1020144, EPI_ISL_1020147, EPI_ISL_1020148, EPI_ISL_1020149, EPI_ISL_1020150 | AZ Klina | AZ Klina | Dr. C. Vael |
| EPI_ISL_1020151 | Laboratorio de Virología HUCA | Laboratorio de Virología HUCA | Sandoval M, Castelló C, Gómez de Oña J, Boga JA, Rojo S, Alvarez-Arguelles ME, Abreu F, Costales I, Perez-Martínez Z, Martín-Rodríguez G, Coto E, Melón S |
| EPI_ISL_1020153 | AZ Klina | AZ Klina | Dr. C. Vael |
| EPI_ISL_1020154 | Laboratorio de Virología HUCA | Laboratorio de Virología HUCA | Sandoval M, Castelló C, Gómez de Oña J, Boga JA, Rojo S, Alvarez-Arguelles ME, Abreu F, Costales I, Perez-Martínez Z, Martín-Rodríguez G, Coto E, Melón S |
| EPI_ISL_1020156, EPI_ISL_1020157, EPI_ISL_1020158, EPI_ISL_1020159 | AZ Klina | AZ Klina | Dr. C. Vael |
| EPI_ISL_1020199 | Balai Besar Rehabilitasi (BNN) | Biosafety Level-3 Laboratory, Indonesian Institute of Sciences (LIPI) | Sri Swasthikawati, Linda Sukmarini, Idris, Masrukhin, Alfi T. Fathurahman, Ruby Setiawan, Ratih Asmana Ningrum, Sugiyono Saputra |

| | | | |
|---|---|---|---|
| EPI_ISL_1020200 | RS Trimitra, Bogor, West java | Biosafety Level-3 Laboratory, Indonesian Institute of Sciences (LIPI) | Sri Swasthikawati, Linda Sukmarini, Idris, Masrukhin, Alfi T. Fathurahman, Ruby Setiawan, Ratih Asmana Ningrum, Sugiyono Saputra |
| EPI_ISL_1020201 | RS Trimitra, Bogor, West java | Biosafety Level-3 Laboratory, Indonesian Institute of Sciences (LIPI) | Isa Nuryana, Syaiful Rizal, Indriawati, Ari S. Wulandari, Syam Budi Iryanto, Anik B. Dharmayanthi, Andri Wardiana, Sugiyono Saputra |
| EPI_ISL_1020202 | RS Trimitra, Bogor, West java | Biosafety Level-3 Laboratory, Indonesian Institute of Sciences (LIPI) | Isa Nuryana, Syaiful Rizal, Indriawati, Ari S. Wulandari, Syam Budi Iryanto, Ratih Asmana Ningrum, Andri Wardiana, Sugiyono Saputra |
| EPI_ISL_1020315, EPI_ISL_1020316 | Department of Laboratory Medicine, National Taiwan University Hospital | Microbial Genomics Core Lab, National Taiwan University Centers of Genomic and Precision Medicine | Shiou-Hwei Yeh, You-Yu Lin, Ya-Yun Lai, Chiao-Ling Li, Shan-Chwen Chang, Pei-Jer Chen, Sui-Yuan Chang |
| EPI_ISL_1034144 | MEDICO COMPETENTE P.O. L'AQUILA | Istituto Zooprofilattico Sperimentale dell'Abruzzo e Molise "G. Caporale" | Lorusso A, Marcacci M, Di Domenico M, Ancora M, Curini V, Mangone I, Rinaldi A, Scialabba S, Di Pasquale A, Cammà C, Puglia I, Calistri P, Savini G |
| EPI_ISL_1034274 | Provincial Sanitary and Epidemiological Inspection in Gdask | Laboratory of Recombinant Vaccines | Lukasz Rabalski, Maciej Kosinski, Maciej Grzybek, Natalia Mazur-Panasiuk, Krysztof Pyr, Boguslaw Szewczyk, Krystyna Bienkowska-Szewczyk |
| EPI_ISL_1034304, EPI_ISL_1034306 | Laboratorio de Ecologia de Doencas Transmissiveis na Amazonia, Instituto Leonidas e Maria Deane - Fiocruz Amazonia | Laboratorio de Ecologia de Doencas Transmissiveis na Amazonia, Instituto Leonidas e Maria Deane - Fiocruz Amazonia | Valdinete Nascimento, Victor Souza, André Corado, Fernanda Nascimento, George Silva, Ágatha Costa, Debora Duarte, Karina Pessoa, Matilde Mejía, Luciana Gonçalves, Maria Júlia Brandão, Michele Jesus, Felipe Naveca on behalf of the Fiocruz COVID-19 Genomic Surveillance Network |
| EPI_ISL_1034793 | Virologia Dipartimento di Scienze Biomediche Università di Sassari Viale San Pietro, 43/B - Sassari | Laboratorio Specialistico di Ematologia, Ospedale "San Francesco", via Mannironi 1, 08100 Nuoro | Giovanna Piras, Tatiana Fancello, Maria Monne, Rosanna Asproni, Caterina Serra, Elena Rimini, Salvatore Rubino |
| EPI_ISL_1034916 | Laboratorio Biologia Molecolare Sars Cov2 - UOC Laboratorio Analisi - Servizio Medicina di Laboratorio, Ospedale "San Francesco" - ATS-ASSL Nuoro | Laboratorio specialistico UOC Ematologia - Ospedale "San Francesco" - ATS-ASSL Nuoro | Piras Giovanna, Asproni Rosanna, Malune Paolo, Fiamma Maura, Monne Maria Itria, Palmas Angelo Domenico, Lo Maglio Iana, Mameli Giuseppe |
| EPI_ISL_1035873, EPI_ISL_1035874, EPI_ISL_1035875, EPI_ISL_1035876, EPI_ISL_1035877, EPI_ISL_1035878, EPI_ISL_1035879, EPI_ISL_1035880, EPI_ISL_1035881, EPI_ISL_1035882, EPI_ISL_1035883, EPI_ISL_1035884, EPI_ISL_1035885, EPI_ISL_1035886, EPI_ISL_1035887, EPI_ISL_1035888 | | | |
| see above | Istituto Zooprofilattico Sperimentale del Mezzogiorno | IZSM-U.O.C. Virologia | Maurizio Viscardi, Lorena Cardillo, Pellegrino Cerino, Massimo Zollo,Giovanna Fusco, Esterina De Carlo, Antonio Limone |
| EPI_ISL_1035893, EPI_ISL_1035894, EPI_ISL_1035895, EPI_ISL_1035896, EPI_ISL_1035897, EPI_ISL_1035898, EPI_ISL_1035899 | Istituto Zooprofilattico Sperimentale del Mezzogiorno | IZSM-U.O.C. Virologia | Maurizio Viscardi, Lorena Cardillo, Sergio Brandi, Pellegrino Cerino, Massimo Zollo,Giovanna Fusco, Esterina De Carlo, Antonio Limone |
| EPI_ISL_1035904, EPI_ISL_1035905, EPI_ISL_1035906, EPI_ISL_1035907, EPI_ISL_1035908, EPI_ISL_1035909, EPI_ISL_1035910, EPI_ISL_1035911, EPI_ISL_1035912, EPI_ISL_1035913, EPI_ISL_1035914, EPI_ISL_1035915, EPI_ISL_1035916 | | | |
| see above | Hopital CH de Troyes | National Reference Center for Viruses of Respiratory Infections, Institut Pasteur, Paris | Marion Barbet, Sylvie Behillil, Méline Bizard, Angela Brisebarre, Camille Capel, Etienne Simon-Lorière, Vincent Enouf, Maud Vanpeene, Sylvie van der Werf,Stoessel-Thouvenin Patricia |
| EPI_ISL_1035917, EPI_ISL_1035918 | hopital Laboratoire de Virologie du CHU de Lille | National Reference Center for Viruses of Respiratory Infections, Institut Pasteur, Paris | Marion Barbet, Sylvie Behillil, Méline Bizard, Angela Brisebarre, Camille Capel, Etienne Simon-Lorière, Vincent Enouf, Maud Vanpeene, Sylvie van der Werf,Guigon Aurélie |
| EPI_ISL_1035919, EPI_ISL_1035920, EPI_ISL_1035921, EPI_ISL_1035922 | Labo analyses med | National Reference Center for Viruses of Respiratory Infections, Institut Pasteur, Paris | Marion Barbet, Sylvie Behillil, Méline Bizard, Angela Brisebarre, Camille Capel, Etienne Simon-Lorière, Vincent Enouf, Maud Vanpeene, Sylvie van der Werf,Goubard Agathe |
| EPI_ISL_1035923 | Outre Mer CH de Mayotte | National Reference Center for Viruses of Respiratory Infections, Institut Pasteur, Paris | Marion Barbet, Sylvie Behillil, Méline Bizard, Angela Brisebarre, Camille Capel, Etienne Simon-Lorière, Vincent Enouf, Maud Vanpeene, Sylvie van der Werf,Combe Patrice |
| EPI_ISL_1035924 | Hopital CH de Troyes | National Reference Center for Viruses of Respiratory Infections, Institut Pasteur, Paris | Marion Barbet, Sylvie Behillil, Méline Bizard, Angela Brisebarre, Camille Capel, Etienne Simon-Lorière, Vincent Enouf, Maud Vanpeene, Sylvie van der Werf,Stoessel-Thouvenin Patricia |
| EPI_ISL_1035925, EPI_ISL_1035926 | Hopital CHRU Tours | National Reference Center for Viruses of Respiratory Infections, Institut Pasteur, Paris | Marion Barbet, Sylvie Behillil, Méline Bizard, Angela Brisebarre, Camille Capel, Etienne Simon-Lorière, Vincent Enouf, Maud Vanpeene, Sylvie van der Werf,Guinoiseau Thibault |
| EPI_ISL_1035927 | Hopital CH de Troyes | National Reference Center for Viruses of Respiratory Infections, Institut Pasteur, Paris | Marion Barbet, Sylvie Behillil, Méline Bizard, Angela Brisebarre, Camille Capel, Etienne Simon-Lorière, Vincent Enouf, Maud Vanpeene, Sylvie van der Werf,Stoessel-Thouvenin Patricia |
| EPI_ISL_1035928 | hopital Laboratoire de Virologie du CHU de Lille | National Reference Center for Viruses of Respiratory Infections, Institut Pasteur, Paris | Marion Barbet, Sylvie Behillil, Méline Bizard, Angela Brisebarre, Camille Capel, Etienne Simon-Lorière, Vincent Enouf, Maud Vanpeene, Sylvie van der Werf,Guigon Aurélie |
| EPI_ISL_1035929 | Hopital CHRU Tours | National Reference Center for Viruses of Respiratory Infections, Institut Pasteur, Paris | Marion Barbet, Sylvie Behillil, Méline Bizard, Angela Brisebarre, Camille Capel, Etienne Simon-Lorière, Vincent Enouf, Maud Vanpeene, Sylvie van der Werf,Guinoiseau Thibault |
| EPI_ISL_1035930 | Hopital Centre Hospitalier d'Arpajon | National Reference Center for Viruses of Respiratory Infections, Institut Pasteur, Paris | Marion Barbet, Sylvie Behillil, Méline Bizard, Angela Brisebarre, Camille Capel, Etienne Simon-Lorière, Vincent Enouf, Maud Vanpeene, Sylvie van der Werf,Barboux |
| EPI_ISL_1035931, EPI_ISL_1035932, EPI_ISL_1035933, EPI_ISL_1035934, EPI_ISL_1035935, EPI_ISL_1035936, EPI_ISL_1035937, EPI_ISL_1035938, EPI_ISL_1035939, EPI_ISL_1035940, EPI_ISL_1035941, EPI_ISL_1035942, EPI_ISL_1035943, EPI_ISL_1035944, EPI_ISL_1035945, EPI_ISL_1035946, EPI_ISL_1035947, EPI_ISL_1035948 | | | |
| see above | Outre Mer CH de Mayotte | National Reference Center for Viruses of Respiratory Infections, Institut Pasteur, Paris | Marion Barbet, Sylvie Behillil, Méline Bizard, Angela Brisebarre, Camille Capel, Etienne Simon-Lorière, Vincent Enouf, Maud Vanpeene, Sylvie van der Werf,Combe Patrice |
| EPI_ISL_1035952, EPI_ISL_1035953, EPI_ISL_1035954, EPI_ISL_1035955, EPI_ISL_1035956, EPI_ISL_1035957 | Hopital CHRU Tours | National Reference Center for Viruses of Respiratory Infections, Institut Pasteur, Paris | Marion Barbet, Sylvie Behillil, Méline Bizard, Angela Brisebarre, Camille Capel, Etienne Simon-Lorière, Vincent Enouf, Maud Vanpeene, Sylvie van der Werf,Guinoiseau Thibault |
| EPI_ISL_1035958 | Hopital CH Jeanne de Navarre Laboratoire de Biologie | National Reference Center for Viruses of Respiratory Infections, Institut Pasteur, Paris | Marion Barbet, Sylvie Behillil, Méline Bizard, Angela Brisebarre, Camille Capel, Etienne Simon-Lorière, Vincent Enouf, Maud Vanpeene, Sylvie van der Werf,Thellier |
| EPI_ISL_1035959 | Outre Mer CH de Mayotte | National Reference Center for Viruses of Respiratory Infections, Institut Pasteur, Paris | Marion Barbet, Sylvie Behillil, Méline Bizard, Angela Brisebarre, Camille Capel, Etienne Simon-Lorière, Vincent Enouf, Maud Vanpeene, Sylvie van der Werf,Combe Patrice |
| EPI_ISL_1035960 | Labo Analyses Med | National Reference Center for Viruses of Respiratory Infections, Institut Pasteur, Paris | Marion Barbet, Sylvie Behillil, Méline Bizard, Angela Brisebarre, Camille Capel, Etienne Simon-Lorière, Vincent Enouf, Maud Vanpeene, Sylvie van der Werf,Le Vicky |
| EPI_ISL_1035961 | Labo analyses med | National Reference Center for Viruses of Respiratory Infections, Institut Pasteur, Paris | Marion Barbet, Sylvie Behillil, Méline Bizard, Angela Brisebarre, Camille Capel, Etienne Simon-Lorière, Vincent Enouf, Maud Vanpeene, Sylvie van der Werf,Goubard Agathe |
| EPI_ISL_1035962 | Labo Analyses Med | National Reference Center for Viruses of Respiratory Infections, Institut Pasteur, Paris | Marion Barbet, Sylvie Behillil, Méline Bizard, Angela Brisebarre, Camille Capel, Etienne Simon-Lorière, Vincent Enouf, Maud Vanpeene, Sylvie van der Werf,Ravaux Jean Jacques |
| EPI_ISL_1035963, EPI_ISL_1035964, EPI_ISL_1035965 | Labo Analyses Med | National Reference Center for Viruses of Respiratory Infections, Institut Pasteur, Paris | Marion Barbet, Sylvie Behillil, Méline Bizard, Angela Brisebarre, Camille Capel, Etienne Simon-Lorière, Vincent Enouf, Maud Vanpeene, Sylvie van der Werf,Coignard Catherine |
| EPI_ISL_1036193, EPI_ISL_1036194, EPI_ISL_1036195, EPI_ISL_1036196, EPI_ISL_1036197 | SIESP CH | Istituto Zooprofilattico Sperimentale dell'Abruzzo e Molise "G. Caporale" | Lorusso A, Marcacci M, Di Domenico M, Ancora M, Curini V, Mangone I, Rinaldi A, Scialabba S, Di Pasquale A, Cammà C, Puglia I, Calistri P, Savini G |
| EPI_ISL_1036198 | ASL AVEZZANO-SULMONA-L'AQUILA DIP PREV SERV IGIENE E SAN PUBBLICA | Istituto Zooprofilattico Sperimentale dell'Abruzzo e Molise "G. Caporale" | Lorusso A, Marcacci M, Di Domenico M, Ancora M, Curini V, Mangone I, Rinaldi A, Scialabba S, Di Pasquale A, Cammà C, Puglia I, Calistri P, Savini G |

| | | | |
|---|---|---|---|
| EPI_ISL_1036199, EPI_ISL_1036200, EPI_ISL_1036201 | SIESP CH | Istituto Zooprofilattico Sperimentale dell'Abruzzo e Molise "G. Caporale" | Lorusso A, Marcacci M, Di Domenico M, Ancora M, Curini V, Mangone I, Rinaldi A, Scialabba S, Di Pasquale A, Cammà C, Puglia I, Calistri P, Savini G |
| EPI_ISL_1036202 | SIESP TERAMO | Istituto Zooprofilattico Sperimentale dell'Abruzzo e Molise "G. Caporale" | Lorusso A, Marcacci M, Di Domenico M, Ancora M, Curini V, Mangone I, Rinaldi A, Scialabba S, Di Pasquale A, Cammà C, Puglia I, Calistri P, Savini G |
| EPI_ISL_1036203, EPI_ISL_1036204 | SIESP CH | Istituto Zooprofilattico Sperimentale dell'Abruzzo e Molise "G. Caporale" | Lorusso A, Marcacci M, Di Domenico M, Ancora M, Curini V, Mangone I, Rinaldi A, Scialabba S, Di Pasquale A, Cammà C, Puglia I, Calistri P, Savini G |
| EPI_ISL_1036205 | PRES TERR CASOLI | Istituto Zooprofilattico Sperimentale dell'Abruzzo e Molise "G. Caporale" | Lorusso A, Marcacci M, Di Domenico M, Ancora M, Curini V, Mangone I, Rinaldi A, Scialabba S, Di Pasquale A, Cammà C, Puglia I, Calistri P, Savini G |
| EPI_ISL_1036206 | SIESP AQ | Istituto Zooprofilattico Sperimentale dell'Abruzzo e Molise "G. Caporale" | Lorusso A, Marcacci M, Di Domenico M, Ancora M, Curini V, Mangone I, Rinaldi A, Scialabba S, Di Pasquale A, Cammà C, Puglia I, Calistri P, Savini G |
| EPI_ISL_1036207, EPI_ISL_1036208 | SIESP TERAMO | Istituto Zooprofilattico Sperimentale dell'Abruzzo e Molise "G. Caporale" | Lorusso A, Marcacci M, Di Domenico M, Ancora M, Curini V, Mangone I, Rinaldi A, Scialabba S, Di Pasquale A, Cammà C, Puglia I, Calistri P, Savini G |
| EPI_ISL_1036209 | SIESP DIP PREV CHIETI | Istituto Zooprofilattico Sperimentale dell'Abruzzo e Molise "G. Caporale" | Lorusso A, Marcacci M, Di Domenico M, Ancora M, Curini V, Mangone I, Rinaldi A, Scialabba S, Di Pasquale A, Cammà C, Puglia I, Calistri P, Savini G |
| EPI_ISL_1036210 | SIESP TERAMO | Istituto Zooprofilattico Sperimentale dell'Abruzzo e Molise "G. Caporale" | Lorusso A, Marcacci M, Di Domenico M, Ancora M, Curini V, Mangone I, Rinaldi A, Scialabba S, Di Pasquale A, Cammà C, Puglia I, Calistri P, Savini G |
| EPI_ISL_1036211 | ASL AVEZZANO-SULMONA-L'AQUILA | Istituto Zooprofilattico Sperimentale dell'Abruzzo e Molise "G. Caporale" | Lorusso A, Marcacci M, Di Domenico M, Ancora M, Curini V, Mangone I, Rinaldi A, Scialabba S, Di Pasquale A, Cammà C, Puglia I, Calistri P, Savini G |
| EPI_ISL_1036212, EPI_ISL_1036215, EPI_ISL_1036216 | SIESP TERAMO | Istituto Zooprofilattico Sperimentale dell'Abruzzo e Molise "G. Caporale" | Lorusso A, Marcacci M, Di Domenico M, Ancora M, Curini V, Mangone I, Rinaldi A, Scialabba S, Di Pasquale A, Cammà C, Puglia I, Calistri P, Savini G |
| EPI_ISL_1036217 | FONDAZIONE PICCOLA OPERA CARITAS | Istituto Zooprofilattico Sperimentale dell'Abruzzo e Molise "G. Caporale" | Lorusso A, Marcacci M, Di Domenico M, Ancora M, Curini V, Mangone I, Rinaldi A, Scialabba S, Di Pasquale A, Cammà C, Puglia I, Calistri P, Savini G |
| EPI_ISL_1036221 | OSP SAN SALVATORE UOC MAL INF | Istituto Zooprofilattico Sperimentale dell'Abruzzo e Molise "G. Caporale" | Lorusso A, Marcacci M, Di Domenico M, Ancora M, Curini V, Mangone I, Rinaldi A, Scialabba S, Di Pasquale A, Cammà C, Puglia I, Calistri P, Savini G |
| EPI_ISL_1036222 | SIESP TERAMO | Istituto Zooprofilattico Sperimentale dell'Abruzzo e Molise "G. Caporale" | Lorusso A, Marcacci M, Di Domenico M, Ancora M, Curini V, Mangone I, Rinaldi A, Scialabba S, Di Pasquale A, Cammà C, Puglia I, Calistri P, Savini G |
| EPI_ISL_1036223 | OSP SAN SALVATORE MED INT | Istituto Zooprofilattico Sperimentale dell'Abruzzo e Molise "G. Caporale" | Lorusso A, Marcacci M, Di Domenico M, Ancora M, Curini V, Mangone I, Rinaldi A, Scialabba S, Di Pasquale A, Cammà C, Puglia I, Calistri P, Savini G |
| EPI_ISL_1036228, EPI_ISL_1036229 | P.O.CARDARELLI | Istituto Zooprofilattico Sperimentale dell'Abruzzo e Molise "G. Caporale" | Scutellà M, Niro G, Lorusso A, Marcacci M, Di Domenico M, Ancora M, Curini V, Mangone I, Rinaldi A, Scialabba S, Di Pasquale A, Cammà C, Puglia I, Calistri P, Savini G |
| EPI_ISL_1036239 | Center of Advanced Studies and Technology, Molecular Genetics Laboratory | Center of Advanced Studies and Technology, Molecular Genetics Laboratory | Ferrante Rossella, Mandatori Domitilla, De Fabritiis Simone, Damiani Verena, Anaclerio Federico |
| EPI_ISL_1036272 | WHO National Influenza Centre Russian Federation | WHO National Influenza Centre Russian Federation | Andrey Komissarov, Artem Fadeev, Anna Ivanova, Kseniya Komissarova, Dmitry Bazhenov, Tamila Musaeva, Maria Timofeeva, Veronika Eder, Maria Pisareva, Daria Danilenko, Ksenia Safina, Elena Nabieva, Georgii Bazykin, Dmitry Lioznov |
| EPI_ISL_1036373, EPI_ISL_1036374, EPI_ISL_1036375 | Labo analyses med | National Reference Center for Viruses of Respiratory Infections, Institut Pasteur, Paris | Marion Barbet, Sylvie Behillil, Méline Bizard, Angela Brisebarre, Camille Capel, Etienne Simon-Lorière, Vincent Enouf, Maud Vanpeene, Sylvie van der Werf,Amzalag Jonas |
| EPI_ISL_1036376, EPI_ISL_1036377, EPI_ISL_1036378, EPI_ISL_1036379, EPI_ISL_1036380, EPI_ISL_1036381, EPI_ISL_1036382, EPI_ISL_1036383, EPI_ISL_1036384, EPI_ISL_1036385, EPI_ISL_1036386, EPI_ISL_1036387, EPI_ISL_1036388, EPI_ISL_1036389, EPI_ISL_1036390, EPI_ISL_1036391, EPI_ISL_1036392, EPI_ISL_1036393, EPI_ISL_1036394, EPI_ISL_1036395, EPI_ISL_1036396, EPI_ISL_1036397 | | | |
| see above | hopital Centre de Biologie humaine CHU Amiens | National Reference Center for Viruses of Respiratory Infections, Institut Pasteur, Paris | Marion Barbet, Sylvie Behillil, Méline Bizard, Angela Brisebarre, Camille Capel, Etienne Simon-Lorière, Vincent Enouf, Maud Vanpeene, Sylvie van der Werf,Castelain Sandrine |
| EPI_ISL_1036398, EPI_ISL_1036399, EPI_ISL_1036400, EPI_ISL_1036401, EPI_ISL_1036402 | Hopital CH René Dubos - Laboratoire de Biologie médicale | National Reference Center for Viruses of Respiratory Infections, Institut Pasteur, Paris | Marion Barbet, Sylvie Behillil, Méline Bizard, Angela Brisebarre, Camille Capel, Etienne Simon-Lorière, Vincent Enouf, Maud Vanpeene, Sylvie van der Werf,Blanchard GenevièVe |
| EPI_ISL_1036403, EPI_ISL_1036404 | Labo Analyses Med | National Reference Center for Viruses of Respiratory Infections, Institut Pasteur, Paris | Marion Barbet, Sylvie Behillil, Méline Bizard, Angela Brisebarre, Camille Capel, Etienne Simon-Lorière, Vincent Enouf, Maud Vanpeene, Sylvie van der Werf,Le Vicky |
| EPI_ISL_1036405, EPI_ISL_1036406 | Hopital Groupe Hospitalier Intercommunal Le Raincy Montfermeil | National Reference Center for Viruses of Respiratory Infections, Institut Pasteur, Paris | Marion Barbet, Sylvie Behillil, Méline Bizard, Angela Brisebarre, Camille Capel, Etienne Simon-Lorière, Vincent Enouf, Maud Vanpeene, Sylvie van der Werf,Luizy Nelly |
| EPI_ISL_1036407 | Hopital HIA PERCY | National Reference Center for Viruses of Respiratory Infections, Institut Pasteur, Paris | Marion Barbet, Sylvie Behillil, Méline Bizard, Angela Brisebarre, Camille Capel, Etienne Simon-Lorière, Vincent Enouf, Maud Vanpeene, Sylvie van der Werf,Foissaud Vincent |
| EPI_ISL_1036408 | hopital Hôpital Necker | National Reference Center for Viruses of Respiratory Infections, Institut Pasteur, Paris | Marion Barbet, Sylvie Behillil, Méline Bizard, Angela Brisebarre, Camille Capel, Etienne Simon-Lorière, Vincent Enouf, Maud Vanpeene, Sylvie van der Werf,Leruez-Ville Marianne |
| EPI_ISL_1036409, EPI_ISL_1036410, EPI_ISL_1036411, EPI_ISL_1036412, EPI_ISL_1036413, EPI_ISL_1036414, EPI_ISL_1036415, EPI_ISL_1036416, EPI_ISL_1036417, EPI_ISL_1036418, EPI_ISL_1036419, EPI_ISL_1036420, EPI_ISL_1036421, EPI_ISL_1036422, EPI_ISL_1036423, EPI_ISL_1036424, EPI_ISL_1036425 | | | |
| see above | Labo analyses med | National Reference Center for Viruses of Respiratory Infections, Institut Pasteur, Paris | Marion Barbet, Sylvie Behillil, Méline Bizard, Angela Brisebarre, Camille Capel, Etienne Simon-Lorière, Vincent Enouf, Maud Vanpeene, Sylvie van der Werf,Goubard Agathe |
| EPI_ISL_1036426 | Hopital Laboratoire CH Laon | National Reference Center for Viruses of Respiratory Infections, Institut Pasteur, Paris | Marion Barbet, Sylvie Behillil, Méline Bizard, Angela Brisebarre, Camille Capel, Etienne Simon-Lorière, Vincent Enouf, Maud Vanpeene, Sylvie van der Werf,Ravaux Jean Jacques |
| EPI_ISL_1036427, EPI_ISL_1036428, EPI_ISL_1036429, EPI_ISL_1036430, EPI_ISL_1036431, EPI_ISL_1036432 | Hopital Laboratoire de Virologie | National Reference Center for Viruses of Respiratory Infections, Institut Pasteur, Paris | Marion Barbet, Sylvie Behillil, Méline Bizard, Angela Brisebarre, Camille Capel, Etienne Simon-Lorière, Vincent Enouf, Maud Vanpeene, Sylvie van der Werf,Griscelli Franck |
| EPI_ISL_1039691, EPI_ISL_1039692, EPI_ISL_1039693, EPI_ISL_1039694, EPI_ISL_1039695 | LACEN do Estado de Goias | Instituto Adolfo Lutz, Interdiciplinary Procedures Center, Strategic Laboratory | Claudio Tavares Sacchi, Claudia Regina Gonçalves, Erica Valessa Ramos Gomes, Karoline Rodrigues Campos |
| EPI_ISL_1040823 | Secretaria Municipal de Saude de Piracaia | Instituto Adolfo Lutz, Interdiciplinary Procedures Center, Strategic Laboratory | Claudio Tavares Sacchi, Claudia Regina Gonçalves, Erica Valessa Ramos Gomes, Karoline Rodrigues Campos |
| EPI_ISL_1040824 | LACEN do Mato Grosso do Sul | Instituto Adolfo Lutz, Interdiciplinary Procedures Center, Strategic Laboratory | Claudio Tavares Sacchi, Claudia Regina Gonçalves, Erica Valessa Ramos Gomes, Karoline Rodrigues Campos |
| EPI_ISL_1041199, EPI_ISL_1041200, EPI_ISL_1041201, EPI_ISL_1041202, EPI_ISL_1041203, EPI_ISL_1041204, EPI_ISL_1041205, EPI_ISL_1041206, EPI_ISL_1041207, EPI_ISL_1041208, EPI_ISL_1041209, EPI_ISL_1041210, EPI_ISL_1041211, EPI_ISL_1041212 | | | |
| see above | Hungarian Defence Forces Military Medical Centre | National Laboratory of Virology, Szentágothai Research Centre | Endre Gábor Tóth, Balázs Somogyi, Ágnes Balázs-Nagy, Csaba Pereszlényi,Ferenc Jakab, Gábor Kemenesi |
| EPI_ISL_1041509 | LACEN do Estado de Goias | Instituto Adolfo Lutz, Interdiciplinary Procedures Center, Strategic Laboratory | Claudio Tavares Sacchi, Claudia Regina Gonçalves, Erica Valessa Ramos Gomes, Karoline Rodrigues Campos |

| | | | |
|---|---|---|---|
| EPI_ISL_1048381 | Farmalab Indoutama Jakarta | National Institute of Health Research and Development | Hana Apsari Pawestri, Subangkit, Kartika Dewi Puspa, Arie Ardiansyah Nugraha, Hartanti Dian Ikawati, Krisna Nur Andriana Pangesti, Nelly Puspandari, Vivi Setiawaty. |
| EPI_ISL_1055037 | Institute for Medical Research, Infectious Disease Research Centre, National Institutes of Health, Ministry of Health Malaysia | Institute for Medical Research, Infectious Disease Research Centre, National Institutes of Health, Ministry of Health Malaysia | Suppiah J, Kamel K, Mohd Zawawi Z, Ramly MN, Robert F, Thayan R |
| EPI_ISL_1055261, EPI_ISL_1055262 | Institute for Medical Research, Infectious Disease Research Centre, National Institutes of Health, Ministry of Health Malaysia | Institute for Medical Research, Infectious Disease Research Centre, National Institutes of Health, Ministry of Health Malaysia | Suppiah J, Kamel K,Moh Zawawi Z, Ramly N, Robert F, Thayan R |
| EPI_ISL_1055264, EPI_ISL_1055361, EPI_ISL_1055362 | Institute for Medical Research, Infectious Disease Research Centre, National Institutes of Health, Ministry of Health Malaysia | Institute for Medical Research, Infectious Disease Research Centre, National Institutes of Health, Ministry of Health Malaysia | Suppiah J, Kamel K, Mohd Zawawi Z, Ramly N, Robert F, Thayan R |
| EPI_ISL_1055363, EPI_ISL_1055364 | Institute for Medical Research, Infectious Disease Research Centre, National Institutes of Health, Ministry of Health Malaysia | Institute for Medical Research, Infectious Disease Research Centre, National Institutes of Health, Ministry of Health Malaysia | Suppiah J, Kamel K, Azizan MA, Thayan R |
| EPI_ISL_1055754 | Microbiology Department. Complexo Hospitalario Universitario de Vigo | Microbiology Department. Complexo Hospitalario Universitario de Vigo | Microbiology Department, Complexo Hospitalario Universitario de Vigo (CHUVI). |
| EPI_ISL_1055761, EPI_ISL_1055762, EPI_ISL_1055764 | Vithas Lab. Hospital Vithas Vigo | Microbiology Department. Complexo Hospitalario Universitario de Vigo | Microbiology Department, Complexo Hospitalario Universitario de Vigo (CHUVI). Vithas Lab, Hospital Vithas Vigo |
| EPI_ISL_1055765 | Virology Unit, Pisa University Hospital | Virology Unit, AOUP | Marialinda Vatteroni, Susi Frateschi |
| EPI_ISL_1055792 | Virology Unit, Pisa University Hospital | Virology Unit, AOUP, Pisa | Marialinda Vatteroni, Susi Frateschi |
| EPI_ISL_1055800 | Virology Unit, AOUP | Virology Unit, AOUP, Pisa | Marialinda Vatteroni, Susi Frateschi |
| EPI_ISL_1055811 | Virology Unit, AOUP | Virolgy Unit, AOUP | Marialinda Vatteroni, Susi Frateschi |
| EPI_ISL_1055815 | Virology Unit, AOUP | Virology Unit, AOUP, Pisa | Marialinda Vatteroni, Susi Frateschi, Mauro Pistello |
| EPI_ISL_1055820 | Virology Unit, AOUP | Virology Unit, AOUP | Marialinda Vatteroni, Susi Frateschi |
| EPI_ISL_1055824 | Virology Unit, AOUP, Pisa, Italy | Virology Unit, AOUP, Pisa | Marialinda Vatteroni, Susi Frateschi, Mauro Pistello |
| EPI_ISL_1058043 | SIESP CHIETI - DRIVE IN CHIETI | Istituto Zooprofilattico Sperimentale dell'Abruzzo e Molise "G. Caporale" | Lorusso A, Marcacci M, Di Domenico M, Ancora M, Curini V, Mangone I, Rinaldi A, Scialabba S, Di Pasquale A, Cammà C, Puglia I, Calistri P, Savini G |
| EPI_ISL_1058044, EPI_ISL_1058045 | SIESP CHIETI - DRIVE IN ORTONA | Istituto Zooprofilattico Sperimentale dell'Abruzzo e Molise "G. Caporale" | Lorusso A, Marcacci M, Di Domenico M, Ancora M, Curini V, Mangone I, Rinaldi A, Scialabba S, Di Pasquale A, Cammà C, Puglia I, Calistri P, Savini G |
| EPI_ISL_1058061 | Baylor Scott & White - Southlake Family Medicine | Baylor Scott & White-Temple | Ari Rao, Linden Morales, Kimberly Walker, Marcus Volz, Shelby Hendrickson |
| EPI_ISL_1060338 | Hopital | National Reference Center for Viruses of Respiratory Infections, Institut Pasteur, Paris | Marion Barbet, Sylvie Behillil, Méline Bizard, Angela Brisebarre, Camille Capel, Etienne Simon-Lorière, Vincent Enouf, Maud Vanpeene, Sylvie van der Werf,Delepine Alain |
| EPI_ISL_1060339 | Hopital | National Reference Center for Viruses of Respiratory Infections, Institut Pasteur, Paris | Marion Barbet, Sylvie Behillil, Méline Bizard, Angela Brisebarre, Camille Capel, Etienne Simon-Lorière, Vincent Enouf, Maud Vanpeene, Sylvie van der Werf,Foissaud Vincent |
| EPI_ISL_1060340, EPI_ISL_1060341, EPI_ISL_1060342, EPI_ISL_1060343, EPI_ISL_1060344, EPI_ISL_1060345, EPI_ISL_1060346, EPI_ISL_1060347, EPI_ISL_1060348, EPI_ISL_1060349, EPI_ISL_1060350, EPI_ISL_1060351, EPI_ISL_1060352, EPI_ISL_1060353, EPI_ISL_1060354, EPI_ISL_1060355, EPI_ISL_1060356, EPI_ISL_1060357, EPI_ISL_1060358 | | | |
| see above | Hopital | National Reference Center for Viruses of Respiratory Infections, Institut Pasteur, Paris | Marion Barbet, Sylvie Behillil, Méline Bizard, Angela Brisebarre, Camille Capel, Etienne Simon-Lorière, Vincent Enouf, Maud Vanpeene, Sylvie van der Werf,Fourgeaud Jacques |
| EPI_ISL_1060368 | Hopital | National Reference Center for Viruses of Respiratory Infections, Institut Pasteur, Paris | Marion Barbet, Sylvie Behillil, Méline Bizard, Angela Brisebarre, Camille Capel, Etienne Simon-Lorière, Vincent Enouf, Maud Vanpeene, Sylvie van der Werf,Griscelli Franck |
| EPI_ISL_1060369, EPI_ISL_1060370 | Labo Analyses Med | National Reference Center for Viruses of Respiratory Infections, Institut Pasteur, Paris | Marion Barbet, Sylvie Behillil, Méline Bizard, Angela Brisebarre, Camille Capel, Etienne Simon-Lorière, Vincent Enouf, Maud Vanpeene, Sylvie van der Werf,Bendayan James |
| EPI_ISL_1060371, EPI_ISL_1060372, EPI_ISL_1060373, EPI_ISL_1060374, EPI_ISL_1060375, EPI_ISL_1060376 | Labo Analyses Med | National Reference Center for Viruses of Respiratory Infections, Institut Pasteur, Paris | Marion Barbet, Sylvie Behillil, Méline Bizard, Angela Brisebarre, Camille Capel, Etienne Simon-Lorière, Vincent Enouf, Maud Vanpeene, Sylvie van der Werf,Fiorina Jc |
| EPI_ISL_1060377, EPI_ISL_1060378 | Labo analyse Med | National Reference Center for Viruses of Respiratory Infections, Institut Pasteur, Paris | Marion Barbet, Sylvie Behillil, Méline Bizard, Angela Brisebarre, Camille Capel, Etienne Simon-Lorière, Vincent Enouf, Maud Vanpeene, Sylvie van der Werf,Goubard Agathe |
| EPI_ISL_1060380, EPI_ISL_1060381, EPI_ISL_1060382, EPI_ISL_1060383, EPI_ISL_1060384, EPI_ISL_1060385, EPI_ISL_1060386, EPI_ISL_1060387, EPI_ISL_1060388 | Hopital | National Reference Center for Viruses of Respiratory Infections, Institut Pasteur, Paris | Marion Barbet, Sylvie Behillil, Méline Bizard, Angela Brisebarre, Camille Capel, Etienne Simon-Lorière, Vincent Enouf, Maud Vanpeene, Sylvie van der Werf,Guinoiseau Thibault |
| EPI_ISL_1060389 | Hopital | National Reference Center for Viruses of Respiratory Infections, Institut Pasteur, Paris | Marion Barbet, Sylvie Behillil, Méline Bizard, Angela Brisebarre, Camille Capel, Etienne Simon-Lorière, Vincent Enouf, Maud Vanpeene, Sylvie van der Werf,Foissaud Vincent |
| EPI_ISL_1060390, EPI_ISL_1060391, EPI_ISL_1060392, EPI_ISL_1060393, EPI_ISL_1060394 | Hopital | National Reference Center for Viruses of Respiratory Infections, Institut Pasteur, Paris | Marion Barbet, Sylvie Behillil, Méline Bizard, Angela Brisebarre, Camille Capel, Etienne Simon-Lorière, Vincent Enouf, Maud Vanpeene, Sylvie van der Werf,Fourgeaud Jacques |
| EPI_ISL_1060396 | Labo Analyses Med | National Reference Center for Viruses of Respiratory Infections, Institut Pasteur, Paris | Marion Barbet, Sylvie Behillil, Méline Bizard, Angela Brisebarre, Camille Capel, Etienne Simon-Lorière, Vincent Enouf, Maud Vanpeene, Sylvie van der Werf,Fiorina Jc |
| EPI_ISL_1060397 | Labo analyse Med | National Reference Center for Viruses of Respiratory Infections, Institut Pasteur, Paris | Marion Barbet, Sylvie Behillil, Méline Bizard, Angela Brisebarre, Camille Capel, Etienne Simon-Lorière, Vincent Enouf, Maud Vanpeene, Sylvie van der Werf,Goubard Agathe |
| EPI_ISL_1060399 | Labo Analyses Med | National Reference Center for Viruses of Respiratory Infections, Institut Pasteur, Paris | Marion Barbet, Sylvie Behillil, Méline Bizard, Angela Brisebarre, Camille Capel, Etienne Simon-Lorière, Vincent Enouf, Maud Vanpeene, Sylvie van der Werf,Fiorina Jc |
| EPI_ISL_1060400, EPI_ISL_1060401 | Labo Analyses Med | National Reference Center for Viruses of Respiratory Infections, Institut Pasteur, Paris | Marion Barbet, Sylvie Behillil, Méline Bizard, Angela Brisebarre, Camille Capel, Etienne Simon-Lorière, Vincent Enouf, Maud Vanpeene, Sylvie van der Werf,Pasquier |
| EPI_ISL_1060402 | Hopital | National Reference Center for Viruses of Respiratory Infections, Institut Pasteur, Paris | Marion Barbet, Sylvie Behillil, Méline Bizard, Angela Brisebarre, Camille Capel, Etienne Simon-Lorière, Vincent Enouf, Maud Vanpeene, Sylvie van der Werf,Guinoiseau Thibault |
| EPI_ISL_1060403 | Labo Analyses Med | National Reference Center for Viruses of Respiratory Infections, Institut Pasteur, Paris | Marion Barbet, Sylvie Behillil, Méline Bizard, Angela Brisebarre, Camille Capel, Etienne Simon-Lorière, Vincent Enouf, Maud Vanpeene, Sylvie van der Werf,Desjardins Nicolas |
| EPI_ISL_1060404, EPI_ISL_1060405, EPI_ISL_1060406, EPI_ISL_1060407, EPI_ISL_1060408, EPI_ISL_1060409, EPI_ISL_1060410, EPI_ISL_1060411, EPI_ISL_1060412, EPI_ISL_1060413, EPI_ISL_1060414, EPI_ISL_1060415, EPI_ISL_1060416, EPI_ISL_1060417, EPI_ISL_1060418, EPI_ISL_1060419, EPI_ISL_1060420, | | | |

| | | | |
|---|---|---|---|
| EPI_ISL_1060421, EPI_ISL_1060422, EPI_ISL_1060423, EPI_ISL_1060424, EPI_ISL_1060425, EPI_ISL_1060426, EPI_ISL_1060427, EPI_ISL_1060428 | | | |
| see above | Labo Analyses Med | National Reference Center for Viruses of Respiratory Infections, Institut Pasteur, Paris | Marion Barbet, Sylvie Behillil, Méline Bizard, Angela Brisebarre, Camille Capel, Etienne Simon-Lorière, Vincent Enouf, Maud Vanpeene, Sylvie van der Werf,Fiorina Jc |
| EPI_ISL_1060429 | Hopital | National Reference Center for Viruses of Respiratory Infections, Institut Pasteur, Paris | Marion Barbet, Sylvie Behillil, Méline Bizard, Angela Brisebarre, Camille Capel, Etienne Simon-Lorière, Vincent Enouf, Maud Vanpeene, Sylvie van der Werf,Guinoiseau Thibault |
| EPI_ISL_1060430, EPI_ISL_1060431, EPI_ISL_1060432 | Hopital | National Reference Center for Viruses of Respiratory Infections, Institut Pasteur, Paris | Marion Barbet, Sylvie Behillil, Méline Bizard, Angela Brisebarre, Camille Capel, Etienne Simon-Lorière, Vincent Enouf, Maud Vanpeene, Sylvie van der Werf,Fourgeaud Jacques |
| EPI_ISL_1060434, EPI_ISL_1060435, EPI_ISL_1060436, EPI_ISL_1060437, EPI_ISL_1060438 | Labo Analyses Med | National Reference Center for Viruses of Respiratory Infections, Institut Pasteur, Paris | Marion Barbet, Sylvie Behillil, Méline Bizard, Angela Brisebarre, Camille Capel, Etienne Simon-Lorière, Vincent Enouf, Maud Vanpeene, Sylvie van der Werf,Desjardins Nicolas |
| EPI_ISL_1060439, EPI_ISL_1060440 | Labo Analyses Med | National Reference Center for Viruses of Respiratory Infections, Institut Pasteur, Paris | Marion Barbet, Sylvie Behillil, Méline Bizard, Angela Brisebarre, Camille Capel, Etienne Simon-Lorière, Vincent Enouf, Maud Vanpeene, Sylvie van der Werf,Le Vicky |
| EPI_ISL_1060442 | Hopital | National Reference Center for Viruses of Respiratory Infections, Institut Pasteur, Paris | Marion Barbet, Sylvie Behillil, Méline Bizard, Angela Brisebarre, Camille Capel, Etienne Simon-Lorière, Vincent Enouf, Maud Vanpeene, Sylvie van der Werf,Guinoiseau Thibault |
| EPI_ISL_1060443 | Labo Analyses Med | National Reference Center for Viruses of Respiratory Infections, Institut Pasteur, Paris | Marion Barbet, Sylvie Behillil, Méline Bizard, Angela Brisebarre, Camille Capel, Etienne Simon-Lorière, Vincent Enouf, Maud Vanpeene, Sylvie van der Werf,Fiorina Jc |
| EPI_ISL_1060446 | Hopital | National Reference Center for Viruses of Respiratory Infections, Institut Pasteur, Paris | Marion Barbet, Sylvie Behillil, Méline Bizard, Angela Brisebarre, Camille Capel, Etienne Simon-Lorière, Vincent Enouf, Maud Vanpeene, Sylvie van der Werf,Pilorge LéA |
| EPI_ISL_1060447, EPI_ISL_1060448, EPI_ISL_1060449, EPI_ISL_1060450, EPI_ISL_1060451, EPI_ISL_1060452, EPI_ISL_1060453 | Hopital | National Reference Center for Viruses of Respiratory Infections, Institut Pasteur, Paris | Marion Barbet, Sylvie Behillil, Méline Bizard, Angela Brisebarre, Camille Capel, Etienne Simon-Lorière, Vincent Enouf, Maud Vanpeene, Sylvie van der Werf,Guinoiseau Thibault |
| EPI_ISL_1060454, EPI_ISL_1060455, EPI_ISL_1060456, EPI_ISL_1060457, EPI_ISL_1060458, EPI_ISL_1060459, EPI_ISL_1060460 | Hopital | National Reference Center for Viruses of Respiratory Infections, Institut Pasteur, Paris | Marion Barbet, Sylvie Behillil, Méline Bizard, Angela Brisebarre, Camille Capel, Etienne Simon-Lorière, Vincent Enouf, Maud Vanpeene, Sylvie van der Werf,Fourgeaud Jacques |
| EPI_ISL_1060463 | Labo Analyses Med | National Reference Center for Viruses of Respiratory Infections, Institut Pasteur, Paris | Marion Barbet, Sylvie Behillil, Méline Bizard, Angela Brisebarre, Camille Capel, Etienne Simon-Lorière, Vincent Enouf, Maud Vanpeene, Sylvie van der Werf,Sarret Benoit |
| EPI_ISL_1060464 | Labo Analyses Med | National Reference Center for Viruses of Respiratory Infections, Institut Pasteur, Paris | Marion Barbet, Sylvie Behillil, Méline Bizard, Angela Brisebarre, Camille Capel, Etienne Simon-Lorière, Vincent Enouf, Maud Vanpeene, Sylvie van der Werf,Chauvet |
| EPI_ISL_1060465, EPI_ISL_1060466, EPI_ISL_1060467, EPI_ISL_1060468, EPI_ISL_1060469, EPI_ISL_1060470, EPI_ISL_1060471, EPI_ISL_1060472, EPI_ISL_1060473, EPI_ISL_1060474, EPI_ISL_1060476, EPI_ISL_1060477, EPI_ISL_1060478, EPI_ISL_1060479, EPI_ISL_1060480, EPI_ISL_1060481, EPI_ISL_1060482, EPI_ISL_1060483, EPI_ISL_1060484, EPI_ISL_1060485, EPI_ISL_1060486, EPI_ISL_1060487, EPI_ISL_1060488, EPI_ISL_1060489 | | | |
| see above | Labo Analyses Med | National Reference Center for Viruses of Respiratory Infections, Institut Pasteur, Paris | Marion Barbet, Sylvie Behillil, Méline Bizard, Angela Brisebarre, Camille Capel, Etienne Simon-Lorière, Vincent Enouf, Maud Vanpeene, Sylvie van der Werf,Fiorina Jc |
| EPI_ISL_1060490 | Labo Analyses Med | National Reference Center for Viruses of Respiratory Infections, Institut Pasteur, Paris | Marion Barbet, Sylvie Behillil, Méline Bizard, Angela Brisebarre, Camille Capel, Etienne Simon-Lorière, Vincent Enouf, Maud Vanpeene, Sylvie van der Werf,Le Vicky |
| EPI_ISL_1060557, EPI_ISL_1060558, EPI_ISL_1060559, EPI_ISL_1060560 | Hopital | National Reference Center for Viruses of Respiratory Infections, Institut Pasteur, Paris | Marion Barbet, Sylvie Behillil, Méline Bizard, Angela Brisebarre, Camille Capel, Etienne Simon-Lorière, Vincent Enouf, Maud Vanpeene, Sylvie van der Werf,Fourgeaud Jacques |
| EPI_ISL_1060562 | Labo Analyses Med | National Reference Center for Viruses of Respiratory Infections, Institut Pasteur, Paris | Marion Barbet, Sylvie Behillil, Méline Bizard, Angela Brisebarre, Camille Capel, Etienne Simon-Lorière, Vincent Enouf, Maud Vanpeene, Sylvie van der Werf,Garand Anne-Laure |
| EPI_ISL_1060567 | Hopital | National Reference Center for Viruses of Respiratory Infections, Institut Pasteur, Paris | Marion Barbet, Sylvie Behillil, Méline Bizard, Angela Brisebarre, Camille Capel, Etienne Simon-Lorière, Vincent Enouf, Maud Vanpeene, Sylvie van der Werf,L'Honneur Anne-Sophie |
| EPI_ISL_1060568, EPI_ISL_1060569 | Labo Analyses Med | National Reference Center for Viruses of Respiratory Infections, Institut Pasteur, Paris | Marion Barbet, Sylvie Behillil, Méline Bizard, Angela Brisebarre, Camille Capel, Etienne Simon-Lorière, Vincent Enouf, Maud Vanpeene, Sylvie van der Werf,Foissaud Vincent |
| EPI_ISL_1060570, EPI_ISL_1060571 | Outre mer | National Reference Center for Viruses of Respiratory Infections, Institut Pasteur, Paris | Marion Barbet, Sylvie Behillil, Méline Bizard, Angela Brisebarre, Camille Capel, Etienne Simon-Lorière, Vincent Enouf, Maud Vanpeene, Sylvie van der Werf,Rousset Dominique |
| EPI_ISL_1060572, EPI_ISL_1060573 | Labo Analyses Med | National Reference Center for Viruses of Respiratory Infections, Institut Pasteur, Paris | Marion Barbet, Sylvie Behillil, Méline Bizard, Angela Brisebarre, Camille Capel, Etienne Simon-Lorière, Vincent Enouf, Maud Vanpeene, Sylvie van der Werf,Garand Anne-Laure |
| EPI_ISL_1060574 | Labo Analyses Med | National Reference Center for Viruses of Respiratory Infections, Institut Pasteur, Paris | Marion Barbet, Sylvie Behillil, Méline Bizard, Angela Brisebarre, Camille Capel, Etienne Simon-Lorière, Vincent Enouf, Maud Vanpeene, Sylvie van der Werf,Le Vicky |
| EPI_ISL_1060575 | Hopital | National Reference Center for Viruses of Respiratory Infections, Institut Pasteur, Paris | Marion Barbet, Sylvie Behillil, Méline Bizard, Angela Brisebarre, Camille Capel, Etienne Simon-Lorière, Vincent Enouf, Maud Vanpeene, Sylvie van der Werf,Cartier Riviere B |
| EPI_ISL_1060576 | Labo Analyses Med | National Reference Center for Viruses of Respiratory Infections, Institut Pasteur, Paris | Marion Barbet, Sylvie Behillil, Méline Bizard, Angela Brisebarre, Camille Capel, Etienne Simon-Lorière, Vincent Enouf, Maud Vanpeene, Sylvie van der Werf,Jouard Charles |
| EPI_ISL_1060577, EPI_ISL_1060578, EPI_ISL_1060579, EPI_ISL_1060580, EPI_ISL_1060581, EPI_ISL_1060582, EPI_ISL_1060583, EPI_ISL_1060584, EPI_ISL_1060585, EPI_ISL_1060586, EPI_ISL_1060587, EPI_ISL_1060588, EPI_ISL_1060589, EPI_ISL_1060590, EPI_ISL_1060591, EPI_ISL_1060592 | | | |
| see above | Outre mer | National Reference Center for Viruses of Respiratory Infections, Institut Pasteur, Paris | Marion Barbet, Sylvie Behillil, Méline Bizard, Angela Brisebarre, Camille Capel, Etienne Simon-Lorière, Vincent Enouf, Maud Vanpeene, Sylvie van der Werf,Rousset Dominique |
| EPI_ISL_1060593 | Labo Analyses Med | National Reference Center for Viruses of Respiratory Infections, Institut Pasteur, Paris | Marion Barbet, Sylvie Behillil, Méline Bizard, Angela Brisebarre, Camille Capel, Etienne Simon-Lorière, Vincent Enouf, Maud Vanpeene, Sylvie van der Werf,Le Vicky |
| EPI_ISL_1060597 | Cerballiance Vauréal | Cerba | Lecorche E, Olivi M, Benazra M, Haim-Boukobza S, Trombert-Paolantoni S, Roquebert B |
| EPI_ISL_1061413 | Istanbul University-Cerrahpasa, Cerrahpasa School of Medicine, COVID-19 Laboratory | Istanbul University-Cerrahpasa, Cerrahpasa School of Medicine, COVID-19 Laboratory | Mert Ahmet Kuskucu, Yesim Tuyji Tok, Okan Kadir Nohut, Zarifa Abullayeva, Ebru Yucebag, Fusun Can, Serap imek Yavuz, Haluk Eraksoy, Kenan Midilli |
| EPI_ISL_1063785, EPI_ISL_1063786, EPI_ISL_1063787 | SIESP CHIETI - DRIVE IN ORTONA | Istituto Zooprofilattico Sperimentale dell'Abruzzo e Molise "G. Caporale" | Lorusso A, Marcacci M, Di Domenico M, Ancora M, Curini V, Mangone I, Rinaldi A, Scialabba S, Di Pasquale A, Cammà C, Puglia I, Calistri P, Savini G |
| EPI_ISL_1063912 | Laboratorio Biologia Molecolare Sars Cov2 - UOC Laboratorio Analisi - Servizio Medicina di Laboratorio, Ospedale "San Francesco" - ATS-ASSL Nuoro | Laboratorio specialistico UOC Ematologia - Ospedale "San Francesco" - ATS-ASSL Nuoro | Piras Giovanna, Asproni Rossana, Malune Paolo, Fiamma Maura, Monne Maria Itria, Palmas Angelo Domenico, Lo Maglio Iana, Mameli Giuseppe |
| EPI_ISL_1067604 | HOSPITAL SAN JUAN DE DIOS | Inciensa, Instituto Costarricense de Investigación y Enseñanza en Nutrición y Salud | Francisco Duarte, Hebleen Porras, Claudio Soto-Garita, Estela Cordero, Adriana Godínez, Melany Calderón & Daniel Cascante-Serrano |
| EPI_ISL_1067608 | unknown | Inciensa, Instituto Costarricense de Investigación y Enseñanza en Nutrición y Salud | Francisco Duarte, Hebleen Porras, Claudio Soto-Garita, Estela Cordero, Adriana Godínez, Melany Calderón, Caterina Guzmán, Nazareth Ruiz & Adriana Bermúdez-Espinoza |

| | | | |
|---|---|---|---|
| EPI_ISL_1068849, EPI_ISL_1068850, EPI_ISL_1068851, EPI_ISL_1068929 | Institute for Medical Research, Infectious Disease Research Centre, National Institutes of Health, Ministry of Health Malaysia | Institute for Medical Research, Infectious Disease Research Centre, National Institutes of Health, Ministry of Health Malaysia | Suppiah J, Kamel K, Azizan MA, Thayan R |
| EPI_ISL_1068996, EPI_ISL_1068997 | UCSI University | Institute for Medical Research, Infectious Disease Research Centre, National Institutes of Health, Ministry of Health Malaysia | Suppiah J, Kamel K, Azizan MA, Sekaran SD, Thayan R |
| EPI_ISL_1068998 | UCSI University | Institute for Medical Research, Infectious Disease Research Centre, National Institutes of Health, Ministry of Health Malaysia | Suppiah J, Kamel K, Azizan MA, Thayan R |
| EPI_ISL_1069076 | UCSI University | Institute for Medical Research, Infectious Disease Research Centre, National Institutes of Health, Ministry of Health Malaysia | Suppiah J, Kamel K, Azizan MA, Sekaran SD, Thayan R |
| EPI_ISL_1073021 | Department of Virology | Department of Virology | Massab Umair, Muhammad Salman, Nazish Badar, Aamer Ikram |
| EPI_ISL_1078991, EPI_ISL_1078993, EPI_ISL_1078995, EPI_ISL_1078996, EPI_ISL_1078997, EPI_ISL_1078999, EPI_ISL_1079000, EPI_ISL_1079002, EPI_ISL_1079003, EPI_ISL_1079004, EPI_ISL_1079006, EPI_ISL_1079007, EPI_ISL_1079008, EPI_ISL_1079158, EPI_ISL_1079159, EPI_ISL_1079161, EPI_ISL_1079162, EPI_ISL_1079163, EPI_ISL_1079165, EPI_ISL_1079166 | | | |
| see above | IAL Regional de Bauru | Instituto Adolfo Lutz, Interdiciplinary Procedures Center, Strategic Laboratory | Claudio Tavares Sacchi, Claudia Regina Gonçalves, Erica Valessa Ramos Gomes, Karoline Rodrigues Campos |
| EPI_ISL_1079696 | Centre Hospitalier de Wallonie Picarde (CHWaPi) | Institut de Pathologie et Génétique (IPG) | Pascale Hilbert, Jérémie Gras |
| EPI_ISL_1080464, EPI_ISL_1080465, EPI_ISL_1080466, EPI_ISL_1080467, EPI_ISL_1080468, EPI_ISL_1080469, EPI_ISL_1080470, EPI_ISL_1080471, EPI_ISL_1080472, EPI_ISL_1080473, EPI_ISL_1080474, EPI_ISL_1080476, EPI_ISL_1080477, EPI_ISL_1080478, EPI_ISL_1080481, EPI_ISL_1080482, EPI_ISL_1080483, EPI_ISL_1080484, EPI_ISL_1080485, EPI_ISL_1080486, EPI_ISL_1080487, EPI_ISL_1080488, EPI_ISL_1080489, EPI_ISL_1080490, EPI_ISL_1080493, EPI_ISL_1080494, EPI_ISL_1080495 | | | |
| see above | Istituto Zooprofilattico Sperimentale del Mezzogiorno | Istituto Zooprofilattico Sperimentale del Mezzogiorno-U.O.C. Virologia | Maurizio Viscardi, Lorena Cardillo, Loredana Cozzolino, Pellegrino Cerino, Massimo Zollo,Giovanna Fusco, Esterina De Carlo, Antonio Limone |
| EPI_ISL_1080853, EPI_ISL_1080855, EPI_ISL_1081136, EPI_ISL_1081173, EPI_ISL_1081176, EPI_ISL_1081177, EPI_ISL_1081178, EPI_ISL_1081179, EPI_ISL_1081181, EPI_ISL_1081182, EPI_ISL_1081183, EPI_ISL_1081184, EPI_ISL_1081189 | | | |
| see above | Baylor Scott & White-Temple | Baylor Scott & White-Temple | Ari Rao, Linden Morales, Kimberly Walker, Marcus Volz, Shelby Hendrickson |
| EPI_ISL_1081312, EPI_ISL_1081313, EPI_ISL_1081315, EPI_ISL_1081318, EPI_ISL_1081319, EPI_ISL_1081320, EPI_ISL_1081321, EPI_ISL_1081322, EPI_ISL_1081323 | Bozeman Health Deaconess Hospital | Wiedenheft lab, Montana State University | Artem Nemudryi, Anna Nemudraia, Tanner Wiegand, Joseph Nichols, Deann T. Snyder, Jodi F. Hedges, Calvin Cicha, Helen Lee, Karl K. Vanderwood, Diane Bimczok, Mark A. Jutila and Blake Wiedenheft |
| EPI_ISL_1081327, EPI_ISL_1081352 | Institute for Medical Research, Infectious Disease Research Centre, National Institutes of Health, Ministry of Health Malaysia | Institute for Medical Research, Infectious Disease Research Centre, National Institutes of Health, Ministry of Health Malaysia | Suppiah J, Kamel K, Azizan MA, Thayan R |
| EPI_ISL_1082253 | Laboratorio Biologia Molecolare Sars Cov2 - UOC Laboratorio Analisi - Servizio Medicina di Laboratorio, Ospedale "San Francesco" - ATS-ASSL Nuoro | Laboratorio specialistico UOC Ematologia - Ospedale "San Francesco" - ATS-ASSL Nuoro | Giovanna Piras, Rosanna Asproni, Paolo Malune, Maura Fiamma, Maria Itria Monne, Angelo Domenico Palmas, Iana Lo Maglio, Giuseppe Mameli |
| EPI_ISL_1082254 | Virologia, Dipartimento di Scienze Biomediche, Università di Sassari, Viale San Pietro, 43/B - Sassari | Laboratorio specialistico UOC Ematologia - Ospedale "San Francesco" - ATS-ASSL Nuoro | Giovanna Piras, Asproni Rosanna, Paolo Malune, Maria Itria Monne, Angelo Domenico Palmas, Caterina Serra, Elena Rimini, Salvatore Rubino |
| EPI_ISL_1082292, EPI_ISL_1082293, EPI_ISL_1082295 | INT Fondazione Pascale | INT Fondazione Pascale | INT Fondazione Pascale |
| EPI_ISL_1084610 | Hospital for Infectious Diseases, Molecular Diagnostics Laboratory, Warsaw, Poland | 8. Laboratory of Recombinant Vaccines, Intercollegiate Faculty of Biotechnology University of Gdansk and Medical University of Gdansk, 2. ViroGenetics - BSL3 Laboratory of Virology, Maopolska Centre of Biotechnology, Jagiellonian University | Lukasz Rabalski, Maciej Kosinski, Natalia Mazur-Panasiuk, Aneta Kopacz, Piotr Zabek, Tomasz Dyda, Andrzej Horban, Krzysztof Pyrc, Krystyna Bienkowska-Szewczyk |
| EPI_ISL_1084611 | Hospital for Infectious Diseases, Molecular Diagnostics Laboratory, Warsaw, Poland | 26. Laboratory of Recombinant Vaccines, Intercollegiate Faculty of Biotechnology University of Gdansk and Medical University of Gdansk, 2. ViroGenetics - BSL3 Laboratory of Virology, Maopolska Centre of Biotechnology, Jagiellonian University | Lukasz Rabalski, Maciej Kosinski, Natalia Mazur-Panasiuk, Aneta Kopacz, Piotr Zabek, Tomasz Dyda, Andrzej Horban, Krzysztof Pyrc, Krystyna Bienkowska-Szewczyk |
| EPI_ISL_1084612 | Hospital for Infectious Diseases, Molecular Diagnostics Laboratory, Warsaw, Poland | 40. Laboratory of Recombinant Vaccines, Intercollegiate Faculty of Biotechnology University of Gdansk and Medical University of Gdansk, 2. ViroGenetics - BSL3 Laboratory of Virology, Maopolska Centre of Biotechnology, Jagiellonian University | Lukasz Rabalski, Maciej Kosinski, Natalia Mazur-Panasiuk, Aneta Kopacz, Piotr Zabek, Tomasz Dyda, Andrzej Horban, Krzysztof Pyrc, Krystyna Bienkowska-Szewczyk |
| EPI_ISL_1084613 | Hospital for Infectious Diseases, Molecular Diagnostics Laboratory, Warsaw, Poland | 63. Laboratory of Recombinant Vaccines, Intercollegiate Faculty of Biotechnology University of Gdansk and Medical University of Gdansk, 2. ViroGenetics - BSL3 Laboratory of Virology, Maopolska Centre of Biotechnology, Jagiellonian University | Lukasz Rabalski, Maciej Kosinski, Natalia Mazur-Panasiuk, Aneta Kopacz, Piotr Zabek, Tomasz Dyda, Andrzej Horban, Krzysztof Pyrc, Krystyna Bienkowska-Szewczyk |
| EPI_ISL_1084614 | Hospital for Infectious Diseases, Molecular Diagnostics Laboratory, Warsaw, Poland | 56. Laboratory of Recombinant Vaccines, Intercollegiate Faculty of Biotechnology University of Gdansk and Medical University of Gdansk, 2. ViroGenetics - BSL3 Laboratory of Virology, Maopolska Centre of Biotechnology, Jagiellonian University | Lukasz Rabalski, Maciej Kosinski, Natalia Mazur-Panasiuk, Aneta Kopacz, Piotr Zabek, Tomasz Dyda, Andrzej Horban, Krzysztof Pyrc, Krystyna Bienkowska-Szewczyk |
| EPI_ISL_1084620 | Hospital for Infectious Diseases, Molecular Diagnostics Laboratory, Warsaw, Poland | 10. Laboratory of Recombinant Vaccines, Intercollegiate Faculty of Biotechnology University of Gdansk and Medical University of Gdansk, 2. ViroGenetics - BSL3 Laboratory of Virology, Maopolska Centre of Biotechnology, Jagiellonian University | Lukasz Rabalski, Maciej Kosinski, Natalia Mazur-Panasiuk, Aneta Kopacz, Piotr Zabek, Tomasz Dyda, Andrzej Horban, Krzysztof Pyrc, Krystyna Bienkowska-Szewczyk |
| EPI_ISL_1084621 | Hospital for Infectious Diseases, Molecular Diagnostics Laboratory, Warsaw, Poland | 18. Laboratory of Recombinant Vaccines, Intercollegiate Faculty of Biotechnology University of Gdansk and Medical University of Gdansk, 2. ViroGenetics - BSL3 Laboratory of Virology, Maopolska Centre of Biotechnology, Jagiellonian University | Lukasz Rabalski, Maciej Kosinski, Natalia Mazur-Panasiuk, Aneta Kopacz, Piotr Zabek, Tomasz Dyda, Andrzej Horban, Krzysztof Pyrc, Krystyna Bienkowska-Szewczyk |
| EPI_ISL_1084622 | Hospital for Infectious Diseases, Molecular Diagnostics | 22. Laboratory of Recombinant Vaccines, Intercollegiate | Lukasz Rabalski, Maciej Kosinski, Natalia Mazur-Panasiuk, Aneta Kopacz, Piotr Zabek, Tomasz Dyda, Andrzej Horban, Krzysztof Pyrc, Krystyna |

| | | | |
|---|---|---|---|
| | Laboratory, Warsaw, Poland | Faculty of Biotechnology University of Gdansk and Medical University of Gdansk, 2. ViroGenetics - BSL3 Laboratory of Virology, Maopolska Centre of Biotechnology, Jagiellonian University | Bienkowska-Szewczyk |
| EPI_ISL_1084623 | Hospital for Infectious Diseases, Molecular Diagnostics Laboratory, Warsaw, Poland | 28. Laboratory of Recombinant Vaccines, Intercollegiate Faculty of Biotechnology University of Gdansk and Medical University of Gdansk, 2. ViroGenetics - BSL3 Laboratory of Virology, Maopolska Centre of Biotechnology, Jagiellonian University | Lukasz Rabalski, Maciej Kosinski, Natalia Mazur-Panasiuk, Aneta Kopacz, Piotr Zabek, Tomasz Dyda, Andrzej Horban, Krzysztof Pyrc, Krystyna Bienkowska-Szewczyk |
| EPI_ISL_1084624 | Hospital for Infectious Diseases, Molecular Diagnostics Laboratory, Warsaw, Poland | 37. Laboratory of Recombinant Vaccines, Intercollegiate Faculty of Biotechnology University of Gdansk and Medical University of Gdansk, 2. ViroGenetics - BSL3 Laboratory of Virology, Maopolska Centre of Biotechnology, Jagiellonian University | Lukasz Rabalski, Maciej Kosinski, Natalia Mazur-Panasiuk, Aneta Kopacz, Piotr Zabek, Tomasz Dyda, Andrzej Horban, Krzysztof Pyrc, Krystyna Bienkowska-Szewczyk |
| EPI_ISL_1084625 | Hospital for Infectious Diseases, Molecular Diagnostics Laboratory, Warsaw, Poland | 38. Laboratory of Recombinant Vaccines, Intercollegiate Faculty of Biotechnology University of Gdansk and Medical University of Gdansk, 2. ViroGenetics - BSL3 Laboratory of Virology, Maopolska Centre of Biotechnology, Jagiellonian University | Lukasz Rabalski, Maciej Kosinski, Natalia Mazur-Panasiuk, Aneta Kopacz, Piotr Zabek, Tomasz Dyda, Andrzej Horban, Krzysztof Pyrc, Krystyna Bienkowska-Szewczyk |
| EPI_ISL_1084626 | Hospital for Infectious Diseases, Molecular Diagnostics Laboratory, Warsaw, Poland | 42. Laboratory of Recombinant Vaccines, Intercollegiate Faculty of Biotechnology University of Gdansk and Medical University of Gdansk, 2. ViroGenetics - BSL3 Laboratory of Virology, Maopolska Centre of Biotechnology, Jagiellonian University | Lukasz Rabalski, Maciej Kosinski, Natalia Mazur-Panasiuk, Aneta Kopacz, Piotr Zabek, Tomasz Dyda, Andrzej Horban, Krzysztof Pyrc, Krystyna Bienkowska-Szewczyk |
| EPI_ISL_1084627 | Hospital for Infectious Diseases, Molecular Diagnostics Laboratory, Warsaw, Poland | 2. Laboratory of Recombinant Vaccines, Intercollegiate Faculty of Biotechnology University of Gdansk and Medical University of Gdansk, 2. ViroGenetics - BSL3 Laboratory of Virology, Maopolska Centre of Biotechnology, Jagiellonian University | Lukasz Rabalski, Maciej Kosinski, Natalia Mazur-Panasiuk, Aneta Kopacz, Piotr Zabek, Tomasz Dyda, Andrzej Horban, Krzysztof Pyrc, Krystyna Bienkowska-Szewczyk |
| EPI_ISL_1084628 | Hospital for Infectious Diseases, Molecular Diagnostics Laboratory, Warsaw, Poland | 7. Laboratory of Recombinant Vaccines, Intercollegiate Faculty of Biotechnology University of Gdansk and Medical University of Gdansk, 2. ViroGenetics - BSL3 Laboratory of Virology, Maopolska Centre of Biotechnology, Jagiellonian University | Lukasz Rabalski, Maciej Kosinski, Natalia Mazur-Panasiuk, Aneta Kopacz, Piotr Zabek, Tomasz Dyda, Andrzej Horban, Krzysztof Pyrc, Krystyna Bienkowska-Szewczyk |
| EPI_ISL_1084629 | Hospital for Infectious Diseases, Molecular Diagnostics Laboratory, Warsaw, Poland | 31. Laboratory of Recombinant Vaccines, Intercollegiate Faculty of Biotechnology University of Gdansk and Medical University of Gdansk, 2. ViroGenetics - BSL3 Laboratory of Virology, Maopolska Centre of Biotechnology, Jagiellonian University | Lukasz Rabalski, Maciej Kosinski, Natalia Mazur-Panasiuk, Aneta Kopacz, Piotr Zabek, Tomasz Dyda, Andrzej Horban, Krzysztof Pyrc, Krystyna Bienkowska-Szewczyk |
| EPI_ISL_1084630 | Hospital for Infectious Diseases, Molecular Diagnostics Laboratory, Warsaw, Poland | 35. Laboratory of Recombinant Vaccines, Intercollegiate Faculty of Biotechnology University of Gdansk and Medical University of Gdansk, 2. ViroGenetics - BSL3 Laboratory of Virology, Maopolska Centre of Biotechnology, Jagiellonian University | Lukasz Rabalski, Maciej Kosinski, Natalia Mazur-Panasiuk, Aneta Kopacz, Piotr Zabek, Tomasz Dyda, Andrzej Horban, Krzysztof Pyrc, Krystyna Bienkowska-Szewczyk |
| EPI_ISL_1084631 | Hospital for Infectious Diseases, Molecular Diagnostics Laboratory, Warsaw, Poland | 7. Laboratory of Recombinant Vaccines, Intercollegiate Faculty of Biotechnology University of Gdansk and Medical University of Gdansk, 2. ViroGenetics - BSL3 Laboratory of Virology, Maopolska Centre of Biotechnology, Jagiellonian University | Lukasz Rabalski, Maciej Kosinski, Natalia Mazur-Panasiuk, Aneta Kopacz, Piotr Zabek, Tomasz Dyda, Andrzej Horban, Krzysztof Pyrc, Krystyna Bienkowska-Szewczyk |
| EPI_ISL_1084632 | Hospital for Infectious Diseases, Molecular Diagnostics Laboratory, Warsaw, Poland | 6. Laboratory of Recombinant Vaccines, Intercollegiate Faculty of Biotechnology University of Gdansk and Medical University of Gdansk, 2. ViroGenetics - BSL3 Laboratory of Virology, Maopolska Centre of Biotechnology, Jagiellonian University | Lukasz Rabalski, Maciej Kosinski, Natalia Mazur-Panasiuk, Aneta Kopacz, Piotr Zabek, Tomasz Dyda, Andrzej Horban, Krzysztof Pyrc, Krystyna Bienkowska-Szewczyk |
| EPI_ISL_1084633 | Hospital for Infectious Diseases, Molecular Diagnostics Laboratory, Warsaw, Poland | 5. Laboratory of Recombinant Vaccines, Intercollegiate Faculty of Biotechnology University of Gdansk and Medical University of Gdansk, 2. ViroGenetics - BSL3 Laboratory of Virology, Maopolska Centre of Biotechnology, Jagiellonian University | Lukasz Rabalski, Maciej Kosinski, Natalia Mazur-Panasiuk, Aneta Kopacz, Piotr Zabek, Tomasz Dyda, Andrzej Horban, Krzysztof Pyrc, Krystyna Bienkowska-Szewczyk |
| EPI_ISL_1084634 | Hospital for Infectious Diseases, Molecular Diagnostics Laboratory, Warsaw, Poland | 3. Laboratory of Recombinant Vaccines, Intercollegiate Faculty of Biotechnology University of Gdansk and Medical University of Gdansk, 2. ViroGenetics - BSL3 Laboratory of Virology, Maopolska Centre of Biotechnology, Jagiellonian University | Lukasz Rabalski, Maciej Kosinski, Natalia Mazur-Panasiuk, Aneta Kopacz, Piotr Zabek, Tomasz Dyda, Andrzej Horban, Krzysztof Pyrc, Krystyna Bienkowska-Szewczyk |
| EPI_ISL_1084635 | Hospital for Infectious Diseases, Molecular Diagnostics Laboratory, Warsaw, Poland | 0. Laboratory of Recombinant Vaccines, Intercollegiate Faculty of Biotechnology University of Gdansk and Medical University of Gdansk, 2. ViroGenetics - BSL3 Laboratory of Virology, Maopolska Centre of Biotechnology, Jagiellonian University | Lukasz Rabalski, Maciej Kosinski, Natalia Mazur-Panasiuk, Aneta Kopacz, Piotr Zabek, Tomasz Dyda, Andrzej Horban, Krzysztof Pyrc, Krystyna Bienkowska-Szewczyk |
| EPI_ISL_1084636 | Hospital for Infectious Diseases, Molecular Diagnostics Laboratory, Warsaw, Poland | 1. Laboratory of Recombinant Vaccines, Intercollegiate Faculty of Biotechnology University of Gdansk and Medical University of Gdansk, 2. ViroGenetics - BSL3 Laboratory of Virology, Maopolska Centre of Biotechnology, Jagiellonian University | Lukasz Rabalski, Maciej Kosinski, Natalia Mazur-Panasiuk, Aneta Kopacz, Piotr Zabek, Tomasz Dyda, Andrzej Horban, Krzysztof Pyrc, Krystyna Bienkowska-Szewczyk |
| EPI_ISL_1084637 | Hospital for Infectious Diseases, Molecular Diagnostics Laboratory, Warsaw, Poland | 3. Laboratory of Recombinant Vaccines, Intercollegiate Faculty of Biotechnology University of Gdansk and Medical University of Gdansk, 2. ViroGenetics - BSL3 Laboratory of Virology, Maopolska Centre of Biotechnology, Jagiellonian University | Lukasz Rabalski, Maciej Kosinski, Natalia Mazur-Panasiuk, Aneta Kopacz, Piotr Zabek, Tomasz Dyda, Andrzej Horban, Krzysztof Pyrc, Krystyna Bienkowska-Szewczyk |

| | | | |
|---|---|---|---|
| EPI_ISL_1084638 | Hospital for Infectious Diseases, Molecular Diagnostics Laboratory, Warsaw, Poland | 4. Laboratory of Recombinant Vaccines, Intercollegiate Faculty of Biotechnology University of Gdansk and Medical University of Gdansk, 2. ViroGenetics - BSL3 Laboratory of Virology, Maopolska Centre of Biotechnology, Jagiellonian University | Lukasz Rabalski, Maciej Kosinski, Natalia Mazur-Panasiuk, Aneta Kopacz, Piotr Zabek, Tomasz Dyda, Andrzej Horban, Krzysztof Pyrc, Krystyna Bienkowska-Szewczyk |
| EPI_ISL_1084639 | Hospital for Infectious Diseases, Molecular Diagnostics Laboratory, Warsaw, Poland | 5. Laboratory of Recombinant Vaccines, Intercollegiate Faculty of Biotechnology University of Gdansk and Medical University of Gdansk, 2. ViroGenetics - BSL3 Laboratory of Virology, Maopolska Centre of Biotechnology, Jagiellonian University | Lukasz Rabalski, Maciej Kosinski, Natalia Mazur-Panasiuk, Aneta Kopacz, Piotr Zabek, Tomasz Dyda, Andrzej Horban, Krzysztof Pyrc, Krystyna Bienkowska-Szewczyk |
| EPI_ISL_1084640 | Hospital for Infectious Diseases, Molecular Diagnostics Laboratory, Warsaw, Poland | 6. Laboratory of Recombinant Vaccines, Intercollegiate Faculty of Biotechnology University of Gdansk and Medical University of Gdansk, 2. ViroGenetics - BSL3 Laboratory of Virology, Maopolska Centre of Biotechnology, Jagiellonian University | Lukasz Rabalski, Maciej Kosinski, Natalia Mazur-Panasiuk, Aneta Kopacz, Piotr Zabek, Tomasz Dyda, Andrzej Horban, Krzysztof Pyrc, Krystyna Bienkowska-Szewczyk |
| EPI_ISL_1084641 | Hospital for Infectious Diseases, Molecular Diagnostics Laboratory, Warsaw, Poland | 9. Laboratory of Recombinant Vaccines, Intercollegiate Faculty of Biotechnology University of Gdansk and Medical University of Gdansk, 2. ViroGenetics - BSL3 Laboratory of Virology, Maopolska Centre of Biotechnology, Jagiellonian University | Lukasz Rabalski, Maciej Kosinski, Natalia Mazur-Panasiuk, Aneta Kopacz, Piotr Zabek, Tomasz Dyda, Andrzej Horban, Krzysztof Pyrc, Krystyna Bienkowska-Szewczyk |
| EPI_ISL_1084642 | Hospital for Infectious Diseases, Molecular Diagnostics Laboratory, Warsaw, Poland | 11. Laboratory of Recombinant Vaccines, Intercollegiate Faculty of Biotechnology University of Gdansk and Medical University of Gdansk, 2. ViroGenetics - BSL3 Laboratory of Virology, Maopolska Centre of Biotechnology, Jagiellonian University | Lukasz Rabalski, Maciej Kosinski, Natalia Mazur-Panasiuk, Aneta Kopacz, Piotr Zabek, Tomasz Dyda, Andrzej Horban, Krzysztof Pyrc, Krystyna Bienkowska-Szewczyk |
| EPI_ISL_1084643 | Hospital for Infectious Diseases, Molecular Diagnostics Laboratory, Warsaw, Poland | 12. Laboratory of Recombinant Vaccines, Intercollegiate Faculty of Biotechnology University of Gdansk and Medical University of Gdansk, 2. ViroGenetics - BSL3 Laboratory of Virology, Maopolska Centre of Biotechnology, Jagiellonian University | Lukasz Rabalski, Maciej Kosinski, Natalia Mazur-Panasiuk, Aneta Kopacz, Piotr Zabek, Tomasz Dyda, Andrzej Horban, Krzysztof Pyrc, Krystyna Bienkowska-Szewczyk |
| EPI_ISL_1084644 | Hospital for Infectious Diseases, Molecular Diagnostics Laboratory, Warsaw, Poland | 14. Laboratory of Recombinant Vaccines, Intercollegiate Faculty of Biotechnology University of Gdansk and Medical University of Gdansk, 2. ViroGenetics - BSL3 Laboratory of Virology, Maopolska Centre of Biotechnology, Jagiellonian University | Lukasz Rabalski, Maciej Kosinski, Natalia Mazur-Panasiuk, Aneta Kopacz, Piotr Zabek, Tomasz Dyda, Andrzej Horban, Krzysztof Pyrc, Krystyna Bienkowska-Szewczyk |
| EPI_ISL_1084645 | Hospital for Infectious Diseases, Molecular Diagnostics Laboratory, Warsaw, Poland | 15. Laboratory of Recombinant Vaccines, Intercollegiate Faculty of Biotechnology University of Gdansk and Medical University of Gdansk, 2. ViroGenetics - BSL3 Laboratory of Virology, Maopolska Centre of Biotechnology, Jagiellonian University | Lukasz Rabalski, Maciej Kosinski, Natalia Mazur-Panasiuk, Aneta Kopacz, Piotr Zabek, Tomasz Dyda, Andrzej Horban, Krzysztof Pyrc, Krystyna Bienkowska-Szewczyk |
| EPI_ISL_1084646 | Hospital for Infectious Diseases, Molecular Diagnostics Laboratory, Warsaw, Poland | 16. Laboratory of Recombinant Vaccines, Intercollegiate Faculty of Biotechnology University of Gdansk and Medical University of Gdansk, 2. ViroGenetics - BSL3 Laboratory of Virology, Maopolska Centre of Biotechnology, Jagiellonian University | Lukasz Rabalski, Maciej Kosinski, Natalia Mazur-Panasiuk, Aneta Kopacz, Piotr Zabek, Tomasz Dyda, Andrzej Horban, Krzysztof Pyrc, Krystyna Bienkowska-Szewczyk |
| EPI_ISL_1084647 | Hospital for Infectious Diseases, Molecular Diagnostics Laboratory, Warsaw, Poland | 17. Laboratory of Recombinant Vaccines, Intercollegiate Faculty of Biotechnology University of Gdansk and Medical University of Gdansk, 2. ViroGenetics - BSL3 Laboratory of Virology, Maopolska Centre of Biotechnology, Jagiellonian University | Lukasz Rabalski, Maciej Kosinski, Natalia Mazur-Panasiuk, Aneta Kopacz, Piotr Zabek, Tomasz Dyda, Andrzej Horban, Krzysztof Pyrc, Krystyna Bienkowska-Szewczyk |
| EPI_ISL_1084648 | Hospital for Infectious Diseases, Molecular Diagnostics Laboratory, Warsaw, Poland | 19. Laboratory of Recombinant Vaccines, Intercollegiate Faculty of Biotechnology University of Gdansk and Medical University of Gdansk, 2. ViroGenetics - BSL3 Laboratory of Virology, Maopolska Centre of Biotechnology, Jagiellonian University | Lukasz Rabalski, Maciej Kosinski, Natalia Mazur-Panasiuk, Aneta Kopacz, Piotr Zabek, Tomasz Dyda, Andrzej Horban, Krzysztof Pyrc, Krystyna Bienkowska-Szewczyk |
| EPI_ISL_1084649 | Hospital for Infectious Diseases, Molecular Diagnostics Laboratory, Warsaw, Poland | 20. Laboratory of Recombinant Vaccines, Intercollegiate Faculty of Biotechnology University of Gdansk and Medical University of Gdansk, 2. ViroGenetics - BSL3 Laboratory of Virology, Maopolska Centre of Biotechnology, Jagiellonian University | Lukasz Rabalski, Maciej Kosinski, Natalia Mazur-Panasiuk, Aneta Kopacz, Piotr Zabek, Tomasz Dyda, Andrzej Horban, Krzysztof Pyrc, Krystyna Bienkowska-Szewczyk |
| EPI_ISL_1084650 | Hospital for Infectious Diseases, Molecular Diagnostics Laboratory, Warsaw, Poland | 21. Laboratory of Recombinant Vaccines, Intercollegiate Faculty of Biotechnology University of Gdansk and Medical University of Gdansk, 2. ViroGenetics - BSL3 Laboratory of Virology, Maopolska Centre of Biotechnology, Jagiellonian University | Lukasz Rabalski, Maciej Kosinski, Natalia Mazur-Panasiuk, Aneta Kopacz, Piotr Zabek, Tomasz Dyda, Andrzej Horban, Krzysztof Pyrc, Krystyna Bienkowska-Szewczyk |
| EPI_ISL_1084651 | Hospital for Infectious Diseases, Molecular Diagnostics Laboratory, Warsaw, Poland | 23. Laboratory of Recombinant Vaccines, Intercollegiate Faculty of Biotechnology University of Gdansk and Medical University of Gdansk, 2. ViroGenetics - BSL3 Laboratory of Virology, Maopolska Centre of Biotechnology, Jagiellonian University | Lukasz Rabalski, Maciej Kosinski, Natalia Mazur-Panasiuk, Aneta Kopacz, Piotr Zabek, Tomasz Dyda, Andrzej Horban, Krzysztof Pyrc, Krystyna Bienkowska-Szewczyk |
| EPI_ISL_1084652 | Hospital for Infectious Diseases, Molecular Diagnostics Laboratory, Warsaw, Poland | 24. Laboratory of Recombinant Vaccines, Intercollegiate Faculty of Biotechnology University of Gdansk and Medical University of Gdansk, 2. ViroGenetics - BSL3 Laboratory of Virology, Maopolska Centre of Biotechnology, Jagiellonian University | Lukasz Rabalski, Maciej Kosinski, Natalia Mazur-Panasiuk, Aneta Kopacz, Piotr Zabek, Tomasz Dyda, Andrzej Horban, Krzysztof Pyrc, Krystyna Bienkowska-Szewczyk |
| EPI_ISL_1084653 | Hospital for Infectious Diseases, Molecular Diagnostics Laboratory, Warsaw, Poland | 25. Laboratory of Recombinant Vaccines, Intercollegiate Faculty of Biotechnology University of Gdansk and Medical University of Gdansk, 2. ViroGenetics - BSL3 Laboratory of Virology, Maopolska Centre of Biotechnology, Jagiellonian University | Lukasz Rabalski, Maciej Kosinski, Natalia Mazur-Panasiuk, Aneta Kopacz, Piotr Zabek, Tomasz Dyda, Andrzej Horban, Krzysztof Pyrc, Krystyna Bienkowska-Szewczyk |

| | | | |
|---|---|---|---|
| EPI_ISL_1084654 | Hospital for Infectious Diseases, Molecular Diagnostics Laboratory, Warsaw, Poland | 29. Laboratory of Recombinant Vaccines, Intercollegiate Faculty of Biotechnology University of Gdansk and Medical University of Gdansk, 2. ViroGenetics - BSL3 Laboratory of Virology, Maopolska Centre of Biotechnology, Jagiellonian University | Lukasz Rabalski, Maciej Kosinski, Natalia Mazur-Panasiuk, Aneta Kopacz, Piotr Zabek, Tomasz Dyda, Andrzej Horban, Krzysztof Pyrc, Krystyna Bienkowska-Szewczyk |
| EPI_ISL_1084655 | Hospital for Infectious Diseases, Molecular Diagnostics Laboratory, Warsaw, Poland | 30. Laboratory of Recombinant Vaccines, Intercollegiate Faculty of Biotechnology University of Gdansk and Medical University of Gdansk, 2. ViroGenetics - BSL3 Laboratory of Virology, Maopolska Centre of Biotechnology, Jagiellonian University | Lukasz Rabalski, Maciej Kosinski, Natalia Mazur-Panasiuk, Aneta Kopacz, Piotr Zabek, Tomasz Dyda, Andrzej Horban, Krzysztof Pyrc, Krystyna Bienkowska-Szewczyk |
| EPI_ISL_1084656 | Hospital for Infectious Diseases, Molecular Diagnostics Laboratory, Warsaw, Poland | 32. Laboratory of Recombinant Vaccines, Intercollegiate Faculty of Biotechnology University of Gdansk and Medical University of Gdansk, 2. ViroGenetics - BSL3 Laboratory of Virology, Maopolska Centre of Biotechnology, Jagiellonian University | Lukasz Rabalski, Maciej Kosinski, Natalia Mazur-Panasiuk, Aneta Kopacz, Piotr Zabek, Tomasz Dyda, Andrzej Horban, Krzysztof Pyrc, Krystyna Bienkowska-Szewczyk |
| EPI_ISL_1084657 | Hospital for Infectious Diseases, Molecular Diagnostics Laboratory, Warsaw, Poland | 33. Laboratory of Recombinant Vaccines, Intercollegiate Faculty of Biotechnology University of Gdansk and Medical University of Gdansk, 2. ViroGenetics - BSL3 Laboratory of Virology, Maopolska Centre of Biotechnology, Jagiellonian University | Lukasz Rabalski, Maciej Kosinski, Natalia Mazur-Panasiuk, Aneta Kopacz, Piotr Zabek, Tomasz Dyda, Andrzej Horban, Krzysztof Pyrc, Krystyna Bienkowska-Szewczyk |
| EPI_ISL_1084658 | Hospital for Infectious Diseases, Molecular Diagnostics Laboratory, Warsaw, Poland | 39. Laboratory of Recombinant Vaccines, Intercollegiate Faculty of Biotechnology University of Gdansk and Medical University of Gdansk, 2. ViroGenetics - BSL3 Laboratory of Virology, Maopolska Centre of Biotechnology, Jagiellonian University | Lukasz Rabalski, Maciej Kosinski, Natalia Mazur-Panasiuk, Aneta Kopacz, Piotr Zabek, Tomasz Dyda, Andrzej Horban, Krzysztof Pyrc, Krystyna Bienkowska-Szewczyk |
| EPI_ISL_1084659 | Hospital for Infectious Diseases, Molecular Diagnostics Laboratory, Warsaw, Poland | 41. Laboratory of Recombinant Vaccines, Intercollegiate Faculty of Biotechnology University of Gdansk and Medical University of Gdansk, 2. ViroGenetics - BSL3 Laboratory of Virology, Maopolska Centre of Biotechnology, Jagiellonian University | Lukasz Rabalski, Maciej Kosinski, Natalia Mazur-Panasiuk, Aneta Kopacz, Piotr Zabek, Tomasz Dyda, Andrzej Horban, Krzysztof Pyrc, Krystyna Bienkowska-Szewczyk |
| EPI_ISL_1084660 | Hospital for Infectious Diseases, Molecular Diagnostics Laboratory, Warsaw, Poland | 44. Laboratory of Recombinant Vaccines, Intercollegiate Faculty of Biotechnology University of Gdansk and Medical University of Gdansk, 2. ViroGenetics - BSL3 Laboratory of Virology, Maopolska Centre of Biotechnology, Jagiellonian University | Lukasz Rabalski, Maciej Kosinski, Natalia Mazur-Panasiuk, Aneta Kopacz, Piotr Zabek, Tomasz Dyda, Andrzej Horban, Krzysztof Pyrc, Krystyna Bienkowska-Szewczyk |
| EPI_ISL_1084661 | Hospital for Infectious Diseases, Molecular Diagnostics Laboratory, Warsaw, Poland | 45. Laboratory of Recombinant Vaccines, Intercollegiate Faculty of Biotechnology University of Gdansk and Medical University of Gdansk, 2. ViroGenetics - BSL3 Laboratory of Virology, Maopolska Centre of Biotechnology, Jagiellonian University | Lukasz Rabalski, Maciej Kosinski, Natalia Mazur-Panasiuk, Aneta Kopacz, Piotr Zabek, Tomasz Dyda, Andrzej Horban, Krzysztof Pyrc, Krystyna Bienkowska-Szewczyk |
| EPI_ISL_1084662 | Hospital for Infectious Diseases, Molecular Diagnostics Laboratory, Warsaw, Poland | 46. Laboratory of Recombinant Vaccines, Intercollegiate Faculty of Biotechnology University of Gdansk and Medical University of Gdansk, 2. ViroGenetics - BSL3 Laboratory of Virology, Maopolska Centre of Biotechnology, Jagiellonian University | Lukasz Rabalski, Maciej Kosinski, Natalia Mazur-Panasiuk, Aneta Kopacz, Piotr Zabek, Tomasz Dyda, Andrzej Horban, Krzysztof Pyrc, Krystyna Bienkowska-Szewczyk |
| EPI_ISL_1084663 | Hospital for Infectious Diseases, Molecular Diagnostics Laboratory, Warsaw, Poland | 47. Laboratory of Recombinant Vaccines, Intercollegiate Faculty of Biotechnology University of Gdansk and Medical University of Gdansk, 2. ViroGenetics - BSL3 Laboratory of Virology, Maopolska Centre of Biotechnology, Jagiellonian University | Lukasz Rabalski, Maciej Kosinski, Natalia Mazur-Panasiuk, Aneta Kopacz, Piotr Zabek, Tomasz Dyda, Andrzej Horban, Krzysztof Pyrc, Krystyna Bienkowska-Szewczyk |
| EPI_ISL_1084664 | Hospital for Infectious Diseases, Molecular Diagnostics Laboratory, Warsaw, Poland | 48. Laboratory of Recombinant Vaccines, Intercollegiate Faculty of Biotechnology University of Gdansk and Medical University of Gdansk, 2. ViroGenetics - BSL3 Laboratory of Virology, Maopolska Centre of Biotechnology, Jagiellonian University | Lukasz Rabalski, Maciej Kosinski, Natalia Mazur-Panasiuk, Aneta Kopacz, Piotr Zabek, Tomasz Dyda, Andrzej Horban, Krzysztof Pyrc, Krystyna Bienkowska-Szewczyk |
| EPI_ISL_1084665 | Hospital for Infectious Diseases, Molecular Diagnostics Laboratory, Warsaw, Poland | 49. Laboratory of Recombinant Vaccines, Intercollegiate Faculty of Biotechnology University of Gdansk and Medical University of Gdansk, 2. ViroGenetics - BSL3 Laboratory of Virology, Maopolska Centre of Biotechnology, Jagiellonian University | Lukasz Rabalski, Maciej Kosinski, Natalia Mazur-Panasiuk, Aneta Kopacz, Piotr Zabek, Tomasz Dyda, Andrzej Horban, Krzysztof Pyrc, Krystyna Bienkowska-Szewczyk |
| EPI_ISL_1084666 | Hospital for Infectious Diseases, Molecular Diagnostics Laboratory, Warsaw, Poland | 50. Laboratory of Recombinant Vaccines, Intercollegiate Faculty of Biotechnology University of Gdansk and Medical University of Gdansk, 2. ViroGenetics - BSL3 Laboratory of Virology, Maopolska Centre of Biotechnology, Jagiellonian University | Lukasz Rabalski, Maciej Kosinski, Natalia Mazur-Panasiuk, Aneta Kopacz, Piotr Zabek, Tomasz Dyda, Andrzej Horban, Krzysztof Pyrc, Krystyna Bienkowska-Szewczyk |
| EPI_ISL_1084667 | Hospital for Infectious Diseases, Molecular Diagnostics Laboratory, Warsaw, Poland | 51. Laboratory of Recombinant Vaccines, Intercollegiate Faculty of Biotechnology University of Gdansk and Medical University of Gdansk, 2. ViroGenetics - BSL3 Laboratory of Virology, Maopolska Centre of Biotechnology, Jagiellonian University | Lukasz Rabalski, Maciej Kosinski, Natalia Mazur-Panasiuk, Aneta Kopacz, Piotr Zabek, Tomasz Dyda, Andrzej Horban, Krzysztof Pyrc, Krystyna Bienkowska-Szewczyk |
| EPI_ISL_1084668 | Hospital for Infectious Diseases, Molecular Diagnostics Laboratory, Warsaw, Poland | 53. Laboratory of Recombinant Vaccines, Intercollegiate Faculty of Biotechnology University of Gdansk and Medical University of Gdansk, 2. ViroGenetics - BSL3 Laboratory of Virology, Maopolska Centre of Biotechnology, Jagiellonian University | Lukasz Rabalski, Maciej Kosinski, Natalia Mazur-Panasiuk, Aneta Kopacz, Piotr Zabek, Tomasz Dyda, Andrzej Horban, Krzysztof Pyrc, Krystyna Bienkowska-Szewczyk |
| EPI_ISL_1084669 | Hospital for Infectious Diseases, Molecular Diagnostics Laboratory, Warsaw, Poland | 54. Laboratory of Recombinant Vaccines, Intercollegiate Faculty of Biotechnology University of Gdansk and Medical University of Gdansk, 2. ViroGenetics - BSL3 Laboratory of Virology, Maopolska Centre of Biotechnology, Jagiellonian University | Lukasz Rabalski, Maciej Kosinski, Natalia Mazur-Panasiuk, Aneta Kopacz, Piotr Zabek, Tomasz Dyda, Andrzej Horban, Krzysztof Pyrc, Krystyna Bienkowska-Szewczyk |

| | | | |
|---|---|---|---|
| EPI_ISL_1084670 | Hospital for Infectious Diseases, Molecular Diagnostics Laboratory, Warsaw, Poland | 58. Laboratory of Recombinant Vaccines, Intercollegiate Faculty of Biotechnology University of Gdansk and Medical University of Gdansk, 2. ViroGenetics - BSL3 Laboratory of Virology, Maopolska Centre of Biotechnology, Jagiellonian University | Lukasz Rabalski, Maciej Kosinski, Natalia Mazur-Panasiuk, Aneta Kopacz, Piotr Zabek, Tomasz Dyda, Andrzej Horban, Krzysztof Pyrc, Krystyna Bienkowska-Szewczyk |
| EPI_ISL_1084671 | Hospital for Infectious Diseases, Molecular Diagnostics Laboratory, Warsaw, Poland | 60. Laboratory of Recombinant Vaccines, Intercollegiate Faculty of Biotechnology University of Gdansk and Medical University of Gdansk, 2. ViroGenetics - BSL3 Laboratory of Virology, Maopolska Centre of Biotechnology, Jagiellonian University | Lukasz Rabalski, Maciej Kosinski, Natalia Mazur-Panasiuk, Aneta Kopacz, Piotr Zabek, Tomasz Dyda, Andrzej Horban, Krzysztof Pyrc, Krystyna Bienkowska-Szewczyk |
| EPI_ISL_1084672 | Hospital for Infectious Diseases, Molecular Diagnostics Laboratory, Warsaw, Poland | 62. Laboratory of Recombinant Vaccines, Intercollegiate Faculty of Biotechnology University of Gdansk and Medical University of Gdansk, 2. ViroGenetics - BSL3 Laboratory of Virology, Maopolska Centre of Biotechnology, Jagiellonian University | Lukasz Rabalski, Maciej Kosinski, Natalia Mazur-Panasiuk, Aneta Kopacz, Piotr Zabek, Tomasz Dyda, Andrzej Horban, Krzysztof Pyrc, Krystyna Bienkowska-Szewczyk |
| EPI_ISL_1085162, EPI_ISL_1085174 | Laboratory of Infectious Diseases, Department of Biomedical and Clinical Sciences L. Sacco, University of Milan | Laboratory of Infectious Diseases, Department of Biomedical and Clinical Sciences L. Sacco, University of Milan | Alessia Lai, Annalisa Bergna, Carla Della Ventura, Claudia Balotta, Massimo Galli, Gianguglielmo Zehender on behalf of SARS-CoV-2 ITALIAN RESEARCH ENTERPRISE-(SCIRE) Collaborative Group |
| EPI_ISL_1085273, EPI_ISL_1085276, EPI_ISL_1085278, EPI_ISL_1085279, EPI_ISL_1085280, EPI_ISL_1085282, EPI_ISL_1085283, EPI_ISL_1085284, EPI_ISL_1085307, EPI_ISL_1085309, EPI_ISL_1085360, EPI_ISL_1085386, EPI_ISL_1085387, EPI_ISL_1085388, EPI_ISL_1085400, EPI_ISL_1085453, EPI_ISL_1085473, EPI_ISL_1085533, EPI_ISL_1085545, EPI_ISL_1085553, EPI_ISL_1085585, EPI_ISL_1085588, EPI_ISL_1085589, EPI_ISL_1085590, EPI_ISL_1085591, EPI_ISL_1085593, EPI_ISL_1085594, EPI_ISL_1085600, EPI_ISL_1085607, EPI_ISL_1085641, EPI_ISL_1085646, EPI_ISL_1085647, EPI_ISL_1085648, EPI_ISL_1085649, EPI_ISL_1085651, EPI_ISL_1085656, EPI_ISL_1085665, EPI_ISL_1085668, EPI_ISL_1085669, EPI_ISL_1085670, EPI_ISL_1085671, EPI_ISL_1085672, EPI_ISL_1085709, EPI_ISL_1085713, EPI_ISL_1085716, EPI_ISL_1085717, EPI_ISL_1085718, EPI_ISL_1085722, EPI_ISL_1085724, EPI_ISL_1085725, EPI_ISL_1085726, EPI_ISL_1085732, EPI_ISL_1085734, EPI_ISL_1085735, EPI_ISL_1085784, EPI_ISL_1085786, EPI_ISL_1085792, EPI_ISL_1085796, EPI_ISL_1085808, EPI_ISL_1085809, EPI_ISL_1085812, EPI_ISL_1085813, EPI_ISL_1085814, EPI_ISL_1085858, EPI_ISL_1085859, EPI_ISL_1085862, EPI_ISL_1085863, EPI_ISL_1085865, EPI_ISL_1085866, EPI_ISL_1085868, EPI_ISL_1085928, EPI_ISL_1085942, EPI_ISL_1085943, EPI_ISL_1085944 | | | |
| see above | Hôpital Henri Mondor | Department of Virology, Henri Mondor University Hospital, Assistance Publique Hôpitaux de Paris, Université Paris-Est Créteil, INSERM U955 | Christophe Rodriguez, Slim Fourati, Vanessa Demontant, Guillaume Gricourt, Melissa N'Debi, Alexandre Soulier, Elisabeth Trawinski, Jean-Michel Pawlotsky |
| EPI_ISL_1086034 | Diagnosticos da America - DASA | Instituto Adolfo Lutz, Interdiciplinary Procedures Center, Strategic Laboratory | Claudio Tavares Sacchi, Claudia Regina Gonçalves, Erica Valessa Ramos Gomes, Karoline Rodrigues Campos, Caio Vinicius Dias Lopes |
| EPI_ISL_1086035, EPI_ISL_1086036 | IAL Regional de Bauru | Instituto Adolfo Lutz, Interdiciplinary Procedures Center, Strategic Laboratory | Claudio Tavares Sacchi, Claudia Regina Gonçalves, Erica Valessa Ramos Gomes, Karoline Rodrigues Campos, Caio Vinicius Dias Lopes |
| EPI_ISL_1086037, EPI_ISL_1086038, EPI_ISL_1086039, EPI_ISL_1086040, EPI_ISL_1086041, EPI_ISL_1086042, EPI_ISL_1086043 | Diagnosticos da America - DASA | Instituto Adolfo Lutz, Interdiciplinary Procedures Center, Strategic Laboratory | Claudio Tavares Sacchi, Claudia Regina Gonçalves, Erica Valessa Ramos Gomes, Karoline Rodrigues Campos, Caio Vinicius Dias Lopes |
| EPI_ISL_1086044, EPI_ISL_1086045, EPI_ISL_1086046, EPI_ISL_1086047, EPI_ISL_1086050, EPI_ISL_1086051, EPI_ISL_1086052, EPI_ISL_1086053, EPI_ISL_1086054, EPI_ISL_1086055, EPI_ISL_1086056, EPI_ISL_1086057 | | | |
| see above | IAL Regional de Bauru | Instituto Adolfo Lutz, Interdiciplinary Procedures Center, Strategic Laboratory | Claudio Tavares Sacchi, Claudia Regina Gonçalves, Erica Valessa Ramos Gomes, Karoline Rodrigues Campos, Caio Vinicius Dias Lopes |
| EPI_ISL_1092360 | IAL Regional de Bauru | Instituto Adolfo Lutz, Interdiciplinary Procedures Center, Strategic Laboratory | Claudio Tavares Sacchi, Claudia Regina Gonçalves, Erica Valessa Ramos Gomes, Karoline Rodrigues Campos |
| EPI_ISL_1092725 | Diagnosticos da America - DASA | Instituto Adolfo Lutz, Interdiciplinary Procedures Center, Strategic Laboratory | Claudio Tavares Sacchi, Claudia Regina Gonçalves, Erica Valessa Ramos Gomes, Karoline Rodrigues Campos |
| EPI_ISL_1093427 | Influenza and respiratory viruses lab. Pasteur Institute in Algeria | Human virology Department | A.Hachid, F.A.Khardine, M.A. Beloufa, F.Derrar |
| EPI_ISL_1093428 | Influenza and respiratory viruses. Pasteur Institute in Algeria | Human virology deprtment | A.Hachid, F.A.Khardine, F.Derrar |
| EPI_ISL_1095384, EPI_ISL_1095449 | Laboratorio Microbiologia e Virologia P.O. Cotugno A.O. dei Colli | Laboratorio Microbiologia e Virologia P.O. Cotugno A.O. dei Colli | Luigi Atripaldi, Claudia Tiberio, Anna Perfetti, Pellegrino Cerino, Biancamaria Pierri, Maria Concetta Cuomo |
| EPI_ISL_1095913 | IAL Regional de Bauru | Instituto Adolfo Lutz, Interdiciplinary Procedures Center, Strategic Laboratory | Claudio Tavares Sacchi, Claudia Regina Gonçalves, Erica Valessa Ramos Gomes, Karoline Rodrigues Campos |
| EPI_ISL_1096121 | Diagnosticos da America - DASA | Instituto Adolfo Lutz, Interdiciplinary Procedures Center, Strategic Laboratory | Claudio Tavares Sacchi, Claudia Regina Gonçalves, Erica Valessa Ramos Gomes, Karoline Rodrigues Campos, Caio Vinicius Dias Lopes |
| EPI_ISL_1096122, EPI_ISL_1096123, EPI_ISL_1096124, EPI_ISL_1096125, EPI_ISL_1096126, EPI_ISL_1096127, EPI_ISL_1096128, EPI_ISL_1096129, EPI_ISL_1096130, EPI_ISL_1096131, EPI_ISL_1096132, EPI_ISL_1096133, EPI_ISL_1096134 | | | |
| see above | IAL Regional de Bauru | Instituto Adolfo Lutz, Interdiciplinary Procedures Center, Strategic Laboratory | Claudio Tavares Sacchi, Claudia Regina Gonçalves, Erica Valessa Ramos Gomes, Karoline Rodrigues Campos, Caio Vinicius Dias Lopes |
| EPI_ISL_1096135 | Diagnosticos da America - DASA | Instituto Adolfo Lutz, Interdiciplinary Procedures Center, Strategic Laboratory | Claudio Tavares Sacchi, Claudia Regina Gonçalves, Erica Valessa Ramos Gomes, Karoline Rodrigues Campos, Caio Vinicius Dias Lopes |
| EPI_ISL_1096136 | IAL Regional de Bauru | Instituto Adolfo Lutz, Interdiciplinary Procedures Center, Strategic Laboratory | Claudio Tavares Sacchi, Claudia Regina Gonçalves, Erica Valessa Ramos Gomes, Karoline Rodrigues Campos, Caio Vinicius Dias Lopes |
| EPI_ISL_1096140 | Cambodian National Public Health Laboratory, National Institute of Public Health | Virology Unit, Institut Pasteur du Cambodge | Sokhoun Yann, Ly Sovann, Kraing Sidonn, Yi Sengdoeurn, Chin Savuth, Chau Darapheak, Veasna Duong, Erik A Karlsson |
| EPI_ISL_1096228, EPI_ISL_1096229, EPI_ISL_1096232, EPI_ISL_1096233 | Laboratorio Microbiologia e Virologia P.O. Cotugno A.O. dei Colli | Laboratorio Microbiologia e Virologia P.O. Cotugno A.O. dei Colli | Luigi Atripaldi, Claudia Tiberio, Anna Perfetti, Pellegrino Cerino, Biancamaria Pierri, Maria Concetta Cuomo |
| EPI_ISL_1096234, EPI_ISL_1096235 | Outre mer | National Reference Center for Viruses of Respiratory Infections, Institut Pasteur, Paris | Marion Barbet, Sylvie Behillil, Méline Bizard, Angela Brisebarre, Camille Capel, Etienne Simon-Lorière, Vincent Enouf, Maud Vanpeene, Sylvie van der Werf,Rousset Dominique |
| EPI_ISL_1096236, EPI_ISL_1096237, EPI_ISL_1096238, EPI_ISL_1096239, EPI_ISL_1096240, EPI_ISL_1096241 | Research | National Reference Center for Viruses of Respiratory Infections, Institut Pasteur, Paris | Marion Barbet, Sylvie Behillil, Méline Bizard, Angela Brisebarre, Camille Capel, Etienne Simon-Lorière, Vincent Enouf, Maud Vanpeene, Sylvie van der Werf,Leruez-Ville Marianne |
| EPI_ISL_1096242 | Hopital | National Reference Center for Viruses of Respiratory Infections, Institut Pasteur, Paris | Marion Barbet, Sylvie Behillil, Méline Bizard, Angela Brisebarre, Camille Capel, Etienne Simon-Lorière, Vincent Enouf, Maud Vanpeene, Sylvie van der Werf,Martres Pascale |
| EPI_ISL_1096243 | Labo Analyses Med | National Reference Center for Viruses of Respiratory Infections, Institut Pasteur, Paris | Marion Barbet, Sylvie Behillil, Méline Bizard, Angela Brisebarre, Camille Capel, Etienne Simon-Lorière, Vincent Enouf, Maud Vanpeene, Sylvie van der Werf,Martres Pascale |
| EPI_ISL_1096244 | Hopital | National Reference Center for Viruses of Respiratory Infections, Institut Pasteur, Paris | Marion Barbet, Sylvie Behillil, Méline Bizard, Angela Brisebarre, Camille Capel, Etienne Simon-Lorière, Vincent Enouf, Maud Vanpeene, Sylvie van der Werf,Martres Pascale |
| EPI_ISL_1096245, EPI_ISL_1096246 | Hopital | National Reference Center for Viruses of Respiratory Infections, Institut Pasteur, Paris | Marion Barbet, Sylvie Behillil, Méline Bizard, Angela Brisebarre, Camille Capel, Etienne Simon-Lorière, Vincent Enouf, Maud Vanpeene, Sylvie van der Werf,Cariou Marie-Estelle |

| | | | |
|---|---|---|---|
| EPI_ISL_1096248 | Hopital | National Reference Center for Viruses of Respiratory Infections, Institut Pasteur, Paris | Marion Barbet, Sylvie Behillil, Méline Bizard, Angela Brisebarre, Camille Capel, Etienne Simon-Lorière, Vincent Enouf, Maud Vanpeene, Sylvie van der Werf,Guinard JéRome |
| EPI_ISL_1096250, EPI_ISL_1096251, EPI_ISL_1096252 | Labo Analyses Med | National Reference Center for Viruses of Respiratory Infections, Institut Pasteur, Paris | Marion Barbet, Sylvie Behillil, Méline Bizard, Angela Brisebarre, Camille Capel, Etienne Simon-Lorière, Vincent Enouf, Maud Vanpeene, Sylvie van der Werf |
| EPI_ISL_1096253 | Hopital | National Reference Center for Viruses of Respiratory Infections, Institut Pasteur, Paris | Marion Barbet, Sylvie Behillil, Méline Bizard, Angela Brisebarre, Camille Capel, Etienne Simon-Lorière, Vincent Enouf, Maud Vanpeene, Sylvie van der Werf,Hochart A |
| EPI_ISL_1096254 | Hopital | National Reference Center for Viruses of Respiratory Infections, Institut Pasteur, Paris | Marion Barbet, Sylvie Behillil, Méline Bizard, Angela Brisebarre, Camille Capel, Etienne Simon-Lorière, Vincent Enouf, Maud Vanpeene, Sylvie van der Werf,Bisiau H |
| EPI_ISL_1096255, EPI_ISL_1096256 | Hopital | National Reference Center for Viruses of Respiratory Infections, Institut Pasteur, Paris | Marion Barbet, Sylvie Behillil, Méline Bizard, Angela Brisebarre, Camille Capel, Etienne Simon-Lorière, Vincent Enouf, Maud Vanpeene, Sylvie van der Werf,Farfour Eric |
| EPI_ISL_1096257 | Hopital | National Reference Center for Viruses of Respiratory Infections, Institut Pasteur, Paris | Marion Barbet, Sylvie Behillil, Méline Bizard, Angela Brisebarre, Camille Capel, Etienne Simon-Lorière, Vincent Enouf, Maud Vanpeene, Sylvie van der Werf,Martres Pascale |
| EPI_ISL_1096258 | Hopital | National Reference Center for Viruses of Respiratory Infections, Institut Pasteur, Paris | Marion Barbet, Sylvie Behillil, Méline Bizard, Angela Brisebarre, Camille Capel, Etienne Simon-Lorière, Vincent Enouf, Maud Vanpeene, Sylvie van der Werf,LefeuvreCaroline |
| EPI_ISL_1096259 | Hopital | National Reference Center for Viruses of Respiratory Infections, Institut Pasteur, Paris | Marion Barbet, Sylvie Behillil, Méline Bizard, Angela Brisebarre, Camille Capel, Etienne Simon-Lorière, Vincent Enouf, Maud Vanpeene, Sylvie van der Werf,Castelain Sandrine |
| EPI_ISL_1096260 | Hopital | National Reference Center for Viruses of Respiratory Infections, Institut Pasteur, Paris | Marion Barbet, Sylvie Behillil, Méline Bizard, Angela Brisebarre, Camille Capel, Etienne Simon-Lorière, Vincent Enouf, Maud Vanpeene, Sylvie van der Werf,Guinard JéRome |
| EPI_ISL_1096262 | Outre mer | National Reference Center for Viruses of Respiratory Infections, Institut Pasteur, Paris | Marion Barbet, Sylvie Behillil, Méline Bizard, Angela Brisebarre, Camille Capel, Etienne Simon-Lorière, Vincent Enouf, Maud Vanpeene, Sylvie van der Werf,Rousset Dominique |
| EPI_ISL_1096263, EPI_ISL_1096267 | Research | National Reference Center for Viruses of Respiratory Infections, Institut Pasteur, Paris | Marion Barbet, Sylvie Behillil, Méline Bizard, Angela Brisebarre, Camille Capel, Etienne Simon-Lorière, Vincent Enouf, Maud Vanpeene, Sylvie van der Werf,Leruez-Ville Marianne |
| EPI_ISL_1096268, EPI_ISL_1096269 | Labo Analyses Med | National Reference Center for Viruses of Respiratory Infections, Institut Pasteur, Paris | Marion Barbet, Sylvie Behillil, Méline Bizard, Angela Brisebarre, Camille Capel, Etienne Simon-Lorière, Vincent Enouf, Maud Vanpeene, Sylvie van der Werf,Coignard Catherine |
| EPI_ISL_1096270, EPI_ISL_1096271 | Hopital | National Reference Center for Viruses of Respiratory Infections, Institut Pasteur, Paris | Marion Barbet, Sylvie Behillil, Méline Bizard, Angela Brisebarre, Camille Capel, Etienne Simon-Lorière, Vincent Enouf, Maud Vanpeene, Sylvie van der Werf,Martres Pascale |
| EPI_ISL_1096272 | Hopital | National Reference Center for Viruses of Respiratory Infections, Institut Pasteur, Paris | Marion Barbet, Sylvie Behillil, Méline Bizard, Angela Brisebarre, Camille Capel, Etienne Simon-Lorière, Vincent Enouf, Maud Vanpeene, Sylvie van der Werf,Guinard JéRome |
| EPI_ISL_1096273 | Hopital | National Reference Center for Viruses of Respiratory Infections, Institut Pasteur, Paris | Marion Barbet, Sylvie Behillil, Méline Bizard, Angela Brisebarre, Camille Capel, Etienne Simon-Lorière, Vincent Enouf, Maud Vanpeene, Sylvie van der Werf,Cariou Marie-Estelle |
| EPI_ISL_1096274 | Research | National Reference Center for Viruses of Respiratory Infections, Institut Pasteur, Paris | Marion Barbet, Sylvie Behillil, Méline Bizard, Angela Brisebarre, Camille Capel, Etienne Simon-Lorière, Vincent Enouf, Maud Vanpeene, Sylvie van der Werf,Leruez-Ville Marianne |
| EPI_ISL_1096275 | Hopital | National Reference Center for Viruses of Respiratory Infections, Institut Pasteur, Paris | Marion Barbet, Sylvie Behillil, Méline Bizard, Angela Brisebarre, Camille Capel, Etienne Simon-Lorière, Vincent Enouf, Maud Vanpeene, Sylvie van der Werf,Lagathu GisèLe |
| EPI_ISL_1096276, EPI_ISL_1096277 | Hopital | National Reference Center for Viruses of Respiratory Infections, Institut Pasteur, Paris | Marion Barbet, Sylvie Behillil, Méline Bizard, Angela Brisebarre, Camille Capel, Etienne Simon-Lorière, Vincent Enouf, Maud Vanpeene, Sylvie van der Werf,Cariou Marie-Estelle |
| EPI_ISL_1096279, EPI_ISL_1096281, EPI_ISL_1096282, EPI_ISL_1096285, EPI_ISL_1096287, EPI_ISL_1096288, EPI_ISL_1096290, EPI_ISL_1096291, EPI_ISL_1096293, EPI_ISL_1096295, EPI_ISL_1096296, EPI_ISL_1096298, EPI_ISL_1096300, EPI_ISL_1096301, EPI_ISL_1096302 | | | |
| see above | Hopital | National Reference Center for Viruses of Respiratory Infections, Institut Pasteur, Paris | Marion Barbet, Sylvie Behillil, Méline Bizard, Angela Brisebarre, Camille Capel, Etienne Simon-Lorière, Vincent Enouf, Maud Vanpeene, Sylvie van der Werf,Guinard JéRome |
| EPI_ISL_1096303 | Labo Analyses Med | National Reference Center for Viruses of Respiratory Infections, Institut Pasteur, Paris | Marion Barbet, Sylvie Behillil, Méline Bizard, Angela Brisebarre, Camille Capel, Etienne Simon-Lorière, Vincent Enouf, Maud Vanpeene, Sylvie van der Werf |
| EPI_ISL_1096304 | Hopital | National Reference Center for Viruses of Respiratory Infections, Institut Pasteur, Paris | Marion Barbet, Sylvie Behillil, Méline Bizard, Angela Brisebarre, Camille Capel, Etienne Simon-Lorière, Vincent Enouf, Maud Vanpeene, Sylvie van der Werf,Guinoiseau Thibault |
| EPI_ISL_1096305, EPI_ISL_1096306, EPI_ISL_1096307, EPI_ISL_1096308 | Hopital | National Reference Center for Viruses of Respiratory Infections, Institut Pasteur, Paris | Marion Barbet, Sylvie Behillil, Méline Bizard, Angela Brisebarre, Camille Capel, Etienne Simon-Lorière, Vincent Enouf, Maud Vanpeene, Sylvie van der Werf,Castelain Sandrine |
| EPI_ISL_1096309, EPI_ISL_1096310, EPI_ISL_1096311, EPI_ISL_1096312 | Hopital | National Reference Center for Viruses of Respiratory Infections, Institut Pasteur, Paris | Marion Barbet, Sylvie Behillil, Méline Bizard, Angela Brisebarre, Camille Capel, Etienne Simon-Lorière, Vincent Enouf, Maud Vanpeene, Sylvie van der Werf,Hochart A |
| EPI_ISL_1096313, EPI_ISL_1096314, EPI_ISL_1096315, EPI_ISL_1096316 | Labo Analyses Med | National Reference Center for Viruses of Respiratory Infections, Institut Pasteur, Paris | Marion Barbet, Sylvie Behillil, Méline Bizard, Angela Brisebarre, Camille Capel, Etienne Simon-Lorière, Vincent Enouf, Maud Vanpeene, Sylvie van der Werf,Fiorina Jc |
| EPI_ISL_1096317, EPI_ISL_1096318, EPI_ISL_1096319, EPI_ISL_1096320 | Hopital | National Reference Center for Viruses of Respiratory Infections, Institut Pasteur, Paris | Marion Barbet, Sylvie Behillil, Méline Bizard, Angela Brisebarre, Camille Capel, Etienne Simon-Lorière, Vincent Enouf, Maud Vanpeene, Sylvie van der Werf,Bisiau H |
| EPI_ISL_1096321, EPI_ISL_1096322, EPI_ISL_1096323, EPI_ISL_1096324, EPI_ISL_1096325, EPI_ISL_1096326, EPI_ISL_1096327, EPI_ISL_1096328, EPI_ISL_1096329, EPI_ISL_1096330, EPI_ISL_1096332, EPI_ISL_1096333 | | | |
| see above | Research | National Reference Center for Viruses of Respiratory Infections, Institut Pasteur, Paris | Marion Barbet, Sylvie Behillil, Méline Bizard, Angela Brisebarre, Camille Capel, Etienne Simon-Lorière, Vincent Enouf, Maud Vanpeene, Sylvie van der Werf,Leruez-Ville Marianne |
| EPI_ISL_1096334 | Hopital | National Reference Center for Viruses of Respiratory Infections, Institut Pasteur, Paris | Marion Barbet, Sylvie Behillil, Méline Bizard, Angela Brisebarre, Camille Capel, Etienne Simon-Lorière, Vincent Enouf, Maud Vanpeene, Sylvie van der Werf,LefeuvreCaroline |
| EPI_ISL_1096347, EPI_ISL_1096348 | Cambodian National Public Health Laboratory, National Institute of Public Health | Virology Unit, Institut Pasteur du Cambodge | Sokhoun Yann, Ly Sovann, Kraing Sidonn, Yi Sengdoeurn, Chin Savuth, Chau Darapheak, Veasna Duong, Erik A Karlsson |
| EPI_ISL_1096982, EPI_ISL_1096986, EPI_ISL_1096989, EPI_ISL_1096991, EPI_ISL_1096993, EPI_ISL_1096994, EPI_ISL_1096996, EPI_ISL_1096998, EPI_ISL_1097000, EPI_ISL_1097004, EPI_ISL_1097005, EPI_ISL_1097008, EPI_ISL_1097009, EPI_ISL_1097010, EPI_ISL_1097011, EPI_ISL_1097012, EPI_ISL_1097013, EPI_ISL_1097014, EPI_ISL_1097015 | | | |
| see above | Laboratorio Microbiologia e Virologia P.O. Cotugno A.O. dei Colli | Laboratorio Microbiologia e Virologia P.O. Cotugno A.O. dei Colli | Luigi Atripaldi, Claudia Tiberio, Anna Perfetti, Pellegrino Cerino, Biancamaria Pierri, Maria Concetta Cuomo |
| EPI_ISL_1097016 | Laboratorio Microbiologia e Virologia P.O. Cotugno A.O. dei Col | Laboratorio Microbiologia e Virologia P.O. Cotugno A.O. dei Colli | Luigi Atripaldi, Claudia Tiberio, Anna Perfetti, Pellegrino Cerino, Biancamaria Pierri, Maria Concetta Cuomo |
| EPI_ISL_1097017 | Laboratorio Microbiologia e Virologia P.O. Cotugno A.O. dei Colli | Laboratorio Microbiologia e Virologia P.O. Cotugno A.O. dei Colli | Luigi Atripaldi, Claudia Tiberio, Anna Perfetti, Pellegrino Cerino, Biancamaria Pierri, Maria Concetta Cuomo |
| EPI_ISL_1097018 | Baylor Scott & White-Temple | Baylor Scott & White-Temple | Ari Rao, Linden Morales, Kimberly Walker, Marcus Volz, Shelby Hendrickson |
| EPI_ISL_1097019, EPI_ISL_1097020, EPI_ISL_1097021, EPI_ISL_1097022 | Laboratorio Microbiologia e Virologia P.O. Cotugno A.O. dei Colli | Laboratorio Microbiologia e Virologia P.O. Cotugno A.O. dei Colli | Luigi Atripaldi, Claudia Tiberio, Anna Perfetti, Pellegrino Cerino, Biancamaria Pierri, Maria Concetta Cuomo |

| | | | |
|---|---|---|---|
| EPI_ISL_1098602, EPI_ISL_1098603, EPI_ISL_1098604, EPI_ISL_1098605 | Cambodian National Public Health Laboratory, National Institute of Public Health | Virology Unit, Institut Pasteur du Cambodge | Sokhoun Yann, Ly Sovann, Kraing Sidonn, Yi Sengdoeurn, Chin Savuth, Chau Darapheak, Veasna Duong, Erik A Karlsson |
| EPI_ISL_1098606, EPI_ISL_1098607, EPI_ISL_1098608 | Virology Unit, Institut Pasteur du Cambodge | Virology Unit, Institut Pasteur du Cambodge | Sokhoun Yann, Ly Sovann, Kraing Sidonn, Yi Sengdoeurn, Chin Savuth, Chau Darapheak, Veasna Duong, Erik A Karlsson |
| EPI_ISL_1103563, EPI_ISL_1103573, EPI_ISL_1103575, EPI_ISL_1103577, EPI_ISL_1103578, EPI_ISL_1103580, EPI_ISL_1103581 | AZ Klina | AZ Klina | Dr. C. Vael |
| EPI_ISL_1104645, EPI_ISL_1104647, EPI_ISL_1104649 | Virologia, Dipartimento di Scienze Biomediche, Università di Sassari, Viale San Pietro, 43/B - Sassari | Laboratorio specialistico UOC Ematologia - Ospedale "San Francesco" - ATS-ASSL Nuoro | Giovanna Piras, Asproni Rosanna, Paolo Malune, Maria Itria Monne, Angelo Domenico Palmas, Caterina Serra, Elena Rimini, Salvatore Rubino |
| EPI_ISL_1104650 | Laboratorio Biologia Molecolare Sars Cov2 - UOC Laboratorio Analisi - Servizio Medicina di Laboratorio, Ospedale "San Francesco" - ATS-ASSL Nuoro | Laboratorio specialistico UOC Ematologia - Ospedale "San Francesco" - ATS-ASSL Nuoro | Giovanna Piras, Rosanna Asproni, Paolo Malune, Maura Fiamma, Maria Itria Monne, Angelo Domenico Palmas, Iana Lo Maglio, Giuseppe Mameli |
| EPI_ISL_1104652, EPI_ISL_1104654, EPI_ISL_1104656 | Virologia, Dipartimento di Scienze Biomediche, Università di Sassari, Viale San Pietro, 43/B - Sassari | Laboratorio specialistico UOC Ematologia - Ospedale "San Francesco" - ATS-ASSL Nuoro | Giovanna Piras, Asproni Rosanna, Paolo Malune, Maria Itria Monne, Angelo Domenico Palmas, Caterina Serra, Elena Rimini, Salvatore Rubino |
| EPI_ISL_1109632 | Instituto de Biotecnologia - UNESP-Botucatu-SP | Instituto de Biotecnologia - UNESP-Botucatu-SP | Leila Sabrina Ullmann; Fábio Sossai Possebon; Cecília Artico Banho; Cíntia Bittar; Guilherme Campos; Helena Lage Ferreira; Jorge A. Petroli Marchesi; Lívia Sacchetto; Maisa C. Pereira Parra; Marília Moraes; Maurício L. Nogueira; Paula Rahal; Paulo Inacio da Costa; João Pessoa Araújo Jr. |
| EPI_ISL_1109638, EPI_ISL_1109639, EPI_ISL_1109640, EPI_ISL_1109641, EPI_ISL_1109642, EPI_ISL_1109643, EPI_ISL_1109644 | Althaia. Xarxa Assistencial Universitària de Manresa | IrsiCaixa - Can Ruti CovidSeq | Marc Noguera-Julian, Mariona Parera, Maria Casadellà, Pilar Armengol, Francesc Catala-Moll, Roger Paredes, Bonaventura Clotet Gloria Trujillo, Rafel Perez Vidal, Jaume Trape Pujol, Carolina Gonzalez Fernandez, Roger Paredes, Eulalia Grau, Bonaventura Clotet |
| EPI_ISL_1109990, EPI_ISL_1110024, EPI_ISL_1110026 | Baylor Scott & White-Irving | Baylor Scott & White-Temple | Ari Rao, Linden Morales, Kimberly Walker, Marcus Volz, Shelby Hendrickson |
| EPI_ISL_1110160, EPI_ISL_1110162, EPI_ISL_1110178 | Hôpital Henri Mondor | Department of Virology, Henri Mondor University Hospital, Assistance Publique Hôpitaux de Paris, Université Paris-Est Créteil, INSERM U955 | Christophe Rodriguez, Slim Fourati, Vanessa Demontant, Guillaume Gricourt, Melissa N'Debi, Alexandre Soulier, Elisabeth Trawinski, Jean-Michel Pawlotsky |
| EPI_ISL_1110290, EPI_ISL_1110292, EPI_ISL_1110294 | Baylor Scott & White-Temple | Baylor Scott & White-Temple | Ari Rao, Linden Morales, Kimberly Walker, Marcus Volz, Shelby Hendrickson |
| EPI_ISL_1110390 | Hopital | National Reference Center for Viruses of Respiratory Infections, Institut Pasteur, Paris | Marion Barbet, Sylvie Behillil, Méline Bizard, Angela Brisebarre, Camille Capel, Etienne Simon-Lorière, Vincent Enouf, Maud Vanpeene, Sylvie van der Werf,Marque Juillet StéPhanie |
| EPI_ISL_1110391, EPI_ISL_1110392 | Hopital | National Reference Center for Viruses of Respiratory Infections, Institut Pasteur, Paris | Marion Barbet, Sylvie Behillil, Méline Bizard, Angela Brisebarre, Camille Capel, Etienne Simon-Lorière, Vincent Enouf, Maud Vanpeene, Sylvie van der Werf,Descamps Dominique |
| EPI_ISL_1110393 | Hopital | National Reference Center for Viruses of Respiratory Infections, Institut Pasteur, Paris | Marion Barbet, Sylvie Behillil, Méline Bizard, Angela Brisebarre, Camille Capel, Etienne Simon-Lorière, Vincent Enouf, Maud Vanpeene, Sylvie van der Werf,Van Agt S. |
| EPI_ISL_1110394 | Hopital | National Reference Center for Viruses of Respiratory Infections, Institut Pasteur, Paris | Marion Barbet, Sylvie Behillil, Méline Bizard, Angela Brisebarre, Camille Capel, Etienne Simon-Lorière, Vincent Enouf, Maud Vanpeene, Sylvie van der Werf,Farfour Eric |
| EPI_ISL_1110395, EPI_ISL_1110396 | Hopital | National Reference Center for Viruses of Respiratory Infections, Institut Pasteur, Paris | Marion Barbet, Sylvie Behillil, Méline Bizard, Angela Brisebarre, Camille Capel, Etienne Simon-Lorière, Vincent Enouf, Maud Vanpeene, Sylvie van der Werf,Gros Isabelle |
| EPI_ISL_1110397, EPI_ISL_1110398 | Hopital | National Reference Center for Viruses of Respiratory Infections, Institut Pasteur, Paris | Marion Barbet, Sylvie Behillil, Méline Bizard, Angela Brisebarre, Camille Capel, Etienne Simon-Lorière, Vincent Enouf, Maud Vanpeene, Sylvie van der Werf,Febreau Christine |
| EPI_ISL_1110399, EPI_ISL_1110400 | Labo Analyses Med | National Reference Center for Viruses of Respiratory Infections, Institut Pasteur, Paris | Marion Barbet, Sylvie Behillil, Méline Bizard, Angela Brisebarre, Camille Capel, Etienne Simon-Lorière, Vincent Enouf, Maud Vanpeene, Sylvie van der Werf,Felloni Claire |
| EPI_ISL_1110401, EPI_ISL_1110402, EPI_ISL_1110403, EPI_ISL_1110404 | Labo Analyses Med | National Reference Center for Viruses of Respiratory Infections, Institut Pasteur, Paris | Marion Barbet, Sylvie Behillil, Méline Bizard, Angela Brisebarre, Camille Capel, Etienne Simon-Lorière, Vincent Enouf, Maud Vanpeene, Sylvie van der Werf, |
| EPI_ISL_1110405 | Labo Analyses Med | National Reference Center for Viruses of Respiratory Infections, Institut Pasteur, Paris | Marion Barbet, Sylvie Behillil, Méline Bizard, Angela Brisebarre, Camille Capel, Etienne Simon-Lorière, Vincent Enouf, Maud Vanpeene, Sylvie van der Werf,Potiron GréGoire |
| EPI_ISL_1110406, EPI_ISL_1110407 | Hopital | National Reference Center for Viruses of Respiratory Infections, Institut Pasteur, Paris | Marion Barbet, Sylvie Behillil, Méline Bizard, Angela Brisebarre, Camille Capel, Etienne Simon-Lorière, Vincent Enouf, Maud Vanpeene, Sylvie van der Werf,Andre-Garnier Elisabeth |
| EPI_ISL_1110408 | Labo Analyses Med | National Reference Center for Viruses of Respiratory Infections, Institut Pasteur, Paris | Marion Barbet, Sylvie Behillil, Méline Bizard, Angela Brisebarre, Camille Capel, Etienne Simon-Lorière, Vincent Enouf, Maud Vanpeene, Sylvie van der Werf,Serazin V |
| EPI_ISL_1110409 | Hopital | National Reference Center for Viruses of Respiratory Infections, Institut Pasteur, Paris | Marion Barbet, Sylvie Behillil, Méline Bizard, Angela Brisebarre, Camille Capel, Etienne Simon-Lorière, Vincent Enouf, Maud Vanpeene, Sylvie van der Werf,Morvan Pauline |
| EPI_ISL_1110414 | Hopital | National Reference Center for Viruses of Respiratory Infections, Institut Pasteur, Paris | Marion Barbet, Sylvie Behillil, Méline Bizard, Angela Brisebarre, Camille Capel, Etienne Simon-Lorière, Vincent Enouf, Maud Vanpeene, Sylvie van der Werf,Marque Juillet StéPhanie |
| EPI_ISL_1110415, EPI_ISL_1110416, EPI_ISL_1110417 | Hopital | National Reference Center for Viruses of Respiratory Infections, Institut Pasteur, Paris | Marion Barbet, Sylvie Behillil, Méline Bizard, Angela Brisebarre, Camille Capel, Etienne Simon-Lorière, Vincent Enouf, Maud Vanpeene, Sylvie van der Werf,Van Agt S. |
| EPI_ISL_1110418, EPI_ISL_1110419 | Hopital | National Reference Center for Viruses of Respiratory Infections, Institut Pasteur, Paris | Marion Barbet, Sylvie Behillil, Méline Bizard, Angela Brisebarre, Camille Capel, Etienne Simon-Lorière, Vincent Enouf, Maud Vanpeene, Sylvie van der Werf,Guinoiseau Thibault |
| EPI_ISL_1110420 | Labo Analyses Med | National Reference Center for Viruses of Respiratory Infections, Institut Pasteur, Paris | Marion Barbet, Sylvie Behillil, Méline Bizard, Angela Brisebarre, Camille Capel, Etienne Simon-Lorière, Vincent Enouf, Maud Vanpeene, Sylvie van der Werf, |
| EPI_ISL_1110421 | Hopital | National Reference Center for Viruses of Respiratory Infections, Institut Pasteur, Paris | Marion Barbet, Sylvie Behillil, Méline Bizard, Angela Brisebarre, Camille Capel, Etienne Simon-Lorière, Vincent Enouf, Maud Vanpeene, Sylvie van der Werf, |
| EPI_ISL_1110422, EPI_ISL_1110423 | Labo Analyses Med | National Reference Center for Viruses of Respiratory Infections, Institut Pasteur, Paris | Marion Barbet, Sylvie Behillil, Méline Bizard, Angela Brisebarre, Camille Capel, Etienne Simon-Lorière, Vincent Enouf, Maud Vanpeene, Sylvie van der Werf, |
| EPI_ISL_1110424, EPI_ISL_1110425, EPI_ISL_1110426, EPI_ISL_1110427 | Labo Analyses Med | National Reference Center for Viruses of Respiratory Infections, Institut Pasteur, Paris | Marion Barbet, Sylvie Behillil, Méline Bizard, Angela Brisebarre, Camille Capel, Etienne Simon-Lorière, Vincent Enouf, Maud Vanpeene, Sylvie van der Werf,Holstein Anne |
| EPI_ISL_1110428 | Hopital | National Reference Center for Viruses of Respiratory Infections, Institut Pasteur, Paris | Marion Barbet, Sylvie Behillil, Méline Bizard, Angela Brisebarre, Camille Capel, Etienne Simon-Lorière, Vincent Enouf, Maud Vanpeene, Sylvie van der Werf,Andre-Garnier Elisabeth |
| EPI_ISL_1110430 | Labo Analyses Med | National Reference Center for Viruses of Respiratory Infections, Institut Pasteur, Paris | Marion Barbet, Sylvie Behillil, Méline Bizard, Angela Brisebarre, Camille Capel, Etienne Simon-Lorière, Vincent Enouf, Maud Vanpeene, Sylvie van der Werf,Serazin V |
| EPI_ISL_1110431 | Hopital | National Reference Center for Viruses of Respiratory Infections, Institut Pasteur, Paris | Marion Barbet, Sylvie Behillil, Méline Bizard, Angela Brisebarre, Camille Capel, Etienne Simon-Lorière, Vincent Enouf, Maud Vanpeene, Sylvie van der Werf,Andre-Garnier Elisabeth |

| | | | |
|---|---|---|---|
| EPI_ISL_1110432, EPI_ISL_1110433, EPI_ISL_1110434 | Labo Analyses Med | National Reference Center for Viruses of Respiratory Infections, Institut Pasteur, Paris | Marion Barbet, Sylvie Behillil, Méline Bizard, Angela Brisebarre, Camille Capel, Etienne Simon-Lorière, Vincent Enouf, Maud Vanpeene, Sylvie van der Werf,Potiron GréGoire |
| EPI_ISL_1110436 | Hopital | National Reference Center for Viruses of Respiratory Infections, Institut Pasteur, Paris | Marion Barbet, Sylvie Behillil, Méline Bizard, Angela Brisebarre, Camille Capel, Etienne Simon-Lorière, Vincent Enouf, Maud Vanpeene, Sylvie van der Werf,Guinoiseau Thibault |
| EPI_ISL_1110441 | Hopital | National Reference Center for Viruses of Respiratory Infections, Institut Pasteur, Paris | Marion Barbet, Sylvie Behillil, Méline Bizard, Angela Brisebarre, Camille Capel, Etienne Simon-Lorière, Vincent Enouf, Maud Vanpeene, Sylvie van der Werf,Ducancelle Alexandra |
| EPI_ISL_1110442 | Labo Analyses Med | National Reference Center for Viruses of Respiratory Infections, Institut Pasteur, Paris | Marion Barbet, Sylvie Behillil, Méline Bizard, Angela Brisebarre, Camille Capel, Etienne Simon-Lorière, Vincent Enouf, Maud Vanpeene, Sylvie van der Werf,Lesimple BéAtrice |
| EPI_ISL_1110443 | Labo Analyses Med | National Reference Center for Viruses of Respiratory Infections, Institut Pasteur, Paris | Marion Barbet, Sylvie Behillil, Méline Bizard, Angela Brisebarre, Camille Capel, Etienne Simon-Lorière, Vincent Enouf, Maud Vanpeene, Sylvie van der Werf,Felloni Claire |
| EPI_ISL_1110444, EPI_ISL_1110445 | Hopital | National Reference Center for Viruses of Respiratory Infections, Institut Pasteur, Paris | Marion Barbet, Sylvie Behillil, Méline Bizard, Angela Brisebarre, Camille Capel, Etienne Simon-Lorière, Vincent Enouf, Maud Vanpeene, Sylvie van der Werf,Marque Juillet StéPhanie |
| EPI_ISL_1110446, EPI_ISL_1110447 | Hopital | National Reference Center for Viruses of Respiratory Infections, Institut Pasteur, Paris | Marion Barbet, Sylvie Behillil, Méline Bizard, Angela Brisebarre, Camille Capel, Etienne Simon-Lorière, Vincent Enouf, Maud Vanpeene, Sylvie van der Werf,Van Agt S. |
| EPI_ISL_1110448, EPI_ISL_1110449, EPI_ISL_1110450, EPI_ISL_1110451, EPI_ISL_1110452 | Hopital | National Reference Center for Viruses of Respiratory Infections, Institut Pasteur, Paris | Marion Barbet, Sylvie Behillil, Méline Bizard, Angela Brisebarre, Camille Capel, Etienne Simon-Lorière, Vincent Enouf, Maud Vanpeene, Sylvie van der Werf,Patoz Pierre |
| EPI_ISL_1110453 | Hopital | National Reference Center for Viruses of Respiratory Infections, Institut Pasteur, Paris | Marion Barbet, Sylvie Behillil, Méline Bizard, Angela Brisebarre, Camille Capel, Etienne Simon-Lorière, Vincent Enouf, Maud Vanpeene, Sylvie van der Werf,Guinoiseau Thibault |
| EPI_ISL_1110454, EPI_ISL_1110455 | Hopital | National Reference Center for Viruses of Respiratory Infections, Institut Pasteur, Paris | Marion Barbet, Sylvie Behillil, Méline Bizard, Angela Brisebarre, Camille Capel, Etienne Simon-Lorière, Vincent Enouf, Maud Vanpeene, Sylvie van der Werf,Gros Isabelle |
| EPI_ISL_1110456 | Hopital | National Reference Center for Viruses of Respiratory Infections, Institut Pasteur, Paris | Marion Barbet, Sylvie Behillil, Méline Bizard, Angela Brisebarre, Camille Capel, Etienne Simon-Lorière, Vincent Enouf, Maud Vanpeene, Sylvie van der Werf,Farfour Eric |
| EPI_ISL_1110457 | Hopital | National Reference Center for Viruses of Respiratory Infections, Institut Pasteur, Paris | Marion Barbet, Sylvie Behillil, Méline Bizard, Angela Brisebarre, Camille Capel, Etienne Simon-Lorière, Vincent Enouf, Maud Vanpeene, Sylvie van der Werf,Febreau Christine |
| EPI_ISL_1110458, EPI_ISL_1110459 | Hopital | National Reference Center for Viruses of Respiratory Infections, Institut Pasteur, Paris | Marion Barbet, Sylvie Behillil, Méline Bizard, Angela Brisebarre, Camille Capel, Etienne Simon-Lorière, Vincent Enouf, Maud Vanpeene, Sylvie van der Werf, |
| EPI_ISL_1110460 | Labo Analyses Med | National Reference Center for Viruses of Respiratory Infections, Institut Pasteur, Paris | Marion Barbet, Sylvie Behillil, Méline Bizard, Angela Brisebarre, Camille Capel, Etienne Simon-Lorière, Vincent Enouf, Maud Vanpeene, Sylvie van der Werf,Felloni Claire |
| EPI_ISL_1110461 | Labo Analyses Med | National Reference Center for Viruses of Respiratory Infections, Institut Pasteur, Paris | Marion Barbet, Sylvie Behillil, Méline Bizard, Angela Brisebarre, Camille Capel, Etienne Simon-Lorière, Vincent Enouf, Maud Vanpeene, Sylvie van der Werf, |
| EPI_ISL_1110462 | Hopital | National Reference Center for Viruses of Respiratory Infections, Institut Pasteur, Paris | Marion Barbet, Sylvie Behillil, Méline Bizard, Angela Brisebarre, Camille Capel, Etienne Simon-Lorière, Vincent Enouf, Maud Vanpeene, Sylvie van der Werf, |
| EPI_ISL_1110464, EPI_ISL_1110465, EPI_ISL_1110466 | Hopital | National Reference Center for Viruses of Respiratory Infections, Institut Pasteur, Paris | Marion Barbet, Sylvie Behillil, Méline Bizard, Angela Brisebarre, Camille Capel, Etienne Simon-Lorière, Vincent Enouf, Maud Vanpeene, Sylvie van der Werf,Ducancelle Alexandra |
| EPI_ISL_1110467 | Labo Analyses Med | National Reference Center for Viruses of Respiratory Infections, Institut Pasteur, Paris | Marion Barbet, Sylvie Behillil, Méline Bizard, Angela Brisebarre, Camille Capel, Etienne Simon-Lorière, Vincent Enouf, Maud Vanpeene, Sylvie van der Werf,Lesimple BéAtrice |
| EPI_ISL_1110468 | Labo Analyses Med | National Reference Center for Viruses of Respiratory Infections, Institut Pasteur, Paris | Marion Barbet, Sylvie Behillil, Méline Bizard, Angela Brisebarre, Camille Capel, Etienne Simon-Lorière, Vincent Enouf, Maud Vanpeene, Sylvie van der Werf,Holstein Anne |
| EPI_ISL_1110469 | Labo Analyses Med | National Reference Center for Viruses of Respiratory Infections, Institut Pasteur, Paris | Marion Barbet, Sylvie Behillil, Méline Bizard, Angela Brisebarre, Camille Capel, Etienne Simon-Lorière, Vincent Enouf, Maud Vanpeene, Sylvie van der Werf,Potiron GréGoire |
| EPI_ISL_1110470, EPI_ISL_1110471, EPI_ISL_1110473 | Labo Analyses Med | National Reference Center for Viruses of Respiratory Infections, Institut Pasteur, Paris | Marion Barbet, Sylvie Behillil, Méline Bizard, Angela Brisebarre, Camille Capel, Etienne Simon-Lorière, Vincent Enouf, Maud Vanpeene, Sylvie van der Werf,Etienne Elodie |
| EPI_ISL_1110474 | Labo Analyses Med | National Reference Center for Viruses of Respiratory Infections, Institut Pasteur, Paris | Marion Barbet, Sylvie Behillil, Méline Bizard, Angela Brisebarre, Camille Capel, Etienne Simon-Lorière, Vincent Enouf, Maud Vanpeene, Sylvie van der Werf,Serazin V |
| EPI_ISL_1110475 | Hopital | National Reference Center for Viruses of Respiratory Infections, Institut Pasteur, Paris | Marion Barbet, Sylvie Behillil, Méline Bizard, Angela Brisebarre, Camille Capel, Etienne Simon-Lorière, Vincent Enouf, Maud Vanpeene, Sylvie van der Werf,Morvan Pauline |
| EPI_ISL_1110476, EPI_ISL_1110477, EPI_ISL_1110478, EPI_ISL_1110479 | Hopital | National Reference Center for Viruses of Respiratory Infections, Institut Pasteur, Paris | Marion Barbet, Sylvie Behillil, Méline Bizard, Angela Brisebarre, Camille Capel, Etienne Simon-Lorière, Vincent Enouf, Maud Vanpeene, Sylvie van der Werf,Pilorge LéA |
| EPI_ISL_1110581, EPI_ISL_1110582, EPI_ISL_1110583, EPI_ISL_1110584, EPI_ISL_1110585, EPI_ISL_1110586, EPI_ISL_1110587, EPI_ISL_1110588, EPI_ISL_1110589, EPI_ISL_1110590, EPI_ISL_1110591, EPI_ISL_1110593, EPI_ISL_1110594, EPI_ISL_1110595 | | | |
| see above | LAB ANALISI PO CARDARELLI CB | Istituto Zooprofilattico Sperimentale dell'Abruzzo e Molise "G. Caporale" | Scutellà M, Niro G, Lorusso A, Marcacci M, Di Domenico M, Ancora M, Curini V, Mangone I, Rinaldi A, Scialabba S, Di Pasquale A, Cammà C, Puglia I, Calistri P, Savini G |
| EPI_ISL_1110598, EPI_ISL_1110599, EPI_ISL_1110600 | SIESP CH-Drive in VASTO | Istituto Zooprofilattico Sperimentale dell'Abruzzo e Molise "G. Caporale" | Lorusso A, Marcacci M, Di Domenico M, Ancora M, Curini V, Mangone I, Rinaldi A, Scialabba S, Di Pasquale A, Cammà C, Puglia I, Calistri P, Savini G |
| EPI_ISL_1110601 | Ospedale Civile Teramo | Istituto Zooprofilattico Sperimentale dell'Abruzzo e Molise "G. Caporale" | Lorusso A, Marcacci M, Di Domenico M, Ancora M, Curini V, Mangone I, Rinaldi A, Scialabba S, Di Pasquale A, Cammà C, Puglia I, Calistri P, Savini G |
| EPI_ISL_1110602 | SIESP Dip. Prev Teramo | Istituto Zooprofilattico Sperimentale dell'Abruzzo e Molise "G. Caporale" | Lorusso A, Marcacci M, Di Domenico M, Ancora M, Curini V, Mangone I, Rinaldi A, Scialabba S, Di Pasquale A, Cammà C, Puglia I, Calistri P, Savini G |
| EPI_ISL_1110881 | Hopital | National Reference Center for Viruses of Respiratory Infections, Institut Pasteur, Paris | Marion Barbet, Sylvie Behillil, Méline Bizard, Angela Brisebarre, Camille Capel, Etienne Simon-Lorière, Vincent Enouf, Maud Vanpeene, Sylvie van der Werf,Fourgeaud Jacques |
| EPI_ISL_1110882, EPI_ISL_1110883 | Hopital | National Reference Center for Viruses of Respiratory Infections, Institut Pasteur, Paris | Marion Barbet, Sylvie Behillil, Méline Bizard, Angela Brisebarre, Camille Capel, Etienne Simon-Lorière, Vincent Enouf, Maud Vanpeene, Sylvie van der Werf,Griscelli Franck |
| EPI_ISL_1110884, EPI_ISL_1110885 | Hopital | National Reference Center for Viruses of Respiratory Infections, Institut Pasteur, Paris | Marion Barbet, Sylvie Behillil, Méline Bizard, Angela Brisebarre, Camille Capel, Etienne Simon-Lorière, Vincent Enouf, Maud Vanpeene, Sylvie van der Werf,Pilorge LéA |
| EPI_ISL_1110886, EPI_ISL_1110887 | Hopital | National Reference Center for Viruses of Respiratory Infections, Institut Pasteur, Paris | Marion Barbet, Sylvie Behillil, Méline Bizard, Angela Brisebarre, Camille Capel, Etienne Simon-Lorière, Vincent Enouf, Maud Vanpeene, Sylvie van der Werf,Ledru Sylvie |
| EPI_ISL_1110888, EPI_ISL_1110889 | Labo Analyses Med | National Reference Center for Viruses of Respiratory Infections, Institut Pasteur, Paris | Marion Barbet, Sylvie Behillil, Méline Bizard, Angela Brisebarre, Camille Capel, Etienne Simon-Lorière, Vincent Enouf, Maud Vanpeene, Sylvie van der Werf,Cerballiance Paris |
| EPI_ISL_1110890, EPI_ISL_1110891 | Hopital | National Reference Center for Viruses of Respiratory Infections, Institut Pasteur, Paris | Marion Barbet, Sylvie Behillil, Méline Bizard, Angela Brisebarre, Camille Capel, Etienne Simon-Lorière, Vincent Enouf, Maud Vanpeene, Sylvie van der Werf,Riverain Emeline |

| | | | |
|---|---|---|---|
| EPI_ISL_1110892, EPI_ISL_1110893, EPI_ISL_1110894 | Labo Analyses Med | National Reference Center for Viruses of Respiratory Infections, Institut Pasteur, Paris | Marion Barbet, Sylvie Behillil, Méline Bizard, Angela Brisebarre, Camille Capel, Etienne Simon-Lorière, Vincent Enouf, Maud Vanpeene, Sylvie van der Werf |
| EPI_ISL_1110895, EPI_ISL_1110896, EPI_ISL_1110897 | Hopital | National Reference Center for Viruses of Respiratory Infections, Institut Pasteur, Paris | Marion Barbet, Sylvie Behillil, Méline Bizard, Angela Brisebarre, Camille Capel, Etienne Simon-Lorière, Vincent Enouf, Maud Vanpeene, Sylvie van der Werf,Cady Anne |
| EPI_ISL_1110898 | Hopital | National Reference Center for Viruses of Respiratory Infections, Institut Pasteur, Paris | Marion Barbet, Sylvie Behillil, Méline Bizard, Angela Brisebarre, Camille Capel, Etienne Simon-Lorière, Vincent Enouf, Maud Vanpeene, Sylvie van der Werf,Fourgeaud Jacques |
| EPI_ISL_1110899, EPI_ISL_1110900, EPI_ISL_1110901 | Hopital | National Reference Center for Viruses of Respiratory Infections, Institut Pasteur, Paris | Marion Barbet, Sylvie Behillil, Méline Bizard, Angela Brisebarre, Camille Capel, Etienne Simon-Lorière, Vincent Enouf, Maud Vanpeene, Sylvie van der Werf,Pilorge LéA |
| EPI_ISL_1110902, EPI_ISL_1110903, EPI_ISL_1110904, EPI_ISL_1110905 | Hopital | National Reference Center for Viruses of Respiratory Infections, Institut Pasteur, Paris | Marion Barbet, Sylvie Behillil, Méline Bizard, Angela Brisebarre, Camille Capel, Etienne Simon-Lorière, Vincent Enouf, Maud Vanpeene, Sylvie van der Werf,Fourgeaud Jacques |
| EPI_ISL_1110906, EPI_ISL_1110907, EPI_ISL_1110908, EPI_ISL_1110909, EPI_ISL_1110910, EPI_ISL_1110911, EPI_ISL_1110912, EPI_ISL_1110913, EPI_ISL_1110914, EPI_ISL_1110915 | Hopital | National Reference Center for Viruses of Respiratory Infections, Institut Pasteur, Paris | Marion Barbet, Sylvie Behillil, Méline Bizard, Angela Brisebarre, Camille Capel, Etienne Simon-Lorière, Vincent Enouf, Maud Vanpeene, Sylvie van der Werf,Gastli Nabil |
| EPI_ISL_1110916 | Hopital | National Reference Center for Viruses of Respiratory Infections, Institut Pasteur, Paris | Marion Barbet, Sylvie Behillil, Méline Bizard, Angela Brisebarre, Camille Capel, Etienne Simon-Lorière, Vincent Enouf, Maud Vanpeene, Sylvie van der Werf,Cady Anne |
| EPI_ISL_1110917 | Armies | National Reference Center for Viruses of Respiratory Infections, Institut Pasteur, Paris | Marion Barbet, Sylvie Behillil, Méline Bizard, Angela Brisebarre, Camille Capel, Etienne Simon-Lorière, Vincent Enouf, Maud Vanpeene, Sylvie van der Werf,Desroches Marine |
| EPI_ISL_1110918 | Hopital | National Reference Center for Viruses of Respiratory Infections, Institut Pasteur, Paris | Marion Barbet, Sylvie Behillil, Méline Bizard, Angela Brisebarre, Camille Capel, Etienne Simon-Lorière, Vincent Enouf, Maud Vanpeene, Sylvie van der Werf,Vachee Anne |
| EPI_ISL_1110919, EPI_ISL_1110920, EPI_ISL_1110921 | Hopital | National Reference Center for Viruses of Respiratory Infections, Institut Pasteur, Paris | Marion Barbet, Sylvie Behillil, Méline Bizard, Angela Brisebarre, Camille Capel, Etienne Simon-Lorière, Vincent Enouf, Maud Vanpeene, Sylvie van der Werf,Engelmann Ilka |
| EPI_ISL_1110922 | Hopital | National Reference Center for Viruses of Respiratory Infections, Institut Pasteur, Paris | Marion Barbet, Sylvie Behillil, Méline Bizard, Angela Brisebarre, Camille Capel, Etienne Simon-Lorière, Vincent Enouf, Maud Vanpeene, Sylvie van der Werf,Bernardi Franck |
| EPI_ISL_1110924, EPI_ISL_1110925, EPI_ISL_1110926 | Labo Analyses Med | National Reference Center for Viruses of Respiratory Infections, Institut Pasteur, Paris | Marion Barbet, Sylvie Behillil, Méline Bizard, Angela Brisebarre, Camille Capel, Etienne Simon-Lorière, Vincent Enouf, Maud Vanpeene, Sylvie van der Werf,Cerballiance Paris |
| EPI_ISL_1110927 | Labo Analyses Med | National Reference Center for Viruses of Respiratory Infections, Institut Pasteur, Paris | Marion Barbet, Sylvie Behillil, Méline Bizard, Angela Brisebarre, Camille Capel, Etienne Simon-Lorière, Vincent Enouf, Maud Vanpeene, Sylvie van der Werf |
| EPI_ISL_1110928 | Hopital | National Reference Center for Viruses of Respiratory Infections, Institut Pasteur, Paris | Marion Barbet, Sylvie Behillil, Méline Bizard, Angela Brisebarre, Camille Capel, Etienne Simon-Lorière, Vincent Enouf, Maud Vanpeene, Sylvie van der Werf,Cady Anne |
| EPI_ISL_1110929, EPI_ISL_1110930, EPI_ISL_1110931, EPI_ISL_1110932 | Hopital | National Reference Center for Viruses of Respiratory Infections, Institut Pasteur, Paris | Marion Barbet, Sylvie Behillil, Méline Bizard, Angela Brisebarre, Camille Capel, Etienne Simon-Lorière, Vincent Enouf, Maud Vanpeene, Sylvie van der Werf,Pilorge LéA |
| EPI_ISL_1110933, EPI_ISL_1110934, EPI_ISL_1110935 | Hopital | National Reference Center for Viruses of Respiratory Infections, Institut Pasteur, Paris | Marion Barbet, Sylvie Behillil, Méline Bizard, Angela Brisebarre, Camille Capel, Etienne Simon-Lorière, Vincent Enouf, Maud Vanpeene, Sylvie van der Werf,Fourgeaud Jacques |
| EPI_ISL_1110936, EPI_ISL_1110937, EPI_ISL_1110938, EPI_ISL_1110939, EPI_ISL_1110940, EPI_ISL_1110941, EPI_ISL_1110942, EPI_ISL_1110943, EPI_ISL_1110944, EPI_ISL_1110945, EPI_ISL_1110946, EPI_ISL_1110947, EPI_ISL_1110948, EPI_ISL_1110949, EPI_ISL_1110950, EPI_ISL_1110951 | | | |
| see above | Hopital | National Reference Center for Viruses of Respiratory Infections, Institut Pasteur, Paris | Marion Barbet, Sylvie Behillil, Méline Bizard, Angela Brisebarre, Camille Capel, Etienne Simon-Lorière, Vincent Enouf, Maud Vanpeene, Sylvie van der Werf,Gastli Nabil |
| EPI_ISL_1110953 | Hopital | National Reference Center for Viruses of Respiratory Infections, Institut Pasteur, Paris | Marion Barbet, Sylvie Behillil, Méline Bizard, Angela Brisebarre, Camille Capel, Etienne Simon-Lorière, Vincent Enouf, Maud Vanpeene, Sylvie van der Werf,Marichal A. |
| EPI_ISL_1110954 | Hopital | National Reference Center for Viruses of Respiratory Infections, Institut Pasteur, Paris | Marion Barbet, Sylvie Behillil, Méline Bizard, Angela Brisebarre, Camille Capel, Etienne Simon-Lorière, Vincent Enouf, Maud Vanpeene, Sylvie van der Werf,Engelmann Ilka |
| EPI_ISL_1110955 | Hopital | National Reference Center for Viruses of Respiratory Infections, Institut Pasteur, Paris | Marion Barbet, Sylvie Behillil, Méline Bizard, Angela Brisebarre, Camille Capel, Etienne Simon-Lorière, Vincent Enouf, Maud Vanpeene, Sylvie van der Werf,Vachee Anne |
| EPI_ISL_1110956 | Hopital | National Reference Center for Viruses of Respiratory Infections, Institut Pasteur, Paris | Marion Barbet, Sylvie Behillil, Méline Bizard, Angela Brisebarre, Camille Capel, Etienne Simon-Lorière, Vincent Enouf, Maud Vanpeene, Sylvie van der Werf,Ledru Sylvie |
| EPI_ISL_1110957, EPI_ISL_1110958, EPI_ISL_1110959, EPI_ISL_1110960, EPI_ISL_1110961, EPI_ISL_1110962, EPI_ISL_1110963 | Hopital | National Reference Center for Viruses of Respiratory Infections, Institut Pasteur, Paris | Marion Barbet, Sylvie Behillil, Méline Bizard, Angela Brisebarre, Camille Capel, Etienne Simon-Lorière, Vincent Enouf, Maud Vanpeene, Sylvie van der Werf,Pilorge LéA |
| EPI_ISL_1110964, EPI_ISL_1110965 | Hopital | National Reference Center for Viruses of Respiratory Infections, Institut Pasteur, Paris | Marion Barbet, Sylvie Behillil, Méline Bizard, Angela Brisebarre, Camille Capel, Etienne Simon-Lorière, Vincent Enouf, Maud Vanpeene, Sylvie van der Werf,Lepiller Quentin |
| EPI_ISL_1110968 | Hopital | National Reference Center for Viruses of Respiratory Infections, Institut Pasteur, Paris | Marion Barbet, Sylvie Behillil, Méline Bizard, Angela Brisebarre, Camille Capel, Etienne Simon-Lorière, Vincent Enouf, Maud Vanpeene, Sylvie van der Werf,Gastli Nabil |
| EPI_ISL_1110969 | Hopital | National Reference Center for Viruses of Respiratory Infections, Institut Pasteur, Paris | Marion Barbet, Sylvie Behillil, Méline Bizard, Angela Brisebarre, Camille Capel, Etienne Simon-Lorière, Vincent Enouf, Maud Vanpeene, Sylvie van der Werf,Ramirez José-Manuel |
| EPI_ISL_1110970, EPI_ISL_1110971 | Hopital | National Reference Center for Viruses of Respiratory Infections, Institut Pasteur, Paris | Marion Barbet, Sylvie Behillil, Méline Bizard, Angela Brisebarre, Camille Capel, Etienne Simon-Lorière, Vincent Enouf, Maud Vanpeene, Sylvie van der Werf,Pilorge LéA |
| EPI_ISL_1110972 | Hopital | National Reference Center for Viruses of Respiratory Infections, Institut Pasteur, Paris | Marion Barbet, Sylvie Behillil, Méline Bizard, Angela Brisebarre, Camille Capel, Etienne Simon-Lorière, Vincent Enouf, Maud Vanpeene, Sylvie van der Werf,Gastli Nabil |
| EPI_ISL_1110973 | Hopital | National Reference Center for Viruses of Respiratory Infections, Institut Pasteur, Paris | Marion Barbet, Sylvie Behillil, Méline Bizard, Angela Brisebarre, Camille Capel, Etienne Simon-Lorière, Vincent Enouf, Maud Vanpeene, Sylvie van der Werf,Fourgeaud Jacques |
| EPI_ISL_1110974 | Hopital | National Reference Center for Viruses of Respiratory Infections, Institut Pasteur, Paris | Marion Barbet, Sylvie Behillil, Méline Bizard, Angela Brisebarre, Camille Capel, Etienne Simon-Lorière, Vincent Enouf, Maud Vanpeene, Sylvie van der Werf,Griscelli Franck |
| EPI_ISL_1110975, EPI_ISL_1110976, EPI_ISL_1110977 | Hopital | National Reference Center for Viruses of Respiratory Infections, Institut Pasteur, Paris | Marion Barbet, Sylvie Behillil, Méline Bizard, Angela Brisebarre, Camille Capel, Etienne Simon-Lorière, Vincent Enouf, Maud Vanpeene, Sylvie van der Werf,Pilorge LéA |
| EPI_ISL_1110978, EPI_ISL_1110979 | Hopital | National Reference Center for Viruses of Respiratory Infections, Institut Pasteur, Paris | Marion Barbet, Sylvie Behillil, Méline Bizard, Angela Brisebarre, Camille Capel, Etienne Simon-Lorière, Vincent Enouf, Maud Vanpeene, Sylvie van der Werf,Farfour Eric |
| EPI_ISL_1110980 | Labo Analyses Med | National Reference Center for Viruses of Respiratory Infections, Institut Pasteur, Paris | Marion Barbet, Sylvie Behillil, Méline Bizard, Angela Brisebarre, Camille Capel, Etienne Simon-Lorière, Vincent Enouf, Maud Vanpeene, Sylvie van der Werf |
| EPI_ISL_1110981, EPI_ISL_1110982, EPI_ISL_1110983, EPI_ISL_1110984, EPI_ISL_1110985, EPI_ISL_1110986, EPI_ISL_1110987, EPI_ISL_1110988, EPI_ISL_1110989, EPI_ISL_1110990, EPI_ISL_1110991, EPI_ISL_1110992, EPI_ISL_1110993, EPI_ISL_1110994, EPI_ISL_1110995, EPI_ISL_1110996, EPI_ISL_1110997, | | | |

| | | | |
|---|---|---|---|
| EPI_ISL_1110998, EPI_ISL_1110999, EPI_ISL_1111000, EPI_ISL_1111001, EPI_ISL_1111002, EPI_ISL_1111003, EPI_ISL_1111004, EPI_ISL_1111005, EPI_ISL_1111006, EPI_ISL_1111007, EPI_ISL_1111008, EPI_ISL_1111009, EPI_ISL_1111010, EPI_ISL_1111011, EPI_ISL_1111012, EPI_ISL_1111013, EPI_ISL_1111014, EPI_ISL_1111015, EPI_ISL_1111016, EPI_ISL_1111017, EPI_ISL_1111018, EPI_ISL_1111019, EPI_ISL_1111020, EPI_ISL_1111021, EPI_ISL_1111022, EPI_ISL_1111023, EPI_ISL_1111024, EPI_ISL_1111025, EPI_ISL_1111026, EPI_ISL_1111027, EPI_ISL_1111028, EPI_ISL_1111029, EPI_ISL_1111030, EPI_ISL_1111031, EPI_ISL_1111032, EPI_ISL_1111033, EPI_ISL_1111034, EPI_ISL_1111035, EPI_ISL_1111036 | | | |
| see above | Hopital | National Reference Center for Viruses of Respiratory Infections, Institut Pasteur, Paris | Marion Barbet, Sylvie Behillil, Méline Bizard, Angela Brisebarre, Camille Capel, Etienne Simon-Lorière, Vincent Enouf, Maud Vanpeene, Sylvie van der Werf,Pilorge LéA |
| EPI_ISL_1111037, EPI_ISL_1111038 | Hopital | National Reference Center for Viruses of Respiratory Infections, Institut Pasteur, Paris | Marion Barbet, Sylvie Behillil, Méline Bizard, Angela Brisebarre, Camille Capel, Etienne Simon-Lorière, Vincent Enouf, Maud Vanpeene, Sylvie van der Werf,Fourgeaud Jacques |
| EPI_ISL_1111039, EPI_ISL_1111040, EPI_ISL_1111041, EPI_ISL_1111042, EPI_ISL_1111043, EPI_ISL_1111044, EPI_ISL_1111045, EPI_ISL_1111046 | Hopital | National Reference Center for Viruses of Respiratory Infections, Institut Pasteur, Paris | Marion Barbet, Sylvie Behillil, Méline Bizard, Angela Brisebarre, Camille Capel, Etienne Simon-Lorière, Vincent Enouf, Maud Vanpeene, Sylvie van der Werf,Gastli Nabil |
| EPI_ISL_1111047 | Armies | National Reference Center for Viruses of Respiratory Infections, Institut Pasteur, Paris | Marion Barbet, Sylvie Behillil, Méline Bizard, Angela Brisebarre, Camille Capel, Etienne Simon-Lorière, Vincent Enouf, Maud Vanpeene, Sylvie van der Werf,Foissaud Vincent |
| EPI_ISL_1111048 | Hopital | National Reference Center for Viruses of Respiratory Infections, Institut Pasteur, Paris | Marion Barbet, Sylvie Behillil, Méline Bizard, Angela Brisebarre, Camille Capel, Etienne Simon-Lorière, Vincent Enouf, Maud Vanpeene, Sylvie van der Werf,Riverain Emeline |
| EPI_ISL_1111049 | Armies | National Reference Center for Viruses of Respiratory Infections, Institut Pasteur, Paris | Marion Barbet, Sylvie Behillil, Méline Bizard, Angela Brisebarre, Camille Capel, Etienne Simon-Lorière, Vincent Enouf, Maud Vanpeene, Sylvie van der Werf,Desroches Marine |
| EPI_ISL_1111050, EPI_ISL_1111051 | Hopital | National Reference Center for Viruses of Respiratory Infections, Institut Pasteur, Paris | Marion Barbet, Sylvie Behillil, Méline Bizard, Angela Brisebarre, Camille Capel, Etienne Simon-Lorière, Vincent Enouf, Maud Vanpeene, Sylvie van der Werf,Bernardi Franck |
| EPI_ISL_1111052, EPI_ISL_1111053, EPI_ISL_1111054 | Hopital | National Reference Center for Viruses of Respiratory Infections, Institut Pasteur, Paris | Marion Barbet, Sylvie Behillil, Méline Bizard, Angela Brisebarre, Camille Capel, Etienne Simon-Lorière, Vincent Enouf, Maud Vanpeene, Sylvie van der Werf,Vachee Anne |
| EPI_ISL_1111055 | Hopital | National Reference Center for Viruses of Respiratory Infections, Institut Pasteur, Paris | Marion Barbet, Sylvie Behillil, Méline Bizard, Angela Brisebarre, Camille Capel, Etienne Simon-Lorière, Vincent Enouf, Maud Vanpeene, Sylvie van der Werf,Engelmann Ilka |
| EPI_ISL_1111056 | Hopital | National Reference Center for Viruses of Respiratory Infections, Institut Pasteur, Paris | Marion Barbet, Sylvie Behillil, Méline Bizard, Angela Brisebarre, Camille Capel, Etienne Simon-Lorière, Vincent Enouf, Maud Vanpeene, Sylvie van der Werf,Bigaillon Christine |
| EPI_ISL_1111059 | Hopital | National Reference Center for Viruses of Respiratory Infections, Institut Pasteur, Paris | Marion Barbet, Sylvie Behillil, Méline Bizard, Angela Brisebarre, Camille Capel, Etienne Simon-Lorière, Vincent Enouf, Maud Vanpeene, Sylvie van der Werf,Gros Isabelle |
| EPI_ISL_1111060 | Labo Analyses Med | National Reference Center for Viruses of Respiratory Infections, Institut Pasteur, Paris | Marion Barbet, Sylvie Behillil, Méline Bizard, Angela Brisebarre, Camille Capel, Etienne Simon-Lorière, Vincent Enouf, Maud Vanpeene, Sylvie van der Werf,Felloni Claire |
| EPI_ISL_1111061 | Hopital | National Reference Center for Viruses of Respiratory Infections, Institut Pasteur, Paris | Marion Barbet, Sylvie Behillil, Méline Bizard, Angela Brisebarre, Camille Capel, Etienne Simon-Lorière, Vincent Enouf, Maud Vanpeene, Sylvie van der Werf,Ducancelle Alexandra |
| EPI_ISL_1111062, EPI_ISL_1111063 | Labo Analyses Med | National Reference Center for Viruses of Respiratory Infections, Institut Pasteur, Paris | Marion Barbet, Sylvie Behillil, Méline Bizard, Angela Brisebarre, Camille Capel, Etienne Simon-Lorière, Vincent Enouf, Maud Vanpeene, Sylvie van der Werf,Serazin V |
| EPI_ISL_1111064 | Labo Analyses Med | National Reference Center for Viruses of Respiratory Infections, Institut Pasteur, Paris | Marion Barbet, Sylvie Behillil, Méline Bizard, Angela Brisebarre, Camille Capel, Etienne Simon-Lorière, Vincent Enouf, Maud Vanpeene, Sylvie van der Werf,Dufour Mj |
| EPI_ISL_1111127, EPI_ISL_1111128, EPI_ISL_1111129, EPI_ISL_1111130, EPI_ISL_1111131, EPI_ISL_1111281, EPI_ISL_1111282, EPI_ISL_1111283, EPI_ISL_1111284, EPI_ISL_1111285, EPI_ISL_1111286, EPI_ISL_1111287, EPI_ISL_1111288, EPI_ISL_1111289, EPI_ISL_1111290, EPI_ISL_1111291, EPI_ISL_1111292, EPI_ISL_1111293, EPI_ISL_1111294, EPI_ISL_1111295, EPI_ISL_1111296, EPI_ISL_1111297, EPI_ISL_1111298, EPI_ISL_1111299, EPI_ISL_1111300, EPI_ISL_1111301, EPI_ISL_1111302, EPI_ISL_1111303, EPI_ISL_1111304, EPI_ISL_1111305, EPI_ISL_1111306, EPI_ISL_1111307, EPI_ISL_1111316, EPI_ISL_1111317, EPI_ISL_1111318, EPI_ISL_1111319, EPI_ISL_1111320, EPI_ISL_1111321, EPI_ISL_1111322, EPI_ISL_1111323, EPI_ISL_1111324, EPI_ISL_1111325, EPI_ISL_1111326, EPI_ISL_1111327, EPI_ISL_1111328, EPI_ISL_1111329, EPI_ISL_1111330, EPI_ISL_1111331, EPI_ISL_1111332, EPI_ISL_1111334, EPI_ISL_1111335, EPI_ISL_1111336, EPI_ISL_1111337, EPI_ISL_1111338, EPI_ISL_1111339, EPI_ISL_1111340, EPI_ISL_1111341, EPI_ISL_1111343, EPI_ISL_1111344, EPI_ISL_1111345, EPI_ISL_1111346, EPI_ISL_1111347, EPI_ISL_1111348, EPI_ISL_1111349, EPI_ISL_1111350, EPI_ISL_1111351, EPI_ISL_1111353, EPI_ISL_1111354, EPI_ISL_1111355, EPI_ISL_1111447, EPI_ISL_1111448, EPI_ISL_1111449, EPI_ISL_1111450, EPI_ISL_1111451, EPI_ISL_1111452, EPI_ISL_1111453, EPI_ISL_1111454, EPI_ISL_1111455, EPI_ISL_1111456, EPI_ISL_1111468 | | | |
| see above | Laboratorio de Referencia Nacional de Virus Respiratorio. Instituto Nacional de Salud Perú | Laboratorio de Referencia Nacional de Enteropatógenos. Instituto Nacional de Salud del Perú | Ronnie Gavilan Chavez, Junior Caro Castro, Willi Quino Sifuentes, Veronica Hurtado Vela, Iris Silva Molina, Fiorella Orellana Peralta |
| EPI_ISL_1112221, EPI_ISL_1112222 | Ospedale Santissima Annunziata | Istituto Zooprofilattico Sperimentale della Puglia e della Basilicata | Parisi A., Bianco A., Capozzi L., Del Sambro L., Simone D., Giannico A., Ridolfi D. |
| EPI_ISL_1112223, EPI_ISL_1112224, EPI_ISL_1112225, EPI_ISL_1112226 | Ospedale Di Venere - Carbonara | Istituto Zooprofilattico Sperimentale della Puglia e della Basilicata | Parisi A., Bianco A., Capozzi L., Del Sambro L., Simone D., Giannico A., Ridolfi D. |
| EPI_ISL_1112228, EPI_ISL_1112232, EPI_ISL_1112234 | Istituto Zooprofilattico Sperimentale della Puglia e della Basilicata | Istituto Zooprofilattico Sperimentale della Puglia e della Basilicata | Parisi A., Bianco A., Capozzi L., Del Sambro L., Simone D., Giannico A., Ridolfi D. |
| EPI_ISL_1112235 | Ospedali Riuniti Azienda Ospedaliera Universitaria - Foggia | Istituto Zooprofilattico Sperimentale della Puglia e della Basilicata | Parisi A., Bianco A., Capozzi L., Del Sambro L., Simone D., Giannico A., Ridolfi D. |
| EPI_ISL_1112236 | Dipartimento di Scienze Biomediche e Oncologia Umana - Azienda Ospedaliero Universitaria Consorziale Policlinico | Istituto Zooprofilattico Sperimentale della Puglia e della Basilicata | Parisi A., Bianco A., Capozzi L., Del Sambro L., Simone D., Giannico A., Ridolfi D. |
| EPI_ISL_1112237, EPI_ISL_1112238, EPI_ISL_1112239, EPI_ISL_1112240, EPI_ISL_1112241, EPI_ISL_1112242, EPI_ISL_1112243, EPI_ISL_1112244, EPI_ISL_1112245, EPI_ISL_1112246, EPI_ISL_1112247, EPI_ISL_1112248, EPI_ISL_1112250, EPI_ISL_1112251, EPI_ISL_1112252, EPI_ISL_1112253, EPI_ISL_1112254, EPI_ISL_1112255, EPI_ISL_1112256, EPI_ISL_1112257, EPI_ISL_1112258, EPI_ISL_1112259, EPI_ISL_1112260, EPI_ISL_1112261, EPI_ISL_1112262, EPI_ISL_1112263, EPI_ISL_1112265, EPI_ISL_1112266, EPI_ISL_1112267, EPI_ISL_1112268, EPI_ISL_1112269, EPI_ISL_1112271, EPI_ISL_1112272, EPI_ISL_1112273, EPI_ISL_1112274, EPI_ISL_1112275, EPI_ISL_1112276, EPI_ISL_1112277, EPI_ISL_1112278, EPI_ISL_1112279, EPI_ISL_1112280, EPI_ISL_1112281, EPI_ISL_1112282, EPI_ISL_1112283, EPI_ISL_1112284, EPI_ISL_1112285, EPI_ISL_1112286, EPI_ISL_1112287, EPI_ISL_1112288, EPI_ISL_1112289, EPI_ISL_1112290, EPI_ISL_1112291 | | | |
| see above | Dipartimento di Scienze Biomediche e Oncologia Umana - Azienda Ospedaliero Universitaria Consorziale Policlinico | Istituto Zooprofilattico Sperimentale della Puglia e della Basilicata | Parisi A., Bianco A., Capozzi L., Del Sambro L., Simone D., Chironna M., Loconsole D., Sallustio A. |
| EPI_ISL_1114717, EPI_ISL_1114718, EPI_ISL_1114719, EPI_ISL_1114720, EPI_ISL_1114721, EPI_ISL_1114722, EPI_ISL_1114723, EPI_ISL_1114724 | Institute for Medical Research, Infectious Disease Research Centre, National Institutes of Health, Ministry of Health Malaysia | Institute for Medical Research, Infectious Disease Research Centre, National Institutes of Health, Ministry of Health Malaysia | Suppiah J, Kamel K, Azizan MA, Thayan R |
| EPI_ISL_1114769 | Virologia, Dipartimento di Scienze Biomediche, Università di Sassari, Viale San Pietro, 43/B - Sassari | Laboratorio specialistico UOC Ematologia - Ospedale "San Francesco" - ATS-ASSL Nuoro | Giovanna Piras, Rosanna Asproni, Paolo Malune, Maria Itria Monne, Angelo Domenico Palmas, Caterina Serra, Elena Rimini, Salvatore Rubino |
| EPI_ISL_1116444, EPI_ISL_1116446, EPI_ISL_1116447, EPI_ISL_1116453, EPI_ISL_1116454, EPI_ISL_1116455, EPI_ISL_1116456, EPI_ISL_1116457 | National Institute of Infectious Diseases-Prof. Dr. Matei Bals Molecular Diagnostics Laboratory | National Institute of Infectious Diseases-Prof. Dr. Matei Bals Molecular Diagnostics Laboratory | Leontina Banica, Marius Surleac, Corina Casangiu, Petre Milu, Andreea Tudor, Simona Paraschiv, Dan Otelea |
| EPI_ISL_1117486, EPI_ISL_1117487 | SIESP DIP PREV CHIETI | Istituto Zooprofilattico Sperimentale dell'Abruzzo e Molise "G. Caporale" | Lorusso A, Marcacci M, Di Domenico M, Ancora M, Curini V, Mangone I, Rinaldi A, Scialabba S, Di Pasquale A, Cammà C, Scialabba S, Puglia I, Calistri P, Savini G |
| EPI_ISL_1117488 | DIP PREV AVEZZANO, SERVIZIO DI IGIENE EPIDEMIOLOGIA E SANITÀ PUBBLICA | Istituto Zooprofilattico Sperimentale dell'Abruzzo e Molise "G. Caporale" | Lorusso A, Marcacci M, Di Domenico M, Ancora M, Curini V, Mangone I, Rinaldi A, Scialabba S, Di Pasquale A, Cammà C, Scialabba S, Puglia I, Calistri P, Savini G |
| EPI_ISL_1117489 | SIESP DIP PREV TERAMO | Istituto Zooprofilattico Sperimentale dell'Abruzzo e Molise "G. | Lorusso A, Marcacci M, Di Domenico M, Ancora M, Curini V, Mangone I, Rinaldi A, Scialabba S, Di Pasquale A, Cammà C, Scialabba S, Puglia I, Calistri P, |

| | | | |
|---|---|---|---|
| | | Caporale" | Savini G |
| EPI_ISL_1117490 | SIESP CHIETI, DRIVE IN VASTO | Istituto Zooprofilattico Sperimentale dell'Abruzzo e Molise "G. Caporale" | Lorusso A, Marcacci M, Di Domenico M, Ancora M, Curini V, Mangone I, Rinaldi A, Scialabba S, Di Pasquale A, Cammà C, Scialabba S, Puglia I, Calistri P, Savini G |
| EPI_ISL_1117491, EPI_ISL_1117492, EPI_ISL_1117494 | SIESP DIP PREV CHIETI | Istituto Zooprofilattico Sperimentale dell'Abruzzo e Molise "G. Caporale" | Lorusso A, Marcacci M, Di Domenico M, Ancora M, Curini V, Mangone I, Rinaldi A, Scialabba S, Di Pasquale A, Cammà C, Scialabba S, Puglia I, Calistri P, Savini G |
| EPI_ISL_1117495 | SIESP DIP PREV TERAMO | Istituto Zooprofilattico Sperimentale dell'Abruzzo e Molise "G. Caporale" | Lorusso A, Marcacci M, Di Domenico M, Ancora M, Curini V, Mangone I, Rinaldi A, Scialabba S, Di Pasquale A, Cammà C, Scialabba S, Puglia I, Calistri P, Savini G |
| EPI_ISL_1117496 | USCA PESCINA | Istituto Zooprofilattico Sperimentale dell'Abruzzo e Molise "G. Caporale" | Lorusso A, Marcacci M, Di Domenico M, Ancora M, Curini V, Mangone I, Rinaldi A, Scialabba S, Di Pasquale A, Cammà C, Scialabba S, Puglia I, Calistri P, Savini G |
| EPI_ISL_1117497 | SIESP CHIETI, DRIVE IN ORTONA | Istituto Zooprofilattico Sperimentale dell'Abruzzo e Molise "G. Caporale" | Lorusso A, Marcacci M, Di Domenico M, Ancora M, Curini V, Mangone I, Rinaldi A, Scialabba S, Di Pasquale A, Cammà C, Scialabba S, Puglia I, Calistri P, Savini G |
| EPI_ISL_1117498 | PRESIDIO TERRITORIALE CASOLI | Istituto Zooprofilattico Sperimentale dell'Abruzzo e Molise "G. Caporale" | Lorusso A, Marcacci M, Di Domenico M, Ancora M, Curini V, Mangone I, Rinaldi A, Scialabba S, Di Pasquale A, Cammà C, Scialabba S, Puglia I, Calistri P, Savini G |
| EPI_ISL_1117499 | SIESP CHIETI, DRIVE IN VASTO | Istituto Zooprofilattico Sperimentale dell'Abruzzo e Molise "G. Caporale" | Lorusso A, Marcacci M, Di Domenico M, Ancora M, Curini V, Mangone I, Rinaldi A, Scialabba S, Di Pasquale A, Cammà C, Scialabba S, Puglia I, Calistri P, Savini G |
| EPI_ISL_1117500 | SIESP CHIETI, DRIVE IN ORTONA | Istituto Zooprofilattico Sperimentale dell'Abruzzo e Molise "G. Caporale" | Lorusso A, Marcacci M, Di Domenico M, Ancora M, Curini V, Mangone I, Rinaldi A, Scialabba S, Di Pasquale A, Cammà C, Scialabba S, Puglia I, Calistri P, Savini G |
| EPI_ISL_1117501 | OSP SAN SALVATORE, MEDICINA INTERNA | Istituto Zooprofilattico Sperimentale dell'Abruzzo e Molise "G. Caporale" | Lorusso A, Marcacci M, Di Domenico M, Ancora M, Curini V, Mangone I, Rinaldi A, Scialabba S, Di Pasquale A, Cammà C, Scialabba S, Puglia I, Calistri P, Savini G |
| EPI_ISL_1117502 | SIESP DIP PREV CHIETI | Istituto Zooprofilattico Sperimentale dell'Abruzzo e Molise "G. Caporale" | Lorusso A, Marcacci M, Di Domenico M, Ancora M, Curini V, Mangone I, Rinaldi A, Scialabba S, Di Pasquale A, Cammà C, Scialabba S, Puglia I, Calistri P, Savini G |
| EPI_ISL_1117503 | SIESP DIP PREV TERAMO | Istituto Zooprofilattico Sperimentale dell'Abruzzo e Molise "G. Caporale" | Lorusso A, Marcacci M, Di Domenico M, Ancora M, Curini V, Mangone I, Rinaldi A, Scialabba S, Di Pasquale A, Cammà C, Scialabba S, Puglia I, Calistri P, Savini G |
| EPI_ISL_1117504, EPI_ISL_1117505 | OSP SAN SALVATORE, MEDICINA INTERNA | Istituto Zooprofilattico Sperimentale dell'Abruzzo e Molise "G. Caporale" | Lorusso A, Marcacci M, Di Domenico M, Ancora M, Curini V, Mangone I, Rinaldi A, Scialabba S, Di Pasquale A, Cammà C, Scialabba S, Puglia I, Calistri P, Savini G |
| EPI_ISL_1117507, EPI_ISL_1117508 | SIESP CHIETI, DRIVE IN LANCIANO | Istituto Zooprofilattico Sperimentale dell'Abruzzo e Molise "G. Caporale" | Lorusso A, Marcacci M, Di Domenico M, Ancora M, Curini V, Mangone I, Rinaldi A, Scialabba S, Di Pasquale A, Cammà C, Scialabba S, Puglia I, Calistri P, Savini G |
| EPI_ISL_1117514 | PRES OSP SAN LIBERATORE | Istituto Zooprofilattico Sperimentale dell'Abruzzo e Molise "G. Caporale" | Lorusso A, Marcacci M, Di Domenico M, Ancora M, Curini V, Mangone I, Rinaldi A, Scialabba S, Di Pasquale A, Cammà C, Scialabba S, Puglia I, Calistri P, Savini G |
| EPI_ISL_1117515 | SIESP DIP PREV TERAMO | Istituto Zooprofilattico Sperimentale dell'Abruzzo e Molise "G. Caporale" | Lorusso A, Marcacci M, Di Domenico M, Ancora M, Curini V, Mangone I, Rinaldi A, Scialabba S, Di Pasquale A, Cammà C, Scialabba S, Puglia I, Calistri P, Savini G |
| EPI_ISL_1117516 | SIESP CHIETI, DRIVE IN LANCIANO | Istituto Zooprofilattico Sperimentale dell'Abruzzo e Molise "G. Caporale" | Lorusso A, Marcacci M, Di Domenico M, Ancora M, Curini V, Mangone I, Rinaldi A, Scialabba S, Di Pasquale A, Cammà C, Scialabba S, Puglia I, Calistri P, Savini G |
| EPI_ISL_1117517 | SIESP CHIETI, DRIVE IN ORTONA | Istituto Zooprofilattico Sperimentale dell'Abruzzo e Molise "G. Caporale" | Lorusso A, Marcacci M, Di Domenico M, Ancora M, Curini V, Mangone I, Rinaldi A, Scialabba S, Di Pasquale A, Cammà C, Scialabba S, Puglia I, Calistri P, Savini G |
| EPI_ISL_1117518 | SIESP DIP PREV TERAMO | Istituto Zooprofilattico Sperimentale dell'Abruzzo e Molise "G. Caporale" | Lorusso A, Marcacci M, Di Domenico M, Ancora M, Curini V, Mangone I, Rinaldi A, Scialabba S, Di Pasquale A, Cammà C, Scialabba S, Puglia I, Calistri P, Savini G |
| EPI_ISL_1117519 | SIESP CHIETI, DRIVE IN CHIETI | Istituto Zooprofilattico Sperimentale dell'Abruzzo e Molise "G. Caporale" | Lorusso A, Marcacci M, Di Domenico M, Ancora M, Curini V, Mangone I, Rinaldi A, Scialabba S, Di Pasquale A, Cammà C, Scialabba S, Puglia I, Calistri P, Savini G |
| EPI_ISL_1117520 | USCA AVEZZANO | Istituto Zooprofilattico Sperimentale dell'Abruzzo e Molise "G. Caporale" | Lorusso A, Marcacci M, Di Domenico M, Ancora M, Curini V, Mangone I, Rinaldi A, Scialabba S, Di Pasquale A, Cammà C, Scialabba S, Puglia I, Calistri P, Savini G |
| EPI_ISL_1117521 | DIP PREV AVEZZANO, SERVIZIO DI IGIENE EPIDEMIOLOGIA E SANITÀ PUBBLICA | Istituto Zooprofilattico Sperimentale dell'Abruzzo e Molise "G. Caporale" | Lorusso A, Marcacci M, Di Domenico M, Ancora M, Curini V, Mangone I, Rinaldi A, Scialabba S, Di Pasquale A, Cammà C, Scialabba S, Puglia I, Calistri P, Savini G |
| EPI_ISL_1117522, EPI_ISL_1117523, EPI_ISL_1117524 | SIESP DIP PREV TERAMO | Istituto Zooprofilattico Sperimentale dell'Abruzzo e Molise "G. Caporale" | Lorusso A, Marcacci M, Di Domenico M, Ancora M, Curini V, Mangone I, Rinaldi A, Scialabba S, Di Pasquale A, Cammà C, Scialabba S, Puglia I, Calistri P, Savini G |
| EPI_ISL_1117525 | OSP SAN SALVATORE, MEDICINA INTERNA | Istituto Zooprofilattico Sperimentale dell'Abruzzo e Molise "G. Caporale" | Lorusso A, Marcacci M, Di Domenico M, Ancora M, Curini V, Mangone I, Rinaldi A, Scialabba S, Di Pasquale A, Cammà C, Scialabba S, Puglia I, Calistri P, Savini G |
| EPI_ISL_1117526 | IZS TERAMO | Istituto Zooprofilattico Sperimentale dell'Abruzzo e Molise "G. Caporale" | Lorusso A, Marcacci M, Di Domenico M, Ancora M, Curini V, Mangone I, Rinaldi A, Scialabba S, Di Pasquale A, Cammà C, Scialabba S, Puglia I, Calistri P, Savini G |
| EPI_ISL_1117673 | SIESP DIP PREV TERAMO | Istituto Zooprofilattico Sperimentale dell'Abruzzo e Molise "G. Caporale" | Lorusso A, Marcacci M, Di Domenico M, Ancora M, Curini V, Mangone I, Rinaldi A, Scialabba S, Di Pasquale A, Cammà C, Puglia I, Calistri P, Savini G |
| EPI_ISL_1117674 | SIESP CHIETI, DRIVE IN CHIETI | Istituto Zooprofilattico Sperimentale dell'Abruzzo e Molise "G. Caporale" | Lorusso A, Marcacci M, Di Domenico M, Ancora M, Curini V, Mangone I, Rinaldi A, Scialabba S, Di Pasquale A, Cammà C, Puglia I, Calistri P, Savini G |
| EPI_ISL_1117675 | DIP PREV AVEZZANO-SERVIZIO DI IGIENE EPIDEMIOLOGIA E SANITÀ PUBBLICA | Istituto Zooprofilattico Sperimentale dell'Abruzzo e Molise "G. Caporale" | Lorusso A, Marcacci M, Di Domenico M, Ancora M, Curini V, Mangone I, Rinaldi A, Scialabba S, Di Pasquale A, Cammà C, Puglia I, Calistri P, Savini G |
| EPI_ISL_1117676 | SIESP CHIETI, DRIVE IN LANCIANO | Istituto Zooprofilattico Sperimentale dell'Abruzzo e Molise "G. Caporale" | Lorusso A, Marcacci M, Di Domenico M, Ancora M, Curini V, Mangone I, Rinaldi A, Scialabba S, Di Pasquale A, Cammà C, Puglia I, Calistri P, Savini G |
| EPI_ISL_1117677 | USCA CASTEL DI SANGRO | Istituto Zooprofilattico Sperimentale dell'Abruzzo e Molise "G. Caporale" | Lorusso A, Marcacci M, Di Domenico M, Ancora M, Curini V, Mangone I, Rinaldi A, Scialabba S, Di Pasquale A, Cammà C, Puglia I, Calistri P, Savini G |
| EPI_ISL_1117678, EPI_ISL_1117679 | SIESP CHIETI, DRIVE IN VASTO | Istituto Zooprofilattico Sperimentale dell'Abruzzo e Molise "G. Caporale" | Lorusso A, Marcacci M, Di Domenico M, Ancora M, Curini V, Mangone I, Rinaldi A, Scialabba S, Di Pasquale A, Cammà C, Puglia I, Calistri P, Savini G |
| EPI_ISL_1117680 | SIESP DIP PREV TERAMO | Istituto Zooprofilattico Sperimentale dell'Abruzzo e Molise "G. Caporale" | Lorusso A, Marcacci M, Di Domenico M, Ancora M, Curini V, Mangone I, Rinaldi A, Scialabba S, Di Pasquale A, Cammà C, Puglia I, Calistri P, Savini G |
| EPI_ISL_1117681 | SIESP CHIETI, DRIVE IN CHIETI | Istituto Zooprofilattico Sperimentale dell'Abruzzo e Molise "G. Caporale" | Lorusso A, Marcacci M, Di Domenico M, Ancora M, Curini V, Mangone I, Rinaldi A, Scialabba S, Di Pasquale A, Cammà C, Puglia I, Calistri P, Savini G |
| EPI_ISL_1117682 | SIESP CHIETI, DRIVE IN LANCIANO | Istituto Zooprofilattico Sperimentale dell'Abruzzo e Molise "G. Caporale" | Lorusso A, Marcacci M, Di Domenico M, Ancora M, Curini V, Mangone I, Rinaldi A, Scialabba S, Di Pasquale A, Cammà C, Puglia I, Calistri P, Savini G |
| EPI_ISL_1117683, EPI_ISL_1117684 | USCA CASTEL DI SANGRO | Istituto Zooprofilattico Sperimentale dell'Abruzzo e Molise "G. Caporale" | Lorusso A, Marcacci M, Di Domenico M, Ancora M, Curini V, Mangone I, Rinaldi A, Scialabba S, Di Pasquale A, Cammà C, Puglia I, Calistri P, Savini G |
| EPI_ISL_1117685 | SIESP DIP PREV TERAMO | Istituto Zooprofilattico Sperimentale dell'Abruzzo e Molise "G. | Lorusso A, Marcacci M, Di Domenico M, Ancora M, Curini V, Mangone I, Rinaldi A, Scialabba S, Di Pasquale A, Cammà C, Puglia I, Calistri P, Savini G |

| | | | |
|---|---|---|---|
| | | Caporale" | |
| EPI_ISL_1117686 | DIP PREV AVEZZANO-SERVIZIO DI IGIENE EPIDEMIOLOGIA E SANITÀ PUBBLICA | Istituto Zooprofilattico Sperimentale dell'Abruzzo e Molise "G. Caporale" | Lorusso A, Marcacci M, Di Domenico M, Ancora M, Curini V, Mangone I, Rinaldi A, Scialabba S, Di Pasquale A, Cammà C, Puglia I, Calistri P, Savini G |
| EPI_ISL_1117687 | SIESP CHIETI, DRIVE IN ORTONA | Istituto Zooprofilattico Sperimentale dell'Abruzzo e Molise "G. Caporale" | Lorusso A, Marcacci M, Di Domenico M, Ancora M, Curini V, Mangone I, Rinaldi A, Scialabba S, Di Pasquale A, Cammà C, Puglia I, Calistri P, Savini G |
| EPI_ISL_1117688 | SIESP DIP PREV CHIETI | Istituto Zooprofilattico Sperimentale dell'Abruzzo e Molise "G. Caporale" | Lorusso A, Marcacci M, Di Domenico M, Ancora M, Curini V, Mangone I, Rinaldi A, Scialabba S, Di Pasquale A, Cammà C, Puglia I, Calistri P, Savini G |
| EPI_ISL_1117689 | SIESP CHIETI, DRIVE IN LANCIANO | Istituto Zooprofilattico Sperimentale dell'Abruzzo e Molise "G. Caporale" | Lorusso A, Marcacci M, Di Domenico M, Ancora M, Curini V, Mangone I, Rinaldi A, Scialabba S, Di Pasquale A, Cammà C, Puglia I, Calistri P, Savini G |
| EPI_ISL_1117691 | SIESP DIP PREV TERAMO | Istituto Zooprofilattico Sperimentale dell'Abruzzo e Molise "G. Caporale" | Lorusso A, Marcacci M, Di Domenico M, Ancora M, Curini V, Mangone I, Rinaldi A, Scialabba S, Di Pasquale A, Cammà C, Puglia I, Calistri P, Savini G |
| EPI_ISL_1117692 | PRESIDIO OSP SAN LIBERATORE | Istituto Zooprofilattico Sperimentale dell'Abruzzo e Molise "G. Caporale" | Lorusso A, Marcacci M, Di Domenico M, Ancora M, Curini V, Mangone I, Rinaldi A, Scialabba S, Di Pasquale A, Cammà C, Puglia I, Calistri P, Savini G |
| EPI_ISL_1117693 | SIESP DIP PREV TERAMO | Istituto Zooprofilattico Sperimentale dell'Abruzzo e Molise "G. Caporale" | Lorusso A, Marcacci M, Di Domenico M, Ancora M, Curini V, Mangone I, Rinaldi A, Scialabba S, Di Pasquale A, Cammà C, Puglia I, Calistri P, Savini G |
| EPI_ISL_1117694, EPI_ISL_1117695 | SIESP CHIETI, DRIVE IN LANCIANO | Istituto Zooprofilattico Sperimentale dell'Abruzzo e Molise "G. Caporale" | Lorusso A, Marcacci M, Di Domenico M, Ancora M, Curini V, Mangone I, Rinaldi A, Scialabba S, Di Pasquale A, Cammà C, Puglia I, Calistri P, Savini G |
| EPI_ISL_1117696 | SIESP CHIETI, DRIVE IN VASTO | Istituto Zooprofilattico Sperimentale dell'Abruzzo e Molise "G. Caporale" | Lorusso A, Marcacci M, Di Domenico M, Ancora M, Curini V, Mangone I, Rinaldi A, Scialabba S, Di Pasquale A, Cammà C, Puglia I, Calistri P, Savini G |
| EPI_ISL_1117697 | DIP PREV AVEZZANO-SERVIZIO DI IGIENE EPIDEMIOLOGIA E SANITÀ PUBBLICA | Istituto Zooprofilattico Sperimentale dell'Abruzzo e Molise "G. Caporale" | Lorusso A, Marcacci M, Di Domenico M, Ancora M, Curini V, Mangone I, Rinaldi A, Scialabba S, Di Pasquale A, Cammà C, Puglia I, Calistri P, Savini G |
| EPI_ISL_1117698 | SIESP CHIETI, DRIVE IN LANCIANO | Istituto Zooprofilattico Sperimentale dell'Abruzzo e Molise "G. Caporale" | Lorusso A, Marcacci M, Di Domenico M, Ancora M, Curini V, Mangone I, Rinaldi A, Scialabba S, Di Pasquale A, Cammà C, Puglia I, Calistri P, Savini G |
| EPI_ISL_1117699 | SIESP CHIETI, DRIVE IN VASTO | Istituto Zooprofilattico Sperimentale dell'Abruzzo e Molise "G. Caporale" | Lorusso A, Marcacci M, Di Domenico M, Ancora M, Curini V, Mangone I, Rinaldi A, Scialabba S, Di Pasquale A, Cammà C, Puglia I, Calistri P, Savini G |
| EPI_ISL_1117810, EPI_ISL_1117811, EPI_ISL_1117812, EPI_ISL_1117813, EPI_ISL_1117814, EPI_ISL_1117815, EPI_ISL_1117816 | Institut für Pathologie, Salzkammergut Klinikum Vöcklabruck | Bergthaler laboratory, CeMM Research Center for Molecular Medicine of the Austrian Academy of Sciences | Lukas Endler, Anna Schedl, Thomas Penz, Benedikt Agerer, Maelle Le Moing, Michael Schuster, Bekir Erguner, Jan Laine, Martin Senekowitsch, Christoph Bock, Andreas Bergthaler |
| EPI_ISL_1117817, EPI_ISL_1117818, EPI_ISL_1117819, EPI_ISL_1117820 | Center for Virology, Medical University of Vienna | Bergthaler laboratory, CeMM Research Center for Molecular Medicine of the Austrian Academy of Sciences | Lukas Endler, Anna Schedl, Thomas Penz, Benedikt Agerer, Maelle Le Moing, Michael Schuster, Bekir Erguner, Jan Laine, Martin Senekowitsch, Christoph Bock, Andreas Bergthaler |
| EPI_ISL_1117821 | Department of Laboratory Medicine, Division of Clinical Virology, University of Medicine, Vienna | Bergthaler laboratory, CeMM Research Center for Molecular Medicine of the Austrian Academy of Sciences | Lukas Endler, Anna Schedl, Thomas Penz, Benedikt Agerer, Maelle Le Moing, Michael Schuster, Bekir Erguner, Jan Laine, Martin Senekowitsch, Christoph Bock, Andreas Bergthaler |
| EPI_ISL_1117822, EPI_ISL_1117823 | Center for Virology, Medical University of Vienna | Bergthaler laboratory, CeMM Research Center for Molecular Medicine of the Austrian Academy of Sciences | Lukas Endler, Anna Schedl, Thomas Penz, Benedikt Agerer, Maelle Le Moing, Michael Schuster, Bekir Erguner, Jan Laine, Martin Senekowitsch, Christoph Bock, Andreas Bergthaler |
| EPI_ISL_1117824, EPI_ISL_1117825, EPI_ISL_1117826, EPI_ISL_1117827, EPI_ISL_1117828, EPI_ISL_1117829, EPI_ISL_1117830 | Austrian Agency for Health and Food Safety (AGES) | Bergthaler laboratory, CeMM Research Center for Molecular Medicine of the Austrian Academy of Sciences | Lukas Endler, Anna Schedl, Thomas Penz, Benedikt Agerer, Maelle Le Moing, Michael Schuster, Bekir Erguner, Jan Laine, Martin Senekowitsch, Christoph Bock, Andreas Bergthaler |
| EPI_ISL_1117831, EPI_ISL_1117832, EPI_ISL_1117833 | HG Pharma GmbH | Bergthaler laboratory, CeMM Research Center for Molecular Medicine of the Austrian Academy of Sciences | Lukas Endler, Anna Schedl, Thomas Penz, Benedikt Agerer, Maelle Le Moing, Michael Schuster, Bekir Erguner, Jan Laine, Martin Senekowitsch, Christoph Bock, Andreas Bergthaler |
| EPI_ISL_1117834, EPI_ISL_1117835 | Klinikum Wels-Grieskirchen | Bergthaler laboratory, CeMM Research Center for Molecular Medicine of the Austrian Academy of Sciences | Lukas Endler, Anna Schedl, Thomas Penz, Benedikt Agerer, Maelle Le Moing, Michael Schuster, Bekir Erguner, Jan Laine, Martin Senekowitsch, Christoph Bock, Andreas Bergthaler |
| EPI_ISL_1117836, EPI_ISL_1117837 | Center for Virology, Medical University of Vienna | Bergthaler laboratory, CeMM Research Center for Molecular Medicine of the Austrian Academy of Sciences | Lukas Endler, Anna Schedl, Thomas Penz, Benedikt Agerer, Maelle Le Moing, Michael Schuster, Bekir Erguner, Jan Laine, Martin Senekowitsch, Christoph Bock, Andreas Bergthaler |
| EPI_ISL_1117838, EPI_ISL_1117839 | Austrian Agency for Health and Food Safety (AGES) | Bergthaler laboratory, CeMM Research Center for Molecular Medicine of the Austrian Academy of Sciences | Lukas Endler, Anna Schedl, Thomas Penz, Benedikt Agerer, Maelle Le Moing, Michael Schuster, Bekir Erguner, Jan Laine, Martin Senekowitsch, Christoph Bock, Andreas Bergthaler |
| EPI_ISL_1117840, EPI_ISL_1117841, EPI_ISL_1117842, EPI_ISL_1117843, EPI_ISL_1117844, EPI_ISL_1117845 | Klinikum Wels-Grieskirchen | Bergthaler laboratory, CeMM Research Center for Molecular Medicine of the Austrian Academy of Sciences | Lukas Endler, Anna Schedl, Thomas Penz, Benedikt Agerer, Maelle Le Moing, Michael Schuster, Bekir Erguner, Jan Laine, Martin Senekowitsch, Christoph Bock, Andreas Bergthaler |
| EPI_ISL_1117846, EPI_ISL_1117847 | Institut für Pathologie, Salzkammergut Klinikum Vöcklabruck | Bergthaler laboratory, CeMM Research Center for Molecular Medicine of the Austrian Academy of Sciences | Lukas Endler, Anna Schedl, Thomas Penz, Benedikt Agerer, Maelle Le Moing, Michael Schuster, Bekir Erguner, Jan Laine, Martin Senekowitsch, Christoph Bock, Andreas Bergthaler |
| EPI_ISL_1117848, EPI_ISL_1117849, EPI_ISL_1117850, EPI_ISL_1117851, EPI_ISL_1117852, EPI_ISL_1117853 | Center for Virology, Medical University of Vienna | Bergthaler laboratory, CeMM Research Center for Molecular Medicine of the Austrian Academy of Sciences | Lukas Endler, Anna Schedl, Thomas Penz, Benedikt Agerer, Maelle Le Moing, Michael Schuster, Bekir Erguner, Jan Laine, Martin Senekowitsch, Christoph Bock, Andreas Bergthaler |
| EPI_ISL_1117854, EPI_ISL_1117855, EPI_ISL_1117856, EPI_ISL_1117857, EPI_ISL_1117858, EPI_ISL_1117859, EPI_ISL_1117860, EPI_ISL_1117861, EPI_ISL_1117862, EPI_ISL_1117863, EPI_ISL_1117864, EPI_ISL_1117865, EPI_ISL_1117866, EPI_ISL_1117867, EPI_ISL_1117868, EPI_ISL_1117869 | | | |
| see above | Austrian Agency for Health and Food Safety (AGES) | Bergthaler laboratory, CeMM Research Center for Molecular Medicine of the Austrian Academy of Sciences | Lukas Endler, Anna Schedl, Thomas Penz, Benedikt Agerer, Maelle Le Moing, Michael Schuster, Bekir Erguner, Jan Laine, Martin Senekowitsch, Christoph Bock, Andreas Bergthaler |
| EPI_ISL_1117870, EPI_ISL_1117871, EPI_ISL_1117872 | HG Pharma GmbH | Bergthaler laboratory, CeMM Research Center for Molecular Medicine of the Austrian Academy of Sciences | Lukas Endler, Anna Schedl, Thomas Penz, Benedikt Agerer, Maelle Le Moing, Michael Schuster, Bekir Erguner, Jan Laine, Martin Senekowitsch, Christoph Bock, Andreas Bergthaler |
| EPI_ISL_1117873, EPI_ISL_1117875 | Tauernklinikum - Standort Zell am See | Bergthaler laboratory, CeMM Research Center for Molecular Medicine of the Austrian Academy of Sciences | Lukas Endler, Anna Schedl, Thomas Penz, Benedikt Agerer, Maelle Le Moing, Michael Schuster, Bekir Erguner, Jan Laine, Martin Senekowitsch, Christoph Bock, Andreas Bergthaler |
| EPI_ISL_1117876, EPI_ISL_1117877, EPI_ISL_1117878, EPI_ISL_1117879, EPI_ISL_1117880, EPI_ISL_1117881 | Center for Virology, Medical University of Vienna | Bergthaler laboratory, CeMM Research Center for Molecular Medicine of the Austrian Academy of Sciences | Lukas Endler, Anna Schedl, Thomas Penz, Benedikt Agerer, Maelle Le Moing, Michael Schuster, Bekir Erguner, Jan Laine, Martin Senekowitsch, Christoph Bock, Andreas Bergthaler |
| EPI_ISL_1117882, EPI_ISL_1117883, EPI_ISL_1117884, EPI_ISL_1117885 | HG Pharma GmbH | Bergthaler laboratory, CeMM Research Center for Molecular Medicine of the Austrian Academy of Sciences | Lukas Endler, Anna Schedl, Thomas Penz, Benedikt Agerer, Maelle Le Moing, Michael Schuster, Bekir Erguner, Jan Laine, Martin Senekowitsch, Christoph Bock, Andreas Bergthaler |
| EPI_ISL_1117886 | Center for Virology, Medical University of Vienna | Bergthaler laboratory, CeMM Research Center for Molecular Medicine of the Austrian Academy of Sciences | Lukas Endler, Anna Schedl, Thomas Penz, Benedikt Agerer, Maelle Le Moing, Michael Schuster, Bekir Erguner, Jan Laine, Martin Senekowitsch, Christoph Bock, Andreas Bergthaler |
| EPI_ISL_1117887 | Austrian Agency for Health and Food Safety (AGES) | Bergthaler laboratory, CeMM Research Center for Molecular Medicine of the Austrian Academy of Sciences | Lukas Endler, Anna Schedl, Thomas Penz, Benedikt Agerer, Maelle Le Moing, Michael Schuster, Bekir Erguner, Jan Laine, Martin Senekowitsch, Christoph Bock, Andreas Bergthaler |

| | | | |
|---|---|---|---|
| EPI_ISL_1117920 | HG Pharma GmbH | Bergthaler laboratory, CeMM Research Center for Molecular Medicine of the Austrian Academy of Sciences | Lukas Endler, Anna Schedl, Thomas Penz, Benedikt Agerer, Maelle Le Moing, Michael Schuster, Bekir Erguner, Jan Laine, Martin Senekowitsch, Christoph Bock, Andreas Bergthaler |
| EPI_ISL_1117921 | Klinikum Wels-Grieskirchen | Bergthaler laboratory, CeMM Research Center for Molecular Medicine of the Austrian Academy of Sciences | Lukas Endler, Anna Schedl, Thomas Penz, Benedikt Agerer, Maelle Le Moing, Michael Schuster, Bekir Erguner, Jan Laine, Martin Senekowitsch, Christoph Bock, Andreas Bergthaler |
| EPI_ISL_1117922 | HG Pharma GmbH | Bergthaler laboratory, CeMM Research Center for Molecular Medicine of the Austrian Academy of Sciences | Lukas Endler, Anna Schedl, Thomas Penz, Benedikt Agerer, Maelle Le Moing, Michael Schuster, Bekir Erguner, Jan Laine, Martin Senekowitsch, Christoph Bock, Andreas Bergthaler |
| EPI_ISL_1117924 | Tauernklinikum - Standort Zell am See | Bergthaler laboratory, CeMM Research Center for Molecular Medicine of the Austrian Academy of Sciences | Lukas Endler, Anna Schedl, Thomas Penz, Benedikt Agerer, Maelle Le Moing, Michael Schuster, Bekir Erguner, Jan Laine, Martin Senekowitsch, Christoph Bock, Andreas Bergthaler |
| EPI_ISL_1117925, EPI_ISL_1117926 | Center for Virology, Medical University of Vienna | Bergthaler laboratory, CeMM Research Center for Molecular Medicine of the Austrian Academy of Sciences | Lukas Endler, Anna Schedl, Thomas Penz, Benedikt Agerer, Maelle Le Moing, Michael Schuster, Bekir Erguner, Jan Laine, Martin Senekowitsch, Christoph Bock, Andreas Bergthaler |
| EPI_ISL_1117927 | Austrian Agency for Health and Food Safety (AGES) | Bergthaler laboratory, CeMM Research Center for Molecular Medicine of the Austrian Academy of Sciences | Lukas Endler, Anna Schedl, Thomas Penz, Benedikt Agerer, Maelle Le Moing, Michael Schuster, Bekir Erguner, Jan Laine, Martin Senekowitsch, Christoph Bock, Andreas Bergthaler |
| EPI_ISL_1117928, EPI_ISL_1117929, EPI_ISL_1117930, EPI_ISL_1117931, EPI_ISL_1117932, EPI_ISL_1117933, EPI_ISL_1117934, EPI_ISL_1117935 | HG Pharma GmbH | Bergthaler laboratory, CeMM Research Center for Molecular Medicine of the Austrian Academy of Sciences | Lukas Endler, Anna Schedl, Thomas Penz, Benedikt Agerer, Maelle Le Moing, Michael Schuster, Bekir Erguner, Jan Laine, Martin Senekowitsch, Christoph Bock, Andreas Bergthaler |
| EPI_ISL_1117949, EPI_ISL_1117950 | Austrian Agency for Health and Food Safety (AGES) | Bergthaler laboratory, CeMM Research Center for Molecular Medicine of the Austrian Academy of Sciences | Lukas Endler, Anna Schedl, Thomas Penz, Benedikt Agerer, Maelle Le Moing, Michael Schuster, Bekir Erguner, Jan Laine, Martin Senekowitsch, Christoph Bock, Andreas Bergthaler |
| EPI_ISL_1117951, EPI_ISL_1117952, EPI_ISL_1117955, EPI_ISL_1117956, EPI_ISL_1117957, EPI_ISL_1117958, EPI_ISL_1117959, EPI_ISL_1117960, EPI_ISL_1117962, EPI_ISL_1117963, EPI_ISL_1117964, EPI_ISL_1117965, EPI_ISL_1117966, EPI_ISL_1117967, EPI_ISL_1117968, EPI_ISL_1117970, EPI_ISL_1117971, EPI_ISL_1117972, EPI_ISL_1117973, EPI_ISL_1117974, EPI_ISL_1117975, EPI_ISL_1117976, EPI_ISL_1117977, EPI_ISL_1117978, EPI_ISL_1117979, EPI_ISL_1117980, EPI_ISL_1117984, EPI_ISL_1117985, EPI_ISL_1117988, EPI_ISL_1117991, EPI_ISL_1117992, EPI_ISL_1117993, EPI_ISL_1117994, EPI_ISL_1117995, EPI_ISL_1117996, EPI_ISL_1117997, EPI_ISL_1117998, EPI_ISL_1117999, EPI_ISL_1118000, EPI_ISL_1118001, EPI_ISL_1118002, EPI_ISL_1118003, EPI_ISL_1118004, EPI_ISL_1118005, EPI_ISL_1118006, EPI_ISL_1118007, EPI_ISL_1118008, EPI_ISL_1118009, EPI_ISL_1118010, EPI_ISL_1118011, EPI_ISL_1118012, EPI_ISL_1118013, EPI_ISL_1118014, EPI_ISL_1118015, EPI_ISL_1118016, EPI_ISL_1118017, EPI_ISL_1118018, EPI_ISL_1118019, EPI_ISL_1118021, EPI_ISL_1118022, EPI_ISL_1118023, EPI_ISL_1118024, EPI_ISL_1118025, EPI_ISL_1118026, EPI_ISL_1118027, EPI_ISL_1118028 | | | |
| see above | HG Pharma GmbH | Bergthaler laboratory, CeMM Research Center for Molecular Medicine of the Austrian Academy of Sciences | Lukas Endler, Anna Schedl, Thomas Penz, Benedikt Agerer, Maelle Le Moing, Michael Schuster, Bekir Erguner, Jan Laine, Martin Senekowitsch, Christoph Bock, Andreas Bergthaler |
| EPI_ISL_1118029, EPI_ISL_1118030, EPI_ISL_1118031, EPI_ISL_1118032, EPI_ISL_1118033, EPI_ISL_1118034, EPI_ISL_1118035, EPI_ISL_1118036, EPI_ISL_1118037, EPI_ISL_1118038, EPI_ISL_1118039, EPI_ISL_1118040, EPI_ISL_1118041, EPI_ISL_1118042, EPI_ISL_1118043, EPI_ISL_1118044, EPI_ISL_1118045, EPI_ISL_1118046, EPI_ISL_1118047, EPI_ISL_1118048, EPI_ISL_1118049 | | | |
| see above | Austrian Agency for Health and Food Safety (AGES) | Bergthaler laboratory, CeMM Research Center for Molecular Medicine of the Austrian Academy of Sciences | Lukas Endler, Anna Schedl, Thomas Penz, Benedikt Agerer, Maelle Le Moing, Michael Schuster, Bekir Erguner, Jan Laine, Martin Senekowitsch, Christoph Bock, Andreas Bergthaler |
| EPI_ISL_1118050, EPI_ISL_1118051, EPI_ISL_1118052, EPI_ISL_1118053, EPI_ISL_1118054, EPI_ISL_1118055, EPI_ISL_1118056, EPI_ISL_1118057, EPI_ISL_1118058, EPI_ISL_1118059, EPI_ISL_1118060, EPI_ISL_1118061 | | | |
| see above | Klinikum Wels-Grieskirchen | Bergthaler laboratory, CeMM Research Center for Molecular Medicine of the Austrian Academy of Sciences | Lukas Endler, Anna Schedl, Thomas Penz, Benedikt Agerer, Maelle Le Moing, Michael Schuster, Bekir Erguner, Jan Laine, Martin Senekowitsch, Christoph Bock, Andreas Bergthaler |
| EPI_ISL_1118062 | Institut für Pathologie, Salzkammergut Klinikum Vöcklabruck | Bergthaler laboratory, CeMM Research Center for Molecular Medicine of the Austrian Academy of Sciences | Lukas Endler, Anna Schedl, Thomas Penz, Benedikt Agerer, Maelle Le Moing, Michael Schuster, Bekir Erguner, Jan Laine, Martin Senekowitsch, Christoph Bock, Andreas Bergthaler |
| EPI_ISL_1118063 | Klinikum Wels-Grieskirchen | Bergthaler laboratory, CeMM Research Center for Molecular Medicine of the Austrian Academy of Sciences | Lukas Endler, Anna Schedl, Thomas Penz, Benedikt Agerer, Maelle Le Moing, Michael Schuster, Bekir Erguner, Jan Laine, Martin Senekowitsch, Christoph Bock, Andreas Bergthaler |
| EPI_ISL_1118064 | 3. Medizinische Abteilung, Hanusch Krankenhaus | Bergthaler laboratory, CeMM Research Center for Molecular Medicine of the Austrian Academy of Sciences | Lukas Endler, Anna Schedl, Thomas Penz, Benedikt Agerer, Maelle Le Moing, Michael Schuster, Bekir Erguner, Jan Laine, Martin Senekowitsch, Christoph Bock, Andreas Bergthaler |
| EPI_ISL_1118065, EPI_ISL_1118066, EPI_ISL_1118067, EPI_ISL_1118068, EPI_ISL_1118069, EPI_ISL_1118070, EPI_ISL_1118071 | Center for Virology, Medical University of Vienna | Bergthaler laboratory, CeMM Research Center for Molecular Medicine of the Austrian Academy of Sciences | Lukas Endler, Anna Schedl, Thomas Penz, Benedikt Agerer, Maelle Le Moing, Michael Schuster, Bekir Erguner, Jan Laine, Martin Senekowitsch, Christoph Bock, Andreas Bergthaler |
| EPI_ISL_1118072 | Elling group, Institute of Molecular Biotechnology (IMBA) | Bergthaler laboratory, CeMM Research Center for Molecular Medicine of the Austrian Academy of Sciences | Lukas Endler, Anna Schedl, Thomas Penz, Benedikt Agerer, Maelle Le Moing, Michael Schuster, Bekir Erguner, Jan Laine, Martin Senekowitsch, Christoph Bock, Andreas Bergthaler |
| EPI_ISL_1118073, EPI_ISL_1118074, EPI_ISL_1118075, EPI_ISL_1118076, EPI_ISL_1118077, EPI_ISL_1118078, EPI_ISL_1118079, EPI_ISL_1118080 | Austrian Agency for Health and Food Safety (AGES) | Bergthaler laboratory, CeMM Research Center for Molecular Medicine of the Austrian Academy of Sciences | Lukas Endler, Anna Schedl, Thomas Penz, Benedikt Agerer, Maelle Le Moing, Michael Schuster, Bekir Erguner, Jan Laine, Martin Senekowitsch, Christoph Bock, Andreas Bergthaler |
| EPI_ISL_1118081, EPI_ISL_1118082 | HG Pharma GmbH | Bergthaler laboratory, CeMM Research Center for Molecular Medicine of the Austrian Academy of Sciences | Lukas Endler, Anna Schedl, Thomas Penz, Benedikt Agerer, Maelle Le Moing, Michael Schuster, Bekir Erguner, Jan Laine, Martin Senekowitsch, Christoph Bock, Andreas Bergthaler |
| EPI_ISL_1118161, EPI_ISL_1118162, EPI_ISL_1118163, EPI_ISL_1118164, EPI_ISL_1118165, EPI_ISL_1118166, EPI_ISL_1118167, EPI_ISL_1118168, EPI_ISL_1118169, EPI_ISL_1118170, EPI_ISL_1118171, EPI_ISL_1118172, EPI_ISL_1118173, EPI_ISL_1118174, EPI_ISL_1118175, EPI_ISL_1118176, EPI_ISL_1118177, EPI_ISL_1118178, EPI_ISL_1118179, EPI_ISL_1118180, EPI_ISL_1118181, EPI_ISL_1118182, EPI_ISL_1118183, EPI_ISL_1118184, EPI_ISL_1118185, EPI_ISL_1118186, EPI_ISL_1118187, EPI_ISL_1118188, EPI_ISL_1118189, EPI_ISL_1118190, EPI_ISL_1118191, EPI_ISL_1118192, EPI_ISL_1118193, EPI_ISL_1118194, EPI_ISL_1118195, EPI_ISL_1118196, EPI_ISL_1118197, EPI_ISL_1118198, EPI_ISL_1118199, EPI_ISL_1118200, EPI_ISL_1118201, EPI_ISL_1118202, EPI_ISL_1118203, EPI_ISL_1118204 | | | |
| see above | LAB ANALISI PO CARDARELLI CAMPOBASSO | Istituto Zooprofilattico Sperimentale dell'Abruzzo e Molise "G. Caporale" | Scutellà M, Niro G, Felice V. Lorusso A, Marcacci M, Di Domenico M, Ancora M, Curini V, Mangone I, Rinaldi A, Scialabba S, Di Pasquale A, Cammà C, Puglia I, Calistri P, Savini G |
| EPI_ISL_1118207, EPI_ISL_1118208, EPI_ISL_1118209, EPI_ISL_1118210, EPI_ISL_1118211, EPI_ISL_1118212, EPI_ISL_1118213, EPI_ISL_1118214, EPI_ISL_1118215, EPI_ISL_1118216 | National Institute of Health Research and Development | National Institute of Health Research and Development | Hana Apsari Pawestri, Subangkit, Kartika Dewi Puspa, Arie Ardiansyah Nugraha, Hartanti Dian Ikawati, Krisna Nur Andriana Pangesti, Yuni Rukminiati, Ririn Ramadhany, Agustiningsih, Kindi Adam, Holy Arif Wibowo, Triyani Soekarso, Ni Ketut Susilarini, Nurika Hariastuti, Ully Alfi Nikmah, Reni Herman, Nike Susanti, Herna, Tati Febriyanti, Natalie Laurencia Kipuw, Fauzul Muna, Irene Lorinda Indalao, Nelly Puspandari, Vivi Setiawaty. |
| EPI_ISL_1118876 | Hopital | National Reference Center for Viruses of Respiratory Infections, Institut Pasteur, Paris | Marion Barbet, Sylvie Behillil, Méline Bizard, Angela Brisebarre, Camille Capel, Etienne Simon-Lorière, Vincent Enouf, Maud Vanpeene, Sylvie van der Werf,Pilorge LéA |
| EPI_ISL_1118877 | Hopital | National Reference Center for Viruses of Respiratory Infections, Institut Pasteur, Paris | Marion Barbet, Sylvie Behillil, Méline Bizard, Angela Brisebarre, Camille Capel, Etienne Simon-Lorière, Vincent Enouf, Maud Vanpeene, Sylvie van der Werf,Gastli Nabil |
| EPI_ISL_1118878 | Hopital | National Reference Center for Viruses of Respiratory Infections, Institut Pasteur, Paris | Marion Barbet, Sylvie Behillil, Méline Bizard, Angela Brisebarre, Camille Capel, Etienne Simon-Lorière, Vincent Enouf, Maud Vanpeene, Sylvie van der Werf,Ledru Sylvie |
| EPI_ISL_1118883 | Hopital | National Reference Center for Viruses of Respiratory Infections, Institut Pasteur, Paris | Marion Barbet, Sylvie Behillil, Méline Bizard, Angela Brisebarre, Camille Capel, Etienne Simon-Lorière, Vincent Enouf, Maud Vanpeene, Sylvie van der Werf,Luizy Nelly |
| EPI_ISL_1118930 | Laboratorio Microbiologia e Virologia P.O. Cotugno A.O. dei Colli | Laboratorio Microbiologia e Virologia P.O. Cotugno A.O. dei Colli | Luigi Atripaldi, Claudia Tiberio, Anna Perfetti, Pellegrino Cerino, Biancamaria Pierri, Maria Concetta Cuomo |
| EPI_ISL_1118931 | National Institute of Health Research and Development | National Institute of Health Research and Development | Hana Apsari Pawestri, Subangkit, Kartika Dewi Puspa, Arie Ardiansyah Nugraha, Hartanti Dian Ikawati, Krisna Nur Andriana Pangesti, Yuni Rukminiati, Ririn Ramadhany, Agustiningsih, Kindi Adam, Holy Arif Wibowo, Triyani Soekarso, Ni Ketut Susilarini, Nurika Hariastuti, Ully Alfi Nikmah, Reni Herman, Nike Susanti, Herna, Tati Febriyanti, Natalie Laurencia Kipuw, Fauzul Muna, Irene Lorinda Indalao, Nelly Puspandari, Vivi Setiawaty. |
| EPI_ISL_1118932 | Laboratorio Microbiologia e Virologia P.O. Cotugno A.O. dei | Laboratorio Microbiologia e Virologia P.O. Cotugno A.O. dei | Luigi Atripaldi, Claudia Tiberio, Anna Perfetti, Pellegrino Cerino, Biancamaria Pierri, Maria Concetta Cuomo |

| | | | |
|---|---|---|---|
| | Colli | Colli | |
| EPI_ISL_1118933 | National Institute of Health Research and Development | National Institute of Health Research and Development | Hana Apsari Pawestri, Subangkit, Kartika Dewi Puspa, Arie Ardiansyah Nugraha, Hartanti Dian Ikawati, Krisna Nur Andriana Pangesti, Yuni Rukminiati, Ririn Ramadhany, Agustiningsih, Kindi Adam, Holy Arif Wibowo, Triyani Soekarso, Ni Ketut Susilarini, Nurika Hariastuti, Ully Alfi Nikmah, Reni Herman, Nike Susanti, Herna, Tati Febriyanti, Natalie Laurencia Kipuw, Fauzul Muna, Irene Lorinda Indalao, Nelly Puspandari, Vivi Setiawaty. |
| EPI_ISL_1118934, EPI_ISL_1119567 | Laboratorio Microbiologia e Virologia P.O. Cotugno A.O. dei Colli | Laboratorio Microbiologia e Virologia P.O. Cotugno A.O. dei Colli | Luigi Atripaldi, Claudia Tiberio, Anna Perfetti, Pellegrino Cerino, Biancamaria Pierri, Maria Concetta Cuomo |
| EPI_ISL_1121033 | USCA CASTEL DI SANGRO | Istituto Zooprofilattico Sperimentale dell'Abruzzo e Molise "G. Caporale" | Lorusso A, Marcacci M, Di Domenico M, Ancora M, Curini V, Mangone I, Rinaldi A, Scialabba S, Di Pasquale A, Cammà C, Puglia I, Calistri P, Savini G |
| EPI_ISL_1121034 | LAB ANALISI PO CARDARELLI CAMPOBASSO | Istituto Zooprofilattico Sperimentale dell'Abruzzo e Molise "G. Caporale" | Scutellà M, Niro G, Felice V. Lorusso A, Marcacci M, Di Domenico M, Ancora M, Curini V, Mangone I, Rinaldi A, Scialabba S, Di Pasquale A, Cammà C, Puglia I, Calistri P, Savini G |
| EPI_ISL_1121307 | Hospital E Antonio Policarpo de Oliveira | Instituto Adolfo Lutz, Interdiciplinary Procedures Center, Strategic Laboratory | Claudio Tavares Sacchi, Claudia Regina Gonçalves, Erica Valessa Ramos Gomes, Karoline Rodrigues Campos, Caio Vinicius Dias Lopes |
| EPI_ISL_1121308 | Hospital e Pronto Socorro Portinari | Instituto Adolfo Lutz, Interdiciplinary Procedures Center, Strategic Laboratory | Claudio Tavares Sacchi, Claudia Regina Gonçalves, Erica Valessa Ramos Gomes, Karoline Rodrigues Campos, Caio Vinicius Dias Lopes |
| EPI_ISL_1121309 | UBS Jose Francisco Rezende | Instituto Adolfo Lutz, Interdiciplinary Procedures Center, Strategic Laboratory | Claudio Tavares Sacchi, Claudia Regina Gonçalves, Erica Valessa Ramos Gomes, Karoline Rodrigues Campos, Caio Vinicius Dias Lopes |
| EPI_ISL_1121310 | IAL Regional de Bauru | Instituto Adolfo Lutz, Interdiciplinary Procedures Center, Strategic Laboratory | Claudio Tavares Sacchi, Claudia Regina Gonçalves, Erica Valessa Ramos Gomes, Karoline Rodrigues Campos, Caio Vinicius Dias Lopes |
| EPI_ISL_1121311 | Centro de Saude II Dr. Jose Paione Mococa | Instituto Adolfo Lutz, Interdiciplinary Procedures Center, Strategic Laboratory | Claudio Tavares Sacchi, Claudia Regina Gonçalves, Erica Valessa Ramos Gomes, Karoline Rodrigues Campos, Caio Vinicius Dias Lopes |
| EPI_ISL_1121312, EPI_ISL_1121313, EPI_ISL_1121314, EPI_ISL_1121315, EPI_ISL_1121317 | IAL Regional de Bauru | Instituto Adolfo Lutz, Interdiciplinary Procedures Center, Strategic Laboratory | Claudio Tavares Sacchi, Claudia Regina Gonçalves, Erica Valessa Ramos Gomes, Karoline Rodrigues Campos, Caio Vinicius Dias Lopes |
| EPI_ISL_1121318, EPI_ISL_1121319 | Hospital de Campanha COVID 19 Caieiras | Instituto Adolfo Lutz, Interdiciplinary Procedures Center, Strategic Laboratory | Claudio Tavares Sacchi, Claudia Regina Gonçalves, Erica Valessa Ramos Gomes, Karoline Rodrigues Campos, Caio Vinicius Dias Lopes |
| EPI_ISL_1121320, EPI_ISL_1121321 | IAL Regional de Bauru | Instituto Adolfo Lutz, Interdiciplinary Procedures Center, Strategic Laboratory | Claudio Tavares Sacchi, Claudia Regina Gonçalves, Erica Valessa Ramos Gomes, Karoline Rodrigues Campos, Caio Vinicius Dias Lopes |
| EPI_ISL_1121322 | Santa Casa de Santa Isabel | Instituto Adolfo Lutz, Interdiciplinary Procedures Center, Strategic Laboratory | Claudio Tavares Sacchi, Claudia Regina Gonçalves, Erica Valessa Ramos Gomes, Karoline Rodrigues Campos, Caio Vinicius Dias Lopes |
| EPI_ISL_1121323, EPI_ISL_1121324 | Complexo Hospitalar Padre Bentode Guarulhos | Instituto Adolfo Lutz, Interdiciplinary Procedures Center, Strategic Laboratory | Claudio Tavares Sacchi, Claudia Regina Gonçalves, Erica Valessa Ramos Gomes, Karoline Rodrigues Campos, Caio Vinicius Dias Lopes |
| EPI_ISL_1121325 | IAL Regional de Santos | Instituto Adolfo Lutz, Interdiciplinary Procedures Center, Strategic Laboratory | Claudio Tavares Sacchi, Claudia Regina Gonçalves, Erica Valessa Ramos Gomes, Karoline Rodrigues Campos, Caio Vinicius Dias Lopes |
| EPI_ISL_1122456 | Manila Doctors Hospital | Philippine Genome Center | Francis A. Tablizo, Kenneth M. Kim, Carlo M. Lapid, Marc Jerrone R. Castro, Maria Sofia L. Yangzon, Benedict A. Maralit, Marc Edsel C. Ayes, Eva Maria Cutiongco-de la Paz, Alethea R. de Guzman, Jan Michael C. Yap, Jo-Hannah S. Llames, Sheila Mae M. Araiza, Kris P. Punayan, Irish Coleen A. Asin, Candice Francheska B. Tambaoan, Asia Louisa U. Chong, Karol Sophia Agape R. Padilla, Rianna Patricia S. Cruz, El King D. Morado, Joshua Gregor A. Dizon, Razel Nikka M. Hao, Arianne A. Zamora, Devon Ray Pacial, Juan Antonio R. Magalang, Marissa Alejandria, Celia Carlos, Anna Ong-Lim, Edsel Maurice Salvaña, John Q. Wong, Jaime C. Montoya, Maria Rosario Singh-Vergeire and Cynthia P. Saloma |
| EPI_ISL_1123297 | Virology Unit, AOUP | Virology Unit, AOUP | Mamrialinda Vatteroni, Susi Frateschi, Mauro Pistello |
| EPI_ISL_1123298, EPI_ISL_1123299 | Virology Unit, AOUP | Virology Unit | Marialinda Vatteroni, Susi Frateschi, Mauro Pistello |
| EPI_ISL_1123325, EPI_ISL_1123332 | Virology Unit, AOUP | Virology Unit, AOUP | Marialinda Vatteroni, Susi Frateschi, Mauro Pistello |
| EPI_ISL_1123344, EPI_ISL_1123346, EPI_ISL_1123347 | Virology Unit, AOUP | Virology Unit | Marialinda Vatteroni, Susi Frateschi, Mauro Pistello |
| EPI_ISL_1123372 | UPA I Santa Isabel | Instituto Adolfo Lutz, Interdiciplinary Procedures Center, Strategic Laboratory | Claudio Tavares Sacchi, Claudia Regina Gonçalves, Erica Valessa Ramos Gomes, Karoline Rodrigues Campos, Caio Vinicius Dias Lopes |
| EPI_ISL_1123373 | Grupo Tecnico de Vigilancia Sanitaria e Epidemiologica | Instituto Adolfo Lutz, Interdiciplinary Procedures Center, Strategic Laboratory | Claudio Tavares Sacchi, Claudia Regina Gonçalves, Erica Valessa Ramos Gomes, Karoline Rodrigues Campos, Caio Vinicius Dias Lopes |
| EPI_ISL_1123374 | IAL Regional de Santos | Instituto Adolfo Lutz, Interdiciplinary Procedures Center, Strategic Laboratory | Claudio Tavares Sacchi, Claudia Regina Gonçalves, Erica Valessa Ramos Gomes, Karoline Rodrigues Campos, Caio Vinicius Dias Lopes |
| EPI_ISL_1123388 | Laboratorio de Virología HUCA | Laboratorio de Virología HUCA | Sandoval M, Castelló C, Gómez de Oña J, Boga JA, Rojo S, Alvarez-Arguelles ME, Abreu F, Costales I, Perez-Martínez Z, Martín-Rodríguez G, Coto E, Melón S |
| EPI_ISL_1133120, EPI_ISL_1133121, EPI_ISL_1133122, EPI_ISL_1133123, EPI_ISL_1133124, EPI_ISL_1133125, EPI_ISL_1133126, EPI_ISL_1133127, EPI_ISL_1133128, EPI_ISL_1133129, EPI_ISL_1133130, EPI_ISL_1133131, EPI_ISL_1133132, EPI_ISL_1133134, EPI_ISL_1133135, EPI_ISL_1133136, EPI_ISL_1133137, EPI_ISL_1133138, EPI_ISL_1133139, EPI_ISL_1133140, EPI_ISL_1133141, EPI_ISL_1133142, EPI_ISL_1133143, EPI_ISL_1133144 | | | |
| see above | LABCOVID_HCPA | LABRESIS_HCPA | Martins AF, Wink PL, Volpato F, Rosset C, de Paris F, Monteiro F, Barth AL |
| EPI_ISL_1138518 | SA Pärnu Hospital Laboratory | 1. Laboratory of Communicable Diseases (Estonia); 2. Eurofins Genomics Europe Sequencing GmbH | Liidia Dotsenko |
| EPI_ISL_1138519, EPI_ISL_1138520, EPI_ISL_1138521, EPI_ISL_1138522, EPI_ISL_1138523 | Laboratory of Communicable Diseases | 1. Laboratory of Communicable Diseases (Estonia); 2. Eurofins Genomics Europe Sequencing GmbH | Liidia Dotsenko |
| EPI_ISL_1138524, EPI_ISL_1138525 | SA Pärnu Hospital Laboratory | 1. Laboratory of Communicable Diseases (Estonia); 2. Eurofins Genomics Europe Sequencing GmbH | Liidia Dotsenko |
| EPI_ISL_1138526, EPI_ISL_1138527, EPI_ISL_1138528, EPI_ISL_1138529, EPI_ISL_1138530, EPI_ISL_1138531, EPI_ISL_1138532, EPI_ISL_1138533, EPI_ISL_1138534, EPI_ISL_1138535, EPI_ISL_1138536 | | | |
| see above | Laboratory of Communicable Diseases | 1. Laboratory of Communicable Diseases (Estonia); 2. Eurofins Genomics Europe Sequencing GmbH | Liidia Dotsenko |
| EPI_ISL_1138537 | SA Pärnu Hospital Laboratory | 1. Laboratory of Communicable Diseases (Estonia); 2. Eurofins Genomics Europe Sequencing GmbH | Liidia Dotsenko |
| EPI_ISL_1138538, EPI_ISL_1138539, EPI_ISL_1138540, EPI_ISL_1138541 | Laboratory of Communicable Diseases | 1. Laboratory of Communicable Diseases (Estonia); 2. Eurofins Genomics Europe Sequencing GmbH | Liidia Dotsenko |
| EPI_ISL_1138542, EPI_ISL_1138543 | SA Pärnu Hospital Laboratory | 1. Laboratory of Communicable Diseases (Estonia); 2. Eurofins Genomics Europe Sequencing GmbH | Liidia Dotsenko |

| | | | |
|---|---|---|---|
| EPI_ISL_1138544, EPI_ISL_1138545, EPI_ISL_1138546 | Laboratory of Communicable Diseases | 1. Laboratory of Communicable Diseases (Estonia); 2. Eurofins Genomics Europe Sequencing GmbH | Liidia Dotsenko |
| EPI_ISL_1138547 | SA Pärnu Hospital Laboratory | 1. Laboratory of Communicable Diseases (Estonia); 2. Eurofins Genomics Europe Sequencing GmbH | Liidia Dotsenko |
| EPI_ISL_1138548, EPI_ISL_1138549, EPI_ISL_1138550, EPI_ISL_1138551, EPI_ISL_1138552 | Laboratory of Communicable Diseases | 1. Laboratory of Communicable Diseases (Estonia); 2. Eurofins Genomics Europe Sequencing GmbH | Liidia Dotsenko |
| EPI_ISL_1138553 | SA Pärnu Hospital Laboratory | 1. Laboratory of Communicable Diseases (Estonia); 2. Eurofins Genomics Europe Sequencing GmbH | Liidia Dotsenko |
| EPI_ISL_1138554, EPI_ISL_1138555, EPI_ISL_1138556, EPI_ISL_1138557, EPI_ISL_1138558, EPI_ISL_1138559, EPI_ISL_1138560, EPI_ISL_1138561, EPI_ISL_1138562, EPI_ISL_1138563, EPI_ISL_1138564, EPI_ISL_1138566 | | | |
| see above | Laboratory of Communicable Diseases | 1. Laboratory of Communicable Diseases (Estonia); 2. Eurofins Genomics Europe Sequencing GmbH | Liidia Dotsenko |
| EPI_ISL_1138567 | SA Pärnu Hospital Laboratory | 1. Laboratory of Communicable Diseases (Estonia); 2. Eurofins Genomics Europe Sequencing GmbH | Liidia Dotsenko |
| EPI_ISL_1138568, EPI_ISL_1138569, EPI_ISL_1138570, EPI_ISL_1138571, EPI_ISL_1138572 | Laboratory of Communicable Diseases | 1. Laboratory of Communicable Diseases (Estonia); 2. Eurofins Genomics Europe Sequencing GmbH | Liidia Dotsenko |
| EPI_ISL_1138573, EPI_ISL_1138574, EPI_ISL_1138575, EPI_ISL_1138576, EPI_ISL_1138577, EPI_ISL_1138579, EPI_ISL_1138580, EPI_ISL_1138581, EPI_ISL_1138582, EPI_ISL_1138583, EPI_ISL_1138584, EPI_ISL_1138585, EPI_ISL_1138586, EPI_ISL_1138587, EPI_ISL_1138588, EPI_ISL_1138589, EPI_ISL_1138590, EPI_ISL_1138592, EPI_ISL_1138593, EPI_ISL_1138594, EPI_ISL_1138595, EPI_ISL_1138596 | | | |
| see above | SYNLAB Eesti OÜ | 1. Laboratory of Communicable Diseases (Estonia); 2. Eurofins Genomics Europe Sequencing GmbH | Liidia Dotsenko |
| EPI_ISL_1138597 | Laboratory of Communicable Diseases | 1. Laboratory of Communicable Diseases (Estonia); 2. Eurofins Genomics Europe Sequencing GmbH | Liidia Dotsenko |
| EPI_ISL_1138745 | Analysis laboratory | National Reference Center for Viruses of Respiratory Infections, Institut Pasteur, Paris | Marion Barbet, Sylvie Behillil, Méline Bizard, Angela Brisebarre, Camille Capel, Etienne Simon-Lorière, Vincent Enouf, Maud Vanpeene, Sylvie van der Werf,J. Besson |
| EPI_ISL_1138969 | Laboratory for HIV and opportunistic infections diagnosis The Republican Research and Practical Center for Epidemiology and Microbiology (RRPCEM) | Laboratory for HIV and opportunistic infections diagnosis The Republican Research and Practical Center for Epidemiology and Microbiology (RRPCEM) | Elena Gasich, Kirill Bulda, Artur Akhremchuk, Leonid Valentovich, Analoly Krasko, Vladimir Gorbunov |
| EPI_ISL_1138983 | Laboratory for HIV and opportunistic infections diagnosis The Republican Research and Practical Center for Epidemiology and Microbiology (RRPCEM) | Laboratory for HIV and opportunistic infections diagnosis The Republican Research and Practical Center for Epidemiology and Microbiology (RRPCEM) | Elena Gasich, Kirill Bulda, Artur Akhremchuk, Leonid Valentovich, Anatoly Krasko, Vladimir Gorbunov |
| EPI_ISL_1139069 | SMS Aruja | Instituto Adolfo Lutz, Interdiciplinary Procedures Center, Strategic Laboratory | Claudio Tavares Sacchi, Claudia Regina Gonçalves, Erica Valessa Ramos Gomes, Karoline Rodrigues Campos, Caio Vinicius Dias Lopes |
| EPI_ISL_1139070 | IAL Regional de Ribeirao Preto | Instituto Adolfo Lutz, Interdiciplinary Procedures Center, Strategic Laboratory | Claudio Tavares Sacchi, Claudia Regina Gonçalves, Erica Valessa Ramos Gomes, Karoline Rodrigues Campos, Caio Vinicius Dias Lopes |
| EPI_ISL_1139071, EPI_ISL_1139072, EPI_ISL_1139073, EPI_ISL_1139074 | Instituto Adolfo Lutz Central | Instituto Adolfo Lutz, Interdiciplinary Procedures Center, Strategic Laboratory | Claudio Tavares Sacchi, Claudia Regina Gonçalves, Erica Valessa Ramos Gomes, Karoline Rodrigues Campos, Caio Vinicius Dias Lopes |
| EPI_ISL_1139075, EPI_ISL_1139077 | Lab Loc - Itapecerica da Serra | Instituto Adolfo Lutz, Interdiciplinary Procedures Center, Strategic Laboratory | Claudio Tavares Sacchi, Claudia Regina Gonçalves, Erica Valessa Ramos Gomes, Karoline Rodrigues Campos, Caio Vinicius Dias Lopes |
| EPI_ISL_1140095, EPI_ISL_1140097 | Lithuanian University of Health Sciences Hospital, Department of Laboratory Medicine | Lithuanian University of Health Sciences Hospital, Department of Genetics and Molecular Medicine | Rasa Ugenskien, Darius ereškeviius, Inga Nasvytien, Živil Žemeckien, Mantas Šarauskas, Marius Šukys, Astra Vitkauskien, Renaldas Jurkeviius |
| EPI_ISL_1159390 | Semen Gresik Hospital | Institute of Tropical Disease, Universitas Airlangga | Rima R Prasetya, Krisnoadi Rahardjo, Aldise M Nastri, Jezzy R Dewantari, Muchdor, Gatot Soegiarto, Laksmi Wulandari, Resti Yudhawati, Soetjipto, Yasuko Mori, Maria I Lusida, Kazufumi Shimizu |
| EPI_ISL_1159391 | Royal Hospital | Institute of Tropical Disease, Universitas Airlangga | Krisnoadi Rahardjo, Aldise M Nastri, Jezzy R Dewantari, Rima R Prasetya, Henny P Margastuti, Gatot Soegiarto, Laksmi Wulandari, Resti Yudhawati, Soetjipto, Yasuko Mori, Maria I Lusida, Kazufumi Shimizu |
| EPI_ISL_1161414, EPI_ISL_1161415, EPI_ISL_1163530 | LABCOVID_HCPA | LABRESIS_HCPA | Martins AF, Wink PL, Volpato F, Rosset C, de Paris F, Monteiro F, Zavascki AP, Barth AL |
| EPI_ISL_1163795, EPI_ISL_1163796, EPI_ISL_1163797 | Azienda USL Umbria 2 | Istituto Zooprofilattico Sperimentale dell'Abruzzo e Molise "G. Caporale" | Proietti A., Pistoni E., Lorusso A, Marcacci M, Di Domenico M, Ancora M, Curini V, Mangone I, Rinaldi A, Scialabba S, Di Pasquale A, Cammà C, Puglia I, Calistri P, Savini G |
| EPI_ISL_1163798, EPI_ISL_1163799, EPI_ISL_1163814, EPI_ISL_1163815, EPI_ISL_1163817, EPI_ISL_1163818, EPI_ISL_1163819, EPI_ISL_1163820, EPI_ISL_1163821, EPI_ISL_1163822, EPI_ISL_1163824, EPI_ISL_1163825, EPI_ISL_1163826, EPI_ISL_1163828 | | | |
| see above | Università degli Studi di Perugia | Istituto Zooprofilattico Sperimentale dell'Abruzzo e Molise "G. Caporale" | Mencacci A., Camilloni B., Lorusso A, Marcacci M, Di Domenico M, Ancora M, Curini V, Mangone I, Rinaldi A, Scialabba S, Di Pasquale A, Cammà C, Puglia I, Calistri P, Savini G |
| EPI_ISL_1163829, EPI_ISL_1163830 | Istituto Zooprofilattico Sperimentale Umbria e Marche "Togo Rosati" | Istituto Zooprofilattico Sperimentale dell'Abruzzo e Molise "G. Caporale" | Biagetti M., Giammarioli M., Lorusso A, Marcacci M, Di Domenico M, Ancora M, Curini V, Mangone I, Rinaldi A, Scialabba S, Di Pasquale A, Cammà C, Puglia I, Calistri P, Savini G |
| EPI_ISL_1163889, EPI_ISL_1163890, EPI_ISL_1163891, EPI_ISL_1163892, EPI_ISL_1163893, EPI_ISL_1163894, EPI_ISL_1163895 | IZSM | IZSM-U.O.C. Virologia | Maurizio Viscardi, Lorena Cardillo, Luigi Atripaldi,Giovanna Fusco, Esterina De Carlo, Antonio Limone |
| EPI_ISL_1164754, EPI_ISL_1164755, EPI_ISL_1164756 | Azienda USL Umbria 2 | Istituto Zooprofilattico Sperimentale dell'Abruzzo e Molise "G. Caporale" | Proietti A, Pistoni E, Lorusso A, Marcacci M, Di Domenico M, Ancora M, Curini V, Mangone I, Rinaldi A, Scialabba S, Di Pasquale A, Cammà C, Puglia I, Calistri P, Savini G |
| EPI_ISL_1164787 | Idaho Bureau of Laboratories | IBL | "R. Beukelman, Matthew C. Burns, Aimee Ceniseros, Robert L. Voermans, Christopher Ball" |
| EPI_ISL_1164950, EPI_ISL_1164951, EPI_ISL_1164953, EPI_ISL_1164960, EPI_ISL_1164961, EPI_ISL_1164963, EPI_ISL_1164964, EPI_ISL_1164966, EPI_ISL_1164967, EPI_ISL_1164968 | Hôpital Georges L. Dumont | National Microbiology Laboratory (NML) | Anna Majer, Shari Tyson, Grace Seo, Philip Mabon, Elsie Grudeski, Rhiannon Huzarewich, Russell Mandes, Anneliese Landgraff, Jennifer Tanner, Natalie Knox, Morag Graham, Gary Van Domselaar, Richard Garceau, Guillaume Desnoyers, Nathalie Bastien, Yan Li, Timothy Booth, Darian Hole, Madison Chapel, Kirsten Biggar, CanCOGeN's metadata curation team, Public Health Agency of Canada CanCOGeN team |
| EPI_ISL_1166147, EPI_ISL_1166148 | Laboratorio Analisi Osp. Città di Castello - Azienda USL Umbria1 | Istituto Zooprofilattico Sperimentale dell'Abruzzo e Molise "G. Caporale" | Malagigi V, Tacconi P, Lorusso A, Marcacci M, Di Domenico M, Ancora M, Curini V, Mangone I, Rinaldi A, Scialabba S, Di Pasquale A, Cammà C, Puglia I, Calistri P, Savini G |
| EPI_ISL_1166149, EPI_ISL_1166151, EPI_ISL_1166152, EPI_ISL_1166153, EPI_ISL_1166154, EPI_ISL_1166155, EPI_ISL_1166156, EPI_ISL_1166157, EPI_ISL_1166159, EPI_ISL_1166160, EPI_ISL_1166162, EPI_ISL_1166163, EPI_ISL_1166164, EPI_ISL_1166165, EPI_ISL_1166166, EPI_ISL_1166167, EPI_ISL_1166168, EPI_ISL_1166169, EPI_ISL_1166170, EPI_ISL_1166171 | | | |
| see above | Università degli Studi di Perugia | Istituto Zooprofilattico Sperimentale dell'Abruzzo e Molise "G. Caporale" | Mencacci A, Camilloni B, Lorusso A, Marcacci M, Di Domenico M, Ancora M, Curini V, Mangone I, Rinaldi A, Scialabba S, Di Pasquale A, Cammà C, Puglia I, Calistri P, Savini G |

| | | | |
|---|---|---|---|
| EPI_ISL_1166172, EPI_ISL_1166173, EPI_ISL_1166174, EPI_ISL_1166175, EPI_ISL_1166176, EPI_ISL_1166177, EPI_ISL_1166178, EPI_ISL_1166179, EPI_ISL_1166180, EPI_ISL_1166181, EPI_ISL_1166182, EPI_ISL_1166183, EPI_ISL_1166184, EPI_ISL_1166185, EPI_ISL_1166186, EPI_ISL_1166187, EPI_ISL_1166188 | | | |
| see above | Istituto Zooprofilattico Sperimentale Umbria e Marche "Togo Rosati" | Istituto Zooprofilattico Sperimentale dell'Abruzzo e Molise "G. Caporale" | Biagetti M, Giammarioli M, Lorusso A, Marcacci M, Di Domenico M, Ancora M, Curini V, Mangone I, Rinaldi A, Scialabba S, Di Pasquale A, Cammà C, Puglia I, Calistri P, Savini G |
| EPI_ISL_1166921, EPI_ISL_1166922, EPI_ISL_1166923, EPI_ISL_1166924, EPI_ISL_1166925, EPI_ISL_1166926, EPI_ISL_1166927, EPI_ISL_1166928, EPI_ISL_1166929, EPI_ISL_1166930, EPI_ISL_1166931, EPI_ISL_1166932, EPI_ISL_1166933, EPI_ISL_1166934, EPI_ISL_1166935, EPI_ISL_1166936, EPI_ISL_1166937, EPI_ISL_1166938, EPI_ISL_1166939, EPI_ISL_1166940, EPI_ISL_1166941, EPI_ISL_1166942, EPI_ISL_1166943, EPI_ISL_1166944, EPI_ISL_1166945, EPI_ISL_1166946, EPI_ISL_1166947, EPI_ISL_1166948, EPI_ISL_1166949, EPI_ISL_1166950, EPI_ISL_1166951, EPI_ISL_1166952, EPI_ISL_1166953, EPI_ISL_1166954, EPI_ISL_1166955, EPI_ISL_1166956, EPI_ISL_1166957, EPI_ISL_1166958, EPI_ISL_1166959, EPI_ISL_1166960, EPI_ISL_1166961, EPI_ISL_1166962, EPI_ISL_1166963, EPI_ISL_1166964, EPI_ISL_1166965, EPI_ISL_1166966, EPI_ISL_1166967, EPI_ISL_1166968, EPI_ISL_1166969, EPI_ISL_1166970, EPI_ISL_1166971, EPI_ISL_1166972, EPI_ISL_1166973, EPI_ISL_1166974, EPI_ISL_1166975, EPI_ISL_1166976, EPI_ISL_1166977, EPI_ISL_1166978, EPI_ISL_1166979, EPI_ISL_1166980, EPI_ISL_1166981, EPI_ISL_1166982, EPI_ISL_1166983, EPI_ISL_1166984, EPI_ISL_1166985, EPI_ISL_1166986, EPI_ISL_1166987, EPI_ISL_1166988, EPI_ISL_1166989, EPI_ISL_1166990, EPI_ISL_1166991, EPI_ISL_1166992, EPI_ISL_1166993, EPI_ISL_1166994, EPI_ISL_1166995, EPI_ISL_1166996, EPI_ISL_1166997, EPI_ISL_1166998, EPI_ISL_1166999, EPI_ISL_1167000, EPI_ISL_1167001, EPI_ISL_1167002, EPI_ISL_1167003, EPI_ISL_1167004, EPI_ISL_1167005, EPI_ISL_1167006, EPI_ISL_1167007, EPI_ISL_1167008, EPI_ISL_1167009, EPI_ISL_1167011, EPI_ISL_1167012, EPI_ISL_1167013, EPI_ISL_1167014, EPI_ISL_1167015, EPI_ISL_1167016, EPI_ISL_1167017, EPI_ISL_1167018, EPI_ISL_1167019, EPI_ISL_1167020, EPI_ISL_1167021 | | | |
| see above | Hopital | National Reference Center for Viruses of Respiratory Infections, Institut Pasteur, Paris | Marion Barbet, Sylvie Behillil, Méline Bizard, Angela Brisebarre, Camille Capel, Etienne Simon-Lorière, Vincent Enouf, Maud Vanpeene, Sylvie van der Werf,Combe Patrice |
| EPI_ISL_1167128, EPI_ISL_1167129, EPI_ISL_1167130, EPI_ISL_1167131, EPI_ISL_1167132, EPI_ISL_1167133, EPI_ISL_1167134, EPI_ISL_1167136, EPI_ISL_1167137, EPI_ISL_1167142, EPI_ISL_1167143, EPI_ISL_1167144, EPI_ISL_1167151, EPI_ISL_1167152, EPI_ISL_1167153, EPI_ISL_1167154, EPI_ISL_1167155, EPI_ISL_1167156, EPI_ISL_1167157, EPI_ISL_1167158, EPI_ISL_1167159, EPI_ISL_1167160, EPI_ISL_1167161, EPI_ISL_1167162, EPI_ISL_1167163, EPI_ISL_1167164, EPI_ISL_1167165, EPI_ISL_1167166, EPI_ISL_1167167, EPI_ISL_1167168, EPI_ISL_1167169, EPI_ISL_1167170, EPI_ISL_1167171, EPI_ISL_1167172, EPI_ISL_1167174, EPI_ISL_1167178, EPI_ISL_1167179, EPI_ISL_1167180, EPI_ISL_1167181, EPI_ISL_1167182, EPI_ISL_1167183, EPI_ISL_1167184 | | | |
| see above | Iressef Genomics lab | L'institut de Recherche en Santé, de Surveillance Épidémiologique et de Formation (IRESSEF) | Souleymane MBOUP, Abdou PADANE, Abdoulie KANTEH ,Abdul Karim SESAY , Khadim GUEYE, Papa Alassane DIAW, Birahim Piere NDIAYE, Barada CISSE, Aminata MBOUP, Moustapha MBOW, Ndeye Coumba Toure KANE, Nafisatou LEYE, Gora LO, Ambroise AHOUIDI , Astou Gaye GAYE, Aminata DIA, Yacine DIA |
| EPI_ISL_1168191 | Hopital | National Reference Center for Viruses of Respiratory Infections, Institut Pasteur, Paris | Marion Barbet, Sylvie Behillil, Méline Bizard, Angela Brisebarre, Camille Capel, Etienne Simon-Lorière, Vincent Enouf, Maud Vanpeene, Sylvie van der Werf |
| EPI_ISL_1168192 | Hopital | National Reference Center for Viruses of Respiratory Infections, Institut Pasteur, Paris | Marion Barbet, Sylvie Behillil, Méline Bizard, Angela Brisebarre, Camille Capel, Etienne Simon-Lorière, Vincent Enouf, Maud Vanpeene, Sylvie van der Werf,Ramirez José-Manuel |
| EPI_ISL_1168193, EPI_ISL_1168194 | Hopital | National Reference Center for Viruses of Respiratory Infections, Institut Pasteur, Paris | Marion Barbet, Sylvie Behillil, Méline Bizard, Angela Brisebarre, Camille Capel, Etienne Simon-Lorière, Vincent Enouf, Maud Vanpeene, Sylvie van der Werf,Hermann CéCile |
| EPI_ISL_1168195, EPI_ISL_1168197, EPI_ISL_1168198, EPI_ISL_1168199 | Labo Analyses Med | National Reference Center for Viruses of Respiratory Infections, Institut Pasteur, Paris | Marion Barbet, Sylvie Behillil, Méline Bizard, Angela Brisebarre, Camille Capel, Etienne Simon-Lorière, Vincent Enouf, Maud Vanpeene, Sylvie van der Werf,Lecorche |
| EPI_ISL_1168200, EPI_ISL_1168201 | Labo Analyses Med | National Reference Center for Viruses of Respiratory Infections, Institut Pasteur, Paris | Marion Barbet, Sylvie Behillil, Méline Bizard, Angela Brisebarre, Camille Capel, Etienne Simon-Lorière, Vincent Enouf, Maud Vanpeene, Sylvie van der Werf,Le Vicky |
| EPI_ISL_1168202 | Hopital | National Reference Center for Viruses of Respiratory Infections, Institut Pasteur, Paris | Marion Barbet, Sylvie Behillil, Méline Bizard, Angela Brisebarre, Camille Capel, Etienne Simon-Lorière, Vincent Enouf, Maud Vanpeene, Sylvie van der Werf,Martres Pascale |
| EPI_ISL_1168203 | Armies | National Reference Center for Viruses of Respiratory Infections, Institut Pasteur, Paris | Marion Barbet, Sylvie Behillil, Méline Bizard, Angela Brisebarre, Camille Capel, Etienne Simon-Lorière, Vincent Enouf, Maud Vanpeene, Sylvie van der Werf,Desroches Marine |
| EPI_ISL_1168209, EPI_ISL_1168210, EPI_ISL_1168211 | Hopital | National Reference Center for Viruses of Respiratory Infections, Institut Pasteur, Paris | Marion Barbet, Sylvie Behillil, Méline Bizard, Angela Brisebarre, Camille Capel, Etienne Simon-Lorière, Vincent Enouf, Maud Vanpeene, Sylvie van der Werf,Martres Pascale |
| EPI_ISL_1168212, EPI_ISL_1168216, EPI_ISL_1168217 | Armies | National Reference Center for Viruses of Respiratory Infections, Institut Pasteur, Paris | Marion Barbet, Sylvie Behillil, Méline Bizard, Angela Brisebarre, Camille Capel, Etienne Simon-Lorière, Vincent Enouf, Maud Vanpeene, Sylvie van der Werf,Desroches Marine |
| EPI_ISL_1168219, EPI_ISL_1168220 | Hopital | National Reference Center for Viruses of Respiratory Infections, Institut Pasteur, Paris | Marion Barbet, Sylvie Behillil, Méline Bizard, Angela Brisebarre, Camille Capel, Etienne Simon-Lorière, Vincent Enouf, Maud Vanpeene, Sylvie van der Werf,Capron Nicolas |
| EPI_ISL_1168452, EPI_ISL_1168453, EPI_ISL_1168454, EPI_ISL_1168455, EPI_ISL_1168456, EPI_ISL_1168457, EPI_ISL_1168458, EPI_ISL_1168459, EPI_ISL_1168460, EPI_ISL_1168462, EPI_ISL_1168463, EPI_ISL_1168464, EPI_ISL_1168465, EPI_ISL_1168466, EPI_ISL_1168467, EPI_ISL_1168468, EPI_ISL_1168469, EPI_ISL_1168470, EPI_ISL_1168471, EPI_ISL_1168472, EPI_ISL_1168473, EPI_ISL_1168474, EPI_ISL_1168475, EPI_ISL_1168476, EPI_ISL_1168477, EPI_ISL_1168478, EPI_ISL_1168479, EPI_ISL_1168480, EPI_ISL_1168482, EPI_ISL_1168483, EPI_ISL_1168484, EPI_ISL_1168485, EPI_ISL_1168486, EPI_ISL_1168487, EPI_ISL_1168489, EPI_ISL_1168490, EPI_ISL_1168491, EPI_ISL_1168492, EPI_ISL_1168493, EPI_ISL_1168495, EPI_ISL_1168496, EPI_ISL_1168498, EPI_ISL_1168499, EPI_ISL_1168500, EPI_ISL_1168501, EPI_ISL_1168502, EPI_ISL_1168503, EPI_ISL_1168504, EPI_ISL_1168505, EPI_ISL_1168506, EPI_ISL_1168507, EPI_ISL_1168509, EPI_ISL_1168510, EPI_ISL_1168511, EPI_ISL_1168512, EPI_ISL_1168513, EPI_ISL_1168514, EPI_ISL_1168515, EPI_ISL_1168516, EPI_ISL_1168517, EPI_ISL_1168518, EPI_ISL_1168519, EPI_ISL_1168520, EPI_ISL_1168521, EPI_ISL_1168522, EPI_ISL_1168523, EPI_ISL_1168524, EPI_ISL_1168525, EPI_ISL_1168526, EPI_ISL_1168527, EPI_ISL_1168528, EPI_ISL_1168530, EPI_ISL_1168531, EPI_ISL_1168537, EPI_ISL_1168538, EPI_ISL_1168539, EPI_ISL_1168540, EPI_ISL_1168541, EPI_ISL_1168542, EPI_ISL_1168543, EPI_ISL_1168544, EPI_ISL_1168545, EPI_ISL_1168546, EPI_ISL_1168547, EPI_ISL_1168549, EPI_ISL_1168550, EPI_ISL_1168551, EPI_ISL_1168553, EPI_ISL_1168557, EPI_ISL_1168558, EPI_ISL_1168560, EPI_ISL_1168561, EPI_ISL_1168562, EPI_ISL_1168563, EPI_ISL_1168565, EPI_ISL_1168566, EPI_ISL_1168567, EPI_ISL_1168569, EPI_ISL_1168570, EPI_ISL_1168571, EPI_ISL_1168572, EPI_ISL_1168573, EPI_ISL_1168574, EPI_ISL_1168583, EPI_ISL_1168590, EPI_ISL_1168596, EPI_ISL_1168597, EPI_ISL_1168599, EPI_ISL_1168604, EPI_ISL_1168607, EPI_ISL_1168608, EPI_ISL_1168610, EPI_ISL_1168612, EPI_ISL_1168613, EPI_ISL_1168614, EPI_ISL_1168616, EPI_ISL_1168617, EPI_ISL_1168618, EPI_ISL_1168624, EPI_ISL_1168625, EPI_ISL_1168626, EPI_ISL_1168627, EPI_ISL_1168630, EPI_ISL_1168631, EPI_ISL_1168632, EPI_ISL_1168633 | | | |
| see above | Instituto de Diagnostico y Referencia Epidemiologicos INDRE_RNLSP | Instituto de Diagnostico y Referencia Epidemiologicos (INDRE) | Claudia Wong-Arambula, Abril Rodriguez-Maldonado, Vanessa Rivero-Arredondo, Ariadna Medina-Benitez, Joaquin Quiroz-Mercado,David Fragoso-Fonseca, Sergio Rangel-Guerrero, Natividad Cruz-Ortiz, Tatiana Nunez-Garcia, Gisela Barrera-Badillo, Lucia Hernandez-Rivas, Irma Lopez-Martinez, Ernesto Ramirez-Gonzalez. |
| EPI_ISL_1168655, EPI_ISL_1168657, EPI_ISL_1168676 | Centers for Disease Control and Prevention, Dengue Branch | Centers for Disease Control and Prevention, Dengue Branch | Gilberto A. Santiago, Glenda Gonzalez, Betzabel Flores, Keyla Charriez, Gabriela Paz-Bailey, Jorge L. Munoz-Jordan |
| EPI_ISL_1169046 | National Institute of Health Research and Development | National Institute of Health Research and Development | Hana Apsari Pawestri, Subangkit, Kartika Dewi Puspa, Arie Ardiansyah Nugraha, Hartanti Dian Ikawati, Krisna Nur Andriana Pangesti, Yuni Rukminiati, Ririn Ramadhany, Agustiningsih, Kindi Adam, Holy Arif Wibowo, Triyani Soekarso, Ni Ketut Susilarini, Nurika Hariastuti, Ully Alfi Nikmah, Reni Herman, Nike Susanti, Herna, Tati Febriyanti, Natalie Laurencia Kipuw, Fauzul Muna, Irene Lorinda Indalao, Nelly Puspandari, Vivi Setiawaty. |
| EPI_ISL_1169049 | RSUP Adam Malik Medan Sumatera Utara | National Institute of Health Research and Development | Vivi Setiawaty, Hana Apsari Pawestri, Subangkit, Kartika Dewi Puspa, Arie Ardiansyah Nugraha, Hartanti Dian Ikawati, Nelly Puspandari |
| EPI_ISL_1169050 | National Institute of Health Research and Development | National Institute of Health Research and Development | Hartanti Dian Ikawati, Hana Apsari Pawestri, Subangkit, Kartika Dewi Puspa, Arie Ardiansyah Nugraha, risna Nur Andriana Pangesti, Yuni Rukminiati, Ririn Ramadhany, Agustiningsih, Kindi Adam, Holy Arif Wibowo, Triyani Soekarso, Ni Ketut Susilarini, Nurika Hariastuti, Ully Alfi Nikmah, Reni Herman, Nike Susanti, Herna, Tati Febriyanti, Natalie Laurencia Kipuw, Fauzul Muna, Irene Lorinda Indalao, Nelly Puspandari, Vivi Setiawaty. |
| EPI_ISL_1169052, EPI_ISL_1169053, EPI_ISL_1169054, EPI_ISL_1169055 | National Institute of Health Research and Development | National Institute of Health Research and Development | Subangkit, Hana Apsari Pawestri, Kartika Dewi Puspa, Arie Ardiansyah Nugraha, Hartanti Dian Ikawati, Krisna Nur Andriana Pangesti, Yuni Rukminiati, Ririn Ramadhany, Agustiningsih, Kindi Adam, Holy Arif Wibowo, Triyani Soekarso, Ni Ketut Susilarini, Nurika Hariastuti, Ully Alfi Nikmah, Reni Herman, Nike Susanti, Herna, Tati Febriyanti, Natalie Laurencia Kipuw, Fauzul Muna, Irene Lorinda Indalao, Nelly Puspandari, Vivi Setiawaty. |
| EPI_ISL_1169056 | National Institute of Health Research and Development | National Institute of Health Research and Development | Kartika Dewi Puspa, Hana Apsari Pawestri, Subangkit, Arie Ardiansyah Nugraha, Hartanti Dian Ikawati, Krisna Nur Andriana Pangesti, Yuni Rukminiati, Ririn Ramadhany, Agustiningsih, Kindi Adam, Holy Arif Wibowo, Triyani Soekarso, Ni Ketut Susilarini, Nurika Hariastuti, Ully Alfi Nikmah, Reni Herman, Nike Susanti, Herna, Tati Febriyanti, Natalie Laurencia Kipuw, Fauzul Muna, Irene Lorinda Indalao, Nelly Puspandari, Vivi Setiawaty. |
| EPI_ISL_1169057, EPI_ISL_1169058 | National Institute of Health Research and Development | National Institute of Health Research and Development | Arie Ardiansyah Nugraha, Kartika Dewi Puspa, Hana Apsari Pawestri, Subangkit, , Hartanti Dian Ikawati, Krisna Nur Andriana Pangesti, Yuni Rukminiati, Ririn Ramadhany, Agustiningsih, Kindi Adam, Holy Arif Wibowo, Triyani Soekarso, Ni Ketut Susilarini, Nurika Hariastuti, Ully Alfi Nikmah, Reni Herman, Nike Susanti, Herna, Tati Febriyanti, Natalie Laurencia Kipuw, Fauzul Muna, Irene Lorinda Indalao, Nelly Puspandari, Vivi Setiawaty. |
| EPI_ISL_1169059, EPI_ISL_1169060 | National Institute of Health Research and Development | National Institute of Health Research and Development | Hartanti Dian Ikawati, Hana Apsari Pawestri, Subangkit, Kartika Dewi Puspa, Arie Ardiansyah Nugraha, risna Nur Andriana Pangesti, Yuni Rukminiati, Ririn Ramadhany, Agustiningsih, Kindi Adam, Holy Arif Wibowo, Triyani Soekarso, Ni Ketut Susilarini, Nurika Hariastuti, Ully Alfi Nikmah, Reni Herman, Nike Susanti, Herna, Tati Febriyanti, Natalie Laurencia Kipuw, Fauzul Muna, Irene Lorinda Indalao, Nelly Puspandari, Vivi Setiawaty. |
| EPI_ISL_1169894, EPI_ISL_1169895 | Istituto Zooprofilattico Sperimentale Umbria e Marche "Togo Rosati" | Istituto Zooprofilattico Sperimentale dell'Abruzzo e Molise "G. Caporale" | Biagetti M., Giammarioli M., Lorusso A, Marcacci M, Di Domenico M, Ancora M, Curini V, Mangone I, Rinaldi A, Scialabba S, Di Pasquale A, Cammà C, Puglia I, Calistri P, Savini G |
| EPI_ISL_1170948 | National Institute of Health Research and Development | National Institute of Health Research and Development | Ririn Ramadhany, Yuni Rukminiati, Agustiningsih, Kindi Adam, Holy Arif Wibowo, Hana Apsari Pawestri, Subangkit; Kartika Dewi Puspa, Arie Ardiansyah Nugraha, Hartanti Dian Ikawati, Krisna Nur Andriana Pangesti, Ni Ketut Susilarini, Nur Ika Hariastuti, Uly Alfi Nikmah, Mursinah, Reni Herman, Nike Susanti, Herna, Tati Febriyanti, Natalie Laurencia Kipuw, Fauzul Muna, Aulia Rizki, Nelly Puspandari, Vivi Setiawaty |

| | | | |
|---|---|---|---|
| EPI_ISL_1171619 | IAL Regional de Presidente Prudente | Instituto Adolfo Lutz, Interdiciplinary Procedures Center, Strategic Laboratory | Claudio Tavares Sacchi, Claudia Regina Gonçalves, Erica Valessa Ramos Gomes, Karoline Rodrigues Campos |
| EPI_ISL_1171626, EPI_ISL_1171628, EPI_ISL_1171634, EPI_ISL_1171638 | IAL Regional de Santos | Instituto Adolfo Lutz, Interdiciplinary Procedures Center, Strategic Laboratory | Claudio Tavares Sacchi, Claudia Regina Gonçalves, Erica Valessa Ramos Gomes, Karoline Rodrigues Campos, Caio Vinicius Dias Lopes |
| EPI_ISL_1171641, EPI_ISL_1171642, EPI_ISL_1171643, EPI_ISL_1171644, EPI_ISL_1171645, EPI_ISL_1171646, EPI_ISL_1171647 | IAL Regional de Marilia | Instituto Adolfo Lutz, Interdiciplinary Procedures Center, Strategic Laboratory | Claudio Tavares Sacchi, Claudia Regina Gonçalves, Erica Valessa Ramos Gomes, Karoline Rodrigues Campos, Caio Vinicius Dias Lopes |
| EPI_ISL_1171651, EPI_ISL_1171652, EPI_ISL_1171653, EPI_ISL_1171654, EPI_ISL_1171655, EPI_ISL_1171656, EPI_ISL_1171657, EPI_ISL_1171658, EPI_ISL_1171659, EPI_ISL_1171660, EPI_ISL_1171661, EPI_ISL_1171662, EPI_ISL_1171663, EPI_ISL_1171664, EPI_ISL_1171665, EPI_ISL_1171666, EPI_ISL_1171667, EPI_ISL_1171671, EPI_ISL_1171672, EPI_ISL_1171673, EPI_ISL_1171674 | | | |
| see above | IAL Regional de Presidente Prudente | Instituto Adolfo Lutz, Interdiciplinary Procedures Center, Strategic Laboratory | Claudio Tavares Sacchi, Claudia Regina Gonçalves, Erica Valessa Ramos Gomes, Karoline Rodrigues Campos, Caio Vinicius Dias Lopes |
| EPI_ISL_1171675, EPI_ISL_1171676, EPI_ISL_1171677, EPI_ISL_1171678, EPI_ISL_1171679, EPI_ISL_1171680, EPI_ISL_1171681, EPI_ISL_1171682, EPI_ISL_1171683, EPI_ISL_1171684, EPI_ISL_1171685, EPI_ISL_1171686, EPI_ISL_1171687, EPI_ISL_1171688, EPI_ISL_1171689, EPI_ISL_1171690, EPI_ISL_1171691, EPI_ISL_1171692, EPI_ISL_1171693, EPI_ISL_1171694, EPI_ISL_1171695, EPI_ISL_1171696, EPI_ISL_1171697, EPI_ISL_1171698, EPI_ISL_1171699, EPI_ISL_1171700, EPI_ISL_1171701, EPI_ISL_1171702, EPI_ISL_1171703, EPI_ISL_1171704, EPI_ISL_1171705 | | | |
| see above | UOC LAB ANALISI PO CARDARELLI CAMPOBASSO | Istituto Zooprofilattico Sperimentale dell'Abruzzo e Molise "G. Caporale" | Scutellà M, Niro G, Felice V. Lorusso A, Marcacci M, Di Domenico M, Ancora M, Curini V, Mangone I, Rinaldi A, Scialabba S, Di Pasquale A, Cammà C, Puglia I, Calistri P, Savini G |
| EPI_ISL_1173508 | Institut de Virologie du CHU de Strasbourg | Institut de Virologie du CHU de Strasbourg | Fafi-Kremer Samira, Gallais Floriane, Gantner Pierre, Laugel Elodie, Solis Morgane, Velay Aurélie, Wendling Maire-Josée |
| EPI_ISL_1173766 | Imelda Hospital | Imelda Hospital | Johan Frans, Hanne Valgaeren, Dagmar Obbels |
| EPI_ISL_1176297, EPI_ISL_1176305, EPI_ISL_1176307 | Institut de Virologie du CHU de Strasbourg | Institut de Virologie du CHU de Strasbourg | Fafi-Kremer Samira, Gallais Floriane, Gantner Pierre, Laugel Elodie, Solis Morgane, Velay Aurélie, Wendling Maire-Josée |
| EPI_ISL_1180210, EPI_ISL_1180211, EPI_ISL_1180213, EPI_ISL_1180214, EPI_ISL_1180216, EPI_ISL_1180217, EPI_ISL_1180219, EPI_ISL_1180221, EPI_ISL_1180223, EPI_ISL_1180225, EPI_ISL_1180227 | | | |
| see above | Althaia. Xarxa Assistencial Universitària de Manresa | IrsiCaixa - Can Ruti CovidSeq | Marc Noguera-Julian, Mariona Parera, Maria Casadellà, Pilar Armengol, Francesc Catala-Moll, Roger Paredes, Bonaventura Clotet Gloria Trujillo, Rafel Perez Vidal, Jaume Trape Pujol, Carolina Gonzalez Fernandez, Roger Paredes, Eulalia Grau, Bonaventura Clotet |
| EPI_ISL_1180715, EPI_ISL_1180716, EPI_ISL_1180718, EPI_ISL_1180719, EPI_ISL_1180720, EPI_ISL_1180722, EPI_ISL_1180724, EPI_ISL_1180725, EPI_ISL_1180726, EPI_ISL_1180727, EPI_ISL_1180728, EPI_ISL_1180729, EPI_ISL_1180730, EPI_ISL_1180731, EPI_ISL_1180732, EPI_ISL_1180733, EPI_ISL_1180734, EPI_ISL_1180735, EPI_ISL_1180736, EPI_ISL_1180737, EPI_ISL_1180738, EPI_ISL_1180739, EPI_ISL_1180740, EPI_ISL_1180741, EPI_ISL_1180742, EPI_ISL_1180743, EPI_ISL_1180744, EPI_ISL_1180747, EPI_ISL_1180748, EPI_ISL_1180749 | | | |
| see above | HG Pharma GmbH | Bergthaler laboratory, CeMM Research Center for Molecular Medicine of the Austrian Academy of Sciences | Lukas Endler, Anna Schedl, Fabian Amman, Thomas Penz, Benedikt Agerer, Maelle Le Moing, Michael Schuster, Bekir Erguner, Jan Laine, Martin Senekowitsch, Christoph Bock, Andreas Bergthaler |
| EPI_ISL_1180752, EPI_ISL_1180753, EPI_ISL_1180754, EPI_ISL_1180755, EPI_ISL_1180756, EPI_ISL_1180757, EPI_ISL_1180758, EPI_ISL_1180759, EPI_ISL_1180760, EPI_ISL_1180763, EPI_ISL_1180764, EPI_ISL_1180765, EPI_ISL_1180766, EPI_ISL_1180768, EPI_ISL_1180769, EPI_ISL_1180771, EPI_ISL_1180772, EPI_ISL_1180773, EPI_ISL_1180774, EPI_ISL_1180777, EPI_ISL_1180778, EPI_ISL_1180779, EPI_ISL_1180781, EPI_ISL_1180782, EPI_ISL_1180783, EPI_ISL_1180784, EPI_ISL_1180785, EPI_ISL_1180786, EPI_ISL_1180787, EPI_ISL_1180788, EPI_ISL_1180789, EPI_ISL_1180790, EPI_ISL_1180791, EPI_ISL_1180792, EPI_ISL_1180794, EPI_ISL_1180795, EPI_ISL_1180796, EPI_ISL_1180797, EPI_ISL_1180798, EPI_ISL_1180799, EPI_ISL_1180800, EPI_ISL_1180801, EPI_ISL_1180802, EPI_ISL_1180803, EPI_ISL_1180804, EPI_ISL_1180805, EPI_ISL_1180806, EPI_ISL_1180807, EPI_ISL_1180808, EPI_ISL_1180809, EPI_ISL_1180810, EPI_ISL_1180811, EPI_ISL_1180812, EPI_ISL_1180813, EPI_ISL_1180814, EPI_ISL_1180818, EPI_ISL_1180819, EPI_ISL_1180820, EPI_ISL_1180821, EPI_ISL_1180822, EPI_ISL_1180823, EPI_ISL_1180824, EPI_ISL_1180825, EPI_ISL_1180826, EPI_ISL_1180827, EPI_ISL_1180828, EPI_ISL_1180830 | | | |
| see above | Austrian Agency for Health and Food Safety (AGES) | Bergthaler laboratory, CeMM Research Center for Molecular Medicine of the Austrian Academy of Sciences | Lukas Endler, Anna Schedl, Fabian Amman, Thomas Penz, Benedikt Agerer, Maelle Le Moing, Michael Schuster, Bekir Erguner, Jan Laine, Martin Senekowitsch, Christoph Bock, Andreas Bergthaler |
| EPI_ISL_1180832, EPI_ISL_1180834, EPI_ISL_1180835, EPI_ISL_1180836, EPI_ISL_1180837 | Klinikum Wels-Grieskirchen | Bergthaler laboratory, CeMM Research Center for Molecular Medicine of the Austrian Academy of Sciences | Lukas Endler, Anna Schedl, Fabian Amman, Thomas Penz, Benedikt Agerer, Maelle Le Moing, Michael Schuster, Bekir Erguner, Jan Laine, Martin Senekowitsch, Christoph Bock, Andreas Bergthaler |
| EPI_ISL_1180840, EPI_ISL_1180841, EPI_ISL_1180842, EPI_ISL_1180843, EPI_ISL_1180844, EPI_ISL_1180845, EPI_ISL_1180846, EPI_ISL_1180847, EPI_ISL_1180848, EPI_ISL_1180849, EPI_ISL_1180850, EPI_ISL_1180851, EPI_ISL_1180852, EPI_ISL_1180853, EPI_ISL_1180854, EPI_ISL_1180855, EPI_ISL_1180857, EPI_ISL_1180858, EPI_ISL_1180859, EPI_ISL_1180860, EPI_ISL_1180861, EPI_ISL_1180862, EPI_ISL_1180863, EPI_ISL_1180864, EPI_ISL_1180865, EPI_ISL_1180866, EPI_ISL_1180868, EPI_ISL_1180869, EPI_ISL_1180870, EPI_ISL_1180871, EPI_ISL_1180872, EPI_ISL_1180873, EPI_ISL_1180874, EPI_ISL_1180875, EPI_ISL_1180876, EPI_ISL_1180877, EPI_ISL_1180878, EPI_ISL_1180879, EPI_ISL_1180880, EPI_ISL_1180885 | | | |
| see above | Department of Laboratory Medicine, Division of Clinical Virology, University of Medicine, Vienna | Bergthaler laboratory, CeMM Research Center for Molecular Medicine of the Austrian Academy of Sciences | Lukas Endler, Anna Schedl, Fabian Amman, Thomas Penz, Benedikt Agerer, Maelle Le Moing, Michael Schuster, Bekir Erguner, Jan Laine, Martin Senekowitsch, Christoph Bock, Andreas Bergthaler |
| EPI_ISL_1180932, EPI_ISL_1180933, EPI_ISL_1180934 | Austrian Agency for Health and Food Safety (AGES) | Bergthaler laboratory, CeMM Research Center for Molecular Medicine of the Austrian Academy of Sciences | Lukas Endler, Anna Schedl, Fabian Amman, Thomas Penz, Benedikt Agerer, Maelle Le Moing, Michael Schuster, Bekir Erguner, Jan Laine, Martin Senekowitsch, Christoph Bock, Andreas Bergthaler |
| EPI_ISL_1181394 | Laboratorio Central de Saude Publica do Estado do Parana (LACEN-PR) | Laboratory of Respiratory Viruses and Measles, Oswaldo Cruz Institute, FIOCRUZ | Paola Resende, Luciana Appolinario, Fernando Motta, Anna Carolina Paixao, Ana Carolina Mendonca, Alice Sampaio Rocha, Renata Serrano Lopes, Maria do Carmo Debur, Irina Nastassja Riediger, Marilda Siqueira on behalf of the Fiocruz COVID-19 Genomic Surveillance Network |
| EPI_ISL_1182098, EPI_ISL_1182099, EPI_ISL_1182100, EPI_ISL_1182102 | Adolfo Lutz Institute Santos Regional Center Brazil | Retrovirus Laboratory Adolfo Lutz Institute | Gabriela Bastos Cabral, Giselle I S Lopez-Lopes, Cintia Ahagon, Audrey Cilli, Paula Morena Guimaraes, Igor Mohamed Hussein, Luis Brigido |
| EPI_ISL_1184113, EPI_ISL_1184114, EPI_ISL_1184135, EPI_ISL_1184136, EPI_ISL_1184137 | Rochester Regional Hospital | Multidrug-Resistant Organism Repository and Surveillance Network (MRSN) | Emil Lesho, Brendan Corey, Francois Lebreton, Ana C. Ong, Brett E. Swierczewski, Jason W. Bennett, Edward E. Walsh, and Patrick Mc Gann |
| EPI_ISL_1186114 | Hospital Queen Elizabeth | Institute for Medical Research, Infectious Disease Research Centre, National Institutes of Health, Ministry of Health Malaysia | Suppiah J, Kamel K, Mohd Zawawi Z, Ramly N, Robert F, Thayan R |
| EPI_ISL_1186115, EPI_ISL_1186199, EPI_ISL_1186205, EPI_ISL_1186206, EPI_ISL_1186207 | Institut de Virologie du CHU de Strasbourg | Institut de Virologie du CHU de Strasbourg | Fafi-Kremer Samira, Gallais Floriane, Gantner Pierre, Laugel Elodie, Solis Morgane, Velay Aurélie, Wendling Marie-Josée |
| EPI_ISL_1190749, EPI_ISL_1190750, EPI_ISL_1190751, EPI_ISL_1190752, EPI_ISL_1190753, EPI_ISL_1190754, EPI_ISL_1190755, EPI_ISL_1190756, EPI_ISL_1190757, EPI_ISL_1190758, EPI_ISL_1190759, EPI_ISL_1190760, EPI_ISL_1190763, EPI_ISL_1190764 | | | |
| see above | CREMER(Centre de Rechercherches sur les Maladies Emergentes et Ré-émergentes) | TransVIHMI(Recherches Translationnelles sur le VIH et les Maladies Infectieuses) | Celestin Godwe, Christelle Butel, Martin Maidadi Foudi, Laetitia Serrano, Dowbiss Meta Djomsi, Nicole Vidal, Esemu Livo, Rodrigue Kamga, Marie Amougou , Eitel Mpoudi Ngole,Ahidjo Ayouba, Marcel Tongo, Martine Peeters, Eric Delaporte |
| EPI_ISL_1190885, EPI_ISL_1190886, EPI_ISL_1190887, EPI_ISL_1190888 | North Estonia Medical Centre laboratory | 1. Laboratory of Communicable Diseases (Estonia); 2. Eurofins Genomics Europe Sequencing GmbH | Liidia Dotsenko et al. |
| EPI_ISL_1190889, EPI_ISL_1190890, EPI_ISL_1190891, EPI_ISL_1190892 | West Tallinn Central Hospital laboratory | 1. Laboratory of Communicable Diseases (Estonia); 2. Eurofins Genomics Europe Sequencing GmbH | Liidia Dotsenko et al. |
| EPI_ISL_1190893, EPI_ISL_1190894 | North Estonia Medical Centre laboratory | 1. Laboratory of Communicable Diseases (Estonia); 2. Eurofins Genomics Europe Sequencing GmbH | Liidia Dotsenko et al. |
| EPI_ISL_1190895, EPI_ISL_1190896 | West Tallinn Central Hospital laboratory | 1. Laboratory of Communicable Diseases (Estonia); 2. Eurofins Genomics Europe Sequencing GmbH | Liidia Dotsenko et al. |
| EPI_ISL_1190897 | North Estonia Medical Centre laboratory | 1. Laboratory of Communicable Diseases (Estonia); 2. Eurofins Genomics Europe Sequencing GmbH | Liidia Dotsenko et al. |
| EPI_ISL_1190898, EPI_ISL_1190899, EPI_ISL_1190900, EPI_ISL_1190901, EPI_ISL_1190902, EPI_ISL_1190903, EPI_ISL_1190904, EPI_ISL_1190905, EPI_ISL_1190906, EPI_ISL_1190907, EPI_ISL_1190908, EPI_ISL_1190909, EPI_ISL_1190910, EPI_ISL_1190911, EPI_ISL_1190912, EPI_ISL_1190913, EPI_ISL_1190914, EPI_ISL_1190915, EPI_ISL_1190916, EPI_ISL_1190917, EPI_ISL_1190918, EPI_ISL_1190919, EPI_ISL_1190920, EPI_ISL_1190921, EPI_ISL_1190922, EPI_ISL_1190923, EPI_ISL_1190924, EPI_ISL_1190925, EPI_ISL_1190926, EPI_ISL_1190927, EPI_ISL_1190928, EPI_ISL_1190929, EPI_ISL_1190930, EPI_ISL_1190931, | | | |

EPI_ISL_1190932, EPI_ISL_1190933

| | | | |
|---|---|---|---|
| see above | Synlab Eesti OÜ | 1. Laboratory of Communicable Diseases (Estonia); 2. Eurofins Genomics Europe Sequencing GmbH | Liidia Dotsenko et al. |
| EPI_ISL_1190934, EPI_ISL_1190935, EPI_ISL_1190936, EPI_ISL_1190937, EPI_ISL_1190938 | Laboratory of Communicable Diseases | 1. Laboratory of Communicable Diseases (Estonia); 2. Eurofins Genomics Europe Sequencing GmbH | Liidia Dotsenko et al. |
| EPI_ISL_1190939 | North Estonia Medical Centre laboratory | 1. Laboratory of Communicable Diseases (Estonia); 2. Eurofins Genomics Europe Sequencing GmbH | Liidia Dotsenko et al. |

EPI_ISL_1190940, EPI_ISL_1190941, EPI_ISL_1190942, EPI_ISL_1190943, EPI_ISL_1190944, EPI_ISL_1190945, EPI_ISL_1190946, EPI_ISL_1190947, EPI_ISL_1190948, EPI_ISL_1190949, EPI_ISL_1190950

| | | | |
|---|---|---|---|
| see above | Laboratory of Communicable Diseases | 1. Laboratory of Communicable Diseases (Estonia); 2. Eurofins Genomics Europe Sequencing GmbH | Liidia Dotsenko et al. |
| EPI_ISL_1190951, EPI_ISL_1190952, EPI_ISL_1190953, EPI_ISL_1190954 | West Tallinn Central Hospital laboratory | 1. Laboratory of Communicable Diseases (Estonia); 2. Eurofins Genomics Europe Sequencing GmbH | Liidia Dotsenko et al. |
| EPI_ISL_1190955 | Laboratory of Communicable Diseases | 1. Laboratory of Communicable Diseases (Estonia); 2. Eurofins Genomics Europe Sequencing GmbH | Liidia Dotsenko et al. |
| EPI_ISL_1190956, EPI_ISL_1190957, EPI_ISL_1190958 | West Tallinn Central Hospital laboratory | 1. Laboratory of Communicable Diseases (Estonia); 2. Eurofins Genomics Europe Sequencing GmbH | Liidia Dotsenko et al. |
| EPI_ISL_1190959, EPI_ISL_1190960, EPI_ISL_1190961, EPI_ISL_1190962, EPI_ISL_1190963 | North Estonia Medical Centre laboratory | 1. Laboratory of Communicable Diseases (Estonia); 2. Eurofins Genomics Europe Sequencing GmbH | Liidia Dotsenko et al. |
| EPI_ISL_1191080, EPI_ISL_1191081, EPI_ISL_1191082, EPI_ISL_1191083 | AZ St.Jan Brugge | AZ SINT-JAN BRUGGE | Marijke Reynders, Patrick Descheemaeker, Laurien Hoornaert, Thomas Van Landschoot, Jorn Hellemans |
| EPI_ISL_1191084, EPI_ISL_1191085 | AZ Zeno | AZ SINT-JAN BRUGGE | Marijke Reynders, Patrick Descheemaeker, Laurien Hoornaert, Thomas Van Landschoot, Jorn Hellemans |
| EPI_ISL_1191086, EPI_ISL_1191087, EPI_ISL_1191088, EPI_ISL_1191089, EPI_ISL_1191090, EPI_ISL_1191091 | AZ St.Jan Brugge | AZ SINT-JAN BRUGGE | Marijke Reynders, Patrick Descheemaeker, Laurien Hoornaert, Thomas Van Landschoot, Jorn Hellemans |
| EPI_ISL_1191092, EPI_ISL_1191093 | AZ Zeno | AZ SINT-JAN BRUGGE | Marijke Reynders, Patrick Descheemaeker, Laurien Hoornaert, Thomas Van Landschoot, Jorn Hellemans |
| EPI_ISL_1191142, EPI_ISL_1191143, EPI_ISL_1191144, EPI_ISL_1191145 | Laboratorio Biologia Molecolare Sars Cov2 - UOC Laboratorio Analisi - Servizio Medicina di Laboratorio, Ospedale "San Francesco" - ATS-ASSL Nuoro | Laboratorio specialistico UOC Ematologia - Ospedale "San Francesco" - ATS-ASSL Nuoro | Piras Giovanna, Asproni Rosanna,,Fiamma Maura, Monne Maria Itria, Palmas Angelo Domenico, Lo Maglio Iana, Mameli Giuseppe |
| EPI_ISL_1191626 | AZ St.Jan | AZ SINT-JAN BRUGGE | Marijke Reynders, Patrick Descheemaeker, Laurien Hoornaert, Thomas Van Landschoot, Jorn Hellemans |
| EPI_ISL_1191627, EPI_ISL_1191628 | AZ Zeno | AZ SINT-JAN BRUGGE | Marijke Reynders, Patrick Descheemaeker, Laurien Hoornaert, Thomas Van Landschoot, Jorn Hellemans |
| EPI_ISL_1194709 | AZ St.jan Brugge | AZ SINT-JAN BRUGGE | Marijke Reynders, Patrick Descheemaeker, Laurien Hoornaert, Thomas Van Landschoot, Jorn Hellemans |
| EPI_ISL_1194710 | AZ Zeno | AZ SINT-JAN BRUGGE | Marijke Reynders, Patrick Descheemaeker, Laurien Hoornaert, Thomas Van Landschoot, Jorn Hellemans |
| EPI_ISL_1194711, EPI_ISL_1194712, EPI_ISL_1194713, EPI_ISL_1194714 | AZ St.jan Brugge | AZ SINT-JAN BRUGGE | Marijke Reynders, Patrick Descheemaeker, Laurien Hoornaert, Thomas Van Landschoot, Jorn Hellemans |
| EPI_ISL_1195962, EPI_ISL_1195963 | Microbiology and Virology Unit, Florence Careggi University Hospital | Microbiology and Virology Unit, Florence Careggi University Hospital | Vincenzo Di Pilato, Marco Coppi, Fabio Morecchiato, Noemi Aiezza, Ilaria Baccani, Nicla Giovacchini, Alberto Antonelli, Emanuele Gori, Gian Maria Rossolini |
| EPI_ISL_1196279, EPI_ISL_1196280 | Universitas Indonesia Hospital | Faculty of Medicine, Universitas Indonesia | Fadilah, Rafika Indah Paramita, Linda Erlina, Khaerunissa Anbar Istiadi, Ardiana Kusumaningrum, Dimas Seto, Astuti Giantini, Doni Usman, Putra Mahanaim Tampubolon, Aulia Reski Widyaningrum, Ferdi Anda Sitepu, Irandi Putra Pratomo, Dwiana Ocviyanti, Budi Wiweko, Anis Karuniawati, Ari Fahrial Syam, Badriul Hegar |
| EPI_ISL_1196295 | UBS Otacilio Firmino Lopes | Instituto Adolfo Lutz, Interdiciplinary Procedures Center, Strategic Laboratory | Claudio Tavares Sacchi, Claudia Regina Gonçalves, Erica Valessa Ramos Gomes, Karoline Rodrigues Campos, Caio Vinicius Dias Lopes |
| EPI_ISL_1196296 | Centro de Saude II Dr. Jose Paione Mococa | Instituto Adolfo Lutz, Interdiciplinary Procedures Center, Strategic Laboratory | Claudio Tavares Sacchi, Claudia Regina Gonçalves, Erica Valessa Ramos Gomes, Karoline Rodrigues Campos, Caio Vinicius Dias Lopes |
| EPI_ISL_1196297, EPI_ISL_1196298, EPI_ISL_1196299, EPI_ISL_1196300, EPI_ISL_1196301, EPI_ISL_1196302 | IAL Regional de Marilia | Instituto Adolfo Lutz, Interdiciplinary Procedures Center, Strategic Laboratory | Claudio Tavares Sacchi, Claudia Regina Gonçalves, Erica Valessa Ramos Gomes, Karoline Rodrigues Campos, Caio Vinicius Dias Lopes |
| EPI_ISL_1196303, EPI_ISL_1196304 | Universitas Indonesia Hospital | Faculty of Medicine, Universitas Indonesia | Fadilah, Rafika Indah Paramita, Linda Erlina, Khaerunissa Anbar Istiadi, Ardiana Kusumaningrum, Dimas Seto, Astuti Giantini, Doni Usman, Putra Mahanaim Tampubolon, Aulia Reski Widyaningrum, Ferdi Anda Sitepu, Irandi Putra Pratomo, Dwiana Ocviyanti, Budi Wiweko, Anis Karuniawati, Ari Fahrial Syam, Badriul Hegar |
| EPI_ISL_1196431 | HOSPITAL MEXICO | Inciensa, Instituto Costarricense de Investigación y Enseñanza en Nutrición y Salud | Francisco Duarte, Hebleen Porras, Claudio Soto-Garita, Estela Cordero, Adriana Godínez, Melany Calderón & Teresita Somogyi |
| EPI_ISL_1196766, EPI_ISL_1196767 | Institut de Virologie du CHU de Strasbourg | Institut de Virologie du CHU de Strasbourg | Fafi-Kremer Samira, Gallais Floriane, Gantner Pierre, Laugel Elodie, Solis Morgane, Velay Aurélie, Wendling Marie-Josée |
| EPI_ISL_1199636, EPI_ISL_1200086 | Unità Operativa di Microbiologia e Virologia, Ospedale Civico e Di Cristina, ARNAS, Palermo, Italy | Tumor Immunology Unit, Department of Health Sciences, University of Palermo School of Medicine; Section of Microbiology, University of Palermo School of Medicine; National Research Council of Italy - High Performance Computing and Networking Institute (CNR-ICAR) of Palermo | Vacca Davide, Fiannaca Antonino, Bonura Celestino, Cancila Valeria, La Paglia Laura, Bonura Floriana, Morello Gaia, Massimo La Rosa, Di Bernardo Francesca, Alfonso Maurizio Urso, Tripodo Claudio, Craxi Antonio, Vitale Francesco, Giammanco Maurizio Giovanni |
| EPI_ISL_1200610 | OCME Office Of Chief Medical Examiner | New York City Public Health Laboratory | Jade Wang, et al. |
| EPI_ISL_1201431, EPI_ISL_1201432, EPI_ISL_1201433, EPI_ISL_1201434, EPI_ISL_1201435, EPI_ISL_1201436 | Laboratory of Genetics and Personalized Medicine, Zan Mitrev Clinic | Laboratory of Genetics and Personalized Medicine, Zan Mitrev Clinic | Kungulovski G. et al. |
| EPI_ISL_1201482 | AZ Zeno | AZ SINT-JAN BRUGGE | Marijke Reynders, Patrick Descheemaeker, Laurien Hoornaert, Thomas Van Landschoot, Jorn Hellemans |
| EPI_ISL_1201511, EPI_ISL_1201512, EPI_ISL_1201513, EPI_ISL_1201514, EPI_ISL_1201515, EPI_ISL_1201516, EPI_ISL_1201517, EPI_ISL_1201518, EPI_ISL_1201519 | Outre mer | National Reference Center for Viruses of Respiratory Infections, Institut Pasteur, Paris | Marion Barbet, Sylvie Behillil, Méline Bizard, Angela Brisebarre, Camille Capel, Etienne Simon-Lorière, Vincent Enouf, Maud Vanpeene, Sylvie van der Werf,Combe Patrice |
| EPI_ISL_1201526, EPI_ISL_1201527, EPI_ISL_1201528, EPI_ISL_1201529, | Outre Mer | National Reference Center for Viruses of Respiratory Infections, Institut Pasteur, Paris | Marion Barbet, Sylvie Behillil, Méline Bizard, Angela Brisebarre, Camille Capel, Etienne Simon-Lorière, Vincent Enouf, Maud Vanpeene, Sylvie van der Werf,Rousset Dominique |

| | | | |
|---|---|---|---|
| EPI_ISL_1201530, EPI_ISL_1201531, EPI_ISL_1201532, EPI_ISL_1201533, EPI_ISL_1201534 | | | |
| EPI_ISL_1201540, EPI_ISL_1201541 | Hopital | National Reference Center for Viruses of Respiratory Infections, Institut Pasteur, Paris | Marion Barbet, Sylvie Behillil, Méline Bizard, Angela Brisebarre, Camille Capel, Etienne Simon-Lorière, Vincent Enouf, Maud Vanpeene, Sylvie van der Werf,Castelain Sandrine |
| EPI_ISL_1201546, EPI_ISL_1201547, EPI_ISL_1201549, EPI_ISL_1201550, EPI_ISL_1201551, EPI_ISL_1201552 | Labo Analyses Med | National Reference Center for Viruses of Respiratory Infections, Institut Pasteur, Paris | Marion Barbet, Sylvie Behillil, Méline Bizard, Angela Brisebarre, Camille Capel, Etienne Simon-Lorière, Vincent Enouf, Maud Vanpeene, Sylvie van der Werf |
| EPI_ISL_1201884, EPI_ISL_1201885, EPI_ISL_1201886, EPI_ISL_1201887, EPI_ISL_1201888 | Aeroporto Internacional de Guarulhos | Instituto Adolfo Lutz, Interdiciplinary Procedures Center, Strategic Laboratory | Claudio Tavares Sacchi, Claudia Regina Gonçalves, Erica Valessa Ramos Gomes, Karoline Rodrigues Campos, Caio Vinicius Dias Lopes |
| EPI_ISL_1201889, EPI_ISL_1201890, EPI_ISL_1201891, EPI_ISL_1201892 | IAL Regional de Marilia | Instituto Adolfo Lutz, Interdiciplinary Procedures Center, Strategic Laboratory | Claudio Tavares Sacchi, Claudia Regina Gonçalves, Erica Valessa Ramos Gomes, Karoline Rodrigues Campos, Caio Vinicius Dias Lopes |
| EPI_ISL_1201893 | IAL Regional de Sorocaba | Instituto Adolfo Lutz, Interdiciplinary Procedures Center, Strategic Laboratory | Claudio Tavares Sacchi, Claudia Regina Gonçalves, Erica Valessa Ramos Gomes, Karoline Rodrigues Campos, Caio Vinicius Dias Lopes |
| EPI_ISL_1201895, EPI_ISL_1201896 | Diagnosticos da America - DASA | Instituto Adolfo Lutz, Interdiciplinary Procedures Center, Strategic Laboratory | Claudio Tavares Sacchi, Claudia Regina Gonçalves, Erica Valessa Ramos Gomes, Karoline Rodrigues Campos, Caio Vinicius Dias Lopes |
| EPI_ISL_1205220, EPI_ISL_1205221, EPI_ISL_1205223, EPI_ISL_1205224, EPI_ISL_1205227, EPI_ISL_1205229, EPI_ISL_1205230, EPI_ISL_1205231, EPI_ISL_1205232, EPI_ISL_1205233, EPI_ISL_1205234, EPI_ISL_1205235, EPI_ISL_1205236, EPI_ISL_1205237, EPI_ISL_1205238, EPI_ISL_1205239, EPI_ISL_1205240, EPI_ISL_1205241, EPI_ISL_1205242, EPI_ISL_1205243, EPI_ISL_1205244, EPI_ISL_1205245, EPI_ISL_1205246, EPI_ISL_1205247, EPI_ISL_1205248, EPI_ISL_1205250, EPI_ISL_1205252, EPI_ISL_1205253, EPI_ISL_1205254, EPI_ISL_1205255, EPI_ISL_1205256, EPI_ISL_1205257, EPI_ISL_1205260, EPI_ISL_1205261, EPI_ISL_1205262, EPI_ISL_1205264, EPI_ISL_1205266, EPI_ISL_1205267, EPI_ISL_1205268, EPI_ISL_1205269, EPI_ISL_1205271, EPI_ISL_1205272, EPI_ISL_1205273, EPI_ISL_1205276, EPI_ISL_1205277, EPI_ISL_1205278, EPI_ISL_1205280, EPI_ISL_1205281, EPI_ISL_1205282, EPI_ISL_1205283, EPI_ISL_1205285, EPI_ISL_1205286, EPI_ISL_1205287, EPI_ISL_1205288, EPI_ISL_1205289, EPI_ISL_1205290, EPI_ISL_1205291, EPI_ISL_1205292, EPI_ISL_1205294, EPI_ISL_1205295, EPI_ISL_1205296, EPI_ISL_1205297, EPI_ISL_1205298, EPI_ISL_1205301, EPI_ISL_1205302 | | | |
| see above | WVU Rapid Development Laboratory | WVU and Marshall University Combined Genomics Core Facilities | James Denvir, Peter Stoilov, Peter Perrotta, Wesley Kimble, Ryan Percifield |
| EPI_ISL_1205462, EPI_ISL_1208665, EPI_ISL_1209015, EPI_ISL_1209016, EPI_ISL_1209019 | Institut de Virologie du CHU de Strasbourg | Institut de Virologie du CHU de Strasbourg | Fafi-Kremer Samira, Gallais Floriane, Gantner Pierre, Laugel Elodie, Solis Morgane, Velay Aurélie, Wendling Marie-Josée |
| EPI_ISL_1209263, EPI_ISL_1209264 | Institute for Laboratory Diagnostics and Microbiology, Klinikum Klagenfurt am Wörthersee | Bergthaler laboratory, CeMM Research Center for Molecular Medicine of the Austrian Academy of Sciences | Lukas Endler, Anna Schedl, Fabian Amman, Thomas Penz, Benedikt Agerer, Maelle Le Moing, Michael Schuster, Bekir Erguner, Jan Laine, Martin Senekowitsch, Christoph Bock, Andreas Bergthaler |
| EPI_ISL_1209269, EPI_ISL_1209270, EPI_ISL_1209271, EPI_ISL_1209272, EPI_ISL_1209273, EPI_ISL_1209275, EPI_ISL_1209276, EPI_ISL_1209277, EPI_ISL_1209278, EPI_ISL_1209279, EPI_ISL_1209280 | | | |
| see above | Department of Laboratory Medicine, Division of Clinical Virology, University of Medicine, Vienna | Bergthaler laboratory, CeMM Research Center for Molecular Medicine of the Austrian Academy of Sciences | Lukas Endler, Anna Schedl, Fabian Amman, Thomas Penz, Benedikt Agerer, Maelle Le Moing, Michael Schuster, Bekir Erguner, Jan Laine, Martin Senekowitsch, Christoph Bock, Andreas Bergthaler |
| EPI_ISL_1209367 | 3. Medizinische Abteilung, Hanusch Krankenhaus | Bergthaler laboratory, CeMM Research Center for Molecular Medicine of the Austrian Academy of Sciences | Lukas Endler, Anna Schedl, Fabian Amman, Thomas Penz, Benedikt Agerer, Maelle Le Moing, Michael Schuster, Bekir Erguner, Jan Laine, Martin Senekowitsch, Christoph Bock, Andreas Bergthaler |
| EPI_ISL_1209371, EPI_ISL_1209372 | CH DOUAI | CHU Lille - Laboratoire de Virologie | AIT YAHYA Emilie, ALIDJINOU Enagnon Kazali, BOCKET Laurence, CREPIN Michel, DEMAY Christophe, ENGELMANN Ilka, GEFFROY Sandrine, GUIGON Aurélie, LAZREK Mouna, NOBILLIAUX Florian, PREVOST Brigitte, THUILLIER Caroline, TINEZ Claire |
| EPI_ISL_1209373 | CH ARRAS | CHU Lille - Laboratoire de Virologie | AIT YAHYA Emilie, ALIDJINOU Enagnon Kazali, BOCKET Laurence, CREPIN Michel, DEMAY Christophe, ENGELMANN Ilka, GEFFROY Sandrine, GUIGON Aurélie, LAZREK Mouna, NOBILLIAUX Florian, PREVOST Brigitte, THUILLIER Caroline, TINEZ Claire |
| EPI_ISL_1209374 | CH SECLIN | CHU Lille - Laboratoire de Virologie | AIT YAHYA Emilie, ALIDJINOU Enagnon Kazali, BOCKET Laurence, CREPIN Michel, DEMAY Christophe, ENGELMANN Ilka, GEFFROY Sandrine, GUIGON Aurélie, LAZREK Mouna, NOBILLIAUX Florian, PREVOST Brigitte, THUILLIER Caroline, TINEZ Claire |
| EPI_ISL_1209375 | CH VALENCIENNES | CHU Lille - Laboratoire de Virologie | AIT YAHYA Emilie, ALIDJINOU Enagnon Kazali, BOCKET Laurence, CREPIN Michel, DEMAY Christophe, ENGELMANN Ilka, GEFFROY Sandrine, GUIGON Aurélie, LAZREK Mouna, NOBILLIAUX Florian, PREVOST Brigitte, THUILLIER Caroline, TINEZ Claire |
| EPI_ISL_1209376 | LAM BIOCENTRE MERCURE | CHU Lille - Laboratoire de Virologie | AIT YAHYA Emilie, ALIDJINOU Enagnon Kazali, BOCKET Laurence, CREPIN Michel, DEMAY Christophe, ENGELMANN Ilka, GEFFROY Sandrine, GUIGON Aurélie, LAZREK Mouna, NOBILLIAUX Florian, PREVOST Brigitte, THUILLIER Caroline, TINEZ Claire |
| EPI_ISL_1209377 | CHU LILLE | CHU Lille - Laboratoire de Virologie | AIT YAHYA Emilie, ALIDJINOU Enagnon Kazali, BOCKET Laurence, CREPIN Michel, DEMAY Christophe, ENGELMANN Ilka, GEFFROY Sandrine, GUIGON Aurélie, LAZREK Mouna, NOBILLIAUX Florian, PREVOST Brigitte, THUILLIER Caroline, TINEZ Claire |
| EPI_ISL_1209378, EPI_ISL_1209379 | LAM DIAGNOVIE RONCHIN | CHU Lille - Laboratoire de Virologie | AIT YAHYA Emilie, ALIDJINOU Enagnon Kazali, BOCKET Laurence, CREPIN Michel, DEMAY Christophe, ENGELMANN Ilka, GEFFROY Sandrine, GUIGON Aurélie, LAZREK Mouna, NOBILLIAUX Florian, PREVOST Brigitte, THUILLIER Caroline, TINEZ Claire |
| EPI_ISL_1209380 | LAM BIOPATH | CHU Lille - Laboratoire de Virologie | AIT YAHYA Emilie, ALIDJINOU Enagnon Kazali, BOCKET Laurence, CREPIN Michel, DEMAY Christophe, ENGELMANN Ilka, GEFFROY Sandrine, GUIGON Aurélie, LAZREK Mouna, NOBILLIAUX Florian, PREVOST Brigitte, THUILLIER Caroline, TINEZ Claire |
| EPI_ISL_1209381 | CH LENS | CHU Lille - Laboratoire de Virologie | AIT YAHYA Emilie, ALIDJINOU Enagnon Kazali, BOCKET Laurence, CREPIN Michel, DEMAY Christophe, ENGELMANN Ilka, GEFFROY Sandrine, GUIGON Aurélie, LAZREK Mouna, NOBILLIAUX Florian, PREVOST Brigitte, THUILLIER Caroline, TINEZ Claire |
| EPI_ISL_1209382 | CH ROUBAIX | CHU Lille - Laboratoire de Virologie | AIT YAHYA Emilie, ALIDJINOU Enagnon Kazali, BOCKET Laurence, CREPIN Michel, DEMAY Christophe, ENGELMANN Ilka, GEFFROY Sandrine, GUIGON Aurélie, LAZREK Mouna, NOBILLIAUX Florian, PREVOST Brigitte, THUILLIER Caroline, TINEZ Claire |
| EPI_ISL_1209383 | CH CALAIS | CHU Lille - Laboratoire de Virologie | AIT YAHYA Emilie, ALIDJINOU Enagnon Kazali, BOCKET Laurence, CREPIN Michel, DEMAY Christophe, ENGELMANN Ilka, GEFFROY Sandrine, GUIGON Aurélie, LAZREK Mouna, NOBILLIAUX Florian, PREVOST Brigitte, THUILLIER Caroline, TINEZ Claire |
| EPI_ISL_1209384 | CH SAINT-OMER | CHU Lille - Laboratoire de Virologie | AIT YAHYA Emilie, ALIDJINOU Enagnon Kazali, BOCKET Laurence, CREPIN Michel, DEMAY Christophe, ENGELMANN Ilka, GEFFROY Sandrine, GUIGON Aurélie, LAZREK Mouna, NOBILLIAUX Florian, PREVOST Brigitte, THUILLIER Caroline, TINEZ Claire |
| EPI_ISL_1209385 | CH ARMENTIERES | CHU Lille - Laboratoire de Virologie | AIT YAHYA Emilie, ALIDJINOU Enagnon Kazali, BOCKET Laurence, CREPIN Michel, DEMAY Christophe, ENGELMANN Ilka, GEFFROY Sandrine, GUIGON Aurélie, LAZREK Mouna, NOBILLIAUX Florian, PREVOST Brigitte, THUILLIER Caroline, TINEZ Claire |
| EPI_ISL_1209386, EPI_ISL_1209387 | CHU LILLE | CHU Lille - Laboratoire de Virologie | AIT YAHYA Emilie, ALIDJINOU Enagnon Kazali, BOCKET Laurence, CREPIN Michel, DEMAY Christophe, ENGELMANN Ilka, GEFFROY Sandrine, GUIGON Aurélie, LAZREK Mouna, NOBILLIAUX Florian, PREVOST Brigitte, THUILLIER Caroline, TINEZ Claire |
| EPI_ISL_1209388 | CH LENS | CHU Lille - Laboratoire de Virologie | AIT YAHYA Emilie, ALIDJINOU Enagnon Kazali, BOCKET Laurence, CREPIN Michel, DEMAY Christophe, ENGELMANN Ilka, GEFFROY Sandrine, GUIGON Aurélie, LAZREK Mouna, NOBILLIAUX Florian, PREVOST Brigitte, THUILLIER Caroline, TINEZ Claire |
| EPI_ISL_1209389 | CH VALENCIENNES | CHU Lille - Laboratoire de Virologie | AIT YAHYA Emilie, ALIDJINOU Enagnon Kazali, BOCKET Laurence, CREPIN Michel, DEMAY Christophe, ENGELMANN Ilka, GEFFROY Sandrine, GUIGON Aurélie, LAZREK Mouna, NOBILLIAUX Florian, PREVOST Brigitte, THUILLIER Caroline, TINEZ Claire |
| EPI_ISL_1209390 | CH BETHUNE | CHU Lille - Laboratoire de Virologie | AIT YAHYA Emilie, ALIDJINOU Enagnon Kazali, BOCKET Laurence, CREPIN Michel, DEMAY Christophe, ENGELMANN Ilka, GEFFROY Sandrine, GUIGON Aurélie, LAZREK Mouna, NOBILLIAUX Florian, PREVOST Brigitte, THUILLIER Caroline, TINEZ Claire |
| EPI_ISL_1209391 | CH MAUBEUGE | CHU Lille - Laboratoire de Virologie | AIT YAHYA Emilie, ALIDJINOU Enagnon Kazali, BOCKET Laurence, CREPIN Michel, DEMAY Christophe, ENGELMANN Ilka, GEFFROY Sandrine, GUIGON Aurélie, LAZREK Mouna, NOBILLIAUX Florian, PREVOST Brigitte, THUILLIER Caroline, TINEZ Claire |
| EPI_ISL_1209392 | CHU LILLE | CHU Lille - Laboratoire de Virologie | AIT YAHYA Emilie, ALIDJINOU Enagnon Kazali, BOCKET Laurence, CREPIN Michel, DEMAY Christophe, ENGELMANN Ilka, GEFFROY Sandrine, GUIGON Aurélie, LAZREK Mouna, NOBILLIAUX Florian, PREVOST Brigitte, THUILLIER Caroline, TINEZ Claire |

| | | | |
|---|---|---|---|
| EPI_ISL_1209393 | CH SAINT-OMER | CHU Lille - Laboratoire de Virologie | AIT YAHYA Emilie, ALIDJINOU Enagnon Kazali, BOCKET Laurence, CREPIN Michel, DEMAY Christophe, ENGELMANN Ilka, GEFFROY Sandrine, GUIGON Aurélie, LAZREK Mouna, NOBILLIAUX Florian, PREVOST Brigitte, THUILLIER Caroline, TINEZ Claire |
| EPI_ISL_1209394 | LAM BIOPATH | CHU Lille - Laboratoire de Virologie | AIT YAHYA Emilie, ALIDJINOU Enagnon Kazali, BOCKET Laurence, CREPIN Michel, DEMAY Christophe, ENGELMANN Ilka, GEFFROY Sandrine, GUIGON Aurélie, LAZREK Mouna, NOBILLIAUX Florian, PREVOST Brigitte, THUILLIER Caroline, TINEZ Claire |
| EPI_ISL_1209395 | CHU LILLE | CHU Lille - Laboratoire de Virologie | AIT YAHYA Emilie, ALIDJINOU Enagnon Kazali, BOCKET Laurence, CREPIN Michel, DEMAY Christophe, ENGELMANN Ilka, GEFFROY Sandrine, GUIGON Aurélie, LAZREK Mouna, NOBILLIAUX Florian, PREVOST Brigitte, THUILLIER Caroline, TINEZ Claire |
| EPI_ISL_1209396 | CH DOUAI | CHU Lille - Laboratoire de Virologie | AIT YAHYA Emilie, ALIDJINOU Enagnon Kazali, BOCKET Laurence, CREPIN Michel, DEMAY Christophe, ENGELMANN Ilka, GEFFROY Sandrine, GUIGON Aurélie, LAZREK Mouna, NOBILLIAUX Florian, PREVOST Brigitte, THUILLIER Caroline, TINEZ Claire |
| EPI_ISL_1209397, EPI_ISL_1209398 | CH ROUBAIX | CHU Lille - Laboratoire de Virologie | AIT YAHYA Emilie, ALIDJINOU Enagnon Kazali, BOCKET Laurence, CREPIN Michel, DEMAY Christophe, ENGELMANN Ilka, GEFFROY Sandrine, GUIGON Aurélie, LAZREK Mouna, NOBILLIAUX Florian, PREVOST Brigitte, THUILLIER Caroline, TINEZ Claire |
| EPI_ISL_1209399 | CH SECLIN | CHU Lille - Laboratoire de Virologie | AIT YAHYA Emilie, ALIDJINOU Enagnon Kazali, BOCKET Laurence, CREPIN Michel, DEMAY Christophe, ENGELMANN Ilka, GEFFROY Sandrine, GUIGON Aurélie, LAZREK Mouna, NOBILLIAUX Florian, PREVOST Brigitte, THUILLIER Caroline, TINEZ Claire |
| EPI_ISL_1209400 | CHU LILLE | CHU Lille - Laboratoire de Virologie | AIT YAHYA Emilie, ALIDJINOU Enagnon Kazali, BOCKET Laurence, CREPIN Michel, DEMAY Christophe, ENGELMANN Ilka, GEFFROY Sandrine, GUIGON Aurélie, LAZREK Mouna, NOBILLIAUX Florian, PREVOST Brigitte, THUILLIER Caroline, TINEZ Claire |
| EPI_ISL_1209401, EPI_ISL_1209402 | CH ARMENTIERES | CHU Lille - Laboratoire de Virologie | AIT YAHYA Emilie, ALIDJINOU Enagnon Kazali, BOCKET Laurence, CREPIN Michel, DEMAY Christophe, ENGELMANN Ilka, GEFFROY Sandrine, GUIGON Aurélie, LAZREK Mouna, NOBILLIAUX Florian, PREVOST Brigitte, THUILLIER Caroline, TINEZ Claire |
| EPI_ISL_1209403 | CH BETHUNE | CHU Lille - Laboratoire de Virologie | AIT YAHYA Emilie, ALIDJINOU Enagnon Kazali, BOCKET Laurence, CREPIN Michel, DEMAY Christophe, ENGELMANN Ilka, GEFFROY Sandrine, GUIGON Aurélie, LAZREK Mouna, NOBILLIAUX Florian, PREVOST Brigitte, THUILLIER Caroline, TINEZ Claire |
| EPI_ISL_1209404 | CH MAUBEUGE | CHU Lille - Laboratoire de Virologie | AIT YAHYA Emilie, ALIDJINOU Enagnon Kazali, BOCKET Laurence, CREPIN Michel, DEMAY Christophe, ENGELMANN Ilka, GEFFROY Sandrine, GUIGON Aurélie, LAZREK Mouna, NOBILLIAUX Florian, PREVOST Brigitte, THUILLIER Caroline, TINEZ Claire |
| EPI_ISL_1209405 | CH BETHUNE | CHU Lille - Laboratoire de Virologie | AIT YAHYA Emilie, ALIDJINOU Enagnon Kazali, BOCKET Laurence, CREPIN Michel, DEMAY Christophe, ENGELMANN Ilka, GEFFROY Sandrine, GUIGON Aurélie, LAZREK Mouna, NOBILLIAUX Florian, PREVOST Brigitte, THUILLIER Caroline, TINEZ Claire |
| EPI_ISL_1209406 | LAM DIAGNOVIE RONCHIN | CHU Lille - Laboratoire de Virologie | AIT YAHYA Emilie, ALIDJINOU Enagnon Kazali, BOCKET Laurence, CREPIN Michel, DEMAY Christophe, ENGELMANN Ilka, GEFFROY Sandrine, GUIGON Aurélie, LAZREK Mouna, NOBILLIAUX Florian, PREVOST Brigitte, THUILLIER Caroline, TINEZ Claire |
| EPI_ISL_1209466, EPI_ISL_1209467, EPI_ISL_1209468 | Unità Operativa di Microbiologia e Virologia, Ospedale Civico e Di Cristina, ARNAS, Palermo, Italy | Tumor Immunology Unit, Department of Health Sciences, University of Palermo School of Medicine; Section of Microbiology, University of Palermo School of Medicine; National Research Council of Italy - High Performance Computing and Networking Institute (CNR-ICAR) of Palermo | Vacca, Davide; Fiannaca, Antonino; Bonura, Celestino; Cancila, Valeria; La Paglia, Laura; Bonura, Floriana; Morello, Gaia; La Rosa, Massimo; Di Bernardo, Francesca; Urso, Alfonso Maurizio; Tripodo, Claudio; Craxi, Antonio; Vitale, Francesco; Giammanco, Maurizio Giovanni. |
| EPI_ISL_1212619, EPI_ISL_1212621, EPI_ISL_1212626, EPI_ISL_1212628, EPI_ISL_1212630, EPI_ISL_1212632, EPI_ISL_1212634, EPI_ISL_1212636, EPI_ISL_1212638, EPI_ISL_1212643, EPI_ISL_1212645, EPI_ISL_1212647, EPI_ISL_1212650, EPI_ISL_1212652, EPI_ISL_1212654, EPI_ISL_1212656, EPI_ISL_1212659, EPI_ISL_1212661, EPI_ISL_1212664, EPI_ISL_1212666, EPI_ISL_1212668, EPI_ISL_1212670 | | | |
| see above | Università degli Studi di Perugia | Istituto Zooprofilattico Sperimentale dell'Abruzzo e Molise "G. Caporale" | Mencacci A., Camilloni B., Lorusso A, Marcacci M, Di Domenico M, Ancora M, Curini V, Mangone I, Rinaldi A, Scialabba S, Di Pasquale A, Cammà C, Puglia I, Calistri P, Savini G |
| EPI_ISL_1212684, EPI_ISL_1212686, EPI_ISL_1212689, EPI_ISL_1212691, EPI_ISL_1212693, EPI_ISL_1212696, EPI_ISL_1212698, EPI_ISL_1212700 | Istituto Zooprofilattico Sperimentale Umbria e Marche "Togo Rosati" | Istituto Zooprofilattico Sperimentale dell'Abruzzo e Molise "G. Caporale" | Biagetti M., Giammarioli M., Lorusso A, Marcacci M, Di Domenico M, Ancora M, Curini V, Mangone I, Rinaldi A, Scialabba S, Di Pasquale A, Cammà C, Puglia I, Calistri P, Savini G |
| EPI_ISL_1213565, EPI_ISL_1213566 | Hi-Precision Diagnostic Center (QC) | Philippine Genome Center | Francis A. Tablizo, Kenneth M. Kim, Carlo M. Lapid, Marc Jerrone R. Castro, Maria Sofia L. Yangzon, Benedict A. Maralit, Marc Edsel C. Ayes, Eva Maria Cutiongco-de la Paz, Alethea R. de Guzman, Jan Michael C. Yap, Jo-Hannah S. Llames, Sheila Mae M. Araiza, Kris P. Punayan, Irish Coleen A. Asin, Candice Francheska B. Tambaoan, Asia Louisa U. Chong, Karol Sophia Agape R. Padilla, Rianna Patricia S. Cruz, El King D. Morado, Joshua Gregor A. Dizon, Razel Nikka M. Hao, Arianne A. Zamora, Devon Ray Pacial, Juan Antonio R. Magalang, Marissa Alejandria, Celia Carlos, Anna Ong-Lim, Edsel Maurice Salvaña, John Q. Wong, Jaime C. Montoya, Maria Rosario Singh-Vergeire and Cynthia P. Saloma |
| EPI_ISL_1213569 | Manila Doctors Hospital | Philippine Genome Center | Francis A. Tablizo, Kenneth M. Kim, Carlo M. Lapid, Marc Jerrone R. Castro, Maria Sofia L. Yangzon, Benedict A. Maralit, Marc Edsel C. Ayes, Eva Maria Cutiongco-de la Paz, Alethea R. de Guzman, Jan Michael C. Yap, Jo-Hannah S. Llames, Sheila Mae M. Araiza, Kris P. Punayan, Irish Coleen A. Asin, Candice Francheska B. Tambaoan, Asia Louisa U. Chong, Karol Sophia Agape R. Padilla, Rianna Patricia S. Cruz, El King D. Morado, Joshua Gregor A. Dizon, Razel Nikka M. Hao, Arianne A. Zamora, Devon Ray Pacial, Juan Antonio R. Magalang, Marissa Alejandria, Celia Carlos, Anna Ong-Lim, Edsel Maurice Salvaña, John Q. Wong, Jaime C. Montoya, Maria Rosario Singh-Vergeire and Cynthia P. Saloma |
| EPI_ISL_1213571 | Negros Oriental Provincial Hospital | Philippine Genome Center | Francis A. Tablizo, Kenneth M. Kim, Carlo M. Lapid, Marc Jerrone R. Castro, Maria Sofia L. Yangzon, Benedict A. Maralit, Marc Edsel C. Ayes, Eva Maria Cutiongco-de la Paz, Alethea R. de Guzman, Jan Michael C. Yap, Jo-Hannah S. Llames, Sheila Mae M. Araiza, Kris P. Punayan, Irish Coleen A. Asin, Candice Francheska B. Tambaoan, Asia Louisa U. Chong, Karol Sophia Agape R. Padilla, Rianna Patricia S. Cruz, El King D. Morado, Joshua Gregor A. Dizon, Razel Nikka M. Hao, Arianne A. Zamora, Devon Ray Pacial, Juan Antonio R. Magalang, Marissa Alejandria, Celia Carlos, Anna Ong-Lim, Edsel Maurice Salvaña, John Q. Wong, Jaime C. Montoya, Maria Rosario Singh-Vergeire and Cynthia P. Saloma |
| EPI_ISL_1213581 | Prime Care Alpha Covid-19 Testing Laboratory | Philippine Genome Center | Francis A. Tablizo, Kenneth M. Kim, Carlo M. Lapid, Marc Jerrone R. Castro, Maria Sofia L. Yangzon, Benedict A. Maralit, Marc Edsel C. Ayes, Eva Maria Cutiongco-de la Paz, Alethea R. de Guzman, Jan Michael C. Yap, Jo-Hannah S. Llames, Sheila Mae M. Araiza, Kris P. Punayan, Irish Coleen A. Asin, Candice Francheska B. Tambaoan, Asia Louisa U. Chong, Karol Sophia Agape R. Padilla, Rianna Patricia S. Cruz, El King D. Morado, Joshua Gregor A. Dizon, Razel Nikka M. Hao, Arianne A. Zamora, Devon Ray Pacial, Juan Antonio R. Magalang, Marissa Alejandria, Celia Carlos, Anna Ong-Lim, Edsel Maurice Salvaña, John Q. Wong, Jaime C. Montoya, Maria Rosario Singh-Vergeire and Cynthia P. Saloma |
| EPI_ISL_1213583 | Teresita Jalandoni Provincial Hospital | Philippine Genome Center | Francis A. Tablizo, Kenneth M. Kim, Carlo M. Lapid, Marc Jerrone R. Castro, Maria Sofia L. Yangzon, Benedict A. Maralit, Marc Edsel C. Ayes, Eva Maria Cutiongco-de la Paz, Alethea R. de Guzman, Jan Michael C. Yap, Jo-Hannah S. Llames, Sheila Mae M. Araiza, Kris P. Punayan, Irish Coleen A. Asin, Candice Francheska B. Tambaoan, Asia Louisa U. Chong, Karol Sophia Agape R. Padilla, Rianna Patricia S. Cruz, El King D. Morado, Joshua Gregor A. Dizon, Razel Nikka M. Hao, Arianne A. Zamora, Devon Ray Pacial, Juan Antonio R. Magalang, Marissa Alejandria, Celia Carlos, Anna Ong-Lim, Edsel Maurice Salvaña, John Q. Wong, Jaime C. Montoya, Maria Rosario Singh-Vergeire and Cynthia P. Saloma |
| EPI_ISL_1213588 | The Medical City - Ortigas | Philippine Genome Center | Francis A. Tablizo, Kenneth M. Kim, Carlo M. Lapid, Marc Jerrone R. Castro, Maria Sofia L. Yangzon, Benedict A. Maralit, Marc Edsel C. Ayes, Eva Maria Cutiongco-de la Paz, Alethea R. de Guzman, Jan Michael C. Yap, Jo-Hannah S. Llames, Sheila Mae M. Araiza, Kris P. Punayan, Irish Coleen A. Asin, Candice Francheska B. Tambaoan, Asia Louisa U. Chong, Karol Sophia Agape R. Padilla, Rianna Patricia S. Cruz, El King D. Morado, Joshua Gregor A. Dizon, Razel Nikka M. Hao, Arianne A. Zamora, Devon Ray Pacial, Juan Antonio R. Magalang, Marissa Alejandria, Celia Carlos, Anna Ong-Lim, Edsel Maurice Salvaña, John Q. Wong, Jaime C. Montoya, Maria Rosario Singh-Vergeire and Cynthia P. Saloma |
| EPI_ISL_1215947, EPI_ISL_1216923, EPI_ISL_1217356, EPI_ISL_1217709 | Department of Sciences for Health Promotion and Mother-Child Care "G. D'Alessandro", University of Palermo | Tumor Immunology Unit, Department of Health Sciences, University of Palermo School of Medicine; Section of Microbiology, University of Palermo School of Medicine; National Research Council of Italy - High Performance Computing and Networking Institute (CNR-ICAR) of Palermo | Vacca, Davide; Fiannaca, Antonino; Bonura, Celestino; Cancila, Valeria; La Paglia, Laura; Bonura, Floriana; Morello, Gaia; La Rosa, Massimo; Di Bernardo, Francesca; Urso, Alfonso Maurizio; Tripodo, Claudio; Craxi, Antonio; Vitale, Francesco; Giammanco, Maurizio Giovanni. |
| EPI_ISL_1219022, EPI_ISL_1219023, EPI_ISL_1219024, EPI_ISL_1219025, EPI_ISL_1219026 | IAL Regional de Sorocaba | Instituto Adolfo Lutz, Interdiciplinary Procedures Center, Strategic Laboratory | Claudio Tavares Sacchi, Claudia Regina Gonçalves, Erica Valessa Ramos Gomes, Karoline Rodrigues Campos, Caio Vinicius Dias Lopes |

| | | | |
|---|---|---|---|
| EPI_ISL_1219027 | IAL Regional de Presidente Prudente | Instituto Adolfo Lutz, Interdiciplinary Procedures Center, Strategic Laboratory | Claudio Tavares Sacchi, Claudia Regina Gonçalves, Erica Valessa Ramos Gomes, Karoline Rodrigues Campos, Caio Vinicius Dias Lopes |
| EPI_ISL_1219028, EPI_ISL_1219029, EPI_ISL_1219030, EPI_ISL_1219031, EPI_ISL_1219032, EPI_ISL_1219033, EPI_ISL_1219034, EPI_ISL_1219035, EPI_ISL_1219036 | Aeroporto Internacional de Guarulhos | Instituto Adolfo Lutz, Interdiciplinary Procedures Center, Strategic Laboratory | Claudio Tavares Sacchi, Claudia Regina Gonçalves, Erica Valessa Ramos Gomes, Karoline Rodrigues Campos, Caio Vinicius Dias Lopes |
| EPI_ISL_1219037 | IAL Regional de Presidente Prudente | Instituto Adolfo Lutz, Interdiciplinary Procedures Center, Strategic Laboratory | Claudio Tavares Sacchi, Claudia Regina Gonçalves, Erica Valessa Ramos Gomes, Karoline Rodrigues Campos, Caio Vinicius Dias Lopes |
| EPI_ISL_1219132, EPI_ISL_1219133 | Laboratorio Central de Saude Publica do Estado do Parana (LACEN-PR) | Laboratory of Respiratory Viruses and Measles, Oswaldo Cruz Institute, FIOCRUZ | Paola Resende, Luciana Appolinario, Fernando Motta, Anna Carolina Paixao, Ana Carolina Mendonca, Alice Sampaio Rocha, Renata Serrano Lopes, Maria do Carmo Debur, Irina Nastassja Riediger, Marilda Siqueira on behalf of the Fiocruz COVID-19 Genomic Surveillance Network |
| EPI_ISL_1219134, EPI_ISL_1219135 | Laboratorio Central de Saude Publica do Estado do Alagoas (LACEN-AL) | Laboratory of Respiratory Viruses and Measles, Oswaldo Cruz Institute, FIOCRUZ | Paola Resende, Luciana Appolinario, Fernando Motta, Anna Carolina Paixao, Ana Carolina Mendonca, Alice Sampaio Rocha, Renata Serrano Lopes, Anderson Brandao Leite, Marilda Siqueira on behalf of the Fiocruz COVID-19 Genomic Surveillance Network |
| EPI_ISL_1219708, EPI_ISL_1219709 | Laboratorio Biologia Molecolare Sars Cov2 - UOC Laboratorio Analisi - Servizio Medicina di Laboratorio, Ospedale "San Francesco" - ATS-ASSL Nuoro | Laboratorio specialistico UOC Ematologia - Ospedale "San Francesco" - ATS-ASSL Nuoro | Piras Giovanna, Asproni Rosanna,,Fiamma Maura, Monne Maria Itria, Palmas Angelo Domenico, Lo Maglio Iana, Mameli Giuseppe |
| EPI_ISL_1219711, EPI_ISL_1219712, EPI_ISL_1219713, EPI_ISL_1219715, EPI_ISL_1219716, EPI_ISL_1219717, EPI_ISL_1219718 | Instituto de Diagnostico y Referencia Epidemiologicos INDRE_RNLSP | Instituto de Diagnostico y Referencia Epidemiologicos (INDRE) | Claudia Wong-Arambula, Abril Rodriguez-Maldonado, Vanessa Rivero-Arredondo, Ariadna Medina-Benitez, Joaquin Quiroz-Mercado,David Fragoso-Fonseca, Sergio Rangel-Guerrero, Natividad Cruz-Ortiz, Tatiana Nunez-Garcia, Gisela Barrera-Badillo, Lucia Hernandez-Rivas, Irma Lopez-Martinez, Ernesto Ramirez-Gonzalez. |
| EPI_ISL_1219938 | Labo Analyses Med | National Reference Center for Viruses of Respiratory Infections, Institut Pasteur, Paris | Marion Barbet, Sylvie Behillil, Méline Bizard, Angela Brisebarre, Camille Capel, Etienne Simon-Lorière, Vincent Enouf, Maud Vanpeene, Sylvie van der Werf |
| EPI_ISL_1219952, EPI_ISL_1219953, EPI_ISL_1219954, EPI_ISL_1219955, EPI_ISL_1219956 | Hopital | National Reference Center for Viruses of Respiratory Infections, Institut Pasteur, Paris | Marion Barbet, Sylvie Behillil, Méline Bizard, Angela Brisebarre, Camille Capel, Etienne Simon-Lorière, Vincent Enouf, Maud Vanpeene, Sylvie van der Werf,Guiard Carine |
| EPI_ISL_1222710 | Cerballiance IDF site Vaureal | Cerba Lab | Lecorche E, Trombert-Paolantoni S, Olivi M, Benasra M, Mallek R, Haïm-Boukobza S, Roquebert B |
| EPI_ISL_1222720 | Cerballiance Maule | Cerba Lab | Lecorche E, Trombert-Paolantoni S, Mallek R, Zimmer S, Olivi M, Besrana M, Haïm-Boukobza S, Roquebert B |
| EPI_ISL_1222721 | Cerballiance Viry Chatillon | Cerba LAb | Trombert-Paolantoni S, Lecorche E, Mallek R, Zimmer S, Olivi M, Lessenne A, Picard B, Haïm-Boukobza S, Roquebert B |
| EPI_ISL_1224909, EPI_ISL_1224911, EPI_ISL_1224912 | Laboratory for Respiratory Viruses, Cantacuzino National Military-Medical Institute for Research and Development | Cantacuzino Institute Virology | Luiza Ustea, Nicoleta Paraschiv, Catalina Pascu, Mihaela Lazar |
| EPI_ISL_1231546, EPI_ISL_1231547, EPI_ISL_1231548, EPI_ISL_1231550, EPI_ISL_1231551, EPI_ISL_1231552, EPI_ISL_1231554, EPI_ISL_1231555, EPI_ISL_1231556, EPI_ISL_1231558, EPI_ISL_1231559, EPI_ISL_1231560, EPI_ISL_1231562, EPI_ISL_1231563, EPI_ISL_1231564, EPI_ISL_1231566, EPI_ISL_1231567, EPI_ISL_1231568, EPI_ISL_1231570, EPI_ISL_1231571, EPI_ISL_1231572, EPI_ISL_1231574, EPI_ISL_1231575, EPI_ISL_1231576, EPI_ISL_1231578, EPI_ISL_1231579, EPI_ISL_1231580, EPI_ISL_1231582, EPI_ISL_1231583, EPI_ISL_1231584, EPI_ISL_1231586, EPI_ISL_1231587, EPI_ISL_1231588, EPI_ISL_1231589, EPI_ISL_1231591, EPI_ISL_1231592, EPI_ISL_1231593, EPI_ISL_1231595, EPI_ISL_1231596, EPI_ISL_1231597, EPI_ISL_1231599, EPI_ISL_1231600, EPI_ISL_1231601, EPI_ISL_1231603, EPI_ISL_1231604, EPI_ISL_1231605, EPI_ISL_1231607, EPI_ISL_1231608, EPI_ISL_1231609, EPI_ISL_1231611, EPI_ISL_1231612, EPI_ISL_1231613, EPI_ISL_1231614, EPI_ISL_1231616, EPI_ISL_1231617, EPI_ISL_1231619, EPI_ISL_1231620, EPI_ISL_1231621, EPI_ISL_1231622, EPI_ISL_1231624, EPI_ISL_1231625, EPI_ISL_1231627, EPI_ISL_1231628, EPI_ISL_1231629, EPI_ISL_1231631, EPI_ISL_1231632, EPI_ISL_1231633, EPI_ISL_1231635, EPI_ISL_1231636, EPI_ISL_1231637, EPI_ISL_1231639, EPI_ISL_1231640, EPI_ISL_1231641, EPI_ISL_1231643, EPI_ISL_1231644, EPI_ISL_1231645, EPI_ISL_1231647, EPI_ISL_1231648, EPI_ISL_1231649, EPI_ISL_1231650, EPI_ISL_1231652, EPI_ISL_1231653, EPI_ISL_1231654, EPI_ISL_1231656, EPI_ISL_1231657, EPI_ISL_1231658, EPI_ISL_1231660, EPI_ISL_1231661, EPI_ISL_1231662, EPI_ISL_1231664, EPI_ISL_1231665, EPI_ISL_1231667, EPI_ISL_1231668, EPI_ISL_1231669, EPI_ISL_1231671, EPI_ISL_1231672, EPI_ISL_1231673, EPI_ISL_1231674, EPI_ISL_1231676, EPI_ISL_1231677, EPI_ISL_1231679, EPI_ISL_1231680, EPI_ISL_1231681, EPI_ISL_1231683, EPI_ISL_1231684, EPI_ISL_1231685, EPI_ISL_1231687, EPI_ISL_1231688, EPI_ISL_1231689, EPI_ISL_1231691, EPI_ISL_1231692, EPI_ISL_1231693, EPI_ISL_1231695, EPI_ISL_1231696, EPI_ISL_1231697, EPI_ISL_1231699, EPI_ISL_1231700, EPI_ISL_1231701, EPI_ISL_1231703, EPI_ISL_1231704, EPI_ISL_1231705, EPI_ISL_1231707, EPI_ISL_1231708, EPI_ISL_1231709, EPI_ISL_1231711, EPI_ISL_1231712, EPI_ISL_1231713, EPI_ISL_1231715, EPI_ISL_1231716, EPI_ISL_1231717, EPI_ISL_1231718, EPI_ISL_1231720, EPI_ISL_1231721, EPI_ISL_1231722, EPI_ISL_1231724, EPI_ISL_1231725, EPI_ISL_1231726, EPI_ISL_1231728, EPI_ISL_1231729, EPI_ISL_1231731, EPI_ISL_1231732, EPI_ISL_1231733, EPI_ISL_1231734, EPI_ISL_1231736, EPI_ISL_1231737, EPI_ISL_1231738, EPI_ISL_1231740, EPI_ISL_1231741, EPI_ISL_1231744, EPI_ISL_1231745, EPI_ISL_1231746, EPI_ISL_1231748, EPI_ISL_1231749, EPI_ISL_1231750, EPI_ISL_1231752, EPI_ISL_1231753, EPI_ISL_1231754, EPI_ISL_1231756, EPI_ISL_1231757, EPI_ISL_1231759, EPI_ISL_1231760, EPI_ISL_1231761, EPI_ISL_1231763, EPI_ISL_1231764, EPI_ISL_1231765, EPI_ISL_1231766, EPI_ISL_1231768, EPI_ISL_1231769, EPI_ISL_1231770, EPI_ISL_1231772, EPI_ISL_1231773 | | | |
| see above | National Center for Infectious and Parasitic Diseases (NCIPD) | National Center for Infectious and Parasitic Diseases (NCIPD) | Alexiev et al |
| EPI_ISL_1233478, EPI_ISL_1233479, EPI_ISL_1233480, EPI_ISL_1233481, EPI_ISL_1233482, EPI_ISL_1233484, EPI_ISL_1233485, EPI_ISL_1233486, EPI_ISL_1233487, EPI_ISL_1233488, EPI_ISL_1233489, EPI_ISL_1233490, EPI_ISL_1233491, EPI_ISL_1233492, EPI_ISL_1233493, EPI_ISL_1233494, EPI_ISL_1233496, EPI_ISL_1233497, EPI_ISL_1233498, EPI_ISL_1233499, EPI_ISL_1233500, EPI_ISL_1233501, EPI_ISL_1233502, EPI_ISL_1233503, EPI_ISL_1233505, EPI_ISL_1233506, EPI_ISL_1233507, EPI_ISL_1233508, EPI_ISL_1233545, EPI_ISL_1233546, EPI_ISL_1233547, EPI_ISL_1233548, EPI_ISL_1233549, EPI_ISL_1233550, EPI_ISL_1233551, EPI_ISL_1233552, EPI_ISL_1233554, EPI_ISL_1233555, EPI_ISL_1233556, EPI_ISL_1233557, EPI_ISL_1233558, EPI_ISL_1233559, EPI_ISL_1233560, EPI_ISL_1233561, EPI_ISL_1233562, EPI_ISL_1233563, EPI_ISL_1233564, EPI_ISL_1233565, EPI_ISL_1233566, EPI_ISL_1233567, EPI_ISL_1233568, EPI_ISL_1233569, EPI_ISL_1233570, EPI_ISL_1233571, EPI_ISL_1233572, EPI_ISL_1233573, EPI_ISL_1233574, EPI_ISL_1233575, EPI_ISL_1233576, EPI_ISL_1233577, EPI_ISL_1233578, EPI_ISL_1233579, EPI_ISL_1233580, EPI_ISL_1233581, EPI_ISL_1233582, EPI_ISL_1233583, EPI_ISL_1233584, EPI_ISL_1233585, EPI_ISL_1233586, EPI_ISL_1233587, EPI_ISL_1233588, EPI_ISL_1233589, EPI_ISL_1233591, EPI_ISL_1233592, EPI_ISL_1233593, EPI_ISL_1233594, EPI_ISL_1233595, EPI_ISL_1233596, EPI_ISL_1233597, EPI_ISL_1233598, EPI_ISL_1233599, EPI_ISL_1233600, EPI_ISL_1233601, EPI_ISL_1233602, EPI_ISL_1233603, EPI_ISL_1233604, EPI_ISL_1233605, EPI_ISL_1233606, EPI_ISL_1233608, EPI_ISL_1233609, EPI_ISL_1233610, EPI_ISL_1233611, EPI_ISL_1233612, EPI_ISL_1233613, EPI_ISL_1233614, EPI_ISL_1233615, EPI_ISL_1233616, EPI_ISL_1233618, EPI_ISL_1233619, EPI_ISL_1233620, EPI_ISL_1233621, EPI_ISL_1233622 | | | |
| see above | National Center for Infectious and Parasitic Diseases (NCIPD) | National Center for Infectious and Parasitic Diseases (NCIPD) | Alexiev et al |
| EPI_ISL_1239143 | National Institute of Health Research and Development | National Institute of Health Research and Development | Agustiningsih, Kindi Adam, Holy Arif Wibowo, Ririn Ramadhany, Yuni Rukminiati, Hana A Pawestri, Subangkit, Kartika Dewi Puspa, Arie Ardiansyah Nugraha, Hartanti Dian Ikawati, Krisna Nur Andriana Pangesti, Triyani Soekarso, Ni Ketut Susilarini, Nur Ika Hariastuti, Uly Alfi Nikmah, Mursinah, Asri Febriyani, Reni Herman, Nike Susanti, Herna, Tati Febriyanti, Natalie Laurencia Kipuw, Fauzul Muna, Aulia Rizki, Nelly Puspandari, Vivi Setiawaty |
| EPI_ISL_1239144 | National Institute of Health Research and Development | National Institute of Health Research and Development | Kindi Adam, Holy Arif Wibowo, Ririn Ramadhany, Yuni Rukminiati, Agustiningsih, Hana A Pawestri, Subangkit, Kartika Dewi Puspa, Arie Ardiansyah Nugraha, Hartanti Dian Ikawati, Krisna Nur Andriana Pangesti, Triyani Soekarso, Ni Ketut Susilarini, Nur Ika Hariastuti, Uly Alfi Nikmah, Mursinah, Asri Febriyani, Reni Herman, Nike Susanti, Herna, Tati Febriyanti, Natalie Laurencia Kipuw, Fauzul Muna, Aulia Rizki, Nelly Puspandari, Vivi Setiawaty |
| EPI_ISL_1242034, EPI_ISL_1242035, EPI_ISL_1242036, EPI_ISL_1242038, EPI_ISL_1242041, EPI_ISL_1242042, EPI_ISL_1242043, EPI_ISL_1242044, EPI_ISL_1242045, EPI_ISL_1242046, EPI_ISL_1242047, EPI_ISL_1246953, EPI_ISL_1246954, EPI_ISL_1246955, EPI_ISL_1246956, EPI_ISL_1246957 | | | |
| see above | Microbiology Department, University Hospital Donostia | Microbiology Department, University Hospital Donostia | Marimon JM, Montes M, Piñeiro L, Sorarrain A, Cilla G. |
| EPI_ISL_1246958, EPI_ISL_1246961, EPI_ISL_1246970 | Institut de Virologie du CHU de Strasbourg | Institut de Virologie du CHU de Strasbourg | Fafi-Kremer Samira, Gallais Floriane, Gantner Pierre, Laugel Elodie, Solis Morgane, Velay Aurélie, Wendling Marie-Josée |
| EPI_ISL_1248934 | Clinical Microbiology Lab RSPTN Universitas Hasanuddin | Clinical Microbiology Lab RSPTN Universitas Hasanuddin | Muhammad Nasrum Massi, Rizalinda Sjahril, Gita Vita Soraya, Triwibawa, Handayani, Muhammad Yusuf Usman, Rufika Shari |
| EPI_ISL_1249610 | Clinical Microbiology Lab RSPTN Universitas Hasanuddin | Clinical Microbiology Lab RSPTN Universitas Hasanuddin | Muhammad Nasrum Massi, Rizalinda Sjahril, Gita Vita Soraya, Triwibawa, Handayani Halik, Muhammad Yusuf Usman, Rufika Shari |
| EPI_ISL_1250612, EPI_ISL_1250626 | Centre Hospitalier Universitaire de Rouen Laboratoire de Virologie | Centre Hospitalier Universitaire de Rouen Laboratoire de Virologie | Alice Moisan, Fabienne De Oliveira, Marie Leoz |
| EPI_ISL_1250859 | Tanjungpura University Hospital | Tanjungpura University Hospital | Puji Astuti, Sofi Siti Shofiyah; Delima Fajar Liana; Mahyarudin; Andriani; Virhan Novianry; Muhammad Ibnu Kahtan; Eka Ardiani Putri; Helmi Sastriawan; Rini Andriani; Ambar Rialita; Hana Apsari Pawestri; Arie Ardiansyah Nugraha; Kartika Dewi Puspa; Hartanti Dian Ikawati; Vivi Setiawaty |
| EPI_ISL_1250956, EPI_ISL_1250966, EPI_ISL_1250977, EPI_ISL_1250983 | Siesp dipartimento di prevenzione | Istituto Zooprofilattico Sperimentale dell'Abruzzo e Molise "G. Caporale" | Lorusso A, Marcacci M, Di Domenico M, Ancora M, Curini V, Mangone I, Rinaldi A, Scialabba S, Di Pasquale A, Cammà C, Puglia I, Calistri P, Savini G |
| EPI_ISL_1250984 | SIESP CHIETI - LANCIANO | Istituto Zooprofilattico Sperimentale dell'Abruzzo e Molise "G. Caporale" | Lorusso A, Marcacci M, Di Domenico M, Ancora M, Curini V, Mangone I, Rinaldi A, Scialabba S, Di Pasquale A, Cammà C, Puglia I, Calistri P, Savini G |
| EPI_ISL_1250985 | Usca castel di sangro | Istituto Zooprofilattico Sperimentale dell'Abruzzo e Molise "G. Caporale" | Lorusso A, Marcacci M, Di Domenico M, Ancora M, Curini V, Mangone I, Rinaldi A, Scialabba S, Di Pasquale A, Cammà C, Puglia I, Calistri P, Savini G |
| EPI_ISL_1250986 | USCA Pescina | Istituto Zooprofilattico Sperimentale dell'Abruzzo e Molise "G. Caporale" | Lorusso A, Marcacci M, Di Domenico M, Ancora M, Curini V, Mangone I, Rinaldi A, Scialabba S, Di Pasquale A, Cammà C, Puglia I, Calistri P, Savini G |

| | | | |
|---|---|---|---|
| EPI_ISL_1250987 | siesp Chieti | Istituto Zooprofilattico Sperimentale dell'Abruzzo e Molise "G. Caporale" | Lorusso A, Marcacci M, Di Domenico M, Ancora M, Curini V, Mangone I, Rinaldi A, Scialabba S, Di Pasquale A, Cammà C, Puglia I, Calistri P, Savini G |
| EPI_ISL_1250988 | Siesp dipartimento di prevenzione | Istituto Zooprofilattico Sperimentale dell'Abruzzo e Molise "G. Caporale" | Lorusso A, Marcacci M, Di Domenico M, Ancora M, Curini V, Mangone I, Rinaldi A, Scialabba S, Di Pasquale A, Cammà C, Puglia I, Calistri P, Savini G |
| EPI_ISL_1250989 | SIESP CHIETI - DRIVE IN ORTONA | Istituto Zooprofilattico Sperimentale dell'Abruzzo e Molise "G. Caporale" | Lorusso A, Marcacci M, Di Domenico M, Ancora M, Curini V, Mangone I, Rinaldi A, Scialabba S, Di Pasquale A, Cammà C, Puglia I, Calistri P, Savini G |
| EPI_ISL_1250990 | SIESP CHIETI - LANCIANO | Istituto Zooprofilattico Sperimentale dell'Abruzzo e Molise "G. Caporale" | Lorusso A, Marcacci M, Di Domenico M, Ancora M, Curini V, Mangone I, Rinaldi A, Scialabba S, Di Pasquale A, Cammà C, Puglia I, Calistri P, Savini G |
| EPI_ISL_1251002 | OSPEDALE SAN SALVATORE L'AQUILA UOC PNEUMOLOGIA | Istituto Zooprofilattico Sperimentale dell'Abruzzo e Molise "G. Caporale" | Lorusso A, Marcacci M, Di Domenico M, Ancora M, Curini V, Mangone I, Rinaldi A, Scialabba S, Di Pasquale A, Cammà C, Puglia I, Calistri P, Savini G |
| EPI_ISL_1251020 | Siesp dipartimento di prevenzione | Istituto Zooprofilattico Sperimentale dell'Abruzzo e Molise "G. Caporale" | Lorusso A, Marcacci M, Di Domenico M, Ancora M, Curini V, Mangone I, Rinaldi A, Scialabba S, Di Pasquale A, Cammà C, Puglia I, Calistri P, Savini G |
| EPI_ISL_1251062 | USCA Pescina | Istituto Zooprofilattico Sperimentale dell'Abruzzo e Molise "G. Caporale" | Lorusso A, Marcacci M, Di Domenico M, Ancora M, Curini V, Mangone I, Rinaldi A, Scialabba S, Di Pasquale A, Cammà C, Puglia I, Calistri P, Savini G |
| EPI_ISL_1251277 | "Stefan S. Nicolau"Institute of Virology | "Stefan S. Nicolau"Institute of Virology | Anca Botezatu, Iulia Virginia Iancu, Coralia Bleotu, Mihaela Economescu, Laura Necula, Denisa Dragu, Lilia Matei, Ioana Pitica, Marinela Bostan, Mirela Mihaila, Laura Grecu, Camelia Sultana, Simona Ruta, Ana Iulia Neagu, Adriana Plesa, Saviana Nedeianu, Alina Nastasie, Cristina Mambet,Gabriela Anton, Carmen Cristina Diaconu |
| EPI_ISL_1254647, EPI_ISL_1254649, EPI_ISL_1254650, EPI_ISL_1254651 | Ospedale di Venere | Istituto Zooprofilattico Sperimentale della Puglia e della Basilicata | Parisi A., Bianco A., Capozzi L., Del Sambro L., Simone D., Iacobellis M., Ridolfi D., Giannico A. |
| EPI_ISL_1254652, EPI_ISL_1254653, EPI_ISL_1254654, EPI_ISL_1254655, EPI_ISL_1254656, EPI_ISL_1254657 | Istituto Zooprofilattico Sperimentale della Puglia e della Basilicata | Istituto Zooprofilattico Sperimentale della Puglia e della Basilicata | Parisi A., Bianco A., Capozzi L., Del Sambro L., Simone D., Manzulli V, Rondinone V., Pace L., Cipolletta D., Galante D. |
| EPI_ISL_1254658, EPI_ISL_1254659 | Presidio di Brindisi Di Summa - Perrino | Istituto Zooprofilattico Sperimentale della Puglia e della Basilicata | Parisi A., Bianco A., Capozzi L., Del Sambro L., Simone D., Difato L., Ridolfi D., Giannico A. |
| EPI_ISL_1254667, EPI_ISL_1254668 | Ospedale di Venere | Istituto Zooprofilattico Sperimentale della Puglia e della Basilicata | Parisi A., Bianco A., Capozzi L., Del Sambro L., Simone D., Iacobellis M., Ridolfi D., Giannico A. |
| EPI_ISL_1254673, EPI_ISL_1254688 | Istituto Zooprofilattico Sperimentale della Puglia e della Basilicata | Istituto Zooprofilattico Sperimentale della Puglia e della Basilicata | Parisi A., Bianco A., Capozzi L., Del Sambro L., Simone D., Manzulli V, Rondinone V., Pace L., Cipolletta D., Galante D. |
| EPI_ISL_1255140, EPI_ISL_1255264, EPI_ISL_1255265, EPI_ISL_1255266, EPI_ISL_1255267, EPI_ISL_1255268, EPI_ISL_1255269, EPI_ISL_1255271, EPI_ISL_1255272, EPI_ISL_1255273, EPI_ISL_1255275, EPI_ISL_1255276, EPI_ISL_1255277 | | | |
| see above | West African Centre for Cell Biology of Infectious Pathogens (WACCBIP), University of Ghana, Accra, Ghana | West African Centre for Cell Biology of Infectious Pathogens (WACCBIP), University of Ghana, Volta Road, Legon-Accra, Ghana | Collins M. Morang'a, Joyce M. Ngoi, Evelyn B. Quansah, Samirah Saiid, Dominic S.Y. Amuzu, Vincent Appiah, Philip M. Soglo, Vanessa Magnussen, Aisha Mohammed, Kesego Tapela, Nelson Kibinge, Abdoulaye B Diallo, Frederick Kumi-Ansah, Theophilus Odoom, Oliver D Boakye5, Emmanuella Amoako4, Abdul-Karim Abass, , Samuel Kaba Akoriyea, Frederick Tei-Maya, Lucas N. Amenga-Etego, Dam Kenneth Mibut, Yaw Bediako, Benjamin Demah Nuertey, Gordon A Awandare, Peter K Quashie, Gordon A Awandare, Yaw Bediako |
| EPI_ISL_1257812, EPI_ISL_1257813 | Institut de Virologie du CHU de Strasbourg | Institut de Virologie du CHU de Strasbourg | Fafi-Kremer Samira, Gallais Floriane, Gantner Pierre, Laugel Elodie, Solis Morgane, Velay Aurélie, Wendling Marie-Josée |
| EPI_ISL_1257814 | WHO National Influenza Centre Russian Federation | WHO National Influenza Centre Russian Federation | Andrey Komissarov, Artem Fadeev, Anna Ivanova, Kseniya Komissarova, Dmitry Bazhenov, Tamila Musaeva, Maria Timofeeva, Veronika Eder, Maria Pisareva, Daria Danilenko, Ksenia Safina, Elena Nabieva, Georgii Bazykin, Dmitry Lioznov |
| EPI_ISL_1257815 | Institut de Virologie du CHU de Strasbourg | Institut de Virologie du CHU de Strasbourg | Fafi-Kremer Samira, Gallais Floriane, Gantner Pierre, Laugel Elodie, Solis Morgane, Velay Aurélie, Wendling Marie-Josée |
| EPI_ISL_1257816, EPI_ISL_1257817 | National Institute of Health Research and Development | National Institute of Health Research and Development | Subangkit, Hana Apsari Pawestri, Kartika Dewi Puspa, Arie Ardiansyah Nugraha, Hartanti Dian Ikawati, Krisna Nur Andriana Pangesti, Yuni Rukminiati, Ririn Ramadhany, Agustiningsih, Kindi Adam, Holy Arif Wibowo, Triyani Soekarso, Ni Ketut Susilarini, Nurika Hariastuti, Ully Alfi Nikmah, Reni Herman, Nike Susanti, Herna, Tati Febriyanti, Natalie Laurencia Kipuw, Fauzul Muna, Irene Lorinda Indalao, Nelly Puspandari, Vivi Setiawaty. |
| EPI_ISL_1257821 | Balai Litbangkes Papua | National Institute of Health Research and Development | Hana Apsari Pawestri, Subangkit, Kartika Dewi Puspa, Arie Ardiansyah Nugraha, Hartanti Dian Ikawati, Krisna Pangesti, Nelly Puspandari, Hotma Sitompul, Antonius Octavian, Vivi Setiawaty |
| EPI_ISL_1257827, EPI_ISL_1257828, EPI_ISL_1257829 | RSUP Dr. Sardjito | National Institute of Health Research and Development | Vivi Setiawaty, Hana Apsari Pawestri, Subangkit, Kartika Dewi Puspa, Arie Ardiansyah Nugraha, Hartanti Dian Ikawati, Nelly Puspandari, Krisna Pangesti |
| EPI_ISL_1257830, EPI_ISL_1257831 | Balai Litbangkes Tanah Bumbu | National Institute of Health Research and Development | Vivi Setiawaty, Hana Apsari Pawestri, Subangkit, Kartika Dewi Puspa, Arie Ardiansyah Nugraha, Hartanti Dian Ikawati, Nelly Puspandari, Juhairiyah, Syarif Hidayat, Krisna Pangesti |
| EPI_ISL_1257833, EPI_ISL_1257835, EPI_ISL_1257836, EPI_ISL_1257837, EPI_ISL_1257838, EPI_ISL_1257839, EPI_ISL_1257840, EPI_ISL_1257841 | RSUP Sanglah | National Institute of Health Research and Development | Arie Ardiansyah Nugraha, Hana Apsari Pawestri, Subangkit, Kartika Dewi Puspa, Hartanti Dian Ikawati, Nelly Puspandari, Krisna Pangesti, I Nengah Tony Rustawan, I Gede Ketut Sajinadiyasa, Vivi Setiawaty |
| EPI_ISL_1257842, EPI_ISL_1257843, EPI_ISL_1257844, EPI_ISL_1257845, EPI_ISL_1257846 | RSUP Fatmawati | National Institute of Health Research and Development | Kartika Dewi Puspa, Hana Apsari Pawestri, Subangkit, Arie Ardiansyah Nugraha, Hartanti Dian Ikawati, Nelly Puspandari, Krisna Pangesti, Lidya Utami, Luci Liana, Lusiya Ningsih, Vivi Setiawaty |
| EPI_ISL_1257847, EPI_ISL_1257848, EPI_ISL_1257849, EPI_ISL_1257850 | RSPI Prof.Dr.Sulianti Saroso | National Institute of Health Research and Development | Hartanti Dian Ikawati, Hana Apsari Pawestri, Subangkit, Kartika Dewi Puspa, Arie Ardiansyah Nugraha, Krisna Pangesti, Nelly Puspandari, Vivi Setiawaty |
| EPI_ISL_1257854 | BBTKLPP Banjarbaru | National Institute of Health Research and Development | Hartanti Dian Ikawati, Hana Apsari Pawestri, Subangkit, Kartika Dewi Puspa, Arie Ardiansyah Nugraha, Krisna Pangesti, Nelly Puspandari, Dewi Hermawati, Mutia Raudah, Vivi Setiawaty |
| EPI_ISL_1257857, EPI_ISL_1257858 | BBTKLPP Yogyakarta | National Institute of Health Research and Development | Kartika Dewi Puspa, Hana Apsari Pawestri, Subangkit, Arie Ardiansyah Nugraha, Hartanti Dian Ikawati, Nelly Puspandari, Krisna Pangesti, Irene, Havid Setyawan, Indaryati, Vivi Setiawaty |
| EPI_ISL_1257859 | KKP Soekarno Hatta | National Institute of Health Research and Development | Hana Apsari Pawestri, Subangkit, Kartika Dewi Puspa, Arie Ardiansyah Nugraha, Hartanti Dian Ikawati, Nelly Puspandari, Krisna Pangesti, Vivi Setiawaty |
| EPI_ISL_1257861 | Farmalab Indoutama | National Institute of Health Research and Development | Hana Apsari Pawestri, Subangkit, Kartika Dewi Puspa, Arie Ardiansyah Nugraha, Hartanti Dian Ikawati, Nelly Puspandari, Krisna Pangesti, Vivi Setiawaty |
| EPI_ISL_1257862 | Farmalab Indoutama | National Institute of Health Research and Development | Vivi Setiawaty, Hana Apsari Pawestri, Subangkit, Kartika Dewi Puspa, Arie Ardiansyah Nugraha, Hartanti Dian Ikawati, Nelly Puspandari, Krisna Pangesti |
| EPI_ISL_1257865 | RSUP Adam Malik | National Institute of Health Research and Development | Kartika Dewi Puspa, Hana Apsari Pawestri, Subangkit, Arie Ardiansyah Nugraha, Hartanti Dian Ikawati, Nelly Puspandari, Krisna Pangesti, Vivi Setiawaty |
| EPI_ISL_1257866 | RSUP Adam Malik | National Institute of Health Research and Development | Vivi Setiawaty, Hana Apsari Pawestri, Subangkit, Kartika Dewi Puspa, Arie Ardiansyah Nugraha, Hartanti Dian Ikawati, Nelly Puspandari, Maliana, Ade Rahmaini, Nelly Elfrida Samosir, Rina Yunita, Krisna Pangesti |
| EPI_ISL_1257881 | Centre Pasteur du Cameroun | Institut Pasteur de Dakar | Njouom Richard, Diagne Moussa Moïse, Dia Ndongo, Diallo Amadou, Sankhe Safietou, Diop Mamadou, Loucoubar Cheikh, Elisabeth Carniel, Faye Ousmane, Sall Amadou Alpha |
| EPI_ISL_1257886 | Hopital Général de Douala | Institut Pasteur de Dakar | Njouom Richard, Diagne Moussa Moïse, Dia Ndongo, Diallo Amadou, Sankhe Safietou, Diop Mamadou, Loucoubar Cheikh, Elisabeth Carniel, Faye Ousmane, Sall Amadou Alpha |
| EPI_ISL_1257891 | Hopital Laquintinie de Douala | Institut Pasteur de Dakar | Njouom Richard, Diagne Moussa Moïse, Dia Ndongo, Diallo Amadou, Sankhe Safietou, Diop Mamadou, Loucoubar Cheikh, Elisabeth Carniel, Faye Ousmane, Sall Amadou Alpha |

| | | | |
|---|---|---|---|
| EPI_ISL_1259027, EPI_ISL_1259028, EPI_ISL_1259029, EPI_ISL_1259030, EPI_ISL_1259031, EPI_ISL_1259032, EPI_ISL_1259033, EPI_ISL_1259034, EPI_ISL_1259035, EPI_ISL_1259036, EPI_ISL_1259037, EPI_ISL_1259039, EPI_ISL_1259040, EPI_ISL_1259041, EPI_ISL_1259042, EPI_ISL_1259043, EPI_ISL_1259044, EPI_ISL_1259045, EPI_ISL_1259046, EPI_ISL_1259047, EPI_ISL_1259048, EPI_ISL_1259049, EPI_ISL_1259050, EPI_ISL_1259051, EPI_ISL_1259052, EPI_ISL_1259053, EPI_ISL_1259054, EPI_ISL_1259055, EPI_ISL_1259056, EPI_ISL_1259057, EPI_ISL_1259058, EPI_ISL_1259059, EPI_ISL_1259060, EPI_ISL_1259061, EPI_ISL_1259062, EPI_ISL_1259063, EPI_ISL_1259064, EPI_ISL_1259065, EPI_ISL_1259066, EPI_ISL_1259067, EPI_ISL_1259068, EPI_ISL_1259070, EPI_ISL_1259071, EPI_ISL_1259073, EPI_ISL_1259074, EPI_ISL_1259075, EPI_ISL_1259076, EPI_ISL_1259077, EPI_ISL_1259078, EPI_ISL_1259079, EPI_ISL_1259080, EPI_ISL_1259081, EPI_ISL_1259082, EPI_ISL_1259083, EPI_ISL_1259084, EPI_ISL_1259085, EPI_ISL_1259087, EPI_ISL_1259088, EPI_ISL_1259089, EPI_ISL_1259090, EPI_ISL_1259091, EPI_ISL_1259092, EPI_ISL_1259093, EPI_ISL_1259094, EPI_ISL_1259095, EPI_ISL_1259096, EPI_ISL_1259097, EPI_ISL_1259098, EPI_ISL_1259099, EPI_ISL_1259100, EPI_ISL_1259101, EPI_ISL_1259102, EPI_ISL_1259103, EPI_ISL_1259104, EPI_ISL_1259105, EPI_ISL_1259106, EPI_ISL_1259107, EPI_ISL_1259108, EPI_ISL_1259109, EPI_ISL_1259110, EPI_ISL_1259111, EPI_ISL_1259113, EPI_ISL_1259114, EPI_ISL_1259115, EPI_ISL_1259116, EPI_ISL_1259117, EPI_ISL_1259118, EPI_ISL_1259119, EPI_ISL_1259120, EPI_ISL_1259121, EPI_ISL_1259123, EPI_ISL_1259124, EPI_ISL_1259125, EPI_ISL_1259126, EPI_ISL_1259127, EPI_ISL_1259128, EPI_ISL_1259129, EPI_ISL_1259130, EPI_ISL_1259131, EPI_ISL_1259132, EPI_ISL_1259133, EPI_ISL_1259134, EPI_ISL_1259135, EPI_ISL_1259136, EPI_ISL_1259137, EPI_ISL_1259138, EPI_ISL_1259139, EPI_ISL_1259140, EPI_ISL_1259141, EPI_ISL_1259142, EPI_ISL_1259143, EPI_ISL_1259145, EPI_ISL_1259146, EPI_ISL_1259147, EPI_ISL_1259148, EPI_ISL_1259149, EPI_ISL_1259150, EPI_ISL_1259151, EPI_ISL_1259152, EPI_ISL_1259153, EPI_ISL_1259154 | | | |
| see above | Synlab Eesti OÜ | 1. Laboratory of Communicable Diseases (Estonia); 2. Eurofins Genomics Europe Sequencing GmbH | Liidia Dotsenko et al. |
| EPI_ISL_1259155, EPI_ISL_1259156, EPI_ISL_1259157, EPI_ISL_1259158, EPI_ISL_1259159 | North Estonia Medical Centre laboratory | 1. Laboratory of Communicable Diseases (Estonia); 2. Eurofins Genomics Europe Sequencing GmbH | Liidia Dotsenko et al. |
| EPI_ISL_1259160, EPI_ISL_1259161, EPI_ISL_1259162, EPI_ISL_1259163, EPI_ISL_1259164, EPI_ISL_1259165, EPI_ISL_1259166, EPI_ISL_1259167, EPI_ISL_1259168, EPI_ISL_1259169, EPI_ISL_1259170, EPI_ISL_1259171, EPI_ISL_1259172, EPI_ISL_1259174, EPI_ISL_1259175, EPI_ISL_1259176, EPI_ISL_1259177, EPI_ISL_1259178, EPI_ISL_1259179, EPI_ISL_1259180 | | | |
| see above | Laboratory of Communicable Diseases | 1. Laboratory of Communicable Diseases (Estonia); 2. Eurofins Genomics Europe Sequencing GmbH | Liidia Dotsenko et al. |
| EPI_ISL_1259182, EPI_ISL_1259184, EPI_ISL_1259185, EPI_ISL_1259188, EPI_ISL_1259189, EPI_ISL_1259191, EPI_ISL_1259193 | Department of Virology, Istituto Zooprofilattico Sperimentale del Lazio e della Toscana (IZSLT) | Department of General Diagnostics; Department of Virology; Istituto Zooprofilattico Sperimentale del Lazio e della Toscana (IZSLT) | Patricia Alba, Giuseppe Manna, Elena L. Diaconu, Fabiola Feltrin, Raffaela Conti, Teresa Scicluna, Virginia Carfora, Antonella Cersini, Alessia Franco, Antonio Battisti. |
| EPI_ISL_1259284, EPI_ISL_1259285, EPI_ISL_1259286, EPI_ISL_1259287, EPI_ISL_1259288, EPI_ISL_1259289, EPI_ISL_1259290, EPI_ISL_1259291, EPI_ISL_1259292, EPI_ISL_1259293, EPI_ISL_1259294, EPI_ISL_1259295 | | | |
| see above | Hopital | National Reference Center for Viruses of Respiratory Infections, Institut Pasteur, Paris | Marion Barbet, Sylvie Behillil, Méline Bizard, Angela Brisebarre, Camille Capel, Etienne Simon-Lorière, Vincent Enouf, Maud Vanpeene, Sylvie van der Werf,Combe Patrice |
| EPI_ISL_1259296 | Hopital | National Reference Center for Viruses of Respiratory Infections, Institut Pasteur, Paris | Marion Barbet, Sylvie Behillil, Méline Bizard, Angela Brisebarre, Camille Capel, Etienne Simon-Lorière, Vincent Enouf, Maud Vanpeene, Sylvie van der Werf,Hermann CéCile |
| EPI_ISL_1259298 | Labo Analyses Med | National Reference Center for Viruses of Respiratory Infections, Institut Pasteur, Paris | Marion Barbet, Sylvie Behillil, Méline Bizard, Angela Brisebarre, Camille Capel, Etienne Simon-Lorière, Vincent Enouf, Maud Vanpeene, Sylvie van der Werf,Ducancelle Alexandra |
| EPI_ISL_1259299, EPI_ISL_1259300 | Hopital | National Reference Center for Viruses of Respiratory Infections, Institut Pasteur, Paris | Marion Barbet, Sylvie Behillil, Méline Bizard, Angela Brisebarre, Camille Capel, Etienne Simon-Lorière, Vincent Enouf, Maud Vanpeene, Sylvie van der Werf,Ducancelle Alexandra |
| EPI_ISL_1259301, EPI_ISL_1259302 | Labo Analyses Med | National Reference Center for Viruses of Respiratory Infections, Institut Pasteur, Paris | Marion Barbet, Sylvie Behillil, Méline Bizard, Angela Brisebarre, Camille Capel, Etienne Simon-Lorière, Vincent Enouf, Maud Vanpeene, Sylvie van der Werf,Coignard Catherine |
| EPI_ISL_1259303 | Labo Analyses Med | National Reference Center for Viruses of Respiratory Infections, Institut Pasteur, Paris | Marion Barbet, Sylvie Behillil, Méline Bizard, Angela Brisebarre, Camille Capel, Etienne Simon-Lorière, Vincent Enouf, Maud Vanpeene, Sylvie van der Werf,Bonnet Cyrile |
| EPI_ISL_1259308 | Outre Mer | National Reference Center for Viruses of Respiratory Infections, Institut Pasteur, Paris | Marion Barbet, Sylvie Behillil, Méline Bizard, Angela Brisebarre, Camille Capel, Etienne Simon-Lorière, Vincent Enouf, Maud Vanpeene, Sylvie van der Werf,Rousset Dominique |
| EPI_ISL_1261868 | Ospedale di Venere | Istituto Zooprofilattico Sperimentale della Puglia e della Basilicata | Parisi A., Bianco A., Capozzi L., Del Sambro L., Simone D., Iacobellis M., Ridolfi D., Giannico A. |
| EPI_ISL_1262740, EPI_ISL_1262741, EPI_ISL_1262742, EPI_ISL_1262743 | Hopital | National Reference Center for Viruses of Respiratory Infections, Institut Pasteur, Paris | Marion Barbet, Sylvie Behillil, Méline Bizard, Angela Brisebarre, Camille Capel, Etienne Simon-Lorière, Vincent Enouf, Maud Vanpeene, Sylvie van der Werf,Castelain Sandrine |
| EPI_ISL_1262909 | Outre Mer | National Reference Center for Viruses of Respiratory Infections, Institut Pasteur, Paris | Marion Barbet, Sylvie Behillil, Méline Bizard, Angela Brisebarre, Camille Capel, Etienne Simon-Lorière, Vincent Enouf, Maud Vanpeene, Sylvie van der Werf,Rousset Dominique |
| EPI_ISL_1263319, EPI_ISL_1263320, EPI_ISL_1263321, EPI_ISL_1263322, EPI_ISL_1263323 | Clinical Microbiology Lab RSPTN Universitas Hasanuddin | Clinical Microbiology Lab RSPTN Universitas Hasanuddin | Muhammad Nasrum Massi, Rizalinda Sjahril, Gita Vita Soraya, Triwibawa, Handayani Halik, Muhammad Yusuf Usman, Rufika Shari |
| EPI_ISL_1263540 | Institute for Medical Research, Infectious Disease Research Centre, National Institutes of Health, Ministry of Health Malaysia | Institute for Medical Research, Infectious Disease Research Centre, National Institutes of Health, Ministry of Health Malaysia | Suppiah J, Kamel K, Mohd Zawawi Z, Azizan MA, Thayan R |
| EPI_ISL_1265144 | AZ Sint-Jan Brugge-Oostende AV | AZ SINT-JAN BRUGGE | Marijke Reynders, Patrick Descheemaeker, Laurien Hoornaert, Thomas Van Landschoot, Jorn Hellemans |
| EPI_ISL_1265161 | Institute for Medical Research, Infectious Disease Research Centre, National Institutes of Health, Ministry of Health Malaysia | Institute for Medical Research, Infectious Disease Research Centre, National Institutes of Health, Ministry of Health Malaysia | Suppiah J, Kamel K, Mohd Zawawi Z, Azizan MA, Thayan R |
| EPI_ISL_1265370 | Department of Biomedical, Surgical and Dental Sciences and Istituto Clinico Città Studi, University of Milan | Department of Biomedical, Surgical and Dental Sciences and Istituto Clinico Città Studi, University of Milan | Delbue Serena, Ferrante Pasquale, Signorini Lucia, Dolci Maria, Ticozzi Rosalia |
| EPI_ISL_1265624, EPI_ISL_1265637 | CHUGA-IBP-laboratoire de Virologie | IBP-laboratoire de virologie | Sylvie Larrat, Patrice Morand, Pascal Poignard, Raphaël Germi, Julien Lupo, Anne Signori -Schmuck , Anne-Karen Faure, Benjamin Nemoz, Aurélie Truffot, Léa Ponderand, Julien Andréani |
| EPI_ISL_1265740 | Klinisch Laboratorium ZNA | Klinisch Laboratorium ZNA | Verstrepen et al. |
| EPI_ISL_1273214 | Oxford University Clinical Research Unit (OUCRU) | Oxford University Clinical Research Unit (OUCRU) | Nguyen Van Vinh Chau, Nguyen Thi Thu Hong, Nghiem My Ngoc, Nguyen To Anh, Huynh Trung Trieu, Le Nguyen Truc Nhu, Lam Minh Yen, Ngo Ngoc Quang Minh, Nguyen Thanh Phong, Nguyen Thanh Truong, le Thi Thu Huong, Tran Nguyen Hoang Tu, Le Manh Hung, Tran Tan Thanh, Nguyen Thanh Dung, Nguyen Tri Dung, Guy Thwaites, Le Van Tan |
| EPI_ISL_1273842 | CH de Lens | CHU Lille - Laboratoire de Virologie | AIT YAHYA Emilie, ALIDJINOU Enagnon Kazali, BOCKET Laurence, CREPIN Michel, DEMAY Christophe, ENGELMANN Ilka, GEFFROY Sandrine, GUIGON Aurélie, LAZREK Mouna, NOBILLIAUX Florian, PREVOST Brigitte, THUILLIER Caroline, TINEZ Claire |
| EPI_ISL_1278103, EPI_ISL_1278110, EPI_ISL_1278119 | NL-Dr. Leonard A. Miller Centre for Health Services | National Microbiology Laboratory (NML) | Anna Majer, Shari Tyson, Grace Seo, Philip Mabon, Elsie Grudeski, Rhiannon Huzarewich, Russell Mandes, Anneliese Landgraff, Jennifer Tanner, Natalie Knox, Morag Graham, Gary Van Domselaar, Robert Needle, Yang Yu, Adel Malek, Laura Gilbert, George Zahariadis, Nathalie Bastien, Yan Li, Timothy Booth, Darian Hole, Madison Chapel, Kirsten Biggar, Kerri Smith, CanCOGeN's metadata curation team, Public Health Agency of Canada CanCOGeN team |
| EPI_ISL_1279263 | Laboratorio Central de Epidemiología (LCE) | Instituto Nacional de Enfermedades Respiratorias (INER): Centro de Investigación en Enfermedades Infecciosas (CIENI) | Consorcio Mexicano de Vigilancia Genómica (CoViGen-Mex). Authors (in alphabetical order): Julio Elias Alvarado-Yaah, Carlos F. Arias, Santiago Ávila-Ríos, Víctor Hugo Borja-Aburto, Celia Boukadida, Juan Bautista Chale-Dzul , José Antonio Enciso-Moreno, Gloria Elena Espinoza-Ayala, Fernando Fontove-Herrera, Concepción Grajales-Muñiz, Ricardo Grande, Alfredo Herrera-Estrella, Carla Ivón Herrera-Najera, Pavel Isa, Brenda Irasema Maldonado-Meza, Bernardo Martínez-Miguel, Margarita Matías-Florentino, María Guadalupe de Jesús Mireles-Rivera, Gloria María Molina-Salinas, Hector Montoya-Fuentes, José Esteban Muñoz-Medina, José de Jesús Nuñez-Contreras, Alicia Ocaña-Mondragón, Luis Alberto Ochoa-Carrera, Hector Esteban Paz-Juárez, Francisco Pulido, Helen Haydee Fernanda Ramírez-Plascencia, Angel Gustavo Salas-Lais, Jorge Ivan Salinal-Nevarez, Alejandro Sanchez-Flores, Clara Esperanza Santacruz-Tinoco, María Guadalupe Santiago-Mauricio , Nelly Sélem-Mojica, Blanca Taboada , Gloria Vazquez |

| | | | |
|---|---|---|---|
| EPI_ISL_1279264 | Centro de Investigación Biomédica del Noreste (CIBIN) | Instituto Nacional de Enfermedades Respiratorias (INER): Centro de Investigación en Enfermedades Infecciosas (CIENI) | Consorcio Mexicano de Vigilancia Genómica (CoViGen-Mex). Authors (in alphabetical order): Julio Elias Alvarado-Yaah, Carlos F. Arias, Santiago Ávila-Ríos, Víctor Hugo Borja-Aburto, Celia Boukadida, Juan Bautista Chale-Dzul , José Antonio Enciso-Moreno, Gloria Elena Espinoza-Ayala, Fernando Fontove-Herrera, Concepción Grajales-Muñiz, Ricardo Grande, Alfredo Herrera-Estrella, Carla Ivón Herrera-Najera, Pavel Isa, Brenda Irasema Maldonado-Meza, Bernardo Martínez-Miguel, Margarita Matías-Florentino, María Guadalupe de Jesús Mireles-Rivera, Gloria María Molina-Salinas, Hector Montoya-Fuentes, José Esteban Muñoz-Medina, José de Jesús Nuñez-Contreras, Alicia Ocaña-Mondragón, Luis Alberto Ochoa-Carrera, Hector Esteban Paz-Juárez, Francisco Pulido, Helen Haydee Fernanda Ramírez-Plascencia, Angel Gustavo Salas-Lais, Jorge Ivan Salinal-Nevarez, Alejandro Sanchez-Flores, Clara Esperanza Santacruz-Tinoco, María Guadalupe Santiago-Mauricio , Nelly Sélem-Mojica, Blanca Taboada , Gloria Vazquez |
| EPI_ISL_1279265 | Unidad de Investigación Biomédica de Zacatecas (UIBZ) | Instituto Nacional de Enfermedades Respiratorias (INER): Centro de Investigación en Enfermedades Infecciosas (CIENI) | Consorcio Mexicano de Vigilancia Genómica (CoViGen-Mex). Authors (in alphabetical order): Julio Elias Alvarado-Yaah, Carlos F. Arias, Santiago Ávila-Ríos, Víctor Hugo Borja-Aburto, Celia Boukadida, Juan Bautista Chale-Dzul , José Antonio Enciso-Moreno, Gloria Elena Espinoza-Ayala, Fernando Fontove-Herrera, Concepción Grajales-Muñiz, Ricardo Grande, Alfredo Herrera-Estrella, Carla Ivón Herrera-Najera, Pavel Isa, Brenda Irasema Maldonado-Meza, Bernardo Martínez-Miguel, Margarita Matías-Florentino, María Guadalupe de Jesús Mireles-Rivera, Gloria María Molina-Salinas, Hector Montoya-Fuentes, José Esteban Muñoz-Medina, José de Jesús Nuñez-Contreras, Alicia Ocaña-Mondragón, Luis Alberto Ochoa-Carrera, Hector Esteban Paz-Juárez, Francisco Pulido, Helen Haydee Fernanda Ramírez-Plascencia, Angel Gustavo Salas-Lais, Jorge Ivan Salinal-Nevarez, Alejandro Sanchez-Flores, Clara Esperanza Santacruz-Tinoco, María Guadalupe Santiago-Mauricio , Nelly Sélem-Mojica, Blanca Taboada , Gloria Vazquez |
| EPI_ISL_1279266, EPI_ISL_1279267 | Centro de Investigación Biomédica del Noreste (CIBIN) | Instituto Nacional de Enfermedades Respiratorias (INER): Centro de Investigación en Enfermedades Infecciosas (CIENI) | Consorcio Mexicano de Vigilancia Genómica (CoViGen-Mex). Authors (in alphabetical order): Julio Elias Alvarado-Yaah, Carlos F. Arias, Santiago Ávila-Ríos, Víctor Hugo Borja-Aburto, Celia Boukadida, Juan Bautista Chale-Dzul , José Antonio Enciso-Moreno, Gloria Elena Espinoza-Ayala, Fernando Fontove-Herrera, Concepción Grajales-Muñiz, Ricardo Grande, Alfredo Herrera-Estrella, Carla Ivón Herrera-Najera, Pavel Isa, Brenda Irasema Maldonado-Meza, Bernardo Martínez-Miguel, Margarita Matías-Florentino, María Guadalupe de Jesús Mireles-Rivera, Gloria María Molina-Salinas, Hector Montoya-Fuentes, José Esteban Muñoz-Medina, José de Jesús Nuñez-Contreras, Alicia Ocaña-Mondragón, Luis Alberto Ochoa-Carrera, Hector Esteban Paz-Juárez, Francisco Pulido, Helen Haydee Fernanda Ramírez-Plascencia, Angel Gustavo Salas-Lais, Jorge Ivan Salinal-Nevarez, Alejandro Sanchez-Flores, Clara Esperanza Santacruz-Tinoco, María Guadalupe Santiago-Mauricio , Nelly Sélem-Mojica, Blanca Taboada , Gloria Vazquez |
| EPI_ISL_1279270, EPI_ISL_1279271, EPI_ISL_1279272 | Unidad de Investigación Biomédica de Zacatecas (UIBZ) | Instituto Nacional de Enfermedades Respiratorias (INER): Centro de Investigación en Enfermedades Infecciosas (CIENI) | Consorcio Mexicano de Vigilancia Genómica (CoViGen-Mex). Authors (in alphabetical order): Julio Elias Alvarado-Yaah, Carlos F. Arias, Santiago Ávila-Ríos, Víctor Hugo Borja-Aburto, Celia Boukadida, Juan Bautista Chale-Dzul , José Antonio Enciso-Moreno, Gloria Elena Espinoza-Ayala, Fernando Fontove-Herrera, Concepción Grajales-Muñiz, Ricardo Grande, Alfredo Herrera-Estrella, Carla Ivón Herrera-Najera, Pavel Isa, Brenda Irasema Maldonado-Meza, Bernardo Martínez-Miguel, Margarita Matías-Florentino, María Guadalupe de Jesús Mireles-Rivera, Gloria María Molina-Salinas, Hector Montoya-Fuentes, José Esteban Muñoz-Medina, José de Jesús Nuñez-Contreras, Alicia Ocaña-Mondragón, Luis Alberto Ochoa-Carrera, Hector Esteban Paz-Juárez, Francisco Pulido, Helen Haydee Fernanda Ramírez-Plascencia, Angel Gustavo Salas-Lais, Jorge Ivan Salinal-Nevarez, Alejandro Sanchez-Flores, Clara Esperanza Santacruz-Tinoco, María Guadalupe Santiago-Mauricio , Nelly Sélem-Mojica, Blanca Taboada , Gloria Vazquez |
| EPI_ISL_1279274 | Laboratorio Central de Epidemiología (LCE) | Instituto Nacional de Enfermedades Respiratorias (INER): Centro de Investigación en Enfermedades Infecciosas (CIENI) | Consorcio Mexicano de Vigilancia Genómica (CoViGen-Mex). Authors (in alphabetical order): Julio Elias Alvarado-Yaah, Carlos F. Arias, Santiago Ávila-Ríos, Víctor Hugo Borja-Aburto, Celia Boukadida, Juan Bautista Chale-Dzul , José Antonio Enciso-Moreno, Gloria Elena Espinoza-Ayala, Fernando Fontove-Herrera, Concepción Grajales-Muñiz, Ricardo Grande, Alfredo Herrera-Estrella, Carla Ivón Herrera-Najera, Pavel Isa, Brenda Irasema Maldonado-Meza, Bernardo Martínez-Miguel, Margarita Matías-Florentino, María Guadalupe de Jesús Mireles-Rivera, Gloria María Molina-Salinas, Hector Montoya-Fuentes, José Esteban Muñoz-Medina, José de Jesús Nuñez-Contreras, Alicia Ocaña-Mondragón, Luis Alberto Ochoa-Carrera, Hector Esteban Paz-Juárez, Francisco Pulido, Helen Haydee Fernanda Ramírez-Plascencia, Angel Gustavo Salas-Lais, Jorge Ivan Salinal-Nevarez, Alejandro Sanchez-Flores, Clara Esperanza Santacruz-Tinoco, María Guadalupe Santiago-Mauricio , Nelly Sélem-Mojica, Blanca Taboada , Gloria Vazquez |
| EPI_ISL_1279275, EPI_ISL_1279276 | Centro de Investigación Biomédica del Noreste (CIBIN) | Instituto Nacional de Enfermedades Respiratorias (INER): Centro de Investigación en Enfermedades Infecciosas (CIENI) | Consorcio Mexicano de Vigilancia Genómica (CoViGen-Mex). Authors (in alphabetical order): Julio Elias Alvarado-Yaah, Carlos F. Arias, Santiago Ávila-Ríos, Víctor Hugo Borja-Aburto, Celia Boukadida, Juan Bautista Chale-Dzul , José Antonio Enciso-Moreno, Gloria Elena Espinoza-Ayala, Fernando Fontove-Herrera, Concepción Grajales-Muñiz, Ricardo Grande, Alfredo Herrera-Estrella, Carla Ivón Herrera-Najera, Pavel Isa, Brenda Irasema Maldonado-Meza, Bernardo Martínez-Miguel, Margarita Matías-Florentino, María Guadalupe de Jesús Mireles-Rivera, Gloria María Molina-Salinas, Hector Montoya-Fuentes, José Esteban Muñoz-Medina, José de Jesús Nuñez-Contreras, Alicia Ocaña-Mondragón, Luis Alberto Ochoa-Carrera, Hector Esteban Paz-Juárez, Francisco Pulido, Helen Haydee Fernanda Ramírez-Plascencia, Angel Gustavo Salas-Lais, Jorge Ivan Salinal-Nevarez, Alejandro Sanchez-Flores, Clara Esperanza Santacruz-Tinoco, María Guadalupe Santiago-Mauricio , Nelly Sélem-Mojica, Blanca Taboada , Gloria Vazquez |
| EPI_ISL_1279279, EPI_ISL_1279280 | Unidad de Investigación Biomédica de Zacatecas (UIBZ) | Instituto Nacional de Enfermedades Respiratorias (INER): Centro de Investigación en Enfermedades Infecciosas (CIENI) | Consorcio Mexicano de Vigilancia Genómica (CoViGen-Mex). Authors (in alphabetical order): Julio Elias Alvarado-Yaah, Carlos F. Arias, Santiago Ávila-Ríos, Víctor Hugo Borja-Aburto, Celia Boukadida, Juan Bautista Chale-Dzul , José Antonio Enciso-Moreno, Gloria Elena Espinoza-Ayala, Fernando Fontove-Herrera, Concepción Grajales-Muñiz, Ricardo Grande, Alfredo Herrera-Estrella, Carla Ivón Herrera-Najera, Pavel Isa, Brenda Irasema Maldonado-Meza, Bernardo Martínez-Miguel, Margarita Matías-Florentino, María Guadalupe de Jesús Mireles-Rivera, Gloria María Molina-Salinas, Hector Montoya-Fuentes, José Esteban Muñoz-Medina, José de Jesús Nuñez-Contreras, Alicia Ocaña-Mondragón, Luis Alberto Ochoa-Carrera, Hector Esteban Paz-Juárez, Francisco Pulido, Helen Haydee Fernanda Ramírez-Plascencia, Angel Gustavo Salas-Lais, Jorge Ivan Salinal-Nevarez, Alejandro Sanchez-Flores, Clara Esperanza Santacruz-Tinoco, María Guadalupe Santiago-Mauricio , Nelly Sélem-Mojica, Blanca Taboada , Gloria Vazquez |
| EPI_ISL_1279282 | Unidad de Investigación Médica de Yucatán (UIMY) | Instituto Nacional de Enfermedades Respiratorias (INER): Centro de Investigación en Enfermedades Infecciosas (CIENI) | Consorcio Mexicano de Vigilancia Genómica (CoViGen-Mex). Authors (in alphabetical order): Julio Elias Alvarado-Yaah, Carlos F. Arias, Santiago Ávila-Ríos, Víctor Hugo Borja-Aburto, Celia Boukadida, Juan Bautista Chale-Dzul , José Antonio Enciso-Moreno, Gloria Elena Espinoza-Ayala, Fernando Fontove-Herrera, Concepción Grajales-Muñiz, Ricardo Grande, Alfredo Herrera-Estrella, Carla Ivón Herrera-Najera, Pavel Isa, Brenda Irasema Maldonado-Meza, Bernardo Martínez-Miguel, Margarita Matías-Florentino, María Guadalupe de Jesús Mireles-Rivera, Gloria María Molina-Salinas, Hector Montoya-Fuentes, José Esteban Muñoz-Medina, José de Jesús Nuñez-Contreras, Alicia Ocaña-Mondragón, Luis Alberto Ochoa-Carrera, Hector Esteban Paz-Juárez, Francisco Pulido, Helen Haydee Fernanda Ramírez-Plascencia, Angel Gustavo Salas-Lais, Jorge Ivan Salinal-Nevarez, Alejandro Sanchez-Flores, Clara Esperanza Santacruz-Tinoco, María Guadalupe Santiago-Mauricio , Nelly Sélem-Mojica, Blanca Taboada , Gloria Vazquez |
| EPI_ISL_1279286 | Laboratorio Central de Epidemiología (LCE) | Instituto Nacional de Enfermedades Respiratorias (INER): Centro de Investigación en Enfermedades Infecciosas (CIENI) | Consorcio Mexicano de Vigilancia Genómica (CoViGen-Mex). Authors (in alphabetical order): Julio Elias Alvarado-Yaah, Carlos F. Arias, Santiago Ávila-Ríos, Víctor Hugo Borja-Aburto, Celia Boukadida, Juan Bautista Chale-Dzul , José Antonio Enciso-Moreno, Gloria Elena Espinoza-Ayala, Fernando Fontove-Herrera, Concepción Grajales-Muñiz, Ricardo Grande, Alfredo Herrera-Estrella, Carla Ivón Herrera-Najera, Pavel Isa, Brenda Irasema Maldonado-Meza, Bernardo Martínez-Miguel, Margarita Matías-Florentino, María Guadalupe de Jesús Mireles-Rivera, Gloria María Molina-Salinas, Hector Montoya-Fuentes, José Esteban Muñoz-Medina, José de Jesús Nuñez-Contreras, Alicia Ocaña-Mondragón, Luis Alberto Ochoa-Carrera, Hector Esteban Paz-Juárez, Francisco Pulido, Helen Haydee Fernanda Ramírez-Plascencia, Angel Gustavo Salas-Lais, Jorge Ivan Salinal-Nevarez, Alejandro Sanchez-Flores, Clara Esperanza Santacruz-Tinoco, María Guadalupe Santiago-Mauricio , Nelly Sélem-Mojica, Blanca Taboada , Gloria Vazquez |
| EPI_ISL_1279287 | Centro de Investigación Biomédica del Noreste (CIBIN) | Instituto Nacional de Enfermedades Respiratorias (INER): Centro de Investigación en Enfermedades Infecciosas (CIENI) | Consorcio Mexicano de Vigilancia Genómica (CoViGen-Mex). Authors (in alphabetical order): Julio Elias Alvarado-Yaah, Carlos F. Arias, Santiago Ávila-Ríos, Víctor Hugo Borja-Aburto, Celia Boukadida, Juan Bautista Chale-Dzul , José Antonio Enciso-Moreno, Gloria Elena Espinoza-Ayala, Fernando Fontove-Herrera, Concepción Grajales-Muñiz, Ricardo Grande, Alfredo Herrera-Estrella, Carla Ivón Herrera-Najera, Pavel Isa, Brenda Irasema Maldonado-Meza, Bernardo Martínez-Miguel, Margarita Matías-Florentino, María Guadalupe de Jesús Mireles-Rivera, Gloria María Molina-Salinas, Hector Montoya-Fuentes, José Esteban Muñoz-Medina, José de Jesús Nuñez-Contreras, Alicia Ocaña-Mondragón, Luis Alberto Ochoa-Carrera, Hector Esteban Paz-Juárez, Francisco Pulido, Helen Haydee Fernanda Ramírez-Plascencia, Angel Gustavo Salas-Lais, Jorge Ivan Salinal-Nevarez, Alejandro Sanchez-Flores, Clara Esperanza Santacruz-Tinoco, María Guadalupe Santiago-Mauricio , Nelly Sélem-Mojica, Blanca Taboada , Gloria Vazquez |
| EPI_ISL_1279288, EPI_ISL_1279289 | Laboratorio Central de Epidemiología (LCE) | Instituto Nacional de Enfermedades Respiratorias (INER): Centro de Investigación en Enfermedades Infecciosas (CIENI) | Consorcio Mexicano de Vigilancia Genómica (CoViGen-Mex). Authors (in alphabetical order): Julio Elias Alvarado-Yaah, Carlos F. Arias, Santiago Ávila-Ríos, Víctor Hugo Borja-Aburto, Celia Boukadida, Juan Bautista Chale-Dzul , José Antonio Enciso-Moreno, Gloria Elena Espinoza-Ayala, Fernando Fontove-Herrera, Concepción Grajales-Muñiz, Ricardo Grande, Alfredo Herrera-Estrella, Carla Ivón Herrera-Najera, Pavel Isa, Brenda Irasema Maldonado-Meza, Bernardo Martínez-Miguel, Margarita Matías-Florentino, María Guadalupe de Jesús Mireles-Rivera, Gloria María Molina-Salinas, Hector Montoya-Fuentes, José Esteban Muñoz-Medina, José de Jesús Nuñez-Contreras, Alicia Ocaña-Mondragón, Luis Alberto Ochoa-Carrera, Hector Esteban Paz-Juárez, Francisco Pulido, Helen Haydee Fernanda Ramírez-Plascencia, Angel Gustavo Salas-Lais, Jorge Ivan Salinal-Nevarez, Alejandro Sanchez-Flores, Clara Esperanza Santacruz-Tinoco, María Guadalupe Santiago-Mauricio , Nelly Sélem-Mojica, Blanca Taboada , Gloria Vazquez |
| EPI_ISL_1279294 | Unidad de Investigación Médica de Yucatán (UIMY) | Instituto Nacional de Enfermedades Respiratorias (INER): Centro de Investigación en Enfermedades Infecciosas (CIENI) | Consorcio Mexicano de Vigilancia Genómica (CoViGen-Mex). Authors (in alphabetical order): Julio Elias Alvarado-Yaah, Carlos F. Arias, Santiago Ávila-Ríos, Víctor Hugo Borja-Aburto, Celia Boukadida, Juan Bautista Chale-Dzul , José Antonio Enciso-Moreno, Gloria Elena Espinoza-Ayala, Fernando Fontove-Herrera, Concepción Grajales-Muñiz, Ricardo Grande, Alfredo Herrera-Estrella, Carla Ivón Herrera-Najera, Pavel Isa, Brenda Irasema Maldonado-Meza, Bernardo Martínez-Miguel, Margarita Matías-Florentino, María Guadalupe de Jesús Mireles-Rivera, Gloria María Molina-Salinas, Hector Montoya-Fuentes, José Esteban Muñoz-Medina, José de Jesús Nuñez-Contreras, Alicia Ocaña-Mondragón, Luis Alberto Ochoa-Carrera, Hector Esteban |

| | | | |
|---|---|---|---|
| | | | Paz-Juárez, Francisco Pulido, Helen Haydee Fernanda Ramírez-Plascencia, Angel Gustavo Salas-Lais, Jorge Ivan Salinal-Nevarez, Alejandro Sanchez-Flores, Clara Esperanza Santacruz-Tinoco, María Guadalupe Santiago-Mauricio , Nelly Sélem-Mojica, Blanca Taboada , Gloria Vazquez |
| EPI_ISL_1279295 | Unidad de Investigación Biomédica de Zacatecas (UIBZ) | Instituto Nacional de Enfermedades Respiratorias (INER): Centro de Investigación en Enfermedades Infecciosas (CIENI) | Consorcio Mexicano de Vigilancia Genómica (CoViGen-Mex). Authors (in alphabetical order): Julio Elias Alvarado-Yaah, Carlos F. Arias, Santiago Ávila-Ríos, Víctor Hugo Borja-Aburto, Celia Boukadida, Juan Bautista Chale-Dzul , José Antonio Enciso-Moreno, Gloria Elena Espinoza-Ayala, Fernando Fontove-Herrera, Concepción Grajales-Muñiz, Ricardo Grande, Alfredo Herrera-Estrella, Carla Ivón Herrera-Najera, Pavel Isa, Brenda Irasema Maldonado-Meza, Bernardo Martínez-Miguel, Margarita Matías-Florentino, María Guadalupe de Jesús Mireles-Rivera, Gloria María Molina-Salinas, Hector Montoya-Fuentes, José Esteban Muñoz-Medina, José de Jesús Nuñez-Contreras, Alicia Ocaña-Mondragón, Luis Alberto Ochoa-Carrera, Hector Esteban Paz-Juárez, Francisco Pulido, Helen Haydee Fernanda Ramírez-Plascencia, Angel Gustavo Salas-Lais, Jorge Ivan Salinal-Nevarez, Alejandro Sanchez-Flores, Clara Esperanza Santacruz-Tinoco, María Guadalupe Santiago-Mauricio , Nelly Sélem-Mojica, Blanca Taboada , Gloria Vazquez |
| EPI_ISL_1279298 | Centro de Investigación Biomédica del Noreste (CIBIN) | Instituto Nacional de Enfermedades Respiratorias (INER): Centro de Investigación en Enfermedades Infecciosas (CIENI) | Consorcio Mexicano de Vigilancia Genómica (CoViGen-Mex). Authors (in alphabetical order): Julio Elias Alvarado-Yaah, Carlos F. Arias, Santiago Ávila-Ríos, Víctor Hugo Borja-Aburto, Celia Boukadida, Juan Bautista Chale-Dzul , José Antonio Enciso-Moreno, Gloria Elena Espinoza-Ayala, Fernando Fontove-Herrera, Concepción Grajales-Muñiz, Ricardo Grande, Alfredo Herrera-Estrella, Carla Ivón Herrera-Najera, Pavel Isa, Brenda Irasema Maldonado-Meza, Bernardo Martínez-Miguel, Margarita Matías-Florentino, María Guadalupe de Jesús Mireles-Rivera, Gloria María Molina-Salinas, Hector Montoya-Fuentes, José Esteban Muñoz-Medina, José de Jesús Nuñez-Contreras, Alicia Ocaña-Mondragón, Luis Alberto Ochoa-Carrera, Hector Esteban Paz-Juárez, Francisco Pulido, Helen Haydee Fernanda Ramírez-Plascencia, Angel Gustavo Salas-Lais, Jorge Ivan Salinal-Nevarez, Alejandro Sanchez-Flores, Clara Esperanza Santacruz-Tinoco, María Guadalupe Santiago-Mauricio , Nelly Sélem-Mojica, Blanca Taboada , Gloria Vazquez |
| EPI_ISL_1279300, EPI_ISL_1279301 | Unidad de Investigación Biomédica de Zacatecas (UIBZ) | Instituto Nacional de Enfermedades Respiratorias (INER): Centro de Investigación en Enfermedades Infecciosas (CIENI) | Consorcio Mexicano de Vigilancia Genómica (CoViGen-Mex). Authors (in alphabetical order): Julio Elias Alvarado-Yaah, Carlos F. Arias, Santiago Ávila-Ríos, Víctor Hugo Borja-Aburto, Celia Boukadida, Juan Bautista Chale-Dzul , José Antonio Enciso-Moreno, Gloria Elena Espinoza-Ayala, Fernando Fontove-Herrera, Concepción Grajales-Muñiz, Ricardo Grande, Alfredo Herrera-Estrella, Carla Ivón Herrera-Najera, Pavel Isa, Brenda Irasema Maldonado-Meza, Bernardo Martínez-Miguel, Margarita Matías-Florentino, María Guadalupe de Jesús Mireles-Rivera, Gloria María Molina-Salinas, Hector Montoya-Fuentes, José Esteban Muñoz-Medina, José de Jesús Nuñez-Contreras, Alicia Ocaña-Mondragón, Luis Alberto Ochoa-Carrera, Hector Esteban Paz-Juárez, Francisco Pulido, Helen Haydee Fernanda Ramírez-Plascencia, Angel Gustavo Salas-Lais, Jorge Ivan Salinal-Nevarez, Alejandro Sanchez-Flores, Clara Esperanza Santacruz-Tinoco, María Guadalupe Santiago-Mauricio , Nelly Sélem-Mojica, Blanca Taboada , Gloria Vazquez |
| EPI_ISL_1279302 | Centro de Investigación Biomédica del Noreste (CIBIN) | Instituto Nacional de Enfermedades Respiratorias (INER): Centro de Investigación en Enfermedades Infecciosas (CIENI) | Consorcio Mexicano de Vigilancia Genómica (CoViGen-Mex). Authors (in alphabetical order): Julio Elias Alvarado-Yaah, Carlos F. Arias, Santiago Ávila-Ríos, Víctor Hugo Borja-Aburto, Celia Boukadida, Juan Bautista Chale-Dzul , José Antonio Enciso-Moreno, Gloria Elena Espinoza-Ayala, Fernando Fontove-Herrera, Concepción Grajales-Muñiz, Ricardo Grande, Alfredo Herrera-Estrella, Carla Ivón Herrera-Najera, Pavel Isa, Brenda Irasema Maldonado-Meza, Bernardo Martínez-Miguel, Margarita Matías-Florentino, María Guadalupe de Jesús Mireles-Rivera, Gloria María Molina-Salinas, Hector Montoya-Fuentes, José Esteban Muñoz-Medina, José de Jesús Nuñez-Contreras, Alicia Ocaña-Mondragón, Luis Alberto Ochoa-Carrera, Hector Esteban Paz-Juárez, Francisco Pulido, Helen Haydee Fernanda Ramírez-Plascencia, Angel Gustavo Salas-Lais, Jorge Ivan Salinal-Nevarez, Alejandro Sanchez-Flores, Clara Esperanza Santacruz-Tinoco, María Guadalupe Santiago-Mauricio , Nelly Sélem-Mojica, Blanca Taboada , Gloria Vazquez |
| EPI_ISL_1279304, EPI_ISL_1279305, EPI_ISL_1279306, EPI_ISL_1279307, EPI_ISL_1279308, EPI_ISL_1279309, EPI_ISL_1279310, EPI_ISL_1279311, EPI_ISL_1279312, EPI_ISL_1279313, EPI_ISL_1279314, EPI_ISL_1279315, EPI_ISL_1279316, EPI_ISL_1279317, EPI_ISL_1279318, EPI_ISL_1279319, EPI_ISL_1279320, EPI_ISL_1279321, EPI_ISL_1279322, EPI_ISL_1279323, EPI_ISL_1279324, EPI_ISL_1279325, EPI_ISL_1279326, EPI_ISL_1279327, EPI_ISL_1279328, EPI_ISL_1279329, EPI_ISL_1279330, EPI_ISL_1279331, EPI_ISL_1279332, EPI_ISL_1279333, EPI_ISL_1279334, EPI_ISL_1279335, EPI_ISL_1279336, EPI_ISL_1279337, EPI_ISL_1279338, EPI_ISL_1279339, EPI_ISL_1279340, EPI_ISL_1279341, EPI_ISL_1279342, EPI_ISL_1279343, EPI_ISL_1279344, EPI_ISL_1279345, EPI_ISL_1279346, EPI_ISL_1279347, EPI_ISL_1279348, EPI_ISL_1279349, EPI_ISL_1279350, EPI_ISL_1279351, EPI_ISL_1279352, EPI_ISL_1279353, EPI_ISL_1279354, EPI_ISL_1279355, EPI_ISL_1279356, EPI_ISL_1279357, EPI_ISL_1279358, EPI_ISL_1279359, EPI_ISL_1279360, EPI_ISL_1279361, EPI_ISL_1279362, EPI_ISL_1279363, EPI_ISL_1279364, EPI_ISL_1279365, EPI_ISL_1279366, EPI_ISL_1279367, EPI_ISL_1279368, EPI_ISL_1279369, EPI_ISL_1279370, EPI_ISL_1279371, EPI_ISL_1279372, EPI_ISL_1279373, EPI_ISL_1279374, EPI_ISL_1279375, EPI_ISL_1279376, EPI_ISL_1279377, EPI_ISL_1279378, EPI_ISL_1279379, EPI_ISL_1279380, EPI_ISL_1279381 | | | |
| see above | Unidad de Investigación Biomédica de Zacatecas (UIBZ) | Instituto Nacional de Enfermedades Respiratorias (INER): Centro de Investigación en Enfermedades Infecciosas (CIENI) | Consorcio Mexicano de Vigilancia Genómica (CoViGen-Mex). Authors (in alphabetical order): Julio Elias Alvarado-Yaah, Carlos F. Arias, Santiago Ávila-Ríos, Víctor Hugo Borja-Aburto, Celia Boukadida, Juan Bautista Chale-Dzul , José Antonio Enciso-Moreno, Gloria Elena Espinoza-Ayala, Fernando Fontove-Herrera, Concepción Grajales-Muñiz, Ricardo Grande, Alfredo Herrera-Estrella, Carla Ivón Herrera-Najera, Pavel Isa, Brenda Irasema Maldonado-Meza, Bernardo Martínez-Miguel, Margarita Matías-Florentino, María Guadalupe de Jesús Mireles-Rivera, Gloria María Molina-Salinas, Hector Montoya-Fuentes, José Esteban Muñoz-Medina, José de Jesús Nuñez-Contreras, Alicia Ocaña-Mondragón, Luis Alberto Ochoa-Carrera, Hector Esteban Paz-Juárez, Francisco Pulido, Helen Haydee Fernanda Ramírez-Plascencia, Angel Gustavo Salas-Lais, Jorge Ivan Salinal-Nevarez, Alejandro Sanchez-Flores, Clara Esperanza Santacruz-Tinoco, María Guadalupe Santiago-Mauricio , Nelly Sélem-Mojica, Blanca Taboada , Gloria Vazquez |
| EPI_ISL_1279403, EPI_ISL_1279404, EPI_ISL_1279405, EPI_ISL_1279406, EPI_ISL_1279407, EPI_ISL_1279411, EPI_ISL_1279412, EPI_ISL_1279413, EPI_ISL_1279419, EPI_ISL_1279420, EPI_ISL_1279421, EPI_ISL_1279422, EPI_ISL_1279423, EPI_ISL_1279424, EPI_ISL_1279425, EPI_ISL_1279426, EPI_ISL_1279427, EPI_ISL_1279430, EPI_ISL_1279431, EPI_ISL_1279432, EPI_ISL_1279439 | | | |
| see above | Laboratorio Central de Epidemiología (LCE) | Instituto Nacional de Enfermedades Respiratorias (INER): Centro de Investigación en Enfermedades Infecciosas (CIENI) | Consorcio Mexicano de Vigilancia Genómica (CoViGen-Mex). Authors (in alphabetical order): Julio Elias Alvarado-Yaah, Carlos F. Arias, Santiago Ávila-Ríos, Víctor Hugo Borja-Aburto, Celia Boukadida, Juan Bautista Chale-Dzul , José Antonio Enciso-Moreno, Gloria Elena Espinoza-Ayala, Fernando Fontove-Herrera, Concepción Grajales-Muñiz, Ricardo Grande, Alfredo Herrera-Estrella, Carla Ivón Herrera-Najera, Pavel Isa, Brenda Irasema Maldonado-Meza, Bernardo Martínez-Miguel, Margarita Matías-Florentino, María Guadalupe de Jesús Mireles-Rivera, Gloria María Molina-Salinas, Hector Montoya-Fuentes, José Esteban Muñoz-Medina, José de Jesús Nuñez-Contreras, Alicia Ocaña-Mondragón, Luis Alberto Ochoa-Carrera, Hector Esteban Paz-Juárez, Francisco Pulido, Helen Haydee Fernanda Ramírez-Plascencia, Angel Gustavo Salas-Lais, Jorge Ivan Salinal-Nevarez, Alejandro Sanchez-Flores, Clara Esperanza Santacruz-Tinoco, María Guadalupe Santiago-Mauricio , Nelly Sélem-Mojica, Blanca Taboada , Gloria Vazquez |
| EPI_ISL_1279440 | Unidad de Investigación Médica de Yucatán (UIMY) | Instituto Nacional de Enfermedades Respiratorias (INER): Centro de Investigación en Enfermedades Infecciosas (CIENI) | Consorcio Mexicano de Vigilancia Genómica (CoViGen-Mex). Authors (in alphabetical order): Julio Elias Alvarado-Yaah, Carlos F. Arias, Santiago Ávila-Ríos, Víctor Hugo Borja-Aburto, Celia Boukadida, Juan Bautista Chale-Dzul , José Antonio Enciso-Moreno, Gloria Elena Espinoza-Ayala, Fernando Fontove-Herrera, Concepción Grajales-Muñiz, Ricardo Grande, Alfredo Herrera-Estrella, Carla Ivón Herrera-Najera, Pavel Isa, Brenda Irasema Maldonado-Meza, Bernardo Martínez-Miguel, Margarita Matías-Florentino, María Guadalupe de Jesús Mireles-Rivera, Gloria María Molina-Salinas, Hector Montoya-Fuentes, José Esteban Muñoz-Medina, José de Jesús Nuñez-Contreras, Alicia Ocaña-Mondragón, Luis Alberto Ochoa-Carrera, Hector Esteban Paz-Juárez, Francisco Pulido, Helen Haydee Fernanda Ramírez-Plascencia, Angel Gustavo Salas-Lais, Jorge Ivan Salinal-Nevarez, Alejandro Sanchez-Flores, Clara Esperanza Santacruz-Tinoco, María Guadalupe Santiago-Mauricio , Nelly Sélem-Mojica, Blanca Taboada , Gloria Vazquez |
| EPI_ISL_1279448, EPI_ISL_1279449, EPI_ISL_1279450, EPI_ISL_1279461 | Laboratorio Central de Epidemiología (LCE) | Instituto Nacional de Enfermedades Respiratorias (INER): Centro de Investigación en Enfermedades Infecciosas (CIENI) | Consorcio Mexicano de Vigilancia Genómica (CoViGen-Mex). Authors (in alphabetical order): Julio Elias Alvarado-Yaah, Carlos F. Arias, Santiago Ávila-Ríos, Víctor Hugo Borja-Aburto, Celia Boukadida, Juan Bautista Chale-Dzul , José Antonio Enciso-Moreno, Gloria Elena Espinoza-Ayala, Fernando Fontove-Herrera, Concepción Grajales-Muñiz, Ricardo Grande, Alfredo Herrera-Estrella, Carla Ivón Herrera-Najera, Pavel Isa, Brenda Irasema Maldonado-Meza, Bernardo Martínez-Miguel, Margarita Matías-Florentino, María Guadalupe de Jesús Mireles-Rivera, Gloria María Molina-Salinas, Hector Montoya-Fuentes, José Esteban Muñoz-Medina, José de Jesús Nuñez-Contreras, Alicia Ocaña-Mondragón, Luis Alberto Ochoa-Carrera, Hector Esteban Paz-Juárez, Francisco Pulido, Helen Haydee Fernanda Ramírez-Plascencia, Angel Gustavo Salas-Lais, Jorge Ivan Salinal-Nevarez, Alejandro Sanchez-Flores, Clara Esperanza Santacruz-Tinoco, María Guadalupe Santiago-Mauricio , Nelly Sélem-Mojica, Blanca Taboada , Gloria Vazquez |
| EPI_ISL_1279462, EPI_ISL_1279463, EPI_ISL_1279464, EPI_ISL_1279465, EPI_ISL_1279466, EPI_ISL_1279467, EPI_ISL_1279468, EPI_ISL_1279469, EPI_ISL_1279470, EPI_ISL_1279471, EPI_ISL_1279472, EPI_ISL_1279473, EPI_ISL_1279474, EPI_ISL_1279475, EPI_ISL_1279476, EPI_ISL_1279477, EPI_ISL_1279478, EPI_ISL_1279479, EPI_ISL_1279480, EPI_ISL_1279481, EPI_ISL_1279482, EPI_ISL_1279483, EPI_ISL_1279484, EPI_ISL_1279485, EPI_ISL_1279486, EPI_ISL_1279487, EPI_ISL_1279488, EPI_ISL_1279489, EPI_ISL_1279490, EPI_ISL_1279491, EPI_ISL_1279492, EPI_ISL_1279493, EPI_ISL_1279494, EPI_ISL_1279495, EPI_ISL_1279496, EPI_ISL_1279497, EPI_ISL_1279498, EPI_ISL_1279499, EPI_ISL_1279500, EPI_ISL_1279501, EPI_ISL_1279502, EPI_ISL_1279503, EPI_ISL_1279504, EPI_ISL_1279505, EPI_ISL_1279506, EPI_ISL_1279507, EPI_ISL_1279508, EPI_ISL_1279509, EPI_ISL_1279510, EPI_ISL_1279511, EPI_ISL_1279512, EPI_ISL_1279513, EPI_ISL_1279514, EPI_ISL_1279515, EPI_ISL_1279516, EPI_ISL_1279517, EPI_ISL_1279518, EPI_ISL_1279519, EPI_ISL_1279520, EPI_ISL_1279521, EPI_ISL_1279522, EPI_ISL_1279523, EPI_ISL_1279524, EPI_ISL_1279525, EPI_ISL_1279526, EPI_ISL_1279527, EPI_ISL_1279528, EPI_ISL_1279529, EPI_ISL_1279530, EPI_ISL_1279531, EPI_ISL_1279532, EPI_ISL_1279533, EPI_ISL_1279534, EPI_ISL_1279535, EPI_ISL_1279536 | | | |
| see above | Centro de Investigación Biomédica del Noreste (CIBIN) | Instituto Nacional de Enfermedades Respiratorias (INER): Centro de Investigación en Enfermedades Infecciosas (CIENI) | Consorcio Mexicano de Vigilancia Genómica (CoViGen-Mex). Authors (in alphabetical order): Julio Elias Alvarado-Yaah, Carlos F. Arias, Santiago Ávila-Ríos, Víctor Hugo Borja-Aburto, Celia Boukadida, Juan Bautista Chale-Dzul , José Antonio Enciso-Moreno, Gloria Elena Espinoza-Ayala, Fernando Fontove-Herrera, Concepción Grajales-Muñiz, Ricardo Grande, Alfredo Herrera-Estrella, Carla Ivón Herrera-Najera, Pavel Isa, Brenda Irasema Maldonado-Meza, Bernardo Martínez-Miguel, Margarita Matías-Florentino, María Guadalupe de Jesús Mireles-Rivera, Gloria María Molina-Salinas, Hector Montoya-Fuentes, José Esteban Muñoz-Medina, José de Jesús Nuñez-Contreras, Alicia Ocaña-Mondragón, Luis Alberto Ochoa-Carrera, Hector Esteban Paz-Juárez, Francisco Pulido, Helen Haydee Fernanda Ramírez-Plascencia, Angel Gustavo Salas-Lais, Jorge Ivan Salinal-Nevarez, Alejandro Sanchez-Flores, Clara Esperanza Santacruz-Tinoco, María Guadalupe Santiago-Mauricio , Nelly Sélem-Mojica, Blanca Taboada , Gloria Vazquez |
| EPI_ISL_1279538, EPI_ISL_1279541, EPI_ISL_1279542, EPI_ISL_1279543, EPI_ISL_1279544, EPI_ISL_1279545, EPI_ISL_1279552, EPI_ISL_1279553, EPI_ISL_1279554, EPI_ISL_1279555, EPI_ISL_1279556, EPI_ISL_1279560, EPI_ISL_1279564, EPI_ISL_1279599, EPI_ISL_1279602, EPI_ISL_1279603 | | | |
| see above | Laboratorio Central de Epidemiología (LCE) | Instituto Nacional de Enfermedades Respiratorias (INER): Centro de Investigación en Enfermedades Infecciosas (CIENI) | Consorcio Mexicano de Vigilancia Genómica (CoViGen-Mex). Authors (in alphabetical order): Julio Elias Alvarado-Yaah, Carlos F. Arias, Santiago Ávila-Ríos, Víctor Hugo Borja-Aburto, Celia Boukadida, Juan Bautista Chale-Dzul , José Antonio Enciso-Moreno, Gloria Elena Espinoza-Ayala, Fernando |

| | | | |
|---|---|---|---|
| | | | Fontove-Herrera, Concepción Grajales-Muñiz, Ricardo Grande, Alfredo Herrera-Estrella, Carla Ivón Herrera-Najera, Pavel Isa, Brenda Irasema Maldonado-Meza, Bernardo Martínez-Miguel, Margarita Matías-Florentino, María Guadalupe de Jesús Mireles-Rivera, Gloria María Molina-Salinas, Hector Montoya-Fuentes, José Esteban Muñoz-Medina, José de Jesús Nuñez-Contreras, Alicia Ocaña-Mondragón, Luis Alberto Ochoa-Carrera, Hector Esteban Paz-Juárez, Francisco Pulido, Helen Haydee Fernanda Ramírez-Plascencia, Angel Gustavo Salas-Lais, Jorge Ivan Salinal-Nevarez, Alejandro Sanchez-Flores, Clara Esperanza Santacruz-Tinoco, María Guadalupe Santiago-Mauricio , Nelly Sélem-Mojica, Blanca Taboada , Gloria Vazquez |
| EPI_ISL_1279604, EPI_ISL_1279605, EPI_ISL_1279606, EPI_ISL_1279607, EPI_ISL_1279608, EPI_ISL_1279609, EPI_ISL_1279610, EPI_ISL_1279611, EPI_ISL_1279612, EPI_ISL_1279613, EPI_ISL_1279614 | | | |
| see above | Unidad de Investigación Médica de Yucatán (UIMY) | Instituto Nacional de Enfermedades Respiratorias (INER): Centro de Investigación en Enfermedades Infecciosas (CIENI) | Consorcio Mexicano de Vigilancia Genómica (CoViGen-Mex). Authors (in alphabetical order): Julio Elias Alvarado-Yaah, Carlos F. Arias, Santiago Ávila-Ríos, Víctor Hugo Borja-Aburto, Celia Boukadida, Juan Bautista Chale-Dzul , José Antonio Enciso-Moreno, Gloria Elena Espinoza-Ayala, Fernando Fontove-Herrera, Concepción Grajales-Muñiz, Ricardo Grande, Alfredo Herrera-Estrella, Carla Ivón Herrera-Najera, Pavel Isa, Brenda Irasema Maldonado-Meza, Bernardo Martínez-Miguel, Margarita Matías-Florentino, María Guadalupe de Jesús Mireles-Rivera, Gloria María Molina-Salinas, Hector Montoya-Fuentes, José Esteban Muñoz-Medina, José de Jesús Nuñez-Contreras, Alicia Ocaña-Mondragón, Luis Alberto Ochoa-Carrera, Hector Esteban Paz-Juárez, Francisco Pulido, Helen Haydee Fernanda Ramírez-Plascencia, Angel Gustavo Salas-Lais, Jorge Ivan Salinal-Nevarez, Alejandro Sanchez-Flores, Clara Esperanza Santacruz-Tinoco, María Guadalupe Santiago-Mauricio , Nelly Sélem-Mojica, Blanca Taboada , Gloria Vazquez |
| EPI_ISL_1284067 | RS Mitra Keluarga Pratama Jatiasih | Eijkman Institute for Molecular Biology, Ministry of Research and Technology/National Agency for Research and Innovation | Frilasita A Yudhaputri, Hidayat Trimarsanto, Iskandar Adnan, Lydia V. Panggalo, Sukma Oktavianthi, Willy Agustine, Edison Johar, Safarina G Malik, Khin Saw Myint, Amin Soebandrio |
| EPI_ISL_1284069, EPI_ISL_1284072 | RS Bina Sehat Mandiri | Eijkman Institute for Molecular Biology, Ministry of Research and Technology/National Agency for Research and Innovation | Frilasita A Yudhaputri, Hidayat Trimarsanto, Iskandar Adnan, Lydia V. Panggalo, Sukma Oktavianthi, Willy Agustine, Edison Johar, Safarina G Malik, Khin Saw Myint, Amin Soebandrio |
| EPI_ISL_1284074, EPI_ISL_1284076 | RSIA PKU Muhammadiyah | Eijkman Institute for Molecular Biology, Ministry of Research and Technology/National Agency for Research and Innovation | Hidayat Trimarsanto, Iskandar Adnan, Lydia V. Panggalo, Sukma Oktavianthi, Willy Agustine, Edison Johar, Frilasita A Yudhaputri, Safarina G Malik, Khin Saw Myint, Amin Soebandrio |
| EPI_ISL_1284078 | RS Rumah Indonesia Sehat | Eijkman Institute for Molecular Biology, Ministry of Research and Technology/National Agency for Research and Innovation | Hidayat Trimarsanto, Iskandar Adnan, Lydia V. Panggalo, Sukma Oktavianthi, Willy Agustine, Edison Johar, Frilasita A Yudhaputri, Safarina G Malik, Khin Saw Myint, Amin Soebandrio |
| EPI_ISL_1284083 | RS Umum Harapan Bunda | Eijkman Institute for Molecular Biology, Ministry of Research and Technology/National Agency for Research and Innovation | Hidayat Trimarsanto, Iskandar Adnan, Lydia V. Panggalo, Sukma Oktavianthi, Willy Agustine, Edison Johar, Frilasita A Yudhaputri, Safarina G Malik, Khin Saw Myint, Amin Soebandrio |
| EPI_ISL_1284087 | RSJ Dr. Soeharto Heerdjan | Eijkman Institute for Molecular Biology, Ministry of Research and Technology/National Agency for Research and Innovation | Hidayat Trimarsanto, Iskandar Adnan, Lydia V. Panggalo, Sukma Oktavianthi, Willy Agustine, Edison Johar, Frilasita A Yudhaputri, Safarina G Malik, Khin Saw Myint, Amin Soebandrio |
| EPI_ISL_1284090 | RSU Sumber Waras | Eijkman Institute for Molecular Biology, Ministry of Research and Technology/National Agency for Research and Innovation | Hidayat Trimarsanto, Iskandar Adnan, Lydia V. Panggalo, Sukma Oktavianthi, Willy Agustine, Edison Johar, Frilasita A Yudhaputri, Safarina G Malik, Khin Saw Myint, Amin Soebandrio |
| EPI_ISL_1284105 | RS Mitra Husada Tangerang | Eijkman Institute for Molecular Biology, Ministry of Research and Technology/National Agency for Research and Innovation | Iskandar Adnan, Lydia V. Panggalo, Sukma Oktavianthi, Willy Agustine, Edison Johar, Hidayat Trimarsanto, Frilasita A Yudhaputri, Safarina G Malik, Khin Saw Myint, Amin Soebandrio |
| EPI_ISL_1284109, EPI_ISL_1284114 | RSU HGA Depok | Eijkman Institute for Molecular Biology, Ministry of Research and Technology/National Agency for Research and Innovation | Iskandar Adnan, Lydia V. Panggalo, Sukma Oktavianthi, Willy Agustine, Edison Johar, Hidayat Trimarsanto, Frilasita A Yudhaputri, Safarina G Malik, Khin Saw Myint, Amin Soebandrio |
| EPI_ISL_1284118 | RSU HGA Depok | Eijkman Institute for Molecular Biology, Ministry of Research and Technology/National Agency for Research and Innovation | Lydia V. Panggalo, Sukma Oktavianthi, Willy Agustine, Edison Johar, Hidayat Trimarsanto, Iskandar Adnan, Frilasita A Yudhaputri, Safarina G Malik, Khin Saw Myint, Amin Soebandrio |
| EPI_ISL_1284133 | RS Mitra Keluarga Gading Serpong | Eijkman Institute for Molecular Biology, Ministry of Research and Technology/National Agency for Research and Innovation | Lydia V. Panggalo, Sukma Oktavianthi, Willy Agustine, Edison Johar, Hidayat Trimarsanto, Iskandar Adnan, Frilasita A Yudhaputri, Safarina G Malik, Khin Saw Myint, Amin Soebandrio |
| EPI_ISL_1284140, EPI_ISL_1284145 | RSAB Harapan Kita | Eijkman Institute for Molecular Biology, Ministry of Research and Technology/National Agency for Research and Innovation | Sukma Oktavianthi, Willy Agustine, Edison Johar, Hidayat Trimarsanto, Iskandar Adnan, Lydia V. Panggalo, Frilasita A Yudhaputri, Safarina G Malik, Khin Saw Myint, Amin Soebandrio |
| EPI_ISL_1284149, EPI_ISL_1284152 | RSIA Bunda Sejahtera | Eijkman Institute for Molecular Biology, Ministry of Research and Technology/National Agency for Research and Innovation | Sukma Oktavianthi, Willy Agustine, Edison Johar, Hidayat Trimarsanto, Iskandar Adnan, Lydia V. Panggalo, Frilasita A Yudhaputri, Safarina G Malik, Khin Saw Myint, Amin Soebandrio |
| EPI_ISL_1284156 | RS Sentra Medika Cikarang | Eijkman Institute for Molecular Biology, Ministry of Research and Technology/National Agency for Research and Innovation | Sukma Oktavianthi, Willy Agustine, Edison Johar, Hidayat Trimarsanto, Iskandar Adnan, Lydia V. Panggalo, Frilasita A Yudhaputri, Safarina G Malik, Khin Saw Myint, Amin Soebandrio |
| EPI_ISL_1284168 | RSU Harapan Bunda | Eijkman Institute for Molecular Biology, Ministry of Research and Technology/National Agency for Research and Innovation | Willy Agustine, Edison Johar, Hidayat Trimarsanto, Iskandar Adnan, Lydia V. Panggalo, Sukma Oktavianthi, Frilasita A Yudhaputri, Safarina G Malik, Khin Saw Myint, Amin Soebandrio |
| EPI_ISL_1284169 | RSU Azra | Eijkman Institute for Molecular Biology, Ministry of Research and Technology/National Agency for Research and Innovation | Willy Agustine, Edison Johar, Hidayat Trimarsanto, Iskandar Adnan, Lydia V. Panggalo, Sukma Oktavianthi, Frilasita A Yudhaputri, Safarina G Malik, Khin Saw Myint, Amin Soebandrio |
| EPI_ISL_1284172 | RSU Azra | Eijkman Institute for Molecular Biology, Ministry of Research and Technology/National Agency for Research and Innovation | Edison Johar, Frilasita A Yudhaputri, Hidayat Trimarsanto, Iskandar Adnan, Lydia V. Panggalo, Sukma Oktavianthi, Willy Agustine, Safarina G Malik, Khin Saw Myint, Amin Soebandrio |
| EPI_ISL_1284176 | RS Harapan Mulia Cibarusah | Eijkman Institute for Molecular Biology, Ministry of Research and Technology/National Agency for Research and Innovation | Edison Johar, Frilasita A Yudhaputri, Hidayat Trimarsanto, Iskandar Adnan, Lydia V. Panggalo, Sukma Oktavianthi, Willy Agustine, Safarina G Malik, Khin Saw Myint, Amin Soebandrio |
| EPI_ISL_1284178 | RSIA Selasih Medika | Eijkman Institute for Molecular Biology, Ministry of Research and Technology/National Agency for Research and Innovation | Edison Johar, Frilasita A Yudhaputri, Hidayat Trimarsanto, Iskandar Adnan, Lydia V. Panggalo, Sukma Oktavianthi, Willy Agustine, Safarina G Malik, Khin Saw Myint, Amin Soebandrio |
| EPI_ISL_1284182 | RSJ Dr. Soeharto Heerdjan | Eijkman Institute for Molecular Biology, Ministry of Research and Technology/National Agency for Research and Innovation | Edison Johar, Frilasita A Yudhaputri, Hidayat Trimarsanto, Iskandar Adnan, Lydia V. Panggalo, Sukma Oktavianthi, Willy Agustine, Safarina G Malik, Khin Saw Myint, Amin Soebandrio |
| EPI_ISL_1284186 | RS Cendana | Eijkman Institute for Molecular Biology, Ministry of Research and Technology/National Agency for Research and Innovation | Edison Johar, Frilasita A Yudhaputri, Hidayat Trimarsanto, Iskandar Adnan, Lydia V. Panggalo, Sukma Oktavianthi, Willy Agustine, Safarina G Malik, Khin Saw Myint, Amin Soebandrio |
| EPI_ISL_1284190 | RS Permata Cibubur | Eijkman Institute for Molecular Biology, Ministry of Research and Technology/National Agency for Research and Innovation | Edison Johar, Frilasita A Yudhaputri, Hidayat Trimarsanto, Iskandar Adnan, Lydia V. Panggalo, Sukma Oktavianthi, Willy Agustine, Safarina G Malik, Khin Saw Myint, Amin Soebandrio |
| EPI_ISL_1284193 | RS Permata Cibubur | Eijkman Institute for Molecular Biology, Ministry of Research and Technology/National Agency for Research and Innovation | Frilasita A Yudhaputri, Hidayat Trimarsanto, Iskandar Adnan, Lydia V. Panggalo, Sukma Oktavianthi, Willy Agustine, Edison Johar, Safarina G Malik, Khin Saw Myint, Amin Soebandrio |
| EPI_ISL_1284201 | RS Bhakti Husada Cikarang | Eijkman Institute for Molecular Biology, Ministry of Research and Technology/National Agency for Research and Innovation | Frilasita A Yudhaputri, Hidayat Trimarsanto, Iskandar Adnan, Lydia V. Panggalo, Sukma Oktavianthi, Willy Agustine, Edison Johar, Safarina G Malik, Khin Saw Myint, Amin Soebandrio |
| EPI_ISL_1284207 | RS Mitra Keluarga Cibubur | Eijkman Institute for Molecular Biology, Ministry of Research and Technology/National Agency for Research and Innovation | Frilasita A Yudhaputri, Hidayat Trimarsanto, Iskandar Adnan, Lydia V. Panggalo, Sukma Oktavianthi, Willy Agustine, Edison Johar, Safarina G Malik, Khin Saw Myint, Amin Soebandrio |
| EPI_ISL_1284210 | RS Mitra Husada Pringsewu | Eijkman Institute for Molecular Biology, Ministry of Research and Technology/National Agency for Research and Innovation | Hidayat Trimarsanto, Iskandar Adnan, Lydia V. Panggalo, Sukma Oktavianthi, Willy Agustine, Edison Johar, Frilasita A Yudhaputri, Safarina G Malik, Khin Saw Myint, Amin Soebandrio |
| EPI_ISL_1284226, EPI_ISL_1284230, EPI_ISL_1284234 | RSU Harapan Bunda | Eijkman Institute for Molecular Biology, Ministry of Research and Technology/National Agency for Research and Innovation | Iskandar Adnan, Lydia V. Panggalo, Sukma Oktavianthi, Willy Agustine, Edison Johar, Hidayat Trimarsanto, Frilasita A Yudhaputri, Safarina G Malik, Khin Saw Myint, Amin Soebandrio |
| EPI_ISL_1284237 | RSU Harapan Bunda | Eijkman Institute for Molecular Biology, Ministry of Research and Technology/National Agency for Research and Innovation | Lydia V. Panggalo, Sukma Oktavianthi, Willy Agustine, Edison Johar, Hidayat Trimarsanto, Iskandar Adnan, Frilasita A Yudhaputri, Safarina G Malik, Khin Saw Myint, Amin Soebandrio |
| EPI_ISL_1284244, EPI_ISL_1284247 | RS Masmitra | Eijkman Institute for Molecular Biology, Ministry of Research and Technology/National Agency for Research and Innovation | Lydia V. Panggalo, Sukma Oktavianthi, Willy Agustine, Edison Johar, Hidayat Trimarsanto, Iskandar Adnan, Frilasita A Yudhaputri, Safarina G Malik, Khin Saw Myint, Amin Soebandrio |
| EPI_ISL_1284251 | RS Mitra Keluarga Gading Serpong | Eijkman Institute for Molecular Biology, Ministry of Research | Lydia V. Panggalo, Sukma Oktavianthi, Willy Agustine, Edison Johar, Hidayat Trimarsanto, Iskandar Adnan, Frilasita A Yudhaputri, Safarina G Malik, Khin |

| | | | |
|---|---|---|---|
| | | and Technology/National Agency for Research and Innovation | Saw Myint, Amin Soebandrio |
| EPI_ISL_1284254 | RS Mitra Keluarga Gading Serpong | Eijkman Institute for Molecular Biology, Ministry of Research and Technology/National Agency for Research and Innovation | Sukma Oktavianthi, Willy Agustine, Edison Johar, Hidayat Trimarsanto, Iskandar Adnan, Lydia V. Panggalo, Frilasita A Yudhaputri, Safarina G Malik, Khin Saw Myint, Amin Soebandrio |
| EPI_ISL_1284269 | RS Mitra Keluarga Bekasi Barat | Eijkman Institute for Molecular Biology, Ministry of Research and Technology/National Agency for Research and Innovation | Willy Agustine, Edison Johar, Hidayat Trimarsanto, Iskandar Adnan, Lydia V. Panggalo, Sukma Oktavianthi, Frilasita A Yudhaputri, Safarina G Malik, Khin Saw Myint, Amin Soebandrio |
| EPI_ISL_1284273 | RS Medirossa Cikarang | Eijkman Institute for Molecular Biology, Ministry of Research and Technology/National Agency for Research and Innovation | Willy Agustine, Edison Johar, Hidayat Trimarsanto, Iskandar Adnan, Lydia V. Panggalo, Sukma Oktavianthi, Frilasita A Yudhaputri, Safarina G Malik, Khin Saw Myint, Amin Soebandrio |
| EPI_ISL_1284288 | RS Mitra Keluarga Kalideres | Eijkman Institute for Molecular Biology, Ministry of Research and Technology/National Agency for Research and Innovation | Willy Agustine, Edison Johar, Hidayat Trimarsanto, Iskandar Adnan, Lydia V. Panggalo, Sukma Oktavianthi, Frilasita A Yudhaputri, Safarina G Malik, Khin Saw Myint, Amin Soebandrio |
| EPI_ISL_1284292 | RS Mitra Keluarga Kalideres | Eijkman Institute for Molecular Biology, Ministry of Research and Technology/National Agency for Research and Innovation | Edison Johar, Frilasita A Yudhaputri, Hidayat Trimarsanto, Iskandar Adnan, Lydia V. Panggalo, Sukma Oktavianthi, Willy Agustine, Safarina G Malik, Khin Saw Myint, Amin Soebandrio |
| EPI_ISL_1284296 | RSJ Dr. Soeharto Heerdjan | Eijkman Institute for Molecular Biology, Ministry of Research and Technology/National Agency for Research and Innovation | Edison Johar, Frilasita A Yudhaputri, Hidayat Trimarsanto, Iskandar Adnan, Lydia V. Panggalo, Sukma Oktavianthi, Willy Agustine, Safarina G Malik, Khin Saw Myint, Amin Soebandrio |
| EPI_ISL_1284307 | RS Mitra Keluarga Gading Serpong | Eijkman Institute for Molecular Biology, Ministry of Research and Technology/National Agency for Research and Innovation | Edison Johar, Frilasita A Yudhaputri, Hidayat Trimarsanto, Iskandar Adnan, Lydia V. Panggalo, Sukma Oktavianthi, Willy Agustine, Safarina G Malik, Khin Saw Myint, Amin Soebandrio |
| EPI_ISL_1284310 | RS Bhakti Husada Cikarang | Eijkman Institute for Molecular Biology, Ministry of Research and Technology/National Agency for Research and Innovation | Edison Johar, Frilasita A Yudhaputri, Hidayat Trimarsanto, Iskandar Adnan, Lydia V. Panggalo, Sukma Oktavianthi, Willy Agustine, Safarina G Malik, Khin Saw Myint, Amin Soebandrio |
| EPI_ISL_1284318 | RS Sentra Medika Cikarang | Eijkman Institute for Molecular Biology, Ministry of Research and Technology/National Agency for Research and Innovation | Edison Johar, Frilasita A Yudhaputri, Hidayat Trimarsanto, Iskandar Adnan, Lydia V. Panggalo, Sukma Oktavianthi, Willy Agustine, Safarina G Malik, Khin Saw Myint, Amin Soebandrio |
| EPI_ISL_1284322 | RS Islam Sari Asih Ar-Rahmah | Eijkman Institute for Molecular Biology, Ministry of Research and Technology/National Agency for Research and Innovation | Frilasita A Yudhaputri, Hidayat Trimarsanto, Iskandar Adnan, Lydia V. Panggalo, Sukma Oktavianthi, Willy Agustine, Edison Johar, Safarina G Malik, Khin Saw Myint, Amin Soebandrio |
| EPI_ISL_1287761, EPI_ISL_1287762, EPI_ISL_1287763, EPI_ISL_1287764, EPI_ISL_1287765, EPI_ISL_1287766, EPI_ISL_1287767 | Unidad de Investigación Biomédica de Zacatecas (UIBZ) | Instituto Nacional de Enfermedades Respiratorias (INER): Centro de Investigación en Enfermedades Infecciosas (CIENI) | Consorcio Mexicano de Vigilancia Genómica (CoViGen-Mex). Authors (in alphabetical order): Julio Elias Alvarado-Yaah, Carlos F. Arias, Santiago Ávila-Ríos, Víctor Hugo Borja-Aburto, Celia Boukadida, Juan Bautista Chale-Dzul , José Antonio Enciso-Moreno, Gloria Elena Espinoza-Ayala, Fernando Fontove-Herrera, Concepción Grajales-Muñiz, Ricardo Grande, Alfredo Herrera-Estrella, Carla Ivón Herrera-Najera, Pavel Isa, Brenda Irasema Maldonado-Meza, Bernardo Martínez-Miguel, Margarita Matías-Florentino, María Guadalupe de Jesús Mireles-Rivera, Gloria María Molina-Salinas, Hector Montoya-Fuentes, José Esteban Muñoz-Medina, José de Jesús Nuñez-Contreras, Alicia Ocaña-Mondragón, Luis Alberto Ochoa-Carrera, Hector Esteban Paz-Juárez, Francisco Pulido, Helen Haydee Fernanda Ramírez-Plascencia, Angel Gustavo Salas-Lais, Jorge Ivan Salinal-Nevarez, Alejandro Sanchez-Flores, Clara Esperanza Santacruz-Tinoco, María Guadalupe Santiago-Mauricio , Nelly Sélem-Mojica, Blanca Taboada , Gloria Vazquez |
| EPI_ISL_1287768, EPI_ISL_1287770 | Laboratorio Central de Epidemiología (LCE) | Instituto Nacional de Enfermedades Respiratorias (INER): Centro de Investigación en Enfermedades Infecciosas (CIENI) | Consorcio Mexicano de Vigilancia Genómica (CoViGen-Mex). Authors (in alphabetical order): Julio Elias Alvarado-Yaah, Carlos F. Arias, Santiago Ávila-Ríos, Víctor Hugo Borja-Aburto, Celia Boukadida, Juan Bautista Chale-Dzul , José Antonio Enciso-Moreno, Gloria Elena Espinoza-Ayala, Fernando Fontove-Herrera, Concepción Grajales-Muñiz, Ricardo Grande, Alfredo Herrera-Estrella, Carla Ivón Herrera-Najera, Pavel Isa, Brenda Irasema Maldonado-Meza, Bernardo Martínez-Miguel, Margarita Matías-Florentino, María Guadalupe de Jesús Mireles-Rivera, Gloria María Molina-Salinas, Hector Montoya-Fuentes, José Esteban Muñoz-Medina, José de Jesús Nuñez-Contreras, Alicia Ocaña-Mondragón, Luis Alberto Ochoa-Carrera, Hector Esteban Paz-Juárez, Francisco Pulido, Helen Haydee Fernanda Ramírez-Plascencia, Angel Gustavo Salas-Lais, Jorge Ivan Salinal-Nevarez, Alejandro Sanchez-Flores, Clara Esperanza Santacruz-Tinoco, María Guadalupe Santiago-Mauricio , Nelly Sélem-Mojica, Blanca Taboada , Gloria Vazquez |
| EPI_ISL_1287771, EPI_ISL_1287772, EPI_ISL_1287773 | Centro de Investigación Biomédica del Noreste (CIBIN) | Instituto Nacional de Enfermedades Respiratorias (INER): Centro de Investigación en Enfermedades Infecciosas (CIENI) | Consorcio Mexicano de Vigilancia Genómica (CoViGen-Mex). Authors (in alphabetical order): Julio Elias Alvarado-Yaah, Carlos F. Arias, Santiago Ávila-Ríos, Víctor Hugo Borja-Aburto, Celia Boukadida, Juan Bautista Chale-Dzul , José Antonio Enciso-Moreno, Gloria Elena Espinoza-Ayala, Fernando Fontove-Herrera, Concepción Grajales-Muñiz, Ricardo Grande, Alfredo Herrera-Estrella, Carla Ivón Herrera-Najera, Pavel Isa, Brenda Irasema Maldonado-Meza, Bernardo Martínez-Miguel, Margarita Matías-Florentino, María Guadalupe de Jesús Mireles-Rivera, Gloria María Molina-Salinas, Hector Montoya-Fuentes, José Esteban Muñoz-Medina, José de Jesús Nuñez-Contreras, Alicia Ocaña-Mondragón, Luis Alberto Ochoa-Carrera, Hector Esteban Paz-Juárez, Francisco Pulido, Helen Haydee Fernanda Ramírez-Plascencia, Angel Gustavo Salas-Lais, Jorge Ivan Salinal-Nevarez, Alejandro Sanchez-Flores, Clara Esperanza Santacruz-Tinoco, María Guadalupe Santiago-Mauricio , Nelly Sélem-Mojica, Blanca Taboada , Gloria Vazquez |
| EPI_ISL_1287776 | Laboratorio Central de Epidemiología (LCE) | Instituto Nacional de Enfermedades Respiratorias (INER): Centro de Investigación en Enfermedades Infecciosas (CIENI) | Consorcio Mexicano de Vigilancia Genómica (CoViGen-Mex). Authors (in alphabetical order): Julio Elias Alvarado-Yaah, Carlos F. Arias, Santiago Ávila-Ríos, Víctor Hugo Borja-Aburto, Celia Boukadida, Juan Bautista Chale-Dzul , José Antonio Enciso-Moreno, Gloria Elena Espinoza-Ayala, Fernando Fontove-Herrera, Concepción Grajales-Muñiz, Ricardo Grande, Alfredo Herrera-Estrella, Carla Ivón Herrera-Najera, Pavel Isa, Brenda Irasema Maldonado-Meza, Bernardo Martínez-Miguel, Margarita Matías-Florentino, María Guadalupe de Jesús Mireles-Rivera, Gloria María Molina-Salinas, Hector Montoya-Fuentes, José Esteban Muñoz-Medina, José de Jesús Nuñez-Contreras, Alicia Ocaña-Mondragón, Luis Alberto Ochoa-Carrera, Hector Esteban Paz-Juárez, Francisco Pulido, Helen Haydee Fernanda Ramírez-Plascencia, Angel Gustavo Salas-Lais, Jorge Ivan Salinal-Nevarez, Alejandro Sanchez-Flores, Clara Esperanza Santacruz-Tinoco, María Guadalupe Santiago-Mauricio , Nelly Sélem-Mojica, Blanca Taboada , Gloria Vazquez |
| EPI_ISL_1288153, EPI_ISL_1288154, EPI_ISL_1288155, EPI_ISL_1288156, EPI_ISL_1288157, EPI_ISL_1288159, EPI_ISL_1288161, EPI_ISL_1288162, EPI_ISL_1288163, EPI_ISL_1288164, EPI_ISL_1288165, EPI_ISL_1288166, EPI_ISL_1288168, EPI_ISL_1288170, EPI_ISL_1288171, EPI_ISL_1288172, EPI_ISL_1288173, EPI_ISL_1288175, EPI_ISL_1288176, EPI_ISL_1288177, EPI_ISL_1288178, EPI_ISL_1288179, EPI_ISL_1288180, EPI_ISL_1288181, EPI_ISL_1288182, EPI_ISL_1288183, EPI_ISL_1288184, EPI_ISL_1288185, EPI_ISL_1288186, EPI_ISL_1288187, EPI_ISL_1288188, EPI_ISL_1288189, EPI_ISL_1288190, EPI_ISL_1288191, EPI_ISL_1288192, EPI_ISL_1288193, EPI_ISL_1288194, EPI_ISL_1288195, EPI_ISL_1288197, EPI_ISL_1288199, EPI_ISL_1288200, EPI_ISL_1288201, EPI_ISL_1288202, EPI_ISL_1288203, EPI_ISL_1288204, EPI_ISL_1288205, EPI_ISL_1288206, EPI_ISL_1288207, EPI_ISL_1288208, EPI_ISL_1288209, EPI_ISL_1288210, EPI_ISL_1288211, EPI_ISL_1288213, EPI_ISL_1288214, EPI_ISL_1288216, EPI_ISL_1288217, EPI_ISL_1288218, EPI_ISL_1288219, EPI_ISL_1288220, EPI_ISL_1288221, EPI_ISL_1288222, EPI_ISL_1288223, EPI_ISL_1288224, EPI_ISL_1288225, EPI_ISL_1288227, EPI_ISL_1288228, EPI_ISL_1288229, EPI_ISL_1288230, EPI_ISL_1288231, EPI_ISL_1288232, EPI_ISL_1288233, EPI_ISL_1288234, EPI_ISL_1288236, EPI_ISL_1288237, EPI_ISL_1288238, EPI_ISL_1288239, EPI_ISL_1288240, EPI_ISL_1288241, EPI_ISL_1288242, EPI_ISL_1288243, EPI_ISL_1288244, EPI_ISL_1288245, EPI_ISL_1288246, EPI_ISL_1288247, EPI_ISL_1288248, EPI_ISL_1288249, EPI_ISL_1288251, EPI_ISL_1288252, EPI_ISL_1288253, EPI_ISL_1288254, EPI_ISL_1288255, EPI_ISL_1288256, EPI_ISL_1288257, EPI_ISL_1288259, EPI_ISL_1288260, EPI_ISL_1288261, EPI_ISL_1288263, EPI_ISL_1288264, EPI_ISL_1288265, EPI_ISL_1288266, EPI_ISL_1288267, EPI_ISL_1288268, EPI_ISL_1288269, EPI_ISL_1288270, EPI_ISL_1288271, EPI_ISL_1288272, EPI_ISL_1288273, EPI_ISL_1288274, EPI_ISL_1288275, EPI_ISL_1288276, EPI_ISL_1288277, EPI_ISL_1288278, EPI_ISL_1288280, EPI_ISL_1288281, EPI_ISL_1288282, EPI_ISL_1288283, EPI_ISL_1288284, EPI_ISL_1288285, EPI_ISL_1288287, EPI_ISL_1288288, EPI_ISL_1288290, EPI_ISL_1288291, EPI_ISL_1288292, EPI_ISL_1288293, EPI_ISL_1288294, EPI_ISL_1288295, EPI_ISL_1288296, EPI_ISL_1288298, EPI_ISL_1288299, EPI_ISL_1288300, EPI_ISL_1288301, EPI_ISL_1288302, EPI_ISL_1288303, EPI_ISL_1288304, EPI_ISL_1288305, EPI_ISL_1288306, EPI_ISL_1288307, EPI_ISL_1288308, EPI_ISL_1288309, EPI_ISL_1288310, EPI_ISL_1288311, EPI_ISL_1288312, EPI_ISL_1288313, EPI_ISL_1288314, EPI_ISL_1288315, EPI_ISL_1288317, EPI_ISL_1288318, EPI_ISL_1288319, EPI_ISL_1288320, EPI_ISL_1288321, EPI_ISL_1288323, EPI_ISL_1288324, EPI_ISL_1288325, EPI_ISL_1288326, EPI_ISL_1288327, EPI_ISL_1288328, EPI_ISL_1288329, EPI_ISL_1288330, EPI_ISL_1288331, EPI_ISL_1288332, EPI_ISL_1288333, EPI_ISL_1288334, EPI_ISL_1288335, EPI_ISL_1288336, EPI_ISL_1288337, EPI_ISL_1288338, EPI_ISL_1288339, EPI_ISL_1288340, EPI_ISL_1288341, EPI_ISL_1288342, EPI_ISL_1288343, EPI_ISL_1288344, EPI_ISL_1288345, EPI_ISL_1288347, EPI_ISL_1288348, EPI_ISL_1288349, EPI_ISL_1288350, EPI_ISL_1288352, EPI_ISL_1288353, EPI_ISL_1288354, EPI_ISL_1288355, EPI_ISL_1288356, EPI_ISL_1288357, EPI_ISL_1288358, EPI_ISL_1288359, EPI_ISL_1288360, EPI_ISL_1288362, EPI_ISL_1288363, EPI_ISL_1288364, EPI_ISL_1288365, EPI_ISL_1288366, EPI_ISL_1288367, EPI_ISL_1288368, EPI_ISL_1288370, EPI_ISL_1288371, EPI_ISL_1288372, EPI_ISL_1288373, EPI_ISL_1288374, EPI_ISL_1288375, EPI_ISL_1288376, EPI_ISL_1288377, EPI_ISL_1288378, EPI_ISL_1288379, EPI_ISL_1288380, EPI_ISL_1288381, EPI_ISL_1288382, EPI_ISL_1288383, EPI_ISL_1288384, EPI_ISL_1288385, EPI_ISL_1288386, EPI_ISL_1288387, EPI_ISL_1288388, EPI_ISL_1288389, EPI_ISL_1288390, EPI_ISL_1288391, EPI_ISL_1288392, EPI_ISL_1288393, EPI_ISL_1288394, EPI_ISL_1288395, EPI_ISL_1288396, EPI_ISL_1288397, EPI_ISL_1288398, EPI_ISL_1288399, EPI_ISL_1288400, EPI_ISL_1288401, EPI_ISL_1288402, EPI_ISL_1288403, EPI_ISL_1288404, EPI_ISL_1288405, EPI_ISL_1288406, EPI_ISL_1288407, EPI_ISL_1288408, EPI_ISL_1288409, EPI_ISL_1288410, EPI_ISL_1288412, EPI_ISL_1288413, EPI_ISL_1288414, EPI_ISL_1288415, EPI_ISL_1288416, EPI_ISL_1288417, EPI_ISL_1288418, EPI_ISL_1288419, EPI_ISL_1288420, EPI_ISL_1288421, EPI_ISL_1288422, EPI_ISL_1288423, EPI_ISL_1288424, EPI_ISL_1288425, EPI_ISL_1288426, EPI_ISL_1288427, EPI_ISL_1288429, EPI_ISL_1288430, EPI_ISL_1288431, EPI_ISL_1288432, EPI_ISL_1288433, EPI_ISL_1288434, EPI_ISL_1288435, EPI_ISL_1288436, EPI_ISL_1288438, EPI_ISL_1288439, EPI_ISL_1288440, EPI_ISL_1288441, EPI_ISL_1288442, EPI_ISL_1288443, EPI_ISL_1288444, EPI_ISL_1288445, EPI_ISL_1288446, EPI_ISL_1288447, EPI_ISL_1288448, EPI_ISL_1288449, EPI_ISL_1288450, EPI_ISL_1288451, EPI_ISL_1288452, EPI_ISL_1288453, EPI_ISL_1288454, EPI_ISL_1288455, EPI_ISL_1288456, EPI_ISL_1288457, EPI_ISL_1288459, EPI_ISL_1288460, EPI_ISL_1288461, EPI_ISL_1288462, EPI_ISL_1288463, EPI_ISL_1288464, EPI_ISL_1288465, EPI_ISL_1288466, EPI_ISL_1288467, EPI_ISL_1288468, EPI_ISL_1288469, EPI_ISL_1288470, EPI_ISL_1288471, EPI_ISL_1288472, EPI_ISL_1288473, EPI_ISL_1288474, EPI_ISL_1288475, EPI_ISL_1288476, EPI_ISL_1288477, EPI_ISL_1288478, EPI_ISL_1288479, EPI_ISL_1288480, EPI_ISL_1288481, EPI_ISL_1288482, EPI_ISL_1288483, EPI_ISL_1288485, EPI_ISL_1288486, EPI_ISL_1288487, EPI_ISL_1288488, EPI_ISL_1288489, EPI_ISL_1288490, EPI_ISL_1288491, EPI_ISL_1288492, EPI_ISL_1288493, EPI_ISL_1288494, EPI_ISL_1288495, EPI_ISL_1288496, EPI_ISL_1288497, EPI_ISL_1288498, EPI_ISL_1288499, EPI_ISL_1288500, EPI_ISL_1288501, EPI_ISL_1288503, EPI_ISL_1288504, EPI_ISL_1288505, EPI_ISL_1288506, EPI_ISL_1288507, EPI_ISL_1288508, EPI_ISL_1288509, EPI_ISL_1288510, EPI_ISL_1288511, EPI_ISL_1288512, EPI_ISL_1288513, EPI_ISL_1288514, EPI_ISL_1288515, EPI_ISL_1288516, EPI_ISL_1288517, EPI_ISL_1288518, EPI_ISL_1288519, EPI_ISL_1288520, EPI_ISL_1288522, EPI_ISL_1288523, EPI_ISL_1288524, EPI_ISL_1288525, EPI_ISL_1288526, EPI_ISL_1288527, EPI_ISL_1288529, EPI_ISL_1288530, EPI_ISL_1288531, EPI_ISL_1288532, EPI_ISL_1288533 | | | |
| see above | Laboratorio Central de Epidemiología (LCE) | Instituto de Biotecnología de la UNAM | Consorcio Mexicano de Vigilancia Genómica (CoViGen-Mex). Authors (in alphabetical order): Julio Elias Alvarado-Yaah, Carlos F. Arias, Santiago Ávila-Ríos, Víctor Hugo Borja-Aburto, Celia Boukadida, Juan Bautista Chale-Dzul , José Antonio Enciso-Moreno, Gloria Elena Espinoza-Ayala, Fernando Fontove-Herrera, Concepción Grajales-Muñiz, Ricardo Grande, Alfredo Herrera-Estrella, Carla Ivón Herrera-Najera, Pavel Isa, Brenda Irasema Maldonado-Meza, Bernardo Martínez-Miguel, Margarita Matías-Florentino, María Guadalupe de Jesús Mireles-Rivera, Gloria María Molina-Salinas, Hector Montoya-Fuentes, José Esteban Muñoz-Medina, José de Jesús Nuñez-Contreras, Alicia Ocaña-Mondragón, Luis Alberto Ochoa-Carrera, Hector Esteban Paz-Juárez, Francisco Pulido, Helen Haydee Fernanda Ramírez-Plascencia, Angel Gustavo Salas-Lais, Jorge Ivan Salinal-Nevarez, Alejandro Sanchez-Flores, Clara Esperanza Santacruz-Tinoco, María Guadalupe Santiago-Mauricio , Nelly Sélem-Mojica, Blanca Taboada , Gloria Vazquez |
| EPI_ISL_1291549, EPI_ISL_1291554, EPI_ISL_1291555, EPI_ISL_1291556 | Microbiology Department, University Hospital Donostia | Microbiology Department, University Hospital Donostia | Marimon JM, Montes M, Piñeiro L, Sorarrain A, Gomez M, Cilla G. |

| | | | |
|---|---|---|---|
| EPI_ISL_1292807 | CHU Pontchaillou | CHU Pontchaillou | GROLHIER Claire, DUFOUR Marie José, DENOUAL Florent, ETCHEVERRY Amandine, SASSI Mohamed, JAGLINE Steven, PRONIER Charlotte, GALIBERT Marie Dominique, DE TAYRAC Marie, THIBAULT Vincent |
| EPI_ISL_1293052, EPI_ISL_1293053, EPI_ISL_1293054, EPI_ISL_1293055 | LACEN de Rondonia | Instituto Adolfo Lutz, Interdiciplinary Procedures Center, Strategic Laboratory | Claudio Tavares Sacchi, Claudia Regina Gonçalves, Erica Valessa Ramos Gomes, Karoline Rodrigues Campos, Caio Vinicius Dias Lopes |
| EPI_ISL_1293056, EPI_ISL_1293057, EPI_ISL_1293058, EPI_ISL_1293059, EPI_ISL_1293060, EPI_ISL_1293061, EPI_ISL_1293062, EPI_ISL_1293063, EPI_ISL_1293064, EPI_ISL_1293065, EPI_ISL_1293067, EPI_ISL_1293068, EPI_ISL_1293069, EPI_ISL_1293070, EPI_ISL_1293071, EPI_ISL_1293072, EPI_ISL_1293073, EPI_ISL_1293074, EPI_ISL_1293075, EPI_ISL_1293076, EPI_ISL_1293077, EPI_ISL_1293078, EPI_ISL_1293079, EPI_ISL_1293080, EPI_ISL_1293081 | | | |
| see above | IAL Regional de Sorocaba | Instituto Adolfo Lutz, Interdiciplinary Procedures Center, Strategic Laboratory | Claudio Tavares Sacchi, Claudia Regina Gonçalves, Erica Valessa Ramos Gomes, Karoline Rodrigues Campos, Caio Vinicius Dias Lopes |
| EPI_ISL_1293338, EPI_ISL_1293343, EPI_ISL_1293346, EPI_ISL_1293349 | Institut de Virologie du CHU de Strasbourg | Institut de Virologie du CHU de Strasbourg | Fafi-Kremer Samira, Gallais Floriane, Gantner Pierre, Laugel Elodie, Solis Morgane, Velay Aurélie, Wendling Marie-Josée |
| EPI_ISL_1293350 | Oxford University Clinical Research Unit (OUCRU) | Oxford University Clinical Research Unit (OUCRU) | Nguyen Van Vinh Chau, Nguyen Thi Thu Hong, Nghiem My Ngoc, Nguyen To Anh, Huynh Trung Trieu, Le Nguyen Truc Nhu, Lam Minh Yen, Ngo Ngoc Quang Minh, Nguyen Thanh Phong, Nguyen Thanh Truong, Le Thi Thu Huong, Tran Nguyen Hoang Tu, Le Manh Hung, Tran Tan Thanh, Nguyen Thanh Dung, Nguyen Tri Dung, Guy Thwaites, le Van Tan |
| EPI_ISL_1293352 | Institut de Virologie du CHU de Strasbourg | Institut de Virologie du CHU de Strasbourg | Fafi-Kremer Samira, Gallais Floriane, Gantner Pierre, Laugel Elodie, Solis Morgane, Velay Aurélie, Wendling Marie-Josée |
| EPI_ISL_1294358 | OLVZ Aalst | OLVZ Aalst | Astrid Holderbeke |
| EPI_ISL_1295178 | Institut de Virologie du CHU de Strasbourg | Institut de Virologie du CHU de Strasbourg | Fafi-Kremer Samira, Gallais Floriane, Gantner Pierre, Laugel Elodie, Solis Morgane, Velay Aurélie, Wendling Marie-Josée |
| EPI_ISL_1296238, EPI_ISL_1296239, EPI_ISL_1296240, EPI_ISL_1296248, EPI_ISL_1296249, EPI_ISL_1296250, EPI_ISL_1296251 | Università degli Studi di Perugia | Istituto Zooprofilattico Sperimentale dell'Abruzzo e Molise "G. Caporale" | Mencacci A, Camilloni B,Lorusso A, Marcacci M, Di Domenico M, Ancora M, Curini V, Mangone I, Rinaldi A, Scialabba S, Di Pasquale A, Cammà C, Puglia I, Calistri P, Savini G |
| EPI_ISL_1296253, EPI_ISL_1296254, EPI_ISL_1296255, EPI_ISL_1296256, EPI_ISL_1296257 | Laboratorio Analisi Osp. Città di Castello - Azienda USL Umbria1 | Istituto Zooprofilattico Sperimentale dell'Abruzzo e Molise "G. Caporale" | Malagigi V, Tacconi P, Mencacci A, Camilloni B,Lorusso A, Marcacci M, Di Domenico M, Ancora M, Curini V, Mangone I, Rinaldi A, Scialabba S, Di Pasquale A, Cammà C, Puglia I, Calistri P, Savini G |
| EPI_ISL_1296258 | Università degli Studi di Perugia | Istituto Zooprofilattico Sperimentale dell'Abruzzo e Molise "G. Caporale" | Mencacci A, Camilloni B,Lorusso A, Marcacci M, Di Domenico M, Ancora M, Curini V, Mangone I, Rinaldi A, Scialabba S, Di Pasquale A, Cammà C, Puglia I, Calistri P, Savini G |
| EPI_ISL_1296259, EPI_ISL_1296260 | Laboratorio Analisi Osp. Città di Castello - Azienda USL Umbria1 | Istituto Zooprofilattico Sperimentale dell'Abruzzo e Molise "G. Caporale" | Malagigi V, Tacconi P, Mencacci A, Camilloni B,Lorusso A, Marcacci M, Di Domenico M, Ancora M, Curini V, Mangone I, Rinaldi A, Scialabba S, Di Pasquale A, Cammà C, Puglia I, Calistri P, Savini G |
| EPI_ISL_1296261 | Istituto Zooprofilattico Sperimentale Umbria e Marche "Togo Rosati" | Istituto Zooprofilattico Sperimentale dell'Abruzzo e Molise "G. Caporale" | Biagetti M,Giammarioli M,Lorusso A, Marcacci M, Di Domenico M, Ancora M, Curini V, Mangone I, Rinaldi A, Scialabba S, Di Pasquale A, Cammà C, Puglia I, Calistri P, Savini G |
| EPI_ISL_1296264, EPI_ISL_1296265, EPI_ISL_1296266, EPI_ISL_1296267 | Azienda Ospedaliera Terni | Istituto Zooprofilattico Sperimentale dell'Abruzzo e Molise "G. Caporale" | Palumbo M, ScaccettiA,Lorusso A, Marcacci M, Di Domenico M, Ancora M, Curini V, Mangone I, Rinaldi A, Scialabba S, Di Pasquale A, Cammà C, Puglia I, Calistri P, Savini G |
| EPI_ISL_1296269, EPI_ISL_1296270, EPI_ISL_1296271, EPI_ISL_1296272, EPI_ISL_1296273, EPI_ISL_1296274, EPI_ISL_1296275, EPI_ISL_1296276, EPI_ISL_1296277, EPI_ISL_1296278 | Azienda Ospedaliera Terni | Istituto Zooprofilattico Sperimentale dell'Abruzzo e Molise "G. Caporale" | Palumbo M, ScaccettiA, Lorusso A, Marcacci M, Di Domenico M, Ancora M, Curini V, Mangone I, Rinaldi A, Scialabba S, Di Pasquale A, Cammà C, Puglia I, Calistri P, Savini G |
| EPI_ISL_1296279 | Università degli Studi di Perugia | Istituto Zooprofilattico Sperimentale dell'Abruzzo e Molise "G. Caporale" | Mencacci A, Camilloni B, Lorusso A, Marcacci M, Di Domenico M, Ancora M, Curini V, Mangone I, Rinaldi A, Scialabba S, Di Pasquale A, Cammà C, Puglia I, Calistri P, Savini G |
| EPI_ISL_1296280 | Azienda Ospedaliera Terni | Istituto Zooprofilattico Sperimentale dell'Abruzzo e Molise "G. Caporale" | Palumbo M, Scaccetti A,,Lorusso A, Marcacci M, Di Domenico M, Ancora M, Curini V, Mangone I, Rinaldi A, Scialabba S, Di Pasquale A, Cammà C, Puglia I, Calistri P, Savini G |
| EPI_ISL_1296281, EPI_ISL_1296282, EPI_ISL_1296285, EPI_ISL_1296286, EPI_ISL_1296287 | Azienda Ospedaliera Terni | Istituto Zooprofilattico Sperimentale dell'Abruzzo e Molise "G. Caporale" | Palumbo M, Scaccetti A, Lorusso A, Marcacci M, Di Domenico M, Ancora M, Curini V, Mangone I, Rinaldi A, Scialabba S, Di Pasquale A, Cammà C, Puglia I, Calistri P, Savini G |
| EPI_ISL_1296288 | Università degli Studi di Perugia | Istituto Zooprofilattico Sperimentale dell'Abruzzo e Molise "G. Caporale" | Mencacci A,Camilloni B,Lorusso A, Marcacci M, Di Domenico M, Ancora M, Curini V, Mangone I, Rinaldi A, Scialabba S, Di Pasquale A, Cammà C, Puglia I, Calistri P, Savini G |
| EPI_ISL_1296289, EPI_ISL_1296290, EPI_ISL_1296291 | Università degli Studi di Perugia | Istituto Zooprofilattico Sperimentale dell'Abruzzo e Molise "G. Caporale" | Mencacci A, Camilloni B,Lorusso A, Marcacci M, Di Domenico M, Ancora M, Curini V, Mangone I, Rinaldi A, Scialabba S, Di Pasquale A, Cammà C, Puglia I, Calistri P, Savini G |
| EPI_ISL_1301806 | Hopital | National Reference Center for Viruses of Respiratory Infections, Institut Pasteur, Paris | Marion Barbet, Sylvie Behillil, Méline Bizard, Angela Brisebarre, Camille Capel, Louise Lefrançois, Etienne Simon-Lorière, Vincent Enouf, Maud Vanpeene, Sylvie van der Werf,Leruez-Ville Marianne |
| EPI_ISL_1301816, EPI_ISL_1301859 | hopital | National Reference Center for Viruses of Respiratory Infections, Institut Pasteur, Paris | Marion Barbet, Sylvie Behillil, Méline Bizard, Angela Brisebarre, Camille Capel, Louise Lefrançois, Etienne Simon-Lorière, Vincent Enouf, Maud Vanpeene, Sylvie van der Werf,Leruez-Ville Marianne |
| EPI_ISL_1301861, EPI_ISL_1301862 | Labo Analyses Med | National Reference Center for Viruses of Respiratory Infections, Institut Pasteur, Paris | Marion Barbet, Sylvie Behillil, Méline Bizard, Angela Brisebarre, Camille Capel, Louise Lefrançois, Etienne Simon-Lorière, Vincent Enouf, Maud Vanpeene, Sylvie van der Werf,Chauvet |
| EPI_ISL_1301884, EPI_ISL_1301885 | Labo Analyses Med | National Reference Center for Viruses of Respiratory Infections, Institut Pasteur, Paris | Marion Barbet, Sylvie Behillil, Méline Bizard, Angela Brisebarre, Camille Capel, Louise Lefrançois, Etienne Simon-Lorière, Vincent Enouf, Maud Vanpeene, Sylvie van der Werf,Lescieux Alexandre |
| EPI_ISL_1301899, EPI_ISL_1301900 | Outre mer | National Reference Center for Viruses of Respiratory Infections, Institut Pasteur, Paris | Marion Barbet, Sylvie Behillil, Méline Bizard, Angela Brisebarre, Camille Capel, Louise Lefrançois, Etienne Simon-Lorière, Vincent Enouf, Maud Vanpeene, Sylvie van der Werf,Combe Patrice |
| EPI_ISL_1301901, EPI_ISL_1301902 | Labo Analyses Med | National Reference Center for Viruses of Respiratory Infections, Institut Pasteur, Paris | Marion Barbet, Sylvie Behillil, Méline Bizard, Angela Brisebarre, Camille Capel, Louise Lefrançois, Etienne Simon-Lorière, Vincent Enouf, Maud Vanpeene, Sylvie van der Werf,Chauvet |
| EPI_ISL_1301930 | Hopital | National Reference Center for Viruses of Respiratory Infections, Institut Pasteur, Paris | Marion Barbet, Sylvie Behillil, Méline Bizard, Angela Brisebarre, Camille Capel, Louise Lefrançois, Etienne Simon-Lorière, Vincent Enouf, Maud Vanpeene, Sylvie van der Werf,Leruez-Ville Marianne |
| EPI_ISL_1301938, EPI_ISL_1301939, EPI_ISL_1301940, EPI_ISL_1301941, EPI_ISL_1301942, EPI_ISL_1301943 | hopital | National Reference Center for Viruses of Respiratory Infections, Institut Pasteur, Paris | Marion Barbet, Sylvie Behillil, Méline Bizard, Angela Brisebarre, Camille Capel, Louise Lefrançois, Etienne Simon-Lorière, Vincent Enouf, Maud Vanpeene, Sylvie van der Werf,Leruez-Ville Marianne |
| EPI_ISL_1301985, EPI_ISL_1301986, EPI_ISL_1301987, EPI_ISL_1301988, EPI_ISL_1301989, EPI_ISL_1301990, EPI_ISL_1301991, EPI_ISL_1301992, EPI_ISL_1301993, EPI_ISL_1301994, EPI_ISL_1301995, EPI_ISL_1301996, EPI_ISL_1301997, EPI_ISL_1301998, EPI_ISL_1301999, EPI_ISL_1302000, EPI_ISL_1302001, EPI_ISL_1302002, EPI_ISL_1302003, EPI_ISL_1302005, EPI_ISL_1302006, EPI_ISL_1302007, EPI_ISL_1302008, EPI_ISL_1302009, EPI_ISL_1302010, EPI_ISL_1302011, EPI_ISL_1302012, EPI_ISL_1302014, EPI_ISL_1302015, EPI_ISL_1302016, EPI_ISL_1302017, EPI_ISL_1302018, EPI_ISL_1302019, EPI_ISL_1302020, EPI_ISL_1302021, EPI_ISL_1302022, EPI_ISL_1302023, EPI_ISL_1302024, EPI_ISL_1302025, EPI_ISL_1302027, EPI_ISL_1302028, EPI_ISL_1302030, EPI_ISL_1302031, EPI_ISL_1302032, EPI_ISL_1302053, EPI_ISL_1302054, EPI_ISL_1302055, EPI_ISL_1302056, EPI_ISL_1302057, EPI_ISL_1302058, EPI_ISL_1302059, EPI_ISL_1302060, EPI_ISL_1302061, EPI_ISL_1302062, EPI_ISL_1302063, EPI_ISL_1302064, EPI_ISL_1302066, EPI_ISL_1302067, EPI_ISL_1302068, EPI_ISL_1302069, EPI_ISL_1302070, EPI_ISL_1302071, EPI_ISL_1302072, EPI_ISL_1302073, EPI_ISL_1302074, EPI_ISL_1302075, EPI_ISL_1302076, EPI_ISL_1302077, EPI_ISL_1302078, EPI_ISL_1302091, EPI_ISL_1302092, EPI_ISL_1302093, EPI_ISL_1302095, EPI_ISL_1302110, EPI_ISL_1302111, EPI_ISL_1302112, EPI_ISL_1302113, EPI_ISL_1302114, EPI_ISL_1302115, EPI_ISL_1302116, EPI_ISL_1302117, EPI_ISL_1302118, EPI_ISL_1302119, EPI_ISL_1302120, EPI_ISL_1302121, EPI_ISL_1302150, EPI_ISL_1302151 | | | |
| see above | National Center of Infectious and Parasitic Diseases | National Center of Infectious and Parasitic Diseases | Alexiev et al |
| EPI_ISL_1302401 | Laboratorio Central de Epidemiología IMSS | Instituto de Biotecnología de la UNAM | Authors from IBT, IMSS, InDRE and INER (in alphabetical order): Carlos F. Arias, Santiago Ávila-Ríos, Gisela Barrera-Badillo, Eduardo Becerril-Vargas, |

| | | | |
|---|---|---|---|
| | | | Celia Boukadida, Natividad Cruz-Ortiz, Larissa Fernandes-Matano, Ricardo Grande, Lucia Hernandez-Rivas, Alejandra Hernández-Terán, Pavel Isa, Irma Lopez-Martinez, José Arturo Martínez-Orozco, Margarita Matías-Florentino, Fidencio Mejía-Nepomuceno, Edgar Mendieta-Condado, Mario Mújica-Sánchez, José Esteban Muñoz-Medina, Tatiana Nunez-Garcia, Luis Alberto Ochoa-Carrera, Hector Esteban Paz-Juárez, Francisco Pulido, José Ernesto Ramírez-González, Alma Rincón-Rubio, Teresita Rojas-Mendoza, Jorge Salas-Hernández, Alejandro Sanchez-Flores, Clara Esperanza Santacruz-Tinoco, Andrea Santos Coy-Arechavaleta, Blanca Taboada, Gloria Vazquez, Joel Armando Vázquez-Pérez, Jerome Jean Verleyen |
| EPI_ISL_1302408, EPI_ISL_1302409, EPI_ISL_1302410, EPI_ISL_1302411, EPI_ISL_1302412, EPI_ISL_1302413, EPI_ISL_1302414, EPI_ISL_1302415, EPI_ISL_1302416, EPI_ISL_1302417, EPI_ISL_1302418, EPI_ISL_1302419, EPI_ISL_1302420, EPI_ISL_1302421, EPI_ISL_1302422, EPI_ISL_1302423, EPI_ISL_1302443 | | | |
| see above | SC Microbiologia e Virologia AOUSS | AMES Centro Polidiagnostico Strumentale S.r.l. | Salvatore Rubino; Flavia Angioj; Laura Firino; Rosalba Govoni; Gabriele Ibba; Vincenzo Lai; Erica Mura; Bianca Paglietti; Claudia Piu; Anna Puggioni; Elena Rimini; Giulia Rocca; Caterina Serra; Sergio Uzzau. |
| EPI_ISL_1302674 | National Center for Infectious and Parasitic Diseases | National Center for Infectious and Parasitic Diseases | Alexiev et al |
| EPI_ISL_1302788, EPI_ISL_1302789, EPI_ISL_1302790, EPI_ISL_1302791, EPI_ISL_1302793, EPI_ISL_1302794, EPI_ISL_1302796, EPI_ISL_1302797, EPI_ISL_1302798, EPI_ISL_1302800, EPI_ISL_1302801, EPI_ISL_1302804, EPI_ISL_1302806, EPI_ISL_1302807, EPI_ISL_1302808, EPI_ISL_1302809, EPI_ISL_1302813, EPI_ISL_1302815, EPI_ISL_1302816, EPI_ISL_1302817, EPI_ISL_1302818, EPI_ISL_1302819, EPI_ISL_1302821, EPI_ISL_1302823, EPI_ISL_1302828, EPI_ISL_1302829, EPI_ISL_1302831, EPI_ISL_1302832, EPI_ISL_1302834, EPI_ISL_1302839, EPI_ISL_1302843, EPI_ISL_1302849, EPI_ISL_1302852, EPI_ISL_1302853, EPI_ISL_1302854, EPI_ISL_1302855, EPI_ISL_1302859, EPI_ISL_1302860, EPI_ISL_1302861, EPI_ISL_1302862, EPI_ISL_1302863, EPI_ISL_1302864, EPI_ISL_1302865, EPI_ISL_1302866, EPI_ISL_1302869, EPI_ISL_1302872 | | | |
| see above | ADMED Microbiologie | Genomics and Transcriptomics, Philip Morris International | Reto Lienhard, Marie-Lise Tritten, Emmanuel Guedj, Nicolas Sierro, Rémi Dulize, David Bornand, Mehdi Auberson, Maxime Berthouzoz, Nikolai Ivanov, Manuel Peitsch |
| EPI_ISL_1303427 | Microbiology Unit, ASST Monza - San Gerardo Hospital | Laboratory of Clinical Microbiology, Virology and Bioemergencies, ASST Fatebenefratelli Sacco - Sacco Hospital | Valeria Micheli, Alberto Rizzo, Alessandro Mancon, Fiorenza Bracchitta, Luca Rizzuto, Sergio Malandrin, Annalisa Cavallero, Maria Rita Gismondo |
| EPI_ISL_1303499, EPI_ISL_1303500, EPI_ISL_1303501, EPI_ISL_1303502, EPI_ISL_1303503, EPI_ISL_1303505 | LACEN de Rondonia | Instituto Adolfo Lutz, Interdiciplinary Procedures Center, Strategic Laboratory | Claudio Tavares Sacchi, Claudia Regina Gonçalves, Erica Valessa Ramos Gomes, Karoline Rodrigues Campos, Caio Vinicius Dias Lopes |
| EPI_ISL_1303506, EPI_ISL_1303507, EPI_ISL_1303508 | LACEN do Distrito Federal | Instituto Adolfo Lutz, Interdiciplinary Procedures Center, Strategic Laboratory | Claudio Tavares Sacchi, Claudia Regina Gonçalves, Erica Valessa Ramos Gomes, Karoline Rodrigues Campos, Caio Vinicius Dias Lopes |
| EPI_ISL_1303509 | LACEN do Estado de Tocantins | Instituto Adolfo Lutz, Interdiciplinary Procedures Center, Strategic Laboratory | Claudio Tavares Sacchi, Claudia Regina Gonçalves, Erica Valessa Ramos Gomes, Karoline Rodrigues Campos, Caio Vinicius Dias Lopes |
| EPI_ISL_1303510, EPI_ISL_1303511, EPI_ISL_1303512, EPI_ISL_1303514, EPI_ISL_1303517 | LACEN do Estado de Goias | Instituto Adolfo Lutz, Interdiciplinary Procedures Center, Strategic Laboratory | Claudio Tavares Sacchi, Claudia Regina Gonçalves, Erica Valessa Ramos Gomes, Karoline Rodrigues Campos, Caio Vinicius Dias Lopes |
| EPI_ISL_1303518, EPI_ISL_1303519, EPI_ISL_1303520, EPI_ISL_1303521, EPI_ISL_1303522, EPI_ISL_1303523, EPI_ISL_1303524, EPI_ISL_1303525, EPI_ISL_1303526, EPI_ISL_1303527, EPI_ISL_1303528, EPI_ISL_1303529, EPI_ISL_1303530, EPI_ISL_1303531, EPI_ISL_1303532, EPI_ISL_1303533, EPI_ISL_1303534 | | | |
| see above | IAL Regional de São Jose do Rio Preto | Instituto Adolfo Lutz, Interdiciplinary Procedures Center, Strategic Laboratory | Claudio Tavares Sacchi, Claudia Regina Gonçalves, Erica Valessa Ramos Gomes, Karoline Rodrigues Campos, Caio Vinicius Dias Lopes |
| EPI_ISL_1303535, EPI_ISL_1303536 | Hospital Heliopolis | Instituto Adolfo Lutz, Interdiciplinary Procedures Center, Strategic Laboratory | Claudio Tavares Sacchi, Claudia Regina Gonçalves, Erica Valessa Ramos Gomes, Karoline Rodrigues Campos, Caio Vinicius Dias Lopes |
| EPI_ISL_1303537 | Hospital Estadual de Vila Alpina | Instituto Adolfo Lutz, Interdiciplinary Procedures Center, Strategic Laboratory | Claudio Tavares Sacchi, Claudia Regina Gonçalves, Erica Valessa Ramos Gomes, Karoline Rodrigues Campos, Caio Vinicius Dias Lopes |
| EPI_ISL_1303538, EPI_ISL_1303539 | Hospital Presidente | Instituto Adolfo Lutz, Interdiciplinary Procedures Center, Strategic Laboratory | Claudio Tavares Sacchi, Claudia Regina Gonçalves, Erica Valessa Ramos Gomes, Karoline Rodrigues Campos, Caio Vinicius Dias Lopes |
| EPI_ISL_1303540, EPI_ISL_1303541 | Hospital Municipal Cidade Tiradentes Carmen Prudente | Instituto Adolfo Lutz, Interdiciplinary Procedures Center, Strategic Laboratory | Claudio Tavares Sacchi, Claudia Regina Gonçalves, Erica Valessa Ramos Gomes, Karoline Rodrigues Campos, Caio Vinicius Dias Lopes |
| EPI_ISL_1303542, EPI_ISL_1303543 | Hospital Estadual de Campanha Barradas | Instituto Adolfo Lutz, Interdiciplinary Procedures Center, Strategic Laboratory | Claudio Tavares Sacchi, Claudia Regina Gonçalves, Erica Valessa Ramos Gomes, Karoline Rodrigues Campos, Caio Vinicius Dias Lopes |
| EPI_ISL_1303544, EPI_ISL_1303545 | Hospital Municipal Cidade Tiradentes Carmen Prudente | Instituto Adolfo Lutz, Interdiciplinary Procedures Center, Strategic Laboratory | Claudio Tavares Sacchi, Claudia Regina Gonçalves, Erica Valessa Ramos Gomes, Karoline Rodrigues Campos, Caio Vinicius Dias Lopes |
| EPI_ISL_1303546, EPI_ISL_1303547, EPI_ISL_1303548, EPI_ISL_1303549 | UPA Vila Santa Catarina | Instituto Adolfo Lutz, Interdiciplinary Procedures Center, Strategic Laboratory | Claudio Tavares Sacchi, Claudia Regina Gonçalves, Erica Valessa Ramos Gomes, Karoline Rodrigues Campos, Caio Vinicius Dias Lopes |
| EPI_ISL_1307692, EPI_ISL_1307696, EPI_ISL_1307700, EPI_ISL_1307712, EPI_ISL_1307714 | Institut de Virologie du CHU de Strasbourg | Institut de Virologie du CHU de Strasbourg | Fafi-Kremer Samira, Gallais Floriane, Gantner Pierre, Laugel Elodie, Solis Morgane, Velay Aurélie, Wendling Marie-Josée |
| EPI_ISL_1310779, EPI_ISL_1310781, EPI_ISL_1315635 | Laboratorio de Virología HUCA | Laboratorio de Virología HUCA | Sandoval M, Castelló C, Gómez de Oña J, Boga JA, Rojo S, Alvarez-Arguelles ME, Abreu F, Costales I, Perez-Martínez Z, Martín-Rodríguez G, Coto E, Melón S |
| EPI_ISL_1318206, EPI_ISL_1318210, EPI_ISL_1318213, EPI_ISL_1318214, EPI_ISL_1318215, EPI_ISL_1318216, EPI_ISL_1318217, EPI_ISL_1318218, EPI_ISL_1318219 | Ministry of Health Hospitals | Institute of Health and Community Medicine | Chan Chia Jui, David Perera, Ooi Mong How, Chua Hock Hin, Tonnii Sia Loong Loong, Wong Jyn Shan, Wong Kiing Aik |
| EPI_ISL_1319254, EPI_ISL_1319255, EPI_ISL_1319258, EPI_ISL_1319259, EPI_ISL_1319260, EPI_ISL_1319262, EPI_ISL_1319263, EPI_ISL_1319265, EPI_ISL_1319266, EPI_ISL_1319267, EPI_ISL_1319269, EPI_ISL_1319270, EPI_ISL_1319271, EPI_ISL_1319272, EPI_ISL_1319274, EPI_ISL_1319275, EPI_ISL_1319276, EPI_ISL_1319277, EPI_ISL_1319279, EPI_ISL_1319280, EPI_ISL_1319281, EPI_ISL_1319283, EPI_ISL_1319287, EPI_ISL_1319288, EPI_ISL_1319290, EPI_ISL_1319291, EPI_ISL_1319292, EPI_ISL_1319295, EPI_ISL_1319297, EPI_ISL_1319298, EPI_ISL_1319300, EPI_ISL_1319302, EPI_ISL_1319303, EPI_ISL_1319305, EPI_ISL_1319306, EPI_ISL_1319308, EPI_ISL_1319309, EPI_ISL_1319310, EPI_ISL_1319313, EPI_ISL_1319314, EPI_ISL_1319316, EPI_ISL_1319317, EPI_ISL_1319318, EPI_ISL_1319319, EPI_ISL_1319322, EPI_ISL_1319324, EPI_ISL_1319325, EPI_ISL_1319327, EPI_ISL_1319328, EPI_ISL_1319329, EPI_ISL_1319331, EPI_ISL_1319332, EPI_ISL_1319335, EPI_ISL_1319336, EPI_ISL_1319337, EPI_ISL_1319339, EPI_ISL_1319340, EPI_ISL_1319341, EPI_ISL_1319343, EPI_ISL_1319344, EPI_ISL_1319345, EPI_ISL_1319347, EPI_ISL_1319348, EPI_ISL_1319350, EPI_ISL_1319351 | | | |
| see above | Synlab Eesti OÜ | 1. Laboratory of Communicable Diseases (Estonia); 2. Eurofins Genomics Europe Sequencing GmbH | Liidia Dotsenko et al. |
| EPI_ISL_1319352, EPI_ISL_1319354, EPI_ISL_1319355, EPI_ISL_1319356, EPI_ISL_1319358 | North Estonia Medical Centre laboratory | 1. Laboratory of Communicable Diseases (Estonia); 2. Eurofins Genomics Europe Sequencing GmbH | Liidia Dotsenko et al. |
| EPI_ISL_1319371, EPI_ISL_1319372, EPI_ISL_1319373, EPI_ISL_1319374, EPI_ISL_1319375, EPI_ISL_1319377, EPI_ISL_1319378, EPI_ISL_1319379, EPI_ISL_1319382, EPI_ISL_1319383, EPI_ISL_1319384, EPI_ISL_1319385, EPI_ISL_1319388, EPI_ISL_1319390, EPI_ISL_1319392, EPI_ISL_1319393, EPI_ISL_1319395, EPI_ISL_1319397, EPI_ISL_1319399, EPI_ISL_1319401, EPI_ISL_1319403, EPI_ISL_1319404, EPI_ISL_1319406, EPI_ISL_1319407, EPI_ISL_1319409, EPI_ISL_1319411, EPI_ISL_1319412, EPI_ISL_1319413, EPI_ISL_1319415, EPI_ISL_1319416, EPI_ISL_1319417, EPI_ISL_1319419, EPI_ISL_1319420, EPI_ISL_1319422, EPI_ISL_1319423, EPI_ISL_1319424, EPI_ISL_1319426, EPI_ISL_1319427, EPI_ISL_1319429, EPI_ISL_1319430, EPI_ISL_1319431, EPI_ISL_1319432, EPI_ISL_1319433, EPI_ISL_1319434, EPI_ISL_1319436, EPI_ISL_1319437, EPI_ISL_1319438 | | | |
| see above | Synlab Eesti OÜ | 1. Laboratory of Communicable Diseases (Estonia); 2. Eurofins Genomics Europe Sequencing GmbH | Liidia Dotsenko et al. |
| EPI_ISL_1319440, EPI_ISL_1319441, EPI_ISL_1319443, EPI_ISL_1319449 | North Estonia Medical Centre laboratory | 1. Laboratory of Communicable Diseases (Estonia); 2. Eurofins Genomics Europe Sequencing GmbH | Liidia Dotsenko et al. |
| EPI_ISL_1322209 | IrsiCaixa | IrsiCaixa | Marc Noguera-Julian, Mariona Parera, Maria Casadellà, Pilar Armengol, Francesc Catala-Moll, Roger Paredes, Bonaventura Clotet Gloria Trujillo, Rafel Perez Vidal, Jaume Trape Pujol, Carolina Gonzalez Fernandez, Roger Paredes, Eulalia Grau, Bonaventura Clotet |
| EPI_ISL_1323555 | CHU LILLE | CHU Lille - Laboratoire de Virologie | AIT YAHYA Emilie, ALIDJINOU Enagnon Kazali, BOCKET Laurence, CREPIN Michel, DEMAY Christophe, ENGELMANN Ilka, GEFFROY Sandrine, |

| | | | |
|---|---|---|---|
| | | | GUIGON Aurélie, LAZREK Mouna, NOBILLIAUX Florian, PREVOST Brigitte, THUILLIER Caroline, TINEZ Claire |
| EPI_ISL_1323556 | CH ROUBAIX | CHU Lille - Laboratoire de Virologie | AIT YAHYA Emilie, ALIDJINOU Enagnon Kazali, BOCKET Laurence, CREPIN Michel, DEMAY Christophe, ENGELMANN Ilka, GEFFROY Sandrine, GUIGON Aurélie, LAZREK Mouna, NOBILLIAUX Florian, PREVOST Brigitte, THUILLIER Caroline, TINEZ Claire |
| EPI_ISL_1323557 | CH TOURCOING | CHU Lille - Laboratoire de Virologie | AIT YAHYA Emilie, ALIDJINOU Enagnon Kazali, BOCKET Laurence, CREPIN Michel, DEMAY Christophe, ENGELMANN Ilka, GEFFROY Sandrine, GUIGON Aurélie, LAZREK Mouna, NOBILLIAUX Florian, PREVOST Brigitte, THUILLIER Caroline, TINEZ Claire |
| EPI_ISL_1323558 | CH CAMBRAI | CHU Lille - Laboratoire de Virologie | AIT YAHYA Emilie, ALIDJINOU Enagnon Kazali, BOCKET Laurence, CREPIN Michel, DEMAY Christophe, ENGELMANN Ilka, GEFFROY Sandrine, GUIGON Aurélie, LAZREK Mouna, NOBILLIAUX Florian, PREVOST Brigitte, THUILLIER Caroline, TINEZ Claire |
| EPI_ISL_1323559 | CH ARRAS | CHU Lille - Laboratoire de Virologie | AIT YAHYA Emilie, ALIDJINOU Enagnon Kazali, BOCKET Laurence, CREPIN Michel, DEMAY Christophe, ENGELMANN Ilka, GEFFROY Sandrine, GUIGON Aurélie, LAZREK Mouna, NOBILLIAUX Florian, PREVOST Brigitte, THUILLIER Caroline, TINEZ Claire |
| EPI_ISL_1323560, EPI_ISL_1323561 | CHU LILLE | CHU Lille - Laboratoire de Virologie | AIT YAHYA Emilie, ALIDJINOU Enagnon Kazali, BOCKET Laurence, CREPIN Michel, DEMAY Christophe, ENGELMANN Ilka, GEFFROY Sandrine, GUIGON Aurélie, LAZREK Mouna, NOBILLIAUX Florian, PREVOST Brigitte, THUILLIER Caroline, TINEZ Claire |
| EPI_ISL_1323562 | CH SECLIN | CHU Lille - Laboratoire de Virologie | AIT YAHYA Emilie, ALIDJINOU Enagnon Kazali, BOCKET Laurence, CREPIN Michel, DEMAY Christophe, ENGELMANN Ilka, GEFFROY Sandrine, GUIGON Aurélie, LAZREK Mouna, NOBILLIAUX Florian, PREVOST Brigitte, THUILLIER Caroline, TINEZ Claire |
| EPI_ISL_1323565 | CHU LILLE | CHU Lille - Laboratoire de Virologie | AIT YAHYA Emilie, ALIDJINOU Enagnon Kazali, BOCKET Laurence, CREPIN Michel, DEMAY Christophe, ENGELMANN Ilka, GEFFROY Sandrine, GUIGON Aurélie, LAZREK Mouna, NOBILLIAUX Florian, PREVOST Brigitte, THUILLIER Caroline, TINEZ Claire |
| EPI_ISL_1323566 | CH VALENCIENNES | CHU Lille - Laboratoire de Virologie | AIT YAHYA Emilie, ALIDJINOU Enagnon Kazali, BOCKET Laurence, CREPIN Michel, DEMAY Christophe, ENGELMANN Ilka, GEFFROY Sandrine, GUIGON Aurélie, LAZREK Mouna, NOBILLIAUX Florian, PREVOST Brigitte, THUILLIER Caroline, TINEZ Claire |
| EPI_ISL_1323567, EPI_ISL_1323568 | CH ARMENTIERES | CHU Lille - Laboratoire de Virologie | AIT YAHYA Emilie, ALIDJINOU Enagnon Kazali, BOCKET Laurence, CREPIN Michel, DEMAY Christophe, ENGELMANN Ilka, GEFFROY Sandrine, GUIGON Aurélie, LAZREK Mouna, NOBILLIAUX Florian, PREVOST Brigitte, THUILLIER Caroline, TINEZ Claire |
| EPI_ISL_1323569 | CH BETHUNE | CHU Lille - Laboratoire de Virologie | AIT YAHYA Emilie, ALIDJINOU Enagnon Kazali, BOCKET Laurence, CREPIN Michel, DEMAY Christophe, ENGELMANN Ilka, GEFFROY Sandrine, GUIGON Aurélie, LAZREK Mouna, NOBILLIAUX Florian, PREVOST Brigitte, THUILLIER Caroline, TINEZ Claire |
| EPI_ISL_1323570 | CH CALAIS | CHU Lille - Laboratoire de Virologie | AIT YAHYA Emilie, ALIDJINOU Enagnon Kazali, BOCKET Laurence, CREPIN Michel, DEMAY Christophe, ENGELMANN Ilka, GEFFROY Sandrine, GUIGON Aurélie, LAZREK Mouna, NOBILLIAUX Florian, PREVOST Brigitte, THUILLIER Caroline, TINEZ Claire |
| EPI_ISL_1323571, EPI_ISL_1323572 | CHU LILLE | CHU Lille - Laboratoire de Virologie | AIT YAHYA Emilie, ALIDJINOU Enagnon Kazali, BOCKET Laurence, CREPIN Michel, DEMAY Christophe, ENGELMANN Ilka, GEFFROY Sandrine, GUIGON Aurélie, LAZREK Mouna, NOBILLIAUX Florian, PREVOST Brigitte, THUILLIER Caroline, TINEZ Claire |
| EPI_ISL_1323575 | CH DOUAI | CHU Lille - Laboratoire de Virologie | AIT YAHYA Emilie, ALIDJINOU Enagnon Kazali, BOCKET Laurence, CREPIN Michel, DEMAY Christophe, ENGELMANN Ilka, GEFFROY Sandrine, GUIGON Aurélie, LAZREK Mouna, NOBILLIAUX Florian, PREVOST Brigitte, THUILLIER Caroline, TINEZ Claire |
| EPI_ISL_1323576 | CHU LILLE | CHU Lille - Laboratoire de Virologie | AIT YAHYA Emilie, ALIDJINOU Enagnon Kazali, BOCKET Laurence, CREPIN Michel, DEMAY Christophe, ENGELMANN Ilka, GEFFROY Sandrine, GUIGON Aurélie, LAZREK Mouna, NOBILLIAUX Florian, PREVOST Brigitte, THUILLIER Caroline, TINEZ Claire |
| EPI_ISL_1323577 | CH ROUBAIX | CHU Lille - Laboratoire de Virologie | AIT YAHYA Emilie, ALIDJINOU Enagnon Kazali, BOCKET Laurence, CREPIN Michel, DEMAY Christophe, ENGELMANN Ilka, GEFFROY Sandrine, GUIGON Aurélie, LAZREK Mouna, NOBILLIAUX Florian, PREVOST Brigitte, THUILLIER Caroline, TINEZ Claire |
| EPI_ISL_1323578 | CH LENS | CHU Lille - Laboratoire de Virologie | AIT YAHYA Emilie, ALIDJINOU Enagnon Kazali, BOCKET Laurence, CREPIN Michel, DEMAY Christophe, ENGELMANN Ilka, GEFFROY Sandrine, GUIGON Aurélie, LAZREK Mouna, NOBILLIAUX Florian, PREVOST Brigitte, THUILLIER Caroline, TINEZ Claire |
| EPI_ISL_1323579, EPI_ISL_1323580 | CH BETHUNE | CHU Lille - Laboratoire de Virologie | AIT YAHYA Emilie, ALIDJINOU Enagnon Kazali, BOCKET Laurence, CREPIN Michel, DEMAY Christophe, ENGELMANN Ilka, GEFFROY Sandrine, GUIGON Aurélie, LAZREK Mouna, NOBILLIAUX Florian, PREVOST Brigitte, THUILLIER Caroline, TINEZ Claire |
| EPI_ISL_1323581 | CH LENS | CHU Lille - Laboratoire de Virologie | AIT YAHYA Emilie, ALIDJINOU Enagnon Kazali, BOCKET Laurence, CREPIN Michel, DEMAY Christophe, ENGELMANN Ilka, GEFFROY Sandrine, GUIGON Aurélie, LAZREK Mouna, NOBILLIAUX Florian, PREVOST Brigitte, THUILLIER Caroline, TINEZ Claire |
| EPI_ISL_1323582 | CH ROUBAIX | CHU Lille - Laboratoire de Virologie | AIT YAHYA Emilie, ALIDJINOU Enagnon Kazali, BOCKET Laurence, CREPIN Michel, DEMAY Christophe, ENGELMANN Ilka, GEFFROY Sandrine, GUIGON Aurélie, LAZREK Mouna, NOBILLIAUX Florian, PREVOST Brigitte, THUILLIER Caroline, TINEZ Claire |
| EPI_ISL_1323583, EPI_ISL_1323584 | CH LENS | CHU Lille - Laboratoire de Virologie | AIT YAHYA Emilie, ALIDJINOU Enagnon Kazali, BOCKET Laurence, CREPIN Michel, DEMAY Christophe, ENGELMANN Ilka, GEFFROY Sandrine, GUIGON Aurélie, LAZREK Mouna, NOBILLIAUX Florian, PREVOST Brigitte, THUILLIER Caroline, TINEZ Claire |
| EPI_ISL_1323585, EPI_ISL_1323586 | CH ROUBAIX | CHU Lille - Laboratoire de Virologie | AIT YAHYA Emilie, ALIDJINOU Enagnon Kazali, BOCKET Laurence, CREPIN Michel, DEMAY Christophe, ENGELMANN Ilka, GEFFROY Sandrine, GUIGON Aurélie, LAZREK Mouna, NOBILLIAUX Florian, PREVOST Brigitte, THUILLIER Caroline, TINEZ Claire |
| EPI_ISL_1323587, EPI_ISL_1323588 | CH BETHUNE | CHU Lille - Laboratoire de Virologie | AIT YAHYA Emilie, ALIDJINOU Enagnon Kazali, BOCKET Laurence, CREPIN Michel, DEMAY Christophe, ENGELMANN Ilka, GEFFROY Sandrine, GUIGON Aurélie, LAZREK Mouna, NOBILLIAUX Florian, PREVOST Brigitte, THUILLIER Caroline, TINEZ Claire |
| EPI_ISL_1323589, EPI_ISL_1323590, EPI_ISL_1323591 | CH ROUBAIX | CHU Lille - Laboratoire de Virologie | AIT YAHYA Emilie, ALIDJINOU Enagnon Kazali, BOCKET Laurence, CREPIN Michel, DEMAY Christophe, ENGELMANN Ilka, GEFFROY Sandrine, GUIGON Aurélie, LAZREK Mouna, NOBILLIAUX Florian, PREVOST Brigitte, THUILLIER Caroline, TINEZ Claire |
| EPI_ISL_1323592 | CH SECLIN | CHU Lille - Laboratoire de Virologie | AIT YAHYA Emilie, ALIDJINOU Enagnon Kazali, BOCKET Laurence, CREPIN Michel, DEMAY Christophe, ENGELMANN Ilka, GEFFROY Sandrine, GUIGON Aurélie, LAZREK Mouna, NOBILLIAUX Florian, PREVOST Brigitte, THUILLIER Caroline, TINEZ Claire |
| EPI_ISL_1323593 | CHU LILLE | CHU Lille - Laboratoire de Virologie | AIT YAHYA Emilie, ALIDJINOU Enagnon Kazali, BOCKET Laurence, CREPIN Michel, DEMAY Christophe, ENGELMANN Ilka, GEFFROY Sandrine, GUIGON Aurélie, LAZREK Mouna, NOBILLIAUX Florian, PREVOST Brigitte, THUILLIER Caroline, TINEZ Claire |
| EPI_ISL_1323594 | CH CAMBRAI | CHU Lille - Laboratoire de Virologie | AIT YAHYA Emilie, ALIDJINOU Enagnon Kazali, BOCKET Laurence, CREPIN Michel, DEMAY Christophe, ENGELMANN Ilka, GEFFROY Sandrine, GUIGON Aurélie, LAZREK Mouna, NOBILLIAUX Florian, PREVOST Brigitte, THUILLIER Caroline, TINEZ Claire |
| EPI_ISL_1323595 | CHU LILLE | CHU Lille - Laboratoire de Virologie | AIT YAHYA Emilie, ALIDJINOU Enagnon Kazali, BOCKET Laurence, CREPIN Michel, DEMAY Christophe, ENGELMANN Ilka, GEFFROY Sandrine, GUIGON Aurélie, LAZREK Mouna, NOBILLIAUX Florian, PREVOST Brigitte, THUILLIER Caroline, TINEZ Claire |
| EPI_ISL_1323596 | CH VALENCIENNES | CHU Lille - Laboratoire de Virologie | AIT YAHYA Emilie, ALIDJINOU Enagnon Kazali, BOCKET Laurence, CREPIN Michel, DEMAY Christophe, ENGELMANN Ilka, GEFFROY Sandrine, GUIGON Aurélie, LAZREK Mouna, NOBILLIAUX Florian, PREVOST Brigitte, THUILLIER Caroline, TINEZ Claire |
| EPI_ISL_1323597 | CHU LILLE | CHU Lille - Laboratoire de Virologie | AIT YAHYA Emilie, ALIDJINOU Enagnon Kazali, BOCKET Laurence, CREPIN Michel, DEMAY Christophe, ENGELMANN Ilka, GEFFROY Sandrine, GUIGON Aurélie, LAZREK Mouna, NOBILLIAUX Florian, PREVOST Brigitte, THUILLIER Caroline, TINEZ Claire |
| EPI_ISL_1323598 | CH DOUAI | CHU Lille - Laboratoire de Virologie | AIT YAHYA Emilie, ALIDJINOU Enagnon Kazali, BOCKET Laurence, CREPIN Michel, DEMAY Christophe, ENGELMANN Ilka, GEFFROY Sandrine, GUIGON Aurélie, LAZREK Mouna, NOBILLIAUX Florian, PREVOST Brigitte, THUILLIER Caroline, TINEZ Claire |
| EPI_ISL_1323599 | CH LENS | CHU Lille - Laboratoire de Virologie | AIT YAHYA Emilie, ALIDJINOU Enagnon Kazali, BOCKET Laurence, CREPIN Michel, DEMAY Christophe, ENGELMANN Ilka, GEFFROY Sandrine, GUIGON Aurélie, LAZREK Mouna, NOBILLIAUX Florian, PREVOST Brigitte, THUILLIER Caroline, TINEZ Claire |
| EPI_ISL_1323600 | CH TOURCOING | CHU Lille - Laboratoire de Virologie | AIT YAHYA Emilie, ALIDJINOU Enagnon Kazali, BOCKET Laurence, CREPIN Michel, DEMAY Christophe, ENGELMANN Ilka, GEFFROY Sandrine, GUIGON Aurélie, LAZREK Mouna, NOBILLIAUX Florian, PREVOST Brigitte, THUILLIER Caroline, TINEZ Claire |
| EPI_ISL_1323601 | CH DOUAI | CHU Lille - Laboratoire de Virologie | AIT YAHYA Emilie, ALIDJINOU Enagnon Kazali, BOCKET Laurence, CREPIN Michel, DEMAY Christophe, ENGELMANN Ilka, GEFFROY Sandrine, GUIGON Aurélie, LAZREK Mouna, NOBILLIAUX Florian, PREVOST Brigitte, THUILLIER Caroline, TINEZ Claire |
| EPI_ISL_1323602, EPI_ISL_1323603 | CH ARMENTIERES | CHU Lille - Laboratoire de Virologie | AIT YAHYA Emilie, ALIDJINOU Enagnon Kazali, BOCKET Laurence, CREPIN Michel, DEMAY Christophe, ENGELMANN Ilka, GEFFROY Sandrine, GUIGON Aurélie, LAZREK Mouna, NOBILLIAUX Florian, PREVOST Brigitte, THUILLIER Caroline, TINEZ Claire |
| EPI_ISL_1323604, EPI_ISL_1323605, | CH BETHUNE | CHU Lille - Laboratoire de Virologie | AIT YAHYA Emilie, ALIDJINOU Enagnon Kazali, BOCKET Laurence, CREPIN Michel, DEMAY Christophe, ENGELMANN Ilka, GEFFROY Sandrine, |

| | | | |
|---|---|---|---|
| EPI_ISL_1323606 | | | GUIGON Aurélie, LAZREK Mouna, NOBILLIAUX Florian, PREVOST Brigitte, THUILLIER Caroline, TINEZ Claire |
| EPI_ISL_1323607, EPI_ISL_1323608 | CH ARRAS | CHU Lille - Laboratoire de Virologie | AIT YAHYA Emilie, ALIDJINOU Enagnon Kazali, BOCKET Laurence, CREPIN Michel, DEMAY Christophe, ENGELMANN Ilka, GEFFROY Sandrine, GUIGON Aurélie, LAZREK Mouna, NOBILLIAUX Florian, PREVOST Brigitte, THUILLIER Caroline, TINEZ Claire |
| EPI_ISL_1323609, EPI_ISL_1323610 | CH CALAIS | CHU Lille - Laboratoire de Virologie | AIT YAHYA Emilie, ALIDJINOU Enagnon Kazali, BOCKET Laurence, CREPIN Michel, DEMAY Christophe, ENGELMANN Ilka, GEFFROY Sandrine, GUIGON Aurélie, LAZREK Mouna, NOBILLIAUX Florian, PREVOST Brigitte, THUILLIER Caroline, TINEZ Claire |
| EPI_ISL_1323611 | CH DOUAI | CHU Lille - Laboratoire de Virologie | AIT YAHYA Emilie, ALIDJINOU Enagnon Kazali, BOCKET Laurence, CREPIN Michel, DEMAY Christophe, ENGELMANN Ilka, GEFFROY Sandrine, GUIGON Aurélie, LAZREK Mouna, NOBILLIAUX Florian, PREVOST Brigitte, THUILLIER Caroline, TINEZ Claire |
| EPI_ISL_1323612, EPI_ISL_1323613 | CH ROUBAIX | CHU Lille - Laboratoire de Virologie | AIT YAHYA Emilie, ALIDJINOU Enagnon Kazali, BOCKET Laurence, CREPIN Michel, DEMAY Christophe, ENGELMANN Ilka, GEFFROY Sandrine, GUIGON Aurélie, LAZREK Mouna, NOBILLIAUX Florian, PREVOST Brigitte, THUILLIER Caroline, TINEZ Claire |
| EPI_ISL_1323614, EPI_ISL_1323615 | CH TOURCOING | CHU Lille - Laboratoire de Virologie | AIT YAHYA Emilie, ALIDJINOU Enagnon Kazali, BOCKET Laurence, CREPIN Michel, DEMAY Christophe, ENGELMANN Ilka, GEFFROY Sandrine, GUIGON Aurélie, LAZREK Mouna, NOBILLIAUX Florian, PREVOST Brigitte, THUILLIER Caroline, TINEZ Claire |
| EPI_ISL_1323616, EPI_ISL_1323617 | CH VALENCIENNES | CHU Lille - Laboratoire de Virologie | AIT YAHYA Emilie, ALIDJINOU Enagnon Kazali, BOCKET Laurence, CREPIN Michel, DEMAY Christophe, ENGELMANN Ilka, GEFFROY Sandrine, GUIGON Aurélie, LAZREK Mouna, NOBILLIAUX Florian, PREVOST Brigitte, THUILLIER Caroline, TINEZ Claire |
| EPI_ISL_1323618 | CH ARMENTIERES | CHU Lille - Laboratoire de Virologie | AIT YAHYA Emilie, ALIDJINOU Enagnon Kazali, BOCKET Laurence, CREPIN Michel, DEMAY Christophe, ENGELMANN Ilka, GEFFROY Sandrine, GUIGON Aurélie, LAZREK Mouna, NOBILLIAUX Florian, PREVOST Brigitte, THUILLIER Caroline, TINEZ Claire |
| EPI_ISL_1323619 | CHU LILLE | CHU Lille - Laboratoire de Virologie | AIT YAHYA Emilie, ALIDJINOU Enagnon Kazali, BOCKET Laurence, CREPIN Michel, DEMAY Christophe, ENGELMANN Ilka, GEFFROY Sandrine, GUIGON Aurélie, LAZREK Mouna, NOBILLIAUX Florian, PREVOST Brigitte, THUILLIER Caroline, TINEZ Claire |
| EPI_ISL_1323965, EPI_ISL_1323966, EPI_ISL_1323967, EPI_ISL_1323969, EPI_ISL_1323970, EPI_ISL_1323971, EPI_ISL_1323972, EPI_ISL_1323973, EPI_ISL_1323974 | Istituto Zooprofilattico Sperimentale del Mezzogiorno (IZSM) | IZSM-U.O.C. Virologia | Maurizio Viscardi, Lorena Cardillo, Pellegrino Cerino, Giovanna Fusco, Esterina De Carlo, Antonio Limone |
| EPI_ISL_1324758, EPI_ISL_1324759, EPI_ISL_1324760, EPI_ISL_1324761, EPI_ISL_1324762, EPI_ISL_1324763, EPI_ISL_1324764, EPI_ISL_1324765 | Laboratorio de Diagnostico de Enfermedades Emergentes y Reemergentes | Instituto de Diagnostico y Referencia Epidemiologicos (INDRE) | Claudia Wong-Arambula, Abril Rodriguez-Maldonado, Vanessa Rivero-Arredondo, Ariadna Medina-Benitez, Joaquin Quiroz-Mercado, Sergio Rangel-Guerrero,Natali Vega-Magana, Natividad Cruz-Ortiz, Tatiana Nunez-Garcia, Gisela Barrera-Badillo, Lucia Hernandez-Rivas, Irma Lopez-Martinez, Ernesto Ramirez-Gonzalez. |
| EPI_ISL_1324771, EPI_ISL_1324782 | Institut de Virologie du CHU de Strasbourg | Institut de Virologie du CHU de Strasbourg | Fafi-Kremer Samira, Gallais Floriane, Gantner Pierre, Laugel Elodie, Solis Morgane, Velay Aurélie, Wendling Marie-Josée |
| EPI_ISL_1334365 | Ministry of Health Hospitals | Institute of Health and Community Medicine | Chan Chia Jui, David Perera, Ooi Mong How, Chua Hock Hin, Tonnii Sia Loong Loong, Wong Jyn Shan, Wong Kiing Aik |
| EPI_ISL_1334380 | Laboratorios Lister | Instituto de Diagnostico y Referencia Epidemiologicos (INDRE) | Claudia Wong-Arambula, Abril Rodriguez-Maldonado, Vanessa Rivero-Arredondo, Ariadna Medina-Benitez, Joaquin Quiroz-Mercado, Sergio Rangel-Guerrero, Natividad Cruz-Ortiz, Tatiana Nunez-Garcia, Gisela Barrera-Badillo, Lucia Hernandez-Rivas, Irma Lopez-Martinez, Ernesto Ramirez-Gonzalez. |
| EPI_ISL_1335757 | LESP Zacatecas | Instituto de diagnóstico y Referencia Epidemiologicos (INDRE) Departamento de Virología | Claudia Wong-Arambula, Abril Rodriguez-Maldonado, Vanessa Rivero-Arredondo, Ariadna Medina-Benitez, Joaquin Quiroz-Mercado, Sergio Rangel-Guerrero, Natividad Cruz-Ortiz, Tatiana Nunez-Garcia, Gisela Barrera-Badillo, Lucia Hernandez-Rivas, Irma Lopez-Martinez, Ernesto Ramirez-Gonzalez. |
| EPI_ISL_1336329 | Hopital | National Reference Center for Viruses of Respiratory Infections, Institut Pasteur, Paris | Marion Barbet, Sylvie Behillil, Méline Bizard, Angela Brisebarre, Camille Capel, Etienne Simon-Lorière, Vincent Enouf, Maud Vanpeene, Sylvie van der Werf,Castelain Sandrine |
| EPI_ISL_1336797 | DIP. PREV. AVEZZANO SERVIZIO DI IGIENE EPIDEMIOLOGIA E SANITA' PUBBLICA | Istituto Zooprofilattico Sperimentale dell'Abruzzo e Molise "G. Caporale" | Lorusso A, Marcacci M, Di Domenico M, Ancora M, Curini V, Di Lollo V, Mangone I, Rinaldi A, Scialabba S, Caporale M, Di Pasquale A, Cammà C, Puglia I, Calistri P, Savini G |
| EPI_ISL_1337360, EPI_ISL_1337362 | LESP Queretaro | Instituto de Diagnostico y Referencia Epidemiologicos (INDRE) | Claudia Wong-Arambula, Abril Rodriguez-Maldonado, Vanessa Rivero-Arredondo, Ariadna Medina-Benitez, Joaquin Quiroz-Mercado, Sergio Rangel-Guerrero, Natividad Cruz-Ortiz, Tatiana Nunez-Garcia, Gisela Barrera-Badillo, Lucia Hernandez-Rivas, Irma Lopez-Martinez, Ernesto Ramirez-Gonzalez. |
| EPI_ISL_1337363, EPI_ISL_1337365, EPI_ISL_1337367 | LESP Guanajuato | Instituto de Diagnostico y Referencia Epidemiologicos (INDRE) | Claudia Wong-Arambula, Abril Rodriguez-Maldonado, Vanessa Rivero-Arredondo, Ariadna Medina-Benitez, Joaquin Quiroz-Mercado, Sergio Rangel-Guerrero, Natividad Cruz-Ortiz, Tatiana Nunez-Garcia, Gisela Barrera-Badillo, Lucia Hernandez-Rivas, Irma Lopez-Martinez, Ernesto Ramirez-Gonzalez. |
| EPI_ISL_1337368 | LESP Nuevo Leon | Instituto de Diagnostico y Referencia Epidemiologicos (INDRE) | Claudia Wong-Arambula, Abril Rodriguez-Maldonado, Vanessa Rivero-Arredondo, Ariadna Medina-Benitez, Joaquin Quiroz-Mercado, Sergio Rangel-Guerrero, Natividad Cruz-Ortiz, Tatiana Nunez-Garcia, Gisela Barrera-Badillo, Lucia Hernandez-Rivas, Irma Lopez-Martinez, Ernesto Ramirez-Gonzalez. |
| EPI_ISL_1337370, EPI_ISL_1337371, EPI_ISL_1337373, EPI_ISL_1337374 | LESP Baja California Sur | Instituto de Diagnostico y Referencia Epidemiologicos (INDRE) | Claudia Wong-Arambula, Abril Rodriguez-Maldonado, Vanessa Rivero-Arredondo, Ariadna Medina-Benitez, Joaquin Quiroz-Mercado, Sergio Rangel-Guerrero, Natividad Cruz-Ortiz, Tatiana Nunez-Garcia, Gisela Barrera-Badillo, Lucia Hernandez-Rivas, Irma Lopez-Martinez, Ernesto Ramirez-Gonzalez. |
| EPI_ISL_1337395 | LESP Colima | Instituto de Diagnostico y Referencia Epidemiologicos (INDRE) | Claudia Wong-Arambula, Abril Rodriguez-Maldonado, Vanessa Rivero-Arredondo, Ariadna Medina-Benitez, Joaquin Quiroz-Mercado, Sergio Rangel-Guerrero, Natividad Cruz-Ortiz, Tatiana Nunez-Garcia, Gisela Barrera-Badillo, Lucia Hernandez-Rivas, Irma Lopez-Martinez, Ernesto Ramirez-Gonzalez. |
| EPI_ISL_1340611 | LESP Guanajuato | Instituto de Diagnostico y Referencia Epidemiologicos (INDRE) | Claudia Wong-Arambula, Abril Rodriguez-Maldonado, Vanessa Rivero-Arredondo, Ariadna Medina-Benitez, Joaquin Quiroz-Mercado, Sergio Rangel-Guerrero, Natividad Cruz-Ortiz, Tatiana Nunez-Garcia, Gisela Barrera-Badillo, Lucia Hernandez-Rivas, Irma Lopez-Martinez, Ernesto Ramirez-Gonzalez. |
| EPI_ISL_1340613 | Laboratorios DIAGNOMOL | Instituto de Diagnostico y Referencia Epidemiologicos (INDRE) | Claudia Wong-Arambula, Abril Rodriguez-Maldonado, Vanessa Rivero-Arredondo, Ariadna Medina-Benitez, Joaquin Quiroz-Mercado, Sergio Rangel-Guerrero, Natividad Cruz-Ortiz, Tatiana Nunez-Garcia, Gisela Barrera-Badillo, Lucia Hernandez-Rivas, Irma Lopez-Martinez, Ernesto Ramirez-Gonzalez. |
| EPI_ISL_1340614, EPI_ISL_1340616, EPI_ISL_1340617 | LESP Guanajuato | Instituto de Diagnostico y Referencia Epidemiologicos (INDRE) | Claudia Wong-Arambula, Abril Rodriguez-Maldonado, Vanessa Rivero-Arredondo, Ariadna Medina-Benitez, Joaquin Quiroz-Mercado, Sergio Rangel-Guerrero, Natividad Cruz-Ortiz, Tatiana Nunez-Garcia, Gisela Barrera-Badillo, Lucia Hernandez-Rivas, Irma Lopez-Martinez, Ernesto Ramirez-Gonzalez. |
| EPI_ISL_1340619 | Laboratorios BIMODI | Instituto de Diagnostico y Referencia Epidemiologicos (INDRE) | Claudia Wong-Arambula, Abril Rodriguez-Maldonado, Vanessa Rivero-Arredondo, Ariadna Medina-Benitez, Joaquin Quiroz-Mercado, Sergio Rangel-Guerrero, Natividad Cruz-Ortiz, Tatiana Nunez-Garcia, Gisela Barrera-Badillo, Lucia Hernandez-Rivas, Irma Lopez-Martinez, Ernesto Ramirez-Gonzalez. |
| EPI_ISL_1340620, EPI_ISL_1340621, EPI_ISL_1340623 | Laboratorio BIOMEDICA | Instituto de Diagnostico y Referencia Epidemiologicos (INDRE) | Claudia Wong-Arambula, Abril Rodriguez-Maldonado, Vanessa Rivero-Arredondo, Ariadna Medina-Benitez, Joaquin Quiroz-Mercado, Sergio Rangel-Guerrero, Natividad Cruz-Ortiz, Tatiana Nunez-Garcia, Gisela Barrera-Badillo, Lucia Hernandez-Rivas, Irma Lopez-Martinez, Ernesto Ramirez-Gonzalez. |
| EPI_ISL_1340624, EPI_ISL_1340626, EPI_ISL_1340627 | LESP Guanajuato | Instituto de Diagnostico y Referencia Epidemiologicos (INDRE) | Claudia Wong-Arambula, Abril Rodriguez-Maldonado, Vanessa Rivero-Arredondo, Ariadna Medina-Benitez, Joaquin Quiroz-Mercado, Sergio Rangel-Guerrero, Natividad Cruz-Ortiz, Tatiana Nunez-Garcia, Gisela Barrera-Badillo, Lucia Hernandez-Rivas, Irma Lopez-Martinez, Ernesto Ramirez-Gonzalez. |
| EPI_ISL_1340629 | INER | Instituto de Diagnostico y Referencia Epidemiologicos (INDRE) | Claudia Wong-Arambula, Abril Rodriguez-Maldonado, Vanessa Rivero-Arredondo, Ariadna Medina-Benitez, Joaquin Quiroz-Mercado, Sergio Rangel-Guerrero, Natividad Cruz-Ortiz, Tatiana Nunez-Garcia, Gisela Barrera-Badillo, Lucia Hernandez-Rivas, Irma Lopez-Martinez, Ernesto |

| | | | |
|---|---|---|---|
| | | | Ramirez-Gonzalez. |
| EPI_ISL_1340630, EPI_ISL_1340631, EPI_ISL_1340633 | LESP Guanajuato | Instituto de Diagnostico y Referencia Epidemiologicos (INDRE) | Claudia Wong-Arambula, Abril Rodriguez-Maldonado, Vanessa Rivero-Arredondo, Ariadna Medina-Benitez, Joaquin Quiroz-Mercado, Sergio Rangel-Guerrero, Natividad Cruz-Ortiz, Tatiana Nunez-Garcia, Gisela Barrera-Badillo, Lucia Hernandez-Rivas, Irma Lopez-Martinez, Ernesto Ramirez-Gonzalez. |
| EPI_ISL_1340634 | Laboratorios BIMODI | Instituto de Diagnostico y Referencia Epidemiologicos (INDRE) | Claudia Wong-Arambula, Abril Rodriguez-Maldonado, Vanessa Rivero-Arredondo, Ariadna Medina-Benitez, Joaquin Quiroz-Mercado, Sergio Rangel-Guerrero, Natividad Cruz-Ortiz, Tatiana Nunez-Garcia, Gisela Barrera-Badillo, Lucia Hernandez-Rivas, Irma Lopez-Martinez, Ernesto Ramirez-Gonzalez. |
| EPI_ISL_1340636, EPI_ISL_1340637, EPI_ISL_1340639 | LESP Queretaro | Instituto de Diagnostico y Referencia Epidemiologicos (INDRE) | Claudia Wong-Arambula, Abril Rodriguez-Maldonado, Vanessa Rivero-Arredondo, Ariadna Medina-Benitez, Joaquin Quiroz-Mercado, Sergio Rangel-Guerrero, Natividad Cruz-Ortiz, Tatiana Nunez-Garcia, Gisela Barrera-Badillo, Lucia Hernandez-Rivas, Irma Lopez-Martinez, Ernesto Ramirez-Gonzalez. |
| EPI_ISL_1340640, EPI_ISL_1340642 | LESP Hidalgo | Instituto de Diagnostico y Referencia Epidemiologicos (INDRE) | Claudia Wong-Arambula, Abril Rodriguez-Maldonado, Vanessa Rivero-Arredondo, Ariadna Medina-Benitez, Joaquin Quiroz-Mercado, Sergio Rangel-Guerrero, Natividad Cruz-Ortiz, Tatiana Nunez-Garcia, Gisela Barrera-Badillo, Lucia Hernandez-Rivas, Irma Lopez-Martinez, Ernesto Ramirez-Gonzalez. |
| EPI_ISL_1340643, EPI_ISL_1340644, EPI_ISL_1340646, EPI_ISL_1340647, EPI_ISL_1340648, EPI_ISL_1340650 | LESP Guanajuato | Instituto de Diagnostico y Referencia Epidemiologicos (INDRE) | Claudia Wong-Arambula, Abril Rodriguez-Maldonado, Vanessa Rivero-Arredondo, Ariadna Medina-Benitez, Joaquin Quiroz-Mercado, Sergio Rangel-Guerrero, Natividad Cruz-Ortiz, Tatiana Nunez-Garcia, Gisela Barrera-Badillo, Lucia Hernandez-Rivas, Irma Lopez-Martinez, Ernesto Ramirez-Gonzalez. |
| EPI_ISL_1340651, EPI_ISL_1340652 | LESP Puebla | Instituto de Diagnostico y Referencia Epidemiologicos (INDRE) | Claudia Wong-Arambula, Abril Rodriguez-Maldonado, Vanessa Rivero-Arredondo, Ariadna Medina-Benitez, Joaquin Quiroz-Mercado, Sergio Rangel-Guerrero, Natividad Cruz-Ortiz, Tatiana Nunez-Garcia, Gisela Barrera-Badillo, Lucia Hernandez-Rivas, Irma Lopez-Martinez, Ernesto Ramirez-Gonzalez. |
| EPI_ISL_1340654 | Laboratorios BIMODI | Instituto de Diagnostico y Referencia Epidemiologicos (INDRE) | Claudia Wong-Arambula, Abril Rodriguez-Maldonado, Vanessa Rivero-Arredondo, Ariadna Medina-Benitez, Joaquin Quiroz-Mercado, Sergio Rangel-Guerrero, Natividad Cruz-Ortiz, Tatiana Nunez-Garcia, Gisela Barrera-Badillo, Lucia Hernandez-Rivas, Irma Lopez-Martinez, Ernesto Ramirez-Gonzalez. |
| EPI_ISL_1340655, EPI_ISL_1340657 | LESP Queretaro | Instituto de Diagnostico y Referencia Epidemiologicos (INDRE) | Claudia Wong-Arambula, Abril Rodriguez-Maldonado, Vanessa Rivero-Arredondo, Ariadna Medina-Benitez, Joaquin Quiroz-Mercado, Sergio Rangel-Guerrero, Natividad Cruz-Ortiz, Tatiana Nunez-Garcia, Gisela Barrera-Badillo, Lucia Hernandez-Rivas, Irma Lopez-Martinez, Ernesto Ramirez-Gonzalez. |
| EPI_ISL_1340658 | LESP Yucatan | Instituto de Diagnostico y Referencia Epidemiologicos (INDRE) | Claudia Wong-Arambula, Abril Rodriguez-Maldonado, Vanessa Rivero-Arredondo, Ariadna Medina-Benitez, Joaquin Quiroz-Mercado, Sergio Rangel-Guerrero, Natividad Cruz-Ortiz, Tatiana Nunez-Garcia, Gisela Barrera-Badillo, Lucia Hernandez-Rivas, Irma Lopez-Martinez, Ernesto Ramirez-Gonzalez. |
| EPI_ISL_1340660 | LESP Zacatecas | Instituto de Diagnostico y Referencia Epidemiologicos (INDRE) | Claudia Wong-Arambula, Abril Rodriguez-Maldonado, Vanessa Rivero-Arredondo, Ariadna Medina-Benitez, Joaquin Quiroz-Mercado, Sergio Rangel-Guerrero, Natividad Cruz-Ortiz, Tatiana Nunez-Garcia, Gisela Barrera-Badillo, Lucia Hernandez-Rivas, Irma Lopez-Martinez, Ernesto Ramirez-Gonzalez. |
| EPI_ISL_1340661 | LESP Yucatan | Instituto de Diagnostico y Referencia Epidemiologicos (INDRE) | Claudia Wong-Arambula, Abril Rodriguez-Maldonado, Vanessa Rivero-Arredondo, Ariadna Medina-Benitez, Joaquin Quiroz-Mercado, Sergio Rangel-Guerrero, Natividad Cruz-Ortiz, Tatiana Nunez-Garcia, Gisela Barrera-Badillo, Lucia Hernandez-Rivas, Irma Lopez-Martinez, Ernesto Ramirez-Gonzalez. |
| EPI_ISL_1340786, EPI_ISL_1340787, EPI_ISL_1340788, EPI_ISL_1340789, EPI_ISL_1340790, EPI_ISL_1340791, EPI_ISL_1340792, EPI_ISL_1340793, EPI_ISL_1340794, EPI_ISL_1340795, EPI_ISL_1340796, EPI_ISL_1340798, EPI_ISL_1340799, EPI_ISL_1340800, EPI_ISL_1340801, EPI_ISL_1340802, EPI_ISL_1340803, EPI_ISL_1340804, EPI_ISL_1340805, EPI_ISL_1340806, EPI_ISL_1340807, EPI_ISL_1340808, EPI_ISL_1340809, EPI_ISL_1340810, EPI_ISL_1340811, EPI_ISL_1340812, EPI_ISL_1340813, EPI_ISL_1340814, EPI_ISL_1340815, EPI_ISL_1340816, EPI_ISL_1340817, EPI_ISL_1340818, EPI_ISL_1340819, EPI_ISL_1340820, EPI_ISL_1340822, EPI_ISL_1340823, EPI_ISL_1340824, EPI_ISL_1340825, EPI_ISL_1340826, EPI_ISL_1340827, EPI_ISL_1340828, EPI_ISL_1340829, EPI_ISL_1340830, EPI_ISL_1340831, EPI_ISL_1340857 | | | |
| see above | Northwestern Memorial Hospital | Northwestern University - Ozer Lab | Ramon Lorenzo-Redondo, Lacy M. Simons, Taylor J. Dean, Chad J. Achenbach, Lawrence J. Jennings, Chao Qi, Michael G. Ison, Judd F. Hultquist, Egon A. Ozer |
| EPI_ISL_1346634, EPI_ISL_1346636 | Center of Excellence in Clinical Virology | Research Unit of Systems Microbiology | Jiratchaya Puenpa, Pattaraporn Nimsamer, Oraphan Mayuramart, Vorthon Sawaswong, Sunchai Payungporn, Anek Mungaomklang, Ritthideach Yorsaeng, Jira Chansaenroj, Kamolthip Atsawawaranunt, Vichan Pawan, Suwanna Petto, Amornmas Kongklieng, Pakkaporn Panwijitkul, Yong Poovowan |
| EPI_ISL_1347201 | CH Arras | CHU Lille - Laboratoire de Virologie | AIT YAHYA Emilie, ALIDJINOU Enagnon Kazali, BOCKET Laurence, CREPIN Michel, DEMAY Christophe, ENGELMANN Ilka, GEFFROY Sandrine, GUIGON Aurélie, LAZREK Mouna, NOBILLIAUX Florian, PREVOST Brigitte, THUILLIER Caroline, TINEZ Claire |
| EPI_ISL_1358291, EPI_ISL_1358292, EPI_ISL_1358293, EPI_ISL_1358294, EPI_ISL_1358295, EPI_ISL_1358296, EPI_ISL_1358297, EPI_ISL_1358298, EPI_ISL_1358299 | IAL Regional de Santo Andre | Instituto Adolfo Lutz, Interdiciplinary Procedures Center, Strategic Laboratory | Claudio Tavares Sacchi, Claudia Regina Gonçalves, Erica Valessa Ramos Gomes, Karoline Rodrigues Campos, Caio Vinicius Dias Lopes |
| EPI_ISL_1358304, EPI_ISL_1358306, EPI_ISL_1358307, EPI_ISL_1358308, EPI_ISL_1358309, EPI_ISL_1358310, EPI_ISL_1358311, EPI_ISL_1358312, EPI_ISL_1358313, EPI_ISL_1358314, EPI_ISL_1358315 | | | |
| see above | LACEN do Mato Grosso do Sul | Instituto Adolfo Lutz, Interdiciplinary Procedures Center, Strategic Laboratory | Claudio Tavares Sacchi, Claudia Regina Gonçalves, Erica Valessa Ramos Gomes, Karoline Rodrigues Campos, Caio Vinicius Dias Lopes |
| EPI_ISL_1358321 | UPA Vila Santa Catarina | Instituto Adolfo Lutz, Interdiciplinary Procedures Center, Strategic Laboratory | Claudio Tavares Sacchi, Claudia Regina Gonçalves, Erica Valessa Ramos Gomes, Karoline Rodrigues Campos, Caio Vinicius Dias Lopes |
| EPI_ISL_1359064, EPI_ISL_1359065 | LESP Queretaro | Instituto de Diagnostico y Referencia Epidemiologicos (INDRE) | Claudia Wong-Arambula, Abril Rodriguez-Maldonado, Vanessa Rivero-Arredondo, Ariadna Medina-Benitez, Joaquin Quiroz-Mercado, Sergio Rangel-Guerrero, Natividad Cruz-Ortiz, Tatiana Nunez-Garcia, Gisela Barrera-Badillo, Lucia Hernandez-Rivas, Irma Lopez-Martinez, Ernesto Ramirez-Gonzalez. |
| EPI_ISL_1359066, EPI_ISL_1359067 | LESP Yucatan | Instituto de Diagnostico y Referencia Epidemiologicos (INDRE) | Claudia Wong-Arambula, Abril Rodriguez-Maldonado, Vanessa Rivero-Arredondo, Ariadna Medina-Benitez, Joaquin Quiroz-Mercado, Sergio Rangel-Guerrero, Natividad Cruz-Ortiz, Tatiana Nunez-Garcia, Gisela Barrera-Badillo, Lucia Hernandez-Rivas, Irma Lopez-Martinez, Ernesto Ramirez-Gonzalez. |
| EPI_ISL_1359068, EPI_ISL_1359069 | LESP Zacatecas | Instituto de Diagnostico y Referencia Epidemiologicos (INDRE) | Claudia Wong-Arambula, Abril Rodriguez-Maldonado, Vanessa Rivero-Arredondo, Ariadna Medina-Benitez, Joaquin Quiroz-Mercado, Sergio Rangel-Guerrero, Natividad Cruz-Ortiz, Tatiana Nunez-Garcia, Gisela Barrera-Badillo, Lucia Hernandez-Rivas, Irma Lopez-Martinez, Ernesto Ramirez-Gonzalez. |
| EPI_ISL_1359070, EPI_ISL_1359072, EPI_ISL_1359073, EPI_ISL_1359077, EPI_ISL_1359078, EPI_ISL_1359079, EPI_ISL_1359080, EPI_ISL_1359081, EPI_ISL_1359082, EPI_ISL_1359083, EPI_ISL_1359085, EPI_ISL_1359086, EPI_ISL_1359087 | | | |
| see above | LESP Guanajuato | Instituto de Diagnostico y Referencia Epidemiologicos (INDRE) | Claudia Wong-Arambula, Abril Rodriguez-Maldonado, Vanessa Rivero-Arredondo, Ariadna Medina-Benitez, Joaquin Quiroz-Mercado, Sergio Rangel-Guerrero, Natividad Cruz-Ortiz, Tatiana Nunez-Garcia, Gisela Barrera-Badillo, Lucia Hernandez-Rivas, Irma Lopez-Martinez, Ernesto Ramirez-Gonzalez. |
| EPI_ISL_1360339, EPI_ISL_1360340 | CSIR-National Environmental Engineering Research Institute | CSIR-Centre for Cellular and Molecular Biology - INSACOG | Ara Sreenivas, Onkar Kulkarni, Lamuk Zaveri, Sofia Banu, Shreekant Verma, Amareshwar Vodapalli , Viswagithe S L, B Himasri, Sharath Chandra Thota, Karthik Bharadwaj Tallapaka, Krishna Khairnar, Rakesh K Mishra,Divya Tej Sowpati |
| EPI_ISL_1360342 | CSIR-National Environmental Engineering Research Institute | CSIR-Centre for Cellular and Molecular Biology - INSACOG | Sofia Banu, Lamuk Zaveri, Ara Sreenivas, Shreekant Verma, Onkar Kulkarni, Amareshwar Vodapalli ,Viswagithe S L, B Himasri, Sharath Chandra Thota, Karthik Bharadwaj Tallapaka, Krishna Khairnar, Rakesh K Mishra,Divya Tej Sowpati |
| EPI_ISL_1360343, EPI_ISL_1360344 | CSIR-National Environmental Engineering Research Institute | CSIR-Centre for Cellular and Molecular Biology - INSACOG | Onkar Kulkarni, Lamuk Zaveri, Ara Sreenivas, Sofia Banu, Shreekant Verma, Amareshwar Vodapalli , Viswagithe S L, B Himasri, Sharath Chandra Thota, Karthik Bharadwaj Tallapaka, Krishna Khairnar, Rakesh K Mishra,Divya Tej Sowpati |
| EPI_ISL_1360345 | CSIR-National Environmental Engineering Research Institute | CSIR-Centre for Cellular and Molecular Biology - INSACOG | Lamuk Zaveri, Ara Sreenivas, Sofia Banu, Onkar Kulkarni, Shreekant Verma, Amareshwar Vodapalli , Viswagithe S L, B Himasri, Sharath Chandra Thota, |

| | | | |
|---|---|---|---|
| | | | Karthik Bharadwaj Tallapaka, Krishna Khairnar, Rakesh K Mishra,Divya Tej Sowpati |
| EPI_ISL_1360346 | CSIR-National Environmental Engineering Research Institute | CSIR-Centre for Cellular and Molecular Biology - INSACOG | Lamuk Zaveri, Ara Sreenivas, Onkar Kulkarni, Sofia Banu, Shreekant Verma,Amareshwar Vodapalli, Viswagithe S L, B Himasri, Sharath Chandra Thota, Karthik Bharadwaj Tallapaka, Krishna Khairnar, Rakesh K Mishra,Divya Tej Sowpati |
| EPI_ISL_1360348 | CSIR-National Environmental Engineering Research Institute | CSIR-Centre for Cellular and Molecular Biology - INSACOG | Ara Sreenivas, Onkar Kulkarni, Lamuk Zaveri, Sofia Banu, Shreekant Verma, Amareshwar Vodapalli , Viswagithe S L, B Himasri, Sharath Chandra Thota, Karthik Bharadwaj Tallapaka, Krishna Khairnar, Rakesh K Mishra,Divya Tej Sowpati |
| EPI_ISL_1360349, EPI_ISL_1360350 | CSIR-National Environmental Engineering Research Institute | CSIR-Centre for Cellular and Molecular Biology - INSACOG | Sofia Banu, Lamuk Zaveri, Ara Sreenivas, Shreekant Verma, Onkar Kulkarni, Amareshwar Vodapalli ,Viswagithe S L, B Himasri, Sharath Chandra Thota, Karthik Bharadwaj Tallapaka, Krishna Khairnar, Rakesh K Mishra,Divya Tej Sowpati |
| EPI_ISL_1360351, EPI_ISL_1360352 | CSIR-National Environmental Engineering Research Institute | CSIR-Centre for Cellular and Molecular Biology - INSACOG | Onkar Kulkarni, Lamuk Zaveri, Ara Sreenivas, Sofia Banu, Shreekant Verma, Amareshwar Vodapalli , Viswagithe S L, B Himasri, Sharath Chandra Thota, Karthik Bharadwaj Tallapaka, Krishna Khairnar, Rakesh K Mishra,Divya Tej Sowpati |
| EPI_ISL_1360353 | CSIR-National Environmental Engineering Research Institute | CSIR-Centre for Cellular and Molecular Biology - INSACOG | Lamuk Zaveri, Ara Sreenivas, Sofia Banu, Onkar Kulkarni, Shreekant Verma, Amareshwar Vodapalli , Viswagithe S L, B Himasri, Sharath Chandra Thota, Karthik Bharadwaj Tallapaka, Krishna Khairnar, Rakesh K Mishra,Divya Tej Sowpati |
| EPI_ISL_1360354, EPI_ISL_1360355 | CSIR-National Environmental Engineering Research Institute | CSIR-Centre for Cellular and Molecular Biology - INSACOG | Ara Sreenivas, Onkar Kulkarni, Lamuk Zaveri, Sofia Banu, Shreekant Verma, Amareshwar Vodapalli , Viswagithe S L, B Himasri, Sharath Chandra Thota, Karthik Bharadwaj Tallapaka, Krishna Khairnar, Rakesh K Mishra,Divya Tej Sowpati |
| EPI_ISL_1360356 | CSIR-National Environmental Engineering Research Institute | CSIR-Centre for Cellular and Molecular Biology - INSACOG | Sofia Banu, Lamuk Zaveri, Ara Sreenivas, Shreekant Verma, Onkar Kulkarni, Amareshwar Vodapalli ,Viswagithe S L, B Himasri, Sharath Chandra Thota, Karthik Bharadwaj Tallapaka, Krishna Khairnar, Rakesh K Mishra,Divya Tej Sowpati |
| EPI_ISL_1360359 | CSIR-National Environmental Engineering Research Institute | CSIR-Centre for Cellular and Molecular Biology - INSACOG | Lamuk Zaveri, Ara Sreenivas, Sofia Banu, Onkar Kulkarni, Shreekant Verma, Amareshwar Vodapalli , Viswagithe S L, B Himasri, Sharath Chandra Thota, Karthik Bharadwaj Tallapaka, Krishna Khairnar, Rakesh K Mishra,Divya Tej Sowpati |
| EPI_ISL_1360360 | CSIR-National Environmental Engineering Research Institute | CSIR-Centre for Cellular and Molecular Biology - INSACOG | Lamuk Zaveri, Ara Sreenivas, Onkar Kulkarni, Sofia Banu, Shreekant Verma,Amareshwar Vodapalli, Viswagithe S L, B Himasri, Sharath Chandra Thota, Karthik Bharadwaj Tallapaka, Krishna Khairnar, Rakesh K Mishra,Divya Tej Sowpati |
| EPI_ISL_1360361, EPI_ISL_1360362 | CSIR-National Environmental Engineering Research Institute | CSIR-Centre for Cellular and Molecular Biology - INSACOG | Ara Sreenivas, Onkar Kulkarni, Lamuk Zaveri, Sofia Banu, Shreekant Verma, Amareshwar Vodapalli , Viswagithe S L, B Himasri, Sharath Chandra Thota, Karthik Bharadwaj Tallapaka, Krishna Khairnar, Rakesh K Mishra,Divya Tej Sowpati |
| EPI_ISL_1360364 | CSIR-National Environmental Engineering Research Institute | CSIR-Centre for Cellular and Molecular Biology - INSACOG | Sofia Banu, Lamuk Zaveri, Ara Sreenivas, Shreekant Verma, Onkar Kulkarni, Amareshwar Vodapalli ,Viswagithe S L, B Himasri, Sharath Chandra Thota, Karthik Bharadwaj Tallapaka, Krishna Khairnar, Rakesh K Mishra,Divya Tej Sowpati |
| EPI_ISL_1360365, EPI_ISL_1360366 | CSIR-National Environmental Engineering Research Institute | CSIR-Centre for Cellular and Molecular Biology - INSACOG | Onkar Kulkarni, Lamuk Zaveri, Ara Sreenivas, Sofia Banu, Shreekant Verma, Amareshwar Vodapalli , Viswagithe S L, B Himasri, Sharath Chandra Thota, Karthik Bharadwaj Tallapaka, Krishna Khairnar, Rakesh K Mishra,Divya Tej Sowpati |
| EPI_ISL_1360367 | CSIR-National Environmental Engineering Research Institute | CSIR-Centre for Cellular and Molecular Biology - INSACOG | Lamuk Zaveri, Ara Sreenivas, Sofia Banu, Onkar Kulkarni, Shreekant Verma, Amareshwar Vodapalli , Viswagithe S L, B Himasri, Sharath Chandra Thota, Karthik Bharadwaj Tallapaka, Krishna Khairnar, Rakesh K Mishra,Divya Tej Sowpati |
| EPI_ISL_1360368 | CSIR-National Environmental Engineering Research Institute | CSIR-Centre for Cellular and Molecular Biology - INSACOG | Lamuk Zaveri, Ara Sreenivas, Onkar Kulkarni, Sofia Banu, Shreekant Verma,Amareshwar Vodapalli, Viswagithe S L, B Himasri, Sharath Chandra Thota, Karthik Bharadwaj Tallapaka, Krishna Khairnar, Rakesh K Mishra,Divya Tej Sowpati |
| EPI_ISL_1360369, EPI_ISL_1360370 | CSIR-National Environmental Engineering Research Institute | CSIR-Centre for Cellular and Molecular Biology - INSACOG | Ara Sreenivas, Onkar Kulkarni, Lamuk Zaveri, Sofia Banu, Shreekant Verma, Amareshwar Vodapalli , Viswagithe S L, B Himasri, Sharath Chandra Thota, Karthik Bharadwaj Tallapaka, Krishna Khairnar, Rakesh K Mishra,Divya Tej Sowpati |
| EPI_ISL_1360408, EPI_ISL_1360409, EPI_ISL_1360410, EPI_ISL_1360411 | Laboratorio de Enfermedades Emergentes y Reemergentes | Laboratorio de Enfermedades Emergentes y Reemergentes | Vega-Magaña Natali, Peña-Rodriguez Marcela, Viera-Segura Oliver, García-Chagollan Mariel, Hernández-Bello Jorge, Muñoz-Valle José Francisco |
| EPI_ISL_1363115, EPI_ISL_1363116 | Laboratory of Polymerase Chain Reaction of Federal Budget Institution of Science "Central Research Institute of Epidemiology" of The Federal Service on Customers' Rights Protection and Human Well-being Surveillance | Group of Genomics and Postgenomic Technologies of Central Research Institute of Epidemiology | Samoilov AE, Kaptelova VV, Korneenko EV, Valdokhina AV, Bulanenko VP, Saenko SS, Golubeva AG, Zotova MI, Berlina YY, Solovyeva ED, Shipulina OY, Speranskaya AS, Tivanova EV, Kondrasheva LY, Akimkin VG, Cherkashina AS |
| EPI_ISL_1363117, EPI_ISL_1363118 | Molecular diagnostic laboratory of Federal Budget Institution of Science "Central Research Institute of Epidemiology" of The Federal Service on Customers' Rights Protection and Human Well-being Surveillance | Group of Genomics and Postgenomic Technologies of Central Research Institute of Epidemiology | Samoilov AE, Kaptelova VV, Korneenko EV, Valdokhina AV, Bulanenko VP, Saenko SS, Golubeva AG, Zotova MI, Berlina YY, Solovyeva ED, Shipulina OY, Speranskaya AS, Tivanova EV, Kondrasheva LY, Akimkin VG, Cherkashina AS |
| EPI_ISL_1365639, EPI_ISL_1365640, EPI_ISL_1365641, EPI_ISL_1365642, EPI_ISL_1365643, EPI_ISL_1365644, EPI_ISL_1365645, EPI_ISL_1365646, EPI_ISL_1365647, EPI_ISL_1365648 | LESP Nuevo Leon | Instituto de Diagnostico y Referencia Epidemiologicos (INDRE) | Claudia Wong-Arambula, Abril Rodriguez-Maldonado, Vanessa Rivero-Arredondo, Ariadna Medina-Benitez, Joaquin Quiroz-Mercado, Sergio Rangel-Guerrero, Natividad Cruz-Ortiz, Tatiana Nunez-Garcia, Gisela Barrera-Badillo, Lucia Hernandez-Rivas, Irma Lopez-Martinez, Ernesto Ramirez-Gonzalez. |
| EPI_ISL_1365650, EPI_ISL_1365651, EPI_ISL_1365652, EPI_ISL_1365653, EPI_ISL_1365654, EPI_ISL_1365655 | LESP Morelos | Instituto de Diagnostico y Referencia Epidemiologicos (INDRE) | Claudia Wong-Arambula, Abril Rodriguez-Maldonado, Vanessa Rivero-Arredondo, Ariadna Medina-Benitez, Joaquin Quiroz-Mercado, Sergio Rangel-Guerrero, Natividad Cruz-Ortiz, Tatiana Nunez-Garcia, Gisela Barrera-Badillo, Lucia Hernandez-Rivas, Irma Lopez-Martinez, Ernesto Ramirez-Gonzalez. |
| EPI_ISL_1365656 | LESP Veracruz | Instituto de Diagnostico y Referencia Epidemiologicos (INDRE) | Claudia Wong-Arambula, Abril Rodriguez-Maldonado, Vanessa Rivero-Arredondo, Ariadna Medina-Benitez, Joaquin Quiroz-Mercado, Sergio Rangel-Guerrero, Natividad Cruz-Ortiz, Tatiana Nunez-Garcia, Gisela Barrera-Badillo, Lucia Hernandez-Rivas, Irma Lopez-Martinez, Ernesto Ramirez-Gonzalez. |
| EPI_ISL_1365657 | LESP Ciudad de Mexico | Instituto de Diagnostico y Referencia Epidemiologicos (INDRE) | Claudia Wong-Arambula, Abril Rodriguez-Maldonado, Vanessa Rivero-Arredondo, Ariadna Medina-Benitez, Joaquin Quiroz-Mercado, Sergio Rangel-Guerrero, Natividad Cruz-Ortiz, Tatiana Nunez-Garcia, Gisela Barrera-Badillo, Lucia Hernandez-Rivas, Irma Lopez-Martinez, Ernesto Ramirez-Gonzalez. |
| EPI_ISL_1365660 | LESP Baja California | Instituto de Diagnostico y Referencia Epidemiologicos (INDRE) | Claudia Wong-Arambula, Abril Rodriguez-Maldonado, Vanessa Rivero-Arredondo, Ariadna Medina-Benitez, Joaquin Quiroz-Mercado, Sergio Rangel-Guerrero, Natividad Cruz-Ortiz, Tatiana Nunez-Garcia, Gisela Barrera-Badillo, Lucia Hernandez-Rivas, Irma Lopez-Martinez, Ernesto Ramirez-Gonzalez. |
| EPI_ISL_1365661 | LESP Quintana Roo | Instituto de Diagnostico y Referencia Epidemiologicos (INDRE) | Claudia Wong-Arambula, Abril Rodriguez-Maldonado, Vanessa Rivero-Arredondo, Ariadna Medina-Benitez, Joaquin Quiroz-Mercado, Sergio Rangel-Guerrero, Natividad Cruz-Ortiz, Tatiana Nunez-Garcia, Gisela Barrera-Badillo, Lucia Hernandez-Rivas, Irma Lopez-Martinez, Ernesto Ramirez-Gonzalez. |
| EPI_ISL_1365663 | LESP Coahuila | Instituto de Diagnostico y Referencia Epidemiologicos (INDRE) | Claudia Wong-Arambula, Abril Rodriguez-Maldonado, Vanessa Rivero-Arredondo, Ariadna Medina-Benitez, Joaquin Quiroz-Mercado, Sergio Rangel-Guerrero, Natividad Cruz-Ortiz, Tatiana Nunez-Garcia, Gisela Barrera-Badillo, Lucia Hernandez-Rivas, Irma Lopez-Martinez, Ernesto Ramirez-Gonzalez. |
| EPI_ISL_1366661, EPI_ISL_1366662 | LESP Jalisco | Instituto de Diagnostico y Referencia Epidemiologicos (INDRE) | Claudia Wong-Arambula, Abril Rodriguez-Maldonado, Vanessa Rivero-Arredondo, Ariadna Medina-Benitez, Joaquin Quiroz-Mercado, Sergio Rangel-Guerrero, Natividad Cruz-Ortiz, Tatiana Nunez-Garcia, Gisela Barrera-Badillo, Lucia Hernandez-Rivas, Irma Lopez-Martinez, Ernesto Ramirez-Gonzalez. |
| EPI_ISL_1366663, EPI_ISL_1366664, EPI_ISL_1366665 | LESP Nuevo Leon | Instituto de Diagnostico y Referencia Epidemiologicos (INDRE) | Claudia Wong-Arambula, Abril Rodriguez-Maldonado, Vanessa Rivero-Arredondo, Ariadna Medina-Benitez, Joaquin Quiroz-Mercado, Sergio Rangel-Guerrero, Natividad Cruz-Ortiz, Tatiana Nunez-Garcia, Gisela Barrera-Badillo, Lucia Hernandez-Rivas, Irma Lopez-Martinez, Ernesto Ramirez-Gonzalez. |
| EPI_ISL_1366666 | LESP Yucatan | Instituto de Diagnostico y Referencia Epidemiologicos (INDRE) | Claudia Wong-Arambula, Abril Rodriguez-Maldonado, Vanessa Rivero-Arredondo, Ariadna Medina-Benitez, Joaquin Quiroz-Mercado, Sergio Rangel-Guerrero, Natividad Cruz-Ortiz, Tatiana Nunez-Garcia, Gisela Barrera-Badillo, Lucia Hernandez-Rivas, Irma Lopez-Martinez, Ernesto Ramirez-Gonzalez. |

| | | | |
|---|---|---|---|
| EPI_ISL_1366667, EPI_ISL_1366668, EPI_ISL_1366669, EPI_ISL_1366670, EPI_ISL_1366671 | LESP Queretaro | Instituto de Diagnostico y Referencia Epidemiologicos (INDRE) | Claudia Wong-Arambula, Abril Rodriguez-Maldonado, Vanessa Rivero-Arredondo, Ariadna Medina-Benitez, Joaquin Quiroz-Mercado, Sergio Rangel-Guerrero, Natividad Cruz-Ortiz, Tatiana Nunez-Garcia, Gisela Barrera-Badillo, Lucia Hernandez-Rivas, Irma Lopez-Martinez, Ernesto Ramirez-Gonzalez. |
| EPI_ISL_1366677, EPI_ISL_1366678, EPI_ISL_1366679, EPI_ISL_1366680, EPI_ISL_1366681 | LESP Morelos | Instituto de Diagnostico y Referencia Epidemiologicos (INDRE) | Claudia Wong-Arambula, Abril Rodriguez-Maldonado, Vanessa Rivero-Arredondo, Ariadna Medina-Benitez, Joaquin Quiroz-Mercado, Sergio Rangel-Guerrero, Natividad Cruz-Ortiz, Tatiana Nunez-Garcia, Gisela Barrera-Badillo, Lucia Hernandez-Rivas, Irma Lopez-Martinez, Ernesto Ramirez-Gonzalez. |
| EPI_ISL_1366682, EPI_ISL_1366685 | LESP Veracruz | Instituto de Diagnostico y Referencia Epidemiologicos (INDRE) | Claudia Wong-Arambula, Abril Rodriguez-Maldonado, Vanessa Rivero-Arredondo, Ariadna Medina-Benitez, Joaquin Quiroz-Mercado, Sergio Rangel-Guerrero, Natividad Cruz-Ortiz, Tatiana Nunez-Garcia, Gisela Barrera-Badillo, Lucia Hernandez-Rivas, Irma Lopez-Martinez, Ernesto Ramirez-Gonzalez. |
| EPI_ISL_1366687, EPI_ISL_1366688, EPI_ISL_1366689 | LESP Ciudad de Mexico | Instituto de Diagnostico y Referencia Epidemiologicos (INDRE) | Claudia Wong-Arambula, Abril Rodriguez-Maldonado, Vanessa Rivero-Arredondo, Ariadna Medina-Benitez, Joaquin Quiroz-Mercado, Sergio Rangel-Guerrero, Natividad Cruz-Ortiz, Tatiana Nunez-Garcia, Gisela Barrera-Badillo, Lucia Hernandez-Rivas, Irma Lopez-Martinez, Ernesto Ramirez-Gonzalez. |
| EPI_ISL_1366742, EPI_ISL_1366743, EPI_ISL_1366744, EPI_ISL_1366745 | OUCRU | OUCRU | Nguyen Van Vinh Chau, Nguyen Thi Thu Hong, Nghiem My Ngoc, Nguyen To Anh, Huynh Trung Trieu, Le Nguyen Truc Nhu, Lam Minh Yen, Ngo Ngoc Quang Minh, Nguyen Thanh Phong, Nguyen Thanh Truong, Le Thi Thu Huong, Tran Nguyen Hoang Tu, Le Manh Hung, Tran Tan Thanh, Nguyen Thanh Dung, Nguyen Tri Dung, Guy Thwaites, Le Van Tan |
| EPI_ISL_1366746 | Laboratori de Referencia de Catalunya | Laboratori de Referencia de Catalunya | Ramirez A., Bellosillo B., Padilla E., Vilas A., Hernandez JJ., Canal M. |
| EPI_ISL_1369223, EPI_ISL_1369224 | Labo Marquise | CHU Lille - Laboratoire de Virologie | AIT YAHYA Emilie, ALIDJINOU Enagnon Kazali, BOCKET Laurence, CREPIN Michel, DEMAY Christophe, ENGELMANN Ilka, GEFFROY Sandrine, GUIGON Aurélie, LAZREK Mouna, NOBILLIAUX Florian, PREVOST Brigitte, THUILLIER Caroline, TINEZ Claire |
| EPI_ISL_1369226, EPI_ISL_1369227 | CH Armentieres | CHU Lille - Laboratoire de Virologie | AIT YAHYA Emilie, ALIDJINOU Enagnon Kazali, BOCKET Laurence, CREPIN Michel, DEMAY Christophe, ENGELMANN Ilka, GEFFROY Sandrine, GUIGON Aurélie, LAZREK Mouna, NOBILLIAUX Florian, PREVOST Brigitte, THUILLIER Caroline, TINEZ Claire |
| EPI_ISL_1369228, EPI_ISL_1369229, EPI_ISL_1369230, EPI_ISL_1369232 | CH St Philibert | CHU Lille - Laboratoire de Virologie | AIT YAHYA Emilie, ALIDJINOU Enagnon Kazali, BOCKET Laurence, CREPIN Michel, DEMAY Christophe, ENGELMANN Ilka, GEFFROY Sandrine, GUIGON Aurélie, LAZREK Mouna, NOBILLIAUX Florian, PREVOST Brigitte, THUILLIER Caroline, TINEZ Claire |
| EPI_ISL_1369233, EPI_ISL_1369234, EPI_ISL_1369235, EPI_ISL_1369236, EPI_ISL_1369237 | CH Maubeuge | CHU Lille - Laboratoire de Virologie | AIT YAHYA Emilie, ALIDJINOU Enagnon Kazali, BOCKET Laurence, CREPIN Michel, DEMAY Christophe, ENGELMANN Ilka, GEFFROY Sandrine, GUIGON Aurélie, LAZREK Mouna, NOBILLIAUX Florian, PREVOST Brigitte, THUILLIER Caroline, TINEZ Claire |
| EPI_ISL_1369238, EPI_ISL_1369239, EPI_ISL_1369240 | CH Dunkerque | CHU Lille - Laboratoire de Virologie | AIT YAHYA Emilie, ALIDJINOU Enagnon Kazali, BOCKET Laurence, CREPIN Michel, DEMAY Christophe, ENGELMANN Ilka, GEFFROY Sandrine, GUIGON Aurélie, LAZREK Mouna, NOBILLIAUX Florian, PREVOST Brigitte, THUILLIER Caroline, TINEZ Claire |
| EPI_ISL_1369241, EPI_ISL_1369242, EPI_ISL_1369243, EPI_ISL_1369244, EPI_ISL_1369245, EPI_ISL_1369246 | CH Tourcoing | CHU Lille - Laboratoire de Virologie | AIT YAHYA Emilie, ALIDJINOU Enagnon Kazali, BOCKET Laurence, CREPIN Michel, DEMAY Christophe, ENGELMANN Ilka, GEFFROY Sandrine, GUIGON Aurélie, LAZREK Mouna, NOBILLIAUX Florian, PREVOST Brigitte, THUILLIER Caroline, TINEZ Claire |
| EPI_ISL_1369249 | CH Calais | CHU Lille - Laboratoire de Virologie | AIT YAHYA Emilie, ALIDJINOU Enagnon Kazali, BOCKET Laurence, CREPIN Michel, DEMAY Christophe, ENGELMANN Ilka, GEFFROY Sandrine, GUIGON Aurélie, LAZREK Mouna, NOBILLIAUX Florian, PREVOST Brigitte, THUILLIER Caroline, TINEZ Claire |
| EPI_ISL_1369250, EPI_ISL_1369251, EPI_ISL_1369252, EPI_ISL_1369253 | CH Roubaix | CHU Lille - Laboratoire de Virologie | AIT YAHYA Emilie, ALIDJINOU Enagnon Kazali, BOCKET Laurence, CREPIN Michel, DEMAY Christophe, ENGELMANN Ilka, GEFFROY Sandrine, GUIGON Aurélie, LAZREK Mouna, NOBILLIAUX Florian, PREVOST Brigitte, THUILLIER Caroline, TINEZ Claire |
| EPI_ISL_1369257 | CH Cambrai | CHU Lille - Laboratoire de Virologie | AIT YAHYA Emilie, ALIDJINOU Enagnon Kazali, BOCKET Laurence, CREPIN Michel, DEMAY Christophe, ENGELMANN Ilka, GEFFROY Sandrine, GUIGON Aurélie, LAZREK Mouna, NOBILLIAUX Florian, PREVOST Brigitte, THUILLIER Caroline, TINEZ Claire |
| EPI_ISL_1369258 | CH Arras | CHU Lille - Laboratoire de Virologie | AIT YAHYA Emilie, ALIDJINOU Enagnon Kazali, BOCKET Laurence, CREPIN Michel, DEMAY Christophe, ENGELMANN Ilka, GEFFROY Sandrine, GUIGON Aurélie, LAZREK Mouna, NOBILLIAUX Florian, PREVOST Brigitte, THUILLIER Caroline, TINEZ Claire |
| EPI_ISL_1369260 | CH Lens | CHU Lille - Laboratoire de Virologie | AIT YAHYA Emilie, ALIDJINOU Enagnon Kazali, BOCKET Laurence, CREPIN Michel, DEMAY Christophe, ENGELMANN Ilka, GEFFROY Sandrine, GUIGON Aurélie, LAZREK Mouna, NOBILLIAUX Florian, PREVOST Brigitte, THUILLIER Caroline, TINEZ Claire |
| EPI_ISL_1369265 | CH Seclin | CHU Lille - Laboratoire de Virologie | AIT YAHYA Emilie, ALIDJINOU Enagnon Kazali, BOCKET Laurence, CREPIN Michel, DEMAY Christophe, ENGELMANN Ilka, GEFFROY Sandrine, GUIGON Aurélie, LAZREK Mouna, NOBILLIAUX Florian, PREVOST Brigitte, THUILLIER Caroline, TINEZ Claire |
| EPI_ISL_1369266, EPI_ISL_1369267, EPI_ISL_1369268, EPI_ISL_1369269, EPI_ISL_1369270 | CH Valenciennes | CHU Lille - Laboratoire de Virologie | AIT YAHYA Emilie, ALIDJINOU Enagnon Kazali, BOCKET Laurence, CREPIN Michel, DEMAY Christophe, ENGELMANN Ilka, GEFFROY Sandrine, GUIGON Aurélie, LAZREK Mouna, NOBILLIAUX Florian, PREVOST Brigitte, THUILLIER Caroline, TINEZ Claire |
| EPI_ISL_1369279 | CHU LILLE | CHU Lille - Laboratoire de Virologie | AIT YAHYA Emilie, ALIDJINOU Enagnon Kazali, BOCKET Laurence, CREPIN Michel, DEMAY Christophe, ENGELMANN Ilka, GEFFROY Sandrine, GUIGON Aurélie, LAZREK Mouna, NOBILLIAUX Florian, PREVOST Brigitte, THUILLIER Caroline, TINEZ Claire |
| EPI_ISL_1369283 | CH Douai | CHU Lille - Laboratoire de Virologie | AIT YAHYA Emilie, ALIDJINOU Enagnon Kazali, BOCKET Laurence, CREPIN Michel, DEMAY Christophe, ENGELMANN Ilka, GEFFROY Sandrine, GUIGON Aurélie, LAZREK Mouna, NOBILLIAUX Florian, PREVOST Brigitte, THUILLIER Caroline, TINEZ Claire |
| EPI_ISL_1369287 | CH Bethune | CHU Lille - Laboratoire de Virologie | AIT YAHYA Emilie, ALIDJINOU Enagnon Kazali, BOCKET Laurence, CREPIN Michel, DEMAY Christophe, ENGELMANN Ilka, GEFFROY Sandrine, GUIGON Aurélie, LAZREK Mouna, NOBILLIAUX Florian, PREVOST Brigitte, THUILLIER Caroline, TINEZ Claire |
| EPI_ISL_1369289, EPI_ISL_1369290, EPI_ISL_1369291 | CH BOULOGNE SUR MER | CHU Lille - Laboratoire de Virologie | AIT YAHYA Emilie, ALIDJINOU Enagnon Kazali, BOCKET Laurence, CREPIN Michel, DEMAY Christophe, ENGELMANN Ilka, GEFFROY Sandrine, GUIGON Aurélie, LAZREK Mouna, NOBILLIAUX Florian, PREVOST Brigitte, THUILLIER Caroline, TINEZ Claire |
| EPI_ISL_1372185, EPI_ISL_1372186 | Cliniques universitaires Saint-Luc | UCLouvain/IREC/MBLG | Jean Ruelle, Lysa Pinsmaye, Eleonore Ngyuvula Mantu, Benoit Kabamba Mukadi |
| EPI_ISL_1372288, EPI_ISL_1372291, EPI_ISL_1372292, EPI_ISL_1372293, EPI_ISL_1372294, EPI_ISL_1372295, EPI_ISL_1372296, EPI_ISL_1372297, EPI_ISL_1372314, EPI_ISL_1372315, EPI_ISL_1372316, EPI_ISL_1372317 | | | |
| see above | WHO National Influenza Centre Russian Federation | WHO National Influenza Centre Russian Federation | Andrey Komissarov, Artem Fadeev, Anna Ivanova, Kseniya Komissarova, Dmitry Bazhenov, Tamila Musaeva, Maria Timofeeva, Veronika Eder, Maria Pisareva, Daria Danilenko, Ksenia Safina, Elena Nabieva, Georgii Bazykin, Dmitry Lioznov |
| EPI_ISL_1372425 | OLVZ Aalst | OLVZ Aalst | Astrid Holderbeke |
| EPI_ISL_1379392 | HOSPITAL CIUDAD NEILY | Inciensa, Instituto Costarricense de Investigación y Enseñanza en Nutrición y Salud | Francisco Duarte, Hebleen Porras, Claudio Soto-Garita, Estela Cordero, Adriana Godínez & Melany Calderón |
| EPI_ISL_1379393 | LABORATORIO CLINICO LABIN | Inciensa, Instituto Costarricense de Investigación y Enseñanza en Nutrición y Salud | Francisco Duarte, Hebleen Porras, Claudio Soto-Garita, Estela Cordero, Adriana Godínez & Melany Calderón |
| EPI_ISL_1379394, EPI_ISL_1379395 | AREA DE SALUD CORREDORES | Inciensa, Instituto Costarricense de Investigación y Enseñanza en Nutrición y Salud | Francisco Duarte, Hebleen Porras, Claudio Soto-Garita, Estela Cordero, Adriana Godínez & Melany Calderón |
| EPI_ISL_1379399 | HOSPITAL SAN VITO DE COTO BRUS | Inciensa, Instituto Costarricense de Investigación y Enseñanza en Nutrición y Salud | Francisco Duarte, Hebleen Porras, Claudio Soto-Garita, Estela Cordero, Adriana Godínez & Melany Calderón |
| EPI_ISL_1379400 | AREA DE SALUD SAN JUAN-SAN DIEGO-CONCEPCION 2 | Inciensa, Instituto Costarricense de Investigación y Enseñanza en Nutrición y Salud | Francisco Duarte, Hebleen Porras, Claudio Soto-Garita, Estela Cordero, Adriana Godínez & Melany Calderón |
| EPI_ISL_1379402 | AREA DE SALUD CATEDRAL NORESTE | Inciensa, Instituto Costarricense de Investigación y Enseñanza en Nutrición y Salud | Francisco Duarte, Hebleen Porras, Claudio Soto-Garita, Estela Cordero, Adriana Godínez & Melany Calderón |
| EPI_ISL_1379403 | AREA DE SALUD GOICOECHEA 2 - CLINICA DR. JIMENEZ | Inciensa, Instituto Costarricense de Investigación y | Francisco Duarte, Hebleen Porras, Claudio Soto-Garita, Estela Cordero, Adriana Godínez & Melany Calderón |

| | | | |
|---|---|---|---|
| | NUÑEZ | Enseñanza en Nutrición y Salud | |
| EPI_ISL_1379404, EPI_ISL_1379405 | CLINICA BIBLICA | Inciensa, Instituto Costarricense de Investigación y Enseñanza en Nutrición y Salud | Francisco Duarte, Hebleen Porras, Claudio Soto-Garita, Estela Cordero, Adriana Godínez, Melany Calderón & Karla Gutiérrez-González |
| EPI_ISL_1379406 | LABORATORIO CLINICO DR. ECHANDI | Inciensa, Instituto Costarricense de Investigación y Enseñanza en Nutrición y Salud | Francisco Duarte, Hebleen Porras, Claudio Soto-Garita, Estela Cordero, Adriana Godínez & Melany Calderón |
| EPI_ISL_1379408 | AREA DE SALUD CATEDRAL NORESTE | Inciensa, Instituto Costarricense de Investigación y Enseñanza en Nutrición y Salud | Francisco Duarte, Hebleen Porras, Claudio Soto-Garita, Estela Cordero, Adriana Godínez & Melany Calderón |
| EPI_ISL_1379409 | AREA DE SALUD CURRIDABAT 2 | Inciensa, Instituto Costarricense de Investigación y Enseñanza en Nutrición y Salud | Francisco Duarte, Hebleen Porras, Claudio Soto-Garita, Estela Cordero, Adriana Godínez & Melany Calderón |
| EPI_ISL_1379410, EPI_ISL_1379411, EPI_ISL_1379412 | AREA DE SALUD CATEDRAL NORESTE | Inciensa, Instituto Costarricense de Investigación y Enseñanza en Nutrición y Salud | Francisco Duarte, Hebleen Porras, Claudio Soto-Garita, Estela Cordero, Adriana Godínez & Melany Calderón |
| EPI_ISL_1379417, EPI_ISL_1379418, EPI_ISL_1379419 | HOSPITAL DR. TONY FACIO CASTRO | Inciensa, Instituto Costarricense de Investigación y Enseñanza en Nutrición y Salud | Francisco Duarte, Hebleen Porras, Claudio Soto-Garita, Estela Cordero, Adriana Godínez, Melany Calderón & Tashana Anglin-Williams |
| EPI_ISL_1379421 | HOSPITAL SAN JUAN DE DIOS | Inciensa, Instituto Costarricense de Investigación y Enseñanza en Nutrición y Salud | Francisco Duarte, Hebleen Porras, Claudio Soto-Garita, Estela Cordero, Adriana Godínez, Melany Calderón & Daniel Cascante-Serrano |
| EPI_ISL_1379423, EPI_ISL_1379424 | AREA DE SALUD MORAVIA | Inciensa, Instituto Costarricense de Investigación y Enseñanza en Nutrición y Salud | Francisco Duarte, Hebleen Porras, Claudio Soto-Garita, Estela Cordero, Adriana Godínez & Melany Calderón |
| EPI_ISL_1379425 | AREA DE SALUD PARAISO-CERVANTES | Inciensa, Instituto Costarricense de Investigación y Enseñanza en Nutrición y Salud | Francisco Duarte, Hebleen Porras, Claudio Soto-Garita, Estela Cordero, Adriana Godínez, Melany Calderón & Mónica Charpentier-Artavia |
| EPI_ISL_1379426 | HOSPITAL DR. MAX PERALTA JIMENEZ | Inciensa, Instituto Costarricense de Investigación y Enseñanza en Nutrición y Salud | Francisco Duarte, Hebleen Porras, Claudio Soto-Garita, Estela Cordero, Adriana Godínez, Melany Calderón & Mónica Charpentier-Artavia |
| EPI_ISL_1379428 | HOSPITAL DR. ENRIQUE BALTODANO BRICEÑO | Inciensa, Instituto Costarricense de Investigación y Enseñanza en Nutrición y Salud | Francisco Duarte, Hebleen Porras, Claudio Soto-Garita, Estela Cordero, Adriana Godínez, Melany Calderón & Karol Matamoros-Montoya |
| EPI_ISL_1379430 | HOSPITAL DR. CARLOS LUIS VALVERDE VEGA | Inciensa, Instituto Costarricense de Investigación y Enseñanza en Nutrición y Salud | Francisco Duarte, Hebleen Porras, Claudio Soto-Garita, Estela Cordero, Adriana Godínez, Melany Calderón & Teresita Somogyi |
| EPI_ISL_1379431 | LABORATORIO CLINICO LABIN | Inciensa, Instituto Costarricense de Investigación y Enseñanza en Nutrición y Salud | Francisco Duarte, Hebleen Porras, Claudio Soto-Garita, Estela Cordero, Adriana Godínez, Melany Calderón & Pei Ling Chan Ma |
| EPI_ISL_1381043, EPI_ISL_1381044, EPI_ISL_1381045, EPI_ISL_1381046, EPI_ISL_1381047, EPI_ISL_1381048, EPI_ISL_1381049, EPI_ISL_1381050, EPI_ISL_1381051, EPI_ISL_1381052, EPI_ISL_1381053, EPI_ISL_1381054, EPI_ISL_1381055, EPI_ISL_1381056, EPI_ISL_1381057, EPI_ISL_1381058, EPI_ISL_1381059, EPI_ISL_1381060, EPI_ISL_1381061, EPI_ISL_1381062, EPI_ISL_1381063, EPI_ISL_1381064, EPI_ISL_1381065 | | | |
| see above | IAL Regional de Santo Andre | Instituto Adolfo Lutz, Interdiciplinary Procedures Center, Strategic Laboratory | Claudio Tavares Sacchi, Claudia Regina Gonçalves, Erica Valessa Ramos Gomes, Karoline Rodrigues Campos, Caio Vinicius Dias Lopes |
| EPI_ISL_1381147, EPI_ISL_1381148, EPI_ISL_1381149, EPI_ISL_1381151, EPI_ISL_1381152, EPI_ISL_1381156 | Hospital | National Reference Center for Viruses of Respiratory Infections, Institut Pasteur, Paris | Marion Barbet, Sylvie Behillil, Méline Bizard, Angela Brisebarre, Camille Capel, Louise Lefrançois, Etienne Simon-Lorière, Vincent Enouf, Maud Vanpeene, Sylvie van der Werf,Hecquet Denise |
| EPI_ISL_1381203, EPI_ISL_1381204, EPI_ISL_1381205, EPI_ISL_1381206, EPI_ISL_1381207, EPI_ISL_1381208 | Hospital | National Reference Center for Viruses of Respiratory Infections, Institut Pasteur, Paris | Marion Barbet, Sylvie Behillil, Méline Bizard, Angela Brisebarre, Camille Capel, Louise Lefrançois, Etienne Simon-Lorière, Vincent Enouf, Maud Vanpeene, Sylvie van der Werf,Bressollette CéLine |
| EPI_ISL_1381306, EPI_ISL_1381307, EPI_ISL_1381308, EPI_ISL_1381309, EPI_ISL_1381314, EPI_ISL_1381315, EPI_ISL_1381316, EPI_ISL_1381317, EPI_ISL_1381318, EPI_ISL_1381319, EPI_ISL_1381320, EPI_ISL_1381321, EPI_ISL_1381322, EPI_ISL_1381323, EPI_ISL_1381326, EPI_ISL_1381332, EPI_ISL_1381333, EPI_ISL_1381334, EPI_ISL_1381335, EPI_ISL_1381336, EPI_ISL_1381338, EPI_ISL_1381340, EPI_ISL_1381341, EPI_ISL_1381342, EPI_ISL_1381343, EPI_ISL_1381344, EPI_ISL_1381346, EPI_ISL_1381347, EPI_ISL_1381348, EPI_ISL_1381349, EPI_ISL_1381350, EPI_ISL_1381351, EPI_ISL_1381353, EPI_ISL_1381354, EPI_ISL_1381356, EPI_ISL_1381385 | | | |
| see above | Laboratory for Respiratory Viruses, Cantacuzino National Military-Medical Institute for Research and Development | Cantacuzino Institute Virology | Luiza Ustea, Nicoleta Paraschiv, Catalina Pascu, Sorin Dinu, Mihaela Lazar |
| EPI_ISL_1391009, EPI_ISL_1391011, EPI_ISL_1391012, EPI_ISL_1391013, EPI_ISL_1391014, EPI_ISL_1391015 | National Institute of Health Research and Development | National Institute of Health Research and Development | Subangkit, Hana Apsari Pawestri, Kartika Dewi Puspa, Arie Ardiansyah Nugraha, Hartanti Dian Ikawati, Krisna Nur Andriana Pangesti, Yuni Rukminiati, Ririn Ramadhany, Agustiningsih, Kindi Adam, Holy Arif Wibowo, Triyani Soekarso, Ni Ketut Susilarini, Nurika Hariastuti, Ully Alfi Nikmah, Reni Herman, Nike Susanti, Herna, Tati Febriyanti, Natalie Laurencia Kipuw, Fauzul Muna, Irene Lorinda Indalao, Nelly Puspandari, Vivi Setiawaty. |
| EPI_ISL_1391279, EPI_ISL_1391285, EPI_ISL_1391288, EPI_ISL_1391297, EPI_ISL_1391298, EPI_ISL_1391305, EPI_ISL_1391306, EPI_ISL_1391309, EPI_ISL_1391312, EPI_ISL_1391313, EPI_ISL_1391315, EPI_ISL_1391323, EPI_ISL_1391324, EPI_ISL_1391325, EPI_ISL_1391326, EPI_ISL_1391334, EPI_ISL_1391335, EPI_ISL_1391337, EPI_ISL_1391341, EPI_ISL_1391342, EPI_ISL_1391348, EPI_ISL_1391349, EPI_ISL_1391350, EPI_ISL_1391354, EPI_ISL_1391355, EPI_ISL_1391362 | | | |
| see above | Motol University Hospital | Institute of Applied Biotechnologies a.s. | Petr Klempt, Ondej Brzo, Martin Kašný, Kateina Kvapilová, Pavel Devínek, Petr Kvapil |
| EPI_ISL_1392647 | CHU Nantes Virology | CHU Nantes Virology | Celine Bressollette-Bodin, Thomas Drumel, Audrey Rodallec, Berthe-Marie Imbert-Marcille |
| EPI_ISL_1392923, EPI_ISL_1392924, EPI_ISL_1392925, EPI_ISL_1392926, EPI_ISL_1392927, EPI_ISL_1392928, EPI_ISL_1392929, EPI_ISL_1392930, EPI_ISL_1392931, EPI_ISL_1392932, EPI_ISL_1392933, EPI_ISL_1392934, EPI_ISL_1392935, EPI_ISL_1392936, EPI_ISL_1392937, EPI_ISL_1392938, EPI_ISL_1392939, EPI_ISL_1392940, EPI_ISL_1392941, EPI_ISL_1392942, EPI_ISL_1392943 | | | |
| see above | SC Microbiologia e Virologia AOUSS | AMES Centro Polidiagnostico Strumentale S.r.l. | Salvatore Rubino; Flavia Angioj; Laura Firino; Rosalba Govoni; Gabriele Ibba; Vincenzo Lai; Erica Mura; Bianca Paglietti; Claudia Piu; Anna Puggioni; Elena Rimini; Giulia Rocca; Caterina Serra; Sergio Uzzau. |
| EPI_ISL_1398266, EPI_ISL_1398332, EPI_ISL_1398358, EPI_ISL_1398368, EPI_ISL_1398373, EPI_ISL_1398926, EPI_ISL_1398933 | Institut de Virologie du CHU de Strasbourg | Institut de Virologie du CHU de Strasbourg | Fafi-Kremer Samira, Gallais Floriane, Gantner Pierre, Laugel Elodie, Solis Morgane, Velay Aurélie, Wendling Marie-Josée |
| EPI_ISL_1398934 | West Bandung Public Health | West Java Health Laboratory | Azzania Fibriani, Ema Rahmawati, Ryan Bayusantika Ristandi, Rifky Waluyajati Rachman, Cut Nur Cinthia Alamanda, Isak Solihin, Rini Robiani, Miftahul Faridl, Karimatu Khoirunnisa, Kamila Tania |
| EPI_ISL_1398935, EPI_ISL_1398938, EPI_ISL_1398940, EPI_ISL_1398982 | Institut de Virologie du CHU de Strasbourg | Institut de Virologie du CHU de Strasbourg | Fafi-Kremer Samira, Gallais Floriane, Gantner Pierre, Laugel Elodie, Solis Morgane, Velay Aurélie, Wendling Marie-Josée |
| EPI_ISL_1399259 | LESP Jalisco | Instituto de Diagnostico y Referencia Epidemiologicos (INDRE) | Claudia Wong-Arambula, Abril Rodriguez-Maldonado, Vanessa Rivero-Arredondo, Ariadna Medina-Benitez, Joaquin Quiroz-Mercado, Sergio Rangel-Guerrero, Natividad Cruz-Ortiz, Tatiana Nunez-Garcia, Gisela Barrera-Badillo, Lucia Hernandez-Rivas, Irma Lopez-Martinez, Ernesto Ramirez-Gonzalez. |
| EPI_ISL_1399260 | LESP Guanajuato | Instituto de Diagnostico y Referencia Epidemiologicos (INDRE) | Claudia Wong-Arambula, Abril Rodriguez-Maldonado, Vanessa Rivero-Arredondo, Ariadna Medina-Benitez, Joaquin Quiroz-Mercado, Sergio Rangel-Guerrero, Natividad Cruz-Ortiz, Tatiana Nunez-Garcia, Gisela Barrera-Badillo, Lucia Hernandez-Rivas, Irma Lopez-Martinez, Ernesto Ramirez-Gonzalez. |
| EPI_ISL_1399262, EPI_ISL_1399263, EPI_ISL_1399264 | LESP Nuevo Leon | Instituto de Diagnostico y Referencia Epidemiologicos (INDRE) | Claudia Wong-Arambula, Abril Rodriguez-Maldonado, Vanessa Rivero-Arredondo, Ariadna Medina-Benitez, Joaquin Quiroz-Mercado, Sergio Rangel-Guerrero, Natividad Cruz-Ortiz, Tatiana Nunez-Garcia, Gisela Barrera-Badillo, Lucia Hernandez-Rivas, Irma Lopez-Martinez, Ernesto Ramirez-Gonzalez. |
| EPI_ISL_1399275, EPI_ISL_1399276, EPI_ISL_1399277 | LESP Tamaulipas | Instituto de Diagnostico y Referencia Epidemiologicos (INDRE) | Claudia Wong-Arambula, Abril Rodriguez-Maldonado, Vanessa Rivero-Arredondo, Ariadna Medina-Benitez, Joaquin Quiroz-Mercado, Sergio Rangel-Guerrero, Natividad Cruz-Ortiz, Tatiana Nunez-Garcia, Gisela Barrera-Badillo, Lucia Hernandez-Rivas, Irma Lopez-Martinez, Ernesto |

| | | | |
|---|---|---|---|
| | | | Ramirez-Gonzalez. |
| EPI_ISL_1400310 | LESP Puebla | Instituto de Diagnostico y Referencia Epidemiologicos (INDRE) | Claudia Wong-Arambula, Abril Rodriguez-Maldonado, Vanessa Rivero-Arredondo, Ariadna Medina-Benitez, Joaquin Quiroz-Mercado, Sergio Rangel-Guerrero, Natividad Cruz-Ortiz, Tatiana Nunez-Garcia, Gisela Barrera-Badillo, Lucia Hernandez-Rivas, Irma Lopez-Martinez, Ernesto Ramirez-Gonzalez. |
| EPI_ISL_1401099, EPI_ISL_1401100, EPI_ISL_1401102, EPI_ISL_1401103, EPI_ISL_1401104, EPI_ISL_1401105, EPI_ISL_1401106, EPI_ISL_1401107, EPI_ISL_1401108, EPI_ISL_1401109, EPI_ISL_1401111, EPI_ISL_1401112, EPI_ISL_1401113, EPI_ISL_1401114, EPI_ISL_1401115, EPI_ISL_1401116, EPI_ISL_1401117, EPI_ISL_1401118, EPI_ISL_1401119, EPI_ISL_1401282, EPI_ISL_1401283, EPI_ISL_1401284, EPI_ISL_1401285, EPI_ISL_1401286, EPI_ISL_1401287, EPI_ISL_1401288, EPI_ISL_1401289, EPI_ISL_1401290, EPI_ISL_1401291, EPI_ISL_1401292, EPI_ISL_1401293, EPI_ISL_1401294 | | | |
| see above | National Center of Infectious and Parasitic Diseases | National Center of Infectious and Parasitic Diseases | Alexiev et al |
| EPI_ISL_1404376, EPI_ISL_1404377, EPI_ISL_1404378, EPI_ISL_1404384, EPI_ISL_1404385, EPI_ISL_1404386, EPI_ISL_1404387 | Klinisch Laboratorium ZNA | Klinisch Laboratorium ZNA | Verstrepen et al. |
| EPI_ISL_1405917 | Hospital Queen Elizabeth | Institute for Medical Research, Infectious Disease Research Centre, National Institutes of Health, Ministry of Health Malaysia | Suppiah J, Kamel K, Azizan MA, Thayan R |
| EPI_ISL_1406153 | Institute for Medical Research, Infectious Disease Research Centre, National Institutes of Health, Ministry of Health Malaysia | Institute for Medical Research, Infectious Disease Research Centre, National Institutes of Health, Ministry of Health Malaysia | Suppiah J, Kamel K, Mohd Zawawi Z, Ramly N, Robert F, Thayan R |
| EPI_ISL_1406540 | Centre Jean Perrin | CHU Clermont-Ferrand, service de virologie | Bisseux Maxime, Mirand Audrey, Combes Patricia, Henquell Cécile |
| EPI_ISL_1406544, EPI_ISL_1406621 | Centre Hospitalier Universitaire Clermont-Ferrand | CHU Clermont-Ferrand, service de virologie | Bisseux Maxime, Mirand Audrey, Combes Patricia, Henquell Cécile |
| EPI_ISL_1406695 | National Center of Infectious and Parasitic Diseases | National Center of Infectious and Parasitic Diseases | Alexiev et al |
| EPI_ISL_1411159 | National Institute of Health Research and Development | National Institute of Health Research and Development | Subangkit, Hana Apsari Pawestri, Kartika Dewi Puspa, Arie Ardiansyah Nugraha, Hartanti Dian Ikawati, Krisna Nur Andriana Pangesti, Yuni Rukminiati, Ririn Ramadhany, Agustiningsih, Kindi Adam, Holy Arif Wibowo, Triyani Soekarso, Ni Ketut Susilarini, Nurika Hariastuti, Ully Alfi Nikmah, Reni Herman, Nike Susanti, Herna, Tati Febriyanti, Natalie Laurencia Kipuw, Fauzul Muna, Irene Lorinda Indalao, Nelly Puspandari, Vivi Setiawaty. |
| EPI_ISL_1415898 | Laboratorium Intibios Karawang | National Institute of Health Research and Development | Vivi Setiawaty, Hana Apsari Pawestri, Subangkit, Kartika Dewi Puspa, Arie Ardiansyah Nugraha, Hartanti Dian Ikawati, Nelly Puspandari, Krisna Nur Andriana Pangesti |
| EPI_ISL_1416330 | Microbiology and Virology Laboratory, "Policlinico Riuniti, Azienda Ospedaliero Universitaria, Foggia" | Microbiology and Virology Laboratory, "Policlinico Riuniti, Azienda Ospedaliero Universitaria, Foggia" | Fabio Arena, Settimia Altamura, Donatella Cedola, Valeria Delli Carri, Annamaria D'Aprile, Maria Rosaria Lipsi, Loris Micelli, Maria Pia Patrizio, Daniela Pisanelli, Tiziana Rollo, Maurizio Margaglione and Rosella De Nittis. |
| EPI_ISL_1416976 | Outre mer | National Reference Center for Viruses of Respiratory Infections, Institut Pasteur, Paris | Marion Barbet, Sylvie Behillil, Méline Bizard,Frédéric Lemoine,Corinne Maufrais,Christophe Malabat, Angela Brisebarre, Camille Capel, Louise Lefrançois, Etienne Simon-Lorière, Vincent Enouf, Maud Vanpeene, Sylvie van der Werf,Esther Gyde |
| EPI_ISL_1424007 | LESP Yucatan | Instituto de Diagnostico y Referencia Epidemiologicos (INDRE) | Claudia Wong-Arambula, Abril Rodriguez-Maldonado, Vanessa Rivero-Arredondo, Ariadna Medina-Benitez, Joaquin Quiroz-Mercado, Sergio Rangel-Guerrero, Natividad Cruz-Ortiz, Tatiana Nunez-Garcia, Gisela Barrera-Badillo, Lucia Hernandez-Rivas, Irma Lopez-Martinez, Ernesto Ramirez-Gonzalez. |
| EPI_ISL_1424009 | LESP Hidalgo | Instituto de Diagnostico y Referencia Epidemiologicos (INDRE) | Claudia Wong-Arambula, Abril Rodriguez-Maldonado, Vanessa Rivero-Arredondo, Ariadna Medina-Benitez, Joaquin Quiroz-Mercado, Sergio Rangel-Guerrero, Natividad Cruz-Ortiz, Tatiana Nunez-Garcia, Gisela Barrera-Badillo, Lucia Hernandez-Rivas, Irma Lopez-Martinez, Ernesto Ramirez-Gonzalez. |
| EPI_ISL_1424010, EPI_ISL_1424011 | LESP Guanajuato | Instituto de Diagnostico y Referencia Epidemiologicos (INDRE) | Claudia Wong-Arambula, Abril Rodriguez-Maldonado, Vanessa Rivero-Arredondo, Ariadna Medina-Benitez, Joaquin Quiroz-Mercado, Sergio Rangel-Guerrero, Natividad Cruz-Ortiz, Tatiana Nunez-Garcia, Gisela Barrera-Badillo, Lucia Hernandez-Rivas, Irma Lopez-Martinez, Ernesto Ramirez-Gonzalez. |
| EPI_ISL_1424013 | LESP Chiapas | Instituto de Diagnostico y Referencia Epidemiologicos (INDRE) | Claudia Wong-Arambula, Abril Rodriguez-Maldonado, Vanessa Rivero-Arredondo, Ariadna Medina-Benitez, Joaquin Quiroz-Mercado, Sergio Rangel-Guerrero, Natividad Cruz-Ortiz, Tatiana Nunez-Garcia, Gisela Barrera-Badillo, Lucia Hernandez-Rivas, Irma Lopez-Martinez, Ernesto Ramirez-Gonzalez. |
| EPI_ISL_1424015 | LESP Campeche | Instituto de Diagnostico y Referencia Epidemiologicos (INDRE) | Claudia Wong-Arambula, Abril Rodriguez-Maldonado, Vanessa Rivero-Arredondo, Ariadna Medina-Benitez, Joaquin Quiroz-Mercado, Sergio Rangel-Guerrero, Natividad Cruz-Ortiz, Tatiana Nunez-Garcia, Gisela Barrera-Badillo, Lucia Hernandez-Rivas, Irma Lopez-Martinez, Ernesto Ramirez-Gonzalez. |
| EPI_ISL_1424016 | LESP Queretaro | Instituto de Diagnostico y Referencia Epidemiologicos (INDRE) | Claudia Wong-Arambula, Abril Rodriguez-Maldonado, Vanessa Rivero-Arredondo, Ariadna Medina-Benitez, Joaquin Quiroz-Mercado, Sergio Rangel-Guerrero, Natividad Cruz-Ortiz, Tatiana Nunez-Garcia, Gisela Barrera-Badillo, Lucia Hernandez-Rivas, Irma Lopez-Martinez, Ernesto Ramirez-Gonzalez. |
| EPI_ISL_1424017, EPI_ISL_1424019 | LESP Morelos | Instituto de Diagnostico y Referencia Epidemiologicos (INDRE) | Claudia Wong-Arambula, Abril Rodriguez-Maldonado, Vanessa Rivero-Arredondo, Ariadna Medina-Benitez, Joaquin Quiroz-Mercado, Sergio Rangel-Guerrero, Natividad Cruz-Ortiz, Tatiana Nunez-Garcia, Gisela Barrera-Badillo, Lucia Hernandez-Rivas, Irma Lopez-Martinez, Ernesto Ramirez-Gonzalez. |
| EPI_ISL_1424039, EPI_ISL_1424040 | LESP Tamaulipas | Instituto de Diagnostico y Referencia Epidemiologicos (INDRE) | Claudia Wong-Arambula, Abril Rodriguez-Maldonado, Vanessa Rivero-Arredondo, Ariadna Medina-Benitez, Joaquin Quiroz-Mercado, Sergio Rangel-Guerrero, Natividad Cruz-Ortiz, Tatiana Nunez-Garcia, Gisela Barrera-Badillo, Lucia Hernandez-Rivas, Irma Lopez-Martinez, Ernesto Ramirez-Gonzalez. |
| EPI_ISL_1424046 | LESP Veracruz | Instituto de Diagnostico y Referencia Epidemiologicos (INDRE) | Claudia Wong-Arambula, Abril Rodriguez-Maldonado, Vanessa Rivero-Arredondo, Ariadna Medina-Benitez, Joaquin Quiroz-Mercado, Sergio Rangel-Guerrero, Natividad Cruz-Ortiz, Tatiana Nunez-Garcia, Gisela Barrera-Badillo, Lucia Hernandez-Rivas, Irma Lopez-Martinez, Ernesto Ramirez-Gonzalez. |
| EPI_ISL_1424064 | Institute for Medical Research, Infectious Disease Research Centre, National Institutes of Health, Ministry of Health Malaysia | Institute for Medical Research, Infectious Disease Research Centre, National Institutes of Health, Ministry of Health Malaysia | Suppiah J, Kamel K, Mohd Zawawi Z, Ramly N, Robert F, Thayan R |
| EPI_ISL_1442121, EPI_ISL_1442143 | Azienda Ospedaliero - Universitaria di Modena Policlinico - Virologia e Microbiologia Molecolare | Istituto Zooprofilattico Sperimentale della Lombardia e dell'Emilia Romagna (IZSLER), Risk Analysis and Genomic Epidemiology Unit | Monica Pecorari, William Gennari, Giulia Fregni Serpini, Marina Morganti, Ilaria Menozzi, Erika Scaltriti, Stefano Pongolini |
| EPI_ISL_1468411 | Centro de Saude II Matao | Instituto Adolfo Lutz, Interdiciplinary Procedures Center, Strategic Laboratory | Claudio Tavares Sacchi, Claudia Regina Gonçalves, Erica Valessa Ramos Gomes, Karoline Rodrigues Campos, Caio Vinicius Dias Lopes |
| EPI_ISL_1468412 | Hospital Estadual de Mirandopolis | Instituto Adolfo Lutz, Interdiciplinary Procedures Center, Strategic Laboratory | Claudio Tavares Sacchi, Claudia Regina Gonçalves, Erica Valessa Ramos Gomes, Karoline Rodrigues Campos, Caio Vinicius Dias Lopes |
| EPI_ISL_1468417 | Santa Casa de Aracatuba Hospital Sagrado Coracao de Jesus | Instituto Adolfo Lutz, Interdiciplinary Procedures Center, Strategic Laboratory | Claudio Tavares Sacchi, Claudia Regina Gonçalves, Erica Valessa Ramos Gomes, Karoline Rodrigues Campos, Caio Vinicius Dias Lopes |
| EPI_ISL_1468422, EPI_ISL_1468423 | Penitenciaria Compacta de Avanhandava | Instituto Adolfo Lutz, Interdiciplinary Procedures Center, Strategic Laboratory | Claudio Tavares Sacchi, Claudia Regina Gonçalves, Erica Valessa Ramos Gomes, Karoline Rodrigues Campos, Caio Vinicius Dias Lopes |
| EPI_ISL_1468425 | Santa Casa de Birigui | Instituto Adolfo Lutz, Interdiciplinary Procedures Center, Strategic Laboratory | Claudio Tavares Sacchi, Claudia Regina Gonçalves, Erica Valessa Ramos Gomes, Karoline Rodrigues Campos, Caio Vinicius Dias Lopes |
| EPI_ISL_1468426, EPI_ISL_1468427 | Centro de Saude II Matao | Instituto Adolfo Lutz, Interdiciplinary Procedures Center, | Claudio Tavares Sacchi, Claudia Regina Gonçalves, Erica Valessa Ramos Gomes, Karoline Rodrigues Campos, Caio Vinicius Dias Lopes |

| | | | |
|---|---|---|---|
| | | Strategic Laboratory | |
| EPI_ISL_1468428 | SAE Servico de Atendimento Especializado | Instituto Adolfo Lutz, Interdiciplinary Procedures Center, Strategic Laboratory | Claudio Tavares Sacchi, Claudia Regina Gonçalves, Erica Valessa Ramos Gomes, Karoline Rodrigues Campos, Caio Vinicius Dias Lopes |
| EPI_ISL_1468430 | Centro de Saude II Matao | Instituto Adolfo Lutz, Interdiciplinary Procedures Center, Strategic Laboratory | Claudio Tavares Sacchi, Claudia Regina Gonçalves, Erica Valessa Ramos Gomes, Karoline Rodrigues Campos, Caio Vinicius Dias Lopes |
| EPI_ISL_1468442 | Santa Casa de Birigui | Instituto Adolfo Lutz, Interdiciplinary Procedures Center, Strategic Laboratory | Claudio Tavares Sacchi, Claudia Regina Gonçalves, Erica Valessa Ramos Gomes, Karoline Rodrigues Campos, Caio Vinicius Dias Lopes |
| EPI_ISL_1468444 | Secretaria Municipal de Saude de Birigui | Instituto Adolfo Lutz, Interdiciplinary Procedures Center, Strategic Laboratory | Claudio Tavares Sacchi, Claudia Regina Gonçalves, Erica Valessa Ramos Gomes, Karoline Rodrigues Campos, Caio Vinicius Dias Lopes |
| EPI_ISL_1468445 | Santa Casa de Misericordia de Pereira Barreto | Instituto Adolfo Lutz, Interdiciplinary Procedures Center, Strategic Laboratory | Claudio Tavares Sacchi, Claudia Regina Gonçalves, Erica Valessa Ramos Gomes, Karoline Rodrigues Campos, Caio Vinicius Dias Lopes |
| EPI_ISL_1468446 | Secretaria Municipal de Saude de Birigui | Instituto Adolfo Lutz, Interdiciplinary Procedures Center, Strategic Laboratory | Claudio Tavares Sacchi, Claudia Regina Gonçalves, Erica Valessa Ramos Gomes, Karoline Rodrigues Campos, Caio Vinicius Dias Lopes |
| EPI_ISL_1468449, EPI_ISL_1468450 | Santa Casa de Aracatuba Hospital Sagrado Coracao de Jesus | Instituto Adolfo Lutz, Interdiciplinary Procedures Center, Strategic Laboratory | Claudio Tavares Sacchi, Claudia Regina Gonçalves, Erica Valessa Ramos Gomes, Karoline Rodrigues Campos, Caio Vinicius Dias Lopes |
| EPI_ISL_1468452 | Centro de Atendimento COVID | Instituto Adolfo Lutz, Interdiciplinary Procedures Center, Strategic Laboratory | Claudio Tavares Sacchi, Claudia Regina Gonçalves, Erica Valessa Ramos Gomes, Karoline Rodrigues Campos, Caio Vinicius Dias Lopes |
| EPI_ISL_1468453 | Santa Casa de Misericordia de Pereira Barreto | Instituto Adolfo Lutz, Interdiciplinary Procedures Center, Strategic Laboratory | Claudio Tavares Sacchi, Claudia Regina Gonçalves, Erica Valessa Ramos Gomes, Karoline Rodrigues Campos, Caio Vinicius Dias Lopes |
| EPI_ISL_1468454 | Santa Casa de Andradina | Instituto Adolfo Lutz, Interdiciplinary Procedures Center, Strategic Laboratory | Claudio Tavares Sacchi, Claudia Regina Gonçalves, Erica Valessa Ramos Gomes, Karoline Rodrigues Campos, Caio Vinicius Dias Lopes |
| EPI_ISL_1468455 | UBS 02 Jardim Toselar Birigui | Instituto Adolfo Lutz, Interdiciplinary Procedures Center, Strategic Laboratory | Claudio Tavares Sacchi, Claudia Regina Gonçalves, Erica Valessa Ramos Gomes, Karoline Rodrigues Campos, Caio Vinicius Dias Lopes |
| EPI_ISL_1468456, EPI_ISL_1468457 | UBS Dr Alfredo Dantas de Souza Umuarama | Instituto Adolfo Lutz, Interdiciplinary Procedures Center, Strategic Laboratory | Claudio Tavares Sacchi, Claudia Regina Gonçalves, Erica Valessa Ramos Gomes, Karoline Rodrigues Campos, Caio Vinicius Dias Lopes |
| EPI_ISL_1468458, EPI_ISL_1468459 | Secretaria Municipal de Saude de Valparaiso SP | Instituto Adolfo Lutz, Interdiciplinary Procedures Center, Strategic Laboratory | Claudio Tavares Sacchi, Claudia Regina Gonçalves, Erica Valessa Ramos Gomes, Karoline Rodrigues Campos, Caio Vinicius Dias Lopes |
| EPI_ISL_1468460 | Santa Casa de Birigui | Instituto Adolfo Lutz, Interdiciplinary Procedures Center, Strategic Laboratory | Claudio Tavares Sacchi, Claudia Regina Gonçalves, Erica Valessa Ramos Gomes, Karoline Rodrigues Campos, Caio Vinicius Dias Lopes |
| EPI_ISL_1468461 | Santa Casa de Aracatuba Hospital Sagrado Coracao de Jesus | Instituto Adolfo Lutz, Interdiciplinary Procedures Center, Strategic Laboratory | Claudio Tavares Sacchi, Claudia Regina Gonçalves, Erica Valessa Ramos Gomes, Karoline Rodrigues Campos, Caio Vinicius Dias Lopes |
| EPI_ISL_1468462 | Servico Especial de Saude de Araraquara Sesa Araraquara | Instituto Adolfo Lutz, Interdiciplinary Procedures Center, Strategic Laboratory | Claudio Tavares Sacchi, Claudia Regina Gonçalves, Erica Valessa Ramos Gomes, Karoline Rodrigues Campos, Caio Vinicius Dias Lopes |
| EPI_ISL_1468463 | Centro de Saude II Matao | Instituto Adolfo Lutz, Interdiciplinary Procedures Center, Strategic Laboratory | Claudio Tavares Sacchi, Claudia Regina Gonçalves, Erica Valessa Ramos Gomes, Karoline Rodrigues Campos, Caio Vinicius Dias Lopes |
| EPI_ISL_1468464, EPI_ISL_1468465 | Sae servico de Atendimento Especializado | Instituto Adolfo Lutz, Interdiciplinary Procedures Center, Strategic Laboratory | Claudio Tavares Sacchi, Claudia Regina Gonçalves, Erica Valessa Ramos Gomes, Karoline Rodrigues Campos, Caio Vinicius Dias Lopes |
| EPI_ISL_1468466 | Santa Casa de Sao Carlos | Instituto Adolfo Lutz, Interdiciplinary Procedures Center, Strategic Laboratory | Claudio Tavares Sacchi, Claudia Regina Gonçalves, Erica Valessa Ramos Gomes, Karoline Rodrigues Campos, Caio Vinicius Dias Lopes |
| EPI_ISL_1468467 | Secretaria Municipal de Saude Descalvado | Instituto Adolfo Lutz, Interdiciplinary Procedures Center, Strategic Laboratory | Claudio Tavares Sacchi, Claudia Regina Gonçalves, Erica Valessa Ramos Gomes, Karoline Rodrigues Campos, Caio Vinicius Dias Lopes |
| EPI_ISL_1468469 | Secretaria Municipal de Saude Porto Ferreira | Instituto Adolfo Lutz, Interdiciplinary Procedures Center, Strategic Laboratory | Claudio Tavares Sacchi, Claudia Regina Gonçalves, Erica Valessa Ramos Gomes, Karoline Rodrigues Campos, Caio Vinicius Dias Lopes |
| EPI_ISL_1468470, EPI_ISL_1468471 | Secretaria Municipal de Saude Descalvado | Instituto Adolfo Lutz, Interdiciplinary Procedures Center, Strategic Laboratory | Claudio Tavares Sacchi, Claudia Regina Gonçalves, Erica Valessa Ramos Gomes, Karoline Rodrigues Campos, Caio Vinicius Dias Lopes |
| EPI_ISL_1468472, EPI_ISL_1468473 | Centro de Saude II Matao | Instituto Adolfo Lutz, Interdiciplinary Procedures Center, Strategic Laboratory | Claudio Tavares Sacchi, Claudia Regina Gonçalves, Erica Valessa Ramos Gomes, Karoline Rodrigues Campos, Caio Vinicius Dias Lopes |
| EPI_ISL_1468474 | Sae servico de Atendimento Especializado | Instituto Adolfo Lutz, Interdiciplinary Procedures Center, Strategic Laboratory | Claudio Tavares Sacchi, Claudia Regina Gonçalves, Erica Valessa Ramos Gomes, Karoline Rodrigues Campos, Caio Vinicius Dias Lopes |
| EPI_ISL_1469108, EPI_ISL_1469109, EPI_ISL_1469110, EPI_ISL_1469111, EPI_ISL_1469112, EPI_ISL_1469113, EPI_ISL_1469114, EPI_ISL_1469115, EPI_ISL_1469116, EPI_ISL_1469117, EPI_ISL_1469118, EPI_ISL_1469119 | | | |
| see above | Centro de Investigacion en Ciencias de la Salud y Biomedicina | CINVESTAV | Sofia Bernal, Andreu Comas |
| EPI_ISL_1469234 | Diagnostic and Research Center of Infectious Diseases, Medical Faculty, Andalas University | Diagnostic and Research Center of Infectious Diseases, Medical Faculty, Andalas University | Andani Eka Putra, Syafrizayanti, Ikhwan R. Sudji, Linosefa, Dede Rahman Agustian, Ayu Novita Trisnawati, Siskalil Fahma, Nia Ayuni Putri, Sekar Asri Tresnaningtyas, SM Rezvi, Fauzul Azhim, Dessy Arisanty, Gestina Aliska, Desmawati, Nita Afriani ,Syandrez Prima Putra, Juane Plantika Menra, Yolani Syaputri, Mutia Lailani |
| EPI_ISL_1469242 | National Institute Health Research and Development | National Institute of Health Research and Development | Subangkit, Hana Apsari Pawestri, Kartika Dewi Puspa, Arie Ardiansyah Nugraha, Hartanti Dian Ikawati, Krisna Nur Andriana Pangesti, Yuni Rukminiati, Ririn Ramadhany, Agustiningsih, Kindi Adam, Holy Arif Wibowo, Triyani Soekarso, Ni Ketut Susilarini, Nurika Hariastuti, Ully Alfi Nikmah, Reni Herman, Nike Susanti, Herna, Tati Febriyanti, Natalie Laurencia Kipuw, Fauzul Muna, Irene Lorinda Indalao, Nelly Puspandari, Vivi Setiawaty. |
| EPI_ISL_1469243 | National Institute Health Research and Development | National Institute of Health Research and Development | Reni Herman, Krisna Pangesti, Hana Apsari Pawestri, Subangkit, Kartika Dewi Puspa, Arie Ardiansyah Nugraha, Hartanti Dian Ikawati, Nelly Puspandari, Vivi Setiawaty |
| EPI_ISL_1469246 | Dinas Kesehatan Kabupaten Ponorogo | National Institute of Health Research and Development | Arie Ardiansyah Nugraha, Hana Apsari Pawestri, Subangkit, Kartika Dewi Puspa, Hartanti Dian Ikawati, Nelly Puspandari, Krisna Pangesti, Vivi Setiawaty |
| EPI_ISL_1469249 | Lab Intibios Karawang | National Institute of Health Research and Development | Hana Apsari Pawestri, Subangkit, Kartika Dewi Puspa, Arie Ardiansyah Nugraha, Hartanti Dian Ikawati, Nelly Puspandari, Krisna Pangesti, Vivi Setiawaty |
| EPI_ISL_1469252, EPI_ISL_1469259 | National Institute Health Research and Development | National Institute of Health Research and Development | Subangkit, Hana Apsari Pawestri, Kartika Dewi Puspa, Arie Ardiansyah Nugraha, Hartanti Dian Ikawati, Krisna Nur Andriana Pangesti, Yuni Rukminiati, Ririn Ramadhany, Agustiningsih, Kindi Adam, Holy Arif Wibowo, Triyani Soekarso, Ni Ketut Susilarini, Nurika Hariastuti, Ully Alfi Nikmah, Reni Herman, Nike Susanti, Herna, Tati Febriyanti, Natalie Laurencia Kipuw, Fauzul Muna, Irene Lorinda Indalao, Nelly Puspandari, Vivi Setiawaty. |
| EPI_ISL_1469271, EPI_ISL_1469272, EPI_ISL_1469273, EPI_ISL_1469274, EPI_ISL_1469275 | Dinas Kesehatan Kabupaten Ponorogo | National Institute of Health Research and Development | Arie Ardiansyah Nugraha, Hana Apsari Pawestri, Subangkit, Kartika Dewi Puspa, Hartanti Dian Ikawati, Nelly Puspandari, Krisna Pangesti, Vivi Setiawaty |
| EPI_ISL_1469280, EPI_ISL_1469281, EPI_ISL_1469282, EPI_ISL_1469283 | RSUD Surakarta | National Institute of Health Research and Development | Kartika Dewi Puspa, Hana Apsari Pawestri, Subangkit, Arie Ardiansyah Nugraha, Hartanti Dian Ikawati, Nelly Puspandari, Krisna Pangesti, Vivi Setiawaty |
| EPI_ISL_1469284 | Dinas Kesehatan Kabupaten Ponorogo | National Institute of Health Research and Development | Kartika Dewi Puspa, Hana Apsari Pawestri, Subangkit, Arie Ardiansyah Nugraha, Hartanti Dian Ikawati, Nelly Puspandari, Krisna Pangesti, Vivi Setiawaty |

| | | | |
|---|---|---|---|
| EPI_ISL_1470001, EPI_ISL_1470002, EPI_ISL_1470003, EPI_ISL_1470004, EPI_ISL_1470005, EPI_ISL_1470006, EPI_ISL_1470008, EPI_ISL_1470009, EPI_ISL_1470010, EPI_ISL_1470011, EPI_ISL_1470012, EPI_ISL_1470013, EPI_ISL_1470014, EPI_ISL_1470015, EPI_ISL_1470017, EPI_ISL_1470018, EPI_ISL_1470019, EPI_ISL_1470021, EPI_ISL_1470022, EPI_ISL_1470024, EPI_ISL_1470025, EPI_ISL_1470026, EPI_ISL_1470027, EPI_ISL_1470028, EPI_ISL_1470029, EPI_ISL_1470030, EPI_ISL_1470031, EPI_ISL_1470032, EPI_ISL_1470033, EPI_ISL_1470036, EPI_ISL_1470037, EPI_ISL_1470038, EPI_ISL_1470039, EPI_ISL_1470040, EPI_ISL_1470041, EPI_ISL_1470042, EPI_ISL_1470043, EPI_ISL_1470044, EPI_ISL_1470045, EPI_ISL_1470046, EPI_ISL_1470047, EPI_ISL_1470048, EPI_ISL_1470049, EPI_ISL_1470050, EPI_ISL_1470051, EPI_ISL_1470052, EPI_ISL_1470053, EPI_ISL_1470054, EPI_ISL_1470055, EPI_ISL_1470056, EPI_ISL_1470057, EPI_ISL_1470058 | | | |
| see above | Synlab Eesti OÜ | 1. Laboratory of Communicable Diseases (Estonia); 2. Eurofins Genomics Europe Sequencing GmbH | Liidia Dotsenko et al. |
| EPI_ISL_1470069 | North Estonia Medical Centre laboratory | 1. Laboratory of Communicable Diseases (Estonia); 2. Eurofins Genomics Europe Sequencing GmbH | Liidia Dotsenko et al. |
| EPI_ISL_1470070, EPI_ISL_1470071, EPI_ISL_1470073, EPI_ISL_1470074, EPI_ISL_1470075, EPI_ISL_1470076, EPI_ISL_1470077 | Laboratory of Communicable Diseases | 1. Laboratory of Communicable Diseases (Estonia); 2. Eurofins Genomics Europe Sequencing GmbH | Liidia Dotsenko et al. |
| EPI_ISL_1470078, EPI_ISL_1470079, EPI_ISL_1470080, EPI_ISL_1470081, EPI_ISL_1470082, EPI_ISL_1470083, EPI_ISL_1470084, EPI_ISL_1470085, EPI_ISL_1470087, EPI_ISL_1470088, EPI_ISL_1470089, EPI_ISL_1470090, EPI_ISL_1470091, EPI_ISL_1470092, EPI_ISL_1470093, EPI_ISL_1470094, EPI_ISL_1470095, EPI_ISL_1470096, EPI_ISL_1470097, EPI_ISL_1470098, EPI_ISL_1470099, EPI_ISL_1470100, EPI_ISL_1470101, EPI_ISL_1470102, EPI_ISL_1470103, EPI_ISL_1470104, EPI_ISL_1470105, EPI_ISL_1470106, EPI_ISL_1470107, EPI_ISL_1470108, EPI_ISL_1470109, EPI_ISL_1470112, EPI_ISL_1470114, EPI_ISL_1470115, EPI_ISL_1470116, EPI_ISL_1470117, EPI_ISL_1470120, EPI_ISL_1470121 | | | |
| see above | Synlab Eesti OÜ | 1. Laboratory of Communicable Diseases (Estonia); 2. Eurofins Genomics Europe Sequencing GmbH | Liidia Dotsenko et al. |
| EPI_ISL_1470122, EPI_ISL_1470123, EPI_ISL_1470124, EPI_ISL_1470125, EPI_ISL_1470129, EPI_ISL_1470130, EPI_ISL_1470131, EPI_ISL_1470132, EPI_ISL_1470133, EPI_ISL_1470135 | North Estonia Medical Centre laboratory | 1. Laboratory of Communicable Diseases (Estonia); 2. Eurofins Genomics Europe Sequencing GmbH | Liidia Dotsenko et al. |
| EPI_ISL_1478755, EPI_ISL_1478756 | Complejo Hospitalario Universitario de Pontevedra | Microbiology Department. Complexo Hospitalario Universitario de Vigo | Campello M, Perez S, Regueiro B, Trigo M |
| EPI_ISL_1478757, EPI_ISL_1478759, EPI_ISL_1478760, EPI_ISL_1478761, EPI_ISL_1478762, EPI_ISL_1478763, EPI_ISL_1478764, EPI_ISL_1478765, EPI_ISL_1478766, EPI_ISL_1478767, EPI_ISL_1478768, EPI_ISL_1478769, EPI_ISL_1478770, EPI_ISL_1478771, EPI_ISL_1478772 | | | |
| see above | Microbiology Department. Complexo Hospitalario Universitario de Vigo | Microbiology Department. Complexo Hospitalario Universitario de Vigo | Alfaia N, Alonso I, Alvarez M, Cabrera JJ, Carballo R, Cores O, Cortizo S, del-Campo V, Martinez L, Mediero G, Perez S, Potel C, Regueiro B, Rey S, Vasallo FJ |
| EPI_ISL_1493572 | Centro de Saude II Dr Alcides Facundo Arroyo | Instituto Adolfo Lutz, Interdiciplinary Procedures Center, Strategic Laboratory | Claudio Tavares Sacchi, Claudia Regina Gonçalves, Erica Valessa Ramos Gomes, Karoline Rodrigues Campos, Caio Vinicius Dias Lopes |
| EPI_ISL_1493580 | CS II Dr Miguel Vitaliano Orlandia | Instituto Adolfo Lutz, Interdiciplinary Procedures Center, Strategic Laboratory | Claudio Tavares Sacchi, Claudia Regina Gonçalves, Erica Valessa Ramos Gomes, Karoline Rodrigues Campos, Caio Vinicius Dias Lopes |
| EPI_ISL_1493581 | Sae servico de Atendimento Especializado | Instituto Adolfo Lutz, Interdiciplinary Procedures Center, Strategic Laboratory | Claudio Tavares Sacchi, Claudia Regina Gonçalves, Erica Valessa Ramos Gomes, Karoline Rodrigues Campos, Caio Vinicius Dias Lopes |
| EPI_ISL_1493582 | CS II Dr Miguel Vitaliano Orlandia | Instituto Adolfo Lutz, Interdiciplinary Procedures Center, Strategic Laboratory | Claudio Tavares Sacchi, Claudia Regina Gonçalves, Erica Valessa Ramos Gomes, Karoline Rodrigues Campos, Caio Vinicius Dias Lopes |
| EPI_ISL_1493584 | LACEN do Estado de Rondonia | Instituto Adolfo Lutz, Interdiciplinary Procedures Center, Strategic Laboratory | Claudio Tavares Sacchi, Claudia Regina Gonçalves, Erica Valessa Ramos Gomes, Karoline Rodrigues Campos, Caio Vinicius Dias Lopes |
| EPI_ISL_1493585 | Unidade de Saude Dr Phebo de Oliveira Roge Ferreira | Instituto Adolfo Lutz, Interdiciplinary Procedures Center, Strategic Laboratory | Claudio Tavares Sacchi, Claudia Regina Gonçalves, Erica Valessa Ramos Gomes, Karoline Rodrigues Campos, Caio Vinicius Dias Lopes |
| EPI_ISL_1493586 | Hospital Sao Marcos da Samamorro Agudo | Instituto Adolfo Lutz, Interdiciplinary Procedures Center, Strategic Laboratory | Claudio Tavares Sacchi, Claudia Regina Gonçalves, Erica Valessa Ramos Gomes, Karoline Rodrigues Campos, Caio Vinicius Dias Lopes |
| EPI_ISL_1493587, EPI_ISL_1493588 | CS III de Patrocinio Paulista | Instituto Adolfo Lutz, Interdiciplinary Procedures Center, Strategic Laboratory | Claudio Tavares Sacchi, Claudia Regina Gonçalves, Erica Valessa Ramos Gomes, Karoline Rodrigues Campos, Caio Vinicius Dias Lopes |
| EPI_ISL_1493589 | CS II Dr Jose Ferreira Telles | Instituto Adolfo Lutz, Interdiciplinary Procedures Center, Strategic Laboratory | Claudio Tavares Sacchi, Claudia Regina Gonçalves, Erica Valessa Ramos Gomes, Karoline Rodrigues Campos, Caio Vinicius Dias Lopes |
| EPI_ISL_1493590 | Hospital Sao Marcos da Samamorro Agudo | Instituto Adolfo Lutz, Interdiciplinary Procedures Center, Strategic Laboratory | Claudio Tavares Sacchi, Claudia Regina Gonçalves, Erica Valessa Ramos Gomes, Karoline Rodrigues Campos, Caio Vinicius Dias Lopes |
| EPI_ISL_1493591 | Centro de Saude II Dr Alcides Facundo Arroyo | Instituto Adolfo Lutz, Interdiciplinary Procedures Center, Strategic Laboratory | Claudio Tavares Sacchi, Claudia Regina Gonçalves, Erica Valessa Ramos Gomes, Karoline Rodrigues Campos, Caio Vinicius Dias Lopes |
| EPI_ISL_1493592 | Santa Casa de Guaira | Instituto Adolfo Lutz, Interdiciplinary Procedures Center, Strategic Laboratory | Claudio Tavares Sacchi, Claudia Regina Gonçalves, Erica Valessa Ramos Gomes, Karoline Rodrigues Campos, Caio Vinicius Dias Lopes |
| EPI_ISL_1493593 | CS II Dr Jahyr de Paula Ribeiro Guara | Instituto Adolfo Lutz, Interdiciplinary Procedures Center, Strategic Laboratory | Claudio Tavares Sacchi, Claudia Regina Gonçalves, Erica Valessa Ramos Gomes, Karoline Rodrigues Campos, Caio Vinicius Dias Lopes |
| EPI_ISL_1493594 | Unidade de Saude Dr Phebo de Oliveira Roge Ferreira | Instituto Adolfo Lutz, Interdiciplinary Procedures Center, Strategic Laboratory | Claudio Tavares Sacchi, Claudia Regina Gonçalves, Erica Valessa Ramos Gomes, Karoline Rodrigues Campos, Caio Vinicius Dias Lopes |
| EPI_ISL_1493595, EPI_ISL_1493596, EPI_ISL_1494924 | LACEN do Estado de Rondonia | Instituto Adolfo Lutz, Interdiciplinary Procedures Center, Strategic Laboratory | Claudio Tavares Sacchi, Claudia Regina Gonçalves, Erica Valessa Ramos Gomes, Karoline Rodrigues Campos, Caio Vinicius Dias Lopes |
| EPI_ISL_1495975, EPI_ISL_1495976, EPI_ISL_1495977, EPI_ISL_1495978, EPI_ISL_1495979, EPI_ISL_1498305 | Microbiology Department, University Hospital Donostia | Microbiology Department, University Hospital Donostia | Marimon JM, Montes M, Piñeiro L, Sorarrain A, Gomez M, Cilla G. |
| EPI_ISL_1498917 | Hospital Estadual de Mirandopolis | Instituto Adolfo Lutz, Interdiciplinary Procedures Center, Strategic Laboratory | Claudio Tavares Sacchi, Claudia Regina Gonçalves, Erica Valessa Ramos Gomes, Karoline Rodrigues Campos, Caio Vinicius Dias Lopes |
| EPI_ISL_1500948 | Institut National d'Hygiène (INH) | Unité Mixte Internationale TransVIHMI (UMI 233 IRD - U1175 INSERM - Université de Montpellier)IRD (Institut de recherche pour le développement) | Mounerou SALOU, Christelle BUTEL, Wembo A. HALATOKO,Issaka Maman, Abla A. KONOU, Amivi EHLAN, Adodo SADJI, Kokou TEGUENI,Sidonie A.M.KAGNISSODE, Akoélé SILIADIN, Alassane OURO-MEDELI, Messanh DOUFFAN,Déléma MABA,Sika DOSSIM, Améyo DORKENOO, Mireille PRINCE-DAVID,Anoumou DAGNRA,Laetitia SERRANO,Ahidjo AYOUBA,Eric DELAPORTE, Martine PEETERS |
| EPI_ISL_1500949, EPI_ISL_1500950, EPI_ISL_1500951, EPI_ISL_1500952, EPI_ISL_1500953, EPI_ISL_1500954, EPI_ISL_1500955, EPI_ISL_1500956, EPI_ISL_1500957, EPI_ISL_1500958, EPI_ISL_1500959, EPI_ISL_1500972, EPI_ISL_1500974, EPI_ISL_1500975, EPI_ISL_1500976, EPI_ISL_1500977, EPI_ISL_1500978, EPI_ISL_1500979, EPI_ISL_1500980, EPI_ISL_1500981, EPI_ISL_1500982, EPI_ISL_1500983, EPI_ISL_1500984, EPI_ISL_1500985, EPI_ISL_1500986, EPI_ISL_1500987, EPI_ISL_1500988, EPI_ISL_1500989, EPI_ISL_1500990, EPI_ISL_1500991, EPI_ISL_1500992, EPI_ISL_1500993, EPI_ISL_1501004, EPI_ISL_1501005, EPI_ISL_1501006, EPI_ISL_1501007, EPI_ISL_1501008, EPI_ISL_1501009, EPI_ISL_1501010, EPI_ISL_1501011, EPI_ISL_1501012, EPI_ISL_1501013, EPI_ISL_1501014, EPI_ISL_1501015, EPI_ISL_1501016, EPI_ISL_1501017, EPI_ISL_1501018, EPI_ISL_1501019, EPI_ISL_1501020, EPI_ISL_1501021, EPI_ISL_1501022, EPI_ISL_1501023, EPI_ISL_1501024, EPI_ISL_1501025, EPI_ISL_1501026, EPI_ISL_1501027, EPI_ISL_1501028, EPI_ISL_1501029, EPI_ISL_1501030, EPI_ISL_1501031, EPI_ISL_1501032, EPI_ISL_1501033, EPI_ISL_1501034, EPI_ISL_1501035, EPI_ISL_1501036, EPI_ISL_1501037, EPI_ISL_1501038, EPI_ISL_1501039, EPI_ISL_1501040, EPI_ISL_1501041, EPI_ISL_1501042, EPI_ISL_1501043, EPI_ISL_1501044, EPI_ISL_1501045, EPI_ISL_1501046, EPI_ISL_1501047, EPI_ISL_1501048, EPI_ISL_1501049, EPI_ISL_1501050, EPI_ISL_1501051, EPI_ISL_1501052, EPI_ISL_1501053, EPI_ISL_1501054, EPI_ISL_1501055, EPI_ISL_1501056, EPI_ISL_1501057, EPI_ISL_1501058, EPI_ISL_1501059, EPI_ISL_1501060, EPI_ISL_1501061, EPI_ISL_1501062, EPI_ISL_1501063, EPI_ISL_1501064, EPI_ISL_1501083, EPI_ISL_1501084, EPI_ISL_1501085, EPI_ISL_1501086, EPI_ISL_1501087, EPI_ISL_1501088, EPI_ISL_1501089, EPI_ISL_1501090, EPI_ISL_1501091, EPI_ISL_1501092, EPI_ISL_1501093, EPI_ISL_1501137 | | | |
| see above | HEGP - Laboratoire de Virologie | HEGP - Laboratoire de Virologie | David Veyer |

| | | | |
|---|---|---|---|
| EPI_ISL_1502113, EPI_ISL_1502114, EPI_ISL_1502115, EPI_ISL_1502116, EPI_ISL_1502117, EPI_ISL_1502118, EPI_ISL_1502119, EPI_ISL_1502120, EPI_ISL_1502121, EPI_ISL_1502122, EPI_ISL_1502123, EPI_ISL_1502124, EPI_ISL_1502125, EPI_ISL_1502126, EPI_ISL_1502127, EPI_ISL_1502128, EPI_ISL_1502199, EPI_ISL_1502200, EPI_ISL_1502201, EPI_ISL_1502202, EPI_ISL_1502203, EPI_ISL_1502204, EPI_ISL_1502205, EPI_ISL_1502206, EPI_ISL_1502207, EPI_ISL_1502208, EPI_ISL_1502209, EPI_ISL_1502210 | | | |
| see above | Microbiology Department, University Hospital Donostia | Microbiology Department, University Hospital Donostia | Marimon JM, Montes M, Piñeiro L, Sorarrain A, Gomez M, Cilla G. |
| EPI_ISL_1502567 | Laboratoire Biolim/FSS/UL | Unité Mixte Internationale TransVIHMI (UMI 233 IRD - U1175 INSERM - Université de Montpellier) IRD (Institut de recherche pour le développement) | Mounerou SALOU, Christelle BUTEL, Wembo A. HALATOKO,Amivi EHLAN,Abla A. KONOU,Issaka Maman,Syntiche DEVATCHAGNI Adodo SADJI, Kokou TEGUENI,Koku AGBODEKA, Sidonie A.M.KAGNISSODE, Akoélé SILIADIN, Alassane OURO-MEDELI, Messanh DOUFFAN,Déléma MABA,Sika DOSSIM, Améyo DORKENOO, Mireille PRINCE-DAVID,Anoumou DAGNRA, Laetitia SERRANO,Ahidjo AYOUBA, Eric DELAPORTE, Martine PEETERS |
| EPI_ISL_1502815 | Infectious Diseases Department , Instituto Nacional de Ciencias Medicas y Nutrición | Instituto Nacional de Ciencias Medicas y Nutricion Infectious Diseases | Guillermo M. Ruiz-Palacios, Pilar Ramos Cervantes, Violeta Ibarra Gonzalez, Fernando Ledesma Barrientos, Miriam Arciniega Fuentes , Fernando Arteaga Cabello, Diana Paola Davalos Martinez, Luis Alberto Garcia Andrade, Luz Elena Cervantes Villar |
| EPI_ISL_1503006, EPI_ISL_1503008, EPI_ISL_1503009, EPI_ISL_1503010, EPI_ISL_1503011, EPI_ISL_1503012, EPI_ISL_1503013, EPI_ISL_1503014, EPI_ISL_1503015, EPI_ISL_1503016, EPI_ISL_1503019, EPI_ISL_1503021, EPI_ISL_1503022, EPI_ISL_1503025, EPI_ISL_1503026, EPI_ISL_1503028, EPI_ISL_1503029, EPI_ISL_1503030, EPI_ISL_1503031, EPI_ISL_1503032, EPI_ISL_1503033, EPI_ISL_1503034, EPI_ISL_1503035, EPI_ISL_1503036, EPI_ISL_1503040, EPI_ISL_1503045, EPI_ISL_1503077, EPI_ISL_1503079, EPI_ISL_1503085, EPI_ISL_1503086, EPI_ISL_1503088, EPI_ISL_1503089, EPI_ISL_1503104 | | | |
| see above | Gorgas Memorial Laboratory of Health Studies | Gorgas Memorial Laboratory of Health Studies | Gonzalez Claudia, Leyda Abrego, Moreno Ambar, Oris Chavarria, Jessica Gondola, Marlenne Castillo, Ortiz Alma, Castillo Jorge, Moreno Brechla, Franco Danilo, Lopez-Verges Sandra, Martinez Alexander |
| EPI_ISL_1503125, EPI_ISL_1503126, EPI_ISL_1503127, EPI_ISL_1503128, EPI_ISL_1503129 | Hospital Regional Anita Moreno Los Santos | Gorgas Memorial Laboratory of Health Studies | Gonzalez Claudia, Best Dayana, Espino Juliee, Leyda Abrego, Moreno Ambar, Oris Chavarria, Jessica Gondola, Marlenne Castillo, Ortiz Alma, Castillo Jorge, Moreno Brechla, Franco Danilo, Lopez-Verges Sandra, Martinez Alexander |
| EPI_ISL_1503130, EPI_ISL_1503131, EPI_ISL_1503132, EPI_ISL_1503133, EPI_ISL_1503134, EPI_ISL_1503135, EPI_ISL_1503139, EPI_ISL_1503140, EPI_ISL_1503141 | Gorgas Memorial Laboratory of Health Studies | Gorgas Memorial Laboratory of Health Studies | Gonzalez Claudia, Leyda Abrego, Moreno Ambar, Oris Chavarria, Jessica Gondola, Marlenne Castillo, Ortiz Alma, Castillo Jorge, Moreno Brechla, Franco Danilo, Lopez-Verges Sandra, Martinez Alexander |
| EPI_ISL_1503143 | Infectious Diseases Department , Instituto Nacional de Ciencias Medicas y Nutrición | Instituto Nacional de Ciencias Medicas y Nutricion Infectious Diseases | Guillermo M. Ruiz-Palacios, Pilar Ramos Cervantes, Violeta Ibarra Gonzalez, Fernando Ledesma Barrientos, Miriam Arciniega Fuentes , Fernando Arteaga Cabello, Diana Paola Davalos Martinez, Luis Alberto Garcia Andrade, Luz Elena Cervantes Villar |
| EPI_ISL_1508826, EPI_ISL_1508829, EPI_ISL_1508830, EPI_ISL_1508832, EPI_ISL_1508838, EPI_ISL_1508845 | Medical Genetics Laboratory, Regional Centre of Medical Genetics, Emergency County Hospital Craiova | Medical Genetics Laboratory, Regional Centre of Medical Genetics, Emergency County Hospital Craiova | Anca-Lelia (Riza) Costache, Ioana Streata, Stefania Dorobantu, Adina Dragos, Mihai Cucu, Ana-Maria Buga, Razvan Plesea, Andrei Pirvu, Elena Plesea, Monica Cara, Mihai Ioana |
| EPI_ISL_1508849 | Laboratoire Biolim/FSS/UL | Unité Mixte Internationale TransVIHMI (UMI 233 IRD - U1175 INSERM - Université de Montpellier) IRD (Institut de recherche pour le développement) | Mounerou SALOU, Christelle BUTEL, Wembo A. HALATOKO, Amivi EHLAN, Abla A. KONOU, Issaka Maman, Syntyche DEVATCHAGNI, Adodo SADJI, Kokou TEGUENI,Koku AGBODEKA, Sidonie A.M.KAGNISSODE, Akoélé SILIADIN, Alassane OURO-MEDELI, Messanh DOUFFAN,Déléma MABA,Sika DOSSIM, Améyo DORKENOO, Mireille PRINCE-DAVID,Anoumou DAGNRA, Laetitia SERRANO,Ahidjo AYOUBA, Eric DELAPORTE, Martine PEETERS |
| EPI_ISL_1508855, EPI_ISL_1508860, EPI_ISL_1508861, EPI_ISL_1508877, EPI_ISL_1508878, EPI_ISL_1508879, EPI_ISL_1508881, EPI_ISL_1508882 | University of Oregon COVID-19 MAP Laboratory | University of Oregon Genomics and Cell Characterization Core Facility (GC3F) | Douglas Turnbull, Ariana White, Jeff Bishop, Jason Carriere |
| EPI_ISL_1508894 | Laboratoire Biolim/FSS/UL | Unité Mixte Internationale TransVIHMI (UMI 233 IRD - U1175 INSERM - Université de Montpellier) IRD (Institut de recherche pour le développement) | Mounerou SALOU, Christelle BUTEL, Wembo A. HALATOKO,Amivi EHLAN, Abla A. KONOU, Issaka Maman,Syntyche DEVATCHAGNI, Adodo SADJI, Kokou TEGUENI,Koku AGBODEKA, Sidonie A.M.KAGNISSODE, Akoélé SILIADIN, Alassane OURO-MEDELI, Messanh DOUFFAN,Déléma MABA,Sika DOSSIM, Améyo DORKENOO, Mireille PRINCE-DAVID,Anoumou DAGNRA, Laetitia SERRANO, Ahidjo AYOUBA, Eric DELAPORTE, Martine PEETERS |
| EPI_ISL_1508896 | Laboratoire Biolim/FSS/UL | Unité Mixte Internationale TransVIHMI (UMI 233 IRD - U1175 INSERM - Université de Montpellier) IRD (Institut de recherche pour le développement) | Mounerou SALOU, Christelle BUTEL, Wembo A. HALATOKO, Amivi EHLAN, Abla A. KONOU, Issaka Maman,Syntyche DEVATCHAGNI, Adodo SADJI, Kokou TEGUENI,Koku AGBODEKA, Sidonie A.M.KAGNISSODE, Akoélé SILIADIN, Alassane OURO-MEDELI, Messanh DOUFFAN,Déléma MABA,Sika DOSSIM, Améyo DORKENOO, Mireille PRINCE-DAVID,Anoumou DAGNRA, Laetitia SERRANO,Ahidjo AYOUBA, Eric DELAPORTE, Martine PEETERS |
| EPI_ISL_1508897 | Laboratoire Biolim/FSS/UL | Unité Mixte Internationale TransVIHMI (UMI 233 IRD - U1175 INSERM - Université de Montpellier) IRD (Institut de recherche pour le développement) | Mounerou SALOU, Christelle BUTEL, Wembo A. HALATOKO, Amivi EHLAN, Abla A. KONOU, Issaka Maman,Syntyche DEVATCHAGNI, Adodo SADJI, Kokou TEGUENI, Koku AGBODEKA, Sidonie A.M.KAGNISSODE, Akoélé SILIADIN, Alassane OURO-MEDELI, Messanh DOUFFAN,Déléma MABA,Sika DOSSIM, Améyo DORKENOO, Mireille PRINCE-DAVID,Anoumou DAGNRA, Laetitia SERRANO, Ahidjo AYOUBA, Eric DELAPORTE, Martine PEETERS |
| EPI_ISL_1508916 | Laboratoire Biolim/FSS/UL | Unité Mixte Internationale TransVIHMI (UMI 233 IRD - U1175 INSERM - Université de Montpellier) IRD (Institut de recherche pour le développement) | Mounerou SALOU, Christelle BUTEL, Wembo A. HALATOKO, Amivi EHLAN, Abla A. KONOU, Issaka Maman, Syntyche DEVATCHAGNI, Adodo SADJI, Kokou TEGUENI,Koku AGBODEKA, Sidonie A.M.KAGNISSODE, Akoélé SILIADIN, Alassane OURO-MEDELI, Messanh DOUFFAN,Déléma MABA,Sika DOSSIM, Améyo DORKENOO, Mireille PRINCE-DAVID,Anoumou DAGNRA, Laetitia SERRANO,Ahidjo AYOUBA, Eric DELAPORTE, Martine PEETERS |
| EPI_ISL_1508941 | Laboratoire Biolim/FSS/UL | Unité Mixte Internationale TransVIHMI (UMI 233 IRD - U1175 INSERM - Université de Montpellier) IRD (Institut de recherche pour le développement) | Mounerou SALOU, Christelle BUTEL, Wembo A. HALATOKO, Amivi EHLAN, Abla A. KONOU, Issaka Maman,Syntyche DEVATCHAGNI, Adodo SADJI, Kokou TEGUENI,Koku AGBODEKA, Sidonie A.M.KAGNISSODE, Akoélé SILIADIN, Alassane OURO-MEDELI, Messanh DOUFFAN,Déléma MABA,Sika DOSSIM, Améyo DORKENOO, Mireille PRINCE-DAVID,Anoumou DAGNRA, Laetitia SERRANO,Ahidjo AYOUBA, Eric DELAPORTE, Martine PEETERS |
| EPI_ISL_1508944 | Laboratoire Biolim/FSS/UL | Unité Mixte Internationale TransVIHMI (UMI 233 IRD - U1175 INSERM - Université de Montpellier) IRD (Institut de recherche pour le développement) | Mounerou SALOU, Christelle BUTEL, Wembo A. HALATOKO, Amivi EHLAN, Abla A. KONOU, Issaka Maman, Syntyche DEVATCHAGNI, Adodo SADJI, Kokou TEGUENI,Koku AGBODEKA, Sidonie A.M.KAGNISSODE, Akoélé SILIADIN, Alassane OURO-MEDELI, Messanh DOUFFAN,Déléma MABA,Sika DOSSIM, Améyo DORKENOO, Mireille PRINCE-DAVID,Anoumou DAGNRA, Laetitia SERRANO,Ahidjo AYOUBA, Eric DELAPORTE, Martine PEETERS |
| EPI_ISL_1508949 | Laboratoire Biolim/FSS/UL | Unité Mixte Internationale TransVIHMI (UMI 233 IRD - U1175 INSERM - Université de Montpellier) IRD (Institut de recherche pour le développement) | Mounerou SALOU, Christelle BUTEL, Wembo A. HALATOKO,Amivi EHLAN, Abla A. KONOU, Issaka Maman,Syntyche DEVATCHAGNI, Adodo SADJI, Kokou TEGUENI, Koku AGBODEKA, Sidonie A.M.KAGNISSODE, Akoélé SILIADIN, Alassane OURO-MEDELI, Messanh DOUFFAN,Déléma MABA,Sika DOSSIM, Améyo DORKENOO, Mireille PRINCE-DAVID,Anoumou DAGNRA, Laetitia SERRANO,Ahidjo AYOUBA, Eric DELAPORTE, Martine PEETERS |
| EPI_ISL_1508957, EPI_ISL_1508959, EPI_ISL_1508962 | Laboratoire Biolim/FSS/UL | Unité Mixte Internationale TransVIHMI (UMI 233 IRD - U1175 INSERM - Université de Montpellier) IRD (Institut de recherche pour le développement) | Mounerou SALOU, Christelle BUTEL, Wembo A. HALATOKO, Amivi EHLAN, Abla A. KONOU, Issaka Maman, Syntyche DEVATCHAGNI, Adodo SADJI, Kokou TEGUENI,Koku AGBODEKA, Sidonie A.M.KAGNISSODE, Akoélé SILIADIN, Alassane OURO-MEDELI, Messanh DOUFFAN,Déléma MABA,Sika DOSSIM, Améyo DORKENOO, Mireille PRINCE-DAVID,Anoumou DAGNRA, Laetitia SERRANO,Ahidjo AYOUBA, Eric DELAPORTE, Martine PEETERS |
| EPI_ISL_1508988 | Institut National d'hygiène | Unité Mixte Internationale TransVIHMI (UMI 233 IRD - U1175 INSERM - Université de Montpellier) IRD (Institut de recherche pour le développement) | Mounerou SALOU, Christelle BUTEL, Wembo A. HALATOKO,Amivi EHLAN, Abla A. KONOU, Issaka Maman, Syntyche DEVATCHAGNI, Adodo SADJI, Kokou TEGUENI,Koku AGBODEKA, Sidonie A.M.KAGNISSODE, Akoélé SILIADIN, Alassane OURO-MEDELI, Messanh DOUFFAN,Déléma MABA,Sika DOSSIM, Améyo DORKENOO, Mireille PRINCE-DAVID,Anoumou DAGNRA, Laetitia SERRANO, Ahidjo AYOUBA, Eric DELAPORTE, Martine PEETERS |
| EPI_ISL_1508989, EPI_ISL_1508990, EPI_ISL_1508991 | Laboratoire Biolim/FSS/UL | Unité Mixte Internationale TransVIHMI (UMI 233 IRD - U1175 INSERM - Université de Montpellier) IRD (Institut de recherche pour le développement) | Mounerou SALOU, Christelle BUTEL, Wembo A. HALATOKO, Amivi EHLAN, Abla A. KONOU, Issaka Maman, Syntyche DEVATCHAGNI, Adodo SADJI, Kokou TEGUENI,Koku AGBODEKA, Sidonie A.M.KAGNISSODE, Akoélé SILIADIN, Alassane OURO-MEDELI, Messanh DOUFFAN,Déléma MABA,Sika DOSSIM, Améyo DORKENOO, Mireille PRINCE-DAVID,Anoumou DAGNRA, Laetitia SERRANO,Ahidjo AYOUBA, Eric DELAPORTE, Martine PEETERS |
| EPI_ISL_1508998, EPI_ISL_1509002 | Institut National d'hygiène | Unité Mixte Internationale TransVIHMI (UMI 233 IRD - U1175 INSERM - Université de Montpellier) IRD (Institut de recherche pour le développement) | Mounerou SALOU, Christelle BUTEL, Wembo A. HALATOKO,Amivi EHLAN, Abla A. KONOU, Issaka Maman,Syntyche DEVATCHAGNI, Adodo SADJI, Kokou TEGUENI,Koku AGBODEKA, Sidonie A.M.KAGNISSODE, Akoélé SILIADIN, Alassane OURO-MEDELI, Messanh DOUFFAN,Déléma MABA,Sika DOSSIM, Améyo DORKENOO, Mireille PRINCE-DAVID,Anoumou DAGNRA, Laetitia SERRANO,Ahidjo AYOUBA, Eric DELAPORTE, Martine PEETERS |
| EPI_ISL_1509095 | Institut National d'hygiène | Unité Mixte Internationale TransVIHMI (UMI 233 IRD - U1175 INSERM - Université de Montpellier) IRD (Institut de recherche pour le développement) | Mounerou SALOU, Christelle BUTEL, Wembo A. HALATOKO, Amivi EHLAN, Abla A. KONOU, Issaka Maman,Syntyche DEVATCHAGNI, Adodo SADJI, Kokou TEGUENI,Koku AGBODEKA, Sidonie A.M.KAGNISSODE, Akoélé SILIADIN, Alassane OURO-MEDELI, Messanh DOUFFAN,Déléma MABA,Sika DOSSIM, Améyo DORKENOO, Mireille PRINCE-DAVID,Anoumou DAGNRA, Laetitia SERRANO,Ahidjo AYOUBA, Eric DELAPORTE, Martine PEETERS |
| EPI_ISL_1509096 | Laboratoire Biolim/FSS/UL | Unité Mixte Internationale TransVIHMI (UMI 233 IRD - U1175 INSERM - Université de Montpellier) IRD (Institut de recherche pour le développement) | Mounerou SALOU, Christelle BUTEL, Wembo A. HALATOKO, Amivi EHLAN, Abla A. KONOU, Issaka Maman, Syntyche DEVATCHAGNI, Adodo SADJI, Kokou TEGUENI,Koku AGBODEKA, Sidonie A.M.KAGNISSODE, Akoélé SILIADIN, Alassane OURO-MEDELI, Messanh DOUFFAN,Déléma MABA,Sika DOSSIM, Améyo DORKENOO, Mireille PRINCE-DAVID,Anoumou DAGNRA, Laetitia SERRANO,Ahidjo AYOUBA, Eric DELAPORTE, Martine PEETERS |
| EPI_ISL_1509097 | Institut National d'hygiène | Unité Mixte Internationale TransVIHMI (UMI 233 IRD - U1175 INSERM - Université de Montpellier) IRD (Institut de recherche pour le développement) | Mounerou SALOU, Christelle BUTEL, Wembo A. HALATOKO,Amivi EHLAN, Abla A. KONOU, Issaka Maman,Syntyche DEVATCHAGNI, Adodo SADJI, Kokou TEGUENI,Koku AGBODEKA, Sidonie A.M.KAGNISSODE, Akoélé SILIADIN, Alassane OURO-MEDELI, Messanh DOUFFAN,Déléma MABA,Sika DOSSIM, Améyo DORKENOO, Mireille PRINCE-DAVID,Anoumou DAGNRA, Laetitia SERRANO,Ahidjo AYOUBA, Eric DELAPORTE, Martine PEETERS |

| | | | |
|---|---|---|---|
| EPI_ISL_1509098 | Laboratoire Biolim/FSS/UL | Unité Mixte Internationale TransVIHMI (UMI 233 IRD - U1175 INSERM - Université de Montpellier) IRD (Institut de recherche pour le développement) | Mounerou SALOU, Christelle BUTEL, Wembo A. HALATOKO, Amivi EHLAN, Abla A. KONOU, Issaka Maman, Syntyche DEVATCHAGNI, Adodo SADJI, Kokou TEGUENI,Koku AGBODEKA, Sidonie A.M.KAGNISSODE, Akoélé SILIADIN, Alassane OURO-MEDELI, Messanh DOUFFAN,Déléma MABA,Sika DOSSIM, Améyo DORKENOO, Mireille PRINCE-DAVID,Anoumou DAGNRA, Laetitia SERRANO,Ahidjo AYOUBA, Eric DELAPORTE, Martine PEETERS |
| EPI_ISL_1509099 | Institut National d'hygiène | Unité Mixte Internationale TransVIHMI (UMI 233 IRD - U1175 INSERM - Université de Montpellier) IRD (Institut de recherche pour le développement) | Mounerou SALOU, Christelle BUTEL, Wembo A. HALATOKO, Amivi EHLAN, Abla A. KONOU,Issaka Maman, Syntyche DEVATCHAGNI, Adodo SADJI, Kokou TEGUENI,Koku AGBODEKA, Sidonie A.M.KAGNISSODE, Akoélé SILIADIN, Alassane OURO-MEDELI, Messanh DOUFFAN,Déléma MABA,Sika DOSSIM, Améyo DORKENOO, Mireille PRINCE-DAVID,Anoumou DAGNRA, Laetitia SERRANO,Ahidjo AYOUBA, Eric DELAPORTE, Martine PEETERS |
| EPI_ISL_1509100 | Laboratoire Biolim/FSS/UL | Unité Mixte Internationale TransVIHMI (UMI 233 IRD - U1175 INSERM - Université de Montpellier) IRD (Institut de recherche pour le développement) | Mounerou SALOU, Christelle BUTEL, Wembo A. HALATOKO, Amivi EHLAN, Abla A. KONOU, Issaka Maman, Syntyche DEVATCHAGNI, Adodo SADJI, Kokou TEGUENI,Koku AGBODEKA, Sidonie A.M.KAGNISSODE, Akoélé SILIADIN, Alassane OURO-MEDELI, Messanh DOUFFAN,Déléma MABA,Sika DOSSIM, Améyo DORKENOO, Mireille PRINCE-DAVID,Anoumou DAGNRA, Laetitia SERRANO,Ahidjo AYOUBA, Eric DELAPORTE, Martine PEETERS |
| EPI_ISL_1509101 | Institut National d'hygiène | Unité Mixte Internationale TransVIHMI (UMI 233 IRD - U1175 INSERM - Université de Montpellier) IRD (Institut de recherche pour le développement) | Mounerou SALOU, Christelle BUTEL, Wembo A. HALATOKO, Amivi EHLAN, Abla A. KONOU, Issaka Maman, Syntyche DEVATCHAGNI, Adodo SADJI, Kokou TEGUENI,Koku AGBODEKA, Sidonie A.M.KAGNISSODE, Akoélé SILIADIN, Alassane OURO-MEDELI, Messanh DOUFFAN,Déléma MABA,Sika DOSSIM, Améyo DORKENOO, Mireille PRINCE-DAVID,Anoumou DAGNRA, Laetitia SERRANO,Ahidjo AYOUBA, Eric DELAPORTE, Martine PEETERS |
| EPI_ISL_1509102, EPI_ISL_1509273, EPI_ISL_1509638 | Laboratoire Biolim/FSS/UL | Unité Mixte Internationale TransVIHMI (UMI 233 IRD - U1175 INSERM - Université de Montpellier) IRD (Institut de recherche pour le développement) | Mounerou SALOU, Christelle BUTEL, Wembo A. HALATOKO, Amivi EHLAN, Abla A. KONOU, Issaka Maman, Syntyche DEVATCHAGNI, Adodo SADJI, Kokou TEGUENI,Koku AGBODEKA, Sidonie A.M.KAGNISSODE, Akoélé SILIADIN, Alassane OURO-MEDELI, Messanh DOUFFAN,Déléma MABA,Sika DOSSIM, Améyo DORKENOO, Mireille PRINCE-DAVID,Anoumou DAGNRA, Laetitia SERRANO,Ahidjo AYOUBA, Eric DELAPORTE, Martine PEETERS |
| EPI_ISL_1509720 | UBS de Auriflama | Instituto Adolfo Lutz, Interdiciplinary Procedures Center, Strategic Laboratory | Claudio Tavares Sacchi, Claudia Regina Gonçalves, Erica Valessa Ramos Gomes, Karoline Rodrigues Campos, Caio Vinicius Dias Lopes |
| EPI_ISL_1509760, EPI_ISL_1509761, EPI_ISL_1509762, EPI_ISL_1509763, EPI_ISL_1509764, EPI_ISL_1509765, EPI_ISL_1509766, EPI_ISL_1509767, EPI_ISL_1509769, EPI_ISL_1509775, EPI_ISL_1509778, EPI_ISL_1509781, EPI_ISL_1509782, EPI_ISL_1509783, EPI_ISL_1509784 | | | |
| see above | National Center of Infectious and Parasitic Diseases | National Center of Infectious and Parasitic Diseases | Alexiev, Ivanov, Korsun, Stoitsova, Philipova, Dimitrova, Grigorova L., Hristova, Donchev, Stoykov, Trifonova, Dobrinov, Grigorova I., Kantardjiev |
| EPI_ISL_1509952, EPI_ISL_1509953, EPI_ISL_1509954, EPI_ISL_1509955 | Microbiology Department, University Hospital Donostia | Microbiology Department, University Hospital Donostia | Marimon JM, Montes M, Piñeiro L, Sorarrain A, Gomez M, Cilla G. |
| EPI_ISL_1510606 | Laboratory of Extremely Dangerous Infections of Federal Budget Health Care Instute "Center of Hygiene and Epidemiology in Sverdlovsk region" | Group of Genomics and Postgenomic Technologies of Central Research Institute of Epidemilology | Samoilov AE, Kaptelova VV, Korneenko EV, Valdokhina AV, Bulanenko VP, Saenko SS, Golubeva AG, Esaulkova AY, Durasova AL, Egorova EA, Snitkovskaya TE, Zotova MI, Berlina YY, Solovyeva ED,Cherkashina AS, Shipulina OY, Speranskaya AS, Tivanova EV, Kondrasheva LY, Akimkin VG |
| EPI_ISL_1510608, EPI_ISL_1510609, EPI_ISL_1510610, EPI_ISL_1510613, EPI_ISL_1510614, EPI_ISL_1510615 | Molecular diagnostic laboratory of Federal Budget Institution of Science "Central Research Institute of Epidemiology" of The Federal Service on Customers' Rights Protection and Human Well-being Surveillance | Group of Genomics and Postgenomic Technologies of Central Research Institute of Epidemilology | Samoilov AE, Kaptelova VV, Korneenko EV, Valdokhina AV, Bulanenko VP, Saenko SS, Golubeva AG, Zotova MI, Berlina YY, Solovyeva ED, Cherkashina AS, Shipulina OY, Speranskaya AS, Tivanova EV, Kondrasheva LY, Akimkin VG |
| EPI_ISL_1511271 | Hospital General de México, Medicina Genómica | Hospital General de México, Medicina Genómica | Ana Laura Sánchez-Sandoval, María Eligia Juárez-Torres, Mariana Villalba-De la Peña, Siddhartha Ríos-Zaragoza, Guadalupe Mercedes Lucía Guerrero-Avendaño, Eira Valeria Barrón-Palma |
| EPI_ISL_1511643 | Genomic and molecular Biology Group, A.C.Camargo Cancer Center | Laboratory of Bioinformatics and Computational Biology, A.C.Camargo Cancer Center | Giovana Torrezan, Dirce Carraro, Israel Tojal, Alexandre Defelicibus |
| EPI_ISL_1516959, EPI_ISL_1516960, EPI_ISL_1517011, EPI_ISL_1517012, EPI_ISL_1517014, EPI_ISL_1517015, EPI_ISL_1517017 | Hospital | National Reference Center for Viruses of Respiratory Infections, Institut Pasteur, Paris | Marion Barbet, Sylvie Behillil, Méline Bizard, Angela Brisebarre, Camille Capel,Frédéric Lemoine,Corinne Maufrais,Christophe Malabat, Louise Lefrançois, Etienne Simon-Lorière, Vincent Enouf, Maud Vanpeene, Sylvie van der Werf,Adeline Baron |
| EPI_ISL_1517408 | Inciensa, Instituto Costarricense de Investigación y Enseñanza en Nutrición y Salud | Inciensa, Instituto Costarricense de Investigación y Enseñanza en Nutrición y Salud | Cristian Peréz-Corrales, Valeria Peralta-Barquero & Centeno-Miranda MJ |
| EPI_ISL_1517409 | Inciensa, Instituto Costarricense de Investigación y Enseñanza en Nutrición y Salud | Inciensa, Instituto Costarricense de Investigación y Enseñanza en Nutrición y Salud | Cristian Peréz-Corrales & Valeria Peralta-Barquero |
| EPI_ISL_1517410 | Inciensa, Instituto Costarricense de Investigación y Enseñanza en Nutrición y Salud | Inciensa, Instituto Costarricense de Investigación y Enseñanza en Nutrición y Salud | Cristian Peréz-Corrales, Valeria Peralta-Barquero & César Cerdas Quesada |
| EPI_ISL_1520110, EPI_ISL_1520111, EPI_ISL_1520112 | Hospital Municipal Reynaldo Guerra Cajati | Instituto Adolfo Lutz, Interdiciplinary Procedures Center, Strategic Laboratory | Claudio Tavares Sacchi, Claudia Regina Gonçalves, Erica Valessa Ramos Gomes, Karoline Rodrigues Campos, Caio Vinicius Dias Lopes |
| EPI_ISL_1520113 | Hospital Santo Antonio de Juquia Juquia | Instituto Adolfo Lutz, Interdiciplinary Procedures Center, Strategic Laboratory | Claudio Tavares Sacchi, Claudia Regina Gonçalves, Erica Valessa Ramos Gomes, Karoline Rodrigues Campos, Caio Vinicius Dias Lopes |
| EPI_ISL_1520114 | UBS III de Pariquera Acu Pariquera Acu | Instituto Adolfo Lutz, Interdiciplinary Procedures Center, Strategic Laboratory | Claudio Tavares Sacchi, Claudia Regina Gonçalves, Erica Valessa Ramos Gomes, Karoline Rodrigues Campos, Caio Vinicius Dias Lopes |
| EPI_ISL_1520115, EPI_ISL_1520116 | Unidade de Pronto Atendimento UPA | Instituto Adolfo Lutz, Interdiciplinary Procedures Center, Strategic Laboratory | Claudio Tavares Sacchi, Claudia Regina Gonçalves, Erica Valessa Ramos Gomes, Karoline Rodrigues Campos, Caio Vinicius Dias Lopes |
| EPI_ISL_1520117, EPI_ISL_1520118, EPI_ISL_1520119, EPI_ISL_1520120, EPI_ISL_1520121, EPI_ISL_1520122, EPI_ISL_1520123, EPI_ISL_1520124, EPI_ISL_1520125, EPI_ISL_1520126, EPI_ISL_1520127, EPI_ISL_1520128 | | | |
| see above | Centro de Saude II Dr Jose de Felipe Espito Santo do Pinhal SP | Instituto Adolfo Lutz, Interdiciplinary Procedures Center, Strategic Laboratory | Claudio Tavares Sacchi, Claudia Regina Gonçalves, Erica Valessa Ramos Gomes, Karoline Rodrigues Campos, Caio Vinicius Dias Lopes |
| EPI_ISL_1520129, EPI_ISL_1520130, EPI_ISL_1520131, EPI_ISL_1520132, EPI_ISL_1520133, EPI_ISL_1520134, EPI_ISL_1520135, EPI_ISL_1520136, EPI_ISL_1520137 | Centro de Saude II Dr Jose Paione Mococa | Instituto Adolfo Lutz, Interdiciplinary Procedures Center, Strategic Laboratory | Claudio Tavares Sacchi, Claudia Regina Gonçalves, Erica Valessa Ramos Gomes, Karoline Rodrigues Campos, Caio Vinicius Dias Lopes |
| EPI_ISL_1524330, EPI_ISL_1524333, EPI_ISL_1524338, EPI_ISL_1524340, EPI_ISL_1524343, EPI_ISL_1524345, EPI_ISL_1524346, EPI_ISL_1524347, EPI_ISL_1524349, EPI_ISL_1524351, EPI_ISL_1524353, EPI_ISL_1524356, EPI_ISL_1524359, EPI_ISL_1524361, EPI_ISL_1524362, EPI_ISL_1524364, EPI_ISL_1524369, EPI_ISL_1524372, EPI_ISL_1524374, EPI_ISL_1524376 | | | |
| see above | Biology Department, College of Science, Al Muthanna University and Public Health Laboratory, Al-Muthanna Health Directorate | Department of Virology, Faculty of Medicine, University of Helsinki, Helsinki, Finland | Nihad Al-Rashedi, Hussein Alburkat, Murad Munahi, Alaa Hameed, Ali Jasim, Olli Vapalahti, Tarja Sironen,Teemu Smura |
| EPI_ISL_1524650, EPI_ISL_1524708, EPI_ISL_1524717, EPI_ISL_1524720, EPI_ISL_1524723 | Laboratoire Biolim/FSS/UL | Unité Mixte Internationale TransVIHMI (UMI 233 IRD - U1175 INSERM - Université de Montpellier) IRD (Institut de recherche pour le développement) | Mounerou SALOU, Christelle BUTEL, Wembo A. HALATOKO, Amivi EHLAN, Abla A. KONOU, Issaka Maman, Syntyche DEVATCHAGNI, Adodo SADJI, Kokou TEGUENI,Koku AGBODEKA, Sidonie A.M.KAGNISSODE, Akoélé SILIADIN, Alassane OURO-MEDELI, Messanh DOUFFAN,Déléma MABA,Sika DOSSIM, Améyo DORKENOO, Mireille PRINCE-DAVID,Anoumou DAGNRA, Laetitia SERRANO,Ahidjo AYOUBA, Eric DELAPORTE, Martine PEETERS |
| EPI_ISL_1526781 | Hospital | National Reference Center for Viruses of Respiratory Infections, Institut Pasteur, Paris | Marion Barbet, Sylvie Behillil, Méline Bizard, Angela Brisebarre, Camille Capel,Frédéric Lemoine,Corinne Maufrais,Christophe Malabat, Louise Lefrançois, Etienne Simon-Lorière, Vincent Enouf, Maud Vanpeene, Sylvie van der Werf,Adeline Baron |
| EPI_ISL_1527011 | unknown | Inciensa, Instituto Costarricense de Investigación y Enseñanza en Nutrición y Salud | Pérez-Corrales C, Barboza-Arguedas E & Aguilar-Monge R |
| EPI_ISL_1527012 | Hospital Guápiles | Inciensa, Instituto Costarricense de Investigación y Enseñanza en Nutrición y Salud | Pérez-Corrales C, Barboza-Arguedas E & Cerdas-Quesada C |
| EPI_ISL_1527013 | COOPESAIN-TIBÁS | Inciensa, Instituto Costarricense de Investigación y | Pérez-Corrales C, Barboza-Arguedas E & Flores-León G |

| | | | |
|---|---|---|---|
| | | Enseñanza en Nutrición y Salud | |
| EPI_ISL_1527014 | COOPESIBA BARVA | Inciensa, Instituto Costarricense de Investigación y Enseñanza en Nutrición y Salud | Pérez-Corrales C, Barboza-Arguedas E & Kellerman-Montero K |
| EPI_ISL_1528239 | Hospital | National Reference Center for Viruses of Respiratory Infections, Institut Pasteur, Paris | Marion Barbet, Sylvie Behillil, Frédéric Lemoine, Corinne Maufrais, Christophe Malabat, Méline Bizard, Angela Brisebarre, Camille Capel, Louise Lefrançois, Etienne Simon-Lorière, Vincent Enouf, Maud Vanpeene, Sylvie van der Werf,Eric Farfour |
| EPI_ISL_1531770, EPI_ISL_1531773, EPI_ISL_1531774, EPI_ISL_1531775 | University of Oregon COVID-19 MAP Laboratory | University of Oregon Genomics and Cell Characterization Core Facility (GC3F) | Douglas Turnbull, Ariana White, Jeff Bishop, Jason Carriere |
| EPI_ISL_1532796 | Microbiology Department. Complexo Hospitalario Universitario de Vigo | Microbiology Department. Complexo Hospitalario Universitario de Vigo | Alfaya N, Alonso I, Alvarez M, Cabrera JJ, Carballo R, Cores O, Cortizo S, del-Campo V, Martinez L, Mediero G, Perez S, Potel C, Regueiro B, Rey S, Vasallo FJ |
| EPI_ISL_1532800, EPI_ISL_1532803 | Cambodian National Public Health Laboratory, National Institute of Public Health | Virology Unit, Institut Pasteur du Cambodge | Sokhoun Yann, Teyputita Ou, Leakhena Pum, Ly Sovann, Kraing Sidonn, Yi Sengdoeurn, Chin Savuth, Chau Darapheak, Veasna Duong, Erik A Karlsson |
| EPI_ISL_1532809 | Virology Unit, Institut Pasteur du Cambodge | Virology Unit, Institut Pasteur du Cambodge | Sokhoun Yann, Teyputita Ou, Leakhena Pum, Ly Sovann, Kraing Sidonn, Yi Sengdoeurn, Chin Savuth, Chau Darapheak, Veasna Duong, Erik A Karlsson |
| EPI_ISL_1532812 | Cambodian National Public Health Laboratory, National Institute of Public Health | Virology Unit, Institut Pasteur du Cambodge | Sokhoun Yann, Teyputita Ou, Leakhena Pum, Ly Sovann, Kraing Sidonn, Yi Sengdoeurn, Chin Savuth, Chau Darapheak, Veasna Duong, Erik A Karlsson |
| EPI_ISL_1533037, EPI_ISL_1533044, EPI_ISL_1533058, EPI_ISL_1533061, EPI_ISL_1533069, EPI_ISL_1533082, EPI_ISL_1533089 | Diagnostic and Research Center of Infectious Diseases, Medical Faculty, Andalas University | Diagnostic and Research Center of Infectious Diseases, Medical Faculty, Andalas University | Andani Eka Putra, Syafrizayanti, Ikhwan R. Sudji, Linosefa, Dede Rahman Agustian, Ayu Novita Trisnawati, Siskalil Fahma, Nia Ayuni Putri, Sekar Asri Tresnaningtyas, SM Rezvi, Fauzul Azhim, Dessy Arisanty, Gestina Aliska, Desmawati, Nita Afriani ,Syandrez Prima Putra, Juane Plantika Menra, Yolani Syaputri, Mutia Lailani |
| EPI_ISL_1533689, EPI_ISL_1533690, EPI_ISL_1533691 | Centro de Saude II Dr. Jose Paione Mococa | Instituto Adolfo Lutz, Interdiciplinary Procedures Center, Strategic Laboratory | Claudio Tavares Sacchi, Claudia Regina Gonçalves, Erica Valessa Ramos Gomes, Karoline Rodrigues Campos, Caio Vinicius Dias Lopes, Leonardo Jose Tadeu de Araujo |
| EPI_ISL_1533692 | Santa Casa de Sao Paulo | Instituto Adolfo Lutz, Interdiciplinary Procedures Center, Strategic Laboratory | Claudio Tavares Sacchi, Claudia Regina Gonçalves, Erica Valessa Ramos Gomes, Karoline Rodrigues Campos, Caio Vinicius Dias Lopes, Leonardo Jose Tadeu de Araujo |
| EPI_ISL_1533693 | Secretaria Municipal de Saude de Ubatuba | Instituto Adolfo Lutz, Interdiciplinary Procedures Center, Strategic Laboratory | Claudio Tavares Sacchi, Claudia Regina Gonçalves, Erica Valessa Ramos Gomes, Karoline Rodrigues Campos, Caio Vinicius Dias Lopes, Leonardo Jose Tadeu de Araujo |
| EPI_ISL_1533694 | Santa Casa de Birigui | Instituto Adolfo Lutz, Interdiciplinary Procedures Center, Strategic Laboratory | Claudio Tavares Sacchi, Claudia Regina Gonçalves, Erica Valessa Ramos Gomes, Karoline Rodrigues Campos, Caio Vinicius Dias Lopes, Leonardo Jose Tadeu de Araujo |
| EPI_ISL_1533695 | Hospital Municipal Dr Mario Gatti | Instituto Adolfo Lutz, Interdiciplinary Procedures Center, Strategic Laboratory | Claudio Tavares Sacchi, Claudia Regina Gonçalves, Erica Valessa Ramos Gomes, Karoline Rodrigues Campos, Caio Vinicius Dias Lopes, Leonardo Jose Tadeu de Araujo |
| EPI_ISL_1533696 | Hospital Municipal de Pedreira | Instituto Adolfo Lutz, Interdiciplinary Procedures Center, Strategic Laboratory | Claudio Tavares Sacchi, Claudia Regina Gonçalves, Erica Valessa Ramos Gomes, Karoline Rodrigues Campos, Caio Vinicius Dias Lopes, Leonardo Jose Tadeu de Araujo |
| EPI_ISL_1533697 | Hospital Estadual de Vila Alpina | Instituto Adolfo Lutz, Interdiciplinary Procedures Center, Strategic Laboratory | Claudio Tavares Sacchi, Claudia Regina Gonçalves, Erica Valessa Ramos Gomes, Karoline Rodrigues Campos, Caio Vinicius Dias Lopes, Leonardo Jose Tadeu de Araujo |
| EPI_ISL_1533698 | Secretaria Municipal de Saude de Birigui | Instituto Adolfo Lutz, Interdiciplinary Procedures Center, Strategic Laboratory | Claudio Tavares Sacchi, Claudia Regina Gonçalves, Erica Valessa Ramos Gomes, Karoline Rodrigues Campos, Caio Vinicius Dias Lopes, Leonardo Jose Tadeu de Araujo |
| EPI_ISL_1533699 | Hospital das Clinicas Luzia de Pinho Melo | Instituto Adolfo Lutz, Interdiciplinary Procedures Center, Strategic Laboratory | Claudio Tavares Sacchi, Claudia Regina Gonçalves, Erica Valessa Ramos Gomes, Karoline Rodrigues Campos, Caio Vinicius Dias Lopes, Leonardo Jose Tadeu de Araujo |
| EPI_ISL_1533700 | Hospital Estadual de Sapopemba Sao Paulo | Instituto Adolfo Lutz, Interdiciplinary Procedures Center, Strategic Laboratory | Claudio Tavares Sacchi, Claudia Regina Gonçalves, Erica Valessa Ramos Gomes, Karoline Rodrigues Campos, Caio Vinicius Dias Lopes, Leonardo Jose Tadeu de Araujo |
| EPI_ISL_1533701 | Hospital Estadual de Vila Alpina Org Social Seconci Sao Paulo | Instituto Adolfo Lutz, Interdiciplinary Procedures Center, Strategic Laboratory | Claudio Tavares Sacchi, Claudia Regina Gonçalves, Erica Valessa Ramos Gomes, Karoline Rodrigues Campos, Caio Vinicius Dias Lopes, Leonardo Jose Tadeu de Araujo |
| EPI_ISL_1533702 | Hospital Universitario da USP Sao Paulo | Instituto Adolfo Lutz, Interdiciplinary Procedures Center, Strategic Laboratory | Claudio Tavares Sacchi, Claudia Regina Gonçalves, Erica Valessa Ramos Gomes, Karoline Rodrigues Campos, Caio Vinicius Dias Lopes, Leonardo Jose Tadeu de Araujo |
| EPI_ISL_1533708 | Hospital Municipal Dr Mario Gatti Campinas | Instituto Adolfo Lutz, Interdiciplinary Procedures Center, Strategic Laboratory | Claudio Tavares Sacchi, Claudia Regina Gonçalves, Erica Valessa Ramos Gomes, Karoline Rodrigues Campos, Caio Vinicius Dias Lopes, Leonardo Jose Tadeu de Araujo |
| EPI_ISL_1534690, EPI_ISL_1534691, EPI_ISL_1534692, EPI_ISL_1534693, EPI_ISL_1534694, EPI_ISL_1534696 | BTKLPP kelas 1 Batam | National Institute of Health Research and Development | Hana Apsari Pawestri, Subangkit, Kartika Dewi Puspa, Arie Ardiansyah Nugraha, Hartanti Dian Ikawati, Nelly Puspandari, Krisna Pangesti, Budi Santosa, Ismail, Ida Neni Haryanti, Vivi Setiawaty |
| EPI_ISL_1534697, EPI_ISL_1534698, EPI_ISL_1534699 | Ciputra Hospital Citra Garden City | National Institute of Health Research and Development | Vivi Setiawaty, Hana Apsari Pawestri, Subangkit, Kartika Dewi Puspa, Arie Ardiansyah Nugraha, Hartanti Dian Ikawati, Nelly Puspandari, Krisna Pangesti |
| EPI_ISL_1534700 | Ciputra Hospital Citra Garden City | National Institute of Health Research and Development | Hartanti Dian Ikawati, Hana Apsari Pawestri, Subangkit, Kartika Dewi Puspa, Arie Ardiansyah Nugraha, Krisna Pangesti, Nelly Puspandari, Vivi Setiawaty |
| EPI_ISL_1534704, EPI_ISL_1534705 | BBLK Surabaya | National Institute of Health Research and Development | Kartika Dewi Puspa, Hana Apsari Pawestri, Subangkit, Arie Ardiansyah Nugraha, Hartanti Dian Ikawati, Nelly Puspandari, Krisna Pangesti, Vivi Setiawaty |
| EPI_ISL_1534707 | National Institute Health Research and Development | National Institute of Health Research and Development | Subangkit, Hana Apsari Pawestri, Kartika Dewi Puspa, Arie Ardiansyah Nugraha, Hartanti Dian Ikawati, Krisna Nur Andriana Pangesti, Yuni Rukminiati, Ririn Ramadhany, Agustiningsih, Kindi Adam, Holy Arif Wibowo, Triyani Soekarso, Ni Ketut Susilarini, Nurika Hariastuti, Ully Alfi Nikmah, Reni Herman, Nike Susanti, Herna, Tati Febriyanti, Natalie Laurencia Kipuw, Fauzul Muna, Irene Lorinda Indalao, Nelly Puspandari, Vivi Setiawaty. |
| EPI_ISL_1544110 | B.J. Medical College and Civil hospital, Ahmedabad | Gujarat Biotechnology Research Centre | Pranay Shah,Zuber Saiyed,Ramesh Pandit,Janvi Raval,Zarna Patel,Nitin Savaliya,Dinesh Kumar,Twinkle Soni,Sonal Sharma ,Kamlesh J Upadhyay,Sanjay Kapadia,Dipa Kinariwala,Umang Mishra,Nitesh Shah,Chaitanya Joshi,Madhvi Joshi |
| EPI_ISL_1544111 | B.J. Medical College and Civil hospital, Ahmedabad | Gujarat Biotechnology Research Centre | Kamlesh J Upadhyay,Ramesh Pandit,Janvi Raval,Zarna Patel,Nitin Savaliya,Dinesh Kumar,Twinkle Soni,Sonal Sharma ,Zuber Saiyed,Pranay Shah,Sanjay Kapadia,Dipa Kinariwala,Umang Mishra,Nitesh Shah,Chaitanya Joshi,Madhvi Joshi |
| EPI_ISL_1548049, EPI_ISL_1548050, EPI_ISL_1548051, EPI_ISL_1548052, EPI_ISL_1548053, EPI_ISL_1548054, EPI_ISL_1548055, EPI_ISL_1548056, EPI_ISL_1548057, EPI_ISL_1548058, EPI_ISL_1548060, EPI_ISL_1548061, EPI_ISL_1548062, EPI_ISL_1548063, EPI_ISL_1548064, EPI_ISL_1548065, EPI_ISL_1548066, EPI_ISL_1548067, EPI_ISL_1548068, EPI_ISL_1548069, EPI_ISL_1548070 | | | |
| see above | University of Rome Tor Vergata: Departm Experim Medicine Chair of Virology | University of Rome Tor Vergata: Departm Experim Medicine Chair of Virology | Maria Concetta Bellocchi, Luca Carioti, Maria Botticelli, Lorenzo Piermatteo, Francesca Ceccherini-Silberstein |
| EPI_ISL_1548504 | Laboratoire Biolim/FSS/UL | Unité Mixte Internationale TransVIHMI (UMI 233 IRD - U1175 INSERM - Université de Montpellier) IRD (Institut de recherche pour le développement) | Mounerou SALOU, Christelle BUTEL, Wembo A. HALATOKO, Amivi EHLAN, Abla A. KONOU, Issaka Maman, Syntyche DEVATCHAGNI, Adodo SADJI, Kokou TEGUENI,Koku AGBODEKA, Sidonie A.M.KAGNISSODE, Akoélé SILIADIN, Alassane OURO-MEDELI, Messanh DOUFFAN,Déléma MABA,Sika DOSSIM, Améyo DORKENOO, Mireille PRINCE-DAVID,Anoumou DAGNRA, Laetitia SERRANO,Ahidjo AYOUBA, Eric DELAPORTE, Martine PEETERS |
| EPI_ISL_1554918 | Hospital | National Reference Center for Viruses of Respiratory Infections, Institut Pasteur, Paris | Marion Barbet, Sylvie Behillil, Méline Bizard, Angela Brisebarre, Camille Capel,Frédéric Lemoine,Corinne Maufrais,Christophe Malabat, Damien Mornico, Louise Lefrançois, Etienne Simon-Lorière, Vincent Enouf, Maud Vanpeene, Sylvie van der Werf,Ducancelle Alexandra |
| EPI_ISL_1554934, EPI_ISL_1554935, EPI_ISL_1554937, EPI_ISL_1554942, EPI_ISL_1554961 | Hospital | National Reference Center for Viruses of Respiratory Infections, Institut Pasteur, Paris | Marion Barbet, Sylvie Behillil, Méline Bizard, Angela Brisebarre, Camille Capel,Frédéric Lemoine,Corinne Maufrais,Christophe Malabat, Damien Mornico, Louise Lefrançois, Etienne Simon-Lorière, Vincent Enouf, Maud Vanpeene, Sylvie van der Werf,Guinard JéRome |
| EPI_ISL_1558831, EPI_ISL_1558832, | LESP Jalisco/UdeG | Instituto de Diagnostico y Referencia Epidemiologicos | Claudia Wong-Arambula, Abril Rodriguez-Maldonado, Vanessa Rivero-Arredondo, Ariadna Medina-Benitez, Joaquin Quiroz-Mercado, Sergio |

| | | | |
|---|---|---|---|
| EPI_ISL_1558833 | | (INDRE) | Rangel-Guerrero, Natividad Cruz-Ortiz, Tatiana Nunez-Garcia, Gisela Barrera-Badillo, Lucia Hernandez-Rivas, Irma Lopez-Martinez, Ernesto Ramirez-Gonzalez. |
| EPI_ISL_1575035 | Cruces University Hospital | Biocruces Bizkaia | Mikel Gallego Rodrigo, Ana Belen De la Hoz Rastrollo |
| EPI_ISL_1577912 | Laboratory for Clinical Immunology and Molecular Genetics - University Clinic Golnik Laboratory for Respiratory Microbiology - University Clinic Golnik | Laboratory for Clinical Immunology and Molecular Genetics -University Clinic Golnik | Matija Rijavec, Julij Šelb, Urška Bidovec Stojkovi, Žan Kogovšek, Nina Rupar, Viktorija Tomi, Peter Korošec |
| EPI_ISL_1586902 | Laboratoire Biolim/FSS/UL | Unité Mixte Internationale TransVIHMI (UMI 233 IRD - U1175 INSERM - Université de Montpellier) IRD (Institut de recherche pour le développement) | Mounerou SALOU, Christelle BUTEL, Wembo A. HALATOKO, Amivi EHLAN, Abla A. KONOU, Issaka Maman, Syntyche DEVATCHAGNI, Adodo SADJI, Kokou TEGUENI,Koku AGBODEKA, Sidonie A.M.KAGNISSODE, Akoélé SILIADIN, Alassane OURO-MEDELI, Messanh DOUFFAN,Déléma MABA,Sika DOSSIM, Améyo DORKENOO, Mireille PRINCE-DAVID,Anoumou DAGNRA, Laetitia SERRANO,Ahidjo AYOUBA, Eric DELAPORTE, Martine PEETERS |
| EPI_ISL_1587822, EPI_ISL_1587840, EPI_ISL_1587910 | NL-Dr. Leonard A. Miller Centre for Health Services | National Microbiology Laboratory (NML) | Anna Majer, Shari Tyson, Grace Seo, Philip Mabon, Elsie Grudeski, Rhiannon Huzarewich, Russell Mandes, Anneliese Landgraff, Jennifer Tanner, Natalie Knox, Morag Graham, Gary Van Domselaar, Robert Needle, Yang Yu, Adel Malek, Laura Gilbert, George Zahariadis, Nathalie Bastien, Yan Li, Timothy Booth, Darian Hole, Madison Chapel, Kirsten Biggar, Kerri Smith, CanCOGeN's metadata curation team, Public Health Agency of Canada CanCOGeN team |
| EPI_ISL_1588069 | NB-Hôpital Georges L. Dumont | National Microbiology Laboratory (NML) | Anna Majer, Shari Tyson, Grace Seo, Philip Mabon, Elsie Grudeski, Rhiannon Huzarewich, Russell Mandes, Anneliese Landgraff, Jennifer Tanner, Natalie Knox, Morag Graham, Gary Van Domselaar, Richard Garceau, Guillaume Desnoyers, Nathalie Bastien, Yan Li, Timothy Booth, Darian Hole, Madison Chapel, Kirsten Biggar, CanCOGeN's metadata curation team, Public Health Agency of Canada CanCOGeN team |
| EPI_ISL_1588833 | Hospital | National Reference Center for Viruses of Respiratory Infections, Institut Pasteur, Paris | Marion Barbet, Sylvie Behillil, Méline Bizard, Angela Brisebarre, Camille Capel,Frédéric Lemoine,Corinne Maufrais,Christophe Malabat, Damien Mornico, Louise Lefrançois, Etienne Simon-Lorière, Vincent Enouf, Maud Vanpeene, Sylvie van der Werf,Guinard JéRome |
| EPI_ISL_1589120, EPI_ISL_1589122, EPI_ISL_1589123, EPI_ISL_1589124, EPI_ISL_1589125, EPI_ISL_1589126 | Department of Medical, Surgical, and Health Sciences, University of Trieste and Institute for Maternal and Child Health "IRCCS Burlo Garofolo" | Department of Biomedical, Surgical and Dental Sciences; University of Milan | Ferrante Pasquale, Delbue Serena, Signorini Lucia, Dolci Maria, Comar Manola, Campisciano Giuseppina, Cason Carolina, Ticozzi Rosalia |
| EPI_ISL_1593826, EPI_ISL_1593828, EPI_ISL_1593829, EPI_ISL_1593830 | National Institute of Public Health | State Veterinary Institute Prague | Nagy,A;Jirincova,H;Suri,T;Trnka,D;Vecerova,J |
| EPI_ISL_1608033, EPI_ISL_1608034, EPI_ISL_1608035, EPI_ISL_1608036, EPI_ISL_1608043 | Cliniques Saint-Pierre Ottignies | UCLouvain/IREC/MBLG | Jean Ruelle, Lysa Pinsmaye, Eleonore Ngyuvula Mantu, Benoit Kabamba Mukadi |
| EPI_ISL_1608103, EPI_ISL_1608104, EPI_ISL_1608105, EPI_ISL_1608106, EPI_ISL_1608107, EPI_ISL_1608108 | Programa de Oncovirologia, Instituto Nacional de Câncer | Programa de Oncovirologia, Instituto Nacional de Câncer | Livia R. Goes, Juliana D. Siqueira,Marianne Garrido, Brunna M. Alves, Ana Cristina Pereira, Claudia Cicala, James Arthos, João P.B. Viola, Marcelo A. Soares |
| EPI_ISL_1608161, EPI_ISL_1608162, EPI_ISL_1608163, EPI_ISL_1608164, EPI_ISL_1608165, EPI_ISL_1608166, EPI_ISL_1608167, EPI_ISL_1608168 | Hospital São Rafael - IDOR | Hospital São Rafael - IDOR | Carolina Kymie Vasques Nonaka, Tiago Gräf, Camila Araújo de Lorenzo Barcia, Vanessa Ferreira Costa, Janderson Lopes de Oliveira, Rogério da Hora Passos, Clarissa Araújo Gurgel Rocha, Iasmin Nogueira Bastos, Maria Clara Brito de Santana, Ian Marinho Santos, Karoline Almeida Felix de Sousa, Isadora Cristina de Siqueira, Thamires Gomes Lopes Weber, Ana Verena Almeida Mendes, Bruno Solano de Freitas Souza |
| EPI_ISL_1608516, EPI_ISL_1608518, EPI_ISL_1608519, EPI_ISL_1608520, EPI_ISL_1608521, EPI_ISL_1608522, EPI_ISL_1608523 | National Center of Infectious and Parasitic Diseases | National Center of Infectious and Parasitic Diseases | Alexiev et al |
| EPI_ISL_1622381 | RS Bhakti Husada Cikarang | Eijkman Institute for Molecular Biology, Ministry of Research and Technology/National Agency for Research and Innovation | Hidayat Trimarsanto, Lydia V. Panggalo, Iskandar Adnan, Sukma Oktavianthi, Edison Johar, Frilasita A Yudhaputri, Muhammad Rezki Rasyak, Willy Agustine, Safarina G Malik, Khin Saw Myint, Amin Soebandrio |
| EPI_ISL_1622382 | RS Medika Dramaga | Eijkman Institute for Molecular Biology, Ministry of Research and Technology/National Agency for Research and Innovation | Hidayat Trimarsanto, Lydia V. Panggalo, Iskandar Adnan, Sukma Oktavianthi, Edison Johar, Frilasita A Yudhaputri, Muhammad Rezki Rasyak, Willy Agustine, Safarina G Malik, Khin Saw Myint, Amin Soebandrio |
| EPI_ISL_1622383 | RS Bina Sehat Mandiri | Eijkman Institute for Molecular Biology, Ministry of Research and Technology/National Agency for Research and Innovation | Hidayat Trimarsanto, Lydia V. Panggalo, Iskandar Adnan, Sukma Oktavianthi, Edison Johar, Frilasita A Yudhaputri, Muhammad Rezki Rasyak, Willy Agustine, Safarina G Malik, Khin Saw Myint, Amin Soebandrio |
| EPI_ISL_1622384, EPI_ISL_1622385 | RS Harapan Bunda | Eijkman Institute for Molecular Biology, Ministry of Research and Technology/National Agency for Research and Innovation | Lydia V. Panggalo, Iskandar Adnan, Sukma Oktavianthi, Edison Johar, Frilasita A Yudhaputri, Muhammad Rezki Rasyak, Willy Agustine, Hidayat Trimarsanto, Safarina G Malik, Khin Saw Myint, Amin Soebandrio |
| EPI_ISL_1622386 | RSJ Dr. Soeharto Heerdjan | Eijkman Institute for Molecular Biology, Ministry of Research and Technology/National Agency for Research and Innovation | Lydia V. Panggalo, Iskandar Adnan, Sukma Oktavianthi, Edison Johar, Frilasita A Yudhaputri, Muhammad Rezki Rasyak, Willy Agustine, Hidayat Trimarsanto, Safarina G Malik, Khin Saw Myint, Amin Soebandrio |
| EPI_ISL_1622387 | RSU HGA Depok | Eijkman Institute for Molecular Biology, Ministry of Research and Technology/National Agency for Research and Innovation | Lydia V. Panggalo, Iskandar Adnan, Sukma Oktavianthi, Edison Johar, Frilasita A Yudhaputri, Muhammad Rezki Rasyak, Willy Agustine, Hidayat Trimarsanto, Safarina G Malik, Khin Saw Myint, Amin Soebandrio |
| EPI_ISL_1622389 | RS Masmitra | Eijkman Institute for Molecular Biology, Ministry of Research and Technology/National Agency for Research and Innovation | Lydia V. Panggalo, Iskandar Adnan, Sukma Oktavianthi, Edison Johar, Frilasita A Yudhaputri, Muhammad Rezki Rasyak, Willy Agustine, Hidayat Trimarsanto, Safarina G Malik, Khin Saw Myint, Amin Soebandrio |
| EPI_ISL_1622390 | RS Sentra Medika Cikarang | Eijkman Institute for Molecular Biology, Ministry of Research and Technology/National Agency for Research and Innovation | Lydia V. Panggalo, Iskandar Adnan, Sukma Oktavianthi, Edison Johar, Frilasita A Yudhaputri, Muhammad Rezki Rasyak, Willy Agustine, Hidayat Trimarsanto, Safarina G Malik, Khin Saw Myint, Amin Soebandrio |
| EPI_ISL_1622391 | RS Medirossa Cikarang | Eijkman Institute for Molecular Biology, Ministry of Research and Technology/National Agency for Research and Innovation | Lydia V. Panggalo, Iskandar Adnan, Sukma Oktavianthi, Edison Johar, Frilasita A Yudhaputri, Muhammad Rezki Rasyak, Willy Agustine, Hidayat Trimarsanto, Safarina G Malik, Khin Saw Myint, Amin Soebandrio |
| EPI_ISL_1622392 | RSIA Kenari Graha Medika | Eijkman Institute for Molecular Biology, Ministry of Research and Technology/National Agency for Research and Innovation | Iskandar Adnan, Sukma Oktavianthi, Edison Johar, Frilasita A Yudhaputri, Muhammad Rezki Rasyak, Willy Agustine, Hidayat Trimarsanto, Lydia V. Panggalo, Safarina G Malik, Khin Saw Myint, Amin Soebandrio |
| EPI_ISL_1622393 | RSJ Dr. Soeharto Heerdjan | Eijkman Institute for Molecular Biology, Ministry of Research and Technology/National Agency for Research and Innovation | Iskandar Adnan, Sukma Oktavianthi, Edison Johar, Frilasita A Yudhaputri, Muhammad Rezki Rasyak, Willy Agustine, Hidayat Trimarsanto, Lydia V. Panggalo, Safarina G Malik, Khin Saw Myint, Amin Soebandrio |
| EPI_ISL_1622394 | RSU Proklamasi | Eijkman Institute for Molecular Biology, Ministry of Research and Technology/National Agency for Research and Innovation | Iskandar Adnan, Sukma Oktavianthi, Edison Johar, Frilasita A Yudhaputri, Muhammad Rezki Rasyak, Willy Agustine, Hidayat Trimarsanto, Lydia V. Panggalo, Safarina G Malik, Khin Saw Myint, Amin Soebandrio |
| EPI_ISL_1622395, EPI_ISL_1622396 | RSUD Cileungsi | Eijkman Institute for Molecular Biology, Ministry of Research and Technology/National Agency for Research and Innovation | Iskandar Adnan, Sukma Oktavianthi, Edison Johar, Frilasita A Yudhaputri, Muhammad Rezki Rasyak, Willy Agustine, Hidayat Trimarsanto, Lydia V. Panggalo, Safarina G Malik, Khin Saw Myint, Amin Soebandrio |
| EPI_ISL_1622397, EPI_ISL_1622398, EPI_ISL_1622399 | RSAB Harapan Kita | Eijkman Institute for Molecular Biology, Ministry of Research and Technology/National Agency for Research and Innovation | Iskandar Adnan, Sukma Oktavianthi, Edison Johar, Frilasita A Yudhaputri, Muhammad Rezki Rasyak, Willy Agustine, Hidayat Trimarsanto, Lydia V. Panggalo, Safarina G Malik, Khin Saw Myint, Amin Soebandrio |
| EPI_ISL_1622400 | RSU HGA Depok | Eijkman Institute for Molecular Biology, Ministry of Research and Technology/National Agency for Research and Innovation | Iskandar Adnan, Sukma Oktavianthi, Edison Johar, Frilasita A Yudhaputri, Muhammad Rezki Rasyak, Willy Agustine, Hidayat Trimarsanto, Lydia V. Panggalo, Safarina G Malik, Khin Saw Myint, Amin Soebandrio |
| EPI_ISL_1622401 | RSU Proklamasi | Eijkman Institute for Molecular Biology, Ministry of Research and Technology/National Agency for Research and Innovation | Iskandar Adnan, Sukma Oktavianthi, Edison Johar, Frilasita A Yudhaputri, Muhammad Rezki Rasyak, Willy Agustine, Hidayat Trimarsanto, Lydia V. Panggalo, Safarina G Malik, Khin Saw Myint, Amin Soebandrio |
| EPI_ISL_1622402 | RSU Proklamasi | Eijkman Institute for Molecular Biology, Ministry of Research and Technology/National Agency for Research and Innovation | Sukma Oktavianthi, Edison Johar, Frilasita A Yudhaputri, Muhammad Rezki Rasyak, Willy Agustine, Hidayat Trimarsanto, Lydia V. Panggalo, Iskandar Adnan, Safarina G Malik, Khin Saw Myint, Amin Soebandrio |
| EPI_ISL_1622403, EPI_ISL_1622404 | RS Bhakti Husada Cikarang | Eijkman Institute for Molecular Biology, Ministry of Research and Technology/National Agency for Research and Innovation | Sukma Oktavianthi, Edison Johar, Frilasita A Yudhaputri, Muhammad Rezki Rasyak, Willy Agustine, Hidayat Trimarsanto, Lydia V. Panggalo, Iskandar Adnan, Safarina G Malik, Khin Saw Myint, Amin Soebandrio |

| | | | |
|---|---|---|---|
| EPI_ISL_1622405, EPI_ISL_1622406 | RSUD Cileungsi | Eijkman Institute for Molecular Biology, Ministry of Research and Technology/National Agency for Research and Innovation | Sukma Oktavianthi, Edison Johar, Frilasita A Yudhaputri, Muhammad Rezki Rasyak, Willy Agustine, Hidayat Trimarsanto, Lydia V. Panggalo, Iskandar Adnan, Safarina G Malik, Khin Saw Myint, Amin Soebandrio |
| EPI_ISL_1622408 | RS Medirossa Cikarang | Eijkman Institute for Molecular Biology, Ministry of Research and Technology/National Agency for Research and Innovation | Edison Johar, Frilasita A Yudhaputri, Muhammad Rezki Rasyak, Willy Agustine, Hidayat Trimarsanto, Lydia V. Panggalo, Iskandar Adnan, Sukma Oktavianthi, Safarina G Malik, Khin Saw Myint, Amin Soebandrio |
| EPI_ISL_1622410 | RS Cendana | Eijkman Institute for Molecular Biology, Ministry of Research and Technology/National Agency for Research and Innovation | Edison Johar, Frilasita A Yudhaputri, Muhammad Rezki Rasyak, Willy Agustine, Hidayat Trimarsanto, Lydia V. Panggalo, Iskandar Adnan, Sukma Oktavianthi, Safarina G Malik, Khin Saw Myint, Amin Soebandrio |
| EPI_ISL_1622412 | RS Medirossa Cikarang | Eijkman Institute for Molecular Biology, Ministry of Research and Technology/National Agency for Research and Innovation | Edison Johar, Frilasita A Yudhaputri, Muhammad Rezki Rasyak, Willy Agustine, Hidayat Trimarsanto, Lydia V. Panggalo, Iskandar Adnan, Sukma Oktavianthi, Safarina G Malik, Khin Saw Myint, Amin Soebandrio |
| EPI_ISL_1622415 | RSU HGA Depok | Eijkman Institute for Molecular Biology, Ministry of Research and Technology/National Agency for Research and Innovation | Edison Johar, Frilasita A Yudhaputri, Muhammad Rezki Rasyak, Willy Agustine, Hidayat Trimarsanto, Lydia V. Panggalo, Iskandar Adnan, Sukma Oktavianthi, Safarina G Malik, Khin Saw Myint, Amin Soebandrio |
| EPI_ISL_1622416 | RS Permata Cibubur | Eijkman Institute for Molecular Biology, Ministry of Research and Technology/National Agency for Research and Innovation | Frilasita A Yudhaputri, Muhammad Rezki Rasyak, Willy Agustine, Hidayat Trimarsanto, Lydia V. Panggalo, Iskandar Adnan, Sukma Oktavianthi, Edison Johar, Safarina G Malik, Khin Saw Myint, Amin Soebandrio |
| EPI_ISL_1624246 | Naval Infectious Diseases Diagnostic Laboratory | Naval Medical Research Center Biological Defense Research Directorate | Logan Voegtly, Catherine Arnold, Bishwo Adhikari, Francisco Malagon Bautista, Andrea Luquette, Gregory Rice, Andrew Bennett, Kyle Long, Lindsay Glang, Michael Deschenes, Megan Schilling, Victor Sugiharto, Regina Cer, Kimberly Bishop-Lilly |
| EPI_ISL_1625977, EPI_ISL_1625978, EPI_ISL_1625979, EPI_ISL_1625980, EPI_ISL_1625981, EPI_ISL_1625982, EPI_ISL_1625983, EPI_ISL_1625984, EPI_ISL_1625985, EPI_ISL_1625986 | Instituto Adolfo Lutz - Regional de Rio Claro | Instituto Adolfo Lutz, Interdiciplinary Procedures Center, Strategic Laboratory | Claudio Tavares Sacchi, Claudia Regina Gonçalves, Erica Valessa Ramos Gomes, Karoline Rodrigues Campos, Caio Vinicius Dias Lopes, Leonardo Jose Tadeu de Araujo, Katia Correa de Oliveira Santos |
| EPI_ISL_1627344 | Ospedale Vito Fazzi | Istituto Zooprofilattico Sperimentale della Puglia e della Basilicata | Parisi A., Bianco A., Capozzi L., Del Sambro L., Simone D., Difato L., Ridolfi D., Giannico A. |
| EPI_ISL_1627345, EPI_ISL_1627346 | Presidio di Brindisi Di Summa - Perrino | Istituto Zooprofilattico Sperimentale della Puglia e della Basilicata | Parisi A., Bianco A., Capozzi L., Del Sambro L., Simone D., Difato L., Santoro A., Ridolfi D., Giannico A. |
| EPI_ISL_1627347 | Ospedale di Venere | Istituto Zooprofilattico Sperimentale della Puglia e della Basilicata | Parisi A., Bianco A., Capozzi L., Del Sambro L., Simone D., Iacobellis M., Ridolfi D., Giannico A. |
| EPI_ISL_1627348, EPI_ISL_1627349, EPI_ISL_1627350, EPI_ISL_1627351 | Presidio di Brindisi Di Summa - Perrino | Istituto Zooprofilattico Sperimentale della Puglia e della Basilicata | Parisi A., Bianco A., Capozzi L., Del Sambro L., Simone D., Difato L., Santoro A., Ridolfi D., Giannico A. |
| EPI_ISL_1627352, EPI_ISL_1627353, EPI_ISL_1627354, EPI_ISL_1627355, EPI_ISL_1627356, EPI_ISL_1627357, EPI_ISL_1627358, EPI_ISL_1627360 | Istituto Zooprofilattico Sperimentale della Puglia e della Basilicata | Istituto Zooprofilattico Sperimentale della Puglia e della Basilicata | Parisi A., Bianco A., Capozzi L., Del Sambro L., Simone D., Galante D., Pace L. |
| EPI_ISL_1628686, EPI_ISL_1628688, EPI_ISL_1628692, EPI_ISL_1628693, EPI_ISL_1628695, EPI_ISL_1628697 | CERBALLIANCE | UMR PIMIT | Dr David A Wilkinson, Dr Patrick Mavingui, Dr Camille Lebarbenchon, Magali Turpin |
| EPI_ISL_1628700, EPI_ISL_1628701, EPI_ISL_1628702, EPI_ISL_1628705, EPI_ISL_1628706, EPI_ISL_1628707 | REUNILAB | UMR PIMIT | Dr David A Wilkinson, Dr Patrick Mavingui, Dr Camille Lebarbenchon, Magali Turpin |
| EPI_ISL_1628710, EPI_ISL_1628713, EPI_ISL_1628716, EPI_ISL_1628719, EPI_ISL_1628720, EPI_ISL_1628721, EPI_ISL_1628722 | CERBALLIANCE | UMR PIMIT | Dr David A Wilkinson, Dr Patrick Mavingui, Dr Camille Lebarbenchon, Magali Turpin |
| EPI_ISL_1628725, EPI_ISL_1628726, EPI_ISL_1628727 | SAINT BENOIT | UMR PIMIT | Dr David A Wilkinson, Dr Patrick Mavingui, Dr Camille Lebarbenchon, Magali Turpin |
| EPI_ISL_1628728 | REUNILAB | UMR PIMIT | Dr David A Wilkinson, Dr Patrick Mavingui, Dr Camille Lebarbenchon, Magali Turpin |
| EPI_ISL_1628730, EPI_ISL_1628734 | SAINT BENOIT | UMR PIMIT | Dr David A Wilkinson, Dr Patrick Mavingui, Dr Camille Lebarbenchon, Magali Turpin |
| EPI_ISL_1628741, EPI_ISL_1628743 | CERBALLIANCE | UMR PIMIT | Dr David A Wilkinson, Dr Patrick Mavingui, Dr Camille Lebarbenchon, Magali Turpin |
| EPI_ISL_1628745 | SAINT BENOIT | UMR PIMIT | Dr David A Wilkinson, Dr Patrick Mavingui, Dr Camille Lebarbenchon, Magali Turpin |
| EPI_ISL_1628747 | CHU NORD | UMR PIMIT | Dr David A Wilkinson, Dr Patrick Mavingui, Dr Camille Lebarbenchon, Magali Turpin |
| EPI_ISL_1628753 | CERBALLIANCE | UMR PIMIT | Dr David A Wilkinson, Dr Patrick Mavingui, Dr Camille Lebarbenchon, Magali Turpin |
| EPI_ISL_1628764, EPI_ISL_1628765, EPI_ISL_1628767 | REUNILAB | UMR PIMIT | Dr David A Wilkinson, Dr Patrick Mavingui, Dr Camille Lebarbenchon, Magali Turpin |
| EPI_ISL_1628770, EPI_ISL_1628771, EPI_ISL_1628772 | CERBALLIANCE | UMR PIMIT | Dr David A Wilkinson, Dr Patrick Mavingui, Dr Camille Lebarbenchon, Magali Turpin |
| EPI_ISL_1628774 | REUNILAB | UMR PIMIT | Dr David A Wilkinson, Dr Patrick Mavingui, Dr Camille Lebarbenchon, Magali Turpin |
| EPI_ISL_1628775 | BIO AUSTRAL | UMR PIMIT | Dr David A Wilkinson, Dr Patrick Mavingui, Dr Camille Lebarbenchon, Magali Turpin |
| EPI_ISL_1628776 | CERBALLIANCE | UMR PIMIT | Dr David A Wilkinson, Dr Patrick Mavingui, Dr Camille Lebarbenchon, Magali Turpin |
| EPI_ISL_1628779, EPI_ISL_1628780, EPI_ISL_1628781 | REUNILAB | UMR PIMIT | Dr David A Wilkinson, Dr Patrick Mavingui, Dr Camille Lebarbenchon, Magali Turpin |
| EPI_ISL_1628785, EPI_ISL_1628786, EPI_ISL_1628787, EPI_ISL_1628788, EPI_ISL_1628789, EPI_ISL_1628798, EPI_ISL_1628800, EPI_ISL_1628801, EPI_ISL_1628802 | BIO AUSTRAL | UMR PIMIT | Dr David A Wilkinson, Dr Patrick Mavingui, Dr Camille Lebarbenchon, Magali Turpin |
| EPI_ISL_1628804, EPI_ISL_1628805, EPI_ISL_1628811 | CHU | UMR PIMIT | Dr David A Wilkinson, Dr Patrick Mavingui, Dr Camille Lebarbenchon, Magali Turpin |
| EPI_ISL_1628812, EPI_ISL_1628814, EPI_ISL_1628815 | CERBALLIANCE | UMR PIMIT | Dr David A Wilkinson, Dr Patrick Mavingui, Dr Camille Lebarbenchon, Magali Turpin |
| EPI_ISL_1628816, EPI_ISL_1628817 | REUNILAB | UMR PIMIT | Dr David A Wilkinson, Dr Patrick Mavingui, Dr Camille Lebarbenchon, Magali Turpin |
| EPI_ISL_1628818, EPI_ISL_1628819, EPI_ISL_1628820, EPI_ISL_1628821, EPI_ISL_1628823 | CERBALLIANCE | UMR PIMIT | Dr David A Wilkinson, Dr Patrick Mavingui, Dr Camille Lebarbenchon, Magali Turpin |

| | | | |
|---|---|---|---|
| EPI_ISL_1628825 | REUNILAB | UMR PIMIT | Dr David A Wilkinson, Dr Patrick Mavingui, Dr Camille Lebarbenchon, Magali Turpin |
| EPI_ISL_1628827, EPI_ISL_1628828, EPI_ISL_1628833 | CHU | UMR PIMIT | Dr David A Wilkinson, Dr Patrick Mavingui, Dr Camille Lebarbenchon, Magali Turpin |
| EPI_ISL_1628835, EPI_ISL_1628836, EPI_ISL_1628837, EPI_ISL_1628838, EPI_ISL_1628840, EPI_ISL_1628841, EPI_ISL_1628842 | CERBALLIANCE | UMR PIMIT | Dr David A Wilkinson, Dr Patrick Mavingui, Dr Camille Lebarbenchon, Magali Turpin |
| EPI_ISL_1628846 | GHER | UMR PIMIT | Dr David A Wilkinson, Dr Patrick Mavingui, Dr Camille Lebarbenchon, Magali Turpin |
| EPI_ISL_1628851, EPI_ISL_1628852, EPI_ISL_1628853 | CERBALLIANCE | UMR PIMIT | Dr David A Wilkinson, Dr Patrick Mavingui, Dr Camille Lebarbenchon, Magali Turpin |
| EPI_ISL_1628857 | GHER | UMR PIMIT | Dr David A Wilkinson, Dr Patrick Mavingui, Dr Camille Lebarbenchon, Magali Turpin |
| EPI_ISL_1628861, EPI_ISL_1628867, EPI_ISL_1628868 | REUNILAB | UMR PIMIT | Dr David A Wilkinson, Dr Patrick Mavingui, Dr Camille Lebarbenchon, Magali Turpin |
| EPI_ISL_1628872, EPI_ISL_1628873, EPI_ISL_1628875, EPI_ISL_1628876, EPI_ISL_1628877, EPI_ISL_1628878, EPI_ISL_1628880, EPI_ISL_1628883, EPI_ISL_1628886, EPI_ISL_1628887, EPI_ISL_1628888 | | | |
| see above | CERBALLIANCE | UMR PIMIT | Dr David A Wilkinson, Dr Patrick Mavingui, Dr Camille Lebarbenchon, Magali Turpin |
| EPI_ISL_1628890 | CHU | UMR PIMIT | Dr David A Wilkinson, Dr Patrick Mavingui, Dr Camille Lebarbenchon, Magali Turpin |
| EPI_ISL_1628895, EPI_ISL_1628902, EPI_ISL_1628904, EPI_ISL_1628908 | REUNILAB | UMR PIMIT | Dr David A Wilkinson, Dr Patrick Mavingui, Dr Camille Lebarbenchon, Magali Turpin |
| EPI_ISL_1628914 | CERBALLIANCE | UMR PIMIT | Dr David A Wilkinson, Dr Patrick Mavingui, Dr Camille Lebarbenchon, Magali Turpin |
| EPI_ISL_1628921, EPI_ISL_1628924 | CHU | UMR PIMIT | Dr David A Wilkinson, Dr Patrick Mavingui, Dr Camille Lebarbenchon, Magali Turpin |
| EPI_ISL_1628932, EPI_ISL_1628946, EPI_ISL_1628948 | GHER | UMR PIMIT | Dr David A Wilkinson, Dr Patrick Mavingui, Dr Camille Lebarbenchon, Magali Turpin |
| EPI_ISL_1628956, EPI_ISL_1628965, EPI_ISL_1628968, EPI_ISL_1628969, EPI_ISL_1628979, EPI_ISL_1628982, EPI_ISL_1628983, EPI_ISL_1628984, EPI_ISL_1628993, EPI_ISL_1628995, EPI_ISL_1628996, EPI_ISL_1628998, EPI_ISL_1628999, EPI_ISL_1629000 | | | |
| see above | CERBALLIANCE | UMR PIMIT | Dr David A Wilkinson, Dr Patrick Mavingui, Dr Camille Lebarbenchon, Magali Turpin |
| EPI_ISL_1629006 | CHU | UMR PIMIT | Dr David A Wilkinson, Dr Patrick Mavingui, Dr Camille Lebarbenchon, Magali Turpin |
| EPI_ISL_1629012 | CERBALLIANCE | UMR PIMIT | Dr David A Wilkinson, Dr Patrick Mavingui, Dr Camille Lebarbenchon, Magali Turpin |
| EPI_ISL_1629015 | GHER | UMR PIMIT | Dr David A Wilkinson, Dr Patrick Mavingui, Dr Camille Lebarbenchon, Magali Turpin |
| EPI_ISL_1629027 | CHU | UMR PIMIT | Dr David A Wilkinson, Dr Patrick Mavingui, Dr Camille Lebarbenchon, Magali Turpin |
| EPI_ISL_1629035, EPI_ISL_1629037, EPI_ISL_1629038, EPI_ISL_1629040, EPI_ISL_1629045, EPI_ISL_1629049 | CERBALLIANCE | UMR PIMIT | Dr David A Wilkinson, Dr Patrick Mavingui, Dr Camille Lebarbenchon, Magali Turpin |
| EPI_ISL_1629062, EPI_ISL_1629063 | CHU | UMR PIMIT | Dr David A Wilkinson, Dr Patrick Mavingui, Dr Camille Lebarbenchon, Magali Turpin |
| EPI_ISL_1629078, EPI_ISL_1629089, EPI_ISL_1629090, EPI_ISL_1629091, EPI_ISL_1629093, EPI_ISL_1629096, EPI_ISL_1629097, EPI_ISL_1629101 | GHER | UMR PIMIT | Dr David A Wilkinson, Dr Patrick Mavingui, Dr Camille Lebarbenchon, Magali Turpin |
| EPI_ISL_1629127, EPI_ISL_1629128, EPI_ISL_1629129, EPI_ISL_1629130, EPI_ISL_1629131, EPI_ISL_1629133, EPI_ISL_1629137 | CERBALLIANCE | UMR PIMIT | Dr David A Wilkinson, Dr Patrick Mavingui, Dr Camille Lebarbenchon, Magali Turpin |
| EPI_ISL_1629187, EPI_ISL_1629188 | CHU | UMR PIMIT | Dr David A Wilkinson, Dr Patrick Mavingui, Dr Camille Lebarbenchon, Magali Turpin |
| EPI_ISL_1629192, EPI_ISL_1629197, EPI_ISL_1629200, EPI_ISL_1629207, EPI_ISL_1629342, EPI_ISL_1629343, EPI_ISL_1629349 | REUNILAB | UMR PIMIT | Dr David A Wilkinson, Dr Patrick Mavingui, Dr Camille Lebarbenchon, Magali Turpin |
| EPI_ISL_1629352, EPI_ISL_1629355 | CERBALLIANCE | UMR PIMIT | Dr David A Wilkinson, Dr Patrick Mavingui, Dr Camille Lebarbenchon, Magali Turpin |
| EPI_ISL_1629398 | SAINT BENOIT | UMR PIMIT | Dr David A Wilkinson, Dr Patrick Mavingui, Dr Camille Lebarbenchon, Magali Turpin |
| EPI_ISL_1629400, EPI_ISL_1629407, EPI_ISL_1629410, EPI_ISL_1629412 | REUNILAB | UMR PIMIT | Dr David A Wilkinson, Dr Patrick Mavingui, Dr Camille Lebarbenchon, Magali Turpin |
| EPI_ISL_1629420 | BIO AUSTRAL | UMR PIMIT | Dr David A Wilkinson, Dr Patrick Mavingui, Dr Camille Lebarbenchon, Magali Turpin |
| EPI_ISL_1629472 | CHU | UMR PIMIT | Dr David A Wilkinson, Dr Patrick Mavingui, Dr Camille Lebarbenchon, Magali Turpin |
| EPI_ISL_1629473 | REUNILAB | UMR PIMIT | Dr David A Wilkinson, Dr Patrick Mavingui, Dr Camille Lebarbenchon, Magali Turpin |
| EPI_ISL_1629711 | Universidad Industrial de Santander (Laboratorio Central de Investigaciones - Clínica Chicamocha). | Universidad Industrial de Santander. | Francisco Martinez-Perez, Cristian E. Cadena-Caballero, Diego Rueda-Plata, Carolina S. Torres-Jiménez, Lizeth J. Forero-Buitrago, Erika Lizarazo-Gutiérrez, Carlos Barrios-Hernández, Lina M. Vera-Cala. |
| EPI_ISL_1629731, EPI_ISL_1629732 | UDEA | Universidad Nacional de Colombia - Laboratorio Genómico One Health | Laura Silvana Perez, Andres F. Cardona-Rios, Daniel O. Maldonado-Perez, Maria Stella López, Simón Villegas Velásquez, Karl A Ciuoderis, Idabely Betancur Ortiz, Sandra Ines Cano, Eliana Patricia Calvo Tapiero, Diego A. Álvarez-Díaz, Carlos Franco-Muñoz, Hector Alejandro Ruiz-Moreno, Marcela Mercado-Reyes, Jorge E. Osorio, Juan P. Hernandez-Ortiz |
| EPI_ISL_1629736, EPI_ISL_1629737 | SURA | Universidad Nacional de Colombia - Laboratorio Genómico One Health | Karl A Ciuoderis, Andres F. Cardona-Rios, Daniel O. Maldonado-Perez, Maria Stella López, Simón Villegas Velásquez, Laura Silvana Perez, Idabely Betancur Ortiz, Sandra Ines Cano, Eliana Patricia Calvo Tapiero, Diego A. Álvarez-Díaz, Carlos Franco-Muñoz, Hector Alejandro Ruiz-Moreno, Marcela Mercado-Reyes, Jorge E. Osorio, Juan P. Hernandez-Ortiz |
| EPI_ISL_1629748 | SYNLAB | Universidad Nacional de Colombia - Laboratorio Genómico One Health | Andres F. Cardona-Rios, Daniel O. Maldonado-Perez, Maria Stella López, Simón Villegas Velásquez, Laura Silvana Perez, Karl A Ciuoderis, Idabely Betancur Ortiz, Sandra Ines Cano, Eliana Patricia Calvo Tapiero, Diego A. Álvarez-Díaz, Carlos Franco-Muñoz, Hector Alejandro Ruiz-Moreno, Marcela Mercado-Reyes, Jorge E. Osorio, Juan P. Hernandez-Ortiz |
| EPI_ISL_1629791 | Instituto de Medicina Tropical Alexander Von Humboldt, Universidad Peruana Cayetano Heredia | Laboratorio de Genómica Microbiana, Universidad Peruana Cayetano Heredia | Lenin Maturrano, Pablo Tsukayama, Alejandra Dávila-Barclay, Guillermo Salvatierra, Luis González, Pedro E. Romero, Diego Cuicapuza, Janet Huancachoque, Pool Marcos |
| EPI_ISL_1630279 | Clinical Microbiology Lab RSPTN Universitas Hasanuddin | Clinical Microbiology Lab RSPTN Universitas Hasanuddin | Muhammad Nasrum Massi, Rizalinda Sjahril, Gita Vita Soraya, Triwibawa, Handayani Halik, Muhammad Yusuf Usman, Rufika Shari |
| EPI_ISL_1632706 | Presidio di Brindisi Di Summa - Perrino | Istituto Zooprofilattico Sperimentale della Puglia e della Basilicata | Parisi A., Bianco A., Capozzi L., Del Sambro L., Simone D., Difato L., Santoro A., Ridolfi D., Giannico A. |

| | | | |
|---|---|---|---|
| EPI_ISL_1647423, EPI_ISL_1647424, EPI_ISL_1647427, EPI_ISL_1647428, EPI_ISL_1647429, EPI_ISL_1647430, EPI_ISL_1647431, EPI_ISL_1647432, EPI_ISL_1647433, EPI_ISL_1647434, EPI_ISL_1647435, EPI_ISL_1647436, EPI_ISL_1647437, EPI_ISL_1647438, EPI_ISL_1647439, EPI_ISL_1647440, EPI_ISL_1647441, EPI_ISL_1647442, EPI_ISL_1647443, EPI_ISL_1647444, EPI_ISL_1647445, EPI_ISL_1647453, EPI_ISL_1647454, EPI_ISL_1647455 | | | |
| see above | Indian Council of Medical Research-National Institute of Virology, Microbial Containment Complex | Indian Council of Medical Research-National Institute of Virology, Microbial Containment Complex | Pragya D Yadav |
| EPI_ISL_1651212 | CH Loire Vendée Océan | CHU Nantes Virology | Celine Bressollette-Bodin, Thomas Drumel, Audrey Rodallec, Berthe-Marie Imbert-Marcille, Chappey Olivier |
| EPI_ISL_1652055, EPI_ISL_1652056, EPI_ISL_1652059, EPI_ISL_1652060, EPI_ISL_1652069, EPI_ISL_1652070 | Integrated Biorepository of H3Africa Uganda - IBRH3AU | Molecular Biology Laboratory | Savannah Mwesigwa, Eric Katagirya, Gerald Mboowa, David Patrick Kateete, Misaki Wayengera, Emmanuel Nasinghe, Ashaba Fred Katabazi, Edgar Kigozi, Samuel Kirimunda, Rogers Kamulegeya, Moses Luutu, Lwanga Newton, Nsubuga Gideon, Bernard Ssentalo Bagaya, Sarah Stanley, Moses Joloba |
| EPI_ISL_1654203 | Research | National Reference Center for Viruses of Respiratory Infections, Institut Pasteur, Paris | Marion Barbet, Sylvie Behillil, Frédéric Lemoine, Corinne Maufrais, Christophe Malabat, Damien Mornico, Méline Bizard, Angela Brisebarre, Camille Capel, Louise Lefrançois, Etienne Simon-Lorière, Vincent Enouf, Maud Vanpeene, Sylvie van der Werf,Marianne Leruez-Ville |
| EPI_ISL_1654906, EPI_ISL_1654907, EPI_ISL_1654909, EPI_ISL_1654988, EPI_ISL_1654989, EPI_ISL_1654990, EPI_ISL_1654992, EPI_ISL_1654993, EPI_ISL_1654994, EPI_ISL_1654995, EPI_ISL_1654996, EPI_ISL_1654997, EPI_ISL_1654998, EPI_ISL_1655000, EPI_ISL_1655001, EPI_ISL_1655002, EPI_ISL_1655003, EPI_ISL_1655004, EPI_ISL_1655005, EPI_ISL_1655007, EPI_ISL_1655008, EPI_ISL_1655009, EPI_ISL_1655010, EPI_ISL_1655011, EPI_ISL_1655013, EPI_ISL_1655014, EPI_ISL_1655015, EPI_ISL_1655016, EPI_ISL_1655017, EPI_ISL_1655018, EPI_ISL_1655019, EPI_ISL_1655020, EPI_ISL_1655021, EPI_ISL_1655022, EPI_ISL_1655023, EPI_ISL_1655024, EPI_ISL_1655025, EPI_ISL_1655026, EPI_ISL_1655027, EPI_ISL_1655028, EPI_ISL_1655029, EPI_ISL_1655030, EPI_ISL_1655032, EPI_ISL_1655033, EPI_ISL_1655034, EPI_ISL_1655035, EPI_ISL_1655036, EPI_ISL_1655037, EPI_ISL_1655038, EPI_ISL_1655039, EPI_ISL_1655040, EPI_ISL_1655041, EPI_ISL_1655042, EPI_ISL_1655043, EPI_ISL_1655044, EPI_ISL_1655046, EPI_ISL_1655047, EPI_ISL_1655049, EPI_ISL_1655050, EPI_ISL_1655051, EPI_ISL_1655052, EPI_ISL_1655054, EPI_ISL_1655055, EPI_ISL_1655056, EPI_ISL_1655057, EPI_ISL_1655058, EPI_ISL_1655059, EPI_ISL_1655060, EPI_ISL_1655061, EPI_ISL_1655062, EPI_ISL_1655063, EPI_ISL_1655064, EPI_ISL_1655065, EPI_ISL_1655066, EPI_ISL_1655067, EPI_ISL_1655068, EPI_ISL_1655069, EPI_ISL_1655070 | | | |
| see above | NMVRVI | National Public Health Surveillance Laboratory | Lukas Zemaitis, Migle Gabrielaite, Jelena Razmuk, Svajune Muralyte, Ana Steponkiene, Lukas Vasionis, Danas Baksa |
| EPI_ISL_1659513, EPI_ISL_1659514, EPI_ISL_1659517, EPI_ISL_1659524, EPI_ISL_1659525, EPI_ISL_1659526, EPI_ISL_1659527, EPI_ISL_1659528, EPI_ISL_1659531, EPI_ISL_1659532, EPI_ISL_1659533, EPI_ISL_1659534, EPI_ISL_1659535, EPI_ISL_1659536, EPI_ISL_1659537, EPI_ISL_1659538, EPI_ISL_1659539, EPI_ISL_1659542, EPI_ISL_1659545, EPI_ISL_1659546 | | | |
| see above | NVSPL | National Public Health Surveillance Laboratory | Lukas Zemaitis, Migle Gabrielaite, Jelena Razmuk, Svajune Muralyte, Ana Steponkiene, Lukas Vasionis, Danas Baksa |
| EPI_ISL_1663583, EPI_ISL_1663584, EPI_ISL_1663585 | IU-Cerrahpasa, Cerrahpasa School of Medicine, COVID-19 Lab | IU-Cerrahpasa, Cerrahpasa School of Medicine, COVID-19 Lab | Mert Kuskucu, Yesim Tuyji Tok, Kenan Midilli |
| EPI_ISL_1663588 | Biolytix AG | Swiss Tropical and Public Health Institute | Salome Hosch, Philipp Wagner, Curdin Decurtins, Anna Henger, Ralf Seyfarth, Adrian Härri, Claudia Daubenberger, Tobias Schindler |
| EPI_ISL_1667469, EPI_ISL_1667471 | Department of Laboratory Medicine, National Taiwan University Hospital | Microbial Genomics Core Lab, National Taiwan University Centers of Genomic and Precision Medicine | Shiou-Hwei Yeh, You-Yu Lin, Ya-Yun Lai, Chiao-Ling Li, Shan-Chwen Chang, Pei-Jer Chen, Sui-Yuan Chang |
| EPI_ISL_1669766, EPI_ISL_1669767, EPI_ISL_1669768, EPI_ISL_1669769, EPI_ISL_1669770, EPI_ISL_1669771, EPI_ISL_1669772, EPI_ISL_1669773, EPI_ISL_1669774, EPI_ISL_1669775, EPI_ISL_1669776, EPI_ISL_1669777, EPI_ISL_1669778 | | | |
| see above | Indian Council of Medical Research-National Institute of Virology, Microbial Containment Complex | Indian Council of Medical Research-National Institute of Virology, Microbial Containment Complex | Pragya D Yadav |
| EPI_ISL_1670883 | Laboratorio Analisi Osp. Città di Castello - Azienda USL Umbria1 | Istituto Zooprofilattico Sperimentale dell'Abruzzo e Molise "G. Caporale" | Malagigi V., Tacconi P., Lorusso A, Marcacci M, Di Domenico M, Ancora M, Curini V, Mangone I, Rinaldi A, Delli Compagni E, Scialabba S, Di Pasquale A, Cammà C, Puglia I, Calistri P, Savini G |
| EPI_ISL_1672718 | Hôpital Paul Brousse | Department of Virology, Henri Mondor University Hospital, Assistance Publique Hôpitaux de Paris, Université Paris-Est Créteil, INSERM U955 | Christophe Rodriguez, Slim Fourati, Vanessa Demontant, Guillaume Gricourt, Melissa N'Debi, Alexandre Soulier, Elisabeth Trawinski, Jean-Michel Pawlotsky |
| EPI_ISL_1672864, EPI_ISL_1672866 | Biologia molecular de enfermedades emergentes y EPOC, Instituto Nacional de Enfermedades Respiratorias | Biologia molecular de enfermedades emergentes y EPOC, Instituto Nacional de Enfermedades Respiratorias | Vazquez-Perez Joel Armando, Mejía-Nepomuceno Fidencio, Hernández-Teran Alejandra, Serna Ricardo, Salas-Hernández Jorge, Mario Mújica-Sánchez, Eduardo Becerril-Vargas, José Arturo Martínez-Orozco, Perez-Padilla Rogelio |
| EPI_ISL_1673259 | LABORATORIO LAS AMERICAS | Universidad Nacional de Colombia - Laboratorio Genómico One Health | Maria Stella López, Andres F. Cardona-Rios, Daniel O. Maldonado-Perez, Simón Villegas Velásquez, Lina M. Hurtado, Laura Silvana Perez, Karl A Ciuoderis, Idabely Betancur Ortiz, Sandra Ines Cano, Eliana Patricia Calvo Tapiero, Diego A. Álvarez-Díaz, Carlos Franco-Muñoz, Hector Alejandro Ruiz-Moreno, Marcela Mercado-Reyes, Jorge E. Osorio, Juan P. Hernandez-Ortiz |
| EPI_ISL_1673302 | DNA Solution Ltd. | Genomic Research Lab, BCSIR | Md. Saddam Hossain, Md. Murshed Hasan Sarkar, Mohammad Samir Uzzaman, Eshrar Osman, Md. Ahasan Habib, Shahina Akter, Tanjina Akhter Banu, Abu Sayeed Mohammad Mahmud, Barna Goswami, Iffat Jahan, Mohammad Mohi Uddin, Mohammad Fazle Alam Rabbi, Md Firoz Kabir, Kazi Nadim Hasan, Md. Mizanur Rahman, Md. Abdul Khaleque, Sharif Akhteruzzamani, Md. Salim Khan |
| EPI_ISL_1685376, EPI_ISL_1685377 | Pärnu Hospital | 1. Laboratory of Communicable Diseases (Estonia); 2. Eurofins Genomics Europe Sequencing GmbH | Liidia Dotsenko et al. |
| EPI_ISL_1708319 | Department of Infectious, Respiratory and Digestive Medicine, Graduate School of Medicine, University of the Ryukyus | Genome Information Research Center, Research Institute for Microbial Diseases, Osaka University | Wakaki Kami, Takeshi Kinjo, Wakako Arakaki, Hiroya Oki, Daisuke Motooka, Shota Nakamura, and Jiro Fujita |
| EPI_ISL_1708517, EPI_ISL_1708518 | State collection of viruses, N.F. Gamaleya National Research Center for Epidemiology and Microbiology of the Ministry of Health of the Russian Federation | Laboratory of genome analysis, N.F. Gamaleya National Research Center for Epidemiology and Microbiology of the Ministry of Health of the Russian Federation | Olga Voronina, Ekaterina Aksenova, Natalia Ryzhova, Marina Kunda, Inna Dolzhikova, Andrey Botikov, Daria Grousova, Anna Kovyrshina, Denis Logunov |
| EPI_ISL_1710855, EPI_ISL_1710856, EPI_ISL_1710857, EPI_ISL_1710858, EPI_ISL_1710859, EPI_ISL_1710860, EPI_ISL_1710861, EPI_ISL_1710862, EPI_ISL_1710863, EPI_ISL_1710864, EPI_ISL_1710865 | | | |
| see above | Pathogenic Microorganisms Variability Laboratory | Pathogenic Microorganisms Variability Laboratory | Alexey Shchetinin, Maria Nikiforova, Andrei Siniavin, Andrey Pochtovyy, Evgeny Usachev, Elena Shidlovskaya, Elizaveta Divisenko, Nadezhda Kuznetsova, Valeria Bacalin, Olga Burgasova, Ludmila Kolobukhina, Svetlana Smetanina, Denis Logunov, Vladimir Gushchin, Alexander Gintsburg |
| EPI_ISL_1752621, EPI_ISL_1752622, EPI_ISL_1752623, EPI_ISL_1752624, EPI_ISL_1752625, EPI_ISL_1752626, EPI_ISL_1752627, EPI_ISL_1752628, EPI_ISL_1752629, EPI_ISL_1752630, EPI_ISL_1752631 | | | |
| see above | Showa University PCR Center | Showa University PCR Center | Fumihiro Ishikawa, Hideto Oyamada, Yuko Udaka, Keiko Ishino, Yuji Kiuchi |
| EPI_ISL_1753848, EPI_ISL_1753852, EPI_ISL_1753867, EPI_ISL_1753871, EPI_ISL_1753877, EPI_ISL_1753881, EPI_ISL_1753885, EPI_ISL_1753890, EPI_ISL_1753894, EPI_ISL_1753895, EPI_ISL_1753896, EPI_ISL_1753897, EPI_ISL_1753914, EPI_ISL_1753916, EPI_ISL_1753917, EPI_ISL_1753918, EPI_ISL_1753920, EPI_ISL_1753924, EPI_ISL_1753929, EPI_ISL_1753930, EPI_ISL_1753931, EPI_ISL_1753935, EPI_ISL_1753937, EPI_ISL_1753938 | | | |
| see above | Servicio Microbiología Hospital La Paz | Servicio Microbiología Hospital La Paz | Fernando Lázaro, Rubén Cáceres, Jesús Mingorance Cruz, Elie Dahdouh |
| EPI_ISL_1760236, EPI_ISL_1760237, EPI_ISL_1760238, EPI_ISL_1760239, EPI_ISL_1760240, EPI_ISL_1760243, EPI_ISL_1760244, EPI_ISL_1760245, EPI_ISL_1760246, EPI_ISL_1760247, EPI_ISL_1760248, EPI_ISL_1760249, EPI_ISL_1760257, EPI_ISL_1760258 | | | |
| see above | Centers for Disease Control and Prevention, Dengue Branch | Centers for Disease Control and Prevention, Dengue Branch | Gilberto A. Santiago, Glenda Gonzalez, Betzabel Flores, Keyla Charriez, Gabriela Paz-Bailey, Jorge L. Munoz-Jordan |
| EPI_ISL_1761341, EPI_ISL_1761342, EPI_ISL_1761343, EPI_ISL_1761344, EPI_ISL_1761345, EPI_ISL_1761346, EPI_ISL_1761347, EPI_ISL_1761348, EPI_ISL_1761349, EPI_ISL_1761350, EPI_ISL_1761351, EPI_ISL_1761352, EPI_ISL_1761353, EPI_ISL_1761354, EPI_ISL_1761355, EPI_ISL_1761356, EPI_ISL_1761357, EPI_ISL_1761358 | | | |
| see above | Tokyo Medical and Dental University | Tokyo Medical and Dental University | Kousuke Tanimoto, Yukie Tanaka, Yoko Nukui, Akinori Kimura, Hiroaki Takeuchi |
| EPI_ISL_1761588 | Hospital de Palamós. Serveis de Salut Integral Baix Empordà (SSIBE) | IrsiCaixa | Marc Noguera-Julian, Mariona Parera, Maria Casadellà, Pilar Armengol, Francesc Catala-Moll, Roger Paredes, Bonaventura Clotet Martin-Urda Diez-Canseco, Anabel; Toda Savall, Mº de la Roca; Marchena Romero, Jose Andres; Rodriguez Jareño, Mari Cruz; Erick Fernando Mayer Arispe; Claver Belver, Natalia |
| EPI_ISL_1789202 | Paru Hospital | Institute of Tropical Disease, Universitas Airlangga | Maria I Lusida, Krisnoadi Rahardjo, Aldise M Nastri, Jezzy R Dewantari, Rima R Prasetya, Dyah Retno, Gatot Soegiarto, Laksmi Wulandari, Resti Yudhawati, Soetjipto, Yasuko Mori, Kazufumi Shimizu |
| EPI_ISL_1790099, EPI_ISL_1790101 | Centre Pasteur Annexe de Garoua | Institut Pasteur de Dakar | Njouom Richard, Diagne Moussa Moïse, Dia Ndongo, Diallo Amadou, Sankhe Safietou, Diop Mamadou, Ndiaye Ndack, Loucoubar Cheikh, Carniel Elisabeth, Faye Ousmane, Sall Amadou Alpha |
| EPI_ISL_1790107, EPI_ISL_1790108, EPI_ISL_1790109, EPI_ISL_1790124, EPI_ISL_1790125, EPI_ISL_1790126 | Centre Pasteur du Cameroun | Institut Pasteur de Dakar | Njouom Richard, Diagne Moussa Moïse, Dia Ndongo, Diallo Amadou, Sankhe Safietou, Diop Mamadou, Ndiaye Ndack, Loucoubar Cheikh, Carniel Elisabeth, Faye Ousmane, Sall Amadou Alpha |

| | | | |
|---|---|---|---|
| EPI_ISL_1791459, EPI_ISL_1791461, EPI_ISL_1791465, EPI_ISL_1791466, EPI_ISL_1791470, EPI_ISL_1791472, EPI_ISL_1791547, EPI_ISL_1791548, EPI_ISL_1791549, EPI_ISL_1791550, EPI_ISL_1791551, EPI_ISL_1791552, EPI_ISL_1791553, EPI_ISL_1791554, EPI_ISL_1791556, EPI_ISL_1791557, EPI_ISL_1791558, EPI_ISL_1791559, EPI_ISL_1791560, EPI_ISL_1791561, EPI_ISL_1791562, EPI_ISL_1791563, EPI_ISL_1791564, EPI_ISL_1791565, EPI_ISL_1791566, EPI_ISL_1791567, EPI_ISL_1791568, EPI_ISL_1791570, EPI_ISL_1791571, EPI_ISL_1791572, EPI_ISL_1791573, EPI_ISL_1791574, EPI_ISL_1791575, EPI_ISL_1791576, EPI_ISL_1791578, EPI_ISL_1791579, EPI_ISL_1791581, EPI_ISL_1791582, EPI_ISL_1791585, EPI_ISL_1791589, EPI_ISL_1791590 | | | |
| see above | University of Oregon COVID-19 MAP Laboratory | University of Oregon Genomics and Cell Characterization Core Facility (GC3F) | Douglas Turnbull, Ariana White, Jeff Bishop, Jason Carriere |
| EPI_ISL_1800014 | Laboratoire Biolim/FSS/UL | Unité Mixte Internationale TransVIHMI (UMI 233 IRD - U1175 INSERM - Université de Montpellier) IRD (Institut de recherche pour le développement) | Mounerou SALOU, Christelle BUTEL, Wembo A. HALATOKO, Amivi EHLAN, Abla A. KONOU, Issaka Maman, Syntyche DEVATCHAGNI, Adodo SADJI, Kokou TEGUENI,Koku AGBODEKA, Sidonie A.M.KAGNISSODE, Akoélé SILIADIN, Alassane OURO-MEDELI, Messanh DOUFFAN,Déléma MABA,Sika DOSSIM, Améyo DORKENOO, Mireille PRINCE-DAVID,Anoumou DAGNRA, Laetitia SERRANO,Ahidjo AYOUBA, Eric DELAPORTE, Martine PEETERS |
| EPI_ISL_1805932, EPI_ISL_1805933, EPI_ISL_1805957, EPI_ISL_1805958 | National Center for Communicable Diseases (NCCD) National Influenza Center | National Center for Communicable Diseases (NCCD) National Influenza Center | Naranzul Ts,Bayasgalan N,Khishigmunkh Ch,Seiichiro F,Hideka M,Mina N,Shinji W,Ankhbayar S,Tsogzolmaa G,Darmaa B,Battur L,Nymadawa P |
| EPI_ISL_1821225, EPI_ISL_1821226, EPI_ISL_1821227, EPI_ISL_1821228, EPI_ISL_1821229, EPI_ISL_1821230, EPI_ISL_1821231, EPI_ISL_1821232, EPI_ISL_1821233 | Instituto Adolfo Lutz - Regional de Marilia | Instituto Adolfo Lutz, Interdiciplinary Procedures Center, Strategic Laboratory | Claudio Tavares Sacchi, Claudia Regina Gonçalves, Erica Valessa Ramos Gomes, Karoline Rodrigues Campos, Caio Vinicius Dias Lopes, Leonardo Jose Tadeu de Araujo |
| EPI_ISL_1823568 | Indiana Animal Disease Diagnostic Laboratory | Carpi Laboratory - Purdue University | Jack Dorman, Ilinca I Ciubotariu, Nicole M Perry, Jobin J Kattoor, Lev Gorenstein, Abebe A Fola, G Kenitra Hendrix, Rebecca P Wilkes, Giovanna Carpi |
| EPI_ISL_1824605 | RSUP Sanglah | National Institute of Health Research and Development | Vivi Setiawaty, Hana Apsari Pawestri, Subangkit, Kartika Dewi Puspa, Arie Ardiansyah Nugraha, Hartanti Dian Ikawati, Krisna Pangesti, Nelly Puspandari, I Nengah Tony Rustawan, I Gede Ketut Sajinadiyasa |
| EPI_ISL_1856464, EPI_ISL_1856466, EPI_ISL_1856469, EPI_ISL_1856472, EPI_ISL_1856476 | UVA Clinical Microbiology Lab | Mathers Lab | Hardik I. Parikh, Katie E. Barry, Amy J. Mathers |
| EPI_ISL_1893390, EPI_ISL_1893392, EPI_ISL_1893394, EPI_ISL_1893396, EPI_ISL_1893399, EPI_ISL_1893401, EPI_ISL_1893408, EPI_ISL_1893409, EPI_ISL_1893411, EPI_ISL_1893413, EPI_ISL_1893415, EPI_ISL_1893418, EPI_ISL_1893419, EPI_ISL_1893427, EPI_ISL_1893435, EPI_ISL_1893457, EPI_ISL_1893511, EPI_ISL_1893513, EPI_ISL_1893515, EPI_ISL_1893517, EPI_ISL_1893525, EPI_ISL_1893527, EPI_ISL_1893529, EPI_ISL_1893559, EPI_ISL_1893561, EPI_ISL_1893563, EPI_ISL_1893565, EPI_ISL_1893567, EPI_ISL_1893570, EPI_ISL_1893572, EPI_ISL_1893573, EPI_ISL_1893576, EPI_ISL_1893584, EPI_ISL_1893585, EPI_ISL_1893588, EPI_ISL_1893590, EPI_ISL_1893594, EPI_ISL_1893595, EPI_ISL_1893598, EPI_ISL_1893618, EPI_ISL_1893626, EPI_ISL_1893628, EPI_ISL_1893630, EPI_ISL_1893632, EPI_ISL_1893634, EPI_ISL_1893636, EPI_ISL_1893638, EPI_ISL_1893645, EPI_ISL_1893652, EPI_ISL_1893658, EPI_ISL_1893660 | | | |
| see above | IPK | Respiratory Infections Laboratory | Lissette Perez, Yahisel Tejero, Yanet Pintos, Daylin Mederos, Odalys Valdes, Mirtha Elisa Aguado, Liena Ponce |
| EPI_ISL_1905205 | AZ St.Jan Brugge-Oostende | AZ SINT-JAN BRUGGE | Marijke Reynders, Patrick Descheemaeker, Laurien Hoornaert, Thomas Van Landschoot, Jorn Hellemans |
| EPI_ISL_1910444 | Centre Hospitalier Universitaire Clermont-Ferrand | CHU Clermont-Ferrand, service de virologie | Bisseux Maxime, Mirand Audrey, Combes Patricia, Henquell Cécile |
| EPI_ISL_1910445 | HOPITAL DE MONTLUCON | CHU Clermont-Ferrand, service de virologie | Bisseux Maxime, Mirand Audrey, Combes Patricia, Henquell Cécile |
| EPI_ISL_1916435, EPI_ISL_1916470 | CSIR-National Environmental Engineering Research Institute | CSIR-Centre for Cellular and Molecular Biology - INSACOG | Ara Sreenivas, Onkar Kulkarni, Lamuk Zaveri, Sofia Banu, Shreekant Verma, Amareshwar Vodapalli , Viswagithe S L, B Himasri, Sharath Chandra Thota, Karthik Bharadwaj Tallapaka, Krishna Khairnar, Rakesh K Mishra,Divya Tej Sowpati |
| EPI_ISL_1936296, EPI_ISL_1936297, EPI_ISL_1936298, EPI_ISL_1936299, EPI_ISL_1936300, EPI_ISL_1936301, EPI_ISL_1936302, EPI_ISL_1936303, EPI_ISL_1936304, EPI_ISL_1936305, EPI_ISL_1936306, EPI_ISL_1936307, EPI_ISL_1936308, EPI_ISL_1936309, EPI_ISL_1936310, EPI_ISL_1936311, EPI_ISL_1936312, EPI_ISL_1936313, EPI_ISL_1936314, EPI_ISL_1936315, EPI_ISL_1936316, EPI_ISL_1936317, EPI_ISL_1936318, EPI_ISL_1936319, EPI_ISL_1936320, EPI_ISL_1936321, EPI_ISL_1936322, EPI_ISL_1936323, EPI_ISL_1936324, EPI_ISL_1936325, EPI_ISL_1936326, EPI_ISL_1936327, EPI_ISL_1936328, EPI_ISL_1936329, EPI_ISL_1936330, EPI_ISL_1936331, EPI_ISL_1936332, EPI_ISL_1936333, EPI_ISL_1936334, EPI_ISL_1936335, EPI_ISL_1936336, EPI_ISL_1936337, EPI_ISL_1936338, EPI_ISL_1936339, EPI_ISL_1936340, EPI_ISL_1936341, EPI_ISL_1936342, EPI_ISL_1936343, EPI_ISL_1936344, EPI_ISL_1936345, EPI_ISL_1936346, EPI_ISL_1936347, EPI_ISL_1936348, EPI_ISL_1936349, EPI_ISL_1936350, EPI_ISL_1936351, EPI_ISL_1936352, EPI_ISL_1936353, EPI_ISL_1936354, EPI_ISL_1936355, EPI_ISL_1936356, EPI_ISL_1936357, EPI_ISL_1936358, EPI_ISL_1936359, EPI_ISL_1936360, EPI_ISL_1936361, EPI_ISL_1936362, EPI_ISL_1936363, EPI_ISL_1936364, EPI_ISL_1936365, EPI_ISL_1936366, EPI_ISL_1936367, EPI_ISL_1936368, EPI_ISL_1936369, EPI_ISL_1936370, EPI_ISL_1936371, EPI_ISL_1936372, EPI_ISL_1936373, EPI_ISL_1936374 | | | |
| see above | Main Chemical Laboratories Egypt Army | Main Chemical Laboratories Egypt Army | Sherine Helmy, Laila Elsawy, Ola Elroby, Mostfa Yakout, Mohamed Seadawy |
| EPI_ISL_1939475, EPI_ISL_1939476, EPI_ISL_1939477, EPI_ISL_1939478, EPI_ISL_1939479, EPI_ISL_1939480, EPI_ISL_1939481, EPI_ISL_1939482, EPI_ISL_1939483, EPI_ISL_1939485, EPI_ISL_1939486, EPI_ISL_1939487, EPI_ISL_1939488, EPI_ISL_1939489, EPI_ISL_1939490, EPI_ISL_1939491, EPI_ISL_1939492, EPI_ISL_1939493, EPI_ISL_1939494, EPI_ISL_1939495, EPI_ISL_1939496, EPI_ISL_1939497, EPI_ISL_1939498, EPI_ISL_1939499, EPI_ISL_1939500, EPI_ISL_1939501, EPI_ISL_1939503, EPI_ISL_1939504, EPI_ISL_1939505, EPI_ISL_1939506, EPI_ISL_1939507, EPI_ISL_1939508, EPI_ISL_1939509, EPI_ISL_1939510, EPI_ISL_1939511, EPI_ISL_1939512, EPI_ISL_1939513, EPI_ISL_1939514, EPI_ISL_1939515, EPI_ISL_1939516, EPI_ISL_1939517, EPI_ISL_1939518, EPI_ISL_1939519, EPI_ISL_1939520, EPI_ISL_1939521, EPI_ISL_1939522, EPI_ISL_1939523, EPI_ISL_1939524, EPI_ISL_1939525, EPI_ISL_1939526, EPI_ISL_1939532, EPI_ISL_1939533, EPI_ISL_1939534, EPI_ISL_1939535, EPI_ISL_1939536, EPI_ISL_1939537, EPI_ISL_1939538, EPI_ISL_1939539, EPI_ISL_1939540, EPI_ISL_1939541, EPI_ISL_1939542, EPI_ISL_1939543 | | | |
| see above | Institute of Tropical Medicine "Pedro Kourí" (IPK) | Respiratory Infections Laboratory | Lissette Perez Santos, Yahisel Tejero Suarez, Yanet Pintos, Dailyn Medero, Odalys Valdes , Mirtha Elisa Aguado , Liena de Regla Ponce |
| EPI_ISL_1972103, EPI_ISL_1972104, EPI_ISL_1972105 | AZ St.Jan Brugge-Oostende | AZ SINT-JAN BRUGGE | Marijke Reynders, Patrick Descheemaeker, Laurien Hoornaert, Thomas Van Landschoot, Jorn Hellemans |
| EPI_ISL_1972282 | Baylor Scott & White-Temple | Baylor Scott & White-Temple | Ari Rao, Linden Morales, Kimberly Walker, Marcus Volz, Shelby Hendrickson |
| EPI_ISL_2001156 | PanGenomics International Pvt. Ltd. | Gujarat Biotechnology Research Centre | Sonal Sharma ,Zuber Saiyed,Ramesh Pandit,Janvi Raval,Zarna Patel,Nitin Savaliya,Dinesh Kumar,Twinkle Soni,Dipali Dhawan,Umang Mishra,Nitesh Shah,Chaitanya Joshi,Madhvi Joshi |
| EPI_ISL_2001157 | PanGenomics International Pvt. Ltd. | Gujarat Biotechnology Research Centre | Zuber Saiyed,Ramesh Pandit,Janvi Raval,Zarna Patel,Nitin Savaliya,Dinesh Kumar,Twinkle Soni,Sonal Sharma ,Dipali Dhawan,Umang Mishra,Nitesh Shah,Chaitanya Joshi,Madhvi Joshi |
| EPI_ISL_2001158 | PanGenomics International Pvt. Ltd. | Gujarat Biotechnology Research Centre | Ramesh Pandit,Janvi Raval,Zarna Patel,Nitin Savaliya,Dinesh Kumar,Twinkle Soni,Sonal Sharma ,Zuber Saiyed,Dipali Dhawan,Umang Mishra,Nitesh Shah,Chaitanya Joshi,Madhvi Joshi |
| EPI_ISL_2001159 | PanGenomics International Pvt. Ltd. | Gujarat Biotechnology Research Centre | Janvi Raval,Zarna Patel,Nitin Savaliya,Dinesh Kumar,Twinkle Soni,Sonal Sharma ,Zuber Saiyed,Ramesh Pandit,Dipali Dhawan,Umang Mishra,Nitesh Shah,Chaitanya Joshi,Madhvi Joshi |
| EPI_ISL_2001160 | PanGenomics International Pvt. Ltd. | Gujarat Biotechnology Research Centre | Zarna Patel,Nitin Savaliya,Dinesh Kumar,Twinkle Soni,Sonal Sharma ,Zuber Saiyed,Ramesh Pandit,Janvi Raval,Dipali Dhawan,Umang Mishra,Nitesh Shah,Chaitanya Joshi,Madhvi Joshi |
| EPI_ISL_2001161 | PanGenomics International Pvt. Ltd. | Gujarat Biotechnology Research Centre | Nitin Savaliya,Dinesh Kumar,Twinkle Soni,Sonal Sharma ,Zuber Saiyed,Ramesh Pandit,Janvi Raval,Zarna Patel,Dipali Dhawan,Umang Mishra,Nitesh Shah,Chaitanya Joshi,Madhvi Joshi |
| EPI_ISL_2001162 | PanGenomics International Pvt. Ltd. | Gujarat Biotechnology Research Centre | Dinesh Kumar,Twinkle Soni,Sonal Sharma ,Zuber Saiyed,Ramesh Pandit,Janvi Raval,Zarna Patel,Nitin Savaliya,Dipali Dhawan,Umang Mishra,Nitesh Shah,Chaitanya Joshi,Madhvi Joshi |
| EPI_ISL_2001163 | PanGenomics International Pvt. Ltd. | Gujarat Biotechnology Research Centre | Twinkle Soni,Sonal Sharma ,Zuber Saiyed,Ramesh Pandit,Janvi Raval,Zarna Patel,Nitin Savaliya,Dinesh Kumar,Dipali Dhawan,Umang Mishra,Nitesh Shah,Chaitanya Joshi,Madhvi Joshi |
| EPI_ISL_2001164 | PanGenomics International Pvt. Ltd. | Gujarat Biotechnology Research Centre | Sonal Sharma ,Zuber Saiyed,Ramesh Pandit,Janvi Raval,Zarna Patel,Nitin Savaliya,Dinesh Kumar,Twinkle Soni,Dipali Dhawan,Umang Mishra,Nitesh Shah,Chaitanya Joshi,Madhvi Joshi |
| EPI_ISL_2001165 | PanGenomics International Pvt. Ltd. | Gujarat Biotechnology Research Centre | Zuber Saiyed,Ramesh Pandit,Janvi Raval,Zarna Patel,Nitin Savaliya,Dinesh Kumar,Twinkle Soni,Sonal Sharma ,Dipali Dhawan,Umang Mishra,Nitesh Shah,Chaitanya Joshi,Madhvi Joshi |
| EPI_ISL_2001167 | PanGenomics International Pvt. Ltd. | Gujarat Biotechnology Research Centre | Janvi Raval,Zarna Patel,Nitin Savaliya,Dinesh Kumar,Twinkle Soni,Sonal Sharma ,Zuber Saiyed,Ramesh Pandit,Dipali Dhawan,Umang Mishra,Nitesh Shah,Chaitanya Joshi,Madhvi Joshi |
| EPI_ISL_2001168 | PanGenomics International Pvt. Ltd. | Gujarat Biotechnology Research Centre | Zarna Patel,Nitin Savaliya,Dinesh Kumar,Twinkle Soni,Sonal Sharma ,Zuber Saiyed,Ramesh Pandit,Janvi Raval,Dipali Dhawan,Umang Mishra,Nitesh Shah,Chaitanya Joshi,Madhvi Joshi |
| EPI_ISL_2001169 | PanGenomics International Pvt. Ltd. | Gujarat Biotechnology Research Centre | Nitin Savaliya,Dinesh Kumar,Twinkle Soni,Sonal Sharma ,Zuber Saiyed,Ramesh Pandit,Janvi Raval,Zarna Patel,Dipali Dhawan,Umang Mishra,Nitesh Shah,Chaitanya Joshi,Madhvi Joshi |
| EPI_ISL_2003459 | Virology Laboratory, IDD, icddr,b | Virology Laboratory, IDD, icddr,b | Muntasir Alam, M Ishrat Jahan, Shafina Jahan, Rubaiyat Yameen, Afruna Rahman, Emily S. Gurley, Shams El Arifeen, Mustafizur Rahman |

| | | | |
|---|---|---|---|
| EPI_ISL_2004089, EPI_ISL_2004098, EPI_ISL_2004099 | IU-Cerrahpasa, Cerrahpasa School of Medicine, COVID-19 Lab | IU-Cerrahpasa, Cerrahpasa School of Medicine, COVID-19 Lab | Mert Kuskucu, Yesim Tuyji Tok, Kenan Midilli |
| EPI_ISL_2038930, EPI_ISL_2038931, EPI_ISL_2038932, EPI_ISL_2038934, EPI_ISL_2038935, EPI_ISL_2038936, EPI_ISL_2038937, EPI_ISL_2038938, EPI_ISL_2038939, EPI_ISL_2038940, EPI_ISL_2038941, EPI_ISL_2038942, EPI_ISL_2038943, EPI_ISL_2038945, EPI_ISL_2038946, EPI_ISL_2038954 | | | |
| see above | Laboratorio Central de Saude Publica do Estado de Santa Catarina (LACEN-SC) | Laboratory of Respiratory Viruses and Measles, Oswaldo Cruz Institute, FIOCRUZ | Paola Resende, Luciana Appolinario, Fernando Motta, Anna Carolina Paixao, Ana Carolina Mendonca, Alice Sampaio Rocha, Taina Venas, Elisa Cavalcante Pereira, Renata Serrano Lopes, Darcita Buerger Rovaris, Sandra Bianchini Fernandes, Marilda Siqueira on behalf of the Fiocruz COVID-19 Genomic Surveillance Network |
| EPI_ISL_2038955 | Laboratorio Central de Saude Publica do Estado do Parana (LACEN-PR) | Laboratory of Respiratory Viruses and Measles, Oswaldo Cruz Institute, FIOCRUZ | Paola Resende, Luciana Appolinario, Fernando Motta, Anna Carolina Paixao, Ana Carolina Mendonca, Alice Sampaio Rocha, Taina Venas, Elisa Cavalcante Pereira, Renata Serrano Lopes, Irina Riediger, Marilda Siqueira on behalf of the Fiocruz COVID-19 Genomic Surveillance Network |
| EPI_ISL_2038957, EPI_ISL_2038958 | Laboratorio Central de Saude Publica do Estado do Rio Grande do Sul (LACEN-RS) | Laboratory of Respiratory Viruses and Measles, Oswaldo Cruz Institute, FIOCRUZ | Paola Resende, Luciana Appolinario, Fernando Motta, Anna Carolina Paixao, Ana Carolina Mendonca, Alice Sampaio Rocha, Taina Venas, Elisa Cavalcante Pereira, Renata Serrano Lopes, Tatiana Schaffer Gregianini, Richard Salvato, Marilda Siqueira on behalf of the Fiocruz COVID-19 Genomic Surveillance Network |
| EPI_ISL_2047853 | Istituto Zooprofilattico Sperimentale Umbria e Marche "Togo Rosati" | Istituto Zooprofilattico Sperimentale dell'Abruzzo e Molise "G. Caporale" | Biagetti M., Giammarioli M., Lorusso A, Marcacci M, Di Domenico M, Ancora M, Curini V, Mangone I, Rinaldi A, Delli Compagni E, Scialabba S, Di Pasquale A, Cammà C, Puglia I, Calistri P, Savini G |
| EPI_ISL_2047854 | Istituto Zooprofilattico Sperimentale Umbria e Marche "Togo Rosati" | Istituto Zooprofilattico Sperimentale dell'Abruzzo e Molise "G. Caporale" | Biagetti M, Giammarioli M, Lorusso A, Marcacci M, Di Domenico M, Ancora M, Curini V, Mangone I, Rinaldi A, Delli Compagni E, Scialabba S, Di Pasquale A, Cammà C, Puglia I, Calistri P, Savini G |
| EPI_ISL_2047856 | Azienda Ospedaliera Terni | Istituto Zooprofilattico Sperimentale dell'Abruzzo e Molise "G. Caporale" | Palumbo M, Scaccetti A, Lorusso A, Marcacci M, Di Domenico M, Ancora M, Curini V, Mangone I, Rinaldi A, Delli Compagni E, Scialabba S, Di Pasquale A, Cammà C, Puglia I, Calistri P, Savini G |
| EPI_ISL_2047857 | Azienda Ospedaliera Terni | Istituto Zooprofilattico Sperimentale dell'Abruzzo e Molise "G. Caporale" | Palumbo A, Scaccetti A, Lorusso A, Marcacci M, Di Domenico M, Ancora M, Curini V, Mangone I, Rinaldi A, Delli Compagni E, Scialabba S, Di Pasquale A, Cammà C, Puglia I, Calistri P, Savini G |
| EPI_ISL_2047873 | Azienda USL Umbria 2 | Istituto Zooprofilattico Sperimentale dell'Abruzzo e Molise "G. Caporale" | Proietti A, Pistoni E, Lorusso A, Marcacci M, Di Domenico M, Ancora M, Curini V, Mangone I, Rinaldi A, Delli Compagni E, Scialabba S, Di Pasquale A, Cammà C, Puglia I, Calistri P, Savini G |
| EPI_ISL_2047879 | Ochsner Health | Bioinfoexperts, LLC | Amy Feehan, David J. Nolan, Rebecca Rose, Sissy Cross, David Moraga Amador, Tong Yang, Luke Caruso, Wayra Navia, Lydia Von Borstel, Xiao Hui Zhou, Julia-Garcia-Diaz, Susanna L. Lamers |
| EPI_ISL_2081198, EPI_ISL_2081199, EPI_ISL_2081200, EPI_ISL_2081201, EPI_ISL_2081202, EPI_ISL_2081203, EPI_ISL_2081204, EPI_ISL_2081205, EPI_ISL_2081206, EPI_ISL_2081207, EPI_ISL_2081208, EPI_ISL_2081209, EPI_ISL_2081210, EPI_ISL_2081211, EPI_ISL_2081212, EPI_ISL_2081213, EPI_ISL_2081214, EPI_ISL_2081215, EPI_ISL_2081216, EPI_ISL_2081217 | | | |
| see above | Premier Laboratory Services | BioInfoExperts LLC | Cheryl Metzler, Valerie Murray, Rebecca Rose, David Nolan, Susanna Lamers |
| EPI_ISL_2081835, EPI_ISL_2081836, EPI_ISL_2081838, EPI_ISL_2081917, EPI_ISL_2081918, EPI_ISL_2081919, EPI_ISL_2081920, EPI_ISL_2081922, EPI_ISL_2081923, EPI_ISL_2081924, EPI_ISL_2081925, EPI_ISL_2081926, EPI_ISL_2081927, EPI_ISL_2081928, EPI_ISL_2081949, EPI_ISL_2081950, EPI_ISL_2081951, EPI_ISL_2081952, EPI_ISL_2081953, EPI_ISL_2081954, EPI_ISL_2081955, EPI_ISL_2081956, EPI_ISL_2081957, EPI_ISL_2081964, EPI_ISL_2081965, EPI_ISL_2081966, EPI_ISL_2081967, EPI_ISL_2081968, EPI_ISL_2081969, EPI_ISL_2081970, EPI_ISL_2081971, EPI_ISL_2081972, EPI_ISL_2081973, EPI_ISL_2081974, EPI_ISL_2081975, EPI_ISL_2081976, EPI_ISL_2081977, EPI_ISL_2081978, EPI_ISL_2081979, EPI_ISL_2081980, EPI_ISL_2081981, EPI_ISL_2081982, EPI_ISL_2081983, EPI_ISL_2081984, EPI_ISL_2081985, EPI_ISL_2081986, EPI_ISL_2081987, EPI_ISL_2081988, EPI_ISL_2081989, EPI_ISL_2081990, EPI_ISL_2081991, EPI_ISL_2081992, EPI_ISL_2081993, EPI_ISL_2081994, EPI_ISL_2081995, EPI_ISL_2081996, EPI_ISL_2081997, EPI_ISL_2081998, EPI_ISL_2081999, EPI_ISL_2082000, EPI_ISL_2082001, EPI_ISL_2082002, EPI_ISL_2082003, EPI_ISL_2082004, EPI_ISL_2082005, EPI_ISL_2082006, EPI_ISL_2082007, EPI_ISL_2082008, EPI_ISL_2082009, EPI_ISL_2082010, EPI_ISL_2082011, EPI_ISL_2082012, EPI_ISL_2082025, EPI_ISL_2082026, EPI_ISL_2082027, EPI_ISL_2082028, EPI_ISL_2082032 | | | |
| see above | National Center of Infectious and Parasitic Diseases | National Center of Infectious and Parasitic Diseases | Alexiev, Ivanov, Korsun, Stoitsova, Philipova, Dimitrova, Grigorova L., Hristova, Donchev, Stoykov, Trifonova, Dobrinov, Grigorova I., Kantardjiev |
| EPI_ISL_2086121 | Microbiology Department. Complexo Hospitalario Universitario de Vigo | Microbiology Department. Complexo Hospitalario Universitario de Vigo | Alfaya N, Alonso I, Alvarez M, Cabrera JJ, Carballo R, Cores O, Cortizo S, del-Campo V, Martinez L, Mediero G, Perez S, Potel C, Regueiro B, Rey S, Vasallo FJ |
| EPI_ISL_2086585, EPI_ISL_2086586, EPI_ISL_2086587, EPI_ISL_2086588, EPI_ISL_2086589, EPI_ISL_2086590, EPI_ISL_2086591, EPI_ISL_2086592, EPI_ISL_2086593, EPI_ISL_2086763, EPI_ISL_2086764 | | | |
| see above | LABCOVID_HCPA | LABRESIS_HCPA | Wink PL, Martins AF, Volpato F, Monteiro F, Zavascki AP, Barth AL |
| EPI_ISL_2090732 | REUNILAB | UMR PIMIT | Dr David A Wilkinson, Dr Patrick Mavingui, Dr Camille Lebarbenchon, Magali Turpin |
| EPI_ISL_2090735 | CERBALLIANCE | UMR PIMIT | Dr David A Wilkinson, Dr Patrick Mavingui, Dr Camille Lebarbenchon, Magali Turpin |
| EPI_ISL_2090739 | REUNILAB | UMR PIMIT | Dr David A Wilkinson, Dr Patrick Mavingui, Dr Camille Lebarbenchon, Magali Turpin |
| EPI_ISL_2090741 | CHU NORD | UMR PIMIT | Dr David A Wilkinson, Dr Patrick Mavingui, Dr Camille Lebarbenchon, Magali Turpin |
| EPI_ISL_2090742 | CHOR | UMR PIMIT | Dr David A Wilkinson, Dr Patrick Mavingui, Dr Camille Lebarbenchon, Magali Turpin |
| EPI_ISL_2090745, EPI_ISL_2090746, EPI_ISL_2090747 | CERBALLIANCE | UMR PIMIT | Dr David A Wilkinson, Dr Patrick Mavingui, Dr Camille Lebarbenchon, Magali Turpin |
| EPI_ISL_2091132 | Unidad de Investigación Biomédica de Zacatecas (UIBZ) | Centro de Investigación en Enfermedades Infecciosas (CIENI), Instituto Nacional de Enfermedades Respiratorias (INER) | Consorcio Mexicano de Vigilancia Genómica (CoViGen-Mex). Authors (in alphabetical order): Julio Elias Alvarado-Yaah, Carlos F. Arias, Santiago Ávila-Ríos, Víctor Hugo Borja-Aburto, Celia Boukadida, Juan Bautista Chale-Dzul, Célida Duque Molina, José Antonio Enciso-Moreno, Gloria Elena Espinosa-Ayala, Fernando Fontove-Herrera, Victor Eduardo Garcia-Arias, Concepción Grajales-Muñiz, Ricardo Grande, Alfredo Herrera-Estrella, Carla Ivón Herrera-Najera, Pavel Isa, Brenda Irasema Maldonado-Meza, Bernardo Martínez-Miguel, Margarita Matías-Florentino, María Guadalupe de Jesús Mireles-Rivera, Gloria María Molina-Salinas, Hector Montoya-Fuentes, José Esteban Muñoz-Medina, José de Jesús Nuñez-Contreras, Alicia Ocaña-Mondragón, Luis Alberto Ochoa-Carrera, Hector Esteban Paz-Juárez, Francisco Pulido, Helen Haydee Fernanda Ramírez-Plascencia, Angel Gustavo Salas-Lais, Alejandro Sanchez-Flores, Clara Esperanza Santacruz-Tinoco, María Guadalupe Santiago-Mauricio, Nelly Sélem-Mojica, Blanca Taboada, Gloria Vazquez |
| EPI_ISL_2091134 | Laboratorio Central de Epidemiología (LCE) | Centro de Investigación en Enfermedades Infecciosas (CIENI), Instituto Nacional de Enfermedades Respiratorias (INER) | Consorcio Mexicano de Vigilancia Genómica (CoViGen-Mex). Authors (in alphabetical order): Julio Elias Alvarado-Yaah, Carlos F. Arias, Santiago Ávila-Ríos, Víctor Hugo Borja-Aburto, Celia Boukadida, Juan Bautista Chale-Dzul, Célida Duque Molina, José Antonio Enciso-Moreno, Gloria Elena Espinosa-Ayala, Fernando Fontove-Herrera, Victor Eduardo Garcia-Arias, Concepción Grajales-Muñiz, Ricardo Grande, Alfredo Herrera-Estrella, Carla Ivón Herrera-Najera, Pavel Isa, Brenda Irasema Maldonado-Meza, Bernardo Martínez-Miguel, Margarita Matías-Florentino, María Guadalupe de Jesús Mireles-Rivera, Gloria María Molina-Salinas, Hector Montoya-Fuentes, José Esteban Muñoz-Medina, José de Jesús Nuñez-Contreras, Alicia Ocaña-Mondragón, Luis Alberto Ochoa-Carrera, Hector Esteban Paz-Juárez, Francisco Pulido, Helen Haydee Fernanda Ramírez-Plascencia, Angel Gustavo Salas-Lais, Alejandro Sanchez-Flores, Clara Esperanza Santacruz-Tinoco, María Guadalupe Santiago-Mauricio, Nelly Sélem-Mojica, Blanca Taboada, Gloria Vazquez |
| EPI_ISL_2091135 | Centro de Investigación Biomédica del Noreste (CIBIN) | Centro de Investigación en Enfermedades Infecciosas (CIENI), Instituto Nacional de Enfermedades Respiratorias (INER) | Consorcio Mexicano de Vigilancia Genómica (CoViGen-Mex). Authors (in alphabetical order): Julio Elias Alvarado-Yaah, Carlos F. Arias, Santiago Ávila-Ríos, Víctor Hugo Borja-Aburto, Celia Boukadida, Juan Bautista Chale-Dzul, Célida Duque Molina, José Antonio Enciso-Moreno, Gloria Elena Espinosa-Ayala, Fernando Fontove-Herrera, Victor Eduardo Garcia-Arias, Concepción Grajales-Muñiz, Ricardo Grande, Alfredo Herrera-Estrella, Carla Ivón Herrera-Najera, Pavel Isa, Brenda Irasema Maldonado-Meza, Bernardo Martínez-Miguel, Margarita Matías-Florentino, María Guadalupe de Jesús Mireles-Rivera, Gloria María Molina-Salinas, Hector Montoya-Fuentes, José Esteban Muñoz-Medina, José de Jesús Nuñez-Contreras, Alicia Ocaña-Mondragón, Luis Alberto Ochoa-Carrera, Hector Esteban Paz-Juárez, Francisco Pulido, Helen Haydee Fernanda Ramírez-Plascencia, Angel Gustavo Salas-Lais, Alejandro Sanchez-Flores, Clara Esperanza Santacruz-Tinoco, María Guadalupe Santiago-Mauricio, Nelly Sélem-Mojica, Blanca Taboada, Gloria Vazquez |
| EPI_ISL_2100891, EPI_ISL_2100894, EPI_ISL_2100895, EPI_ISL_2100896, EPI_ISL_2100898, EPI_ISL_2100899, EPI_ISL_2100900, EPI_ISL_2100901, EPI_ISL_2100902, EPI_ISL_2100903, EPI_ISL_2100905, EPI_ISL_2100906, EPI_ISL_2100907, EPI_ISL_2100908, EPI_ISL_2100909, EPI_ISL_2100910, EPI_ISL_2100911, EPI_ISL_2100912, EPI_ISL_2100913, EPI_ISL_2100914, EPI_ISL_2100917, EPI_ISL_2100918, EPI_ISL_2100919, EPI_ISL_2100921, EPI_ISL_2100922, EPI_ISL_2100923, EPI_ISL_2100924, EPI_ISL_2100926, EPI_ISL_2100928, EPI_ISL_2100929, EPI_ISL_2100930, EPI_ISL_2100931, EPI_ISL_2100932, EPI_ISL_2100933, EPI_ISL_2100934, EPI_ISL_2100936, EPI_ISL_2100940, EPI_ISL_2100941, EPI_ISL_2100942, EPI_ISL_2100944, EPI_ISL_2100945, EPI_ISL_2100946, EPI_ISL_2100947, EPI_ISL_2100948, EPI_ISL_2100949, EPI_ISL_2100951, EPI_ISL_2100952, EPI_ISL_2100953, EPI_ISL_2100954, EPI_ISL_2100955, EPI_ISL_2100956, EPI_ISL_2100957, EPI_ISL_2100958, EPI_ISL_2100960, EPI_ISL_2100961, EPI_ISL_2100962, EPI_ISL_2100963 | | | |
| see above | The Ohio State University Applied Microbiology Services Laboratory | The Ohio State University Applied Microbiology Services Laboratory | Seth A. Faith PhD |

| | | | |
|---|---|---|---|
| EPI_ISL_2101062 | Molecular Biology Laboratory, Faculty Medicine and Health Sciences, Warmadewa University | Eijkman Institute for Molecular Biology, National Agency for Research and Innovation; Molecular Biology Laboratory, Faculty Medicine and Health Sciences, Warmadewa University | Sukma Oktavianthi, Lidwina Priliani, Edison Johar, Frilasita A Yudhaputri, Muhammad Rezki Rasyak, Willy Agustine, Hidayat Trimarsanto, Lydia V. Panggalo, Iskandar Adnan, Sri Masyeni, Erly Sintya, Safarina G Malik, Khin Saw Myint, Amin Soebandrio |
| EPI_ISL_2101063 | Molecular Biology Laboratory, Faculty Medicine and Health Sciences, Warmadewa University | Eijkman Institute for Molecular Biology, National Agency for Research and Innovation; Molecular Biology Laboratory, Faculty Medicine and Health Sciences, Warmadewa University | Lidwina Priliani, Edison Johar, Frilasita A Yudhaputri, Muhammad Rezki Rasyak, Willy Agustine, Hidayat Trimarsanto, Lydia V. Panggalo, Iskandar Adnan, Sukma Oktavianthi, Sri Masyeni, Erly Sintya, Safarina G Malik, Khin Saw Myint, Amin Soebandrio |
| EPI_ISL_2101064 | Molecular Biology Laboratory, Faculty Medicine and Health Sciences, Warmadewa University | Eijkman Institute for Molecular Biology, National Agency for Research and Innovation; Molecular Biology Laboratory, Faculty Medicine and Health Sciences, Warmadewa University | Edison Johar, Frilasita A Yudhaputri, Muhammad Rezki Rasyak, Willy Agustine, Hidayat Trimarsanto, Lydia V. Panggalo, Iskandar Adnan, Sukma Oktavianthi, Lidwina Priliani, Sri Masyeni, Erly Sintya, Safarina G Malik, Khin Saw Myint, Amin Soebandrio |
| EPI_ISL_2101065 | Molecular Biology Laboratory, Faculty Medicine and Health Sciences, Warmadewa University | Eijkman Institute for Molecular Biology, National Agency for Research and Innovation; Molecular Biology Laboratory, Faculty Medicine and Health Sciences, Warmadewa University | Lydia V. Panggalo, Iskandar Adnan, Sukma Oktavianthi, Lidwina Priliani, Edison Johar, Frilasita A Yudhaputri, Muhammad Rezki Rasyak, Willy Agustine, Hidayat Trimarsanto, Sri Masyeni, Erly Sintya, Safarina G Malik, Khin Saw Myint, Amin Soebandrio |
| EPI_ISL_2101066 | Molecular Biology Laboratory, Faculty Medicine and Health Sciences, Warmadewa University | Eijkman Institute for Molecular Biology, National Agency for Research and Innovation; Molecular Biology Laboratory, Faculty Medicine and Health Sciences, Warmadewa University | Iskandar Adnan, Sukma Oktavianthi, Lidwina Priliani, Edison Johar, Frilasita A Yudhaputri, Muhammad Rezki Rasyak, Willy Agustine, Hidayat Trimarsanto, Lydia V. Panggalo, Sri Masyeni, Erly Sintya, Safarina G Malik, Khin Saw Myint, Amin Soebandrio |
| EPI_ISL_2101067 | Molecular Biology Laboratory, Faculty Medicine and Health Sciences, Warmadewa University | Eijkman Institute for Molecular Biology, National Agency for Research and Innovation; Molecular Biology Laboratory, Faculty Medicine and Health Sciences, Warmadewa University | Sukma Oktavianthi, Lidwina Priliani, Edison Johar, Frilasita A Yudhaputri, Muhammad Rezki Rasyak, Willy Agustine, Hidayat Trimarsanto, Lydia V. Panggalo, Iskandar Adnan, Sri Masyeni, Erly Sintya, Safarina G Malik, Khin Saw Myint, Amin Soebandrio |
| EPI_ISL_2101068 | Molecular Biology Laboratory, Faculty Medicine and Health Sciences, Warmadewa University | Eijkman Institute for Molecular Biology, National Agency for Research and Innovation; Molecular Biology Laboratory, Faculty Medicine and Health Sciences, Warmadewa University | Lidwina Priliani, Edison Johar, Frilasita A Yudhaputri, Muhammad Rezki Rasyak, Willy Agustine, Hidayat Trimarsanto, Lydia V. Panggalo, Iskandar Adnan, Sukma Oktavianthi, Sri Masyeni, Erly Sintya, Safarina G Malik, Khin Saw Myint, Amin Soebandrio |
| EPI_ISL_2101069 | Molecular Biology Laboratory, Faculty Medicine and Health Sciences, Warmadewa University | Eijkman Institute for Molecular Biology, National Agency for Research and Innovation; Molecular Biology Laboratory, Faculty Medicine and Health Sciences, Warmadewa University | Edison Johar, Frilasita A Yudhaputri, Muhammad Rezki Rasyak, Willy Agustine, Hidayat Trimarsanto, Lydia V. Panggalo, Iskandar Adnan, Sukma Oktavianthi, Lidwina Priliani, Sri Masyeni, Erly Sintya, Safarina G Malik, Khin Saw Myint, Amin Soebandrio |
| EPI_ISL_2101070 | Molecular Biology Laboratory, Faculty Medicine and Health Sciences, Warmadewa University | Eijkman Institute for Molecular Biology, National Agency for Research and Innovation; Molecular Biology Laboratory, Faculty Medicine and Health Sciences, Warmadewa University | Frilasita A Yudhaputri, Muhammad Rezki Rasyak, Willy Agustine, Hidayat Trimarsanto, Lydia V. Panggalo, Iskandar Adnan, Sukma Oktavianthi, Lidwina Priliani, Edison Johar, Sri Masyeni, Erly Sintya, Safarina G Malik, Khin Saw Myint, Amin Soebandrio |
| EPI_ISL_2101071 | Molecular Biology Laboratory, Faculty Medicine and Health Sciences, Warmadewa University | Eijkman Institute for Molecular Biology, National Agency for Research and Innovation; Molecular Biology Laboratory, Faculty Medicine and Health Sciences, Warmadewa University | Hidayat Trimarsanto, Lydia V. Panggalo, Iskandar Adnan, Sukma Oktavianthi, Lidwina Priliani, Edison Johar, Frilasita A Yudhaputri, Muhammad Rezki Rasyak, Willy Agustine, Sri Masyeni, Erly Sintya, Safarina G Malik, Khin Saw Myint, Amin Soebandrio |
| EPI_ISL_2101072 | Molecular Biology Laboratory, Faculty Medicine and Health Sciences, Warmadewa University | Eijkman Institute for Molecular Biology, National Agency for Research and Innovation; Molecular Biology Laboratory, Faculty Medicine and Health Sciences, Warmadewa University | Lydia V. Panggalo, Iskandar Adnan, Sukma Oktavianthi, Lidwina Priliani, Edison Johar, Frilasita A Yudhaputri, Muhammad Rezki Rasyak, Willy Agustine, Hidayat Trimarsanto, Sri Masyeni, Erly Sintya, Safarina G Malik, Khin Saw Myint, Amin Soebandrio |
| EPI_ISL_2101073 | Molecular Biology Laboratory, Faculty Medicine and Health Sciences, Warmadewa University | Eijkman Institute for Molecular Biology, National Agency for Research and Innovation; Molecular Biology Laboratory, Faculty Medicine and Health Sciences, Warmadewa University | Iskandar Adnan, Sukma Oktavianthi, Lidwina Priliani, Edison Johar, Frilasita A Yudhaputri, Muhammad Rezki Rasyak, Willy Agustine, Hidayat Trimarsanto, Lydia V. Panggalo, Sri Masyeni, Erly Sintya, Safarina G Malik, Khin Saw Myint, Amin Soebandrio |
| EPI_ISL_2101229 | RSUD Klungkung | Eijkman Institute for Molecular Biology, National Agency for Research and Innovation; Molecular Biology Laboratory, Faculty Medicine and Health Sciences, Warmadewa University | Iskandar Adnan, Sukma Oktavianthi, Lidwina Priliani, Edison Johar, Frilasita A Yudhaputri, Muhammad Rezki Rasyak, Willy Agustine, Hidayat Trimarsanto, Lydia V. Panggalo, Sri Masyeni, Erly Sintya, I Komang Parwata, Safarina G Malik, Khin Saw Myint, Amin Soebandrio |
| EPI_ISL_2101230 | RSUD Klungkung | Eijkman Institute for Molecular Biology, National Agency for Research and Innovation; Molecular Biology Laboratory, Faculty Medicine and Health Sciences, Warmadewa University | Lidwina Priliani, Edison Johar, Frilasita A Yudhaputri, Muhammad Rezki Rasyak, Willy Agustine, Hidayat Trimarsanto, Lydia V. Panggalo, Iskandar Adnan, Sukma Oktavianthi, Sri Masyeni, Erly Sintya, I Komang Parwata, Safarina G Malik, Khin Saw Myint, Amin Soebandrio |
| EPI_ISL_2101231 | RSUD Klungkung | Eijkman Institute for Molecular Biology, National Agency for Research and Innovation; Molecular Biology Laboratory, Faculty Medicine and Health Sciences, Warmadewa University | Willy Agustine, Hidayat Trimarsanto, Lydia V. Panggalo, Iskandar Adnan, Sukma Oktavianthi, Lidwina Priliani, Edison Johar, Frilasita A Yudhaputri, Muhammad Rezki Rasyak, Sri Masyeni, Erly Sintya, I Komang Parwata, Safarina G Malik, Khin Saw Myint, Amin Soebandrio |
| EPI_ISL_2101232 | RSUD Klungkung | Eijkman Institute for Molecular Biology, National Agency for Research and Innovation; Molecular Biology Laboratory, Faculty Medicine and Health Sciences, Warmadewa University | Lidwina Priliani, Edison Johar, Frilasita A Yudhaputri, Muhammad Rezki Rasyak, Willy Agustine, Hidayat Trimarsanto, Lydia V. Panggalo, Iskandar Adnan, Sukma Oktavianthi, Sri Masyeni, Erly Sintya, I Komang Parwata, Safarina G Malik, Khin Saw Myint, Amin Soebandrio |
| EPI_ISL_2101233 | RSUD Klungkung | Eijkman Institute for Molecular Biology, National Agency for Research and Innovation; Molecular Biology Laboratory, Faculty Medicine and Health Sciences, Warmadewa University | Frilasita A Yudhaputri, Muhammad Rezki Rasyak, Willy Agustine, Hidayat Trimarsanto, Lydia V. Panggalo, Iskandar Adnan, Sukma Oktavianthi, Lidwina Priliani, Edison Johar, Sri Masyeni, Erly Sintya, I Komang Parwata, Safarina G Malik, Khin Saw Myint, Amin Soebandrio |
| EPI_ISL_2101234 | RSUD Klungkung | Eijkman Institute for Molecular Biology, National Agency for Research and Innovation; Molecular Biology Laboratory, Faculty Medicine and Health Sciences, Warmadewa University | Muhammad Rezki Rasyak, Willy Agustine, Hidayat Trimarsanto, Lydia V. Panggalo, Iskandar Adnan, Sukma Oktavianthi, Lidwina Priliani, Edison Johar, Frilasita A Yudhaputri, Sri Masyeni, Erly Sintya, I Komang Parwata, Safarina G Malik, Khin Saw Myint, Amin Soebandrio |
| EPI_ISL_2101235 | RSUD Klungkung | Eijkman Institute for Molecular Biology, National Agency for Research and Innovation; Molecular Biology Laboratory, Faculty Medicine and Health Sciences, Warmadewa University | Hidayat Trimarsanto, Lydia V. Panggalo, Iskandar Adnan, Sukma Oktavianthi, Lidwina Priliani, Edison Johar, Frilasita A Yudhaputri, Muhammad Rezki Rasyak, Willy Agustine, Sri Masyeni, Erly Sintya, I Komang Parwata, Safarina G Malik, Khin Saw Myint, Amin Soebandrio |
| EPI_ISL_2101292 | RSUD Mangusada | Eijkman Institute for Molecular Biology, National Agency for Research and Innovation; Molecular Biology Laboratory, | Frilasita A Yudhaputri, Muhammad Rezki Rasyak, Willy Agustine, Hidayat Trimarsanto, Lydia V. Panggalo, Iskandar Adnan, Sukma Oktavianthi, Lidwina Priliani, Edison Johar, Sri Masyeni, Erly Sintya, Ida Ayu Wayan Mahayani, Safarina G Malik, Khin Saw Myint, Amin Soebandrio |

| | | | |
|---|---|---|---|
| | | Faculty Medicine and Health Sciences, Warmadewa University | |
| EPI_ISL_2101293 | RSUD Mangusada | Eijkman Institute for Molecular Biology, National Agency for Research and Innovation; Molecular Biology Laboratory, Faculty Medicine and Health Sciences, Warmadewa University | Muhammad Rezki Rasyak, Willy Agustine, Hidayat Trimarsanto, Lydia V. Panggalo, Iskandar Adnan, Sukma Oktavianthi, Lidwina Priliani, Edison Johar, Frilasita A Yudhaputri, Sri Masyeni, Erly Sintya, Ida Ayu Wayan Mahayani, Safarina G Malik, Khin Saw Myint, Amin Soebandrio |
| EPI_ISL_2101294 | RSUD Mangusada | Eijkman Institute for Molecular Biology, National Agency for Research and Innovation; Molecular Biology Laboratory, Faculty Medicine and Health Sciences, Warmadewa University | Willy Agustine, Hidayat Trimarsanto, Lydia V. Panggalo, Iskandar Adnan, Sukma Oktavianthi, Lidwina Priliani, Edison Johar, Frilasita A Yudhaputri, Muhammad Rezki Rasyak, Sri Masyeni, Erly Sintya, Ida Ayu Wayan Mahayani, Safarina G Malik, Khin Saw Myint, Amin Soebandrio |
| EPI_ISL_2101295 | RSUD Mangusada | Eijkman Institute for Molecular Biology, National Agency for Research and Innovation; Molecular Biology Laboratory, Faculty Medicine and Health Sciences, Warmadewa University | Hidayat Trimarsanto, Lydia V. Panggalo, Iskandar Adnan, Sukma Oktavianthi, Lidwina Priliani, Edison Johar, Frilasita A Yudhaputri, Muhammad Rezki Rasyak, Willy Agustine, Sri Masyeni, Erly Sintya, Ida Ayu Wayan Mahayani, Safarina G Malik, Khin Saw Myint, Amin Soebandrio |
| EPI_ISL_2101296 | RSUD Mangusada | Eijkman Institute for Molecular Biology, National Agency for Research and Innovation; Molecular Biology Laboratory, Faculty Medicine and Health Sciences, Warmadewa University | Lydia V. Panggalo, Iskandar Adnan, Sukma Oktavianthi, Lidwina Priliani, Edison Johar, Frilasita A Yudhaputri, Muhammad Rezki Rasyak, Willy Agustine, Hidayat Trimarsanto, Sri Masyeni, Erly Sintya, Ida Ayu Wayan Mahayani, Safarina G Malik, Khin Saw Myint, Amin Soebandrio |
| EPI_ISL_2101297 | RSUD Mangusada | Eijkman Institute for Molecular Biology, National Agency for Research and Innovation; Molecular Biology Laboratory, Faculty Medicine and Health Sciences, Warmadewa University | Edison Johar, Frilasita A Yudhaputri, Muhammad Rezki Rasyak, Willy Agustine, Hidayat Trimarsanto, Lydia V. Panggalo, Iskandar Adnan, Sukma Oktavianthi, Lidwina Priliani, Sri Masyeni, Erly Sintya, Ida Ayu Wayan Mahayani, Safarina G Malik, Khin Saw Myint, Amin Soebandrio |
| EPI_ISL_2101298 | RSUD Mangusada | Eijkman Institute for Molecular Biology, National Agency for Research and Innovation; Molecular Biology Laboratory, Faculty Medicine and Health Sciences, Warmadewa University | Muhammad Rezki Rasyak, Willy Agustine, Hidayat Trimarsanto, Lydia V. Panggalo, Iskandar Adnan, Sukma Oktavianthi, Lidwina Priliani, Edison Johar, Frilasita A Yudhaputri, Sri Masyeni, Erly Sintya, Ida Ayu Wayan Mahayani, Safarina G Malik, Khin Saw Myint, Amin Soebandrio |
| EPI_ISL_2101299 | RSUD Mangusada | Eijkman Institute for Molecular Biology, National Agency for Research and Innovation; Molecular Biology Laboratory, Faculty Medicine and Health Sciences, Warmadewa University | Willy Agustine, Hidayat Trimarsanto, Lydia V. Panggalo, Iskandar Adnan, Sukma Oktavianthi, Lidwina Priliani, Edison Johar, Frilasita A Yudhaputri, Muhammad Rezki Rasyak, Sri Masyeni, Erly Sintya, Ida Ayu Wayan Mahayani, Safarina G Malik, Khin Saw Myint, Amin Soebandrio |
| EPI_ISL_2101300 | RSUD Mangusada | Eijkman Institute for Molecular Biology, National Agency for Research and Innovation; Molecular Biology Laboratory, Faculty Medicine and Health Sciences, Warmadewa University | Hidayat Trimarsanto, Lydia V. Panggalo, Iskandar Adnan, Sukma Oktavianthi, Lidwina Priliani, Edison Johar, Frilasita A Yudhaputri, Muhammad Rezki Rasyak, Willy Agustine, Sri Masyeni, Erly Sintya, Ida Ayu Wayan Mahayani, Safarina G Malik, Khin Saw Myint, Amin Soebandrio |
| EPI_ISL_2101301 | RSUD Mangusada | Eijkman Institute for Molecular Biology, National Agency for Research and Innovation; Molecular Biology Laboratory, Faculty Medicine and Health Sciences, Warmadewa University | Lydia V. Panggalo, Iskandar Adnan, Sukma Oktavianthi, Lidwina Priliani, Edison Johar, Frilasita A Yudhaputri, Muhammad Rezki Rasyak, Willy Agustine, Hidayat Trimarsanto, Sri Masyeni, Erly Sintya, Ida Ayu Wayan Mahayani, Safarina G Malik, Khin Saw Myint, Amin Soebandrio |
| EPI_ISL_2101302 | RSUD Mangusada | Eijkman Institute for Molecular Biology, National Agency for Research and Innovation; Molecular Biology Laboratory, Faculty Medicine and Health Sciences, Warmadewa University | Iskandar Adnan, Sukma Oktavianthi, Lidwina Priliani, Edison Johar, Frilasita A Yudhaputri, Muhammad Rezki Rasyak, Willy Agustine, Hidayat Trimarsanto, Lydia V. Panggalo, Sri Masyeni, Erly Sintya, Ida Ayu Wayan Mahayani, Safarina G Malik, Khin Saw Myint, Amin Soebandrio |
| EPI_ISL_2101303 | RSUD Mangusada | Eijkman Institute for Molecular Biology, National Agency for Research and Innovation; Molecular Biology Laboratory, Faculty Medicine and Health Sciences, Warmadewa University | Sukma Oktavianthi, Lidwina Priliani, Edison Johar, Frilasita A Yudhaputri, Muhammad Rezki Rasyak, Willy Agustine, Hidayat Trimarsanto, Lydia V. Panggalo, Iskandar Adnan, Sri Masyeni, Erly Sintya, Ida Ayu Wayan Mahayani, Safarina G Malik, Khin Saw Myint, Amin Soebandrio |
| EPI_ISL_2101304 | RSUD Mangusada | Eijkman Institute for Molecular Biology, National Agency for Research and Innovation; Molecular Biology Laboratory, Faculty Medicine and Health Sciences, Warmadewa University | Lidwina Priliani, Edison Johar, Frilasita A Yudhaputri, Muhammad Rezki Rasyak, Willy Agustine, Hidayat Trimarsanto, Lydia V. Panggalo, Iskandar Adnan, Sukma Oktavianthi, Sri Masyeni, Erly Sintya, Ida Ayu Wayan Mahayani, Safarina G Malik, Khin Saw Myint, Amin Soebandrio |
| EPI_ISL_2101305 | RSUD Mangusada | Eijkman Institute for Molecular Biology, National Agency for Research and Innovation; Molecular Biology Laboratory, Faculty Medicine and Health Sciences, Warmadewa University | Edison Johar, Frilasita A Yudhaputri, Muhammad Rezki Rasyak, Willy Agustine, Hidayat Trimarsanto, Lydia V. Panggalo, Iskandar Adnan, Sukma Oktavianthi, Lidwina Priliani, Sri Masyeni, Erly Sintya, Ida Ayu Wayan Mahayani, Safarina G Malik, Khin Saw Myint, Amin Soebandrio |
| EPI_ISL_2101306 | RSUD Mangusada | Eijkman Institute for Molecular Biology, National Agency for Research and Innovation; Molecular Biology Laboratory, Faculty Medicine and Health Sciences, Warmadewa University | Frilasita A Yudhaputri, Muhammad Rezki Rasyak, Willy Agustine, Hidayat Trimarsanto, Lydia V. Panggalo, Iskandar Adnan, Sukma Oktavianthi, Lidwina Priliani, Edison Johar, Sri Masyeni, Erly Sintya, Ida Ayu Wayan Mahayani, Safarina G Malik, Khin Saw Myint, Amin Soebandrio |
| EPI_ISL_2101307 | RSUD Mangusada | Eijkman Institute for Molecular Biology, National Agency for Research and Innovation; Molecular Biology Laboratory, Faculty Medicine and Health Sciences, Warmadewa University | Muhammad Rezki Rasyak, Willy Agustine, Hidayat Trimarsanto, Lydia V. Panggalo, Iskandar Adnan, Sukma Oktavianthi, Lidwina Priliani, Edison Johar, Frilasita A Yudhaputri, Sri Masyeni, Erly Sintya, Ida Ayu Wayan Mahayani, Safarina G Malik, Khin Saw Myint, Amin Soebandrio |
| EPI_ISL_2101308 | RSUD Mangusada | Eijkman Institute for Molecular Biology, National Agency for Research and Innovation; Molecular Biology Laboratory, Faculty Medicine and Health Sciences, Warmadewa University | Willy Agustine, Hidayat Trimarsanto, Lydia V. Panggalo, Iskandar Adnan, Sukma Oktavianthi, Lidwina Priliani, Edison Johar, Frilasita A Yudhaputri, Muhammad Rezki Rasyak, Sri Masyeni, Erly Sintya, Ida Ayu Wayan Mahayani, Safarina G Malik, Khin Saw Myint, Amin Soebandrio |
| EPI_ISL_2101309 | RSUD Mangusada | Eijkman Institute for Molecular Biology, National Agency for Research and Innovation; Molecular Biology Laboratory, Faculty Medicine and Health Sciences, Warmadewa University | Hidayat Trimarsanto, Lydia V. Panggalo, Iskandar Adnan, Sukma Oktavianthi, Lidwina Priliani, Edison Johar, Frilasita A Yudhaputri, Muhammad Rezki Rasyak, Willy Agustine, Sri Masyeni, Erly Sintya, Ida Ayu Wayan Mahayani, Safarina G Malik, Khin Saw Myint, Amin Soebandrio |
| EPI_ISL_2101310 | RSUD Mangusada | Eijkman Institute for Molecular Biology, National Agency for Research and Innovation; Molecular Biology Laboratory, Faculty Medicine and Health Sciences, Warmadewa University | Lydia V. Panggalo, Iskandar Adnan, Sukma Oktavianthi, Lidwina Priliani, Edison Johar, Frilasita A Yudhaputri, Muhammad Rezki Rasyak, Willy Agustine, Hidayat Trimarsanto, Sri Masyeni, Erly Sintya, Ida Ayu Wayan Mahayani, Safarina G Malik, Khin Saw Myint, Amin Soebandrio |
| EPI_ISL_2101311 | RSUD Mangusada | Eijkman Institute for Molecular Biology, National Agency for Research and Innovation; Molecular Biology Laboratory, Faculty Medicine and Health Sciences, Warmadewa University | Iskandar Adnan, Sukma Oktavianthi, Lidwina Priliani, Edison Johar, Frilasita A Yudhaputri, Muhammad Rezki Rasyak, Willy Agustine, Hidayat Trimarsanto, Lydia V. Panggalo, Sri Masyeni, Erly Sintya, Ida Ayu Wayan Mahayani, Safarina G Malik, Khin Saw Myint, Amin Soebandrio |

| | | | |
|---|---|---|---|
| EPI_ISL_2101312 | RSUD Mangusada | Eijkman Institute for Molecular Biology, National Agency for Research and Innovation; Molecular Biology Laboratory, Faculty Medicine and Health Sciences, Warmadewa University | Sukma Oktavianthi, Lidwina Priliani, Edison Johar, Frilasita A Yudhaputri, Muhammad Rezki Rasyak, Willy Agustine, Hidayat Trimarsanto, Lydia V. Panggalo, Iskandar Adnan, Sri Masyeni, Erly Sintya, Ida Ayu Wayan Mahayani, Safarina G Malik, Khin Saw Myint, Amin Soebandrio |
| EPI_ISL_2101313 | RSUD Mangusada | Eijkman Institute for Molecular Biology, National Agency for Research and Innovation; Molecular Biology Laboratory, Faculty Medicine and Health Sciences, Warmadewa University | Edison Johar, Frilasita A Yudhaputri, Muhammad Rezki Rasyak, Willy Agustine, Hidayat Trimarsanto, Lydia V. Panggalo, Iskandar Adnan, Sukma Oktavianthi, Lidwina Priliani, Sri Masyeni, Erly Sintya, Ida Ayu Wayan Mahayani, Safarina G Malik, Khin Saw Myint, Amin Soebandrio |
| EPI_ISL_2101314 | RSUD Mangusada | Eijkman Institute for Molecular Biology, National Agency for Research and Innovation; Molecular Biology Laboratory, Faculty Medicine and Health Sciences, Warmadewa University | Frilasita A Yudhaputri, Muhammad Rezki Rasyak, Willy Agustine, Hidayat Trimarsanto, Lydia V. Panggalo, Iskandar Adnan, Sukma Oktavianthi, Lidwina Priliani, Edison Johar, Sri Masyeni, Erly Sintya, Ida Ayu Wayan Mahayani, Safarina G Malik, Khin Saw Myint, Amin Soebandrio |
| EPI_ISL_2101315 | RSUD Mangusada | Eijkman Institute for Molecular Biology, National Agency for Research and Innovation; Molecular Biology Laboratory, Faculty Medicine and Health Sciences, Warmadewa University | Muhammad Rezki Rasyak, Willy Agustine, Hidayat Trimarsanto, Lydia V. Panggalo, Iskandar Adnan, Sukma Oktavianthi, Lidwina Priliani, Edison Johar, Frilasita A Yudhaputri, Sri Masyeni, Erly Sintya, Ida Ayu Wayan Mahayani, Safarina G Malik, Khin Saw Myint, Amin Soebandrio |
| EPI_ISL_2101316 | RSUD Mangusada | Eijkman Institute for Molecular Biology, National Agency for Research and Innovation; Molecular Biology Laboratory, Faculty Medicine and Health Sciences, Warmadewa University | Willy Agustine, Hidayat Trimarsanto, Lydia V. Panggalo, Iskandar Adnan, Sukma Oktavianthi, Lidwina Priliani, Edison Johar, Frilasita A Yudhaputri, Muhammad Rezki Rasyak, Sri Masyeni, Erly Sintya, Ida Ayu Wayan Mahayani, Safarina G Malik, Khin Saw Myint, Amin Soebandrio |
| EPI_ISL_2101317 | RSUD Mangusada | Eijkman Institute for Molecular Biology, National Agency for Research and Innovation; Molecular Biology Laboratory, Faculty Medicine and Health Sciences, Warmadewa University | Hidayat Trimarsanto, Lydia V. Panggalo, Iskandar Adnan, Sukma Oktavianthi, Lidwina Priliani, Edison Johar, Frilasita A Yudhaputri, Muhammad Rezki Rasyak, Willy Agustine, Sri Masyeni, Erly Sintya, Ida Ayu Wayan Mahayani, Safarina G Malik, Khin Saw Myint, Amin Soebandrio |
| EPI_ISL_2101318 | RSUD Mangusada | Eijkman Institute for Molecular Biology, National Agency for Research and Innovation; Molecular Biology Laboratory, Faculty Medicine and Health Sciences, Warmadewa University | Iskandar Adnan, Sukma Oktavianthi, Lidwina Priliani, Edison Johar, Frilasita A Yudhaputri, Muhammad Rezki Rasyak, Willy Agustine, Hidayat Trimarsanto, Lydia V. Panggalo, Sri Masyeni, Erly Sintya, Ida Ayu Wayan Mahayani, Safarina G Malik, Khin Saw Myint, Amin Soebandrio |
| EPI_ISL_2101319 | RSUD Mangusada | Eijkman Institute for Molecular Biology, National Agency for Research and Innovation; Molecular Biology Laboratory, Faculty Medicine and Health Sciences, Warmadewa University | Lidwina Priliani, Edison Johar, Frilasita A Yudhaputri, Muhammad Rezki Rasyak, Willy Agustine, Hidayat Trimarsanto, Lydia V. Panggalo, Iskandar Adnan, Sukma Oktavianthi, Sri Masyeni, Erly Sintya, Ida Ayu Wayan Mahayani, Safarina G Malik, Khin Saw Myint, Amin Soebandrio |
| EPI_ISL_2101320 | RSUD Mangusada | Eijkman Institute for Molecular Biology, National Agency for Research and Innovation; Molecular Biology Laboratory, Faculty Medicine and Health Sciences, Warmadewa University | Edison Johar, Frilasita A Yudhaputri, Muhammad Rezki Rasyak, Willy Agustine, Hidayat Trimarsanto, Lydia V. Panggalo, Iskandar Adnan, Sukma Oktavianthi, Lidwina Priliani, Sri Masyeni, Erly Sintya, Ida Ayu Wayan Mahayani, Safarina G Malik, Khin Saw Myint, Amin Soebandrio |
| EPI_ISL_2101321 | RSUD Mangusada | Eijkman Institute for Molecular Biology, National Agency for Research and Innovation; Molecular Biology Laboratory, Faculty Medicine and Health Sciences, Warmadewa University | Frilasita A Yudhaputri, Muhammad Rezki Rasyak, Willy Agustine, Hidayat Trimarsanto, Lydia V. Panggalo, Iskandar Adnan, Sukma Oktavianthi, Lidwina Priliani, Edison Johar, Sri Masyeni, Erly Sintya, Ida Ayu Wayan Mahayani, Safarina G Malik, Khin Saw Myint, Amin Soebandrio |
| EPI_ISL_2101322 | RSUD Mangusada | Eijkman Institute for Molecular Biology, National Agency for Research and Innovation; Molecular Biology Laboratory, Faculty Medicine and Health Sciences, Warmadewa University | Muhammad Rezki Rasyak, Willy Agustine, Hidayat Trimarsanto, Lydia V. Panggalo, Iskandar Adnan, Sukma Oktavianthi, Lidwina Priliani, Edison Johar, Frilasita A Yudhaputri, Sri Masyeni, Erly Sintya, Ida Ayu Wayan Mahayani, Safarina G Malik, Khin Saw Myint, Amin Soebandrio |
| EPI_ISL_2101323 | RSUD Mangusada | Eijkman Institute for Molecular Biology, National Agency for Research and Innovation; Molecular Biology Laboratory, Faculty Medicine and Health Sciences, Warmadewa University | Willy Agustine, Hidayat Trimarsanto, Lydia V. Panggalo, Iskandar Adnan, Sukma Oktavianthi, Lidwina Priliani, Edison Johar, Frilasita A Yudhaputri, Muhammad Rezki Rasyak, Sri Masyeni, Erly Sintya, Ida Ayu Wayan Mahayani, Safarina G Malik, Khin Saw Myint, Amin Soebandrio |
| EPI_ISL_2105287 | San Antonio City of Ilagan Hospital | Philippine Genome Center | Francis A. Tablizo, Kenneth M. Kim, Carlo M. Lapid, Marc Jerrone R. Castro, Maria Sofia L. Yangzon, Benedict A. Maralit, Marc Edsel C. Ayes, Eva Maria Cutiongco-de la Paz, Alethea R. de Guzman, Jan Michael C. Yap, Jo-Hannah S. Llames, Sheila Mae M. Araiza, Kris P. Punayan, Irish Coleen A. Asin, Candice Francheska B. Tambaoan, Asia Louisa U. Chong, Karol Sophia Agape R. Padilla, Rianna Patricia S. Cruz, El King D. Morado, Joshua Gregor A. Dizon, Razel Nikka M. Hao, Arianne A. Zamora, Devon Ray Pacial, Juan Antonio R. Magalang, Marissa Alejandria, Celia Carlos, Anna Ong-Lim, Edsel Maurice Salvaña, John Q. Wong, Jaime C. Montoya, Maria Rosario Singh-Vergeire and Cynthia P. Saloma |
| EPI_ISL_2105290 | Southern Philippines Medical Center (SPMC-SNL-MBL) | Philippine Genome Center | Francis A. Tablizo, Kenneth M. Kim, Carlo M. Lapid, Marc Jerrone R. Castro, Maria Sofia L. Yangzon, Benedict A. Maralit, Marc Edsel C. Ayes, Eva Maria Cutiongco-de la Paz, Alethea R. de Guzman, Jan Michael C. Yap, Jo-Hannah S. Llames, Sheila Mae M. Araiza, Kris P. Punayan, Irish Coleen A. Asin, Candice Francheska B. Tambaoan, Asia Louisa U. Chong, Karol Sophia Agape R. Padilla, Rianna Patricia S. Cruz, El King D. Morado, Joshua Gregor A. Dizon, Razel Nikka M. Hao, Arianne A. Zamora, Devon Ray Pacial, Juan Antonio R. Magalang, Marissa Alejandria, Celia Carlos, Anna Ong-Lim, Edsel Maurice Salvaña, John Q. Wong, Jaime C. Montoya, Maria Rosario Singh-Vergeire and Cynthia P. Saloma |
| EPI_ISL_2105292 | The Medical City -Â Ortigas | Philippine Genome Center | Francis A. Tablizo, Kenneth M. Kim, Carlo M. Lapid, Marc Jerrone R. Castro, Maria Sofia L. Yangzon, Benedict A. Maralit, Marc Edsel C. Ayes, Eva Maria Cutiongco-de la Paz, Alethea R. de Guzman, Jan Michael C. Yap, Jo-Hannah S. Llames, Sheila Mae M. Araiza, Kris P. Punayan, Irish Coleen A. Asin, Candice Francheska B. Tambaoan, Asia Louisa U. Chong, Karol Sophia Agape R. Padilla, Rianna Patricia S. Cruz, El King D. Morado, Joshua Gregor A. Dizon, Razel Nikka M. Hao, Arianne A. Zamora, Devon Ray Pacial, Juan Antonio R. Magalang, Marissa Alejandria, Celia Carlos, Anna Ong-Lim, Edsel Maurice Salvaña, John Q. Wong, Jaime C. Montoya, Maria Rosario Singh-Vergeire and Cynthia P. Saloma |
| EPI_ISL_2105293 | Southern Philippines Medical Center (SPMC-SNL-MBL) | Philippine Genome Center | Francis A. Tablizo, Kenneth M. Kim, Carlo M. Lapid, Marc Jerrone R. Castro, Maria Sofia L. Yangzon, Benedict A. Maralit, Marc Edsel C. Ayes, Eva Maria Cutiongco-de la Paz, Alethea R. de Guzman, Jan Michael C. Yap, Jo-Hannah S. Llames, Sheila Mae M. Araiza, Kris P. Punayan, Irish Coleen A. Asin, Candice Francheska B. Tambaoan, Asia Louisa U. Chong, Karol Sophia Agape R. Padilla, Rianna Patricia S. Cruz, El King D. Morado, Joshua Gregor A. Dizon, Razel Nikka M. Hao, Arianne A. Zamora, Devon Ray Pacial, Juan Antonio R. Magalang, Marissa Alejandria, Celia Carlos, Anna Ong-Lim, Edsel Maurice Salvaña, John Q. Wong, Jaime C. Montoya, Maria Rosario Singh-Vergeire and Cynthia P. Saloma |
| EPI_ISL_2105299 | Cagayan Valley Medical Center Molecular Laboratory | Philippine Genome Center | Francis A. Tablizo, Kenneth M. Kim, Carlo M. Lapid, Marc Jerrone R. Castro, Maria Sofia L. Yangzon, Benedict A. Maralit, Marc Edsel C. Ayes, Eva Maria Cutiongco-de la Paz, Alethea R. de Guzman, Jan Michael C. Yap, Jo-Hannah S. Llames, Sheila Mae M. Araiza, Kris P. Punayan, Irish Coleen A. Asin, Candice Francheska B. Tambaoan, Asia Louisa U. Chong, Karol Sophia Agape R. Padilla, Rianna Patricia S. Cruz, El King D. Morado, Joshua Gregor A. Dizon, Razel Nikka M. Hao, Arianne A. Zamora, Devon Ray Pacial, Juan Antonio R. Magalang, Marissa Alejandria, Celia Carlos, Anna Ong-Lim, Edsel Maurice Salvaña, John Q. Wong, Jaime C. Montoya, Maria Rosario Singh-Vergeire and Cynthia P. Saloma |
| EPI_ISL_2105301, EPI_ISL_2105305, EPI_ISL_2105307, EPI_ISL_2105308, EPI_ISL_2105314, EPI_ISL_2105319 | Southern Philippines Medical Center (SPMC-SNL-MBL) | Philippine Genome Center | Francis A. Tablizo, Kenneth M. Kim, Carlo M. Lapid, Marc Jerrone R. Castro, Maria Sofia L. Yangzon, Benedict A. Maralit, Marc Edsel C. Ayes, Eva Maria Cutiongco-de la Paz, Alethea R. de Guzman, Jan Michael C. Yap, Jo-Hannah S. Llames, Sheila Mae M. Araiza, Kris P. Punayan, Irish Coleen A. Asin, Candice Francheska B. Tambaoan, Asia Louisa U. Chong, Karol Sophia Agape R. Padilla, Rianna Patricia S. Cruz, El King D. Morado, Joshua Gregor A. Dizon, Razel Nikka M. Hao, Arianne A. Zamora, Devon Ray Pacial, Juan Antonio R. Magalang, Marissa Alejandria, Celia Carlos, Anna Ong-Lim, Edsel Maurice Salvaña, John Q. Wong, Jaime C. Montoya, Maria Rosario Singh-Vergeire and Cynthia P. Saloma |

| | | | |
|---|---|---|---|
| EPI_ISL_2105322 | Cagayan Valley Medical Center Molecular Laboratory | Philippine Genome Center | Francis A. Tablizo, Kenneth M. Kim, Carlo M. Lapid, Marc Jerrone R. Castro, Maria Sofia L. Yangzon, Benedict A. Maralit, Marc Edsel C. Ayes, Eva Maria Cutiongco-de la Paz, Alethea R. de Guzman, Jan Michael C. Yap, Jo-Hannah S. Llames, Sheila Mae M. Araiza, Kris P. Punayan, Irish Coleen A. Asin, Candice Franceska B. Tambaoan, Asia Louisa U. Chong, Karol Sophia Agape R. Padilla, Rianna Patricia S. Cruz, El King D. Morado, Joshua Gregor A. Dizon, Razel Nikka M. Hao, Arianne A. Zamora, Devon Ray Pacial, Juan Antonio R. Magalang, Marissa Alejandria, Celia Carlos, Anna Ong-Lim, Edsel Maurice Salvaña, John Q. Wong, Jaime C. Montoya, Maria Rosario Singh-Vergeire and Cynthia P. Saloma |
| EPI_ISL_2105330 | Southern Philippines Medical Center (SPMC-SNL-MBL) | Philippine Genome Center | Francis A. Tablizo, Kenneth M. Kim, Carlo M. Lapid, Marc Jerrone R. Castro, Maria Sofia L. Yangzon, Benedict A. Maralit, Marc Edsel C. Ayes, Eva Maria Cutiongco-de la Paz, Alethea R. de Guzman, Jan Michael C. Yap, Jo-Hannah S. Llames, Sheila Mae M. Araiza, Kris P. Punayan, Irish Coleen A. Asin, Candice Franceska B. Tambaoan, Asia Louisa U. Chong, Karol Sophia Agape R. Padilla, Rianna Patricia S. Cruz, El King D. Morado, Joshua Gregor A. Dizon, Razel Nikka M. Hao, Arianne A. Zamora, Devon Ray Pacial, Juan Antonio R. Magalang, Marissa Alejandria, Celia Carlos, Anna Ong-Lim, Edsel Maurice Salvaña, John Q. Wong, Jaime C. Montoya, Maria Rosario Singh-Vergeire and Cynthia P. Saloma |
| EPI_ISL_2105331 | Davao One World Diagnostic Center Incorporated | Philippine Genome Center | Francis A. Tablizo, Kenneth M. Kim, Carlo M. Lapid, Marc Jerrone R. Castro, Maria Sofia L. Yangzon, Benedict A. Maralit, Marc Edsel C. Ayes, Eva Maria Cutiongco-de la Paz, Alethea R. de Guzman, Jan Michael C. Yap, Jo-Hannah S. Llames, Sheila Mae M. Araiza, Kris P. Punayan, Irish Coleen A. Asin, Candice Franceska B. Tambaoan, Asia Louisa U. Chong, Karol Sophia Agape R. Padilla, Rianna Patricia S. Cruz, El King D. Morado, Joshua Gregor A. Dizon, Razel Nikka M. Hao, Arianne A. Zamora, Devon Ray Pacial, Juan Antonio R. Magalang, Marissa Alejandria, Celia Carlos, Anna Ong-Lim, Edsel Maurice Salvaña, John Q. Wong, Jaime C. Montoya, Maria Rosario Singh-Vergeire and Cynthia P. Saloma |
| EPI_ISL_2105334 | Southern Philippines Medical Center (SPMC-SNL-MBL) | Philippine Genome Center | Francis A. Tablizo, Kenneth M. Kim, Carlo M. Lapid, Marc Jerrone R. Castro, Maria Sofia L. Yangzon, Benedict A. Maralit, Marc Edsel C. Ayes, Eva Maria Cutiongco-de la Paz, Alethea R. de Guzman, Jan Michael C. Yap, Jo-Hannah S. Llames, Sheila Mae M. Araiza, Kris P. Punayan, Irish Coleen A. Asin, Candice Franceska B. Tambaoan, Asia Louisa U. Chong, Karol Sophia Agape R. Padilla, Rianna Patricia S. Cruz, El King D. Morado, Joshua Gregor A. Dizon, Razel Nikka M. Hao, Arianne A. Zamora, Devon Ray Pacial, Juan Antonio R. Magalang, Marissa Alejandria, Celia Carlos, Anna Ong-Lim, Edsel Maurice Salvaña, John Q. Wong, Jaime C. Montoya, Maria Rosario Singh-Vergeire and Cynthia P. Saloma |
| EPI_ISL_2105335 | The Medical City -Â Ortigas | Philippine Genome Center | Francis A. Tablizo, Kenneth M. Kim, Carlo M. Lapid, Marc Jerrone R. Castro, Maria Sofia L. Yangzon, Benedict A. Maralit, Marc Edsel C. Ayes, Eva Maria Cutiongco-de la Paz, Alethea R. de Guzman, Jan Michael C. Yap, Jo-Hannah S. Llames, Sheila Mae M. Araiza, Kris P. Punayan, Irish Coleen A. Asin, Candice Franceska B. Tambaoan, Asia Louisa U. Chong, Karol Sophia Agape R. Padilla, Rianna Patricia S. Cruz, El King D. Morado, Joshua Gregor A. Dizon, Razel Nikka M. Hao, Arianne A. Zamora, Devon Ray Pacial, Juan Antonio R. Magalang, Marissa Alejandria, Celia Carlos, Anna Ong-Lim, Edsel Maurice Salvaña, John Q. Wong, Jaime C. Montoya, Maria Rosario Singh-Vergeire and Cynthia P. Saloma |
| EPI_ISL_2105338 | Southern Philippines Medical Center (SPMC-SNL-MBL) | Philippine Genome Center | Francis A. Tablizo, Kenneth M. Kim, Carlo M. Lapid, Marc Jerrone R. Castro, Maria Sofia L. Yangzon, Benedict A. Maralit, Marc Edsel C. Ayes, Eva Maria Cutiongco-de la Paz, Alethea R. de Guzman, Jan Michael C. Yap, Jo-Hannah S. Llames, Sheila Mae M. Araiza, Kris P. Punayan, Irish Coleen A. Asin, Candice Franceska B. Tambaoan, Asia Louisa U. Chong, Karol Sophia Agape R. Padilla, Rianna Patricia S. Cruz, El King D. Morado, Joshua Gregor A. Dizon, Razel Nikka M. Hao, Arianne A. Zamora, Devon Ray Pacial, Juan Antonio R. Magalang, Marissa Alejandria, Celia Carlos, Anna Ong-Lim, Edsel Maurice Salvaña, John Q. Wong, Jaime C. Montoya, Maria Rosario Singh-Vergeire and Cynthia P. Saloma |
| EPI_ISL_2105340 | Safeguard DNA Diagnostic, Inc. | Philippine Genome Center | Francis A. Tablizo, Kenneth M. Kim, Carlo M. Lapid, Marc Jerrone R. Castro, Maria Sofia L. Yangzon, Benedict A. Maralit, Marc Edsel C. Ayes, Eva Maria Cutiongco-de la Paz, Alethea R. de Guzman, Jan Michael C. Yap, Jo-Hannah S. Llames, Sheila Mae M. Araiza, Kris P. Punayan, Irish Coleen A. Asin, Candice Franceska B. Tambaoan, Asia Louisa U. Chong, Karol Sophia Agape R. Padilla, Rianna Patricia S. Cruz, El King D. Morado, Joshua Gregor A. Dizon, Razel Nikka M. Hao, Arianne A. Zamora, Devon Ray Pacial, Juan Antonio R. Magalang, Marissa Alejandria, Celia Carlos, Anna Ong-Lim, Edsel Maurice Salvaña, John Q. Wong, Jaime C. Montoya, Maria Rosario Singh-Vergeire and Cynthia P. Saloma |
| EPI_ISL_2105341 | Valenzuela Hope Molecular Laboratory | Philippine Genome Center | Francis A. Tablizo, Kenneth M. Kim, Carlo M. Lapid, Marc Jerrone R. Castro, Maria Sofia L. Yangzon, Benedict A. Maralit, Marc Edsel C. Ayes, Eva Maria Cutiongco-de la Paz, Alethea R. de Guzman, Jan Michael C. Yap, Jo-Hannah S. Llames, Sheila Mae M. Araiza, Kris P. Punayan, Irish Coleen A. Asin, Candice Franceska B. Tambaoan, Asia Louisa U. Chong, Karol Sophia Agape R. Padilla, Rianna Patricia S. Cruz, El King D. Morado, Joshua Gregor A. Dizon, Razel Nikka M. Hao, Arianne A. Zamora, Devon Ray Pacial, Juan Antonio R. Magalang, Marissa Alejandria, Celia Carlos, Anna Ong-Lim, Edsel Maurice Salvaña, John Q. Wong, Jaime C. Montoya, Maria Rosario Singh-Vergeire and Cynthia P. Saloma |
| EPI_ISL_2105342, EPI_ISL_2105343, EPI_ISL_2105345, EPI_ISL_2105350, EPI_ISL_2105354, EPI_ISL_2105357, EPI_ISL_2105359, EPI_ISL_2105364, EPI_ISL_2105368 | Southern Philippines Medical Center (SPMC-SNL-MBL) | Philippine Genome Center | Francis A. Tablizo, Kenneth M. Kim, Carlo M. Lapid, Marc Jerrone R. Castro, Maria Sofia L. Yangzon, Benedict A. Maralit, Marc Edsel C. Ayes, Eva Maria Cutiongco-de la Paz, Alethea R. de Guzman, Jan Michael C. Yap, Jo-Hannah S. Llames, Sheila Mae M. Araiza, Kris P. Punayan, Irish Coleen A. Asin, Candice Franceska B. Tambaoan, Asia Louisa U. Chong, Karol Sophia Agape R. Padilla, Rianna Patricia S. Cruz, El King D. Morado, Joshua Gregor A. Dizon, Razel Nikka M. Hao, Arianne A. Zamora, Devon Ray Pacial, Juan Antonio R. Magalang, Marissa Alejandria, Celia Carlos, Anna Ong-Lim, Edsel Maurice Salvaña, John Q. Wong, Jaime C. Montoya, Maria Rosario Singh-Vergeire and Cynthia P. Saloma |
| EPI_ISL_2105369 | The Medical City -Â Ortigas | Philippine Genome Center | Francis A. Tablizo, Kenneth M. Kim, Carlo M. Lapid, Marc Jerrone R. Castro, Maria Sofia L. Yangzon, Benedict A. Maralit, Marc Edsel C. Ayes, Eva Maria Cutiongco-de la Paz, Alethea R. de Guzman, Jan Michael C. Yap, Jo-Hannah S. Llames, Sheila Mae M. Araiza, Kris P. Punayan, Irish Coleen A. Asin, Candice Franceska B. Tambaoan, Asia Louisa U. Chong, Karol Sophia Agape R. Padilla, Rianna Patricia S. Cruz, El King D. Morado, Joshua Gregor A. Dizon, Razel Nikka M. Hao, Arianne A. Zamora, Devon Ray Pacial, Juan Antonio R. Magalang, Marissa Alejandria, Celia Carlos, Anna Ong-Lim, Edsel Maurice Salvaña, John Q. Wong, Jaime C. Montoya, Maria Rosario Singh-Vergeire and Cynthia P. Saloma |
| EPI_ISL_2105372, EPI_ISL_2105376 | Southern Philippines Medical Center (SPMC-SNL-MBL) | Philippine Genome Center | Francis A. Tablizo, Kenneth M. Kim, Carlo M. Lapid, Marc Jerrone R. Castro, Maria Sofia L. Yangzon, Benedict A. Maralit, Marc Edsel C. Ayes, Eva Maria Cutiongco-de la Paz, Alethea R. de Guzman, Jan Michael C. Yap, Jo-Hannah S. Llames, Sheila Mae M. Araiza, Kris P. Punayan, Irish Coleen A. Asin, Candice Franceska B. Tambaoan, Asia Louisa U. Chong, Karol Sophia Agape R. Padilla, Rianna Patricia S. Cruz, El King D. Morado, Joshua Gregor A. Dizon, Razel Nikka M. Hao, Arianne A. Zamora, Devon Ray Pacial, Juan Antonio R. Magalang, Marissa Alejandria, Celia Carlos, Anna Ong-Lim, Edsel Maurice Salvaña, John Q. Wong, Jaime C. Montoya, Maria Rosario Singh-Vergeire and Cynthia P. Saloma |
| EPI_ISL_2105383 | Cebu TB Reference Laboratory | Philippine Genome Center | Francis A. Tablizo, Kenneth M. Kim, Carlo M. Lapid, Marc Jerrone R. Castro, Maria Sofia L. Yangzon, Benedict A. Maralit, Marc Edsel C. Ayes, Eva Maria Cutiongco-de la Paz, Alethea R. de Guzman, Jan Michael C. Yap, Jo-Hannah S. Llames, Sheila Mae M. Araiza, Kris P. Punayan, Irish Coleen A. Asin, Candice Franceska B. Tambaoan, Asia Louisa U. Chong, Karol Sophia Agape R. Padilla, Rianna Patricia S. Cruz, El King D. Morado, Joshua Gregor A. Dizon, Razel Nikka M. Hao, Arianne A. Zamora, Devon Ray Pacial, Juan Antonio R. Magalang, Marissa Alejandria, Celia Carlos, Anna Ong-Lim, Edsel Maurice Salvaña, John Q. Wong, Jaime C. Montoya, Maria Rosario Singh-Vergeire and Cynthia P. Saloma |
| EPI_ISL_2105385 | Southern Philippines Medical Center (SPMC-SNL-MBL) | Philippine Genome Center | Francis A. Tablizo, Kenneth M. Kim, Carlo M. Lapid, Marc Jerrone R. Castro, Maria Sofia L. Yangzon, Benedict A. Maralit, Marc Edsel C. Ayes, Eva Maria Cutiongco-de la Paz, Alethea R. de Guzman, Jan Michael C. Yap, Jo-Hannah S. Llames, Sheila Mae M. Araiza, Kris P. Punayan, Irish Coleen A. Asin, Candice Franceska B. Tambaoan, Asia Louisa U. Chong, Karol Sophia Agape R. Padilla, Rianna Patricia S. Cruz, El King D. Morado, Joshua Gregor A. Dizon, Razel Nikka M. Hao, Arianne A. Zamora, Devon Ray Pacial, Juan Antonio R. Magalang, Marissa Alejandria, Celia Carlos, Anna Ong-Lim, Edsel Maurice Salvaña, John Q. Wong, Jaime C. Montoya, Maria Rosario Singh-Vergeire and Cynthia P. Saloma |
| EPI_ISL_2105391 | University of the Philippines National Institutes of Health (UP NIH) | Philippine Genome Center | Francis A. Tablizo, Kenneth M. Kim, Carlo M. Lapid, Marc Jerrone R. Castro, Maria Sofia L. Yangzon, Benedict A. Maralit, Marc Edsel C. Ayes, Eva Maria Cutiongco-de la Paz, Alethea R. de Guzman, Jan Michael C. Yap, Jo-Hannah S. Llames, Sheila Mae M. Araiza, Kris P. Punayan, Irish Coleen A. Asin, Candice Franceska B. Tambaoan, Asia Louisa U. Chong, Karol Sophia Agape R. Padilla, Rianna Patricia S. Cruz, El King D. Morado, Joshua Gregor A. Dizon, Razel Nikka M. Hao, Arianne A. Zamora, Devon Ray Pacial, Juan Antonio R. Magalang, Marissa Alejandria, Celia Carlos, Anna Ong-Lim, Edsel Maurice Salvaña, John Q. Wong, Jaime C. Montoya, Maria Rosario Singh-Vergeire and Cynthia P. Saloma |
| EPI_ISL_2105394 | Safeguard DNA Diagnostic, Inc. | Philippine Genome Center | Francis A. Tablizo, Kenneth M. Kim, Carlo M. Lapid, Marc Jerrone R. Castro, Maria Sofia L. Yangzon, Benedict A. Maralit, Marc Edsel C. Ayes, Eva Maria Cutiongco-de la Paz, Alethea R. de Guzman, Jan Michael C. Yap, Jo-Hannah S. Llames, Sheila Mae M. Araiza, Kris P. Punayan, Irish Coleen A. Asin, Candice Franceska B. Tambaoan, Asia Louisa U. Chong, Karol Sophia Agape R. Padilla, Rianna Patricia S. Cruz, El King D. Morado, Joshua Gregor A. Dizon, Razel Nikka M. Hao, Arianne A. Zamora, Devon Ray Pacial, Juan Antonio R. Magalang, Marissa Alejandria, Celia Carlos, Anna Ong-Lim, Edsel Maurice Salvaña, John Q. Wong, Jaime C. Montoya, Maria Rosario Singh-Vergeire and Cynthia P. Saloma |
| EPI_ISL_2105395, EPI_ISL_2105396 | Davao One World Diagnostic Center Incorporated | Philippine Genome Center | Francis A. Tablizo, Kenneth M. Kim, Carlo M. Lapid, Marc Jerrone R. Castro, Maria Sofia L. Yangzon, Benedict A. Maralit, Marc Edsel C. Ayes, Eva Maria Cutiongco-de la Paz, Alethea R. de Guzman, Jan Michael C. Yap, Jo-Hannah S. Llames, Sheila Mae M. Araiza, Kris P. Punayan, Irish Coleen A. Asin, Candice Franceska B. Tambaoan, Asia Louisa U. Chong, Karol Sophia Agape R. Padilla, Rianna Patricia S. Cruz, El King D. Morado, Joshua Gregor A. Dizon, Razel Nikka M. Hao, Arianne A. Zamora, Devon Ray Pacial, Juan Antonio R. Magalang, Marissa Alejandria, Celia Carlos, Anna Ong-Lim, Edsel Maurice Salvaña, John Q. Wong, Jaime C. Montoya, Maria Rosario Singh-Vergeire and Cynthia P. Saloma |

| | | | |
|---|---|---|---|
| EPI_ISL_2105401 | Northern Mindanao TB Regional Center | Philippine Genome Center | Francis A. Tablizo, Kenneth M. Kim, Carlo M. Lapid, Marc Jerrone R. Castro, Maria Sofia L. Yangzon, Benedict A. Maralit, Marc Edsel C. Ayes, Eva Maria Cutiongco-de la Paz, Alethea R. de Guzman, Jan Michael C. Yap, Jo-Hannah S. Llames, Sheila Mae M. Araiza, Kris P. Punayan, Irish Coleen A. Asin, Candice Francheska B. Tambaoan, Asia Louisa U. Chong, Karol Sophia Agape R. Padilla, Rianna Patricia S. Cruz, El King D. Morado, Joshua Gregor A. Dizon, Razel Nikka M. Hao, Arianne A. Zamora, Devon Ray Pacial, Juan Antonio R. Magalang, Marissa Alejandria, Celia Carlos, Anna Ong-Lim, Edsel Maurice Salvaña, John Q. Wong, Jaime C. Montoya, Maria Rosario Singh-Vergeire and Cynthia P. Saloma |
| EPI_ISL_2105405 | Philippine Red Cross - Misamis Oriental | Philippine Genome Center | Francis A. Tablizo, Kenneth M. Kim, Carlo M. Lapid, Marc Jerrone R. Castro, Maria Sofia L. Yangzon, Benedict A. Maralit, Marc Edsel C. Ayes, Eva Maria Cutiongco-de la Paz, Alethea R. de Guzman, Jan Michael C. Yap, Jo-Hannah S. Llames, Sheila Mae M. Araiza, Kris P. Punayan, Irish Coleen A. Asin, Candice Francheska B. Tambaoan, Asia Louisa U. Chong, Karol Sophia Agape R. Padilla, Rianna Patricia S. Cruz, El King D. Morado, Joshua Gregor A. Dizon, Razel Nikka M. Hao, Arianne A. Zamora, Devon Ray Pacial, Juan Antonio R. Magalang, Marissa Alejandria, Celia Carlos, Anna Ong-Lim, Edsel Maurice Salvaña, John Q. Wong, Jaime C. Montoya, Maria Rosario Singh-Vergeire and Cynthia P. Saloma |
| EPI_ISL_2105410 | Negros Oriental Provincial Hospital | Philippine Genome Center | Francis A. Tablizo, Kenneth M. Kim, Carlo M. Lapid, Marc Jerrone R. Castro, Maria Sofia L. Yangzon, Benedict A. Maralit, Marc Edsel C. Ayes, Eva Maria Cutiongco-de la Paz, Alethea R. de Guzman, Jan Michael C. Yap, Jo-Hannah S. Llames, Sheila Mae M. Araiza, Kris P. Punayan, Irish Coleen A. Asin, Candice Francheska B. Tambaoan, Asia Louisa U. Chong, Karol Sophia Agape R. Padilla, Rianna Patricia S. Cruz, El King D. Morado, Joshua Gregor A. Dizon, Razel Nikka M. Hao, Arianne A. Zamora, Devon Ray Pacial, Juan Antonio R. Magalang, Marissa Alejandria, Celia Carlos, Anna Ong-Lim, Edsel Maurice Salvaña, John Q. Wong, Jaime C. Montoya, Maria Rosario Singh-Vergeire and Cynthia P. Saloma |
| EPI_ISL_2105411 | The Medical City -Â Ortigas | Philippine Genome Center | Francis A. Tablizo, Kenneth M. Kim, Carlo M. Lapid, Marc Jerrone R. Castro, Maria Sofia L. Yangzon, Benedict A. Maralit, Marc Edsel C. Ayes, Eva Maria Cutiongco-de la Paz, Alethea R. de Guzman, Jan Michael C. Yap, Jo-Hannah S. Llames, Sheila Mae M. Araiza, Kris P. Punayan, Irish Coleen A. Asin, Candice Francheska B. Tambaoan, Asia Louisa U. Chong, Karol Sophia Agape R. Padilla, Rianna Patricia S. Cruz, El King D. Morado, Joshua Gregor A. Dizon, Razel Nikka M. Hao, Arianne A. Zamora, Devon Ray Pacial, Juan Antonio R. Magalang, Marissa Alejandria, Celia Carlos, Anna Ong-Lim, Edsel Maurice Salvaña, John Q. Wong, Jaime C. Montoya, Maria Rosario Singh-Vergeire and Cynthia P. Saloma |
| EPI_ISL_2105414 | Butuan Medical Center | Philippine Genome Center | Francis A. Tablizo, Kenneth M. Kim, Carlo M. Lapid, Marc Jerrone R. Castro, Maria Sofia L. Yangzon, Benedict A. Maralit, Marc Edsel C. Ayes, Eva Maria Cutiongco-de la Paz, Alethea R. de Guzman, Jan Michael C. Yap, Jo-Hannah S. Llames, Sheila Mae M. Araiza, Kris P. Punayan, Irish Coleen A. Asin, Candice Francheska B. Tambaoan, Asia Louisa U. Chong, Karol Sophia Agape R. Padilla, Rianna Patricia S. Cruz, El King D. Morado, Joshua Gregor A. Dizon, Razel Nikka M. Hao, Arianne A. Zamora, Devon Ray Pacial, Juan Antonio R. Magalang, Marissa Alejandria, Celia Carlos, Anna Ong-Lim, Edsel Maurice Salvaña, John Q. Wong, Jaime C. Montoya, Maria Rosario Singh-Vergeire and Cynthia P. Saloma |
| EPI_ISL_2105420 | Philippine Red Cross - Misamis Oriental | Philippine Genome Center | Francis A. Tablizo, Kenneth M. Kim, Carlo M. Lapid, Marc Jerrone R. Castro, Maria Sofia L. Yangzon, Benedict A. Maralit, Marc Edsel C. Ayes, Eva Maria Cutiongco-de la Paz, Alethea R. de Guzman, Jan Michael C. Yap, Jo-Hannah S. Llames, Sheila Mae M. Araiza, Kris P. Punayan, Irish Coleen A. Asin, Candice Francheska B. Tambaoan, Asia Louisa U. Chong, Karol Sophia Agape R. Padilla, Rianna Patricia S. Cruz, El King D. Morado, Joshua Gregor A. Dizon, Razel Nikka M. Hao, Arianne A. Zamora, Devon Ray Pacial, Juan Antonio R. Magalang, Marissa Alejandria, Celia Carlos, Anna Ong-Lim, Edsel Maurice Salvaña, John Q. Wong, Jaime C. Montoya, Maria Rosario Singh-Vergeire and Cynthia P. Saloma |
| EPI_ISL_2105421 | Baguio General Hospital Medical Center (BGHMC) | Philippine Genome Center | Francis A. Tablizo, Kenneth M. Kim, Carlo M. Lapid, Marc Jerrone R. Castro, Maria Sofia L. Yangzon, Benedict A. Maralit, Marc Edsel C. Ayes, Eva Maria Cutiongco-de la Paz, Alethea R. de Guzman, Jan Michael C. Yap, Jo-Hannah S. Llames, Sheila Mae M. Araiza, Kris P. Punayan, Irish Coleen A. Asin, Candice Francheska B. Tambaoan, Asia Louisa U. Chong, Karol Sophia Agape R. Padilla, Rianna Patricia S. Cruz, El King D. Morado, Joshua Gregor A. Dizon, Razel Nikka M. Hao, Arianne A. Zamora, Devon Ray Pacial, Juan Antonio R. Magalang, Marissa Alejandria, Celia Carlos, Anna Ong-Lim, Edsel Maurice Salvaña, John Q. Wong, Jaime C. Montoya, Maria Rosario Singh-Vergeire and Cynthia P. Saloma |
| EPI_ISL_2105424 | Philippine Red Cross Logistics and Multipurpose Center | Philippine Genome Center | Francis A. Tablizo, Kenneth M. Kim, Carlo M. Lapid, Marc Jerrone R. Castro, Maria Sofia L. Yangzon, Benedict A. Maralit, Marc Edsel C. Ayes, Eva Maria Cutiongco-de la Paz, Alethea R. de Guzman, Jan Michael C. Yap, Jo-Hannah S. Llames, Sheila Mae M. Araiza, Kris P. Punayan, Irish Coleen A. Asin, Candice Francheska B. Tambaoan, Asia Louisa U. Chong, Karol Sophia Agape R. Padilla, Rianna Patricia S. Cruz, El King D. Morado, Joshua Gregor A. Dizon, Razel Nikka M. Hao, Arianne A. Zamora, Devon Ray Pacial, Juan Antonio R. Magalang, Marissa Alejandria, Celia Carlos, Anna Ong-Lim, Edsel Maurice Salvaña, John Q. Wong, Jaime C. Montoya, Maria Rosario Singh-Vergeire and Cynthia P. Saloma |
| EPI_ISL_2105428 | Southern Philippines Medical Center (SPMC-SNL-MBL) | Philippine Genome Center | Francis A. Tablizo, Kenneth M. Kim, Carlo M. Lapid, Marc Jerrone R. Castro, Maria Sofia L. Yangzon, Benedict A. Maralit, Marc Edsel C. Ayes, Eva Maria Cutiongco-de la Paz, Alethea R. de Guzman, Jan Michael C. Yap, Jo-Hannah S. Llames, Sheila Mae M. Araiza, Kris P. Punayan, Irish Coleen A. Asin, Candice Francheska B. Tambaoan, Asia Louisa U. Chong, Karol Sophia Agape R. Padilla, Rianna Patricia S. Cruz, El King D. Morado, Joshua Gregor A. Dizon, Razel Nikka M. Hao, Arianne A. Zamora, Devon Ray Pacial, Juan Antonio R. Magalang, Marissa Alejandria, Celia Carlos, Anna Ong-Lim, Edsel Maurice Salvaña, John Q. Wong, Jaime C. Montoya, Maria Rosario Singh-Vergeire and Cynthia P. Saloma |
| EPI_ISL_2105432 | Philippine Red Cross - Misamis Oriental | Philippine Genome Center | Francis A. Tablizo, Kenneth M. Kim, Carlo M. Lapid, Marc Jerrone R. Castro, Maria Sofia L. Yangzon, Benedict A. Maralit, Marc Edsel C. Ayes, Eva Maria Cutiongco-de la Paz, Alethea R. de Guzman, Jan Michael C. Yap, Jo-Hannah S. Llames, Sheila Mae M. Araiza, Kris P. Punayan, Irish Coleen A. Asin, Candice Francheska B. Tambaoan, Asia Louisa U. Chong, Karol Sophia Agape R. Padilla, Rianna Patricia S. Cruz, El King D. Morado, Joshua Gregor A. Dizon, Razel Nikka M. Hao, Arianne A. Zamora, Devon Ray Pacial, Juan Antonio R. Magalang, Marissa Alejandria, Celia Carlos, Anna Ong-Lim, Edsel Maurice Salvaña, John Q. Wong, Jaime C. Montoya, Maria Rosario Singh-Vergeire and Cynthia P. Saloma |
| EPI_ISL_2105434 | Philippine Red Cross Logistics and Multipurpose Center | Philippine Genome Center | Francis A. Tablizo, Kenneth M. Kim, Carlo M. Lapid, Marc Jerrone R. Castro, Maria Sofia L. Yangzon, Benedict A. Maralit, Marc Edsel C. Ayes, Eva Maria Cutiongco-de la Paz, Alethea R. de Guzman, Jan Michael C. Yap, Jo-Hannah S. Llames, Sheila Mae M. Araiza, Kris P. Punayan, Irish Coleen A. Asin, Candice Francheska B. Tambaoan, Asia Louisa U. Chong, Karol Sophia Agape R. Padilla, Rianna Patricia S. Cruz, El King D. Morado, Joshua Gregor A. Dizon, Razel Nikka M. Hao, Arianne A. Zamora, Devon Ray Pacial, Juan Antonio R. Magalang, Marissa Alejandria, Celia Carlos, Anna Ong-Lim, Edsel Maurice Salvaña, John Q. Wong, Jaime C. Montoya, Maria Rosario Singh-Vergeire and Cynthia P. Saloma |
| EPI_ISL_2105439 | Negros Oriental Provincial Hospital | Philippine Genome Center | Francis A. Tablizo, Kenneth M. Kim, Carlo M. Lapid, Marc Jerrone R. Castro, Maria Sofia L. Yangzon, Benedict A. Maralit, Marc Edsel C. Ayes, Eva Maria Cutiongco-de la Paz, Alethea R. de Guzman, Jan Michael C. Yap, Jo-Hannah S. Llames, Sheila Mae M. Araiza, Kris P. Punayan, Irish Coleen A. Asin, Candice Francheska B. Tambaoan, Asia Louisa U. Chong, Karol Sophia Agape R. Padilla, Rianna Patricia S. Cruz, El King D. Morado, Joshua Gregor A. Dizon, Razel Nikka M. Hao, Arianne A. Zamora, Devon Ray Pacial, Juan Antonio R. Magalang, Marissa Alejandria, Celia Carlos, Anna Ong-Lim, Edsel Maurice Salvaña, John Q. Wong, Jaime C. Montoya, Maria Rosario Singh-Vergeire and Cynthia P. Saloma |
| EPI_ISL_2105446 | Philippine Red Cross - Misamis Oriental | Philippine Genome Center | Francis A. Tablizo, Kenneth M. Kim, Carlo M. Lapid, Marc Jerrone R. Castro, Maria Sofia L. Yangzon, Benedict A. Maralit, Marc Edsel C. Ayes, Eva Maria Cutiongco-de la Paz, Alethea R. de Guzman, Jan Michael C. Yap, Jo-Hannah S. Llames, Sheila Mae M. Araiza, Kris P. Punayan, Irish Coleen A. Asin, Candice Francheska B. Tambaoan, Asia Louisa U. Chong, Karol Sophia Agape R. Padilla, Rianna Patricia S. Cruz, El King D. Morado, Joshua Gregor A. Dizon, Razel Nikka M. Hao, Arianne A. Zamora, Devon Ray Pacial, Juan Antonio R. Magalang, Marissa Alejandria, Celia Carlos, Anna Ong-Lim, Edsel Maurice Salvaña, John Q. Wong, Jaime C. Montoya, Maria Rosario Singh-Vergeire and Cynthia P. Saloma |
| EPI_ISL_2105456 | Butuan Medical Center | Philippine Genome Center | Francis A. Tablizo, Kenneth M. Kim, Carlo M. Lapid, Marc Jerrone R. Castro, Maria Sofia L. Yangzon, Benedict A. Maralit, Marc Edsel C. Ayes, Eva Maria Cutiongco-de la Paz, Alethea R. de Guzman, Jan Michael C. Yap, Jo-Hannah S. Llames, Sheila Mae M. Araiza, Kris P. Punayan, Irish Coleen A. Asin, Candice Francheska B. Tambaoan, Asia Louisa U. Chong, Karol Sophia Agape R. Padilla, Rianna Patricia S. Cruz, El King D. Morado, Joshua Gregor A. Dizon, Razel Nikka M. Hao, Arianne A. Zamora, Devon Ray Pacial, Juan Antonio R. Magalang, Marissa Alejandria, Celia Carlos, Anna Ong-Lim, Edsel Maurice Salvaña, John Q. Wong, Jaime C. Montoya, Maria Rosario Singh-Vergeire and Cynthia P. Saloma |
| EPI_ISL_2105459 | Philippine Red Cross - Misamis Oriental | Philippine Genome Center | Francis A. Tablizo, Kenneth M. Kim, Carlo M. Lapid, Marc Jerrone R. Castro, Maria Sofia L. Yangzon, Benedict A. Maralit, Marc Edsel C. Ayes, Eva Maria Cutiongco-de la Paz, Alethea R. de Guzman, Jan Michael C. Yap, Jo-Hannah S. Llames, Sheila Mae M. Araiza, Kris P. Punayan, Irish Coleen A. Asin, Candice Francheska B. Tambaoan, Asia Louisa U. Chong, Karol Sophia Agape R. Padilla, Rianna Patricia S. Cruz, El King D. Morado, Joshua Gregor A. Dizon, Razel Nikka M. Hao, Arianne A. Zamora, Devon Ray Pacial, Juan Antonio R. Magalang, Marissa Alejandria, Celia Carlos, Anna Ong-Lim, Edsel Maurice Salvaña, John Q. Wong, Jaime C. Montoya, Maria Rosario Singh-Vergeire and Cynthia P. Saloma |
| EPI_ISL_2105461 | Research Institute for Tropical Medicine, Inc. (RITM) | Philippine Genome Center | Francis A. Tablizo, Kenneth M. Kim, Carlo M. Lapid, Marc Jerrone R. Castro, Maria Sofia L. Yangzon, Benedict A. Maralit, Marc Edsel C. Ayes, Eva Maria Cutiongco-de la Paz, Alethea R. de Guzman, Jan Michael C. Yap, Jo-Hannah S. Llames, Sheila Mae M. Araiza, Kris P. Punayan, Irish Coleen A. Asin, Candice Francheska B. Tambaoan, Asia Louisa U. Chong, Karol Sophia Agape R. Padilla, Rianna Patricia S. Cruz, El King D. Morado, Joshua Gregor A. Dizon, Razel Nikka M. Hao, Arianne A. Zamora, Devon Ray Pacial, Juan Antonio R. Magalang, Marissa Alejandria, Celia Carlos, Anna Ong-Lim, Edsel Maurice Salvaña, John Q. Wong, Jaime C. Montoya, Maria Rosario Singh-Vergeire and Cynthia P. Saloma |

| | | | |
|---|---|---|---|
| EPI_ISL_2105463, EPI_ISL_2105465, EPI_ISL_2105467, EPI_ISL_2105480, EPI_ISL_2105486, EPI_ISL_2105507 | Bohol Containerized PCR Laboratory | Philippine Genome Center | Francis A. Tablizo, Kenneth M. Kim, Carlo M. Lapid, Marc Jerrone R. Castro, Maria Sofia L. Yangzon, Benedict A. Maralit, Marc Edsel C. Ayes, Eva Maria Cutiongco-de la Paz, Alethea R. de Guzman, Jan Michael C. Yap, Jo-Hannah S. Llames, Sheila Mae M. Araiza, Kris P. Punayan, Irish Coleen A. Asin, Candice Francheska B. Tambaoan, Asia Louisa U. Chong, Karol Sophia Agape R. Padilla, Rianna Patricia S. Cruz, El King D. Morado, Joshua Gregor A. Dizon, Razel Nikka M. Hao, Arianne A. Zamora, Devon Ray Pacial, Juan Antonio R. Magalang, Marissa Alejandria, Celia Carlos, Anna Ong-Lim, Edsel Maurice Salvaña, John Q. Wong, Jaime C. Montoya, Maria Rosario Singh-Vergeire and Cynthia P. Saloma |
| EPI_ISL_2105522, EPI_ISL_2105524 | Butuan Medical Center | Philippine Genome Center | Francis A. Tablizo, Kenneth M. Kim, Carlo M. Lapid, Marc Jerrone R. Castro, Maria Sofia L. Yangzon, Benedict A. Maralit, Marc Edsel C. Ayes, Eva Maria Cutiongco-de la Paz, Alethea R. de Guzman, Jan Michael C. Yap, Jo-Hannah S. Llames, Sheila Mae M. Araiza, Kris P. Punayan, Irish Coleen A. Asin, Candice Francheska B. Tambaoan, Asia Louisa U. Chong, Karol Sophia Agape R. Padilla, Rianna Patricia S. Cruz, El King D. Morado, Joshua Gregor A. Dizon, Razel Nikka M. Hao, Arianne A. Zamora, Devon Ray Pacial, Juan Antonio R. Magalang, Marissa Alejandria, Celia Carlos, Anna Ong-Lim, Edsel Maurice Salvaña, John Q. Wong, Jaime C. Montoya, Maria Rosario Singh-Vergeire and Cynthia P. Saloma |
| EPI_ISL_2105529 | Cagayan Valley Medical Center Molecular Laboratory | Philippine Genome Center | Francis A. Tablizo, Kenneth M. Kim, Carlo M. Lapid, Marc Jerrone R. Castro, Maria Sofia L. Yangzon, Benedict A. Maralit, Marc Edsel C. Ayes, Eva Maria Cutiongco-de la Paz, Alethea R. de Guzman, Jan Michael C. Yap, Jo-Hannah S. Llames, Sheila Mae M. Araiza, Kris P. Punayan, Irish Coleen A. Asin, Candice Francheska B. Tambaoan, Asia Louisa U. Chong, Karol Sophia Agape R. Padilla, Rianna Patricia S. Cruz, El King D. Morado, Joshua Gregor A. Dizon, Razel Nikka M. Hao, Arianne A. Zamora, Devon Ray Pacial, Juan Antonio R. Magalang, Marissa Alejandria, Celia Carlos, Anna Ong-Lim, Edsel Maurice Salvaña, John Q. Wong, Jaime C. Montoya, Maria Rosario Singh-Vergeire and Cynthia P. Saloma |
| EPI_ISL_2105531 | Butuan Medical Center | Philippine Genome Center | Francis A. Tablizo, Kenneth M. Kim, Carlo M. Lapid, Marc Jerrone R. Castro, Maria Sofia L. Yangzon, Benedict A. Maralit, Marc Edsel C. Ayes, Eva Maria Cutiongco-de la Paz, Alethea R. de Guzman, Jan Michael C. Yap, Jo-Hannah S. Llames, Sheila Mae M. Araiza, Kris P. Punayan, Irish Coleen A. Asin, Candice Francheska B. Tambaoan, Asia Louisa U. Chong, Karol Sophia Agape R. Padilla, Rianna Patricia S. Cruz, El King D. Morado, Joshua Gregor A. Dizon, Razel Nikka M. Hao, Arianne A. Zamora, Devon Ray Pacial, Juan Antonio R. Magalang, Marissa Alejandria, Celia Carlos, Anna Ong-Lim, Edsel Maurice Salvaña, John Q. Wong, Jaime C. Montoya, Maria Rosario Singh-Vergeire and Cynthia P. Saloma |
| EPI_ISL_2105532 | Cagayan Valley Medical Center Molecular Laboratory | Philippine Genome Center | Francis A. Tablizo, Kenneth M. Kim, Carlo M. Lapid, Marc Jerrone R. Castro, Maria Sofia L. Yangzon, Benedict A. Maralit, Marc Edsel C. Ayes, Eva Maria Cutiongco-de la Paz, Alethea R. de Guzman, Jan Michael C. Yap, Jo-Hannah S. Llames, Sheila Mae M. Araiza, Kris P. Punayan, Irish Coleen A. Asin, Candice Francheska B. Tambaoan, Asia Louisa U. Chong, Karol Sophia Agape R. Padilla, Rianna Patricia S. Cruz, El King D. Morado, Joshua Gregor A. Dizon, Razel Nikka M. Hao, Arianne A. Zamora, Devon Ray Pacial, Juan Antonio R. Magalang, Marissa Alejandria, Celia Carlos, Anna Ong-Lim, Edsel Maurice Salvaña, John Q. Wong, Jaime C. Montoya, Maria Rosario Singh-Vergeire and Cynthia P. Saloma |
| EPI_ISL_2105541 | Dr. Jorge P. Royeca Hospital | Philippine Genome Center | Francis A. Tablizo, Kenneth M. Kim, Carlo M. Lapid, Marc Jerrone R. Castro, Maria Sofia L. Yangzon, Benedict A. Maralit, Marc Edsel C. Ayes, Eva Maria Cutiongco-de la Paz, Alethea R. de Guzman, Jan Michael C. Yap, Jo-Hannah S. Llames, Sheila Mae M. Araiza, Kris P. Punayan, Irish Coleen A. Asin, Candice Francheska B. Tambaoan, Asia Louisa U. Chong, Karol Sophia Agape R. Padilla, Rianna Patricia S. Cruz, El King D. Morado, Joshua Gregor A. Dizon, Razel Nikka M. Hao, Arianne A. Zamora, Devon Ray Pacial, Juan Antonio R. Magalang, Marissa Alejandria, Celia Carlos, Anna Ong-Lim, Edsel Maurice Salvaña, John Q. Wong, Jaime C. Montoya, Maria Rosario Singh-Vergeire and Cynthia P. Saloma |
| EPI_ISL_2105543 | Cotabato Regional and Medical Center | Philippine Genome Center | Francis A. Tablizo, Kenneth M. Kim, Carlo M. Lapid, Marc Jerrone R. Castro, Maria Sofia L. Yangzon, Benedict A. Maralit, Marc Edsel C. Ayes, Eva Maria Cutiongco-de la Paz, Alethea R. de Guzman, Jan Michael C. Yap, Jo-Hannah S. Llames, Sheila Mae M. Araiza, Kris P. Punayan, Irish Coleen A. Asin, Candice Francheska B. Tambaoan, Asia Louisa U. Chong, Karol Sophia Agape R. Padilla, Rianna Patricia S. Cruz, El King D. Morado, Joshua Gregor A. Dizon, Razel Nikka M. Hao, Arianne A. Zamora, Devon Ray Pacial, Juan Antonio R. Magalang, Marissa Alejandria, Celia Carlos, Anna Ong-Lim, Edsel Maurice Salvaña, John Q. Wong, Jaime C. Montoya, Maria Rosario Singh-Vergeire and Cynthia P. Saloma |
| EPI_ISL_2105544 | Butuan Medical Center | Philippine Genome Center | Francis A. Tablizo, Kenneth M. Kim, Carlo M. Lapid, Marc Jerrone R. Castro, Maria Sofia L. Yangzon, Benedict A. Maralit, Marc Edsel C. Ayes, Eva Maria Cutiongco-de la Paz, Alethea R. de Guzman, Jan Michael C. Yap, Jo-Hannah S. Llames, Sheila Mae M. Araiza, Kris P. Punayan, Irish Coleen A. Asin, Candice Francheska B. Tambaoan, Asia Louisa U. Chong, Karol Sophia Agape R. Padilla, Rianna Patricia S. Cruz, El King D. Morado, Joshua Gregor A. Dizon, Razel Nikka M. Hao, Arianne A. Zamora, Devon Ray Pacial, Juan Antonio R. Magalang, Marissa Alejandria, Celia Carlos, Anna Ong-Lim, Edsel Maurice Salvaña, John Q. Wong, Jaime C. Montoya, Maria Rosario Singh-Vergeire and Cynthia P. Saloma |
| EPI_ISL_2105546 | The Lord's Grace Medical and Industrial Clinic | Philippine Genome Center | Francis A. Tablizo, Kenneth M. Kim, Carlo M. Lapid, Marc Jerrone R. Castro, Maria Sofia L. Yangzon, Benedict A. Maralit, Marc Edsel C. Ayes, Eva Maria Cutiongco-de la Paz, Alethea R. de Guzman, Jan Michael C. Yap, Jo-Hannah S. Llames, Sheila Mae M. Araiza, Kris P. Punayan, Irish Coleen A. Asin, Candice Francheska B. Tambaoan, Asia Louisa U. Chong, Karol Sophia Agape R. Padilla, Rianna Patricia S. Cruz, El King D. Morado, Joshua Gregor A. Dizon, Razel Nikka M. Hao, Arianne A. Zamora, Devon Ray Pacial, Juan Antonio R. Magalang, Marissa Alejandria, Celia Carlos, Anna Ong-Lim, Edsel Maurice Salvaña, John Q. Wong, Jaime C. Montoya, Maria Rosario Singh-Vergeire and Cynthia P. Saloma |
| EPI_ISL_2105548, EPI_ISL_2105552 | Butuan Medical Center | Philippine Genome Center | Francis A. Tablizo, Kenneth M. Kim, Carlo M. Lapid, Marc Jerrone R. Castro, Maria Sofia L. Yangzon, Benedict A. Maralit, Marc Edsel C. Ayes, Eva Maria Cutiongco-de la Paz, Alethea R. de Guzman, Jan Michael C. Yap, Jo-Hannah S. Llames, Sheila Mae M. Araiza, Kris P. Punayan, Irish Coleen A. Asin, Candice Francheska B. Tambaoan, Asia Louisa U. Chong, Karol Sophia Agape R. Padilla, Rianna Patricia S. Cruz, El King D. Morado, Joshua Gregor A. Dizon, Razel Nikka M. Hao, Arianne A. Zamora, Devon Ray Pacial, Juan Antonio R. Magalang, Marissa Alejandria, Celia Carlos, Anna Ong-Lim, Edsel Maurice Salvaña, John Q. Wong, Jaime C. Montoya, Maria Rosario Singh-Vergeire and Cynthia P. Saloma |
| EPI_ISL_2105553 | First Aide Diagnostic Center | Philippine Genome Center | Francis A. Tablizo, Kenneth M. Kim, Carlo M. Lapid, Marc Jerrone R. Castro, Maria Sofia L. Yangzon, Benedict A. Maralit, Marc Edsel C. Ayes, Eva Maria Cutiongco-de la Paz, Alethea R. de Guzman, Jan Michael C. Yap, Jo-Hannah S. Llames, Sheila Mae M. Araiza, Kris P. Punayan, Irish Coleen A. Asin, Candice Francheska B. Tambaoan, Asia Louisa U. Chong, Karol Sophia Agape R. Padilla, Rianna Patricia S. Cruz, El King D. Morado, Joshua Gregor A. Dizon, Razel Nikka M. Hao, Arianne A. Zamora, Devon Ray Pacial, Juan Antonio R. Magalang, Marissa Alejandria, Celia Carlos, Anna Ong-Lim, Edsel Maurice Salvaña, John Q. Wong, Jaime C. Montoya, Maria Rosario Singh-Vergeire and Cynthia P. Saloma |
| EPI_ISL_2105556 | Northern Mindanao TB Regional Center | Philippine Genome Center | Francis A. Tablizo, Kenneth M. Kim, Carlo M. Lapid, Marc Jerrone R. Castro, Maria Sofia L. Yangzon, Benedict A. Maralit, Marc Edsel C. Ayes, Eva Maria Cutiongco-de la Paz, Alethea R. de Guzman, Jan Michael C. Yap, Jo-Hannah S. Llames, Sheila Mae M. Araiza, Kris P. Punayan, Irish Coleen A. Asin, Candice Francheska B. Tambaoan, Asia Louisa U. Chong, Karol Sophia Agape R. Padilla, Rianna Patricia S. Cruz, El King D. Morado, Joshua Gregor A. Dizon, Razel Nikka M. Hao, Arianne A. Zamora, Devon Ray Pacial, Juan Antonio R. Magalang, Marissa Alejandria, Celia Carlos, Anna Ong-Lim, Edsel Maurice Salvaña, John Q. Wong, Jaime C. Montoya, Maria Rosario Singh-Vergeire and Cynthia P. Saloma |
| EPI_ISL_2105557 | PROVIDERS MULTI-PURPOSE COOPERATIVE MEDICAL CENTER | Philippine Genome Center | Francis A. Tablizo, Kenneth M. Kim, Carlo M. Lapid, Marc Jerrone R. Castro, Maria Sofia L. Yangzon, Benedict A. Maralit, Marc Edsel C. Ayes, Eva Maria Cutiongco-de la Paz, Alethea R. de Guzman, Jan Michael C. Yap, Jo-Hannah S. Llames, Sheila Mae M. Araiza, Kris P. Punayan, Irish Coleen A. Asin, Candice Francheska B. Tambaoan, Asia Louisa U. Chong, Karol Sophia Agape R. Padilla, Rianna Patricia S. Cruz, El King D. Morado, Joshua Gregor A. Dizon, Razel Nikka M. Hao, Arianne A. Zamora, Devon Ray Pacial, Juan Antonio R. Magalang, Marissa Alejandria, Celia Carlos, Anna Ong-Lim, Edsel Maurice Salvaña, John Q. Wong, Jaime C. Montoya, Maria Rosario Singh-Vergeire and Cynthia P. Saloma |
| EPI_ISL_2107804 | National Center of Infectious and Parasitic Diseases | National Center of Infectious and Parasitic Diseases | Alexiev, Ivanov, Korsun, Stoitsova, Philipova, Dimitrova, Grigorova L., Hristova, Donchev, Stoykov, Trifonova, Dobrinov, Grigorova I., Kantardjiev |
| EPI_ISL_2110847, EPI_ISL_2110856 | Reitor da Universidade Jean Piaget Guine-Bissau | MRCG at LSHTM, Genomics lab | Aladje Balde, Abdul Karim Sesay, Abdoulie Kanteh, Bakary Sanyang, Simão Tchuda Bióté, Rui Inndi, Adul Candé, Faatu Cassama, Milanca Agostinho Cá, Rei José Pereira, Erica Luis Maria Magalhães, Aicha Balde, Bubacar Delgado Pinto Embaló, Edmira Maria da Costa, Paulina Joãozinho da Costa Jarra Manneh, Mariama Kujabi, Dabiri Damilari, Sainabou Laye Ndure |
| EPI_ISL_2131268 | OLVZ Aalst | OLVZ Aalst | Astrid Holderbeke |
| EPI_ISL_2134822 | HospitalIsraelita Albert Eistein | Virology Laboratory- Institute of Tropical Medicine SP | Fernando Lucas Melo, Camila Malta Romano |
| EPI_ISL_2140800, EPI_ISL_2140801, EPI_ISL_2140802, EPI_ISL_2140804, EPI_ISL_2140805, EPI_ISL_2140807, EPI_ISL_2140808, EPI_ISL_2140809, EPI_ISL_2140810, EPI_ISL_2140811, EPI_ISL_2140812, EPI_ISL_2140814, EPI_ISL_2140815, EPI_ISL_2140816, EPI_ISL_2140817, EPI_ISL_2140819, EPI_ISL_2140821, EPI_ISL_2140822, EPI_ISL_2140823, EPI_ISL_2140824, EPI_ISL_2140825, EPI_ISL_2140827, EPI_ISL_2140828, EPI_ISL_2140829, EPI_ISL_2140830, EPI_ISL_2140831, EPI_ISL_2140832, EPI_ISL_2140833, EPI_ISL_2140835, EPI_ISL_2140836, EPI_ISL_2140837, EPI_ISL_2140838, EPI_ISL_2140839, EPI_ISL_2140840, EPI_ISL_2140841, EPI_ISL_2140842, EPI_ISL_2140843, EPI_ISL_2140845, EPI_ISL_2140846, EPI_ISL_2140847, EPI_ISL_2140849, EPI_ISL_2140850, EPI_ISL_2140855, EPI_ISL_2140856, EPI_ISL_2140857, EPI_ISL_2140861, EPI_ISL_2140862, EPI_ISL_2140863, EPI_ISL_2140864, EPI_ISL_2140865, EPI_ISL_2140866, EPI_ISL_2140867, EPI_ISL_2140868, EPI_ISL_2140870, EPI_ISL_2140871, EPI_ISL_2140874, EPI_ISL_2140875, EPI_ISL_2153737, EPI_ISL_2153748, EPI_ISL_2153756, EPI_ISL_2153760, EPI_ISL_2153763, EPI_ISL_2153764, EPI_ISL_2153766, EPI_ISL_2153768, EPI_ISL_2153770, EPI_ISL_2153780, EPI_ISL_2153785, EPI_ISL_2153787, EPI_ISL_2153792, EPI_ISL_2153796, EPI_ISL_2153799, EPI_ISL_2153802 | | | |
| see above | The Ohio State University Applied Microbiology Services Laboratory | The Ohio State University Applied Microbiology Services Laboratory | Seth A. Faith PhD |

| | | | |
|---|---|---|---|
| EPI_ISL_2154995 | Parkway Medical and Diagnostic Center | Philippine Genome Center | Francis A. Tablizo, Kenneth M. Kim, Carlo M. Lapid, Marc Jerrone R. Castro, Maria Sofia L. Yangzon, Benedict A. Maralit, Marc Edsel C. Ayes, Eva Maria Cutiongco-de la Paz, Alethea R. de Guzman, Jan Michael C. Yap, Jo-Hannah S. Llames, Sheila Mae M. Araiza, Kris P. Punayan, Irish Coleen A. Asin, Candice Francheska B. Tambaoan, Asia Louisa U. Chong, Karol Sophia Agape R. Padilla, Rianna Patricia S. Cruz, El King D. Morado, Joshua Gregor A. Dizon, Razel Nikka M. Hao, Arianne A. Zamora, Devon Ray Pacial, Juan Antonio R. Magalang, Marissa Alejandria, Celia Carlos, Anna Ong-Lim, Edsel Maurice Salvaña, John Q. Wong, Jaime C. Montoya, Maria Rosario Singh-Vergeire and Cynthia P. Saloma |
| EPI_ISL_2155001 | Ilocos Training and Regional Medical Center | Philippine Genome Center | Francis A. Tablizo, Kenneth M. Kim, Carlo M. Lapid, Marc Jerrone R. Castro, Maria Sofia L. Yangzon, Benedict A. Maralit, Marc Edsel C. Ayes, Eva Maria Cutiongco-de la Paz, Alethea R. de Guzman, Jan Michael C. Yap, Jo-Hannah S. Llames, Sheila Mae M. Araiza, Kris P. Punayan, Irish Coleen A. Asin, Candice Francheska B. Tambaoan, Asia Louisa U. Chong, Karol Sophia Agape R. Padilla, Rianna Patricia S. Cruz, El King D. Morado, Joshua Gregor A. Dizon, Razel Nikka M. Hao, Arianne A. Zamora, Devon Ray Pacial, Juan Antonio R. Magalang, Marissa Alejandria, Celia Carlos, Anna Ong-Lim, Edsel Maurice Salvaña, John Q. Wong, Jaime C. Montoya, Maria Rosario Singh-Vergeire and Cynthia P. Saloma |
| EPI_ISL_2155002, EPI_ISL_2155003, EPI_ISL_2155004, EPI_ISL_2155005 | Baguio General Hospital Medical Center (BGHMC) | Philippine Genome Center | Francis A. Tablizo, Kenneth M. Kim, Carlo M. Lapid, Marc Jerrone R. Castro, Maria Sofia L. Yangzon, Benedict A. Maralit, Marc Edsel C. Ayes, Eva Maria Cutiongco-de la Paz, Alethea R. de Guzman, Jan Michael C. Yap, Jo-Hannah S. Llames, Sheila Mae M. Araiza, Kris P. Punayan, Irish Coleen A. Asin, Candice Francheska B. Tambaoan, Asia Louisa U. Chong, Karol Sophia Agape R. Padilla, Rianna Patricia S. Cruz, El King D. Morado, Joshua Gregor A. Dizon, Razel Nikka M. Hao, Arianne A. Zamora, Devon Ray Pacial, Juan Antonio R. Magalang, Marissa Alejandria, Celia Carlos, Anna Ong-Lim, Edsel Maurice Salvaña, John Q. Wong, Jaime C. Montoya, Maria Rosario Singh-Vergeire and Cynthia P. Saloma |
| EPI_ISL_2155007 | First Aide Diagnostic Center | Philippine Genome Center | Francis A. Tablizo, Kenneth M. Kim, Carlo M. Lapid, Marc Jerrone R. Castro, Maria Sofia L. Yangzon, Benedict A. Maralit, Marc Edsel C. Ayes, Eva Maria Cutiongco-de la Paz, Alethea R. de Guzman, Jan Michael C. Yap, Jo-Hannah S. Llames, Sheila Mae M. Araiza, Kris P. Punayan, Irish Coleen A. Asin, Candice Francheska B. Tambaoan, Asia Louisa U. Chong, Karol Sophia Agape R. Padilla, Rianna Patricia S. Cruz, El King D. Morado, Joshua Gregor A. Dizon, Razel Nikka M. Hao, Arianne A. Zamora, Devon Ray Pacial, Juan Antonio R. Magalang, Marissa Alejandria, Celia Carlos, Anna Ong-Lim, Edsel Maurice Salvaña, John Q. Wong, Jaime C. Montoya, Maria Rosario Singh-Vergeire and Cynthia P. Saloma |
| EPI_ISL_2155008 | Baguio General Hospital Medical Center (BGHMC) | Philippine Genome Center | Francis A. Tablizo, Kenneth M. Kim, Carlo M. Lapid, Marc Jerrone R. Castro, Maria Sofia L. Yangzon, Benedict A. Maralit, Marc Edsel C. Ayes, Eva Maria Cutiongco-de la Paz, Alethea R. de Guzman, Jan Michael C. Yap, Jo-Hannah S. Llames, Sheila Mae M. Araiza, Kris P. Punayan, Irish Coleen A. Asin, Candice Francheska B. Tambaoan, Asia Louisa U. Chong, Karol Sophia Agape R. Padilla, Rianna Patricia S. Cruz, El King D. Morado, Joshua Gregor A. Dizon, Razel Nikka M. Hao, Arianne A. Zamora, Devon Ray Pacial, Juan Antonio R. Magalang, Marissa Alejandria, Celia Carlos, Anna Ong-Lim, Edsel Maurice Salvaña, John Q. Wong, Jaime C. Montoya, Maria Rosario Singh-Vergeire and Cynthia P. Saloma |
| EPI_ISL_2155009 | Parkway Medical and Diagnostic Center | Philippine Genome Center | Francis A. Tablizo, Kenneth M. Kim, Carlo M. Lapid, Marc Jerrone R. Castro, Maria Sofia L. Yangzon, Benedict A. Maralit, Marc Edsel C. Ayes, Eva Maria Cutiongco-de la Paz, Alethea R. de Guzman, Jan Michael C. Yap, Jo-Hannah S. Llames, Sheila Mae M. Araiza, Kris P. Punayan, Irish Coleen A. Asin, Candice Francheska B. Tambaoan, Asia Louisa U. Chong, Karol Sophia Agape R. Padilla, Rianna Patricia S. Cruz, El King D. Morado, Joshua Gregor A. Dizon, Razel Nikka M. Hao, Arianne A. Zamora, Devon Ray Pacial, Juan Antonio R. Magalang, Marissa Alejandria, Celia Carlos, Anna Ong-Lim, Edsel Maurice Salvaña, John Q. Wong, Jaime C. Montoya, Maria Rosario Singh-Vergeire and Cynthia P. Saloma |
| EPI_ISL_2155010 | Manila Doctors Hospital | Philippine Genome Center | Francis A. Tablizo, Kenneth M. Kim, Carlo M. Lapid, Marc Jerrone R. Castro, Maria Sofia L. Yangzon, Benedict A. Maralit, Marc Edsel C. Ayes, Eva Maria Cutiongco-de la Paz, Alethea R. de Guzman, Jan Michael C. Yap, Jo-Hannah S. Llames, Sheila Mae M. Araiza, Kris P. Punayan, Irish Coleen A. Asin, Candice Francheska B. Tambaoan, Asia Louisa U. Chong, Karol Sophia Agape R. Padilla, Rianna Patricia S. Cruz, El King D. Morado, Joshua Gregor A. Dizon, Razel Nikka M. Hao, Arianne A. Zamora, Devon Ray Pacial, Juan Antonio R. Magalang, Marissa Alejandria, Celia Carlos, Anna Ong-Lim, Edsel Maurice Salvaña, John Q. Wong, Jaime C. Montoya, Maria Rosario Singh-Vergeire and Cynthia P. Saloma |
| EPI_ISL_2155011, EPI_ISL_2155012, EPI_ISL_2155014 | Baguio General Hospital Medical Center (BGHMC) | Philippine Genome Center | Francis A. Tablizo, Kenneth M. Kim, Carlo M. Lapid, Marc Jerrone R. Castro, Maria Sofia L. Yangzon, Benedict A. Maralit, Marc Edsel C. Ayes, Eva Maria Cutiongco-de la Paz, Alethea R. de Guzman, Jan Michael C. Yap, Jo-Hannah S. Llames, Sheila Mae M. Araiza, Kris P. Punayan, Irish Coleen A. Asin, Candice Francheska B. Tambaoan, Asia Louisa U. Chong, Karol Sophia Agape R. Padilla, Rianna Patricia S. Cruz, El King D. Morado, Joshua Gregor A. Dizon, Razel Nikka M. Hao, Arianne A. Zamora, Devon Ray Pacial, Juan Antonio R. Magalang, Marissa Alejandria, Celia Carlos, Anna Ong-Lim, Edsel Maurice Salvaña, John Q. Wong, Jaime C. Montoya, Maria Rosario Singh-Vergeire and Cynthia P. Saloma |
| EPI_ISL_2155015, EPI_ISL_2155052 | Philippine Red Cross - Port Area | Philippine Genome Center | Francis A. Tablizo, Kenneth M. Kim, Carlo M. Lapid, Marc Jerrone R. Castro, Maria Sofia L. Yangzon, Benedict A. Maralit, Marc Edsel C. Ayes, Eva Maria Cutiongco-de la Paz, Alethea R. de Guzman, Jan Michael C. Yap, Jo-Hannah S. Llames, Sheila Mae M. Araiza, Kris P. Punayan, Irish Coleen A. Asin, Candice Francheska B. Tambaoan, Asia Louisa U. Chong, Karol Sophia Agape R. Padilla, Rianna Patricia S. Cruz, El King D. Morado, Joshua Gregor A. Dizon, Razel Nikka M. Hao, Arianne A. Zamora, Devon Ray Pacial, Juan Antonio R. Magalang, Marissa Alejandria, Celia Carlos, Anna Ong-Lim, Edsel Maurice Salvaña, John Q. Wong, Jaime C. Montoya, Maria Rosario Singh-Vergeire and Cynthia P. Saloma |
| EPI_ISL_2155053 | Baguio General Hospital Medical Center (BGHMC) | Philippine Genome Center | Francis A. Tablizo, Kenneth M. Kim, Carlo M. Lapid, Marc Jerrone R. Castro, Maria Sofia L. Yangzon, Benedict A. Maralit, Marc Edsel C. Ayes, Eva Maria Cutiongco-de la Paz, Alethea R. de Guzman, Jan Michael C. Yap, Jo-Hannah S. Llames, Sheila Mae M. Araiza, Kris P. Punayan, Irish Coleen A. Asin, Candice Francheska B. Tambaoan, Asia Louisa U. Chong, Karol Sophia Agape R. Padilla, Rianna Patricia S. Cruz, El King D. Morado, Joshua Gregor A. Dizon, Razel Nikka M. Hao, Arianne A. Zamora, Devon Ray Pacial, Juan Antonio R. Magalang, Marissa Alejandria, Celia Carlos, Anna Ong-Lim, Edsel Maurice Salvaña, John Q. Wong, Jaime C. Montoya, Maria Rosario Singh-Vergeire and Cynthia P. Saloma |
| EPI_ISL_2155057, EPI_ISL_2155071, EPI_ISL_2155082 | Philippine Red Cross - Port Area | Philippine Genome Center | Francis A. Tablizo, Kenneth M. Kim, Carlo M. Lapid, Marc Jerrone R. Castro, Maria Sofia L. Yangzon, Benedict A. Maralit, Marc Edsel C. Ayes, Eva Maria Cutiongco-de la Paz, Alethea R. de Guzman, Jan Michael C. Yap, Jo-Hannah S. Llames, Sheila Mae M. Araiza, Kris P. Punayan, Irish Coleen A. Asin, Candice Francheska B. Tambaoan, Asia Louisa U. Chong, Karol Sophia Agape R. Padilla, Rianna Patricia S. Cruz, El King D. Morado, Joshua Gregor A. Dizon, Razel Nikka M. Hao, Arianne A. Zamora, Devon Ray Pacial, Juan Antonio R. Magalang, Marissa Alejandria, Celia Carlos, Anna Ong-Lim, Edsel Maurice Salvaña, John Q. Wong, Jaime C. Montoya, Maria Rosario Singh-Vergeire and Cynthia P. Saloma |
| EPI_ISL_2155098 | San Lazaro Hospital (SLH) | Philippine Genome Center | Francis A. Tablizo, Kenneth M. Kim, Carlo M. Lapid, Marc Jerrone R. Castro, Maria Sofia L. Yangzon, Benedict A. Maralit, Marc Edsel C. Ayes, Eva Maria Cutiongco-de la Paz, Alethea R. de Guzman, Jan Michael C. Yap, Jo-Hannah S. Llames, Sheila Mae M. Araiza, Kris P. Punayan, Irish Coleen A. Asin, Candice Francheska B. Tambaoan, Asia Louisa U. Chong, Karol Sophia Agape R. Padilla, Rianna Patricia S. Cruz, El King D. Morado, Joshua Gregor A. Dizon, Razel Nikka M. Hao, Arianne A. Zamora, Devon Ray Pacial, Juan Antonio R. Magalang, Marissa Alejandria, Celia Carlos, Anna Ong-Lim, Edsel Maurice Salvaña, John Q. Wong, Jaime C. Montoya, Maria Rosario Singh-Vergeire and Cynthia P. Saloma |
| EPI_ISL_2155113 | Philippine Red Cross - Port Area | Philippine Genome Center | Francis A. Tablizo, Kenneth M. Kim, Carlo M. Lapid, Marc Jerrone R. Castro, Maria Sofia L. Yangzon, Benedict A. Maralit, Marc Edsel C. Ayes, Eva Maria Cutiongco-de la Paz, Alethea R. de Guzman, Jan Michael C. Yap, Jo-Hannah S. Llames, Sheila Mae M. Araiza, Kris P. Punayan, Irish Coleen A. Asin, Candice Francheska B. Tambaoan, Asia Louisa U. Chong, Karol Sophia Agape R. Padilla, Rianna Patricia S. Cruz, El King D. Morado, Joshua Gregor A. Dizon, Razel Nikka M. Hao, Arianne A. Zamora, Devon Ray Pacial, Juan Antonio R. Magalang, Marissa Alejandria, Celia Carlos, Anna Ong-Lim, Edsel Maurice Salvaña, John Q. Wong, Jaime C. Montoya, Maria Rosario Singh-Vergeire and Cynthia P. Saloma |
| EPI_ISL_2155125 | Philippine Red Cross - National Blood Center | Philippine Genome Center | Francis A. Tablizo, Kenneth M. Kim, Carlo M. Lapid, Marc Jerrone R. Castro, Maria Sofia L. Yangzon, Benedict A. Maralit, Marc Edsel C. Ayes, Eva Maria Cutiongco-de la Paz, Alethea R. de Guzman, Jan Michael C. Yap, Jo-Hannah S. Llames, Sheila Mae M. Araiza, Kris P. Punayan, Irish Coleen A. Asin, Candice Francheska B. Tambaoan, Asia Louisa U. Chong, Karol Sophia Agape R. Padilla, Rianna Patricia S. Cruz, El King D. Morado, Joshua Gregor A. Dizon, Razel Nikka M. Hao, Arianne A. Zamora, Devon Ray Pacial, Juan Antonio R. Magalang, Marissa Alejandria, Celia Carlos, Anna Ong-Lim, Edsel Maurice Salvaña, John Q. Wong, Jaime C. Montoya, Maria Rosario Singh-Vergeire and Cynthia P. Saloma |
| EPI_ISL_2155150, EPI_ISL_2155151 | Baguio General Hospital Medical Center (BGHMC) | Philippine Genome Center | Francis A. Tablizo, Kenneth M. Kim, Carlo M. Lapid, Marc Jerrone R. Castro, Maria Sofia L. Yangzon, Benedict A. Maralit, Marc Edsel C. Ayes, Eva Maria Cutiongco-de la Paz, Alethea R. de Guzman, Jan Michael C. Yap, Jo-Hannah S. Llames, Sheila Mae M. Araiza, Kris P. Punayan, Irish Coleen A. Asin, Candice Francheska B. Tambaoan, Asia Louisa U. Chong, Karol Sophia Agape R. Padilla, Rianna Patricia S. Cruz, El King D. Morado, Joshua Gregor A. Dizon, Razel Nikka M. Hao, Arianne A. Zamora, Devon Ray Pacial, Juan Antonio R. Magalang, Marissa Alejandria, Celia Carlos, Anna Ong-Lim, Edsel Maurice Salvaña, John Q. Wong, Jaime C. Montoya, Maria Rosario Singh-Vergeire and Cynthia P. Saloma |
| EPI_ISL_2155157 | Philippine Red Cross - Port Area | Philippine Genome Center | Francis A. Tablizo, Kenneth M. Kim, Carlo M. Lapid, Marc Jerrone R. Castro, Maria Sofia L. Yangzon, Benedict A. Maralit, Marc Edsel C. Ayes, Eva Maria Cutiongco-de la Paz, Alethea R. de Guzman, Jan Michael C. Yap, Jo-Hannah S. Llames, Sheila Mae M. Araiza, Kris P. Punayan, Irish Coleen A. Asin, Candice Francheska B. Tambaoan, Asia Louisa U. Chong, Karol Sophia Agape R. Padilla, Rianna Patricia S. Cruz, El King D. Morado, Joshua Gregor A. Dizon, Razel Nikka M. Hao, Arianne A. Zamora, Devon Ray Pacial, Juan Antonio R. Magalang, Marissa Alejandria, Celia Carlos, Anna Ong-Lim, Edsel Maurice Salvaña, John Q. Wong, Jaime C. Montoya, Maria Rosario Singh-Vergeire and Cynthia P. Saloma |

| | | | |
|---|---|---|---|
| EPI_ISL_2155169 | Pasig City Children's Hospital - Child's Hope | Philippine Genome Center | Francis A. Tablizo, Kenneth M. Kim, Carlo M. Lapid, Marc Jerrone R. Castro, Maria Sofia L. Yangzon, Benedict A. Maralit, Marc Edsel C. Ayes, Eva Maria Cutiongco-de la Paz, Alethea R. de Guzman, Jan Michael C. Yap, Jo-Hannah S. Llames, Sheila Mae M. Araiza, Kris P. Punayan, Irish Coleen A. Asin, Candice Francheska B. Tambaoan, Asia Louisa U. Chong, Karol Sophia Agape R. Padilla, Rianna Patricia S. Cruz, El King D. Morado, Joshua Gregor A. Dizon, Razel Nikka M. Hao, Arianne A. Zamora, Devon Ray Pacial, Juan Antonio R. Magalang, Marissa Alejandria, Celia Carlos, Anna Ong-Lim, Edsel Maurice Salvaña, John Q. Wong, Jaime C. Montoya, Maria Rosario Singh-Vergeire and Cynthia P. Saloma |
| EPI_ISL_2155171, EPI_ISL_2155179 | Philippine Red Cross - Port Area | Philippine Genome Center | Francis A. Tablizo, Kenneth M. Kim, Carlo M. Lapid, Marc Jerrone R. Castro, Maria Sofia L. Yangzon, Benedict A. Maralit, Marc Edsel C. Ayes, Eva Maria Cutiongco-de la Paz, Alethea R. de Guzman, Jan Michael C. Yap, Jo-Hannah S. Llames, Sheila Mae M. Araiza, Kris P. Punayan, Irish Coleen A. Asin, Candice Francheska B. Tambaoan, Asia Louisa U. Chong, Karol Sophia Agape R. Padilla, Rianna Patricia S. Cruz, El King D. Morado, Joshua Gregor A. Dizon, Razel Nikka M. Hao, Arianne A. Zamora, Devon Ray Pacial, Juan Antonio R. Magalang, Marissa Alejandria, Celia Carlos, Anna Ong-Lim, Edsel Maurice Salvaña, John Q. Wong, Jaime C. Montoya, Maria Rosario Singh-Vergeire and Cynthia P. Saloma |
| EPI_ISL_2155184 | Hi-Precision Diagnostic Center (QC) | Philippine Genome Center | Francis A. Tablizo, Kenneth M. Kim, Carlo M. Lapid, Marc Jerrone R. Castro, Maria Sofia L. Yangzon, Benedict A. Maralit, Marc Edsel C. Ayes, Eva Maria Cutiongco-de la Paz, Alethea R. de Guzman, Jan Michael C. Yap, Jo-Hannah S. Llames, Sheila Mae M. Araiza, Kris P. Punayan, Irish Coleen A. Asin, Candice Francheska B. Tambaoan, Asia Louisa U. Chong, Karol Sophia Agape R. Padilla, Rianna Patricia S. Cruz, El King D. Morado, Joshua Gregor A. Dizon, Razel Nikka M. Hao, Arianne A. Zamora, Devon Ray Pacial, Juan Antonio R. Magalang, Marissa Alejandria, Celia Carlos, Anna Ong-Lim, Edsel Maurice Salvaña, John Q. Wong, Jaime C. Montoya, Maria Rosario Singh-Vergeire and Cynthia P. Saloma |
| EPI_ISL_2155204 | Baguio General Hospital Medical Center (BGHMC) | Philippine Genome Center | Francis A. Tablizo, Kenneth M. Kim, Carlo M. Lapid, Marc Jerrone R. Castro, Maria Sofia L. Yangzon, Benedict A. Maralit, Marc Edsel C. Ayes, Eva Maria Cutiongco-de la Paz, Alethea R. de Guzman, Jan Michael C. Yap, Jo-Hannah S. Llames, Sheila Mae M. Araiza, Kris P. Punayan, Irish Coleen A. Asin, Candice Francheska B. Tambaoan, Asia Louisa U. Chong, Karol Sophia Agape R. Padilla, Rianna Patricia S. Cruz, El King D. Morado, Joshua Gregor A. Dizon, Razel Nikka M. Hao, Arianne A. Zamora, Devon Ray Pacial, Juan Antonio R. Magalang, Marissa Alejandria, Celia Carlos, Anna Ong-Lim, Edsel Maurice Salvaña, John Q. Wong, Jaime C. Montoya, Maria Rosario Singh-Vergeire and Cynthia P. Saloma |
| EPI_ISL_2155205 | Philippine Red Cross - National Blood Center | Philippine Genome Center | Francis A. Tablizo, Kenneth M. Kim, Carlo M. Lapid, Marc Jerrone R. Castro, Maria Sofia L. Yangzon, Benedict A. Maralit, Marc Edsel C. Ayes, Eva Maria Cutiongco-de la Paz, Alethea R. de Guzman, Jan Michael C. Yap, Jo-Hannah S. Llames, Sheila Mae M. Araiza, Kris P. Punayan, Irish Coleen A. Asin, Candice Francheska B. Tambaoan, Asia Louisa U. Chong, Karol Sophia Agape R. Padilla, Rianna Patricia S. Cruz, El King D. Morado, Joshua Gregor A. Dizon, Razel Nikka M. Hao, Arianne A. Zamora, Devon Ray Pacial, Juan Antonio R. Magalang, Marissa Alejandria, Celia Carlos, Anna Ong-Lim, Edsel Maurice Salvaña, John Q. Wong, Jaime C. Montoya, Maria Rosario Singh-Vergeire and Cynthia P. Saloma |
| EPI_ISL_2155209 | Baguio General Hospital Medical Center (BGHMC) | Philippine Genome Center | Francis A. Tablizo, Kenneth M. Kim, Carlo M. Lapid, Marc Jerrone R. Castro, Maria Sofia L. Yangzon, Benedict A. Maralit, Marc Edsel C. Ayes, Eva Maria Cutiongco-de la Paz, Alethea R. de Guzman, Jan Michael C. Yap, Jo-Hannah S. Llames, Sheila Mae M. Araiza, Kris P. Punayan, Irish Coleen A. Asin, Candice Francheska B. Tambaoan, Asia Louisa U. Chong, Karol Sophia Agape R. Padilla, Rianna Patricia S. Cruz, El King D. Morado, Joshua Gregor A. Dizon, Razel Nikka M. Hao, Arianne A. Zamora, Devon Ray Pacial, Juan Antonio R. Magalang, Marissa Alejandria, Celia Carlos, Anna Ong-Lim, Edsel Maurice Salvaña, John Q. Wong, Jaime C. Montoya, Maria Rosario Singh-Vergeire and Cynthia P. Saloma |
| EPI_ISL_2155211 | Parkway Medical and Diagnostic Center | Philippine Genome Center | Francis A. Tablizo, Kenneth M. Kim, Carlo M. Lapid, Marc Jerrone R. Castro, Maria Sofia L. Yangzon, Benedict A. Maralit, Marc Edsel C. Ayes, Eva Maria Cutiongco-de la Paz, Alethea R. de Guzman, Jan Michael C. Yap, Jo-Hannah S. Llames, Sheila Mae M. Araiza, Kris P. Punayan, Irish Coleen A. Asin, Candice Francheska B. Tambaoan, Asia Louisa U. Chong, Karol Sophia Agape R. Padilla, Rianna Patricia S. Cruz, El King D. Morado, Joshua Gregor A. Dizon, Razel Nikka M. Hao, Arianne A. Zamora, Devon Ray Pacial, Juan Antonio R. Magalang, Marissa Alejandria, Celia Carlos, Anna Ong-Lim, Edsel Maurice Salvaña, John Q. Wong, Jaime C. Montoya, Maria Rosario Singh-Vergeire and Cynthia P. Saloma |
| EPI_ISL_2155215 | Hi-Precision Diagnostic Center (QC) | Philippine Genome Center | Francis A. Tablizo, Kenneth M. Kim, Carlo M. Lapid, Marc Jerrone R. Castro, Maria Sofia L. Yangzon, Benedict A. Maralit, Marc Edsel C. Ayes, Eva Maria Cutiongco-de la Paz, Alethea R. de Guzman, Jan Michael C. Yap, Jo-Hannah S. Llames, Sheila Mae M. Araiza, Kris P. Punayan, Irish Coleen A. Asin, Candice Francheska B. Tambaoan, Asia Louisa U. Chong, Karol Sophia Agape R. Padilla, Rianna Patricia S. Cruz, El King D. Morado, Joshua Gregor A. Dizon, Razel Nikka M. Hao, Arianne A. Zamora, Devon Ray Pacial, Juan Antonio R. Magalang, Marissa Alejandria, Celia Carlos, Anna Ong-Lim, Edsel Maurice Salvaña, John Q. Wong, Jaime C. Montoya, Maria Rosario Singh-Vergeire and Cynthia P. Saloma |
| EPI_ISL_2155220 | Philippine Red Cross - Port Area | Philippine Genome Center | Francis A. Tablizo, Kenneth M. Kim, Carlo M. Lapid, Marc Jerrone R. Castro, Maria Sofia L. Yangzon, Benedict A. Maralit, Marc Edsel C. Ayes, Eva Maria Cutiongco-de la Paz, Alethea R. de Guzman, Jan Michael C. Yap, Jo-Hannah S. Llames, Sheila Mae M. Araiza, Kris P. Punayan, Irish Coleen A. Asin, Candice Francheska B. Tambaoan, Asia Louisa U. Chong, Karol Sophia Agape R. Padilla, Rianna Patricia S. Cruz, El King D. Morado, Joshua Gregor A. Dizon, Razel Nikka M. Hao, Arianne A. Zamora, Devon Ray Pacial, Juan Antonio R. Magalang, Marissa Alejandria, Celia Carlos, Anna Ong-Lim, Edsel Maurice Salvaña, John Q. Wong, Jaime C. Montoya, Maria Rosario Singh-Vergeire and Cynthia P. Saloma |
| EPI_ISL_2155251, EPI_ISL_2155252 | Bataan General Hospital and Medical Center | Philippine Genome Center | Francis A. Tablizo, Kenneth M. Kim, Carlo M. Lapid, Marc Jerrone R. Castro, Maria Sofia L. Yangzon, Benedict A. Maralit, Marc Edsel C. Ayes, Eva Maria Cutiongco-de la Paz, Alethea R. de Guzman, Jan Michael C. Yap, Jo-Hannah S. Llames, Sheila Mae M. Araiza, Kris P. Punayan, Irish Coleen A. Asin, Candice Francheska B. Tambaoan, Asia Louisa U. Chong, Karol Sophia Agape R. Padilla, Rianna Patricia S. Cruz, El King D. Morado, Joshua Gregor A. Dizon, Razel Nikka M. Hao, Arianne A. Zamora, Devon Ray Pacial, Juan Antonio R. Magalang, Marissa Alejandria, Celia Carlos, Anna Ong-Lim, Edsel Maurice Salvaña, John Q. Wong, Jaime C. Montoya, Maria Rosario Singh-Vergeire and Cynthia P. Saloma |
| EPI_ISL_2155256 | Philippine Red Cross- Batangas Chapter Molecular Laboratory | Philippine Genome Center | Francis A. Tablizo, Kenneth M. Kim, Carlo M. Lapid, Marc Jerrone R. Castro, Maria Sofia L. Yangzon, Benedict A. Maralit, Marc Edsel C. Ayes, Eva Maria Cutiongco-de la Paz, Alethea R. de Guzman, Jan Michael C. Yap, Jo-Hannah S. Llames, Sheila Mae M. Araiza, Kris P. Punayan, Irish Coleen A. Asin, Candice Francheska B. Tambaoan, Asia Louisa U. Chong, Karol Sophia Agape R. Padilla, Rianna Patricia S. Cruz, El King D. Morado, Joshua Gregor A. Dizon, Razel Nikka M. Hao, Arianne A. Zamora, Devon Ray Pacial, Juan Antonio R. Magalang, Marissa Alejandria, Celia Carlos, Anna Ong-Lim, Edsel Maurice Salvaña, John Q. Wong, Jaime C. Montoya, Maria Rosario Singh-Vergeire and Cynthia P. Saloma |
| EPI_ISL_2155257 | The Medical City -Â Ortigas | Philippine Genome Center | Francis A. Tablizo, Kenneth M. Kim, Carlo M. Lapid, Marc Jerrone R. Castro, Maria Sofia L. Yangzon, Benedict A. Maralit, Marc Edsel C. Ayes, Eva Maria Cutiongco-de la Paz, Alethea R. de Guzman, Jan Michael C. Yap, Jo-Hannah S. Llames, Sheila Mae M. Araiza, Kris P. Punayan, Irish Coleen A. Asin, Candice Francheska B. Tambaoan, Asia Louisa U. Chong, Karol Sophia Agape R. Padilla, Rianna Patricia S. Cruz, El King D. Morado, Joshua Gregor A. Dizon, Razel Nikka M. Hao, Arianne A. Zamora, Devon Ray Pacial, Juan Antonio R. Magalang, Marissa Alejandria, Celia Carlos, Anna Ong-Lim, Edsel Maurice Salvaña, John Q. Wong, Jaime C. Montoya, Maria Rosario Singh-Vergeire and Cynthia P. Saloma |
| EPI_ISL_2155258 | Philippine Red Cross - Port Area | Philippine Genome Center | Francis A. Tablizo, Kenneth M. Kim, Carlo M. Lapid, Marc Jerrone R. Castro, Maria Sofia L. Yangzon, Benedict A. Maralit, Marc Edsel C. Ayes, Eva Maria Cutiongco-de la Paz, Alethea R. de Guzman, Jan Michael C. Yap, Jo-Hannah S. Llames, Sheila Mae M. Araiza, Kris P. Punayan, Irish Coleen A. Asin, Candice Francheska B. Tambaoan, Asia Louisa U. Chong, Karol Sophia Agape R. Padilla, Rianna Patricia S. Cruz, El King D. Morado, Joshua Gregor A. Dizon, Razel Nikka M. Hao, Arianne A. Zamora, Devon Ray Pacial, Juan Antonio R. Magalang, Marissa Alejandria, Celia Carlos, Anna Ong-Lim, Edsel Maurice Salvaña, John Q. Wong, Jaime C. Montoya, Maria Rosario Singh-Vergeire and Cynthia P. Saloma |
| EPI_ISL_2155264 | Baguio General Hospital Medical Center (BGHMC) | Philippine Genome Center | Francis A. Tablizo, Kenneth M. Kim, Carlo M. Lapid, Marc Jerrone R. Castro, Maria Sofia L. Yangzon, Benedict A. Maralit, Marc Edsel C. Ayes, Eva Maria Cutiongco-de la Paz, Alethea R. de Guzman, Jan Michael C. Yap, Jo-Hannah S. Llames, Sheila Mae M. Araiza, Kris P. Punayan, Irish Coleen A. Asin, Candice Francheska B. Tambaoan, Asia Louisa U. Chong, Karol Sophia Agape R. Padilla, Rianna Patricia S. Cruz, El King D. Morado, Joshua Gregor A. Dizon, Razel Nikka M. Hao, Arianne A. Zamora, Devon Ray Pacial, Juan Antonio R. Magalang, Marissa Alejandria, Celia Carlos, Anna Ong-Lim, Edsel Maurice Salvaña, John Q. Wong, Jaime C. Montoya, Maria Rosario Singh-Vergeire and Cynthia P. Saloma |
| EPI_ISL_2155267 | Butuan Medical Center | Philippine Genome Center | Francis A. Tablizo, Kenneth M. Kim, Carlo M. Lapid, Marc Jerrone R. Castro, Maria Sofia L. Yangzon, Benedict A. Maralit, Marc Edsel C. Ayes, Eva Maria Cutiongco-de la Paz, Alethea R. de Guzman, Jan Michael C. Yap, Jo-Hannah S. Llames, Sheila Mae M. Araiza, Kris P. Punayan, Irish Coleen A. Asin, Candice Francheska B. Tambaoan, Asia Louisa U. Chong, Karol Sophia Agape R. Padilla, Rianna Patricia S. Cruz, El King D. Morado, Joshua Gregor A. Dizon, Razel Nikka M. Hao, Arianne A. Zamora, Devon Ray Pacial, Juan Antonio R. Magalang, Marissa Alejandria, Celia Carlos, Anna Ong-Lim, Edsel Maurice Salvaña, John Q. Wong, Jaime C. Montoya, Maria Rosario Singh-Vergeire and Cynthia P. Saloma |
| EPI_ISL_2155278 | Philippine Red Cross - National Blood Center | Philippine Genome Center | Francis A. Tablizo, Kenneth M. Kim, Carlo M. Lapid, Marc Jerrone R. Castro, Maria Sofia L. Yangzon, Benedict A. Maralit, Marc Edsel C. Ayes, Eva Maria Cutiongco-de la Paz, Alethea R. de Guzman, Jan Michael C. Yap, Jo-Hannah S. Llames, Sheila Mae M. Araiza, Kris P. Punayan, Irish Coleen A. Asin, Candice Francheska B. Tambaoan, Asia Louisa U. Chong, Karol Sophia Agape R. Padilla, Rianna Patricia S. Cruz, El King D. Morado, Joshua Gregor A. Dizon, Razel Nikka M. Hao, Arianne A. Zamora, Devon Ray Pacial, Juan Antonio R. Magalang, Marissa Alejandria, Celia Carlos, Anna Ong-Lim, Edsel Maurice Salvaña, John Q. Wong, Jaime C. Montoya, Maria Rosario Singh-Vergeire and Cynthia P. Saloma |

| | | | |
|---|---|---|---|
| EPI_ISL_2155290 | Philippine Red Cross - Port Area | Philippine Genome Center | Francis A. Tablizo, Kenneth M. Kim, Carlo M. Lapid, Marc Jerrone R. Castro, Maria Sofia L. Yangzon, Benedict A. Maralit, Marc Edsel C. Ayes, Eva Maria Cutiongco-de la Paz, Alethea R. de Guzman, Jan Michael C. Yap, Jo-Hannah S. Llames, Sheila Mae M. Araiza, Kris P. Punayan, Irish Coleen A. Asin, Candice Francheska B. Tambaoan, Asia Louisa U. Chong, Karol Sophia Agape R. Padilla, Rianna Patricia S. Cruz, El King D. Morado, Joshua Gregor A. Dizon, Razel Nikka M. Hao, Arianne A. Zamora, Devon Ray Pacial, Juan Antonio R. Magalang, Marissa Alejandria, Celia Carlos, Anna Ong-Lim, Edsel Maurice Salvaña, John Q. Wong, Jaime C. Montoya, Maria Rosario Singh-Vergeire and Cynthia P. Saloma |
| EPI_ISL_2155308 | Pasig City Children's Hospital - Child's Hope | Philippine Genome Center | Francis A. Tablizo, Kenneth M. Kim, Carlo M. Lapid, Marc Jerrone R. Castro, Maria Sofia L. Yangzon, Benedict A. Maralit, Marc Edsel C. Ayes, Eva Maria Cutiongco-de la Paz, Alethea R. de Guzman, Jan Michael C. Yap, Jo-Hannah S. Llames, Sheila Mae M. Araiza, Kris P. Punayan, Irish Coleen A. Asin, Candice Francheska B. Tambaoan, Asia Louisa U. Chong, Karol Sophia Agape R. Padilla, Rianna Patricia S. Cruz, El King D. Morado, Joshua Gregor A. Dizon, Razel Nikka M. Hao, Arianne A. Zamora, Devon Ray Pacial, Juan Antonio R. Magalang, Marissa Alejandria, Celia Carlos, Anna Ong-Lim, Edsel Maurice Salvaña, John Q. Wong, Jaime C. Montoya, Maria Rosario Singh-Vergeire and Cynthia P. Saloma |
| EPI_ISL_2155319 | RURAL HEALTH UNIT - RIZAL | Philippine Genome Center | Francis A. Tablizo, Kenneth M. Kim, Carlo M. Lapid, Marc Jerrone R. Castro, Maria Sofia L. Yangzon, Benedict A. Maralit, Marc Edsel C. Ayes, Eva Maria Cutiongco-de la Paz, Alethea R. de Guzman, Jan Michael C. Yap, Jo-Hannah S. Llames, Sheila Mae M. Araiza, Kris P. Punayan, Irish Coleen A. Asin, Candice Francheska B. Tambaoan, Asia Louisa U. Chong, Karol Sophia Agape R. Padilla, Rianna Patricia S. Cruz, El King D. Morado, Joshua Gregor A. Dizon, Razel Nikka M. Hao, Arianne A. Zamora, Devon Ray Pacial, Juan Antonio R. Magalang, Marissa Alejandria, Celia Carlos, Anna Ong-Lim, Edsel Maurice Salvaña, John Q. Wong, Jaime C. Montoya, Maria Rosario Singh-Vergeire and Cynthia P. Saloma |
| EPI_ISL_2155323 | Ilocos Training and Regional Medical Center | Philippine Genome Center | Francis A. Tablizo, Kenneth M. Kim, Carlo M. Lapid, Marc Jerrone R. Castro, Maria Sofia L. Yangzon, Benedict A. Maralit, Marc Edsel C. Ayes, Eva Maria Cutiongco-de la Paz, Alethea R. de Guzman, Jan Michael C. Yap, Jo-Hannah S. Llames, Sheila Mae M. Araiza, Kris P. Punayan, Irish Coleen A. Asin, Candice Francheska B. Tambaoan, Asia Louisa U. Chong, Karol Sophia Agape R. Padilla, Rianna Patricia S. Cruz, El King D. Morado, Joshua Gregor A. Dizon, Razel Nikka M. Hao, Arianne A. Zamora, Devon Ray Pacial, Juan Antonio R. Magalang, Marissa Alejandria, Celia Carlos, Anna Ong-Lim, Edsel Maurice Salvaña, John Q. Wong, Jaime C. Montoya, Maria Rosario Singh-Vergeire and Cynthia P. Saloma |
| EPI_ISL_2155330 | Parkway Medical and Diagnostic Center | Philippine Genome Center | Francis A. Tablizo, Kenneth M. Kim, Carlo M. Lapid, Marc Jerrone R. Castro, Maria Sofia L. Yangzon, Benedict A. Maralit, Marc Edsel C. Ayes, Eva Maria Cutiongco-de la Paz, Alethea R. de Guzman, Jan Michael C. Yap, Jo-Hannah S. Llames, Sheila Mae M. Araiza, Kris P. Punayan, Irish Coleen A. Asin, Candice Francheska B. Tambaoan, Asia Louisa U. Chong, Karol Sophia Agape R. Padilla, Rianna Patricia S. Cruz, El King D. Morado, Joshua Gregor A. Dizon, Razel Nikka M. Hao, Arianne A. Zamora, Devon Ray Pacial, Juan Antonio R. Magalang, Marissa Alejandria, Celia Carlos, Anna Ong-Lim, Edsel Maurice Salvaña, John Q. Wong, Jaime C. Montoya, Maria Rosario Singh-Vergeire and Cynthia P. Saloma |
| EPI_ISL_2155338 | Philippine Red Cross - Port Area | Philippine Genome Center | Francis A. Tablizo, Kenneth M. Kim, Carlo M. Lapid, Marc Jerrone R. Castro, Maria Sofia L. Yangzon, Benedict A. Maralit, Marc Edsel C. Ayes, Eva Maria Cutiongco-de la Paz, Alethea R. de Guzman, Jan Michael C. Yap, Jo-Hannah S. Llames, Sheila Mae M. Araiza, Kris P. Punayan, Irish Coleen A. Asin, Candice Francheska B. Tambaoan, Asia Louisa U. Chong, Karol Sophia Agape R. Padilla, Rianna Patricia S. Cruz, El King D. Morado, Joshua Gregor A. Dizon, Razel Nikka M. Hao, Arianne A. Zamora, Devon Ray Pacial, Juan Antonio R. Magalang, Marissa Alejandria, Celia Carlos, Anna Ong-Lim, Edsel Maurice Salvaña, John Q. Wong, Jaime C. Montoya, Maria Rosario Singh-Vergeire and Cynthia P. Saloma |
| EPI_ISL_2155339 | Baguio General Hospital Medical Center (BGHMC) | Philippine Genome Center | Francis A. Tablizo, Kenneth M. Kim, Carlo M. Lapid, Marc Jerrone R. Castro, Maria Sofia L. Yangzon, Benedict A. Maralit, Marc Edsel C. Ayes, Eva Maria Cutiongco-de la Paz, Alethea R. de Guzman, Jan Michael C. Yap, Jo-Hannah S. Llames, Sheila Mae M. Araiza, Kris P. Punayan, Irish Coleen A. Asin, Candice Francheska B. Tambaoan, Asia Louisa U. Chong, Karol Sophia Agape R. Padilla, Rianna Patricia S. Cruz, El King D. Morado, Joshua Gregor A. Dizon, Razel Nikka M. Hao, Arianne A. Zamora, Devon Ray Pacial, Juan Antonio R. Magalang, Marissa Alejandria, Celia Carlos, Anna Ong-Lim, Edsel Maurice Salvaña, John Q. Wong, Jaime C. Montoya, Maria Rosario Singh-Vergeire and Cynthia P. Saloma |
| EPI_ISL_2155352 | The Lord's Grace Medical and Industrial Clinic | Philippine Genome Center | Francis A. Tablizo, Kenneth M. Kim, Carlo M. Lapid, Marc Jerrone R. Castro, Maria Sofia L. Yangzon, Benedict A. Maralit, Marc Edsel C. Ayes, Eva Maria Cutiongco-de la Paz, Alethea R. de Guzman, Jan Michael C. Yap, Jo-Hannah S. Llames, Sheila Mae M. Araiza, Kris P. Punayan, Irish Coleen A. Asin, Candice Francheska B. Tambaoan, Asia Louisa U. Chong, Karol Sophia Agape R. Padilla, Rianna Patricia S. Cruz, El King D. Morado, Joshua Gregor A. Dizon, Razel Nikka M. Hao, Arianne A. Zamora, Devon Ray Pacial, Juan Antonio R. Magalang, Marissa Alejandria, Celia Carlos, Anna Ong-Lim, Edsel Maurice Salvaña, John Q. Wong, Jaime C. Montoya, Maria Rosario Singh-Vergeire and Cynthia P. Saloma |
| EPI_ISL_2155354 | Baguio General Hospital Medical Center (BGHMC) | Philippine Genome Center | Francis A. Tablizo, Kenneth M. Kim, Carlo M. Lapid, Marc Jerrone R. Castro, Maria Sofia L. Yangzon, Benedict A. Maralit, Marc Edsel C. Ayes, Eva Maria Cutiongco-de la Paz, Alethea R. de Guzman, Jan Michael C. Yap, Jo-Hannah S. Llames, Sheila Mae M. Araiza, Kris P. Punayan, Irish Coleen A. Asin, Candice Francheska B. Tambaoan, Asia Louisa U. Chong, Karol Sophia Agape R. Padilla, Rianna Patricia S. Cruz, El King D. Morado, Joshua Gregor A. Dizon, Razel Nikka M. Hao, Arianne A. Zamora, Devon Ray Pacial, Juan Antonio R. Magalang, Marissa Alejandria, Celia Carlos, Anna Ong-Lim, Edsel Maurice Salvaña, John Q. Wong, Jaime C. Montoya, Maria Rosario Singh-Vergeire and Cynthia P. Saloma |
| EPI_ISL_2155356 | The Lord's Grace Medical and Industrial Clinic | Philippine Genome Center | Francis A. Tablizo, Kenneth M. Kim, Carlo M. Lapid, Marc Jerrone R. Castro, Maria Sofia L. Yangzon, Benedict A. Maralit, Marc Edsel C. Ayes, Eva Maria Cutiongco-de la Paz, Alethea R. de Guzman, Jan Michael C. Yap, Jo-Hannah S. Llames, Sheila Mae M. Araiza, Kris P. Punayan, Irish Coleen A. Asin, Candice Francheska B. Tambaoan, Asia Louisa U. Chong, Karol Sophia Agape R. Padilla, Rianna Patricia S. Cruz, El King D. Morado, Joshua Gregor A. Dizon, Razel Nikka M. Hao, Arianne A. Zamora, Devon Ray Pacial, Juan Antonio R. Magalang, Marissa Alejandria, Celia Carlos, Anna Ong-Lim, Edsel Maurice Salvaña, John Q. Wong, Jaime C. Montoya, Maria Rosario Singh-Vergeire and Cynthia P. Saloma |
| EPI_ISL_2155378, EPI_ISL_2155379, EPI_ISL_2155381, EPI_ISL_2155383, EPI_ISL_2155384, EPI_ISL_2155385 | Mariano Marcos Memorial Hospital Medical Center | Philippine Genome Center | Francis A. Tablizo, Kenneth M. Kim, Carlo M. Lapid, Marc Jerrone R. Castro, Maria Sofia L. Yangzon, Benedict A. Maralit, Marc Edsel C. Ayes, Eva Maria Cutiongco-de la Paz, Alethea R. de Guzman, Jan Michael C. Yap, Jo-Hannah S. Llames, Sheila Mae M. Araiza, Kris P. Punayan, Irish Coleen A. Asin, Candice Francheska B. Tambaoan, Asia Louisa U. Chong, Karol Sophia Agape R. Padilla, Rianna Patricia S. Cruz, El King D. Morado, Joshua Gregor A. Dizon, Razel Nikka M. Hao, Arianne A. Zamora, Devon Ray Pacial, Juan Antonio R. Magalang, Marissa Alejandria, Celia Carlos, Anna Ong-Lim, Edsel Maurice Salvaña, John Q. Wong, Jaime C. Montoya, Maria Rosario Singh-Vergeire and Cynthia P. Saloma |
| EPI_ISL_2155398, EPI_ISL_2155399, EPI_ISL_2155401, EPI_ISL_2155403, EPI_ISL_2155404 | Pasig City Children's Hospital - Child's Hope | Philippine Genome Center | Francis A. Tablizo, Kenneth M. Kim, Carlo M. Lapid, Marc Jerrone R. Castro, Maria Sofia L. Yangzon, Benedict A. Maralit, Marc Edsel C. Ayes, Eva Maria Cutiongco-de la Paz, Alethea R. de Guzman, Jan Michael C. Yap, Jo-Hannah S. Llames, Sheila Mae M. Araiza, Kris P. Punayan, Irish Coleen A. Asin, Candice Francheska B. Tambaoan, Asia Louisa U. Chong, Karol Sophia Agape R. Padilla, Rianna Patricia S. Cruz, El King D. Morado, Joshua Gregor A. Dizon, Razel Nikka M. Hao, Arianne A. Zamora, Devon Ray Pacial, Juan Antonio R. Magalang, Marissa Alejandria, Celia Carlos, Anna Ong-Lim, Edsel Maurice Salvaña, John Q. Wong, Jaime C. Montoya, Maria Rosario Singh-Vergeire and Cynthia P. Saloma |
| EPI_ISL_2155407, EPI_ISL_2155410 | Makati Medical Center | Philippine Genome Center | Francis A. Tablizo, Kenneth M. Kim, Carlo M. Lapid, Marc Jerrone R. Castro, Maria Sofia L. Yangzon, Benedict A. Maralit, Marc Edsel C. Ayes, Eva Maria Cutiongco-de la Paz, Alethea R. de Guzman, Jan Michael C. Yap, Jo-Hannah S. Llames, Sheila Mae M. Araiza, Kris P. Punayan, Irish Coleen A. Asin, Candice Francheska B. Tambaoan, Asia Louisa U. Chong, Karol Sophia Agape R. Padilla, Rianna Patricia S. Cruz, El King D. Morado, Joshua Gregor A. Dizon, Razel Nikka M. Hao, Arianne A. Zamora, Devon Ray Pacial, Juan Antonio R. Magalang, Marissa Alejandria, Celia Carlos, Anna Ong-Lim, Edsel Maurice Salvaña, John Q. Wong, Jaime C. Montoya, Maria Rosario Singh-Vergeire and Cynthia P. Saloma |
| EPI_ISL_2155415, EPI_ISL_2155416, EPI_ISL_2155418, EPI_ISL_2155419, EPI_ISL_2155420, EPI_ISL_2155422, EPI_ISL_2155423 | Parkway Medical and Diagnostic Center | Philippine Genome Center | Francis A. Tablizo, Kenneth M. Kim, Carlo M. Lapid, Marc Jerrone R. Castro, Maria Sofia L. Yangzon, Benedict A. Maralit, Marc Edsel C. Ayes, Eva Maria Cutiongco-de la Paz, Alethea R. de Guzman, Jan Michael C. Yap, Jo-Hannah S. Llames, Sheila Mae M. Araiza, Kris P. Punayan, Irish Coleen A. Asin, Candice Francheska B. Tambaoan, Asia Louisa U. Chong, Karol Sophia Agape R. Padilla, Rianna Patricia S. Cruz, El King D. Morado, Joshua Gregor A. Dizon, Razel Nikka M. Hao, Arianne A. Zamora, Devon Ray Pacial, Juan Antonio R. Magalang, Marissa Alejandria, Celia Carlos, Anna Ong-Lim, Edsel Maurice Salvaña, John Q. Wong, Jaime C. Montoya, Maria Rosario Singh-Vergeire and Cynthia P. Saloma |
| EPI_ISL_2155424, EPI_ISL_2155426, EPI_ISL_2155427, EPI_ISL_2155428, EPI_ISL_2155430, EPI_ISL_2155431, EPI_ISL_2155432, EPI_ISL_2155433, EPI_ISL_2155434 | Valenzuela Hope Molecular Laboratory | Philippine Genome Center | Francis A. Tablizo, Kenneth M. Kim, Carlo M. Lapid, Marc Jerrone R. Castro, Maria Sofia L. Yangzon, Benedict A. Maralit, Marc Edsel C. Ayes, Eva Maria Cutiongco-de la Paz, Alethea R. de Guzman, Jan Michael C. Yap, Jo-Hannah S. Llames, Sheila Mae M. Araiza, Kris P. Punayan, Irish Coleen A. Asin, Candice Francheska B. Tambaoan, Asia Louisa U. Chong, Karol Sophia Agape R. Padilla, Rianna Patricia S. Cruz, El King D. Morado, Joshua Gregor A. Dizon, Razel Nikka M. Hao, Arianne A. Zamora, Devon Ray Pacial, Juan Antonio R. Magalang, Marissa Alejandria, Celia Carlos, Anna Ong-Lim, Edsel Maurice Salvaña, John Q. Wong, Jaime C. Montoya, Maria Rosario Singh-Vergeire and Cynthia P. Saloma |
| EPI_ISL_2155436, EPI_ISL_2155437, EPI_ISL_2155441, EPI_ISL_2155442, EPI_ISL_2155444, EPI_ISL_2155445, EPI_ISL_2155448, EPI_ISL_2155449, EPI_ISL_2155451, EPI_ISL_2155453, EPI_ISL_2155456, EPI_ISL_2155459, EPI_ISL_2155460 | | | |
| see above | Philippine Red Cross - National Blood Center | Philippine Genome Center | Francis A. Tablizo, Kenneth M. Kim, Carlo M. Lapid, Marc Jerrone R. Castro, Maria Sofia L. Yangzon, Benedict A. Maralit, Marc Edsel C. Ayes, Eva Maria Cutiongco-de la Paz, Alethea R. de Guzman, Jan Michael C. Yap, Jo-Hannah S. Llames, Sheila Mae M. Araiza, Kris P. Punayan, Irish Coleen A. Asin, Candice Francheska B. Tambaoan, Asia Louisa U. Chong, Karol Sophia Agape R. Padilla, Rianna Patricia S. Cruz, El King D. Morado, Joshua Gregor A. |

| | | | |
|---|---|---|---|
| | | | Dizon, Razel Nikka M. Hao, Arianne A. Zamora, Devon Ray Pacial, Juan Antonio R. Magalang, Marissa Alejandria, Celia Carlos, Anna Ong-Lim, Edsel Maurice Salvaña, John Q. Wong, Jaime C. Montoya, Maria Rosario Singh-Vergeire and Cynthia P. Saloma |
| EPI_ISL_2155462, EPI_ISL_2155464, EPI_ISL_2155467, EPI_ISL_2155469, EPI_ISL_2155471, EPI_ISL_2155473, EPI_ISL_2155474 | Hi-Precision Diagnostic Center (QC) | Philippine Genome Center | Francis A. Tablizo, Kenneth M. Kim, Carlo M. Lapid, Marc Jerrone R. Castro, Maria Sofia L. Yangzon, Benedict A. Maralit, Marc Edsel C. Ayes, Eva Maria Cutiongco-de la Paz, Alethea R. de Guzman, Jan Michael C. Yap, Jo-Hannah S. Llames, Sheila Mae M. Araiza, Kris P. Punayan, Irish Coleen A. Asin, Candice Francheska B. Tambaoan, Asia Louisa U. Chong, Karol Sophia Agape R. Padilla, Rianna Patricia S. Cruz, El King D. Morado, Joshua Gregor A. Dizon, Razel Nikka M. Hao, Arianne A. Zamora, Devon Ray Pacial, Juan Antonio R. Magalang, Marissa Alejandria, Celia Carlos, Anna Ong-Lim, Edsel Maurice Salvaña, John Q. Wong, Jaime C. Montoya, Maria Rosario Singh-Vergeire and Cynthia P. Saloma |
| EPI_ISL_2155477, EPI_ISL_2155480, EPI_ISL_2155481, EPI_ISL_2155482, EPI_ISL_2155483, EPI_ISL_2155484 | University of the Philippines National Institutes of Health (UP NIH) | Philippine Genome Center | Francis A. Tablizo, Kenneth M. Kim, Carlo M. Lapid, Marc Jerrone R. Castro, Maria Sofia L. Yangzon, Benedict A. Maralit, Marc Edsel C. Ayes, Eva Maria Cutiongco-de la Paz, Alethea R. de Guzman, Jan Michael C. Yap, Jo-Hannah S. Llames, Sheila Mae M. Araiza, Kris P. Punayan, Irish Coleen A. Asin, Candice Francheska B. Tambaoan, Asia Louisa U. Chong, Karol Sophia Agape R. Padilla, Rianna Patricia S. Cruz, El King D. Morado, Joshua Gregor A. Dizon, Razel Nikka M. Hao, Arianne A. Zamora, Devon Ray Pacial, Juan Antonio R. Magalang, Marissa Alejandria, Celia Carlos, Anna Ong-Lim, Edsel Maurice Salvaña, John Q. Wong, Jaime C. Montoya, Maria Rosario Singh-Vergeire and Cynthia P. Saloma |
| EPI_ISL_2155499, EPI_ISL_2155504, EPI_ISL_2155512 | Manila Doctors Hospital | Philippine Genome Center | Francis A. Tablizo, Kenneth M. Kim, Carlo M. Lapid, Marc Jerrone R. Castro, Maria Sofia L. Yangzon, Benedict A. Maralit, Marc Edsel C. Ayes, Eva Maria Cutiongco-de la Paz, Alethea R. de Guzman, Jan Michael C. Yap, Jo-Hannah S. Llames, Sheila Mae M. Araiza, Kris P. Punayan, Irish Coleen A. Asin, Candice Francheska B. Tambaoan, Asia Louisa U. Chong, Karol Sophia Agape R. Padilla, Rianna Patricia S. Cruz, El King D. Morado, Joshua Gregor A. Dizon, Razel Nikka M. Hao, Arianne A. Zamora, Devon Ray Pacial, Juan Antonio R. Magalang, Marissa Alejandria, Celia Carlos, Anna Ong-Lim, Edsel Maurice Salvaña, John Q. Wong, Jaime C. Montoya, Maria Rosario Singh-Vergeire and Cynthia P. Saloma |
| EPI_ISL_2155530, EPI_ISL_2155533, EPI_ISL_2155537, EPI_ISL_2155538, EPI_ISL_2155539, EPI_ISL_2155540, EPI_ISL_2155541, EPI_ISL_2155542, EPI_ISL_2155546, EPI_ISL_2155547, EPI_ISL_2155548, EPI_ISL_2155549, EPI_ISL_2155550, EPI_ISL_2155551, EPI_ISL_2155552, EPI_ISL_2155558, EPI_ISL_2155560, EPI_ISL_2155563, EPI_ISL_2155565, EPI_ISL_2155568, EPI_ISL_2155572, EPI_ISL_2155573, EPI_ISL_2155574, EPI_ISL_2155578, EPI_ISL_2155584, EPI_ISL_2155586 | | | |
| see above | UPLB Covid-19 Molecular Diagnostic Laboratory | Philippine Genome Center | Francis A. Tablizo, Kenneth M. Kim, Carlo M. Lapid, Marc Jerrone R. Castro, Maria Sofia L. Yangzon, Benedict A. Maralit, Marc Edsel C. Ayes, Eva Maria Cutiongco-de la Paz, Alethea R. de Guzman, Jan Michael C. Yap, Jo-Hannah S. Llames, Sheila Mae M. Araiza, Kris P. Punayan, Irish Coleen A. Asin, Candice Francheska B. Tambaoan, Asia Louisa U. Chong, Karol Sophia Agape R. Padilla, Rianna Patricia S. Cruz, El King D. Morado, Joshua Gregor A. Dizon, Razel Nikka M. Hao, Arianne A. Zamora, Devon Ray Pacial, Juan Antonio R. Magalang, Marissa Alejandria, Celia Carlos, Anna Ong-Lim, Edsel Maurice Salvaña, John Q. Wong, Jaime C. Montoya, Maria Rosario Singh-Vergeire and Cynthia P. Saloma |
| EPI_ISL_2155592, EPI_ISL_2155593, EPI_ISL_2155596, EPI_ISL_2155598 | Bataan General Hospital and Medical Center | Philippine Genome Center | Francis A. Tablizo, Kenneth M. Kim, Carlo M. Lapid, Marc Jerrone R. Castro, Maria Sofia L. Yangzon, Benedict A. Maralit, Marc Edsel C. Ayes, Eva Maria Cutiongco-de la Paz, Alethea R. de Guzman, Jan Michael C. Yap, Jo-Hannah S. Llames, Sheila Mae M. Araiza, Kris P. Punayan, Irish Coleen A. Asin, Candice Francheska B. Tambaoan, Asia Louisa U. Chong, Karol Sophia Agape R. Padilla, Rianna Patricia S. Cruz, El King D. Morado, Joshua Gregor A. Dizon, Razel Nikka M. Hao, Arianne A. Zamora, Devon Ray Pacial, Juan Antonio R. Magalang, Marissa Alejandria, Celia Carlos, Anna Ong-Lim, Edsel Maurice Salvaña, John Q. Wong, Jaime C. Montoya, Maria Rosario Singh-Vergeire and Cynthia P. Saloma |
| EPI_ISL_2155600 | SAN PABLO CITY GENERAL HOSPITAL | Philippine Genome Center | Francis A. Tablizo, Kenneth M. Kim, Carlo M. Lapid, Marc Jerrone R. Castro, Maria Sofia L. Yangzon, Benedict A. Maralit, Marc Edsel C. Ayes, Eva Maria Cutiongco-de la Paz, Alethea R. de Guzman, Jan Michael C. Yap, Jo-Hannah S. Llames, Sheila Mae M. Araiza, Kris P. Punayan, Irish Coleen A. Asin, Candice Francheska B. Tambaoan, Asia Louisa U. Chong, Karol Sophia Agape R. Padilla, Rianna Patricia S. Cruz, El King D. Morado, Joshua Gregor A. Dizon, Razel Nikka M. Hao, Arianne A. Zamora, Devon Ray Pacial, Juan Antonio R. Magalang, Marissa Alejandria, Celia Carlos, Anna Ong-Lim, Edsel Maurice Salvaña, John Q. Wong, Jaime C. Montoya, Maria Rosario Singh-Vergeire and Cynthia P. Saloma |
| EPI_ISL_2155603, EPI_ISL_2155604 | Philippine Red Cross- Batangas Chapter Molecular Laboratory | Philippine Genome Center | Francis A. Tablizo, Kenneth M. Kim, Carlo M. Lapid, Marc Jerrone R. Castro, Maria Sofia L. Yangzon, Benedict A. Maralit, Marc Edsel C. Ayes, Eva Maria Cutiongco-de la Paz, Alethea R. de Guzman, Jan Michael C. Yap, Jo-Hannah S. Llames, Sheila Mae M. Araiza, Kris P. Punayan, Irish Coleen A. Asin, Candice Francheska B. Tambaoan, Asia Louisa U. Chong, Karol Sophia Agape R. Padilla, Rianna Patricia S. Cruz, El King D. Morado, Joshua Gregor A. Dizon, Razel Nikka M. Hao, Arianne A. Zamora, Devon Ray Pacial, Juan Antonio R. Magalang, Marissa Alejandria, Celia Carlos, Anna Ong-Lim, Edsel Maurice Salvaña, John Q. Wong, Jaime C. Montoya, Maria Rosario Singh-Vergeire and Cynthia P. Saloma |
| EPI_ISL_2155617 | Research Institute for Tropical Medicine, Inc. (RITM) | Philippine Genome Center | Francis A. Tablizo, Kenneth M. Kim, Carlo M. Lapid, Marc Jerrone R. Castro, Maria Sofia L. Yangzon, Benedict A. Maralit, Marc Edsel C. Ayes, Eva Maria Cutiongco-de la Paz, Alethea R. de Guzman, Jan Michael C. Yap, Jo-Hannah S. Llames, Sheila Mae M. Araiza, Kris P. Punayan, Irish Coleen A. Asin, Candice Francheska B. Tambaoan, Asia Louisa U. Chong, Karol Sophia Agape R. Padilla, Rianna Patricia S. Cruz, El King D. Morado, Joshua Gregor A. Dizon, Razel Nikka M. Hao, Arianne A. Zamora, Devon Ray Pacial, Juan Antonio R. Magalang, Marissa Alejandria, Celia Carlos, Anna Ong-Lim, Edsel Maurice Salvaña, John Q. Wong, Jaime C. Montoya, Maria Rosario Singh-Vergeire and Cynthia P. Saloma |
| EPI_ISL_2155645, EPI_ISL_2155647, EPI_ISL_2155648, EPI_ISL_2155649, EPI_ISL_2155650, EPI_ISL_2155651, EPI_ISL_2155652, EPI_ISL_2155654, EPI_ISL_2155655, EPI_ISL_2155656 | Ilocos Training and Regional Medical Center | Philippine Genome Center | Francis A. Tablizo, Kenneth M. Kim, Carlo M. Lapid, Marc Jerrone R. Castro, Maria Sofia L. Yangzon, Benedict A. Maralit, Marc Edsel C. Ayes, Eva Maria Cutiongco-de la Paz, Alethea R. de Guzman, Jan Michael C. Yap, Jo-Hannah S. Llames, Sheila Mae M. Araiza, Kris P. Punayan, Irish Coleen A. Asin, Candice Francheska B. Tambaoan, Asia Louisa U. Chong, Karol Sophia Agape R. Padilla, Rianna Patricia S. Cruz, El King D. Morado, Joshua Gregor A. Dizon, Razel Nikka M. Hao, Arianne A. Zamora, Devon Ray Pacial, Juan Antonio R. Magalang, Marissa Alejandria, Celia Carlos, Anna Ong-Lim, Edsel Maurice Salvaña, John Q. Wong, Jaime C. Montoya, Maria Rosario Singh-Vergeire and Cynthia P. Saloma |
| EPI_ISL_2155659, EPI_ISL_2155660, EPI_ISL_2155661, EPI_ISL_2155662 | Mary Mediatrix Medical Center | Philippine Genome Center | Francis A. Tablizo, Kenneth M. Kim, Carlo M. Lapid, Marc Jerrone R. Castro, Maria Sofia L. Yangzon, Benedict A. Maralit, Marc Edsel C. Ayes, Eva Maria Cutiongco-de la Paz, Alethea R. de Guzman, Jan Michael C. Yap, Jo-Hannah S. Llames, Sheila Mae M. Araiza, Kris P. Punayan, Irish Coleen A. Asin, Candice Francheska B. Tambaoan, Asia Louisa U. Chong, Karol Sophia Agape R. Padilla, Rianna Patricia S. Cruz, El King D. Morado, Joshua Gregor A. Dizon, Razel Nikka M. Hao, Arianne A. Zamora, Devon Ray Pacial, Juan Antonio R. Magalang, Marissa Alejandria, Celia Carlos, Anna Ong-Lim, Edsel Maurice Salvaña, John Q. Wong, Jaime C. Montoya, Maria Rosario Singh-Vergeire and Cynthia P. Saloma |
| EPI_ISL_2155664, EPI_ISL_2155665, EPI_ISL_2155666, EPI_ISL_2155667, EPI_ISL_2155668, EPI_ISL_2155669, EPI_ISL_2155670, EPI_ISL_2155671, EPI_ISL_2155672, EPI_ISL_2155673, EPI_ISL_2155674, EPI_ISL_2155675 | | | |
| see above | San Antonio City of Ilagan Hospital | Philippine Genome Center | Francis A. Tablizo, Kenneth M. Kim, Carlo M. Lapid, Marc Jerrone R. Castro, Maria Sofia L. Yangzon, Benedict A. Maralit, Marc Edsel C. Ayes, Eva Maria Cutiongco-de la Paz, Alethea R. de Guzman, Jan Michael C. Yap, Jo-Hannah S. Llames, Sheila Mae M. Araiza, Kris P. Punayan, Irish Coleen A. Asin, Candice Francheska B. Tambaoan, Asia Louisa U. Chong, Karol Sophia Agape R. Padilla, Rianna Patricia S. Cruz, El King D. Morado, Joshua Gregor A. Dizon, Razel Nikka M. Hao, Arianne A. Zamora, Devon Ray Pacial, Juan Antonio R. Magalang, Marissa Alejandria, Celia Carlos, Anna Ong-Lim, Edsel Maurice Salvaña, John Q. Wong, Jaime C. Montoya, Maria Rosario Singh-Vergeire and Cynthia P. Saloma |
| EPI_ISL_2155676, EPI_ISL_2155677 | Philippine Red Cross - Clark Molecular Laboratory | Philippine Genome Center | Francis A. Tablizo, Kenneth M. Kim, Carlo M. Lapid, Marc Jerrone R. Castro, Maria Sofia L. Yangzon, Benedict A. Maralit, Marc Edsel C. Ayes, Eva Maria Cutiongco-de la Paz, Alethea R. de Guzman, Jan Michael C. Yap, Jo-Hannah S. Llames, Sheila Mae M. Araiza, Kris P. Punayan, Irish Coleen A. Asin, Candice Francheska B. Tambaoan, Asia Louisa U. Chong, Karol Sophia Agape R. Padilla, Rianna Patricia S. Cruz, El King D. Morado, Joshua Gregor A. Dizon, Razel Nikka M. Hao, Arianne A. Zamora, Devon Ray Pacial, Juan Antonio R. Magalang, Marissa Alejandria, Celia Carlos, Anna Ong-Lim, Edsel Maurice Salvaña, John Q. Wong, Jaime C. Montoya, Maria Rosario Singh-Vergeire and Cynthia P. Saloma |
| EPI_ISL_2155679, EPI_ISL_2155680, EPI_ISL_2155682, EPI_ISL_2155684, EPI_ISL_2155686, EPI_ISL_2155687, EPI_ISL_2155688, EPI_ISL_2155689, EPI_ISL_2155690, EPI_ISL_2155691, EPI_ISL_2155694, EPI_ISL_2155695, EPI_ISL_2155696, EPI_ISL_2155697, EPI_ISL_2155698, EPI_ISL_2155699, EPI_ISL_2155700 | | | |
| see above | The Medical City -Â Ortigas | Philippine Genome Center | Francis A. Tablizo, Kenneth M. Kim, Carlo M. Lapid, Marc Jerrone R. Castro, Maria Sofia L. Yangzon, Benedict A. Maralit, Marc Edsel C. Ayes, Eva Maria Cutiongco-de la Paz, Alethea R. de Guzman, Jan Michael C. Yap, Jo-Hannah S. Llames, Sheila Mae M. Araiza, Kris P. Punayan, Irish Coleen A. Asin, Candice Francheska B. Tambaoan, Asia Louisa U. Chong, Karol Sophia Agape R. Padilla, Rianna Patricia S. Cruz, El King D. Morado, Joshua Gregor A. Dizon, Razel Nikka M. Hao, Arianne A. Zamora, Devon Ray Pacial, Juan Antonio R. Magalang, Marissa Alejandria, Celia Carlos, Anna Ong-Lim, Edsel Maurice Salvaña, John Q. Wong, Jaime C. Montoya, Maria Rosario Singh-Vergeire and Cynthia P. Saloma |
| EPI_ISL_2155703, EPI_ISL_2155709, EPI_ISL_2155716, EPI_ISL_2155722, EPI_ISL_2155723 | Cagayan Valley Medical Center Molecular Laboratory | Philippine Genome Center | Francis A. Tablizo, Kenneth M. Kim, Carlo M. Lapid, Marc Jerrone R. Castro, Maria Sofia L. Yangzon, Benedict A. Maralit, Marc Edsel C. Ayes, Eva Maria Cutiongco-de la Paz, Alethea R. de Guzman, Jan Michael C. Yap, Jo-Hannah S. Llames, Sheila Mae M. Araiza, Kris P. Punayan, Irish Coleen A. Asin, Candice Francheska B. Tambaoan, Asia Louisa U. Chong, Karol Sophia Agape R. Padilla, Rianna Patricia S. Cruz, El King D. Morado, Joshua Gregor A. Dizon, Razel Nikka M. Hao, Arianne A. Zamora, Devon Ray Pacial, Juan Antonio R. Magalang, Marissa Alejandria, Celia Carlos, Anna Ong-Lim, Edsel Maurice Salvaña, John Q. Wong, Jaime C. Montoya, Maria Rosario Singh-Vergeire and Cynthia P. Saloma |
| EPI_ISL_2155729, EPI_ISL_2155730 | Laguna Holy Family Hospital, Inc. | Philippine Genome Center | Francis A. Tablizo, Kenneth M. Kim, Carlo M. Lapid, Marc Jerrone R. Castro, Maria Sofia L. Yangzon, Benedict A. Maralit, Marc Edsel C. Ayes, Eva Maria Cutiongco-de la Paz, Alethea R. de Guzman, Jan Michael C. Yap, Jo-Hannah S. Llames, Sheila Mae M. Araiza, Kris P. Punayan, Irish Coleen A. Asin, |

| | | | |
|---|---|---|---|
| | | | Candice Francheska B. Tambaoan, Asia Louisa U. Chong, Karol Sophia Agape R. Padilla, Rianna Patricia S. Cruz, El King D. Morado, Joshua Gregor A. Dizon, Razel Nikka M. Hao, Arianne A. Zamora, Devon Ray Pacial, Juan Antonio R. Magalang, Marissa Alejandria, Celia Carlos, Anna Ong-Lim, Edsel Maurice Salvaña, John Q. Wong, Jaime C. Montoya, Maria Rosario Singh-Vergeire and Cynthia P. Saloma |
| EPI_ISL_2155731, EPI_ISL_2155732, EPI_ISL_2155734, EPI_ISL_2155735, EPI_ISL_2155737, EPI_ISL_2155739, EPI_ISL_2155740, EPI_ISL_2155742, EPI_ISL_2155743, EPI_ISL_2155744, EPI_ISL_2155745, EPI_ISL_2155746, EPI_ISL_2155747, EPI_ISL_2155749, EPI_ISL_2155750, EPI_ISL_2155751, EPI_ISL_2155752 | | | |
| see above | Philippine Red Cross Logistics and Multipurpose Center | Philippine Genome Center | Francis A. Tablizo, Kenneth M. Kim, Carlo M. Lapid, Marc Jerrone R. Castro, Maria Sofia L. Yangzon, Benedict A. Maralit, Marc Edsel C. Ayes, Eva Maria Cutiongco-de la Paz, Alethea R. de Guzman, Jan Michael C. Yap, Jo-Hannah S. Llames, Sheila Mae M. Araiza, Kris P. Punayan, Irish Coleen A. Asin, Candice Francheska B. Tambaoan, Asia Louisa U. Chong, Karol Sophia Agape R. Padilla, Rianna Patricia S. Cruz, El King D. Morado, Joshua Gregor A. Dizon, Razel Nikka M. Hao, Arianne A. Zamora, Devon Ray Pacial, Juan Antonio R. Magalang, Marissa Alejandria, Celia Carlos, Anna Ong-Lim, Edsel Maurice Salvaña, John Q. Wong, Jaime C. Montoya, Maria Rosario Singh-Vergeire and Cynthia P. Saloma |
| EPI_ISL_2155757 | First Aide Diagnostic Center | Philippine Genome Center | Francis A. Tablizo, Kenneth M. Kim, Carlo M. Lapid, Marc Jerrone R. Castro, Maria Sofia L. Yangzon, Benedict A. Maralit, Marc Edsel C. Ayes, Eva Maria Cutiongco-de la Paz, Alethea R. de Guzman, Jan Michael C. Yap, Jo-Hannah S. Llames, Sheila Mae M. Araiza, Kris P. Punayan, Irish Coleen A. Asin, Candice Francheska B. Tambaoan, Asia Louisa U. Chong, Karol Sophia Agape R. Padilla, Rianna Patricia S. Cruz, El King D. Morado, Joshua Gregor A. Dizon, Razel Nikka M. Hao, Arianne A. Zamora, Devon Ray Pacial, Juan Antonio R. Magalang, Marissa Alejandria, Celia Carlos, Anna Ong-Lim, Edsel Maurice Salvaña, John Q. Wong, Jaime C. Montoya, Maria Rosario Singh-Vergeire and Cynthia P. Saloma |
| EPI_ISL_2155763, EPI_ISL_2155764, EPI_ISL_2155765 | Lung Center of the Philippines (LCP) | Philippine Genome Center | Francis A. Tablizo, Kenneth M. Kim, Carlo M. Lapid, Marc Jerrone R. Castro, Maria Sofia L. Yangzon, Benedict A. Maralit, Marc Edsel C. Ayes, Eva Maria Cutiongco-de la Paz, Alethea R. de Guzman, Jan Michael C. Yap, Jo-Hannah S. Llames, Sheila Mae M. Araiza, Kris P. Punayan, Irish Coleen A. Asin, Candice Francheska B. Tambaoan, Asia Louisa U. Chong, Karol Sophia Agape R. Padilla, Rianna Patricia S. Cruz, El King D. Morado, Joshua Gregor A. Dizon, Razel Nikka M. Hao, Arianne A. Zamora, Devon Ray Pacial, Juan Antonio R. Magalang, Marissa Alejandria, Celia Carlos, Anna Ong-Lim, Edsel Maurice Salvaña, John Q. Wong, Jaime C. Montoya, Maria Rosario Singh-Vergeire and Cynthia P. Saloma |
| EPI_ISL_2155772, EPI_ISL_2155781 | Philippine Red Cross - Port Area | Philippine Genome Center | Francis A. Tablizo, Kenneth M. Kim, Carlo M. Lapid, Marc Jerrone R. Castro, Maria Sofia L. Yangzon, Benedict A. Maralit, Marc Edsel C. Ayes, Eva Maria Cutiongco-de la Paz, Alethea R. de Guzman, Jan Michael C. Yap, Jo-Hannah S. Llames, Sheila Mae M. Araiza, Kris P. Punayan, Irish Coleen A. Asin, Candice Francheska B. Tambaoan, Asia Louisa U. Chong, Karol Sophia Agape R. Padilla, Rianna Patricia S. Cruz, El King D. Morado, Joshua Gregor A. Dizon, Razel Nikka M. Hao, Arianne A. Zamora, Devon Ray Pacial, Juan Antonio R. Magalang, Marissa Alejandria, Celia Carlos, Anna Ong-Lim, Edsel Maurice Salvaña, John Q. Wong, Jaime C. Montoya, Maria Rosario Singh-Vergeire and Cynthia P. Saloma |
| EPI_ISL_2155788 | Southern Philippines Medical Center (SPMC-SNL-MBL) | Philippine Genome Center | Francis A. Tablizo, Kenneth M. Kim, Carlo M. Lapid, Marc Jerrone R. Castro, Maria Sofia L. Yangzon, Benedict A. Maralit, Marc Edsel C. Ayes, Eva Maria Cutiongco-de la Paz, Alethea R. de Guzman, Jan Michael C. Yap, Jo-Hannah S. Llames, Sheila Mae M. Araiza, Kris P. Punayan, Irish Coleen A. Asin, Candice Francheska B. Tambaoan, Asia Louisa U. Chong, Karol Sophia Agape R. Padilla, Rianna Patricia S. Cruz, El King D. Morado, Joshua Gregor A. Dizon, Razel Nikka M. Hao, Arianne A. Zamora, Devon Ray Pacial, Juan Antonio R. Magalang, Marissa Alejandria, Celia Carlos, Anna Ong-Lim, Edsel Maurice Salvaña, John Q. Wong, Jaime C. Montoya, Maria Rosario Singh-Vergeire and Cynthia P. Saloma |
| EPI_ISL_2155791, EPI_ISL_2155792, EPI_ISL_2155793, EPI_ISL_2155794 | Philippine General Hospital (PGH) | Philippine Genome Center | Francis A. Tablizo, Kenneth M. Kim, Carlo M. Lapid, Marc Jerrone R. Castro, Maria Sofia L. Yangzon, Benedict A. Maralit, Marc Edsel C. Ayes, Eva Maria Cutiongco-de la Paz, Alethea R. de Guzman, Jan Michael C. Yap, Jo-Hannah S. Llames, Sheila Mae M. Araiza, Kris P. Punayan, Irish Coleen A. Asin, Candice Francheska B. Tambaoan, Asia Louisa U. Chong, Karol Sophia Agape R. Padilla, Rianna Patricia S. Cruz, El King D. Morado, Joshua Gregor A. Dizon, Razel Nikka M. Hao, Arianne A. Zamora, Devon Ray Pacial, Juan Antonio R. Magalang, Marissa Alejandria, Celia Carlos, Anna Ong-Lim, Edsel Maurice Salvaña, John Q. Wong, Jaime C. Montoya, Maria Rosario Singh-Vergeire and Cynthia P. Saloma |
| EPI_ISL_2155799, EPI_ISL_2155800, EPI_ISL_2155801, EPI_ISL_2155802, EPI_ISL_2155804, EPI_ISL_2155806, EPI_ISL_2155807, EPI_ISL_2155809, EPI_ISL_2155810, EPI_ISL_2155811 | Safeguard DNA Diagnostic, Inc. | Philippine Genome Center | Francis A. Tablizo, Kenneth M. Kim, Carlo M. Lapid, Marc Jerrone R. Castro, Maria Sofia L. Yangzon, Benedict A. Maralit, Marc Edsel C. Ayes, Eva Maria Cutiongco-de la Paz, Alethea R. de Guzman, Jan Michael C. Yap, Jo-Hannah S. Llames, Sheila Mae M. Araiza, Kris P. Punayan, Irish Coleen A. Asin, Candice Francheska B. Tambaoan, Asia Louisa U. Chong, Karol Sophia Agape R. Padilla, Rianna Patricia S. Cruz, El King D. Morado, Joshua Gregor A. Dizon, Razel Nikka M. Hao, Arianne A. Zamora, Devon Ray Pacial, Juan Antonio R. Magalang, Marissa Alejandria, Celia Carlos, Anna Ong-Lim, Edsel Maurice Salvaña, John Q. Wong, Jaime C. Montoya, Maria Rosario Singh-Vergeire and Cynthia P. Saloma |
| EPI_ISL_2155813, EPI_ISL_2155814, EPI_ISL_2155815, EPI_ISL_2155819, EPI_ISL_2155820, EPI_ISL_2155821 | Southern Isabela Medical Center Molecular Diagnostic & Research Laboratory (SIMC - MDRL) | Philippine Genome Center | Francis A. Tablizo, Kenneth M. Kim, Carlo M. Lapid, Marc Jerrone R. Castro, Maria Sofia L. Yangzon, Benedict A. Maralit, Marc Edsel C. Ayes, Eva Maria Cutiongco-de la Paz, Alethea R. de Guzman, Jan Michael C. Yap, Jo-Hannah S. Llames, Sheila Mae M. Araiza, Kris P. Punayan, Irish Coleen A. Asin, Candice Francheska B. Tambaoan, Asia Louisa U. Chong, Karol Sophia Agape R. Padilla, Rianna Patricia S. Cruz, El King D. Morado, Joshua Gregor A. Dizon, Razel Nikka M. Hao, Arianne A. Zamora, Devon Ray Pacial, Juan Antonio R. Magalang, Marissa Alejandria, Celia Carlos, Anna Ong-Lim, Edsel Maurice Salvaña, John Q. Wong, Jaime C. Montoya, Maria Rosario Singh-Vergeire and Cynthia P. Saloma |
| EPI_ISL_2155843, EPI_ISL_2155844 | San Lazaro Hospital | Philippine Genome Center | Francis A. Tablizo, Kenneth M. Kim, Carlo M. Lapid, Marc Jerrone R. Castro, Maria Sofia L. Yangzon, Benedict A. Maralit, Marc Edsel C. Ayes, Eva Maria Cutiongco-de la Paz, Alethea R. de Guzman, Jan Michael C. Yap, Jo-Hannah S. Llames, Sheila Mae M. Araiza, Kris P. Punayan, Irish Coleen A. Asin, Candice Francheska B. Tambaoan, Asia Louisa U. Chong, Karol Sophia Agape R. Padilla, Rianna Patricia S. Cruz, El King D. Morado, Joshua Gregor A. Dizon, Razel Nikka M. Hao, Arianne A. Zamora, Devon Ray Pacial, Juan Antonio R. Magalang, Marissa Alejandria, Celia Carlos, Anna Ong-Lim, Edsel Maurice Salvaña, John Q. Wong, Jaime C. Montoya, Maria Rosario Singh-Vergeire and Cynthia P. Saloma |
| EPI_ISL_2155892, EPI_ISL_2155899, EPI_ISL_2155906, EPI_ISL_2155908 | Philippine Red Cross - National Blood Center | Philippine Genome Center | Francis A. Tablizo, Kenneth M. Kim, Carlo M. Lapid, Marc Jerrone R. Castro, Maria Sofia L. Yangzon, Benedict A. Maralit, Marc Edsel C. Ayes, Eva Maria Cutiongco-de la Paz, Alethea R. de Guzman, Jan Michael C. Yap, Jo-Hannah S. Llames, Sheila Mae M. Araiza, Kris P. Punayan, Irish Coleen A. Asin, Candice Francheska B. Tambaoan, Asia Louisa U. Chong, Karol Sophia Agape R. Padilla, Rianna Patricia S. Cruz, El King D. Morado, Joshua Gregor A. Dizon, Razel Nikka M. Hao, Arianne A. Zamora, Devon Ray Pacial, Juan Antonio R. Magalang, Marissa Alejandria, Celia Carlos, Anna Ong-Lim, Edsel Maurice Salvaña, John Q. Wong, Jaime C. Montoya, Maria Rosario Singh-Vergeire and Cynthia P. Saloma |
| EPI_ISL_2155931 | Cebu TB Reference Laboratory | Philippine Genome Center | Francis A. Tablizo, Kenneth M. Kim, Carlo M. Lapid, Marc Jerrone R. Castro, Maria Sofia L. Yangzon, Benedict A. Maralit, Marc Edsel C. Ayes, Eva Maria Cutiongco-de la Paz, Alethea R. de Guzman, Jan Michael C. Yap, Jo-Hannah S. Llames, Sheila Mae M. Araiza, Kris P. Punayan, Irish Coleen A. Asin, Candice Francheska B. Tambaoan, Asia Louisa U. Chong, Karol Sophia Agape R. Padilla, Rianna Patricia S. Cruz, El King D. Morado, Joshua Gregor A. Dizon, Razel Nikka M. Hao, Arianne A. Zamora, Devon Ray Pacial, Juan Antonio R. Magalang, Marissa Alejandria, Celia Carlos, Anna Ong-Lim, Edsel Maurice Salvaña, John Q. Wong, Jaime C. Montoya, Maria Rosario Singh-Vergeire and Cynthia P. Saloma |
| EPI_ISL_2155951 | PHILIPPINE AIRPORT DIAGNOSTIC LABORATORY | Philippine Genome Center | Francis A. Tablizo, Kenneth M. Kim, Carlo M. Lapid, Marc Jerrone R. Castro, Maria Sofia L. Yangzon, Benedict A. Maralit, Marc Edsel C. Ayes, Eva Maria Cutiongco-de la Paz, Alethea R. de Guzman, Jan Michael C. Yap, Jo-Hannah S. Llames, Sheila Mae M. Araiza, Kris P. Punayan, Irish Coleen A. Asin, Candice Francheska B. Tambaoan, Asia Louisa U. Chong, Karol Sophia Agape R. Padilla, Rianna Patricia S. Cruz, El King D. Morado, Joshua Gregor A. Dizon, Razel Nikka M. Hao, Arianne A. Zamora, Devon Ray Pacial, Juan Antonio R. Magalang, Marissa Alejandria, Celia Carlos, Anna Ong-Lim, Edsel Maurice Salvaña, John Q. Wong, Jaime C. Montoya, Maria Rosario Singh-Vergeire and Cynthia P. Saloma |
| EPI_ISL_2155979, EPI_ISL_2155980, EPI_ISL_2155981 | Philippine Red Cross - Clark Molecular Laboratory | Philippine Genome Center | Francis A. Tablizo, Kenneth M. Kim, Carlo M. Lapid, Marc Jerrone R. Castro, Maria Sofia L. Yangzon, Benedict A. Maralit, Marc Edsel C. Ayes, Eva Maria Cutiongco-de la Paz, Alethea R. de Guzman, Jan Michael C. Yap, Jo-Hannah S. Llames, Sheila Mae M. Araiza, Kris P. Punayan, Irish Coleen A. Asin, Candice Francheska B. Tambaoan, Asia Louisa U. Chong, Karol Sophia Agape R. Padilla, Rianna Patricia S. Cruz, El King D. Morado, Joshua Gregor A. Dizon, Razel Nikka M. Hao, Arianne A. Zamora, Devon Ray Pacial, Juan Antonio R. Magalang, Marissa Alejandria, Celia Carlos, Anna Ong-Lim, Edsel Maurice Salvaña, John Q. Wong, Jaime C. Montoya, Maria Rosario Singh-Vergeire and Cynthia P. Saloma |
| EPI_ISL_2155984, EPI_ISL_2155986, EPI_ISL_2155987, EPI_ISL_2155989, EPI_ISL_2155990, EPI_ISL_2155992, EPI_ISL_2155996, EPI_ISL_2155997 | The Medical City -Â Ortigas | Philippine Genome Center | Francis A. Tablizo, Kenneth M. Kim, Carlo M. Lapid, Marc Jerrone R. Castro, Maria Sofia L. Yangzon, Benedict A. Maralit, Marc Edsel C. Ayes, Eva Maria Cutiongco-de la Paz, Alethea R. de Guzman, Jan Michael C. Yap, Jo-Hannah S. Llames, Sheila Mae M. Araiza, Kris P. Punayan, Irish Coleen A. Asin, Candice Francheska B. Tambaoan, Asia Louisa U. Chong, Karol Sophia Agape R. Padilla, Rianna Patricia S. Cruz, El King D. Morado, Joshua Gregor A. Dizon, Razel Nikka M. Hao, Arianne A. Zamora, Devon Ray Pacial, Juan Antonio R. Magalang, Marissa Alejandria, Celia Carlos, Anna Ong-Lim, Edsel Maurice Salvaña, John Q. Wong, Jaime C. Montoya, Maria Rosario Singh-Vergeire and Cynthia P. Saloma |
| EPI_ISL_2156001, EPI_ISL_2156002, EPI_ISL_2156006, EPI_ISL_2156007, EPI_ISL_2156008, EPI_ISL_2156009 | Philippine Red Cross Logistics and Multipurpose Center | Philippine Genome Center | Francis A. Tablizo, Kenneth M. Kim, Carlo M. Lapid, Marc Jerrone R. Castro, Maria Sofia L. Yangzon, Benedict A. Maralit, Marc Edsel C. Ayes, Eva Maria Cutiongco-de la Paz, Alethea R. de Guzman, Jan Michael C. Yap, Jo-Hannah S. Llames, Sheila Mae M. Araiza, Kris P. Punayan, Irish Coleen A. Asin, Candice Francheska B. Tambaoan, Asia Louisa U. Chong, Karol Sophia Agape R. Padilla, Rianna Patricia S. Cruz, El King D. Morado, Joshua Gregor A. Dizon, Razel Nikka M. Hao, Arianne A. Zamora, Devon Ray Pacial, Juan Antonio R. Magalang, Marissa Alejandria, Celia Carlos, Anna Ong-Lim, Edsel Maurice Salvaña, John Q. Wong, Jaime C. Montoya, Maria Rosario Singh-Vergeire and Cynthia P. Saloma |

| | | | |
|---|---|---|---|
| EPI_ISL_2156021 | Philippine General Hospital (PGH) | Philippine Genome Center | Francis A. Tablizo, Kenneth M. Kim, Carlo M. Lapid, Marc Jerrone R. Castro, Maria Sofia L. Yangzon, Benedict A. Maralit, Marc Edsel C. Ayes, Eva Maria Cutiongco-de la Paz, Alethea R. de Guzman, Jan Michael C. Yap, Jo-Hannah S. Llames, Sheila Mae M. Araiza, Kris P. Punayan, Irish Coleen A. Asin, Candice Francheska B. Tambaoan, Asia Louisa U. Chong, Karol Sophia Agape R. Padilla, Rianna Patricia S. Cruz, El King D. Morado, Joshua Gregor A. Dizon, Razel Nikka M. Hao, Arianne A. Zamora, Devon Ray Pacial, Juan Antonio R. Magalang, Marissa Alejandria, Celia Carlos, Anna Ong-Lim, Edsel Maurice Salvaña, John Q. Wong, Jaime C. Montoya, Maria Rosario Singh-Vergeire and Cynthia P. Saloma |
| EPI_ISL_2156022 | Bohol Containerized PCR Laboratory | Philippine Genome Center | Francis A. Tablizo, Kenneth M. Kim, Carlo M. Lapid, Marc Jerrone R. Castro, Maria Sofia L. Yangzon, Benedict A. Maralit, Marc Edsel C. Ayes, Eva Maria Cutiongco-de la Paz, Alethea R. de Guzman, Jan Michael C. Yap, Jo-Hannah S. Llames, Sheila Mae M. Araiza, Kris P. Punayan, Irish Coleen A. Asin, Candice Francheska B. Tambaoan, Asia Louisa U. Chong, Karol Sophia Agape R. Padilla, Rianna Patricia S. Cruz, El King D. Morado, Joshua Gregor A. Dizon, Razel Nikka M. Hao, Arianne A. Zamora, Devon Ray Pacial, Juan Antonio R. Magalang, Marissa Alejandria, Celia Carlos, Anna Ong-Lim, Edsel Maurice Salvaña, John Q. Wong, Jaime C. Montoya, Maria Rosario Singh-Vergeire and Cynthia P. Saloma |
| EPI_ISL_2156099, EPI_ISL_2156101, EPI_ISL_2156103, EPI_ISL_2156104, EPI_ISL_2156106, EPI_ISL_2156107, EPI_ISL_2156108 | The Lord's Grace Medical and Industrial Clinic | Philippine Genome Center | Francis A. Tablizo, Kenneth M. Kim, Carlo M. Lapid, Marc Jerrone R. Castro, Maria Sofia L. Yangzon, Benedict A. Maralit, Marc Edsel C. Ayes, Eva Maria Cutiongco-de la Paz, Alethea R. de Guzman, Jan Michael C. Yap, Jo-Hannah S. Llames, Sheila Mae M. Araiza, Kris P. Punayan, Irish Coleen A. Asin, Candice Francheska B. Tambaoan, Asia Louisa U. Chong, Karol Sophia Agape R. Padilla, Rianna Patricia S. Cruz, El King D. Morado, Joshua Gregor A. Dizon, Razel Nikka M. Hao, Arianne A. Zamora, Devon Ray Pacial, Juan Antonio R. Magalang, Marissa Alejandria, Celia Carlos, Anna Ong-Lim, Edsel Maurice Salvaña, John Q. Wong, Jaime C. Montoya, Maria Rosario Singh-Vergeire and Cynthia P. Saloma |
| EPI_ISL_2156109 | Pasig City Children's Hospital - Child's Hope | Philippine Genome Center | Francis A. Tablizo, Kenneth M. Kim, Carlo M. Lapid, Marc Jerrone R. Castro, Maria Sofia L. Yangzon, Benedict A. Maralit, Marc Edsel C. Ayes, Eva Maria Cutiongco-de la Paz, Alethea R. de Guzman, Jan Michael C. Yap, Jo-Hannah S. Llames, Sheila Mae M. Araiza, Kris P. Punayan, Irish Coleen A. Asin, Candice Francheska B. Tambaoan, Asia Louisa U. Chong, Karol Sophia Agape R. Padilla, Rianna Patricia S. Cruz, El King D. Morado, Joshua Gregor A. Dizon, Razel Nikka M. Hao, Arianne A. Zamora, Devon Ray Pacial, Juan Antonio R. Magalang, Marissa Alejandria, Celia Carlos, Anna Ong-Lim, Edsel Maurice Salvaña, John Q. Wong, Jaime C. Montoya, Maria Rosario Singh-Vergeire and Cynthia P. Saloma |
| EPI_ISL_2156111 | Parkway Medical and Diagnostic Center | Philippine Genome Center | Francis A. Tablizo, Kenneth M. Kim, Carlo M. Lapid, Marc Jerrone R. Castro, Maria Sofia L. Yangzon, Benedict A. Maralit, Marc Edsel C. Ayes, Eva Maria Cutiongco-de la Paz, Alethea R. de Guzman, Jan Michael C. Yap, Jo-Hannah S. Llames, Sheila Mae M. Araiza, Kris P. Punayan, Irish Coleen A. Asin, Candice Francheska B. Tambaoan, Asia Louisa U. Chong, Karol Sophia Agape R. Padilla, Rianna Patricia S. Cruz, El King D. Morado, Joshua Gregor A. Dizon, Razel Nikka M. Hao, Arianne A. Zamora, Devon Ray Pacial, Juan Antonio R. Magalang, Marissa Alejandria, Celia Carlos, Anna Ong-Lim, Edsel Maurice Salvaña, John Q. Wong, Jaime C. Montoya, Maria Rosario Singh-Vergeire and Cynthia P. Saloma |
| EPI_ISL_2156116 | Hi-Precision Diagnostic Center (QC) | Philippine Genome Center | Francis A. Tablizo, Kenneth M. Kim, Carlo M. Lapid, Marc Jerrone R. Castro, Maria Sofia L. Yangzon, Benedict A. Maralit, Marc Edsel C. Ayes, Eva Maria Cutiongco-de la Paz, Alethea R. de Guzman, Jan Michael C. Yap, Jo-Hannah S. Llames, Sheila Mae M. Araiza, Kris P. Punayan, Irish Coleen A. Asin, Candice Francheska B. Tambaoan, Asia Louisa U. Chong, Karol Sophia Agape R. Padilla, Rianna Patricia S. Cruz, El King D. Morado, Joshua Gregor A. Dizon, Razel Nikka M. Hao, Arianne A. Zamora, Devon Ray Pacial, Juan Antonio R. Magalang, Marissa Alejandria, Celia Carlos, Anna Ong-Lim, Edsel Maurice Salvaña, John Q. Wong, Jaime C. Montoya, Maria Rosario Singh-Vergeire and Cynthia P. Saloma |
| EPI_ISL_2156117 | University of the East Ramon Magsaysay Memorial Medical Center (UERM) | Philippine Genome Center | Francis A. Tablizo, Kenneth M. Kim, Carlo M. Lapid, Marc Jerrone R. Castro, Maria Sofia L. Yangzon, Benedict A. Maralit, Marc Edsel C. Ayes, Eva Maria Cutiongco-de la Paz, Alethea R. de Guzman, Jan Michael C. Yap, Jo-Hannah S. Llames, Sheila Mae M. Araiza, Kris P. Punayan, Irish Coleen A. Asin, Candice Francheska B. Tambaoan, Asia Louisa U. Chong, Karol Sophia Agape R. Padilla, Rianna Patricia S. Cruz, El King D. Morado, Joshua Gregor A. Dizon, Razel Nikka M. Hao, Arianne A. Zamora, Devon Ray Pacial, Juan Antonio R. Magalang, Marissa Alejandria, Celia Carlos, Anna Ong-Lim, Edsel Maurice Salvaña, John Q. Wong, Jaime C. Montoya, Maria Rosario Singh-Vergeire and Cynthia P. Saloma |
| EPI_ISL_2156129, EPI_ISL_2156130, EPI_ISL_2156133, EPI_ISL_2156134, EPI_ISL_2156136, EPI_ISL_2156139, EPI_ISL_2156140 | Baguio General Hospital Medical Center (BGHMC) | Philippine Genome Center | Francis A. Tablizo, Kenneth M. Kim, Carlo M. Lapid, Marc Jerrone R. Castro, Maria Sofia L. Yangzon, Benedict A. Maralit, Marc Edsel C. Ayes, Eva Maria Cutiongco-de la Paz, Alethea R. de Guzman, Jan Michael C. Yap, Jo-Hannah S. Llames, Sheila Mae M. Araiza, Kris P. Punayan, Irish Coleen A. Asin, Candice Francheska B. Tambaoan, Asia Louisa U. Chong, Karol Sophia Agape R. Padilla, Rianna Patricia S. Cruz, El King D. Morado, Joshua Gregor A. Dizon, Razel Nikka M. Hao, Arianne A. Zamora, Devon Ray Pacial, Juan Antonio R. Magalang, Marissa Alejandria, Celia Carlos, Anna Ong-Lim, Edsel Maurice Salvaña, John Q. Wong, Jaime C. Montoya, Maria Rosario Singh-Vergeire and Cynthia P. Saloma |
| EPI_ISL_2156146 | Bataan General Hospital and Medical Center | Philippine Genome Center | Francis A. Tablizo, Kenneth M. Kim, Carlo M. Lapid, Marc Jerrone R. Castro, Maria Sofia L. Yangzon, Benedict A. Maralit, Marc Edsel C. Ayes, Eva Maria Cutiongco-de la Paz, Alethea R. de Guzman, Jan Michael C. Yap, Jo-Hannah S. Llames, Sheila Mae M. Araiza, Kris P. Punayan, Irish Coleen A. Asin, Candice Francheska B. Tambaoan, Asia Louisa U. Chong, Karol Sophia Agape R. Padilla, Rianna Patricia S. Cruz, El King D. Morado, Joshua Gregor A. Dizon, Razel Nikka M. Hao, Arianne A. Zamora, Devon Ray Pacial, Juan Antonio R. Magalang, Marissa Alejandria, Celia Carlos, Anna Ong-Lim, Edsel Maurice Salvaña, John Q. Wong, Jaime C. Montoya, Maria Rosario Singh-Vergeire and Cynthia P. Saloma |
| EPI_ISL_2156159, EPI_ISL_2156162 | Philippine Airport Diagnostic Laboratory | Philippine Genome Center | Francis A. Tablizo, Kenneth M. Kim, Carlo M. Lapid, Marc Jerrone R. Castro, Maria Sofia L. Yangzon, Benedict A. Maralit, Marc Edsel C. Ayes, Eva Maria Cutiongco-de la Paz, Alethea R. de Guzman, Jan Michael C. Yap, Jo-Hannah S. Llames, Sheila Mae M. Araiza, Kris P. Punayan, Irish Coleen A. Asin, Candice Francheska B. Tambaoan, Asia Louisa U. Chong, Karol Sophia Agape R. Padilla, Rianna Patricia S. Cruz, El King D. Morado, Joshua Gregor A. Dizon, Razel Nikka M. Hao, Arianne A. Zamora, Devon Ray Pacial, Juan Antonio R. Magalang, Marissa Alejandria, Celia Carlos, Anna Ong-Lim, Edsel Maurice Salvaña, John Q. Wong, Jaime C. Montoya, Maria Rosario Singh-Vergeire and Cynthia P. Saloma |
| EPI_ISL_2156168 | Dr. Jorge P. Royeca Hospital | Philippine Genome Center | Francis A. Tablizo, Kenneth M. Kim, Carlo M. Lapid, Marc Jerrone R. Castro, Maria Sofia L. Yangzon, Benedict A. Maralit, Marc Edsel C. Ayes, Eva Maria Cutiongco-de la Paz, Alethea R. de Guzman, Jan Michael C. Yap, Jo-Hannah S. Llames, Sheila Mae M. Araiza, Kris P. Punayan, Irish Coleen A. Asin, Candice Francheska B. Tambaoan, Asia Louisa U. Chong, Karol Sophia Agape R. Padilla, Rianna Patricia S. Cruz, El King D. Morado, Joshua Gregor A. Dizon, Razel Nikka M. Hao, Arianne A. Zamora, Devon Ray Pacial, Juan Antonio R. Magalang, Marissa Alejandria, Celia Carlos, Anna Ong-Lim, Edsel Maurice Salvaña, John Q. Wong, Jaime C. Montoya, Maria Rosario Singh-Vergeire and Cynthia P. Saloma |
| EPI_ISL_2156187 | Northern Mindanao TB Regional Center | Philippine Genome Center | Francis A. Tablizo, Kenneth M. Kim, Carlo M. Lapid, Marc Jerrone R. Castro, Maria Sofia L. Yangzon, Benedict A. Maralit, Marc Edsel C. Ayes, Eva Maria Cutiongco-de la Paz, Alethea R. de Guzman, Jan Michael C. Yap, Jo-Hannah S. Llames, Sheila Mae M. Araiza, Kris P. Punayan, Irish Coleen A. Asin, Candice Francheska B. Tambaoan, Asia Louisa U. Chong, Karol Sophia Agape R. Padilla, Rianna Patricia S. Cruz, El King D. Morado, Joshua Gregor A. Dizon, Razel Nikka M. Hao, Arianne A. Zamora, Devon Ray Pacial, Juan Antonio R. Magalang, Marissa Alejandria, Celia Carlos, Anna Ong-Lim, Edsel Maurice Salvaña, John Q. Wong, Jaime C. Montoya, Maria Rosario Singh-Vergeire and Cynthia P. Saloma |
| EPI_ISL_2156189 | Graman Medical & Maternity Hospital Inc Molecular Laboratory | Philippine Genome Center | Francis A. Tablizo, Kenneth M. Kim, Carlo M. Lapid, Marc Jerrone R. Castro, Maria Sofia L. Yangzon, Benedict A. Maralit, Marc Edsel C. Ayes, Eva Maria Cutiongco-de la Paz, Alethea R. de Guzman, Jan Michael C. Yap, Jo-Hannah S. Llames, Sheila Mae M. Araiza, Kris P. Punayan, Irish Coleen A. Asin, Candice Francheska B. Tambaoan, Asia Louisa U. Chong, Karol Sophia Agape R. Padilla, Rianna Patricia S. Cruz, El King D. Morado, Joshua Gregor A. Dizon, Razel Nikka M. Hao, Arianne A. Zamora, Devon Ray Pacial, Juan Antonio R. Magalang, Marissa Alejandria, Celia Carlos, Anna Ong-Lim, Edsel Maurice Salvaña, John Q. Wong, Jaime C. Montoya, Maria Rosario Singh-Vergeire and Cynthia P. Saloma |
| EPI_ISL_2156190, EPI_ISL_2156191, EPI_ISL_2156194, EPI_ISL_2156196 | Ilocos Training and Regional Medical Center | Philippine Genome Center | Francis A. Tablizo, Kenneth M. Kim, Carlo M. Lapid, Marc Jerrone R. Castro, Maria Sofia L. Yangzon, Benedict A. Maralit, Marc Edsel C. Ayes, Eva Maria Cutiongco-de la Paz, Alethea R. de Guzman, Jan Michael C. Yap, Jo-Hannah S. Llames, Sheila Mae M. Araiza, Kris P. Punayan, Irish Coleen A. Asin, Candice Francheska B. Tambaoan, Asia Louisa U. Chong, Karol Sophia Agape R. Padilla, Rianna Patricia S. Cruz, El King D. Morado, Joshua Gregor A. Dizon, Razel Nikka M. Hao, Arianne A. Zamora, Devon Ray Pacial, Juan Antonio R. Magalang, Marissa Alejandria, Celia Carlos, Anna Ong-Lim, Edsel Maurice Salvaña, John Q. Wong, Jaime C. Montoya, Maria Rosario Singh-Vergeire and Cynthia P. Saloma |
| EPI_ISL_2156198 | Prime Care Alpha Covid-19 Testing Laboratory | Philippine Genome Center | Francis A. Tablizo, Kenneth M. Kim, Carlo M. Lapid, Marc Jerrone R. Castro, Maria Sofia L. Yangzon, Benedict A. Maralit, Marc Edsel C. Ayes, Eva Maria Cutiongco-de la Paz, Alethea R. de Guzman, Jan Michael C. Yap, Jo-Hannah S. Llames, Sheila Mae M. Araiza, Kris P. Punayan, Irish Coleen A. Asin, Candice Francheska B. Tambaoan, Asia Louisa U. Chong, Karol Sophia Agape R. Padilla, Rianna Patricia S. Cruz, El King D. Morado, Joshua Gregor A. Dizon, Razel Nikka M. Hao, Arianne A. Zamora, Devon Ray Pacial, Juan Antonio R. Magalang, Marissa Alejandria, Celia Carlos, Anna Ong-Lim, Edsel Maurice Salvaña, John Q. Wong, Jaime C. Montoya, Maria Rosario Singh-Vergeire and Cynthia P. Saloma |
| EPI_ISL_2156201 | Philippine Red Cross - Clark Molecular Laboratory | Philippine Genome Center | Francis A. Tablizo, Kenneth M. Kim, Carlo M. Lapid, Marc Jerrone R. Castro, Maria Sofia L. Yangzon, Benedict A. Maralit, Marc Edsel C. Ayes, Eva Maria Cutiongco-de la Paz, Alethea R. de Guzman, Jan Michael C. Yap, Jo-Hannah S. Llames, Sheila Mae M. Araiza, Kris P. Punayan, Irish Coleen A. Asin, Candice Francheska B. Tambaoan, Asia Louisa U. Chong, Karol Sophia Agape R. Padilla, Rianna Patricia S. Cruz, El King D. Morado, Joshua Gregor A. Dizon, Razel Nikka M. Hao, Arianne A. Zamora, Devon Ray Pacial, Juan Antonio R. Magalang, Marissa Alejandria, Celia Carlos, Anna Ong-Lim, Edsel Maurice Salvaña, John Q. Wong, Jaime C. Montoya, Maria Rosario Singh-Vergeire and Cynthia P. Saloma |

| | | | |
|---|---|---|---|
| EPI_ISL_2156202, EPI_ISL_2156203, EPI_ISL_2156205 | The Medical City | Philippine Genome Center | Francis A. Tablizo, Kenneth M. Kim, Carlo M. Lapid, Marc Jerrone R. Castro, Maria Sofia L. Yangzon, Benedict A. Maralit, Marc Edsel C. Ayes, Eva Maria Cutiongco-de la Paz, Alethea R. de Guzman, Jan Michael C. Yap, Jo-Hannah S. Llames, Sheila Mae M. Araiza, Kris P. Punayan, Irish Coleen A. Asin, Candice Francheska B. Tambaoan, Asia Louisa U. Chong, Karol Sophia Agape R. Padilla, Rianna Patricia S. Cruz, El King D. Morado, Joshua Gregor A. Dizon, Razel Nikka M. Hao, Arianne A. Zamora, Devon Ray Pacial, Juan Antonio R. Magalang, Marissa Alejandria, Celia Carlos, Anna Ong-Lim, Edsel Maurice Salvaña, John Q. Wong, Jaime C. Montoya, Maria Rosario Singh-Vergeire and Cynthia P. Saloma |
| EPI_ISL_2156206, EPI_ISL_2156210 | Cagayan Valley Medical Center Molecular Laboratory | Philippine Genome Center | Francis A. Tablizo, Kenneth M. Kim, Carlo M. Lapid, Marc Jerrone R. Castro, Maria Sofia L. Yangzon, Benedict A. Maralit, Marc Edsel C. Ayes, Eva Maria Cutiongco-de la Paz, Alethea R. de Guzman, Jan Michael C. Yap, Jo-Hannah S. Llames, Sheila Mae M. Araiza, Kris P. Punayan, Irish Coleen A. Asin, Candice Francheska B. Tambaoan, Asia Louisa U. Chong, Karol Sophia Agape R. Padilla, Rianna Patricia S. Cruz, El King D. Morado, Joshua Gregor A. Dizon, Razel Nikka M. Hao, Arianne A. Zamora, Devon Ray Pacial, Juan Antonio R. Magalang, Marissa Alejandria, Celia Carlos, Anna Ong-Lim, Edsel Maurice Salvaña, John Q. Wong, Jaime C. Montoya, Maria Rosario Singh-Vergeire and Cynthia P. Saloma |
| EPI_ISL_2156220 | Philippine Red Cross Logistics and Multipurpose Center | Philippine Genome Center | Francis A. Tablizo, Kenneth M. Kim, Carlo M. Lapid, Marc Jerrone R. Castro, Maria Sofia L. Yangzon, Benedict A. Maralit, Marc Edsel C. Ayes, Eva Maria Cutiongco-de la Paz, Alethea R. de Guzman, Jan Michael C. Yap, Jo-Hannah S. Llames, Sheila Mae M. Araiza, Kris P. Punayan, Irish Coleen A. Asin, Candice Francheska B. Tambaoan, Asia Louisa U. Chong, Karol Sophia Agape R. Padilla, Rianna Patricia S. Cruz, El King D. Morado, Joshua Gregor A. Dizon, Razel Nikka M. Hao, Arianne A. Zamora, Devon Ray Pacial, Juan Antonio R. Magalang, Marissa Alejandria, Celia Carlos, Anna Ong-Lim, Edsel Maurice Salvaña, John Q. Wong, Jaime C. Montoya, Maria Rosario Singh-Vergeire and Cynthia P. Saloma |
| EPI_ISL_2156222 | First Aide Diagnostic Center | Philippine Genome Center | Francis A. Tablizo, Kenneth M. Kim, Carlo M. Lapid, Marc Jerrone R. Castro, Maria Sofia L. Yangzon, Benedict A. Maralit, Marc Edsel C. Ayes, Eva Maria Cutiongco-de la Paz, Alethea R. de Guzman, Jan Michael C. Yap, Jo-Hannah S. Llames, Sheila Mae M. Araiza, Kris P. Punayan, Irish Coleen A. Asin, Candice Francheska B. Tambaoan, Asia Louisa U. Chong, Karol Sophia Agape R. Padilla, Rianna Patricia S. Cruz, El King D. Morado, Joshua Gregor A. Dizon, Razel Nikka M. Hao, Arianne A. Zamora, Devon Ray Pacial, Juan Antonio R. Magalang, Marissa Alejandria, Celia Carlos, Anna Ong-Lim, Edsel Maurice Salvaña, John Q. Wong, Jaime C. Montoya, Maria Rosario Singh-Vergeire and Cynthia P. Saloma |
| EPI_ISL_2156223, EPI_ISL_2156225 | Lung Center of the Philippines (LCP) | Philippine Genome Center | Francis A. Tablizo, Kenneth M. Kim, Carlo M. Lapid, Marc Jerrone R. Castro, Maria Sofia L. Yangzon, Benedict A. Maralit, Marc Edsel C. Ayes, Eva Maria Cutiongco-de la Paz, Alethea R. de Guzman, Jan Michael C. Yap, Jo-Hannah S. Llames, Sheila Mae M. Araiza, Kris P. Punayan, Irish Coleen A. Asin, Candice Francheska B. Tambaoan, Asia Louisa U. Chong, Karol Sophia Agape R. Padilla, Rianna Patricia S. Cruz, El King D. Morado, Joshua Gregor A. Dizon, Razel Nikka M. Hao, Arianne A. Zamora, Devon Ray Pacial, Juan Antonio R. Magalang, Marissa Alejandria, Celia Carlos, Anna Ong-Lim, Edsel Maurice Salvaña, John Q. Wong, Jaime C. Montoya, Maria Rosario Singh-Vergeire and Cynthia P. Saloma |
| EPI_ISL_2156238, EPI_ISL_2156239, EPI_ISL_2156243 | Philippine Red Cross - Port Area | Philippine Genome Center | Francis A. Tablizo, Kenneth M. Kim, Carlo M. Lapid, Marc Jerrone R. Castro, Maria Sofia L. Yangzon, Benedict A. Maralit, Marc Edsel C. Ayes, Eva Maria Cutiongco-de la Paz, Alethea R. de Guzman, Jan Michael C. Yap, Jo-Hannah S. Llames, Sheila Mae M. Araiza, Kris P. Punayan, Irish Coleen A. Asin, Candice Francheska B. Tambaoan, Asia Louisa U. Chong, Karol Sophia Agape R. Padilla, Rianna Patricia S. Cruz, El King D. Morado, Joshua Gregor A. Dizon, Razel Nikka M. Hao, Arianne A. Zamora, Devon Ray Pacial, Juan Antonio R. Magalang, Marissa Alejandria, Celia Carlos, Anna Ong-Lim, Edsel Maurice Salvaña, John Q. Wong, Jaime C. Montoya, Maria Rosario Singh-Vergeire and Cynthia P. Saloma |
| EPI_ISL_2156250, EPI_ISL_2156251 | Safeguard DNA Diagnostic, Inc. | Philippine Genome Center | Francis A. Tablizo, Kenneth M. Kim, Carlo M. Lapid, Marc Jerrone R. Castro, Maria Sofia L. Yangzon, Benedict A. Maralit, Marc Edsel C. Ayes, Eva Maria Cutiongco-de la Paz, Alethea R. de Guzman, Jan Michael C. Yap, Jo-Hannah S. Llames, Sheila Mae M. Araiza, Kris P. Punayan, Irish Coleen A. Asin, Candice Francheska B. Tambaoan, Asia Louisa U. Chong, Karol Sophia Agape R. Padilla, Rianna Patricia S. Cruz, El King D. Morado, Joshua Gregor A. Dizon, Razel Nikka M. Hao, Arianne A. Zamora, Devon Ray Pacial, Juan Antonio R. Magalang, Marissa Alejandria, Celia Carlos, Anna Ong-Lim, Edsel Maurice Salvaña, John Q. Wong, Jaime C. Montoya, Maria Rosario Singh-Vergeire and Cynthia P. Saloma |
| EPI_ISL_2156315 | Philippine Red Cross - Port Area | Philippine Genome Center | Francis A. Tablizo, Kenneth M. Kim, Carlo M. Lapid, Marc Jerrone R. Castro, Maria Sofia L. Yangzon, Elcid Aaron R. Pangilinan, Benedict A. Maralit, Marc Edsel C. Ayes, Eva Maria Cutiongco-de la Paz, Alethea R. de Guzman, Jan Michael C. Yap, Jo-Hannah S. Llames, Sheila Mae M. Araiza, Kris P. Punayan, Irish Coleen A. Asin, Candice Francheska B. Tambaoan, Asia Louisa U. Chong, Karol Sophia Agape R. Padilla, Rianna Patricia S. Cruz, El King D. Morado, Joshua Gregor A. Dizon, Razel Nikka M. Hao, Arianne A. Zamora, Devon Ray Pacial, Juan Antonio R. Magalang, Marissa Alejandria, Celia Carlos, Anna Ong-Lim, Edsel Maurice Salvaña, John Q. Wong, Jaime C. Montoya, Maria Rosario Singh-Vergeire and Cynthia P. Saloma |
| EPI_ISL_2156319 | BULACAN MEDICAL CENTER | Philippine Genome Center | Francis A. Tablizo, Kenneth M. Kim, Carlo M. Lapid, Marc Jerrone R. Castro, Maria Sofia L. Yangzon, Elcid Aaron R. Pangilinan, Benedict A. Maralit, Marc Edsel C. Ayes, Eva Maria Cutiongco-de la Paz, Alethea R. de Guzman, Jan Michael C. Yap, Jo-Hannah S. Llames, Sheila Mae M. Araiza, Kris P. Punayan, Irish Coleen A. Asin, Candice Francheska B. Tambaoan, Asia Louisa U. Chong, Karol Sophia Agape R. Padilla, Rianna Patricia S. Cruz, El King D. Morado, Joshua Gregor A. Dizon, Razel Nikka M. Hao, Arianne A. Zamora, Devon Ray Pacial, Juan Antonio R. Magalang, Marissa Alejandria, Celia Carlos, Anna Ong-Lim, Edsel Maurice Salvaña, John Q. Wong, Jaime C. Montoya, Maria Rosario Singh-Vergeire and Cynthia P. Saloma |
| EPI_ISL_2156320 | Philippine Red Cross - National Blood Center | Philippine Genome Center | Francis A. Tablizo, Kenneth M. Kim, Carlo M. Lapid, Marc Jerrone R. Castro, Maria Sofia L. Yangzon, Elcid Aaron R. Pangilinan, Benedict A. Maralit, Marc Edsel C. Ayes, Eva Maria Cutiongco-de la Paz, Alethea R. de Guzman, Jan Michael C. Yap, Jo-Hannah S. Llames, Sheila Mae M. Araiza, Kris P. Punayan, Irish Coleen A. Asin, Candice Francheska B. Tambaoan, Asia Louisa U. Chong, Karol Sophia Agape R. Padilla, Rianna Patricia S. Cruz, El King D. Morado, Joshua Gregor A. Dizon, Razel Nikka M. Hao, Arianne A. Zamora, Devon Ray Pacial, Juan Antonio R. Magalang, Marissa Alejandria, Celia Carlos, Anna Ong-Lim, Edsel Maurice Salvaña, John Q. Wong, Jaime C. Montoya, Maria Rosario Singh-Vergeire and Cynthia P. Saloma |
| EPI_ISL_2156323 | Philippine Red Cross Logistics and Multipurpose Center | Philippine Genome Center | Francis A. Tablizo, Kenneth M. Kim, Carlo M. Lapid, Marc Jerrone R. Castro, Maria Sofia L. Yangzon, Elcid Aaron R. Pangilinan, Benedict A. Maralit, Marc Edsel C. Ayes, Eva Maria Cutiongco-de la Paz, Alethea R. de Guzman, Jan Michael C. Yap, Jo-Hannah S. Llames, Sheila Mae M. Araiza, Kris P. Punayan, Irish Coleen A. Asin, Candice Francheska B. Tambaoan, Asia Louisa U. Chong, Karol Sophia Agape R. Padilla, Rianna Patricia S. Cruz, El King D. Morado, Joshua Gregor A. Dizon, Razel Nikka M. Hao, Arianne A. Zamora, Devon Ray Pacial, Juan Antonio R. Magalang, Marissa Alejandria, Celia Carlos, Anna Ong-Lim, Edsel Maurice Salvaña, John Q. Wong, Jaime C. Montoya, Maria Rosario Singh-Vergeire and Cynthia P. Saloma |
| EPI_ISL_2156325 | Philippine Red Cross - Clark Molecular Laboratory | Philippine Genome Center | Francis A. Tablizo, Kenneth M. Kim, Carlo M. Lapid, Marc Jerrone R. Castro, Maria Sofia L. Yangzon, Elcid Aaron R. Pangilinan, Benedict A. Maralit, Marc Edsel C. Ayes, Eva Maria Cutiongco-de la Paz, Alethea R. de Guzman, Jan Michael C. Yap, Jo-Hannah S. Llames, Sheila Mae M. Araiza, Kris P. Punayan, Irish Coleen A. Asin, Candice Francheska B. Tambaoan, Asia Louisa U. Chong, Karol Sophia Agape R. Padilla, Rianna Patricia S. Cruz, El King D. Morado, Joshua Gregor A. Dizon, Razel Nikka M. Hao, Arianne A. Zamora, Devon Ray Pacial, Juan Antonio R. Magalang, Marissa Alejandria, Celia Carlos, Anna Ong-Lim, Edsel Maurice Salvaña, John Q. Wong, Jaime C. Montoya, Maria Rosario Singh-Vergeire and Cynthia P. Saloma |
| EPI_ISL_2156328 | PHILIPPINE AIRPORT DIAGNOSTIC LABORATORY | Philippine Genome Center | Francis A. Tablizo, Kenneth M. Kim, Carlo M. Lapid, Marc Jerrone R. Castro, Maria Sofia L. Yangzon, Elcid Aaron R. Pangilinan, Benedict A. Maralit, Marc Edsel C. Ayes, Eva Maria Cutiongco-de la Paz, Alethea R. de Guzman, Jan Michael C. Yap, Jo-Hannah S. Llames, Sheila Mae M. Araiza, Kris P. Punayan, Irish Coleen A. Asin, Candice Francheska B. Tambaoan, Asia Louisa U. Chong, Karol Sophia Agape R. Padilla, Rianna Patricia S. Cruz, El King D. Morado, Joshua Gregor A. Dizon, Razel Nikka M. Hao, Arianne A. Zamora, Devon Ray Pacial, Juan Antonio R. Magalang, Marissa Alejandria, Celia Carlos, Anna Ong-Lim, Edsel Maurice Salvaña, John Q. Wong, Jaime C. Montoya, Maria Rosario Singh-Vergeire and Cynthia P. Saloma |
| EPI_ISL_2156329 | Baguio General Hospital Medical Center (BGHMC) | Philippine Genome Center | Francis A. Tablizo, Kenneth M. Kim, Carlo M. Lapid, Marc Jerrone R. Castro, Maria Sofia L. Yangzon, Elcid Aaron R. Pangilinan, Benedict A. Maralit, Marc Edsel C. Ayes, Eva Maria Cutiongco-de la Paz, Alethea R. de Guzman, Jan Michael C. Yap, Jo-Hannah S. Llames, Sheila Mae M. Araiza, Kris P. Punayan, Irish Coleen A. Asin, Candice Francheska B. Tambaoan, Asia Louisa U. Chong, Karol Sophia Agape R. Padilla, Rianna Patricia S. Cruz, El King D. Morado, Joshua Gregor A. Dizon, Razel Nikka M. Hao, Arianne A. Zamora, Devon Ray Pacial, Juan Antonio R. Magalang, Marissa Alejandria, Celia Carlos, Anna Ong-Lim, Edsel Maurice Salvaña, John Q. Wong, Jaime C. Montoya, Maria Rosario Singh-Vergeire and Cynthia P. Saloma |
| EPI_ISL_2156338, EPI_ISL_2156339 | Philippine Red Cross - Clark Molecular Laboratory | Philippine Genome Center | Francis A. Tablizo, Kenneth M. Kim, Carlo M. Lapid, Marc Jerrone R. Castro, Maria Sofia L. Yangzon, Elcid Aaron R. Pangilinan, Benedict A. Maralit, Marc Edsel C. Ayes, Eva Maria Cutiongco-de la Paz, Alethea R. de Guzman, Jan Michael C. Yap, Jo-Hannah S. Llames, Sheila Mae M. Araiza, Kris P. Punayan, Irish Coleen A. Asin, Candice Francheska B. Tambaoan, Asia Louisa U. Chong, Karol Sophia Agape R. Padilla, Rianna Patricia S. Cruz, El King D. Morado, Joshua Gregor A. Dizon, Razel Nikka M. Hao, Arianne A. Zamora, Devon Ray Pacial, Juan Antonio R. Magalang, Marissa Alejandria, Celia Carlos, Anna Ong-Lim, Edsel Maurice Salvaña, John Q. Wong, Jaime C. Montoya, Maria Rosario Singh-Vergeire and Cynthia P. Saloma |
| EPI_ISL_2156340, EPI_ISL_2156343 | Butuan Medical Center | Philippine Genome Center | Francis A. Tablizo, Kenneth M. Kim, Carlo M. Lapid, Marc Jerrone R. Castro, Maria Sofia L. Yangzon, Elcid Aaron R. Pangilinan, Benedict A. Maralit, Marc Edsel C. Ayes, Eva Maria Cutiongco-de la Paz, Alethea R. de Guzman, Jan Michael C. Yap, Jo-Hannah S. Llames, Sheila Mae M. Araiza, Kris P. Punayan, Irish Coleen A. Asin, Candice Francheska B. Tambaoan, Asia Louisa U. Chong, Karol Sophia Agape R. Padilla, Rianna Patricia S. Cruz, El King D. Morado, Joshua Gregor A. Dizon, Razel Nikka M. Hao, Arianne A. Zamora, Devon Ray Pacial, Juan Antonio R. Magalang, Marissa Alejandria, Celia Carlos, Anna Ong-Lim, Edsel Maurice Salvaña, John Q. Wong, Jaime C. Montoya, Maria Rosario Singh-Vergeire and Cynthia P. Saloma |

| | | | |
|---|---|---|---|
| EPI_ISL_2156344, EPI_ISL_2156346 | Baguio General Hospital Medical Center (BGHMC) | Philippine Genome Center | Francis A. Tablizo, Kenneth M. Kim, Carlo M. Lapid, Marc Jerrone R. Castro, Maria Sofia L. Yangzon, Elcid Aaron R. Pangilinan, Benedict A. Maralit, Marc Edsel C. Ayes, Eva Maria Cutiongco-de la Paz, Alethea R. de Guzman, Jan Michael C. Yap, Jo-Hannah S. Llames, Sheila Mae M. Araiza, Kris P. Punayan, Irish Coleen A. Asin, Candice Francheska B. Tambaoan, Asia Louisa U. Chong, Karol Sophia Agape R. Padilla, Rianna Patricia S. Cruz, El King D. Morado, Joshua Gregor A. Dizon, Razel Nikka M. Hao, Arianne A. Zamora, Devon Ray Pacial, Juan Antonio R. Magalang, Marissa Alejandria, Celia Carlos, Anna Ong-Lim, Edsel Maurice Salvaña, John Q. Wong, Jaime C. Montoya, Maria Rosario Singh-Vergeire and Cynthia P. Saloma |
| EPI_ISL_2156347, EPI_ISL_2156348 | Philippine Red Cross - Port Area | Philippine Genome Center | Francis A. Tablizo, Kenneth M. Kim, Carlo M. Lapid, Marc Jerrone R. Castro, Maria Sofia L. Yangzon, Elcid Aaron R. Pangilinan, Benedict A. Maralit, Marc Edsel C. Ayes, Eva Maria Cutiongco-de la Paz, Alethea R. de Guzman, Jan Michael C. Yap, Jo-Hannah S. Llames, Sheila Mae M. Araiza, Kris P. Punayan, Irish Coleen A. Asin, Candice Francheska B. Tambaoan, Asia Louisa U. Chong, Karol Sophia Agape R. Padilla, Rianna Patricia S. Cruz, El King D. Morado, Joshua Gregor A. Dizon, Razel Nikka M. Hao, Arianne A. Zamora, Devon Ray Pacial, Juan Antonio R. Magalang, Marissa Alejandria, Celia Carlos, Anna Ong-Lim, Edsel Maurice Salvaña, John Q. Wong, Jaime C. Montoya, Maria Rosario Singh-Vergeire and Cynthia P. Saloma |
| EPI_ISL_2156349 | The Medical City -Â Ortigas | Philippine Genome Center | Francis A. Tablizo, Kenneth M. Kim, Carlo M. Lapid, Marc Jerrone R. Castro, Maria Sofia L. Yangzon, Elcid Aaron R. Pangilinan, Benedict A. Maralit, Marc Edsel C. Ayes, Eva Maria Cutiongco-de la Paz, Alethea R. de Guzman, Jan Michael C. Yap, Jo-Hannah S. Llames, Sheila Mae M. Araiza, Kris P. Punayan, Irish Coleen A. Asin, Candice Francheska B. Tambaoan, Asia Louisa U. Chong, Karol Sophia Agape R. Padilla, Rianna Patricia S. Cruz, El King D. Morado, Joshua Gregor A. Dizon, Razel Nikka M. Hao, Arianne A. Zamora, Devon Ray Pacial, Juan Antonio R. Magalang, Marissa Alejandria, Celia Carlos, Anna Ong-Lim, Edsel Maurice Salvaña, John Q. Wong, Jaime C. Montoya, Maria Rosario Singh-Vergeire and Cynthia P. Saloma |
| EPI_ISL_2156350 | Philippine Red Cross - Clark Molecular Laboratory | Philippine Genome Center | Francis A. Tablizo, Kenneth M. Kim, Carlo M. Lapid, Marc Jerrone R. Castro, Maria Sofia L. Yangzon, Elcid Aaron R. Pangilinan, Benedict A. Maralit, Marc Edsel C. Ayes, Eva Maria Cutiongco-de la Paz, Alethea R. de Guzman, Jan Michael C. Yap, Jo-Hannah S. Llames, Sheila Mae M. Araiza, Kris P. Punayan, Irish Coleen A. Asin, Candice Francheska B. Tambaoan, Asia Louisa U. Chong, Karol Sophia Agape R. Padilla, Rianna Patricia S. Cruz, El King D. Morado, Joshua Gregor A. Dizon, Razel Nikka M. Hao, Arianne A. Zamora, Devon Ray Pacial, Juan Antonio R. Magalang, Marissa Alejandria, Celia Carlos, Anna Ong-Lim, Edsel Maurice Salvaña, John Q. Wong, Jaime C. Montoya, Maria Rosario Singh-Vergeire and Cynthia P. Saloma |
| EPI_ISL_2156351, EPI_ISL_2156352, EPI_ISL_2156353, EPI_ISL_2156354, EPI_ISL_2156356, EPI_ISL_2156360, EPI_ISL_2156361 | Research Institute for Tropical Medicine, Inc. (RITM) | Philippine Genome Center | Francis A. Tablizo, Kenneth M. Kim, Carlo M. Lapid, Marc Jerrone R. Castro, Maria Sofia L. Yangzon, Elcid Aaron R. Pangilinan, Benedict A. Maralit, Marc Edsel C. Ayes, Eva Maria Cutiongco-de la Paz, Alethea R. de Guzman, Jan Michael C. Yap, Jo-Hannah S. Llames, Sheila Mae M. Araiza, Kris P. Punayan, Irish Coleen A. Asin, Candice Francheska B. Tambaoan, Asia Louisa U. Chong, Karol Sophia Agape R. Padilla, Rianna Patricia S. Cruz, El King D. Morado, Joshua Gregor A. Dizon, Razel Nikka M. Hao, Arianne A. Zamora, Devon Ray Pacial, Juan Antonio R. Magalang, Marissa Alejandria, Celia Carlos, Anna Ong-Lim, Edsel Maurice Salvaña, John Q. Wong, Jaime C. Montoya, Maria Rosario Singh-Vergeire and Cynthia P. Saloma |
| EPI_ISL_2156364 | Pasig City Children's Hospital - Child's Hope | Philippine Genome Center | Francis A. Tablizo, Kenneth M. Kim, Carlo M. Lapid, Marc Jerrone R. Castro, Maria Sofia L. Yangzon, Elcid Aaron R. Pangilinan, Benedict A. Maralit, Marc Edsel C. Ayes, Eva Maria Cutiongco-de la Paz, Alethea R. de Guzman, Jan Michael C. Yap, Jo-Hannah S. Llames, Sheila Mae M. Araiza, Kris P. Punayan, Irish Coleen A. Asin, Candice Francheska B. Tambaoan, Asia Louisa U. Chong, Karol Sophia Agape R. Padilla, Rianna Patricia S. Cruz, El King D. Morado, Joshua Gregor A. Dizon, Razel Nikka M. Hao, Arianne A. Zamora, Devon Ray Pacial, Juan Antonio R. Magalang, Marissa Alejandria, Celia Carlos, Anna Ong-Lim, Edsel Maurice Salvaña, John Q. Wong, Jaime C. Montoya, Maria Rosario Singh-Vergeire and Cynthia P. Saloma |
| EPI_ISL_2156366 | Philippine Red Cross - Port Area | Philippine Genome Center | Francis A. Tablizo, Kenneth M. Kim, Carlo M. Lapid, Marc Jerrone R. Castro, Maria Sofia L. Yangzon, Elcid Aaron R. Pangilinan, Benedict A. Maralit, Marc Edsel C. Ayes, Eva Maria Cutiongco-de la Paz, Alethea R. de Guzman, Jan Michael C. Yap, Jo-Hannah S. Llames, Sheila Mae M. Araiza, Kris P. Punayan, Irish Coleen A. Asin, Candice Francheska B. Tambaoan, Asia Louisa U. Chong, Karol Sophia Agape R. Padilla, Rianna Patricia S. Cruz, El King D. Morado, Joshua Gregor A. Dizon, Razel Nikka M. Hao, Arianne A. Zamora, Devon Ray Pacial, Juan Antonio R. Magalang, Marissa Alejandria, Celia Carlos, Anna Ong-Lim, Edsel Maurice Salvaña, John Q. Wong, Jaime C. Montoya, Maria Rosario Singh-Vergeire and Cynthia P. Saloma |
| EPI_ISL_2156367 | Cagayan Valley Medical Center Molecular Laboratory | Philippine Genome Center | Francis A. Tablizo, Kenneth M. Kim, Carlo M. Lapid, Marc Jerrone R. Castro, Maria Sofia L. Yangzon, Elcid Aaron R. Pangilinan, Benedict A. Maralit, Marc Edsel C. Ayes, Eva Maria Cutiongco-de la Paz, Alethea R. de Guzman, Jan Michael C. Yap, Jo-Hannah S. Llames, Sheila Mae M. Araiza, Kris P. Punayan, Irish Coleen A. Asin, Candice Francheska B. Tambaoan, Asia Louisa U. Chong, Karol Sophia Agape R. Padilla, Rianna Patricia S. Cruz, El King D. Morado, Joshua Gregor A. Dizon, Razel Nikka M. Hao, Arianne A. Zamora, Devon Ray Pacial, Juan Antonio R. Magalang, Marissa Alejandria, Celia Carlos, Anna Ong-Lim, Edsel Maurice Salvaña, John Q. Wong, Jaime C. Montoya, Maria Rosario Singh-Vergeire and Cynthia P. Saloma |
| EPI_ISL_2156369, EPI_ISL_2156370 | The Medical City -Â Ortigas | Philippine Genome Center | Francis A. Tablizo, Kenneth M. Kim, Carlo M. Lapid, Marc Jerrone R. Castro, Maria Sofia L. Yangzon, Elcid Aaron R. Pangilinan, Benedict A. Maralit, Marc Edsel C. Ayes, Eva Maria Cutiongco-de la Paz, Alethea R. de Guzman, Jan Michael C. Yap, Jo-Hannah S. Llames, Sheila Mae M. Araiza, Kris P. Punayan, Irish Coleen A. Asin, Candice Francheska B. Tambaoan, Asia Louisa U. Chong, Karol Sophia Agape R. Padilla, Rianna Patricia S. Cruz, El King D. Morado, Joshua Gregor A. Dizon, Razel Nikka M. Hao, Arianne A. Zamora, Devon Ray Pacial, Juan Antonio R. Magalang, Marissa Alejandria, Celia Carlos, Anna Ong-Lim, Edsel Maurice Salvaña, John Q. Wong, Jaime C. Montoya, Maria Rosario Singh-Vergeire and Cynthia P. Saloma |
| EPI_ISL_2156371 | Davao One World Diagnostic Center Incorporated | Philippine Genome Center | Francis A. Tablizo, Kenneth M. Kim, Carlo M. Lapid, Marc Jerrone R. Castro, Maria Sofia L. Yangzon, Elcid Aaron R. Pangilinan, Benedict A. Maralit, Marc Edsel C. Ayes, Eva Maria Cutiongco-de la Paz, Alethea R. de Guzman, Jan Michael C. Yap, Jo-Hannah S. Llames, Sheila Mae M. Araiza, Kris P. Punayan, Irish Coleen A. Asin, Candice Francheska B. Tambaoan, Asia Louisa U. Chong, Karol Sophia Agape R. Padilla, Rianna Patricia S. Cruz, El King D. Morado, Joshua Gregor A. Dizon, Razel Nikka M. Hao, Arianne A. Zamora, Devon Ray Pacial, Juan Antonio R. Magalang, Marissa Alejandria, Celia Carlos, Anna Ong-Lim, Edsel Maurice Salvaña, John Q. Wong, Jaime C. Montoya, Maria Rosario Singh-Vergeire and Cynthia P. Saloma |
| EPI_ISL_2156372, EPI_ISL_2156375, EPI_ISL_2156378 | Research Institute for Tropical Medicine, Inc. (RITM) | Philippine Genome Center | Francis A. Tablizo, Kenneth M. Kim, Carlo M. Lapid, Marc Jerrone R. Castro, Maria Sofia L. Yangzon, Elcid Aaron R. Pangilinan, Benedict A. Maralit, Marc Edsel C. Ayes, Eva Maria Cutiongco-de la Paz, Alethea R. de Guzman, Jan Michael C. Yap, Jo-Hannah S. Llames, Sheila Mae M. Araiza, Kris P. Punayan, Irish Coleen A. Asin, Candice Francheska B. Tambaoan, Asia Louisa U. Chong, Karol Sophia Agape R. Padilla, Rianna Patricia S. Cruz, El King D. Morado, Joshua Gregor A. Dizon, Razel Nikka M. Hao, Arianne A. Zamora, Devon Ray Pacial, Juan Antonio R. Magalang, Marissa Alejandria, Celia Carlos, Anna Ong-Lim, Edsel Maurice Salvaña, John Q. Wong, Jaime C. Montoya, Maria Rosario Singh-Vergeire and Cynthia P. Saloma |
| EPI_ISL_2156380 | Butuan Medical Center | Philippine Genome Center | Francis A. Tablizo, Kenneth M. Kim, Carlo M. Lapid, Marc Jerrone R. Castro, Maria Sofia L. Yangzon, Elcid Aaron R. Pangilinan, Benedict A. Maralit, Marc Edsel C. Ayes, Eva Maria Cutiongco-de la Paz, Alethea R. de Guzman, Jan Michael C. Yap, Jo-Hannah S. Llames, Sheila Mae M. Araiza, Kris P. Punayan, Irish Coleen A. Asin, Candice Francheska B. Tambaoan, Asia Louisa U. Chong, Karol Sophia Agape R. Padilla, Rianna Patricia S. Cruz, El King D. Morado, Joshua Gregor A. Dizon, Razel Nikka M. Hao, Arianne A. Zamora, Devon Ray Pacial, Juan Antonio R. Magalang, Marissa Alejandria, Celia Carlos, Anna Ong-Lim, Edsel Maurice Salvaña, John Q. Wong, Jaime C. Montoya, Maria Rosario Singh-Vergeire and Cynthia P. Saloma |
| EPI_ISL_2156383, EPI_ISL_2156385, EPI_ISL_2156386, EPI_ISL_2156387 | Baguio General Hospital Medical Center (BGHMC) | Philippine Genome Center | Francis A. Tablizo, Kenneth M. Kim, Carlo M. Lapid, Marc Jerrone R. Castro, Maria Sofia L. Yangzon, Elcid Aaron R. Pangilinan, Benedict A. Maralit, Marc Edsel C. Ayes, Eva Maria Cutiongco-de la Paz, Alethea R. de Guzman, Jan Michael C. Yap, Jo-Hannah S. Llames, Sheila Mae M. Araiza, Kris P. Punayan, Irish Coleen A. Asin, Candice Francheska B. Tambaoan, Asia Louisa U. Chong, Karol Sophia Agape R. Padilla, Rianna Patricia S. Cruz, El King D. Morado, Joshua Gregor A. Dizon, Razel Nikka M. Hao, Arianne A. Zamora, Devon Ray Pacial, Juan Antonio R. Magalang, Marissa Alejandria, Celia Carlos, Anna Ong-Lim, Edsel Maurice Salvaña, John Q. Wong, Jaime C. Montoya, Maria Rosario Singh-Vergeire and Cynthia P. Saloma |
| EPI_ISL_2156389 | San Lazaro Hospital | Philippine Genome Center | Francis A. Tablizo, Kenneth M. Kim, Carlo M. Lapid, Marc Jerrone R. Castro, Maria Sofia L. Yangzon, Elcid Aaron R. Pangilinan, Benedict A. Maralit, Marc Edsel C. Ayes, Eva Maria Cutiongco-de la Paz, Alethea R. de Guzman, Jan Michael C. Yap, Jo-Hannah S. Llames, Sheila Mae M. Araiza, Kris P. Punayan, Irish Coleen A. Asin, Candice Francheska B. Tambaoan, Asia Louisa U. Chong, Karol Sophia Agape R. Padilla, Rianna Patricia S. Cruz, El King D. Morado, Joshua Gregor A. Dizon, Razel Nikka M. Hao, Arianne A. Zamora, Devon Ray Pacial, Juan Antonio R. Magalang, Marissa Alejandria, Celia Carlos, Anna Ong-Lim, Edsel Maurice Salvaña, John Q. Wong, Jaime C. Montoya, Maria Rosario Singh-Vergeire and Cynthia P. Saloma |
| EPI_ISL_2156390 | Safeguard DNA Diagnostic, Inc. | Philippine Genome Center | Francis A. Tablizo, Kenneth M. Kim, Carlo M. Lapid, Marc Jerrone R. Castro, Maria Sofia L. Yangzon, Elcid Aaron R. Pangilinan, Benedict A. Maralit, Marc Edsel C. Ayes, Eva Maria Cutiongco-de la Paz, Alethea R. de Guzman, Jan Michael C. Yap, Jo-Hannah S. Llames, Sheila Mae M. Araiza, Kris P. Punayan, Irish Coleen A. Asin, Candice Francheska B. Tambaoan, Asia Louisa U. Chong, Karol Sophia Agape R. Padilla, Rianna Patricia S. Cruz, El King D. Morado, Joshua Gregor A. Dizon, Razel Nikka M. Hao, Arianne A. Zamora, Devon Ray Pacial, Juan Antonio R. Magalang, Marissa Alejandria, Celia Carlos, Anna Ong-Lim, Edsel Maurice Salvaña, John Q. Wong, Jaime C. Montoya, Maria Rosario Singh-Vergeire and Cynthia P. Saloma |
| EPI_ISL_2156392, EPI_ISL_2156393 | Philippine Red Cross - Port Area | Philippine Genome Center | Francis A. Tablizo, Kenneth M. Kim, Carlo M. Lapid, Marc Jerrone R. Castro, Maria Sofia L. Yangzon, Elcid Aaron R. Pangilinan, Benedict A. Maralit, Marc Edsel C. Ayes, Eva Maria Cutiongco-de la Paz, Alethea R. de Guzman, Jan Michael C. Yap, Jo-Hannah S. Llames, Sheila Mae M. Araiza, Kris P. Punayan, Irish Coleen A. Asin, Candice Francheska B. Tambaoan, Asia Louisa U. Chong, Karol Sophia Agape R. Padilla, Rianna Patricia S. Cruz, El King D. Morado, Joshua Gregor A. Dizon, Razel Nikka M. Hao, Arianne A. Zamora, Devon Ray Pacial, Juan Antonio R. Magalang, Marissa Alejandria, Celia Carlos, Anna Ong-Lim, Edsel Maurice Salvaña, John Q. Wong, Jaime C. Montoya, Maria Rosario Singh-Vergeire and Cynthia P. Saloma |

| | | | |
|---|---|---|---|
| EPI_ISL_2156394 | Ospital ng Imus | Philippine Genome Center | Francis A. Tablizo, Kenneth M. Kim, Carlo M. Lapid, Marc Jerrone R. Castro, Maria Sofia L. Yangzon, Elcid Aaron R. Pangilinan, Benedict A. Maralit, Marc Edsel C. Ayes, Eva Maria Cutiongco-de la Paz, Alethea R. de Guzman, Jan Michael C. Yap, Jo-Hannah S. Llames, Sheila Mae M. Araiza, Kris P. Punayan, Irish Coleen A. Asin, Candice Francheska B. Tambaoan, Asia Louisa U. Chong, Karol Sophia Agape R. Padilla, Rianna Patricia S. Cruz, El King D. Morado, Joshua Gregor A. Dizon, Razel Nikka M. Hao, Arianne A. Zamora, Devon Ray Pacial, Juan Antonio R. Magalang, Marissa Alejandria, Celia Carlos, Anna Ong-Lim, Edsel Maurice Salvaña, John Q. Wong, Jaime C. Montoya, Maria Rosario Singh-Vergeire and Cynthia P. Saloma |
| EPI_ISL_2156395 | The Medical City -Â Ortigas | Philippine Genome Center | Francis A. Tablizo, Kenneth M. Kim, Carlo M. Lapid, Marc Jerrone R. Castro, Maria Sofia L. Yangzon, Elcid Aaron R. Pangilinan, Benedict A. Maralit, Marc Edsel C. Ayes, Eva Maria Cutiongco-de la Paz, Alethea R. de Guzman, Jan Michael C. Yap, Jo-Hannah S. Llames, Sheila Mae M. Araiza, Kris P. Punayan, Irish Coleen A. Asin, Candice Francheska B. Tambaoan, Asia Louisa U. Chong, Karol Sophia Agape R. Padilla, Rianna Patricia S. Cruz, El King D. Morado, Joshua Gregor A. Dizon, Razel Nikka M. Hao, Arianne A. Zamora, Devon Ray Pacial, Juan Antonio R. Magalang, Marissa Alejandria, Celia Carlos, Anna Ong-Lim, Edsel Maurice Salvaña, John Q. Wong, Jaime C. Montoya, Maria Rosario Singh-Vergeire and Cynthia P. Saloma |
| EPI_ISL_2156397, EPI_ISL_2156398 | Butuan Medical Center | Philippine Genome Center | Francis A. Tablizo, Kenneth M. Kim, Carlo M. Lapid, Marc Jerrone R. Castro, Maria Sofia L. Yangzon, Elcid Aaron R. Pangilinan, Benedict A. Maralit, Marc Edsel C. Ayes, Eva Maria Cutiongco-de la Paz, Alethea R. de Guzman, Jan Michael C. Yap, Jo-Hannah S. Llames, Sheila Mae M. Araiza, Kris P. Punayan, Irish Coleen A. Asin, Candice Francheska B. Tambaoan, Asia Louisa U. Chong, Karol Sophia Agape R. Padilla, Rianna Patricia S. Cruz, El King D. Morado, Joshua Gregor A. Dizon, Razel Nikka M. Hao, Arianne A. Zamora, Devon Ray Pacial, Juan Antonio R. Magalang, Marissa Alejandria, Celia Carlos, Anna Ong-Lim, Edsel Maurice Salvaña, John Q. Wong, Jaime C. Montoya, Maria Rosario Singh-Vergeire and Cynthia P. Saloma |
| EPI_ISL_2156401 | First Aide Diagnostic Center | Philippine Genome Center | Francis A. Tablizo, Kenneth M. Kim, Carlo M. Lapid, Marc Jerrone R. Castro, Maria Sofia L. Yangzon, Elcid Aaron R. Pangilinan, Benedict A. Maralit, Marc Edsel C. Ayes, Eva Maria Cutiongco-de la Paz, Alethea R. de Guzman, Jan Michael C. Yap, Jo-Hannah S. Llames, Sheila Mae M. Araiza, Kris P. Punayan, Irish Coleen A. Asin, Candice Francheska B. Tambaoan, Asia Louisa U. Chong, Karol Sophia Agape R. Padilla, Rianna Patricia S. Cruz, El King D. Morado, Joshua Gregor A. Dizon, Razel Nikka M. Hao, Arianne A. Zamora, Devon Ray Pacial, Juan Antonio R. Magalang, Marissa Alejandria, Celia Carlos, Anna Ong-Lim, Edsel Maurice Salvaña, John Q. Wong, Jaime C. Montoya, Maria Rosario Singh-Vergeire and Cynthia P. Saloma |
| EPI_ISL_2156403, EPI_ISL_2156405, EPI_ISL_2156406 | Davao One World Diagnostic Center Incorporated | Philippine Genome Center | Francis A. Tablizo, Kenneth M. Kim, Carlo M. Lapid, Marc Jerrone R. Castro, Maria Sofia L. Yangzon, Elcid Aaron R. Pangilinan, Benedict A. Maralit, Marc Edsel C. Ayes, Eva Maria Cutiongco-de la Paz, Alethea R. de Guzman, Jan Michael C. Yap, Jo-Hannah S. Llames, Sheila Mae M. Araiza, Kris P. Punayan, Irish Coleen A. Asin, Candice Francheska B. Tambaoan, Asia Louisa U. Chong, Karol Sophia Agape R. Padilla, Rianna Patricia S. Cruz, El King D. Morado, Joshua Gregor A. Dizon, Razel Nikka M. Hao, Arianne A. Zamora, Devon Ray Pacial, Juan Antonio R. Magalang, Marissa Alejandria, Celia Carlos, Anna Ong-Lim, Edsel Maurice Salvaña, John Q. Wong, Jaime C. Montoya, Maria Rosario Singh-Vergeire and Cynthia P. Saloma |
| EPI_ISL_2156409 | Baguio General Hospital Medical Center (BGHMC) | Philippine Genome Center | Francis A. Tablizo, Kenneth M. Kim, Carlo M. Lapid, Marc Jerrone R. Castro, Maria Sofia L. Yangzon, Elcid Aaron R. Pangilinan, Benedict A. Maralit, Marc Edsel C. Ayes, Eva Maria Cutiongco-de la Paz, Alethea R. de Guzman, Jan Michael C. Yap, Jo-Hannah S. Llames, Sheila Mae M. Araiza, Kris P. Punayan, Irish Coleen A. Asin, Candice Francheska B. Tambaoan, Asia Louisa U. Chong, Karol Sophia Agape R. Padilla, Rianna Patricia S. Cruz, El King D. Morado, Joshua Gregor A. Dizon, Razel Nikka M. Hao, Arianne A. Zamora, Devon Ray Pacial, Juan Antonio R. Magalang, Marissa Alejandria, Celia Carlos, Anna Ong-Lim, Edsel Maurice Salvaña, John Q. Wong, Jaime C. Montoya, Maria Rosario Singh-Vergeire and Cynthia P. Saloma |
| EPI_ISL_2156410, EPI_ISL_2156411 | Hi-Precision Diagnostic Center (QC) | Philippine Genome Center | Francis A. Tablizo, Kenneth M. Kim, Carlo M. Lapid, Marc Jerrone R. Castro, Maria Sofia L. Yangzon, Elcid Aaron R. Pangilinan, Benedict A. Maralit, Marc Edsel C. Ayes, Eva Maria Cutiongco-de la Paz, Alethea R. de Guzman, Jan Michael C. Yap, Jo-Hannah S. Llames, Sheila Mae M. Araiza, Kris P. Punayan, Irish Coleen A. Asin, Candice Francheska B. Tambaoan, Asia Louisa U. Chong, Karol Sophia Agape R. Padilla, Rianna Patricia S. Cruz, El King D. Morado, Joshua Gregor A. Dizon, Razel Nikka M. Hao, Arianne A. Zamora, Devon Ray Pacial, Juan Antonio R. Magalang, Marissa Alejandria, Celia Carlos, Anna Ong-Lim, Edsel Maurice Salvaña, John Q. Wong, Jaime C. Montoya, Maria Rosario Singh-Vergeire and Cynthia P. Saloma |
| EPI_ISL_2156412 | Pasig City Children's Hospital - Child's Hope | Philippine Genome Center | Francis A. Tablizo, Kenneth M. Kim, Carlo M. Lapid, Marc Jerrone R. Castro, Maria Sofia L. Yangzon, Elcid Aaron R. Pangilinan, Benedict A. Maralit, Marc Edsel C. Ayes, Eva Maria Cutiongco-de la Paz, Alethea R. de Guzman, Jan Michael C. Yap, Jo-Hannah S. Llames, Sheila Mae M. Araiza, Kris P. Punayan, Irish Coleen A. Asin, Candice Francheska B. Tambaoan, Asia Louisa U. Chong, Karol Sophia Agape R. Padilla, Rianna Patricia S. Cruz, El King D. Morado, Joshua Gregor A. Dizon, Razel Nikka M. Hao, Arianne A. Zamora, Devon Ray Pacial, Juan Antonio R. Magalang, Marissa Alejandria, Celia Carlos, Anna Ong-Lim, Edsel Maurice Salvaña, John Q. Wong, Jaime C. Montoya, Maria Rosario Singh-Vergeire and Cynthia P. Saloma |
| EPI_ISL_2156413 | Safeguard DNA Diagnostic, Inc. | Philippine Genome Center | Francis A. Tablizo, Kenneth M. Kim, Carlo M. Lapid, Marc Jerrone R. Castro, Maria Sofia L. Yangzon, Elcid Aaron R. Pangilinan, Benedict A. Maralit, Marc Edsel C. Ayes, Eva Maria Cutiongco-de la Paz, Alethea R. de Guzman, Jan Michael C. Yap, Jo-Hannah S. Llames, Sheila Mae M. Araiza, Kris P. Punayan, Irish Coleen A. Asin, Candice Francheska B. Tambaoan, Asia Louisa U. Chong, Karol Sophia Agape R. Padilla, Rianna Patricia S. Cruz, El King D. Morado, Joshua Gregor A. Dizon, Razel Nikka M. Hao, Arianne A. Zamora, Devon Ray Pacial, Juan Antonio R. Magalang, Marissa Alejandria, Celia Carlos, Anna Ong-Lim, Edsel Maurice Salvaña, John Q. Wong, Jaime C. Montoya, Maria Rosario Singh-Vergeire and Cynthia P. Saloma |
| EPI_ISL_2156415 | Cagayan Valley Medical Center Molecular Laboratory | Philippine Genome Center | Francis A. Tablizo, Kenneth M. Kim, Carlo M. Lapid, Marc Jerrone R. Castro, Maria Sofia L. Yangzon, Elcid Aaron R. Pangilinan, Benedict A. Maralit, Marc Edsel C. Ayes, Eva Maria Cutiongco-de la Paz, Alethea R. de Guzman, Jan Michael C. Yap, Jo-Hannah S. Llames, Sheila Mae M. Araiza, Kris P. Punayan, Irish Coleen A. Asin, Candice Francheska B. Tambaoan, Asia Louisa U. Chong, Karol Sophia Agape R. Padilla, Rianna Patricia S. Cruz, El King D. Morado, Joshua Gregor A. Dizon, Razel Nikka M. Hao, Arianne A. Zamora, Devon Ray Pacial, Juan Antonio R. Magalang, Marissa Alejandria, Celia Carlos, Anna Ong-Lim, Edsel Maurice Salvaña, John Q. Wong, Jaime C. Montoya, Maria Rosario Singh-Vergeire and Cynthia P. Saloma |
| EPI_ISL_2156416 | The Medical City -Â Ortigas | Philippine Genome Center | Francis A. Tablizo, Kenneth M. Kim, Carlo M. Lapid, Marc Jerrone R. Castro, Maria Sofia L. Yangzon, Elcid Aaron R. Pangilinan, Benedict A. Maralit, Marc Edsel C. Ayes, Eva Maria Cutiongco-de la Paz, Alethea R. de Guzman, Jan Michael C. Yap, Jo-Hannah S. Llames, Sheila Mae M. Araiza, Kris P. Punayan, Irish Coleen A. Asin, Candice Francheska B. Tambaoan, Asia Louisa U. Chong, Karol Sophia Agape R. Padilla, Rianna Patricia S. Cruz, El King D. Morado, Joshua Gregor A. Dizon, Razel Nikka M. Hao, Arianne A. Zamora, Devon Ray Pacial, Juan Antonio R. Magalang, Marissa Alejandria, Celia Carlos, Anna Ong-Lim, Edsel Maurice Salvaña, John Q. Wong, Jaime C. Montoya, Maria Rosario Singh-Vergeire and Cynthia P. Saloma |
| EPI_ISL_2156419 | Davao One World Diagnostic Center Incorporated | Philippine Genome Center | Francis A. Tablizo, Kenneth M. Kim, Carlo M. Lapid, Marc Jerrone R. Castro, Maria Sofia L. Yangzon, Elcid Aaron R. Pangilinan, Benedict A. Maralit, Marc Edsel C. Ayes, Eva Maria Cutiongco-de la Paz, Alethea R. de Guzman, Jan Michael C. Yap, Jo-Hannah S. Llames, Sheila Mae M. Araiza, Kris P. Punayan, Irish Coleen A. Asin, Candice Francheska B. Tambaoan, Asia Louisa U. Chong, Karol Sophia Agape R. Padilla, Rianna Patricia S. Cruz, El King D. Morado, Joshua Gregor A. Dizon, Razel Nikka M. Hao, Arianne A. Zamora, Devon Ray Pacial, Juan Antonio R. Magalang, Marissa Alejandria, Celia Carlos, Anna Ong-Lim, Edsel Maurice Salvaña, John Q. Wong, Jaime C. Montoya, Maria Rosario Singh-Vergeire and Cynthia P. Saloma |
| EPI_ISL_2156421, EPI_ISL_2156422 | Butuan Medical Center | Philippine Genome Center | Francis A. Tablizo, Kenneth M. Kim, Carlo M. Lapid, Marc Jerrone R. Castro, Maria Sofia L. Yangzon, Elcid Aaron R. Pangilinan, Benedict A. Maralit, Marc Edsel C. Ayes, Eva Maria Cutiongco-de la Paz, Alethea R. de Guzman, Jan Michael C. Yap, Jo-Hannah S. Llames, Sheila Mae M. Araiza, Kris P. Punayan, Irish Coleen A. Asin, Candice Francheska B. Tambaoan, Asia Louisa U. Chong, Karol Sophia Agape R. Padilla, Rianna Patricia S. Cruz, El King D. Morado, Joshua Gregor A. Dizon, Razel Nikka M. Hao, Arianne A. Zamora, Devon Ray Pacial, Juan Antonio R. Magalang, Marissa Alejandria, Celia Carlos, Anna Ong-Lim, Edsel Maurice Salvaña, John Q. Wong, Jaime C. Montoya, Maria Rosario Singh-Vergeire and Cynthia P. Saloma |
| EPI_ISL_2156423, EPI_ISL_2156424, EPI_ISL_2156425 | Baguio General Hospital Medical Center (BGHMC) | Philippine Genome Center | Francis A. Tablizo, Kenneth M. Kim, Carlo M. Lapid, Marc Jerrone R. Castro, Maria Sofia L. Yangzon, Elcid Aaron R. Pangilinan, Benedict A. Maralit, Marc Edsel C. Ayes, Eva Maria Cutiongco-de la Paz, Alethea R. de Guzman, Jan Michael C. Yap, Jo-Hannah S. Llames, Sheila Mae M. Araiza, Kris P. Punayan, Irish Coleen A. Asin, Candice Francheska B. Tambaoan, Asia Louisa U. Chong, Karol Sophia Agape R. Padilla, Rianna Patricia S. Cruz, El King D. Morado, Joshua Gregor A. Dizon, Razel Nikka M. Hao, Arianne A. Zamora, Devon Ray Pacial, Juan Antonio R. Magalang, Marissa Alejandria, Celia Carlos, Anna Ong-Lim, Edsel Maurice Salvaña, John Q. Wong, Jaime C. Montoya, Maria Rosario Singh-Vergeire and Cynthia P. Saloma |
| EPI_ISL_2156427, EPI_ISL_2156428, EPI_ISL_2156429, EPI_ISL_2156431, EPI_ISL_2156432, EPI_ISL_2156433 | Cagayan Valley Medical Center Molecular Laboratory | Philippine Genome Center | Francis A. Tablizo, Kenneth M. Kim, Carlo M. Lapid, Marc Jerrone R. Castro, Maria Sofia L. Yangzon, Elcid Aaron R. Pangilinan, Benedict A. Maralit, Marc Edsel C. Ayes, Eva Maria Cutiongco-de la Paz, Alethea R. de Guzman, Jan Michael C. Yap, Jo-Hannah S. Llames, Sheila Mae M. Araiza, Kris P. Punayan, Irish Coleen A. Asin, Candice Francheska B. Tambaoan, Asia Louisa U. Chong, Karol Sophia Agape R. Padilla, Rianna Patricia S. Cruz, El King D. Morado, Joshua Gregor A. Dizon, Razel Nikka M. Hao, Arianne A. Zamora, Devon Ray Pacial, Juan Antonio R. Magalang, Marissa Alejandria, Celia Carlos, Anna Ong-Lim, Edsel Maurice Salvaña, John Q. Wong, Jaime C. Montoya, Maria Rosario Singh-Vergeire and Cynthia P. Saloma |
| EPI_ISL_2156435, EPI_ISL_2156436, EPI_ISL_2156438, EPI_ISL_2156440, EPI_ISL_2156441, EPI_ISL_2156442, EPI_ISL_2156443 | Research Institute for Tropical Medicine, Inc. (RITM) | Philippine Genome Center | Francis A. Tablizo, Kenneth M. Kim, Carlo M. Lapid, Marc Jerrone R. Castro, Maria Sofia L. Yangzon, Elcid Aaron R. Pangilinan, Benedict A. Maralit, Marc Edsel C. Ayes, Eva Maria Cutiongco-de la Paz, Alethea R. de Guzman, Jan Michael C. Yap, Jo-Hannah S. Llames, Sheila Mae M. Araiza, Kris P. Punayan, Irish Coleen A. Asin, Candice Francheska B. Tambaoan, Asia Louisa U. Chong, Karol Sophia Agape R. Padilla, Rianna Patricia S. Cruz, El King D. Morado, Joshua Gregor A. Dizon, Razel Nikka M. Hao, Arianne A. Zamora, Devon Ray Pacial, Juan Antonio R. Magalang, Marissa Alejandria, Celia Carlos, Anna Ong-Lim, Edsel Maurice Salvaña, John Q. Wong, Jaime C. Montoya, Maria Rosario Singh-Vergeire and Cynthia P. Saloma |

| | | | |
|---|---|---|---|
| EPI_ISL_2156445, EPI_ISL_2156447 | Butuan Medical Center | Philippine Genome Center | Francis A. Tablizo, Kenneth M. Kim, Carlo M. Lapid, Marc Jerrone R. Castro, Maria Sofia L. Yangzon, Elcid Aaron R. Pangilinan, Benedict A. Maralit, Marc Edsel C. Ayes, Eva Maria Cutiongco-de la Paz, Alethea R. de Guzman, Jan Michael C. Yap, Jo-Hannah S. Llames, Sheila Mae M. Araiza, Kris P. Punayan, Irish Coleen A. Asin, Candice Francheska B. Tambaoan, Asia Louisa U. Chong, Karol Sophia Agape R. Padilla, Rianna Patricia S. Cruz, El King D. Morado, Joshua Gregor A. Dizon, Razel Nikka M. Hao, Arianne A. Zamora, Devon Ray Pacial, Juan Antonio R. Magalang, Marissa Alejandria, Celia Carlos, Anna Ong-Lim, Edsel Maurice Salvaña, John Q. Wong, Jaime C. Montoya, Maria Rosario Singh-Vergeire and Cynthia P. Saloma |
| EPI_ISL_2156449, EPI_ISL_2156450 | Baguio General Hospital Medical Center (BGHMC) | Philippine Genome Center | Francis A. Tablizo, Kenneth M. Kim, Carlo M. Lapid, Marc Jerrone R. Castro, Maria Sofia L. Yangzon, Elcid Aaron R. Pangilinan, Benedict A. Maralit, Marc Edsel C. Ayes, Eva Maria Cutiongco-de la Paz, Alethea R. de Guzman, Jan Michael C. Yap, Jo-Hannah S. Llames, Sheila Mae M. Araiza, Kris P. Punayan, Irish Coleen A. Asin, Candice Francheska B. Tambaoan, Asia Louisa U. Chong, Karol Sophia Agape R. Padilla, Rianna Patricia S. Cruz, El King D. Morado, Joshua Gregor A. Dizon, Razel Nikka M. Hao, Arianne A. Zamora, Devon Ray Pacial, Juan Antonio R. Magalang, Marissa Alejandria, Celia Carlos, Anna Ong-Lim, Edsel Maurice Salvaña, John Q. Wong, Jaime C. Montoya, Maria Rosario Singh-Vergeire and Cynthia P. Saloma |
| EPI_ISL_2156454, EPI_ISL_2156455 | Philippine Red Cross - National Blood Center | Philippine Genome Center | Francis A. Tablizo, Kenneth M. Kim, Carlo M. Lapid, Marc Jerrone R. Castro, Maria Sofia L. Yangzon, Elcid Aaron R. Pangilinan, Benedict A. Maralit, Marc Edsel C. Ayes, Eva Maria Cutiongco-de la Paz, Alethea R. de Guzman, Jan Michael C. Yap, Jo-Hannah S. Llames, Sheila Mae M. Araiza, Kris P. Punayan, Irish Coleen A. Asin, Candice Francheska B. Tambaoan, Asia Louisa U. Chong, Karol Sophia Agape R. Padilla, Rianna Patricia S. Cruz, El King D. Morado, Joshua Gregor A. Dizon, Razel Nikka M. Hao, Arianne A. Zamora, Devon Ray Pacial, Juan Antonio R. Magalang, Marissa Alejandria, Celia Carlos, Anna Ong-Lim, Edsel Maurice Salvaña, John Q. Wong, Jaime C. Montoya, Maria Rosario Singh-Vergeire and Cynthia P. Saloma |
| EPI_ISL_2156457 | St. Luke's Medical Center - BGC (HB) | Philippine Genome Center | Francis A. Tablizo, Kenneth M. Kim, Carlo M. Lapid, Marc Jerrone R. Castro, Maria Sofia L. Yangzon, Elcid Aaron R. Pangilinan, Benedict A. Maralit, Marc Edsel C. Ayes, Eva Maria Cutiongco-de la Paz, Alethea R. de Guzman, Jan Michael C. Yap, Jo-Hannah S. Llames, Sheila Mae M. Araiza, Kris P. Punayan, Irish Coleen A. Asin, Candice Francheska B. Tambaoan, Asia Louisa U. Chong, Karol Sophia Agape R. Padilla, Rianna Patricia S. Cruz, El King D. Morado, Joshua Gregor A. Dizon, Razel Nikka M. Hao, Arianne A. Zamora, Devon Ray Pacial, Juan Antonio R. Magalang, Marissa Alejandria, Celia Carlos, Anna Ong-Lim, Edsel Maurice Salvaña, John Q. Wong, Jaime C. Montoya, Maria Rosario Singh-Vergeire and Cynthia P. Saloma |
| EPI_ISL_2156458 | First Aide Diagnostic Center | Philippine Genome Center | Francis A. Tablizo, Kenneth M. Kim, Carlo M. Lapid, Marc Jerrone R. Castro, Maria Sofia L. Yangzon, Elcid Aaron R. Pangilinan, Benedict A. Maralit, Marc Edsel C. Ayes, Eva Maria Cutiongco-de la Paz, Alethea R. de Guzman, Jan Michael C. Yap, Jo-Hannah S. Llames, Sheila Mae M. Araiza, Kris P. Punayan, Irish Coleen A. Asin, Candice Francheska B. Tambaoan, Asia Louisa U. Chong, Karol Sophia Agape R. Padilla, Rianna Patricia S. Cruz, El King D. Morado, Joshua Gregor A. Dizon, Razel Nikka M. Hao, Arianne A. Zamora, Devon Ray Pacial, Juan Antonio R. Magalang, Marissa Alejandria, Celia Carlos, Anna Ong-Lim, Edsel Maurice Salvaña, John Q. Wong, Jaime C. Montoya, Maria Rosario Singh-Vergeire and Cynthia P. Saloma |
| EPI_ISL_2156459 | Philippine Red Cross Logistics and Multipurpose Center | Philippine Genome Center | Francis A. Tablizo, Kenneth M. Kim, Carlo M. Lapid, Marc Jerrone R. Castro, Maria Sofia L. Yangzon, Elcid Aaron R. Pangilinan, Benedict A. Maralit, Marc Edsel C. Ayes, Eva Maria Cutiongco-de la Paz, Alethea R. de Guzman, Jan Michael C. Yap, Jo-Hannah S. Llames, Sheila Mae M. Araiza, Kris P. Punayan, Irish Coleen A. Asin, Candice Francheska B. Tambaoan, Asia Louisa U. Chong, Karol Sophia Agape R. Padilla, Rianna Patricia S. Cruz, El King D. Morado, Joshua Gregor A. Dizon, Razel Nikka M. Hao, Arianne A. Zamora, Devon Ray Pacial, Juan Antonio R. Magalang, Marissa Alejandria, Celia Carlos, Anna Ong-Lim, Edsel Maurice Salvaña, John Q. Wong, Jaime C. Montoya, Maria Rosario Singh-Vergeire and Cynthia P. Saloma |
| EPI_ISL_2156460 | Cagayan Valley Medical Center Molecular Laboratory | Philippine Genome Center | Francis A. Tablizo, Kenneth M. Kim, Carlo M. Lapid, Marc Jerrone R. Castro, Maria Sofia L. Yangzon, Elcid Aaron R. Pangilinan, Benedict A. Maralit, Marc Edsel C. Ayes, Eva Maria Cutiongco-de la Paz, Alethea R. de Guzman, Jan Michael C. Yap, Jo-Hannah S. Llames, Sheila Mae M. Araiza, Kris P. Punayan, Irish Coleen A. Asin, Candice Francheska B. Tambaoan, Asia Louisa U. Chong, Karol Sophia Agape R. Padilla, Rianna Patricia S. Cruz, El King D. Morado, Joshua Gregor A. Dizon, Razel Nikka M. Hao, Arianne A. Zamora, Devon Ray Pacial, Juan Antonio R. Magalang, Marissa Alejandria, Celia Carlos, Anna Ong-Lim, Edsel Maurice Salvaña, John Q. Wong, Jaime C. Montoya, Maria Rosario Singh-Vergeire and Cynthia P. Saloma |
| EPI_ISL_2156461, EPI_ISL_2156462 | The Medical City -Â Ortigas | Philippine Genome Center | Francis A. Tablizo, Kenneth M. Kim, Carlo M. Lapid, Marc Jerrone R. Castro, Maria Sofia L. Yangzon, Elcid Aaron R. Pangilinan, Benedict A. Maralit, Marc Edsel C. Ayes, Eva Maria Cutiongco-de la Paz, Alethea R. de Guzman, Jan Michael C. Yap, Jo-Hannah S. Llames, Sheila Mae M. Araiza, Kris P. Punayan, Irish Coleen A. Asin, Candice Francheska B. Tambaoan, Asia Louisa U. Chong, Karol Sophia Agape R. Padilla, Rianna Patricia S. Cruz, El King D. Morado, Joshua Gregor A. Dizon, Razel Nikka M. Hao, Arianne A. Zamora, Devon Ray Pacial, Juan Antonio R. Magalang, Marissa Alejandria, Celia Carlos, Anna Ong-Lim, Edsel Maurice Salvaña, John Q. Wong, Jaime C. Montoya, Maria Rosario Singh-Vergeire and Cynthia P. Saloma |
| EPI_ISL_2156464 | Butuan Medical Center | Philippine Genome Center | Francis A. Tablizo, Kenneth M. Kim, Carlo M. Lapid, Marc Jerrone R. Castro, Maria Sofia L. Yangzon, Elcid Aaron R. Pangilinan, Benedict A. Maralit, Marc Edsel C. Ayes, Eva Maria Cutiongco-de la Paz, Alethea R. de Guzman, Jan Michael C. Yap, Jo-Hannah S. Llames, Sheila Mae M. Araiza, Kris P. Punayan, Irish Coleen A. Asin, Candice Francheska B. Tambaoan, Asia Louisa U. Chong, Karol Sophia Agape R. Padilla, Rianna Patricia S. Cruz, El King D. Morado, Joshua Gregor A. Dizon, Razel Nikka M. Hao, Arianne A. Zamora, Devon Ray Pacial, Juan Antonio R. Magalang, Marissa Alejandria, Celia Carlos, Anna Ong-Lim, Edsel Maurice Salvaña, John Q. Wong, Jaime C. Montoya, Maria Rosario Singh-Vergeire and Cynthia P. Saloma |
| EPI_ISL_2156469, EPI_ISL_2156470, EPI_ISL_2156472, EPI_ISL_2156473 | Baguio General Hospital Medical Center (BGHMC) | Philippine Genome Center | Francis A. Tablizo, Kenneth M. Kim, Carlo M. Lapid, Marc Jerrone R. Castro, Maria Sofia L. Yangzon, Elcid Aaron R. Pangilinan, Benedict A. Maralit, Marc Edsel C. Ayes, Eva Maria Cutiongco-de la Paz, Alethea R. de Guzman, Jan Michael C. Yap, Jo-Hannah S. Llames, Sheila Mae M. Araiza, Kris P. Punayan, Irish Coleen A. Asin, Candice Francheska B. Tambaoan, Asia Louisa U. Chong, Karol Sophia Agape R. Padilla, Rianna Patricia S. Cruz, El King D. Morado, Joshua Gregor A. Dizon, Razel Nikka M. Hao, Arianne A. Zamora, Devon Ray Pacial, Juan Antonio R. Magalang, Marissa Alejandria, Celia Carlos, Anna Ong-Lim, Edsel Maurice Salvaña, John Q. Wong, Jaime C. Montoya, Maria Rosario Singh-Vergeire and Cynthia P. Saloma |
| EPI_ISL_2156474 | Philippine Red Cross - National Blood Center | Philippine Genome Center | Francis A. Tablizo, Kenneth M. Kim, Carlo M. Lapid, Marc Jerrone R. Castro, Maria Sofia L. Yangzon, Elcid Aaron R. Pangilinan, Benedict A. Maralit, Marc Edsel C. Ayes, Eva Maria Cutiongco-de la Paz, Alethea R. de Guzman, Jan Michael C. Yap, Jo-Hannah S. Llames, Sheila Mae M. Araiza, Kris P. Punayan, Irish Coleen A. Asin, Candice Francheska B. Tambaoan, Asia Louisa U. Chong, Karol Sophia Agape R. Padilla, Rianna Patricia S. Cruz, El King D. Morado, Joshua Gregor A. Dizon, Razel Nikka M. Hao, Arianne A. Zamora, Devon Ray Pacial, Juan Antonio R. Magalang, Marissa Alejandria, Celia Carlos, Anna Ong-Lim, Edsel Maurice Salvaña, John Q. Wong, Jaime C. Montoya, Maria Rosario Singh-Vergeire and Cynthia P. Saloma |
| EPI_ISL_2156476 | Philippine Red Cross Logistics and Multipurpose Center | Philippine Genome Center | Francis A. Tablizo, Kenneth M. Kim, Carlo M. Lapid, Marc Jerrone R. Castro, Maria Sofia L. Yangzon, Elcid Aaron R. Pangilinan, Benedict A. Maralit, Marc Edsel C. Ayes, Eva Maria Cutiongco-de la Paz, Alethea R. de Guzman, Jan Michael C. Yap, Jo-Hannah S. Llames, Sheila Mae M. Araiza, Kris P. Punayan, Irish Coleen A. Asin, Candice Francheska B. Tambaoan, Asia Louisa U. Chong, Karol Sophia Agape R. Padilla, Rianna Patricia S. Cruz, El King D. Morado, Joshua Gregor A. Dizon, Razel Nikka M. Hao, Arianne A. Zamora, Devon Ray Pacial, Juan Antonio R. Magalang, Marissa Alejandria, Celia Carlos, Anna Ong-Lim, Edsel Maurice Salvaña, John Q. Wong, Jaime C. Montoya, Maria Rosario Singh-Vergeire and Cynthia P. Saloma |
| EPI_ISL_2156480 | The Lord's Grace Medical and Industrial Clinic | Philippine Genome Center | Francis A. Tablizo, Kenneth M. Kim, Carlo M. Lapid, Marc Jerrone R. Castro, Maria Sofia L. Yangzon, Elcid Aaron R. Pangilinan, Benedict A. Maralit, Marc Edsel C. Ayes, Eva Maria Cutiongco-de la Paz, Alethea R. de Guzman, Jan Michael C. Yap, Jo-Hannah S. Llames, Sheila Mae M. Araiza, Kris P. Punayan, Irish Coleen A. Asin, Candice Francheska B. Tambaoan, Asia Louisa U. Chong, Karol Sophia Agape R. Padilla, Rianna Patricia S. Cruz, El King D. Morado, Joshua Gregor A. Dizon, Razel Nikka M. Hao, Arianne A. Zamora, Devon Ray Pacial, Juan Antonio R. Magalang, Marissa Alejandria, Celia Carlos, Anna Ong-Lim, Edsel Maurice Salvaña, John Q. Wong, Jaime C. Montoya, Maria Rosario Singh-Vergeire and Cynthia P. Saloma |
| EPI_ISL_2156483 | First Aide Diagnostic Center | Philippine Genome Center | Francis A. Tablizo, Kenneth M. Kim, Carlo M. Lapid, Marc Jerrone R. Castro, Maria Sofia L. Yangzon, Elcid Aaron R. Pangilinan, Benedict A. Maralit, Marc Edsel C. Ayes, Eva Maria Cutiongco-de la Paz, Alethea R. de Guzman, Jan Michael C. Yap, Jo-Hannah S. Llames, Sheila Mae M. Araiza, Kris P. Punayan, Irish Coleen A. Asin, Candice Francheska B. Tambaoan, Asia Louisa U. Chong, Karol Sophia Agape R. Padilla, Rianna Patricia S. Cruz, El King D. Morado, Joshua Gregor A. Dizon, Razel Nikka M. Hao, Arianne A. Zamora, Devon Ray Pacial, Juan Antonio R. Magalang, Marissa Alejandria, Celia Carlos, Anna Ong-Lim, Edsel Maurice Salvaña, John Q. Wong, Jaime C. Montoya, Maria Rosario Singh-Vergeire and Cynthia P. Saloma |
| EPI_ISL_2156484 | Philippine Red Cross Logistics and Multipurpose Center | Philippine Genome Center | Francis A. Tablizo, Kenneth M. Kim, Carlo M. Lapid, Marc Jerrone R. Castro, Maria Sofia L. Yangzon, Elcid Aaron R. Pangilinan, Benedict A. Maralit, Marc Edsel C. Ayes, Eva Maria Cutiongco-de la Paz, Alethea R. de Guzman, Jan Michael C. Yap, Jo-Hannah S. Llames, Sheila Mae M. Araiza, Kris P. Punayan, Irish Coleen A. Asin, Candice Francheska B. Tambaoan, Asia Louisa U. Chong, Karol Sophia Agape R. Padilla, Rianna Patricia S. Cruz, El King D. Morado, Joshua Gregor A. Dizon, Razel Nikka M. Hao, Arianne A. Zamora, Devon Ray Pacial, Juan Antonio R. Magalang, Marissa Alejandria, Celia Carlos, Anna Ong-Lim, Edsel Maurice Salvaña, John Q. Wong, Jaime C. Montoya, Maria Rosario Singh-Vergeire and Cynthia P. Saloma |
| EPI_ISL_2156485 | Northern Mindanao TB Regional Center | Philippine Genome Center | Francis A. Tablizo, Kenneth M. Kim, Carlo M. Lapid, Marc Jerrone R. Castro, Maria Sofia L. Yangzon, Elcid Aaron R. Pangilinan, Benedict A. Maralit, Marc Edsel C. Ayes, Eva Maria Cutiongco-de la Paz, Alethea R. de Guzman, Jan Michael C. Yap, Jo-Hannah S. Llames, Sheila Mae M. Araiza, Kris P. Punayan, Irish Coleen A. Asin, Candice Francheska B. Tambaoan, Asia Louisa U. Chong, Karol Sophia Agape R. Padilla, Rianna Patricia S. Cruz, El King D. Morado, Joshua Gregor A. Dizon, Razel Nikka M. Hao, Arianne A. Zamora, Devon Ray Pacial, Juan Antonio R. Magalang, Marissa Alejandria, Celia Carlos, Anna Ong-Lim, Edsel Maurice Salvaña, John Q. Wong, Jaime C. Montoya, Maria Rosario Singh-Vergeire and Cynthia P. Saloma |

| | | | |
|---|---|---|---|
| EPI_ISL_2156489 | Laguna Provincial Hospital - San Pablo District Hospital | Philippine Genome Center | Francis A. Tablizo, Kenneth M. Kim, Carlo M. Lapid, Marc Jerrone R. Castro, Maria Sofia L. Yangzon, Elcid Aaron R. Pangilinan, Benedict A. Maralit, Marc Edsel C. Ayes, Eva Maria Cutiongco-de la Paz, Alethea R. de Guzman, Jan Michael C. Yap, Jo-Hannah S. Llames, Sheila Mae M. Araiza, Kris P. Punayan, Irish Coleen A. Asin, Candice Francheska B. Tambaoan, Asia Louisa U. Chong, Karol Sophia Agape R. Padilla, Rianna Patricia S. Cruz, El King D. Morado, Joshua Gregor A. Dizon, Razel Nikka M. Hao, Arianne A. Zamora, Devon Ray Pacial, Juan Antonio R. Magalang, Marissa Alejandria, Celia Carlos, Anna Ong-Lim, Edsel Maurice Salvaña, John Q. Wong, Jaime C. Montoya, Maria Rosario Singh-Vergeire and Cynthia P. Saloma |
| EPI_ISL_2156491 | The Lord's Grace Medical and Industrial Clinic | Philippine Genome Center | Francis A. Tablizo, Kenneth M. Kim, Carlo M. Lapid, Marc Jerrone R. Castro, Maria Sofia L. Yangzon, Elcid Aaron R. Pangilinan, Benedict A. Maralit, Marc Edsel C. Ayes, Eva Maria Cutiongco-de la Paz, Alethea R. de Guzman, Jan Michael C. Yap, Jo-Hannah S. Llames, Sheila Mae M. Araiza, Kris P. Punayan, Irish Coleen A. Asin, Candice Francheska B. Tambaoan, Asia Louisa U. Chong, Karol Sophia Agape R. Padilla, Rianna Patricia S. Cruz, El King D. Morado, Joshua Gregor A. Dizon, Razel Nikka M. Hao, Arianne A. Zamora, Devon Ray Pacial, Juan Antonio R. Magalang, Marissa Alejandria, Celia Carlos, Anna Ong-Lim, Edsel Maurice Salvaña, John Q. Wong, Jaime C. Montoya, Maria Rosario Singh-Vergeire and Cynthia P. Saloma |
| EPI_ISL_2156492 | The Medical City -Â Ortigas | Philippine Genome Center | Francis A. Tablizo, Kenneth M. Kim, Carlo M. Lapid, Marc Jerrone R. Castro, Maria Sofia L. Yangzon, Elcid Aaron R. Pangilinan, Benedict A. Maralit, Marc Edsel C. Ayes, Eva Maria Cutiongco-de la Paz, Alethea R. de Guzman, Jan Michael C. Yap, Jo-Hannah S. Llames, Sheila Mae M. Araiza, Kris P. Punayan, Irish Coleen A. Asin, Candice Francheska B. Tambaoan, Asia Louisa U. Chong, Karol Sophia Agape R. Padilla, Rianna Patricia S. Cruz, El King D. Morado, Joshua Gregor A. Dizon, Razel Nikka M. Hao, Arianne A. Zamora, Devon Ray Pacial, Juan Antonio R. Magalang, Marissa Alejandria, Celia Carlos, Anna Ong-Lim, Edsel Maurice Salvaña, John Q. Wong, Jaime C. Montoya, Maria Rosario Singh-Vergeire and Cynthia P. Saloma |
| EPI_ISL_2156493 | Research Institute for Tropical Medicine, Inc. (RITM) | Philippine Genome Center | Francis A. Tablizo, Kenneth M. Kim, Carlo M. Lapid, Marc Jerrone R. Castro, Maria Sofia L. Yangzon, Elcid Aaron R. Pangilinan, Benedict A. Maralit, Marc Edsel C. Ayes, Eva Maria Cutiongco-de la Paz, Alethea R. de Guzman, Jan Michael C. Yap, Jo-Hannah S. Llames, Sheila Mae M. Araiza, Kris P. Punayan, Irish Coleen A. Asin, Candice Francheska B. Tambaoan, Asia Louisa U. Chong, Karol Sophia Agape R. Padilla, Rianna Patricia S. Cruz, El King D. Morado, Joshua Gregor A. Dizon, Razel Nikka M. Hao, Arianne A. Zamora, Devon Ray Pacial, Juan Antonio R. Magalang, Marissa Alejandria, Celia Carlos, Anna Ong-Lim, Edsel Maurice Salvaña, John Q. Wong, Jaime C. Montoya, Maria Rosario Singh-Vergeire and Cynthia P. Saloma |
| EPI_ISL_2156496 | Philippine General Hospital (PGH) | Philippine Genome Center | Francis A. Tablizo, Kenneth M. Kim, Carlo M. Lapid, Marc Jerrone R. Castro, Maria Sofia L. Yangzon, Elcid Aaron R. Pangilinan, Benedict A. Maralit, Marc Edsel C. Ayes, Eva Maria Cutiongco-de la Paz, Alethea R. de Guzman, Jan Michael C. Yap, Jo-Hannah S. Llames, Sheila Mae M. Araiza, Kris P. Punayan, Irish Coleen A. Asin, Candice Francheska B. Tambaoan, Asia Louisa U. Chong, Karol Sophia Agape R. Padilla, Rianna Patricia S. Cruz, El King D. Morado, Joshua Gregor A. Dizon, Razel Nikka M. Hao, Arianne A. Zamora, Devon Ray Pacial, Juan Antonio R. Magalang, Marissa Alejandria, Celia Carlos, Anna Ong-Lim, Edsel Maurice Salvaña, John Q. Wong, Jaime C. Montoya, Maria Rosario Singh-Vergeire and Cynthia P. Saloma |
| EPI_ISL_2156497 | San Antonio City of Ilagan Hospital | Philippine Genome Center | Francis A. Tablizo, Kenneth M. Kim, Carlo M. Lapid, Marc Jerrone R. Castro, Maria Sofia L. Yangzon, Elcid Aaron R. Pangilinan, Benedict A. Maralit, Marc Edsel C. Ayes, Eva Maria Cutiongco-de la Paz, Alethea R. de Guzman, Jan Michael C. Yap, Jo-Hannah S. Llames, Sheila Mae M. Araiza, Kris P. Punayan, Irish Coleen A. Asin, Candice Francheska B. Tambaoan, Asia Louisa U. Chong, Karol Sophia Agape R. Padilla, Rianna Patricia S. Cruz, El King D. Morado, Joshua Gregor A. Dizon, Razel Nikka M. Hao, Arianne A. Zamora, Devon Ray Pacial, Juan Antonio R. Magalang, Marissa Alejandria, Celia Carlos, Anna Ong-Lim, Edsel Maurice Salvaña, John Q. Wong, Jaime C. Montoya, Maria Rosario Singh-Vergeire and Cynthia P. Saloma |
| EPI_ISL_2156500 | Detoxicare Molecular Diagnostics Laboratory | Philippine Genome Center | Francis A. Tablizo, Kenneth M. Kim, Carlo M. Lapid, Marc Jerrone R. Castro, Maria Sofia L. Yangzon, Elcid Aaron R. Pangilinan, Benedict A. Maralit, Marc Edsel C. Ayes, Eva Maria Cutiongco-de la Paz, Alethea R. de Guzman, Jan Michael C. Yap, Jo-Hannah S. Llames, Sheila Mae M. Araiza, Kris P. Punayan, Irish Coleen A. Asin, Candice Francheska B. Tambaoan, Asia Louisa U. Chong, Karol Sophia Agape R. Padilla, Rianna Patricia S. Cruz, El King D. Morado, Joshua Gregor A. Dizon, Razel Nikka M. Hao, Arianne A. Zamora, Devon Ray Pacial, Juan Antonio R. Magalang, Marissa Alejandria, Celia Carlos, Anna Ong-Lim, Edsel Maurice Salvaña, John Q. Wong, Jaime C. Montoya, Maria Rosario Singh-Vergeire and Cynthia P. Saloma |
| EPI_ISL_2156501 | Valenzuela Hope Molecular Laboratory | Philippine Genome Center | Francis A. Tablizo, Kenneth M. Kim, Carlo M. Lapid, Marc Jerrone R. Castro, Maria Sofia L. Yangzon, Elcid Aaron R. Pangilinan, Benedict A. Maralit, Marc Edsel C. Ayes, Eva Maria Cutiongco-de la Paz, Alethea R. de Guzman, Jan Michael C. Yap, Jo-Hannah S. Llames, Sheila Mae M. Araiza, Kris P. Punayan, Irish Coleen A. Asin, Candice Francheska B. Tambaoan, Asia Louisa U. Chong, Karol Sophia Agape R. Padilla, Rianna Patricia S. Cruz, El King D. Morado, Joshua Gregor A. Dizon, Razel Nikka M. Hao, Arianne A. Zamora, Devon Ray Pacial, Juan Antonio R. Magalang, Marissa Alejandria, Celia Carlos, Anna Ong-Lim, Edsel Maurice Salvaña, John Q. Wong, Jaime C. Montoya, Maria Rosario Singh-Vergeire and Cynthia P. Saloma |
| EPI_ISL_2156504 | Philippine Red Cross Logistics and Multipurpose Center | Philippine Genome Center | Francis A. Tablizo, Kenneth M. Kim, Carlo M. Lapid, Marc Jerrone R. Castro, Maria Sofia L. Yangzon, Elcid Aaron R. Pangilinan, Benedict A. Maralit, Marc Edsel C. Ayes, Eva Maria Cutiongco-de la Paz, Alethea R. de Guzman, Jan Michael C. Yap, Jo-Hannah S. Llames, Sheila Mae M. Araiza, Kris P. Punayan, Irish Coleen A. Asin, Candice Francheska B. Tambaoan, Asia Louisa U. Chong, Karol Sophia Agape R. Padilla, Rianna Patricia S. Cruz, El King D. Morado, Joshua Gregor A. Dizon, Razel Nikka M. Hao, Arianne A. Zamora, Devon Ray Pacial, Juan Antonio R. Magalang, Marissa Alejandria, Celia Carlos, Anna Ong-Lim, Edsel Maurice Salvaña, John Q. Wong, Jaime C. Montoya, Maria Rosario Singh-Vergeire and Cynthia P. Saloma |
| EPI_ISL_2156511, EPI_ISL_2156512 | The Lord's Grace Medical and Industrial Clinic | Philippine Genome Center | Francis A. Tablizo, Kenneth M. Kim, Carlo M. Lapid, Marc Jerrone R. Castro, Maria Sofia L. Yangzon, Elcid Aaron R. Pangilinan, Benedict A. Maralit, Marc Edsel C. Ayes, Eva Maria Cutiongco-de la Paz, Alethea R. de Guzman, Jan Michael C. Yap, Jo-Hannah S. Llames, Sheila Mae M. Araiza, Kris P. Punayan, Irish Coleen A. Asin, Candice Francheska B. Tambaoan, Asia Louisa U. Chong, Karol Sophia Agape R. Padilla, Rianna Patricia S. Cruz, El King D. Morado, Joshua Gregor A. Dizon, Razel Nikka M. Hao, Arianne A. Zamora, Devon Ray Pacial, Juan Antonio R. Magalang, Marissa Alejandria, Celia Carlos, Anna Ong-Lim, Edsel Maurice Salvaña, John Q. Wong, Jaime C. Montoya, Maria Rosario Singh-Vergeire and Cynthia P. Saloma |
| EPI_ISL_2156513, EPI_ISL_2156514 | Lung Center of the Philippines (LCP) | Philippine Genome Center | Francis A. Tablizo, Kenneth M. Kim, Carlo M. Lapid, Marc Jerrone R. Castro, Maria Sofia L. Yangzon, Elcid Aaron R. Pangilinan, Benedict A. Maralit, Marc Edsel C. Ayes, Eva Maria Cutiongco-de la Paz, Alethea R. de Guzman, Jan Michael C. Yap, Jo-Hannah S. Llames, Sheila Mae M. Araiza, Kris P. Punayan, Irish Coleen A. Asin, Candice Francheska B. Tambaoan, Asia Louisa U. Chong, Karol Sophia Agape R. Padilla, Rianna Patricia S. Cruz, El King D. Morado, Joshua Gregor A. Dizon, Razel Nikka M. Hao, Arianne A. Zamora, Devon Ray Pacial, Juan Antonio R. Magalang, Marissa Alejandria, Celia Carlos, Anna Ong-Lim, Edsel Maurice Salvaña, John Q. Wong, Jaime C. Montoya, Maria Rosario Singh-Vergeire and Cynthia P. Saloma |
| EPI_ISL_2156515 | Cagayan Valley Medical Center Molecular Laboratory | Philippine Genome Center | Francis A. Tablizo, Kenneth M. Kim, Carlo M. Lapid, Marc Jerrone R. Castro, Maria Sofia L. Yangzon, Elcid Aaron R. Pangilinan, Benedict A. Maralit, Marc Edsel C. Ayes, Eva Maria Cutiongco-de la Paz, Alethea R. de Guzman, Jan Michael C. Yap, Jo-Hannah S. Llames, Sheila Mae M. Araiza, Kris P. Punayan, Irish Coleen A. Asin, Candice Francheska B. Tambaoan, Asia Louisa U. Chong, Karol Sophia Agape R. Padilla, Rianna Patricia S. Cruz, El King D. Morado, Joshua Gregor A. Dizon, Razel Nikka M. Hao, Arianne A. Zamora, Devon Ray Pacial, Juan Antonio R. Magalang, Marissa Alejandria, Celia Carlos, Anna Ong-Lim, Edsel Maurice Salvaña, John Q. Wong, Jaime C. Montoya, Maria Rosario Singh-Vergeire and Cynthia P. Saloma |
| EPI_ISL_2156516 | The Lord's Grace Medical and Industrial Clinic | Philippine Genome Center | Francis A. Tablizo, Kenneth M. Kim, Carlo M. Lapid, Marc Jerrone R. Castro, Maria Sofia L. Yangzon, Elcid Aaron R. Pangilinan, Benedict A. Maralit, Marc Edsel C. Ayes, Eva Maria Cutiongco-de la Paz, Alethea R. de Guzman, Jan Michael C. Yap, Jo-Hannah S. Llames, Sheila Mae M. Araiza, Kris P. Punayan, Irish Coleen A. Asin, Candice Francheska B. Tambaoan, Asia Louisa U. Chong, Karol Sophia Agape R. Padilla, Rianna Patricia S. Cruz, El King D. Morado, Joshua Gregor A. Dizon, Razel Nikka M. Hao, Arianne A. Zamora, Devon Ray Pacial, Juan Antonio R. Magalang, Marissa Alejandria, Celia Carlos, Anna Ong-Lim, Edsel Maurice Salvaña, John Q. Wong, Jaime C. Montoya, Maria Rosario Singh-Vergeire and Cynthia P. Saloma |
| EPI_ISL_2156523 | Lung Center of the Philippines (LCP) | Philippine Genome Center | Francis A. Tablizo, Kenneth M. Kim, Carlo M. Lapid, Marc Jerrone R. Castro, Maria Sofia L. Yangzon, Elcid Aaron R. Pangilinan, Benedict A. Maralit, Marc Edsel C. Ayes, Eva Maria Cutiongco-de la Paz, Alethea R. de Guzman, Jan Michael C. Yap, Jo-Hannah S. Llames, Sheila Mae M. Araiza, Kris P. Punayan, Irish Coleen A. Asin, Candice Francheska B. Tambaoan, Asia Louisa U. Chong, Karol Sophia Agape R. Padilla, Rianna Patricia S. Cruz, El King D. Morado, Joshua Gregor A. Dizon, Razel Nikka M. Hao, Arianne A. Zamora, Devon Ray Pacial, Juan Antonio R. Magalang, Marissa Alejandria, Celia Carlos, Anna Ong-Lim, Edsel Maurice Salvaña, John Q. Wong, Jaime C. Montoya, Maria Rosario Singh-Vergeire and Cynthia P. Saloma |
| EPI_ISL_2156526 | Parkway Medical and Diagnostic Center | Philippine Genome Center | Francis A. Tablizo, Kenneth M. Kim, Carlo M. Lapid, Marc Jerrone R. Castro, Maria Sofia L. Yangzon, Elcid Aaron R. Pangilinan, Benedict A. Maralit, Marc Edsel C. Ayes, Eva Maria Cutiongco-de la Paz, Alethea R. de Guzman, Jan Michael C. Yap, Jo-Hannah S. Llames, Sheila Mae M. Araiza, Kris P. Punayan, Irish Coleen A. Asin, Candice Francheska B. Tambaoan, Asia Louisa U. Chong, Karol Sophia Agape R. Padilla, Rianna Patricia S. Cruz, El King D. Morado, Joshua Gregor A. Dizon, Razel Nikka M. Hao, Arianne A. Zamora, Devon Ray Pacial, Juan Antonio R. Magalang, Marissa Alejandria, Celia Carlos, Anna Ong-Lim, Edsel Maurice Salvaña, John Q. Wong, Jaime C. Montoya, Maria Rosario Singh-Vergeire and Cynthia P. Saloma |
| EPI_ISL_2156527 | Lung Center of the Philippines (LCP) | Philippine Genome Center | Francis A. Tablizo, Kenneth M. Kim, Carlo M. Lapid, Marc Jerrone R. Castro, Maria Sofia L. Yangzon, Elcid Aaron R. Pangilinan, Benedict A. Maralit, Marc Edsel C. Ayes, Eva Maria Cutiongco-de la Paz, Alethea R. de Guzman, Jan Michael C. Yap, Jo-Hannah S. Llames, Sheila Mae M. Araiza, Kris P. Punayan, Irish Coleen A. Asin, Candice Francheska B. Tambaoan, Asia Louisa U. Chong, Karol Sophia Agape R. Padilla, Rianna Patricia S. Cruz, El King D. Morado, Joshua Gregor A. Dizon, Razel Nikka M. Hao, Arianne A. Zamora, Devon Ray Pacial, Juan Antonio R. Magalang, Marissa Alejandria, Celia Carlos, Anna Ong-Lim, Edsel Maurice Salvaña, John Q. Wong, Jaime C. Montoya, Maria Rosario Singh-Vergeire and Cynthia P. Saloma |

| | | | |
|---|---|---|---|
| EPI_ISL_2156529 | Baguio General Hospital Medical Center (BGHMC) | Philippine Genome Center | Francis A. Tablizo, Kenneth M. Kim, Carlo M. Lapid, Marc Jerrone R. Castro, Maria Sofia L. Yangzon, Elcid Aaron R. Pangilinan, Benedict A. Maralit, Marc Edsel C. Ayes, Eva Maria Cutiongco-de la Paz, Alethea R. de Guzman, Jan Michael C. Yap, Jo-Hannah S. Llames, Sheila Mae M. Araiza, Kris P. Punayan, Irish Coleen A. Asin, Candice Francheska B. Tambaoan, Asia Louisa U. Chong, Karol Sophia Agape R. Padilla, Rianna Patricia S. Cruz, El King D. Morado, Joshua Gregor A. Dizon, Razel Nikka M. Hao, Arianne A. Zamora, Devon Ray Pacial, Juan Antonio R. Magalang, Marissa Alejandria, Celia Carlos, Anna Ong-Lim, Edsel Maurice Salvaña, John Q. Wong, Jaime C. Montoya, Maria Rosario Singh-Vergeire and Cynthia P. Saloma |
| EPI_ISL_2156531 | Qualimed Hospital Sta. Rosa | Philippine Genome Center | Francis A. Tablizo, Kenneth M. Kim, Carlo M. Lapid, Marc Jerrone R. Castro, Maria Sofia L. Yangzon, Elcid Aaron R. Pangilinan, Benedict A. Maralit, Marc Edsel C. Ayes, Eva Maria Cutiongco-de la Paz, Alethea R. de Guzman, Jan Michael C. Yap, Jo-Hannah S. Llames, Sheila Mae M. Araiza, Kris P. Punayan, Irish Coleen A. Asin, Candice Francheska B. Tambaoan, Asia Louisa U. Chong, Karol Sophia Agape R. Padilla, Rianna Patricia S. Cruz, El King D. Morado, Joshua Gregor A. Dizon, Razel Nikka M. Hao, Arianne A. Zamora, Devon Ray Pacial, Juan Antonio R. Magalang, Marissa Alejandria, Celia Carlos, Anna Ong-Lim, Edsel Maurice Salvaña, John Q. Wong, Jaime C. Montoya, Maria Rosario Singh-Vergeire and Cynthia P. Saloma |
| EPI_ISL_2156533 | University of the East Ramon Magsaysay Memorial Medical Center (UERM) | Philippine Genome Center | Francis A. Tablizo, Kenneth M. Kim, Carlo M. Lapid, Marc Jerrone R. Castro, Maria Sofia L. Yangzon, Elcid Aaron R. Pangilinan, Benedict A. Maralit, Marc Edsel C. Ayes, Eva Maria Cutiongco-de la Paz, Alethea R. de Guzman, Jan Michael C. Yap, Jo-Hannah S. Llames, Sheila Mae M. Araiza, Kris P. Punayan, Irish Coleen A. Asin, Candice Francheska B. Tambaoan, Asia Louisa U. Chong, Karol Sophia Agape R. Padilla, Rianna Patricia S. Cruz, El King D. Morado, Joshua Gregor A. Dizon, Razel Nikka M. Hao, Arianne A. Zamora, Devon Ray Pacial, Juan Antonio R. Magalang, Marissa Alejandria, Celia Carlos, Anna Ong-Lim, Edsel Maurice Salvaña, John Q. Wong, Jaime C. Montoya, Maria Rosario Singh-Vergeire and Cynthia P. Saloma |
| EPI_ISL_2156534 | The Lord's Grace Medical and Industrial Clinic | Philippine Genome Center | Francis A. Tablizo, Kenneth M. Kim, Carlo M. Lapid, Marc Jerrone R. Castro, Maria Sofia L. Yangzon, Elcid Aaron R. Pangilinan, Benedict A. Maralit, Marc Edsel C. Ayes, Eva Maria Cutiongco-de la Paz, Alethea R. de Guzman, Jan Michael C. Yap, Jo-Hannah S. Llames, Sheila Mae M. Araiza, Kris P. Punayan, Irish Coleen A. Asin, Candice Francheska B. Tambaoan, Asia Louisa U. Chong, Karol Sophia Agape R. Padilla, Rianna Patricia S. Cruz, El King D. Morado, Joshua Gregor A. Dizon, Razel Nikka M. Hao, Arianne A. Zamora, Devon Ray Pacial, Juan Antonio R. Magalang, Marissa Alejandria, Celia Carlos, Anna Ong-Lim, Edsel Maurice Salvaña, John Q. Wong, Jaime C. Montoya, Maria Rosario Singh-Vergeire and Cynthia P. Saloma |
| EPI_ISL_2156535 | Philippine Red Cross - Port Area | Philippine Genome Center | Francis A. Tablizo, Kenneth M. Kim, Carlo M. Lapid, Marc Jerrone R. Castro, Maria Sofia L. Yangzon, Elcid Aaron R. Pangilinan, Benedict A. Maralit, Marc Edsel C. Ayes, Eva Maria Cutiongco-de la Paz, Alethea R. de Guzman, Jan Michael C. Yap, Jo-Hannah S. Llames, Sheila Mae M. Araiza, Kris P. Punayan, Irish Coleen A. Asin, Candice Francheska B. Tambaoan, Asia Louisa U. Chong, Karol Sophia Agape R. Padilla, Rianna Patricia S. Cruz, El King D. Morado, Joshua Gregor A. Dizon, Razel Nikka M. Hao, Arianne A. Zamora, Devon Ray Pacial, Juan Antonio R. Magalang, Marissa Alejandria, Celia Carlos, Anna Ong-Lim, Edsel Maurice Salvaña, John Q. Wong, Jaime C. Montoya, Maria Rosario Singh-Vergeire and Cynthia P. Saloma |
| EPI_ISL_2156536 | Philippine Airport Diagnostic Laboratory | Philippine Genome Center | Francis A. Tablizo, Kenneth M. Kim, Carlo M. Lapid, Marc Jerrone R. Castro, Maria Sofia L. Yangzon, Elcid Aaron R. Pangilinan, Benedict A. Maralit, Marc Edsel C. Ayes, Eva Maria Cutiongco-de la Paz, Alethea R. de Guzman, Jan Michael C. Yap, Jo-Hannah S. Llames, Sheila Mae M. Araiza, Kris P. Punayan, Irish Coleen A. Asin, Candice Francheska B. Tambaoan, Asia Louisa U. Chong, Karol Sophia Agape R. Padilla, Rianna Patricia S. Cruz, El King D. Morado, Joshua Gregor A. Dizon, Razel Nikka M. Hao, Arianne A. Zamora, Devon Ray Pacial, Juan Antonio R. Magalang, Marissa Alejandria, Celia Carlos, Anna Ong-Lim, Edsel Maurice Salvaña, John Q. Wong, Jaime C. Montoya, Maria Rosario Singh-Vergeire and Cynthia P. Saloma |
| EPI_ISL_2156546 | Baguio General Hospital Medical Center (BGHMC) | Philippine Genome Center | Francis A. Tablizo, Kenneth M. Kim, Carlo M. Lapid, Marc Jerrone R. Castro, Maria Sofia L. Yangzon, Elcid Aaron R. Pangilinan, Benedict A. Maralit, Marc Edsel C. Ayes, Eva Maria Cutiongco-de la Paz, Alethea R. de Guzman, Jan Michael C. Yap, Jo-Hannah S. Llames, Sheila Mae M. Araiza, Kris P. Punayan, Irish Coleen A. Asin, Candice Francheska B. Tambaoan, Asia Louisa U. Chong, Karol Sophia Agape R. Padilla, Rianna Patricia S. Cruz, El King D. Morado, Joshua Gregor A. Dizon, Razel Nikka M. Hao, Arianne A. Zamora, Devon Ray Pacial, Juan Antonio R. Magalang, Marissa Alejandria, Celia Carlos, Anna Ong-Lim, Edsel Maurice Salvaña, John Q. Wong, Jaime C. Montoya, Maria Rosario Singh-Vergeire and Cynthia P. Saloma |
| EPI_ISL_2156547, EPI_ISL_2156548 | Philippine Red Cross - National Blood Center | Philippine Genome Center | Francis A. Tablizo, Kenneth M. Kim, Carlo M. Lapid, Marc Jerrone R. Castro, Maria Sofia L. Yangzon, Elcid Aaron R. Pangilinan, Benedict A. Maralit, Marc Edsel C. Ayes, Eva Maria Cutiongco-de la Paz, Alethea R. de Guzman, Jan Michael C. Yap, Jo-Hannah S. Llames, Sheila Mae M. Araiza, Kris P. Punayan, Irish Coleen A. Asin, Candice Francheska B. Tambaoan, Asia Louisa U. Chong, Karol Sophia Agape R. Padilla, Rianna Patricia S. Cruz, El King D. Morado, Joshua Gregor A. Dizon, Razel Nikka M. Hao, Arianne A. Zamora, Devon Ray Pacial, Juan Antonio R. Magalang, Marissa Alejandria, Celia Carlos, Anna Ong-Lim, Edsel Maurice Salvaña, John Q. Wong, Jaime C. Montoya, Maria Rosario Singh-Vergeire and Cynthia P. Saloma |
| EPI_ISL_2156549 | Valenzuela Hope Molecular Laboratory | Philippine Genome Center | Francis A. Tablizo, Kenneth M. Kim, Carlo M. Lapid, Marc Jerrone R. Castro, Maria Sofia L. Yangzon, Elcid Aaron R. Pangilinan, Benedict A. Maralit, Marc Edsel C. Ayes, Eva Maria Cutiongco-de la Paz, Alethea R. de Guzman, Jan Michael C. Yap, Jo-Hannah S. Llames, Sheila Mae M. Araiza, Kris P. Punayan, Irish Coleen A. Asin, Candice Francheska B. Tambaoan, Asia Louisa U. Chong, Karol Sophia Agape R. Padilla, Rianna Patricia S. Cruz, El King D. Morado, Joshua Gregor A. Dizon, Razel Nikka M. Hao, Arianne A. Zamora, Devon Ray Pacial, Juan Antonio R. Magalang, Marissa Alejandria, Celia Carlos, Anna Ong-Lim, Edsel Maurice Salvaña, John Q. Wong, Jaime C. Montoya, Maria Rosario Singh-Vergeire and Cynthia P. Saloma |
| EPI_ISL_2156550 | The Lord's Grace Medical and Industrial Clinic | Philippine Genome Center | Francis A. Tablizo, Kenneth M. Kim, Carlo M. Lapid, Marc Jerrone R. Castro, Maria Sofia L. Yangzon, Elcid Aaron R. Pangilinan, Benedict A. Maralit, Marc Edsel C. Ayes, Eva Maria Cutiongco-de la Paz, Alethea R. de Guzman, Jan Michael C. Yap, Jo-Hannah S. Llames, Sheila Mae M. Araiza, Kris P. Punayan, Irish Coleen A. Asin, Candice Francheska B. Tambaoan, Asia Louisa U. Chong, Karol Sophia Agape R. Padilla, Rianna Patricia S. Cruz, El King D. Morado, Joshua Gregor A. Dizon, Razel Nikka M. Hao, Arianne A. Zamora, Devon Ray Pacial, Juan Antonio R. Magalang, Marissa Alejandria, Celia Carlos, Anna Ong-Lim, Edsel Maurice Salvaña, John Q. Wong, Jaime C. Montoya, Maria Rosario Singh-Vergeire and Cynthia P. Saloma |
| EPI_ISL_2156551, EPI_ISL_2156552 | Amosup Seamen's Hospital | Philippine Genome Center | Francis A. Tablizo, Kenneth M. Kim, Carlo M. Lapid, Marc Jerrone R. Castro, Maria Sofia L. Yangzon, Elcid Aaron R. Pangilinan, Benedict A. Maralit, Marc Edsel C. Ayes, Eva Maria Cutiongco-de la Paz, Alethea R. de Guzman, Jan Michael C. Yap, Jo-Hannah S. Llames, Sheila Mae M. Araiza, Kris P. Punayan, Irish Coleen A. Asin, Candice Francheska B. Tambaoan, Asia Louisa U. Chong, Karol Sophia Agape R. Padilla, Rianna Patricia S. Cruz, El King D. Morado, Joshua Gregor A. Dizon, Razel Nikka M. Hao, Arianne A. Zamora, Devon Ray Pacial, Juan Antonio R. Magalang, Marissa Alejandria, Celia Carlos, Anna Ong-Lim, Edsel Maurice Salvaña, John Q. Wong, Jaime C. Montoya, Maria Rosario Singh-Vergeire and Cynthia P. Saloma |
| EPI_ISL_2156553 | Cagayan Valley Medical Center Molecular Laboratory | Philippine Genome Center | Francis A. Tablizo, Kenneth M. Kim, Carlo M. Lapid, Marc Jerrone R. Castro, Maria Sofia L. Yangzon, Elcid Aaron R. Pangilinan, Benedict A. Maralit, Marc Edsel C. Ayes, Eva Maria Cutiongco-de la Paz, Alethea R. de Guzman, Jan Michael C. Yap, Jo-Hannah S. Llames, Sheila Mae M. Araiza, Kris P. Punayan, Irish Coleen A. Asin, Candice Francheska B. Tambaoan, Asia Louisa U. Chong, Karol Sophia Agape R. Padilla, Rianna Patricia S. Cruz, El King D. Morado, Joshua Gregor A. Dizon, Razel Nikka M. Hao, Arianne A. Zamora, Devon Ray Pacial, Juan Antonio R. Magalang, Marissa Alejandria, Celia Carlos, Anna Ong-Lim, Edsel Maurice Salvaña, John Q. Wong, Jaime C. Montoya, Maria Rosario Singh-Vergeire and Cynthia P. Saloma |
| EPI_ISL_2156556 | Philippine Red Cross - National Blood Center | Philippine Genome Center | Francis A. Tablizo, Kenneth M. Kim, Carlo M. Lapid, Marc Jerrone R. Castro, Maria Sofia L. Yangzon, Elcid Aaron R. Pangilinan, Benedict A. Maralit, Marc Edsel C. Ayes, Eva Maria Cutiongco-de la Paz, Alethea R. de Guzman, Jan Michael C. Yap, Jo-Hannah S. Llames, Sheila Mae M. Araiza, Kris P. Punayan, Irish Coleen A. Asin, Candice Francheska B. Tambaoan, Asia Louisa U. Chong, Karol Sophia Agape R. Padilla, Rianna Patricia S. Cruz, El King D. Morado, Joshua Gregor A. Dizon, Razel Nikka M. Hao, Arianne A. Zamora, Devon Ray Pacial, Juan Antonio R. Magalang, Marissa Alejandria, Celia Carlos, Anna Ong-Lim, Edsel Maurice Salvaña, John Q. Wong, Jaime C. Montoya, Maria Rosario Singh-Vergeire and Cynthia P. Saloma |
| EPI_ISL_2156557 | Qualimed Hospital Sta. Rosa | Philippine Genome Center | Francis A. Tablizo, Kenneth M. Kim, Carlo M. Lapid, Marc Jerrone R. Castro, Maria Sofia L. Yangzon, Elcid Aaron R. Pangilinan, Benedict A. Maralit, Marc Edsel C. Ayes, Eva Maria Cutiongco-de la Paz, Alethea R. de Guzman, Jan Michael C. Yap, Jo-Hannah S. Llames, Sheila Mae M. Araiza, Kris P. Punayan, Irish Coleen A. Asin, Candice Francheska B. Tambaoan, Asia Louisa U. Chong, Karol Sophia Agape R. Padilla, Rianna Patricia S. Cruz, El King D. Morado, Joshua Gregor A. Dizon, Razel Nikka M. Hao, Arianne A. Zamora, Devon Ray Pacial, Juan Antonio R. Magalang, Marissa Alejandria, Celia Carlos, Anna Ong-Lim, Edsel Maurice Salvaña, John Q. Wong, Jaime C. Montoya, Maria Rosario Singh-Vergeire and Cynthia P. Saloma |
| EPI_ISL_2156558, EPI_ISL_2156561 | Research Institute for Tropical Medicine, Inc. (RITM) | Philippine Genome Center | Francis A. Tablizo, Kenneth M. Kim, Carlo M. Lapid, Marc Jerrone R. Castro, Maria Sofia L. Yangzon, Elcid Aaron R. Pangilinan, Benedict A. Maralit, Marc Edsel C. Ayes, Eva Maria Cutiongco-de la Paz, Alethea R. de Guzman, Jan Michael C. Yap, Jo-Hannah S. Llames, Sheila Mae M. Araiza, Kris P. Punayan, Irish Coleen A. Asin, Candice Francheska B. Tambaoan, Asia Louisa U. Chong, Karol Sophia Agape R. Padilla, Rianna Patricia S. Cruz, El King D. Morado, Joshua Gregor A. Dizon, Razel Nikka M. Hao, Arianne A. Zamora, Devon Ray Pacial, Juan Antonio R. Magalang, Marissa Alejandria, Celia Carlos, Anna Ong-Lim, Edsel Maurice Salvaña, John Q. Wong, Jaime C. Montoya, Maria Rosario Singh-Vergeire and Cynthia P. Saloma |
| EPI_ISL_2156564 | Daniel O. Mercado Medical Center | Philippine Genome Center | Francis A. Tablizo, Kenneth M. Kim, Carlo M. Lapid, Marc Jerrone R. Castro, Maria Sofia L. Yangzon, Elcid Aaron R. Pangilinan, Benedict A. Maralit, Marc Edsel C. Ayes, Eva Maria Cutiongco-de la Paz, Alethea R. de Guzman, Jan Michael C. Yap, Jo-Hannah S. Llames, Sheila Mae M. Araiza, Kris P. Punayan, |

| | | | |
|---|---|---|---|
| | | | Irish Coleen A. Asin, Candice Francheska B. Tambaoan, Asia Louisa U. Chong, Karol Sophia Agape R. Padilla, Rianna Patricia S. Cruz, El King D. Morado, Joshua Gregor A. Dizon, Razel Nikka M. Hao, Arianne A. Zamora, Devon Ray Pacial, Juan Antonio R. Magalang, Marissa Alejandria, Celia Carlos, Anna Ong-Lim, Edsel Maurice Salvaña, John Q. Wong, Jaime C. Montoya, Maria Rosario Singh-Vergeire and Cynthia P. Saloma |
| EPI_ISL_2156569 | Research Institute for Tropical Medicine, Inc. (RITM) | Philippine Genome Center | Francis A. Tablizo, Kenneth M. Kim, Carlo M. Lapid, Marc Jerrone R. Castro, Maria Sofia L. Yangzon, Elcid Aaron R. Pangilinan, Benedict A. Maralit, Marc Edsel C. Ayes, Eva Maria Cutiongco-de la Paz, Alethea R. de Guzman, Jan Michael C. Yap, Jo-Hannah S. Llames, Sheila Mae M. Araiza, Kris P. Punayan, Irish Coleen A. Asin, Candice Francheska B. Tambaoan, Asia Louisa U. Chong, Karol Sophia Agape R. Padilla, Rianna Patricia S. Cruz, El King D. Morado, Joshua Gregor A. Dizon, Razel Nikka M. Hao, Arianne A. Zamora, Devon Ray Pacial, Juan Antonio R. Magalang, Marissa Alejandria, Celia Carlos, Anna Ong-Lim, Edsel Maurice Salvaña, John Q. Wong, Jaime C. Montoya, Maria Rosario Singh-Vergeire and Cynthia P. Saloma |
| EPI_ISL_2156571, EPI_ISL_2156573 | Baguio General Hospital Medical Center (BGHMC) | Philippine Genome Center | Francis A. Tablizo, Kenneth M. Kim, Carlo M. Lapid, Marc Jerrone R. Castro, Maria Sofia L. Yangzon, Elcid Aaron R. Pangilinan, Benedict A. Maralit, Marc Edsel C. Ayes, Eva Maria Cutiongco-de la Paz, Alethea R. de Guzman, Jan Michael C. Yap, Jo-Hannah S. Llames, Sheila Mae M. Araiza, Kris P. Punayan, Irish Coleen A. Asin, Candice Francheska B. Tambaoan, Asia Louisa U. Chong, Karol Sophia Agape R. Padilla, Rianna Patricia S. Cruz, El King D. Morado, Joshua Gregor A. Dizon, Razel Nikka M. Hao, Arianne A. Zamora, Devon Ray Pacial, Juan Antonio R. Magalang, Marissa Alejandria, Celia Carlos, Anna Ong-Lim, Edsel Maurice Salvaña, John Q. Wong, Jaime C. Montoya, Maria Rosario Singh-Vergeire and Cynthia P. Saloma |
| EPI_ISL_2156574 | Research Institute for Tropical Medicine, Inc. (RITM) | Philippine Genome Center | Francis A. Tablizo, Kenneth M. Kim, Carlo M. Lapid, Marc Jerrone R. Castro, Maria Sofia L. Yangzon, Elcid Aaron R. Pangilinan, Benedict A. Maralit, Marc Edsel C. Ayes, Eva Maria Cutiongco-de la Paz, Alethea R. de Guzman, Jan Michael C. Yap, Jo-Hannah S. Llames, Sheila Mae M. Araiza, Kris P. Punayan, Irish Coleen A. Asin, Candice Francheska B. Tambaoan, Asia Louisa U. Chong, Karol Sophia Agape R. Padilla, Rianna Patricia S. Cruz, El King D. Morado, Joshua Gregor A. Dizon, Razel Nikka M. Hao, Arianne A. Zamora, Devon Ray Pacial, Juan Antonio R. Magalang, Marissa Alejandria, Celia Carlos, Anna Ong-Lim, Edsel Maurice Salvaña, John Q. Wong, Jaime C. Montoya, Maria Rosario Singh-Vergeire and Cynthia P. Saloma |
| EPI_ISL_2156577 | Bataan General Hospital and Medical Center | Philippine Genome Center | Francis A. Tablizo, Kenneth M. Kim, Carlo M. Lapid, Marc Jerrone R. Castro, Maria Sofia L. Yangzon, Elcid Aaron R. Pangilinan, Benedict A. Maralit, Marc Edsel C. Ayes, Eva Maria Cutiongco-de la Paz, Alethea R. de Guzman, Jan Michael C. Yap, Jo-Hannah S. Llames, Sheila Mae M. Araiza, Kris P. Punayan, Irish Coleen A. Asin, Candice Francheska B. Tambaoan, Asia Louisa U. Chong, Karol Sophia Agape R. Padilla, Rianna Patricia S. Cruz, El King D. Morado, Joshua Gregor A. Dizon, Razel Nikka M. Hao, Arianne A. Zamora, Devon Ray Pacial, Juan Antonio R. Magalang, Marissa Alejandria, Celia Carlos, Anna Ong-Lim, Edsel Maurice Salvaña, John Q. Wong, Jaime C. Montoya, Maria Rosario Singh-Vergeire and Cynthia P. Saloma |
| EPI_ISL_2156580, EPI_ISL_2156581 | Cagayan Valley Medical Center Molecular Laboratory | Philippine Genome Center | Francis A. Tablizo, Kenneth M. Kim, Carlo M. Lapid, Marc Jerrone R. Castro, Maria Sofia L. Yangzon, Elcid Aaron R. Pangilinan, Benedict A. Maralit, Marc Edsel C. Ayes, Eva Maria Cutiongco-de la Paz, Alethea R. de Guzman, Jan Michael C. Yap, Jo-Hannah S. Llames, Sheila Mae M. Araiza, Kris P. Punayan, Irish Coleen A. Asin, Candice Francheska B. Tambaoan, Asia Louisa U. Chong, Karol Sophia Agape R. Padilla, Rianna Patricia S. Cruz, El King D. Morado, Joshua Gregor A. Dizon, Razel Nikka M. Hao, Arianne A. Zamora, Devon Ray Pacial, Juan Antonio R. Magalang, Marissa Alejandria, Celia Carlos, Anna Ong-Lim, Edsel Maurice Salvaña, John Q. Wong, Jaime C. Montoya, Maria Rosario Singh-Vergeire and Cynthia P. Saloma |
| EPI_ISL_2156583 | San Antonio City of Ilagan Hospital | Philippine Genome Center | Francis A. Tablizo, Kenneth M. Kim, Carlo M. Lapid, Marc Jerrone R. Castro, Maria Sofia L. Yangzon, Elcid Aaron R. Pangilinan, Benedict A. Maralit, Marc Edsel C. Ayes, Eva Maria Cutiongco-de la Paz, Alethea R. de Guzman, Jan Michael C. Yap, Jo-Hannah S. Llames, Sheila Mae M. Araiza, Kris P. Punayan, Irish Coleen A. Asin, Candice Francheska B. Tambaoan, Asia Louisa U. Chong, Karol Sophia Agape R. Padilla, Rianna Patricia S. Cruz, El King D. Morado, Joshua Gregor A. Dizon, Razel Nikka M. Hao, Arianne A. Zamora, Devon Ray Pacial, Juan Antonio R. Magalang, Marissa Alejandria, Celia Carlos, Anna Ong-Lim, Edsel Maurice Salvaña, John Q. Wong, Jaime C. Montoya, Maria Rosario Singh-Vergeire and Cynthia P. Saloma |
| EPI_ISL_2156584 | Northern Mindanao TB Regional Center | Philippine Genome Center | Francis A. Tablizo, Kenneth M. Kim, Carlo M. Lapid, Marc Jerrone R. Castro, Maria Sofia L. Yangzon, Elcid Aaron R. Pangilinan, Benedict A. Maralit, Marc Edsel C. Ayes, Eva Maria Cutiongco-de la Paz, Alethea R. de Guzman, Jan Michael C. Yap, Jo-Hannah S. Llames, Sheila Mae M. Araiza, Kris P. Punayan, Irish Coleen A. Asin, Candice Francheska B. Tambaoan, Asia Louisa U. Chong, Karol Sophia Agape R. Padilla, Rianna Patricia S. Cruz, El King D. Morado, Joshua Gregor A. Dizon, Razel Nikka M. Hao, Arianne A. Zamora, Devon Ray Pacial, Juan Antonio R. Magalang, Marissa Alejandria, Celia Carlos, Anna Ong-Lim, Edsel Maurice Salvaña, John Q. Wong, Jaime C. Montoya, Maria Rosario Singh-Vergeire and Cynthia P. Saloma |
| EPI_ISL_2156588 | Baguio General Hospital Medical Center (BGHMC) | Philippine Genome Center | Francis A. Tablizo, Kenneth M. Kim, Carlo M. Lapid, Marc Jerrone R. Castro, Maria Sofia L. Yangzon, Elcid Aaron R. Pangilinan, Benedict A. Maralit, Marc Edsel C. Ayes, Eva Maria Cutiongco-de la Paz, Alethea R. de Guzman, Jan Michael C. Yap, Jo-Hannah S. Llames, Sheila Mae M. Araiza, Kris P. Punayan, Irish Coleen A. Asin, Candice Francheska B. Tambaoan, Asia Louisa U. Chong, Karol Sophia Agape R. Padilla, Rianna Patricia S. Cruz, El King D. Morado, Joshua Gregor A. Dizon, Razel Nikka M. Hao, Arianne A. Zamora, Devon Ray Pacial, Juan Antonio R. Magalang, Marissa Alejandria, Celia Carlos, Anna Ong-Lim, Edsel Maurice Salvaña, John Q. Wong, Jaime C. Montoya, Maria Rosario Singh-Vergeire and Cynthia P. Saloma |
| EPI_ISL_2156591, EPI_ISL_2156592, EPI_ISL_2156593 | Northern Mindanao TB Regional Center | Philippine Genome Center | Francis A. Tablizo, Kenneth M. Kim, Carlo M. Lapid, Marc Jerrone R. Castro, Maria Sofia L. Yangzon, Elcid Aaron R. Pangilinan, Benedict A. Maralit, Marc Edsel C. Ayes, Eva Maria Cutiongco-de la Paz, Alethea R. de Guzman, Jan Michael C. Yap, Jo-Hannah S. Llames, Sheila Mae M. Araiza, Kris P. Punayan, Irish Coleen A. Asin, Candice Francheska B. Tambaoan, Asia Louisa U. Chong, Karol Sophia Agape R. Padilla, Rianna Patricia S. Cruz, El King D. Morado, Joshua Gregor A. Dizon, Razel Nikka M. Hao, Arianne A. Zamora, Devon Ray Pacial, Juan Antonio R. Magalang, Marissa Alejandria, Celia Carlos, Anna Ong-Lim, Edsel Maurice Salvaña, John Q. Wong, Jaime C. Montoya, Maria Rosario Singh-Vergeire and Cynthia P. Saloma |
| EPI_ISL_2156596 | Manila Doctors Hospital | Philippine Genome Center | Francis A. Tablizo, Kenneth M. Kim, Carlo M. Lapid, Marc Jerrone R. Castro, Maria Sofia L. Yangzon, Elcid Aaron R. Pangilinan, Benedict A. Maralit, Marc Edsel C. Ayes, Eva Maria Cutiongco-de la Paz, Alethea R. de Guzman, Jan Michael C. Yap, Jo-Hannah S. Llames, Sheila Mae M. Araiza, Kris P. Punayan, Irish Coleen A. Asin, Candice Francheska B. Tambaoan, Asia Louisa U. Chong, Karol Sophia Agape R. Padilla, Rianna Patricia S. Cruz, El King D. Morado, Joshua Gregor A. Dizon, Razel Nikka M. Hao, Arianne A. Zamora, Devon Ray Pacial, Juan Antonio R. Magalang, Marissa Alejandria, Celia Carlos, Anna Ong-Lim, Edsel Maurice Salvaña, John Q. Wong, Jaime C. Montoya, Maria Rosario Singh-Vergeire and Cynthia P. Saloma |
| EPI_ISL_2156598, EPI_ISL_2156599 | Baguio General Hospital Medical Center (BGHMC) | Philippine Genome Center | Francis A. Tablizo, Kenneth M. Kim, Carlo M. Lapid, Marc Jerrone R. Castro, Maria Sofia L. Yangzon, Elcid Aaron R. Pangilinan, Benedict A. Maralit, Marc Edsel C. Ayes, Eva Maria Cutiongco-de la Paz, Alethea R. de Guzman, Jan Michael C. Yap, Jo-Hannah S. Llames, Sheila Mae M. Araiza, Kris P. Punayan, Irish Coleen A. Asin, Candice Francheska B. Tambaoan, Asia Louisa U. Chong, Karol Sophia Agape R. Padilla, Rianna Patricia S. Cruz, El King D. Morado, Joshua Gregor A. Dizon, Razel Nikka M. Hao, Arianne A. Zamora, Devon Ray Pacial, Juan Antonio R. Magalang, Marissa Alejandria, Celia Carlos, Anna Ong-Lim, Edsel Maurice Salvaña, John Q. Wong, Jaime C. Montoya, Maria Rosario Singh-Vergeire and Cynthia P. Saloma |
| EPI_ISL_2156600 | Research Institute for Tropical Medicine, Inc. (RITM) | Philippine Genome Center | Francis A. Tablizo, Kenneth M. Kim, Carlo M. Lapid, Marc Jerrone R. Castro, Maria Sofia L. Yangzon, Elcid Aaron R. Pangilinan, Benedict A. Maralit, Marc Edsel C. Ayes, Eva Maria Cutiongco-de la Paz, Alethea R. de Guzman, Jan Michael C. Yap, Jo-Hannah S. Llames, Sheila Mae M. Araiza, Kris P. Punayan, Irish Coleen A. Asin, Candice Francheska B. Tambaoan, Asia Louisa U. Chong, Karol Sophia Agape R. Padilla, Rianna Patricia S. Cruz, El King D. Morado, Joshua Gregor A. Dizon, Razel Nikka M. Hao, Arianne A. Zamora, Devon Ray Pacial, Juan Antonio R. Magalang, Marissa Alejandria, Celia Carlos, Anna Ong-Lim, Edsel Maurice Salvaña, John Q. Wong, Jaime C. Montoya, Maria Rosario Singh-Vergeire and Cynthia P. Saloma |
| EPI_ISL_2156604 | Baguio General Hospital Medical Center (BGHMC) | Philippine Genome Center | Francis A. Tablizo, Kenneth M. Kim, Carlo M. Lapid, Marc Jerrone R. Castro, Maria Sofia L. Yangzon, Elcid Aaron R. Pangilinan, Benedict A. Maralit, Marc Edsel C. Ayes, Eva Maria Cutiongco-de la Paz, Alethea R. de Guzman, Jan Michael C. Yap, Jo-Hannah S. Llames, Sheila Mae M. Araiza, Kris P. Punayan, Irish Coleen A. Asin, Candice Francheska B. Tambaoan, Asia Louisa U. Chong, Karol Sophia Agape R. Padilla, Rianna Patricia S. Cruz, El King D. Morado, Joshua Gregor A. Dizon, Razel Nikka M. Hao, Arianne A. Zamora, Devon Ray Pacial, Juan Antonio R. Magalang, Marissa Alejandria, Celia Carlos, Anna Ong-Lim, Edsel Maurice Salvaña, John Q. Wong, Jaime C. Montoya, Maria Rosario Singh-Vergeire and Cynthia P. Saloma |
| EPI_ISL_2156606 | National Kidney and Transplant Institute (NKTI) | Philippine Genome Center | Francis A. Tablizo, Kenneth M. Kim, Carlo M. Lapid, Marc Jerrone R. Castro, Maria Sofia L. Yangzon, Elcid Aaron R. Pangilinan, Benedict A. Maralit, Marc Edsel C. Ayes, Eva Maria Cutiongco-de la Paz, Alethea R. de Guzman, Jan Michael C. Yap, Jo-Hannah S. Llames, Sheila Mae M. Araiza, Kris P. Punayan, Irish Coleen A. Asin, Candice Francheska B. Tambaoan, Asia Louisa U. Chong, Karol Sophia Agape R. Padilla, Rianna Patricia S. Cruz, El King D. Morado, Joshua Gregor A. Dizon, Razel Nikka M. Hao, Arianne A. Zamora, Devon Ray Pacial, Juan Antonio R. Magalang, Marissa Alejandria, Celia Carlos, Anna Ong-Lim, Edsel Maurice Salvaña, John Q. Wong, Jaime C. Montoya, Maria Rosario Singh-Vergeire and Cynthia P. Saloma |
| EPI_ISL_2156612 | Philippine Red Cross- Batangas Chapter Molecular Laboratory | Philippine Genome Center | Francis A. Tablizo, Kenneth M. Kim, Carlo M. Lapid, Marc Jerrone R. Castro, Maria Sofia L. Yangzon, Elcid Aaron R. Pangilinan, Benedict A. Maralit, Marc Edsel C. Ayes, Eva Maria Cutiongco-de la Paz, Alethea R. de Guzman, Jan Michael C. Yap, Jo-Hannah S. Llames, Sheila Mae M. Araiza, Kris P. Punayan, Irish Coleen A. Asin, Candice Francheska B. Tambaoan, Asia Louisa U. Chong, Karol Sophia Agape R. Padilla, Rianna Patricia S. Cruz, El King D. Morado, Joshua Gregor A. Dizon, Razel Nikka M. Hao, Arianne A. Zamora, Devon Ray Pacial, Juan Antonio R. Magalang, Marissa Alejandria, Celia Carlos, Anna Ong-Lim, Edsel Maurice Salvaña, John Q. Wong, Jaime C. Montoya, Maria Rosario Singh-Vergeire and Cynthia P. Saloma |
| EPI_ISL_2157344, EPI_ISL_2157347, EPI_ISL_2157348, EPI_ISL_2157349, EPI_ISL_2157350, EPI_ISL_2157351, EPI_ISL_2157352, EPI_ISL_2157353, EPI_ISL_2157354, EPI_ISL_2157355, EPI_ISL_2157356, EPI_ISL_2157357, EPI_ISL_2157358, EPI_ISL_2157359, EPI_ISL_2157360, EPI_ISL_2157378, EPI_ISL_2157551, | | | |

| | | | |
|---|---|---|---|
| EPI_ISL_2157552, EPI_ISL_2157553, EPI_ISL_2157554, EPI_ISL_2157555, EPI_ISL_2157579, EPI_ISL_2157580, EPI_ISL_2157583, EPI_ISL_2157584, EPI_ISL_2157585 | | | |
| see above | Red - Regional de Vigilancia Genómica del COVID-19 | Laboratory of Respiratory Viruses and Measles, Oswaldo Cruz Institute, FIOCRUZ | Paola Resende, Mitzi Castro, Claudia Díaz, Sandra Paola Paz, Luciana Appolinario, Fernando Motta, Anna Carolina Paixao, Ana Carolina Mendonca, Alice Sampaio Rocha, Taina Venas, Eilisa Cavalcante Pereira, Renata Serrano Lopes, Marilda Siqueira on behalf of the Fiocruz COVID-19 Genomic Surveillance Network |
| EPI_ISL_2161030 | National Hematology and Transfusiology Center | National Hematology and Transfusiology Center | Aghayev AR |
| EPI_ISL_2178453 | Labo Analyses Med | National Reference Center for Viruses of Respiratory Infections, Institut Pasteur, Paris | Marion Barbet, Sylvie Behillil, Frédéric Lemoine, Corinne Maufrais, Christophe Malabat, Damien Mornico, Méline Bizard, Angela Brisebarre, Camille Capel, Louise Lefrançois, Etienne Simon-Lorière, Vincent Enouf, Maud Vanpeene, Sylvie van der Werf,Le Vicky |
| EPI_ISL_2178454 | Labo Analyses Med | National Reference Center for Viruses of Respiratory Infections, Institut Pasteur, Paris | Marion Barbet, Sylvie Behillil, Frédéric Lemoine, Corinne Maufrais, Christophe Malabat, Damien Mornico, Méline Bizard, Angela Brisebarre, Camille Capel, Louise Lefrançois, Etienne Simon-Lorière, Vincent Enouf, Maud Vanpeene, Sylvie van der Werf,Mallet C |
| EPI_ISL_2178456 | Labo Analyses Med | National Reference Center for Viruses of Respiratory Infections, Institut Pasteur, Paris | Marion Barbet, Sylvie Behillil, Frédéric Lemoine, Corinne Maufrais, Christophe Malabat, Damien Mornico, Méline Bizard, Angela Brisebarre, Camille Capel, Louise Lefrançois, Etienne Simon-Lorière, Vincent Enouf, Maud Vanpeene, Sylvie van der Werf,Le Vicky |
| EPI_ISL_2178457, EPI_ISL_2178458, EPI_ISL_2178459 | Labo Analyses Med | National Reference Center for Viruses of Respiratory Infections, Institut Pasteur, Paris | Marion Barbet, Sylvie Behillil, Frédéric Lemoine, Corinne Maufrais, Christophe Malabat, Damien Mornico, Méline Bizard, Angela Brisebarre, Camille Capel, Louise Lefrançois, Etienne Simon-Lorière, Vincent Enouf, Maud Vanpeene, Sylvie van der Werf,Mallet C |
| EPI_ISL_2178460 | Labo Analyses Med | National Reference Center for Viruses of Respiratory Infections, Institut Pasteur, Paris | Marion Barbet, Sylvie Behillil, Frédéric Lemoine, Corinne Maufrais, Christophe Malabat, Damien Mornico, Méline Bizard, Angela Brisebarre, Camille Capel, Louise Lefrançois, Etienne Simon-Lorière, Vincent Enouf, Maud Vanpeene, Sylvie van der Werf,Le Vicky |
| EPI_ISL_2178461, EPI_ISL_2178462, EPI_ISL_2178463 | Labo Analyses Med | National Reference Center for Viruses of Respiratory Infections, Institut Pasteur, Paris | Marion Barbet, Sylvie Behillil, Frédéric Lemoine, Corinne Maufrais, Christophe Malabat, Damien Mornico, Méline Bizard, Angela Brisebarre, Camille Capel, Louise Lefrançois, Etienne Simon-Lorière, Vincent Enouf, Maud Vanpeene, Sylvie van der Werf,Mallet C |
| EPI_ISL_2200078, EPI_ISL_2200081, EPI_ISL_2200083, EPI_ISL_2200086, EPI_ISL_2200091, EPI_ISL_2200092, EPI_ISL_2200093, EPI_ISL_2200097, EPI_ISL_2200102, EPI_ISL_2200104, EPI_ISL_2200105, EPI_ISL_2200106, EPI_ISL_2200108, EPI_ISL_2200111, EPI_ISL_2200114, EPI_ISL_2200116, EPI_ISL_2200117, EPI_ISL_2200121, EPI_ISL_2200122, EPI_ISL_2200126, EPI_ISL_2200127, EPI_ISL_2200129, EPI_ISL_2200130, EPI_ISL_2200132, EPI_ISL_2200136, EPI_ISL_2200137, EPI_ISL_2200139, EPI_ISL_2200140, EPI_ISL_2200148, EPI_ISL_2200151, EPI_ISL_2200154 | | | |
| see above | The Ohio State University Applied Microbiology Services Laboratory | The Ohio State University Applied Microbiology Services Laboratory | Seth A. Faith PhD |
| EPI_ISL_2214920, EPI_ISL_2214925, EPI_ISL_2214932, EPI_ISL_2214936, EPI_ISL_2214942, EPI_ISL_2214948, EPI_ISL_2214953, EPI_ISL_2214957, EPI_ISL_2214963 | Centre Hospitalier Universitaire Clermont-Ferrand | CHU Clermont-Ferrand, service de virologie | Bisseux Maxime, Mirand Audrey, Combes Patricia, Henquell Cécile |
| EPI_ISL_2226319 | MEDILAB | CHU Poitiers | Caroline MICHAUD, Maxime PICHON, Manon PRAT, Valentin BON-BARET, Birama N'DIAYE, Magali GARCIA, Agnes BEBY-DEFAUX, Nicolas LEVEQUE |
| EPI_ISL_2226326 | BIO86 CHAUMONT | CHU Poitiers | Caroline MICHAUD, Maxime PICHON, Manon PRAT, Valentin BON-BARET, Birama N'DIAYE, Magali GARCIA, Agnes BEBY-DEFAUX, Nicolas LEVEQUE |
| EPI_ISL_2226331 | BIO86 GENCAY | CHU Poitiers | Caroline MICHAUD, Maxime PICHON, Manon PRAT, Valentin BON-BARET, Birama N'DIAYE, Magali GARCIA, Agnes BEBY-DEFAUX, Nicolas LEVEQUE |
| EPI_ISL_2226333 | CHU POITIERS | CHU Poitiers | Caroline MICHAUD, Maxime PICHON, Manon PRAT, Valentin BON-BARET, Birama N'DIAYE, Magali GARCIA, Agnes BEBY-DEFAUX, Nicolas LEVEQUE |
| EPI_ISL_2226334 | MEDILAB | CHU Poitiers | Caroline MICHAUD, Maxime PICHON, Manon PRAT, Valentin BON-BARET, Birama N'DIAYE, Magali GARCIA, Agnes BEBY-DEFAUX, Nicolas LEVEQUE |
| EPI_ISL_2226338 | BIO86 CHAUMONT | CHU Poitiers | Caroline MICHAUD, Maxime PICHON, Manon PRAT, Valentin BON-BARET, Birama N'DIAYE, Magali GARCIA, Agnes BEBY-DEFAUX, Nicolas LEVEQUE |
| EPI_ISL_2231466, EPI_ISL_2231472, EPI_ISL_2231476, EPI_ISL_2231479, EPI_ISL_2231486, EPI_ISL_2231494, EPI_ISL_2231498, EPI_ISL_2231499, EPI_ISL_2231502, EPI_ISL_2231503, EPI_ISL_2231515, EPI_ISL_2231516, EPI_ISL_2231523, EPI_ISL_2231524 | | | |
| see above | Centro de Diagnostico COVID-19 UABC Tijuana | Andersen lab at Scripps Research | SEARCH Alliance San Diego with Idanya Rubi Serafin Higuera, Manuel Sanchez Alavez, Jorge Luis Jimenez Niebla, German Ibarra, Jonathan Vincent Baena, Oscar Efren Zazueta Fierro |
| EPI_ISL_2254131 | Department of Virology, School of Medicine, Shiraz University of Medical Sciences, Shiraz, Iran | Genetics Research Center, University of Social Welfare and Rehabilitation Sciences | Zohreh Fattahi, Marzieh Mohseni, Kimia Kahrizi, Moattari A, moghadami M, Hossein Najmabadi |
| EPI_ISL_2254500 | Veterinary institute in Zvolen Slovakia | Veterinary institute in Zvolen Zvolen Slovakia | Dirbáková Z., Sujová S., Mokryšová S., Tinák M., Mojžiš M |
| EPI_ISL_2254670 | Veterinary institute in Zvolen Slovakia | Veterinary institute in Zvolen Slovakia | Dirbáková Z., Sujová S., Mokryšová S., Tinák M., Mojžiš M |
| EPI_ISL_2254718, EPI_ISL_2254719 | Department of Virology, School of Medicine, Shiraz University of Medical Sciences, shiraz, Iran | Genetics Research Center, University of Social Welfare and Rehabilitation Sciences | Zohreh Fattahi, Marzieh Mohseni, Kimia Kahrizi, Moattari A, moghadami M, Hossein Najmabadi. |
| EPI_ISL_2271906, EPI_ISL_2271907, EPI_ISL_2271908, EPI_ISL_2271909 | Hospital General de México, Medicina Genómica | Hospital General de México, Medicina Genómica | Ana Laura Sánchez-Sandoval, María Eligia Juárez-Torres, Mariana Villalba-De la Peña, Siddhartha Ríos-Zaragoza, Guadalupe Mercedes Lucía Guerrero-Avendaño, Erika Anabel Alvarado-Silva, Eira Valeria Barrón-Palma |
| EPI_ISL_2279586, EPI_ISL_2279587, EPI_ISL_2279588, EPI_ISL_2279590, EPI_ISL_2279591, EPI_ISL_2279598 | Hospital | National Reference Center for Viruses of Respiratory Infections, Institut Pasteur, Paris | Marion Barbet, Sylvie Behillil, Méline Bizard, Angela Brisebarre, Camille Capel, Vincent Enouf, Louise Lefrançois, Frédéric Lemoine, Christophe Malabat, Corinne Maufrais, Etienne Simon-Lorière, Maud Vanpeene, Sylvie Van der Werf ,StéPhanie Marque Juillet |
| EPI_ISL_2280284 | Institut Pasteur de Guyane | Institut Pasteur | Marion Barbet, Sylvie Behillil, Méline Bizard, Angela Brisebarre, Camille Capel, Vincent Enouf, Louise Lefrançois, Frédéric Lemoine, Christophe Malabat, Corinne Maufrais, Emmanuelle Permal, Etienne Simon-Lorière, Maud Vanpeene, Sylvie Van der Werf ,Dominique Rousset |
| EPI_ISL_2284875 | Darus Syifa' Islamic Hospital | Institute of Tropical Disease, Universitas Airlangga | Krisnoadi Rahardjo, Aldise M Nastri, Jezzy R Dewantari, Rima R Prasetya, HM Faiz, Gatot Soegiarto, Laksmi Wulandari, Resti Yudhawati, Yasuko Mori, Soetjipto, Kazufumi Shimizu, Maria I Lusida |
| EPI_ISL_2284879 | Siti Khodijah Hospital | Institute of Tropical Disease, Universitas Airlangga | Kazufumi Shimizu, Krisnoadi Rahardjo, Aldise M Nastri, Jezzy R Dewantari, Rima R Prasetya, Muhammad Hamdan, Gatot Soegiarto, Laksmi Wulandari, Resti Yudhawati, Yasuko Mori, Soetjipto, Maria I Lusida |
| EPI_ISL_2285107, EPI_ISL_2285123 | Microbiology Department. Complexo Hospitalario Universitario de Vigo | Microbiology Department. Complexo Hospitalario Universitario de Vigo | Alfaya N, Alonso I, Alvarez M, Cabrera JJ, Carballo R, Cores O, Cortizo S, del-Campo V, Martinez L, Mediero G, Perez S, Potel C, Regueiro B, Rey S, Vasallo FJ |
| EPI_ISL_2290144, EPI_ISL_2290145, EPI_ISL_2290147, EPI_ISL_2290148, EPI_ISL_2290150, EPI_ISL_2290151, EPI_ISL_2290154, EPI_ISL_2290155, EPI_ISL_2290156, EPI_ISL_2290157, EPI_ISL_2290158, EPI_ISL_2290159, EPI_ISL_2290160, EPI_ISL_2290161 | | | |
| see above | Turkish Society of Internal Medicine | Gen Era Diagnostics Corporation Life Science Research and Molecular Diagnostics | Serhat Ünal |
| EPI_ISL_2307329, EPI_ISL_2307330, EPI_ISL_2307342, EPI_ISL_2307343 | Genome Analysis Center, Yamanashi Central Hospital | Genome Analysis Center, Yamanashi Central Hospital | Yosuke Hirotsu |
| EPI_ISL_2312656 | Centre Hospitalier Universitaire Clermont-Ferrand | CHU Clermont-Ferrand, service de virologie | Bisseux Maxime, Mirand Audrey, Combes Patricia, Henquell Cécile |
| EPI_ISL_2312729, EPI_ISL_2312730, EPI_ISL_2312731, EPI_ISL_2312732, EPI_ISL_2312733, EPI_ISL_2312734, EPI_ISL_2312735, EPI_ISL_2312736, EPI_ISL_2312737, EPI_ISL_2312738, EPI_ISL_2312739, EPI_ISL_2312740, EPI_ISL_2312741, EPI_ISL_2312742, EPI_ISL_2312743, EPI_ISL_2312744, EPI_ISL_2312745, EPI_ISL_2312746, EPI_ISL_2312747, EPI_ISL_2312748, EPI_ISL_2312749, EPI_ISL_2312750, EPI_ISL_2312751, EPI_ISL_2312752, EPI_ISL_2312753, EPI_ISL_2312754, EPI_ISL_2312755, EPI_ISL_2312756, EPI_ISL_2312757, EPI_ISL_2312758, EPI_ISL_2312759, EPI_ISL_2312760, EPI_ISL_2312761, EPI_ISL_2312762, EPI_ISL_2312763, EPI_ISL_2312764, EPI_ISL_2312765, EPI_ISL_2312766, EPI_ISL_2312767, EPI_ISL_2312768, EPI_ISL_2312769, EPI_ISL_2312771, EPI_ISL_2312772, EPI_ISL_2312773, EPI_ISL_2312774, EPI_ISL_2312775, EPI_ISL_2312776, EPI_ISL_2312777, EPI_ISL_2312779, EPI_ISL_2312781, EPI_ISL_2312782, EPI_ISL_2312783, EPI_ISL_2312784, EPI_ISL_2312785, EPI_ISL_2312786, EPI_ISL_2312791, EPI_ISL_2312792, EPI_ISL_2312793, EPI_ISL_2312794, EPI_ISL_2312796, EPI_ISL_2312797, EPI_ISL_2312798, EPI_ISL_2312800, EPI_ISL_2312801, EPI_ISL_2312802, EPI_ISL_2312803, EPI_ISL_2312806, EPI_ISL_2312807, EPI_ISL_2312809, EPI_ISL_2312810, EPI_ISL_2312811, EPI_ISL_2312812, EPI_ISL_2312813, EPI_ISL_2312814, EPI_ISL_2312815, EPI_ISL_2312816, EPI_ISL_2312817, EPI_ISL_2312818, EPI_ISL_2312820, EPI_ISL_2312821, EPI_ISL_2312822, EPI_ISL_2312823, EPI_ISL_2312824, EPI_ISL_2312826, EPI_ISL_2312827, EPI_ISL_2312828, EPI_ISL_2312829, EPI_ISL_2312830, EPI_ISL_2312831, EPI_ISL_2312832, EPI_ISL_2312833, EPI_ISL_2312834, EPI_ISL_2312836, EPI_ISL_2312837, EPI_ISL_2312840, EPI_ISL_2312841, EPI_ISL_2312842, EPI_ISL_2312843, EPI_ISL_2312844, EPI_ISL_2312846, EPI_ISL_2312847, EPI_ISL_2312848, EPI_ISL_2312849, EPI_ISL_2312850, EPI_ISL_2312851, EPI_ISL_2312852, EPI_ISL_2312855, EPI_ISL_2312856, EPI_ISL_2312858, EPI_ISL_2312859, EPI_ISL_2312860, EPI_ISL_2312861, EPI_ISL_2312862, EPI_ISL_2312864, EPI_ISL_2312865, EPI_ISL_2312866, EPI_ISL_2312867, EPI_ISL_2312869, EPI_ISL_2312870, EPI_ISL_2312871, EPI_ISL_2312872, EPI_ISL_2312873, EPI_ISL_2312874, EPI_ISL_2312875, EPI_ISL_2312876, EPI_ISL_2312877, EPI_ISL_2312878, EPI_ISL_2312879, EPI_ISL_2312881, EPI_ISL_2312882, EPI_ISL_2312884, EPI_ISL_2312885, EPI_ISL_2312886, EPI_ISL_2312887, EPI_ISL_2312888, EPI_ISL_2312891, EPI_ISL_2312892, EPI_ISL_2312894, EPI_ISL_2312896, EPI_ISL_2312897, EPI_ISL_2312898, EPI_ISL_2312899, EPI_ISL_2312903, EPI_ISL_2312904, EPI_ISL_2312905, EPI_ISL_2312906, EPI_ISL_2312908, EPI_ISL_2312909, EPI_ISL_2312911, EPI_ISL_2312912, EPI_ISL_2312913, EPI_ISL_2312914, EPI_ISL_2312915, | | | |

| | | | |
|---|---|---|---|
| EPI_ISL_2312916, EPI_ISL_2312917, EPI_ISL_2312918, EPI_ISL_2312919, EPI_ISL_2312920, EPI_ISL_2312922, EPI_ISL_2312923, EPI_ISL_2312924, EPI_ISL_2312925, EPI_ISL_2312926, EPI_ISL_2312927, EPI_ISL_2312928, EPI_ISL_2312929, EPI_ISL_2312930, EPI_ISL_2312931, EPI_ISL_2312933, EPI_ISL_2312934, EPI_ISL_2312935, EPI_ISL_2312936, EPI_ISL_2312937, EPI_ISL_2312938, EPI_ISL_2312939, EPI_ISL_2312940, EPI_ISL_2312941, EPI_ISL_2312942, EPI_ISL_2312943, EPI_ISL_2312944, EPI_ISL_2312945, EPI_ISL_2312946, EPI_ISL_2312947, EPI_ISL_2312949, EPI_ISL_2312950, EPI_ISL_2312951, EPI_ISL_2312952, EPI_ISL_2312954, EPI_ISL_2312955, EPI_ISL_2312956, EPI_ISL_2312957, EPI_ISL_2312958, EPI_ISL_2312959, EPI_ISL_2312960 | | | |
| see above | The Ohio State University Applied Microbiology Services Laboratory | The Ohio State University Applied Microbiology Services Laboratory | Seth A. Faith PhD |
| EPI_ISL_2321227, EPI_ISL_2321228, EPI_ISL_2322468 | Genome Analysis Center, Yamanashi Central Hospital | Genome Analysis Center, Yamanashi Central Hospital | Yosuke Hirotsu |
| EPI_ISL_2322745, EPI_ISL_2322746 | AZ Sint-Lucas Gent | AZ Sint-Lucas Gent | Jos Van Acker, Charlotte Verfaillie, Elke Vanlaere, Klara De Rauw |
| EPI_ISL_2322762 | Collection of WHO National Influenza Center, N.F. Gamaleya National Research Center for Epidemiology and Microbiology of the Ministry of Health of the Russian Federation | Laboratory of genome analysis, N.F. Gamaleya National Research Center for Epidemiology and Microbiology of the Ministry of Health of the Russian Federation | Olga Voronina, Ekaterina Aksenova, Natalia Ryzhova, Marina Kunda, Kirill Krasnoslobodtsev, Elena Burtseva, Denis Logunov |
| EPI_ISL_2348648, EPI_ISL_2348659 | Gyncentrum | Gyncentrum | Emilia Morawiec, Pawel Czerwinski, Adam Pudelko, Aleksandra Skubis-Sikora, Magdalena Samul, Agnieszka Polak, Celina Kruszniewska-Rajs, Tomasz Wasik, Robert Wojtyczka, Jolanta Bartosiewicz- Wasik, Maria Maklasinska-Majdanik |
| EPI_ISL_2361908, EPI_ISL_2361910, EPI_ISL_2361911, EPI_ISL_2361912 | Noguchi Memorial Institute for Medical Research, University of Ghana, Legon, Ghana | Institute of Tropical Medicine, Universitätsklinikum Tübingen, Germany | Bright Adu, Quaneeta Mohktar, Le Thi Kieu Linh, Sivaramakrishna Rachakonda, Hilda Opoku Frempong, Keren Okyerebea Attiku, Joyce Appiah-Kubi, Joseph Humphrey Kofi Bonney, John Kofi Odoom, Abraham Kwabena Anang, Srinivas-reddy Pallerla, Dorothy Yeboah-Manu, Thirumalaisamy P Velavan |
| EPI_ISL_2364488, EPI_ISL_2364504, EPI_ISL_2364510 | Genome Analysis Center, Yamanashi Central Hospital | Genome Analysis Center, Yamanashi Central Hospital | Yosuke Hirotsu |
| EPI_ISL_2365885 | Microbiology Department. Complexo Hospitalario Universitario de Vigo | Microbiology Department. Complexo Hospitalario Universitario de Vigo | Alfaya N, Alonso I, Alvarez M, Cabrera JJ, Carballo R, Cores O, Cortizo S, del-Campo V, Martinez L, Mediero G, Perez S, Potel C, Regueiro B, Rey S, Vasallo FJ |
| EPI_ISL_2375850, EPI_ISL_2375851, EPI_ISL_2375852, EPI_ISL_2375853, EPI_ISL_2375854, EPI_ISL_2375855, EPI_ISL_2375856, EPI_ISL_2375857, EPI_ISL_2375858, EPI_ISL_2375859, EPI_ISL_2375860, EPI_ISL_2375861, EPI_ISL_2375862, EPI_ISL_2375863, EPI_ISL_2375864, EPI_ISL_2375865, EPI_ISL_2375866, EPI_ISL_2375867, EPI_ISL_2375868, EPI_ISL_2375869, EPI_ISL_2375870, EPI_ISL_2375871, EPI_ISL_2375872, EPI_ISL_2375874, EPI_ISL_2375875, EPI_ISL_2375876 | | | |
| see above | Programa de Oncovirologia, Instituto Nacional de Câncer | Programa de Oncovirologia, Instituto Nacional de Câncer | Livia R. Goes, Juliana D. Siqueira, Marianne M. Garrido, Brunna M. Alves, Ana Cristina P. M. Pereira, Claudia Cicala, James Arthos, João P.B. Viola, Marcelo A. Soares |
| EPI_ISL_2376383 | Noguchi Memorial Institute for Medical Research, University of Ghana, Legon, Ghana | Institute of Tropical Medicine, Universitätsklinikum Tübingen, Germany | Bright Adu, Quaneeta Mohktar, Le Thi Kieu Linh, Sivaramakrishna Rachakonda, Hilda Opoku Frempong, Keren Okyerebea Attiku, Joyce Appiah-Kubi, Joseph Humphrey Kofi Bonney, John Kofi Odoom, Abraham Kwabena Anang, Srinivas-reddy Pallerla, Dorothy Yeboah-Manu, Thirumalaisamy P Velavan |
| EPI_ISL_2381993 | AZ Zeno Knokke-Blankenberge | AZ Sint Jan | Marijke Reynders, Patrick Descheemaeker, Laurien Hoornaert, Thomas Van Landschoot, Marie Madeleine Chabert-Consen, Jorn Hellemans |
| EPI_ISL_2382010, EPI_ISL_2382013, EPI_ISL_2382014, EPI_ISL_2382015, EPI_ISL_2382017, EPI_ISL_2382018, EPI_ISL_2382019 | HG Pharma GmbH | Bergthaler laboratory, CeMM Research Center for Molecular Medicine of the Austrian Academy of Sciences | Lukas Endler, Anna Schedl, Fabian Amman, Petr Triska, Thomas Penz, Benedikt Agerer, Maelle Le Moing, Michael Schuster, Bekir Erguner, Jan Laine, Martin Senekowitsch, Christoph Bock, Andreas Bergthaler |
| EPI_ISL_2382452, EPI_ISL_2382454, EPI_ISL_2382455, EPI_ISL_2382456, EPI_ISL_2382457, EPI_ISL_2382458, EPI_ISL_2382459, EPI_ISL_2382460, EPI_ISL_2382461, EPI_ISL_2382462, EPI_ISL_2382464, EPI_ISL_2382465 | | | |
| see above | The Ohio State University Applied Microbiology Services Laboratory | The Ohio State University Applied Microbiology Services Laboratory | Seth A. Faith PhD |
| EPI_ISL_2383206 | The teaching Baghdad hospital | PCR lab | Abdulhussein Thair, Fadhil Hula |
| EPI_ISL_2399743, EPI_ISL_2399745, EPI_ISL_2399746, EPI_ISL_2399747, EPI_ISL_2399749, EPI_ISL_2399751, EPI_ISL_2399752, EPI_ISL_2399753, EPI_ISL_2399754, EPI_ISL_2399756 | The Ohio State University Applied Microbiology Services Laboratory | The Ohio State University Applied Microbiology Services Laboratory | Seth A. Faith PhD |
| EPI_ISL_2422725, EPI_ISL_2422732, EPI_ISL_2422735, EPI_ISL_2422741, EPI_ISL_2422746, EPI_ISL_2422756, EPI_ISL_2422761, EPI_ISL_2422762, EPI_ISL_2422773, EPI_ISL_2422777 | San Diego County Public Health Laboratory | Andersen lab at Scripps Research | SEARCH Alliance San Diego with Ashleigh Murphy, Jovan Shephard, Brett Austin |
| EPI_ISL_2426941 | Centers for Disease Control and Prevention, Dengue Branch | Centers for Disease Control and Prevention, Dengue Branch | Gilberto A. Santiago, Glenda Gonzalez, Betzabel Flores, Keyla Charriez, Gabriela Paz-Bailey, Jorge L. Munoz-Jordan |
| EPI_ISL_2431843 | Laboratório de Microbiologia Molecular - Universidade FEEVALE | Molecular Microbiology Laboratory | Alana Witt Hansen, Fágner Henrique Heldt, Fernando Rosado Spilki, Flávio Silveira, Juliana Schons Gularte, Juliane Deise Fleck, Mariana Soares da Silva, Meriane Demoliner, Matheus Nunes Weber, Paula Rodrigues de Almeida, Micheli Filippi. |
| EPI_ISL_2443781 | The Ohio State University Applied Microbiology Services Laboratory | The Ohio State University Applied Microbiology Services Laboratory | Seth A. Faith PhD |
| EPI_ISL_2458067, EPI_ISL_2458068, EPI_ISL_2458069, EPI_ISL_2458070, EPI_ISL_2458071, EPI_ISL_2458072, EPI_ISL_2458073, EPI_ISL_2458074, EPI_ISL_2458075, EPI_ISL_2458076, EPI_ISL_2458077, EPI_ISL_2458078, EPI_ISL_2458079, EPI_ISL_2458080, EPI_ISL_2458081, EPI_ISL_2458082, EPI_ISL_2458083, EPI_ISL_2458084, EPI_ISL_2458085, EPI_ISL_2458086, EPI_ISL_2458087, EPI_ISL_2458088, EPI_ISL_2458089, EPI_ISL_2458090, EPI_ISL_2458091, EPI_ISL_2458092, EPI_ISL_2458093, EPI_ISL_2458094, EPI_ISL_2458095, EPI_ISL_2458096, EPI_ISL_2458097, EPI_ISL_2458098, EPI_ISL_2458099, EPI_ISL_2458100, EPI_ISL_2458101, EPI_ISL_2458102, EPI_ISL_2458103, EPI_ISL_2458104, EPI_ISL_2458105 | | | |
| see above | Adolfo Lutz Institute Santos Regional Center | Retrovirus Laboratory Adolfo Lutz Institute | Gabriela Bastos Cabral, Giselle I S Lopez-Lopes, Cintia Ahagon, Audrey Cilli, Paula Morena Guimaraes, Igor Mohamed Hussein, Luis Brigido |
| EPI_ISL_2458106 | Adolfo Lutz Institute Santos Regional Center | Dr Arnaldo 355 Sao Paulo City, State of Sao Paulo CEP 01246-902, Brazil | Gabriela Bastos Cabral, Giselle I S Lopez-Lopes, Cintia Ahagon, Audrey Cilli, Paula Morena Guimaraes, Igor Mohamed Hussein, Luis Brigido |
| EPI_ISL_2467837, EPI_ISL_2467838, EPI_ISL_2467839, EPI_ISL_2467840, EPI_ISL_2467841, EPI_ISL_2467842, EPI_ISL_2467843, EPI_ISL_2467845, EPI_ISL_2467846, EPI_ISL_2467847, EPI_ISL_2467848, EPI_ISL_2467849, EPI_ISL_2467850, EPI_ISL_2467851, EPI_ISL_2467852, EPI_ISL_2467853, EPI_ISL_2467854, EPI_ISL_2467855, EPI_ISL_2467856, EPI_ISL_2467857, EPI_ISL_2467858, EPI_ISL_2467859, EPI_ISL_2467860, EPI_ISL_2467861, EPI_ISL_2467862, EPI_ISL_2467863, EPI_ISL_2467864 | | | |
| see above | Adolfo Lutz Institute Sao Paulo Virology Center | Retrovirus Laboratory Adolfo Lutz Institute | Gabriela Bastos Cabral, Giselle I S Lopez-Lopes, Cintia Ahagon, Audrey Cilli, Paula Morena Guimaraes, Igor Mohamed Hussein, Luis Brigido |
| EPI_ISL_2467909, EPI_ISL_2467911, EPI_ISL_2467912, EPI_ISL_2467913, EPI_ISL_2467919, EPI_ISL_2467924, EPI_ISL_2467929 | Biology Department, College of Science, Al Muthanna University and Public Health Laboratory, Al-Muthanna Health Directorate | Department of Virology, Faculty of Medicine, University of Helsinki, Helsinki, Finland | Nihad Al-Rashedi, Hussein Alburkat, Murad Munahi, Alaa Hameed, Ali Jasim, Olli Vapalahti, Tarja Sironen,Teemu Smura |
| EPI_ISL_2474178, EPI_ISL_2474179, EPI_ISL_2474183, EPI_ISL_2474184, EPI_ISL_2474185 | Adolfo Lutz Institute Santos Regional Center | Retrovirus Laboratory Adolfo Lutz Institute | Gabriela Bastos Cabral, Giselle I S Lopez-Lopes, Cintia Ahagon, Audrey Cilli, Paula Morena Guimaraes, Igor Mohamed Hussein, Luis Brigido |
| EPI_ISL_2484759 | Queen Elizabeth Hospital | Hong Kong Department of Health | Alan K.L. Tsang, Gannon C.K. Mak, Peter K.C. Cheng, Peter C.W. Yip, Ken H.L. Ng, Edman T.K. Lam, Rickjason C.W. Chan, Dominic N.C. Tsang |
| EPI_ISL_2485170, EPI_ISL_2485175, EPI_ISL_2485177, EPI_ISL_2485178, EPI_ISL_2485179, EPI_ISL_2485185, EPI_ISL_2485193, EPI_ISL_2485199, EPI_ISL_2485200, EPI_ISL_2485205, EPI_ISL_2485211, EPI_ISL_2485212, EPI_ISL_2485213, EPI_ISL_2485218, EPI_ISL_2485219 | | | |
| see above | ILV Kärnten | Bergthaler laboratory, CeMM Research Center for Molecular Medicine of the Austrian Academy of Sciences | Lukas Endler, Anna Schedl, Fabian Amman, Petr Triska, Thomas Penz, Benedikt Agerer, Maelle Le Moing, Michael Schuster, Bekir Erguner, Jan Laine, Martin Senekowitsch, Christoph Bock, Andreas Bergthaler |
| EPI_ISL_2491667 | HOPITAL DE MONTLUCON | CHU Clermont-Ferrand, service de virologie | Bisseux Maxime, Mirand Audrey, Combes Patricia, Henquell Cécile |
| EPI_ISL_2516944, EPI_ISL_2516965 | Microbiology Department. Complexo Hospitalario Universitario | Microbiology Department. Complexo Hospitalario Universitario | Alfaya N, Alonso I, Alvarez M, Cabrera JJ, Carballo R, Cores O, Cortizo S, del-Campo V, Martinez L, Mediero G, Perez S, Potel C, Regueiro B, Rey S, |

| | | | |
|---|---|---|---|
| | de Vigo | de Vigo | Vasallo FJ |
| EPI_ISL_2534057 | S.C. Laboratorio Analisi, Ospedale di Lavagna, ASL 4 Liguria | U.O. Igiene, Ospedale Policlinico San Martino | Bruzzone Bianca, Bandettini Roberto, Caligiuri Patrizia, De Pace Vanessa, Domnich Alexander, Orsi Andrea, Ricucci Valentina, Icardi Giancarlo |
| EPI_ISL_2534058 | U.O. Igiene, Ospedale Policlinico San Martino | U.O. Igiene, Ospedale Policlinico San Martino | Bruzzone Bianca, Caligiuri Patrizia, De Pace Vanessa, Domnich Alexander, Orsi Andrea, Ricucci Valentina, Icardi Giancarlo |
| EPI_ISL_2535802, EPI_ISL_2535803, EPI_ISL_2535804, EPI_ISL_2535805, EPI_ISL_2535806, EPI_ISL_2535807, EPI_ISL_2535808, EPI_ISL_2535809, EPI_ISL_2535810, EPI_ISL_2535811, EPI_ISL_2535812, EPI_ISL_2535813, EPI_ISL_2535815, EPI_ISL_2535817, EPI_ISL_2535818, EPI_ISL_2535819, EPI_ISL_2535823, EPI_ISL_2535824 | | | |
| see above | National Influenza Centre | National Influenza Centre | William K. Ampofo, Michael Marks, Ivy A. Asante, Sharon Hsu, Benjamiin B. Lindsey, Benjamin H. Foulkes, Mildred Adusei-Poku, Linda Boatemaa, Lorreta Kwasah, Joseph Oliver-Commey, Ernest Asiedu, Franklin Asiedu-Bekoe, Gordon Awandare, Joyce Ngoi, Dennis Laryea, Mathew D. Parker, Thushan I de Silva, |
| EPI_ISL_2536303, EPI_ISL_2536305, EPI_ISL_2536352, EPI_ISL_2536353 | Laboratorio Central de Saude Publica do Estado da Paraiba (LACEN-PB) | Laboratory of Respiratory Viruses and Measles, Oswaldo Cruz Institute, FIOCRUZ | Paola Resende, Luciana Appolinario, Fernando Motta, Anna Carolina Paixao, Ana Carolina Mendonca, Alice Sampaio Rocha, Taina Venas, Elisa Cavalcante Pereira, Renata Serrano Lopes, Joao Felipe Bezerra, Dalane Loudal Florentino Teixeira, Marilda Siqueira on behalf of the Fiocruz COVID-19 Genomic Surveillance Network |
| EPI_ISL_2544508 | Provincial Public Health Reference Laboratory (PPHRL) P& SHD | Provincial Public Health Reference Laboratory (PPHRL) P& SHD | ANDLEEB HANIF |
| EPI_ISL_2544837, EPI_ISL_2544838, EPI_ISL_2544839, EPI_ISL_2544840, EPI_ISL_2544841, EPI_ISL_2544844, EPI_ISL_2544847, EPI_ISL_2544848, EPI_ISL_2544849, EPI_ISL_2544850, EPI_ISL_2544851, EPI_ISL_2544852, EPI_ISL_2544857, EPI_ISL_2544858, EPI_ISL_2544859, EPI_ISL_2544860, EPI_ISL_2544861, EPI_ISL_2544862, EPI_ISL_2544864, EPI_ISL_2544865, EPI_ISL_2544866, EPI_ISL_2544867, EPI_ISL_2544868, EPI_ISL_2544869, EPI_ISL_2544870, EPI_ISL_2544872, EPI_ISL_2544873, EPI_ISL_2544874, EPI_ISL_2544875, EPI_ISL_2544876 | | | |
| see above | Laboratorio de Pesquisa em Virologia, FAMERP, SJRP | Laboratorio de Pesquisa em Virologia, FAMERP, SJRP | Cecília Artico Banho; Lívia Sacchetto; Guilherme Campos; Fábio Sossai Possebon; Leila Sabrina Ullmann; Cíntia Bittar; Helena Lage Ferreira; Jorge A. Petroli Marchesi; Maisa C. Pereira Parra; Marília Moraes; Paula Rahal; Paulo Inacio da Costa; João Pessoa Araújo Jr.; Maurício L. Nogueira. |
| EPI_ISL_2548847 | Department of Medical Virology, Faculty of Medicine, Hamadan University of Medical Sciences, Hamadan, Iran | Genetics Research Center, University of Social Welfare and Rehabilitation Sciences | Zohreh Fattahi, Marzieh Mohseni, Maryam Beheshtian, Kimia Kahrizi, Farid Azizi Jalilian, Nastaran Ansari, Mehdi Norouzi, Azam Khamseh, Milad Zandi, Vadat Poortahmasebi, Hossein Najmabadi. |
| EPI_ISL_2566167, EPI_ISL_2566168, EPI_ISL_2566182, EPI_ISL_2566184, EPI_ISL_2566192, EPI_ISL_2566194, EPI_ISL_2566195, EPI_ISL_2566196, EPI_ISL_2566197, EPI_ISL_2566198 | Microbiology Department. Complexo Hospitalario Universitario de Vigo | Microbiology Department. Complexo Hospitalario Universitario de Vigo | Alfaya N, Alonso I, Alvarez M, Cabrera JJ, Carballo R, Cores O, Cortizo S, del-Campo V, Martinez L, Mediero G, Perez S, Potel C, Regueiro B, Rey S, Vasallo FJ |
| EPI_ISL_2566445 | Singapore General Hospital | Department of Microbiology | Nurdyana Abdul Rahman, Kenneth Xin Long Chan, Chayaporn Suphavilai, Kun Lee Lim, Sui Sin Goh, Kian Sing Chan, Lynette Oon, Kern Rei Chng, James Sim Heng Chiak, Niranjan Nagarajan, Karrie Ko |
| EPI_ISL_2603243, EPI_ISL_2603244, EPI_ISL_2603254 | Microbiology Department. Complexo Hospitalario Universitario de Vigo | Microbiology Department. Complexo Hospitalario Universitario de Vigo | Alfaya N, Alonso I, Alvarez M, Cabrera JJ, Carballo R, Cores O, Cortizo S, del-Campo V, Martinez L, Mediero G, Perez S, Potel C, Regueiro B, Rey S, Vasallo FJ |
| EPI_ISL_2614353 | Labortorio Central de Saude Publica do Estado do Rio de Janeiro (LACEN/RJ) | Laboratory of Respiratory Viruses and Measles, Oswaldo Cruz Institute, FIOCRUZ | Paola Resende, Luciana Appolinario, Fernando Motta, Anna Carolina Paixao, Ana Carolina Mendonca, Alice Sampaio Rocha, Taina Venas, Elisa Cavalcante Pereira, Renata Serrano Lopes, Andrea Cony Cavalcanti, Marilda Siqueira on behalf of the Fiocruz COVID-19 Genomic Surveillance Network |
| EPI_ISL_2614385, EPI_ISL_2614386 | Adolfo Lutz Institute Sao Paulo Virology Center | Retrovirus Laboratory Adolfo Lutz Institute | Gabriela Bastos Cabral, Giselle I S Lopez-Lopes, Cintia Ahagon, Audrey Cilli, Paula Morena Guimaraes, Igor Mohamed Hussein, Luis Brigido |
| EPI_ISL_2614519 | Instituto Adolfo Lutz - Regional de Campinas | Instituto Adolfo Lutz, Interdiciplinary Procedures Center, Strategic Laboratory | Claudio Tavares Sacchi, Claudia Regina Gonçalves, Erica Valessa Ramos Gomes, Karoline Rodrigues Campos, Caio Vinicius Dias Lopes, Leonardo Jose Tadeu de Araujo |
| EPI_ISL_2614542 | LACEN do Estado de Goias | Instituto Adolfo Lutz, Interdiciplinary Procedures Center, Strategic Laboratory | Claudio Tavares Sacchi, Claudia Regina Gonçalves, Erica Valessa Ramos Gomes, Karoline Rodrigues Campos, Caio Vinicius Dias Lopes, Leonardo Jose Tadeu de Araujo |
| EPI_ISL_2617088, EPI_ISL_2617089, EPI_ISL_2617091 | Africa_CDC - Malawi P6 S2 | CERI, Centre for Epidemic Response and Innvoation, Stellenbosch University and KRISP, KZN Research Innovation and Sequencing Platform, UKZN. | Mvula B, Chilima B, Chiwaula M, Mwangomba W, Panja L, Kasambara W, Auld A, Kim L, Kampira E, Kaba M, Wadonda N, Maida A, Giandhari J, Pillay S, Naidoo Y, Lessells R, Emmanuel SJ, Tegally H, Wilkinson E, de Oliveira T |
| EPI_ISL_2645661, EPI_ISL_2645662, EPI_ISL_2645663 | Laboratorio Central de Saude Publica do Estado de Alagoas (LACEN/AL) | Laboratory of Respiratory Viruses and Measles, Oswaldo Cruz Institute, FIOCRUZ | Paola Resende, Luciana Appolinario, Fernando Motta, Anna Carolina Paixao, Ana Carolina Mendonca, Alice Sampaio Rocha, Taina Venas, Elisa Cavalcante Pereira, Renata Serrano Lopes, Anderson Brandao Leite, Marilda Siqueira on behalf of the Fiocruz COVID-19 Genomic Surveillance Network |
| EPI_ISL_2652118 | AZ Klina | AZ Klina | Carl Vael - Lynsey Berckmans |
| EPI_ISL_2657676 | CLINICA BIBLICA | Inciensa, Instituto Costarricense de Investigación y Enseñanza en Nutrición y Salud | Francisco Duarte, Hebleen Porras, Claudio Soto-Garita, Estela Cordero, Adriana Godínez, Melany Calderón & Karla Gutiérrez-González |
| EPI_ISL_2661829, EPI_ISL_2661831, EPI_ISL_2661835, EPI_ISL_2661841, EPI_ISL_2661842, EPI_ISL_2661844, EPI_ISL_2661845, EPI_ISL_2661847, EPI_ISL_2661852, EPI_ISL_2661853, EPI_ISL_2661855, EPI_ISL_2661856, EPI_ISL_2661861, EPI_ISL_2661862, EPI_ISL_2661863 | | | |
| see above | Laboratorio Central de Saude Publica do Estado do Rio Grande do Sul (LACEN-RS) | Laboratory of Respiratory Viruses and Measles, Oswaldo Cruz Institute, FIOCRUZ | Paola Resende, Luciana Appolinario, Fernando Motta, Anna Carolina Paixao, Ana Carolina Mendonca, Alice Sampaio Rocha, Taina Venas, Elisa Cavalcante Pereira, Renata Serrano Lopes, Tatiana Schaffer Gregianini, Richard Salvato, Marilda Siqueira on behalf of the Fiocruz COVID-19 Genomic Surveillance Network |
| EPI_ISL_2663148, EPI_ISL_2663149, EPI_ISL_2663151, EPI_ISL_2663152, EPI_ISL_2663154, EPI_ISL_2663155, EPI_ISL_2663157, EPI_ISL_2663158, EPI_ISL_2663169, EPI_ISL_2663177 | Ochsner Health | Bioinfoexperts, LLC | Judy Crabtree, Amy Feehan, Rebecca Rose, David J. Nolan, Tessa LaFleur, Julia-Garcia-Diaz, Lucio Miele, Susanna L. Lamers |
| EPI_ISL_2663376 | LESP Queretaro | Instituto de Diagnostico y Referencia Epidemiologicos (INDRE) | Claudia Wong-Arambula, Abril Rodriguez-Maldonado, Vanessa Rivero-Arredondo, Ariadna Medina-Benitez, Joaquin Quiroz-Mercado, Sergio Rangel-Guerrero, Natividad Cruz-Ortiz, Tatiana Nunez-Garcia, Gisela Barrera-Badillo, Lucia Hernandez-Rivas, Irma Lopez-Martinez, Ernesto Ramirez-Gonzalez. |
| EPI_ISL_2677080, EPI_ISL_2677081, EPI_ISL_2677160 | Labortorio Central de Saude Publica do Estado de Santa Catarina (LACEN/SC) | Laboratory of Respiratory Viruses and Measles, Oswaldo Cruz Institute, FIOCRUZ | Paola Resende, Luciana Appolinario, Fernando Motta, Anna Carolina Paixao, Ana Carolina Mendonca, Alice Sampaio Rocha, Taina Venas, Elisa Cavalcante Pereira, Renata Serrano Lopes, Darcita Buerger Rovaris, Sandra Bianchini Fernandes, Marilda Siqueira on behalf of the Fiocruz COVID-19 Genomic Surveillance Network |
| EPI_ISL_2688670 | Centre Hospitalier Universitaire de Rouen Laboratoire de Virologie | Centre Hospitalier Universitaire de Rouen Laboratoire de Virologie | Alice Moisan, Fabienne De Oliveira, Marie Leoz |
| EPI_ISL_2689144, EPI_ISL_2689146 | MSK Microbiology Lab | MSK Microbiology Lab | Esther Babady, Tracy McMillen, Krupa Jani |
| EPI_ISL_2690138, EPI_ISL_2690139, EPI_ISL_2690140, EPI_ISL_2690141, EPI_ISL_2690142, EPI_ISL_2690143, EPI_ISL_2690144, EPI_ISL_2690145, EPI_ISL_2690168, EPI_ISL_2690169, EPI_ISL_2690170, EPI_ISL_2690171, EPI_ISL_2690178 | | | |
| see above | Ochsner Health | Bioinfoexperts, LLC | Judy Crabtree, Amy Feehan, Rebecca Rose, David J. Nolan, Tessa LaFleur, Julia-Garcia-Diaz, Lucio Miele, Susanna L. Lamers |
| EPI_ISL_2690310, EPI_ISL_2690404, EPI_ISL_2690405, EPI_ISL_2690406, EPI_ISL_2690407, EPI_ISL_2690426, EPI_ISL_2690428, EPI_ISL_2690436, EPI_ISL_2690437, EPI_ISL_2690438, EPI_ISL_2690440, EPI_ISL_2690441, EPI_ISL_2690658, EPI_ISL_2691009, EPI_ISL_2691013, EPI_ISL_2691016, EPI_ISL_2691017, EPI_ISL_2691019, EPI_ISL_2691020, EPI_ISL_2691022, EPI_ISL_2691023, EPI_ISL_2691026, EPI_ISL_2691027, EPI_ISL_2691028, EPI_ISL_2691029, EPI_ISL_2691030, EPI_ISL_2691035, EPI_ISL_2691041, EPI_ISL_2691042, EPI_ISL_2691046, EPI_ISL_2691048, EPI_ISL_2691049, EPI_ISL_2691051, EPI_ISL_2691052, EPI_ISL_2691053 | | | |
| see above | MSK Microbiology Lab | MSK Microbiology Lab | Esther Babady, Tracy McMillen, Krupa Jani |
| EPI_ISL_2691204, EPI_ISL_2691205 | LACEN do Estado de Goias | Instituto Adolfo Lutz, Interdiciplinary Procedures Center, Strategic Laboratory | Claudio Tavares Sacchi, Claudia Regina Gonçalves, Erica Valessa Ramos Gomes, Karoline Rodrigues Campos, Caio Vinicius Dias Lopes, Leonardo Jose Tadeu de Araujo |
| EPI_ISL_2692143, EPI_ISL_2692144, EPI_ISL_2692169, EPI_ISL_2692170, | Clinical Microbiology Lab RSPTN Universitas Hasanuddin | Clinical Microbiology Lab RSPTN Universitas Hasanuddin | Muhammad Nasrum Massi, Handayani Halik, Rizalinda Sjahril, Gita Vita Soraya, Triwibawa, Muhammad Yusuf Usman, Rufika Shari |

| | | | |
|---|---|---|---|
| EPI_ISL_2692171, EPI_ISL_2692179, EPI_ISL_2692205, EPI_ISL_2692889, EPI_ISL_2692891, EPI_ISL_2692892 | | | |
| EPI_ISL_2692992 | Clinical Microbiology Laboratory, Faculty of Medicine, Universitas Indonesia | Clinical Microbiology Laboratory, Faculty of Medicine, Universitas Indonesia | Ibrahim F, Natasha A, Yasmon A, Fitrhiyah, Saharman YR, Karuniawati A, Sudarmono P. |
| EPI_ISL_2710020 | Ochsner Health | Bioinfoexperts, LLC | Judy Crabtree, Amy Feehan, Rebecca Rose, David J. Nolan, Tessa LaFleur, Julia-Garcia-Diaz, Lucio Miele, Susanna L. Lamers |
| EPI_ISL_2713091, EPI_ISL_2713095, EPI_ISL_2713096, EPI_ISL_2713097, EPI_ISL_2713100, EPI_ISL_2713101, EPI_ISL_2713102, EPI_ISL_2713103, EPI_ISL_2713104, EPI_ISL_2713105, EPI_ISL_2713106, EPI_ISL_2713107, EPI_ISL_2713108, EPI_ISL_2713109, EPI_ISL_2713110, EPI_ISL_2713111, EPI_ISL_2713112, EPI_ISL_2713113, EPI_ISL_2713121, EPI_ISL_2713122, EPI_ISL_2713124, EPI_ISL_2713125, EPI_ISL_2713127, EPI_ISL_2713128, EPI_ISL_2713129, EPI_ISL_2713130, EPI_ISL_2713131, EPI_ISL_2713132, EPI_ISL_2713133, EPI_ISL_2713134, EPI_ISL_2713135, EPI_ISL_2713136, EPI_ISL_2713137, EPI_ISL_2713138, EPI_ISL_2713139, EPI_ISL_2713140, EPI_ISL_2713141, EPI_ISL_2713142, EPI_ISL_2713143, EPI_ISL_2713144, EPI_ISL_2713151, EPI_ISL_2713153, EPI_ISL_2713154, EPI_ISL_2713155, EPI_ISL_2713156, EPI_ISL_2713158, EPI_ISL_2713159, EPI_ISL_2713160, EPI_ISL_2713161, EPI_ISL_2713162, EPI_ISL_2713163, EPI_ISL_2713164, EPI_ISL_2713165, EPI_ISL_2713171, EPI_ISL_2713172, EPI_ISL_2713173, EPI_ISL_2713174, EPI_ISL_2713175, EPI_ISL_2713176, EPI_ISL_2713177, EPI_ISL_2713179, EPI_ISL_2713181, EPI_ISL_2713182, EPI_ISL_2713183, EPI_ISL_2713184, EPI_ISL_2713185, EPI_ISL_2713186, EPI_ISL_2713187, EPI_ISL_2713188, EPI_ISL_2713189, EPI_ISL_2713190, EPI_ISL_2713191, EPI_ISL_2713192, EPI_ISL_2713193, EPI_ISL_2713217, EPI_ISL_2713218, EPI_ISL_2713220 | | | |
| see above | Department of Health Technology and Informatics, The Hong Kong Polytechnic University | Department of Health Technology and Informatics, The Hong Kong Polytechnic University | Gilman Kit-Hang Siu, Lam-Kwong Lee, Kenneth Siu-Sing Leung, Jake Siu-Lun Leung, Wing-Hei Lo, Timothy Ting-Leung Ng, Chloe Toi-Mei Chan, Hoi-Ching Jim, Ivan Tak-Fai Wong, Kingsley King-Gee Tam, Hiu-Yin Lao, Denise Sze-Hang Wong, Alan Ka-Lun Wu, Miranda Chong-Yee Yau, Yvette Wai-Man Lai, Kitty Sau-Chun Fung, Sandy Ka-Yee Chau, Barry Kin-Chung Wong, Wing-Kin To, Kristine Luk, Alex Yat-Man Ho, Tak-Lun Que, Kam-Tong Yip, Wing Cheong Yam, David Ho-Keung Shum, Shea Ping Yip |
| EPI_ISL_2718547 | Klaus Überla | Armin Ensser | Antonia Sophia Peter, Klaus Überla, Armin Ensser |
| EPI_ISL_2727270, EPI_ISL_2727272 | ZARV/NHLS, Department Medical Virology, University of Pretoria | KRISP, KZN Research Innovation and Sequencing Platform | Amy Strydom, Adriano Mendes, Micheala Davids, Sim Mayaphi and Marietjie Venter, Giandhari J, Pillay S, Naidoo Y, Ramphal U, Lessells R, Emmanuel SJ, Tegally H, Wilkinson E, de Oliveira T |
| EPI_ISL_2728581, EPI_ISL_2728582, EPI_ISL_2728583, EPI_ISL_2728584, EPI_ISL_2728585, EPI_ISL_2728586, EPI_ISL_2728588, EPI_ISL_2728589, EPI_ISL_2728591, EPI_ISL_2728597 | Laboratorio de Infectologia y Virologia Molecular | Laboratory of Molecular Virology, School of Medicine, Pontificia Universidad Catolica de Chile | Catalina Pardo-Roa, Constanza Martinez-Valdevenito, Jennifer Angulo, Tamara Garcia-Salum, Estefany Poblete, Maria Jose Avendano, Leonardo I. Almonacid, Ana Maria Contreras, Carlos Palma, Jorge Levican, Eileen Serrano, Constanza Maldonado, M. Belen Leyton, Erick Salinas, Andres E. Munoz-Marcos, Francisco Melo, Marcela Ferres, Rafael A. Medina |
| EPI_ISL_2756432, EPI_ISL_2756437, EPI_ISL_2756438, EPI_ISL_2756439, EPI_ISL_2756441, EPI_ISL_2756448, EPI_ISL_2756449, EPI_ISL_2756450, EPI_ISL_2756451, EPI_ISL_2756452, EPI_ISL_2756454, EPI_ISL_2756455, EPI_ISL_2756456, EPI_ISL_2756480, EPI_ISL_2756481 | | | |
| see above | Instituto Adolfo Lutz Central | Instituto Adolfo Lutz, Interdiciplinary Procedures Center, Strategic Laboratory | Claudio Tavares Sacchi, Claudia Regina Gonçalves, Erica Valessa Ramos Gomes, Karoline Rodrigues Campos, Caio Vinicius Dias Lopes, Leonardo Jose Tadeu de Araujo |
| EPI_ISL_2779290 | Busia border point | USAMRD-A, Basic Science Laboratory | Gathii Kimita, Josphat Nyataya, Esther Omuseni, Faith Sigei, Alan Lemtudo, Eric Muthanje, Brian Andika, Rehema Liyai, Rachel Githii, Clement Masakwe, Stephen Ochola, George Awinda, Carol Kifude, Beth Mutai, John Waitumbi |
| EPI_ISL_2779301 | KEMRI/DEID Surveillance Site (Lamu County Hospital) | USAMRD-A, Basic Science Laboratory | Gathii Kimita, Josphat Nyataya, Esther Omuseni, Faith Sigei, Alan Lemtudo, Eric Muthanje, Brian Andika, Rehema Liyai, Rachel Githii, Clement Masakwe, Stephen Ochola, George Awinda, Carol Kifude, Beth Mutai, John Waitumbi |
| EPI_ISL_2779317 | Port Health Clinic, Busia border point | USAMRD-A, Basic Science Laboratory | Gathii Kimita, Josphat Nyataya, Esther Omuseni, Faith Sigei, Alan Lemtudo, Eric Muthanje, Brian Andika, Rehema Liyai, Rachel Githii, Clement Masakwe, Stephen Ochola, George Awinda, Carol Kifude, Beth Mutai, John Waitumbi |
| EPI_ISL_2779318 | Port Health Clinic, Malaba border point | USAMRD-A, Basic Science Laboratory | Gathii Kimita, Josphat Nyataya, Esther Omuseni, Faith Sigei, Alan Lemtudo, Eric Muthanje, Brian Andika, Rehema Liyai, Rachel Githii, Clement Masakwe, Stephen Ochola, George Awinda, Carol Kifude, Beth Mutai, John Waitumbi |
| EPI_ISL_2779389 | GK Prison Kiambu | USAMRD-A, Basic Science Laboratory | Gathii Kimita, Josphat Nyataya, Esther Omuseni, Faith Sigei, Alan Lemtudo, Eric Muthanje, Brian Andika, Rehema Liyai, Rachel Githii, Clement Masakwe, Stephen Ochola, George Awinda, Carol Kifude, Beth Mutai, John Waitumbi |
| EPI_ISL_2779390, EPI_ISL_2779394, EPI_ISL_2779400, EPI_ISL_2779401, EPI_ISL_2779402, EPI_ISL_2779403, EPI_ISL_2779404, EPI_ISL_2779405, EPI_ISL_2779406, EPI_ISL_2779407, EPI_ISL_2779408, EPI_ISL_2779410, EPI_ISL_2779412, EPI_ISL_2779413, EPI_ISL_2779414, EPI_ISL_2779416, EPI_ISL_2779417 | | | |
| see above | BATUK | USAMRD-A, Basic Science Laboratory | Gathii Kimita, Josphat Nyataya, Esther Omuseni, Faith Sigei, Alan Lemtudo, Eric Muthanje, Brian Andika, Rehema Liyai, Rachel Githii, Clement Masakwe, Stephen Ochola, George Awinda, Carol Kifude, Beth Mutai, John Waitumbi |
| EPI_ISL_2779466 | KCRH KEMRI/DEID Surveillance Site | USAMRD-A, Basic Science Laboratory | Gathii Kimita, Josphat Nyataya, Esther Omuseni, Faith Sigei, Alan Lemtudo, Eric Muthanje, Brian Andika, Rehema Liyai, Rachel Githii, Clement Masakwe, Stephen Ochola, George Awinda, Carol Kifude, Beth Mutai, John Waitumbi |
| EPI_ISL_2779488 | GK Prison Kitengela | USAMRD-A, Basic Science Laboratory | Gathii Kimita, Josphat Nyataya, Esther Omuseni, Faith Sigei, Alan Lemtudo, Eric Muthanje, Brian Andika, Rehema Liyai, Rachel Githii, Clement Masakwe, Stephen Ochola, George Awinda, Carol Kifude, Beth Mutai, John Waitumbi |
| EPI_ISL_2779510, EPI_ISL_2779512, EPI_ISL_2779513, EPI_ISL_2779514, EPI_ISL_2779516, EPI_ISL_2779517, EPI_ISL_2779518, EPI_ISL_2779519, EPI_ISL_2779520, EPI_ISL_2779521 | BATUK | USAMRD-A, Basic Science Laboratory | Gathii Kimita, Josphat Nyataya, Esther Omuseni, Faith Sigei, Alan Lemtudo, Eric Muthanje, Brian Andika, Rehema Liyai, Rachel Githii, Clement Masakwe, Stephen Ochola, George Awinda, Carol Kifude, Beth Mutai, John Waitumbi |
| EPI_ISL_2789814, EPI_ISL_2789823, EPI_ISL_2789824, EPI_ISL_2789825, EPI_ISL_2789826, EPI_ISL_2789827, EPI_ISL_2789828, EPI_ISL_2789829, EPI_ISL_2789830, EPI_ISL_2789831, EPI_ISL_2789832, EPI_ISL_2789833, EPI_ISL_2789834, EPI_ISL_2789876, EPI_ISL_2789877, EPI_ISL_2789878 | | | |
| see above | Centre Hospitalier Universitaire Clermont-Ferrand | CHU Clermont-Ferrand, service de virologie | Bisseux Maxime, Mirand Audrey, Combes Patricia, Henquell Cécile |
| EPI_ISL_2790906, EPI_ISL_2790907 | Labo analyses Med | National Reference Center for Viruses of Respiratory Infections, Institut Pasteur, Paris | Marion Barbet, Sylvie Behillil, Méline Bizard, Angela Brisebarre, Camille Capel, Vincent Enouf, Louise Lefrançois, Frédéric Lemoine, Christophe Malabat, Corinne Maufrais, Emmanuelle Permal, Etienne Simon-Lorière, Maud Vanpeene, Sylvie Van der Werf ,GréGoire Potiron |
| EPI_ISL_2790908 | Medecine sentinel network | National Reference Center for Viruses of Respiratory Infections, Institut Pasteur, Paris | Marion Barbet, Sylvie Behillil, Méline Bizard, Angela Brisebarre, Camille Capel, Vincent Enouf, Louise Lefrançois, Frédéric Lemoine, Christophe Malabat, Corinne Maufrais, Emmanuelle Permal, Etienne Simon-Lorière, Maud Vanpeene, Sylvie Van der Werf ,Ghatfan Hassan |
| EPI_ISL_2795653, EPI_ISL_2795654, EPI_ISL_2795656 | Infectious Diseases & Tropical Medicine Research Center, Health Research Institute, Babol University of Medical Sciences, Babol, Iran | Genetics Research Center, University of Social Welfare and Rehabilitation Sciences | Zohreh Fattahi, Marzieh Mohseni, Kimia Kahrizi, Yousef Yahyapour, Masomeh Bayani, Farzin Sadeghi, Hossein Najmabadi. |
| EPI_ISL_2828704, EPI_ISL_2828705, EPI_ISL_2828706 | LACEN do Estado de Goias | Instituto Adolfo Lutz, Interdiciplinary Procedures Center, Strategic Laboratory | Claudio Tavares Sacchi, Claudia Regina Gonçalves, Erica Valessa Ramos Gomes, Karoline Rodrigues Campos, Caio Vinicius Dias Lopes |
| EPI_ISL_2835511, EPI_ISL_2835512, EPI_ISL_2835513, EPI_ISL_2835514, EPI_ISL_2835515, EPI_ISL_2835516, EPI_ISL_2835517, EPI_ISL_2835518, EPI_ISL_2835519, EPI_ISL_2835520, EPI_ISL_2835521, EPI_ISL_2835522, EPI_ISL_2835523, EPI_ISL_2835524, EPI_ISL_2835525, EPI_ISL_2835526, EPI_ISL_2835527, EPI_ISL_2835528, EPI_ISL_2835529, EPI_ISL_2835530, EPI_ISL_2835531, EPI_ISL_2835532, EPI_ISL_2835533, EPI_ISL_2835534, EPI_ISL_2835535, EPI_ISL_2835536, EPI_ISL_2835537, EPI_ISL_2835538, EPI_ISL_2835539, EPI_ISL_2835540, EPI_ISL_2835541, EPI_ISL_2835542, EPI_ISL_2835543, EPI_ISL_2835544, EPI_ISL_2835545, EPI_ISL_2835546, EPI_ISL_2835547, EPI_ISL_2835548, EPI_ISL_2835549, EPI_ISL_2835550, EPI_ISL_2835551 | | | |
| see above | Sharp HealthCare Laboratory | Andersen lab at Scripps Research | SEARCH Alliance San Diego with Aaron Harding, Jacquelyn Berumen, Cathy Woerle, Liam McGinnis, Art Mendoza, Omid Bakhtar |
| EPI_ISL_2860784, EPI_ISL_2860785, EPI_ISL_2860786 | Department of Health Technology and Informatics, The Hong Kong Polytechnic University | Department of Health Technology and Informatics, The Hong Kong Polytechnic University | Gilman Kit-Hang Siu, Lam-Kwong Lee, Kenneth Siu-Sing Leung, Jake Siu-Lun Leung, Wing-Hei Lo, Timothy Ting-Leung Ng, Chloe Toi-Mei Chan, Hoi-Ching Jim, Ivan Tak-Fai Wong, Kingsley King-Gee Tam, Hiu-Yin Lao, Denise Sze-Hang Wong, Alan Ka-Lun Wu, Miranda Chong-Yee Yau, Yvette Wai-Man Lai, Kitty Sau-Chun Fung, Sandy Ka-Yee Chau, Barry Kin-Chung Wong, Wing-Kin To, Kristine Luk, Alex Yat-Man Ho, Tak-Lun Que, Kam-Tong Yip, Wing Cheong Yam, David Ho-Keung Shum, Shea Ping Yip |
| EPI_ISL_861488 | Cerballiance Côte d'Azur | CERBA LAB | Prots.L; Pichon R; Barrieu-Moussat S. |
| EPI_ISL_861489 | Cerballiance Côte d'Azur | CERBA LAB | Adjtoutah Z; Mangiarotti T; Prots L. |
| EPI_ISL_861490 | Cerballiance Côte d'Azur | CERBA LAB | De La Chapelle A; Prots.L; |
| EPI_ISL_861491 | Cerballiance Côte d'Azur | CERBA LAB | Chobert F; Prots.L |

| | | | |
|---|---|---|---|
| EPI_ISL_861492 | Cerballiance Côte d'Azur | CERBA LAB | Barrieu-Moussat S;Adoul O;Boudjahem Z; Roman E; Chaudet L; Martin L; Aankour N; Jaugin P; Gastaud S; Prots L. |
| EPI_ISL_861493 | Cerballiance Côte d'Azur | CERBA LAB | Marius M; Barsanti M; Maltese S; Francois J; Sinsoulieu T; Van Der Rhyn I; Morello V; Marcadal M; Gandillet A; Prots L. |
| EPI_ISL_861494 | Cerballiance Côte d'Azur | CERBA LAB | Prots L. Raimondi V. |
| EPI_ISL_861495 | Cerballiance Côte d'Azur | CERBA LAB | Pichon R. Quintle S; Sanza O; Mangiarotti T; Viltard B; Prots L. |
| EPI_ISL_861497 | Cerballiance Centre Est | CERBA LAB | Chanard C; Gazzano V; Munier C; Prots L. |
| EPI_ISL_861500, EPI_ISL_861501, EPI_ISL_861502 | Cerballiance Centre Est | CERBA LAB | Chanard E; Gazzano V; Munier C; Prots L. |
| EPI_ISL_862835, EPI_ISL_862836 | National Institute of Infectious Diseases-Prof. Dr. Matei Bals Molecular Diagnostics Laboratory | National Institute of Infectious Diseases-Prof. Dr. Matei Bals Molecular Diagnostics Laboratory | Leontina Banica, Marius Surleac, Corina Casangiu, Petre Milu, Andreea Tudor, Simona Paraschiv, Dan Otelea |
| EPI_ISL_869166 | Laboratory of Microbiology,ASST Settelaghi, Varese, Italy | Laboratory of Microbiology,ASST Settelaghi, Varese, Italy | Maggi,F., Novazzi,F., Genoni,A., Baj,A., Spezia,P.G., Focosi,D., Zago,C., Colombo,A., Cassani,G., Pasciuta,R., Tamborini,A.,Rossi,A., Prestia,M., Capuano,R., Azzi,L., Donadini,A., Catanoso,G., Grossi,P., Maffioli,L. and Bonelli,G. |
| EPI_ISL_869240 | Laboratory of Microbiology, ASST Settelaghi, Varese, Italy | Laboratory of Microbiology, ASST Settelaghi, Varese, Italy | Novazzi,F., Genoni,A., Focosi,D., Baj,A., Spezia,P.G., Zago,C., Colombo,A., Cassani,G., Pasciuta,R., Tamborini,A.,Rossi,A., Prestia,M., Capuano,R., Azzi,L., Donadini,A.,Catanoso,G., Maggi,F. |
| EPI_ISL_871872 | Botswana Harvard HIV Reference Laboratory | Botswana Harvard HIV Reference Laboratory | Sikhulile Moyo, Dorcas Maruapula, Wonderful Choga, Botshelo Radibe, Boitumelo Zuze, David Lawrence, Roger Shapiro, Shahin Lockman, Mosepele Mosepele, Joseph, Makhema, Simani Gaseitsiwe |
| EPI_ISL_872192 | Conjunto Hospitalar do Mandaqui de Sao Paulo | Instituto Adolfo Lutz, Interdiciplinary Procedures Center, Strategic Laboratory | Claudio Tavares Sacchi, Claudia Regina Gonçalves, Erica Valessa Ramos Gomes, Karoline Rodrigues Campos, Katia Correa de Oliveira Santos, Ana Lucia de Carvalho Avelino, Fabiana Cristina Pereira dos Santos |
| EPI_ISL_875566, EPI_ISL_875567, EPI_ISL_875568 | SIESP L'AQUILA | Istituto Zooprofilattico Sperimentale dell'Abruzzo e Molise "G. Caporale" | Lorusso A, Marcacci M, Di Domenico M, Ancora M, Curini V, Mangone I, Rinaldi A, Scialabba S, Di Pasquale A, Cammà C, Puglia I, Calistri P, Savini G |
| EPI_ISL_883155 | San Donato Arezzo Analysis Laboratory - Clinical Molecular Pathology sector | San Donato Arezzo Analysis Laboratory - Clinical Molecular Pathology sector | Alessandro Pancrazzi and Alice Moncada |
| EPI_ISL_883325, EPI_ISL_883344, EPI_ISL_883345, EPI_ISL_883406, EPI_ISL_883407, EPI_ISL_883408 | OCME Office Of Chief Medical Examiner | New York City Public Health Laboratory | Jade Wang, et al. |
| EPI_ISL_885143 | Central public health laboratory | Molecular Diagnostics Department, Central public health laboratory | Dler,H., Dlshad,H., Furat,S., Sharmeen,F.-A., Dalia,F., Mohsen,A.,Hemdad,A., Fahmi,A., Hemn,M. and Idrees,H. |
| EPI_ISL_891249, EPI_ISL_891250 | AZ Delta | AZ Delta | Geert Martens, Dieter De Smet, Brigitte Maes |
| EPI_ISL_891251 | Singapore General Hospital | Department of Microbiology | Nurdyana Abdul Rahman, Kun Lee Lim, Chenhao Li, Sui Sin Goh, Kenneth Xin Long Chan, Kian Sing Chan, Lynette Oon, Kern Rei Chng, Niranjan Nagarajan, Karrie Ko |
| EPI_ISL_906053 | Tilia Laboratories s.r.o. | Tilia Laboratories s.r.o. | Sona Pekova, MD, PhD. |
| EPI_ISL_906070 | UPA Dr. Akira Tada | Instituto Adolfo Lutz, Interdiciplinary Procedures Center, Strategic Laboratory | Claudio Tavares Sacchi, Claudia Regina Gonçalves, Erica Valessa Ramos Gomes, Karoline Rodrigues Campos |
| EPI_ISL_906071 | LACEN-PI DR. Costa Alvarenga | Instituto Adolfo Lutz, Interdiciplinary Procedures Center, Strategic Laboratory | Claudio Tavares Sacchi, Claudia Regina Gonçalves, Erica Valessa Ramos Gomes, Karoline Rodrigues Campos |
| EPI_ISL_906072 | UPA Dr. Akira Tada | Instituto Adolfo Lutz, Interdiciplinary Procedures Center, Strategic Laboratory | Claudio Tavares Sacchi, Claudia Regina Gonçalves, Erica Valessa Ramos Gomes, Karoline Rodrigues Campos |
| EPI_ISL_906073 | Hospital Vila Lobos | Instituto Adolfo Lutz, Interdiciplinary Procedures Center, Strategic Laboratory | Claudio Tavares Sacchi, Claudia Regina Gonçalves, Erica Valessa Ramos Gomes, Karoline Rodrigues Campos |
| EPI_ISL_906074 | Searom Laboratorio Diagnostico | Instituto Adolfo Lutz, Interdiciplinary Procedures Center, Strategic Laboratory | Claudio Tavares Sacchi, Claudia Regina Gonçalves, Erica Valessa Ramos Gomes, Karoline Rodrigues Campos |
| EPI_ISL_906075 | Hospital Geral de Vila Penteado Dr Jose Pangella Sao Paulo | Instituto Adolfo Lutz, Interdiciplinary Procedures Center, Strategic Laboratory | Claudio Tavares Sacchi, Claudia Regina Gonçalves, Erica Valessa Ramos Gomes, Karoline Rodrigues Campos |
| EPI_ISL_906076, EPI_ISL_906077 | Hospital Sao Luiz Sao Caetano | Instituto Adolfo Lutz, Interdiciplinary Procedures Center, Strategic Laboratory | Claudio Tavares Sacchi, Claudia Regina Gonçalves, Erica Valessa Ramos Gomes, Karoline Rodrigues Campos |
| EPI_ISL_906078 | Hospital Carlos Chagas | Instituto Adolfo Lutz, Interdiciplinary Procedures Center, Strategic Laboratory | Claudio Tavares Sacchi, Claudia Regina Gonçalves, Erica Valessa Ramos Gomes, Karoline Rodrigues Campos |
| EPI_ISL_906079 | Hospital Paulistano Paulista | Instituto Adolfo Lutz, Interdiciplinary Procedures Center, Strategic Laboratory | Claudio Tavares Sacchi, Claudia Regina Gonçalves, Erica Valessa Ramos Gomes, Karoline Rodrigues Campos |
| EPI_ISL_906080, EPI_ISL_906081 | Hospital Beneficiencia Portuguesa | Instituto Adolfo Lutz, Interdiciplinary Procedures Center, Strategic Laboratory | Claudio Tavares Sacchi, Claudia Regina Gonçalves, Erica Valessa Ramos Gomes, Karoline Rodrigues Campos |
| EPI_ISL_909953 | AZ Delta | AZ Delta | Geert Martens, Dieter De Smet |
| EPI_ISL_909954 | AZ Delta | AZ Delta | Geert Marten; Dieter De Smet |
| EPI_ISL_909963 | AZ Delta | AZ Delta | Geert Martens; Dieter De Smet |
| EPI_ISL_910011 | AZ Delta | AZ Delta | Geert Martens; Dieter De Smet |
| EPI_ISL_910012, EPI_ISL_910013, EPI_ISL_910015 | AZ Delta | AZ Delta | Geert Martens; Dieter De Smet |
| EPI_ISL_911386, EPI_ISL_911387 | AZDelta | AZDelta | Geert Martens; Dieter De Smet |
| EPI_ISL_911525 | Microbiology and Virology Unit, Florence Careggi University Hospital | Microbiology and Virology Unit, Florence Careggi University Hospital | Vincenzo Di Pilato, Marco Coppi, Fabio Morecchiato, Noemi Aiezza, Ilaria Baccani, Alberto Antonelli, Emanuele Gori, Gian Maria Rossolini |
| EPI_ISL_911548 | ARUP laboratories | ARUP Laboratories | Hymas W, Slechta ES, Pyne MT, Mallory MA, Simmon KE, Shakir SM, Hillyard DR, Barker AP |
| EPI_ISL_912579 | Service Biologie -CH de Versailles | National Reference Center for Viruses of Respiratory Infections, Institut Pasteur, Paris | Marion Barbet, Sylvie Behillil, Méline Bizard, Angela Brisebarre, Camille Capel, Etienne Simon-Lorière, Vincent Enouf, Maud Vanpeene, Sylvie van der Werf,Marque Juillet StéPhanie |
| EPI_ISL_912598 | Hopital Avicenne Laboratoire central de bactériologie virologie hygiène | National Reference Center for Viruses of Respiratory Infections, Institut Pasteur, Paris | Marion Barbet, Sylvie Behillil, Méline Bizard, Angela Brisebarre, Camille Capel, Etienne Simon-Lorière, Vincent Enouf, Maud Vanpeene, Sylvie van der Werf,Brichler Ségolène |
| EPI_ISL_912601 | Labo Analyses Med | National Reference Center for Viruses of Respiratory Infections, Institut Pasteur, Paris | Marion Barbet, Sylvie Behillil, Méline Bizard, Angela Brisebarre, Camille Capel, Etienne Simon-Lorière, Vincent Enouf, Maud Vanpeene, Sylvie van der Werf |
| EPI_ISL_912614 | CENTRE HOSPITALIER JACQUES LACARIN LABORATOIRE | National Reference Center for Viruses of Respiratory Infections, Institut Pasteur, Paris | Marion Barbet, Sylvie Behillil, Méline Bizard, Angela Brisebarre, Camille Capel, Etienne Simon-Lorière, Vincent Enouf, Maud Vanpeene, Sylvie van der Werf |

| | | | |
|---|---|---|---|
| EPI_ISL_912623 | Labo Analyses Med | National Reference Center for Viruses of Respiratory Infections, Institut Pasteur, Paris | Marion Barbet, Sylvie Behillil, Méline Bizard, Angela Brisebarre, Camille Capel, Etienne Simon-Lorière, Vincent Enouf, Maud Vanpeene, Sylvie van der Werf |
| EPI_ISL_912670, EPI_ISL_912671, EPI_ISL_912678, EPI_ISL_912682, EPI_ISL_912684, EPI_ISL_912751, EPI_ISL_912755, EPI_ISL_912757, EPI_ISL_912758, EPI_ISL_912759, EPI_ISL_912762, EPI_ISL_912766, EPI_ISL_912767, EPI_ISL_912768, EPI_ISL_912769, EPI_ISL_912770, EPI_ISL_912772, EPI_ISL_912773, EPI_ISL_912776, EPI_ISL_912777, EPI_ISL_912779, EPI_ISL_912797, EPI_ISL_912798, EPI_ISL_912802, EPI_ISL_912831, EPI_ISL_912834, EPI_ISL_912835, EPI_ISL_912836, EPI_ISL_912843, EPI_ISL_912847, EPI_ISL_912848, EPI_ISL_912849, EPI_ISL_912867, EPI_ISL_912872, EPI_ISL_912875, EPI_ISL_912877, EPI_ISL_912882, EPI_ISL_912883, EPI_ISL_912884, EPI_ISL_912890, EPI_ISL_912960 | | | |
| see above | Hôpital Henri Mondor | Department of Virology, Henri Mondor University Hospital, Assistance Publique Hôpitaux de Paris, Université Paris-Est Créteil, INSERM U955 | Christophe Rodriguez, Slim Fourati, Vanessa Demontant, Guillaume Gricourt, Melissa N'Debi, Alexandre Soulier, Elisabeth Trawinski, Jean-Michel Pawlotsky |
| EPI_ISL_913025, EPI_ISL_913029 | Hospital los Arcos | Instituto de Salud Carlos III | Iglesias-Caballero, M. Camarero, S. Sandonís,V. Vázquez, S. Pozo, F. Casas, I. Jiménez, P. Zaballos, A. Monzón, S. Varona, S. Cuesta, I. Cámara, M. |
| EPI_ISL_913496, EPI_ISL_913497, EPI_ISL_913498, EPI_ISL_913499, EPI_ISL_913500, EPI_ISL_913501, EPI_ISL_913502, EPI_ISL_913503, EPI_ISL_913504, EPI_ISL_913505, EPI_ISL_913506, EPI_ISL_913507, EPI_ISL_913508, EPI_ISL_913509, EPI_ISL_913510, EPI_ISL_913511, EPI_ISL_913512, EPI_ISL_913513, EPI_ISL_913594, EPI_ISL_913595, EPI_ISL_913596, EPI_ISL_913597, EPI_ISL_913598, EPI_ISL_913599, EPI_ISL_913600, EPI_ISL_913601, EPI_ISL_913602, EPI_ISL_913603, EPI_ISL_913604, EPI_ISL_913605 | | | |
| see above | AZDelta | AZDelta | Geert Martens; Dieter De Smet |
| EPI_ISL_918269 | SIESP DIPARTIMENTO DI PREVENZIONE TERAMO | Istituto Zooprofilattico Sperimentale dell'Abruzzo e Molise "G. Caporale" | Lorusso A, Marcacci M, Di Domenico M, Ancora M, Curini V, Mangone I, Rinaldi A, Scialabba S, Di Pasquale A, Cammà C, Puglia I, Calistri P, Savini G |
| EPI_ISL_918410 | Ospedale Di Venere - Carbonara | Istituto Zooprofilattico Sperimentale della Puglia e della Basilicata | Parisi A., Bianco A., Capozzi L., Del Sambro L., Simone D., Manzulli V, Rondinone V., Pace L., Cipolletta D., Galante D. |
| EPI_ISL_918483 | Laboratory of Microbiology, ASST Settelaghi, Varese, Italy | Laboratory of Microbiology, ASST Settelaghi, Varese, Italy | Novazzi,F., Genoni,A., Baj,A., Focosi,D., Spezia,P.G., Zago,C., Colombo,A., Cassani,G., Pasciuta,R., Tamborini,A., Rossi,A., Prestia,M., Capuano,R., Maggi,F. |
| EPI_ISL_918496 | BSWMC-Marble Falls | BSWMC-Temple Molecular | Ari Rao, Linden Morales, Kimberly Walker, Shelby Hendrickson |
| EPI_ISL_925173 | Laboratorio de Referencia Nacional de Enteropatógenos. Instituto Nacional de Salud del Perú | Laboratorio de Referencia Nacional de Enteropatógenos. Instituto Nacional de Salud del Perú | Ronnie Gavilan Chavez, Junior Caro Castro, Willi Quino Sifuentes, Veronica Hurtado Vela, Iris Silva Molina, Fiorella Orellana Peralta, |
| EPI_ISL_925208 | Baylor Scott & White-Temple | Baylor Scott & White-Temple | Ari Rao, Linden Morales, Kimberly Walker, Marcus Volz, Shelby Hendrickson |
| EPI_ISL_934424 | Institute for Medical Research, Infectious Disease Research Centre, National Institutes of Health, Ministry of Health Malaysia | Institute for Medical Research, Infectious Disease Research Centre, National Institutes of Health, Ministry of Health Malaysia | Suppiah J, Kamel K, Azizan MA, Thayan R |
| EPI_ISL_934992, EPI_ISL_934993, EPI_ISL_934994, EPI_ISL_934998, EPI_ISL_934999 | ADMED Microbiologie | Genomics and Transcriptomics, Philip Morris International | Reto Lienhard, Marie-Lise Tritten, Emmanuel Guedj, Nicolas Sierro, Rémi Dulize, David Bornand, Mehdi Auberson, Maxime Berthouzoz, Nikolai Ivanov, Manuel Peitsch |
| EPI_ISL_935017, EPI_ISL_935021 | Laboratory for Respiratory Viruses, Cantacuzino National Military-Medical Institute for Research and Development | Cantacuzino Institute Virology | Luiza Ustea, Nicoleta Paraschiv, Mihaela Lazar |
| EPI_ISL_935170 | Molecular Diagnostics Pathology Department Mater Dei Hospital | Molecular Diagnostics Pathology Department Mater Dei Hospital | Graziella Zahra |
| EPI_ISL_935533 | Labo Analyses Med | National Reference Center for Viruses of Respiratory Infections, Institut Pasteur, Paris | Marion Barbet, Sylvie Behillil, Méline Bizard, Angela Brisebarre, Camille Capel, Etienne Simon-Lorière, Vincent Enouf, Maud Vanpeene, Sylvie van der Werf,Jacques Alexandra |
| EPI_ISL_935534 | Labo Analyses Med | National Reference Center for Viruses of Respiratory Infections, Institut Pasteur, Paris | Marion Barbet, Sylvie Behillil, Méline Bizard, Angela Brisebarre, Camille Capel, Etienne Simon-Lorière, Vincent Enouf, Maud Vanpeene, Sylvie van der Werf,Garel FréDéRick |
| EPI_ISL_935546 | Labo Analyses Med | National Reference Center for Viruses of Respiratory Infections, Institut Pasteur, Paris | Marion Barbet, Sylvie Behillil, Méline Bizard, Angela Brisebarre, Camille Capel, Etienne Simon-Lorière, Vincent Enouf, Maud Vanpeene, Sylvie van der Werf,Besson J |
| EPI_ISL_935549 | Labo Analyses Med | National Reference Center for Viruses of Respiratory Infections, Institut Pasteur, Paris | Marion Barbet, Sylvie Behillil, Méline Bizard, Angela Brisebarre, Camille Capel, Etienne Simon-Lorière, Vincent Enouf, Maud Vanpeene, Sylvie van der Werf |
| EPI_ISL_935550 | Labo Analyses Med | National Reference Center for Viruses of Respiratory Infections, Institut Pasteur, Paris | Marion Barbet, Sylvie Behillil, Méline Bizard, Angela Brisebarre, Camille Capel, Etienne Simon-Lorière, Vincent Enouf, Maud Vanpeene, Sylvie van der Werf,Lefaure Brieuc |
| EPI_ISL_935558, EPI_ISL_935559 | Hopital | National Reference Center for Viruses of Respiratory Infections, Institut Pasteur, Paris | Marion Barbet, Sylvie Behillil, Méline Bizard, Angela Brisebarre, Camille Capel, Etienne Simon-Lorière, Vincent Enouf, Maud Vanpeene, Sylvie van der Werf |
| EPI_ISL_935561 | Hopital | National Reference Center for Viruses of Respiratory Infections, Institut Pasteur, Paris | Marion Barbet, Sylvie Behillil, Méline Bizard, Angela Brisebarre, Camille Capel, Etienne Simon-Lorière, Vincent Enouf, Maud Vanpeene, Sylvie van der Werf,Gaschet Anne |
| EPI_ISL_935563 | Labo Analyses Med | National Reference Center for Viruses of Respiratory Infections, Institut Pasteur, Paris | Marion Barbet, Sylvie Behillil, Méline Bizard, Angela Brisebarre, Camille Capel, Etienne Simon-Lorière, Vincent Enouf, Maud Vanpeene, Sylvie van der Werf |
| EPI_ISL_935565 | Hopital | National Reference Center for Viruses of Respiratory Infections, Institut Pasteur, Paris | Marion Barbet, Sylvie Behillil, Méline Bizard, Angela Brisebarre, Camille Capel, Etienne Simon-Lorière, Vincent Enouf, Maud Vanpeene, Sylvie van der Werf,Bret Laurent |
| EPI_ISL_935570 | Hopital | National Reference Center for Viruses of Respiratory Infections, Institut Pasteur, Paris | Marion Barbet, Sylvie Behillil, Méline Bizard, Angela Brisebarre, Camille Capel, Etienne Simon-Lorière, Vincent Enouf, Maud Vanpeene, Sylvie van der Werf,Castelain Sandrine |
| EPI_ISL_935576 | Labo Analyses Med | National Reference Center for Viruses of Respiratory Infections, Institut Pasteur, Paris | Marion Barbet, Sylvie Behillil, Méline Bizard, Angela Brisebarre, Camille Capel, Etienne Simon-Lorière, Vincent Enouf, Maud Vanpeene, Sylvie van der Werf,Breton-Cazaux JéRôMe |
| EPI_ISL_935577, EPI_ISL_935578, EPI_ISL_935579, EPI_ISL_935580 | Hopital | National Reference Center for Viruses of Respiratory Infections, Institut Pasteur, Paris | Marion Barbet, Sylvie Behillil, Méline Bizard, Angela Brisebarre, Camille Capel, Etienne Simon-Lorière, Vincent Enouf, Maud Vanpeene, Sylvie van der Werf,Brichler SéGolèNe |
| EPI_ISL_935582 | Hopital | National Reference Center for Viruses of Respiratory Infections, Institut Pasteur, Paris | Marion Barbet, Sylvie Behillil, Méline Bizard, Angela Brisebarre, Camille Capel, Etienne Simon-Lorière, Vincent Enouf, Maud Vanpeene, Sylvie van der Werf,Blazejewski C. |
| EPI_ISL_935588 | Labo Analyses med | National Reference Center for Viruses of Respiratory Infections, Institut Pasteur, Paris | Marion Barbet, Sylvie Behillil, Méline Bizard, Angela Brisebarre, Camille Capel, Etienne Simon-Lorière, Vincent Enouf, Maud Vanpeene, Sylvie van der Werf,Amzalag Jonas |
| EPI_ISL_935590 | Labo Analyses Med | National Reference Center for Viruses of Respiratory Infections, Institut Pasteur, Paris | Marion Barbet, Sylvie Behillil, Méline Bizard, Angela Brisebarre, Camille Capel, Etienne Simon-Lorière, Vincent Enouf, Maud Vanpeene, Sylvie van der Werf,Amzalag Jonas |
| EPI_ISL_935591, EPI_ISL_935592, EPI_ISL_935593, EPI_ISL_935594, EPI_ISL_935595, EPI_ISL_935602, EPI_ISL_935603, EPI_ISL_935604, EPI_ISL_935607, EPI_ISL_935608, EPI_ISL_935611, EPI_ISL_935612, EPI_ISL_935613, EPI_ISL_935615, EPI_ISL_935616, EPI_ISL_935617, EPI_ISL_935618, EPI_ISL_935619, EPI_ISL_935620, EPI_ISL_935621, EPI_ISL_935622, EPI_ISL_935623, EPI_ISL_935624, EPI_ISL_935625, EPI_ISL_935628, EPI_ISL_935629, EPI_ISL_935630, EPI_ISL_935631, EPI_ISL_935634, EPI_ISL_935635, EPI_ISL_935636 | | | |
| see above | Labo Analyses med | National Reference Center for Viruses of Respiratory Infections, Institut Pasteur, Paris | Marion Barbet, Sylvie Behillil, Méline Bizard, Angela Brisebarre, Camille Capel, Etienne Simon-Lorière, Vincent Enouf, Maud Vanpeene, Sylvie van der Werf,Amzalag Jonas |
| EPI_ISL_935637 | Labo Analyses Med | National Reference Center for Viruses of Respiratory Infections, Institut Pasteur, Paris | Marion Barbet, Sylvie Behillil, Méline Bizard, Angela Brisebarre, Camille Capel, Etienne Simon-Lorière, Vincent Enouf, Maud Vanpeene, Sylvie van der Werf,Jacques Alexandra |
| EPI_ISL_935642 | Labo Analyses Med | National Reference Center for Viruses of Respiratory Infections, Institut Pasteur, Paris | Marion Barbet, Sylvie Behillil, Méline Bizard, Angela Brisebarre, Camille Capel, Etienne Simon-Lorière, Vincent Enouf, Maud Vanpeene, Sylvie van der Werf |

| | | | |
|---|---|---|---|
| EPI_ISL_935643 | Labo Analyses Med | National Reference Center for Viruses of Respiratory Infections, Institut Pasteur, Paris | Marion Barbet, Sylvie Behillil, Méline Bizard, Angela Brisebarre, Camille Capel, Etienne Simon-Lorière, Vincent Enouf, Maud Vanpeene, Sylvie van der Werf,Le Berre (D) David |
| EPI_ISL_935645, EPI_ISL_935646, EPI_ISL_935647, EPI_ISL_935648, EPI_ISL_935649 | Labo Analyses Med | National Reference Center for Viruses of Respiratory Infections, Institut Pasteur, Paris | Marion Barbet, Sylvie Behillil, Méline Bizard, Angela Brisebarre, Camille Capel, Etienne Simon-Lorière, Vincent Enouf, Maud Vanpeene, Sylvie van der Werf |
| EPI_ISL_937186, EPI_ISL_937187, EPI_ISL_937188, EPI_ISL_937189, EPI_ISL_937195, EPI_ISL_937196, EPI_ISL_937198, EPI_ISL_937199, EPI_ISL_937240, EPI_ISL_937241, EPI_ISL_937242, EPI_ISL_937243, EPI_ISL_937244, EPI_ISL_937245, EPI_ISL_937246, EPI_ISL_937247, EPI_ISL_937252, EPI_ISL_937253, EPI_ISL_937254, EPI_ISL_937255 | | | |
| see above | OCME Office Of Chief Medical Examiner | New York City Public Health Laboratory | Jade Wang, et al. |
| EPI_ISL_939629, EPI_ISL_939643 | Laboratory of Virology and Molecular Diagnostics | Institute of Public Health of Republic of North Macedonia Laboratory of Virology and Molecular Diagnostics | Maja Kuzmanovska, Golubinka Boshevska, Elizabeta Janchevska |
| EPI_ISL_939647, EPI_ISL_940144, EPI_ISL_940554 | Laboratory for Respiratory Viruses, Cantacuzino National Military-Medical Institute for Research and Development | Cantacuzino Institute Virology | Luiza Ustea, Nicoleta Paraschiv, Mihaela Lazar |
| EPI_ISL_940564 | Servizo de Microbioloxía. Complexo Hospitalario Universitario de Santiago de Compostela | Servizo de Microbioloxía. Complexo Hospitalario Universitario de Santiago de Compostela | Antonio Aguilera, Gema Barbeito, Amparo Coira, Rocio Trastoy, María Luisa Pérez del Molino |
| EPI_ISL_940565 | University of Bari Biomedical Sciences and Human Oncology | University of Bari Biomedical Sciences and Human Oncology | Chironna M., Sallustio A., Loconsole D., Accogli M. |
| EPI_ISL_940609 | Hospital e Maternidade Celso Pierro | Instituto Adolfo Lutz, Interdiciplinary Procedures Center, Strategic Laboratory | Claudio Tavares Sacchi, Claudia Regina Gonçalves, Erica Valessa Ramos Gomes, Karoline Rodrigues Campos |
| EPI_ISL_940610, EPI_ISL_940611 | Hospital Paulistano Paulista | Instituto Adolfo Lutz, Interdiciplinary Procedures Center, Strategic Laboratory | Claudio Tavares Sacchi, Claudia Regina Gonçalves, Erica Valessa Ramos Gomes, Karoline Rodrigues Campos |
| EPI_ISL_940613, EPI_ISL_940614, EPI_ISL_940615, EPI_ISL_940616, EPI_ISL_940617, EPI_ISL_940618 | LACEN-PI DR. Costa Alvarenga | Instituto Adolfo Lutz, Interdiciplinary Procedures Center, Strategic Laboratory | Claudio Tavares Sacchi, Claudia Regina Gonçalves, Erica Valessa Ramos Gomes, Karoline Rodrigues Campos |
| EPI_ISL_940625 | Hospital Sao Joaquim - Beneficiencia Portuguesa | Instituto Adolfo Lutz, Interdiciplinary Procedures Center, Strategic Laboratory | Claudio Tavares Sacchi, Claudia Regina Gonçalves, Erica Valessa Ramos Gomes, Karoline Rodrigues Campos |
| EPI_ISL_940626, EPI_ISL_940627 | Hospital Central Sao Caetano do Sul | Instituto Adolfo Lutz, Interdiciplinary Procedures Center, Strategic Laboratory | Claudio Tavares Sacchi, Claudia Regina Gonçalves, Erica Valessa Ramos Gomes, Karoline Rodrigues Campos |
| EPI_ISL_940628 | Unidade Mista de Iguape | Instituto Adolfo Lutz, Interdiciplinary Procedures Center, Strategic Laboratory | Claudio Tavares Sacchi, Claudia Regina Gonçalves, Erica Valessa Ramos Gomes, Karoline Rodrigues Campos |
| EPI_ISL_940629 | Hospital Municipal Josanias Castanha Braga | Instituto Adolfo Lutz, Interdiciplinary Procedures Center, Strategic Laboratory | Claudio Tavares Sacchi, Claudia Regina Gonçalves, Erica Valessa Ramos Gomes, Karoline Rodrigues Campos |
| EPI_ISL_940630 | Hospital Geral de Sao Paulo | Instituto Adolfo Lutz, Interdiciplinary Procedures Center, Strategic Laboratory | Claudio Tavares Sacchi, Claudia Regina Gonçalves, Erica Valessa Ramos Gomes, Karoline Rodrigues Campos |
| EPI_ISL_940631, EPI_ISL_940632 | University of Bari Biomedical Sciences and Human Oncology | University of Bari Biomedical Sciences and Human Oncology | Chironna M., Sallustio A., Loconsole D., Accogli M. |
| EPI_ISL_940683 | Servizo de Microbioloxía. Complexo Hospitalario Universitario de Santiago de Compostela | Servizo de Microbioloxía. Complexo Hospitalario Universitario de Santiago de Compostela | Antonio Aguilera, Gema Barbeito, Amparo Coira, Rocio Trastoy, María Luisa Pérez del Molino |
| EPI_ISL_940739 | University of Bari Biomedical Sciences and Human Oncology | University of Bari Biomedical Sciences and Human Oncology | Chironna M., Sallustio A., Loconsole D., Accogli M. |
| EPI_ISL_940890, EPI_ISL_940891 | Vaccines and Infectious Diseases Analytics Research Unit (VIDA) | KRISP, KZN Research Innovation and Sequencing Platform | Baillie Vicky, du Plessis Jeanine, Giandhari Jennifer, Pillay Sureshnee, Naidoo Yeshnee, Tegally Houriiyah, de Oliveira Tulio, Madhi Shabir |
| EPI_ISL_940996 | Sentinelles Idf | National Reference Center for Viruses of Respiratory Infections, Institut Pasteur, Paris | Marion Barbet, Sylvie Behillil, Méline Bizard, Angela Brisebarre, Camille Capel, Etienne Simon-Lorière, Vincent Enouf, Maud Vanpeene, Sylvie van der Werf, Garrigues Anne |
| EPI_ISL_941026 | Labo Analyses Med | National Reference Center for Viruses of Respiratory Infections, Institut Pasteur, Paris | Marion Barbet, Sylvie Behillil, Méline Bizard, Angela Brisebarre, Camille Capel, Etienne Simon-Lorière, Vincent Enouf, Maud Vanpeene, Sylvie van der Werf, Griscelli Franck |
| EPI_ISL_941027 | Hopital | National Reference Center for Viruses of Respiratory Infections, Institut Pasteur, Paris | Marion Barbet, Sylvie Behillil, Méline Bizard, Angela Brisebarre, Camille Capel, Etienne Simon-Lorière, Vincent Enouf, Maud Vanpeene, Sylvie van der Werf, Marque Juillet StéPhanie |
| EPI_ISL_941030, EPI_ISL_941031, EPI_ISL_941033, EPI_ISL_941034, EPI_ISL_941035, EPI_ISL_941036, EPI_ISL_941037, EPI_ISL_941038, EPI_ISL_941040, EPI_ISL_941041, EPI_ISL_941044, EPI_ISL_941045, EPI_ISL_941046, EPI_ISL_941048, EPI_ISL_941049, EPI_ISL_941050, EPI_ISL_941051, EPI_ISL_941053, EPI_ISL_941056, EPI_ISL_941057, EPI_ISL_941058, EPI_ISL_941060, EPI_ISL_941061, EPI_ISL_941063, EPI_ISL_941066, EPI_ISL_941069, EPI_ISL_941071, EPI_ISL_941073, EPI_ISL_941074, EPI_ISL_941075, EPI_ISL_941076 | | | |
| see above | Labo Analyses Med | National Reference Center for Viruses of Respiratory Infections, Institut Pasteur, Paris | Marion Barbet, Sylvie Behillil, Méline Bizard, Angela Brisebarre, Camille Capel, Etienne Simon-Lorière, Vincent Enouf, Maud Vanpeene, Sylvie van der Werf, Merah Kader |
| EPI_ISL_942927 | LESP Nuevo Leon/Grupo de Diagnostico ARIES | Instituto de Diagnostico y Referencia Epidemiologicos (INDRE) | Claudia Wong-Arambula, Abril Rodriguez-Maldonado, Fabiola Garces-Ayala, Natividad Cruz-Ortiz, Tatiana Nunez-Garcia, Gisela Barrera-Badillo, Lucia Hernandez-Rivas, Irma Lopez-Martinez, Ernesto Ramirez-Gonzalez. |
| EPI_ISL_942972, EPI_ISL_942973, EPI_ISL_942974 | Servizo de Microbioloxía. Complexo Hospitalario Universitario de Santiago de Compostela | Servizo de Microbioloxía. Complexo Hospitalario Universitario de Santiago de Compostela | Antonio Aguilera, Gema Barbeito, Amparo Coira, José Costa, Rocio Trastoy, María Luisa Pérez del Molino. |
| EPI_ISL_943550 | Baylor Scott & White-Temple | Baylor Scott & White-Temple | Ari Rao, Linden Morales, Kimberly Walker, Marcus Volz, Shelby Hendrickson |
| EPI_ISL_943556, EPI_ISL_943558, EPI_ISL_943560, EPI_ISL_943562 | Servizo de Microbioloxía. Complexo Hospitalario Universitario de Santiago de Compostela | Servizo de Microbioloxía. Complexo Hospitalario Universitario de Santiago de Compostela | Antonio Aguilera, Gema Barbeito, Amparo Coira, José Costa, Rocio Trastoy, María Luisa Pérez del Molino. |
| EPI_ISL_943570 | Laboratorio de Referencia Nacional de Virus Respiratorio. Instituto Nacional de Salud Perú | Laboratorio de Referencia Nacional de Biotecnología y Biología Molecular. Instituto Nacional de Salud Perú | Carlos Padilla Rojas, Karolyn Vega Chozo, Luis Barcena, Priscila Lope Pari, Omar Caceres Rey, Marco Galarza Perez, Maribel Huaringa Nuñez, Johanna Balbuena Torrez, Henri Bailon Calderon, Nancy Rojas Serrano |
| EPI_ISL_943967, EPI_ISL_943968, EPI_ISL_943969, EPI_ISL_943970, EPI_ISL_943971 | Hospital Geral de Sao Paulo | Instituto Adolfo Lutz, Interdiciplinary Procedures Center, Strategic Laboratory | Claudio Tavares Sacchi, Claudia Regina Gonçalves, Erica Valessa Ramos Gomes, Karoline Rodrigues Campos |
| EPI_ISL_943987 | LACEN do Estado de Tocantins | Instituto Adolfo Lutz, Interdiciplinary Procedures Center, Strategic Laboratory | Claudio Tavares Sacchi, Claudia Regina Gonçalves, Erica Valessa Ramos Gomes, Karoline Rodrigues Campos |
| EPI_ISL_943990 | LACEN do Estado de Goias | Instituto Adolfo Lutz, Interdiciplinary Procedures Center, Strategic Laboratory | Claudio Tavares Sacchi, Claudia Regina Gonçalves, Erica Valessa Ramos Gomes, Karoline Rodrigues Campos |
| EPI_ISL_944101 | Institute for Medical Research, Infectious Disease Research Centre, National Institutes of Health, Ministry of Health Malaysia | Institute for Medical Research, Infectious Disease Research Centre, National Institutes of Health, Ministry of Health Malaysia | Suppiah J, Kamel K, Azizan MA, Thayan R |
| EPI_ISL_944764, EPI_ISL_944778 | National Institute of Infectious Diseases-Prof. Dr. Matei Bals Molecular Diagnostics Laboratory | National Institute of Infectious Diseases-Prof. Dr. Matei Bals Molecular Diagnostics Laboratory | Leontina Banica, Marius Surleac, Corina Casangiu, Petre Milu, Andreea Tudor, Simona Paraschiv, Dan Otelea |
| EPI_ISL_949182 | University of Bari Biomedical Sciences and Human Oncology | University of Bari Biomedical Sciences and Human Oncology | Chironna M., Sallustio A., Loconsole D., Accogli M. |

| | | | |
|---|---|---|---|
| EPI_ISL_949184 | University of Bari Biomedical Sciences and Human Oncology | University of Bari Biomedical Sciences and Human Oncology | Chironna M., Sallustio A., Loconsole D., Accogli A. |
| EPI_ISL_949185 | University of Bari Biomedical Sciences and Human Oncology | University of Bari Biomedical Sciences and Human Oncology | Chironna M., Sallustio A., Loconsole D., Accogli M. |
| EPI_ISL_949188 | University of Bari Biomedical Sciences and Human Oncology | University of Bari Biomedical Sciences and Human Oncology | Chironna A., Sallustio A., Loconsole D., Accogli M. |
| EPI_ISL_949191 | University of Bari Biomedical Sciences and Human Oncology | University of Bari Biomedical Sciences and Human Oncology | Chironna M., Sallustio A., Loconsole D., Accogli M. |
| EPI_ISL_953942, EPI_ISL_953943, EPI_ISL_953944, EPI_ISL_953945, EPI_ISL_953946, EPI_ISL_953947, EPI_ISL_953948, EPI_ISL_953949, EPI_ISL_953950 | Hopital | National Reference Center for Viruses of Respiratory Infections, Institut Pasteur, Paris | Marion Barbet, Sylvie Behillil, Méline Bizard, Angela Brisebarre, Camille Capel, Etienne Simon-Lorière, Vincent Enouf, Maud Vanpeene, Sylvie van der Werf,Cady Anne |
| EPI_ISL_953959 | Hopital | National Reference Center for Viruses of Respiratory Infections, Institut Pasteur, Paris | Marion Barbet, Sylvie Behillil, Méline Bizard, Angela Brisebarre, Camille Capel, Etienne Simon-Lorière, Vincent Enouf, Maud Vanpeene, Sylvie van der Werf,Ramirez José-Manuel |
| EPI_ISL_953960 | Hopital | National Reference Center for Viruses of Respiratory Infections, Institut Pasteur, Paris | Marion Barbet, Sylvie Behillil, Méline Bizard, Angela Brisebarre, Camille Capel, Etienne Simon-Lorière, Vincent Enouf, Maud Vanpeene, Sylvie van der Werf,Griscelli Franck |
| EPI_ISL_953961, EPI_ISL_953962 | Hopital | National Reference Center for Viruses of Respiratory Infections, Institut Pasteur, Paris | Marion Barbet, Sylvie Behillil, Méline Bizard, Angela Brisebarre, Camille Capel, Etienne Simon-Lorière, Vincent Enouf, Maud Vanpeene, Sylvie van der Werf,Martres Pascale |
| EPI_ISL_953963 | Recherche | National Reference Center for Viruses of Respiratory Infections, Institut Pasteur, Paris | Marion Barbet, Sylvie Behillil, Méline Bizard, Angela Brisebarre, Camille Capel, Etienne Simon-Lorière, Vincent Enouf, Maud Vanpeene, Sylvie van der Werf,Leruez-Ville Marianne |
| EPI_ISL_953964 | Hopital | National Reference Center for Viruses of Respiratory Infections, Institut Pasteur, Paris | Marion Barbet, Sylvie Behillil, Méline Bizard, Angela Brisebarre, Camille Capel, Etienne Simon-Lorière, Vincent Enouf, Maud Vanpeene, Sylvie van der Werf,Ramirez José-Manuel |
| EPI_ISL_953965 | Hopital | National Reference Center for Viruses of Respiratory Infections, Institut Pasteur, Paris | Marion Barbet, Sylvie Behillil, Méline Bizard, Angela Brisebarre, Camille Capel, Etienne Simon-Lorière, Vincent Enouf, Maud Vanpeene, Sylvie van der Werf,Martres Pascale |
| EPI_ISL_953966 | Hopital | National Reference Center for Viruses of Respiratory Infections, Institut Pasteur, Paris | Marion Barbet, Sylvie Behillil, Méline Bizard, Angela Brisebarre, Camille Capel, Etienne Simon-Lorière, Vincent Enouf, Maud Vanpeene, Sylvie van der Werf,Louvetl |
| EPI_ISL_953967 | Hopital | National Reference Center for Viruses of Respiratory Infections, Institut Pasteur, Paris | Marion Barbet, Sylvie Behillil, Méline Bizard, Angela Brisebarre, Camille Capel, Etienne Simon-Lorière, Vincent Enouf, Maud Vanpeene, Sylvie van der Werf,Guinard JéRome |
| EPI_ISL_953968 | Labo Analyses Med | National Reference Center for Viruses of Respiratory Infections, Institut Pasteur, Paris | Marion Barbet, Sylvie Behillil, Méline Bizard, Angela Brisebarre, Camille Capel, Etienne Simon-Lorière, Vincent Enouf, Maud Vanpeene, Sylvie van der Werf,Le Berre (D) David |
| EPI_ISL_953969 | Hopital | National Reference Center for Viruses of Respiratory Infections, Institut Pasteur, Paris | Marion Barbet, Sylvie Behillil, Méline Bizard, Angela Brisebarre, Camille Capel, Etienne Simon-Lorière, Vincent Enouf, Maud Vanpeene, Sylvie van der Werf,Maneglie Benjamin |
| EPI_ISL_953970 | Hopital | National Reference Center for Viruses of Respiratory Infections, Institut Pasteur, Paris | Marion Barbet, Sylvie Behillil, Méline Bizard, Angela Brisebarre, Camille Capel, Etienne Simon-Lorière, Vincent Enouf, Maud Vanpeene, Sylvie van der Werf,Fourgeaud Jacques |
| EPI_ISL_953971 | Hopital | National Reference Center for Viruses of Respiratory Infections, Institut Pasteur, Paris | Marion Barbet, Sylvie Behillil, Méline Bizard, Angela Brisebarre, Camille Capel, Etienne Simon-Lorière, Vincent Enouf, Maud Vanpeene, Sylvie van der Werf,Lesimple BéAtrice |
| EPI_ISL_953972, EPI_ISL_953973 | Hopital | National Reference Center for Viruses of Respiratory Infections, Institut Pasteur, Paris | Marion Barbet, Sylvie Behillil, Méline Bizard, Angela Brisebarre, Camille Capel, Etienne Simon-Lorière, Vincent Enouf, Maud Vanpeene, Sylvie van der Werf,Eloy Clarence |
| EPI_ISL_953974, EPI_ISL_953975 | Hopital | National Reference Center for Viruses of Respiratory Infections, Institut Pasteur, Paris | Marion Barbet, Sylvie Behillil, Méline Bizard, Angela Brisebarre, Camille Capel, Etienne Simon-Lorière, Vincent Enouf, Maud Vanpeene, Sylvie van der Werf,Ducancelle Alexandra |
| EPI_ISL_953976 | Hopital | National Reference Center for Viruses of Respiratory Infections, Institut Pasteur, Paris | Marion Barbet, Sylvie Behillil, Méline Bizard, Angela Brisebarre, Camille Capel, Etienne Simon-Lorière, Vincent Enouf, Maud Vanpeene, Sylvie van der Werf,Louvetl |
| EPI_ISL_953977 | Hopital | National Reference Center for Viruses of Respiratory Infections, Institut Pasteur, Paris | Marion Barbet, Sylvie Behillil, Méline Bizard, Angela Brisebarre, Camille Capel, Etienne Simon-Lorière, Vincent Enouf, Maud Vanpeene, Sylvie van der Werf,Brichler SéGolèNe |
| EPI_ISL_953978 | Hopital | National Reference Center for Viruses of Respiratory Infections, Institut Pasteur, Paris | Marion Barbet, Sylvie Behillil, Méline Bizard, Angela Brisebarre, Camille Capel, Etienne Simon-Lorière, Vincent Enouf, Maud Vanpeene, Sylvie van der Werf,Fourgeaud Jacques |
| EPI_ISL_953979 | Hopital | National Reference Center for Viruses of Respiratory Infections, Institut Pasteur, Paris | Marion Barbet, Sylvie Behillil, Méline Bizard, Angela Brisebarre, Camille Capel, Etienne Simon-Lorière, Vincent Enouf, Maud Vanpeene, Sylvie van der Werf,Lesimple BéAtrice |
| EPI_ISL_953980, EPI_ISL_953981, EPI_ISL_953982, EPI_ISL_953983, EPI_ISL_953984 | Hopital | National Reference Center for Viruses of Respiratory Infections, Institut Pasteur, Paris | Marion Barbet, Sylvie Behillil, Méline Bizard, Angela Brisebarre, Camille Capel, Etienne Simon-Lorière, Vincent Enouf, Maud Vanpeene, Sylvie van der Werf,Ducancelle Alexandra |
| EPI_ISL_953985 | Hopital | National Reference Center for Viruses of Respiratory Infections, Institut Pasteur, Paris | Marion Barbet, Sylvie Behillil, Méline Bizard, Angela Brisebarre, Camille Capel, Etienne Simon-Lorière, Vincent Enouf, Maud Vanpeene, Sylvie van der Werf,Lagathu GisèLe |
| EPI_ISL_953986 | Labo Analyses Med | National Reference Center for Viruses of Respiratory Infections, Institut Pasteur, Paris | Marion Barbet, Sylvie Behillil, Méline Bizard, Angela Brisebarre, Camille Capel, Etienne Simon-Lorière, Vincent Enouf, Maud Vanpeene, Sylvie van der Werf |
| EPI_ISL_953987 | Hopital | National Reference Center for Viruses of Respiratory Infections, Institut Pasteur, Paris | Marion Barbet, Sylvie Behillil, Méline Bizard, Angela Brisebarre, Camille Capel, Etienne Simon-Lorière, Vincent Enouf, Maud Vanpeene, Sylvie van der Werf,Lagathu GisèLe |
| EPI_ISL_953988 | Labo Analyses Med | National Reference Center for Viruses of Respiratory Infections, Institut Pasteur, Paris | Marion Barbet, Sylvie Behillil, Méline Bizard, Angela Brisebarre, Camille Capel, Etienne Simon-Lorière, Vincent Enouf, Maud Vanpeene, Sylvie van der Werf,Selas Labomaine |
| EPI_ISL_953989 | Hopital | National Reference Center for Viruses of Respiratory Infections, Institut Pasteur, Paris | Marion Barbet, Sylvie Behillil, Méline Bizard, Angela Brisebarre, Camille Capel, Etienne Simon-Lorière, Vincent Enouf, Maud Vanpeene, Sylvie van der Werf,Lesimple BéAtrice |
| EPI_ISL_953990 | Hopital | National Reference Center for Viruses of Respiratory Infections, Institut Pasteur, Paris | Marion Barbet, Sylvie Behillil, Méline Bizard, Angela Brisebarre, Camille Capel, Etienne Simon-Lorière, Vincent Enouf, Maud Vanpeene, Sylvie van der Werf,Eloy Clarence |
| EPI_ISL_953991, EPI_ISL_953992, EPI_ISL_953993 | Hopital | National Reference Center for Viruses of Respiratory Infections, Institut Pasteur, Paris | Marion Barbet, Sylvie Behillil, Méline Bizard, Angela Brisebarre, Camille Capel, Etienne Simon-Lorière, Vincent Enouf, Maud Vanpeene, Sylvie van der Werf,Lagathu GisèLe |
| EPI_ISL_954009, EPI_ISL_954010 | Hopital | National Reference Center for Viruses of Respiratory Infections, Institut Pasteur, Paris | Marion Barbet, Sylvie Behillil, Méline Bizard, Angela Brisebarre, Camille Capel, Etienne Simon-Lorière, Vincent Enouf, Maud Vanpeene, Sylvie van der Werf,Fourgeaud Jacques |
| EPI_ISL_954011 | Hopital | National Reference Center for Viruses of Respiratory Infections, Institut Pasteur, Paris | Marion Barbet, Sylvie Behillil, Méline Bizard, Angela Brisebarre, Camille Capel, Etienne Simon-Lorière, Vincent Enouf, Maud Vanpeene, Sylvie van der Werf,Bigaillon Christine |
| EPI_ISL_954012 | Hopital | National Reference Center for Viruses of Respiratory Infections, Institut Pasteur, Paris | Marion Barbet, Sylvie Behillil, Méline Bizard, Angela Brisebarre, Camille Capel, Etienne Simon-Lorière, Vincent Enouf, Maud Vanpeene, Sylvie van der Werf,Breuil J. |
| EPI_ISL_954013 | Recherche | National Reference Center for Viruses of Respiratory Infections, Institut Pasteur, Paris | Marion Barbet, Sylvie Behillil, Méline Bizard, Angela Brisebarre, Camille Capel, Etienne Simon-Lorière, Vincent Enouf, Maud Vanpeene, Sylvie van der Werf,Leruez-Ville Marianne |

| | | | |
|---|---|---|---|
| EPI_ISL_954014 | Hopital | National Reference Center for Viruses of Respiratory Infections, Institut Pasteur, Paris | Marion Barbet, Sylvie Behillil, Méline Bizard, Angela Brisebarre, Camille Capel, Etienne Simon-Lorière, Vincent Enouf, Maud Vanpeene, Sylvie van der Werf,Farfour Eric |
| EPI_ISL_954015 | Hopital | National Reference Center for Viruses of Respiratory Infections, Institut Pasteur, Paris | Marion Barbet, Sylvie Behillil, Méline Bizard, Angela Brisebarre, Camille Capel, Etienne Simon-Lorière, Vincent Enouf, Maud Vanpeene, Sylvie van der Werf,Fourgeaud Jacques |
| EPI_ISL_954016 | Hopital | National Reference Center for Viruses of Respiratory Infections, Institut Pasteur, Paris | Marion Barbet, Sylvie Behillil, Méline Bizard, Angela Brisebarre, Camille Capel, Etienne Simon-Lorière, Vincent Enouf, Maud Vanpeene, Sylvie van der Werf,Larreche |
| EPI_ISL_954017 | Hopital | National Reference Center for Viruses of Respiratory Infections, Institut Pasteur, Paris | Marion Barbet, Sylvie Behillil, Méline Bizard, Angela Brisebarre, Camille Capel, Etienne Simon-Lorière, Vincent Enouf, Maud Vanpeene, Sylvie van der Werf,Griscelli Franck |
| EPI_ISL_954018 | Hopital | National Reference Center for Viruses of Respiratory Infections, Institut Pasteur, Paris | Marion Barbet, Sylvie Behillil, Méline Bizard, Angela Brisebarre, Camille Capel, Etienne Simon-Lorière, Vincent Enouf, Maud Vanpeene, Sylvie van der Werf,Ducancelle Alexandra |
| EPI_ISL_954019 | Hopital | National Reference Center for Viruses of Respiratory Infections, Institut Pasteur, Paris | Marion Barbet, Sylvie Behillil, Méline Bizard, Angela Brisebarre, Camille Capel, Etienne Simon-Lorière, Vincent Enouf, Maud Vanpeene, Sylvie van der Werf,Louvetl |
| EPI_ISL_954020 | Hopital | National Reference Center for Viruses of Respiratory Infections, Institut Pasteur, Paris | Marion Barbet, Sylvie Behillil, Méline Bizard, Angela Brisebarre, Camille Capel, Etienne Simon-Lorière, Vincent Enouf, Maud Vanpeene, Sylvie van der Werf,Breuil J. |
| EPI_ISL_954021 | hopital | National Reference Center for Viruses of Respiratory Infections, Institut Pasteur, Paris | Marion Barbet, Sylvie Behillil, Méline Bizard, Angela Brisebarre, Camille Capel, Etienne Simon-Lorière, Vincent Enouf, Maud Vanpeene, Sylvie van der Werf,Isabelle Joly |
| EPI_ISL_954030 | Labo Analyses Med | National Reference Center for Viruses of Respiratory Infections, Institut Pasteur, Paris | Marion Barbet, Sylvie Behillil, Méline Bizard, Angela Brisebarre, Camille Capel, Etienne Simon-Lorière, Vincent Enouf, Maud Vanpeene, Sylvie van der Werf |
| EPI_ISL_954032 | Hopital | National Reference Center for Viruses of Respiratory Infections, Institut Pasteur, Paris | Marion Barbet, Sylvie Behillil, Méline Bizard, Angela Brisebarre, Camille Capel, Etienne Simon-Lorière, Vincent Enouf, Maud Vanpeene, Sylvie van der Werf,Griscelli Franck |
| EPI_ISL_954033 | Hopital | National Reference Center for Viruses of Respiratory Infections, Institut Pasteur, Paris | Marion Barbet, Sylvie Behillil, Méline Bizard, Angela Brisebarre, Camille Capel, Etienne Simon-Lorière, Vincent Enouf, Maud Vanpeene, Sylvie van der Werf,Brichler SéGolèNe |
| EPI_ISL_954034 | Recherche | National Reference Center for Viruses of Respiratory Infections, Institut Pasteur, Paris | Marion Barbet, Sylvie Behillil, Méline Bizard, Angela Brisebarre, Camille Capel, Etienne Simon-Lorière, Vincent Enouf, Maud Vanpeene, Sylvie van der Werf,Leruez-Ville Marianne |
| EPI_ISL_954036 | Labo Analyses Med | National Reference Center for Viruses of Respiratory Infections, Institut Pasteur, Paris | Marion Barbet, Sylvie Behillil, Méline Bizard, Angela Brisebarre, Camille Capel, Etienne Simon-Lorière, Vincent Enouf, Maud Vanpeene, Sylvie van der Werf,Besson J |
| EPI_ISL_954037, EPI_ISL_954038 | Labo Analyses Med | National Reference Center for Viruses of Respiratory Infections, Institut Pasteur, Paris | Marion Barbet, Sylvie Behillil, Méline Bizard, Angela Brisebarre, Camille Capel, Etienne Simon-Lorière, Vincent Enouf, Maud Vanpeene, Sylvie van der Werf |
| EPI_ISL_954039 | Recherche | National Reference Center for Viruses of Respiratory Infections, Institut Pasteur, Paris | Marion Barbet, Sylvie Behillil, Méline Bizard, Angela Brisebarre, Camille Capel, Etienne Simon-Lorière, Vincent Enouf, Maud Vanpeene, Sylvie van der Werf,Leruez-Ville Marianne |
| EPI_ISL_954040 | Labo Analyses Med | National Reference Center for Viruses of Respiratory Infections, Institut Pasteur, Paris | Marion Barbet, Sylvie Behillil, Méline Bizard, Angela Brisebarre, Camille Capel, Etienne Simon-Lorière, Vincent Enouf, Maud Vanpeene, Sylvie van der Werf,Labm Hignard |
| EPI_ISL_954041 | Labo Analyses Med | National Reference Center for Viruses of Respiratory Infections, Institut Pasteur, Paris | Marion Barbet, Sylvie Behillil, Méline Bizard, Angela Brisebarre, Camille Capel, Etienne Simon-Lorière, Vincent Enouf, Maud Vanpeene, Sylvie van der Werf |
| EPI_ISL_954042 | Recherche | National Reference Center for Viruses of Respiratory Infections, Institut Pasteur, Paris | Marion Barbet, Sylvie Behillil, Méline Bizard, Angela Brisebarre, Camille Capel, Etienne Simon-Lorière, Vincent Enouf, Maud Vanpeene, Sylvie van der Werf,Leruez-Ville Marianne |
| EPI_ISL_954043, EPI_ISL_954044, EPI_ISL_954045, EPI_ISL_954046 | Outre Mer | National Reference Center for Viruses of Respiratory Infections, Institut Pasteur, Paris | Marion Barbet, Sylvie Behillil, Méline Bizard, Angela Brisebarre, Camille Capel, Etienne Simon-Lorière, Vincent Enouf, Maud Vanpeene, Sylvie van der Werf,Rousset Dominique |
| EPI_ISL_954047 | Hopital | National Reference Center for Viruses of Respiratory Infections, Institut Pasteur, Paris | Marion Barbet, Sylvie Behillil, Méline Bizard, Angela Brisebarre, Camille Capel, Etienne Simon-Lorière, Vincent Enouf, Maud Vanpeene, Sylvie van der Werf,Centre Hospitalier De Manosque |
| EPI_ISL_954048 | Hopital | National Reference Center for Viruses of Respiratory Infections, Institut Pasteur, Paris | Marion Barbet, Sylvie Behillil, Méline Bizard, Angela Brisebarre, Camille Capel, Etienne Simon-Lorière, Vincent Enouf, Maud Vanpeene, Sylvie van der Werf,Guinard JéRome |
| EPI_ISL_954049 | Labo Analyses Med | National Reference Center for Viruses of Respiratory Infections, Institut Pasteur, Paris | Marion Barbet, Sylvie Behillil, Méline Bizard, Angela Brisebarre, Camille Capel, Etienne Simon-Lorière, Vincent Enouf, Maud Vanpeene, Sylvie van der Werf,Nativelle |
| EPI_ISL_954050 | Labo Analyses Med | National Reference Center for Viruses of Respiratory Infections, Institut Pasteur, Paris | Marion Barbet, Sylvie Behillil, Méline Bizard, Angela Brisebarre, Camille Capel, Etienne Simon-Lorière, Vincent Enouf, Maud Vanpeene, Sylvie van der Werf,Dornic Catherine |
| EPI_ISL_954051, EPI_ISL_954052 | Labo Analyses Med | National Reference Center for Viruses of Respiratory Infections, Institut Pasteur, Paris | Marion Barbet, Sylvie Behillil, Méline Bizard, Angela Brisebarre, Camille Capel, Etienne Simon-Lorière, Vincent Enouf, Maud Vanpeene, Sylvie van der Werf |
| EPI_ISL_954053 | Recherche | National Reference Center for Viruses of Respiratory Infections, Institut Pasteur, Paris | Marion Barbet, Sylvie Behillil, Méline Bizard, Angela Brisebarre, Camille Capel, Etienne Simon-Lorière, Vincent Enouf, Maud Vanpeene, Sylvie van der Werf,Leruez-Ville Marianne |
| EPI_ISL_954054 | Labo Analyses Med | National Reference Center for Viruses of Respiratory Infections, Institut Pasteur, Paris | Marion Barbet, Sylvie Behillil, Méline Bizard, Angela Brisebarre, Camille Capel, Etienne Simon-Lorière, Vincent Enouf, Maud Vanpeene, Sylvie van der Werf,Holstein Anne |
| EPI_ISL_954055, EPI_ISL_954056, EPI_ISL_954057, EPI_ISL_954058 | Outre Mer | National Reference Center for Viruses of Respiratory Infections, Institut Pasteur, Paris | Marion Barbet, Sylvie Behillil, Méline Bizard, Angela Brisebarre, Camille Capel, Etienne Simon-Lorière, Vincent Enouf, Maud Vanpeene, Sylvie van der Werf,Rousset Dominique |
| EPI_ISL_954059 | Labo Analyses Med | National Reference Center for Viruses of Respiratory Infections, Institut Pasteur, Paris | Marion Barbet, Sylvie Behillil, Méline Bizard, Angela Brisebarre, Camille Capel, Etienne Simon-Lorière, Vincent Enouf, Maud Vanpeene, Sylvie van der Werf,Holstein Anne |
| EPI_ISL_954060, EPI_ISL_954061, EPI_ISL_954062, EPI_ISL_954063, EPI_ISL_954064, EPI_ISL_954065, EPI_ISL_954066, EPI_ISL_954067 | Labo Analyses Med | National Reference Center for Viruses of Respiratory Infections, Institut Pasteur, Paris | Marion Barbet, Sylvie Behillil, Méline Bizard, Angela Brisebarre, Camille Capel, Etienne Simon-Lorière, Vincent Enouf, Maud Vanpeene, Sylvie van der Werf,Breton-Cazaux JéRôMe |
| EPI_ISL_954068, EPI_ISL_954069 | Labo Analyses Med | National Reference Center for Viruses of Respiratory Infections, Institut Pasteur, Paris | Marion Barbet, Sylvie Behillil, Méline Bizard, Angela Brisebarre, Camille Capel, Etienne Simon-Lorière, Vincent Enouf, Maud Vanpeene, Sylvie van der Werf |
| EPI_ISL_954070 | Labo Analyses Med | National Reference Center for Viruses of Respiratory Infections, Institut Pasteur, Paris | Marion Barbet, Sylvie Behillil, Méline Bizard, Angela Brisebarre, Camille Capel, Etienne Simon-Lorière, Vincent Enouf, Maud Vanpeene, Sylvie van der Werf,Michez Karine |
| EPI_ISL_954071, EPI_ISL_954072 | Hopital | National Reference Center for Viruses of Respiratory Infections, Institut Pasteur, Paris | Marion Barbet, Sylvie Behillil, Méline Bizard, Angela Brisebarre, Camille Capel, Etienne Simon-Lorière, Vincent Enouf, Maud Vanpeene, Sylvie van der Werf,Lagathu GisèLe |
| EPI_ISL_954073, EPI_ISL_954074, EPI_ISL_954075 | Outre Mer | National Reference Center for Viruses of Respiratory Infections, Institut Pasteur, Paris | Marion Barbet, Sylvie Behillil, Méline Bizard, Angela Brisebarre, Camille Capel, Etienne Simon-Lorière, Vincent Enouf, Maud Vanpeene, Sylvie van der Werf,Rousset Dominique |
| EPI_ISL_954076, EPI_ISL_954077 | Hopital | National Reference Center for Viruses of Respiratory Infections, Institut Pasteur, Paris | Marion Barbet, Sylvie Behillil, Méline Bizard, Angela Brisebarre, Camille Capel, Etienne Simon-Lorière, Vincent Enouf, Maud Vanpeene, Sylvie van der Werf,Ramanantsoa CéLine |
| EPI_ISL_954078, EPI_ISL_954079, | Labo Analyses Med | National Reference Center for Viruses of Respiratory | Marion Barbet, Sylvie Behillil, Méline Bizard, Angela Brisebarre, Camille Capel, Etienne Simon-Lorière, Vincent Enouf, Maud Vanpeene, Sylvie van der |

| | | | |
|---|---|---|---|
| EPI_ISL_954080 | | Infections, Institut Pasteur, Paris | Werf,Takoudju Eve-Marie |
| EPI_ISL_954081 | Hopital | National Reference Center for Viruses of Respiratory Infections, Institut Pasteur, Paris | Marion Barbet, Sylvie Behillil, Méline Bizard, Angela Brisebarre, Camille Capel, Etienne Simon-Lorière, Vincent Enouf, Maud Vanpeene, Sylvie van der Werf,Cady Anne |
| EPI_ISL_954082, EPI_ISL_954083, EPI_ISL_954084 | Recherche | National Reference Center for Viruses of Respiratory Infections, Institut Pasteur, Paris | Marion Barbet, Sylvie Behillil, Méline Bizard, Angela Brisebarre, Camille Capel, Etienne Simon-Lorière, Vincent Enouf, Maud Vanpeene, Sylvie van der Werf,Leruez-Ville Marianne |
| EPI_ISL_954085, EPI_ISL_954086, EPI_ISL_954087, EPI_ISL_954088, EPI_ISL_954089, EPI_ISL_954090 | Outre Mer | National Reference Center for Viruses of Respiratory Infections, Institut Pasteur, Paris | Marion Barbet, Sylvie Behillil, Méline Bizard, Angela Brisebarre, Camille Capel, Etienne Simon-Lorière, Vincent Enouf, Maud Vanpeene, Sylvie van der Werf,Rousset Dominique |
| EPI_ISL_954091 | Labo Analyses Med | National Reference Center for Viruses of Respiratory Infections, Institut Pasteur, Paris | Marion Barbet, Sylvie Behillil, Méline Bizard, Angela Brisebarre, Camille Capel, Etienne Simon-Lorière, Vincent Enouf, Maud Vanpeene, Sylvie van der Werf,Besson J |
| EPI_ISL_954092 | Labo Analyses Med | National Reference Center for Viruses of Respiratory Infections, Institut Pasteur, Paris | Marion Barbet, Sylvie Behillil, Méline Bizard, Angela Brisebarre, Camille Capel, Etienne Simon-Lorière, Vincent Enouf, Maud Vanpeene, Sylvie van der Werf,Dlab Dieppe |
| EPI_ISL_954093 | Labo Analyses Med | National Reference Center for Viruses of Respiratory Infections, Institut Pasteur, Paris | Marion Barbet, Sylvie Behillil, Méline Bizard, Angela Brisebarre, Camille Capel, Etienne Simon-Lorière, Vincent Enouf, Maud Vanpeene, Sylvie van der Werf,Holstein Anne |
| EPI_ISL_954094, EPI_ISL_954095, EPI_ISL_954096, EPI_ISL_954097, EPI_ISL_954098, EPI_ISL_954099, EPI_ISL_954100, EPI_ISL_954101, EPI_ISL_954102 | Outre Mer | National Reference Center for Viruses of Respiratory Infections, Institut Pasteur, Paris | Marion Barbet, Sylvie Behillil, Méline Bizard, Angela Brisebarre, Camille Capel, Etienne Simon-Lorière, Vincent Enouf, Maud Vanpeene, Sylvie van der Werf,Rousset Dominique |
| EPI_ISL_954103, EPI_ISL_954104 | Hopital | National Reference Center for Viruses of Respiratory Infections, Institut Pasteur, Paris | Marion Barbet, Sylvie Behillil, Méline Bizard, Angela Brisebarre, Camille Capel, Etienne Simon-Lorière, Vincent Enouf, Maud Vanpeene, Sylvie van der Werf,Raulin Olivia |
| EPI_ISL_954105 | Labo Analyses Med | National Reference Center for Viruses of Respiratory Infections, Institut Pasteur, Paris | Marion Barbet, Sylvie Behillil, Méline Bizard, Angela Brisebarre, Camille Capel, Etienne Simon-Lorière, Vincent Enouf, Maud Vanpeene, Sylvie van der Werf,Takoudju Eve-Marie |
| EPI_ISL_954106, EPI_ISL_954107 | Hopital | National Reference Center for Viruses of Respiratory Infections, Institut Pasteur, Paris | Marion Barbet, Sylvie Behillil, Méline Bizard, Angela Brisebarre, Camille Capel, Etienne Simon-Lorière, Vincent Enouf, Maud Vanpeene, Sylvie van der Werf,Lesimple BéAtrice |
| EPI_ISL_954108, EPI_ISL_954109, EPI_ISL_954110, EPI_ISL_954111 | Outre Mer | National Reference Center for Viruses of Respiratory Infections, Institut Pasteur, Paris | Marion Barbet, Sylvie Behillil, Méline Bizard, Angela Brisebarre, Camille Capel, Etienne Simon-Lorière, Vincent Enouf, Maud Vanpeene, Sylvie van der Werf,Rousset Dominique |
| EPI_ISL_954112 | Hopital | National Reference Center for Viruses of Respiratory Infections, Institut Pasteur, Paris | Marion Barbet, Sylvie Behillil, Méline Bizard, Angela Brisebarre, Camille Capel, Etienne Simon-Lorière, Vincent Enouf, Maud Vanpeene, Sylvie van der Werf,Lesimple BéAtrice |
| EPI_ISL_954113 | Hopital | National Reference Center for Viruses of Respiratory Infections, Institut Pasteur, Paris | Marion Barbet, Sylvie Behillil, Méline Bizard, Angela Brisebarre, Camille Capel, Etienne Simon-Lorière, Vincent Enouf, Maud Vanpeene, Sylvie van der Werf,Lagathu GisèLe |
| EPI_ISL_954755, EPI_ISL_954756, EPI_ISL_954757, EPI_ISL_954758, EPI_ISL_954759, EPI_ISL_954760, EPI_ISL_954761, EPI_ISL_954762, EPI_ISL_954763, EPI_ISL_954764, EPI_ISL_954765, EPI_ISL_954766, EPI_ISL_954767, EPI_ISL_954768, EPI_ISL_954769, EPI_ISL_954771, EPI_ISL_954772, EPI_ISL_954773, EPI_ISL_954775, EPI_ISL_954777, EPI_ISL_954778, EPI_ISL_954779 | | | |
| see above | AZDelta | AZDelta | Geert Martens; Dieter De Smet |
| EPI_ISL_954782, EPI_ISL_954783 | Hospital Universitario Virgen de la Arrixaca | Instituto de Salud Carlos III | Iglesias-Caballero, M. Camarero, S. Sandonís,V. Vázquez, S. Pozo, F. Casas, I. Jiménez, P. Zaballos, A. Monzón, S. Varona, S. Cuesta, I. Moreno, L. |
| EPI_ISL_954785 | Complejo Asistencial Universitario de Salamanca | Instituto de Salud Carlos III | Iglesias-Caballero, M. Camarero, S. Sandonís,V. Vázquez, S. Pozo, F. Casas, I. Jiménez, P. Zaballos, A. Monzón, S. Varona, S. Cuesta, I. Ávila,A. |
| EPI_ISL_954787 | Hospital J.M. Morales Meseguer | Instituto de Salud Carlos III | Iglesias-Caballero, M. Camarero, S. Sandonís,V. Vázquez, S. Pozo, F. Casas, I. Jiménez, P. Zaballos, A. Monzón, S. Varona, S. Cuesta, I. Navarro, D. |
| EPI_ISL_954789 | Complejo Hospitalario Universitario La Coruña | Instituto de Salud Carlos III | Iglesias-Caballero, M. Camarero, S. Sandonís,V. Vázquez, S. Pozo, F. Casas, I. Jiménez, P. Zaballos, A. Monzón, S. Varona, S. Cuesta, I. Peña, F. |
| EPI_ISL_954791 | Complejo Hospitalario de Navarra | Instituto de Salud Carlos III | Iglesias-Caballero, M. Camarero, S. Sandonís,V. Vázquez, S. Pozo, F. Casas, I. Jiménez, P. Zaballos, A. Monzón, S. Varona, S. Cuesta, I. Ezpeleta,C. |
| EPI_ISL_954794 | Complejo Hospitalario Universitario La Coruña | Instituto de Salud Carlos III | Iglesias-Caballero, M. Camarero, S. Sandonís,V. Vázquez, S. Pozo, F. Casas, I. Jiménez, P. Zaballos, A. Monzón, S. Varona, S. Cuesta, I. Peña, F. |
| EPI_ISL_954795 | HOSPITAL GENERAL RIO CARRION | Instituto de Salud Carlos III | Iglesias-Caballero, M. Camarero, S. Sandonís,V. Vázquez, S. Pozo, F. Casas, I. Jiménez, P. Zaballos, A. Monzón, S. Varona, S. Cuesta, I. García, A. |
| EPI_ISL_954797 | Hospital Los Arcos | Instituto de Salud Carlos III | Iglesias-Caballero, M. Camarero, S. Sandonís,V. Vázquez, S. Pozo, F. Casas, I. Jiménez, P. Zaballos, A. Monzón, S. Varona, S. Cuesta, I. Camara, M. |
| EPI_ISL_954799 | Hospital Universitario Virgen de la Arrixaca | Instituto de Salud Carlos III | Iglesias-Caballero, M. Camarero, S. Sandonís,V. Vázquez, S. Pozo, F. Casas, I. Jiménez, P. Zaballos, A. Monzón, S. Varona, S. Cuesta, I. Moreno, L. |
| EPI_ISL_954800 | Complejo Hospitalario Universitario La Coruña | Instituto de Salud Carlos III | Iglesias-Caballero, M. Camarero, S. Sandonís,V. Vázquez, S. Pozo, F. Casas, I. Jiménez, P. Zaballos, A. Monzón, S. Varona, S. Cuesta, I. Peña, F. |
| EPI_ISL_954801 | Hospital Los Arcos | Instituto de Salud Carlos III | Iglesias-Caballero, M. Camarero, S. Sandonís,V. Vázquez, S. Pozo, F. Casas, I. Jiménez, P. Zaballos, A. Monzón, S. Varona, S. Cuesta, I. Camara, M. |
| EPI_ISL_954802 | Complejo Hospitalario de Navarra | Instituto de Salud Carlos III | Iglesias-Caballero, M. Camarero, S. Sandonís,V. Vázquez, S. Pozo, F. Casas, I. Jiménez, P. Zaballos, A. Monzón, S. Varona, S. Cuesta, I. Ezpeleta,C. |
| EPI_ISL_954803 | Hospital Los Arcos | Instituto de Salud Carlos III | Iglesias-Caballero, M. Camarero, S. Sandonís,V. Vázquez, S. Pozo, F. Casas, I. Jiménez, P. Zaballos, A. Monzón, S. Varona, S. Cuesta, I. Camara, M. |
| EPI_ISL_954812 | Hospital Universitario Virgen de la Arrixaca | Instituto de Salud Carlos III | Iglesias-Caballero, M. Camarero, S. Sandonís,V. Vázquez, S. Pozo, F. Casas, I. Jiménez, P. Zaballos, A. Monzón, S. Varona, S. Cuesta, I. Moreno, L. |
| EPI_ISL_954814, EPI_ISL_954815, EPI_ISL_954816, EPI_ISL_954817 | AZDelta | AZDelta | Geert Martens; Dieter De Smet |
| EPI_ISL_955061 | Hopital | National Reference Center for Viruses of Respiratory Infections, Institut Pasteur, Paris | Marion Barbet, Sylvie Behillil, Méline Bizard, Angela Brisebarre, Camille Capel, Etienne Simon-Lorière, Vincent Enouf, Maud Vanpeene, Sylvie van der Werf,Fourgeaud Jacques |
| EPI_ISL_955062, EPI_ISL_955063 | Labo analyses Med | National Reference Center for Viruses of Respiratory Infections, Institut Pasteur, Paris | Marion Barbet, Sylvie Behillil, Méline Bizard, Angela Brisebarre, Camille Capel, Etienne Simon-Lorière, Vincent Enouf, Maud Vanpeene, Sylvie van der Werf,Potiron GréGoire |
| EPI_ISL_955064 | Hopital | National Reference Center for Viruses of Respiratory Infections, Institut Pasteur, Paris | Marion Barbet, Sylvie Behillil, Méline Bizard, Angela Brisebarre, Camille Capel, Etienne Simon-Lorière, Vincent Enouf, Maud Vanpeene, Sylvie van der Werf,Guinoiseau Thibault |
| EPI_ISL_955065 | Hopital | National Reference Center for Viruses of Respiratory Infections, Institut Pasteur, Paris | Marion Barbet, Sylvie Behillil, Méline Bizard, Angela Brisebarre, Camille Capel, Etienne Simon-Lorière, Vincent Enouf, Maud Vanpeene, Sylvie van der Werf,Fourgeaud Jacques |
| EPI_ISL_955066, EPI_ISL_955067 | Labo analyses med | National Reference Center for Viruses of Respiratory Infections, Institut Pasteur, Paris | Marion Barbet, Sylvie Behillil, Méline Bizard, Angela Brisebarre, Camille Capel, Etienne Simon-Lorière, Vincent Enouf, Maud Vanpeene, Sylvie van der Werf,Amzalag Jonas |
| EPI_ISL_955068, EPI_ISL_955069, EPI_ISL_955070, EPI_ISL_955071, EPI_ISL_955072 | Labo analyses Med | National Reference Center for Viruses of Respiratory Infections, Institut Pasteur, Paris | Marion Barbet, Sylvie Behillil, Méline Bizard, Angela Brisebarre, Camille Capel, Etienne Simon-Lorière, Vincent Enouf, Maud Vanpeene, Sylvie van der Werf,Potiron GréGoire |
| EPI_ISL_955073 | Labo Analyses Med | National Reference Center for Viruses of Respiratory Infections, Institut Pasteur, Paris | Marion Barbet, Sylvie Behillil, Méline Bizard, Angela Brisebarre, Camille Capel, Etienne Simon-Lorière, Vincent Enouf, Maud Vanpeene, Sylvie van der Werf,Jacques Alexandra |
| EPI_ISL_955074, EPI_ISL_955075 | Hopital | National Reference Center for Viruses of Respiratory Infections, Institut Pasteur, Paris | Marion Barbet, Sylvie Behillil, Méline Bizard, Angela Brisebarre, Camille Capel, Etienne Simon-Lorière, Vincent Enouf, Maud Vanpeene, Sylvie van der Werf,Guinoiseau Thibault |
| EPI_ISL_955076 | Labo analyses Med | National Reference Center for Viruses of Respiratory | Marion Barbet, Sylvie Behillil, Méline Bizard, Angela Brisebarre, Camille Capel, Etienne Simon-Lorière, Vincent Enouf, Maud Vanpeene, Sylvie van der |

| | | | |
|---|---|---|---|
| | | Infections, Institut Pasteur, Paris | Werf,Potiron GréGoire |
| EPI_ISL_955077 | Hopital | National Reference Center for Viruses of Respiratory Infections, Institut Pasteur, Paris | Marion Barbet, Sylvie Behillil, Méline Bizard, Angela Brisebarre, Camille Capel, Etienne Simon-Lorière, Vincent Enouf, Maud Vanpeene, Sylvie van der Werf,Guinoiseau Thibault |
| EPI_ISL_955078 | Hopital | National Reference Center for Viruses of Respiratory Infections, Institut Pasteur, Paris | Marion Barbet, Sylvie Behillil, Méline Bizard, Angela Brisebarre, Camille Capel, Etienne Simon-Lorière, Vincent Enouf, Maud Vanpeene, Sylvie van der Werf,Fourgeaud Jacques |
| EPI_ISL_955079, EPI_ISL_955080 | Labo analyses Med | National Reference Center for Viruses of Respiratory Infections, Institut Pasteur, Paris | Marion Barbet, Sylvie Behillil, Méline Bizard, Angela Brisebarre, Camille Capel, Etienne Simon-Lorière, Vincent Enouf, Maud Vanpeene, Sylvie van der Werf,Potiron GréGoire |
| EPI_ISL_955081 | Hopital | National Reference Center for Viruses of Respiratory Infections, Institut Pasteur, Paris | Marion Barbet, Sylvie Behillil, Méline Bizard, Angela Brisebarre, Camille Capel, Etienne Simon-Lorière, Vincent Enouf, Maud Vanpeene, Sylvie van der Werf,Guinoiseau Thibault |
| EPI_ISL_955082 | Labo analyses Med | National Reference Center for Viruses of Respiratory Infections, Institut Pasteur, Paris | Marion Barbet, Sylvie Behillil, Méline Bizard, Angela Brisebarre, Camille Capel, Etienne Simon-Lorière, Vincent Enouf, Maud Vanpeene, Sylvie van der Werf,Potiron GréGoire |
| EPI_ISL_955083, EPI_ISL_955084 | Hopital | National Reference Center for Viruses of Respiratory Infections, Institut Pasteur, Paris | Marion Barbet, Sylvie Behillil, Méline Bizard, Angela Brisebarre, Camille Capel, Etienne Simon-Lorière, Vincent Enouf, Maud Vanpeene, Sylvie van der Werf,Fourgeaud Jacques |
| EPI_ISL_955085 | Hopital | National Reference Center for Viruses of Respiratory Infections, Institut Pasteur, Paris | Marion Barbet, Sylvie Behillil, Méline Bizard, Angela Brisebarre, Camille Capel, Etienne Simon-Lorière, Vincent Enouf, Maud Vanpeene, Sylvie van der Werf,Joly Isabelle |
| EPI_ISL_955086, EPI_ISL_955087, EPI_ISL_955088, EPI_ISL_955089, EPI_ISL_955090, EPI_ISL_955091, EPI_ISL_955092, EPI_ISL_955093 | Labo analyses Med | National Reference Center for Viruses of Respiratory Infections, Institut Pasteur, Paris | Marion Barbet, Sylvie Behillil, Méline Bizard, Angela Brisebarre, Camille Capel, Etienne Simon-Lorière, Vincent Enouf, Maud Vanpeene, Sylvie van der Werf,Potiron GréGoire |
| EPI_ISL_955094 | Labo Analyses Med | National Reference Center for Viruses of Respiratory Infections, Institut Pasteur, Paris | Marion Barbet, Sylvie Behillil, Méline Bizard, Angela Brisebarre, Camille Capel, Etienne Simon-Lorière, Vincent Enouf, Maud Vanpeene, Sylvie van der Werf,Potiron GréGoire |
| EPI_ISL_955095 | Hopital | National Reference Center for Viruses of Respiratory Infections, Institut Pasteur, Paris | Marion Barbet, Sylvie Behillil, Méline Bizard, Angela Brisebarre, Camille Capel, Etienne Simon-Lorière, Vincent Enouf, Maud Vanpeene, Sylvie van der Werf,Joly Isabelle |
| EPI_ISL_955096, EPI_ISL_955097, EPI_ISL_955098, EPI_ISL_955099, EPI_ISL_955100 | Hopital | National Reference Center for Viruses of Respiratory Infections, Institut Pasteur, Paris | Marion Barbet, Sylvie Behillil, Méline Bizard, Angela Brisebarre, Camille Capel, Etienne Simon-Lorière, Vincent Enouf, Maud Vanpeene, Sylvie van der Werf,Guinoiseau Thibault |
| EPI_ISL_955176 | Hopital | National Reference Center for Viruses of Respiratory Infections, Institut Pasteur, Paris | Marion Barbet, Sylvie Behillil, Méline Bizard, Angela Brisebarre, Camille Capel, Etienne Simon-Lorière, Vincent Enouf, Maud Vanpeene, Sylvie van der Werf,Brichler SéGolèNe |
| EPI_ISL_955177 | Hopital | National Reference Center for Viruses of Respiratory Infections, Institut Pasteur, Paris | Marion Barbet, Sylvie Behillil, Méline Bizard, Angela Brisebarre, Camille Capel, Etienne Simon-Lorière, Vincent Enouf, Maud Vanpeene, Sylvie van der Werf,Fourgeaud Jacques |
| EPI_ISL_955181 | Labo Analyses Med | National Reference Center for Viruses of Respiratory Infections, Institut Pasteur, Paris | Marion Barbet, Sylvie Behillil, Méline Bizard, Angela Brisebarre, Camille Capel, Etienne Simon-Lorière, Vincent Enouf, Maud Vanpeene, Sylvie van der Werf,Amzalag Jonas |
| EPI_ISL_955183 | Labo Analyses Med | National Reference Center for Viruses of Respiratory Infections, Institut Pasteur, Paris | Marion Barbet, Sylvie Behillil, Méline Bizard, Angela Brisebarre, Camille Capel, Etienne Simon-Lorière, Vincent Enouf, Maud Vanpeene, Sylvie van der Werf,Le Berre (D) David |
| EPI_ISL_955187 | University of Sarajevo, Veterinary Faculty, Laboratory for Molecular Diagnostic and Research Laboratory | University of Sarajevo, Veterinary Faculty, Laboratory for Molecular Diagnostic and Research Laboratory | Goleti Š., Goleti T., Ali-Šeho A., Softi A., Hodži A., Terzi I., Šabi E., Jaži A., Nicevi M. |
| EPI_ISL_955188 | University of Sarajevo, Veterinary Faculty, Laboratory for Molecular Diagnostic and Research Laboratory | University of Sarajevo, Veterinary Faculty, Laboratory for Molecular Diagnostic and Research Laboratory | Goleti Š., Goleti T., Ali-Šeho A., Softi A., Jaži A., Terzi I., Nicevi M., Šabi E., Hodži A. |
| EPI_ISL_955210 | University of Sarajevo, Veterinary Faculty, Laboratory for Molecular Diagnostic and Research Laboratory | University of Sarajevo, Veterinary Faculty, Laboratory for Molecular Diagnostic and Research Laboratory | Goleti Š., Goleti T., Ali-Šeho A., Softi A., Hodži A., Terzi I., Jaži A., Nicevi M., Šabi E. |
| EPI_ISL_955229 | University of Sarajevo, Veterinary Faculty, Laboratory for Molecular Diagnostic and Research Laboratory | University of Sarajevo, Veterinary Faculty, Laboratory for Molecular Diagnostic and Research Laboratory | Goleti Š., Goleti T., Ali-Šeho A., Softi A., Nicevi M., Terzi I., Šabi E., Hodži A., Jaži A. |
| EPI_ISL_956285 | Instituto Nacional de Salud- Dirección de Redes de Laboratorios de Salud Pública | Instituto Nacional de Salud- Dirección de Investigación en Salud Pública | Katherine Laiton-Donato, Diego A. Álvarez-Díaz, Carlos Franco-Muñoz, Mauricio Pacheco-Montealegre, Hector Alejandro Ruiz-Moreno, Maria T. Herrera-Sepúlveda, Diego Andrés Prada, Jhonnatan Reales-González, Sheryll Corchuelo, Julian Naizaque, Gerardo Santamaría, Magdalena Wiesner, Martha Lucia Ospina Martinez, Marcela Mercado-Reyes |
| EPI_ISL_960406, EPI_ISL_960420, EPI_ISL_960421, EPI_ISL_960422, EPI_ISL_960427, EPI_ISL_960428 | The National Institute of Public Health | State Veterinary Institute Prague | Nagy,A;Vecerova,J;Cernikova,L;Stara,M;Jirincova,H;Trnka,D |
| EPI_ISL_961066 | SIESP CHIETI - DRIVE IN LANCIANO | Istituto Zooprofilattico Sperimentale dell'Abruzzo e Molise "G. Caporale" | Lorusso A, Marcacci M, Di Domenico M, Ancora M, Curini V, Mangone I, Rinaldi A, Scialabba S, Di Pasquale A, Cammà C, Puglia I, Calistri P, Savini G |
| EPI_ISL_961067, EPI_ISL_961068 | SIESP DIPARTIMENTO DI PREVENZIONE CHIETI | Istituto Zooprofilattico Sperimentale dell'Abruzzo e Molise "G. Caporale" | Lorusso A, Marcacci M, Di Domenico M, Ancora M, Curini V, Mangone I, Rinaldi A, Scialabba S, Di Pasquale A, Cammà C, Puglia I, Calistri P, Savini G |
| EPI_ISL_961069 | SIESP CHIETI - DRIVE IN CHIETI | Istituto Zooprofilattico Sperimentale dell'Abruzzo e Molise "G. Caporale" | Lorusso A, Marcacci M, Di Domenico M, Ancora M, Curini V, Mangone I, Rinaldi A, Scialabba S, Di Pasquale A, Cammà C, Puglia I, Calistri P, Savini G |
| EPI_ISL_961070 | SIESP DIPARTIMENTO DI PREVENZIONE CHIETI | Istituto Zooprofilattico Sperimentale dell'Abruzzo e Molise "G. Caporale" | Lorusso A, Marcacci M, Di Domenico M, Ancora M, Curini V, Mangone I, Rinaldi A, Scialabba S, Di Pasquale A, Cammà C, Puglia I, Calistri P, Savini G |
| EPI_ISL_961071, EPI_ISL_961072, EPI_ISL_961073 | SIESP CHIETI - DRIVE IN CHIETI | Istituto Zooprofilattico Sperimentale dell'Abruzzo e Molise "G. Caporale" | Lorusso A, Marcacci M, Di Domenico M, Ancora M, Curini V, Mangone I, Rinaldi A, Scialabba S, Di Pasquale A, Cammà C, Puglia I, Calistri P, Savini G |
| EPI_ISL_961074 | SIESP CHIETI - DRIVE IN ORTONA | Istituto Zooprofilattico Sperimentale dell'Abruzzo e Molise "G. Caporale" | Lorusso A, Marcacci M, Di Domenico M, Ancora M, Curini V, Mangone I, Rinaldi A, Scialabba S, Di Pasquale A, Cammà C, Puglia I, Calistri P, Savini G |
| EPI_ISL_961075 | SIESP DIPARTIMENTO DI PREVENZIONE CHIETI | Istituto Zooprofilattico Sperimentale dell'Abruzzo e Molise "G. Caporale" | Lorusso A, Marcacci M, Di Domenico M, Ancora M, Curini V, Mangone I, Rinaldi A, Scialabba S, Di Pasquale A, Cammà C, Puglia I, Calistri P, Savini G |
| EPI_ISL_961076 | SIESP CHIETI - DRIVE IN ORTONA | Istituto Zooprofilattico Sperimentale dell'Abruzzo e Molise "G. Caporale" | Lorusso A, Marcacci M, Di Domenico M, Ancora M, Curini V, Mangone I, Rinaldi A, Scialabba S, Di Pasquale A, Cammà C, Puglia I, Calistri P, Savini G |
| EPI_ISL_961077 | SIESP TERAMO | Istituto Zooprofilattico Sperimentale dell'Abruzzo e Molise "G. Caporale" | Lorusso A, Marcacci M, Di Domenico M, Ancora M, Curini V, Mangone I, Rinaldi A, Scialabba S, Di Pasquale A, Cammà C, Puglia I, Calistri P, Savini G |
| EPI_ISL_961078, EPI_ISL_961079, EPI_ISL_961080 | DIP. PREV. AVEZZANO SERVIZIO DI IGIENE EPIDEMIOLOGIAE SANITA' PUBBLICA | Istituto Zooprofilattico Sperimentale dell'Abruzzo e Molise "G. Caporale" | Lorusso A, Marcacci M, Di Domenico M, Ancora M, Curini V, Mangone I, Rinaldi A, Scialabba S, Di Pasquale A, Cammà C, Puglia I, Calistri P, Savini G |
| EPI_ISL_961081 | SIESP CHIETI - DRIVE IN CHIETI | Istituto Zooprofilattico Sperimentale dell'Abruzzo e Molise "G. Caporale" | Lorusso A, Marcacci M, Di Domenico M, Ancora M, Curini V, Mangone I, Rinaldi A, Scialabba S, Di Pasquale A, Cammà C, Puglia I, Calistri P, Savini G |
| EPI_ISL_961082 | SIESP DIPARTIMENTO DI PREVENZIONE CHIETI | Istituto Zooprofilattico Sperimentale dell'Abruzzo e Molise "G. | Lorusso A, Marcacci M, Di Domenico M, Ancora M, Curini V, Mangone I, Rinaldi A, Scialabba S, Di Pasquale A, Cammà C, Puglia I, Calistri P, Savini G |

| | | | |
|---|---|---|---|
| | | Caporale" | |
| EPI_ISL_961083 | DIP. PREV. AVEZZANO SERVIZIO DI IGIENE EPIDEMIOLOGIAE SANITA' PUBBLICA | Istituto Zooprofilattico Sperimentale dell'Abruzzo e Molise "G. Caporale" | Lorusso A, Marcacci M, Di Domenico M, Ancora M, Curini V, Mangone I, Rinaldi A, Scialabba S, Di Pasquale A, Cammà C, Puglia I, Calistri P, Savini G |
| EPI_ISL_961084 | SIESP TERAMO | Istituto Zooprofilattico Sperimentale dell'Abruzzo e Molise "G. Caporale" | Lorusso A, Marcacci M, Di Domenico M, Ancora M, Curini V, Mangone I, Rinaldi A, Scialabba S, Di Pasquale A, Cammà C, Puglia I, Calistri P, Savini G |
| EPI_ISL_961085 | SIESP CHIETI - DRIVE IN ORTONA | Istituto Zooprofilattico Sperimentale dell'Abruzzo e Molise "G. Caporale" | Lorusso A, Marcacci M, Di Domenico M, Ancora M, Curini V, Mangone I, Rinaldi A, Scialabba S, Di Pasquale A, Cammà C, Puglia I, Calistri P, Savini G |
| EPI_ISL_961086 | SIESP CHIETI - DRIVE IN CHIETI | Istituto Zooprofilattico Sperimentale dell'Abruzzo e Molise "G. Caporale" | Lorusso A, Marcacci M, Di Domenico M, Ancora M, Curini V, Mangone I, Rinaldi A, Scialabba S, Di Pasquale A, Cammà C, Puglia I, Calistri P, Savini G |
| EPI_ISL_961087, EPI_ISL_961088, EPI_ISL_961089 | SIESP DIPARTIMENTO DI PREVENZIONE CHIETI | Istituto Zooprofilattico Sperimentale dell'Abruzzo e Molise "G. Caporale" | Lorusso A, Marcacci M, Di Domenico M, Ancora M, Curini V, Mangone I, Rinaldi A, Scialabba S, Di Pasquale A, Cammà C, Puglia I, Calistri P, Savini G |
| EPI_ISL_961090, EPI_ISL_961091, EPI_ISL_961092 | SIESP CHIETI - DRIVE IN CHIETI | Istituto Zooprofilattico Sperimentale dell'Abruzzo e Molise "G. Caporale" | Lorusso A, Marcacci M, Di Domenico M, Ancora M, Curini V, Mangone I, Rinaldi A, Scialabba S, Di Pasquale A, Cammà C, Puglia I, Calistri P, Savini G |
| EPI_ISL_961093 | Ospedale SS Annunziata | Istituto Zooprofilattico Sperimentale dell'Abruzzo e Molise "G. Caporale" | Lorusso A, Marcacci M, Di Domenico M, Ancora M, Curini V, Mangone I, Rinaldi A, Scialabba S, Di Pasquale A, Cammà C, Puglia I, Calistri P, Savini G |
| EPI_ISL_961094, EPI_ISL_961095 | DIP. PREV. AVEZZANO SERVIZIO DI IGIENE EPIDEMIOLOGIAE SANITA' PUBBLICA | Istituto Zooprofilattico Sperimentale dell'Abruzzo e Molise "G. Caporale" | Lorusso A, Marcacci M, Di Domenico M, Ancora M, Curini V, Mangone I, Rinaldi A, Scialabba S, Di Pasquale A, Cammà C, Puglia I, Calistri P, Savini G |
| EPI_ISL_961097, EPI_ISL_961098, EPI_ISL_961099, EPI_ISL_961100, EPI_ISL_961101, EPI_ISL_961102, EPI_ISL_961103 | SIESP TERAMO | Istituto Zooprofilattico Sperimentale dell'Abruzzo e Molise "G. Caporale" | Lorusso A, Marcacci M, Di Domenico M, Ancora M, Curini V, Mangone I, Rinaldi A, Scialabba S, Di Pasquale A, Cammà C, Puglia I, Calistri P, Savini G |
| EPI_ISL_961467, EPI_ISL_961468, EPI_ISL_961469 | LESP Nuevo Leon/Grupo de Diagnostico ARIES | Instituto de Diagnostico y Referencia Epidemiologicos (INDRE) | Claudia Wong-Arambula, Abril Rodriguez-Maldonado, Fabiola Garces-Ayala, Natividad Cruz-Ortiz, Tatiana Nunez-Garcia, Gisela Barrera-Badillo, Lucia Hernandez-Rivas, Irma Lopez-Martinez, Ernesto Ramirez-Gonzalez. |
| EPI_ISL_961710, EPI_ISL_961711 | SIESP CHIETI - DRIVE IN CHIETI | Istituto Zooprofilattico Sperimentale dell'Abruzzo e Molise "G. Caporale" | Lorusso A, Marcacci M, Di Domenico M, Ancora M, Curini V, Mangone I, Rinaldi A, Scialabba S, Di Pasquale A, Cammà C, Puglia I, Calistri P, Savini G |
| EPI_ISL_961712, EPI_ISL_961713 | SIESP CHIETI DRIVE IN LANCIANO | Istituto Zooprofilattico Sperimentale dell'Abruzzo e Molise "G. Caporale" | Lorusso A, Marcacci M, Di Domenico M, Ancora M, Curini V, Mangone I, Rinaldi A, Scialabba S, Di Pasquale A, Cammà C, Puglia I, Calistri P, Savini G |
| EPI_ISL_961714 | SIESP CHIETI - DRIVE IN CHIETI | Istituto Zooprofilattico Sperimentale dell'Abruzzo e Molise "G. Caporale" | Lorusso A, Marcacci M, Di Domenico M, Ancora M, Curini V, Mangone I, Rinaldi A, Scialabba S, Di Pasquale A, Cammà C, Puglia I, Calistri P, Savini G |
| EPI_ISL_961715, EPI_ISL_961716 | SIESP CHIETI - DRIVE IN ORTONA | Istituto Zooprofilattico Sperimentale dell'Abruzzo e Molise "G. Caporale" | Lorusso A, Marcacci M, Di Domenico M, Ancora M, Curini V, Mangone I, Rinaldi A, Scialabba S, Di Pasquale A, Cammà C, Puglia I, Calistri P, Savini G |
| EPI_ISL_961717 | SIESP CHIETI - DRIVE IN CHIETI | Istituto Zooprofilattico Sperimentale dell'Abruzzo e Molise "G. Caporale" | Lorusso A, Marcacci M, Di Domenico M, Ancora M, Curini V, Mangone I, Rinaldi A, Scialabba S, Di Pasquale A, Cammà C, Puglia I, Calistri P, Savini G |
| EPI_ISL_961718, EPI_ISL_961719, EPI_ISL_961720, EPI_ISL_961721, EPI_ISL_961722 | SIESP CHIETI - DRIVE IN ORTONA | Istituto Zooprofilattico Sperimentale dell'Abruzzo e Molise "G. Caporale" | Lorusso A, Marcacci M, Di Domenico M, Ancora M, Curini V, Mangone I, Rinaldi A, Scialabba S, Di Pasquale A, Cammà C, Puglia I, Calistri P, Savini G |
| EPI_ISL_961723 | SIESP CHIETI DRIVE IN LANCIANO | Istituto Zooprofilattico Sperimentale dell'Abruzzo e Molise "G. Caporale" | Lorusso A, Marcacci M, Di Domenico M, Ancora M, Curini V, Mangone I, Rinaldi A, Scialabba S, Di Pasquale A, Cammà C, Puglia I, Calistri P, Savini G |
| EPI_ISL_961724 | SIESP CHIETI - DRIVE IN ORTONA | Istituto Zooprofilattico Sperimentale dell'Abruzzo e Molise "G. Caporale" | Lorusso A, Marcacci M, Di Domenico M, Ancora M, Curini V, Mangone I, Rinaldi A, Scialabba S, Di Pasquale A, Cammà C, Puglia I, Calistri P, Savini G |
| EPI_ISL_961725 | SIESP CHIETI DRIVE IN GISSI | Istituto Zooprofilattico Sperimentale dell'Abruzzo e Molise "G. Caporale" | Lorusso A, Marcacci M, Di Domenico M, Ancora M, Curini V, Mangone I, Rinaldi A, Scialabba S, Di Pasquale A, Cammà C, Puglia I, Calistri P, Savini G |
| EPI_ISL_961726, EPI_ISL_961727 | SIESP CHIETI - DRIVE IN CHIETI | Istituto Zooprofilattico Sperimentale dell'Abruzzo e Molise "G. Caporale" | Lorusso A, Marcacci M, Di Domenico M, Ancora M, Curini V, Mangone I, Rinaldi A, Scialabba S, Di Pasquale A, Cammà C, Puglia I, Calistri P, Savini G |
| EPI_ISL_961728 | SIESP DIPARTIMENTO DI PREVENZIONE CHIETI | Istituto Zooprofilattico Sperimentale dell'Abruzzo e Molise "G. Caporale" | Lorusso A, Marcacci M, Di Domenico M, Ancora M, Curini V, Mangone I, Rinaldi A, Scialabba S, Di Pasquale A, Cammà C, Puglia I, Calistri P, Savini G |
| EPI_ISL_961729 | SIESP CHIETI - DRIVE IN CHIETI | Istituto Zooprofilattico Sperimentale dell'Abruzzo e Molise "G. Caporale" | Lorusso A, Marcacci M, Di Domenico M, Ancora M, Curini V, Mangone I, Rinaldi A, Scialabba S, Di Pasquale A, Cammà C, Puglia I, Calistri P, Savini G |
| EPI_ISL_961730 | Dipartimento Prevenzione Avezzano-Servizio Igiene epidemiologia Sanità Pubblica | Istituto Zooprofilattico Sperimentale dell'Abruzzo e Molise "G. Caporale" | Lorusso A, Marcacci M, Di Domenico M, Ancora M, Curini V, Mangone I, Rinaldi A, Scialabba S, Di Pasquale A, Cammà C, Puglia I, Calistri P, Savini G |
| EPI_ISL_961731, EPI_ISL_961732 | SIESP CHIETI - DRIVE IN CHIETI | Istituto Zooprofilattico Sperimentale dell'Abruzzo e Molise "G. Caporale" | Lorusso A, Marcacci M, Di Domenico M, Ancora M, Curini V, Mangone I, Rinaldi A, Scialabba S, Di Pasquale A, Cammà C, Puglia I, Calistri P, Savini G |
| EPI_ISL_961733 | SIESP CHIETI - DRIVE IN ORTONA | Istituto Zooprofilattico Sperimentale dell'Abruzzo e Molise "G. Caporale" | Lorusso A, Marcacci M, Di Domenico M, Ancora M, Curini V, Mangone I, Rinaldi A, Scialabba S, Di Pasquale A, Cammà C, Puglia I, Calistri P, Savini G |
| EPI_ISL_961734 | Dipartimento Prevenzione Avezzano-Servizio Igiene epidemiologia Sanità Pubblica | Istituto Zooprofilattico Sperimentale dell'Abruzzo e Molise "G. Caporale" | Lorusso A, Marcacci M, Di Domenico M, Ancora M, Curini V, Mangone I, Rinaldi A, Scialabba S, Di Pasquale A, Cammà C, Puglia I, Calistri P, Savini G |
| EPI_ISL_961735, EPI_ISL_961736 | SIESP CHIETI - DRIVE IN CHIETI | Istituto Zooprofilattico Sperimentale dell'Abruzzo e Molise "G. Caporale" | Lorusso A, Marcacci M, Di Domenico M, Ancora M, Curini V, Mangone I, Rinaldi A, Scialabba S, Di Pasquale A, Cammà C, Puglia I, Calistri P, Savini G |
| EPI_ISL_961737 | SIESP CHIETI- DRIVE IN VASTO | Istituto Zooprofilattico Sperimentale dell'Abruzzo e Molise "G. Caporale" | Lorusso A, Marcacci M, Di Domenico M, Ancora M, Curini V, Mangone I, Rinaldi A, Scialabba S, Di Pasquale A, Cammà C, Puglia I, Calistri P, Savini G |
| EPI_ISL_961738, EPI_ISL_961739 | SIESP CHIETI DRIVE IN LANCIANO | Istituto Zooprofilattico Sperimentale dell'Abruzzo e Molise "G. Caporale" | Lorusso A, Marcacci M, Di Domenico M, Ancora M, Curini V, Mangone I, Rinaldi A, Scialabba S, Di Pasquale A, Cammà C, Puglia I, Calistri P, Savini G |
| EPI_ISL_961740 | SIESP CHIETI - DRIVE IN CHIETI | Istituto Zooprofilattico Sperimentale dell'Abruzzo e Molise "G. Caporale" | Lorusso A, Marcacci M, Di Domenico M, Ancora M, Curini V, Mangone I, Rinaldi A, Scialabba S, Di Pasquale A, Cammà C, Puglia I, Calistri P, Savini G |
| EPI_ISL_961741 | SIESP CHIETI- DRIVE IN VASTO | Istituto Zooprofilattico Sperimentale dell'Abruzzo e Molise "G. Caporale" | Lorusso A, Marcacci M, Di Domenico M, Ancora M, Curini V, Mangone I, Rinaldi A, Scialabba S, Di Pasquale A, Cammà C, Puglia I, Calistri P, Savini G |
| EPI_ISL_961742 | SIESP CHIETI - DRIVE IN CHIETI | Istituto Zooprofilattico Sperimentale dell'Abruzzo e Molise "G. Caporale" | Lorusso A, Marcacci M, Di Domenico M, Ancora M, Curini V, Mangone I, Rinaldi A, Scialabba S, Di Pasquale A, Cammà C, Puglia I, Calistri P, Savini G |
| EPI_ISL_961743 | SIESP CHIETI DRIVE IN GISSI | Istituto Zooprofilattico Sperimentale dell'Abruzzo e Molise "G. Caporale" | Lorusso A, Marcacci M, Di Domenico M, Ancora M, Curini V, Mangone I, Rinaldi A, Scialabba S, Di Pasquale A, Cammà C, Puglia I, Calistri P, Savini G |
| EPI_ISL_961744 | SIESP CHIETI - DRIVE IN ORTONA | Istituto Zooprofilattico Sperimentale dell'Abruzzo e Molise "G. | Lorusso A, Marcacci M, Di Domenico M, Ancora M, Curini V, Mangone I, Rinaldi A, Scialabba S, Di Pasquale A, Cammà C, Puglia I, Calistri P, Savini G |

| | | | |
|---|---|---|---|
| | | Caporale" | |
| EPI_ISL_961745 | SIESP CHIETI DRIVE IN GISSI | Istituto Zooprofilattico Sperimentale dell'Abruzzo e Molise "G. Caporale" | Lorusso A, Marcacci M, Di Domenico M, Ancora M, Curini V, Mangone I, Rinaldi A, Scialabba S, Di Pasquale A, Cammà C, Puglia I, Calistri P, Savini G |
| EPI_ISL_961746 | SIESP CHIETI - DRIVE IN CHIETI | Istituto Zooprofilattico Sperimentale dell'Abruzzo e Molise "G. Caporale" | Lorusso A, Marcacci M, Di Domenico M, Ancora M, Curini V, Mangone I, Rinaldi A, Scialabba S, Di Pasquale A, Cammà C, Puglia I, Calistri P, Savini G |
| EPI_ISL_961747 | Dipartimento Prevenzione Avezzano-Servizio Igiene epidemiologia Sanità Pubblica | Istituto Zooprofilattico Sperimentale dell'Abruzzo e Molise "G. Caporale" | Lorusso A, Marcacci M, Di Domenico M, Ancora M, Curini V, Mangone I, Rinaldi A, Scialabba S, Di Pasquale A, Cammà C, Puglia I, Calistri P, Savini G |
| EPI_ISL_961748 | SIESP CHIETI DRIVE IN LANCIANO | Istituto Zooprofilattico Sperimentale dell'Abruzzo e Molise "G. Caporale" | Lorusso A, Marcacci M, Di Domenico M, Ancora M, Curini V, Mangone I, Rinaldi A, Scialabba S, Di Pasquale A, Cammà C, Puglia I, Calistri P, Savini G |
| EPI_ISL_961749 | SIESP DIPARTIMENTO DI PREVENZIONE CHIETI | Istituto Zooprofilattico Sperimentale dell'Abruzzo e Molise "G. Caporale" | Lorusso A, Marcacci M, Di Domenico M, Ancora M, Curini V, Mangone I, Rinaldi A, Scialabba S, Di Pasquale A, Cammà C, Puglia I, Calistri P, Savini G |
| EPI_ISL_961750 | Ospedale Cvile Atri Med. Interna | Istituto Zooprofilattico Sperimentale dell'Abruzzo e Molise "G. Caporale" | Lorusso A, Marcacci M, Di Domenico M, Ancora M, Curini V, Mangone I, Rinaldi A, Scialabba S, Di Pasquale A, Cammà C, Puglia I, Calistri P, Savini G |
| EPI_ISL_961751 | SIESP DIPARTIMENTO DI PREVENZIONE CHIETI | Istituto Zooprofilattico Sperimentale dell'Abruzzo e Molise "G. Caporale" | Lorusso A, Marcacci M, Di Domenico M, Ancora M, Curini V, Mangone I, Rinaldi A, Scialabba S, Di Pasquale A, Cammà C, Puglia I, Calistri P, Savini G |
| EPI_ISL_961752, EPI_ISL_961753 | SIESP SULMONA | Istituto Zooprofilattico Sperimentale dell'Abruzzo e Molise "G. Caporale" | Lorusso A, Marcacci M, Di Domenico M, Ancora M, Curini V, Mangone I, Rinaldi A, Scialabba S, Di Pasquale A, Cammà C, Puglia I, Calistri P, Savini G |
| EPI_ISL_961754 | Presidio Ospedaliero Sulmona | Istituto Zooprofilattico Sperimentale dell'Abruzzo e Molise "G. Caporale" | Lorusso A, Marcacci M, Di Domenico M, Ancora M, Curini V, Mangone I, Rinaldi A, Scialabba S, Di Pasquale A, Cammà C, Puglia I, Calistri P, Savini G |
| EPI_ISL_961755, EPI_ISL_961756, EPI_ISL_961757, EPI_ISL_961758 | SIESP SULMONA | Istituto Zooprofilattico Sperimentale dell'Abruzzo e Molise "G. Caporale" | Lorusso A, Marcacci M, Di Domenico M, Ancora M, Curini V, Mangone I, Rinaldi A, Scialabba S, Di Pasquale A, Cammà C, Puglia I, Calistri P, Savini G |
| EPI_ISL_962205, EPI_ISL_962207, EPI_ISL_962523, EPI_ISL_962524, EPI_ISL_962525, EPI_ISL_962526, EPI_ISL_962527 | Institute for Medical Research, Infectious Disease Research Centre, National Institutes of Health, Ministry of Health Malaysia | Institute for Medical Research, Infectious Disease Research Centre, National Institutes of Health, Ministry of Health Malaysia | Suppiah J, Kamel K, Azizan MA, Thayan R |
| EPI_ISL_965028, EPI_ISL_965031, EPI_ISL_965114 | Laboratorio Biologia Molecolare Sars Cov2 - UOC Laboratorio Analisi - Servizio Medicina di Laboratorio, Ospedale "San Francesco" - ATS-ASSL Nuoro | Laboratorio specialistico UOC Ematologia - Ospedale "San Francesco" - ATS-ASSL Nuoro | Piras Giovanna, Asproni Rosanna, Malune Paolo, Fiamma Maura, Monne Maria Itria, Palmas Angelo Domenico, Lo Maglio Iana, Mameli Giuseppe |
| EPI_ISL_965253 | SIESP CHIETI | Istituto Zooprofilattico Sperimentale dell'Abruzzo e Molise "G. Caporale" | Lorusso A, Marcacci M, Di Domenico M, Ancora M, Curini V, Mangone I, Rinaldi A, Scialabba S, Di Pasquale A, Cammà C, Puglia I, Calistri P, Savini G |
| EPI_ISL_965254, EPI_ISL_965255 | SIESP CHIETI - DRIVE IN CHIETI | Istituto Zooprofilattico Sperimentale dell'Abruzzo e Molise "G. Caporale" | Lorusso A, Marcacci M, Di Domenico M, Ancora M, Curini V, Mangone I, Rinaldi A, Scialabba S, Di Pasquale A, Cammà C, Puglia I, Calistri P, Savini G |
| EPI_ISL_965256 | SIESP CHIETI-DRIVE IN VASTO | Istituto Zooprofilattico Sperimentale dell'Abruzzo e Molise "G. Caporale" | Lorusso A, Marcacci M, Di Domenico M, Ancora M, Curini V, Mangone I, Rinaldi A, Scialabba S, Di Pasquale A, Cammà C, Puglia I, Calistri P, Savini G |
| EPI_ISL_965257 | SIESP CHIETI-DRIVE IN GISSI | Istituto Zooprofilattico Sperimentale dell'Abruzzo e Molise "G. Caporale" | Lorusso A, Marcacci M, Di Domenico M, Ancora M, Curini V, Mangone I, Rinaldi A, Scialabba S, Di Pasquale A, Cammà C, Puglia I, Calistri P, Savini G |
| EPI_ISL_965258, EPI_ISL_965259 | SIESP CHIETI-DRIVE IN VASTO | Istituto Zooprofilattico Sperimentale dell'Abruzzo e Molise "G. Caporale" | Lorusso A, Marcacci M, Di Domenico M, Ancora M, Curini V, Mangone I, Rinaldi A, Scialabba S, Di Pasquale A, Cammà C, Puglia I, Calistri P, Savini G |
| EPI_ISL_965260 | SIESP CHIETI-DRIVE IN GISSI | Istituto Zooprofilattico Sperimentale dell'Abruzzo e Molise "G. Caporale" | Lorusso A, Marcacci M, Di Domenico M, Ancora M, Curini V, Mangone I, Rinaldi A, Scialabba S, Di Pasquale A, Cammà C, Puglia I, Calistri P, Savini G |
| EPI_ISL_965261 | SIESP CHIETI | Istituto Zooprofilattico Sperimentale dell'Abruzzo e Molise "G. Caporale" | Lorusso A, Marcacci M, Di Domenico M, Ancora M, Curini V, Mangone I, Rinaldi A, Scialabba S, Di Pasquale A, Cammà C, Puglia I, Calistri P, Savini G |
| EPI_ISL_965262 | SIESP CHIETI, DISTRETTO SAN CHIETI | Istituto Zooprofilattico Sperimentale dell'Abruzzo e Molise "G. Caporale" | Lorusso A, Marcacci M, Di Domenico M, Ancora M, Curini V, Mangone I, Rinaldi A, Scialabba S, Di Pasquale A, Cammà C, Puglia I, Calistri P, Savini G |
| EPI_ISL_965263 | SIESP CHIETI-DRIVE IN ORTONA | Istituto Zooprofilattico Sperimentale dell'Abruzzo e Molise "G. Caporale" | Lorusso A, Marcacci M, Di Domenico M, Ancora M, Curini V, Mangone I, Rinaldi A, Scialabba S, Di Pasquale A, Cammà C, Puglia I, Calistri P, Savini G |
| EPI_ISL_965265 | SIESP CHIETI - DRIVE IN CHIETI | Istituto Zooprofilattico Sperimentale dell'Abruzzo e Molise "G. Caporale" | Lorusso A, Marcacci M, Di Domenico M, Ancora M, Curini V, Mangone I, Rinaldi A, Scialabba S, Di Pasquale A, Cammà C, Puglia I, Calistri P, Savini G |
| EPI_ISL_965266 | SIESP CHIETI | Istituto Zooprofilattico Sperimentale dell'Abruzzo e Molise "G. Caporale" | Lorusso A, Marcacci M, Di Domenico M, Ancora M, Curini V, Mangone I, Rinaldi A, Scialabba S, Di Pasquale A, Cammà C, Puglia I, Calistri P, Savini G |
| EPI_ISL_965267 | SIESP CHIETI, DISTRETTON SAN CHIETI | Istituto Zooprofilattico Sperimentale dell'Abruzzo e Molise "G. Caporale" | Lorusso A, Marcacci M, Di Domenico M, Ancora M, Curini V, Mangone I, Rinaldi A, Scialabba S, Di Pasquale A, Cammà C, Puglia I, Calistri P, Savini G |
| EPI_ISL_965268, EPI_ISL_965269 | SIESP CHIETI, DISTRETTO SAN CHIETI | Istituto Zooprofilattico Sperimentale dell'Abruzzo e Molise "G. Caporale" | Lorusso A, Marcacci M, Di Domenico M, Ancora M, Curini V, Mangone I, Rinaldi A, Scialabba S, Di Pasquale A, Cammà C, Puglia I, Calistri P, Savini G |
| EPI_ISL_965270 | SIESP CHIETI | Istituto Zooprofilattico Sperimentale dell'Abruzzo e Molise "G. Caporale" | Lorusso A, Marcacci M, Di Domenico M, Ancora M, Curini V, Mangone I, Rinaldi A, Scialabba S, Di Pasquale A, Cammà C, Puglia I, Calistri P, Savini G |
| EPI_ISL_965271 | SIESP CHIETI, DISTRETTO SAN CHIETI | Istituto Zooprofilattico Sperimentale dell'Abruzzo e Molise "G. Caporale" | Lorusso A, Marcacci M, Di Domenico M, Ancora M, Curini V, Mangone I, Rinaldi A, Scialabba S, Di Pasquale A, Cammà C, Puglia I, Calistri P, Savini G |
| EPI_ISL_965272 | SIESP CHIETI | Istituto Zooprofilattico Sperimentale dell'Abruzzo e Molise "G. Caporale" | Lorusso A, Marcacci M, Di Domenico M, Ancora M, Curini V, Mangone I, Rinaldi A, Scialabba S, Di Pasquale A, Cammà C, Puglia I, Calistri P, Savini G |
| EPI_ISL_965273, EPI_ISL_965274, EPI_ISL_965275 | SIESP CHIETI, DISTRETTO SAN CHIETI | Istituto Zooprofilattico Sperimentale dell'Abruzzo e Molise "G. Caporale" | Lorusso A, Marcacci M, Di Domenico M, Ancora M, Curini V, Mangone I, Rinaldi A, Scialabba S, Di Pasquale A, Cammà C, Puglia I, Calistri P, Savini G |
| EPI_ISL_965276 | OSP CIV ATRI | Istituto Zooprofilattico Sperimentale dell'Abruzzo e Molise "G. Caporale" | Lorusso A, Marcacci M, Di Domenico M, Ancora M, Curini V, Mangone I, Rinaldi A, Scialabba S, Di Pasquale A, Cammà C, Puglia I, Calistri P, Savini G |
| EPI_ISL_965280, EPI_ISL_965281, EPI_ISL_965282, EPI_ISL_965283, EPI_ISL_965284, EPI_ISL_965285, EPI_ISL_965286, EPI_ISL_965287, EPI_ISL_965288, EPI_ISL_965289, EPI_ISL_965290, EPI_ISL_965291, EPI_ISL_965292, EPI_ISL_965293, EPI_ISL_965294, EPI_ISL_965295, EPI_ISL_965296, EPI_ISL_965297, EPI_ISL_965298, EPI_ISL_965299, EPI_ISL_965300 | | | |
| see above | P.O.CARDARELLI | Istituto Zooprofilattico Sperimentale dell'Abruzzo e Molise "G. Caporale" | Scutellà M, Niro G, Lorusso A, Marcacci M, Di Domenico M, Ancora M, Curini V, Mangone I, Rinaldi A, Scialabba S, Di Pasquale A, Cammà C, Puglia I, Calistri P, Savini G |
| EPI_ISL_965911, EPI_ISL_965913 | Hospital Los Arcos del Mar Menor | Instituto de Salud Carlos III | Vázquez, S. Iglesias-Caballero, M. Sandonís,V. Camarero, S. Pozo, F. Casas, I. Jiménez, P. Zaballos, A. Monzón, S. Varona, S. Cuesta, I. Cámara, M. |
| EPI_ISL_965914, EPI_ISL_965915 | Hospital General Universitario de Ciudad Real | Instituto de Salud Carlos III | Vázquez, S. Iglesias-Caballero, M. Sandonís,V. Camarero, S. Pozo, F. Casas, I. Jiménez, P. Zaballos, A. Monzón, S. Varona, S. Cuesta, I. Illescas, S. |

| | | | |
|---|---|---|---|
| EPI_ISL_965916 | Servicio Murciano de Salud | Instituto de Salud Carlos III | Vázquez, S. Iglesias-Caballero, M. Sandonís,V. Camarero, S. Pozo, F. Casas, I. Jiménez, P. Zaballos, A. Monzón, S. Varona, S. Cuesta, I. Blázquez, A. |
| EPI_ISL_965917, EPI_ISL_965918, EPI_ISL_965919 | Hospital Los Arcos del Mar Menor | Instituto de Salud Carlos III | Vázquez, S. Iglesias-Caballero, M. Sandonís,V. Camarero, S. Pozo, F. Casas, I. Jiménez, P. Zaballos, A. Monzón, S. Varona, S. Cuesta, I. Cámara, M. |
| EPI_ISL_965920 | Servicio Murciano de Salud | Instituto de Salud Carlos III | Vázquez, S. Iglesias-Caballero, M. Sandonís,V. Camarero, S. Pozo, F. Casas, I. Jiménez, P. Zaballos, A. Monzón, S. Varona, S. Cuesta, I. Blázquez, A. |
| EPI_ISL_965921 | Consejería de Sanidad y Asuntos Sociales de Castilla La Mancha | Instituto de Salud Carlos III | Vázquez, S. Iglesias-Caballero, M. Sandonís,V. Camarero, S. Pozo, F. Casas, I. Jiménez, P. Zaballos, A. Monzón, S. Varona, S. Cuesta, I. Gutiérrez, G. |
| EPI_ISL_965922 | Servicio Murciano de Salud | Instituto de Salud Carlos III | Vázquez, S. Iglesias-Caballero, M. Sandonís,V. Camarero, S. Pozo, F. Casas, I. Jiménez, P. Zaballos, A. Monzón, S. Varona, S. Cuesta, I. Blázquez, A. |
| EPI_ISL_965923 | Hospital Los Arcos del Mar Menor | Instituto de Salud Carlos III | Vázquez, S. Iglesias-Caballero, M. Sandonís,V. Camarero, S. Pozo, F. Casas, I. Jiménez, P. Zaballos, A. Monzón, S. Varona, S. Cuesta, I. Cámara, M. |
| EPI_ISL_965924, EPI_ISL_965925 | Servicio Murciano de Salud | Instituto de Salud Carlos III | Vázquez, S. Iglesias-Caballero, M. Sandonís,V. Camarero, S. Pozo, F. Casas, I. Jiménez, P. Zaballos, A. Monzón, S. Varona, S. Cuesta, I. Blázquez, A. |
| EPI_ISL_965926 | Hospital General Universitario de Ciudad Real | Instituto de Salud Carlos III | Sandonís,V. Vázquez, S. Iglesias-Caballero, M. Camarero, S. Pozo, F. Casas, I. Jiménez, P. Zaballos, A. Monzón, S. Varona, S. Cuesta, I. Illescas, S. |
| EPI_ISL_965927 | Servicio Murciano de Salud | Instituto de Salud Carlos III | Sandonís,V. Vázquez, S. Iglesias-Caballero, M. Camarero, S. Pozo, F. Casas, I. Jiménez, P. Zaballos, A. Monzón, S. Varona, S. Cuesta, I. Blázquez, A. |
| EPI_ISL_965928, EPI_ISL_965929 | Hospital Los Arcos del Mar Menor | Instituto de Salud Carlos III | Sandonís,V. Vázquez, S. Iglesias-Caballero, M. Camarero, S. Pozo, F. Casas, I. Jiménez, P. Zaballos, A. Monzón, S. Varona, S. Cuesta, I. Cámara, M. |
| EPI_ISL_965930 | Hospital Comarcal de Melilla | Instituto de Salud Carlos III | Sandonís,V. Vázquez, S. Iglesias-Caballero, M. Camarero, S. Pozo, F. Casas, I. Jiménez, P. Zaballos, A. Monzón, S. Varona, S. Cuesta, I. Román, S. |
| EPI_ISL_965931 | Servicio Murciano de Salud | Instituto de Salud Carlos III | Sandonís,V. Vázquez, S. Iglesias-Caballero, M. Camarero, S. Pozo, F. Casas, I. Jiménez, P. Zaballos, A. Monzón, S. Varona, S. Cuesta, I. Blázquez, A. |
| EPI_ISL_965932 | Hospital Los Arcos del Mar Menor | Instituto de Salud Carlos III | Sandonís,V. Vázquez, S. Iglesias-Caballero, M. Camarero, S. Pozo, F. Casas, I. Jiménez, P. Zaballos, A. Monzón, S. Varona, S. Cuesta, I. Cámara, M. |
| EPI_ISL_965933 | Complejo Hospitalario de Navarra | Instituto de Salud Carlos III | Sandonís,V. Vázquez, S. Iglesias-Caballero, M. Camarero, S. Pozo, F. Casas, I. Jiménez, P. Zaballos, A. Monzón, S. Varona, S. Cuesta, I. Navascués, A. |
| EPI_ISL_965935 | Hospital Los Arcos del Mar Menor | Instituto de Salud Carlos III | Sandonís,V. Vázquez, S. Iglesias-Caballero, M. Camarero, S. Pozo, F. Casas, I. Jiménez, P. Zaballos, A. Monzón, S. Varona, S. Cuesta, I. Cámara, M. |
| EPI_ISL_965936 | Hospital Comarcal de Melilla | Instituto de Salud Carlos III | Sandonís,V. Vázquez, S. Iglesias-Caballero, M. Camarero, S. Pozo, F. Casas, I. Jiménez, P. Zaballos, A. Monzón, S. Varona, S. Cuesta, I. Román, S. |
| EPI_ISL_965937 | Complejo Hospitalario de Navarra | Instituto de Salud Carlos III | Sandonís,V. Vázquez, S. Iglesias-Caballero, M. Camarero, S. Pozo, F. Casas, I. Jiménez, P. Zaballos, A. Monzón, S. Varona, S. Cuesta, I. Navascués, A. |
| EPI_ISL_965939 | Hospital Los Arcos del Mar Menor | Instituto de Salud Carlos III | Sandonís,V. Vázquez, S. Iglesias-Caballero, M. Camarero, S. Pozo, F. Casas, I. Jiménez, P. Zaballos, A. Monzón, S. Varona, S. Cuesta, I. Cámara, M. |
| EPI_ISL_965940 | Hospital General Universitario de Ciudad Real | Instituto de Salud Carlos III | Sandonís,V. Vázquez, S. Iglesias-Caballero, M. Camarero, S. Pozo, F. Casas, I. Jiménez, P. Zaballos, A. Monzón, S. Varona, S. Cuesta, I. Illescas, S. |
| EPI_ISL_965941 | Hospital Los Arcos del Mar Menor | Instituto de Salud Carlos III | Sandonís,V. Vázquez, S. Iglesias-Caballero, M. Camarero, S. Pozo, F. Casas, I. Jiménez, P. Zaballos, A. Monzón, S. Varona, S. Cuesta, I. Cámara, M. |
| EPI_ISL_965942, EPI_ISL_965943, EPI_ISL_965944 | Complejo Hospitalario de Navarra | Instituto de Salud Carlos III | Iglesias-Caballero, M. Sandonís,V. Vázquez, S. Camarero, S. Pozo, F. Casas, I. Jiménez, P. Zaballos, A. Monzón, S. Varona, S. Cuesta, I. Navascués, A. |
| EPI_ISL_965945 | Hospital Los Arcos del Mar Menor | Instituto de Salud Carlos III | Iglesias-Caballero, M. Sandonís,V. Vázquez, S. Camarero, S. Pozo, F. Casas, I. Jiménez, P. Zaballos, A. Monzón, S. Varona, S. Cuesta, I. Cámara, M. |
| EPI_ISL_965946 | Hospital General Universitario de Ciudad Real | Instituto de Salud Carlos III | Iglesias-Caballero, M. Sandonís,V. Vázquez, S. Camarero, S. Pozo, F. Casas, I. Jiménez, P. Zaballos, A. Monzón, S. Varona, S. Cuesta, I. Illescas, S. |
| EPI_ISL_965947 | Hospital Los Arcos del Mar Menor | Instituto de Salud Carlos III | Iglesias-Caballero, M. Sandonís,V. Vázquez, S. Camarero, S. Pozo, F. Casas, I. Jiménez, P. Zaballos, A. Monzón, S. Varona, S. Cuesta, I. Cámara, M. |
| EPI_ISL_965949 | Hospital Comarcal de Melilla | Instituto de Salud Carlos III | Iglesias-Caballero, M. Sandonís,V. Vázquez, S. Camarero, S. Pozo, F. Casas, I. Jiménez, P. Zaballos, A. Monzón, S. Varona, S. Cuesta, I. Román, S. |
| EPI_ISL_965950 | Complejo Hospitalario de Navarra | Instituto de Salud Carlos III | Iglesias-Caballero, M. Sandonís,V. Vázquez, S. Camarero, S. Pozo, F. Casas, I. Jiménez, P. Zaballos, A. Monzón, S. Varona, S. Cuesta, I. Navascués, A. |
| EPI_ISL_965951 | Hospital Los Arcos del Mar Menor | Instituto de Salud Carlos III | Iglesias-Caballero, M. Sandonís,V. Vázquez, S. Camarero, S. Pozo, F. Casas, I. Jiménez, P. Zaballos, A. Monzón, S. Varona, S. Cuesta, I. Cámara, M. |
| EPI_ISL_965953, EPI_ISL_965954 | Servicio Murciano de Salud | Instituto de Salud Carlos III | Iglesias-Caballero, M. Sandonís,V. Vázquez, S. Camarero, S. Pozo, F. Casas, I. Jiménez, P. Zaballos, A. Monzón, S. Varona, S. Cuesta, I. Blázquez, A. |
| EPI_ISL_965955 | Hospital Los Arcos del Mar Menor | Instituto de Salud Carlos III | Iglesias-Caballero, M. Sandonís,V. Vázquez, S. Camarero, S. Pozo, F. Casas, I. Jiménez, P. Zaballos, A. Monzón, S. Varona, S. Cuesta, I. Cámara, M. |
| EPI_ISL_965956 | Complejo Hospitalario de Navarra | Instituto de Salud Carlos III | Iglesias-Caballero, M. Sandonís,V. Vázquez, S. Camarero, S. Pozo, F. Casas, I. Jiménez, P. Zaballos, A. Monzón, S. Varona, S. Cuesta, I. Navascués, A. |
| EPI_ISL_965957 | Consejería de Sanidad y Asuntos Sociales de Castilla La Mancha | Instituto de Salud Carlos III | Iglesias-Caballero, M. Sandonís,V. Vázquez, S. Camarero, S. Pozo, F. Casas, I. Jiménez, P. Zaballos, A. Monzón, S. Varona, S. Cuesta, I. Gutiérrez, G. |
| EPI_ISL_965958 | Hospital Los Arcos del Mar Menor | Instituto de Salud Carlos III | Iglesias-Caballero, M. Sandonís,V. Vázquez, S. Camarero, S. Pozo, F. Casas, I. Jiménez, P. Zaballos, A. Monzón, S. Varona, S. Cuesta, I. Cámara, M. |
| EPI_ISL_965960 | Servicio Murciano de Salud | Instituto de Salud Carlos III | Iglesias-Caballero, M. Sandonís,V. Vázquez, S. Camarero, S. Pozo, F. Casas, I. Jiménez, P. Zaballos, A. Monzón, S. Varona, S. Cuesta, I. Blázquez, A. |
| EPI_ISL_966285 | Baylor Scott & White - Irving | Baylor Scott & White - Temple | Ari Rao, Linden Morales, Kimberly Walker, Marcus Volz, Shelby Hendrickson |
| EPI_ISL_966310, EPI_ISL_966313 | Baylor Scott & White - Temple | Baylor Scott & White - Temple | Ari Rao, Linden Morales, Kimberly Walker, Marcus Volz, Shelby Hendrickson |
| EPI_ISL_968089 | Institute for Medical Research, Infectious Disease Research Centre, National Institutes of Health, Ministry of Health Malaysia | Institute for Medical Research, Infectious Disease Research Centre, National Institutes of Health, Ministry of Health Malaysia | Suppiah J, Kamel K, Azizan MA, Thayan R |
| EPI_ISL_969131 | Laboratorio Microbiologia e Virologia P.O. Cotugno A.O. dei Colli | Laboratorio Microbiologia e Virologia P.O. Cotugno A.O. dei Colli | Luigi Atripaldi, Claudia Tiberio, Anna Perfetti, |
| EPI_ISL_969227 | Laboratorio Microbiologia e Virologia P.O. Cotugno A.O. dei Colli | Laboratorio Microbiologia e Virologia P.O. Cotugno A.O. dei Colli | Luigi Atripaldi, Claudia Tiberio, Anna Perfetti |
| EPI_ISL_969297 | Laboratorio Microbiologia e Virologia, P.O. Cotugno, A.O. dei Colli | Laboratorio Microbiologia e Virologia, P.O. Cotugno, A.O. dei Colli | Luigi Atripaldi, Claudia Tiberio, Anna Perfetti |
| EPI_ISL_969884 | Laboratorio Microbiologia e Virologia P.O. Cotugno A.O. dei Colli | Laboratorio Microbiologia e Virologia P.O. Cotugno A.O. dei Colli | Luigi Atripaldi,Claudia Tiberio, Anna Perfetti |
| EPI_ISL_970647 | Laboratorio Microbiologia e Virologia P.O. Cotugno A.O. dei Colli | Laboratorio Microbiologia e Virologia P.O. Cotugno A.O. dei Colli | Luigi Atripaldi, Claudia Tiberio,Anna Perfetti |
| EPI_ISL_974745 | Laboratorio Microbiologia e Virologia P.O. Cotugno A.O. dei Colli | Laboratorio Microbiologia e Virologia P.O. Cotugno A.O. dei Colli | Luigi Atripaldi, Claudia Tiberio, Anna Perfetti, |
| EPI_ISL_974853 | Laboratorio de Virología / Hospital Universitario Central de Asturias (HUCA) | Laboratorio de Virología / Hospital Universitario Central de Asturias (HUCA) | Castelló C, Gómez de Oña J, Boga JA, Rojo S, Alvarez-Arguelles ME, Abreu F, Costales I, Sandoval M, Perez-Martínez Z, Martín-Rodríguez G, Coto E, Melón S |
| EPI_ISL_977417 | Hospital de Cabueñes | Laboratorio de Virología / Hospital Universitario Central de Asturias (HUCA) | Castelló C, Gómez de Oña J, Boga JA, Rojo S, Alvarez-Arguelles ME, Abreu F, Costales I, Sandoval M, Perez-Martínez Z, Martín-Rodríguez G, Coto E, Melón S |
| EPI_ISL_977489 | UPA Dr. Akira Tada | Instituto Adolfo Lutz, Interdiciplinary Procedures Center, Strategic Laboratory | Claudio Tavares Sacchi, Claudia Regina Gonçalves, Erica Valessa Ramos Gomes, Karoline Rodrigues Campos |
| EPI_ISL_977490 | Hospital Municipal Guido Guida | Instituto Adolfo Lutz, Interdiciplinary Procedures Center, Strategic Laboratory | Claudio Tavares Sacchi, Claudia Regina Gonçalves, Erica Valessa Ramos Gomes, Karoline Rodrigues Campos |
| EPI_ISL_977495, EPI_ISL_977496, EPI_ISL_977497, EPI_ISL_977498 | University of Bari Biomedical Sciences and Human Oncology | University of Bari Biomedical Sciences and Human Oncology | Chironna M., Sallustio A., Loconsole D., Accogli M. |

| | | | |
|---|---|---|---|
| EPI_ISL_977505, EPI_ISL_977506 | Laboratorio de Virología HUCA | Laboratorio de Virología HUCA | Sandoval M, Castelló C, Gómez de Oña J, Boga JA, Rojo S, Alvarez-Arguelles ME, Abreu F, Costales I, Perez-Martínez Z, Martín-Rodríguez G, Coto E, Melón S |
| EPI_ISL_977507 | Hospital Valle del Nalon | Laboratorio de Virología HUCA | Sandoval M, Castelló C, Gómez de Oña J, Boga JA, Rojo S, Alvarez-Arguelles ME, Abreu F, Costales I, Perez-Martínez Z, Martín-Rodríguez G, Coto E, Melón S |
| EPI_ISL_977511, EPI_ISL_977512 | Laboratorio de Virología HUCA | Laboratorio de Virología HUCA | Sandoval M, Castelló C, Gómez de Oña J, Boga JA, Rojo S, Alvarez-Arguelles ME, Abreu F, Costales I, Perez-Martínez Z, Martín-Rodríguez G, Coto E, Melón S |
| EPI_ISL_977597 | Servicio Murciano de Salud | Instituto de Salud Carlos III | Vázquez, S. Iglesias-Caballero, M. Sandonís,V. Camarero, S. Pozo, F. Casas, I. Jiménez, P. Zaballos, A. Monzón, S. Varona, S. Cuesta, I. Blázquez, A. |
| EPI_ISL_977598 | Laboratorio Microbiologia e Virologia P.O. Cotugno A.O. dei Colli | Laboratorio Microbiologia e Virologia P.O. Cotugno A.O. dei Colli | Luigi Atripaldi, Claudia Tiberio, Anna Perfetti |
| EPI_ISL_977601, EPI_ISL_977603 | Laboratorio Microbiologia e Virologia P.O. Cotugno A.O. dei Colli | Laboratorio Microbiologia e Virologia P.O. Cotugno A.O. dei Colli | Luigi Atripaldi, Claudia Tiberio, Anna Perfetti, |
| EPI_ISL_977605, EPI_ISL_977606, EPI_ISL_977607 | SC (UCO) Igiene e Sanità Pubblica (funzione integrata con SC Microbiologia e Virologia) e Laboratory of Molecular Virology of the International Centre for Genetic Engineering and Biotechnology (ICGEB) | ARGO Laboratorio Genomica ed Epigenomica | Licastro D, Dal Monego S, Degasperi M, Marcello A, D'Agaro P, De Rosa R |
| EPI_ISL_977622, EPI_ISL_977623, EPI_ISL_977626, EPI_ISL_977627, EPI_ISL_977630, EPI_ISL_977631, EPI_ISL_977632, EPI_ISL_977635, EPI_ISL_977636 | SC (UCO) Igiene e Sanità Pubblica (funzione integrata con SC Microbiologia e Virologia) e Laboratory of Molecular Virology of the International Centre for Genetic Engineering and Biotechnology (ICGEB) | ARGO Laboratorio Genomica ed Epigenomica | Licastro D, Dal Monego S, Degasperi M, Marcello A, Segat L, Piscianz E, D'Agaro P |
| EPI_ISL_977643, EPI_ISL_977644, EPI_ISL_977645, EPI_ISL_977646, EPI_ISL_977647, EPI_ISL_977649 | SC (UCO) Igiene e Sanità Pubblica (funzione integrata con SC Microbiologia e Virologia) e Laboratory of Molecular Virology of the International Centre for Genetic Engineering and Biotechnology (ICGEB) | ARGO Laboratorio Genomica ed Epigenomica | Licastro D, Dal Monego S, Degasperi M, Marcello A, D'Agaro P, Lombardo F |
| EPI_ISL_977651 | Laboratorio Microbiologia e Virologia P.O. Cotugno A.O. dei Colli | Laboratorio Microbiologia e Virologia P.O. Cotugno A.O. dei Colli | Luigi Atripaldi, Claudia Tiberio, Anna Perfetti, |
| EPI_ISL_979544, EPI_ISL_979798, EPI_ISL_979800, EPI_ISL_979801, EPI_ISL_979802, EPI_ISL_979803, EPI_ISL_979804, EPI_ISL_979893, EPI_ISL_979959, EPI_ISL_979960, EPI_ISL_979961, EPI_ISL_979963, EPI_ISL_979966, EPI_ISL_979967, EPI_ISL_979968, EPI_ISL_979969 | | | |
| see above | National Institute of Infectious Diseases-Prof. Dr. Matei Bals Molecular Diagnostics Laboratory | National Institute of Infectious Diseases-Prof. Dr. Matei Bals Molecular Diagnostics Laboratory | Leontina Banica, Marius Surleac, Corina Casangiu, Petre Milu, Andreea Tudor, Simona Paraschiv, Dan Otelea |
| EPI_ISL_981371 | AZ Klina | AZ Klina | Dr. C. Vael |
| EPI_ISL_981372 | Laborarory for Respiratory Viruses, "Cantacuzino" National Military-Medical Institute for Resararch and Development | Cantacuzino Institute Virology | Luiza Ustea, Nicoleta Paraschiv, Catalina Pascu, Mihaela Lazar |
| EPI_ISL_981373 | AZ Klina | AZ Klina | Dr. C. Vael |
| EPI_ISL_981374, EPI_ISL_981375 | Laboratory for Respiratory Viruses, Cantacuzino National Military-Medical Institute for Research and Development | Cantacuzino Institute Virology | Luiza Ustea, Nicoleta Paraschiv, Catalina Pascu, Mihaela Lazar |
| EPI_ISL_981376 | AZ Klina | AZ Klina | Dr. C. Vael |
| EPI_ISL_981377 | Laboratory for Respiratory Viruses, Cantacuzino National Military-Medical Institute for Research and Development | Cantacuzino Institute Virology | Luiza Ustea, Nicoleta Paraschiv, Catalina Pascu, Mihaela Lazar |
| EPI_ISL_981378, EPI_ISL_981379 | AZ Klina | AZ Klina | Dr. C. Vael |
| EPI_ISL_981383, EPI_ISL_981385, EPI_ISL_981387 | IAL Regional de Bauru | Instituto Adolfo Lutz, Interdiciplinary Procedures Center, Strategic Laboratory | Claudio Tavares Sacchi, Claudia Regina Gonçalves, Erica Valessa Ramos Gomes, Karoline Rodrigues Campos |
| EPI_ISL_981468 | CHI VILLENEUVE ST GEORGES | Department of Virology, Henri Mondor University Hospital, Assistance Publique Hôpitaux de Paris, Université Paris-Est Créteil, INSERM U955 | Christophe Rodriguez, Slim Fourati, Vanessa Demontant, Guillaume Gricourt, Melissa N'Debi, Alexandre Soulier, Elisabeth Trawinski, Jean-Michel Pawlotsky |
| EPI_ISL_982139, EPI_ISL_982170, EPI_ISL_982180, EPI_ISL_982204, EPI_ISL_982205, EPI_ISL_982206, EPI_ISL_982219, EPI_ISL_982221, EPI_ISL_982222, EPI_ISL_982224, EPI_ISL_982225 | | | |
| see above | Hôpital Henri Mondor | Department of Virology, Henri Mondor University Hospital, Assistance Publique Hôpitaux de Paris, Université Paris-Est Créteil, INSERM U955 | Christophe Rodriguez, Slim Fourati, Vanessa Demontant, Guillaume Gricourt, Melissa N'Debi, Alexandre Soulier, Elisabeth Trawinski, Jean-Michel Pawlotsky |
| EPI_ISL_983097, EPI_ISL_983098 | Microbiology and Virology Unit, Florence Careggi University Hospital | Microbiology and Virology Unit, Florence Careggi University Hospital | Vincenzo Di Pilato, Marco Coppi, Fabio Morecchiato, Noemi Aiezza, Ilaria Baccani, Alberto Antonelli, Emanuele Gori, Gian Maria Rossolini |
| EPI_ISL_983865, EPI_ISL_984619, EPI_ISL_984620, EPI_ISL_984621 | Central Laboratory of Public Health of Rio Grande do Sul (Lacen-RS) | State Center for Health Surveillance of the Health Department of the State of Rio Grande do Sul (CEVS/SES-RS) | Aline Campos, Cynthia Molina, Lara Crescente, Leticia Garay, Ludmila Fiorenzano Baethgen, Richard Salvato, Tatiana Gregianini |
| EPI_ISL_984678, EPI_ISL_984683 | Laboratorio de Virología HUCA | Laboratorio de Virología HUCA | Sandoval M, Castelló C, Gómez de Oña J, Boga JA, Rojo S, Alvarez-Arguelles ME, Abreu F, Costales I, Perez-Martínez Z, Martín-Rodríguez G, Coto E, Melón S |
| EPI_ISL_984996 | U.O. Igiene, Ospedale Policlinico San Martino | U.O. Igiene, Ospedale Policlinico San Martino | Bruzzone Bianca, Caligiuri Patrizia, De Pace Vanessa, Domnich Alexander, Orsi Andrea, Ricucci Valentina, Icardi Giancarlo |
| EPI_ISL_984997, EPI_ISL_984998, EPI_ISL_984999 | S.C. Laboratorio Analisi, ASL 3 Liguria | U.O. Igiene, Ospedale Policlinico San Martino | Bruzzone Bianca, Caligiuri Patrizia, De Pace Vanessa, Domnich Alexander, Orsi Andrea, Ricucci Valentina, Spitaleri Antonino, Icardi Giancarlo |
| EPI_ISL_985000 | U.O. Igiene, Ospedale Policlinico San Martino | U.O. Igiene, Ospedale Policlinico San Martino | Bruzzone Bianca, Caligiuri Patrizia, De Pace Vanessa, Domnich Alexander, Orsi Andrea, Ricucci Valentina, Spitaleri Antonino, Icardi Giancarlo |
| EPI_ISL_985001, EPI_ISL_985002, EPI_ISL_985003, EPI_ISL_985004, EPI_ISL_985005, EPI_ISL_985006, EPI_ISL_985007, EPI_ISL_985008, EPI_ISL_985009, EPI_ISL_985010, EPI_ISL_985011, EPI_ISL_985012, EPI_ISL_985013, EPI_ISL_985014, EPI_ISL_985015, EPI_ISL_985016 | | | |
| see above | S.C. Laboratorio Analisi, ASL 3 Liguria | U.O. Igiene, Ospedale Policlinico San Martino | Bruzzone Bianca, Caligiuri Patrizia, De Pace Vanessa, Domnich Alexander, Orsi Andrea, Ricucci Valentina, Spitaleri Antonino, Icardi Giancarlo |
| EPI_ISL_985017 | S.C. Patologia Clinica, Ospedale Sant'Andrea, ASL 5 | U.O. Igiene, Ospedale Policlinico San Martino | Bruzzone Bianca, Battolla Enrico, Caligiuri Patrizia, De Pace Vanessa, Domnich Alexander, Orsi Andrea, Ricucci Valentina, Icardi Giancarlo |
| EPI_ISL_985018, EPI_ISL_985019, EPI_ISL_985020, EPI_ISL_985021, EPI_ISL_985022, EPI_ISL_985023, EPI_ISL_985024 | S.S.D. Microbiologia, Stabilimento ospedaliero di Sanremo, ASL 1 Liguria | U.O. Igiene, Ospedale Policlinico San Martino | Bruzzone Bianca, Caligiuri Patrizia, De Pace Vanessa, Domnich Alexander, Dusi Pier Andrea, Orsi Andrea, Ricucci Valentina, Icardi Giancarlo |
| EPI_ISL_985025, EPI_ISL_985026 | S.S.D. Microbiologia, Ospedale Santa Corona di Pietra Ligure, ASL 2 Liguria | U.O. Igiene, Ospedale Policlinico San Martino | Bruzzone Bianca, Caligiuri Patrizia, De Pace Vanessa, Domnich Alexander, Orsi Andrea, Ricucci Valentina, Valle Caterina, Icardi Giancarlo |
| EPI_ISL_985027, EPI_ISL_985028, EPI_ISL_985029, EPI_ISL_985030, EPI_ISL_985031 | Laboratorio di Patologia Clinica, Ospedale San Paolo in Valloria, ASL 2 Liguria | U.O. Igiene, Ospedale Policlinico San Martino | Bruzzone Bianca, Caligiuri Patrizia, De Pace Vanessa, Domnich Alexander, Lillo Flavia, Orsi Andrea, Ricucci Valentina, Icardi Giancarlo |

| | | | |
|---|---|---|---|
| EPI_ISL_985032 | U.O. Igiene, Ospedale Policlinico San Martino | U.O. Igiene, Ospedale Policlinico San Martino | Bruzzone Bianca, Caligiuri Patrizia, De Pace Vanessa, Domnich Alexander, Orsi Andrea, Ricucci Valentina, Icardi Giancarlo |
| EPI_ISL_985303, EPI_ISL_985304, EPI_ISL_985305, EPI_ISL_985306, EPI_ISL_985307, EPI_ISL_985308, EPI_ISL_985309, EPI_ISL_985310, EPI_ISL_985311, EPI_ISL_985312, EPI_ISL_985313, EPI_ISL_985314, EPI_ISL_985315, EPI_ISL_985316, EPI_ISL_985317 | | | |
| see above | LACEN do Estado de Goias | Instituto Adolfo Lutz, Interdiciplinary Procedures Center, Strategic Laboratory | Claudio Tavares Sacchi, Claudia Regina Gonçalves, Erica Valessa Ramos Gomes, Karoline Rodrigues Campos |
| EPI_ISL_995331, EPI_ISL_995332, EPI_ISL_995333, EPI_ISL_995334, EPI_ISL_995335, EPI_ISL_995336, EPI_ISL_995337, EPI_ISL_995338, EPI_ISL_995339, EPI_ISL_995340, EPI_ISL_995341, EPI_ISL_995342, EPI_ISL_995344, EPI_ISL_995345, EPI_ISL_995346, EPI_ISL_995347, EPI_ISL_995348, EPI_ISL_995349, EPI_ISL_995350, EPI_ISL_995351, EPI_ISL_995353, EPI_ISL_995354, EPI_ISL_995355, EPI_ISL_995356, EPI_ISL_995357, EPI_ISL_995358, EPI_ISL_995359, EPI_ISL_995360, EPI_ISL_995361, EPI_ISL_995362, EPI_ISL_995363, EPI_ISL_995364, EPI_ISL_995365, EPI_ISL_995366, EPI_ISL_995367, EPI_ISL_995368, EPI_ISL_995370, EPI_ISL_995371, EPI_ISL_995372, EPI_ISL_995373, EPI_ISL_995374, EPI_ISL_995375, EPI_ISL_995376, EPI_ISL_995378, EPI_ISL_995379, EPI_ISL_995380, EPI_ISL_995381, EPI_ISL_995382, EPI_ISL_995383, EPI_ISL_995384, EPI_ISL_995385, EPI_ISL_995386, EPI_ISL_995387, EPI_ISL_995388, EPI_ISL_995389, EPI_ISL_995390, EPI_ISL_995392, EPI_ISL_995393, EPI_ISL_995394, EPI_ISL_995395, EPI_ISL_995396, EPI_ISL_995397, EPI_ISL_995398, EPI_ISL_995399, EPI_ISL_995400, EPI_ISL_995401, EPI_ISL_995402, EPI_ISL_995403, EPI_ISL_995404, EPI_ISL_995405, EPI_ISL_995406, EPI_ISL_995407, EPI_ISL_995409, EPI_ISL_995410, EPI_ISL_995411, EPI_ISL_995412, EPI_ISL_995413, EPI_ISL_995414, EPI_ISL_995416, EPI_ISL_995418, EPI_ISL_995419, EPI_ISL_995420, EPI_ISL_995421, EPI_ISL_995422, EPI_ISL_995423, EPI_ISL_995424, EPI_ISL_995425, EPI_ISL_995426, EPI_ISL_995427, EPI_ISL_995428, EPI_ISL_995429, EPI_ISL_995430, EPI_ISL_995431, EPI_ISL_995432, EPI_ISL_995433, EPI_ISL_995434, EPI_ISL_995435, EPI_ISL_995436, EPI_ISL_995437, EPI_ISL_995438, EPI_ISL_995439, EPI_ISL_995440, EPI_ISL_995441, EPI_ISL_995442, EPI_ISL_995443, EPI_ISL_995444, EPI_ISL_995445 | | | |
| see above | Outre mer | National Reference Center for Viruses of Respiratory Infections, Institut Pasteur, Paris | Marion Barbet, Sylvie Behillil, Méline Bizard, Angela Brisebarre, Camille Capel, Etienne Simon-Lorière, Vincent Enouf, Maud Vanpeene, Sylvie van der Werf,Combe Patrice |
| EPI_ISL_995449 | Hopital | National Reference Center for Viruses of Respiratory Infections, Institut Pasteur, Paris | Marion Barbet, Sylvie Behillil, Méline Bizard, Angela Brisebarre, Camille Capel, Etienne Simon-Lorière, Vincent Enouf, Maud Vanpeene, Sylvie van der Werf,Marque Juillet StéPhanie |
| EPI_ISL_995450, EPI_ISL_995451, EPI_ISL_995452, EPI_ISL_995453, EPI_ISL_995454, EPI_ISL_995455, EPI_ISL_995456, EPI_ISL_995457, EPI_ISL_995458, EPI_ISL_995459, EPI_ISL_995460, EPI_ISL_995461 | | | |
| see above | Hopital | National Reference Center for Viruses of Respiratory Infections, Institut Pasteur, Paris | Marion Barbet, Sylvie Behillil, Méline Bizard, Angela Brisebarre, Camille Capel, Etienne Simon-Lorière, Vincent Enouf, Maud Vanpeene, Sylvie van der Werf,Fourgeaud Jacques |
| EPI_ISL_995462, EPI_ISL_995463, EPI_ISL_995464, EPI_ISL_995465, EPI_ISL_995466, EPI_ISL_995467 | Hopital | National Reference Center for Viruses of Respiratory Infections, Institut Pasteur, Paris | Marion Barbet, Sylvie Behillil, Méline Bizard, Angela Brisebarre, Camille Capel, Etienne Simon-Lorière, Vincent Enouf, Maud Vanpeene, Sylvie van der Werf,Guinoiseau Thibault |
| EPI_ISL_995468, EPI_ISL_995469, EPI_ISL_995470 | Labo Analyses Med | National Reference Center for Viruses of Respiratory Infections, Institut Pasteur, Paris | Marion Barbet, Sylvie Behillil, Méline Bizard, Angela Brisebarre, Camille Capel, Etienne Simon-Lorière, Vincent Enouf, Maud Vanpeene, Sylvie van der Werf,Garand Anne-Laure |
| EPI_ISL_995471, EPI_ISL_995472 | Hopital | National Reference Center for Viruses of Respiratory Infections, Institut Pasteur, Paris | Marion Barbet, Sylvie Behillil, Méline Bizard, Angela Brisebarre, Camille Capel, Etienne Simon-Lorière, Vincent Enouf, Maud Vanpeene, Sylvie van der Werf,Merens Audrey |
| EPI_ISL_995473 | Recherche | National Reference Center for Viruses of Respiratory Infections, Institut Pasteur, Paris | Marion Barbet, Sylvie Behillil, Méline Bizard, Angela Brisebarre, Camille Capel, Etienne Simon-Lorière, Vincent Enouf, Maud Vanpeene, Sylvie van der Werf,Leruez-Ville Marianne |
| EPI_ISL_995474, EPI_ISL_995475, EPI_ISL_995476, EPI_ISL_995477, EPI_ISL_995478, EPI_ISL_995479, EPI_ISL_995480, EPI_ISL_995481, EPI_ISL_995482, EPI_ISL_995483, EPI_ISL_995484, EPI_ISL_995485, EPI_ISL_995486, EPI_ISL_995488, EPI_ISL_995489, EPI_ISL_995490 | | | |
| see above | Hopital | National Reference Center for Viruses of Respiratory Infections, Institut Pasteur, Paris | Marion Barbet, Sylvie Behillil, Méline Bizard, Angela Brisebarre, Camille Capel, Etienne Simon-Lorière, Vincent Enouf, Maud Vanpeene, Sylvie van der Werf,Delamare Catherine |
| EPI_ISL_995491, EPI_ISL_995492, EPI_ISL_995493, EPI_ISL_995494 | Labo Analyses Med | National Reference Center for Viruses of Respiratory Infections, Institut Pasteur, Paris | Marion Barbet, Sylvie Behillil, Méline Bizard, Angela Brisebarre, Camille Capel, Etienne Simon-Lorière, Vincent Enouf, Maud Vanpeene, Sylvie van der Werf,Delamare Catherine |

We gratefully acknowledge the following Authors from the Originating laboratories responsible for obtaining the specimens, as well as the Submitting laboratories where the genome data were generated and shared via GISAID, on which this research is based.

All Submitters of data may be contacted directly via www.gisaid.org

Authors are sorted alphabetically.

| Accession ID | Originating Laboratory | Submitting Laboratory | Authors |
|---|---|---|---|
| EPI_ISL_1042001 | Istinye university genetic diseases assessment center | Istinye university genetic diseases assessment center | Zeynep Ocak MD,Prof Dr , Mursel Caliskan ,Msc |
| EPI_ISL_1063399 | Istinye university genetic diseases assessment center | Istinye university genetic diseases assessment center | Zeynep Ocak MD,Prof Dr , Mursel Caliskan ,Msc |
| EPI_ISL_1081350 | Institute for Medical Research, Infectious Disease Research Centre, National Institutes of Health, Ministry of Health Malaysia | Institute for Medical Research, Infectious Disease Research Centre, National Institutes of Health, Ministry of Health Malaysia | Suppiah J, Kamel K, Azizan MA, Thayan R |
| EPI_ISL_1082449 | National Laboratory for Health, Environment and Food | CISLD (Clinical Institute of Special Laboratory Diagnostics), University Children's Hospital, University Medical Center Ljubljana | Jernej Kova, Barbara Jenko Bizjan, Tine Tesovnik, Robert Šket, Ema Lovšin, Katarina Kozmos, Ana Grom, Maruša Debeljak, Marko Pokorn, Tadej Battelino |
| EPI_ISL_1085267, EPI_ISL_1085268, EPI_ISL_1085269, EPI_ISL_1085270, EPI_ISL_1085306, EPI_ISL_1085384, EPI_ISL_1085385 | Hôpital Henri Mondor | Department of Virology, Henri Mondor University Hospital, Assistance Publique Hôpitaux de Paris, Université Paris-Est Créteil, INSERM U955 | Christophe Rodriguez, Slim Fourati, Vanessa Demontant, Guillaume Gricourt, Melissa N'Debi, Alexandre Soulier, Elisabeth Trawinski, Jean-Michel Pawlotsky |
| EPI_ISL_1103583 | AZ Klina | AZ Klina | Dr. C. Vael |
| EPI_ISL_1108017 | AZ Klina | AZ Klina | Dr.C. Vael |
| EPI_ISL_1108255, EPI_ISL_1108503, EPI_ISL_1108770, EPI_ISL_1109036, EPI_ISL_1109283, EPI_ISL_1109392, EPI_ISL_1109393, EPI_ISL_1109483, EPI_ISL_1109631 | AZ Klina | AZ Klina | Dr. C. Vael |
| EPI_ISL_1110204 | Hôpital Henri Mondor | Department of Virology, Henri Mondor University Hospital, Assistance Publique Hôpitaux de Paris, Université Paris-Est Créteil, INSERM U955 | Christophe Rodriguez, Slim Fourati, Vanessa Demontant, Guillaume Gricourt, Melissa N'Debi, Alexandre Soulier, Elisabeth Trawinski, Jean-Michel Pawlotsky |
| EPI_ISL_1110480, EPI_ISL_1110481, EPI_ISL_1110482, EPI_ISL_1110483, EPI_ISL_1110484, EPI_ISL_1110485, EPI_ISL_1110486, EPI_ISL_1110487, EPI_ISL_1110488, EPI_ISL_1110489, EPI_ISL_1110490, EPI_ISL_1110491 | | | |
| see above | Hopital | National Reference Center for Viruses of Respiratory Infections, Institut Pasteur, Paris | Marion Barbet, Sylvie Behillil, Méline Bizard, Angela Brisebarre, Camille Capel, Etienne Simon-Lorière, Vincent Enouf, Maud Vanpeene, Sylvie van der Werf,Pilorge LéA |
| EPI_ISL_1110596 | SIESP L'Aquila | Istituto Zooprofilattico Sperimentale dell'Abruzzo e Molise "G. Caporale" | Lorusso A, Marcacci M, Di Domenico M, Ancora M, Curini V, Mangone I, Rinaldi A, Scialabba S, Di Pasquale A, Cammà C, Puglia I, Calistri P, Savini G |
| EPI_ISL_1110597 | SIESP CHIETI-Drive in Chieti | Istituto Zooprofilattico Sperimentale dell'Abruzzo e Molise "G. Caporale" | Lorusso A, Marcacci M, Di Domenico M, Ancora M, Curini V, Mangone I, Rinaldi A, Scialabba S, Di Pasquale A, Cammà C, Puglia I, Calistri P, Savini G |
| EPI_ISL_1114767, EPI_ISL_1114768, EPI_ISL_1114771, EPI_ISL_1116429, EPI_ISL_1116430, EPI_ISL_1116432, EPI_ISL_1116433, EPI_ISL_1116434, EPI_ISL_1116435, EPI_ISL_1116436, EPI_ISL_1116437, EPI_ISL_1116438, EPI_ISL_1116439, EPI_ISL_1116441, EPI_ISL_1116442, EPI_ISL_1116443, EPI_ISL_1116462 | | | |
| see above | National Institute of Infectious Diseases-Prof. Dr. Matei Bals Molecular Diagnostics Laboratory | National Institute of Infectious Diseases-Prof. Dr. Matei Bals Molecular Diagnostics Laboratory | Leontina Banica, Marius Surleac, Corina Casangiu, Petre Milu, Andreea Tudor, Simona Paraschiv, Dan Otelea |
| EPI_ISL_1117493 | OSP SAN SALVATORE, MEDICINA INTERNA | Istituto Zooprofilattico Sperimentale dell'Abruzzo e Molise "G. Caporale" | Lorusso A, Marcacci M, Di Domenico M, Ancora M, Curini V, Mangone I, Rinaldi A, Scialabba S, Di Pasquale A, Cammà C, Scialabba S, Puglia I, Calistri P, Savini G |
| EPI_ISL_1117506, EPI_ISL_1117510, EPI_ISL_1117511, EPI_ISL_1117512, EPI_ISL_1117513 | SIESP DIP PREV CHIETI | Istituto Zooprofilattico Sperimentale dell'Abruzzo e Molise "G. Caporale" | Lorusso A, Marcacci M, Di Domenico M, Ancora M, Curini V, Mangone I, Rinaldi A, Scialabba S, Di Pasquale A, Cammà C, Scialabba S, Puglia I, Calistri P, Savini G |
| EPI_ISL_1163910 | IZSM | IZSM-U.O.C. Virologia | Maurizio Viscardi, Lorena Cardillo, Giuseppe Prota, Vincenzo Petrella, Luigi Atripaldi, Giovanna Fusco, Esterina De Carlo, Antonio Limone |
| EPI_ISL_1163911, EPI_ISL_1163912, EPI_ISL_1163913 | IZSM | IZSM-U.O.C. Virologia | Maurizio Viscardi, Lorena Cardillo, Luigi Atripaldi,Giovanna Fusco, Esterina De Carlo, Antonio Limone |
| EPI_ISL_1164855 | Diagnostic Genomics Lab and Functional Genomics Core University of South Carolina | Diagnostic Genomics Lab and Functional Genomics Core, University of South Carolina | Carolyn Banister, Hao Ji, Diego Altomare, Herr Doktor Professor Genomiklaborsdirektor Universitätkovidenpandemieretter Phillip Buckhaults, Michael Shtutman, Helmut Albrecht |
| EPI_ISL_1166877, EPI_ISL_1166878, EPI_ISL_1166879, EPI_ISL_1166880, EPI_ISL_1166881 | IZSM | IZSM-U.O.C. Virologia | Maurizio Viscardi, Lorena Cardillo, Luigi Atripaldi,Giovanna Fusco, Esterina De Carlo, Antonio Limone |
| EPI_ISL_1168206, EPI_ISL_1168207, EPI_ISL_1168208 | Hopital | National Reference Center for Viruses of Respiratory Infections, Institut Pasteur, Paris | Marion Barbet, Sylvie Behillil, Méline Bizard, Angela Brisebarre, Camille Capel, Etienne Simon-Lorière, Vincent Enouf, Maud Vanpeene, Sylvie van der Werf,Labourdette Isabelle |
| EPI_ISL_1168213, EPI_ISL_1168214, EPI_ISL_1168215 | Armies | National Reference Center for Viruses of Respiratory Infections, Institut Pasteur, Paris | Marion Barbet, Sylvie Behillil, Méline Bizard, Angela Brisebarre, Camille Capel, Etienne Simon-Lorière, Vincent Enouf, Maud Vanpeene, Sylvie van der Werf,Desroches Marine |
| EPI_ISL_1168218 | Hopital | National Reference Center for Viruses of Respiratory Infections, Institut Pasteur, Paris | Marion Barbet, Sylvie Behillil, Méline Bizard, Angela Brisebarre, Camille Capel, Etienne Simon-Lorière, Vincent Enouf, Maud Vanpeene, Sylvie van der Werf,Guinard JéRome |
| EPI_ISL_1169901, EPI_ISL_1169902, EPI_ISL_1169903 | Imelda Hospital | Imelda Hospital | Johan Frans, Dagmar Obbels, Hanne Valgaeren |
| EPI_ISL_1169920 | Galdakao University Hospital | Cruces University Hospital | Ana Gual-de-Torrella, Pilar Berdonces, Mikel Gallego, Ana Belén de la Oz |
| EPI_ISL_1171707 | Baylor Scott & White-Temple | Baylor Scott & White-Temple | ri Rao, Linden Morales, Kimberly Walker, Marcus Volz, Shelby Hendrickson, Caitlin Maloney |
| EPI_ISL_1171711, EPI_ISL_1172018, EPI_ISL_1172019 | Baylor Scott & White-Temple | Baylor Scott & White-Temple | Ari Rao, Linden Morales, Kimberly Walker, Marcus Volz, Shelby Hendrickson, Caitlin Maloney |
| EPI_ISL_1172020 | Baylor Scott & White-Irving | Baylor Scott & White-Temple | Ari Rao, Linden Morales, Kimberly Walker, Marcus Volz, Shelby Hendrickson, Caitlin Maloney |
| EPI_ISL_1173205 | Istituto Zooprofilattico Sperimentale del Mezzogiorno | Istituto Zooprofilattico Sperimentale del Mezzogiorno-U.O.C. Virologia | Maurizio Viscardi, Lorena Cardillo, Luigi Atripaldi,Giovanna Fusco, Esterina De Carlo, Antonio Limone |

| | | | |
|---|---|---|---|
| EPI_ISL_1173768, EPI_ISL_1176298, EPI_ISL_1176300 | Imelda Hospital | Imelda Hospital | Johan Frans, Hanne Valgaeren, Dagmar Obbels |
| EPI_ISL_1176301, EPI_ISL_1176302, EPI_ISL_1176304, EPI_ISL_1176306, EPI_ISL_1176308, EPI_ISL_1176309, EPI_ISL_1176310, EPI_ISL_1176312, EPI_ISL_1176314, EPI_ISL_1176315, EPI_ISL_1176317, EPI_ISL_1176319, EPI_ISL_1176321, EPI_ISL_1176322, EPI_ISL_1176323, EPI_ISL_1176324, EPI_ISL_1176325, EPI_ISL_1176326, EPI_ISL_1176327, EPI_ISL_1176328, EPI_ISL_1176329, EPI_ISL_1176330, EPI_ISL_1176331, EPI_ISL_1176332 | | | |
| see above | AZ Klina | AZ Klina | Dr. C. Vael |
| EPI_ISL_1180141, EPI_ISL_1180142, EPI_ISL_1180143, EPI_ISL_1180144 | Laboratorio Biologia Molecolare Sars Cov2 - UOC Laboratorio Analisi - Servizio Medicina di Laboratorio, Ospedale "San Francesco" - ATS-ASSL Nuoro | Laboratorio specialistico UOC Ematologia - Ospedale "San Francesco" - ATS-ASSL Nuoro | Giovanna Piras, Rosanna Asproni, Paolo Malune, Maria Itria Monne, Maura Fiamma, Angelo Domenico Palmas, Iana Lo Maglio, Giuseppe Mameli |
| EPI_ISL_1180212, EPI_ISL_1180215, EPI_ISL_1180218, EPI_ISL_1180220, EPI_ISL_1180224, EPI_ISL_1180226, EPI_ISL_1180228 | Althaia. Xarxa Assistencial Universitària de Manresa | IrsiCaixa - Can Ruti CovidSeq | Marc Noguera-Julian, Mariona Parera, Maria Casadellà, Pilar Armengol, Francesc Catala-Moll, Roger Paredes, Bonaventura Clotet Gloria Trujillo, Rafel Perez Vidal, Jaume Trape Pujol, Carolina Gonzalez Fernandez, Roger Paredes, Eulalia Grau, Bonaventura Clotet |
| EPI_ISL_1180229, EPI_ISL_1180231, EPI_ISL_1180235, EPI_ISL_1180237, EPI_ISL_1180238, EPI_ISL_1180255, EPI_ISL_1180256, EPI_ISL_1180258, EPI_ISL_1180279, EPI_ISL_1180669, EPI_ISL_1180670, EPI_ISL_1180958 | | | |
| see above | Baylor Scott & White-Temple | Baylor Scott & White-Temple | Ari Rao, Linden Morales, Kimberly Walker, Marcus Volz, Shelby Hendrickson, Caitlin Maloney |
| EPI_ISL_1184825, EPI_ISL_1185852, EPI_ISL_1186023, EPI_ISL_1186024, EPI_ISL_1186069, EPI_ISL_1186070, EPI_ISL_1186113, EPI_ISL_1186203, EPI_ISL_1186213, EPI_ISL_1186215, EPI_ISL_1186217 | | | |
| see above | National Institute of Infectious Diseases-Prof. Dr. Matei Bals Molecular Diagnostics Laboratory | National Institute of Infectious Diseases-Prof. Dr. Matei Bals Molecular Diagnostics Laboratory | Leontina Banica, Marius Surleac, Corina Casangiu, Petre Milu, Andreea Tudor, Simona Paraschiv, Dan Otelea |
| EPI_ISL_1191728 | CHWAPI - SITE NOTRE DAME | Institut de Pathologie et Génétique (IPG) | Pascale Hilbert, Jérémie Gras |
| EPI_ISL_1195444, EPI_ISL_1195445, EPI_ISL_1195446, EPI_ISL_1195447, EPI_ISL_1195448, EPI_ISL_1195449, EPI_ISL_1195450, EPI_ISL_1195451, EPI_ISL_1195452, EPI_ISL_1195453, EPI_ISL_1195454, EPI_ISL_1195455, EPI_ISL_1195456, EPI_ISL_1195457, EPI_ISL_1195458, EPI_ISL_1195459, EPI_ISL_1195460, EPI_ISL_1195461, EPI_ISL_1195462, EPI_ISL_1195463, EPI_ISL_1195464, EPI_ISL_1195465, EPI_ISL_1195466, EPI_ISL_1195467, EPI_ISL_1195468, EPI_ISL_1195469, EPI_ISL_1195475, EPI_ISL_1195476 | | | |
| see above | SC Microbiologia e Virologia AOUSS | AMES Centro Polidiagnostico Strumentale S.r.l. | Gabriele Ibba, Rosangela Sau, Flavia Angioj, Marcello Abbondio, Salvatore Rubino, Sergio Uzzau. |
| EPI_ISL_1195961 | Microbiology and Virology Unit, Florence Careggi University Hospital | Microbiology and Virology Unit, Florence Careggi University Hospital | Vincenzo Di Pilato, Marco Coppi, Fabio Morecchiato, Noemi Aiezza, Ilaria Baccani, Nicla Giovacchini, Alberto Antonelli, Emanuele Gori, Gian Maria Rossolini |
| EPI_ISL_1196281 | Universitas Indonesia Hospital | Faculty of Medicine, Universitas Indonesia | Fadilah, Rafika Indah Paramita, Linda Erlina, Khaerunissa Anbar Istiadi, Ardiana Kusumaningrum, Dimas Seto, Astuti Giantini, Doni Usman, Putra Mahanaim Tampubolon, Aulia Reski Widyaningrum, Ferdi Anda Sitepu, Irandi Putra Pratomo, Dwiana Ocviyanti, Budi Wiweko, Anis Karuniawati, Ari Fahrial Syam, Badriul Hegar |
| EPI_ISL_1197037 | Laboratoire de virologie clinique - Institut Pasteur de Tunis | 1-Laboratory of Microbiology, National Reference Lab, Charles Nicolle Hospital; 2-University of Tunis ElManar, Faculty of Medicine of Tunis, LR99ES09, Tunis, Tunisia | Sameh Trabelsi, Nissaf Ben Alaya, Mouna Ben Sassi, Sana Ferjani, Mouna Safer, Salma Abid, Fares Wasfi, Mariem Gdoura, Sondes Haddad, Anissa Chouikha, Roua Ben Othman, Sara Chammam, Ines Mdini, Manel Ben Sassi, Imen Kacem, Maher Kharrat, Alia BenKahla, Jalila Ben Khelil, Riadh Daghfous, Riadh Gouider, Henda Triki, Ilhem Boutiba-Ben Boubaker. |
| EPI_ISL_1198698 | INMI Lazzaro Spallanzani IRCCS | INMI Lazzaro Spallanzani IRCCS | E Giombini, F Messina, M Rueca, O Butera, CEM Gruber, B Bartolini, A Di Caro, MR Capobianchi |
| EPI_ISL_1200549, EPI_ISL_1200555, EPI_ISL_1200576, EPI_ISL_1200607, EPI_ISL_1200608, EPI_ISL_1200609, EPI_ISL_1200627 | OCME Office Of Chief Medical Examiner | New York City Public Health Laboratory | Jade Wang, et al. |
| EPI_ISL_1201507 | Hopital | National Reference Center for Viruses of Respiratory Infections, Institut Pasteur, Paris | Marion Barbet, Sylvie Behillil, Méline Bizard, Angela Brisebarre, Camille Capel, Etienne Simon-Lorière, Vincent Enouf, Maud Vanpeene, Sylvie van der Werf,Durivault JéRôMe |
| EPI_ISL_1201508, EPI_ISL_1201509, EPI_ISL_1201510 | Hopital | National Reference Center for Viruses of Respiratory Infections, Institut Pasteur, Paris | Marion Barbet, Sylvie Behillil, Méline Bizard, Angela Brisebarre, Camille Capel, Etienne Simon-Lorière, Vincent Enouf, Maud Vanpeene, Sylvie van der Werf,Pham StéPhanie |
| EPI_ISL_1201520 | Hopital | National Reference Center for Viruses of Respiratory Infections, Institut Pasteur, Paris | Marion Barbet, Sylvie Behillil, Méline Bizard, Angela Brisebarre, Camille Capel, Etienne Simon-Lorière, Vincent Enouf, Maud Vanpeene, Sylvie van der Werf,Woercel Isabelle |
| EPI_ISL_1201521 | Labo Analyses Med | National Reference Center for Viruses of Respiratory Infections, Institut Pasteur, Paris | Marion Barbet, Sylvie Behillil, Méline Bizard, Angela Brisebarre, Camille Capel, Etienne Simon-Lorière, Vincent Enouf, Maud Vanpeene, Sylvie van der Werf |
| EPI_ISL_1201522, EPI_ISL_1201523, EPI_ISL_1201524, EPI_ISL_1201525 | Hopital | National Reference Center for Viruses of Respiratory Infections, Institut Pasteur, Paris | Marion Barbet, Sylvie Behillil, Méline Bizard, Angela Brisebarre, Camille Capel, Etienne Simon-Lorière, Vincent Enouf, Maud Vanpeene, Sylvie van der Werf,Desroches Marine |
| EPI_ISL_1201537 | Armies | National Reference Center for Viruses of Respiratory Infections, Institut Pasteur, Paris | Marion Barbet, Sylvie Behillil, Méline Bizard, Angela Brisebarre, Camille Capel, Etienne Simon-Lorière, Vincent Enouf, Maud Vanpeene, Sylvie van der Werf,Rousset Dominique |
| EPI_ISL_1201538 | Armies | National Reference Center for Viruses of Respiratory Infections, Institut Pasteur, Paris | Marion Barbet, Sylvie Behillil, Méline Bizard, Angela Brisebarre, Camille Capel, Etienne Simon-Lorière, Vincent Enouf, Maud Vanpeene, Sylvie van der Werf,Desroches Marine |
| EPI_ISL_1201539 | Hopital | National Reference Center for Viruses of Respiratory Infections, Institut Pasteur, Paris | Marion Barbet, Sylvie Behillil, Méline Bizard, Angela Brisebarre, Camille Capel, Etienne Simon-Lorière, Vincent Enouf, Maud Vanpeene, Sylvie van der Werf,Castelain Sandrine |
| EPI_ISL_1201542, EPI_ISL_1201543 | Hopital | National Reference Center for Viruses of Respiratory Infections, Institut Pasteur, Paris | Marion Barbet, Sylvie Behillil, Méline Bizard, Angela Brisebarre, Camille Capel, Etienne Simon-Lorière, Vincent Enouf, Maud Vanpeene, Sylvie van der Werf,Amade (W) |
| EPI_ISL_1201544, EPI_ISL_1201545, EPI_ISL_1201548 | Labo Analyses Med | National Reference Center for Viruses of Respiratory Infections, Institut Pasteur, Paris | Marion Barbet, Sylvie Behillil, Méline Bizard, Angela Brisebarre, Camille Capel, Etienne Simon-Lorière, Vincent Enouf, Maud Vanpeene, Sylvie van der Werf |
| EPI_ISL_1201553 | Labo Analyses Med | National Reference Center for Viruses of Respiratory Infections, Institut Pasteur, Paris | Marion Barbet, Sylvie Behillil, Méline Bizard, Angela Brisebarre, Camille Capel, Etienne Simon-Lorière, Vincent Enouf, Maud Vanpeene, Sylvie van der Werf,Le Vicky |
| EPI_ISL_1209265, EPI_ISL_1209266 | Institute for Laboratory Diagnostics and Microbiology, Klinikum Klagenfurt am Wörthersee | Bergthaler laboratory, CeMM Research Center for Molecular Medicine of the Austrian Academy of Sciences | Lukas Endler, Anna Schedl, Fabian Amman, Thomas Penz, Benedikt Agerer, Maelle Le Moing, Michael Schuster, Bekir Erguner, Jan Laine, Martin Senekowitsch, Christoph Bock, Andreas Bergthaler |
| EPI_ISL_1209281, EPI_ISL_1209282, EPI_ISL_1209283, EPI_ISL_1209285, EPI_ISL_1209286, EPI_ISL_1209287, EPI_ISL_1209288, EPI_ISL_1209289, EPI_ISL_1209290, EPI_ISL_1209291, EPI_ISL_1209292, EPI_ISL_1209293, EPI_ISL_1209294, EPI_ISL_1209295, EPI_ISL_1209296, EPI_ISL_1209297, EPI_ISL_1209298 | | | |
| see above | Department of Laboratory Medicine, Division of Clinical Virology, University of Medicine, Vienna | Bergthaler laboratory, CeMM Research Center for Molecular Medicine of the Austrian Academy of Sciences | Lukas Endler, Anna Schedl, Fabian Amman, Thomas Penz, Benedikt Agerer, Maelle Le Moing, Michael Schuster, Bekir Erguner, Jan Laine, Martin Senekowitsch, Christoph Bock, Andreas Bergthaler |
| EPI_ISL_1209365, EPI_ISL_1209366, EPI_ISL_1209368 | 3. Medizinische Abteilung, Hanusch Krankenhaus | Bergthaler laboratory, CeMM Research Center for Molecular Medicine of the Austrian Academy of Sciences | Lukas Endler, Anna Schedl, Fabian Amman, Thomas Penz, Benedikt Agerer, Maelle Le Moing, Michael Schuster, Bekir Erguner, Jan Laine, Martin Senekowitsch, Christoph Bock, Andreas Bergthaler |
| EPI_ISL_1209407 | Laboratory for HIV and opportunistic infections diagnosis The Republican Research and Practical Center for Epidemiology and Microbiology (RRPCEM) | Laboratory for HIV and opportunistic infections diagnosis The Republican Research and Practical Center for Epidemiology and Microbiology (RRPCEM) | Elena Gasich, Kirill Bulda, Artur Akhremchuk, Leonid Valentovich, Anatoly Krasko, Vladimir Gorbunov |
| EPI_ISL_1213580 | Philippine Red Cross Logistics and Multipurpose Center | Philippine Genome Center | Francis A. Tablizo, Kenneth M. Kim, Carlo M. Lapid, Marc Jerrone R. Castro, Maria Sofia L. Yangzon, Benedict A. Maralit, Marc Edsel C. Ayes, Eva Maria Cutiongco-de la Paz, Alethea R. de Guzman, Jan Michael C. Yap, Jo-Hannah S. Llames, Sheila Mae M. Araiza, Kris P. Punayan, Irish Coleen A. Asin, Candice Francheska B. Tambaoan, Asia Louisa U. Chong, Karol Sophia Agape R. Padilla, Rianna Patricia S. Cruz, El King D. Morado, Joshua Gregor A. Dizon, Razel Nikka M. Hao, Arianne A. Zamora, Devon Ray Pacial, Juan Antonio R. Magalang, Marissa Alejandria, Celia Carlos, Anna Ong-Lim, Edsel Maurice Salvaña, John Q. Wong, Jaime C. Montoya, Maria Rosario Singh-Vergeire and Cynthia P. Saloma |

| | | | |
|---|---|---|---|
| EPI_ISL_1219011 | CHWAPI - SITE NOTRE DAME | Institut de Pathologie et Génétique (IPG) | Pascale Hilbert, Jérémie Gras |
| EPI_ISL_1219021 | Hospital Aviccena | Instituto Adolfo Lutz, Interdiciplinary Procedures Center, Strategic Laboratory | Claudio Tavares Sacchi, Claudia Regina Gonçalves, Erica Valessa Ramos Gomes, Karoline Rodrigues Campos, Caio Vinicius Dias Lopes |
| EPI_ISL_1219927, EPI_ISL_1219928, EPI_ISL_1219929 | Hopital | National Reference Center for Viruses of Respiratory Infections, Institut Pasteur, Paris | Marion Barbet, Sylvie Behillil, Méline Bizard, Angela Brisebarre, Camille Capel, Etienne Simon-Lorière, Vincent Enouf, Maud Vanpeene, Sylvie van der Werf,Serazin V |
| EPI_ISL_1219932 | Hopital | National Reference Center for Viruses of Respiratory Infections, Institut Pasteur, Paris | Marion Barbet, Sylvie Behillil, Méline Bizard, Angela Brisebarre, Camille Capel, Etienne Simon-Lorière, Vincent Enouf, Maud Vanpeene, Sylvie van der Werf,Delamare Catherine |
| EPI_ISL_1219933, EPI_ISL_1219934 | Hopital | National Reference Center for Viruses of Respiratory Infections, Institut Pasteur, Paris | Marion Barbet, Sylvie Behillil, Méline Bizard, Angela Brisebarre, Camille Capel, Etienne Simon-Lorière, Vincent Enouf, Maud Vanpeene, Sylvie van der Werf,Marque Juillet StéPhanie |
| EPI_ISL_1219935, EPI_ISL_1219936 | Labo Analyses Med | National Reference Center for Viruses of Respiratory Infections, Institut Pasteur, Paris | Marion Barbet, Sylvie Behillil, Méline Bizard, Angela Brisebarre, Camille Capel, Etienne Simon-Lorière, Vincent Enouf, Maud Vanpeene, Sylvie van der Werf |
| EPI_ISL_1219937 | Hopital | National Reference Center for Viruses of Respiratory Infections, Institut Pasteur, Paris | Marion Barbet, Sylvie Behillil, Méline Bizard, Angela Brisebarre, Camille Capel, Etienne Simon-Lorière, Vincent Enouf, Maud Vanpeene, Sylvie van der Werf,Serazin V |
| EPI_ISL_1219939 | Hopital | National Reference Center for Viruses of Respiratory Infections, Institut Pasteur, Paris | Marion Barbet, Sylvie Behillil, Méline Bizard, Angela Brisebarre, Camille Capel, Etienne Simon-Lorière, Vincent Enouf, Maud Vanpeene, Sylvie van der Werf,Delamare Catherine |
| EPI_ISL_1219940, EPI_ISL_1219941 | Hopital | National Reference Center for Viruses of Respiratory Infections, Institut Pasteur, Paris | Marion Barbet, Sylvie Behillil, Méline Bizard, Angela Brisebarre, Camille Capel, Etienne Simon-Lorière, Vincent Enouf, Maud Vanpeene, Sylvie van der Werf,Stoessel-Thouvenin Patricia |
| EPI_ISL_1219943, EPI_ISL_1219944, EPI_ISL_1219945, EPI_ISL_1219946 | Hopital | National Reference Center for Viruses of Respiratory Infections, Institut Pasteur, Paris | Marion Barbet, Sylvie Behillil, Méline Bizard, Angela Brisebarre, Camille Capel, Etienne Simon-Lorière, Vincent Enouf, Maud Vanpeene, Sylvie van der Werf,Delamare Catherine |
| EPI_ISL_1219947, EPI_ISL_1219948 | Hopital | National Reference Center for Viruses of Respiratory Infections, Institut Pasteur, Paris | Marion Barbet, Sylvie Behillil, Méline Bizard, Angela Brisebarre, Camille Capel, Etienne Simon-Lorière, Vincent Enouf, Maud Vanpeene, Sylvie van der Werf,Lepiller Quentin |
| EPI_ISL_1219949 | Labo Analyses Med | National Reference Center for Viruses of Respiratory Infections, Institut Pasteur, Paris | Marion Barbet, Sylvie Behillil, Méline Bizard, Angela Brisebarre, Camille Capel, Etienne Simon-Lorière, Vincent Enouf, Maud Vanpeene, Sylvie van der Werf,Durivault JéRôMe |
| EPI_ISL_1219950 | Labo Analyses Med | National Reference Center for Viruses of Respiratory Infections, Institut Pasteur, Paris | Marion Barbet, Sylvie Behillil, Méline Bizard, Angela Brisebarre, Camille Capel, Etienne Simon-Lorière, Vincent Enouf, Maud Vanpeene, Sylvie van der Werf,Ebel Anne |
| EPI_ISL_1219951 | Hopital | National Reference Center for Viruses of Respiratory Infections, Institut Pasteur, Paris | Marion Barbet, Sylvie Behillil, Méline Bizard, Angela Brisebarre, Camille Capel, Etienne Simon-Lorière, Vincent Enouf, Maud Vanpeene, Sylvie van der Werf,Serazin V |
| EPI_ISL_1219957, EPI_ISL_1219958 | Hopital | National Reference Center for Viruses of Respiratory Infections, Institut Pasteur, Paris | Marion Barbet, Sylvie Behillil, Méline Bizard, Angela Brisebarre, Camille Capel, Etienne Simon-Lorière, Vincent Enouf, Maud Vanpeene, Sylvie van der Werf,Delamare Catherine |
| EPI_ISL_1219959, EPI_ISL_1219960 | Labo Analyses Med | National Reference Center for Viruses of Respiratory Infections, Institut Pasteur, Paris | Marion Barbet, Sylvie Behillil, Méline Bizard, Angela Brisebarre, Camille Capel, Etienne Simon-Lorière, Vincent Enouf, Maud Vanpeene, Sylvie van der Werf,Oliel Jonathan |
| EPI_ISL_1219962, EPI_ISL_1219963 | Labo Analyses Med | National Reference Center for Viruses of Respiratory Infections, Institut Pasteur, Paris | Marion Barbet, Sylvie Behillil, Méline Bizard, Angela Brisebarre, Camille Capel, Etienne Simon-Lorière, Vincent Enouf, Maud Vanpeene, Sylvie van der Werf,Durivault JéRôMe |
| EPI_ISL_1220862 | Laboratoire de microbiologie CHU NIMES | Laboratoire de microbiologie CHU NIMES | Stephan Robin, Marie-josée Carles, Sophie Bravo, Agathe Boudet |
| EPI_ISL_1222766 | Central Scientific Research Department, Gomel State Medical University, | Laboratory of Genomics and Bioinformatics of the Forest Research Institute of the NAS of Belarus | Stoma,I.O., Baranov,O.Y., Voropaev,E.V., Osipkina,O.V.,Zyatskov,A.A., Shaforost,A.S., Padutov,V.E., Panteleev,S.V., Kiryanov,P.S. and Mozharovskaya,L.V. |
| EPI_ISL_1228692, EPI_ISL_1228908, EPI_ISL_1228909 | Laboratory for Respiratory Viruses, Cantacuzino National Military-Medical Institute for Research and Development | Cantacuzino Institute Virology | Luiza Ustea, Nicoleta Paraschiv, Catalina Pascu, Mihaela Lazar |
| EPI_ISL_1231436 | Baylor Esoteric + Molecular Lab | Baylor Esoteric + Molecular Lab | Elizabeth Forrester and Dawn Richards |
| EPI_ISL_1233224, EPI_ISL_1233225, EPI_ISL_1233226, EPI_ISL_1233227, EPI_ISL_1233228, EPI_ISL_1233229, EPI_ISL_1233230 | Hopital | National Reference Center for Viruses of Respiratory Infections, Institut Pasteur, Paris | Marion Barbet, Sylvie Behillil, Méline Bizard, Angela Brisebarre, Camille Capel, Etienne Simon-Lorière, Vincent Enouf, Maud Vanpeene, Sylvie van der Werf,Djamdjian Laura |
| EPI_ISL_1233231, EPI_ISL_1233232 | Hopital | National Reference Center for Viruses of Respiratory Infections, Institut Pasteur, Paris | Marion Barbet, Sylvie Behillil, Méline Bizard, Angela Brisebarre, Camille Capel, Etienne Simon-Lorière, Vincent Enouf, Maud Vanpeene, Sylvie van der Werf,Descamps Dominique |
| EPI_ISL_1233234, EPI_ISL_1233235 | Hopital | National Reference Center for Viruses of Respiratory Infections, Institut Pasteur, Paris | Marion Barbet, Sylvie Behillil, Méline Bizard, Angela Brisebarre, Camille Capel, Etienne Simon-Lorière, Vincent Enouf, Maud Vanpeene, Sylvie van der Werf,Vachee Anne |
| EPI_ISL_1233236 | Hopital | National Reference Center for Viruses of Respiratory Infections, Institut Pasteur, Paris | Marion Barbet, Sylvie Behillil, Méline Bizard, Angela Brisebarre, Camille Capel, Etienne Simon-Lorière, Vincent Enouf, Maud Vanpeene, Sylvie van der Werf,Stoessel-Thouvenin Patricia |
| EPI_ISL_1233237, EPI_ISL_1233238, EPI_ISL_1233240, EPI_ISL_1233241, EPI_ISL_1233242, EPI_ISL_1233243, EPI_ISL_1233244, EPI_ISL_1233245, EPI_ISL_1233246 | Labo Analyses Med | National Reference Center for Viruses of Respiratory Infections, Institut Pasteur, Paris | Marion Barbet, Sylvie Behillil, Méline Bizard, Angela Brisebarre, Camille Capel, Etienne Simon-Lorière, Vincent Enouf, Maud Vanpeene, Sylvie van der Werf |
| EPI_ISL_1233247, EPI_ISL_1233248, EPI_ISL_1233249, EPI_ISL_1233250, EPI_ISL_1233251, EPI_ISL_1233252 | Hopital | National Reference Center for Viruses of Respiratory Infections, Institut Pasteur, Paris | Marion Barbet, Sylvie Behillil, Méline Bizard, Angela Brisebarre, Camille Capel, Etienne Simon-Lorière, Vincent Enouf, Maud Vanpeene, Sylvie van der Werf,Ducancelle Alexandra |
| EPI_ISL_1233253 | Hopital | National Reference Center for Viruses of Respiratory Infections, Institut Pasteur, Paris | Marion Barbet, Sylvie Behillil, Méline Bizard, Angela Brisebarre, Camille Capel, Etienne Simon-Lorière, Vincent Enouf, Maud Vanpeene, Sylvie van der Werf,Cady Anne |
| EPI_ISL_1233820, EPI_ISL_1233821, EPI_ISL_1233822, EPI_ISL_1233823, EPI_ISL_1233824, EPI_ISL_1233825, EPI_ISL_1233826, EPI_ISL_1233827, EPI_ISL_1233828, EPI_ISL_1233830, EPI_ISL_1233831, EPI_ISL_1233832, EPI_ISL_1233833, EPI_ISL_1233834, EPI_ISL_1233835, EPI_ISL_1233836, EPI_ISL_1233837, EPI_ISL_1233838, EPI_ISL_1233839, EPI_ISL_1233840, EPI_ISL_1233842, EPI_ISL_1233843, EPI_ISL_1233844, EPI_ISL_1233845, EPI_ISL_1233846, EPI_ISL_1233847, EPI_ISL_1233848, EPI_ISL_1233849, EPI_ISL_1233850, EPI_ISL_1233851, EPI_ISL_1233852, EPI_ISL_1233853, EPI_ISL_1233854, EPI_ISL_1233856, EPI_ISL_1233857, EPI_ISL_1233858, EPI_ISL_1233859, EPI_ISL_1233860, EPI_ISL_1233861, EPI_ISL_1233862, EPI_ISL_1233863, EPI_ISL_1233864, EPI_ISL_1233865, EPI_ISL_1233866, EPI_ISL_1233867 | | | |
| see above | Institute of Molecular and Translational Medicine / Laboratory of Experimental Medicine, Faculty of Medicine and Dentistry, Palacky University | Institute of Molecular and Translational Medicine / Laboratory of Experimental Medicine | Rastislav Slavkovský, Hana Jaworek, Vladimíra Koudeláková, Marián Hajdúch |
| EPI_ISL_1235679 | Baylor Scott & White-Temple | Baylor Scott & White-Temple | Ari Rao, Linden Morales, Kimberly Walker, Marcus Volz, Shelby Hendrickson, Caitlin Maloney |
| EPI_ISL_1235680 | Baylor Scott & White-Irving | Baylor Scott & White-Temple | Ari Rao, Linden Morales, Kimberly Walker, Marcus Volz, Shelby Hendrickson, Caitlin Maloney |
| EPI_ISL_1235685 | Baylor Scott & White-Temple | Baylor Scott & White-Temple | Ari Rao, Linden Morales, Kimberly Walker, Marcus Volz, Shelby Hendrickson, Caitlin Maloney |
| EPI_ISL_1239142 | Ohio State University Wexner Medical Cente | Polaris Molecular Laboratory | Chang Y-S, Ru P, Chappell D, Tu H, Snyder P, Pancholi P, Koenig S, Corcoran S, Jones D |
| EPI_ISL_1239369 | Hopital | National Reference Center for Viruses of Respiratory Infections, Institut Pasteur, Paris | Marion Barbet, Sylvie Behillil, Méline Bizard, Angela Brisebarre, Camille Capel, Etienne Simon-Lorière, Vincent Enouf, Maud Vanpeene, Sylvie van der Werf,Descamps Dominique |

| | | | |
|---|---|---|---|
| EPI_ISL_1239370 | Hopital | National Reference Center for Viruses of Respiratory Infections, Institut Pasteur, Paris | Marion Barbet, Sylvie Behillil, Méline Bizard, Angela Brisebarre, Camille Capel, Etienne Simon-Lorière, Vincent Enouf, Maud Vanpeene, Sylvie van der Werf,Dufour Mj |
| EPI_ISL_1239371 | Hopital | National Reference Center for Viruses of Respiratory Infections, Institut Pasteur, Paris | Marion Barbet, Sylvie Behillil, Méline Bizard, Angela Brisebarre, Camille Capel, Etienne Simon-Lorière, Vincent Enouf, Maud Vanpeene, Sylvie van der Werf,Ducancelle Alexandra |
| EPI_ISL_1240071, EPI_ISL_1240072, EPI_ISL_1240073, EPI_ISL_1240074, EPI_ISL_1240075, EPI_ISL_1240076 | Clinical Center, University of Sarajevo; Unit for Clinical Microbiology | Clinical Center, University of Sarajevo; Unit for Clinical Microbiology | Irma Salimovi-Beši, Amela Dedei-Ljubovi, Edina Zahirovi, Suzana Arapi, Sebija Izetbegovi, Sandra Vegar-Zubovi |
| EPI_ISL_1240719, EPI_ISL_1240720, EPI_ISL_1240721, EPI_ISL_1240722, EPI_ISL_1240723, EPI_ISL_1240724, EPI_ISL_1240725 | Influenza and respiratory viruses | Human virology Department | A.Hachid, F.Khardine, M.A.Beloufa, A.Bensalem, F.Derrar |
| EPI_ISL_1250946 | Usca castel di sangro | Istituto Zooprofilattico Sperimentale dell'Abruzzo e Molise "G. Caporale" | Lorusso A, Marcacci M, Di Domenico M, Ancora M, Curini V, Mangone I, Rinaldi A, Scialabba S, Di Pasquale A, Cammà C, Puglia I, Calistri P, Savini G |
| EPI_ISL_1250947 | siesp Chieti | Istituto Zooprofilattico Sperimentale dell'Abruzzo e Molise "G. Caporale" | Lorusso A, Marcacci M, Di Domenico M, Ancora M, Curini V, Mangone I, Rinaldi A, Scialabba S, Di Pasquale A, Cammà C, Puglia I, Calistri P, Savini G |
| EPI_ISL_1250948 | Usca castel di sangro | Istituto Zooprofilattico Sperimentale dell'Abruzzo e Molise "G. Caporale" | Lorusso A, Marcacci M, Di Domenico M, Ancora M, Curini V, Mangone I, Rinaldi A, Scialabba S, Di Pasquale A, Cammà C, Puglia I, Calistri P, Savini G |
| EPI_ISL_1250949 | Siesp dipartimento di prevenzione | Istituto Zooprofilattico Sperimentale dell'Abruzzo e Molise "G. Caporale" | Lorusso A, Marcacci M, Di Domenico M, Ancora M, Curini V, Mangone I, Rinaldi A, Scialabba S, Di Pasquale A, Cammà C, Puglia I, Calistri P, Savini G |
| EPI_ISL_1250950, EPI_ISL_1250951 | siesp dipartimento di prevenzione | Istituto Zooprofilattico Sperimentale dell'Abruzzo e Molise "G. Caporale" | Lorusso A, Marcacci M, Di Domenico M, Ancora M, Curini V, Mangone I, Rinaldi A, Scialabba S, Di Pasquale A, Cammà C, Puglia I, Calistri P, Savini G |
| EPI_ISL_1250952 | ospedale civile teramo | Istituto Zooprofilattico Sperimentale dell'Abruzzo e Molise "G. Caporale" | Lorusso A, Marcacci M, Di Domenico M, Ancora M, Curini V, Mangone I, Rinaldi A, Scialabba S, Di Pasquale A, Cammà C, Puglia I, Calistri P, Savini G |
| EPI_ISL_1250953 | dip prev avezzano | Istituto Zooprofilattico Sperimentale dell'Abruzzo e Molise "G. Caporale" | Lorusso A, Marcacci M, Di Domenico M, Ancora M, Curini V, Mangone I, Rinaldi A, Scialabba S, Di Pasquale A, Cammà C, Puglia I, Calistri P, Savini G |
| EPI_ISL_1250954 | siesp Chieti | Istituto Zooprofilattico Sperimentale dell'Abruzzo e Molise "G. Caporale" | Lorusso A, Marcacci M, Di Domenico M, Ancora M, Curini V, Mangone I, Rinaldi A, Scialabba S, Di Pasquale A, Cammà C, Puglia I, Calistri P, Savini G |
| EPI_ISL_1250957 | Siesp dipartimento di prevenzione | Istituto Zooprofilattico Sperimentale dell'Abruzzo e Molise "G. Caporale" | Lorusso A, Marcacci M, Di Domenico M, Ancora M, Curini V, Mangone I, Rinaldi A, Scialabba S, Di Pasquale A, Cammà C, Puglia I, Calistri P, Savini G |
| EPI_ISL_1250958 | dip prev avezzano | Istituto Zooprofilattico Sperimentale dell'Abruzzo e Molise "G. Caporale" | Lorusso A, Marcacci M, Di Domenico M, Ancora M, Curini V, Mangone I, Rinaldi A, Scialabba S, Di Pasquale A, Cammà C, Puglia I, Calistri P, Savini G |
| EPI_ISL_1250959, EPI_ISL_1250961 | siesp Chieti | Istituto Zooprofilattico Sperimentale dell'Abruzzo e Molise "G. Caporale" | Lorusso A, Marcacci M, Di Domenico M, Ancora M, Curini V, Mangone I, Rinaldi A, Scialabba S, Di Pasquale A, Cammà C, Puglia I, Calistri P, Savini G |
| EPI_ISL_1250962, EPI_ISL_1250965 | Siesp dipartimento di prevenzione | Istituto Zooprofilattico Sperimentale dell'Abruzzo e Molise "G. Caporale" | Lorusso A, Marcacci M, Di Domenico M, Ancora M, Curini V, Mangone I, Rinaldi A, Scialabba S, Di Pasquale A, Cammà C, Puglia I, Calistri P, Savini G |
| EPI_ISL_1250967, EPI_ISL_1250968 | SIESP CHIETI - LANCIANO | Istituto Zooprofilattico Sperimentale dell'Abruzzo e Molise "G. Caporale" | Lorusso A, Marcacci M, Di Domenico M, Ancora M, Curini V, Mangone I, Rinaldi A, Scialabba S, Di Pasquale A, Cammà C, Puglia I, Calistri P, Savini G |
| EPI_ISL_1250969 | Siesp dipartimento di prevenzione | Istituto Zooprofilattico Sperimentale dell'Abruzzo e Molise "G. Caporale" | Lorusso A, Marcacci M, Di Domenico M, Ancora M, Curini V, Mangone I, Rinaldi A, Scialabba S, Di Pasquale A, Cammà C, Puglia I, Calistri P, Savini G |
| EPI_ISL_1250970, EPI_ISL_1250971 | SIESP CHIETI - LANCIANO | Istituto Zooprofilattico Sperimentale dell'Abruzzo e Molise "G. Caporale" | Lorusso A, Marcacci M, Di Domenico M, Ancora M, Curini V, Mangone I, Rinaldi A, Scialabba S, Di Pasquale A, Cammà C, Puglia I, Calistri P, Savini G |
| EPI_ISL_1250972 | OSPEDALE SANT'OMERO | Istituto Zooprofilattico Sperimentale dell'Abruzzo e Molise "G. Caporale" | Lorusso A, Marcacci M, Di Domenico M, Ancora M, Curini V, Mangone I, Rinaldi A, Scialabba S, Di Pasquale A, Cammà C, Puglia I, Calistri P, Savini G |
| EPI_ISL_1250974 | SIESP CHIETI - LANCIANO | Istituto Zooprofilattico Sperimentale dell'Abruzzo e Molise "G. Caporale" | Lorusso A, Marcacci M, Di Domenico M, Ancora M, Curini V, Mangone I, Rinaldi A, Scialabba S, Di Pasquale A, Cammà C, Puglia I, Calistri P, Savini G |
| EPI_ISL_1250975, EPI_ISL_1250976, EPI_ISL_1250978 | Siesp dipartimento di prevenzione | Istituto Zooprofilattico Sperimentale dell'Abruzzo e Molise "G. Caporale" | Lorusso A, Marcacci M, Di Domenico M, Ancora M, Curini V, Mangone I, Rinaldi A, Scialabba S, Di Pasquale A, Cammà C, Puglia I, Calistri P, Savini G |
| EPI_ISL_1250981, EPI_ISL_1250982 | DIP. PREV. AVEZZANO | Istituto Zooprofilattico Sperimentale dell'Abruzzo e Molise "G. Caporale" | Lorusso A, Marcacci M, Di Domenico M, Ancora M, Curini V, Mangone I, Rinaldi A, Scialabba S, Di Pasquale A, Cammà C, Puglia I, Calistri P, Savini G |
| EPI_ISL_1250991, EPI_ISL_1250992, EPI_ISL_1250994, EPI_ISL_1250995, EPI_ISL_1250996, EPI_ISL_1250997 | Siesp dipartimento di prevenzione | Istituto Zooprofilattico Sperimentale dell'Abruzzo e Molise "G. Caporale" | Lorusso A, Marcacci M, Di Domenico M, Ancora M, Curini V, Mangone I, Rinaldi A, Scialabba S, Di Pasquale A, Cammà C, Puglia I, Calistri P, Savini G |
| EPI_ISL_1251000 | SIESP CHIETI - LANCIANO | Istituto Zooprofilattico Sperimentale dell'Abruzzo e Molise "G. Caporale" | Lorusso A, Marcacci M, Di Domenico M, Ancora M, Curini V, Mangone I, Rinaldi A, Scialabba S, Di Pasquale A, Cammà C, Puglia I, Calistri P, Savini G |
| EPI_ISL_1251001 | SIESP CHIETI - DRIVE IN ORTONA | Istituto Zooprofilattico Sperimentale dell'Abruzzo e Molise "G. Caporale" | Lorusso A, Marcacci M, Di Domenico M, Ancora M, Curini V, Mangone I, Rinaldi A, Scialabba S, Di Pasquale A, Cammà C, Puglia I, Calistri P, Savini G |
| EPI_ISL_1251003 | SIESP CHIETI - DRIVE IN CHIET | Istituto Zooprofilattico Sperimentale dell'Abruzzo e Molise "G. Caporale" | Lorusso A, Marcacci M, Di Domenico M, Ancora M, Curini V, Mangone I, Rinaldi A, Scialabba S, Di Pasquale A, Cammà C, Puglia I, Calistri P, Savini G |
| EPI_ISL_1251004 | siesp Chieti | Istituto Zooprofilattico Sperimentale dell'Abruzzo e Molise "G. Caporale" | Lorusso A, Marcacci M, Di Domenico M, Ancora M, Curini V, Mangone I, Rinaldi A, Scialabba S, Di Pasquale A, Cammà C, Puglia I, Calistri P, Savini G |
| EPI_ISL_1251005, EPI_ISL_1251006, EPI_ISL_1251007 | OSPEDALE SAN SALVATORE | Istituto Zooprofilattico Sperimentale dell'Abruzzo e Molise "G. Caporale" | Lorusso A, Marcacci M, Di Domenico M, Ancora M, Curini V, Mangone I, Rinaldi A, Scialabba S, Di Pasquale A, Cammà C, Puglia I, Calistri P, Savini G |
| EPI_ISL_1251008, EPI_ISL_1251009, EPI_ISL_1251011, EPI_ISL_1251014, EPI_ISL_1251015, EPI_ISL_1251016 | Siesp dipartimento di prevenzione | Istituto Zooprofilattico Sperimentale dell'Abruzzo e Molise "G. Caporale" | Lorusso A, Marcacci M, Di Domenico M, Ancora M, Curini V, Mangone I, Rinaldi A, Scialabba S, Di Pasquale A, Cammà C, Puglia I, Calistri P, Savini G |
| EPI_ISL_1251018 | "DIP. PREV. AVEZZANO SERVIZIO DI IGIENE EPIDEMIOLOGIA E SANITA' PUBBLICA | Istituto Zooprofilattico Sperimentale dell'Abruzzo e Molise "G. Caporale" | Lorusso A, Marcacci M, Di Domenico M, Ancora M, Curini V, Mangone I, Rinaldi A, Scialabba S, Di Pasquale A, Cammà C, Puglia I, Calistri P, Savini G |
| EPI_ISL_1251019, EPI_ISL_1251021, EPI_ISL_1251022 | Usca castel di sangro | Istituto Zooprofilattico Sperimentale dell'Abruzzo e Molise "G. Caporale" | Lorusso A, Marcacci M, Di Domenico M, Ancora M, Curini V, Mangone I, Rinaldi A, Scialabba S, Di Pasquale A, Cammà C, Puglia I, Calistri P, Savini G |
| EPI_ISL_1251023 | USCA Pescina | Istituto Zooprofilattico Sperimentale dell'Abruzzo e Molise "G. Caporale" | Lorusso A, Marcacci M, Di Domenico M, Ancora M, Curini V, Mangone I, Rinaldi A, Scialabba S, Di Pasquale A, Cammà C, Puglia I, Calistri P, Savini G |

| | | | |
|---|---|---|---|
| EPI_ISL_1251024 | OSPEDALE CIVILE GIULIANOVA | Istituto Zooprofilattico Sperimentale dell'Abruzzo e Molise "G. Caporale" | Lorusso A, Marcacci M, Di Domenico M, Ancora M, Curini V, Mangone I, Rinaldi A, Scialabba S, Di Pasquale A, Cammà C, Puglia I, Calistri P, Savini G |
| EPI_ISL_1251025 | PRESIDIO TERRITORIALE CASOLI | Istituto Zooprofilattico Sperimentale dell'Abruzzo e Molise "G. Caporale" | Lorusso A, Marcacci M, Di Domenico M, Ancora M, Curini V, Mangone I, Rinaldi A, Scialabba S, Di Pasquale A, Cammà C, Puglia I, Calistri P, Savini G |
| EPI_ISL_1251026 | siesp Chieti | Istituto Zooprofilattico Sperimentale dell'Abruzzo e Molise "G. Caporale" | Lorusso A, Marcacci M, Di Domenico M, Ancora M, Curini V, Mangone I, Rinaldi A, Scialabba S, Di Pasquale A, Cammà C, Puglia I, Calistri P, Savini G |
| EPI_ISL_1251027 | USCA Tagliacozzo | Istituto Zooprofilattico Sperimentale dell'Abruzzo e Molise "G. Caporale" | Lorusso A, Marcacci M, Di Domenico M, Ancora M, Curini V, Mangone I, Rinaldi A, Scialabba S, Di Pasquale A, Cammà C, Puglia I, Calistri P, Savini G |
| EPI_ISL_1251028 | siesp Chieti | Istituto Zooprofilattico Sperimentale dell'Abruzzo e Molise "G. Caporale" | Lorusso A, Marcacci M, Di Domenico M, Ancora M, Curini V, Mangone I, Rinaldi A, Scialabba S, Di Pasquale A, Cammà C, Puglia I, Calistri P, Savini G |
| EPI_ISL_1251029, EPI_ISL_1251030, EPI_ISL_1251032 | SIESP CHIETI - DRIVE IN VASTO | Istituto Zooprofilattico Sperimentale dell'Abruzzo e Molise "G. Caporale" | Lorusso A, Marcacci M, Di Domenico M, Ancora M, Curini V, Mangone I, Rinaldi A, Scialabba S, Di Pasquale A, Cammà C, Puglia I, Calistri P, Savini G |
| EPI_ISL_1251033, EPI_ISL_1251034 | Siesp dipartimento di prevenzione | Istituto Zooprofilattico Sperimentale dell'Abruzzo e Molise "G. Caporale" | Lorusso A, Marcacci M, Di Domenico M, Ancora M, Curini V, Mangone I, Rinaldi A, Scialabba S, Di Pasquale A, Cammà C, Puglia I, Calistri P, Savini G |
| EPI_ISL_1251037 | SIESP SULMONA | Istituto Zooprofilattico Sperimentale dell'Abruzzo e Molise "G. Caporale" | Lorusso A, Marcacci M, Di Domenico M, Ancora M, Curini V, Mangone I, Rinaldi A, Scialabba S, Di Pasquale A, Cammà C, Puglia I, Calistri P, Savini G |
| EPI_ISL_1251038 | SIESP CHIETI - DRIVE IN ORTONA | Istituto Zooprofilattico Sperimentale dell'Abruzzo e Molise "G. Caporale" | Lorusso A, Marcacci M, Di Domenico M, Ancora M, Curini V, Mangone I, Rinaldi A, Scialabba S, Di Pasquale A, Cammà C, Puglia I, Calistri P, Savini G |
| EPI_ISL_1251039 | DIP. PREV. AVEZZANO SERVIZIO DI IGIENE EPIDEMIOLOGIA E SANITA' PUBBLICA | Istituto Zooprofilattico Sperimentale dell'Abruzzo e Molise "G. Caporale" | Lorusso A, Marcacci M, Di Domenico M, Ancora M, Curini V, Mangone I, Rinaldi A, Scialabba S, Di Pasquale A, Cammà C, Puglia I, Calistri P, Savini G |
| EPI_ISL_1251040 | SIESP L'AQUILA | Istituto Zooprofilattico Sperimentale dell'Abruzzo e Molise "G. Caporale" | Lorusso A, Marcacci M, Di Domenico M, Ancora M, Curini V, Mangone I, Rinaldi A, Scialabba S, Di Pasquale A, Cammà C, Puglia I, Calistri P, Savini G |
| EPI_ISL_1251043 | SIESP CHIETI - LANCIANO | Istituto Zooprofilattico Sperimentale dell'Abruzzo e Molise "G. Caporale" | Lorusso A, Marcacci M, Di Domenico M, Ancora M, Curini V, Mangone I, Rinaldi A, Scialabba S, Di Pasquale A, Cammà C, Puglia I, Calistri P, Savini G |
| EPI_ISL_1251044 | USCA Avezzano | Istituto Zooprofilattico Sperimentale dell'Abruzzo e Molise "G. Caporale" | Lorusso A, Marcacci M, Di Domenico M, Ancora M, Curini V, Mangone I, Rinaldi A, Scialabba S, Di Pasquale A, Cammà C, Puglia I, Calistri P, Savini G |
| EPI_ISL_1251045 | OSPEDALE CIVILE GIULIANOVA | Istituto Zooprofilattico Sperimentale dell'Abruzzo e Molise "G. Caporale" | Lorusso A, Marcacci M, Di Domenico M, Ancora M, Curini V, Mangone I, Rinaldi A, Scialabba S, Di Pasquale A, Cammà C, Puglia I, Calistri P, Savini G |
| EPI_ISL_1251046, EPI_ISL_1251047, EPI_ISL_1251048, EPI_ISL_1251049, EPI_ISL_1251050 | Siesp dipartimento di prevenzione | Istituto Zooprofilattico Sperimentale dell'Abruzzo e Molise "G. Caporale" | Lorusso A, Marcacci M, Di Domenico M, Ancora M, Curini V, Mangone I, Rinaldi A, Scialabba S, Di Pasquale A, Cammà C, Puglia I, Calistri P, Savini G |
| EPI_ISL_1251055 | OSPEDALE CIVILE GIULIANOVA - MEDICINA INTERNA | Istituto Zooprofilattico Sperimentale dell'Abruzzo e Molise "G. Caporale" | Lorusso A, Marcacci M, Di Domenico M, Ancora M, Curini V, Mangone I, Rinaldi A, Scialabba S, Di Pasquale A, Cammà C, Puglia I, Calistri P, Savini G |
| EPI_ISL_1251057, EPI_ISL_1251059, EPI_ISL_1251060 | Siesp dipartimento di prevenzione | Istituto Zooprofilattico Sperimentale dell'Abruzzo e Molise "G. Caporale" | Lorusso A, Marcacci M, Di Domenico M, Ancora M, Curini V, Mangone I, Rinaldi A, Scialabba S, Di Pasquale A, Cammà C, Puglia I, Calistri P, Savini G |
| EPI_ISL_1251061 | DIP. PREV. AVEZZANO SERVIZIO DI IGIENE EPIDEMIOLOGIA E SANITA' PUBBLICA | Istituto Zooprofilattico Sperimentale dell'Abruzzo e Molise "G. Caporale" | Lorusso A, Marcacci M, Di Domenico M, Ancora M, Curini V, Mangone I, Rinaldi A, Scialabba S, Di Pasquale A, Cammà C, Puglia I, Calistri P, Savini G |
| EPI_ISL_1251063, EPI_ISL_1251064 | Siesp dipartimento di prevenzione | Istituto Zooprofilattico Sperimentale dell'Abruzzo e Molise "G. Caporale" | Lorusso A, Marcacci M, Di Domenico M, Ancora M, Curini V, Mangone I, Rinaldi A, Scialabba S, Di Pasquale A, Cammà C, Puglia I, Calistri P, Savini G |
| EPI_ISL_1251066, EPI_ISL_1251067 | USCA Castel di Sangro | Istituto Zooprofilattico Sperimentale dell'Abruzzo e Molise "G. Caporale" | Lorusso A, Marcacci M, Di Domenico M, Ancora M, Curini V, Mangone I, Rinaldi A, Scialabba S, Di Pasquale A, Cammà C, Puglia I, Calistri P, Savini G |
| EPI_ISL_1251068, EPI_ISL_1251069, EPI_ISL_1251071 | SIESP CHIETI - DRIVE IN ORTONA | Istituto Zooprofilattico Sperimentale dell'Abruzzo e Molise "G. Caporale" | Lorusso A, Marcacci M, Di Domenico M, Ancora M, Curini V, Mangone I, Rinaldi A, Scialabba S, Di Pasquale A, Cammà C, Puglia I, Calistri P, Savini G |
| EPI_ISL_1251072 | USCA Castel di Sangro | Istituto Zooprofilattico Sperimentale dell'Abruzzo e Molise "G. Caporale" | Lorusso A, Marcacci M, Di Domenico M, Ancora M, Curini V, Mangone I, Rinaldi A, Scialabba S, Di Pasquale A, Cammà C, Puglia I, Calistri P, Savini G |
| EPI_ISL_1251077 | siesp Chieti | Istituto Zooprofilattico Sperimentale dell'Abruzzo e Molise "G. Caporale" | Lorusso A, Marcacci M, Di Domenico M, Ancora M, Curini V, Mangone I, Rinaldi A, Scialabba S, Di Pasquale A, Cammà C, Puglia I, Calistri P, Savini G |
| EPI_ISL_1251078 | "Presidio Ospedaliero "San Liberatore" Atri | Istituto Zooprofilattico Sperimentale dell'Abruzzo e Molise "G. Caporale" | Lorusso A, Marcacci M, Di Domenico M, Ancora M, Curini V, Mangone I, Rinaldi A, Scialabba S, Di Pasquale A, Cammà C, Puglia I, Calistri P, Savini G |
| EPI_ISL_1251079 | RSA GIULIANOVA | Istituto Zooprofilattico Sperimentale dell'Abruzzo e Molise "G. Caporale" | Lorusso A, Marcacci M, Di Domenico M, Ancora M, Curini V, Mangone I, Rinaldi A, Scialabba S, Di Pasquale A, Cammà C, Puglia I, Calistri P, Savini G |
| EPI_ISL_1251080, EPI_ISL_1251082 | Siesp dipartimento di prevenzione | Istituto Zooprofilattico Sperimentale dell'Abruzzo e Molise "G. Caporale" | Lorusso A, Marcacci M, Di Domenico M, Ancora M, Curini V, Mangone I, Rinaldi A, Scialabba S, Di Pasquale A, Cammà C, Puglia I, Calistri P, Savini G |
| EPI_ISL_1251083 | SIESP CHIETI - DRIVE IN CHIETI | Istituto Zooprofilattico Sperimentale dell'Abruzzo e Molise "G. Caporale" | Lorusso A, Marcacci M, Di Domenico M, Ancora M, Curini V, Mangone I, Rinaldi A, Scialabba S, Di Pasquale A, Cammà C, Puglia I, Calistri P, Savini G |
| EPI_ISL_1251086, EPI_ISL_1251087, EPI_ISL_1251088, EPI_ISL_1251090, EPI_ISL_1251091 | Siesp dipartimento di prevenzione | Istituto Zooprofilattico Sperimentale dell'Abruzzo e Molise "G. Caporale" | Lorusso A, Marcacci M, Di Domenico M, Ancora M, Curini V, Mangone I, Rinaldi A, Scialabba S, Di Pasquale A, Cammà C, Puglia I, Calistri P, Savini G |
| EPI_ISL_1251094 | SIESP CHIETI - LANCIANO | Istituto Zooprofilattico Sperimentale dell'Abruzzo e Molise "G. Caporale" | Lorusso A, Marcacci M, Di Domenico M, Ancora M, Curini V, Mangone I, Rinaldi A, Scialabba S, Di Pasquale A, Cammà C, Puglia I, Calistri P, Savini G |
| EPI_ISL_1251096 | Presidio Ospedaliero "San Liberatore" Atri | Istituto Zooprofilattico Sperimentale dell'Abruzzo e Molise "G. Caporale" | Lorusso A, Marcacci M, Di Domenico M, Ancora M, Curini V, Mangone I, Rinaldi A, Scialabba S, Di Pasquale A, Cammà C, Puglia I, Calistri P, Savini G |
| EPI_ISL_1251098, EPI_ISL_1251099, EPI_ISL_1251100, EPI_ISL_1251104, EPI_ISL_1251106 | Siesp dipartimento di prevenzione | Istituto Zooprofilattico Sperimentale dell'Abruzzo e Molise "G. Caporale" | Lorusso A, Marcacci M, Di Domenico M, Ancora M, Curini V, Mangone I, Rinaldi A, Scialabba S, Di Pasquale A, Cammà C, Puglia I, Calistri P, Savini G |
| EPI_ISL_1251108 | SIESP CHIETI - DRIVE IN ORTONA | Istituto Zooprofilattico Sperimentale dell'Abruzzo e Molise "G. Caporale" | Lorusso A, Marcacci M, Di Domenico M, Ancora M, Curini V, Mangone I, Rinaldi A, Scialabba S, Di Pasquale A, Cammà C, Puglia I, Calistri P, Savini G |
| EPI_ISL_1251109, EPI_ISL_1251112 | SIESP CHIETI - LANCIANO | Istituto Zooprofilattico Sperimentale dell'Abruzzo e Molise "G. Caporale" | Lorusso A, Marcacci M, Di Domenico M, Ancora M, Curini V, Mangone I, Rinaldi A, Scialabba S, Di Pasquale A, Cammà C, Puglia I, Calistri P, Savini G |
| EPI_ISL_1251114 | RSA GIULIANOVA | Istituto Zooprofilattico Sperimentale dell'Abruzzo e Molise "G. Caporale" | Lorusso A, Marcacci M, Di Domenico M, Ancora M, Curini V, Mangone I, Rinaldi A, Scialabba S, Di Pasquale A, Cammà C, Puglia I, Calistri P, Savini G |

| | | | |
|---|---|---|---|
| EPI_ISL_1251116 | SIESP L'AQUILA | Istituto Zooprofilattico Sperimentale dell'Abruzzo e Molise "G. Caporale" | Lorusso A, Marcacci M, Di Domenico M, Ancora M, Curini V, Mangone I, Rinaldi A, Scialabba S, Di Pasquale A, Cammà C, Puglia I, Calistri P, Savini G |
| EPI_ISL_1251244, EPI_ISL_1251246 | SC (UCO) Igiene e Sanità Pubblica | ARGO Laboratorio Genomica ed Epigenomica | Licastro D, Dal Monego S, Degasperi M, Segat L, Piscianz E, D'Agaro P |
| EPI_ISL_1254646 | Diagnostic Genomics Lab and Functional Genomics Core University of South Carolina | Diagnostic Genomics Lab and Functional Genomics Core, University of South Carolina | Carolyn Banister, Hao Ji, Diego Altomare, Herr Doktor Professor Genomiklaborsdirektor Universitätkovidenpandemieretter Phillip Buckhaults, Michael Shtutman, Helmut Albrecht |
| EPI_ISL_1254660, EPI_ISL_1254661, EPI_ISL_1254662, EPI_ISL_1254663, EPI_ISL_1254664 | Presidio di Brindisi Di Summa - Perrino | Istituto Zooprofilattico Sperimentale della Puglia e della Basilicata | Parisi A., Bianco A., Capozzi L., Del Sambro L., Simone D., Difato L., Ridolfi D., Giannico A. |
| EPI_ISL_1254665 | Ospedale Santa Caterina Novella | Istituto Zooprofilattico Sperimentale della Puglia e della Basilicata | Parisi A., Bianco A., Capozzi L., Del Sambro L., Simone D., Difato L., Ridolfi D., Giannico A. |
| EPI_ISL_1254666, EPI_ISL_1254669, EPI_ISL_1254670 | Ospedale di Venere | Istituto Zooprofilattico Sperimentale della Puglia e della Basilicata | Parisi A., Bianco A., Capozzi L., Del Sambro L., Simone D., Iacobellis M., Ridolfi D., Giannico A. |
| EPI_ISL_1254671, EPI_ISL_1254672 | Istituto Zooprofilattico Sperimentale della Puglia e della Basilicata | Istituto Zooprofilattico Sperimentale della Puglia e della Basilicata | Parisi A., Bianco A., Capozzi L., Del Sambro L., Simone D., Manzulli V, Rondinone V., Pace L., Cipolletta D., Galante D. |
| EPI_ISL_1254675, EPI_ISL_1254676, EPI_ISL_1254677, EPI_ISL_1254678, EPI_ISL_1254679, EPI_ISL_1254680 | Presidio di Brindisi Di Summa - Perrino | Istituto Zooprofilattico Sperimentale della Puglia e della Basilicata | Parisi A., Bianco A., Capozzi L., Del Sambro L., Simone D., Difato L., Ridolfi D., Giannico A. |
| EPI_ISL_1254682, EPI_ISL_1254683, EPI_ISL_1254684, EPI_ISL_1254685, EPI_ISL_1254686, EPI_ISL_1254687 | Ospedale Generale Regionale F. Miulli | Istituto Zooprofilattico Sperimentale della Puglia e della Basilicata | Parisi A., Bianco A., Capozzi L., Del Sambro L., Simone D., Difato L., Ridolfi D., Giannico A. |
| EPI_ISL_1254690, EPI_ISL_1254691, EPI_ISL_1254692, EPI_ISL_1254693, EPI_ISL_1254695 | Istituto Zooprofilattico Sperimentale della Puglia e della Basilicata | Istituto Zooprofilattico Sperimentale della Puglia e della Basilicata | Parisi A., Bianco A., Capozzi L., Del Sambro L., Simone D., Manzulli V, Rondinone V., Pace L., Cipolletta D., Galante D. |
| EPI_ISL_1254696, EPI_ISL_1254697, EPI_ISL_1254698, EPI_ISL_1254699, EPI_ISL_1254700 | Ospedale Santissima Annunziata | Istituto Zooprofilattico Sperimentale della Puglia e della Basilicata | Parisi A., Bianco A., Capozzi L., Del Sambro L., Simone D., Tampoia M., Difato L., Ridolfi D., Giannico A. |
| EPI_ISL_1254702, EPI_ISL_1254703 | Ospedale Casa Sollievo della Sofferenza | Istituto Zooprofilattico Sperimentale della Puglia e della Basilicata | Parisi A., Bianco A., Capozzi L., Del Sambro L., Simone D., Difato L., Ridolfi D., Giannico A. |
| EPI_ISL_1257856 | BBTKLPP Yogyakarta | National Institute of Health Research and Development | Kartika Dewi Puspa, Hana Apsari Pawestri, Subangkit, Arie Ardiansyah Nugraha, Hartanti Dian Ikawati, Nelly Puspandari, Krisna Pangesti, Irene, Havid Setyawan, Indaryati, Vivi Setiawaty |
| EPI_ISL_1257898 | National Medicines Institute | DNA Sequencing and Synthesis Facility (oligo.pl), Institute of Biochemistry and Biophysics PAS | Gawor Jan, Baraniak Anna |
| EPI_ISL_1258426, EPI_ISL_1258427, EPI_ISL_1258428, EPI_ISL_1258431, EPI_ISL_1258432, EPI_ISL_1258433, EPI_ISL_1258434, EPI_ISL_1258435, EPI_ISL_1258437, EPI_ISL_1258438, EPI_ISL_1258439, EPI_ISL_1258440, EPI_ISL_1258441, EPI_ISL_1258442 | | | |
| see above | National Institute of Infectious Diseases-Prof. Dr. Matei Bals Molecular Diagnostics Laboratory | National Institute of Infectious Diseases-Prof. Dr. Matei Bals Molecular Diagnostics Laboratory | Corina Casangiu, Leontina Banica, Marius Surleac, Ovidiu Vlaicu, Ionelia Nicolae, Simona Paraschiv, Dan Otelea |
| EPI_ISL_1258443 | Elias Emergency University Hospital, Bucharest | National Institute of Infectious Diseases-Prof. Dr. Matei Bals Molecular Diagnostics Laboratory | Corina Casangiu, Leontina Banica, Marius Surleac, Ovidiu Vlaicu, Ionelia Nicolae, Simona Paraschiv, Dan Otelea |
| EPI_ISL_1258445, EPI_ISL_1258446, EPI_ISL_1258455, EPI_ISL_1258456, EPI_ISL_1258457, EPI_ISL_1258458 | National Institute of Infectious Diseases-Prof. Dr. Matei Bals Molecular Diagnostics Laboratory | National Institute of Infectious Diseases-Prof. Dr. Matei Bals Molecular Diagnostics Laboratory | Corina Casangiu, Leontina Banica, Marius Surleac, Ovidiu Vlaicu, Ionelia Nicolae, Simona Paraschiv, Dan Otelea |
| EPI_ISL_1258460, EPI_ISL_1258461 | Gorj County Directorate of Public Health | National Institute of Infectious Diseases-Prof. Dr. Matei Bals Molecular Diagnostics Laboratory | Corina Casangiu, Leontina Banica, Marius Surleac, Ovidiu Vlaicu, Ionelia Nicolae, Simona Paraschiv, Dan Otelea |
| EPI_ISL_1259194, EPI_ISL_1259195, EPI_ISL_1259196, EPI_ISL_1259197, EPI_ISL_1259198, EPI_ISL_1259199, EPI_ISL_1259201, EPI_ISL_1259202, EPI_ISL_1259203, EPI_ISL_1259204, EPI_ISL_1259205, EPI_ISL_1259206, EPI_ISL_1259207 | | | |
| see above | Department of Virology, Istituto Zooprofilattico Sperimentale del Lazio e della Toscana (IZSLT) | Department of General Diagnostics; Department of Virology; Istituto Zooprofilattico Sperimentale del Lazio e della Toscana (IZSLT) | Patricia Alba, Giuseppe Manna, Elena L. Diaconu, Fabiola Feltrin, Raffaela Conti, Teresa Scicluna, Virginia Carfora, Antonella Cersini, Alessia Franco, Antonio Battisti. |
| EPI_ISL_1259307 | Hopital | National Reference Center for Viruses of Respiratory Infections, Institut Pasteur, Paris | Marion Barbet, Sylvie Behillil, Méline Bizard, Angela Brisebarre, Camille Capel, Etienne Simon-Lorière, Vincent Enouf, Maud Vanpeene, Sylvie van der Werf,Dufour Mj |
| EPI_ISL_1259310 | Hopital | National Reference Center for Viruses of Respiratory Infections, Institut Pasteur, Paris | Marion Barbet, Sylvie Behillil, Méline Bizard, Angela Brisebarre, Camille Capel, Etienne Simon-Lorière, Vincent Enouf, Maud Vanpeene, Sylvie van der Werf,Marque Juillet StéPhanie |
| EPI_ISL_1259313 | Klinisch Laboratorium ZNA | Klinisch Laboratorium ZNA | Verstrepen et al. |
| EPI_ISL_1261136 | National Institute of Infectious Diseases-Prof. Dr. Matei Bals Molecular Diagnostics Laboratory | National Institute of Infectious Diseases-Prof. Dr. Matei Bals Molecular Diagnostics Laboratory | Corina Casangiu, Leontina Banica, Marius Surleac, Ovidiu Vlaicu, Ionelia Nicolae, Simona Paraschiv, Dan Otelea |
| EPI_ISL_1261811, EPI_ISL_1261827, EPI_ISL_1261828, EPI_ISL_1261829, EPI_ISL_1261846, EPI_ISL_1261854, EPI_ISL_1261865 | OCME Office Of Chief Medical Examiner | New York City Public Health Laboratory | Jade Wang, et al. |
| EPI_ISL_1261869 | Ospedale Generale Regionale F. Miulli | Istituto Zooprofilattico Sperimentale della Puglia e della Basilicata | Parisi A., Bianco A., Capozzi L., Del Sambro L., Simone D., Difato L., Ridolfi D., Giannico A. |
| EPI_ISL_1261870 | Istituto Zooprofilattico Sperimentale della Puglia e della Basilicata | Istituto Zooprofilattico Sperimentale della Puglia e della Basilicata | Parisi A., Bianco A., Capozzi L., Del Sambro L., Simone D., Manzulli V, Rondinone V., Pace L., Cipolletta D., Galante D. |
| EPI_ISL_1262718, EPI_ISL_1262719, EPI_ISL_1262720, EPI_ISL_1262721 | Hopital | National Reference Center for Viruses of Respiratory Infections, Institut Pasteur, Paris | Marion Barbet, Sylvie Behillil, Méline Bizard, Angela Brisebarre, Camille Capel, Etienne Simon-Lorière, Vincent Enouf, Maud Vanpeene, Sylvie van der Werf,Marque Juillet StéPhanie |
| EPI_ISL_1262722, EPI_ISL_1262723, EPI_ISL_1262724, EPI_ISL_1262725, EPI_ISL_1262726, EPI_ISL_1262727, EPI_ISL_1262728 | Hopital | National Reference Center for Viruses of Respiratory Infections, Institut Pasteur, Paris | Marion Barbet, Sylvie Behillil, Méline Bizard, Angela Brisebarre, Camille Capel, Etienne Simon-Lorière, Vincent Enouf, Maud Vanpeene, Sylvie van der Werf,Martres Pascale |
| EPI_ISL_1262729, EPI_ISL_1262730, EPI_ISL_1262732 | Hopital | National Reference Center for Viruses of Respiratory Infections, Institut Pasteur, Paris | Marion Barbet, Sylvie Behillil, Méline Bizard, Angela Brisebarre, Camille Capel, Etienne Simon-Lorière, Vincent Enouf, Maud Vanpeene, Sylvie van der Werf,Farfour Eric |
| EPI_ISL_1262733, EPI_ISL_1262734, EPI_ISL_1262735, EPI_ISL_1262736, EPI_ISL_1262737, EPI_ISL_1262738, | Hopital | National Reference Center for Viruses of Respiratory Infections, Institut Pasteur, Paris | Marion Barbet, Sylvie Behillil, Méline Bizard, Angela Brisebarre, Camille Capel, Etienne Simon-Lorière, Vincent Enouf, Maud Vanpeene, Sylvie van der Werf,Castelain Sandrine |

| | | | |
|---|---|---|---|
| EPI_ISL_1262739, EPI_ISL_1262744, EPI_ISL_1262745, EPI_ISL_1262746 | | | |
| EPI_ISL_1262747, EPI_ISL_1262748, EPI_ISL_1262749, EPI_ISL_1262750 | Armies | National Reference Center for Viruses of Respiratory Infections, Institut Pasteur, Paris | Marion Barbet, Sylvie Behillil, Méline Bizard, Angela Brisebarre, Camille Capel, Etienne Simon-Lorière, Vincent Enouf, Maud Vanpeene, Sylvie van der Werf,Desroches Marine |
| EPI_ISL_1262751, EPI_ISL_1262752, EPI_ISL_1262753, EPI_ISL_1262754 | Hopital | National Reference Center for Viruses of Respiratory Infections, Institut Pasteur, Paris | Marion Barbet, Sylvie Behillil, Méline Bizard, Angela Brisebarre, Camille Capel, Etienne Simon-Lorière, Vincent Enouf, Maud Vanpeene, Sylvie van der Werf,Djamdjian Laura |
| EPI_ISL_1262755, EPI_ISL_1262756, EPI_ISL_1262757, EPI_ISL_1262758, EPI_ISL_1262759 | Hopital | National Reference Center for Viruses of Respiratory Infections, Institut Pasteur, Paris | Marion Barbet, Sylvie Behillil, Méline Bizard, Angela Brisebarre, Camille Capel, Etienne Simon-Lorière, Vincent Enouf, Maud Vanpeene, Sylvie van der Werf,Cady Anne |
| EPI_ISL_1262763, EPI_ISL_1262764, EPI_ISL_1262765, EPI_ISL_1262766, EPI_ISL_1262767, EPI_ISL_1262768, EPI_ISL_1262769, EPI_ISL_1262770, EPI_ISL_1262771, EPI_ISL_1262772, EPI_ISL_1262773, EPI_ISL_1262774, EPI_ISL_1262775 | | | |
| see above | Labo Analyses Med | National Reference Center for Viruses of Respiratory Infections, Institut Pasteur, Paris | Marion Barbet, Sylvie Behillil, Méline Bizard, Angela Brisebarre, Camille Capel, Etienne Simon-Lorière, Vincent Enouf, Maud Vanpeene, Sylvie van der Werf,Gestin Brieuc |
| EPI_ISL_1262776, EPI_ISL_1262778, EPI_ISL_1262779, EPI_ISL_1262780, EPI_ISL_1262781, EPI_ISL_1262782, EPI_ISL_1262783, EPI_ISL_1262784, EPI_ISL_1262786, EPI_ISL_1262787, EPI_ISL_1262788, EPI_ISL_1262789, EPI_ISL_1262790, EPI_ISL_1262791, EPI_ISL_1262792, EPI_ISL_1262793, EPI_ISL_1262794, EPI_ISL_1262795, EPI_ISL_1262796, EPI_ISL_1262797, EPI_ISL_1262798, EPI_ISL_1262799, EPI_ISL_1262800, EPI_ISL_1262801, EPI_ISL_1262802, EPI_ISL_1262803, EPI_ISL_1262804, EPI_ISL_1262805 | | | |
| see above | Labo Analyses Med | National Reference Center for Viruses of Respiratory Infections, Institut Pasteur, Paris | Marion Barbet, Sylvie Behillil, Méline Bizard, Angela Brisebarre, Camille Capel, Etienne Simon-Lorière, Vincent Enouf, Maud Vanpeene, Sylvie van der Werf,Holstein Anne |
| EPI_ISL_1262806, EPI_ISL_1262807, EPI_ISL_1262809, EPI_ISL_1262810, EPI_ISL_1262811, EPI_ISL_1262812 | Labo Analyses Med | National Reference Center for Viruses of Respiratory Infections, Institut Pasteur, Paris | Marion Barbet, Sylvie Behillil, Méline Bizard, Angela Brisebarre, Camille Capel, Etienne Simon-Lorière, Vincent Enouf, Maud Vanpeene, Sylvie van der Werf,Potiron GréGoire |
| EPI_ISL_1262813 | hopital | National Reference Center for Viruses of Respiratory Infections, Institut Pasteur, Paris | Marion Barbet, Sylvie Behillil, Méline Bizard, Angela Brisebarre, Camille Capel, Etienne Simon-Lorière, Vincent Enouf, Maud Vanpeene, Sylvie van der Werf,Leruez-Ville Marianne |
| EPI_ISL_1262815 | Hopital | National Reference Center for Viruses of Respiratory Infections, Institut Pasteur, Paris | Marion Barbet, Sylvie Behillil, Méline Bizard, Angela Brisebarre, Camille Capel, Etienne Simon-Lorière, Vincent Enouf, Maud Vanpeene, Sylvie van der Werf,Woercel Isabelle |
| EPI_ISL_1262819, EPI_ISL_1262820, EPI_ISL_1262822, EPI_ISL_1262823, EPI_ISL_1262825, EPI_ISL_1262826, EPI_ISL_1262827, EPI_ISL_1262828, EPI_ISL_1262829, EPI_ISL_1262830, EPI_ISL_1262831, EPI_ISL_1262832, EPI_ISL_1262833, EPI_ISL_1262834, EPI_ISL_1262835, EPI_ISL_1262836, EPI_ISL_1262837, EPI_ISL_1262838, EPI_ISL_1262839, EPI_ISL_1262840, EPI_ISL_1262841, EPI_ISL_1262842, EPI_ISL_1262843, EPI_ISL_1262844, EPI_ISL_1262845, EPI_ISL_1262846, EPI_ISL_1262847, EPI_ISL_1262848, EPI_ISL_1262849, EPI_ISL_1262850, EPI_ISL_1262851, EPI_ISL_1262852, EPI_ISL_1262853, EPI_ISL_1262854, EPI_ISL_1262855, EPI_ISL_1262856, EPI_ISL_1262857, EPI_ISL_1262858, EPI_ISL_1262859, EPI_ISL_1262860, EPI_ISL_1262861, EPI_ISL_1262862, EPI_ISL_1262863, EPI_ISL_1262864, EPI_ISL_1262865, EPI_ISL_1262866, EPI_ISL_1262867, EPI_ISL_1262868, EPI_ISL_1262869, EPI_ISL_1262870, EPI_ISL_1262872, EPI_ISL_1262874, EPI_ISL_1262875, EPI_ISL_1262876, EPI_ISL_1262877, EPI_ISL_1262878, EPI_ISL_1262879 | | | |
| see above | hopital | National Reference Center for Viruses of Respiratory Infections, Institut Pasteur, Paris | Marion Barbet, Sylvie Behillil, Méline Bizard, Angela Brisebarre, Camille Capel, Etienne Simon-Lorière, Vincent Enouf, Maud Vanpeene, Sylvie van der Werf,Gastli Nabil |
| EPI_ISL_1262881, EPI_ISL_1262882, EPI_ISL_1262883, EPI_ISL_1262884, EPI_ISL_1262885, EPI_ISL_1262886, EPI_ISL_1262887, EPI_ISL_1262888, EPI_ISL_1262889, EPI_ISL_1262890, EPI_ISL_1262891, EPI_ISL_1262892, EPI_ISL_1262893, EPI_ISL_1262894, EPI_ISL_1262895, EPI_ISL_1262896, EPI_ISL_1262897 | | | |
| see above | Labo Analyses Med | National Reference Center for Viruses of Respiratory Infections, Institut Pasteur, Paris | Marion Barbet, Sylvie Behillil, Méline Bizard, Angela Brisebarre, Camille Capel, Etienne Simon-Lorière, Vincent Enouf, Maud Vanpeene, Sylvie van der Werf,Felloni Claire |
| EPI_ISL_1262898, EPI_ISL_1262899, EPI_ISL_1262900, EPI_ISL_1262901, EPI_ISL_1262902, EPI_ISL_1262903 | Hopital | National Reference Center for Viruses of Respiratory Infections, Institut Pasteur, Paris | Marion Barbet, Sylvie Behillil, Méline Bizard, Angela Brisebarre, Camille Capel, Etienne Simon-Lorière, Vincent Enouf, Maud Vanpeene, Sylvie van der Werf,Billy Pierre Alain |
| EPI_ISL_1262904 | Hopital | National Reference Center for Viruses of Respiratory Infections, Institut Pasteur, Paris | Marion Barbet, Sylvie Behillil, Méline Bizard, Angela Brisebarre, Camille Capel, Etienne Simon-Lorière, Vincent Enouf, Maud Vanpeene, Sylvie van der Werf,Riverain Emeline |
| EPI_ISL_1262905, EPI_ISL_1262906, EPI_ISL_1262907 | Hopital | National Reference Center for Viruses of Respiratory Infections, Institut Pasteur, Paris | Marion Barbet, Sylvie Behillil, Méline Bizard, Angela Brisebarre, Camille Capel, Etienne Simon-Lorière, Vincent Enouf, Maud Vanpeene, Sylvie van der Werf,Bernardi Franck |
| EPI_ISL_1262910, EPI_ISL_1262911, EPI_ISL_1262912, EPI_ISL_1262913 | Outre Mer | National Reference Center for Viruses of Respiratory Infections, Institut Pasteur, Paris | Marion Barbet, Sylvie Behillil, Méline Bizard, Angela Brisebarre, Camille Capel, Etienne Simon-Lorière, Vincent Enouf, Maud Vanpeene, Sylvie van der Werf,Rousset Dominique |
| EPI_ISL_1262914 | Hopital | National Reference Center for Viruses of Respiratory Infections, Institut Pasteur, Paris | Marion Barbet, Sylvie Behillil, Méline Bizard, Angela Brisebarre, Camille Capel, Etienne Simon-Lorière, Vincent Enouf, Maud Vanpeene, Sylvie van der Werf,Irimia Alix |
| EPI_ISL_1262915, EPI_ISL_1262916 | Labo Analyses Med | National Reference Center for Viruses of Respiratory Infections, Institut Pasteur, Paris | Marion Barbet, Sylvie Behillil, Méline Bizard, Angela Brisebarre, Camille Capel, Etienne Simon-Lorière, Vincent Enouf, Maud Vanpeene, Sylvie van der Werf,Laroche Christine |
| EPI_ISL_1262917, EPI_ISL_1262918, EPI_ISL_1262919, EPI_ISL_1262920 | Hopital | National Reference Center for Viruses of Respiratory Infections, Institut Pasteur, Paris | Marion Barbet, Sylvie Behillil, Méline Bizard, Angela Brisebarre, Camille Capel, Etienne Simon-Lorière, Vincent Enouf, Maud Vanpeene, Sylvie van der Werf,Guillotel Eleha |
| EPI_ISL_1262921, EPI_ISL_1262922, EPI_ISL_1262923, EPI_ISL_1262924, EPI_ISL_1262925, EPI_ISL_1262926, EPI_ISL_1262927, EPI_ISL_1262928, EPI_ISL_1262929, EPI_ISL_1262930, EPI_ISL_1262931, EPI_ISL_1262932, EPI_ISL_1262933 | | | |
| see above | Hopital | National Reference Center for Viruses of Respiratory Infections, Institut Pasteur, Paris | Marion Barbet, Sylvie Behillil, Méline Bizard, Angela Brisebarre, Camille Capel, Etienne Simon-Lorière, Vincent Enouf, Maud Vanpeene, Sylvie van der Werf,Thibault Vincent |
| EPI_ISL_1262934, EPI_ISL_1262935, EPI_ISL_1262936, EPI_ISL_1262937, EPI_ISL_1262938, EPI_ISL_1262939, EPI_ISL_1262940, EPI_ISL_1262941, EPI_ISL_1262942, EPI_ISL_1262943, EPI_ISL_1262944, EPI_ISL_1262945, EPI_ISL_1262946, EPI_ISL_1262947, EPI_ISL_1262948, EPI_ISL_1262949, EPI_ISL_1262950, EPI_ISL_1262951, EPI_ISL_1262952, EPI_ISL_1262953, EPI_ISL_1262954, EPI_ISL_1262955, EPI_ISL_1262956, EPI_ISL_1262957, EPI_ISL_1262958, EPI_ISL_1262959, EPI_ISL_1262960, EPI_ISL_1262961, EPI_ISL_1262962, EPI_ISL_1262963, EPI_ISL_1262964, EPI_ISL_1262965, EPI_ISL_1262966, EPI_ISL_1262967, EPI_ISL_1262968, EPI_ISL_1262969, EPI_ISL_1262970, EPI_ISL_1262971, EPI_ISL_1262972, EPI_ISL_1262973, EPI_ISL_1262974, EPI_ISL_1262975, EPI_ISL_1262976, EPI_ISL_1262977, EPI_ISL_1262978, EPI_ISL_1262980 | | | |
| see above | Labo Analyses Med | National Reference Center for Viruses of Respiratory Infections, Institut Pasteur, Paris | Marion Barbet, Sylvie Behillil, Méline Bizard, Angela Brisebarre, Camille Capel, Etienne Simon-Lorière, Vincent Enouf, Maud Vanpeene, Sylvie van der Werf,Leonard Pierre-Yves |
| EPI_ISL_1262981, EPI_ISL_1262982, EPI_ISL_1262983, EPI_ISL_1262984, EPI_ISL_1262985, EPI_ISL_1262986 | Hopital | National Reference Center for Viruses of Respiratory Infections, Institut Pasteur, Paris | Marion Barbet, Sylvie Behillil, Méline Bizard, Angela Brisebarre, Camille Capel, Etienne Simon-Lorière, Vincent Enouf, Maud Vanpeene, Sylvie van der Werf,Guillaume CléMence |
| EPI_ISL_1262987, EPI_ISL_1262988 | Hopital | National Reference Center for Viruses of Respiratory Infections, Institut Pasteur, Paris | Marion Barbet, Sylvie Behillil, Méline Bizard, Angela Brisebarre, Camille Capel, Etienne Simon-Lorière, Vincent Enouf, Maud Vanpeene, Sylvie van der Werf,Woercel Isabelle |
| EPI_ISL_1262989 | Hopital | National Reference Center for Viruses of Respiratory Infections, Institut Pasteur, Paris | Marion Barbet, Sylvie Behillil, Méline Bizard, Angela Brisebarre, Camille Capel, Etienne Simon-Lorière, Vincent Enouf, Maud Vanpeene, Sylvie van der Werf,Bour Jean Baptiste |
| EPI_ISL_1262990, EPI_ISL_1262991, EPI_ISL_1262992, EPI_ISL_1262993, EPI_ISL_1262994, EPI_ISL_1262995, EPI_ISL_1262996, EPI_ISL_1262997, EPI_ISL_1262998, EPI_ISL_1262999, EPI_ISL_1263000, EPI_ISL_1263001, EPI_ISL_1263002, EPI_ISL_1263004, EPI_ISL_1263005, EPI_ISL_1263006, EPI_ISL_1263007, EPI_ISL_1263008, EPI_ISL_1263009, EPI_ISL_1263010, EPI_ISL_1263011, EPI_ISL_1263012, EPI_ISL_1263013, EPI_ISL_1263014, EPI_ISL_1263015, EPI_ISL_1263016, EPI_ISL_1263017, EPI_ISL_1263018, EPI_ISL_1263019, EPI_ISL_1263020, EPI_ISL_1263021, EPI_ISL_1263022, EPI_ISL_1263023, EPI_ISL_1263024, EPI_ISL_1263025, EPI_ISL_1263026, EPI_ISL_1263027, EPI_ISL_1263028, EPI_ISL_1263029, EPI_ISL_1263030, EPI_ISL_1263031, EPI_ISL_1263032, EPI_ISL_1263033, EPI_ISL_1263034, EPI_ISL_1263035, EPI_ISL_1263036, EPI_ISL_1263037, EPI_ISL_1263038, EPI_ISL_1263039 | | | |
| see above | Labo Analyses Med | National Reference Center for Viruses of Respiratory Infections, Institut Pasteur, Paris | Marion Barbet, Sylvie Behillil, Méline Bizard, Angela Brisebarre, Camille Capel, Etienne Simon-Lorière, Vincent Enouf, Maud Vanpeene, Sylvie van der Werf,Said-Delattre OphéLie |
| EPI_ISL_1263040, EPI_ISL_1263041, EPI_ISL_1263042, EPI_ISL_1263044, EPI_ISL_1263045, EPI_ISL_1263046, EPI_ISL_1263047, EPI_ISL_1263048, EPI_ISL_1263049, EPI_ISL_1263050, EPI_ISL_1263051, EPI_ISL_1263052, EPI_ISL_1263053, EPI_ISL_1263054, EPI_ISL_1263055, EPI_ISL_1263056, EPI_ISL_1263057, EPI_ISL_1263058, EPI_ISL_1263059, EPI_ISL_1263060, EPI_ISL_1263061, EPI_ISL_1263062, EPI_ISL_1263064, EPI_ISL_1263065, EPI_ISL_1263066, EPI_ISL_1263067, EPI_ISL_1263068, EPI_ISL_1263069, EPI_ISL_1263070, EPI_ISL_1263071, EPI_ISL_1263073, EPI_ISL_1263074, EPI_ISL_1263075, EPI_ISL_1263076, EPI_ISL_1263077, EPI_ISL_1263078, EPI_ISL_1263079, EPI_ISL_1263080, EPI_ISL_1263081, EPI_ISL_1263082, EPI_ISL_1263083, EPI_ISL_1263084, EPI_ISL_1263086, EPI_ISL_1263087, EPI_ISL_1263088, EPI_ISL_1263089 | | | |
| see above | Labo Analyses Med | National Reference Center for Viruses of Respiratory Infections, Institut Pasteur, Paris | Marion Barbet, Sylvie Behillil, Méline Bizard, Angela Brisebarre, Camille Capel, Etienne Simon-Lorière, Vincent Enouf, Maud Vanpeene, Sylvie van der Werf,Roquebert BéNéDicte |

| | | | |
|---|---|---|---|
| EPI_ISL_1264762 | Institute for Medical Research, Infectious Disease Research Centre, National Institutes of Health, Ministry of Health Malaysia | Institute for Medical Research, Infectious Disease Research Centre, National Institutes of Health, Ministry of Health Malaysia | Suppiah J, Kamel K, Mohd Zawawi Z, Azizan MA, Thayan R |
| EPI_ISL_1265143 | AZ Zeno | AZ SINT-JAN BRUGGE | Marijke Reynders, Patrick Descheemaeker, Laurien Hoornaert, Thomas Van Landschoot, Jorn Hellemans |
| EPI_ISL_1265148, EPI_ISL_1265155 | AZ Sint-Jan Brugge-Oostende AV | AZ SINT-JAN BRUGGE | Marijke Reynders, Patrick Descheemaeker, Laurien Hoornaert, Thomas Van Landschoot, Jorn Hellemans |
| EPI_ISL_1265156 | AZ Zeno | AZ SINT-JAN BRUGGE | Marijke Reynders, Patrick Descheemaeker, Laurien Hoornaert, Thomas Van Landschoot, Jorn Hellemans |
| EPI_ISL_1265158, EPI_ISL_1265160 | AZ Sint-Jan Brugge-Oostende AV | AZ SINT-JAN BRUGGE | Marijke Reynders, Patrick Descheemaeker, Laurien Hoornaert, Thomas Van Landschoot, Jorn Hellemans |
| EPI_ISL_1265368, EPI_ISL_1265372 | "Stefan S. Nicolau"Institute of Virology | "Stefan S. Nicolau"Institute of Virology | Anca Botezatu, Iulia Virginia Iancu, Coralia Bleotu, Mihaela Economescu, Laura Necula, Denisa Dragu, Lilia Matei, Ioana Pitica, Marinela Bostan, Mirela Mihaila, Laura Grecu, Camelia Sultana, Simona Ruta, Ana Iulia Neagu, Adriana Plesa, Saviana Nedeianu, Alina Nastasie, Cristina Mambet,Gabriela Anton, Carmen Cristina Diaconu |
| EPI_ISL_1265441, EPI_ISL_1265442, EPI_ISL_1265443 | BTKLPP Kelas I Makassar | Eijkman Institute for Molecular Biology, Ministry of Research and Technology/National Agency for Research and Innovation; National Institute of Health Research and Development | Edison Johar, Hidayat Trimarsanto, Frilasita A Yudhaputri, Iskandar Adnan, Willy Agustine, Sukma Oktavianthi, Lydia V. Panggalo, Slamet, Vivi Setiawaty, Hana Apsari Pawestri, Safarina G Malik, Khin Saw Myint, Amin Soebandrio |
| EPI_ISL_1265444, EPI_ISL_1265445, EPI_ISL_1265446 | BTKLPP Kelas I Makassar | Eijkman Institute for Molecular Biology, Ministry of Research and Technology/National Agency for Research and Innovation; National Institute of Health Research and Development | Hidayat Trimarsanto, Frilasita A Yudhaputri, Iskandar Adnan, Willy Agustine, Sukma Oktavianthi, Lydia V. Panggalo, Edison Johar, Slamet, Vivi Setiawaty, Hana Apsari Pawestri, Safarina G Malik, Khin Saw Myint, Amin Soebandrio |
| EPI_ISL_1265447, EPI_ISL_1265448, EPI_ISL_1265449 | BTKLPP Kelas I Makassar | Eijkman Institute for Molecular Biology, Ministry of Research and Technology/National Agency for Research and Innovation; National Institute of Health Research and Development | Frilasita A Yudhaputri, Iskandar Adnan, Willy Agustine, Sukma Oktavianthi, Lydia V. Panggalo, Edison Johar, Hidayat Trimarsanto, Slamet, Vivi Setiawaty, Hana Apsari Pawestri, Safarina G Malik, Khin Saw Myint, Amin Soebandrio |
| EPI_ISL_1265450, EPI_ISL_1265451, EPI_ISL_1265452 | BTKLPP Kelas I Makassar | Eijkman Institute for Molecular Biology, Ministry of Research and Technology/National Agency for Research and Innovation; National Institute of Health Research and Development | Iskandar Adnan, Willy Agustine, Sukma Oktavianthi, Lydia V. Panggalo, Edison Johar, Hidayat Trimarsanto, Frilasita A Yudhaputri, Slamet, Vivi Setiawaty, Hana Apsari Pawestri, Safarina G Malik, Khin Saw Myint, Amin Soebandrio |
| EPI_ISL_1265453, EPI_ISL_1265454, EPI_ISL_1265455 | BTKLPP Kelas I Makassar | Eijkman Institute for Molecular Biology, Ministry of Research and Technology/National Agency for Research and Innovation; National Institute of Health Research and Development | Willy Agustine, Sukma Oktavianthi, Lydia V. Panggalo, Edison Johar, Hidayat Trimarsanto, Frilasita A Yudhaputri, Iskandar Adnan, Slamet, Vivi Setiawaty, Hana Apsari Pawestri, Safarina G Malik, Khin Saw Myint, Amin Soebandrio |
| EPI_ISL_1265456, EPI_ISL_1265457, EPI_ISL_1265458, EPI_ISL_1265459 | BTKLPP Kelas I Makassar | Eijkman Institute for Molecular Biology, Ministry of Research and Technology/National Agency for Research and Innovation; National Institute of Health Research and Development | Sukma Oktavianthi, Lydia V. Panggalo, Edison Johar, Hidayat Trimarsanto, Frilasita A Yudhaputri, Iskandar Adnan, Willy Agustine, Slamet, Vivi Setiawaty, Hana Apsari Pawestri, Safarina G Malik, Khin Saw Myint, Amin Soebandrio |
| EPI_ISL_1265460, EPI_ISL_1265461, EPI_ISL_1265462 | BTKLPP Kelas I Makassar | Eijkman Institute for Molecular Biology, Ministry of Research and Technology/National Agency for Research and Innovation; National Institute of Health Research and Development | Lydia V. Panggalo, Edison Johar, Hidayat Trimarsanto, Frilasita A Yudhaputri, Iskandar Adnan, Willy Agustine, Sukma Oktavianthi, Slamet, Vivi Setiawaty, Hana Apsari Pawestri, Safarina G Malik, Khin Saw Myint, Amin Soebandrio |
| EPI_ISL_1265741, EPI_ISL_1265742, EPI_ISL_1265743, EPI_ISL_1265744, EPI_ISL_1265745, EPI_ISL_1265746, EPI_ISL_1265747, EPI_ISL_1265748, EPI_ISL_1265749, EPI_ISL_1265750 | Klinisch Laboratorium ZNA | Klinisch Laboratorium ZNA | Verstrepen et al. |
| EPI_ISL_1271520 | Cerballiance Vauréal | Cerba Lab | Roquebert B, Lecorche E, Trombert-Paolantoni S, Olivi M, Costa JM, Benazra M, Zimmer S, Verdurme L, Haïm-Boukobza S |
| EPI_ISL_1279268, EPI_ISL_1279269, EPI_ISL_1279277, EPI_ISL_1279278, EPI_ISL_1279290, EPI_ISL_1279291, EPI_ISL_1279292, EPI_ISL_1279293, EPI_ISL_1279296, EPI_ISL_1279297, EPI_ISL_1279299, EPI_ISL_1279303 | | | |
| see above | Laboratorio Central de Epidemiología (LCE) | Instituto Nacional de Enfermedades Respiratorias (INER): Centro de Investigación en Enfermedades Infecciosas (CIENI) | Consorcio Mexicano de Vigilancia Genómica (CoViGen-Mex). Authors (in alphabetical order): Julio Elias Alvarado-Yaah, Carlos F. Arias, Santiago Ávila-Ríos, Víctor Hugo Borja-Aburto, Celia Boukadida, Juan Bautista Chale-Dzul , José Antonio Enciso-Moreno, Gloria Elena Espinoza-Ayala, Fernando Fontove-Herrera, Concepción Grajales-Muñiz, Ricardo Grande, Alfredo Herrera-Estrella, Carla Ivón Herrera-Najera, Pavel Isa, Brenda Irasema Maldonado-Meza, Bernardo Martínez-Miguel, Margarita Matías-Florentino, María Guadalupe de Jesús Mireles-Rivera, Gloria María Molina-Salinas, Hector Montoya-Fuentes, José Esteban Muñoz-Medina, José de Jesús Nuñez-Contreras, Alicia Ocaña-Mondragón, Luis Alberto Ochoa-Carrera, Hector Esteban Paz-Juárez, Francisco Pulido, Helen Haydee Fernanda Ramírez-Plascencia, Angel Gustavo Salas-Lais, Jorge Ivan Salinal-Nevarez, Alejandro Sanchez-Flores, Clara Esperanza Santacruz-Tinoco, María Guadalupe Santiago-Mauricio , Nelly Sélem-Mojica, Blanca Taboada , Gloria Vazquez |
| EPI_ISL_1279382 | Unidad de Investigación Biomédica de Zacatecas (UIBZ) | Instituto Nacional de Enfermedades Respiratorias (INER): Centro de Investigación en Enfermedades Infecciosas (CIENI) | Consorcio Mexicano de Vigilancia Genómica (CoViGen-Mex). Authors (in alphabetical order): Julio Elias Alvarado-Yaah, Carlos F. Arias, Santiago Ávila-Ríos, Víctor Hugo Borja-Aburto, Celia Boukadida, Juan Bautista Chale-Dzul , José Antonio Enciso-Moreno, Gloria Elena Espinoza-Ayala, Fernando Fontove-Herrera, Concepción Grajales-Muñiz, Ricardo Grande, Alfredo Herrera-Estrella, Carla Ivón Herrera-Najera, Pavel Isa, Brenda Irasema Maldonado-Meza, Bernardo Martínez-Miguel, Margarita Matías-Florentino, María Guadalupe de Jesús Mireles-Rivera, Gloria María Molina-Salinas, Hector Montoya-Fuentes, José Esteban Muñoz-Medina, José de Jesús Nuñez-Contreras, Alicia Ocaña-Mondragón, Luis Alberto Ochoa-Carrera, Hector Esteban Paz-Juárez, Francisco Pulido, Helen Haydee Fernanda Ramírez-Plascencia, Angel Gustavo Salas-Lais, Jorge Ivan Salinal-Nevarez, Alejandro Sanchez-Flores, Clara Esperanza Santacruz-Tinoco, María Guadalupe Santiago-Mauricio , Nelly Sélem-Mojica, Blanca Taboada , Gloria Vazquez |
| EPI_ISL_1279383, EPI_ISL_1279384, EPI_ISL_1279385, EPI_ISL_1279386, EPI_ISL_1279387, EPI_ISL_1279388, EPI_ISL_1279389, EPI_ISL_1279390, EPI_ISL_1279391, EPI_ISL_1279392, EPI_ISL_1279393, EPI_ISL_1279394, EPI_ISL_1279395, EPI_ISL_1279396, EPI_ISL_1279397, EPI_ISL_1279398, EPI_ISL_1279399, EPI_ISL_1279400, EPI_ISL_1279401, EPI_ISL_1279402, EPI_ISL_1279408, EPI_ISL_1279409, EPI_ISL_1279410, EPI_ISL_1279414, EPI_ISL_1279415, EPI_ISL_1279416, EPI_ISL_1279417, EPI_ISL_1279418, EPI_ISL_1279428, EPI_ISL_1279429, EPI_ISL_1279433, EPI_ISL_1279434, EPI_ISL_1279435, EPI_ISL_1279436, EPI_ISL_1279437, EPI_ISL_1279438, EPI_ISL_1279441, EPI_ISL_1279442, EPI_ISL_1279443, EPI_ISL_1279444, EPI_ISL_1279445, EPI_ISL_1279446, EPI_ISL_1279447, EPI_ISL_1279451, EPI_ISL_1279452, EPI_ISL_1279453, EPI_ISL_1279454, EPI_ISL_1279455, EPI_ISL_1279456, EPI_ISL_1279457, EPI_ISL_1279458, EPI_ISL_1279459, EPI_ISL_1279460, EPI_ISL_1279537, EPI_ISL_1279539, EPI_ISL_1279540, EPI_ISL_1279546, EPI_ISL_1279547, EPI_ISL_1279548, EPI_ISL_1279549, EPI_ISL_1279550, EPI_ISL_1279551, EPI_ISL_1279557, EPI_ISL_1279558, EPI_ISL_1279559, EPI_ISL_1279561, EPI_ISL_1279562, EPI_ISL_1279563, EPI_ISL_1279565, EPI_ISL_1279566, EPI_ISL_1279567, EPI_ISL_1279568, EPI_ISL_1279569, EPI_ISL_1279570, EPI_ISL_1279571, EPI_ISL_1279572, EPI_ISL_1279573, EPI_ISL_1279574, EPI_ISL_1279575, EPI_ISL_1279576, EPI_ISL_1279577, EPI_ISL_1279578, EPI_ISL_1279579, EPI_ISL_1279580, EPI_ISL_1279581, EPI_ISL_1279582, EPI_ISL_1279583, EPI_ISL_1279584, EPI_ISL_1279585, EPI_ISL_1279586, EPI_ISL_1279587, EPI_ISL_1279588, EPI_ISL_1279589, EPI_ISL_1279590, EPI_ISL_1279591, EPI_ISL_1279592, EPI_ISL_1279593, EPI_ISL_1279594, EPI_ISL_1279595, EPI_ISL_1279596, EPI_ISL_1279597, EPI_ISL_1279598, EPI_ISL_1279600, EPI_ISL_1279601 | | | |
| see above | Laboratorio Central de Epidemiología (LCE) | Instituto Nacional de Enfermedades Respiratorias (INER): Centro de Investigación en Enfermedades Infecciosas (CIENI) | Consorcio Mexicano de Vigilancia Genómica (CoViGen-Mex). Authors (in alphabetical order): Julio Elias Alvarado-Yaah, Carlos F. Arias, Santiago Ávila-Ríos, Víctor Hugo Borja-Aburto, Celia Boukadida, Juan Bautista Chale-Dzul , José Antonio Enciso-Moreno, Gloria Elena Espinoza-Ayala, Fernando Fontove-Herrera, Concepción Grajales-Muñiz, Ricardo Grande, Alfredo Herrera-Estrella, Carla Ivón Herrera-Najera, Pavel Isa, Brenda Irasema Maldonado-Meza, Bernardo Martínez-Miguel, Margarita Matías-Florentino, María Guadalupe de Jesús Mireles-Rivera, Gloria María Molina-Salinas, Hector Montoya-Fuentes, José Esteban Muñoz-Medina, José de Jesús Nuñez-Contreras, Alicia Ocaña-Mondragón, Luis Alberto Ochoa-Carrera, Hector Esteban Paz-Juárez, Francisco Pulido, Helen Haydee Fernanda Ramírez-Plascencia, Angel Gustavo Salas-Lais, Jorge Ivan Salinal-Nevarez, Alejandro Sanchez-Flores, Clara Esperanza Santacruz-Tinoco, María Guadalupe Santiago-Mauricio , Nelly Sélem-Mojica, Blanca Taboada , Gloria Vazquez |
| EPI_ISL_1281683 | Labo Somedi | Imelda hospital Bonheiden | Johan Frans, Dagmar Obbels, Hanne Valgaeren |
| EPI_ISL_1281692 | Imelda hospital Bonheiden | Imelda hospital Bonheiden | Johan Frans, Dagmar Obbels, Hanne Valgaeren |
| EPI_ISL_1281693 | Heilig hart Lier | Imelda hospital Bonheiden | Johan Frans, Dagmar Obbels, Hanne Valgaeren |

| | | | |
|---|---|---|---|
| EPI_ISL_1284327, EPI_ISL_1284335 | RS Mitra Keluarga Kalideres | Eijkman Institute for Molecular Biology, Ministry of Research and Technology/National Agency for Research and Innovation | Frilasita A Yudhaputri, Hidayat Trimarsanto, Iskandar Adnan, Lydia V. Panggalo, Sukma Oktavianthi, Willy Agustine, Edison Johar, Safarina G Malik, Khin Saw Myint, Amin Soebandrio |
| EPI_ISL_1287349 | Botswana Harvard HIV Reference Laboratory | Botswana Harvard HIV Reference Laboratory | Sikhulile Moyo, Wonderful T. Choga, Dorcas Maruapula,Thongbotho Mphoyakgosi, Boitumelo Zuze, Botshelo Radibe, Legodile Kooepile, David Lawrence, Roger Shapiro, Shahin Lockman, Mosepele Mosepele, Joseph Makhema, Simani Gaseitsiwe |
| EPI_ISL_1287769, EPI_ISL_1287774, EPI_ISL_1287775, EPI_ISL_1287777 | Laboratorio Central de Epidemiología (LCE) | Instituto Nacional de Enfermedades Respiratorias (INER): Centro de Investigación en Enfermedades Infecciosas (CIENI) | Consorcio Mexicano de Vigilancia Genómica (CoViGen-Mex). Authors (in alphabetical order): Julio Elias Alvarado-Yaah, Carlos F. Arias, Santiago Ávila-Ríos, Víctor Hugo Borja-Aburto, Celia Boukadida, Juan Bautista Chale-Dzul , José Antonio Enciso-Moreno, Gloria Elena Espinoza-Ayala, Fernando Fontove-Herrera, Concepción Grajales-Muñiz, Ricardo Grande, Alfredo Herrera-Estrella, Carla Ivón Herrera-Najera, Pavel Isa, Brenda Irasema Maldonado-Meza, Bernardo Martínez-Miguel, Margarita Matías-Florentino, María Guadalupe de Jesús Mireles-Rivera, Gloria María Molina-Salinas, Hector Montoya-Fuentes, José Esteban Muñoz-Medina, José de Jesús Nuñez-Contreras, Alicia Ocaña-Mondragón, Luis Alberto Ochoa-Carrera, Hector Esteban Paz-Juárez, Francisco Pulido, Helen Haydee Fernanda Ramírez-Plascencia, Angel Gustavo Salas-Lais, Jorge Ivan Salinal-Nevarez, Alejandro Sanchez-Flores, Clara Esperanza Santacruz-Tinoco, María Guadalupe Santiago-Mauricio , Nelly Sélem-Mojica, Blanca Taboada , Gloria Vazquez |
| EPI_ISL_1288351, EPI_ISL_1288369 | Laboratorio Central de Epidemiología (LCE) | Instituto de Biotecnología de la UNAM | Consorcio Mexicano de Vigilancia Genómica (CoViGen-Mex). Authors (in alphabetical order): Julio Elias Alvarado-Yaah, Carlos F. Arias, Santiago Ávila-Ríos, Víctor Hugo Borja-Aburto, Celia Boukadida, Juan Bautista Chale-Dzul , José Antonio Enciso-Moreno, Gloria Elena Espinoza-Ayala, Fernando Fontove-Herrera, Concepción Grajales-Muñiz, Ricardo Grande, Alfredo Herrera-Estrella, Carla Ivón Herrera-Najera, Pavel Isa, Brenda Irasema Maldonado-Meza, Bernardo Martínez-Miguel, Margarita Matías-Florentino, María Guadalupe de Jesús Mireles-Rivera, Gloria María Molina-Salinas, Hector Montoya-Fuentes, José Esteban Muñoz-Medina, José de Jesús Nuñez-Contreras, Alicia Ocaña-Mondragón, Luis Alberto Ochoa-Carrera, Hector Esteban Paz-Juárez, Francisco Pulido, Helen Haydee Fernanda Ramírez-Plascencia, Angel Gustavo Salas-Lais, Jorge Ivan Salinal-Nevarez, Alejandro Sanchez-Flores, Clara Esperanza Santacruz-Tinoco, María Guadalupe Santiago-Mauricio , Nelly Sélem-Mojica, Blanca Taboada , Gloria Vazquez |
| EPI_ISL_1290805 | DIP. PREV. AVEZZANO SERVIZIO DI IGIENE EPIDEMIOLOGIA E SANITA' PUBBLICA | Istituto Zooprofilattico Sperimentale dell'Abruzzo e Molise "G. Caporale" | Lorusso A, Marcacci M, Di Domenico M, Ancora M, Curini V, Mangone I, Rinaldi A, Scialabba S, Di Pasquale A, Cammà C, Puglia I, Calistri P, Savini G |
| EPI_ISL_1290807 | Siesp dipartimento di prevenzione | Istituto Zooprofilattico Sperimentale dell'Abruzzo e Molise "G. Caporale" | Lorusso A, Marcacci M, Di Domenico M, Ancora M, Curini V, Mangone I, Rinaldi A, Scialabba S, Di Pasquale A, Cammà C, Puglia I, Calistri P, Savini G |
| EPI_ISL_1290808 | SIESP L'AQUILA | Istituto Zooprofilattico Sperimentale dell'Abruzzo e Molise "G. Caporale" | Lorusso A, Marcacci M, Di Domenico M, Ancora M, Curini V, Mangone I, Rinaldi A, Scialabba S, Di Pasquale A, Cammà C, Puglia I, Calistri P, Savini G |
| EPI_ISL_1291550, EPI_ISL_1291551, EPI_ISL_1291552, EPI_ISL_1291553, EPI_ISL_1291557 | Microbiology Department, University Hospital Donostia | Microbiology Department, University Hospital Donostia | Marimon JM, Montes M, Piñeiro L, Sorarrain A, Gomez M, Cilla G. |
| EPI_ISL_1292089 | Baylor Scott & White-Irving | Baylor Scott & White-Temple | Ari Rao, Linden Morales, Kimberly Walker, Marcus Volz, Shelby Hendrickson, Caitlin Maloney |
| EPI_ISL_1292808 | CHU Pontchaillou | CHU Pontchaillou | GROLHIER Claire, DUFOUR Marie José, DENOUAL Florent, ETCHEVERRY Amandine, SASSI Mohamed, JAGLINE Steven, PRONIER Charlotte, GALIBERT Marie Dominique, DE TAYRAC Marie, THIBAULT Vincent |
| EPI_ISL_1295641, EPI_ISL_1295643, EPI_ISL_1295644, EPI_ISL_1295645 | Laboratory for Respiratory Viruses, Cantacuzino National Military-Medical Institute for Research and Development | Cantacuzino Institute Virology | Luiza Ustea, Nicoleta Paraschiv, Catalina Pascu, Mihaela Lazar |
| EPI_ISL_1296241, EPI_ISL_1296242 | Università degli Studi di Perugia | Istituto Zooprofilattico Sperimentale dell'Abruzzo e Molise "G. Caporale" | Mencacci A, Camilloni B,Lorusso A, Marcacci M, Di Domenico M, Ancora M, Curini V, Mangone I, Rinaldi A, Scialabba S, Di Pasquale A, Cammà C, Puglia I, Calistri P, Savini G |
| EPI_ISL_1296262 | Istituto Zooprofilattico Sperimentale Umbria e Marche "Togo Rosati" | Istituto Zooprofilattico Sperimentale dell'Abruzzo e Molise "G. Caporale" | Biagetti M,Giammarioli M,Lorusso A, Marcacci M, Di Domenico M, Ancora M, Curini V, Mangone I, Rinaldi A, Scialabba S, Di Pasquale A, Cammà C, Puglia I, Calistri P, Savini G |
| EPI_ISL_1296268 | Università degli Studi di Perugia | Istituto Zooprofilattico Sperimentale dell'Abruzzo e Molise "G. Caporale" | Mencacci A, Camilloni B, Lorusso A, Marcacci M, Di Domenico M, Ancora M, Curini V, Mangone I, Rinaldi A, Scialabba S, Di Pasquale A, Cammà C, Puglia I, Calistri P, Savini G |
| EPI_ISL_1296283, EPI_ISL_1296284 | Azienda Ospedaliera Terni | Istituto Zooprofilattico Sperimentale dell'Abruzzo e Molise "G. Caporale" | Palumbo M, Scaccetti A, Lorusso A, Marcacci M, Di Domenico M, Ancora M, Curini V, Mangone I, Rinaldi A, Scialabba S, Di Pasquale A, Cammà C, Puglia I, Calistri P, Savini G |
| EPI_ISL_1298474 | Burshtyn CCH | The Institute of Molecular Biology and Genetics of NASU | M.Tukalo et al. |
| EPI_ISL_1298475 | Kosov CRH | The Institute of Molecular Biology and Genetics of NASU | M.Tukalo et al. |
| EPI_ISL_1298476, EPI_ISL_1298477 | Ivano-Frankivsk CCH#1 | The Institute of Molecular Biology and Genetics of NASU | M.Tukalo et al. |
| EPI_ISL_1298478, EPI_ISL_1298479 | Kosov CRH | The Institute of Molecular Biology and Genetics of NASU | M.Tukalo et al. |
| EPI_ISL_1298480 | Nadvirna CRH | The Institute of Molecular Biology and Genetics of NASU | M.Tukalo et al. |
| EPI_ISL_1298481 | Ivano-Frankivsk CCH#1 | The Institute of Molecular Biology and Genetics of NASU | M.Tukalo et al. |
| EPI_ISL_1298482 | Tysmenytsia CH | The Institute of Molecular Biology and Genetics of NASU | M.Tukalo et al. |
| EPI_ISL_1298483 | Bogorodchany CRH | The Institute of Molecular Biology and Genetics of NASU | M.Tukalo et al. |
| EPI_ISL_1298484 | Nadvirna CRH | The Institute of Molecular Biology and Genetics of NASU | M.Tukalo et al. |
| EPI_ISL_1298485, EPI_ISL_1298486 | Kosov CRH | The Institute of Molecular Biology and Genetics of NASU | M.Tukalo et al. |
| EPI_ISL_1298487 | Kalush CRH | The Institute of Molecular Biology and Genetics of NASU | M.Tukalo et al. |
| EPI_ISL_1298488 | Bogorodchany CRH | The Institute of Molecular Biology and Genetics of NASU | M.Tukalo et al. |
| EPI_ISL_1298492 | National Platform bis UMONS/Jolimont | National Platform bis UMONS/Jolimont | François Dufrasne, Gautier Detry, Ruddy Wattiez |
| EPI_ISL_1299229, EPI_ISL_1299231, EPI_ISL_1299236, EPI_ISL_1299238 | OLVZ Aalst | OLVZ Aalst | Astrid Holderbeke |
| EPI_ISL_1299245, EPI_ISL_1299247, EPI_ISL_1299251 | CHWAPI - SITE NOTRE DAME | Institut de Pathologie et Genetique (IPG) | Pascale Hilbert, Jérémie Gras |
| EPI_ISL_1299500 | Azienda USL Umbria 2 | Istituto Zooprofilattico Sperimentale dell'Abruzzo e Molise "G. Caporale" | Proietti A, Pistoni E, Lorusso A, Marcacci M, Di Domenico M, Ancora M, Curini V, Mangone I, Rinaldi A, Scialabba S, Di Pasquale A, Cammà C, Puglia I, Calistri P, Savini G |
| EPI_ISL_1299501, EPI_ISL_1299502, EPI_ISL_1299503, EPI_ISL_1299504, EPI_ISL_1299505, EPI_ISL_1299507, EPI_ISL_1299508, EPI_ISL_1299509, EPI_ISL_1299510 | Azienda Ospedaliera Terni | Istituto Zooprofilattico Sperimentale dell'Abruzzo e Molise "G. Caporale" | Palumbo M, ScaccettiA, Lorusso A, Marcacci M, Di Domenico M, Ancora M, Curini V, Mangone I, Rinaldi A, Scialabba S, Di Pasquale A, Cammà C, Puglia I, Calistri P, Savini G |
| EPI_ISL_1299511, EPI_ISL_1299512, EPI_ISL_1299513, EPI_ISL_1299514, EPI_ISL_1299515, EPI_ISL_1299516, EPI_ISL_1299517, EPI_ISL_1299518, EPI_ISL_1299519, EPI_ISL_1299520 | Azienda Ospedaliera Terni | Istituto Zooprofilattico Sperimentale dell'Abruzzo e Molise "G. Caporale" | Palumbo m, Scaccetti A, Lorusso A, Marcacci M, Di Domenico M, Ancora M, Curini V, Mangone I, Rinaldi A, Scialabba S, Di Pasquale A, Cammà C, Puglia I, Calistri P, Savini G |
| EPI_ISL_1299521 | Azienda USL Umbria 2 | Istituto Zooprofilattico Sperimentale dell'Abruzzo e Molise "G. Caporale" | Proietti A, Pistoni E, Lorusso A, Marcacci M, Di Domenico M, Ancora M, Curini V, Mangone I, Rinaldi A, Scialabba S, Di Pasquale A, Cammà C, Puglia I, Calistri P, Savini G |

| | | | |
|---|---|---|---|
| EPI_ISL_1299522, EPI_ISL_1299523, EPI_ISL_1299524, EPI_ISL_1299526 | Azienda Ospedaliera Terni | Istituto Zooprofilattico Sperimentale dell'Abruzzo e Molise "G. Caporale" | Palumbo m, Scaccetti A, Lorusso A, Marcacci M, Di Domenico M, Ancora M, Curini V, Mangone I, Rinaldi A, Scialabba S, Di Pasquale A, Cammà C, Puglia I, Calistri P, Savini G |
| EPI_ISL_1299527, EPI_ISL_1299528, EPI_ISL_1299529, EPI_ISL_1299530 | Istituto Zooprofilattico Sperimentale Umbria e Marche "Togo Rosati" | Istituto Zooprofilattico Sperimentale dell'Abruzzo e Molise "G. Caporale" | Biagetti m, Giammarioli M, Lorusso A, Marcacci M, Di Domenico M, Ancora M, Curini V, Mangone I, Rinaldi A, Scialabba S, Di Pasquale A, Cammà C, Puglia I, Calistri P, Savini G |
| EPI_ISL_1299532 | Azienda USL Umbria 2 | Istituto Zooprofilattico Sperimentale dell'Abruzzo e Molise "G. Caporale" | Proietti A, Pistoni E, Lorusso A, Marcacci M, Di Domenico M, Ancora M, Curini V, Mangone I, Rinaldi A, Scialabba S, Di Pasquale A, Cammà C, Puglia I, Calistri P, Savini G |
| EPI_ISL_1299533, EPI_ISL_1299534, EPI_ISL_1299535, EPI_ISL_1299536, EPI_ISL_1299537, EPI_ISL_1299538, EPI_ISL_1299539, EPI_ISL_1299540, EPI_ISL_1299541, EPI_ISL_1299542 | Istituto Zooprofilattico Sperimentale Umbria e Marche "Togo Rosati" | Istituto Zooprofilattico Sperimentale dell'Abruzzo e Molise "G. Caporale" | Biagetti M, Giammarioli M, Lorusso A, Marcacci M, Di Domenico M, Ancora M, Curini V, Mangone I, Rinaldi A, Scialabba S, Di Pasquale A, Cammà C, Puglia I, Calistri P, Savini G |
| EPI_ISL_1299543 | Azienda USL Umbria 2 | Istituto Zooprofilattico Sperimentale dell'Abruzzo e Molise "G. Caporale" | Proietti A, Pistoni E, Lorusso A, Marcacci M, Di Domenico M, Ancora M, Curini V, Mangone I, Rinaldi A, Scialabba S, Di Pasquale A, Cammà C, Puglia I, Calistri P, Savini G |
| EPI_ISL_1299544 | Istituto Zooprofilattico Sperimentale Umbria e Marche "Togo Rosati" | Istituto Zooprofilattico Sperimentale dell'Abruzzo e Molise "G. Caporale" | Biagetti M, Giammarioli M,Lorusso A, Marcacci M, Di Domenico M, Ancora M, Curini V, Mangone I, Rinaldi A, Scialabba S, Di Pasquale A, Cammà C, Puglia I, Calistri P, Savini G |
| EPI_ISL_1299545, EPI_ISL_1299546, EPI_ISL_1299547, EPI_ISL_1299548, EPI_ISL_1299549, EPI_ISL_1299550, EPI_ISL_1299551, EPI_ISL_1299552, EPI_ISL_1299553 | Laboratorio Analisi Osp. Città di Castello - Azienda USL Umbria1 | Istituto Zooprofilattico Sperimentale dell'Abruzzo e Molise "G. Caporale" | Malagigi V, Tacconi P,Lorusso A, Marcacci M, Di Domenico M, Ancora M, Curini V, Mangone I, Rinaldi A, Scialabba S, Di Pasquale A, Cammà C, Puglia I, Calistri P, Savini G |
| EPI_ISL_1299554 | Azienda USL Umbria 2 | Istituto Zooprofilattico Sperimentale dell'Abruzzo e Molise "G. Caporale" | Proietti A, Pistoni E, Lorusso A, Marcacci M, Di Domenico M, Ancora M, Curini V, Mangone I, Rinaldi A, Scialabba S, Di Pasquale A, Cammà C, Puglia I, Calistri P, Savini G |
| EPI_ISL_1299555 | Laboratorio Analisi Osp. Città di Castello - Azienda USL Umbria1 | Istituto Zooprofilattico Sperimentale dell'Abruzzo e Molise "G. Caporale" | Malagigi V, Tacconi P,Lorusso A, Marcacci M, Di Domenico M, Ancora M, Curini V, Mangone I, Rinaldi A, Scialabba S, Di Pasquale A, Cammà C, Puglia I, Calistri P, Savini G |
| EPI_ISL_1299556, EPI_ISL_1299557, EPI_ISL_1299558, EPI_ISL_1299559, EPI_ISL_1299560, EPI_ISL_1299561, EPI_ISL_1299562 | Università degli Studi di Perugia | Istituto Zooprofilattico Sperimentale dell'Abruzzo e Molise "G. Caporale" | Mencacci A, Camilloni B,Lorusso A, Marcacci M, Di Domenico M, Ancora M, Curini V, Mangone I, Rinaldi A, Scialabba S, Di Pasquale A, Cammà C, Puglia I, Calistri P, Savini G |
| EPI_ISL_1299563 | Azienda Ospedaliera Terni | Istituto Zooprofilattico Sperimentale dell'Abruzzo e Molise "G. Caporale" | Palumbo m, Scaccetti A, Lorusso A, Marcacci M, Di Domenico M, Ancora M, Curini V, Mangone I, Rinaldi A, Scialabba S, Di Pasquale A, Cammà C, Puglia I, Calistri P, Savini G |
| EPI_ISL_1299564, EPI_ISL_1299565, EPI_ISL_1299566, EPI_ISL_1299567, EPI_ISL_1299568, EPI_ISL_1299569 | Università degli Studi di Perugia | Istituto Zooprofilattico Sperimentale dell'Abruzzo e Molise "G. Caporale" | Mencacci A, Camilloni B,Lorusso A, Marcacci M, Di Domenico M, Ancora M, Curini V, Mangone I, Rinaldi A, Scialabba S, Di Pasquale A, Cammà C, Puglia I, Calistri P, Savini G |
| EPI_ISL_1299570 | Azienda Ospedaliera Terni | Istituto Zooprofilattico Sperimentale dell'Abruzzo e Molise "G. Caporale" | Palumbo M, Scaccetti A, Lorusso A, Marcacci M, Di Domenico M, Ancora M, Curini V, Mangone I, Rinaldi A, Scialabba S, Di Pasquale A, Cammà C, Puglia I, Calistri P, Savini G |
| EPI_ISL_1300645 | Clinical Center, University of Sarajevo; Unit for Clinical Microbiology | Clinical Center, University of Sarajevo; Unit for Clinical Microbiology | Irma Salimovi-Beši, Amela Dedei-Ljubovi, Edina Zahirovi, Suzana Arapi, Sebija Izetbegovi, Sandra Vegar-Zubovi, Maja Kuzmanovska, Golubinka Boshevska |
| EPI_ISL_1300646 | Cantonal Hospital Zenica; Department of Microbiological Diagnostics | Clinical Center, University of Sarajevo; Unit for Clinical Microbiology | Irma Salimovi-Beši, Amela Dedei-Ljubovi, Edina Zahirovi, Suzana Arapi, Sebija Izetbegovi, Sandra Vegar-Zubovi, Maja Kuzmanovska, Golubinka Boshevska |
| EPI_ISL_1300647, EPI_ISL_1300648, EPI_ISL_1300649, EPI_ISL_1300650, EPI_ISL_1300651 | Clinical Center, University of Sarajevo; Unit for Clinical Microbiology | Clinical Center, University of Sarajevo; Unit for Clinical Microbiology | Irma Salimovi-Beši, Amela Dedei-Ljubovi, Edina Zahirovi, Suzana Arapi, Sebija Izetbegovi, Sandra Vegar-Zubovi, Maja Kuzmanovska, Golubinka Boshevska |
| EPI_ISL_1300652 | Institute of Public Health of Central Bosnia Canton, Travnik | Clinical Center, University of Sarajevo; Unit for Clinical Microbiology | Irma Salimovi-Beši, Amela Dedei-Ljubovi, Edina Zahirovi, Suzana Arapi, Sebija Izetbegovi, Sandra Vegar-Zubovi, Maja Kuzmanovska, Golubinka Boshevska |
| EPI_ISL_1300653 | Clinical Center, University of Sarajevo; Unit for Clinical Microbiology | Clinical Center, University of Sarajevo; Unit for Clinical Microbiology | Irma Salimovi-Beši, Amela Dedei-Ljubovi, Edina Zahirovi, Suzana Arapi, Sebija Izetbegovi, Sandra Vegar-Zubovi, Maja Kuzmanovska, Golubinka Boshevska |
| EPI_ISL_1300654, EPI_ISL_1300655 | Cantonal Hospital Zenica; Department of Microbiological Diagnostics | Clinical Center, University of Sarajevo; Unit for Clinical Microbiology | Irma Salimovi-Beši, Amela Dedei-Ljubovi, Edina Zahirovi, Suzana Arapi, Sebija Izetbegovi, Sandra Vegar-Zubovi, Maja Kuzmanovska, Golubinka Boshevska |
| EPI_ISL_1300656, EPI_ISL_1300657, EPI_ISL_1300658, EPI_ISL_1300659 | Clinical Center, University of Sarajevo; Unit for Clinical Microbiology | Clinical Center, University of Sarajevo; Unit for Clinical Microbiology | Irma Salimovi-Beši, Amela Dedei-Ljubovi, Edina Zahirovi, Suzana Arapi, Sebija Izetbegovi, Sandra Vegar-Zubovi, Maja Kuzmanovska, Golubinka Boshevska |
| EPI_ISL_1301805 | Hopital | National Reference Center for Viruses of Respiratory Infections, Institut Pasteur, Paris | Marion Barbet, Sylvie Behillil, Méline Bizard, Angela Brisebarre, Camille Capel, Louise Lefrançois, Etienne Simon-Lorière, Vincent Enouf, Maud Vanpeene, Sylvie van der Werf,Griscelli Franck |
| EPI_ISL_1301807, EPI_ISL_1301808 | Hopital | National Reference Center for Viruses of Respiratory Infections, Institut Pasteur, Paris | Marion Barbet, Sylvie Behillil, Méline Bizard, Angela Brisebarre, Camille Capel, Louise Lefrançois, Etienne Simon-Lorière, Vincent Enouf, Maud Vanpeene, Sylvie van der Werf,Guinoiseau Thibault |
| EPI_ISL_1301809 | Hopital | National Reference Center for Viruses of Respiratory Infections, Institut Pasteur, Paris | Marion Barbet, Sylvie Behillil, Méline Bizard, Angela Brisebarre, Camille Capel, Louise Lefrançois, Etienne Simon-Lorière, Vincent Enouf, Maud Vanpeene, Sylvie van der Werf,Ramanantsoa CéLine |
| EPI_ISL_1301810 | Hopital | National Reference Center for Viruses of Respiratory Infections, Institut Pasteur, Paris | Marion Barbet, Sylvie Behillil, Méline Bizard, Angela Brisebarre, Camille Capel, Louise Lefrançois, Etienne Simon-Lorière, Vincent Enouf, Maud Vanpeene, Sylvie van der Werf,Coste-Burel Marianne |
| EPI_ISL_1301811 | Hopital | National Reference Center for Viruses of Respiratory Infections, Institut Pasteur, Paris | Marion Barbet, Sylvie Behillil, Méline Bizard, Angela Brisebarre, Camille Capel, Louise Lefrançois, Etienne Simon-Lorière, Vincent Enouf, Maud Vanpeene, Sylvie van der Werf,Leruez-Ville Marianne |
| EPI_ISL_1301812, EPI_ISL_1301813, EPI_ISL_1301814, EPI_ISL_1301815 | Hopital | National Reference Center for Viruses of Respiratory Infections, Institut Pasteur, Paris | Marion Barbet, Sylvie Behillil, Méline Bizard, Angela Brisebarre, Camille Capel, Louise Lefrançois, Etienne Simon-Lorière, Vincent Enouf, Maud Vanpeene, Sylvie van der Werf,Griscelli Franck |
| EPI_ISL_1301817, EPI_ISL_1301818, EPI_ISL_1301819 | Research | National Reference Center for Viruses of Respiratory Infections, Institut Pasteur, Paris | Marion Barbet, Sylvie Behillil, Méline Bizard, Angela Brisebarre, Camille Capel, Louise Lefrançois, Etienne Simon-Lorière, Vincent Enouf, Maud Vanpeene, Sylvie van der Werf,Leruez-Ville Marianne |
| EPI_ISL_1301820 | hopital | National Reference Center for Viruses of Respiratory Infections, Institut Pasteur, Paris | Marion Barbet, Sylvie Behillil, Méline Bizard, Angela Brisebarre, Camille Capel, Louise Lefrançois, Etienne Simon-Lorière, Vincent Enouf, Maud Vanpeene, Sylvie van der Werf,Leruez-Ville Marianne |
| EPI_ISL_1301821, EPI_ISL_1301822 | Labo Analyses Med | National Reference Center for Viruses of Respiratory Infections, Institut Pasteur, Paris | Marion Barbet, Sylvie Behillil, Méline Bizard, Angela Brisebarre, Camille Capel, Louise Lefrançois, Etienne Simon-Lorière, Vincent Enouf, Maud Vanpeene, Sylvie van der Werf,Roquebert BéNéDicte |
| EPI_ISL_1301823, EPI_ISL_1301824, EPI_ISL_1301825, EPI_ISL_1301826, EPI_ISL_1301827 | hopital | National Reference Center for Viruses of Respiratory Infections, Institut Pasteur, Paris | Marion Barbet, Sylvie Behillil, Méline Bizard, Angela Brisebarre, Camille Capel, Louise Lefrançois, Etienne Simon-Lorière, Vincent Enouf, Maud Vanpeene, Sylvie van der Werf,Guigon AuréLie |
| EPI_ISL_1301828 | hopital | National Reference Center for Viruses of Respiratory Infections, Institut Pasteur, Paris | Marion Barbet, Sylvie Behillil, Méline Bizard, Angela Brisebarre, Camille Capel, Louise Lefrançois, Etienne Simon-Lorière, Vincent Enouf, Maud Vanpeene, Sylvie van der Werf,Paluch Maxime |

| | | | |
|---|---|---|---|
| EPI_ISL_1301829 | Armies | National Reference Center for Viruses of Respiratory Infections, Institut Pasteur, Paris | Marion Barbet, Sylvie Behillil, Méline Bizard, Angela Brisebarre, Camille Capel, Louise Lefrançois, Etienne Simon-Lorière, Vincent Enouf, Maud Vanpeene, Sylvie van der Werf,Desroches Marine |
| EPI_ISL_1301831 | Hopital | National Reference Center for Viruses of Respiratory Infections, Institut Pasteur, Paris | Marion Barbet, Sylvie Behillil, Méline Bizard, Angela Brisebarre, Camille Capel, Louise Lefrançois, Etienne Simon-Lorière, Vincent Enouf, Maud Vanpeene, Sylvie van der Werf,Ramanantsoa CéLine |
| EPI_ISL_1301833 | hopital | National Reference Center for Viruses of Respiratory Infections, Institut Pasteur, Paris | Marion Barbet, Sylvie Behillil, Méline Bizard, Angela Brisebarre, Camille Capel, Louise Lefrançois, Etienne Simon-Lorière, Vincent Enouf, Maud Vanpeene, Sylvie van der Werf,Guigon AuréLie |
| EPI_ISL_1301838 | Hopital | National Reference Center for Viruses of Respiratory Infections, Institut Pasteur, Paris | Marion Barbet, Sylvie Behillil, Méline Bizard, Angela Brisebarre, Camille Capel, Louise Lefrançois, Etienne Simon-Lorière, Vincent Enouf, Maud Vanpeene, Sylvie van der Werf,Leruez-Ville Marianne |
| EPI_ISL_1301839, EPI_ISL_1301840, EPI_ISL_1301841 | Hopital | National Reference Center for Viruses of Respiratory Infections, Institut Pasteur, Paris | Marion Barbet, Sylvie Behillil, Méline Bizard, Angela Brisebarre, Camille Capel, Louise Lefrançois, Etienne Simon-Lorière, Vincent Enouf, Maud Vanpeene, Sylvie van der Werf,Guinoiseau Thibault |
| EPI_ISL_1301842 | Hopital | National Reference Center for Viruses of Respiratory Infections, Institut Pasteur, Paris | Marion Barbet, Sylvie Behillil, Méline Bizard, Angela Brisebarre, Camille Capel, Louise Lefrançois, Etienne Simon-Lorière, Vincent Enouf, Maud Vanpeene, Sylvie van der Werf,Ramanantsoa CéLine |
| EPI_ISL_1301843, EPI_ISL_1301844, EPI_ISL_1301845, EPI_ISL_1301846 | Hopital | National Reference Center for Viruses of Respiratory Infections, Institut Pasteur, Paris | Marion Barbet, Sylvie Behillil, Méline Bizard, Angela Brisebarre, Camille Capel, Louise Lefrançois, Etienne Simon-Lorière, Vincent Enouf, Maud Vanpeene, Sylvie van der Werf,Coste-Burel Marianne |
| EPI_ISL_1301847, EPI_ISL_1301848, EPI_ISL_1301849, EPI_ISL_1301850 | Hopital | National Reference Center for Viruses of Respiratory Infections, Institut Pasteur, Paris | Marion Barbet, Sylvie Behillil, Méline Bizard, Angela Brisebarre, Camille Capel, Louise Lefrançois, Etienne Simon-Lorière, Vincent Enouf, Maud Vanpeene, Sylvie van der Werf,Leruez-Ville Marianne |
| EPI_ISL_1301851 | Hopital | National Reference Center for Viruses of Respiratory Infections, Institut Pasteur, Paris | Marion Barbet, Sylvie Behillil, Méline Bizard, Angela Brisebarre, Camille Capel, Louise Lefrançois, Etienne Simon-Lorière, Vincent Enouf, Maud Vanpeene, Sylvie van der Werf,Ramanantsoa CéLine |
| EPI_ISL_1301853, EPI_ISL_1301854, EPI_ISL_1301855, EPI_ISL_1301856, EPI_ISL_1301857, EPI_ISL_1301858 | Hopital | National Reference Center for Viruses of Respiratory Infections, Institut Pasteur, Paris | Marion Barbet, Sylvie Behillil, Méline Bizard, Angela Brisebarre, Camille Capel, Louise Lefrançois, Etienne Simon-Lorière, Vincent Enouf, Maud Vanpeene, Sylvie van der Werf,Griscelli Franck |
| EPI_ISL_1301860 | Armies | National Reference Center for Viruses of Respiratory Infections, Institut Pasteur, Paris | Marion Barbet, Sylvie Behillil, Méline Bizard, Angela Brisebarre, Camille Capel, Louise Lefrançois, Etienne Simon-Lorière, Vincent Enouf, Maud Vanpeene, Sylvie van der Werf,Desroches Marine |
| EPI_ISL_1301863, EPI_ISL_1301864 | Research | National Reference Center for Viruses of Respiratory Infections, Institut Pasteur, Paris | Marion Barbet, Sylvie Behillil, Méline Bizard, Angela Brisebarre, Camille Capel, Louise Lefrançois, Etienne Simon-Lorière, Vincent Enouf, Maud Vanpeene, Sylvie van der Werf,Leruez-Ville Marianne |
| EPI_ISL_1301865 | Labo Analyses Med | National Reference Center for Viruses of Respiratory Infections, Institut Pasteur, Paris | Marion Barbet, Sylvie Behillil, Méline Bizard, Angela Brisebarre, Camille Capel, Louise Lefrançois, Etienne Simon-Lorière, Vincent Enouf, Maud Vanpeene, Sylvie van der Werf,Roquebert BéNéDicte |
| EPI_ISL_1301866, EPI_ISL_1301867, EPI_ISL_1301868, EPI_ISL_1301869, EPI_ISL_1301870, EPI_ISL_1301871, EPI_ISL_1301872, EPI_ISL_1301873 | hopital | National Reference Center for Viruses of Respiratory Infections, Institut Pasteur, Paris | Marion Barbet, Sylvie Behillil, Méline Bizard, Angela Brisebarre, Camille Capel, Louise Lefrançois, Etienne Simon-Lorière, Vincent Enouf, Maud Vanpeene, Sylvie van der Werf,Guigon AuréLie |
| EPI_ISL_1301874 | hopital | National Reference Center for Viruses of Respiratory Infections, Institut Pasteur, Paris | Marion Barbet, Sylvie Behillil, Méline Bizard, Angela Brisebarre, Camille Capel, Louise Lefrançois, Etienne Simon-Lorière, Vincent Enouf, Maud Vanpeene, Sylvie van der Werf,Descamps Dominique |
| EPI_ISL_1301875, EPI_ISL_1301876 | hopital | National Reference Center for Viruses of Respiratory Infections, Institut Pasteur, Paris | Marion Barbet, Sylvie Behillil, Méline Bizard, Angela Brisebarre, Camille Capel, Louise Lefrançois, Etienne Simon-Lorière, Vincent Enouf, Maud Vanpeene, Sylvie van der Werf,Paluch Maxime |
| EPI_ISL_1301877, EPI_ISL_1301878, EPI_ISL_1301879, EPI_ISL_1301880, EPI_ISL_1301881 | Armies | National Reference Center for Viruses of Respiratory Infections, Institut Pasteur, Paris | Marion Barbet, Sylvie Behillil, Méline Bizard, Angela Brisebarre, Camille Capel, Louise Lefrançois, Etienne Simon-Lorière, Vincent Enouf, Maud Vanpeene, Sylvie van der Werf,Desroches Marine |
| EPI_ISL_1301882 | Hopital | National Reference Center for Viruses of Respiratory Infections, Institut Pasteur, Paris | Marion Barbet, Sylvie Behillil, Méline Bizard, Angela Brisebarre, Camille Capel, Louise Lefrançois, Etienne Simon-Lorière, Vincent Enouf, Maud Vanpeene, Sylvie van der Werf,Coste-Burel Marianne |
| EPI_ISL_1301883 | Research | National Reference Center for Viruses of Respiratory Infections, Institut Pasteur, Paris | Marion Barbet, Sylvie Behillil, Méline Bizard, Angela Brisebarre, Camille Capel, Louise Lefrançois, Etienne Simon-Lorière, Vincent Enouf, Maud Vanpeene, Sylvie van der Werf,Leruez-Ville Marianne |
| EPI_ISL_1301886 | Labo Analyses Med | National Reference Center for Viruses of Respiratory Infections, Institut Pasteur, Paris | Marion Barbet, Sylvie Behillil, Méline Bizard, Angela Brisebarre, Camille Capel, Louise Lefrançois, Etienne Simon-Lorière, Vincent Enouf, Maud Vanpeene, Sylvie van der Werf,Potiron GréGoire |
| EPI_ISL_1301888 | hopital | National Reference Center for Viruses of Respiratory Infections, Institut Pasteur, Paris | Marion Barbet, Sylvie Behillil, Méline Bizard, Angela Brisebarre, Camille Capel, Louise Lefrançois, Etienne Simon-Lorière, Vincent Enouf, Maud Vanpeene, Sylvie van der Werf,Paluch Maxime |
| EPI_ISL_1301890, EPI_ISL_1301891, EPI_ISL_1301892, EPI_ISL_1301893, EPI_ISL_1301894 | Hopital | National Reference Center for Viruses of Respiratory Infections, Institut Pasteur, Paris | Marion Barbet, Sylvie Behillil, Méline Bizard, Angela Brisebarre, Camille Capel, Louise Lefrançois, Etienne Simon-Lorière, Vincent Enouf, Maud Vanpeene, Sylvie van der Werf,Ramanantsoa CéLine |
| EPI_ISL_1301896, EPI_ISL_1301897, EPI_ISL_1301898 | Hopital | National Reference Center for Viruses of Respiratory Infections, Institut Pasteur, Paris | Marion Barbet, Sylvie Behillil, Méline Bizard, Angela Brisebarre, Camille Capel, Louise Lefrançois, Etienne Simon-Lorière, Vincent Enouf, Maud Vanpeene, Sylvie van der Werf,Pilorge LéA |
| EPI_ISL_1301916, EPI_ISL_1301917, EPI_ISL_1301918 | Labo Analyses Med | National Reference Center for Viruses of Respiratory Infections, Institut Pasteur, Paris | Marion Barbet, Sylvie Behillil, Méline Bizard, Angela Brisebarre, Camille Capel, Louise Lefrançois, Etienne Simon-Lorière, Vincent Enouf, Maud Vanpeene, Sylvie van der Werf,Le Vicky |
| EPI_ISL_1301919 | Labo Analyses Med | National Reference Center for Viruses of Respiratory Infections, Institut Pasteur, Paris | Marion Barbet, Sylvie Behillil, Méline Bizard, Angela Brisebarre, Camille Capel, Louise Lefrançois, Etienne Simon-Lorière, Vincent Enouf, Maud Vanpeene, Sylvie van der Werf,Lebihan David |
| EPI_ISL_1301922 | Hopital | National Reference Center for Viruses of Respiratory Infections, Institut Pasteur, Paris | Marion Barbet, Sylvie Behillil, Méline Bizard, Angela Brisebarre, Camille Capel, Louise Lefrançois, Etienne Simon-Lorière, Vincent Enouf, Maud Vanpeene, Sylvie van der Werf,Ramanantsoa CéLine |
| EPI_ISL_1301923 | Labo Analyses Med | National Reference Center for Viruses of Respiratory Infections, Institut Pasteur, Paris | Marion Barbet, Sylvie Behillil, Méline Bizard, Angela Brisebarre, Camille Capel, Louise Lefrançois, Etienne Simon-Lorière, Vincent Enouf, Maud Vanpeene, Sylvie van der Werf,Le Vicky |
| EPI_ISL_1301931 | Hopital | National Reference Center for Viruses of Respiratory Infections, Institut Pasteur, Paris | Marion Barbet, Sylvie Behillil, Méline Bizard, Angela Brisebarre, Camille Capel, Louise Lefrançois, Etienne Simon-Lorière, Vincent Enouf, Maud Vanpeene, Sylvie van der Werf,Coste-Burel Marianne |
| EPI_ISL_1301932, EPI_ISL_1301933 | Hopital | National Reference Center for Viruses of Respiratory Infections, Institut Pasteur, Paris | Marion Barbet, Sylvie Behillil, Méline Bizard, Angela Brisebarre, Camille Capel, Louise Lefrançois, Etienne Simon-Lorière, Vincent Enouf, Maud Vanpeene, Sylvie van der Werf,Ramanantsoa CéLine |
| EPI_ISL_1301934 | Hopital | National Reference Center for Viruses of Respiratory Infections, Institut Pasteur, Paris | Marion Barbet, Sylvie Behillil, Méline Bizard, Angela Brisebarre, Camille Capel, Louise Lefrançois, Etienne Simon-Lorière, Vincent Enouf, Maud Vanpeene, Sylvie van der Werf,Marty Quinternet Solene |
| EPI_ISL_1301935, EPI_ISL_1301936, EPI_ISL_1301937 | Hopital | National Reference Center for Viruses of Respiratory Infections, Institut Pasteur, Paris | Marion Barbet, Sylvie Behillil, Méline Bizard, Angela Brisebarre, Camille Capel, Louise Lefrançois, Etienne Simon-Lorière, Vincent Enouf, Maud Vanpeene, Sylvie van der Werf,Ramanantsoa CéLine |
| EPI_ISL_1301944 | Research | National Reference Center for Viruses of Respiratory Infections, Institut Pasteur, Paris | Marion Barbet, Sylvie Behillil, Méline Bizard, Angela Brisebarre, Camille Capel, Louise Lefrançois, Etienne Simon-Lorière, Vincent Enouf, Maud Vanpeene, Sylvie van der Werf,Leruez-Ville Marianne |
| EPI_ISL_1301945, EPI_ISL_1301946, EPI_ISL_1301947 | Labo Analyses Med | National Reference Center for Viruses of Respiratory Infections, Institut Pasteur, Paris | Marion Barbet, Sylvie Behillil, Méline Bizard, Angela Brisebarre, Camille Capel, Louise Lefrançois, Etienne Simon-Lorière, Vincent Enouf, Maud Vanpeene, Sylvie van der Werf,Roquebert BéNéDicte |

| | | | |
|---|---|---|---|
| EPI_ISL_1301948, EPI_ISL_1301949, EPI_ISL_1301950 | hopital | National Reference Center for Viruses of Respiratory Infections, Institut Pasteur, Paris | Marion Barbet, Sylvie Behillil, Méline Bizard, Angela Brisebarre, Camille Capel, Louise Lefrançois, Etienne Simon-Lorière, Vincent Enouf, Maud Vanpeene, Sylvie van der Werf,Guigon AuréLie |
| EPI_ISL_1301951 | hopital | National Reference Center for Viruses of Respiratory Infections, Institut Pasteur, Paris | Marion Barbet, Sylvie Behillil, Méline Bizard, Angela Brisebarre, Camille Capel, Louise Lefrançois, Etienne Simon-Lorière, Vincent Enouf, Maud Vanpeene, Sylvie van der Werf,Descamps Dominique |
| EPI_ISL_1301952, EPI_ISL_1301953, EPI_ISL_1301954, EPI_ISL_1301955, EPI_ISL_1301956 | hopital | National Reference Center for Viruses of Respiratory Infections, Institut Pasteur, Paris | Marion Barbet, Sylvie Behillil, Méline Bizard, Angela Brisebarre, Camille Capel, Louise Lefrançois, Etienne Simon-Lorière, Vincent Enouf, Maud Vanpeene, Sylvie van der Werf,Paluch Maxime |
| EPI_ISL_1301957, EPI_ISL_1301958 | Research | National Reference Center for Viruses of Respiratory Infections, Institut Pasteur, Paris | Marion Barbet, Sylvie Behillil, Méline Bizard, Angela Brisebarre, Camille Capel, Louise Lefrançois, Etienne Simon-Lorière, Vincent Enouf, Maud Vanpeene, Sylvie van der Werf,Leruez-Ville Marianne |
| EPI_ISL_1301959 | Labo Analyses Med | National Reference Center for Viruses of Respiratory Infections, Institut Pasteur, Paris | Marion Barbet, Sylvie Behillil, Méline Bizard, Angela Brisebarre, Camille Capel, Louise Lefrançois, Etienne Simon-Lorière, Vincent Enouf, Maud Vanpeene, Sylvie van der Werf,Girard Sophie |
| EPI_ISL_1302004, EPI_ISL_1302013, EPI_ISL_1302026, EPI_ISL_1302029, EPI_ISL_1302033, EPI_ISL_1302034, EPI_ISL_1302035, EPI_ISL_1302036, EPI_ISL_1302037, EPI_ISL_1302038, EPI_ISL_1302039, EPI_ISL_1302040, EPI_ISL_1302041, EPI_ISL_1302042, EPI_ISL_1302043, EPI_ISL_1302044, EPI_ISL_1302045, EPI_ISL_1302046, EPI_ISL_1302047, EPI_ISL_1302048, EPI_ISL_1302049, EPI_ISL_1302050, EPI_ISL_1302051, EPI_ISL_1302052, EPI_ISL_1302079, EPI_ISL_1302080, EPI_ISL_1302081, EPI_ISL_1302082, EPI_ISL_1302083, EPI_ISL_1302084, EPI_ISL_1302085, EPI_ISL_1302086, EPI_ISL_1302087, EPI_ISL_1302088, EPI_ISL_1302089, EPI_ISL_1302096, EPI_ISL_1302149 | | | |
| see above | National Center of Infectious and Parasitic Diseases | National Center of Infectious and Parasitic Diseases | Alexiev et al |
| EPI_ISL_1302873, EPI_ISL_1302875, EPI_ISL_1302876, EPI_ISL_1302877, EPI_ISL_1302878, EPI_ISL_1302881, EPI_ISL_1302882, EPI_ISL_1302883, EPI_ISL_1302888, EPI_ISL_1302889, EPI_ISL_1302890, EPI_ISL_1302891, EPI_ISL_1302896, EPI_ISL_1302897, EPI_ISL_1302898, EPI_ISL_1302899, EPI_ISL_1302900, EPI_ISL_1302902, EPI_ISL_1302903, EPI_ISL_1302904, EPI_ISL_1302905, EPI_ISL_1302906, EPI_ISL_1302907, EPI_ISL_1302909, EPI_ISL_1302910, EPI_ISL_1302912, EPI_ISL_1302913, EPI_ISL_1302915, EPI_ISL_1302917, EPI_ISL_1302920, EPI_ISL_1302921, EPI_ISL_1302922, EPI_ISL_1302923, EPI_ISL_1302924, EPI_ISL_1302925, EPI_ISL_1302926, EPI_ISL_1302927, EPI_ISL_1302928, EPI_ISL_1302929, EPI_ISL_1302930, EPI_ISL_1302934, EPI_ISL_1302935, EPI_ISL_1302936, EPI_ISL_1302937, EPI_ISL_1302938, EPI_ISL_1302939, EPI_ISL_1302940, EPI_ISL_1302943, EPI_ISL_1302945, EPI_ISL_1302946, EPI_ISL_1302950, EPI_ISL_1302953, EPI_ISL_1302954, EPI_ISL_1302956, EPI_ISL_1302957, EPI_ISL_1302958, EPI_ISL_1302959, EPI_ISL_1302960, EPI_ISL_1302961, EPI_ISL_1302963, EPI_ISL_1302965, EPI_ISL_1302966, EPI_ISL_1302969, EPI_ISL_1302970, EPI_ISL_1302974, EPI_ISL_1302975, EPI_ISL_1302976, EPI_ISL_1302977, EPI_ISL_1302978, EPI_ISL_1302979, EPI_ISL_1302980, EPI_ISL_1302981, EPI_ISL_1302984, EPI_ISL_1302985, EPI_ISL_1302986, EPI_ISL_1302992, EPI_ISL_1302993, EPI_ISL_1302994, EPI_ISL_1302996, EPI_ISL_1302997, EPI_ISL_1302999, EPI_ISL_1303000, EPI_ISL_1303001, EPI_ISL_1303002, EPI_ISL_1303003, EPI_ISL_1303004, EPI_ISL_1303005, EPI_ISL_1303006, EPI_ISL_1303007, EPI_ISL_1303008, EPI_ISL_1303009, EPI_ISL_1303013, EPI_ISL_1303014, EPI_ISL_1303015, EPI_ISL_1303016, EPI_ISL_1303017, EPI_ISL_1303018, EPI_ISL_1303019, EPI_ISL_1303020, EPI_ISL_1303021, EPI_ISL_1303025, EPI_ISL_1303026, EPI_ISL_1303027, EPI_ISL_1303029 | | | |
| see above | ADMED Microbiologie | Genomics and Transcriptomics, Philip Morris International | Reto Lienhard, Marie-Lise Tritten, Emmanuel Guedj, Nicolas Sierro, Rémi Dulize, David Bornand, Mehdi Auberson, Maxime Berthouzoz, Nikolai Ivanov, Manuel Peitsch |
| EPI_ISL_1307711 | Institut de Virologie du CHU de Strasbourg | Institut de Virologie du CHU de Strasbourg | Fafi-Kremer Samira, Gallais Floriane, Gantner Pierre, Laugel Elodie, Solis Morgane, Velay Aurélie, Wendling Marie-Josée |
| EPI_ISL_1311862 | Laboratorio Biologia Molecolare Sars Cov2 - UOC Laboratorio Analisi - Servizio Medicina di Laboratorio, Ospedale "San Francesco" - ATS-ASSL Nuoro | Laboratorio specialistico UOC Ematologia - Ospedale "San Francesco" - ATS-ASSL Nuoro | Piras Giovanna, Asproni Rosanna,Fiamma Maura, Monne Maria Itria, Malune Paolo, Palmas Angelo Domenico, Lo Maglio Iana, Mameli Giuseppe |
| EPI_ISL_1311965 | The Ohio State University Applied Microbiology Services Laboratory | The Ohio State University Applied Microbiology Services Laboratory | Seth A. Faith PhD |
| EPI_ISL_1311970, EPI_ISL_1311972 | CHWAPI - SITE NOTRE DAME | Institut de Pathologie et Genetique (IPG) | Pascale Hilbert, Jérémie Gras |
| EPI_ISL_1315327, EPI_ISL_1315421, EPI_ISL_1315422, EPI_ISL_1315423, EPI_ISL_1315424, EPI_ISL_1315425, EPI_ISL_1315426 | Laboratory for Respiratory Viruses, Cantacuzino National Military-Medical Institute for Research and Development | Cantacuzino Institute Virology | Luiza Ustea, Nicoleta Paraschiv, Catalina Pascu, Mihaela Lazar |
| EPI_ISL_1315578, EPI_ISL_1315579, EPI_ISL_1315581, EPI_ISL_1315605, EPI_ISL_1315607, EPI_ISL_1315608, EPI_ISL_1315610, EPI_ISL_1315611, EPI_ISL_1315612, EPI_ISL_1315616, EPI_ISL_1315618, EPI_ISL_1315619, EPI_ISL_1315621, EPI_ISL_1315622, EPI_ISL_1315624, EPI_ISL_1315625, EPI_ISL_1315630, EPI_ISL_1315632, EPI_ISL_1315633, EPI_ISL_1315637, EPI_ISL_1315639, EPI_ISL_1315640, EPI_ISL_1315641, EPI_ISL_1315642 | | | |
| see above | AZ Klina | AZ Klina | Carl Vael, Lynsey Berckmans |
| EPI_ISL_1317214 | Laboratorio de Virología HUCA | Laboratorio de Virología HUCA | Sandoval M, Castelló C, Gómez de Oña J, Boga JA, Rojo S, Alvarez-Arguelles ME, Abreu F, Costales I, Perez-Martínez Z, Martín-Rodríguez G, Coto E, Melón S |
| EPI_ISL_1318207 | Ministry of Health Hospitals | Institute of Health and Community Medicine | Chan Chia Jui, David Perera, Ooi Mong How, Chua Hock Hin, Tonnii Sia Loong Loong, Wong Jyn Shan, Wong Kiing Aik |
| EPI_ISL_1319359, EPI_ISL_1319360, EPI_ISL_1319362, EPI_ISL_1319363, EPI_ISL_1319365, EPI_ISL_1319366, EPI_ISL_1319367, EPI_ISL_1319369, EPI_ISL_1319370 | Laboratory of Communicable Diseases | 1. Laboratory of Communicable Diseases (Estonia); 2. Eurofins Genomics Europe Sequencing GmbH | Liidia Dotsenko et al. |
| EPI_ISL_1319445, EPI_ISL_1319447, EPI_ISL_1319454, EPI_ISL_1319455, EPI_ISL_1319457 | North Estonia Medical Centre laboratory | 1. Laboratory of Communicable Diseases (Estonia); 2. Eurofins Genomics Europe Sequencing GmbH | Liidia Dotsenko et al. |
| EPI_ISL_1320174, EPI_ISL_1320180 | IZSM | IZSM-U.O.C. Virologia | Maurizio Viscardi, Lorena Cardillo, Giovanna Fusco, Esterina De Carlo, Antonio Limone |
| EPI_ISL_1320193 | IZSM | IZSM-U.O.C. Virologia | Maurizio Viscardi, Lorena Cardillo, Claudio de Martinis, Giovanna Fusco, Esterina De Carlo, Antonio Limone |
| EPI_ISL_1320197 | IZSM | IZSM-U.O.C. Virologia | Maurizio Viscardi, Lorena Cardillo, Giovanna Fusco, Esterina De Carlo, Antonio Limone |
| EPI_ISL_1321734, EPI_ISL_1321735, EPI_ISL_1321736, EPI_ISL_1321737, EPI_ISL_1321738, EPI_ISL_1321739, EPI_ISL_1321740, EPI_ISL_1321741, EPI_ISL_1321742, EPI_ISL_1321743, EPI_ISL_1321744, EPI_ISL_1321745, EPI_ISL_1321746, EPI_ISL_1321747, EPI_ISL_1321748, EPI_ISL_1321749, EPI_ISL_1321750, EPI_ISL_1321751, EPI_ISL_1321752, EPI_ISL_1321753, EPI_ISL_1321754, EPI_ISL_1321755 | | | |
| see above | Università degli Studi di Perugia | Istituto Zooprofilattico Sperimentale dell'Abruzzo e Molise "G. Caporale" | Mencacci A, Camilloni B, Lorusso A, Marcacci M, Di Domenico M, Ancora M, Curini V, Mangone I, Rinaldi A, Scialabba S, Di Pasquale A, Cammà C, Puglia I, Calistri P, Savini G |
| EPI_ISL_1321756, EPI_ISL_1321757, EPI_ISL_1321758, EPI_ISL_1321759, EPI_ISL_1321760, EPI_ISL_1321761, EPI_ISL_1321762, EPI_ISL_1321763, EPI_ISL_1321765, EPI_ISL_1321766, EPI_ISL_1321767, EPI_ISL_1321768, EPI_ISL_1321769, EPI_ISL_1321770, EPI_ISL_1321771, EPI_ISL_1321772, EPI_ISL_1321773 | | | |
| see above | Azienda Ospedaliera Terni | Istituto Zooprofilattico Sperimentale dell'Abruzzo e Molise "G. Caporale" | Palumbo M, Scaccetti A, Lorusso A, Marcacci M, Di Domenico M, Ancora M, Curini V, Mangone I, Rinaldi A, Scialabba S, Di Pasquale A, Cammà C, Puglia I, Calistri P, Savini G |
| EPI_ISL_1321774 | Università degli Studi di Perugia | Istituto Zooprofilattico Sperimentale dell'Abruzzo e Molise "G. Caporale" | Mencacci A, Camilloni B, Lorusso A, Marcacci M, Di Domenico M, Ancora M, Curini V, Mangone I, Rinaldi A, Scialabba S, Di Pasquale A, Cammà C, Puglia I, Calistri P, Savini G |
| EPI_ISL_1321775, EPI_ISL_1321776, EPI_ISL_1321777, EPI_ISL_1321778, EPI_ISL_1321779, EPI_ISL_1321780, EPI_ISL_1321781, EPI_ISL_1321782, EPI_ISL_1321783, EPI_ISL_1321784 | Azienda Ospedaliera Terni | Istituto Zooprofilattico Sperimentale dell'Abruzzo e Molise "G. Caporale" | Palumbo M, Scaccetti A, Lorusso A, Marcacci M, Di Domenico M, Ancora M, Curini V, Mangone I, Rinaldi A, Scialabba S, Di Pasquale A, Cammà C, Puglia I, Calistri P, Savini G |
| EPI_ISL_1321785 | Università degli Studi di Perugia | Istituto Zooprofilattico Sperimentale dell'Abruzzo e Molise "G. Caporale" | Mencacci A, Camilloni B, Lorusso A, Marcacci M, Di Domenico M, Ancora M, Curini V, Mangone I, Rinaldi A, Scialabba S, Di Pasquale A, Cammà C, Puglia I, Calistri P, Savini G |
| EPI_ISL_1321786, EPI_ISL_1321787, EPI_ISL_1321788, EPI_ISL_1321789, EPI_ISL_1321790, EPI_ISL_1321791, EPI_ISL_1321792, EPI_ISL_1321793, EPI_ISL_1321794 | Azienda Ospedaliera Terni | Istituto Zooprofilattico Sperimentale dell'Abruzzo e Molise "G. Caporale" | Palumbo M, Scaccetti A, Lorusso A, Marcacci M, Di Domenico M, Ancora M, Curini V, Mangone I, Rinaldi A, Scialabba S, Di Pasquale A, Cammà C, Puglia I, Calistri P, Savini G |

| | | | |
|---|---|---|---|
| EPI_ISL_1321795 | Istituto Zooprofilattico Sperimentale Umbria e Marche "Togo Rosati" | Istituto Zooprofilattico Sperimentale dell'Abruzzo e Molise "G. Caporale" | Biagetti M, Giammarioli M,Lorusso A, Marcacci M, Di Domenico M, Ancora M, Curini V, Mangone I, Rinaldi A, Scialabba S, Di Pasquale A, Cammà C, Puglia I, Calistri P, Savini G |
| EPI_ISL_1321796 | Università degli Studi di Perugia | Istituto Zooprofilattico Sperimentale dell'Abruzzo e Molise "G. Caporale" | Mencacci A, Camilloni B, Lorusso A, Marcacci M, Di Domenico M, Ancora M, Curini V, Mangone I, Rinaldi A, Scialabba S, Di Pasquale A, Cammà C, Puglia I, Calistri P, Savini G |
| EPI_ISL_1321797, EPI_ISL_1321798, EPI_ISL_1321799, EPI_ISL_1321800, EPI_ISL_1321801, EPI_ISL_1321802, EPI_ISL_1321803 | Istituto Zooprofilattico Sperimentale Umbria e Marche "Togo Rosati" | Istituto Zooprofilattico Sperimentale dell'Abruzzo e Molise "G. Caporale" | Biagetti M, Giammarioli M,Lorusso A, Marcacci M, Di Domenico M, Ancora M, Curini V, Mangone I, Rinaldi A, Scialabba S, Di Pasquale A, Cammà C, Puglia I, Calistri P, Savini G |
| EPI_ISL_1321804, EPI_ISL_1321805, EPI_ISL_1321806 | Università degli Studi di Perugia | Istituto Zooprofilattico Sperimentale dell'Abruzzo e Molise "G. Caporale" | Mencacci A, Camilloni B, Lorusso A, Marcacci M, Di Domenico M, Ancora M, Curini V, Mangone I, Rinaldi A, Scialabba S, Di Pasquale A, Cammà C, Puglia I, Calistri P, Savini G |
| EPI_ISL_1322110, EPI_ISL_1322111, EPI_ISL_1322112, EPI_ISL_1322113, EPI_ISL_1322114, EPI_ISL_1322115, EPI_ISL_1322116, EPI_ISL_1322117, EPI_ISL_1322118, EPI_ISL_1322119, EPI_ISL_1322121, EPI_ISL_1322123, EPI_ISL_1322125, EPI_ISL_1322127, EPI_ISL_1322128, EPI_ISL_1322129, EPI_ISL_1322132, EPI_ISL_1322133, EPI_ISL_1322134, EPI_ISL_1322135, EPI_ISL_1322136, EPI_ISL_1322138, EPI_ISL_1322139, EPI_ISL_1322142, EPI_ISL_1322143, EPI_ISL_1322144, EPI_ISL_1322146, EPI_ISL_1322147, EPI_ISL_1322148, EPI_ISL_1322149, EPI_ISL_1322150, EPI_ISL_1322151, EPI_ISL_1322153, EPI_ISL_1322154, EPI_ISL_1322155, EPI_ISL_1322156, EPI_ISL_1322158, EPI_ISL_1322161, EPI_ISL_1322164, EPI_ISL_1322165, EPI_ISL_1322166, EPI_ISL_1322167, EPI_ISL_1322168, EPI_ISL_1322169, EPI_ISL_1322170, EPI_ISL_1322171, EPI_ISL_1322172, EPI_ISL_1322173, EPI_ISL_1322174, EPI_ISL_1322175, EPI_ISL_1322177, EPI_ISL_1322178, EPI_ISL_1322179, EPI_ISL_1322184, EPI_ISL_1322186, EPI_ISL_1322187, EPI_ISL_1322188, EPI_ISL_1322189, EPI_ISL_1322190, EPI_ISL_1322191, EPI_ISL_1322192, EPI_ISL_1322193, EPI_ISL_1322194, EPI_ISL_1322196, EPI_ISL_1322197, EPI_ISL_1322198, EPI_ISL_1322199, EPI_ISL_1322200, EPI_ISL_1322201, EPI_ISL_1322203 | | | |
| see above | Microbiology Department, Laboratori Clínic Metropolitana Nord. Hospital Universitari Germans Trias i Pujol. | Can Ruti SARS-CoV-2 Sequencing Hub (HUGTiP/IrsiCaixa/IGTP) | Marc Noguera-Julian, Pilar Armengol, Ignacio Blanco, Antoni E Bordoy, Francesc Catala-Moll, Pere-Joan Cardona, Julia G Prado, Carol Galvez Maria Casadellà, Cristina Casañ, Gemma Clara, Irina Pey, Jordi Barretina, Bonaventura Clotet, Cristina Esteban, Montserrat Giménez, Mercedes Guerrero, Anna Not, Roger Paredes, Mariona Parera, Verónica Saludes, Alba Sánchez, and Elisa Martró on behalf of the Can Ruti SARS-CoV-2 Sequencing Hub. |
| EPI_ISL_1322204, EPI_ISL_1322205, EPI_ISL_1322207, EPI_ISL_1322208, EPI_ISL_1322211, EPI_ISL_1322212, EPI_ISL_1322213, EPI_ISL_1322214, EPI_ISL_1322215, EPI_ISL_1322216, EPI_ISL_1322217, EPI_ISL_1322218 | | | |
| see above | IrsiCaixa | IrsiCaixa | Marc Noguera-Julian, Mariona Parera, Maria Casadellà, Pilar Armengol, Francesc Catala-Moll, Roger Paredes, Bonaventura Clotet Gloria Trujillo, Rafel Perez Vidal, Jaume Trape Pujol, Carolina Gonzalez Fernandez, Roger Paredes, Eulalia Grau, Bonaventura Clotet |
| EPI_ISL_1323552, EPI_ISL_1323553 | CHU LILLE | CHU Lille - Laboratoire de Virologie | AIT YAHYA Emilie, ALIDJINOU Enagnon Kazali, BOCKET Laurence, CREPIN Michel, DEMAY Christophe, ENGELMANN Ilka, GEFFROY Sandrine, GUIGON Aurélie, LAZREK Mouna, NOBILLIAUX Florian, PREVOST Brigitte, THUILLIER Caroline, TINEZ Claire |
| EPI_ISL_1323554 | CH VALENCIENNES | CHU Lille - Laboratoire de Virologie | AIT YAHYA Emilie, ALIDJINOU Enagnon Kazali, BOCKET Laurence, CREPIN Michel, DEMAY Christophe, ENGELMANN Ilka, GEFFROY Sandrine, GUIGON Aurélie, LAZREK Mouna, NOBILLIAUX Florian, PREVOST Brigitte, THUILLIER Caroline, TINEZ Claire |
| EPI_ISL_1323563, EPI_ISL_1323564, EPI_ISL_1323574 | CHU LILLE | CHU Lille - Laboratoire de Virologie | AIT YAHYA Emilie, ALIDJINOU Enagnon Kazali, BOCKET Laurence, CREPIN Michel, DEMAY Christophe, ENGELMANN Ilka, GEFFROY Sandrine, GUIGON Aurélie, LAZREK Mouna, NOBILLIAUX Florian, PREVOST Brigitte, THUILLIER Caroline, TINEZ Claire |
| EPI_ISL_1323980 | Istituto Zooprofilattico Sperimentale del Mezzogiorno (IZSM) | IZSM-U.O.C. Virologia | Maurizio Viscardi, Lorena Cardillo, Claudio de Martinis, Giovanna Fusco, Esterina De Carlo, Antonio Limone |
| EPI_ISL_1324467, EPI_ISL_1324469, EPI_ISL_1324470, EPI_ISL_1324471, EPI_ISL_1324472, EPI_ISL_1324473, EPI_ISL_1324477 | National Platform bis UMONS/Jolimont | National Platform bis UMONS/Jolimont | François Dufrasne, Gautier Detry, Ruddy Wattiez |
| EPI_ISL_1324655 | Laboratorio de Virología HUCA | Laboratorio de Virología HUCA | Sandoval M, Castelló C, Gómez de Oña J, Boga JA, Rojo S, Alvarez-Arguelles ME, Abreu F, Costales I, Perez-Martínez Z, Martín-Rodríguez G, Coto E, Melón S |
| EPI_ISL_1334382, EPI_ISL_1334383, EPI_ISL_1334384, EPI_ISL_1334385 | Laboratorios Lister | Instituto de Diagnostico y Referencia Epidemiologicos (INDRE) | Claudia Wong-Arambula, Abril Rodriguez-Maldonado, Vanessa Rivero-Arredondo, Ariadna Medina-Benitez, Joaquin Quiroz-Mercado, Sergio Rangel-Guerrero, Natividad Cruz-Ortiz, Tatiana Nunez-Garcia, Gisela Barrera-Badillo, Lucia Hernandez-Rivas, Irma Lopez-Martinez, Ernesto Ramirez-Gonzalez. |
| EPI_ISL_1334386 | LESP Guanajuato | Instituto de Diagnostico y Referencia Epidemiologicos (INDRE) | Claudia Wong-Arambula, Abril Rodriguez-Maldonado, Vanessa Rivero-Arredondo, Ariadna Medina-Benitez, Joaquin Quiroz-Mercado, Sergio Rangel-Guerrero, Natividad Cruz-Ortiz, Tatiana Nunez-Garcia, Gisela Barrera-Badillo, Lucia Hernandez-Rivas, Irma Lopez-Martinez, Ernesto Ramirez-Gonzalez. |
| EPI_ISL_1335873, EPI_ISL_1335875 | AZ Sint-Jan | AZ SINT-JAN BRUGGE | Marijke Reynders, Patrick Descheemaeker, Laurien Hoornaert, Thomas Van Landschoot, Jorn Hellemans |
| EPI_ISL_1335876, EPI_ISL_1335877, EPI_ISL_1335880, EPI_ISL_1335881 | AZ Zeno | AZ SINT-JAN BRUGGE | Marijke Reynders, Patrick Descheemaeker, Laurien Hoornaert, Thomas Van Landschoot, Jorn Hellemans |
| EPI_ISL_1336083, EPI_ISL_1336084 | SIESP DIPARTIMENTO DI PREVENZIONE TERAMO | Istituto Zooprofilattico Sperimentale dell'Abruzzo e Molise "G. Caporale" | Lorusso A, Marcacci M, Di Domenico M, Ancora M, Curini V, Mangone I, Rinaldi A, Scialabba S, Di Pasquale A, Cammà C, Puglia I, Calistri P, Savini G |
| EPI_ISL_1336178 | Azienda Ospedaliera Terni | Istituto Zooprofilattico Sperimentale dell'Abruzzo e Molise "G. Caporale" | Palumbo M, Scaccetti A, Lorusso A, Marcacci M, Di Domenico M, Ancora M, Curini V, Mangone I, Rinaldi A, Scialabba S, Di Pasquale A, Cammà C, Puglia I, Calistri P, Savini G |
| EPI_ISL_1336179 | Istituto Zooprofilattico Sperimentale Umbria e Marche "Togo Rosati" | Istituto Zooprofilattico Sperimentale dell'Abruzzo e Molise "G. Caporale" | Biagetti M, Giammarioli M,Lorusso A, Marcacci M, Di Domenico M, Ancora M, Curini V, Mangone I, Rinaldi A, Scialabba S, Di Pasquale A, Cammà C, Puglia I, Calistri P, Savini G |
| EPI_ISL_1336180 | Virology Center Adolfo Lutz Institute Sao Paulo Brazil | Retrovirus Laboratory Adolfo Lutz Institute | Gabriela Bastos Cabral, Giselle I S Lopez-Lopes, Cintia Ahagon, Audrey Cilli, Paula Morena Guimaraes, Igor Mohamed Hussein,Junior Pereira de Souza , Roberta Shiavon, Luis Brigido |
| EPI_ISL_1336181 | Virology Center Adolfo Lutz Institute Sao Paulo Brazil | Retrovirus Laboratory Adolfo Lutz Institute | Gabriela Bastos Cabral, Giselle I S Lopez-Lopes, Cintia Ahagon, Audrey Cilli, Paula Morena Guimaraes, Igor Mohamed Hussein, Junior Pereira de Souza , Roberta Shiavon, Luis Brigido |
| EPI_ISL_1336330 | Labo Analyses Med | National Reference Center for Viruses of Respiratory Infections, Institut Pasteur, Paris | Marion Barbet, Sylvie Behillil, Méline Bizard, Angela Brisebarre, Camille Capel, Etienne Simon-Lorière, Vincent Enouf, Maud Vanpeene, Sylvie van der Werf,Gestin Brieuc |
| EPI_ISL_1336331, EPI_ISL_1336332, EPI_ISL_1336333 | Labo Analyses Med | National Reference Center for Viruses of Respiratory Infections, Institut Pasteur, Paris | Marion Barbet, Sylvie Behillil, Méline Bizard, Angela Brisebarre, Camille Capel, Etienne Simon-Lorière, Vincent Enouf, Maud Vanpeene, Sylvie van der Werf,Holstein Anne |
| EPI_ISL_1336334 | Hopital | National Reference Center for Viruses of Respiratory Infections, Institut Pasteur, Paris | Marion Barbet, Sylvie Behillil, Méline Bizard, Angela Brisebarre, Camille Capel, Etienne Simon-Lorière, Vincent Enouf, Maud Vanpeene, Sylvie van der Werf,Martres Pascale |
| EPI_ISL_1336335, EPI_ISL_1336336, EPI_ISL_1336337, EPI_ISL_1336338, EPI_ISL_1336339, EPI_ISL_1336340 | hopital | National Reference Center for Viruses of Respiratory Infections, Institut Pasteur, Paris | Marion Barbet, Sylvie Behillil, Méline Bizard, Angela Brisebarre, Camille Capel, Etienne Simon-Lorière, Vincent Enouf, Maud Vanpeene, Sylvie van der Werf,Gastli Nabil |
| EPI_ISL_1336341 | Hopital | National Reference Center for Viruses of Respiratory Infections, Institut Pasteur, Paris | Marion Barbet, Sylvie Behillil, Méline Bizard, Angela Brisebarre, Camille Capel, Etienne Simon-Lorière, Vincent Enouf, Maud Vanpeene, Sylvie van der Werf,Thibault Vincent |
| EPI_ISL_1336342, EPI_ISL_1336343, EPI_ISL_1336344 | Labo Analyses Med | National Reference Center for Viruses of Respiratory Infections, Institut Pasteur, Paris | Marion Barbet, Sylvie Behillil, Méline Bizard, Angela Brisebarre, Camille Capel, Etienne Simon-Lorière, Vincent Enouf, Maud Vanpeene, Sylvie van der Werf,Leonard Pierre-Yves |
| EPI_ISL_1336345, EPI_ISL_1336346, EPI_ISL_1336347, EPI_ISL_1336348, EPI_ISL_1336349 | Labo Analyses Med | National Reference Center for Viruses of Respiratory Infections, Institut Pasteur, Paris | Marion Barbet, Sylvie Behillil, Méline Bizard, Angela Brisebarre, Camille Capel, Etienne Simon-Lorière, Vincent Enouf, Maud Vanpeene, Sylvie van der Werf,Said-Delattre OphéLie |
| EPI_ISL_1336350, EPI_ISL_1336351 | Labo Analyses Med | National Reference Center for Viruses of Respiratory Infections, Institut Pasteur, Paris | Marion Barbet, Sylvie Behillil, Méline Bizard, Angela Brisebarre, Camille Capel, Etienne Simon-Lorière, Vincent Enouf, Maud Vanpeene, Sylvie van der Werf,Roquebert BéNéDicte |
| EPI_ISL_1336491, EPI_ISL_1336492, EPI_ISL_1336493, EPI_ISL_1336494, EPI_ISL_1336495, EPI_ISL_1336496, EPI_ISL_1336497, EPI_ISL_1336498, EPI_ISL_1336499, EPI_ISL_1336500, EPI_ISL_1336501, EPI_ISL_1336502, EPI_ISL_1336503, EPI_ISL_1336504, EPI_ISL_1336505, EPI_ISL_1336506, EPI_ISL_1336507, | | | |

EPI_ISL_1336508, EPI_ISL_1336509, EPI_ISL_1336510

| | | | |
|---|---|---|---|
| see above | Microbiology Department, University Hospital Donostia | Microbiology Department, University Hospital Donostia | Marimon JM, Montes M, Piñeiro L, Sorarrain A, Gomez M, Cilla G. |
| EPI_ISL_1336650 | Università degli Studi di Perugia | Istituto Zooprofilattico Sperimentale dell'Abruzzo e Molise "G. Caporale" | Mencacci A, Camilloni B, Lorusso A, Marcacci M, Di Domenico M, Ancora M, Curini V, Mangone I, Rinaldi A, Scialabba S, Di Pasquale A, Cammà C, Puglia I, Calistri P, Savini G |
| EPI_ISL_1336651 | Azienda Ospedaliera Terni | Istituto Zooprofilattico Sperimentale dell'Abruzzo e Molise "G. Caporale" | Palumbo M, Scaccetti A, Lorusso A, Marcacci M, Di Domenico M, Ancora M, Curini V, Mangone I, Rinaldi A, Scialabba S, Di Pasquale A, Cammà C, Puglia I, Calistri P, Savini G |
| EPI_ISL_1336661 | USCA Tagliacozzo | Istituto Zooprofilattico Sperimentale dell'Abruzzo e Molise "G. Caporale" | Lorusso A, Marcacci M, Di Domenico M, Ancora M, Curini V, Di Lollo V, Mangone I, Rinaldi A, Scialabba S, Caporale M, Di Pasquale A, Cammà C, Puglia I, Calistri P, Savini G |
| EPI_ISL_1336706 | DIP. PREV. AVEZZANO SERVIZIO DI IGIENE EPIDEMIOLOGIA E SANITA' PUBBLICA | Istituto Zooprofilattico Sperimentale dell'Abruzzo e Molise "G. Caporale" | Lorusso A, Marcacci M, Di Domenico M, Ancora M, Curini V, Di Lollo V, Mangone I, Rinaldi A, Scialabba S, Caporale M, Di Pasquale A, Cammà C, Puglia I, Calistri P, Savini G |
| EPI_ISL_1336707 | SIESP CHIETI - DRIVE IN ORTONA | Istituto Zooprofilattico Sperimentale dell'Abruzzo e Molise "G. Caporale" | Lorusso A, Marcacci M, Di Domenico M, Ancora M, Curini V, Di Lollo V, Mangone I, Rinaldi A, Scialabba S, Caporale M, Di Pasquale A, Cammà C, Puglia I, Calistri P, Savini G |
| EPI_ISL_1336708 | USCA Pescina | Istituto Zooprofilattico Sperimentale dell'Abruzzo e Molise "G. Caporale" | Lorusso A, Marcacci M, Di Domenico M, Ancora M, Curini V, Di Lollo V, Mangone I, Rinaldi A, Scialabba S, Caporale M, Di Pasquale A, Cammà C, Puglia I, Calistri P, Savini G |
| EPI_ISL_1336709, EPI_ISL_1336710, EPI_ISL_1336711 | SIESP DIPARTIMENTO DI PREVENZIONE TERAMO | Istituto Zooprofilattico Sperimentale dell'Abruzzo e Molise "G. Caporale" | Lorusso A, Marcacci M, Di Domenico M, Ancora M, Curini V, Di Lollo V, Mangone I, Rinaldi A, Scialabba S, Caporale M, Di Pasquale A, Cammà C, Puglia I, Calistri P, Savini G |
| EPI_ISL_1336712 | SIESP CHIETI - DRIVE IN CHIETI | Istituto Zooprofilattico Sperimentale dell'Abruzzo e Molise "G. Caporale" | Lorusso A, Marcacci M, Di Domenico M, Ancora M, Curini V, Di Lollo V, Mangone I, Rinaldi A, Scialabba S, Caporale M, Di Pasquale A, Cammà C, Puglia I, Calistri P, Savini G |
| EPI_ISL_1336713 | SIESP DIPARTIMENTO DI PREVENZIONE TERAMO | Istituto Zooprofilattico Sperimentale dell'Abruzzo e Molise "G. Caporale" | Lorusso A, Marcacci M, Di Domenico M, Ancora M, Curini V, Di Lollo V, Mangone I, Rinaldi A, Scialabba S, Caporale M, Di Pasquale A, Cammà C, Puglia I, Calistri P, Savini G |
| EPI_ISL_1336714 | SIESP CHIETI - DRIVE IN CHIETI | Istituto Zooprofilattico Sperimentale dell'Abruzzo e Molise "G. Caporale" | Lorusso A, Marcacci M, Di Domenico M, Ancora M, Curini V, Di Lollo V, Mangone I, Rinaldi A, Scialabba S, Caporale M, Di Pasquale A, Cammà C, Puglia I, Calistri P, Savini G |
| EPI_ISL_1336715, EPI_ISL_1336716, EPI_ISL_1336717 | USCA Castel di Sangro | Istituto Zooprofilattico Sperimentale dell'Abruzzo e Molise "G. Caporale" | Lorusso A, Marcacci M, Di Domenico M, Ancora M, Curini V, Di Lollo V, Mangone I, Rinaldi A, Scialabba S, Caporale M, Di Pasquale A, Cammà C, Puglia I, Calistri P, Savini G |
| EPI_ISL_1336718 | SIESP DIPARTIMENTO DI PREVENZIONE TERAMO | Istituto Zooprofilattico Sperimentale dell'Abruzzo e Molise "G. Caporale" | Lorusso A, Marcacci M, Di Domenico M, Ancora M, Curini V, Di Lollo V, Mangone I, Rinaldi A, Scialabba S, Caporale M, Di Pasquale A, Cammà C, Puglia I, Calistri P, Savini G |
| EPI_ISL_1336719, EPI_ISL_1336720 | SIESP CHIETI - DRIVE IN ORTONA | Istituto Zooprofilattico Sperimentale dell'Abruzzo e Molise "G. Caporale" | Lorusso A, Marcacci M, Di Domenico M, Ancora M, Curini V, Di Lollo V, Mangone I, Rinaldi A, Scialabba S, Caporale M, Di Pasquale A, Cammà C, Puglia I, Calistri P, Savini G |
| EPI_ISL_1336721 | USCA Pescina | Istituto Zooprofilattico Sperimentale dell'Abruzzo e Molise "G. Caporale" | Lorusso A, Marcacci M, Di Domenico M, Ancora M, Curini V, Di Lollo V, Mangone I, Rinaldi A, Scialabba S, Caporale M, Di Pasquale A, Cammà C, Puglia I, Calistri P, Savini G |
| EPI_ISL_1336722 | USCA Castel di Sangro | Istituto Zooprofilattico Sperimentale dell'Abruzzo e Molise "G. Caporale" | Lorusso A, Marcacci M, Di Domenico M, Ancora M, Curini V, Di Lollo V, Mangone I, Rinaldi A, Scialabba S, Caporale M, Di Pasquale A, Cammà C, Puglia I, Calistri P, Savini G |
| EPI_ISL_1336723 | SIESP DIPARTIMENTO DI PREVENZIONE TERAMO | Istituto Zooprofilattico Sperimentale dell'Abruzzo e Molise "G. Caporale" | Lorusso A, Marcacci M, Di Domenico M, Ancora M, Curini V, Di Lollo V, Mangone I, Rinaldi A, Scialabba S, Caporale M, Di Pasquale A, Cammà C, Puglia I, Calistri P, Savini G |
| EPI_ISL_1336724 | USCA Castel di Sangro | Istituto Zooprofilattico Sperimentale dell'Abruzzo e Molise "G. Caporale" | Lorusso A, Marcacci M, Di Domenico M, Ancora M, Curini V, Di Lollo V, Mangone I, Rinaldi A, Scialabba S, Caporale M, Di Pasquale A, Cammà C, Puglia I, Calistri P, Savini G |
| EPI_ISL_1336725 | SIESP CHIETI - DRIVE IN VASTO | Istituto Zooprofilattico Sperimentale dell'Abruzzo e Molise "G. Caporale" | Lorusso A, Marcacci M, Di Domenico M, Ancora M, Curini V, Di Lollo V, Mangone I, Rinaldi A, Scialabba S, Caporale M, Di Pasquale A, Cammà C, Puglia I, Calistri P, Savini G |
| EPI_ISL_1336726 | DIP. PREV. AVEZZANO SERVIZIO DI IGIENE EPIDEMIOLOGIA E SANITA' PUBBLICA | Istituto Zooprofilattico Sperimentale dell'Abruzzo e Molise "G. Caporale" | Lorusso A, Marcacci M, Di Domenico M, Ancora M, Curini V, Di Lollo V, Mangone I, Rinaldi A, Scialabba S, Caporale M, Di Pasquale A, Cammà C, Puglia I, Calistri P, Savini G |
| EPI_ISL_1336727 | USCA Tagliacozzo | Istituto Zooprofilattico Sperimentale dell'Abruzzo e Molise "G. Caporale" | Lorusso A, Marcacci M, Di Domenico M, Ancora M, Curini V, Di Lollo V, Mangone I, Rinaldi A, Scialabba S, Caporale M, Di Pasquale A, Cammà C, Puglia I, Calistri P, Savini G |
| EPI_ISL_1336728, EPI_ISL_1336729 | SIESP DIPARTIMENTO DI PREVENZIONE TERAMO | Istituto Zooprofilattico Sperimentale dell'Abruzzo e Molise "G. Caporale" | Lorusso A, Marcacci M, Di Domenico M, Ancora M, Curini V, Di Lollo V, Mangone I, Rinaldi A, Scialabba S, Caporale M, Di Pasquale A, Cammà C, Puglia I, Calistri P, Savini G |
| EPI_ISL_1336730, EPI_ISL_1336731 | Simeoni - La Casa del Sole | Istituto Zooprofilattico Sperimentale dell'Abruzzo e Molise "G. Caporale" | Lorusso A, Marcacci M, Di Domenico M, Ancora M, Curini V, Di Lollo V, Mangone I, Rinaldi A, Scialabba S, Caporale M, Di Pasquale A, Cammà C, Puglia I, Calistri P, Savini G |
| EPI_ISL_1336732 | OSPEDALE CIVILE GIULIANOVA - PRONTO SOCCORSO | Istituto Zooprofilattico Sperimentale dell'Abruzzo e Molise "G. Caporale" | Lorusso A, Marcacci M, Di Domenico M, Ancora M, Curini V, Di Lollo V, Mangone I, Rinaldi A, Scialabba S, Caporale M, Di Pasquale A, Cammà C, Puglia I, Calistri P, Savini G |
| EPI_ISL_1336733, EPI_ISL_1336734 | SIESP DIPARTIMENTO DI PREVENZIONE TERAMO | Istituto Zooprofilattico Sperimentale dell'Abruzzo e Molise "G. Caporale" | Lorusso A, Marcacci M, Di Domenico M, Ancora M, Curini V, Di Lollo V, Mangone I, Rinaldi A, Scialabba S, Caporale M, Di Pasquale A, Cammà C, Puglia I, Calistri P, Savini G |
| EPI_ISL_1336735 | DIP. PREV. AVEZZANO SERVIZIO DI IGIENE EPIDEMIOLOGIA E SANITA' PUBBLICA | Istituto Zooprofilattico Sperimentale dell'Abruzzo e Molise "G. Caporale" | Lorusso A, Marcacci M, Di Domenico M, Ancora M, Curini V, Di Lollo V, Mangone I, Rinaldi A, Scialabba S, Caporale M, Di Pasquale A, Cammà C, Puglia I, Calistri P, Savini G |
| EPI_ISL_1336736 | SIESP CHIETI - DRIVE IN CHIETI | Istituto Zooprofilattico Sperimentale dell'Abruzzo e Molise "G. Caporale" | Lorusso A, Marcacci M, Di Domenico M, Ancora M, Curini V, Di Lollo V, Mangone I, Rinaldi A, Scialabba S, Caporale M, Di Pasquale A, Cammà C, Puglia I, Calistri P, Savini G |
| EPI_ISL_1336737 | USCA Tagliacozzo | Istituto Zooprofilattico Sperimentale dell'Abruzzo e Molise "G. Caporale" | Lorusso A, Marcacci M, Di Domenico M, Ancora M, Curini V, Di Lollo V, Mangone I, Rinaldi A, Scialabba S, Caporale M, Di Pasquale A, Cammà C, Puglia I, Calistri P, Savini G |
| EPI_ISL_1336738, EPI_ISL_1336739 | SIESP DIPARTIMENTO DI PREVENZIONE TERAMO | Istituto Zooprofilattico Sperimentale dell'Abruzzo e Molise "G. Caporale" | Lorusso A, Marcacci M, Di Domenico M, Ancora M, Curini V, Di Lollo V, Mangone I, Rinaldi A, Scialabba S, Caporale M, Di Pasquale A, Cammà C, Puglia I, Calistri P, Savini G |
| EPI_ISL_1336740, EPI_ISL_1336741 | SIESP CHIETI - DRIVE IN CHIETI | Istituto Zooprofilattico Sperimentale dell'Abruzzo e Molise "G. Caporale" | Lorusso A, Marcacci M, Di Domenico M, Ancora M, Curini V, Di Lollo V, Mangone I, Rinaldi A, Scialabba S, Caporale M, Di Pasquale A, Cammà C, Puglia I, Calistri P, Savini G |
| EPI_ISL_1336742 | USCA Castel di Sangro | Istituto Zooprofilattico Sperimentale dell'Abruzzo e Molise "G. Caporale" | Lorusso A, Marcacci M, Di Domenico M, Ancora M, Curini V, Di Lollo V, Mangone I, Rinaldi A, Scialabba S, Caporale M, Di Pasquale A, Cammà C, Puglia I, Calistri P, Savini G |
| EPI_ISL_1336743 | RSA GIULIANOVA | Istituto Zooprofilattico Sperimentale dell'Abruzzo e Molise "G. Caporale" | Lorusso A, Marcacci M, Di Domenico M, Ancora M, Curini V, Di Lollo V, Mangone I, Rinaldi A, Scialabba S, Caporale M, Di Pasquale A, Cammà C, Puglia I, Calistri P, Savini G |
| EPI_ISL_1336744, EPI_ISL_1336745 | USCA Castel di Sangro | Istituto Zooprofilattico Sperimentale dell'Abruzzo e Molise "G. Caporale" | Lorusso A, Marcacci M, Di Domenico M, Ancora M, Curini V, Di Lollo V, Mangone I, Rinaldi A, Scialabba S, Caporale M, Di Pasquale A, Cammà C, Puglia I, Calistri P, Savini G |
| EPI_ISL_1336746 | SIESP DIPARTIMENTO DI PREVENZIONE TERAMO | Istituto Zooprofilattico Sperimentale dell'Abruzzo e Molise "G. Caporale" | Lorusso A, Marcacci M, Di Domenico M, Ancora M, Curini V, Di Lollo V, Mangone I, Rinaldi A, Scialabba S, Caporale M, Di Pasquale A, Cammà C, Puglia I, Calistri P, Savini G |
| EPI_ISL_1336747, EPI_ISL_1336748 | SIESP CHIETI - DRIVE IN ORTONA | Istituto Zooprofilattico Sperimentale dell'Abruzzo e Molise "G. Caporale" | Lorusso A, Marcacci M, Di Domenico M, Ancora M, Curini V, Di Lollo V, Mangone I, Rinaldi A, Scialabba S, Caporale M, Di Pasquale A, Cammà C, Puglia I, Calistri P, Savini G |

| | | | |
|---|---|---|---|
| EPI_ISL_1336749 | DIP. PREV. AVEZZANO SERVIZIO DI IGIENE EPIDEMIOLOGIA E SANITA' PUBBLICA | Istituto Zooprofilattico Sperimentale dell'Abruzzo e Molise "G. Caporale" | Lorusso A, Marcacci M, Di Domenico M, Ancora M, Curini V, Di Lollo V, Mangone I, Rinaldi A, Scialabba S, Caporale M, Di Pasquale A, Cammà C, Puglia I, Calistri P, Savini G |
| EPI_ISL_1336750 | USCA Pescina | Istituto Zooprofilattico Sperimentale dell'Abruzzo e Molise "G. Caporale" | Lorusso A, Marcacci M, Di Domenico M, Ancora M, Curini V, Di Lollo V, Mangone I, Rinaldi A, Scialabba S, Caporale M, Di Pasquale A, Cammà C, Puglia I, Calistri P, Savini G |
| EPI_ISL_1336751 | SIESP CHIETI - DRIVE IN CHIETI | Istituto Zooprofilattico Sperimentale dell'Abruzzo e Molise "G. Caporale" | Lorusso A, Marcacci M, Di Domenico M, Ancora M, Curini V, Di Lollo V, Mangone I, Rinaldi A, Scialabba S, Caporale M, Di Pasquale A, Cammà C, Puglia I, Calistri P, Savini G |
| EPI_ISL_1336752, EPI_ISL_1336753 | USCA Castel di Sangro | Istituto Zooprofilattico Sperimentale dell'Abruzzo e Molise "G. Caporale" | Lorusso A, Marcacci M, Di Domenico M, Ancora M, Curini V, Di Lollo V, Mangone I, Rinaldi A, Scialabba S, Caporale M, Di Pasquale A, Cammà C, Puglia I, Calistri P, Savini G |
| EPI_ISL_1336754 | SIESP CHIETI - DRIVE IN VASTO | Istituto Zooprofilattico Sperimentale dell'Abruzzo e Molise "G. Caporale" | Lorusso A, Marcacci M, Di Domenico M, Ancora M, Curini V, Di Lollo V, Mangone I, Rinaldi A, Scialabba S, Caporale M, Di Pasquale A, Cammà C, Puglia I, Calistri P, Savini G |
| EPI_ISL_1336755 | SIESP DIPARTIMENTO DI PREVENZIONE TERAMO | Istituto Zooprofilattico Sperimentale dell'Abruzzo e Molise "G. Caporale" | Lorusso A, Marcacci M, Di Domenico M, Ancora M, Curini V, Di Lollo V, Mangone I, Rinaldi A, Scialabba S, Caporale M, Di Pasquale A, Cammà C, Puglia I, Calistri P, Savini G |
| EPI_ISL_1336756 | Simeoni - La Casa del Sole | Istituto Zooprofilattico Sperimentale dell'Abruzzo e Molise "G. Caporale" | Lorusso A, Marcacci M, Di Domenico M, Ancora M, Curini V, Di Lollo V, Mangone I, Rinaldi A, Scialabba S, Caporale M, Di Pasquale A, Cammà C, Puglia I, Calistri P, Savini G |
| EPI_ISL_1336757, EPI_ISL_1336758, EPI_ISL_1336760 | SIESP DIPARTIMENTO DI PREVENZIONE TERAMO | Istituto Zooprofilattico Sperimentale dell'Abruzzo e Molise "G. Caporale" | Lorusso A, Marcacci M, Di Domenico M, Ancora M, Curini V, Di Lollo V, Mangone I, Rinaldi A, Scialabba S, Caporale M, Di Pasquale A, Cammà C, Puglia I, Calistri P, Savini G |
| EPI_ISL_1336761 | DIP. PREV. AVEZZANO SERVIZIO DI IGIENE EPIDEMIOLOGIA E SANITA' PUBBLICA | Istituto Zooprofilattico Sperimentale dell'Abruzzo e Molise "G. Caporale" | Lorusso A, Marcacci M, Di Domenico M, Ancora M, Curini V, Di Lollo V, Mangone I, Rinaldi A, Scialabba S, Caporale M, Di Pasquale A, Cammà C, Puglia I, Calistri P, Savini G |
| EPI_ISL_1336762, EPI_ISL_1336763, EPI_ISL_1336764, EPI_ISL_1336765 | SIESP CHIETI - DRIVE IN LANCIANO | Istituto Zooprofilattico Sperimentale dell'Abruzzo e Molise "G. Caporale" | Lorusso A, Marcacci M, Di Domenico M, Ancora M, Curini V, Di Lollo V, Mangone I, Rinaldi A, Scialabba S, Caporale M, Di Pasquale A, Cammà C, Puglia I, Calistri P, Savini G |
| EPI_ISL_1336766 | SIESP DIPARTIMENTO DI PREVENZIONE TERAMO | Istituto Zooprofilattico Sperimentale dell'Abruzzo e Molise "G. Caporale" | Lorusso A, Marcacci M, Di Domenico M, Ancora M, Curini V, Di Lollo V, Mangone I, Rinaldi A, Scialabba S, Caporale M, Di Pasquale A, Cammà C, Puglia I, Calistri P, Savini G |
| EPI_ISL_1336767 | PRESIDIO TERRITORIALE CASOLI | Istituto Zooprofilattico Sperimentale dell'Abruzzo e Molise "G. Caporale" | Lorusso A, Marcacci M, Di Domenico M, Ancora M, Curini V, Di Lollo V, Mangone I, Rinaldi A, Scialabba S, Caporale M, Di Pasquale A, Cammà C, Puglia I, Calistri P, Savini G |
| EPI_ISL_1336768, EPI_ISL_1336769, EPI_ISL_1336770, EPI_ISL_1336771 | SIESP DIPARTIMENTO DI PREVENZIONE TERAMO | Istituto Zooprofilattico Sperimentale dell'Abruzzo e Molise "G. Caporale" | Lorusso A, Marcacci M, Di Domenico M, Ancora M, Curini V, Di Lollo V, Mangone I, Rinaldi A, Scialabba S, Caporale M, Di Pasquale A, Cammà C, Puglia I, Calistri P, Savini G |
| EPI_ISL_1336772, EPI_ISL_1336773 | SIESP CHIETI - DRIVE IN LANCIANO | Istituto Zooprofilattico Sperimentale dell'Abruzzo e Molise "G. Caporale" | Lorusso A, Marcacci M, Di Domenico M, Ancora M, Curini V, Di Lollo V, Mangone I, Rinaldi A, Scialabba S, Caporale M, Di Pasquale A, Cammà C, Puglia I, Calistri P, Savini G |
| EPI_ISL_1336774, EPI_ISL_1336775 | SIESP DIPARTIMENTO DI PREVENZIONE TERAMO | Istituto Zooprofilattico Sperimentale dell'Abruzzo e Molise "G. Caporale" | Lorusso A, Marcacci M, Di Domenico M, Ancora M, Curini V, Di Lollo V, Mangone I, Rinaldi A, Scialabba S, Caporale M, Di Pasquale A, Cammà C, Puglia I, Calistri P, Savini G |
| EPI_ISL_1336776, EPI_ISL_1336777 | RSA GIULIANOVA | Istituto Zooprofilattico Sperimentale dell'Abruzzo e Molise "G. Caporale" | Lorusso A, Marcacci M, Di Domenico M, Ancora M, Curini V, Di Lollo V, Mangone I, Rinaldi A, Scialabba S, Caporale M, Di Pasquale A, Cammà C, Puglia I, Calistri P, Savini G |
| EPI_ISL_1336778, EPI_ISL_1336779 | SIESP DIPARTIMENTO DI PREVENZIONE TERAMO | Istituto Zooprofilattico Sperimentale dell'Abruzzo e Molise "G. Caporale" | Lorusso A, Marcacci M, Di Domenico M, Ancora M, Curini V, Di Lollo V, Mangone I, Rinaldi A, Scialabba S, Caporale M, Di Pasquale A, Cammà C, Puglia I, Calistri P, Savini G |
| EPI_ISL_1336780 | SIESP CHIETI - DRIVE IN LANCIANO | Istituto Zooprofilattico Sperimentale dell'Abruzzo e Molise "G. Caporale" | Lorusso A, Marcacci M, Di Domenico M, Ancora M, Curini V, Di Lollo V, Mangone I, Rinaldi A, Scialabba S, Caporale M, Di Pasquale A, Cammà C, Puglia I, Calistri P, Savini G |
| EPI_ISL_1336781 | SIESP DIPARTIMENTO DI PREVENZIONE TERAMO | Istituto Zooprofilattico Sperimentale dell'Abruzzo e Molise "G. Caporale" | Lorusso A, Marcacci M, Di Domenico M, Ancora M, Curini V, Di Lollo V, Mangone I, Rinaldi A, Scialabba S, Caporale M, Di Pasquale A, Cammà C, Puglia I, Calistri P, Savini G |
| EPI_ISL_1336782 | SIESP DIPARTIMENTO DI PREVENZIONE CHIETI | Istituto Zooprofilattico Sperimentale dell'Abruzzo e Molise "G. Caporale" | Lorusso A, Marcacci M, Di Domenico M, Ancora M, Curini V, Di Lollo V, Mangone I, Rinaldi A, Scialabba S, Caporale M, Di Pasquale A, Cammà C, Puglia I, Calistri P, Savini G |
| EPI_ISL_1336783, EPI_ISL_1336784 | SIESP DIPARTIMENTO DI PREVENZIONE TERAMO | Istituto Zooprofilattico Sperimentale dell'Abruzzo e Molise "G. Caporale" | Lorusso A, Marcacci M, Di Domenico M, Ancora M, Curini V, Di Lollo V, Mangone I, Rinaldi A, Scialabba S, Caporale M, Di Pasquale A, Cammà C, Puglia I, Calistri P, Savini G |
| EPI_ISL_1336785 | DIP. PREV. AVEZZANO SERVIZIO DI IGIENE EPIDEMIOLOGIA E SANITA' PUBBLICA | Istituto Zooprofilattico Sperimentale dell'Abruzzo e Molise "G. Caporale" | Lorusso A, Marcacci M, Di Domenico M, Ancora M, Curini V, Di Lollo V, Mangone I, Rinaldi A, Scialabba S, Caporale M, Di Pasquale A, Cammà C, Puglia I, Calistri P, Savini G |
| EPI_ISL_1336786 | USCA Avezzano | Istituto Zooprofilattico Sperimentale dell'Abruzzo e Molise "G. Caporale" | Lorusso A, Marcacci M, Di Domenico M, Ancora M, Curini V, Di Lollo V, Mangone I, Rinaldi A, Scialabba S, Caporale M, Di Pasquale A, Cammà C, Puglia I, Calistri P, Savini G |
| EPI_ISL_1336787 | SIESP CHIETI - DRIVE IN ORTONA | Istituto Zooprofilattico Sperimentale dell'Abruzzo e Molise "G. Caporale" | Lorusso A, Marcacci M, Di Domenico M, Ancora M, Curini V, Di Lollo V, Mangone I, Rinaldi A, Scialabba S, Caporale M, Di Pasquale A, Cammà C, Puglia I, Calistri P, Savini G |
| EPI_ISL_1336788 | OSPEDALE CIVILE SANT'OMERO - PRONTO SOCCORSO | Istituto Zooprofilattico Sperimentale dell'Abruzzo e Molise "G. Caporale" | Lorusso A, Marcacci M, Di Domenico M, Ancora M, Curini V, Di Lollo V, Mangone I, Rinaldi A, Scialabba S, Caporale M, Di Pasquale A, Cammà C, Puglia I, Calistri P, Savini G |
| EPI_ISL_1336789, EPI_ISL_1336790, EPI_ISL_1336791 | SIESP DIPARTIMENTO DI PREVENZIONE TERAMO | Istituto Zooprofilattico Sperimentale dell'Abruzzo e Molise "G. Caporale" | Lorusso A, Marcacci M, Di Domenico M, Ancora M, Curini V, Di Lollo V, Mangone I, Rinaldi A, Scialabba S, Caporale M, Di Pasquale A, Cammà C, Puglia I, Calistri P, Savini G |
| EPI_ISL_1336792 | SIESP CHIETI - DRIVE IN LANCIANO | Istituto Zooprofilattico Sperimentale dell'Abruzzo e Molise "G. Caporale" | Lorusso A, Marcacci M, Di Domenico M, Ancora M, Curini V, Di Lollo V, Mangone I, Rinaldi A, Scialabba S, Caporale M, Di Pasquale A, Cammà C, Puglia I, Calistri P, Savini G |
| EPI_ISL_1336793 | SIESP CHIETI - DRIVE IN VASTO | Istituto Zooprofilattico Sperimentale dell'Abruzzo e Molise "G. Caporale" | Lorusso A, Marcacci M, Di Domenico M, Ancora M, Curini V, Di Lollo V, Mangone I, Rinaldi A, Scialabba S, Caporale M, Di Pasquale A, Cammà C, Puglia I, Calistri P, Savini G |
| EPI_ISL_1336794, EPI_ISL_1336795 | SIESP CHIETI - DRIVE IN LANCIANO | Istituto Zooprofilattico Sperimentale dell'Abruzzo e Molise "G. Caporale" | Lorusso A, Marcacci M, Di Domenico M, Ancora M, Curini V, Di Lollo V, Mangone I, Rinaldi A, Scialabba S, Caporale M, Di Pasquale A, Cammà C, Puglia I, Calistri P, Savini G |
| EPI_ISL_1336796 | SIESP DIPARTIMENTO DI PREVENZIONE TERAMO | Istituto Zooprofilattico Sperimentale dell'Abruzzo e Molise "G. Caporale" | Lorusso A, Marcacci M, Di Domenico M, Ancora M, Curini V, Di Lollo V, Mangone I, Rinaldi A, Scialabba S, Caporale M, Di Pasquale A, Cammà C, Puglia I, Calistri P, Savini G |
| EPI_ISL_1336800, EPI_ISL_1336801, EPI_ISL_1336802, EPI_ISL_1336803, EPI_ISL_1336804, EPI_ISL_1336805, EPI_ISL_1336806, EPI_ISL_1336807, EPI_ISL_1336808, EPI_ISL_1336809, EPI_ISL_1336810, EPI_ISL_1336811, EPI_ISL_1336812, EPI_ISL_1336813, EPI_ISL_1336814, EPI_ISL_1336816, EPI_ISL_1336817, EPI_ISL_1336818, EPI_ISL_1336819, EPI_ISL_1336820 | | | |
| see above | Azienda Ospedaliera Terni | Istituto Zooprofilattico Sperimentale dell'Abruzzo e Molise "G. Caporale" | Palumbo M, Scaccetti A, Lorusso A, Marcacci M, Di Domenico M, Ancora M, Curini V, Mangone I, Rinaldi A, Scialabba S, Caporale M, Di Pasquale A, Cammà C, Puglia I, Calistri P, Savini G |
| EPI_ISL_1336821 | Istituto Zooprofilattico Sperimentale Umbria e Marche "Togo Rosati" | Istituto Zooprofilattico Sperimentale dell'Abruzzo e Molise "G. Caporale" | Biagetti M, Giammarioli M, Lorusso A, Marcacci M, Di Domenico M, Ancora M, Curini V, Mangone I, Rinaldi A, Scialabba S, Caporale M, Di Pasquale A, Cammà C, Puglia I, Calistri P, Savini G |
| EPI_ISL_1336822, EPI_ISL_1336823, EPI_ISL_1336824, EPI_ISL_1336825, EPI_ISL_1336826, EPI_ISL_1336827, EPI_ISL_1336828, EPI_ISL_1336829, EPI_ISL_1336830, EPI_ISL_1336831 | Azienda Ospedaliera Terni | Istituto Zooprofilattico Sperimentale dell'Abruzzo e Molise "G. Caporale" | Palumbo M, Scaccetti A, Lorusso A, Marcacci M, Di Domenico M, Ancora M, Curini V, Mangone I, Rinaldi A, Scialabba S, Caporale M, Di Pasquale A, Cammà C, Puglia I, Calistri P, Savini G |

| | | | |
|---|---|---|---|
| EPI_ISL_1336832 | Istituto Zooprofilattico Sperimentale Umbria e Marche "Togo Rosati" | Istituto Zooprofilattico Sperimentale dell'Abruzzo e Molise "G. Caporale" | Biagetti M, Giammarioli M, Lorusso A, Marcacci M, Di Domenico M, Ancora M, Curini V, Mangone I, Rinaldi A, Scialabba S, Caporale M, Di Pasquale A, Cammà C, Puglia I, Calistri P, Savini G |
| EPI_ISL_1336833, EPI_ISL_1336834, EPI_ISL_1336835, EPI_ISL_1336836, EPI_ISL_1336837, EPI_ISL_1336838, EPI_ISL_1336839, EPI_ISL_1336840 | Azienda Ospedaliera Terni | Istituto Zooprofilattico Sperimentale dell'Abruzzo e Molise "G. Caporale" | Palumbo M, Scaccetti A, Lorusso A, Marcacci M, Di Domenico M, Ancora M, Curini V, Mangone I, Rinaldi A, Scialabba S, Caporale M, Di Pasquale A, Cammà C, Puglia I, Calistri P, Savini G |
| EPI_ISL_1336842 | Istituto Zooprofilattico Sperimentale Umbria e Marche "Togo Rosati" | Istituto Zooprofilattico Sperimentale dell'Abruzzo e Molise "G. Caporale" | Biagetti M, Giammarioli M, Lorusso A, Marcacci M, Di Domenico M, Ancora M, Curini V, Mangone I, Rinaldi A, Scialabba S, Caporale M, Di Pasquale A, Cammà C, Puglia I, Calistri P, Savini G |
| EPI_ISL_1336843, EPI_ISL_1336844, EPI_ISL_1336846, EPI_ISL_1336847, EPI_ISL_1336848, EPI_ISL_1336849, EPI_ISL_1336850, EPI_ISL_1336851, EPI_ISL_1336852 | Azienda Ospedaliera Terni | Istituto Zooprofilattico Sperimentale dell'Abruzzo e Molise "G. Caporale" | Palumbo M, Scaccetti A, Lorusso A, Marcacci M, Di Domenico M, Ancora M, Curini V, Mangone I, Rinaldi A, Scialabba S, Caporale M, Di Pasquale A, Cammà C, Puglia I, Calistri P, Savini G |
| EPI_ISL_1336853 | Istituto Zooprofilattico Sperimentale Umbria e Marche "Togo Rosati" | Istituto Zooprofilattico Sperimentale dell'Abruzzo e Molise "G. Caporale" | Biagetti M, Giammarioli M, Lorusso A, Marcacci M, Di Domenico M, Ancora M, Curini V, Mangone I, Rinaldi A, Scialabba S, Caporale M, Di Pasquale A, Cammà C, Puglia I, Calistri P, Savini G |
| EPI_ISL_1336854, EPI_ISL_1336855, EPI_ISL_1336856, EPI_ISL_1336857, EPI_ISL_1336858, EPI_ISL_1336859 | Azienda Ospedaliera Terni | Istituto Zooprofilattico Sperimentale dell'Abruzzo e Molise "G. Caporale" | Palumbo M, Scaccetti A, Lorusso A, Marcacci M, Di Domenico M, Ancora M, Curini V, Mangone I, Rinaldi A, Scialabba S, Caporale M, Di Pasquale A, Cammà C, Puglia I, Calistri P, Savini G |
| EPI_ISL_1336860 | Azienda USL Umbria 2 | Istituto Zooprofilattico Sperimentale dell'Abruzzo e Molise "G. Caporale" | Proietti A, Pistoni E, Lorusso A, Marcacci M, Di Domenico M, Ancora M, Curini V, Mangone I, Rinaldi A, Scialabba S, Caporale M, Di Pasquale A, Cammà C, Puglia I, Calistri P, Savini G |
| EPI_ISL_1336861 | Azienda USL Umbria 3 | Istituto Zooprofilattico Sperimentale dell'Abruzzo e Molise "G. Caporale" | Proietti A, Pistoni E, Lorusso A, Marcacci M, Di Domenico M, Ancora M, Curini V, Mangone I, Rinaldi A, Scialabba S, Caporale M, Di Pasquale A, Cammà C, Puglia I, Calistri P, Savini G |
| EPI_ISL_1336862 | Azienda Ospedaliera Terni | Istituto Zooprofilattico Sperimentale dell'Abruzzo e Molise "G. Caporale" | Palumbo M, Scaccetti A, Lorusso A, Marcacci M, Di Domenico M, Ancora M, Curini V, Mangone I, Rinaldi A, Scialabba S, Caporale M, Di Pasquale A, Cammà C, Puglia I, Calistri P, Savini G |
| EPI_ISL_1336863 | Laboratorio Analisi Osp. Città di Castello - Azienda USL Umbria1 | Istituto Zooprofilattico Sperimentale dell'Abruzzo e Molise "G. Caporale" | Malagigi V, Tacconi P, Lorusso A, Marcacci M, Di Domenico M, Ancora M, Curini V, Mangone I, Rinaldi A, Scialabba S, Caporale M, Di Pasquale A, Cammà C, Puglia I, Calistri P, Savini G |
| EPI_ISL_1336864 | Laboratorio Analisi Osp. Città di Castello - Azienda USL Umbria2 | Istituto Zooprofilattico Sperimentale dell'Abruzzo e Molise "G. Caporale" | Malagigi V, Tacconi P, Lorusso A, Marcacci M, Di Domenico M, Ancora M, Curini V, Mangone I, Rinaldi A, Scialabba S, Caporale M, Di Pasquale A, Cammà C, Puglia I, Calistri P, Savini G |
| EPI_ISL_1336865 | Laboratorio Analisi Osp. Città di Castello - Azienda USL Umbria3 | Istituto Zooprofilattico Sperimentale dell'Abruzzo e Molise "G. Caporale" | Malagigi V, Tacconi P, Lorusso A, Marcacci M, Di Domenico M, Ancora M, Curini V, Mangone I, Rinaldi A, Scialabba S, Caporale M, Di Pasquale A, Cammà C, Puglia I, Calistri P, Savini G |
| EPI_ISL_1336866 | Laboratorio Analisi Osp. Città di Castello - Azienda USL Umbria4 | Istituto Zooprofilattico Sperimentale dell'Abruzzo e Molise "G. Caporale" | Malagigi V, Tacconi P, Lorusso A, Marcacci M, Di Domenico M, Ancora M, Curini V, Mangone I, Rinaldi A, Scialabba S, Caporale M, Di Pasquale A, Cammà C, Puglia I, Calistri P, Savini G |
| EPI_ISL_1336867 | Laboratorio Analisi Osp. Città di Castello - Azienda USL Umbria5 | Istituto Zooprofilattico Sperimentale dell'Abruzzo e Molise "G. Caporale" | Malagigi V, Tacconi P, Lorusso A, Marcacci M, Di Domenico M, Ancora M, Curini V, Mangone I, Rinaldi A, Scialabba S, Caporale M, Di Pasquale A, Cammà C, Puglia I, Calistri P, Savini G |
| EPI_ISL_1336868 | Laboratorio Analisi Osp. Città di Castello - Azienda USL Umbria6 | Istituto Zooprofilattico Sperimentale dell'Abruzzo e Molise "G. Caporale" | Malagigi V, Tacconi P, Lorusso A, Marcacci M, Di Domenico M, Ancora M, Curini V, Mangone I, Rinaldi A, Scialabba S, Caporale M, Di Pasquale A, Cammà C, Puglia I, Calistri P, Savini G |
| EPI_ISL_1336869 | Laboratorio Analisi Osp. Città di Castello - Azienda USL Umbria7 | Istituto Zooprofilattico Sperimentale dell'Abruzzo e Molise "G. Caporale" | Malagigi V, Tacconi P, Lorusso A, Marcacci M, Di Domenico M, Ancora M, Curini V, Mangone I, Rinaldi A, Scialabba S, Caporale M, Di Pasquale A, Cammà C, Puglia I, Calistri P, Savini G |
| EPI_ISL_1336870 | Laboratorio Analisi Osp. Città di Castello - Azienda USL Umbria8 | Istituto Zooprofilattico Sperimentale dell'Abruzzo e Molise "G. Caporale" | Malagigi V, Tacconi P, Lorusso A, Marcacci M, Di Domenico M, Ancora M, Curini V, Mangone I, Rinaldi A, Scialabba S, Caporale M, Di Pasquale A, Cammà C, Puglia I, Calistri P, Savini G |
| EPI_ISL_1336871, EPI_ISL_1336872 | Università degli Studi di Perugia | Istituto Zooprofilattico Sperimentale dell'Abruzzo e Molise "G. Caporale" | Mencacci A, Camilloni B, Lorusso A, Marcacci M, Di Domenico M, Ancora M, Curini V, Mangone I, Rinaldi A, Scialabba S, Caporale M, Di Pasquale A, Cammà C, Puglia I, Calistri P, Savini G |
| EPI_ISL_1336873 | Istituto Zooprofilattico Sperimentale Umbria e Marche "Togo Rosati" | Istituto Zooprofilattico Sperimentale dell'Abruzzo e Molise "G. Caporale" | Biagetti M, Giammarioli M, Lorusso A, Marcacci M, Di Domenico M, Ancora M, Curini V, Mangone I, Rinaldi A, Scialabba S, Caporale M, Di Pasquale A, Cammà C, Puglia I, Calistri P, Savini G |
| EPI_ISL_1336874, EPI_ISL_1336875, EPI_ISL_1336876, EPI_ISL_1336877, EPI_ISL_1336878, EPI_ISL_1336879 | Università degli Studi di Perugia | Istituto Zooprofilattico Sperimentale dell'Abruzzo e Molise "G. Caporale" | Mencacci A, Camilloni B, Lorusso A, Marcacci M, Di Domenico M, Ancora M, Curini V, Mangone I, Rinaldi A, Scialabba S, Caporale M, Di Pasquale A, Cammà C, Puglia I, Calistri P, Savini G |
| EPI_ISL_1336880 | Istituto Zooprofilattico Sperimentale Umbria e Marche "Togo Rosati" | Istituto Zooprofilattico Sperimentale dell'Abruzzo e Molise "G. Caporale" | Biagetti M, Giammarioli M, Lorusso A, Marcacci M, Di Domenico M, Ancora M, Curini V, Mangone I, Rinaldi A, Scialabba S, Caporale M, Di Pasquale A, Cammà C, Puglia I, Calistri P, Savini G |
| EPI_ISL_1336884 | SIESP CHIETI - DRIVE IN LANCIANO | Istituto Zooprofilattico Sperimentale dell'Abruzzo e Molise "G. Caporale" | Lorusso A, Marcacci M, Di Domenico M, Ancora M, Curini V, Di Lollo Valeria, Mangone I, Rinaldi A, Scialabba S, Caporale M, Di Pasquale A, Cammà C, Puglia I, Calistri P, Savini G |
| EPI_ISL_1336885 | SIESP DIPARTIMENTO DI PREVENZIONE TERAMO | Istituto Zooprofilattico Sperimentale dell'Abruzzo e Molise "G. Caporale" | Lorusso A, Marcacci M, Di Domenico M, Ancora M, Curini V, Di Lollo Valeria, Mangone I, Rinaldi A, Scialabba S, Caporale M, Di Pasquale A, Cammà C, Puglia I, Calistri P, Savini G |
| EPI_ISL_1336886 | SIESP SULMONA | Istituto Zooprofilattico Sperimentale dell'Abruzzo e Molise "G. Caporale" | Lorusso A, Marcacci M, Di Domenico M, Ancora M, Curini V, Di Lollo Valeria, Mangone I, Rinaldi A, Scialabba S, Caporale M, Di Pasquale A, Cammà C, Puglia I, Calistri P, Savini G |
| EPI_ISL_1336887 | SIESP CHIETI - DRIVE IN LANCIANO | Istituto Zooprofilattico Sperimentale dell'Abruzzo e Molise "G. Caporale" | Lorusso A, Marcacci M, Di Domenico M, Ancora M, Curini V, Di Lollo Valeria, Mangone I, Rinaldi A, Scialabba S, Caporale M, Di Pasquale A, Cammà C, Puglia I, Calistri P, Savini G |
| EPI_ISL_1336888 | DIP. PREV. AVEZZANO SERVIZIO DI IGIENE EPIDEMIOLOGIA E SANITA' PUBBLICA | Istituto Zooprofilattico Sperimentale dell'Abruzzo e Molise "G. Caporale" | Lorusso A, Marcacci M, Di Domenico M, Ancora M, Curini V, Di Lollo Valeria, Mangone I, Rinaldi A, Scialabba S, Caporale M, Di Pasquale A, Cammà C, Puglia I, Calistri P, Savini G |
| EPI_ISL_1336889, EPI_ISL_1336890, EPI_ISL_1336891 | SIESP DIPARTIMENTO DI PREVENZIONE TERAMO | Istituto Zooprofilattico Sperimentale dell'Abruzzo e Molise "G. Caporale" | Lorusso A, Marcacci M, Di Domenico M, Ancora M, Curini V, Di Lollo Valeria, Mangone I, Rinaldi A, Scialabba S, Caporale M, Di Pasquale A, Cammà C, Puglia I, Calistri P, Savini G |
| EPI_ISL_1336892 | SIESP CHIETI - DRIVE IN ATESSA | Istituto Zooprofilattico Sperimentale dell'Abruzzo e Molise "G. Caporale" | Lorusso A, Marcacci M, Di Domenico M, Ancora M, Curini V, Di Lollo Valeria, Mangone I, Rinaldi A, Scialabba S, Caporale M, Di Pasquale A, Cammà C, Puglia I, Calistri P, Savini G |
| EPI_ISL_1336893, EPI_ISL_1336894 | SIESP DIPARTIMENTO DI PREVENZIONE TERAMO | Istituto Zooprofilattico Sperimentale dell'Abruzzo e Molise "G. Caporale" | Lorusso A, Marcacci M, Di Domenico M, Ancora M, Curini V, Di Lollo Valeria, Mangone I, Rinaldi A, Scialabba S, Caporale M, Di Pasquale A, Cammà C, Puglia I, Calistri P, Savini G |
| EPI_ISL_1336895 | USCA Pescina | Istituto Zooprofilattico Sperimentale dell'Abruzzo e Molise "G. Caporale" | Lorusso A, Marcacci M, Di Domenico M, Ancora M, Curini V, Di Lollo Valeria, Mangone I, Rinaldi A, Scialabba S, Caporale M, Di Pasquale A, Cammà C, Puglia I, Calistri P, Savini G |
| EPI_ISL_1336896 | USCA Castel di Sangro | Istituto Zooprofilattico Sperimentale dell'Abruzzo e Molise "G. Caporale" | Lorusso A, Marcacci M, Di Domenico M, Ancora M, Curini V, Di Lollo Valeria, Mangone I, Rinaldi A, Scialabba S, Caporale M, Di Pasquale A, Cammà C, Puglia I, Calistri P, Savini G |
| EPI_ISL_1336897 | SIESP CHIETI - DRIVE IN GISSI | Istituto Zooprofilattico Sperimentale dell'Abruzzo e Molise "G. Caporale" | Lorusso A, Marcacci M, Di Domenico M, Ancora M, Curini V, Di Lollo Valeria, Mangone I, Rinaldi A, Scialabba S, Caporale M, Di Pasquale A, Cammà C, Puglia I, Calistri P, Savini G |
| EPI_ISL_1336898 | USCA Sulmona | Istituto Zooprofilattico Sperimentale dell'Abruzzo e Molise "G. | Lorusso A, Marcacci M, Di Domenico M, Ancora M, Curini V, Di Lollo Valeria, Mangone I, Rinaldi A, Scialabba S, Caporale M, Di Pasquale A, Cammà C, |

| | | | |
|---|---|---|---|
| | | Caporale" | Puglia I, Calistri P, Savini G |
| EPI_ISL_1336899 | SIESP DIPARTIMENTO DI PREVENZIONE TERAMO | Istituto Zooprofilattico Sperimentale dell'Abruzzo e Molise "G. Caporale" | Lorusso A, Marcacci M, Di Domenico M, Ancora M, Curini V, Di Lollo Valeria, Mangone I, Rinaldi A, Scialabba S, Caporale M, Di Pasquale A, Cammà C, Puglia I, Calistri P, Savini G |
| EPI_ISL_1336900 | OSPEDALE "SAN CAMILLO" ATESSA - MEDICINA E LUNGODEGENZA | Istituto Zooprofilattico Sperimentale dell'Abruzzo e Molise "G. Caporale" | Lorusso A, Marcacci M, Di Domenico M, Ancora M, Curini V, Di Lollo Valeria, Mangone I, Rinaldi A, Scialabba S, Caporale M, Di Pasquale A, Cammà C, Puglia I, Calistri P, Savini G |
| EPI_ISL_1336901 | DIP. PREV. AVEZZANO SERVIZIO DI IGIENE EPIDEMIOLOGIA E SANITA' PUBBLICA | Istituto Zooprofilattico Sperimentale dell'Abruzzo e Molise "G. Caporale" | Lorusso A, Marcacci M, Di Domenico M, Ancora M, Curini V, Di Lollo Valeria, Mangone I, Rinaldi A, Scialabba S, Caporale M, Di Pasquale A, Cammà C, Puglia I, Calistri P, Savini G |
| EPI_ISL_1336902 | SIESP CHIETI - DRIVE IN GISSI | Istituto Zooprofilattico Sperimentale dell'Abruzzo e Molise "G. Caporale" | Lorusso A, Marcacci M, Di Domenico M, Ancora M, Curini V, Di Lollo Valeria, Mangone I, Rinaldi A, Scialabba S, Caporale M, Di Pasquale A, Cammà C, Puglia I, Calistri P, Savini G |
| EPI_ISL_1336903 | SIESP CHIETI - DRIVE IN LANCIANO | Istituto Zooprofilattico Sperimentale dell'Abruzzo e Molise "G. Caporale" | Lorusso A, Marcacci M, Di Domenico M, Ancora M, Curini V, Di Lollo Valeria, Mangone I, Rinaldi A, Scialabba S, Caporale M, Di Pasquale A, Cammà C, Puglia I, Calistri P, Savini G |
| EPI_ISL_1336904 | SIESP CHIETI - DRIVE IN ORTONA | Istituto Zooprofilattico Sperimentale dell'Abruzzo e Molise "G. Caporale" | Lorusso A, Marcacci M, Di Domenico M, Ancora M, Curini V, Di Lollo Valeria, Mangone I, Rinaldi A, Scialabba S, Caporale M, Di Pasquale A, Cammà C, Puglia I, Calistri P, Savini G |
| EPI_ISL_1336905 | SIESP DIPARTIMENTO DI PREVENZIONE TERAMO | Istituto Zooprofilattico Sperimentale dell'Abruzzo e Molise "G. Caporale" | Lorusso A, Marcacci M, Di Domenico M, Ancora M, Curini V, Di Lollo Valeria, Mangone I, Rinaldi A, Scialabba S, Caporale M, Di Pasquale A, Cammà C, Puglia I, Calistri P, Savini G |
| EPI_ISL_1336906 | SIESP DIPARTIMENTO DI PREVENZIONE CHIETI | Istituto Zooprofilattico Sperimentale dell'Abruzzo e Molise "G. Caporale" | Lorusso A, Marcacci M, Di Domenico M, Ancora M, Curini V, Di Lollo Valeria, Mangone I, Rinaldi A, Scialabba S, Caporale M, Di Pasquale A, Cammà C, Puglia I, Calistri P, Savini G |
| EPI_ISL_1336907, EPI_ISL_1336908, EPI_ISL_1336909 | SIESP SULMONA | Istituto Zooprofilattico Sperimentale dell'Abruzzo e Molise "G. Caporale" | Lorusso A, Marcacci M, Di Domenico M, Ancora M, Curini V, Di Lollo Valeria, Mangone I, Rinaldi A, Scialabba S, Caporale M, Di Pasquale A, Cammà C, Puglia I, Calistri P, Savini G |
| EPI_ISL_1336910 | SIESP CHIETI - DRIVE IN CHIETI | Istituto Zooprofilattico Sperimentale dell'Abruzzo e Molise "G. Caporale" | Lorusso A, Marcacci M, Di Domenico M, Ancora M, Curini V, Di Lollo Valeria, Mangone I, Rinaldi A, Scialabba S, Caporale M, Di Pasquale A, Cammà C, Puglia I, Calistri P, Savini G |
| EPI_ISL_1336911 | L'AQUILONE | Istituto Zooprofilattico Sperimentale dell'Abruzzo e Molise "G. Caporale" | Lorusso A, Marcacci M, Di Domenico M, Ancora M, Curini V, Di Lollo Valeria, Mangone I, Rinaldi A, Scialabba S, Caporale M, Di Pasquale A, Cammà C, Puglia I, Calistri P, Savini G |
| EPI_ISL_1336912 | SIESP DIPARTIMENTO DI PREVENZIONE TERAMO | Istituto Zooprofilattico Sperimentale dell'Abruzzo e Molise "G. Caporale" | Lorusso A, Marcacci M, Di Domenico M, Ancora M, Curini V, Di Lollo Valeria, Mangone I, Rinaldi A, Scialabba S, Caporale M, Di Pasquale A, Cammà C, Puglia I, Calistri P, Savini G |
| EPI_ISL_1336913 | OSPEDALE CIVILE SANT'OMERO - PRONTO SOCCORSO | Istituto Zooprofilattico Sperimentale dell'Abruzzo e Molise "G. Caporale" | Lorusso A, Marcacci M, Di Domenico M, Ancora M, Curini V, Di Lollo Valeria, Mangone I, Rinaldi A, Scialabba S, Caporale M, Di Pasquale A, Cammà C, Puglia I, Calistri P, Savini G |
| EPI_ISL_1336914 | SIESP DIPARTIMENTO DI PREVENZIONE TERAMO | Istituto Zooprofilattico Sperimentale dell'Abruzzo e Molise "G. Caporale" | Lorusso A, Marcacci M, Di Domenico M, Ancora M, Curini V, Di Lollo Valeria, Mangone I, Rinaldi A, Scialabba S, Caporale M, Di Pasquale A, Cammà C, Puglia I, Calistri P, Savini G |
| EPI_ISL_1336915 | SIESP CHIETI - DRIVE IN LANCIANO | Istituto Zooprofilattico Sperimentale dell'Abruzzo e Molise "G. Caporale" | Lorusso A, Marcacci M, Di Domenico M, Ancora M, Curini V, Di Lollo Valeria, Mangone I, Rinaldi A, Scialabba S, Caporale M, Di Pasquale A, Cammà C, Puglia I, Calistri P, Savini G |
| EPI_ISL_1336916, EPI_ISL_1336917, EPI_ISL_1336918, EPI_ISL_1336919 | SIESP DIPARTIMENTO DI PREVENZIONE TERAMO | Istituto Zooprofilattico Sperimentale dell'Abruzzo e Molise "G. Caporale" | Lorusso A, Marcacci M, Di Domenico M, Ancora M, Curini V, Di Lollo Valeria, Mangone I, Rinaldi A, Scialabba S, Caporale M, Di Pasquale A, Cammà C, Puglia I, Calistri P, Savini G |
| EPI_ISL_1336920 | SIESP CHIETI - DRIVE IN VASTO | Istituto Zooprofilattico Sperimentale dell'Abruzzo e Molise "G. Caporale" | Lorusso A, Marcacci M, Di Domenico M, Ancora M, Curini V, Di Lollo Valeria, Mangone I, Rinaldi A, Scialabba S, Caporale M, Di Pasquale A, Cammà C, Puglia I, Calistri P, Savini G |
| EPI_ISL_1336921 | SIESP DIPARTIMENTO DI PREVENZIONE TERAMO | Istituto Zooprofilattico Sperimentale dell'Abruzzo e Molise "G. Caporale" | Lorusso A, Marcacci M, Di Domenico M, Ancora M, Curini V, Di Lollo Valeria, Mangone I, Rinaldi A, Scialabba S, Caporale M, Di Pasquale A, Cammà C, Puglia I, Calistri P, Savini G |
| EPI_ISL_1336922 | DIP. PREV. AVEZZANO SERVIZIO DI IGIENE EPIDEMIOLOGIA E SANITA' PUBBLICA | Istituto Zooprofilattico Sperimentale dell'Abruzzo e Molise "G. Caporale" | Lorusso A, Marcacci M, Di Domenico M, Ancora M, Curini V, Di Lollo Valeria, Mangone I, Rinaldi A, Scialabba S, Caporale M, Di Pasquale A, Cammà C, Puglia I, Calistri P, Savini G |
| EPI_ISL_1336923 | Casa di Cura Di Lorenzo | Istituto Zooprofilattico Sperimentale dell'Abruzzo e Molise "G. Caporale" | Lorusso A, Marcacci M, Di Domenico M, Ancora M, Curini V, Di Lollo Valeria, Mangone I, Rinaldi A, Scialabba S, Caporale M, Di Pasquale A, Cammà C, Puglia I, Calistri P, Savini G |
| EPI_ISL_1336924 | SIESP SULMONA | Istituto Zooprofilattico Sperimentale dell'Abruzzo e Molise "G. Caporale" | Lorusso A, Marcacci M, Di Domenico M, Ancora M, Curini V, Di Lollo Valeria, Mangone I, Rinaldi A, Scialabba S, Caporale M, Di Pasquale A, Cammà C, Puglia I, Calistri P, Savini G |
| EPI_ISL_1336925 | DIP. PREV. AVEZZANO SERVIZIO DI IGIENE EPIDEMIOLOGIA E SANITA' PUBBLICA | Istituto Zooprofilattico Sperimentale dell'Abruzzo e Molise "G. Caporale" | Lorusso A, Marcacci M, Di Domenico M, Ancora M, Curini V, Di Lollo Valeria, Mangone I, Rinaldi A, Scialabba S, Caporale M, Di Pasquale A, Cammà C, Puglia I, Calistri P, Savini G |
| EPI_ISL_1336926 | SIESP CHIETI - DRIVE IN CHIETI | Istituto Zooprofilattico Sperimentale dell'Abruzzo e Molise "G. Caporale" | Lorusso A, Marcacci M, Di Domenico M, Ancora M, Curini V, Di Lollo Valeria, Mangone I, Rinaldi A, Scialabba S, Caporale M, Di Pasquale A, Cammà C, Puglia I, Calistri P, Savini G |
| EPI_ISL_1336927 | USCA Castel di Sangro | Istituto Zooprofilattico Sperimentale dell'Abruzzo e Molise "G. Caporale" | Lorusso A, Marcacci M, Di Domenico M, Ancora M, Curini V, Di Lollo Valeria, Mangone I, Rinaldi A, Scialabba S, Caporale M, Di Pasquale A, Cammà C, Puglia I, Calistri P, Savini G |
| EPI_ISL_1336928 | SIESP CHIETI - DRIVE IN VASTO | Istituto Zooprofilattico Sperimentale dell'Abruzzo e Molise "G. Caporale" | Lorusso A, Marcacci M, Di Domenico M, Ancora M, Curini V, Di Lollo Valeria, Mangone I, Rinaldi A, Scialabba S, Caporale M, Di Pasquale A, Cammà C, Puglia I, Calistri P, Savini G |
| EPI_ISL_1336929, EPI_ISL_1336930 | SIESP DIPARTIMENTO DI PREVENZIONE CHIETI | Istituto Zooprofilattico Sperimentale dell'Abruzzo e Molise "G. Caporale" | Lorusso A, Marcacci M, Di Domenico M, Ancora M, Curini V, Di Lollo Valeria, Mangone I, Rinaldi A, Scialabba S, Caporale M, Di Pasquale A, Cammà C, Puglia I, Calistri P, Savini G |
| EPI_ISL_1336931 | USCA Sulmona | Istituto Zooprofilattico Sperimentale dell'Abruzzo e Molise "G. Caporale" | Lorusso A, Marcacci M, Di Domenico M, Ancora M, Curini V, Di Lollo Valeria, Mangone I, Rinaldi A, Scialabba S, Caporale M, Di Pasquale A, Cammà C, Puglia I, Calistri P, Savini G |
| EPI_ISL_1336932 | DIP. PREV. AVEZZANO SERVIZIO DI IGIENE EPIDEMIOLOGIA E SANITA' PUBBLICA | Istituto Zooprofilattico Sperimentale dell'Abruzzo e Molise "G. Caporale" | Lorusso A, Marcacci M, Di Domenico M, Ancora M, Curini V, Di Lollo Valeria, Mangone I, Rinaldi A, Scialabba S, Caporale M, Di Pasquale A, Cammà C, Puglia I, Calistri P, Savini G |
| EPI_ISL_1336933 | SIESP CHIETI - DRIVE IN CHIETI | Istituto Zooprofilattico Sperimentale dell'Abruzzo e Molise "G. Caporale" | Lorusso A, Marcacci M, Di Domenico M, Ancora M, Curini V, Di Lollo Valeria, Mangone I, Rinaldi A, Scialabba S, Caporale M, Di Pasquale A, Cammà C, Puglia I, Calistri P, Savini G |
| EPI_ISL_1336934 | SIESP CHIETI - DRIVE IN VASTO | Istituto Zooprofilattico Sperimentale dell'Abruzzo e Molise "G. Caporale" | Lorusso A, Marcacci M, Di Domenico M, Ancora M, Curini V, Di Lollo Valeria, Mangone I, Rinaldi A, Scialabba S, Caporale M, Di Pasquale A, Cammà C, Puglia I, Calistri P, Savini G |
| EPI_ISL_1336935 | SIESP CHIETI - DRIVE IN LANCIANO | Istituto Zooprofilattico Sperimentale dell'Abruzzo e Molise "G. Caporale" | Lorusso A, Marcacci M, Di Domenico M, Ancora M, Curini V, Di Lollo Valeria, Mangone I, Rinaldi A, Scialabba S, Caporale M, Di Pasquale A, Cammà C, Puglia I, Calistri P, Savini G |
| EPI_ISL_1336936 | SIESP CHIETI - DRIVE IN ORTONA | Istituto Zooprofilattico Sperimentale dell'Abruzzo e Molise "G. Caporale" | Lorusso A, Marcacci M, Di Domenico M, Ancora M, Curini V, Di Lollo Valeria, Mangone I, Rinaldi A, Scialabba S, Caporale M, Di Pasquale A, Cammà C, Puglia I, Calistri P, Savini G |
| EPI_ISL_1336937 | SIESP CHIETI - DRIVE IN GISSI | Istituto Zooprofilattico Sperimentale dell'Abruzzo e Molise "G. Caporale" | Lorusso A, Marcacci M, Di Domenico M, Ancora M, Curini V, Di Lollo Valeria, Mangone I, Rinaldi A, Scialabba S, Caporale M, Di Pasquale A, Cammà C, Puglia I, Calistri P, Savini G |
| EPI_ISL_1336938 | SIESP CHIETI - DRIVE IN LANCIANO | Istituto Zooprofilattico Sperimentale dell'Abruzzo e Molise "G. Caporale" | Lorusso A, Marcacci M, Di Domenico M, Ancora M, Curini V, Di Lollo Valeria, Mangone I, Rinaldi A, Scialabba S, Caporale M, Di Pasquale A, Cammà C, Puglia I, Calistri P, Savini G |
| EPI_ISL_1336939, EPI_ISL_1336940 | USCA Castel di Sangro | Istituto Zooprofilattico Sperimentale dell'Abruzzo e Molise "G. | Lorusso A, Marcacci M, Di Domenico M, Ancora M, Curini V, Di Lollo Valeria, Mangone I, Rinaldi A, Scialabba S, Caporale M, Di Pasquale A, Cammà C, |

| | | | |
|---|---|---|---|
| | | Caporale" | Puglia I, Calistri P, Savini G |
| EPI_ISL_1336941 | USCA Avezzano | Istituto Zooprofilattico Sperimentale dell'Abruzzo e Molise "G. Caporale" | Lorusso A, Marcacci M, Di Domenico M, Ancora M, Curini V, Di Lollo Valeria, Mangone I, Rinaldi A, Scialabba S, Caporale M, Di Pasquale A, Cammà C, Puglia I, Calistri P, Savini G |
| EPI_ISL_1336942 | SIESP DIPARTIMENTO DI PREVENZIONE TERAMO | Istituto Zooprofilattico Sperimentale dell'Abruzzo e Molise "G. Caporale" | Lorusso A, Marcacci M, Di Domenico M, Ancora M, Curini V, Di Lollo Valeria, Mangone I, Rinaldi A, Scialabba S, Caporale M, Di Pasquale A, Cammà C, Puglia I, Calistri P, Savini G |
| EPI_ISL_1336943 | SIESP CHIETI - DRIVE IN VASTO | Istituto Zooprofilattico Sperimentale dell'Abruzzo e Molise "G. Caporale" | Lorusso A, Marcacci M, Di Domenico M, Ancora M, Curini V, Di Lollo Valeria, Mangone I, Rinaldi A, Scialabba S, Caporale M, Di Pasquale A, Cammà C, Puglia I, Calistri P, Savini G |
| EPI_ISL_1336944 | USCA Tagliacozzo | Istituto Zooprofilattico Sperimentale dell'Abruzzo e Molise "G. Caporale" | Lorusso A, Marcacci M, Di Domenico M, Ancora M, Curini V, Di Lollo Valeria, Mangone I, Rinaldi A, Scialabba S, Caporale M, Di Pasquale A, Cammà C, Puglia I, Calistri P, Savini G |
| EPI_ISL_1336945 | SIESP DIPARTIMENTO DI PREVENZIONE CHIETI | Istituto Zooprofilattico Sperimentale dell'Abruzzo e Molise "G. Caporale" | Lorusso A, Marcacci M, Di Domenico M, Ancora M, Curini V, Di Lollo Valeria, Mangone I, Rinaldi A, Scialabba S, Caporale M, Di Pasquale A, Cammà C, Puglia I, Calistri P, Savini G |
| EPI_ISL_1336946 | SIESP DIPARTIMENTO DI PREVENZIONE TERAMO | Istituto Zooprofilattico Sperimentale dell'Abruzzo e Molise "G. Caporale" | Lorusso A, Marcacci M, Di Domenico M, Ancora M, Curini V, Di Lollo Valeria, Mangone I, Rinaldi A, Scialabba S, Caporale M, Di Pasquale A, Cammà C, Puglia I, Calistri P, Savini G |
| EPI_ISL_1336947 | USCA Pescina | Istituto Zooprofilattico Sperimentale dell'Abruzzo e Molise "G. Caporale" | Lorusso A, Marcacci M, Di Domenico M, Ancora M, Curini V, Di Lollo Valeria, Mangone I, Rinaldi A, Scialabba S, Caporale M, Di Pasquale A, Cammà C, Puglia I, Calistri P, Savini G |
| EPI_ISL_1336948, EPI_ISL_1336949 | SIESP DIPARTIMENTO DI PREVENZIONE TERAMO | Istituto Zooprofilattico Sperimentale dell'Abruzzo e Molise "G. Caporale" | Lorusso A, Marcacci M, Di Domenico M, Ancora M, Curini V, Di Lollo Valeria, Mangone I, Rinaldi A, Scialabba S, Caporale M, Di Pasquale A, Cammà C, Puglia I, Calistri P, Savini G |
| EPI_ISL_1336951 | OSPEDALE CIVILE TERAMO - CENTRO TRASFUSIONALE | Istituto Zooprofilattico Sperimentale dell'Abruzzo e Molise "G. Caporale" | Lorusso A, Marcacci M, Di Domenico M, Ancora M, Curini V, Di Lollo Valeria, Mangone I, Rinaldi A, Scialabba S, Caporale M, Di Pasquale A, Cammà C, Puglia I, Calistri P, Savini G |
| EPI_ISL_1336952, EPI_ISL_1336953, EPI_ISL_1336954 | SIESP DIPARTIMENTO DI PREVENZIONE TERAMO | Istituto Zooprofilattico Sperimentale dell'Abruzzo e Molise "G. Caporale" | Lorusso A, Marcacci M, Di Domenico M, Ancora M, Curini V, Di Lollo Valeria, Mangone I, Rinaldi A, Scialabba S, Caporale M, Di Pasquale A, Cammà C, Puglia I, Calistri P, Savini G |
| EPI_ISL_1336955 | SIESP CHIETI - DRIVE IN CHIETI | Istituto Zooprofilattico Sperimentale dell'Abruzzo e Molise "G. Caporale" | Lorusso A, Marcacci M, Di Domenico M, Ancora M, Curini V, Di Lollo Valeria, Mangone I, Rinaldi A, Scialabba S, Caporale M, Di Pasquale A, Cammà C, Puglia I, Calistri P, Savini G |
| EPI_ISL_1336956 | SIESP DIPARTIMENTO DI PREVENZIONE TERAMO | Istituto Zooprofilattico Sperimentale dell'Abruzzo e Molise "G. Caporale" | Lorusso A, Marcacci M, Di Domenico M, Ancora M, Curini V, Di Lollo Valeria, Mangone I, Rinaldi A, Scialabba S, Caporale M, Di Pasquale A, Cammà C, Puglia I, Calistri P, Savini G |
| EPI_ISL_1336957 | SIESP CHIETI - DRIVE IN VASTO | Istituto Zooprofilattico Sperimentale dell'Abruzzo e Molise "G. Caporale" | Lorusso A, Marcacci M, Di Domenico M, Ancora M, Curini V, Di Lollo Valeria, Mangone I, Rinaldi A, Scialabba S, Caporale M, Di Pasquale A, Cammà C, Puglia I, Calistri P, Savini G |
| EPI_ISL_1336958 | SIESP CHIETI - DRIVE IN ORTONA | Istituto Zooprofilattico Sperimentale dell'Abruzzo e Molise "G. Caporale" | Lorusso A, Marcacci M, Di Domenico M, Ancora M, Curini V, Di Lollo Valeria, Mangone I, Rinaldi A, Scialabba S, Caporale M, Di Pasquale A, Cammà C, Puglia I, Calistri P, Savini G |
| EPI_ISL_1336959 | SIESP CHIETI - DRIVE IN VASTO | Istituto Zooprofilattico Sperimentale dell'Abruzzo e Molise "G. Caporale" | Lorusso A, Marcacci M, Di Domenico M, Ancora M, Curini V, Di Lollo Valeria, Mangone I, Rinaldi A, Scialabba S, Caporale M, Di Pasquale A, Cammà C, Puglia I, Calistri P, Savini G |
| EPI_ISL_1336960 | SIESP DIPARTIMENTO DI PREVENZIONE TERAMO | Istituto Zooprofilattico Sperimentale dell'Abruzzo e Molise "G. Caporale" | Lorusso A, Marcacci M, Di Domenico M, Ancora M, Curini V, Di Lollo Valeria, Mangone I, Rinaldi A, Scialabba S, Caporale M, Di Pasquale A, Cammà C, Puglia I, Calistri P, Savini G |
| EPI_ISL_1336961 | SIESP CHIETI - DRIVE IN CHIETI | Istituto Zooprofilattico Sperimentale dell'Abruzzo e Molise "G. Caporale" | Lorusso A, Marcacci M, Di Domenico M, Ancora M, Curini V, Di Lollo Valeria, Mangone I, Rinaldi A, Scialabba S, Caporale M, Di Pasquale A, Cammà C, Puglia I, Calistri P, Savini G |
| EPI_ISL_1336962 | OSPEDALE CIVILE GIULIANOVA - PRONTO SOCCORSO | Istituto Zooprofilattico Sperimentale dell'Abruzzo e Molise "G. Caporale" | Lorusso A, Marcacci M, Di Domenico M, Ancora M, Curini V, Di Lollo Valeria, Mangone I, Rinaldi A, Scialabba S, Caporale M, Di Pasquale A, Cammà C, Puglia I, Calistri P, Savini G |
| EPI_ISL_1336963 | SIESP CHIETI - DRIVE IN ORTONA | Istituto Zooprofilattico Sperimentale dell'Abruzzo e Molise "G. Caporale" | Lorusso A, Marcacci M, Di Domenico M, Ancora M, Curini V, Di Lollo Valeria, Mangone I, Rinaldi A, Scialabba S, Caporale M, Di Pasquale A, Cammà C, Puglia I, Calistri P, Savini G |
| EPI_ISL_1337376, EPI_ISL_1337377, EPI_ISL_1337379, EPI_ISL_1337380, EPI_ISL_1337382, EPI_ISL_1337383, EPI_ISL_1337384, EPI_ISL_1337386, EPI_ISL_1337387 | LESP Guanajuato | Instituto de Diagnostico y Referencia Epidemiologicos (INDRE) | Claudia Wong-Arambula, Abril Rodriguez-Maldonado, Vanessa Rivero-Arredondo, Ariadna Medina-Benitez, Joaquin Quiroz-Mercado, Sergio Rangel-Guerrero, Natividad Cruz-Ortiz, Tatiana Nunez-Garcia, Gisela Barrera-Badillo, Lucia Hernandez-Rivas, Irma Lopez-Martinez, Ernesto Ramirez-Gonzalez. |
| EPI_ISL_1337389 | LESP Sinaloa | Instituto de Diagnostico y Referencia Epidemiologicos (INDRE) | Claudia Wong-Arambula, Abril Rodriguez-Maldonado, Vanessa Rivero-Arredondo, Ariadna Medina-Benitez, Joaquin Quiroz-Mercado, Sergio Rangel-Guerrero, Natividad Cruz-Ortiz, Tatiana Nunez-Garcia, Gisela Barrera-Badillo, Lucia Hernandez-Rivas, Irma Lopez-Martinez, Ernesto Ramirez-Gonzalez. |
| EPI_ISL_1337390 | LESP Baja California | Instituto de Diagnostico y Referencia Epidemiologicos (INDRE) | Claudia Wong-Arambula, Abril Rodriguez-Maldonado, Vanessa Rivero-Arredondo, Ariadna Medina-Benitez, Joaquin Quiroz-Mercado, Sergio Rangel-Guerrero, Natividad Cruz-Ortiz, Tatiana Nunez-Garcia, Gisela Barrera-Badillo, Lucia Hernandez-Rivas, Irma Lopez-Martinez, Ernesto Ramirez-Gonzalez. |
| EPI_ISL_1337392, EPI_ISL_1337393 | LESP Baja California Sur | Instituto de Diagnostico y Referencia Epidemiologicos (INDRE) | Claudia Wong-Arambula, Abril Rodriguez-Maldonado, Vanessa Rivero-Arredondo, Ariadna Medina-Benitez, Joaquin Quiroz-Mercado, Sergio Rangel-Guerrero, Natividad Cruz-Ortiz, Tatiana Nunez-Garcia, Gisela Barrera-Badillo, Lucia Hernandez-Rivas, Irma Lopez-Martinez, Ernesto Ramirez-Gonzalez. |
| EPI_ISL_1337518, EPI_ISL_1337519, EPI_ISL_1337524, EPI_ISL_1337527, EPI_ISL_1337529, EPI_ISL_1337539, EPI_ISL_1337541, EPI_ISL_1337546 | OCME Office Of Chief Medical Examiner | New York City Public Health Laboratory | Jade Wang, et al. |
| EPI_ISL_1340608, EPI_ISL_1340610 | LESP Guanajuato | Instituto de Diagnostico y Referencia Epidemiologicos (INDRE) | Claudia Wong-Arambula, Abril Rodriguez-Maldonado, Vanessa Rivero-Arredondo, Ariadna Medina-Benitez, Joaquin Quiroz-Mercado, Sergio Rangel-Guerrero, Natividad Cruz-Ortiz, Tatiana Nunez-Garcia, Gisela Barrera-Badillo, Lucia Hernandez-Rivas, Irma Lopez-Martinez, Ernesto Ramirez-Gonzalez. |
| EPI_ISL_1340832, EPI_ISL_1340833, EPI_ISL_1340835, EPI_ISL_1340836, EPI_ISL_1340837, EPI_ISL_1340838, EPI_ISL_1340839, EPI_ISL_1340840, EPI_ISL_1340841, EPI_ISL_1340842, EPI_ISL_1340843, EPI_ISL_1340844, EPI_ISL_1340845, EPI_ISL_1340846, EPI_ISL_1340847, EPI_ISL_1340848, EPI_ISL_1340850, EPI_ISL_1340851, EPI_ISL_1340852, EPI_ISL_1340853, EPI_ISL_1340854, EPI_ISL_1340855, EPI_ISL_1340856, EPI_ISL_1340860, EPI_ISL_1340861 | | | |
| see above | Northwestern Memorial Hospital | Northwestern University - Ozer Lab | Ramon Lorenzo-Redondo, Lacy M. Simons, Taylor J. Dean, Chad J. Achenbach, Lawrence J. Jennings, Chao Qi, Michael G. Ison, Judd F. Hultquist, Egon A. Ozer |
| EPI_ISL_1346639, EPI_ISL_1346640, EPI_ISL_1346641, EPI_ISL_1346751, EPI_ISL_1346804, EPI_ISL_1346856 | Institut de Virologie du CHU de Strasbourg | Institut de Virologie du CHU de Strasbourg | Fafi-Kremer Samira, Gallais Floriane, Gantner Pierre, Laugel Elodie, Solis Morgane, Velay Aurélie, Wendling Marie-Josée |
| EPI_ISL_1347245 | INT Fondazione Pascale | INT Fondazione Pascale | INT Fondazione Pascale |
| EPI_ISL_1351418, EPI_ISL_1351422, EPI_ISL_1351428, EPI_ISL_1351430, EPI_ISL_1351435, EPI_ISL_1351454, EPI_ISL_1351460, EPI_ISL_1351462 | Centro de InvestigaciÃn BiomÃ©dica del Noreste (CIBIN) | Unidad de Genomica Avanzada | Consorcio Mexicano de Vigilancia Genomica (CoViGen-Mex). Authors (in alphabetical order): Julio Elias Alvarado-Yaah, Carlos F. Arias, Santiago Avila-Rios, Victor Hugo Borja-Aburto, Celia Boukadida, Juan Bautista Chale-Dzul , Jose Antonio Enciso-Moreno, Gloria Elena Espinoza-Ayala, Fernando Fontove-Herrera, Concepcion Grajales-Muniz, Ricardo Grande, Alfredo Herrera-Estrella, Carla Ivon Herrera-Najera, Pavel Isa, Brenda Irasema Maldonado-Meza, Bernardo Martinez-Miguel, Margarita Matias-Florentino, Maria Guadalupe de Jesus Mireles-Rivera, Gloria Maria Molina-Salinas, Hector Montoya-Fuentes, Jose Esteban Munoz-Medina, Jose de Jesus Nunez-Contreras, Alicia Ocana-Mondragon, Luis Alberto Ochoa-Carrera, Hector Esteban |

| | | | |
|---|---|---|---|
| | | | Paz-Juarez, Francisco Pulido, Helen Haydee Fernanda Ramirez-Plascencia, Angel Gustavo Salas-Lais, Jorge Ivan Salinal-Nevarez, Alejandro Sanchez-Flores, Clara Esperanza Santacruz-Tinoco, Maria Guadalupe Santiago-Mauricio, Nelly Selem-Mojica, Blanca Taboada, Gloria Vazquez |
| EPI_ISL_1351477, EPI_ISL_1351486, EPI_ISL_1351489, EPI_ISL_1351494, EPI_ISL_1351498, EPI_ISL_1351500, EPI_ISL_1351506, EPI_ISL_1351509 | Unidad de InvestigaciÃn MÃ©dica de YucatÃ¡n (UIMY) | Unidad de Genomica Avanzada | Consorcio Mexicano de Vigilancia Genomica (CoViGen-Mex). Authors (in alphabetical order): Julio Elias Alvarado-Yaah, Carlos F. Arias, Santiago Avila-Rios, Victor Hugo Borja-Aburto, Celia Boukadida, Juan Bautista Chale-Dzul , Jose Antonio Enciso-Moreno, Gloria Elena Espinoza-Ayala, Fernando Fontove-Herrera, Concepcion Grajales-Muniz, Ricardo Grande, Alfredo Herrera-Estrella, Carla Ivon Herrera-Najera, Pavel Isa, Brenda Irasema Maldonado-Meza, Bernardo Martinez-Miguel, Margarita Matias-Florentino, Maria Guadalupe de Jesus Mireles-Rivera, Gloria Maria Molina-Salinas, Hector Montoya-Fuentes, Jose Esteban Munoz-Medina, Jose de Jesus Nunez-Contreras, Alicia Ocana-Mondragon, Luis Alberto Ochoa-Carrera, Hector Esteban Paz-Juarez, Francisco Pulido, Helen Haydee Fernanda Ramirez-Plascencia, Angel Gustavo Salas-Lais, Jorge Ivan Salinal-Nevarez, Alejandro Sanchez-Flores, Clara Esperanza Santacruz-Tinoco, Maria Guadalupe Santiago-Mauricio, Nelly Selem-Mojica, Blanca Taboada, Gloria Vazquez |
| EPI_ISL_1351511, EPI_ISL_1351514, EPI_ISL_1351517, EPI_ISL_1351520, EPI_ISL_1351523, EPI_ISL_1351526, EPI_ISL_1351529, EPI_ISL_1351532, EPI_ISL_1351535, EPI_ISL_1351538, EPI_ISL_1351540, EPI_ISL_1351543, EPI_ISL_1351547, EPI_ISL_1351552, EPI_ISL_1351555, EPI_ISL_1351558, EPI_ISL_1351561, EPI_ISL_1351567, EPI_ISL_1351571, EPI_ISL_1351574, EPI_ISL_1351577, EPI_ISL_1351580, EPI_ISL_1351583, EPI_ISL_1351585, EPI_ISL_1351592, EPI_ISL_1351595, EPI_ISL_1351601, EPI_ISL_1351605, EPI_ISL_1351608, EPI_ISL_1351611, EPI_ISL_1351614, EPI_ISL_1351620, EPI_ISL_1351626, EPI_ISL_1351629, EPI_ISL_1351635, EPI_ISL_1351638, EPI_ISL_1351651, EPI_ISL_1351654, EPI_ISL_1351656, EPI_ISL_1351659, EPI_ISL_1351668, EPI_ISL_1351670, EPI_ISL_1351672, EPI_ISL_1351675, EPI_ISL_1351688, EPI_ISL_1351691, EPI_ISL_1351700, EPI_ISL_1351703, EPI_ISL_1351706, EPI_ISL_1351709 | | | |
| see above | Centro de InvestigaciÃn BiomÃ©dica de Occidente (CIBO) | Unidad de Genomica Avanzada | Consorcio Mexicano de Vigilancia Genomica (CoViGen-Mex). Authors (in alphabetical order): Julio Elias Alvarado-Yaah, Carlos F. Arias, Santiago Avila-Rios, Victor Hugo Borja-Aburto, Celia Boukadida, Juan Bautista Chale-Dzul , Jose Antonio Enciso-Moreno, Gloria Elena Espinoza-Ayala, Fernando Fontove-Herrera, Concepcion Grajales-Muniz, Ricardo Grande, Alfredo Herrera-Estrella, Carla Ivon Herrera-Najera, Pavel Isa, Brenda Irasema Maldonado-Meza, Bernardo Martinez-Miguel, Margarita Matias-Florentino, Maria Guadalupe de Jesus Mireles-Rivera, Gloria Maria Molina-Salinas, Hector Montoya-Fuentes, Jose Esteban Munoz-Medina, Jose de Jesus Nunez-Contreras, Alicia Ocana-Mondragon, Luis Alberto Ochoa-Carrera, Hector Esteban Paz-Juarez, Francisco Pulido, Helen Haydee Fernanda Ramirez-Plascencia, Angel Gustavo Salas-Lais, Jorge Ivan Salinal-Nevarez, Alejandro Sanchez-Flores, Clara Esperanza Santacruz-Tinoco, Maria Guadalupe Santiago-Mauricio, Nelly Selem-Mojica, Blanca Taboada, Gloria Vazquez |
| EPI_ISL_1351713, EPI_ISL_1351716, EPI_ISL_1351725, EPI_ISL_1351731, EPI_ISL_1351734, EPI_ISL_1351739, EPI_ISL_1351743, EPI_ISL_1351749, EPI_ISL_1351752, EPI_ISL_1351764, EPI_ISL_1351770, EPI_ISL_1351787, EPI_ISL_1351790, EPI_ISL_1351793, EPI_ISL_1351799, EPI_ISL_1351802, EPI_ISL_1351805, EPI_ISL_1351809, EPI_ISL_1351811, EPI_ISL_1351815, EPI_ISL_1351818, EPI_ISL_1351824, EPI_ISL_1351827, EPI_ISL_1351833, EPI_ISL_1351836, EPI_ISL_1351842, EPI_ISL_1351848, EPI_ISL_1351854, EPI_ISL_1351857, EPI_ISL_1351866, EPI_ISL_1351869, EPI_ISL_1351875, EPI_ISL_1351878, EPI_ISL_1351880, EPI_ISL_1351886, EPI_ISL_1351890, EPI_ISL_1351902, EPI_ISL_1351905, EPI_ISL_1351908, EPI_ISL_1351911, EPI_ISL_1351914, EPI_ISL_1351917, EPI_ISL_1351921, EPI_ISL_1351924, EPI_ISL_1351927, EPI_ISL_1351930, EPI_ISL_1351933, EPI_ISL_1351935, EPI_ISL_1351937, EPI_ISL_1351941, EPI_ISL_1351944, EPI_ISL_1351950, EPI_ISL_1351963, EPI_ISL_1351966, EPI_ISL_1351969, EPI_ISL_1351972, EPI_ISL_1351975, EPI_ISL_1351978, EPI_ISL_1351981, EPI_ISL_1351984, EPI_ISL_1351987, EPI_ISL_1351990, EPI_ISL_1351993, EPI_ISL_1351997, EPI_ISL_1352012, EPI_ISL_1352018, EPI_ISL_1352032, EPI_ISL_1352038, EPI_ISL_1352051, EPI_ISL_1352055, EPI_ISL_1352057, EPI_ISL_1352060, EPI_ISL_1352066, EPI_ISL_1352070 | | | |
| see above | Laboratorio Central de EpidemiologÃ-a (LCE) | Unidad de Genomica Avanzada | Consorcio Mexicano de Vigilancia Genomica (CoViGen-Mex). Authors (in alphabetical order): Julio Elias Alvarado-Yaah, Carlos F. Arias, Santiago Avila-Rios, Victor Hugo Borja-Aburto, Celia Boukadida, Juan Bautista Chale-Dzul , Jose Antonio Enciso-Moreno, Gloria Elena Espinoza-Ayala, Fernando Fontove-Herrera, Concepcion Grajales-Muniz, Ricardo Grande, Alfredo Herrera-Estrella, Carla Ivon Herrera-Najera, Pavel Isa, Brenda Irasema Maldonado-Meza, Bernardo Martinez-Miguel, Margarita Matias-Florentino, Maria Guadalupe de Jesus Mireles-Rivera, Gloria Maria Molina-Salinas, Hector Montoya-Fuentes, Jose Esteban Munoz-Medina, Jose de Jesus Nunez-Contreras, Alicia Ocana-Mondragon, Luis Alberto Ochoa-Carrera, Hector Esteban Paz-Juarez, Francisco Pulido, Helen Haydee Fernanda Ramirez-Plascencia, Angel Gustavo Salas-Lais, Jorge Ivan Salinal-Nevarez, Alejandro Sanchez-Flores, Clara Esperanza Santacruz-Tinoco, Maria Guadalupe Santiago-Mauricio, Nelly Selem-Mojica, Blanca Taboada, Gloria Vazquez |
| EPI_ISL_1352083 | Unidad de InvestigaciÃn BiomÃ©dica de Zacatecas (UIBZ) | Unidad de Genomica Avanzada | Consorcio Mexicano de Vigilancia Genomica (CoViGen-Mex). Authors (in alphabetical order): Julio Elias Alvarado-Yaah, Carlos F. Arias, Santiago Avila-Rios, Victor Hugo Borja-Aburto, Celia Boukadida, Juan Bautista Chale-Dzul , Jose Antonio Enciso-Moreno, Gloria Elena Espinoza-Ayala, Fernando Fontove-Herrera, Concepcion Grajales-Muniz, Ricardo Grande, Alfredo Herrera-Estrella, Carla Ivon Herrera-Najera, Pavel Isa, Brenda Irasema Maldonado-Meza, Bernardo Martinez-Miguel, Margarita Matias-Florentino, Maria Guadalupe de Jesus Mireles-Rivera, Gloria Maria Molina-Salinas, Hector Montoya-Fuentes, Jose Esteban Munoz-Medina, Jose de Jesus Nunez-Contreras, Alicia Ocana-Mondragon, Luis Alberto Ochoa-Carrera, Hector Esteban Paz-Juarez, Francisco Pulido, Helen Haydee Fernanda Ramirez-Plascencia, Angel Gustavo Salas-Lais, Jorge Ivan Salinal-Nevarez, Alejandro Sanchez-Flores, Clara Esperanza Santacruz-Tinoco, Maria Guadalupe Santiago-Mauricio, Nelly Selem-Mojica, Blanca Taboada, Gloria Vazquez |
| EPI_ISL_1358285 | Centro de Treinamento e Referencia DST AIDS | Instituto Adolfo Lutz, Interdiciplinary Procedures Center, Strategic Laboratory | Claudio Tavares Sacchi, Claudia Regina Gonçalves, Erica Valessa Ramos Gomes, Karoline Rodrigues Campos, Caio Vinicius Dias Lopes |
| EPI_ISL_1358286 | Hospital Municipal Josanias Castanha Braga | Instituto Adolfo Lutz, Interdiciplinary Procedures Center, Strategic Laboratory | Claudio Tavares Sacchi, Claudia Regina Gonçalves, Erica Valessa Ramos Gomes, Karoline Rodrigues Campos, Caio Vinicius Dias Lopes |
| EPI_ISL_1358287 | Hospital Nipo Brasileiro | Instituto Adolfo Lutz, Interdiciplinary Procedures Center, Strategic Laboratory | Claudio Tavares Sacchi, Claudia Regina Gonçalves, Erica Valessa Ramos Gomes, Karoline Rodrigues Campos, Caio Vinicius Dias Lopes |
| EPI_ISL_1358288, EPI_ISL_1358289, EPI_ISL_1358290 | IAL Regional de Aracatuba | Instituto Adolfo Lutz, Interdiciplinary Procedures Center, Strategic Laboratory | Claudio Tavares Sacchi, Claudia Regina Gonçalves, Erica Valessa Ramos Gomes, Karoline Rodrigues Campos, Caio Vinicius Dias Lopes |
| EPI_ISL_1358300, EPI_ISL_1358301, EPI_ISL_1358302, EPI_ISL_1358303 | Lacen de Tocantins | Instituto Adolfo Lutz, Interdiciplinary Procedures Center, Strategic Laboratory | Claudio Tavares Sacchi, Claudia Regina Gonçalves, Erica Valessa Ramos Gomes, Karoline Rodrigues Campos, Caio Vinicius Dias Lopes |
| EPI_ISL_1358305 | LACEN do Mato Grosso do Sul | Instituto Adolfo Lutz, Interdiciplinary Procedures Center, Strategic Laboratory | Claudio Tavares Sacchi, Claudia Regina Gonçalves, Erica Valessa Ramos Gomes, Karoline Rodrigues Campos, Caio Vinicius Dias Lopes |
| EPI_ISL_1358318, EPI_ISL_1358319, EPI_ISL_1358320, EPI_ISL_1358322 | UPA Vila Santa Catarina | Instituto Adolfo Lutz, Interdiciplinary Procedures Center, Strategic Laboratory | Claudio Tavares Sacchi, Claudia Regina Gonçalves, Erica Valessa Ramos Gomes, Karoline Rodrigues Campos, Caio Vinicius Dias Lopes |
| EPI_ISL_1359062 | Laboratorios LISTER | Instituto de Diagnostico y Referencia Epidemiologicos (INDRE) | Claudia Wong-Arambula, Abril Rodriguez-Maldonado, Vanessa Rivero-Arredondo, Ariadna Medina-Benitez, Joaquin Quiroz-Mercado, Sergio Rangel-Guerrero, Natividad Cruz-Ortiz, Tatiana Nunez-Garcia, Gisela Barrera-Badillo, Lucia Hernandez-Rivas, Irma Lopez-Martinez, Ernesto Ramirez-Gonzalez. |
| EPI_ISL_1359063 | Laboratorio BIOMEDICA | Instituto de Diagnostico y Referencia Epidemiologicos (INDRE) | Claudia Wong-Arambula, Abril Rodriguez-Maldonado, Vanessa Rivero-Arredondo, Ariadna Medina-Benitez, Joaquin Quiroz-Mercado, Sergio Rangel-Guerrero, Natividad Cruz-Ortiz, Tatiana Nunez-Garcia, Gisela Barrera-Badillo, Lucia Hernandez-Rivas, Irma Lopez-Martinez, Ernesto Ramirez-Gonzalez. |
| EPI_ISL_1359075, EPI_ISL_1359076 | LESP Guanajuato | Instituto de Diagnostico y Referencia Epidemiologicos (INDRE) | Claudia Wong-Arambula, Abril Rodriguez-Maldonado, Vanessa Rivero-Arredondo, Ariadna Medina-Benitez, Joaquin Quiroz-Mercado, Sergio Rangel-Guerrero, Natividad Cruz-Ortiz, Tatiana Nunez-Garcia, Gisela Barrera-Badillo, Lucia Hernandez-Rivas, Irma Lopez-Martinez, Ernesto Ramirez-Gonzalez. |
| EPI_ISL_1360031, EPI_ISL_1360032, EPI_ISL_1360033, EPI_ISL_1360034, EPI_ISL_1360035, EPI_ISL_1360036, EPI_ISL_1360037, EPI_ISL_1360038, EPI_ISL_1360039, EPI_ISL_1360040, EPI_ISL_1360041, EPI_ISL_1360042, EPI_ISL_1360043, EPI_ISL_1360044, EPI_ISL_1360045, EPI_ISL_1360046, EPI_ISL_1360047, EPI_ISL_1360048, EPI_ISL_1360049, EPI_ISL_1360050, EPI_ISL_1360051, EPI_ISL_1360052, EPI_ISL_1360053, EPI_ISL_1360054 | | | |
| see above | Klinisch Laboratorium ZNA | Klinisch Laboratorium ZNA | Verstrepen et al. |
| EPI_ISL_1360371, EPI_ISL_1360372 | CSIR-National Environmental Engineering Research Institute | CSIR-Centre for Cellular and Molecular Biology - INSACOG | Sofia Banu, Lamuk Zaveri, Ara Sreenivas, Shreekant Verma, Onkar Kulkarni, Amareshwar Vodapalli ,Viswagithe S L, B Himasri, Sharath Chandra Thota, Karthik Bharadwaj Tallapaka, Krishna Khairnar, Rakesh K Mishra,Divya Tej Sowpati |
| EPI_ISL_1360373 | CSIR-National Environmental Engineering Research Institute | CSIR-Centre for Cellular and Molecular Biology - INSACOG | Onkar Kulkarni, Lamuk Zaveri, Ara Sreenivas, Sofia Banu, Shreekant Verma, Amareshwar Vodapalli , Viswagithe S L, B Himasri, Sharath Chandra Thota, Karthik Bharadwaj Tallapaka, Krishna Khairnar, Rakesh K Mishra,Divya Tej Sowpati |
| EPI_ISL_1360375 | CSIR-National Environmental Engineering Research Institute | CSIR-Centre for Cellular and Molecular Biology - INSACOG | Lamuk Zaveri, Ara Sreenivas, Sofia Banu, Onkar Kulkarni, Shreekant Verma, Amareshwar Vodapalli , Viswagithe S L, B Himasri, Sharath Chandra Thota, Karthik Bharadwaj Tallapaka, Krishna Khairnar, Rakesh K Mishra,Divya Tej Sowpati |
| EPI_ISL_1360377, EPI_ISL_1360378 | CSIR-National Environmental Engineering Research Institute | CSIR-Centre for Cellular and Molecular Biology - INSACOG | Ara Sreenivas, Onkar Kulkarni, Lamuk Zaveri, Sofia Banu, Shreekant Verma, Amareshwar Vodapalli , Viswagithe S L, B Himasri, Sharath Chandra Thota, Karthik Bharadwaj Tallapaka, Krishna Khairnar, Rakesh K Mishra,Divya Tej Sowpati |
| EPI_ISL_1360381, EPI_ISL_1360382 | CSIR-National Environmental Engineering Research Institute | CSIR-Centre for Cellular and Molecular Biology - INSACOG | Onkar Kulkarni, Lamuk Zaveri, Ara Sreenivas, Sofia Banu, Shreekant Verma, Amareshwar Vodapalli , Viswagithe S L, B Himasri, Sharath Chandra Thota, Karthik Bharadwaj Tallapaka, Krishna Khairnar, Rakesh K Mishra,Divya Tej Sowpati |
| EPI_ISL_1360383 | CSIR-National Environmental Engineering Research Institute | CSIR-Centre for Cellular and Molecular Biology - INSACOG | Lamuk Zaveri, Ara Sreenivas, Sofia Banu, Onkar Kulkarni, Shreekant Verma, Amareshwar Vodapalli , Viswagithe S L, B Himasri, Sharath Chandra Thota, |

| | | | |
|---|---|---|---|
| | | | Karthik Bharadwaj Tallapaka, Krishna Khairnar, Rakesh K Mishra,Divya Tej Sowpati |
| EPI_ISL_1360384 | CSIR-National Environmental Engineering Research Institute | CSIR-Centre for Cellular and Molecular Biology - INSACOG | Lamuk Zaveri, Ara Sreenivas, Onkar Kulkarni, Sofia Banu, Shreekant Verma,Amareshwar Vodapalli, Viswagithe S L, B Himasri, Sharath Chandra Thota, Karthik Bharadwaj Tallapaka, Krishna Khairnar, Rakesh K Mishra,Divya Tej Sowpati |
| EPI_ISL_1360385 | CSIR-National Environmental Engineering Research Institute | CSIR-Centre for Cellular and Molecular Biology - INSACOG | Ara Sreenivas, Onkar Kulkarni, Lamuk Zaveri, Sofia Banu, Shreekant Verma, Amareshwar Vodapalli , Viswagithe S L, B Himasri, Sharath Chandra Thota, Karthik Bharadwaj Tallapaka, Krishna Khairnar, Rakesh K Mishra,Divya Tej Sowpati |
| EPI_ISL_1360387, EPI_ISL_1360388 | CSIR-National Environmental Engineering Research Institute | CSIR-Centre for Cellular and Molecular Biology - INSACOG | Sofia Banu, Lamuk Zaveri, Ara Sreenivas, Shreekant Verma, Onkar Kulkarni, Amareshwar Vodapalli ,Viswagithe S L, B Himasri, Sharath Chandra Thota, Karthik Bharadwaj Tallapaka, Krishna Khairnar, Rakesh K Mishra,Divya Tej Sowpati |
| EPI_ISL_1365662, EPI_ISL_1365664, EPI_ISL_1365665, EPI_ISL_1365666, EPI_ISL_1365667 | LESP Coahuila | Instituto de Diagnostico y Referencia Epidemiologicos (INDRE) | Claudia Wong-Arambula, Abril Rodriguez-Maldonado, Vanessa Rivero-Arredondo, Ariadna Medina-Benitez, Joaquin Quiroz-Mercado, Sergio Rangel-Guerrero, Natividad Cruz-Ortiz, Tatiana Nunez-Garcia, Gisela Barrera-Badillo, Lucia Hernandez-Rivas, Irma Lopez-Martinez, Ernesto Ramirez-Gonzalez. |
| EPI_ISL_1365668, EPI_ISL_1365669, EPI_ISL_1365670, EPI_ISL_1365671, EPI_ISL_1365672, EPI_ISL_1365673, EPI_ISL_1365674 | LESP Nuevo Leon | Instituto de Diagnostico y Referencia Epidemiologicos (INDRE) | Claudia Wong-Arambula, Abril Rodriguez-Maldonado, Vanessa Rivero-Arredondo, Ariadna Medina-Benitez, Joaquin Quiroz-Mercado, Sergio Rangel-Guerrero, Natividad Cruz-Ortiz, Tatiana Nunez-Garcia, Gisela Barrera-Badillo, Lucia Hernandez-Rivas, Irma Lopez-Martinez, Ernesto Ramirez-Gonzalez. |
| EPI_ISL_1365675 | LESP Guanajuato | Instituto de Diagnostico y Referencia Epidemiologicos (INDRE) | Claudia Wong-Arambula, Abril Rodriguez-Maldonado, Vanessa Rivero-Arredondo, Ariadna Medina-Benitez, Joaquin Quiroz-Mercado, Sergio Rangel-Guerrero, Natividad Cruz-Ortiz, Tatiana Nunez-Garcia, Gisela Barrera-Badillo, Lucia Hernandez-Rivas, Irma Lopez-Martinez, Ernesto Ramirez-Gonzalez. |
| EPI_ISL_1365747 | Associação Fundo de Incentivo a Pesquisa | Associação Fundo de Incentivo a Pesquisa | Priscila Farias Tempaku, Juliana Nogueira Martins Rodrigues, Erika Rodrigues de Oliveira, Soraya Sgambatti de Andrade, Debora Ribeiro Ramadan, Sergio Tufik |
| EPI_ISL_1366655 | Usansolo-Galdakao University Hospital | Cruces University Hospital | Ana Gual-de-Torrella, Izaskun Alejo-Cancho, mikel Gallego; Ana Belén de la Hoz |
| EPI_ISL_1366657 | Usansolo-Galdakao University Hospital | Cruces University Hospital | Ana Gual-de-Torrella, Izaskun Alejo-Cancho, Mikel Gallego, Ana Belén de la Oz |
| EPI_ISL_1366660 | LESP Puebla | Instituto de Diagnostico y Referencia Epidemiologicos (INDRE) | Claudia Wong-Arambula, Abril Rodriguez-Maldonado, Vanessa Rivero-Arredondo, Ariadna Medina-Benitez, Joaquin Quiroz-Mercado, Sergio Rangel-Guerrero, Natividad Cruz-Ortiz, Tatiana Nunez-Garcia, Gisela Barrera-Badillo, Lucia Hernandez-Rivas, Irma Lopez-Martinez, Ernesto Ramirez-Gonzalez. |
| EPI_ISL_1366672, EPI_ISL_1366673, EPI_ISL_1366674, EPI_ISL_1366675, EPI_ISL_1366676 | LESP Queretaro | Instituto de Diagnostico y Referencia Epidemiologicos (INDRE) | Claudia Wong-Arambula, Abril Rodriguez-Maldonado, Vanessa Rivero-Arredondo, Ariadna Medina-Benitez, Joaquin Quiroz-Mercado, Sergio Rangel-Guerrero, Natividad Cruz-Ortiz, Tatiana Nunez-Garcia, Gisela Barrera-Badillo, Lucia Hernandez-Rivas, Irma Lopez-Martinez, Ernesto Ramirez-Gonzalez. |
| EPI_ISL_1366683, EPI_ISL_1366684, EPI_ISL_1366686 | LESP Veracruz | Instituto de Diagnostico y Referencia Epidemiologicos (INDRE) | Claudia Wong-Arambula, Abril Rodriguez-Maldonado, Vanessa Rivero-Arredondo, Ariadna Medina-Benitez, Joaquin Quiroz-Mercado, Sergio Rangel-Guerrero, Natividad Cruz-Ortiz, Tatiana Nunez-Garcia, Gisela Barrera-Badillo, Lucia Hernandez-Rivas, Irma Lopez-Martinez, Ernesto Ramirez-Gonzalez. |
| EPI_ISL_1366690, EPI_ISL_1366691, EPI_ISL_1366692 | LESP Hidalgo | Instituto de Diagnostico y Referencia Epidemiologicos (INDRE) | Claudia Wong-Arambula, Abril Rodriguez-Maldonado, Vanessa Rivero-Arredondo, Ariadna Medina-Benitez, Joaquin Quiroz-Mercado, Sergio Rangel-Guerrero, Natividad Cruz-Ortiz, Tatiana Nunez-Garcia, Gisela Barrera-Badillo, Lucia Hernandez-Rivas, Irma Lopez-Martinez, Ernesto Ramirez-Gonzalez. |
| EPI_ISL_1366693, EPI_ISL_1366694, EPI_ISL_1366695, EPI_ISL_1366696, EPI_ISL_1366697, EPI_ISL_1366698, EPI_ISL_1366699 | LESP Nuevo Leon | Instituto de Diagnostico y Referencia Epidemiologicos (INDRE) | Claudia Wong-Arambula, Abril Rodriguez-Maldonado, Vanessa Rivero-Arredondo, Ariadna Medina-Benitez, Joaquin Quiroz-Mercado, Sergio Rangel-Guerrero, Natividad Cruz-Ortiz, Tatiana Nunez-Garcia, Gisela Barrera-Badillo, Lucia Hernandez-Rivas, Irma Lopez-Martinez, Ernesto Ramirez-Gonzalez. |
| EPI_ISL_1366700 | LESP Guanajuato | Instituto de Diagnostico y Referencia Epidemiologicos (INDRE) | Claudia Wong-Arambula, Abril Rodriguez-Maldonado, Vanessa Rivero-Arredondo, Ariadna Medina-Benitez, Joaquin Quiroz-Mercado, Sergio Rangel-Guerrero, Natividad Cruz-Ortiz, Tatiana Nunez-Garcia, Gisela Barrera-Badillo, Lucia Hernandez-Rivas, Irma Lopez-Martinez, Ernesto Ramirez-Gonzalez. |
| EPI_ISL_1369225 | Labo Marquise | CHU Lille - Laboratoire de Virologie | AIT YAHYA Emilie, ALIDJINOU Enagnon Kazali, BOCKET Laurence, CREPIN Michel, DEMAY Christophe, ENGELMANN Ilka, GEFFROY Sandrine, GUIGON Aurélie, LAZREK Mouna, NOBILLIAUX Florian, PREVOST Brigitte, THUILLIER Caroline, TINEZ Claire |
| EPI_ISL_1369247, EPI_ISL_1369248 | CH Tourcoing | CHU Lille - Laboratoire de Virologie | AIT YAHYA Emilie, ALIDJINOU Enagnon Kazali, BOCKET Laurence, CREPIN Michel, DEMAY Christophe, ENGELMANN Ilka, GEFFROY Sandrine, GUIGON Aurélie, LAZREK Mouna, NOBILLIAUX Florian, PREVOST Brigitte, THUILLIER Caroline, TINEZ Claire |
| EPI_ISL_1369254 | CH Roubaix | CHU Lille - Laboratoire de Virologie | AIT YAHYA Emilie, ALIDJINOU Enagnon Kazali, BOCKET Laurence, CREPIN Michel, DEMAY Christophe, ENGELMANN Ilka, GEFFROY Sandrine, GUIGON Aurélie, LAZREK Mouna, NOBILLIAUX Florian, PREVOST Brigitte, THUILLIER Caroline, TINEZ Claire |
| EPI_ISL_1369255 | Synlab | CHU Lille - Laboratoire de Virologie | AIT YAHYA Emilie, ALIDJINOU Enagnon Kazali, BOCKET Laurence, CREPIN Michel, DEMAY Christophe, ENGELMANN Ilka, GEFFROY Sandrine, GUIGON Aurélie, LAZREK Mouna, NOBILLIAUX Florian, PREVOST Brigitte, THUILLIER Caroline, TINEZ Claire |
| EPI_ISL_1369259 | CH Arras | CHU Lille - Laboratoire de Virologie | AIT YAHYA Emilie, ALIDJINOU Enagnon Kazali, BOCKET Laurence, CREPIN Michel, DEMAY Christophe, ENGELMANN Ilka, GEFFROY Sandrine, GUIGON Aurélie, LAZREK Mouna, NOBILLIAUX Florian, PREVOST Brigitte, THUILLIER Caroline, TINEZ Claire |
| EPI_ISL_1369261, EPI_ISL_1369262, EPI_ISL_1369263, EPI_ISL_1369264 | CH Lens | CHU Lille - Laboratoire de Virologie | AIT YAHYA Emilie, ALIDJINOU Enagnon Kazali, BOCKET Laurence, CREPIN Michel, DEMAY Christophe, ENGELMANN Ilka, GEFFROY Sandrine, GUIGON Aurélie, LAZREK Mouna, NOBILLIAUX Florian, PREVOST Brigitte, THUILLIER Caroline, TINEZ Claire |
| EPI_ISL_1369271 | CH Valenciennes | CHU Lille - Laboratoire de Virologie | AIT YAHYA Emilie, ALIDJINOU Enagnon Kazali, BOCKET Laurence, CREPIN Michel, DEMAY Christophe, ENGELMANN Ilka, GEFFROY Sandrine, GUIGON Aurélie, LAZREK Mouna, NOBILLIAUX Florian, PREVOST Brigitte, THUILLIER Caroline, TINEZ Claire |
| EPI_ISL_1369272, EPI_ISL_1369273, EPI_ISL_1369274, EPI_ISL_1369276, EPI_ISL_1369277 | CHU LILLE | CHU Lille - Laboratoire de Virologie | AIT YAHYA Emilie, ALIDJINOU Enagnon Kazali, BOCKET Laurence, CREPIN Michel, DEMAY Christophe, ENGELMANN Ilka, GEFFROY Sandrine, GUIGON Aurélie, LAZREK Mouna, NOBILLIAUX Florian, PREVOST Brigitte, THUILLIER Caroline, TINEZ Claire |
| EPI_ISL_1369278 | CH St Omer | CHU Lille - Laboratoire de Virologie | AIT YAHYA Emilie, ALIDJINOU Enagnon Kazali, BOCKET Laurence, CREPIN Michel, DEMAY Christophe, ENGELMANN Ilka, GEFFROY Sandrine, GUIGON Aurélie, LAZREK Mouna, NOBILLIAUX Florian, PREVOST Brigitte, THUILLIER Caroline, TINEZ Claire |
| EPI_ISL_1369280, EPI_ISL_1369281, EPI_ISL_1369282 | CHU LILLE | CHU Lille - Laboratoire de Virologie | AIT YAHYA Emilie, ALIDJINOU Enagnon Kazali, BOCKET Laurence, CREPIN Michel, DEMAY Christophe, ENGELMANN Ilka, GEFFROY Sandrine, GUIGON Aurélie, LAZREK Mouna, NOBILLIAUX Florian, PREVOST Brigitte, THUILLIER Caroline, TINEZ Claire |
| EPI_ISL_1369284, EPI_ISL_1369285, EPI_ISL_1369286 | CH Douai | CHU Lille - Laboratoire de Virologie | AIT YAHYA Emilie, ALIDJINOU Enagnon Kazali, BOCKET Laurence, CREPIN Michel, DEMAY Christophe, ENGELMANN Ilka, GEFFROY Sandrine, GUIGON Aurélie, LAZREK Mouna, NOBILLIAUX Florian, PREVOST Brigitte, THUILLIER Caroline, TINEZ Claire |
| EPI_ISL_1369288 | CH Bethune | CHU Lille - Laboratoire de Virologie | AIT YAHYA Emilie, ALIDJINOU Enagnon Kazali, BOCKET Laurence, CREPIN Michel, DEMAY Christophe, ENGELMANN Ilka, GEFFROY Sandrine, GUIGON Aurélie, LAZREK Mouna, NOBILLIAUX Florian, PREVOST Brigitte, THUILLIER Caroline, TINEZ Claire |
| EPI_ISL_1369292 | CH BOULOGNE SUR MER | CHU Lille - Laboratoire de Virologie | AIT YAHYA Emilie, ALIDJINOU Enagnon Kazali, BOCKET Laurence, CREPIN Michel, DEMAY Christophe, ENGELMANN Ilka, GEFFROY Sandrine, GUIGON Aurélie, LAZREK Mouna, NOBILLIAUX Florian, PREVOST Brigitte, THUILLIER Caroline, TINEZ Claire |
| EPI_ISL_1372183, EPI_ISL_1372184 | Cliniques universitaires Saint-Luc | UCLouvain/IREC/MBLG | Jean Ruelle, Lysa Pinsmaye, Eleonore Ngyuvula Mantu, Benoit Kabamba Mukadi |
| EPI_ISL_1372420, EPI_ISL_1372421, EPI_ISL_1372424 | OLVZ Aalst | OLVZ Aalst | Astrid Holderbeke |
| EPI_ISL_1373369 | Somedi laboratories | Imelda Hospital | Johan Frans, Dagmar Obbels, Hanne Valgaeren |
| EPI_ISL_1373370 | Imelda Hospital | Imelda Hospital | Johan Frans, Dagmar Obbels, Hanne Valgaeren |

| | | | |
|---|---|---|---|
| EPI_ISL_1379413, EPI_ISL_1379414 | AREA DE SALUD CORONADO | Inciensa, Instituto Costarricense de Investigación y Enseñanza en Nutrición y Salud | Francisco Duarte, Hebleen Porras, Claudio Soto-Garita, Estela Cordero, Adriana Godínez & Melany Calderón |
| EPI_ISL_1379415 | AREA DE SALUD PAVAS (COOPESALUD) | Inciensa, Instituto Costarricense de Investigación y Enseñanza en Nutrición y Salud | Francisco Duarte, Hebleen Porras, Claudio Soto-Garita, Estela Cordero, Adriana Godínez & Melany Calderón |
| EPI_ISL_1379416 | AREA DE SALUD GOICOECHEA 2 - CLINICA DR. JIMENEZ NUÑEZ | Inciensa, Instituto Costarricense de Investigación y Enseñanza en Nutrición y Salud | Francisco Duarte, Hebleen Porras, Claudio Soto-Garita, Estela Cordero, Adriana Godínez & Melany Calderón |
| EPI_ISL_1379420 | AREA DE SALUD DESAMPARADOS 2 (COOPESALUD) | Inciensa, Instituto Costarricense de Investigación y Enseñanza en Nutrición y Salud | Francisco Duarte, Hebleen Porras, Claudio Soto-Garita, Estela Cordero, Adriana Godínez, Melany Calderón & Tashana Anglin-Williams |
| EPI_ISL_1379422 | HOSPITAL SAN JUAN DE DIOS | Inciensa, Instituto Costarricense de Investigación y Enseñanza en Nutrición y Salud | Francisco Duarte, Hebleen Porras, Claudio Soto-Garita, Estela Cordero, Adriana Godínez, Melany Calderón & Daniel Cascante-Serrano |
| EPI_ISL_1379427 | LABORATORIO CLINICO LABIN | Inciensa, Instituto Costarricense de Investigación y Enseñanza en Nutrición y Salud | Francisco Duarte, Hebleen Porras, Claudio Soto-Garita, Estela Cordero, Adriana Godínez, Melany Calderón & Pei Ling Chan Ma |
| EPI_ISL_1379979, EPI_ISL_1379980, EPI_ISL_1379983, EPI_ISL_1379984, EPI_ISL_1379985, EPI_ISL_1379987, EPI_ISL_1379997, EPI_ISL_1379998, EPI_ISL_1379999, EPI_ISL_1380003, EPI_ISL_1380008, EPI_ISL_1380012, EPI_ISL_1380014, EPI_ISL_1380015, EPI_ISL_1380018, EPI_ISL_1380019, EPI_ISL_1380020, EPI_ISL_1380024, EPI_ISL_1380025, EPI_ISL_1380026, EPI_ISL_1380028, EPI_ISL_1380029, EPI_ISL_1380030, EPI_ISL_1380035, EPI_ISL_1380036, EPI_ISL_1380044, EPI_ISL_1380045 | | | |
| see above | SC Microbiologia e Virologia AOUSS | AMES Centro Polidiagnostico Strumentale S.r.l. | Salvatore Rubino; Flavia Angioj; Laura Firino; Rosalba Govoni; Gabriele Ibba; Vincenzo Lai; Erica Mura; Bianca Paglietti; Claudia Piu; Anna Puggioni; Elena Rimini; Giulia Rocca; Caterina Serra; Sergio Uzzau. |
| EPI_ISL_1381068 | Conjunto Hospitalar do Mandaqui de Sao Paulo | Instituto Adolfo Lutz, Interdiciplinary Procedures Center, Strategic Laboratory | Claudio Tavares Sacchi, Claudia Regina Gonçalves, Erica Valessa Ramos Gomes, Karoline Rodrigues Campos, Caio Vinicius Dias Lopes |
| EPI_ISL_1381069 | Hospital Heliopolis | Instituto Adolfo Lutz, Interdiciplinary Procedures Center, Strategic Laboratory | Claudio Tavares Sacchi, Claudia Regina Gonçalves, Erica Valessa Ramos Gomes, Karoline Rodrigues Campos, Caio Vinicius Dias Lopes |
| EPI_ISL_1381070, EPI_ISL_1381071 | Hospital Municipal Cidade Tiradentes Carmem Prudente | Instituto Adolfo Lutz, Interdiciplinary Procedures Center, Strategic Laboratory | Claudio Tavares Sacchi, Claudia Regina Gonçalves, Erica Valessa Ramos Gomes, Karoline Rodrigues Campos, Caio Vinicius Dias Lopes |
| EPI_ISL_1381072, EPI_ISL_1381073, EPI_ISL_1381074 | Hospital | National Reference Center for Viruses of Respiratory Infections, Institut Pasteur, Paris | Marion Barbet, Sylvie Behillil, Méline Bizard, Angela Brisebarre, Camille Capel, Louise Lefrançois, Etienne Simon-Lorière, Vincent Enouf, Maud Vanpeene, Sylvie van der Werf,Leruez-Ville Marianne |
| EPI_ISL_1381075, EPI_ISL_1381076, EPI_ISL_1381077, EPI_ISL_1381078, EPI_ISL_1381079, EPI_ISL_1381080, EPI_ISL_1381081, EPI_ISL_1381082, EPI_ISL_1381083, EPI_ISL_1381084, EPI_ISL_1381085, EPI_ISL_1381086, EPI_ISL_1381087, EPI_ISL_1381088, EPI_ISL_1381089, EPI_ISL_1381090, EPI_ISL_1381091, EPI_ISL_1381092, EPI_ISL_1381093, EPI_ISL_1381094, EPI_ISL_1381095, EPI_ISL_1381097, EPI_ISL_1381098, EPI_ISL_1381099, EPI_ISL_1381100, EPI_ISL_1381101, EPI_ISL_1381102, EPI_ISL_1381103, EPI_ISL_1381104, EPI_ISL_1381105, EPI_ISL_1381106, EPI_ISL_1381107, EPI_ISL_1381108, EPI_ISL_1381109, EPI_ISL_1381110, EPI_ISL_1381111, EPI_ISL_1381112, EPI_ISL_1381113, EPI_ISL_1381114, EPI_ISL_1381115, EPI_ISL_1381116, EPI_ISL_1381117, EPI_ISL_1381118, EPI_ISL_1381119, EPI_ISL_1381120, EPI_ISL_1381121, EPI_ISL_1381122, EPI_ISL_1381123, EPI_ISL_1381124, EPI_ISL_1381125, EPI_ISL_1381126, EPI_ISL_1381127, EPI_ISL_1381129, EPI_ISL_1381130, EPI_ISL_1381131, EPI_ISL_1381132, EPI_ISL_1381133, EPI_ISL_1381134, EPI_ISL_1381135, EPI_ISL_1381136, EPI_ISL_1381137, EPI_ISL_1381138, EPI_ISL_1381139, EPI_ISL_1381140, EPI_ISL_1381141, EPI_ISL_1381142, EPI_ISL_1381143, EPI_ISL_1381144 | | | |
| see above | Hospital | National Reference Center for Viruses of Respiratory Infections, Institut Pasteur, Paris | Marion Barbet, Sylvie Behillil, Méline Bizard, Angela Brisebarre, Camille Capel, Louise Lefrançois, Etienne Simon-Lorière, Vincent Enouf, Maud Vanpeene, Sylvie van der Werf,Gastli Nabil |
| EPI_ISL_1381153, EPI_ISL_1381154, EPI_ISL_1381155, EPI_ISL_1381157, EPI_ISL_1381158, EPI_ISL_1381159 | Hospital | National Reference Center for Viruses of Respiratory Infections, Institut Pasteur, Paris | Marion Barbet, Sylvie Behillil, Méline Bizard, Angela Brisebarre, Camille Capel, Louise Lefrançois, Etienne Simon-Lorière, Vincent Enouf, Maud Vanpeene, Sylvie van der Werf,Hecquet Denise |
| EPI_ISL_1381160, EPI_ISL_1381162 | Labo Analyses Med | National Reference Center for Viruses of Respiratory Infections, Institut Pasteur, Paris | Marion Barbet, Sylvie Behillil, Méline Bizard, Angela Brisebarre, Camille Capel, Louise Lefrançois, Etienne Simon-Lorière, Vincent Enouf, Maud Vanpeene, Sylvie van der Werf,Besson J |
| EPI_ISL_1381163, EPI_ISL_1381164 | Hospital | National Reference Center for Viruses of Respiratory Infections, Institut Pasteur, Paris | Marion Barbet, Sylvie Behillil, Méline Bizard, Angela Brisebarre, Camille Capel, Louise Lefrançois, Etienne Simon-Lorière, Vincent Enouf, Maud Vanpeene, Sylvie van der Werf,Bastian Sylvaine |
| EPI_ISL_1381200 | Hospital | National Reference Center for Viruses of Respiratory Infections, Institut Pasteur, Paris | Marion Barbet, Sylvie Behillil, Méline Bizard, Angela Brisebarre, Camille Capel, Louise Lefrançois, Etienne Simon-Lorière, Vincent Enouf, Maud Vanpeene, Sylvie van der Werf,Bressollette CéLine |
| EPI_ISL_1381209 | Sentinelles Province | National Reference Center for Viruses of Respiratory Infections, Institut Pasteur, Paris | Marion Barbet, Sylvie Behillil, Méline Bizard, Angela Brisebarre, Camille Capel, Louise Lefrançois, Etienne Simon-Lorière, Vincent Enouf, Maud Vanpeene, Sylvie van der Werf,Lefevre StéPhane |
| EPI_ISL_1381211, EPI_ISL_1381212 | Armies | National Reference Center for Viruses of Respiratory Infections, Institut Pasteur, Paris | Marion Barbet, Sylvie Behillil, Méline Bizard, Angela Brisebarre, Camille Capel, Louise Lefrançois, Etienne Simon-Lorière, Vincent Enouf, Maud Vanpeene, Sylvie van der Werf,Desroches Marine |
| EPI_ISL_1381213 | Hospital | National Reference Center for Viruses of Respiratory Infections, Institut Pasteur, Paris | Marion Barbet, Sylvie Behillil, Méline Bizard, Angela Brisebarre, Camille Capel, Louise Lefrançois, Etienne Simon-Lorière, Vincent Enouf, Maud Vanpeene, Sylvie van der Werf,Louvet Laurence |
| EPI_ISL_1381214 | Sentinelles Paris | National Reference Center for Viruses of Respiratory Infections, Institut Pasteur, Paris | Marion Barbet, Sylvie Behillil, Méline Bizard, Angela Brisebarre, Camille Capel, Louise Lefrançois, Etienne Simon-Lorière, Vincent Enouf, Maud Vanpeene, Sylvie van der Werf,Rousset Dominique |
| EPI_ISL_1381215, EPI_ISL_1381216, EPI_ISL_1381217 | Hospital | National Reference Center for Viruses of Respiratory Infections, Institut Pasteur, Paris | Marion Barbet, Sylvie Behillil, Méline Bizard, Angela Brisebarre, Camille Capel, Louise Lefrançois, Etienne Simon-Lorière, Vincent Enouf, Maud Vanpeene, Sylvie van der Werf,Rousset Dominique |
| EPI_ISL_1381218, EPI_ISL_1381219, EPI_ISL_1381220, EPI_ISL_1381221, EPI_ISL_1381222, EPI_ISL_1381223 | Labo Analyses Med | National Reference Center for Viruses of Respiratory Infections, Institut Pasteur, Paris | Marion Barbet, Sylvie Behillil, Méline Bizard, Angela Brisebarre, Camille Capel, Louise Lefrançois, Etienne Simon-Lorière, Vincent Enouf, Maud Vanpeene, Sylvie van der Werf,Rousset Dominique |
| EPI_ISL_1381224 | Labo Analyses Med | National Reference Center for Viruses of Respiratory Infections, Institut Pasteur, Paris | Marion Barbet, Sylvie Behillil, Méline Bizard, Angela Brisebarre, Camille Capel, Louise Lefrançois, Etienne Simon-Lorière, Vincent Enouf, Maud Vanpeene, Sylvie van der Werf,Le Vicky |
| EPI_ISL_1381225, EPI_ISL_1381226, EPI_ISL_1381227, EPI_ISL_1381228, EPI_ISL_1381229, EPI_ISL_1381230 | Hospital | National Reference Center for Viruses of Respiratory Infections, Institut Pasteur, Paris | Marion Barbet, Sylvie Behillil, Méline Bizard, Angela Brisebarre, Camille Capel, Louise Lefrançois, Etienne Simon-Lorière, Vincent Enouf, Maud Vanpeene, Sylvie van der Werf,Desroches Marine |
| EPI_ISL_1381233 | Hospital | National Reference Center for Viruses of Respiratory Infections, Institut Pasteur, Paris | Marion Barbet, Sylvie Behillil, Méline Bizard, Angela Brisebarre, Camille Capel, Louise Lefrançois, Etienne Simon-Lorière, Vincent Enouf, Maud Vanpeene, Sylvie van der Werf,Hervo CéLine |
| EPI_ISL_1381234, EPI_ISL_1381235, EPI_ISL_1381236, EPI_ISL_1381237, EPI_ISL_1381239, EPI_ISL_1381240, EPI_ISL_1381241, EPI_ISL_1381243, EPI_ISL_1381245, EPI_ISL_1381246, EPI_ISL_1381248, EPI_ISL_1381249, EPI_ISL_1381250, EPI_ISL_1381252, EPI_ISL_1381253, EPI_ISL_1381255, EPI_ISL_1381256, EPI_ISL_1381257 | | | |
| see above | Centro de Investigacion Biomedica del Noreste (CIBIN) | Unidad de Genomica Avanzada | Consorcio Mexicano de Vigilancia Genomica (CoViGen-Mex). Authors (in alphabetical order): Julio Elias Alvarado-Yaah, Carlos F. Arias, Santiago Avila-Rios, Victor Hugo Borja-Aburto, Celia Boukadida, Juan Bautista Chale-Dzul , Jose Antonio Enciso-Moreno, Gloria Elena Espinoza-Ayala, Fernando Fontove-Herrera, Concepcion Grajales-Muniz, Ricardo Grande, Alfredo Herrera-Estrella, Carla Ivon Herrera-Najera, Pavel Isa, Brenda Irasema Maldonado-Meza, Bernardo Martinez-Miguel, Margarita Matias-Florentino, Maria Guadalupe de Jesus Mireles-Rivera, Gloria Maria Molina-Salinas, Hector Montoya-Fuentes, Jose Esteban Munoz-Medina, Jose de Jesus Nunez-Contreras, Alicia Ocana-Mondragon, Luis Alberto Ochoa-Carrera, Hector Esteban Paz-Juarez, Francisco Pulido, Helen Haydee Fernanda Ramirez-Plascencia, Angel Gustavo Salas-Lais, Jorge Ivan Salinal-Nevarez, Alejandro Sanchez-Flores, Clara Esperanza Santacruz-Tinoco, Maria Guadalupe Santiago-Mauricio, Nelly Selem-Mojica, Blanca Taboada, Gloria Vazquez |
| EPI_ISL_1381310, EPI_ISL_1381311, EPI_ISL_1381312, EPI_ISL_1381313, EPI_ISL_1381324, EPI_ISL_1381325, EPI_ISL_1381327, EPI_ISL_1381328, EPI_ISL_1381329, EPI_ISL_1381330, EPI_ISL_1381331, EPI_ISL_1381358, EPI_ISL_1381359, EPI_ISL_1381360, EPI_ISL_1381361, EPI_ISL_1381362, EPI_ISL_1381363, EPI_ISL_1381364, EPI_ISL_1381365, EPI_ISL_1381366, EPI_ISL_1381368, EPI_ISL_1381369, EPI_ISL_1381370, EPI_ISL_1381371, EPI_ISL_1381372, EPI_ISL_1381373, EPI_ISL_1381374, EPI_ISL_1381375, EPI_ISL_1381376, EPI_ISL_1381380, EPI_ISL_1381382, EPI_ISL_1381384 | | | |
| see above | Laboratory for Respiratory Viruses, Cantacuzino National Military-Medical Institute for Research and Development | Cantacuzino Institute Virology | Luiza Ustea, Nicoleta Paraschiv, Catalina Pascu, Sorin Dinu, Mihaela Lazar |

| | | | |
|---|---|---|---|
| EPI_ISL_1381816, EPI_ISL_1381817 | Hospital | National Reference Center for Viruses of Respiratory Infections, Institut Pasteur, Paris | Marion Barbet, Sylvie Behillil, Méline Bizard, Angela Brisebarre, Camille Capel, Louise Lefrançois, Etienne Simon-Lorière, Vincent Enouf, Maud Vanpeene, Sylvie van der Werf,Leruez-Ville Marianne |
| EPI_ISL_1381818, EPI_ISL_1381819, EPI_ISL_1381820, EPI_ISL_1381821 | Hospital | National Reference Center for Viruses of Respiratory Infections, Institut Pasteur, Paris | Marion Barbet, Sylvie Behillil, Méline Bizard, Angela Brisebarre, Camille Capel, Louise Lefrançois, Etienne Simon-Lorière, Vincent Enouf, Maud Vanpeene, Sylvie van der Werf,Gastli Nabil |
| EPI_ISL_1381828 | Hospital | National Reference Center for Viruses of Respiratory Infections, Institut Pasteur, Paris | Marion Barbet, Sylvie Behillil, Méline Bizard, Angela Brisebarre, Camille Capel, Louise Lefrançois, Etienne Simon-Lorière, Vincent Enouf, Maud Vanpeene, Sylvie van der Werf,Louvet Laurence |
| EPI_ISL_1381829 | Hospital | National Reference Center for Viruses of Respiratory Infections, Institut Pasteur, Paris | Marion Barbet, Sylvie Behillil, Méline Bizard, Angela Brisebarre, Camille Capel, Louise Lefrançois, Etienne Simon-Lorière, Vincent Enouf, Maud Vanpeene, Sylvie van der Werf,Dufour Mj |
| EPI_ISL_1381830 | Hospital | National Reference Center for Viruses of Respiratory Infections, Institut Pasteur, Paris | Marion Barbet, Sylvie Behillil, Méline Bizard, Angela Brisebarre, Camille Capel, Louise Lefrançois, Etienne Simon-Lorière, Vincent Enouf, Maud Vanpeene, Sylvie van der Werf,Pilorge LéA |
| EPI_ISL_1382117 | Clinical Microbiology Laboratory Basurto University Hospital | BioCruces Bizkaia | Mikel Urrutikoetxea-Gutierrez, Mª Carmen Nieto Toboso, Estibaliz Ugalde Zarraga, Mikel Gallego Rodrigo, Ana Belén de la Hoz Rastrollo, Jose Luis Díaz de Tuesta Del Arco |
| EPI_ISL_1385841, EPI_ISL_1386108, EPI_ISL_1386109, EPI_ISL_1386111, EPI_ISL_1386113 | Institut de Virologie du CHU de Strasbourg | Institut de Virologie du CHU de Strasbourg | Fafi-Kremer Samira, Gallais Floriane, Gantner Pierre, Laugel Elodie, Solis Morgane, Velay Aurélie, Wendling Marie-Josée |
| EPI_ISL_1386926, EPI_ISL_1387178, EPI_ISL_1387637, EPI_ISL_1388503, EPI_ISL_1389156, EPI_ISL_1389157 | National Platform bis UMONS/Jolimont | National Platform bis UMONS/Jolimont | François Dufrasne, Guillaume Bayon-Vicente, Gautier Detry, Ruddy Wattiez |
| EPI_ISL_1389164 | Hospital | National Reference Center for Viruses of Respiratory Infections, Institut Pasteur, Paris | Marion Barbet, Sylvie Behillil, Méline Bizard, Angela Brisebarre, Camille Capel, Louise Lefrançois, Etienne Simon-Lorière, Vincent Enouf, Maud Vanpeene, Sylvie van der Werf,Pilorge LéA |
| EPI_ISL_1389165, EPI_ISL_1389166, EPI_ISL_1389167, EPI_ISL_1389168, EPI_ISL_1389169, EPI_ISL_1389170, EPI_ISL_1389171, EPI_ISL_1389172, EPI_ISL_1389173, EPI_ISL_1389174, EPI_ISL_1389175, EPI_ISL_1389176, EPI_ISL_1389177, EPI_ISL_1389178, EPI_ISL_1389179, EPI_ISL_1389180, EPI_ISL_1389181 | | | |
| see above | Labo Analyses Med | National Reference Center for Viruses of Respiratory Infections, Institut Pasteur, Paris | Marion Barbet, Sylvie Behillil, Méline Bizard, Angela Brisebarre, Camille Capel, Louise Lefrançois, Etienne Simon-Lorière, Vincent Enouf, Maud Vanpeene, Sylvie van der Werf,Pichard Ronan |
| EPI_ISL_1389182, EPI_ISL_1389183, EPI_ISL_1389184, EPI_ISL_1389185, EPI_ISL_1389186, EPI_ISL_1389187, EPI_ISL_1389188 | Hospital | National Reference Center for Viruses of Respiratory Infections, Institut Pasteur, Paris | Marion Barbet, Sylvie Behillil, Méline Bizard, Angela Brisebarre, Camille Capel, Louise Lefrançois, Etienne Simon-Lorière, Vincent Enouf, Maud Vanpeene, Sylvie van der Werf,Pilorge LéA |
| EPI_ISL_1389189, EPI_ISL_1389190, EPI_ISL_1389191, EPI_ISL_1389192, EPI_ISL_1389193, EPI_ISL_1389194, EPI_ISL_1389195 | Hospital | National Reference Center for Viruses of Respiratory Infections, Institut Pasteur, Paris | Marion Barbet, Sylvie Behillil, Méline Bizard, Angela Brisebarre, Camille Capel, Louise Lefrançois, Etienne Simon-Lorière, Vincent Enouf, Maud Vanpeene, Sylvie van der Werf,Febreau Christine |
| EPI_ISL_1389196 | Labo Analyses Med | National Reference Center for Viruses of Respiratory Infections, Institut Pasteur, Paris | Marion Barbet, Sylvie Behillil, Méline Bizard, Angela Brisebarre, Camille Capel, Louise Lefrançois, Etienne Simon-Lorière, Vincent Enouf, Maud Vanpeene, Sylvie van der Werf,Gentilhomme Hervé |
| EPI_ISL_1389197, EPI_ISL_1389198, EPI_ISL_1389199, EPI_ISL_1389200, EPI_ISL_1389201, EPI_ISL_1389202, EPI_ISL_1389203, EPI_ISL_1389204, EPI_ISL_1389205, EPI_ISL_1389206, EPI_ISL_1389207, EPI_ISL_1389208, EPI_ISL_1389209, EPI_ISL_1389210, EPI_ISL_1389211, EPI_ISL_1389212, EPI_ISL_1389213, EPI_ISL_1389214, EPI_ISL_1389215, EPI_ISL_1389216, EPI_ISL_1389217, EPI_ISL_1389218, EPI_ISL_1389219, EPI_ISL_1389220, EPI_ISL_1389221, EPI_ISL_1389222, EPI_ISL_1389223, EPI_ISL_1389224, EPI_ISL_1389225, EPI_ISL_1389226, EPI_ISL_1389227, EPI_ISL_1389228, EPI_ISL_1389229 | | | |
| see above | Labo Analyses Med | National Reference Center for Viruses of Respiratory Infections, Institut Pasteur, Paris | Marion Barbet, Sylvie Behillil, Méline Bizard, Angela Brisebarre, Camille Capel, Louise Lefrançois, Etienne Simon-Lorière, Vincent Enouf, Maud Vanpeene, Sylvie van der Werf,Pichard Ronan |
| EPI_ISL_1389230, EPI_ISL_1389231, EPI_ISL_1389232, EPI_ISL_1389233, EPI_ISL_1389234, EPI_ISL_1389235, EPI_ISL_1389236, EPI_ISL_1389237, EPI_ISL_1389238, EPI_ISL_1389239, EPI_ISL_1389240, EPI_ISL_1389241, EPI_ISL_1389242, EPI_ISL_1389243, EPI_ISL_1389244, EPI_ISL_1389245, EPI_ISL_1389246, EPI_ISL_1389247, EPI_ISL_1389248, EPI_ISL_1389249, EPI_ISL_1389250, EPI_ISL_1389251, EPI_ISL_1389252, EPI_ISL_1389253 | | | |
| see above | Labo Analyses Med | National Reference Center for Viruses of Respiratory Infections, Institut Pasteur, Paris | Marion Barbet, Sylvie Behillil, Méline Bizard, Angela Brisebarre, Camille Capel, Louise Lefrançois, Etienne Simon-Lorière, Vincent Enouf, Maud Vanpeene, Sylvie van der Werf,Gestin Brieuc |
| EPI_ISL_1389254, EPI_ISL_1389255 | Hospital | National Reference Center for Viruses of Respiratory Infections, Institut Pasteur, Paris | Marion Barbet, Sylvie Behillil, Méline Bizard, Angela Brisebarre, Camille Capel, Louise Lefrançois, Etienne Simon-Lorière, Vincent Enouf, Maud Vanpeene, Sylvie van der Werf,Guillotel Eleha |
| EPI_ISL_1389256 | Hospital | National Reference Center for Viruses of Respiratory Infections, Institut Pasteur, Paris | Marion Barbet, Sylvie Behillil, Méline Bizard, Angela Brisebarre, Camille Capel, Louise Lefrançois, Etienne Simon-Lorière, Vincent Enouf, Maud Vanpeene, Sylvie van der Werf,Martres Pascale |
| EPI_ISL_1389257, EPI_ISL_1389258, EPI_ISL_1389259, EPI_ISL_1389260, EPI_ISL_1389261 | Labo Analyses Med | National Reference Center for Viruses of Respiratory Infections, Institut Pasteur, Paris | Marion Barbet, Sylvie Behillil, Méline Bizard, Angela Brisebarre, Camille Capel, Louise Lefrançois, Etienne Simon-Lorière, Vincent Enouf, Maud Vanpeene, Sylvie van der Werf,Felloni Claire |
| EPI_ISL_1389262, EPI_ISL_1389263 | Labo Analyses Med | National Reference Center for Viruses of Respiratory Infections, Institut Pasteur, Paris | Marion Barbet, Sylvie Behillil, Méline Bizard, Angela Brisebarre, Camille Capel, Louise Lefrançois, Etienne Simon-Lorière, Vincent Enouf, Maud Vanpeene, Sylvie van der Werf,Pichard Ronan |
| EPI_ISL_1389264, EPI_ISL_1389265 | Labo Analyses Med | National Reference Center for Viruses of Respiratory Infections, Institut Pasteur, Paris | Marion Barbet, Sylvie Behillil, Méline Bizard, Angela Brisebarre, Camille Capel, Louise Lefrançois, Etienne Simon-Lorière, Vincent Enouf, Maud Vanpeene, Sylvie van der Werf,Gestin Brieuc |
| EPI_ISL_1389266 | Labo Analyses Med | National Reference Center for Viruses of Respiratory Infections, Institut Pasteur, Paris | Marion Barbet, Sylvie Behillil, Méline Bizard, Angela Brisebarre, Camille Capel, Louise Lefrançois, Etienne Simon-Lorière, Vincent Enouf, Maud Vanpeene, Sylvie van der Werf,Felloni Claire |
| EPI_ISL_1389267 | Hospital | National Reference Center for Viruses of Respiratory Infections, Institut Pasteur, Paris | Marion Barbet, Sylvie Behillil, Méline Bizard, Angela Brisebarre, Camille Capel, Louise Lefrançois, Etienne Simon-Lorière, Vincent Enouf, Maud Vanpeene, Sylvie van der Werf,Guillotel Eleha |
| EPI_ISL_1389268, EPI_ISL_1389269 | Labo Analyses Med | National Reference Center for Viruses of Respiratory Infections, Institut Pasteur, Paris | Marion Barbet, Sylvie Behillil, Méline Bizard, Angela Brisebarre, Camille Capel, Louise Lefrançois, Etienne Simon-Lorière, Vincent Enouf, Maud Vanpeene, Sylvie van der Werf,Felloni Claire |
| EPI_ISL_1389270, EPI_ISL_1389271, EPI_ISL_1389272, EPI_ISL_1389273, EPI_ISL_1389274, EPI_ISL_1389276, EPI_ISL_1389277, EPI_ISL_1389278, EPI_ISL_1389279, EPI_ISL_1389280, EPI_ISL_1389281, EPI_ISL_1389282, EPI_ISL_1389283, EPI_ISL_1389284, EPI_ISL_1389285, EPI_ISL_1389286 | | | |
| see above | Labo Analyses Med | National Reference Center for Viruses of Respiratory Infections, Institut Pasteur, Paris | Marion Barbet, Sylvie Behillil, Méline Bizard, Angela Brisebarre, Camille Capel, Louise Lefrançois, Etienne Simon-Lorière, Vincent Enouf, Maud Vanpeene, Sylvie van der Werf,Pichard Ronan |
| EPI_ISL_1389287, EPI_ISL_1389288 | Hospital | National Reference Center for Viruses of Respiratory Infections, Institut Pasteur, Paris | Marion Barbet, Sylvie Behillil, Méline Bizard, Angela Brisebarre, Camille Capel, Louise Lefrançois, Etienne Simon-Lorière, Vincent Enouf, Maud Vanpeene, Sylvie van der Werf,Pilorge LéA |
| EPI_ISL_1389289, EPI_ISL_1389290, EPI_ISL_1389291, EPI_ISL_1389292, EPI_ISL_1389293, EPI_ISL_1389294, EPI_ISL_1389295, EPI_ISL_1389296 | Labo Analyses Med | National Reference Center for Viruses of Respiratory Infections, Institut Pasteur, Paris | Marion Barbet, Sylvie Behillil, Méline Bizard, Angela Brisebarre, Camille Capel, Louise Lefrançois, Etienne Simon-Lorière, Vincent Enouf, Maud Vanpeene, Sylvie van der Werf,Pichard Ronan |
| EPI_ISL_1389297, EPI_ISL_1389298, EPI_ISL_1389299, EPI_ISL_1389300, EPI_ISL_1389301, EPI_ISL_1389302, EPI_ISL_1389303, EPI_ISL_1389304, EPI_ISL_1389305, EPI_ISL_1389306 | Labo Analyses Med | National Reference Center for Viruses of Respiratory Infections, Institut Pasteur, Paris | Marion Barbet, Sylvie Behillil, Méline Bizard, Angela Brisebarre, Camille Capel, Louise Lefrançois, Etienne Simon-Lorière, Vincent Enouf, Maud Vanpeene, Sylvie van der Werf,Gestin Brieuc |

| | | | |
|---|---|---|---|
| EPI_ISL_1389307, EPI_ISL_1389308, EPI_ISL_1389309, EPI_ISL_1389310 | Hospital | National Reference Center for Viruses of Respiratory Infections, Institut Pasteur, Paris | Marion Barbet, Sylvie Behillil, Méline Bizard, Angela Brisebarre, Camille Capel, Louise Lefrançois, Etienne Simon-Lorière, Vincent Enouf, Maud Vanpeene, Sylvie van der Werf,Guillotel Eleha |
| EPI_ISL_1389311 | Armies | National Reference Center for Viruses of Respiratory Infections, Institut Pasteur, Paris | Marion Barbet, Sylvie Behillil, Méline Bizard, Angela Brisebarre, Camille Capel, Louise Lefrançois, Etienne Simon-Lorière, Vincent Enouf, Maud Vanpeene, Sylvie van der Werf,Desroches Marine |
| EPI_ISL_1389312, EPI_ISL_1389313, EPI_ISL_1389314 | Labo Analyses Med | National Reference Center for Viruses of Respiratory Infections, Institut Pasteur, Paris | Marion Barbet, Sylvie Behillil, Méline Bizard, Angela Brisebarre, Camille Capel, Louise Lefrançois, Etienne Simon-Lorière, Vincent Enouf, Maud Vanpeene, Sylvie van der Werf,Felloni Claire |
| EPI_ISL_1389315, EPI_ISL_1389316, EPI_ISL_1389317 | Labo Analyses Med | National Reference Center for Viruses of Respiratory Infections, Institut Pasteur, Paris | Marion Barbet, Sylvie Behillil, Méline Bizard, Angela Brisebarre, Camille Capel, Louise Lefrançois, Etienne Simon-Lorière, Vincent Enouf, Maud Vanpeene, Sylvie van der Werf,Pichard Ronan |
| EPI_ISL_1389318 | Labo Analyses Med | National Reference Center for Viruses of Respiratory Infections, Institut Pasteur, Paris | Marion Barbet, Sylvie Behillil, Méline Bizard, Angela Brisebarre, Camille Capel, Louise Lefrançois, Etienne Simon-Lorière, Vincent Enouf, Maud Vanpeene, Sylvie van der Werf,Gestin Brieuc |
| EPI_ISL_1389319 | Hospital | National Reference Center for Viruses of Respiratory Infections, Institut Pasteur, Paris | Marion Barbet, Sylvie Behillil, Méline Bizard, Angela Brisebarre, Camille Capel, Louise Lefrançois, Etienne Simon-Lorière, Vincent Enouf, Maud Vanpeene, Sylvie van der Werf,Febreau Christine |
| EPI_ISL_1389320 | Labo Analyses Med | National Reference Center for Viruses of Respiratory Infections, Institut Pasteur, Paris | Marion Barbet, Sylvie Behillil, Méline Bizard, Angela Brisebarre, Camille Capel, Louise Lefrançois, Etienne Simon-Lorière, Vincent Enouf, Maud Vanpeene, Sylvie van der Werf,Felloni Claire |
| EPI_ISL_1389321, EPI_ISL_1389322, EPI_ISL_1389323, EPI_ISL_1389324 | Labo Analyses Med | National Reference Center for Viruses of Respiratory Infections, Institut Pasteur, Paris | Marion Barbet, Sylvie Behillil, Méline Bizard, Angela Brisebarre, Camille Capel, Louise Lefrançois, Etienne Simon-Lorière, Vincent Enouf, Maud Vanpeene, Sylvie van der Werf,Pichard Ronan |
| EPI_ISL_1389326, EPI_ISL_1389327 | Labo Analyses Med | National Reference Center for Viruses of Respiratory Infections, Institut Pasteur, Paris | Marion Barbet, Sylvie Behillil, Méline Bizard, Angela Brisebarre, Camille Capel, Louise Lefrançois, Etienne Simon-Lorière, Vincent Enouf, Maud Vanpeene, Sylvie van der Werf,Gestin Brieuc |
| EPI_ISL_1389328, EPI_ISL_1389329 | Labo Analyses Med | National Reference Center for Viruses of Respiratory Infections, Institut Pasteur, Paris | Marion Barbet, Sylvie Behillil, Méline Bizard, Angela Brisebarre, Camille Capel, Louise Lefrançois, Etienne Simon-Lorière, Vincent Enouf, Maud Vanpeene, Sylvie van der Werf,Pichard Ronan |
| EPI_ISL_1389330, EPI_ISL_1389331, EPI_ISL_1389332 | Labo Analyses Med | National Reference Center for Viruses of Respiratory Infections, Institut Pasteur, Paris | Marion Barbet, Sylvie Behillil, Méline Bizard, Angela Brisebarre, Camille Capel, Louise Lefrançois, Etienne Simon-Lorière, Vincent Enouf, Maud Vanpeene, Sylvie van der Werf,Gestin Brieuc |
| EPI_ISL_1389333 | Labo Analyses Med | National Reference Center for Viruses of Respiratory Infections, Institut Pasteur, Paris | Marion Barbet, Sylvie Behillil, Méline Bizard, Angela Brisebarre, Camille Capel, Louise Lefrançois, Etienne Simon-Lorière, Vincent Enouf, Maud Vanpeene, Sylvie van der Werf |
| EPI_ISL_1389334 | Labo Analyses Med | National Reference Center for Viruses of Respiratory Infections, Institut Pasteur, Paris | Marion Barbet, Sylvie Behillil, Méline Bizard, Angela Brisebarre, Camille Capel, Louise Lefrançois, Etienne Simon-Lorière, Vincent Enouf, Maud Vanpeene, Sylvie van der Werf,Pichard Ronan |
| EPI_ISL_1389335 | Labo Analyses Med | National Reference Center for Viruses of Respiratory Infections, Institut Pasteur, Paris | Marion Barbet, Sylvie Behillil, Méline Bizard, Angela Brisebarre, Camille Capel, Louise Lefrançois, Etienne Simon-Lorière, Vincent Enouf, Maud Vanpeene, Sylvie van der Werf,Gestin Brieuc |
| EPI_ISL_1389336 | Labo Analyses Med | National Reference Center for Viruses of Respiratory Infections, Institut Pasteur, Paris | Marion Barbet, Sylvie Behillil, Méline Bizard, Angela Brisebarre, Camille Capel, Louise Lefrançois, Etienne Simon-Lorière, Vincent Enouf, Maud Vanpeene, Sylvie van der Werf,Felloni Claire |
| EPI_ISL_1389337, EPI_ISL_1389338, EPI_ISL_1389339, EPI_ISL_1389340, EPI_ISL_1389341, EPI_ISL_1389342, EPI_ISL_1389343 | Labo Analyses Med | National Reference Center for Viruses of Respiratory Infections, Institut Pasteur, Paris | Marion Barbet, Sylvie Behillil, Méline Bizard, Angela Brisebarre, Camille Capel, Louise Lefrançois, Etienne Simon-Lorière, Vincent Enouf, Maud Vanpeene, Sylvie van der Werf,Pichard Ronan |
| EPI_ISL_1389344 | Hospital | National Reference Center for Viruses of Respiratory Infections, Institut Pasteur, Paris | Marion Barbet, Sylvie Behillil, Méline Bizard, Angela Brisebarre, Camille Capel, Louise Lefrançois, Etienne Simon-Lorière, Vincent Enouf, Maud Vanpeene, Sylvie van der Werf,Febreau Christine |
| EPI_ISL_1389345, EPI_ISL_1389346, EPI_ISL_1389347, EPI_ISL_1389348 | Labo Analyses Med | National Reference Center for Viruses of Respiratory Infections, Institut Pasteur, Paris | Marion Barbet, Sylvie Behillil, Méline Bizard, Angela Brisebarre, Camille Capel, Louise Lefrançois, Etienne Simon-Lorière, Vincent Enouf, Maud Vanpeene, Sylvie van der Werf,Pichard Ronan |
| EPI_ISL_1389349, EPI_ISL_1389350, EPI_ISL_1389351, EPI_ISL_1389352 | Labo Analyses Med | National Reference Center for Viruses of Respiratory Infections, Institut Pasteur, Paris | Marion Barbet, Sylvie Behillil, Méline Bizard, Angela Brisebarre, Camille Capel, Louise Lefrançois, Etienne Simon-Lorière, Vincent Enouf, Maud Vanpeene, Sylvie van der Werf,Gestin Brieuc |
| EPI_ISL_1389353 | Hospital | National Reference Center for Viruses of Respiratory Infections, Institut Pasteur, Paris | Marion Barbet, Sylvie Behillil, Méline Bizard, Angela Brisebarre, Camille Capel, Louise Lefrançois, Etienne Simon-Lorière, Vincent Enouf, Maud Vanpeene, Sylvie van der Werf,Martres Pascale |
| EPI_ISL_1390769 | Azienda Ospedaliera Terni | Istituto Zooprofilattico Sperimentale dell'Abruzzo e Molise "G. Caporale" | Palumbo M, Scaccetti A, Lorusso A, Marcacci M, Di Domenico M, Ancora M, Curini V, Mangone I, Rinaldi A, Scialabba S, Caporale M, Di Pasquale A, Cammà C, Puglia I, Calistri P, Savini G |
| EPI_ISL_1391016, EPI_ISL_1391017, EPI_ISL_1391018 | National Institute of Health Research and Development | National Institute of Health Research and Development | Subangkit, Hana Apsari Pawestri, Kartika Dewi Puspa, Arie Ardiansyah Nugraha, Hartanti Dian Ikawati, Krisna Nur Andriana Pangesti, Yuni Rukminiati, Ririn Ramadhany, Agustiningsih, Kindi Adam, Holy Arif Wibowo, Triyani Soekarso, Ni Ketut Susilarini, Nurika Hariastuti, Ully Alfi Nikmah, Reni Herman, Nike Susanti, Herna, Tati Febriyanti, Natalie Laurencia Kipuw, Fauzul Muna, Irene Lorinda Indalao, Nelly Puspandari, Vivi Setiawaty. |
| EPI_ISL_1391020, EPI_ISL_1391021, EPI_ISL_1391023 | Ospedale Generale Regionale F. Miulli | Istituto Zooprofilattico Sperimentale della Puglia e della Basilicata | Parisi A., Bianco A., Capozzi L., Del Sambro L., Simone D., Ceci E. |
| EPI_ISL_1391077, EPI_ISL_1391083, EPI_ISL_1391097 | Ospedale Santissima Annunziata | Istituto Zooprofilattico Sperimentale della Puglia e della Basilicata | Parisi A., Bianco A., Capozzi L., Del Sambro L., Simone D., Difato L., Ridolfi D., Giannico A, Tampoia M. |
| EPI_ISL_1391393 | SIESP DIPARTIMENTO DI PREVENZIONE CHIETI | Istituto Zooprofilattico Sperimentale dell'Abruzzo e Molise "G. Caporale" | Lorusso A, Marcacci M, Di Domenico M, Ancora M, Curini V, Di Lollo Valeria, Mangone I, Rinaldi A, Scialabba S, Caporale M, Di Pasquale A, Cammà C, Puglia I, Calistri P, Savini G |
| EPI_ISL_1391394 | SIESP DIPARTIMENTO DI PREVENZIONE TERAMO | Istituto Zooprofilattico Sperimentale dell'Abruzzo e Molise "G. Caporale" | Lorusso A, Marcacci M, Di Domenico M, Ancora M, Curini V, Di Lollo Valeria, Mangone I, Rinaldi A, Scialabba S, Caporale M, Di Pasquale A, Cammà C, Puglia I, Calistri P, Savini G |
| EPI_ISL_1391395 | OSPEDALE S.S. ANNUNZIATA CHIETI | Istituto Zooprofilattico Sperimentale dell'Abruzzo e Molise "G. Caporale" | Lorusso A, Marcacci M, Di Domenico M, Ancora M, Curini V, Di Lollo Valeria, Mangone I, Rinaldi A, Scialabba S, Caporale M, Di Pasquale A, Cammà C, Puglia I, Calistri P, Savini G |
| EPI_ISL_1391396 | SIESP CHIETI - DRIVE IN CHIETI | Istituto Zooprofilattico Sperimentale dell'Abruzzo e Molise "G. Caporale" | Lorusso A, Marcacci M, Di Domenico M, Ancora M, Curini V, Di Lollo Valeria, Mangone I, Rinaldi A, Scialabba S, Caporale M, Di Pasquale A, Cammà C, Puglia I, Calistri P, Savini G |
| EPI_ISL_1391397 | SIESP CHIETI - DRIVE IN ORTONA | Istituto Zooprofilattico Sperimentale dell'Abruzzo e Molise "G. Caporale" | Lorusso A, Marcacci M, Di Domenico M, Ancora M, Curini V, Di Lollo Valeria, Mangone I, Rinaldi A, Scialabba S, Caporale M, Di Pasquale A, Cammà C, Puglia I, Calistri P, Savini G |
| EPI_ISL_1391398 | SIESP SULMONA | Istituto Zooprofilattico Sperimentale dell'Abruzzo e Molise "G. Caporale" | Lorusso A, Marcacci M, Di Domenico M, Ancora M, Curini V, Di Lollo Valeria, Mangone I, Rinaldi A, Scialabba S, Caporale M, Di Pasquale A, Cammà C, Puglia I, Calistri P, Savini G |
| EPI_ISL_1391399, EPI_ISL_1391400 | SIESP CHIETI - DRIVE IN ORTONA | Istituto Zooprofilattico Sperimentale dell'Abruzzo e Molise "G. Caporale" | Lorusso A, Marcacci M, Di Domenico M, Ancora M, Curini V, Di Lollo Valeria, Mangone I, Rinaldi A, Scialabba S, Caporale M, Di Pasquale A, Cammà C, Puglia I, Calistri P, Savini G |
| EPI_ISL_1391401, EPI_ISL_1391402 | SIESP DIPARTIMENTO DI PREVENZIONE TERAMO | Istituto Zooprofilattico Sperimentale dell'Abruzzo e Molise "G. Caporale" | Lorusso A, Marcacci M, Di Domenico M, Ancora M, Curini V, Di Lollo Valeria, Mangone I, Rinaldi A, Scialabba S, Caporale M, Di Pasquale A, Cammà C, Puglia I, Calistri P, Savini G |
| EPI_ISL_1391403 | DIP. PREV. AVEZZANO SERVIZIO DI IGIENE EPIDEMIOLOGIA E SANITA' PUBBLICA | Istituto Zooprofilattico Sperimentale dell'Abruzzo e Molise "G. Caporale" | Lorusso A, Marcacci M, Di Domenico M, Ancora M, Curini V, Di Lollo Valeria, Mangone I, Rinaldi A, Scialabba S, Caporale M, Di Pasquale A, Cammà C, Puglia I, Calistri P, Savini G |
| EPI_ISL_1391404 | SIESP DIPARTIMENTO DI PREVENZIONE CHIETI | Istituto Zooprofilattico Sperimentale dell'Abruzzo e Molise "G. Caporale" | Lorusso A, Marcacci M, Di Domenico M, Ancora M, Curini V, Di Lollo Valeria, Mangone I, Rinaldi A, Scialabba S, Caporale M, Di Pasquale A, Cammà C, Puglia I, Calistri P, Savini G |

| | | | |
|---|---|---|---|
| EPI_ISL_1391405 | SIESP CHIETI - DRIVE IN CHIETI | Istituto Zooprofilattico Sperimentale dell'Abruzzo e Molise "G. Caporale" | Lorusso A, Marcacci M, Di Domenico M, Ancora M, Curini V, Di Lollo Valeria, Mangone I, Rinaldi A, Scialabba S, Caporale M, Di Pasquale A, Cammà C, Puglia I, Calistri P, Savini G |
| EPI_ISL_1391406 | SIESP SULMONA | Istituto Zooprofilattico Sperimentale dell'Abruzzo e Molise "G. Caporale" | Lorusso A, Marcacci M, Di Domenico M, Ancora M, Curini V, Di Lollo Valeria, Mangone I, Rinaldi A, Scialabba S, Caporale M, Di Pasquale A, Cammà C, Puglia I, Calistri P, Savini G |
| EPI_ISL_1391407 | SIESP DIPARTIMENTO DI PREVENZIONE TERAMO | Istituto Zooprofilattico Sperimentale dell'Abruzzo e Molise "G. Caporale" | Lorusso A, Marcacci M, Di Domenico M, Ancora M, Curini V, Di Lollo Valeria, Mangone I, Rinaldi A, Scialabba S, Caporale M, Di Pasquale A, Cammà C, Puglia I, Calistri P, Savini G |
| EPI_ISL_1391408 | SIESP CHIETI - DRIVE IN ORTONA | Istituto Zooprofilattico Sperimentale dell'Abruzzo e Molise "G. Caporale" | Lorusso A, Marcacci M, Di Domenico M, Ancora M, Curini V, Di Lollo Valeria, Mangone I, Rinaldi A, Scialabba S, Caporale M, Di Pasquale A, Cammà C, Puglia I, Calistri P, Savini G |
| EPI_ISL_1391413 | Baylor Scott & White-Irving | Baylor Scott & White-Temple | Ari Rao, Linden Morales, Kimberly Walker, Marcus Volz, Shelby Hendrickson, Caitlin Maloney |
| EPI_ISL_1391415 | Baylor Scott & White-Waxahachie | Baylor Scott & White-Temple | Ari Rao, Linden Morales, Kimberly Walker, Marcus Volz, Shelby Hendrickson, Caitlin Maloney |
| EPI_ISL_1392650, EPI_ISL_1392651 | OSPEDALE CIVILE GIULIANOVA - PRONTO SOCCORSO GIULIANOVA(TERAMO) | Istituto Zooprofilattico Sperimentale dell'Abruzzo e Molise "G. Caporale" | Lorusso A, Marcacci M, Di Domenico M, Ancora M, Curini V, Di Lollo Valeria, Mangone I, Rinaldi A, Delli Compagni E, Scialabba S, Caporale M, Di Pasquale A, Cammà C, Puglia I, Calistri P, Savini G |
| EPI_ISL_1392652, EPI_ISL_1392653, EPI_ISL_1392654, EPI_ISL_1392655, EPI_ISL_1392656, EPI_ISL_1392657, EPI_ISL_1392658, EPI_ISL_1392659, EPI_ISL_1392660, EPI_ISL_1392661, EPI_ISL_1392662 | | | |
| see above | SIESP DIPARTIMENTO DI PREVENZIONE TERAMO TERAMO(TERAMO) | Istituto Zooprofilattico Sperimentale dell'Abruzzo e Molise "G. Caporale" | Lorusso A, Marcacci M, Di Domenico M, Ancora M, Curini V, Di Lollo Valeria, Mangone I, Rinaldi A, Delli Compagni E, Scialabba S, Caporale M, Di Pasquale A, Cammà C, Puglia I, Calistri P, Savini G |
| EPI_ISL_1392663, EPI_ISL_1392665 | RSA GIULIANOVA GIULIANOVA(TERAMO) | Istituto Zooprofilattico Sperimentale dell'Abruzzo e Molise "G. Caporale" | Lorusso A, Marcacci M, Di Domenico M, Ancora M, Curini V, Di Lollo Valeria, Mangone I, Rinaldi A, Delli Compagni E, Scialabba S, Caporale M, Di Pasquale A, Cammà C, Puglia I, Calistri P, Savini G |
| EPI_ISL_1392666 | SIESP DIPARTIMENTO DI PREVENZIONE CHIETI CHIETI(CHIETI) | Istituto Zooprofilattico Sperimentale dell'Abruzzo e Molise "G. Caporale" | Lorusso A, Marcacci M, Di Domenico M, Ancora M, Curini V, Di Lollo Valeria, Mangone I, Rinaldi A, Delli Compagni E, Scialabba S, Caporale M, Di Pasquale A, Cammà C, Puglia I, Calistri P, Savini G |
| EPI_ISL_1392667, EPI_ISL_1392668, EPI_ISL_1392669 | DIP. PREV. AVEZZANO SERVIZIO DI IGIENE EPIDEMIOLOGIA E SANITA' PUBBLICA AVEZZANO(L'AQUILA) | Istituto Zooprofilattico Sperimentale dell'Abruzzo e Molise "G. Caporale" | Lorusso A, Marcacci M, Di Domenico M, Ancora M, Curini V, Di Lollo Valeria, Mangone I, Rinaldi A, Delli Compagni E, Scialabba S, Caporale M, Di Pasquale A, Cammà C, Puglia I, Calistri P, Savini G |
| EPI_ISL_1392670, EPI_ISL_1392671 | SIESP SULMONA SULMONA(L'AQUILA) | Istituto Zooprofilattico Sperimentale dell'Abruzzo e Molise "G. Caporale" | Lorusso A, Marcacci M, Di Domenico M, Ancora M, Curini V, Di Lollo Valeria, Mangone I, Rinaldi A, Delli Compagni E, Scialabba S, Caporale M, Di Pasquale A, Cammà C, Puglia I, Calistri P, Savini G |
| EPI_ISL_1392672, EPI_ISL_1392673 | DIP. PREV. AVEZZANO SERVIZIO DI IGIENE EPIDEMIOLOGIA E SANITA' PUBBLICA AVEZZANO(L'AQUILA) | Istituto Zooprofilattico Sperimentale dell'Abruzzo e Molise "G. Caporale" | Lorusso A, Marcacci M, Di Domenico M, Ancora M, Curini V, Di Lollo Valeria, Mangone I, Rinaldi A, Delli Compagni E, Scialabba S, Caporale M, Di Pasquale A, Cammà C, Puglia I, Calistri P, Savini G |
| EPI_ISL_1392674 | SIESP SULMONA SULMONA(L'AQUILA) | Istituto Zooprofilattico Sperimentale dell'Abruzzo e Molise "G. Caporale" | Lorusso A, Marcacci M, Di Domenico M, Ancora M, Curini V, Di Lollo Valeria, Mangone I, Rinaldi A, Delli Compagni E, Scialabba S, Caporale M, Di Pasquale A, Cammà C, Puglia I, Calistri P, Savini G |
| EPI_ISL_1392676, EPI_ISL_1392677, EPI_ISL_1392678, EPI_ISL_1392679 | DIP. PREV. AVEZZANO SERVIZIO DI IGIENE EPIDEMIOLOGIA E SANITA' PUBBLICA AVEZZANO(L'AQUILA) | Istituto Zooprofilattico Sperimentale dell'Abruzzo e Molise "G. Caporale" | Lorusso A, Marcacci M, Di Domenico M, Ancora M, Curini V, Di Lollo Valeria, Mangone I, Rinaldi A, Delli Compagni E, Scialabba S, Caporale M, Di Pasquale A, Cammà C, Puglia I, Calistri P, Savini G |
| EPI_ISL_1392680 | SIESP CHIETI - DRIVE IN CHIETI CHIETI(CHIETI) | Istituto Zooprofilattico Sperimentale dell'Abruzzo e Molise "G. Caporale" | Lorusso A, Marcacci M, Di Domenico M, Ancora M, Curini V, Di Lollo Valeria, Mangone I, Rinaldi A, Delli Compagni E, Scialabba S, Caporale M, Di Pasquale A, Cammà C, Puglia I, Calistri P, Savini G |
| EPI_ISL_1392681 | SIESP CHIETI - DRIVE IN LANCIANO LANCIANO(CHIETI) | Istituto Zooprofilattico Sperimentale dell'Abruzzo e Molise "G. Caporale" | Lorusso A, Marcacci M, Di Domenico M, Ancora M, Curini V, Di Lollo Valeria, Mangone I, Rinaldi A, Delli Compagni E, Scialabba S, Caporale M, Di Pasquale A, Cammà C, Puglia I, Calistri P, Savini G |
| EPI_ISL_1392682 | SIESP DIPARTIMENTO DI PREVENZIONE TERAMO TERAMO(TERAMO) | Istituto Zooprofilattico Sperimentale dell'Abruzzo e Molise "G. Caporale" | Lorusso A, Marcacci M, Di Domenico M, Ancora M, Curini V, Di Lollo Valeria, Mangone I, Rinaldi A, Delli Compagni E, Scialabba S, Caporale M, Di Pasquale A, Cammà C, Puglia I, Calistri P, Savini G |
| EPI_ISL_1392683, EPI_ISL_1392684 | USCA Tagliacozzo TAGLIACOZZO(L'AQUILA) | Istituto Zooprofilattico Sperimentale dell'Abruzzo e Molise "G. Caporale" | Lorusso A, Marcacci M, Di Domenico M, Ancora M, Curini V, Di Lollo Valeria, Mangone I, Rinaldi A, Delli Compagni E, Scialabba S, Caporale M, Di Pasquale A, Cammà C, Puglia I, Calistri P, Savini G |
| EPI_ISL_1392685, EPI_ISL_1392686, EPI_ISL_1392687 | USCA Pescina PESCINA(L'AQUILA) | Istituto Zooprofilattico Sperimentale dell'Abruzzo e Molise "G. Caporale" | Lorusso A, Marcacci M, Di Domenico M, Ancora M, Curini V, Di Lollo Valeria, Mangone I, Rinaldi A, Delli Compagni E, Scialabba S, Caporale M, Di Pasquale A, Cammà C, Puglia I, Calistri P, Savini G |
| EPI_ISL_1392688, EPI_ISL_1392689 | SIESP CHIETI - DRIVE IN VASTO VASTO(CHIETI) | Istituto Zooprofilattico Sperimentale dell'Abruzzo e Molise "G. Caporale" | Lorusso A, Marcacci M, Di Domenico M, Ancora M, Curini V, Di Lollo Valeria, Mangone I, Rinaldi A, Delli Compagni E, Scialabba S, Caporale M, Di Pasquale A, Cammà C, Puglia I, Calistri P, Savini G |
| EPI_ISL_1392690 | SIESP DIPARTIMENTO DI PREVENZIONE CHIETI CHIETI(CHIETI) | Istituto Zooprofilattico Sperimentale dell'Abruzzo e Molise "G. Caporale" | Lorusso A, Marcacci M, Di Domenico M, Ancora M, Curini V, Di Lollo Valeria, Mangone I, Rinaldi A, Delli Compagni E, Scialabba S, Caporale M, Di Pasquale A, Cammà C, Puglia I, Calistri P, Savini G |
| EPI_ISL_1392691, EPI_ISL_1392693 | SIESP CHIETI - DRIVE IN GISSI GISSI(CHIETI) | Istituto Zooprofilattico Sperimentale dell'Abruzzo e Molise "G. Caporale" | Lorusso A, Marcacci M, Di Domenico M, Ancora M, Curini V, Di Lollo Valeria, Mangone I, Rinaldi A, Delli Compagni E, Scialabba S, Caporale M, Di Pasquale A, Cammà C, Puglia I, Calistri P, Savini G |
| EPI_ISL_1392694 | SIESP CHIETI - DRIVE IN VASTO VASTO(CHIETI) | Istituto Zooprofilattico Sperimentale dell'Abruzzo e Molise "G. Caporale" | Lorusso A, Marcacci M, Di Domenico M, Ancora M, Curini V, Di Lollo Valeria, Mangone I, Rinaldi A, Delli Compagni E, Scialabba S, Caporale M, Di Pasquale A, Cammà C, Puglia I, Calistri P, Savini G |
| EPI_ISL_1392695, EPI_ISL_1392696, EPI_ISL_1392697 | SIESP CHIETI - DRIVE IN LANCIANO LANCIANO(CHIETI) | Istituto Zooprofilattico Sperimentale dell'Abruzzo e Molise "G. Caporale" | Lorusso A, Marcacci M, Di Domenico M, Ancora M, Curini V, Di Lollo Valeria, Mangone I, Rinaldi A, Delli Compagni E, Scialabba S, Caporale M, Di Pasquale A, Cammà C, Puglia I, Calistri P, Savini G |
| EPI_ISL_1392698, EPI_ISL_1392699, EPI_ISL_1392700, EPI_ISL_1392701 | SIESP CHIETI - DRIVE IN CHIETI CHIETI(CHIETI) | Istituto Zooprofilattico Sperimentale dell'Abruzzo e Molise "G. Caporale" | Lorusso A, Marcacci M, Di Domenico M, Ancora M, Curini V, Di Lollo Valeria, Mangone I, Rinaldi A, Delli Compagni E, Scialabba S, Caporale M, Di Pasquale A, Cammà C, Puglia I, Calistri P, Savini G |
| EPI_ISL_1392702 | SIESP CHIETI - DRIVE IN ORTONA ORTONA(CHIETI) | Istituto Zooprofilattico Sperimentale dell'Abruzzo e Molise "G. Caporale" | Lorusso A, Marcacci M, Di Domenico M, Ancora M, Curini V, Di Lollo Valeria, Mangone I, Rinaldi A, Delli Compagni E, Scialabba S, Caporale M, Di Pasquale A, Cammà C, Puglia I, Calistri P, Savini G |
| EPI_ISL_1392703, EPI_ISL_1392704, EPI_ISL_1392705, EPI_ISL_1392706, EPI_ISL_1392707, EPI_ISL_1392708, EPI_ISL_1392709 | SIESP DIPARTIMENTO DI PREVENZIONE TERAMO TERAMO(TERAMO) | Istituto Zooprofilattico Sperimentale dell'Abruzzo e Molise "G. Caporale" | Lorusso A, Marcacci M, Di Domenico M, Ancora M, Curini V, Di Lollo Valeria, Mangone I, Rinaldi A, Delli Compagni E, Scialabba S, Caporale M, Di Pasquale A, Cammà C, Puglia I, Calistri P, Savini G |
| EPI_ISL_1392710 | SIESP CHIETI - DRIVE IN CHIETI CHIETI(CHIETI) | Istituto Zooprofilattico Sperimentale dell'Abruzzo e Molise "G. Caporale" | Lorusso A, Marcacci M, Di Domenico M, Ancora M, Curini V, Di Lollo Valeria, Mangone I, Rinaldi A, Delli Compagni E, Scialabba S, Caporale M, Di Pasquale A, Cammà C, Puglia I, Calistri P, Savini G |
| EPI_ISL_1392711, EPI_ISL_1392712 | DIP. PREV. AVEZZANO SERVIZIO DI IGIENE EPIDEMIOLOGIA E SANITA' PUBBLICA AVEZZANO(L'AQUILA) | Istituto Zooprofilattico Sperimentale dell'Abruzzo e Molise "G. Caporale" | Lorusso A, Marcacci M, Di Domenico M, Ancora M, Curini V, Di Lollo Valeria, Mangone I, Rinaldi A, Delli Compagni E, Scialabba S, Caporale M, Di Pasquale A, Cammà C, Puglia I, Calistri P, Savini G |
| EPI_ISL_1392713 | SIESP DIPARTIMENTO DI PREVENZIONE TERAMO TERAMO(TERAMO) | Istituto Zooprofilattico Sperimentale dell'Abruzzo e Molise "G. Caporale" | Lorusso A, Marcacci M, Di Domenico M, Ancora M, Curini V, Di Lollo Valeria, Mangone I, Rinaldi A, Delli Compagni E, Scialabba S, Caporale M, Di Pasquale A, Cammà C, Puglia I, Calistri P, Savini G |
| EPI_ISL_1394290, EPI_ISL_1394291, EPI_ISL_1394292, EPI_ISL_1394293, EPI_ISL_1394294, EPI_ISL_1394295, EPI_ISL_1394296, EPI_ISL_1394297, EPI_ISL_1394298, EPI_ISL_1394299, EPI_ISL_1394300, EPI_ISL_1394301, EPI_ISL_1394302, EPI_ISL_1394303, EPI_ISL_1394304, EPI_ISL_1394305, EPI_ISL_1394306, EPI_ISL_1394307, EPI_ISL_1394308, EPI_ISL_1394310, EPI_ISL_1394311, EPI_ISL_1394312, EPI_ISL_1394313 | | | |
| see above | Hospital for Infectious Diseases, Molecular Diagnostics Laboratory, Warsaw, Poland | 1. Virogenetics Laboratory of Virology, Maopolska Centre of Biotechnology, Jagiellonian University. 2. Intercollegiate | Lukasz Rabalski, Maciej Kosinski, Natalia Mazur-Panasiuk, Aneta Kopacz, Piotr Zabek, Tomasz Dyda, Andrzej Horban, Krzysztof Pyrc, Krystyna Bienkowska-Szewczyk |

| | | | |
|---|---|---|---|
| | | Faculty of Biotechnology University of Gdansk and Medical University of Gdansk | |
| EPI_ISL_1394314, EPI_ISL_1394315, EPI_ISL_1394316, EPI_ISL_1394317, EPI_ISL_1394318, EPI_ISL_1394319, EPI_ISL_1394320, EPI_ISL_1394322, EPI_ISL_1394323, EPI_ISL_1394324, EPI_ISL_1394325, EPI_ISL_1394326, EPI_ISL_1394327, EPI_ISL_1394328, EPI_ISL_1394329, EPI_ISL_1394330, EPI_ISL_1394331, EPI_ISL_1394332, EPI_ISL_1394333, EPI_ISL_1394335, EPI_ISL_1394336, EPI_ISL_1394337 | | | |
| see above | Hospital for Infectious Diseases, Molecular Diagnostics Laboratory, Warsaw, Poland | 1. Virogenetics Laboratory of Virology, Maopolska Centre of Biotechnology, Jagiellonian University. 2. Intercollegiate Faculty of Biotechnology University of Gdansk and Medical University of Gdansk | Lukasz Rabalski, Maciej Kosinski, Karolina Gackowska, Natalia Mazur-Panasiuk, Aneta Kopacz, Piotr Zabek, Tomasz Dyda, Andrzej Horban, Krzysztof Pyrc, Krystyna Bienkowska-Szewczyk |
| EPI_ISL_1396651 | SIESP DIPARTIMENTO DI PREVENZIONE TERAMO | Istituto Zooprofilattico Sperimentale dell'Abruzzo e Molise "G. Caporale" | Lorusso A, Marcacci M, Di Domenico M, Ancora M, Curini V, Di Lollo Valeria, Mangone I, Rinaldi A, Scialabba S, Caporale M, Di Pasquale A, Cammà C, Puglia I, Calistri P, Savini G |
| EPI_ISL_1396653 | SIESP DIPARTIMENTO DI PREVENZIONE CHIETI | Istituto Zooprofilattico Sperimentale dell'Abruzzo e Molise "G. Caporale" | Lorusso A, Marcacci M, Di Domenico M, Ancora M, Curini V, Di Lollo Valeria, Mangone I, Rinaldi A, Scialabba S, Caporale M, Di Pasquale A, Cammà C, Puglia I, Calistri P, Savini G |
| EPI_ISL_1396667, EPI_ISL_1396669, EPI_ISL_1396670, EPI_ISL_1396671, EPI_ISL_1396672, EPI_ISL_1396673, EPI_ISL_1396674, EPI_ISL_1396686, EPI_ISL_1396687, EPI_ISL_1396688, EPI_ISL_1396694, EPI_ISL_1396695, EPI_ISL_1396696, EPI_ISL_1396697, EPI_ISL_1396698, EPI_ISL_1396699, EPI_ISL_1396700, EPI_ISL_1396701, EPI_ISL_1396702, EPI_ISL_1396705, EPI_ISL_1396706, EPI_ISL_1396707, EPI_ISL_1396714, EPI_ISL_1396715, EPI_ISL_1396716, EPI_ISL_1396717, EPI_ISL_1396719, EPI_ISL_1396722, EPI_ISL_1396723, EPI_ISL_1396724, EPI_ISL_1396725, EPI_ISL_1396726, EPI_ISL_1396727, EPI_ISL_1396728, EPI_ISL_1396729, EPI_ISL_1396730, EPI_ISL_1396731, EPI_ISL_1396734 | | | |
| see above | CHU Lille | CHU Lille - Laboratoire de Virologie | AIT YAHYA Emilie, ALIDJINOU Enagnon Kazali, BOCKET Laurence, CREPIN Michel, DEMAY Christophe, ENGELMANN Ilka, GEFFROY Sandrine, GUIGON Aurélie, LAMBERT Valérie, LAZREK Mouna, NOBILLIAUX Florian, PREVOST Brigitte, THUILLIER Caroline, TINEZ Claire ,TCHANTCHOU NJOSSE YANICK |
| EPI_ISL_1396756, EPI_ISL_1396759, EPI_ISL_1396761, EPI_ISL_1396762, EPI_ISL_1396764, EPI_ISL_1396765, EPI_ISL_1396770, EPI_ISL_1396771, EPI_ISL_1396772, EPI_ISL_1396773, EPI_ISL_1396776, EPI_ISL_1396799, EPI_ISL_1396803 | | | |
| see above | Hôpital Henri Mondor | Department of Virology, Henri Mondor University Hospital, Assistance Publique Hôpitaux de Paris, Université Paris-Est Créteil, INSERM U955 | Christophe Rodriguez, Slim Fourati, Vanessa Demontant, Guillaume Gricourt, Melissa N'Debi, Alexandre Soulier, Elisabeth Trawinski, Jean-Michel Pawlotsky |
| EPI_ISL_1399261 | LESP Guanajuato | Instituto de Diagnostico y Referencia Epidemiologicos (INDRE) | Claudia Wong-Arambula, Abril Rodriguez-Maldonado, Vanessa Rivero-Arredondo, Ariadna Medina-Benitez, Joaquin Quiroz-Mercado, Sergio Rangel-Guerrero, Natividad Cruz-Ortiz, Tatiana Nunez-Garcia, Gisela Barrera-Badillo, Lucia Hernandez-Rivas, Irma Lopez-Martinez, Ernesto Ramirez-Gonzalez. |
| EPI_ISL_1399265, EPI_ISL_1399266, EPI_ISL_1399267, EPI_ISL_1399268 | LESP Nuevo Leon | Instituto de Diagnostico y Referencia Epidemiologicos (INDRE) | Claudia Wong-Arambula, Abril Rodriguez-Maldonado, Vanessa Rivero-Arredondo, Ariadna Medina-Benitez, Joaquin Quiroz-Mercado, Sergio Rangel-Guerrero, Natividad Cruz-Ortiz, Tatiana Nunez-Garcia, Gisela Barrera-Badillo, Lucia Hernandez-Rivas, Irma Lopez-Martinez, Ernesto Ramirez-Gonzalez. |
| EPI_ISL_1399271 | LESP Chihuahua | Instituto de Diagnostico y Referencia Epidemiologicos (INDRE) | Claudia Wong-Arambula, Abril Rodriguez-Maldonado, Vanessa Rivero-Arredondo, Ariadna Medina-Benitez, Joaquin Quiroz-Mercado, Sergio Rangel-Guerrero, Natividad Cruz-Ortiz, Tatiana Nunez-Garcia, Gisela Barrera-Badillo, Lucia Hernandez-Rivas, Irma Lopez-Martinez, Ernesto Ramirez-Gonzalez. |
| EPI_ISL_1399272 | LESP Queretaro | Instituto de Diagnostico y Referencia Epidemiologicos (INDRE) | Claudia Wong-Arambula, Abril Rodriguez-Maldonado, Vanessa Rivero-Arredondo, Ariadna Medina-Benitez, Joaquin Quiroz-Mercado, Sergio Rangel-Guerrero, Natividad Cruz-Ortiz, Tatiana Nunez-Garcia, Gisela Barrera-Badillo, Lucia Hernandez-Rivas, Irma Lopez-Martinez, Ernesto Ramirez-Gonzalez. |
| EPI_ISL_1399273 | LESP Nayarit | Instituto de Diagnostico y Referencia Epidemiologicos (INDRE) | Claudia Wong-Arambula, Abril Rodriguez-Maldonado, Vanessa Rivero-Arredondo, Ariadna Medina-Benitez, Joaquin Quiroz-Mercado, Sergio Rangel-Guerrero, Natividad Cruz-Ortiz, Tatiana Nunez-Garcia, Gisela Barrera-Badillo, Lucia Hernandez-Rivas, Irma Lopez-Martinez, Ernesto Ramirez-Gonzalez. |
| EPI_ISL_1399278 | LESP Coahuila | Instituto de Diagnostico y Referencia Epidemiologicos (INDRE) | Claudia Wong-Arambula, Abril Rodriguez-Maldonado, Vanessa Rivero-Arredondo, Ariadna Medina-Benitez, Joaquin Quiroz-Mercado, Sergio Rangel-Guerrero, Natividad Cruz-Ortiz, Tatiana Nunez-Garcia, Gisela Barrera-Badillo, Lucia Hernandez-Rivas, Irma Lopez-Martinez, Ernesto Ramirez-Gonzalez. |
| EPI_ISL_1399279 | LESP Veracruz | Instituto de Diagnostico y Referencia Epidemiologicos (INDRE) | Claudia Wong-Arambula, Abril Rodriguez-Maldonado, Vanessa Rivero-Arredondo, Ariadna Medina-Benitez, Joaquin Quiroz-Mercado, Sergio Rangel-Guerrero, Natividad Cruz-Ortiz, Tatiana Nunez-Garcia, Gisela Barrera-Badillo, Lucia Hernandez-Rivas, Irma Lopez-Martinez, Ernesto Ramirez-Gonzalez. |
| EPI_ISL_1399280 | LESP Baja California | Instituto de Diagnostico y Referencia Epidemiologicos (INDRE) | Claudia Wong-Arambula, Abril Rodriguez-Maldonado, Vanessa Rivero-Arredondo, Ariadna Medina-Benitez, Joaquin Quiroz-Mercado, Sergio Rangel-Guerrero, Natividad Cruz-Ortiz, Tatiana Nunez-Garcia, Gisela Barrera-Badillo, Lucia Hernandez-Rivas, Irma Lopez-Martinez, Ernesto Ramirez-Gonzalez. |
| EPI_ISL_1399281 | LESP Ciudad de Mexico | Instituto de Diagnostico y Referencia Epidemiologicos (INDRE) | Claudia Wong-Arambula, Abril Rodriguez-Maldonado, Vanessa Rivero-Arredondo, Ariadna Medina-Benitez, Joaquin Quiroz-Mercado, Sergio Rangel-Guerrero, Natividad Cruz-Ortiz, Tatiana Nunez-Garcia, Gisela Barrera-Badillo, Lucia Hernandez-Rivas, Irma Lopez-Martinez, Ernesto Ramirez-Gonzalez. |
| EPI_ISL_1400298 | LESP Jalisco | Instituto de Diagnostico y Referencia Epidemiologicos (INDRE) | Claudia Wong-Arambula, Abril Rodriguez-Maldonado, Vanessa Rivero-Arredondo, Ariadna Medina-Benitez, Joaquin Quiroz-Mercado, Sergio Rangel-Guerrero, Natividad Cruz-Ortiz, Tatiana Nunez-Garcia, Gisela Barrera-Badillo, Lucia Hernandez-Rivas, Irma Lopez-Martinez, Ernesto Ramirez-Gonzalez. |
| EPI_ISL_1400304 | LESP Guanajuato | Instituto de Diagnostico y Referencia Epidemiologicos (INDRE) | Claudia Wong-Arambula, Abril Rodriguez-Maldonado, Vanessa Rivero-Arredondo, Ariadna Medina-Benitez, Joaquin Quiroz-Mercado, Sergio Rangel-Guerrero, Natividad Cruz-Ortiz, Tatiana Nunez-Garcia, Gisela Barrera-Badillo, Lucia Hernandez-Rivas, Irma Lopez-Martinez, Ernesto Ramirez-Gonzalez. |
| EPI_ISL_1400305, EPI_ISL_1400306, EPI_ISL_1400307, EPI_ISL_1400308, EPI_ISL_1400309 | LESP Morelos | Instituto de Diagnostico y Referencia Epidemiologicos (INDRE) | Claudia Wong-Arambula, Abril Rodriguez-Maldonado, Vanessa Rivero-Arredondo, Ariadna Medina-Benitez, Joaquin Quiroz-Mercado, Sergio Rangel-Guerrero, Natividad Cruz-Ortiz, Tatiana Nunez-Garcia, Gisela Barrera-Badillo, Lucia Hernandez-Rivas, Irma Lopez-Martinez, Ernesto Ramirez-Gonzalez. |
| EPI_ISL_1400311, EPI_ISL_1400312, EPI_ISL_1400313 | LESP Puebla | Instituto de Diagnostico y Referencia Epidemiologicos (INDRE) | Claudia Wong-Arambula, Abril Rodriguez-Maldonado, Vanessa Rivero-Arredondo, Ariadna Medina-Benitez, Joaquin Quiroz-Mercado, Sergio Rangel-Guerrero, Natividad Cruz-Ortiz, Tatiana Nunez-Garcia, Gisela Barrera-Badillo, Lucia Hernandez-Rivas, Irma Lopez-Martinez, Ernesto Ramirez-Gonzalez. |
| EPI_ISL_1400314, EPI_ISL_1400315, EPI_ISL_1400317, EPI_ISL_1400318, EPI_ISL_1400319, EPI_ISL_1400320, EPI_ISL_1400321, EPI_ISL_1400322, EPI_ISL_1400323, EPI_ISL_1400324, EPI_ISL_1400326, EPI_ISL_1400327, EPI_ISL_1400328, EPI_ISL_1400329, EPI_ISL_1400331, EPI_ISL_1400332, EPI_ISL_1400333, EPI_ISL_1400334, EPI_ISL_1400335, EPI_ISL_1400336, EPI_ISL_1400337, EPI_ISL_1400338, EPI_ISL_1400339, EPI_ISL_1400341, EPI_ISL_1400342, EPI_ISL_1400343, EPI_ISL_1400344, EPI_ISL_1400345, EPI_ISL_1400346, EPI_ISL_1400347, EPI_ISL_1400348, EPI_ISL_1400349 | | | |
| see above | LESP Guanajuato | Instituto de Diagnostico y Referencia Epidemiologicos (INDRE) | Claudia Wong-Arambula, Abril Rodriguez-Maldonado, Vanessa Rivero-Arredondo, Ariadna Medina-Benitez, Joaquin Quiroz-Mercado, Sergio Rangel-Guerrero, Natividad Cruz-Ortiz, Tatiana Nunez-Garcia, Gisela Barrera-Badillo, Lucia Hernandez-Rivas, Irma Lopez-Martinez, Ernesto Ramirez-Gonzalez. |
| EPI_ISL_1401101, EPI_ISL_1401120, EPI_ISL_1401121, EPI_ISL_1401122, EPI_ISL_1401123, EPI_ISL_1401124, EPI_ISL_1401125, EPI_ISL_1401126, EPI_ISL_1401127, EPI_ISL_1401128, EPI_ISL_1401129, EPI_ISL_1401130, EPI_ISL_1401131, EPI_ISL_1401132, EPI_ISL_1401133, EPI_ISL_1401134, EPI_ISL_1401135, EPI_ISL_1401136, EPI_ISL_1401137, EPI_ISL_1401139, EPI_ISL_1401140, EPI_ISL_1401141, EPI_ISL_1401142, EPI_ISL_1401143, EPI_ISL_1401144, EPI_ISL_1401145, EPI_ISL_1401146, EPI_ISL_1401147, EPI_ISL_1401148, EPI_ISL_1401149, EPI_ISL_1401150, EPI_ISL_1401151, EPI_ISL_1401152, EPI_ISL_1401153, EPI_ISL_1401154, EPI_ISL_1401155, EPI_ISL_1401156, EPI_ISL_1401157, EPI_ISL_1401158, EPI_ISL_1401159, EPI_ISL_1401161, EPI_ISL_1401162, EPI_ISL_1401163, EPI_ISL_1401164, EPI_ISL_1401165, EPI_ISL_1401166, EPI_ISL_1401167, EPI_ISL_1401168, EPI_ISL_1401169, EPI_ISL_1401170, EPI_ISL_1401171, EPI_ISL_1401172, EPI_ISL_1401173, EPI_ISL_1401174, EPI_ISL_1401175, EPI_ISL_1401176, EPI_ISL_1401177, EPI_ISL_1401178, EPI_ISL_1401179, EPI_ISL_1401180, EPI_ISL_1401182, EPI_ISL_1401183, EPI_ISL_1401184, EPI_ISL_1401185, EPI_ISL_1401186, EPI_ISL_1401187, EPI_ISL_1401188, EPI_ISL_1401189, EPI_ISL_1401190, EPI_ISL_1401191, EPI_ISL_1401192, EPI_ISL_1401197, EPI_ISL_1401199, EPI_ISL_1401200, EPI_ISL_1401201, EPI_ISL_1401202, EPI_ISL_1401203, EPI_ISL_1401204, EPI_ISL_1401205, EPI_ISL_1401206, EPI_ISL_1401207, EPI_ISL_1401209, EPI_ISL_1401210, EPI_ISL_1401211, EPI_ISL_1401212, EPI_ISL_1401213, EPI_ISL_1401214, EPI_ISL_1401215, EPI_ISL_1401216, EPI_ISL_1401217, EPI_ISL_1401218, EPI_ISL_1401219, EPI_ISL_1401220, EPI_ISL_1401221, EPI_ISL_1401222, EPI_ISL_1401223, EPI_ISL_1401224, EPI_ISL_1401225, EPI_ISL_1401226, EPI_ISL_1401227, EPI_ISL_1401228, EPI_ISL_1401229, EPI_ISL_1401231, EPI_ISL_1401232, EPI_ISL_1401233, EPI_ISL_1401234, EPI_ISL_1401235, EPI_ISL_1401236, EPI_ISL_1401237, EPI_ISL_1401238, EPI_ISL_1401240, EPI_ISL_1401241, EPI_ISL_1401242, EPI_ISL_1401243, EPI_ISL_1401244, EPI_ISL_1401245, EPI_ISL_1401246, EPI_ISL_1401247, EPI_ISL_1401248, EPI_ISL_1401249, EPI_ISL_1401250, EPI_ISL_1401251, EPI_ISL_1401252, EPI_ISL_1401253, EPI_ISL_1401254, EPI_ISL_1401255, EPI_ISL_1401256, EPI_ISL_1401257, EPI_ISL_1401258, EPI_ISL_1401259, EPI_ISL_1401261, EPI_ISL_1401262, EPI_ISL_1401263, EPI_ISL_1401264, EPI_ISL_1401265, EPI_ISL_1401266, EPI_ISL_1401267, EPI_ISL_1401268, EPI_ISL_1401269, EPI_ISL_1401270, EPI_ISL_1401271, EPI_ISL_1401272, EPI_ISL_1401273, EPI_ISL_1401274, EPI_ISL_1401275, EPI_ISL_1401276, EPI_ISL_1401277, EPI_ISL_1401278, EPI_ISL_1401279, EPI_ISL_1401280, EPI_ISL_1401281, EPI_ISL_1401295, EPI_ISL_1401296, | | | |

| | | | |
|---|---|---|---|
| EPI_ISL_1401297, EPI_ISL_1401298, EPI_ISL_1401299, EPI_ISL_1401300, EPI_ISL_1401301, EPI_ISL_1401302, EPI_ISL_1401303, EPI_ISL_1401304, EPI_ISL_1401305, EPI_ISL_1401306, EPI_ISL_1401307, EPI_ISL_1401308, EPI_ISL_1401309, EPI_ISL_1401310, EPI_ISL_1401311, EPI_ISL_1401312, EPI_ISL_1401313, EPI_ISL_1401314, EPI_ISL_1401315, EPI_ISL_1401316, EPI_ISL_1401317, EPI_ISL_1401318, EPI_ISL_1401319, EPI_ISL_1401320, EPI_ISL_1401321, EPI_ISL_1401322, EPI_ISL_1401323, EPI_ISL_1401324, EPI_ISL_1401325, EPI_ISL_1401326, EPI_ISL_1401327, EPI_ISL_1401328, EPI_ISL_1401331, EPI_ISL_1401334, EPI_ISL_1401335, EPI_ISL_1401336, EPI_ISL_1401337, EPI_ISL_1401338, EPI_ISL_1401339, EPI_ISL_1401340, EPI_ISL_1401341, EPI_ISL_1401342, EPI_ISL_1401343, EPI_ISL_1401344, EPI_ISL_1401345, EPI_ISL_1401346, EPI_ISL_1401347, EPI_ISL_1401348, EPI_ISL_1401349, EPI_ISL_1401350, EPI_ISL_1401351, EPI_ISL_1401352, EPI_ISL_1401353, EPI_ISL_1401354, EPI_ISL_1401355, EPI_ISL_1401356, EPI_ISL_1401360, EPI_ISL_1401361, EPI_ISL_1401363, EPI_ISL_1401364, EPI_ISL_1401365, EPI_ISL_1401366, EPI_ISL_1401367, EPI_ISL_1401368, EPI_ISL_1401369, EPI_ISL_1401370, EPI_ISL_1401372, EPI_ISL_1401373, EPI_ISL_1401374, EPI_ISL_1401375, EPI_ISL_1401376, EPI_ISL_1401377, EPI_ISL_1401378, EPI_ISL_1401379, EPI_ISL_1401380, EPI_ISL_1401381, EPI_ISL_1401382, EPI_ISL_1401383, EPI_ISL_1401384, EPI_ISL_1401385, EPI_ISL_1401386, EPI_ISL_1401389, EPI_ISL_1401390, EPI_ISL_1401391, EPI_ISL_1401392, EPI_ISL_1401393, EPI_ISL_1401395, EPI_ISL_1401396, EPI_ISL_1401397, EPI_ISL_1401398, EPI_ISL_1401399, EPI_ISL_1401400, EPI_ISL_1401401, EPI_ISL_1401402, EPI_ISL_1401403, EPI_ISL_1401404, EPI_ISL_1401405, EPI_ISL_1401406, EPI_ISL_1401407, EPI_ISL_1401408, EPI_ISL_1401409, EPI_ISL_1401410, EPI_ISL_1401411, EPI_ISL_1401412, EPI_ISL_1401413, EPI_ISL_1401414, EPI_ISL_1401415, EPI_ISL_1401416, EPI_ISL_1401417, EPI_ISL_1401418, EPI_ISL_1401420, EPI_ISL_1401421, EPI_ISL_1401422, EPI_ISL_1401424, EPI_ISL_1401425, EPI_ISL_1401426, EPI_ISL_1401427, EPI_ISL_1401428, EPI_ISL_1401430, EPI_ISL_1401431, EPI_ISL_1401432, EPI_ISL_1401433, EPI_ISL_1401434, EPI_ISL_1401435, EPI_ISL_1401443, EPI_ISL_1401444, EPI_ISL_1401445, EPI_ISL_1401446, EPI_ISL_1401447, EPI_ISL_1401448, EPI_ISL_1401449, EPI_ISL_1401450, EPI_ISL_1401451, EPI_ISL_1401452, EPI_ISL_1401453, EPI_ISL_1401454, EPI_ISL_1401455, EPI_ISL_1401456, EPI_ISL_1401457, EPI_ISL_1401458, EPI_ISL_1401459, EPI_ISL_1401460, EPI_ISL_1401461, EPI_ISL_1401462, EPI_ISL_1401463, EPI_ISL_1401464 | | | |
| see above | National Center of Infectious and Parasitic Diseases | National Center of Infectious and Parasitic Diseases | Alexiev et al |
| EPI_ISL_1402408, EPI_ISL_1402409, EPI_ISL_1402410 | AZ Zeno | AZ SINT-JAN BRUGGE | Marijke Reynders, Patrick Descheemaeker, Laurien Hoornaert, Thomas Van Landschoot, Jorn Hellemans |
| EPI_ISL_1402411, EPI_ISL_1402412, EPI_ISL_1402413, EPI_ISL_1402417 | AZ Sint-Jan | AZ SINT-JAN BRUGGE | Marijke Reynders, Patrick Descheemaeker, Laurien Hoornaert, Thomas Van Landschoot, Jorn Hellemans |
| EPI_ISL_1403384 | AZ Klina | AZ Klina | Carl Vael, Lynsey Berckmans |
| EPI_ISL_1403422, EPI_ISL_1403423, EPI_ISL_1403424, EPI_ISL_1403425, EPI_ISL_1403426 | AZ Klina | AZ Klina | Carl Vael - Lynsey Berckmans |
| EPI_ISL_1403502 | SIESP DIPARTIMENTO DI PREVENZIONE TERAMO TERAMO(TERAMO) | Istituto Zooprofilattico Sperimentale dell'Abruzzo e Molise "G. Caporale" | Lorusso A, Marcacci M, Di Domenico M, Ancora M, Curini V, Mangone I, Delli Compagni E, Rinaldi A, Scialabba S, Di Pasquale A, Cammà C, Puglia I, Calistri P, Savini G |
| EPI_ISL_1403514, EPI_ISL_1403523, EPI_ISL_1403524 | SIESP SULMONA SULMONA(L'AQUILA) | Istituto Zooprofilattico Sperimentale dell'Abruzzo e Molise "G. Caporale" | Lorusso A, Marcacci M, Di Domenico M, Ancora M, Curini V, Mangone I, Delli Compagni E, Rinaldi A, Scialabba S, Di Pasquale A, Cammà C, Puglia I, Calistri P, Savini G |
| EPI_ISL_1403549, EPI_ISL_1403554, EPI_ISL_1403556, EPI_ISL_1403557, EPI_ISL_1403558 | SIESP DIPARTIMENTO DI PREVENZIONE TERAMO TERAMO(TERAMO) | Istituto Zooprofilattico Sperimentale dell'Abruzzo e Molise "G. Caporale" | Lorusso A, Marcacci M, Di Domenico M, Ancora M, Curini V, Mangone I, Delli Compagni E, Rinaldi A, Scialabba S, Di Pasquale A, Cammà C, Puglia I, Calistri P, Savini G |
| EPI_ISL_1403560 | SIESP CHIETI - DRIVE IN LANCIANO LANCIANO(CHIETI) | Istituto Zooprofilattico Sperimentale dell'Abruzzo e Molise "G. Caporale" | Lorusso A, Marcacci M, Di Domenico M, Ancora M, Curini V, Mangone I, Delli Compagni E, Rinaldi A, Scialabba S, Di Pasquale A, Cammà C, Puglia I, Calistri P, Savini G |
| EPI_ISL_1403564, EPI_ISL_1403572, EPI_ISL_1403573, EPI_ISL_1403582, EPI_ISL_1403602, EPI_ISL_1403603 | SIESP DIPARTIMENTO DI PREVENZIONE TERAMO TERAMO(TERAMO) | Istituto Zooprofilattico Sperimentale dell'Abruzzo e Molise "G. Caporale" | Lorusso A, Marcacci M, Di Domenico M, Ancora M, Curini V, Mangone I, Delli Compagni E, Rinaldi A, Scialabba S, Di Pasquale A, Cammà C, Puglia I, Calistri P, Savini G |
| EPI_ISL_1403606 | SIESP CHIETI - DRIVE IN LANCIANO LANCIANO(CHIETI) | Istituto Zooprofilattico Sperimentale dell'Abruzzo e Molise "G. Caporale" | Lorusso A, Marcacci M, Di Domenico M, Ancora M, Curini V, Mangone I, Delli Compagni E, Rinaldi A, Scialabba S, Di Pasquale A, Cammà C, Puglia I, Calistri P, Savini G |
| EPI_ISL_1404374 | Cerballliance Les Clayes | Cerba Lab | Lecorche E, Trombert-Paolantoni S, Olivi M, Benazra M, Zimmer S, Haim-Boukobza S |
| EPI_ISL_1404379, EPI_ISL_1404380, EPI_ISL_1404381, EPI_ISL_1404382, EPI_ISL_1404388, EPI_ISL_1404389, EPI_ISL_1404390, EPI_ISL_1404391, EPI_ISL_1404392, EPI_ISL_1404393, EPI_ISL_1404394 | | | |
| see above | Klinisch Laboratorium ZNA | Klinisch Laboratorium ZNA | Verstrepen et al. |
| EPI_ISL_1405900, EPI_ISL_1405901, EPI_ISL_1405902, EPI_ISL_1405903 | LESP Guanajuato | Instituto de Diagnostico y Referencia Epidemiologicos (INDRE) | Claudia Wong-Arambula, Abril Rodriguez-Maldonado, Vanessa Rivero-Arredondo, Ariadna Medina-Benitez, Joaquin Quiroz-Mercado, Sergio Rangel-Guerrero, Natividad Cruz-Ortiz, Tatiana Nunez-Garcia, Gisela Barrera-Badillo, Lucia Hernandez-Rivas, Irma Lopez-Martinez, Ernesto Ramirez-Gonzalez. |
| EPI_ISL_1405905 | LESP Chihuahua | Instituto de Diagnostico y Referencia Epidemiologicos (INDRE) | Claudia Wong-Arambula, Abril Rodriguez-Maldonado, Vanessa Rivero-Arredondo, Ariadna Medina-Benitez, Joaquin Quiroz-Mercado, Sergio Rangel-Guerrero, Natividad Cruz-Ortiz, Tatiana Nunez-Garcia, Gisela Barrera-Badillo, Lucia Hernandez-Rivas, Irma Lopez-Martinez, Ernesto Ramirez-Gonzalez. |
| EPI_ISL_1405906 | LESP Guanajuato | Instituto de Diagnostico y Referencia Epidemiologicos (INDRE) | Claudia Wong-Arambula, Abril Rodriguez-Maldonado, Vanessa Rivero-Arredondo, Ariadna Medina-Benitez, Joaquin Quiroz-Mercado, Sergio Rangel-Guerrero, Natividad Cruz-Ortiz, Tatiana Nunez-Garcia, Gisela Barrera-Badillo, Lucia Hernandez-Rivas, Irma Lopez-Martinez, Ernesto Ramirez-Gonzalez. |
| EPI_ISL_1405908 | LESP Jalisco | Instituto de Diagnostico y Referencia Epidemiologicos (INDRE) | Claudia Wong-Arambula, Abril Rodriguez-Maldonado, Vanessa Rivero-Arredondo, Ariadna Medina-Benitez, Joaquin Quiroz-Mercado, Sergio Rangel-Guerrero, Natividad Cruz-Ortiz, Tatiana Nunez-Garcia, Gisela Barrera-Badillo, Lucia Hernandez-Rivas, Irma Lopez-Martinez, Ernesto Ramirez-Gonzalez. |
| EPI_ISL_1405909, EPI_ISL_1405910 | LESP Guanajuato | Instituto de Diagnostico y Referencia Epidemiologicos (INDRE) | Claudia Wong-Arambula, Abril Rodriguez-Maldonado, Vanessa Rivero-Arredondo, Ariadna Medina-Benitez, Joaquin Quiroz-Mercado, Sergio Rangel-Guerrero, Natividad Cruz-Ortiz, Tatiana Nunez-Garcia, Gisela Barrera-Badillo, Lucia Hernandez-Rivas, Irma Lopez-Martinez, Ernesto Ramirez-Gonzalez. |
| EPI_ISL_1405911 | LESP Jalisco | Instituto de Diagnostico y Referencia Epidemiologicos (INDRE) | Claudia Wong-Arambula, Abril Rodriguez-Maldonado, Vanessa Rivero-Arredondo, Ariadna Medina-Benitez, Joaquin Quiroz-Mercado, Sergio Rangel-Guerrero, Natividad Cruz-Ortiz, Tatiana Nunez-Garcia, Gisela Barrera-Badillo, Lucia Hernandez-Rivas, Irma Lopez-Martinez, Ernesto Ramirez-Gonzalez. |
| EPI_ISL_1405912, EPI_ISL_1405915 | LESP Guanajuato | Instituto de Diagnostico y Referencia Epidemiologicos (INDRE) | Claudia Wong-Arambula, Abril Rodriguez-Maldonado, Vanessa Rivero-Arredondo, Ariadna Medina-Benitez, Joaquin Quiroz-Mercado, Sergio Rangel-Guerrero, Natividad Cruz-Ortiz, Tatiana Nunez-Garcia, Gisela Barrera-Badillo, Lucia Hernandez-Rivas, Irma Lopez-Martinez, Ernesto Ramirez-Gonzalez. |
| EPI_ISL_1405966, EPI_ISL_1406078 | Institute for Medical Research, Infectious Disease Research Centre, National Institutes of Health, Ministry of Health Malaysia | Institute for Medical Research, Infectious Disease Research Centre, National Institutes of Health, Ministry of Health Malaysia | Suppiah J, Kamel K, Mohd Zawawi Z, Ramly N, Robert F, Thayan R |
| EPI_ISL_1406145, EPI_ISL_1406147 | Institut de Virologie du CHU de Strasbourg | Institut de Virologie du CHU de Strasbourg | Fafi-Kremer Samira, Gallais Floriane, Gantner Pierre, Laugel Elodie, Solis Morgane, Velay Aurélie, Wendling Marie-Josée |
| EPI_ISL_1406148, EPI_ISL_1406187, EPI_ISL_1406192, EPI_ISL_1406194, EPI_ISL_1406197, EPI_ISL_1406199, EPI_ISL_1406201, EPI_ISL_1406204, EPI_ISL_1406242, EPI_ISL_1406244 | Institute for Medical Research, Infectious Disease Research Centre, National Institutes of Health, Ministry of Health Malaysia | Institute for Medical Research, Infectious Disease Research Centre, National Institutes of Health, Ministry of Health Malaysia | Suppiah J, Kamel K, Mohd Zawawi Z, Ramly N, Robert F, Thayan R |
| EPI_ISL_1406245 | Hospital Monte Naranco | Laboratorio de Virología HUCA | Sandoval M, Castelló C, Gómez de Oña J, Boga JA, Rojo S, Alvarez-Arguelles ME, Abreu F, Costales I, Perez-Martínez Z, Martín-Rodríguez G, Coto E, Melón S |
| EPI_ISL_1406247, EPI_ISL_1406250 | Institute for Medical Research, Infectious Disease Research Centre, National Institutes of Health, Ministry of Health Malaysia | Institute for Medical Research, Infectious Disease Research Centre, National Institutes of Health, Ministry of Health Malaysia | Suppiah J, Kamel K, Mohd Zawawi Z, Ramly N, Robert F, Thayan R |
| EPI_ISL_1406274, EPI_ISL_1406277, EPI_ISL_1406281, EPI_ISL_1406286, EPI_ISL_1406292, EPI_ISL_1406294 | Institute for Medical Research, Infectious Disease Research Centre, National Institutes of Health, Ministry of Health Malaysia | Institute for Medical Research, Infectious Disease Research Centre, National Institutes of Health, Ministry of Health Malaysia | Suppiah J, Kamel K, Thayan R |

| | | | |
|---|---|---|---|
| EPI_ISL_1406390 | Hospital Monte Naranco | Laboratorio de Virología HUCA | Sandoval M, Castelló C, Gómez de Oña J, Boga JA, Rojo S, Alvarez-Arguelles ME, Abreu F, Costales I, Perez-Martínez Z, Martín-Rodríguez G, Coto E, Melón S |
| EPI_ISL_1406518 | Centre De Prélévement COVID VICHY | CHU Clermont-Ferrand, service de virologie | Bisseux Maxime, Mirand Audrey, Combes Patricia, Henquell Cécile |
| EPI_ISL_1406522, EPI_ISL_1406523, EPI_ISL_1406524, EPI_ISL_1406525, EPI_ISL_1406526, EPI_ISL_1406527 | Centre Hospitalier Universitaire Clermont-Ferrand | CHU Clermont-Ferrand, service de virologie | Bisseux Maxime, Mirand Audrey, Combes Patricia, Henquell Cécile |
| EPI_ISL_1406528, EPI_ISL_1406529 | Centre Hospitalier RIOM | CHU Clermont-Ferrand, service de virologie | Bisseux Maxime, Mirand Audrey, Combes Patricia, Henquell Cécile |
| EPI_ISL_1406530, EPI_ISL_1406531 | HOPITAL DU MONT DORE | CHU Clermont-Ferrand, service de virologie | Bisseux Maxime, Mirand Audrey, Combes Patricia, Henquell Cécile |
| EPI_ISL_1406532 | Centre Hospitalier Universitaire Clermont-Ferrand | CHU Clermont-Ferrand, service de virologie | Bisseux Maxime, Mirand Audrey, Combes Patricia, Henquell Cécile |
| EPI_ISL_1406533 | Centre hospitalier D'AURILLAC | CHU Clermont-Ferrand, service de virologie | Bisseux Maxime, Mirand Audrey, Combes Patricia, Henquell Cécile |
| EPI_ISL_1406534 | Centre Hospitalier Universitaire Clermont-Ferrand | CHU Clermont-Ferrand, service de virologie | Bisseux Maxime, Mirand Audrey, Combes Patricia, Henquell Cécile |
| EPI_ISL_1406535 | HOPITAL DU MONT DORE | CHU Clermont-Ferrand, service de virologie | Bisseux Maxime, Mirand Audrey, Combes Patricia, Henquell Cécile |
| EPI_ISL_1406536, EPI_ISL_1406537, EPI_ISL_1406538, EPI_ISL_1406539, EPI_ISL_1406548, EPI_ISL_1406550, EPI_ISL_1406551, EPI_ISL_1406552, EPI_ISL_1406553, EPI_ISL_1406554, EPI_ISL_1406555, EPI_ISL_1406556, EPI_ISL_1406557, EPI_ISL_1406558, EPI_ISL_1406559, EPI_ISL_1406560, EPI_ISL_1406561, EPI_ISL_1406564, EPI_ISL_1406566, EPI_ISL_1406568, EPI_ISL_1406569, EPI_ISL_1406570 | | | |
| see above | Centre Hospitalier Universitaire Clermont-Ferrand | CHU Clermont-Ferrand, service de virologie | Bisseux Maxime, Mirand Audrey, Combes Patricia, Henquell Cécile |
| EPI_ISL_1406571 | HOPITAL DU MONT DORE | CHU Clermont-Ferrand, service de virologie | Bisseux Maxime, Mirand Audrey, Combes Patricia, Henquell Cécile |
| EPI_ISL_1406572, EPI_ISL_1406573 | Centre Hospitalier Universitaire Clermont-Ferrand | CHU Clermont-Ferrand, service de virologie | Bisseux Maxime, Mirand Audrey, Combes Patricia, Henquell Cécile |
| EPI_ISL_1406574 | HOPITAL DU MONT DORE | CHU Clermont-Ferrand, service de virologie | Bisseux Maxime, Mirand Audrey, Combes Patricia, Henquell Cécile |
| EPI_ISL_1406575 | Centre Hospitalier Paul Ardier | CHU Clermont-Ferrand, service de virologie | Bisseux Maxime, Mirand Audrey, Combes Patricia, Henquell Cécile |
| EPI_ISL_1406576, EPI_ISL_1406577, EPI_ISL_1406578, EPI_ISL_1406579, EPI_ISL_1406580, EPI_ISL_1406581, EPI_ISL_1406582, EPI_ISL_1406583, EPI_ISL_1406584 | Centre Hospitalier Universitaire Clermont-Ferrand | CHU Clermont-Ferrand, service de virologie | Bisseux Maxime, Mirand Audrey, Combes Patricia, Henquell Cécile |
| EPI_ISL_1406585 | Centre De Prelevement COVID RIOM | CHU Clermont-Ferrand, service de virologie | Bisseux Maxime, Mirand Audrey, Combes Patricia, Henquell Cécile |
| EPI_ISL_1406586 | Centre Hospitalier Universitaire Clermont-Ferrand | CHU Clermont-Ferrand, service de virologie | Bisseux Maxime, Mirand Audrey, Combes Patricia, Henquell Cécile |
| EPI_ISL_1406587 | HOPITAL DE MONTLUCON | CHU Clermont-Ferrand, service de virologie | Bisseux Maxime, Mirand Audrey, Combes Patricia, Henquell Cécile |
| EPI_ISL_1406588 | Centre Hospitalier RIOM | CHU Clermont-Ferrand, service de virologie | Bisseux Maxime, Mirand Audrey, Combes Patricia, Henquell Cécile |
| EPI_ISL_1406589 | CDP COVID MONTLUCON | CHU Clermont-Ferrand, service de virologie | Bisseux Maxime, Mirand Audrey, Combes Patricia, Henquell Cécile |
| EPI_ISL_1406590 | Centre De Prelevement COVID RIOM | CHU Clermont-Ferrand, service de virologie | Bisseux Maxime, Mirand Audrey, Combes Patricia, Henquell Cécile |
| EPI_ISL_1406591 | Centre Hospitalier Universitaire Clermont-Ferrand | CHU Clermont-Ferrand, service de virologie | Bisseux Maxime, Mirand Audrey, Combes Patricia, Henquell Cécile |
| EPI_ISL_1406592 | Centre hospitalier D'AURILLAC | CHU Clermont-Ferrand, service de virologie | Bisseux Maxime, Mirand Audrey, Combes Patricia, Henquell Cécile |
| EPI_ISL_1406593 | HOPITAL DU MONT DORE | CHU Clermont-Ferrand, service de virologie | Bisseux Maxime, Mirand Audrey, Combes Patricia, Henquell Cécile |
| EPI_ISL_1406594 | Centre De Prelevement COVID RIOM | CHU Clermont-Ferrand, service de virologie | Bisseux Maxime, Mirand Audrey, Combes Patricia, Henquell Cécile |
| EPI_ISL_1406595 | Centre Hospitalier Universitaire Clermont-Ferrand | CHU Clermont-Ferrand, service de virologie | Bisseux Maxime, Mirand Audrey, Combes Patricia, Henquell Cécile |
| EPI_ISL_1406596 | HOPITAL DE VICHY | CHU Clermont-Ferrand, service de virologie | Bisseux Maxime, Mirand Audrey, Combes Patricia, Henquell Cécile |
| EPI_ISL_1406597 | Centre De Prélévement COVID VICHY | CHU Clermont-Ferrand, service de virologie | Bisseux Maxime, Mirand Audrey, Combes Patricia, Henquell Cécile |
| EPI_ISL_1406599 | Centre De Prelevement COVID RIOM | CHU Clermont-Ferrand, service de virologie | Bisseux Maxime, Mirand Audrey, Combes Patricia, Henquell Cécile |
| EPI_ISL_1406600 | Centre Hospitalier Universitaire Clermont-Ferrand | CHU Clermont-Ferrand, service de virologie | Bisseux Maxime, Mirand Audrey, Combes Patricia, Henquell Cécile |
| EPI_ISL_1406601 | Centre hospitalier D'AURILLAC | CHU Clermont-Ferrand, service de virologie | Bisseux Maxime, Mirand Audrey, Combes Patricia, Henquell Cécile |
| EPI_ISL_1406602 | Centre Hospitalier Universitaire Clermont-Ferrand | CHU Clermont-Ferrand, service de virologie | Bisseux Maxime, Mirand Audrey, Combes Patricia, Henquell Cécile |
| EPI_ISL_1406603 | Centre Hospitalier RIOM | CHU Clermont-Ferrand, service de virologie | Bisseux Maxime, Mirand Audrey, Combes Patricia, Henquell Cécile |
| EPI_ISL_1406604, EPI_ISL_1406605 | Centre Hospitalier Universitaire Clermont-Ferrand | CHU Clermont-Ferrand, service de virologie | Bisseux Maxime, Mirand Audrey, Combes Patricia, Henquell Cécile |
| EPI_ISL_1406606 | Centre De Prelevement COVID RIOM | CHU Clermont-Ferrand, service de virologie | Bisseux Maxime, Mirand Audrey, Combes Patricia, Henquell Cécile |
| EPI_ISL_1406607 | Centre Hospitalier Universitaire Clermont-Ferrand | CHU Clermont-Ferrand, service de virologie | Bisseux Maxime, Mirand Audrey, Combes Patricia, Henquell Cécile |
| EPI_ISL_1406608 | Centre hospitalier D'AURILLAC | CHU Clermont-Ferrand, service de virologie | Bisseux Maxime, Mirand Audrey, Combes Patricia, Henquell Cécile |
| EPI_ISL_1406609, EPI_ISL_1406610 | Centre Hospitalier Universitaire Clermont-Ferrand | CHU Clermont-Ferrand, service de virologie | Bisseux Maxime, Mirand Audrey, Combes Patricia, Henquell Cécile |
| EPI_ISL_1406611 | HOPITAL DE VICHY | CHU Clermont-Ferrand, service de virologie | Bisseux Maxime, Mirand Audrey, Combes Patricia, Henquell Cécile |
| EPI_ISL_1406612, EPI_ISL_1406613, EPI_ISL_1406614, EPI_ISL_1406615 | Centre Hospitalier Universitaire Clermont-Ferrand | CHU Clermont-Ferrand, service de virologie | Bisseux Maxime, Mirand Audrey, Combes Patricia, Henquell Cécile |
| EPI_ISL_1406616 | Centre De Prelevement COVID RIOM | CHU Clermont-Ferrand, service de virologie | Bisseux Maxime, Mirand Audrey, Combes Patricia, Henquell Cécile |
| EPI_ISL_1406617 | Centre Hospitalier Universitaire Clermont-Ferrand | CHU Clermont-Ferrand, service de virologie | Bisseux Maxime, Mirand Audrey, Combes Patricia, Henquell Cécile |
| EPI_ISL_1406618 | Centre hospitalier D'AURILLAC | CHU Clermont-Ferrand, service de virologie | Bisseux Maxime, Mirand Audrey, Combes Patricia, Henquell Cécile |
| EPI_ISL_1406619 | Centre De Prelevement COVID RIOM | CHU Clermont-Ferrand, service de virologie | Bisseux Maxime, Mirand Audrey, Combes Patricia, Henquell Cécile |
| EPI_ISL_1406620, EPI_ISL_1406622 | Centre Hospitalier Universitaire Clermont-Ferrand | CHU Clermont-Ferrand, service de virologie | Bisseux Maxime, Mirand Audrey, Combes Patricia, Henquell Cécile |
| EPI_ISL_1406623 | HOPITAL DE VICHY | CHU Clermont-Ferrand, service de virologie | Bisseux Maxime, Mirand Audrey, Combes Patricia, Henquell Cécile |
| EPI_ISL_1406639, EPI_ISL_1406640 | Centre De Prelevement COVID RIOM | CHU Clermont-Ferrand, service de virologie | Bisseux Maxime, Mirand Audrey, Combes Patricia, Henquell Cécile |
| EPI_ISL_1406644 | Centre Hospitalier Universitaire Clermont-Ferrand | CHU Clermont-Ferrand, service de virologie | Bisseux Maxime, Mirand Audrey, Combes Patricia, Henquell Cécile |
| EPI_ISL_1406649 | CDP COVID MONTLUCON | CHU Clermont-Ferrand, service de virologie | Bisseux Maxime, Mirand Audrey, Combes Patricia, Henquell Cécile |
| EPI_ISL_1406652 | Centre De Prélévement COVID VICHY | CHU Clermont-Ferrand, service de virologie | Bisseux Maxime, Mirand Audrey, Combes Patricia, Henquell Cécile |
| EPI_ISL_1406655 | Centre Hospitalier Universitaire Clermont-Ferrand | CHU Clermont-Ferrand, service de virologie | Bisseux Maxime, Mirand Audrey, Combes Patricia, Henquell Cécile |
| EPI_ISL_1406656 | Centre De Prelevement COVID RIOM | CHU Clermont-Ferrand, service de virologie | Bisseux Maxime, Mirand Audrey, Combes Patricia, Henquell Cécile |

| | | | |
|---|---|---|---|
| EPI_ISL_1406658, EPI_ISL_1406659 | Centre Hospitalier Universitaire Clermont-Ferrand | CHU Clermont-Ferrand, service de virologie | Bisseux Maxime, Mirand Audrey, Combes Patricia, Henquell Cécile |
| EPI_ISL_1406660, EPI_ISL_1406661, EPI_ISL_1406662 | HOPITAL DE VICHY | CHU Clermont-Ferrand, service de virologie | Bisseux Maxime, Mirand Audrey, Combes Patricia, Henquell Cécile |
| EPI_ISL_1406663 | Centre Hospitalier Universitaire Clermont-Ferrand | CHU Clermont-Ferrand, service de virologie | Bisseux Maxime, Mirand Audrey, Combes Patricia, Henquell Cécile |
| EPI_ISL_1406664 | Centre De Prelevement COVID RIOM | CHU Clermont-Ferrand, service de virologie | Bisseux Maxime, Mirand Audrey, Combes Patricia, Henquell Cécile |
| EPI_ISL_1406665 | HOPITAL DE VICHY | CHU Clermont-Ferrand, service de virologie | Bisseux Maxime, Mirand Audrey, Combes Patricia, Henquell Cécile |
| EPI_ISL_1406666, EPI_ISL_1406667, EPI_ISL_1406668 | Centre Hospitalier Universitaire Clermont-Ferrand | CHU Clermont-Ferrand, service de virologie | Bisseux Maxime, Mirand Audrey, Combes Patricia, Henquell Cécile |
| EPI_ISL_1406669 | HOPITAL DE VICHY | CHU Clermont-Ferrand, service de virologie | Bisseux Maxime, Mirand Audrey, Combes Patricia, Henquell Cécile |
| EPI_ISL_1406670 | Centre Hospitalier RIOM | CHU Clermont-Ferrand, service de virologie | Bisseux Maxime, Mirand Audrey, Combes Patricia, Henquell Cécile |
| EPI_ISL_1406671 | HOPITAL DE VICHY | CHU Clermont-Ferrand, service de virologie | Bisseux Maxime, Mirand Audrey, Combes Patricia, Henquell Cécile |
| EPI_ISL_1406672 | HOPITAL DU MONT DORE | CHU Clermont-Ferrand, service de virologie | Bisseux Maxime, Mirand Audrey, Combes Patricia, Henquell Cécile |
| EPI_ISL_1406673 | HOPITAL DE VICHY | CHU Clermont-Ferrand, service de virologie | Bisseux Maxime, Mirand Audrey, Combes Patricia, Henquell Cécile |
| EPI_ISL_1406674 | EHPAD | CHU Clermont-Ferrand, service de virologie | Bisseux Maxime, Mirand Audrey, Combes Patricia, Henquell Cécile |
| EPI_ISL_1406675 | Centre Hospitalier Universitaire Clermont-Ferrand | CHU Clermont-Ferrand, service de virologie | Bisseux Maxime, Mirand Audrey, Combes Patricia, Henquell Cécile |
| EPI_ISL_1406676, EPI_ISL_1406677, EPI_ISL_1406678 | HOPITAL DE VICHY | CHU Clermont-Ferrand, service de virologie | Bisseux Maxime, Mirand Audrey, Combes Patricia, Henquell Cécile |
| EPI_ISL_1406679 | HOPITAL DE MONTLUCON | CHU Clermont-Ferrand, service de virologie | Bisseux Maxime, Mirand Audrey, Combes Patricia, Henquell Cécile |
| EPI_ISL_1406680 | Centre Hospitalier Universitaire Clermont-Ferrand | CHU Clermont-Ferrand, service de virologie | Bisseux Maxime, Mirand Audrey, Combes Patricia, Henquell Cécile |
| EPI_ISL_1406682 | HOPITAL DE VICHY | CHU Clermont-Ferrand, service de virologie | Bisseux Maxime, Mirand Audrey, Combes Patricia, Henquell Cécile |
| EPI_ISL_1406683 | Centre Hospitalier Universitaire Clermont-Ferrand | CHU Clermont-Ferrand, service de virologie | Bisseux Maxime, Mirand Audrey, Combes Patricia, Henquell Cécile |
| EPI_ISL_1406690, EPI_ISL_1406691, EPI_ISL_1406692, EPI_ISL_1406693 | National Center of Infectious and Parasitic Diseases | National Center of Infectious and Parasitic Diseases | Alexiev et al |
| EPI_ISL_1408900, EPI_ISL_1408902, EPI_ISL_1408904 | AZ Zeno | AZ SINT-JAN BRUGGE | Marijke Reynders, Patrick Descheemaeker, Laurien Hoornaert, Thomas Van Landschoot, Jorn Hellemans |
| EPI_ISL_1413427 | National Institute of Health Research and Development | National Institute of Health Research and Development | Subangkit, Hana Apsari Pawestri, Kartika Dewi Puspa, Arie Ardiansyah Nugraha, Hartanti Dian Ikawati, Krisna Nur Andriana Pangesti, Yuni Rukminiati, Ririn Ramadhany, Agustiningsih, Kindi Adam, Holy Arif Wibowo, Triyani Soekarso, Ni Ketut Susilarini, Nurika Hariastuti, Ully Alfi Nikmah, Reni Herman, Nike Susanti, Herna, Tati Febriyanti, Natalie Laurencia Kipuw, Fauzul Muna, Irene Lorinda Indalao, Nelly Puspandari, Vivi Setiawaty. |
| EPI_ISL_1415420 | SIESP DIPARTIMENTO DI PREVENZIONE TERAMO TERAMO(TERAMO) | Istituto Zooprofilattico Sperimentale dell'Abruzzo e Molise "G. Caporale" | Lorusso A, Marcacci M, Di Domenico M, Ancora M, Curini V, Di Lollo Valeria, Mangone I, Rinaldi A, Delli Compagni E, Scialabba S, Caporale M, Di Pasquale A, Cammà C, Puglia I, Calistri P, Savini G |
| EPI_ISL_1415421 | OSPEDALE CIVILE SANT'OMERO - PRONTO SOCCORSO SANT'OMERO(TERAMO) | Istituto Zooprofilattico Sperimentale dell'Abruzzo e Molise "G. Caporale" | Lorusso A, Marcacci M, Di Domenico M, Ancora M, Curini V, Di Lollo Valeria, Mangone I, Rinaldi A, Delli Compagni E, Scialabba S, Caporale M, Di Pasquale A, Cammà C, Puglia I, Calistri P, Savini G |
| EPI_ISL_1415422 | USCA Avezzano AVEZZANO(L'AQUILA) | Istituto Zooprofilattico Sperimentale dell'Abruzzo e Molise "G. Caporale" | Lorusso A, Marcacci M, Di Domenico M, Ancora M, Curini V, Di Lollo Valeria, Mangone I, Rinaldi A, Delli Compagni E, Scialabba S, Caporale M, Di Pasquale A, Cammà C, Puglia I, Calistri P, Savini G |
| EPI_ISL_1415423 | SIESP CHIETI - DRIVE IN ORTONA ORTONA(CHIETI) | Istituto Zooprofilattico Sperimentale dell'Abruzzo e Molise "G. Caporale" | Lorusso A, Marcacci M, Di Domenico M, Ancora M, Curini V, Di Lollo Valeria, Mangone I, Rinaldi A, Delli Compagni E, Scialabba S, Caporale M, Di Pasquale A, Cammà C, Puglia I, Calistri P, Savini G |
| EPI_ISL_1415424 | SIESP DIPARTIMENTO DI PREVENZIONE TERAMO TERAMO(TERAMO) | Istituto Zooprofilattico Sperimentale dell'Abruzzo e Molise "G. Caporale" | Lorusso A, Marcacci M, Di Domenico M, Ancora M, Curini V, Di Lollo Valeria, Mangone I, Rinaldi A, Delli Compagni E, Scialabba S, Caporale M, Di Pasquale A, Cammà C, Puglia I, Calistri P, Savini G |
| EPI_ISL_1415426 | SIESP CHIETI - DRIVE IN LANCIANO | Istituto Zooprofilattico Sperimentale dell'Abruzzo e Molise "G. Caporale" | Lorusso A, Marcacci M, Di Domenico M, Ancora M, Curini V, Di Lollo Valeria, Mangone I, Rinaldi A, Delli Compagni E, Scialabba S, Caporale M, Di Pasquale A, Cammà C, Puglia I, Calistri P, Savini G |
| EPI_ISL_1415427 | National Institute of Health Research and Development | National Institute of Health Research and Development | Subangkit, Hana Apsari Pawestri, Kartika Dewi Puspa, Arie Ardiansyah Nugraha, Hartanti Dian Ikawati, Krisna Nur Andriana Pangesti, Yuni Rukminiati, Ririn Ramadhany, Agustiningsih, Kindi Adam, Holy Arif Wibowo, Triyani Soekarso, Ni Ketut Susilarini, Nurika Hariastuti, Ully Alfi Nikmah, Reni Herman, Nike Susanti, Herna, Tati Febriyanti, Natalie Laurencia Kipuw, Fauzul Muna, Irene Lorinda Indalao, Nelly Puspandari, Vivi Setiawaty. |
| EPI_ISL_1416191 | Fakultas Kedokteran Universitas Sumatera Utara | National Institute of Health Research and Development | Vivi Setiawaty, Hana Apsari Pawestri, Subangkit, Kartika Dewi Puspa, Arie Ardiansyah Nugraha, Hartanti Dian Ikawati, Nelly Puspandari, Krisna Nur Andriana Pangesti |
| EPI_ISL_1416332, EPI_ISL_1416337, EPI_ISL_1416339, EPI_ISL_1416341, EPI_ISL_1416342, EPI_ISL_1416343, EPI_ISL_1416344, EPI_ISL_1416346, EPI_ISL_1416347, EPI_ISL_1416348, EPI_ISL_1416349, EPI_ISL_1416350, EPI_ISL_1416352, EPI_ISL_1416355, EPI_ISL_1416357, EPI_ISL_1416359, EPI_ISL_1416361, EPI_ISL_1416362, EPI_ISL_1416363, EPI_ISL_1416364, EPI_ISL_1416365, EPI_ISL_1416366, EPI_ISL_1416367, EPI_ISL_1416368, EPI_ISL_1416369, EPI_ISL_1416370, EPI_ISL_1416371, EPI_ISL_1416374, EPI_ISL_1416376, EPI_ISL_1416379, EPI_ISL_1416380, EPI_ISL_1416381, EPI_ISL_1416384, EPI_ISL_1416385, EPI_ISL_1416390, EPI_ISL_1416393, EPI_ISL_1416394, EPI_ISL_1416395, EPI_ISL_1416397, EPI_ISL_1416398, EPI_ISL_1416399, EPI_ISL_1416402, EPI_ISL_1416403, EPI_ISL_1416405, EPI_ISL_1416408, EPI_ISL_1416409, EPI_ISL_1416412, EPI_ISL_1416416, EPI_ISL_1416417, EPI_ISL_1416418, EPI_ISL_1416419, EPI_ISL_1416420, EPI_ISL_1416421, EPI_ISL_1416424, EPI_ISL_1416425, EPI_ISL_1416426, EPI_ISL_1416427, EPI_ISL_1416428, EPI_ISL_1416429, EPI_ISL_1416430, EPI_ISL_1416432, EPI_ISL_1416433, EPI_ISL_1416434, EPI_ISL_1416435, EPI_ISL_1416436, EPI_ISL_1416438, EPI_ISL_1416439, EPI_ISL_1416440, EPI_ISL_1416441, EPI_ISL_1416442, EPI_ISL_1416443, EPI_ISL_1416444, EPI_ISL_1416445, EPI_ISL_1416447, EPI_ISL_1416448, EPI_ISL_1416449, EPI_ISL_1416452, EPI_ISL_1416453, EPI_ISL_1416454, EPI_ISL_1416460, EPI_ISL_1416461, EPI_ISL_1416462, EPI_ISL_1416463, EPI_ISL_1416464, EPI_ISL_1416466, EPI_ISL_1416467, EPI_ISL_1416469, EPI_ISL_1416470, EPI_ISL_1416474, EPI_ISL_1416475, EPI_ISL_1416476, EPI_ISL_1416477, EPI_ISL_1416478, EPI_ISL_1416480, EPI_ISL_1416481, EPI_ISL_1416482, EPI_ISL_1416483, EPI_ISL_1416484, EPI_ISL_1416487, EPI_ISL_1416488, EPI_ISL_1416496, EPI_ISL_1416497, EPI_ISL_1416498, EPI_ISL_1416501, EPI_ISL_1416502, EPI_ISL_1416504, EPI_ISL_1416505, EPI_ISL_1416507, EPI_ISL_1416508, EPI_ISL_1416509, EPI_ISL_1416511, EPI_ISL_1416512, EPI_ISL_1416514, EPI_ISL_1416516, EPI_ISL_1416518, EPI_ISL_1416519, EPI_ISL_1416521, EPI_ISL_1416522, EPI_ISL_1416523, EPI_ISL_1416524, EPI_ISL_1416528, EPI_ISL_1416529, EPI_ISL_1416530, EPI_ISL_1416531, EPI_ISL_1416532, EPI_ISL_1416533, EPI_ISL_1416535, EPI_ISL_1416536, EPI_ISL_1416538, EPI_ISL_1416539, EPI_ISL_1416540, EPI_ISL_1416541, EPI_ISL_1416542, EPI_ISL_1416543, EPI_ISL_1416544, EPI_ISL_1416545, EPI_ISL_1416547, EPI_ISL_1416548, EPI_ISL_1416550, EPI_ISL_1416551, EPI_ISL_1416553, EPI_ISL_1416557, EPI_ISL_1416558, EPI_ISL_1416559, EPI_ISL_1416560, EPI_ISL_1416561, EPI_ISL_1416562, EPI_ISL_1416566, EPI_ISL_1416567, EPI_ISL_1416569, EPI_ISL_1416571, EPI_ISL_1416572, EPI_ISL_1416574, EPI_ISL_1416575, EPI_ISL_1416576, EPI_ISL_1416578, EPI_ISL_1416579, EPI_ISL_1416580, EPI_ISL_1416581, EPI_ISL_1416582, EPI_ISL_1416583, EPI_ISL_1416584, EPI_ISL_1416585, EPI_ISL_1416586, EPI_ISL_1416587, EPI_ISL_1416589, EPI_ISL_1416591, EPI_ISL_1416592, EPI_ISL_1416595, EPI_ISL_1416597, EPI_ISL_1416598, EPI_ISL_1416600, EPI_ISL_1416602, EPI_ISL_1416603, EPI_ISL_1416604, EPI_ISL_1416605, EPI_ISL_1416606, EPI_ISL_1416607, EPI_ISL_1416609, EPI_ISL_1416610, EPI_ISL_1416611, EPI_ISL_1416612, EPI_ISL_1416613, EPI_ISL_1416614, EPI_ISL_1416615, EPI_ISL_1416617, EPI_ISL_1416618, EPI_ISL_1416619, EPI_ISL_1416620, EPI_ISL_1416621, EPI_ISL_1416622, EPI_ISL_1416623, EPI_ISL_1416624, EPI_ISL_1416625, EPI_ISL_1416626, EPI_ISL_1416627, EPI_ISL_1416628, EPI_ISL_1416629, EPI_ISL_1416632, EPI_ISL_1416634, EPI_ISL_1416636, EPI_ISL_1416638, EPI_ISL_1416640, EPI_ISL_1416644, EPI_ISL_1416645, EPI_ISL_1416646, EPI_ISL_1416647, EPI_ISL_1416648, EPI_ISL_1416649, EPI_ISL_1416650, EPI_ISL_1416651, EPI_ISL_1416652, EPI_ISL_1416653, EPI_ISL_1416654, EPI_ISL_1416655, EPI_ISL_1416656, EPI_ISL_1416658, EPI_ISL_1416659, EPI_ISL_1416660, EPI_ISL_1416661, EPI_ISL_1416663, EPI_ISL_1416664, EPI_ISL_1416665, EPI_ISL_1416666, EPI_ISL_1416667, EPI_ISL_1416668, EPI_ISL_1416670, EPI_ISL_1416671, EPI_ISL_1416672, EPI_ISL_1416673, EPI_ISL_1416674, EPI_ISL_1416675, EPI_ISL_1416676, EPI_ISL_1416678, EPI_ISL_1416679, EPI_ISL_1416680, EPI_ISL_1416681, EPI_ISL_1416682 | | | |
| see above | Laboratorio Central de Epidemiología (LCE) | Instituto de Biotecnología de la UNAM | Consorcio Mexicano de Vigilancia Genómica (CoViGen-Mex). Authors (in alphabetical order): Julio Elias Alvarado-Yaah, Carlos F. Arias, Santiago Ávila-Ríos, Víctor Hugo Borja-Aburto, Celia Boukadida, Juan Bautista Chale-Dzul , José Antonio Enciso-Moreno, Gloria Elena Espinoza-Ayala, Fernando Fontove-Herrera, Concepción Grajales-Muñiz, Ricardo Grande, Alfredo Herrera-Estrella, Carla Ivón Herrera-Najera, Pavel Isa, Brenda Irasema Maldonado-Meza, Bernardo Martínez-Miguel, Margarita Matías-Florentino, María Guadalupe de Jesús Mireles-Rivera, Gloria María Molina-Salinas, Hector Montoya-Fuentes, José Esteban Muñoz-Medina, José de Jesús Nuñez-Contreras, Alicia Ocaña-Mondragón, Luis Alberto Ochoa-Carrera, Hector Esteban Paz-Juárez, Francisco Pulido, Helen Haydee Fernanda Ramírez-Plascencia, Angel Gustavo Salas-Lais, Jorge Ivan Salinal-Nevarez, Alejandro Sanchez-Flores, Clara Esperanza Santacruz-Tinoco, María Guadalupe Santiago-Mauricio , Nelly Sélem-Mojica, Blanca Taboada , Gloria Vazquez |
| EPI_ISL_1416811 | Hospital | National Reference Center for Viruses of Respiratory Infections, Institut Pasteur, Paris | Marion Barbet, Sylvie Behillil, Méline Bizard,Frédéric Lemoine,Corinne Maufrais,Christophe Malabat, Angela Brisebarre, Camille Capel, Louise Lefrançois, Etienne Simon-Lorière, Vincent Enouf, Maud Vanpeene, Sylvie van der Werf,Mj Dufour |
| EPI_ISL_1416812 | Labo Analyses Med | National Reference Center for Viruses of Respiratory Infections, Institut Pasteur, Paris | Marion Barbet, Sylvie Behillil, Méline Bizard,Frédéric Lemoine,Corinne Maufrais,Christophe Malabat, Angela Brisebarre, Camille Capel, Louise Lefrançois, Etienne Simon-Lorière, Vincent Enouf, Maud Vanpeene, Sylvie van der Werf,Ilka Engelmann |
| EPI_ISL_1416813, EPI_ISL_1416815, | Labo Analyses Med | National Reference Center for Viruses of Respiratory | Marion Barbet, Sylvie Behillil, Méline Bizard,Frédéric Lemoine,Corinne Maufrais,Christophe Malabat, Angela Brisebarre, Camille Capel, Louise Lefrançois, |

| | | | |
|---|---|---|---|
| EPI_ISL_1416816, EPI_ISL_1416817, EPI_ISL_1416818, EPI_ISL_1416819 | | Infections, Institut Pasteur, Paris | Etienne Simon-Lorière, Vincent Enouf, Maud Vanpeene, Sylvie van der Werf,Ronan Pichard |
| EPI_ISL_1416820 | Labo Analyses Med | National Reference Center for Viruses of Respiratory Infections, Institut Pasteur, Paris | Marion Barbet, Sylvie Behillil, Méline Bizard,Frédéric Lemoine,Corinne Maufrais,Christophe Malabat, Angela Brisebarre, Camille Capel, Louise Lefrançois, Etienne Simon-Lorière, Vincent Enouf, Maud Vanpeene, Sylvie van der Werf,Anne Cady |
| EPI_ISL_1416821, EPI_ISL_1416822 | Labo Analyses Med | National Reference Center for Viruses of Respiratory Infections, Institut Pasteur, Paris | Marion Barbet, Sylvie Behillil, Méline Bizard,Frédéric Lemoine,Corinne Maufrais,Christophe Malabat, Angela Brisebarre, Camille Capel, Louise Lefrançois, Etienne Simon-Lorière, Vincent Enouf, Maud Vanpeene, Sylvie van der Werf,CéLine Ramanantsoa |
| EPI_ISL_1416823, EPI_ISL_1416824, EPI_ISL_1416825, EPI_ISL_1416826, EPI_ISL_1416827, EPI_ISL_1416828 | Hospital | National Reference Center for Viruses of Respiratory Infections, Institut Pasteur, Paris | Marion Barbet, Sylvie Behillil, Méline Bizard,Frédéric Lemoine,Corinne Maufrais,Christophe Malabat, Angela Brisebarre, Camille Capel, Louise Lefrançois, Etienne Simon-Lorière, Vincent Enouf, Maud Vanpeene, Sylvie van der Werf,Jean Baptiste Bour |
| EPI_ISL_1416829 | Hospital | National Reference Center for Viruses of Respiratory Infections, Institut Pasteur, Paris | Marion Barbet, Sylvie Behillil, Méline Bizard,Frédéric Lemoine,Corinne Maufrais,Christophe Malabat, Angela Brisebarre, Camille Capel, Louise Lefrançois, Etienne Simon-Lorière, Vincent Enouf, Maud Vanpeene, Sylvie van der Werf,Anne Cady |
| EPI_ISL_1416830 | Hospital | National Reference Center for Viruses of Respiratory Infections, Institut Pasteur, Paris | Marion Barbet, Sylvie Behillil, Méline Bizard,Frédéric Lemoine,Corinne Maufrais,Christophe Malabat, Angela Brisebarre, Camille Capel, Louise Lefrançois, Etienne Simon-Lorière, Vincent Enouf, Maud Vanpeene, Sylvie van der Werf,Jean Baptiste Bour |
| EPI_ISL_1416831 | Hospital | National Reference Center for Viruses of Respiratory Infections, Institut Pasteur, Paris | Marion Barbet, Sylvie Behillil, Méline Bizard,Frédéric Lemoine,Corinne Maufrais,Christophe Malabat, Angela Brisebarre, Camille Capel, Louise Lefrançois, Etienne Simon-Lorière, Vincent Enouf, Maud Vanpeene, Sylvie van der Werf,Ronan Pichard |
| EPI_ISL_1416832 | Hospital | National Reference Center for Viruses of Respiratory Infections, Institut Pasteur, Paris | Marion Barbet, Sylvie Behillil, Méline Bizard,Frédéric Lemoine,Corinne Maufrais,Christophe Malabat, Angela Brisebarre, Camille Capel, Louise Lefrançois, Etienne Simon-Lorière, Vincent Enouf, Maud Vanpeene, Sylvie van der Werf,Sophie Chalmin |
| EPI_ISL_1416833, EPI_ISL_1416834, EPI_ISL_1416835, EPI_ISL_1416836, EPI_ISL_1416837, EPI_ISL_1416838 | Labo Analyses Med | National Reference Center for Viruses of Respiratory Infections, Institut Pasteur, Paris | Marion Barbet, Sylvie Behillil, Méline Bizard,Frédéric Lemoine,Corinne Maufrais,Christophe Malabat, Angela Brisebarre, Camille Capel, Louise Lefrançois, Etienne Simon-Lorière, Vincent Enouf, Maud Vanpeene, Sylvie van der Werf,Sophie Chalmin |
| EPI_ISL_1416839 | Labo Analyses Med | National Reference Center for Viruses of Respiratory Infections, Institut Pasteur, Paris | Marion Barbet, Sylvie Behillil, Méline Bizard,Frédéric Lemoine,Corinne Maufrais,Christophe Malabat, Angela Brisebarre, Camille Capel, Louise Lefrançois, Etienne Simon-Lorière, Vincent Enouf, Maud Vanpeene, Sylvie van der Werf,Dominique Descamps |
| EPI_ISL_1416840, EPI_ISL_1416841, EPI_ISL_1416842, EPI_ISL_1416843 | Labo Analyses Med | National Reference Center for Viruses of Respiratory Infections, Institut Pasteur, Paris | Marion Barbet, Sylvie Behillil, Méline Bizard,Frédéric Lemoine,Corinne Maufrais,Christophe Malabat, Angela Brisebarre, Camille Capel, Louise Lefrançois, Etienne Simon-Lorière, Vincent Enouf, Maud Vanpeene, Sylvie van der Werf,Sophie Chalmin |
| EPI_ISL_1416846, EPI_ISL_1416847, EPI_ISL_1416848 | Labo Analyses Med | National Reference Center for Viruses of Respiratory Infections, Institut Pasteur, Paris | Marion Barbet, Sylvie Behillil, Méline Bizard,Frédéric Lemoine,Corinne Maufrais,Christophe Malabat, Angela Brisebarre, Camille Capel, Louise Lefrançois, Etienne Simon-Lorière, Vincent Enouf, Maud Vanpeene, Sylvie van der Werf,StéPhanie Guyomard-Rabenirina |
| EPI_ISL_1416849 | Labo Analyses Med | National Reference Center for Viruses of Respiratory Infections, Institut Pasteur, Paris | Marion Barbet, Sylvie Behillil, Méline Bizard,Frédéric Lemoine,Corinne Maufrais,Christophe Malabat, Angela Brisebarre, Camille Capel, Louise Lefrançois, Etienne Simon-Lorière, Vincent Enouf, Maud Vanpeene, Sylvie van der Werf,Claire Felloni |
| EPI_ISL_1416850, EPI_ISL_1416851, EPI_ISL_1416852, EPI_ISL_1416853, EPI_ISL_1416854, EPI_ISL_1416855, EPI_ISL_1416856 | Hospital | National Reference Center for Viruses of Respiratory Infections, Institut Pasteur, Paris | Marion Barbet, Sylvie Behillil, Méline Bizard,Frédéric Lemoine,Corinne Maufrais,Christophe Malabat, Angela Brisebarre, Camille Capel, Louise Lefrançois, Etienne Simon-Lorière, Vincent Enouf, Maud Vanpeene, Sylvie van der Werf,StéPhanie Guyomard-Rabenirina |
| EPI_ISL_1416859 | Labo analyses med | National Reference Center for Viruses of Respiratory Infections, Institut Pasteur, Paris | Marion Barbet, Sylvie Behillil, Méline Bizard,Frédéric Lemoine,Corinne Maufrais,Christophe Malabat, Angela Brisebarre, Camille Capel, Louise Lefrançois, Etienne Simon-Lorière, Vincent Enouf, Maud Vanpeene, Sylvie van der Werf,JéRome Guinard |
| EPI_ISL_1416860, EPI_ISL_1416861, EPI_ISL_1416862, EPI_ISL_1416863, EPI_ISL_1416864, EPI_ISL_1416865, EPI_ISL_1416866, EPI_ISL_1416867, EPI_ISL_1416868 | Labo analyses med | National Reference Center for Viruses of Respiratory Infections, Institut Pasteur, Paris | Marion Barbet, Sylvie Behillil, Méline Bizard,Frédéric Lemoine,Corinne Maufrais,Christophe Malabat, Angela Brisebarre, Camille Capel, Louise Lefrançois, Etienne Simon-Lorière, Vincent Enouf, Maud Vanpeene, Sylvie van der Werf,StéPhanie Guyomard-Rabenirina |
| EPI_ISL_1416869 | Labo analyses med | National Reference Center for Viruses of Respiratory Infections, Institut Pasteur, Paris | Marion Barbet, Sylvie Behillil, Méline Bizard,Frédéric Lemoine,Corinne Maufrais,Christophe Malabat, Angela Brisebarre, Camille Capel, Louise Lefrançois, Etienne Simon-Lorière, Vincent Enouf, Maud Vanpeene, Sylvie van der Werf,GréGoire Potiron |
| EPI_ISL_1416870, EPI_ISL_1416871, EPI_ISL_1416872, EPI_ISL_1416874, EPI_ISL_1416875, EPI_ISL_1416876 | Labo analyses med | National Reference Center for Viruses of Respiratory Infections, Institut Pasteur, Paris | Marion Barbet, Sylvie Behillil, Méline Bizard,Frédéric Lemoine,Corinne Maufrais,Christophe Malabat, Angela Brisebarre, Camille Capel, Louise Lefrançois, Etienne Simon-Lorière, Vincent Enouf, Maud Vanpeene, Sylvie van der Werf,StéPhanie Guyomard-Rabenirina |
| EPI_ISL_1416877, EPI_ISL_1416878 | Labo analyses med | National Reference Center for Viruses of Respiratory Infections, Institut Pasteur, Paris | Marion Barbet, Sylvie Behillil, Méline Bizard,Frédéric Lemoine,Corinne Maufrais,Christophe Malabat, Angela Brisebarre, Camille Capel, Louise Lefrançois, Etienne Simon-Lorière, Vincent Enouf, Maud Vanpeene, Sylvie van der Werf,FréDéRique Canis |
| EPI_ISL_1416879 | Labo analyses med | National Reference Center for Viruses of Respiratory Infections, Institut Pasteur, Paris | Marion Barbet, Sylvie Behillil, Méline Bizard,Frédéric Lemoine,Corinne Maufrais,Christophe Malabat, Angela Brisebarre, Camille Capel, Louise Lefrançois, Etienne Simon-Lorière, Vincent Enouf, Maud Vanpeene, Sylvie van der Werf,Ronan Pichard |
| EPI_ISL_1416880, EPI_ISL_1416881, EPI_ISL_1416882, EPI_ISL_1416883, EPI_ISL_1416884, EPI_ISL_1416885, EPI_ISL_1416886, EPI_ISL_1416887, EPI_ISL_1416888, EPI_ISL_1416889, EPI_ISL_1416890, EPI_ISL_1416891, EPI_ISL_1416892, EPI_ISL_1416893, EPI_ISL_1416894, EPI_ISL_1416895, EPI_ISL_1416896, EPI_ISL_1416897, EPI_ISL_1416898, EPI_ISL_1416899, EPI_ISL_1416900 | | | |
| see above | Labo analyses med | National Reference Center for Viruses of Respiratory Infections, Institut Pasteur, Paris | Marion Barbet, Sylvie Behillil, Méline Bizard,Frédéric Lemoine,Corinne Maufrais,Christophe Malabat, Angela Brisebarre, Camille Capel, Louise Lefrançois, Etienne Simon-Lorière, Vincent Enouf, Maud Vanpeene, Sylvie van der Werf,Anne Holstein |
| EPI_ISL_1416901 | Labo analyses med | National Reference Center for Viruses of Respiratory Infections, Institut Pasteur, Paris | Marion Barbet, Sylvie Behillil, Méline Bizard,Frédéric Lemoine,Corinne Maufrais,Christophe Malabat, Angela Brisebarre, Camille Capel, Louise Lefrançois, Etienne Simon-Lorière, Vincent Enouf, Maud Vanpeene, Sylvie van der Werf,Nathalie Coppe-Dron |
| EPI_ISL_1416903, EPI_ISL_1416904 | Labo analyses med | National Reference Center for Viruses of Respiratory Infections, Institut Pasteur, Paris | Marion Barbet, Sylvie Behillil, Méline Bizard,Frédéric Lemoine,Corinne Maufrais,Christophe Malabat, Angela Brisebarre, Camille Capel, Louise Lefrançois, Etienne Simon-Lorière, Vincent Enouf, Maud Vanpeene, Sylvie van der Werf,Anne Holstein |
| EPI_ISL_1416905 | Labo analyses med | National Reference Center for Viruses of Respiratory Infections, Institut Pasteur, Paris | Marion Barbet, Sylvie Behillil, Méline Bizard,Frédéric Lemoine,Corinne Maufrais,Christophe Malabat, Angela Brisebarre, Camille Capel, Louise Lefrançois, Etienne Simon-Lorière, Vincent Enouf, Maud Vanpeene, Sylvie van der Werf,Thibault Guinoiseau |
| EPI_ISL_1416906 | Labo analyses med | National Reference Center for Viruses of Respiratory Infections, Institut Pasteur, Paris | Marion Barbet, Sylvie Behillil, Méline Bizard,Frédéric Lemoine,Corinne Maufrais,Christophe Malabat, Angela Brisebarre, Camille Capel, Louise Lefrançois, Etienne Simon-Lorière, Vincent Enouf, Maud Vanpeene, Sylvie van der Werf,Anne Holstein |
| EPI_ISL_1416907 | Hospital | National Reference Center for Viruses of Respiratory Infections, Institut Pasteur, Paris | Marion Barbet, Sylvie Behillil, Méline Bizard,Frédéric Lemoine,Corinne Maufrais,Christophe Malabat, Angela Brisebarre, Camille Capel, Louise Lefrançois, Etienne Simon-Lorière, Vincent Enouf, Maud Vanpeene, Sylvie van der Werf,Anne Holstein |
| EPI_ISL_1416908 | Labo Analyses Med | National Reference Center for Viruses of Respiratory Infections, Institut Pasteur, Paris | Marion Barbet, Sylvie Behillil, Méline Bizard,Frédéric Lemoine,Corinne Maufrais,Christophe Malabat, Angela Brisebarre, Camille Capel, Louise Lefrançois, Etienne Simon-Lorière, Vincent Enouf, Maud Vanpeene, Sylvie van der Werf,BéAtrice Lesimple |
| EPI_ISL_1416909, EPI_ISL_1416910, EPI_ISL_1416911 | Hospital | National Reference Center for Viruses of Respiratory Infections, Institut Pasteur, Paris | Marion Barbet, Sylvie Behillil, Méline Bizard,Frédéric Lemoine,Corinne Maufrais,Christophe Malabat, Angela Brisebarre, Camille Capel, Louise Lefrançois, Etienne Simon-Lorière, Vincent Enouf, Maud Vanpeene, Sylvie van der Werf,Thibault Guinoiseau |
| EPI_ISL_1416912 | Hospital | National Reference Center for Viruses of Respiratory Infections, Institut Pasteur, Paris | Marion Barbet, Sylvie Behillil, Méline Bizard,Frédéric Lemoine,Corinne Maufrais,Christophe Malabat, Angela Brisebarre, Camille Capel, Louise Lefrançois, Etienne Simon-Lorière, Vincent Enouf, Maud Vanpeene, Sylvie van der Werf,Dominique Descamps |
| EPI_ISL_1416914, EPI_ISL_1416915 | Hospital | National Reference Center for Viruses of Respiratory Infections, Institut Pasteur, Paris | Marion Barbet, Sylvie Behillil, Méline Bizard,Frédéric Lemoine,Corinne Maufrais,Christophe Malabat, Angela Brisebarre, Camille Capel, Louise Lefrançois, Etienne Simon-Lorière, Vincent Enouf, Maud Vanpeene, Sylvie van der Werf,Thibault Guinoiseau |
| EPI_ISL_1416916 | Hospital | National Reference Center for Viruses of Respiratory Infections, Institut Pasteur, Paris | Marion Barbet, Sylvie Behillil, Méline Bizard,Frédéric Lemoine,Corinne Maufrais,Christophe Malabat, Angela Brisebarre, Camille Capel, Louise Lefrançois, Etienne Simon-Lorière, Vincent Enouf, Maud Vanpeene, Sylvie van der Werf,Catherine Coignard |

| | | | |
|---|---|---|---|
| EPI_ISL_1416917 | Hospital | National Reference Center for Viruses of Respiratory Infections, Institut Pasteur, Paris | Marion Barbet, Sylvie Behillil, Méline Bizard,Frédéric Lemoine,Corinne Maufrais,Christophe Malabat, Angela Brisebarre, Camille Capel, Louise Lefrançois, Etienne Simon-Lorière, Vincent Enouf, Maud Vanpeene, Sylvie van der Werf,Thibault Guinoiseau |
| EPI_ISL_1416918 | Hospital | National Reference Center for Viruses of Respiratory Infections, Institut Pasteur, Paris | Marion Barbet, Sylvie Behillil, Méline Bizard,Frédéric Lemoine,Corinne Maufrais,Christophe Malabat, Angela Brisebarre, Camille Capel, Louise Lefrançois, Etienne Simon-Lorière, Vincent Enouf, Maud Vanpeene, Sylvie van der Werf,Patricia Stoessel-Thouvenin |
| EPI_ISL_1416919 | Hospital | National Reference Center for Viruses of Respiratory Infections, Institut Pasteur, Paris | Marion Barbet, Sylvie Behillil, Méline Bizard,Frédéric Lemoine,Corinne Maufrais,Christophe Malabat, Angela Brisebarre, Camille Capel, Louise Lefrançois, Etienne Simon-Lorière, Vincent Enouf, Maud Vanpeene, Sylvie van der Werf,Dominique Descamps |
| EPI_ISL_1416920 | Hospital | National Reference Center for Viruses of Respiratory Infections, Institut Pasteur, Paris | Marion Barbet, Sylvie Behillil, Méline Bizard,Frédéric Lemoine,Corinne Maufrais,Christophe Malabat, Angela Brisebarre, Camille Capel, Louise Lefrançois, Etienne Simon-Lorière, Vincent Enouf, Maud Vanpeene, Sylvie van der Werf,Ilka Engelmann |
| EPI_ISL_1416921, EPI_ISL_1416922 | Hospital | National Reference Center for Viruses of Respiratory Infections, Institut Pasteur, Paris | Marion Barbet, Sylvie Behillil, Méline Bizard,Frédéric Lemoine,Corinne Maufrais,Christophe Malabat, Angela Brisebarre, Camille Capel, Louise Lefrançois, Etienne Simon-Lorière, Vincent Enouf, Maud Vanpeene, Sylvie van der Werf,Dominique Descamps |
| EPI_ISL_1416923 | Hospital | National Reference Center for Viruses of Respiratory Infections, Institut Pasteur, Paris | Marion Barbet, Sylvie Behillil, Méline Bizard,Frédéric Lemoine,Corinne Maufrais,Christophe Malabat, Angela Brisebarre, Camille Capel, Louise Lefrançois, Etienne Simon-Lorière, Vincent Enouf, Maud Vanpeene, Sylvie van der Werf,Ilka Engelmann |
| EPI_ISL_1416924 | Hospital | National Reference Center for Viruses of Respiratory Infections, Institut Pasteur, Paris | Marion Barbet, Sylvie Behillil, Méline Bizard,Frédéric Lemoine,Corinne Maufrais,Christophe Malabat, Angela Brisebarre, Camille Capel, Louise Lefrançois, Etienne Simon-Lorière, Vincent Enouf, Maud Vanpeene, Sylvie van der Werf,Ben Hadj Yahia |
| EPI_ISL_1416925, EPI_ISL_1416926 | Hospital | National Reference Center for Viruses of Respiratory Infections, Institut Pasteur, Paris | Marion Barbet, Sylvie Behillil, Méline Bizard,Frédéric Lemoine,Corinne Maufrais,Christophe Malabat, Angela Brisebarre, Camille Capel, Louise Lefrançois, Etienne Simon-Lorière, Vincent Enouf, Maud Vanpeene, Sylvie van der Werf,Ilka Engelmann |
| EPI_ISL_1416927 | Hospital | National Reference Center for Viruses of Respiratory Infections, Institut Pasteur, Paris | Marion Barbet, Sylvie Behillil, Méline Bizard,Frédéric Lemoine,Corinne Maufrais,Christophe Malabat, Angela Brisebarre, Camille Capel, Louise Lefrançois, Etienne Simon-Lorière, Vincent Enouf, Maud Vanpeene, Sylvie van der Werf,S. Van Agt |
| EPI_ISL_1416928, EPI_ISL_1416929, EPI_ISL_1416930, EPI_ISL_1416931 | Hospital | National Reference Center for Viruses of Respiratory Infections, Institut Pasteur, Paris | Marion Barbet, Sylvie Behillil, Méline Bizard,Frédéric Lemoine,Corinne Maufrais,Christophe Malabat, Angela Brisebarre, Camille Capel, Louise Lefrançois, Etienne Simon-Lorière, Vincent Enouf, Maud Vanpeene, Sylvie van der Werf,Ilka Engelmann |
| EPI_ISL_1416932 | Hospital | National Reference Center for Viruses of Respiratory Infections, Institut Pasteur, Paris | Marion Barbet, Sylvie Behillil, Méline Bizard,Frédéric Lemoine,Corinne Maufrais,Christophe Malabat, Angela Brisebarre, Camille Capel, Louise Lefrançois, Etienne Simon-Lorière, Vincent Enouf, Maud Vanpeene, Sylvie van der Werf,Mj Dufour |
| EPI_ISL_1416933 | Hospital | National Reference Center for Viruses of Respiratory Infections, Institut Pasteur, Paris | Marion Barbet, Sylvie Behillil, Méline Bizard,Frédéric Lemoine,Corinne Maufrais,Christophe Malabat, Angela Brisebarre, Camille Capel, Louise Lefrançois, Etienne Simon-Lorière, Vincent Enouf, Maud Vanpeene, Sylvie van der Werf,Ronan Pichard |
| EPI_ISL_1416934, EPI_ISL_1416935 | Hospital | National Reference Center for Viruses of Respiratory Infections, Institut Pasteur, Paris | Marion Barbet, Sylvie Behillil, Méline Bizard,Frédéric Lemoine,Corinne Maufrais,Christophe Malabat, Angela Brisebarre, Camille Capel, Louise Lefrançois, Etienne Simon-Lorière, Vincent Enouf, Maud Vanpeene, Sylvie van der Werf,S. Van Agt |
| EPI_ISL_1416936, EPI_ISL_1416937, EPI_ISL_1416938, EPI_ISL_1416939 | Hospital | National Reference Center for Viruses of Respiratory Infections, Institut Pasteur, Paris | Marion Barbet, Sylvie Behillil, Méline Bizard,Frédéric Lemoine,Corinne Maufrais,Christophe Malabat, Angela Brisebarre, Camille Capel, Louise Lefrançois, Etienne Simon-Lorière, Vincent Enouf, Maud Vanpeene, Sylvie van der Werf,Pascale Martres |
| EPI_ISL_1416940 | Hospital | National Reference Center for Viruses of Respiratory Infections, Institut Pasteur, Paris | Marion Barbet, Sylvie Behillil, Méline Bizard,Frédéric Lemoine,Corinne Maufrais,Christophe Malabat, Angela Brisebarre, Camille Capel, Louise Lefrançois, Etienne Simon-Lorière, Vincent Enouf, Maud Vanpeene, Sylvie van der Werf,Anne Cady |
| EPI_ISL_1416941, EPI_ISL_1416942, EPI_ISL_1416943 | Hospital | National Reference Center for Viruses of Respiratory Infections, Institut Pasteur, Paris | Marion Barbet, Sylvie Behillil, Méline Bizard,Frédéric Lemoine,Corinne Maufrais,Christophe Malabat, Angela Brisebarre, Camille Capel, Louise Lefrançois, Etienne Simon-Lorière, Vincent Enouf, Maud Vanpeene, Sylvie van der Werf,Ronan Pichard |
| EPI_ISL_1416944 | Hospital | National Reference Center for Viruses of Respiratory Infections, Institut Pasteur, Paris | Marion Barbet, Sylvie Behillil, Méline Bizard,Frédéric Lemoine,Corinne Maufrais,Christophe Malabat, Angela Brisebarre, Camille Capel, Louise Lefrançois, Etienne Simon-Lorière, Vincent Enouf, Maud Vanpeene, Sylvie van der Werf,Jean Baptiste Bour |
| EPI_ISL_1416945, EPI_ISL_1416946, EPI_ISL_1416947, EPI_ISL_1416948 | Hospital | National Reference Center for Viruses of Respiratory Infections, Institut Pasteur, Paris | Marion Barbet, Sylvie Behillil, Méline Bizard,Frédéric Lemoine,Corinne Maufrais,Christophe Malabat, Angela Brisebarre, Camille Capel, Louise Lefrançois, Etienne Simon-Lorière, Vincent Enouf, Maud Vanpeene, Sylvie van der Werf,Anne Cady |
| EPI_ISL_1416949 | Hospital | National Reference Center for Viruses of Respiratory Infections, Institut Pasteur, Paris | Marion Barbet, Sylvie Behillil, Méline Bizard,Frédéric Lemoine,Corinne Maufrais,Christophe Malabat, Angela Brisebarre, Camille Capel, Louise Lefrançois, Etienne Simon-Lorière, Vincent Enouf, Maud Vanpeene, Sylvie van der Werf,Jean Baptiste Bour |
| EPI_ISL_1416950 | Hospital | National Reference Center for Viruses of Respiratory Infections, Institut Pasteur, Paris | Marion Barbet, Sylvie Behillil, Méline Bizard,Frédéric Lemoine,Corinne Maufrais,Christophe Malabat, Angela Brisebarre, Camille Capel, Louise Lefrançois, Etienne Simon-Lorière, Vincent Enouf, Maud Vanpeene, Sylvie van der Werf,Ronan Pichard |
| EPI_ISL_1416951 | Hospital | National Reference Center for Viruses of Respiratory Infections, Institut Pasteur, Paris | Marion Barbet, Sylvie Behillil, Méline Bizard,Frédéric Lemoine,Corinne Maufrais,Christophe Malabat, Angela Brisebarre, Camille Capel, Louise Lefrançois, Etienne Simon-Lorière, Vincent Enouf, Maud Vanpeene, Sylvie van der Werf,Anne Cady |
| EPI_ISL_1416952, EPI_ISL_1416953 | Labo Analyses Med | National Reference Center for Viruses of Respiratory Infections, Institut Pasteur, Paris | Marion Barbet, Sylvie Behillil, Méline Bizard,Frédéric Lemoine,Corinne Maufrais,Christophe Malabat, Angela Brisebarre, Camille Capel, Louise Lefrançois, Etienne Simon-Lorière, Vincent Enouf, Maud Vanpeene, Sylvie van der Werf,Anne Cady |
| EPI_ISL_1416954 | Labo Analyses Med | National Reference Center for Viruses of Respiratory Infections, Institut Pasteur, Paris | Marion Barbet, Sylvie Behillil, Méline Bizard,Frédéric Lemoine,Corinne Maufrais,Christophe Malabat, Angela Brisebarre, Camille Capel, Louise Lefrançois, Etienne Simon-Lorière, Vincent Enouf, Maud Vanpeene, Sylvie van der Werf,Ronan Pichard |
| EPI_ISL_1416955 | Labo Analyses Med | National Reference Center for Viruses of Respiratory Infections, Institut Pasteur, Paris | Marion Barbet, Sylvie Behillil, Méline Bizard,Frédéric Lemoine,Corinne Maufrais,Christophe Malabat, Angela Brisebarre, Camille Capel, Louise Lefrançois, Etienne Simon-Lorière, Vincent Enouf, Maud Vanpeene, Sylvie van der Werf,Sophie Chalmin |
| EPI_ISL_1416956, EPI_ISL_1416957, EPI_ISL_1416958, EPI_ISL_1416959 | Labo Analyses Med | National Reference Center for Viruses of Respiratory Infections, Institut Pasteur, Paris | Marion Barbet, Sylvie Behillil, Méline Bizard,Frédéric Lemoine,Corinne Maufrais,Christophe Malabat, Angela Brisebarre, Camille Capel, Louise Lefrançois, Etienne Simon-Lorière, Vincent Enouf, Maud Vanpeene, Sylvie van der Werf,Ronan Pichard |
| EPI_ISL_1416960, EPI_ISL_1416963 | Labo Analyses Med | National Reference Center for Viruses of Respiratory Infections, Institut Pasteur, Paris | Marion Barbet, Sylvie Behillil, Méline Bizard,Frédéric Lemoine,Corinne Maufrais,Christophe Malabat, Angela Brisebarre, Camille Capel, Louise Lefrançois, Etienne Simon-Lorière, Vincent Enouf, Maud Vanpeene, Sylvie van der Werf,Claire Felloni |
| EPI_ISL_1416964, EPI_ISL_1416965 | Labo Analyses Med | National Reference Center for Viruses of Respiratory Infections, Institut Pasteur, Paris | Marion Barbet, Sylvie Behillil, Méline Bizard,Frédéric Lemoine,Corinne Maufrais,Christophe Malabat, Angela Brisebarre, Camille Capel, Louise Lefrançois, Etienne Simon-Lorière, Vincent Enouf, Maud Vanpeene, Sylvie van der Werf,Ronan Pichard |
| EPI_ISL_1416966 | Outre mer | National Reference Center for Viruses of Respiratory Infections, Institut Pasteur, Paris | Marion Barbet, Sylvie Behillil, Méline Bizard,Frédéric Lemoine,Corinne Maufrais,Christophe Malabat, Angela Brisebarre, Camille Capel, Louise Lefrançois, Etienne Simon-Lorière, Vincent Enouf, Maud Vanpeene, Sylvie van der Werf,Claire Felloni |
| EPI_ISL_1416967 | Outre mer | National Reference Center for Viruses of Respiratory Infections, Institut Pasteur, Paris | Marion Barbet, Sylvie Behillil, Méline Bizard,Frédéric Lemoine,Corinne Maufrais,Christophe Malabat, Angela Brisebarre, Camille Capel, Louise Lefrançois, Etienne Simon-Lorière, Vincent Enouf, Maud Vanpeene, Sylvie van der Werf,Esther Gyde |
| EPI_ISL_1416968, EPI_ISL_1416969 | Outre mer | National Reference Center for Viruses of Respiratory Infections, Institut Pasteur, Paris | Marion Barbet, Sylvie Behillil, Méline Bizard,Frédéric Lemoine,Corinne Maufrais,Christophe Malabat, Angela Brisebarre, Camille Capel, Louise Lefrançois, Etienne Simon-Lorière, Vincent Enouf, Maud Vanpeene, Sylvie van der Werf,Claire Felloni |
| EPI_ISL_1416970 | Outre mer | National Reference Center for Viruses of Respiratory Infections, Institut Pasteur, Paris | Marion Barbet, Sylvie Behillil, Méline Bizard,Frédéric Lemoine,Corinne Maufrais,Christophe Malabat, Angela Brisebarre, Camille Capel, Louise Lefrançois, Etienne Simon-Lorière, Vincent Enouf, Maud Vanpeene, Sylvie van der Werf,Esther Gyde |
| EPI_ISL_1416971 | Outre mer | National Reference Center for Viruses of Respiratory Infections, Institut Pasteur, Paris | Marion Barbet, Sylvie Behillil, Méline Bizard,Frédéric Lemoine,Corinne Maufrais,Christophe Malabat, Angela Brisebarre, Camille Capel, Louise Lefrançois, Etienne Simon-Lorière, Vincent Enouf, Maud Vanpeene, Sylvie van der Werf,StéPhanie Guyomard-Rabenirina |
| EPI_ISL_1416972, EPI_ISL_1416973, EPI_ISL_1416975, EPI_ISL_1416977 | Outre mer | National Reference Center for Viruses of Respiratory Infections, Institut Pasteur, Paris | Marion Barbet, Sylvie Behillil, Méline Bizard,Frédéric Lemoine,Corinne Maufrais,Christophe Malabat, Angela Brisebarre, Camille Capel, Louise Lefrançois, Etienne Simon-Lorière, Vincent Enouf, Maud Vanpeene, Sylvie van der Werf,Esther Gyde |
| EPI_ISL_1416978 | Outre mer | National Reference Center for Viruses of Respiratory Infections, Institut Pasteur, Paris | Marion Barbet, Sylvie Behillil, Méline Bizard,Frédéric Lemoine,Corinne Maufrais,Christophe Malabat, Angela Brisebarre, Camille Capel, Louise Lefrançois, Etienne Simon-Lorière, Vincent Enouf, Maud Vanpeene, Sylvie van der Werf,CéLine Ramanantsoa |
| EPI_ISL_1416979 | Outre mer | National Reference Center for Viruses of Respiratory Infections, Institut Pasteur, Paris | Marion Barbet, Sylvie Behillil, Méline Bizard,Frédéric Lemoine,Corinne Maufrais,Christophe Malabat, Angela Brisebarre, Camille Capel, Louise Lefrançois, Etienne Simon-Lorière, Vincent Enouf, Maud Vanpeene, Sylvie van der Werf,JéRome Guinard |

| | | | |
|---|---|---|---|
| EPI_ISL_1416980, EPI_ISL_1416981, EPI_ISL_1416982, EPI_ISL_1416983, EPI_ISL_1416984, EPI_ISL_1416985, EPI_ISL_1416986, EPI_ISL_1416987, EPI_ISL_1416988 | Outre mer | National Reference Center for Viruses of Respiratory Infections, Institut Pasteur, Paris | Marion Barbet, Sylvie Behillil, Méline Bizard,Frédéric Lemoine,Corinne Maufrais,Christophe Malabat, Angela Brisebarre, Camille Capel, Louise Lefrançois, Etienne Simon-Lorière, Vincent Enouf, Maud Vanpeene, Sylvie van der Werf,GréGoire Potiron |
| EPI_ISL_1416990 | Outre mer | National Reference Center for Viruses of Respiratory Infections, Institut Pasteur, Paris | Marion Barbet, Sylvie Behillil, Méline Bizard,Frédéric Lemoine,Corinne Maufrais,Christophe Malabat, Angela Brisebarre, Camille Capel, Louise Lefrançois, Etienne Simon-Lorière, Vincent Enouf, Maud Vanpeene, Sylvie van der Werf,Anne Holstein |
| EPI_ISL_1416998, EPI_ISL_1416999 | Hospital | National Reference Center for Viruses of Respiratory Infections, Institut Pasteur, Paris | Marion Barbet, Sylvie Behillil, Méline Bizard,Frédéric Lemoine,Corinne Maufrais,Christophe Malabat, Angela Brisebarre, Camille Capel, Louise Lefrançois, Etienne Simon-Lorière, Vincent Enouf, Maud Vanpeene, Sylvie van der Werf,Anne Holstein |
| EPI_ISL_1417000, EPI_ISL_1417001, EPI_ISL_1417002, EPI_ISL_1417003, EPI_ISL_1417005, EPI_ISL_1417006, EPI_ISL_1417007, EPI_ISL_1417008, EPI_ISL_1417009, EPI_ISL_1417010, EPI_ISL_1417012, EPI_ISL_1417013, EPI_ISL_1417014 | | | |
| see above | Hospital | National Reference Center for Viruses of Respiratory Infections, Institut Pasteur, Paris | Marion Barbet, Sylvie Behillil, Méline Bizard,Frédéric Lemoine,Corinne Maufrais,Christophe Malabat, Angela Brisebarre, Camille Capel, Louise Lefrançois, Etienne Simon-Lorière, Vincent Enouf, Maud Vanpeene, Sylvie van der Werf,FréDéRique Canis |
| EPI_ISL_1417015, EPI_ISL_1417016, EPI_ISL_1417017, EPI_ISL_1417018, EPI_ISL_1417020, EPI_ISL_1417021, EPI_ISL_1417022, EPI_ISL_1417023 | Labo Analyses Med | National Reference Center for Viruses of Respiratory Infections, Institut Pasteur, Paris | Marion Barbet, Sylvie Behillil, Méline Bizard,Frédéric Lemoine,Corinne Maufrais,Christophe Malabat, Angela Brisebarre, Camille Capel, Louise Lefrançois, Etienne Simon-Lorière, Vincent Enouf, Maud Vanpeene, Sylvie van der Werf,Catherine Coignard |
| EPI_ISL_1417024, EPI_ISL_1417025 | Hospital | National Reference Center for Viruses of Respiratory Infections, Institut Pasteur, Paris | Marion Barbet, Sylvie Behillil, Méline Bizard,Frédéric Lemoine,Corinne Maufrais,Christophe Malabat, Angela Brisebarre, Camille Capel, Louise Lefrançois, Etienne Simon-Lorière, Vincent Enouf, Maud Vanpeene, Sylvie van der Werf,FrançOise Lunel Fabiani |
| EPI_ISL_1417026, EPI_ISL_1417027, EPI_ISL_1417028, EPI_ISL_1417029, EPI_ISL_1417030, EPI_ISL_1417031, EPI_ISL_1417032 | Labo Analyses Med | National Reference Center for Viruses of Respiratory Infections, Institut Pasteur, Paris | Marion Barbet, Sylvie Behillil, Méline Bizard,Frédéric Lemoine,Corinne Maufrais,Christophe Malabat, Angela Brisebarre, Camille Capel, Louise Lefrançois, Etienne Simon-Lorière, Vincent Enouf, Maud Vanpeene, Sylvie van der Werf,Ronan Pichard |
| EPI_ISL_1417033, EPI_ISL_1417034, EPI_ISL_1417035, EPI_ISL_1417036, EPI_ISL_1417037, EPI_ISL_1417038, EPI_ISL_1417039, EPI_ISL_1417041, EPI_ISL_1417042, EPI_ISL_1417043 | Labo Analyses Med | National Reference Center for Viruses of Respiratory Infections, Institut Pasteur, Paris | Marion Barbet, Sylvie Behillil, Méline Bizard,Frédéric Lemoine,Corinne Maufrais,Christophe Malabat, Angela Brisebarre, Camille Capel, Louise Lefrançois, Etienne Simon-Lorière, Vincent Enouf, Maud Vanpeene, Sylvie van der Werf,Pierre-Yves Leonard |
| EPI_ISL_1417044, EPI_ISL_1417045, EPI_ISL_1417046, EPI_ISL_1417048, EPI_ISL_1417049 | Labo Analyses Med | National Reference Center for Viruses of Respiratory Infections, Institut Pasteur, Paris | Marion Barbet, Sylvie Behillil, Méline Bizard,Frédéric Lemoine,Corinne Maufrais,Christophe Malabat, Angela Brisebarre, Camille Capel, Louise Lefrançois, Etienne Simon-Lorière, Vincent Enouf, Maud Vanpeene, Sylvie van der Werf,Sophie Chalmin |
| EPI_ISL_1417050 | Labo Analyses Med | National Reference Center for Viruses of Respiratory Infections, Institut Pasteur, Paris | Marion Barbet, Sylvie Behillil, Méline Bizard,Frédéric Lemoine,Corinne Maufrais,Christophe Malabat, Angela Brisebarre, Camille Capel, Louise Lefrançois, Etienne Simon-Lorière, Vincent Enouf, Maud Vanpeene, Sylvie van der Werf,Jean-FrançOis Comes |
| EPI_ISL_1417052 | Hospital | National Reference Center for Viruses of Respiratory Infections, Institut Pasteur, Paris | Marion Barbet, Sylvie Behillil, Méline Bizard,Frédéric Lemoine,Corinne Maufrais,Christophe Malabat, Angela Brisebarre, Camille Capel, Louise Lefrançois, Etienne Simon-Lorière, Vincent Enouf, Maud Vanpeene, Sylvie van der Werf,Jean-FrançOis Comes |
| EPI_ISL_1417053, EPI_ISL_1417054, EPI_ISL_1417055 | Hospital | National Reference Center for Viruses of Respiratory Infections, Institut Pasteur, Paris | Marion Barbet, Sylvie Behillil, Méline Bizard,Frédéric Lemoine,Corinne Maufrais,Christophe Malabat, Angela Brisebarre, Camille Capel, Louise Lefrançois, Etienne Simon-Lorière, Vincent Enouf, Maud Vanpeene, Sylvie van der Werf,OphéLie Said-Delattre |
| EPI_ISL_1417056 | Hospital | National Reference Center for Viruses of Respiratory Infections, Institut Pasteur, Paris | Marion Barbet, Sylvie Behillil, Méline Bizard,Frédéric Lemoine,Corinne Maufrais,Christophe Malabat, Angela Brisebarre, Camille Capel, Louise Lefrançois, Etienne Simon-Lorière, Vincent Enouf, Maud Vanpeene, Sylvie van der Werf,CéLine Bressollette |
| EPI_ISL_1417057 | Hospital | National Reference Center for Viruses of Respiratory Infections, Institut Pasteur, Paris | Marion Barbet, Sylvie Behillil, Méline Bizard,Frédéric Lemoine,Corinne Maufrais,Christophe Malabat, Angela Brisebarre, Camille Capel, Louise Lefrançois, Etienne Simon-Lorière, Vincent Enouf, Maud Vanpeene, Sylvie van der Werf,OphéLie Said-Delattre |
| EPI_ISL_1417058 | Hospital | National Reference Center for Viruses of Respiratory Infections, Institut Pasteur, Paris | Marion Barbet, Sylvie Behillil, Méline Bizard,Frédéric Lemoine,Corinne Maufrais,Christophe Malabat, Angela Brisebarre, Camille Capel, Louise Lefrançois, Etienne Simon-Lorière, Vincent Enouf, Maud Vanpeene, Sylvie van der Werf,CéLine Bressollette |
| EPI_ISL_1417060, EPI_ISL_1417061, EPI_ISL_1417062, EPI_ISL_1417063, EPI_ISL_1417064, EPI_ISL_1417065 | Hospital | National Reference Center for Viruses of Respiratory Infections, Institut Pasteur, Paris | Marion Barbet, Sylvie Behillil, Méline Bizard,Frédéric Lemoine,Corinne Maufrais,Christophe Malabat, Angela Brisebarre, Camille Capel, Louise Lefrançois, Etienne Simon-Lorière, Vincent Enouf, Maud Vanpeene, Sylvie van der Werf,OphéLie Said-Delattre |
| EPI_ISL_1417066 | Hospital | National Reference Center for Viruses of Respiratory Infections, Institut Pasteur, Paris | Marion Barbet, Sylvie Behillil, Méline Bizard,Frédéric Lemoine,Corinne Maufrais,Christophe Malabat, Angela Brisebarre, Camille Capel, Louise Lefrançois, Etienne Simon-Lorière, Vincent Enouf, Maud Vanpeene, Sylvie van der Werf,CéLine Bressollette |
| EPI_ISL_1417071, EPI_ISL_1417072, EPI_ISL_1417073, EPI_ISL_1417074, EPI_ISL_1417075, EPI_ISL_1417076, EPI_ISL_1417077, EPI_ISL_1417078 | Labo Analyses Med | National Reference Center for Viruses of Respiratory Infections, Institut Pasteur, Paris | Marion Barbet, Sylvie Behillil, Méline Bizard,Frédéric Lemoine,Corinne Maufrais,Christophe Malabat, Angela Brisebarre, Camille Capel, Louise Lefrançois, Etienne Simon-Lorière, Vincent Enouf, Maud Vanpeene, Sylvie van der Werf,OphéLie Said-Delattre |
| EPI_ISL_1417079, EPI_ISL_1417080, EPI_ISL_1417081, EPI_ISL_1417082, EPI_ISL_1417083, EPI_ISL_1417084, EPI_ISL_1417085, EPI_ISL_1417086 | Labo Analyses Med | National Reference Center for Viruses of Respiratory Infections, Institut Pasteur, Paris | Marion Barbet, Sylvie Behillil, Méline Bizard,Frédéric Lemoine,Corinne Maufrais,Christophe Malabat, Angela Brisebarre, Camille Capel, Louise Lefrançois, Etienne Simon-Lorière, Vincent Enouf, Maud Vanpeene, Sylvie van der Werf,Catherine Coignard |
| EPI_ISL_1417087 | Labo Analyses Med | National Reference Center for Viruses of Respiratory Infections, Institut Pasteur, Paris | Marion Barbet, Sylvie Behillil, Méline Bizard,Frédéric Lemoine,Corinne Maufrais,Christophe Malabat, Angela Brisebarre, Camille Capel, Louise Lefrançois, Etienne Simon-Lorière, Vincent Enouf, Maud Vanpeene, Sylvie van der Werf,Jean-FrançOis Comes |
| EPI_ISL_1417088, EPI_ISL_1417089 | Labo Analyses Med | National Reference Center for Viruses of Respiratory Infections, Institut Pasteur, Paris | Marion Barbet, Sylvie Behillil, Méline Bizard,Frédéric Lemoine,Corinne Maufrais,Christophe Malabat, Angela Brisebarre, Camille Capel, Louise Lefrançois, Etienne Simon-Lorière, Vincent Enouf, Maud Vanpeene, Sylvie van der Werf,Catherine Coignard |
| EPI_ISL_1417090 | Labo Analyses Med | National Reference Center for Viruses of Respiratory Infections, Institut Pasteur, Paris | Marion Barbet, Sylvie Behillil, Méline Bizard,Frédéric Lemoine,Corinne Maufrais,Christophe Malabat, Angela Brisebarre, Camille Capel, Louise Lefrançois, Etienne Simon-Lorière, Vincent Enouf, Maud Vanpeene, Sylvie van der Werf,Sophie Chalmin |
| EPI_ISL_1417091, EPI_ISL_1417092, EPI_ISL_1417093 | Labo Analyses Med | National Reference Center for Viruses of Respiratory Infections, Institut Pasteur, Paris | Marion Barbet, Sylvie Behillil, Méline Bizard,Frédéric Lemoine,Corinne Maufrais,Christophe Malabat, Angela Brisebarre, Camille Capel, Louise Lefrançois, Etienne Simon-Lorière, Vincent Enouf, Maud Vanpeene, Sylvie van der Werf,Catherine Coignard |
| EPI_ISL_1417094 | Labo Analyses Med | National Reference Center for Viruses of Respiratory Infections, Institut Pasteur, Paris | Marion Barbet, Sylvie Behillil, Méline Bizard,Frédéric Lemoine,Corinne Maufrais,Christophe Malabat, Angela Brisebarre, Camille Capel, Louise Lefrançois, Etienne Simon-Lorière, Vincent Enouf, Maud Vanpeene, Sylvie van der Werf,Sophie Chalmin |
| EPI_ISL_1417095 | Hospital | National Reference Center for Viruses of Respiratory Infections, Institut Pasteur, Paris | Marion Barbet, Sylvie Behillil, Méline Bizard,Frédéric Lemoine,Corinne Maufrais,Christophe Malabat, Angela Brisebarre, Camille Capel, Louise Lefrançois, Etienne Simon-Lorière, Vincent Enouf, Maud Vanpeene, Sylvie van der Werf,Catherine Coignard |
| EPI_ISL_1417096, EPI_ISL_1417097 | Hospital | National Reference Center for Viruses of Respiratory Infections, Institut Pasteur, Paris | Marion Barbet, Sylvie Behillil, Méline Bizard,Frédéric Lemoine,Corinne Maufrais,Christophe Malabat, Angela Brisebarre, Camille Capel, Louise Lefrançois, Etienne Simon-Lorière, Vincent Enouf, Maud Vanpeene, Sylvie van der Werf,Sophie Chalmin |
| EPI_ISL_1417098 | Hospital | National Reference Center for Viruses of Respiratory Infections, Institut Pasteur, Paris | Marion Barbet, Sylvie Behillil, Méline Bizard,Frédéric Lemoine,Corinne Maufrais,Christophe Malabat, Angela Brisebarre, Camille Capel, Louise Lefrançois, Etienne Simon-Lorière, Vincent Enouf, Maud Vanpeene, Sylvie van der Werf,Catherine Coignard |
| EPI_ISL_1417099, EPI_ISL_1417100, EPI_ISL_1417101, EPI_ISL_1417102, EPI_ISL_1417103, EPI_ISL_1417104, EPI_ISL_1417105, EPI_ISL_1417106, EPI_ISL_1417107, EPI_ISL_1417109, EPI_ISL_1417110, EPI_ISL_1417111 | | | |
| see above | Hospital | National Reference Center for Viruses of Respiratory | Marion Barbet, Sylvie Behillil, Méline Bizard,Frédéric Lemoine,Corinne Maufrais,Christophe Malabat, Angela Brisebarre, Camille Capel, Louise Lefrançois, |

| | | | |
|---|---|---|---|
| | | Infections, Institut Pasteur, Paris | Etienne Simon-Lorière, Vincent Enouf, Maud Vanpeene, Sylvie van der Werf,Sophie Chalmin |
| EPI_ISL_1417112, EPI_ISL_1417113, EPI_ISL_1417114, EPI_ISL_1417115 | Labo Analyses Med | National Reference Center for Viruses of Respiratory Infections, Institut Pasteur, Paris | Marion Barbet, Sylvie Behillil, Méline Bizard,Frédéric Lemoine,Corinne Maufrais,Christophe Malabat, Angela Brisebarre, Camille Capel, Louise Lefrançois, Etienne Simon-Lorière, Vincent Enouf, Maud Vanpeene, Sylvie van der Werf,Sophie Chalmin |
| EPI_ISL_1417116, EPI_ISL_1417117, EPI_ISL_1417118, EPI_ISL_1417119, EPI_ISL_1417120, EPI_ISL_1417121, EPI_ISL_1417122, EPI_ISL_1417123, EPI_ISL_1417124, EPI_ISL_1417125 | Labo Analyses Med | National Reference Center for Viruses of Respiratory Infections, Institut Pasteur, Paris | Marion Barbet, Sylvie Behillil, Méline Bizard,Frédéric Lemoine,Corinne Maufrais,Christophe Malabat, Angela Brisebarre, Camille Capel, Louise Lefrançois, Etienne Simon-Lorière, Vincent Enouf, Maud Vanpeene, Sylvie van der Werf,Catherine Coignard |
| EPI_ISL_1417126, EPI_ISL_1417127, EPI_ISL_1417128, EPI_ISL_1417129, EPI_ISL_1417131, EPI_ISL_1417132, EPI_ISL_1417133, EPI_ISL_1417134, EPI_ISL_1417137, EPI_ISL_1417138, EPI_ISL_1417139, EPI_ISL_1417140, EPI_ISL_1417142, EPI_ISL_1417143 | | | |
| see above | Labo Analyses Med | National Reference Center for Viruses of Respiratory Infections, Institut Pasteur, Paris | Marion Barbet, Sylvie Behillil, Méline Bizard,Frédéric Lemoine,Corinne Maufrais,Christophe Malabat, Angela Brisebarre, Camille Capel, Louise Lefrançois, Etienne Simon-Lorière, Vincent Enouf, Maud Vanpeene, Sylvie van der Werf,Jean-FrançOis Comes |
| EPI_ISL_1417145, EPI_ISL_1417146, EPI_ISL_1417147, EPI_ISL_1417148, EPI_ISL_1417149, EPI_ISL_1417150, EPI_ISL_1417151, EPI_ISL_1417152 | Labo Analyses Med | National Reference Center for Viruses of Respiratory Infections, Institut Pasteur, Paris | Marion Barbet, Sylvie Behillil, Méline Bizard,Frédéric Lemoine,Corinne Maufrais,Christophe Malabat, Angela Brisebarre, Camille Capel, Louise Lefrançois, Etienne Simon-Lorière, Vincent Enouf, Maud Vanpeene, Sylvie van der Werf,Catherine Coignard |
| EPI_ISL_1417153, EPI_ISL_1417154 | Labo Analyses Med | National Reference Center for Viruses of Respiratory Infections, Institut Pasteur, Paris | Marion Barbet, Sylvie Behillil, Méline Bizard,Frédéric Lemoine,Corinne Maufrais,Christophe Malabat, Angela Brisebarre, Camille Capel, Louise Lefrançois, Etienne Simon-Lorière, Vincent Enouf, Maud Vanpeene, Sylvie van der Werf,M.-J. Blanc-Galiby |
| EPI_ISL_1417156, EPI_ISL_1417157, EPI_ISL_1417158, EPI_ISL_1417159, EPI_ISL_1417160, EPI_ISL_1417161, EPI_ISL_1417162, EPI_ISL_1417163, EPI_ISL_1417164, EPI_ISL_1417165, EPI_ISL_1417166, EPI_ISL_1417167, EPI_ISL_1417168, EPI_ISL_1417169 | | | |
| see above | Labo Analyses Med | National Reference Center for Viruses of Respiratory Infections, Institut Pasteur, Paris | Marion Barbet, Sylvie Behillil, Méline Bizard,Frédéric Lemoine,Corinne Maufrais,Christophe Malabat, Angela Brisebarre, Camille Capel, Louise Lefrançois, Etienne Simon-Lorière, Vincent Enouf, Maud Vanpeene, Sylvie van der Werf,Jean-FrançOis Comes |
| EPI_ISL_1417170 | Labo Analyses Med | National Reference Center for Viruses of Respiratory Infections, Institut Pasteur, Paris | Marion Barbet, Sylvie Behillil, Méline Bizard,Frédéric Lemoine,Corinne Maufrais,Christophe Malabat, Angela Brisebarre, Camille Capel, Louise Lefrançois, Etienne Simon-Lorière, Vincent Enouf, Maud Vanpeene, Sylvie van der Werf,CéLine Bressollette |
| EPI_ISL_1417171 | Labo Analyses Med | National Reference Center for Viruses of Respiratory Infections, Institut Pasteur, Paris | Marion Barbet, Sylvie Behillil, Méline Bizard,Frédéric Lemoine,Corinne Maufrais,Christophe Malabat, Angela Brisebarre, Camille Capel, Louise Lefrançois, Etienne Simon-Lorière, Vincent Enouf, Maud Vanpeene, Sylvie van der Werf,OphéLie Said-Delattre |
| EPI_ISL_1417172, EPI_ISL_1417173 | Labo Analyses Med | National Reference Center for Viruses of Respiratory Infections, Institut Pasteur, Paris | Marion Barbet, Sylvie Behillil, Méline Bizard,Frédéric Lemoine,Corinne Maufrais,Christophe Malabat, Angela Brisebarre, Camille Capel, Louise Lefrançois, Etienne Simon-Lorière, Vincent Enouf, Maud Vanpeene, Sylvie van der Werf,Sophie Chalmin |
| EPI_ISL_1417174, EPI_ISL_1417175, EPI_ISL_1417176, EPI_ISL_1417177, EPI_ISL_1417178, EPI_ISL_1417179, EPI_ISL_1417180 | Labo Analyses Med | National Reference Center for Viruses of Respiratory Infections, Institut Pasteur, Paris | Marion Barbet, Sylvie Behillil, Méline Bizard,Frédéric Lemoine,Corinne Maufrais,Christophe Malabat, Angela Brisebarre, Camille Capel, Louise Lefrançois, Etienne Simon-Lorière, Vincent Enouf, Maud Vanpeene, Sylvie van der Werf,CéLine Bressollette |
| EPI_ISL_1417181, EPI_ISL_1417182 | Labo Analyses Med | National Reference Center for Viruses of Respiratory Infections, Institut Pasteur, Paris | Marion Barbet, Sylvie Behillil, Méline Bizard,Frédéric Lemoine,Corinne Maufrais,Christophe Malabat, Angela Brisebarre, Camille Capel, Louise Lefrançois, Etienne Simon-Lorière, Vincent Enouf, Maud Vanpeene, Sylvie van der Werf,OphéLie Said-Delattre |
| EPI_ISL_1417183, EPI_ISL_1417184, EPI_ISL_1417185, EPI_ISL_1417186, EPI_ISL_1417187, EPI_ISL_1417188, EPI_ISL_1417189 | Labo Analyses Med | National Reference Center for Viruses of Respiratory Infections, Institut Pasteur, Paris | Marion Barbet, Sylvie Behillil, Méline Bizard,Frédéric Lemoine,Corinne Maufrais,Christophe Malabat, Angela Brisebarre, Camille Capel, Louise Lefrançois, Etienne Simon-Lorière, Vincent Enouf, Maud Vanpeene, Sylvie van der Werf,CéLine Bressollette |
| EPI_ISL_1417190 | Labo Analyses Med | National Reference Center for Viruses of Respiratory Infections, Institut Pasteur, Paris | Marion Barbet, Sylvie Behillil, Méline Bizard,Frédéric Lemoine,Corinne Maufrais,Christophe Malabat, Angela Brisebarre, Camille Capel, Louise Lefrançois, Etienne Simon-Lorière, Vincent Enouf, Maud Vanpeene, Sylvie van der Werf,Catherine Coignard |
| EPI_ISL_1417191 | Labo Analyses Med | National Reference Center for Viruses of Respiratory Infections, Institut Pasteur, Paris | Marion Barbet, Sylvie Behillil, Méline Bizard,Frédéric Lemoine,Corinne Maufrais,Christophe Malabat, Angela Brisebarre, Camille Capel, Louise Lefrançois, Etienne Simon-Lorière, Vincent Enouf, Maud Vanpeene, Sylvie van der Werf,OphéLie Said-Delattre |
| EPI_ISL_1417192 | Labo Analyses Med | National Reference Center for Viruses of Respiratory Infections, Institut Pasteur, Paris | Marion Barbet, Sylvie Behillil, Méline Bizard,Frédéric Lemoine,Corinne Maufrais,Christophe Malabat, Angela Brisebarre, Camille Capel, Louise Lefrançois, Etienne Simon-Lorière, Vincent Enouf, Maud Vanpeene, Sylvie van der Werf,CéLine Bressollette |
| EPI_ISL_1417193, EPI_ISL_1417194, EPI_ISL_1417195, EPI_ISL_1417196, EPI_ISL_1417197, EPI_ISL_1417199 | Labo Analyses Med | National Reference Center for Viruses of Respiratory Infections, Institut Pasteur, Paris | Marion Barbet, Sylvie Behillil, Méline Bizard,Frédéric Lemoine,Corinne Maufrais,Christophe Malabat, Angela Brisebarre, Camille Capel, Louise Lefrançois, Etienne Simon-Lorière, Vincent Enouf, Maud Vanpeene, Sylvie van der Werf,Catherine Coignard |
| EPI_ISL_1417200, EPI_ISL_1417201, EPI_ISL_1417202, EPI_ISL_1417203, EPI_ISL_1417204, EPI_ISL_1417205, EPI_ISL_1417206, EPI_ISL_1417207, EPI_ISL_1417208, EPI_ISL_1417209, EPI_ISL_1417210, EPI_ISL_1417211, EPI_ISL_1417212, EPI_ISL_1417213, EPI_ISL_1417214, EPI_ISL_1417215 | | | |
| see above | Labo Analyses Med | National Reference Center for Viruses of Respiratory Infections, Institut Pasteur, Paris | Marion Barbet, Sylvie Behillil, Méline Bizard,Frédéric Lemoine,Corinne Maufrais,Christophe Malabat, Angela Brisebarre, Camille Capel, Louise Lefrançois, Etienne Simon-Lorière, Vincent Enouf, Maud Vanpeene, Sylvie van der Werf,OphéLie Said-Delattre |
| EPI_ISL_1418319, EPI_ISL_1418320, EPI_ISL_1418321, EPI_ISL_1418322, EPI_ISL_1418326, EPI_ISL_1418327, EPI_ISL_1418328, EPI_ISL_1418329 | National Institute of Infectious Diseases-Prof. Dr. Matei Bals Molecular Diagnostics Laboratory | National Institute of Infectious Diseases-Prof. Dr. Matei Bals Molecular Diagnostics Laboratory | Corina Casangiu, Leontina Banica, Marius Surleac, Ovidiu Vlaicu, Ionelia Nicolae, Simona Paraschiv, Dan Otelea |
| EPI_ISL_1423505 | Unidad de Patologia Clinica | Instituto de Diagnostico y Referencia Epidemiologicos (INDRE) | Claudia Wong-Arambula, Abril Rodriguez-Maldonado, Vanessa Rivero-Arredondo, Ariadna Medina-Benitez, Joaquin Quiroz-Mercado, Sergio Rangel-Guerrero, Natividad Cruz-Ortiz, Tatiana Nunez-Garcia, Gisela Barrera-Badillo, Lucia Hernandez-Rivas, Irma Lopez-Martinez, Ernesto Ramirez-Gonzalez. |
| EPI_ISL_1423507 | LESP Chihuahua | Instituto de Diagnostico y Referencia Epidemiologicos (INDRE) | Claudia Wong-Arambula, Abril Rodriguez-Maldonado, Vanessa Rivero-Arredondo, Ariadna Medina-Benitez, Joaquin Quiroz-Mercado, Sergio Rangel-Guerrero, Natividad Cruz-Ortiz, Tatiana Nunez-Garcia, Gisela Barrera-Badillo, Lucia Hernandez-Rivas, Irma Lopez-Martinez, Ernesto Ramirez-Gonzalez. |
| EPI_ISL_1423511 | LESP Tamaulipas | Instituto de Diagnostico y Referencia Epidemiologicos (INDRE) | Claudia Wong-Arambula, Abril Rodriguez-Maldonado, Vanessa Rivero-Arredondo, Ariadna Medina-Benitez, Joaquin Quiroz-Mercado, Sergio Rangel-Guerrero, Natividad Cruz-Ortiz, Tatiana Nunez-Garcia, Gisela Barrera-Badillo, Lucia Hernandez-Rivas, Irma Lopez-Martinez, Ernesto Ramirez-Gonzalez. |
| EPI_ISL_1423512 | LESP Chihuahua | Instituto de Diagnostico y Referencia Epidemiologicos (INDRE) | Claudia Wong-Arambula, Abril Rodriguez-Maldonado, Vanessa Rivero-Arredondo, Ariadna Medina-Benitez, Joaquin Quiroz-Mercado, Sergio Rangel-Guerrero, Natividad Cruz-Ortiz, Tatiana Nunez-Garcia, Gisela Barrera-Badillo, Lucia Hernandez-Rivas, Irma Lopez-Martinez, Ernesto Ramirez-Gonzalez. |
| EPI_ISL_1423513 | LESP Queretaro | Instituto de Diagnostico y Referencia Epidemiologicos (INDRE) | Claudia Wong-Arambula, Abril Rodriguez-Maldonado, Vanessa Rivero-Arredondo, Ariadna Medina-Benitez, Joaquin Quiroz-Mercado, Sergio Rangel-Guerrero, Natividad Cruz-Ortiz, Tatiana Nunez-Garcia, Gisela Barrera-Badillo, Lucia Hernandez-Rivas, Irma Lopez-Martinez, Ernesto Ramirez-Gonzalez. |
| EPI_ISL_1423514 | LESP Yucatan | Instituto de Diagnostico y Referencia Epidemiologicos (INDRE) | Claudia Wong-Arambula, Abril Rodriguez-Maldonado, Vanessa Rivero-Arredondo, Ariadna Medina-Benitez, Joaquin Quiroz-Mercado, Sergio Rangel-Guerrero, Natividad Cruz-Ortiz, Tatiana Nunez-Garcia, Gisela Barrera-Badillo, Lucia Hernandez-Rivas, Irma Lopez-Martinez, Ernesto Ramirez-Gonzalez. |
| EPI_ISL_1424008 | LESP Hidalgo | Instituto de Diagnostico y Referencia Epidemiologicos (INDRE) | Claudia Wong-Arambula, Abril Rodriguez-Maldonado, Vanessa Rivero-Arredondo, Ariadna Medina-Benitez, Joaquin Quiroz-Mercado, Sergio Rangel-Guerrero, Natividad Cruz-Ortiz, Tatiana Nunez-Garcia, Gisela Barrera-Badillo, Lucia Hernandez-Rivas, Irma Lopez-Martinez, Ernesto Ramirez-Gonzalez. |

| | | | |
|---|---|---|---|
| EPI_ISL_1424014 | LESP Yucatan | Instituto de Diagnostico y Referencia Epidemiologicos (INDRE) | Claudia Wong-Arambula, Abril Rodriguez-Maldonado, Vanessa Rivero-Arredondo, Ariadna Medina-Benitez, Joaquin Quiroz-Mercado, Sergio Rangel-Guerrero, Natividad Cruz-Ortiz, Tatiana Nunez-Garcia, Gisela Barrera-Badillo, Lucia Hernandez-Rivas, Irma Lopez-Martinez, Ernesto Ramirez-Gonzalez. |
| EPI_ISL_1424018 | LESP Veracruz | Instituto de Diagnostico y Referencia Epidemiologicos (INDRE) | Claudia Wong-Arambula, Abril Rodriguez-Maldonado, Vanessa Rivero-Arredondo, Ariadna Medina-Benitez, Joaquin Quiroz-Mercado, Sergio Rangel-Guerrero, Natividad Cruz-Ortiz, Tatiana Nunez-Garcia, Gisela Barrera-Badillo, Lucia Hernandez-Rivas, Irma Lopez-Martinez, Ernesto Ramirez-Gonzalez. |
| EPI_ISL_1424020 | LESP Chiapas | Instituto de Diagnostico y Referencia Epidemiologicos (INDRE) | Claudia Wong-Arambula, Abril Rodriguez-Maldonado, Vanessa Rivero-Arredondo, Ariadna Medina-Benitez, Joaquin Quiroz-Mercado, Sergio Rangel-Guerrero, Natividad Cruz-Ortiz, Tatiana Nunez-Garcia, Gisela Barrera-Badillo, Lucia Hernandez-Rivas, Irma Lopez-Martinez, Ernesto Ramirez-Gonzalez. |
| EPI_ISL_1424021, EPI_ISL_1424022, EPI_ISL_1424024, EPI_ISL_1424026, EPI_ISL_1424029, EPI_ISL_1424030, EPI_ISL_1424031, EPI_ISL_1424032, EPI_ISL_1424033 | LESP Queretaro | Instituto de Diagnostico y Referencia Epidemiologicos (INDRE) | Claudia Wong-Arambula, Abril Rodriguez-Maldonado, Vanessa Rivero-Arredondo, Ariadna Medina-Benitez, Joaquin Quiroz-Mercado, Sergio Rangel-Guerrero, Natividad Cruz-Ortiz, Tatiana Nunez-Garcia, Gisela Barrera-Badillo, Lucia Hernandez-Rivas, Irma Lopez-Martinez, Ernesto Ramirez-Gonzalez. |
| EPI_ISL_1424034 | LESP Guanajuato | Instituto de Diagnostico y Referencia Epidemiologicos (INDRE) | Claudia Wong-Arambula, Abril Rodriguez-Maldonado, Vanessa Rivero-Arredondo, Ariadna Medina-Benitez, Joaquin Quiroz-Mercado, Sergio Rangel-Guerrero, Natividad Cruz-Ortiz, Tatiana Nunez-Garcia, Gisela Barrera-Badillo, Lucia Hernandez-Rivas, Irma Lopez-Martinez, Ernesto Ramirez-Gonzalez. |
| EPI_ISL_1424035, EPI_ISL_1424037, EPI_ISL_1424038 | LESP Yucatan | Instituto de Diagnostico y Referencia Epidemiologicos (INDRE) | Claudia Wong-Arambula, Abril Rodriguez-Maldonado, Vanessa Rivero-Arredondo, Ariadna Medina-Benitez, Joaquin Quiroz-Mercado, Sergio Rangel-Guerrero, Natividad Cruz-Ortiz, Tatiana Nunez-Garcia, Gisela Barrera-Badillo, Lucia Hernandez-Rivas, Irma Lopez-Martinez, Ernesto Ramirez-Gonzalez. |
| EPI_ISL_1424043 | LESP Guerrero | Instituto de Diagnostico y Referencia Epidemiologicos (INDRE) | Claudia Wong-Arambula, Abril Rodriguez-Maldonado, Vanessa Rivero-Arredondo, Ariadna Medina-Benitez, Joaquin Quiroz-Mercado, Sergio Rangel-Guerrero, Natividad Cruz-Ortiz, Tatiana Nunez-Garcia, Gisela Barrera-Badillo, Lucia Hernandez-Rivas, Irma Lopez-Martinez, Ernesto Ramirez-Gonzalez. |
| EPI_ISL_1424044, EPI_ISL_1424045 | LESP Coahuila | Instituto de Diagnostico y Referencia Epidemiologicos (INDRE) | Claudia Wong-Arambula, Abril Rodriguez-Maldonado, Vanessa Rivero-Arredondo, Ariadna Medina-Benitez, Joaquin Quiroz-Mercado, Sergio Rangel-Guerrero, Natividad Cruz-Ortiz, Tatiana Nunez-Garcia, Gisela Barrera-Badillo, Lucia Hernandez-Rivas, Irma Lopez-Martinez, Ernesto Ramirez-Gonzalez. |
| EPI_ISL_1424047, EPI_ISL_1424048, EPI_ISL_1424049 | LESP Veracruz | Instituto de Diagnostico y Referencia Epidemiologicos (INDRE) | Claudia Wong-Arambula, Abril Rodriguez-Maldonado, Vanessa Rivero-Arredondo, Ariadna Medina-Benitez, Joaquin Quiroz-Mercado, Sergio Rangel-Guerrero, Natividad Cruz-Ortiz, Tatiana Nunez-Garcia, Gisela Barrera-Badillo, Lucia Hernandez-Rivas, Irma Lopez-Martinez, Ernesto Ramirez-Gonzalez. |
| EPI_ISL_1424050 | LESP Baja California | Instituto de Diagnostico y Referencia Epidemiologicos (INDRE) | Claudia Wong-Arambula, Abril Rodriguez-Maldonado, Vanessa Rivero-Arredondo, Ariadna Medina-Benitez, Joaquin Quiroz-Mercado, Sergio Rangel-Guerrero, Natividad Cruz-Ortiz, Tatiana Nunez-Garcia, Gisela Barrera-Badillo, Lucia Hernandez-Rivas, Irma Lopez-Martinez, Ernesto Ramirez-Gonzalez. |
| EPI_ISL_1424051, EPI_ISL_1424052, EPI_ISL_1424053 | LESP Ciudad de Mexico | Instituto de Diagnostico y Referencia Epidemiologicos (INDRE) | Claudia Wong-Arambula, Abril Rodriguez-Maldonado, Vanessa Rivero-Arredondo, Ariadna Medina-Benitez, Joaquin Quiroz-Mercado, Sergio Rangel-Guerrero, Natividad Cruz-Ortiz, Tatiana Nunez-Garcia, Gisela Barrera-Badillo, Lucia Hernandez-Rivas, Irma Lopez-Martinez, Ernesto Ramirez-Gonzalez. |
| EPI_ISL_1424130, EPI_ISL_1424143, EPI_ISL_1424466, EPI_ISL_1424467, EPI_ISL_1424468, EPI_ISL_1424470, EPI_ISL_1424471 | Institute for Medical Research, Infectious Disease Research Centre, National Institutes of Health, Ministry of Health Malaysia | Institute for Medical Research, Infectious Disease Research Centre, National Institutes of Health, Ministry of Health Malaysia | Suppiah J, Kamel K, Thayan R |
| EPI_ISL_1424503, EPI_ISL_1424509 | Institut de Virologie du CHU de Strasbourg | Institut de Virologie du CHU de Strasbourg | Fafi-Kremer Samira, Gallais Floriane, Gantner Pierre, Laugel Elodie, Solis Morgane, Velay Aurélie, Wendling Marie-Josée |
| EPI_ISL_1424510, EPI_ISL_1424518 | Institute for Medical Research, Infectious Disease Research Centre, National Institutes of Health, Ministry of Health Malaysia | Institute for Medical Research, Infectious Disease Research Centre, National Institutes of Health, Ministry of Health Malaysia | Suppiah J, Kamel K, Thayan R |
| EPI_ISL_1442249, EPI_ISL_1442254 | Azienda Ospedaliero - Universitaria di Modena Policlinico - Virologia e Microbiologia Molecolare | Istituto Zooprofilattico Sperimentale della Lombardia e dell'Emilia Romagna (IZSLER), Risk Analysis and Genomic Epidemiology Unit | Monica Pecorari, William Gennari, Giulia Fregni Serpini, Marina Morganti, Ilaria Menozzi, Erika Scaltriti, Stefano Pongolini |
| EPI_ISL_1442622, EPI_ISL_1442626, EPI_ISL_1442631, EPI_ISL_1442636, EPI_ISL_1442639, EPI_ISL_1442650, EPI_ISL_1442654, EPI_ISL_1442658, EPI_ISL_1442660, EPI_ISL_1442662, EPI_ISL_1442664, EPI_ISL_1442671, EPI_ISL_1442676, EPI_ISL_1442678, EPI_ISL_1442680 | | | |
| see above | SC Microbiologia e Virologia AOUSS | AMES Centro Polidiagnostico Strumentale S.r.l. | Salvatore Rubino; Flavia Angioj; Laura Firino; Rosalba Govoni; Gabriele Ibba; Vincenzo Lai; Erica Mura; Bianca Paglietti; Claudia Piu; Anna Puggioni; Elena Rimini; Giulia Rocca; Caterina Serra; Sergio Uzzau. |
| EPI_ISL_1443024 | Arcispedale Santa Maria Nuova, Autoimmunità, Allergologia e Biotecnologie Innovative | Istituto Zooprofilattico Sperimentale della Lombardia e dell'Emilia Romagna (IZSLER), Risk Analysis and Genomic Epidemiology Unit | Alessandro Zerbini, Lucia Belloni, Stefania Croci, Marina Morganti, Ilaria Menozzi, Erika Scaltriti, Stefano Pongolini |
| EPI_ISL_1443869, EPI_ISL_1443870, EPI_ISL_1443871 | Hospital | National Reference Center for Viruses of Respiratory Infections, Institut Pasteur, Paris | Marion Barbet, Sylvie Behillil, Méline Bizard,Frédéric Lemoine,Corinne Maufrais,Christophe Malabat, Angela Brisebarre, Camille Capel, Louise Lefrançois, Etienne Simon-Lorière, Vincent Enouf, Maud Vanpeene, Sylvie van der Werf,Anne Cady |
| EPI_ISL_1443872 | Hospital | National Reference Center for Viruses of Respiratory Infections, Institut Pasteur, Paris | Marion Barbet, Sylvie Behillil, Méline Bizard,Frédéric Lemoine,Corinne Maufrais,Christophe Malabat, Angela Brisebarre, Camille Capel, Louise Lefrançois, Etienne Simon-Lorière, Vincent Enouf, Maud Vanpeene, Sylvie van der Werf,Esther Gyde |
| EPI_ISL_1443873, EPI_ISL_1443874 | Labo analyses med | National Reference Center for Viruses of Respiratory Infections, Institut Pasteur, Paris | Marion Barbet, Sylvie Behillil, Méline Bizard,Frédéric Lemoine,Corinne Maufrais,Christophe Malabat, Angela Brisebarre, Camille Capel, Louise Lefrançois, Etienne Simon-Lorière, Vincent Enouf, Maud Vanpeene, Sylvie van der Werf,GréGoire Potiron |
| EPI_ISL_1443875, EPI_ISL_1443876, EPI_ISL_1443877, EPI_ISL_1443878, EPI_ISL_1443879 | Labo analyses med | National Reference Center for Viruses of Respiratory Infections, Institut Pasteur, Paris | Marion Barbet, Sylvie Behillil, Méline Bizard,Frédéric Lemoine,Corinne Maufrais,Christophe Malabat, Angela Brisebarre, Camille Capel, Louise Lefrançois, Etienne Simon-Lorière, Vincent Enouf, Maud Vanpeene, Sylvie van der Werf,Anne Holstein |
| EPI_ISL_1443880 | Labo Analyses Med | National Reference Center for Viruses of Respiratory Infections, Institut Pasteur, Paris | Marion Barbet, Sylvie Behillil, Méline Bizard,Frédéric Lemoine,Corinne Maufrais,Christophe Malabat, Angela Brisebarre, Camille Capel, Louise Lefrançois, Etienne Simon-Lorière, Vincent Enouf, Maud Vanpeene, Sylvie van der Werf,Mj Dufour |
| EPI_ISL_1443881, EPI_ISL_1443882, EPI_ISL_1443883, EPI_ISL_1443884 | Hospital | National Reference Center for Viruses of Respiratory Infections, Institut Pasteur, Paris | Marion Barbet, Sylvie Behillil, Méline Bizard,Frédéric Lemoine,Corinne Maufrais,Christophe Malabat, Angela Brisebarre, Camille Capel, Louise Lefrançois, Etienne Simon-Lorière, Vincent Enouf, Maud Vanpeene, Sylvie van der Werf,Thibault Guinoiseau |
| EPI_ISL_1443885, EPI_ISL_1443886 | Hospital | National Reference Center for Viruses of Respiratory Infections, Institut Pasteur, Paris | Marion Barbet, Sylvie Behillil, Méline Bizard,Frédéric Lemoine,Corinne Maufrais,Christophe Malabat, Angela Brisebarre, Camille Capel, Louise Lefrançois, Etienne Simon-Lorière, Vincent Enouf, Maud Vanpeene, Sylvie van der Werf,Ilka Engelmann |
| EPI_ISL_1443887, EPI_ISL_1443888 | Hospital | National Reference Center for Viruses of Respiratory Infections, Institut Pasteur, Paris | Marion Barbet, Sylvie Behillil, Méline Bizard,Frédéric Lemoine,Corinne Maufrais,Christophe Malabat, Angela Brisebarre, Camille Capel, Louise Lefrançois, Etienne Simon-Lorière, Vincent Enouf, Maud Vanpeene, Sylvie van der Werf,CéLine Ramanantsoa |
| EPI_ISL_1443889 | Hospital | National Reference Center for Viruses of Respiratory Infections, Institut Pasteur, Paris | Marion Barbet, Sylvie Behillil, Méline Bizard,Frédéric Lemoine,Corinne Maufrais,Christophe Malabat, Angela Brisebarre, Camille Capel, Louise Lefrançois, Etienne Simon-Lorière, Vincent Enouf, Maud Vanpeene, Sylvie van der Werf,Jean Baptiste Bour |
| EPI_ISL_1443890 | Labo Analyses Med | National Reference Center for Viruses of Respiratory Infections, Institut Pasteur, Paris | Marion Barbet, Sylvie Behillil, Méline Bizard,Frédéric Lemoine,Corinne Maufrais,Christophe Malabat, Angela Brisebarre, Camille Capel, Louise Lefrançois, Etienne Simon-Lorière, Vincent Enouf, Maud Vanpeene, Sylvie van der Werf,Sophie Chalmin |

| | | | |
|---|---|---|---|
| EPI_ISL_1443891, EPI_ISL_1443892, EPI_ISL_1443893, EPI_ISL_1443894 | Outre mer | National Reference Center for Viruses of Respiratory Infections, Institut Pasteur, Paris | Marion Barbet, Sylvie Behillil, Méline Bizard,Frédéric Lemoine,Corinne Maufrais,Christophe Malabat, Angela Brisebarre, Camille Capel, Louise Lefrançois, Etienne Simon-Lorière, Vincent Enouf, Maud Vanpeene, Sylvie van der Werf,StéPhanie Guyomard-Rabenirina |
| EPI_ISL_1443895 | Hospital | National Reference Center for Viruses of Respiratory Infections, Institut Pasteur, Paris | Marion Barbet, Sylvie Behillil, Méline Bizard,Frédéric Lemoine,Corinne Maufrais,Christophe Malabat, Angela Brisebarre, Camille Capel, Louise Lefrançois, Etienne Simon-Lorière, Vincent Enouf, Maud Vanpeene, Sylvie van der Werf,FréDéRique Canis |
| EPI_ISL_1443896 | Labo Analyses Med | National Reference Center for Viruses of Respiratory Infections, Institut Pasteur, Paris | Marion Barbet, Sylvie Behillil, Méline Bizard,Frédéric Lemoine,Corinne Maufrais,Christophe Malabat, Angela Brisebarre, Camille Capel, Louise Lefrançois, Etienne Simon-Lorière, Vincent Enouf, Maud Vanpeene, Sylvie van der Werf,Sophie Chalmin |
| EPI_ISL_1443897, EPI_ISL_1443898, EPI_ISL_1443899, EPI_ISL_1443900 | Hospital | National Reference Center for Viruses of Respiratory Infections, Institut Pasteur, Paris | Marion Barbet, Sylvie Behillil, Méline Bizard,Frédéric Lemoine,Corinne Maufrais,Christophe Malabat, Angela Brisebarre, Camille Capel, Louise Lefrançois, Etienne Simon-Lorière, Vincent Enouf, Maud Vanpeene, Sylvie van der Werf,CéLine Bressollette |
| EPI_ISL_1443903, EPI_ISL_1443904 | Labo Analyses Med | National Reference Center for Viruses of Respiratory Infections, Institut Pasteur, Paris | Marion Barbet, Sylvie Behillil, Méline Bizard,Frédéric Lemoine,Corinne Maufrais,Christophe Malabat, Angela Brisebarre, Camille Capel, Louise Lefrançois, Etienne Simon-Lorière, Vincent Enouf, Maud Vanpeene, Sylvie van der Werf,Catherine Coignard |
| EPI_ISL_1443905, EPI_ISL_1443906, EPI_ISL_1443907 | Hospital | National Reference Center for Viruses of Respiratory Infections, Institut Pasteur, Paris | Marion Barbet, Sylvie Behillil, Méline Bizard,Frédéric Lemoine,Corinne Maufrais,Christophe Malabat, Angela Brisebarre, Camille Capel, Louise Lefrançois, Etienne Simon-Lorière, Vincent Enouf, Maud Vanpeene, Sylvie van der Werf,Sophie Chalmin |
| EPI_ISL_1443908 | Labo Analyses Med | National Reference Center for Viruses of Respiratory Infections, Institut Pasteur, Paris | Marion Barbet, Sylvie Behillil, Méline Bizard,Frédéric Lemoine,Corinne Maufrais,Christophe Malabat, Angela Brisebarre, Camille Capel, Louise Lefrançois, Etienne Simon-Lorière, Vincent Enouf, Maud Vanpeene, Sylvie van der Werf,Jean-FrançOis Comes |
| EPI_ISL_1443909, EPI_ISL_1443910 | Labo Analyses Med | National Reference Center for Viruses of Respiratory Infections, Institut Pasteur, Paris | Marion Barbet, Sylvie Behillil, Méline Bizard,Frédéric Lemoine,Corinne Maufrais,Christophe Malabat, Angela Brisebarre, Camille Capel, Louise Lefrançois, Etienne Simon-Lorière, Vincent Enouf, Maud Vanpeene, Sylvie van der Werf,OphéLie Said-Delattre |
| EPI_ISL_1447351, EPI_ISL_1447352 | Diagnostic and Research Center of Infectious Diseases, Medical Faculty, Andalas University | Diagnostic and Research Center of Infectious Diseases, Medical Faculty, Andalas University | Andani Eka Putra, Syafrizayanti, Ikhwan R. Sudji, Linosefa, Dede Rahman Agustian, Ayu Novita Trisnawati, Siskalil Fahma, Nia Ayuni Putri, Sekar Asri Tresnaningtyas, SM Rezvi, Fauzul Azhim, Dessy Arisanty, Gestina Aliska, Desmawati, Nita Afriani ,Syandrez Prima Putra, Juane Plantika Menra, Yolani Syaputri, Mutia Lailani |
| EPI_ISL_1455670 | Marius Nasta Institute of Pneumology | "Stefan S. Nicolau"Institute of Virology | Anca Botezatu, Iulia Virginia Iancu, Coralia Bleotu, Mihaela Economescu, Laura Necula, Denisa Dragu, Lilia Matei, Ioana Pitica, Marinela Bostan, Mirela Mihaila, Laura Grecu, Camelia Sultana, Simona Ruta, Ana Iulia Neagu, Adriana Plesa, Saviana Nedeianu, Alina Nastasie, Cristina Mambet,Gabriela Anton,Tudor Constantinescu, Carmen Cristina Diaconu |
| EPI_ISL_1457773 | Dr. Carol Davila Central Military Emergency University Hospital | "Stefan S. Nicolau"Institute of Virology | Anca Botezatu, Iulia Virginia Iancu, Coralia Bleotu, Mihaela Economescu, Laura Necula, Denisa Dragu, Lilia Matei, Ioana Pitica, Marinela Bostan, Mirela Mihaila, Laura Grecu, Camelia Sultana, Simona Ruta, Ana Iulia Neagu, Adriana Plesa, Saviana Nedeianu, Alina Nastasie, Cristina Mambet,Gabriela Anton,Codruta-Georgiana Carp, Carmen Cristina Diaconu |
| EPI_ISL_1461640, EPI_ISL_1461641, EPI_ISL_1461643, EPI_ISL_1461646, EPI_ISL_1461648 | CHU NIMES | CHU NIMES | Stephan Robin, Marie-josée Carles, Sophie Bravo, Agathe Boudet |
| EPI_ISL_1463448, EPI_ISL_1463451, EPI_ISL_1463452, EPI_ISL_1463453, EPI_ISL_1463454, EPI_ISL_1463455, EPI_ISL_1463457, EPI_ISL_1463458, EPI_ISL_1463459, EPI_ISL_1463460, EPI_ISL_1463461, EPI_ISL_1463462, EPI_ISL_1463463, EPI_ISL_1463464, EPI_ISL_1463465, EPI_ISL_1463466 | | | |
| see above | Labo Analyses Med | National Reference Center for Viruses of Respiratory Infections, Institut Pasteur, Paris | Marion Barbet, Sylvie Behillil,Frédéric Lemoine,Corinne Maufrais,Christophe Malabat, Méline Bizard, Angela Brisebarre, Camille Capel, Louise Lefrançois, Etienne Simon-Lorière, Vincent Enouf, Maud Vanpeene, Sylvie van der Werf,Nabil Gastli |
| EPI_ISL_1463467 | Labo Analyses Med | National Reference Center for Viruses of Respiratory Infections, Institut Pasteur, Paris | Marion Barbet, Sylvie Behillil,Frédéric Lemoine,Corinne Maufrais,Christophe Malabat,Adrien Pain, Méline Bizard, Angela Brisebarre, Camille Capel, Louise Lefrançois, Etienne Simon-Lorière, Vincent Enouf, Maud Vanpeene, Sylvie van der Werf,Nabil Gastli |
| EPI_ISL_1463468 | Labo Analyses Med | National Reference Center for Viruses of Respiratory Infections, Institut Pasteur, Paris | Marion Barbet, Sylvie Behillil,Frédéric Lemoine,Corinne Maufrais,Christophe Malabat, Méline Bizard, Angela Brisebarre, Camille Capel, Louise Lefrançois, Etienne Simon-Lorière, Vincent Enouf, Maud Vanpeene, Sylvie van der Werf,Nabil Gastli |
| EPI_ISL_1463469 | Labo Analyses Med | National Reference Center for Viruses of Respiratory Infections, Institut Pasteur, Paris | Marion Barbet, Sylvie Behillil,Frédéric Lemoine,Corinne Maufrais,Christophe Malabat,Amaury Vaysse, Méline Bizard, Angela Brisebarre, Camille Capel, Louise Lefrançois, Etienne Simon-Lorière, Vincent Enouf, Maud Vanpeene, Sylvie van der Werf,Nabil Gastli |
| EPI_ISL_1463470, EPI_ISL_1463471 | Labo Analyses Med | National Reference Center for Viruses of Respiratory Infections, Institut Pasteur, Paris | Marion Barbet, Sylvie Behillil,Frédéric Lemoine,Corinne Maufrais,Christophe Malabat, Méline Bizard, Angela Brisebarre, Camille Capel, Louise Lefrançois, Etienne Simon-Lorière, Vincent Enouf, Maud Vanpeene, Sylvie van der Werf,Nabil Gastli |
| EPI_ISL_1463473 | Labo Analyses Med | National Reference Center for Viruses of Respiratory Infections, Institut Pasteur, Paris | Marion Barbet, Sylvie Behillil,Frédéric Lemoine,Corinne Maufrais,Christophe Malabat,Amaury Vaysse, Méline Bizard, Angela Brisebarre, Camille Capel, Louise Lefrançois, Etienne Simon-Lorière, Vincent Enouf, Maud Vanpeene, Sylvie van der Werf,Nabil Gastli |
| EPI_ISL_1463474, EPI_ISL_1463475, EPI_ISL_1463476 | Labo Analyses Med | National Reference Center for Viruses of Respiratory Infections, Institut Pasteur, Paris | Marion Barbet, Sylvie Behillil,Frédéric Lemoine,Corinne Maufrais,Christophe Malabat, Méline Bizard, Angela Brisebarre, Camille Capel, Louise Lefrançois, Etienne Simon-Lorière, Vincent Enouf, Maud Vanpeene, Sylvie van der Werf,Nabil Gastli |
| EPI_ISL_1463477, EPI_ISL_1463478 | Labo Analyses Med | National Reference Center for Viruses of Respiratory Infections, Institut Pasteur, Paris | Marion Barbet, Sylvie Behillil,Frédéric Lemoine,Corinne Maufrais,Christophe Malabat,Amaury Vaysse, Méline Bizard, Angela Brisebarre, Camille Capel, Louise Lefrançois, Etienne Simon-Lorière, Vincent Enouf, Maud Vanpeene, Sylvie van der Werf,Nabil Gastli |
| EPI_ISL_1463479 | Labo Analyses Med | National Reference Center for Viruses of Respiratory Infections, Institut Pasteur, Paris | Marion Barbet, Sylvie Behillil,Frédéric Lemoine,Corinne Maufrais,Christophe Malabat,Adrien Pain, Méline Bizard, Angela Brisebarre, Camille Capel, Louise Lefrançois, Etienne Simon-Lorière, Vincent Enouf, Maud Vanpeene, Sylvie van der Werf,Nabil Gastli |
| EPI_ISL_1463480, EPI_ISL_1463481 | Labo Analyses Med | National Reference Center for Viruses of Respiratory Infections, Institut Pasteur, Paris | Marion Barbet, Sylvie Behillil,Frédéric Lemoine,Corinne Maufrais,Christophe Malabat, Méline Bizard, Angela Brisebarre, Camille Capel, Louise Lefrançois, Etienne Simon-Lorière, Vincent Enouf, Maud Vanpeene, Sylvie van der Werf,Nabil Gastli |
| EPI_ISL_1463482 | Labo Analyses Med | National Reference Center for Viruses of Respiratory Infections, Institut Pasteur, Paris | Marion Barbet, Sylvie Behillil,Frédéric Lemoine,Corinne Maufrais,Christophe Malabat,Amaury Vaysse, Méline Bizard, Angela Brisebarre, Camille Capel, Louise Lefrançois, Etienne Simon-Lorière, Vincent Enouf, Maud Vanpeene, Sylvie van der Werf,Nabil Gastli |
| EPI_ISL_1463483, EPI_ISL_1463484, EPI_ISL_1463485 | Labo Analyses Med | National Reference Center for Viruses of Respiratory Infections, Institut Pasteur, Paris | Marion Barbet, Sylvie Behillil,Frédéric Lemoine,Corinne Maufrais,Christophe Malabat, Méline Bizard, Angela Brisebarre, Camille Capel, Louise Lefrançois, Etienne Simon-Lorière, Vincent Enouf, Maud Vanpeene, Sylvie van der Werf,Nabil Gastli |
| EPI_ISL_1463486 | Labo Analyses Med | National Reference Center for Viruses of Respiratory Infections, Institut Pasteur, Paris | Marion Barbet, Sylvie Behillil,Frédéric Lemoine,Corinne Maufrais,Christophe Malabat,Amaury Vaysse, Méline Bizard, Angela Brisebarre, Camille Capel, Louise Lefrançois, Etienne Simon-Lorière, Vincent Enouf, Maud Vanpeene, Sylvie van der Werf,Nabil Gastli |
| EPI_ISL_1463487, EPI_ISL_1463488, EPI_ISL_1463490, EPI_ISL_1463491, EPI_ISL_1463492, EPI_ISL_1463493, EPI_ISL_1463494, EPI_ISL_1463495, EPI_ISL_1463496 | Labo Analyses Med | National Reference Center for Viruses of Respiratory Infections, Institut Pasteur, Paris | Marion Barbet, Sylvie Behillil,Frédéric Lemoine,Corinne Maufrais,Christophe Malabat, Méline Bizard, Angela Brisebarre, Camille Capel, Louise Lefrançois, Etienne Simon-Lorière, Vincent Enouf, Maud Vanpeene, Sylvie van der Werf,Nabil Gastli |
| EPI_ISL_1463497 | Labo Analyses Med | National Reference Center for Viruses of Respiratory Infections, Institut Pasteur, Paris | Marion Barbet, Sylvie Behillil,Frédéric Lemoine,Corinne Maufrais,Christophe Malabat,Amaury Vaysse, Méline Bizard, Angela Brisebarre, Camille Capel, Louise Lefrançois, Etienne Simon-Lorière, Vincent Enouf, Maud Vanpeene, Sylvie van der Werf,Nabil Gastli |
| EPI_ISL_1463498 | Labo Analyses Med | National Reference Center for Viruses of Respiratory Infections, Institut Pasteur, Paris | Marion Barbet, Sylvie Behillil,Frédéric Lemoine,Corinne Maufrais,Christophe Malabat, Méline Bizard, Angela Brisebarre, Camille Capel, Louise Lefrançois, Etienne Simon-Lorière, Vincent Enouf, Maud Vanpeene, Sylvie van der Werf,Nabil Gastli |
| EPI_ISL_1463499 | Labo Analyses Med | National Reference Center for Viruses of Respiratory Infections, Institut Pasteur, Paris | Marion Barbet, Sylvie Behillil,Frédéric Lemoine,Corinne Maufrais,Christophe Malabat,Amaury Vaysse, Méline Bizard, Angela Brisebarre, Camille Capel, Louise Lefrançois, Etienne Simon-Lorière, Vincent Enouf, Maud Vanpeene, Sylvie van der Werf,Nabil Gastli |
| EPI_ISL_1463500, EPI_ISL_1463501, EPI_ISL_1463502 | Labo Analyses Med | National Reference Center for Viruses of Respiratory Infections, Institut Pasteur, Paris | Marion Barbet, Sylvie Behillil,Frédéric Lemoine,Corinne Maufrais,Christophe Malabat, Méline Bizard, Angela Brisebarre, Camille Capel, Louise Lefrançois, Etienne Simon-Lorière, Vincent Enouf, Maud Vanpeene, Sylvie van der Werf,Nabil Gastli |
| EPI_ISL_1463504 | Labo Analyses Med | National Reference Center for Viruses of Respiratory Infections, Institut Pasteur, Paris | Marion Barbet, Sylvie Behillil,Frédéric Lemoine,Corinne Maufrais,Christophe Malabat,Amaury Vaysse, Méline Bizard, Angela Brisebarre, Camille Capel, Louise Lefrançois, Etienne Simon-Lorière, Vincent Enouf, Maud Vanpeene, Sylvie van der Werf,Nabil Gastli |
| EPI_ISL_1463505, EPI_ISL_1463506, EPI_ISL_1463507, EPI_ISL_1463508, EPI_ISL_1463509, EPI_ISL_1463510, EPI_ISL_1463511, EPI_ISL_1463512, EPI_ISL_1463513, EPI_ISL_1463515, EPI_ISL_1463517 | | | |

| | | | |
|---|---|---|---|
| see above | Labo Analyses Med | National Reference Center for Viruses of Respiratory Infections, Institut Pasteur, Paris | Marion Barbet, Sylvie Behillil,Frédéric Lemoine,Corinne Maufrais,Christophe Malabat, Méline Bizard, Angela Brisebarre, Camille Capel, Louise Lefrançois, Etienne Simon-Lorière, Vincent Enouf, Maud Vanpeene, Sylvie van der Werf,Nabil Gastli |
| EPI_ISL_1463518 | Labo Analyses Med | National Reference Center for Viruses of Respiratory Infections, Institut Pasteur, Paris | Marion Barbet, Sylvie Behillil,Frédéric Lemoine,Corinne Maufrais,Christophe Malabat,Emmanuelle Permal, Méline Bizard, Angela Brisebarre, Camille Capel, Louise Lefrançois, Etienne Simon-Lorière, Vincent Enouf, Maud Vanpeene, Sylvie van der Werf,Nabil Gastli |
| EPI_ISL_1463519, EPI_ISL_1463520, EPI_ISL_1463521, EPI_ISL_1463522, EPI_ISL_1463523, EPI_ISL_1463524, EPI_ISL_1463525, EPI_ISL_1463526, EPI_ISL_1463527, EPI_ISL_1463528 | Labo Analyses Med | National Reference Center for Viruses of Respiratory Infections, Institut Pasteur, Paris | Marion Barbet, Sylvie Behillil,Frédéric Lemoine,Corinne Maufrais,Christophe Malabat, Méline Bizard, Angela Brisebarre, Camille Capel, Louise Lefrançois, Etienne Simon-Lorière, Vincent Enouf, Maud Vanpeene, Sylvie van der Werf,Nabil Gastli |
| EPI_ISL_1463529 | Labo Analyses Med | National Reference Center for Viruses of Respiratory Infections, Institut Pasteur, Paris | Marion Barbet, Sylvie Behillil,Frédéric Lemoine,Corinne Maufrais,Christophe Malabat,Emmanuelle Permal, Méline Bizard, Angela Brisebarre, Camille Capel, Louise Lefrançois, Etienne Simon-Lorière, Vincent Enouf, Maud Vanpeene, Sylvie van der Werf,Nabil Gastli |
| EPI_ISL_1463530, EPI_ISL_1463531 | Labo Analyses Med | National Reference Center for Viruses of Respiratory Infections, Institut Pasteur, Paris | Marion Barbet, Sylvie Behillil,Frédéric Lemoine,Corinne Maufrais,Christophe Malabat, Méline Bizard, Angela Brisebarre, Camille Capel, Louise Lefrançois, Etienne Simon-Lorière, Vincent Enouf, Maud Vanpeene, Sylvie van der Werf,Nabil Gastli |
| EPI_ISL_1463532 | Labo Analyses Med | National Reference Center for Viruses of Respiratory Infections, Institut Pasteur, Paris | Marion Barbet, Sylvie Behillil,Frédéric Lemoine,Corinne Maufrais,Christophe Malabat,Emmanuelle Permal, Méline Bizard, Angela Brisebarre, Camille Capel, Louise Lefrançois, Etienne Simon-Lorière, Vincent Enouf, Maud Vanpeene, Sylvie van der Werf,Nabil Gastli |
| EPI_ISL_1463533 | Labo Analyses Med | National Reference Center for Viruses of Respiratory Infections, Institut Pasteur, Paris | Marion Barbet, Sylvie Behillil,Frédéric Lemoine,Corinne Maufrais,Christophe Malabat, Méline Bizard, Angela Brisebarre, Camille Capel, Louise Lefrançois, Etienne Simon-Lorière, Vincent Enouf, Maud Vanpeene, Sylvie van der Werf,Nabil Gastli |
| EPI_ISL_1463534 | Labo Analyses Med | National Reference Center for Viruses of Respiratory Infections, Institut Pasteur, Paris | Marion Barbet, Sylvie Behillil,Frédéric Lemoine,Corinne Maufrais,Christophe Malabat,Emmanuelle Permal, Méline Bizard, Angela Brisebarre, Camille Capel, Louise Lefrançois, Etienne Simon-Lorière, Vincent Enouf, Maud Vanpeene, Sylvie van der Werf,Nabil Gastli |
| EPI_ISL_1463535, EPI_ISL_1463536, EPI_ISL_1463537, EPI_ISL_1463538, EPI_ISL_1463539 | Labo Analyses Med | National Reference Center for Viruses of Respiratory Infections, Institut Pasteur, Paris | Marion Barbet, Sylvie Behillil,Frédéric Lemoine,Corinne Maufrais,Christophe Malabat, Méline Bizard, Angela Brisebarre, Camille Capel, Louise Lefrançois, Etienne Simon-Lorière, Vincent Enouf, Maud Vanpeene, Sylvie van der Werf,Nabil Gastli |
| EPI_ISL_1463540 | Labo Analyses Med | National Reference Center for Viruses of Respiratory Infections, Institut Pasteur, Paris | Marion Barbet, Sylvie Behillil,Frédéric Lemoine,Corinne Maufrais,Christophe Malabat,Emmanuelle Permal, Méline Bizard, Angela Brisebarre, Camille Capel, Louise Lefrançois, Etienne Simon-Lorière, Vincent Enouf, Maud Vanpeene, Sylvie van der Werf,Karine Michez |
| EPI_ISL_1465449, EPI_ISL_1465450 | Institute of Molecular and Translational Medicine / Laboratory of Experimental Medicine, Faculty of Medicine and Dentistry, Palacky University and University Hospital Olomouc | Institute of Molecular and Translational Medicine / Laboratory of Experimental Medicine, Faculty of Medicine and Dentistry, Palacky University | Rastislav Slavkovský, Hana Jaworek, Vladimíra Koudeláková, Barbora Blumová, Tomáš Pospíšil, Marián Hajdúch |
| EPI_ISL_1465451 | Institute of Molecular and Translational Medicine / Laboratory of Experimental Medicine, Faculty of Medicine and Dentistry, Palacky University and University Hospital Olomouc | Institute of Molecular and Translational Medicine / Laboratory of Experimental Medicine, Faculty of Medicine and Dentistry, Palacky University | Rastislav Slavkovský, Hana Jaworek, Vladimíra Koudeláková, Barbora Blumová, Tomáš Pospíšil, Marián Hajdúch |
| EPI_ISL_1465452, EPI_ISL_1465453, EPI_ISL_1465454, EPI_ISL_1465455, EPI_ISL_1465456, EPI_ISL_1465457, EPI_ISL_1465458, EPI_ISL_1465459, EPI_ISL_1465460, EPI_ISL_1465461, EPI_ISL_1465462, EPI_ISL_1465463, EPI_ISL_1465464, EPI_ISL_1465465, EPI_ISL_1465466, EPI_ISL_1465467, EPI_ISL_1465468, EPI_ISL_1465469, EPI_ISL_1465470, EPI_ISL_1465471, EPI_ISL_1465472, EPI_ISL_1465473, EPI_ISL_1465476, EPI_ISL_1465477, EPI_ISL_1465478, EPI_ISL_1465479 | | | |
| see above | Institute of Molecular and Translational Medicine / Laboratory of Experimental Medicine, Faculty of Medicine and Dentistry, Palacky University and University Hospital Olomouc | Institute of Molecular and Translational Medicine / Laboratory of Experimental Medicine, Faculty of Medicine and Dentistry, Palacky University | Rastislav Slavkovský, Hana Jaworek, Vladimíra Koudeláková, Barbora Blumová, Tomáš Pospíšil, Marián Hajdúch |
| EPI_ISL_1465482 | Medicine sentinel network Province | National Reference Center for Viruses of Respiratory Infections, Institut Pasteur, Paris | Marion Barbet, Sylvie Behillil,Frédéric Lemoine,Corinne Maufrais,Christophe Malabat,Damien Mornico, Méline Bizard, Angela Brisebarre, Camille Capel, Louise Lefrançois, Etienne Simon-Lorière, Vincent Enouf, Maud Vanpeene, Sylvie van der Werf,Nicolas Le Duff |
| EPI_ISL_1465483, EPI_ISL_1465484, EPI_ISL_1465485, EPI_ISL_1465486, EPI_ISL_1465487, EPI_ISL_1465488, EPI_ISL_1465489, EPI_ISL_1465490, EPI_ISL_1465491 | Labo Analyses Med | National Reference Center for Viruses of Respiratory Infections, Institut Pasteur, Paris | Marion Barbet, Sylvie Behillil,Frédéric Lemoine,Corinne Maufrais,Christophe Malabat, Méline Bizard, Angela Brisebarre, Camille Capel, Louise Lefrançois, Etienne Simon-Lorière, Vincent Enouf, Maud Vanpeene, Sylvie van der Werf,OphéLie Said-Delattre |
| EPI_ISL_1465492 | Labo Analyses Med | National Reference Center for Viruses of Respiratory Infections, Institut Pasteur, Paris | Marion Barbet, Sylvie Behillil,Frédéric Lemoine,Corinne Maufrais,Christophe Malabat,Damien Mornico, Méline Bizard, Angela Brisebarre, Camille Capel, Louise Lefrançois, Etienne Simon-Lorière, Vincent Enouf, Maud Vanpeene, Sylvie van der Werf,OphéLie Said-Delattre |
| EPI_ISL_1465493, EPI_ISL_1465494, EPI_ISL_1465495, EPI_ISL_1465496 | Labo Analyses Med | National Reference Center for Viruses of Respiratory Infections, Institut Pasteur, Paris | Marion Barbet, Sylvie Behillil,Frédéric Lemoine,Corinne Maufrais,Christophe Malabat, Méline Bizard, Angela Brisebarre, Camille Capel, Louise Lefrançois, Etienne Simon-Lorière, Vincent Enouf, Maud Vanpeene, Sylvie van der Werf,OphéLie Said-Delattre |
| EPI_ISL_1465497 | Labo Analyses Med | National Reference Center for Viruses of Respiratory Infections, Institut Pasteur, Paris | Marion Barbet, Sylvie Behillil,Frédéric Lemoine,Corinne Maufrais,Christophe Malabat,Damien Mornico, Méline Bizard, Angela Brisebarre, Camille Capel, Louise Lefrançois, Etienne Simon-Lorière, Vincent Enouf, Maud Vanpeene, Sylvie van der Werf,OphéLie Said-Delattre |
| EPI_ISL_1465498, EPI_ISL_1465499, EPI_ISL_1465500, EPI_ISL_1465501, EPI_ISL_1465502, EPI_ISL_1465503, EPI_ISL_1465504, EPI_ISL_1465505, EPI_ISL_1465506, EPI_ISL_1465507, EPI_ISL_1465508, EPI_ISL_1465509, EPI_ISL_1465510, EPI_ISL_1465511, EPI_ISL_1465513, EPI_ISL_1465514, EPI_ISL_1465515, EPI_ISL_1465516 | | | |
| see above | Labo Analyses Med | National Reference Center for Viruses of Respiratory Infections, Institut Pasteur, Paris | Marion Barbet, Sylvie Behillil,Frédéric Lemoine,Corinne Maufrais,Christophe Malabat, Méline Bizard, Angela Brisebarre, Camille Capel, Louise Lefrançois, Etienne Simon-Lorière, Vincent Enouf, Maud Vanpeene, Sylvie van der Werf,OphéLie Said-Delattre |
| EPI_ISL_1465517 | Labo Analyses Med | National Reference Center for Viruses of Respiratory Infections, Institut Pasteur, Paris | Marion Barbet, Sylvie Behillil,Frédéric Lemoine,Corinne Maufrais,Christophe Malabat,Amaury Vaysse, Méline Bizard, Angela Brisebarre, Camille Capel, Louise Lefrançois, Etienne Simon-Lorière, Vincent Enouf, Maud Vanpeene, Sylvie van der Werf,OphéLie Said-Delattre |
| EPI_ISL_1465518 | Labo Analyses Med | National Reference Center for Viruses of Respiratory Infections, Institut Pasteur, Paris | Marion Barbet, Sylvie Behillil,Frédéric Lemoine,Corinne Maufrais,Christophe Malabat, Méline Bizard, Angela Brisebarre, Camille Capel, Louise Lefrançois, Etienne Simon-Lorière, Vincent Enouf, Maud Vanpeene, Sylvie van der Werf,OphéLie Said-Delattre |
| EPI_ISL_1465519 | Labo Analyses Med | National Reference Center for Viruses of Respiratory Infections, Institut Pasteur, Paris | Marion Barbet, Sylvie Behillil,Frédéric Lemoine,Corinne Maufrais,Christophe Malabat,Amaury Vaysse, Méline Bizard, Angela Brisebarre, Camille Capel, Louise Lefrançois, Etienne Simon-Lorière, Vincent Enouf, Maud Vanpeene, Sylvie van der Werf,OphéLie Said-Delattre |
| EPI_ISL_1465520 | Labo Analyses Med | National Reference Center for Viruses of Respiratory Infections, Institut Pasteur, Paris | Marion Barbet, Sylvie Behillil,Frédéric Lemoine,Corinne Maufrais,Christophe Malabat, Méline Bizard, Angela Brisebarre, Camille Capel, Louise Lefrançois, Etienne Simon-Lorière, Vincent Enouf, Maud Vanpeene, Sylvie van der Werf,OphéLie Said-Delattre |
| EPI_ISL_1465521 | Labo Analyses Med | National Reference Center for Viruses of Respiratory Infections, Institut Pasteur, Paris | Marion Barbet, Sylvie Behillil,Frédéric Lemoine,Corinne Maufrais,Christophe Malabat,Amaury Vaysse, Méline Bizard, Angela Brisebarre, Camille Capel, Louise Lefrançois, Etienne Simon-Lorière, Vincent Enouf, Maud Vanpeene, Sylvie van der Werf,OphéLie Said-Delattre |
| EPI_ISL_1465522, EPI_ISL_1465523, EPI_ISL_1465524, EPI_ISL_1465525, EPI_ISL_1465526, EPI_ISL_1465527, EPI_ISL_1465528, EPI_ISL_1465529, EPI_ISL_1465530, EPI_ISL_1465531, EPI_ISL_1465532, EPI_ISL_1465533 | | | |
| see above | Labo Analyses Med | National Reference Center for Viruses of Respiratory Infections, Institut Pasteur, Paris | Marion Barbet, Sylvie Behillil,Frédéric Lemoine,Corinne Maufrais,Christophe Malabat, Méline Bizard, Angela Brisebarre, Camille Capel, Louise Lefrançois, Etienne Simon-Lorière, Vincent Enouf, Maud Vanpeene, Sylvie van der Werf,OphéLie Said-Delattre |
| EPI_ISL_1465534 | Labo Analyses Med | National Reference Center for Viruses of Respiratory Infections, Institut Pasteur, Paris | Marion Barbet, Sylvie Behillil,Frédéric Lemoine,Corinne Maufrais,Christophe Malabat,Adrien Pain, Méline Bizard, Angela Brisebarre, Camille Capel, Louise Lefrançois, Etienne Simon-Lorière, Vincent Enouf, Maud Vanpeene, Sylvie van der Werf,OphéLie Said-Delattre |
| EPI_ISL_1465536 | Hospital | National Reference Center for Viruses of Respiratory Infections, Institut Pasteur, Paris | Marion Barbet, Sylvie Behillil,Frédéric Lemoine,Corinne Maufrais,Christophe Malabat, Méline Bizard, Angela Brisebarre, Camille Capel, Louise Lefrançois, Etienne Simon-Lorière, Vincent Enouf, Maud Vanpeene, Sylvie van der Werf,SéVerine Haouisee |
| EPI_ISL_1465537 | Labo Analyses Med | National Reference Center for Viruses of Respiratory Infections, Institut Pasteur, Paris | Marion Barbet, Sylvie Behillil,Frédéric Lemoine,Corinne Maufrais,Christophe Malabat, Méline Bizard, Angela Brisebarre, Camille Capel, Louise Lefrançois, Etienne Simon-Lorière, Vincent Enouf, Maud Vanpeene, Sylvie van der Werf,SéVerine Haouisee |
| EPI_ISL_1465538, EPI_ISL_1465540 | Hospital | National Reference Center for Viruses of Respiratory | Marion Barbet, Sylvie Behillil,Frédéric Lemoine,Corinne Maufrais,Christophe Malabat, Méline Bizard, Angela Brisebarre, Camille Capel, Louise Lefrançois, |

| | | | |
|---|---|---|---|
| | | Infections, Institut Pasteur, Paris | Etienne Simon-Lorière, Vincent Enouf, Maud Vanpeene, Sylvie van der Werf,SéVerine Haouisee |
| EPI_ISL_1465541, EPI_ISL_1465542, EPI_ISL_1465543 | Labo Analyses Med | National Reference Center for Viruses of Respiratory Infections, Institut Pasteur, Paris | Marion Barbet, Sylvie Behillil,Frédéric Lemoine,Corinne Maufrais,Christophe Malabat,Damien Mornico, Méline Bizard, Angela Brisebarre, Camille Capel, Louise Lefrançois, Etienne Simon-Lorière, Vincent Enouf, Maud Vanpeene, Sylvie van der Werf,Jean-FrançOis Comes |
| EPI_ISL_1465544, EPI_ISL_1465545 | Labo Analyses Med | National Reference Center for Viruses of Respiratory Infections, Institut Pasteur, Paris | Marion Barbet, Sylvie Behillil,Frédéric Lemoine,Corinne Maufrais,Christophe Malabat, Méline Bizard, Angela Brisebarre, Camille Capel, Louise Lefrançois, Etienne Simon-Lorière, Vincent Enouf, Maud Vanpeene, Sylvie van der Werf,Jean-FrançOis Comes |
| EPI_ISL_1465546, EPI_ISL_1465547 | Labo Analyses Med | National Reference Center for Viruses of Respiratory Infections, Institut Pasteur, Paris | Marion Barbet, Sylvie Behillil,Frédéric Lemoine,Corinne Maufrais,Christophe Malabat,Damien Mornico, Méline Bizard, Angela Brisebarre, Camille Capel, Louise Lefrançois, Etienne Simon-Lorière, Vincent Enouf, Maud Vanpeene, Sylvie van der Werf,Jean-FrançOis Comes |
| EPI_ISL_1465548, EPI_ISL_1465549, EPI_ISL_1465550 | Labo Analyses Med | National Reference Center for Viruses of Respiratory Infections, Institut Pasteur, Paris | Marion Barbet, Sylvie Behillil,Frédéric Lemoine,Corinne Maufrais,Christophe Malabat, Méline Bizard, Angela Brisebarre, Camille Capel, Louise Lefrançois, Etienne Simon-Lorière, Vincent Enouf, Maud Vanpeene, Sylvie van der Werf,Jean-FrançOis Comes |
| EPI_ISL_1465551 | Labo Analyses Med | National Reference Center for Viruses of Respiratory Infections, Institut Pasteur, Paris | Marion Barbet, Sylvie Behillil,Frédéric Lemoine,Corinne Maufrais,Christophe Malabat,Damien Mornico, Méline Bizard, Angela Brisebarre, Camille Capel, Louise Lefrançois, Etienne Simon-Lorière, Vincent Enouf, Maud Vanpeene, Sylvie van der Werf,Jean-FrançOis Comes |
| EPI_ISL_1465552, EPI_ISL_1465553, EPI_ISL_1465554, EPI_ISL_1465555, EPI_ISL_1465557, EPI_ISL_1465558, EPI_ISL_1465559, EPI_ISL_1465560, EPI_ISL_1465561, EPI_ISL_1465562, EPI_ISL_1465563, EPI_ISL_1465564, EPI_ISL_1465565 | | | |
| see above | Labo Analyses Med | National Reference Center for Viruses of Respiratory Infections, Institut Pasteur, Paris | Marion Barbet, Sylvie Behillil,Frédéric Lemoine,Corinne Maufrais,Christophe Malabat, Méline Bizard, Angela Brisebarre, Camille Capel, Louise Lefrançois, Etienne Simon-Lorière, Vincent Enouf, Maud Vanpeene, Sylvie van der Werf,Jean-FrançOis Comes |
| EPI_ISL_1465566 | Labo Analyses Med | National Reference Center for Viruses of Respiratory Infections, Institut Pasteur, Paris | Marion Barbet, Sylvie Behillil,Frédéric Lemoine,Corinne Maufrais,Christophe Malabat,Damien Mornico, Méline Bizard, Angela Brisebarre, Camille Capel, Louise Lefrançois, Etienne Simon-Lorière, Vincent Enouf, Maud Vanpeene, Sylvie van der Werf,Jean-FrançOis Comes |
| EPI_ISL_1465567, EPI_ISL_1465568 | Labo Analyses Med | National Reference Center for Viruses of Respiratory Infections, Institut Pasteur, Paris | Marion Barbet, Sylvie Behillil,Frédéric Lemoine,Corinne Maufrais,Christophe Malabat, Méline Bizard, Angela Brisebarre, Camille Capel, Louise Lefrançois, Etienne Simon-Lorière, Vincent Enouf, Maud Vanpeene, Sylvie van der Werf,Jean-FrançOis Comes |
| EPI_ISL_1465569, EPI_ISL_1465570 | Labo Analyses Med | National Reference Center for Viruses of Respiratory Infections, Institut Pasteur, Paris | Marion Barbet, Sylvie Behillil,Frédéric Lemoine,Corinne Maufrais,Christophe Malabat,Damien Mornico, Méline Bizard, Angela Brisebarre, Camille Capel, Louise Lefrançois, Etienne Simon-Lorière, Vincent Enouf, Maud Vanpeene, Sylvie van der Werf,Jean-FrançOis Comes |
| EPI_ISL_1465571, EPI_ISL_1465572 | Labo Analyses Med | National Reference Center for Viruses of Respiratory Infections, Institut Pasteur, Paris | Marion Barbet, Sylvie Behillil,Frédéric Lemoine,Corinne Maufrais,Christophe Malabat, Méline Bizard, Angela Brisebarre, Camille Capel, Louise Lefrançois, Etienne Simon-Lorière, Vincent Enouf, Maud Vanpeene, Sylvie van der Werf,Jean-FrançOis Comes |
| EPI_ISL_1465573, EPI_ISL_1465574, EPI_ISL_1465575 | Labo Analyses Med | National Reference Center for Viruses of Respiratory Infections, Institut Pasteur, Paris | Marion Barbet, Sylvie Behillil,Frédéric Lemoine,Corinne Maufrais,Christophe Malabat,Damien Mornico, Méline Bizard, Angela Brisebarre, Camille Capel, Louise Lefrançois, Etienne Simon-Lorière, Vincent Enouf, Maud Vanpeene, Sylvie van der Werf,Jean-FrançOis Comes |
| EPI_ISL_1465576, EPI_ISL_1465577, EPI_ISL_1465578, EPI_ISL_1465579, EPI_ISL_1465580, EPI_ISL_1465581, EPI_ISL_1465582, EPI_ISL_1465583 | Labo Analyses Med | National Reference Center for Viruses of Respiratory Infections, Institut Pasteur, Paris | Marion Barbet, Sylvie Behillil,Frédéric Lemoine,Corinne Maufrais,Christophe Malabat, Méline Bizard, Angela Brisebarre, Camille Capel, Louise Lefrançois, Etienne Simon-Lorière, Vincent Enouf, Maud Vanpeene, Sylvie van der Werf,Jean-FrançOis Comes |
| EPI_ISL_1465584 | Labo Analyses Med | National Reference Center for Viruses of Respiratory Infections, Institut Pasteur, Paris | Marion Barbet, Sylvie Behillil,Frédéric Lemoine,Corinne Maufrais,Christophe Malabat,Emmanuelle Permal, Méline Bizard, Angela Brisebarre, Camille Capel, Louise Lefrançois, Etienne Simon-Lorière, Vincent Enouf, Maud Vanpeene, Sylvie van der Werf,Jean-FrançOis Comes |
| EPI_ISL_1465585, EPI_ISL_1465586, EPI_ISL_1465587 | Labo Analyses Med | National Reference Center for Viruses of Respiratory Infections, Institut Pasteur, Paris | Marion Barbet, Sylvie Behillil,Frédéric Lemoine,Corinne Maufrais,Christophe Malabat, Méline Bizard, Angela Brisebarre, Camille Capel, Louise Lefrançois, Etienne Simon-Lorière, Vincent Enouf, Maud Vanpeene, Sylvie van der Werf,Jean-FrançOis Comes |
| EPI_ISL_1465588 | Labo Analyses Med | National Reference Center for Viruses of Respiratory Infections, Institut Pasteur, Paris | Marion Barbet, Sylvie Behillil,Frédéric Lemoine,Corinne Maufrais,Christophe Malabat,Emmanuelle Permal, Méline Bizard, Angela Brisebarre, Camille Capel, Louise Lefrançois, Etienne Simon-Lorière, Vincent Enouf, Maud Vanpeene, Sylvie van der Werf,Jean-FrançOis Comes |
| EPI_ISL_1465589, EPI_ISL_1465590, EPI_ISL_1465591 | Labo Analyses Med | National Reference Center for Viruses of Respiratory Infections, Institut Pasteur, Paris | Marion Barbet, Sylvie Behillil,Frédéric Lemoine,Corinne Maufrais,Christophe Malabat, Méline Bizard, Angela Brisebarre, Camille Capel, Louise Lefrançois, Etienne Simon-Lorière, Vincent Enouf, Maud Vanpeene, Sylvie van der Werf,Jean-FrançOis Comes |
| EPI_ISL_1465592, EPI_ISL_1465593 | Labo Analyses Med | National Reference Center for Viruses of Respiratory Infections, Institut Pasteur, Paris | Marion Barbet, Sylvie Behillil,Frédéric Lemoine,Corinne Maufrais,Christophe Malabat,Emmanuelle Permal, Méline Bizard, Angela Brisebarre, Camille Capel, Louise Lefrançois, Etienne Simon-Lorière, Vincent Enouf, Maud Vanpeene, Sylvie van der Werf,Jean-FrançOis Comes |
| EPI_ISL_1465594, EPI_ISL_1465595, EPI_ISL_1465596, EPI_ISL_1465597, EPI_ISL_1465599 | Labo Analyses Med | National Reference Center for Viruses of Respiratory Infections, Institut Pasteur, Paris | Marion Barbet, Sylvie Behillil,Frédéric Lemoine,Corinne Maufrais,Christophe Malabat, Méline Bizard, Angela Brisebarre, Camille Capel, Louise Lefrançois, Etienne Simon-Lorière, Vincent Enouf, Maud Vanpeene, Sylvie van der Werf,Jean-FrançOis Comes |
| EPI_ISL_1465601 | Labo Analyses Med | National Reference Center for Viruses of Respiratory Infections, Institut Pasteur, Paris | Marion Barbet, Sylvie Behillil,Frédéric Lemoine,Corinne Maufrais,Christophe Malabat,Damien Mornico, Méline Bizard, Angela Brisebarre, Camille Capel, Louise Lefrançois, Etienne Simon-Lorière, Vincent Enouf, Maud Vanpeene, Sylvie van der Werf,Jean-FrançOis Comes |
| EPI_ISL_1465602 | Labo Analyses Med | National Reference Center for Viruses of Respiratory Infections, Institut Pasteur, Paris | Marion Barbet, Sylvie Behillil,Frédéric Lemoine,Corinne Maufrais,Christophe Malabat, Méline Bizard, Angela Brisebarre, Camille Capel, Louise Lefrançois, Etienne Simon-Lorière, Vincent Enouf, Maud Vanpeene, Sylvie van der Werf,Jean-FrançOis Comes |
| EPI_ISL_1465603 | Labo Analyses Med | National Reference Center for Viruses of Respiratory Infections, Institut Pasteur, Paris | Marion Barbet, Sylvie Behillil,Frédéric Lemoine,Corinne Maufrais,Christophe Malabat,Damien Mornico, Méline Bizard, Angela Brisebarre, Camille Capel, Louise Lefrançois, Etienne Simon-Lorière, Vincent Enouf, Maud Vanpeene, Sylvie van der Werf,Jean-FrançOis Comes |
| EPI_ISL_1465604, EPI_ISL_1465606, EPI_ISL_1465607 | Labo Analyses Med | National Reference Center for Viruses of Respiratory Infections, Institut Pasteur, Paris | Marion Barbet, Sylvie Behillil,Frédéric Lemoine,Corinne Maufrais,Christophe Malabat, Méline Bizard, Angela Brisebarre, Camille Capel, Louise Lefrançois, Etienne Simon-Lorière, Vincent Enouf, Maud Vanpeene, Sylvie van der Werf,Catherine Delamare |
| EPI_ISL_1465608 | Labo Analyses Med | National Reference Center for Viruses of Respiratory Infections, Institut Pasteur, Paris | Marion Barbet, Sylvie Behillil,Frédéric Lemoine,Corinne Maufrais,Christophe Malabat,Damien Mornico, Méline Bizard, Angela Brisebarre, Camille Capel, Louise Lefrançois, Etienne Simon-Lorière, Vincent Enouf, Maud Vanpeene, Sylvie van der Werf,Catherine Delamare |
| EPI_ISL_1465609 | Labo Analyses Med | National Reference Center for Viruses of Respiratory Infections, Institut Pasteur, Paris | Marion Barbet, Sylvie Behillil,Frédéric Lemoine,Corinne Maufrais,Christophe Malabat, Méline Bizard, Angela Brisebarre, Camille Capel, Louise Lefrançois, Etienne Simon-Lorière, Vincent Enouf, Maud Vanpeene, Sylvie van der Werf,Catherine Delamare |
| EPI_ISL_1465610 | Hospital | National Reference Center for Viruses of Respiratory Infections, Institut Pasteur, Paris | Marion Barbet, Sylvie Behillil,Frédéric Lemoine,Corinne Maufrais,Christophe Malabat,Damien Mornico, Méline Bizard, Angela Brisebarre, Camille Capel, Louise Lefrançois, Etienne Simon-Lorière, Vincent Enouf, Maud Vanpeene, Sylvie van der Werf,Isabelle Woercel |
| EPI_ISL_1465611 | Armies | National Reference Center for Viruses of Respiratory Infections, Institut Pasteur, Paris | Marion Barbet, Sylvie Behillil,Frédéric Lemoine,Corinne Maufrais,Christophe Malabat, Méline Bizard, Angela Brisebarre, Camille Capel, Louise Lefrançois, Etienne Simon-Lorière, Vincent Enouf, Maud Vanpeene, Sylvie van der Werf,Marine Desroches |
| EPI_ISL_1468413, EPI_ISL_1468414, EPI_ISL_1468415 | LACEN do Estado de Goias | Instituto Adolfo Lutz, Interdiciplinary Procedures Center, Strategic Laboratory | Claudio Tavares Sacchi, Claudia Regina Gonçalves, Erica Valessa Ramos Gomes, Karoline Rodrigues Campos, Caio Vinicius Dias Lopes |
| EPI_ISL_1468416 | Secretaria Municipal de Saude de Andradina | Instituto Adolfo Lutz, Interdiciplinary Procedures Center, Strategic Laboratory | Claudio Tavares Sacchi, Claudia Regina Gonçalves, Erica Valessa Ramos Gomes, Karoline Rodrigues Campos, Caio Vinicius Dias Lopes |
| EPI_ISL_1468419 | Santa Casa de Aracatuba Hospital Sagrado Coracao de Jesus | Instituto Adolfo Lutz, Interdiciplinary Procedures Center, Strategic Laboratory | Claudio Tavares Sacchi, Claudia Regina Gonçalves, Erica Valessa Ramos Gomes, Karoline Rodrigues Campos, Caio Vinicius Dias Lopes |
| EPI_ISL_1468420, EPI_ISL_1468421 | Santa Casa de Birigui | Instituto Adolfo Lutz, Interdiciplinary Procedures Center, Strategic Laboratory | Claudio Tavares Sacchi, Claudia Regina Gonçalves, Erica Valessa Ramos Gomes, Karoline Rodrigues Campos, Caio Vinicius Dias Lopes |
| EPI_ISL_1468424 | Santa Casa de Aracatuba Hospital Sagrado Coracao de Jesus | Instituto Adolfo Lutz, Interdiciplinary Procedures Center, Strategic Laboratory | Claudio Tavares Sacchi, Claudia Regina Gonçalves, Erica Valessa Ramos Gomes, Karoline Rodrigues Campos, Caio Vinicius Dias Lopes |
| EPI_ISL_1468429 | Centro de Saude II Matao | Instituto Adolfo Lutz, Interdiciplinary Procedures Center, Strategic Laboratory | Claudio Tavares Sacchi, Claudia Regina Gonçalves, Erica Valessa Ramos Gomes, Karoline Rodrigues Campos, Caio Vinicius Dias Lopes |
| EPI_ISL_1468431 | LACEN do Estado de Goias | Instituto Adolfo Lutz, Interdiciplinary Procedures Center, Strategic Laboratory | Claudio Tavares Sacchi, Claudia Regina Gonçalves, Erica Valessa Ramos Gomes, Karoline Rodrigues Campos, Caio Vinicius Dias Lopes |

| | | | |
|---|---|---|---|
| EPI_ISL_1468443 | Hospital Estadual de Mirandopolis | Instituto Adolfo Lutz, Interdiciplinary Procedures Center, Strategic Laboratory | Claudio Tavares Sacchi, Claudia Regina Gonçalves, Erica Valessa Ramos Gomes, Karoline Rodrigues Campos, Caio Vinicius Dias Lopes |
| EPI_ISL_1468447 | Secretaria Municipal de Saude de Andradina | Instituto Adolfo Lutz, Interdiciplinary Procedures Center, Strategic Laboratory | Claudio Tavares Sacchi, Claudia Regina Gonçalves, Erica Valessa Ramos Gomes, Karoline Rodrigues Campos, Caio Vinicius Dias Lopes |
| EPI_ISL_1468448 | Santa Casa de Birigui | Instituto Adolfo Lutz, Interdiciplinary Procedures Center, Strategic Laboratory | Claudio Tavares Sacchi, Claudia Regina Gonçalves, Erica Valessa Ramos Gomes, Karoline Rodrigues Campos, Caio Vinicius Dias Lopes |
| EPI_ISL_1468451 | UBS IV Guararapes | Instituto Adolfo Lutz, Interdiciplinary Procedures Center, Strategic Laboratory | Claudio Tavares Sacchi, Claudia Regina Gonçalves, Erica Valessa Ramos Gomes, Karoline Rodrigues Campos, Caio Vinicius Dias Lopes |
| EPI_ISL_1468468 | Secretaria municipal de saude de Itapolis | Instituto Adolfo Lutz, Interdiciplinary Procedures Center, Strategic Laboratory | Claudio Tavares Sacchi, Claudia Regina Gonçalves, Erica Valessa Ramos Gomes, Karoline Rodrigues Campos, Caio Vinicius Dias Lopes |
| EPI_ISL_1469244 | National Institute Health Research and Development | National Institute of Health Research and Development | Reni Herman, Krisna Pangesti, Hana Apsari Pawestri, Subangkit, Kartika Dewi Puspa, Arie Ardiansyah Nugraha, Hartanti Dian Ikawati, Nelly Puspandari, Vivi Setiawaty |
| EPI_ISL_1469245 | National Institute Health Research and Development | National Institute of Health Research and Development | Subangkit, Hana Apsari Pawestri, Kartika Dewi Puspa, Arie Ardiansyah Nugraha, Hartanti Dian Ikawati, Krisna Nur Andriana Pangesti, Yuni Rukminiati, Ririn Ramadhany, Agustiningsih, Kindi Adam, Holy Arif Wibowo, Triyani Soekarso, Ni Ketut Susilarini, Nurika Hariastuti, Ully Alfi Nikmah, Reni Herman, Nike Susanti, Herna, Tati Febriyanti, Natalie Laurencia Kipuw, Fauzul Muna, Irene Lorinda Indalao, Nelly Puspandari, Vivi Setiawaty. |
| EPI_ISL_1469247 | RS Khusus Paru Karawang | National Institute of Health Research and Development | Hartanti Dian Ikawati, Hana Apsari Pawestri, Subangkit, Kartika Dewi Puspa, Arie Ardiansyah Nugraha, Krisna Pangesti, Nelly Puspandari, Vivi Setiawaty |
| EPI_ISL_1469248 | RS Mandaya Karawang | National Institute of Health Research and Development | Vivi Setiawaty, Hana Apsari Pawestri, Subangkit, Kartika Dewi Puspa, Arie Ardiansyah Nugraha, Hartanti Dian Ikawati, Nelly Puspandari, Krisna Pangesti |
| EPI_ISL_1469250 | Dinas Kesehatan Blitar | National Institute of Health Research and Development | Hana Apsari Pawestri, Subangkit, Kartika Dewi Puspa, Arie Ardiansyah Nugraha, Hartanti Dian Ikawati, Nelly Puspandari, Krisna Pangesti, Vivi Setiawaty |
| EPI_ISL_1469251 | Lab Intibios Karawang | National Institute of Health Research and Development | Hana Apsari Pawestri, Subangkit, Kartika Dewi Puspa, Arie Ardiansyah Nugraha, Hartanti Dian Ikawati, Nelly Puspandari, Krisna Pangesti, Vivi Setiawaty |
| EPI_ISL_1469253 | RS Mandaya Karawang | National Institute of Health Research and Development | Vivi Setiawaty, Hana Apsari Pawestri, Subangkit, Kartika Dewi Puspa, Arie Ardiansyah Nugraha, Hartanti Dian Ikawati, Nelly Puspandari, Krisna Pangesti |
| EPI_ISL_1469254 | RS Lira Medika Karawang | National Institute of Health Research and Development | Vivi Setiawaty, Hana Apsari Pawestri, Subangkit, Kartika Dewi Puspa, Arie Ardiansyah Nugraha, Hartanti Dian Ikawati, Nelly Puspandari, Krisna Pangesti |
| EPI_ISL_1469258 | PKM TEMPURAN KARAWANG | National Institute of Health Research and Development | Subangkit, Hana Apsari Pawestri, Kartika Dewi Puspa, Arie Ardiansyah Nugraha, Hartanti Dian Ikawati, Krisna Nur Andriana Pangesti, Yuni Rukminiati, Ririn Ramadhany, Agustiningsih, Kindi Adam, Holy Arif Wibowo, Triyani Soekarso, Ni Ketut Susilarini, Nurika Hariastuti, Ully Alfi Nikmah, Reni Herman, Nike Susanti, Herna, Tati Febriyanti, Natalie Laurencia Kipuw, Fauzul Muna, Irene Lorinda Indalao, Nelly Puspandari, Vivi Setiawaty. |
| EPI_ISL_1469260, EPI_ISL_1469261, EPI_ISL_1469262, EPI_ISL_1469263, EPI_ISL_1469264, EPI_ISL_1469265, EPI_ISL_1469266, EPI_ISL_1469267, EPI_ISL_1469268, EPI_ISL_1469269, EPI_ISL_1469270, EPI_ISL_1469276, EPI_ISL_1469277, EPI_ISL_1469278 | | | |
| see above | National Institute Health Research and Development | National Institute of Health Research and Development | Subangkit, Hana Apsari Pawestri, Kartika Dewi Puspa, Arie Ardiansyah Nugraha, Hartanti Dian Ikawati, Krisna Nur Andriana Pangesti, Yuni Rukminiati, Ririn Ramadhany, Agustiningsih, Kindi Adam, Holy Arif Wibowo, Triyani Soekarso, Ni Ketut Susilarini, Nurika Hariastuti, Ully Alfi Nikmah, Reni Herman, Nike Susanti, Herna, Tati Febriyanti, Natalie Laurencia Kipuw, Fauzul Muna, Irene Lorinda Indalao, Nelly Puspandari, Vivi Setiawaty. |
| EPI_ISL_1469285 | RS Khusus Paru Karawang | National Institute of Health Research and Development | Hartanti Dian Ikawati, Hana Apsari Pawestri, Subangkit, Kartika Dewi Puspa, Arie Ardiansyah Nugraha, Krisna Pangesti, Nelly Puspandari, Vivi Setiawaty |
| EPI_ISL_1469286 | RS Khusus Paru Karawang | National Institute of Health Research and Development | Vivi Setiawaty, Hana Apsari Pawestri, Subangkit, Kartika Dewi Puspa, Arie Ardiansyah Nugraha, Hartanti Dian Ikawati, Nelly Puspandari, Krisna Pangesti |
| EPI_ISL_1469287 | RS Mandaya Karawang | National Institute of Health Research and Development | Vivi Setiawaty, Hana Apsari Pawestri, Subangkit, Kartika Dewi Puspa, Arie Ardiansyah Nugraha, Hartanti Dian Ikawati, Nelly Puspandari, Krisna Pangesti |
| EPI_ISL_1469288, EPI_ISL_1469289 | Labkesda Karawang | National Institute of Health Research and Development | Hana Apsari Pawestri, Subangkit, Kartika Dewi Puspa, Arie Ardiansyah Nugraha, Hartanti Dian Ikawati, Nelly Puspandari, Krisna Pangesti, Vivi Setiawaty |
| EPI_ISL_1469938, EPI_ISL_1469939 | LESP Nuevo Leon | Instituto de Diagnostico y Referencia Epidemiologicos (INDRE) | Claudia Wong-Arambula, Abril Rodriguez-Maldonado, Vanessa Rivero-Arredondo, Ariadna Medina-Benitez, Joaquin Quiroz-Mercado, Sergio Rangel-Guerrero, Natividad Cruz-Ortiz, Tatiana Nunez-Garcia, Gisela Barrera-Badillo, Lucia Hernandez-Rivas, Irma Lopez-Martinez, Ernesto Ramirez-Gonzalez. |
| EPI_ISL_1469940 | LESP Sinaloa | Instituto de Diagnostico y Referencia Epidemiologicos (INDRE) | Claudia Wong-Arambula, Abril Rodriguez-Maldonado, Vanessa Rivero-Arredondo, Ariadna Medina-Benitez, Joaquin Quiroz-Mercado, Sergio Rangel-Guerrero, Natividad Cruz-Ortiz, Tatiana Nunez-Garcia, Gisela Barrera-Badillo, Lucia Hernandez-Rivas, Irma Lopez-Martinez, Ernesto Ramirez-Gonzalez. |
| EPI_ISL_1469941 | LESP Ciudad de Mexico | Instituto de Diagnostico y Referencia Epidemiologicos (INDRE) | Claudia Wong-Arambula, Abril Rodriguez-Maldonado, Vanessa Rivero-Arredondo, Ariadna Medina-Benitez, Joaquin Quiroz-Mercado, Sergio Rangel-Guerrero, Natividad Cruz-Ortiz, Tatiana Nunez-Garcia, Gisela Barrera-Badillo, Lucia Hernandez-Rivas, Irma Lopez-Martinez, Ernesto Ramirez-Gonzalez. |
| EPI_ISL_1469942, EPI_ISL_1469943, EPI_ISL_1469944 | LESP Guanajuato | Instituto de Diagnostico y Referencia Epidemiologicos (INDRE) | Claudia Wong-Arambula, Abril Rodriguez-Maldonado, Vanessa Rivero-Arredondo, Ariadna Medina-Benitez, Joaquin Quiroz-Mercado, Sergio Rangel-Guerrero, Natividad Cruz-Ortiz, Tatiana Nunez-Garcia, Gisela Barrera-Badillo, Lucia Hernandez-Rivas, Irma Lopez-Martinez, Ernesto Ramirez-Gonzalez. |
| EPI_ISL_1469989, EPI_ISL_1469990, EPI_ISL_1469991, EPI_ISL_1469992, EPI_ISL_1469993, EPI_ISL_1469994, EPI_ISL_1469995, EPI_ISL_1469996, EPI_ISL_1469997, EPI_ISL_1469998, EPI_ISL_1469999, EPI_ISL_1470000 | | | |
| see above | Laboratory of Communicable Diseases | 1. Laboratory of Communicable Diseases (Estonia); 2. Eurofins Genomics Europe Sequencing GmbH | Liidia Dotsenko et al. |
| EPI_ISL_1470059, EPI_ISL_1470060, EPI_ISL_1470061, EPI_ISL_1470062, EPI_ISL_1470064, EPI_ISL_1470065, EPI_ISL_1470066, EPI_ISL_1470136 | North Estonia Medical Centre laboratory | 1. Laboratory of Communicable Diseases (Estonia); 2. Eurofins Genomics Europe Sequencing GmbH | Liidia Dotsenko et al. |
| EPI_ISL_1470137, EPI_ISL_1470138, EPI_ISL_1470139, EPI_ISL_1470140, EPI_ISL_1470142, EPI_ISL_1470143, EPI_ISL_1470144, EPI_ISL_1470145, EPI_ISL_1470146, EPI_ISL_1470147, EPI_ISL_1470148, EPI_ISL_1470149, EPI_ISL_1470150, EPI_ISL_1470152, EPI_ISL_1470153, EPI_ISL_1470154, EPI_ISL_1470155, EPI_ISL_1470156, EPI_ISL_1470157, EPI_ISL_1470158, EPI_ISL_1470159, EPI_ISL_1470160, EPI_ISL_1470161, EPI_ISL_1470162, EPI_ISL_1470163, EPI_ISL_1470165, EPI_ISL_1470167, EPI_ISL_1470168, EPI_ISL_1470169, EPI_ISL_1470170, EPI_ISL_1470171, EPI_ISL_1470172, EPI_ISL_1470173, EPI_ISL_1470174, EPI_ISL_1470175, EPI_ISL_1470176, EPI_ISL_1470177, EPI_ISL_1470178, EPI_ISL_1470179, EPI_ISL_1470180, EPI_ISL_1470181, EPI_ISL_1470182, EPI_ISL_1470183, EPI_ISL_1470184, EPI_ISL_1470185, EPI_ISL_1470186, EPI_ISL_1470187, EPI_ISL_1470188, EPI_ISL_1470190, EPI_ISL_1470191, EPI_ISL_1470192, EPI_ISL_1470193, EPI_ISL_1470194, EPI_ISL_1470195, EPI_ISL_1470197, EPI_ISL_1470198, EPI_ISL_1470199, EPI_ISL_1470200, EPI_ISL_1470201, EPI_ISL_1470202, EPI_ISL_1470203, EPI_ISL_1470204, EPI_ISL_1470205, EPI_ISL_1470206, EPI_ISL_1470207, EPI_ISL_1470208, EPI_ISL_1470209, EPI_ISL_1470210, EPI_ISL_1470211, EPI_ISL_1470212, EPI_ISL_1470214, EPI_ISL_1470215, EPI_ISL_1470216, EPI_ISL_1470217, EPI_ISL_1470218, EPI_ISL_1470219, EPI_ISL_1470220, EPI_ISL_1470222, EPI_ISL_1470223, EPI_ISL_1470224, EPI_ISL_1470225, EPI_ISL_1470226, EPI_ISL_1470228, EPI_ISL_1470229, EPI_ISL_1470230, EPI_ISL_1470231, EPI_ISL_1470232, EPI_ISL_1470233, EPI_ISL_1470234, EPI_ISL_1470235, EPI_ISL_1470236, EPI_ISL_1470237, EPI_ISL_1470238, EPI_ISL_1470239, EPI_ISL_1470240, EPI_ISL_1470241, EPI_ISL_1470242, EPI_ISL_1470244, EPI_ISL_1470245, EPI_ISL_1470246, EPI_ISL_1470247, EPI_ISL_1470248, EPI_ISL_1470249, EPI_ISL_1470250, EPI_ISL_1470251, EPI_ISL_1470252, EPI_ISL_1470253, EPI_ISL_1470254, EPI_ISL_1470255, EPI_ISL_1470256, EPI_ISL_1470257, EPI_ISL_1470258, EPI_ISL_1470259, EPI_ISL_1470260, EPI_ISL_1470261, EPI_ISL_1470262, EPI_ISL_1470263, EPI_ISL_1470264, EPI_ISL_1470265, EPI_ISL_1470266, EPI_ISL_1470267, EPI_ISL_1470268, EPI_ISL_1470270, EPI_ISL_1470271, EPI_ISL_1470275, EPI_ISL_1470276, EPI_ISL_1470277, EPI_ISL_1470278, EPI_ISL_1470280, EPI_ISL_1470281, EPI_ISL_1470282, EPI_ISL_1470283, EPI_ISL_1470284, EPI_ISL_1470285, EPI_ISL_1470286, EPI_ISL_1470287, EPI_ISL_1470288, EPI_ISL_1470289, EPI_ISL_1470290, EPI_ISL_1470291, EPI_ISL_1470292, EPI_ISL_1470294, EPI_ISL_1470295, EPI_ISL_1470296 | | | |
| see above | Synlab Eesti OÜ | 1. Laboratory of Communicable Diseases (Estonia); 2. Eurofins Genomics Europe Sequencing GmbH | Liidia Dotsenko et al. |
| EPI_ISL_1478773, EPI_ISL_1478774, EPI_ISL_1478775, EPI_ISL_1478776, EPI_ISL_1478777, EPI_ISL_1478778, EPI_ISL_1478779, EPI_ISL_1478780, EPI_ISL_1478781, EPI_ISL_1478782, EPI_ISL_1478783, EPI_ISL_1478784, EPI_ISL_1478785, EPI_ISL_1478786, EPI_ISL_1478787, EPI_ISL_1478788, EPI_ISL_1478789, EPI_ISL_1478790, EPI_ISL_1478791, EPI_ISL_1478792, EPI_ISL_1478793, EPI_ISL_1478794, EPI_ISL_1478795, EPI_ISL_1478796, EPI_ISL_1478797, EPI_ISL_1478798, EPI_ISL_1478799, EPI_ISL_1478800, EPI_ISL_1478801, EPI_ISL_1478802, EPI_ISL_1478803, EPI_ISL_1478804, EPI_ISL_1478805, EPI_ISL_1478806, EPI_ISL_1478807, EPI_ISL_1478808, EPI_ISL_1478809, EPI_ISL_1478810, EPI_ISL_1478811, EPI_ISL_1478812, EPI_ISL_1478813, EPI_ISL_1478814, EPI_ISL_1478815, EPI_ISL_1478816 | | | |
| see above | Microbiology Department. Complexo Hospitalario Universitario de Vigo | Microbiology Department. Complexo Hospitalario Universitario de Vigo | Alfaia N, Alonso I, Alvarez M, Cabrera JJ, Carballo R, Cores O, Cortizo S, del-Campo V, Martinez L, Mediero G, Perez S, Potel C, Regueiro B, Rey S, Vasallo FJ |
| EPI_ISL_1483032, EPI_ISL_1483033, EPI_ISL_1483035, EPI_ISL_1483036, EPI_ISL_1483037 | LESP Nuevo Leon | Instituto de Diagnostico y Referencia Epidemiologicos (INDRE) | Claudia Wong-Arambula, Abril Rodriguez-Maldonado, Vanessa Rivero-Arredondo, Ariadna Medina-Benitez, Joaquin Quiroz-Mercado, Sergio Rangel-Guerrero, Natividad Cruz-Ortiz, Tatiana Nunez-Garcia, Gisela Barrera-Badillo, Lucia Hernandez-Rivas, Irma Lopez-Martinez, Ernesto Ramirez-Gonzalez. |
| EPI_ISL_1483041 | LESP Nayarit | Instituto de Diagnostico y Referencia Epidemiologicos | Claudia Wong-Arambula, Abril Rodriguez-Maldonado, Vanessa Rivero-Arredondo, Ariadna Medina-Benitez, Joaquin Quiroz-Mercado, Sergio |

| | | | |
|---|---|---|---|
| | | (INDRE) | Rangel-Guerrero, Natividad Cruz-Ortiz, Tatiana Nunez-Garcia, Gisela Barrera-Badillo, Lucia Hernandez-Rivas, Irma Lopez-Martinez, Ernesto Ramirez-Gonzalez. |
| EPI_ISL_1483042, EPI_ISL_1483043 | LESP Sinaloa | Instituto de Diagnostico y Referencia Epidemiologicos (INDRE) | Claudia Wong-Arambula, Abril Rodriguez-Maldonado, Vanessa Rivero-Arredondo, Ariadna Medina-Benitez, Joaquin Quiroz-Mercado, Sergio Rangel-Guerrero, Natividad Cruz-Ortiz, Tatiana Nunez-Garcia, Gisela Barrera-Badillo, Lucia Hernandez-Rivas, Irma Lopez-Martinez, Ernesto Ramirez-Gonzalez. |
| EPI_ISL_1483044, EPI_ISL_1483045, EPI_ISL_1483047, EPI_ISL_1483049, EPI_ISL_1483050 | LESP Queretaro | Instituto de Diagnostico y Referencia Epidemiologicos (INDRE) | Claudia Wong-Arambula, Abril Rodriguez-Maldonado, Vanessa Rivero-Arredondo, Ariadna Medina-Benitez, Joaquin Quiroz-Mercado, Sergio Rangel-Guerrero, Natividad Cruz-Ortiz, Tatiana Nunez-Garcia, Gisela Barrera-Badillo, Lucia Hernandez-Rivas, Irma Lopez-Martinez, Ernesto Ramirez-Gonzalez. |
| EPI_ISL_1483056, EPI_ISL_1483057 | LESP Veracruz | Instituto de Diagnostico y Referencia Epidemiologicos (INDRE) | Claudia Wong-Arambula, Abril Rodriguez-Maldonado, Vanessa Rivero-Arredondo, Ariadna Medina-Benitez, Joaquin Quiroz-Mercado, Sergio Rangel-Guerrero, Natividad Cruz-Ortiz, Tatiana Nunez-Garcia, Gisela Barrera-Badillo, Lucia Hernandez-Rivas, Irma Lopez-Martinez, Ernesto Ramirez-Gonzalez. |
| EPI_ISL_1483058 | LESP Hidalgo | Instituto de Diagnostico y Referencia Epidemiologicos (INDRE) | Claudia Wong-Arambula, Abril Rodriguez-Maldonado, Vanessa Rivero-Arredondo, Ariadna Medina-Benitez, Joaquin Quiroz-Mercado, Sergio Rangel-Guerrero, Natividad Cruz-Ortiz, Tatiana Nunez-Garcia, Gisela Barrera-Badillo, Lucia Hernandez-Rivas, Irma Lopez-Martinez, Ernesto Ramirez-Gonzalez. |
| EPI_ISL_1483059, EPI_ISL_1483060, EPI_ISL_1483061, EPI_ISL_1483062, EPI_ISL_1483063, EPI_ISL_1483064, EPI_ISL_1483065 | LESP Puebla | Instituto de Diagnostico y Referencia Epidemiologicos (INDRE) | Claudia Wong-Arambula, Abril Rodriguez-Maldonado, Vanessa Rivero-Arredondo, Ariadna Medina-Benitez, Joaquin Quiroz-Mercado, Sergio Rangel-Guerrero, Natividad Cruz-Ortiz, Tatiana Nunez-Garcia, Gisela Barrera-Badillo, Lucia Hernandez-Rivas, Irma Lopez-Martinez, Ernesto Ramirez-Gonzalez. |
| EPI_ISL_1483066, EPI_ISL_1483067 | LESP Campeche | Instituto de Diagnostico y Referencia Epidemiologicos (INDRE) | Claudia Wong-Arambula, Abril Rodriguez-Maldonado, Vanessa Rivero-Arredondo, Ariadna Medina-Benitez, Joaquin Quiroz-Mercado, Sergio Rangel-Guerrero, Natividad Cruz-Ortiz, Tatiana Nunez-Garcia, Gisela Barrera-Badillo, Lucia Hernandez-Rivas, Irma Lopez-Martinez, Ernesto Ramirez-Gonzalez. |
| EPI_ISL_1483068, EPI_ISL_1483069, EPI_ISL_1483070, EPI_ISL_1483071, EPI_ISL_1483072 | LESP Jalisco | Instituto de Diagnostico y Referencia Epidemiologicos (INDRE) | Claudia Wong-Arambula, Abril Rodriguez-Maldonado, Vanessa Rivero-Arredondo, Ariadna Medina-Benitez, Joaquin Quiroz-Mercado, Sergio Rangel-Guerrero, Natividad Cruz-Ortiz, Tatiana Nunez-Garcia, Gisela Barrera-Badillo, Lucia Hernandez-Rivas, Irma Lopez-Martinez, Ernesto Ramirez-Gonzalez. |
| EPI_ISL_1483073 | LESP Guerrero | Instituto de Diagnostico y Referencia Epidemiologicos (INDRE) | Claudia Wong-Arambula, Abril Rodriguez-Maldonado, Vanessa Rivero-Arredondo, Ariadna Medina-Benitez, Joaquin Quiroz-Mercado, Sergio Rangel-Guerrero, Natividad Cruz-Ortiz, Tatiana Nunez-Garcia, Gisela Barrera-Badillo, Lucia Hernandez-Rivas, Irma Lopez-Martinez, Ernesto Ramirez-Gonzalez. |
| EPI_ISL_1483074 | LESP Chihuahua | Instituto de Diagnostico y Referencia Epidemiologicos (INDRE) | Claudia Wong-Arambula, Abril Rodriguez-Maldonado, Vanessa Rivero-Arredondo, Ariadna Medina-Benitez, Joaquin Quiroz-Mercado, Sergio Rangel-Guerrero, Natividad Cruz-Ortiz, Tatiana Nunez-Garcia, Gisela Barrera-Badillo, Lucia Hernandez-Rivas, Irma Lopez-Martinez, Ernesto Ramirez-Gonzalez. |
| EPI_ISL_1489729 | Hospital | National Reference Center for Viruses of Respiratory Infections, Institut Pasteur, Paris | Marion Barbet, Sylvie Behillil,Frédéric Lemoine,Corinne Maufrais,Christophe Malabat,Damien Mornico, Méline Bizard, Angela Brisebarre, Camille Capel, Louise Lefrançois, Etienne Simon-Lorière, Vincent Enouf, Maud Vanpeene, Sylvie van der Werf,Mj Dufour |
| EPI_ISL_1489730 | Labo Analyses Med | National Reference Center for Viruses of Respiratory Infections, Institut Pasteur, Paris | Marion Barbet, Sylvie Behillil,Frédéric Lemoine,Corinne Maufrais,Christophe Malabat,Damien Mornico, Méline Bizard, Angela Brisebarre, Camille Capel, Louise Lefrançois, Etienne Simon-Lorière, Vincent Enouf, Maud Vanpeene, Sylvie van der Werf,OphéLie Said-Delattre |
| EPI_ISL_1489731, EPI_ISL_1489732, EPI_ISL_1489733, EPI_ISL_1489734 | Labo Analyses Med | National Reference Center for Viruses of Respiratory Infections, Institut Pasteur, Paris | Marion Barbet, Sylvie Behillil,Frédéric Lemoine,Corinne Maufrais,Christophe Malabat,Amaury Vaysse, Méline Bizard, Angela Brisebarre, Camille Capel, Louise Lefrançois, Etienne Simon-Lorière, Vincent Enouf, Maud Vanpeene, Sylvie van der Werf,OphéLie Said-Delattre |
| EPI_ISL_1489735 | Labo Analyses Med | National Reference Center for Viruses of Respiratory Infections, Institut Pasteur, Paris | Marion Barbet, Sylvie Behillil,Frédéric Lemoine,Corinne Maufrais,Christophe Malabat,Adrien Pain, Méline Bizard, Angela Brisebarre, Camille Capel, Louise Lefrançois, Etienne Simon-Lorière, Vincent Enouf, Maud Vanpeene, Sylvie van der Werf,OphéLie Said-Delattre |
| EPI_ISL_1489736 | Labo Analyses Med | National Reference Center for Viruses of Respiratory Infections, Institut Pasteur, Paris | Marion Barbet, Sylvie Behillil,Frédéric Lemoine,Corinne Maufrais,Christophe Malabat,Amaury Vaysse, Méline Bizard, Angela Brisebarre, Camille Capel, Louise Lefrançois, Etienne Simon-Lorière, Vincent Enouf, Maud Vanpeene, Sylvie van der Werf,OphéLie Said-Delattre |
| EPI_ISL_1489738 | Hospital | National Reference Center for Viruses of Respiratory Infections, Institut Pasteur, Paris | Marion Barbet, Sylvie Behillil,Frédéric Lemoine,Corinne Maufrais,Christophe Malabat,Damien Mornico, Méline Bizard, Angela Brisebarre, Camille Capel, Louise Lefrançois, Etienne Simon-Lorière, Vincent Enouf, Maud Vanpeene, Sylvie van der Werf,SéVerine Haouisee |
| EPI_ISL_1489739 | Labo Analyses Med | National Reference Center for Viruses of Respiratory Infections, Institut Pasteur, Paris | Marion Barbet, Sylvie Behillil,Frédéric Lemoine,Corinne Maufrais,Christophe Malabat, Méline Bizard, Angela Brisebarre, Camille Capel, Louise Lefrançois, Etienne Simon-Lorière, Vincent Enouf, Maud Vanpeene, Sylvie van der Werf,Catherine Delamare |
| EPI_ISL_1489835, EPI_ISL_1489836 | Labo Analyses Med | National Reference Center for Viruses of Respiratory Infections, Institut Pasteur, Paris | Marion Barbet, Sylvie Behillil,Frédéric Lemoine,Corinne Maufrais,Christophe Malabat,Adrien Pain, Méline Bizard, Angela Brisebarre, Camille Capel, Louise Lefrançois, Etienne Simon-Lorière, Vincent Enouf, Maud Vanpeene, Sylvie van der Werf,Nabil Gastli |
| EPI_ISL_1489837 | Labo Analyses Med | National Reference Center for Viruses of Respiratory Infections, Institut Pasteur, Paris | Marion Barbet, Sylvie Behillil,Frédéric Lemoine,Corinne Maufrais,Christophe Malabat,Amaury Vaysse, Méline Bizard, Angela Brisebarre, Camille Capel, Louise Lefrançois, Etienne Simon-Lorière, Vincent Enouf, Maud Vanpeene, Sylvie van der Werf,Nabil Gastli |
| EPI_ISL_1489839 | Labo Analyses Med | National Reference Center for Viruses of Respiratory Infections, Institut Pasteur, Paris | Marion Barbet, Sylvie Behillil,Frédéric Lemoine,Corinne Maufrais,Christophe Malabat,Adrien Pain, Méline Bizard, Angela Brisebarre, Camille Capel, Louise Lefrançois, Etienne Simon-Lorière, Vincent Enouf, Maud Vanpeene, Sylvie van der Werf,Nabil Gastli |
| EPI_ISL_1489840, EPI_ISL_1489842 | Labo Analyses Med | National Reference Center for Viruses of Respiratory Infections, Institut Pasteur, Paris | Marion Barbet, Sylvie Behillil,Frédéric Lemoine,Corinne Maufrais,Christophe Malabat,Emmanuelle Permal, Méline Bizard, Angela Brisebarre, Camille Capel, Louise Lefrançois, Etienne Simon-Lorière, Vincent Enouf, Maud Vanpeene, Sylvie van der Werf,Nabil Gastli |
| EPI_ISL_1490234 | HOPITAL DE VICHY | CHU Clermont-Ferrand, service de virologie | Bisseux Maxime, Mirand Audrey, Combes Patricia, Henquell Cécile |
| EPI_ISL_1490236 | Centre Hospitalier Universitaire Clermont-Ferrand | CHU Clermont-Ferrand, service de virologie | Bisseux Maxime, Mirand Audrey, Combes Patricia, Henquell Cécile |
| EPI_ISL_1490237 | Centre Jean Perrin | CHU Clermont-Ferrand, service de virologie | Bisseux Maxime, Mirand Audrey, Combes Patricia, Henquell Cécile |
| EPI_ISL_1490238 | Centre Hospitalier Universitaire Clermont-Ferrand | CHU Clermont-Ferrand, service de virologie | Bisseux Maxime, Mirand Audrey, Combes Patricia, Henquell Cécile |
| EPI_ISL_1492456 | LESP Jalisco | Instituto de diagnóstico y Referencia Epidemiologicos (INDRE) | Claudia Wong-Arambula, Abril Rodriguez-Maldonado, Vanessa Rivero-Arredondo, Ariadna Medina-Benitez, Joaquin Quiroz-Mercado, Sergio Rangel-Guerrero, Natividad Cruz-Ortiz, Tatiana Nunez-Garcia, Gisela Barrera-Badillo, Lucia Hernandez-Rivas, Irma Lopez-Martinez, Ernesto Ramirez-Gonzalez. |
| EPI_ISL_1492561 | HOPITAL DE VICHY | CHU Clermont-Ferrand, service de virologie | Bisseux Maxime, Mirand Audrey, Combes Patricia, Henquell Cécile |
| EPI_ISL_1492562, EPI_ISL_1492563 | Centre Hospitalier Universitaire Clermont-Ferrand | CHU Clermont-Ferrand, service de virologie | Bisseux Maxime, Mirand Audrey, Combes Patricia, Henquell Cécile |
| EPI_ISL_1493573, EPI_ISL_1493574, EPI_ISL_1493575, EPI_ISL_1493576, EPI_ISL_1493577 | LACEN do Estado de Goias | Instituto Adolfo Lutz, Interdiciplinary Procedures Center, Strategic Laboratory | Claudio Tavares Sacchi, Claudia Regina Gonçalves, Erica Valessa Ramos Gomes, Karoline Rodrigues Campos, Caio Vinicius Dias Lopes |
| EPI_ISL_1493578, EPI_ISL_1493579 | LACEN do Estado de Rondonia | Instituto Adolfo Lutz, Interdiciplinary Procedures Center, Strategic Laboratory | Claudio Tavares Sacchi, Claudia Regina Gonçalves, Erica Valessa Ramos Gomes, Karoline Rodrigues Campos, Caio Vinicius Dias Lopes |
| EPI_ISL_1494626, EPI_ISL_1494630 | LESP Chihuahua | Instituto de Diagnostico y Referencia Epidemiologicos (INDRE) | Claudia Wong-Arambula, Abril Rodriguez-Maldonado, Vanessa Rivero-Arredondo, Ariadna Medina-Benitez, Joaquin Quiroz-Mercado, Sergio Rangel-Guerrero, Natividad Cruz-Ortiz, Tatiana Nunez-Garcia, Gisela Barrera-Badillo, Lucia Hernandez-Rivas, Irma Lopez-Martinez, Ernesto Ramirez-Gonzalez. |
| EPI_ISL_1494631, EPI_ISL_1494633, EPI_ISL_1494634, EPI_ISL_1494635 | LESP Nuevo Leon | Instituto de Diagnostico y Referencia Epidemiologicos (INDRE) | Claudia Wong-Arambula, Abril Rodriguez-Maldonado, Vanessa Rivero-Arredondo, Ariadna Medina-Benitez, Joaquin Quiroz-Mercado, Sergio Rangel-Guerrero, Natividad Cruz-Ortiz, Tatiana Nunez-Garcia, Gisela Barrera-Badillo, Lucia Hernandez-Rivas, Irma Lopez-Martinez, Ernesto Ramirez-Gonzalez. |

| | | | |
|---|---|---|---|
| EPI_ISL_1494636, EPI_ISL_1494639, EPI_ISL_1494640, EPI_ISL_1494643, EPI_ISL_1494644, EPI_ISL_1494645 | LESP Ciudad de Mexico | Instituto de Diagnostico y Referencia Epidemiologicos (INDRE) | Claudia Wong-Arambula, Abril Rodriguez-Maldonado, Vanessa Rivero-Arredondo, Ariadna Medina-Benitez, Joaquin Quiroz-Mercado, Sergio Rangel-Guerrero, Natividad Cruz-Ortiz, Tatiana Nunez-Garcia, Gisela Barrera-Badillo, Lucia Hernandez-Rivas, Irma Lopez-Martinez, Ernesto Ramirez-Gonzalez. |
| EPI_ISL_1494648, EPI_ISL_1494649 | LESP Nuevo Leon | Instituto de Diagnostico y Referencia Epidemiologicos (INDRE) | Claudia Wong-Arambula, Abril Rodriguez-Maldonado, Vanessa Rivero-Arredondo, Ariadna Medina-Benitez, Joaquin Quiroz-Mercado, Sergio Rangel-Guerrero, Natividad Cruz-Ortiz, Tatiana Nunez-Garcia, Gisela Barrera-Badillo, Lucia Hernandez-Rivas, Irma Lopez-Martinez, Ernesto Ramirez-Gonzalez. |
| EPI_ISL_1494650, EPI_ISL_1494651, EPI_ISL_1494652, EPI_ISL_1494653, EPI_ISL_1494654 | LESP Tamaulipas | Instituto de Diagnostico y Referencia Epidemiologicos (INDRE) | Claudia Wong-Arambula, Abril Rodriguez-Maldonado, Vanessa Rivero-Arredondo, Ariadna Medina-Benitez, Joaquin Quiroz-Mercado, Sergio Rangel-Guerrero, Natividad Cruz-Ortiz, Tatiana Nunez-Garcia, Gisela Barrera-Badillo, Lucia Hernandez-Rivas, Irma Lopez-Martinez, Ernesto Ramirez-Gonzalez. |
| EPI_ISL_1494655, EPI_ISL_1494656 | LESP Hidalgo | Instituto de Diagnostico y Referencia Epidemiologicos (INDRE) | Claudia Wong-Arambula, Abril Rodriguez-Maldonado, Vanessa Rivero-Arredondo, Ariadna Medina-Benitez, Joaquin Quiroz-Mercado, Sergio Rangel-Guerrero, Natividad Cruz-Ortiz, Tatiana Nunez-Garcia, Gisela Barrera-Badillo, Lucia Hernandez-Rivas, Irma Lopez-Martinez, Ernesto Ramirez-Gonzalez. |
| EPI_ISL_1494657, EPI_ISL_1494658, EPI_ISL_1494659, EPI_ISL_1494660, EPI_ISL_1494661, EPI_ISL_1494662, EPI_ISL_1494663, EPI_ISL_1494664, EPI_ISL_1494665, EPI_ISL_1494666, EPI_ISL_1494667, EPI_ISL_1494668, EPI_ISL_1494669 | | | |
| see above | LESP Guanajuato | Instituto de Diagnostico y Referencia Epidemiologicos (INDRE) | Claudia Wong-Arambula, Abril Rodriguez-Maldonado, Vanessa Rivero-Arredondo, Ariadna Medina-Benitez, Joaquin Quiroz-Mercado, Sergio Rangel-Guerrero, Natividad Cruz-Ortiz, Tatiana Nunez-Garcia, Gisela Barrera-Badillo, Lucia Hernandez-Rivas, Irma Lopez-Martinez, Ernesto Ramirez-Gonzalez. |
| EPI_ISL_1494670, EPI_ISL_1494671, EPI_ISL_1494672 | LESP Sinaloa | Instituto de Diagnostico y Referencia Epidemiologicos (INDRE) | Claudia Wong-Arambula, Abril Rodriguez-Maldonado, Vanessa Rivero-Arredondo, Ariadna Medina-Benitez, Joaquin Quiroz-Mercado, Sergio Rangel-Guerrero, Natividad Cruz-Ortiz, Tatiana Nunez-Garcia, Gisela Barrera-Badillo, Lucia Hernandez-Rivas, Irma Lopez-Martinez, Ernesto Ramirez-Gonzalez. |
| EPI_ISL_1494673 | LESP Nuevo Leon | Instituto de Diagnostico y Referencia Epidemiologicos (INDRE) | Claudia Wong-Arambula, Abril Rodriguez-Maldonado, Vanessa Rivero-Arredondo, Ariadna Medina-Benitez, Joaquin Quiroz-Mercado, Sergio Rangel-Guerrero, Natividad Cruz-Ortiz, Tatiana Nunez-Garcia, Gisela Barrera-Badillo, Lucia Hernandez-Rivas, Irma Lopez-Martinez, Ernesto Ramirez-Gonzalez. |
| EPI_ISL_1494923 | CS II Dr Jose Ferreira Telles | Instituto Adolfo Lutz, Interdiciplinary Procedures Center, Strategic Laboratory | Claudio Tavares Sacchi, Claudia Regina Gonçalves, Erica Valessa Ramos Gomes, Karoline Rodrigues Campos, Caio Vinicius Dias Lopes |
| EPI_ISL_1495044, EPI_ISL_1495055 | Institut de Virologie du CHU de Strasbourg | Institut de Virologie du CHU de Strasbourg | Fafi-Kremer Samira, Gallais Floriane, Gantner Pierre, Laugel Elodie, Solis Morgane, Velay Aurélie, Wendling Marie-Josée |
| EPI_ISL_1498306, EPI_ISL_1498307, EPI_ISL_1498308, EPI_ISL_1498309, EPI_ISL_1498310, EPI_ISL_1498311, EPI_ISL_1498312 | Microbiology Department, University Hospital Donostia | Microbiology Department, University Hospital Donostia | Marimon JM, Montes M, Piñeiro L, Sorarrain A, Gomez M, Cilla G. |
| EPI_ISL_1499474, EPI_ISL_1499475, EPI_ISL_1499476, EPI_ISL_1499477, EPI_ISL_1499478, EPI_ISL_1499479, EPI_ISL_1499480, EPI_ISL_1499481, EPI_ISL_1499483, EPI_ISL_1499484, EPI_ISL_1499485, EPI_ISL_1499486, EPI_ISL_1499487, EPI_ISL_1499488, EPI_ISL_1499489, EPI_ISL_1499490, EPI_ISL_1499491, EPI_ISL_1499492, EPI_ISL_1499493, EPI_ISL_1499495, EPI_ISL_1499499, EPI_ISL_1499500, EPI_ISL_1499501, EPI_ISL_1499502, EPI_ISL_1499503 | | | |
| see above | Azienda Ospedaliera Terni | Istituto Zooprofilattico Sperimentale dell'Abruzzo e Molise "G. Caporale" | Palumbo M, Scaccetti A, Lorusso A, Marcacci M, Di Domenico M, Ancora M, Curini V, Di Lollo V, Delli Compagni E, Mangone I, Rinaldi A, Scialabba S, Caporale M, Di Pasquale A, Cammà C, Puglia I, Calistri P, Savini G |
| EPI_ISL_1499625, EPI_ISL_1499626 | Azienda Ospedaliera Terni | Istituto Zooprofilattico Sperimentale dell'Abruzzo e Molise "G. Caporale" | Palumbo M, Scaccetti A, Lorusso A, Marcacci M, Di Domenico M, Ancora M, Curini V, Di Lollo Valeria, Mangone I, Rinaldi A, Delli Compagni E, Scialabba S, Caporale M, Di Pasquale A, Cammà C, Puglia I, Calistri P, Savini G |
| EPI_ISL_1499645 | Laboratorio Analisi Osp. Città di Castello - Azienda USL Umbria1 | Istituto Zooprofilattico Sperimentale dell'Abruzzo e Molise "G. Caporale" | Malagigi V, Tacconi P,Lorusso A, Marcacci M, Di Domenico M, Ancora M, Curini V, Di Lollo Valeria, Mangone I, Rinaldi A, Delli Compagni E, Scialabba S, Caporale M, Di Pasquale A, Cammà C, Puglia I, Calistri P, Savini G |
| EPI_ISL_1501065, EPI_ISL_1501066, EPI_ISL_1501067, EPI_ISL_1501068, EPI_ISL_1501069, EPI_ISL_1501070, EPI_ISL_1501071, EPI_ISL_1501072, EPI_ISL_1501073, EPI_ISL_1501074, EPI_ISL_1501075, EPI_ISL_1501076, EPI_ISL_1501077, EPI_ISL_1501078, EPI_ISL_1501079, EPI_ISL_1501080, EPI_ISL_1501081, EPI_ISL_1501082, EPI_ISL_1501094, EPI_ISL_1501095, EPI_ISL_1501096, EPI_ISL_1501097, EPI_ISL_1501098, EPI_ISL_1501100, EPI_ISL_1501101, EPI_ISL_1501102, EPI_ISL_1501103, EPI_ISL_1501104, EPI_ISL_1501105, EPI_ISL_1501106, EPI_ISL_1501107, EPI_ISL_1501108, EPI_ISL_1501109, EPI_ISL_1501110, EPI_ISL_1501111, EPI_ISL_1501112, EPI_ISL_1501113, EPI_ISL_1501114, EPI_ISL_1501115, EPI_ISL_1501116, EPI_ISL_1501117, EPI_ISL_1501118, EPI_ISL_1501119, EPI_ISL_1501120, EPI_ISL_1501121, EPI_ISL_1501122, EPI_ISL_1501123, EPI_ISL_1501124, EPI_ISL_1501125, EPI_ISL_1501126, EPI_ISL_1501127 | | | |
| see above | HEGP - Laboratoire de Virologie | HEGP - Laboratoire de Virologie | David Veyer |
| EPI_ISL_1501129, EPI_ISL_1501130, EPI_ISL_1501131, EPI_ISL_1501132, EPI_ISL_1501133, EPI_ISL_1501134, EPI_ISL_1501135, EPI_ISL_1501136 | Broussais | HEGP - Laboratoire de Virologie | David Veyer |
| EPI_ISL_1501138, EPI_ISL_1501139, EPI_ISL_1501140, EPI_ISL_1501141, EPI_ISL_1501142, EPI_ISL_1501143, EPI_ISL_1501144, EPI_ISL_1501145, EPI_ISL_1501146, EPI_ISL_1501148, EPI_ISL_1501149, EPI_ISL_1501150, EPI_ISL_1501151, EPI_ISL_1501152, EPI_ISL_1501153, EPI_ISL_1501155, EPI_ISL_1501156, EPI_ISL_1501157, EPI_ISL_1501158, EPI_ISL_1501159, EPI_ISL_1501160, EPI_ISL_1501161, EPI_ISL_1501162, EPI_ISL_1501163, EPI_ISL_1501164, EPI_ISL_1501165, EPI_ISL_1501166, EPI_ISL_1501167, EPI_ISL_1501168, EPI_ISL_1501169 | | | |
| see above | HEGP - Laboratoire de Virologie | HEGP - Laboratoire de Virologie | David Veyer |
| EPI_ISL_1501170, EPI_ISL_1501171, EPI_ISL_1501172, EPI_ISL_1501173 | Broussais | HEGP - Laboratoire de Virologie | David Veyer |
| EPI_ISL_1501174, EPI_ISL_1501175, EPI_ISL_1501176, EPI_ISL_1501177, EPI_ISL_1501178, EPI_ISL_1501179, EPI_ISL_1501180, EPI_ISL_1501181, EPI_ISL_1501182, EPI_ISL_1501183, EPI_ISL_1501184, EPI_ISL_1501185, EPI_ISL_1501186, EPI_ISL_1501187, EPI_ISL_1501188, EPI_ISL_1501189, EPI_ISL_1501190, EPI_ISL_1501191, EPI_ISL_1501192, EPI_ISL_1501193, EPI_ISL_1501194, EPI_ISL_1501195, EPI_ISL_1501196, EPI_ISL_1501197, EPI_ISL_1501198, EPI_ISL_1501199, EPI_ISL_1501200, EPI_ISL_1501201, EPI_ISL_1501202, EPI_ISL_1501203, EPI_ISL_1501204, EPI_ISL_1501205, EPI_ISL_1501206, EPI_ISL_1501207, EPI_ISL_1501208, EPI_ISL_1501209, EPI_ISL_1501210 | | | |
| see above | HEGP - Laboratoire de Virologie | HEGP - Laboratoire de Virologie | David Veyer |
| EPI_ISL_1501211, EPI_ISL_1501212, EPI_ISL_1501216 | Broussais | HEGP - Laboratoire de Virologie | David Veyer |
| EPI_ISL_1501220 | HEGP - Laboratoire de Virologie | HEGP - Laboratoire de Virologie | David Veyer |
| EPI_ISL_1501259, EPI_ISL_1501260, EPI_ISL_1501261, EPI_ISL_1501262, EPI_ISL_1501263, EPI_ISL_1501264 | Broussais | HEGP - Laboratoire de Virologie | David Veyer |
| EPI_ISL_1502540, EPI_ISL_1502541, EPI_ISL_1502542, EPI_ISL_1502543, EPI_ISL_1502544, EPI_ISL_1502545 | OSPEDALE SAN SALVATORE - MEDICINA DI LABORATORIO L'AQUILA(L'AQUILA) | Istituto Zooprofilattico Sperimentale dell'Abruzzo e Molise "G. Caporale" | Lorusso A, Marcacci M, Di Domenico M, Ancora M, Curini V, Di Lollo Valeria, Mangone I, Rinaldi A, Delli Compagni E, Scialabba S, Caporale M, Di Pasquale A, Cammà C, Puglia I, Calistri P, Savini G |
| EPI_ISL_1503112, EPI_ISL_1503114, EPI_ISL_1503115, EPI_ISL_1503119 | Gorgas Memorial Laboratory of Health Studies | Gorgas Memorial Laboratory of Health Studies | Gonzalez Claudia, Leyda Abrego, Moreno Ambar, Oris Chavarria, Jessica Gondola, Marlenne Castillo, Ortiz Alma, Castillo Jorge, Moreno Brechla, Franco Danilo, Lopez-Verges Sandra, Martinez Alexander |
| EPI_ISL_1503980, EPI_ISL_1503982 | Institut de Virologie du CHU de Strasbourg | Institut de Virologie du CHU de Strasbourg | Fafi-Kremer Samira, Gallais Floriane, Gantner Pierre, Laugel Elodie, Solis Morgane, Velay Aurélie, Wendling Marie-Josée |
| EPI_ISL_1504058, EPI_ISL_1504059, EPI_ISL_1504060, EPI_ISL_1504061, EPI_ISL_1504062, EPI_ISL_1504063, EPI_ISL_1504064, EPI_ISL_1504065, EPI_ISL_1504066, EPI_ISL_1504067 | LESP Nuevo Leon | Instituto de Diagnostico y Referencia Epidemiologicos (INDRE) | Claudia Wong-Arambula, Abril Rodriguez-Maldonado, Vanessa Rivero-Arredondo, Ariadna Medina-Benitez, Joaquin Quiroz-Mercado, Sergio Rangel-Guerrero, Natividad Cruz-Ortiz, Tatiana Nunez-Garcia, Gisela Barrera-Badillo, Lucia Hernandez-Rivas, Irma Lopez-Martinez, Ernesto Ramirez-Gonzalez. |
| EPI_ISL_1504068, EPI_ISL_1504069, EPI_ISL_1504070, EPI_ISL_1504072, EPI_ISL_1504073, EPI_ISL_1504074, EPI_ISL_1504075, EPI_ISL_1504076, EPI_ISL_1504077, EPI_ISL_1504078, EPI_ISL_1504080 | | | |

| | | | |
|---|---|---|---|
| see above | LESP Morelos | Instituto de Diagnostico y Referencia Epidemiologicos (INDRE) | Claudia Wong-Arambula, Abril Rodriguez-Maldonado, Vanessa Rivero-Arredondo, Ariadna Medina-Benitez, Joaquin Quiroz-Mercado, Sergio Rangel-Guerrero, Natividad Cruz-Ortiz, Tatiana Nunez-Garcia, Gisela Barrera-Badillo, Lucia Hernandez-Rivas, Irma Lopez-Martinez, Ernesto Ramirez-Gonzalez. |
| EPI_ISL_1504083 | LESP Ciudad de Mexico | Instituto de Diagnostico y Referencia Epidemiologicos (INDRE) | Claudia Wong-Arambula, Abril Rodriguez-Maldonado, Vanessa Rivero-Arredondo, Ariadna Medina-Benitez, Joaquin Quiroz-Mercado, Sergio Rangel-Guerrero, Natividad Cruz-Ortiz, Tatiana Nunez-Garcia, Gisela Barrera-Badillo, Lucia Hernandez-Rivas, Irma Lopez-Martinez, Ernesto Ramirez-Gonzalez. |
| EPI_ISL_1504084, EPI_ISL_1504085, EPI_ISL_1504086, EPI_ISL_1504087, EPI_ISL_1504088 | LESP Chihuahua | Instituto de Diagnostico y Referencia Epidemiologicos (INDRE) | Claudia Wong-Arambula, Abril Rodriguez-Maldonado, Vanessa Rivero-Arredondo, Ariadna Medina-Benitez, Joaquin Quiroz-Mercado, Sergio Rangel-Guerrero, Natividad Cruz-Ortiz, Tatiana Nunez-Garcia, Gisela Barrera-Badillo, Lucia Hernandez-Rivas, Irma Lopez-Martinez, Ernesto Ramirez-Gonzalez. |
| EPI_ISL_1504089, EPI_ISL_1504091, EPI_ISL_1504092, EPI_ISL_1504093, EPI_ISL_1504094, EPI_ISL_1504095, EPI_ISL_1504096, EPI_ISL_1504097, EPI_ISL_1504098, EPI_ISL_1504099, EPI_ISL_1504100, EPI_ISL_1504101, EPI_ISL_1504102 | | | |
| see above | LESP Queretaro | Instituto de Diagnostico y Referencia Epidemiologicos (INDRE) | Claudia Wong-Arambula, Abril Rodriguez-Maldonado, Vanessa Rivero-Arredondo, Ariadna Medina-Benitez, Joaquin Quiroz-Mercado, Sergio Rangel-Guerrero, Natividad Cruz-Ortiz, Tatiana Nunez-Garcia, Gisela Barrera-Badillo, Lucia Hernandez-Rivas, Irma Lopez-Martinez, Ernesto Ramirez-Gonzalez. |
| EPI_ISL_1504104 | LESP Guerrero | Instituto de Diagnostico y Referencia Epidemiologicos (INDRE) | Claudia Wong-Arambula, Abril Rodriguez-Maldonado, Vanessa Rivero-Arredondo, Ariadna Medina-Benitez, Joaquin Quiroz-Mercado, Sergio Rangel-Guerrero, Natividad Cruz-Ortiz, Tatiana Nunez-Garcia, Gisela Barrera-Badillo, Lucia Hernandez-Rivas, Irma Lopez-Martinez, Ernesto Ramirez-Gonzalez. |
| EPI_ISL_1504105 | LESP Baja California Sur | Instituto de Diagnostico y Referencia Epidemiologicos (INDRE) | Claudia Wong-Arambula, Abril Rodriguez-Maldonado, Vanessa Rivero-Arredondo, Ariadna Medina-Benitez, Joaquin Quiroz-Mercado, Sergio Rangel-Guerrero, Natividad Cruz-Ortiz, Tatiana Nunez-Garcia, Gisela Barrera-Badillo, Lucia Hernandez-Rivas, Irma Lopez-Martinez, Ernesto Ramirez-Gonzalez. |
| EPI_ISL_1504106, EPI_ISL_1504107 | LESPZacatecas | Instituto de Diagnostico y Referencia Epidemiologicos (INDRE) | Claudia Wong-Arambula, Abril Rodriguez-Maldonado, Vanessa Rivero-Arredondo, Ariadna Medina-Benitez, Joaquin Quiroz-Mercado, Sergio Rangel-Guerrero, Natividad Cruz-Ortiz, Tatiana Nunez-Garcia, Gisela Barrera-Badillo, Lucia Hernandez-Rivas, Irma Lopez-Martinez, Ernesto Ramirez-Gonzalez. |
| EPI_ISL_1504108 | LESP Baja California | Instituto de Diagnostico y Referencia Epidemiologicos (INDRE) | Claudia Wong-Arambula, Abril Rodriguez-Maldonado, Vanessa Rivero-Arredondo, Ariadna Medina-Benitez, Joaquin Quiroz-Mercado, Sergio Rangel-Guerrero, Natividad Cruz-Ortiz, Tatiana Nunez-Garcia, Gisela Barrera-Badillo, Lucia Hernandez-Rivas, Irma Lopez-Martinez, Ernesto Ramirez-Gonzalez. |
| EPI_ISL_1504109 | LESP Jalisco | Instituto de Diagnostico y Referencia Epidemiologicos (INDRE) | Claudia Wong-Arambula, Abril Rodriguez-Maldonado, Vanessa Rivero-Arredondo, Ariadna Medina-Benitez, Joaquin Quiroz-Mercado, Sergio Rangel-Guerrero, Natividad Cruz-Ortiz, Tatiana Nunez-Garcia, Gisela Barrera-Badillo, Lucia Hernandez-Rivas, Irma Lopez-Martinez, Ernesto Ramirez-Gonzalez. |
| EPI_ISL_1507483 | Medical Genetics Laboratory, Regional Centre of Medical Genetics, Emergency County Hospital Craiova | Medical Genetics Laboratory, Regional Centre of Medical Genetics, Emergency County Hospital Craiova | Anca-Lelia (Riza) Costache, Ioana Streata, Stefania Dorobantu, Adina Dragos, Mihai Cucu, Ana-Maria Buga, Razvan Plesea, Elena Plesea, Monica Cara, Mihai Ioana |
| EPI_ISL_1507877 | Institut de Virologie du CHU de Strasbourg | Institut de Virologie du CHU de Strasbourg | Fafi-Kremer Samira, Gallais Floriane, Gantner Pierre, Laugel Elodie, Solis Morgane, Velay Aurélie, Wendling Marie-Josée |
| EPI_ISL_1508711 | Medical Genetics Laboratory, Regional Centre of Medical Genetics, Emergency County Hospital Craiova | Medical Genetics Laboratory, Regional Centre of Medical Genetics, Emergency County Hospital Craiova | Anca-Lelia (Riza) Costache, Ioana Streata, Stefania Dorobantu, Adina Dragos, Mihai Cucu, Ana-Maria Buga, Razvan Plesea, Elena Plesea, Monica Cara, Mihai Ioana |
| EPI_ISL_1508820, EPI_ISL_1508827 | Institut de Virologie du CHU de Strasbourg | Institut de Virologie du CHU de Strasbourg | Fafi-Kremer Samira, Gallais Floriane, Gantner Pierre, Laugel Elodie, Solis Morgane, Velay Aurélie, Wendling Marie-Josée |
| EPI_ISL_1508831 | Medical Genetics Laboratory, Regional Centre of Medical Genetics, Emergency County Hospital Craiova | Medical Genetics Laboratory, Regional Centre of Medical Genetics, Emergency County Hospital Craiova | Anca-Lelia (Riza) Costache, Ioana Streata, Stefania Dorobantu, Adina Dragos, Mihai Cucu, Ana-Maria Buga, Razvan Plesea, Andrei Pirvu, Elena Plesea, Monica Cara, Mihai Ioana |
| EPI_ISL_1508834 | Institut de Virologie du CHU de Strasbourg | Institut de Virologie du CHU de Strasbourg | Fafi-Kremer Samira, Gallais Floriane, Gantner Pierre, Laugel Elodie, Solis Morgane, Velay Aurélie, Wendling Marie-Josée |
| EPI_ISL_1508835, EPI_ISL_1508836, EPI_ISL_1508840, EPI_ISL_1508843, EPI_ISL_1508846, EPI_ISL_1508847, EPI_ISL_1508850, EPI_ISL_1508851 | Medical Genetics Laboratory, Regional Centre of Medical Genetics, Emergency County Hospital Craiova | Medical Genetics Laboratory, Regional Centre of Medical Genetics, Emergency County Hospital Craiova | Anca-Lelia (Riza) Costache, Ioana Streata, Stefania Dorobantu, Adina Dragos, Mihai Cucu, Ana-Maria Buga, Razvan Plesea, Andrei Pirvu, Elena Plesea, Monica Cara, Mihai Ioana |
| EPI_ISL_1508853, EPI_ISL_1508854, EPI_ISL_1508856, EPI_ISL_1508857, EPI_ISL_1508858, EPI_ISL_1508859, EPI_ISL_1508862, EPI_ISL_1508863, EPI_ISL_1508864, EPI_ISL_1508865, EPI_ISL_1508866, EPI_ISL_1508867, EPI_ISL_1508868, EPI_ISL_1508869, EPI_ISL_1508870, EPI_ISL_1508871, EPI_ISL_1508872, EPI_ISL_1508873, EPI_ISL_1508874, EPI_ISL_1508875, EPI_ISL_1508876, EPI_ISL_1508880, EPI_ISL_1508883, EPI_ISL_1508884, EPI_ISL_1508885, EPI_ISL_1508886, EPI_ISL_1508888, EPI_ISL_1508889 | | | |
| see above | University of Oregon COVID-19 MAP Laboratory | University of Oregon Genomics and Cell Characterization Core Facility (GC3F) | Douglas Turnbull, Ariana White, Jeff Bishop, Jason Carriere |
| EPI_ISL_1509639 | UBS de Auriflama | Instituto Adolfo Lutz, Interdiciplinary Procedures Center, Strategic Laboratory | Claudio Tavares Sacchi, Claudia Regina Gonçalves, Erica Valessa Ramos Gomes, Karoline Rodrigues Campos, Caio Vinicius Dias Lopes, Leonardo Jose Tadeu de Araujo |
| EPI_ISL_1509768, EPI_ISL_1509770, EPI_ISL_1509771, EPI_ISL_1509772, EPI_ISL_1509773, EPI_ISL_1509774, EPI_ISL_1509785, EPI_ISL_1509788, EPI_ISL_1509790, EPI_ISL_1509792, EPI_ISL_1509794, EPI_ISL_1509799, EPI_ISL_1509800, EPI_ISL_1509801, EPI_ISL_1509802, EPI_ISL_1509803, EPI_ISL_1509804, EPI_ISL_1509805, EPI_ISL_1509806, EPI_ISL_1509807, EPI_ISL_1509808, EPI_ISL_1509809, EPI_ISL_1509810, EPI_ISL_1509812, EPI_ISL_1509813, EPI_ISL_1509814, EPI_ISL_1509815, EPI_ISL_1509816, EPI_ISL_1509817, EPI_ISL_1509818, EPI_ISL_1509819, EPI_ISL_1509820, EPI_ISL_1509821, EPI_ISL_1509822, EPI_ISL_1509823, EPI_ISL_1509824, EPI_ISL_1509825, EPI_ISL_1509826, EPI_ISL_1509828, EPI_ISL_1509829, EPI_ISL_1509830, EPI_ISL_1509831, EPI_ISL_1509832, EPI_ISL_1509833, EPI_ISL_1509834, EPI_ISL_1509836, EPI_ISL_1509837, EPI_ISL_1509838, EPI_ISL_1509839, EPI_ISL_1509840, EPI_ISL_1509880, EPI_ISL_1509881 | | | |
| see above | National Center of Infectious and Parasitic Diseases | National Center of Infectious and Parasitic Diseases | Alexiev, Ivanov, Korsun, Stoitsova, Philipova, Dimitrova, Grigorova L., Hristova, Donchev, Stoykov, Trifonova, Dobrinov, Grigorova I., Kantardjiev |
| EPI_ISL_1509923 | Dipartimento di Medicina e Chirurgia Universita' degli Studi dell'Insubria Laboratorio di Microbiologia Medica Varese Italia | Laboratorio di Microbiologia e Virologia, Università Vita-Salute San Raffaele, Milano | Fabrizio Maggi, Matteo Castelli, Roberta Antonia Diotti, Michela Sampaolo, Enzo Boeri, Elena Criscuolo, Roberto Ferrarese, Massimo Clementi, Nicasio Mancini and Nicola Clementi |
| EPI_ISL_1509956, EPI_ISL_1509957, EPI_ISL_1509958, EPI_ISL_1509959 | Microbiology Department, University Hospital Donostia | Microbiology Department, University Hospital Donostia | Marimon JM, Montes M, Piñeiro L, Sorarrain A, Gomez M, Cilla G. |
| EPI_ISL_1510605 | Università di Parma, Laboratorio di Igiene e Sanità Pubblica | Istituto Zooprofilattico Sperimentale della Lombardia e dell'Emilia Romagna (IZSLER), Risk Analysis and Genomic Epidemiology Unit | Maria Eugenia Colucci, Licia Veronesi, Paola Affanni, Marina Morganti, Ilaria Menozzi, Erika Scaltriti, Stefano Pongolini |
| EPI_ISL_1510611, EPI_ISL_1510617, EPI_ISL_1510618, EPI_ISL_1510619 | Molecular diagnostic laboratory of Federal Budget Institution of Science "Central Research Institute of Epidemiology" of The Federal Service on Customers' Rights Protection and Human Well-being Surveillance | Group of Genomics and Postgenomic Technologies of Central Research Institute of Epidemiology | Samoilov AE, Kaptelova VV, Korneenko EV, Valdokhina AV, Bulanenko VP, Saenko SS, Golubeva AG, Zotova MI, Berlina YY, Solovyeva ED, Cherkashina AS, Shipulina OY, Speranskaya AS, Tivanova EV, Kondrasheva LY, Akimkin VG |
| EPI_ISL_1510632 | Azienda Sanitaria Locale di Piacenza - Presidio Ospedaliero - Laboratorio di Microbiologia | Istituto Zooprofilattico Sperimentale della Lombardia e dell'Emilia Romagna (IZSLER), Risk Analysis and Genomic Epidemiology Unit | Lo Cascio Giuliana, Schiavo Roberta, Peroni Irene, Marina Morganti, Ilaria Menozzi, Erika Scaltriti, Stefano Pongolini |
| EPI_ISL_1516763 | LESP Hidalgo | Instituto de Diagnostico y Referencia Epidemiologicos (INDRE) | Claudia Wong-Arambula, Abril Rodriguez-Maldonado, Vanessa Rivero-Arredondo, Ariadna Medina-Benitez, Joaquin Quiroz-Mercado, Sergio Rangel-Guerrero, Natividad Cruz-Ortiz, Tatiana Nunez-Garcia, Gisela Barrera-Badillo, Lucia Hernandez-Rivas, Irma Lopez-Martinez, Ernesto Ramirez-Gonzalez. |
| EPI_ISL_1516764 | LESP Puebla | Instituto de Diagnostico y Referencia Epidemiologicos (INDRE) | Claudia Wong-Arambula, Abril Rodriguez-Maldonado, Vanessa Rivero-Arredondo, Ariadna Medina-Benitez, Joaquin Quiroz-Mercado, Sergio Rangel-Guerrero, Natividad Cruz-Ortiz, Tatiana Nunez-Garcia, Gisela Barrera-Badillo, Lucia Hernandez-Rivas, Irma Lopez-Martinez, Ernesto Ramirez-Gonzalez. |
| EPI_ISL_1516767, EPI_ISL_1516768 | LESP Chihuahua | Instituto de Diagnostico y Referencia Epidemiologicos | Claudia Wong-Arambula, Abril Rodriguez-Maldonado, Vanessa Rivero-Arredondo, Ariadna Medina-Benitez, Joaquin Quiroz-Mercado, Sergio |

| | | | |
|---|---|---|---|
| | | (INDRE) | Rangel-Guerrero, Natividad Cruz-Ortiz, Tatiana Nunez-Garcia, Gisela Barrera-Badillo, Lucia Hernandez-Rivas, Irma Lopez-Martinez, Ernesto Ramirez-Gonzalez. |
| EPI_ISL_1516769, EPI_ISL_1516770, EPI_ISL_1516773 | LESP Nuevo Leon | Instituto de Diagnostico y Referencia Epidemiologicos (INDRE) | Claudia Wong-Arambula, Abril Rodriguez-Maldonado, Vanessa Rivero-Arredondo, Ariadna Medina-Benitez, Joaquin Quiroz-Mercado, Sergio Rangel-Guerrero, Natividad Cruz-Ortiz, Tatiana Nunez-Garcia, Gisela Barrera-Badillo, Lucia Hernandez-Rivas, Irma Lopez-Martinez, Ernesto Ramirez-Gonzalez. |
| EPI_ISL_1516776 | LESP Tamaulipas | Instituto de Diagnostico y Referencia Epidemiologicos (INDRE) | Claudia Wong-Arambula, Abril Rodriguez-Maldonado, Vanessa Rivero-Arredondo, Ariadna Medina-Benitez, Joaquin Quiroz-Mercado, Sergio Rangel-Guerrero, Natividad Cruz-Ortiz, Tatiana Nunez-Garcia, Gisela Barrera-Badillo, Lucia Hernandez-Rivas, Irma Lopez-Martinez, Ernesto Ramirez-Gonzalez. |
| EPI_ISL_1516787 | LESP Ciudad de Mexico | Instituto de Diagnostico y Referencia Epidemiologicos (INDRE) | Claudia Wong-Arambula, Abril Rodriguez-Maldonado, Vanessa Rivero-Arredondo, Ariadna Medina-Benitez, Joaquin Quiroz-Mercado, Sergio Rangel-Guerrero, Natividad Cruz-Ortiz, Tatiana Nunez-Garcia, Gisela Barrera-Badillo, Lucia Hernandez-Rivas, Irma Lopez-Martinez, Ernesto Ramirez-Gonzalez. |
| EPI_ISL_1516789, EPI_ISL_1516790 | LESP Tamaulipas | Instituto de Diagnostico y Referencia Epidemiologicos (INDRE) | Claudia Wong-Arambula, Abril Rodriguez-Maldonado, Vanessa Rivero-Arredondo, Ariadna Medina-Benitez, Joaquin Quiroz-Mercado, Sergio Rangel-Guerrero, Natividad Cruz-Ortiz, Tatiana Nunez-Garcia, Gisela Barrera-Badillo, Lucia Hernandez-Rivas, Irma Lopez-Martinez, Ernesto Ramirez-Gonzalez. |
| EPI_ISL_1516791, EPI_ISL_1516792, EPI_ISL_1516793 | LESP Coahuila | Instituto de Diagnostico y Referencia Epidemiologicos (INDRE) | Claudia Wong-Arambula, Abril Rodriguez-Maldonado, Vanessa Rivero-Arredondo, Ariadna Medina-Benitez, Joaquin Quiroz-Mercado, Sergio Rangel-Guerrero, Natividad Cruz-Ortiz, Tatiana Nunez-Garcia, Gisela Barrera-Badillo, Lucia Hernandez-Rivas, Irma Lopez-Martinez, Ernesto Ramirez-Gonzalez. |
| EPI_ISL_1516794 | LESP Guanajuato | Instituto de Diagnostico y Referencia Epidemiologicos (INDRE) | Claudia Wong-Arambula, Abril Rodriguez-Maldonado, Vanessa Rivero-Arredondo, Ariadna Medina-Benitez, Joaquin Quiroz-Mercado, Sergio Rangel-Guerrero, Natividad Cruz-Ortiz, Tatiana Nunez-Garcia, Gisela Barrera-Badillo, Lucia Hernandez-Rivas, Irma Lopez-Martinez, Ernesto Ramirez-Gonzalez. |
| EPI_ISL_1516795 | LESP Jalisco | Instituto de Diagnostico y Referencia Epidemiologicos (INDRE) | Claudia Wong-Arambula, Abril Rodriguez-Maldonado, Vanessa Rivero-Arredondo, Ariadna Medina-Benitez, Joaquin Quiroz-Mercado, Sergio Rangel-Guerrero, Natividad Cruz-Ortiz, Tatiana Nunez-Garcia, Gisela Barrera-Badillo, Lucia Hernandez-Rivas, Irma Lopez-Martinez, Ernesto Ramirez-Gonzalez. |
| EPI_ISL_1516796 | LESP Guanajuato | Instituto de Diagnostico y Referencia Epidemiologicos (INDRE) | Claudia Wong-Arambula, Abril Rodriguez-Maldonado, Vanessa Rivero-Arredondo, Ariadna Medina-Benitez, Joaquin Quiroz-Mercado, Sergio Rangel-Guerrero, Natividad Cruz-Ortiz, Tatiana Nunez-Garcia, Gisela Barrera-Badillo, Lucia Hernandez-Rivas, Irma Lopez-Martinez, Ernesto Ramirez-Gonzalez. |
| EPI_ISL_1516797, EPI_ISL_1516799, EPI_ISL_1516801, EPI_ISL_1516802, EPI_ISL_1516803, EPI_ISL_1516804, EPI_ISL_1516805, EPI_ISL_1516806, EPI_ISL_1516807 | LESP Nuevo Leon | Instituto de Diagnostico y Referencia Epidemiologicos (INDRE) | Claudia Wong-Arambula, Abril Rodriguez-Maldonado, Vanessa Rivero-Arredondo, Ariadna Medina-Benitez, Joaquin Quiroz-Mercado, Sergio Rangel-Guerrero, Natividad Cruz-Ortiz, Tatiana Nunez-Garcia, Gisela Barrera-Badillo, Lucia Hernandez-Rivas, Irma Lopez-Martinez, Ernesto Ramirez-Gonzalez. |
| EPI_ISL_1516808, EPI_ISL_1516810, EPI_ISL_1516811, EPI_ISL_1516812, EPI_ISL_1516813, EPI_ISL_1516815 | LESP Morelos | Instituto de Diagnostico y Referencia Epidemiologicos (INDRE) | Claudia Wong-Arambula, Abril Rodriguez-Maldonado, Vanessa Rivero-Arredondo, Ariadna Medina-Benitez, Joaquin Quiroz-Mercado, Sergio Rangel-Guerrero, Natividad Cruz-Ortiz, Tatiana Nunez-Garcia, Gisela Barrera-Badillo, Lucia Hernandez-Rivas, Irma Lopez-Martinez, Ernesto Ramirez-Gonzalez. |
| EPI_ISL_1516818, EPI_ISL_1516819, EPI_ISL_1516820, EPI_ISL_1516821, EPI_ISL_1516823 | LESP Chihuahua | Instituto de Diagnostico y Referencia Epidemiologicos (INDRE) | Claudia Wong-Arambula, Abril Rodriguez-Maldonado, Vanessa Rivero-Arredondo, Ariadna Medina-Benitez, Joaquin Quiroz-Mercado, Sergio Rangel-Guerrero, Natividad Cruz-Ortiz, Tatiana Nunez-Garcia, Gisela Barrera-Badillo, Lucia Hernandez-Rivas, Irma Lopez-Martinez, Ernesto Ramirez-Gonzalez. |
| EPI_ISL_1516824, EPI_ISL_1516825 | LESP Queretaro | Instituto de Diagnostico y Referencia Epidemiologicos (INDRE) | Claudia Wong-Arambula, Abril Rodriguez-Maldonado, Vanessa Rivero-Arredondo, Ariadna Medina-Benitez, Joaquin Quiroz-Mercado, Sergio Rangel-Guerrero, Natividad Cruz-Ortiz, Tatiana Nunez-Garcia, Gisela Barrera-Badillo, Lucia Hernandez-Rivas, Irma Lopez-Martinez, Ernesto Ramirez-Gonzalez. |
| EPI_ISL_1516891, EPI_ISL_1516892, EPI_ISL_1516893, EPI_ISL_1516894, EPI_ISL_1516895, EPI_ISL_1516896, EPI_ISL_1516897 | Hospital | National Reference Center for Viruses of Respiratory Infections, Institut Pasteur, Paris | Marion Barbet, Sylvie Behillil, Méline Bizard, Angela Brisebarre, Camille Capel,Frédéric Lemoine,Corinne Maufrais,Christophe Malabat,Pierre Lechat, Louise Lefrançois, Etienne Simon-Lorière, Vincent Enouf, Maud Vanpeene, Sylvie van der Werf,FrançOise Jouble |
| EPI_ISL_1516907 | Hospital | National Reference Center for Viruses of Respiratory Infections, Institut Pasteur, Paris | Marion Barbet, Sylvie Behillil, Méline Bizard, Angela Brisebarre, Camille Capel,Frédéric Lemoine,Corinne Maufrais,Christophe Malabat,Gael Millot, Louise Lefrançois, Etienne Simon-Lorière, Vincent Enouf, Maud Vanpeene, Sylvie van der Werf,Dominique Descamps |
| EPI_ISL_1516908 | Hospital | National Reference Center for Viruses of Respiratory Infections, Institut Pasteur, Paris | Marion Barbet, Sylvie Behillil, Méline Bizard, Angela Brisebarre, Camille Capel,Frédéric Lemoine,Corinne Maufrais,Christophe Malabat,Pierre Lechat, Louise Lefrançois, Etienne Simon-Lorière, Vincent Enouf, Maud Vanpeene, Sylvie van der Werf,Dominique Descamps |
| EPI_ISL_1516909, EPI_ISL_1516910 | Hospital | National Reference Center for Viruses of Respiratory Infections, Institut Pasteur, Paris | Marion Barbet, Sylvie Behillil, Méline Bizard, Angela Brisebarre, Camille Capel,Frédéric Lemoine,Corinne Maufrais,Christophe Malabat, Louise Lefrançois, Etienne Simon-Lorière, Vincent Enouf, Maud Vanpeene, Sylvie van der Werf,Dominique Descamps |
| EPI_ISL_1516911, EPI_ISL_1516912 | Hospital | National Reference Center for Viruses of Respiratory Infections, Institut Pasteur, Paris | Marion Barbet, Sylvie Behillil, Méline Bizard, Angela Brisebarre, Camille Capel,Frédéric Lemoine,Corinne Maufrais,Christophe Malabat,Pierre Lechat, Louise Lefrançois, Etienne Simon-Lorière, Vincent Enouf, Maud Vanpeene, Sylvie van der Werf,Dominique Descamps |
| EPI_ISL_1516913, EPI_ISL_1516914 | Hospital | National Reference Center for Viruses of Respiratory Infections, Institut Pasteur, Paris | Marion Barbet, Sylvie Behillil, Méline Bizard, Angela Brisebarre, Camille Capel,Frédéric Lemoine,Corinne Maufrais,Christophe Malabat, Louise Lefrançois, Etienne Simon-Lorière, Vincent Enouf, Maud Vanpeene, Sylvie van der Werf,Dominique Descamps |
| EPI_ISL_1516915 | Hospital | National Reference Center for Viruses of Respiratory Infections, Institut Pasteur, Paris | Marion Barbet, Sylvie Behillil, Méline Bizard, Angela Brisebarre, Camille Capel,Frédéric Lemoine,Corinne Maufrais,Christophe Malabat,Pierre Lechat, Louise Lefrançois, Etienne Simon-Lorière, Vincent Enouf, Maud Vanpeene, Sylvie van der Werf,Dominique Descamps |
| EPI_ISL_1516916 | Hospital | National Reference Center for Viruses of Respiratory Infections, Institut Pasteur, Paris | Marion Barbet, Sylvie Behillil, Méline Bizard, Angela Brisebarre, Camille Capel,Frédéric Lemoine,Corinne Maufrais,Christophe Malabat, Louise Lefrançois, Etienne Simon-Lorière, Vincent Enouf, Maud Vanpeene, Sylvie van der Werf,Dominique Descamps |
| EPI_ISL_1516918, EPI_ISL_1516919 | Labo Analyses Med | National Reference Center for Viruses of Respiratory Infections, Institut Pasteur, Paris | Marion Barbet, Sylvie Behillil, Méline Bizard, Angela Brisebarre, Camille Capel,Frédéric Lemoine,Corinne Maufrais,Christophe Malabat, Louise Lefrançois, Etienne Simon-Lorière, Vincent Enouf, Maud Vanpeene, Sylvie van der Werf |
| EPI_ISL_1516926, EPI_ISL_1516927, EPI_ISL_1516928, EPI_ISL_1516929, EPI_ISL_1516930, EPI_ISL_1516931, EPI_ISL_1516932, EPI_ISL_1516933, EPI_ISL_1516934, EPI_ISL_1516936, EPI_ISL_1516937, EPI_ISL_1516938, EPI_ISL_1516939 | | | |
| see above | Hospital | National Reference Center for Viruses of Respiratory Infections, Institut Pasteur, Paris | Marion Barbet, Sylvie Behillil, Méline Bizard, Angela Brisebarre, Camille Capel,Frédéric Lemoine,Corinne Maufrais,Christophe Malabat, Louise Lefrançois, Etienne Simon-Lorière, Vincent Enouf, Maud Vanpeene, Sylvie van der Werf,Odile Morvan |
| EPI_ISL_1516941 | Hospital | National Reference Center for Viruses of Respiratory Infections, Institut Pasteur, Paris | Marion Barbet, Sylvie Behillil, Méline Bizard, Angela Brisebarre, Camille Capel,Frédéric Lemoine,Corinne Maufrais,Christophe Malabat,Pierre Lechat, Louise Lefrançois, Etienne Simon-Lorière, Vincent Enouf, Maud Vanpeene, Sylvie van der Werf,Odile Morvan |
| EPI_ISL_1516942, EPI_ISL_1516943 | Labo Analyses Med | National Reference Center for Viruses of Respiratory Infections, Institut Pasteur, Paris | Marion Barbet, Sylvie Behillil, Méline Bizard, Angela Brisebarre, Camille Capel,Frédéric Lemoine,Corinne Maufrais,Christophe Malabat, Louise Lefrançois, Etienne Simon-Lorière, Vincent Enouf, Maud Vanpeene, Sylvie van der Werf,Fabienne Huet |
| EPI_ISL_1516944 | Labo Analyses Med | National Reference Center for Viruses of Respiratory Infections, Institut Pasteur, Paris | Marion Barbet, Sylvie Behillil, Méline Bizard, Angela Brisebarre, Camille Capel,Frédéric Lemoine,Corinne Maufrais,Christophe Malabat,Pierre Lechat, Louise Lefrançois, Etienne Simon-Lorière, Vincent Enouf, Maud Vanpeene, Sylvie van der Werf,Fabienne Huet |
| EPI_ISL_1516945, EPI_ISL_1516946 | Labo Analyses Med | National Reference Center for Viruses of Respiratory Infections, Institut Pasteur, Paris | Marion Barbet, Sylvie Behillil, Méline Bizard, Angela Brisebarre, Camille Capel,Frédéric Lemoine,Corinne Maufrais,Christophe Malabat, Louise Lefrançois, Etienne Simon-Lorière, Vincent Enouf, Maud Vanpeene, Sylvie van der Werf,Fabienne Huet |
| EPI_ISL_1516947 | Labo Analyses Med | National Reference Center for Viruses of Respiratory Infections, Institut Pasteur, Paris | Marion Barbet, Sylvie Behillil, Méline Bizard, Angela Brisebarre, Camille Capel,Frédéric Lemoine,Corinne Maufrais,Christophe Malabat,Pierre Lechat, Louise Lefrançois, Etienne Simon-Lorière, Vincent Enouf, Maud Vanpeene, Sylvie van der Werf,Fabienne Huet |

| | | | |
|---|---|---|---|
| EPI_ISL_1516948, EPI_ISL_1516949, EPI_ISL_1516950, EPI_ISL_1516951, EPI_ISL_1516952, EPI_ISL_1516953, EPI_ISL_1516954, EPI_ISL_1516955, EPI_ISL_1516956 | Labo Analyses Med | National Reference Center for Viruses of Respiratory Infections, Institut Pasteur, Paris | Marion Barbet, Sylvie Behillil, Méline Bizard, Angela Brisebarre, Camille Capel,Frédéric Lemoine,Corinne Maufrais,Christophe Malabat, Louise Lefrançois, Etienne Simon-Lorière, Vincent Enouf, Maud Vanpeene, Sylvie van der Werf,Fabienne Huet |
| EPI_ISL_1516957 | Labo Analyses Med | National Reference Center for Viruses of Respiratory Infections, Institut Pasteur, Paris | Marion Barbet, Sylvie Behillil, Méline Bizard, Angela Brisebarre, Camille Capel,Frédéric Lemoine,Corinne Maufrais,Christophe Malabat,Gael Millot, Louise Lefrançois, Etienne Simon-Lorière, Vincent Enouf, Maud Vanpeene, Sylvie van der Werf,Fabienne Huet |
| EPI_ISL_1516983, EPI_ISL_1516984, EPI_ISL_1516986, EPI_ISL_1516987, EPI_ISL_1516988, EPI_ISL_1516990, EPI_ISL_1516991, EPI_ISL_1516993, EPI_ISL_1516994, EPI_ISL_1516996, EPI_ISL_1516997, EPI_ISL_1516999, EPI_ISL_1517000, EPI_ISL_1517001 | | | |
| see above | Labo Analyses Med | National Reference Center for Viruses of Respiratory Infections, Institut Pasteur, Paris | Marion Barbet, Sylvie Behillil, Méline Bizard, Angela Brisebarre, Camille Capel,Frédéric Lemoine,Corinne Maufrais,Christophe Malabat, Louise Lefrançois, Etienne Simon-Lorière, Vincent Enouf, Maud Vanpeene, Sylvie van der Werf,Jean-FrançOis Comes |
| EPI_ISL_1517009, EPI_ISL_1517018, EPI_ISL_1517019 | Hospital | National Reference Center for Viruses of Respiratory Infections, Institut Pasteur, Paris | Marion Barbet, Sylvie Behillil, Méline Bizard, Angela Brisebarre, Camille Capel,Frédéric Lemoine,Corinne Maufrais,Christophe Malabat, Louise Lefrançois, Etienne Simon-Lorière, Vincent Enouf, Maud Vanpeene, Sylvie van der Werf,Adeline Baron |
| EPI_ISL_1517023, EPI_ISL_1517025, EPI_ISL_1517026 | Hospital | National Reference Center for Viruses of Respiratory Infections, Institut Pasteur, Paris | Marion Barbet, Sylvie Behillil, Méline Bizard, Angela Brisebarre, Camille Capel,Frédéric Lemoine,Corinne Maufrais,Christophe Malabat, Louise Lefrançois, Etienne Simon-Lorière, Vincent Enouf, Maud Vanpeene, Sylvie van der Werf,JéRome Guinard |
| EPI_ISL_1517054, EPI_ISL_1517055, EPI_ISL_1517057, EPI_ISL_1517058, EPI_ISL_1517059, EPI_ISL_1517061, EPI_ISL_1517062, EPI_ISL_1517064, EPI_ISL_1517065, EPI_ISL_1517067, EPI_ISL_1517068 | | | |
| see above | Hospital | National Reference Center for Viruses of Respiratory Infections, Institut Pasteur, Paris | Marion Barbet, Sylvie Behillil, Méline Bizard, Angela Brisebarre, Camille Capel,Frédéric Lemoine,Corinne Maufrais,Christophe Malabat, Louise Lefrançois, Etienne Simon-Lorière, Vincent Enouf, Maud Vanpeene, Sylvie van der Werf,LéA Pilorge |
| EPI_ISL_1517071, EPI_ISL_1517073, EPI_ISL_1517074, EPI_ISL_1517076 | Labo Analyses Med | National Reference Center for Viruses of Respiratory Infections, Institut Pasteur, Paris | Marion Barbet, Sylvie Behillil, Méline Bizard, Angela Brisebarre, Camille Capel,Frédéric Lemoine,Corinne Maufrais,Christophe Malabat, Louise Lefrançois, Etienne Simon-Lorière, Vincent Enouf, Maud Vanpeene, Sylvie van der Werf,Vicky Le |
| EPI_ISL_1517108, EPI_ISL_1517109, EPI_ISL_1517110, EPI_ISL_1517115 | Labo Analyses Med | National Reference Center for Viruses of Respiratory Infections, Institut Pasteur, Paris | Marion Barbet, Sylvie Behillil, Méline Bizard, Angela Brisebarre, Camille Capel,Frédéric Lemoine,Corinne Maufrais,Christophe Malabat, Louise Lefrançois, Etienne Simon-Lorière, Vincent Enouf, Maud Vanpeene, Sylvie van der Werf,Jean-FrançOis Comes |
| EPI_ISL_1517190, EPI_ISL_1517385, EPI_ISL_1517386 | Institut de Virologie du CHU de Strasbourg | Institut de Virologie du CHU de Strasbourg | Fafi-Kremer Samira, Gallais Floriane, Gantner Pierre, Laugel Elodie, Solis Morgane, Velay Aurélie, Wendling Marie-Josée |
| EPI_ISL_1517429 | Inciensa, Instituto Costarricense de Investigación y Enseñanza en Nutrición y Salud | Inciensa, Instituto Costarricense de Investigación y Enseñanza en Nutrición y Salud | Cristian Peréz-Corrales, Barboza-Arguedas E & Blanco-Delgado C |
| EPI_ISL_1517430 | Inciensa, Instituto Costarricense de Investigación y Enseñanza en Nutrición y Salud | Inciensa, Instituto Costarricense de Investigación y Enseñanza en Nutrición y Salud | Cristian Peréz-Corrales, Barboza-Arguedas E & Kellerman-Montero K |
| EPI_ISL_1517431 | Inciensa, Instituto Costarricense de Investigación y Enseñanza en Nutrición y Salud | Inciensa, Instituto Costarricense de Investigación y Enseñanza en Nutrición y Salud | Cristian Peréz-Corrales, Barboza-Arguedas E & Blanco-Delgado C |
| EPI_ISL_1517435, EPI_ISL_1517436, EPI_ISL_1517437, EPI_ISL_1517438, EPI_ISL_1517439, EPI_ISL_1517440, EPI_ISL_1517441, EPI_ISL_1517442, EPI_ISL_1517443, EPI_ISL_1517444, EPI_ISL_1517445, EPI_ISL_1517446, EPI_ISL_1517447, EPI_ISL_1517448, EPI_ISL_1517449, EPI_ISL_1517450, EPI_ISL_1517451, EPI_ISL_1517452, EPI_ISL_1517453, EPI_ISL_1517454, EPI_ISL_1517455, EPI_ISL_1517456 | | | |
| see above | Labo Analyses Med | National Reference Center for Viruses of Respiratory Infections, Institut Pasteur, Paris | Marion Barbet, Sylvie Behillil, Méline Bizard, Angela Brisebarre, Camille Capel,Frédéric Lemoine,Corinne Maufrais,Christophe Malabat, Louise Lefrançois, Etienne Simon-Lorière, Vincent Enouf, Maud Vanpeene, Sylvie van der Werf,Karine Michez |
| EPI_ISL_1517459, EPI_ISL_1517460, EPI_ISL_1517461, EPI_ISL_1517462 | Labo Analyses Med | National Reference Center for Viruses of Respiratory Infections, Institut Pasteur, Paris | Marion Barbet, Sylvie Behillil, Méline Bizard, Angela Brisebarre, Camille Capel,Frédéric Lemoine,Corinne Maufrais,Christophe Malabat, Louise Lefrançois, Etienne Simon-Lorière, Vincent Enouf, Maud Vanpeene, Sylvie van der Werf,Jean-FrançOis Comes |
| EPI_ISL_1517463 | Labo Analyses Med | National Reference Center for Viruses of Respiratory Infections, Institut Pasteur, Paris | Marion Barbet, Sylvie Behillil, Méline Bizard, Angela Brisebarre, Camille Capel,Frédéric Lemoine,Corinne Maufrais,Christophe Malabat, Louise Lefrançois, Etienne Simon-Lorière, Vincent Enouf, Maud Vanpeene, Sylvie van der Werf,Hue |
| EPI_ISL_1517468 | Labo Analyses Med | National Reference Center for Viruses of Respiratory Infections, Institut Pasteur, Paris | Marion Barbet, Sylvie Behillil, Méline Bizard, Angela Brisebarre, Camille Capel,Frédéric Lemoine,Corinne Maufrais,Christophe Malabat, Louise Lefrançois, Etienne Simon-Lorière, Vincent Enouf, Maud Vanpeene, Sylvie van der Werf,Alain Delepine |
| EPI_ISL_1517469 | Labo Analyses Med | National Reference Center for Viruses of Respiratory Infections, Institut Pasteur, Paris | Marion Barbet, Sylvie Behillil, Méline Bizard, Angela Brisebarre, Camille Capel,Frédéric Lemoine,Corinne Maufrais,Christophe Malabat, Louise Lefrançois, Etienne Simon-Lorière, Vincent Enouf, Maud Vanpeene, Sylvie van der Werf,GréGoire Potiron |
| EPI_ISL_1517491, EPI_ISL_1517492, EPI_ISL_1517493 | Hospital | National Reference Center for Viruses of Respiratory Infections, Institut Pasteur, Paris | Marion Barbet, Sylvie Behillil, Méline Bizard, Angela Brisebarre, Camille Capel,Frédéric Lemoine,Corinne Maufrais,Christophe Malabat, Louise Lefrançois, Etienne Simon-Lorière, Vincent Enouf, Maud Vanpeene, Sylvie van der Werf,Sylvaine Bastian |
| EPI_ISL_1520108, EPI_ISL_1520109 | LACEN do Estado de Rondonia | Instituto Adolfo Lutz, Interdiciplinary Procedures Center, Strategic Laboratory | Claudio Tavares Sacchi, Claudia Regina Gonçalves, Erica Valessa Ramos Gomes, Karoline Rodrigues Campos, Caio Vinicius Dias Lopes |
| EPI_ISL_1520141, EPI_ISL_1520178, EPI_ISL_1520367 | Institut de Virologie du CHU de Strasbourg | Institut de Virologie du CHU de Strasbourg | Fafi-Kremer Samira, Gallais Floriane, Gantner Pierre, Laugel Elodie, Solis Morgane, Velay Aurélie, Wendling Marie-Josée |
| EPI_ISL_1524121, EPI_ISL_1524122, EPI_ISL_1524142, EPI_ISL_1524146, EPI_ISL_1524147, EPI_ISL_1524148, EPI_ISL_1524149, EPI_ISL_1524150 | Public Health Authority of the Slovak Republic | Laboratory of Genomics and Bioinformatics, Comenius University Science Park | Tatiana Sedláková, Diana Rusáková, Miroslav Böhmer, Anna Giová, Jaroslav Budiš, Tomáš Szemes |
| EPI_ISL_1524331, EPI_ISL_1524334, EPI_ISL_1524335, EPI_ISL_1524336, EPI_ISL_1524337, EPI_ISL_1524339, EPI_ISL_1524341, EPI_ISL_1524342, EPI_ISL_1524348, EPI_ISL_1524350, EPI_ISL_1524352, EPI_ISL_1524354, EPI_ISL_1524355, EPI_ISL_1524357, EPI_ISL_1524358, EPI_ISL_1524360, EPI_ISL_1524363, EPI_ISL_1524365, EPI_ISL_1524366, EPI_ISL_1524367, EPI_ISL_1524368, EPI_ISL_1524370, EPI_ISL_1524371, EPI_ISL_1524375, EPI_ISL_1524377, EPI_ISL_1524378, EPI_ISL_1524379, EPI_ISL_1524380, EPI_ISL_1524381 | | | |
| see above | Biology Department, College of Science, Al Muthanna University and Public Health Laboratory, Al-Muthanna Health Directorate | Department of Virology, Faculty of Medicine, University of Helsinki, Helsinki, Finland | Nihad Al-Rashedi, Hussein Alburkat, Murad Munahi, Alaa Hameed, Ali Jasim, Olli Vapalahti, Tarja Sironen,Teemu Smura |
| EPI_ISL_1524709, EPI_ISL_1524710 | Azienda Ospedaliera Terni | Istituto Zooprofilattico Sperimentale dell'Abruzzo e Molise "G. Caporale" | Palumbo M, Scaccetti A, Lorusso A, Marcacci M, Di Domenico M, Ancora M, Curini V, Di Lollo V, Delli Compagni E, Mangone I, Rinaldi A, Scialabba S, Caporale M, Di Pasquale A, Cammà C, Puglia I, Calistri P, Savini G |
| EPI_ISL_1525261, EPI_ISL_1525262 | Hospital | National Reference Center for Viruses of Respiratory Infections, Institut Pasteur, Paris | Marion Barbet, Sylvie Behillil, Méline Bizard, Angela Brisebarre, Camille Capel,Frédéric Lemoine,Corinne Maufrais,Christophe Malabat,Pierre Lechat, Louise Lefrançois, Etienne Simon-Lorière, Vincent Enouf, Maud Vanpeene, Sylvie van der Werf,Dominique Descamps |
| EPI_ISL_1525263, EPI_ISL_1525265 | Hospital | National Reference Center for Viruses of Respiratory Infections, Institut Pasteur, Paris | Marion Barbet, Sylvie Behillil, Méline Bizard, Angela Brisebarre, Camille Capel,Frédéric Lemoine,Corinne Maufrais,Christophe Malabat,Gael Millot, Louise Lefrançois, Etienne Simon-Lorière, Vincent Enouf, Maud Vanpeene, Sylvie van der Werf,Odile Morvan |
| EPI_ISL_1525266 | Hospital | National Reference Center for Viruses of Respiratory Infections, Institut Pasteur, Paris | Marion Barbet, Sylvie Behillil, Méline Bizard, Angela Brisebarre, Camille Capel,Frédéric Lemoine,Corinne Maufrais,Christophe Malabat,Pierre Lechat, Louise Lefrançois, Etienne Simon-Lorière, Vincent Enouf, Maud Vanpeene, Sylvie van der Werf,Odile Morvan |
| EPI_ISL_1526792 | Labo Analyses Med | National Reference Center for Viruses of Respiratory Infections, Institut Pasteur, Paris | Marion Barbet, Sylvie Behillil, Méline Bizard, Angela Brisebarre, Camille Capel,Frédéric Lemoine,Corinne Maufrais,Christophe Malabat, Louise Lefrançois, Etienne Simon-Lorière, Vincent Enouf, Maud Vanpeene, Sylvie van der Werf,Karine Michez |
| EPI_ISL_1527015 | Hospital Guápiles | Inciensa, Instituto Costarricense de Investigación y Enseñanza en Nutrición y Salud | Pérez-Corrales C, Barboza-Arguedas E & Cerdas-Quesada C |
| EPI_ISL_1527016 | Clínica Clorito Picado | Inciensa, Instituto Costarricense de Investigación y Enseñanza en Nutrición y Salud | Pérez-Corrales C, Barboza-Arguedas E & Salas-Abarca P |
| EPI_ISL_1527017, EPI_ISL_1527018 | unknown | Inciensa, Instituto Costarricense de Investigación y | Pérez-Corrales C, Barboza-Arguedas E & Murillo-Bustos H |

| | | | |
|---|---|---|---|
| | | Enseñanza en Nutrición y Salud | |
| EPI_ISL_1527019 | unknown | Inciensa, Instituto Costarricense de Investigación y Enseñanza en Nutrición y Salud | Pérez-Corrales C, Barboza-Arguedas E & Blanco-Delgado C |
| EPI_ISL_1527020 | unknown | Inciensa, Instituto Costarricense de Investigación y Enseñanza en Nutrición y Salud | Pérez-Corrales C, Barboza-Arguedas E & Ramírez-Pacheco J |
| EPI_ISL_1527021 | Hospital Guápiles | Inciensa, Instituto Costarricense de Investigación y Enseñanza en Nutrición y Salud | Pérez-Corrales C, Barboza-Arguedas E & Cerdas-Quesada C |
| EPI_ISL_1527022 | Clínica Clorito Picado | Inciensa, Instituto Costarricense de Investigación y Enseñanza en Nutrición y Salud | Pérez-Corrales C, Barboza-Arguedas E & Salas-Abarca P |
| EPI_ISL_1527023 | unknown | Inciensa, Instituto Costarricense de Investigación y Enseñanza en Nutrición y Salud | Pérez-Corrales C, Barboza-Arguedas E & Murillo-Bustos H |
| EPI_ISL_1527024 | AS Belén Flores | Inciensa, Instituto Costarricense de Investigación y Enseñanza en Nutrición y Salud | Pérez-Corrales C, Barboza-Arguedas E & Centeno-Miranda M |
| EPI_ISL_1527025 | unknown | Inciensa, Instituto Costarricense de Investigación y Enseñanza en Nutrición y Salud | Pérez-Corrales C & Barboza-Arguedas E |
| EPI_ISL_1527026 | unknown | Inciensa, Instituto Costarricense de Investigación y Enseñanza en Nutrición y Salud | Pérez-Corrales C, Barboza-Arguedas E & Murillo-Bustos H |
| EPI_ISL_1527027 | unknown | Inciensa, Instituto Costarricense de Investigación y Enseñanza en Nutrición y Salud | Pérez-Corrales C, Barboza-Arguedas E & Cerdas-Quesada C |
| EPI_ISL_1527028 | unknown | Inciensa, Instituto Costarricense de Investigación y Enseñanza en Nutrición y Salud | Pérez-Corrales C, Barboza-Arguedas E & Ramírez-Pacheco J |
| EPI_ISL_1528182 | Hospital | National Reference Center for Viruses of Respiratory Infections, Institut Pasteur, Paris | Marion Barbet, Sylvie Behillil, Frédéric Lemoine, Corinne Maufrais, Christophe Malabat, Méline Bizard, Angela Brisebarre, Camille Capel, Louise Lefrançois, Etienne Simon-Lorière, Vincent Enouf, Maud Vanpeene, Sylvie van der Werf,Mj Dufour |
| EPI_ISL_1528195, EPI_ISL_1528196 | Hospital | National Reference Center for Viruses of Respiratory Infections, Institut Pasteur, Paris | Marion Barbet, Sylvie Behillil, Frédéric Lemoine, Corinne Maufrais, Christophe Malabat, Méline Bizard, Angela Brisebarre, Camille Capel, Louise Lefrançois, Etienne Simon-Lorière, Vincent Enouf, Maud Vanpeene, Sylvie van der Werf,Sylvaine Bastian |
| EPI_ISL_1528197 | Hospital | National Reference Center for Viruses of Respiratory Infections, Institut Pasteur, Paris | Marion Barbet, Sylvie Behillil, Frédéric Lemoine, Corinne Maufrais, Christophe Malabat,Adrien Pain, Méline Bizard, Angela Brisebarre, Camille Capel, Louise Lefrançois, Etienne Simon-Lorière, Vincent Enouf, Maud Vanpeene, Sylvie van der Werf,Sylvaine Bastian |
| EPI_ISL_1528236 | Hospital | National Reference Center for Viruses of Respiratory Infections, Institut Pasteur, Paris | Marion Barbet, Sylvie Behillil, Frédéric Lemoine, Corinne Maufrais, Christophe Malabat,Amaury Vaysse, Méline Bizard, Angela Brisebarre, Camille Capel, Louise Lefrançois, Etienne Simon-Lorière, Vincent Enouf, Maud Vanpeene, Sylvie van der Werf,Alexandra Ducancelle |
| EPI_ISL_1529535 | Global Health Research Complex (GHRC) | Texas A&M Institute for Genomic Sciences and Society (TIGSS) | Benjamin W. Neuman, Melissa M. Kahl-Mcdonagh, Kurt A. Zuelke, David Threadgill, Allison Ficht, Sankar P. Chaki, Sierra Guidry, Drew Hillhouse, Wesley Brashear |
| EPI_ISL_1531768, EPI_ISL_1531772 | University of Oregon COVID-19 MAP Laboratory | University of Oregon Genomics and Cell Characterization Core Facility (GC3F) | Douglas Turnbull, Ariana White, Jeff Bishop, Jason Carriere |
| EPI_ISL_1532236 | LESP Queretaro | Instituto de Diagnostico y Referencia Epidemiologicos (INDRE) | Claudia Wong-Arambula, Abril Rodriguez-Maldonado, Vanessa Rivero-Arredondo, Ariadna Medina-Benitez, Joaquin Quiroz-Mercado, Sergio Rangel-Guerrero, Natividad Cruz-Ortiz, Tatiana Nunez-Garcia, Gisela Barrera-Badillo, Lucia Hernandez-Rivas, Irma Lopez-Martinez, Ernesto Ramirez-Gonzalez. |
| EPI_ISL_1532238 | LESP Guanajuato | Instituto de Diagnostico y Referencia Epidemiologicos (INDRE) | Claudia Wong-Arambula, Abril Rodriguez-Maldonado, Vanessa Rivero-Arredondo, Ariadna Medina-Benitez, Joaquin Quiroz-Mercado, Sergio Rangel-Guerrero, Natividad Cruz-Ortiz, Tatiana Nunez-Garcia, Gisela Barrera-Badillo, Lucia Hernandez-Rivas, Irma Lopez-Martinez, Ernesto Ramirez-Gonzalez. |
| EPI_ISL_1532239 | LESP Queretaro | Instituto de Diagnostico y Referencia Epidemiologicos (INDRE) | Claudia Wong-Arambula, Abril Rodriguez-Maldonado, Vanessa Rivero-Arredondo, Ariadna Medina-Benitez, Joaquin Quiroz-Mercado, Sergio Rangel-Guerrero, Natividad Cruz-Ortiz, Tatiana Nunez-Garcia, Gisela Barrera-Badillo, Lucia Hernandez-Rivas, Irma Lopez-Martinez, Ernesto Ramirez-Gonzalez. |
| EPI_ISL_1532241, EPI_ISL_1532242 | LESP Guerrero | Instituto de Diagnostico y Referencia Epidemiologicos (INDRE) | Claudia Wong-Arambula, Abril Rodriguez-Maldonado, Vanessa Rivero-Arredondo, Ariadna Medina-Benitez, Joaquin Quiroz-Mercado, Sergio Rangel-Guerrero, Natividad Cruz-Ortiz, Tatiana Nunez-Garcia, Gisela Barrera-Badillo, Lucia Hernandez-Rivas, Irma Lopez-Martinez, Ernesto Ramirez-Gonzalez. |
| EPI_ISL_1532243, EPI_ISL_1532244, EPI_ISL_1532245 | LESP Baja California Sur | Instituto de Diagnostico y Referencia Epidemiologicos (INDRE) | Claudia Wong-Arambula, Abril Rodriguez-Maldonado, Vanessa Rivero-Arredondo, Ariadna Medina-Benitez, Joaquin Quiroz-Mercado, Sergio Rangel-Guerrero, Natividad Cruz-Ortiz, Tatiana Nunez-Garcia, Gisela Barrera-Badillo, Lucia Hernandez-Rivas, Irma Lopez-Martinez, Ernesto Ramirez-Gonzalez. |
| EPI_ISL_1532246, EPI_ISL_1532247 | LESP Chiapas | Instituto de Diagnostico y Referencia Epidemiologicos (INDRE) | Claudia Wong-Arambula, Abril Rodriguez-Maldonado, Vanessa Rivero-Arredondo, Ariadna Medina-Benitez, Joaquin Quiroz-Mercado, Sergio Rangel-Guerrero, Natividad Cruz-Ortiz, Tatiana Nunez-Garcia, Gisela Barrera-Badillo, Lucia Hernandez-Rivas, Irma Lopez-Martinez, Ernesto Ramirez-Gonzalez. |
| EPI_ISL_1532248 | LESP Yucatan | Instituto de Diagnostico y Referencia Epidemiologicos (INDRE) | Claudia Wong-Arambula, Abril Rodriguez-Maldonado, Vanessa Rivero-Arredondo, Ariadna Medina-Benitez, Joaquin Quiroz-Mercado, Sergio Rangel-Guerrero, Natividad Cruz-Ortiz, Tatiana Nunez-Garcia, Gisela Barrera-Badillo, Lucia Hernandez-Rivas, Irma Lopez-Martinez, Ernesto Ramirez-Gonzalez. |
| EPI_ISL_1532249 | LESP Estado de Mexico | Instituto de Diagnostico y Referencia Epidemiologicos (INDRE) | Claudia Wong-Arambula, Abril Rodriguez-Maldonado, Vanessa Rivero-Arredondo, Ariadna Medina-Benitez, Joaquin Quiroz-Mercado, Sergio Rangel-Guerrero, Natividad Cruz-Ortiz, Tatiana Nunez-Garcia, Gisela Barrera-Badillo, Lucia Hernandez-Rivas, Irma Lopez-Martinez, Ernesto Ramirez-Gonzalez. |
| EPI_ISL_1532250, EPI_ISL_1532252, EPI_ISL_1532253, EPI_ISL_1532254, EPI_ISL_1532255, EPI_ISL_1532256, EPI_ISL_1532257, EPI_ISL_1532258, EPI_ISL_1532259, EPI_ISL_1532260, EPI_ISL_1532261, EPI_ISL_1532262, EPI_ISL_1532263 | | | |
| see above | LESPZacatecas | Instituto de Diagnostico y Referencia Epidemiologicos (INDRE) | Claudia Wong-Arambula, Abril Rodriguez-Maldonado, Vanessa Rivero-Arredondo, Ariadna Medina-Benitez, Joaquin Quiroz-Mercado, Sergio Rangel-Guerrero, Natividad Cruz-Ortiz, Tatiana Nunez-Garcia, Gisela Barrera-Badillo, Lucia Hernandez-Rivas, Irma Lopez-Martinez, Ernesto Ramirez-Gonzalez. |
| EPI_ISL_1532264 | LESP Baja California | Instituto de Diagnostico y Referencia Epidemiologicos (INDRE) | Claudia Wong-Arambula, Abril Rodriguez-Maldonado, Vanessa Rivero-Arredondo, Ariadna Medina-Benitez, Joaquin Quiroz-Mercado, Sergio Rangel-Guerrero, Natividad Cruz-Ortiz, Tatiana Nunez-Garcia, Gisela Barrera-Badillo, Lucia Hernandez-Rivas, Irma Lopez-Martinez, Ernesto Ramirez-Gonzalez. |
| EPI_ISL_1532265 | LESP Ciudad de Mexico | Instituto de Diagnostico y Referencia Epidemiologicos (INDRE) | Claudia Wong-Arambula, Abril Rodriguez-Maldonado, Vanessa Rivero-Arredondo, Ariadna Medina-Benitez, Joaquin Quiroz-Mercado, Sergio Rangel-Guerrero, Natividad Cruz-Ortiz, Tatiana Nunez-Garcia, Gisela Barrera-Badillo, Lucia Hernandez-Rivas, Irma Lopez-Martinez, Ernesto Ramirez-Gonzalez. |
| EPI_ISL_1532266, EPI_ISL_1532269 | LESP Jalisco | Instituto de Diagnostico y Referencia Epidemiologicos (INDRE) | Claudia Wong-Arambula, Abril Rodriguez-Maldonado, Vanessa Rivero-Arredondo, Ariadna Medina-Benitez, Joaquin Quiroz-Mercado, Sergio Rangel-Guerrero, Natividad Cruz-Ortiz, Tatiana Nunez-Garcia, Gisela Barrera-Badillo, Lucia Hernandez-Rivas, Irma Lopez-Martinez, Ernesto Ramirez-Gonzalez. |
| EPI_ISL_1532273, EPI_ISL_1532274, EPI_ISL_1532277 | LESP Queretaro | Instituto de Diagnostico y Referencia Epidemiologicos (INDRE) | Claudia Wong-Arambula, Abril Rodriguez-Maldonado, Vanessa Rivero-Arredondo, Ariadna Medina-Benitez, Joaquin Quiroz-Mercado, Sergio Rangel-Guerrero, Natividad Cruz-Ortiz, Tatiana Nunez-Garcia, Gisela Barrera-Badillo, Lucia Hernandez-Rivas, Irma Lopez-Martinez, Ernesto Ramirez-Gonzalez. |

| | | | |
|---|---|---|---|
| EPI_ISL_1532279 | LESP Ciudad de Mexico | Instituto de Diagnostico y Referencia Epidemiologicos (INDRE) | Claudia Wong-Arambula, Abril Rodriguez-Maldonado, Vanessa Rivero-Arredondo, Ariadna Medina-Benitez, Joaquin Quiroz-Mercado, Sergio Rangel-Guerrero, Natividad Cruz-Ortiz, Tatiana Nunez-Garcia, Gisela Barrera-Badillo, Lucia Hernandez-Rivas, Irma Lopez-Martinez, Ernesto Ramirez-Gonzalez. |
| EPI_ISL_1532281 | LESP Quintana Roo | Instituto de Diagnostico y Referencia Epidemiologicos (INDRE) | Claudia Wong-Arambula, Abril Rodriguez-Maldonado, Vanessa Rivero-Arredondo, Ariadna Medina-Benitez, Joaquin Quiroz-Mercado, Sergio Rangel-Guerrero, Natividad Cruz-Ortiz, Tatiana Nunez-Garcia, Gisela Barrera-Badillo, Lucia Hernandez-Rivas, Irma Lopez-Martinez, Ernesto Ramirez-Gonzalez. |
| EPI_ISL_1532801, EPI_ISL_1532802, EPI_ISL_1532804, EPI_ISL_1532805, EPI_ISL_1532806, EPI_ISL_1532807, EPI_ISL_1532808, EPI_ISL_1532810, EPI_ISL_1532811, EPI_ISL_1532813, EPI_ISL_1532814, EPI_ISL_1532815 | | | |
| see above | Virology Unit, Institut Pasteur du Cambodge | Virology Unit, Institut Pasteur du Cambodge | Sokhoun Yann, Teyputita Ou, Leakhena Pum, Ly Sovann, Kraing Sidonn, Yi Sengdoeurn, Chin Savuth, Chau Darapheak, Veasna Duong, Erik A Karlsson |
| EPI_ISL_1533036, EPI_ISL_1533040, EPI_ISL_1533046 | Diagnostic and Research Center of Infectious Diseases, Medical Faculty, Andalas University | Diagnostic and Research Center of Infectious Diseases, Medical Faculty, Andalas University | Andani Eka Putra, Syafrizayanti, Ikhwan R. Sudji, Linosefa, Dede Rahman Agustian, Ayu Novita Trisnawati, Siskalil Fahma, Nia Ayuni Putri, Sekar Asri Tresnaningtyas, SM Rezvi, Fauzul Azhim, Dessy Arisanty, Gestina Aliska, Desmawati, Nita Afriani ,Syandrez Prima Putra, Juane Plantika Menra, Yolani Syaputri, Mutia Lailani |
| EPI_ISL_1533703 | Pronto Atendimento Sao Jose | Instituto Adolfo Lutz, Interdiciplinary Procedures Center, Strategic Laboratory | Claudio Tavares Sacchi, Claudia Regina Gonçalves, Erica Valessa Ramos Gomes, Karoline Rodrigues Campos, Caio Vinicius Dias Lopes, Leonardo Jose Tadeu de Araujo |
| EPI_ISL_1533704 | Santa Casa de Atibaia Pro Saude | Instituto Adolfo Lutz, Interdiciplinary Procedures Center, Strategic Laboratory | Claudio Tavares Sacchi, Claudia Regina Gonçalves, Erica Valessa Ramos Gomes, Karoline Rodrigues Campos, Caio Vinicius Dias Lopes, Leonardo Jose Tadeu de Araujo |
| EPI_ISL_1533705 | Vigilancia em Saude | Instituto Adolfo Lutz, Interdiciplinary Procedures Center, Strategic Laboratory | Claudio Tavares Sacchi, Claudia Regina Gonçalves, Erica Valessa Ramos Gomes, Karoline Rodrigues Campos, Caio Vinicius Dias Lopes, Leonardo Jose Tadeu de Araujo |
| EPI_ISL_1533706 | PS Mun Santana Lauro Ribas Braga | Instituto Adolfo Lutz, Interdiciplinary Procedures Center, Strategic Laboratory | Claudio Tavares Sacchi, Claudia Regina Gonçalves, Erica Valessa Ramos Gomes, Karoline Rodrigues Campos, Caio Vinicius Dias Lopes, Leonardo Jose Tadeu de Araujo |
| EPI_ISL_1533707 | Hosp Mun Planalto Waldomiro de Paula | Instituto Adolfo Lutz, Interdiciplinary Procedures Center, Strategic Laboratory | Claudio Tavares Sacchi, Claudia Regina Gonçalves, Erica Valessa Ramos Gomes, Karoline Rodrigues Campos, Caio Vinicius Dias Lopes, Leonardo Jose Tadeu de Araujo |
| EPI_ISL_1533709, EPI_ISL_1533710 | Hospital Estadual de Vila Alpina Org Social Seconci Sao Paulo | Instituto Adolfo Lutz, Interdiciplinary Procedures Center, Strategic Laboratory | Claudio Tavares Sacchi, Claudia Regina Gonçalves, Erica Valessa Ramos Gomes, Karoline Rodrigues Campos, Caio Vinicius Dias Lopes, Leonardo Jose Tadeu de Araujo |
| EPI_ISL_1533711 | Hospital Santa Clara | Instituto Adolfo Lutz, Interdiciplinary Procedures Center, Strategic Laboratory | Claudio Tavares Sacchi, Claudia Regina Gonçalves, Erica Valessa Ramos Gomes, Karoline Rodrigues Campos, Caio Vinicius Dias Lopes, Leonardo Jose Tadeu de Araujo |
| EPI_ISL_1533712 | Hospital Universitario da USP Sao Paulo | Instituto Adolfo Lutz, Interdiciplinary Procedures Center, Strategic Laboratory | Claudio Tavares Sacchi, Claudia Regina Gonçalves, Erica Valessa Ramos Gomes, Karoline Rodrigues Campos, Caio Vinicius Dias Lopes, Leonardo Jose Tadeu de Araujo |
| EPI_ISL_1533713 | Unidade de Pronto Atendimento Jd Amanda | Instituto Adolfo Lutz, Interdiciplinary Procedures Center, Strategic Laboratory | Claudio Tavares Sacchi, Claudia Regina Gonçalves, Erica Valessa Ramos Gomes, Karoline Rodrigues Campos, Caio Vinicius Dias Lopes, Leonardo Jose Tadeu de Araujo |
| EPI_ISL_1533714 | Pronto Socorro Dr Osmar Mesquita | Instituto Adolfo Lutz, Interdiciplinary Procedures Center, Strategic Laboratory | Claudio Tavares Sacchi, Claudia Regina Gonçalves, Erica Valessa Ramos Gomes, Karoline Rodrigues Campos, Caio Vinicius Dias Lopes, Leonardo Jose Tadeu de Araujo |
| EPI_ISL_1533715 | Santa Casa de Aracatuba Hospital Sagrado Coracao de Jesus | Instituto Adolfo Lutz, Interdiciplinary Procedures Center, Strategic Laboratory | Claudio Tavares Sacchi, Claudia Regina Gonçalves, Erica Valessa Ramos Gomes, Karoline Rodrigues Campos, Caio Vinicius Dias Lopes, Leonardo Jose Tadeu de Araujo |
| EPI_ISL_1533716 | Vigilancia Em Saude | Instituto Adolfo Lutz, Interdiciplinary Procedures Center, Strategic Laboratory | Claudio Tavares Sacchi, Claudia Regina Gonçalves, Erica Valessa Ramos Gomes, Karoline Rodrigues Campos, Caio Vinicius Dias Lopes, Leonardo Jose Tadeu de Araujo |
| EPI_ISL_1533717 | Unidade de Pronto Atendimento Jd Amanda | Instituto Adolfo Lutz, Interdiciplinary Procedures Center, Strategic Laboratory | Claudio Tavares Sacchi, Claudia Regina Gonçalves, Erica Valessa Ramos Gomes, Karoline Rodrigues Campos, Caio Vinicius Dias Lopes, Leonardo Jose Tadeu de Araujo |
| EPI_ISL_1533718 | Servico de Verificacao de Obitos Svo Guarulhos | Instituto Adolfo Lutz, Interdiciplinary Procedures Center, Strategic Laboratory | Claudio Tavares Sacchi, Claudia Regina Gonçalves, Erica Valessa Ramos Gomes, Karoline Rodrigues Campos, Caio Vinicius Dias Lopes, Leonardo Jose Tadeu de Araujo |
| EPI_ISL_1533719 | Policlinica Munic da Est Turistica de Holambra | Instituto Adolfo Lutz, Interdiciplinary Procedures Center, Strategic Laboratory | Claudio Tavares Sacchi, Claudia Regina Gonçalves, Erica Valessa Ramos Gomes, Karoline Rodrigues Campos, Caio Vinicius Dias Lopes, Leonardo Jose Tadeu de Araujo |
| EPI_ISL_1533720 | Santa Casa de Penapolis | Instituto Adolfo Lutz, Interdiciplinary Procedures Center, Strategic Laboratory | Claudio Tavares Sacchi, Claudia Regina Gonçalves, Erica Valessa Ramos Gomes, Karoline Rodrigues Campos, Caio Vinicius Dias Lopes, Leonardo Jose Tadeu de Araujo |
| EPI_ISL_1533721 | Santa Casa de Aracatuba Hospital Sagrado Coracao De Jesus | Instituto Adolfo Lutz, Interdiciplinary Procedures Center, Strategic Laboratory | Claudio Tavares Sacchi, Claudia Regina Gonçalves, Erica Valessa Ramos Gomes, Karoline Rodrigues Campos, Caio Vinicius Dias Lopes, Leonardo Jose Tadeu de Araujo |
| EPI_ISL_1533722 | Secretaria Municipal de Saude De Piracaia | Instituto Adolfo Lutz, Interdiciplinary Procedures Center, Strategic Laboratory | Claudio Tavares Sacchi, Claudia Regina Gonçalves, Erica Valessa Ramos Gomes, Karoline Rodrigues Campos, Caio Vinicius Dias Lopes, Leonardo Jose Tadeu de Araujo |
| EPI_ISL_1533723 | Hospital Geral de Itaquaquecetuba | Instituto Adolfo Lutz, Interdiciplinary Procedures Center, Strategic Laboratory | Claudio Tavares Sacchi, Claudia Regina Gonçalves, Erica Valessa Ramos Gomes, Karoline Rodrigues Campos, Caio Vinicius Dias Lopes, Leonardo Jose Tadeu de Araujo |
| EPI_ISL_1533724 | Diretoria Municipal de Saude | Instituto Adolfo Lutz, Interdiciplinary Procedures Center, Strategic Laboratory | Claudio Tavares Sacchi, Claudia Regina Gonçalves, Erica Valessa Ramos Gomes, Karoline Rodrigues Campos, Caio Vinicius Dias Lopes, Leonardo Jose Tadeu de Araujo |
| EPI_ISL_1533725 | Hospital Geral de Guarulhos | Instituto Adolfo Lutz, Interdiciplinary Procedures Center, Strategic Laboratory | Claudio Tavares Sacchi, Claudia Regina Gonçalves, Erica Valessa Ramos Gomes, Karoline Rodrigues Campos, Caio Vinicius Dias Lopes, Leonardo Jose Tadeu de Araujo |
| EPI_ISL_1533726 | Hospital Estadual de Campanha Covid 19 Barradas | Instituto Adolfo Lutz, Interdiciplinary Procedures Center, Strategic Laboratory | Claudio Tavares Sacchi, Claudia Regina Gonçalves, Erica Valessa Ramos Gomes, Karoline Rodrigues Campos, Caio Vinicius Dias Lopes, Leonardo Jose Tadeu de Araujo |
| EPI_ISL_1533727 | Hospital Universitario da USP Sao Paulo | Instituto Adolfo Lutz, Interdiciplinary Procedures Center, Strategic Laboratory | Claudio Tavares Sacchi, Claudia Regina Gonçalves, Erica Valessa Ramos Gomes, Karoline Rodrigues Campos, Caio Vinicius Dias Lopes, Leonardo Jose Tadeu de Araujo |
| EPI_ISL_1534002 | Laboratorio Central de Saude Publica do Estado de Santa Catarina (LACEN-SC) | Laboratory of Respiratory Viruses and Measles, Oswaldo Cruz Institute, FIOCRUZ | Paola Resende, Luciana Appolinario, Fernando Motta, Anna Carolina Paixao, Ana Carolina Mendonca, Alice Sampaio Rocha, Renata Serrano Lopes, Darcita Buerger Rovaris, Sandra Bianchini Fernandes, Marilda Siqueira on behalf of the Fiocruz COVID-19 Genomic Surveillance Network |
| EPI_ISL_1534528, EPI_ISL_1534529, EPI_ISL_1534530, EPI_ISL_1534531, EPI_ISL_1534532, EPI_ISL_1534533, EPI_ISL_1534534, EPI_ISL_1534535, EPI_ISL_1534536, EPI_ISL_1534537, EPI_ISL_1534538, EPI_ISL_1534539, EPI_ISL_1534540, EPI_ISL_1534541, EPI_ISL_1534542, EPI_ISL_1534543 | | | |
| see above | Virology Unit, Institut Pasteur du Cambodge | Virology Unit, Institut Pasteur du Cambodge | Sokhoun Yann, Teyputita Ou, Leakhena Pum, Ly Sovann, Kraing Sidonn, Yi Sengdoeurn, Chin Savuth, Chau Darapheak, Veasna Duong, Erik A Karlsson |
| EPI_ISL_1534701, EPI_ISL_1534702 | Fakultas kedokteran Universitas Sumatra Utara | National Institute of Health Research and Development | Hartanti Dian Ikawati, Hana Apsari Pawestri, Subangkit, Kartika Dewi Puspa, Arie Ardiansyah Nugraha, Krisna Pangesti, Nelly Puspandari, Vivi Setiawaty |
| EPI_ISL_1534703 | Fakultas kedokteran Universitas Sumatra Utara | National Institute of Health Research and Development | Kartika Dewi Puspa, Hana Apsari Pawestri, Subangkit, Arie Ardiansyah Nugraha, Hartanti Dian Ikawati, Nelly Puspandari, Krisna Pangesti, Vivi Setiawaty |
| EPI_ISL_1534708 | Labkesda Karawang | National Institute of Health Research and Development | Arie Ardiansyah Nugraha, Hana Apsari Pawestri, Subangkit, Kartika Dewi Puspa, Hartanti Dian Ikawati, Nelly Puspandari, Krisna Pangesti, Vivi Setiawaty |
| EPI_ISL_1538412 | Hospital Universitario Basurto Clinical Microbiology Service | Biocruces Bizkaia | Mikel Joseba Urrutikoetxea-Gutierrez, Mª Carmen Nieto Toboso, Estibaliza Ugalde Zarraga, Mikel Gallego Rodrigo, Ana Belen de la Hoz, Jose Luis Díaz de Tuesta del Arco |
| EPI_ISL_1538413 | Basurto University Hospital Microbiology Lab | Biocruces | Mikel Urrutikoetxea-Gutierrez, Mª Carmen Nieto Toboso, Estibalez Ugalde Zarrraga, Mikel Gallego Rodrigo, Ana de la Hoz, Jose Luis Díaz de Tuesta del Arco |
| EPI_ISL_1538414 | Basurto University Hospital | Biocruces | Mikel Urrutikoetxea-Gutierrez, Mª Carmen Nieto Toboso, Estibaliz Ugalde Zarraga, Mikel Gallego Rodrigo, Ana de la Hoz, José Luís Díaz de Tuesta del |

| | | | |
|---|---|---|---|
| | | | Arco |
| EPI_ISL_1540447 | Hospital | National Reference Center for Viruses of Respiratory Infections, Institut Pasteur, Paris | Marion Barbet, Sylvie Behillil, Frédéric Lemoine, Corinne Maufrais, Christophe Malabat, Méline Bizard, Angela Brisebarre, Camille Capel, Louise Lefrançois, Etienne Simon-Lorière, Vincent Enouf, Maud Vanpeene, Sylvie van der Werf,Sylvaine Bastian |
| EPI_ISL_1543932, EPI_ISL_1544109 | Institut de Virologie du CHU de Strasbourg | Institut de Virologie du CHU de Strasbourg | Fafi-Kremer Samira, Gallais Floriane, Gantner Pierre, Laugel Elodie, Solis Morgane, Velay Aurélie, Wendling Marie-Josée |
| EPI_ISL_1544112 | B.J. Medical College and Civil hospital, Ahmedabad | Gujarat Biotechnology Research Centre | Sanjay Kapadia,Janvi Raval,Zarna Patel,Nitin Savaliya,Dinesh Kumar,Twinkle Soni,Sonal Sharma ,Zuber Saiyed,Ramesh Pandit,Pranay Shah,Kamlesh J Upadhyay,Dipa Kinariwala,Umang Mishra,Nitesh Shah,Chaitanya Joshi,Madhvi Joshi |
| EPI_ISL_1544113 | B.J. Medical College and Civil hospital, Ahmedabad | Gujarat Biotechnology Research Centre | Dipa Kinariwala,Zarna Patel,Nitin Savaliya,Dinesh Kumar,Twinkle Soni,Sonal Sharma ,Zuber Saiyed,Ramesh Pandit,Janvi Raval,Pranay Shah,Kamlesh J Upadhyay,Sanjay Kapadia,Umang Mishra,Nitesh Shah,Chaitanya Joshi,Madhvi Joshi |
| EPI_ISL_1544114 | B.J. Medical College and Civil hospital, Ahmedabad | Gujarat Biotechnology Research Centre | Umang Mishra,Nitin Savaliya,Dinesh Kumar,Twinkle Soni,Sonal Sharma ,Zuber Saiyed,Ramesh Pandit,Janvi Raval,Zarna Patel,Pranay Shah,Kamlesh J Upadhyay,Sanjay Kapadia,Dipa Kinariwala,Nitesh Shah,Chaitanya Joshi,Madhvi Joshi |
| EPI_ISL_1544115 | B.J. Medical College and Civil hospital, Ahmedabad | Gujarat Biotechnology Research Centre | Umang Mishra,Dinesh Kumar,Twinkle Soni,Sonal Sharma ,Zuber Saiyed,Ramesh Pandit,Janvi Raval,Zarna Patel,Nitin Savaliya,Pranay Shah,Kamlesh J Upadhyay,Sanjay Kapadia,Dipa Kinariwala,Nitesh Shah,Chaitanya Joshi,Madhvi Joshi |
| EPI_ISL_1544116 | B.J. Medical College and Civil hospital, Ahmedabad | Gujarat Biotechnology Research Centre | Nitesh Shah,Twinkle Soni,Sonal Sharma ,Zuber Saiyed,Ramesh Pandit,Janvi Raval,Zarna Patel,Nitin Savaliya,Dinesh Kumar,Pranay Shah,Kamlesh J Upadhyay,Sanjay Kapadia,Dipa Kinariwala,Umang Mishra,Chaitanya Joshi,Madhvi Joshi |
| EPI_ISL_1544117 | B.J. Medical College and Civil hospital, Ahmedabad | Gujarat Biotechnology Research Centre | Nitesh Shah,Sonal Sharma ,Zuber Saiyed,Ramesh Pandit,Janvi Raval,Zarna Patel,Nitin Savaliya,Dinesh Kumar,Twinkle Soni,Pranay Shah,Kamlesh J Upadhyay,Sanjay Kapadia,Dipa Kinariwala,Umang Mishra,Chaitanya Joshi,Madhvi Joshi |
| EPI_ISL_1544133 | GMERS, Medical College, Gandhinagar | Gujarat Biotechnology Research Centre | Gaurishankar Shrimali,Sonal Sharma ,Zuber Saiyed,Ramesh Pandit,Janvi Raval,Zarna Patel,Nitin Savaliya,Dinesh Kumar,Twinkle Soni,Umang Mishra,Nitesh Shah,Chaitanya Joshi,Madhvi Joshi |
| EPI_ISL_1544134 | GMERS, Medical College, Gandhinagar | Gujarat Biotechnology Research Centre | Zuber Saiyed,Ramesh Pandit,Janvi Raval,Zarna Patel,Nitin Savaliya,Dinesh Kumar,Twinkle Soni,Sonal Sharma ,Gaurishankar Shrimali,Umang Mishra,Nitesh Shah,Chaitanya Joshi,Madhvi Joshi |
| EPI_ISL_1547802, EPI_ISL_1547803, EPI_ISL_1547804, EPI_ISL_1547805 | CSIR-National Environmental Engineering Research Institute | CSIR-Centre for Cellular and Molecular Biology - INSACOG | Onkar Kulkarni, Lamuk Zaveri, Ara Sreenivas, Sofia Banu, Shreekant Verma, Amareshwar Vodapalli , Viswagithe S L, B Himasri, Sharath Chandra Thota, Karthik Bharadwaj Tallapaka, Krishna Khairnar, Rakesh K Mishra,Divya Tej Sowpati |
| EPI_ISL_1548071 | University of Rome Tor Vergata: Departm Experim Medicine Chair of Virology | University of Rome Tor Vergata: Departm Experim Medicine Chair of Virology | Maria Concetta Bellocchi, Luca Carioti, Maria Botticelli, Lorenzo Piermatteo, Francesca Ceccherini-Silberstein |
| EPI_ISL_1554887 | Sentinelles Idf | National Reference Center for Viruses of Respiratory Infections, Institut Pasteur, Paris | Marion Barbet, Sylvie Behillil, Méline Bizard, Angela Brisebarre, Camille Capel,Frédéric Lemoine,Corinne Maufrais,Christophe Malabat, Damien Mornico, Louise Lefrançois, Etienne Simon-Lorière, Vincent Enouf, Maud Vanpeene, Sylvie van der Werf,Askin Dilek |
| EPI_ISL_1554888, EPI_ISL_1554889, EPI_ISL_1554890, EPI_ISL_1554893, EPI_ISL_1554894, EPI_ISL_1554895, EPI_ISL_1554896, EPI_ISL_1554897 | Hospital | National Reference Center for Viruses of Respiratory Infections, Institut Pasteur, Paris | Marion Barbet, Sylvie Behillil, Méline Bizard, Angela Brisebarre, Camille Capel,Frédéric Lemoine,Corinne Maufrais,Christophe Malabat, Damien Mornico, Louise Lefrançois, Etienne Simon-Lorière, Vincent Enouf, Maud Vanpeene, Sylvie van der Werf,Leruez-Ville Marianne |
| EPI_ISL_1554898 | Hospital | National Reference Center for Viruses of Respiratory Infections, Institut Pasteur, Paris | Marion Barbet, Sylvie Behillil, Méline Bizard, Angela Brisebarre, Camille Capel,Frédéric Lemoine,Corinne Maufrais,Christophe Malabat, Damien Mornico, Louise Lefrançois, Etienne Simon-Lorière, Vincent Enouf, Maud Vanpeene, Sylvie van der Werf,Farfour Eric |
| EPI_ISL_1554900, EPI_ISL_1554901 | Hospital | National Reference Center for Viruses of Respiratory Infections, Institut Pasteur, Paris | Marion Barbet, Sylvie Behillil, Méline Bizard, Angela Brisebarre, Camille Capel,Frédéric Lemoine,Corinne Maufrais,Christophe Malabat, Damien Mornico, Louise Lefrançois, Etienne Simon-Lorière, Vincent Enouf, Maud Vanpeene, Sylvie van der Werf,Dorval I. |
| EPI_ISL_1554902, EPI_ISL_1554903 | Hospital | National Reference Center for Viruses of Respiratory Infections, Institut Pasteur, Paris | Marion Barbet, Sylvie Behillil, Méline Bizard, Angela Brisebarre, Camille Capel,Frédéric Lemoine,Corinne Maufrais,Christophe Malabat, Damien Mornico, Louise Lefrançois, Etienne Simon-Lorière, Vincent Enouf, Maud Vanpeene, Sylvie van der Werf,Hochart A |
| EPI_ISL_1554904 | Labo Analyses Med | National Reference Center for Viruses of Respiratory Infections, Institut Pasteur, Paris | Marion Barbet, Sylvie Behillil, Méline Bizard, Angela Brisebarre, Camille Capel,Frédéric Lemoine,Corinne Maufrais,Christophe Malabat, Damien Mornico, Louise Lefrançois, Etienne Simon-Lorière, Vincent Enouf, Maud Vanpeene, Sylvie van der Werf,Selas Medilys |
| EPI_ISL_1554905, EPI_ISL_1554906, EPI_ISL_1554907, EPI_ISL_1554908, EPI_ISL_1554909, EPI_ISL_1554910, EPI_ISL_1554911, EPI_ISL_1554912, EPI_ISL_1554913, EPI_ISL_1554914, EPI_ISL_1554915, EPI_ISL_1554916 | | | |
| see above | Hospital | National Reference Center for Viruses of Respiratory Infections, Institut Pasteur, Paris | Marion Barbet, Sylvie Behillil, Méline Bizard, Angela Brisebarre, Camille Capel,Frédéric Lemoine,Corinne Maufrais,Christophe Malabat, Damien Mornico, Louise Lefrançois, Etienne Simon-Lorière, Vincent Enouf, Maud Vanpeene, Sylvie van der Werf,Serazin ValéRie |
| EPI_ISL_1554917, EPI_ISL_1554919, EPI_ISL_1554920, EPI_ISL_1554922, EPI_ISL_1554923 | Hospital | National Reference Center for Viruses of Respiratory Infections, Institut Pasteur, Paris | Marion Barbet, Sylvie Behillil, Méline Bizard, Angela Brisebarre, Camille Capel,Frédéric Lemoine,Corinne Maufrais,Christophe Malabat, Damien Mornico, Louise Lefrançois, Etienne Simon-Lorière, Vincent Enouf, Maud Vanpeene, Sylvie van der Werf,Ducancelle Alexandra |
| EPI_ISL_1554924, EPI_ISL_1554925, EPI_ISL_1554926, EPI_ISL_1554927, EPI_ISL_1554928, EPI_ISL_1554929, EPI_ISL_1554930, EPI_ISL_1554931, EPI_ISL_1554932, EPI_ISL_1554933, EPI_ISL_1554936, EPI_ISL_1554938, EPI_ISL_1554939, EPI_ISL_1554940, EPI_ISL_1554941, EPI_ISL_1554943, EPI_ISL_1554944, EPI_ISL_1554945, EPI_ISL_1554946, EPI_ISL_1554947, EPI_ISL_1554948, EPI_ISL_1554949, EPI_ISL_1554952, EPI_ISL_1554955, EPI_ISL_1554958 | | | |
| see above | Hospital | National Reference Center for Viruses of Respiratory Infections, Institut Pasteur, Paris | Marion Barbet, Sylvie Behillil, Méline Bizard, Angela Brisebarre, Camille Capel,Frédéric Lemoine,Corinne Maufrais,Christophe Malabat, Damien Mornico, Louise Lefrançois, Etienne Simon-Lorière, Vincent Enouf, Maud Vanpeene, Sylvie van der Werf,Guinard JéRome |
| EPI_ISL_1558595, EPI_ISL_1558596, EPI_ISL_1558597, EPI_ISL_1558599, EPI_ISL_1558600, EPI_ISL_1558601, EPI_ISL_1558603, EPI_ISL_1558604, EPI_ISL_1558605, EPI_ISL_1558606, EPI_ISL_1558607, EPI_ISL_1558608, EPI_ISL_1558609, EPI_ISL_1558610, EPI_ISL_1558611, EPI_ISL_1558613, EPI_ISL_1558614 | | | |
| see above | Department of Virology, Istituto Zooprofilattico Sperimentale del Lazio e della Toscana (IZSLT) | Department of General Diagnostics; Department of Virology; Istituto Zooprofilattico Sperimentale del Lazio e della Toscana (IZSLT) | Patricia Alba, Giuseppe Manna, Elena L. Diaconu, Fabiola Feltrin, Raffaela Conti, Teresa Scicluna, Virginia Carfora, Antonella Cersini, Alessia Franco, Antonio Battisti. |
| EPI_ISL_1558791, EPI_ISL_1558792, EPI_ISL_1558793 | LESP Chihuahua | Instituto de Diagnostico y Referencia Epidemiologicos (INDRE) | Claudia Wong-Arambula, Abril Rodriguez-Maldonado, Vanessa Rivero-Arredondo, Ariadna Medina-Benitez, Joaquin Quiroz-Mercado, Sergio Rangel-Guerrero, Natividad Cruz-Ortiz, Tatiana Nunez-Garcia, Gisela Barrera-Badillo, Lucia Hernandez-Rivas, Irma Lopez-Martinez, Ernesto Ramirez-Gonzalez. |
| EPI_ISL_1558795 | LESP Queretaro | Instituto de Diagnostico y Referencia Epidemiologicos (INDRE) | Claudia Wong-Arambula, Abril Rodriguez-Maldonado, Vanessa Rivero-Arredondo, Ariadna Medina-Benitez, Joaquin Quiroz-Mercado, Sergio Rangel-Guerrero, Natividad Cruz-Ortiz, Tatiana Nunez-Garcia, Gisela Barrera-Badillo, Lucia Hernandez-Rivas, Irma Lopez-Martinez, Ernesto Ramirez-Gonzalez. |
| EPI_ISL_1558796, EPI_ISL_1558797 | LESPZacatecas | Instituto de Diagnostico y Referencia Epidemiologicos (INDRE) | Claudia Wong-Arambula, Abril Rodriguez-Maldonado, Vanessa Rivero-Arredondo, Ariadna Medina-Benitez, Joaquin Quiroz-Mercado, Sergio Rangel-Guerrero, Natividad Cruz-Ortiz, Tatiana Nunez-Garcia, Gisela Barrera-Badillo, Lucia Hernandez-Rivas, Irma Lopez-Martinez, Ernesto Ramirez-Gonzalez. |
| EPI_ISL_1558798 | LESP Baja California | Instituto de Diagnostico y Referencia Epidemiologicos (INDRE) | Claudia Wong-Arambula, Abril Rodriguez-Maldonado, Vanessa Rivero-Arredondo, Ariadna Medina-Benitez, Joaquin Quiroz-Mercado, Sergio Rangel-Guerrero, Natividad Cruz-Ortiz, Tatiana Nunez-Garcia, Gisela Barrera-Badillo, Lucia Hernandez-Rivas, Irma Lopez-Martinez, Ernesto Ramirez-Gonzalez. |
| EPI_ISL_1558799 | LESP Jalisco | Instituto de Diagnostico y Referencia Epidemiologicos (INDRE) | Claudia Wong-Arambula, Abril Rodriguez-Maldonado, Vanessa Rivero-Arredondo, Ariadna Medina-Benitez, Joaquin Quiroz-Mercado, Sergio Rangel-Guerrero, Natividad Cruz-Ortiz, Tatiana Nunez-Garcia, Gisela Barrera-Badillo, Lucia Hernandez-Rivas, Irma Lopez-Martinez, Ernesto Ramirez-Gonzalez. |
| EPI_ISL_1558801 | LESP Baja California Sur | Instituto de Diagnostico y Referencia Epidemiologicos (INDRE) | Claudia Wong-Arambula, Abril Rodriguez-Maldonado, Vanessa Rivero-Arredondo, Ariadna Medina-Benitez, Joaquin Quiroz-Mercado, Sergio Rangel-Guerrero, Natividad Cruz-Ortiz, Tatiana Nunez-Garcia, Gisela Barrera-Badillo, Lucia Hernandez-Rivas, Irma Lopez-Martinez, Ernesto Ramirez-Gonzalez. |
| EPI_ISL_1558828, EPI_ISL_1558830 | LESP Jalisco/UdeG | Instituto de Diagnostico y Referencia Epidemiologicos (INDRE) | Claudia Wong-Arambula, Abril Rodriguez-Maldonado, Vanessa Rivero-Arredondo, Ariadna Medina-Benitez, Joaquin Quiroz-Mercado, Sergio Rangel-Guerrero, Natividad Cruz-Ortiz, Tatiana Nunez-Garcia, Gisela Barrera-Badillo, Lucia Hernandez-Rivas, Irma Lopez-Martinez, Ernesto |

| | | | |
|---|---|---|---|
| | | | Ramirez-Gonzalez. |
| EPI_ISL_1563738, EPI_ISL_1563739 | Institut de Virologie du CHU de Strasbourg | Institut de Virologie du CHU de Strasbourg | Fafi-Kremer Samira, Gallais Floriane, Gantner Pierre, Laugel Elodie, Solis Morgane, Velay Aurélie, Wendling Marie-Josée |
| EPI_ISL_1577331 | CH Lens | CHU Lille - Laboratoire de Virologie | AIT YAHYA Emilie, ALIDJINOU Enagnon Kazali, BOCKET Laurence, CREPIN Michel, DEMAY Christophe, ENGELMANN Ilka, GEFFROY Sandrine, GUIGON Aurélie, LAMBERT Valérie, LAZREK Mouna, NOBILLIAUX Florian, PREVOST Brigitte, THUILLIER Caroline, TINEZ Claire ,TCHANTCHOU NJOSSE YANICK |
| EPI_ISL_1577332 | CH Boulogne sur mer | CHU Lille - Laboratoire de Virologie | AIT YAHYA Emilie, ALIDJINOU Enagnon Kazali, BOCKET Laurence, CREPIN Michel, DEMAY Christophe, ENGELMANN Ilka, GEFFROY Sandrine, GUIGON Aurélie, LAMBERT Valérie, LAZREK Mouna, NOBILLIAUX Florian, PREVOST Brigitte, THUILLIER Caroline, TINEZ Claire ,TCHANTCHOU NJOSSE YANICK |
| EPI_ISL_1577334 | CH Douai | CHU Lille - Laboratoire de Virologie | AIT YAHYA Emilie, ALIDJINOU Enagnon Kazali, BOCKET Laurence, CREPIN Michel, DEMAY Christophe, ENGELMANN Ilka, GEFFROY Sandrine, GUIGON Aurélie, LAMBERT Valérie, LAZREK Mouna, NOBILLIAUX Florian, PREVOST Brigitte, THUILLIER Caroline, TINEZ Claire ,TCHANTCHOU NJOSSE YANICK |
| EPI_ISL_1577335, EPI_ISL_1577336, EPI_ISL_1577337 | CHU Lille | CHU Lille - Laboratoire de Virologie | AIT YAHYA Emilie, ALIDJINOU Enagnon Kazali, BOCKET Laurence, CREPIN Michel, DEMAY Christophe, ENGELMANN Ilka, GEFFROY Sandrine, GUIGON Aurélie, LAMBERT Valérie, LAZREK Mouna, NOBILLIAUX Florian, PREVOST Brigitte, THUILLIER Caroline, TINEZ Claire ,TCHANTCHOU NJOSSE YANICK |
| EPI_ISL_1577338 | CH Cambrai | CHU Lille - Laboratoire de Virologie | AIT YAHYA Emilie, ALIDJINOU Enagnon Kazali, BOCKET Laurence, CREPIN Michel, DEMAY Christophe, ENGELMANN Ilka, GEFFROY Sandrine, GUIGON Aurélie, LAMBERT Valérie, LAZREK Mouna, NOBILLIAUX Florian, PREVOST Brigitte, THUILLIER Caroline, TINEZ Claire ,TCHANTCHOU NJOSSE YANICK |
| EPI_ISL_1577340, EPI_ISL_1577341, EPI_ISL_1577342, EPI_ISL_1577343, EPI_ISL_1577344, EPI_ISL_1577345, EPI_ISL_1577346, EPI_ISL_1577347, EPI_ISL_1577348, EPI_ISL_1577349 | CHU Lille | CHU Lille - Laboratoire de Virologie | AIT YAHYA Emilie, ALIDJINOU Enagnon Kazali, BOCKET Laurence, CREPIN Michel, DEMAY Christophe, ENGELMANN Ilka, GEFFROY Sandrine, GUIGON Aurélie, LAMBERT Valérie, LAZREK Mouna, NOBILLIAUX Florian, PREVOST Brigitte, THUILLIER Caroline, TINEZ Claire ,TCHANTCHOU NJOSSE YANICK |
| EPI_ISL_1577351 | CH SAINT PHILIBERT | CHU Lille - Laboratoire de Virologie | AIT YAHYA Emilie, ALIDJINOU Enagnon Kazali, BOCKET Laurence, CREPIN Michel, DEMAY Christophe, ENGELMANN Ilka, GEFFROY Sandrine, GUIGON Aurélie, LAMBERT Valérie, LAZREK Mouna, NOBILLIAUX Florian, PREVOST Brigitte, THUILLIER Caroline, TINEZ Claire ,TCHANTCHOU NJOSSE YANICK |
| EPI_ISL_1577352 | CHU Lille | CHU Lille - Laboratoire de Virologie | AIT YAHYA Emilie, ALIDJINOU Enagnon Kazali, BOCKET Laurence, CREPIN Michel, DEMAY Christophe, ENGELMANN Ilka, GEFFROY Sandrine, GUIGON Aurélie, LAMBERT Valérie, LAZREK Mouna, NOBILLIAUX Florian, PREVOST Brigitte, THUILLIER Caroline, TINEZ Claire ,TCHANTCHOU NJOSSE YANICK |
| EPI_ISL_1577353 | CH Bethune | CHU Lille - Laboratoire de Virologie | AIT YAHYA Emilie, ALIDJINOU Enagnon Kazali, BOCKET Laurence, CREPIN Michel, DEMAY Christophe, ENGELMANN Ilka, GEFFROY Sandrine, GUIGON Aurélie, LAMBERT Valérie, LAZREK Mouna, NOBILLIAUX Florian, PREVOST Brigitte, THUILLIER Caroline, TINEZ Claire ,TCHANTCHOU NJOSSE YANICK |
| EPI_ISL_1577355 | CHU Lille | CHU Lille - Laboratoire de Virologie | AIT YAHYA Emilie, ALIDJINOU Enagnon Kazali, BOCKET Laurence, CREPIN Michel, DEMAY Christophe, ENGELMANN Ilka, GEFFROY Sandrine, GUIGON Aurélie, LAMBERT Valérie, LAZREK Mouna, NOBILLIAUX Florian, PREVOST Brigitte, THUILLIER Caroline, TINEZ Claire ,TCHANTCHOU NJOSSE YANICK |
| EPI_ISL_1577360, EPI_ISL_1577361 | CH Arras | CHU Lille - Laboratoire de Virologie | AIT YAHYA Emilie, ALIDJINOU Enagnon Kazali, BOCKET Laurence, CREPIN Michel, DEMAY Christophe, ENGELMANN Ilka, GEFFROY Sandrine, GUIGON Aurélie, LAMBERT Valérie, LAZREK Mouna, NOBILLIAUX Florian, PREVOST Brigitte, THUILLIER Caroline, TINEZ Claire ,TCHANTCHOU NJOSSE YANICK |
| EPI_ISL_1577362 | CH Tourcoing | CHU Lille - Laboratoire de Virologie | AIT YAHYA Emilie, ALIDJINOU Enagnon Kazali, BOCKET Laurence, CREPIN Michel, DEMAY Christophe, ENGELMANN Ilka, GEFFROY Sandrine, GUIGON Aurélie, LAMBERT Valérie, LAZREK Mouna, NOBILLIAUX Florian, PREVOST Brigitte, THUILLIER Caroline, TINEZ Claire ,TCHANTCHOU NJOSSE YANICK |
| EPI_ISL_1577363 | CH Roubaix | CHU Lille - Laboratoire de Virologie | AIT YAHYA Emilie, ALIDJINOU Enagnon Kazali, BOCKET Laurence, CREPIN Michel, DEMAY Christophe, ENGELMANN Ilka, GEFFROY Sandrine, GUIGON Aurélie, LAMBERT Valérie, LAZREK Mouna, NOBILLIAUX Florian, PREVOST Brigitte, THUILLIER Caroline, TINEZ Claire ,TCHANTCHOU NJOSSE YANICK |
| EPI_ISL_1577365, EPI_ISL_1577366, EPI_ISL_1577367, EPI_ISL_1577368 | CH Bethune | CHU Lille - Laboratoire de Virologie | AIT YAHYA Emilie, ALIDJINOU Enagnon Kazali, BOCKET Laurence, CREPIN Michel, DEMAY Christophe, ENGELMANN Ilka, GEFFROY Sandrine, GUIGON Aurélie, LAMBERT Valérie, LAZREK Mouna, NOBILLIAUX Florian, PREVOST Brigitte, THUILLIER Caroline, TINEZ Claire ,TCHANTCHOU NJOSSE YANICK |
| EPI_ISL_1577375 | CH Arras | CHU Lille - Laboratoire de Virologie | AIT YAHYA Emilie, ALIDJINOU Enagnon Kazali, BOCKET Laurence, CREPIN Michel, DEMAY Christophe, ENGELMANN Ilka, GEFFROY Sandrine, GUIGON Aurélie, LAMBERT Valérie, LAZREK Mouna, NOBILLIAUX Florian, PREVOST Brigitte, THUILLIER Caroline, TINEZ Claire ,TCHANTCHOU NJOSSE YANICK |
| EPI_ISL_1577560, EPI_ISL_1577561, EPI_ISL_1577562, EPI_ISL_1577563, EPI_ISL_1577564, EPI_ISL_1577565, EPI_ISL_1577566, EPI_ISL_1577567, EPI_ISL_1577568, EPI_ISL_1577569, EPI_ISL_1577570, EPI_ISL_1577571, EPI_ISL_1577572, EPI_ISL_1577573, EPI_ISL_1577574, EPI_ISL_1577575, EPI_ISL_1577576, EPI_ISL_1577577, EPI_ISL_1577578, EPI_ISL_1577579, EPI_ISL_1577580, EPI_ISL_1577581, EPI_ISL_1577582, EPI_ISL_1577583, EPI_ISL_1577584, EPI_ISL_1577585 | | | |
| see above | University of Rome Tor Vergata: Departm Experim Medicine Chair of Virology | University of Rome Tor Vergata: Departm Experim Medicine Chair of Virology | Maria Concetta Bellocchi, Luca Carioti, Maria Botticelli, Lorenzo Piermatteo, Marco Iannetta, Loredana Sarmati, Massimo Andreoni, Francesca Ceccherini-Silberstein |
| EPI_ISL_1577908, EPI_ISL_1577913 | Laboratory for Clinical Immunology and Molecular Genetics - University Clinic Golnik Laboratory for Respiratory Microbiology - University Clinic Golnik | Laboratory for Clinical Immunology and Molecular Genetics -University Clinic Golnik | Matija Rijavec, Julij Šelb, Urška Bidovec Stojkovi, Žan Kogovšek, Nina Rupar, Viktorija Tomi, Peter Korošec |
| EPI_ISL_1579823, EPI_ISL_1580175, EPI_ISL_1580275, EPI_ISL_1580316, EPI_ISL_1580340 | NMVRVI | National Public Health Surveillance Laboratory | Lukas Zemaitis, Migle Gabrielaite, Jelena Razmuk, Svajune Muralyte, Ana Steponkiene, Lukas Vasionis, Danas Baksa |
| EPI_ISL_1583095, EPI_ISL_1583096, EPI_ISL_1583157 | Hospital | National Reference Center for Viruses of Respiratory Infections, Institut Pasteur, Paris | Marion Barbet, Sylvie Behillil, Frédéric Lemoine, Corinne Maufrais, Christophe Malabat, Méline Bizard, Angela Brisebarre, Camille Capel, Louise Lefrançois, Etienne Simon-Lorière, Vincent Enouf, Maud Vanpeene, Sylvie van der Werf,Sandrine Castelain |
| EPI_ISL_1583632 | Institut de Virologie du CHU de Strasbourg | Institut de Virologie du CHU de Strasbourg | Fafi-Kremer Samira, Gallais Floriane, Gantner Pierre, Laugel Elodie, Solis Morgane, Velay Aurélie, Wendling Marie-Josée |
| EPI_ISL_1585281, EPI_ISL_1585282 | Laboratorio di Riferimento Regionale della Sicilia Occidentale per l'Emergenza COVID-19 | Laboratorio di Riferimento Regionale della Sicilia Occidentale per l'Emergenza COVID-19 | Fabio Tramuto, Carmelo Massimo Maida, Daniela Di Naro, Giulia Randazzo, Walter Mazzucco, Giorgio Graziano, Vincenzo Restivo, Claudio Costantino, Francesco Vitale |
| EPI_ISL_1585284 | Virology Unit, AOUP | Virology Unit, AOUP | Marialinda Vatteroni, Susi Frateschi |
| EPI_ISL_1585285 | Virology Unit | Virology Unit | Marialinda Vatteroni, Susi Frateschi |
| EPI_ISL_1585286 | Virology Unit, AOUP | Virology Unit, AOUP | Marialinda Vatteroni, Susi Frateschi |
| EPI_ISL_1585287 | Virology Unit, AOUP | Virology Unit, AOUP | Maralinda Vatteroni, Susi Frateschi |
| EPI_ISL_1585384, EPI_ISL_1585385, EPI_ISL_1585386, EPI_ISL_1585387, EPI_ISL_1585388, EPI_ISL_1585389, EPI_ISL_1585390, EPI_ISL_1585391, EPI_ISL_1585392, EPI_ISL_1585393, EPI_ISL_1585394, EPI_ISL_1585395, EPI_ISL_1585396, EPI_ISL_1585397, EPI_ISL_1585398, EPI_ISL_1585399, EPI_ISL_1585400, EPI_ISL_1585401, EPI_ISL_1585402, EPI_ISL_1585403, EPI_ISL_1585404, EPI_ISL_1585405, EPI_ISL_1585406, EPI_ISL_1585407, EPI_ISL_1585409, EPI_ISL_1585410, EPI_ISL_1585411, EPI_ISL_1585412, EPI_ISL_1585413, EPI_ISL_1585414, EPI_ISL_1585415, EPI_ISL_1585416, EPI_ISL_1585417, EPI_ISL_1585419, EPI_ISL_1585420, EPI_ISL_1585421, EPI_ISL_1585422, EPI_ISL_1585424, EPI_ISL_1585425, EPI_ISL_1585426, EPI_ISL_1585427 | | | |
| see above | Centro de Investigación Biomédica del Noreste (CIBIN) | Instituto Nacional de Enfermedades Respiratorias (INER): Centro de Investigación en Enfermedades Infecciosas (CIENI) | Consorcio Mexicano de Vigilancia Genómica (CoViGen-Mex). Authors (in alphabetical order): Julio Elias Alvarado-Yaah, Carlos F. Arias, Santiago Ávila-Ríos, Víctor Hugo Borja-Aburto, Celia Boukadida, Juan Bautista Chale-Dzul, Célida Duque Molina, José Antonio Enciso-Moreno, Gloria Elena |

| | | | |
|---|---|---|---|
| | | | Espinosa-Ayala, Fernando Fontove-Herrera, Victor Eduardo Garcia-Arias, Concepción Grajales-Muñiz, Ricardo Grande, Alfredo Herrera-Estrella, Carla Ivón Herrera-Najera, Pavel Isa, Brenda Irasema Maldonado-Meza, Bernardo Martínez-Miguel, Margarita Matías-Florentino, María Guadalupe de Jesús Mireles-Rivera, Gloria María Molina-Salinas, Hector Montoya-Fuentes, José Esteban Muñoz-Medina, José de Jesús Nuñez-Contreras, Alicia Ocaña-Mondragón, Luis Alberto Ochoa-Carrera, Hector Esteban Paz-Juárez, Francisco Pulido, Helen Haydee Fernanda Ramírez-Plascencia, Angel Gustavo Salas-Lais, Alejandro Sanchez-Flores, Clara Esperanza Santacruz-Tinoco, María Guadalupe Santiago-Mauricio, Nelly Sélem-Mojica, Blanca Taboada, Gloria Vazquez |
| EPI_ISL_1585428, EPI_ISL_1585429, EPI_ISL_1585430, EPI_ISL_1585431, EPI_ISL_1585432, EPI_ISL_1585433, EPI_ISL_1585434, EPI_ISL_1585435, EPI_ISL_1585436, EPI_ISL_1585437, EPI_ISL_1585438, EPI_ISL_1585439, EPI_ISL_1585440, EPI_ISL_1585441, EPI_ISL_1585442, EPI_ISL_1585443, EPI_ISL_1585444, EPI_ISL_1585445, EPI_ISL_1585447, EPI_ISL_1585449, EPI_ISL_1585450, EPI_ISL_1585451, EPI_ISL_1585452, EPI_ISL_1585453, EPI_ISL_1585454 | | | |
| see above | Unidad de Investigación Médica de Yucatán (UIMY) | Instituto Nacional de Enfermedades Respiratorias (INER): Centro de Investigación en Enfermedades Infecciosas (CIENI) | Consorcio Mexicano de Vigilancia Genómica (CoViGen-Mex). Authors (in alphabetical order): Julio Elias Alvarado-Yaah, Carlos F. Arias, Santiago Ávila-Ríos, Víctor Hugo Borja-Aburto, Celia Boukadida, Juan Bautista Chale-Dzul, Célida Duque Molina, José Antonio Enciso-Moreno, Gloria Elena Espinosa-Ayala, Fernando Fontove-Herrera, Victor Eduardo Garcia-Arias, Concepción Grajales-Muñiz, Ricardo Grande, Alfredo Herrera-Estrella, Carla Ivón Herrera-Najera, Pavel Isa, Brenda Irasema Maldonado-Meza, Bernardo Martínez-Miguel, Margarita Matías-Florentino, María Guadalupe de Jesús Mireles-Rivera, Gloria María Molina-Salinas, Hector Montoya-Fuentes, José Esteban Muñoz-Medina, José de Jesús Nuñez-Contreras, Alicia Ocaña-Mondragón, Luis Alberto Ochoa-Carrera, Hector Esteban Paz-Juárez, Francisco Pulido, Helen Haydee Fernanda Ramírez-Plascencia, Angel Gustavo Salas-Lais, Alejandro Sanchez-Flores, Clara Esperanza Santacruz-Tinoco, María Guadalupe Santiago-Mauricio, Nelly Sélem-Mojica, Blanca Taboada, Gloria Vazquez |
| EPI_ISL_1585455, EPI_ISL_1585456, EPI_ISL_1585457, EPI_ISL_1585458, EPI_ISL_1585459, EPI_ISL_1585460, EPI_ISL_1585461, EPI_ISL_1585462, EPI_ISL_1585463, EPI_ISL_1585464, EPI_ISL_1585465, EPI_ISL_1585466, EPI_ISL_1585467, EPI_ISL_1585468, EPI_ISL_1585469, EPI_ISL_1585470, EPI_ISL_1585471, EPI_ISL_1585472, EPI_ISL_1585473, EPI_ISL_1585474, EPI_ISL_1585476, EPI_ISL_1585477 | | | |
| see above | Unidad de Investigación Biomédica de Zacatecas (UIBZ) | Instituto Nacional de Enfermedades Respiratorias (INER): Centro de Investigación en Enfermedades Infecciosas (CIENI) | Consorcio Mexicano de Vigilancia Genómica (CoViGen-Mex). Authors (in alphabetical order): Julio Elias Alvarado-Yaah, Carlos F. Arias, Santiago Ávila-Ríos, Víctor Hugo Borja-Aburto, Celia Boukadida, Juan Bautista Chale-Dzul, Célida Duque Molina, José Antonio Enciso-Moreno, Gloria Elena Espinosa-Ayala, Fernando Fontove-Herrera, Victor Eduardo Garcia-Arias, Concepción Grajales-Muñiz, Ricardo Grande, Alfredo Herrera-Estrella, Carla Ivón Herrera-Najera, Pavel Isa, Brenda Irasema Maldonado-Meza, Bernardo Martínez-Miguel, Margarita Matías-Florentino, María Guadalupe de Jesús Mireles-Rivera, Gloria María Molina-Salinas, Hector Montoya-Fuentes, José Esteban Muñoz-Medina, José de Jesús Nuñez-Contreras, Alicia Ocaña-Mondragón, Luis Alberto Ochoa-Carrera, Hector Esteban Paz-Juárez, Francisco Pulido, Helen Haydee Fernanda Ramírez-Plascencia, Angel Gustavo Salas-Lais, Alejandro Sanchez-Flores, Clara Esperanza Santacruz-Tinoco, María Guadalupe Santiago-Mauricio, Nelly Sélem-Mojica, Blanca Taboada, Gloria Vazquez |
| EPI_ISL_1585480, EPI_ISL_1585481, EPI_ISL_1585482, EPI_ISL_1585483, EPI_ISL_1585484, EPI_ISL_1585485, EPI_ISL_1585486, EPI_ISL_1585487, EPI_ISL_1585488, EPI_ISL_1585489, EPI_ISL_1585490, EPI_ISL_1585491, EPI_ISL_1585492, EPI_ISL_1585493, EPI_ISL_1585494, EPI_ISL_1585495, EPI_ISL_1585496, EPI_ISL_1585497, EPI_ISL_1585499, EPI_ISL_1585500, EPI_ISL_1585501, EPI_ISL_1585502, EPI_ISL_1585504, EPI_ISL_1585505, EPI_ISL_1585506, EPI_ISL_1585507, EPI_ISL_1585508, EPI_ISL_1585509, EPI_ISL_1585511, EPI_ISL_1585512, EPI_ISL_1585513, EPI_ISL_1585514, EPI_ISL_1585515, EPI_ISL_1585516, EPI_ISL_1585517, EPI_ISL_1585519, EPI_ISL_1585520, EPI_ISL_1585522, EPI_ISL_1585523, EPI_ISL_1585524, EPI_ISL_1585525, EPI_ISL_1585526, EPI_ISL_1585527, EPI_ISL_1585528, EPI_ISL_1585529, EPI_ISL_1585530, EPI_ISL_1585531, EPI_ISL_1585532, EPI_ISL_1585533, EPI_ISL_1585534, EPI_ISL_1585535, EPI_ISL_1585536, EPI_ISL_1585537, EPI_ISL_1585538, EPI_ISL_1585539, EPI_ISL_1585543, EPI_ISL_1585544, EPI_ISL_1585545, EPI_ISL_1585547, EPI_ISL_1585548, EPI_ISL_1585549, EPI_ISL_1585550, EPI_ISL_1585552, EPI_ISL_1585553, EPI_ISL_1585554 | | | |
| see above | Centro de Investigación Biomédica de Occidente (CIBO) | Instituto Nacional de Enfermedades Respiratorias (INER): Centro de Investigación en Enfermedades Infecciosas (CIENI) | Consorcio Mexicano de Vigilancia Genómica (CoViGen-Mex). Authors (in alphabetical order): Julio Elias Alvarado-Yaah, Carlos F. Arias, Santiago Ávila-Ríos, Víctor Hugo Borja-Aburto, Celia Boukadida, Juan Bautista Chale-Dzul, Célida Duque Molina, José Antonio Enciso-Moreno, Gloria Elena Espinosa-Ayala, Fernando Fontove-Herrera, Victor Eduardo Garcia-Arias, Concepción Grajales-Muñiz, Ricardo Grande, Alfredo Herrera-Estrella, Carla Ivón Herrera-Najera, Pavel Isa, Brenda Irasema Maldonado-Meza, Bernardo Martínez-Miguel, Margarita Matías-Florentino, María Guadalupe de Jesús Mireles-Rivera, Gloria María Molina-Salinas, Hector Montoya-Fuentes, José Esteban Muñoz-Medina, José de Jesús Nuñez-Contreras, Alicia Ocaña-Mondragón, Luis Alberto Ochoa-Carrera, Hector Esteban Paz-Juárez, Francisco Pulido, Helen Haydee Fernanda Ramírez-Plascencia, Angel Gustavo Salas-Lais, Alejandro Sanchez-Flores, Clara Esperanza Santacruz-Tinoco, María Guadalupe Santiago-Mauricio, Nelly Sélem-Mojica, Blanca Taboada, Gloria Vazquez |
| EPI_ISL_1585564, EPI_ISL_1585565, EPI_ISL_1585566, EPI_ISL_1585567, EPI_ISL_1585568, EPI_ISL_1585569, EPI_ISL_1585570, EPI_ISL_1585571, EPI_ISL_1585572, EPI_ISL_1585573, EPI_ISL_1585574, EPI_ISL_1585575, EPI_ISL_1585576, EPI_ISL_1585577, EPI_ISL_1585578, EPI_ISL_1585579, EPI_ISL_1585580, EPI_ISL_1585581, EPI_ISL_1585582, EPI_ISL_1585583, EPI_ISL_1585584, EPI_ISL_1585585, EPI_ISL_1585586, EPI_ISL_1585587, EPI_ISL_1585588, EPI_ISL_1585589, EPI_ISL_1585590, EPI_ISL_1585591, EPI_ISL_1585592, EPI_ISL_1585593, EPI_ISL_1585594, EPI_ISL_1585595, EPI_ISL_1585597, EPI_ISL_1585599, EPI_ISL_1585600, EPI_ISL_1585601, EPI_ISL_1585602, EPI_ISL_1585604, EPI_ISL_1585605, EPI_ISL_1585606, EPI_ISL_1585610, EPI_ISL_1585613, EPI_ISL_1585615, EPI_ISL_1585617, EPI_ISL_1585619, EPI_ISL_1585624, EPI_ISL_1585625, EPI_ISL_1585641, EPI_ISL_1585642, EPI_ISL_1585658 | | | |
| see above | Laboratorio Central de Epidemiología (LCE) | Instituto Nacional de Enfermedades Respiratorias (INER): Centro de Investigación en Enfermedades Infecciosas (CIENI) | Consorcio Mexicano de Vigilancia Genómica (CoViGen-Mex). Authors (in alphabetical order): Julio Elias Alvarado-Yaah, Carlos F. Arias, Santiago Ávila-Ríos, Víctor Hugo Borja-Aburto, Celia Boukadida, Juan Bautista Chale-Dzul, Célida Duque Molina, José Antonio Enciso-Moreno, Gloria Elena Espinosa-Ayala, Fernando Fontove-Herrera, Victor Eduardo Garcia-Arias, Concepción Grajales-Muñiz, Ricardo Grande, Alfredo Herrera-Estrella, Carla Ivón Herrera-Najera, Pavel Isa, Brenda Irasema Maldonado-Meza, Bernardo Martínez-Miguel, Margarita Matías-Florentino, María Guadalupe de Jesús Mireles-Rivera, Gloria María Molina-Salinas, Hector Montoya-Fuentes, José Esteban Muñoz-Medina, José de Jesús Nuñez-Contreras, Alicia Ocaña-Mondragón, Luis Alberto Ochoa-Carrera, Hector Esteban Paz-Juárez, Francisco Pulido, Helen Haydee Fernanda Ramírez-Plascencia, Angel Gustavo Salas-Lais, Alejandro Sanchez-Flores, Clara Esperanza Santacruz-Tinoco, María Guadalupe Santiago-Mauricio, Nelly Sélem-Mojica, Blanca Taboada, Gloria Vazquez |
| EPI_ISL_1585890 | Alphabio Laboratory | LBM ALPHABIO, Marseille | Vincent GARCIA |
| EPI_ISL_1587827, EPI_ISL_1587831, EPI_ISL_1587833, EPI_ISL_1587882, EPI_ISL_1587884, EPI_ISL_1587893, EPI_ISL_1587897, EPI_ISL_1587899, EPI_ISL_1587905, EPI_ISL_1587912 | NL-Dr. Leonard A. Miller Centre for Health Services | National Microbiology Laboratory (NML) | Anna Majer, Shari Tyson, Grace Seo, Philip Mabon, Elsie Grudeski, Rhiannon Huzarewich, Russell Mandes, Anneliese Landgraff, Jennifer Tanner, Natalie Knox, Morag Graham, Gary Van Domselaar, Robert Needle, Yang Yu, Adel Malek, Laura Gilbert, George Zahariadis, Nathalie Bastien, Yan Li, Timothy Booth, Darian Hole, Madison Chapel, Kirsten Biggar, Kerri Smith, CanCOGeN's metadata curation team, Public Health Agency of Canada CanCOGeN team |
| EPI_ISL_1588078, EPI_ISL_1588100, EPI_ISL_1588101, EPI_ISL_1588104, EPI_ISL_1588106, EPI_ISL_1588107, EPI_ISL_1588110, EPI_ISL_1588111, EPI_ISL_1588113, EPI_ISL_1588114, EPI_ISL_1588116, EPI_ISL_1588117, EPI_ISL_1588119, EPI_ISL_1588120, EPI_ISL_1588125, EPI_ISL_1588126, EPI_ISL_1588135 | | | |
| see above | NB-Hôpital Georges L. Dumont | National Microbiology Laboratory (NML) | Anna Majer, Shari Tyson, Grace Seo, Philip Mabon, Elsie Grudeski, Rhiannon Huzarewich, Russell Mandes, Anneliese Landgraff, Jennifer Tanner, Natalie Knox, Morag Graham, Gary Van Domselaar, Richard Garceau, Guillaume Desnoyers, Nathalie Bastien, Yan Li, Timothy Booth, Darian Hole, Madison Chapel, Kirsten Biggar, CanCOGeN's metadata curation team, Public Health Agency of Canada CanCOGeN team |
| EPI_ISL_1588443, EPI_ISL_1588444, EPI_ISL_1588445, EPI_ISL_1588479, EPI_ISL_1588480, EPI_ISL_1588481, EPI_ISL_1588487, EPI_ISL_1588488, EPI_ISL_1588493, EPI_ISL_1588494, EPI_ISL_1588495, EPI_ISL_1588496 | | | |
| see above | National Institute of Public Health | National Institute of Public Health | Helena Jirincova, Jaromira Vecerova, Timotej Suri, Dusan Trnka, Alexander Nagy |
| EPI_ISL_1588710 | Bioptická laborato s.r.o. | Bioptická laborato s.r.o. | Petr Šteiner, Tomáš Vanek, Nikola Bílá, Martina Putzová, Michaela íhová, Silva Vondráková |
| EPI_ISL_1588829 | Hospital | National Reference Center for Viruses of Respiratory Infections, Institut Pasteur, Paris | Marion Barbet, Sylvie Behillil, Méline Bizard, Angela Brisebarre, Camille Capel,Frédéric Lemoine,Corinne Maufrais,Christophe Malabat, Damien Mornico, Louise Lefrançois, Etienne Simon-Lorière, Vincent Enouf, Maud Vanpeene, Sylvie van der Werf,Hochart A |
| EPI_ISL_1588830, EPI_ISL_1588831 | Hospital | National Reference Center for Viruses of Respiratory Infections, Institut Pasteur, Paris | Marion Barbet, Sylvie Behillil, Méline Bizard, Angela Brisebarre, Camille Capel,Frédéric Lemoine,Corinne Maufrais,Christophe Malabat, Damien Mornico, Louise Lefrançois, Etienne Simon-Lorière, Vincent Enouf, Maud Vanpeene, Sylvie van der Werf,Serazin ValéRie |
| EPI_ISL_1588832, EPI_ISL_1588834 | Hospital | National Reference Center for Viruses of Respiratory Infections, Institut Pasteur, Paris | Marion Barbet, Sylvie Behillil, Méline Bizard, Angela Brisebarre, Camille Capel,Frédéric Lemoine,Corinne Maufrais,Christophe Malabat, Damien Mornico, Louise Lefrançois, Etienne Simon-Lorière, Vincent Enouf, Maud Vanpeene, Sylvie van der Werf,Guinard JéRome |
| EPI_ISL_1591105, EPI_ISL_1591106, EPI_ISL_1591107 | "St. Parascheva" Infectious Diseases Clinical Hospital | Cantacuzino Institute Virology | Luiza Ustea, Nicoleta Paraschiv, Cherciu Carmen, Catalina Pascu, Mihaela Oprea, Sorin Dinu, Mihaela Lazar |
| EPI_ISL_1591108, EPI_ISL_1591109, EPI_ISL_1591110, EPI_ISL_1591111, EPI_ISL_1591112, EPI_ISL_1591113, EPI_ISL_1591114, EPI_ISL_1591116, EPI_ISL_1591117, EPI_ISL_1591118 | Laboratory for Respiratory Viruses, Cantacuzino National Military-Medical Institute for Research and Development | Cantacuzino Institute Virology | Luiza Ustea, Nicoleta Paraschiv, Cherciu Carmen, Catalina Pascu, Mihaela Oprea, Sorin Dinu, Mihaela Lazar |
| EPI_ISL_1591119, EPI_ISL_1591123 | "St. Parascheva" Infectious Diseases Clinical Hospital | Cantacuzino Institute Virology | Luiza Ustea, Nicoleta Paraschiv, Cherciu Carmen, Catalina Pascu, Mihaela Oprea, Sorin Dinu, Mihaela Lazar |

| | | | |
|---|---|---|---|
| EPI_ISL_1591124, EPI_ISL_1591125 | Laboratory for Respiratory Viruses, Cantacuzino National Military-Medical Institute for Research and Development | Cantacuzino Institute Virology | Luiza Ustea, Nicoleta Paraschiv, Cherciu Carmen, Catalina Pascu, Mihaela Oprea, Sorin Dinu, Mihaela Lazar |
| EPI_ISL_1591126 | "St. Parascheva" Infectious Diseases Clinical Hospital | Cantacuzino Institute Virology | Luiza Ustea, Nicoleta Paraschiv, Cherciu Carmen, Catalina Pascu, Mihaela Oprea, Sorin Dinu, Mihaela Lazar |
| EPI_ISL_1591127, EPI_ISL_1591128, EPI_ISL_1591129, EPI_ISL_1591130, EPI_ISL_1591131, EPI_ISL_1591132, EPI_ISL_1591134, EPI_ISL_1591135, EPI_ISL_1591137, EPI_ISL_1591138, EPI_ISL_1591139, EPI_ISL_1591141, EPI_ISL_1591142, EPI_ISL_1591143, EPI_ISL_1591144, EPI_ISL_1591145 | | | |
| see above | Laboratory for Respiratory Viruses, Cantacuzino National Military-Medical Institute for Research and Development | Cantacuzino Institute Virology | Luiza Ustea, Nicoleta Paraschiv, Cherciu Carmen, Catalina Pascu, Mihaela Oprea, Sorin Dinu, Mihaela Lazar |
| EPI_ISL_1591252 | Department of Virology, Istituto Zooprofilattico Sperimentale del Lazio e della Toscana (IZSLT) | Department of General Diagnostics; Department of Virology; Istituto Zooprofilattico Sperimentale del Lazio e della Toscana (IZSLT) | Patricia Alba, Giuseppe Manna, Elena L. Diaconu, Fabiola Feltrin, Raffaela Conti, Teresa Scicluna, Virginia Carfora, Antonella Cersini, Alessia Franco, Antonio Battisti. |
| EPI_ISL_1593832, EPI_ISL_1593833, EPI_ISL_1593834, EPI_ISL_1593835 | National Institute of Public Health | State Veterinary Institute Prague | Nagy,A;Jirincova,H;Suri,T;Trnka,D;Vecerova,J |
| EPI_ISL_1593977, EPI_ISL_1593978, EPI_ISL_1593979, EPI_ISL_1593980 | CHC Andrée Rosemon | Institut Pasteur de la Guyane | Anne Lavergne, Dominique Rousset |
| EPI_ISL_1593982 | CNR Institut Pasteur de la Guyane | Institut Pasteur de la Guyane | Anne Lavergne, Dominique Rousset |
| EPI_ISL_1593983, EPI_ISL_1593984, EPI_ISL_1593985, EPI_ISL_1593986, EPI_ISL_1593987 | Labo Carage | Institut Pasteur de la Guyane | Anne Lavergne, Dominique Rousset |
| EPI_ISL_1595478 | Labo Analyses Med | National Reference Center for Viruses of Respiratory Infections, Institut Pasteur, Paris | Marion Barbet, Sylvie Behillil, Frédéric Lemoine, Corinne Maufrais, Christophe Malabat, Victoire Baillet, Méline Bizard, Angela Brisebarre, Camille Capel, Louise Lefrançois, Etienne Simon-Lorière, Vincent Enouf, Maud Vanpeene, Sylvie van der Werf,Nelly Luizy |
| EPI_ISL_1595595, EPI_ISL_1595596 | Centro de Investigación Biomédica del Noreste (CIBIN) | Instituto Nacional de Enfermedades Respiratorias (INER): Centro de Investigación en Enfermedades Infecciosas (CIENI) | Consorcio Mexicano de Vigilancia Genómica (CoViGen-Mex). Authors (in alphabetical order): Julio Elias Alvarado-Yaah, Carlos F. Arias, Santiago Ávila-Ríos, Víctor Hugo Borja-Aburto, Celia Boukadida, Juan Bautista Chale-Dzul, Célida Duque Molina, José Antonio Enciso-Moreno, Gloria Elena Espinosa-Ayala, Fernando Fontove-Herrera, Victor Eduardo Garcia-Arias, Concepción Grajales-Muñiz, Ricardo Grande, Alfredo Herrera-Estrella, Carla Ivón Herrera-Najera, Pavel Isa, Brenda Irasema Maldonado-Meza, Bernardo Martínez-Miguel, Margarita Matías-Florentino, María Guadalupe de Jesús Mireles-Rivera, Gloria María Molina-Salinas, Hector Montoya-Fuentes, José Esteban Muñoz-Medina, José de Jesús Nuñez-Contreras, Alicia Ocaña-Mondragón, Luis Alberto Ochoa-Carrera, Hector Esteban Paz-Juárez, Francisco Pulido, Helen Haydee Fernanda Ramírez-Plascencia, Angel Gustavo Salas-Lais, Alejandro Sanchez-Flores, Clara Esperanza Santacruz-Tinoco, María Guadalupe Santiago-Mauricio, Nelly Sélem-Mojica, Blanca Taboada, Gloria Vazquez |
| EPI_ISL_1595599 | Unidad de Investigación Médica de Yucatán (UIMY) | Instituto Nacional de Enfermedades Respiratorias (INER): Centro de Investigación en Enfermedades Infecciosas (CIENI) | Consorcio Mexicano de Vigilancia Genómica (CoViGen-Mex). Authors (in alphabetical order): Julio Elias Alvarado-Yaah, Carlos F. Arias, Santiago Ávila-Ríos, Víctor Hugo Borja-Aburto, Celia Boukadida, Juan Bautista Chale-Dzul, Célida Duque Molina, José Antonio Enciso-Moreno, Gloria Elena Espinosa-Ayala, Fernando Fontove-Herrera, Victor Eduardo Garcia-Arias, Concepción Grajales-Muñiz, Ricardo Grande, Alfredo Herrera-Estrella, Carla Ivón Herrera-Najera, Pavel Isa, Brenda Irasema Maldonado-Meza, Bernardo Martínez-Miguel, Margarita Matías-Florentino, María Guadalupe de Jesús Mireles-Rivera, Gloria María Molina-Salinas, Hector Montoya-Fuentes, José Esteban Muñoz-Medina, José de Jesús Nuñez-Contreras, Alicia Ocaña-Mondragón, Luis Alberto Ochoa-Carrera, Hector Esteban Paz-Juárez, Francisco Pulido, Helen Haydee Fernanda Ramírez-Plascencia, Angel Gustavo Salas-Lais, Alejandro Sanchez-Flores, Clara Esperanza Santacruz-Tinoco, María Guadalupe Santiago-Mauricio, Nelly Sélem-Mojica, Blanca Taboada, Gloria Vazquez |
| EPI_ISL_1595602 | Centro de Investigación Biomédica de Occidente (CIBO) | Instituto Nacional de Enfermedades Respiratorias (INER): Centro de Investigación en Enfermedades Infecciosas (CIENI) | Consorcio Mexicano de Vigilancia Genómica (CoViGen-Mex). Authors (in alphabetical order): Julio Elias Alvarado-Yaah, Carlos F. Arias, Santiago Ávila-Ríos, Víctor Hugo Borja-Aburto, Celia Boukadida, Juan Bautista Chale-Dzul, Célida Duque Molina, José Antonio Enciso-Moreno, Gloria Elena Espinosa-Ayala, Fernando Fontove-Herrera, Victor Eduardo Garcia-Arias, Concepción Grajales-Muñiz, Ricardo Grande, Alfredo Herrera-Estrella, Carla Ivón Herrera-Najera, Pavel Isa, Brenda Irasema Maldonado-Meza, Bernardo Martínez-Miguel, Margarita Matías-Florentino, María Guadalupe de Jesús Mireles-Rivera, Gloria María Molina-Salinas, Hector Montoya-Fuentes, José Esteban Muñoz-Medina, José de Jesús Nuñez-Contreras, Alicia Ocaña-Mondragón, Luis Alberto Ochoa-Carrera, Hector Esteban Paz-Juárez, Francisco Pulido, Helen Haydee Fernanda Ramírez-Plascencia, Angel Gustavo Salas-Lais, Alejandro Sanchez-Flores, Clara Esperanza Santacruz-Tinoco, María Guadalupe Santiago-Mauricio, Nelly Sélem-Mojica, Blanca Taboada, Gloria Vazquez |
| EPI_ISL_1595612 | Institut de Virologie du CHU de Strasbourg | Institut de Virologie du CHU de Strasbourg | Fafi-Kremer Samira, Gallais Floriane, Gantner Pierre, Laugel Elodie, Solis Morgane, Velay Aurélie, Wendling Marie-Josée |
| EPI_ISL_1599541, EPI_ISL_1599542, EPI_ISL_1599543, EPI_ISL_1599544 | Klinisch Laboratorium GZA | Klinisch Laboratorium ZNA | Verstrepen et al. |
| EPI_ISL_1608032, EPI_ISL_1608037, EPI_ISL_1608038, EPI_ISL_1608039, EPI_ISL_1608040, EPI_ISL_1608041, EPI_ISL_1608042, EPI_ISL_1608044, EPI_ISL_1608045 | Cliniques Saint-Pierre Ottignies | UCLouvain/IREC/MBLG | Jean Ruelle, Lysa Pinsmaye, Eleonore Ngyuvula Mantu, Benoit Kabamba Mukadi |
| EPI_ISL_1608109, EPI_ISL_1608110, EPI_ISL_1608111, EPI_ISL_1608112, EPI_ISL_1608113, EPI_ISL_1608114, EPI_ISL_1608115, EPI_ISL_1608116, EPI_ISL_1608117, EPI_ISL_1608118, EPI_ISL_1608119, EPI_ISL_1608120, EPI_ISL_1608121, EPI_ISL_1608122 | | | |
| see above | Programa de Oncovirologia, Instituto Nacional de Câncer | Programa de Oncovirologia, Instituto Nacional de Câncer | Livia R. Goes, Juliana D. Siqueira, Marianne M. Garrido, Brunna M. Alves, Ana Cristina P. M. Pereira, Claudia Cicala, James Arthos, João P.B. Viola, Marcelo A. Soares |
| EPI_ISL_1608169, EPI_ISL_1608170, EPI_ISL_1608171 | Hospital São Rafael - IDOR | Hospital São Rafael - IDOR | Carolina Kymie Vasques Nonaka, Tiago Gräf, Camila Araújo de Lorenzo Barcia, Vanessa Ferreira Costa, Janderson Lopes de Oliveira, Rogério da Hora Passos, Clarissa Araújo Gurgel Rocha, Iasmin Nogueira Bastos, Maria Clara Brito de Santana, Ian Marinho Santos, Karoline Almeida Felix de Sousa, Isadora Cristina de Siqueira, Thamires Gomes Lopes Weber, Ana Verena Almeida Mendes, Bruno Solano de Freitas Souza |
| EPI_ISL_1608224, EPI_ISL_1608225, EPI_ISL_1608226, EPI_ISL_1608227, EPI_ISL_1608228, EPI_ISL_1608231, EPI_ISL_1608232, EPI_ISL_1608233, EPI_ISL_1608234, EPI_ISL_1608235, EPI_ISL_1608236, EPI_ISL_1608237, EPI_ISL_1608239, EPI_ISL_1608240, EPI_ISL_1608242, EPI_ISL_1608243, EPI_ISL_1608246, EPI_ISL_1608247, EPI_ISL_1608248, EPI_ISL_1608249, EPI_ISL_1608250, EPI_ISL_1608251, EPI_ISL_1608253, EPI_ISL_1608254, EPI_ISL_1608255, EPI_ISL_1608256, EPI_ISL_1608257, EPI_ISL_1608258, EPI_ISL_1608259, EPI_ISL_1608261, EPI_ISL_1608262, EPI_ISL_1608263, EPI_ISL_1608264, EPI_ISL_1608265, EPI_ISL_1608266, EPI_ISL_1608267, EPI_ISL_1608268, EPI_ISL_1608269, EPI_ISL_1608270, EPI_ISL_1608271, EPI_ISL_1608272, EPI_ISL_1608273, EPI_ISL_1608275, EPI_ISL_1608277, EPI_ISL_1608278, EPI_ISL_1608279, EPI_ISL_1608280, EPI_ISL_1608281, EPI_ISL_1608282, EPI_ISL_1608283, EPI_ISL_1608284, EPI_ISL_1608285, EPI_ISL_1608287, EPI_ISL_1608288, EPI_ISL_1608289, EPI_ISL_1608290, EPI_ISL_1608291, EPI_ISL_1608292, EPI_ISL_1608293, EPI_ISL_1608294, EPI_ISL_1608295, EPI_ISL_1608296, EPI_ISL_1608297, EPI_ISL_1608298, EPI_ISL_1608299, EPI_ISL_1608300, EPI_ISL_1608302, EPI_ISL_1608303, EPI_ISL_1608304, EPI_ISL_1608305, EPI_ISL_1608306, EPI_ISL_1608307, EPI_ISL_1608308, EPI_ISL_1608309, EPI_ISL_1608310, EPI_ISL_1608311, EPI_ISL_1608312, EPI_ISL_1608313, EPI_ISL_1608314, EPI_ISL_1608315, EPI_ISL_1608316, EPI_ISL_1608317, EPI_ISL_1608318, EPI_ISL_1608319, EPI_ISL_1608320, EPI_ISL_1608321, EPI_ISL_1608322, EPI_ISL_1608324, EPI_ISL_1608325, EPI_ISL_1608326, EPI_ISL_1608329, EPI_ISL_1608332, EPI_ISL_1608333, EPI_ISL_1608334, EPI_ISL_1608335, EPI_ISL_1608336, EPI_ISL_1608337, EPI_ISL_1608341, EPI_ISL_1608342, EPI_ISL_1608343, EPI_ISL_1608344, EPI_ISL_1608345, EPI_ISL_1608350, EPI_ISL_1608353, EPI_ISL_1608357, EPI_ISL_1608358, EPI_ISL_1608359, EPI_ISL_1608360, EPI_ISL_1608363, EPI_ISL_1608364, EPI_ISL_1608366, EPI_ISL_1608367, EPI_ISL_1608369, EPI_ISL_1608372, EPI_ISL_1608373, EPI_ISL_1608374, EPI_ISL_1608376, EPI_ISL_1608377, EPI_ISL_1608378, EPI_ISL_1608381, EPI_ISL_1608382, EPI_ISL_1608383, EPI_ISL_1608384, EPI_ISL_1608385, EPI_ISL_1608386, EPI_ISL_1608388, EPI_ISL_1608390, EPI_ISL_1608391, EPI_ISL_1608392, EPI_ISL_1608394, EPI_ISL_1608395, EPI_ISL_1608396, EPI_ISL_1608397, EPI_ISL_1608398, EPI_ISL_1608399, EPI_ISL_1608400, EPI_ISL_1608401, EPI_ISL_1608402, EPI_ISL_1608403, EPI_ISL_1608404, EPI_ISL_1608405, EPI_ISL_1608406, EPI_ISL_1608407, EPI_ISL_1608408, EPI_ISL_1608410, EPI_ISL_1608411, EPI_ISL_1608412, EPI_ISL_1608413, EPI_ISL_1608414, EPI_ISL_1608415, EPI_ISL_1608416, EPI_ISL_1608417, EPI_ISL_1608418, EPI_ISL_1608419, EPI_ISL_1608420, EPI_ISL_1608421, EPI_ISL_1608422, EPI_ISL_1608424, EPI_ISL_1608425, EPI_ISL_1608426, EPI_ISL_1608427, EPI_ISL_1608428, EPI_ISL_1608430, EPI_ISL_1608431, EPI_ISL_1608432, EPI_ISL_1608433, EPI_ISL_1608434, EPI_ISL_1608435, EPI_ISL_1608436, EPI_ISL_1608437, EPI_ISL_1608438, EPI_ISL_1608439, EPI_ISL_1608440, EPI_ISL_1608442, EPI_ISL_1608444, EPI_ISL_1608445, EPI_ISL_1608447, EPI_ISL_1608448, EPI_ISL_1608449, EPI_ISL_1608450, EPI_ISL_1608451, EPI_ISL_1608452, EPI_ISL_1608453, EPI_ISL_1608454, EPI_ISL_1608455, EPI_ISL_1608456, EPI_ISL_1608457, EPI_ISL_1608458, EPI_ISL_1608459, EPI_ISL_1608460, EPI_ISL_1608461, EPI_ISL_1608462, EPI_ISL_1608463, EPI_ISL_1608464, EPI_ISL_1608465, EPI_ISL_1608466, EPI_ISL_1608468, EPI_ISL_1608470, EPI_ISL_1608471, EPI_ISL_1608472, EPI_ISL_1608473, EPI_ISL_1608474, EPI_ISL_1608475, EPI_ISL_1608476, EPI_ISL_1608477, EPI_ISL_1608478, EPI_ISL_1608479, EPI_ISL_1608480, EPI_ISL_1608481, EPI_ISL_1608482, EPI_ISL_1608483, EPI_ISL_1608484, EPI_ISL_1608485, EPI_ISL_1608487, EPI_ISL_1608488, EPI_ISL_1608489, EPI_ISL_1608490, EPI_ISL_1608492, EPI_ISL_1608493, EPI_ISL_1608494, EPI_ISL_1608495, EPI_ISL_1608496, EPI_ISL_1608497, EPI_ISL_1608499, EPI_ISL_1608500, EPI_ISL_1608502, EPI_ISL_1608503, EPI_ISL_1608507, EPI_ISL_1608508, EPI_ISL_1608509, EPI_ISL_1608511, EPI_ISL_1608512, EPI_ISL_1608513, EPI_ISL_1608514, EPI_ISL_1608515, EPI_ISL_1608524, EPI_ISL_1608525, EPI_ISL_1608526, EPI_ISL_1608527, EPI_ISL_1608528, EPI_ISL_1608529, EPI_ISL_1608530, EPI_ISL_1608531, EPI_ISL_1608533, EPI_ISL_1608534, EPI_ISL_1608535, EPI_ISL_1608536, EPI_ISL_1608537, EPI_ISL_1608538, EPI_ISL_1608539, EPI_ISL_1608540, EPI_ISL_1608541, EPI_ISL_1608542, EPI_ISL_1608543, EPI_ISL_1608545, EPI_ISL_1608547, EPI_ISL_1608548, EPI_ISL_1608549, EPI_ISL_1608550, EPI_ISL_1608551, EPI_ISL_1608552, EPI_ISL_1608553, EPI_ISL_1608556, EPI_ISL_1608557, EPI_ISL_1608558, EPI_ISL_1608559, EPI_ISL_1608560, EPI_ISL_1608561, EPI_ISL_1608562, EPI_ISL_1608563, EPI_ISL_1608564, EPI_ISL_1608566, EPI_ISL_1608567 | | | |
| see above | National Center of Infectious and Parasitic Diseases | National Center of Infectious and Parasitic Diseases | Alexiev et al |
| EPI_ISL_1615556 | Basurto University Hospital: Clinical Microbiology Laboratory | Biocruces Bizkaia | Mikel Urrutikoetxea-Gutierrez, Mª Carmen Nieto Toboso, Estibaliz Ugalde Zarraga, Mikel Gallego Rodrigo, Ana de la Hoz, José Luis Díaz de Tuesta del Arco. |

| | | | |
|---|---|---|---|
| EPI_ISL_1615557 | Basurto University Hospital: Clinical Microbiology Laboratory | Biocruces Bizkaia | Mikel Urrutikoetxea-Gutierrez, Mª Carmen Nieto Toboso, Estibaliz Ugalde Zarraga, Mikel Gallego Rodrigo, Ana de la Hoz, José Luis Díaz de Tuesta del Arco |
| EPI_ISL_1620175 | MRCG at LSHTM Genomics lab | MRCG at LSHTM Genomics lab | Abdul Karim sesay, Abdoulie Kanteh, Jarra Manneh, Mariama Kujabi, Bakary Sanyang |
| EPI_ISL_1620536, EPI_ISL_1620537, EPI_ISL_1620546, EPI_ISL_1620547, EPI_ISL_1620548, EPI_ISL_1620549, EPI_ISL_1620550, EPI_ISL_1620551, EPI_ISL_1620552, EPI_ISL_1620553, EPI_ISL_1620580, EPI_ISL_1620594, EPI_ISL_1620595, EPI_ISL_1620596, EPI_ISL_1620597, EPI_ISL_1620598, EPI_ISL_1620599, EPI_ISL_1620600, EPI_ISL_1620601, EPI_ISL_1620602, EPI_ISL_1620603, EPI_ISL_1620604, EPI_ISL_1620605, EPI_ISL_1620606, EPI_ISL_1620609, EPI_ISL_1620610, EPI_ISL_1620611, EPI_ISL_1620612, EPI_ISL_1620613, EPI_ISL_1620614, EPI_ISL_1620615, EPI_ISL_1620616, EPI_ISL_1620617, EPI_ISL_1620618, EPI_ISL_1620619, EPI_ISL_1620620, EPI_ISL_1620621, EPI_ISL_1620622, EPI_ISL_1620623, EPI_ISL_1620624, EPI_ISL_1620625, EPI_ISL_1620626, EPI_ISL_1620627 | | | |
| see above | Puerto Rico Department of Health | Centers for Disease Control and Prevention, Dengue Branch | Gilberto A. Santiago, Glenda Gonzalez, Betzabel Flores, Keyla Charriez, Gabriela Paz-Bailey, Jorge L. Munoz-Jordan |
| EPI_ISL_1621485 | Hospital | National Reference Center for Viruses of Respiratory Infections, Institut Pasteur, Paris | Marion Barbet, Sylvie Behillil, Frédéric Lemoine, Corinne Maufrais, Christophe Malabat, Méline Bizard, Angela Brisebarre, Camille Capel, Louise Lefrançois, Etienne Simon-Lorière, Vincent Enouf, Maud Vanpeene, Sylvie van der Werf,JéRome Guinard |
| EPI_ISL_1621486 | Hospital | National Reference Center for Viruses of Respiratory Infections, Institut Pasteur, Paris | Marion Barbet, Sylvie Behillil, Frédéric Lemoine, Corinne Maufrais, Christophe Malabat, Emmanuelle Permal, Méline Bizard, Angela Brisebarre, Camille Capel, Louise Lefrançois, Etienne Simon-Lorière, Vincent Enouf, Maud Vanpeene, Sylvie van der Werf,JéRome Guinard |
| EPI_ISL_1621487, EPI_ISL_1621488, EPI_ISL_1621489 | Hospital | National Reference Center for Viruses of Respiratory Infections, Institut Pasteur, Paris | Marion Barbet, Sylvie Behillil, Frédéric Lemoine, Corinne Maufrais, Christophe Malabat, Méline Bizard, Angela Brisebarre, Camille Capel, Louise Lefrançois, Etienne Simon-Lorière, Vincent Enouf, Maud Vanpeene, Sylvie van der Werf,JéRome Guinard |
| EPI_ISL_1621491 | Hospital | National Reference Center for Viruses of Respiratory Infections, Institut Pasteur, Paris | Marion Barbet, Sylvie Behillil, Frédéric Lemoine, Corinne Maufrais, Christophe Malabat, Amaury Vaysse, Méline Bizard, Angela Brisebarre, Camille Capel, Louise Lefrançois, Etienne Simon-Lorière, Vincent Enouf, Maud Vanpeene, Sylvie van der Werf,Pascale Martres |
| EPI_ISL_1621492, EPI_ISL_1621493, EPI_ISL_1621495, EPI_ISL_1621496, EPI_ISL_1621497 | Hospital | National Reference Center for Viruses of Respiratory Infections, Institut Pasteur, Paris | Marion Barbet, Sylvie Behillil, Frédéric Lemoine, Corinne Maufrais, Christophe Malabat, Méline Bizard, Angela Brisebarre, Camille Capel, Louise Lefrançois, Etienne Simon-Lorière, Vincent Enouf, Maud Vanpeene, Sylvie van der Werf,Pascale Martres |
| EPI_ISL_1621498, EPI_ISL_1621499, EPI_ISL_1621500 | Hospital | National Reference Center for Viruses of Respiratory Infections, Institut Pasteur, Paris | Marion Barbet, Sylvie Behillil, Frédéric Lemoine, Corinne Maufrais, Christophe Malabat, Méline Bizard, Angela Brisebarre, Camille Capel, Louise Lefrançois, Etienne Simon-Lorière, Vincent Enouf, Maud Vanpeene, Sylvie van der Werf,Marie-Pierre Weillaert |
| EPI_ISL_1621501 | Hospital | National Reference Center for Viruses of Respiratory Infections, Institut Pasteur, Paris | Marion Barbet, Sylvie Behillil, Frédéric Lemoine, Corinne Maufrais, Christophe Malabat, Méline Bizard, Angela Brisebarre, Camille Capel, Louise Lefrançois, Etienne Simon-Lorière, Vincent Enouf, Maud Vanpeene, Sylvie van der Werf,Sandrine Castelain |
| EPI_ISL_1621502, EPI_ISL_1621503 | Hospital | National Reference Center for Viruses of Respiratory Infections, Institut Pasteur, Paris | Marion Barbet, Sylvie Behillil, Frédéric Lemoine, Corinne Maufrais, Christophe Malabat, Amaury Vaysse, Méline Bizard, Angela Brisebarre, Camille Capel, Louise Lefrançois, Etienne Simon-Lorière, Vincent Enouf, Maud Vanpeene, Sylvie van der Werf,Sandrine Castelain |
| EPI_ISL_1621504, EPI_ISL_1621505, EPI_ISL_1621506, EPI_ISL_1621507, EPI_ISL_1621509 | Hospital | National Reference Center for Viruses of Respiratory Infections, Institut Pasteur, Paris | Marion Barbet, Sylvie Behillil, Frédéric Lemoine, Corinne Maufrais, Christophe Malabat, Méline Bizard, Angela Brisebarre, Camille Capel, Louise Lefrançois, Etienne Simon-Lorière, Vincent Enouf, Maud Vanpeene, Sylvie van der Werf,Sandrine Castelain |
| EPI_ISL_1621510, EPI_ISL_1621511 | Hospital | National Reference Center for Viruses of Respiratory Infections, Institut Pasteur, Paris | Marion Barbet, Sylvie Behillil, Frédéric Lemoine, Corinne Maufrais, Christophe Malabat, Amaury Vaysse, Méline Bizard, Angela Brisebarre, Camille Capel, Louise Lefrançois, Etienne Simon-Lorière, Vincent Enouf, Maud Vanpeene, Sylvie van der Werf,Isabelle Woercel |
| EPI_ISL_1621512 | Hospital | National Reference Center for Viruses of Respiratory Infections, Institut Pasteur, Paris | Marion Barbet, Sylvie Behillil, Frédéric Lemoine, Corinne Maufrais, Christophe Malabat, Amaury Vaysse, Méline Bizard, Angela Brisebarre, Camille Capel, Louise Lefrançois, Etienne Simon-Lorière, Vincent Enouf, Maud Vanpeene, Sylvie van der Werf,Dominique Descamps |
| EPI_ISL_1621513, EPI_ISL_1621514, EPI_ISL_1621515 | Hospital | National Reference Center for Viruses of Respiratory Infections, Institut Pasteur, Paris | Marion Barbet, Sylvie Behillil, Frédéric Lemoine, Corinne Maufrais, Christophe Malabat, Méline Bizard, Angela Brisebarre, Camille Capel, Louise Lefrançois, Etienne Simon-Lorière, Vincent Enouf, Maud Vanpeene, Sylvie van der Werf,Dominique Descamps |
| EPI_ISL_1621516 | Hospital | National Reference Center for Viruses of Respiratory Infections, Institut Pasteur, Paris | Marion Barbet, Sylvie Behillil, Frédéric Lemoine, Corinne Maufrais, Christophe Malabat, Amaury Vaysse, Méline Bizard, Angela Brisebarre, Camille Capel, Louise Lefrançois, Etienne Simon-Lorière, Vincent Enouf, Maud Vanpeene, Sylvie van der Werf,Dominique Descamps |
| EPI_ISL_1621517 | Hospital | National Reference Center for Viruses of Respiratory Infections, Institut Pasteur, Paris | Marion Barbet, Sylvie Behillil, Frédéric Lemoine, Corinne Maufrais, Christophe Malabat, Emmanuelle Permal, Méline Bizard, Angela Brisebarre, Camille Capel, Louise Lefrançois, Etienne Simon-Lorière, Vincent Enouf, Maud Vanpeene, Sylvie van der Werf,Dominique Descamps |
| EPI_ISL_1621518 | Hospital | National Reference Center for Viruses of Respiratory Infections, Institut Pasteur, Paris | Marion Barbet, Sylvie Behillil, Frédéric Lemoine, Corinne Maufrais, Christophe Malabat, Méline Bizard, Angela Brisebarre, Camille Capel, Louise Lefrançois, Etienne Simon-Lorière, Vincent Enouf, Maud Vanpeene, Sylvie van der Werf,Eric Farfour |
| EPI_ISL_1621519 | Hospital | National Reference Center for Viruses of Respiratory Infections, Institut Pasteur, Paris | Marion Barbet, Sylvie Behillil, Frédéric Lemoine, Corinne Maufrais, Christophe Malabat, Amaury Vaysse, Méline Bizard, Angela Brisebarre, Camille Capel, Louise Lefrançois, Etienne Simon-Lorière, Vincent Enouf, Maud Vanpeene, Sylvie van der Werf,Eric Farfour |
| EPI_ISL_1621520, EPI_ISL_1621521, EPI_ISL_1621522 | Hospital | National Reference Center for Viruses of Respiratory Infections, Institut Pasteur, Paris | Marion Barbet, Sylvie Behillil, Frédéric Lemoine, Corinne Maufrais, Christophe Malabat, Méline Bizard, Angela Brisebarre, Camille Capel, Louise Lefrançois, Etienne Simon-Lorière, Vincent Enouf, Maud Vanpeene, Sylvie van der Werf,Eric Farfour |
| EPI_ISL_1621523, EPI_ISL_1621524 | Hospital | National Reference Center for Viruses of Respiratory Infections, Institut Pasteur, Paris | Marion Barbet, Sylvie Behillil, Frédéric Lemoine, Corinne Maufrais, Christophe Malabat, Méline Bizard, Angela Brisebarre, Camille Capel, Louise Lefrançois, Etienne Simon-Lorière, Vincent Enouf, Maud Vanpeene, Sylvie van der Werf,Pierre Alain Billy |
| EPI_ISL_1621525, EPI_ISL_1621526, EPI_ISL_1621529, EPI_ISL_1621530 | Hospital | National Reference Center for Viruses of Respiratory Infections, Institut Pasteur, Paris | Marion Barbet, Sylvie Behillil, Frédéric Lemoine, Corinne Maufrais, Christophe Malabat, Méline Bizard, Angela Brisebarre, Camille Capel, Louise Lefrançois, Etienne Simon-Lorière, Vincent Enouf, Maud Vanpeene, Sylvie van der Werf,Marine Desroches |
| EPI_ISL_1621531, EPI_ISL_1621532, EPI_ISL_1621533 | Hospital | National Reference Center for Viruses of Respiratory Infections, Institut Pasteur, Paris | Marion Barbet, Sylvie Behillil, Frédéric Lemoine, Corinne Maufrais, Christophe Malabat, Méline Bizard, Angela Brisebarre, Camille Capel, Louise Lefrançois, Etienne Simon-Lorière, Vincent Enouf, Maud Vanpeene, Sylvie van der Werf,CéCile Farrugia |
| EPI_ISL_1621534, EPI_ISL_1621535, EPI_ISL_1621536, EPI_ISL_1621537 | Hospital | National Reference Center for Viruses of Respiratory Infections, Institut Pasteur, Paris | Marion Barbet, Sylvie Behillil, Frédéric Lemoine, Corinne Maufrais, Christophe Malabat, Méline Bizard, Angela Brisebarre, Camille Capel, Louise Lefrançois, Etienne Simon-Lorière, Vincent Enouf, Maud Vanpeene, Sylvie van der Werf,StéPhanie Marque Juillet |
| EPI_ISL_1621538 | Hospital | National Reference Center for Viruses of Respiratory Infections, Institut Pasteur, Paris | Marion Barbet, Sylvie Behillil, Frédéric Lemoine, Corinne Maufrais, Christophe Malabat, Méline Bizard, Angela Brisebarre, Camille Capel, Louise Lefrançois, Etienne Simon-Lorière, Vincent Enouf, Maud Vanpeene, Sylvie van der Werf,CéCile Farrugia |
| EPI_ISL_1621539, EPI_ISL_1621540 | Hospital | National Reference Center for Viruses of Respiratory Infections, Institut Pasteur, Paris | Marion Barbet, Sylvie Behillil, Frédéric Lemoine, Corinne Maufrais, Christophe Malabat, Méline Bizard, Angela Brisebarre, Camille Capel, Louise Lefrançois, Etienne Simon-Lorière, Vincent Enouf, Maud Vanpeene, Sylvie van der Werf,AuréLie Emirian |
| EPI_ISL_1621541, EPI_ISL_1621542, EPI_ISL_1621543, EPI_ISL_1621544, EPI_ISL_1621545 | Hospital | National Reference Center for Viruses of Respiratory Infections, Institut Pasteur, Paris | Marion Barbet, Sylvie Behillil, Frédéric Lemoine, Corinne Maufrais, Christophe Malabat, Méline Bizard, Angela Brisebarre, Camille Capel, Louise Lefrançois, Etienne Simon-Lorière, Vincent Enouf, Maud Vanpeene, Sylvie van der Werf,Sylvie Ledru |
| EPI_ISL_1621546 | Hospital | National Reference Center for Viruses of Respiratory Infections, Institut Pasteur, Paris | Marion Barbet, Sylvie Behillil, Frédéric Lemoine, Corinne Maufrais, Christophe Malabat, Méline Bizard, Angela Brisebarre, Camille Capel, Louise Lefrançois, Etienne Simon-Lorière, Vincent Enouf, Maud Vanpeene, Sylvie van der Werf,Jp Gangneux |
| EPI_ISL_1621547 | Hospital | National Reference Center for Viruses of Respiratory Infections, Institut Pasteur, Paris | Marion Barbet, Sylvie Behillil, Frédéric Lemoine, Corinne Maufrais, Christophe Malabat, Méline Bizard, Angela Brisebarre, Camille Capel, Louise Lefrançois, Etienne Simon-Lorière, Vincent Enouf, Maud Vanpeene, Sylvie van der Werf,Franck Bernardi |
| EPI_ISL_1621548 | Hospital | National Reference Center for Viruses of Respiratory Infections, Institut Pasteur, Paris | Marion Barbet, Sylvie Behillil, Frédéric Lemoine, Corinne Maufrais, Christophe Malabat, Méline Bizard, Angela Brisebarre, Camille Capel, Louise Lefrançois, Etienne Simon-Lorière, Vincent Enouf, Maud Vanpeene, Sylvie van der Werf,Pierre-Yves Meudec |
| EPI_ISL_1621549, EPI_ISL_1621550, EPI_ISL_1621551, EPI_ISL_1621552, EPI_ISL_1621553 | Labo Analyses Med | National Reference Center for Viruses of Respiratory Infections, Institut Pasteur, Paris | Marion Barbet, Sylvie Behillil, Frédéric Lemoine, Corinne Maufrais, Christophe Malabat, Méline Bizard, Angela Brisebarre, Camille Capel, Louise Lefrançois, Etienne Simon-Lorière, Vincent Enouf, Maud Vanpeene, Sylvie van der Werf,Jean-Philippe Memain |
| EPI_ISL_1621554 | Armees | National Reference Center for Viruses of Respiratory Infections, Institut Pasteur, Paris | Marion Barbet, Sylvie Behillil, Frédéric Lemoine, Corinne Maufrais, Christophe Malabat, Méline Bizard, Angela Brisebarre, Camille Capel, Louise Lefrançois, Etienne Simon-Lorière, Vincent Enouf, Maud Vanpeene, Sylvie van der Werf,Janvier |
| EPI_ISL_1621555, EPI_ISL_1621556, EPI_ISL_1621557, EPI_ISL_1621558 | Hospital | National Reference Center for Viruses of Respiratory Infections, Institut Pasteur, Paris | Marion Barbet, Sylvie Behillil, Frédéric Lemoine, Corinne Maufrais, Christophe Malabat, Méline Bizard, Angela Brisebarre, Camille Capel, Louise Lefrançois, Etienne Simon-Lorière, Vincent Enouf, Maud Vanpeene, Sylvie van der Werf,Mj Dufour |

| | | | |
|---|---|---|---|
| EPI_ISL_1621559, EPI_ISL_1621560 | Labo Analyses Med | National Reference Center for Viruses of Respiratory Infections, Institut Pasteur, Paris | Marion Barbet, Sylvie Behillil, Frédéric Lemoine, Corinne Maufrais, Christophe Malabat, Méline Bizard, Angela Brisebarre, Camille Capel, Louise Lefrançois, Etienne Simon-Lorière, Vincent Enouf, Maud Vanpeene, Sylvie van der Werf,Ronan Pichard |
| EPI_ISL_1621561 | Labo Analyses Med | National Reference Center for Viruses of Respiratory Infections, Institut Pasteur, Paris | Marion Barbet, Sylvie Behillil, Frédéric Lemoine, Corinne Maufrais, Christophe Malabat, Méline Bizard, Angela Brisebarre, Camille Capel, Louise Lefrançois, Etienne Simon-Lorière, Vincent Enouf, Maud Vanpeene, Sylvie van der Werf |
| EPI_ISL_1621564, EPI_ISL_1621565, EPI_ISL_1621566, EPI_ISL_1621567, EPI_ISL_1621568, EPI_ISL_1621569, EPI_ISL_1621571, EPI_ISL_1621572, EPI_ISL_1621573, EPI_ISL_1621574 | Labo Analyses Med | National Reference Center for Viruses of Respiratory Infections, Institut Pasteur, Paris | Marion Barbet, Sylvie Behillil, Frédéric Lemoine, Corinne Maufrais, Christophe Malabat, Méline Bizard, Angela Brisebarre, Camille Capel, Louise Lefrançois, Etienne Simon-Lorière, Vincent Enouf, Maud Vanpeene, Sylvie van der Werf,Ronan Pichard |
| EPI_ISL_1621575 | Labo Analyses Med | National Reference Center for Viruses of Respiratory Infections, Institut Pasteur, Paris | Marion Barbet, Sylvie Behillil, Frédéric Lemoine, Corinne Maufrais, Christophe Malabat, Amaury Vaysse, Méline Bizard, Angela Brisebarre, Camille Capel, Louise Lefrançois, Etienne Simon-Lorière, Vincent Enouf, Maud Vanpeene, Sylvie van der Werf,Ronan Pichard |
| EPI_ISL_1621576, EPI_ISL_1621577, EPI_ISL_1621578, EPI_ISL_1621579, EPI_ISL_1621580, EPI_ISL_1621581, EPI_ISL_1621582, EPI_ISL_1621583, EPI_ISL_1621584, EPI_ISL_1621585, EPI_ISL_1621586, EPI_ISL_1621587, EPI_ISL_1621588, EPI_ISL_1621589, EPI_ISL_1621590 | | | |
| see above | Labo Analyses Med | National Reference Center for Viruses of Respiratory Infections, Institut Pasteur, Paris | Marion Barbet, Sylvie Behillil, Frédéric Lemoine, Corinne Maufrais, Christophe Malabat, Méline Bizard, Angela Brisebarre, Camille Capel, Louise Lefrançois, Etienne Simon-Lorière, Vincent Enouf, Maud Vanpeene, Sylvie van der Werf,Ronan Pichard |
| EPI_ISL_1622417 | RS Permata Cibubur | Eijkman Institute for Molecular Biology, Ministry of Research and Technology/National Agency for Research and Innovation | Frilasita A Yudhaputri, Muhammad Rezki Rasyak, Willy Agustine, Hidayat Trimarsanto, Lydia V. Panggalo, Iskandar Adnan, Sukma Oktavianthi, Edison Johar, Safarina G Malik, Khin Saw Myint, Amin Soebandrio |
| EPI_ISL_1622418 | RS Qadr | Eijkman Institute for Molecular Biology, Ministry of Research and Technology/National Agency for Research and Innovation | Frilasita A Yudhaputri, Muhammad Rezki Rasyak, Willy Agustine, Hidayat Trimarsanto, Lydia V. Panggalo, Iskandar Adnan, Sukma Oktavianthi, Edison Johar, Safarina G Malik, Khin Saw Myint, Amin Soebandrio |
| EPI_ISL_1622420 | RS Sentra Medika Cikarang | Eijkman Institute for Molecular Biology, Ministry of Research and Technology/National Agency for Research and Innovation | Frilasita A Yudhaputri, Muhammad Rezki Rasyak, Willy Agustine, Hidayat Trimarsanto, Lydia V. Panggalo, Iskandar Adnan, Sukma Oktavianthi, Edison Johar, Safarina G Malik, Khin Saw Myint, Amin Soebandrio |
| EPI_ISL_1622421, EPI_ISL_1622422 | RS Medirossa Cikarang | Eijkman Institute for Molecular Biology, Ministry of Research and Technology/National Agency for Research and Innovation | Frilasita A Yudhaputri, Muhammad Rezki Rasyak, Willy Agustine, Hidayat Trimarsanto, Lydia V. Panggalo, Iskandar Adnan, Sukma Oktavianthi, Edison Johar, Safarina G Malik, Khin Saw Myint, Amin Soebandrio |
| EPI_ISL_1622423 | RSJ Dr. Soeharto Heerdjan | Eijkman Institute for Molecular Biology, Ministry of Research and Technology/National Agency for Research and Innovation | Frilasita A Yudhaputri, Muhammad Rezki Rasyak, Willy Agustine, Hidayat Trimarsanto, Lydia V. Panggalo, Iskandar Adnan, Sukma Oktavianthi, Edison Johar, Safarina G Malik, Khin Saw Myint, Amin Soebandrio |
| EPI_ISL_1622424 | RS Sumber Waras | Eijkman Institute for Molecular Biology, Ministry of Research and Technology/National Agency for Research and Innovation | Muhammad Rezki Rasyak, Willy Agustine, Hidayat Trimarsanto, Lydia V. Panggalo, Iskandar Adnan, Sukma Oktavianthi, Edison Johar, Frilasita A Yudhaputri, Safarina G Malik, Khin Saw Myint, Amin Soebandrio |
| EPI_ISL_1622425 | RS Mitra Keluarga Kelapa Gading | Eijkman Institute for Molecular Biology, Ministry of Research and Technology/National Agency for Research and Innovation | Muhammad Rezki Rasyak, Willy Agustine, Hidayat Trimarsanto, Lydia V. Panggalo, Iskandar Adnan, Sukma Oktavianthi, Edison Johar, Frilasita A Yudhaputri, Safarina G Malik, Khin Saw Myint, Amin Soebandrio |
| EPI_ISL_1622426 | RS Mary Cileungsi Hijau | Eijkman Institute for Molecular Biology, Ministry of Research and Technology/National Agency for Research and Innovation | Muhammad Rezki Rasyak, Willy Agustine, Hidayat Trimarsanto, Lydia V. Panggalo, Iskandar Adnan, Sukma Oktavianthi, Edison Johar, Frilasita A Yudhaputri, Safarina G Malik, Khin Saw Myint, Amin Soebandrio |
| EPI_ISL_1622427 | RS OMNI Pulomas | Eijkman Institute for Molecular Biology, Ministry of Research and Technology/National Agency for Research and Innovation | Muhammad Rezki Rasyak, Willy Agustine, Hidayat Trimarsanto, Lydia V. Panggalo, Iskandar Adnan, Sukma Oktavianthi, Edison Johar, Frilasita A Yudhaputri, Safarina G Malik, Khin Saw Myint, Amin Soebandrio |
| EPI_ISL_1622430 | RS Mitra Keluarga Gading Serpong | Eijkman Institute for Molecular Biology, Ministry of Research and Technology/National Agency for Research and Innovation | Muhammad Rezki Rasyak, Willy Agustine, Hidayat Trimarsanto, Lydia V. Panggalo, Iskandar Adnan, Sukma Oktavianthi, Edison Johar, Frilasita A Yudhaputri, Safarina G Malik, Khin Saw Myint, Amin Soebandrio |
| EPI_ISL_1622432 | RS Sentra Medika Cikarang | Eijkman Institute for Molecular Biology, Ministry of Research and Technology/National Agency for Research and Innovation | Muhammad Rezki Rasyak, Willy Agustine, Hidayat Trimarsanto, Lydia V. Panggalo, Iskandar Adnan, Sukma Oktavianthi, Edison Johar, Frilasita A Yudhaputri, Safarina G Malik, Khin Saw Myint, Amin Soebandrio |
| EPI_ISL_1622433 | RSJ Dr. Soeharto Heerdjan | Eijkman Institute for Molecular Biology, Ministry of Research and Technology/National Agency for Research and Innovation | Muhammad Rezki Rasyak, Willy Agustine, Hidayat Trimarsanto, Lydia V. Panggalo, Iskandar Adnan, Sukma Oktavianthi, Edison Johar, Frilasita A Yudhaputri, Safarina G Malik, Khin Saw Myint, Amin Soebandrio |
| EPI_ISL_1622435 | RSIA Selasih Medika | Eijkman Institute for Molecular Biology, Ministry of Research and Technology/National Agency for Research and Innovation | Lydia V. Panggalo, Iskandar Adnan, Sukma Oktavianthi, Edison Johar, Frilasita A Yudhaputri, Muhammad Rezki Rasyak, Willy Agustine, Hidayat Trimarsanto, Safarina G Malik, Khin Saw Myint, Amin Soebandrio |
| EPI_ISL_1623666, EPI_ISL_1623680, EPI_ISL_1623689 | National Platform bis UMONS/Jolimont | National Platform bis UMONS/Jolimont | François Dufrasne, Guillaume Bayon-Vicente, Gautier Detry, Ruddy Wattiez |
| EPI_ISL_1623763, EPI_ISL_1623767, EPI_ISL_1623768, EPI_ISL_1623769, EPI_ISL_1623770, EPI_ISL_1623773, EPI_ISL_1623774, EPI_ISL_1623775, EPI_ISL_1623776, EPI_ISL_1623777, EPI_ISL_1623778, EPI_ISL_1623779, EPI_ISL_1623780, EPI_ISL_1623781, EPI_ISL_1623782, EPI_ISL_1623784, EPI_ISL_1623785, EPI_ISL_1623786, EPI_ISL_1623787, EPI_ISL_1623789, EPI_ISL_1623790, EPI_ISL_1623791, EPI_ISL_1623792, EPI_ISL_1623793, EPI_ISL_1623794, EPI_ISL_1623795, EPI_ISL_1623796, EPI_ISL_1623797, EPI_ISL_1623798, EPI_ISL_1623799, EPI_ISL_1623800, EPI_ISL_1623801, EPI_ISL_1623802, EPI_ISL_1623804, EPI_ISL_1623805, EPI_ISL_1623806, EPI_ISL_1623807, EPI_ISL_1623808, EPI_ISL_1623809, EPI_ISL_1623811, EPI_ISL_1623812, EPI_ISL_1623813, EPI_ISL_1623814, EPI_ISL_1623815 | | | |
| see above | Baylor Scott & White-Temple | Baylor Scott & White-Temple | Ari Rao, Linden Morales, Kimberly Walker, Marcus Volz, Shelby Hendrickson, Caitlin Maloney |
| EPI_ISL_1625972 | Ama J Angela | Instituto Adolfo Lutz, Interdiciplinary Procedures Center, Strategic Laboratory | Claudio Tavares Sacchi, Claudia Regina Gonçalves, Erica Valessa Ramos Gomes, Karoline Rodrigues Campos, Caio Vinicius Dias Lopes, Leonardo Jose Tadeu de Araujo, Katia Correa de Oliveira Santos |
| EPI_ISL_1625973, EPI_ISL_1625974 | Santa Casa de Aracatuba Hospital Sagrado Coracao de Jesus | Instituto Adolfo Lutz, Interdiciplinary Procedures Center, Strategic Laboratory | Claudio Tavares Sacchi, Claudia Regina Gonçalves, Erica Valessa Ramos Gomes, Karoline Rodrigues Campos, Caio Vinicius Dias Lopes, Leonardo Jose Tadeu de Araujo, Katia Correa de Oliveira Santos |
| EPI_ISL_1625975 | CS II Dr Antonio Vicoso Moreira De Rezende Sumare | Instituto Adolfo Lutz, Interdiciplinary Procedures Center, Strategic Laboratory | Claudio Tavares Sacchi, Claudia Regina Gonçalves, Erica Valessa Ramos Gomes, Karoline Rodrigues Campos, Caio Vinicius Dias Lopes, Leonardo Jose Tadeu de Araujo, Katia Correa de Oliveira Santos |
| EPI_ISL_1625976 | Centro de Saude III de Divinolandia | Instituto Adolfo Lutz, Interdiciplinary Procedures Center, Strategic Laboratory | Claudio Tavares Sacchi, Claudia Regina Gonçalves, Erica Valessa Ramos Gomes, Karoline Rodrigues Campos, Caio Vinicius Dias Lopes, Leonardo Jose Tadeu de Araujo, Katia Correa de Oliveira Santos |
| EPI_ISL_1625987, EPI_ISL_1625988, EPI_ISL_1625989, EPI_ISL_1625990, EPI_ISL_1625991, EPI_ISL_1625992, EPI_ISL_1625993, EPI_ISL_1625994, EPI_ISL_1625995, EPI_ISL_1625996, EPI_ISL_1625997, EPI_ISL_1625998, EPI_ISL_1625999, EPI_ISL_1626000, EPI_ISL_1626001, EPI_ISL_1626002, EPI_ISL_1626003, EPI_ISL_1626004, EPI_ISL_1626005, EPI_ISL_1626006, EPI_ISL_1626007, EPI_ISL_1626008, EPI_ISL_1626606 | | | |
| see above | Instituto Adolfo Lutz - Regional de Rio Claro | Instituto Adolfo Lutz, Interdiciplinary Procedures Center, Strategic Laboratory | Claudio Tavares Sacchi, Claudia Regina Gonçalves, Erica Valessa Ramos Gomes, Karoline Rodrigues Campos, Caio Vinicius Dias Lopes, Leonardo Jose Tadeu de Araujo, Katia Correa de Oliveira Santos |
| EPI_ISL_1626794, EPI_ISL_1626795, EPI_ISL_1626796, EPI_ISL_1626797, EPI_ISL_1626798, EPI_ISL_1626799, EPI_ISL_1626800, EPI_ISL_1626801 | LESP Chihuahua | Instituto de Diagnostico y Referencia Epidemiologicos (INDRE) | Claudia Wong-Arambula, Abril Rodriguez-Maldonado, Vanessa Rivero-Arredondo, Ariadna Medina-Benitez, Joaquin Quiroz-Mercado, Sergio Rangel-Guerrero, Natividad Cruz-Ortiz, Tatiana Nunez-Garcia, Gisela Barrera-Badillo, Lucia Hernandez-Rivas, Irma Lopez-Martinez, Ernesto Ramirez-Gonzalez. |
| EPI_ISL_1626806 | LESP Nuevo Leon | Instituto de Diagnostico y Referencia Epidemiologicos (INDRE) | Claudia Wong-Arambula, Abril Rodriguez-Maldonado, Vanessa Rivero-Arredondo, Ariadna Medina-Benitez, Joaquin Quiroz-Mercado, Sergio Rangel-Guerrero, Natividad Cruz-Ortiz, Tatiana Nunez-Garcia, Gisela Barrera-Badillo, Lucia Hernandez-Rivas, Irma Lopez-Martinez, Ernesto Ramirez-Gonzalez. |
| EPI_ISL_1626814 | LESP Baja California Sur | Instituto de Diagnostico y Referencia Epidemiologicos (INDRE) | Claudia Wong-Arambula, Abril Rodriguez-Maldonado, Vanessa Rivero-Arredondo, Ariadna Medina-Benitez, Joaquin Quiroz-Mercado, Sergio Rangel-Guerrero, Natividad Cruz-Ortiz, Tatiana Nunez-Garcia, Gisela Barrera-Badillo, Lucia Hernandez-Rivas, Irma Lopez-Martinez, Ernesto Ramirez-Gonzalez. |
| EPI_ISL_1627343 | Presidio di Brindisi Di Summa - Perrino | Istituto Zooprofilattico Sperimentale della Puglia e della Basilicata | Parisi A., Bianco A., Capozzi L., Del Sambro L., Simone D., Difato L., Santoro A., Ridolfi D., Giannico A. |
| EPI_ISL_1627361 | Istituto Zooprofilattico Sperimentale della Puglia e della Basilicata | Istituto Zooprofilattico Sperimentale della Puglia e della Basilicata | Parisi A., Bianco A., Capozzi L., Del Sambro L., Simone D., Galante D., Pace L. |

| | | | |
|---|---|---|---|
| EPI_ISL_1628344 | Hospital Geral de Pedreira | Instituto Adolfo Lutz, Interdiciplinary Procedures Center, Strategic Laboratory | Claudio Tavares Sacchi, Claudia Regina Gonçalves, Erica Valessa Ramos Gomes, Karoline Rodrigues Campos, Caio Vinicius Dias Lopes, Leonardo Jose Tadeu de Araujo, Katia Correa de Oliveira Santos |
| EPI_ISL_1628347, EPI_ISL_1628348, EPI_ISL_1628349, EPI_ISL_1628350, EPI_ISL_1628351, EPI_ISL_1628352, EPI_ISL_1628353, EPI_ISL_1628354, EPI_ISL_1628355, EPI_ISL_1628356, EPI_ISL_1628357, EPI_ISL_1628358, EPI_ISL_1628359, EPI_ISL_1628360, EPI_ISL_1628361, EPI_ISL_1628362 | | | |
| see above | Instituto Adolfo Lutz Central | Instituto Adolfo Lutz, Interdiciplinary Procedures Center, Strategic Laboratory | Claudio Tavares Sacchi, Claudia Regina Gonçalves, Erica Valessa Ramos Gomes, Karoline Rodrigues Campos, Caio Vinicius Dias Lopes, Leonardo Jose Tadeu de Araujo, Katia Correa de Oliveira Santos |
| EPI_ISL_1628363, EPI_ISL_1628364, EPI_ISL_1628365 | LACEN do Estado de Goias | Instituto Adolfo Lutz, Interdiciplinary Procedures Center, Strategic Laboratory | Claudio Tavares Sacchi, Claudia Regina Gonçalves, Erica Valessa Ramos Gomes, Karoline Rodrigues Campos, Caio Vinicius Dias Lopes, Leonardo Jose Tadeu de Araujo, Katia Correa de Oliveira Santos |
| EPI_ISL_1628368 | Centro Medico Social Comunitario Januario Teodoro de Souza | Instituto Adolfo Lutz, Interdiciplinary Procedures Center, Strategic Laboratory | Claudio Tavares Sacchi, Claudia Regina Gonçalves, Erica Valessa Ramos Gomes, Karoline Rodrigues Campos, Caio Vinicius Dias Lopes, Leonardo Jose Tadeu de Araujo, Katia Correa de Oliveira Santos |
| EPI_ISL_1628369 | UPA Dr Luis Atilio Losi Viana Ribeirao Preto | Instituto Adolfo Lutz, Interdiciplinary Procedures Center, Strategic Laboratory | Claudio Tavares Sacchi, Claudia Regina Gonçalves, Erica Valessa Ramos Gomes, Karoline Rodrigues Campos, Caio Vinicius Dias Lopes, Leonardo Jose Tadeu de Araujo, Katia Correa de Oliveira Santos |
| EPI_ISL_1628370 | Sae Servico De Atendimento Especializado | Instituto Adolfo Lutz, Interdiciplinary Procedures Center, Strategic Laboratory | Claudio Tavares Sacchi, Claudia Regina Gonçalves, Erica Valessa Ramos Gomes, Karoline Rodrigues Campos, Caio Vinicius Dias Lopes, Leonardo Jose Tadeu de Araujo, Katia Correa de Oliveira Santos |
| EPI_ISL_1628371 | Centro De Saude II Ibitinga | Instituto Adolfo Lutz, Interdiciplinary Procedures Center, Strategic Laboratory | Claudio Tavares Sacchi, Claudia Regina Gonçalves, Erica Valessa Ramos Gomes, Karoline Rodrigues Campos, Caio Vinicius Dias Lopes, Leonardo Jose Tadeu de Araujo, Katia Correa de Oliveira Santos |
| EPI_ISL_1628372 | Hospital Sao Marcos da Sama Morro Agudo | Instituto Adolfo Lutz, Interdiciplinary Procedures Center, Strategic Laboratory | Claudio Tavares Sacchi, Claudia Regina Gonçalves, Erica Valessa Ramos Gomes, Karoline Rodrigues Campos, Caio Vinicius Dias Lopes, Leonardo Jose Tadeu de Araujo, Katia Correa de Oliveira Santos |
| EPI_ISL_1628374 | Santa Casa De Cravinhos | Instituto Adolfo Lutz, Interdiciplinary Procedures Center, Strategic Laboratory | Claudio Tavares Sacchi, Claudia Regina Gonçalves, Erica Valessa Ramos Gomes, Karoline Rodrigues Campos, Caio Vinicius Dias Lopes, Leonardo Jose Tadeu de Araujo, Katia Correa de Oliveira Santos |
| EPI_ISL_1628377 | Centro De Saude II Ibitinga | Instituto Adolfo Lutz, Interdiciplinary Procedures Center, Strategic Laboratory | Claudio Tavares Sacchi, Claudia Regina Gonçalves, Erica Valessa Ramos Gomes, Karoline Rodrigues Campos, Caio Vinicius Dias Lopes, Leonardo Jose Tadeu de Araujo, Katia Correa de Oliveira Santos |
| EPI_ISL_1628379 | UPA Dr Luis Atilio Losi Viana Ribeirao Preto | Instituto Adolfo Lutz, Interdiciplinary Procedures Center, Strategic Laboratory | Claudio Tavares Sacchi, Claudia Regina Gonçalves, Erica Valessa Ramos Gomes, Karoline Rodrigues Campos, Caio Vinicius Dias Lopes, Leonardo Jose Tadeu de Araujo, Katia Correa de Oliveira Santos |
| EPI_ISL_1628501, EPI_ISL_1628502, EPI_ISL_1628503, EPI_ISL_1628504, EPI_ISL_1628505, EPI_ISL_1628506, EPI_ISL_1628507, EPI_ISL_1628508, EPI_ISL_1628509, EPI_ISL_1628510, EPI_ISL_1628511, EPI_ISL_1628512, EPI_ISL_1628513, EPI_ISL_1628514, EPI_ISL_1628515, EPI_ISL_1628516, EPI_ISL_1628517, EPI_ISL_1628518 | | | |
| see above | National Center of Infectious and Parasitic Diseases | National Center of Infectious and Parasitic Diseases | Alexiev et al |
| EPI_ISL_1628933, EPI_ISL_1628938, EPI_ISL_1628940, EPI_ISL_1628942, EPI_ISL_1628943, EPI_ISL_1628944, EPI_ISL_1628945, EPI_ISL_1628947, EPI_ISL_1629014, EPI_ISL_1629030, EPI_ISL_1629031 | | | |
| see above | GHER | UMR PIMIT | Dr David A Wilkinson, Dr Patrick Mavingui, Dr Camille Lebarbenchon, Magali Turpin |
| EPI_ISL_1629054, EPI_ISL_1629102 | CHU | UMR PIMIT | Dr David A Wilkinson, Dr Patrick Mavingui, Dr Camille Lebarbenchon, Magali Turpin |
| EPI_ISL_1629117, EPI_ISL_1629121, EPI_ISL_1629124, EPI_ISL_1629125 | GHER | UMR PIMIT | Dr David A Wilkinson, Dr Patrick Mavingui, Dr Camille Lebarbenchon, Magali Turpin |
| EPI_ISL_1629145, EPI_ISL_1629151, EPI_ISL_1629152, EPI_ISL_1629153, EPI_ISL_1629154, EPI_ISL_1629155, EPI_ISL_1629160, EPI_ISL_1629163 | CERBALLIANCE | UMR PIMIT | Dr David A Wilkinson, Dr Patrick Mavingui, Dr Camille Lebarbenchon, Magali Turpin |
| EPI_ISL_1629166 | GHER | UMR PIMIT | Dr David A Wilkinson, Dr Patrick Mavingui, Dr Camille Lebarbenchon, Magali Turpin |
| EPI_ISL_1629175 | CHU | UMR PIMIT | Dr David A Wilkinson, Dr Patrick Mavingui, Dr Camille Lebarbenchon, Magali Turpin |
| EPI_ISL_1629178 | GHER | UMR PIMIT | Dr David A Wilkinson, Dr Patrick Mavingui, Dr Camille Lebarbenchon, Magali Turpin |
| EPI_ISL_1629184 | CHU | UMR PIMIT | Dr David A Wilkinson, Dr Patrick Mavingui, Dr Camille Lebarbenchon, Magali Turpin |
| EPI_ISL_1629209, EPI_ISL_1629210, EPI_ISL_1629211, EPI_ISL_1629213, EPI_ISL_1629214, EPI_ISL_1629216, EPI_ISL_1629218, EPI_ISL_1629222, EPI_ISL_1629223, EPI_ISL_1629224, EPI_ISL_1629229 | | | |
| see above | REUNILAB | UMR PIMIT | Dr David A Wilkinson, Dr Patrick Mavingui, Dr Camille Lebarbenchon, Magali Turpin |
| EPI_ISL_1629243 | CHU | UMR PIMIT | Dr David A Wilkinson, Dr Patrick Mavingui, Dr Camille Lebarbenchon, Magali Turpin |
| EPI_ISL_1629249 | CHOR | UMR PIMIT | Dr David A Wilkinson, Dr Patrick Mavingui, Dr Camille Lebarbenchon, Magali Turpin |
| EPI_ISL_1629251 | CHU NORD | UMR PIMIT | Dr David A Wilkinson, Dr Patrick Mavingui, Dr Camille Lebarbenchon, Magali Turpin |
| EPI_ISL_1629267, EPI_ISL_1629269, EPI_ISL_1629271, EPI_ISL_1629272, EPI_ISL_1629276, EPI_ISL_1629279 | REUNILAB | UMR PIMIT | Dr David A Wilkinson, Dr Patrick Mavingui, Dr Camille Lebarbenchon, Magali Turpin |
| EPI_ISL_1629291 | CERBALLIANCE | UMR PIMIT | Dr David A Wilkinson, Dr Patrick Mavingui, Dr Camille Lebarbenchon, Magali Turpin |
| EPI_ISL_1629304, EPI_ISL_1629305, EPI_ISL_1629308, EPI_ISL_1629311, EPI_ISL_1629312, EPI_ISL_1629314, EPI_ISL_1629318, EPI_ISL_1629322, EPI_ISL_1629329, EPI_ISL_1629331, EPI_ISL_1629332, EPI_ISL_1629335, EPI_ISL_1629340, EPI_ISL_1629425, EPI_ISL_1629429, EPI_ISL_1629431, EPI_ISL_1629436, EPI_ISL_1629439, EPI_ISL_1629441, EPI_ISL_1629443 | | | |
| see above | REUNILAB | UMR PIMIT | Dr David A Wilkinson, Dr Patrick Mavingui, Dr Camille Lebarbenchon, Magali Turpin |
| EPI_ISL_1629458 | CHU | UMR PIMIT | Dr David A Wilkinson, Dr Patrick Mavingui, Dr Camille Lebarbenchon, Magali Turpin |
| EPI_ISL_1629468, EPI_ISL_1629480 | REUNILAB | UMR PIMIT | Dr David A Wilkinson, Dr Patrick Mavingui, Dr Camille Lebarbenchon, Magali Turpin |
| EPI_ISL_1629809 | Laboratório de Microbiologia Molecular - Universidade FEEVALE | Molecular Microbiology Laboratory | Alana Witt Hansen, Fágner Henrique Heldt, Fernando Rosado Spilki, Flávio Silveira, Juliana Schons Gulart, Juliane Deise Fleck, Mariana Soares da Silva, Meriane Demoliner, Matheus Nunes Weber, Paula Rodrigues de Almeida, Michele Filippi. |
| EPI_ISL_1630278 | Clinical Microbiology Lab RSPTN Universitas Hasanuddin | Clinical Microbiology Lab RSPTN Universitas Hasanuddin | Muhammad Nasrum Massi, Rizalinda Sjahril, Gita Vita Soraya, Triwibawa, Handayani Halik, Muhammad Yusuf Usman, Rufika Shari |
| EPI_ISL_1651214, EPI_ISL_1651215 | CH La Roche sur Yon | CHU Nantes Virology | Celine Bressollette-Bodin, Thomas Drumel, Audrey Rodallec, Berthe-Marie Imbert-Marcille, Cecile Le Boteff |
| EPI_ISL_1651229, EPI_ISL_1651230 | LESP Chiapas | Instituto de Diagnostico y Referencia Epidemiologicos (INDRE) | Claudia Wong-Arambula, Abril Rodriguez-Maldonado, Vanessa Rivero-Arredondo, Ariadna Medina-Benitez, Joaquin Quiroz-Mercado, Sergio Rangel-Guerrero, Natividad Cruz-Ortiz, Tatiana Nunez-Garcia, Gisela Barrera-Badillo, Lucia Hernandez-Rivas, Irma Lopez-Martinez, Ernesto Ramirez-Gonzalez. |
| EPI_ISL_1651231, EPI_ISL_1651232, EPI_ISL_1651233, EPI_ISL_1651234 | LESP Puebla | Instituto de Diagnostico y Referencia Epidemiologicos (INDRE) | Claudia Wong-Arambula, Abril Rodriguez-Maldonado, Vanessa Rivero-Arredondo, Ariadna Medina-Benitez, Joaquin Quiroz-Mercado, Sergio Rangel-Guerrero, Natividad Cruz-Ortiz, Tatiana Nunez-Garcia, Gisela Barrera-Badillo, Lucia Hernandez-Rivas, Irma Lopez-Martinez, Ernesto Ramirez-Gonzalez. |
| EPI_ISL_1651235, EPI_ISL_1651236, EPI_ISL_1651237, EPI_ISL_1651238, EPI_ISL_1651239, EPI_ISL_1651240, EPI_ISL_1651241, EPI_ISL_1651242, EPI_ISL_1651243, EPI_ISL_1651244, EPI_ISL_1651245, EPI_ISL_1651246, EPI_ISL_1651247, EPI_ISL_1651248, EPI_ISL_1651249 | | | |
| see above | LESP Nuevo Leon | Instituto de Diagnostico y Referencia Epidemiologicos (INDRE) | Claudia Wong-Arambula, Abril Rodriguez-Maldonado, Vanessa Rivero-Arredondo, Ariadna Medina-Benitez, Joaquin Quiroz-Mercado, Sergio Rangel-Guerrero, Natividad Cruz-Ortiz, Tatiana Nunez-Garcia, Gisela Barrera-Badillo, Lucia Hernandez-Rivas, Irma Lopez-Martinez, Ernesto Ramirez-Gonzalez. |

| | | | |
|---|---|---|---|
| EPI_ISL_1651252 | LESP Oaxaca | Instituto de Diagnostico y Referencia Epidemiologicos (INDRE) | Claudia Wong-Arambula, Abril Rodriguez-Maldonado, Vanessa Rivero-Arredondo, Ariadna Medina-Benitez, Joaquin Quiroz-Mercado, Sergio Rangel-Guerrero, Natividad Cruz-Ortiz, Tatiana Nunez-Garcia, Gisela Barrera-Badillo, Lucia Hernandez-Rivas, Irma Lopez-Martinez, Ernesto Ramirez-Gonzalez. |
| EPI_ISL_1651253 | LESP Sinaloa | Instituto de Diagnostico y Referencia Epidemiologicos (INDRE) | Claudia Wong-Arambula, Abril Rodriguez-Maldonado, Vanessa Rivero-Arredondo, Ariadna Medina-Benitez, Joaquin Quiroz-Mercado, Sergio Rangel-Guerrero, Natividad Cruz-Ortiz, Tatiana Nunez-Garcia, Gisela Barrera-Badillo, Lucia Hernandez-Rivas, Irma Lopez-Martinez, Ernesto Ramirez-Gonzalez. |
| EPI_ISL_1651254 | LESP Sonora | Instituto de Diagnostico y Referencia Epidemiologicos (INDRE) | Claudia Wong-Arambula, Abril Rodriguez-Maldonado, Vanessa Rivero-Arredondo, Ariadna Medina-Benitez, Joaquin Quiroz-Mercado, Sergio Rangel-Guerrero, Natividad Cruz-Ortiz, Tatiana Nunez-Garcia, Gisela Barrera-Badillo, Lucia Hernandez-Rivas, Irma Lopez-Martinez, Ernesto Ramirez-Gonzalez. |
| EPI_ISL_1651255, EPI_ISL_1651256, EPI_ISL_1651257, EPI_ISL_1651258, EPI_ISL_1651259, EPI_ISL_1651260, EPI_ISL_1651261 | LESP Morelos | Instituto de Diagnostico y Referencia Epidemiologicos (INDRE) | Claudia Wong-Arambula, Abril Rodriguez-Maldonado, Vanessa Rivero-Arredondo, Ariadna Medina-Benitez, Joaquin Quiroz-Mercado, Sergio Rangel-Guerrero, Natividad Cruz-Ortiz, Tatiana Nunez-Garcia, Gisela Barrera-Badillo, Lucia Hernandez-Rivas, Irma Lopez-Martinez, Ernesto Ramirez-Gonzalez. |
| EPI_ISL_1651277, EPI_ISL_1651278 | LESP Queretaro | Instituto de Diagnostico y Referencia Epidemiologicos (INDRE) | Claudia Wong-Arambula, Abril Rodriguez-Maldonado, Vanessa Rivero-Arredondo, Ariadna Medina-Benitez, Joaquin Quiroz-Mercado, Sergio Rangel-Guerrero, Natividad Cruz-Ortiz, Tatiana Nunez-Garcia, Gisela Barrera-Badillo, Lucia Hernandez-Rivas, Irma Lopez-Martinez, Ernesto Ramirez-Gonzalez. |
| EPI_ISL_1651308, EPI_ISL_1651309, EPI_ISL_1651310 | CHU Nantes Virology | CHU Nantes Virology | Celine Bressollette-Bodin, Thomas Drumel, Audrey Rodallec, Berthe-Marie Imbert-Marcille |
| EPI_ISL_1651877, EPI_ISL_1651878, EPI_ISL_1651879, EPI_ISL_1651880 | LESP Chihuahua | Instituto de Diagnostico y Referencia Epidemiologicos (INDRE) | Claudia Wong-Arambula, Abril Rodriguez-Maldonado, Vanessa Rivero-Arredondo, Ariadna Medina-Benitez, Joaquin Quiroz-Mercado, Sergio Rangel-Guerrero, Natividad Cruz-Ortiz, Tatiana Nunez-Garcia, Gisela Barrera-Badillo, Lucia Hernandez-Rivas, Irma Lopez-Martinez, Ernesto Ramirez-Gonzalez. |
| EPI_ISL_1651881, EPI_ISL_1651882 | LESP Veracruz | Instituto de Diagnostico y Referencia Epidemiologicos (INDRE) | Claudia Wong-Arambula, Abril Rodriguez-Maldonado, Vanessa Rivero-Arredondo, Ariadna Medina-Benitez, Joaquin Quiroz-Mercado, Sergio Rangel-Guerrero, Natividad Cruz-Ortiz, Tatiana Nunez-Garcia, Gisela Barrera-Badillo, Lucia Hernandez-Rivas, Irma Lopez-Martinez, Ernesto Ramirez-Gonzalez. |
| EPI_ISL_1651887 | LESP Tamaulipas | Instituto de Diagnostico y Referencia Epidemiologicos (INDRE) | Claudia Wong-Arambula, Abril Rodriguez-Maldonado, Vanessa Rivero-Arredondo, Ariadna Medina-Benitez, Joaquin Quiroz-Mercado, Sergio Rangel-Guerrero, Natividad Cruz-Ortiz, Tatiana Nunez-Garcia, Gisela Barrera-Badillo, Lucia Hernandez-Rivas, Irma Lopez-Martinez, Ernesto Ramirez-Gonzalez. |
| EPI_ISL_1651888, EPI_ISL_1651899, EPI_ISL_1651902 | LESP Ciudad de Mexico | Instituto de Diagnostico y Referencia Epidemiologicos (INDRE) | Claudia Wong-Arambula, Abril Rodriguez-Maldonado, Vanessa Rivero-Arredondo, Ariadna Medina-Benitez, Joaquin Quiroz-Mercado, Sergio Rangel-Guerrero, Natividad Cruz-Ortiz, Tatiana Nunez-Garcia, Gisela Barrera-Badillo, Lucia Hernandez-Rivas, Irma Lopez-Martinez, Ernesto Ramirez-Gonzalez. |
| EPI_ISL_1653935 | Alea Genetic Center | Alea Genetic Center | Rijad Konjhodzic, Lana Salihefendic, Dino Pecar, Lejla Pojskic, Adis Kandic |
| EPI_ISL_1654206, EPI_ISL_1654207, EPI_ISL_1654208 | Research | National Reference Center for Viruses of Respiratory Infections, Institut Pasteur, Paris | Marion Barbet, Sylvie Behillil, Frédéric Lemoine, Corinne Maufrais, Christophe Malabat, Damien Mornico, Méline Bizard, Angela Brisebarre, Camille Capel, Louise Lefrançois, Etienne Simon-Lorière, Vincent Enouf, Maud Vanpeene, Sylvie van der Werf,Marianne Leruez-Ville |
| EPI_ISL_1654899, EPI_ISL_1654901, EPI_ISL_1654902, EPI_ISL_1654903, EPI_ISL_1654904, EPI_ISL_1654905, EPI_ISL_1654910, EPI_ISL_1654911, EPI_ISL_1654912, EPI_ISL_1654913, EPI_ISL_1654914, EPI_ISL_1654917, EPI_ISL_1654918, EPI_ISL_1654919, EPI_ISL_1654920, EPI_ISL_1654921, EPI_ISL_1654922, EPI_ISL_1654923, EPI_ISL_1654924, EPI_ISL_1654925, EPI_ISL_1654926, EPI_ISL_1654927, EPI_ISL_1654928, EPI_ISL_1654929, EPI_ISL_1654930, EPI_ISL_1654932, EPI_ISL_1654933, EPI_ISL_1654934, EPI_ISL_1654935, EPI_ISL_1654936, EPI_ISL_1654937, EPI_ISL_1654938, EPI_ISL_1654939, EPI_ISL_1654940, EPI_ISL_1654941, EPI_ISL_1654943, EPI_ISL_1654944, EPI_ISL_1654945, EPI_ISL_1654946, EPI_ISL_1654948, EPI_ISL_1654951, EPI_ISL_1654952, EPI_ISL_1654953, EPI_ISL_1654954, EPI_ISL_1654956, EPI_ISL_1654958, EPI_ISL_1654959, EPI_ISL_1654961, EPI_ISL_1654962, EPI_ISL_1654963, EPI_ISL_1654965, EPI_ISL_1654966, EPI_ISL_1654967, EPI_ISL_1654968, EPI_ISL_1654969, EPI_ISL_1654970, EPI_ISL_1654971, EPI_ISL_1654972, EPI_ISL_1654973, EPI_ISL_1654974, EPI_ISL_1654977, EPI_ISL_1654978, EPI_ISL_1654979, EPI_ISL_1654980, EPI_ISL_1654981, EPI_ISL_1654982, EPI_ISL_1654983, EPI_ISL_1654984, EPI_ISL_1654985, EPI_ISL_1654986 | | | |
| see above | NMVRVI | National Public Health Surveillance Laboratory | Lukas Zemaitis, Migle Gabrielaite, Jelena Razmuk, Svajune Muralyte, Ana Steponkiene, Lukas Vasionis, Danas Baksa |
| EPI_ISL_1655072 | NVSPL | National Public Health Surveillance Laboratory | Lukas Zemaitis, Migle Gabrielaite, Jelena Razmuk, Svajune Muralyte, Ana Steponkiene, Lukas Vasionis, Danas Baksa |
| EPI_ISL_1655076 | Hospital | National Reference Center for Viruses of Respiratory Infections, Institut Pasteur, Paris | Marion Barbet, Sylvie Behillil, Frédéric Lemoine, Corinne Maufrais, Christophe Malabat, Amaury Vaysse, Méline Bizard, Angela Brisebarre, Camille Capel, Louise Lefrançois, Etienne Simon-Lorière, Vincent Enouf, Maud Vanpeene, Sylvie van der Werf,Pascale Martres |
| EPI_ISL_1655077 | Hospital | National Reference Center for Viruses of Respiratory Infections, Institut Pasteur, Paris | Marion Barbet, Sylvie Behillil, Frédéric Lemoine, Corinne Maufrais, Christophe Malabat, Emmanuelle Permal, Méline Bizard, Angela Brisebarre, Camille Capel, Louise Lefrançois, Etienne Simon-Lorière, Vincent Enouf, Maud Vanpeene, Sylvie van der Werf,Dominique Descamps |
| EPI_ISL_1655078 | Hospital | National Reference Center for Viruses of Respiratory Infections, Institut Pasteur, Paris | Marion Barbet, Sylvie Behillil, Frédéric Lemoine, Corinne Maufrais, Christophe Malabat, Amaury Vaysse, Méline Bizard, Angela Brisebarre, Camille Capel, Louise Lefrançois, Etienne Simon-Lorière, Vincent Enouf, Maud Vanpeene, Sylvie van der Werf,Dominique Descamps |
| EPI_ISL_1655079 | Hospital | National Reference Center for Viruses of Respiratory Infections, Institut Pasteur, Paris | Marion Barbet, Sylvie Behillil, Frédéric Lemoine, Corinne Maufrais, Christophe Malabat, Méline Bizard, Angela Brisebarre, Camille Capel, Louise Lefrançois, Etienne Simon-Lorière, Vincent Enouf, Maud Vanpeene, Sylvie van der Werf,Eric Farfour |
| EPI_ISL_1655080 | Hospital | National Reference Center for Viruses of Respiratory Infections, Institut Pasteur, Paris | Marion Barbet, Sylvie Behillil, Frédéric Lemoine, Corinne Maufrais, Christophe Malabat, Méline Bizard, Angela Brisebarre, Camille Capel, Louise Lefrançois, Etienne Simon-Lorière, Vincent Enouf, Maud Vanpeene, Sylvie van der Werf,Pierre Alain Billy |
| EPI_ISL_1655081 | Hospital | National Reference Center for Viruses of Respiratory Infections, Institut Pasteur, Paris | Marion Barbet, Sylvie Behillil, Frédéric Lemoine, Corinne Maufrais, Christophe Malabat, Méline Bizard, Angela Brisebarre, Camille Capel, Louise Lefrançois, Etienne Simon-Lorière, Vincent Enouf, Maud Vanpeene, Sylvie van der Werf,Franck Bernardi |
| EPI_ISL_1655965, EPI_ISL_1655966, EPI_ISL_1655967, EPI_ISL_1655968, EPI_ISL_1655969, EPI_ISL_1655970, EPI_ISL_1655971, EPI_ISL_1655972, EPI_ISL_1655973, EPI_ISL_1655974, EPI_ISL_1655975, EPI_ISL_1655976, EPI_ISL_1655977, EPI_ISL_1655978, EPI_ISL_1655979, EPI_ISL_1655980, EPI_ISL_1655982, EPI_ISL_1655983, EPI_ISL_1655984, EPI_ISL_1655985, EPI_ISL_1655986, EPI_ISL_1655987, EPI_ISL_1655988, EPI_ISL_1655989, EPI_ISL_1655990, EPI_ISL_1655991, EPI_ISL_1655992, EPI_ISL_1655993, EPI_ISL_1655994, EPI_ISL_1655995, EPI_ISL_1655996, EPI_ISL_1655997, EPI_ISL_1655998, EPI_ISL_1655999, EPI_ISL_1656000, EPI_ISL_1656001, EPI_ISL_1656002, EPI_ISL_1656003, EPI_ISL_1656004, EPI_ISL_1656005, EPI_ISL_1656006, EPI_ISL_1656007, EPI_ISL_1656008, EPI_ISL_1656009, EPI_ISL_1656010, EPI_ISL_1656011, EPI_ISL_1656012, EPI_ISL_1656013, EPI_ISL_1656014, EPI_ISL_1656015, EPI_ISL_1656016, EPI_ISL_1656017, EPI_ISL_1656018, EPI_ISL_1656019, EPI_ISL_1656020, EPI_ISL_1656021, EPI_ISL_1656022, EPI_ISL_1656023, EPI_ISL_1656024, EPI_ISL_1656025, EPI_ISL_1656026, EPI_ISL_1656027, EPI_ISL_1656028, EPI_ISL_1656029, EPI_ISL_1656030, EPI_ISL_1656031, EPI_ISL_1656032, EPI_ISL_1656033, EPI_ISL_1656034, EPI_ISL_1656035, EPI_ISL_1656036, EPI_ISL_1656037, EPI_ISL_1656038, EPI_ISL_1656039, EPI_ISL_1656040, EPI_ISL_1656041, EPI_ISL_1656042, EPI_ISL_1656043, EPI_ISL_1656044, EPI_ISL_1656045, EPI_ISL_1656046, EPI_ISL_1656047, EPI_ISL_1656048, EPI_ISL_1656049, EPI_ISL_1656050, EPI_ISL_1656051, EPI_ISL_1656052, EPI_ISL_1656053, EPI_ISL_1656054, EPI_ISL_1656056, EPI_ISL_1656057, EPI_ISL_1656058, EPI_ISL_1656059, EPI_ISL_1656060, EPI_ISL_1656061, EPI_ISL_1656062, EPI_ISL_1656063, EPI_ISL_1656065, EPI_ISL_1656066, EPI_ISL_1656067, EPI_ISL_1656069, EPI_ISL_1656070, EPI_ISL_1656071, EPI_ISL_1656072, EPI_ISL_1656073, EPI_ISL_1656074, EPI_ISL_1656075, EPI_ISL_1656076, EPI_ISL_1656077, EPI_ISL_1656078, EPI_ISL_1656079, EPI_ISL_1656080, EPI_ISL_1656081, EPI_ISL_1656082, EPI_ISL_1656083, EPI_ISL_1656084, EPI_ISL_1656085, EPI_ISL_1656086, EPI_ISL_1656087, EPI_ISL_1656088, EPI_ISL_1656089, EPI_ISL_1656090, EPI_ISL_1656091, EPI_ISL_1656092, EPI_ISL_1656093, EPI_ISL_1656094, EPI_ISL_1656095, EPI_ISL_1656096, EPI_ISL_1656097, EPI_ISL_1656098, EPI_ISL_1656099, EPI_ISL_1656100, EPI_ISL_1656101, EPI_ISL_1656102, EPI_ISL_1656103, EPI_ISL_1656104, EPI_ISL_1656105, EPI_ISL_1656106, EPI_ISL_1656107, EPI_ISL_1656108, EPI_ISL_1656109, EPI_ISL_1656110, EPI_ISL_1656111, EPI_ISL_1656112, EPI_ISL_1656113, EPI_ISL_1656114, EPI_ISL_1656115, EPI_ISL_1656116, EPI_ISL_1656117, EPI_ISL_1656118, EPI_ISL_1656119, EPI_ISL_1656120, EPI_ISL_1656121, EPI_ISL_1656122, EPI_ISL_1656123, EPI_ISL_1656124, EPI_ISL_1656125, EPI_ISL_1656126, EPI_ISL_1656127, EPI_ISL_1656128, EPI_ISL_1656129, EPI_ISL_1656130, EPI_ISL_1656131, EPI_ISL_1656132, EPI_ISL_1656133, EPI_ISL_1656134, EPI_ISL_1656135, EPI_ISL_1656136, EPI_ISL_1656137, EPI_ISL_1656138, EPI_ISL_1656139, EPI_ISL_1656140, EPI_ISL_1656141, EPI_ISL_1656142, EPI_ISL_1656143, EPI_ISL_1656144, EPI_ISL_1656145 | | | |
| see above | NMVRVI | National Public Health Surveillance Laboratory | Lukas Zemaitis, Migle Gabrielaite, Jelena Razmuk, Svajune Muralyte, Ana Steponkiene, Lukas Vasionis, Danas Baksa |
| EPI_ISL_1656833 | Public Health Authority of the Slovak Republic | Laboratory of Genomics and Bioinformatics, Comenius University Science Park | Tatiana Sedláková, Diana Rusáková, Miroslav Böhmer, Anna Giová, Jaroslav Budiš, Tomáš Szemes |
| EPI_ISL_1657113, EPI_ISL_1657114, EPI_ISL_1657115 | National Institute of Infectious Diseases-Prof. Dr. Matei Bals Molecular Diagnostics Laboratory | National Institute of Infectious Diseases-Prof. Dr. Matei Bals Molecular Diagnostics Laboratory | Corina Casangiu, Leontina Banica, Marius Surleac, Ovidiu Vlaicu, Andreea Tudor, Simona Paraschiv, Dan Otelea |
| EPI_ISL_1661490 | LESP Ciudad de Mexico | Instituto de Diagnostico y Referencia Epidemiologicos (INDRE) | Claudia Wong-Arambula, Abril Rodriguez-Maldonado, Vanessa Rivero-Arredondo, Ariadna Medina-Benitez, Joaquin Quiroz-Mercado, Sergio Rangel-Guerrero, Natividad Cruz-Ortiz, Tatiana Nunez-Garcia, Gisela Barrera-Badillo, Lucia Hernandez-Rivas, Irma Lopez-Martinez, Ernesto Ramirez-Gonzalez. |
| EPI_ISL_1661492 | LESP Estado de Mexico | Instituto de Diagnostico y Referencia Epidemiologicos (INDRE) | Claudia Wong-Arambula, Abril Rodriguez-Maldonado, Vanessa Rivero-Arredondo, Ariadna Medina-Benitez, Joaquin Quiroz-Mercado, Sergio Rangel-Guerrero, Natividad Cruz-Ortiz, Tatiana Nunez-Garcia, Gisela Barrera-Badillo, Lucia Hernandez-Rivas, Irma Lopez-Martinez, Ernesto Ramirez-Gonzalez. |

| | | | |
|---|---|---|---|
| EPI_ISL_1661493, EPI_ISL_1661494 | LESP Ciudad de Mexico | Instituto de Diagnostico y Referencia Epidemiologicos (INDRE) | Claudia Wong-Arambula, Abril Rodriguez-Maldonado, Vanessa Rivero-Arredondo, Ariadna Medina-Benitez, Joaquin Quiroz-Mercado, Sergio Rangel-Guerrero, Natividad Cruz-Ortiz, Tatiana Nunez-Garcia, Gisela Barrera-Badillo, Lucia Hernandez-Rivas, Irma Lopez-Martinez, Ernesto Ramirez-Gonzalez. |
| EPI_ISL_1661497, EPI_ISL_1661498, EPI_ISL_1661501, EPI_ISL_1661502, EPI_ISL_1661503 | LESP Guanajuato | Instituto de Diagnostico y Referencia Epidemiologicos (INDRE) | Claudia Wong-Arambula, Abril Rodriguez-Maldonado, Vanessa Rivero-Arredondo, Ariadna Medina-Benitez, Joaquin Quiroz-Mercado, Sergio Rangel-Guerrero, Natividad Cruz-Ortiz, Tatiana Nunez-Garcia, Gisela Barrera-Badillo, Lucia Hernandez-Rivas, Irma Lopez-Martinez, Ernesto Ramirez-Gonzalez. |
| EPI_ISL_1661548 | LESP Michoacan | Instituto de Diagnostico y Referencia Epidemiologicos (INDRE) | Claudia Wong-Arambula, Abril Rodriguez-Maldonado, Vanessa Rivero-Arredondo, Ariadna Medina-Benitez, Joaquin Quiroz-Mercado, Sergio Rangel-Guerrero, Natividad Cruz-Ortiz, Tatiana Nunez-Garcia, Gisela Barrera-Badillo, Lucia Hernandez-Rivas, Irma Lopez-Martinez, Ernesto Ramirez-Gonzalez. |
| EPI_ISL_1662062, EPI_ISL_1662064 | Centro de Investigacion Biomedica de Occidente (CIBO) | Unidad de Genomica Avanzada | Consorcio Mexicano de Vigilancia Genomica (CoViGen-Mex). Authors (in alphabetical order): Julio Elias Alvarado-Yaah, Carlos F. Arias, Santiago Ávila-Rios, Victor Hugo Borja-Aburto, Celia Boukadida, Juan Bautista Chale-Dzul, Celida Duque Molina, Jose Antonio Enciso-Moreno, Gloria Elena Espinosa-Ayala, Fernando Fontove-Herrera, Victor Eduardo Garcia-Arias,Concepcion Grajales-Muniz, Ricardo Grande, Alfredo Herrera-Estrella, Carla Ivon Herrera-Najera, Pavel Isa, Brenda Irasema Maldonado-Meza, Bernardo Martinez-Miguel, Margarita Matias-Florentino, Maria Guadalupe de Jesus Mireles-Rivera, Gloria Maria Molina-Salinas, Hector Montoya-Fuentes, Jose Esteban Munoz-Medina, Jose de Jesus Nunez-Contreras, Alicia Ocana-Mondragon, Luis Alberto Ochoa-Carrera, Hector Esteban Paz-Juarez, Francisco Pulido, Helen Haydee Fernanda Ramirez-Plascencia, Angel Gustavo Salas-Lais, Alejandro Sanchez-Flores, Clara Esperanza Santacruz-Tinoco, Maria Guadalupe Santiago-Mauricio, Nelly Selem-Mojica, Blanca Taboada, Gloria Vazquez |
| EPI_ISL_1662072 | Laboratorio Central de Epidemiologia (LCE) | Unidad de Genomica Avanzada | Consorcio Mexicano de Vigilancia Genomica (CoViGen-Mex). Authors (in alphabetical order): Julio Elias Alvarado-Yaah, Carlos F. Arias, Santiago Ávila-Rios, Victor Hugo Borja-Aburto, Celia Boukadida, Juan Bautista Chale-Dzul, Celida Duque Molina, Jose Antonio Enciso-Moreno, Gloria Elena Espinosa-Ayala, Fernando Fontove-Herrera, Victor Eduardo Garcia-Arias,Concepcion Grajales-Muniz, Ricardo Grande, Alfredo Herrera-Estrella, Carla Ivon Herrera-Najera, Pavel Isa, Brenda Irasema Maldonado-Meza, Bernardo Martinez-Miguel, Margarita Matias-Florentino, Maria Guadalupe de Jesus Mireles-Rivera, Gloria Maria Molina-Salinas, Hector Montoya-Fuentes, Jose Esteban Munoz-Medina, Jose de Jesus Nunez-Contreras, Alicia Ocana-Mondragon, Luis Alberto Ochoa-Carrera, Hector Esteban Paz-Juarez, Francisco Pulido, Helen Haydee Fernanda Ramirez-Plascencia, Angel Gustavo Salas-Lais, Alejandro Sanchez-Flores, Clara Esperanza Santacruz-Tinoco, Maria Guadalupe Santiago-Mauricio, Nelly Selem-Mojica, Blanca Taboada, Gloria Vazquez |
| EPI_ISL_1662076, EPI_ISL_1662123 | Centro de Investigacion Biomedica de Occidente (CIBO) | Unidad de Genomica Avanzada | Consorcio Mexicano de Vigilancia Genomica (CoViGen-Mex). Authors (in alphabetical order): Julio Elias Alvarado-Yaah, Carlos F. Arias, Santiago Ávila-Rios, Victor Hugo Borja-Aburto, Celia Boukadida, Juan Bautista Chale-Dzul, Celida Duque Molina, Jose Antonio Enciso-Moreno, Gloria Elena Espinosa-Ayala, Fernando Fontove-Herrera, Victor Eduardo Garcia-Arias,Concepcion Grajales-Muniz, Ricardo Grande, Alfredo Herrera-Estrella, Carla Ivon Herrera-Najera, Pavel Isa, Brenda Irasema Maldonado-Meza, Bernardo Martinez-Miguel, Margarita Matias-Florentino, Maria Guadalupe de Jesus Mireles-Rivera, Gloria Maria Molina-Salinas, Hector Montoya-Fuentes, Jose Esteban Munoz-Medina, Jose de Jesus Nunez-Contreras, Alicia Ocana-Mondragon, Luis Alberto Ochoa-Carrera, Hector Esteban Paz-Juarez, Francisco Pulido, Helen Haydee Fernanda Ramirez-Plascencia, Angel Gustavo Salas-Lais, Alejandro Sanchez-Flores, Clara Esperanza Santacruz-Tinoco, Maria Guadalupe Santiago-Mauricio, Nelly Selem-Mojica, Blanca Taboada, Gloria Vazquez |
| EPI_ISL_1662184 | Unidad de Investigacion Medica de Yucatan (UIMY) | Unidad de Genomica Avanzada | Consorcio Mexicano de Vigilancia Genomica (CoViGen-Mex). Authors (in alphabetical order): Julio Elias Alvarado-Yaah, Carlos F. Arias, Santiago Ávila-Rios, Victor Hugo Borja-Aburto, Celia Boukadida, Juan Bautista Chale-Dzul, Celida Duque Molina, Jose Antonio Enciso-Moreno, Gloria Elena Espinosa-Ayala, Fernando Fontove-Herrera, Victor Eduardo Garcia-Arias,Concepcion Grajales-Muniz, Ricardo Grande, Alfredo Herrera-Estrella, Carla Ivon Herrera-Najera, Pavel Isa, Brenda Irasema Maldonado-Meza, Bernardo Martinez-Miguel, Margarita Matias-Florentino, Maria Guadalupe de Jesus Mireles-Rivera, Gloria Maria Molina-Salinas, Hector Montoya-Fuentes, Jose Esteban Munoz-Medina, Jose de Jesus Nunez-Contreras, Alicia Ocana-Mondragon, Luis Alberto Ochoa-Carrera, Hector Esteban Paz-Juarez, Francisco Pulido, Helen Haydee Fernanda Ramirez-Plascencia, Angel Gustavo Salas-Lais, Alejandro Sanchez-Flores, Clara Esperanza Santacruz-Tinoco, Maria Guadalupe Santiago-Mauricio, Nelly Selem-Mojica, Blanca Taboada, Gloria Vazquez |
| EPI_ISL_1662186 | Centro de Investigacion Biomedica de Occidente (CIBO) | Unidad de Genomica Avanzada | Consorcio Mexicano de Vigilancia Genomica (CoViGen-Mex). Authors (in alphabetical order): Julio Elias Alvarado-Yaah, Carlos F. Arias, Santiago Ávila-Rios, Victor Hugo Borja-Aburto, Celia Boukadida, Juan Bautista Chale-Dzul, Celida Duque Molina, Jose Antonio Enciso-Moreno, Gloria Elena Espinosa-Ayala, Fernando Fontove-Herrera, Victor Eduardo Garcia-Arias,Concepcion Grajales-Muniz, Ricardo Grande, Alfredo Herrera-Estrella, Carla Ivon Herrera-Najera, Pavel Isa, Brenda Irasema Maldonado-Meza, Bernardo Martinez-Miguel, Margarita Matias-Florentino, Maria Guadalupe de Jesus Mireles-Rivera, Gloria Maria Molina-Salinas, Hector Montoya-Fuentes, Jose Esteban Munoz-Medina, Jose de Jesus Nunez-Contreras, Alicia Ocana-Mondragon, Luis Alberto Ochoa-Carrera, Hector Esteban Paz-Juarez, Francisco Pulido, Helen Haydee Fernanda Ramirez-Plascencia, Angel Gustavo Salas-Lais, Alejandro Sanchez-Flores, Clara Esperanza Santacruz-Tinoco, Maria Guadalupe Santiago-Mauricio, Nelly Selem-Mojica, Blanca Taboada, Gloria Vazquez |
| EPI_ISL_1662469, EPI_ISL_1663586, EPI_ISL_1663589, EPI_ISL_1663595, EPI_ISL_1663596, EPI_ISL_1663597, EPI_ISL_1663598 | IU-Cerrahpasa, Cerrahpasa School of Medicine, COVID-19 Lab | IU-Cerrahpasa, Cerrahpasa School of Medicine, COVID-19 Lab | Mert Kuskucu, Yesim Tuyji Tok, Kenan Midilli |
| EPI_ISL_1667472 | Department of Laboratory Medicine, National Taiwan University Hospital | Microbial Genomics Core Lab, National Taiwan University Centers of Genomic and Precision Medicine | Shiou-Hwei Yeh, You-Yu Lin, Ya-Yun Lai, Chiao-Ling Li, Shan-Chwen Chang, Pei-Jer Chen, Sui-Yuan Chang |
| EPI_ISL_1669789 | Istituto Zooprofilattico Sperimentale della Lombardia e dell'Emilia Romagna (IZSLER), Sede Territoriale di Modena | Istituto Zooprofilattico Sperimentale della Lombardia e dell'Emilia Romagna (IZSLER), Risk Analysis and Genomic Epidemiology Unit | Carra Elena, Gianluca Rugna, Marina Morganti, Ilaria Menozzi, Erika Scaltriti, Stefano Pongolini |
| EPI_ISL_1669804, EPI_ISL_1669805, EPI_ISL_1669806, EPI_ISL_1669807, EPI_ISL_1669808, EPI_ISL_1669809, EPI_ISL_1669811, EPI_ISL_1669812, EPI_ISL_1669813, EPI_ISL_1669814, EPI_ISL_1669815, EPI_ISL_1669817, EPI_ISL_1669818, EPI_ISL_1669819, EPI_ISL_1669820, EPI_ISL_1669822, EPI_ISL_1669823, EPI_ISL_1669824, EPI_ISL_1669825, EPI_ISL_1669827, EPI_ISL_1669828, EPI_ISL_1669829, EPI_ISL_1669830, EPI_ISL_1669831, EPI_ISL_1669832, EPI_ISL_1669833, EPI_ISL_1669834, EPI_ISL_1669835, EPI_ISL_1669836, EPI_ISL_1669837, EPI_ISL_1669838 | | | |
| see above | Microbiology Department, University Hospital Donostia | Microbiology Department, University Hospital Donostia | Marimon JM, Montes M, Piñeiro L, Sorarrain A, Gomez M, Cilla G. |
| EPI_ISL_1669960 | General Hospital - Prilep | Laboratory of virology and molecular diagnostics, Institute of Public Health | Kuzmanovska M, Boshevska G, Janchevska E. |
| EPI_ISL_1669966 | Laboratory of virology and molecular diagnostics, Institute of Public Health | Laboratory of virology and molecular diagnostics, Institute of Public Health | Kuzmanovska M, Boshevska G, Janchevska E. |
| EPI_ISL_1669967 | General Hospital - Prilep | Laboratory of virology and molecular diagnostics, Institute of Public Health | Kuzmanovska M, Boshevska G, Janchevska E. |
| EPI_ISL_1669968, EPI_ISL_1669976 | General Hospital - Stip | Laboratory of virology and molecular diagnostics, Institute of Public Health | Kuzmanovska M, Boshevska G, Janchevska E. |
| EPI_ISL_1669977 | General Hospital - Prilep | Laboratory of virology and molecular diagnostics, Institute of Public Health | Kuzmanovska M, Boshevska G, Janchevska E. |
| EPI_ISL_1669978 | General Hospital - Kochani | Laboratory of virology and molecular diagnostics, Institute of Public Health | Kuzmanovska M, Boshevska G, Janchevska E. |
| EPI_ISL_1669979, EPI_ISL_1669980 | General Hospital - Prilep | Laboratory of virology and molecular diagnostics, Institute of Public Health | Kuzmanovska M, Boshevska G, Janchevska E. |
| EPI_ISL_1669983 | General Hospital - Stip | Laboratory of virology and molecular diagnostics, Institute of Public Health | Kuzmanovska M, Boshevska G, Janchevska E. |

| | | | |
|---|---|---|---|
| EPI_ISL_1669984 | General Hospital - Ohrid | Laboratory of virology and molecular diagnostics, Institute of Public Health | Kuzmanovska M, Boshevska G, Janchevska E. |
| EPI_ISL_1670694, EPI_ISL_1670697, EPI_ISL_1670699, EPI_ISL_1670701 | Azienda Sanitaria Locale di Piacenza - Presidio Ospedaliero - Laboratorio di Microbiologia | Istituto Zooprofilattico Sperimentale della Lombardia e dell'Emilia Romagna (IZSLER), Risk Analysis and Genomic Epidemiology Unit | Lo Cascio Giuliana, Schiavo Roberta, Peroni Irene, Marina Morganti, Ilaria Menozzi, Erika Scaltriti, Stefano Pongolini |
| EPI_ISL_1672416 | GH A.CHENEVIER-H.MONDOR | Department of Virology, Henri Mondor University Hospital, Assistance Publique Hôpitaux de Paris, Université Paris-Est Créteil, INSERM U955 | Christophe Rodriguez, Slim Fourati, Vanessa Demontant, Guillaume Gricourt, Melissa N'Debi, Alexandre Soulier, Elisabeth Trawinski, Jean-Michel Pawlotsky |
| EPI_ISL_1672418 | GH JOFFRE DUPUYTREN | Department of Virology, Henri Mondor University Hospital, Assistance Publique Hôpitaux de Paris, Université Paris-Est Créteil, INSERM U955 | Christophe Rodriguez, Slim Fourati, Vanessa Demontant, Guillaume Gricourt, Melissa N'Debi, Alexandre Soulier, Elisabeth Trawinski, Jean-Michel Pawlotsky |
| EPI_ISL_1672422, EPI_ISL_1672423, EPI_ISL_1672424 | GH A.CHENEVIER-H.MONDOR | Department of Virology, Henri Mondor University Hospital, Assistance Publique Hôpitaux de Paris, Université Paris-Est Créteil, INSERM U955 | Christophe Rodriguez, Slim Fourati, Vanessa Demontant, Guillaume Gricourt, Melissa N'Debi, Alexandre Soulier, Elisabeth Trawinski, Jean-Michel Pawlotsky |
| EPI_ISL_1672426, EPI_ISL_1672463, EPI_ISL_1672471 | GH JOFFRE DUPUYTREN | Department of Virology, Henri Mondor University Hospital, Assistance Publique Hôpitaux de Paris, Université Paris-Est Créteil, INSERM U955 | Christophe Rodriguez, Slim Fourati, Vanessa Demontant, Guillaume Gricourt, Melissa N'Debi, Alexandre Soulier, Elisabeth Trawinski, Jean-Michel Pawlotsky |
| EPI_ISL_1672474, EPI_ISL_1672475, EPI_ISL_1672477, EPI_ISL_1672487, EPI_ISL_1672491, EPI_ISL_1672494, EPI_ISL_1672495, EPI_ISL_1672507, EPI_ISL_1672510, EPI_ISL_1672521, EPI_ISL_1672523, EPI_ISL_1672531, EPI_ISL_1672533, EPI_ISL_1672535, EPI_ISL_1672537, EPI_ISL_1672539, EPI_ISL_1672542, EPI_ISL_1672545, EPI_ISL_1672549 | | | |
| see above | GH A.CHENEVIER-H.MONDOR | Department of Virology, Henri Mondor University Hospital, Assistance Publique Hôpitaux de Paris, Université Paris-Est Créteil, INSERM U955 | Christophe Rodriguez, Slim Fourati, Vanessa Demontant, Guillaume Gricourt, Melissa N'Debi, Alexandre Soulier, Elisabeth Trawinski, Jean-Michel Pawlotsky |
| EPI_ISL_1672757 | HOPITAL EMILE ROUX | Department of Virology, Henri Mondor University Hospital, Assistance Publique Hôpitaux de Paris, Université Paris-Est Créteil, INSERM U955 | Christophe Rodriguez, Slim Fourati, Vanessa Demontant, Guillaume Gricourt, Melissa N'Debi, Alexandre Soulier, Elisabeth Trawinski, Jean-Michel Pawlotsky |
| EPI_ISL_1672762 | GH JOFFRE DUPUYTREN | Department of Virology, Henri Mondor University Hospital, Assistance Publique Hôpitaux de Paris, Université Paris-Est Créteil, INSERM U955 | Christophe Rodriguez, Slim Fourati, Vanessa Demontant, Guillaume Gricourt, Melissa N'Debi, Alexandre Soulier, Elisabeth Trawinski, Jean-Michel Pawlotsky |
| EPI_ISL_1672766, EPI_ISL_1672768 | GH A.CHENEVIER-H.MONDOR | Department of Virology, Henri Mondor University Hospital, Assistance Publique Hôpitaux de Paris, Université Paris-Est Créteil, INSERM U955 | Christophe Rodriguez, Slim Fourati, Vanessa Demontant, Guillaume Gricourt, Melissa N'Debi, Alexandre Soulier, Elisabeth Trawinski, Jean-Michel Pawlotsky |
| EPI_ISL_1672772 | GH JOFFRE DUPUYTREN | Department of Virology, Henri Mondor University Hospital, Assistance Publique Hôpitaux de Paris, Université Paris-Est Créteil, INSERM U955 | Christophe Rodriguez, Slim Fourati, Vanessa Demontant, Guillaume Gricourt, Melissa N'Debi, Alexandre Soulier, Elisabeth Trawinski, Jean-Michel Pawlotsky |
| EPI_ISL_1673263 | LESP Sinaloa | Instituto de Diagnostico y Referencia Epidemiologicos (INDRE) | Claudia Wong-Arambula, Abril Rodriguez-Maldonado, Vanessa Rivero-Arredondo, Ariadna Medina-Benitez, Joaquin Quiroz-Mercado, Sergio Rangel-Guerrero, Natividad Cruz-Ortiz, Tatiana Nunez-Garcia, Gisela Barrera-Badillo, Lucia Hernandez-Rivas, Irma Lopez-Martinez, Ernesto Ramirez-Gonzalez. |
| EPI_ISL_1673265 | LESP Guerrero | Instituto de Diagnostico y Referencia Epidemiologicos (INDRE) | Claudia Wong-Arambula, Abril Rodriguez-Maldonado, Vanessa Rivero-Arredondo, Ariadna Medina-Benitez, Joaquin Quiroz-Mercado, Sergio Rangel-Guerrero, Natividad Cruz-Ortiz, Tatiana Nunez-Garcia, Gisela Barrera-Badillo, Lucia Hernandez-Rivas, Irma Lopez-Martinez, Ernesto Ramirez-Gonzalez. |
| EPI_ISL_1673270, EPI_ISL_1673271, EPI_ISL_1673272, EPI_ISL_1673273 | LESP Guanajuato | Instituto de Diagnostico y Referencia Epidemiologicos (INDRE) | Claudia Wong-Arambula, Abril Rodriguez-Maldonado, Vanessa Rivero-Arredondo, Ariadna Medina-Benitez, Joaquin Quiroz-Mercado, Sergio Rangel-Guerrero, Natividad Cruz-Ortiz, Tatiana Nunez-Garcia, Gisela Barrera-Badillo, Lucia Hernandez-Rivas, Irma Lopez-Martinez, Ernesto Ramirez-Gonzalez. |
| EPI_ISL_1673307, EPI_ISL_1673332 | Institute for Medical Research, Infectious Disease Research Centre, National Institutes of Health, Ministry of Health Malaysia | Institute for Medical Research, Infectious Disease Research Centre, National Institutes of Health, Ministry of Health Malaysia | Suppiah J, Kamel K, Thayan R |
| EPI_ISL_1673683, EPI_ISL_1673735 | Institute for Medical Research, Infectious Disease Research Centre, National Institutes of Health, Ministry of Health Malaysia | Institute for Medical Research, Infectious Disease Research Centre, National Institutes of Health, Ministry of Health Malaysia | Suppiah J, Kamel K, Mohd Zawawi Z, Thayan R |
| EPI_ISL_1675010 | Labo Analyses Med | National Reference Center for Viruses of Respiratory Infections, Institut Pasteur, Paris | Marion Barbet, Sylvie Behillil, Méline Bizard, Angela Brisebarre, Camille Capel, Vincent Enouf, Louise Lefrançois, Frédéric Lemoine, Christophe Malabat, Corinne Maufrais, Pierre Lechat, Etienne Simon-Lorière, Maud Vanpeene, Sylvie Van der Werf ,Gilles Desvaux |
| EPI_ISL_1678002 | CQRC_QUALITY CONTROL CHEMICAL BIOLOGICAL RISK_AOOR Villa Sofia Cervello Palermo | CQRC_QUALITY CONTROL CHEMICAL BIOLOGICAL RISK_AOOR Villa Sofia Cervello Palermo | Di Gaudio, F., Brunacci, G., Contino, F., |
| EPI_ISL_1678003 | CQRC_QUALITY CONTROL CHEMICAL BIOLOGICAL RISK_AOOR Villa Sofia Cervello Palermo | CQRC_QUALITY CONTROL CHEMICAL BIOLOGICAL RISK_AOOR Villa Sofia Cervello Palermo | Di Gaudio, F., Contino, F., Brunacci, G., |
| EPI_ISL_1678006 | CQRC_QUALITY CONTROL CHEMICAL BIOLOGICAL RISK_AOOR Villa Sofia Cervello Palermo | CQRC_QUALITY CONTROL CHEMICAL BIOLOGICAL RISK_AOOR Villa Sofia Cervello Palermo | Di Gaudio, F., Brunacci, G., Contino, F., |
| EPI_ISL_1678007 | CQRC_QUALITY CONTROL CHEMICAL BIOLOGICAL RISK_AOOR Villa Sofia Cervello Palermo | CQRC_QUALITY CONTROL CHEMICAL BIOLOGICAL RISK_AOOR Villa Sofia Cervello Palermo | Di Gaudio, F., Contino, F., Brunacci, G., |
| EPI_ISL_1678008 | CQRC_QUALITY CONTROL CHEMICAL BIOLOGICAL RISK_AOOR Villa Sofia Cervello Palermo | CQRC_QUALITY CONTROL CHEMICAL BIOLOGICAL RISK_AOOR Villa Sofia Cervello Palermo | Di Gaudio, F., Brunacci, G., Contino, F., |
| EPI_ISL_1678009 | CQRC_QUALITY CONTROL CHEMICAL BIOLOGICAL RISK_AOOR Villa Sofia Cervello Palermo | CQRC_QUALITY CONTROL CHEMICAL BIOLOGICAL RISK_AOOR Villa Sofia Cervello Palermo | Di Gaudio, F., Contino, F., Brunacci, G., |
| EPI_ISL_1678010, EPI_ISL_1678014 | CQRC_QUALITY CONTROL CHEMICAL BIOLOGICAL RISK_AOOR Villa Sofia Cervello Palermo | CQRC_QUALITY CONTROL CHEMICAL BIOLOGICAL RISK_AOOR Villa Sofia Cervello Palermo | Di Gaudio, F., Brunacci, G., Contino, F., |
| EPI_ISL_1678017 | CQRC_QUALITY CONTROL CHEMICAL BIOLOGICAL RISK_AOOR Villa Sofia Cervello Palermo | CQRC_QUALITY CONTROL CHEMICAL BIOLOGICAL RISK_AOOR Villa Sofia Cervello Palermo | Di Gaudio, F., Contino, F., Brunacci, G., |
| EPI_ISL_1678018 | CQRC_QUALITY CONTROL CHEMICAL BIOLOGICAL RISK_AOOR Villa Sofia Cervello Palermo | CQRC_QUALITY CONTROL CHEMICAL BIOLOGICAL RISK_AOOR Villa Sofia Cervello Palermo | Di Gaudio, F., Contino, F., Orlando, V., Buffa, V., Bonvissuto, M., Di Chiara, V., Todaro, I., Seidita, G., Di Sclafani, A., Lungari, A., Brunacci, G. |
| EPI_ISL_1678019 | CQRC_QUALITY CONTROL CHEMICAL BIOLOGICAL RISK_AOOR Villa Sofia Cervello Palermo | CQRC_QUALITY CONTROL CHEMICAL BIOLOGICAL RISK_AOOR Villa Sofia Cervello Palermo | Di Gaudio, F., Contino, F., Brunacci, G. |
| EPI_ISL_1678020 | CQRC_QUALITY CONTROL CHEMICAL BIOLOGICAL RISK_AOOR Villa Sofia Cervello Palermo | CQRC_QUALITY CONTROL CHEMICAL BIOLOGICAL RISK_AOOR Villa Sofia Cervello Palermo | Di Gaudio, F., Brunacci, G., Contino, F., |

| | | | |
|---|---|---|---|
| EPI_ISL_1678021 | CQRC_QUALITY CONTROL CHEMICAL BIOLOGICAL RISK_AOOR Villa Sofia Cervello Palermo | CQRC_QUALITY CONTROL CHEMICAL BIOLOGICAL RISK_AOOR Villa Sofia Cervello Palermo | Di Gaudio, F., Contino, F., Brunacci, G., |
| EPI_ISL_1678387, EPI_ISL_1678388 | Laboratory for Clinical Immunology and Molecular Genetics - University Clinic Golnik Laboratory for Respiratory Microbiology - University Clinic Golnik | Laboratory for Clinical Immunology and Molecular Genetics -University Clinic Golnik | Matija Rijavec, Julij Šelb, Urška Bidovec Stojkovi, Žan Kogovšek, Nina Rupar, Viktorija Tomi, Peter Korošec |
| EPI_ISL_1678389, EPI_ISL_1678391, EPI_ISL_1678392, EPI_ISL_1678393, EPI_ISL_1678395, EPI_ISL_1678396, EPI_ISL_1678398, EPI_ISL_1678399, EPI_ISL_1678400, EPI_ISL_1678401, EPI_ISL_1678402, EPI_ISL_1678403, EPI_ISL_1678404, EPI_ISL_1678405, EPI_ISL_1678406, EPI_ISL_1678407, EPI_ISL_1678408, EPI_ISL_1678414, EPI_ISL_1678415, EPI_ISL_1678416, EPI_ISL_1678417, EPI_ISL_1678418, EPI_ISL_1678420, EPI_ISL_1678421, EPI_ISL_1678422, EPI_ISL_1678423, EPI_ISL_1678425, EPI_ISL_1678426, EPI_ISL_1678427, EPI_ISL_1678428, EPI_ISL_1678430, EPI_ISL_1678431, EPI_ISL_1678432, EPI_ISL_1678433, EPI_ISL_1678434, EPI_ISL_1678435, EPI_ISL_1678436, EPI_ISL_1678437, EPI_ISL_1678438, EPI_ISL_1678439, EPI_ISL_1678440, EPI_ISL_1678441, EPI_ISL_1678442, EPI_ISL_1678443, EPI_ISL_1678444, EPI_ISL_1678445, EPI_ISL_1678446, EPI_ISL_1678447, EPI_ISL_1678448, EPI_ISL_1678449, EPI_ISL_1678450, EPI_ISL_1678451, EPI_ISL_1678452, EPI_ISL_1678453, EPI_ISL_1678454, EPI_ISL_1678455, EPI_ISL_1678456, EPI_ISL_1678457, EPI_ISL_1678458, EPI_ISL_1678460, EPI_ISL_1678461, EPI_ISL_1678462, EPI_ISL_1678463, EPI_ISL_1678464, EPI_ISL_1678465, EPI_ISL_1678466, EPI_ISL_1678467, EPI_ISL_1678468, EPI_ISL_1678469, EPI_ISL_1678470, EPI_ISL_1678471, EPI_ISL_1678472, EPI_ISL_1678473, EPI_ISL_1678477, EPI_ISL_1678479, EPI_ISL_1678480, EPI_ISL_1678482, EPI_ISL_1678483, EPI_ISL_1678577, EPI_ISL_1678578, EPI_ISL_1678579, EPI_ISL_1678580, EPI_ISL_1678581, EPI_ISL_1678582 | | | |
| see above | HG Pharma GmbH | Bergthaler laboratory, CeMM Research Center for Molecular Medicine of the Austrian Academy of Sciences | Lukas Endler, Anna Schedl, Fabian Amman, Petr Triska, Thomas Penz, Benedikt Agerer, Maelle Le Moing, Michael Schuster, Bekir Erguner, Jan Laine, Martin Senekowitsch, Christoph Bock, Andreas Bergthaler |
| EPI_ISL_1685229, EPI_ISL_1685230, EPI_ISL_1685231, EPI_ISL_1685232, EPI_ISL_1685233, EPI_ISL_1685234, EPI_ISL_1685235, EPI_ISL_1685236, EPI_ISL_1685237, EPI_ISL_1685238, EPI_ISL_1685239, EPI_ISL_1685241, EPI_ISL_1685242, EPI_ISL_1685244, EPI_ISL_1685246, EPI_ISL_1685247, EPI_ISL_1685248, EPI_ISL_1685249, EPI_ISL_1685251, EPI_ISL_1685253, EPI_ISL_1685254, EPI_ISL_1685255, EPI_ISL_1685256, EPI_ISL_1685257, EPI_ISL_1685258, EPI_ISL_1685259, EPI_ISL_1685260, EPI_ISL_1685261, EPI_ISL_1685262, EPI_ISL_1685263, EPI_ISL_1685264, EPI_ISL_1685265, EPI_ISL_1685266, EPI_ISL_1685268, EPI_ISL_1685269, EPI_ISL_1685270, EPI_ISL_1685271, EPI_ISL_1685272, EPI_ISL_1685273, EPI_ISL_1685274, EPI_ISL_1685275, EPI_ISL_1685276, EPI_ISL_1685277, EPI_ISL_1685278, EPI_ISL_1685279, EPI_ISL_1685280, EPI_ISL_1685281, EPI_ISL_1685282, EPI_ISL_1685283, EPI_ISL_1685284, EPI_ISL_1685285, EPI_ISL_1685288, EPI_ISL_1685290, EPI_ISL_1685291, EPI_ISL_1685292, EPI_ISL_1685293, EPI_ISL_1685294, EPI_ISL_1685295, EPI_ISL_1685297, EPI_ISL_1685298, EPI_ISL_1685299, EPI_ISL_1685300, EPI_ISL_1685301, EPI_ISL_1685302, EPI_ISL_1685303, EPI_ISL_1685304, EPI_ISL_1685305, EPI_ISL_1685306, EPI_ISL_1685307, EPI_ISL_1685308, EPI_ISL_1685309, EPI_ISL_1685310 | | | |
| see above | Synlab Eesti OÜ | 1. Laboratory of Communicable Diseases (Estonia); 2. Eurofins Genomics Europe Sequencing GmbH | Liidia Dotsenko et al. |
| EPI_ISL_1685311, EPI_ISL_1685312, EPI_ISL_1685313, EPI_ISL_1685314, EPI_ISL_1685315, EPI_ISL_1685316, EPI_ISL_1685317, EPI_ISL_1685318, EPI_ISL_1685319, EPI_ISL_1685320, EPI_ISL_1685321, EPI_ISL_1685322, EPI_ISL_1685324, EPI_ISL_1685325, EPI_ISL_1685326, EPI_ISL_1685328, EPI_ISL_1685329, EPI_ISL_1685330, EPI_ISL_1685331, EPI_ISL_1685332, EPI_ISL_1685334, EPI_ISL_1685335, EPI_ISL_1685336, EPI_ISL_1685337, EPI_ISL_1685338, EPI_ISL_1685339, EPI_ISL_1685340, EPI_ISL_1685341, EPI_ISL_1685344, EPI_ISL_1685345, EPI_ISL_1685346, EPI_ISL_1685347, EPI_ISL_1685349, EPI_ISL_1685350, EPI_ISL_1685351, EPI_ISL_1685352, EPI_ISL_1685353, EPI_ISL_1685355, EPI_ISL_1685356, EPI_ISL_1685358, EPI_ISL_1685359, EPI_ISL_1685360, EPI_ISL_1685362, EPI_ISL_1685363, EPI_ISL_1685364, EPI_ISL_1685365, EPI_ISL_1685366, EPI_ISL_1685367, EPI_ISL_1685368 | | | |
| see above | Noth Estonia Medical Centre | 1. Laboratory of Communicable Diseases (Estonia); 2. Eurofins Genomics Europe Sequencing GmbH | Liidia Dotsenko et al. |
| EPI_ISL_1685375, EPI_ISL_1685378, EPI_ISL_1685380, EPI_ISL_1685381, EPI_ISL_1685382 | Pärnu Hospital | 1. Laboratory of Communicable Diseases (Estonia); 2. Eurofins Genomics Europe Sequencing GmbH | Liidia Dotsenko et al. |
| EPI_ISL_1685383, EPI_ISL_1685384, EPI_ISL_1685385, EPI_ISL_1685386, EPI_ISL_1685387, EPI_ISL_1685389, EPI_ISL_1685391, EPI_ISL_1685392, EPI_ISL_1685393, EPI_ISL_1685395, EPI_ISL_1685396, EPI_ISL_1685397, EPI_ISL_1685398, EPI_ISL_1685399, EPI_ISL_1685400, EPI_ISL_1685401, EPI_ISL_1685403, EPI_ISL_1685404, EPI_ISL_1685405, EPI_ISL_1685406, EPI_ISL_1685409, EPI_ISL_1685410, EPI_ISL_1685411, EPI_ISL_1685413, EPI_ISL_1685414, EPI_ISL_1685415, EPI_ISL_1685416, EPI_ISL_1685417, EPI_ISL_1685418, EPI_ISL_1685419, EPI_ISL_1685420, EPI_ISL_1685421, EPI_ISL_1685422, EPI_ISL_1685423, EPI_ISL_1685424, EPI_ISL_1685425, EPI_ISL_1685426, EPI_ISL_1685427, EPI_ISL_1685428, EPI_ISL_1685429, EPI_ISL_1685430, EPI_ISL_1685431, EPI_ISL_1685432, EPI_ISL_1685433, EPI_ISL_1685434, EPI_ISL_1685435, EPI_ISL_1685436, EPI_ISL_1685437, EPI_ISL_1685438, EPI_ISL_1685439, EPI_ISL_1685440, EPI_ISL_1685441, EPI_ISL_1685442, EPI_ISL_1685443, EPI_ISL_1685444, EPI_ISL_1685445, EPI_ISL_1685446, EPI_ISL_1685447, EPI_ISL_1685448, EPI_ISL_1685450, EPI_ISL_1685451, EPI_ISL_1685452, EPI_ISL_1685453, EPI_ISL_1685454, EPI_ISL_1685455, EPI_ISL_1685456, EPI_ISL_1685457, EPI_ISL_1685458, EPI_ISL_1685459, EPI_ISL_1685460, EPI_ISL_1685461, EPI_ISL_1685462, EPI_ISL_1685463, EPI_ISL_1685464, EPI_ISL_1685465, EPI_ISL_1685994 | | | |
| see above | Synlab Eesti OÜ | 1. Laboratory of Communicable Diseases (Estonia); 2. Eurofins Genomics Europe Sequencing GmbH | Liidia Dotsenko et al. |
| EPI_ISL_1695922 | Hôpital Avicenne | Department of Virology, Henri Mondor University Hospital, Assistance Publique Hôpitaux de Paris, Université Paris-Est Créteil, INSERM U955 | Christophe Rodriguez, Slim Fourati, Vanessa Demontant, Guillaume Gricourt, Melissa N'Debi, Alexandre Soulier, Elisabeth Trawinski, Jean-Michel Pawlotsky |
| EPI_ISL_1696746, EPI_ISL_1696747, EPI_ISL_1696748, EPI_ISL_1696749 | Institute for Medical Research, Infectious Disease Research Centre, National Institutes of Health, Ministry of Health Malaysia | Institute for Medical Research, Infectious Disease Research Centre, National Institutes of Health, Ministry of Health Malaysia | Suppiah J, Kamel K, Thayan R |
| EPI_ISL_1696750, EPI_ISL_1696751, EPI_ISL_1696752, EPI_ISL_1696753, EPI_ISL_1696754, EPI_ISL_1696755 | Institute for Medical Research, Infectious Disease Research Centre, National Institutes of Health, Ministry of Health Malaysia | Institute for Medical Research, Infectious Disease Research Centre, National Institutes of Health, Ministry of Health Malaysia | Suppiah J, Kamel K, Mohd Zawawi Z, Thayan R |
| EPI_ISL_1700563 | Research | National Reference Center for Viruses of Respiratory Infections, Institut Pasteur, Paris | Marion Barbet, Sylvie Behillil, Méline Bizard, Angela Brisebarre, Camille Capel, Vincent Enouf, Louise Lefrançois, Frédéric Lemoine, Christophe Malabat, Corinne Maufrais, Etienne Simon-Lorière, Maud Vanpeene, Sylvie Van der Werf, Marianne Leruez-Ville |
| EPI_ISL_1700783 | HG Pharma GmbH | Bergthaler laboratory, CeMM Research Center for Molecular Medicine of the Austrian Academy of Sciences | Lukas Endler, Anna Schedl, Fabian Amman, Petr Triska, Thomas Penz, Benedikt Agerer, Maelle Le Moing, Michael Schuster, Bekir Erguner, Jan Laine, Martin Senekowitsch, Christoph Bock, Andreas Bergthaler |
| EPI_ISL_1700796, EPI_ISL_1700799 | LESP Nuevo Leon | Instituto de Diagnostico y Referencia Epidemiologicos (INDRE) | Claudia Wong-Arambula, Abril Rodriguez-Maldonado, Vanessa Rivero-Arredondo, Ariadna Medina-Benitez, Joaquin Quiroz-Mercado, Sergio Rangel-Guerrero, Natividad Cruz-Ortiz, Tatiana Nunez-Garcia, Gisela Barrera-Badillo, Lucia Hernandez-Rivas, Irma Lopez-Martinez, Ernesto Ramirez-Gonzalez. |
| EPI_ISL_1700801 | LESP Sonora | Instituto de Diagnostico y Referencia Epidemiologicos (INDRE) | Claudia Wong-Arambula, Abril Rodriguez-Maldonado, Vanessa Rivero-Arredondo, Ariadna Medina-Benitez, Joaquin Quiroz-Mercado, Sergio Rangel-Guerrero, Natividad Cruz-Ortiz, Tatiana Nunez-Garcia, Gisela Barrera-Badillo, Lucia Hernandez-Rivas, Irma Lopez-Martinez, Ernesto Ramirez-Gonzalez. |
| EPI_ISL_1700802 | LESP Morelos | Instituto de Diagnostico y Referencia Epidemiologicos (INDRE) | Claudia Wong-Arambula, Abril Rodriguez-Maldonado, Vanessa Rivero-Arredondo, Ariadna Medina-Benitez, Joaquin Quiroz-Mercado, Sergio Rangel-Guerrero, Natividad Cruz-Ortiz, Tatiana Nunez-Garcia, Gisela Barrera-Badillo, Lucia Hernandez-Rivas, Irma Lopez-Martinez, Ernesto Ramirez-Gonzalez. |
| EPI_ISL_1700807, EPI_ISL_1700808, EPI_ISL_1700809 | LESP Chihuahua | Instituto de Diagnostico y Referencia Epidemiologicos (INDRE) | Claudia Wong-Arambula, Abril Rodriguez-Maldonado, Vanessa Rivero-Arredondo, Ariadna Medina-Benitez, Joaquin Quiroz-Mercado, Sergio Rangel-Guerrero, Natividad Cruz-Ortiz, Tatiana Nunez-Garcia, Gisela Barrera-Badillo, Lucia Hernandez-Rivas, Irma Lopez-Martinez, Ernesto Ramirez-Gonzalez. |
| EPI_ISL_1700810 | LESP Guerrero | Instituto de Diagnostico y Referencia Epidemiologicos (INDRE) | Claudia Wong-Arambula, Abril Rodriguez-Maldonado, Vanessa Rivero-Arredondo, Ariadna Medina-Benitez, Joaquin Quiroz-Mercado, Sergio Rangel-Guerrero, Natividad Cruz-Ortiz, Tatiana Nunez-Garcia, Gisela Barrera-Badillo, Lucia Hernandez-Rivas, Irma Lopez-Martinez, Ernesto Ramirez-Gonzalez. |
| EPI_ISL_1700811, EPI_ISL_1700812, EPI_ISL_1700813, EPI_ISL_1700815 | LESP Veracruz | Instituto de Diagnostico y Referencia Epidemiologicos (INDRE) | Claudia Wong-Arambula, Abril Rodriguez-Maldonado, Vanessa Rivero-Arredondo, Ariadna Medina-Benitez, Joaquin Quiroz-Mercado, Sergio Rangel-Guerrero, Natividad Cruz-Ortiz, Tatiana Nunez-Garcia, Gisela Barrera-Badillo, Lucia Hernandez-Rivas, Irma Lopez-Martinez, Ernesto Ramirez-Gonzalez. |
| EPI_ISL_1700816 | LESP Campeche | Instituto de Diagnostico y Referencia Epidemiologicos (INDRE) | Claudia Wong-Arambula, Abril Rodriguez-Maldonado, Vanessa Rivero-Arredondo, Ariadna Medina-Benitez, Joaquin Quiroz-Mercado, Sergio Rangel-Guerrero, Natividad Cruz-Ortiz, Tatiana Nunez-Garcia, Gisela Barrera-Badillo, Lucia Hernandez-Rivas, Irma Lopez-Martinez, Ernesto Ramirez-Gonzalez. |
| EPI_ISL_1700817 | LESP Tamaulipas | Instituto de Diagnostico y Referencia Epidemiologicos (INDRE) | Claudia Wong-Arambula, Abril Rodriguez-Maldonado, Vanessa Rivero-Arredondo, Ariadna Medina-Benitez, Joaquin Quiroz-Mercado, Sergio Rangel-Guerrero, Natividad Cruz-Ortiz, Tatiana Nunez-Garcia, Gisela Barrera-Badillo, Lucia Hernandez-Rivas, Irma Lopez-Martinez, Ernesto Ramirez-Gonzalez. |
| EPI_ISL_1700829 | LESP Guanajuato | Instituto de Diagnostico y Referencia Epidemiologicos (INDRE) | Claudia Wong-Arambula, Abril Rodriguez-Maldonado, Vanessa Rivero-Arredondo, Ariadna Medina-Benitez, Joaquin Quiroz-Mercado, Sergio Rangel-Guerrero, Natividad Cruz-Ortiz, Tatiana Nunez-Garcia, Gisela Barrera-Badillo, Lucia Hernandez-Rivas, Irma Lopez-Martinez, Ernesto Ramirez-Gonzalez. |
| EPI_ISL_1706476, EPI_ISL_1706477, | Centre Hospitalier Universitaire de Rouen Laboratoire de | Centre Hospitalier Universitaire de Rouen Laboratoire de | Alice Moisan, Fabienne De Oliveira, Marie Leoz |

| | | | |
|---|---|---|---|
| EPI_ISL_1706479 | Virologie | Virologie | |
| EPI_ISL_1706774, EPI_ISL_1706776, EPI_ISL_1706777 | GH A.CHENEVIER-H.MONDOR | Department of Virology, Henri Mondor University Hospital, Assistance Publique Hôpitaux de Paris, Université Paris-Est Créteil, INSERM U955 | Christophe Rodriguez, Slim Fourati, Vanessa Demontant, Guillaume Gricourt, Melissa N'Debi, Alexandre Soulier, Elisabeth Trawinski, Jean-Michel Pawlotsky |
| EPI_ISL_1706780 | GH JOFFRE DUPUYTREN | Department of Virology, Henri Mondor University Hospital, Assistance Publique Hôpitaux de Paris, Université Paris-Est Créteil, INSERM U955 | Christophe Rodriguez, Slim Fourati, Vanessa Demontant, Guillaume Gricourt, Melissa N'Debi, Alexandre Soulier, Elisabeth Trawinski, Jean-Michel Pawlotsky |
| EPI_ISL_1706782, EPI_ISL_1706784, EPI_ISL_1706789 | GH A.CHENEVIER-H.MONDOR | Department of Virology, Henri Mondor University Hospital, Assistance Publique Hôpitaux de Paris, Université Paris-Est Créteil, INSERM U955 | Christophe Rodriguez, Slim Fourati, Vanessa Demontant, Guillaume Gricourt, Melissa N'Debi, Alexandre Soulier, Elisabeth Trawinski, Jean-Michel Pawlotsky |
| EPI_ISL_1706794 | GH JOFFRE DUPUYTREN | Department of Virology, Henri Mondor University Hospital, Assistance Publique Hôpitaux de Paris, Université Paris-Est Créteil, INSERM U955 | Christophe Rodriguez, Slim Fourati, Vanessa Demontant, Guillaume Gricourt, Melissa N'Debi, Alexandre Soulier, Elisabeth Trawinski, Jean-Michel Pawlotsky |
| EPI_ISL_1706796, EPI_ISL_1706797, EPI_ISL_1706799, EPI_ISL_1706807 | GH A.CHENEVIER-H.MONDOR | Department of Virology, Henri Mondor University Hospital, Assistance Publique Hôpitaux de Paris, Université Paris-Est Créteil, INSERM U955 | Christophe Rodriguez, Slim Fourati, Vanessa Demontant, Guillaume Gricourt, Melissa N'Debi, Alexandre Soulier, Elisabeth Trawinski, Jean-Michel Pawlotsky |
| EPI_ISL_1706811 | HOPITAL G. CLEMENCEAU | Department of Virology, Henri Mondor University Hospital, Assistance Publique Hôpitaux de Paris, Université Paris-Est Créteil, INSERM U955 | Christophe Rodriguez, Slim Fourati, Vanessa Demontant, Guillaume Gricourt, Melissa N'Debi, Alexandre Soulier, Elisabeth Trawinski, Jean-Michel Pawlotsky |
| EPI_ISL_1706812, EPI_ISL_1706817, EPI_ISL_1706820, EPI_ISL_1706823, EPI_ISL_1706824, EPI_ISL_1706825, EPI_ISL_1706826, EPI_ISL_1706860, EPI_ISL_1706861 | GH A.CHENEVIER-H.MONDOR | Department of Virology, Henri Mondor University Hospital, Assistance Publique Hôpitaux de Paris, Université Paris-Est Créteil, INSERM U955 | Christophe Rodriguez, Slim Fourati, Vanessa Demontant, Guillaume Gricourt, Melissa N'Debi, Alexandre Soulier, Elisabeth Trawinski, Jean-Michel Pawlotsky |
| EPI_ISL_1707073, EPI_ISL_1707074, EPI_ISL_1707075 | Klinisch Laboratorium ZNA | Klinisch Laboratorium ZNA | Verstrepen et al. |
| EPI_ISL_1707688 | UBDS Dr Marco Antonio Sahao Vila Virginia | Instituto Adolfo Lutz, Interdiciplinary Procedures Center, Strategic Laboratory | Claudio Tavares Sacchi, Claudia Regina Gonçalves, Erica Valessa Ramos Gomes, Karoline Rodrigues Campos, Caio Vinicius Dias Lopes, Leonardo Jose Tadeu de Araujo, Katia Correa de Oliveira Santos |
| EPI_ISL_1708320, EPI_ISL_1708321 | Department of Infectious, Respiratory and Digestive Medicine, Graduate School of Medicine, University of the Ryukyus | Genome Information Research Center, Research Institute for Microbial Diseases, Osaka University | Wakaki Kami, Takeshi Kinjo, Wakako Arakaki, Hiroya Oki, Daisuke Motooka, Shota Nakamura, and Jiro Fujita |
| EPI_ISL_1708607 | LESP Puebla | Instituto de Diagnostico y Referencia Epidemiologicos (INDRE) | Claudia Wong-Arambula, Abril Rodriguez-Maldonado, Vanessa Rivero-Arredondo, Ariadna Medina-Benitez, Joaquin Quiroz-Mercado, Sergio Rangel-Guerrero, Natividad Cruz-Ortiz, Tatiana Nunez-Garcia, Gisela Barrera-Badillo, Lucia Hernandez-Rivas, Irma Lopez-Martinez, Ernesto Ramirez-Gonzalez. |
| EPI_ISL_1710866 | Pathogenic Microorganisms Variability Laboratory | Pathogenic Microorganisms Variability Laboratory | Alexey Shchetinin, Maria Nikiforova, Andrei Siniavin, Andrey Pochtovyy, Evgeny Usachev, Elena Shidlovskaya, Elizaveta Divisenko, Nadezhda Kuznetsova, Valeria Bacalin, Olga Burgasova, Ludmila Kolobukhina, Svetlana Smetanina, Denis Logunov, Vladimir Gushchin, Alexander Gintsburg |
| EPI_ISL_1710966 | LESP Ciudad de Mexico | Instituto de Diagnostico y Referencia Epidemiologicos (INDRE) | Claudia Wong-Arambula, Abril Rodriguez-Maldonado, Vanessa Rivero-Arredondo, Ariadna Medina-Benitez, Joaquin Quiroz-Mercado, Sergio Rangel-Guerrero, Natividad Cruz-Ortiz, Tatiana Nunez-Garcia, Gisela Barrera-Badillo, Lucia Hernandez-Rivas, Irma Lopez-Martinez, Ernesto Ramirez-Gonzalez. |
| EPI_ISL_1712378 | AREA DE SALUD OSA | Inciensa, Instituto Costarricense de Investigación y Enseñanza en Nutrición y Salud | Francisco Duarte, Hebleen Porras, Claudio Soto-Garita, Estela Cordero, Adriana Godínez, Melany Calderón, José Luis Vargas, Mariela Gutiérrez, Joselyn Prado & María Fernanda Matamoros-Madrigal |
| EPI_ISL_1712379 | HOSPITAL SAN JUAN DE DIOS | Inciensa, Instituto Costarricense de Investigación y Enseñanza en Nutrición y Salud | Francisco Duarte, Hebleen Porras, Claudio Soto-Garita, Estela Cordero, Adriana Godínez, Melany Calderón, José Luis Vargas, Mariela Gutiérrez, Joselyn Prado & Alberto Quirós Hidalgo |
| EPI_ISL_1712380 | HOSPITAL GUAPILES | Inciensa, Instituto Costarricense de Investigación y Enseñanza en Nutrición y Salud | Francisco Duarte, Hebleen Porras, Claudio Soto-Garita, Estela Cordero, Adriana Godínez, Melany Calderón, José Luis Vargas, Mariela Gutiérrez, Joselyn Prado & César Cerdas-Quesada |
| EPI_ISL_1712381 | AREA DE SALUD ALAJUELITA | Inciensa, Instituto Costarricense de Investigación y Enseñanza en Nutrición y Salud | Francisco Duarte, Hebleen Porras, Claudio Soto-Garita, Estela Cordero, Adriana Godínez, Melany Calderón, José Luis Vargas, Mariela Gutiérrez, Joselyn Prado & Jonathan Rojas |
| EPI_ISL_1712382, EPI_ISL_1712383, EPI_ISL_1712384, EPI_ISL_1712385 | HOSPITAL MEXICO | Inciensa, Instituto Costarricense de Investigación y Enseñanza en Nutrición y Salud | Francisco Duarte, Hebleen Porras, Claudio Soto-Garita, Estela Cordero, Adriana Godínez, Melany Calderón, José Luis Vargas, Mariela Gutiérrez, Joselyn Prado & Teresita Somogyi |
| EPI_ISL_1712386 | AREA DE SALUD OREAMUNO-PACAYAS-TIERRA BLANCA | Inciensa, Instituto Costarricense de Investigación y Enseñanza en Nutrición y Salud | Francisco Duarte, Hebleen Porras, Claudio Soto-Garita, Estela Cordero, Adriana Godínez, Melany Calderón, José Luis Vargas, Mariela Gutiérrez, Joselyn Prado & Carolina Loría Acosta |
| EPI_ISL_1712387 | AREA DE SALUD CORONADO | Inciensa, Instituto Costarricense de Investigación y Enseñanza en Nutrición y Salud | Francisco Duarte, Hebleen Porras, Claudio Soto-Garita, Estela Cordero, Adriana Godínez, Melany Calderón, José Luis Vargas, Mariela Gutiérrez & Joselyn Prado |
| EPI_ISL_1712388 | AREA DE SALUD LOS SANTOS | Inciensa, Instituto Costarricense de Investigación y Enseñanza en Nutrición y Salud | Francisco Duarte, Hebleen Porras, Claudio Soto-Garita, Estela Cordero, Adriana Godínez, Melany Calderón, José Luis Vargas, Mariela Gutiérrez, Joselyn Prado & Mónica Charpentier-Artavia |
| EPI_ISL_1712389 | AREA DE SALUD TURRIALBA-JIMENEZ | Inciensa, Instituto Costarricense de Investigación y Enseñanza en Nutrición y Salud | Francisco Duarte, Hebleen Porras, Claudio Soto-Garita, Estela Cordero, Adriana Godínez, Melany Calderón, José Luis Vargas, Mariela Gutiérrez, Joselyn Prado & Mónica Charpentier-Artavia |
| EPI_ISL_1712390 | HOSPITAL SAN VICENTE DE PAUL | Inciensa, Instituto Costarricense de Investigación y Enseñanza en Nutrición y Salud | Francisco Duarte, Hebleen Porras, Claudio Soto-Garita, Estela Cordero, Adriana Godínez, Melany Calderón, José Luis Vargas, Mariela Gutiérrez, Joselyn Prado & Silvia Sáenz-García |
| EPI_ISL_1712391, EPI_ISL_1712392, EPI_ISL_1712393, EPI_ISL_1712394, EPI_ISL_1712395 | HOSPITAL DR. TONY FACIO CASTRO | Inciensa, Instituto Costarricense de Investigación y Enseñanza en Nutrición y Salud | Francisco Duarte, Hebleen Porras, Claudio Soto-Garita, Estela Cordero, Adriana Godínez, Melany Calderón, José Luis Vargas, Mariela Gutiérrez, Joselyn Prado & Marcela Rodríguez-Gómez |
| EPI_ISL_1712396 | HOSPITAL DR. FERNANDO ESCALANTE PRADILLA | Inciensa, Instituto Costarricense de Investigación y Enseñanza en Nutrición y Salud | Francisco Duarte, Hebleen Porras, Claudio Soto-Garita, Estela Cordero, Adriana Godínez, Melany Calderón, José Luis Vargas, Mariela Gutiérrez, Joselyn Prado & Fabricio Aguilar |
| EPI_ISL_1712397 | AREA DE SALUD SAN JUAN-SAN DIEGO-CONCEPCION 2 | Inciensa, Instituto Costarricense de Investigación y Enseñanza en Nutrición y Salud | Francisco Duarte, Hebleen Porras, Claudio Soto-Garita, Estela Cordero, Adriana Godínez, Melany Calderón, José Luis Vargas, Mariela Gutiérrez & Joselyn Prado |
| EPI_ISL_1712398 | AREA DE SALUD CORONADO | Inciensa, Instituto Costarricense de Investigación y Enseñanza en Nutrición y Salud | Francisco Duarte, Hebleen Porras, Claudio Soto-Garita, Estela Cordero, Adriana Godínez, Melany Calderón, José Luis Vargas, Mariela Gutiérrez & Joselyn Prado |
| EPI_ISL_1712399 | AREA DE SALUD CATEDRAL NORESTE | Inciensa, Instituto Costarricense de Investigación y Enseñanza en Nutrición y Salud | Francisco Duarte, Hebleen Porras, Claudio Soto-Garita, Estela Cordero, Adriana Godínez, Melany Calderón, José Luis Vargas, Mariela Gutiérrez & Joselyn Prado |
| EPI_ISL_1712400, EPI_ISL_1712401 | HOSPITAL CIUDAD NEILY | Inciensa, Instituto Costarricense de Investigación y Enseñanza en Nutrición y Salud | Francisco Duarte, Hebleen Porras, Claudio Soto-Garita, Estela Cordero, Adriana Godínez, Melany Calderón, José Luis Vargas, Mariela Gutiérrez, Joselyn Prado & Raúl Zeledón-Mayorga |
| EPI_ISL_1712402, EPI_ISL_1712403, EPI_ISL_1712404, EPI_ISL_1712405, | HOSPITAL DR. ENRIQUE BALTODANO BRICEÑO | Inciensa, Instituto Costarricense de Investigación y Enseñanza en Nutrición y Salud | Francisco Duarte, Hebleen Porras, Claudio Soto-Garita, Estela Cordero, Adriana Godínez, Melany Calderón, José Luis Vargas, Mariela Gutiérrez, Joselyn Prado & Adriana Bermúdez-Espinoza |

| | | | |
|---|---|---|---|
| EPI_ISL_1712406 | | | |
| EPI_ISL_1712407 | HOSPITAL SAN VICENTE DE PAUL | Inciensa, Instituto Costarricense de Investigación y Enseñanza en Nutrición y Salud | Francisco Duarte, Hebleen Porras, Claudio Soto-Garita, Estela Cordero, Adriana Godínez, Melany Calderón, José Luis Vargas, Mariela Gutiérrez, Joselyn Prado & Silvia Sáenz-García |
| EPI_ISL_1712408 | AREA DE SALUD BARRANCA - CLINICA DR. ROBERTO SOTO | Inciensa, Instituto Costarricense de Investigación y Enseñanza en Nutrición y Salud | Francisco Duarte, Hebleen Porras, Claudio Soto-Garita, Estela Cordero, Adriana Godínez, Melany Calderón, José Luis Vargas, Mariela Gutiérrez, Joselyn Prado & Andrea Moreno-Carvajal |
| EPI_ISL_1712409 | AREA DE SALUD GARABITO | Inciensa, Instituto Costarricense de Investigación y Enseñanza en Nutrición y Salud | Francisco Duarte, Hebleen Porras, Claudio Soto-Garita, Estela Cordero, Adriana Godínez, Melany Calderón, José Luis Vargas, Mariela Gutiérrez, Joselyn Prado & Andrea Moreno-Carvajal |
| EPI_ISL_1712410 | AREA DE SALUD ALAJUELITA | Inciensa, Instituto Costarricense de Investigación y Enseñanza en Nutrición y Salud | Francisco Duarte, Hebleen Porras, Claudio Soto-Garita, Estela Cordero, Adriana Godínez, Melany Calderón, José Luis Vargas, Mariela Gutiérrez, Joselyn Prado & Jonathan Rojas |
| EPI_ISL_1712411 | AREA DE SALUD CORONADO | Inciensa, Instituto Costarricense de Investigación y Enseñanza en Nutrición y Salud | Francisco Duarte, Hebleen Porras, Claudio Soto-Garita, Estela Cordero, Adriana Godínez, Melany Calderón, José Luis Vargas, Mariela Gutiérrez & Joselyn Prado. |
| EPI_ISL_1712412, EPI_ISL_1712413 | LABORATORIO CLINICO LABIN | Inciensa, Instituto Costarricense de Investigación y Enseñanza en Nutrición y Salud | Francisco Duarte, Hebleen Porras, Claudio Soto-Garita, Estela Cordero, Adriana Godínez, Melany Calderón, José Luis Vargas, Mariela Gutiérrez, Joselyn Prado & Pei Ling Chan Ma |
| EPI_ISL_1712414 | AREA DE SALUD DESAMPARADOS 1 - CLINICA DR. MARCIAL FALLAS | Inciensa, Instituto Costarricense de Investigación y Enseñanza en Nutrición y Salud | Francisco Duarte, Hebleen Porras, Claudio Soto-Garita, Estela Cordero, Adriana Godínez, Melany Calderón, José Luis Vargas, Mariela Gutiérrez, Joselyn Prado & Katherine Alfaro |
| EPI_ISL_1712415 | AREA DE SALUD ASERRI | Inciensa, Instituto Costarricense de Investigación y Enseñanza en Nutrición y Salud | Francisco Duarte, Hebleen Porras, Claudio Soto-Garita, Estela Cordero, Adriana Godínez, Melany Calderón, José Luis Vargas, Mariela Gutiérrez, Joselyn Prado & Alberto Quirós-Hidalgo |
| EPI_ISL_1712416 | HOSPITAL DR. CARLOS LUIS VALVERDE VEGA | Inciensa, Instituto Costarricense de Investigación y Enseñanza en Nutrición y Salud | Francisco Duarte, Hebleen Porras, Claudio Soto-Garita, Estela Cordero, Adriana Godínez, Melany Calderón, José Luis Vargas, Mariela Gutiérrez, Joselyn Prado & Teresita Somogyi |
| EPI_ISL_1712417 | HOSPITAL SAN FRANCISCO DE ASIS | Inciensa, Instituto Costarricense de Investigación y Enseñanza en Nutrición y Salud | Francisco Duarte, Hebleen Porras, Claudio Soto-Garita, Estela Cordero, Adriana Godínez, Melany Calderón, José Luis Vargas, Mariela Gutiérrez, Joselyn Prado & Teresita Somogyi |
| EPI_ISL_1712418 | AREA DE SALUD NARANJO | Inciensa, Instituto Costarricense de Investigación y Enseñanza en Nutrición y Salud | Francisco Duarte, Hebleen Porras, Claudio Soto-Garita, Estela Cordero, Adriana Godínez, Melany Calderón, José Luis Vargas, Mariela Gutiérrez, Joselyn Prado & Teresita Somogyi |
| EPI_ISL_1712419 | AREA DE SALUD HEREDIA-CUBUJUQUI - CLINICA DR. FRANCISCO BOLAÑOS | Inciensa, Instituto Costarricense de Investigación y Enseñanza en Nutrición y Salud | Francisco Duarte, Hebleen Porras, Claudio Soto-Garita, Estela Cordero, Adriana Godínez, Melany Calderón, José Luis Vargas, Mariela Gutiérrez, Joselyn Prado & Teresita Somogyi |
| EPI_ISL_1712420, EPI_ISL_1712421 | LABORATORIO CLINICO LABIN | Inciensa, Instituto Costarricense de Investigación y Enseñanza en Nutrición y Salud | Francisco Duarte, Hebleen Porras, Claudio Soto-Garita, Estela Cordero, Adriana Godínez, Melany Calderón, José Luis Vargas, Mariela Gutiérrez, Joselyn Prado & David Rodríguez-Masis |
| EPI_ISL_1714796, EPI_ISL_1714797 | Institute for Medical Research, Infectious Disease Research Centre, National Institutes of Health, Ministry of Health Malaysia | Institute for Medical Research, Infectious Disease Research Centre, National Institutes of Health, Ministry of Health Malaysia | Suppiah J, Kamel K, Mohd Zawawi Z, Thayan R |
| EPI_ISL_1715134 | Centro de Saude II Ibitinga | Instituto Adolfo Lutz, Interdiciplinary Procedures Center, Strategic Laboratory | Claudio Tavares Sacchi, Claudia Regina Gonçalves, Erica Valessa Ramos Gomes, Karoline Rodrigues Campos, Caio Vinicius Dias Lopes, Leonardo Jose Tadeu de Araujo, Katia Correa de Oliveira Santos |
| EPI_ISL_1715135, EPI_ISL_1715136 | CS II Dr Jose Ferreira Telles | Instituto Adolfo Lutz, Interdiciplinary Procedures Center, Strategic Laboratory | Claudio Tavares Sacchi, Claudia Regina Gonçalves, Erica Valessa Ramos Gomes, Karoline Rodrigues Campos, Caio Vinicius Dias Lopes, Leonardo Jose Tadeu de Araujo, Katia Correa de Oliveira Santos |
| EPI_ISL_1715137 | Secretaria Municipal De Saude | Instituto Adolfo Lutz, Interdiciplinary Procedures Center, Strategic Laboratory | Claudio Tavares Sacchi, Claudia Regina Gonçalves, Erica Valessa Ramos Gomes, Karoline Rodrigues Campos, Caio Vinicius Dias Lopes, Leonardo Jose Tadeu de Araujo, Katia Correa de Oliveira Santos |
| EPI_ISL_1715138 | Santa Casa De Cravinhos | Instituto Adolfo Lutz, Interdiciplinary Procedures Center, Strategic Laboratory | Claudio Tavares Sacchi, Claudia Regina Gonçalves, Erica Valessa Ramos Gomes, Karoline Rodrigues Campos, Caio Vinicius Dias Lopes, Leonardo Jose Tadeu de Araujo, Katia Correa de Oliveira Santos |
| EPI_ISL_1715139 | Sae Servico De Atendimento Especializado | Instituto Adolfo Lutz, Interdiciplinary Procedures Center, Strategic Laboratory | Claudio Tavares Sacchi, Claudia Regina Gonçalves, Erica Valessa Ramos Gomes, Karoline Rodrigues Campos, Caio Vinicius Dias Lopes, Leonardo Jose Tadeu de Araujo, Katia Correa de Oliveira Santos |
| EPI_ISL_1715141 | Centro De Saude II Ibitinga | Instituto Adolfo Lutz, Interdiciplinary Procedures Center, Strategic Laboratory | Claudio Tavares Sacchi, Claudia Regina Gonçalves, Erica Valessa Ramos Gomes, Karoline Rodrigues Campos, Caio Vinicius Dias Lopes, Leonardo Jose Tadeu de Araujo, Katia Correa de Oliveira Santos |
| EPI_ISL_1715143 | Secretaria Municipal de Saude De Guariba | Instituto Adolfo Lutz, Interdiciplinary Procedures Center, Strategic Laboratory | Claudio Tavares Sacchi, Claudia Regina Gonçalves, Erica Valessa Ramos Gomes, Karoline Rodrigues Campos, Caio Vinicius Dias Lopes, Leonardo Jose Tadeu de Araujo, Katia Correa de Oliveira Santos |
| EPI_ISL_1715144 | UBDS Dr Joao Baptista Quartim Central | Instituto Adolfo Lutz, Interdiciplinary Procedures Center, Strategic Laboratory | Claudio Tavares Sacchi, Claudia Regina Gonçalves, Erica Valessa Ramos Gomes, Karoline Rodrigues Campos, Caio Vinicius Dias Lopes, Leonardo Jose Tadeu de Araujo, Katia Correa de Oliveira Santos |
| EPI_ISL_1715145 | UPA de Bebedouro | Instituto Adolfo Lutz, Interdiciplinary Procedures Center, Strategic Laboratory | Claudio Tavares Sacchi, Claudia Regina Gonçalves, Erica Valessa Ramos Gomes, Karoline Rodrigues Campos, Caio Vinicius Dias Lopes, Leonardo Jose Tadeu de Araujo, Katia Correa de Oliveira Santos |
| EPI_ISL_1715159 | Instituto Adolfo Lutz Central | Instituto Adolfo Lutz, Interdiciplinary Procedures Center, Strategic Laboratory | Claudio Tavares Sacchi, Claudia Regina Gonçalves, Erica Valessa Ramos Gomes, Karoline Rodrigues Campos, Caio Vinicius Dias Lopes, Leonardo Jose Tadeu de Araujo, Katia Correa de Oliveira Santos |
| EPI_ISL_1715418, EPI_ISL_1715420, EPI_ISL_1715421, EPI_ISL_1715422, EPI_ISL_1715423, EPI_ISL_1715424 | University of Rome Tor Vergata: Departm Experim Medicine Chair of Virology | University of Rome Tor Vergata: Departm Experim Medicine Chair of Virology | Maria Concetta Bellocchi, Luca Carioti, Maria Botticelli, Lorenzo Piermatteo, Marco Iannetta, Loredana Sarmati, Massimo Andreoni, Francesca Ceccherini-Silberstein |
| EPI_ISL_1731562, EPI_ISL_1731563, EPI_ISL_1731564, EPI_ISL_1731565, EPI_ISL_1731566 | MRCG at LSHTM Genomics lab | MRCG at LSHTM Genomics lab | Abdul Karim sesay, Abdoulie Kanteh, Jarra Manneh, Mariama Kujabi, Bakary Sanyang |
| EPI_ISL_1731574 | UBDS Dr Marco Antonio Sahao Vila Virginia | Instituto Adolfo Lutz, Interdiciplinary Procedures Center, Strategic Laboratory | Claudio Tavares Sacchi, Claudia Regina Gonçalves, Erica Valessa Ramos Gomes, Karoline Rodrigues Campos, Caio Vinicius Dias Lopes, Leonardo Jose Tadeu de Araujo, Katia Correa de Oliveira Santos |
| EPI_ISL_1731576 | UPA Dr Luis Atilio Losi Viana Ribeirao Preto | Instituto Adolfo Lutz, Interdiciplinary Procedures Center, Strategic Laboratory | Claudio Tavares Sacchi, Claudia Regina Gonçalves, Erica Valessa Ramos Gomes, Karoline Rodrigues Campos, Caio Vinicius Dias Lopes, Leonardo Jose Tadeu de Araujo, Katia Correa de Oliveira Santos |
| EPI_ISL_1732020 | North Estonia Medical Centre | 1. Laboratory of Communicable Diseases (Estonia); 2. Eurofins Genomics Europe Sequencing GmbH | Liidia Dotsenko et al. |
| EPI_ISL_1743715 | Usansolo-Galdakao University Hospital | Cruces University Hospital | Ana Gual-de-Torrella, Izaskun Alejo-Cancho, Mikel Gallego, Ana Belén de la Hoz |
| EPI_ISL_1749433 | Singapore General Hospital | Department of Microbiology | Nurdyana Abdul Rahman, Kun Lee Lim, Chenhao Li, Sui Sin Goh, Kenneth Xin Long Chan, Kian Sing Chan, Lynette Oon, Kern Rei Chng, Niranjan Nagarajan, Karrie Ko |
| EPI_ISL_1752649, EPI_ISL_1752650, EPI_ISL_1752651, EPI_ISL_1752652, EPI_ISL_1752653, EPI_ISL_1752654, EPI_ISL_1752657, EPI_ISL_1752662 | Instituto Adolfo Lutz Central | Instituto Adolfo Lutz, Interdiciplinary Procedures Center, Strategic Laboratory | Claudio Tavares Sacchi, Claudia Regina Gonçalves, Erica Valessa Ramos Gomes, Karoline Rodrigues Campos, Caio Vinicius Dias Lopes, Leonardo Jose Tadeu de Araujo, Katia Correa de Oliveira Santos |

EPI_ISL_1753942, EPI_ISL_1753944, EPI_ISL_1753945, EPI_ISL_1753952, EPI_ISL_1753953, EPI_ISL_1753956, EPI_ISL_1753960, EPI_ISL_1753964, EPI_ISL_1753983, EPI_ISL_1753986, EPI_ISL_1753988, EPI_ISL_1753989, EPI_ISL_1753992, EPI_ISL_1753993, EPI_ISL_1753995, EPI_ISL_1753996, EPI_ISL_1753998, EPI_ISL_1754001, EPI_ISL_1754005, EPI_ISL_1754009, EPI_ISL_1754010, EPI_ISL_1754017, EPI_ISL_1754018, EPI_ISL_1754021, EPI_ISL_1754025, EPI_ISL_1754030, EPI_ISL_1754033, EPI_ISL_1754035, EPI_ISL_1754036, EPI_ISL_1754037, EPI_ISL_1754039, EPI_ISL_1754043, EPI_ISL_1754048, EPI_ISL_1754056,

| | | | |
|---|---|---|---|
| EPI_ISL_1754057, EPI_ISL_1754058, EPI_ISL_1754060, EPI_ISL_1754063, EPI_ISL_1754064, EPI_ISL_1754066, EPI_ISL_1754067, EPI_ISL_1754069, EPI_ISL_1754070, EPI_ISL_1754072, EPI_ISL_1754073, EPI_ISL_1754074, EPI_ISL_1754078, EPI_ISL_1754079, EPI_ISL_1754082, EPI_ISL_1754083, EPI_ISL_1754086, EPI_ISL_1754087, EPI_ISL_1754090, EPI_ISL_1754091, EPI_ISL_1754095, EPI_ISL_1754096, EPI_ISL_1754098, EPI_ISL_1754099 | | | |
| see above | Servicio Microbiología Hospital La Paz | Servicio Microbiología Hospital La Paz | Fernando Lázaro, Rubén Cáceres, Jesús Mingorance Cruz, Elie Dahdouh |
| EPI_ISL_1754220, EPI_ISL_1754223, EPI_ISL_1754224, EPI_ISL_1754227, EPI_ISL_1754251 | GH A.CHENEVIER-H.MONDOR | Department of Virology, Henri Mondor University Hospital, Assistance Publique Hôpitaux de Paris, Université Paris-Est Créteil, INSERM U955 | Christophe Rodriguez, Slim Fourati, Vanessa Demontant, Guillaume Gricourt, Melissa N'Debi, Alexandre Soulier, Elisabeth Trawinski, Jean-Michel Pawlotsky |
| EPI_ISL_1760250, EPI_ISL_1760252, EPI_ISL_1760256, EPI_ISL_1760259, EPI_ISL_1760260, EPI_ISL_1760292, EPI_ISL_1760293, EPI_ISL_1760294, EPI_ISL_1760295 | Centers for Disease Control and Prevention, Dengue Branch | Centers for Disease Control and Prevention, Dengue Branch | Gilberto A. Santiago, Glenda Gonzalez, Betzabel Flores, Keyla Charriez, Gabriela Paz-Bailey, Jorge L. Munoz-Jordan |
| EPI_ISL_1760296, EPI_ISL_1760297, EPI_ISL_1760298, EPI_ISL_1760299 | Puerto Rico Department of Health | Centers for Disease Control and Prevention, Dengue Branch | Gilberto A. Santiago, Glenda Gonzalez, Betzabel Flores, Keyla Charriez, Gabriela Paz-Bailey, Jorge L. Munoz-Jordan |
| EPI_ISL_1761359, EPI_ISL_1761360, EPI_ISL_1761361, EPI_ISL_1761362, EPI_ISL_1761364, EPI_ISL_1761365 | Tokyo Medical and Dental University | Tokyo Medical and Dental University | Kousuke Tanimoto, Yukie Tanaka, Yoko Nukui, Akinori Kimura, Hiroaki Takeuchi |
| EPI_ISL_1761589, EPI_ISL_1761590, EPI_ISL_1761592, EPI_ISL_1761593, EPI_ISL_1761596, EPI_ISL_1761600, EPI_ISL_1761601, EPI_ISL_1761603, EPI_ISL_1761604, EPI_ISL_1761606, EPI_ISL_1761607, EPI_ISL_1761608 | | | |
| see above | Althaia. Xarxa Assistencial Universitària de Manresa | IrsiCaixa | Marc Noguera-Julian, Mariona Parera, Maria Casadellà, Pilar Armengol, Francesc Catala-Moll, Roger Paredes, Bonaventura Clotet Gloria Trujillo, Rafel Perez Vidal, Jaume Trape Pujol, Carolina Gonzalez Fernandez, Miquel Micó, Antonia Flor, Eulalia Grau |
| EPI_ISL_1788762 | Research | National Reference Center for Viruses of Respiratory Infections, Institut Pasteur, Paris | Marion Barbet, Sylvie Behillil, Méline Bizard, Angela Brisebarre, Camille Capel, Vincent Enouf, Louise Lefrançois, Frédéric Lemoine, Christophe Malabat, Corinne Maufrais, Etienne Simon-Lorière, Maud Vanpeene, Sylvie Van der Werf ,Marianne Leruez-Ville |
| EPI_ISL_1805632 | LDSP TOLIMA | Instituto Nacional de Salud- Dirección de Investigación en Salud Pública | Katherine Laiton-Donato, Diego A. Álvarez-Díaz, Carlos Franco-Muñoz, Hector Alejandro Ruiz-Moreno, Paola Rojas, Maria T. Herrera-Sepúlveda, Diego Andrés Prada, Jhonnatan Reales-González, Sheryll Corchuelo, Julian Naizaque, Jorge Rivera, Gerardo Santamaría, Sergio Gomez, Lisseth Pardo, Juan Camilo Martinez, Marta Lopez Blanco, Ángela Alarcon Cruz, Diana Malo, Carmen Osorio, Magdalena Wiesner, Martha Lucia Ospina Martinez, Marcela Mercado-Reyes |
| EPI_ISL_1805959, EPI_ISL_1805960, EPI_ISL_1805961, EPI_ISL_1805962 | National Center for Communicable Diseases (NCCD) National Influenza Center | National Center for Communicable Diseases (NCCD) National Influenza Center | Naranzul Ts,Bayasgalan N,Khishigmunkh Ch,Seiichiro F,Hideka M,Mina N,Shinji W,Ankhbayar S,Tsogzolmaa G,Darmaa B,Battur L,Nymadawa P |
| EPI_ISL_1817584 | Servicio Microbiología Hospital La Paz | Servicio Microbiología Hospital La Paz | Fernando Lázaro, Rubén Cáceres, Jesús Mingorance Cruz, Elie Dahdouh |
| EPI_ISL_1819132, EPI_ISL_1819133, EPI_ISL_1819134, EPI_ISL_1819135 | Laboratori de Referencia de Catalunya | Laboratori de Referencia de Catalunya | Ramirez A., Bellosillo B., Padilla E., Vilas A., Hernandez JJ., Canal M. |
| EPI_ISL_1821204 | Santa Casa de Cravinhos | Instituto Adolfo Lutz, Interdiciplinary Procedures Center, Strategic Laboratory | Claudio Tavares Sacchi, Claudia Regina Gonçalves, Erica Valessa Ramos Gomes, Karoline Rodrigues Campos, Caio Vinicius Dias Lopes, Leonardo Jose Tadeu de Araujo |
| EPI_ISL_1821206 | Sae Servico de Atendimento Especializado | Instituto Adolfo Lutz, Interdiciplinary Procedures Center, Strategic Laboratory | Claudio Tavares Sacchi, Claudia Regina Gonçalves, Erica Valessa Ramos Gomes, Karoline Rodrigues Campos, Caio Vinicius Dias Lopes, Leonardo Jose Tadeu de Araujo |
| EPI_ISL_1821207 | Secretaria Municipal de Saúde de Lins | Instituto Adolfo Lutz, Interdiciplinary Procedures Center, Strategic Laboratory | Claudio Tavares Sacchi, Claudia Regina Gonçalves, Erica Valessa Ramos Gomes, Karoline Rodrigues Campos, Caio Vinicius Dias Lopes, Leonardo Jose Tadeu de Araujo |
| EPI_ISL_1821210 | Hospital Universitario de Marilia | Instituto Adolfo Lutz, Interdiciplinary Procedures Center, Strategic Laboratory | Claudio Tavares Sacchi, Claudia Regina Gonçalves, Erica Valessa Ramos Gomes, Karoline Rodrigues Campos, Caio Vinicius Dias Lopes, Leonardo Jose Tadeu de Araujo |
| EPI_ISL_1821211, EPI_ISL_1821212, EPI_ISL_1821213 | Diagnóstico da America S/A | Instituto Adolfo Lutz, Interdiciplinary Procedures Center, Strategic Laboratory | Claudio Tavares Sacchi, Claudia Regina Gonçalves, Erica Valessa Ramos Gomes, Karoline Rodrigues Campos, Caio Vinicius Dias Lopes, Leonardo Jose Tadeu de Araujo |
| EPI_ISL_1821234, EPI_ISL_1821235, EPI_ISL_1821236, EPI_ISL_1821237, EPI_ISL_1821238, EPI_ISL_1821243, EPI_ISL_1821244, EPI_ISL_1821245 | Instituto Adolfo Lutz - Regional de Marilia | Instituto Adolfo Lutz, Interdiciplinary Procedures Center, Strategic Laboratory | Claudio Tavares Sacchi, Claudia Regina Gonçalves, Erica Valessa Ramos Gomes, Karoline Rodrigues Campos, Caio Vinicius Dias Lopes, Leonardo Jose Tadeu de Araujo |
| EPI_ISL_1823404, EPI_ISL_1823406, EPI_ISL_1823410, EPI_ISL_1823414, EPI_ISL_1823415, EPI_ISL_1823419, EPI_ISL_1823421, EPI_ISL_1823423, EPI_ISL_1823425, EPI_ISL_1823428, EPI_ISL_1823430, EPI_ISL_1823432, EPI_ISL_1823433, EPI_ISL_1823437, EPI_ISL_1823440, EPI_ISL_1823441, EPI_ISL_1823442, EPI_ISL_1823444, EPI_ISL_1823445, EPI_ISL_1823446, EPI_ISL_1823450 | | | |
| see above | Indiana Animal Disease Diagnostic Laboratory | Carpi Laboratory - Purdue University | Jack Dorman, Ilinca I Ciubotariu, Nicole M Perry, Jobin J Kattoor, Lev Gorenstein, Abebe A Fola, G Kenitra Hendrix, Rebecca P Wilkes, Giovanna Carpi |
| EPI_ISL_1824418 | AZ Klina | Augustijnslei 100 2930 Brasschaat | Carl Vael - Lynsey Berckmans |
| EPI_ISL_1824601, EPI_ISL_1824602, EPI_ISL_1824603, EPI_ISL_1824610, EPI_ISL_1824611, EPI_ISL_1824613, EPI_ISL_1824614 | National Institute of Health Research and Development | National Institute of Health Research and Development | Subangkit, Hana Apsari Pawestri, Kartika Dewi Puspa, Arie Ardiansyah Nugraha, Hartanti Dian Ikawati, Krisna Nur Andriana Pangesti, Yuni Rukminiati, Ririn Ramadhany, Agustiningsih, Kindi Adam, Holy Arif Wibowo, Triyani Soekarso, Ni Ketut Susilarini, Nurika Hariastuti, Ully Alfi Nikmah, Reni Herman, Nike Susanti, Herna, Tati Febriyanti, Natalie Laurencia Kipuw, Fauzul Muna, Irene Lorinda Indalao, Nelly Puspandari, Vivi Setiawaty. |
| EPI_ISL_1824618 | RSUD Bahteramas Kendari | National Institute of Health Research and Development | Hana Apsari Pawestri, Subangkit, Kartika Dewi Puspa, Arie Ardiansyah Nugraha, Hartanti Dian Ikawati, Krisna Pangesti, Nelly Puspandari, Vivi Setiawaty |
| EPI_ISL_1824636 | COVID Laboratory AOU Messina | Molecular Hepatology Lab AOU Messina | Giuseppina Raffa, Cristina Musolino, Valeria Chines, Daniele Lombardo, Domenico Giosa, Teresa Pollicino. |
| EPI_ISL_1824719 | National Institute of Health Research and Development | National Institute of Health Research and Development | Subangkit, Hana Apsari Pawestri, Kartika Dewi Puspa, Arie Ardiansyah Nugraha, Hartanti Dian Ikawati, Krisna Nur Andriana Pangesti, Yuni Rukminiati, Ririn Ramadhany, Agustiningsih, Kindi Adam, Holy Arif Wibowo, Triyani Soekarso, Ni Ketut Susilarini, Nurika Hariastuti, Ully Alfi Nikmah, Reni Herman, Nike Susanti, Herna, Tati Febriyanti, Natalie Laurencia Kipuw, Fauzul Muna, Irene Lorinda Indalao, Nelly Puspandari, Vivi Setiawaty. |
| EPI_ISL_1827503 | AREA DE SALUD CATEDRAL NORESTE | Inciensa, Instituto Costarricense de Investigación y Enseñanza en Nutrición y Salud | Francisco Duarte, Hebleen Porras, Claudio Soto-Garita, Estela Cordero, Adriana Godínez, Melany Calderón, José Luis Vargas, Mariela Gutiérrez & Joselyn Prado |
| EPI_ISL_1827504 | AREA DE SALUD COTO BRUS | Inciensa, Instituto Costarricense de Investigación y Enseñanza en Nutrición y Salud | Francisco Duarte, Hebleen Porras, Claudio Soto-Garita, Estela Cordero, Adriana Godínez, Melany Calderón, José Luis Vargas, Mariela Gutiérrez, Joselyn Prado & María José Ramírez |
| EPI_ISL_1827505, EPI_ISL_1827506 | HOSPITAL METROPOLITANO | Inciensa, Instituto Costarricense de Investigación y Enseñanza en Nutrición y Salud | Francisco Duarte, Hebleen Porras, Claudio Soto-Garita, Estela Cordero, Adriana Godínez, Melany Calderón, José Luis Vargas, Mariela Gutiérrez, Joselyn Prado & Margarita Lee-Lui |
| EPI_ISL_1827507 | CENTRO PENAL LUIS PAULINO MORA | Inciensa, Instituto Costarricense de Investigación y Enseñanza en Nutrición y Salud | Francisco Duarte, Hebleen Porras, Claudio Soto-Garita, Estela Cordero, Adriana Godínez, Melany Calderón, José Luis Vargas, Mariela Gutiérrez & Joselyn Prado |
| EPI_ISL_1827508 | AREA DE SALUD LOS SANTOS | Inciensa, Instituto Costarricense de Investigación y Enseñanza en Nutrición y Salud | Francisco Duarte, Hebleen Porras, Claudio Soto-Garita, Estela Cordero, Adriana Godínez, Melany Calderón, José Luis Vargas, Mariela Gutiérrez & Joselyn Prado |
| EPI_ISL_1827509 | AREA DE SALUD GUATUSO | Inciensa, Instituto Costarricense de Investigación y Enseñanza en Nutrición y Salud | Francisco Duarte, Hebleen Porras, Claudio Soto-Garita, Estela Cordero, Adriana Godínez, Melany Calderón, José Luis Vargas, Mariela Gutiérrez, Joselyn Prado & Francisco Chacón-Valverde |
| EPI_ISL_1827510 | HOSPITAL CIUDAD NEILY | Inciensa, Instituto Costarricense de Investigación y Enseñanza en Nutrición y Salud | Francisco Duarte, Hebleen Porras, Claudio Soto-Garita, Estela Cordero, Adriana Godínez, Melany Calderón, José Luis Vargas, Mariela Gutiérrez, Joselyn Prado & Nelly Castro-Bonilla |

| | | | |
|---|---|---|---|
| EPI_ISL_1827511 | AREA DE SALUD ALAJUELITA | Inciensa, Instituto Costarricense de Investigación y Enseñanza en Nutrición y Salud | Francisco Duarte, Hebleen Porras, Claudio Soto-Garita, Estela Cordero, Adriana Godínez, Melany Calderón, José Luis Vargas, Mariela Gutiérrez, Joselyn Prado & Jonathan Rojas |
| EPI_ISL_1827512, EPI_ISL_1827513 | HOSPITAL SAN VICENTE DE PAUL | Inciensa, Instituto Costarricense de Investigación y Enseñanza en Nutrición y Salud | Francisco Duarte, Hebleen Porras, Claudio Soto-Garita, Estela Cordero, Adriana Godínez, Melany Calderón, José Luis Vargas, Mariela Gutiérrez, Joselyn Prado & Silvia Sáenz-García |
| EPI_ISL_1827514 | HOSPITAL GUAPILES | Inciensa, Instituto Costarricense de Investigación y Enseñanza en Nutrición y Salud | Francisco Duarte, Hebleen Porras, Claudio Soto-Garita, Estela Cordero, Adriana Godínez, Melany Calderón, José Luis Vargas, Mariela Gutiérrez, Joselyn Prado & César Cerdas-Quesada |
| EPI_ISL_1827515, EPI_ISL_1827516, EPI_ISL_1827517 | HOSPITAL MONSEÑOR SANABRIA | Inciensa, Instituto Costarricense de Investigación y Enseñanza en Nutrición y Salud | Francisco Duarte, Hebleen Porras, Claudio Soto-Garita, Estela Cordero, Adriana Godínez, Melany Calderón, José Luis Vargas, Mariela Gutiérrez, Joselyn Prado & Andrea Moreno-Carvajal |
| EPI_ISL_1827518 | AREA DE SALUD PURISCAL-TURRUBARES | Inciensa, Instituto Costarricense de Investigación y Enseñanza en Nutrición y Salud | Francisco Duarte, Hebleen Porras, Claudio Soto-Garita, Estela Cordero, Adriana Godínez, Melany Calderón, José Luis Vargas, Mariela Gutiérrez, Joselyn Prado & Maribel Chavarría Azofeifa |
| EPI_ISL_1827519 | LABORATORIO CLINICO LABIN | Inciensa, Instituto Costarricense de Investigación y Enseñanza en Nutrición y Salud | Francisco Duarte, Hebleen Porras, Claudio Soto-Garita, Estela Cordero, Adriana Godínez, Melany Calderón, José Luis Vargas, Mariela Gutiérrez, Joselyn Prado & Pei Ling Chan Ma |
| EPI_ISL_1827520 | AREA DE SALUD SIQUIRRES | Inciensa, Instituto Costarricense de Investigación y Enseñanza en Nutrición y Salud | Francisco Duarte, Hebleen Porras, Claudio Soto-Garita, Estela Cordero, Adriana Godínez, Melany Calderón, José Luis Vargas, Mariela Gutiérrez, Joselyn Prado & Ileana Chaves-Peraza |
| EPI_ISL_1827521 | AREA DE SALUD DESAMPARADOS 2 (COOPESALUD) | Inciensa, Instituto Costarricense de Investigación y Enseñanza en Nutrición y Salud | Francisco Duarte, Hebleen Porras, Claudio Soto-Garita, Estela Cordero, Adriana Godínez, Melany Calderón, José Luis Vargas, Mariela Gutiérrez, Joselyn Prado & Nicole Vargas-Víquez |
| EPI_ISL_1827522 | AREA DE SALUD DESAMPARADOS 1 - CLINICA DR. MARCIAL FALLAS | Inciensa, Instituto Costarricense de Investigación y Enseñanza en Nutrición y Salud | Francisco Duarte, Hebleen Porras, Claudio Soto-Garita, Estela Cordero, Adriana Godínez, Melany Calderón, José Luis Vargas, Mariela Gutiérrez, Joselyn Prado & Nicole Vargas-Víquez |
| EPI_ISL_1827523 | AREA DE SALUD CURRIDABAT 2 | Inciensa, Instituto Costarricense de Investigación y Enseñanza en Nutrición y Salud | Francisco Duarte, Hebleen Porras, Claudio Soto-Garita, Estela Cordero, Adriana Godínez, Melany Calderón, José Luis Vargas, Mariela Gutiérrez & Joselyn Prado |
| EPI_ISL_1827524 | HOSPITAL DR. CARLOS LUIS VALVERDE VEGA | Inciensa, Instituto Costarricense de Investigación y Enseñanza en Nutrición y Salud | Francisco Duarte, Hebleen Porras, Claudio Soto-Garita, Estela Cordero, Adriana Godínez, Melany Calderón, José Luis Vargas, Mariela Gutiérrez, Joselyn Prado & Teresita Somogyi |
| EPI_ISL_1827525 | AREA DE SALUD LOS CHILES | Inciensa, Instituto Costarricense de Investigación y Enseñanza en Nutrición y Salud | Francisco Duarte, Hebleen Porras, Claudio Soto-Garita, Estela Cordero, Adriana Godínez, Melany Calderón, José Luis Vargas, Mariela Gutiérrez, Joselyn Prado & Teresita Somogyi |
| EPI_ISL_1827526 | AREA DE SALUD ALAJUELA OESTE | Inciensa, Instituto Costarricense de Investigación y Enseñanza en Nutrición y Salud | Francisco Duarte, Hebleen Porras, Claudio Soto-Garita, Estela Cordero, Adriana Godínez, Melany Calderón, José Luis Vargas, Mariela Gutiérrez, Joselyn Prado & Teresita Somogyi |
| EPI_ISL_1827530 | HOSPITAL METROPOLITANO | Inciensa, Instituto Costarricense de Investigación y Enseñanza en Nutrición y Salud | Francisco Duarte, Hebleen Porras, Claudio Soto-Garita, Estela Cordero, Adriana Godínez, Melany Calderón, José Luis Vargas, Mariela Gutiérrez, Joselyn Prado & Margarita Lee-Lui |
| EPI_ISL_1827531 | AREA DE SALUD LIMON | Inciensa, Instituto Costarricense de Investigación y Enseñanza en Nutrición y Salud | Francisco Duarte, Hebleen Porras, Claudio Soto-Garita, Estela Cordero, Adriana Godínez, Melany Calderón, José Luis Vargas, Mariela Gutiérrez, Joselyn Prado & Marcela Rodríguez-Gómez |
| EPI_ISL_1827533 | AREA DE SALUD SIQUIRRES | Inciensa, Instituto Costarricense de Investigación y Enseñanza en Nutrición y Salud | Francisco Duarte, Hebleen Porras, Claudio Soto-Garita, Estela Cordero, Adriana Godínez, Melany Calderón, José Luis Vargas, Mariela Gutiérrez, Joselyn Prado & Ileana Chaves-Peraza |
| EPI_ISL_1827536 | AREA DE SALUD COTO BRUS | Inciensa, Instituto Costarricense de Investigación y Enseñanza en Nutrición y Salud | Francisco Duarte, Hebleen Porras, Claudio Soto-Garita, Estela Cordero, Adriana Godínez, Melany Calderón, José Luis Vargas, Mariela Gutiérrez, Joselyn Prado & Fabricio Aguilar |
| EPI_ISL_1827537 | AREA DE SALUD GOLFITO | Inciensa, Instituto Costarricense de Investigación y Enseñanza en Nutrición y Salud | Francisco Duarte, Hebleen Porras, Claudio Soto-Garita, Estela Cordero, Adriana Godínez, Melany Calderón, José Luis Vargas, Mariela Gutiérrez, Joselyn Prado & Lizzeth Blanco |
| EPI_ISL_1827542 | AREA DE SALUD SANTA CRUZ | Inciensa, Instituto Costarricense de Investigación y Enseñanza en Nutrición y Salud | Francisco Duarte, Hebleen Porras, Claudio Soto-Garita, Estela Cordero, Adriana Godínez, Melany Calderón, José Luis Vargas, Mariela Gutiérrez, Joselyn Prado & Arnoldo Zamora-Villegas |
| EPI_ISL_1827545 | AREA DE SALUD ESPARZA | Inciensa, Instituto Costarricense de Investigación y Enseñanza en Nutrición y Salud | Francisco Duarte, Hebleen Porras, Claudio Soto-Garita, Estela Cordero, Adriana Godínez, Melany Calderón, José Luis Vargas, Mariela Gutiérrez, Joselyn Prado & Andrea Moreno-Carvajal |
| EPI_ISL_1827546 | AREA DE SALUD PARRITA | Inciensa, Instituto Costarricense de Investigación y Enseñanza en Nutrición y Salud | Francisco Duarte, Hebleen Porras, Claudio Soto-Garita, Estela Cordero, Adriana Godínez, Melany Calderón, José Luis Vargas, Mariela Gutiérrez, Joselyn Prado & Andrea Moreno-Carvajal |
| EPI_ISL_1827547 | HOSPITAL SAN VICENTE DE PAUL | Inciensa, Instituto Costarricense de Investigación y Enseñanza en Nutrición y Salud | Francisco Duarte, Hebleen Porras, Claudio Soto-Garita, Estela Cordero, Adriana Godínez, Melany Calderón, José Luis Vargas, Mariela Gutiérrez, Joselyn Prado & Silvia Sáenz-García |
| EPI_ISL_1856445, EPI_ISL_1856447, EPI_ISL_1856450, EPI_ISL_1856453, EPI_ISL_1856456, EPI_ISL_1856457, EPI_ISL_1856460, EPI_ISL_1856504, EPI_ISL_1856509, EPI_ISL_1856510, EPI_ISL_1856511, EPI_ISL_1856512, EPI_ISL_1856513, EPI_ISL_1856514, EPI_ISL_1856515, EPI_ISL_1856516, EPI_ISL_1856518, EPI_ISL_1856539, EPI_ISL_1856540, EPI_ISL_1856541, EPI_ISL_1856544, EPI_ISL_1856545, EPI_ISL_1856546, EPI_ISL_1856547, EPI_ISL_1856550, EPI_ISL_1856552, EPI_ISL_1856553, EPI_ISL_1856554, EPI_ISL_1856555, EPI_ISL_1856556, EPI_ISL_1856557, EPI_ISL_1856558, EPI_ISL_1856560, EPI_ISL_1856562, EPI_ISL_1856563, EPI_ISL_1856564, EPI_ISL_1856565, EPI_ISL_1856566, EPI_ISL_1856567, EPI_ISL_1856568, EPI_ISL_1856569, EPI_ISL_1856570, EPI_ISL_1856571, EPI_ISL_1856572, EPI_ISL_1856574, EPI_ISL_1856576, EPI_ISL_1856577, EPI_ISL_1856578, EPI_ISL_1856579, EPI_ISL_1856580, EPI_ISL_1856581, EPI_ISL_1856583, EPI_ISL_1856584, EPI_ISL_1856586, EPI_ISL_1856587, EPI_ISL_1856588, EPI_ISL_1856590, EPI_ISL_1856591, EPI_ISL_1856592, EPI_ISL_1856593, EPI_ISL_1856595, EPI_ISL_1856597, EPI_ISL_1856598, EPI_ISL_1856599, EPI_ISL_1856600, EPI_ISL_1856601, EPI_ISL_1856602, EPI_ISL_1856605, EPI_ISL_1856606, EPI_ISL_1856608, EPI_ISL_1856609, EPI_ISL_1856610, EPI_ISL_1856611, EPI_ISL_1856612, EPI_ISL_1856613, EPI_ISL_1856616, EPI_ISL_1856617, EPI_ISL_1856618, EPI_ISL_1856619, EPI_ISL_1856620, EPI_ISL_1856624, EPI_ISL_1856638, EPI_ISL_1856639, EPI_ISL_1856640, EPI_ISL_1856642, EPI_ISL_1856643, EPI_ISL_1856644, EPI_ISL_1856645, EPI_ISL_1856650, EPI_ISL_1856656, EPI_ISL_1856659, EPI_ISL_1856664, EPI_ISL_1856724, EPI_ISL_1856820, EPI_ISL_1856821, EPI_ISL_1856839 | | | |
| see above | UVA Clinical Microbiology Lab | Mathers Lab | Hardik I. Parikh, Katie E. Barry, Amy J. Mathers |
| EPI_ISL_1905207 | AZ St.Jan Brugge-Oostende | AZ SINT-JAN BRUGGE | Marijke Reynders, Patrick Descheemaeker, Laurien Hoornaert, Thomas Van Landschoot, Jorn Hellemans |
| EPI_ISL_1908981 | Provincial Public Health Reference Laboratory | Provincial Public Health Reference Laboratory | ANDLEEB HANIF |
| EPI_ISL_1909780 | Institute of Molecular and Translational Medicine / Laboratory of Experimental Medicine, Faculty of Medicine and Dentistry, Palacky University and University Hospital Olomouc | Institute of Molecular and Translational Medicine / Laboratory of Experimental Medicine, Faculty of Medicine and Dentistry, Palacky University | Rastislav Slavkovský, Hana Jaworek, Vladimíra Koudeláková, Barbora Blumová, Tomáš Pospíšil, Marián Hajdúch |
| EPI_ISL_1909831 | Provincial Public Health Reference Laboratory | Provincial Public Health Reference Laboratory | ANDLEEB HANIF, Asim Altaf, Shahida Hussain, Mariam Javed |
| EPI_ISL_1909924, EPI_ISL_1910016, EPI_ISL_1910106 | Provincial Public Health Reference Laboratory | Provincial Public Health Reference Laboratory | ANDLEEB HANIF |
| EPI_ISL_1910861 | CH Tourcoing | CHU Lille | AIT YAHYA Emilie, ALIDJINOU Enagnon Kazali, BOCKET Laurence, CREPIN Michel, DEMAY Christophe, ENGELMANN Ilka, GEFFROY Sandrine, GUIGON Aurélie, LAMBERT Valérie, LAZREK Mouna, NOBILLIAUX Florian, PREVOST Brigitte, THUILLIER Caroline, TINEZ Claire |
| EPI_ISL_1913070 | Centre de Recherches Médicales de Lambaréné (CERMEL) | Centre de Recherches Médicales de Lambaréné (CERMEL) | Gédéon Prince Manouana, Anicet Mouity Matoumba, Michel Ngonga Dikongo, Georgelin Nguema Ondo, Rodrigue Bikangui, Samira Zoa Assoumou, Srinivas reddy Pallerla, Jean Bernard Lekana-Douki, Joël-Fleury Djoba Siawaya, Steffen Borrmann, Thirumalaisamy P. Velavan, Bertrand Lell and Ayola Akim Adegnika |
| EPI_ISL_1914581, EPI_ISL_1914584 | State Virus Research and Diagnostic Laboratory (VRDL), AIIMS Raipur | State Virus Research and Diagnostic Laboratory (VRDL), AIIMS Raipur | Pushpendra Singh, Kuldeep Sharma, Priyanka Singh, Somya Sharma, Sanjay Singh Negi, Anudita Bhargava |
| EPI_ISL_1915515 | National Institute of Health Research and Development | National Institute of Health Research and Development | Subangkit, Hana Apsari Pawestri, Kartika Dewi Puspa, Arie Ardiansyah Nugraha, Hartanti Dian Ikawati, Krisna Nur Andriana Pangesti, Yuni Rukminiati, Ririn Ramadhany, Agustiningsih, Kindi Adam, Holy Arif Wibowo, Triyani Soekarso, Ni Ketut Susilarini, Nurika Hariastuti, Ully Alfi Nikmah, Reni Herman, Nike Susanti, Herna, Tati Febriyanti, Natalie Laurencia Kipuw, Fauzul Muna, Irene Lorinda Indalao, Nelly Puspandari, Vivi Setiawaty. |
| EPI_ISL_1916430, EPI_ISL_1916432, EPI_ISL_1916440, EPI_ISL_1916441, | CSIR-National Environmental Engineering Research Institute | CSIR-Centre for Cellular and Molecular Biology - INSACOG | Ara Sreenivas, Onkar Kulkarni, Lamuk Zaveri, Sofia Banu, Shreekant Verma, Amareshwar Vodapalli , Viswagithe S L, B Himasri, Sharath Chandra Thota, Karthik Bharadwaj Tallapaka, Krishna Khairnar, Rakesh K Mishra,Divya Tej Sowpati |

| | | | |
|---|---|---|---|
| EPI_ISL_1916462, EPI_ISL_1916473, EPI_ISL_1916474, EPI_ISL_1916475 | | | |
| EPI_ISL_1935579, EPI_ISL_1935580 | UVA Clinical Microbiology Lab | Mathers Lab | Hardik I. Parikh, Katie E. Barry, Amy J. Mathers |
| EPI_ISL_1939544, EPI_ISL_1939545, EPI_ISL_1939546, EPI_ISL_1939547, EPI_ISL_1939548, EPI_ISL_1939550, EPI_ISL_1939551, EPI_ISL_1939552, EPI_ISL_1939553, EPI_ISL_1939554, EPI_ISL_1939555, EPI_ISL_1939556, EPI_ISL_1939557, EPI_ISL_1939558, EPI_ISL_1939559, EPI_ISL_1939560, EPI_ISL_1939561, EPI_ISL_1939562, EPI_ISL_1939563, EPI_ISL_1939564, EPI_ISL_1939565, EPI_ISL_1939566, EPI_ISL_1939567, EPI_ISL_1939568, EPI_ISL_1939569, EPI_ISL_1939570, EPI_ISL_1939571, EPI_ISL_1939572, EPI_ISL_1939573, EPI_ISL_1939574, EPI_ISL_1939575, EPI_ISL_1939576, EPI_ISL_1939577, EPI_ISL_1939578, EPI_ISL_1939579, EPI_ISL_1939580, EPI_ISL_1939581, EPI_ISL_1939582, EPI_ISL_1939583, EPI_ISL_1939584, EPI_ISL_1939586, EPI_ISL_1939588, EPI_ISL_1939589, EPI_ISL_1939591, EPI_ISL_1939592, EPI_ISL_1939597, EPI_ISL_1939600, EPI_ISL_1939601, EPI_ISL_1939602, EPI_ISL_1939604, EPI_ISL_1939605, EPI_ISL_1939606, EPI_ISL_1939607, EPI_ISL_1939608, EPI_ISL_1939609, EPI_ISL_1939610, EPI_ISL_1939611, EPI_ISL_1939613, EPI_ISL_1939614, EPI_ISL_1939615, EPI_ISL_1939616 | | | |
| see above | Institute of Tropical Medicine "Pedro Kourí" (IPK) | Respiratory Infections Laboratory | Lissette Perez Santos, Yahisel Tejero Suarez, Yanet Pintos, Dailyn Medero, Odalys Valdes , Mirtha Elisa Aguado , Liena de Regla Ponce |
| EPI_ISL_1942246 | SVFI, Veterinary institute in Zvolen | SVFI, Veterinary institute in Zvolen | Dirbáková Z., Sujová S., Mokryšová S., Tinák M., Mojžiš M., |
| EPI_ISL_1965121, EPI_ISL_1965122, EPI_ISL_1965123, EPI_ISL_1965124, EPI_ISL_1965125, EPI_ISL_1965126, EPI_ISL_1965127, EPI_ISL_1965128, EPI_ISL_1965129, EPI_ISL_1965130, EPI_ISL_1965131, EPI_ISL_1965132, EPI_ISL_1965133, EPI_ISL_1965134, EPI_ISL_1965135, EPI_ISL_1965137, EPI_ISL_1965138, EPI_ISL_1965139, EPI_ISL_1965140, EPI_ISL_1965141, EPI_ISL_1965142, EPI_ISL_1965143, EPI_ISL_1965144, EPI_ISL_1965145, EPI_ISL_1965146, EPI_ISL_1965147, EPI_ISL_1965148, EPI_ISL_1965149, EPI_ISL_1965150, EPI_ISL_1965151, EPI_ISL_1965152, EPI_ISL_1965153, EPI_ISL_1965154, EPI_ISL_1965155, EPI_ISL_1965156, EPI_ISL_1965157, EPI_ISL_1965158, EPI_ISL_1965159, EPI_ISL_1965160, EPI_ISL_1965161, EPI_ISL_1965162, EPI_ISL_1965163, EPI_ISL_1965164, EPI_ISL_1965165, EPI_ISL_1965166, EPI_ISL_1965167, EPI_ISL_1965168, EPI_ISL_1965169, EPI_ISL_1965172, EPI_ISL_1965173, EPI_ISL_1965174, EPI_ISL_1965175, EPI_ISL_1965176, EPI_ISL_1965177, EPI_ISL_1965178, EPI_ISL_1965179, EPI_ISL_1965180, EPI_ISL_1965181, EPI_ISL_1965182, EPI_ISL_1965183, EPI_ISL_1965184, EPI_ISL_1965185, EPI_ISL_1965186, EPI_ISL_1965187, EPI_ISL_1965188, EPI_ISL_1965189, EPI_ISL_1965190, EPI_ISL_1965191, EPI_ISL_1965192, EPI_ISL_1965193, EPI_ISL_1965194, EPI_ISL_1965196, EPI_ISL_1965197, EPI_ISL_1965198, EPI_ISL_1965199, EPI_ISL_1965200, EPI_ISL_1965201, EPI_ISL_1965202, EPI_ISL_1965203, EPI_ISL_1965204, EPI_ISL_1965205, EPI_ISL_1965206, EPI_ISL_1965207, EPI_ISL_1965208, EPI_ISL_1965209, EPI_ISL_1965210, EPI_ISL_1965211, EPI_ISL_1965212, EPI_ISL_1965213, EPI_ISL_1965214, EPI_ISL_1965215, EPI_ISL_1965216, EPI_ISL_1965279, EPI_ISL_1965280, EPI_ISL_1965281 | | | |
| see above | IPK | Respiratory Infections Laboratory | Lissette Perez , Yahisel Tejero , Yanet Pintos, Dailyn Medero,Odalys Valdes , Mirtha Elisa Aguado , Liena de Regla Ponce |
| EPI_ISL_1965535, EPI_ISL_1965537, EPI_ISL_1965539, EPI_ISL_1965540, EPI_ISL_1965541, EPI_ISL_1965545, EPI_ISL_1965546, EPI_ISL_1965547, EPI_ISL_1965550, EPI_ISL_1965558, EPI_ISL_1965562, EPI_ISL_1965563, EPI_ISL_1965565, EPI_ISL_1965567, EPI_ISL_1965568, EPI_ISL_1965571, EPI_ISL_1965573, EPI_ISL_1965574, EPI_ISL_1965575, EPI_ISL_1965577, EPI_ISL_1965578, EPI_ISL_1965579, EPI_ISL_1965580, EPI_ISL_1965588, EPI_ISL_1965596, EPI_ISL_1965597, EPI_ISL_1965600, EPI_ISL_1965601, EPI_ISL_1965602, EPI_ISL_1965603, EPI_ISL_1965604, EPI_ISL_1965605, EPI_ISL_1965609, EPI_ISL_1965610, EPI_ISL_1965618, EPI_ISL_1965619, EPI_ISL_1965620, EPI_ISL_1965627, EPI_ISL_1965635 | | | |
| see above | HG Pharma GmbH | Bergthaler laboratory, CeMM Research Center for Molecular Medicine of the Austrian Academy of Sciences | Lukas Endler, Anna Schedl, Fabian Amman, Petr Triska, Thomas Penz, Benedikt Agerer, Maelle Le Moing, Michael Schuster, Bekir Erguner, Jan Laine, Martin Senekowitsch, Christoph Bock, Andreas Bergthaler |
| EPI_ISL_1972106 | AZ Zeno | AZ SINT-JAN BRUGGE | Marijke Reynders, Patrick Descheemaeker, Laurien Hoornaert, Thomas Van Landschoot, Jorn Hellemans |
| EPI_ISL_1972283, EPI_ISL_1972284 | Baylor Scott & White-Temple | Baylor Scott & White-Temple | Ari Rao, Linden Morales, Kimberly Walker, Marcus Volz, Shelby Hendrickson |
| EPI_ISL_2001123 | GMERS, Government Medical College, Gandhinagar | Gujarat Biotechnology Research Centre | Nitin Savaliya,Dinesh Kumar,Twinkle Soni,Sonal Sharma ,Zuber Saiyed,Ramesh Pandit,Janvi Raval,Zarna Patel,Gaurishankar Shrimali,Umang Mishra,Nitesh Shah,Chaitanya Joshi,Madhvi Joshi |
| EPI_ISL_2001124 | GMERS, Government Medical College, Gandhinagar | Gujarat Biotechnology Research Centre | Dinesh Kumar,Twinkle Soni,Sonal Sharma ,Zuber Saiyed,Ramesh Pandit,Janvi Raval,Zarna Patel,Nitin Savaliya,Gaurishankar Shrimali,Umang Mishra,Nitesh Shah,Chaitanya Joshi,Madhvi Joshi |
| EPI_ISL_2001125 | GMERS, Government Medical College, Gandhinagar | Gujarat Biotechnology Research Centre | Twinkle Soni,Sonal Sharma ,Zuber Saiyed,Ramesh Pandit,Janvi Raval,Zarna Patel,Nitin Savaliya,Dinesh Kumar,Gaurishankar Shrimali,Umang Mishra,Nitesh Shah,Chaitanya Joshi,Madhvi Joshi |
| EPI_ISL_2001126 | GMERS, Government Medical College, Gandhinagar | Gujarat Biotechnology Research Centre | Sonal Sharma ,Zuber Saiyed,Ramesh Pandit,Janvi Raval,Zarna Patel,Nitin Savaliya,Dinesh Kumar,Twinkle Soni,Gaurishankar Shrimali,Umang Mishra,Nitesh Shah,Chaitanya Joshi,Madhvi Joshi |
| EPI_ISL_2001170 | PanGenomics International Pvt. Ltd. | Gujarat Biotechnology Research Centre | Dinesh Kumar,Twinkle Soni,Sonal Sharma ,Zuber Saiyed,Ramesh Pandit,Janvi Raval,Zarna Patel,Nitin Savaliya,Dipali Dhawan,Umang Mishra,Nitesh Shah,Chaitanya Joshi,Madhvi Joshi |
| EPI_ISL_2001171 | PanGenomics International Pvt. Ltd. | Gujarat Biotechnology Research Centre | Twinkle Soni,Sonal Sharma ,Zuber Saiyed,Ramesh Pandit,Janvi Raval,Zarna Patel,Nitin Savaliya,Dinesh Kumar,Dipali Dhawan,Umang Mishra,Nitesh Shah,Chaitanya Joshi,Madhvi Joshi |
| EPI_ISL_2001172 | PanGenomics International Pvt. Ltd. | Gujarat Biotechnology Research Centre | Sonal Sharma ,Zuber Saiyed,Ramesh Pandit,Janvi Raval,Zarna Patel,Nitin Savaliya,Dinesh Kumar,Twinkle Soni,Dipali Dhawan,Umang Mishra,Nitesh Shah,Chaitanya Joshi,Madhvi Joshi |
| EPI_ISL_2001173 | PanGenomics International Pvt. Ltd. | Gujarat Biotechnology Research Centre | Zuber Saiyed,Ramesh Pandit,Janvi Raval,Zarna Patel,Nitin Savaliya,Dinesh Kumar,Twinkle Soni,Sonal Sharma ,Dipali Dhawan,Umang Mishra,Nitesh Shah,Chaitanya Joshi,Madhvi Joshi |
| EPI_ISL_2001174 | PanGenomics International Pvt. Ltd. | Gujarat Biotechnology Research Centre | Ramesh Pandit,Janvi Raval,Zarna Patel,Nitin Savaliya,Dinesh Kumar,Twinkle Soni,Sonal Sharma ,Zuber Saiyed,Dipali Dhawan,Umang Mishra,Nitesh Shah,Chaitanya Joshi,Madhvi Joshi |
| EPI_ISL_2001175 | PanGenomics International Pvt. Ltd. | Gujarat Biotechnology Research Centre | Janvi Raval,Zarna Patel,Nitin Savaliya,Dinesh Kumar,Twinkle Soni,Sonal Sharma ,Zuber Saiyed,Ramesh Pandit,Dipali Dhawan,Umang Mishra,Nitesh Shah,Chaitanya Joshi,Madhvi Joshi |
| EPI_ISL_2001176 | PanGenomics International Pvt. Ltd. | Gujarat Biotechnology Research Centre | Zarna Patel,Nitin Savaliya,Dinesh Kumar,Twinkle Soni,Sonal Sharma ,Zuber Saiyed,Ramesh Pandit,Janvi Raval,Dipali Dhawan,Umang Mishra,Nitesh Shah,Chaitanya Joshi,Madhvi Joshi |
| EPI_ISL_2001177 | PanGenomics International Pvt. Ltd. | Gujarat Biotechnology Research Centre | Dinesh Kumar,Twinkle Soni,Sonal Sharma ,Zuber Saiyed,Ramesh Pandit,Janvi Raval,Zarna Patel,Nitin Savaliya,Dipali Dhawan,Umang Mishra,Nitesh Shah,Chaitanya Joshi,Madhvi Joshi |
| EPI_ISL_2003111 | Pronto Socorro Central de Bauru | Instituto Adolfo Lutz, Interdiciplinary Procedures Center, Strategic Laboratory | Claudio Tavares Sacchi, Claudia Regina Gonçalves, Erica Valessa Ramos Gomes, Karoline Rodrigues Campos, Caio Vinicius Dias Lopes, Leonardo Jose Tadeu de Araujo |
| EPI_ISL_2003112 | CS de Paulo de Faria | Instituto Adolfo Lutz, Interdiciplinary Procedures Center, Strategic Laboratory | Claudio Tavares Sacchi, Claudia Regina Gonçalves, Erica Valessa Ramos Gomes, Karoline Rodrigues Campos, Caio Vinicius Dias Lopes, Leonardo Jose Tadeu de Araujo |
| EPI_ISL_2003134, EPI_ISL_2003135, EPI_ISL_2003136, EPI_ISL_2003137, EPI_ISL_2003138 | Instituto Adolfo Lutz - Regional de Santos | Instituto Adolfo Lutz, Interdiciplinary Procedures Center, Strategic Laboratory | Claudio Tavares Sacchi, Claudia Regina Gonçalves, Erica Valessa Ramos Gomes, Karoline Rodrigues Campos, Caio Vinicius Dias Lopes, Leonardo Jose Tadeu de Araujo |
| EPI_ISL_2003150 | Instituto Adolfo Lutz - Regional de Bauru | Instituto Adolfo Lutz, Interdiciplinary Procedures Center, Strategic Laboratory | Claudio Tavares Sacchi, Claudia Regina Gonçalves, Erica Valessa Ramos Gomes, Karoline Rodrigues Campos, Caio Vinicius Dias Lopes, Leonardo Jose Tadeu de Araujo |
| EPI_ISL_2007588 | Dipartimento di Medicina e Chirurgia Universita' degli Studi dell'Insubria Laboratorio di Microbiologia Medica Varese Italia | Laboratorio di Microbiologia e Virologia, Università Vita-Salute San Raffaele, Milano | Matteo Castelli, Roberto Ferrarese, Elena Criscuolo, Roberta Antonia Diotti, Massimo Clementi, Nicasio Mancini, Nicola Clementi |
| EPI_ISL_2013598 | PanGenomics International Pvt. Ltd. | Gujarat Biotechnology Research Centre | Nitin Savaliya,Dinesh Kumar,Twinkle Soni,Sonal Sharma ,Zuber Saiyed,Ramesh Pandit,Janvi Raval,Zarna Patel,Dipali Dhawan,Umang Mishra,Nitesh Shah,Chaitanya Joshi,Madhvi Joshi |
| EPI_ISL_2029991 | Hospital | National Reference Center for Viruses of Respiratory Infections, Institut Pasteur, Paris | Marion Barbet, Sylvie Behillil, Méline Bizard, Angela Brisebarre, Camille Capel, Vincent Enouf, Louise Lefrançois, Frédéric Lemoine, Christophe Malabat, Corinne Maufrais, Amaury Vaysse, Etienne Simon-Lorière, Maud Vanpeene, Sylvie Van der Werf ,CéLine Bressollette |
| EPI_ISL_2029997, EPI_ISL_2030057, EPI_ISL_2030096 | Hospital | National Reference Center for Viruses of Respiratory Infections, Institut Pasteur, Paris | Marion Barbet, Sylvie Behillil, Méline Bizard, Angela Brisebarre, Camille Capel, Vincent Enouf, Louise Lefrançois, Frédéric Lemoine, Christophe Malabat, Corinne Maufrais, Etienne Simon-Lorière, Maud Vanpeene, Sylvie Van der Werf ,CéLine Bressollette |
| EPI_ISL_2030123 | Hospital | National Reference Center for Viruses of Respiratory Infections, Institut Pasteur, Paris | Marion Barbet, Sylvie Behillil, Méline Bizard, Angela Brisebarre, Camille Capel, Vincent Enouf, Louise Lefrançois, Frédéric Lemoine, Christophe Malabat, Corinne Maufrais, Amaury Vaysse, Etienne Simon-Lorière, Maud Vanpeene, Sylvie Van der Werf ,CéLine Bressollette |
| EPI_ISL_2037442 | Instituto de Virologia "Dr. J. M. Vanella", Facultad de Ciencias Médicas, Universidad Nacional de Córdoba. | Centro de Investigaciones Agropecuarias (CIAP), Instituto Nacional de Tecnologia Agropecuaria (INTA) Cordoba, Argentina, on behalf of 'Proyecto Argentino Interinstitucional de Genomica de SARS-CoV-2 (PAIS Consortium) | Brenda Konigheim, Lorena Spinsanti, Adrian Diaz, Javier Aguilar, Sebastian Blanco, Mauricio Beranek, Maria Elisa Rivarola, Maria Belen Pisano, Viviana Re, Gonzalo Castro, Gabriela Barbas, Franco Fernandez, Humberto Debat, Nathalie Marquez, Sandra Gallego. |
| EPI_ISL_2040580 | CHU Nantes Virology | CHU Nantes Virology | Celine Bressollette-Bodin, Thomas Drumel, Audrey Rodallec, Berthe-Marie Imbert-Marcille |

| | | | |
|---|---|---|---|
| EPI_ISL_2047859 | Azienda Ospedaliera Terni | Istituto Zooprofilattico Sperimentale dell'Abruzzo e Molise "G. Caporale" | Palumbo M, Scaccetti A, Lorusso A, Marcacci M, Di Domenico M, Ancora M, Curini V, Mangone I, Rinaldi A, Delli Compagni E, Scialabba S, Di Pasquale A, Cammà C, Puglia I, Calistri P, Savini G |
| EPI_ISL_2081194, EPI_ISL_2081195, EPI_ISL_2081196, EPI_ISL_2081197, EPI_ISL_2081218, EPI_ISL_2081219, EPI_ISL_2081220, EPI_ISL_2081223 | Premier Laboratory Services | BioInfoExperts LLC | Cheryl Metzler, Valerie Murray, Rebecca Rose, David Nolan, Susanna Lamers |
| EPI_ISL_2081834, EPI_ISL_2081837, EPI_ISL_2081840, EPI_ISL_2081841, EPI_ISL_2081842, EPI_ISL_2081843, EPI_ISL_2081844, EPI_ISL_2081845, EPI_ISL_2081846, EPI_ISL_2081847, EPI_ISL_2081848, EPI_ISL_2081849, EPI_ISL_2081850, EPI_ISL_2081851, EPI_ISL_2081852, EPI_ISL_2081853, EPI_ISL_2081854, EPI_ISL_2081855, EPI_ISL_2081856, EPI_ISL_2081857, EPI_ISL_2081858, EPI_ISL_2081859, EPI_ISL_2081860, EPI_ISL_2081861, EPI_ISL_2081864, EPI_ISL_2081865, EPI_ISL_2081866, EPI_ISL_2081867, EPI_ISL_2081868, EPI_ISL_2081869, EPI_ISL_2081870, EPI_ISL_2081871, EPI_ISL_2081872, EPI_ISL_2081873, EPI_ISL_2081874 | | | |
| see above | National Center of Infectious and Parasitic Diseases | National Center of Infectious and Parasitic Diseases | Alexiev, Ivanov, Korsun, Stoitsova, Philipova, Dimitrova, Grigorova L., Hristova, Donchev, Stoykov, Trifonova, Dobrinov, Grigorova I., Kantardjiev |
| EPI_ISL_2083044 | Baylor Scott & White-Temple | Baylor Scott & White-Temple | Ari Rao, Linden Morales, Kimberly Walker, Marcus Volz, Shelby Hendrickson |
| EPI_ISL_2086594, EPI_ISL_2086595, EPI_ISL_2086596, EPI_ISL_2086597, EPI_ISL_2086598, EPI_ISL_2086599, EPI_ISL_2086600, EPI_ISL_2086601, EPI_ISL_2086602, EPI_ISL_2086603, EPI_ISL_2086604, EPI_ISL_2086605, EPI_ISL_2086606, EPI_ISL_2086607, EPI_ISL_2086608, EPI_ISL_2086609 | | | |
| see above | LABCOVID_HCPA | LABRESIS_HCPA | Wink PL, Martins AF, Volpato F, Monteiro F, Zavascki AP, Barth AL |
| EPI_ISL_2090758 | REUNILAB | UMR PIMIT | Dr David A Wilkinson, Dr Patrick Mavingui, Dr Camille Lebarbenchon, Magali Turpin |
| EPI_ISL_2091133, EPI_ISL_2091137 | Laboratorio Central de Epidemiología (LCE) | Centro de Investigación en Enfermedades Infecciosas (CIENI), Instituto Nacional de Enfermedades Respiratorias (INER) | Consorcio Mexicano de Vigilancia Genómica (CoViGen-Mex). Authors (in alphabetical order): Julio Elias Alvarado-Yaah, Carlos F. Arias, Santiago Ávila-Ríos, Víctor Hugo Borja-Aburto, Celia Boukadida, Juan Bautista Chale-Dzul, Célida Duque Molina, José Antonio Enciso-Moreno, Gloria Elena Espinosa-Ayala, Fernando Fontove-Herrera, Victor Eduardo Garcia-Arias, Concepción Grajales-Muñiz, Ricardo Grande, Alfredo Herrera-Estrella, Carla Ivón Herrera-Najera, Pavel Isa, Brenda Irasema Maldonado-Meza, Bernardo Martínez-Miguel, Margarita Matías-Florentino, María Guadalupe de Jesús Mireles-Rivera, Gloria María Molina-Salinas, Hector Montoya-Fuentes, José Esteban Muñoz-Medina, José de Jesús Nuñez-Contreras, Alicia Ocaña-Mondragón, Luis Alberto Ochoa-Carrera, Hector Esteban Paz-Juárez, Francisco Pulido, Helen Haydee Fernanda Ramírez-Plascencia, Angel Gustavo Salas-Lais, Alejandro Sanchez-Flores, Clara Esperanza Santacruz-Tinoco, María Guadalupe Santiago-Mauricio, Nelly Sélem-Mojica, Blanca Taboada, Gloria Vazquez |
| EPI_ISL_2091138 | Centro de Investigación Biomédica del Noreste (CIBIN) | Centro de Investigación en Enfermedades Infecciosas (CIENI), Instituto Nacional de Enfermedades Respiratorias (INER) | Consorcio Mexicano de Vigilancia Genómica (CoViGen-Mex). Authors (in alphabetical order): Julio Elias Alvarado-Yaah, Carlos F. Arias, Santiago Ávila-Ríos, Víctor Hugo Borja-Aburto, Celia Boukadida, Juan Bautista Chale-Dzul, Célida Duque Molina, José Antonio Enciso-Moreno, Gloria Elena Espinosa-Ayala, Fernando Fontove-Herrera, Victor Eduardo Garcia-Arias, Concepción Grajales-Muñiz, Ricardo Grande, Alfredo Herrera-Estrella, Carla Ivón Herrera-Najera, Pavel Isa, Brenda Irasema Maldonado-Meza, Bernardo Martínez-Miguel, Margarita Matías-Florentino, María Guadalupe de Jesús Mireles-Rivera, Gloria María Molina-Salinas, Hector Montoya-Fuentes, José Esteban Muñoz-Medina, José de Jesús Nuñez-Contreras, Alicia Ocaña-Mondragón, Luis Alberto Ochoa-Carrera, Hector Esteban Paz-Juárez, Francisco Pulido, Helen Haydee Fernanda Ramírez-Plascencia, Angel Gustavo Salas-Lais, Alejandro Sanchez-Flores, Clara Esperanza Santacruz-Tinoco, María Guadalupe Santiago-Mauricio, Nelly Sélem-Mojica, Blanca Taboada, Gloria Vazquez |
| EPI_ISL_2091140, EPI_ISL_2091141, EPI_ISL_2091142 | Centro de Investigación Biomédica de Occidente (CIBO) | Centro de Investigación en Enfermedades Infecciosas (CIENI), Instituto Nacional de Enfermedades Respiratorias (INER) | Consorcio Mexicano de Vigilancia Genómica (CoViGen-Mex). Authors (in alphabetical order): Julio Elias Alvarado-Yaah, Carlos F. Arias, Santiago Ávila-Ríos, Víctor Hugo Borja-Aburto, Celia Boukadida, Juan Bautista Chale-Dzul, Célida Duque Molina, José Antonio Enciso-Moreno, Gloria Elena Espinosa-Ayala, Fernando Fontove-Herrera, Victor Eduardo Garcia-Arias, Concepción Grajales-Muñiz, Ricardo Grande, Alfredo Herrera-Estrella, Carla Ivón Herrera-Najera, Pavel Isa, Brenda Irasema Maldonado-Meza, Bernardo Martínez-Miguel, Margarita Matías-Florentino, María Guadalupe de Jesús Mireles-Rivera, Gloria María Molina-Salinas, Hector Montoya-Fuentes, José Esteban Muñoz-Medina, José de Jesús Nuñez-Contreras, Alicia Ocaña-Mondragón, Luis Alberto Ochoa-Carrera, Hector Esteban Paz-Juárez, Francisco Pulido, Helen Haydee Fernanda Ramírez-Plascencia, Angel Gustavo Salas-Lais, Alejandro Sanchez-Flores, Clara Esperanza Santacruz-Tinoco, María Guadalupe Santiago-Mauricio, Nelly Sélem-Mojica, Blanca Taboada, Gloria Vazquez |
| EPI_ISL_2091143 | Laboratorio Central de Epidemiología (LCE) | Centro de Investigación en Enfermedades Infecciosas (CIENI), Instituto Nacional de Enfermedades Respiratorias (INER) | Consorcio Mexicano de Vigilancia Genómica (CoViGen-Mex). Authors (in alphabetical order): Julio Elias Alvarado-Yaah, Carlos F. Arias, Santiago Ávila-Ríos, Víctor Hugo Borja-Aburto, Celia Boukadida, Juan Bautista Chale-Dzul, Célida Duque Molina, José Antonio Enciso-Moreno, Gloria Elena Espinosa-Ayala, Fernando Fontove-Herrera, Victor Eduardo Garcia-Arias, Concepción Grajales-Muñiz, Ricardo Grande, Alfredo Herrera-Estrella, Carla Ivón Herrera-Najera, Pavel Isa, Brenda Irasema Maldonado-Meza, Bernardo Martínez-Miguel, Margarita Matías-Florentino, María Guadalupe de Jesús Mireles-Rivera, Gloria María Molina-Salinas, Hector Montoya-Fuentes, José Esteban Muñoz-Medina, José de Jesús Nuñez-Contreras, Alicia Ocaña-Mondragón, Luis Alberto Ochoa-Carrera, Hector Esteban Paz-Juárez, Francisco Pulido, Helen Haydee Fernanda Ramírez-Plascencia, Angel Gustavo Salas-Lais, Alejandro Sanchez-Flores, Clara Esperanza Santacruz-Tinoco, María Guadalupe Santiago-Mauricio, Nelly Sélem-Mojica, Blanca Taboada, Gloria Vazquez |
| EPI_ISL_2091145 | Centro de Investigación Biomédica de Occidente (CIBO) | Centro de Investigación en Enfermedades Infecciosas (CIENI), Instituto Nacional de Enfermedades Respiratorias (INER) | Consorcio Mexicano de Vigilancia Genómica (CoViGen-Mex). Authors (in alphabetical order): Julio Elias Alvarado-Yaah, Carlos F. Arias, Santiago Ávila-Ríos, Víctor Hugo Borja-Aburto, Celia Boukadida, Juan Bautista Chale-Dzul, Célida Duque Molina, José Antonio Enciso-Moreno, Gloria Elena Espinosa-Ayala, Fernando Fontove-Herrera, Victor Eduardo Garcia-Arias, Concepción Grajales-Muñiz, Ricardo Grande, Alfredo Herrera-Estrella, Carla Ivón Herrera-Najera, Pavel Isa, Brenda Irasema Maldonado-Meza, Bernardo Martínez-Miguel, Margarita Matías-Florentino, María Guadalupe de Jesús Mireles-Rivera, Gloria María Molina-Salinas, Hector Montoya-Fuentes, José Esteban Muñoz-Medina, José de Jesús Nuñez-Contreras, Alicia Ocaña-Mondragón, Luis Alberto Ochoa-Carrera, Hector Esteban Paz-Juárez, Francisco Pulido, Helen Haydee Fernanda Ramírez-Plascencia, Angel Gustavo Salas-Lais, Alejandro Sanchez-Flores, Clara Esperanza Santacruz-Tinoco, María Guadalupe Santiago-Mauricio, Nelly Sélem-Mojica, Blanca Taboada, Gloria Vazquez |
| EPI_ISL_2092866, EPI_ISL_2092867, EPI_ISL_2092868 | CQRC_QUALITY CONTROL CHEMICAL BIOLOGICAL RISK_AOOR Villa Sofia Cervello Palermo | CQRC_QUALITY CONTROL CHEMICAL BIOLOGICAL RISK_AOOR Villa Sofia Cervello Palermo | Di Gaudio, F., Contino, F., Brunacci, G., |
| EPI_ISL_2092869, EPI_ISL_2092870 | CQRC_QUALITY CONTROL CHEMICAL BIOLOGICAL RISK_AOOR Villa Sofia Cervello Palermo | CQRC_QUALITY CONTROL CHEMICAL BIOLOGICAL RISK_AOOR Villa Sofia Cervello Palermo | Di Gaudio, F., Brunacci, G., Contino, F., |
| EPI_ISL_2092871 | CQRC_QUALITY CONTROL CHEMICAL BIOLOGICAL RISK_AOOR Villa Sofia Cervello Palermo | CQRC_QUALITY CONTROL CHEMICAL BIOLOGICAL RISK_AOOR Villa Sofia Cervello Palermo | Di Gaudio, F., Contino, F., Brunacci, G., |
| EPI_ISL_2096005, EPI_ISL_2096006, EPI_ISL_2096007, EPI_ISL_2096008, EPI_ISL_2096009, EPI_ISL_2096010, EPI_ISL_2096011, EPI_ISL_2096012, EPI_ISL_2096013, EPI_ISL_2096014, EPI_ISL_2096015, EPI_ISL_2096016, EPI_ISL_2096017, EPI_ISL_2096018, EPI_ISL_2096019, EPI_ISL_2096020, EPI_ISL_2096021, EPI_ISL_2096022, EPI_ISL_2096023, EPI_ISL_2096024, EPI_ISL_2096025, EPI_ISL_2096026, EPI_ISL_2096027, EPI_ISL_2096028, EPI_ISL_2096029, EPI_ISL_2096030, EPI_ISL_2096031, EPI_ISL_2096032, EPI_ISL_2096033, EPI_ISL_2096034, EPI_ISL_2096035, EPI_ISL_2096036, EPI_ISL_2096037, EPI_ISL_2096038, EPI_ISL_2096039, EPI_ISL_2096040, EPI_ISL_2096041, EPI_ISL_2096042, EPI_ISL_2096043, EPI_ISL_2096044, EPI_ISL_2096045, EPI_ISL_2096046, EPI_ISL_2096047, EPI_ISL_2096048, EPI_ISL_2096049, EPI_ISL_2096050, EPI_ISL_2096051 | | | |
| see above | National Food and Veterinary Risk Assessment Institute | National Public Health Surveillance Laboratory | Lukas Zemaitis, Migle Gabrielaite, Jelena Razmuk, Svajune Muralyte, Ana Steponkiene, Lukas Vasionis, Danas Baksa |
| EPI_ISL_2099351, EPI_ISL_2099352, EPI_ISL_2099353, EPI_ISL_2099354, EPI_ISL_2099355, EPI_ISL_2099356, EPI_ISL_2099357, EPI_ISL_2099455, EPI_ISL_2099456, EPI_ISL_2099457 | National Center of Infectious and Parasitic Diseases | National Center of Infectious and Parasitic Diseases | Alexiev, Ivanov, Korsun, Stoitsova, Philipova, Dimitrova, Grigorova L., Hristova, Donchev, Stoykov, Trifonova, Dobrinov, Grigorova I., Kantardjiev |
| EPI_ISL_2101074 | Molecular Biology Laboratory, Faculty Medicine and Health Sciences, Warmadewa University | Eijkman Institute for Molecular Biology, National Agency for Research and Innovation; Molecular Biology Laboratory, Faculty Medicine and Health Sciences, Warmadewa University | Sukma Oktavianthi, Lidwina Priliani, Edison Johar, Frilasita A Yudhaputri, Muhammad Rezki Rasyak, Willy Agustine, Hidayat Trimarsanto, Lydia V. Panggalo, Iskandar Adnan, Sri Masyeni, Erly Sintya, Safarina G Malik, Khin Saw Myint, Amin Soebandrio |
| EPI_ISL_2101075 | Molecular Biology Laboratory, Faculty Medicine and Health Sciences, Warmadewa University | Eijkman Institute for Molecular Biology, National Agency for Research and Innovation; Molecular Biology Laboratory, Faculty Medicine and Health Sciences, Warmadewa University | Lidwina Priliani, Edison Johar, Frilasita A Yudhaputri, Muhammad Rezki Rasyak, Willy Agustine, Hidayat Trimarsanto, Lydia V. Panggalo, Iskandar Adnan, Sukma Oktavianthi, Sri Masyeni, Erly Sintya, Safarina G Malik, Khin Saw Myint, Amin Soebandrio |

| | | | |
|---|---|---|---|
| EPI_ISL_2101076 | Molecular Biology Laboratory, Faculty Medicine and Health Sciences, Warmadewa University | Eijkman Institute for Molecular Biology, National Agency for Research and Innovation; Molecular Biology Laboratory, Faculty Medicine and Health Sciences, Warmadewa University | Edison Johar, Frilasita A Yudhaputri, Muhammad Rezki Rasyak, Willy Agustine, Hidayat Trimarsanto, Lydia V. Panggalo, Iskandar Adnan, Sukma Oktavianthi, Lidwina Priliani, Sri Masyeni, Erly Sintya, Safarina G Malik, Khin Saw Myint, Amin Soebandrio |
| EPI_ISL_2101077 | Molecular Biology Laboratory, Faculty Medicine and Health Sciences, Warmadewa University | Eijkman Institute for Molecular Biology, National Agency for Research and Innovation; Molecular Biology Laboratory, Faculty Medicine and Health Sciences, Warmadewa University | Frilasita A Yudhaputri, Muhammad Rezki Rasyak, Willy Agustine, Hidayat Trimarsanto, Lydia V. Panggalo, Iskandar Adnan, Sukma Oktavianthi, Lidwina Priliani, Edison Johar, Sri Masyeni, Erly Sintya, Safarina G Malik, Khin Saw Myint, Amin Soebandrio |
| EPI_ISL_2101078 | Molecular Biology Laboratory, Faculty Medicine and Health Sciences, Warmadewa University | Eijkman Institute for Molecular Biology, National Agency for Research and Innovation; Molecular Biology Laboratory, Faculty Medicine and Health Sciences, Warmadewa University | Willy Agustine, Hidayat Trimarsanto, Lydia V. Panggalo, Iskandar Adnan, Sukma Oktavianthi, Lidwina Priliani, Edison Johar, Frilasita A Yudhaputri, Muhammad Rezki Rasyak, Sri Masyeni, Erly Sintya, Safarina G Malik, Khin Saw Myint, Amin Soebandrio |
| EPI_ISL_2101079 | Molecular Biology Laboratory, Faculty Medicine and Health Sciences, Warmadewa University | Eijkman Institute for Molecular Biology, National Agency for Research and Innovation; Molecular Biology Laboratory, Faculty Medicine and Health Sciences, Warmadewa University | Hidayat Trimarsanto, Lydia V. Panggalo, Iskandar Adnan, Sukma Oktavianthi, Lidwina Priliani, Edison Johar, Frilasita A Yudhaputri, Muhammad Rezki Rasyak, Willy Agustine, Sri Masyeni, Erly Sintya, Safarina G Malik, Khin Saw Myint, Amin Soebandrio |
| EPI_ISL_2101080 | Molecular Biology Laboratory, Faculty Medicine and Health Sciences, Warmadewa University | Eijkman Institute for Molecular Biology, National Agency for Research and Innovation; Molecular Biology Laboratory, Faculty Medicine and Health Sciences, Warmadewa University | Lydia V. Panggalo, Iskandar Adnan, Sukma Oktavianthi, Lidwina Priliani, Edison Johar, Frilasita A Yudhaputri, Muhammad Rezki Rasyak, Willy Agustine, Hidayat Trimarsanto, Sri Masyeni, Erly Sintya, Safarina G Malik, Khin Saw Myint, Amin Soebandrio |
| EPI_ISL_2101081 | Molecular Biology Laboratory, Faculty Medicine and Health Sciences, Warmadewa University | Eijkman Institute for Molecular Biology, National Agency for Research and Innovation; Molecular Biology Laboratory, Faculty Medicine and Health Sciences, Warmadewa University | Sukma Oktavianthi, Lidwina Priliani, Edison Johar, Frilasita A Yudhaputri, Muhammad Rezki Rasyak, Willy Agustine, Hidayat Trimarsanto, Lydia V. Panggalo, Iskandar Adnan, Sri Masyeni, Erly Sintya, Safarina G Malik, Khin Saw Myint, Amin Soebandrio |
| EPI_ISL_2101082 | Molecular Biology Laboratory, Faculty Medicine and Health Sciences, Warmadewa University | Eijkman Institute for Molecular Biology, National Agency for Research and Innovation; Molecular Biology Laboratory, Faculty Medicine and Health Sciences, Warmadewa University | Lidwina Priliani, Edison Johar, Frilasita A Yudhaputri, Muhammad Rezki Rasyak, Willy Agustine, Hidayat Trimarsanto, Lydia V. Panggalo, Iskandar Adnan, Sukma Oktavianthi, Sri Masyeni, Erly Sintya, Safarina G Malik, Khin Saw Myint, Amin Soebandrio |
| EPI_ISL_2101083 | Molecular Biology Laboratory, Faculty Medicine and Health Sciences, Warmadewa University | Eijkman Institute for Molecular Biology, National Agency for Research and Innovation; Molecular Biology Laboratory, Faculty Medicine and Health Sciences, Warmadewa University | Edison Johar, Frilasita A Yudhaputri, Muhammad Rezki Rasyak, Willy Agustine, Hidayat Trimarsanto, Lydia V. Panggalo, Iskandar Adnan, Sukma Oktavianthi, Lidwina Priliani, Sri Masyeni, Erly Sintya, Safarina G Malik, Khin Saw Myint, Amin Soebandrio |
| EPI_ISL_2101084 | Molecular Biology Laboratory, Faculty Medicine and Health Sciences, Warmadewa University | Eijkman Institute for Molecular Biology, National Agency for Research and Innovation; Molecular Biology Laboratory, Faculty Medicine and Health Sciences, Warmadewa University | Frilasita A Yudhaputri, Muhammad Rezki Rasyak, Willy Agustine, Hidayat Trimarsanto, Lydia V. Panggalo, Iskandar Adnan, Sukma Oktavianthi, Lidwina Priliani, Edison Johar, Sri Masyeni, Erly Sintya, Safarina G Malik, Khin Saw Myint, Amin Soebandrio |
| EPI_ISL_2101085 | Molecular Biology Laboratory, Faculty Medicine and Health Sciences, Warmadewa University | Eijkman Institute for Molecular Biology, National Agency for Research and Innovation; Molecular Biology Laboratory, Faculty Medicine and Health Sciences, Warmadewa University | Willy Agustine, Hidayat Trimarsanto, Lydia V. Panggalo, Iskandar Adnan, Sukma Oktavianthi, Lidwina Priliani, Edison Johar, Frilasita A Yudhaputri, Muhammad Rezki Rasyak, Sri Masyeni, Erly Sintya, Safarina G Malik, Khin Saw Myint, Amin Soebandrio |
| EPI_ISL_2101086 | Molecular Biology Laboratory, Faculty Medicine and Health Sciences, Warmadewa University | Eijkman Institute for Molecular Biology, National Agency for Research and Innovation; Molecular Biology Laboratory, Faculty Medicine and Health Sciences, Warmadewa University | Hidayat Trimarsanto, Lydia V. Panggalo, Iskandar Adnan, Sukma Oktavianthi, Lidwina Priliani, Edison Johar, Frilasita A Yudhaputri, Muhammad Rezki Rasyak, Willy Agustine, Sri Masyeni, Erly Sintya, Safarina G Malik, Khin Saw Myint, Amin Soebandrio |
| EPI_ISL_2101087 | Molecular Biology Laboratory, Faculty Medicine and Health Sciences, Warmadewa University | Eijkman Institute for Molecular Biology, National Agency for Research and Innovation; Molecular Biology Laboratory, Faculty Medicine and Health Sciences, Warmadewa University | Lydia V. Panggalo, Iskandar Adnan, Sukma Oktavianthi, Lidwina Priliani, Edison Johar, Frilasita A Yudhaputri, Muhammad Rezki Rasyak, Willy Agustine, Hidayat Trimarsanto, Sri Masyeni, Erly Sintya, Safarina G Malik, Khin Saw Myint, Amin Soebandrio |
| EPI_ISL_2101088 | Molecular Biology Laboratory, Faculty Medicine and Health Sciences, Warmadewa University | Eijkman Institute for Molecular Biology, National Agency for Research and Innovation; Molecular Biology Laboratory, Faculty Medicine and Health Sciences, Warmadewa University | Iskandar Adnan, Sukma Oktavianthi, Lidwina Priliani, Edison Johar, Frilasita A Yudhaputri, Muhammad Rezki Rasyak, Willy Agustine, Hidayat Trimarsanto, Lydia V. Panggalo, Sri Masyeni, Erly Sintya, Safarina G Malik, Khin Saw Myint, Amin Soebandrio |
| EPI_ISL_2101089 | Molecular Biology Laboratory, Faculty Medicine and Health Sciences, Warmadewa University | Eijkman Institute for Molecular Biology, National Agency for Research and Innovation; Molecular Biology Laboratory, Faculty Medicine and Health Sciences, Warmadewa University | Sukma Oktavianthi, Lidwina Priliani, Edison Johar, Frilasita A Yudhaputri, Muhammad Rezki Rasyak, Willy Agustine, Hidayat Trimarsanto, Lydia V. Panggalo, Iskandar Adnan, Sri Masyeni, Erly Sintya, Safarina G Malik, Khin Saw Myint, Amin Soebandrio |
| EPI_ISL_2101090 | Molecular Biology Laboratory, Faculty Medicine and Health Sciences, Warmadewa University | Eijkman Institute for Molecular Biology, National Agency for Research and Innovation; Molecular Biology Laboratory, Faculty Medicine and Health Sciences, Warmadewa University | Lidwina Priliani, Edison Johar, Frilasita A Yudhaputri, Muhammad Rezki Rasyak, Willy Agustine, Hidayat Trimarsanto, Lydia V. Panggalo, Iskandar Adnan, Sukma Oktavianthi, Sri Masyeni, Erly Sintya, Safarina G Malik, Khin Saw Myint, Amin Soebandrio |
| EPI_ISL_2101091 | Molecular Biology Laboratory, Faculty Medicine and Health Sciences, Warmadewa University | Eijkman Institute for Molecular Biology, National Agency for Research and Innovation; Molecular Biology Laboratory, Faculty Medicine and Health Sciences, Warmadewa University | Frilasita A Yudhaputri, Muhammad Rezki Rasyak, Willy Agustine, Hidayat Trimarsanto, Lydia V. Panggalo, Iskandar Adnan, Sukma Oktavianthi, Lidwina Priliani, Edison Johar, Sri Masyeni, Erly Sintya, Safarina G Malik, Khin Saw Myint, Amin Soebandrio |
| EPI_ISL_2101092 | Molecular Biology Laboratory, Faculty Medicine and Health Sciences, Warmadewa University | Eijkman Institute for Molecular Biology, National Agency for Research and Innovation; Molecular Biology Laboratory, Faculty Medicine and Health Sciences, Warmadewa University | Willy Agustine, Hidayat Trimarsanto, Lydia V. Panggalo, Iskandar Adnan, Sukma Oktavianthi, Lidwina Priliani, Edison Johar, Frilasita A Yudhaputri, Muhammad Rezki Rasyak, Sri Masyeni, Erly Sintya, Safarina G Malik, Khin Saw Myint, Amin Soebandrio |
| EPI_ISL_2101093 | Molecular Biology Laboratory, Faculty Medicine and Health Sciences, Warmadewa University | Eijkman Institute for Molecular Biology, National Agency for Research and Innovation; Molecular Biology Laboratory, Faculty Medicine and Health Sciences, Warmadewa University | Hidayat Trimarsanto, Lydia V. Panggalo, Iskandar Adnan, Sukma Oktavianthi, Lidwina Priliani, Edison Johar, Frilasita A Yudhaputri, Muhammad Rezki Rasyak, Willy Agustine, Sri Masyeni, Erly Sintya, Safarina G Malik, Khin Saw Myint, Amin Soebandrio |
| EPI_ISL_2101094 | Molecular Biology Laboratory, Faculty Medicine and Health Sciences, Warmadewa University | Eijkman Institute for Molecular Biology, National Agency for Research and Innovation; Molecular Biology Laboratory, Faculty Medicine and Health Sciences, Warmadewa University | Lydia V. Panggalo, Iskandar Adnan, Sukma Oktavianthi, Lidwina Priliani, Edison Johar, Frilasita A Yudhaputri, Muhammad Rezki Rasyak, Willy Agustine, Hidayat Trimarsanto, Sri Masyeni, Erly Sintya, Safarina G Malik, Khin Saw Myint, Amin Soebandrio |
| EPI_ISL_2101095 | Molecular Biology Laboratory, Faculty Medicine and Health Sciences, Warmadewa University | Eijkman Institute for Molecular Biology, National Agency for Research and Innovation; Molecular Biology Laboratory, | Iskandar Adnan, Sukma Oktavianthi, Lidwina Priliani, Edison Johar, Frilasita A Yudhaputri, Muhammad Rezki Rasyak, Willy Agustine, Hidayat Trimarsanto, Lydia V. Panggalo, Sri Masyeni, Erly Sintya, Safarina G Malik, Khin Saw Myint, Amin Soebandrio |

| | | | |
|---|---|---|---|
| | | Faculty Medicine and Health Sciences, Warmadewa University | |
| EPI_ISL_2101096 | Molecular Biology Laboratory, Faculty Medicine and Health Sciences, Warmadewa University | Eijkman Institute for Molecular Biology, National Agency for Research and Innovation; Molecular Biology Laboratory, Faculty Medicine and Health Sciences, Warmadewa University | Lidwina Priliani, Edison Johar, Frilasita A Yudhaputri, Muhammad Rezki Rasyak, Willy Agustine, Hidayat Trimarsanto, Lydia V. Panggalo, Iskandar Adnan, Sukma Oktavianthi, Sri Masyeni, Erly Sintya, Safarina G Malik, Khin Saw Myint, Amin Soebandrio |
| EPI_ISL_2101097 | Molecular Biology Laboratory, Faculty Medicine and Health Sciences, Warmadewa University | Eijkman Institute for Molecular Biology, National Agency for Research and Innovation; Molecular Biology Laboratory, Faculty Medicine and Health Sciences, Warmadewa University | Edison Johar, Frilasita A Yudhaputri, Muhammad Rezki Rasyak, Willy Agustine, Hidayat Trimarsanto, Lydia V. Panggalo, Iskandar Adnan, Sukma Oktavianthi, Lidwina Priliani, Sri Masyeni, Erly Sintya, Safarina G Malik, Khin Saw Myint, Amin Soebandrio |
| EPI_ISL_2101098 | Molecular Biology Laboratory, Faculty Medicine and Health Sciences, Warmadewa University | Eijkman Institute for Molecular Biology, National Agency for Research and Innovation; Molecular Biology Laboratory, Faculty Medicine and Health Sciences, Warmadewa University | Frilasita A Yudhaputri, Muhammad Rezki Rasyak, Willy Agustine, Hidayat Trimarsanto, Lydia V. Panggalo, Iskandar Adnan, Sukma Oktavianthi, Lidwina Priliani, Edison Johar, Sri Masyeni, Erly Sintya, Safarina G Malik, Khin Saw Myint, Amin Soebandrio |
| EPI_ISL_2101099 | Molecular Biology Laboratory, Faculty Medicine and Health Sciences, Warmadewa University | Eijkman Institute for Molecular Biology, National Agency for Research and Innovation; Molecular Biology Laboratory, Faculty Medicine and Health Sciences, Warmadewa University | Willy Agustine, Hidayat Trimarsanto, Lydia V. Panggalo, Iskandar Adnan, Sukma Oktavianthi, Lidwina Priliani, Edison Johar, Frilasita A Yudhaputri, Muhammad Rezki Rasyak, Sri Masyeni, Erly Sintya, Safarina G Malik, Khin Saw Myint, Amin Soebandrio |
| EPI_ISL_2101100 | Molecular Biology Laboratory, Faculty Medicine and Health Sciences, Warmadewa University | Eijkman Institute for Molecular Biology, National Agency for Research and Innovation; Molecular Biology Laboratory, Faculty Medicine and Health Sciences, Warmadewa University | Lydia V. Panggalo, Iskandar Adnan, Sukma Oktavianthi, Lidwina Priliani, Edison Johar, Frilasita A Yudhaputri, Muhammad Rezki Rasyak, Willy Agustine, Hidayat Trimarsanto, Sri Masyeni, Erly Sintya, Safarina G Malik, Khin Saw Myint, Amin Soebandrio |
| EPI_ISL_2101102 | Molecular Biology Laboratory, Faculty Medicine and Health Sciences, Warmadewa University | Eijkman Institute for Molecular Biology, National Agency for Research and Innovation; Molecular Biology Laboratory, Faculty Medicine and Health Sciences, Warmadewa University | Sukma Oktavianthi, Lidwina Priliani, Edison Johar, Frilasita A Yudhaputri, Muhammad Rezki Rasyak, Willy Agustine, Hidayat Trimarsanto, Lydia V. Panggalo, Iskandar Adnan, Sri Masyeni, Erly Sintya, Safarina G Malik, Khin Saw Myint, Amin Soebandrio |
| EPI_ISL_2101105 | Molecular Biology Laboratory, Faculty Medicine and Health Sciences, Warmadewa University | Eijkman Institute for Molecular Biology, National Agency for Research and Innovation; Molecular Biology Laboratory, Faculty Medicine and Health Sciences, Warmadewa University | Frilasita A Yudhaputri, Muhammad Rezki Rasyak, Willy Agustine, Hidayat Trimarsanto, Lydia V. Panggalo, Iskandar Adnan, Sukma Oktavianthi, Lidwina Priliani, Edison Johar, Sri Masyeni, Erly Sintya, Safarina G Malik, Khin Saw Myint, Amin Soebandrio |
| EPI_ISL_2101106 | Molecular Biology Laboratory, Faculty Medicine and Health Sciences, Warmadewa University | Eijkman Institute for Molecular Biology, National Agency for Research and Innovation; Molecular Biology Laboratory, Faculty Medicine and Health Sciences, Warmadewa University | Willy Agustine, Hidayat Trimarsanto, Lydia V. Panggalo, Iskandar Adnan, Sukma Oktavianthi, Lidwina Priliani, Edison Johar, Frilasita A Yudhaputri, Muhammad Rezki Rasyak, Sri Masyeni, Erly Sintya, Safarina G Malik, Khin Saw Myint, Amin Soebandrio |
| EPI_ISL_2101107 | Molecular Biology Laboratory, Faculty Medicine and Health Sciences, Warmadewa University | Eijkman Institute for Molecular Biology, National Agency for Research and Innovation; Molecular Biology Laboratory, Faculty Medicine and Health Sciences, Warmadewa University | Hidayat Trimarsanto, Lydia V. Panggalo, Iskandar Adnan, Sukma Oktavianthi, Lidwina Priliani, Edison Johar, Frilasita A Yudhaputri, Muhammad Rezki Rasyak, Willy Agustine, Sri Masyeni, Erly Sintya, Safarina G Malik, Khin Saw Myint, Amin Soebandrio |
| EPI_ISL_2101108 | Molecular Biology Laboratory, Faculty Medicine and Health Sciences, Warmadewa University | Eijkman Institute for Molecular Biology, National Agency for Research and Innovation; Molecular Biology Laboratory, Faculty Medicine and Health Sciences, Warmadewa University | Lydia V. Panggalo, Iskandar Adnan, Sukma Oktavianthi, Lidwina Priliani, Edison Johar, Frilasita A Yudhaputri, Muhammad Rezki Rasyak, Willy Agustine, Hidayat Trimarsanto, Sri Masyeni, Erly Sintya, Safarina G Malik, Khin Saw Myint, Amin Soebandrio |
| EPI_ISL_2101109 | Molecular Biology Laboratory, Faculty Medicine and Health Sciences, Warmadewa University | Eijkman Institute for Molecular Biology, National Agency for Research and Innovation; Molecular Biology Laboratory, Faculty Medicine and Health Sciences, Warmadewa University | Iskandar Adnan, Sukma Oktavianthi, Lidwina Priliani, Edison Johar, Frilasita A Yudhaputri, Muhammad Rezki Rasyak, Willy Agustine, Hidayat Trimarsanto, Lydia V. Panggalo, Sri Masyeni, Erly Sintya, Safarina G Malik, Khin Saw Myint, Amin Soebandrio |
| EPI_ISL_2101110 | Molecular Biology Laboratory, Faculty Medicine and Health Sciences, Warmadewa University | Eijkman Institute for Molecular Biology, National Agency for Research and Innovation; Molecular Biology Laboratory, Faculty Medicine and Health Sciences, Warmadewa University | Lidwina Priliani, Edison Johar, Frilasita A Yudhaputri, Muhammad Rezki Rasyak, Willy Agustine, Hidayat Trimarsanto, Lydia V. Panggalo, Iskandar Adnan, Sukma Oktavianthi, Sri Masyeni, Erly Sintya, Safarina G Malik, Khin Saw Myint, Amin Soebandrio |
| EPI_ISL_2101111 | Molecular Biology Laboratory, Faculty Medicine and Health Sciences, Warmadewa University | Eijkman Institute for Molecular Biology, National Agency for Research and Innovation; Molecular Biology Laboratory, Faculty Medicine and Health Sciences, Warmadewa University | Edison Johar, Frilasita A Yudhaputri, Muhammad Rezki Rasyak, Willy Agustine, Hidayat Trimarsanto, Lydia V. Panggalo, Iskandar Adnan, Sukma Oktavianthi, Lidwina Priliani, Sri Masyeni, Erly Sintya, Safarina G Malik, Khin Saw Myint, Amin Soebandrio |
| EPI_ISL_2101113 | Molecular Biology Laboratory, Faculty Medicine and Health Sciences, Warmadewa University | Eijkman Institute for Molecular Biology, National Agency for Research and Innovation; Molecular Biology Laboratory, Faculty Medicine and Health Sciences, Warmadewa University | Frilasita A Yudhaputri, Muhammad Rezki Rasyak, Willy Agustine, Hidayat Trimarsanto, Lydia V. Panggalo, Iskandar Adnan, Sukma Oktavianthi, Lidwina Priliani, Edison Johar, Sri Masyeni, Erly Sintya, Safarina G Malik, Khin Saw Myint, Amin Soebandrio |
| EPI_ISL_2101114 | Molecular Biology Laboratory, Faculty Medicine and Health Sciences, Warmadewa University | Eijkman Institute for Molecular Biology, National Agency for Research and Innovation; Molecular Biology Laboratory, Faculty Medicine and Health Sciences, Warmadewa University | Willy Agustine, Hidayat Trimarsanto, Lydia V. Panggalo, Iskandar Adnan, Sukma Oktavianthi, Lidwina Priliani, Edison Johar, Frilasita A Yudhaputri, Muhammad Rezki Rasyak, Sri Masyeni, Erly Sintya, Safarina G Malik, Khin Saw Myint, Amin Soebandrio |
| EPI_ISL_2101115 | Molecular Biology Laboratory, Faculty Medicine and Health Sciences, Warmadewa University | Eijkman Institute for Molecular Biology, National Agency for Research and Innovation; Molecular Biology Laboratory, Faculty Medicine and Health Sciences, Warmadewa University | Hidayat Trimarsanto, Lydia V. Panggalo, Iskandar Adnan, Sukma Oktavianthi, Lidwina Priliani, Edison Johar, Frilasita A Yudhaputri, Muhammad Rezki Rasyak, Willy Agustine, Sri Masyeni, Erly Sintya, Safarina G Malik, Khin Saw Myint, Amin Soebandrio |
| EPI_ISL_2101116 | Molecular Biology Laboratory, Faculty Medicine and Health Sciences, Warmadewa University | Eijkman Institute for Molecular Biology, National Agency for Research and Innovation; Molecular Biology Laboratory, Faculty Medicine and Health Sciences, Warmadewa University | Lydia V. Panggalo, Iskandar Adnan, Sukma Oktavianthi, Lidwina Priliani, Edison Johar, Frilasita A Yudhaputri, Muhammad Rezki Rasyak, Willy Agustine, Hidayat Trimarsanto, Sri Masyeni, Erly Sintya, Safarina G Malik, Khin Saw Myint, Amin Soebandrio |
| EPI_ISL_2101117 | Molecular Biology Laboratory, Faculty Medicine and Health Sciences, Warmadewa University | Eijkman Institute for Molecular Biology, National Agency for Research and Innovation; Molecular Biology Laboratory, Faculty Medicine and Health Sciences, Warmadewa University | Iskandar Adnan, Sukma Oktavianthi, Lidwina Priliani, Edison Johar, Frilasita A Yudhaputri, Muhammad Rezki Rasyak, Willy Agustine, Hidayat Trimarsanto, Lydia V. Panggalo, Sri Masyeni, Erly Sintya, Safarina G Malik, Khin Saw Myint, Amin Soebandrio |
| EPI_ISL_2101118 | Molecular Biology Laboratory, Faculty Medicine and Health Sciences, Warmadewa University | Eijkman Institute for Molecular Biology, National Agency for Research and Innovation; Molecular Biology Laboratory, Faculty Medicine and Health Sciences, Warmadewa University | Sukma Oktavianthi, Lidwina Priliani, Edison Johar, Frilasita A Yudhaputri, Muhammad Rezki Rasyak, Willy Agustine, Hidayat Trimarsanto, Lydia V. Panggalo, Iskandar Adnan, Sri Masyeni, Erly Sintya, Safarina G Malik, Khin Saw Myint, Amin Soebandrio |

| | | | |
|---|---|---|---|
| EPI_ISL_2101119 | Molecular Biology Laboratory, Faculty Medicine and Health Sciences, Warmadewa University | Eijkman Institute for Molecular Biology, National Agency for Research and Innovation; Molecular Biology Laboratory, Faculty Medicine and Health Sciences, Warmadewa University | Lidwina Priliani, Edison Johar, Frilasita A Yudhaputri, Muhammad Rezki Rasyak, Willy Agustine, Hidayat Trimarsanto, Lydia V. Panggalo, Iskandar Adnan, Sukma Oktavianthi, Sri Masyeni, Erly Sintya, Safarina G Malik, Khin Saw Myint, Amin Soebandrio |
| EPI_ISL_2101120 | Molecular Biology Laboratory, Faculty Medicine and Health Sciences, Warmadewa University | Eijkman Institute for Molecular Biology, National Agency for Research and Innovation; Molecular Biology Laboratory, Faculty Medicine and Health Sciences, Warmadewa University | Edison Johar, Frilasita A Yudhaputri, Muhammad Rezki Rasyak, Willy Agustine, Hidayat Trimarsanto, Lydia V. Panggalo, Iskandar Adnan, Sukma Oktavianthi, Lidwina Priliani, Sri Masyeni, Erly Sintya, Safarina G Malik, Khin Saw Myint, Amin Soebandrio |
| EPI_ISL_2101121 | Molecular Biology Laboratory, Faculty Medicine and Health Sciences, Warmadewa University | Eijkman Institute for Molecular Biology, National Agency for Research and Innovation; Molecular Biology Laboratory, Faculty Medicine and Health Sciences, Warmadewa University | Willy Agustine, Hidayat Trimarsanto, Lydia V. Panggalo, Iskandar Adnan, Sukma Oktavianthi, Lidwina Priliani, Edison Johar, Frilasita A Yudhaputri, Muhammad Rezki Rasyak, Sri Masyeni, Erly Sintya, Safarina G Malik, Khin Saw Myint, Amin Soebandrio |
| EPI_ISL_2101122 | Molecular Biology Laboratory, Faculty Medicine and Health Sciences, Warmadewa University | Eijkman Institute for Molecular Biology, National Agency for Research and Innovation; Molecular Biology Laboratory, Faculty Medicine and Health Sciences, Warmadewa University | Hidayat Trimarsanto, Lydia V. Panggalo, Iskandar Adnan, Sukma Oktavianthi, Lidwina Priliani, Edison Johar, Frilasita A Yudhaputri, Muhammad Rezki Rasyak, Willy Agustine, Sri Masyeni, Erly Sintya, Safarina G Malik, Khin Saw Myint, Amin Soebandrio |
| EPI_ISL_2101123 | Molecular Biology Laboratory, Faculty Medicine and Health Sciences, Warmadewa University | Eijkman Institute for Molecular Biology, National Agency for Research and Innovation; Molecular Biology Laboratory, Faculty Medicine and Health Sciences, Warmadewa University | Lydia V. Panggalo, Iskandar Adnan, Sukma Oktavianthi, Lidwina Priliani, Edison Johar, Frilasita A Yudhaputri, Muhammad Rezki Rasyak, Willy Agustine, Hidayat Trimarsanto, Sri Masyeni, Erly Sintya, Safarina G Malik, Khin Saw Myint, Amin Soebandrio |
| EPI_ISL_2101125 | Molecular Biology Laboratory, Faculty Medicine and Health Sciences, Warmadewa University | Eijkman Institute for Molecular Biology, National Agency for Research and Innovation; Molecular Biology Laboratory, Faculty Medicine and Health Sciences, Warmadewa University | Willy Agustine, Hidayat Trimarsanto, Lydia V. Panggalo, Iskandar Adnan, Sukma Oktavianthi, Lidwina Priliani, Edison Johar, Frilasita A Yudhaputri, Muhammad Rezki Rasyak, Sri Masyeni, Erly Sintya, Safarina G Malik, Khin Saw Myint, Amin Soebandrio |
| EPI_ISL_2101126 | Molecular Biology Laboratory, Faculty Medicine and Health Sciences, Warmadewa University | Eijkman Institute for Molecular Biology, National Agency for Research and Innovation; Molecular Biology Laboratory, Faculty Medicine and Health Sciences, Warmadewa University | Hidayat Trimarsanto, Lydia V. Panggalo, Iskandar Adnan, Sukma Oktavianthi, Lidwina Priliani, Edison Johar, Frilasita A Yudhaputri, Muhammad Rezki Rasyak, Willy Agustine, Sri Masyeni, Erly Sintya, Safarina G Malik, Khin Saw Myint, Amin Soebandrio |
| EPI_ISL_2101127 | Molecular Biology Laboratory, Faculty Medicine and Health Sciences, Warmadewa University | Eijkman Institute for Molecular Biology, National Agency for Research and Innovation; Molecular Biology Laboratory, Faculty Medicine and Health Sciences, Warmadewa University | Iskandar Adnan, Sukma Oktavianthi, Lidwina Priliani, Edison Johar, Frilasita A Yudhaputri, Muhammad Rezki Rasyak, Willy Agustine, Hidayat Trimarsanto, Lydia V. Panggalo, Sri Masyeni, Erly Sintya, Safarina G Malik, Khin Saw Myint, Amin Soebandrio |
| EPI_ISL_2101128 | Molecular Biology Laboratory, Faculty Medicine and Health Sciences, Warmadewa University | Eijkman Institute for Molecular Biology, National Agency for Research and Innovation; Molecular Biology Laboratory, Faculty Medicine and Health Sciences, Warmadewa University | Sukma Oktavianthi, Lidwina Priliani, Edison Johar, Frilasita A Yudhaputri, Muhammad Rezki Rasyak, Willy Agustine, Hidayat Trimarsanto, Lydia V. Panggalo, Iskandar Adnan, Sri Masyeni, Erly Sintya, Safarina G Malik, Khin Saw Myint, Amin Soebandrio |
| EPI_ISL_2101129 | Molecular Biology Laboratory, Faculty Medicine and Health Sciences, Warmadewa University | Eijkman Institute for Molecular Biology, National Agency for Research and Innovation; Molecular Biology Laboratory, Faculty Medicine and Health Sciences, Warmadewa University | Edison Johar, Frilasita A Yudhaputri, Muhammad Rezki Rasyak, Willy Agustine, Hidayat Trimarsanto, Lydia V. Panggalo, Iskandar Adnan, Sukma Oktavianthi, Lidwina Priliani, Sri Masyeni, Erly Sintya, Safarina G Malik, Khin Saw Myint, Amin Soebandrio |
| EPI_ISL_2101131 | Molecular Biology Laboratory, Faculty Medicine and Health Sciences, Warmadewa University | Eijkman Institute for Molecular Biology, National Agency for Research and Innovation; Molecular Biology Laboratory, Faculty Medicine and Health Sciences, Warmadewa University | Willy Agustine, Hidayat Trimarsanto, Lydia V. Panggalo, Iskandar Adnan, Sukma Oktavianthi, Lidwina Priliani, Edison Johar, Frilasita A Yudhaputri, Muhammad Rezki Rasyak, Sri Masyeni, Erly Sintya, Safarina G Malik, Khin Saw Myint, Amin Soebandrio |
| EPI_ISL_2101132 | Molecular Biology Laboratory, Faculty Medicine and Health Sciences, Warmadewa University | Eijkman Institute for Molecular Biology, National Agency for Research and Innovation; Molecular Biology Laboratory, Faculty Medicine and Health Sciences, Warmadewa University | Hidayat Trimarsanto, Lydia V. Panggalo, Iskandar Adnan, Sukma Oktavianthi, Lidwina Priliani, Edison Johar, Frilasita A Yudhaputri, Muhammad Rezki Rasyak, Willy Agustine, Sri Masyeni, Erly Sintya, Safarina G Malik, Khin Saw Myint, Amin Soebandrio |
| EPI_ISL_2101133 | Molecular Biology Laboratory, Faculty Medicine and Health Sciences, Warmadewa University | Eijkman Institute for Molecular Biology, National Agency for Research and Innovation; Molecular Biology Laboratory, Faculty Medicine and Health Sciences, Warmadewa University | Lydia V. Panggalo, Iskandar Adnan, Sukma Oktavianthi, Lidwina Priliani, Edison Johar, Frilasita A Yudhaputri, Muhammad Rezki Rasyak, Willy Agustine, Hidayat Trimarsanto, Sri Masyeni, Erly Sintya, Safarina G Malik, Khin Saw Myint, Amin Soebandrio |
| EPI_ISL_2101134 | Molecular Biology Laboratory, Faculty Medicine and Health Sciences, Warmadewa University | Eijkman Institute for Molecular Biology, National Agency for Research and Innovation; Molecular Biology Laboratory, Faculty Medicine and Health Sciences, Warmadewa University | Iskandar Adnan, Sukma Oktavianthi, Lidwina Priliani, Edison Johar, Frilasita A Yudhaputri, Muhammad Rezki Rasyak, Willy Agustine, Hidayat Trimarsanto, Lydia V. Panggalo, Sri Masyeni, Erly Sintya, Safarina G Malik, Khin Saw Myint, Amin Soebandrio |
| EPI_ISL_2101135 | Molecular Biology Laboratory, Faculty Medicine and Health Sciences, Warmadewa University | Eijkman Institute for Molecular Biology, National Agency for Research and Innovation; Molecular Biology Laboratory, Faculty Medicine and Health Sciences, Warmadewa University | Sukma Oktavianthi, Lidwina Priliani, Edison Johar, Frilasita A Yudhaputri, Muhammad Rezki Rasyak, Willy Agustine, Hidayat Trimarsanto, Lydia V. Panggalo, Iskandar Adnan, Sri Masyeni, Erly Sintya, Safarina G Malik, Khin Saw Myint, Amin Soebandrio |
| EPI_ISL_2101136 | Molecular Biology Laboratory, Faculty Medicine and Health Sciences, Warmadewa University | Eijkman Institute for Molecular Biology, National Agency for Research and Innovation; Molecular Biology Laboratory, Faculty Medicine and Health Sciences, Warmadewa University | Edison Johar, Frilasita A Yudhaputri, Muhammad Rezki Rasyak, Willy Agustine, Hidayat Trimarsanto, Lydia V. Panggalo, Iskandar Adnan, Sukma Oktavianthi, Lidwina Priliani, Sri Masyeni, Erly Sintya, Safarina G Malik, Khin Saw Myint, Amin Soebandrio |
| EPI_ISL_2101137 | Molecular Biology Laboratory, Faculty Medicine and Health Sciences, Warmadewa University | Eijkman Institute for Molecular Biology, National Agency for Research and Innovation; Molecular Biology Laboratory, Faculty Medicine and Health Sciences, Warmadewa University | Frilasita A Yudhaputri, Muhammad Rezki Rasyak, Willy Agustine, Hidayat Trimarsanto, Lydia V. Panggalo, Iskandar Adnan, Sukma Oktavianthi, Lidwina Priliani, Edison Johar, Sri Masyeni, Erly Sintya, Safarina G Malik, Khin Saw Myint, Amin Soebandrio |
| EPI_ISL_2101138 | Molecular Biology Laboratory, Faculty Medicine and Health Sciences, Warmadewa University | Eijkman Institute for Molecular Biology, National Agency for Research and Innovation; Molecular Biology Laboratory, Faculty Medicine and Health Sciences, Warmadewa University | Willy Agustine, Hidayat Trimarsanto, Lydia V. Panggalo, Iskandar Adnan, Sukma Oktavianthi, Lidwina Priliani, Edison Johar, Frilasita A Yudhaputri, Muhammad Rezki Rasyak, Sri Masyeni, Erly Sintya, Safarina G Malik, Khin Saw Myint, Amin Soebandrio |
| EPI_ISL_2101139 | Molecular Biology Laboratory, Faculty Medicine and Health Sciences, Warmadewa University | Eijkman Institute for Molecular Biology, National Agency for Research and Innovation; Molecular Biology Laboratory, Faculty Medicine and Health Sciences, Warmadewa University | Hidayat Trimarsanto, Lydia V. Panggalo, Iskandar Adnan, Sukma Oktavianthi, Lidwina Priliani, Edison Johar, Frilasita A Yudhaputri, Muhammad Rezki Rasyak, Willy Agustine, Sri Masyeni, Erly Sintya, Safarina G Malik, Khin Saw Myint, Amin Soebandrio |
| EPI_ISL_2101140 | Molecular Biology Laboratory, Faculty Medicine and Health Sciences, Warmadewa University | Eijkman Institute for Molecular Biology, National Agency for Research and Innovation; Molecular Biology Laboratory, | Lydia V. Panggalo, Iskandar Adnan, Sukma Oktavianthi, Lidwina Priliani, Edison Johar, Frilasita A Yudhaputri, Muhammad Rezki Rasyak, Willy Agustine, Hidayat Trimarsanto, Sri Masyeni, Erly Sintya, Safarina G Malik, Khin Saw Myint, Amin Soebandrio |

| | | | |
|---|---|---|---|
| | | Faculty Medicine and Health Sciences, Warmadewa University | |
| EPI_ISL_2101141 | Molecular Biology Laboratory, Faculty Medicine and Health Sciences, Warmadewa University | Eijkman Institute for Molecular Biology, National Agency for Research and Innovation; Molecular Biology Laboratory, Faculty Medicine and Health Sciences, Warmadewa University | Iskandar Adnan, Sukma Oktavianthi, Lidwina Priliani, Edison Johar, Frilasita A Yudhaputri, Muhammad Rezki Rasyak, Willy Agustine, Hidayat Trimarsanto, Lydia V. Panggalo, Sri Masyeni, Erly Sintya, Safarina G Malik, Khin Saw Myint, Amin Soebandrio |
| EPI_ISL_2101142 | Molecular Biology Laboratory, Faculty Medicine and Health Sciences, Warmadewa University | Eijkman Institute for Molecular Biology, National Agency for Research and Innovation; Molecular Biology Laboratory, Faculty Medicine and Health Sciences, Warmadewa University | Lidwina Priliani, Edison Johar, Frilasita A Yudhaputri, Muhammad Rezki Rasyak, Willy Agustine, Hidayat Trimarsanto, Lydia V. Panggalo, Iskandar Adnan, Sukma Oktavianthi, Sri Masyeni, Erly Sintya, Safarina G Malik, Khin Saw Myint, Amin Soebandrio |
| EPI_ISL_2101143 | Molecular Biology Laboratory, Faculty Medicine and Health Sciences, Warmadewa University | Eijkman Institute for Molecular Biology, National Agency for Research and Innovation; Molecular Biology Laboratory, Faculty Medicine and Health Sciences, Warmadewa University | Frilasita A Yudhaputri, Muhammad Rezki Rasyak, Willy Agustine, Hidayat Trimarsanto, Lydia V. Panggalo, Iskandar Adnan, Sukma Oktavianthi, Lidwina Priliani, Edison Johar, Sri Masyeni, Erly Sintya, Safarina G Malik, Khin Saw Myint, Amin Soebandrio |
| EPI_ISL_2101144 | Molecular Biology Laboratory, Faculty Medicine and Health Sciences, Warmadewa University | Eijkman Institute for Molecular Biology, National Agency for Research and Innovation; Molecular Biology Laboratory, Faculty Medicine and Health Sciences, Warmadewa University | Hidayat Trimarsanto, Lydia V. Panggalo, Iskandar Adnan, Sukma Oktavianthi, Lidwina Priliani, Edison Johar, Frilasita A Yudhaputri, Muhammad Rezki Rasyak, Willy Agustine, Sri Masyeni, Erly Sintya, Safarina G Malik, Khin Saw Myint, Amin Soebandrio |
| EPI_ISL_2101145 | Molecular Biology Laboratory, Faculty Medicine and Health Sciences, Warmadewa University | Eijkman Institute for Molecular Biology, National Agency for Research and Innovation; Molecular Biology Laboratory, Faculty Medicine and Health Sciences, Warmadewa University | Lydia V. Panggalo, Iskandar Adnan, Sukma Oktavianthi, Lidwina Priliani, Edison Johar, Frilasita A Yudhaputri, Muhammad Rezki Rasyak, Willy Agustine, Hidayat Trimarsanto, Sri Masyeni, Erly Sintya, Safarina G Malik, Khin Saw Myint, Amin Soebandrio |
| EPI_ISL_2101146 | Molecular Biology Laboratory, Faculty Medicine and Health Sciences, Warmadewa University | Eijkman Institute for Molecular Biology, National Agency for Research and Innovation; Molecular Biology Laboratory, Faculty Medicine and Health Sciences, Warmadewa University | Iskandar Adnan, Sukma Oktavianthi, Lidwina Priliani, Edison Johar, Frilasita A Yudhaputri, Muhammad Rezki Rasyak, Willy Agustine, Hidayat Trimarsanto, Lydia V. Panggalo, Sri Masyeni, Erly Sintya, Safarina G Malik, Khin Saw Myint, Amin Soebandrio |
| EPI_ISL_2101216 | BRSU Tabanan | Eijkman Institute for Molecular Biology, National Agency for Research and Innovation; Molecular Biology Laboratory, Faculty Medicine and Health Sciences, Warmadewa University | Frilasita A Yudhaputri, Muhammad Rezki Rasyak, Willy Agustine, Hidayat Trimarsanto, Lydia V. Panggalo, Iskandar Adnan, Sukma Oktavianthi, Lidwina Priliani, Edison Johar, Sri Masyeni, Erly Sintya, I Wayan Duta Krisna, Safarina G Malik, Khin Saw Myint, Amin Soebandrio |
| EPI_ISL_2101217 | BRSU Tabanan | Eijkman Institute for Molecular Biology, National Agency for Research and Innovation; Molecular Biology Laboratory, Faculty Medicine and Health Sciences, Warmadewa University | Muhammad Rezki Rasyak, Willy Agustine, Hidayat Trimarsanto, Lydia V. Panggalo, Iskandar Adnan, Sukma Oktavianthi, Lidwina Priliani, Edison Johar, Frilasita A Yudhaputri, Sri Masyeni, Erly Sintya, I Wayan Duta Krisna, Safarina G Malik, Khin Saw Myint, Amin Soebandrio |
| EPI_ISL_2101218 | BRSU Tabanan | Eijkman Institute for Molecular Biology, National Agency for Research and Innovation; Molecular Biology Laboratory, Faculty Medicine and Health Sciences, Warmadewa University | Willy Agustine, Hidayat Trimarsanto, Lydia V. Panggalo, Iskandar Adnan, Sukma Oktavianthi, Lidwina Priliani, Edison Johar, Frilasita A Yudhaputri, Muhammad Rezki Rasyak, Sri Masyeni, Erly Sintya, I Wayan Duta Krisna, Safarina G Malik, Khin Saw Myint, Amin Soebandrio |
| EPI_ISL_2101219 | BRSU Tabanan | Eijkman Institute for Molecular Biology, National Agency for Research and Innovation; Molecular Biology Laboratory, Faculty Medicine and Health Sciences, Warmadewa University | Hidayat Trimarsanto, Lydia V. Panggalo, Iskandar Adnan, Sukma Oktavianthi, Lidwina Priliani, Edison Johar, Frilasita A Yudhaputri, Muhammad Rezki Rasyak, Willy Agustine, Sri Masyeni, Erly Sintya, I Wayan Duta Krisna, Safarina G Malik, Khin Saw Myint, Amin Soebandrio |
| EPI_ISL_2101220 | BRSU Tabanan | Eijkman Institute for Molecular Biology, National Agency for Research and Innovation; Molecular Biology Laboratory, Faculty Medicine and Health Sciences, Warmadewa University | Lydia V. Panggalo, Iskandar Adnan, Sukma Oktavianthi, Lidwina Priliani, Edison Johar, Frilasita A Yudhaputri, Muhammad Rezki Rasyak, Willy Agustine, Hidayat Trimarsanto, Sri Masyeni, Erly Sintya, I Wayan Duta Krisna, Safarina G Malik, Khin Saw Myint, Amin Soebandrio |
| EPI_ISL_2101221 | BRSU Tabanan | Eijkman Institute for Molecular Biology, National Agency for Research and Innovation; Molecular Biology Laboratory, Faculty Medicine and Health Sciences, Warmadewa University | Iskandar Adnan, Sukma Oktavianthi, Lidwina Priliani, Edison Johar, Frilasita A Yudhaputri, Muhammad Rezki Rasyak, Willy Agustine, Hidayat Trimarsanto, Lydia V. Panggalo, Sri Masyeni, Erly Sintya, I Wayan Duta Krisna, Safarina G Malik, Khin Saw Myint, Amin Soebandrio |
| EPI_ISL_2101222 | BRSU Tabanan | Eijkman Institute for Molecular Biology, National Agency for Research and Innovation; Molecular Biology Laboratory, Faculty Medicine and Health Sciences, Warmadewa University | Sukma Oktavianthi, Lidwina Priliani, Edison Johar, Frilasita A Yudhaputri, Muhammad Rezki Rasyak, Willy Agustine, Hidayat Trimarsanto, Lydia V. Panggalo, Iskandar Adnan, Sri Masyeni, Erly Sintya, I Wayan Duta Krisna, Safarina G Malik, Khin Saw Myint, Amin Soebandrio |
| EPI_ISL_2101223 | BRSU Tabanan | Eijkman Institute for Molecular Biology, National Agency for Research and Innovation; Molecular Biology Laboratory, Faculty Medicine and Health Sciences, Warmadewa University | Lidwina Priliani, Edison Johar, Frilasita A Yudhaputri, Muhammad Rezki Rasyak, Willy Agustine, Hidayat Trimarsanto, Lydia V. Panggalo, Iskandar Adnan, Sukma Oktavianthi, Sri Masyeni, Erly Sintya, I Wayan Duta Krisna, Safarina G Malik, Khin Saw Myint, Amin Soebandrio |
| EPI_ISL_2101224 | BRSU Tabanan | Eijkman Institute for Molecular Biology, National Agency for Research and Innovation; Molecular Biology Laboratory, Faculty Medicine and Health Sciences, Warmadewa University | Frilasita A Yudhaputri, Muhammad Rezki Rasyak, Willy Agustine, Hidayat Trimarsanto, Lydia V. Panggalo, Iskandar Adnan, Sukma Oktavianthi, Lidwina Priliani, Edison Johar, Sri Masyeni, Erly Sintya, I Wayan Duta Krisna, Safarina G Malik, Khin Saw Myint, Amin Soebandrio |
| EPI_ISL_2101225 | BRSU Tabanan | Eijkman Institute for Molecular Biology, National Agency for Research and Innovation; Molecular Biology Laboratory, Faculty Medicine and Health Sciences, Warmadewa University | Muhammad Rezki Rasyak, Willy Agustine, Hidayat Trimarsanto, Lydia V. Panggalo, Iskandar Adnan, Sukma Oktavianthi, Lidwina Priliani, Edison Johar, Frilasita A Yudhaputri, Sri Masyeni, Erly Sintya, I Wayan Duta Krisna, Safarina G Malik, Khin Saw Myint, Amin Soebandrio |
| EPI_ISL_2101226 | BRSU Tabanan | Eijkman Institute for Molecular Biology, National Agency for Research and Innovation; Molecular Biology Laboratory, Faculty Medicine and Health Sciences, Warmadewa University | Willy Agustine, Hidayat Trimarsanto, Lydia V. Panggalo, Iskandar Adnan, Sukma Oktavianthi, Lidwina Priliani, Edison Johar, Frilasita A Yudhaputri, Muhammad Rezki Rasyak, Sri Masyeni, Erly Sintya, I Wayan Duta Krisna, Safarina G Malik, Khin Saw Myint, Amin Soebandrio |
| EPI_ISL_2101228 | BRSU Tabanan | Eijkman Institute for Molecular Biology, National Agency for Research and Innovation; Molecular Biology Laboratory, Faculty Medicine and Health Sciences, Warmadewa University | Lydia V. Panggalo, Iskandar Adnan, Sukma Oktavianthi, Lidwina Priliani, Edison Johar, Frilasita A Yudhaputri, Muhammad Rezki Rasyak, Willy Agustine, Hidayat Trimarsanto, Sri Masyeni, Erly Sintya, I Wayan Duta Krisna, Safarina G Malik, Khin Saw Myint, Amin Soebandrio |
| EPI_ISL_2101246 | RSUD Klungkung | Eijkman Institute for Molecular Biology, National Agency for Research and Innovation; Molecular Biology Laboratory, Faculty Medicine and Health Sciences, Warmadewa University | Sukma Oktavianthi, Lidwina Priliani, Edison Johar, Frilasita A Yudhaputri, Muhammad Rezki Rasyak, Willy Agustine, Hidayat Trimarsanto, Lydia V. Panggalo, Iskandar Adnan, Sri Masyeni, Erly Sintya, I Komang Parwata, Safarina G Malik, Khin Saw Myint, Amin Soebandrio |

| | | | |
|---|---|---|---|
| EPI_ISL_2101247 | RSUD Klungkung | Eijkman Institute for Molecular Biology, National Agency for Research and Innovation; Molecular Biology Laboratory, Faculty Medicine and Health Sciences, Warmadewa University | Lidwina Priliani, Edison Johar, Frilasita A Yudhaputri, Muhammad Rezki Rasyak, Willy Agustine, Hidayat Trimarsanto, Lydia V. Panggalo, Iskandar Adnan, Sukma Oktavianthi, Sri Masyeni, Erly Sintya, I Komang Parwata, Safarina G Malik, Khin Saw Myint, Amin Soebandrio |
| EPI_ISL_2101248 | RSUD Klungkung | Eijkman Institute for Molecular Biology, National Agency for Research and Innovation; Molecular Biology Laboratory, Faculty Medicine and Health Sciences, Warmadewa University | Frilasita A Yudhaputri, Muhammad Rezki Rasyak, Willy Agustine, Hidayat Trimarsanto, Lydia V. Panggalo, Iskandar Adnan, Sukma Oktavianthi, Lidwina Priliani, Edison Johar, Sri Masyeni, Erly Sintya, I Komang Parwata, Safarina G Malik, Khin Saw Myint, Amin Soebandrio |
| EPI_ISL_2101249 | RSUD Klungkung | Eijkman Institute for Molecular Biology, National Agency for Research and Innovation; Molecular Biology Laboratory, Faculty Medicine and Health Sciences, Warmadewa University | Muhammad Rezki Rasyak, Willy Agustine, Hidayat Trimarsanto, Lydia V. Panggalo, Iskandar Adnan, Sukma Oktavianthi, Lidwina Priliani, Edison Johar, Frilasita A Yudhaputri, Sri Masyeni, Erly Sintya, I Komang Parwata, Safarina G Malik, Khin Saw Myint, Amin Soebandrio |
| EPI_ISL_2101250 | RSUD Klungkung | Eijkman Institute for Molecular Biology, National Agency for Research and Innovation; Molecular Biology Laboratory, Faculty Medicine and Health Sciences, Warmadewa University | Willy Agustine, Hidayat Trimarsanto, Lydia V. Panggalo, Iskandar Adnan, Sukma Oktavianthi, Lidwina Priliani, Edison Johar, Frilasita A Yudhaputri, Muhammad Rezki Rasyak, Sri Masyeni, Erly Sintya, I Komang Parwata, Safarina G Malik, Khin Saw Myint, Amin Soebandrio |
| EPI_ISL_2101251 | RSUD Klungkung | Eijkman Institute for Molecular Biology, National Agency for Research and Innovation; Molecular Biology Laboratory, Faculty Medicine and Health Sciences, Warmadewa University | Hidayat Trimarsanto, Lydia V. Panggalo, Iskandar Adnan, Sukma Oktavianthi, Lidwina Priliani, Edison Johar, Frilasita A Yudhaputri, Muhammad Rezki Rasyak, Willy Agustine, Sri Masyeni, Erly Sintya, I Komang Parwata, Safarina G Malik, Khin Saw Myint, Amin Soebandrio |
| EPI_ISL_2101252 | RSUD Klungkung | Eijkman Institute for Molecular Biology, National Agency for Research and Innovation; Molecular Biology Laboratory, Faculty Medicine and Health Sciences, Warmadewa University | Lydia V. Panggalo, Iskandar Adnan, Sukma Oktavianthi, Lidwina Priliani, Edison Johar, Frilasita A Yudhaputri, Muhammad Rezki Rasyak, Willy Agustine, Hidayat Trimarsanto, Sri Masyeni, Erly Sintya, I Komang Parwata, Safarina G Malik, Khin Saw Myint, Amin Soebandrio |
| EPI_ISL_2101253 | RSUD Klungkung | Eijkman Institute for Molecular Biology, National Agency for Research and Innovation; Molecular Biology Laboratory, Faculty Medicine and Health Sciences, Warmadewa University | Edison Johar, Frilasita A Yudhaputri, Muhammad Rezki Rasyak, Willy Agustine, Hidayat Trimarsanto, Lydia V. Panggalo, Iskandar Adnan, Sukma Oktavianthi, Lidwina Priliani, Sri Masyeni, Erly Sintya, I Komang Parwata, Safarina G Malik, Khin Saw Myint, Amin Soebandrio |
| EPI_ISL_2101254 | RSUD Klungkung | Eijkman Institute for Molecular Biology, National Agency for Research and Innovation; Molecular Biology Laboratory, Faculty Medicine and Health Sciences, Warmadewa University | Hidayat Trimarsanto, Lydia V. Panggalo, Iskandar Adnan, Sukma Oktavianthi, Lidwina Priliani, Edison Johar, Frilasita A Yudhaputri, Muhammad Rezki Rasyak, Willy Agustine, Sri Masyeni, Erly Sintya, I Komang Parwata, Safarina G Malik, Khin Saw Myint, Amin Soebandrio |
| EPI_ISL_2101255 | RSUD Klungkung | Eijkman Institute for Molecular Biology, National Agency for Research and Innovation; Molecular Biology Laboratory, Faculty Medicine and Health Sciences, Warmadewa University | Lydia V. Panggalo, Iskandar Adnan, Sukma Oktavianthi, Lidwina Priliani, Edison Johar, Frilasita A Yudhaputri, Muhammad Rezki Rasyak, Willy Agustine, Hidayat Trimarsanto, Sri Masyeni, Erly Sintya, I Komang Parwata, Safarina G Malik, Khin Saw Myint, Amin Soebandrio |
| EPI_ISL_2101324 | RSUD Mangusada | Eijkman Institute for Molecular Biology, National Agency for Research and Innovation; Molecular Biology Laboratory, Faculty Medicine and Health Sciences, Warmadewa University | Hidayat Trimarsanto, Lydia V. Panggalo, Iskandar Adnan, Sukma Oktavianthi, Lidwina Priliani, Edison Johar, Frilasita A Yudhaputri, Muhammad Rezki Rasyak, Willy Agustine, Sri Masyeni, Erly Sintya, Ida Ayu Wayan Mahayani, Safarina G Malik, Khin Saw Myint, Amin Soebandrio |
| EPI_ISL_2101325 | RSUD Mangusada | Eijkman Institute for Molecular Biology, National Agency for Research and Innovation; Molecular Biology Laboratory, Faculty Medicine and Health Sciences, Warmadewa University | Lydia V. Panggalo, Iskandar Adnan, Sukma Oktavianthi, Lidwina Priliani, Edison Johar, Frilasita A Yudhaputri, Muhammad Rezki Rasyak, Willy Agustine, Hidayat Trimarsanto, Sri Masyeni, Erly Sintya, Ida Ayu Wayan Mahayani, Safarina G Malik, Khin Saw Myint, Amin Soebandrio |
| EPI_ISL_2101326 | RSUD Mangusada | Eijkman Institute for Molecular Biology, National Agency for Research and Innovation; Molecular Biology Laboratory, Faculty Medicine and Health Sciences, Warmadewa University | Iskandar Adnan, Sukma Oktavianthi, Lidwina Priliani, Edison Johar, Frilasita A Yudhaputri, Muhammad Rezki Rasyak, Willy Agustine, Hidayat Trimarsanto, Lydia V. Panggalo, Sri Masyeni, Erly Sintya, Ida Ayu Wayan Mahayani, Safarina G Malik, Khin Saw Myint, Amin Soebandrio |
| EPI_ISL_2101327 | RSUD Mangusada | Eijkman Institute for Molecular Biology, National Agency for Research and Innovation; Molecular Biology Laboratory, Faculty Medicine and Health Sciences, Warmadewa University | Sukma Oktavianthi, Lidwina Priliani, Edison Johar, Frilasita A Yudhaputri, Muhammad Rezki Rasyak, Willy Agustine, Hidayat Trimarsanto, Lydia V. Panggalo, Iskandar Adnan, Sri Masyeni, Erly Sintya, Ida Ayu Wayan Mahayani, Safarina G Malik, Khin Saw Myint, Amin Soebandrio |
| EPI_ISL_2101328 | RSUD Mangusada | Eijkman Institute for Molecular Biology, National Agency for Research and Innovation; Molecular Biology Laboratory, Faculty Medicine and Health Sciences, Warmadewa University | Lidwina Priliani, Edison Johar, Frilasita A Yudhaputri, Muhammad Rezki Rasyak, Willy Agustine, Hidayat Trimarsanto, Lydia V. Panggalo, Iskandar Adnan, Sukma Oktavianthi, Sri Masyeni, Erly Sintya, Ida Ayu Wayan Mahayani, Safarina G Malik, Khin Saw Myint, Amin Soebandrio |
| EPI_ISL_2101329 | RSUD Mangusada | Eijkman Institute for Molecular Biology, National Agency for Research and Innovation; Molecular Biology Laboratory, Faculty Medicine and Health Sciences, Warmadewa University | Edison Johar, Frilasita A Yudhaputri, Muhammad Rezki Rasyak, Willy Agustine, Hidayat Trimarsanto, Lydia V. Panggalo, Iskandar Adnan, Sukma Oktavianthi, Lidwina Priliani, Sri Masyeni, Erly Sintya, Ida Ayu Wayan Mahayani, Safarina G Malik, Khin Saw Myint, Amin Soebandrio |
| EPI_ISL_2101330 | RSUD Mangusada | Eijkman Institute for Molecular Biology, National Agency for Research and Innovation; Molecular Biology Laboratory, Faculty Medicine and Health Sciences, Warmadewa University | Frilasita A Yudhaputri, Muhammad Rezki Rasyak, Willy Agustine, Hidayat Trimarsanto, Lydia V. Panggalo, Iskandar Adnan, Sukma Oktavianthi, Lidwina Priliani, Edison Johar, Sri Masyeni, Erly Sintya, Ida Ayu Wayan Mahayani, Safarina G Malik, Khin Saw Myint, Amin Soebandrio |
| EPI_ISL_2101331 | RSUD Mangusada | Eijkman Institute for Molecular Biology, National Agency for Research and Innovation; Molecular Biology Laboratory, Faculty Medicine and Health Sciences, Warmadewa University | Muhammad Rezki Rasyak, Willy Agustine, Hidayat Trimarsanto, Lydia V. Panggalo, Iskandar Adnan, Sukma Oktavianthi, Lidwina Priliani, Edison Johar, Frilasita A Yudhaputri, Sri Masyeni, Erly Sintya, Ida Ayu Wayan Mahayani, Safarina G Malik, Khin Saw Myint, Amin Soebandrio |
| EPI_ISL_2101332 | RSUD Mangusada | Eijkman Institute for Molecular Biology, National Agency for Research and Innovation; Molecular Biology Laboratory, Faculty Medicine and Health Sciences, Warmadewa University | Hidayat Trimarsanto, Lydia V. Panggalo, Iskandar Adnan, Sukma Oktavianthi, Lidwina Priliani, Edison Johar, Frilasita A Yudhaputri, Muhammad Rezki Rasyak, Willy Agustine, Sri Masyeni, Erly Sintya, Ida Ayu Wayan Mahayani, Safarina G Malik, Khin Saw Myint, Amin Soebandrio |
| EPI_ISL_2101333 | RSUD Mangusada | Eijkman Institute for Molecular Biology, National Agency for Research and Innovation; Molecular Biology Laboratory, Faculty Medicine and Health Sciences, Warmadewa University | Lydia V. Panggalo, Iskandar Adnan, Sukma Oktavianthi, Lidwina Priliani, Edison Johar, Frilasita A Yudhaputri, Muhammad Rezki Rasyak, Willy Agustine, Hidayat Trimarsanto, Sri Masyeni, Erly Sintya, Ida Ayu Wayan Mahayani, Safarina G Malik, Khin Saw Myint, Amin Soebandrio |
| EPI_ISL_2101334 | RSUD Mangusada | Eijkman Institute for Molecular Biology, National Agency for Research and Innovation; Molecular Biology Laboratory, | Iskandar Adnan, Sukma Oktavianthi, Lidwina Priliani, Edison Johar, Frilasita A Yudhaputri, Muhammad Rezki Rasyak, Willy Agustine, Hidayat Trimarsanto, Lydia V. Panggalo, Sri Masyeni, Erly Sintya, Ida Ayu Wayan Mahayani, Safarina G Malik, Khin Saw Myint, Amin Soebandrio |

| | | | |
|---|---|---|---|
| | | Faculty Medicine and Health Sciences, Warmadewa University | |
| EPI_ISL_2101335 | RSUD Mangusada | Eijkman Institute for Molecular Biology, National Agency for Research and Innovation; Molecular Biology Laboratory, Faculty Medicine and Health Sciences, Warmadewa University | Sukma Oktavianthi, Lidwina Priliani, Edison Johar, Frilasita A Yudhaputri, Muhammad Rezki Rasyak, Willy Agustine, Hidayat Trimarsanto, Lydia V. Panggalo, Iskandar Adnan, Sri Masyeni, Erly Sintya, Ida Ayu Wayan Mahayani, Safarina G Malik, Khin Saw Myint, Amin Soebandrio |
| EPI_ISL_2101336 | RSUD Mangusada | Eijkman Institute for Molecular Biology, National Agency for Research and Innovation; Molecular Biology Laboratory, Faculty Medicine and Health Sciences, Warmadewa University | Edison Johar, Frilasita A Yudhaputri, Muhammad Rezki Rasyak, Willy Agustine, Hidayat Trimarsanto, Lydia V. Panggalo, Iskandar Adnan, Sukma Oktavianthi, Lidwina Priliani, Sri Masyeni, Erly Sintya, Ida Ayu Wayan Mahayani, Safarina G Malik, Khin Saw Myint, Amin Soebandrio |
| EPI_ISL_2101337 | RSUD Mangusada | Eijkman Institute for Molecular Biology, National Agency for Research and Innovation; Molecular Biology Laboratory, Faculty Medicine and Health Sciences, Warmadewa University | Frilasita A Yudhaputri, Muhammad Rezki Rasyak, Willy Agustine, Hidayat Trimarsanto, Lydia V. Panggalo, Iskandar Adnan, Sukma Oktavianthi, Lidwina Priliani, Edison Johar, Sri Masyeni, Erly Sintya, Ida Ayu Wayan Mahayani, Safarina G Malik, Khin Saw Myint, Amin Soebandrio |
| EPI_ISL_2101338 | RSUD Mangusada | Eijkman Institute for Molecular Biology, National Agency for Research and Innovation; Molecular Biology Laboratory, Faculty Medicine and Health Sciences, Warmadewa University | Muhammad Rezki Rasyak, Willy Agustine, Hidayat Trimarsanto, Lydia V. Panggalo, Iskandar Adnan, Sukma Oktavianthi, Lidwina Priliani, Edison Johar, Frilasita A Yudhaputri, Sri Masyeni, Erly Sintya, Ida Ayu Wayan Mahayani, Safarina G Malik, Khin Saw Myint, Amin Soebandrio |
| EPI_ISL_2101339 | RSUD Mangusada | Eijkman Institute for Molecular Biology, National Agency for Research and Innovation; Molecular Biology Laboratory, Faculty Medicine and Health Sciences, Warmadewa University | Willy Agustine, Hidayat Trimarsanto, Lydia V. Panggalo, Iskandar Adnan, Sukma Oktavianthi, Lidwina Priliani, Edison Johar, Frilasita A Yudhaputri, Muhammad Rezki Rasyak, Sri Masyeni, Erly Sintya, Ida Ayu Wayan Mahayani, Safarina G Malik, Khin Saw Myint, Amin Soebandrio |
| EPI_ISL_2101340 | RSUD Mangusada | Eijkman Institute for Molecular Biology, National Agency for Research and Innovation; Molecular Biology Laboratory, Faculty Medicine and Health Sciences, Warmadewa University | Hidayat Trimarsanto, Lydia V. Panggalo, Iskandar Adnan, Sukma Oktavianthi, Lidwina Priliani, Edison Johar, Frilasita A Yudhaputri, Muhammad Rezki Rasyak, Willy Agustine, Sri Masyeni, Erly Sintya, Ida Ayu Wayan Mahayani, Safarina G Malik, Khin Saw Myint, Amin Soebandrio |
| EPI_ISL_2101341 | RSUD Mangusada | Eijkman Institute for Molecular Biology, National Agency for Research and Innovation; Molecular Biology Laboratory, Faculty Medicine and Health Sciences, Warmadewa University | Lydia V. Panggalo, Iskandar Adnan, Sukma Oktavianthi, Lidwina Priliani, Edison Johar, Frilasita A Yudhaputri, Muhammad Rezki Rasyak, Willy Agustine, Hidayat Trimarsanto, Sri Masyeni, Erly Sintya, Ida Ayu Wayan Mahayani, Safarina G Malik, Khin Saw Myint, Amin Soebandrio |
| EPI_ISL_2101342 | RSUD Mangusada | Eijkman Institute for Molecular Biology, National Agency for Research and Innovation; Molecular Biology Laboratory, Faculty Medicine and Health Sciences, Warmadewa University | Iskandar Adnan, Sukma Oktavianthi, Lidwina Priliani, Edison Johar, Frilasita A Yudhaputri, Muhammad Rezki Rasyak, Willy Agustine, Hidayat Trimarsanto, Lydia V. Panggalo, Sri Masyeni, Erly Sintya, Ida Ayu Wayan Mahayani, Safarina G Malik, Khin Saw Myint, Amin Soebandrio |
| EPI_ISL_2103416 | AREA DE SALUD TURRIALBA-JIMENEZ | Inciensa, Instituto Costarricense de Investigación y Enseñanza en Nutrición y Salud | Francisco Duarte, Hebleen Porras, Claudio Soto-Garita, Estela Cordero, Adriana Godínez, Melany Calderón, José Luis Vargas, Mariela Gutiérrez & Joselyn Prado |
| EPI_ISL_2103417 | AREA DE SALUD OROTINA-SAN MATEO | Inciensa, Instituto Costarricense de Investigación y Enseñanza en Nutrición y Salud | Francisco Duarte, Hebleen Porras, Claudio Soto-Garita, Estela Cordero, Adriana Godínez, Melany Calderón, José Luis Vargas, Mariela Gutiérrez, Joselyn Prado & Andrea Moreno-Carvajal |
| EPI_ISL_2103418 | HOSPITAL MONSEÑOR SANABRIA | Inciensa, Instituto Costarricense de Investigación y Enseñanza en Nutrición y Salud | Francisco Duarte, Hebleen Porras, Claudio Soto-Garita, Estela Cordero, Adriana Godínez, Melany Calderón, José Luis Vargas, Mariela Gutiérrez, Joselyn Prado & Jason Vieto-Carranza |
| EPI_ISL_2105487 | Philippine Red Cross Logistics and Multipurpose Center | Philippine Genome Center | Francis A. Tablizo, Kenneth M. Kim, Carlo M. Lapid, Marc Jerrone R. Castro, Maria Sofia L. Yangzon, Benedict A. Maralit, Marc Edsel C. Ayes, Eva Maria Cutiongco-de la Paz, Alethea R. de Guzman, Jan Michael C. Yap, Jo-Hannah S. Llames, Sheila Mae M. Araiza, Kris P. Punayan, Irish Coleen A. Asin, Candice Francheska B. Tambaoan, Asia Louisa U. Chong, Karol Sophia Agape R. Padilla, Rianna Patricia S. Cruz, El King D. Morado, Joshua Gregor A. Dizon, Razel Nikka M. Hao, Arianne A. Zamora, Devon Ray Pacial, Juan Antonio R. Magalang, Marissa Alejandria, Celia Carlos, Anna Ong-Lim, Edsel Maurice Salvaña, John Q. Wong, Jaime C. Montoya, Maria Rosario Singh-Vergeire and Cynthia P. Saloma |
| EPI_ISL_2107436 | Dr. Yoshitaka Tamura Department of Clinical Laboratory, Osaka Habikino Medical Center | Department of Developmental Medicine, Research Institute, Osaka Women's and Children's Hospital | Itaru Yanagihara, Yukiko Nakura |
| EPI_ISL_2139494, EPI_ISL_2139495, EPI_ISL_2139496, EPI_ISL_2139497, EPI_ISL_2139498, EPI_ISL_2139499, EPI_ISL_2139500, EPI_ISL_2139501, EPI_ISL_2139502, EPI_ISL_2139504, EPI_ISL_2139505, EPI_ISL_2139506, EPI_ISL_2139507, EPI_ISL_2139508, EPI_ISL_2139509, EPI_ISL_2139510, EPI_ISL_2139511, EPI_ISL_2139512, EPI_ISL_2139513, EPI_ISL_2139515, EPI_ISL_2139516, EPI_ISL_2139518, EPI_ISL_2139519, EPI_ISL_2139520, EPI_ISL_2139521, EPI_ISL_2139522, EPI_ISL_2139523, EPI_ISL_2139525, EPI_ISL_2139526, EPI_ISL_2139527, EPI_ISL_2139528, EPI_ISL_2139529, EPI_ISL_2139530, EPI_ISL_2139531, EPI_ISL_2139532, EPI_ISL_2139534, EPI_ISL_2139535, EPI_ISL_2139536, EPI_ISL_2139538, EPI_ISL_2139539, EPI_ISL_2139540, EPI_ISL_2139541, EPI_ISL_2139542, EPI_ISL_2139543, EPI_ISL_2139545, EPI_ISL_2139546, EPI_ISL_2139547, EPI_ISL_2139548, EPI_ISL_2139549 | | | |
| see above | Laboratorio Exame | Universidade Federal de Ciencias da Saude de Porto Alegre | Vinícius Bonetti Franceschi, Gabriel Dickin Caldana et al. |
| EPI_ISL_2142744, EPI_ISL_2142746 | MRCG at LSHTM Genomics lab | MRCG at LSHTM Genomics lab | Abdul Karim sesay, Abdoulie Kanteh, Jarra Manneh, Mariama Kujabi, Bakary Sanyang, Damiri Damilari, Sainabou laye Ndure |
| EPI_ISL_2155845, EPI_ISL_2155846, EPI_ISL_2155849 | Hi-Precision Diagnostic Center | Philippine Genome Center | Francis A. Tablizo, Kenneth M. Kim, Carlo M. Lapid, Marc Jerrone R. Castro, Maria Sofia L. Yangzon, Benedict A. Maralit, Marc Edsel C. Ayes, Eva Maria Cutiongco-de la Paz, Alethea R. de Guzman, Jan Michael C. Yap, Jo-Hannah S. Llames, Sheila Mae M. Araiza, Kris P. Punayan, Irish Coleen A. Asin, Candice Francheska B. Tambaoan, Asia Louisa U. Chong, Karol Sophia Agape R. Padilla, Rianna Patricia S. Cruz, El King D. Morado, Joshua Gregor A. Dizon, Razel Nikka M. Hao, Arianne A. Zamora, Devon Ray Pacial, Juan Antonio R. Magalang, Marissa Alejandria, Celia Carlos, Anna Ong-Lim, Edsel Maurice Salvaña, John Q. Wong, Jaime C. Montoya, Maria Rosario Singh-Vergeire and Cynthia P. Saloma |
| EPI_ISL_2155850, EPI_ISL_2155855, EPI_ISL_2155856, EPI_ISL_2155860, EPI_ISL_2155865, EPI_ISL_2155866, EPI_ISL_2155869, EPI_ISL_2155870, EPI_ISL_2155871, EPI_ISL_2155872, EPI_ISL_2155873, EPI_ISL_2155874, EPI_ISL_2155876, EPI_ISL_2155881, EPI_ISL_2155882 | | | |
| see above | THE LORD'S GRACE MEDICAL AND INDUSTRIAL CLINIC | Philippine Genome Center | Francis A. Tablizo, Kenneth M. Kim, Carlo M. Lapid, Marc Jerrone R. Castro, Maria Sofia L. Yangzon, Benedict A. Maralit, Marc Edsel C. Ayes, Eva Maria Cutiongco-de la Paz, Alethea R. de Guzman, Jan Michael C. Yap, Jo-Hannah S. Llames, Sheila Mae M. Araiza, Kris P. Punayan, Irish Coleen A. Asin, Candice Francheska B. Tambaoan, Asia Louisa U. Chong, Karol Sophia Agape R. Padilla, Rianna Patricia S. Cruz, El King D. Morado, Joshua Gregor A. Dizon, Razel Nikka M. Hao, Arianne A. Zamora, Devon Ray Pacial, Juan Antonio R. Magalang, Marissa Alejandria, Celia Carlos, Anna Ong-Lim, Edsel Maurice Salvaña, John Q. Wong, Jaime C. Montoya, Maria Rosario Singh-Vergeire and Cynthia P. Saloma |
| EPI_ISL_2155889, EPI_ISL_2155890, EPI_ISL_2155891, EPI_ISL_2155904, EPI_ISL_2155905, EPI_ISL_2155911, EPI_ISL_2155916 | Philippine Red Cross - National Blood Center | Philippine Genome Center | Francis A. Tablizo, Kenneth M. Kim, Carlo M. Lapid, Marc Jerrone R. Castro, Maria Sofia L. Yangzon, Benedict A. Maralit, Marc Edsel C. Ayes, Eva Maria Cutiongco-de la Paz, Alethea R. de Guzman, Jan Michael C. Yap, Jo-Hannah S. Llames, Sheila Mae M. Araiza, Kris P. Punayan, Irish Coleen A. Asin, Candice Francheska B. Tambaoan, Asia Louisa U. Chong, Karol Sophia Agape R. Padilla, Rianna Patricia S. Cruz, El King D. Morado, Joshua Gregor A. Dizon, Razel Nikka M. Hao, Arianne A. Zamora, Devon Ray Pacial, Juan Antonio R. Magalang, Marissa Alejandria, Celia Carlos, Anna Ong-Lim, Edsel Maurice Salvaña, John Q. Wong, Jaime C. Montoya, Maria Rosario Singh-Vergeire and Cynthia P. Saloma |
| EPI_ISL_2155943, EPI_ISL_2155946, EPI_ISL_2155948, EPI_ISL_2155955, EPI_ISL_2155958, EPI_ISL_2155963 | PHILIPPINE AIRPORT DIAGNOSTIC LABORATORY | Philippine Genome Center | Francis A. Tablizo, Kenneth M. Kim, Carlo M. Lapid, Marc Jerrone R. Castro, Maria Sofia L. Yangzon, Benedict A. Maralit, Marc Edsel C. Ayes, Eva Maria Cutiongco-de la Paz, Alethea R. de Guzman, Jan Michael C. Yap, Jo-Hannah S. Llames, Sheila Mae M. Araiza, Kris P. Punayan, Irish Coleen A. Asin, Candice Francheska B. Tambaoan, Asia Louisa U. Chong, Karol Sophia Agape R. Padilla, Rianna Patricia S. Cruz, El King D. Morado, Joshua Gregor A. Dizon, Razel Nikka M. Hao, Arianne A. Zamora, Devon Ray Pacial, Juan Antonio R. Magalang, Marissa Alejandria, Celia Carlos, Anna Ong-Lim, Edsel Maurice Salvaña, John Q. Wong, Jaime C. Montoya, Maria Rosario Singh-Vergeire and Cynthia P. Saloma |
| EPI_ISL_2155985, EPI_ISL_2155988, EPI_ISL_2155994 | The Medical City -Â Ortigas | Philippine Genome Center | Francis A. Tablizo, Kenneth M. Kim, Carlo M. Lapid, Marc Jerrone R. Castro, Maria Sofia L. Yangzon, Benedict A. Maralit, Marc Edsel C. Ayes, Eva Maria Cutiongco-de la Paz, Alethea R. de Guzman, Jan Michael C. Yap, Jo-Hannah S. Llames, Sheila Mae M. Araiza, Kris P. Punayan, Irish Coleen A. Asin, Candice Francheska B. Tambaoan, Asia Louisa U. Chong, Karol Sophia Agape R. Padilla, Rianna Patricia S. Cruz, El King D. Morado, Joshua Gregor A. |

| | | | |
|---|---|---|---|
| | | | Dizon, Razel Nikka M. Hao, Arianne A. Zamora, Devon Ray Pacial, Juan Antonio R. Magalang, Marissa Alejandria, Celia Carlos, Anna Ong-Lim, Edsel Maurice Salvaña, John Q. Wong, Jaime C. Montoya, Maria Rosario Singh-Vergeire and Cynthia P. Saloma |
| EPI_ISL_2156010 | Qualimed Hospital Sta. Rosa | Philippine Genome Center | Francis A. Tablizo, Kenneth M. Kim, Carlo M. Lapid, Marc Jerrone R. Castro, Maria Sofia L. Yangzon, Benedict A. Maralit, Marc Edsel C. Ayes, Eva Maria Cutiongco-de la Paz, Alethea R. de Guzman, Jan Michael C. Yap, Jo-Hannah S. Llames, Sheila Mae M. Araiza, Kris P. Punayan, Irish Coleen A. Asin, Candice Francheska B. Tambaoan, Asia Louisa U. Chong, Karol Sophia Agape R. Padilla, Rianna Patricia S. Cruz, El King D. Morado, Joshua Gregor A. Dizon, Razel Nikka M. Hao, Arianne A. Zamora, Devon Ray Pacial, Juan Antonio R. Magalang, Marissa Alejandria, Celia Carlos, Anna Ong-Lim, Edsel Maurice Salvaña, John Q. Wong, Jaime C. Montoya, Maria Rosario Singh-Vergeire and Cynthia P. Saloma |
| EPI_ISL_2156258, EPI_ISL_2156259, EPI_ISL_2156262, EPI_ISL_2156263, EPI_ISL_2156264 | THE LORD'S GRACE MEDICAL AND INDUSTRIAL CLINIC | Philippine Genome Center | Francis A. Tablizo, Kenneth M. Kim, Carlo M. Lapid, Marc Jerrone R. Castro, Maria Sofia L. Yangzon, Elcid Aaron R. Pangilinan, Benedict A. Maralit, Marc Edsel C. Ayes, Eva Maria Cutiongco-de la Paz, Alethea R. de Guzman, Jan Michael C. Yap, Jo-Hannah S. Llames, Sheila Mae M. Araiza, Kris P. Punayan, Irish Coleen A. Asin, Candice Francheska B. Tambaoan, Asia Louisa U. Chong, Karol Sophia Agape R. Padilla, Rianna Patricia S. Cruz, El King D. Morado, Joshua Gregor A. Dizon, Razel Nikka M. Hao, Arianne A. Zamora, Devon Ray Pacial, Juan Antonio R. Magalang, Marissa Alejandria, Celia Carlos, Anna Ong-Lim, Edsel Maurice Salvaña, John Q. Wong, Jaime C. Montoya, Maria Rosario Singh-Vergeire and Cynthia P. Saloma |
| EPI_ISL_2156265, EPI_ISL_2156269, EPI_ISL_2156274 | Tarlac Provincial Hospital - Molecular Pathology Laboratory | Philippine Genome Center | Francis A. Tablizo, Kenneth M. Kim, Carlo M. Lapid, Marc Jerrone R. Castro, Maria Sofia L. Yangzon, Elcid Aaron R. Pangilinan, Benedict A. Maralit, Marc Edsel C. Ayes, Eva Maria Cutiongco-de la Paz, Alethea R. de Guzman, Jan Michael C. Yap, Jo-Hannah S. Llames, Sheila Mae M. Araiza, Kris P. Punayan, Irish Coleen A. Asin, Candice Francheska B. Tambaoan, Asia Louisa U. Chong, Karol Sophia Agape R. Padilla, Rianna Patricia S. Cruz, El King D. Morado, Joshua Gregor A. Dizon, Razel Nikka M. Hao, Arianne A. Zamora, Devon Ray Pacial, Juan Antonio R. Magalang, Marissa Alejandria, Celia Carlos, Anna Ong-Lim, Edsel Maurice Salvaña, John Q. Wong, Jaime C. Montoya, Maria Rosario Singh-Vergeire and Cynthia P. Saloma |
| EPI_ISL_2156277 | The Medical City -Â Ortigas | Philippine Genome Center | Francis A. Tablizo, Kenneth M. Kim, Carlo M. Lapid, Marc Jerrone R. Castro, Maria Sofia L. Yangzon, Elcid Aaron R. Pangilinan, Benedict A. Maralit, Marc Edsel C. Ayes, Eva Maria Cutiongco-de la Paz, Alethea R. de Guzman, Jan Michael C. Yap, Jo-Hannah S. Llames, Sheila Mae M. Araiza, Kris P. Punayan, Irish Coleen A. Asin, Candice Francheska B. Tambaoan, Asia Louisa U. Chong, Karol Sophia Agape R. Padilla, Rianna Patricia S. Cruz, El King D. Morado, Joshua Gregor A. Dizon, Razel Nikka M. Hao, Arianne A. Zamora, Devon Ray Pacial, Juan Antonio R. Magalang, Marissa Alejandria, Celia Carlos, Anna Ong-Lim, Edsel Maurice Salvaña, John Q. Wong, Jaime C. Montoya, Maria Rosario Singh-Vergeire and Cynthia P. Saloma |
| EPI_ISL_2156282, EPI_ISL_2156283 | Cotabato Regional and Medical Center | Philippine Genome Center | Francis A. Tablizo, Kenneth M. Kim, Carlo M. Lapid, Marc Jerrone R. Castro, Maria Sofia L. Yangzon, Elcid Aaron R. Pangilinan, Benedict A. Maralit, Marc Edsel C. Ayes, Eva Maria Cutiongco-de la Paz, Alethea R. de Guzman, Jan Michael C. Yap, Jo-Hannah S. Llames, Sheila Mae M. Araiza, Kris P. Punayan, Irish Coleen A. Asin, Candice Francheska B. Tambaoan, Asia Louisa U. Chong, Karol Sophia Agape R. Padilla, Rianna Patricia S. Cruz, El King D. Morado, Joshua Gregor A. Dizon, Razel Nikka M. Hao, Arianne A. Zamora, Devon Ray Pacial, Juan Antonio R. Magalang, Marissa Alejandria, Celia Carlos, Anna Ong-Lim, Edsel Maurice Salvaña, John Q. Wong, Jaime C. Montoya, Maria Rosario Singh-Vergeire and Cynthia P. Saloma |
| EPI_ISL_2156285 | Philippine Red Cross - National Blood Center | Philippine Genome Center | Francis A. Tablizo, Kenneth M. Kim, Carlo M. Lapid, Marc Jerrone R. Castro, Maria Sofia L. Yangzon, Elcid Aaron R. Pangilinan, Benedict A. Maralit, Marc Edsel C. Ayes, Eva Maria Cutiongco-de la Paz, Alethea R. de Guzman, Jan Michael C. Yap, Jo-Hannah S. Llames, Sheila Mae M. Araiza, Kris P. Punayan, Irish Coleen A. Asin, Candice Francheska B. Tambaoan, Asia Louisa U. Chong, Karol Sophia Agape R. Padilla, Rianna Patricia S. Cruz, El King D. Morado, Joshua Gregor A. Dizon, Razel Nikka M. Hao, Arianne A. Zamora, Devon Ray Pacial, Juan Antonio R. Magalang, Marissa Alejandria, Celia Carlos, Anna Ong-Lim, Edsel Maurice Salvaña, John Q. Wong, Jaime C. Montoya, Maria Rosario Singh-Vergeire and Cynthia P. Saloma |
| EPI_ISL_2156286, EPI_ISL_2156287, EPI_ISL_2156288, EPI_ISL_2156289, EPI_ISL_2156290 | THE LORD'S GRACE MEDICAL AND INDUSTRIAL CLINIC | Philippine Genome Center | Francis A. Tablizo, Kenneth M. Kim, Carlo M. Lapid, Marc Jerrone R. Castro, Maria Sofia L. Yangzon, Elcid Aaron R. Pangilinan, Benedict A. Maralit, Marc Edsel C. Ayes, Eva Maria Cutiongco-de la Paz, Alethea R. de Guzman, Jan Michael C. Yap, Jo-Hannah S. Llames, Sheila Mae M. Araiza, Kris P. Punayan, Irish Coleen A. Asin, Candice Francheska B. Tambaoan, Asia Louisa U. Chong, Karol Sophia Agape R. Padilla, Rianna Patricia S. Cruz, El King D. Morado, Joshua Gregor A. Dizon, Razel Nikka M. Hao, Arianne A. Zamora, Devon Ray Pacial, Juan Antonio R. Magalang, Marissa Alejandria, Celia Carlos, Anna Ong-Lim, Edsel Maurice Salvaña, John Q. Wong, Jaime C. Montoya, Maria Rosario Singh-Vergeire and Cynthia P. Saloma |
| EPI_ISL_2156293 | First Aide Diagnostic Center | Philippine Genome Center | Francis A. Tablizo, Kenneth M. Kim, Carlo M. Lapid, Marc Jerrone R. Castro, Maria Sofia L. Yangzon, Elcid Aaron R. Pangilinan, Benedict A. Maralit, Marc Edsel C. Ayes, Eva Maria Cutiongco-de la Paz, Alethea R. de Guzman, Jan Michael C. Yap, Jo-Hannah S. Llames, Sheila Mae M. Araiza, Kris P. Punayan, Irish Coleen A. Asin, Candice Francheska B. Tambaoan, Asia Louisa U. Chong, Karol Sophia Agape R. Padilla, Rianna Patricia S. Cruz, El King D. Morado, Joshua Gregor A. Dizon, Razel Nikka M. Hao, Arianne A. Zamora, Devon Ray Pacial, Juan Antonio R. Magalang, Marissa Alejandria, Celia Carlos, Anna Ong-Lim, Edsel Maurice Salvaña, John Q. Wong, Jaime C. Montoya, Maria Rosario Singh-Vergeire and Cynthia P. Saloma |
| EPI_ISL_2156298, EPI_ISL_2156299 | Cotabato Regional and Medical Center | Philippine Genome Center | Francis A. Tablizo, Kenneth M. Kim, Carlo M. Lapid, Marc Jerrone R. Castro, Maria Sofia L. Yangzon, Elcid Aaron R. Pangilinan, Benedict A. Maralit, Marc Edsel C. Ayes, Eva Maria Cutiongco-de la Paz, Alethea R. de Guzman, Jan Michael C. Yap, Jo-Hannah S. Llames, Sheila Mae M. Araiza, Kris P. Punayan, Irish Coleen A. Asin, Candice Francheska B. Tambaoan, Asia Louisa U. Chong, Karol Sophia Agape R. Padilla, Rianna Patricia S. Cruz, El King D. Morado, Joshua Gregor A. Dizon, Razel Nikka M. Hao, Arianne A. Zamora, Devon Ray Pacial, Juan Antonio R. Magalang, Marissa Alejandria, Celia Carlos, Anna Ong-Lim, Edsel Maurice Salvaña, John Q. Wong, Jaime C. Montoya, Maria Rosario Singh-Vergeire and Cynthia P. Saloma |
| EPI_ISL_2156300 | PHILIPPINE AIRPORT DIAGNOSTIC LABORATORY | Philippine Genome Center | Francis A. Tablizo, Kenneth M. Kim, Carlo M. Lapid, Marc Jerrone R. Castro, Maria Sofia L. Yangzon, Elcid Aaron R. Pangilinan, Benedict A. Maralit, Marc Edsel C. Ayes, Eva Maria Cutiongco-de la Paz, Alethea R. de Guzman, Jan Michael C. Yap, Jo-Hannah S. Llames, Sheila Mae M. Araiza, Kris P. Punayan, Irish Coleen A. Asin, Candice Francheska B. Tambaoan, Asia Louisa U. Chong, Karol Sophia Agape R. Padilla, Rianna Patricia S. Cruz, El King D. Morado, Joshua Gregor A. Dizon, Razel Nikka M. Hao, Arianne A. Zamora, Devon Ray Pacial, Juan Antonio R. Magalang, Marissa Alejandria, Celia Carlos, Anna Ong-Lim, Edsel Maurice Salvaña, John Q. Wong, Jaime C. Montoya, Maria Rosario Singh-Vergeire and Cynthia P. Saloma |
| EPI_ISL_2156302 | THE LORD'S GRACE MEDICAL AND INDUSTRIAL CLINIC | Philippine Genome Center | Francis A. Tablizo, Kenneth M. Kim, Carlo M. Lapid, Marc Jerrone R. Castro, Maria Sofia L. Yangzon, Elcid Aaron R. Pangilinan, Benedict A. Maralit, Marc Edsel C. Ayes, Eva Maria Cutiongco-de la Paz, Alethea R. de Guzman, Jan Michael C. Yap, Jo-Hannah S. Llames, Sheila Mae M. Araiza, Kris P. Punayan, Irish Coleen A. Asin, Candice Francheska B. Tambaoan, Asia Louisa U. Chong, Karol Sophia Agape R. Padilla, Rianna Patricia S. Cruz, El King D. Morado, Joshua Gregor A. Dizon, Razel Nikka M. Hao, Arianne A. Zamora, Devon Ray Pacial, Juan Antonio R. Magalang, Marissa Alejandria, Celia Carlos, Anna Ong-Lim, Edsel Maurice Salvaña, John Q. Wong, Jaime C. Montoya, Maria Rosario Singh-Vergeire and Cynthia P. Saloma |
| EPI_ISL_2156303, EPI_ISL_2156304, EPI_ISL_2156305, EPI_ISL_2156306, EPI_ISL_2156307, EPI_ISL_2156308, EPI_ISL_2156311, EPI_ISL_2156312 | Philippine Red Cross - Port Area | Philippine Genome Center | Francis A. Tablizo, Kenneth M. Kim, Carlo M. Lapid, Marc Jerrone R. Castro, Maria Sofia L. Yangzon, Elcid Aaron R. Pangilinan, Benedict A. Maralit, Marc Edsel C. Ayes, Eva Maria Cutiongco-de la Paz, Alethea R. de Guzman, Jan Michael C. Yap, Jo-Hannah S. Llames, Sheila Mae M. Araiza, Kris P. Punayan, Irish Coleen A. Asin, Candice Francheska B. Tambaoan, Asia Louisa U. Chong, Karol Sophia Agape R. Padilla, Rianna Patricia S. Cruz, El King D. Morado, Joshua Gregor A. Dizon, Razel Nikka M. Hao, Arianne A. Zamora, Devon Ray Pacial, Juan Antonio R. Magalang, Marissa Alejandria, Celia Carlos, Anna Ong-Lim, Edsel Maurice Salvaña, John Q. Wong, Jaime C. Montoya, Maria Rosario Singh-Vergeire and Cynthia P. Saloma |
| EPI_ISL_2156313 | THE LORD'S GRACE MEDICAL AND INDUSTRIAL CLINIC | Philippine Genome Center | Francis A. Tablizo, Kenneth M. Kim, Carlo M. Lapid, Marc Jerrone R. Castro, Maria Sofia L. Yangzon, Elcid Aaron R. Pangilinan, Benedict A. Maralit, Marc Edsel C. Ayes, Eva Maria Cutiongco-de la Paz, Alethea R. de Guzman, Jan Michael C. Yap, Jo-Hannah S. Llames, Sheila Mae M. Araiza, Kris P. Punayan, Irish Coleen A. Asin, Candice Francheska B. Tambaoan, Asia Louisa U. Chong, Karol Sophia Agape R. Padilla, Rianna Patricia S. Cruz, El King D. Morado, Joshua Gregor A. Dizon, Razel Nikka M. Hao, Arianne A. Zamora, Devon Ray Pacial, Juan Antonio R. Magalang, Marissa Alejandria, Celia Carlos, Anna Ong-Lim, Edsel Maurice Salvaña, John Q. Wong, Jaime C. Montoya, Maria Rosario Singh-Vergeire and Cynthia P. Saloma |
| EPI_ISL_2156314 | San Lazaro Hospital | Philippine Genome Center | Francis A. Tablizo, Kenneth M. Kim, Carlo M. Lapid, Marc Jerrone R. Castro, Maria Sofia L. Yangzon, Elcid Aaron R. Pangilinan, Benedict A. Maralit, Marc Edsel C. Ayes, Eva Maria Cutiongco-de la Paz, Alethea R. de Guzman, Jan Michael C. Yap, Jo-Hannah S. Llames, Sheila Mae M. Araiza, Kris P. Punayan, Irish Coleen A. Asin, Candice Francheska B. Tambaoan, Asia Louisa U. Chong, Karol Sophia Agape R. Padilla, Rianna Patricia S. Cruz, El King D. Morado, Joshua Gregor A. Dizon, Razel Nikka M. Hao, Arianne A. Zamora, Devon Ray Pacial, Juan Antonio R. Magalang, Marissa Alejandria, Celia Carlos, Anna Ong-Lim, Edsel Maurice Salvaña, John Q. Wong, Jaime C. Montoya, Maria Rosario Singh-Vergeire and Cynthia P. Saloma |
| EPI_ISL_2157343 | Lboratorio Central de Saude Publica do Estado do Parana (LACEN/PR) | Laboratory of Respiratory Viruses and Measles, Oswaldo Cruz Institute, FIOCRUZ | Paola Resende, Luciana Appolinario, Fernando Motta, Anna Carolina Paixao, Ana Carolina Mendonca, Alice Sampaio Rocha, Taina Venas, Elisa Cavalcante Pereira, Renata Serrano Lopes, Irina Riediger, Marilda Siqueira on behalf of the Fiocruz COVID-19 Genomic Surveillance Network |
| EPI_ISL_2157376 | Laboratorio Central de Saude Publica do Estado do Espirito Santo (LACEN/ES) | Laboratory of Respiratory Viruses and Measles, Oswaldo Cruz Institute, FIOCRUZ | Paola Resende, Luciana Appolinario, Fernando Motta, Anna Carolina Paixao, Ana Carolina Mendonca, Alice Sampaio Rocha, Taina Venas, Eilisa Cavalcante Pereira, Renata Serrano Lopes, Rodrigo Ribeiro Rodrigues, Marilda Siqueira on behalf of the Fiocruz COVID-19 Genomic Surveillance Network |
| EPI_ISL_2157386, EPI_ISL_2157388, EPI_ISL_2157397, EPI_ISL_2157402, EPI_ISL_2157403, EPI_ISL_2157404, EPI_ISL_2157408, EPI_ISL_2157413, EPI_ISL_2157416, EPI_ISL_2157417, EPI_ISL_2157420, EPI_ISL_2157424, EPI_ISL_2157431, EPI_ISL_2157452, EPI_ISL_2157479 | | | |

| | | | |
|---|---|---|---|
| see above | Lboratorio Central de Saude Publica do Estado do Parana (LACEN/PR) | Laboratory of Respiratory Viruses and Measles, Oswaldo Cruz Institute, FIOCRUZ | Paola Resende, Luciana Appolinario, Fernando Motta, Anna Carolina Paixao, Ana Carolina Mendonca, Alice Sampaio Rocha, Taina Venas, Elisa Cavalcante Pereira, Renata Serrano Lopes, Irina Riediger, Marilda Siqueira on behalf of the Fiocruz COVID-19 Genomic Surveillance Network |
| EPI_ISL_2157480, EPI_ISL_2157481 | Laboratorio Central de Saude Publica do Estado de Minas Gerais (LACEN/MG) | Laboratory of Respiratory Viruses and Measles, Oswaldo Cruz Institute, FIOCRUZ | Paola Resende, Luciana Appolinario, Fernando Motta, Anna Carolina Paixao, Ana Carolina Mendonca, Alice Sampaio Rocha, Taina Venas, Elisa Cavalcante Pereira, Renata Serrano Lopes, Andre Felipe Leal Bernardes, Marilda Siqueira on behalf of the Fiocruz COVID-19 Genomic Surveillance Network |
| EPI_ISL_2157482 | Laboratorio Central de Saude Publica do Estado do Parana (LACEN/PR) | Laboratory of Respiratory Viruses and Measles, Oswaldo Cruz Institute, FIOCRUZ | Paola Resende, Luciana Appolinario, Fernando Motta, Anna Carolina Paixao, Ana Carolina Mendonca, Alice Sampaio Rocha, Taina Venas, Elisa Cavalcante Pereira, Renata Serrano Lopes, Irina Riediger, Marilda Siqueira on behalf of the Fiocruz COVID-19 Genomic Surveillance Network |
| EPI_ISL_2157485, EPI_ISL_2157486, EPI_ISL_2157487, EPI_ISL_2157488 | Lboratorio Central de Saude Publica do Estado do Parana (LACEN/PR) | Laboratory of Respiratory Viruses and Measles, Oswaldo Cruz Institute, FIOCRUZ | Paola Resende, Luciana Appolinario, Fernando Motta, Anna Carolina Paixao, Ana Carolina Mendonca, Alice Sampaio Rocha, Taina Venas, Elisa Cavalcante Pereira, Renata Serrano Lopes, Irina Riediger, Marilda Siqueira on behalf of the Fiocruz COVID-19 Genomic Surveillance Network |
| EPI_ISL_2157491 | Laboratorio Central de Saude Publica do Estado de Minas Gerais (LACEN/MG) | Laboratory of Respiratory Viruses and Measles, Oswaldo Cruz Institute, FIOCRUZ | Paola Resende, Luciana Appolinario, Fernando Motta, Anna Carolina Paixao, Ana Carolina Mendonca, Alice Sampaio Rocha, Taina Venas, Elisa Cavalcante Pereira, Renata Serrano Lopes, Andre Felipe Leal Bernardes, Marilda Siqueira on behalf of the Fiocruz COVID-19 Genomic Surveillance Network |
| EPI_ISL_2157492, EPI_ISL_2157496, EPI_ISL_2157595, EPI_ISL_2157597 | Lboratorio Central de Saude Publica do Estado do Parana (LACEN/PR) | Laboratory of Respiratory Viruses and Measles, Oswaldo Cruz Institute, FIOCRUZ | Paola Resende, Luciana Appolinario, Fernando Motta, Anna Carolina Paixao, Ana Carolina Mendonca, Alice Sampaio Rocha, Taina Venas, Elisa Cavalcante Pereira, Renata Serrano Lopes, Irina Riediger, Marilda Siqueira on behalf of the Fiocruz COVID-19 Genomic Surveillance Network |
| EPI_ISL_2171090 | Philippine Red Cross - National Blood Center | Philippine Genome Center | Francis A. Tablizo, Kenneth M. Kim, Carlo M. Lapid, Marc Jerrone R. Castro, Maria Sofia L. Yangzon, Elcid Aaron R. Pangilinan, Benedict A. Maralit, Marc Edsel C. Ayes, Eva Maria Cutiongco-de la Paz, Alethea R. de Guzman, Jan Michael C. Yap, Jo-Hannah S. Llames, Sheila Mae M. Araiza, Kris P. Punayan, Irish Coleen A. Asin, Candice Francheska B. Tambaoan, Asia Louisa U. Chong, Karol Sophia Agape R. Padilla, Rianna Patricia S. Cruz, El King D. Morado, Joshua Gregor A. Dizon, Razel Nikka M. Hao, Arianne A. Zamora, Devon Ray Pacial, Juan Antonio R. Magalang, Marissa Alejandria, Celia Carlos, Anna Ong-Lim, Edsel Maurice Salvaña, John Q. Wong, Jaime C. Montoya, Maria Rosario Singh-Vergeire and Cynthia P. Saloma |
| EPI_ISL_2171091, EPI_ISL_2171092, EPI_ISL_2171093 | Philippine Red Cross - Port Area | Philippine Genome Center | Francis A. Tablizo, Kenneth M. Kim, Carlo M. Lapid, Marc Jerrone R. Castro, Maria Sofia L. Yangzon, Elcid Aaron R. Pangilinan, Benedict A. Maralit, Marc Edsel C. Ayes, Eva Maria Cutiongco-de la Paz, Alethea R. de Guzman, Jan Michael C. Yap, Jo-Hannah S. Llames, Sheila Mae M. Araiza, Kris P. Punayan, Irish Coleen A. Asin, Candice Francheska B. Tambaoan, Asia Louisa U. Chong, Karol Sophia Agape R. Padilla, Rianna Patricia S. Cruz, El King D. Morado, Joshua Gregor A. Dizon, Razel Nikka M. Hao, Arianne A. Zamora, Devon Ray Pacial, Juan Antonio R. Magalang, Marissa Alejandria, Celia Carlos, Anna Ong-Lim, Edsel Maurice Salvaña, John Q. Wong, Jaime C. Montoya, Maria Rosario Singh-Vergeire and Cynthia P. Saloma |
| EPI_ISL_2171094, EPI_ISL_2171095 | Pasig City Children's Hospital - Child's Hope | Philippine Genome Center | Francis A. Tablizo, Kenneth M. Kim, Carlo M. Lapid, Marc Jerrone R. Castro, Maria Sofia L. Yangzon, Elcid Aaron R. Pangilinan, Benedict A. Maralit, Marc Edsel C. Ayes, Eva Maria Cutiongco-de la Paz, Alethea R. de Guzman, Jan Michael C. Yap, Jo-Hannah S. Llames, Sheila Mae M. Araiza, Kris P. Punayan, Irish Coleen A. Asin, Candice Francheska B. Tambaoan, Asia Louisa U. Chong, Karol Sophia Agape R. Padilla, Rianna Patricia S. Cruz, El King D. Morado, Joshua Gregor A. Dizon, Razel Nikka M. Hao, Arianne A. Zamora, Devon Ray Pacial, Juan Antonio R. Magalang, Marissa Alejandria, Celia Carlos, Anna Ong-Lim, Edsel Maurice Salvaña, John Q. Wong, Jaime C. Montoya, Maria Rosario Singh-Vergeire and Cynthia P. Saloma |
| EPI_ISL_2171096 | The Lord's Grace Medical and Industrial Clinic | Philippine Genome Center | Francis A. Tablizo, Kenneth M. Kim, Carlo M. Lapid, Marc Jerrone R. Castro, Maria Sofia L. Yangzon, Elcid Aaron R. Pangilinan, Benedict A. Maralit, Marc Edsel C. Ayes, Eva Maria Cutiongco-de la Paz, Alethea R. de Guzman, Jan Michael C. Yap, Jo-Hannah S. Llames, Sheila Mae M. Araiza, Kris P. Punayan, Irish Coleen A. Asin, Candice Francheska B. Tambaoan, Asia Louisa U. Chong, Karol Sophia Agape R. Padilla, Rianna Patricia S. Cruz, El King D. Morado, Joshua Gregor A. Dizon, Razel Nikka M. Hao, Arianne A. Zamora, Devon Ray Pacial, Juan Antonio R. Magalang, Marissa Alejandria, Celia Carlos, Anna Ong-Lim, Edsel Maurice Salvaña, John Q. Wong, Jaime C. Montoya, Maria Rosario Singh-Vergeire and Cynthia P. Saloma |
| EPI_ISL_2171097, EPI_ISL_2171101, EPI_ISL_2171102 | Philippine Red Cross - Port Area | Philippine Genome Center | Francis A. Tablizo, Kenneth M. Kim, Carlo M. Lapid, Marc Jerrone R. Castro, Maria Sofia L. Yangzon, Elcid Aaron R. Pangilinan, Benedict A. Maralit, Marc Edsel C. Ayes, Eva Maria Cutiongco-de la Paz, Alethea R. de Guzman, Jan Michael C. Yap, Jo-Hannah S. Llames, Sheila Mae M. Araiza, Kris P. Punayan, Irish Coleen A. Asin, Candice Francheska B. Tambaoan, Asia Louisa U. Chong, Karol Sophia Agape R. Padilla, Rianna Patricia S. Cruz, El King D. Morado, Joshua Gregor A. Dizon, Razel Nikka M. Hao, Arianne A. Zamora, Devon Ray Pacial, Juan Antonio R. Magalang, Marissa Alejandria, Celia Carlos, Anna Ong-Lim, Edsel Maurice Salvaña, John Q. Wong, Jaime C. Montoya, Maria Rosario Singh-Vergeire and Cynthia P. Saloma |
| EPI_ISL_2171104 | STA. ROSA CHO - GENEXPERT LABORATORY | Philippine Genome Center | Francis A. Tablizo, Kenneth M. Kim, Carlo M. Lapid, Marc Jerrone R. Castro, Maria Sofia L. Yangzon, Elcid Aaron R. Pangilinan, Benedict A. Maralit, Marc Edsel C. Ayes, Eva Maria Cutiongco-de la Paz, Alethea R. de Guzman, Jan Michael C. Yap, Jo-Hannah S. Llames, Sheila Mae M. Araiza, Kris P. Punayan, Irish Coleen A. Asin, Candice Francheska B. Tambaoan, Asia Louisa U. Chong, Karol Sophia Agape R. Padilla, Rianna Patricia S. Cruz, El King D. Morado, Joshua Gregor A. Dizon, Razel Nikka M. Hao, Arianne A. Zamora, Devon Ray Pacial, Juan Antonio R. Magalang, Marissa Alejandria, Celia Carlos, Anna Ong-Lim, Edsel Maurice Salvaña, John Q. Wong, Jaime C. Montoya, Maria Rosario Singh-Vergeire and Cynthia P. Saloma |
| EPI_ISL_2171105 | Philippine Red Cross - Port Area | Philippine Genome Center | Francis A. Tablizo, Kenneth M. Kim, Carlo M. Lapid, Marc Jerrone R. Castro, Maria Sofia L. Yangzon, Elcid Aaron R. Pangilinan, Benedict A. Maralit, Marc Edsel C. Ayes, Eva Maria Cutiongco-de la Paz, Alethea R. de Guzman, Jan Michael C. Yap, Jo-Hannah S. Llames, Sheila Mae M. Araiza, Kris P. Punayan, Irish Coleen A. Asin, Candice Francheska B. Tambaoan, Asia Louisa U. Chong, Karol Sophia Agape R. Padilla, Rianna Patricia S. Cruz, El King D. Morado, Joshua Gregor A. Dizon, Razel Nikka M. Hao, Arianne A. Zamora, Devon Ray Pacial, Juan Antonio R. Magalang, Marissa Alejandria, Celia Carlos, Anna Ong-Lim, Edsel Maurice Salvaña, John Q. Wong, Jaime C. Montoya, Maria Rosario Singh-Vergeire and Cynthia P. Saloma |
| EPI_ISL_2171106 | Pasig City Children's Hospital - Child's Hope | Philippine Genome Center | Francis A. Tablizo, Kenneth M. Kim, Carlo M. Lapid, Marc Jerrone R. Castro, Maria Sofia L. Yangzon, Elcid Aaron R. Pangilinan, Benedict A. Maralit, Marc Edsel C. Ayes, Eva Maria Cutiongco-de la Paz, Alethea R. de Guzman, Jan Michael C. Yap, Jo-Hannah S. Llames, Sheila Mae M. Araiza, Kris P. Punayan, Irish Coleen A. Asin, Candice Francheska B. Tambaoan, Asia Louisa U. Chong, Karol Sophia Agape R. Padilla, Rianna Patricia S. Cruz, El King D. Morado, Joshua Gregor A. Dizon, Razel Nikka M. Hao, Arianne A. Zamora, Devon Ray Pacial, Juan Antonio R. Magalang, Marissa Alejandria, Celia Carlos, Anna Ong-Lim, Edsel Maurice Salvaña, John Q. Wong, Jaime C. Montoya, Maria Rosario Singh-Vergeire and Cynthia P. Saloma |
| EPI_ISL_2171107 | The Lord's Grace Medical and Industrial Clinic | Philippine Genome Center | Francis A. Tablizo, Kenneth M. Kim, Carlo M. Lapid, Marc Jerrone R. Castro, Maria Sofia L. Yangzon, Elcid Aaron R. Pangilinan, Benedict A. Maralit, Marc Edsel C. Ayes, Eva Maria Cutiongco-de la Paz, Alethea R. de Guzman, Jan Michael C. Yap, Jo-Hannah S. Llames, Sheila Mae M. Araiza, Kris P. Punayan, Irish Coleen A. Asin, Candice Francheska B. Tambaoan, Asia Louisa U. Chong, Karol Sophia Agape R. Padilla, Rianna Patricia S. Cruz, El King D. Morado, Joshua Gregor A. Dizon, Razel Nikka M. Hao, Arianne A. Zamora, Devon Ray Pacial, Juan Antonio R. Magalang, Marissa Alejandria, Celia Carlos, Anna Ong-Lim, Edsel Maurice Salvaña, John Q. Wong, Jaime C. Montoya, Maria Rosario Singh-Vergeire and Cynthia P. Saloma |
| EPI_ISL_2171108, EPI_ISL_2171109, EPI_ISL_2171110 | Philippine Red Cross - Port Area | Philippine Genome Center | Francis A. Tablizo, Kenneth M. Kim, Carlo M. Lapid, Marc Jerrone R. Castro, Maria Sofia L. Yangzon, Elcid Aaron R. Pangilinan, Benedict A. Maralit, Marc Edsel C. Ayes, Eva Maria Cutiongco-de la Paz, Alethea R. de Guzman, Jan Michael C. Yap, Jo-Hannah S. Llames, Sheila Mae M. Araiza, Kris P. Punayan, Irish Coleen A. Asin, Candice Francheska B. Tambaoan, Asia Louisa U. Chong, Karol Sophia Agape R. Padilla, Rianna Patricia S. Cruz, El King D. Morado, Joshua Gregor A. Dizon, Razel Nikka M. Hao, Arianne A. Zamora, Devon Ray Pacial, Juan Antonio R. Magalang, Marissa Alejandria, Celia Carlos, Anna Ong-Lim, Edsel Maurice Salvaña, John Q. Wong, Jaime C. Montoya, Maria Rosario Singh-Vergeire and Cynthia P. Saloma |
| EPI_ISL_2171111 | The Lord's Grace Medical and Industrial Clinic | Philippine Genome Center | Francis A. Tablizo, Kenneth M. Kim, Carlo M. Lapid, Marc Jerrone R. Castro, Maria Sofia L. Yangzon, Elcid Aaron R. Pangilinan, Benedict A. Maralit, Marc Edsel C. Ayes, Eva Maria Cutiongco-de la Paz, Alethea R. de Guzman, Jan Michael C. Yap, Jo-Hannah S. Llames, Sheila Mae M. Araiza, Kris P. Punayan, Irish Coleen A. Asin, Candice Francheska B. Tambaoan, Asia Louisa U. Chong, Karol Sophia Agape R. Padilla, Rianna Patricia S. Cruz, El King D. Morado, Joshua Gregor A. Dizon, Razel Nikka M. Hao, Arianne A. Zamora, Devon Ray Pacial, Juan Antonio R. Magalang, Marissa Alejandria, Celia Carlos, Anna Ong-Lim, Edsel Maurice Salvaña, John Q. Wong, Jaime C. Montoya, Maria Rosario Singh-Vergeire and Cynthia P. Saloma |
| EPI_ISL_2171112, EPI_ISL_2171114 | Philippine Red Cross - Port Area | Philippine Genome Center | Francis A. Tablizo, Kenneth M. Kim, Carlo M. Lapid, Marc Jerrone R. Castro, Maria Sofia L. Yangzon, Elcid Aaron R. Pangilinan, Benedict A. Maralit, Marc Edsel C. Ayes, Eva Maria Cutiongco-de la Paz, Alethea R. de Guzman, Jan Michael C. Yap, Jo-Hannah S. Llames, Sheila Mae M. Araiza, Kris P. Punayan, Irish Coleen A. Asin, Candice Francheska B. Tambaoan, Asia Louisa U. Chong, Karol Sophia Agape R. Padilla, Rianna Patricia S. Cruz, El King D. Morado, Joshua Gregor A. Dizon, Razel Nikka M. Hao, Arianne A. Zamora, Devon Ray Pacial, Juan Antonio R. Magalang, Marissa Alejandria, Celia Carlos, Anna Ong-Lim, Edsel Maurice Salvaña, John Q. Wong, Jaime C. Montoya, Maria Rosario Singh-Vergeire and Cynthia P. Saloma |
| EPI_ISL_2171115, EPI_ISL_2171116 | The Lord's Grace Medical and Industrial Clinic | Philippine Genome Center | Francis A. Tablizo, Kenneth M. Kim, Carlo M. Lapid, Marc Jerrone R. Castro, Maria Sofia L. Yangzon, Elcid Aaron R. Pangilinan, Benedict A. Maralit, Marc Edsel C. Ayes, Eva Maria Cutiongco-de la Paz, Alethea R. de Guzman, Jan Michael C. Yap, Jo-Hannah S. Llames, Sheila Mae M. Araiza, Kris P. |

| | | | |
|---|---|---|---|
| | | | Punayan, Irish Coleen A. Asin, Candice Francheska B. Tambaoan, Asia Louisa U. Chong, Karol Sophia Agape R. Padilla, Rianna Patricia S. Cruz, El King D. Morado, Joshua Gregor A. Dizon, Razel Nikka M. Hao, Arianne A. Zamora, Devon Ray Pacial, Juan Antonio R. Magalang, Marissa Alejandria, Celia Carlos, Anna Ong-Lim, Edsel Maurice Salvaña, John Q. Wong, Jaime C. Montoya, Maria Rosario Singh-Vergeire and Cynthia P. Saloma |
| EPI_ISL_2171117 | University of the Philippines National Institutes of Health (UP NIH) | Philippine Genome Center | Francis A. Tablizo, Kenneth M. Kim, Carlo M. Lapid, Marc Jerrone R. Castro, Maria Sofia L. Yangzon, Elcid Aaron R. Pangilinan, Benedict A. Maralit, Marc Edsel C. Ayes, Eva Maria Cutiongco-de la Paz, Alethea R. de Guzman, Jan Michael C. Yap, Jo-Hannah S. Llames, Sheila Mae M. Araiza, Kris P. Punayan, Irish Coleen A. Asin, Candice Francheska B. Tambaoan, Asia Louisa U. Chong, Karol Sophia Agape R. Padilla, Rianna Patricia S. Cruz, El King D. Morado, Joshua Gregor A. Dizon, Razel Nikka M. Hao, Arianne A. Zamora, Devon Ray Pacial, Juan Antonio R. Magalang, Marissa Alejandria, Celia Carlos, Anna Ong-Lim, Edsel Maurice Salvaña, John Q. Wong, Jaime C. Montoya, Maria Rosario Singh-Vergeire and Cynthia P. Saloma |
| EPI_ISL_2171118 | The Lord's Grace Medical and Industrial Clinic | Philippine Genome Center | Francis A. Tablizo, Kenneth M. Kim, Carlo M. Lapid, Marc Jerrone R. Castro, Maria Sofia L. Yangzon, Elcid Aaron R. Pangilinan, Benedict A. Maralit, Marc Edsel C. Ayes, Eva Maria Cutiongco-de la Paz, Alethea R. de Guzman, Jan Michael C. Yap, Jo-Hannah S. Llames, Sheila Mae M. Araiza, Kris P. Punayan, Irish Coleen A. Asin, Candice Francheska B. Tambaoan, Asia Louisa U. Chong, Karol Sophia Agape R. Padilla, Rianna Patricia S. Cruz, El King D. Morado, Joshua Gregor A. Dizon, Razel Nikka M. Hao, Arianne A. Zamora, Devon Ray Pacial, Juan Antonio R. Magalang, Marissa Alejandria, Celia Carlos, Anna Ong-Lim, Edsel Maurice Salvaña, John Q. Wong, Jaime C. Montoya, Maria Rosario Singh-Vergeire and Cynthia P. Saloma |
| EPI_ISL_2171119 | Philippine Red Cross - Port Area | Philippine Genome Center | Francis A. Tablizo, Kenneth M. Kim, Carlo M. Lapid, Marc Jerrone R. Castro, Maria Sofia L. Yangzon, Elcid Aaron R. Pangilinan, Benedict A. Maralit, Marc Edsel C. Ayes, Eva Maria Cutiongco-de la Paz, Alethea R. de Guzman, Jan Michael C. Yap, Jo-Hannah S. Llames, Sheila Mae M. Araiza, Kris P. Punayan, Irish Coleen A. Asin, Candice Francheska B. Tambaoan, Asia Louisa U. Chong, Karol Sophia Agape R. Padilla, Rianna Patricia S. Cruz, El King D. Morado, Joshua Gregor A. Dizon, Razel Nikka M. Hao, Arianne A. Zamora, Devon Ray Pacial, Juan Antonio R. Magalang, Marissa Alejandria, Celia Carlos, Anna Ong-Lim, Edsel Maurice Salvaña, John Q. Wong, Jaime C. Montoya, Maria Rosario Singh-Vergeire and Cynthia P. Saloma |
| EPI_ISL_2171120 | The Lord's Grace Medical and Industrial Clinic | Philippine Genome Center | Francis A. Tablizo, Kenneth M. Kim, Carlo M. Lapid, Marc Jerrone R. Castro, Maria Sofia L. Yangzon, Elcid Aaron R. Pangilinan, Benedict A. Maralit, Marc Edsel C. Ayes, Eva Maria Cutiongco-de la Paz, Alethea R. de Guzman, Jan Michael C. Yap, Jo-Hannah S. Llames, Sheila Mae M. Araiza, Kris P. Punayan, Irish Coleen A. Asin, Candice Francheska B. Tambaoan, Asia Louisa U. Chong, Karol Sophia Agape R. Padilla, Rianna Patricia S. Cruz, El King D. Morado, Joshua Gregor A. Dizon, Razel Nikka M. Hao, Arianne A. Zamora, Devon Ray Pacial, Juan Antonio R. Magalang, Marissa Alejandria, Celia Carlos, Anna Ong-Lim, Edsel Maurice Salvaña, John Q. Wong, Jaime C. Montoya, Maria Rosario Singh-Vergeire and Cynthia P. Saloma |
| EPI_ISL_2171121 | South Super Highway Molecular Diagnostic Laboratory | Philippine Genome Center | Francis A. Tablizo, Kenneth M. Kim, Carlo M. Lapid, Marc Jerrone R. Castro, Maria Sofia L. Yangzon, Elcid Aaron R. Pangilinan, Benedict A. Maralit, Marc Edsel C. Ayes, Eva Maria Cutiongco-de la Paz, Alethea R. de Guzman, Jan Michael C. Yap, Jo-Hannah S. Llames, Sheila Mae M. Araiza, Kris P. Punayan, Irish Coleen A. Asin, Candice Francheska B. Tambaoan, Asia Louisa U. Chong, Karol Sophia Agape R. Padilla, Rianna Patricia S. Cruz, El King D. Morado, Joshua Gregor A. Dizon, Razel Nikka M. Hao, Arianne A. Zamora, Devon Ray Pacial, Juan Antonio R. Magalang, Marissa Alejandria, Celia Carlos, Anna Ong-Lim, Edsel Maurice Salvaña, John Q. Wong, Jaime C. Montoya, Maria Rosario Singh-Vergeire and Cynthia P. Saloma |
| EPI_ISL_2171122 | Philippine Red Cross - Port Area | Philippine Genome Center | Francis A. Tablizo, Kenneth M. Kim, Carlo M. Lapid, Marc Jerrone R. Castro, Maria Sofia L. Yangzon, Elcid Aaron R. Pangilinan, Benedict A. Maralit, Marc Edsel C. Ayes, Eva Maria Cutiongco-de la Paz, Alethea R. de Guzman, Jan Michael C. Yap, Jo-Hannah S. Llames, Sheila Mae M. Araiza, Kris P. Punayan, Irish Coleen A. Asin, Candice Francheska B. Tambaoan, Asia Louisa U. Chong, Karol Sophia Agape R. Padilla, Rianna Patricia S. Cruz, El King D. Morado, Joshua Gregor A. Dizon, Razel Nikka M. Hao, Arianne A. Zamora, Devon Ray Pacial, Juan Antonio R. Magalang, Marissa Alejandria, Celia Carlos, Anna Ong-Lim, Edsel Maurice Salvaña, John Q. Wong, Jaime C. Montoya, Maria Rosario Singh-Vergeire and Cynthia P. Saloma |
| EPI_ISL_2171123 | Research Institute for Tropical Medicine, Inc. (RITM) | Philippine Genome Center | Francis A. Tablizo, Kenneth M. Kim, Carlo M. Lapid, Marc Jerrone R. Castro, Maria Sofia L. Yangzon, Elcid Aaron R. Pangilinan, Benedict A. Maralit, Marc Edsel C. Ayes, Eva Maria Cutiongco-de la Paz, Alethea R. de Guzman, Jan Michael C. Yap, Jo-Hannah S. Llames, Sheila Mae M. Araiza, Kris P. Punayan, Irish Coleen A. Asin, Candice Francheska B. Tambaoan, Asia Louisa U. Chong, Karol Sophia Agape R. Padilla, Rianna Patricia S. Cruz, El King D. Morado, Joshua Gregor A. Dizon, Razel Nikka M. Hao, Arianne A. Zamora, Devon Ray Pacial, Juan Antonio R. Magalang, Marissa Alejandria, Celia Carlos, Anna Ong-Lim, Edsel Maurice Salvaña, John Q. Wong, Jaime C. Montoya, Maria Rosario Singh-Vergeire and Cynthia P. Saloma |
| EPI_ISL_2171124 | PCR MED Molecular Laboratory | Philippine Genome Center | Francis A. Tablizo, Kenneth M. Kim, Carlo M. Lapid, Marc Jerrone R. Castro, Maria Sofia L. Yangzon, Elcid Aaron R. Pangilinan, Benedict A. Maralit, Marc Edsel C. Ayes, Eva Maria Cutiongco-de la Paz, Alethea R. de Guzman, Jan Michael C. Yap, Jo-Hannah S. Llames, Sheila Mae M. Araiza, Kris P. Punayan, Irish Coleen A. Asin, Candice Francheska B. Tambaoan, Asia Louisa U. Chong, Karol Sophia Agape R. Padilla, Rianna Patricia S. Cruz, El King D. Morado, Joshua Gregor A. Dizon, Razel Nikka M. Hao, Arianne A. Zamora, Devon Ray Pacial, Juan Antonio R. Magalang, Marissa Alejandria, Celia Carlos, Anna Ong-Lim, Edsel Maurice Salvaña, John Q. Wong, Jaime C. Montoya, Maria Rosario Singh-Vergeire and Cynthia P. Saloma |
| EPI_ISL_2171125, EPI_ISL_2171126 | The Lord's Grace Medical and Industrial Clinic | Philippine Genome Center | Francis A. Tablizo, Kenneth M. Kim, Carlo M. Lapid, Marc Jerrone R. Castro, Maria Sofia L. Yangzon, Elcid Aaron R. Pangilinan, Benedict A. Maralit, Marc Edsel C. Ayes, Eva Maria Cutiongco-de la Paz, Alethea R. de Guzman, Jan Michael C. Yap, Jo-Hannah S. Llames, Sheila Mae M. Araiza, Kris P. Punayan, Irish Coleen A. Asin, Candice Francheska B. Tambaoan, Asia Louisa U. Chong, Karol Sophia Agape R. Padilla, Rianna Patricia S. Cruz, El King D. Morado, Joshua Gregor A. Dizon, Razel Nikka M. Hao, Arianne A. Zamora, Devon Ray Pacial, Juan Antonio R. Magalang, Marissa Alejandria, Celia Carlos, Anna Ong-Lim, Edsel Maurice Salvaña, John Q. Wong, Jaime C. Montoya, Maria Rosario Singh-Vergeire and Cynthia P. Saloma |
| EPI_ISL_2171127, EPI_ISL_2171128 | Philippine Red Cross - Port Area | Philippine Genome Center | Francis A. Tablizo, Kenneth M. Kim, Carlo M. Lapid, Marc Jerrone R. Castro, Maria Sofia L. Yangzon, Elcid Aaron R. Pangilinan, Benedict A. Maralit, Marc Edsel C. Ayes, Eva Maria Cutiongco-de la Paz, Alethea R. de Guzman, Jan Michael C. Yap, Jo-Hannah S. Llames, Sheila Mae M. Araiza, Kris P. Punayan, Irish Coleen A. Asin, Candice Francheska B. Tambaoan, Asia Louisa U. Chong, Karol Sophia Agape R. Padilla, Rianna Patricia S. Cruz, El King D. Morado, Joshua Gregor A. Dizon, Razel Nikka M. Hao, Arianne A. Zamora, Devon Ray Pacial, Juan Antonio R. Magalang, Marissa Alejandria, Celia Carlos, Anna Ong-Lim, Edsel Maurice Salvaña, John Q. Wong, Jaime C. Montoya, Maria Rosario Singh-Vergeire and Cynthia P. Saloma |
| EPI_ISL_2171129, EPI_ISL_2171130 | The Lord's Grace Medical and Industrial Clinic | Philippine Genome Center | Francis A. Tablizo, Kenneth M. Kim, Carlo M. Lapid, Marc Jerrone R. Castro, Maria Sofia L. Yangzon, Elcid Aaron R. Pangilinan, Benedict A. Maralit, Marc Edsel C. Ayes, Eva Maria Cutiongco-de la Paz, Alethea R. de Guzman, Jan Michael C. Yap, Jo-Hannah S. Llames, Sheila Mae M. Araiza, Kris P. Punayan, Irish Coleen A. Asin, Candice Francheska B. Tambaoan, Asia Louisa U. Chong, Karol Sophia Agape R. Padilla, Rianna Patricia S. Cruz, El King D. Morado, Joshua Gregor A. Dizon, Razel Nikka M. Hao, Arianne A. Zamora, Devon Ray Pacial, Juan Antonio R. Magalang, Marissa Alejandria, Celia Carlos, Anna Ong-Lim, Edsel Maurice Salvaña, John Q. Wong, Jaime C. Montoya, Maria Rosario Singh-Vergeire and Cynthia P. Saloma |
| EPI_ISL_2171131 | Pasig City Children's Hospital - Child's Hope | Philippine Genome Center | Francis A. Tablizo, Kenneth M. Kim, Carlo M. Lapid, Marc Jerrone R. Castro, Maria Sofia L. Yangzon, Elcid Aaron R. Pangilinan, Benedict A. Maralit, Marc Edsel C. Ayes, Eva Maria Cutiongco-de la Paz, Alethea R. de Guzman, Jan Michael C. Yap, Jo-Hannah S. Llames, Sheila Mae M. Araiza, Kris P. Punayan, Irish Coleen A. Asin, Candice Francheska B. Tambaoan, Asia Louisa U. Chong, Karol Sophia Agape R. Padilla, Rianna Patricia S. Cruz, El King D. Morado, Joshua Gregor A. Dizon, Razel Nikka M. Hao, Arianne A. Zamora, Devon Ray Pacial, Juan Antonio R. Magalang, Marissa Alejandria, Celia Carlos, Anna Ong-Lim, Edsel Maurice Salvaña, John Q. Wong, Jaime C. Montoya, Maria Rosario Singh-Vergeire and Cynthia P. Saloma |
| EPI_ISL_2171132 | The Lord's Grace Medical and Industrial Clinic | Philippine Genome Center | Francis A. Tablizo, Kenneth M. Kim, Carlo M. Lapid, Marc Jerrone R. Castro, Maria Sofia L. Yangzon, Elcid Aaron R. Pangilinan, Benedict A. Maralit, Marc Edsel C. Ayes, Eva Maria Cutiongco-de la Paz, Alethea R. de Guzman, Jan Michael C. Yap, Jo-Hannah S. Llames, Sheila Mae M. Araiza, Kris P. Punayan, Irish Coleen A. Asin, Candice Francheska B. Tambaoan, Asia Louisa U. Chong, Karol Sophia Agape R. Padilla, Rianna Patricia S. Cruz, El King D. Morado, Joshua Gregor A. Dizon, Razel Nikka M. Hao, Arianne A. Zamora, Devon Ray Pacial, Juan Antonio R. Magalang, Marissa Alejandria, Celia Carlos, Anna Ong-Lim, Edsel Maurice Salvaña, John Q. Wong, Jaime C. Montoya, Maria Rosario Singh-Vergeire and Cynthia P. Saloma |
| EPI_ISL_2171133 | South Super Highway Molecular Diagnostic Laboratory | Philippine Genome Center | Francis A. Tablizo, Kenneth M. Kim, Carlo M. Lapid, Marc Jerrone R. Castro, Maria Sofia L. Yangzon, Elcid Aaron R. Pangilinan, Benedict A. Maralit, Marc Edsel C. Ayes, Eva Maria Cutiongco-de la Paz, Alethea R. de Guzman, Jan Michael C. Yap, Jo-Hannah S. Llames, Sheila Mae M. Araiza, Kris P. Punayan, Irish Coleen A. Asin, Candice Francheska B. Tambaoan, Asia Louisa U. Chong, Karol Sophia Agape R. Padilla, Rianna Patricia S. Cruz, El King D. Morado, Joshua Gregor A. Dizon, Razel Nikka M. Hao, Arianne A. Zamora, Devon Ray Pacial, Juan Antonio R. Magalang, Marissa Alejandria, Celia Carlos, Anna Ong-Lim, Edsel Maurice Salvaña, John Q. Wong, Jaime C. Montoya, Maria Rosario Singh-Vergeire and Cynthia P. Saloma |
| EPI_ISL_2171134, EPI_ISL_2171136 | Philippine Red Cross - Port Area | Philippine Genome Center | Francis A. Tablizo, Kenneth M. Kim, Carlo M. Lapid, Marc Jerrone R. Castro, Maria Sofia L. Yangzon, Elcid Aaron R. Pangilinan, Benedict A. Maralit, Marc Edsel C. Ayes, Eva Maria Cutiongco-de la Paz, Alethea R. de Guzman, Jan Michael C. Yap, Jo-Hannah S. Llames, Sheila Mae M. Araiza, Kris P. Punayan, Irish Coleen A. Asin, Candice Francheska B. Tambaoan, Asia Louisa U. Chong, Karol Sophia Agape R. Padilla, Rianna Patricia S. Cruz, El King D. Morado, Joshua Gregor A. Dizon, Razel Nikka M. Hao, Arianne A. Zamora, Devon Ray Pacial, Juan Antonio R. Magalang, Marissa Alejandria, Celia Carlos, Anna Ong-Lim, Edsel Maurice Salvaña, John Q. Wong, Jaime C. Montoya, Maria Rosario Singh-Vergeire and Cynthia P. Saloma |
| EPI_ISL_2171137 | First Aide Diagnostic Center | Philippine Genome Center | Francis A. Tablizo, Kenneth M. Kim, Carlo M. Lapid, Marc Jerrone R. Castro, Maria Sofia L. Yangzon, Elcid Aaron R. Pangilinan, Benedict A. Maralit, Marc |

| | | | |
|---|---|---|---|
| | | | Edsel C. Ayes, Eva Maria Cutiongco-de la Paz, Alethea R. de Guzman, Jan Michael C. Yap, Jo-Hannah S. Llames, Sheila Mae M. Araiza, Kris P. Punayan, Irish Coleen A. Asin, Candice Francheska B. Tambaoan, Asia Louisa U. Chong, Karol Sophia Agape R. Padilla, Rianna Patricia S. Cruz, El King D. Morado, Joshua Gregor A. Dizon, Razel Nikka M. Hao, Arianne A. Zamora, Devon Ray Pacial, Juan Antonio R. Magalang, Marissa Alejandria, Celia Carlos, Anna Ong-Lim, Edsel Maurice Salvaña, John Q. Wong, Jaime C. Montoya, Maria Rosario Singh-Vergeire and Cynthia P. Saloma |
| EPI_ISL_2171139 | Safeguard DNA Diagnostic, Inc. | Philippine Genome Center | Francis A. Tablizo, Kenneth M. Kim, Carlo M. Lapid, Marc Jerrone R. Castro, Maria Sofia L. Yangzon, Elcid Aaron R. Pangilinan, Benedict A. Maralit, Marc Edsel C. Ayes, Eva Maria Cutiongco-de la Paz, Alethea R. de Guzman, Jan Michael C. Yap, Jo-Hannah S. Llames, Sheila Mae M. Araiza, Kris P. Punayan, Irish Coleen A. Asin, Candice Francheska B. Tambaoan, Asia Louisa U. Chong, Karol Sophia Agape R. Padilla, Rianna Patricia S. Cruz, El King D. Morado, Joshua Gregor A. Dizon, Razel Nikka M. Hao, Arianne A. Zamora, Devon Ray Pacial, Juan Antonio R. Magalang, Marissa Alejandria, Celia Carlos, Anna Ong-Lim, Edsel Maurice Salvaña, John Q. Wong, Jaime C. Montoya, Maria Rosario Singh-Vergeire and Cynthia P. Saloma |
| EPI_ISL_2171140, EPI_ISL_2171141, EPI_ISL_2171142 | Philippine Red Cross - Port Area | Philippine Genome Center | Francis A. Tablizo, Kenneth M. Kim, Carlo M. Lapid, Marc Jerrone R. Castro, Maria Sofia L. Yangzon, Elcid Aaron R. Pangilinan, Benedict A. Maralit, Marc Edsel C. Ayes, Eva Maria Cutiongco-de la Paz, Alethea R. de Guzman, Jan Michael C. Yap, Jo-Hannah S. Llames, Sheila Mae M. Araiza, Kris P. Punayan, Irish Coleen A. Asin, Candice Francheska B. Tambaoan, Asia Louisa U. Chong, Karol Sophia Agape R. Padilla, Rianna Patricia S. Cruz, El King D. Morado, Joshua Gregor A. Dizon, Razel Nikka M. Hao, Arianne A. Zamora, Devon Ray Pacial, Juan Antonio R. Magalang, Marissa Alejandria, Celia Carlos, Anna Ong-Lim, Edsel Maurice Salvaña, John Q. Wong, Jaime C. Montoya, Maria Rosario Singh-Vergeire and Cynthia P. Saloma |
| EPI_ISL_2171143 | Philippine Red Cross Logistics and Multipurpose Center | Philippine Genome Center | Francis A. Tablizo, Kenneth M. Kim, Carlo M. Lapid, Marc Jerrone R. Castro, Maria Sofia L. Yangzon, Elcid Aaron R. Pangilinan, Benedict A. Maralit, Marc Edsel C. Ayes, Eva Maria Cutiongco-de la Paz, Alethea R. de Guzman, Jan Michael C. Yap, Jo-Hannah S. Llames, Sheila Mae M. Araiza, Kris P. Punayan, Irish Coleen A. Asin, Candice Francheska B. Tambaoan, Asia Louisa U. Chong, Karol Sophia Agape R. Padilla, Rianna Patricia S. Cruz, El King D. Morado, Joshua Gregor A. Dizon, Razel Nikka M. Hao, Arianne A. Zamora, Devon Ray Pacial, Juan Antonio R. Magalang, Marissa Alejandria, Celia Carlos, Anna Ong-Lim, Edsel Maurice Salvaña, John Q. Wong, Jaime C. Montoya, Maria Rosario Singh-Vergeire and Cynthia P. Saloma |
| EPI_ISL_2171146 | Philippine Red Cross - Port Area | Philippine Genome Center | Francis A. Tablizo, Kenneth M. Kim, Carlo M. Lapid, Marc Jerrone R. Castro, Maria Sofia L. Yangzon, Elcid Aaron R. Pangilinan, Benedict A. Maralit, Marc Edsel C. Ayes, Eva Maria Cutiongco-de la Paz, Alethea R. de Guzman, Jan Michael C. Yap, Jo-Hannah S. Llames, Sheila Mae M. Araiza, Kris P. Punayan, Irish Coleen A. Asin, Candice Francheska B. Tambaoan, Asia Louisa U. Chong, Karol Sophia Agape R. Padilla, Rianna Patricia S. Cruz, El King D. Morado, Joshua Gregor A. Dizon, Razel Nikka M. Hao, Arianne A. Zamora, Devon Ray Pacial, Juan Antonio R. Magalang, Marissa Alejandria, Celia Carlos, Anna Ong-Lim, Edsel Maurice Salvaña, John Q. Wong, Jaime C. Montoya, Maria Rosario Singh-Vergeire and Cynthia P. Saloma |
| EPI_ISL_2171147 | Philippine Red Cross - National Blood Center | Philippine Genome Center | Francis A. Tablizo, Kenneth M. Kim, Carlo M. Lapid, Marc Jerrone R. Castro, Maria Sofia L. Yangzon, Elcid Aaron R. Pangilinan, Benedict A. Maralit, Marc Edsel C. Ayes, Eva Maria Cutiongco-de la Paz, Alethea R. de Guzman, Jan Michael C. Yap, Jo-Hannah S. Llames, Sheila Mae M. Araiza, Kris P. Punayan, Irish Coleen A. Asin, Candice Francheska B. Tambaoan, Asia Louisa U. Chong, Karol Sophia Agape R. Padilla, Rianna Patricia S. Cruz, El King D. Morado, Joshua Gregor A. Dizon, Razel Nikka M. Hao, Arianne A. Zamora, Devon Ray Pacial, Juan Antonio R. Magalang, Marissa Alejandria, Celia Carlos, Anna Ong-Lim, Edsel Maurice Salvaña, John Q. Wong, Jaime C. Montoya, Maria Rosario Singh-Vergeire and Cynthia P. Saloma |
| EPI_ISL_2171148 | Philippine Red Cross - Port Area | Philippine Genome Center | Francis A. Tablizo, Kenneth M. Kim, Carlo M. Lapid, Marc Jerrone R. Castro, Maria Sofia L. Yangzon, Elcid Aaron R. Pangilinan, Benedict A. Maralit, Marc Edsel C. Ayes, Eva Maria Cutiongco-de la Paz, Alethea R. de Guzman, Jan Michael C. Yap, Jo-Hannah S. Llames, Sheila Mae M. Araiza, Kris P. Punayan, Irish Coleen A. Asin, Candice Francheska B. Tambaoan, Asia Louisa U. Chong, Karol Sophia Agape R. Padilla, Rianna Patricia S. Cruz, El King D. Morado, Joshua Gregor A. Dizon, Razel Nikka M. Hao, Arianne A. Zamora, Devon Ray Pacial, Juan Antonio R. Magalang, Marissa Alejandria, Celia Carlos, Anna Ong-Lim, Edsel Maurice Salvaña, John Q. Wong, Jaime C. Montoya, Maria Rosario Singh-Vergeire and Cynthia P. Saloma |
| EPI_ISL_2171149 | Philippine Red Cross Logistics and Multipurpose Center | Philippine Genome Center | Francis A. Tablizo, Kenneth M. Kim, Carlo M. Lapid, Marc Jerrone R. Castro, Maria Sofia L. Yangzon, Elcid Aaron R. Pangilinan, Benedict A. Maralit, Marc Edsel C. Ayes, Eva Maria Cutiongco-de la Paz, Alethea R. de Guzman, Jan Michael C. Yap, Jo-Hannah S. Llames, Sheila Mae M. Araiza, Kris P. Punayan, Irish Coleen A. Asin, Candice Francheska B. Tambaoan, Asia Louisa U. Chong, Karol Sophia Agape R. Padilla, Rianna Patricia S. Cruz, El King D. Morado, Joshua Gregor A. Dizon, Razel Nikka M. Hao, Arianne A. Zamora, Devon Ray Pacial, Juan Antonio R. Magalang, Marissa Alejandria, Celia Carlos, Anna Ong-Lim, Edsel Maurice Salvaña, John Q. Wong, Jaime C. Montoya, Maria Rosario Singh-Vergeire and Cynthia P. Saloma |
| EPI_ISL_2171150 | Ospital ng ParaÃ±aque II | Philippine Genome Center | Francis A. Tablizo, Kenneth M. Kim, Carlo M. Lapid, Marc Jerrone R. Castro, Maria Sofia L. Yangzon, Elcid Aaron R. Pangilinan, Benedict A. Maralit, Marc Edsel C. Ayes, Eva Maria Cutiongco-de la Paz, Alethea R. de Guzman, Jan Michael C. Yap, Jo-Hannah S. Llames, Sheila Mae M. Araiza, Kris P. Punayan, Irish Coleen A. Asin, Candice Francheska B. Tambaoan, Asia Louisa U. Chong, Karol Sophia Agape R. Padilla, Rianna Patricia S. Cruz, El King D. Morado, Joshua Gregor A. Dizon, Razel Nikka M. Hao, Arianne A. Zamora, Devon Ray Pacial, Juan Antonio R. Magalang, Marissa Alejandria, Celia Carlos, Anna Ong-Lim, Edsel Maurice Salvaña, John Q. Wong, Jaime C. Montoya, Maria Rosario Singh-Vergeire and Cynthia P. Saloma |
| EPI_ISL_2171151, EPI_ISL_2171152 | Philippine Red Cross - Port Area | Philippine Genome Center | Francis A. Tablizo, Kenneth M. Kim, Carlo M. Lapid, Marc Jerrone R. Castro, Maria Sofia L. Yangzon, Elcid Aaron R. Pangilinan, Benedict A. Maralit, Marc Edsel C. Ayes, Eva Maria Cutiongco-de la Paz, Alethea R. de Guzman, Jan Michael C. Yap, Jo-Hannah S. Llames, Sheila Mae M. Araiza, Kris P. Punayan, Irish Coleen A. Asin, Candice Francheska B. Tambaoan, Asia Louisa U. Chong, Karol Sophia Agape R. Padilla, Rianna Patricia S. Cruz, El King D. Morado, Joshua Gregor A. Dizon, Razel Nikka M. Hao, Arianne A. Zamora, Devon Ray Pacial, Juan Antonio R. Magalang, Marissa Alejandria, Celia Carlos, Anna Ong-Lim, Edsel Maurice Salvaña, John Q. Wong, Jaime C. Montoya, Maria Rosario Singh-Vergeire and Cynthia P. Saloma |
| EPI_ISL_2171154 | Prime Care Alpha Covid-19 Testing Laboratory | Philippine Genome Center | Francis A. Tablizo, Kenneth M. Kim, Carlo M. Lapid, Marc Jerrone R. Castro, Maria Sofia L. Yangzon, Elcid Aaron R. Pangilinan, Benedict A. Maralit, Marc Edsel C. Ayes, Eva Maria Cutiongco-de la Paz, Alethea R. de Guzman, Jan Michael C. Yap, Jo-Hannah S. Llames, Sheila Mae M. Araiza, Kris P. Punayan, Irish Coleen A. Asin, Candice Francheska B. Tambaoan, Asia Louisa U. Chong, Karol Sophia Agape R. Padilla, Rianna Patricia S. Cruz, El King D. Morado, Joshua Gregor A. Dizon, Razel Nikka M. Hao, Arianne A. Zamora, Devon Ray Pacial, Juan Antonio R. Magalang, Marissa Alejandria, Celia Carlos, Anna Ong-Lim, Edsel Maurice Salvaña, John Q. Wong, Jaime C. Montoya, Maria Rosario Singh-Vergeire and Cynthia P. Saloma |
| EPI_ISL_2171155, EPI_ISL_2171156, EPI_ISL_2171157 | Philippine Red Cross - Port Area | Philippine Genome Center | Francis A. Tablizo, Kenneth M. Kim, Carlo M. Lapid, Marc Jerrone R. Castro, Maria Sofia L. Yangzon, Elcid Aaron R. Pangilinan, Benedict A. Maralit, Marc Edsel C. Ayes, Eva Maria Cutiongco-de la Paz, Alethea R. de Guzman, Jan Michael C. Yap, Jo-Hannah S. Llames, Sheila Mae M. Araiza, Kris P. Punayan, Irish Coleen A. Asin, Candice Francheska B. Tambaoan, Asia Louisa U. Chong, Karol Sophia Agape R. Padilla, Rianna Patricia S. Cruz, El King D. Morado, Joshua Gregor A. Dizon, Razel Nikka M. Hao, Arianne A. Zamora, Devon Ray Pacial, Juan Antonio R. Magalang, Marissa Alejandria, Celia Carlos, Anna Ong-Lim, Edsel Maurice Salvaña, John Q. Wong, Jaime C. Montoya, Maria Rosario Singh-Vergeire and Cynthia P. Saloma |
| EPI_ISL_2171159 | Philippine Red Cross Logistics and Multipurpose Center | Philippine Genome Center | Francis A. Tablizo, Kenneth M. Kim, Carlo M. Lapid, Marc Jerrone R. Castro, Maria Sofia L. Yangzon, Elcid Aaron R. Pangilinan, Benedict A. Maralit, Marc Edsel C. Ayes, Eva Maria Cutiongco-de la Paz, Alethea R. de Guzman, Jan Michael C. Yap, Jo-Hannah S. Llames, Sheila Mae M. Araiza, Kris P. Punayan, Irish Coleen A. Asin, Candice Francheska B. Tambaoan, Asia Louisa U. Chong, Karol Sophia Agape R. Padilla, Rianna Patricia S. Cruz, El King D. Morado, Joshua Gregor A. Dizon, Razel Nikka M. Hao, Arianne A. Zamora, Devon Ray Pacial, Juan Antonio R. Magalang, Marissa Alejandria, Celia Carlos, Anna Ong-Lim, Edsel Maurice Salvaña, John Q. Wong, Jaime C. Montoya, Maria Rosario Singh-Vergeire and Cynthia P. Saloma |
| EPI_ISL_2171160 | Philippine Red Cross - Port Area | Philippine Genome Center | Francis A. Tablizo, Kenneth M. Kim, Carlo M. Lapid, Marc Jerrone R. Castro, Maria Sofia L. Yangzon, Elcid Aaron R. Pangilinan, Benedict A. Maralit, Marc Edsel C. Ayes, Eva Maria Cutiongco-de la Paz, Alethea R. de Guzman, Jan Michael C. Yap, Jo-Hannah S. Llames, Sheila Mae M. Araiza, Kris P. Punayan, Irish Coleen A. Asin, Candice Francheska B. Tambaoan, Asia Louisa U. Chong, Karol Sophia Agape R. Padilla, Rianna Patricia S. Cruz, El King D. Morado, Joshua Gregor A. Dizon, Razel Nikka M. Hao, Arianne A. Zamora, Devon Ray Pacial, Juan Antonio R. Magalang, Marissa Alejandria, Celia Carlos, Anna Ong-Lim, Edsel Maurice Salvaña, John Q. Wong, Jaime C. Montoya, Maria Rosario Singh-Vergeire and Cynthia P. Saloma |
| EPI_ISL_2171161 | Philippine Red Cross Logistics and Multipurpose Center | Philippine Genome Center | Francis A. Tablizo, Kenneth M. Kim, Carlo M. Lapid, Marc Jerrone R. Castro, Maria Sofia L. Yangzon, Elcid Aaron R. Pangilinan, Benedict A. Maralit, Marc Edsel C. Ayes, Eva Maria Cutiongco-de la Paz, Alethea R. de Guzman, Jan Michael C. Yap, Jo-Hannah S. Llames, Sheila Mae M. Araiza, Kris P. Punayan, Irish Coleen A. Asin, Candice Francheska B. Tambaoan, Asia Louisa U. Chong, Karol Sophia Agape R. Padilla, Rianna Patricia S. Cruz, El King D. Morado, Joshua Gregor A. Dizon, Razel Nikka M. Hao, Arianne A. Zamora, Devon Ray Pacial, Juan Antonio R. Magalang, Marissa Alejandria, Celia Carlos, Anna Ong-Lim, Edsel Maurice Salvaña, John Q. Wong, Jaime C. Montoya, Maria Rosario Singh-Vergeire and Cynthia P. Saloma |
| EPI_ISL_2171162 | Ospital ng ParaÃ±aque II | Philippine Genome Center | Francis A. Tablizo, Kenneth M. Kim, Carlo M. Lapid, Marc Jerrone R. Castro, Maria Sofia L. Yangzon, Elcid Aaron R. Pangilinan, Benedict A. Maralit, Marc Edsel C. Ayes, Eva Maria Cutiongco-de la Paz, Alethea R. de Guzman, Jan Michael C. Yap, Jo-Hannah S. Llames, Sheila Mae M. Araiza, Kris P. Punayan, Irish Coleen A. Asin, Candice Francheska B. Tambaoan, Asia Louisa U. Chong, Karol Sophia Agape R. Padilla, Rianna Patricia S. Cruz, El King D. Morado, Joshua Gregor A. Dizon, Razel Nikka M. Hao, Arianne A. Zamora, Devon Ray Pacial, Juan Antonio R. Magalang, Marissa Alejandria, Celia Carlos, Anna Ong-Lim, Edsel Maurice Salvaña, John Q. Wong, Jaime C. Montoya, Maria Rosario Singh-Vergeire and Cynthia P. Saloma |

| | | | |
|---|---|---|---|
| EPI_ISL_2171163 | Cotabato Regional and Medical Center | Philippine Genome Center | Francis A. Tablizo, Kenneth M. Kim, Carlo M. Lapid, Marc Jerrone R. Castro, Maria Sofia L. Yangzon, Elcid Aaron R. Pangilinan, Benedict A. Maralit, Marc Edsel C. Ayes, Eva Maria Cutiongco-de la Paz, Alethea R. de Guzman, Jan Michael C. Yap, Jo-Hannah S. Llames, Sheila Mae M. Araiza, Kris P. Punayan, Irish Coleen A. Asin, Candice Francheska B. Tambaoan, Asia Louisa U. Chong, Karol Sophia Agape R. Padilla, Rianna Patricia S. Cruz, El King D. Morado, Joshua Gregor A. Dizon, Razel Nikka M. Hao, Arianne A. Zamora, Devon Ray Pacial, Juan Antonio R. Magalang, Marissa Alejandria, Celia Carlos, Anna Ong-Lim, Edsel Maurice Salvaña, John Q. Wong, Jaime C. Montoya, Maria Rosario Singh-Vergeire and Cynthia P. Saloma |
| EPI_ISL_2171164 | Philippine Red Cross - Port Area | Philippine Genome Center | Francis A. Tablizo, Kenneth M. Kim, Carlo M. Lapid, Marc Jerrone R. Castro, Maria Sofia L. Yangzon, Elcid Aaron R. Pangilinan, Benedict A. Maralit, Marc Edsel C. Ayes, Eva Maria Cutiongco-de la Paz, Alethea R. de Guzman, Jan Michael C. Yap, Jo-Hannah S. Llames, Sheila Mae M. Araiza, Kris P. Punayan, Irish Coleen A. Asin, Candice Francheska B. Tambaoan, Asia Louisa U. Chong, Karol Sophia Agape R. Padilla, Rianna Patricia S. Cruz, El King D. Morado, Joshua Gregor A. Dizon, Razel Nikka M. Hao, Arianne A. Zamora, Devon Ray Pacial, Juan Antonio R. Magalang, Marissa Alejandria, Celia Carlos, Anna Ong-Lim, Edsel Maurice Salvaña, John Q. Wong, Jaime C. Montoya, Maria Rosario Singh-Vergeire and Cynthia P. Saloma |
| EPI_ISL_2171165 | Pasig City Children's Hospital - Child's Hope | Philippine Genome Center | Francis A. Tablizo, Kenneth M. Kim, Carlo M. Lapid, Marc Jerrone R. Castro, Maria Sofia L. Yangzon, Elcid Aaron R. Pangilinan, Benedict A. Maralit, Marc Edsel C. Ayes, Eva Maria Cutiongco-de la Paz, Alethea R. de Guzman, Jan Michael C. Yap, Jo-Hannah S. Llames, Sheila Mae M. Araiza, Kris P. Punayan, Irish Coleen A. Asin, Candice Francheska B. Tambaoan, Asia Louisa U. Chong, Karol Sophia Agape R. Padilla, Rianna Patricia S. Cruz, El King D. Morado, Joshua Gregor A. Dizon, Razel Nikka M. Hao, Arianne A. Zamora, Devon Ray Pacial, Juan Antonio R. Magalang, Marissa Alejandria, Celia Carlos, Anna Ong-Lim, Edsel Maurice Salvaña, John Q. Wong, Jaime C. Montoya, Maria Rosario Singh-Vergeire and Cynthia P. Saloma |
| EPI_ISL_2171166 | Philippine Red Cross - Port Area | Philippine Genome Center | Francis A. Tablizo, Kenneth M. Kim, Carlo M. Lapid, Marc Jerrone R. Castro, Maria Sofia L. Yangzon, Elcid Aaron R. Pangilinan, Benedict A. Maralit, Marc Edsel C. Ayes, Eva Maria Cutiongco-de la Paz, Alethea R. de Guzman, Jan Michael C. Yap, Jo-Hannah S. Llames, Sheila Mae M. Araiza, Kris P. Punayan, Irish Coleen A. Asin, Candice Francheska B. Tambaoan, Asia Louisa U. Chong, Karol Sophia Agape R. Padilla, Rianna Patricia S. Cruz, El King D. Morado, Joshua Gregor A. Dizon, Razel Nikka M. Hao, Arianne A. Zamora, Devon Ray Pacial, Juan Antonio R. Magalang, Marissa Alejandria, Celia Carlos, Anna Ong-Lim, Edsel Maurice Salvaña, John Q. Wong, Jaime C. Montoya, Maria Rosario Singh-Vergeire and Cynthia P. Saloma |
| EPI_ISL_2171168 | Pasig City Children's Hospital - Child's Hope | Philippine Genome Center | Francis A. Tablizo, Kenneth M. Kim, Carlo M. Lapid, Marc Jerrone R. Castro, Maria Sofia L. Yangzon, Elcid Aaron R. Pangilinan, Benedict A. Maralit, Marc Edsel C. Ayes, Eva Maria Cutiongco-de la Paz, Alethea R. de Guzman, Jan Michael C. Yap, Jo-Hannah S. Llames, Sheila Mae M. Araiza, Kris P. Punayan, Irish Coleen A. Asin, Candice Francheska B. Tambaoan, Asia Louisa U. Chong, Karol Sophia Agape R. Padilla, Rianna Patricia S. Cruz, El King D. Morado, Joshua Gregor A. Dizon, Razel Nikka M. Hao, Arianne A. Zamora, Devon Ray Pacial, Juan Antonio R. Magalang, Marissa Alejandria, Celia Carlos, Anna Ong-Lim, Edsel Maurice Salvaña, John Q. Wong, Jaime C. Montoya, Maria Rosario Singh-Vergeire and Cynthia P. Saloma |
| EPI_ISL_2171170 | Ospital ng ParaÃ±aque II | Philippine Genome Center | Francis A. Tablizo, Kenneth M. Kim, Carlo M. Lapid, Marc Jerrone R. Castro, Maria Sofia L. Yangzon, Elcid Aaron R. Pangilinan, Benedict A. Maralit, Marc Edsel C. Ayes, Eva Maria Cutiongco-de la Paz, Alethea R. de Guzman, Jan Michael C. Yap, Jo-Hannah S. Llames, Sheila Mae M. Araiza, Kris P. Punayan, Irish Coleen A. Asin, Candice Francheska B. Tambaoan, Asia Louisa U. Chong, Karol Sophia Agape R. Padilla, Rianna Patricia S. Cruz, El King D. Morado, Joshua Gregor A. Dizon, Razel Nikka M. Hao, Arianne A. Zamora, Devon Ray Pacial, Juan Antonio R. Magalang, Marissa Alejandria, Celia Carlos, Anna Ong-Lim, Edsel Maurice Salvaña, John Q. Wong, Jaime C. Montoya, Maria Rosario Singh-Vergeire and Cynthia P. Saloma |
| EPI_ISL_2171171 | South Super Highway Molecular Diagnostic Laboratory | Philippine Genome Center | Francis A. Tablizo, Kenneth M. Kim, Carlo M. Lapid, Marc Jerrone R. Castro, Maria Sofia L. Yangzon, Elcid Aaron R. Pangilinan, Benedict A. Maralit, Marc Edsel C. Ayes, Eva Maria Cutiongco-de la Paz, Alethea R. de Guzman, Jan Michael C. Yap, Jo-Hannah S. Llames, Sheila Mae M. Araiza, Kris P. Punayan, Irish Coleen A. Asin, Candice Francheska B. Tambaoan, Asia Louisa U. Chong, Karol Sophia Agape R. Padilla, Rianna Patricia S. Cruz, El King D. Morado, Joshua Gregor A. Dizon, Razel Nikka M. Hao, Arianne A. Zamora, Devon Ray Pacial, Juan Antonio R. Magalang, Marissa Alejandria, Celia Carlos, Anna Ong-Lim, Edsel Maurice Salvaña, John Q. Wong, Jaime C. Montoya, Maria Rosario Singh-Vergeire and Cynthia P. Saloma |
| EPI_ISL_2171172, EPI_ISL_2171173 | The Lord's Grace Medical and Industrial Clinic | Philippine Genome Center | Francis A. Tablizo, Kenneth M. Kim, Carlo M. Lapid, Marc Jerrone R. Castro, Maria Sofia L. Yangzon, Elcid Aaron R. Pangilinan, Benedict A. Maralit, Marc Edsel C. Ayes, Eva Maria Cutiongco-de la Paz, Alethea R. de Guzman, Jan Michael C. Yap, Jo-Hannah S. Llames, Sheila Mae M. Araiza, Kris P. Punayan, Irish Coleen A. Asin, Candice Francheska B. Tambaoan, Asia Louisa U. Chong, Karol Sophia Agape R. Padilla, Rianna Patricia S. Cruz, El King D. Morado, Joshua Gregor A. Dizon, Razel Nikka M. Hao, Arianne A. Zamora, Devon Ray Pacial, Juan Antonio R. Magalang, Marissa Alejandria, Celia Carlos, Anna Ong-Lim, Edsel Maurice Salvaña, John Q. Wong, Jaime C. Montoya, Maria Rosario Singh-Vergeire and Cynthia P. Saloma |
| EPI_ISL_2171174 | Philippine Red Cross - Port Area | Philippine Genome Center | Francis A. Tablizo, Kenneth M. Kim, Carlo M. Lapid, Marc Jerrone R. Castro, Maria Sofia L. Yangzon, Elcid Aaron R. Pangilinan, Benedict A. Maralit, Marc Edsel C. Ayes, Eva Maria Cutiongco-de la Paz, Alethea R. de Guzman, Jan Michael C. Yap, Jo-Hannah S. Llames, Sheila Mae M. Araiza, Kris P. Punayan, Irish Coleen A. Asin, Candice Francheska B. Tambaoan, Asia Louisa U. Chong, Karol Sophia Agape R. Padilla, Rianna Patricia S. Cruz, El King D. Morado, Joshua Gregor A. Dizon, Razel Nikka M. Hao, Arianne A. Zamora, Devon Ray Pacial, Juan Antonio R. Magalang, Marissa Alejandria, Celia Carlos, Anna Ong-Lim, Edsel Maurice Salvaña, John Q. Wong, Jaime C. Montoya, Maria Rosario Singh-Vergeire and Cynthia P. Saloma |
| EPI_ISL_2171175, EPI_ISL_2171177 | UP-PGH Molecular Laboratory | Philippine Genome Center | Francis A. Tablizo, Kenneth M. Kim, Carlo M. Lapid, Marc Jerrone R. Castro, Maria Sofia L. Yangzon, Elcid Aaron R. Pangilinan, Benedict A. Maralit, Marc Edsel C. Ayes, Eva Maria Cutiongco-de la Paz, Alethea R. de Guzman, Jan Michael C. Yap, Jo-Hannah S. Llames, Sheila Mae M. Araiza, Kris P. Punayan, Irish Coleen A. Asin, Candice Francheska B. Tambaoan, Asia Louisa U. Chong, Karol Sophia Agape R. Padilla, Rianna Patricia S. Cruz, El King D. Morado, Joshua Gregor A. Dizon, Razel Nikka M. Hao, Arianne A. Zamora, Devon Ray Pacial, Juan Antonio R. Magalang, Marissa Alejandria, Celia Carlos, Anna Ong-Lim, Edsel Maurice Salvaña, John Q. Wong, Jaime C. Montoya, Maria Rosario Singh-Vergeire and Cynthia P. Saloma |
| EPI_ISL_2171178 | Philippine Red Cross - National Blood Center | Philippine Genome Center | Francis A. Tablizo, Kenneth M. Kim, Carlo M. Lapid, Marc Jerrone R. Castro, Maria Sofia L. Yangzon, Elcid Aaron R. Pangilinan, Benedict A. Maralit, Marc Edsel C. Ayes, Eva Maria Cutiongco-de la Paz, Alethea R. de Guzman, Jan Michael C. Yap, Jo-Hannah S. Llames, Sheila Mae M. Araiza, Kris P. Punayan, Irish Coleen A. Asin, Candice Francheska B. Tambaoan, Asia Louisa U. Chong, Karol Sophia Agape R. Padilla, Rianna Patricia S. Cruz, El King D. Morado, Joshua Gregor A. Dizon, Razel Nikka M. Hao, Arianne A. Zamora, Devon Ray Pacial, Juan Antonio R. Magalang, Marissa Alejandria, Celia Carlos, Anna Ong-Lim, Edsel Maurice Salvaña, John Q. Wong, Jaime C. Montoya, Maria Rosario Singh-Vergeire and Cynthia P. Saloma |
| EPI_ISL_2171179 | UP-PGH Molecular Laboratory | Philippine Genome Center | Francis A. Tablizo, Kenneth M. Kim, Carlo M. Lapid, Marc Jerrone R. Castro, Maria Sofia L. Yangzon, Elcid Aaron R. Pangilinan, Benedict A. Maralit, Marc Edsel C. Ayes, Eva Maria Cutiongco-de la Paz, Alethea R. de Guzman, Jan Michael C. Yap, Jo-Hannah S. Llames, Sheila Mae M. Araiza, Kris P. Punayan, Irish Coleen A. Asin, Candice Francheska B. Tambaoan, Asia Louisa U. Chong, Karol Sophia Agape R. Padilla, Rianna Patricia S. Cruz, El King D. Morado, Joshua Gregor A. Dizon, Razel Nikka M. Hao, Arianne A. Zamora, Devon Ray Pacial, Juan Antonio R. Magalang, Marissa Alejandria, Celia Carlos, Anna Ong-Lim, Edsel Maurice Salvaña, John Q. Wong, Jaime C. Montoya, Maria Rosario Singh-Vergeire and Cynthia P. Saloma |
| EPI_ISL_2171180 | Pasig City Children's Hospital - Child's Hope | Philippine Genome Center | Francis A. Tablizo, Kenneth M. Kim, Carlo M. Lapid, Marc Jerrone R. Castro, Maria Sofia L. Yangzon, Elcid Aaron R. Pangilinan, Benedict A. Maralit, Marc Edsel C. Ayes, Eva Maria Cutiongco-de la Paz, Alethea R. de Guzman, Jan Michael C. Yap, Jo-Hannah S. Llames, Sheila Mae M. Araiza, Kris P. Punayan, Irish Coleen A. Asin, Candice Francheska B. Tambaoan, Asia Louisa U. Chong, Karol Sophia Agape R. Padilla, Rianna Patricia S. Cruz, El King D. Morado, Joshua Gregor A. Dizon, Razel Nikka M. Hao, Arianne A. Zamora, Devon Ray Pacial, Juan Antonio R. Magalang, Marissa Alejandria, Celia Carlos, Anna Ong-Lim, Edsel Maurice Salvaña, John Q. Wong, Jaime C. Montoya, Maria Rosario Singh-Vergeire and Cynthia P. Saloma |
| EPI_ISL_2171181 | The Lord's Grace Medical and Industrial Clinic | Philippine Genome Center | Francis A. Tablizo, Kenneth M. Kim, Carlo M. Lapid, Marc Jerrone R. Castro, Maria Sofia L. Yangzon, Elcid Aaron R. Pangilinan, Benedict A. Maralit, Marc Edsel C. Ayes, Eva Maria Cutiongco-de la Paz, Alethea R. de Guzman, Jan Michael C. Yap, Jo-Hannah S. Llames, Sheila Mae M. Araiza, Kris P. Punayan, Irish Coleen A. Asin, Candice Francheska B. Tambaoan, Asia Louisa U. Chong, Karol Sophia Agape R. Padilla, Rianna Patricia S. Cruz, El King D. Morado, Joshua Gregor A. Dizon, Razel Nikka M. Hao, Arianne A. Zamora, Devon Ray Pacial, Juan Antonio R. Magalang, Marissa Alejandria, Celia Carlos, Anna Ong-Lim, Edsel Maurice Salvaña, John Q. Wong, Jaime C. Montoya, Maria Rosario Singh-Vergeire and Cynthia P. Saloma |
| EPI_ISL_2171183 | Ospital ng ParaÃ±aque II | Philippine Genome Center | Francis A. Tablizo, Kenneth M. Kim, Carlo M. Lapid, Marc Jerrone R. Castro, Maria Sofia L. Yangzon, Elcid Aaron R. Pangilinan, Benedict A. Maralit, Marc Edsel C. Ayes, Eva Maria Cutiongco-de la Paz, Alethea R. de Guzman, Jan Michael C. Yap, Jo-Hannah S. Llames, Sheila Mae M. Araiza, Kris P. Punayan, Irish Coleen A. Asin, Candice Francheska B. Tambaoan, Asia Louisa U. Chong, Karol Sophia Agape R. Padilla, Rianna Patricia S. Cruz, El King D. Morado, Joshua Gregor A. Dizon, Razel Nikka M. Hao, Arianne A. Zamora, Devon Ray Pacial, Juan Antonio R. Magalang, Marissa Alejandria, Celia Carlos, Anna Ong-Lim, Edsel Maurice Salvaña, John Q. Wong, Jaime C. Montoya, Maria Rosario Singh-Vergeire and Cynthia P. Saloma |
| EPI_ISL_2171185 | The Lord's Grace Medical and Industrial Clinic | Philippine Genome Center | Francis A. Tablizo, Kenneth M. Kim, Carlo M. Lapid, Marc Jerrone R. Castro, Maria Sofia L. Yangzon, Elcid Aaron R. Pangilinan, Benedict A. Maralit, Marc Edsel C. Ayes, Eva Maria Cutiongco-de la Paz, Alethea R. de Guzman, Jan Michael C. Yap, Jo-Hannah S. Llames, Sheila Mae M. Araiza, Kris P. Punayan, Irish Coleen A. Asin, Candice Francheska B. Tambaoan, Asia Louisa U. Chong, Karol Sophia Agape R. Padilla, Rianna Patricia S. Cruz, El King D. Morado, Joshua Gregor A. Dizon, Razel Nikka M. Hao, Arianne A. Zamora, Devon Ray Pacial, Juan Antonio R. Magalang, Marissa Alejandria, Celia Carlos, Anna Ong-Lim, Edsel Maurice Salvaña, John Q. Wong, Jaime C. Montoya, Maria Rosario Singh-Vergeire and Cynthia P. Saloma |

| | | | |
|---|---|---|---|
| EPI_ISL_2171186 | Philippine Red Cross - National Blood Center | Philippine Genome Center | Francis A. Tablizo, Kenneth M. Kim, Carlo M. Lapid, Marc Jerrone R. Castro, Maria Sofia L. Yangzon, Elcid Aaron R. Pangilinan, Benedict A. Maralit, Marc Edsel C. Ayes, Eva Maria Cutiongco-de la Paz, Alethea R. de Guzman, Jan Michael C. Yap, Jo-Hannah S. Llames, Sheila Mae M. Araiza, Kris P. Punayan, Irish Coleen A. Asin, Candice Francheska B. Tambaoan, Asia Louisa U. Chong, Karol Sophia Agape R. Padilla, Rianna Patricia S. Cruz, El King D. Morado, Joshua Gregor A. Dizon, Razel Nikka M. Hao, Arianne A. Zamora, Devon Ray Pacial, Juan Antonio R. Magalang, Marissa Alejandria, Celia Carlos, Anna Ong-Lim, Edsel Maurice Salvaña, John Q. Wong, Jaime C. Montoya, Maria Rosario Singh-Vergeire and Cynthia P. Saloma |
| EPI_ISL_2171187, EPI_ISL_2171188 | The Lord's Grace Medical and Industrial Clinic | Philippine Genome Center | Francis A. Tablizo, Kenneth M. Kim, Carlo M. Lapid, Marc Jerrone R. Castro, Maria Sofia L. Yangzon, Elcid Aaron R. Pangilinan, Benedict A. Maralit, Marc Edsel C. Ayes, Eva Maria Cutiongco-de la Paz, Alethea R. de Guzman, Jan Michael C. Yap, Jo-Hannah S. Llames, Sheila Mae M. Araiza, Kris P. Punayan, Irish Coleen A. Asin, Candice Francheska B. Tambaoan, Asia Louisa U. Chong, Karol Sophia Agape R. Padilla, Rianna Patricia S. Cruz, El King D. Morado, Joshua Gregor A. Dizon, Razel Nikka M. Hao, Arianne A. Zamora, Devon Ray Pacial, Juan Antonio R. Magalang, Marissa Alejandria, Celia Carlos, Anna Ong-Lim, Edsel Maurice Salvaña, John Q. Wong, Jaime C. Montoya, Maria Rosario Singh-Vergeire and Cynthia P. Saloma |
| EPI_ISL_2171189 | Philippine Red Cross - Port Area | Philippine Genome Center | Francis A. Tablizo, Kenneth M. Kim, Carlo M. Lapid, Marc Jerrone R. Castro, Maria Sofia L. Yangzon, Elcid Aaron R. Pangilinan, Benedict A. Maralit, Marc Edsel C. Ayes, Eva Maria Cutiongco-de la Paz, Alethea R. de Guzman, Jan Michael C. Yap, Jo-Hannah S. Llames, Sheila Mae M. Araiza, Kris P. Punayan, Irish Coleen A. Asin, Candice Francheska B. Tambaoan, Asia Louisa U. Chong, Karol Sophia Agape R. Padilla, Rianna Patricia S. Cruz, El King D. Morado, Joshua Gregor A. Dizon, Razel Nikka M. Hao, Arianne A. Zamora, Devon Ray Pacial, Juan Antonio R. Magalang, Marissa Alejandria, Celia Carlos, Anna Ong-Lim, Edsel Maurice Salvaña, John Q. Wong, Jaime C. Montoya, Maria Rosario Singh-Vergeire and Cynthia P. Saloma |
| EPI_ISL_2171190 | Safeguard DNA Diagnostic, Inc. | Philippine Genome Center | Francis A. Tablizo, Kenneth M. Kim, Carlo M. Lapid, Marc Jerrone R. Castro, Maria Sofia L. Yangzon, Elcid Aaron R. Pangilinan, Benedict A. Maralit, Marc Edsel C. Ayes, Eva Maria Cutiongco-de la Paz, Alethea R. de Guzman, Jan Michael C. Yap, Jo-Hannah S. Llames, Sheila Mae M. Araiza, Kris P. Punayan, Irish Coleen A. Asin, Candice Francheska B. Tambaoan, Asia Louisa U. Chong, Karol Sophia Agape R. Padilla, Rianna Patricia S. Cruz, El King D. Morado, Joshua Gregor A. Dizon, Razel Nikka M. Hao, Arianne A. Zamora, Devon Ray Pacial, Juan Antonio R. Magalang, Marissa Alejandria, Celia Carlos, Anna Ong-Lim, Edsel Maurice Salvaña, John Q. Wong, Jaime C. Montoya, Maria Rosario Singh-Vergeire and Cynthia P. Saloma |
| EPI_ISL_2171191, EPI_ISL_2171192 | Philippine Red Cross - Port Area | Philippine Genome Center | Francis A. Tablizo, Kenneth M. Kim, Carlo M. Lapid, Marc Jerrone R. Castro, Maria Sofia L. Yangzon, Elcid Aaron R. Pangilinan, Benedict A. Maralit, Marc Edsel C. Ayes, Eva Maria Cutiongco-de la Paz, Alethea R. de Guzman, Jan Michael C. Yap, Jo-Hannah S. Llames, Sheila Mae M. Araiza, Kris P. Punayan, Irish Coleen A. Asin, Candice Francheska B. Tambaoan, Asia Louisa U. Chong, Karol Sophia Agape R. Padilla, Rianna Patricia S. Cruz, El King D. Morado, Joshua Gregor A. Dizon, Razel Nikka M. Hao, Arianne A. Zamora, Devon Ray Pacial, Juan Antonio R. Magalang, Marissa Alejandria, Celia Carlos, Anna Ong-Lim, Edsel Maurice Salvaña, John Q. Wong, Jaime C. Montoya, Maria Rosario Singh-Vergeire and Cynthia P. Saloma |
| EPI_ISL_2171193 | The Lord's Grace Medical and Industrial Clinic | Philippine Genome Center | Francis A. Tablizo, Kenneth M. Kim, Carlo M. Lapid, Marc Jerrone R. Castro, Maria Sofia L. Yangzon, Elcid Aaron R. Pangilinan, Benedict A. Maralit, Marc Edsel C. Ayes, Eva Maria Cutiongco-de la Paz, Alethea R. de Guzman, Jan Michael C. Yap, Jo-Hannah S. Llames, Sheila Mae M. Araiza, Kris P. Punayan, Irish Coleen A. Asin, Candice Francheska B. Tambaoan, Asia Louisa U. Chong, Karol Sophia Agape R. Padilla, Rianna Patricia S. Cruz, El King D. Morado, Joshua Gregor A. Dizon, Razel Nikka M. Hao, Arianne A. Zamora, Devon Ray Pacial, Juan Antonio R. Magalang, Marissa Alejandria, Celia Carlos, Anna Ong-Lim, Edsel Maurice Salvaña, John Q. Wong, Jaime C. Montoya, Maria Rosario Singh-Vergeire and Cynthia P. Saloma |
| EPI_ISL_2171195 | Philippine Red Cross - Port Area | Philippine Genome Center | Francis A. Tablizo, Kenneth M. Kim, Carlo M. Lapid, Marc Jerrone R. Castro, Maria Sofia L. Yangzon, Elcid Aaron R. Pangilinan, Benedict A. Maralit, Marc Edsel C. Ayes, Eva Maria Cutiongco-de la Paz, Alethea R. de Guzman, Jan Michael C. Yap, Jo-Hannah S. Llames, Sheila Mae M. Araiza, Kris P. Punayan, Irish Coleen A. Asin, Candice Francheska B. Tambaoan, Asia Louisa U. Chong, Karol Sophia Agape R. Padilla, Rianna Patricia S. Cruz, El King D. Morado, Joshua Gregor A. Dizon, Razel Nikka M. Hao, Arianne A. Zamora, Devon Ray Pacial, Juan Antonio R. Magalang, Marissa Alejandria, Celia Carlos, Anna Ong-Lim, Edsel Maurice Salvaña, John Q. Wong, Jaime C. Montoya, Maria Rosario Singh-Vergeire and Cynthia P. Saloma |
| EPI_ISL_2171196 | Bulacan Medical Center | Philippine Genome Center | Francis A. Tablizo, Kenneth M. Kim, Carlo M. Lapid, Marc Jerrone R. Castro, Maria Sofia L. Yangzon, Elcid Aaron R. Pangilinan, Benedict A. Maralit, Marc Edsel C. Ayes, Eva Maria Cutiongco-de la Paz, Alethea R. de Guzman, Jan Michael C. Yap, Jo-Hannah S. Llames, Sheila Mae M. Araiza, Kris P. Punayan, Irish Coleen A. Asin, Candice Francheska B. Tambaoan, Asia Louisa U. Chong, Karol Sophia Agape R. Padilla, Rianna Patricia S. Cruz, El King D. Morado, Joshua Gregor A. Dizon, Razel Nikka M. Hao, Arianne A. Zamora, Devon Ray Pacial, Juan Antonio R. Magalang, Marissa Alejandria, Celia Carlos, Anna Ong-Lim, Edsel Maurice Salvaña, John Q. Wong, Jaime C. Montoya, Maria Rosario Singh-Vergeire and Cynthia P. Saloma |
| EPI_ISL_2171198 | Philippine Red Cross - Clark Molecular Laboratory | Philippine Genome Center | Francis A. Tablizo, Kenneth M. Kim, Carlo M. Lapid, Marc Jerrone R. Castro, Maria Sofia L. Yangzon, Elcid Aaron R. Pangilinan, Benedict A. Maralit, Marc Edsel C. Ayes, Eva Maria Cutiongco-de la Paz, Alethea R. de Guzman, Jan Michael C. Yap, Jo-Hannah S. Llames, Sheila Mae M. Araiza, Kris P. Punayan, Irish Coleen A. Asin, Candice Francheska B. Tambaoan, Asia Louisa U. Chong, Karol Sophia Agape R. Padilla, Rianna Patricia S. Cruz, El King D. Morado, Joshua Gregor A. Dizon, Razel Nikka M. Hao, Arianne A. Zamora, Devon Ray Pacial, Juan Antonio R. Magalang, Marissa Alejandria, Celia Carlos, Anna Ong-Lim, Edsel Maurice Salvaña, John Q. Wong, Jaime C. Montoya, Maria Rosario Singh-Vergeire and Cynthia P. Saloma |
| EPI_ISL_2171199 | UP-PGH Molecular Laboratory | Philippine Genome Center | Francis A. Tablizo, Kenneth M. Kim, Carlo M. Lapid, Marc Jerrone R. Castro, Maria Sofia L. Yangzon, Elcid Aaron R. Pangilinan, Benedict A. Maralit, Marc Edsel C. Ayes, Eva Maria Cutiongco-de la Paz, Alethea R. de Guzman, Jan Michael C. Yap, Jo-Hannah S. Llames, Sheila Mae M. Araiza, Kris P. Punayan, Irish Coleen A. Asin, Candice Francheska B. Tambaoan, Asia Louisa U. Chong, Karol Sophia Agape R. Padilla, Rianna Patricia S. Cruz, El King D. Morado, Joshua Gregor A. Dizon, Razel Nikka M. Hao, Arianne A. Zamora, Devon Ray Pacial, Juan Antonio R. Magalang, Marissa Alejandria, Celia Carlos, Anna Ong-Lim, Edsel Maurice Salvaña, John Q. Wong, Jaime C. Montoya, Maria Rosario Singh-Vergeire and Cynthia P. Saloma |
| EPI_ISL_2171200 | Philippine Red Cross - Port Area | Philippine Genome Center | Francis A. Tablizo, Kenneth M. Kim, Carlo M. Lapid, Marc Jerrone R. Castro, Maria Sofia L. Yangzon, Elcid Aaron R. Pangilinan, Benedict A. Maralit, Marc Edsel C. Ayes, Eva Maria Cutiongco-de la Paz, Alethea R. de Guzman, Jan Michael C. Yap, Jo-Hannah S. Llames, Sheila Mae M. Araiza, Kris P. Punayan, Irish Coleen A. Asin, Candice Francheska B. Tambaoan, Asia Louisa U. Chong, Karol Sophia Agape R. Padilla, Rianna Patricia S. Cruz, El King D. Morado, Joshua Gregor A. Dizon, Razel Nikka M. Hao, Arianne A. Zamora, Devon Ray Pacial, Juan Antonio R. Magalang, Marissa Alejandria, Celia Carlos, Anna Ong-Lim, Edsel Maurice Salvaña, John Q. Wong, Jaime C. Montoya, Maria Rosario Singh-Vergeire and Cynthia P. Saloma |
| EPI_ISL_2171201 | Philippine Red Cross - National Blood Center | Philippine Genome Center | Francis A. Tablizo, Kenneth M. Kim, Carlo M. Lapid, Marc Jerrone R. Castro, Maria Sofia L. Yangzon, Elcid Aaron R. Pangilinan, Benedict A. Maralit, Marc Edsel C. Ayes, Eva Maria Cutiongco-de la Paz, Alethea R. de Guzman, Jan Michael C. Yap, Jo-Hannah S. Llames, Sheila Mae M. Araiza, Kris P. Punayan, Irish Coleen A. Asin, Candice Francheska B. Tambaoan, Asia Louisa U. Chong, Karol Sophia Agape R. Padilla, Rianna Patricia S. Cruz, El King D. Morado, Joshua Gregor A. Dizon, Razel Nikka M. Hao, Arianne A. Zamora, Devon Ray Pacial, Juan Antonio R. Magalang, Marissa Alejandria, Celia Carlos, Anna Ong-Lim, Edsel Maurice Salvaña, John Q. Wong, Jaime C. Montoya, Maria Rosario Singh-Vergeire and Cynthia P. Saloma |
| EPI_ISL_2171203 | Philippine Red Cross Logistics and Multipurpose Center | Philippine Genome Center | Francis A. Tablizo, Kenneth M. Kim, Carlo M. Lapid, Marc Jerrone R. Castro, Maria Sofia L. Yangzon, Elcid Aaron R. Pangilinan, Benedict A. Maralit, Marc Edsel C. Ayes, Eva Maria Cutiongco-de la Paz, Alethea R. de Guzman, Jan Michael C. Yap, Jo-Hannah S. Llames, Sheila Mae M. Araiza, Kris P. Punayan, Irish Coleen A. Asin, Candice Francheska B. Tambaoan, Asia Louisa U. Chong, Karol Sophia Agape R. Padilla, Rianna Patricia S. Cruz, El King D. Morado, Joshua Gregor A. Dizon, Razel Nikka M. Hao, Arianne A. Zamora, Devon Ray Pacial, Juan Antonio R. Magalang, Marissa Alejandria, Celia Carlos, Anna Ong-Lim, Edsel Maurice Salvaña, John Q. Wong, Jaime C. Montoya, Maria Rosario Singh-Vergeire and Cynthia P. Saloma |
| EPI_ISL_2171205 | Philippine Red Cross - Port Area | Philippine Genome Center | Francis A. Tablizo, Kenneth M. Kim, Carlo M. Lapid, Marc Jerrone R. Castro, Maria Sofia L. Yangzon, Elcid Aaron R. Pangilinan, Benedict A. Maralit, Marc Edsel C. Ayes, Eva Maria Cutiongco-de la Paz, Alethea R. de Guzman, Jan Michael C. Yap, Jo-Hannah S. Llames, Sheila Mae M. Araiza, Kris P. Punayan, Irish Coleen A. Asin, Candice Francheska B. Tambaoan, Asia Louisa U. Chong, Karol Sophia Agape R. Padilla, Rianna Patricia S. Cruz, El King D. Morado, Joshua Gregor A. Dizon, Razel Nikka M. Hao, Arianne A. Zamora, Devon Ray Pacial, Juan Antonio R. Magalang, Marissa Alejandria, Celia Carlos, Anna Ong-Lim, Edsel Maurice Salvaña, John Q. Wong, Jaime C. Montoya, Maria Rosario Singh-Vergeire and Cynthia P. Saloma |
| EPI_ISL_2171206 | South Super Highway Molecular Diagnostic Laboratory | Philippine Genome Center | Francis A. Tablizo, Kenneth M. Kim, Carlo M. Lapid, Marc Jerrone R. Castro, Maria Sofia L. Yangzon, Elcid Aaron R. Pangilinan, Benedict A. Maralit, Marc Edsel C. Ayes, Eva Maria Cutiongco-de la Paz, Alethea R. de Guzman, Jan Michael C. Yap, Jo-Hannah S. Llames, Sheila Mae M. Araiza, Kris P. Punayan, Irish Coleen A. Asin, Candice Francheska B. Tambaoan, Asia Louisa U. Chong, Karol Sophia Agape R. Padilla, Rianna Patricia S. Cruz, El King D. Morado, Joshua Gregor A. Dizon, Razel Nikka M. Hao, Arianne A. Zamora, Devon Ray Pacial, Juan Antonio R. Magalang, Marissa Alejandria, Celia Carlos, Anna Ong-Lim, Edsel Maurice Salvaña, John Q. Wong, Jaime C. Montoya, Maria Rosario Singh-Vergeire and Cynthia P. Saloma |
| EPI_ISL_2171207 | Philippine Red Cross - Port Area | Philippine Genome Center | Francis A. Tablizo, Kenneth M. Kim, Carlo M. Lapid, Marc Jerrone R. Castro, Maria Sofia L. Yangzon, Elcid Aaron R. Pangilinan, Benedict A. Maralit, Marc Edsel C. Ayes, Eva Maria Cutiongco-de la Paz, Alethea R. de Guzman, Jan Michael C. Yap, Jo-Hannah S. Llames, Sheila Mae M. Araiza, Kris P. Punayan, Irish Coleen A. Asin, Candice Francheska B. Tambaoan, Asia Louisa U. Chong, Karol Sophia Agape R. Padilla, Rianna Patricia S. Cruz, El King D. Morado, Joshua Gregor A. Dizon, Razel Nikka M. Hao, Arianne A. Zamora, Devon Ray Pacial, Juan Antonio R. Magalang, Marissa Alejandria, Celia Carlos, Anna Ong-Lim, Edsel Maurice Salvaña, John Q. Wong, Jaime C. Montoya, Maria Rosario Singh-Vergeire and Cynthia P. Saloma |

| | | | |
|---|---|---|---|
| EPI_ISL_2171210 | The Lord's Grace Medical and Industrial Clinic | Philippine Genome Center | Francis A. Tablizo, Kenneth M. Kim, Carlo M. Lapid, Marc Jerrone R. Castro, Maria Sofia L. Yangzon, Elcid Aaron R. Pangilinan, Benedict A. Maralit, Marc Edsel C. Ayes, Eva Maria Cutiongco-de la Paz, Alethea R. de Guzman, Jan Michael C. Yap, Jo-Hannah S. Llames, Sheila Mae M. Araiza, Kris P. Punayan, Irish Coleen A. Asin, Candice Francheska B. Tambaoan, Asia Louisa U. Chong, Karol Sophia Agape R. Padilla, Rianna Patricia S. Cruz, El King D. Morado, Joshua Gregor A. Dizon, Razel Nikka M. Hao, Arianne A. Zamora, Devon Ray Pacial, Juan Antonio R. Magalang, Marissa Alejandria, Celia Carlos, Anna Ong-Lim, Edsel Maurice Salvaña, John Q. Wong, Jaime C. Montoya, Maria Rosario Singh-Vergeire and Cynthia P. Saloma |
| EPI_ISL_2171211 | Philippine Red Cross - National Blood Center | Philippine Genome Center | Francis A. Tablizo, Kenneth M. Kim, Carlo M. Lapid, Marc Jerrone R. Castro, Maria Sofia L. Yangzon, Elcid Aaron R. Pangilinan, Benedict A. Maralit, Marc Edsel C. Ayes, Eva Maria Cutiongco-de la Paz, Alethea R. de Guzman, Jan Michael C. Yap, Jo-Hannah S. Llames, Sheila Mae M. Araiza, Kris P. Punayan, Irish Coleen A. Asin, Candice Francheska B. Tambaoan, Asia Louisa U. Chong, Karol Sophia Agape R. Padilla, Rianna Patricia S. Cruz, El King D. Morado, Joshua Gregor A. Dizon, Razel Nikka M. Hao, Arianne A. Zamora, Devon Ray Pacial, Juan Antonio R. Magalang, Marissa Alejandria, Celia Carlos, Anna Ong-Lim, Edsel Maurice Salvaña, John Q. Wong, Jaime C. Montoya, Maria Rosario Singh-Vergeire and Cynthia P. Saloma |
| EPI_ISL_2171213 | The Lord's Grace Medical and Industrial Clinic | Philippine Genome Center | Francis A. Tablizo, Kenneth M. Kim, Carlo M. Lapid, Marc Jerrone R. Castro, Maria Sofia L. Yangzon, Elcid Aaron R. Pangilinan, Benedict A. Maralit, Marc Edsel C. Ayes, Eva Maria Cutiongco-de la Paz, Alethea R. de Guzman, Jan Michael C. Yap, Jo-Hannah S. Llames, Sheila Mae M. Araiza, Kris P. Punayan, Irish Coleen A. Asin, Candice Francheska B. Tambaoan, Asia Louisa U. Chong, Karol Sophia Agape R. Padilla, Rianna Patricia S. Cruz, El King D. Morado, Joshua Gregor A. Dizon, Razel Nikka M. Hao, Arianne A. Zamora, Devon Ray Pacial, Juan Antonio R. Magalang, Marissa Alejandria, Celia Carlos, Anna Ong-Lim, Edsel Maurice Salvaña, John Q. Wong, Jaime C. Montoya, Maria Rosario Singh-Vergeire and Cynthia P. Saloma |
| EPI_ISL_2171214 | Philippine Red Cross - Port Area | Philippine Genome Center | Francis A. Tablizo, Kenneth M. Kim, Carlo M. Lapid, Marc Jerrone R. Castro, Maria Sofia L. Yangzon, Elcid Aaron R. Pangilinan, Benedict A. Maralit, Marc Edsel C. Ayes, Eva Maria Cutiongco-de la Paz, Alethea R. de Guzman, Jan Michael C. Yap, Jo-Hannah S. Llames, Sheila Mae M. Araiza, Kris P. Punayan, Irish Coleen A. Asin, Candice Francheska B. Tambaoan, Asia Louisa U. Chong, Karol Sophia Agape R. Padilla, Rianna Patricia S. Cruz, El King D. Morado, Joshua Gregor A. Dizon, Razel Nikka M. Hao, Arianne A. Zamora, Devon Ray Pacial, Juan Antonio R. Magalang, Marissa Alejandria, Celia Carlos, Anna Ong-Lim, Edsel Maurice Salvaña, John Q. Wong, Jaime C. Montoya, Maria Rosario Singh-Vergeire and Cynthia P. Saloma |
| EPI_ISL_2171215 | Research Institute for Tropical Medicine, Inc. (RITM) | Philippine Genome Center | Francis A. Tablizo, Kenneth M. Kim, Carlo M. Lapid, Marc Jerrone R. Castro, Maria Sofia L. Yangzon, Elcid Aaron R. Pangilinan, Benedict A. Maralit, Marc Edsel C. Ayes, Eva Maria Cutiongco-de la Paz, Alethea R. de Guzman, Jan Michael C. Yap, Jo-Hannah S. Llames, Sheila Mae M. Araiza, Kris P. Punayan, Irish Coleen A. Asin, Candice Francheska B. Tambaoan, Asia Louisa U. Chong, Karol Sophia Agape R. Padilla, Rianna Patricia S. Cruz, El King D. Morado, Joshua Gregor A. Dizon, Razel Nikka M. Hao, Arianne A. Zamora, Devon Ray Pacial, Juan Antonio R. Magalang, Marissa Alejandria, Celia Carlos, Anna Ong-Lim, Edsel Maurice Salvaña, John Q. Wong, Jaime C. Montoya, Maria Rosario Singh-Vergeire and Cynthia P. Saloma |
| EPI_ISL_2171216 | The Lord's Grace Medical and Industrial Clinic | Philippine Genome Center | Francis A. Tablizo, Kenneth M. Kim, Carlo M. Lapid, Marc Jerrone R. Castro, Maria Sofia L. Yangzon, Elcid Aaron R. Pangilinan, Benedict A. Maralit, Marc Edsel C. Ayes, Eva Maria Cutiongco-de la Paz, Alethea R. de Guzman, Jan Michael C. Yap, Jo-Hannah S. Llames, Sheila Mae M. Araiza, Kris P. Punayan, Irish Coleen A. Asin, Candice Francheska B. Tambaoan, Asia Louisa U. Chong, Karol Sophia Agape R. Padilla, Rianna Patricia S. Cruz, El King D. Morado, Joshua Gregor A. Dizon, Razel Nikka M. Hao, Arianne A. Zamora, Devon Ray Pacial, Juan Antonio R. Magalang, Marissa Alejandria, Celia Carlos, Anna Ong-Lim, Edsel Maurice Salvaña, John Q. Wong, Jaime C. Montoya, Maria Rosario Singh-Vergeire and Cynthia P. Saloma |
| EPI_ISL_2171217 | Philippine Red Cross - Port Area | Philippine Genome Center | Francis A. Tablizo, Kenneth M. Kim, Carlo M. Lapid, Marc Jerrone R. Castro, Maria Sofia L. Yangzon, Elcid Aaron R. Pangilinan, Benedict A. Maralit, Marc Edsel C. Ayes, Eva Maria Cutiongco-de la Paz, Alethea R. de Guzman, Jan Michael C. Yap, Jo-Hannah S. Llames, Sheila Mae M. Araiza, Kris P. Punayan, Irish Coleen A. Asin, Candice Francheska B. Tambaoan, Asia Louisa U. Chong, Karol Sophia Agape R. Padilla, Rianna Patricia S. Cruz, El King D. Morado, Joshua Gregor A. Dizon, Razel Nikka M. Hao, Arianne A. Zamora, Devon Ray Pacial, Juan Antonio R. Magalang, Marissa Alejandria, Celia Carlos, Anna Ong-Lim, Edsel Maurice Salvaña, John Q. Wong, Jaime C. Montoya, Maria Rosario Singh-Vergeire and Cynthia P. Saloma |
| EPI_ISL_2171218 | South Super Highway Molecular Diagnostic Laboratory | Philippine Genome Center | Francis A. Tablizo, Kenneth M. Kim, Carlo M. Lapid, Marc Jerrone R. Castro, Maria Sofia L. Yangzon, Elcid Aaron R. Pangilinan, Benedict A. Maralit, Marc Edsel C. Ayes, Eva Maria Cutiongco-de la Paz, Alethea R. de Guzman, Jan Michael C. Yap, Jo-Hannah S. Llames, Sheila Mae M. Araiza, Kris P. Punayan, Irish Coleen A. Asin, Candice Francheska B. Tambaoan, Asia Louisa U. Chong, Karol Sophia Agape R. Padilla, Rianna Patricia S. Cruz, El King D. Morado, Joshua Gregor A. Dizon, Razel Nikka M. Hao, Arianne A. Zamora, Devon Ray Pacial, Juan Antonio R. Magalang, Marissa Alejandria, Celia Carlos, Anna Ong-Lim, Edsel Maurice Salvaña, John Q. Wong, Jaime C. Montoya, Maria Rosario Singh-Vergeire and Cynthia P. Saloma |
| EPI_ISL_2171222, EPI_ISL_2171223 | Philippine Red Cross - Port Area | Philippine Genome Center | Francis A. Tablizo, Kenneth M. Kim, Carlo M. Lapid, Marc Jerrone R. Castro, Maria Sofia L. Yangzon, Elcid Aaron R. Pangilinan, Benedict A. Maralit, Marc Edsel C. Ayes, Eva Maria Cutiongco-de la Paz, Alethea R. de Guzman, Jan Michael C. Yap, Jo-Hannah S. Llames, Sheila Mae M. Araiza, Kris P. Punayan, Irish Coleen A. Asin, Candice Francheska B. Tambaoan, Asia Louisa U. Chong, Karol Sophia Agape R. Padilla, Rianna Patricia S. Cruz, El King D. Morado, Joshua Gregor A. Dizon, Razel Nikka M. Hao, Arianne A. Zamora, Devon Ray Pacial, Juan Antonio R. Magalang, Marissa Alejandria, Celia Carlos, Anna Ong-Lim, Edsel Maurice Salvaña, John Q. Wong, Jaime C. Montoya, Maria Rosario Singh-Vergeire and Cynthia P. Saloma |
| EPI_ISL_2171224 | Ospital ng Imus | Philippine Genome Center | Francis A. Tablizo, Kenneth M. Kim, Carlo M. Lapid, Marc Jerrone R. Castro, Maria Sofia L. Yangzon, Elcid Aaron R. Pangilinan, Benedict A. Maralit, Marc Edsel C. Ayes, Eva Maria Cutiongco-de la Paz, Alethea R. de Guzman, Jan Michael C. Yap, Jo-Hannah S. Llames, Sheila Mae M. Araiza, Kris P. Punayan, Irish Coleen A. Asin, Candice Francheska B. Tambaoan, Asia Louisa U. Chong, Karol Sophia Agape R. Padilla, Rianna Patricia S. Cruz, El King D. Morado, Joshua Gregor A. Dizon, Razel Nikka M. Hao, Arianne A. Zamora, Devon Ray Pacial, Juan Antonio R. Magalang, Marissa Alejandria, Celia Carlos, Anna Ong-Lim, Edsel Maurice Salvaña, John Q. Wong, Jaime C. Montoya, Maria Rosario Singh-Vergeire and Cynthia P. Saloma |
| EPI_ISL_2171225, EPI_ISL_2171226 | Philippine Red Cross - Port Area | Philippine Genome Center | Francis A. Tablizo, Kenneth M. Kim, Carlo M. Lapid, Marc Jerrone R. Castro, Maria Sofia L. Yangzon, Elcid Aaron R. Pangilinan, Benedict A. Maralit, Marc Edsel C. Ayes, Eva Maria Cutiongco-de la Paz, Alethea R. de Guzman, Jan Michael C. Yap, Jo-Hannah S. Llames, Sheila Mae M. Araiza, Kris P. Punayan, Irish Coleen A. Asin, Candice Francheska B. Tambaoan, Asia Louisa U. Chong, Karol Sophia Agape R. Padilla, Rianna Patricia S. Cruz, El King D. Morado, Joshua Gregor A. Dizon, Razel Nikka M. Hao, Arianne A. Zamora, Devon Ray Pacial, Juan Antonio R. Magalang, Marissa Alejandria, Celia Carlos, Anna Ong-Lim, Edsel Maurice Salvaña, John Q. Wong, Jaime C. Montoya, Maria Rosario Singh-Vergeire and Cynthia P. Saloma |
| EPI_ISL_2171229 | Cotabato Regional and Medical Center | Philippine Genome Center | Francis A. Tablizo, Kenneth M. Kim, Carlo M. Lapid, Marc Jerrone R. Castro, Maria Sofia L. Yangzon, Elcid Aaron R. Pangilinan, Benedict A. Maralit, Marc Edsel C. Ayes, Eva Maria Cutiongco-de la Paz, Alethea R. de Guzman, Jan Michael C. Yap, Jo-Hannah S. Llames, Sheila Mae M. Araiza, Kris P. Punayan, Irish Coleen A. Asin, Candice Francheska B. Tambaoan, Asia Louisa U. Chong, Karol Sophia Agape R. Padilla, Rianna Patricia S. Cruz, El King D. Morado, Joshua Gregor A. Dizon, Razel Nikka M. Hao, Arianne A. Zamora, Devon Ray Pacial, Juan Antonio R. Magalang, Marissa Alejandria, Celia Carlos, Anna Ong-Lim, Edsel Maurice Salvaña, John Q. Wong, Jaime C. Montoya, Maria Rosario Singh-Vergeire and Cynthia P. Saloma |
| EPI_ISL_2171230 | The Lord's Grace Medical and Industrial Clinic | Philippine Genome Center | Francis A. Tablizo, Kenneth M. Kim, Carlo M. Lapid, Marc Jerrone R. Castro, Maria Sofia L. Yangzon, Elcid Aaron R. Pangilinan, Benedict A. Maralit, Marc Edsel C. Ayes, Eva Maria Cutiongco-de la Paz, Alethea R. de Guzman, Jan Michael C. Yap, Jo-Hannah S. Llames, Sheila Mae M. Araiza, Kris P. Punayan, Irish Coleen A. Asin, Candice Francheska B. Tambaoan, Asia Louisa U. Chong, Karol Sophia Agape R. Padilla, Rianna Patricia S. Cruz, El King D. Morado, Joshua Gregor A. Dizon, Razel Nikka M. Hao, Arianne A. Zamora, Devon Ray Pacial, Juan Antonio R. Magalang, Marissa Alejandria, Celia Carlos, Anna Ong-Lim, Edsel Maurice Salvaña, John Q. Wong, Jaime C. Montoya, Maria Rosario Singh-Vergeire and Cynthia P. Saloma |
| EPI_ISL_2171231, EPI_ISL_2171232 | Philippine Red Cross - Port Area | Philippine Genome Center | Francis A. Tablizo, Kenneth M. Kim, Carlo M. Lapid, Marc Jerrone R. Castro, Maria Sofia L. Yangzon, Elcid Aaron R. Pangilinan, Benedict A. Maralit, Marc Edsel C. Ayes, Eva Maria Cutiongco-de la Paz, Alethea R. de Guzman, Jan Michael C. Yap, Jo-Hannah S. Llames, Sheila Mae M. Araiza, Kris P. Punayan, Irish Coleen A. Asin, Candice Francheska B. Tambaoan, Asia Louisa U. Chong, Karol Sophia Agape R. Padilla, Rianna Patricia S. Cruz, El King D. Morado, Joshua Gregor A. Dizon, Razel Nikka M. Hao, Arianne A. Zamora, Devon Ray Pacial, Juan Antonio R. Magalang, Marissa Alejandria, Celia Carlos, Anna Ong-Lim, Edsel Maurice Salvaña, John Q. Wong, Jaime C. Montoya, Maria Rosario Singh-Vergeire and Cynthia P. Saloma |
| EPI_ISL_2171233, EPI_ISL_2171234 | The Lord's Grace Medical and Industrial Clinic | Philippine Genome Center | Francis A. Tablizo, Kenneth M. Kim, Carlo M. Lapid, Marc Jerrone R. Castro, Maria Sofia L. Yangzon, Elcid Aaron R. Pangilinan, Benedict A. Maralit, Marc Edsel C. Ayes, Eva Maria Cutiongco-de la Paz, Alethea R. de Guzman, Jan Michael C. Yap, Jo-Hannah S. Llames, Sheila Mae M. Araiza, Kris P. Punayan, Irish Coleen A. Asin, Candice Francheska B. Tambaoan, Asia Louisa U. Chong, Karol Sophia Agape R. Padilla, Rianna Patricia S. Cruz, El King D. Morado, Joshua Gregor A. Dizon, Razel Nikka M. Hao, Arianne A. Zamora, Devon Ray Pacial, Juan Antonio R. Magalang, Marissa Alejandria, Celia Carlos, Anna Ong-Lim, Edsel Maurice Salvaña, John Q. Wong, Jaime C. Montoya, Maria Rosario Singh-Vergeire and Cynthia P. Saloma |
| EPI_ISL_2171235, EPI_ISL_2171237, EPI_ISL_2171238, EPI_ISL_2171239 | Philippine Red Cross - Port Area | Philippine Genome Center | Francis A. Tablizo, Kenneth M. Kim, Carlo M. Lapid, Marc Jerrone R. Castro, Maria Sofia L. Yangzon, Elcid Aaron R. Pangilinan, Benedict A. Maralit, Marc Edsel C. Ayes, Eva Maria Cutiongco-de la Paz, Alethea R. de Guzman, Jan Michael C. Yap, Jo-Hannah S. Llames, Sheila Mae M. Araiza, Kris P. Punayan, Irish Coleen A. Asin, Candice Francheska B. Tambaoan, Asia Louisa U. Chong, Karol Sophia Agape R. Padilla, Rianna Patricia S. Cruz, El King D. Morado, Joshua Gregor A. Dizon, Razel Nikka M. Hao, Arianne A. Zamora, Devon Ray Pacial, Juan Antonio R. Magalang, Marissa Alejandria, Celia Carlos, Anna Ong-Lim, Edsel Maurice Salvaña, John Q. Wong, Jaime C. Montoya, Maria Rosario Singh-Vergeire and Cynthia P. Saloma |

| | | | |
|---|---|---|---|
| EPI_ISL_2171244, EPI_ISL_2171247, EPI_ISL_2171252, EPI_ISL_2171253, EPI_ISL_2171256, EPI_ISL_2171257, EPI_ISL_2171260, EPI_ISL_2171262, EPI_ISL_2171263, EPI_ISL_2171264 | Philippine General Hospital (PGH) | Philippine Genome Center | Francis A. Tablizo, Kenneth M. Kim, Carlo M. Lapid, Marc Jerrone R. Castro, Maria Sofia L. Yangzon, Elcid Aaron R. Pangilinan, Benedict A. Maralit, Marc Edsel C. Ayes, Eva Maria Cutiongco-de la Paz, Alethea R. de Guzman, Jan Michael C. Yap, Jo-Hannah S. Llames, Sheila Mae M. Araiza, Kris P. Punayan, Irish Coleen A. Asin, Candice Francheska B. Tambaoan, Asia Louisa U. Chong, Karol Sophia Agape R. Padilla, Rianna Patricia S. Cruz, El King D. Morado, Joshua Gregor A. Dizon, Razel Nikka M. Hao, Arianne A. Zamora, Devon Ray Pacial, Juan Antonio R. Magalang, Marissa Alejandria, Celia Carlos, Anna Ong-Lim, Edsel Maurice Salvaña, John Q. Wong, Jaime C. Montoya, Maria Rosario Singh-Vergeire and Cynthia P. Saloma |
| EPI_ISL_2178452 | Labo Analyses Med | National Reference Center for Viruses of Respiratory Infections, Institut Pasteur, Paris | Marion Barbet, Sylvie Behillil, Frédéric Lemoine, Corinne Maufrais, Christophe Malabat, Damien Mornico, Méline Bizard, Angela Brisebarre, Camille Capel, Louise Lefrançois, Etienne Simon-Lorière, Vincent Enouf, Maud Vanpeene, Sylvie van der Werf,Le Berre David |
| EPI_ISL_2178455 | Labo Analyses Med | National Reference Center for Viruses of Respiratory Infections, Institut Pasteur, Paris | Marion Barbet, Sylvie Behillil, Frédéric Lemoine, Corinne Maufrais, Christophe Malabat, Damien Mornico, Méline Bizard, Angela Brisebarre, Camille Capel, Louise Lefrançois, Etienne Simon-Lorière, Vincent Enouf, Maud Vanpeene, Sylvie van der Werf,Mallet C |
| EPI_ISL_2178464, EPI_ISL_2178465 | Hopital | National Reference Center for Viruses of Respiratory Infections, Institut Pasteur, Paris | Marion Barbet, Sylvie Behillil, Frédéric Lemoine, Corinne Maufrais, Christophe Malabat, Damien Mornico, Méline Bizard, Angela Brisebarre, Camille Capel, Louise Lefrançois, Etienne Simon-Lorière, Vincent Enouf, Maud Vanpeene, Sylvie van der Werf,Scanvic AgnèS |
| EPI_ISL_2178468 | Outre mer | National Reference Center for Viruses of Respiratory Infections, Institut Pasteur, Paris | Marion Barbet, Sylvie Behillil, Méline Bizard, Angela Brisebarre, Camille Capel, Vincent Enouf, Louise Lefrançois, Frédéric Lemoine, Christophe Malabat, Corinne Maufrais, Amaury Vaysse, Etienne Simon-Lorière, Maud Vanpeene, Sylvie Van der Werf ,Patrice Combe |
| EPI_ISL_2178508 | Hospital | National Reference Center for Viruses of Respiratory Infections, Institut Pasteur, Paris | Marion Barbet, Sylvie Behillil, Méline Bizard, Angela Brisebarre, Camille Capel, Vincent Enouf, Louise Lefrançois, Frédéric Lemoine, Christophe Malabat, Corinne Maufrais, Etienne Simon-Lorière, Maud Vanpeene, Sylvie Van der Werf ,CéLine Bressollette |
| EPI_ISL_2179312 | GH JOFFRE DUPUYTREN | Department of Virology, Henri Mondor University Hospital, Assistance Publique Hôpitaux de Paris, Université Paris-Est Créteil, INSERM U955 | Christophe Rodriguez, Slim Fourati, Vanessa Demontant, Guillaume Gricourt, Melissa N'Debi, Alexandre Soulier, Elisabeth Trawinski, Jean-Michel Pawlotsky |
| EPI_ISL_2179313 | GH A.CHENEVIER-H.MONDOR | Department of Virology, Henri Mondor University Hospital, Assistance Publique Hôpitaux de Paris, Université Paris-Est Créteil, INSERM U955 | Christophe Rodriguez, Slim Fourati, Vanessa Demontant, Guillaume Gricourt, Melissa N'Debi, Alexandre Soulier, Elisabeth Trawinski, Jean-Michel Pawlotsky |
| EPI_ISL_2179325, EPI_ISL_2179333, EPI_ISL_2179336, EPI_ISL_2179338, EPI_ISL_2179339, EPI_ISL_2179341 | Hôpital Henri Mondor | Department of Virology, Henri Mondor University Hospital, Assistance Publique Hôpitaux de Paris, Université Paris-Est Créteil, INSERM U955 | Christophe Rodriguez, Slim Fourati, Vanessa Demontant, Guillaume Gricourt, Melissa N'Debi, Alexandre Soulier, Elisabeth Trawinski, Jean-Michel Pawlotsky |
| EPI_ISL_2179378 | GH JOFFRE DUPUYTREN | Department of Virology, Henri Mondor University Hospital, Assistance Publique Hôpitaux de Paris, Université Paris-Est Créteil, INSERM U955 | Christophe Rodriguez, Slim Fourati, Vanessa Demontant, Guillaume Gricourt, Melissa N'Debi, Alexandre Soulier, Elisabeth Trawinski, Jean-Michel Pawlotsky |
| EPI_ISL_2179690 | Servicio Microbiología Hospital La Paz | Servicio Microbiología Hospital La Paz | Fernando Lázaro, Rubén Cáceres, Jesús Mingorance Cruz, Elie Dahdouh |
| EPI_ISL_2180329, EPI_ISL_2180330, EPI_ISL_2180332, EPI_ISL_2180333, EPI_ISL_2180334, EPI_ISL_2180335, EPI_ISL_2180336, EPI_ISL_2180337 | National Center of Infectious and Parasitic Diseases | National Center of Infectious and Parasitic Diseases | Alexiev et al |
| EPI_ISL_2187824 | HLAGYN - Laboratorio de Imunologia de Transplantes de Goias | HLAGYN - Laboratorio de Imunologia de Transplantes de Goias | Fernando Antonio Vinhal dos Santos, Erika Lopes Rocha Batista, Alessandro Leonardo Alvares Magalhaes, Frederico Rodrigues Vinhal, Sabrina Sara Moreira Duarte, Lucas Carlos Gomes Pereira, Daniel Ferreira de Sousa |
| EPI_ISL_2188037 | Qualimed Hospital Sta. Rosa | Philippine Genome Center | Francis A. Tablizo, Kenneth M. Kim, Carlo M. Lapid, Marc Jerrone R. Castro, Maria Sofia L. Yangzon, Elcid Aaron R. Pangilinan, Benedict A. Maralit, Marc Edsel C. Ayes, Eva Maria Cutiongco-de la Paz, Alethea R. de Guzman, Jan Michael C. Yap, Jo-Hannah S. Llames, Sheila Mae M. Araiza, Kris P. Punayan, Irish Coleen A. Asin, Candice Francheska B. Tambaoan, Asia Louisa U. Chong, Karol Sophia Agape R. Padilla, Rianna Patricia S. Cruz, El King D. Morado, Joshua Gregor A. Dizon, Razel Nikka M. Hao, Arianne A. Zamora, Devon Ray Pacial, Juan Antonio R. Magalang, Marissa Alejandria, Celia Carlos, Anna Ong-Lim, Edsel Maurice Salvaña, John Q. Wong, Jaime C. Montoya, Maria Rosario Singh-Vergeire and Cynthia P. Saloma |
| EPI_ISL_2188038 | Ospital ng Imus | Philippine Genome Center | Francis A. Tablizo, Kenneth M. Kim, Carlo M. Lapid, Marc Jerrone R. Castro, Maria Sofia L. Yangzon, Elcid Aaron R. Pangilinan, Benedict A. Maralit, Marc Edsel C. Ayes, Eva Maria Cutiongco-de la Paz, Alethea R. de Guzman, Jan Michael C. Yap, Jo-Hannah S. Llames, Sheila Mae M. Araiza, Kris P. Punayan, Irish Coleen A. Asin, Candice Francheska B. Tambaoan, Asia Louisa U. Chong, Karol Sophia Agape R. Padilla, Rianna Patricia S. Cruz, El King D. Morado, Joshua Gregor A. Dizon, Razel Nikka M. Hao, Arianne A. Zamora, Devon Ray Pacial, Juan Antonio R. Magalang, Marissa Alejandria, Celia Carlos, Anna Ong-Lim, Edsel Maurice Salvaña, John Q. Wong, Jaime C. Montoya, Maria Rosario Singh-Vergeire and Cynthia P. Saloma |
| EPI_ISL_2188041 | The Lord's Grace Medical and Industrial Clinic | Philippine Genome Center | Francis A. Tablizo, Kenneth M. Kim, Carlo M. Lapid, Marc Jerrone R. Castro, Maria Sofia L. Yangzon, Elcid Aaron R. Pangilinan, Benedict A. Maralit, Marc Edsel C. Ayes, Eva Maria Cutiongco-de la Paz, Alethea R. de Guzman, Jan Michael C. Yap, Jo-Hannah S. Llames, Sheila Mae M. Araiza, Kris P. Punayan, Irish Coleen A. Asin, Candice Francheska B. Tambaoan, Asia Louisa U. Chong, Karol Sophia Agape R. Padilla, Rianna Patricia S. Cruz, El King D. Morado, Joshua Gregor A. Dizon, Razel Nikka M. Hao, Arianne A. Zamora, Devon Ray Pacial, Juan Antonio R. Magalang, Marissa Alejandria, Celia Carlos, Anna Ong-Lim, Edsel Maurice Salvaña, John Q. Wong, Jaime C. Montoya, Maria Rosario Singh-Vergeire and Cynthia P. Saloma |
| EPI_ISL_2188042 | Ospital ng Imus | Philippine Genome Center | Francis A. Tablizo, Kenneth M. Kim, Carlo M. Lapid, Marc Jerrone R. Castro, Maria Sofia L. Yangzon, Elcid Aaron R. Pangilinan, Benedict A. Maralit, Marc Edsel C. Ayes, Eva Maria Cutiongco-de la Paz, Alethea R. de Guzman, Jan Michael C. Yap, Jo-Hannah S. Llames, Sheila Mae M. Araiza, Kris P. Punayan, Irish Coleen A. Asin, Candice Francheska B. Tambaoan, Asia Louisa U. Chong, Karol Sophia Agape R. Padilla, Rianna Patricia S. Cruz, El King D. Morado, Joshua Gregor A. Dizon, Razel Nikka M. Hao, Arianne A. Zamora, Devon Ray Pacial, Juan Antonio R. Magalang, Marissa Alejandria, Celia Carlos, Anna Ong-Lim, Edsel Maurice Salvaña, John Q. Wong, Jaime C. Montoya, Maria Rosario Singh-Vergeire and Cynthia P. Saloma |
| EPI_ISL_2188043 | Research Institute for Tropical Medicine, Inc. (RITM) | Philippine Genome Center | Francis A. Tablizo, Kenneth M. Kim, Carlo M. Lapid, Marc Jerrone R. Castro, Maria Sofia L. Yangzon, Elcid Aaron R. Pangilinan, Benedict A. Maralit, Marc Edsel C. Ayes, Eva Maria Cutiongco-de la Paz, Alethea R. de Guzman, Jan Michael C. Yap, Jo-Hannah S. Llames, Sheila Mae M. Araiza, Kris P. Punayan, Irish Coleen A. Asin, Candice Francheska B. Tambaoan, Asia Louisa U. Chong, Karol Sophia Agape R. Padilla, Rianna Patricia S. Cruz, El King D. Morado, Joshua Gregor A. Dizon, Razel Nikka M. Hao, Arianne A. Zamora, Devon Ray Pacial, Juan Antonio R. Magalang, Marissa Alejandria, Celia Carlos, Anna Ong-Lim, Edsel Maurice Salvaña, John Q. Wong, Jaime C. Montoya, Maria Rosario Singh-Vergeire and Cynthia P. Saloma |
| EPI_ISL_2188044 | The Medical City -Â Ortigas | Philippine Genome Center | Francis A. Tablizo, Kenneth M. Kim, Carlo M. Lapid, Marc Jerrone R. Castro, Maria Sofia L. Yangzon, Elcid Aaron R. Pangilinan, Benedict A. Maralit, Marc Edsel C. Ayes, Eva Maria Cutiongco-de la Paz, Alethea R. de Guzman, Jan Michael C. Yap, Jo-Hannah S. Llames, Sheila Mae M. Araiza, Kris P. Punayan, Irish Coleen A. Asin, Candice Francheska B. Tambaoan, Asia Louisa U. Chong, Karol Sophia Agape R. Padilla, Rianna Patricia S. Cruz, El King D. Morado, Joshua Gregor A. Dizon, Razel Nikka M. Hao, Arianne A. Zamora, Devon Ray Pacial, Juan Antonio R. Magalang, Marissa Alejandria, Celia Carlos, Anna Ong-Lim, Edsel Maurice Salvaña, John Q. Wong, Jaime C. Montoya, Maria Rosario Singh-Vergeire and Cynthia P. Saloma |
| EPI_ISL_2188045 | Philippine Red Cross - Port Area | Philippine Genome Center | Francis A. Tablizo, Kenneth M. Kim, Carlo M. Lapid, Marc Jerrone R. Castro, Maria Sofia L. Yangzon, Elcid Aaron R. Pangilinan, Benedict A. Maralit, Marc Edsel C. Ayes, Eva Maria Cutiongco-de la Paz, Alethea R. de Guzman, Jan Michael C. Yap, Jo-Hannah S. Llames, Sheila Mae M. Araiza, Kris P. Punayan, Irish Coleen A. Asin, Candice Francheska B. Tambaoan, Asia Louisa U. Chong, Karol Sophia Agape R. Padilla, Rianna Patricia S. Cruz, El King D. Morado, Joshua Gregor A. Dizon, Razel Nikka M. Hao, Arianne A. Zamora, Devon Ray Pacial, Juan Antonio R. Magalang, Marissa Alejandria, Celia Carlos, Anna Ong-Lim, Edsel Maurice Salvaña, John Q. Wong, Jaime C. Montoya, Maria Rosario Singh-Vergeire and Cynthia P. Saloma |
| EPI_ISL_2188046 | Laguna Provincial Hospital - San Pablo District Hospital | Philippine Genome Center | Francis A. Tablizo, Kenneth M. Kim, Carlo M. Lapid, Marc Jerrone R. Castro, Maria Sofia L. Yangzon, Elcid Aaron R. Pangilinan, Benedict A. Maralit, Marc Edsel C. Ayes, Eva Maria Cutiongco-de la Paz, Alethea R. de Guzman, Jan Michael C. Yap, Jo-Hannah S. Llames, Sheila Mae M. Araiza, Kris P. Punayan, Irish Coleen A. Asin, Candice Francheska B. Tambaoan, Asia Louisa U. Chong, Karol Sophia Agape R. Padilla, Rianna Patricia S. Cruz, El King D. Morado, Joshua Gregor A. Dizon, Razel Nikka M. Hao, Arianne A. Zamora, Devon Ray Pacial, Juan Antonio R. Magalang, Marissa Alejandria, Celia Carlos, Anna Ong-Lim, Edsel Maurice Salvaña, John Q. Wong, Jaime C. Montoya, Maria Rosario Singh-Vergeire and Cynthia P. Saloma |
| EPI_ISL_2188047 | The Medical City -Â Ortigas | Philippine Genome Center | Francis A. Tablizo, Kenneth M. Kim, Carlo M. Lapid, Marc Jerrone R. Castro, Maria Sofia L. Yangzon, Elcid Aaron R. Pangilinan, Benedict A. Maralit, Marc Edsel C. Ayes, Eva Maria Cutiongco-de la Paz, Alethea R. de Guzman, Jan Michael C. Yap, Jo-Hannah S. Llames, Sheila Mae M. Araiza, Kris P. Punayan, Irish Coleen A. Asin, Candice Francheska B. Tambaoan, Asia Louisa U. Chong, Karol Sophia Agape R. Padilla, Rianna Patricia S. Cruz, El King |

| | | | |
|---|---|---|---|
| | | | D. Morado, Joshua Gregor A. Dizon, Razel Nikka M. Hao, Arianne A. Zamora, Devon Ray Pacial, Juan Antonio R. Magalang, Marissa Alejandria, Celia Carlos, Anna Ong-Lim, Edsel Maurice Salvaña, John Q. Wong, Jaime C. Montoya, Maria Rosario Singh-Vergeire and Cynthia P. Saloma |
| EPI_ISL_2188050 | Baguio General Hospital Medical Center (BGHMC) | Philippine Genome Center | Francis A. Tablizo, Kenneth M. Kim, Carlo M. Lapid, Marc Jerrone R. Castro, Maria Sofia L. Yangzon, Elcid Aaron R. Pangilinan, Benedict A. Maralit, Marc Edsel C. Ayes, Eva Maria Cutiongco-de la Paz, Alethea R. de Guzman, Jan Michael C. Yap, Jo-Hannah S. Llames, Sheila Mae M. Araiza, Kris P. Punayan, Irish Coleen A. Asin, Candice Francheska B. Tambaoan, Asia Louisa U. Chong, Karol Sophia Agape R. Padilla, Rianna Patricia S. Cruz, El King D. Morado, Joshua Gregor A. Dizon, Razel Nikka M. Hao, Arianne A. Zamora, Devon Ray Pacial, Juan Antonio R. Magalang, Marissa Alejandria, Celia Carlos, Anna Ong-Lim, Edsel Maurice Salvaña, John Q. Wong, Jaime C. Montoya, Maria Rosario Singh-Vergeire and Cynthia P. Saloma |
| EPI_ISL_2188051 | The Medical City -Â Ortigas | Philippine Genome Center | Francis A. Tablizo, Kenneth M. Kim, Carlo M. Lapid, Marc Jerrone R. Castro, Maria Sofia L. Yangzon, Elcid Aaron R. Pangilinan, Benedict A. Maralit, Marc Edsel C. Ayes, Eva Maria Cutiongco-de la Paz, Alethea R. de Guzman, Jan Michael C. Yap, Jo-Hannah S. Llames, Sheila Mae M. Araiza, Kris P. Punayan, Irish Coleen A. Asin, Candice Francheska B. Tambaoan, Asia Louisa U. Chong, Karol Sophia Agape R. Padilla, Rianna Patricia S. Cruz, El King D. Morado, Joshua Gregor A. Dizon, Razel Nikka M. Hao, Arianne A. Zamora, Devon Ray Pacial, Juan Antonio R. Magalang, Marissa Alejandria, Celia Carlos, Anna Ong-Lim, Edsel Maurice Salvaña, John Q. Wong, Jaime C. Montoya, Maria Rosario Singh-Vergeire and Cynthia P. Saloma |
| EPI_ISL_2188054 | The Lord's Grace Medical and Industrial Clinic | Philippine Genome Center | Francis A. Tablizo, Kenneth M. Kim, Carlo M. Lapid, Marc Jerrone R. Castro, Maria Sofia L. Yangzon, Elcid Aaron R. Pangilinan, Benedict A. Maralit, Marc Edsel C. Ayes, Eva Maria Cutiongco-de la Paz, Alethea R. de Guzman, Jan Michael C. Yap, Jo-Hannah S. Llames, Sheila Mae M. Araiza, Kris P. Punayan, Irish Coleen A. Asin, Candice Francheska B. Tambaoan, Asia Louisa U. Chong, Karol Sophia Agape R. Padilla, Rianna Patricia S. Cruz, El King D. Morado, Joshua Gregor A. Dizon, Razel Nikka M. Hao, Arianne A. Zamora, Devon Ray Pacial, Juan Antonio R. Magalang, Marissa Alejandria, Celia Carlos, Anna Ong-Lim, Edsel Maurice Salvaña, John Q. Wong, Jaime C. Montoya, Maria Rosario Singh-Vergeire and Cynthia P. Saloma |
| EPI_ISL_2188055 | The Medical City -Â Ortigas | Philippine Genome Center | Francis A. Tablizo, Kenneth M. Kim, Carlo M. Lapid, Marc Jerrone R. Castro, Maria Sofia L. Yangzon, Elcid Aaron R. Pangilinan, Benedict A. Maralit, Marc Edsel C. Ayes, Eva Maria Cutiongco-de la Paz, Alethea R. de Guzman, Jan Michael C. Yap, Jo-Hannah S. Llames, Sheila Mae M. Araiza, Kris P. Punayan, Irish Coleen A. Asin, Candice Francheska B. Tambaoan, Asia Louisa U. Chong, Karol Sophia Agape R. Padilla, Rianna Patricia S. Cruz, El King D. Morado, Joshua Gregor A. Dizon, Razel Nikka M. Hao, Arianne A. Zamora, Devon Ray Pacial, Juan Antonio R. Magalang, Marissa Alejandria, Celia Carlos, Anna Ong-Lim, Edsel Maurice Salvaña, John Q. Wong, Jaime C. Montoya, Maria Rosario Singh-Vergeire and Cynthia P. Saloma |
| EPI_ISL_2188060 | SAN ANTONIO CITY OF ILAGAN HOSPITAL | Philippine Genome Center | Francis A. Tablizo, Kenneth M. Kim, Carlo M. Lapid, Marc Jerrone R. Castro, Maria Sofia L. Yangzon, Elcid Aaron R. Pangilinan, Benedict A. Maralit, Marc Edsel C. Ayes, Eva Maria Cutiongco-de la Paz, Alethea R. de Guzman, Jan Michael C. Yap, Jo-Hannah S. Llames, Sheila Mae M. Araiza, Kris P. Punayan, Irish Coleen A. Asin, Candice Francheska B. Tambaoan, Asia Louisa U. Chong, Karol Sophia Agape R. Padilla, Rianna Patricia S. Cruz, El King D. Morado, Joshua Gregor A. Dizon, Razel Nikka M. Hao, Arianne A. Zamora, Devon Ray Pacial, Juan Antonio R. Magalang, Marissa Alejandria, Celia Carlos, Anna Ong-Lim, Edsel Maurice Salvaña, John Q. Wong, Jaime C. Montoya, Maria Rosario Singh-Vergeire and Cynthia P. Saloma |
| EPI_ISL_2188061 | Cagayan Valley Medical Center Molecular Laboratory | Philippine Genome Center | Francis A. Tablizo, Kenneth M. Kim, Carlo M. Lapid, Marc Jerrone R. Castro, Maria Sofia L. Yangzon, Elcid Aaron R. Pangilinan, Benedict A. Maralit, Marc Edsel C. Ayes, Eva Maria Cutiongco-de la Paz, Alethea R. de Guzman, Jan Michael C. Yap, Jo-Hannah S. Llames, Sheila Mae M. Araiza, Kris P. Punayan, Irish Coleen A. Asin, Candice Francheska B. Tambaoan, Asia Louisa U. Chong, Karol Sophia Agape R. Padilla, Rianna Patricia S. Cruz, El King D. Morado, Joshua Gregor A. Dizon, Razel Nikka M. Hao, Arianne A. Zamora, Devon Ray Pacial, Juan Antonio R. Magalang, Marissa Alejandria, Celia Carlos, Anna Ong-Lim, Edsel Maurice Salvaña, John Q. Wong, Jaime C. Montoya, Maria Rosario Singh-Vergeire and Cynthia P. Saloma |
| EPI_ISL_2188063, EPI_ISL_2188064 | The Medical City -Â Ortigas | Philippine Genome Center | Francis A. Tablizo, Kenneth M. Kim, Carlo M. Lapid, Marc Jerrone R. Castro, Maria Sofia L. Yangzon, Elcid Aaron R. Pangilinan, Benedict A. Maralit, Marc Edsel C. Ayes, Eva Maria Cutiongco-de la Paz, Alethea R. de Guzman, Jan Michael C. Yap, Jo-Hannah S. Llames, Sheila Mae M. Araiza, Kris P. Punayan, Irish Coleen A. Asin, Candice Francheska B. Tambaoan, Asia Louisa U. Chong, Karol Sophia Agape R. Padilla, Rianna Patricia S. Cruz, El King D. Morado, Joshua Gregor A. Dizon, Razel Nikka M. Hao, Arianne A. Zamora, Devon Ray Pacial, Juan Antonio R. Magalang, Marissa Alejandria, Celia Carlos, Anna Ong-Lim, Edsel Maurice Salvaña, John Q. Wong, Jaime C. Montoya, Maria Rosario Singh-Vergeire and Cynthia P. Saloma |
| EPI_ISL_2188065 | Qualimed Hospital Sta. Rosa | Philippine Genome Center | Francis A. Tablizo, Kenneth M. Kim, Carlo M. Lapid, Marc Jerrone R. Castro, Maria Sofia L. Yangzon, Elcid Aaron R. Pangilinan, Benedict A. Maralit, Marc Edsel C. Ayes, Eva Maria Cutiongco-de la Paz, Alethea R. de Guzman, Jan Michael C. Yap, Jo-Hannah S. Llames, Sheila Mae M. Araiza, Kris P. Punayan, Irish Coleen A. Asin, Candice Francheska B. Tambaoan, Asia Louisa U. Chong, Karol Sophia Agape R. Padilla, Rianna Patricia S. Cruz, El King D. Morado, Joshua Gregor A. Dizon, Razel Nikka M. Hao, Arianne A. Zamora, Devon Ray Pacial, Juan Antonio R. Magalang, Marissa Alejandria, Celia Carlos, Anna Ong-Lim, Edsel Maurice Salvaña, John Q. Wong, Jaime C. Montoya, Maria Rosario Singh-Vergeire and Cynthia P. Saloma |
| EPI_ISL_2188066 | Sta. Rosa CHO | Philippine Genome Center | Francis A. Tablizo, Kenneth M. Kim, Carlo M. Lapid, Marc Jerrone R. Castro, Maria Sofia L. Yangzon, Elcid Aaron R. Pangilinan, Benedict A. Maralit, Marc Edsel C. Ayes, Eva Maria Cutiongco-de la Paz, Alethea R. de Guzman, Jan Michael C. Yap, Jo-Hannah S. Llames, Sheila Mae M. Araiza, Kris P. Punayan, Irish Coleen A. Asin, Candice Francheska B. Tambaoan, Asia Louisa U. Chong, Karol Sophia Agape R. Padilla, Rianna Patricia S. Cruz, El King D. Morado, Joshua Gregor A. Dizon, Razel Nikka M. Hao, Arianne A. Zamora, Devon Ray Pacial, Juan Antonio R. Magalang, Marissa Alejandria, Celia Carlos, Anna Ong-Lim, Edsel Maurice Salvaña, John Q. Wong, Jaime C. Montoya, Maria Rosario Singh-Vergeire and Cynthia P. Saloma |
| EPI_ISL_2188067 | Ospital ng ParaÃ±aque II | Philippine Genome Center | Francis A. Tablizo, Kenneth M. Kim, Carlo M. Lapid, Marc Jerrone R. Castro, Maria Sofia L. Yangzon, Elcid Aaron R. Pangilinan, Benedict A. Maralit, Marc Edsel C. Ayes, Eva Maria Cutiongco-de la Paz, Alethea R. de Guzman, Jan Michael C. Yap, Jo-Hannah S. Llames, Sheila Mae M. Araiza, Kris P. Punayan, Irish Coleen A. Asin, Candice Francheska B. Tambaoan, Asia Louisa U. Chong, Karol Sophia Agape R. Padilla, Rianna Patricia S. Cruz, El King D. Morado, Joshua Gregor A. Dizon, Razel Nikka M. Hao, Arianne A. Zamora, Devon Ray Pacial, Juan Antonio R. Magalang, Marissa Alejandria, Celia Carlos, Anna Ong-Lim, Edsel Maurice Salvaña, John Q. Wong, Jaime C. Montoya, Maria Rosario Singh-Vergeire and Cynthia P. Saloma |
| EPI_ISL_2188068 | The Medical City -Â Ortigas | Philippine Genome Center | Francis A. Tablizo, Kenneth M. Kim, Carlo M. Lapid, Marc Jerrone R. Castro, Maria Sofia L. Yangzon, Elcid Aaron R. Pangilinan, Benedict A. Maralit, Marc Edsel C. Ayes, Eva Maria Cutiongco-de la Paz, Alethea R. de Guzman, Jan Michael C. Yap, Jo-Hannah S. Llames, Sheila Mae M. Araiza, Kris P. Punayan, Irish Coleen A. Asin, Candice Francheska B. Tambaoan, Asia Louisa U. Chong, Karol Sophia Agape R. Padilla, Rianna Patricia S. Cruz, El King D. Morado, Joshua Gregor A. Dizon, Razel Nikka M. Hao, Arianne A. Zamora, Devon Ray Pacial, Juan Antonio R. Magalang, Marissa Alejandria, Celia Carlos, Anna Ong-Lim, Edsel Maurice Salvaña, John Q. Wong, Jaime C. Montoya, Maria Rosario Singh-Vergeire and Cynthia P. Saloma |
| EPI_ISL_2188069 | The Lord's Grace Medical and Industrial Clinic | Philippine Genome Center | Francis A. Tablizo, Kenneth M. Kim, Carlo M. Lapid, Marc Jerrone R. Castro, Maria Sofia L. Yangzon, Elcid Aaron R. Pangilinan, Benedict A. Maralit, Marc Edsel C. Ayes, Eva Maria Cutiongco-de la Paz, Alethea R. de Guzman, Jan Michael C. Yap, Jo-Hannah S. Llames, Sheila Mae M. Araiza, Kris P. Punayan, Irish Coleen A. Asin, Candice Francheska B. Tambaoan, Asia Louisa U. Chong, Karol Sophia Agape R. Padilla, Rianna Patricia S. Cruz, El King D. Morado, Joshua Gregor A. Dizon, Razel Nikka M. Hao, Arianne A. Zamora, Devon Ray Pacial, Juan Antonio R. Magalang, Marissa Alejandria, Celia Carlos, Anna Ong-Lim, Edsel Maurice Salvaña, John Q. Wong, Jaime C. Montoya, Maria Rosario Singh-Vergeire and Cynthia P. Saloma |
| EPI_ISL_2188072 | The Medical City -Â Ortigas | Philippine Genome Center | Francis A. Tablizo, Kenneth M. Kim, Carlo M. Lapid, Marc Jerrone R. Castro, Maria Sofia L. Yangzon, Elcid Aaron R. Pangilinan, Benedict A. Maralit, Marc Edsel C. Ayes, Eva Maria Cutiongco-de la Paz, Alethea R. de Guzman, Jan Michael C. Yap, Jo-Hannah S. Llames, Sheila Mae M. Araiza, Kris P. Punayan, Irish Coleen A. Asin, Candice Francheska B. Tambaoan, Asia Louisa U. Chong, Karol Sophia Agape R. Padilla, Rianna Patricia S. Cruz, El King D. Morado, Joshua Gregor A. Dizon, Razel Nikka M. Hao, Arianne A. Zamora, Devon Ray Pacial, Juan Antonio R. Magalang, Marissa Alejandria, Celia Carlos, Anna Ong-Lim, Edsel Maurice Salvaña, John Q. Wong, Jaime C. Montoya, Maria Rosario Singh-Vergeire and Cynthia P. Saloma |
| EPI_ISL_2188073, EPI_ISL_2188074 | The Lord's Grace Medical and Industrial Clinic | Philippine Genome Center | Francis A. Tablizo, Kenneth M. Kim, Carlo M. Lapid, Marc Jerrone R. Castro, Maria Sofia L. Yangzon, Elcid Aaron R. Pangilinan, Benedict A. Maralit, Marc Edsel C. Ayes, Eva Maria Cutiongco-de la Paz, Alethea R. de Guzman, Jan Michael C. Yap, Jo-Hannah S. Llames, Sheila Mae M. Araiza, Kris P. Punayan, Irish Coleen A. Asin, Candice Francheska B. Tambaoan, Asia Louisa U. Chong, Karol Sophia Agape R. Padilla, Rianna Patricia S. Cruz, El King D. Morado, Joshua Gregor A. Dizon, Razel Nikka M. Hao, Arianne A. Zamora, Devon Ray Pacial, Juan Antonio R. Magalang, Marissa Alejandria, Celia Carlos, Anna Ong-Lim, Edsel Maurice Salvaña, John Q. Wong, Jaime C. Montoya, Maria Rosario Singh-Vergeire and Cynthia P. Saloma |
| EPI_ISL_2188075 | Philippine Red Cross - Port Area | Philippine Genome Center | Francis A. Tablizo, Kenneth M. Kim, Carlo M. Lapid, Marc Jerrone R. Castro, Maria Sofia L. Yangzon, Elcid Aaron R. Pangilinan, Benedict A. Maralit, Marc Edsel C. Ayes, Eva Maria Cutiongco-de la Paz, Alethea R. de Guzman, Jan Michael C. Yap, Jo-Hannah S. Llames, Sheila Mae M. Araiza, Kris P. Punayan, Irish Coleen A. Asin, Candice Francheska B. Tambaoan, Asia Louisa U. Chong, Karol Sophia Agape R. Padilla, Rianna Patricia S. Cruz, El King D. Morado, Joshua Gregor A. Dizon, Razel Nikka M. Hao, Arianne A. Zamora, Devon Ray Pacial, Juan Antonio R. Magalang, Marissa Alejandria, Celia Carlos, Anna Ong-Lim, Edsel Maurice Salvaña, John Q. Wong, Jaime C. Montoya, Maria Rosario Singh-Vergeire and Cynthia P. Saloma |
| EPI_ISL_2188076 | Ospital ng Imus | Philippine Genome Center | Francis A. Tablizo, Kenneth M. Kim, Carlo M. Lapid, Marc Jerrone R. Castro, Maria Sofia L. Yangzon, Elcid Aaron R. Pangilinan, Benedict A. Maralit, Marc Edsel C. Ayes, Eva Maria Cutiongco-de la Paz, Alethea R. de Guzman, Jan Michael C. Yap, Jo-Hannah S. Llames, Sheila Mae M. Araiza, Kris P. |

| | | | |
|---|---|---|---|
| | | | Punayan, Irish Coleen A. Asin, Candice Francheska B. Tambaoan, Asia Louisa U. Chong, Karol Sophia Agape R. Padilla, Rianna Patricia S. Cruz, El King D. Morado, Joshua Gregor A. Dizon, Razel Nikka M. Hao, Arianne A. Zamora, Devon Ray Pacial, Juan Antonio R. Magalang, Marissa Alejandria, Celia Carlos, Anna Ong-Lim, Edsel Maurice Salvaña, John Q. Wong, Jaime C. Montoya, Maria Rosario Singh-Vergeire and Cynthia P. Saloma |
| EPI_ISL_2188078 | Philippine Red Cross Logistics and Multipurpose Center | Philippine Genome Center | Francis A. Tablizo, Kenneth M. Kim, Carlo M. Lapid, Marc Jerrone R. Castro, Maria Sofia L. Yangzon, Elcid Aaron R. Pangilinan, Benedict A. Maralit, Marc Edsel C. Ayes, Eva Maria Cutiongco-de la Paz, Alethea R. de Guzman, Jan Michael C. Yap, Jo-Hannah S. Llames, Sheila Mae M. Araiza, Kris P. Punayan, Irish Coleen A. Asin, Candice Francheska B. Tambaoan, Asia Louisa U. Chong, Karol Sophia Agape R. Padilla, Rianna Patricia S. Cruz, El King D. Morado, Joshua Gregor A. Dizon, Razel Nikka M. Hao, Arianne A. Zamora, Devon Ray Pacial, Juan Antonio R. Magalang, Marissa Alejandria, Celia Carlos, Anna Ong-Lim, Edsel Maurice Salvaña, John Q. Wong, Jaime C. Montoya, Maria Rosario Singh-Vergeire and Cynthia P. Saloma |
| EPI_ISL_2188079 | Research Institute for Tropical Medicine, Inc. (RITM) | Philippine Genome Center | Francis A. Tablizo, Kenneth M. Kim, Carlo M. Lapid, Marc Jerrone R. Castro, Maria Sofia L. Yangzon, Elcid Aaron R. Pangilinan, Benedict A. Maralit, Marc Edsel C. Ayes, Eva Maria Cutiongco-de la Paz, Alethea R. de Guzman, Jan Michael C. Yap, Jo-Hannah S. Llames, Sheila Mae M. Araiza, Kris P. Punayan, Irish Coleen A. Asin, Candice Francheska B. Tambaoan, Asia Louisa U. Chong, Karol Sophia Agape R. Padilla, Rianna Patricia S. Cruz, El King D. Morado, Joshua Gregor A. Dizon, Razel Nikka M. Hao, Arianne A. Zamora, Devon Ray Pacial, Juan Antonio R. Magalang, Marissa Alejandria, Celia Carlos, Anna Ong-Lim, Edsel Maurice Salvaña, John Q. Wong, Jaime C. Montoya, Maria Rosario Singh-Vergeire and Cynthia P. Saloma |
| EPI_ISL_2188081 | University of the Philippines National Institutes of Health (UP NIH) | Philippine Genome Center | Francis A. Tablizo, Kenneth M. Kim, Carlo M. Lapid, Marc Jerrone R. Castro, Maria Sofia L. Yangzon, Elcid Aaron R. Pangilinan, Benedict A. Maralit, Marc Edsel C. Ayes, Eva Maria Cutiongco-de la Paz, Alethea R. de Guzman, Jan Michael C. Yap, Jo-Hannah S. Llames, Sheila Mae M. Araiza, Kris P. Punayan, Irish Coleen A. Asin, Candice Francheska B. Tambaoan, Asia Louisa U. Chong, Karol Sophia Agape R. Padilla, Rianna Patricia S. Cruz, El King D. Morado, Joshua Gregor A. Dizon, Razel Nikka M. Hao, Arianne A. Zamora, Devon Ray Pacial, Juan Antonio R. Magalang, Marissa Alejandria, Celia Carlos, Anna Ong-Lim, Edsel Maurice Salvaña, John Q. Wong, Jaime C. Montoya, Maria Rosario Singh-Vergeire and Cynthia P. Saloma |
| EPI_ISL_2188082, EPI_ISL_2188083 | The Medical City -Â Ortigas | Philippine Genome Center | Francis A. Tablizo, Kenneth M. Kim, Carlo M. Lapid, Marc Jerrone R. Castro, Maria Sofia L. Yangzon, Elcid Aaron R. Pangilinan, Benedict A. Maralit, Marc Edsel C. Ayes, Eva Maria Cutiongco-de la Paz, Alethea R. de Guzman, Jan Michael C. Yap, Jo-Hannah S. Llames, Sheila Mae M. Araiza, Kris P. Punayan, Irish Coleen A. Asin, Candice Francheska B. Tambaoan, Asia Louisa U. Chong, Karol Sophia Agape R. Padilla, Rianna Patricia S. Cruz, El King D. Morado, Joshua Gregor A. Dizon, Razel Nikka M. Hao, Arianne A. Zamora, Devon Ray Pacial, Juan Antonio R. Magalang, Marissa Alejandria, Celia Carlos, Anna Ong-Lim, Edsel Maurice Salvaña, John Q. Wong, Jaime C. Montoya, Maria Rosario Singh-Vergeire and Cynthia P. Saloma |
| EPI_ISL_2188084 | Amang Rodriguez Memorial Medical Center | Philippine Genome Center | Francis A. Tablizo, Kenneth M. Kim, Carlo M. Lapid, Marc Jerrone R. Castro, Maria Sofia L. Yangzon, Elcid Aaron R. Pangilinan, Benedict A. Maralit, Marc Edsel C. Ayes, Eva Maria Cutiongco-de la Paz, Alethea R. de Guzman, Jan Michael C. Yap, Jo-Hannah S. Llames, Sheila Mae M. Araiza, Kris P. Punayan, Irish Coleen A. Asin, Candice Francheska B. Tambaoan, Asia Louisa U. Chong, Karol Sophia Agape R. Padilla, Rianna Patricia S. Cruz, El King D. Morado, Joshua Gregor A. Dizon, Razel Nikka M. Hao, Arianne A. Zamora, Devon Ray Pacial, Juan Antonio R. Magalang, Marissa Alejandria, Celia Carlos, Anna Ong-Lim, Edsel Maurice Salvaña, John Q. Wong, Jaime C. Montoya, Maria Rosario Singh-Vergeire and Cynthia P. Saloma |
| EPI_ISL_2188085, EPI_ISL_2188087 | The Medical City -Â Ortigas | Philippine Genome Center | Francis A. Tablizo, Kenneth M. Kim, Carlo M. Lapid, Marc Jerrone R. Castro, Maria Sofia L. Yangzon, Elcid Aaron R. Pangilinan, Benedict A. Maralit, Marc Edsel C. Ayes, Eva Maria Cutiongco-de la Paz, Alethea R. de Guzman, Jan Michael C. Yap, Jo-Hannah S. Llames, Sheila Mae M. Araiza, Kris P. Punayan, Irish Coleen A. Asin, Candice Francheska B. Tambaoan, Asia Louisa U. Chong, Karol Sophia Agape R. Padilla, Rianna Patricia S. Cruz, El King D. Morado, Joshua Gregor A. Dizon, Razel Nikka M. Hao, Arianne A. Zamora, Devon Ray Pacial, Juan Antonio R. Magalang, Marissa Alejandria, Celia Carlos, Anna Ong-Lim, Edsel Maurice Salvaña, John Q. Wong, Jaime C. Montoya, Maria Rosario Singh-Vergeire and Cynthia P. Saloma |
| EPI_ISL_2188089 | Philippine Red Cross - Port Area | Philippine Genome Center | Francis A. Tablizo, Kenneth M. Kim, Carlo M. Lapid, Marc Jerrone R. Castro, Maria Sofia L. Yangzon, Elcid Aaron R. Pangilinan, Benedict A. Maralit, Marc Edsel C. Ayes, Eva Maria Cutiongco-de la Paz, Alethea R. de Guzman, Jan Michael C. Yap, Jo-Hannah S. Llames, Sheila Mae M. Araiza, Kris P. Punayan, Irish Coleen A. Asin, Candice Francheska B. Tambaoan, Asia Louisa U. Chong, Karol Sophia Agape R. Padilla, Rianna Patricia S. Cruz, El King D. Morado, Joshua Gregor A. Dizon, Razel Nikka M. Hao, Arianne A. Zamora, Devon Ray Pacial, Juan Antonio R. Magalang, Marissa Alejandria, Celia Carlos, Anna Ong-Lim, Edsel Maurice Salvaña, John Q. Wong, Jaime C. Montoya, Maria Rosario Singh-Vergeire and Cynthia P. Saloma |
| EPI_ISL_2188090, EPI_ISL_2188091, EPI_ISL_2188092 | The Medical City -Â Ortigas | Philippine Genome Center | Francis A. Tablizo, Kenneth M. Kim, Carlo M. Lapid, Marc Jerrone R. Castro, Maria Sofia L. Yangzon, Elcid Aaron R. Pangilinan, Benedict A. Maralit, Marc Edsel C. Ayes, Eva Maria Cutiongco-de la Paz, Alethea R. de Guzman, Jan Michael C. Yap, Jo-Hannah S. Llames, Sheila Mae M. Araiza, Kris P. Punayan, Irish Coleen A. Asin, Candice Francheska B. Tambaoan, Asia Louisa U. Chong, Karol Sophia Agape R. Padilla, Rianna Patricia S. Cruz, El King D. Morado, Joshua Gregor A. Dizon, Razel Nikka M. Hao, Arianne A. Zamora, Devon Ray Pacial, Juan Antonio R. Magalang, Marissa Alejandria, Celia Carlos, Anna Ong-Lim, Edsel Maurice Salvaña, John Q. Wong, Jaime C. Montoya, Maria Rosario Singh-Vergeire and Cynthia P. Saloma |
| EPI_ISL_2188093 | Ospital ng Imus | Philippine Genome Center | Francis A. Tablizo, Kenneth M. Kim, Carlo M. Lapid, Marc Jerrone R. Castro, Maria Sofia L. Yangzon, Elcid Aaron R. Pangilinan, Benedict A. Maralit, Marc Edsel C. Ayes, Eva Maria Cutiongco-de la Paz, Alethea R. de Guzman, Jan Michael C. Yap, Jo-Hannah S. Llames, Sheila Mae M. Araiza, Kris P. Punayan, Irish Coleen A. Asin, Candice Francheska B. Tambaoan, Asia Louisa U. Chong, Karol Sophia Agape R. Padilla, Rianna Patricia S. Cruz, El King D. Morado, Joshua Gregor A. Dizon, Razel Nikka M. Hao, Arianne A. Zamora, Devon Ray Pacial, Juan Antonio R. Magalang, Marissa Alejandria, Celia Carlos, Anna Ong-Lim, Edsel Maurice Salvaña, John Q. Wong, Jaime C. Montoya, Maria Rosario Singh-Vergeire and Cynthia P. Saloma |
| EPI_ISL_2188095 | The Medical City -Â Ortigas | Philippine Genome Center | Francis A. Tablizo, Kenneth M. Kim, Carlo M. Lapid, Marc Jerrone R. Castro, Maria Sofia L. Yangzon, Elcid Aaron R. Pangilinan, Benedict A. Maralit, Marc Edsel C. Ayes, Eva Maria Cutiongco-de la Paz, Alethea R. de Guzman, Jan Michael C. Yap, Jo-Hannah S. Llames, Sheila Mae M. Araiza, Kris P. Punayan, Irish Coleen A. Asin, Candice Francheska B. Tambaoan, Asia Louisa U. Chong, Karol Sophia Agape R. Padilla, Rianna Patricia S. Cruz, El King D. Morado, Joshua Gregor A. Dizon, Razel Nikka M. Hao, Arianne A. Zamora, Devon Ray Pacial, Juan Antonio R. Magalang, Marissa Alejandria, Celia Carlos, Anna Ong-Lim, Edsel Maurice Salvaña, John Q. Wong, Jaime C. Montoya, Maria Rosario Singh-Vergeire and Cynthia P. Saloma |
| EPI_ISL_2188096 | The Lord's Grace Medical and Industrial Clinic | Philippine Genome Center | Francis A. Tablizo, Kenneth M. Kim, Carlo M. Lapid, Marc Jerrone R. Castro, Maria Sofia L. Yangzon, Elcid Aaron R. Pangilinan, Benedict A. Maralit, Marc Edsel C. Ayes, Eva Maria Cutiongco-de la Paz, Alethea R. de Guzman, Jan Michael C. Yap, Jo-Hannah S. Llames, Sheila Mae M. Araiza, Kris P. Punayan, Irish Coleen A. Asin, Candice Francheska B. Tambaoan, Asia Louisa U. Chong, Karol Sophia Agape R. Padilla, Rianna Patricia S. Cruz, El King D. Morado, Joshua Gregor A. Dizon, Razel Nikka M. Hao, Arianne A. Zamora, Devon Ray Pacial, Juan Antonio R. Magalang, Marissa Alejandria, Celia Carlos, Anna Ong-Lim, Edsel Maurice Salvaña, John Q. Wong, Jaime C. Montoya, Maria Rosario Singh-Vergeire and Cynthia P. Saloma |
| EPI_ISL_2188097 | Butuan Medical Center | Philippine Genome Center | Francis A. Tablizo, Kenneth M. Kim, Carlo M. Lapid, Marc Jerrone R. Castro, Maria Sofia L. Yangzon, Elcid Aaron R. Pangilinan, Benedict A. Maralit, Marc Edsel C. Ayes, Eva Maria Cutiongco-de la Paz, Alethea R. de Guzman, Jan Michael C. Yap, Jo-Hannah S. Llames, Sheila Mae M. Araiza, Kris P. Punayan, Irish Coleen A. Asin, Candice Francheska B. Tambaoan, Asia Louisa U. Chong, Karol Sophia Agape R. Padilla, Rianna Patricia S. Cruz, El King D. Morado, Joshua Gregor A. Dizon, Razel Nikka M. Hao, Arianne A. Zamora, Devon Ray Pacial, Juan Antonio R. Magalang, Marissa Alejandria, Celia Carlos, Anna Ong-Lim, Edsel Maurice Salvaña, John Q. Wong, Jaime C. Montoya, Maria Rosario Singh-Vergeire and Cynthia P. Saloma |
| EPI_ISL_2188098 | The Medical City -Â Ortigas | Philippine Genome Center | Francis A. Tablizo, Kenneth M. Kim, Carlo M. Lapid, Marc Jerrone R. Castro, Maria Sofia L. Yangzon, Elcid Aaron R. Pangilinan, Benedict A. Maralit, Marc Edsel C. Ayes, Eva Maria Cutiongco-de la Paz, Alethea R. de Guzman, Jan Michael C. Yap, Jo-Hannah S. Llames, Sheila Mae M. Araiza, Kris P. Punayan, Irish Coleen A. Asin, Candice Francheska B. Tambaoan, Asia Louisa U. Chong, Karol Sophia Agape R. Padilla, Rianna Patricia S. Cruz, El King D. Morado, Joshua Gregor A. Dizon, Razel Nikka M. Hao, Arianne A. Zamora, Devon Ray Pacial, Juan Antonio R. Magalang, Marissa Alejandria, Celia Carlos, Anna Ong-Lim, Edsel Maurice Salvaña, John Q. Wong, Jaime C. Montoya, Maria Rosario Singh-Vergeire and Cynthia P. Saloma |
| EPI_ISL_2188099 | Philippine Red Cross - Port Area | Philippine Genome Center | Francis A. Tablizo, Kenneth M. Kim, Carlo M. Lapid, Marc Jerrone R. Castro, Maria Sofia L. Yangzon, Elcid Aaron R. Pangilinan, Benedict A. Maralit, Marc Edsel C. Ayes, Eva Maria Cutiongco-de la Paz, Alethea R. de Guzman, Jan Michael C. Yap, Jo-Hannah S. Llames, Sheila Mae M. Araiza, Kris P. Punayan, Irish Coleen A. Asin, Candice Francheska B. Tambaoan, Asia Louisa U. Chong, Karol Sophia Agape R. Padilla, Rianna Patricia S. Cruz, El King D. Morado, Joshua Gregor A. Dizon, Razel Nikka M. Hao, Arianne A. Zamora, Devon Ray Pacial, Juan Antonio R. Magalang, Marissa Alejandria, Celia Carlos, Anna Ong-Lim, Edsel Maurice Salvaña, John Q. Wong, Jaime C. Montoya, Maria Rosario Singh-Vergeire and Cynthia P. Saloma |
| EPI_ISL_2188100 | PASIG CITY CHILDREN'S HOSPITAL | Philippine Genome Center | Francis A. Tablizo, Kenneth M. Kim, Carlo M. Lapid, Marc Jerrone R. Castro, Maria Sofia L. Yangzon, Elcid Aaron R. Pangilinan, Benedict A. Maralit, Marc Edsel C. Ayes, Eva Maria Cutiongco-de la Paz, Alethea R. de Guzman, Jan Michael C. Yap, Jo-Hannah S. Llames, Sheila Mae M. Araiza, Kris P. Punayan, Irish Coleen A. Asin, Candice Francheska B. Tambaoan, Asia Louisa U. Chong, Karol Sophia Agape R. Padilla, Rianna Patricia S. Cruz, El King D. Morado, Joshua Gregor A. Dizon, Razel Nikka M. Hao, Arianne A. Zamora, Devon Ray Pacial, Juan Antonio R. Magalang, Marissa Alejandria, Celia Carlos, Anna Ong-Lim, Edsel Maurice Salvaña, John Q. Wong, Jaime C. Montoya, Maria Rosario Singh-Vergeire and Cynthia P. Saloma |
| EPI_ISL_2188101 | Ospital ng Imus | Philippine Genome Center | Francis A. Tablizo, Kenneth M. Kim, Carlo M. Lapid, Marc Jerrone R. Castro, Maria Sofia L. Yangzon, Elcid Aaron R. Pangilinan, Benedict A. Maralit, Marc |

| | | | |
|---|---|---|---|
| | | | Edsel C. Ayes, Eva Maria Cutiongco-de la Paz, Alethea R. de Guzman, Jan Michael C. Yap, Jo-Hannah S. Llames, Sheila Mae M. Araiza, Kris P. Punayan, Irish Coleen A. Asin, Candice Francheska B. Tambaoan, Asia Louisa U. Chong, Karol Sophia Agape R. Padilla, Rianna Patricia S. Cruz, El King D. Morado, Joshua Gregor A. Dizon, Razel Nikka M. Hao, Arianne A. Zamora, Devon Ray Pacial, Juan Antonio R. Magalang, Marissa Alejandria, Celia Carlos, Anna Ong-Lim, Edsel Maurice Salvaña, John Q. Wong, Jaime C. Montoya, Maria Rosario Singh-Vergeire and Cynthia P. Saloma |
| EPI_ISL_2188102 | The Lord's Grace Medical and Industrial Clinic | Philippine Genome Center | Francis A. Tablizo, Kenneth M. Kim, Carlo M. Lapid, Marc Jerrone R. Castro, Maria Sofia L. Yangzon, Elcid Aaron R. Pangilinan, Benedict A. Maralit, Marc Edsel C. Ayes, Eva Maria Cutiongco-de la Paz, Alethea R. de Guzman, Jan Michael C. Yap, Jo-Hannah S. Llames, Sheila Mae M. Araiza, Kris P. Punayan, Irish Coleen A. Asin, Candice Francheska B. Tambaoan, Asia Louisa U. Chong, Karol Sophia Agape R. Padilla, Rianna Patricia S. Cruz, El King D. Morado, Joshua Gregor A. Dizon, Razel Nikka M. Hao, Arianne A. Zamora, Devon Ray Pacial, Juan Antonio R. Magalang, Marissa Alejandria, Celia Carlos, Anna Ong-Lim, Edsel Maurice Salvaña, John Q. Wong, Jaime C. Montoya, Maria Rosario Singh-Vergeire and Cynthia P. Saloma |
| EPI_ISL_2188103 | University of the Philippines National Institutes of Health (UP NIH) | Philippine Genome Center | Francis A. Tablizo, Kenneth M. Kim, Carlo M. Lapid, Marc Jerrone R. Castro, Maria Sofia L. Yangzon, Elcid Aaron R. Pangilinan, Benedict A. Maralit, Marc Edsel C. Ayes, Eva Maria Cutiongco-de la Paz, Alethea R. de Guzman, Jan Michael C. Yap, Jo-Hannah S. Llames, Sheila Mae M. Araiza, Kris P. Punayan, Irish Coleen A. Asin, Candice Francheska B. Tambaoan, Asia Louisa U. Chong, Karol Sophia Agape R. Padilla, Rianna Patricia S. Cruz, El King D. Morado, Joshua Gregor A. Dizon, Razel Nikka M. Hao, Arianne A. Zamora, Devon Ray Pacial, Juan Antonio R. Magalang, Marissa Alejandria, Celia Carlos, Anna Ong-Lim, Edsel Maurice Salvaña, John Q. Wong, Jaime C. Montoya, Maria Rosario Singh-Vergeire and Cynthia P. Saloma |
| EPI_ISL_2188104 | The Lord's Grace Medical and Industrial Clinic | Philippine Genome Center | Francis A. Tablizo, Kenneth M. Kim, Carlo M. Lapid, Marc Jerrone R. Castro, Maria Sofia L. Yangzon, Elcid Aaron R. Pangilinan, Benedict A. Maralit, Marc Edsel C. Ayes, Eva Maria Cutiongco-de la Paz, Alethea R. de Guzman, Jan Michael C. Yap, Jo-Hannah S. Llames, Sheila Mae M. Araiza, Kris P. Punayan, Irish Coleen A. Asin, Candice Francheska B. Tambaoan, Asia Louisa U. Chong, Karol Sophia Agape R. Padilla, Rianna Patricia S. Cruz, El King D. Morado, Joshua Gregor A. Dizon, Razel Nikka M. Hao, Arianne A. Zamora, Devon Ray Pacial, Juan Antonio R. Magalang, Marissa Alejandria, Celia Carlos, Anna Ong-Lim, Edsel Maurice Salvaña, John Q. Wong, Jaime C. Montoya, Maria Rosario Singh-Vergeire and Cynthia P. Saloma |
| EPI_ISL_2188105 | Qualimed Hospital Sta. Rosa | Philippine Genome Center | Francis A. Tablizo, Kenneth M. Kim, Carlo M. Lapid, Marc Jerrone R. Castro, Maria Sofia L. Yangzon, Elcid Aaron R. Pangilinan, Benedict A. Maralit, Marc Edsel C. Ayes, Eva Maria Cutiongco-de la Paz, Alethea R. de Guzman, Jan Michael C. Yap, Jo-Hannah S. Llames, Sheila Mae M. Araiza, Kris P. Punayan, Irish Coleen A. Asin, Candice Francheska B. Tambaoan, Asia Louisa U. Chong, Karol Sophia Agape R. Padilla, Rianna Patricia S. Cruz, El King D. Morado, Joshua Gregor A. Dizon, Razel Nikka M. Hao, Arianne A. Zamora, Devon Ray Pacial, Juan Antonio R. Magalang, Marissa Alejandria, Celia Carlos, Anna Ong-Lim, Edsel Maurice Salvaña, John Q. Wong, Jaime C. Montoya, Maria Rosario Singh-Vergeire and Cynthia P. Saloma |
| EPI_ISL_2188106 | Ospital ng ParaÃ±aque II | Philippine Genome Center | Francis A. Tablizo, Kenneth M. Kim, Carlo M. Lapid, Marc Jerrone R. Castro, Maria Sofia L. Yangzon, Elcid Aaron R. Pangilinan, Benedict A. Maralit, Marc Edsel C. Ayes, Eva Maria Cutiongco-de la Paz, Alethea R. de Guzman, Jan Michael C. Yap, Jo-Hannah S. Llames, Sheila Mae M. Araiza, Kris P. Punayan, Irish Coleen A. Asin, Candice Francheska B. Tambaoan, Asia Louisa U. Chong, Karol Sophia Agape R. Padilla, Rianna Patricia S. Cruz, El King D. Morado, Joshua Gregor A. Dizon, Razel Nikka M. Hao, Arianne A. Zamora, Devon Ray Pacial, Juan Antonio R. Magalang, Marissa Alejandria, Celia Carlos, Anna Ong-Lim, Edsel Maurice Salvaña, John Q. Wong, Jaime C. Montoya, Maria Rosario Singh-Vergeire and Cynthia P. Saloma |
| EPI_ISL_2188107 | The Lord's Grace Medical and Industrial Clinic | Philippine Genome Center | Francis A. Tablizo, Kenneth M. Kim, Carlo M. Lapid, Marc Jerrone R. Castro, Maria Sofia L. Yangzon, Elcid Aaron R. Pangilinan, Benedict A. Maralit, Marc Edsel C. Ayes, Eva Maria Cutiongco-de la Paz, Alethea R. de Guzman, Jan Michael C. Yap, Jo-Hannah S. Llames, Sheila Mae M. Araiza, Kris P. Punayan, Irish Coleen A. Asin, Candice Francheska B. Tambaoan, Asia Louisa U. Chong, Karol Sophia Agape R. Padilla, Rianna Patricia S. Cruz, El King D. Morado, Joshua Gregor A. Dizon, Razel Nikka M. Hao, Arianne A. Zamora, Devon Ray Pacial, Juan Antonio R. Magalang, Marissa Alejandria, Celia Carlos, Anna Ong-Lim, Edsel Maurice Salvaña, John Q. Wong, Jaime C. Montoya, Maria Rosario Singh-Vergeire and Cynthia P. Saloma |
| EPI_ISL_2188108 | SAN ANTONIO CITY OF ILAGAN HOSPITAL | Philippine Genome Center | Francis A. Tablizo, Kenneth M. Kim, Carlo M. Lapid, Marc Jerrone R. Castro, Maria Sofia L. Yangzon, Elcid Aaron R. Pangilinan, Benedict A. Maralit, Marc Edsel C. Ayes, Eva Maria Cutiongco-de la Paz, Alethea R. de Guzman, Jan Michael C. Yap, Jo-Hannah S. Llames, Sheila Mae M. Araiza, Kris P. Punayan, Irish Coleen A. Asin, Candice Francheska B. Tambaoan, Asia Louisa U. Chong, Karol Sophia Agape R. Padilla, Rianna Patricia S. Cruz, El King D. Morado, Joshua Gregor A. Dizon, Razel Nikka M. Hao, Arianne A. Zamora, Devon Ray Pacial, Juan Antonio R. Magalang, Marissa Alejandria, Celia Carlos, Anna Ong-Lim, Edsel Maurice Salvaña, John Q. Wong, Jaime C. Montoya, Maria Rosario Singh-Vergeire and Cynthia P. Saloma |
| EPI_ISL_2188109 | Mary Mediatrix Medical Center | Philippine Genome Center | Francis A. Tablizo, Kenneth M. Kim, Carlo M. Lapid, Marc Jerrone R. Castro, Maria Sofia L. Yangzon, Elcid Aaron R. Pangilinan, Benedict A. Maralit, Marc Edsel C. Ayes, Eva Maria Cutiongco-de la Paz, Alethea R. de Guzman, Jan Michael C. Yap, Jo-Hannah S. Llames, Sheila Mae M. Araiza, Kris P. Punayan, Irish Coleen A. Asin, Candice Francheska B. Tambaoan, Asia Louisa U. Chong, Karol Sophia Agape R. Padilla, Rianna Patricia S. Cruz, El King D. Morado, Joshua Gregor A. Dizon, Razel Nikka M. Hao, Arianne A. Zamora, Devon Ray Pacial, Juan Antonio R. Magalang, Marissa Alejandria, Celia Carlos, Anna Ong-Lim, Edsel Maurice Salvaña, John Q. Wong, Jaime C. Montoya, Maria Rosario Singh-Vergeire and Cynthia P. Saloma |
| EPI_ISL_2188110 | Philippine Red Cross - National Blood Center | Philippine Genome Center | Francis A. Tablizo, Kenneth M. Kim, Carlo M. Lapid, Marc Jerrone R. Castro, Maria Sofia L. Yangzon, Elcid Aaron R. Pangilinan, Benedict A. Maralit, Marc Edsel C. Ayes, Eva Maria Cutiongco-de la Paz, Alethea R. de Guzman, Jan Michael C. Yap, Jo-Hannah S. Llames, Sheila Mae M. Araiza, Kris P. Punayan, Irish Coleen A. Asin, Candice Francheska B. Tambaoan, Asia Louisa U. Chong, Karol Sophia Agape R. Padilla, Rianna Patricia S. Cruz, El King D. Morado, Joshua Gregor A. Dizon, Razel Nikka M. Hao, Arianne A. Zamora, Devon Ray Pacial, Juan Antonio R. Magalang, Marissa Alejandria, Celia Carlos, Anna Ong-Lim, Edsel Maurice Salvaña, John Q. Wong, Jaime C. Montoya, Maria Rosario Singh-Vergeire and Cynthia P. Saloma |
| EPI_ISL_2188111 | The Lord's Grace Medical and Industrial Clinic | Philippine Genome Center | Francis A. Tablizo, Kenneth M. Kim, Carlo M. Lapid, Marc Jerrone R. Castro, Maria Sofia L. Yangzon, Elcid Aaron R. Pangilinan, Benedict A. Maralit, Marc Edsel C. Ayes, Eva Maria Cutiongco-de la Paz, Alethea R. de Guzman, Jan Michael C. Yap, Jo-Hannah S. Llames, Sheila Mae M. Araiza, Kris P. Punayan, Irish Coleen A. Asin, Candice Francheska B. Tambaoan, Asia Louisa U. Chong, Karol Sophia Agape R. Padilla, Rianna Patricia S. Cruz, El King D. Morado, Joshua Gregor A. Dizon, Razel Nikka M. Hao, Arianne A. Zamora, Devon Ray Pacial, Juan Antonio R. Magalang, Marissa Alejandria, Celia Carlos, Anna Ong-Lim, Edsel Maurice Salvaña, John Q. Wong, Jaime C. Montoya, Maria Rosario Singh-Vergeire and Cynthia P. Saloma |
| EPI_ISL_2188112 | Mariano Marcos Memorial Hospital Medical Center | Philippine Genome Center | Francis A. Tablizo, Kenneth M. Kim, Carlo M. Lapid, Marc Jerrone R. Castro, Maria Sofia L. Yangzon, Elcid Aaron R. Pangilinan, Benedict A. Maralit, Marc Edsel C. Ayes, Eva Maria Cutiongco-de la Paz, Alethea R. de Guzman, Jan Michael C. Yap, Jo-Hannah S. Llames, Sheila Mae M. Araiza, Kris P. Punayan, Irish Coleen A. Asin, Candice Francheska B. Tambaoan, Asia Louisa U. Chong, Karol Sophia Agape R. Padilla, Rianna Patricia S. Cruz, El King D. Morado, Joshua Gregor A. Dizon, Razel Nikka M. Hao, Arianne A. Zamora, Devon Ray Pacial, Juan Antonio R. Magalang, Marissa Alejandria, Celia Carlos, Anna Ong-Lim, Edsel Maurice Salvaña, John Q. Wong, Jaime C. Montoya, Maria Rosario Singh-Vergeire and Cynthia P. Saloma |
| EPI_ISL_2188114 | The Lord's Grace Medical and Industrial Clinic | Philippine Genome Center | Francis A. Tablizo, Kenneth M. Kim, Carlo M. Lapid, Marc Jerrone R. Castro, Maria Sofia L. Yangzon, Elcid Aaron R. Pangilinan, Benedict A. Maralit, Marc Edsel C. Ayes, Eva Maria Cutiongco-de la Paz, Alethea R. de Guzman, Jan Michael C. Yap, Jo-Hannah S. Llames, Sheila Mae M. Araiza, Kris P. Punayan, Irish Coleen A. Asin, Candice Francheska B. Tambaoan, Asia Louisa U. Chong, Karol Sophia Agape R. Padilla, Rianna Patricia S. Cruz, El King D. Morado, Joshua Gregor A. Dizon, Razel Nikka M. Hao, Arianne A. Zamora, Devon Ray Pacial, Juan Antonio R. Magalang, Marissa Alejandria, Celia Carlos, Anna Ong-Lim, Edsel Maurice Salvaña, John Q. Wong, Jaime C. Montoya, Maria Rosario Singh-Vergeire and Cynthia P. Saloma |
| EPI_ISL_2188117 | Ospital ng ParaÃ±aque II | Philippine Genome Center | Francis A. Tablizo, Kenneth M. Kim, Carlo M. Lapid, Marc Jerrone R. Castro, Maria Sofia L. Yangzon, Elcid Aaron R. Pangilinan, Benedict A. Maralit, Marc Edsel C. Ayes, Eva Maria Cutiongco-de la Paz, Alethea R. de Guzman, Jan Michael C. Yap, Jo-Hannah S. Llames, Sheila Mae M. Araiza, Kris P. Punayan, Irish Coleen A. Asin, Candice Francheska B. Tambaoan, Asia Louisa U. Chong, Karol Sophia Agape R. Padilla, Rianna Patricia S. Cruz, El King D. Morado, Joshua Gregor A. Dizon, Razel Nikka M. Hao, Arianne A. Zamora, Devon Ray Pacial, Juan Antonio R. Magalang, Marissa Alejandria, Celia Carlos, Anna Ong-Lim, Edsel Maurice Salvaña, John Q. Wong, Jaime C. Montoya, Maria Rosario Singh-Vergeire and Cynthia P. Saloma |
| EPI_ISL_2188118 | The Lord's Grace Medical and Industrial Clinic | Philippine Genome Center | Francis A. Tablizo, Kenneth M. Kim, Carlo M. Lapid, Marc Jerrone R. Castro, Maria Sofia L. Yangzon, Elcid Aaron R. Pangilinan, Benedict A. Maralit, Marc Edsel C. Ayes, Eva Maria Cutiongco-de la Paz, Alethea R. de Guzman, Jan Michael C. Yap, Jo-Hannah S. Llames, Sheila Mae M. Araiza, Kris P. Punayan, Irish Coleen A. Asin, Candice Francheska B. Tambaoan, Asia Louisa U. Chong, Karol Sophia Agape R. Padilla, Rianna Patricia S. Cruz, El King D. Morado, Joshua Gregor A. Dizon, Razel Nikka M. Hao, Arianne A. Zamora, Devon Ray Pacial, Juan Antonio R. Magalang, Marissa Alejandria, Celia Carlos, Anna Ong-Lim, Edsel Maurice Salvaña, John Q. Wong, Jaime C. Montoya, Maria Rosario Singh-Vergeire and Cynthia P. Saloma |
| EPI_ISL_2188119 | The Medical City -Â Ortigas | Philippine Genome Center | Francis A. Tablizo, Kenneth M. Kim, Carlo M. Lapid, Marc Jerrone R. Castro, Maria Sofia L. Yangzon, Elcid Aaron R. Pangilinan, Benedict A. Maralit, Marc Edsel C. Ayes, Eva Maria Cutiongco-de la Paz, Alethea R. de Guzman, Jan Michael C. Yap, Jo-Hannah S. Llames, Sheila Mae M. Araiza, Kris P. Punayan, Irish Coleen A. Asin, Candice Francheska B. Tambaoan, Asia Louisa U. Chong, Karol Sophia Agape R. Padilla, Rianna Patricia S. Cruz, El King D. Morado, Joshua Gregor A. Dizon, Razel Nikka M. Hao, Arianne A. Zamora, Devon Ray Pacial, Juan Antonio R. Magalang, Marissa Alejandria, Celia Carlos, Anna Ong-Lim, Edsel Maurice Salvaña, John Q. Wong, Jaime C. Montoya, Maria Rosario Singh-Vergeire and Cynthia P. Saloma |

| | | | |
|---|---|---|---|
| EPI_ISL_2188121 | Dr. Jorge P. Royeca Hospital | Philippine Genome Center | Francis A. Tablizo, Kenneth M. Kim, Carlo M. Lapid, Marc Jerrone R. Castro, Maria Sofia L. Yangzon, Elcid Aaron R. Pangilinan, Benedict A. Maralit, Marc Edsel C. Ayes, Eva Maria Cutiongco-de la Paz, Alethea R. de Guzman, Jan Michael C. Yap, Jo-Hannah S. Llames, Sheila Mae M. Araiza, Kris P. Punayan, Irish Coleen A. Asin, Candice Francheska B. Tambaoan, Asia Louisa U. Chong, Karol Sophia Agape R. Padilla, Rianna Patricia S. Cruz, El King D. Morado, Joshua Gregor A. Dizon, Razel Nikka M. Hao, Arianne A. Zamora, Devon Ray Pacial, Juan Antonio R. Magalang, Marissa Alejandria, Celia Carlos, Anna Ong-Lim, Edsel Maurice Salvaña, John Q. Wong, Jaime C. Montoya, Maria Rosario Singh-Vergeire and Cynthia P. Saloma |
| EPI_ISL_2188122 | Philippine Red Cross Logistics and Multipurpose Center | Philippine Genome Center | Francis A. Tablizo, Kenneth M. Kim, Carlo M. Lapid, Marc Jerrone R. Castro, Maria Sofia L. Yangzon, Elcid Aaron R. Pangilinan, Benedict A. Maralit, Marc Edsel C. Ayes, Eva Maria Cutiongco-de la Paz, Alethea R. de Guzman, Jan Michael C. Yap, Jo-Hannah S. Llames, Sheila Mae M. Araiza, Kris P. Punayan, Irish Coleen A. Asin, Candice Francheska B. Tambaoan, Asia Louisa U. Chong, Karol Sophia Agape R. Padilla, Rianna Patricia S. Cruz, El King D. Morado, Joshua Gregor A. Dizon, Razel Nikka M. Hao, Arianne A. Zamora, Devon Ray Pacial, Juan Antonio R. Magalang, Marissa Alejandria, Celia Carlos, Anna Ong-Lim, Edsel Maurice Salvaña, John Q. Wong, Jaime C. Montoya, Maria Rosario Singh-Vergeire and Cynthia P. Saloma |
| EPI_ISL_2188123 | University of the Philippines National Institutes of Health (UP NIH) | Philippine Genome Center | Francis A. Tablizo, Kenneth M. Kim, Carlo M. Lapid, Marc Jerrone R. Castro, Maria Sofia L. Yangzon, Elcid Aaron R. Pangilinan, Benedict A. Maralit, Marc Edsel C. Ayes, Eva Maria Cutiongco-de la Paz, Alethea R. de Guzman, Jan Michael C. Yap, Jo-Hannah S. Llames, Sheila Mae M. Araiza, Kris P. Punayan, Irish Coleen A. Asin, Candice Francheska B. Tambaoan, Asia Louisa U. Chong, Karol Sophia Agape R. Padilla, Rianna Patricia S. Cruz, El King D. Morado, Joshua Gregor A. Dizon, Razel Nikka M. Hao, Arianne A. Zamora, Devon Ray Pacial, Juan Antonio R. Magalang, Marissa Alejandria, Celia Carlos, Anna Ong-Lim, Edsel Maurice Salvaña, John Q. Wong, Jaime C. Montoya, Maria Rosario Singh-Vergeire and Cynthia P. Saloma |
| EPI_ISL_2188126 | SAN ANTONIO CITY OF ILAGAN HOSPITAL | Philippine Genome Center | Francis A. Tablizo, Kenneth M. Kim, Carlo M. Lapid, Marc Jerrone R. Castro, Maria Sofia L. Yangzon, Elcid Aaron R. Pangilinan, Benedict A. Maralit, Marc Edsel C. Ayes, Eva Maria Cutiongco-de la Paz, Alethea R. de Guzman, Jan Michael C. Yap, Jo-Hannah S. Llames, Sheila Mae M. Araiza, Kris P. Punayan, Irish Coleen A. Asin, Candice Francheska B. Tambaoan, Asia Louisa U. Chong, Karol Sophia Agape R. Padilla, Rianna Patricia S. Cruz, El King D. Morado, Joshua Gregor A. Dizon, Razel Nikka M. Hao, Arianne A. Zamora, Devon Ray Pacial, Juan Antonio R. Magalang, Marissa Alejandria, Celia Carlos, Anna Ong-Lim, Edsel Maurice Salvaña, John Q. Wong, Jaime C. Montoya, Maria Rosario Singh-Vergeire and Cynthia P. Saloma |
| EPI_ISL_2188127 | Qualimed Hospital Sta. Rosa | Philippine Genome Center | Francis A. Tablizo, Kenneth M. Kim, Carlo M. Lapid, Marc Jerrone R. Castro, Maria Sofia L. Yangzon, Elcid Aaron R. Pangilinan, Benedict A. Maralit, Marc Edsel C. Ayes, Eva Maria Cutiongco-de la Paz, Alethea R. de Guzman, Jan Michael C. Yap, Jo-Hannah S. Llames, Sheila Mae M. Araiza, Kris P. Punayan, Irish Coleen A. Asin, Candice Francheska B. Tambaoan, Asia Louisa U. Chong, Karol Sophia Agape R. Padilla, Rianna Patricia S. Cruz, El King D. Morado, Joshua Gregor A. Dizon, Razel Nikka M. Hao, Arianne A. Zamora, Devon Ray Pacial, Juan Antonio R. Magalang, Marissa Alejandria, Celia Carlos, Anna Ong-Lim, Edsel Maurice Salvaña, John Q. Wong, Jaime C. Montoya, Maria Rosario Singh-Vergeire and Cynthia P. Saloma |
| EPI_ISL_2188129 | PNP General Hospital | Philippine Genome Center | Francis A. Tablizo, Kenneth M. Kim, Carlo M. Lapid, Marc Jerrone R. Castro, Maria Sofia L. Yangzon, Elcid Aaron R. Pangilinan, Benedict A. Maralit, Marc Edsel C. Ayes, Eva Maria Cutiongco-de la Paz, Alethea R. de Guzman, Jan Michael C. Yap, Jo-Hannah S. Llames, Sheila Mae M. Araiza, Kris P. Punayan, Irish Coleen A. Asin, Candice Francheska B. Tambaoan, Asia Louisa U. Chong, Karol Sophia Agape R. Padilla, Rianna Patricia S. Cruz, El King D. Morado, Joshua Gregor A. Dizon, Razel Nikka M. Hao, Arianne A. Zamora, Devon Ray Pacial, Juan Antonio R. Magalang, Marissa Alejandria, Celia Carlos, Anna Ong-Lim, Edsel Maurice Salvaña, John Q. Wong, Jaime C. Montoya, Maria Rosario Singh-Vergeire and Cynthia P. Saloma |
| EPI_ISL_2188130, EPI_ISL_2188131, EPI_ISL_2188132 | The Medical City -Â Ortigas | Philippine Genome Center | Francis A. Tablizo, Kenneth M. Kim, Carlo M. Lapid, Marc Jerrone R. Castro, Maria Sofia L. Yangzon, Elcid Aaron R. Pangilinan, Benedict A. Maralit, Marc Edsel C. Ayes, Eva Maria Cutiongco-de la Paz, Alethea R. de Guzman, Jan Michael C. Yap, Jo-Hannah S. Llames, Sheila Mae M. Araiza, Kris P. Punayan, Irish Coleen A. Asin, Candice Francheska B. Tambaoan, Asia Louisa U. Chong, Karol Sophia Agape R. Padilla, Rianna Patricia S. Cruz, El King D. Morado, Joshua Gregor A. Dizon, Razel Nikka M. Hao, Arianne A. Zamora, Devon Ray Pacial, Juan Antonio R. Magalang, Marissa Alejandria, Celia Carlos, Anna Ong-Lim, Edsel Maurice Salvaña, John Q. Wong, Jaime C. Montoya, Maria Rosario Singh-Vergeire and Cynthia P. Saloma |
| EPI_ISL_2188133 | Philippine Red Cross Logistics and Multipurpose Center | Philippine Genome Center | Francis A. Tablizo, Kenneth M. Kim, Carlo M. Lapid, Marc Jerrone R. Castro, Maria Sofia L. Yangzon, Elcid Aaron R. Pangilinan, Benedict A. Maralit, Marc Edsel C. Ayes, Eva Maria Cutiongco-de la Paz, Alethea R. de Guzman, Jan Michael C. Yap, Jo-Hannah S. Llames, Sheila Mae M. Araiza, Kris P. Punayan, Irish Coleen A. Asin, Candice Francheska B. Tambaoan, Asia Louisa U. Chong, Karol Sophia Agape R. Padilla, Rianna Patricia S. Cruz, El King D. Morado, Joshua Gregor A. Dizon, Razel Nikka M. Hao, Arianne A. Zamora, Devon Ray Pacial, Juan Antonio R. Magalang, Marissa Alejandria, Celia Carlos, Anna Ong-Lim, Edsel Maurice Salvaña, John Q. Wong, Jaime C. Montoya, Maria Rosario Singh-Vergeire and Cynthia P. Saloma |
| EPI_ISL_2188134 | The Medical City -Â Ortigas | Philippine Genome Center | Francis A. Tablizo, Kenneth M. Kim, Carlo M. Lapid, Marc Jerrone R. Castro, Maria Sofia L. Yangzon, Elcid Aaron R. Pangilinan, Benedict A. Maralit, Marc Edsel C. Ayes, Eva Maria Cutiongco-de la Paz, Alethea R. de Guzman, Jan Michael C. Yap, Jo-Hannah S. Llames, Sheila Mae M. Araiza, Kris P. Punayan, Irish Coleen A. Asin, Candice Francheska B. Tambaoan, Asia Louisa U. Chong, Karol Sophia Agape R. Padilla, Rianna Patricia S. Cruz, El King D. Morado, Joshua Gregor A. Dizon, Razel Nikka M. Hao, Arianne A. Zamora, Devon Ray Pacial, Juan Antonio R. Magalang, Marissa Alejandria, Celia Carlos, Anna Ong-Lim, Edsel Maurice Salvaña, John Q. Wong, Jaime C. Montoya, Maria Rosario Singh-Vergeire and Cynthia P. Saloma |
| EPI_ISL_2188135 | SAN ANTONIO CITY OF ILAGAN HOSPITAL | Philippine Genome Center | Francis A. Tablizo, Kenneth M. Kim, Carlo M. Lapid, Marc Jerrone R. Castro, Maria Sofia L. Yangzon, Elcid Aaron R. Pangilinan, Benedict A. Maralit, Marc Edsel C. Ayes, Eva Maria Cutiongco-de la Paz, Alethea R. de Guzman, Jan Michael C. Yap, Jo-Hannah S. Llames, Sheila Mae M. Araiza, Kris P. Punayan, Irish Coleen A. Asin, Candice Francheska B. Tambaoan, Asia Louisa U. Chong, Karol Sophia Agape R. Padilla, Rianna Patricia S. Cruz, El King D. Morado, Joshua Gregor A. Dizon, Razel Nikka M. Hao, Arianne A. Zamora, Devon Ray Pacial, Juan Antonio R. Magalang, Marissa Alejandria, Celia Carlos, Anna Ong-Lim, Edsel Maurice Salvaña, John Q. Wong, Jaime C. Montoya, Maria Rosario Singh-Vergeire and Cynthia P. Saloma |
| EPI_ISL_2188136, EPI_ISL_2188137 | The Medical City -Â Ortigas | Philippine Genome Center | Francis A. Tablizo, Kenneth M. Kim, Carlo M. Lapid, Marc Jerrone R. Castro, Maria Sofia L. Yangzon, Elcid Aaron R. Pangilinan, Benedict A. Maralit, Marc Edsel C. Ayes, Eva Maria Cutiongco-de la Paz, Alethea R. de Guzman, Jan Michael C. Yap, Jo-Hannah S. Llames, Sheila Mae M. Araiza, Kris P. Punayan, Irish Coleen A. Asin, Candice Francheska B. Tambaoan, Asia Louisa U. Chong, Karol Sophia Agape R. Padilla, Rianna Patricia S. Cruz, El King D. Morado, Joshua Gregor A. Dizon, Razel Nikka M. Hao, Arianne A. Zamora, Devon Ray Pacial, Juan Antonio R. Magalang, Marissa Alejandria, Celia Carlos, Anna Ong-Lim, Edsel Maurice Salvaña, John Q. Wong, Jaime C. Montoya, Maria Rosario Singh-Vergeire and Cynthia P. Saloma |
| EPI_ISL_2188138 | Philippine Red Cross Logistics and Multipurpose Center | Philippine Genome Center | Francis A. Tablizo, Kenneth M. Kim, Carlo M. Lapid, Marc Jerrone R. Castro, Maria Sofia L. Yangzon, Elcid Aaron R. Pangilinan, Benedict A. Maralit, Marc Edsel C. Ayes, Eva Maria Cutiongco-de la Paz, Alethea R. de Guzman, Jan Michael C. Yap, Jo-Hannah S. Llames, Sheila Mae M. Araiza, Kris P. Punayan, Irish Coleen A. Asin, Candice Francheska B. Tambaoan, Asia Louisa U. Chong, Karol Sophia Agape R. Padilla, Rianna Patricia S. Cruz, El King D. Morado, Joshua Gregor A. Dizon, Razel Nikka M. Hao, Arianne A. Zamora, Devon Ray Pacial, Juan Antonio R. Magalang, Marissa Alejandria, Celia Carlos, Anna Ong-Lim, Edsel Maurice Salvaña, John Q. Wong, Jaime C. Montoya, Maria Rosario Singh-Vergeire and Cynthia P. Saloma |
| EPI_ISL_2188139 | The Lord's Grace Medical and Industrial Clinic | Philippine Genome Center | Francis A. Tablizo, Kenneth M. Kim, Carlo M. Lapid, Marc Jerrone R. Castro, Maria Sofia L. Yangzon, Elcid Aaron R. Pangilinan, Benedict A. Maralit, Marc Edsel C. Ayes, Eva Maria Cutiongco-de la Paz, Alethea R. de Guzman, Jan Michael C. Yap, Jo-Hannah S. Llames, Sheila Mae M. Araiza, Kris P. Punayan, Irish Coleen A. Asin, Candice Francheska B. Tambaoan, Asia Louisa U. Chong, Karol Sophia Agape R. Padilla, Rianna Patricia S. Cruz, El King D. Morado, Joshua Gregor A. Dizon, Razel Nikka M. Hao, Arianne A. Zamora, Devon Ray Pacial, Juan Antonio R. Magalang, Marissa Alejandria, Celia Carlos, Anna Ong-Lim, Edsel Maurice Salvaña, John Q. Wong, Jaime C. Montoya, Maria Rosario Singh-Vergeire and Cynthia P. Saloma |
| EPI_ISL_2188140 | PASIG CITY CHILDREN'S HOSPITAL | Philippine Genome Center | Francis A. Tablizo, Kenneth M. Kim, Carlo M. Lapid, Marc Jerrone R. Castro, Maria Sofia L. Yangzon, Elcid Aaron R. Pangilinan, Benedict A. Maralit, Marc Edsel C. Ayes, Eva Maria Cutiongco-de la Paz, Alethea R. de Guzman, Jan Michael C. Yap, Jo-Hannah S. Llames, Sheila Mae M. Araiza, Kris P. Punayan, Irish Coleen A. Asin, Candice Francheska B. Tambaoan, Asia Louisa U. Chong, Karol Sophia Agape R. Padilla, Rianna Patricia S. Cruz, El King D. Morado, Joshua Gregor A. Dizon, Razel Nikka M. Hao, Arianne A. Zamora, Devon Ray Pacial, Juan Antonio R. Magalang, Marissa Alejandria, Celia Carlos, Anna Ong-Lim, Edsel Maurice Salvaña, John Q. Wong, Jaime C. Montoya, Maria Rosario Singh-Vergeire and Cynthia P. Saloma |
| EPI_ISL_2188141 | The Medical City -Â Ortigas | Philippine Genome Center | Francis A. Tablizo, Kenneth M. Kim, Carlo M. Lapid, Marc Jerrone R. Castro, Maria Sofia L. Yangzon, Elcid Aaron R. Pangilinan, Benedict A. Maralit, Marc Edsel C. Ayes, Eva Maria Cutiongco-de la Paz, Alethea R. de Guzman, Jan Michael C. Yap, Jo-Hannah S. Llames, Sheila Mae M. Araiza, Kris P. Punayan, Irish Coleen A. Asin, Candice Francheska B. Tambaoan, Asia Louisa U. Chong, Karol Sophia Agape R. Padilla, Rianna Patricia S. Cruz, El King D. Morado, Joshua Gregor A. Dizon, Razel Nikka M. Hao, Arianne A. Zamora, Devon Ray Pacial, Juan Antonio R. Magalang, Marissa Alejandria, Celia Carlos, Anna Ong-Lim, Edsel Maurice Salvaña, John Q. Wong, Jaime C. Montoya, Maria Rosario Singh-Vergeire and Cynthia P. Saloma |
| EPI_ISL_2188142 | Baguio General Hospital Medical Center (BGHMC) | Philippine Genome Center | Francis A. Tablizo, Kenneth M. Kim, Carlo M. Lapid, Marc Jerrone R. Castro, Maria Sofia L. Yangzon, Elcid Aaron R. Pangilinan, Benedict A. Maralit, Marc Edsel C. Ayes, Eva Maria Cutiongco-de la Paz, Alethea R. de Guzman, Jan Michael C. Yap, Jo-Hannah S. Llames, Sheila Mae M. Araiza, Kris P. Punayan, Irish Coleen A. Asin, Candice Francheska B. Tambaoan, Asia Louisa U. Chong, Karol Sophia Agape R. Padilla, Rianna Patricia S. Cruz, El King D. Morado, Joshua Gregor A. Dizon, Razel Nikka M. Hao, Arianne A. Zamora, Devon Ray Pacial, Juan Antonio R. Magalang, Marissa Alejandria, Celia Carlos, Anna Ong-Lim, Edsel Maurice Salvaña, John Q. Wong, Jaime C. Montoya, Maria Rosario Singh-Vergeire and Cynthia P. Saloma |

| | | | |
|---|---|---|---|
| EPI_ISL_2188143 | Sta. Rosa CHO | Philippine Genome Center | Francis A. Tablizo, Kenneth M. Kim, Carlo M. Lapid, Marc Jerrone R. Castro, Maria Sofia L. Yangzon, Elcid Aaron R. Pangilinan, Benedict A. Maralit, Marc Edsel C. Ayes, Eva Maria Cutiongco-de la Paz, Alethea R. de Guzman, Jan Michael C. Yap, Jo-Hannah S. Llames, Sheila Mae M. Araiza, Kris P. Punayan, Irish Coleen A. Asin, Candice Francheska B. Tambaoan, Asia Louisa U. Chong, Karol Sophia Agape R. Padilla, Rianna Patricia S. Cruz, El King D. Morado, Joshua Gregor A. Dizon, Razel Nikka M. Hao, Arianne A. Zamora, Devon Ray Pacial, Juan Antonio R. Magalang, Marissa Alejandria, Celia Carlos, Anna Ong-Lim, Edsel Maurice Salvaña, John Q. Wong, Jaime C. Montoya, Maria Rosario Singh-Vergeire and Cynthia P. Saloma |
| EPI_ISL_2188144, EPI_ISL_2188145, EPI_ISL_2188146 | The Medical City -Â Ortigas | Philippine Genome Center | Francis A. Tablizo, Kenneth M. Kim, Carlo M. Lapid, Marc Jerrone R. Castro, Maria Sofia L. Yangzon, Elcid Aaron R. Pangilinan, Benedict A. Maralit, Marc Edsel C. Ayes, Eva Maria Cutiongco-de la Paz, Alethea R. de Guzman, Jan Michael C. Yap, Jo-Hannah S. Llames, Sheila Mae M. Araiza, Kris P. Punayan, Irish Coleen A. Asin, Candice Francheska B. Tambaoan, Asia Louisa U. Chong, Karol Sophia Agape R. Padilla, Rianna Patricia S. Cruz, El King D. Morado, Joshua Gregor A. Dizon, Razel Nikka M. Hao, Arianne A. Zamora, Devon Ray Pacial, Juan Antonio R. Magalang, Marissa Alejandria, Celia Carlos, Anna Ong-Lim, Edsel Maurice Salvaña, John Q. Wong, Jaime C. Montoya, Maria Rosario Singh-Vergeire and Cynthia P. Saloma |
| EPI_ISL_2188147 | Philippine Red Cross - Port Area | Philippine Genome Center | Francis A. Tablizo, Kenneth M. Kim, Carlo M. Lapid, Marc Jerrone R. Castro, Maria Sofia L. Yangzon, Elcid Aaron R. Pangilinan, Benedict A. Maralit, Marc Edsel C. Ayes, Eva Maria Cutiongco-de la Paz, Alethea R. de Guzman, Jan Michael C. Yap, Jo-Hannah S. Llames, Sheila Mae M. Araiza, Kris P. Punayan, Irish Coleen A. Asin, Candice Francheska B. Tambaoan, Asia Louisa U. Chong, Karol Sophia Agape R. Padilla, Rianna Patricia S. Cruz, El King D. Morado, Joshua Gregor A. Dizon, Razel Nikka M. Hao, Arianne A. Zamora, Devon Ray Pacial, Juan Antonio R. Magalang, Marissa Alejandria, Celia Carlos, Anna Ong-Lim, Edsel Maurice Salvaña, John Q. Wong, Jaime C. Montoya, Maria Rosario Singh-Vergeire and Cynthia P. Saloma |
| EPI_ISL_2188148 | Ospital ng Imus | Philippine Genome Center | Francis A. Tablizo, Kenneth M. Kim, Carlo M. Lapid, Marc Jerrone R. Castro, Maria Sofia L. Yangzon, Elcid Aaron R. Pangilinan, Benedict A. Maralit, Marc Edsel C. Ayes, Eva Maria Cutiongco-de la Paz, Alethea R. de Guzman, Jan Michael C. Yap, Jo-Hannah S. Llames, Sheila Mae M. Araiza, Kris P. Punayan, Irish Coleen A. Asin, Candice Francheska B. Tambaoan, Asia Louisa U. Chong, Karol Sophia Agape R. Padilla, Rianna Patricia S. Cruz, El King D. Morado, Joshua Gregor A. Dizon, Razel Nikka M. Hao, Arianne A. Zamora, Devon Ray Pacial, Juan Antonio R. Magalang, Marissa Alejandria, Celia Carlos, Anna Ong-Lim, Edsel Maurice Salvaña, John Q. Wong, Jaime C. Montoya, Maria Rosario Singh-Vergeire and Cynthia P. Saloma |
| EPI_ISL_2188149, EPI_ISL_2188150, EPI_ISL_2188151, EPI_ISL_2188152, EPI_ISL_2188153 | The Medical City -Â Ortigas | Philippine Genome Center | Francis A. Tablizo, Kenneth M. Kim, Carlo M. Lapid, Marc Jerrone R. Castro, Maria Sofia L. Yangzon, Elcid Aaron R. Pangilinan, Benedict A. Maralit, Marc Edsel C. Ayes, Eva Maria Cutiongco-de la Paz, Alethea R. de Guzman, Jan Michael C. Yap, Jo-Hannah S. Llames, Sheila Mae M. Araiza, Kris P. Punayan, Irish Coleen A. Asin, Candice Francheska B. Tambaoan, Asia Louisa U. Chong, Karol Sophia Agape R. Padilla, Rianna Patricia S. Cruz, El King D. Morado, Joshua Gregor A. Dizon, Razel Nikka M. Hao, Arianne A. Zamora, Devon Ray Pacial, Juan Antonio R. Magalang, Marissa Alejandria, Celia Carlos, Anna Ong-Lim, Edsel Maurice Salvaña, John Q. Wong, Jaime C. Montoya, Maria Rosario Singh-Vergeire and Cynthia P. Saloma |
| EPI_ISL_2188154 | Philippine Red Cross - National Blood Center | Philippine Genome Center | Francis A. Tablizo, Kenneth M. Kim, Carlo M. Lapid, Marc Jerrone R. Castro, Maria Sofia L. Yangzon, Elcid Aaron R. Pangilinan, Benedict A. Maralit, Marc Edsel C. Ayes, Eva Maria Cutiongco-de la Paz, Alethea R. de Guzman, Jan Michael C. Yap, Jo-Hannah S. Llames, Sheila Mae M. Araiza, Kris P. Punayan, Irish Coleen A. Asin, Candice Francheska B. Tambaoan, Asia Louisa U. Chong, Karol Sophia Agape R. Padilla, Rianna Patricia S. Cruz, El King D. Morado, Joshua Gregor A. Dizon, Razel Nikka M. Hao, Arianne A. Zamora, Devon Ray Pacial, Juan Antonio R. Magalang, Marissa Alejandria, Celia Carlos, Anna Ong-Lim, Edsel Maurice Salvaña, John Q. Wong, Jaime C. Montoya, Maria Rosario Singh-Vergeire and Cynthia P. Saloma |
| EPI_ISL_2188155 | The Lord's Grace Medical and Industrial Clinic | Philippine Genome Center | Francis A. Tablizo, Kenneth M. Kim, Carlo M. Lapid, Marc Jerrone R. Castro, Maria Sofia L. Yangzon, Elcid Aaron R. Pangilinan, Benedict A. Maralit, Marc Edsel C. Ayes, Eva Maria Cutiongco-de la Paz, Alethea R. de Guzman, Jan Michael C. Yap, Jo-Hannah S. Llames, Sheila Mae M. Araiza, Kris P. Punayan, Irish Coleen A. Asin, Candice Francheska B. Tambaoan, Asia Louisa U. Chong, Karol Sophia Agape R. Padilla, Rianna Patricia S. Cruz, El King D. Morado, Joshua Gregor A. Dizon, Razel Nikka M. Hao, Arianne A. Zamora, Devon Ray Pacial, Juan Antonio R. Magalang, Marissa Alejandria, Celia Carlos, Anna Ong-Lim, Edsel Maurice Salvaña, John Q. Wong, Jaime C. Montoya, Maria Rosario Singh-Vergeire and Cynthia P. Saloma |
| EPI_ISL_2188156, EPI_ISL_2188157 | The Medical City -Â Ortigas | Philippine Genome Center | Francis A. Tablizo, Kenneth M. Kim, Carlo M. Lapid, Marc Jerrone R. Castro, Maria Sofia L. Yangzon, Elcid Aaron R. Pangilinan, Benedict A. Maralit, Marc Edsel C. Ayes, Eva Maria Cutiongco-de la Paz, Alethea R. de Guzman, Jan Michael C. Yap, Jo-Hannah S. Llames, Sheila Mae M. Araiza, Kris P. Punayan, Irish Coleen A. Asin, Candice Francheska B. Tambaoan, Asia Louisa U. Chong, Karol Sophia Agape R. Padilla, Rianna Patricia S. Cruz, El King D. Morado, Joshua Gregor A. Dizon, Razel Nikka M. Hao, Arianne A. Zamora, Devon Ray Pacial, Juan Antonio R. Magalang, Marissa Alejandria, Celia Carlos, Anna Ong-Lim, Edsel Maurice Salvaña, John Q. Wong, Jaime C. Montoya, Maria Rosario Singh-Vergeire and Cynthia P. Saloma |
| EPI_ISL_2188158 | PASIG CITY CHILDREN'S HOSPITAL | Philippine Genome Center | Francis A. Tablizo, Kenneth M. Kim, Carlo M. Lapid, Marc Jerrone R. Castro, Maria Sofia L. Yangzon, Elcid Aaron R. Pangilinan, Benedict A. Maralit, Marc Edsel C. Ayes, Eva Maria Cutiongco-de la Paz, Alethea R. de Guzman, Jan Michael C. Yap, Jo-Hannah S. Llames, Sheila Mae M. Araiza, Kris P. Punayan, Irish Coleen A. Asin, Candice Francheska B. Tambaoan, Asia Louisa U. Chong, Karol Sophia Agape R. Padilla, Rianna Patricia S. Cruz, El King D. Morado, Joshua Gregor A. Dizon, Razel Nikka M. Hao, Arianne A. Zamora, Devon Ray Pacial, Juan Antonio R. Magalang, Marissa Alejandria, Celia Carlos, Anna Ong-Lim, Edsel Maurice Salvaña, John Q. Wong, Jaime C. Montoya, Maria Rosario Singh-Vergeire and Cynthia P. Saloma |
| EPI_ISL_2188159 | National Kidney and Transplant Institute (NKTI) | Philippine Genome Center | Francis A. Tablizo, Kenneth M. Kim, Carlo M. Lapid, Marc Jerrone R. Castro, Maria Sofia L. Yangzon, Elcid Aaron R. Pangilinan, Benedict A. Maralit, Marc Edsel C. Ayes, Eva Maria Cutiongco-de la Paz, Alethea R. de Guzman, Jan Michael C. Yap, Jo-Hannah S. Llames, Sheila Mae M. Araiza, Kris P. Punayan, Irish Coleen A. Asin, Candice Francheska B. Tambaoan, Asia Louisa U. Chong, Karol Sophia Agape R. Padilla, Rianna Patricia S. Cruz, El King D. Morado, Joshua Gregor A. Dizon, Razel Nikka M. Hao, Arianne A. Zamora, Devon Ray Pacial, Juan Antonio R. Magalang, Marissa Alejandria, Celia Carlos, Anna Ong-Lim, Edsel Maurice Salvaña, John Q. Wong, Jaime C. Montoya, Maria Rosario Singh-Vergeire and Cynthia P. Saloma |
| EPI_ISL_2188161, EPI_ISL_2188162 | The Medical City -Â Ortigas | Philippine Genome Center | Francis A. Tablizo, Kenneth M. Kim, Carlo M. Lapid, Marc Jerrone R. Castro, Maria Sofia L. Yangzon, Elcid Aaron R. Pangilinan, Benedict A. Maralit, Marc Edsel C. Ayes, Eva Maria Cutiongco-de la Paz, Alethea R. de Guzman, Jan Michael C. Yap, Jo-Hannah S. Llames, Sheila Mae M. Araiza, Kris P. Punayan, Irish Coleen A. Asin, Candice Francheska B. Tambaoan, Asia Louisa U. Chong, Karol Sophia Agape R. Padilla, Rianna Patricia S. Cruz, El King D. Morado, Joshua Gregor A. Dizon, Razel Nikka M. Hao, Arianne A. Zamora, Devon Ray Pacial, Juan Antonio R. Magalang, Marissa Alejandria, Celia Carlos, Anna Ong-Lim, Edsel Maurice Salvaña, John Q. Wong, Jaime C. Montoya, Maria Rosario Singh-Vergeire and Cynthia P. Saloma |
| EPI_ISL_2188163 | Qualimed Hospital Sta. Rosa | Philippine Genome Center | Francis A. Tablizo, Kenneth M. Kim, Carlo M. Lapid, Marc Jerrone R. Castro, Maria Sofia L. Yangzon, Elcid Aaron R. Pangilinan, Benedict A. Maralit, Marc Edsel C. Ayes, Eva Maria Cutiongco-de la Paz, Alethea R. de Guzman, Jan Michael C. Yap, Jo-Hannah S. Llames, Sheila Mae M. Araiza, Kris P. Punayan, Irish Coleen A. Asin, Candice Francheska B. Tambaoan, Asia Louisa U. Chong, Karol Sophia Agape R. Padilla, Rianna Patricia S. Cruz, El King D. Morado, Joshua Gregor A. Dizon, Razel Nikka M. Hao, Arianne A. Zamora, Devon Ray Pacial, Juan Antonio R. Magalang, Marissa Alejandria, Celia Carlos, Anna Ong-Lim, Edsel Maurice Salvaña, John Q. Wong, Jaime C. Montoya, Maria Rosario Singh-Vergeire and Cynthia P. Saloma |
| EPI_ISL_2188164, EPI_ISL_2188165, EPI_ISL_2188166, EPI_ISL_2188167 | The Medical City -Â Ortigas | Philippine Genome Center | Francis A. Tablizo, Kenneth M. Kim, Carlo M. Lapid, Marc Jerrone R. Castro, Maria Sofia L. Yangzon, Elcid Aaron R. Pangilinan, Benedict A. Maralit, Marc Edsel C. Ayes, Eva Maria Cutiongco-de la Paz, Alethea R. de Guzman, Jan Michael C. Yap, Jo-Hannah S. Llames, Sheila Mae M. Araiza, Kris P. Punayan, Irish Coleen A. Asin, Candice Francheska B. Tambaoan, Asia Louisa U. Chong, Karol Sophia Agape R. Padilla, Rianna Patricia S. Cruz, El King D. Morado, Joshua Gregor A. Dizon, Razel Nikka M. Hao, Arianne A. Zamora, Devon Ray Pacial, Juan Antonio R. Magalang, Marissa Alejandria, Celia Carlos, Anna Ong-Lim, Edsel Maurice Salvaña, John Q. Wong, Jaime C. Montoya, Maria Rosario Singh-Vergeire and Cynthia P. Saloma |
| EPI_ISL_2188168, EPI_ISL_2188169 | The Lord's Grace Medical and Industrial Clinic | Philippine Genome Center | Francis A. Tablizo, Kenneth M. Kim, Carlo M. Lapid, Marc Jerrone R. Castro, Maria Sofia L. Yangzon, Elcid Aaron R. Pangilinan, Benedict A. Maralit, Marc Edsel C. Ayes, Eva Maria Cutiongco-de la Paz, Alethea R. de Guzman, Jan Michael C. Yap, Jo-Hannah S. Llames, Sheila Mae M. Araiza, Kris P. Punayan, Irish Coleen A. Asin, Candice Francheska B. Tambaoan, Asia Louisa U. Chong, Karol Sophia Agape R. Padilla, Rianna Patricia S. Cruz, El King D. Morado, Joshua Gregor A. Dizon, Razel Nikka M. Hao, Arianne A. Zamora, Devon Ray Pacial, Juan Antonio R. Magalang, Marissa Alejandria, Celia Carlos, Anna Ong-Lim, Edsel Maurice Salvaña, John Q. Wong, Jaime C. Montoya, Maria Rosario Singh-Vergeire and Cynthia P. Saloma |
| EPI_ISL_2188171, EPI_ISL_2188173 | The Medical City -Â Ortigas | Philippine Genome Center | Francis A. Tablizo, Kenneth M. Kim, Carlo M. Lapid, Marc Jerrone R. Castro, Maria Sofia L. Yangzon, Elcid Aaron R. Pangilinan, Benedict A. Maralit, Marc Edsel C. Ayes, Eva Maria Cutiongco-de la Paz, Alethea R. de Guzman, Jan Michael C. Yap, Jo-Hannah S. Llames, Sheila Mae M. Araiza, Kris P. Punayan, Irish Coleen A. Asin, Candice Francheska B. Tambaoan, Asia Louisa U. Chong, Karol Sophia Agape R. Padilla, Rianna Patricia S. Cruz, El King D. Morado, Joshua Gregor A. Dizon, Razel Nikka M. Hao, Arianne A. Zamora, Devon Ray Pacial, Juan Antonio R. Magalang, Marissa Alejandria, Celia Carlos, Anna Ong-Lim, Edsel Maurice Salvaña, John Q. Wong, Jaime C. Montoya, Maria Rosario Singh-Vergeire and Cynthia P. Saloma |
| EPI_ISL_2188174 | The Lord's Grace Medical and Industrial Clinic | Philippine Genome Center | Francis A. Tablizo, Kenneth M. Kim, Carlo M. Lapid, Marc Jerrone R. Castro, Maria Sofia L. Yangzon, Elcid Aaron R. Pangilinan, Benedict A. Maralit, Marc Edsel C. Ayes, Eva Maria Cutiongco-de la Paz, Alethea R. de Guzman, Jan Michael C. Yap, Jo-Hannah S. Llames, Sheila Mae M. Araiza, Kris P. Punayan, Irish Coleen A. Asin, Candice Francheska B. Tambaoan, Asia Louisa U. Chong, Karol Sophia Agape R. Padilla, Rianna Patricia S. Cruz, El King D. Morado, Joshua Gregor A. Dizon, Razel Nikka M. Hao, Arianne A. Zamora, Devon Ray Pacial, Juan Antonio R. Magalang, Marissa Alejandria, Celia Carlos, Anna Ong-Lim, Edsel Maurice Salvaña, John Q. Wong, Jaime C. Montoya, Maria Rosario Singh-Vergeire and Cynthia P. Saloma |

| | | | |
|---|---|---|---|
| EPI_ISL_2188175, EPI_ISL_2188176 | The Medical City -Â Ortigas | Philippine Genome Center | Francis A. Tablizo, Kenneth M. Kim, Carlo M. Lapid, Marc Jerrone R. Castro, Maria Sofia L. Yangzon, Elcid Aaron R. Pangilinan, Benedict A. Maralit, Marc Edsel C. Ayes, Eva Maria Cutiongco-de la Paz, Alethea R. de Guzman, Jan Michael C. Yap, Jo-Hannah S. Llames, Sheila Mae M. Araiza, Kris P. Punayan, Irish Coleen A. Asin, Candice Francheska B. Tambaoan, Asia Louisa U. Chong, Karol Sophia Agape R. Padilla, Rianna Patricia S. Cruz, El King D. Morado, Joshua Gregor A. Dizon, Razel Nikka M. Hao, Arianne A. Zamora, Devon Ray Pacial, Juan Antonio R. Magalang, Marissa Alejandria, Celia Carlos, Anna Ong-Lim, Edsel Maurice Salvaña, John Q. Wong, Jaime C. Montoya, Maria Rosario Singh-Vergeire and Cynthia P. Saloma |
| EPI_ISL_2188177 | Ilocos Training and Regional Medical Center | Philippine Genome Center | Francis A. Tablizo, Kenneth M. Kim, Carlo M. Lapid, Marc Jerrone R. Castro, Maria Sofia L. Yangzon, Elcid Aaron R. Pangilinan, Benedict A. Maralit, Marc Edsel C. Ayes, Eva Maria Cutiongco-de la Paz, Alethea R. de Guzman, Jan Michael C. Yap, Jo-Hannah S. Llames, Sheila Mae M. Araiza, Kris P. Punayan, Irish Coleen A. Asin, Candice Francheska B. Tambaoan, Asia Louisa U. Chong, Karol Sophia Agape R. Padilla, Rianna Patricia S. Cruz, El King D. Morado, Joshua Gregor A. Dizon, Razel Nikka M. Hao, Arianne A. Zamora, Devon Ray Pacial, Juan Antonio R. Magalang, Marissa Alejandria, Celia Carlos, Anna Ong-Lim, Edsel Maurice Salvaña, John Q. Wong, Jaime C. Montoya, Maria Rosario Singh-Vergeire and Cynthia P. Saloma |
| EPI_ISL_2188178 | Philippine Red Cross - Port Area | Philippine Genome Center | Francis A. Tablizo, Kenneth M. Kim, Carlo M. Lapid, Marc Jerrone R. Castro, Maria Sofia L. Yangzon, Elcid Aaron R. Pangilinan, Benedict A. Maralit, Marc Edsel C. Ayes, Eva Maria Cutiongco-de la Paz, Alethea R. de Guzman, Jan Michael C. Yap, Jo-Hannah S. Llames, Sheila Mae M. Araiza, Kris P. Punayan, Irish Coleen A. Asin, Candice Francheska B. Tambaoan, Asia Louisa U. Chong, Karol Sophia Agape R. Padilla, Rianna Patricia S. Cruz, El King D. Morado, Joshua Gregor A. Dizon, Razel Nikka M. Hao, Arianne A. Zamora, Devon Ray Pacial, Juan Antonio R. Magalang, Marissa Alejandria, Celia Carlos, Anna Ong-Lim, Edsel Maurice Salvaña, John Q. Wong, Jaime C. Montoya, Maria Rosario Singh-Vergeire and Cynthia P. Saloma |
| EPI_ISL_2188179 | The Lord's Grace Medical and Industrial Clinic | Philippine Genome Center | Francis A. Tablizo, Kenneth M. Kim, Carlo M. Lapid, Marc Jerrone R. Castro, Maria Sofia L. Yangzon, Elcid Aaron R. Pangilinan, Benedict A. Maralit, Marc Edsel C. Ayes, Eva Maria Cutiongco-de la Paz, Alethea R. de Guzman, Jan Michael C. Yap, Jo-Hannah S. Llames, Sheila Mae M. Araiza, Kris P. Punayan, Irish Coleen A. Asin, Candice Francheska B. Tambaoan, Asia Louisa U. Chong, Karol Sophia Agape R. Padilla, Rianna Patricia S. Cruz, El King D. Morado, Joshua Gregor A. Dizon, Razel Nikka M. Hao, Arianne A. Zamora, Devon Ray Pacial, Juan Antonio R. Magalang, Marissa Alejandria, Celia Carlos, Anna Ong-Lim, Edsel Maurice Salvaña, John Q. Wong, Jaime C. Montoya, Maria Rosario Singh-Vergeire and Cynthia P. Saloma |
| EPI_ISL_2188180 | Mary Mediatrix Medical Center | Philippine Genome Center | Francis A. Tablizo, Kenneth M. Kim, Carlo M. Lapid, Marc Jerrone R. Castro, Maria Sofia L. Yangzon, Elcid Aaron R. Pangilinan, Benedict A. Maralit, Marc Edsel C. Ayes, Eva Maria Cutiongco-de la Paz, Alethea R. de Guzman, Jan Michael C. Yap, Jo-Hannah S. Llames, Sheila Mae M. Araiza, Kris P. Punayan, Irish Coleen A. Asin, Candice Francheska B. Tambaoan, Asia Louisa U. Chong, Karol Sophia Agape R. Padilla, Rianna Patricia S. Cruz, El King D. Morado, Joshua Gregor A. Dizon, Razel Nikka M. Hao, Arianne A. Zamora, Devon Ray Pacial, Juan Antonio R. Magalang, Marissa Alejandria, Celia Carlos, Anna Ong-Lim, Edsel Maurice Salvaña, John Q. Wong, Jaime C. Montoya, Maria Rosario Singh-Vergeire and Cynthia P. Saloma |
| EPI_ISL_2188181 | The Medical City -Â Ortigas | Philippine Genome Center | Francis A. Tablizo, Kenneth M. Kim, Carlo M. Lapid, Marc Jerrone R. Castro, Maria Sofia L. Yangzon, Elcid Aaron R. Pangilinan, Benedict A. Maralit, Marc Edsel C. Ayes, Eva Maria Cutiongco-de la Paz, Alethea R. de Guzman, Jan Michael C. Yap, Jo-Hannah S. Llames, Sheila Mae M. Araiza, Kris P. Punayan, Irish Coleen A. Asin, Candice Francheska B. Tambaoan, Asia Louisa U. Chong, Karol Sophia Agape R. Padilla, Rianna Patricia S. Cruz, El King D. Morado, Joshua Gregor A. Dizon, Razel Nikka M. Hao, Arianne A. Zamora, Devon Ray Pacial, Juan Antonio R. Magalang, Marissa Alejandria, Celia Carlos, Anna Ong-Lim, Edsel Maurice Salvaña, John Q. Wong, Jaime C. Montoya, Maria Rosario Singh-Vergeire and Cynthia P. Saloma |
| EPI_ISL_2188182 | Las PiÃ±as General Hospital and Salellite Trauma Center | Philippine Genome Center | Francis A. Tablizo, Kenneth M. Kim, Carlo M. Lapid, Marc Jerrone R. Castro, Maria Sofia L. Yangzon, Elcid Aaron R. Pangilinan, Benedict A. Maralit, Marc Edsel C. Ayes, Eva Maria Cutiongco-de la Paz, Alethea R. de Guzman, Jan Michael C. Yap, Jo-Hannah S. Llames, Sheila Mae M. Araiza, Kris P. Punayan, Irish Coleen A. Asin, Candice Francheska B. Tambaoan, Asia Louisa U. Chong, Karol Sophia Agape R. Padilla, Rianna Patricia S. Cruz, El King D. Morado, Joshua Gregor A. Dizon, Razel Nikka M. Hao, Arianne A. Zamora, Devon Ray Pacial, Juan Antonio R. Magalang, Marissa Alejandria, Celia Carlos, Anna Ong-Lim, Edsel Maurice Salvaña, John Q. Wong, Jaime C. Montoya, Maria Rosario Singh-Vergeire and Cynthia P. Saloma |
| EPI_ISL_2188183 | The Medical City -Â Ortigas | Philippine Genome Center | Francis A. Tablizo, Kenneth M. Kim, Carlo M. Lapid, Marc Jerrone R. Castro, Maria Sofia L. Yangzon, Elcid Aaron R. Pangilinan, Benedict A. Maralit, Marc Edsel C. Ayes, Eva Maria Cutiongco-de la Paz, Alethea R. de Guzman, Jan Michael C. Yap, Jo-Hannah S. Llames, Sheila Mae M. Araiza, Kris P. Punayan, Irish Coleen A. Asin, Candice Francheska B. Tambaoan, Asia Louisa U. Chong, Karol Sophia Agape R. Padilla, Rianna Patricia S. Cruz, El King D. Morado, Joshua Gregor A. Dizon, Razel Nikka M. Hao, Arianne A. Zamora, Devon Ray Pacial, Juan Antonio R. Magalang, Marissa Alejandria, Celia Carlos, Anna Ong-Lim, Edsel Maurice Salvaña, John Q. Wong, Jaime C. Montoya, Maria Rosario Singh-Vergeire and Cynthia P. Saloma |
| EPI_ISL_2188184 | The Lord's Grace Medical and Industrial Clinic | Philippine Genome Center | Francis A. Tablizo, Kenneth M. Kim, Carlo M. Lapid, Marc Jerrone R. Castro, Maria Sofia L. Yangzon, Elcid Aaron R. Pangilinan, Benedict A. Maralit, Marc Edsel C. Ayes, Eva Maria Cutiongco-de la Paz, Alethea R. de Guzman, Jan Michael C. Yap, Jo-Hannah S. Llames, Sheila Mae M. Araiza, Kris P. Punayan, Irish Coleen A. Asin, Candice Francheska B. Tambaoan, Asia Louisa U. Chong, Karol Sophia Agape R. Padilla, Rianna Patricia S. Cruz, El King D. Morado, Joshua Gregor A. Dizon, Razel Nikka M. Hao, Arianne A. Zamora, Devon Ray Pacial, Juan Antonio R. Magalang, Marissa Alejandria, Celia Carlos, Anna Ong-Lim, Edsel Maurice Salvaña, John Q. Wong, Jaime C. Montoya, Maria Rosario Singh-Vergeire and Cynthia P. Saloma |
| EPI_ISL_2188185 | Baguio General Hospital Medical Center (BGHMC) | Philippine Genome Center | Francis A. Tablizo, Kenneth M. Kim, Carlo M. Lapid, Marc Jerrone R. Castro, Maria Sofia L. Yangzon, Elcid Aaron R. Pangilinan, Benedict A. Maralit, Marc Edsel C. Ayes, Eva Maria Cutiongco-de la Paz, Alethea R. de Guzman, Jan Michael C. Yap, Jo-Hannah S. Llames, Sheila Mae M. Araiza, Kris P. Punayan, Irish Coleen A. Asin, Candice Francheska B. Tambaoan, Asia Louisa U. Chong, Karol Sophia Agape R. Padilla, Rianna Patricia S. Cruz, El King D. Morado, Joshua Gregor A. Dizon, Razel Nikka M. Hao, Arianne A. Zamora, Devon Ray Pacial, Juan Antonio R. Magalang, Marissa Alejandria, Celia Carlos, Anna Ong-Lim, Edsel Maurice Salvaña, John Q. Wong, Jaime C. Montoya, Maria Rosario Singh-Vergeire and Cynthia P. Saloma |
| EPI_ISL_2188186, EPI_ISL_2188187 | The Medical City -Â Ortigas | Philippine Genome Center | Francis A. Tablizo, Kenneth M. Kim, Carlo M. Lapid, Marc Jerrone R. Castro, Maria Sofia L. Yangzon, Elcid Aaron R. Pangilinan, Benedict A. Maralit, Marc Edsel C. Ayes, Eva Maria Cutiongco-de la Paz, Alethea R. de Guzman, Jan Michael C. Yap, Jo-Hannah S. Llames, Sheila Mae M. Araiza, Kris P. Punayan, Irish Coleen A. Asin, Candice Francheska B. Tambaoan, Asia Louisa U. Chong, Karol Sophia Agape R. Padilla, Rianna Patricia S. Cruz, El King D. Morado, Joshua Gregor A. Dizon, Razel Nikka M. Hao, Arianne A. Zamora, Devon Ray Pacial, Juan Antonio R. Magalang, Marissa Alejandria, Celia Carlos, Anna Ong-Lim, Edsel Maurice Salvaña, John Q. Wong, Jaime C. Montoya, Maria Rosario Singh-Vergeire and Cynthia P. Saloma |
| EPI_ISL_2188188 | Philippine Red Cross- Batangas Chapter Molecular Laboratory | Philippine Genome Center | Francis A. Tablizo, Kenneth M. Kim, Carlo M. Lapid, Marc Jerrone R. Castro, Maria Sofia L. Yangzon, Elcid Aaron R. Pangilinan, Benedict A. Maralit, Marc Edsel C. Ayes, Eva Maria Cutiongco-de la Paz, Alethea R. de Guzman, Jan Michael C. Yap, Jo-Hannah S. Llames, Sheila Mae M. Araiza, Kris P. Punayan, Irish Coleen A. Asin, Candice Francheska B. Tambaoan, Asia Louisa U. Chong, Karol Sophia Agape R. Padilla, Rianna Patricia S. Cruz, El King D. Morado, Joshua Gregor A. Dizon, Razel Nikka M. Hao, Arianne A. Zamora, Devon Ray Pacial, Juan Antonio R. Magalang, Marissa Alejandria, Celia Carlos, Anna Ong-Lim, Edsel Maurice Salvaña, John Q. Wong, Jaime C. Montoya, Maria Rosario Singh-Vergeire and Cynthia P. Saloma |
| EPI_ISL_2188189 | Qualimed Hospital Sta. Rosa | Philippine Genome Center | Francis A. Tablizo, Kenneth M. Kim, Carlo M. Lapid, Marc Jerrone R. Castro, Maria Sofia L. Yangzon, Elcid Aaron R. Pangilinan, Benedict A. Maralit, Marc Edsel C. Ayes, Eva Maria Cutiongco-de la Paz, Alethea R. de Guzman, Jan Michael C. Yap, Jo-Hannah S. Llames, Sheila Mae M. Araiza, Kris P. Punayan, Irish Coleen A. Asin, Candice Francheska B. Tambaoan, Asia Louisa U. Chong, Karol Sophia Agape R. Padilla, Rianna Patricia S. Cruz, El King D. Morado, Joshua Gregor A. Dizon, Razel Nikka M. Hao, Arianne A. Zamora, Devon Ray Pacial, Juan Antonio R. Magalang, Marissa Alejandria, Celia Carlos, Anna Ong-Lim, Edsel Maurice Salvaña, John Q. Wong, Jaime C. Montoya, Maria Rosario Singh-Vergeire and Cynthia P. Saloma |
| EPI_ISL_2188190 | The Lord's Grace Medical and Industrial Clinic | Philippine Genome Center | Francis A. Tablizo, Kenneth M. Kim, Carlo M. Lapid, Marc Jerrone R. Castro, Maria Sofia L. Yangzon, Elcid Aaron R. Pangilinan, Benedict A. Maralit, Marc Edsel C. Ayes, Eva Maria Cutiongco-de la Paz, Alethea R. de Guzman, Jan Michael C. Yap, Jo-Hannah S. Llames, Sheila Mae M. Araiza, Kris P. Punayan, Irish Coleen A. Asin, Candice Francheska B. Tambaoan, Asia Louisa U. Chong, Karol Sophia Agape R. Padilla, Rianna Patricia S. Cruz, El King D. Morado, Joshua Gregor A. Dizon, Razel Nikka M. Hao, Arianne A. Zamora, Devon Ray Pacial, Juan Antonio R. Magalang, Marissa Alejandria, Celia Carlos, Anna Ong-Lim, Edsel Maurice Salvaña, John Q. Wong, Jaime C. Montoya, Maria Rosario Singh-Vergeire and Cynthia P. Saloma |
| EPI_ISL_2188191 | Ospital ng ParaÃ±aque II | Philippine Genome Center | Francis A. Tablizo, Kenneth M. Kim, Carlo M. Lapid, Marc Jerrone R. Castro, Maria Sofia L. Yangzon, Elcid Aaron R. Pangilinan, Benedict A. Maralit, Marc Edsel C. Ayes, Eva Maria Cutiongco-de la Paz, Alethea R. de Guzman, Jan Michael C. Yap, Jo-Hannah S. Llames, Sheila Mae M. Araiza, Kris P. Punayan, Irish Coleen A. Asin, Candice Francheska B. Tambaoan, Asia Louisa U. Chong, Karol Sophia Agape R. Padilla, Rianna Patricia S. Cruz, El King D. Morado, Joshua Gregor A. Dizon, Razel Nikka M. Hao, Arianne A. Zamora, Devon Ray Pacial, Juan Antonio R. Magalang, Marissa Alejandria, Celia Carlos, Anna Ong-Lim, Edsel Maurice Salvaña, John Q. Wong, Jaime C. Montoya, Maria Rosario Singh-Vergeire and Cynthia P. Saloma |
| EPI_ISL_2188193 | The Lord's Grace Medical and Industrial Clinic | Philippine Genome Center | Francis A. Tablizo, Kenneth M. Kim, Carlo M. Lapid, Marc Jerrone R. Castro, Maria Sofia L. Yangzon, Elcid Aaron R. Pangilinan, Benedict A. Maralit, Marc Edsel C. Ayes, Eva Maria Cutiongco-de la Paz, Alethea R. de Guzman, Jan Michael C. Yap, Jo-Hannah S. Llames, Sheila Mae M. Araiza, Kris P. Punayan, Irish Coleen A. Asin, Candice Francheska B. Tambaoan, Asia Louisa U. Chong, Karol Sophia Agape R. Padilla, Rianna Patricia S. Cruz, El King D. Morado, Joshua Gregor A. Dizon, Razel Nikka M. Hao, Arianne A. Zamora, Devon Ray Pacial, Juan Antonio R. Magalang, Marissa Alejandria, Celia Carlos, Anna Ong-Lim, Edsel Maurice Salvaña, John Q. Wong, Jaime C. Montoya, Maria Rosario Singh-Vergeire and Cynthia P. Saloma |

| | | | |
|---|---|---|---|
| EPI_ISL_2188194 | The Medical City -Â Ortigas | Philippine Genome Center | Francis A. Tablizo, Kenneth M. Kim, Carlo M. Lapid, Marc Jerrone R. Castro, Maria Sofia L. Yangzon, Elcid Aaron R. Pangilinan, Benedict A. Maralit, Marc Edsel C. Ayes, Eva Maria Cutiongco-de la Paz, Alethea R. de Guzman, Jan Michael C. Yap, Jo-Hannah S. Llames, Sheila Mae M. Araiza, Kris P. Punayan, Irish Coleen A. Asin, Candice Francheska B. Tambaoan, Asia Louisa U. Chong, Karol Sophia Agape R. Padilla, Rianna Patricia S. Cruz, El King D. Morado, Joshua Gregor A. Dizon, Razel Nikka M. Hao, Arianne A. Zamora, Devon Ray Pacial, Juan Antonio R. Magalang, Marissa Alejandria, Celia Carlos, Anna Ong-Lim, Edsel Maurice Salvaña, John Q. Wong, Jaime C. Montoya, Maria Rosario Singh-Vergeire and Cynthia P. Saloma |
| EPI_ISL_2188195 | The Lord's Grace Medical and Industrial Clinic | Philippine Genome Center | Francis A. Tablizo, Kenneth M. Kim, Carlo M. Lapid, Marc Jerrone R. Castro, Maria Sofia L. Yangzon, Elcid Aaron R. Pangilinan, Benedict A. Maralit, Marc Edsel C. Ayes, Eva Maria Cutiongco-de la Paz, Alethea R. de Guzman, Jan Michael C. Yap, Jo-Hannah S. Llames, Sheila Mae M. Araiza, Kris P. Punayan, Irish Coleen A. Asin, Candice Francheska B. Tambaoan, Asia Louisa U. Chong, Karol Sophia Agape R. Padilla, Rianna Patricia S. Cruz, El King D. Morado, Joshua Gregor A. Dizon, Razel Nikka M. Hao, Arianne A. Zamora, Devon Ray Pacial, Juan Antonio R. Magalang, Marissa Alejandria, Celia Carlos, Anna Ong-Lim, Edsel Maurice Salvaña, John Q. Wong, Jaime C. Montoya, Maria Rosario Singh-Vergeire and Cynthia P. Saloma |
| EPI_ISL_2188196 | The Medical City -Â Ortigas | Philippine Genome Center | Francis A. Tablizo, Kenneth M. Kim, Carlo M. Lapid, Marc Jerrone R. Castro, Maria Sofia L. Yangzon, Elcid Aaron R. Pangilinan, Benedict A. Maralit, Marc Edsel C. Ayes, Eva Maria Cutiongco-de la Paz, Alethea R. de Guzman, Jan Michael C. Yap, Jo-Hannah S. Llames, Sheila Mae M. Araiza, Kris P. Punayan, Irish Coleen A. Asin, Candice Francheska B. Tambaoan, Asia Louisa U. Chong, Karol Sophia Agape R. Padilla, Rianna Patricia S. Cruz, El King D. Morado, Joshua Gregor A. Dizon, Razel Nikka M. Hao, Arianne A. Zamora, Devon Ray Pacial, Juan Antonio R. Magalang, Marissa Alejandria, Celia Carlos, Anna Ong-Lim, Edsel Maurice Salvaña, John Q. Wong, Jaime C. Montoya, Maria Rosario Singh-Vergeire and Cynthia P. Saloma |
| EPI_ISL_2188197 | Baguio General Hospital Medical Center (BGHMC) | Philippine Genome Center | Francis A. Tablizo, Kenneth M. Kim, Carlo M. Lapid, Marc Jerrone R. Castro, Maria Sofia L. Yangzon, Elcid Aaron R. Pangilinan, Benedict A. Maralit, Marc Edsel C. Ayes, Eva Maria Cutiongco-de la Paz, Alethea R. de Guzman, Jan Michael C. Yap, Jo-Hannah S. Llames, Sheila Mae M. Araiza, Kris P. Punayan, Irish Coleen A. Asin, Candice Francheska B. Tambaoan, Asia Louisa U. Chong, Karol Sophia Agape R. Padilla, Rianna Patricia S. Cruz, El King D. Morado, Joshua Gregor A. Dizon, Razel Nikka M. Hao, Arianne A. Zamora, Devon Ray Pacial, Juan Antonio R. Magalang, Marissa Alejandria, Celia Carlos, Anna Ong-Lim, Edsel Maurice Salvaña, John Q. Wong, Jaime C. Montoya, Maria Rosario Singh-Vergeire and Cynthia P. Saloma |
| EPI_ISL_2188198, EPI_ISL_2188202, EPI_ISL_2188203 | The Medical City -Â Ortigas | Philippine Genome Center | Francis A. Tablizo, Kenneth M. Kim, Carlo M. Lapid, Marc Jerrone R. Castro, Maria Sofia L. Yangzon, Elcid Aaron R. Pangilinan, Benedict A. Maralit, Marc Edsel C. Ayes, Eva Maria Cutiongco-de la Paz, Alethea R. de Guzman, Jan Michael C. Yap, Jo-Hannah S. Llames, Sheila Mae M. Araiza, Kris P. Punayan, Irish Coleen A. Asin, Candice Francheska B. Tambaoan, Asia Louisa U. Chong, Karol Sophia Agape R. Padilla, Rianna Patricia S. Cruz, El King D. Morado, Joshua Gregor A. Dizon, Razel Nikka M. Hao, Arianne A. Zamora, Devon Ray Pacial, Juan Antonio R. Magalang, Marissa Alejandria, Celia Carlos, Anna Ong-Lim, Edsel Maurice Salvaña, John Q. Wong, Jaime C. Montoya, Maria Rosario Singh-Vergeire and Cynthia P. Saloma |
| EPI_ISL_2188204 | Baguio General Hospital Medical Center (BGHMC) | Philippine Genome Center | Francis A. Tablizo, Kenneth M. Kim, Carlo M. Lapid, Marc Jerrone R. Castro, Maria Sofia L. Yangzon, Elcid Aaron R. Pangilinan, Benedict A. Maralit, Marc Edsel C. Ayes, Eva Maria Cutiongco-de la Paz, Alethea R. de Guzman, Jan Michael C. Yap, Jo-Hannah S. Llames, Sheila Mae M. Araiza, Kris P. Punayan, Irish Coleen A. Asin, Candice Francheska B. Tambaoan, Asia Louisa U. Chong, Karol Sophia Agape R. Padilla, Rianna Patricia S. Cruz, El King D. Morado, Joshua Gregor A. Dizon, Razel Nikka M. Hao, Arianne A. Zamora, Devon Ray Pacial, Juan Antonio R. Magalang, Marissa Alejandria, Celia Carlos, Anna Ong-Lim, Edsel Maurice Salvaña, John Q. Wong, Jaime C. Montoya, Maria Rosario Singh-Vergeire and Cynthia P. Saloma |
| EPI_ISL_2188205, EPI_ISL_2188206 | Laguna Provincial Hospital - San Pablo District Hospital | Philippine Genome Center | Francis A. Tablizo, Kenneth M. Kim, Carlo M. Lapid, Marc Jerrone R. Castro, Maria Sofia L. Yangzon, Elcid Aaron R. Pangilinan, Benedict A. Maralit, Marc Edsel C. Ayes, Eva Maria Cutiongco-de la Paz, Alethea R. de Guzman, Jan Michael C. Yap, Jo-Hannah S. Llames, Sheila Mae M. Araiza, Kris P. Punayan, Irish Coleen A. Asin, Candice Francheska B. Tambaoan, Asia Louisa U. Chong, Karol Sophia Agape R. Padilla, Rianna Patricia S. Cruz, El King D. Morado, Joshua Gregor A. Dizon, Razel Nikka M. Hao, Arianne A. Zamora, Devon Ray Pacial, Juan Antonio R. Magalang, Marissa Alejandria, Celia Carlos, Anna Ong-Lim, Edsel Maurice Salvaña, John Q. Wong, Jaime C. Montoya, Maria Rosario Singh-Vergeire and Cynthia P. Saloma |
| EPI_ISL_2188207, EPI_ISL_2188208 | The Medical City -Â Ortigas | Philippine Genome Center | Francis A. Tablizo, Kenneth M. Kim, Carlo M. Lapid, Marc Jerrone R. Castro, Maria Sofia L. Yangzon, Elcid Aaron R. Pangilinan, Benedict A. Maralit, Marc Edsel C. Ayes, Eva Maria Cutiongco-de la Paz, Alethea R. de Guzman, Jan Michael C. Yap, Jo-Hannah S. Llames, Sheila Mae M. Araiza, Kris P. Punayan, Irish Coleen A. Asin, Candice Francheska B. Tambaoan, Asia Louisa U. Chong, Karol Sophia Agape R. Padilla, Rianna Patricia S. Cruz, El King D. Morado, Joshua Gregor A. Dizon, Razel Nikka M. Hao, Arianne A. Zamora, Devon Ray Pacial, Juan Antonio R. Magalang, Marissa Alejandria, Celia Carlos, Anna Ong-Lim, Edsel Maurice Salvaña, John Q. Wong, Jaime C. Montoya, Maria Rosario Singh-Vergeire and Cynthia P. Saloma |
| EPI_ISL_2188209 | Prime Care Alpha Covid-19 Testing Laboratory | Philippine Genome Center | Francis A. Tablizo, Kenneth M. Kim, Carlo M. Lapid, Marc Jerrone R. Castro, Maria Sofia L. Yangzon, Elcid Aaron R. Pangilinan, Benedict A. Maralit, Marc Edsel C. Ayes, Eva Maria Cutiongco-de la Paz, Alethea R. de Guzman, Jan Michael C. Yap, Jo-Hannah S. Llames, Sheila Mae M. Araiza, Kris P. Punayan, Irish Coleen A. Asin, Candice Francheska B. Tambaoan, Asia Louisa U. Chong, Karol Sophia Agape R. Padilla, Rianna Patricia S. Cruz, El King D. Morado, Joshua Gregor A. Dizon, Razel Nikka M. Hao, Arianne A. Zamora, Devon Ray Pacial, Juan Antonio R. Magalang, Marissa Alejandria, Celia Carlos, Anna Ong-Lim, Edsel Maurice Salvaña, John Q. Wong, Jaime C. Montoya, Maria Rosario Singh-Vergeire and Cynthia P. Saloma |
| EPI_ISL_2188210 | The Medical City -Â Ortigas | Philippine Genome Center | Francis A. Tablizo, Kenneth M. Kim, Carlo M. Lapid, Marc Jerrone R. Castro, Maria Sofia L. Yangzon, Elcid Aaron R. Pangilinan, Benedict A. Maralit, Marc Edsel C. Ayes, Eva Maria Cutiongco-de la Paz, Alethea R. de Guzman, Jan Michael C. Yap, Jo-Hannah S. Llames, Sheila Mae M. Araiza, Kris P. Punayan, Irish Coleen A. Asin, Candice Francheska B. Tambaoan, Asia Louisa U. Chong, Karol Sophia Agape R. Padilla, Rianna Patricia S. Cruz, El King D. Morado, Joshua Gregor A. Dizon, Razel Nikka M. Hao, Arianne A. Zamora, Devon Ray Pacial, Juan Antonio R. Magalang, Marissa Alejandria, Celia Carlos, Anna Ong-Lim, Edsel Maurice Salvaña, John Q. Wong, Jaime C. Montoya, Maria Rosario Singh-Vergeire and Cynthia P. Saloma |
| EPI_ISL_2188211 | Baguio General Hospital Medical Center (BGHMC) | Philippine Genome Center | Francis A. Tablizo, Kenneth M. Kim, Carlo M. Lapid, Marc Jerrone R. Castro, Maria Sofia L. Yangzon, Elcid Aaron R. Pangilinan, Benedict A. Maralit, Marc Edsel C. Ayes, Eva Maria Cutiongco-de la Paz, Alethea R. de Guzman, Jan Michael C. Yap, Jo-Hannah S. Llames, Sheila Mae M. Araiza, Kris P. Punayan, Irish Coleen A. Asin, Candice Francheska B. Tambaoan, Asia Louisa U. Chong, Karol Sophia Agape R. Padilla, Rianna Patricia S. Cruz, El King D. Morado, Joshua Gregor A. Dizon, Razel Nikka M. Hao, Arianne A. Zamora, Devon Ray Pacial, Juan Antonio R. Magalang, Marissa Alejandria, Celia Carlos, Anna Ong-Lim, Edsel Maurice Salvaña, John Q. Wong, Jaime C. Montoya, Maria Rosario Singh-Vergeire and Cynthia P. Saloma |
| EPI_ISL_2188213 | The Lord's Grace Medical and Industrial Clinic | Philippine Genome Center | Francis A. Tablizo, Kenneth M. Kim, Carlo M. Lapid, Marc Jerrone R. Castro, Maria Sofia L. Yangzon, Elcid Aaron R. Pangilinan, Benedict A. Maralit, Marc Edsel C. Ayes, Eva Maria Cutiongco-de la Paz, Alethea R. de Guzman, Jan Michael C. Yap, Jo-Hannah S. Llames, Sheila Mae M. Araiza, Kris P. Punayan, Irish Coleen A. Asin, Candice Francheska B. Tambaoan, Asia Louisa U. Chong, Karol Sophia Agape R. Padilla, Rianna Patricia S. Cruz, El King D. Morado, Joshua Gregor A. Dizon, Razel Nikka M. Hao, Arianne A. Zamora, Devon Ray Pacial, Juan Antonio R. Magalang, Marissa Alejandria, Celia Carlos, Anna Ong-Lim, Edsel Maurice Salvaña, John Q. Wong, Jaime C. Montoya, Maria Rosario Singh-Vergeire and Cynthia P. Saloma |
| EPI_ISL_2188214, EPI_ISL_2188215 | The Medical City -Â Ortigas | Philippine Genome Center | Francis A. Tablizo, Kenneth M. Kim, Carlo M. Lapid, Marc Jerrone R. Castro, Maria Sofia L. Yangzon, Elcid Aaron R. Pangilinan, Benedict A. Maralit, Marc Edsel C. Ayes, Eva Maria Cutiongco-de la Paz, Alethea R. de Guzman, Jan Michael C. Yap, Jo-Hannah S. Llames, Sheila Mae M. Araiza, Kris P. Punayan, Irish Coleen A. Asin, Candice Francheska B. Tambaoan, Asia Louisa U. Chong, Karol Sophia Agape R. Padilla, Rianna Patricia S. Cruz, El King D. Morado, Joshua Gregor A. Dizon, Razel Nikka M. Hao, Arianne A. Zamora, Devon Ray Pacial, Juan Antonio R. Magalang, Marissa Alejandria, Celia Carlos, Anna Ong-Lim, Edsel Maurice Salvaña, John Q. Wong, Jaime C. Montoya, Maria Rosario Singh-Vergeire and Cynthia P. Saloma |
| EPI_ISL_2188217, EPI_ISL_2188218 | The Lord's Grace Medical and Industrial Clinic | Philippine Genome Center | Francis A. Tablizo, Kenneth M. Kim, Carlo M. Lapid, Marc Jerrone R. Castro, Maria Sofia L. Yangzon, Elcid Aaron R. Pangilinan, Benedict A. Maralit, Marc Edsel C. Ayes, Eva Maria Cutiongco-de la Paz, Alethea R. de Guzman, Jan Michael C. Yap, Jo-Hannah S. Llames, Sheila Mae M. Araiza, Kris P. Punayan, Irish Coleen A. Asin, Candice Francheska B. Tambaoan, Asia Louisa U. Chong, Karol Sophia Agape R. Padilla, Rianna Patricia S. Cruz, El King D. Morado, Joshua Gregor A. Dizon, Razel Nikka M. Hao, Arianne A. Zamora, Devon Ray Pacial, Juan Antonio R. Magalang, Marissa Alejandria, Celia Carlos, Anna Ong-Lim, Edsel Maurice Salvaña, John Q. Wong, Jaime C. Montoya, Maria Rosario Singh-Vergeire and Cynthia P. Saloma |
| EPI_ISL_2188219 | Ospital ng Imus | Philippine Genome Center | Francis A. Tablizo, Kenneth M. Kim, Carlo M. Lapid, Marc Jerrone R. Castro, Maria Sofia L. Yangzon, Elcid Aaron R. Pangilinan, Benedict A. Maralit, Marc Edsel C. Ayes, Eva Maria Cutiongco-de la Paz, Alethea R. de Guzman, Jan Michael C. Yap, Jo-Hannah S. Llames, Sheila Mae M. Araiza, Kris P. Punayan, Irish Coleen A. Asin, Candice Francheska B. Tambaoan, Asia Louisa U. Chong, Karol Sophia Agape R. Padilla, Rianna Patricia S. Cruz, El King D. Morado, Joshua Gregor A. Dizon, Razel Nikka M. Hao, Arianne A. Zamora, Devon Ray Pacial, Juan Antonio R. Magalang, Marissa Alejandria, Celia Carlos, Anna Ong-Lim, Edsel Maurice Salvaña, John Q. Wong, Jaime C. Montoya, Maria Rosario Singh-Vergeire and Cynthia P. Saloma |
| EPI_ISL_2188220 | PASIG CITY CHILDREN'S HOSPITAL | Philippine Genome Center | Francis A. Tablizo, Kenneth M. Kim, Carlo M. Lapid, Marc Jerrone R. Castro, Maria Sofia L. Yangzon, Elcid Aaron R. Pangilinan, Benedict A. Maralit, Marc Edsel C. Ayes, Eva Maria Cutiongco-de la Paz, Alethea R. de Guzman, Jan Michael C. Yap, Jo-Hannah S. Llames, Sheila Mae M. Araiza, Kris P. Punayan, Irish Coleen A. Asin, Candice Francheska B. Tambaoan, Asia Louisa U. Chong, Karol Sophia Agape R. Padilla, Rianna Patricia S. Cruz, El King D. Morado, Joshua Gregor A. Dizon, Razel Nikka M. Hao, Arianne A. Zamora, Devon Ray Pacial, Juan Antonio R. Magalang, Marissa Alejandria, Celia Carlos, Anna Ong-Lim, Edsel Maurice Salvaña, John Q. Wong, Jaime C. Montoya, Maria Rosario Singh-Vergeire and Cynthia P. Saloma |

| | | | |
|---|---|---|---|
| EPI_ISL_2188221 | Philippine Red Cross - National Blood Center | Philippine Genome Center | Francis A. Tablizo, Kenneth M. Kim, Carlo M. Lapid, Marc Jerrone R. Castro, Maria Sofia L. Yangzon, Elcid Aaron R. Pangilinan, Benedict A. Maralit, Marc Edsel C. Ayes, Eva Maria Cutiongco-de la Paz, Alethea R. de Guzman, Jan Michael C. Yap, Jo-Hannah S. Llames, Sheila Mae M. Araiza, Kris P. Punayan, Irish Coleen A. Asin, Candice Francheska B. Tambaoan, Asia Louisa U. Chong, Karol Sophia Agape R. Padilla, Rianna Patricia S. Cruz, El King D. Morado, Joshua Gregor A. Dizon, Razel Nikka M. Hao, Arianne A. Zamora, Devon Ray Pacial, Juan Antonio R. Magalang, Marissa Alejandria, Celia Carlos, Anna Ong-Lim, Edsel Maurice Salvaña, John Q. Wong, Jaime C. Montoya, Maria Rosario Singh-Vergeire and Cynthia P. Saloma |
| EPI_ISL_2188222, EPI_ISL_2188223 | The Medical City -Â Ortigas | Philippine Genome Center | Francis A. Tablizo, Kenneth M. Kim, Carlo M. Lapid, Marc Jerrone R. Castro, Maria Sofia L. Yangzon, Elcid Aaron R. Pangilinan, Benedict A. Maralit, Marc Edsel C. Ayes, Eva Maria Cutiongco-de la Paz, Alethea R. de Guzman, Jan Michael C. Yap, Jo-Hannah S. Llames, Sheila Mae M. Araiza, Kris P. Punayan, Irish Coleen A. Asin, Candice Francheska B. Tambaoan, Asia Louisa U. Chong, Karol Sophia Agape R. Padilla, Rianna Patricia S. Cruz, El King D. Morado, Joshua Gregor A. Dizon, Razel Nikka M. Hao, Arianne A. Zamora, Devon Ray Pacial, Juan Antonio R. Magalang, Marissa Alejandria, Celia Carlos, Anna Ong-Lim, Edsel Maurice Salvaña, John Q. Wong, Jaime C. Montoya, Maria Rosario Singh-Vergeire and Cynthia P. Saloma |
| EPI_ISL_2188225 | Philippine Red Cross - Port Area | Philippine Genome Center | Francis A. Tablizo, Kenneth M. Kim, Carlo M. Lapid, Marc Jerrone R. Castro, Maria Sofia L. Yangzon, Elcid Aaron R. Pangilinan, Benedict A. Maralit, Marc Edsel C. Ayes, Eva Maria Cutiongco-de la Paz, Alethea R. de Guzman, Jan Michael C. Yap, Jo-Hannah S. Llames, Sheila Mae M. Araiza, Kris P. Punayan, Irish Coleen A. Asin, Candice Francheska B. Tambaoan, Asia Louisa U. Chong, Karol Sophia Agape R. Padilla, Rianna Patricia S. Cruz, El King D. Morado, Joshua Gregor A. Dizon, Razel Nikka M. Hao, Arianne A. Zamora, Devon Ray Pacial, Juan Antonio R. Magalang, Marissa Alejandria, Celia Carlos, Anna Ong-Lim, Edsel Maurice Salvaña, John Q. Wong, Jaime C. Montoya, Maria Rosario Singh-Vergeire and Cynthia P. Saloma |
| EPI_ISL_2188226 | Philippine Red Cross Logistics and Multipurpose Center | Philippine Genome Center | Francis A. Tablizo, Kenneth M. Kim, Carlo M. Lapid, Marc Jerrone R. Castro, Maria Sofia L. Yangzon, Elcid Aaron R. Pangilinan, Benedict A. Maralit, Marc Edsel C. Ayes, Eva Maria Cutiongco-de la Paz, Alethea R. de Guzman, Jan Michael C. Yap, Jo-Hannah S. Llames, Sheila Mae M. Araiza, Kris P. Punayan, Irish Coleen A. Asin, Candice Francheska B. Tambaoan, Asia Louisa U. Chong, Karol Sophia Agape R. Padilla, Rianna Patricia S. Cruz, El King D. Morado, Joshua Gregor A. Dizon, Razel Nikka M. Hao, Arianne A. Zamora, Devon Ray Pacial, Juan Antonio R. Magalang, Marissa Alejandria, Celia Carlos, Anna Ong-Lim, Edsel Maurice Salvaña, John Q. Wong, Jaime C. Montoya, Maria Rosario Singh-Vergeire and Cynthia P. Saloma |
| EPI_ISL_2188227, EPI_ISL_2188229, EPI_ISL_2188230, EPI_ISL_2188232, EPI_ISL_2188233, EPI_ISL_2188236 | The Medical City -Â Ortigas | Philippine Genome Center | Francis A. Tablizo, Kenneth M. Kim, Carlo M. Lapid, Marc Jerrone R. Castro, Maria Sofia L. Yangzon, Elcid Aaron R. Pangilinan, Benedict A. Maralit, Marc Edsel C. Ayes, Eva Maria Cutiongco-de la Paz, Alethea R. de Guzman, Jan Michael C. Yap, Jo-Hannah S. Llames, Sheila Mae M. Araiza, Kris P. Punayan, Irish Coleen A. Asin, Candice Francheska B. Tambaoan, Asia Louisa U. Chong, Karol Sophia Agape R. Padilla, Rianna Patricia S. Cruz, El King D. Morado, Joshua Gregor A. Dizon, Razel Nikka M. Hao, Arianne A. Zamora, Devon Ray Pacial, Juan Antonio R. Magalang, Marissa Alejandria, Celia Carlos, Anna Ong-Lim, Edsel Maurice Salvaña, John Q. Wong, Jaime C. Montoya, Maria Rosario Singh-Vergeire and Cynthia P. Saloma |
| EPI_ISL_2188237 | Parkway Medical and Diagnostic Center | Philippine Genome Center | Francis A. Tablizo, Kenneth M. Kim, Carlo M. Lapid, Marc Jerrone R. Castro, Maria Sofia L. Yangzon, Elcid Aaron R. Pangilinan, Benedict A. Maralit, Marc Edsel C. Ayes, Eva Maria Cutiongco-de la Paz, Alethea R. de Guzman, Jan Michael C. Yap, Jo-Hannah S. Llames, Sheila Mae M. Araiza, Kris P. Punayan, Irish Coleen A. Asin, Candice Francheska B. Tambaoan, Asia Louisa U. Chong, Karol Sophia Agape R. Padilla, Rianna Patricia S. Cruz, El King D. Morado, Joshua Gregor A. Dizon, Razel Nikka M. Hao, Arianne A. Zamora, Devon Ray Pacial, Juan Antonio R. Magalang, Marissa Alejandria, Celia Carlos, Anna Ong-Lim, Edsel Maurice Salvaña, John Q. Wong, Jaime C. Montoya, Maria Rosario Singh-Vergeire and Cynthia P. Saloma |
| EPI_ISL_2188238 | Sta. Rosa CHO | Philippine Genome Center | Francis A. Tablizo, Kenneth M. Kim, Carlo M. Lapid, Marc Jerrone R. Castro, Maria Sofia L. Yangzon, Elcid Aaron R. Pangilinan, Benedict A. Maralit, Marc Edsel C. Ayes, Eva Maria Cutiongco-de la Paz, Alethea R. de Guzman, Jan Michael C. Yap, Jo-Hannah S. Llames, Sheila Mae M. Araiza, Kris P. Punayan, Irish Coleen A. Asin, Candice Francheska B. Tambaoan, Asia Louisa U. Chong, Karol Sophia Agape R. Padilla, Rianna Patricia S. Cruz, El King D. Morado, Joshua Gregor A. Dizon, Razel Nikka M. Hao, Arianne A. Zamora, Devon Ray Pacial, Juan Antonio R. Magalang, Marissa Alejandria, Celia Carlos, Anna Ong-Lim, Edsel Maurice Salvaña, John Q. Wong, Jaime C. Montoya, Maria Rosario Singh-Vergeire and Cynthia P. Saloma |
| EPI_ISL_2188239 | Ilocos Training and Regional Medical Center | Philippine Genome Center | Francis A. Tablizo, Kenneth M. Kim, Carlo M. Lapid, Marc Jerrone R. Castro, Maria Sofia L. Yangzon, Elcid Aaron R. Pangilinan, Benedict A. Maralit, Marc Edsel C. Ayes, Eva Maria Cutiongco-de la Paz, Alethea R. de Guzman, Jan Michael C. Yap, Jo-Hannah S. Llames, Sheila Mae M. Araiza, Kris P. Punayan, Irish Coleen A. Asin, Candice Francheska B. Tambaoan, Asia Louisa U. Chong, Karol Sophia Agape R. Padilla, Rianna Patricia S. Cruz, El King D. Morado, Joshua Gregor A. Dizon, Razel Nikka M. Hao, Arianne A. Zamora, Devon Ray Pacial, Juan Antonio R. Magalang, Marissa Alejandria, Celia Carlos, Anna Ong-Lim, Edsel Maurice Salvaña, John Q. Wong, Jaime C. Montoya, Maria Rosario Singh-Vergeire and Cynthia P. Saloma |
| EPI_ISL_2188240, EPI_ISL_2188241 | Mary Mediatrix Medical Center | Philippine Genome Center | Francis A. Tablizo, Kenneth M. Kim, Carlo M. Lapid, Marc Jerrone R. Castro, Maria Sofia L. Yangzon, Elcid Aaron R. Pangilinan, Benedict A. Maralit, Marc Edsel C. Ayes, Eva Maria Cutiongco-de la Paz, Alethea R. de Guzman, Jan Michael C. Yap, Jo-Hannah S. Llames, Sheila Mae M. Araiza, Kris P. Punayan, Irish Coleen A. Asin, Candice Francheska B. Tambaoan, Asia Louisa U. Chong, Karol Sophia Agape R. Padilla, Rianna Patricia S. Cruz, El King D. Morado, Joshua Gregor A. Dizon, Razel Nikka M. Hao, Arianne A. Zamora, Devon Ray Pacial, Juan Antonio R. Magalang, Marissa Alejandria, Celia Carlos, Anna Ong-Lim, Edsel Maurice Salvaña, John Q. Wong, Jaime C. Montoya, Maria Rosario Singh-Vergeire and Cynthia P. Saloma |
| EPI_ISL_2188242 | The Medical City -Â Ortigas | Philippine Genome Center | Francis A. Tablizo, Kenneth M. Kim, Carlo M. Lapid, Marc Jerrone R. Castro, Maria Sofia L. Yangzon, Elcid Aaron R. Pangilinan, Benedict A. Maralit, Marc Edsel C. Ayes, Eva Maria Cutiongco-de la Paz, Alethea R. de Guzman, Jan Michael C. Yap, Jo-Hannah S. Llames, Sheila Mae M. Araiza, Kris P. Punayan, Irish Coleen A. Asin, Candice Francheska B. Tambaoan, Asia Louisa U. Chong, Karol Sophia Agape R. Padilla, Rianna Patricia S. Cruz, El King D. Morado, Joshua Gregor A. Dizon, Razel Nikka M. Hao, Arianne A. Zamora, Devon Ray Pacial, Juan Antonio R. Magalang, Marissa Alejandria, Celia Carlos, Anna Ong-Lim, Edsel Maurice Salvaña, John Q. Wong, Jaime C. Montoya, Maria Rosario Singh-Vergeire and Cynthia P. Saloma |
| EPI_ISL_2188243, EPI_ISL_2188245, EPI_ISL_2188246 | The Lord's Grace Medical and Industrial Clinic | Philippine Genome Center | Francis A. Tablizo, Kenneth M. Kim, Carlo M. Lapid, Marc Jerrone R. Castro, Maria Sofia L. Yangzon, Elcid Aaron R. Pangilinan, Benedict A. Maralit, Marc Edsel C. Ayes, Eva Maria Cutiongco-de la Paz, Alethea R. de Guzman, Jan Michael C. Yap, Jo-Hannah S. Llames, Sheila Mae M. Araiza, Kris P. Punayan, Irish Coleen A. Asin, Candice Francheska B. Tambaoan, Asia Louisa U. Chong, Karol Sophia Agape R. Padilla, Rianna Patricia S. Cruz, El King D. Morado, Joshua Gregor A. Dizon, Razel Nikka M. Hao, Arianne A. Zamora, Devon Ray Pacial, Juan Antonio R. Magalang, Marissa Alejandria, Celia Carlos, Anna Ong-Lim, Edsel Maurice Salvaña, John Q. Wong, Jaime C. Montoya, Maria Rosario Singh-Vergeire and Cynthia P. Saloma |
| EPI_ISL_2188247 | Philippine Red Cross - Port Area | Philippine Genome Center | Francis A. Tablizo, Kenneth M. Kim, Carlo M. Lapid, Marc Jerrone R. Castro, Maria Sofia L. Yangzon, Elcid Aaron R. Pangilinan, Benedict A. Maralit, Marc Edsel C. Ayes, Eva Maria Cutiongco-de la Paz, Alethea R. de Guzman, Jan Michael C. Yap, Jo-Hannah S. Llames, Sheila Mae M. Araiza, Kris P. Punayan, Irish Coleen A. Asin, Candice Francheska B. Tambaoan, Asia Louisa U. Chong, Karol Sophia Agape R. Padilla, Rianna Patricia S. Cruz, El King D. Morado, Joshua Gregor A. Dizon, Razel Nikka M. Hao, Arianne A. Zamora, Devon Ray Pacial, Juan Antonio R. Magalang, Marissa Alejandria, Celia Carlos, Anna Ong-Lim, Edsel Maurice Salvaña, John Q. Wong, Jaime C. Montoya, Maria Rosario Singh-Vergeire and Cynthia P. Saloma |
| EPI_ISL_2188251 | Qualimed Hospital Sta. Rosa | Philippine Genome Center | Francis A. Tablizo, Kenneth M. Kim, Carlo M. Lapid, Marc Jerrone R. Castro, Maria Sofia L. Yangzon, Elcid Aaron R. Pangilinan, Benedict A. Maralit, Marc Edsel C. Ayes, Eva Maria Cutiongco-de la Paz, Alethea R. de Guzman, Jan Michael C. Yap, Jo-Hannah S. Llames, Sheila Mae M. Araiza, Kris P. Punayan, Irish Coleen A. Asin, Candice Francheska B. Tambaoan, Asia Louisa U. Chong, Karol Sophia Agape R. Padilla, Rianna Patricia S. Cruz, El King D. Morado, Joshua Gregor A. Dizon, Razel Nikka M. Hao, Arianne A. Zamora, Devon Ray Pacial, Juan Antonio R. Magalang, Marissa Alejandria, Celia Carlos, Anna Ong-Lim, Edsel Maurice Salvaña, John Q. Wong, Jaime C. Montoya, Maria Rosario Singh-Vergeire and Cynthia P. Saloma |
| EPI_ISL_2188252, EPI_ISL_2188253 | The Lord's Grace Medical and Industrial Clinic | Philippine Genome Center | Francis A. Tablizo, Kenneth M. Kim, Carlo M. Lapid, Marc Jerrone R. Castro, Maria Sofia L. Yangzon, Elcid Aaron R. Pangilinan, Benedict A. Maralit, Marc Edsel C. Ayes, Eva Maria Cutiongco-de la Paz, Alethea R. de Guzman, Jan Michael C. Yap, Jo-Hannah S. Llames, Sheila Mae M. Araiza, Kris P. Punayan, Irish Coleen A. Asin, Candice Francheska B. Tambaoan, Asia Louisa U. Chong, Karol Sophia Agape R. Padilla, Rianna Patricia S. Cruz, El King D. Morado, Joshua Gregor A. Dizon, Razel Nikka M. Hao, Arianne A. Zamora, Devon Ray Pacial, Juan Antonio R. Magalang, Marissa Alejandria, Celia Carlos, Anna Ong-Lim, Edsel Maurice Salvaña, John Q. Wong, Jaime C. Montoya, Maria Rosario Singh-Vergeire and Cynthia P. Saloma |
| EPI_ISL_2188254, EPI_ISL_2188255, EPI_ISL_2188257, EPI_ISL_2188258, EPI_ISL_2188259 | The Medical City -Â Ortigas | Philippine Genome Center | Francis A. Tablizo, Kenneth M. Kim, Carlo M. Lapid, Marc Jerrone R. Castro, Maria Sofia L. Yangzon, Elcid Aaron R. Pangilinan, Benedict A. Maralit, Marc Edsel C. Ayes, Eva Maria Cutiongco-de la Paz, Alethea R. de Guzman, Jan Michael C. Yap, Jo-Hannah S. Llames, Sheila Mae M. Araiza, Kris P. Punayan, Irish Coleen A. Asin, Candice Francheska B. Tambaoan, Asia Louisa U. Chong, Karol Sophia Agape R. Padilla, Rianna Patricia S. Cruz, El King D. Morado, Joshua Gregor A. Dizon, Razel Nikka M. Hao, Arianne A. Zamora, Devon Ray Pacial, Juan Antonio R. Magalang, Marissa Alejandria, Celia Carlos, Anna Ong-Lim, Edsel Maurice Salvaña, John Q. Wong, Jaime C. Montoya, Maria Rosario Singh-Vergeire and Cynthia P. Saloma |
| EPI_ISL_2188260 | Philippine Red Cross - Port Area | Philippine Genome Center | Francis A. Tablizo, Kenneth M. Kim, Carlo M. Lapid, Marc Jerrone R. Castro, Maria Sofia L. Yangzon, Elcid Aaron R. Pangilinan, Benedict A. Maralit, Marc Edsel C. Ayes, Eva Maria Cutiongco-de la Paz, Alethea R. de Guzman, Jan Michael C. Yap, Jo-Hannah S. Llames, Sheila Mae M. Araiza, Kris P. Punayan, Irish Coleen A. Asin, Candice Francheska B. Tambaoan, Asia Louisa U. Chong, Karol Sophia Agape R. Padilla, Rianna Patricia S. Cruz, El King D. Morado, Joshua Gregor A. Dizon, Razel Nikka M. Hao, Arianne A. Zamora, Devon Ray Pacial, Juan Antonio R. Magalang, Marissa Alejandria, Celia Carlos, Anna Ong-Lim, Edsel Maurice Salvaña, John Q. Wong, Jaime C. Montoya, Maria Rosario Singh-Vergeire and Cynthia P. Saloma |

| | | | |
|---|---|---|---|
| EPI_ISL_2188262 | The Medical City -Â Ortigas | Philippine Genome Center | Francis A. Tablizo, Kenneth M. Kim, Carlo M. Lapid, Marc Jerrone R. Castro, Maria Sofia L. Yangzon, Elcid Aaron R. Pangilinan, Benedict A. Maralit, Marc Edsel C. Ayes, Eva Maria Cutiongco-de la Paz, Alethea R. de Guzman, Jan Michael C. Yap, Jo-Hannah S. Llames, Sheila Mae M. Araiza, Kris P. Punayan, Irish Coleen A. Asin, Candice Francheska B. Tambaoan, Asia Louisa U. Chong, Karol Sophia Agape R. Padilla, Rianna Patricia S. Cruz, El King D. Morado, Joshua Gregor A. Dizon, Razel Nikka M. Hao, Arianne A. Zamora, Devon Ray Pacial, Juan Antonio R. Magalang, Marissa Alejandria, Celia Carlos, Anna Ong-Lim, Edsel Maurice Salvaña, John Q. Wong, Jaime C. Montoya, Maria Rosario Singh-Vergeire and Cynthia P. Saloma |
| EPI_ISL_2188263 | Qualimed Hospital Sta. Rosa | Philippine Genome Center | Francis A. Tablizo, Kenneth M. Kim, Carlo M. Lapid, Marc Jerrone R. Castro, Maria Sofia L. Yangzon, Elcid Aaron R. Pangilinan, Benedict A. Maralit, Marc Edsel C. Ayes, Eva Maria Cutiongco-de la Paz, Alethea R. de Guzman, Jan Michael C. Yap, Jo-Hannah S. Llames, Sheila Mae M. Araiza, Kris P. Punayan, Irish Coleen A. Asin, Candice Francheska B. Tambaoan, Asia Louisa U. Chong, Karol Sophia Agape R. Padilla, Rianna Patricia S. Cruz, El King D. Morado, Joshua Gregor A. Dizon, Razel Nikka M. Hao, Arianne A. Zamora, Devon Ray Pacial, Juan Antonio R. Magalang, Marissa Alejandria, Celia Carlos, Anna Ong-Lim, Edsel Maurice Salvaña, John Q. Wong, Jaime C. Montoya, Maria Rosario Singh-Vergeire and Cynthia P. Saloma |
| EPI_ISL_2188264, EPI_ISL_2188265 | The Medical City -Â Ortigas | Philippine Genome Center | Francis A. Tablizo, Kenneth M. Kim, Carlo M. Lapid, Marc Jerrone R. Castro, Maria Sofia L. Yangzon, Elcid Aaron R. Pangilinan, Benedict A. Maralit, Marc Edsel C. Ayes, Eva Maria Cutiongco-de la Paz, Alethea R. de Guzman, Jan Michael C. Yap, Jo-Hannah S. Llames, Sheila Mae M. Araiza, Kris P. Punayan, Irish Coleen A. Asin, Candice Francheska B. Tambaoan, Asia Louisa U. Chong, Karol Sophia Agape R. Padilla, Rianna Patricia S. Cruz, El King D. Morado, Joshua Gregor A. Dizon, Razel Nikka M. Hao, Arianne A. Zamora, Devon Ray Pacial, Juan Antonio R. Magalang, Marissa Alejandria, Celia Carlos, Anna Ong-Lim, Edsel Maurice Salvaña, John Q. Wong, Jaime C. Montoya, Maria Rosario Singh-Vergeire and Cynthia P. Saloma |
| EPI_ISL_2188267 | The Lord's Grace Medical and Industrial Clinic | Philippine Genome Center | Francis A. Tablizo, Kenneth M. Kim, Carlo M. Lapid, Marc Jerrone R. Castro, Maria Sofia L. Yangzon, Elcid Aaron R. Pangilinan, Benedict A. Maralit, Marc Edsel C. Ayes, Eva Maria Cutiongco-de la Paz, Alethea R. de Guzman, Jan Michael C. Yap, Jo-Hannah S. Llames, Sheila Mae M. Araiza, Kris P. Punayan, Irish Coleen A. Asin, Candice Francheska B. Tambaoan, Asia Louisa U. Chong, Karol Sophia Agape R. Padilla, Rianna Patricia S. Cruz, El King D. Morado, Joshua Gregor A. Dizon, Razel Nikka M. Hao, Arianne A. Zamora, Devon Ray Pacial, Juan Antonio R. Magalang, Marissa Alejandria, Celia Carlos, Anna Ong-Lim, Edsel Maurice Salvaña, John Q. Wong, Jaime C. Montoya, Maria Rosario Singh-Vergeire and Cynthia P. Saloma |
| EPI_ISL_2188268 | The Medical City -Â Ortigas | Philippine Genome Center | Francis A. Tablizo, Kenneth M. Kim, Carlo M. Lapid, Marc Jerrone R. Castro, Maria Sofia L. Yangzon, Elcid Aaron R. Pangilinan, Benedict A. Maralit, Marc Edsel C. Ayes, Eva Maria Cutiongco-de la Paz, Alethea R. de Guzman, Jan Michael C. Yap, Jo-Hannah S. Llames, Sheila Mae M. Araiza, Kris P. Punayan, Irish Coleen A. Asin, Candice Francheska B. Tambaoan, Asia Louisa U. Chong, Karol Sophia Agape R. Padilla, Rianna Patricia S. Cruz, El King D. Morado, Joshua Gregor A. Dizon, Razel Nikka M. Hao, Arianne A. Zamora, Devon Ray Pacial, Juan Antonio R. Magalang, Marissa Alejandria, Celia Carlos, Anna Ong-Lim, Edsel Maurice Salvaña, John Q. Wong, Jaime C. Montoya, Maria Rosario Singh-Vergeire and Cynthia P. Saloma |
| EPI_ISL_2188269 | Philippine Red Cross - Port Area | Philippine Genome Center | Francis A. Tablizo, Kenneth M. Kim, Carlo M. Lapid, Marc Jerrone R. Castro, Maria Sofia L. Yangzon, Elcid Aaron R. Pangilinan, Benedict A. Maralit, Marc Edsel C. Ayes, Eva Maria Cutiongco-de la Paz, Alethea R. de Guzman, Jan Michael C. Yap, Jo-Hannah S. Llames, Sheila Mae M. Araiza, Kris P. Punayan, Irish Coleen A. Asin, Candice Francheska B. Tambaoan, Asia Louisa U. Chong, Karol Sophia Agape R. Padilla, Rianna Patricia S. Cruz, El King D. Morado, Joshua Gregor A. Dizon, Razel Nikka M. Hao, Arianne A. Zamora, Devon Ray Pacial, Juan Antonio R. Magalang, Marissa Alejandria, Celia Carlos, Anna Ong-Lim, Edsel Maurice Salvaña, John Q. Wong, Jaime C. Montoya, Maria Rosario Singh-Vergeire and Cynthia P. Saloma |
| EPI_ISL_2188270 | The Medical City -Â Ortigas | Philippine Genome Center | Francis A. Tablizo, Kenneth M. Kim, Carlo M. Lapid, Marc Jerrone R. Castro, Maria Sofia L. Yangzon, Elcid Aaron R. Pangilinan, Benedict A. Maralit, Marc Edsel C. Ayes, Eva Maria Cutiongco-de la Paz, Alethea R. de Guzman, Jan Michael C. Yap, Jo-Hannah S. Llames, Sheila Mae M. Araiza, Kris P. Punayan, Irish Coleen A. Asin, Candice Francheska B. Tambaoan, Asia Louisa U. Chong, Karol Sophia Agape R. Padilla, Rianna Patricia S. Cruz, El King D. Morado, Joshua Gregor A. Dizon, Razel Nikka M. Hao, Arianne A. Zamora, Devon Ray Pacial, Juan Antonio R. Magalang, Marissa Alejandria, Celia Carlos, Anna Ong-Lim, Edsel Maurice Salvaña, John Q. Wong, Jaime C. Montoya, Maria Rosario Singh-Vergeire and Cynthia P. Saloma |
| EPI_ISL_2188271 | Philippine Red Cross - Port Area | Philippine Genome Center | Francis A. Tablizo, Kenneth M. Kim, Carlo M. Lapid, Marc Jerrone R. Castro, Maria Sofia L. Yangzon, Elcid Aaron R. Pangilinan, Benedict A. Maralit, Marc Edsel C. Ayes, Eva Maria Cutiongco-de la Paz, Alethea R. de Guzman, Jan Michael C. Yap, Jo-Hannah S. Llames, Sheila Mae M. Araiza, Kris P. Punayan, Irish Coleen A. Asin, Candice Francheska B. Tambaoan, Asia Louisa U. Chong, Karol Sophia Agape R. Padilla, Rianna Patricia S. Cruz, El King D. Morado, Joshua Gregor A. Dizon, Razel Nikka M. Hao, Arianne A. Zamora, Devon Ray Pacial, Juan Antonio R. Magalang, Marissa Alejandria, Celia Carlos, Anna Ong-Lim, Edsel Maurice Salvaña, John Q. Wong, Jaime C. Montoya, Maria Rosario Singh-Vergeire and Cynthia P. Saloma |
| EPI_ISL_2188272 | The Medical City -Â Ortigas | Philippine Genome Center | Francis A. Tablizo, Kenneth M. Kim, Carlo M. Lapid, Marc Jerrone R. Castro, Maria Sofia L. Yangzon, Elcid Aaron R. Pangilinan, Benedict A. Maralit, Marc Edsel C. Ayes, Eva Maria Cutiongco-de la Paz, Alethea R. de Guzman, Jan Michael C. Yap, Jo-Hannah S. Llames, Sheila Mae M. Araiza, Kris P. Punayan, Irish Coleen A. Asin, Candice Francheska B. Tambaoan, Asia Louisa U. Chong, Karol Sophia Agape R. Padilla, Rianna Patricia S. Cruz, El King D. Morado, Joshua Gregor A. Dizon, Razel Nikka M. Hao, Arianne A. Zamora, Devon Ray Pacial, Juan Antonio R. Magalang, Marissa Alejandria, Celia Carlos, Anna Ong-Lim, Edsel Maurice Salvaña, John Q. Wong, Jaime C. Montoya, Maria Rosario Singh-Vergeire and Cynthia P. Saloma |
| EPI_ISL_2188274 | Research Institute for Tropical Medicine, Inc. (RITM) | Philippine Genome Center | Francis A. Tablizo, Kenneth M. Kim, Carlo M. Lapid, Marc Jerrone R. Castro, Maria Sofia L. Yangzon, Elcid Aaron R. Pangilinan, Benedict A. Maralit, Marc Edsel C. Ayes, Eva Maria Cutiongco-de la Paz, Alethea R. de Guzman, Jan Michael C. Yap, Jo-Hannah S. Llames, Sheila Mae M. Araiza, Kris P. Punayan, Irish Coleen A. Asin, Candice Francheska B. Tambaoan, Asia Louisa U. Chong, Karol Sophia Agape R. Padilla, Rianna Patricia S. Cruz, El King D. Morado, Joshua Gregor A. Dizon, Razel Nikka M. Hao, Arianne A. Zamora, Devon Ray Pacial, Juan Antonio R. Magalang, Marissa Alejandria, Celia Carlos, Anna Ong-Lim, Edsel Maurice Salvaña, John Q. Wong, Jaime C. Montoya, Maria Rosario Singh-Vergeire and Cynthia P. Saloma |
| EPI_ISL_2188275 | The Lord's Grace Medical and Industrial Clinic | Philippine Genome Center | Francis A. Tablizo, Kenneth M. Kim, Carlo M. Lapid, Marc Jerrone R. Castro, Maria Sofia L. Yangzon, Elcid Aaron R. Pangilinan, Benedict A. Maralit, Marc Edsel C. Ayes, Eva Maria Cutiongco-de la Paz, Alethea R. de Guzman, Jan Michael C. Yap, Jo-Hannah S. Llames, Sheila Mae M. Araiza, Kris P. Punayan, Irish Coleen A. Asin, Candice Francheska B. Tambaoan, Asia Louisa U. Chong, Karol Sophia Agape R. Padilla, Rianna Patricia S. Cruz, El King D. Morado, Joshua Gregor A. Dizon, Razel Nikka M. Hao, Arianne A. Zamora, Devon Ray Pacial, Juan Antonio R. Magalang, Marissa Alejandria, Celia Carlos, Anna Ong-Lim, Edsel Maurice Salvaña, John Q. Wong, Jaime C. Montoya, Maria Rosario Singh-Vergeire and Cynthia P. Saloma |
| EPI_ISL_2188276 | SAN ANTONIO CITY OF ILAGAN HOSPITAL | Philippine Genome Center | Francis A. Tablizo, Kenneth M. Kim, Carlo M. Lapid, Marc Jerrone R. Castro, Maria Sofia L. Yangzon, Elcid Aaron R. Pangilinan, Benedict A. Maralit, Marc Edsel C. Ayes, Eva Maria Cutiongco-de la Paz, Alethea R. de Guzman, Jan Michael C. Yap, Jo-Hannah S. Llames, Sheila Mae M. Araiza, Kris P. Punayan, Irish Coleen A. Asin, Candice Francheska B. Tambaoan, Asia Louisa U. Chong, Karol Sophia Agape R. Padilla, Rianna Patricia S. Cruz, El King D. Morado, Joshua Gregor A. Dizon, Razel Nikka M. Hao, Arianne A. Zamora, Devon Ray Pacial, Juan Antonio R. Magalang, Marissa Alejandria, Celia Carlos, Anna Ong-Lim, Edsel Maurice Salvaña, John Q. Wong, Jaime C. Montoya, Maria Rosario Singh-Vergeire and Cynthia P. Saloma |
| EPI_ISL_2188277 | The Medical City -Â Ortigas | Philippine Genome Center | Francis A. Tablizo, Kenneth M. Kim, Carlo M. Lapid, Marc Jerrone R. Castro, Maria Sofia L. Yangzon, Elcid Aaron R. Pangilinan, Benedict A. Maralit, Marc Edsel C. Ayes, Eva Maria Cutiongco-de la Paz, Alethea R. de Guzman, Jan Michael C. Yap, Jo-Hannah S. Llames, Sheila Mae M. Araiza, Kris P. Punayan, Irish Coleen A. Asin, Candice Francheska B. Tambaoan, Asia Louisa U. Chong, Karol Sophia Agape R. Padilla, Rianna Patricia S. Cruz, El King D. Morado, Joshua Gregor A. Dizon, Razel Nikka M. Hao, Arianne A. Zamora, Devon Ray Pacial, Juan Antonio R. Magalang, Marissa Alejandria, Celia Carlos, Anna Ong-Lim, Edsel Maurice Salvaña, John Q. Wong, Jaime C. Montoya, Maria Rosario Singh-Vergeire and Cynthia P. Saloma |
| EPI_ISL_2188278 | Dr. Jorge P. Royeca Hospital | Philippine Genome Center | Francis A. Tablizo, Kenneth M. Kim, Carlo M. Lapid, Marc Jerrone R. Castro, Maria Sofia L. Yangzon, Elcid Aaron R. Pangilinan, Benedict A. Maralit, Marc Edsel C. Ayes, Eva Maria Cutiongco-de la Paz, Alethea R. de Guzman, Jan Michael C. Yap, Jo-Hannah S. Llames, Sheila Mae M. Araiza, Kris P. Punayan, Irish Coleen A. Asin, Candice Francheska B. Tambaoan, Asia Louisa U. Chong, Karol Sophia Agape R. Padilla, Rianna Patricia S. Cruz, El King D. Morado, Joshua Gregor A. Dizon, Razel Nikka M. Hao, Arianne A. Zamora, Devon Ray Pacial, Juan Antonio R. Magalang, Marissa Alejandria, Celia Carlos, Anna Ong-Lim, Edsel Maurice Salvaña, John Q. Wong, Jaime C. Montoya, Maria Rosario Singh-Vergeire and Cynthia P. Saloma |
| EPI_ISL_2188439 | Hospital | National Reference Center for Viruses of Respiratory Infections, Institut Pasteur, Paris | Marion Barbet, Sylvie Behillil, Méline Bizard, Angela Brisebarre, Camille Capel, Vincent Enouf, Louise Lefrançois, Frédéric Lemoine, Christophe Malabat, Corinne Maufrais, Etienne Simon-Lorière, Maud Vanpeene, Sylvie Van der Werf ,Patricia Stoessel-Thouvenin |
| EPI_ISL_2188676 | Ospital ng Imus | Philippine Genome Center | Francis A. Tablizo, Kenneth M. Kim, Maria Sofia L. Yangzon, Elcid Aaron R. Pangilinan, Renato Jacinto Q. Mantaring, Benedict A. Maralit, Marc Edsel C. Ayes, Eva Maria Cutiongco-de la Paz, Alethea R. de Guzman, Jan Michael C. Yap, Jo-Hannah S. Llames, Sheila Mae M. Araiza, Kris P. Punayan, Irish Coleen A. Asin, Candice Francheska B. Tambaoan, Asia Louisa U. Chong, Karol Sophia Agape R. Padilla, Rianna Patricia S. Cruz, Carlo M. Lapid, El King D. Morado, Joshua Gregor A. Dizon, Razel Nikka M. Hao, Arianne A. Zamora, Devon Ray Pacial, Juan Antonio R. Magalang, Marissa Alejandria, Celia Carlos, Anna Ong-Lim, Edsel Maurice Salvaña, John Q. Wong, Jaime C. Montoya, Maria Rosario Singh-Vergeire and Cynthia P. Saloma |
| EPI_ISL_2188677 | Cotabato Regional and Medical Center | Philippine Genome Center | Francis A. Tablizo, Kenneth M. Kim, Maria Sofia L. Yangzon, Elcid Aaron R. Pangilinan, Renato Jacinto Q. Mantaring, Benedict A. Maralit, Marc Edsel C. Ayes, Eva Maria Cutiongco-de la Paz, Alethea R. de Guzman, Jan Michael C. Yap, Jo-Hannah S. Llames, Sheila Mae M. Araiza, Kris P. Punayan, Irish |

| | | | |
|---|---|---|---|
| | | | Coleen A. Asin, Candice Francheska B. Tambaoan, Asia Louisa U. Chong, Karol Sophia Agape R. Padilla, Rianna Patricia S. Cruz, Carlo M. Lapid, El King D. Morado, Joshua Gregor A. Dizon, Razel Nikka M. Hao, Arianne A. Zamora, Devon Ray Pacial, Juan Antonio R. Magalang, Marissa Alejandria, Celia Carlos, Anna Ong-Lim, Edsel Maurice Salvaña, John Q. Wong, Jaime C. Montoya, Maria Rosario Singh-Vergeire and Cynthia P. Saloma |
| EPI_ISL_2188678 | Philippine Red Cross - National Blood Center | Philippine Genome Center | Francis A. Tablizo, Kenneth M. Kim, Maria Sofia L. Yangzon, Elcid Aaron R. Pangilinan, Renato Jacinto Q. Mantaring, Benedict A. Maralit, Marc Edsel C. Ayes, Eva Maria Cutiongco-de la Paz, Alethea R. de Guzman, Jan Michael C. Yap, Jo-Hannah S. Llames, Sheila Mae M. Araiza, Kris P. Punayan, Irish Coleen A. Asin, Candice Francheska B. Tambaoan, Asia Louisa U. Chong, Karol Sophia Agape R. Padilla, Rianna Patricia S. Cruz, Carlo M. Lapid, El King D. Morado, Joshua Gregor A. Dizon, Razel Nikka M. Hao, Arianne A. Zamora, Devon Ray Pacial, Juan Antonio R. Magalang, Marissa Alejandria, Celia Carlos, Anna Ong-Lim, Edsel Maurice Salvaña, John Q. Wong, Jaime C. Montoya, Maria Rosario Singh-Vergeire and Cynthia P. Saloma |
| EPI_ISL_2188680 | Philippine Red Cross Logistics and Multipurpose Center | Philippine Genome Center | Francis A. Tablizo, Kenneth M. Kim, Maria Sofia L. Yangzon, Elcid Aaron R. Pangilinan, Renato Jacinto Q. Mantaring, Benedict A. Maralit, Marc Edsel C. Ayes, Eva Maria Cutiongco-de la Paz, Alethea R. de Guzman, Jan Michael C. Yap, Jo-Hannah S. Llames, Sheila Mae M. Araiza, Kris P. Punayan, Irish Coleen A. Asin, Candice Francheska B. Tambaoan, Asia Louisa U. Chong, Karol Sophia Agape R. Padilla, Rianna Patricia S. Cruz, Carlo M. Lapid, El King D. Morado, Joshua Gregor A. Dizon, Razel Nikka M. Hao, Arianne A. Zamora, Devon Ray Pacial, Juan Antonio R. Magalang, Marissa Alejandria, Celia Carlos, Anna Ong-Lim, Edsel Maurice Salvaña, John Q. Wong, Jaime C. Montoya, Maria Rosario Singh-Vergeire and Cynthia P. Saloma |
| EPI_ISL_2188682, EPI_ISL_2188687 | The Medical City -Â Ortigas | Philippine Genome Center | Francis A. Tablizo, Kenneth M. Kim, Maria Sofia L. Yangzon, Elcid Aaron R. Pangilinan, Renato Jacinto Q. Mantaring, Benedict A. Maralit, Marc Edsel C. Ayes, Eva Maria Cutiongco-de la Paz, Alethea R. de Guzman, Jan Michael C. Yap, Jo-Hannah S. Llames, Sheila Mae M. Araiza, Kris P. Punayan, Irish Coleen A. Asin, Candice Francheska B. Tambaoan, Asia Louisa U. Chong, Karol Sophia Agape R. Padilla, Rianna Patricia S. Cruz, Carlo M. Lapid, El King D. Morado, Joshua Gregor A. Dizon, Razel Nikka M. Hao, Arianne A. Zamora, Devon Ray Pacial, Juan Antonio R. Magalang, Marissa Alejandria, Celia Carlos, Anna Ong-Lim, Edsel Maurice Salvaña, John Q. Wong, Jaime C. Montoya, Maria Rosario Singh-Vergeire and Cynthia P. Saloma |
| EPI_ISL_2188688 | The Lord's Grace Medical and Industrial Clinic | Philippine Genome Center | Francis A. Tablizo, Kenneth M. Kim, Maria Sofia L. Yangzon, Elcid Aaron R. Pangilinan, Renato Jacinto Q. Mantaring, Benedict A. Maralit, Marc Edsel C. Ayes, Eva Maria Cutiongco-de la Paz, Alethea R. de Guzman, Jan Michael C. Yap, Jo-Hannah S. Llames, Sheila Mae M. Araiza, Kris P. Punayan, Irish Coleen A. Asin, Candice Francheska B. Tambaoan, Asia Louisa U. Chong, Karol Sophia Agape R. Padilla, Rianna Patricia S. Cruz, Carlo M. Lapid, El King D. Morado, Joshua Gregor A. Dizon, Razel Nikka M. Hao, Arianne A. Zamora, Devon Ray Pacial, Juan Antonio R. Magalang, Marissa Alejandria, Celia Carlos, Anna Ong-Lim, Edsel Maurice Salvaña, John Q. Wong, Jaime C. Montoya, Maria Rosario Singh-Vergeire and Cynthia P. Saloma |
| EPI_ISL_2188689 | Philippine Red Cross - National Blood Center | Philippine Genome Center | Francis A. Tablizo, Kenneth M. Kim, Maria Sofia L. Yangzon, Elcid Aaron R. Pangilinan, Renato Jacinto Q. Mantaring, Benedict A. Maralit, Marc Edsel C. Ayes, Eva Maria Cutiongco-de la Paz, Alethea R. de Guzman, Jan Michael C. Yap, Jo-Hannah S. Llames, Sheila Mae M. Araiza, Kris P. Punayan, Irish Coleen A. Asin, Candice Francheska B. Tambaoan, Asia Louisa U. Chong, Karol Sophia Agape R. Padilla, Rianna Patricia S. Cruz, Carlo M. Lapid, El King D. Morado, Joshua Gregor A. Dizon, Razel Nikka M. Hao, Arianne A. Zamora, Devon Ray Pacial, Juan Antonio R. Magalang, Marissa Alejandria, Celia Carlos, Anna Ong-Lim, Edsel Maurice Salvaña, John Q. Wong, Jaime C. Montoya, Maria Rosario Singh-Vergeire and Cynthia P. Saloma |
| EPI_ISL_2188696 | The Lord's Grace Medical and Industrial Clinic | Philippine Genome Center | Francis A. Tablizo, Kenneth M. Kim, Maria Sofia L. Yangzon, Elcid Aaron R. Pangilinan, Renato Jacinto Q. Mantaring, Benedict A. Maralit, Marc Edsel C. Ayes, Eva Maria Cutiongco-de la Paz, Alethea R. de Guzman, Jan Michael C. Yap, Jo-Hannah S. Llames, Sheila Mae M. Araiza, Kris P. Punayan, Irish Coleen A. Asin, Candice Francheska B. Tambaoan, Asia Louisa U. Chong, Karol Sophia Agape R. Padilla, Rianna Patricia S. Cruz, Carlo M. Lapid, El King D. Morado, Joshua Gregor A. Dizon, Razel Nikka M. Hao, Arianne A. Zamora, Devon Ray Pacial, Juan Antonio R. Magalang, Marissa Alejandria, Celia Carlos, Anna Ong-Lim, Edsel Maurice Salvaña, John Q. Wong, Jaime C. Montoya, Maria Rosario Singh-Vergeire and Cynthia P. Saloma |
| EPI_ISL_2188699 | Philippine Red Cross - Port Area | Philippine Genome Center | Francis A. Tablizo, Kenneth M. Kim, Maria Sofia L. Yangzon, Elcid Aaron R. Pangilinan, Renato Jacinto Q. Mantaring, Benedict A. Maralit, Marc Edsel C. Ayes, Eva Maria Cutiongco-de la Paz, Alethea R. de Guzman, Jan Michael C. Yap, Jo-Hannah S. Llames, Sheila Mae M. Araiza, Kris P. Punayan, Irish Coleen A. Asin, Candice Francheska B. Tambaoan, Asia Louisa U. Chong, Karol Sophia Agape R. Padilla, Rianna Patricia S. Cruz, Carlo M. Lapid, El King D. Morado, Joshua Gregor A. Dizon, Razel Nikka M. Hao, Arianne A. Zamora, Devon Ray Pacial, Juan Antonio R. Magalang, Marissa Alejandria, Celia Carlos, Anna Ong-Lim, Edsel Maurice Salvaña, John Q. Wong, Jaime C. Montoya, Maria Rosario Singh-Vergeire and Cynthia P. Saloma |
| EPI_ISL_2188700 | The Lord's Grace Medical and Industrial Clinic | Philippine Genome Center | Francis A. Tablizo, Kenneth M. Kim, Maria Sofia L. Yangzon, Elcid Aaron R. Pangilinan, Renato Jacinto Q. Mantaring, Benedict A. Maralit, Marc Edsel C. Ayes, Eva Maria Cutiongco-de la Paz, Alethea R. de Guzman, Jan Michael C. Yap, Jo-Hannah S. Llames, Sheila Mae M. Araiza, Kris P. Punayan, Irish Coleen A. Asin, Candice Francheska B. Tambaoan, Asia Louisa U. Chong, Karol Sophia Agape R. Padilla, Rianna Patricia S. Cruz, Carlo M. Lapid, El King D. Morado, Joshua Gregor A. Dizon, Razel Nikka M. Hao, Arianne A. Zamora, Devon Ray Pacial, Juan Antonio R. Magalang, Marissa Alejandria, Celia Carlos, Anna Ong-Lim, Edsel Maurice Salvaña, John Q. Wong, Jaime C. Montoya, Maria Rosario Singh-Vergeire and Cynthia P. Saloma |
| EPI_ISL_2188701 | Research Institute for Tropical Medicine, Inc. (RITM) | Philippine Genome Center | Francis A. Tablizo, Kenneth M. Kim, Maria Sofia L. Yangzon, Elcid Aaron R. Pangilinan, Renato Jacinto Q. Mantaring, Benedict A. Maralit, Marc Edsel C. Ayes, Eva Maria Cutiongco-de la Paz, Alethea R. de Guzman, Jan Michael C. Yap, Jo-Hannah S. Llames, Sheila Mae M. Araiza, Kris P. Punayan, Irish Coleen A. Asin, Candice Francheska B. Tambaoan, Asia Louisa U. Chong, Karol Sophia Agape R. Padilla, Rianna Patricia S. Cruz, Carlo M. Lapid, El King D. Morado, Joshua Gregor A. Dizon, Razel Nikka M. Hao, Arianne A. Zamora, Devon Ray Pacial, Juan Antonio R. Magalang, Marissa Alejandria, Celia Carlos, Anna Ong-Lim, Edsel Maurice Salvaña, John Q. Wong, Jaime C. Montoya, Maria Rosario Singh-Vergeire and Cynthia P. Saloma |
| EPI_ISL_2188705 | The Medical City -Â Ortigas | Philippine Genome Center | Francis A. Tablizo, Kenneth M. Kim, Maria Sofia L. Yangzon, Elcid Aaron R. Pangilinan, Renato Jacinto Q. Mantaring, Benedict A. Maralit, Marc Edsel C. Ayes, Eva Maria Cutiongco-de la Paz, Alethea R. de Guzman, Jan Michael C. Yap, Jo-Hannah S. Llames, Sheila Mae M. Araiza, Kris P. Punayan, Irish Coleen A. Asin, Candice Francheska B. Tambaoan, Asia Louisa U. Chong, Karol Sophia Agape R. Padilla, Rianna Patricia S. Cruz, Carlo M. Lapid, El King D. Morado, Joshua Gregor A. Dizon, Razel Nikka M. Hao, Arianne A. Zamora, Devon Ray Pacial, Juan Antonio R. Magalang, Marissa Alejandria, Celia Carlos, Anna Ong-Lim, Edsel Maurice Salvaña, John Q. Wong, Jaime C. Montoya, Maria Rosario Singh-Vergeire and Cynthia P. Saloma |
| EPI_ISL_2188706 | The Lord's Grace Medical and Industrial Clinic | Philippine Genome Center | Francis A. Tablizo, Kenneth M. Kim, Maria Sofia L. Yangzon, Elcid Aaron R. Pangilinan, Renato Jacinto Q. Mantaring, Benedict A. Maralit, Marc Edsel C. Ayes, Eva Maria Cutiongco-de la Paz, Alethea R. de Guzman, Jan Michael C. Yap, Jo-Hannah S. Llames, Sheila Mae M. Araiza, Kris P. Punayan, Irish Coleen A. Asin, Candice Francheska B. Tambaoan, Asia Louisa U. Chong, Karol Sophia Agape R. Padilla, Rianna Patricia S. Cruz, Carlo M. Lapid, El King D. Morado, Joshua Gregor A. Dizon, Razel Nikka M. Hao, Arianne A. Zamora, Devon Ray Pacial, Juan Antonio R. Magalang, Marissa Alejandria, Celia Carlos, Anna Ong-Lim, Edsel Maurice Salvaña, John Q. Wong, Jaime C. Montoya, Maria Rosario Singh-Vergeire and Cynthia P. Saloma |
| EPI_ISL_2188713, EPI_ISL_2188714 | The Medical City -Â Ortigas | Philippine Genome Center | Francis A. Tablizo, Kenneth M. Kim, Maria Sofia L. Yangzon, Elcid Aaron R. Pangilinan, Renato Jacinto Q. Mantaring, Benedict A. Maralit, Marc Edsel C. Ayes, Eva Maria Cutiongco-de la Paz, Alethea R. de Guzman, Jan Michael C. Yap, Jo-Hannah S. Llames, Sheila Mae M. Araiza, Kris P. Punayan, Irish Coleen A. Asin, Candice Francheska B. Tambaoan, Asia Louisa U. Chong, Karol Sophia Agape R. Padilla, Rianna Patricia S. Cruz, Carlo M. Lapid, El King D. Morado, Joshua Gregor A. Dizon, Razel Nikka M. Hao, Arianne A. Zamora, Devon Ray Pacial, Juan Antonio R. Magalang, Marissa Alejandria, Celia Carlos, Anna Ong-Lim, Edsel Maurice Salvaña, John Q. Wong, Jaime C. Montoya, Maria Rosario Singh-Vergeire and Cynthia P. Saloma |
| EPI_ISL_2188716 | Mariano Marcos Memorial Hospital Medical Center | Philippine Genome Center | Francis A. Tablizo, Kenneth M. Kim, Maria Sofia L. Yangzon, Elcid Aaron R. Pangilinan, Renato Jacinto Q. Mantaring, Benedict A. Maralit, Marc Edsel C. Ayes, Eva Maria Cutiongco-de la Paz, Alethea R. de Guzman, Jan Michael C. Yap, Jo-Hannah S. Llames, Sheila Mae M. Araiza, Kris P. Punayan, Irish Coleen A. Asin, Candice Francheska B. Tambaoan, Asia Louisa U. Chong, Karol Sophia Agape R. Padilla, Rianna Patricia S. Cruz, Carlo M. Lapid, El King D. Morado, Joshua Gregor A. Dizon, Razel Nikka M. Hao, Arianne A. Zamora, Devon Ray Pacial, Juan Antonio R. Magalang, Marissa Alejandria, Celia Carlos, Anna Ong-Lim, Edsel Maurice Salvaña, John Q. Wong, Jaime C. Montoya, Maria Rosario Singh-Vergeire and Cynthia P. Saloma |
| EPI_ISL_2188720 | Cagayan Valley Medical Center Molecular Laboratory | Philippine Genome Center | Francis A. Tablizo, Kenneth M. Kim, Maria Sofia L. Yangzon, Elcid Aaron R. Pangilinan, Renato Jacinto Q. Mantaring, Benedict A. Maralit, Marc Edsel C. Ayes, Eva Maria Cutiongco-de la Paz, Alethea R. de Guzman, Jan Michael C. Yap, Jo-Hannah S. Llames, Sheila Mae M. Araiza, Kris P. Punayan, Irish Coleen A. Asin, Candice Francheska B. Tambaoan, Asia Louisa U. Chong, Karol Sophia Agape R. Padilla, Rianna Patricia S. Cruz, Carlo M. Lapid, El King D. Morado, Joshua Gregor A. Dizon, Razel Nikka M. Hao, Arianne A. Zamora, Devon Ray Pacial, Juan Antonio R. Magalang, Marissa Alejandria, Celia Carlos, Anna Ong-Lim, Edsel Maurice Salvaña, John Q. Wong, Jaime C. Montoya, Maria Rosario Singh-Vergeire and Cynthia P. Saloma |
| EPI_ISL_2188722 | The Lord's Grace Medical and Industrial Clinic | Philippine Genome Center | Francis A. Tablizo, Kenneth M. Kim, Maria Sofia L. Yangzon, Elcid Aaron R. Pangilinan, Renato Jacinto Q. Mantaring, Benedict A. Maralit, Marc Edsel C. Ayes, Eva Maria Cutiongco-de la Paz, Alethea R. de Guzman, Jan Michael C. Yap, Jo-Hannah S. Llames, Sheila Mae M. Araiza, Kris P. Punayan, Irish Coleen A. Asin, Candice Francheska B. Tambaoan, Asia Louisa U. Chong, Karol Sophia Agape R. Padilla, Rianna Patricia S. Cruz, Carlo M. Lapid, El King D. Morado, Joshua Gregor A. Dizon, Razel Nikka M. Hao, Arianne A. Zamora, Devon Ray Pacial, Juan Antonio R. Magalang, Marissa Alejandria, Celia Carlos, Anna Ong-Lim, Edsel Maurice Salvaña, John Q. Wong, Jaime C. Montoya, Maria Rosario Singh-Vergeire and Cynthia P. Saloma |
| EPI_ISL_2188724 | Baguio General Hospital Medical Center (BGHMC) | Philippine Genome Center | Francis A. Tablizo, Kenneth M. Kim, Maria Sofia L. Yangzon, Elcid Aaron R. Pangilinan, Renato Jacinto Q. Mantaring, Benedict A. Maralit, Marc Edsel C. |

| | | | |
|---|---|---|---|
| | | | Ayes, Eva Maria Cutiongco-de la Paz, Alethea R. de Guzman, Jan Michael C. Yap, Jo-Hannah S. Llames, Sheila Mae M. Araiza, Kris P. Punayan, Irish Coleen A. Asin, Candice Francheska B. Tambaoan, Asia Louisa U. Chong, Karol Sophia Agape R. Padilla, Rianna Patricia S. Cruz, Carlo M. Lapid, El King D. Morado, Joshua Gregor A. Dizon, Razel Nikka M. Hao, Arianne A. Zamora, Devon Ray Pacial, Juan Antonio R. Magalang, Marissa Alejandria, Celia Carlos, Anna Ong-Lim, Edsel Maurice Salvaña, John Q. Wong, Jaime C. Montoya, Maria Rosario Singh-Vergeire and Cynthia P. Saloma |
| EPI_ISL_2188725 | The Lord's Grace Medical and Industrial Clinic | Philippine Genome Center | Francis A. Tablizo, Kenneth M. Kim, Maria Sofia L. Yangzon, Elcid Aaron R. Pangilinan, Renato Jacinto Q. Mantaring, Benedict A. Maralit, Marc Edsel C. Ayes, Eva Maria Cutiongco-de la Paz, Alethea R. de Guzman, Jan Michael C. Yap, Jo-Hannah S. Llames, Sheila Mae M. Araiza, Kris P. Punayan, Irish Coleen A. Asin, Candice Francheska B. Tambaoan, Asia Louisa U. Chong, Karol Sophia Agape R. Padilla, Rianna Patricia S. Cruz, Carlo M. Lapid, El King D. Morado, Joshua Gregor A. Dizon, Razel Nikka M. Hao, Arianne A. Zamora, Devon Ray Pacial, Juan Antonio R. Magalang, Marissa Alejandria, Celia Carlos, Anna Ong-Lim, Edsel Maurice Salvaña, John Q. Wong, Jaime C. Montoya, Maria Rosario Singh-Vergeire and Cynthia P. Saloma |
| EPI_ISL_2188726 | Philippine Red Cross Logistics and Multipurpose Center | Philippine Genome Center | Francis A. Tablizo, Kenneth M. Kim, Maria Sofia L. Yangzon, Elcid Aaron R. Pangilinan, Renato Jacinto Q. Mantaring, Benedict A. Maralit, Marc Edsel C. Ayes, Eva Maria Cutiongco-de la Paz, Alethea R. de Guzman, Jan Michael C. Yap, Jo-Hannah S. Llames, Sheila Mae M. Araiza, Kris P. Punayan, Irish Coleen A. Asin, Candice Francheska B. Tambaoan, Asia Louisa U. Chong, Karol Sophia Agape R. Padilla, Rianna Patricia S. Cruz, Carlo M. Lapid, El King D. Morado, Joshua Gregor A. Dizon, Razel Nikka M. Hao, Arianne A. Zamora, Devon Ray Pacial, Juan Antonio R. Magalang, Marissa Alejandria, Celia Carlos, Anna Ong-Lim, Edsel Maurice Salvaña, John Q. Wong, Jaime C. Montoya, Maria Rosario Singh-Vergeire and Cynthia P. Saloma |
| EPI_ISL_2188727 | The Medical City -Â Ortigas | Philippine Genome Center | Francis A. Tablizo, Kenneth M. Kim, Maria Sofia L. Yangzon, Elcid Aaron R. Pangilinan, Renato Jacinto Q. Mantaring, Benedict A. Maralit, Marc Edsel C. Ayes, Eva Maria Cutiongco-de la Paz, Alethea R. de Guzman, Jan Michael C. Yap, Jo-Hannah S. Llames, Sheila Mae M. Araiza, Kris P. Punayan, Irish Coleen A. Asin, Candice Francheska B. Tambaoan, Asia Louisa U. Chong, Karol Sophia Agape R. Padilla, Rianna Patricia S. Cruz, Carlo M. Lapid, El King D. Morado, Joshua Gregor A. Dizon, Razel Nikka M. Hao, Arianne A. Zamora, Devon Ray Pacial, Juan Antonio R. Magalang, Marissa Alejandria, Celia Carlos, Anna Ong-Lim, Edsel Maurice Salvaña, John Q. Wong, Jaime C. Montoya, Maria Rosario Singh-Vergeire and Cynthia P. Saloma |
| EPI_ISL_2188728 | Cotabato Regional and Medical Center | Philippine Genome Center | Francis A. Tablizo, Kenneth M. Kim, Maria Sofia L. Yangzon, Elcid Aaron R. Pangilinan, Renato Jacinto Q. Mantaring, Benedict A. Maralit, Marc Edsel C. Ayes, Eva Maria Cutiongco-de la Paz, Alethea R. de Guzman, Jan Michael C. Yap, Jo-Hannah S. Llames, Sheila Mae M. Araiza, Kris P. Punayan, Irish Coleen A. Asin, Candice Francheska B. Tambaoan, Asia Louisa U. Chong, Karol Sophia Agape R. Padilla, Rianna Patricia S. Cruz, Carlo M. Lapid, El King D. Morado, Joshua Gregor A. Dizon, Razel Nikka M. Hao, Arianne A. Zamora, Devon Ray Pacial, Juan Antonio R. Magalang, Marissa Alejandria, Celia Carlos, Anna Ong-Lim, Edsel Maurice Salvaña, John Q. Wong, Jaime C. Montoya, Maria Rosario Singh-Vergeire and Cynthia P. Saloma |
| EPI_ISL_2188729 | Detoxicare Molecular Diagnostics Laboratory | Philippine Genome Center | Francis A. Tablizo, Kenneth M. Kim, Maria Sofia L. Yangzon, Elcid Aaron R. Pangilinan, Renato Jacinto Q. Mantaring, Benedict A. Maralit, Marc Edsel C. Ayes, Eva Maria Cutiongco-de la Paz, Alethea R. de Guzman, Jan Michael C. Yap, Jo-Hannah S. Llames, Sheila Mae M. Araiza, Kris P. Punayan, Irish Coleen A. Asin, Candice Francheska B. Tambaoan, Asia Louisa U. Chong, Karol Sophia Agape R. Padilla, Rianna Patricia S. Cruz, Carlo M. Lapid, El King D. Morado, Joshua Gregor A. Dizon, Razel Nikka M. Hao, Arianne A. Zamora, Devon Ray Pacial, Juan Antonio R. Magalang, Marissa Alejandria, Celia Carlos, Anna Ong-Lim, Edsel Maurice Salvaña, John Q. Wong, Jaime C. Montoya, Maria Rosario Singh-Vergeire and Cynthia P. Saloma |
| EPI_ISL_2188730, EPI_ISL_2188731 | The Medical City -Â Ortigas | Philippine Genome Center | Francis A. Tablizo, Kenneth M. Kim, Maria Sofia L. Yangzon, Elcid Aaron R. Pangilinan, Renato Jacinto Q. Mantaring, Benedict A. Maralit, Marc Edsel C. Ayes, Eva Maria Cutiongco-de la Paz, Alethea R. de Guzman, Jan Michael C. Yap, Jo-Hannah S. Llames, Sheila Mae M. Araiza, Kris P. Punayan, Irish Coleen A. Asin, Candice Francheska B. Tambaoan, Asia Louisa U. Chong, Karol Sophia Agape R. Padilla, Rianna Patricia S. Cruz, Carlo M. Lapid, El King D. Morado, Joshua Gregor A. Dizon, Razel Nikka M. Hao, Arianne A. Zamora, Devon Ray Pacial, Juan Antonio R. Magalang, Marissa Alejandria, Celia Carlos, Anna Ong-Lim, Edsel Maurice Salvaña, John Q. Wong, Jaime C. Montoya, Maria Rosario Singh-Vergeire and Cynthia P. Saloma |
| EPI_ISL_2188732 | The Lord's Grace Medical and Industrial Clinic | Philippine Genome Center | Francis A. Tablizo, Kenneth M. Kim, Maria Sofia L. Yangzon, Elcid Aaron R. Pangilinan, Renato Jacinto Q. Mantaring, Benedict A. Maralit, Marc Edsel C. Ayes, Eva Maria Cutiongco-de la Paz, Alethea R. de Guzman, Jan Michael C. Yap, Jo-Hannah S. Llames, Sheila Mae M. Araiza, Kris P. Punayan, Irish Coleen A. Asin, Candice Francheska B. Tambaoan, Asia Louisa U. Chong, Karol Sophia Agape R. Padilla, Rianna Patricia S. Cruz, Carlo M. Lapid, El King D. Morado, Joshua Gregor A. Dizon, Razel Nikka M. Hao, Arianne A. Zamora, Devon Ray Pacial, Juan Antonio R. Magalang, Marissa Alejandria, Celia Carlos, Anna Ong-Lim, Edsel Maurice Salvaña, John Q. Wong, Jaime C. Montoya, Maria Rosario Singh-Vergeire and Cynthia P. Saloma |
| EPI_ISL_2188735 | Mary Mediatrix Medical Center | Philippine Genome Center | Francis A. Tablizo, Kenneth M. Kim, Maria Sofia L. Yangzon, Elcid Aaron R. Pangilinan, Renato Jacinto Q. Mantaring, Benedict A. Maralit, Marc Edsel C. Ayes, Eva Maria Cutiongco-de la Paz, Alethea R. de Guzman, Jan Michael C. Yap, Jo-Hannah S. Llames, Sheila Mae M. Araiza, Kris P. Punayan, Irish Coleen A. Asin, Candice Francheska B. Tambaoan, Asia Louisa U. Chong, Karol Sophia Agape R. Padilla, Rianna Patricia S. Cruz, Carlo M. Lapid, El King D. Morado, Joshua Gregor A. Dizon, Razel Nikka M. Hao, Arianne A. Zamora, Devon Ray Pacial, Juan Antonio R. Magalang, Marissa Alejandria, Celia Carlos, Anna Ong-Lim, Edsel Maurice Salvaña, John Q. Wong, Jaime C. Montoya, Maria Rosario Singh-Vergeire and Cynthia P. Saloma |
| EPI_ISL_2188736 | Philippine Red Cross - Port Area | Philippine Genome Center | Francis A. Tablizo, Kenneth M. Kim, Maria Sofia L. Yangzon, Elcid Aaron R. Pangilinan, Renato Jacinto Q. Mantaring, Benedict A. Maralit, Marc Edsel C. Ayes, Eva Maria Cutiongco-de la Paz, Alethea R. de Guzman, Jan Michael C. Yap, Jo-Hannah S. Llames, Sheila Mae M. Araiza, Kris P. Punayan, Irish Coleen A. Asin, Candice Francheska B. Tambaoan, Asia Louisa U. Chong, Karol Sophia Agape R. Padilla, Rianna Patricia S. Cruz, Carlo M. Lapid, El King D. Morado, Joshua Gregor A. Dizon, Razel Nikka M. Hao, Arianne A. Zamora, Devon Ray Pacial, Juan Antonio R. Magalang, Marissa Alejandria, Celia Carlos, Anna Ong-Lim, Edsel Maurice Salvaña, John Q. Wong, Jaime C. Montoya, Maria Rosario Singh-Vergeire and Cynthia P. Saloma |
| EPI_ISL_2188737 | The Lord's Grace Medical and Industrial Clinic | Philippine Genome Center | Francis A. Tablizo, Kenneth M. Kim, Maria Sofia L. Yangzon, Elcid Aaron R. Pangilinan, Renato Jacinto Q. Mantaring, Benedict A. Maralit, Marc Edsel C. Ayes, Eva Maria Cutiongco-de la Paz, Alethea R. de Guzman, Jan Michael C. Yap, Jo-Hannah S. Llames, Sheila Mae M. Araiza, Kris P. Punayan, Irish Coleen A. Asin, Candice Francheska B. Tambaoan, Asia Louisa U. Chong, Karol Sophia Agape R. Padilla, Rianna Patricia S. Cruz, Carlo M. Lapid, El King D. Morado, Joshua Gregor A. Dizon, Razel Nikka M. Hao, Arianne A. Zamora, Devon Ray Pacial, Juan Antonio R. Magalang, Marissa Alejandria, Celia Carlos, Anna Ong-Lim, Edsel Maurice Salvaña, John Q. Wong, Jaime C. Montoya, Maria Rosario Singh-Vergeire and Cynthia P. Saloma |
| EPI_ISL_2188739 | Philippine Red Cross- Batangas Chapter Molecular Laboratory | Philippine Genome Center | Francis A. Tablizo, Kenneth M. Kim, Maria Sofia L. Yangzon, Elcid Aaron R. Pangilinan, Renato Jacinto Q. Mantaring, Benedict A. Maralit, Marc Edsel C. Ayes, Eva Maria Cutiongco-de la Paz, Alethea R. de Guzman, Jan Michael C. Yap, Jo-Hannah S. Llames, Sheila Mae M. Araiza, Kris P. Punayan, Irish Coleen A. Asin, Candice Francheska B. Tambaoan, Asia Louisa U. Chong, Karol Sophia Agape R. Padilla, Rianna Patricia S. Cruz, Carlo M. Lapid, El King D. Morado, Joshua Gregor A. Dizon, Razel Nikka M. Hao, Arianne A. Zamora, Devon Ray Pacial, Juan Antonio R. Magalang, Marissa Alejandria, Celia Carlos, Anna Ong-Lim, Edsel Maurice Salvaña, John Q. Wong, Jaime C. Montoya, Maria Rosario Singh-Vergeire and Cynthia P. Saloma |
| EPI_ISL_2188745 | Ilocos Training and Regional Medical Center | Philippine Genome Center | Francis A. Tablizo, Kenneth M. Kim, Maria Sofia L. Yangzon, Elcid Aaron R. Pangilinan, Renato Jacinto Q. Mantaring, Benedict A. Maralit, Marc Edsel C. Ayes, Eva Maria Cutiongco-de la Paz, Alethea R. de Guzman, Jan Michael C. Yap, Jo-Hannah S. Llames, Sheila Mae M. Araiza, Kris P. Punayan, Irish Coleen A. Asin, Candice Francheska B. Tambaoan, Asia Louisa U. Chong, Karol Sophia Agape R. Padilla, Rianna Patricia S. Cruz, Carlo M. Lapid, El King D. Morado, Joshua Gregor A. Dizon, Razel Nikka M. Hao, Arianne A. Zamora, Devon Ray Pacial, Juan Antonio R. Magalang, Marissa Alejandria, Celia Carlos, Anna Ong-Lim, Edsel Maurice Salvaña, John Q. Wong, Jaime C. Montoya, Maria Rosario Singh-Vergeire and Cynthia P. Saloma |
| EPI_ISL_2188746 | The Medical City -Â Ortigas | Philippine Genome Center | Francis A. Tablizo, Kenneth M. Kim, Maria Sofia L. Yangzon, Elcid Aaron R. Pangilinan, Renato Jacinto Q. Mantaring, Benedict A. Maralit, Marc Edsel C. Ayes, Eva Maria Cutiongco-de la Paz, Alethea R. de Guzman, Jan Michael C. Yap, Jo-Hannah S. Llames, Sheila Mae M. Araiza, Kris P. Punayan, Irish Coleen A. Asin, Candice Francheska B. Tambaoan, Asia Louisa U. Chong, Karol Sophia Agape R. Padilla, Rianna Patricia S. Cruz, Carlo M. Lapid, El King D. Morado, Joshua Gregor A. Dizon, Razel Nikka M. Hao, Arianne A. Zamora, Devon Ray Pacial, Juan Antonio R. Magalang, Marissa Alejandria, Celia Carlos, Anna Ong-Lim, Edsel Maurice Salvaña, John Q. Wong, Jaime C. Montoya, Maria Rosario Singh-Vergeire and Cynthia P. Saloma |
| EPI_ISL_2188748 | Ospital ng Imus | Philippine Genome Center | Francis A. Tablizo, Kenneth M. Kim, Maria Sofia L. Yangzon, Elcid Aaron R. Pangilinan, Renato Jacinto Q. Mantaring, Benedict A. Maralit, Marc Edsel C. Ayes, Eva Maria Cutiongco-de la Paz, Alethea R. de Guzman, Jan Michael C. Yap, Jo-Hannah S. Llames, Sheila Mae M. Araiza, Kris P. Punayan, Irish Coleen A. Asin, Candice Francheska B. Tambaoan, Asia Louisa U. Chong, Karol Sophia Agape R. Padilla, Rianna Patricia S. Cruz, Carlo M. Lapid, El King D. Morado, Joshua Gregor A. Dizon, Razel Nikka M. Hao, Arianne A. Zamora, Devon Ray Pacial, Juan Antonio R. Magalang, Marissa Alejandria, Celia Carlos, Anna Ong-Lim, Edsel Maurice Salvaña, John Q. Wong, Jaime C. Montoya, Maria Rosario Singh-Vergeire and Cynthia P. Saloma |
| EPI_ISL_2188750 | Research Institute for Tropical Medicine, Inc. (RITM) | Philippine Genome Center | Francis A. Tablizo, Kenneth M. Kim, Maria Sofia L. Yangzon, Elcid Aaron R. Pangilinan, Renato Jacinto Q. Mantaring, Benedict A. Maralit, Marc Edsel C. Ayes, Eva Maria Cutiongco-de la Paz, Alethea R. de Guzman, Jan Michael C. Yap, Jo-Hannah S. Llames, Sheila Mae M. Araiza, Kris P. Punayan, Irish Coleen A. Asin, Candice Francheska B. Tambaoan, Asia Louisa U. Chong, Karol Sophia Agape R. Padilla, Rianna Patricia S. Cruz, Carlo M. Lapid, El King D. Morado, Joshua Gregor A. Dizon, Razel Nikka M. Hao, Arianne A. Zamora, Devon Ray Pacial, Juan Antonio R. Magalang, Marissa Alejandria, Celia Carlos, Anna Ong-Lim, Edsel Maurice Salvaña, John Q. Wong, Jaime C. Montoya, Maria Rosario Singh-Vergeire and Cynthia P. Saloma |

| | | | |
|---|---|---|---|
| EPI_ISL_2188751 | Philippine Red Cross - National Blood Center | Philippine Genome Center | Francis A. Tablizo, Kenneth M. Kim, Maria Sofia L. Yangzon, Elcid Aaron R. Pangilinan, Renato Jacinto Q. Mantaring, Benedict A. Maralit, Marc Edsel C. Ayes, Eva Maria Cutiongco-de la Paz, Alethea R. de Guzman, Jan Michael C. Yap, Jo-Hannah S. Llames, Sheila Mae M. Araiza, Kris P. Punayan, Irish Coleen A. Asin, Candice Francheska B. Tambaoan, Asia Louisa U. Chong, Karol Sophia Agape R. Padilla, Rianna Patricia S. Cruz, Carlo M. Lapid, El King D. Morado, Joshua Gregor A. Dizon, Razel Nikka M. Hao, Arianne A. Zamora, Devon Ray Pacial, Juan Antonio R. Magalang, Marissa Alejandria, Celia Carlos, Anna Ong-Lim, Edsel Maurice Salvaña, John Q. Wong, Jaime C. Montoya, Maria Rosario Singh-Vergeire and Cynthia P. Saloma |
| EPI_ISL_2188753 | National Kidney and Transplant Institute (NKTI) | Philippine Genome Center | Francis A. Tablizo, Kenneth M. Kim, Maria Sofia L. Yangzon, Elcid Aaron R. Pangilinan, Renato Jacinto Q. Mantaring, Benedict A. Maralit, Marc Edsel C. Ayes, Eva Maria Cutiongco-de la Paz, Alethea R. de Guzman, Jan Michael C. Yap, Jo-Hannah S. Llames, Sheila Mae M. Araiza, Kris P. Punayan, Irish Coleen A. Asin, Candice Francheska B. Tambaoan, Asia Louisa U. Chong, Karol Sophia Agape R. Padilla, Rianna Patricia S. Cruz, Carlo M. Lapid, El King D. Morado, Joshua Gregor A. Dizon, Razel Nikka M. Hao, Arianne A. Zamora, Devon Ray Pacial, Juan Antonio R. Magalang, Marissa Alejandria, Celia Carlos, Anna Ong-Lim, Edsel Maurice Salvaña, John Q. Wong, Jaime C. Montoya, Maria Rosario Singh-Vergeire and Cynthia P. Saloma |
| EPI_ISL_2188754 | Qualimed Hospital Sta. Rosa | Philippine Genome Center | Francis A. Tablizo, Kenneth M. Kim, Maria Sofia L. Yangzon, Elcid Aaron R. Pangilinan, Renato Jacinto Q. Mantaring, Benedict A. Maralit, Marc Edsel C. Ayes, Eva Maria Cutiongco-de la Paz, Alethea R. de Guzman, Jan Michael C. Yap, Jo-Hannah S. Llames, Sheila Mae M. Araiza, Kris P. Punayan, Irish Coleen A. Asin, Candice Francheska B. Tambaoan, Asia Louisa U. Chong, Karol Sophia Agape R. Padilla, Rianna Patricia S. Cruz, Carlo M. Lapid, El King D. Morado, Joshua Gregor A. Dizon, Razel Nikka M. Hao, Arianne A. Zamora, Devon Ray Pacial, Juan Antonio R. Magalang, Marissa Alejandria, Celia Carlos, Anna Ong-Lim, Edsel Maurice Salvaña, John Q. Wong, Jaime C. Montoya, Maria Rosario Singh-Vergeire and Cynthia P. Saloma |
| EPI_ISL_2188758 | Philippine Red Cross - Port Area | Philippine Genome Center | Francis A. Tablizo, Kenneth M. Kim, Maria Sofia L. Yangzon, Elcid Aaron R. Pangilinan, Renato Jacinto Q. Mantaring, Benedict A. Maralit, Marc Edsel C. Ayes, Eva Maria Cutiongco-de la Paz, Alethea R. de Guzman, Jan Michael C. Yap, Jo-Hannah S. Llames, Sheila Mae M. Araiza, Kris P. Punayan, Irish Coleen A. Asin, Candice Francheska B. Tambaoan, Asia Louisa U. Chong, Karol Sophia Agape R. Padilla, Rianna Patricia S. Cruz, Carlo M. Lapid, El King D. Morado, Joshua Gregor A. Dizon, Razel Nikka M. Hao, Arianne A. Zamora, Devon Ray Pacial, Juan Antonio R. Magalang, Marissa Alejandria, Celia Carlos, Anna Ong-Lim, Edsel Maurice Salvaña, John Q. Wong, Jaime C. Montoya, Maria Rosario Singh-Vergeire and Cynthia P. Saloma |
| EPI_ISL_2188759 | Philippine Red Cross Logistics and Multipurpose Center | Philippine Genome Center | Francis A. Tablizo, Kenneth M. Kim, Maria Sofia L. Yangzon, Elcid Aaron R. Pangilinan, Renato Jacinto Q. Mantaring, Benedict A. Maralit, Marc Edsel C. Ayes, Eva Maria Cutiongco-de la Paz, Alethea R. de Guzman, Jan Michael C. Yap, Jo-Hannah S. Llames, Sheila Mae M. Araiza, Kris P. Punayan, Irish Coleen A. Asin, Candice Francheska B. Tambaoan, Asia Louisa U. Chong, Karol Sophia Agape R. Padilla, Rianna Patricia S. Cruz, Carlo M. Lapid, El King D. Morado, Joshua Gregor A. Dizon, Razel Nikka M. Hao, Arianne A. Zamora, Devon Ray Pacial, Juan Antonio R. Magalang, Marissa Alejandria, Celia Carlos, Anna Ong-Lim, Edsel Maurice Salvaña, John Q. Wong, Jaime C. Montoya, Maria Rosario Singh-Vergeire and Cynthia P. Saloma |
| EPI_ISL_2188763 | Philippine Red Cross - Port Area | Philippine Genome Center | Francis A. Tablizo, Kenneth M. Kim, Maria Sofia L. Yangzon, Elcid Aaron R. Pangilinan, Renato Jacinto Q. Mantaring, Benedict A. Maralit, Marc Edsel C. Ayes, Eva Maria Cutiongco-de la Paz, Alethea R. de Guzman, Jan Michael C. Yap, Jo-Hannah S. Llames, Sheila Mae M. Araiza, Kris P. Punayan, Irish Coleen A. Asin, Candice Francheska B. Tambaoan, Asia Louisa U. Chong, Karol Sophia Agape R. Padilla, Rianna Patricia S. Cruz, Carlo M. Lapid, El King D. Morado, Joshua Gregor A. Dizon, Razel Nikka M. Hao, Arianne A. Zamora, Devon Ray Pacial, Juan Antonio R. Magalang, Marissa Alejandria, Celia Carlos, Anna Ong-Lim, Edsel Maurice Salvaña, John Q. Wong, Jaime C. Montoya, Maria Rosario Singh-Vergeire and Cynthia P. Saloma |
| EPI_ISL_2188764 | Mary Mediatrix Medical Center | Philippine Genome Center | Francis A. Tablizo, Kenneth M. Kim, Maria Sofia L. Yangzon, Elcid Aaron R. Pangilinan, Renato Jacinto Q. Mantaring, Benedict A. Maralit, Marc Edsel C. Ayes, Eva Maria Cutiongco-de la Paz, Alethea R. de Guzman, Jan Michael C. Yap, Jo-Hannah S. Llames, Sheila Mae M. Araiza, Kris P. Punayan, Irish Coleen A. Asin, Candice Francheska B. Tambaoan, Asia Louisa U. Chong, Karol Sophia Agape R. Padilla, Rianna Patricia S. Cruz, Carlo M. Lapid, El King D. Morado, Joshua Gregor A. Dizon, Razel Nikka M. Hao, Arianne A. Zamora, Devon Ray Pacial, Juan Antonio R. Magalang, Marissa Alejandria, Celia Carlos, Anna Ong-Lim, Edsel Maurice Salvaña, John Q. Wong, Jaime C. Montoya, Maria Rosario Singh-Vergeire and Cynthia P. Saloma |
| EPI_ISL_2188765 | The Medical City -Â Ortigas | Philippine Genome Center | Francis A. Tablizo, Kenneth M. Kim, Maria Sofia L. Yangzon, Elcid Aaron R. Pangilinan, Renato Jacinto Q. Mantaring, Benedict A. Maralit, Marc Edsel C. Ayes, Eva Maria Cutiongco-de la Paz, Alethea R. de Guzman, Jan Michael C. Yap, Jo-Hannah S. Llames, Sheila Mae M. Araiza, Kris P. Punayan, Irish Coleen A. Asin, Candice Francheska B. Tambaoan, Asia Louisa U. Chong, Karol Sophia Agape R. Padilla, Rianna Patricia S. Cruz, Carlo M. Lapid, El King D. Morado, Joshua Gregor A. Dizon, Razel Nikka M. Hao, Arianne A. Zamora, Devon Ray Pacial, Juan Antonio R. Magalang, Marissa Alejandria, Celia Carlos, Anna Ong-Lim, Edsel Maurice Salvaña, John Q. Wong, Jaime C. Montoya, Maria Rosario Singh-Vergeire and Cynthia P. Saloma |
| EPI_ISL_2188770 | Philippine Red Cross - Port Area | Philippine Genome Center | Francis A. Tablizo, Kenneth M. Kim, Maria Sofia L. Yangzon, Elcid Aaron R. Pangilinan, Renato Jacinto Q. Mantaring, Benedict A. Maralit, Marc Edsel C. Ayes, Eva Maria Cutiongco-de la Paz, Alethea R. de Guzman, Jan Michael C. Yap, Jo-Hannah S. Llames, Sheila Mae M. Araiza, Kris P. Punayan, Irish Coleen A. Asin, Candice Francheska B. Tambaoan, Asia Louisa U. Chong, Karol Sophia Agape R. Padilla, Rianna Patricia S. Cruz, Carlo M. Lapid, El King D. Morado, Joshua Gregor A. Dizon, Razel Nikka M. Hao, Arianne A. Zamora, Devon Ray Pacial, Juan Antonio R. Magalang, Marissa Alejandria, Celia Carlos, Anna Ong-Lim, Edsel Maurice Salvaña, John Q. Wong, Jaime C. Montoya, Maria Rosario Singh-Vergeire and Cynthia P. Saloma |
| EPI_ISL_2188771 | Mariano Marcos Memorial Hospital Medical Center | Philippine Genome Center | Francis A. Tablizo, Kenneth M. Kim, Maria Sofia L. Yangzon, Elcid Aaron R. Pangilinan, Renato Jacinto Q. Mantaring, Benedict A. Maralit, Marc Edsel C. Ayes, Eva Maria Cutiongco-de la Paz, Alethea R. de Guzman, Jan Michael C. Yap, Jo-Hannah S. Llames, Sheila Mae M. Araiza, Kris P. Punayan, Irish Coleen A. Asin, Candice Francheska B. Tambaoan, Asia Louisa U. Chong, Karol Sophia Agape R. Padilla, Rianna Patricia S. Cruz, Carlo M. Lapid, El King D. Morado, Joshua Gregor A. Dizon, Razel Nikka M. Hao, Arianne A. Zamora, Devon Ray Pacial, Juan Antonio R. Magalang, Marissa Alejandria, Celia Carlos, Anna Ong-Lim, Edsel Maurice Salvaña, John Q. Wong, Jaime C. Montoya, Maria Rosario Singh-Vergeire and Cynthia P. Saloma |
| EPI_ISL_2188772 | First Aide Diagnostic Center | Philippine Genome Center | Francis A. Tablizo, Kenneth M. Kim, Maria Sofia L. Yangzon, Elcid Aaron R. Pangilinan, Renato Jacinto Q. Mantaring, Benedict A. Maralit, Marc Edsel C. Ayes, Eva Maria Cutiongco-de la Paz, Alethea R. de Guzman, Jan Michael C. Yap, Jo-Hannah S. Llames, Sheila Mae M. Araiza, Kris P. Punayan, Irish Coleen A. Asin, Candice Francheska B. Tambaoan, Asia Louisa U. Chong, Karol Sophia Agape R. Padilla, Rianna Patricia S. Cruz, Carlo M. Lapid, El King D. Morado, Joshua Gregor A. Dizon, Razel Nikka M. Hao, Arianne A. Zamora, Devon Ray Pacial, Juan Antonio R. Magalang, Marissa Alejandria, Celia Carlos, Anna Ong-Lim, Edsel Maurice Salvaña, John Q. Wong, Jaime C. Montoya, Maria Rosario Singh-Vergeire and Cynthia P. Saloma |
| EPI_ISL_2188774 | The Lord's Grace Medical and Industrial Clinic | Philippine Genome Center | Francis A. Tablizo, Kenneth M. Kim, Maria Sofia L. Yangzon, Elcid Aaron R. Pangilinan, Renato Jacinto Q. Mantaring, Benedict A. Maralit, Marc Edsel C. Ayes, Eva Maria Cutiongco-de la Paz, Alethea R. de Guzman, Jan Michael C. Yap, Jo-Hannah S. Llames, Sheila Mae M. Araiza, Kris P. Punayan, Irish Coleen A. Asin, Candice Francheska B. Tambaoan, Asia Louisa U. Chong, Karol Sophia Agape R. Padilla, Rianna Patricia S. Cruz, Carlo M. Lapid, El King D. Morado, Joshua Gregor A. Dizon, Razel Nikka M. Hao, Arianne A. Zamora, Devon Ray Pacial, Juan Antonio R. Magalang, Marissa Alejandria, Celia Carlos, Anna Ong-Lim, Edsel Maurice Salvaña, John Q. Wong, Jaime C. Montoya, Maria Rosario Singh-Vergeire and Cynthia P. Saloma |
| EPI_ISL_2188775 | Cotabato Regional and Medical Center | Philippine Genome Center | Francis A. Tablizo, Kenneth M. Kim, Maria Sofia L. Yangzon, Elcid Aaron R. Pangilinan, Renato Jacinto Q. Mantaring, Benedict A. Maralit, Marc Edsel C. Ayes, Eva Maria Cutiongco-de la Paz, Alethea R. de Guzman, Jan Michael C. Yap, Jo-Hannah S. Llames, Sheila Mae M. Araiza, Kris P. Punayan, Irish Coleen A. Asin, Candice Francheska B. Tambaoan, Asia Louisa U. Chong, Karol Sophia Agape R. Padilla, Rianna Patricia S. Cruz, Carlo M. Lapid, El King D. Morado, Joshua Gregor A. Dizon, Razel Nikka M. Hao, Arianne A. Zamora, Devon Ray Pacial, Juan Antonio R. Magalang, Marissa Alejandria, Celia Carlos, Anna Ong-Lim, Edsel Maurice Salvaña, John Q. Wong, Jaime C. Montoya, Maria Rosario Singh-Vergeire and Cynthia P. Saloma |
| EPI_ISL_2188777 | The Medical City -Â Ortigas | Philippine Genome Center | Francis A. Tablizo, Kenneth M. Kim, Maria Sofia L. Yangzon, Elcid Aaron R. Pangilinan, Renato Jacinto Q. Mantaring, Benedict A. Maralit, Marc Edsel C. Ayes, Eva Maria Cutiongco-de la Paz, Alethea R. de Guzman, Jan Michael C. Yap, Jo-Hannah S. Llames, Sheila Mae M. Araiza, Kris P. Punayan, Irish Coleen A. Asin, Candice Francheska B. Tambaoan, Asia Louisa U. Chong, Karol Sophia Agape R. Padilla, Rianna Patricia S. Cruz, Carlo M. Lapid, El King D. Morado, Joshua Gregor A. Dizon, Razel Nikka M. Hao, Arianne A. Zamora, Devon Ray Pacial, Juan Antonio R. Magalang, Marissa Alejandria, Celia Carlos, Anna Ong-Lim, Edsel Maurice Salvaña, John Q. Wong, Jaime C. Montoya, Maria Rosario Singh-Vergeire and Cynthia P. Saloma |
| EPI_ISL_2188778 | Detoxicare Molecular Diagnostics Laboratory | Philippine Genome Center | Francis A. Tablizo, Kenneth M. Kim, Maria Sofia L. Yangzon, Elcid Aaron R. Pangilinan, Renato Jacinto Q. Mantaring, Benedict A. Maralit, Marc Edsel C. Ayes, Eva Maria Cutiongco-de la Paz, Alethea R. de Guzman, Jan Michael C. Yap, Jo-Hannah S. Llames, Sheila Mae M. Araiza, Kris P. Punayan, Irish Coleen A. Asin, Candice Francheska B. Tambaoan, Asia Louisa U. Chong, Karol Sophia Agape R. Padilla, Rianna Patricia S. Cruz, Carlo M. Lapid, El King D. Morado, Joshua Gregor A. Dizon, Razel Nikka M. Hao, Arianne A. Zamora, Devon Ray Pacial, Juan Antonio R. Magalang, Marissa Alejandria, Celia Carlos, Anna Ong-Lim, Edsel Maurice Salvaña, John Q. Wong, Jaime C. Montoya, Maria Rosario Singh-Vergeire and Cynthia P. Saloma |
| EPI_ISL_2188782 | Research Institute for Tropical Medicine, Inc. (RITM) | Philippine Genome Center | Francis A. Tablizo, Kenneth M. Kim, Maria Sofia L. Yangzon, Elcid Aaron R. Pangilinan, Renato Jacinto Q. Mantaring, Benedict A. Maralit, Marc Edsel C. Ayes, Eva Maria Cutiongco-de la Paz, Alethea R. de Guzman, Jan Michael C. Yap, Jo-Hannah S. Llames, Sheila Mae M. Araiza, Kris P. Punayan, Irish Coleen A. Asin, Candice Francheska B. Tambaoan, Asia Louisa U. Chong, Karol Sophia Agape R. Padilla, Rianna Patricia S. Cruz, Carlo M. Lapid, El King D. Morado, Joshua Gregor A. Dizon, Razel Nikka M. Hao, Arianne A. Zamora, Devon Ray Pacial, Juan Antonio R. Magalang, Marissa Alejandria, Celia Carlos, Anna Ong-Lim, Edsel Maurice Salvaña, John Q. Wong, Jaime C. Montoya, Maria Rosario Singh-Vergeire and Cynthia P. Saloma |

| | | | |
|---|---|---|---|
| EPI_ISL_2188784 | Qualimed Hospital Sta. Rosa | Philippine Genome Center | Francis A. Tablizo, Kenneth M. Kim, Maria Sofia L. Yangzon, Elcid Aaron R. Pangilinan, Renato Jacinto Q. Mantaring, Benedict A. Maralit, Marc Edsel C. Ayes, Eva Maria Cutiongco-de la Paz, Alethea R. de Guzman, Jan Michael C. Yap, Jo-Hannah S. Llames, Sheila Mae M. Araiza, Kris P. Punayan, Irish Coleen A. Asin, Candice Francheska B. Tambaoan, Asia Louisa U. Chong, Karol Sophia Agape R. Padilla, Rianna Patricia S. Cruz, Carlo M. Lapid, El King D. Morado, Joshua Gregor A. Dizon, Razel Nikka M. Hao, Arianne A. Zamora, Devon Ray Pacial, Juan Antonio R. Magalang, Marissa Alejandria, Celia Carlos, Anna Ong-Lim, Edsel Maurice Salvaña, John Q. Wong, Jaime C. Montoya, Maria Rosario Singh-Vergeire and Cynthia P. Saloma |
| EPI_ISL_2188786 | Parkway Medical and Diagnostic Center | Philippine Genome Center | Francis A. Tablizo, Kenneth M. Kim, Maria Sofia L. Yangzon, Elcid Aaron R. Pangilinan, Renato Jacinto Q. Mantaring, Benedict A. Maralit, Marc Edsel C. Ayes, Eva Maria Cutiongco-de la Paz, Alethea R. de Guzman, Jan Michael C. Yap, Jo-Hannah S. Llames, Sheila Mae M. Araiza, Kris P. Punayan, Irish Coleen A. Asin, Candice Francheska B. Tambaoan, Asia Louisa U. Chong, Karol Sophia Agape R. Padilla, Rianna Patricia S. Cruz, Carlo M. Lapid, El King D. Morado, Joshua Gregor A. Dizon, Razel Nikka M. Hao, Arianne A. Zamora, Devon Ray Pacial, Juan Antonio R. Magalang, Marissa Alejandria, Celia Carlos, Anna Ong-Lim, Edsel Maurice Salvaña, John Q. Wong, Jaime C. Montoya, Maria Rosario Singh-Vergeire and Cynthia P. Saloma |
| EPI_ISL_2188788 | Philippine Red Cross - National Blood Center | Philippine Genome Center | Francis A. Tablizo, Kenneth M. Kim, Maria Sofia L. Yangzon, Elcid Aaron R. Pangilinan, Renato Jacinto Q. Mantaring, Benedict A. Maralit, Marc Edsel C. Ayes, Eva Maria Cutiongco-de la Paz, Alethea R. de Guzman, Jan Michael C. Yap, Jo-Hannah S. Llames, Sheila Mae M. Araiza, Kris P. Punayan, Irish Coleen A. Asin, Candice Francheska B. Tambaoan, Asia Louisa U. Chong, Karol Sophia Agape R. Padilla, Rianna Patricia S. Cruz, Carlo M. Lapid, El King D. Morado, Joshua Gregor A. Dizon, Razel Nikka M. Hao, Arianne A. Zamora, Devon Ray Pacial, Juan Antonio R. Magalang, Marissa Alejandria, Celia Carlos, Anna Ong-Lim, Edsel Maurice Salvaña, John Q. Wong, Jaime C. Montoya, Maria Rosario Singh-Vergeire and Cynthia P. Saloma |
| EPI_ISL_2188789 | Mary Mediatrix Medical Center | Philippine Genome Center | Francis A. Tablizo, Kenneth M. Kim, Maria Sofia L. Yangzon, Elcid Aaron R. Pangilinan, Renato Jacinto Q. Mantaring, Benedict A. Maralit, Marc Edsel C. Ayes, Eva Maria Cutiongco-de la Paz, Alethea R. de Guzman, Jan Michael C. Yap, Jo-Hannah S. Llames, Sheila Mae M. Araiza, Kris P. Punayan, Irish Coleen A. Asin, Candice Francheska B. Tambaoan, Asia Louisa U. Chong, Karol Sophia Agape R. Padilla, Rianna Patricia S. Cruz, Carlo M. Lapid, El King D. Morado, Joshua Gregor A. Dizon, Razel Nikka M. Hao, Arianne A. Zamora, Devon Ray Pacial, Juan Antonio R. Magalang, Marissa Alejandria, Celia Carlos, Anna Ong-Lim, Edsel Maurice Salvaña, John Q. Wong, Jaime C. Montoya, Maria Rosario Singh-Vergeire and Cynthia P. Saloma |
| EPI_ISL_2188790 | The Medical City -Â Ortigas | Philippine Genome Center | Francis A. Tablizo, Kenneth M. Kim, Maria Sofia L. Yangzon, Elcid Aaron R. Pangilinan, Renato Jacinto Q. Mantaring, Benedict A. Maralit, Marc Edsel C. Ayes, Eva Maria Cutiongco-de la Paz, Alethea R. de Guzman, Jan Michael C. Yap, Jo-Hannah S. Llames, Sheila Mae M. Araiza, Kris P. Punayan, Irish Coleen A. Asin, Candice Francheska B. Tambaoan, Asia Louisa U. Chong, Karol Sophia Agape R. Padilla, Rianna Patricia S. Cruz, Carlo M. Lapid, El King D. Morado, Joshua Gregor A. Dizon, Razel Nikka M. Hao, Arianne A. Zamora, Devon Ray Pacial, Juan Antonio R. Magalang, Marissa Alejandria, Celia Carlos, Anna Ong-Lim, Edsel Maurice Salvaña, John Q. Wong, Jaime C. Montoya, Maria Rosario Singh-Vergeire and Cynthia P. Saloma |
| EPI_ISL_2188791 | The Lord's Grace Medical and Industrial Clinic | Philippine Genome Center | Francis A. Tablizo, Kenneth M. Kim, Maria Sofia L. Yangzon, Elcid Aaron R. Pangilinan, Renato Jacinto Q. Mantaring, Benedict A. Maralit, Marc Edsel C. Ayes, Eva Maria Cutiongco-de la Paz, Alethea R. de Guzman, Jan Michael C. Yap, Jo-Hannah S. Llames, Sheila Mae M. Araiza, Kris P. Punayan, Irish Coleen A. Asin, Candice Francheska B. Tambaoan, Asia Louisa U. Chong, Karol Sophia Agape R. Padilla, Rianna Patricia S. Cruz, Carlo M. Lapid, El King D. Morado, Joshua Gregor A. Dizon, Razel Nikka M. Hao, Arianne A. Zamora, Devon Ray Pacial, Juan Antonio R. Magalang, Marissa Alejandria, Celia Carlos, Anna Ong-Lim, Edsel Maurice Salvaña, John Q. Wong, Jaime C. Montoya, Maria Rosario Singh-Vergeire and Cynthia P. Saloma |
| EPI_ISL_2188792 | The Medical City -Â Ortigas | Philippine Genome Center | Francis A. Tablizo, Kenneth M. Kim, Maria Sofia L. Yangzon, Elcid Aaron R. Pangilinan, Renato Jacinto Q. Mantaring, Benedict A. Maralit, Marc Edsel C. Ayes, Eva Maria Cutiongco-de la Paz, Alethea R. de Guzman, Jan Michael C. Yap, Jo-Hannah S. Llames, Sheila Mae M. Araiza, Kris P. Punayan, Irish Coleen A. Asin, Candice Francheska B. Tambaoan, Asia Louisa U. Chong, Karol Sophia Agape R. Padilla, Rianna Patricia S. Cruz, Carlo M. Lapid, El King D. Morado, Joshua Gregor A. Dizon, Razel Nikka M. Hao, Arianne A. Zamora, Devon Ray Pacial, Juan Antonio R. Magalang, Marissa Alejandria, Celia Carlos, Anna Ong-Lim, Edsel Maurice Salvaña, John Q. Wong, Jaime C. Montoya, Maria Rosario Singh-Vergeire and Cynthia P. Saloma |
| EPI_ISL_2188794 | Philippine Red Cross Logistics and Multipurpose Center | Philippine Genome Center | Francis A. Tablizo, Kenneth M. Kim, Maria Sofia L. Yangzon, Elcid Aaron R. Pangilinan, Renato Jacinto Q. Mantaring, Benedict A. Maralit, Marc Edsel C. Ayes, Eva Maria Cutiongco-de la Paz, Alethea R. de Guzman, Jan Michael C. Yap, Jo-Hannah S. Llames, Sheila Mae M. Araiza, Kris P. Punayan, Irish Coleen A. Asin, Candice Francheska B. Tambaoan, Asia Louisa U. Chong, Karol Sophia Agape R. Padilla, Rianna Patricia S. Cruz, Carlo M. Lapid, El King D. Morado, Joshua Gregor A. Dizon, Razel Nikka M. Hao, Arianne A. Zamora, Devon Ray Pacial, Juan Antonio R. Magalang, Marissa Alejandria, Celia Carlos, Anna Ong-Lim, Edsel Maurice Salvaña, John Q. Wong, Jaime C. Montoya, Maria Rosario Singh-Vergeire and Cynthia P. Saloma |
| EPI_ISL_2188798 | Philippine Red Cross - Port Area | Philippine Genome Center | Francis A. Tablizo, Kenneth M. Kim, Maria Sofia L. Yangzon, Elcid Aaron R. Pangilinan, Renato Jacinto Q. Mantaring, Benedict A. Maralit, Marc Edsel C. Ayes, Eva Maria Cutiongco-de la Paz, Alethea R. de Guzman, Jan Michael C. Yap, Jo-Hannah S. Llames, Sheila Mae M. Araiza, Kris P. Punayan, Irish Coleen A. Asin, Candice Francheska B. Tambaoan, Asia Louisa U. Chong, Karol Sophia Agape R. Padilla, Rianna Patricia S. Cruz, Carlo M. Lapid, El King D. Morado, Joshua Gregor A. Dizon, Razel Nikka M. Hao, Arianne A. Zamora, Devon Ray Pacial, Juan Antonio R. Magalang, Marissa Alejandria, Celia Carlos, Anna Ong-Lim, Edsel Maurice Salvaña, John Q. Wong, Jaime C. Montoya, Maria Rosario Singh-Vergeire and Cynthia P. Saloma |
| EPI_ISL_2188799 | Prime Care Alpha Covid-19 Testing Laboratory | Philippine Genome Center | Francis A. Tablizo, Kenneth M. Kim, Maria Sofia L. Yangzon, Elcid Aaron R. Pangilinan, Renato Jacinto Q. Mantaring, Benedict A. Maralit, Marc Edsel C. Ayes, Eva Maria Cutiongco-de la Paz, Alethea R. de Guzman, Jan Michael C. Yap, Jo-Hannah S. Llames, Sheila Mae M. Araiza, Kris P. Punayan, Irish Coleen A. Asin, Candice Francheska B. Tambaoan, Asia Louisa U. Chong, Karol Sophia Agape R. Padilla, Rianna Patricia S. Cruz, Carlo M. Lapid, El King D. Morado, Joshua Gregor A. Dizon, Razel Nikka M. Hao, Arianne A. Zamora, Devon Ray Pacial, Juan Antonio R. Magalang, Marissa Alejandria, Celia Carlos, Anna Ong-Lim, Edsel Maurice Salvaña, John Q. Wong, Jaime C. Montoya, Maria Rosario Singh-Vergeire and Cynthia P. Saloma |
| EPI_ISL_2188802 | Philippine Red Cross Logistics and Multipurpose Center | Philippine Genome Center | Francis A. Tablizo, Kenneth M. Kim, Maria Sofia L. Yangzon, Elcid Aaron R. Pangilinan, Renato Jacinto Q. Mantaring, Benedict A. Maralit, Marc Edsel C. Ayes, Eva Maria Cutiongco-de la Paz, Alethea R. de Guzman, Jan Michael C. Yap, Jo-Hannah S. Llames, Sheila Mae M. Araiza, Kris P. Punayan, Irish Coleen A. Asin, Candice Francheska B. Tambaoan, Asia Louisa U. Chong, Karol Sophia Agape R. Padilla, Rianna Patricia S. Cruz, Carlo M. Lapid, El King D. Morado, Joshua Gregor A. Dizon, Razel Nikka M. Hao, Arianne A. Zamora, Devon Ray Pacial, Juan Antonio R. Magalang, Marissa Alejandria, Celia Carlos, Anna Ong-Lim, Edsel Maurice Salvaña, John Q. Wong, Jaime C. Montoya, Maria Rosario Singh-Vergeire and Cynthia P. Saloma |
| EPI_ISL_2188803 | Philippine Red Cross - Port Area | Philippine Genome Center | Francis A. Tablizo, Kenneth M. Kim, Maria Sofia L. Yangzon, Elcid Aaron R. Pangilinan, Renato Jacinto Q. Mantaring, Benedict A. Maralit, Marc Edsel C. Ayes, Eva Maria Cutiongco-de la Paz, Alethea R. de Guzman, Jan Michael C. Yap, Jo-Hannah S. Llames, Sheila Mae M. Araiza, Kris P. Punayan, Irish Coleen A. Asin, Candice Francheska B. Tambaoan, Asia Louisa U. Chong, Karol Sophia Agape R. Padilla, Rianna Patricia S. Cruz, Carlo M. Lapid, El King D. Morado, Joshua Gregor A. Dizon, Razel Nikka M. Hao, Arianne A. Zamora, Devon Ray Pacial, Juan Antonio R. Magalang, Marissa Alejandria, Celia Carlos, Anna Ong-Lim, Edsel Maurice Salvaña, John Q. Wong, Jaime C. Montoya, Maria Rosario Singh-Vergeire and Cynthia P. Saloma |
| EPI_ISL_2188804 | PASIG CITY CHILDREN'S HOSPITAL | Philippine Genome Center | Francis A. Tablizo, Kenneth M. Kim, Maria Sofia L. Yangzon, Elcid Aaron R. Pangilinan, Renato Jacinto Q. Mantaring, Benedict A. Maralit, Marc Edsel C. Ayes, Eva Maria Cutiongco-de la Paz, Alethea R. de Guzman, Jan Michael C. Yap, Jo-Hannah S. Llames, Sheila Mae M. Araiza, Kris P. Punayan, Irish Coleen A. Asin, Candice Francheska B. Tambaoan, Asia Louisa U. Chong, Karol Sophia Agape R. Padilla, Rianna Patricia S. Cruz, Carlo M. Lapid, El King D. Morado, Joshua Gregor A. Dizon, Razel Nikka M. Hao, Arianne A. Zamora, Devon Ray Pacial, Juan Antonio R. Magalang, Marissa Alejandria, Celia Carlos, Anna Ong-Lim, Edsel Maurice Salvaña, John Q. Wong, Jaime C. Montoya, Maria Rosario Singh-Vergeire and Cynthia P. Saloma |
| EPI_ISL_2188806 | Philippine Red Cross- Batangas Chapter Molecular Laboratory | Philippine Genome Center | Francis A. Tablizo, Kenneth M. Kim, Maria Sofia L. Yangzon, Elcid Aaron R. Pangilinan, Renato Jacinto Q. Mantaring, Benedict A. Maralit, Marc Edsel C. Ayes, Eva Maria Cutiongco-de la Paz, Alethea R. de Guzman, Jan Michael C. Yap, Jo-Hannah S. Llames, Sheila Mae M. Araiza, Kris P. Punayan, Irish Coleen A. Asin, Candice Francheska B. Tambaoan, Asia Louisa U. Chong, Karol Sophia Agape R. Padilla, Rianna Patricia S. Cruz, Carlo M. Lapid, El King D. Morado, Joshua Gregor A. Dizon, Razel Nikka M. Hao, Arianne A. Zamora, Devon Ray Pacial, Juan Antonio R. Magalang, Marissa Alejandria, Celia Carlos, Anna Ong-Lim, Edsel Maurice Salvaña, John Q. Wong, Jaime C. Montoya, Maria Rosario Singh-Vergeire and Cynthia P. Saloma |
| EPI_ISL_2188808 | The Lord's Grace Medical and Industrial Clinic | Philippine Genome Center | Francis A. Tablizo, Kenneth M. Kim, Maria Sofia L. Yangzon, Elcid Aaron R. Pangilinan, Renato Jacinto Q. Mantaring, Benedict A. Maralit, Marc Edsel C. Ayes, Eva Maria Cutiongco-de la Paz, Alethea R. de Guzman, Jan Michael C. Yap, Jo-Hannah S. Llames, Sheila Mae M. Araiza, Kris P. Punayan, Irish Coleen A. Asin, Candice Francheska B. Tambaoan, Asia Louisa U. Chong, Karol Sophia Agape R. Padilla, Rianna Patricia S. Cruz, Carlo M. Lapid, El King D. Morado, Joshua Gregor A. Dizon, Razel Nikka M. Hao, Arianne A. Zamora, Devon Ray Pacial, Juan Antonio R. Magalang, Marissa Alejandria, Celia Carlos, Anna Ong-Lim, Edsel Maurice Salvaña, John Q. Wong, Jaime C. Montoya, Maria Rosario Singh-Vergeire and Cynthia P. Saloma |
| EPI_ISL_2188810 | The Medical City -Â Ortigas | Philippine Genome Center | Francis A. Tablizo, Kenneth M. Kim, Maria Sofia L. Yangzon, Elcid Aaron R. Pangilinan, Renato Jacinto Q. Mantaring, Benedict A. Maralit, Marc Edsel C. Ayes, Eva Maria Cutiongco-de la Paz, Alethea R. de Guzman, Jan Michael C. Yap, Jo-Hannah S. Llames, Sheila Mae M. Araiza, Kris P. Punayan, Irish Coleen A. Asin, Candice Francheska B. Tambaoan, Asia Louisa U. Chong, Karol Sophia Agape R. Padilla, Rianna Patricia S. Cruz, Carlo M. Lapid, El King D. Morado, Joshua Gregor A. Dizon, Razel Nikka M. Hao, Arianne A. Zamora, Devon Ray Pacial, Juan Antonio R. Magalang, Marissa Alejandria, Celia Carlos, Anna Ong-Lim, Edsel Maurice Salvaña, John Q. Wong, Jaime C. Montoya, Maria Rosario Singh-Vergeire and Cynthia P. Saloma |

| | | | |
|---|---|---|---|
| EPI_ISL_2188811 | Philippine Red Cross - Port Area | Philippine Genome Center | Francis A. Tablizo, Kenneth M. Kim, Maria Sofia L. Yangzon, Elcid Aaron R. Pangilinan, Renato Jacinto Q. Mantaring, Benedict A. Maralit, Marc Edsel C. Ayes, Eva Maria Cutiongco-de la Paz, Alethea R. de Guzman, Jan Michael C. Yap, Jo-Hannah S. Llames, Sheila Mae M. Araiza, Kris P. Punayan, Irish Coleen A. Asin, Candice Francheska B. Tambaoan, Asia Louisa U. Chong, Karol Sophia Agape R. Padilla, Rianna Patricia S. Cruz, Carlo M. Lapid, El King D. Morado, Joshua Gregor A. Dizon, Razel Nikka M. Hao, Arianne A. Zamora, Devon Ray Pacial, Juan Antonio R. Magalang, Marissa Alejandria, Celia Carlos, Anna Ong-Lim, Edsel Maurice Salvaña, John Q. Wong, Jaime C. Montoya, Maria Rosario Singh-Vergeire and Cynthia P. Saloma |
| EPI_ISL_2188813 | Mary Mediatrix Medical Center | Philippine Genome Center | Francis A. Tablizo, Kenneth M. Kim, Maria Sofia L. Yangzon, Elcid Aaron R. Pangilinan, Renato Jacinto Q. Mantaring, Benedict A. Maralit, Marc Edsel C. Ayes, Eva Maria Cutiongco-de la Paz, Alethea R. de Guzman, Jan Michael C. Yap, Jo-Hannah S. Llames, Sheila Mae M. Araiza, Kris P. Punayan, Irish Coleen A. Asin, Candice Francheska B. Tambaoan, Asia Louisa U. Chong, Karol Sophia Agape R. Padilla, Rianna Patricia S. Cruz, Carlo M. Lapid, El King D. Morado, Joshua Gregor A. Dizon, Razel Nikka M. Hao, Arianne A. Zamora, Devon Ray Pacial, Juan Antonio R. Magalang, Marissa Alejandria, Celia Carlos, Anna Ong-Lim, Edsel Maurice Salvaña, John Q. Wong, Jaime C. Montoya, Maria Rosario Singh-Vergeire and Cynthia P. Saloma |
| EPI_ISL_2188814 | The Medical City -Â Ortigas | Philippine Genome Center | Francis A. Tablizo, Kenneth M. Kim, Maria Sofia L. Yangzon, Elcid Aaron R. Pangilinan, Renato Jacinto Q. Mantaring, Benedict A. Maralit, Marc Edsel C. Ayes, Eva Maria Cutiongco-de la Paz, Alethea R. de Guzman, Jan Michael C. Yap, Jo-Hannah S. Llames, Sheila Mae M. Araiza, Kris P. Punayan, Irish Coleen A. Asin, Candice Francheska B. Tambaoan, Asia Louisa U. Chong, Karol Sophia Agape R. Padilla, Rianna Patricia S. Cruz, Carlo M. Lapid, El King D. Morado, Joshua Gregor A. Dizon, Razel Nikka M. Hao, Arianne A. Zamora, Devon Ray Pacial, Juan Antonio R. Magalang, Marissa Alejandria, Celia Carlos, Anna Ong-Lim, Edsel Maurice Salvaña, John Q. Wong, Jaime C. Montoya, Maria Rosario Singh-Vergeire and Cynthia P. Saloma |
| EPI_ISL_2188815 | Ospital ng Imus | Philippine Genome Center | Francis A. Tablizo, Kenneth M. Kim, Maria Sofia L. Yangzon, Elcid Aaron R. Pangilinan, Renato Jacinto Q. Mantaring, Benedict A. Maralit, Marc Edsel C. Ayes, Eva Maria Cutiongco-de la Paz, Alethea R. de Guzman, Jan Michael C. Yap, Jo-Hannah S. Llames, Sheila Mae M. Araiza, Kris P. Punayan, Irish Coleen A. Asin, Candice Francheska B. Tambaoan, Asia Louisa U. Chong, Karol Sophia Agape R. Padilla, Rianna Patricia S. Cruz, Carlo M. Lapid, El King D. Morado, Joshua Gregor A. Dizon, Razel Nikka M. Hao, Arianne A. Zamora, Devon Ray Pacial, Juan Antonio R. Magalang, Marissa Alejandria, Celia Carlos, Anna Ong-Lim, Edsel Maurice Salvaña, John Q. Wong, Jaime C. Montoya, Maria Rosario Singh-Vergeire and Cynthia P. Saloma |
| EPI_ISL_2188818 | Prime Care Alpha Covid-19 Testing Laboratory | Philippine Genome Center | Francis A. Tablizo, Kenneth M. Kim, Maria Sofia L. Yangzon, Elcid Aaron R. Pangilinan, Renato Jacinto Q. Mantaring, Benedict A. Maralit, Marc Edsel C. Ayes, Eva Maria Cutiongco-de la Paz, Alethea R. de Guzman, Jan Michael C. Yap, Jo-Hannah S. Llames, Sheila Mae M. Araiza, Kris P. Punayan, Irish Coleen A. Asin, Candice Francheska B. Tambaoan, Asia Louisa U. Chong, Karol Sophia Agape R. Padilla, Rianna Patricia S. Cruz, Carlo M. Lapid, El King D. Morado, Joshua Gregor A. Dizon, Razel Nikka M. Hao, Arianne A. Zamora, Devon Ray Pacial, Juan Antonio R. Magalang, Marissa Alejandria, Celia Carlos, Anna Ong-Lim, Edsel Maurice Salvaña, John Q. Wong, Jaime C. Montoya, Maria Rosario Singh-Vergeire and Cynthia P. Saloma |
| EPI_ISL_2188821 | The Medical City -Â Ortigas | Philippine Genome Center | Francis A. Tablizo, Kenneth M. Kim, Maria Sofia L. Yangzon, Elcid Aaron R. Pangilinan, Renato Jacinto Q. Mantaring, Benedict A. Maralit, Marc Edsel C. Ayes, Eva Maria Cutiongco-de la Paz, Alethea R. de Guzman, Jan Michael C. Yap, Jo-Hannah S. Llames, Sheila Mae M. Araiza, Kris P. Punayan, Irish Coleen A. Asin, Candice Francheska B. Tambaoan, Asia Louisa U. Chong, Karol Sophia Agape R. Padilla, Rianna Patricia S. Cruz, Carlo M. Lapid, El King D. Morado, Joshua Gregor A. Dizon, Razel Nikka M. Hao, Arianne A. Zamora, Devon Ray Pacial, Juan Antonio R. Magalang, Marissa Alejandria, Celia Carlos, Anna Ong-Lim, Edsel Maurice Salvaña, John Q. Wong, Jaime C. Montoya, Maria Rosario Singh-Vergeire and Cynthia P. Saloma |
| EPI_ISL_2188824 | Philippine Red Cross - National Blood Center | Philippine Genome Center | Francis A. Tablizo, Kenneth M. Kim, Maria Sofia L. Yangzon, Elcid Aaron R. Pangilinan, Renato Jacinto Q. Mantaring, Benedict A. Maralit, Marc Edsel C. Ayes, Eva Maria Cutiongco-de la Paz, Alethea R. de Guzman, Jan Michael C. Yap, Jo-Hannah S. Llames, Sheila Mae M. Araiza, Kris P. Punayan, Irish Coleen A. Asin, Candice Francheska B. Tambaoan, Asia Louisa U. Chong, Karol Sophia Agape R. Padilla, Rianna Patricia S. Cruz, Carlo M. Lapid, El King D. Morado, Joshua Gregor A. Dizon, Razel Nikka M. Hao, Arianne A. Zamora, Devon Ray Pacial, Juan Antonio R. Magalang, Marissa Alejandria, Celia Carlos, Anna Ong-Lim, Edsel Maurice Salvaña, John Q. Wong, Jaime C. Montoya, Maria Rosario Singh-Vergeire and Cynthia P. Saloma |
| EPI_ISL_2188828 | The Lord's Grace Medical and Industrial Clinic | Philippine Genome Center | Francis A. Tablizo, Kenneth M. Kim, Maria Sofia L. Yangzon, Elcid Aaron R. Pangilinan, Renato Jacinto Q. Mantaring, Benedict A. Maralit, Marc Edsel C. Ayes, Eva Maria Cutiongco-de la Paz, Alethea R. de Guzman, Jan Michael C. Yap, Jo-Hannah S. Llames, Sheila Mae M. Araiza, Kris P. Punayan, Irish Coleen A. Asin, Candice Francheska B. Tambaoan, Asia Louisa U. Chong, Karol Sophia Agape R. Padilla, Rianna Patricia S. Cruz, Carlo M. Lapid, El King D. Morado, Joshua Gregor A. Dizon, Razel Nikka M. Hao, Arianne A. Zamora, Devon Ray Pacial, Juan Antonio R. Magalang, Marissa Alejandria, Celia Carlos, Anna Ong-Lim, Edsel Maurice Salvaña, John Q. Wong, Jaime C. Montoya, Maria Rosario Singh-Vergeire and Cynthia P. Saloma |
| EPI_ISL_2188831, EPI_ISL_2188832 | The Medical City -Â Ortigas | Philippine Genome Center | Francis A. Tablizo, Kenneth M. Kim, Maria Sofia L. Yangzon, Elcid Aaron R. Pangilinan, Renato Jacinto Q. Mantaring, Benedict A. Maralit, Marc Edsel C. Ayes, Eva Maria Cutiongco-de la Paz, Alethea R. de Guzman, Jan Michael C. Yap, Jo-Hannah S. Llames, Sheila Mae M. Araiza, Kris P. Punayan, Irish Coleen A. Asin, Candice Francheska B. Tambaoan, Asia Louisa U. Chong, Karol Sophia Agape R. Padilla, Rianna Patricia S. Cruz, Carlo M. Lapid, El King D. Morado, Joshua Gregor A. Dizon, Razel Nikka M. Hao, Arianne A. Zamora, Devon Ray Pacial, Juan Antonio R. Magalang, Marissa Alejandria, Celia Carlos, Anna Ong-Lim, Edsel Maurice Salvaña, John Q. Wong, Jaime C. Montoya, Maria Rosario Singh-Vergeire and Cynthia P. Saloma |
| EPI_ISL_2188836, EPI_ISL_2188841 | Research Institute for Tropical Medicine, Inc. (RITM) | Philippine Genome Center | Francis A. Tablizo, Kenneth M. Kim, Maria Sofia L. Yangzon, Elcid Aaron R. Pangilinan, Renato Jacinto Q. Mantaring, Benedict A. Maralit, Marc Edsel C. Ayes, Eva Maria Cutiongco-de la Paz, Alethea R. de Guzman, Jan Michael C. Yap, Jo-Hannah S. Llames, Sheila Mae M. Araiza, Kris P. Punayan, Irish Coleen A. Asin, Candice Francheska B. Tambaoan, Asia Louisa U. Chong, Karol Sophia Agape R. Padilla, Rianna Patricia S. Cruz, Carlo M. Lapid, El King D. Morado, Joshua Gregor A. Dizon, Razel Nikka M. Hao, Arianne A. Zamora, Devon Ray Pacial, Juan Antonio R. Magalang, Marissa Alejandria, Celia Carlos, Anna Ong-Lim, Edsel Maurice Salvaña, John Q. Wong, Jaime C. Montoya, Maria Rosario Singh-Vergeire and Cynthia P. Saloma |
| EPI_ISL_2188843 | Philippine Red Cross - Port Area | Philippine Genome Center | Francis A. Tablizo, Kenneth M. Kim, Maria Sofia L. Yangzon, Elcid Aaron R. Pangilinan, Renato Jacinto Q. Mantaring, Benedict A. Maralit, Marc Edsel C. Ayes, Eva Maria Cutiongco-de la Paz, Alethea R. de Guzman, Jan Michael C. Yap, Jo-Hannah S. Llames, Sheila Mae M. Araiza, Kris P. Punayan, Irish Coleen A. Asin, Candice Francheska B. Tambaoan, Asia Louisa U. Chong, Karol Sophia Agape R. Padilla, Rianna Patricia S. Cruz, Carlo M. Lapid, El King D. Morado, Joshua Gregor A. Dizon, Razel Nikka M. Hao, Arianne A. Zamora, Devon Ray Pacial, Juan Antonio R. Magalang, Marissa Alejandria, Celia Carlos, Anna Ong-Lim, Edsel Maurice Salvaña, John Q. Wong, Jaime C. Montoya, Maria Rosario Singh-Vergeire and Cynthia P. Saloma |
| EPI_ISL_2188844 | The Medical City -Â Ortigas | Philippine Genome Center | Francis A. Tablizo, Kenneth M. Kim, Maria Sofia L. Yangzon, Elcid Aaron R. Pangilinan, Renato Jacinto Q. Mantaring, Benedict A. Maralit, Marc Edsel C. Ayes, Eva Maria Cutiongco-de la Paz, Alethea R. de Guzman, Jan Michael C. Yap, Jo-Hannah S. Llames, Sheila Mae M. Araiza, Kris P. Punayan, Irish Coleen A. Asin, Candice Francheska B. Tambaoan, Asia Louisa U. Chong, Karol Sophia Agape R. Padilla, Rianna Patricia S. Cruz, Carlo M. Lapid, El King D. Morado, Joshua Gregor A. Dizon, Razel Nikka M. Hao, Arianne A. Zamora, Devon Ray Pacial, Juan Antonio R. Magalang, Marissa Alejandria, Celia Carlos, Anna Ong-Lim, Edsel Maurice Salvaña, John Q. Wong, Jaime C. Montoya, Maria Rosario Singh-Vergeire and Cynthia P. Saloma |
| EPI_ISL_2188845 | National Kidney and Transplant Institute (NKTI) | Philippine Genome Center | Francis A. Tablizo, Kenneth M. Kim, Maria Sofia L. Yangzon, Elcid Aaron R. Pangilinan, Renato Jacinto Q. Mantaring, Benedict A. Maralit, Marc Edsel C. Ayes, Eva Maria Cutiongco-de la Paz, Alethea R. de Guzman, Jan Michael C. Yap, Jo-Hannah S. Llames, Sheila Mae M. Araiza, Kris P. Punayan, Irish Coleen A. Asin, Candice Francheska B. Tambaoan, Asia Louisa U. Chong, Karol Sophia Agape R. Padilla, Rianna Patricia S. Cruz, Carlo M. Lapid, El King D. Morado, Joshua Gregor A. Dizon, Razel Nikka M. Hao, Arianne A. Zamora, Devon Ray Pacial, Juan Antonio R. Magalang, Marissa Alejandria, Celia Carlos, Anna Ong-Lim, Edsel Maurice Salvaña, John Q. Wong, Jaime C. Montoya, Maria Rosario Singh-Vergeire and Cynthia P. Saloma |
| EPI_ISL_2188846 | Research Institute for Tropical Medicine, Inc. (RITM) | Philippine Genome Center | Francis A. Tablizo, Kenneth M. Kim, Maria Sofia L. Yangzon, Elcid Aaron R. Pangilinan, Renato Jacinto Q. Mantaring, Benedict A. Maralit, Marc Edsel C. Ayes, Eva Maria Cutiongco-de la Paz, Alethea R. de Guzman, Jan Michael C. Yap, Jo-Hannah S. Llames, Sheila Mae M. Araiza, Kris P. Punayan, Irish Coleen A. Asin, Candice Francheska B. Tambaoan, Asia Louisa U. Chong, Karol Sophia Agape R. Padilla, Rianna Patricia S. Cruz, Carlo M. Lapid, El King D. Morado, Joshua Gregor A. Dizon, Razel Nikka M. Hao, Arianne A. Zamora, Devon Ray Pacial, Juan Antonio R. Magalang, Marissa Alejandria, Celia Carlos, Anna Ong-Lim, Edsel Maurice Salvaña, John Q. Wong, Jaime C. Montoya, Maria Rosario Singh-Vergeire and Cynthia P. Saloma |
| EPI_ISL_2188854 | Philippine Red Cross - Port Area | Philippine Genome Center | Francis A. Tablizo, Kenneth M. Kim, Maria Sofia L. Yangzon, Elcid Aaron R. Pangilinan, Renato Jacinto Q. Mantaring, Benedict A. Maralit, Marc Edsel C. Ayes, Eva Maria Cutiongco-de la Paz, Alethea R. de Guzman, Jan Michael C. Yap, Jo-Hannah S. Llames, Sheila Mae M. Araiza, Kris P. Punayan, Irish Coleen A. Asin, Candice Francheska B. Tambaoan, Asia Louisa U. Chong, Karol Sophia Agape R. Padilla, Rianna Patricia S. Cruz, Carlo M. Lapid, El King D. Morado, Joshua Gregor A. Dizon, Razel Nikka M. Hao, Arianne A. Zamora, Devon Ray Pacial, Juan Antonio R. Magalang, Marissa Alejandria, Celia Carlos, Anna Ong-Lim, Edsel Maurice Salvaña, John Q. Wong, Jaime C. Montoya, Maria Rosario Singh-Vergeire and Cynthia P. Saloma |
| EPI_ISL_2188855 | Philippine Red Cross Logistics and Multipurpose Center | Philippine Genome Center | Francis A. Tablizo, Kenneth M. Kim, Maria Sofia L. Yangzon, Elcid Aaron R. Pangilinan, Renato Jacinto Q. Mantaring, Benedict A. Maralit, Marc Edsel C. Ayes, Eva Maria Cutiongco-de la Paz, Alethea R. de Guzman, Jan Michael C. Yap, Jo-Hannah S. Llames, Sheila Mae M. Araiza, Kris P. Punayan, Irish Coleen A. Asin, Candice Francheska B. Tambaoan, Asia Louisa U. Chong, Karol Sophia Agape R. Padilla, Rianna Patricia S. Cruz, Carlo M. Lapid, El King D. Morado, Joshua Gregor A. Dizon, Razel Nikka M. Hao, Arianne A. Zamora, Devon Ray Pacial, Juan Antonio R. Magalang, Marissa Alejandria, Celia Carlos, Anna Ong-Lim, Edsel Maurice Salvaña, John Q. Wong, Jaime C. Montoya, Maria Rosario Singh-Vergeire and Cynthia P. Saloma |

| | | | |
|---|---|---|---|
| EPI_ISL_2188856 | Philippine Red Cross - Port Area | Philippine Genome Center | Francis A. Tablizo, Kenneth M. Kim, Maria Sofia L. Yangzon, Elcid Aaron R. Pangilinan, Renato Jacinto Q. Mantaring, Benedict A. Maralit, Marc Edsel C. Ayes, Eva Maria Cutiongco-de la Paz, Alethea R. de Guzman, Jan Michael C. Yap, Jo-Hannah S. Llames, Sheila Mae M. Araiza, Kris P. Punayan, Irish Coleen A. Asin, Candice Francheska B. Tambaoan, Asia Louisa U. Chong, Karol Sophia Agape R. Padilla, Rianna Patricia S. Cruz, Carlo M. Lapid, El King D. Morado, Joshua Gregor A. Dizon, Razel Nikka M. Hao, Arianne A. Zamora, Devon Ray Pacial, Juan Antonio R. Magalang, Marissa Alejandria, Celia Carlos, Anna Ong-Lim, Edsel Maurice Salvaña, John Q. Wong, Jaime C. Montoya, Maria Rosario Singh-Vergeire and Cynthia P. Saloma |
| EPI_ISL_2188857 | Prime Care Alpha Covid-19 Testing Laboratory | Philippine Genome Center | Francis A. Tablizo, Kenneth M. Kim, Maria Sofia L. Yangzon, Elcid Aaron R. Pangilinan, Renato Jacinto Q. Mantaring, Benedict A. Maralit, Marc Edsel C. Ayes, Eva Maria Cutiongco-de la Paz, Alethea R. de Guzman, Jan Michael C. Yap, Jo-Hannah S. Llames, Sheila Mae M. Araiza, Kris P. Punayan, Irish Coleen A. Asin, Candice Francheska B. Tambaoan, Asia Louisa U. Chong, Karol Sophia Agape R. Padilla, Rianna Patricia S. Cruz, Carlo M. Lapid, El King D. Morado, Joshua Gregor A. Dizon, Razel Nikka M. Hao, Arianne A. Zamora, Devon Ray Pacial, Juan Antonio R. Magalang, Marissa Alejandria, Celia Carlos, Anna Ong-Lim, Edsel Maurice Salvaña, John Q. Wong, Jaime C. Montoya, Maria Rosario Singh-Vergeire and Cynthia P. Saloma |
| EPI_ISL_2188858 | Philippine Red Cross - Port Area | Philippine Genome Center | Francis A. Tablizo, Kenneth M. Kim, Maria Sofia L. Yangzon, Elcid Aaron R. Pangilinan, Renato Jacinto Q. Mantaring, Benedict A. Maralit, Marc Edsel C. Ayes, Eva Maria Cutiongco-de la Paz, Alethea R. de Guzman, Jan Michael C. Yap, Jo-Hannah S. Llames, Sheila Mae M. Araiza, Kris P. Punayan, Irish Coleen A. Asin, Candice Francheska B. Tambaoan, Asia Louisa U. Chong, Karol Sophia Agape R. Padilla, Rianna Patricia S. Cruz, Carlo M. Lapid, El King D. Morado, Joshua Gregor A. Dizon, Razel Nikka M. Hao, Arianne A. Zamora, Devon Ray Pacial, Juan Antonio R. Magalang, Marissa Alejandria, Celia Carlos, Anna Ong-Lim, Edsel Maurice Salvaña, John Q. Wong, Jaime C. Montoya, Maria Rosario Singh-Vergeire and Cynthia P. Saloma |
| EPI_ISL_2188859 | Philippine Red Cross - National Blood Center | Philippine Genome Center | Francis A. Tablizo, Kenneth M. Kim, Maria Sofia L. Yangzon, Elcid Aaron R. Pangilinan, Renato Jacinto Q. Mantaring, Benedict A. Maralit, Marc Edsel C. Ayes, Eva Maria Cutiongco-de la Paz, Alethea R. de Guzman, Jan Michael C. Yap, Jo-Hannah S. Llames, Sheila Mae M. Araiza, Kris P. Punayan, Irish Coleen A. Asin, Candice Francheska B. Tambaoan, Asia Louisa U. Chong, Karol Sophia Agape R. Padilla, Rianna Patricia S. Cruz, Carlo M. Lapid, El King D. Morado, Joshua Gregor A. Dizon, Razel Nikka M. Hao, Arianne A. Zamora, Devon Ray Pacial, Juan Antonio R. Magalang, Marissa Alejandria, Celia Carlos, Anna Ong-Lim, Edsel Maurice Salvaña, John Q. Wong, Jaime C. Montoya, Maria Rosario Singh-Vergeire and Cynthia P. Saloma |
| EPI_ISL_2188861 | The Medical City -Â Ortigas | Philippine Genome Center | Francis A. Tablizo, Kenneth M. Kim, Maria Sofia L. Yangzon, Elcid Aaron R. Pangilinan, Renato Jacinto Q. Mantaring, Benedict A. Maralit, Marc Edsel C. Ayes, Eva Maria Cutiongco-de la Paz, Alethea R. de Guzman, Jan Michael C. Yap, Jo-Hannah S. Llames, Sheila Mae M. Araiza, Kris P. Punayan, Irish Coleen A. Asin, Candice Francheska B. Tambaoan, Asia Louisa U. Chong, Karol Sophia Agape R. Padilla, Rianna Patricia S. Cruz, Carlo M. Lapid, El King D. Morado, Joshua Gregor A. Dizon, Razel Nikka M. Hao, Arianne A. Zamora, Devon Ray Pacial, Juan Antonio R. Magalang, Marissa Alejandria, Celia Carlos, Anna Ong-Lim, Edsel Maurice Salvaña, John Q. Wong, Jaime C. Montoya, Maria Rosario Singh-Vergeire and Cynthia P. Saloma |
| EPI_ISL_2188863 | Philippine Red Cross Logistics and Multipurpose Center | Philippine Genome Center | Francis A. Tablizo, Kenneth M. Kim, Maria Sofia L. Yangzon, Elcid Aaron R. Pangilinan, Renato Jacinto Q. Mantaring, Benedict A. Maralit, Marc Edsel C. Ayes, Eva Maria Cutiongco-de la Paz, Alethea R. de Guzman, Jan Michael C. Yap, Jo-Hannah S. Llames, Sheila Mae M. Araiza, Kris P. Punayan, Irish Coleen A. Asin, Candice Francheska B. Tambaoan, Asia Louisa U. Chong, Karol Sophia Agape R. Padilla, Rianna Patricia S. Cruz, Carlo M. Lapid, El King D. Morado, Joshua Gregor A. Dizon, Razel Nikka M. Hao, Arianne A. Zamora, Devon Ray Pacial, Juan Antonio R. Magalang, Marissa Alejandria, Celia Carlos, Anna Ong-Lim, Edsel Maurice Salvaña, John Q. Wong, Jaime C. Montoya, Maria Rosario Singh-Vergeire and Cynthia P. Saloma |
| EPI_ISL_2188864 | Prime Care Alpha Covid-19 Testing Laboratory | Philippine Genome Center | Francis A. Tablizo, Kenneth M. Kim, Maria Sofia L. Yangzon, Elcid Aaron R. Pangilinan, Renato Jacinto Q. Mantaring, Benedict A. Maralit, Marc Edsel C. Ayes, Eva Maria Cutiongco-de la Paz, Alethea R. de Guzman, Jan Michael C. Yap, Jo-Hannah S. Llames, Sheila Mae M. Araiza, Kris P. Punayan, Irish Coleen A. Asin, Candice Francheska B. Tambaoan, Asia Louisa U. Chong, Karol Sophia Agape R. Padilla, Rianna Patricia S. Cruz, Carlo M. Lapid, El King D. Morado, Joshua Gregor A. Dizon, Razel Nikka M. Hao, Arianne A. Zamora, Devon Ray Pacial, Juan Antonio R. Magalang, Marissa Alejandria, Celia Carlos, Anna Ong-Lim, Edsel Maurice Salvaña, John Q. Wong, Jaime C. Montoya, Maria Rosario Singh-Vergeire and Cynthia P. Saloma |
| EPI_ISL_2188866 | The Medical City -Â Ortigas | Philippine Genome Center | Francis A. Tablizo, Kenneth M. Kim, Maria Sofia L. Yangzon, Elcid Aaron R. Pangilinan, Renato Jacinto Q. Mantaring, Benedict A. Maralit, Marc Edsel C. Ayes, Eva Maria Cutiongco-de la Paz, Alethea R. de Guzman, Jan Michael C. Yap, Jo-Hannah S. Llames, Sheila Mae M. Araiza, Kris P. Punayan, Irish Coleen A. Asin, Candice Francheska B. Tambaoan, Asia Louisa U. Chong, Karol Sophia Agape R. Padilla, Rianna Patricia S. Cruz, Carlo M. Lapid, El King D. Morado, Joshua Gregor A. Dizon, Razel Nikka M. Hao, Arianne A. Zamora, Devon Ray Pacial, Juan Antonio R. Magalang, Marissa Alejandria, Celia Carlos, Anna Ong-Lim, Edsel Maurice Salvaña, John Q. Wong, Jaime C. Montoya, Maria Rosario Singh-Vergeire and Cynthia P. Saloma |
| EPI_ISL_2188894, EPI_ISL_2189018, EPI_ISL_2189030, EPI_ISL_2189068 | The Medical City | Philippine Genome Center | Francis A. Tablizo, Kenneth M. Kim, Maria Sofia L. Yangzon, Elcid Aaron R. Pangilinan, Renato Jacinto Q. Mantaring, Benedict A. Maralit, Marc Edsel C. Ayes, Eva Maria Cutiongco-de la Paz, Alethea R. de Guzman, Jan Michael C. Yap, Jo-Hannah S. Llames, Sheila Mae M. Araiza, Kris P. Punayan, Irish Coleen A. Asin, Candice Francheska B. Tambaoan, Asia Louisa U. Chong, Karol Sophia Agape R. Padilla, Rianna Patricia S. Cruz, Carlo M. Lapid, El King D. Morado, Joshua Gregor A. Dizon, Razel Nikka M. Hao, Arianne A. Zamora, Devon Ray Pacial, Juan Antonio R. Magalang, Marissa Alejandria, Celia Carlos, Anna Ong-Lim, Edsel Maurice Salvaña, John Q. Wong, Jaime C. Montoya, Maria Rosario Singh-Vergeire and Cynthia P. Saloma |
| EPI_ISL_2189075 | Qualimed Hospital Sta. Rosa | Philippine Genome Center | Francis A. Tablizo, Kenneth M. Kim, Maria Sofia L. Yangzon, Elcid Aaron R. Pangilinan, Renato Jacinto Q. Mantaring, Benedict A. Maralit, Marc Edsel C. Ayes, Eva Maria Cutiongco-de la Paz, Alethea R. de Guzman, Jan Michael C. Yap, Jo-Hannah S. Llames, Sheila Mae M. Araiza, Kris P. Punayan, Irish Coleen A. Asin, Candice Francheska B. Tambaoan, Asia Louisa U. Chong, Karol Sophia Agape R. Padilla, Rianna Patricia S. Cruz, Carlo M. Lapid, El King D. Morado, Joshua Gregor A. Dizon, Razel Nikka M. Hao, Arianne A. Zamora, Devon Ray Pacial, Juan Antonio R. Magalang, Marissa Alejandria, Celia Carlos, Anna Ong-Lim, Edsel Maurice Salvaña, John Q. Wong, Jaime C. Montoya, Maria Rosario Singh-Vergeire and Cynthia P. Saloma |
| EPI_ISL_2189096, EPI_ISL_2189374, EPI_ISL_2189446, EPI_ISL_2189488 | The Medical City | Philippine Genome Center | Francis A. Tablizo, Kenneth M. Kim, Maria Sofia L. Yangzon, Elcid Aaron R. Pangilinan, Renato Jacinto Q. Mantaring, Benedict A. Maralit, Marc Edsel C. Ayes, Eva Maria Cutiongco-de la Paz, Alethea R. de Guzman, Jan Michael C. Yap, Jo-Hannah S. Llames, Sheila Mae M. Araiza, Kris P. Punayan, Irish Coleen A. Asin, Candice Francheska B. Tambaoan, Asia Louisa U. Chong, Karol Sophia Agape R. Padilla, Rianna Patricia S. Cruz, Carlo M. Lapid, El King D. Morado, Joshua Gregor A. Dizon, Razel Nikka M. Hao, Arianne A. Zamora, Devon Ray Pacial, Juan Antonio R. Magalang, Marissa Alejandria, Celia Carlos, Anna Ong-Lim, Edsel Maurice Salvaña, John Q. Wong, Jaime C. Montoya, Maria Rosario Singh-Vergeire and Cynthia P. Saloma |
| EPI_ISL_2196185, EPI_ISL_2196187, EPI_ISL_2196190, EPI_ISL_2196191, EPI_ISL_2196192, EPI_ISL_2196193, EPI_ISL_2196194 | Laboratory of Respiratory Viruses and Measles, Oswaldo Cruz Institute, FIOCRUZ | Laboratory of Respiratory Viruses and Measles, Oswaldo Cruz Institute, FIOCRUZ | Paola Resende, Luciana Appolinario, Fernando Motta, Ana Carolina Paixao, Ana Carolina Mendonca, Alice Sampaio Rocha, Taina Venas, Elisa Cavalcante Pereira, Renata Serrano Lopes, Marilda Siqueira on behalf of the Fiocruz COVID-19 Genomic Surveillance Network |
| EPI_ISL_2196214 | Laboratorio Central de Saude Publica do Estado de Sergipe (LACEN/SE) | Laboratory of Respiratory Viruses and Measles, Oswaldo Cruz Institute, FIOCRUZ | Paola Resende, Luciana Appolinario, Fernando Motta, Anna Carolina Paixao, Ana Carolina Mendonca, Alice Sampaio Rocha, Tainá Moreira Martins Venas, Elisa Cavalcante Pereira, Renata Serrano Lopes, Cliomar Alves dos Santos, Marilda Siqueira on behalf of the Fiocruz COVID-19 Genomic Surveillance Network |
| EPI_ISL_2196244 | Lboratorio Central de Saude Publica do Estado do Parana (LACEN/PR) | Laboratory of Respiratory Viruses and Measles, Oswaldo Cruz Institute, FIOCRUZ | Paola Resende, Luciana Appolinario, Fernando Motta, Anna Carolina Paixao, Ana Carolina Mendonca, Alice Sampaio Rocha, Taina Venas, Elisa Cavalcante Pereira, Renata Serrano Lopes, Irina Riediger, Marilda Siqueira on behalf of the Fiocruz COVID-19 Genomic Surveillance Network |
| EPI_ISL_2196246 | Laboratorio Central de Saude Publica do Estado de Minas Gerais (LACEN/MG) | Laboratory of Respiratory Viruses and Measles, Oswaldo Cruz Institute, FIOCRUZ | Paola Resende, Luciana Appolinario, Fernando Motta, Anna Carolina Paixao, Ana Carolina Mendonca, Alice Sampaio Rocha, Taina Venas, Elisa Cavalcante Pereira, Renata Serrano Lopes, Andre Felipe Leal Bernardes, Marilda Siqueira on behalf of the Fiocruz COVID-19 Genomic Surveillance Network |
| EPI_ISL_2196254, EPI_ISL_2196255, EPI_ISL_2196256 | Labortorio Central de Saude Publica do Estado de Santa Catarina (LACEN/SC) | Laboratory of Respiratory Viruses and Measles, Oswaldo Cruz Institute, FIOCRUZ | Paola Resende, Luciana Appolinario, Fernando Motta, Anna Carolina Paixao, Ana Carolina Mendonca, Alice Sampaio Rocha, Taina Venas, Elisa Cavalcante Pereira, Renata Serrano Lopes, Darcita Buerger Rovaris, Sandra Bianchini Fernandes, Marilda Siqueira on behalf of the Fiocruz COVID-19 Genomic Surveillance Network |
| EPI_ISL_2196258 | Laboratorio Central de Saude Publica do Estado de Minas Gerais (LACEN/MG) | Laboratory of Respiratory Viruses and Measles, Oswaldo Cruz Institute, FIOCRUZ | Paola Resende, Luciana Appolinario, Fernando Motta, Anna Carolina Paixao, Ana Carolina Mendonca, Alice Sampaio Rocha, Taina Venas, Elisa Cavalcante Pereira, Renata Serrano Lopes, Andre Felipe Leal Bernardes, Marilda Siqueira on behalf of the Fiocruz COVID-19 Genomic Surveillance Network |
| EPI_ISL_2214968 | Centre Hospitalier Universitaire Clermont-Ferrand | CHU Clermont-Ferrand, service de virologie | Bisseux Maxime, Mirand Audrey, Combes Patricia, Henquell Cécile |
| EPI_ISL_2225609 | CHU Pontchaillou | CHU Pontchaillou | GROLHIER Claire, DENOUAL Florent, ETCHEVERRY Amandine, SASSI Mohamed, JAGLINE Steven, FEBREAU Christine, PRONIER Charlotte, GALIBERT Marie Dominique, DE TAYRAC Marie, THIBAULT Vincent |
| EPI_ISL_2226281 | CHNDS | CHU Poitiers | Caroline MICHAUD, Maxime PICHON, Manon PRAT, Valentin BON-BARET, Birama N'DIAYE, Magali GARCIA, Agnes BEBY-DEFAUX, Nicolas LEVEQUE |

| | | | |
|---|---|---|---|
| EPI_ISL_2226283 | MEDILAB | CHU Poitiers | Caroline MICHAUD, Maxime PICHON, Manon PRAT, Valentin BON-BARET, Birama N'DIAYE, Magali GARCIA, Agnes BEBY-DEFAUX, Nicolas LEVEQUE |
| EPI_ISL_2226297 | BIO86 CHAUMONT | CHU Poitiers | Caroline MICHAUD, Maxime PICHON, Manon PRAT, Valentin BON-BARET, Birama N'DIAYE, Magali GARCIA, Agnes BEBY-DEFAUX, Nicolas LEVEQUE |
| EPI_ISL_2226302, EPI_ISL_2226305 | CHU POITIERS | CHU Poitiers | Caroline MICHAUD, Maxime PICHON, Manon PRAT, Valentin BON-BARET, Birama N'DIAYE, Magali GARCIA, Agnes BEBY-DEFAUX, Nicolas LEVEQUE |
| EPI_ISL_2226651, EPI_ISL_2226653, EPI_ISL_2226654, EPI_ISL_2226656, EPI_ISL_2226657, EPI_ISL_2226658, EPI_ISL_2226660, EPI_ISL_2226662, EPI_ISL_2226664, EPI_ISL_2226665, EPI_ISL_2226666, EPI_ISL_2226667, EPI_ISL_2226668, EPI_ISL_2226669, EPI_ISL_2226670, EPI_ISL_2226672, EPI_ISL_2226673, EPI_ISL_2226674 | | | |
| see above | ADMED Microbiologie | Genomics and Transcriptomics, Philip Morris International | Reto Lienhard, Marie-Lise Tritten, Emmanuel Guedj, Nicolas Sierro, Rémi Dulize, David Bornand, Mehdi Auberson, Maxime Berthouzoz, Nikolai Ivanov, Manuel Peitsch |
| EPI_ISL_2231509, EPI_ISL_2231517, EPI_ISL_2231525 | Centro de Diagnostico COVID-19 UABC Tijuana | Andersen lab at Scripps Research | SEARCH Alliance San Diego with Idanya Rubi Serafin Higuera, Manuel Sanchez Alavez, Jorge Luis Jimenez Niebla, German Ibarra, Jonathan Vincent Baena, Oscar Efren Zazueta Fierro |
| EPI_ISL_2232698, EPI_ISL_2232699, EPI_ISL_2232700, EPI_ISL_2232701, EPI_ISL_2232702, EPI_ISL_2232703, EPI_ISL_2232704, EPI_ISL_2232705, EPI_ISL_2232706, EPI_ISL_2232707, EPI_ISL_2232708, EPI_ISL_2232709, EPI_ISL_2232710, EPI_ISL_2232712, EPI_ISL_2232713, EPI_ISL_2232714, EPI_ISL_2232715, EPI_ISL_2232716, EPI_ISL_2232717, EPI_ISL_2232718, EPI_ISL_2232719, EPI_ISL_2232720, EPI_ISL_2232721, EPI_ISL_2232722, EPI_ISL_2232723, EPI_ISL_2232724, EPI_ISL_2232725, EPI_ISL_2232726, EPI_ISL_2232727 | | | |
| see above | Klinikum Wels-Grieskirchen | Bergthaler laboratory, CeMM Research Center for Molecular Medicine of the Austrian Academy of Sciences | Lukas Endler, Anna Schedl, Fabian Amman, Petr Triska, Thomas Penz, Benedikt Agerer, Maelle Le Moing, Michael Schuster, Bekir Erguner, Jan Laine, Martin Senekowitsch, Christoph Bock, Andreas Bergthaler |
| EPI_ISL_2254717, EPI_ISL_2254720, EPI_ISL_2254722 | Veterinary institute in Zvolen Slovakia | Veterinary institute in Zvolen Slovakia | Dirbáková Z., Sujová S., Mokryšová S., Tinák M., Mojžiš M |
| EPI_ISL_2259254 | National Center of Infectious and Parasitic Diseases | National Center of Infectious and Parasitic Diseases | Alexiev et al |
| EPI_ISL_2274094, EPI_ISL_2274095 | Lboratorio Central de Saude Publica do Estado do Parana (LACEN/PR) | Laboratory of Respiratory Viruses and Measles, Oswaldo Cruz Institute, FIOCRUZ | Paola Resende, Luciana Appolinario, Fernando Motta, Anna Carolina Paixao, Ana Carolina Mendonca, Alice Sampaio Rocha, Taina Venas, Elisa Cavalcante Pereira, Renata Serrano Lopes, Irina Riediger, Marilda Siqueira on behalf of the Fiocruz COVID-19 Genomic Surveillance Network |
| EPI_ISL_2274097, EPI_ISL_2274100, EPI_ISL_2274101, EPI_ISL_2274102, EPI_ISL_2274103, EPI_ISL_2274104, EPI_ISL_2274105, EPI_ISL_2274106, EPI_ISL_2274107 | Laboratory of Respiratory Viruses and Measles, Oswaldo Cruz Institute, FIOCRUZ | Laboratory of Respiratory Viruses and Measles, Oswaldo Cruz Institute, FIOCRUZ | Paola Resende, Luciana Appolinario, Fernando Motta, Anna Carolina Paixao, Ana Carolina Mendonca, Alice Sampaio Rocha, Taina Venas, Elisa Cavalcante Pereira, Renata Serrano Lopes, Marilda Siqueira on behalf of the Fiocruz COVID-19 Genomic Surveillance Network |
| EPI_ISL_2274119 | Laboratorio Central de Saude Publica do Estado do Rio Grande do Sul (LACEN-RS) | Laboratory of Respiratory Viruses and Measles, Oswaldo Cruz Institute, FIOCRUZ | Paola Resende, Luciana Appolinario, Fernando Motta, Anna Carolina Paixao, Ana Carolina Mendonca, Alice Sampaio Rocha, Taina Venas, Elisa Cavalcante Pereira, Renata Serrano Lopes, Tatiana Schaffer Gregianini, Richard Salvato, Marilda Siqueira on behalf of the Fiocruz COVID-19 Genomic Surveillance Network |
| EPI_ISL_2274133 | Laboratório Central de Saude Publica do Estado de Santa Catarina (LACEN/SC) | Laboratory of Respiratory Viruses and Measles, Oswaldo Cruz Institute, FIOCRUZ | Paola Resende, Luciana Appolinario, Fernando Motta, Anna Carolina Paixao, Ana Carolina Mendonca, Alice Sampaio Rocha, Taina Venas, Elisa Cavalcante Pereira, Renata Serrano Lopes, Darcita Buerger Rovaris, Sandra Bianchini Fernandes, Marilda Siqueira on behalf of the Fiocruz COVID-19 Genomic Surveillance Network |
| EPI_ISL_2274437 | RSUD Dr. Soetomo | Institute of Tropical Disease, Universitas Airlangga | Kazufumi Shimizu, Krisnoadi Rahardjo, Aldise M Nastri, Jezzy R Dewantari, Rima R Prasetya, Joni Wahyuhadi, Neneng D Kurniati, Gatot Soegiarto, Laksmi Wulandari, Resti Yudhawati, Yasuko Mori, Soetjipto, Maria I Lusida |
| EPI_ISL_2279589 | Hospital | National Reference Center for Viruses of Respiratory Infections, Institut Pasteur, Paris | Marion Barbet, Sylvie Behillil, Méline Bizard, Angela Brisebarre, Camille Capel, Vincent Enouf, Louise Lefrançois, Frédéric Lemoine, Christophe Malabat, Corinne Maufrais, Etienne Simon-Lorière, Maud Vanpeene, Sylvie Van der Werf ,StéPhanie Marque Juillet |
| EPI_ISL_2279592, EPI_ISL_2279593 | Hospital | National Reference Center for Viruses of Respiratory Infections, Institut Pasteur, Paris | Marion Barbet, Sylvie Behillil, Méline Bizard, Angela Brisebarre, Camille Capel, Vincent Enouf, Louise Lefrançois, Frédéric Lemoine, Christophe Malabat, Corinne Maufrais, Amaury Vaysse, Etienne Simon-Lorière, Maud Vanpeene, Sylvie Van der Werf ,StéPhanie Marque Juillet |
| EPI_ISL_2279594, EPI_ISL_2279595, EPI_ISL_2279596, EPI_ISL_2279597, EPI_ISL_2279599, EPI_ISL_2279600 | Hospital | National Reference Center for Viruses of Respiratory Infections, Institut Pasteur, Paris | Marion Barbet, Sylvie Behillil, Méline Bizard, Angela Brisebarre, Camille Capel, Vincent Enouf, Louise Lefrançois, Frédéric Lemoine, Christophe Malabat, Corinne Maufrais, Etienne Simon-Lorière, Maud Vanpeene, Sylvie Van der Werf ,StéPhanie Marque Juillet |
| EPI_ISL_2279601 | Hospital | National Reference Center for Viruses of Respiratory Infections, Institut Pasteur, Paris | Marion Barbet, Sylvie Behillil, Méline Bizard, Angela Brisebarre, Camille Capel, Vincent Enouf, Louise Lefrançois, Frédéric Lemoine, Christophe Malabat, Corinne Maufrais, Amaury Vaysse, Etienne Simon-Lorière, Maud Vanpeene, Sylvie Van der Werf ,StéPhanie Marque Juillet |
| EPI_ISL_2279602, EPI_ISL_2279603, EPI_ISL_2279604, EPI_ISL_2279605, EPI_ISL_2279606, EPI_ISL_2279607 | Hospital | National Reference Center for Viruses of Respiratory Infections, Institut Pasteur, Paris | Marion Barbet, Sylvie Behillil, Méline Bizard, Angela Brisebarre, Camille Capel, Vincent Enouf, Louise Lefrançois, Frédéric Lemoine, Christophe Malabat, Corinne Maufrais, Etienne Simon-Lorière, Maud Vanpeene, Sylvie Van der Werf ,StéPhanie Marque Juillet |
| EPI_ISL_2279608 | Hospital | National Reference Center for Viruses of Respiratory Infections, Institut Pasteur, Paris | Marion Barbet, Sylvie Behillil, Méline Bizard, Angela Brisebarre, Camille Capel, Vincent Enouf, Louise Lefrançois, Frédéric Lemoine, Christophe Malabat, Corinne Maufrais, Ammanuelle Permal, Etienne Simon-Lorière, Maud Vanpeene, Sylvie Van der Werf ,StéPhanie Marque Juillet |
| EPI_ISL_2279609 | Hospital | National Reference Center for Viruses of Respiratory Infections, Institut Pasteur, Paris | Marion Barbet, Sylvie Behillil, Méline Bizard, Angela Brisebarre, Camille Capel, Vincent Enouf, Louise Lefrançois, Frédéric Lemoine, Christophe Malabat, Corinne Maufrais, Etienne Simon-Lorière, Maud Vanpeene, Sylvie Van der Werf ,StéPhanie Marque Juillet |
| EPI_ISL_2279614 | Hospital | National Reference Center for Viruses of Respiratory Infections, Institut Pasteur, Paris | Marion Barbet, Sylvie Behillil, Méline Bizard, Angela Brisebarre, Camille Capel, Vincent Enouf, Louise Lefrançois, Frédéric Lemoine, Christophe Malabat, Corinne Maufrais, Amaury Vaysse, Etienne Simon-Lorière, Maud Vanpeene, Sylvie Van der Werf ,StéPhanie Marque Juillet |
| EPI_ISL_2281746 | Laboratoire du CH de Lannion | Institut Pasteur | Marion Barbet, Sylvie Behillil, Méline Bizard, Angela Brisebarre, Camille Capel, Vincent Enouf, Louise Lefrançois, Frédéric Lemoine, Christophe Malabat, Corinne Maufrais, Emmanuelle Permal, Etienne Simon-Lorière, Maud Vanpeene, Sylvie Van der Werf ,Marie-josé Dufour |
| EPI_ISL_2283704 | LESP Ciudad de Mexico | Instituto de Diagnostico y Referencia Epidemiologicos (INDRE) | Claudia Wong-Arambula, Abril Rodriguez-Maldonado, Vanessa Rivero-Arredondo, Ariadna Medina-Benitez, Joaquin Quiroz-Mercado, Sergio Rangel-Guerrero, Natividad Cruz-Ortiz, Tatiana Nunez-Garcia, Gisela Barrera-Badillo, Lucia Hernandez-Rivas, Irma Lopez-Martinez, Ernesto Ramirez-Gonzalez. |
| EPI_ISL_2284869 | RSUD Dr. Soetomo | Institute of Tropical Disease, Universitas Airlangga | Rima R Prasetya, Krisnoadi Rahardjo, Aldise M Nastri, Jezzy R Dewantari, Joni Wahyuhadi, Neneng D Kurniati, Gatot Soegiarto, Laksmi Wulandari, Resti Yudhawati, Yasuko Mori, Soetjipto, Kazufumi Shimizu, Maria I Lusida |
| EPI_ISL_2284870 | RSUD Dr. Soetomo | Institute of Tropical Disease, Universitas Airlangga | Kazufumi Shimizu, Krisnoadi Rahardjo, Aldise M Nastri, Jezzy R Dewantari, Rima R Prasetya, Joni Wahyuhadi, Neneng D Kurniati, Gatot Soegiarto, Laksmi Wulandari, Resti Yudhawati, Yasuko Mori, Soetjipto, Maria I Lusida |
| EPI_ISL_2284871 | RSUD Dr. Soetomo | Institute of Tropical Disease, Universitas Airlangga | Krisnoadi Rahardjo, Aldise M Nastri, Jezzy R Dewantari, Rima R Prasetya, Joni Wahyuhadi, Neneng D Kurniati, Gatot Soegiarto, Laksmi Wulandari, Resti Yudhawati, Yasuko Mori, Soetjipto, Kazufumi Shimizu, Maria I Lusida |
| EPI_ISL_2284872 | RSUD Dr. Soetomo | Institute of Tropical Disease, Universitas Airlangga | Aldise M Nastri, Krisnoadi Rahardjo, Jezzy R Dewantari, Rima R Prasetya, Joni Wahyuhadi, Neneng D Kurniati, Gatot Soegiarto, Laksmi Wulandari, Resti Yudhawati, Yasuko Mori, Soetjipto, Kazufumi Shimizu, Maria I Lusida |
| EPI_ISL_2284882 | Primasatya Husada Citra Hospital | Institute of Tropical Disease, Universitas Airlangga | Jezzy R Dewantari, Rima R Prasetya, Krisnoadi Rahardjo, Aldise M Nastri, Pudji Djanuartono, Gatot Soegiarto, Laksmi Wulandari, Resti Yudhawati, Yasuko Mori, Soetjipto, Kazufumi Shimizu, Maria I Lusida |
| EPI_ISL_2284883 | Siti Khodijah Hospital | Institute of Tropical Disease, Universitas Airlangga | Rima R Prasetya, Krisnoadi Rahardjo, Aldise M Nastri, Jezzy R Dewantari, Muhammad Hamdan, Gatot Soegiarto, Laksmi Wulandari, Resti Yudhawati, Yasuko Mori, Soetjipto, Kazufumi Shimizu, Maria I Lusida |
| EPI_ISL_2284884 | RSUD Dr. Soetomo | Institute of Tropical Disease, Universitas Airlangga | Krisnoadi Rahardjo, Aldise M Nastri, Jezzy R Dewantari, Rima R Prasetya, Joni Wahyuhadi, Neneng D Kurniati, Gatot Soegiarto, Laksmi Wulandari, Resti Yudhawati, Yasuko Mori, Soetjipto, Kazufumi Shimizu, Maria I Lusida |

| | | | |
|---|---|---|---|
| EPI_ISL_2290152, EPI_ISL_2290153 | Turkish Society of Internal Medicine | Gen Era Diagnostics Corporation Life Science Research and Molecular Diagnostics | Serhat Ünal |
| EPI_ISL_2293423 | Research | National Reference Center for Viruses of Respiratory Infections, Institut Pasteur, Paris | Marion Barbet, Sylvie Behillil, Méline Bizard, Angela Brisebarre, Camille Capel, Vincent Enouf, Louise Lefrançois, Frédéric Lemoine, Christophe Malabat, Corinne Maufrais, Etienne Simon-Lorière, Maud Vanpeene, Sylvie Van der Werf ,Marianne Leruez-Ville |
| EPI_ISL_2312657 | Centre Hospitalier Universitaire Clermont-Ferrand | CHU Clermont-Ferrand, service de virologie | Bisseux Maxime, Mirand Audrey, Combes Patricia, Henquell Cécile |
| EPI_ISL_2321388 | Tripler Army Medical Center | Tripler Army Medical Center | Jonathan A. D'Ambrozio, Rachael Downham, Timothy S. Horseman, Lindsey L. O'Neal, Zakariae Bram, and Catherine F. Uyehara |
| EPI_ISL_2322747, EPI_ISL_2322748, EPI_ISL_2322749, EPI_ISL_2322750, EPI_ISL_2322751 | AZ Sint-Lucas Gent | AZ Sint-Lucas Gent | Jos Van Acker, Charlotte Verfaillie, Elke Vanlaere, Klara De Rauw |
| EPI_ISL_2324169, EPI_ISL_2324170, EPI_ISL_2324171, EPI_ISL_2324172, EPI_ISL_2324173, EPI_ISL_2324174, EPI_ISL_2324175, EPI_ISL_2324176, EPI_ISL_2324177, EPI_ISL_2324178 | Klinikum Wels-Grieskirchen | Bergthaler laboratory, CeMM Research Center for Molecular Medicine of the Austrian Academy of Sciences | Lukas Endler, Anna Schedl, Fabian Amman, Petr Triska, Thomas Penz, Benedikt Agerer, Maelle Le Moing, Michael Schuster, Bekir Erguner, Jan Laine, Martin Senekowitsch, Christoph Bock, Andreas Bergthaler |
| EPI_ISL_2324179, EPI_ISL_2324181, EPI_ISL_2324183, EPI_ISL_2324185, EPI_ISL_2324187, EPI_ISL_2324189, EPI_ISL_2324190, EPI_ISL_2324192, EPI_ISL_2324193, EPI_ISL_2324195, EPI_ISL_2324196, EPI_ISL_2324197, EPI_ISL_2324198, EPI_ISL_2324199, EPI_ISL_2324200, EPI_ISL_2324201, EPI_ISL_2324202, EPI_ISL_2324203, EPI_ISL_2324204, EPI_ISL_2324205, EPI_ISL_2324206, EPI_ISL_2324208, EPI_ISL_2324209, EPI_ISL_2324210, EPI_ISL_2324211, EPI_ISL_2324212, EPI_ISL_2324213, EPI_ISL_2324214, EPI_ISL_2324215, EPI_ISL_2324216, EPI_ISL_2324217, EPI_ISL_2324219, EPI_ISL_2324220, EPI_ISL_2324221, EPI_ISL_2324222, EPI_ISL_2324224, EPI_ISL_2324225, EPI_ISL_2324226, EPI_ISL_2324227, EPI_ISL_2324228, EPI_ISL_2324229, EPI_ISL_2324230, EPI_ISL_2324231, EPI_ISL_2324232, EPI_ISL_2324233, EPI_ISL_2324234, EPI_ISL_2324235, EPI_ISL_2324236, EPI_ISL_2324237, EPI_ISL_2324238, EPI_ISL_2324239, EPI_ISL_2324240, EPI_ISL_2324241, EPI_ISL_2324243, EPI_ISL_2324245, EPI_ISL_2324246, EPI_ISL_2324247, EPI_ISL_2324248, EPI_ISL_2324249, EPI_ISL_2324250, EPI_ISL_2324251, EPI_ISL_2324253, EPI_ISL_2324254, EPI_ISL_2324255, EPI_ISL_2324256, EPI_ISL_2324257 | | | |
| see above | Austrian Agency for Health and Food Safety (AGES) | Bergthaler laboratory, CeMM Research Center for Molecular Medicine of the Austrian Academy of Sciences | Lukas Endler, Anna Schedl, Fabian Amman, Petr Triska, Thomas Penz, Benedikt Agerer, Maelle Le Moing, Michael Schuster, Bekir Erguner, Jan Laine, Martin Senekowitsch, Christoph Bock, Andreas Bergthaler |
| EPI_ISL_2340903, EPI_ISL_2340904, EPI_ISL_2340905, EPI_ISL_2340906, EPI_ISL_2340907, EPI_ISL_2340908, EPI_ISL_2340909, EPI_ISL_2340910, EPI_ISL_2340911, EPI_ISL_2340912, EPI_ISL_2340913, EPI_ISL_2340914, EPI_ISL_2342995, EPI_ISL_2343000, EPI_ISL_2343003 | | | |
| see above | LESP Yucatan | Instituto de Diagnostico y Referencia Epidemiologicos (INDRE) | Claudia Wong-Arambula, Abril Rodriguez-Maldonado, Vanessa Rivero-Arredondo, Ariadna Medina-Benitez, Joaquin Quiroz-Mercado, Sergio Rangel-Guerrero, Natividad Cruz-Ortiz, Tatiana Nunez-Garcia, Gisela Barrera-Badillo, Lucia Hernandez-Rivas, Irma Lopez-Martinez, Ernesto Ramirez-Gonzalez. |
| EPI_ISL_2348501, EPI_ISL_2348502 | Gyncentrum | Gyncentrum | Emilia Morawiec, Pawe Czerwiski, Adam Pudeko, Aleksandra Skubis-Sikora, Magdalena Samul, Agnieszka Polak, Celina Kruszniewska-Rajs, Tomasz Wsik, Robert Wojtyczka, Jolanta Bartosiewicz- Wsik, Maria Maklasiska-Majdanik |
| EPI_ISL_2348503, EPI_ISL_2348504, EPI_ISL_2348505, EPI_ISL_2348506 | Gyncentrum | Gyncentrum | Emilia Morawiec, Pawe? Czerwi?ski, Adam Pude?ko, Aleksandra Skubis-Sikora, Magdalena Samul, Agnieszka Polak, Celina Kruszniewska-Rajs, Tomasz W?sik, Robert Wojtyczka, Jolanta Bartosiewicz- W?sik, Maria Maklasi?ska-Majdanik |
| EPI_ISL_2348507, EPI_ISL_2348508, EPI_ISL_2348509, EPI_ISL_2348510, EPI_ISL_2348618, EPI_ISL_2348619, EPI_ISL_2348620, EPI_ISL_2348621, EPI_ISL_2348665, EPI_ISL_2348668, EPI_ISL_2348669, EPI_ISL_2348670, EPI_ISL_2348677, EPI_ISL_2348679, EPI_ISL_2348680, EPI_ISL_2348682, EPI_ISL_2348684, EPI_ISL_2348686, EPI_ISL_2348687, EPI_ISL_2348689, EPI_ISL_2348690, EPI_ISL_2348691, EPI_ISL_2348693, EPI_ISL_2348694, EPI_ISL_2348695, EPI_ISL_2348696, EPI_ISL_2348698, EPI_ISL_2348699, EPI_ISL_2348700, EPI_ISL_2348701, EPI_ISL_2348702, EPI_ISL_2348703, EPI_ISL_2348707, EPI_ISL_2348708, EPI_ISL_2348709, EPI_ISL_2348711, EPI_ISL_2348712, EPI_ISL_2348713, EPI_ISL_2348714, EPI_ISL_2348715, EPI_ISL_2348717, EPI_ISL_2348718, EPI_ISL_2348720, EPI_ISL_2348721, EPI_ISL_2348722, EPI_ISL_2348723, EPI_ISL_2348724, EPI_ISL_2348725, EPI_ISL_2348726, EPI_ISL_2348728 | | | |
| see above | Gyncentrum | Gyncentrum | Emilia Morawiec, Pawel Czerwinski, Adam Pudelko, Aleksandra Skubis-Sikora, Magdalena Samul, Agnieszka Polak, Celina Kruszniewska-Rajs, Tomasz Wasik, Robert Wojtyczka, Jolanta Bartosiewicz- Wasik, Maria Maklasinska-Majdanik |
| EPI_ISL_2351221, EPI_ISL_2352031 | Dr. Gernot Walder GmbH | Dr. Gernot Walder GmbH | Sissy T. Sonnleitner |
| EPI_ISL_2356912 | Collection of WHO National Influenza Center, N.F. Gamaleya National Research Center for Epidemiology and Microbiology of the Ministry of Health of the Russian Federation | N.F. Gamaleya Research Center for Epidemiology and Microbiology | Olga Voronina, Ekaterina Aksenova, Natalia Ryzhova, Marina Kunda, Kirill Krasnoslobodtsev, Elena Burtseva, Denis Logunov |
| EPI_ISL_2361904, EPI_ISL_2361906, EPI_ISL_2361907, EPI_ISL_2361913, EPI_ISL_2361914, EPI_ISL_2361915, EPI_ISL_2361916, EPI_ISL_2361917, EPI_ISL_2361919, EPI_ISL_2361920, EPI_ISL_2361921, EPI_ISL_2361922, EPI_ISL_2361923 | | | |
| see above | Noguchi Memorial Institute for Medical Research, University of Ghana, Legon, Ghana | Institute of Tropical Medicine, Universitätsklinikum Tübingen, Germany | Bright Adu, Quaneeta Mohktar, Le Thi Kieu Linh, Sivaramakrishna Rachakonda, Hilda Opoku Frempong, Keren Okyerebea Attiku, Joyce Appiah-Kubi, Joseph Humphrey Kofi Bonney, John Kofi Odoom, Abraham Kwabena Anang, Srinivas-reddy Pallerla, Dorothy Yeboah-Manu, Thirumalaisamy P Velavan |
| EPI_ISL_2364519, EPI_ISL_2364520, EPI_ISL_2364521, EPI_ISL_2364522 | Genome Analysis Center, Yamanashi Central Hospital | Genome Analysis Center, Yamanashi Central Hospital | Yosuke Hirotsu |
| EPI_ISL_2365881 | Bezmialem Vakif University Hospital Microbiology Laboratory | Bezmialem Vakif University Hospital Microbiology Laboratory | Mehmet Z. Doymaz, Elif Karaaslan, Merve Yazici-Kalkan, Nesibe Selma Çetin, Bilge Sumbul |
| EPI_ISL_2365970 | University of Bari Biomedical Sciences and Human Oncology, Policlinico | University of Bari Biomedical Sciences and Human Oncology | Maria Chironna, Anna Sallustio, Daniela Loconsole, Marisa Accogli |
| EPI_ISL_2374391 | Hospital | National Reference Center for Viruses of Respiratory Infections, Institut Pasteur, Paris | Marion Barbet, Sylvie Behillil, Méline Bizard, Angela Brisebarre, Camille Capel, Vincent Enouf, Louise Lefrançois, Frédéric Lemoine, Christophe Malabat, Corinne Maufrais, Emmanuelle Permal, Etienne Simon-Lorière, Maud Vanpeene, Sylvie Van der Werf ,Alexandra Ducancelle |
| EPI_ISL_2375877, EPI_ISL_2375878, EPI_ISL_2375879, EPI_ISL_2375880, EPI_ISL_2375881, EPI_ISL_2375882, EPI_ISL_2375883, EPI_ISL_2375884, EPI_ISL_2375885, EPI_ISL_2375886, EPI_ISL_2375887, EPI_ISL_2375888, EPI_ISL_2375889, EPI_ISL_2375890, EPI_ISL_2375891, EPI_ISL_2375892, EPI_ISL_2375893, EPI_ISL_2375894, EPI_ISL_2375895, EPI_ISL_2375896, EPI_ISL_2375897, EPI_ISL_2375898 | | | |
| see above | Programa de Oncovirologia, Instituto Nacional de Câncer | Programa de Oncovirologia, Instituto Nacional de Câncer | Livia R. Goes, Juliana D. Siqueira, Marianne M. Garrido, Brunna M. Alves, Ana Cristina P. M. Pereira, Claudia Cicala, James Arthos, João P.B. Viola, Marcelo A. Soares |
| EPI_ISL_2376384, EPI_ISL_2376385, EPI_ISL_2376386, EPI_ISL_2376387, EPI_ISL_2376388, EPI_ISL_2376389 | Noguchi Memorial Institute for Medical Research, University of Ghana, Legon, Ghana | Institute of Tropical Medicine, Universitätsklinikum Tübingen, Germany | Bright Adu, Quaneeta Mohktar, Le Thi Kieu Linh, Sivaramakrishna Rachakonda, Hilda Opoku Frempong, Keren Okyerebea Attiku, Joyce Appiah-Kubi, Joseph Humphrey Kofi Bonney, John Kofi Odoom, Abraham Kwabena Anang, Srinivas-reddy Pallerla, Dorothy Yeboah-Manu, Thirumalaisamy P Velavan |
| EPI_ISL_2378984, EPI_ISL_2378985, EPI_ISL_2378986, EPI_ISL_2378987, EPI_ISL_2378988, EPI_ISL_2378989, EPI_ISL_2378990, EPI_ISL_2378991, EPI_ISL_2378992, EPI_ISL_2378995, EPI_ISL_2378996, EPI_ISL_2378997, EPI_ISL_2378998, EPI_ISL_2378999, EPI_ISL_2379000, EPI_ISL_2379002, EPI_ISL_2379003, EPI_ISL_2379004, EPI_ISL_2379005, EPI_ISL_2379008, EPI_ISL_2379009, EPI_ISL_2379011, EPI_ISL_2379014, EPI_ISL_2379015, EPI_ISL_2379018, EPI_ISL_2379020, EPI_ISL_2379021, EPI_ISL_2379022, EPI_ISL_2379023, EPI_ISL_2379024, EPI_ISL_2379026, EPI_ISL_2379027, EPI_ISL_2379028, EPI_ISL_2379030, EPI_ISL_2379032, EPI_ISL_2379037, EPI_ISL_2379038, EPI_ISL_2379039, EPI_ISL_2379128, EPI_ISL_2379130, EPI_ISL_2379131, EPI_ISL_2379132, EPI_ISL_2379133, EPI_ISL_2379135, EPI_ISL_2379136, EPI_ISL_2379137, EPI_ISL_2379138, EPI_ISL_2379139, EPI_ISL_2379142, EPI_ISL_2379143, EPI_ISL_2379144 | | | |
| see above | National Center of Infectious and Parasitic Diseases | National Center of Infectious and Parasitic Diseases | Alexiev et al |
| EPI_ISL_2379893 | The teaching Baghdad hospital | PCR lab | Abdulhussein Thair, Fadhil Hula |
| EPI_ISL_2380369 | National Institute of Public Health | Charles University, Faculty of Science, BIOCEV, OMICS Genomics | Vladimír Hampl, Štpánka Hrdá, Ruth Tachezy, Blanka Hamplová, Sebastian Cristian Treitli, Zoltán Füssy, Magdalena Janáová, Ingrid Poláková, Jana Šmahelová, Jií Novák |
| EPI_ISL_2381994 | AZ Zeno Knokke-Blankenberge | AZ Sint Jan | Marijke Reynders, Patrick Descheemaeker, Laurien Hoornaert, Thomas Van Landschoot, Marie Madeleine Chabert-Consen, Jorn Hellemans |
| EPI_ISL_2382466, EPI_ISL_2382467, EPI_ISL_2382469, EPI_ISL_2382470, EPI_ISL_2382471, EPI_ISL_2382472, EPI_ISL_2382473, EPI_ISL_2382474, EPI_ISL_2382475, EPI_ISL_2382476, EPI_ISL_2382477, EPI_ISL_2382478, EPI_ISL_2382479, EPI_ISL_2382481, EPI_ISL_2382482, EPI_ISL_2382483, EPI_ISL_2382484, EPI_ISL_2382486, EPI_ISL_2382487, EPI_ISL_2382488, EPI_ISL_2382490, EPI_ISL_2382491, EPI_ISL_2382492, EPI_ISL_2382493, EPI_ISL_2382494, EPI_ISL_2382495, EPI_ISL_2382496, EPI_ISL_2382497, EPI_ISL_2382498, EPI_ISL_2382499, EPI_ISL_2382500, EPI_ISL_2382502, EPI_ISL_2382503, EPI_ISL_2382505, EPI_ISL_2382506, EPI_ISL_2382508, EPI_ISL_2382509, EPI_ISL_2382510, EPI_ISL_2382512, EPI_ISL_2382513, EPI_ISL_2382514, EPI_ISL_2382515, EPI_ISL_2382518, EPI_ISL_2382519, EPI_ISL_2382520, EPI_ISL_2382521, EPI_ISL_2382749, EPI_ISL_2382750, EPI_ISL_2382751, EPI_ISL_2382752 | | | |
| see above | The Ohio State University Applied Microbiology Services Laboratory | The Ohio State University Applied Microbiology Services Laboratory | Seth A. Faith PhD |
| EPI_ISL_2389618 | B.J. Medical College and Civil Hospital, Ahmedabad | Gujarat Biotechnology Research Centre | Ramesh Pandit,Janvi Raval,Zarna Patel,Nitin Savaliya,Dinesh Kumar,Twinkle Soni,Sonal Sharma ,Zuber Saiyed,Pranay Shah,Umang Mishra,Nitesh Shah,Chaitanya Joshi,Madhvi Joshi |
| EPI_ISL_2389620 | B.J. Medical College and Civil Hospital, Ahmedabad | Gujarat Biotechnology Research Centre | Janvi Raval,Zarna Patel,Nitin Savaliya,Dinesh Kumar,Twinkle Soni,Sonal Sharma ,Zuber Saiyed,Ramesh Pandit,Pranay Shah,Umang Mishra,Nitesh Shah,Chaitanya Joshi,Madhvi Joshi |

| | | | |
|---|---|---|---|
| EPI_ISL_2389622 | B.J. Medical College and Civil Hospital, Ahmedabad | Gujarat Biotechnology Research Centre | Zarna Patel,Nitin Savaliya,Dinesh Kumar,Twinkle Soni,Sonal Sharma ,Zuber Saiyed,Ramesh Pandit,Janvi Raval,Pranay Shah,Umang Mishra,Nitesh Shah,Chaitanya Joshi,Madhvi Joshi |
| EPI_ISL_2389624 | B.J. Medical College and Civil Hospital, Ahmedabad | Gujarat Biotechnology Research Centre | Nitin Savaliya,Dinesh Kumar,Twinkle Soni,Sonal Sharma ,Zuber Saiyed,Ramesh Pandit,Janvi Raval,Zarna Patel,Pranay Shah,Umang Mishra,Nitesh Shah,Chaitanya Joshi,Madhvi Joshi |
| EPI_ISL_2389626 | B.J. Medical College and Civil Hospital, Ahmedabad | Gujarat Biotechnology Research Centre | Dinesh Kumar,Twinkle Soni,Sonal Sharma ,Zuber Saiyed,Ramesh Pandit,Janvi Raval,Zarna Patel,Nitin Savaliya,Pranay Shah,Umang Mishra,Nitesh Shah,Chaitanya Joshi,Madhvi Joshi |
| EPI_ISL_2389628 | B.J. Medical College and Civil Hospital, Ahmedabad | Gujarat Biotechnology Research Centre | Twinkle Soni,Sonal Sharma ,Zuber Saiyed,Ramesh Pandit,Janvi Raval,Zarna Patel,Nitin Savaliya,Dinesh Kumar,Pranay Shah,Umang Mishra,Nitesh Shah,Chaitanya Joshi,Madhvi Joshi |
| EPI_ISL_2389630 | B.J. Medical College and Civil Hospital, Ahmedabad | Gujarat Biotechnology Research Centre | Sonal Sharma ,Zuber Saiyed,Ramesh Pandit,Janvi Raval,Zarna Patel,Nitin Savaliya,Dinesh Kumar,Twinkle Soni,Pranay Shah,Umang Mishra,Nitesh Shah,Chaitanya Joshi,Madhvi Joshi |
| EPI_ISL_2391533 | Centro de Investigación Biomédica del Noreste (CIBIN) | Instituto de Biotecnología de la UNAM | Consorcio Mexicano de Vigilancia Genómica (CoViGen-Mex). Authors (in alphabetical order): Julio Elias Alvarado-Yaah, Carlos F. Arias, Santiago Ávila-Ríos, Eduardo Becerril-Vargas, Víctor Hugo Borja-Aburto, Celia Boukadida, Cristóbal Cháidez-Quiróz, Juan Bautista Chale-Dzul, Ricardo Ciria Merce, Andreu Comas-García, Célida Duque Molina, Julissa Enciso-Ibarra, José Antonio Enciso-Moreno, Gloria Elena Espinosa-Ayala, Fernando Fontove-Herrera, Daniel Fregoso-Rueda, Victor Eduardo Garcia-Arias, Alejandra Garcia-Gasca, Bruno Gomez-Gil, Jean Pierre González, Irvin González-López, Concepción Grajales-Muñiz, Ricardo Grande, Rosa María Gutierrez Rios, Alejandra Hernández-Terán, Alfredo Herrera-Estrella, Carla Ivón Herrera-Najera, Pavel Isa, Daniel Lira Morales, Susana Lopez, Antonio Loza Román, Brenda Irasema Maldonado-Meza, Bernardo Martínez-Miguel, José Arturo Martínez-Orozco, Célida Martínez- Rodríguez, Margarita Matías-Florentino, Fidencio Mejía-Nepomuceno, María Guadalupe de Jesús Mireles-Rivera, Gloria María Molina-Salinas, Hector Montoya-Fuentes, , Mario Mújica-Sánchez, José Esteban Muñoz-Medina, José de Jesús Nuñez-Contreras, Alicia Ocaña-Mondragón, Luis Alberto Ochoa-Carrera, Hector Esteban Paz-Juárez, Francisco Pulido, Helen Haydee Fernanda Ramírez-Plascencia, Jorge Salas-Hernández, Angel Gustavo Salas-Lais, Alejandro Sanchez-Flores, Clara Esperanza Santacruz-Tinoco, María Guadalupe Santiago-Mauricio, Selene Zárate, Nelly Sélem-Mojica, Blanca Taboada, Gloria Vazquez, Joel Armando Vázquez-Pérez. |
| EPI_ISL_2391540 | Unidad de Investigación Biomédica de Zacatecas (UIBZ) | Instituto de Biotecnología de la UNAM | Consorcio Mexicano de Vigilancia Genómica (CoViGen-Mex). Authors (in alphabetical order): Julio Elias Alvarado-Yaah, Carlos F. Arias, Santiago Ávila-Ríos, Eduardo Becerril-Vargas, Víctor Hugo Borja-Aburto, Celia Boukadida, Cristóbal Cháidez-Quiróz, Juan Bautista Chale-Dzul, Ricardo Ciria Merce, Andreu Comas-García, Célida Duque Molina, Julissa Enciso-Ibarra, José Antonio Enciso-Moreno, Gloria Elena Espinosa-Ayala, Fernando Fontove-Herrera, Daniel Fregoso-Rueda, Victor Eduardo Garcia-Arias, Alejandra Garcia-Gasca, Bruno Gomez-Gil, Jean Pierre González, Irvin González-López, Concepción Grajales-Muñiz, Ricardo Grande, Rosa María Gutierrez Rios, Alejandra Hernández-Terán, Alfredo Herrera-Estrella, Carla Ivón Herrera-Najera, Pavel Isa, Daniel Lira Morales, Susana Lopez, Antonio Loza Román, Brenda Irasema Maldonado-Meza, Bernardo Martínez-Miguel, José Arturo Martínez-Orozco, Célida Martínez- Rodríguez, Margarita Matías-Florentino, Fidencio Mejía-Nepomuceno, María Guadalupe de Jesús Mireles-Rivera, Gloria María Molina-Salinas, Hector Montoya-Fuentes, , Mario Mújica-Sánchez, José Esteban Muñoz-Medina, José de Jesús Nuñez-Contreras, Alicia Ocaña-Mondragón, Luis Alberto Ochoa-Carrera, Hector Esteban Paz-Juárez, Francisco Pulido, Helen Haydee Fernanda Ramírez-Plascencia, Jorge Salas-Hernández, Angel Gustavo Salas-Lais, Alejandro Sanchez-Flores, Clara Esperanza Santacruz-Tinoco, María Guadalupe Santiago-Mauricio, Selene Zárate, Nelly Sélem-Mojica, Blanca Taboada, Gloria Vazquez, Joel Armando Vázquez-Pérez. |
| EPI_ISL_2391575, EPI_ISL_2391598, EPI_ISL_2391655 | Centro de Investigación Biomédica del Noreste (CIBIN) | Instituto de Biotecnología de la UNAM | Consorcio Mexicano de Vigilancia Genómica (CoViGen-Mex). Authors (in alphabetical order): Julio Elias Alvarado-Yaah, Carlos F. Arias, Santiago Ávila-Ríos, Eduardo Becerril-Vargas, Víctor Hugo Borja-Aburto, Celia Boukadida, Cristóbal Cháidez-Quiróz, Juan Bautista Chale-Dzul, Ricardo Ciria Merce, Andreu Comas-García, Célida Duque Molina, Julissa Enciso-Ibarra, José Antonio Enciso-Moreno, Gloria Elena Espinosa-Ayala, Fernando Fontove-Herrera, Daniel Fregoso-Rueda, Victor Eduardo Garcia-Arias, Alejandra Garcia-Gasca, Bruno Gomez-Gil, Jean Pierre González, Irvin González-López, Concepción Grajales-Muñiz, Ricardo Grande, Rosa María Gutierrez Rios, Alejandra Hernández-Terán, Alfredo Herrera-Estrella, Carla Ivón Herrera-Najera, Pavel Isa, Daniel Lira Morales, Susana Lopez, Antonio Loza Román, Brenda Irasema Maldonado-Meza, Bernardo Martínez-Miguel, José Arturo Martínez-Orozco, Célida Martínez- Rodríguez, Margarita Matías-Florentino, Fidencio Mejía-Nepomuceno, María Guadalupe de Jesús Mireles-Rivera, Gloria María Molina-Salinas, Hector Montoya-Fuentes, , Mario Mújica-Sánchez, José Esteban Muñoz-Medina, José de Jesús Nuñez-Contreras, Alicia Ocaña-Mondragón, Luis Alberto Ochoa-Carrera, Hector Esteban Paz-Juárez, Francisco Pulido, Helen Haydee Fernanda Ramírez-Plascencia, Jorge Salas-Hernández, Angel Gustavo Salas-Lais, Alejandro Sanchez-Flores, Clara Esperanza Santacruz-Tinoco, María Guadalupe Santiago-Mauricio, Selene Zárate, Nelly Sélem-Mojica, Blanca Taboada, Gloria Vazquez, Joel Armando Vázquez-Pérez. |
| EPI_ISL_2399547, EPI_ISL_2399548, EPI_ISL_2399549, EPI_ISL_2399551, EPI_ISL_2399552, EPI_ISL_2399553, EPI_ISL_2399554, EPI_ISL_2399556, EPI_ISL_2399557, EPI_ISL_2399558, EPI_ISL_2399559, EPI_ISL_2399561, EPI_ISL_2399562, EPI_ISL_2399563, EPI_ISL_2399564, EPI_ISL_2399565, EPI_ISL_2399566, EPI_ISL_2399567, EPI_ISL_2399568, EPI_ISL_2399569, EPI_ISL_2399570, EPI_ISL_2399571, EPI_ISL_2399572, EPI_ISL_2399573, EPI_ISL_2399575, EPI_ISL_2399576, EPI_ISL_2399577, EPI_ISL_2399578, EPI_ISL_2399579, EPI_ISL_2399580, EPI_ISL_2399581, EPI_ISL_2399582, EPI_ISL_2399583, EPI_ISL_2399584, EPI_ISL_2399585, EPI_ISL_2399586, EPI_ISL_2399587, EPI_ISL_2399588, EPI_ISL_2399589, EPI_ISL_2399590, EPI_ISL_2399591, EPI_ISL_2399592, EPI_ISL_2399593, EPI_ISL_2399594, EPI_ISL_2399595, EPI_ISL_2399596, EPI_ISL_2399597, EPI_ISL_2399598, EPI_ISL_2399599, EPI_ISL_2399600, EPI_ISL_2399601, EPI_ISL_2399603, EPI_ISL_2399604, EPI_ISL_2399605, EPI_ISL_2399606, EPI_ISL_2399607, EPI_ISL_2399608, EPI_ISL_2399609, EPI_ISL_2399610, EPI_ISL_2399621, EPI_ISL_2399623, EPI_ISL_2399624, EPI_ISL_2399625, EPI_ISL_2399626, EPI_ISL_2399627, EPI_ISL_2399628, EPI_ISL_2399629, EPI_ISL_2399630, EPI_ISL_2399631, EPI_ISL_2399632, EPI_ISL_2399633, EPI_ISL_2399634, EPI_ISL_2399635, EPI_ISL_2399636, EPI_ISL_2399637, EPI_ISL_2399638, EPI_ISL_2399639, EPI_ISL_2399640, EPI_ISL_2399641, EPI_ISL_2399642, EPI_ISL_2399643, EPI_ISL_2399644, EPI_ISL_2399645, EPI_ISL_2399646, EPI_ISL_2399647, EPI_ISL_2399648, EPI_ISL_2399649, EPI_ISL_2399650, EPI_ISL_2399651, EPI_ISL_2399652, EPI_ISL_2399653, EPI_ISL_2399654, EPI_ISL_2399655, EPI_ISL_2399656, EPI_ISL_2399657, EPI_ISL_2399658, EPI_ISL_2399659, EPI_ISL_2399660, EPI_ISL_2399661, EPI_ISL_2399662, EPI_ISL_2399663, EPI_ISL_2399664, EPI_ISL_2399665, EPI_ISL_2399666, EPI_ISL_2399667, EPI_ISL_2399668, EPI_ISL_2399669, EPI_ISL_2399670, EPI_ISL_2399671, EPI_ISL_2399672, EPI_ISL_2399673, EPI_ISL_2399674, EPI_ISL_2399675, EPI_ISL_2399676, EPI_ISL_2399677, EPI_ISL_2399678, EPI_ISL_2399679, EPI_ISL_2399680, EPI_ISL_2399681, EPI_ISL_2399682, EPI_ISL_2399683, EPI_ISL_2399684, EPI_ISL_2399685, EPI_ISL_2399686, EPI_ISL_2399687, EPI_ISL_2399689, EPI_ISL_2399690, EPI_ISL_2399691, EPI_ISL_2399692, EPI_ISL_2399693, EPI_ISL_2399694, EPI_ISL_2399695, EPI_ISL_2399696, EPI_ISL_2399697, EPI_ISL_2399698, EPI_ISL_2399699, EPI_ISL_2399700, EPI_ISL_2399701, EPI_ISL_2399702, EPI_ISL_2399703, EPI_ISL_2399704, EPI_ISL_2399705, EPI_ISL_2399706, EPI_ISL_2399707, EPI_ISL_2399708, EPI_ISL_2399709, EPI_ISL_2399710, EPI_ISL_2399711, EPI_ISL_2399712, EPI_ISL_2399713, EPI_ISL_2399714, EPI_ISL_2399715, EPI_ISL_2399716, EPI_ISL_2399717, EPI_ISL_2399718, EPI_ISL_2399719, EPI_ISL_2399720, EPI_ISL_2399721, EPI_ISL_2399722, EPI_ISL_2399723, EPI_ISL_2399724, EPI_ISL_2399725, EPI_ISL_2399726, EPI_ISL_2399727, EPI_ISL_2399728, EPI_ISL_2399729, EPI_ISL_2399730, EPI_ISL_2399731, EPI_ISL_2399732, EPI_ISL_2399733, EPI_ISL_2399734, EPI_ISL_2399735, EPI_ISL_2399736, EPI_ISL_2399737, EPI_ISL_2399738, EPI_ISL_2399739, EPI_ISL_2399740, EPI_ISL_2399741, EPI_ISL_2399742, EPI_ISL_2399757 | | | |
| see above | The Ohio State University Applied Microbiology Services Laboratory | The Ohio State University Applied Microbiology Services Laboratory | Seth A. Faith PhD |
| EPI_ISL_2426146 | Clinical Virology Laboratory, Institute of Liver and Biliary Sciences | ILBS - IGIB | Ekta Gupta, Sheetalnath Rooge, Pratibha Kale, Amit Pandey, Chhagan Bihari Sharma, Partha Chattopadhyay, Priti Devi, Ranjeet Maurya, Neha Jha, Priyanka Mehta, Manish Kumar, Pooja Sharma, Sheeba Saifi, Aparna S, Sarfaraz Alam, Bharathram Uppili, Mohammed Faruq, Anurag Agrawal, Rajesh Pandey, Shiv Kumar Sarin |
| EPI_ISL_2429144 | National Institute of Health Research and Development | National Institute of Health Research and Development | Vivi Setiawaty, Hana Apsari Pawestri, Hartanti Dian Ikawati, Subangkit, Kartika Dewi Puspa, Arie Ardiansyah Nugraha, Triyani Soekarso, Krisna Pangesti, Nelly Puspandari |
| EPI_ISL_2434989, EPI_ISL_2436469, EPI_ISL_2437300 | The Ohio State University Applied Microbiology Services Laboratory | The Ohio State University Applied Microbiology Services Laboratory | Seth A. Faith PhD |
| EPI_ISL_2438907, EPI_ISL_2438908, EPI_ISL_2438909, EPI_ISL_2438910, EPI_ISL_2438911, EPI_ISL_2438912, EPI_ISL_2438913, EPI_ISL_2438914, EPI_ISL_2438938, EPI_ISL_2438939, EPI_ISL_2438944, EPI_ISL_2438945, EPI_ISL_2438946, EPI_ISL_2438958, EPI_ISL_2438959, EPI_ISL_2438960, EPI_ISL_2438961, EPI_ISL_2438967, EPI_ISL_2438968 | | | |
| see above | National Institute of Public Health | Charles University, Faculty of Science, BIOCEV, OMICS Genomics | Vladimír Hampl, Štpánka Hrdá, Ruth Tachezy, Blanka Hamplová, Sebastian Cristian Treitli, Zoltán Füssy, Magdalena Janáová, Ingrid Poláková, Jana Šmahelová, Jí Novák |
| EPI_ISL_2448641 | CH Niort | CHU Poitiers | Caroline MICHAUD, Maxime PICHON, Manon PRAT, Valentin BON-BARET, Birama N'DIAYE, Magali GARCIA, Agnes BEBY-DEFAUX, Nicolas LEVEQUE |
| EPI_ISL_2462404 | SUNY Upstate Medical University | SUNY Upstate Medical University | Frank A Middleton |
| EPI_ISL_2462860, EPI_ISL_2462861 | AZ Sint-Lucas Gent | AZ Sint-Lucas Gent | Jos Van Acker, Charlotte Verfaillie, Elke Vanlaere, Klara De Rauw |
| EPI_ISL_2467844 | Adolfo Lutz Institute Sao Paulo Virology Center | Retrovirus Laboratory Adolfo Lutz Institute | Gabriela Bastos Cabral, Giselle I S Lopez-Lopes, Cintia Ahagon, Audrey Cilli, Paula Morena Guimaraes, Igor Mohamed Hussein, Luis Brigido |
| EPI_ISL_2467906, EPI_ISL_2467914, EPI_ISL_2467915, EPI_ISL_2467916 | Biology Department, College of Science, Al Muthanna University and Public Health Laboratory, Al-Muthanna Health Directorate | Department of Virology, Faculty of Medicine, University of Helsinki, Helsinki, Finland | Nihad Al-Rashedi, Hussein Alburkat, Murad Munahi, Alaa Hameed, Ali Jasim, Olli Vapalahti, Tarja Sironen,Teemu Smura |
| EPI_ISL_2484760 | HOME QUARANTINE TASKFORCE | Hong Kong Department of Health | Alan K.L. Tsang, Gannon C.K. Mak, Peter K.C. Cheng, Peter C.W. Yip, Ken H.L. Ng, Edman T.K. Lam, Rickjason C.W. Chan, Dominic N.C. Tsang |
| EPI_ISL_2484761 | Queen Mary Hospital | Hong Kong Department of Health | Alan K.L. Tsang, Gannon C.K. Mak, Peter K.C. Cheng, Peter C.W. Yip, Ken H.L. Ng, Edman T.K. Lam, Rickjason C.W. Chan, Dominic N.C. Tsang |

| | | | |
|---|---|---|---|
| EPI_ISL_2484762 | Temporary Specimen Collection Centre at the AsiaWorld-Expo | Hong Kong Department of Health | Alan K.L. Tsang, Gannon C.K. Mak, Peter K.C. Cheng, Peter C.W. Yip, Ken H.L. Ng, Edman T.K. Lam, Rickjason C.W. Chan, Dominic N.C. Tsang |
| EPI_ISL_2484763 | HOME QUARANTINE TASKFORCE | Hong Kong Department of Health | Alan K.L. Tsang, Gannon C.K. Mak, Peter K.C. Cheng, Peter C.W. Yip, Ken H.L. Ng, Edman T.K. Lam, Rickjason C.W. Chan, Dominic N.C. Tsang |
| EPI_ISL_2484764, EPI_ISL_2484765, EPI_ISL_2484766, EPI_ISL_2484767 | Temporary Specimen Collection Centre at the AsiaWorld-Expo | Hong Kong Department of Health | Alan K.L. Tsang, Gannon C.K. Mak, Peter K.C. Cheng, Peter C.W. Yip, Ken H.L. Ng, Edman T.K. Lam, Rickjason C.W. Chan, Dominic N.C. Tsang |
| EPI_ISL_2488799 | LACEN - Laboratório Central de Saúde Pública do Ceará | Evandro Chagas Institute | Santos, M.C.; Silva, A.M.; Junior, W.D.C.; Barbagelata, L.S.; Ferreira, J.A.; Sousa, E.M.A.; da Silva, P.S.; Pinheiro, K.C.; L.C.; Sousa Junior, E.C. |
| EPI_ISL_2491824, EPI_ISL_2491825, EPI_ISL_2491826, EPI_ISL_2491827, EPI_ISL_2491828, EPI_ISL_2491829, EPI_ISL_2491830 | National Institute of Public Health | Charles University, Faculty of Science, BIOCEV, OMICS Genomics | Vladimír Hampl, Štpánka Hrdá, Ruth Tachezy, Blanka Hamplová, Sebastian Cristian Treitli, Zoltán Füssy, Magdalena Janáová, Ingrid Poláková, Jana Šmahelová, Jií Novák |
| EPI_ISL_2495437, EPI_ISL_2495438, EPI_ISL_2495439, EPI_ISL_2495440, EPI_ISL_2495441, EPI_ISL_2495442, EPI_ISL_2495443, EPI_ISL_2495444, EPI_ISL_2495445, EPI_ISL_2495446, EPI_ISL_2495447, EPI_ISL_2495474, EPI_ISL_2495492, EPI_ISL_2495493, EPI_ISL_2495494, EPI_ISL_2495495, EPI_ISL_2495496, EPI_ISL_2495497, EPI_ISL_2495498, EPI_ISL_2495499, EPI_ISL_2495500, EPI_ISL_2495501, EPI_ISL_2495502, EPI_ISL_2495503, EPI_ISL_2495504, EPI_ISL_2495505, EPI_ISL_2495506, EPI_ISL_2495507, EPI_ISL_2495508, EPI_ISL_2495509, EPI_ISL_2495510, EPI_ISL_2495554, EPI_ISL_2495619, EPI_ISL_2495713, EPI_ISL_2495714, EPI_ISL_2495715, EPI_ISL_2495716, EPI_ISL_2495721 | | | |
| see above | MB-Cadham Provincial laboratory | National Microbiology Laboratory (NML) | Anna Majer, Shari Tyson, Grace Seo, Philip Mabon, Elsie Grudeski, Rhiannon Huzarewich, Russell Mandes, Anneliese Landgraff, Jennifer Tanner, Natalie Knox, Morag Graham, Gary Van Domselaar, Paul Van Caeseele, Jared Bullard, David Alexander, Kerry Dust, Nathalie Bastien, Yan Li, Timothy Booth, Darian Hole, Madison Chapel, Kirsten Biggar, CanCOGeN's metadata curation team, Public Health Agency of Canada CanCOGeN team |
| EPI_ISL_2516949, EPI_ISL_2516950, EPI_ISL_2516951, EPI_ISL_2516955 | Microbiology Department. Complexo Hospitalario Universitario de Vigo | Microbiology Department. Complexo Hospitalario Universitario de Vigo | Alfaya N, Alonso I, Alvarez M, Cabrera JJ, Carballo R, Cores O, Cortizo S, del-Campo V, Martinez L, Mediero G, Perez S, Potel C, Regueiro B, Rey S, Vasallo FJ |
| EPI_ISL_2530313 | Hematopathology Laboratory, ACTREC, TMC | Hematopathology Laboratory, ACTREC, TMC | Hematopathology Laboratory, ACTREC |
| EPI_ISL_2532673, EPI_ISL_2532674, EPI_ISL_2532675 | IZSM | IZSM-U.O.C. Virologia | Maurizio Viscardi, Lorena Cardillo, Claudio de Martinis, Giovanna Fusco, Esterina De Carlo, Antonio Limone |
| EPI_ISL_2534059, EPI_ISL_2534060, EPI_ISL_2534061, EPI_ISL_2534062, EPI_ISL_2534063 | S.C. Laboratorio Analisi, Ospedale di Lavagna, ASL 4 Liguria | U.O. Igiene, Ospedale Policlinico San Martino | Bruzzone Bianca, Bandettini Roberto, Caligiuri Patrizia, De Pace Vanessa, Domnich Alexander, Orsi Andrea, Ricucci Valentina, Icardi Giancarlo |
| EPI_ISL_2534064, EPI_ISL_2534065, EPI_ISL_2534066 | S.C. Patologia Clinica, Ospedale Sant'Andrea, ASL 5 | U.O. Igiene, Ospedale Policlinico San Martino | Bruzzone Bianca, Battolla Enrico, Caligiuri Patrizia, De Pace Vanessa, Domnich Alexander, Orsi Andrea, Ricucci Valentina, Icardi Giancarlo |
| EPI_ISL_2534067, EPI_ISL_2534068 | U.O. Igiene, Ospedale Policlinico San Martino | U.O. Igiene, Ospedale Policlinico San Martino | Bruzzone Bianca, Caligiuri Patrizia, De Pace Vanessa, Domnich Alexander, Orsi Andrea, Ricucci Valentina, Icardi Giancarlo |
| EPI_ISL_2534069, EPI_ISL_2534070, EPI_ISL_2534071 | S.C. Laboratorio Analisi, ASL 3 Liguria | U.O. Igiene, Ospedale Policlinico San Martino | Bruzzone Bianca, Caligiuri Patrizia, De Pace Vanessa, Domnich Alexander, Orsi Andrea, Ricucci Valentina, Spitaleri Antonino, Icardi Giancarlo |
| EPI_ISL_2534072, EPI_ISL_2534073, EPI_ISL_2534074, EPI_ISL_2534075, EPI_ISL_2534076 | U.O. Igiene, Ospedale Policlinico San Martino | U.O. Igiene, Ospedale Policlinico San Martino | Bruzzone Bianca, Caligiuri Patrizia, De Pace Vanessa, Domnich Alexander, Orsi Andrea, Ricucci Valentina, Icardi Giancarlo |
| EPI_ISL_2535784, EPI_ISL_2535785, EPI_ISL_2535787, EPI_ISL_2535789, EPI_ISL_2535790, EPI_ISL_2535830, EPI_ISL_2535831, EPI_ISL_2535832 | National Influenza Centre | National Influenza Centre | William K. Ampofo, Michael Marks, Ivy A. Asante, Sharon Hsu, Benjamiin B. Lindsey, Benjamin H. Foulkes, Mildred Adusei-Poku, Linda Boatemaa, Lorreta Kwasah, Joseph Oliver-Commey, Ernest Asiedu, Franklin Asiedu-Bekoe, Gordon Awandare, Joyce Ngoi, Dennis Laryea, Mathew D. Parker, Thushan I de Silva, |
| EPI_ISL_2536155 | Department of Molecular Diagnostic Emam Ali Educational and Therapeutic Center , Alborz University of Medical Sciences, Karaj, Iran | Genetics Research Center, University of Social Welfare and Rehabilitation Sciences | Zohreh Fattahi, Marzieh Mohseni, Maryam Beheshtian, Kimia Kahrizi, Zakiye Mokhames, Alireza Soleimani , Shima Sadeghi , Saber Soltani, Razieh Dowran, Mohammad Foad Abazari, Hossein Najmabadi. |
| EPI_ISL_2536156, EPI_ISL_2536157, EPI_ISL_2536158, EPI_ISL_2536160, EPI_ISL_2536181 | Department of Molecular Diagnostic Emam Ali Educational and Therapeutic Center , Alborz University of Medical Sciences, Karaj, Iran | Genetics Research Center, University of Social Welfare and Rehabilitation Sciences | Zohreh Fattahi, Marzieh Mohseni, Maryam Beheshtian, Kimia Kahrizi, Zakiye Mokhames, Alireza Soleimani, Shima Sadeghi, Saber Soltani, Razieh Dowran, Mohammad Foad Abazari, Hossein Najmabadi. |
| EPI_ISL_2536304, EPI_ISL_2536306, EPI_ISL_2536307, EPI_ISL_2536308, EPI_ISL_2536309, EPI_ISL_2536310, EPI_ISL_2536312, EPI_ISL_2536314, EPI_ISL_2536315, EPI_ISL_2536316, EPI_ISL_2536317, EPI_ISL_2536318, EPI_ISL_2536319, EPI_ISL_2536320, EPI_ISL_2536324, EPI_ISL_2536330 | | | |
| see above | Laboratorio Central de Saude Publica do Estado da Paraiba (LACEN-PB) | Laboratory of Respiratory Viruses and Measles, Oswaldo Cruz Institute, FIOCRUZ | Paola Resende, Luciana Appolinario, Fernando Motta, Anna Carolina Paixao, Ana Carolina Mendonca, Alice Sampaio Rocha, Taina Venas, Elisa Cavalcante Pereira, Renata Serrano Lopes, Joao Felipe Bezerra, Dalane Loudal Florentino Teixeira, Marilda Siqueira on behalf of the Fiocruz COVID-19 Genomic Surveillance Network |
| EPI_ISL_2537078 | Department of Molecular Diagnostic Emam Ali Educational and Therapeutic Center , Alborz University of Medical Sciences, Karaj, Iran | Genetics Research Center, University of Social Welfare and Rehabilitation Sciences | Zohreh Fattahi, Marzieh Mohseni, Maryam Beheshtian, Kimia Kahrizi, Zakiye Mokhames, Alireza Soleimani, Shima Sadeghi, Saber Soltani, Razieh Dowran, Mohammad Foad Abazari, Hossein Najmabadi. |
| EPI_ISL_2537387 | Department of Medical Virology, School of Medicine Ahvaz Jundishapur University of Medical sciences | Genetics Research Center, University of Social Welfare and Rehabilitation Sciences | Zohreh Fattahi, Marzieh Mohseni, Maryam Beheshtian, Kimia Kahrizi, Azarakhsh Azaran, Farid yousefi, Mohsen Savaie, Mohammad Farahmand, Ahmad Tavakoli, Seyed Mohammad Jazayeri, Hossein Najmabadi. |
| EPI_ISL_2544194 | Laboratorio di Microbiologia e Virologia, Università Vita-Salute San Raffaele, Milano | Laboratorio di Microbiologia e Virologia, Università Vita-Salute San Raffaele, Milano | Nicola Clementi, Elena Criscuolo, Matteo Castelli, Roberto Ferrarese, Michela Sampaolo, Roberta Antonia Diotti, Enzo Boeri, Massimo Locatelli, Massimo Clementi, Nicasio Mancini |
| EPI_ISL_2547336 | Department of Medical Virology, School of Medicine Ahvaz Jundishapur University of Medical sciences | Genetics Research Center, University of Social Welfare and Rehabilitation Sciences | Zohreh Fattahi, Marzieh Mohseni, Maryam Beheshtian, Kimia Kahrizi, Azarakhsh Azaran, Farid yousefi, Mohsen Savaie, Mohammad Farahmand, Ahmad Tavakoli, Seyed Mohammad Jazayeri, Hossein Najmabadi. |
| EPI_ISL_2547337 | Department of Medical Virology, School of Medicine Ahvaz Jundishapur University of Medical sciences | Genetics Research Center, University of Social Welfare and Rehabilitation Sciences | Zohreh Fattahi, Marzieh Mohseni, Maryam Beheshtian, Kimia Kahrizi, Azarakhsh Azaran, Farid yousefi, Mohsen Savaie, Mohammad Farahmand, Ahmad Tavakoli, Seyed Mohammad Jazayeri, Hossein Najmbadi. |
| EPI_ISL_2547339, EPI_ISL_2547428, EPI_ISL_2547429, EPI_ISL_2547522 | Department of Medical Virology, School of Medicine Ahvaz Jundishapur University of Medical sciences | Genetics Research Center, University of Social Welfare and Rehabilitation Sciences | Zohreh Fattahi, Marzieh Mohseni, Maryam Beheshtian, Kimia Kahrizi, Azarakhsh Azaran, Farid yousefi, Mohsen Savaie, Mohammad Farahmand, Ahmad Tavakoli, Seyed Mohammad Jazayeri, Hossein Najmabadi. |
| EPI_ISL_2557250 | Provincial Public Health Reference Laboratory (PPHRL) P& SHD | Provincial Public Health Reference Laboratory (PPHRL) P& SHD | ANDLEEB HANIF |
| EPI_ISL_2557347 | Laboratorio Central de Saude Publica do Estado de Minas Gerais (LACEN/MG) | Laboratory of Respiratory Viruses and Measles, Oswaldo Cruz Institute, FIOCRUZ | Paola Resende, Luciana Appolinario, Fernando Motta, Anna Carolina Paixao, Ana Carolina Mendonca, Alice Sampaio Rocha, Taina Venas, Elisa Cavalcante Pereira, Renata Serrano Lopes, Andre Felipe Leal Bernardes, Marilda Siqueira on behalf of the Fiocruz COVID-19 Genomic Surveillance Network |
| EPI_ISL_2565861 | U.O. Igiene, Ospedale Policlinico San Martino | U.O. Igiene, Ospedale Policlinico San Martino | Bruzzone Bianca, Caligiuri Patrizia, De Pace Vanessa, Domnich Alexander, Orsi Andrea, Ricucci Valentina, Icardi Giancarlo |
| EPI_ISL_2566918 | AZ Klina | AZ Klina | Carl Vael - Lynsey Berckmans |
| EPI_ISL_2600360 | Temporary Specimen Collection Centre at the AsiaWorld-Expo | Hong Kong Department of Health | Alan K.L. Tsang, Gannon C.K. Mak, Peter K.C. Cheng, Peter C.W. Yip, Ken H.L. Ng, Edman T.K. Lam, Rickjason C.W. Chan, Dominic N.C. Tsang |
| EPI_ISL_2600361 | HOME QUARANTINE TASKFORCE | Hong Kong Department of Health | Alan K.L. Tsang, Gannon C.K. Mak, Peter K.C. Cheng, Peter C.W. Yip, Ken H.L. Ng, Edman T.K. Lam, Rickjason C.W. Chan, Dominic N.C. Tsang |
| EPI_ISL_2601578, EPI_ISL_2601579, EPI_ISL_2601580, EPI_ISL_2601581, | Laboratory of Communicable Diseases | 1. Laboratory of Communicable Diseases (Estonia); 2. Eurofins Genomics Europe Sequencing GmbH | Liidia Dotsenko et al. |

| | | | |
|---|---|---|---|
| EPI_ISL_2601582, EPI_ISL_2601584, EPI_ISL_2601585, EPI_ISL_2601586, EPI_ISL_2601587, EPI_ISL_2601588 | | | |
| EPI_ISL_2601628 | North Estonia Medical Centre | 1. Laboratory of Communicable Diseases (Estonia); 2. Eurofins Genomics Europe Sequencing GmbH | Liidia Dotsenko et al. |
| EPI_ISL_2603424 | Lboratorio Central de Saude Publica do Estado do Parana (LACEN/PR) | Laboratory of Respiratory Viruses and Measles, Oswaldo Cruz Institute, FIOCRUZ | Paola Resende, Luciana Appolinario, Fernando Motta, Anna Carolina Paixao, Ana Carolina Mendonca, Alice Sampaio Rocha, Taina Venas, Elisa Cavalcante Pereira, Renata Serrano Lopes, Irina Riediger, Marilda Siqueira on behalf of the Fiocruz COVID-19 Genomic Surveillance Network |
| EPI_ISL_2614518 | Instituto Adolfo Lutz - Regional de Campinas | Instituto Adolfo Lutz, Interdiciplinary Procedures Center, Strategic Laboratory | Claudio Tavares Sacchi, Claudia Regina Gonçalves, Erica Valessa Ramos Gomes, Karoline Rodrigues Campos, Caio Vinicius Dias Lopes, Leonardo Jose Tadeu de Araujo |
| EPI_ISL_2614557, EPI_ISL_2614558, EPI_ISL_2614559, EPI_ISL_2614560, EPI_ISL_2614561 | IAL Presidente Prudente | Instituto Adolfo Lutz, Interdiciplinary Procedures Center, Strategic Laboratory | Claudio Tavares Sacchi, Claudia Regina Gonçalves, Erica Valessa Ramos Gomes, Karoline Rodrigues Campos, Caio Vinicius Dias Lopes, Leonardo Jose Tadeu de Araujo |
| EPI_ISL_2614603, EPI_ISL_2614604, EPI_ISL_2614605, EPI_ISL_2614606, EPI_ISL_2614607, EPI_ISL_2614608, EPI_ISL_2614609, EPI_ISL_2614610 | Instituto Biologico | Instituto Adolfo Lutz, Interdiciplinary Procedures Center, Strategic Laboratory | Claudio Tavares Sacchi, Claudia Regina Gonçalves, Erica Valessa Ramos Gomes, Karoline Rodrigues Campos, Caio Vinicius Dias Lopes, Leonardo Jose Tadeu de Araujo |
| EPI_ISL_2614646 | RS Sumber Waras | Eijkman Institute for Molecular Biology, National Research and Innovation Agency | Lydia V. Panggalo, Iskandar Adnan, Sukma Oktavianthi, Lidwina Priliani, Edison Johar, Frilasita A Yudhaputri, Muhammad Rezki Rasyak, Willy Agustine, Hidayat Trimarsanto, Safarina G Malik, Khin Saw Myint, Amin Soebandrio |
| EPI_ISL_2614647 | RS Sumber Waras | Eijkman Institute for Molecular Biology, National Research and Innovation Agency | Hidayat Trimarsanto, Lydia V. Panggalo, Iskandar Adnan, Sukma Oktavianthi, Lidwina Priliani, Edison Johar, Frilasita A Yudhaputri, Muhammad Rezki Rasyak, Willy Agustine, Safarina G Malik, Khin Saw Myint, Amin Soebandrio |
| EPI_ISL_2614650 | RS Mitra Keluarga Gading Serpong | Eijkman Institute for Molecular Biology, National Research and Innovation Agency | Frilasita A Yudhaputri, Muhammad Rezki Rasyak, Willy Agustine, Hidayat Trimarsanto, Lydia V. Panggalo, Iskandar Adnan, Sukma Oktavianthi, Lidwina Priliani, Edison Johar, Safarina G Malik, Khin Saw Myint, Amin Soebandrio |
| EPI_ISL_2614651 | RS Mitra Keluarga Gading Serpong | Eijkman Institute for Molecular Biology, National Research and Innovation Agency | Edison Johar, Frilasita A Yudhaputri, Muhammad Rezki Rasyak, Willy Agustine, Hidayat Trimarsanto, Lydia V. Panggalo, Iskandar Adnan, Sukma Oktavianthi, Lidwina Priliani, Safarina G Malik, Khin Saw Myint, Amin Soebandrio |
| EPI_ISL_2614652 | RS Bhakti Mulia | Eijkman Institute for Molecular Biology, National Research and Innovation Agency | Lidwina Priliani, Edison Johar, Frilasita A Yudhaputri, Muhammad Rezki Rasyak, Willy Agustine, Hidayat Trimarsanto, Lydia V. Panggalo, Iskandar Adnan, Sukma Oktavianthi, Safarina G Malik, Khin Saw Myint, Amin Soebandrio |
| EPI_ISL_2614653 | RS Bhakti Mulia | Eijkman Institute for Molecular Biology, National Research and Innovation Agency | Sukma Oktavianthi, Lidwina Priliani, Edison Johar, Frilasita A Yudhaputri, Muhammad Rezki Rasyak, Willy Agustine, Hidayat Trimarsanto, Lydia V. Panggalo, Iskandar Adnan, Safarina G Malik, Khin Saw Myint, Amin Soebandrio |
| EPI_ISL_2614654 | RS Bhakti Mulia | Eijkman Institute for Molecular Biology, National Research and Innovation Agency | Iskandar Adnan, Sukma Oktavianthi, Lidwina Priliani, Edison Johar, Frilasita A Yudhaputri, Muhammad Rezki Rasyak, Willy Agustine, Hidayat Trimarsanto, Lydia V. Panggalo, Safarina G Malik, Khin Saw Myint, Amin Soebandrio |
| EPI_ISL_2614655 | RS Harapan Bunda | Eijkman Institute for Molecular Biology, National Research and Innovation Agency | Lydia V. Panggalo, Iskandar Adnan, Sukma Oktavianthi, Lidwina Priliani, Edison Johar, Frilasita A Yudhaputri, Muhammad Rezki Rasyak, Willy Agustine, Hidayat Trimarsanto, Safarina G Malik, Khin Saw Myint, Amin Soebandrio |
| EPI_ISL_2614656 | RS Mitra Keluarga Kelapa Gading | Eijkman Institute for Molecular Biology, National Research and Innovation Agency | Hidayat Trimarsanto, Lydia V. Panggalo, Iskandar Adnan, Sukma Oktavianthi, Lidwina Priliani, Edison Johar, Frilasita A Yudhaputri, Muhammad Rezki Rasyak, Willy Agustine, Safarina G Malik, Khin Saw Myint, Amin Soebandrio |
| EPI_ISL_2614657 | RS Mitra Keluarga Kelapa Gading | Eijkman Institute for Molecular Biology, National Research and Innovation Agency | Willy Agustine, Hidayat Trimarsanto, Lydia V. Panggalo, Iskandar Adnan, Sukma Oktavianthi, Lidwina Priliani, Edison Johar, Frilasita A Yudhaputri, Muhammad Rezki Rasyak, Safarina G Malik, Khin Saw Myint, Amin Soebandrio |
| EPI_ISL_2614658 | RSJ Dr. Soeharto Heerdjan | Eijkman Institute for Molecular Biology, National Research and Innovation Agency | Muhammad Rezki Rasyak, Willy Agustine, Hidayat Trimarsanto, Lydia V. Panggalo, Iskandar Adnan, Sukma Oktavianthi, Lidwina Priliani, Edison Johar, Frilasita A Yudhaputri, Safarina G Malik, Khin Saw Myint, Amin Soebandrio |
| EPI_ISL_2614659 | RSJ Dr. Soeharto Heerdjan | Eijkman Institute for Molecular Biology, National Research and Innovation Agency | Frilasita A Yudhaputri, Muhammad Rezki Rasyak, Willy Agustine, Hidayat Trimarsanto, Lydia V. Panggalo, Iskandar Adnan, Sukma Oktavianthi, Lidwina Priliani, Edison Johar, Safarina G Malik, Khin Saw Myint, Amin Soebandrio |
| EPI_ISL_2614660 | RS Sentra Medika Cikarang | Eijkman Institute for Molecular Biology, National Research and Innovation Agency | Edison Johar, Frilasita A Yudhaputri, Muhammad Rezki Rasyak, Willy Agustine, Hidayat Trimarsanto, Lydia V. Panggalo, Iskandar Adnan, Sukma Oktavianthi, Lidwina Priliani, Safarina G Malik, Khin Saw Myint, Amin Soebandrio |
| EPI_ISL_2614661 | RS Mitra Keluarga Gading Serpong | Eijkman Institute for Molecular Biology, National Research and Innovation Agency | Lidwina Priliani, Edison Johar, Frilasita A Yudhaputri, Muhammad Rezki Rasyak, Willy Agustine, Hidayat Trimarsanto, Lydia V. Panggalo, Iskandar Adnan, Sukma Oktavianthi, Safarina G Malik, Khin Saw Myint, Amin Soebandrio |
| EPI_ISL_2614662 | RS Mitra Keluarga Gading Serpong | Eijkman Institute for Molecular Biology, National Research and Innovation Agency | Sukma Oktavianthi, Lidwina Priliani, Edison Johar, Frilasita A Yudhaputri, Muhammad Rezki Rasyak, Willy Agustine, Hidayat Trimarsanto, Lydia V. Panggalo, Iskandar Adnan, Safarina G Malik, Khin Saw Myint, Amin Soebandrio |
| EPI_ISL_2614663 | RSJ Dr. Soeharto Heerdjan | Eijkman Institute for Molecular Biology, National Research and Innovation Agency | Iskandar Adnan, Sukma Oktavianthi, Lidwina Priliani, Edison Johar, Frilasita A Yudhaputri, Muhammad Rezki Rasyak, Willy Agustine, Hidayat Trimarsanto, Lydia V. Panggalo, Safarina G Malik, Khin Saw Myint, Amin Soebandrio |
| EPI_ISL_2614665 | RS Bhakti Mulia | Eijkman Institute for Molecular Biology, National Research and Innovation Agency | Hidayat Trimarsanto, Lydia V. Panggalo, Iskandar Adnan, Sukma Oktavianthi, Lidwina Priliani, Edison Johar, Frilasita A Yudhaputri, Muhammad Rezki Rasyak, Willy Agustine, Safarina G Malik, Khin Saw Myint, Amin Soebandrio |
| EPI_ISL_2614666 | RS Mitra Keluarga Kelapa Gading | Eijkman Institute for Molecular Biology, National Research and Innovation Agency | Willy Agustine, Hidayat Trimarsanto, Lydia V. Panggalo, Iskandar Adnan, Sukma Oktavianthi, Lidwina Priliani, Edison Johar, Frilasita A Yudhaputri, Muhammad Rezki Rasyak, Safarina G Malik, Khin Saw Myint, Amin Soebandrio |
| EPI_ISL_2614667 | RS Medirossa Cikarang | Eijkman Institute for Molecular Biology, National Research and Innovation Agency | Muhammad Rezki Rasyak, Willy Agustine, Hidayat Trimarsanto, Lydia V. Panggalo, Iskandar Adnan, Sukma Oktavianthi, Lidwina Priliani, Edison Johar, Frilasita A Yudhaputri, Safarina G Malik, Khin Saw Myint, Amin Soebandrio |
| EPI_ISL_2614668 | RS Mitra Keluarga Gading Serpong | Eijkman Institute for Molecular Biology, National Research and Innovation Agency | Frilasita A Yudhaputri, Muhammad Rezki Rasyak, Willy Agustine, Hidayat Trimarsanto, Lydia V. Panggalo, Iskandar Adnan, Sukma Oktavianthi, Lidwina Priliani, Edison Johar, Safarina G Malik, Khin Saw Myint, Amin Soebandrio |
| EPI_ISL_2614669 | RS Sumber Waras | Eijkman Institute for Molecular Biology, National Research and Innovation Agency | Edison Johar, Frilasita A Yudhaputri, Muhammad Rezki Rasyak, Willy Agustine, Hidayat Trimarsanto, Lydia V. Panggalo, Iskandar Adnan, Sukma Oktavianthi, Lidwina Priliani, Safarina G Malik, Khin Saw Myint, Amin Soebandrio |
| EPI_ISL_2614670 | RS Sumber Waras | Eijkman Institute for Molecular Biology, National Research and Innovation Agency | Lidwina Priliani, Edison Johar, Frilasita A Yudhaputri, Muhammad Rezki Rasyak, Willy Agustine, Hidayat Trimarsanto, Lydia V. Panggalo, Iskandar Adnan, Sukma Oktavianthi, Safarina G Malik, Khin Saw Myint, Amin Soebandrio |
| EPI_ISL_2614671 | RS Medirossa Cikarang | Eijkman Institute for Molecular Biology, National Research and Innovation Agency | Sukma Oktavianthi, Lidwina Priliani, Edison Johar, Frilasita A Yudhaputri, Muhammad Rezki Rasyak, Willy Agustine, Hidayat Trimarsanto, Lydia V. Panggalo, Iskandar Adnan, Safarina G Malik, Khin Saw Myint, Amin Soebandrio |
| EPI_ISL_2614672 | RS Sentra Medika Cikarang | Eijkman Institute for Molecular Biology, National Research and Innovation Agency | Iskandar Adnan, Sukma Oktavianthi, Lidwina Priliani, Edison Johar, Frilasita A Yudhaputri, Muhammad Rezki Rasyak, Willy Agustine, Hidayat Trimarsanto, Lydia V. Panggalo, Safarina G Malik, Khin Saw Myint, Amin Soebandrio |
| EPI_ISL_2614673 | RS Sentra Medika Cikarang | Eijkman Institute for Molecular Biology, National Research and Innovation Agency | Lydia V. Panggalo, Iskandar Adnan, Sukma Oktavianthi, Lidwina Priliani, Edison Johar, Frilasita A Yudhaputri, Muhammad Rezki Rasyak, Willy Agustine, Hidayat Trimarsanto, Safarina G Malik, Khin Saw Myint, Amin Soebandrio |
| EPI_ISL_2614674 | RS Sentra Medika Cikarang | Eijkman Institute for Molecular Biology, National Research and Innovation Agency | Hidayat Trimarsanto, Lydia V. Panggalo, Iskandar Adnan, Sukma Oktavianthi, Lidwina Priliani, Edison Johar, Frilasita A Yudhaputri, Muhammad Rezki Rasyak, Willy Agustine, Safarina G Malik, Khin Saw Myint, Amin Soebandrio |
| EPI_ISL_2614676 | RS Mitra Keluarga Kelapa Gading | Eijkman Institute for Molecular Biology, National Research and Innovation Agency | Muhammad Rezki Rasyak, Willy Agustine, Hidayat Trimarsanto, Lydia V. Panggalo, Iskandar Adnan, Sukma Oktavianthi, Lidwina Priliani, Edison Johar, Frilasita A Yudhaputri, Safarina G Malik, Khin Saw Myint, Amin Soebandrio |
| EPI_ISL_2614677 | RSJ Dr. Soeharto Heerdjan | Eijkman Institute for Molecular Biology, National Research | Frilasita A Yudhaputri, Muhammad Rezki Rasyak, Willy Agustine, Hidayat Trimarsanto, Lydia V. Panggalo, Iskandar Adnan, Sukma Oktavianthi, Lidwina |

| | | | |
|---|---|---|---|
| | | and Innovation Agency | Priliani, Edison Johar, Safarina G Malik, Khin Saw Myint, Amin Soebandrio |
| EPI_ISL_2614678 | RSAB Harapan Kita | Eijkman Institute for Molecular Biology, National Research and Innovation Agency | Lidwina Priliani, Edison Johar, Frilasita A Yudhaputri, Muhammad Rezki Rasyak, Willy Agustine, Hidayat Trimarsanto, Lydia V. Panggalo, Iskandar Adnan, Sukma Oktavianthi, Safarina G Malik, Khin Saw Myint, Amin Soebandrio |
| EPI_ISL_2614679 | Klinik Lamina | Eijkman Institute for Molecular Biology, National Research and Innovation Agency | Sukma Oktavianthi, Lidwina Priliani, Edison Johar, Frilasita A Yudhaputri, Muhammad Rezki Rasyak, Willy Agustine, Hidayat Trimarsanto, Lydia V. Panggalo, Iskandar Adnan, Safarina G Malik, Khin Saw Myint, Amin Soebandrio |
| EPI_ISL_2614680 | RSIA Selasih Medika | Eijkman Institute for Molecular Biology, National Research and Innovation Agency | Iskandar Adnan, Sukma Oktavianthi, Lidwina Priliani, Edison Johar, Frilasita A Yudhaputri, Muhammad Rezki Rasyak, Willy Agustine, Hidayat Trimarsanto, Lydia V. Panggalo, Safarina G Malik, Khin Saw Myint, Amin Soebandrio |
| EPI_ISL_2614681 | RS Permata Cibubur | Eijkman Institute for Molecular Biology, National Research and Innovation Agency | Lydia V. Panggalo, Iskandar Adnan, Sukma Oktavianthi, Lidwina Priliani, Edison Johar, Frilasita A Yudhaputri, Muhammad Rezki Rasyak, Willy Agustine, Hidayat Trimarsanto, Safarina G Malik, Khin Saw Myint, Amin Soebandrio |
| EPI_ISL_2614682 | RS Permata Cibubur | Eijkman Institute for Molecular Biology, National Research and Innovation Agency | Hidayat Trimarsanto, Lydia V. Panggalo, Iskandar Adnan, Sukma Oktavianthi, Lidwina Priliani, Edison Johar, Frilasita A Yudhaputri, Muhammad Rezki Rasyak, Willy Agustine, Safarina G Malik, Khin Saw Myint, Amin Soebandrio |
| EPI_ISL_2614683 | RS Sentra Medika Cikarang | Eijkman Institute for Molecular Biology, National Research and Innovation Agency | Willy Agustine, Hidayat Trimarsanto, Lydia V. Panggalo, Iskandar Adnan, Sukma Oktavianthi, Lidwina Priliani, Edison Johar, Frilasita A Yudhaputri, Muhammad Rezki Rasyak, Safarina G Malik, Khin Saw Myint, Amin Soebandrio |
| EPI_ISL_2614684 | RS Sentra Medika Cikarang | Eijkman Institute for Molecular Biology, National Research and Innovation Agency | Muhammad Rezki Rasyak, Willy Agustine, Hidayat Trimarsanto, Lydia V. Panggalo, Iskandar Adnan, Sukma Oktavianthi, Lidwina Priliani, Edison Johar, Frilasita A Yudhaputri, Safarina G Malik, Khin Saw Myint, Amin Soebandrio |
| EPI_ISL_2614686 | RS Sentra Medika Cikarang | Eijkman Institute for Molecular Biology, National Research and Innovation Agency | Edison Johar, Frilasita A Yudhaputri, Muhammad Rezki Rasyak, Willy Agustine, Hidayat Trimarsanto, Lydia V. Panggalo, Iskandar Adnan, Sukma Oktavianthi, Lidwina Priliani, Safarina G Malik, Khin Saw Myint, Amin Soebandrio |
| EPI_ISL_2614687 | RS Sentra Medika Cikarang | Eijkman Institute for Molecular Biology, National Research and Innovation Agency | Lidwina Priliani, Edison Johar, Frilasita A Yudhaputri, Muhammad Rezki Rasyak, Willy Agustine, Hidayat Trimarsanto, Lydia V. Panggalo, Iskandar Adnan, Sukma Oktavianthi, Safarina G Malik, Khin Saw Myint, Amin Soebandrio |
| EPI_ISL_2614688 | RS Medirossa Cikarang | Eijkman Institute for Molecular Biology, National Research and Innovation Agency | Sukma Oktavianthi, Lidwina Priliani, Edison Johar, Frilasita A Yudhaputri, Muhammad Rezki Rasyak, Willy Agustine, Hidayat Trimarsanto, Lydia V. Panggalo, Iskandar Adnan, Safarina G Malik, Khin Saw Myint, Amin Soebandrio |
| EPI_ISL_2614689 | RS Medirossa Cikarang | Eijkman Institute for Molecular Biology, National Research and Innovation Agency | Iskandar Adnan, Sukma Oktavianthi, Lidwina Priliani, Edison Johar, Frilasita A Yudhaputri, Muhammad Rezki Rasyak, Willy Agustine, Hidayat Trimarsanto, Lydia V. Panggalo, Safarina G Malik, Khin Saw Myint, Amin Soebandrio |
| EPI_ISL_2614690 | RS Harapan Bunda | Eijkman Institute for Molecular Biology, National Research and Innovation Agency | Lydia V. Panggalo, Iskandar Adnan, Sukma Oktavianthi, Lidwina Priliani, Edison Johar, Frilasita A Yudhaputri, Muhammad Rezki Rasyak, Willy Agustine, Hidayat Trimarsanto, Safarina G Malik, Khin Saw Myint, Amin Soebandrio |
| EPI_ISL_2614691 | RSJ Dr. Soeharto Heerdjan | Eijkman Institute for Molecular Biology, National Research and Innovation Agency | Hidayat Trimarsanto, Lydia V. Panggalo, Iskandar Adnan, Sukma Oktavianthi, Lidwina Priliani, Edison Johar, Frilasita A Yudhaputri, Muhammad Rezki Rasyak, Willy Agustine, Safarina G Malik, Khin Saw Myint, Amin Soebandrio |
| EPI_ISL_2614692 | RSU HGA Depok | Eijkman Institute for Molecular Biology, National Research and Innovation Agency | Willy Agustine, Hidayat Trimarsanto, Lydia V. Panggalo, Iskandar Adnan, Sukma Oktavianthi, Lidwina Priliani, Edison Johar, Frilasita A Yudhaputri, Muhammad Rezki Rasyak, Safarina G Malik, Khin Saw Myint, Amin Soebandrio |
| EPI_ISL_2614693 | RS Mary Cileungsi Hijau | Eijkman Institute for Molecular Biology, National Research and Innovation Agency | Muhammad Rezki Rasyak, Willy Agustine, Hidayat Trimarsanto, Lydia V. Panggalo, Iskandar Adnan, Sukma Oktavianthi, Lidwina Priliani, Edison Johar, Frilasita A Yudhaputri, Safarina G Malik, Khin Saw Myint, Amin Soebandrio |
| EPI_ISL_2614694 | RS Permata Cibubur | Eijkman Institute for Molecular Biology, National Research and Innovation Agency | Frilasita A Yudhaputri, Muhammad Rezki Rasyak, Willy Agustine, Hidayat Trimarsanto, Lydia V. Panggalo, Iskandar Adnan, Sukma Oktavianthi, Lidwina Priliani, Edison Johar, Safarina G Malik, Khin Saw Myint, Amin Soebandrio |
| EPI_ISL_2614695 | RS Sentra Medika Cikarang | Eijkman Institute for Molecular Biology, National Research and Innovation Agency | Edison Johar, Frilasita A Yudhaputri, Muhammad Rezki Rasyak, Willy Agustine, Hidayat Trimarsanto, Lydia V. Panggalo, Iskandar Adnan, Sukma Oktavianthi, Lidwina Priliani, Safarina G Malik, Khin Saw Myint, Amin Soebandrio |
| EPI_ISL_2614696 | RS Sentra Medika Cikarang | Eijkman Institute for Molecular Biology, National Research and Innovation Agency | Lidwina Priliani, Edison Johar, Frilasita A Yudhaputri, Muhammad Rezki Rasyak, Willy Agustine, Hidayat Trimarsanto, Lydia V. Panggalo, Iskandar Adnan, Sukma Oktavianthi, Safarina G Malik, Khin Saw Myint, Amin Soebandrio |
| EPI_ISL_2614697 | RS Harapan Bunda | Eijkman Institute for Molecular Biology, National Research and Innovation Agency | Sukma Oktavianthi, Lidwina Priliani, Edison Johar, Frilasita A Yudhaputri, Muhammad Rezki Rasyak, Willy Agustine, Hidayat Trimarsanto, Lydia V. Panggalo, Iskandar Adnan, Safarina G Malik, Khin Saw Myint, Amin Soebandrio |
| EPI_ISL_2614698 | RS Harapan Bunda | Eijkman Institute for Molecular Biology, National Research and Innovation Agency | Iskandar Adnan, Sukma Oktavianthi, Lidwina Priliani, Edison Johar, Frilasita A Yudhaputri, Muhammad Rezki Rasyak, Willy Agustine, Hidayat Trimarsanto, Lydia V. Panggalo, Safarina G Malik, Khin Saw Myint, Amin Soebandrio |
| EPI_ISL_2614700 | RSU HGA Depok | Eijkman Institute for Molecular Biology, National Research and Innovation Agency | Hidayat Trimarsanto, Lydia V. Panggalo, Iskandar Adnan, Sukma Oktavianthi, Lidwina Priliani, Edison Johar, Frilasita A Yudhaputri, Muhammad Rezki Rasyak, Willy Agustine, Safarina G Malik, Khin Saw Myint, Amin Soebandrio |
| EPI_ISL_2614701 | RS Mitra Keluarga Gading Serpong | Eijkman Institute for Molecular Biology, National Research and Innovation Agency | Willy Agustine, Hidayat Trimarsanto, Lydia V. Panggalo, Iskandar Adnan, Sukma Oktavianthi, Lidwina Priliani, Edison Johar, Frilasita A Yudhaputri, Muhammad Rezki Rasyak, Safarina G Malik, Khin Saw Myint, Amin Soebandrio |
| EPI_ISL_2614702 | RS Mitra Keluarga Gading Serpong | Eijkman Institute for Molecular Biology, National Research and Innovation Agency | Muhammad Rezki Rasyak, Willy Agustine, Hidayat Trimarsanto, Lydia V. Panggalo, Iskandar Adnan, Sukma Oktavianthi, Lidwina Priliani, Edison Johar, Frilasita A Yudhaputri, Safarina G Malik, Khin Saw Myint, Amin Soebandrio |
| EPI_ISL_2614703 | RS Mitra Keluarga Gading Serpong | Eijkman Institute for Molecular Biology, National Research and Innovation Agency | Frilasita A Yudhaputri, Muhammad Rezki Rasyak, Willy Agustine, Hidayat Trimarsanto, Lydia V. Panggalo, Iskandar Adnan, Sukma Oktavianthi, Lidwina Priliani, Edison Johar, Safarina G Malik, Khin Saw Myint, Amin Soebandrio |
| EPI_ISL_2614705 | RSJ Dr. Soeharto Heerdjan | Eijkman Institute for Molecular Biology, National Research and Innovation Agency | Lidwina Priliani, Edison Johar, Frilasita A Yudhaputri, Muhammad Rezki Rasyak, Willy Agustine, Hidayat Trimarsanto, Lydia V. Panggalo, Iskandar Adnan, Sukma Oktavianthi, Safarina G Malik, Khin Saw Myint, Amin Soebandrio |
| EPI_ISL_2614707 | RS Sentra Medika Cikarang | Eijkman Institute for Molecular Biology, National Research and Innovation Agency | Iskandar Adnan, Sukma Oktavianthi, Lidwina Priliani, Edison Johar, Frilasita A Yudhaputri, Muhammad Rezki Rasyak, Willy Agustine, Hidayat Trimarsanto, Lydia V. Panggalo, Safarina G Malik, Khin Saw Myint, Amin Soebandrio |
| EPI_ISL_2614708 | RS Bhakti Husada Cikarang | Eijkman Institute for Molecular Biology, National Research and Innovation Agency | Lydia V. Panggalo, Iskandar Adnan, Sukma Oktavianthi, Lidwina Priliani, Edison Johar, Frilasita A Yudhaputri, Muhammad Rezki Rasyak, Willy Agustine, Hidayat Trimarsanto, Safarina G Malik, Khin Saw Myint, Amin Soebandrio |
| EPI_ISL_2614709 | RSJ Dr. Soeharto Heerdjan | Eijkman Institute for Molecular Biology, National Research and Innovation Agency | Hidayat Trimarsanto, Lydia V. Panggalo, Iskandar Adnan, Sukma Oktavianthi, Lidwina Priliani, Edison Johar, Frilasita A Yudhaputri, Muhammad Rezki Rasyak, Willy Agustine, Safarina G Malik, Khin Saw Myint, Amin Soebandrio |
| EPI_ISL_2614710 | RS Mitra Husada Tangerang | Eijkman Institute for Molecular Biology, National Research and Innovation Agency | Willy Agustine, Hidayat Trimarsanto, Lydia V. Panggalo, Iskandar Adnan, Sukma Oktavianthi, Lidwina Priliani, Edison Johar, Frilasita A Yudhaputri, Muhammad Rezki Rasyak, Safarina G Malik, Khin Saw Myint, Amin Soebandrio |
| EPI_ISL_2614711 | RS Medirossa Cikarang | Eijkman Institute for Molecular Biology, National Research and Innovation Agency | Muhammad Rezki Rasyak, Willy Agustine, Hidayat Trimarsanto, Lydia V. Panggalo, Iskandar Adnan, Sukma Oktavianthi, Lidwina Priliani, Edison Johar, Frilasita A Yudhaputri, Safarina G Malik, Khin Saw Myint, Amin Soebandrio |
| EPI_ISL_2614712 | RS Mitra Keluarga Gading Serpong | Eijkman Institute for Molecular Biology, National Research and Innovation Agency | Frilasita A Yudhaputri, Muhammad Rezki Rasyak, Willy Agustine, Hidayat Trimarsanto, Lydia V. Panggalo, Iskandar Adnan, Sukma Oktavianthi, Lidwina Priliani, Edison Johar, Safarina G Malik, Khin Saw Myint, Amin Soebandrio |
| EPI_ISL_2614714 | RS Harapan Bunda | Eijkman Institute for Molecular Biology, National Research and Innovation Agency | Iskandar Adnan, Sukma Oktavianthi, Lidwina Priliani, Edison Johar, Frilasita A Yudhaputri, Muhammad Rezki Rasyak, Willy Agustine, Hidayat Trimarsanto, Lydia V. Panggalo, Safarina G Malik, Khin Saw Myint, Amin Soebandrio |
| EPI_ISL_2614715 | RS Sumber Waras | Eijkman Institute for Molecular Biology, National Research and Innovation Agency | Sukma Oktavianthi, Lidwina Priliani, Edison Johar, Frilasita A Yudhaputri, Muhammad Rezki Rasyak, Willy Agustine, Hidayat Trimarsanto, Lydia V. Panggalo, Iskandar Adnan, Safarina G Malik, Khin Saw Myint, Amin Soebandrio |
| EPI_ISL_2614716 | RSIA Bunda Sejati | Eijkman Institute for Molecular Biology, National Research and Innovation Agency | Muhammad Rezki Rasyak, Willy Agustine, Hidayat Trimarsanto, Lydia V. Panggalo, Iskandar Adnan, Sukma Oktavianthi, Lidwina Priliani, Edison Johar, Frilasita A Yudhaputri, Safarina G Malik, Khin Saw Myint, Amin Soebandrio |
| EPI_ISL_2614717 | RS Mitra Keluarga Gading Serpong | Eijkman Institute for Molecular Biology, National Research | Lydia V. Panggalo, Iskandar Adnan, Sukma Oktavianthi, Lidwina Priliani, Edison Johar, Frilasita A Yudhaputri, Muhammad Rezki Rasyak, Willy Agustine, |

| | | | |
|---|---|---|---|
| | | and Innovation Agency | Hidayat Trimarsanto, Safarina G Malik, Khin Saw Myint, Amin Soebandrio |
| EPI_ISL_2614718 | RS Mitra Keluarga Gading Serpong | Eijkman Institute for Molecular Biology, National Research and Innovation Agency | Edison Johar, Frilasita A Yudhaputri, Muhammad Rezki Rasyak, Willy Agustine, Hidayat Trimarsanto, Lydia V. Panggalo, Iskandar Adnan, Sukma Oktavianthi, Lidwina Priliani, Safarina G Malik, Khin Saw Myint, Amin Soebandrio |
| EPI_ISL_2614719 | RS OMNI Pulomas | Eijkman Institute for Molecular Biology, National Research and Innovation Agency | Frilasita A Yudhaputri, Muhammad Rezki Rasyak, Willy Agustine, Hidayat Trimarsanto, Lydia V. Panggalo, Iskandar Adnan, Sukma Oktavianthi, Lidwina Priliani, Edison Johar, Safarina G Malik, Khin Saw Myint, Amin Soebandrio |
| EPI_ISL_2614720 | RS Sentra Medika Cikarang | Eijkman Institute for Molecular Biology, National Research and Innovation Agency | Willy Agustine, Hidayat Trimarsanto, Lydia V. Panggalo, Iskandar Adnan, Sukma Oktavianthi, Lidwina Priliani, Edison Johar, Frilasita A Yudhaputri, Muhammad Rezki Rasyak, Safarina G Malik, Khin Saw Myint, Amin Soebandrio |
| EPI_ISL_2615400 | IAL Presidente Prudente | Instituto Adolfo Lutz, Interdiciplinary Procedures Center, Strategic Laboratory | Claudio Tavares Sacchi, Claudia Regina Gonçalves, Erica Valessa Ramos Gomes, Karoline Rodrigues Campos, Caio Vinicius Dias Lopes, Leonardo Jose Tadeu de Araujo |
| EPI_ISL_2617054 | Africa_CDC - Malawi P6 S2 | CERI, Centre for Epidemic Response and Innvoation, Stellenbosch University and KRISP, KZN Research Innovation and Sequencing Platform, UKZN. | Mvula B, Chilima B, Chiwaula M, Mwangomba W, Panja L, Kasambara W, Auld A, Kim L, Kampira E, Kaba M, Wadonda N, Maida A, Giandhari J, Pillay S, Naidoo Y, Lessells R, Emmanuel SJ, Tegally H, Wilkinson E, de Oliveira T |
| EPI_ISL_2617084, EPI_ISL_2617085 | Africa_CDC - Malawi P4 S2 | CERI, Centre for Epidemic Response and Innvoation, Stellenbosch University and KRISP, KZN Research Innovation and Sequencing Platform, UKZN. | Mvula B, Chilima B, Chiwaula M, Mwangomba W, Panja L, Kasambara W, Auld A, Kim L, Kampira E, Kaba M, Wadonda N, Maida A, Giandhari J, Pillay S, Naidoo Y, Lessells R, Emmanuel SJ, Tegally H, Wilkinson E, de Oliveira T |
| EPI_ISL_2617087 | Africa_CDC - Malawi P6 S2 | CERI, Centre for Epidemic Response and Innvoation, Stellenbosch University and KRISP, KZN Research Innovation and Sequencing Platform, UKZN. | Mvula B, Chilima B, Chiwaula M, Mwangomba W, Panja L, Kasambara W, Auld A, Kim L, Kampira E, Kaba M, Wadonda N, Maida A, Giandhari J, Pillay S, Naidoo Y, Lessells R, Emmanuel SJ, Tegally H, Wilkinson E, de Oliveira T |
| EPI_ISL_2627737 | Belo Horizonte center-south emergency care unit - UPA-BH | Laboratório de Virologia Clínica e Molecular | Erick Gustavo Dorlass, Karine Lima Lourenço, Rubens Daniel Miserani Magalhães, Hugo Sato, Alex Fiorini, Renata Peixoto, Helena Perez Coelho, Ana Paula Salles Fernandes, Bruna Larotonda Telezynski, Guilherme Pereira Scagion, Tatiana Ometto, Luciano Matsumiya Thomazelli, Danielle Bruna Leal Oliveira, Edison Luiz Durigon, Flavio Fonseca e Santuza Teixeira |
| EPI_ISL_2629074, EPI_ISL_2629080 | HEGP - Laboratoire de Virologie | HEGP - Laboratoire de Virologie | David Veyer, Sandrine Imbeaud, Julien Puech, Hélène Péré, Maxime Wack, Nicolas Robillard |
| EPI_ISL_2629165, EPI_ISL_2629171, EPI_ISL_2629182 | Turkish Society of Internal Medicine | Gen Era Diagnostics Corporation Life Science Research and Molecular Diagnostics | Serhat Ünal |
| EPI_ISL_2645516 | Laboratorio Central de Saude Publica do Estado do Espirito Santo (LACEN/ES) | Laboratory of Respiratory Viruses and Measles, Oswaldo Cruz Institute, FIOCRUZ | Paola Resende, Luciana Appolinario, Fernando Motta, Anna Carolina Paixao, Ana Carolina Mendonca, Alice Sampaio Rocha, Taina Venas, Eilisa Cavalcante Pereira, Renata Serrano Lopes, Rodrigo Ribeiro Rodrigues, Marilda Siqueira on behalf of the Fiocruz COVID-19 Genomic Surveillance Network |
| EPI_ISL_2657512 | Dr. Gernot Walder GmbH | Bergthaler laboratory, CeMM Research Center for Molecular Medicine of the Austrian Academy of Sciences | Lukas Endler, Anna Schedl, Fabian Amman, Petr Triska, Matthew Thornton, Thomas Penz, Benedikt Agerer, Maelle Le Moing, Michael Schuster, Bekir Erguner, Jan Laine, Martin Senekowitsch, Christoph Bock, Andreas Bergthaler |
| EPI_ISL_2657677 | HOSPITAL DR. ENRIQUE BALTODANO BRICEÑO | Inciensa, Instituto Costarricense de Investigación y Enseñanza en Nutrición y Salud | Francisco Duarte, Hebleen Porras, Claudio Soto-Garita, Estela Cordero, Adriana Godínez, Melany Calderón & Karol Matamoros-Montoya |
| EPI_ISL_2657678, EPI_ISL_2657679, EPI_ISL_2657680 | HOSPITAL MONSEÑOR SANABRIA | Inciensa, Instituto Costarricense de Investigación y Enseñanza en Nutrición y Salud | Francisco Duarte, Hebleen Porras, Claudio Soto-Garita, Estela Cordero, Adriana Godínez, Melany Calderón, José Luis Vargas, Mariela Gutiérrez, Joselyn Prado & Andrea Moreno-Carvajal |
| EPI_ISL_2660649 | Laboratorio Central de Saude Publica do Estado de Sergipe (LACEN/SE) | Laboratory of Respiratory Viruses and Measles, Oswaldo Cruz Institute, FIOCRUZ | Paola Resende, Luciana Appolinario, Fernando Motta, Anna Carolina Paixao, Ana Carolina Mendonca, Alice Sampaio Rocha, Tainá Moreira Martins Venas, Elisa Cavalcante Pereira, Renata Serrano Lopes, Cliomar Alves dos Santos, Marilda Siqueira on behalf of the Fiocruz COVID-19 Genomic Surveillance Network |
| EPI_ISL_2661749, EPI_ISL_2661755, EPI_ISL_2661765, EPI_ISL_2661768, EPI_ISL_2661771, EPI_ISL_2661776, EPI_ISL_2661780 | Laboratorio Central de Saude Publica do Estado do Rio Grande do Sul (LACEN-RS) | Laboratory of Respiratory Viruses and Measles, Oswaldo Cruz Institute, FIOCRUZ | Paola Resende, Luciana Appolinario, Fernando Motta, Anna Carolina Paixao, Ana Carolina Mendonca, Alice Sampaio Rocha, Taina Venas, Elisa Cavalcante Pereira, Renata Serrano Lopes, Tatiana Schaffer Gregianini, Richard Salvato, Marilda Siqueira on behalf of the Fiocruz COVID-19 Genomic Surveillance Network |
| EPI_ISL_2671894, EPI_ISL_2671899 | Juntendo University Hospital | Juntendo University Hospital | Koji Tuchiya, Yoshie Hosaka, Yoko Tabe |
| EPI_ISL_2677149, EPI_ISL_2677162, EPI_ISL_2677219 | Labortorio Central de Saude Publica do Estado de Santa Catarina (LACEN/SC) | Laboratory of Respiratory Viruses and Measles, Oswaldo Cruz Institute, FIOCRUZ | Paola Resende, Luciana Appolinario, Fernando Motta, Anna Carolina Paixao, Ana Carolina Mendonca, Alice Sampaio Rocha, Taina Venas, Elisa Cavalcante Pereira, Renata Serrano Lopes, Darcita Buerger Rovaris, Sandra Bianchini Fernandes, Marilda Siqueira on behalf of the Fiocruz COVID-19 Genomic Surveillance Network |
| EPI_ISL_2690193 | Ochsner Health | Bioinfoexperts, LLC | Judy Crabtree, Amy Feehan, Rebecca Rose, David J. Nolan, Tessa LaFleur, Julia-Garcia-Diaz, Lucio Miele, Susanna L. Lamers |
| EPI_ISL_2690303, EPI_ISL_2690662, EPI_ISL_2691057, EPI_ISL_2691058, EPI_ISL_2691060, EPI_ISL_2691061, EPI_ISL_2691063, EPI_ISL_2691066, EPI_ISL_2691067, EPI_ISL_2691072 | MSK Microbiology Lab | MSK Microbiology Lab | Esther Babady, Tracy McMillen, Krupa Jani |
| EPI_ISL_2691096, EPI_ISL_2691097, EPI_ISL_2691098, EPI_ISL_2691099, EPI_ISL_2691100, EPI_ISL_2691101 | Instituto Adolfo Lutz - Regional de Marilia | Instituto Adolfo Lutz, Interdiciplinary Procedures Center, Strategic Laboratory | Claudio Tavares Sacchi, Claudia Regina Gonçalves, Erica Valessa Ramos Gomes, Karoline Rodrigues Campos, Caio Vinicius Dias Lopes, Leonardo Jose Tadeu de Araujo |
| EPI_ISL_2692206, EPI_ISL_2692531, EPI_ISL_2692706, EPI_ISL_2692880, EPI_ISL_2692882, EPI_ISL_2692896, EPI_ISL_2692897, EPI_ISL_2692899, EPI_ISL_2692902, EPI_ISL_2692903 | Clinical Microbiology Lab RSPTN Universitas Hasanuddin | Clinical Microbiology Lab RSPTN Universitas Hasanuddin | Muhammad Nasrum Massi, Handayani Halik, Rizalinda Sjahril, Gita Vita Soraya, Triwibawa, Muhammad Yusuf Usman, Rufika Shari |
| EPI_ISL_2713145, EPI_ISL_2713146, EPI_ISL_2713147, EPI_ISL_2713148, EPI_ISL_2713149, EPI_ISL_2713150, EPI_ISL_2713166, EPI_ISL_2713167, EPI_ISL_2713168, EPI_ISL_2713169, EPI_ISL_2713194, EPI_ISL_2713195, EPI_ISL_2713196, EPI_ISL_2713197, EPI_ISL_2713198, EPI_ISL_2713199, EPI_ISL_2713200, EPI_ISL_2713201, EPI_ISL_2713202, EPI_ISL_2713203, EPI_ISL_2713204, EPI_ISL_2713205, EPI_ISL_2713206, EPI_ISL_2713207, EPI_ISL_2713208, EPI_ISL_2713209, EPI_ISL_2713210, EPI_ISL_2713212, EPI_ISL_2713221, EPI_ISL_2713222, EPI_ISL_2713223, EPI_ISL_2713225, EPI_ISL_2713226 | | | |
| see above | Department of Health Technology and Informatics, The Hong Kong Polytechnic University | Department of Health Technology and Informatics, The Hong Kong Polytechnic University | Gilman Kit-Hang Siu, Lam-Kwong Lee, Kenneth Siu-Sing Leung, Jake Siu-Lun Leung, Wing-Hei Lo, Timothy Ting-Leung Ng, Chloe Toi-Mei Chan, Hoi-Ching Jim, Ivan Tak-Fai Wong, Kingsley King-Gee Tam, Hiu-Yin Lao, Denise Sze-Hang Wong, Alan Ka-Lun Wu, Miranda Chong-Yee Yau, Yvette Wai-Man Lai, Kitty Sau-Chun Fung, Sandy Ka-Yee Chau, Barry Kin-Chung Wong, Wing-Kin To, Kristine Luk, Alex Yat-Man Ho, Tak-Lun Que, Kam-Tong Yip, Wing Cheong Yam, David Ho-Keung Shum, Shea Ping Yip |
| EPI_ISL_2727288, EPI_ISL_2727289 | ZARV/NHLS, Department Medical Virology, University of Pretoria | KRISP, KZN Research Innovation and Sequencing Platform | Amy Strydom, Adriano Mendes, Micheala Davids, Sim Mayaphi and Marietjie Venter, Giandhari J, Pillay S, Naidoo Y, Ramphal U, Lessells R, Emmanuel SJ, Tegally H, Wilkinson E, de Oliveira T |
| EPI_ISL_2728531, EPI_ISL_2728532, EPI_ISL_2728534, EPI_ISL_2728563 | Laboratorio de Infectologia y Virologia Molecular | Laboratory of Molecular Virology, School of Medicine, Pontificia Universidad Catolica de Chile | Catalina Pardo-Roa, Constanza Martinez-Valdevenito, Jennifer Angulo, Tamara Garcia-Salum, Estefany Poblete, Maria Jose Avendano, Leonardo I. Almonacid, Ana Maria Contreras, Carlos Palma, Jorge Levican, Eileen Serrano, Constanza Maldonado, M. Belen Leyton, Erick Salinas, Andres E. Munoz-Marcos, Francisco Melo, Marcela Ferres, Alejandro Bhrun, Magdalena Vera, Rafael A. Medina |
| EPI_ISL_2728590, EPI_ISL_2728593, EPI_ISL_2728595, EPI_ISL_2728598, EPI_ISL_2728599, EPI_ISL_2728600, EPI_ISL_2728604, EPI_ISL_2728605 | Laboratorio de Infectologia y Virologia Molecular | Laboratory of Molecular Virology, School of Medicine, Pontificia Universidad Catolica de Chile | Catalina Pardo-Roa, Constanza Martinez-Valdevenito, Jennifer Angulo, Tamara Garcia-Salum, Estefany Poblete, Maria Jose Avendano, Leonardo I. Almonacid, Ana Maria Contreras, Carlos Palma, Jorge Levican, Eileen Serrano, Constanza Maldonado, M. Belen Leyton, Erick Salinas, Andres E. Munoz-Marcos, Francisco Melo, Marcela Ferres, Rafael A. Medina |

| | | | |
|---|---|---|---|
| EPI_ISL_2756436, EPI_ISL_2756440, EPI_ISL_2756442, EPI_ISL_2756443, EPI_ISL_2756458, EPI_ISL_2756467, EPI_ISL_2756468, EPI_ISL_2756474, EPI_ISL_2756475, EPI_ISL_2756477, EPI_ISL_2756478, EPI_ISL_2756479, EPI_ISL_2756482, EPI_ISL_2756483, EPI_ISL_2756484, EPI_ISL_2756485, EPI_ISL_2756486, EPI_ISL_2756487 | | | |
| see above | Instituto Adolfo Lutz Central | Instituto Adolfo Lutz, Interdiciplinary Procedures Center, Strategic Laboratory | Claudio Tavares Sacchi, Claudia Regina Gonçalves, Erica Valessa Ramos Gomes, Karoline Rodrigues Campos, Caio Vinicius Dias Lopes, Leonardo Jose Tadeu de Araujo |
| EPI_ISL_2761936 | LACEN do Mato Grosso do Sul | Instituto Adolfo Lutz, Interdiciplinary Procedures Center, Strategic Laboratory | Claudio Tavares Sacchi, Claudia Regina Gonçalves, Erica Valessa Ramos Gomes, Karoline Rodrigues Campos, Caio Vinicius Dias Lopes, Leonardo Jose Tadeu de Araujo |
| EPI_ISL_2779282 | Trans Nzoia hospital | USAMRD-A, Basic Science Laboratory | Gathii Kimita, Josphat Nyataya, Esther Omuseni, Faith Sigei, Alan Lemtudo, Eric Muthanje, Brian Andika, Rehema Liyai, Rachel Githii, Clement Masakwe, Stephen Ochola, George Awinda, Carol Kifude, Beth Mutai, John Waitumbi |
| EPI_ISL_2779285 | GK Prison Busia (Matayo)Matayos Prison | USAMRD-A, Basic Science Laboratory | Gathii Kimita, Josphat Nyataya, Esther Omuseni, Faith Sigei, Alan Lemtudo, Eric Muthanje, Brian Andika, Rehema Liyai, Rachel Githii, Clement Masakwe, Stephen Ochola, George Awinda, Carol Kifude, Beth Mutai, John Waitumbi |
| EPI_ISL_2779292 | Malaba border point | USAMRD-A, Basic Science Laboratory | Gathii Kimita, Josphat Nyataya, Esther Omuseni, Faith Sigei, Alan Lemtudo, Eric Muthanje, Brian Andika, Rehema Liyai, Rachel Githii, Clement Masakwe, Stephen Ochola, George Awinda, Carol Kifude, Beth Mutai, John Waitumbi |
| EPI_ISL_2779293, EPI_ISL_2779294, EPI_ISL_2779295, EPI_ISL_2779296, EPI_ISL_2779297, EPI_ISL_2779298, EPI_ISL_2779299, EPI_ISL_2779300 | Trans Nzoia hospital | USAMRD-A, Basic Science Laboratory | Gathii Kimita, Josphat Nyataya, Esther Omuseni, Faith Sigei, Alan Lemtudo, Eric Muthanje, Brian Andika, Rehema Liyai, Rachel Githii, Clement Masakwe, Stephen Ochola, George Awinda, Carol Kifude, Beth Mutai, John Waitumbi |
| EPI_ISL_2779302 | KEMRI/DEID Surveillance Site (Lamu County Hospital) | USAMRD-A, Basic Science Laboratory | Gathii Kimita, Josphat Nyataya, Esther Omuseni, Faith Sigei, Alan Lemtudo, Eric Muthanje, Brian Andika, Rehema Liyai, Rachel Githii, Clement Masakwe, Stephen Ochola, George Awinda, Carol Kifude, Beth Mutai, John Waitumbi |
| EPI_ISL_2779303, EPI_ISL_2779304, EPI_ISL_2779305, EPI_ISL_2779306, EPI_ISL_2779307 | GK Prison Busia (Matayo)Matayos Prison | USAMRD-A, Basic Science Laboratory | Gathii Kimita, Josphat Nyataya, Esther Omuseni, Faith Sigei, Alan Lemtudo, Eric Muthanje, Brian Andika, Rehema Liyai, Rachel Githii, Clement Masakwe, Stephen Ochola, George Awinda, Carol Kifude, Beth Mutai, John Waitumbi |
| EPI_ISL_2779308, EPI_ISL_2779309 | KCRH KEMRI/DEID Surveillance Site | USAMRD-A, Basic Science Laboratory | Gathii Kimita, Josphat Nyataya, Esther Omuseni, Faith Sigei, Alan Lemtudo, Eric Muthanje, Brian Andika, Rehema Liyai, Rachel Githii, Clement Masakwe, Stephen Ochola, George Awinda, Carol Kifude, Beth Mutai, John Waitumbi |
| EPI_ISL_2779313, EPI_ISL_2779314, EPI_ISL_2779315, EPI_ISL_2779316 | GK Prisons Dispensary Kapsabet | USAMRD-A, Basic Science Laboratory | Gathii Kimita, Josphat Nyataya, Esther Omuseni, Faith Sigei, Alan Lemtudo, Eric Muthanje, Brian Andika, Rehema Liyai, Rachel Githii, Clement Masakwe, Stephen Ochola, George Awinda, Carol Kifude, Beth Mutai, John Waitumbi |
| EPI_ISL_2779322 | KEMRI/DEID Surveillance Site (Isebani Sub-County Hospital) | USAMRD-A, Basic Science Laboratory | Gathii Kimita, Josphat Nyataya, Esther Omuseni, Faith Sigei, Alan Lemtudo, Eric Muthanje, Brian Andika, Rehema Liyai, Rachel Githii, Clement Masakwe, Stephen Ochola, George Awinda, Carol Kifude, Beth Mutai, John Waitumbi |
| EPI_ISL_2789819, EPI_ISL_2789837 | Centre Hospitalier Universitaire Clermont-Ferrand | CHU Clermont-Ferrand, service de virologie | Bisseux Maxime, Mirand Audrey, Combes Patricia, Henquell Cécile |
| EPI_ISL_2790909 | Hospital | National Reference Center for Viruses of Respiratory Infections, Institut Pasteur, Paris | Marion Barbet, Sylvie Behillil, Méline Bizard, Angela Brisebarre, Camille Capel, Vincent Enouf, Louise Lefrançois, Frédéric Lemoine, Christophe Malabat, Corinne Maufrais, Emmanuelle Permal, Etienne Simon-Lorière, Maud Vanpeene, Sylvie Van der Werf ,Isabelle Woercel |
| EPI_ISL_2820476, EPI_ISL_2820477, EPI_ISL_2820478 | The Hong Kong Polytechnic University - Molecular Diagnostic Laboratory | The Hong Kong Polytechnic University - Molecular Diagnostic Laboratory | Hin Fung Tsang, Allen Chi Shing Yu, Aldrin Kay Yuen Yim, Wai Ming Stanley Leung, Jiachi Chiou, Yin Kwan Evelyn Wong, Dominic Ngai Chong Tsang, Peter and SO, Samuel Chun Lap Lo,Wing Tak Wong, Sze Chuen Cesar Wong |
| EPI_ISL_2841607, EPI_ISL_2841608, EPI_ISL_2841609, EPI_ISL_2841610, EPI_ISL_2841611, EPI_ISL_2841612 | Instituto Biologico | Instituto Adolfo Lutz, Interdiciplinary Procedures Center, Strategic Laboratory | Claudio Tavares Sacchi, Claudia Regina Gonçalves, Erica Valessa Ramos Gomes, Karoline Rodrigues Campos, Caio Vinicius Dias Lopes |
| EPI_ISL_2841923, EPI_ISL_2841924, EPI_ISL_2841926, EPI_ISL_2841927, EPI_ISL_2841928, EPI_ISL_2841929, EPI_ISL_2841930, EPI_ISL_2841931, EPI_ISL_2841932, EPI_ISL_2841933, EPI_ISL_2841934, EPI_ISL_2841935, EPI_ISL_2841936, EPI_ISL_2841937, EPI_ISL_2841938, EPI_ISL_2841939, EPI_ISL_2841940, EPI_ISL_2841941 | | | |
| see above | National Center of Infectious and Parasitic Diseases | National Center of Infectious and Parasitic Diseases | Alexiev et al |
| EPI_ISL_2860789, EPI_ISL_2860790, EPI_ISL_2860791, EPI_ISL_2860792, EPI_ISL_2860793, EPI_ISL_2860794, EPI_ISL_2860795, EPI_ISL_2860797, EPI_ISL_2860798, EPI_ISL_2860799, EPI_ISL_2860801, EPI_ISL_2860811, EPI_ISL_2860813, EPI_ISL_2860814, EPI_ISL_2860815, EPI_ISL_2860816, EPI_ISL_2860817, EPI_ISL_2860818 | | | |
| see above | Department of Health Technology and Informatics, The Hong Kong Polytechnic University | Department of Health Technology and Informatics, The Hong Kong Polytechnic University | Gilman Kit-Hang Siu, Lam-Kwong Lee, Kenneth Siu-Sing Leung, Jake Siu-Lun Leung, Wing-Hei Lo, Timothy Ting-Leung Ng, Chloe Toi-Mei Chan, Hoi-Ching Jim, Ivan Tak-Fai Wong, Kingsley King-Gee Tam, Hiu-Yin Lao, Denise Sze-Hang Wong, Alan Ka-Lun Wu, Miranda Chong-Yee Yau, Yvette Wai-Man Lai, Kitty Sau-Chun Fung, Sandy Ka-Yee Chau, Barry Kin-Chung Wong, Wing-Kin To, Kristine Luk, Alex Yat-Man Ho, Tak-Lun Que, Kam-Tong Yip, Wing Cheong Yam, David Ho-Keung Shum, Shea Ping Yip |
| EPI_ISL_2863796 | Laboratorio Central de Saude Publica do Estado do Mato Grosso do Sul (LACEN/MS) | Laboratory of Respiratory Viruses and Measles, Oswaldo Cruz Institute, FIOCRUZ | Paola Resende, Luciana Appolinario, Fernando Motta, Anna Carolina Paixao, Ana Carolina Mendonca, Alice Sampaio Rocha, Taina Venas, Elisa Cavalcante Pereira, Renata Serrano Lopes, Alex Pauvolid-Corrêa, Mia Ferreira de Araujo, Ana Beatriz Machado Lima, Natália Valente da Silva, Thayssa Keren Santos da Silva Neves, Agatha Cristinne Prudencio Soares, Ighor Leonardo Arantes Gomes, Gislene Lichs, Marilda Siqueira on behalf of the Fiocruz COVID-19 Genomic Surveillance Network |
| EPI_ISL_2886170 | Tanjungpura University Hospital | Tanjungpura University Hospital | Andriani; Mahyarudin; Virhan Novianry; Delima Fajar Liana; Sofi Siti Shofiyah; Puji Astuti; Hana Apsari Pawestri; Arie Ardiansyah Nugraha; Kartika Dewi Puspa; Hartanti Dian Ikawati; Vivi Setiawaty |
| EPI_ISL_2886171 | Tanjungpura University Hospital | Tanjungpura University Hospital | Mahyarudin; Andriani; Virhan Novianry; Delima Fajar Liana; Sofi Siti Shofiyah; Puji Astuti; Hana Apsari Pawestri; Arie Ardiansyah Nugraha; Kartika Dewi Puspa; Hartanti Dian Ikawati; Vivi Setiawaty |
| EPI_ISL_2886174 | Tanjungpura University Hospital | Tanjungpura University Hospital | Delima Fajar Liana; Andriani; Mahyarudin; Virhan Novianry; Sofi Siti Shofiyah; Puji Astuti; Hana Apsari Pawestri; Arie Ardiansyah Nugraha; Kartika Dewi Puspa; Hartanti Dian Ikawati; Vivi Setiawaty |
| EPI_ISL_2886177 | Tanjungpura University Hospital | Tanjungpura University Hospital | Virhan Novianry; Andriani; Mahyarudin; Delima Fajar Liana; Sofi Siti Shofiyah; Puji Astuti; Hana Apsari Pawestri; Arie Ardiansyah Nugraha; Kartika Dewi Puspa; Hartanti Dian Ikawati; Vivi Setiawaty |
| EPI_ISL_2886178 | Tanjungpura University Hospital | Tanjungpura University Hospital | Sofi Siti Shofiyah; Andriani; Mahyarudin; Virhan Novianry; Delima Fajar Liana; Puji Astuti; Hana Apsari Pawestri; Arie Ardiansyah Nugraha; Kartika Dewi Puspa; Hartanti Dian Ikawati; Vivi Setiawaty |
| EPI_ISL_2886179 | Tanjungpura University Hospital | Tanjungpura University Hospital | Puji Astuti; Andriani; Mahyarudin; Virhan Novianry; Delima Fajar Liana; Sofi Siti Shofiyah; Hana Apsari Pawestri; Arie Ardiansyah Nugraha; Kartika Dewi Puspa; Hartanti Dian Ikawati; Vivi Setiawaty |

We gratefully acknowledge the following Authors from the Originating laboratories responsible for obtaining the specimens, as well as the Submitting laboratories where the genome data were generated and shared via GISAID, on which this research is based.

All Submitters of data may be contacted directly via www.gisaid.org

Authors are sorted alphabetically.

| Accession ID | Originating Laboratory | Submitting Laboratory | Authors |
|---|---|---|---|
| EPI_ISL_1301852 | Hopital | National Reference Center for Viruses of Respiratory Infections, Institut Pasteur, Paris | Marion Barbet, Sylvie Behillil, Méline Bizard, Angela Brisebarre, Camille Capel, Louise Lefrançois, Etienne Simon-Lorière, Vincent Enouf, Maud Vanpeene, Sylvie van der Werf,Griscelli Franck |
| EPI_ISL_1305774 | QUALITY CONTROL CHEMICAL BIOLOGICAL RISK, AOOR Villa Sofia Cervello Palermo | QUALITY CONTROL CHEMICAL BIOLOGICAL RISK, AOOR Villa Sofia Cervello Palermo | Di Gaudio,F., Seidita,G., Contino,F., and Brunacci,G. |
| EPI_ISL_1310791 | QUALITY CONTROL CHEMICAL BIOLOGICAL RISK, AOOR Villa Sofia Cervello Palermo | QUALITY CONTROL CHEMICAL BIOLOGICAL RISK, AOOR Villa Sofia Cervello Palermo | Di Gaudio,F., Contino,F., Brunacci,G., Orlando, V., Buffa, V., Di Chiara, V. and Seidita,G. |
| EPI_ISL_1311860, EPI_ISL_1311861 | Laboratorio Biologia Molecolare Sars Cov2 - UOC Laboratorio Analisi - Servizio Medicina di Laboratorio, Ospedale "San Francesco" - ATS-ASSL Nuoro | Laboratorio specialistico UOC Ematologia - Ospedale "San Francesco" - ATS-ASSL Nuoro | Piras Giovanna, Asproni Rosanna,Fiamma Maura, Monne Maria Itria, Malune Paolo, Palmas Angelo Domenico, Lo Maglio Iana, Mameli Giuseppe |
| EPI_ISL_1311964 | QUALITY CONTROL CHEMICAL BIOLOGICAL RISK, AOOR Villa Sofia Cervello Palermo | QUALITY CONTROL CHEMICAL BIOLOGICAL RISK, AOOR Villa Sofia Cervello Palermo | Di Gaudio,F., Contino,F. and Brunacci,G. |
| EPI_ISL_1311973 | QUALITY CONTROL CHEMICAL BIOLOGICAL RISK, AOOR Villa Sofia Cervello Palermo | QUALITY CONTROL CHEMICAL BIOLOGICAL RISK, AOOR Villa Sofia Cervello Palermo | Di Gaudio,F., Brunacci, G.Seidita,G., and Contino,F. |
| EPI_ISL_1312063 | QUALITY CONTROL CHEMICAL BIOLOGICAL RISK, AOOR Villa Sofia Cervello Palermo | QUALITY CONTROL CHEMICAL BIOLOGICAL RISK, AOOR Villa Sofia Cervello Palermo | Di Gaudio,F., Seidita,G., Contino,F., Orlando, V., Di Chiara, V., Buffa, V., Todaro, I., Vasto, S., Pace, E., Cangemi, G. and Brunacci,G. |
| EPI_ISL_1312389 | QUALITY CONTROL CHEMICAL BIOLOGICAL RISK, AOOR Villa Sofia Cervello Palermo | QUALITY CONTROL CHEMICAL BIOLOGICAL RISK, AOOR Villa Sofia Cervello Palermo | Di Gaudio,F., Contino,F., Seidita,G. and Brunacci,G. |
| EPI_ISL_1320185, EPI_ISL_1320187, EPI_ISL_1320188, EPI_ISL_1320190 | IZSM | IZSM-U.O.C. Virologia | Maurizio Viscardi, Lorena Cardillo, Giovanna Fusco, Esterina De Carlo, Antonio Limone |
| EPI_ISL_1357596 | QUALITY CONTROL CHEMICAL BIOLOGICAL RISK, AOOR Villa Sofia Cervello Palermo | QUALITY CONTROL CHEMICAL BIOLOGICAL RISK, AOOR Villa Sofia Cervello Palermo | Di Gaudio,F., Contino,F., and Brunacci,G. |
| EPI_ISL_1357598 | QUALITY CONTROL CHEMICAL BIOLOGICAL RISK, AOOR Villa Sofia Cervello Palermo | QUALITY CONTROL CHEMICAL BIOLOGICAL RISK, AOOR Villa Sofia Cervello Palermo | Di Gaudio, F., Contino, F., Brunacci, G., Ciaccio, M. |
| EPI_ISL_1357600 | QUALITY CONTROL CHEMICAL BIOLOGICAL RISK, AOOR Villa Sofia Cervello Palermo | QUALITY CONTROL CHEMICAL BIOLOGICAL RISK, AOOR Villa Sofia Cervello Palermo | Di Gaudio, F., Contino, F., Orlando, V., Buffa, V., Bonvissuto, M., Di Chiara, V., Todaro, I., Seidita, G., Brunacci, G. |
| EPI_ISL_1357602 | QUALITY CONTROL CHEMICAL BIOLOGICAL RISK, AOOR Villa Sofia Cervello Palermo | QUALITY CONTROL CHEMICAL BIOLOGICAL RISK, AOOR Villa Sofia Cervello Palermo | Di Gaudio, F., Contino, F., Gallo, A., Cuscino, N., Brunacci, G |
| EPI_ISL_1357606 | QUALITY CONTROL CHEMICAL BIOLOGICAL RISK, AOOR Villa Sofia Cervello Palermo | QUALITY CONTROL CHEMICAL BIOLOGICAL RISK, AOOR Villa Sofia Cervello Palermo | Di Gaudio, F., Contino, F., Brunacci, G. |
| EPI_ISL_1357607 | QUALITY CONTROL CHEMICAL BIOLOGICAL RISK, AOOR Villa Sofia Cervello Palermo | QUALITY CONTROL CHEMICAL BIOLOGICAL RISK, AOOR Villa Sofia Cervello Palermo | Di Gaudio, F., Brunacci, G., Gallo, A., Cuscino, N., Contino, F. |
| EPI_ISL_1357644 | QUALITY CONTROL CHEMICAL BIOLOGICAL RISK, AOOR Villa Sofia Cervello Palermo | QUALITY CONTROL CHEMICAL BIOLOGICAL RISK, AOOR Villa Sofia Cervello Palermo | Di Gaudio, F., Contino, F., Orlando, V., Buffa, V., Seidita, G., Bonvissuto, M., Di Chiara, V., Todaro, I., Brunacci, G. |
| EPI_ISL_1357645 | QUALITY CONTROL CHEMICAL BIOLOGICAL RISK, AOOR Villa Sofia Cervello Palermo | QUALITY CONTROL CHEMICAL BIOLOGICAL RISK, AOOR Villa Sofia Cervello Palermo | Di Gaudio, F., Brunacci, G., Gallo, A., Cuscino, N., Contino, F. |
| EPI_ISL_1365746 | Laboratorio di Riferimento Regionale della Sicilia Occidentale per l'Emergenza COVID-19 | Laboratorio di Riferimento Regionale della Sicilia Occidentale per l'Emergenza COVID-19 | Fabio Tramuto, Carmelo Massimo Maida, Daniela Di Naro, Giulia Randazzo, Walter Mazzucco, Giorgio Graziano, Vincenzo Restivo, Claudio Costantino, Francesco Vitale |
| EPI_ISL_1366272 | QUALITY CONTROL CHEMICAL BIOLOGICAL RISK, AOOR Villa Sofia Cervello Palermo | QUALITY CONTROL CHEMICAL BIOLOGICAL RISK, AOOR Villa Sofia Cervello Palermo | Di Gaudio, F., Contino, F., Brunacci, G. |
| EPI_ISL_1366274 | QUALITY CONTROL CHEMICAL BIOLOGICAL RISK, AOOR Villa Sofia Cervello Palermo | QUALITY CONTROL CHEMICAL BIOLOGICAL RISK, AOOR Villa Sofia Cervello Palermo | Di Gaudio, F., Contino, F., Orlando, V., Buffa, V., Seidita, G., Bonvissuto, M., Di Chiara, V., Todaro, I., Brunacci, G |
| EPI_ISL_1366497 | QUALITY CONTROL CHEMICAL BIOLOGICAL RISK, AOOR Villa Sofia Cervello Palermo | QUALITY CONTROL CHEMICAL BIOLOGICAL RISK, AOOR Villa Sofia Cervello Palermo | Di Gaudio, F., Brunacci, G., Contino, F. |
| EPI_ISL_1366506 | QUALITY CONTROL CHEMICAL BIOLOGICAL RISK, AOOR Villa Sofia Cervello Palermo | QUALITY CONTROL CHEMICAL BIOLOGICAL RISK, AOOR Villa Sofia Cervello Palermo | Di Gaudio, F., Contino, F., Brunacci, G. |
| EPI_ISL_1366612 | QUALITY CONTROL CHEMICAL BIOLOGICAL RISK, AOOR Villa Sofia Cervello Palermo | QUALITY CONTROL CHEMICAL BIOLOGICAL RISK, AOOR Villa Sofia Cervello Palermo | Di Gaudio, F., Contino, F., Orlando, V., Buffa, V., Bonvissuto, M., Di Chiara, V., Todaro, I., Seidita, G., Brunacci, G. |
| EPI_ISL_1366654, EPI_ISL_1366656 | QUALITY CONTROL CHEMICAL BIOLOGICAL RISK, AOOR Villa Sofia Cervello Palermo | QUALITY CONTROL CHEMICAL BIOLOGICAL RISK, AOOR Villa Sofia Cervello Palermo | Di Gaudio, F., Contino, F., Brunacci, G. |
| EPI_ISL_1366658 | QUALITY CONTROL CHEMICAL BIOLOGICAL RISK, AOOR Villa Sofia Cervello Palermo | QUALITY CONTROL CHEMICAL BIOLOGICAL RISK, AOOR Villa Sofia Cervello Palermo | Di Gaudio, F., Brunacci, G., Contino, F. |
| EPI_ISL_1366659 | QUALITY CONTROL CHEMICAL BIOLOGICAL RISK, AOOR Villa Sofia Cervello Palermo | QUALITY CONTROL CHEMICAL BIOLOGICAL RISK, AOOR Villa Sofia Cervello Palermo | Di Gaudio, F., Contino, F., Orlando, V., Buffa, V., Seidita, G., Bonvissuto, M., Di Chiara, V., Todaro, I., Brunacci, G |
| EPI_ISL_1366752, EPI_ISL_1366753 | QUALITY CONTROL CHEMICAL BIOLOGICAL RISK, AOOR Villa Sofia Cervello Palermo | QUALITY CONTROL CHEMICAL BIOLOGICAL RISK, AOOR Villa Sofia Cervello Palermo | Di Gaudio, F., Contino, F., Brunacci, G. |
| EPI_ISL_1366889 | QUALITY CONTROL CHEMICAL BIOLOGICAL RISK, AOOR Villa Sofia Cervello Palermo | QUALITY CONTROL CHEMICAL BIOLOGICAL RISK, AOOR Villa Sofia Cervello Palermo | Di Gaudio, F., Brunacci, G., Contino, F. |
| EPI_ISL_1366921 | QUALITY CONTROL CHEMICAL BIOLOGICAL RISK, AOOR Villa Sofia Cervello Palermo | QUALITY CONTROL CHEMICAL BIOLOGICAL RISK, AOOR Villa Sofia Cervello Palermo | Di Gaudio, F., Contino, F., Brunacci, G. |
| EPI_ISL_1372422 | OLVZ Aalst | OLVZ Aalst | Astrid Holderbeke |

| | | | |
|---|---|---|---|
| EPI_ISL_1372540 | Laboratorio Biologia Molecolare Sars Cov2 - UOC Laboratorio Analisi - Servizio Medicina di Laboratorio, Ospedale "San Francesco" - ATS-ASSL Nuoro | Laboratorio specialistico UOC Ematologia - Ospedale "San Francesco" - ATS-ASSL Nuoro | Piras Giovanna, Asproni Rosanna,Fiamma Maura, Monne Maria Itria, Malune Paolo, Palmas Angelo Domenico, Lo Maglio Iana, Mameli Giuseppe |
| EPI_ISL_1372541, EPI_ISL_1372542, EPI_ISL_1372544 | SC Microbiologia e Virologia AOUSS | Laboratorio specialistico UOC Ematologia - Ospedale "San Francesco" - ATS-ASSL Nuoro | Piras Giovanna, Asproni Rosanna,Fiamma Maura, Monne Maria Itria, Malune Paolo, Palmas Angelo Domenico, Lo Maglio Iana, Mameli Giuseppe Salvatore Rubino; Marcello Abbondio; Flavia Angioj; Laura Firino; Rosalba Govoni; Gabriele Ibba; Vincenzo Lai; Erica Mura; Bianca Paglietti; Claudia Piu; Anna Puggioni; Elena Rimini; Giulia Rocca; Rosangela Sau; Caterina Serra; Sergio Uzzau. |
| EPI_ISL_1372546 | Laboratorio Biologia Molecolare Sars Cov2 - UOC Laboratorio Analisi - Servizio Medicina di Laboratorio, Ospedale "San Francesco" - ATS-ASSL Nuoro | Laboratorio specialistico UOC Ematologia - Ospedale "San Francesco" - ATS-ASSL Nuoro | Piras Giovanna, Asproni Rosanna,Fiamma Maura, Monne Maria Itria, Malune Paolo, Palmas Angelo Domenico, Lo Maglio Iana, Mameli Giuseppe |
| EPI_ISL_1372547, EPI_ISL_1372548, EPI_ISL_1372549 | SC Microbiologia e Virologia AOUSS | Laboratorio specialistico UOC Ematologia - Ospedale "San Francesco" - ATS-ASSL Nuoro | Piras Giovanna, Asproni Rosanna,Fiamma Maura, Monne Maria Itria, Malune Paolo, Palmas Angelo Domenico, Lo Maglio Iana, Mameli Giuseppe Salvatore Rubino; Marcello Abbondio; Flavia Angioj; Laura Firino; Rosalba Govoni; Gabriele Ibba; Vincenzo Lai; Erica Mura; Bianca Paglietti; Claudia Piu; Anna Puggioni; Elena Rimini; Giulia Rocca; Rosangela Sau; Caterina Serra; Sergio Uzzau. |
| EPI_ISL_1372564 | Somedi laboratories | Imelda Hospital | Johan Frans, Dagmar Obbels, Hanne Valgaeren |
| EPI_ISL_1373371 | Imelda Hospital | Imelda Hospital | Johan Frans, Dagmar Obbels, Hanne Valgaeren |
| EPI_ISL_1385803 | Laboratorio Microbiologia P.O.Cardarelli | Laboratorio Microbiologia P.O. Cardarelli | Valentina Felice, Giovanna Niro, Scutellà Massimiliano |
| EPI_ISL_1385805 | Laboratorio Microbiologia P.O. Cardarelli | Laboratorio Microbiologia P.O. Cardarelli | Valentina Felice, Giovanna Niro, Massimiliano Scutellà |
| EPI_ISL_1385959 | Institut de Virologie du CHU de Strasbourg | Institut de Virologie du CHU de Strasbourg | Fafi-Kremer Samira, Gallais Floriane, Gantner Pierre, Laugel Elodie, Solis Morgane, Velay Aurélie, Wendling Marie-Josée |
| EPI_ISL_1386119 | COVID Laboratory AOU Messina | Molecular Hepatology Lab AOU Messina | Giuseppina Raffa, Cristina Musolino, Valeria Chines, Daniele Lombardo, Domenico Giosa, Teresa Pollicino. |
| EPI_ISL_1386120, EPI_ISL_1386122 | COVID Laboratory AOU Messina | Molecular Hepatology AOU Messina | Giuseppina Raffa, Cristina Musolino, Valeria Chines, Daniele Lombardo, Domenico Giosa, Teresa Pollicino. |
| EPI_ISL_1391022 | Ospedale Generale Regionale F. Miulli | Istituto Zooprofilattico Sperimentale della Puglia e della Basilicata | Parisi A., Bianco A., Capozzi L., Del Sambro L., Simone D., Ceci E. |
| EPI_ISL_1391025, EPI_ISL_1391026, EPI_ISL_1391028 | I.R.C.C.S. "S. De Bellis" - Ente Ospedaliero | Istituto Zooprofilattico Sperimentale della Puglia e della Basilicata | Parisi A., Bianco A., Capozzi L., Del Sambro L., Simone D., Lippolis A., Notarnicola M. |
| EPI_ISL_1391029, EPI_ISL_1391030, EPI_ISL_1391031, EPI_ISL_1391032, EPI_ISL_1391033, EPI_ISL_1391034, EPI_ISL_1391035, EPI_ISL_1391036, EPI_ISL_1391037 | Ospedale Generale Regionale F. Miulli | Istituto Zooprofilattico Sperimentale della Puglia e della Basilicata | Parisi A., Bianco A., Capozzi L., Del Sambro L., Simone D., Ceci E. |
| EPI_ISL_1391041, EPI_ISL_1391042, EPI_ISL_1391044, EPI_ISL_1391045, EPI_ISL_1391046, EPI_ISL_1391047, EPI_ISL_1391048, EPI_ISL_1391049, EPI_ISL_1391052 | Ospedale di Venere | Istituto Zooprofilattico Sperimentale della Puglia e della Basilicata | Parisi A., Bianco A., Capozzi L., Del Sambro L., Simone D., Iacobellis M., Ridolfi D., Giannico A. |
| EPI_ISL_1391053, EPI_ISL_1391054, EPI_ISL_1391055, EPI_ISL_1391056, EPI_ISL_1391057 | Presidio di Brindisi Di Summa - Perrino | Istituto Zooprofilattico Sperimentale della Puglia e della Basilicata | Parisi A., Bianco A., Capozzi L., Del Sambro L., Simone D., Difato L., Santoro A., Ridolfi D., Giannico A. |
| EPI_ISL_1391058, EPI_ISL_1391059, EPI_ISL_1391060, EPI_ISL_1391061, EPI_ISL_1391062, EPI_ISL_1391063, EPI_ISL_1391064 | Ospedale Vito Fazzi | Istituto Zooprofilattico Sperimentale della Puglia e della Basilicata | Parisi A., Bianco A., Capozzi L., Del Sambro L., Simone D., Difato L., Lobreglio G. |
| EPI_ISL_1391065, EPI_ISL_1391066, EPI_ISL_1391067, EPI_ISL_1391068, EPI_ISL_1391069, EPI_ISL_1391070, EPI_ISL_1391071, EPI_ISL_1391072 | Ospedale Santa Caterina Novella | Istituto Zooprofilattico Sperimentale della Puglia e della Basilicata | Parisi A., Bianco A., Capozzi L., Del Sambro L., Simone D., Difato L., Bruno A. R. |
| EPI_ISL_1391073, EPI_ISL_1391074, EPI_ISL_1391075, EPI_ISL_1391076 | Ospedale Casa Sollievo della Sofferenza | Istituto Zooprofilattico Sperimentale della Puglia e della Basilicata | Parisi A., Bianco A., Capozzi L., Del Sambro L., Simone D., Difato L., Ridolfi D., Giannico A, Labonia M. |
| EPI_ISL_1391078, EPI_ISL_1391079, EPI_ISL_1391080, EPI_ISL_1391081, EPI_ISL_1391082, EPI_ISL_1391084, EPI_ISL_1391085, EPI_ISL_1391086, EPI_ISL_1391087, EPI_ISL_1391088, EPI_ISL_1391089, EPI_ISL_1391090, EPI_ISL_1391091, EPI_ISL_1391092, EPI_ISL_1391093, EPI_ISL_1391094, EPI_ISL_1391095, EPI_ISL_1391096 | | | |
| see above | Ospedale Santissima Annunziata | Istituto Zooprofilattico Sperimentale della Puglia e della Basilicata | Parisi A., Bianco A., Capozzi L., Del Sambro L., Simone D., Difato L., Ridolfi D., Giannico A, Tampoia M. |
| EPI_ISL_1391204, EPI_ISL_1391205, EPI_ISL_1391206, EPI_ISL_1391207, EPI_ISL_1391208, EPI_ISL_1391209, EPI_ISL_1391210, EPI_ISL_1391211, EPI_ISL_1391212, EPI_ISL_1391213, EPI_ISL_1391214, EPI_ISL_1391215, EPI_ISL_1391216, EPI_ISL_1391217, EPI_ISL_1391218, EPI_ISL_1391219, EPI_ISL_1391220, EPI_ISL_1391221, EPI_ISL_1391222, EPI_ISL_1391223, EPI_ISL_1391224, EPI_ISL_1391225, EPI_ISL_1391226, EPI_ISL_1391227, EPI_ISL_1391228, EPI_ISL_1391229 | | | |
| see above | SIESP DIPARTIMENTO DI PREVENZIONE TERAMO TERAMO(TERAMO) | Istituto Zooprofilattico Sperimentale dell'Abruzzo e Molise "G. Caporale" | Lorusso A, Marcacci M, Di Domenico M, Ancora M, Curini V, Di Lollo Valeria, Mangone I, Rinaldi A, Delli Compagni E, Scialabba S, Caporale M, Di Pasquale A, Cammà C, Puglia I, Calistri P, Savini G |
| EPI_ISL_1391230, EPI_ISL_1391231, EPI_ISL_1391232 | SIESP CHIETI - DRIVE IN LANCIANO LANCIANO(CHIETI) | Istituto Zooprofilattico Sperimentale dell'Abruzzo e Molise "G. Caporale" | Lorusso A, Marcacci M, Di Domenico M, Ancora M, Curini V, Di Lollo Valeria, Mangone I, Rinaldi A, Delli Compagni E, Scialabba S, Caporale M, Di Pasquale A, Cammà C, Puglia I, Calistri P, Savini G |
| EPI_ISL_1391234, EPI_ISL_1391235, EPI_ISL_1391236, EPI_ISL_1391237 | SIESP SULMONA SULMONA(L'AQUILA) | Istituto Zooprofilattico Sperimentale dell'Abruzzo e Molise "G. Caporale" | Lorusso A, Marcacci M, Di Domenico M, Ancora M, Curini V, Di Lollo Valeria, Mangone I, Rinaldi A, Delli Compagni E, Scialabba S, Caporale M, Di Pasquale A, Cammà C, Puglia I, Calistri P, Savini G |
| EPI_ISL_1391238 | DIP. PREV. AVEZZANO SERVIZIO DI IGIENE EPIDEMIOLOGIA E SANITA' PUBBLICA AVEZZANO(L'AQUILA) | Istituto Zooprofilattico Sperimentale dell'Abruzzo e Molise "G. Caporale" | Lorusso A, Marcacci M, Di Domenico M, Ancora M, Curini V, Di Lollo Valeria, Mangone I, Rinaldi A, Delli Compagni E, Scialabba S, Caporale M, Di Pasquale A, Cammà C, Puglia I, Calistri P, Savini G |
| EPI_ISL_1391239, EPI_ISL_1391240 | SIESP SULMONA SULMONA(L'AQUILA) | Istituto Zooprofilattico Sperimentale dell'Abruzzo e Molise "G. Caporale" | Lorusso A, Marcacci M, Di Domenico M, Ancora M, Curini V, Di Lollo Valeria, Mangone I, Rinaldi A, Delli Compagni E, Scialabba S, Caporale M, Di Pasquale A, Cammà C, Puglia I, Calistri P, Savini G |
| EPI_ISL_1391241, EPI_ISL_1391242, EPI_ISL_1391243, EPI_ISL_1391244, EPI_ISL_1391245, EPI_ISL_1391246, EPI_ISL_1391247 | DIP. PREV. AVEZZANO SERVIZIO DI IGIENE EPIDEMIOLOGIA E SANITA' PUBBLICA AVEZZANO(L'AQUILA) | Istituto Zooprofilattico Sperimentale dell'Abruzzo e Molise "G. Caporale" | Lorusso A, Marcacci M, Di Domenico M, Ancora M, Curini V, Di Lollo Valeria, Mangone I, Rinaldi A, Delli Compagni E, Scialabba S, Caporale M, Di Pasquale A, Cammà C, Puglia I, Calistri P, Savini G |
| EPI_ISL_1391248 | SIESP SULMONA SULMONA(L'AQUILA) | Istituto Zooprofilattico Sperimentale dell'Abruzzo e Molise "G. Caporale" | Lorusso A, Marcacci M, Di Domenico M, Ancora M, Curini V, Di Lollo Valeria, Mangone I, Rinaldi A, Delli Compagni E, Scialabba S, Caporale M, Di Pasquale A, Cammà C, Puglia I, Calistri P, Savini G |
| EPI_ISL_1391249, EPI_ISL_1391250, EPI_ISL_1391251 | SIESP DIPARTIMENTO DI PREVENZIONE CHIETI CHIETI(CHIETI) | Istituto Zooprofilattico Sperimentale dell'Abruzzo e Molise "G. Caporale" | Lorusso A, Marcacci M, Di Domenico M, Ancora M, Curini V, Di Lollo Valeria, Mangone I, Rinaldi A, Delli Compagni E, Scialabba S, Caporale M, Di Pasquale A, Cammà C, Puglia I, Calistri P, Savini G |
| EPI_ISL_1391252 | SIESP CHIETI - DRIVE IN LANCIANO LANCIANO(CHIETI) | Istituto Zooprofilattico Sperimentale dell'Abruzzo e Molise "G. Caporale" | Lorusso A, Marcacci M, Di Domenico M, Ancora M, Curini V, Di Lollo Valeria, Mangone I, Rinaldi A, Delli Compagni E, Scialabba S, Caporale M, Di Pasquale A, Cammà C, Puglia I, Calistri P, Savini G |

| | | | |
|---|---|---|---|
| EPI_ISL_1391253, EPI_ISL_1391254, EPI_ISL_1391255, EPI_ISL_1391256, EPI_ISL_1391257, EPI_ISL_1391258, EPI_ISL_1391259 | SIESP DIPARTIMENTO DI PREVENZIONE CHIETI CHIETI(CHIETI) | Istituto Zooprofilattico Sperimentale dell'Abruzzo e Molise "G. Caporale" | Lorusso A, Marcacci M, Di Domenico M, Ancora M, Curini V, Di Lollo Valeria, Mangone I, Rinaldi A, Delli Compagni E, Scialabba S, Caporale M, Di Pasquale A, Cammà C, Puglia I, Calistri P, Savini G |
| EPI_ISL_1391260, EPI_ISL_1391261, EPI_ISL_1391262 | SIESP CHIETI - DRIVE IN LANCIANO LANCIANO(CHIETI) | Istituto Zooprofilattico Sperimentale dell'Abruzzo e Molise "G. Caporale" | Lorusso A, Marcacci M, Di Domenico M, Ancora M, Curini V, Di Lollo Valeria, Mangone I, Rinaldi A, Delli Compagni E, Scialabba S, Caporale M, Di Pasquale A, Cammà C, Puglia I, Calistri P, Savini G |
| EPI_ISL_1391263, EPI_ISL_1391264 | SIESP DIPARTIMENTO DI PREVENZIONE CHIETI CHIETI(CHIETI) | Istituto Zooprofilattico Sperimentale dell'Abruzzo e Molise "G. Caporale" | Lorusso A, Marcacci M, Di Domenico M, Ancora M, Curini V, Di Lollo Valeria, Mangone I, Rinaldi A, Delli Compagni E, Scialabba S, Caporale M, Di Pasquale A, Cammà C, Puglia I, Calistri P, Savini G |
| EPI_ISL_1391265 | SIESP CHIETI - DRIVE IN LANCIANO LANCIANO(CHIETI) | Istituto Zooprofilattico Sperimentale dell'Abruzzo e Molise "G. Caporale" | Lorusso A, Marcacci M, Di Domenico M, Ancora M, Curini V, Di Lollo Valeria, Mangone I, Rinaldi A, Delli Compagni E, Scialabba S, Caporale M, Di Pasquale A, Cammà C, Puglia I, Calistri P, Savini G |
| EPI_ISL_1391266, EPI_ISL_1391267, EPI_ISL_1391269, EPI_ISL_1391270 | SIESP CHIETI - DRIVE IN CHIETI CHIETI(CHIETI) | Istituto Zooprofilattico Sperimentale dell'Abruzzo e Molise "G. Caporale" | Lorusso A, Marcacci M, Di Domenico M, Ancora M, Curini V, Di Lollo Valeria, Mangone I, Rinaldi A, Delli Compagni E, Scialabba S, Caporale M, Di Pasquale A, Cammà C, Puglia I, Calistri P, Savini G |
| EPI_ISL_1391271, EPI_ISL_1391272, EPI_ISL_1391273 | SIESP CHIETI - DRIVE IN ORTONA ORTONA(CHIETI) | Istituto Zooprofilattico Sperimentale dell'Abruzzo e Molise "G. Caporale" | Lorusso A, Marcacci M, Di Domenico M, Ancora M, Curini V, Di Lollo Valeria, Mangone I, Rinaldi A, Delli Compagni E, Scialabba S, Caporale M, Di Pasquale A, Cammà C, Puglia I, Calistri P, Savini G |
| EPI_ISL_1391274 | SIESP DIPARTIMENTO DI PREVENZIONE TERAMO TERAMO(TERAMO) | Istituto Zooprofilattico Sperimentale dell'Abruzzo e Molise "G. Caporale" | Lorusso A, Marcacci M, Di Domenico M, Ancora M, Curini V, Di Lollo Valeria, Mangone I, Rinaldi A, Delli Compagni E, Scialabba S, Caporale M, Di Pasquale A, Cammà C, Puglia I, Calistri P, Savini G |
| EPI_ISL_1391275 | SIESP CHIETI - DRIVE IN GISSI GISSI(CHIETI) | Istituto Zooprofilattico Sperimentale dell'Abruzzo e Molise "G. Caporale" | Lorusso A, Marcacci M, Di Domenico M, Ancora M, Curini V, Di Lollo Valeria, Mangone I, Rinaldi A, Delli Compagni E, Scialabba S, Caporale M, Di Pasquale A, Cammà C, Puglia I, Calistri P, Savini G |
| EPI_ISL_1391416, EPI_ISL_1391421, EPI_ISL_1391621 | Baylor Scott & White-Temple | Baylor Scott & White-Temple | Ari Rao, Linden Morales, Kimberly Walker, Marcus Volz, Shelby Hendrickson, Caitlin Maloney |
| EPI_ISL_1394738 | Laboratorio Analisi Osp. Città di Castello - Azienda USL Umbria1 | Istituto Zooprofilattico Sperimentale dell'Abruzzo e Molise "G. Caporale" | Malagigi V, Tacconi P, Lorusso A, Marcacci M, Di Domenico M, Ancora M, Curini V, Di Lollo Valeria, Mangone I, Rinaldi A, Scialabba S, Caporale M, Di Pasquale A, Cammà C, Puglia I, Calistri P, Savini G |
| EPI_ISL_1394739, EPI_ISL_1394740, EPI_ISL_1394741, EPI_ISL_1394742, EPI_ISL_1394743 | Azienda USL Umbria 2 | Istituto Zooprofilattico Sperimentale dell'Abruzzo e Molise "G. Caporale" | Proietti A, Pistoni E, Lorusso A, Marcacci M, Di Domenico M, Ancora M, Curini V, Di Lollo Valeria, Mangone I, Rinaldi A, Scialabba S, Caporale M, Di Pasquale A, Cammà C, Puglia I, Calistri P, Savini G |
| EPI_ISL_1394744, EPI_ISL_1394745, EPI_ISL_1394746, EPI_ISL_1394748 | Azienda Ospedaliera Terni | Istituto Zooprofilattico Sperimentale dell'Abruzzo e Molise "G. Caporale" | Palumbo M, Scaccetti A, Lorusso A, Marcacci M, Di Domenico M, Ancora M, Curini V, Di Lollo Valeria, Mangone I, Rinaldi A, Scialabba S, Caporale M, Di Pasquale A, Cammà C, Puglia I, Calistri P, Savini G |
| EPI_ISL_1394749, EPI_ISL_1394750, EPI_ISL_1394752 | Istituto Zooprofilattico Sperimentale Umbria e Marche "Togo Rosati" | Istituto Zooprofilattico Sperimentale dell'Abruzzo e Molise "G. Caporale" | Biagetti M, Giammarioli M, Lorusso A, Marcacci M, Di Domenico M, Ancora M, Curini V, Di Lollo Valeria, Mangone I, Rinaldi A, Scialabba S, Caporale M, Di Pasquale A, Cammà C, Puglia I, Calistri P, Savini G |
| EPI_ISL_1394754 | Università degli Studi di Perugia | Istituto Zooprofilattico Sperimentale dell'Abruzzo e Molise "G. Caporale" | Mencacci A, Camilloni B, Lorusso A, Marcacci M, Di Domenico M, Ancora M, Curini V, Di Lollo Valeria, Mangone I, Rinaldi A, Scialabba S, Caporale M, Di Pasquale A, Cammà C, Puglia I, Calistri P, Savini G |
| EPI_ISL_1394756, EPI_ISL_1394757 | Laboratorio Analisi Osp. Città di Castello - Azienda USL Umbria1 | Istituto Zooprofilattico Sperimentale dell'Abruzzo e Molise "G. Caporale" | Malagigi V, Tacconi P, Lorusso A, Marcacci M, Di Domenico M, Ancora M, Curini V, Di Lollo Valeria, Mangone I, Rinaldi A, Scialabba S, Caporale M, Di Pasquale A, Cammà C, Puglia I, Calistri P, Savini G |
| EPI_ISL_1394758 | Università degli Studi di Perugia | Istituto Zooprofilattico Sperimentale dell'Abruzzo e Molise "G. Caporale" | Mencacci A, Camilloni B, Lorusso A, Marcacci M, Di Domenico M, Ancora M, Curini V, Di Lollo Valeria, Mangone I, Rinaldi A, Scialabba S, Caporale M, Di Pasquale A, Cammà C, Puglia I, Calistri P, Savini G |
| EPI_ISL_1396406 | OLVZ Aalst | OLVZ Aalst | Astrid Holderbeke |
| EPI_ISL_1399230 | INT Fondazione Pascale | INT Fondazione Pascale | INT Fondazione Pascale |
| EPI_ISL_1402414, EPI_ISL_1402415, EPI_ISL_1402416, EPI_ISL_1402419 | AZ Sint-Jan | AZ SINT-JAN BRUGGE | Marijke Reynders, Patrick Descheemaeker, Laurien Hoornaert, Thomas Van Landschoot, Jorn Hellemans |
| EPI_ISL_1402420 | AZ Zeno | AZ SINT-JAN BRUGGE | Marijke Reynders, Patrick Descheemaeker, Laurien Hoornaert, Thomas Van Landschoot, Jorn Hellemans |
| EPI_ISL_1402421 | AZ Sint-Jan | AZ SINT-JAN BRUGGE | Marijke Reynders, Patrick Descheemaeker, Laurien Hoornaert, Thomas Van Landschoot, Jorn Hellemans |
| EPI_ISL_1403497 | AZ Klina | AZ Klina | Carl Vael- Lynsey Berckmans |
| EPI_ISL_1403499 | AZ Klina | AZ Klina | Carl Vael - Lynsey Berckmans |
| EPI_ISL_1403503 | SIESP CHIETI - DRIVE IN GISSI GISSI(CHIETI) | Istituto Zooprofilattico Sperimentale dell'Abruzzo e Molise "G. Caporale" | Lorusso A, Marcacci M, Di Domenico M, Ancora M, Curini V, Mangone I, Delli Compagni E, Rinaldi A, Scialabba S, Di Pasquale A, Cammà C, Puglia I, Calistri P, Savini G |
| EPI_ISL_1403504 | USCA Tagliacozzo TAGLIACOZZO(L'AQUILA) | Istituto Zooprofilattico Sperimentale dell'Abruzzo e Molise "G. Caporale" | Lorusso A, Marcacci M, Di Domenico M, Ancora M, Curini V, Mangone I, Delli Compagni E, Rinaldi A, Scialabba S, Di Pasquale A, Cammà C, Puglia I, Calistri P, Savini G |
| EPI_ISL_1403505, EPI_ISL_1403506, EPI_ISL_1403507 | SIESP CHIETI - DRIVE IN GISSI GISSI(CHIETI) | Istituto Zooprofilattico Sperimentale dell'Abruzzo e Molise "G. Caporale" | Lorusso A, Marcacci M, Di Domenico M, Ancora M, Curini V, Mangone I, Delli Compagni E, Rinaldi A, Scialabba S, Di Pasquale A, Cammà C, Puglia I, Calistri P, Savini G |
| EPI_ISL_1403508 | OSPEDALE CIVILE GIULIANOVA - PRONTO SOCCORSO GIULIANOVA(TERAMO) | Istituto Zooprofilattico Sperimentale dell'Abruzzo e Molise "G. Caporale" | Lorusso A, Marcacci M, Di Domenico M, Ancora M, Curini V, Mangone I, Delli Compagni E, Rinaldi A, Scialabba S, Di Pasquale A, Cammà C, Puglia I, Calistri P, Savini G |
| EPI_ISL_1403509 | SIESP CHIETI - DRIVE IN ORTONA ORTONA(CHIETI) | Istituto Zooprofilattico Sperimentale dell'Abruzzo e Molise "G. Caporale" | Lorusso A, Marcacci M, Di Domenico M, Ancora M, Curini V, Mangone I, Delli Compagni E, Rinaldi A, Scialabba S, Di Pasquale A, Cammà C, Puglia I, Calistri P, Savini G |
| EPI_ISL_1403510 | MEDICO COMPETENTE P.O. SULMONA - CASTEL DI SANGRO SULMONA(L'AQUILA) | Istituto Zooprofilattico Sperimentale dell'Abruzzo e Molise "G. Caporale" | Lorusso A, Marcacci M, Di Domenico M, Ancora M, Curini V, Mangone I, Delli Compagni E, Rinaldi A, Scialabba S, Di Pasquale A, Cammà C, Puglia I, Calistri P, Savini G |
| EPI_ISL_1403511 | SIESP SULMONA SULMONA(L'AQUILA) | Istituto Zooprofilattico Sperimentale dell'Abruzzo e Molise "G. Caporale" | Lorusso A, Marcacci M, Di Domenico M, Ancora M, Curini V, Mangone I, Delli Compagni E, Rinaldi A, Scialabba S, Di Pasquale A, Cammà C, Puglia I, Calistri P, Savini G |
| EPI_ISL_1403512, EPI_ISL_1403513 | DIP. PREV. AVEZZANO SERVIZIO DI IGIENE EPIDEMIOLOGIA E SANITA' PUBBLICA AVEZZANO(L'AQUILA) | Istituto Zooprofilattico Sperimentale dell'Abruzzo e Molise "G. Caporale" | Lorusso A, Marcacci M, Di Domenico M, Ancora M, Curini V, Mangone I, Delli Compagni E, Rinaldi A, Scialabba S, Di Pasquale A, Cammà C, Puglia I, Calistri P, Savini G |
| EPI_ISL_1403515, EPI_ISL_1403516, EPI_ISL_1403517 | USCA Castel di Sangro CASTEL DI SANGRO(L'AQUILA) | Istituto Zooprofilattico Sperimentale dell'Abruzzo e Molise "G. Caporale" | Lorusso A, Marcacci M, Di Domenico M, Ancora M, Curini V, Mangone I, Delli Compagni E, Rinaldi A, Scialabba S, Di Pasquale A, Cammà C, Puglia I, Calistri P, Savini G |
| EPI_ISL_1403518 | SIESP DIPARTIMENTO DI PREVENZIONE TERAMO TERAMO(TERAMO) | Istituto Zooprofilattico Sperimentale dell'Abruzzo e Molise "G. Caporale" | Lorusso A, Marcacci M, Di Domenico M, Ancora M, Curini V, Mangone I, Delli Compagni E, Rinaldi A, Scialabba S, Di Pasquale A, Cammà C, Puglia I, Calistri P, Savini G |
| EPI_ISL_1403519 | SIESP CHIETI - DRIVE IN CHIETI CHIETI(CHIETI) | Istituto Zooprofilattico Sperimentale dell'Abruzzo e Molise "G. Caporale" | Lorusso A, Marcacci M, Di Domenico M, Ancora M, Curini V, Mangone I, Delli Compagni E, Rinaldi A, Scialabba S, Di Pasquale A, Cammà C, Puglia I, Calistri P, Savini G |
| EPI_ISL_1403520 | SIESP DIPARTIMENTO DI PREVENZIONE TERAMO TERAMO(TERAMO) | Istituto Zooprofilattico Sperimentale dell'Abruzzo e Molise "G. Caporale" | Lorusso A, Marcacci M, Di Domenico M, Ancora M, Curini V, Mangone I, Delli Compagni E, Rinaldi A, Scialabba S, Di Pasquale A, Cammà C, Puglia I, Calistri P, Savini G |
| EPI_ISL_1403521 | SIESP CHIETI - DRIVE IN ORTONA ORTONA(CHIETI) | Istituto Zooprofilattico Sperimentale dell'Abruzzo e Molise "G. Caporale" | Lorusso A, Marcacci M, Di Domenico M, Ancora M, Curini V, Mangone I, Delli Compagni E, Rinaldi A, Scialabba S, Di Pasquale A, Cammà C, Puglia I, Calistri P, Savini G |

| | | | |
|---|---|---|---|
| EPI_ISL_1403522 | SIESP DIPARTIMENTO DI PREVENZIONE TERAMO TERAMO(TERAMO) | Istituto Zooprofilattico Sperimentale dell'Abruzzo e Molise "G. Caporale" | Lorusso A, Marcacci M, Di Domenico M, Ancora M, Curini V, Mangone I, Delli Compagni E, Rinaldi A, Scialabba S, Di Pasquale A, Cammà C, Puglia I, Calistri P, Savini G |
| EPI_ISL_1403525 | SIESP CHIETI - DRIVE IN CHIETI CHIETI(CHIETI) | Istituto Zooprofilattico Sperimentale dell'Abruzzo e Molise "G. Caporale" | Lorusso A, Marcacci M, Di Domenico M, Ancora M, Curini V, Mangone I, Delli Compagni E, Rinaldi A, Scialabba S, Di Pasquale A, Cammà C, Puglia I, Calistri P, Savini G |
| EPI_ISL_1403526 | USCA Pescina PESCINA(L'AQUILA) | Istituto Zooprofilattico Sperimentale dell'Abruzzo e Molise "G. Caporale" | Lorusso A, Marcacci M, Di Domenico M, Ancora M, Curini V, Mangone I, Delli Compagni E, Rinaldi A, Scialabba S, Di Pasquale A, Cammà C, Puglia I, Calistri P, Savini G |
| EPI_ISL_1403527 | SIESP SULMONA SULMONA(L'AQUILA) | Istituto Zooprofilattico Sperimentale dell'Abruzzo e Molise "G. Caporale" | Lorusso A, Marcacci M, Di Domenico M, Ancora M, Curini V, Mangone I, Delli Compagni E, Rinaldi A, Scialabba S, Di Pasquale A, Cammà C, Puglia I, Calistri P, Savini G |
| EPI_ISL_1403528 | SIESP CHIETI - DRIVE IN CHIETI CHIETI(CHIETI) | Istituto Zooprofilattico Sperimentale dell'Abruzzo e Molise "G. Caporale" | Lorusso A, Marcacci M, Di Domenico M, Ancora M, Curini V, Mangone I, Delli Compagni E, Rinaldi A, Scialabba S, Di Pasquale A, Cammà C, Puglia I, Calistri P, Savini G |
| EPI_ISL_1403529 | USCA Sulmona SULMONA(L'AQUILA) | Istituto Zooprofilattico Sperimentale dell'Abruzzo e Molise "G. Caporale" | Lorusso A, Marcacci M, Di Domenico M, Ancora M, Curini V, Mangone I, Delli Compagni E, Rinaldi A, Scialabba S, Di Pasquale A, Cammà C, Puglia I, Calistri P, Savini G |
| EPI_ISL_1403530, EPI_ISL_1403531 | MEDICO COMPETENTE P.O. SULMONA - CASTEL DI SANGRO SULMONA(L'AQUILA) | Istituto Zooprofilattico Sperimentale dell'Abruzzo e Molise "G. Caporale" | Lorusso A, Marcacci M, Di Domenico M, Ancora M, Curini V, Mangone I, Delli Compagni E, Rinaldi A, Scialabba S, Di Pasquale A, Cammà C, Puglia I, Calistri P, Savini G |
| EPI_ISL_1403532 | SIESP CHIETI - DRIVE IN LANCIANO LANCIANO(CHIETI) | Istituto Zooprofilattico Sperimentale dell'Abruzzo e Molise "G. Caporale" | Lorusso A, Marcacci M, Di Domenico M, Ancora M, Curini V, Mangone I, Delli Compagni E, Rinaldi A, Scialabba S, Di Pasquale A, Cammà C, Puglia I, Calistri P, Savini G |
| EPI_ISL_1403533 | SIESP DIPARTIMENTO DI PREVENZIONE CHIETI CHIETI(CHIETI) | Istituto Zooprofilattico Sperimentale dell'Abruzzo e Molise "G. Caporale" | Lorusso A, Marcacci M, Di Domenico M, Ancora M, Curini V, Mangone I, Delli Compagni E, Rinaldi A, Scialabba S, Di Pasquale A, Cammà C, Puglia I, Calistri P, Savini G |
| EPI_ISL_1403534 | SIESP CHIETI - DRIVE IN ATESSA ATESSA(CHIETI) | Istituto Zooprofilattico Sperimentale dell'Abruzzo e Molise "G. Caporale" | Lorusso A, Marcacci M, Di Domenico M, Ancora M, Curini V, Mangone I, Delli Compagni E, Rinaldi A, Scialabba S, Di Pasquale A, Cammà C, Puglia I, Calistri P, Savini G |
| EPI_ISL_1403535 | SIESP CHIETI - DRIVE IN VASTO VASTO(CHIETI) | Istituto Zooprofilattico Sperimentale dell'Abruzzo e Molise "G. Caporale" | Lorusso A, Marcacci M, Di Domenico M, Ancora M, Curini V, Mangone I, Delli Compagni E, Rinaldi A, Scialabba S, Di Pasquale A, Cammà C, Puglia I, Calistri P, Savini G |
| EPI_ISL_1403536 | SIESP SULMONA SULMONA(L'AQUILA) | Istituto Zooprofilattico Sperimentale dell'Abruzzo e Molise "G. Caporale" | Lorusso A, Marcacci M, Di Domenico M, Ancora M, Curini V, Mangone I, Delli Compagni E, Rinaldi A, Scialabba S, Di Pasquale A, Cammà C, Puglia I, Calistri P, Savini G |
| EPI_ISL_1403537 | USCA Sulmona SULMONA(L'AQUILA) | Istituto Zooprofilattico Sperimentale dell'Abruzzo e Molise "G. Caporale" | Lorusso A, Marcacci M, Di Domenico M, Ancora M, Curini V, Mangone I, Delli Compagni E, Rinaldi A, Scialabba S, Di Pasquale A, Cammà C, Puglia I, Calistri P, Savini G |
| EPI_ISL_1403538, EPI_ISL_1403539 | SIESP SULMONA SULMONA(L'AQUILA) | Istituto Zooprofilattico Sperimentale dell'Abruzzo e Molise "G. Caporale" | Lorusso A, Marcacci M, Di Domenico M, Ancora M, Curini V, Mangone I, Delli Compagni E, Rinaldi A, Scialabba S, Di Pasquale A, Cammà C, Puglia I, Calistri P, Savini G |
| EPI_ISL_1403540 | SIESP DIPARTIMENTO DI PREVENZIONE TERAMO TERAMO(TERAMO) | Istituto Zooprofilattico Sperimentale dell'Abruzzo e Molise "G. Caporale" | Lorusso A, Marcacci M, Di Domenico M, Ancora M, Curini V, Mangone I, Delli Compagni E, Rinaldi A, Scialabba S, Di Pasquale A, Cammà C, Puglia I, Calistri P, Savini G |
| EPI_ISL_1403541 | DIP. PREV. AVEZZANO SERVIZIO DI IGIENE EPIDEMIOLOGIA E SANITA' PUBBLICA AVEZZANO(L'AQUILA) | Istituto Zooprofilattico Sperimentale dell'Abruzzo e Molise "G. Caporale" | Lorusso A, Marcacci M, Di Domenico M, Ancora M, Curini V, Mangone I, Delli Compagni E, Rinaldi A, Scialabba S, Di Pasquale A, Cammà C, Puglia I, Calistri P, Savini G |
| EPI_ISL_1403542 | USCA Pescina PESCINA(L'AQUILA) | Istituto Zooprofilattico Sperimentale dell'Abruzzo e Molise "G. Caporale" | Lorusso A, Marcacci M, Di Domenico M, Ancora M, Curini V, Mangone I, Delli Compagni E, Rinaldi A, Scialabba S, Di Pasquale A, Cammà C, Puglia I, Calistri P, Savini G |
| EPI_ISL_1403544 | DIP. PREV. AVEZZANO SERVIZIO DI IGIENE EPIDEMIOLOGIA E SANITA' PUBBLICA AVEZZANO(L'AQUILA) | Istituto Zooprofilattico Sperimentale dell'Abruzzo e Molise "G. Caporale" | Lorusso A, Marcacci M, Di Domenico M, Ancora M, Curini V, Mangone I, Delli Compagni E, Rinaldi A, Scialabba S, Di Pasquale A, Cammà C, Puglia I, Calistri P, Savini G |
| EPI_ISL_1403545 | USCA Pescina PESCINA(L'AQUILA) | Istituto Zooprofilattico Sperimentale dell'Abruzzo e Molise "G. Caporale" | Lorusso A, Marcacci M, Di Domenico M, Ancora M, Curini V, Mangone I, Delli Compagni E, Rinaldi A, Scialabba S, Di Pasquale A, Cammà C, Puglia I, Calistri P, Savini G |
| EPI_ISL_1403546 | DIP. PREV. AVEZZANO SERVIZIO DI IGIENE EPIDEMIOLOGIA E SANITA' PUBBLICA AVEZZANO(L'AQUILA) | Istituto Zooprofilattico Sperimentale dell'Abruzzo e Molise "G. Caporale" | Lorusso A, Marcacci M, Di Domenico M, Ancora M, Curini V, Mangone I, Delli Compagni E, Rinaldi A, Scialabba S, Di Pasquale A, Cammà C, Puglia I, Calistri P, Savini G |
| EPI_ISL_1403547 | USCA Pescina PESCINA(L'AQUILA) | Istituto Zooprofilattico Sperimentale dell'Abruzzo e Molise "G. Caporale" | Lorusso A, Marcacci M, Di Domenico M, Ancora M, Curini V, Mangone I, Delli Compagni E, Rinaldi A, Scialabba S, Di Pasquale A, Cammà C, Puglia I, Calistri P, Savini G |
| EPI_ISL_1403548 | OSPEDALE CIVILE SANT'OMERO - PRONTO SOCCORSO SANT'OMERO(TERAMO) | Istituto Zooprofilattico Sperimentale dell'Abruzzo e Molise "G. Caporale" | Lorusso A, Marcacci M, Di Domenico M, Ancora M, Curini V, Mangone I, Delli Compagni E, Rinaldi A, Scialabba S, Di Pasquale A, Cammà C, Puglia I, Calistri P, Savini G |
| EPI_ISL_1403550, EPI_ISL_1403551 | SIESP DIPARTIMENTO DI PREVENZIONE TERAMO TERAMO(TERAMO) | Istituto Zooprofilattico Sperimentale dell'Abruzzo e Molise "G. Caporale" | Lorusso A, Marcacci M, Di Domenico M, Ancora M, Curini V, Mangone I, Delli Compagni E, Rinaldi A, Scialabba S, Di Pasquale A, Cammà C, Puglia I, Calistri P, Savini G |
| EPI_ISL_1403552 | OSPEDALE CIVILE GIULIANOVA - MEDICINA INTERNA GIULIANOVA(TERAMO) | Istituto Zooprofilattico Sperimentale dell'Abruzzo e Molise "G. Caporale" | Lorusso A, Marcacci M, Di Domenico M, Ancora M, Curini V, Mangone I, Delli Compagni E, Rinaldi A, Scialabba S, Di Pasquale A, Cammà C, Puglia I, Calistri P, Savini G |
| EPI_ISL_1403553 | SIESP DIPARTIMENTO DI PREVENZIONE TERAMO TERAMO(TERAMO) | Istituto Zooprofilattico Sperimentale dell'Abruzzo e Molise "G. Caporale" | Lorusso A, Marcacci M, Di Domenico M, Ancora M, Curini V, Mangone I, Delli Compagni E, Rinaldi A, Scialabba S, Di Pasquale A, Cammà C, Puglia I, Calistri P, Savini G |
| EPI_ISL_1403555 | OSPEDALE CIVILE SANT'OMERO - PRONTO SOCCORSO SANT'OMERO(TERAMO) | Istituto Zooprofilattico Sperimentale dell'Abruzzo e Molise "G. Caporale" | Lorusso A, Marcacci M, Di Domenico M, Ancora M, Curini V, Mangone I, Delli Compagni E, Rinaldi A, Scialabba S, Di Pasquale A, Cammà C, Puglia I, Calistri P, Savini G |
| EPI_ISL_1403559 | SIESP CHIETI - DRIVE IN VASTO VASTO(CHIETI) | Istituto Zooprofilattico Sperimentale dell'Abruzzo e Molise "G. Caporale" | Lorusso A, Marcacci M, Di Domenico M, Ancora M, Curini V, Mangone I, Delli Compagni E, Rinaldi A, Scialabba S, Di Pasquale A, Cammà C, Puglia I, Calistri P, Savini G |
| EPI_ISL_1403561, EPI_ISL_1403562, EPI_ISL_1403563, EPI_ISL_1403565 | SIESP DIPARTIMENTO DI PREVENZIONE TERAMO TERAMO(TERAMO) | Istituto Zooprofilattico Sperimentale dell'Abruzzo e Molise "G. Caporale" | Lorusso A, Marcacci M, Di Domenico M, Ancora M, Curini V, Mangone I, Delli Compagni E, Rinaldi A, Scialabba S, Di Pasquale A, Cammà C, Puglia I, Calistri P, Savini G |
| EPI_ISL_1403566 | SIESP CHIETI - DRIVE IN CHIETI CHIETI(CHIETI) | Istituto Zooprofilattico Sperimentale dell'Abruzzo e Molise "G. Caporale" | Lorusso A, Marcacci M, Di Domenico M, Ancora M, Curini V, Mangone I, Delli Compagni E, Rinaldi A, Scialabba S, Di Pasquale A, Cammà C, Puglia I, Calistri P, Savini G |
| EPI_ISL_1403567 | SIESP DIPARTIMENTO DI PREVENZIONE TERAMO TERAMO(TERAMO) | Istituto Zooprofilattico Sperimentale dell'Abruzzo e Molise "G. Caporale" | Lorusso A, Marcacci M, Di Domenico M, Ancora M, Curini V, Mangone I, Delli Compagni E, Rinaldi A, Scialabba S, Di Pasquale A, Cammà C, Puglia I, Calistri P, Savini G |
| EPI_ISL_1403568 | SIESP CHIETI - DRIVE IN ORTONA ORTONA(CHIETI) | Istituto Zooprofilattico Sperimentale dell'Abruzzo e Molise "G. Caporale" | Lorusso A, Marcacci M, Di Domenico M, Ancora M, Curini V, Mangone I, Delli Compagni E, Rinaldi A, Scialabba S, Di Pasquale A, Cammà C, Puglia I, Calistri P, Savini G |
| EPI_ISL_1403569, EPI_ISL_1403570, EPI_ISL_1403571, EPI_ISL_1403574, EPI_ISL_1403575 | SIESP DIPARTIMENTO DI PREVENZIONE TERAMO TERAMO(TERAMO) | Istituto Zooprofilattico Sperimentale dell'Abruzzo e Molise "G. Caporale" | Lorusso A, Marcacci M, Di Domenico M, Ancora M, Curini V, Mangone I, Delli Compagni E, Rinaldi A, Scialabba S, Di Pasquale A, Cammà C, Puglia I, Calistri P, Savini G |
| EPI_ISL_1403576 | SIESP SULMONA SULMONA(L'AQUILA) | Istituto Zooprofilattico Sperimentale dell'Abruzzo e Molise "G. Caporale" | Lorusso A, Marcacci M, Di Domenico M, Ancora M, Curini V, Mangone I, Delli Compagni E, Rinaldi A, Scialabba S, Di Pasquale A, Cammà C, Puglia I, Calistri P, Savini G |
| EPI_ISL_1403577 | SIESP DIPARTIMENTO DI PREVENZIONE CHIETI | Istituto Zooprofilattico Sperimentale dell'Abruzzo e Molise "G. | Lorusso A, Marcacci M, Di Domenico M, Ancora M, Curini V, Mangone I, Delli Compagni E, Rinaldi A, Scialabba S, Di Pasquale A, Cammà C, Puglia I, |

| | | | |
|---|---|---|---|
| | CHIETI(CHIETI) | Caporale" | Calistri P, Savini G |
| EPI_ISL_1403578, EPI_ISL_1403579, EPI_ISL_1403580 | SIESP DIPARTIMENTO DI PREVENZIONE TERAMO TERAMO(TERAMO) | Istituto Zooprofilattico Sperimentale dell'Abruzzo e Molise "G. Caporale" | Lorusso A, Marcacci M, Di Domenico M, Ancora M, Curini V, Mangone I, Delli Compagni E, Rinaldi A, Scialabba S, Di Pasquale A, Cammà C, Puglia I, Calistri P, Savini G |
| EPI_ISL_1403581 | SIESP CHIETI - DRIVE IN ORTONA ORTONA(CHIETI) | Istituto Zooprofilattico Sperimentale dell'Abruzzo e Molise "G. Caporale" | Lorusso A, Marcacci M, Di Domenico M, Ancora M, Curini V, Mangone I, Delli Compagni E, Rinaldi A, Scialabba S, Di Pasquale A, Cammà C, Puglia I, Calistri P, Savini G |
| EPI_ISL_1403583 | SIESP DIPARTIMENTO DI PREVENZIONE TERAMO TERAMO(TERAMO) | Istituto Zooprofilattico Sperimentale dell'Abruzzo e Molise "G. Caporale" | Lorusso A, Marcacci M, Di Domenico M, Ancora M, Curini V, Mangone I, Delli Compagni E, Rinaldi A, Scialabba S, Di Pasquale A, Cammà C, Puglia I, Calistri P, Savini G |
| EPI_ISL_1403584 | DIP. PREV. AVEZZANO SERVIZIO DI IGIENE EPIDEMIOLOGIA E SANITA' PUBBLICA AVEZZANO(L'AQUILA) | Istituto Zooprofilattico Sperimentale dell'Abruzzo e Molise "G. Caporale" | Lorusso A, Marcacci M, Di Domenico M, Ancora M, Curini V, Mangone I, Delli Compagni E, Rinaldi A, Scialabba S, Di Pasquale A, Cammà C, Puglia I, Calistri P, Savini G |
| EPI_ISL_1403585, EPI_ISL_1403586, EPI_ISL_1403587, EPI_ISL_1403588, EPI_ISL_1403589, EPI_ISL_1403590, EPI_ISL_1403591 | PRESIDIO TERRITORIALE CASOLI CASOLI(CHIETI) | Istituto Zooprofilattico Sperimentale dell'Abruzzo e Molise "G. Caporale" | Lorusso A, Marcacci M, Di Domenico M, Ancora M, Curini V, Mangone I, Delli Compagni E, Rinaldi A, Scialabba S, Di Pasquale A, Cammà C, Puglia I, Calistri P, Savini G |
| EPI_ISL_1403592 | SIESP DIPARTIMENTO DI PREVENZIONE TERAMO TERAMO(TERAMO) | Istituto Zooprofilattico Sperimentale dell'Abruzzo e Molise "G. Caporale" | Lorusso A, Marcacci M, Di Domenico M, Ancora M, Curini V, Mangone I, Delli Compagni E, Rinaldi A, Scialabba S, Di Pasquale A, Cammà C, Puglia I, Calistri P, Savini G |
| EPI_ISL_1403593, EPI_ISL_1403594, EPI_ISL_1403595, EPI_ISL_1403596, EPI_ISL_1403597, EPI_ISL_1403598, EPI_ISL_1403599, EPI_ISL_1403600 | PRESIDIO TERRITORIALE CASOLI CASOLI(CHIETI) | Istituto Zooprofilattico Sperimentale dell'Abruzzo e Molise "G. Caporale" | Lorusso A, Marcacci M, Di Domenico M, Ancora M, Curini V, Mangone I, Delli Compagni E, Rinaldi A, Scialabba S, Di Pasquale A, Cammà C, Puglia I, Calistri P, Savini G |
| EPI_ISL_1403601 | DIP. PREV. AVEZZANO SERVIZIO DI IGIENE EPIDEMIOLOGIA E SANITA' PUBBLICA AVEZZANO(L'AQUILA) | Istituto Zooprofilattico Sperimentale dell'Abruzzo e Molise "G. Caporale" | Lorusso A, Marcacci M, Di Domenico M, Ancora M, Curini V, Mangone I, Delli Compagni E, Rinaldi A, Scialabba S, Di Pasquale A, Cammà C, Puglia I, Calistri P, Savini G |
| EPI_ISL_1403604 | SIESP CHIETI - DRIVE IN LANCIANO LANCIANO(CHIETI) | Istituto Zooprofilattico Sperimentale dell'Abruzzo e Molise "G. Caporale" | Lorusso A, Marcacci M, Di Domenico M, Ancora M, Curini V, Mangone I, Delli Compagni E, Rinaldi A, Scialabba S, Di Pasquale A, Cammà C, Puglia I, Calistri P, Savini G |
| EPI_ISL_1403605 | SIESP DIPARTIMENTO DI PREVENZIONE TERAMO TERAMO(TERAMO) | Istituto Zooprofilattico Sperimentale dell'Abruzzo e Molise "G. Caporale" | Lorusso A, Marcacci M, Di Domenico M, Ancora M, Curini V, Mangone I, Delli Compagni E, Rinaldi A, Scialabba S, Di Pasquale A, Cammà C, Puglia I, Calistri P, Savini G |
| EPI_ISL_1403607, EPI_ISL_1403608, EPI_ISL_1403612, EPI_ISL_1403615, EPI_ISL_1403616, EPI_ISL_1403617, EPI_ISL_1403618, EPI_ISL_1403619, EPI_ISL_1403620, EPI_ISL_1403621, EPI_ISL_1403622, EPI_ISL_1403623, EPI_ISL_1403643, EPI_ISL_1403644, EPI_ISL_1403645, EPI_ISL_1403665 | | | |
| see above | AZ Klina | AZ Klina | Carl Vael - Lynsey Berckmans |
| EPI_ISL_1404383 | Klinisch Laboratorium ZNA | Klinisch Laboratorium ZNA | Verstrepen et al. |
| EPI_ISL_1406249 | INT Fondazione Pascale | INT Fondazione Pascale | INT Fondazione Pascale |
| EPI_ISL_1406519, EPI_ISL_1406520, EPI_ISL_1406521, EPI_ISL_1406541, EPI_ISL_1406542 | Centre Hospitalier Universitaire Clermont-Ferrand | CHU Clermont-Ferrand, service de virologie | Bisseux Maxime, Mirand Audrey, Combes Patricia, Henquell Cécile |
| EPI_ISL_1406543, EPI_ISL_1406545 | HOPITAL DU MONT DORE | CHU Clermont-Ferrand, service de virologie | Bisseux Maxime, Mirand Audrey, Combes Patricia, Henquell Cécile |
| EPI_ISL_1406546, EPI_ISL_1406547, EPI_ISL_1406562, EPI_ISL_1406563, EPI_ISL_1406565, EPI_ISL_1406567, EPI_ISL_1406624, EPI_ISL_1406625, EPI_ISL_1406626, EPI_ISL_1406627, EPI_ISL_1406628, EPI_ISL_1406629 | | | |
| see above | Centre Hospitalier Universitaire Clermont-Ferrand | CHU Clermont-Ferrand, service de virologie | Bisseux Maxime, Mirand Audrey, Combes Patricia, Henquell Cécile |
| EPI_ISL_1406630 | HOPITAL DE MONTLUCON | CHU Clermont-Ferrand, service de virologie | Bisseux Maxime, Mirand Audrey, Combes Patricia, Henquell Cécile |
| EPI_ISL_1406631, EPI_ISL_1406632, EPI_ISL_1406633 | Centre Hospitalier Universitaire Clermont-Ferrand | CHU Clermont-Ferrand, service de virologie | Bisseux Maxime, Mirand Audrey, Combes Patricia, Henquell Cécile |
| EPI_ISL_1406634 | Centre Hospitalier RIOM | CHU Clermont-Ferrand, service de virologie | Bisseux Maxime, Mirand Audrey, Combes Patricia, Henquell Cécile |
| EPI_ISL_1406635, EPI_ISL_1406636 | Centre Hospitalier Universitaire Clermont-Ferrand | CHU Clermont-Ferrand, service de virologie | Bisseux Maxime, Mirand Audrey, Combes Patricia, Henquell Cécile |
| EPI_ISL_1406637 | Centre Jean Perrin | CHU Clermont-Ferrand, service de virologie | Bisseux Maxime, Mirand Audrey, Combes Patricia, Henquell Cécile |
| EPI_ISL_1406638, EPI_ISL_1406641, EPI_ISL_1406642, EPI_ISL_1406643, EPI_ISL_1406645, EPI_ISL_1406646, EPI_ISL_1406647, EPI_ISL_1406648, EPI_ISL_1406654 | Centre Hospitalier Universitaire Clermont-Ferrand | CHU Clermont-Ferrand, service de virologie | Bisseux Maxime, Mirand Audrey, Combes Patricia, Henquell Cécile |
| EPI_ISL_1406657 | Centre De Prélévement COVID VICHY | CHU Clermont-Ferrand, service de virologie | Bisseux Maxime, Mirand Audrey, Combes Patricia, Henquell Cécile |
| EPI_ISL_1406684, EPI_ISL_1406685, EPI_ISL_1406686, EPI_ISL_1406687 | Centre Hospitalier Universitaire Clermont-Ferrand | CHU Clermont-Ferrand, service de virologie | Bisseux Maxime, Mirand Audrey, Combes Patricia, Henquell Cécile |
| EPI_ISL_1406688 | HOPITAL DE VICHY | CHU Clermont-Ferrand, service de virologie | Bisseux Maxime, Mirand Audrey, Combes Patricia, Henquell Cécile |
| EPI_ISL_1406778 | Laboratorio Microbiologia PO Cardarelli | laboratorio Microbiologia PO Cardarelli | Felice Valentina,Giovanna Niro, Massimiliano Scutellà |
| EPI_ISL_1406779 | Laboratorio microbiologia PO Cardarelli | laboratorio microbiologia PO Cardarelli | Valentina Felice, Giovanna Niro, Massimiliano Scutellà |
| EPI_ISL_1406780 | Laboratorio microbiologia PO Cardarelli | Laboratorio microbiologia PO Cardarelli | Valentina Felice, Giovanna Niro, Massimiliano Scutellà |
| EPI_ISL_1406781 | Laboratorio microbiologia PO Cardarelli | laboratorio microbiologia PO Cardarelli | Valentina Felice, Giovanna Niro, Massimiliano Scutellà |
| EPI_ISL_1412932 | SIESP CHIETI - DRIVE IN LANCIANO LANCIANO(CHIETI) | Istituto Zooprofilattico Sperimentale dell'Abruzzo e Molise "G. Caporale" | Lorusso A, Marcacci M, Di Domenico M, Ancora M, Curini V, Di Lollo Valeria, Mangone I, Rinaldi A, Delli Compagni E, Scialabba S, Caporale M, Di Pasquale A, Cammà C, Puglia I, Calistri P, Savini G |
| EPI_ISL_1412935 | SIESP CHIETI - DRIVE IN CHIETI CHIETI(CHIETI) | Istituto Zooprofilattico Sperimentale dell'Abruzzo e Molise "G. Caporale" | Lorusso A, Marcacci M, Di Domenico M, Ancora M, Curini V, Di Lollo Valeria, Mangone I, Rinaldi A, Delli Compagni E, Scialabba S, Caporale M, Di Pasquale A, Cammà C, Puglia I, Calistri P, Savini G |
| EPI_ISL_1418309, EPI_ISL_1418315, EPI_ISL_1418317, EPI_ISL_1418318 | National Institute of Infectious Diseases-Prof. Dr. Matei Bals Molecular Diagnostics Laboratory | National Institute of Infectious Diseases-Prof. Dr. Matei Bals Molecular Diagnostics Laboratory | Corina Casangiu, Leontina Banica, Marius Surleac, Ovidiu Vlaicu, Ionelia Nicolae, Simona Paraschiv, Dan Otelea |
| EPI_ISL_1420829, EPI_ISL_1420830 | SIESP CHIETI - DRIVE IN ORTONA ORTONA(CHIETI) | Istituto Zooprofilattico Sperimentale dell'Abruzzo e Molise "G. Caporale" | Lorusso A, Marcacci M, Di Domenico M, Ancora M, Curini V, Mangone I, Delli Compagni E, Rinaldi A, Scialabba S, Di Pasquale A, Cammà C, Puglia I, Calistri P, Savini G |
| EPI_ISL_1420831 | PRESIDIO TERRITORIALE CASOLI CASOLI(CHIETI) | Istituto Zooprofilattico Sperimentale dell'Abruzzo e Molise "G. Caporale" | Lorusso A, Marcacci M, Di Domenico M, Ancora M, Curini V, Mangone I, Delli Compagni E, Rinaldi A, Scialabba S, Di Pasquale A, Cammà C, Puglia I, Calistri P, Savini G |
| EPI_ISL_1423510 | LESP Ciudad de Mexico | Instituto de Diagnostico y Referencia Epidemiologicos (INDRE) | Claudia Wong-Arambula, Abril Rodriguez-Maldonado, Vanessa Rivero-Arredondo, Ariadna Medina-Benitez, Joaquin Quiroz-Mercado, Sergio Rangel-Guerrero, Natividad Cruz-Ortiz, Tatiana Nunez-Garcia, Gisela Barrera-Badillo, Lucia Hernandez-Rivas, Irma Lopez-Martinez, Ernesto Ramirez-Gonzalez. |

| | | | |
|---|---|---|---|
| EPI_ISL_1427568, EPI_ISL_1428569, EPI_ISL_1428644, EPI_ISL_1429341, EPI_ISL_1429572, EPI_ISL_1429573 | COVID Laboratory AOU Messina | Molecular Hepatology Lab AOU Messina | Giuseppina Raffa, Cristina Musolino, Valeria Chines, Daniele Lombardo, Domenico Giosa, Teresa Pollicino. |
| EPI_ISL_1442994, EPI_ISL_1442995, EPI_ISL_1442996, EPI_ISL_1442997, EPI_ISL_1442998, EPI_ISL_1442999 | Laboratorio di Genetica Medica Matera, Italy | Laboratorio di Genetica Medica Matera, Italy | Domenico Dell'Edera, Arianna Allegretti, Angela Mitidieri, Simone Francesca |
| EPI_ISL_1456121, EPI_ISL_1456450, EPI_ISL_1457196, EPI_ISL_1457409, EPI_ISL_1457771, EPI_ISL_1457772, EPI_ISL_1457774, EPI_ISL_1457775, EPI_ISL_1457776, EPI_ISL_1457777, EPI_ISL_1457778, EPI_ISL_1457779, EPI_ISL_1457780, EPI_ISL_1457781, EPI_ISL_1457782, EPI_ISL_1457783, EPI_ISL_1457784, EPI_ISL_1457785, EPI_ISL_1457786, EPI_ISL_1457787, EPI_ISL_1457788, EPI_ISL_1457789, EPI_ISL_1457790, EPI_ISL_1457791 | | | |
| see above | AZ Klina | AZ Klina | Carl Vael - Lynsey Berckmans |
| EPI_ISL_1464602, EPI_ISL_1464606, EPI_ISL_1464610, EPI_ISL_1464616, EPI_ISL_1464617 | Imeda Hospital | Imelda Hospital | Johan Frans, Dagmar Obbels, Hanne Valgaeren |
| EPI_ISL_1465474, EPI_ISL_1465475, EPI_ISL_1465480, EPI_ISL_1465481 | Institute of Molecular and Translational Medicine / Laboratory of Experimental Medicine, Faculty of Medicine and Dentistry, Palacky University and University Hospital Olomouc | Institute of Molecular and Translational Medicine / Laboratory of Experimental Medicine, Faculty of Medicine and Dentistry, Palacky University | Rastislav Slavkovský, Hana Jaworek, Vladimíra Koudeláková, Barbora Blumová, Tomáš Pospíšil, Marián Hajdúch |
| EPI_ISL_1465613 | Armies | National Reference Center for Viruses of Respiratory Infections, Institut Pasteur, Paris | Marion Barbet, Sylvie Behillil,Frédéric Lemoine,Corinne Maufrais,Christophe Malabat, Méline Bizard, Angela Brisebarre, Camille Capel, Louise Lefrançois, Etienne Simon-Lorière, Vincent Enouf, Maud Vanpeene, Sylvie van der Werf,Marine Desroches |
| EPI_ISL_1465614 | Armies | National Reference Center for Viruses of Respiratory Infections, Institut Pasteur, Paris | Marion Barbet, Sylvie Behillil,Frédéric Lemoine,Corinne Maufrais,Christophe Malabat,Damien Mornico, Méline Bizard, Angela Brisebarre, Camille Capel, Louise Lefrançois, Etienne Simon-Lorière, Vincent Enouf, Maud Vanpeene, Sylvie van der Werf,Marine Desroches |
| EPI_ISL_1469945, EPI_ISL_1469946, EPI_ISL_1469947, EPI_ISL_1469948, EPI_ISL_1469949, EPI_ISL_1469950, EPI_ISL_1469951, EPI_ISL_1469952, EPI_ISL_1469953, EPI_ISL_1469954, EPI_ISL_1469955, EPI_ISL_1469956, EPI_ISL_1469957, EPI_ISL_1469960, EPI_ISL_1469962, EPI_ISL_1469963 | | | |
| see above | University of Bari Biomedical Sciences and Human Oncology | University of Bari Biomedical Sciences and Human Oncology | Chironna M., Sallustio A., Loconsole D., Accogli M. |
| EPI_ISL_1478817, EPI_ISL_1478818, EPI_ISL_1478819, EPI_ISL_1478820, EPI_ISL_1478821, EPI_ISL_1478822, EPI_ISL_1478823, EPI_ISL_1478824, EPI_ISL_1478825, EPI_ISL_1478826, EPI_ISL_1478827, EPI_ISL_1478828, EPI_ISL_1478829, EPI_ISL_1478830, EPI_ISL_1478831, EPI_ISL_1478832, EPI_ISL_1478834, EPI_ISL_1478835, EPI_ISL_1478836, EPI_ISL_1478838, EPI_ISL_1478839, EPI_ISL_1478840, EPI_ISL_1478841, EPI_ISL_1478842, EPI_ISL_1478843 | | | |
| see above | Microbiology Department. Complexo Hospitalario Universitario de Vigo | Microbiology Department. Complexo Hospitalario Universitario de Vigo | Alfaia N, Alonso I, Alvarez M, Cabrera JJ, Carballo R, Cores O, Cortizo S, del-Campo V, Martinez L, Mediero G, Perez S, Potel C, Regueiro B, Rey S, Vasallo FJ |
| EPI_ISL_1483701, EPI_ISL_1483702 | laboratorio microbiologia PO Cardarelli | laboratorio microbiologia PO Cardarelli | Valentina Felice, Massimiliano Scutella. Giovanna Niro |
| EPI_ISL_1483703, EPI_ISL_1483704 | laboratorio micorbiologia PO Cardarelli | laboratorio micorbiologia PO Cardarelli | Valentina Felice, Massimiliano Scutella, Giovanna Niro |
| EPI_ISL_1483705, EPI_ISL_1483706 | laboratorio microbiologia PO Cardarelli | laboratorio microbiologia PO Cardarelli | Valentina Felice, Massimiliano Scutella, Giovanna Niro |
| EPI_ISL_1483707 | HOPITAL DE MONTLUCON | CHU Clermont-Ferrand, service de virologie | Bisseux Maxime, Mirand Audrey, Combes Patricia, Henquell Cécile |
| EPI_ISL_1483708, EPI_ISL_1483709 | Centre Hospitalier Universitaire Clermont-Ferrand | CHU Clermont-Ferrand, service de virologie | Bisseux Maxime, Mirand Audrey, Combes Patricia, Henquell Cécile |
| EPI_ISL_1483710 | HOPITAL DE VICHY | CHU Clermont-Ferrand, service de virologie | Bisseux Maxime, Mirand Audrey, Combes Patricia, Henquell Cécile |
| EPI_ISL_1483711, EPI_ISL_1483712, EPI_ISL_1483713, EPI_ISL_1483714, EPI_ISL_1483715, EPI_ISL_1483716, EPI_ISL_1483717, EPI_ISL_1483718 | Centre Hospitalier Universitaire Clermont-Ferrand | CHU Clermont-Ferrand, service de virologie | Bisseux Maxime, Mirand Audrey, Combes Patricia, Henquell Cécile |
| EPI_ISL_1483719, EPI_ISL_1483720 | Centre hospitalier D'AURILLAC | CHU Clermont-Ferrand, service de virologie | Bisseux Maxime, Mirand Audrey, Combes Patricia, Henquell Cécile |
| EPI_ISL_1483721, EPI_ISL_1483722, EPI_ISL_1483723 | Centre Hospitalier Universitaire Clermont-Ferrand | CHU Clermont-Ferrand, service de virologie | Bisseux Maxime, Mirand Audrey, Combes Patricia, Henquell Cécile |
| EPI_ISL_1483724 | HOPITAL DE VICHY | CHU Clermont-Ferrand, service de virologie | Bisseux Maxime, Mirand Audrey, Combes Patricia, Henquell Cécile |
| EPI_ISL_1483725 | Centre Hospitalier Universitaire Clermont-Ferrand | CHU Clermont-Ferrand, service de virologie | Bisseux Maxime, Mirand Audrey, Combes Patricia, Henquell Cécile |
| EPI_ISL_1483726 | Centre hospitalier D'AURILLAC | CHU Clermont-Ferrand, service de virologie | Bisseux Maxime, Mirand Audrey, Combes Patricia, Henquell Cécile |
| EPI_ISL_1483727, EPI_ISL_1483728 | Centre Hospitalier Universitaire Clermont-Ferrand | CHU Clermont-Ferrand, service de virologie | Bisseux Maxime, Mirand Audrey, Combes Patricia, Henquell Cécile |
| EPI_ISL_1483729 | Centre Hospitalier RIOM | CHU Clermont-Ferrand, service de virologie | Bisseux Maxime, Mirand Audrey, Combes Patricia, Henquell Cécile |
| EPI_ISL_1483730, EPI_ISL_1483731, EPI_ISL_1483732, EPI_ISL_1483733, EPI_ISL_1483734, EPI_ISL_1483735, EPI_ISL_1483736, EPI_ISL_1483737 | Centre Hospitalier Universitaire Clermont-Ferrand | CHU Clermont-Ferrand, service de virologie | Bisseux Maxime, Mirand Audrey, Combes Patricia, Henquell Cécile |
| EPI_ISL_1483738 | HOPITAL DE MONTLUCON | CHU Clermont-Ferrand, service de virologie | Bisseux Maxime, Mirand Audrey, Combes Patricia, Henquell Cécile |
| EPI_ISL_1483739 | Centre Hospitalier Universitaire Clermont-Ferrand | CHU Clermont-Ferrand, service de virologie | Bisseux Maxime, Mirand Audrey, Combes Patricia, Henquell Cécile |
| EPI_ISL_1483740, EPI_ISL_1483741 | HOPITAL DE VICHY | CHU Clermont-Ferrand, service de virologie | Bisseux Maxime, Mirand Audrey, Combes Patricia, Henquell Cécile |
| EPI_ISL_1483742, EPI_ISL_1483743, EPI_ISL_1483744, EPI_ISL_1483745, EPI_ISL_1483746, EPI_ISL_1483747 | Centre Hospitalier Universitaire Clermont-Ferrand | CHU Clermont-Ferrand, service de virologie | Bisseux Maxime, Mirand Audrey, Combes Patricia, Henquell Cécile |
| EPI_ISL_1483748 | PRESCRIPTIONS EXTERIEURES | CHU Clermont-Ferrand, service de virologie | Bisseux Maxime, Mirand Audrey, Combes Patricia, Henquell Cécile |
| EPI_ISL_1483749, EPI_ISL_1483750 | Centre Hospitalier Universitaire Clermont-Ferrand | CHU Clermont-Ferrand, service de virologie | Bisseux Maxime, Mirand Audrey, Combes Patricia, Henquell Cécile |
| EPI_ISL_1483752 | Centre Hospitalier RIOM | CHU Clermont-Ferrand, service de virologie | Bisseux Maxime, Mirand Audrey, Combes Patricia, Henquell Cécile |
| EPI_ISL_1483753 | Centre Hospitalier Universitaire Clermont-Ferrand | CHU Clermont-Ferrand, service de virologie | Bisseux Maxime, Mirand Audrey, Combes Patricia, Henquell Cécile |
| EPI_ISL_1483754 | Centre Hospitalier Paul Ardier | CHU Clermont-Ferrand, service de virologie | Bisseux Maxime, Mirand Audrey, Combes Patricia, Henquell Cécile |
| EPI_ISL_1483755 | Centre Hospitalier Universitaire Clermont-Ferrand | CHU Clermont-Ferrand, service de virologie | Bisseux Maxime, Mirand Audrey, Combes Patricia, Henquell Cécile |
| EPI_ISL_1483756 | Centre Hospitalier RIOM | CHU Clermont-Ferrand, service de virologie | Bisseux Maxime, Mirand Audrey, Combes Patricia, Henquell Cécile |
| EPI_ISL_1483757, EPI_ISL_1483758 | Centre Hospitalier Universitaire Clermont-Ferrand | CHU Clermont-Ferrand, service de virologie | Bisseux Maxime, Mirand Audrey, Combes Patricia, Henquell Cécile |
| EPI_ISL_1483759 | HOPITAL DU MONT DORE | CHU Clermont-Ferrand, service de virologie | Bisseux Maxime, Mirand Audrey, Combes Patricia, Henquell Cécile |
| EPI_ISL_1483760 | HOPITAL DE VICHY | CHU Clermont-Ferrand, service de virologie | Bisseux Maxime, Mirand Audrey, Combes Patricia, Henquell Cécile |
| EPI_ISL_1483761 | HOPITAL DU MONT DORE | CHU Clermont-Ferrand, service de virologie | Bisseux Maxime, Mirand Audrey, Combes Patricia, Henquell Cécile |
| EPI_ISL_1483762, EPI_ISL_1483763, EPI_ISL_1483764, EPI_ISL_1483765, EPI_ISL_1483766, EPI_ISL_1483767, | Centre Hospitalier Universitaire Clermont-Ferrand | CHU Clermont-Ferrand, service de virologie | Bisseux Maxime, Mirand Audrey, Combes Patricia, Henquell Cécile |

| | | | |
|---|---|---|---|
| EPI_ISL_1483768 | | | |
| EPI_ISL_1483769, EPI_ISL_1483770 | HOPITAL DE VICHY | CHU Clermont-Ferrand, service de virologie | Bisseux Maxime, Mirand Audrey, Combes Patricia, Henquell Cécile |
| EPI_ISL_1483771 | Centre De Prélévement COVID VICHY | CHU Clermont-Ferrand, service de virologie | Bisseux Maxime, Mirand Audrey, Combes Patricia, Henquell Cécile |
| EPI_ISL_1483772 | CDP COVID MONTLUCON | CHU Clermont-Ferrand, service de virologie | Bisseux Maxime, Mirand Audrey, Combes Patricia, Henquell Cécile |
| EPI_ISL_1483773, EPI_ISL_1483775 | Centre Hospitalier Universitaire Clermont-Ferrand | CHU Clermont-Ferrand, service de virologie | Bisseux Maxime, Mirand Audrey, Combes Patricia, Henquell Cécile |
| EPI_ISL_1483777, EPI_ISL_1483778, EPI_ISL_1483779 | HOPITAL DU MONT DORE | CHU Clermont-Ferrand, service de virologie | Bisseux Maxime, Mirand Audrey, Combes Patricia, Henquell Cécile |
| EPI_ISL_1483781, EPI_ISL_1483782 | Centre De Prelevement COVID RIOM | CHU Clermont-Ferrand, service de virologie | Bisseux Maxime, Mirand Audrey, Combes Patricia, Henquell Cécile |
| EPI_ISL_1483783, EPI_ISL_1483784, EPI_ISL_1483785, EPI_ISL_1483787, EPI_ISL_1490239, EPI_ISL_1490240, EPI_ISL_1490241, EPI_ISL_1490242, EPI_ISL_1490243 | Centre Hospitalier Universitaire Clermont-Ferrand | CHU Clermont-Ferrand, service de virologie | Bisseux Maxime, Mirand Audrey, Combes Patricia, Henquell Cécile |
| EPI_ISL_1490244 | HOPITAL DE MONTLUCON | CHU Clermont-Ferrand, service de virologie | Bisseux Maxime, Mirand Audrey, Combes Patricia, Henquell Cécile |
| EPI_ISL_1494637, EPI_ISL_1494638, EPI_ISL_1494641, EPI_ISL_1494642 | LESP Ciudad de Mexico | Instituto de Diagnostico y Referencia Epidemiologicos (INDRE) | Claudia Wong-Arambula, Abril Rodriguez-Maldonado, Vanessa Rivero-Arredondo, Ariadna Medina-Benitez, Joaquin Quiroz-Mercado, Sergio Rangel-Guerrero, Natividad Cruz-Ortiz, Tatiana Nunez-Garcia, Gisela Barrera-Badillo, Lucia Hernandez-Rivas, Irma Lopez-Martinez, Ernesto Ramirez-Gonzalez. |
| EPI_ISL_1495145 | Laboratorio microbiologia PO Cardarelli | Laboratorio microbiologia PO Cardarelli | Valentna Felice, Massimiliano Scutella, Giovanna niro |
| EPI_ISL_1495146 | Laboratorio microbiologia PO Cardarelli | Laboratorio microbiologia PO Cardarelli | Valentina Felice, Massimiliano Scutella, Giovanna Niro |
| EPI_ISL_1495147 | Laboratorio microbiologia PO Cardarelli | Laboratorio microbiologia PO Cardarelli | Valentina felice, Massimiliano scutella, Giovanna Niro |
| EPI_ISL_1495148 | Laboratorio microbiologia PO Cardarelli | Laboratorio microbiologia PO Cardarelli | Valentina Felice, Massimiliano Scutella,Giovanna Niro |
| EPI_ISL_1495149 | Laboratorio microbiologia PO Cardarelli | Laboratorio microbiologia PO Cardarelli | Valentina Felice, Massimiliano Scutella, giovanna Niro |
| EPI_ISL_1495151 | Laboratorio microbiologia PO Cardarelli | Laboratorio microbiologia PO Cardarelli | Valentina Felice, Massimiliano Scutella, Giovanna Niro |
| EPI_ISL_1495972 | laboratorio microbiologia PO Cardarelli | laboratorio microbiologia PO Cardarelli | Valentina Felice, Massimiliano Scutella, Giovanna Niro |
| EPI_ISL_1497610 | Ospedale di Venere | Istituto Zooprofilattico Sperimentale della Puglia e della Basilicata | Parisi A., Bianco A., Capozzi L., Del Sambro L., Simone D., Iacobellis M., Ridolfi D., Giannico A. |
| EPI_ISL_1497611, EPI_ISL_1497612 | INT Fondazione Pascale | INT Fondazione Pascale | INT Fondazione Pascale |
| EPI_ISL_1499494, EPI_ISL_1499496, EPI_ISL_1499497, EPI_ISL_1499498 | Azienda Ospedaliera Terni | Istituto Zooprofilattico Sperimentale dell'Abruzzo e Molise "G. Caporale" | Palumbo M, Scaccetti A, Lorusso A, Marcacci M, Di Domenico M, Ancora M, Curini V, Di Lollo V, Delli Compagni E, Mangone I, Rinaldi A, Scialabba S, Caporale M, Di Pasquale A, Cammà C, Puglia I, Calistri P, Savini G |
| EPI_ISL_1499621, EPI_ISL_1499622, EPI_ISL_1499623, EPI_ISL_1499627 | Azienda Ospedaliera Terni | Istituto Zooprofilattico Sperimentale dell'Abruzzo e Molise "G. Caporale" | Palumbo M, Scaccetti A, Lorusso A, Marcacci M, Di Domenico M, Ancora M, Curini V, Di Lollo Valeria, Mangone I, Rinaldi A, Delli Compagni E, Scialabba S, Caporale M, Di Pasquale A, Cammà C, Puglia I, Calistri P, Savini G |
| EPI_ISL_1499628, EPI_ISL_1499629, EPI_ISL_1499630, EPI_ISL_1499631, EPI_ISL_1499632, EPI_ISL_1499633, EPI_ISL_1499634, EPI_ISL_1499635, EPI_ISL_1499636, EPI_ISL_1499637, EPI_ISL_1499638, EPI_ISL_1499639, EPI_ISL_1499640, EPI_ISL_1499641, EPI_ISL_1499642, EPI_ISL_1499643, EPI_ISL_1499644 | | | |
| see above | Università degli Studi di Perugia | Istituto Zooprofilattico Sperimentale dell'Abruzzo e Molise "G. Caporale" | Mencacci A, Camilloni B, Lorusso A, Marcacci M, Di Domenico M, Ancora M, Curini V, Di Lollo Valeria, Mangone I, Rinaldi A, Delli Compagni E, Scialabba S, Caporale M, Di Pasquale A, Cammà C, Puglia I, Calistri P, Savini G |
| EPI_ISL_1499646, EPI_ISL_1499647, EPI_ISL_1499648, EPI_ISL_1499649, EPI_ISL_1499650, EPI_ISL_1499651 | Laboratorio Analisi Osp. Città di Castello - Azienda USL Umbria1 | Istituto Zooprofilattico Sperimentale dell'Abruzzo e Molise "G. Caporale" | Malagigi V, Tacconi P,Lorusso A, Marcacci M, Di Domenico M, Ancora M, Curini V, Di Lollo Valeria, Mangone I, Rinaldi A, Delli Compagni E, Scialabba S, Caporale M, Di Pasquale A, Cammà C, Puglia I, Calistri P, Savini G |
| EPI_ISL_1501219, EPI_ISL_1501221, EPI_ISL_1501222, EPI_ISL_1501223, EPI_ISL_1501224, EPI_ISL_1501225, EPI_ISL_1501226, EPI_ISL_1501227, EPI_ISL_1501228, EPI_ISL_1501230, EPI_ISL_1501231, EPI_ISL_1501232, EPI_ISL_1501233, EPI_ISL_1501234, EPI_ISL_1501235, EPI_ISL_1501236, EPI_ISL_1501237, EPI_ISL_1501238, EPI_ISL_1501239, EPI_ISL_1501240, EPI_ISL_1501241, EPI_ISL_1501242, EPI_ISL_1501243, EPI_ISL_1501244, EPI_ISL_1501245, EPI_ISL_1501246, EPI_ISL_1501247, EPI_ISL_1501248, EPI_ISL_1501249, EPI_ISL_1501250, EPI_ISL_1501251, EPI_ISL_1501253, EPI_ISL_1501254, EPI_ISL_1501255, EPI_ISL_1501256, EPI_ISL_1501257 | | | |
| see above | HEGP - Laboratoire de Virologie | HEGP - Laboratoire de Virologie | David Veyer |
| EPI_ISL_1501265, EPI_ISL_1501267 | Broussais | HEGP - Laboratoire de Virologie | David Veyer |
| EPI_ISL_1502435, EPI_ISL_1502436, EPI_ISL_1502437, EPI_ISL_1502438 | SIESP DIPARTIMENTO DI PREVENZIONE TERAMO TERAMO(TERAMO) | Istituto Zooprofilattico Sperimentale dell'Abruzzo e Molise "G. Caporale" | Lorusso A, Marcacci M, Di Domenico M, Ancora M, Curini V, Di Lollo Valeria, Mangone I, Rinaldi A, Delli Compagni E, Scialabba S, Caporale M, Di Pasquale A, Cammà C, Puglia I, Calistri P, Savini G |
| EPI_ISL_1502439 | SIESP DIPARTIMENTO DI PREVENZIONE CHIETI CHIETI(CHIETI) | Istituto Zooprofilattico Sperimentale dell'Abruzzo e Molise "G. Caporale" | Lorusso A, Marcacci M, Di Domenico M, Ancora M, Curini V, Di Lollo Valeria, Mangone I, Rinaldi A, Delli Compagni E, Scialabba S, Caporale M, Di Pasquale A, Cammà C, Puglia I, Calistri P, Savini G |
| EPI_ISL_1502440 | SIESP CHIETI - DRIVE IN LANCIANO LANCIANO(CHIETI) | Istituto Zooprofilattico Sperimentale dell'Abruzzo e Molise "G. Caporale" | Lorusso A, Marcacci M, Di Domenico M, Ancora M, Curini V, Di Lollo Valeria, Mangone I, Rinaldi A, Delli Compagni E, Scialabba S, Caporale M, Di Pasquale A, Cammà C, Puglia I, Calistri P, Savini G |
| EPI_ISL_1502441 | SIESP CHIETI - DRIVE IN ATESSA ATESSA(CHIETI) | Istituto Zooprofilattico Sperimentale dell'Abruzzo e Molise "G. Caporale" | Lorusso A, Marcacci M, Di Domenico M, Ancora M, Curini V, Di Lollo Valeria, Mangone I, Rinaldi A, Delli Compagni E, Scialabba S, Caporale M, Di Pasquale A, Cammà C, Puglia I, Calistri P, Savini G |
| EPI_ISL_1502442, EPI_ISL_1502443, EPI_ISL_1502444 | SIESP CHIETI - DRIVE IN LANCIANO LANCIANO(CHIETI) | Istituto Zooprofilattico Sperimentale dell'Abruzzo e Molise "G. Caporale" | Lorusso A, Marcacci M, Di Domenico M, Ancora M, Curini V, Di Lollo Valeria, Mangone I, Rinaldi A, Delli Compagni E, Scialabba S, Caporale M, Di Pasquale A, Cammà C, Puglia I, Calistri P, Savini G |
| EPI_ISL_1502445, EPI_ISL_1502446 | SIESP CHIETI - DRIVE IN VASTO VASTO(CHIETI) | Istituto Zooprofilattico Sperimentale dell'Abruzzo e Molise "G. Caporale" | Lorusso A, Marcacci M, Di Domenico M, Ancora M, Curini V, Di Lollo Valeria, Mangone I, Rinaldi A, Delli Compagni E, Scialabba S, Caporale M, Di Pasquale A, Cammà C, Puglia I, Calistri P, Savini G |
| EPI_ISL_1502447, EPI_ISL_1502448 | SIESP CHIETI - DRIVE IN ORTONA ORTONA(CHIETI) | Istituto Zooprofilattico Sperimentale dell'Abruzzo e Molise "G. Caporale" | Lorusso A, Marcacci M, Di Domenico M, Ancora M, Curini V, Di Lollo Valeria, Mangone I, Rinaldi A, Delli Compagni E, Scialabba S, Caporale M, Di Pasquale A, Cammà C, Puglia I, Calistri P, Savini G |
| EPI_ISL_1502449 | OSPEDALE CIVILE GIULIANOVA - PRONTO SOCCORSO GIULIANOVA(TERAMO) | Istituto Zooprofilattico Sperimentale dell'Abruzzo e Molise "G. Caporale" | Lorusso A, Marcacci M, Di Domenico M, Ancora M, Curini V, Di Lollo Valeria, Mangone I, Rinaldi A, Delli Compagni E, Scialabba S, Caporale M, Di Pasquale A, Cammà C, Puglia I, Calistri P, Savini G |
| EPI_ISL_1502450, EPI_ISL_1502451 | SIESP CHIETI - DRIVE IN GISSI GISSI(CHIETI) | Istituto Zooprofilattico Sperimentale dell'Abruzzo e Molise "G. Caporale" | Lorusso A, Marcacci M, Di Domenico M, Ancora M, Curini V, Di Lollo Valeria, Mangone I, Rinaldi A, Delli Compagni E, Scialabba S, Caporale M, Di Pasquale A, Cammà C, Puglia I, Calistri P, Savini G |
| EPI_ISL_1502452, EPI_ISL_1502453, EPI_ISL_1502454, EPI_ISL_1502455, EPI_ISL_1502456, EPI_ISL_1502457, EPI_ISL_1502458, EPI_ISL_1502459 | SIESP DIPARTIMENTO DI PREVENZIONE TERAMO TERAMO(TERAMO) | Istituto Zooprofilattico Sperimentale dell'Abruzzo e Molise "G. Caporale" | Lorusso A, Marcacci M, Di Domenico M, Ancora M, Curini V, Di Lollo Valeria, Mangone I, Rinaldi A, Delli Compagni E, Scialabba S, Caporale M, Di Pasquale A, Cammà C, Puglia I, Calistri P, Savini G |
| EPI_ISL_1502460 | RSA GIULIANOVA GIULIANOVA(TERAMO) | Istituto Zooprofilattico Sperimentale dell'Abruzzo e Molise "G. Caporale" | Lorusso A, Marcacci M, Di Domenico M, Ancora M, Curini V, Di Lollo Valeria, Mangone I, Rinaldi A, Delli Compagni E, Scialabba S, Caporale M, Di Pasquale A, Cammà C, Puglia I, Calistri P, Savini G |
| EPI_ISL_1502461, EPI_ISL_1502462, EPI_ISL_1502463, EPI_ISL_1502464, EPI_ISL_1502465 | SIESP DIPARTIMENTO DI PREVENZIONE TERAMO TERAMO(TERAMO) | Istituto Zooprofilattico Sperimentale dell'Abruzzo e Molise "G. Caporale" | Lorusso A, Marcacci M, Di Domenico M, Ancora M, Curini V, Di Lollo Valeria, Mangone I, Rinaldi A, Delli Compagni E, Scialabba S, Caporale M, Di Pasquale A, Cammà C, Puglia I, Calistri P, Savini G |

| | | | |
|---|---|---|---|
| EPI_ISL_1502466 | SIESP CHIETI - DRIVE IN CHIETI CHIETI(CHIETI) | Istituto Zooprofilattico Sperimentale dell'Abruzzo e Molise "G. Caporale" | Lorusso A, Marcacci M, Di Domenico M, Ancora M, Curini V, Di Lollo Valeria, Mangone I, Rinaldi A, Delli Compagni E, Scialabba S, Caporale M, Di Pasquale A, Cammà C, Puglia I, Calistri P, Savini G |
| EPI_ISL_1502467, EPI_ISL_1502468, EPI_ISL_1502469, EPI_ISL_1502470 | SIESP DIPARTIMENTO DI PREVENZIONE TERAMO TERAMO(TERAMO) | Istituto Zooprofilattico Sperimentale dell'Abruzzo e Molise "G. Caporale" | Lorusso A, Marcacci M, Di Domenico M, Ancora M, Curini V, Di Lollo Valeria, Mangone I, Rinaldi A, Delli Compagni E, Scialabba S, Caporale M, Di Pasquale A, Cammà C, Puglia I, Calistri P, Savini G |
| EPI_ISL_1502471 | SIESP CHIETI - DRIVE IN ORTONA ORTONA(CHIETI) | Istituto Zooprofilattico Sperimentale dell'Abruzzo e Molise "G. Caporale" | Lorusso A, Marcacci M, Di Domenico M, Ancora M, Curini V, Di Lollo Valeria, Mangone I, Rinaldi A, Delli Compagni E, Scialabba S, Caporale M, Di Pasquale A, Cammà C, Puglia I, Calistri P, Savini G |
| EPI_ISL_1502472 | OSPEDALE "F. RENZETTI" LANCIANO - MEDICINA INTERNA LANCIANO(CHIETI) | Istituto Zooprofilattico Sperimentale dell'Abruzzo e Molise "G. Caporale" | Lorusso A, Marcacci M, Di Domenico M, Ancora M, Curini V, Di Lollo Valeria, Mangone I, Rinaldi A, Delli Compagni E, Scialabba S, Caporale M, Di Pasquale A, Cammà C, Puglia I, Calistri P, Savini G |
| EPI_ISL_1502473 | SIESP CHIETI - DRIVE IN LANCIANO LANCIANO(CHIETI) | Istituto Zooprofilattico Sperimentale dell'Abruzzo e Molise "G. Caporale" | Lorusso A, Marcacci M, Di Domenico M, Ancora M, Curini V, Di Lollo Valeria, Mangone I, Rinaldi A, Delli Compagni E, Scialabba S, Caporale M, Di Pasquale A, Cammà C, Puglia I, Calistri P, Savini G |
| EPI_ISL_1502474 | SIESP SULMONA SULMONA(L'AQUILA) | Istituto Zooprofilattico Sperimentale dell'Abruzzo e Molise "G. Caporale" | Lorusso A, Marcacci M, Di Domenico M, Ancora M, Curini V, Di Lollo Valeria, Mangone I, Rinaldi A, Delli Compagni E, Scialabba S, Caporale M, Di Pasquale A, Cammà C, Puglia I, Calistri P, Savini G |
| EPI_ISL_1502475, EPI_ISL_1502476, EPI_ISL_1502477, EPI_ISL_1502478, EPI_ISL_1502479 | DIP. PREV. AVEZZANO SERVIZIO DI IGIENE EPIDEMIOLOGIA E SANITA' PUBBLICA AVEZZANO(L'AQUILA) | Istituto Zooprofilattico Sperimentale dell'Abruzzo e Molise "G. Caporale" | Lorusso A, Marcacci M, Di Domenico M, Ancora M, Curini V, Di Lollo Valeria, Mangone I, Rinaldi A, Delli Compagni E, Scialabba S, Caporale M, Di Pasquale A, Cammà C, Puglia I, Calistri P, Savini G |
| EPI_ISL_1502480 | SIESP DIPARTIMENTO DI PREVENZIONE TERAMO TERAMO(TERAMO) | Istituto Zooprofilattico Sperimentale dell'Abruzzo e Molise "G. Caporale" | Lorusso A, Marcacci M, Di Domenico M, Ancora M, Curini V, Di Lollo Valeria, Mangone I, Rinaldi A, Delli Compagni E, Scialabba S, Caporale M, Di Pasquale A, Cammà C, Puglia I, Calistri P, Savini G |
| EPI_ISL_1502481 | DIP. PREV. AVEZZANO SERVIZIO DI IGIENE EPIDEMIOLOGIA E SANITA' PUBBLICA AVEZZANO(L'AQUILA) | Istituto Zooprofilattico Sperimentale dell'Abruzzo e Molise "G. Caporale" | Lorusso A, Marcacci M, Di Domenico M, Ancora M, Curini V, Di Lollo Valeria, Mangone I, Rinaldi A, Delli Compagni E, Scialabba S, Caporale M, Di Pasquale A, Cammà C, Puglia I, Calistri P, Savini G |
| EPI_ISL_1502482, EPI_ISL_1502483 | SIESP CHIETI - DRIVE IN LANCIANO LANCIANO(CHIETI) | Istituto Zooprofilattico Sperimentale dell'Abruzzo e Molise "G. Caporale" | Lorusso A, Marcacci M, Di Domenico M, Ancora M, Curini V, Di Lollo Valeria, Mangone I, Rinaldi A, Delli Compagni E, Scialabba S, Caporale M, Di Pasquale A, Cammà C, Puglia I, Calistri P, Savini G |
| EPI_ISL_1502484 | SIESP CHIETI - DRIVE IN VASTO VASTO(CHIETI) | Istituto Zooprofilattico Sperimentale dell'Abruzzo e Molise "G. Caporale" | Lorusso A, Marcacci M, Di Domenico M, Ancora M, Curini V, Di Lollo Valeria, Mangone I, Rinaldi A, Delli Compagni E, Scialabba S, Caporale M, Di Pasquale A, Cammà C, Puglia I, Calistri P, Savini G |
| EPI_ISL_1502485, EPI_ISL_1502486 | SIESP CHIETI - DRIVE IN ORTONA ORTONA(CHIETI) | Istituto Zooprofilattico Sperimentale dell'Abruzzo e Molise "G. Caporale" | Lorusso A, Marcacci M, Di Domenico M, Ancora M, Curini V, Di Lollo Valeria, Mangone I, Rinaldi A, Delli Compagni E, Scialabba S, Caporale M, Di Pasquale A, Cammà C, Puglia I, Calistri P, Savini G |
| EPI_ISL_1502487, EPI_ISL_1502488 | SIESP CHIETI - DRIVE IN GISSI GISSI(CHIETI) | Istituto Zooprofilattico Sperimentale dell'Abruzzo e Molise "G. Caporale" | Lorusso A, Marcacci M, Di Domenico M, Ancora M, Curini V, Di Lollo Valeria, Mangone I, Rinaldi A, Delli Compagni E, Scialabba S, Caporale M, Di Pasquale A, Cammà C, Puglia I, Calistri P, Savini G |
| EPI_ISL_1502489, EPI_ISL_1502490 | SIESP CHIETI - DRIVE IN CHIETI CHIETI(CHIETI) | Istituto Zooprofilattico Sperimentale dell'Abruzzo e Molise "G. Caporale" | Lorusso A, Marcacci M, Di Domenico M, Ancora M, Curini V, Di Lollo Valeria, Mangone I, Rinaldi A, Delli Compagni E, Scialabba S, Caporale M, Di Pasquale A, Cammà C, Puglia I, Calistri P, Savini G |
| EPI_ISL_1502491 | SIESP DIPARTIMENTO DI PREVENZIONE CHIETI CHIETI(CHIETI) | Istituto Zooprofilattico Sperimentale dell'Abruzzo e Molise "G. Caporale" | Lorusso A, Marcacci M, Di Domenico M, Ancora M, Curini V, Di Lollo Valeria, Mangone I, Rinaldi A, Delli Compagni E, Scialabba S, Caporale M, Di Pasquale A, Cammà C, Puglia I, Calistri P, Savini G |
| EPI_ISL_1502492, EPI_ISL_1502493, EPI_ISL_1502494, EPI_ISL_1502495, EPI_ISL_1502496, EPI_ISL_1502497 | SIESP DIPARTIMENTO DI PREVENZIONE TERAMO TERAMO(TERAMO) | Istituto Zooprofilattico Sperimentale dell'Abruzzo e Molise "G. Caporale" | Lorusso A, Marcacci M, Di Domenico M, Ancora M, Curini V, Di Lollo Valeria, Mangone I, Rinaldi A, Delli Compagni E, Scialabba S, Caporale M, Di Pasquale A, Cammà C, Puglia I, Calistri P, Savini G |
| EPI_ISL_1502498 | RSA GIULIANOVA GIULIANOVA(TERAMO) | Istituto Zooprofilattico Sperimentale dell'Abruzzo e Molise "G. Caporale" | Lorusso A, Marcacci M, Di Domenico M, Ancora M, Curini V, Di Lollo Valeria, Mangone I, Rinaldi A, Delli Compagni E, Scialabba S, Caporale M, Di Pasquale A, Cammà C, Puglia I, Calistri P, Savini G |
| EPI_ISL_1502499 | SIESP DIPARTIMENTO DI PREVENZIONE TERAMO TERAMO(TERAMO) | Istituto Zooprofilattico Sperimentale dell'Abruzzo e Molise "G. Caporale" | Lorusso A, Marcacci M, Di Domenico M, Ancora M, Curini V, Di Lollo Valeria, Mangone I, Rinaldi A, Delli Compagni E, Scialabba S, Caporale M, Di Pasquale A, Cammà C, Puglia I, Calistri P, Savini G |
| EPI_ISL_1502500, EPI_ISL_1502501, EPI_ISL_1502502 | SIESP DIPARTIMENTO DI PREVENZIONE CHIETI CHIETI(CHIETI) | Istituto Zooprofilattico Sperimentale dell'Abruzzo e Molise "G. Caporale" | Lorusso A, Marcacci M, Di Domenico M, Ancora M, Curini V, Di Lollo Valeria, Mangone I, Rinaldi A, Delli Compagni E, Scialabba S, Caporale M, Di Pasquale A, Cammà C, Puglia I, Calistri P, Savini G |
| EPI_ISL_1502503, EPI_ISL_1502504 | SIESP DIPARTIMENTO DI PREVENZIONE TERAMO TERAMO(TERAMO) | Istituto Zooprofilattico Sperimentale dell'Abruzzo e Molise "G. Caporale" | Lorusso A, Marcacci M, Di Domenico M, Ancora M, Curini V, Di Lollo Valeria, Mangone I, Rinaldi A, Delli Compagni E, Scialabba S, Caporale M, Di Pasquale A, Cammà C, Puglia I, Calistri P, Savini G |
| EPI_ISL_1502506 | SIESP CHIETI - DRIVE IN CHIETI CHIETI(CHIETI) | Istituto Zooprofilattico Sperimentale dell'Abruzzo e Molise "G. Caporale" | Lorusso A, Marcacci M, Di Domenico M, Ancora M, Curini V, Di Lollo Valeria, Mangone I, Rinaldi A, Delli Compagni E, Scialabba S, Caporale M, Di Pasquale A, Cammà C, Puglia I, Calistri P, Savini G |
| EPI_ISL_1502507 | SIESP CHIETI - DRIVE IN ORTONA ORTONA(CHIETI) | Istituto Zooprofilattico Sperimentale dell'Abruzzo e Molise "G. Caporale" | Lorusso A, Marcacci M, Di Domenico M, Ancora M, Curini V, Di Lollo Valeria, Mangone I, Rinaldi A, Delli Compagni E, Scialabba S, Caporale M, Di Pasquale A, Cammà C, Puglia I, Calistri P, Savini G |
| EPI_ISL_1502508, EPI_ISL_1502509, EPI_ISL_1502510, EPI_ISL_1502511 | SIESP SULMONA SULMONA(L'AQUILA) | Istituto Zooprofilattico Sperimentale dell'Abruzzo e Molise "G. Caporale" | Lorusso A, Marcacci M, Di Domenico M, Ancora M, Curini V, Di Lollo Valeria, Mangone I, Rinaldi A, Delli Compagni E, Scialabba S, Caporale M, Di Pasquale A, Cammà C, Puglia I, Calistri P, Savini G |
| EPI_ISL_1502512 | DIP. PREV. AVEZZANO SERVIZIO DI IGIENE EPIDEMIOLOGIA E SANITA' PUBBLICA AVEZZANO(L'AQUILA) | Istituto Zooprofilattico Sperimentale dell'Abruzzo e Molise "G. Caporale" | Lorusso A, Marcacci M, Di Domenico M, Ancora M, Curini V, Di Lollo Valeria, Mangone I, Rinaldi A, Delli Compagni E, Scialabba S, Caporale M, Di Pasquale A, Cammà C, Puglia I, Calistri P, Savini G |
| EPI_ISL_1502513 | SIESP SULMONA SULMONA(L'AQUILA) | Istituto Zooprofilattico Sperimentale dell'Abruzzo e Molise "G. Caporale" | Lorusso A, Marcacci M, Di Domenico M, Ancora M, Curini V, Di Lollo Valeria, Mangone I, Rinaldi A, Delli Compagni E, Scialabba S, Caporale M, Di Pasquale A, Cammà C, Puglia I, Calistri P, Savini G |
| EPI_ISL_1502514, EPI_ISL_1502515, EPI_ISL_1502516 | DIP. PREV. AVEZZANO SERVIZIO DI IGIENE EPIDEMIOLOGIA E SANITA' PUBBLICA AVEZZANO(L'AQUILA) | Istituto Zooprofilattico Sperimentale dell'Abruzzo e Molise "G. Caporale" | Lorusso A, Marcacci M, Di Domenico M, Ancora M, Curini V, Di Lollo Valeria, Mangone I, Rinaldi A, Delli Compagni E, Scialabba S, Caporale M, Di Pasquale A, Cammà C, Puglia I, Calistri P, Savini G |
| EPI_ISL_1502517 | OSPEDALE CIVILE GIULIANOVA - PRONTO SOCCORSO GIULIANOVA(TERAMO) | Istituto Zooprofilattico Sperimentale dell'Abruzzo e Molise "G. Caporale" | Lorusso A, Marcacci M, Di Domenico M, Ancora M, Curini V, Di Lollo Valeria, Mangone I, Rinaldi A, Delli Compagni E, Scialabba S, Caporale M, Di Pasquale A, Cammà C, Puglia I, Calistri P, Savini G |
| EPI_ISL_1502518 | SIESP DIPARTIMENTO DI PREVENZIONE TERAMO TERAMO(TERAMO) | Istituto Zooprofilattico Sperimentale dell'Abruzzo e Molise "G. Caporale" | Lorusso A, Marcacci M, Di Domenico M, Ancora M, Curini V, Di Lollo Valeria, Mangone I, Rinaldi A, Delli Compagni E, Scialabba S, Caporale M, Di Pasquale A, Cammà C, Puglia I, Calistri P, Savini G |
| EPI_ISL_1502519, EPI_ISL_1502520, EPI_ISL_1502521 | DIP. PREV. AVEZZANO SERVIZIO DI IGIENE EPIDEMIOLOGIA E SANITA' PUBBLICA AVEZZANO(L'AQUILA) | Istituto Zooprofilattico Sperimentale dell'Abruzzo e Molise "G. Caporale" | Lorusso A, Marcacci M, Di Domenico M, Ancora M, Curini V, Di Lollo Valeria, Mangone I, Rinaldi A, Delli Compagni E, Scialabba S, Caporale M, Di Pasquale A, Cammà C, Puglia I, Calistri P, Savini G |
| EPI_ISL_1502522 | OSPEDALE "F. RENZETTI" LANCIANO - MEDICINA INTERNA LANCIANO(CHIETI) | Istituto Zooprofilattico Sperimentale dell'Abruzzo e Molise "G. Caporale" | Lorusso A, Marcacci M, Di Domenico M, Ancora M, Curini V, Di Lollo Valeria, Mangone I, Rinaldi A, Delli Compagni E, Scialabba S, Caporale M, Di Pasquale A, Cammà C, Puglia I, Calistri P, Savini G |
| EPI_ISL_1502523 | PRESIDIO TERRITORIALE CASOLI CASOLI(CHIETI) | Istituto Zooprofilattico Sperimentale dell'Abruzzo e Molise "G. Caporale" | Lorusso A, Marcacci M, Di Domenico M, Ancora M, Curini V, Di Lollo Valeria, Mangone I, Rinaldi A, Delli Compagni E, Scialabba S, Caporale M, Di Pasquale A, Cammà C, Puglia I, Calistri P, Savini G |
| EPI_ISL_1502524, EPI_ISL_1502525 | OSPEDALE "F. RENZETTI" LANCIANO - MEDICINA INTERNA LANCIANO(CHIETI) | Istituto Zooprofilattico Sperimentale dell'Abruzzo e Molise "G. Caporale" | Lorusso A, Marcacci M, Di Domenico M, Ancora M, Curini V, Di Lollo Valeria, Mangone I, Rinaldi A, Delli Compagni E, Scialabba S, Caporale M, Di Pasquale A, Cammà C, Puglia I, Calistri P, Savini G |
| EPI_ISL_1502526, EPI_ISL_1502527 | PRESIDIO TERRITORIALE CASOLI CASOLI(CHIETI) | Istituto Zooprofilattico Sperimentale dell'Abruzzo e Molise "G. | Lorusso A, Marcacci M, Di Domenico M, Ancora M, Curini V, Di Lollo Valeria, Mangone I, Rinaldi A, Delli Compagni E, Scialabba S, Caporale M, Di |

| | | | |
|---|---|---|---|
| | | Caporale" | Pasquale A, Cammà C, Puglia I, Calistri P, Savini G |
| EPI_ISL_1502528, EPI_ISL_1502529 | USCA Pescina PESCINA(L'AQUILA) | Istituto Zooprofilattico Sperimentale dell'Abruzzo e Molise "G. Caporale" | Lorusso A, Marcacci M, Di Domenico M, Ancora M, Curini V, Di Lollo Valeria, Mangone I, Rinaldi A, Delli Compagni E, Scialabba S, Caporale M, Di Pasquale A, Cammà C, Puglia I, Calistri P, Savini G |
| EPI_ISL_1502530 | SIESP DIPARTIMENTO DI PREVENZIONE CHIETI CHIETI(CHIETI) | Istituto Zooprofilattico Sperimentale dell'Abruzzo e Molise "G. Caporale" | Lorusso A, Marcacci M, Di Domenico M, Ancora M, Curini V, Di Lollo Valeria, Mangone I, Rinaldi A, Delli Compagni E, Scialabba S, Caporale M, Di Pasquale A, Cammà C, Puglia I, Calistri P, Savini G |
| EPI_ISL_1502531 | SIESP CHIETI - DRIVE IN GISSI GISSI(CHIETI) | Istituto Zooprofilattico Sperimentale dell'Abruzzo e Molise "G. Caporale" | Lorusso A, Marcacci M, Di Domenico M, Ancora M, Curini V, Di Lollo Valeria, Mangone I, Rinaldi A, Delli Compagni E, Scialabba S, Caporale M, Di Pasquale A, Cammà C, Puglia I, Calistri P, Savini G |
| EPI_ISL_1502532, EPI_ISL_1502533 | SIESP DIPARTIMENTO DI PREVENZIONE TERAMO TERAMO(TERAMO) | Istituto Zooprofilattico Sperimentale dell'Abruzzo e Molise "G. Caporale" | Lorusso A, Marcacci M, Di Domenico M, Ancora M, Curini V, Di Lollo Valeria, Mangone I, Rinaldi A, Delli Compagni E, Scialabba S, Caporale M, Di Pasquale A, Cammà C, Puglia I, Calistri P, Savini G |
| EPI_ISL_1502534, EPI_ISL_1502535, EPI_ISL_1502536, EPI_ISL_1502537 | VILLA NOVECENTO S.R.L LANCIANO(CHIETI) | Istituto Zooprofilattico Sperimentale dell'Abruzzo e Molise "G. Caporale" | Lorusso A, Marcacci M, Di Domenico M, Ancora M, Curini V, Di Lollo Valeria, Mangone I, Rinaldi A, Delli Compagni E, Scialabba S, Caporale M, Di Pasquale A, Cammà C, Puglia I, Calistri P, Savini G |
| EPI_ISL_1502538 | SIESP CHIETI - DRIVE IN ORTONA ORTONA(CHIETI) | Istituto Zooprofilattico Sperimentale dell'Abruzzo e Molise "G. Caporale" | Lorusso A, Marcacci M, Di Domenico M, Ancora M, Curini V, Di Lollo Valeria, Mangone I, Rinaldi A, Delli Compagni E, Scialabba S, Caporale M, Di Pasquale A, Cammà C, Puglia I, Calistri P, Savini G |
| EPI_ISL_1502539 | SIESP CHIETI - DRIVE IN CHIETI CHIETI(CHIETI) | Istituto Zooprofilattico Sperimentale dell'Abruzzo e Molise "G. Caporale" | Lorusso A, Marcacci M, Di Domenico M, Ancora M, Curini V, Di Lollo Valeria, Mangone I, Rinaldi A, Delli Compagni E, Scialabba S, Caporale M, Di Pasquale A, Cammà C, Puglia I, Calistri P, Savini G |

EPI_ISL_1502546, EPI_ISL_1502547, EPI_ISL_1502548, EPI_ISL_1502549, EPI_ISL_1502550, EPI_ISL_1502551, EPI_ISL_1502552, EPI_ISL_1502553, EPI_ISL_1502554, EPI_ISL_1502555, EPI_ISL_1502556, EPI_ISL_1502557, EPI_ISL_1502558, EPI_ISL_1502559, EPI_ISL_1502560, EPI_ISL_1502561, EPI_ISL_1502562, EPI_ISL_1502563, EPI_ISL_1502564, EPI_ISL_1502565, EPI_ISL_1502566

| | | | |
|---|---|---|---|
| see above | OSPEDALE SAN SALVATORE - MEDICINA DI LABORATORIO L'AQUILA(L'AQUILA) | Istituto Zooprofilattico Sperimentale dell'Abruzzo e Molise "G. Caporale" | Lorusso A, Marcacci M, Di Domenico M, Ancora M, Curini V, Di Lollo Valeria, Mangone I, Rinaldi A, Delli Compagni E, Scialabba S, Caporale M, Di Pasquale A, Cammà C, Puglia I, Calistri P, Savini G |
| EPI_ISL_1503981, EPI_ISL_1503983, EPI_ISL_1503988, EPI_ISL_1504024 | Virology Unit, AOUP | Virology Unit, AOUP | Marialinda Vatteroni, Susi Frateschi, Mauro Pistello |

EPI_ISL_1509841, EPI_ISL_1509842, EPI_ISL_1509843, EPI_ISL_1509844, EPI_ISL_1509845, EPI_ISL_1509846, EPI_ISL_1509847, EPI_ISL_1509848, EPI_ISL_1509850, EPI_ISL_1509851, EPI_ISL_1509852, EPI_ISL_1509853, EPI_ISL_1509854, EPI_ISL_1509855, EPI_ISL_1509856, EPI_ISL_1509857, EPI_ISL_1509858, EPI_ISL_1509859, EPI_ISL_1509860, EPI_ISL_1509861, EPI_ISL_1509862, EPI_ISL_1509863, EPI_ISL_1509864, EPI_ISL_1509865, EPI_ISL_1509868, EPI_ISL_1509869, EPI_ISL_1509870, EPI_ISL_1509871, EPI_ISL_1509872, EPI_ISL_1509873, EPI_ISL_1509874, EPI_ISL_1509875, EPI_ISL_1509876, EPI_ISL_1509877, EPI_ISL_1509878, EPI_ISL_1509879, EPI_ISL_1509882, EPI_ISL_1509883, EPI_ISL_1509884, EPI_ISL_1509885, EPI_ISL_1509887

| | | | |
|---|---|---|---|
| see above | National Center of Infectious and Parasitic Diseases | National Center of Infectious and Parasitic Diseases | Alexiev, Ivanov, Korsun, Stoitsova, Philipova, Dimitrova, Grigorova L., Hristova, Donchev, Stoykov, Trifonova, Dobrinov, Grigorova I., Kantardjiev |
| EPI_ISL_1516901, EPI_ISL_1516902 | Hospital | National Reference Center for Viruses of Respiratory Infections, Institut Pasteur, Paris | Marion Barbet, Sylvie Behillil, Méline Bizard, Angela Brisebarre, Camille Capel,Frédéric Lemoine,Corinne Maufrais,Christophe Malabat, Louise Lefrançois, Etienne Simon-Lorière, Vincent Enouf, Maud Vanpeene, Sylvie van der Werf,Pascale Martres |
| EPI_ISL_1516903 | Hospital | National Reference Center for Viruses of Respiratory Infections, Institut Pasteur, Paris | Marion Barbet, Sylvie Behillil, Méline Bizard, Angela Brisebarre, Camille Capel,Frédéric Lemoine,Corinne Maufrais,Christophe Malabat,Pierre Lechat, Louise Lefrançois, Etienne Simon-Lorière, Vincent Enouf, Maud Vanpeene, Sylvie van der Werf,Pascale Martres |
| EPI_ISL_1516904, EPI_ISL_1516906 | Hospital | National Reference Center for Viruses of Respiratory Infections, Institut Pasteur, Paris | Marion Barbet, Sylvie Behillil, Méline Bizard, Angela Brisebarre, Camille Capel,Frédéric Lemoine,Corinne Maufrais,Christophe Malabat, Louise Lefrançois, Etienne Simon-Lorière, Vincent Enouf, Maud Vanpeene, Sylvie van der Werf,Pascale Martres |
| EPI_ISL_1516917 | Hospital | National Reference Center for Viruses of Respiratory Infections, Institut Pasteur, Paris | Marion Barbet, Sylvie Behillil, Méline Bizard, Angela Brisebarre, Camille Capel,Frédéric Lemoine,Corinne Maufrais,Christophe Malabat, Louise Lefrançois, Etienne Simon-Lorière, Vincent Enouf, Maud Vanpeene, Sylvie van der Werf,Alain Delepine |
| EPI_ISL_1516920 | Hospital | National Reference Center for Viruses of Respiratory Infections, Institut Pasteur, Paris | Marion Barbet, Sylvie Behillil, Méline Bizard, Angela Brisebarre, Camille Capel,Frédéric Lemoine,Corinne Maufrais,Christophe Malabat,Pierre Lechat, Louise Lefrançois, Etienne Simon-Lorière, Vincent Enouf, Maud Vanpeene, Sylvie van der Werf,Pascale Martres |
| EPI_ISL_1516921 | Hospital | National Reference Center for Viruses of Respiratory Infections, Institut Pasteur, Paris | Marion Barbet, Sylvie Behillil, Méline Bizard, Angela Brisebarre, Camille Capel,Frédéric Lemoine,Corinne Maufrais,Christophe Malabat,Gael Millot, Louise Lefrançois, Etienne Simon-Lorière, Vincent Enouf, Maud Vanpeene, Sylvie van der Werf,Pascale Martres |
| EPI_ISL_1516923, EPI_ISL_1516924 | Armies | National Reference Center for Viruses of Respiratory Infections, Institut Pasteur, Paris | Marion Barbet, Sylvie Behillil, Méline Bizard, Angela Brisebarre, Camille Capel,Frédéric Lemoine,Corinne Maufrais,Christophe Malabat, Louise Lefrançois, Etienne Simon-Lorière, Vincent Enouf, Maud Vanpeene, Sylvie van der Werf,Marine Desroches |
| EPI_ISL_1516981 | Labo Analyses Med | National Reference Center for Viruses of Respiratory Infections, Institut Pasteur, Paris | Marion Barbet, Sylvie Behillil, Méline Bizard, Angela Brisebarre, Camille Capel,Frédéric Lemoine,Corinne Maufrais,Christophe Malabat, Louise Lefrançois, Etienne Simon-Lorière, Vincent Enouf, Maud Vanpeene, Sylvie van der Werf,Jean-FrançOis Comes |
| EPI_ISL_1517005, EPI_ISL_1517007, EPI_ISL_1517008 | Hospital | National Reference Center for Viruses of Respiratory Infections, Institut Pasteur, Paris | Marion Barbet, Sylvie Behillil, Méline Bizard, Angela Brisebarre, Camille Capel,Frédéric Lemoine,Corinne Maufrais,Christophe Malabat, Louise Lefrançois, Etienne Simon-Lorière, Vincent Enouf, Maud Vanpeene, Sylvie van der Werf,Odile Petsaris |
| EPI_ISL_1517022 | Hospital | National Reference Center for Viruses of Respiratory Infections, Institut Pasteur, Paris | Marion Barbet, Sylvie Behillil, Méline Bizard, Angela Brisebarre, Camille Capel,Frédéric Lemoine,Corinne Maufrais,Christophe Malabat, Louise Lefrançois, Etienne Simon-Lorière, Vincent Enouf, Maud Vanpeene, Sylvie van der Werf,H Bisiau |
| EPI_ISL_1517027 | Hospital | National Reference Center for Viruses of Respiratory Infections, Institut Pasteur, Paris | Marion Barbet, Sylvie Behillil, Méline Bizard, Angela Brisebarre, Camille Capel,Frédéric Lemoine,Corinne Maufrais,Christophe Malabat, Louise Lefrançois, Etienne Simon-Lorière, Vincent Enouf, Maud Vanpeene, Sylvie van der Werf,JéRome Guinard |
| EPI_ISL_1517034, EPI_ISL_1517035 | Hospital | National Reference Center for Viruses of Respiratory Infections, Institut Pasteur, Paris | Marion Barbet, Sylvie Behillil, Méline Bizard, Angela Brisebarre, Camille Capel,Frédéric Lemoine,Corinne Maufrais,Christophe Malabat, Louise Lefrançois, Etienne Simon-Lorière, Vincent Enouf, Maud Vanpeene, Sylvie van der Werf,Mj Dufour |
| EPI_ISL_1517070 | Hospital | National Reference Center for Viruses of Respiratory Infections, Institut Pasteur, Paris | Marion Barbet, Sylvie Behillil, Méline Bizard, Angela Brisebarre, Camille Capel,Frédéric Lemoine,Corinne Maufrais,Christophe Malabat, Louise Lefrançois, Etienne Simon-Lorière, Vincent Enouf, Maud Vanpeene, Sylvie van der Werf,Alain Delepine |
| EPI_ISL_1517079, EPI_ISL_1517080 | Hospital | National Reference Center for Viruses of Respiratory Infections, Institut Pasteur, Paris | Marion Barbet, Sylvie Behillil, Méline Bizard, Angela Brisebarre, Camille Capel,Frédéric Lemoine,Corinne Maufrais,Christophe Malabat, Louise Lefrançois, Etienne Simon-Lorière, Vincent Enouf, Maud Vanpeene, Sylvie van der Werf,Pascale Martres |
| EPI_ISL_1517082, EPI_ISL_1517084 | Hospital | National Reference Center for Viruses of Respiratory Infections, Institut Pasteur, Paris | Marion Barbet, Sylvie Behillil, Méline Bizard, Angela Brisebarre, Camille Capel,Frédéric Lemoine,Corinne Maufrais,Christophe Malabat, Louise Lefrançois, Etienne Simon-Lorière, Vincent Enouf, Maud Vanpeene, Sylvie van der Werf,Isabelle Woercel |
| EPI_ISL_1517089, EPI_ISL_1517091, EPI_ISL_1517092, EPI_ISL_1517094 | Armies | National Reference Center for Viruses of Respiratory Infections, Institut Pasteur, Paris | Marion Barbet, Sylvie Behillil, Méline Bizard, Angela Brisebarre, Camille Capel,Frédéric Lemoine,Corinne Maufrais,Christophe Malabat, Louise Lefrançois, Etienne Simon-Lorière, Vincent Enouf, Maud Vanpeene, Sylvie van der Werf,Marine Desroches |
| EPI_ISL_1517095 | Hospital | National Reference Center for Viruses of Respiratory Infections, Institut Pasteur, Paris | Marion Barbet, Sylvie Behillil, Méline Bizard, Angela Brisebarre, Camille Capel,Frédéric Lemoine,Corinne Maufrais,Christophe Malabat, Louise Lefrançois, Etienne Simon-Lorière, Vincent Enouf, Maud Vanpeene, Sylvie van der Werf,H Bisiau |
| EPI_ISL_1517097, EPI_ISL_1517098 | Hospital | National Reference Center for Viruses of Respiratory Infections, Institut Pasteur, Paris | Marion Barbet, Sylvie Behillil, Méline Bizard, Angela Brisebarre, Camille Capel,Frédéric Lemoine,Corinne Maufrais,Christophe Malabat, Louise Lefrançois, Etienne Simon-Lorière, Vincent Enouf, Maud Vanpeene, Sylvie van der Werf,Mj Dufour |
| EPI_ISL_1517099 | Hospital | National Reference Center for Viruses of Respiratory Infections, Institut Pasteur, Paris | Marion Barbet, Sylvie Behillil, Méline Bizard, Angela Brisebarre, Camille Capel,Frédéric Lemoine,Corinne Maufrais,Christophe Malabat, Louise Lefrançois, Etienne Simon-Lorière, Vincent Enouf, Maud Vanpeene, Sylvie van der Werf,Lechat |
| EPI_ISL_1517101 | Hospital | National Reference Center for Viruses of Respiratory Infections, Institut Pasteur, Paris | Marion Barbet, Sylvie Behillil, Méline Bizard, Angela Brisebarre, Camille Capel,Frédéric Lemoine,Corinne Maufrais,Christophe Malabat, Louise Lefrançois, Etienne Simon-Lorière, Vincent Enouf, Maud Vanpeene, Sylvie van der Werf,Sellam |
| EPI_ISL_1517102, EPI_ISL_1517112, EPI_ISL_1517117 | Labo Analyses Med | National Reference Center for Viruses of Respiratory Infections, Institut Pasteur, Paris | Marion Barbet, Sylvie Behillil, Méline Bizard, Angela Brisebarre, Camille Capel,Frédéric Lemoine,Corinne Maufrais,Christophe Malabat, Louise Lefrançois, Etienne Simon-Lorière, Vincent Enouf, Maud Vanpeene, Sylvie van der Werf,Jean-FrançOis Comes |
| EPI_ISL_1517121 | Hospital | National Reference Center for Viruses of Respiratory Infections, Institut Pasteur, Paris | Marion Barbet, Sylvie Behillil, Méline Bizard, Angela Brisebarre, Camille Capel,Frédéric Lemoine,Corinne Maufrais,Christophe Malabat, Louise Lefrançois, Etienne Simon-Lorière, Vincent Enouf, Maud Vanpeene, Sylvie van der Werf,Sellam |
| EPI_ISL_1517124 | Hospital | National Reference Center for Viruses of Respiratory | Marion Barbet, Sylvie Behillil, Méline Bizard, Angela Brisebarre, Camille Capel,Frédéric Lemoine,Corinne Maufrais,Christophe Malabat, Louise Lefrançois, |

| | | | |
|---|---|---|---|
| | | Infections, Institut Pasteur, Paris | Etienne Simon-Lorière, Vincent Enouf, Maud Vanpeene, Sylvie van der Werf,Pascale Martres |
| EPI_ISL_1517126, EPI_ISL_1517127, EPI_ISL_1517129, EPI_ISL_1517130, EPI_ISL_1517132, EPI_ISL_1517135, EPI_ISL_1517136, EPI_ISL_1517138, EPI_ISL_1517140, EPI_ISL_1517142, EPI_ISL_1517143, EPI_ISL_1517145, EPI_ISL_1517148, EPI_ISL_1517149, EPI_ISL_1517152, EPI_ISL_1517153, EPI_ISL_1517156, EPI_ISL_1517158, EPI_ISL_1517159 | | | |
| see above | Hospital | National Reference Center for Viruses of Respiratory Infections, Institut Pasteur, Paris | Marion Barbet, Sylvie Behillil, Méline Bizard, Angela Brisebarre, Camille Capel,Frédéric Lemoine,Corinne Maufrais,Christophe Malabat, Louise Lefrançois, Etienne Simon-Lorière, Vincent Enouf, Maud Vanpeene, Sylvie van der Werf,Nabil Gastli |
| EPI_ISL_1517163, EPI_ISL_1517165 | Hospital | National Reference Center for Viruses of Respiratory Infections, Institut Pasteur, Paris | Marion Barbet, Sylvie Behillil, Méline Bizard, Angela Brisebarre, Camille Capel,Frédéric Lemoine,Corinne Maufrais,Christophe Malabat,Victoire, Louise Lefrançois, Etienne Simon-Lorière, Vincent Enouf, Maud Vanpeene, Sylvie van der Werf,Nabil Gastli |
| EPI_ISL_1517166, EPI_ISL_1517168, EPI_ISL_1517172 | Hospital | National Reference Center for Viruses of Respiratory Infections, Institut Pasteur, Paris | Marion Barbet, Sylvie Behillil, Méline Bizard, Angela Brisebarre, Camille Capel,Frédéric Lemoine,Corinne Maufrais,Christophe Malabat, Louise Lefrançois, Etienne Simon-Lorière, Vincent Enouf, Maud Vanpeene, Sylvie van der Werf,Nabil Gastli |
| EPI_ISL_1517173 | Hospital | National Reference Center for Viruses of Respiratory Infections, Institut Pasteur, Paris | Marion Barbet, Sylvie Behillil, Méline Bizard, Angela Brisebarre, Camille Capel,Frédéric Lemoine,Corinne Maufrais,Christophe Malabat,Victoire, Louise Lefrançois, Etienne Simon-Lorière, Vincent Enouf, Maud Vanpeene, Sylvie van der Werf,Nabil Gastli |
| EPI_ISL_1517174, EPI_ISL_1517176, EPI_ISL_1517178 | Hospital | National Reference Center for Viruses of Respiratory Infections, Institut Pasteur, Paris | Marion Barbet, Sylvie Behillil, Méline Bizard, Angela Brisebarre, Camille Capel,Frédéric Lemoine,Corinne Maufrais,Christophe Malabat, Louise Lefrançois, Etienne Simon-Lorière, Vincent Enouf, Maud Vanpeene, Sylvie van der Werf,Nabil Gastli |
| EPI_ISL_1517194, EPI_ISL_1517195, EPI_ISL_1517196, EPI_ISL_1517197, EPI_ISL_1517199, EPI_ISL_1517200, EPI_ISL_1517201, EPI_ISL_1517202, EPI_ISL_1517203, EPI_ISL_1517204, EPI_ISL_1517205, EPI_ISL_1517206, EPI_ISL_1517207, EPI_ISL_1517208, EPI_ISL_1517210, EPI_ISL_1517212, EPI_ISL_1517213, EPI_ISL_1517215, EPI_ISL_1517216, EPI_ISL_1517217, EPI_ISL_1517218, EPI_ISL_1517219, EPI_ISL_1517220, EPI_ISL_1517221, EPI_ISL_1517222, EPI_ISL_1517223, EPI_ISL_1517224, EPI_ISL_1517227, EPI_ISL_1517228, EPI_ISL_1517229, EPI_ISL_1517230, EPI_ISL_1517231, EPI_ISL_1517232, EPI_ISL_1517233, EPI_ISL_1517234, EPI_ISL_1517235, EPI_ISL_1517236, EPI_ISL_1517237, EPI_ISL_1517238 | | | |
| see above | Labo Analyses Med | National Reference Center for Viruses of Respiratory Infections, Institut Pasteur, Paris | Marion Barbet, Sylvie Behillil, Méline Bizard, Angela Brisebarre, Camille Capel,Frédéric Lemoine,Corinne Maufrais,Christophe Malabat, Louise Lefrançois, Etienne Simon-Lorière, Vincent Enouf, Maud Vanpeene, Sylvie van der Werf,Nabil Gastli |
| EPI_ISL_1517239 | Labo Analyses Med | National Reference Center for Viruses of Respiratory Infections, Institut Pasteur, Paris | Marion Barbet, Sylvie Behillil, Méline Bizard, Angela Brisebarre, Camille Capel,Frédéric Lemoine,Corinne Maufrais,Christophe Malabat, Louise Lefrançois, Etienne Simon-Lorière, Vincent Enouf, Maud Vanpeene, Sylvie van der Werf,Geoffroy Lorin De La Grandmaison |
| EPI_ISL_1517240, EPI_ISL_1517241, EPI_ISL_1517242, EPI_ISL_1517243, EPI_ISL_1517244, EPI_ISL_1517245, EPI_ISL_1517248, EPI_ISL_1517249, EPI_ISL_1517250, EPI_ISL_1517251, EPI_ISL_1517252, EPI_ISL_1517254, EPI_ISL_1517255, EPI_ISL_1517256, EPI_ISL_1517257, EPI_ISL_1517258, EPI_ISL_1517259, EPI_ISL_1517260, EPI_ISL_1517261, EPI_ISL_1517262, EPI_ISL_1517263, EPI_ISL_1517265, EPI_ISL_1517266, EPI_ISL_1517267, EPI_ISL_1517268, EPI_ISL_1517269, EPI_ISL_1517270, EPI_ISL_1517271, EPI_ISL_1517272, EPI_ISL_1517273, EPI_ISL_1517275, EPI_ISL_1517276, EPI_ISL_1517277, EPI_ISL_1517279, EPI_ISL_1517280, EPI_ISL_1517281, EPI_ISL_1517282, EPI_ISL_1517283, EPI_ISL_1517284, EPI_ISL_1517285, EPI_ISL_1517286, EPI_ISL_1517288, EPI_ISL_1517289, EPI_ISL_1517290, EPI_ISL_1517291, EPI_ISL_1517292, EPI_ISL_1517293 | | | |
| see above | Labo Analyses Med | National Reference Center for Viruses of Respiratory Infections, Institut Pasteur, Paris | Marion Barbet, Sylvie Behillil, Méline Bizard, Angela Brisebarre, Camille Capel,Frédéric Lemoine,Corinne Maufrais,Christophe Malabat, Louise Lefrançois, Etienne Simon-Lorière, Vincent Enouf, Maud Vanpeene, Sylvie van der Werf,Nabil Gastli |
| EPI_ISL_1517432 | Inciensa, Instituto Costarricense de Investigación y Enseñanza en Nutrición y Salud | Inciensa, Instituto Costarricense de Investigación y Enseñanza en Nutrición y Salud | Cristian Peréz-Corrales & Barboza-Arguedas E |
| EPI_ISL_1517433 | Inciensa, Instituto Costarricense de Investigación y Enseñanza en Nutrición y Salud | Inciensa, Instituto Costarricense de Investigación y Enseñanza en Nutrición y Salud | Cristian Peréz-Corrales, Barboza-Arguedas E & Centeno-Miranda M |
| EPI_ISL_1517434 | Inciensa, Instituto Costarricense de Investigación y Enseñanza en Nutrición y Salud | Inciensa, Instituto Costarricense de Investigación y Enseñanza en Nutrición y Salud | Cristian Peréz-Corrales, Barboza-Arguedas E & Jimenez-Mora A |
| EPI_ISL_1517457, EPI_ISL_1517458 | Labo Analyses Med | National Reference Center for Viruses of Respiratory Infections, Institut Pasteur, Paris | Marion Barbet, Sylvie Behillil, Méline Bizard, Angela Brisebarre, Camille Capel,Frédéric Lemoine,Corinne Maufrais,Christophe Malabat, Louise Lefrançois, Etienne Simon-Lorière, Vincent Enouf, Maud Vanpeene, Sylvie van der Werf,Jean-FrançOis Comes |
| EPI_ISL_1517464, EPI_ISL_1517465, EPI_ISL_1517466, EPI_ISL_1517467 | Armies | National Reference Center for Viruses of Respiratory Infections, Institut Pasteur, Paris | Marion Barbet, Sylvie Behillil, Méline Bizard, Angela Brisebarre, Camille Capel,Frédéric Lemoine,Corinne Maufrais,Christophe Malabat, Louise Lefrançois, Etienne Simon-Lorière, Vincent Enouf, Maud Vanpeene, Sylvie van der Werf,Marine Desroches |
| EPI_ISL_1517470 | Labo Analyses Med | National Reference Center for Viruses of Respiratory Infections, Institut Pasteur, Paris | Marion Barbet, Sylvie Behillil, Méline Bizard, Angela Brisebarre, Camille Capel,Frédéric Lemoine,Corinne Maufrais,Christophe Malabat, Louise Lefrançois, Etienne Simon-Lorière, Vincent Enouf, Maud Vanpeene, Sylvie van der Werf,GréGoire Potiron |
| EPI_ISL_1517471 | Labo Analyses Med | National Reference Center for Viruses of Respiratory Infections, Institut Pasteur, Paris | Marion Barbet, Sylvie Behillil, Méline Bizard, Angela Brisebarre, Camille Capel,Frédéric Lemoine,Corinne Maufrais,Christophe Malabat, Louise Lefrançois, Etienne Simon-Lorière, Vincent Enouf, Maud Vanpeene, Sylvie van der Werf,Le (V) |
| EPI_ISL_1517472, EPI_ISL_1517473, EPI_ISL_1517474, EPI_ISL_1517475, EPI_ISL_1517476 | Hospital | National Reference Center for Viruses of Respiratory Infections, Institut Pasteur, Paris | Marion Barbet, Sylvie Behillil, Méline Bizard, Angela Brisebarre, Camille Capel,Frédéric Lemoine,Corinne Maufrais,Christophe Malabat, Louise Lefrançois, Etienne Simon-Lorière, Vincent Enouf, Maud Vanpeene, Sylvie van der Werf,Nabil Gastli |
| EPI_ISL_1517479, EPI_ISL_1517480, EPI_ISL_1517481, EPI_ISL_1517482 | Armies | National Reference Center for Viruses of Respiratory Infections, Institut Pasteur, Paris | Marion Barbet, Sylvie Behillil, Méline Bizard, Angela Brisebarre, Camille Capel,Frédéric Lemoine,Corinne Maufrais,Christophe Malabat, Louise Lefrançois, Etienne Simon-Lorière, Vincent Enouf, Maud Vanpeene, Sylvie van der Werf,Marine Desroches |
| EPI_ISL_1517483 | Hospital | National Reference Center for Viruses of Respiratory Infections, Institut Pasteur, Paris | Marion Barbet, Sylvie Behillil, Méline Bizard, Angela Brisebarre, Camille Capel,Frédéric Lemoine,Corinne Maufrais,Christophe Malabat, Louise Lefrançois, Etienne Simon-Lorière, Vincent Enouf, Maud Vanpeene, Sylvie van der Werf,Isabelle Woercel |
| EPI_ISL_1517487, EPI_ISL_1517488 | Hospital | National Reference Center for Viruses of Respiratory Infections, Institut Pasteur, Paris | Marion Barbet, Sylvie Behillil, Méline Bizard, Angela Brisebarre, Camille Capel,Frédéric Lemoine,Corinne Maufrais,Christophe Malabat, Louise Lefrançois, Etienne Simon-Lorière, Vincent Enouf, Maud Vanpeene, Sylvie van der Werf,Jean-Baptiste Bour |
| EPI_ISL_1517489 | Labo Analyses Med | National Reference Center for Viruses of Respiratory Infections, Institut Pasteur, Paris | Marion Barbet, Sylvie Behillil, Méline Bizard, Angela Brisebarre, Camille Capel,Frédéric Lemoine,Corinne Maufrais,Christophe Malabat, Louise Lefrançois, Etienne Simon-Lorière, Vincent Enouf, Maud Vanpeene, Sylvie van der Werf,GréGoire Potiron |
| EPI_ISL_1517490 | Hospital | National Reference Center for Viruses of Respiratory Infections, Institut Pasteur, Paris | Marion Barbet, Sylvie Behillil, Méline Bizard, Angela Brisebarre, Camille Capel,Frédéric Lemoine,Corinne Maufrais,Christophe Malabat, Louise Lefrançois, Etienne Simon-Lorière, Vincent Enouf, Maud Vanpeene, Sylvie van der Werf,Isabelle Woercel |
| EPI_ISL_1517494 | Labo Analyses Med | National Reference Center for Viruses of Respiratory Infections, Institut Pasteur, Paris | Marion Barbet, Sylvie Behillil, Méline Bizard, Angela Brisebarre, Camille Capel,Frédéric Lemoine,Corinne Maufrais,Christophe Malabat, Louise Lefrançois, Etienne Simon-Lorière, Vincent Enouf, Maud Vanpeene, Sylvie van der Werf,Jean-FrançOis Comes |
| EPI_ISL_1517495 | Hospital | National Reference Center for Viruses of Respiratory Infections, Institut Pasteur, Paris | Marion Barbet, Sylvie Behillil, Méline Bizard, Angela Brisebarre, Camille Capel,Frédéric Lemoine,Corinne Maufrais,Christophe Malabat, Louise Lefrançois, Etienne Simon-Lorière, Vincent Enouf, Maud Vanpeene, Sylvie van der Werf,Mj Dufour |
| EPI_ISL_1520098 | Università degli Studi di Perugia | Istituto Zooprofilattico Sperimentale dell'Abruzzo e Molise "G. Caporale" | Mencacci A, Camilloni B, Lorusso A, Marcacci M, Di Domenico M, Ancora M, Curini V, Di Lollo Valeria, Mangone I, Rinaldi A, Delli Compagni E, Scialabba S, Caporale M, Di Pasquale A, Cammà C, Puglia I, Calistri P, Savini G |
| EPI_ISL_1520140, EPI_ISL_1520169, EPI_ISL_1520172, EPI_ISL_1520174, EPI_ISL_1520177, EPI_ISL_1520180, EPI_ISL_1520366, EPI_ISL_1520368, EPI_ISL_1520370, EPI_ISL_1520373, EPI_ISL_1520375 | | | |
| see above | University of Bari Biomedical Sciences and Human Oncology | University of Bari Biomedical Sciences and Human Oncology | Chironna M., Sallustio A., Loconsole D., Accogli M. |
| EPI_ISL_1520377 | University of Bari Biomedical Sciences and Human Oncology | University of Bari Biomedical Sciences and Human Oncology | Chironna M., sallustio A., Loconsole D., Accogli M. |
| EPI_ISL_1520378, EPI_ISL_1520379, EPI_ISL_1520380 | University of Bari Biomedical Sciences and Human Oncology | University of Bari Biomedical Sciences and Human Oncology | Chironna M., Sallustio A., Loconsole D., Accogli M. |
| EPI_ISL_1524753, EPI_ISL_1524755, EPI_ISL_1524773, EPI_ISL_1524774 | National Platform bis UMONS/Jolimont | National Platform bis UMONS/Jolimont | François Dufrasne, Guillaume Bayon-Vicente, Gautier Detry, Ruddy Wattiez |
| EPI_ISL_1524851 | USCA Avezzano AVEZZANO(L'AQUILA) | Istituto Zooprofilattico Sperimentale dell'Abruzzo e Molise "G. Caporale" | Lorusso A, Marcacci M, Di Domenico M, Ancora M, Curini V, Di Lollo Valeria, Mangone I, Rinaldi A, Delli Compagni E, Scialabba S, Caporale M, Di Pasquale A, Cammà C, Puglia I, Calistri P, Savini G |
| EPI_ISL_1524852, EPI_ISL_1524853, EPI_ISL_1524854, EPI_ISL_1524855, EPI_ISL_1524856 | SIESP DIPARTIMENTO DI PREVENZIONE TERAMO TERAMO(TERAMO) | Istituto Zooprofilattico Sperimentale dell'Abruzzo e Molise "G. Caporale" | Lorusso A, Marcacci M, Di Domenico M, Ancora M, Curini V, Di Lollo Valeria, Mangone I, Rinaldi A, Delli Compagni E, Scialabba S, Caporale M, Di Pasquale A, Cammà C, Puglia I, Calistri P, Savini G |

| | | | |
|---|---|---|---|
| EPI_ISL_1524857 | SIESP CHIETI - DRIVE IN CHIETI CHIETI(CHIETI) | Istituto Zooprofilattico Sperimentale dell'Abruzzo e Molise "G. Caporale" | Lorusso A, Marcacci M, Di Domenico M, Ancora M, Curini V, Di Lollo Valeria, Mangone I, Rinaldi A, Delli Compagni E, Scialabba S, Caporale M, Di Pasquale A, Cammà C, Puglia I, Calistri P, Savini G |
| EPI_ISL_1524858 | SIESP CHIETI - DRIVE IN VASTO VASTO(CHIETI) | Istituto Zooprofilattico Sperimentale dell'Abruzzo e Molise "G. Caporale" | Lorusso A, Marcacci M, Di Domenico M, Ancora M, Curini V, Di Lollo Valeria, Mangone I, Rinaldi A, Delli Compagni E, Scialabba S, Caporale M, Di Pasquale A, Cammà C, Puglia I, Calistri P, Savini G |
| EPI_ISL_1524859 | SIESP CHIETI - DRIVE IN GISSI GISSI(CHIETI) | Istituto Zooprofilattico Sperimentale dell'Abruzzo e Molise "G. Caporale" | Lorusso A, Marcacci M, Di Domenico M, Ancora M, Curini V, Di Lollo Valeria, Mangone I, Rinaldi A, Delli Compagni E, Scialabba S, Caporale M, Di Pasquale A, Cammà C, Puglia I, Calistri P, Savini G |
| EPI_ISL_1524860 | SIESP CHIETI - DRIVE IN CHIETI CHIETI(CHIETI) | Istituto Zooprofilattico Sperimentale dell'Abruzzo e Molise "G. Caporale" | Lorusso A, Marcacci M, Di Domenico M, Ancora M, Curini V, Di Lollo Valeria, Mangone I, Rinaldi A, Delli Compagni E, Scialabba S, Caporale M, Di Pasquale A, Cammà C, Puglia I, Calistri P, Savini G |
| EPI_ISL_1524861, EPI_ISL_1524862 | SIESP DIPARTIMENTO DI PREVENZIONE TERAMO TERAMO(TERAMO) | Istituto Zooprofilattico Sperimentale dell'Abruzzo e Molise "G. Caporale" | Lorusso A, Marcacci M, Di Domenico M, Ancora M, Curini V, Di Lollo Valeria, Mangone I, Rinaldi A, Delli Compagni E, Scialabba S, Caporale M, Di Pasquale A, Cammà C, Puglia I, Calistri P, Savini G |
| EPI_ISL_1524863, EPI_ISL_1524864 | SIESP CHIETI - DRIVE IN LANCIANO LANCIANO(CHIETI) | Istituto Zooprofilattico Sperimentale dell'Abruzzo e Molise "G. Caporale" | Lorusso A, Marcacci M, Di Domenico M, Ancora M, Curini V, Di Lollo Valeria, Mangone I, Rinaldi A, Delli Compagni E, Scialabba S, Caporale M, Di Pasquale A, Cammà C, Puglia I, Calistri P, Savini G |
| EPI_ISL_1524865, EPI_ISL_1524866 | SIESP SULMONA SULMONA(L'AQUILA) | Istituto Zooprofilattico Sperimentale dell'Abruzzo e Molise "G. Caporale" | Lorusso A, Marcacci M, Di Domenico M, Ancora M, Curini V, Di Lollo Valeria, Mangone I, Rinaldi A, Delli Compagni E, Scialabba S, Caporale M, Di Pasquale A, Cammà C, Puglia I, Calistri P, Savini G |
| EPI_ISL_1524867 | SIESP CHIETI - DRIVE IN LANCIANO LANCIANO(CHIETI) | Istituto Zooprofilattico Sperimentale dell'Abruzzo e Molise "G. Caporale" | Lorusso A, Marcacci M, Di Domenico M, Ancora M, Curini V, Di Lollo Valeria, Mangone I, Rinaldi A, Delli Compagni E, Scialabba S, Caporale M, Di Pasquale A, Cammà C, Puglia I, Calistri P, Savini G |
| EPI_ISL_1524868 | Assistenza Domiciliare Integrata (ADI) TERAMO TERAMO(TERAMO) | Istituto Zooprofilattico Sperimentale dell'Abruzzo e Molise "G. Caporale" | Lorusso A, Marcacci M, Di Domenico M, Ancora M, Curini V, Di Lollo Valeria, Mangone I, Rinaldi A, Delli Compagni E, Scialabba S, Caporale M, Di Pasquale A, Cammà C, Puglia I, Calistri P, Savini G |
| EPI_ISL_1524869, EPI_ISL_1524870 | SIESP DIPARTIMENTO DI PREVENZIONE TERAMO TERAMO(TERAMO) | Istituto Zooprofilattico Sperimentale dell'Abruzzo e Molise "G. Caporale" | Lorusso A, Marcacci M, Di Domenico M, Ancora M, Curini V, Di Lollo Valeria, Mangone I, Rinaldi A, Delli Compagni E, Scialabba S, Caporale M, Di Pasquale A, Cammà C, Puglia I, Calistri P, Savini G |
| EPI_ISL_1524871 | SIESP SULMONA SULMONA(L'AQUILA) | Istituto Zooprofilattico Sperimentale dell'Abruzzo e Molise "G. Caporale" | Lorusso A, Marcacci M, Di Domenico M, Ancora M, Curini V, Di Lollo Valeria, Mangone I, Rinaldi A, Delli Compagni E, Scialabba S, Caporale M, Di Pasquale A, Cammà C, Puglia I, Calistri P, Savini G |
| EPI_ISL_1524907 | CHU NIMES | CHU NIMES | Stephan Robin, Marie-josée Carles, Sophie Bravo, Agathe Boudet |
| EPI_ISL_1525253 | Somedi laboratories | Imelda Hospital | Johan Frans, Hanne Valgaeren, Dagmar Obbels |
| EPI_ISL_1525254 | Imeda Hospital | Imelda Hospital | Johan Frans, Hanne Valgaeren, Dagmar Obbels |
| EPI_ISL_1525255, EPI_ISL_1525256, EPI_ISL_1525258 | Somedi laboratories | Imelda Hospital | Johan Frans, Hanne Valgaeren, Dagmar Obbels |
| EPI_ISL_1525264 | Hospital | National Reference Center for Viruses of Respiratory Infections, Institut Pasteur, Paris | Marion Barbet, Sylvie Behillil, Méline Bizard, Angela Brisebarre, Camille Capel,Frédéric Lemoine,Corinne Maufrais,Christophe Malabat,Pierre Lechat, Louise Lefrançois, Etienne Simon-Lorière, Vincent Enouf, Maud Vanpeene, Sylvie van der Werf,H Bisiau |
| EPI_ISL_1526735 | Public Health Authority of the Slovak Republic | Laboratory of Genomics and Bioinformatics, Comenius University Science Park | Tatiana Sedláková, Diana Rusáková, Miroslav Böhmer, Anna Giová, Jaroslav Budiš, Tomáš Szemes |
| EPI_ISL_1526778, EPI_ISL_1526779, EPI_ISL_1526780 | Hospital | National Reference Center for Viruses of Respiratory Infections, Institut Pasteur, Paris | Marion Barbet, Sylvie Behillil, Méline Bizard, Angela Brisebarre, Camille Capel,Frédéric Lemoine,Corinne Maufrais,Christophe Malabat, Louise Lefrançois, Etienne Simon-Lorière, Vincent Enouf, Maud Vanpeene, Sylvie van der Werf,Odile Petsaris |
| EPI_ISL_1526782 | Hospital | National Reference Center for Viruses of Respiratory Infections, Institut Pasteur, Paris | Marion Barbet, Sylvie Behillil, Méline Bizard, Angela Brisebarre, Camille Capel,Frédéric Lemoine,Corinne Maufrais,Christophe Malabat, Louise Lefrançois, Etienne Simon-Lorière, Vincent Enouf, Maud Vanpeene, Sylvie van der Werf,H Bisiau |
| EPI_ISL_1526783 | Hospital | National Reference Center for Viruses of Respiratory Infections, Institut Pasteur, Paris | Marion Barbet, Sylvie Behillil, Méline Bizard, Angela Brisebarre, Camille Capel,Frédéric Lemoine,Corinne Maufrais,Christophe Malabat, Louise Lefrançois, Etienne Simon-Lorière, Vincent Enouf, Maud Vanpeene, Sylvie van der Werf,Mj Dufour |
| EPI_ISL_1526785 | Hospital | National Reference Center for Viruses of Respiratory Infections, Institut Pasteur, Paris | Marion Barbet, Sylvie Behillil, Méline Bizard, Angela Brisebarre, Camille Capel,Frédéric Lemoine,Corinne Maufrais,Christophe Malabat, Louise Lefrançois, Etienne Simon-Lorière, Vincent Enouf, Maud Vanpeene, Sylvie van der Werf,Alexandra Ducancelle |
| EPI_ISL_1526786, EPI_ISL_1526787 | Hospital | National Reference Center for Viruses of Respiratory Infections, Institut Pasteur, Paris | Marion Barbet, Sylvie Behillil, Méline Bizard, Angela Brisebarre, Camille Capel,Frédéric Lemoine,Corinne Maufrais,Christophe Malabat, Louise Lefrançois, Etienne Simon-Lorière, Vincent Enouf, Maud Vanpeene, Sylvie van der Werf,Odile Petsaris |
| EPI_ISL_1526788 | Hospital | National Reference Center for Viruses of Respiratory Infections, Institut Pasteur, Paris | Marion Barbet, Sylvie Behillil, Méline Bizard, Angela Brisebarre, Camille Capel,Frédéric Lemoine,Corinne Maufrais,Christophe Malabat, Louise Lefrançois, Etienne Simon-Lorière, Vincent Enouf, Maud Vanpeene, Sylvie van der Werf,Nabil Gastli |
| EPI_ISL_1526789, EPI_ISL_1526790, EPI_ISL_1526791 | Labo Analyses Med | National Reference Center for Viruses of Respiratory Infections, Institut Pasteur, Paris | Marion Barbet, Sylvie Behillil, Méline Bizard, Angela Brisebarre, Camille Capel,Frédéric Lemoine,Corinne Maufrais,Christophe Malabat, Louise Lefrançois, Etienne Simon-Lorière, Vincent Enouf, Maud Vanpeene, Sylvie van der Werf,Nabil Gastli |
| EPI_ISL_1526793 | Hospital | National Reference Center for Viruses of Respiratory Infections, Institut Pasteur, Paris | Marion Barbet, Sylvie Behillil, Méline Bizard, Angela Brisebarre, Camille Capel,Frédéric Lemoine,Corinne Maufrais,Christophe Malabat, Louise Lefrançois, Etienne Simon-Lorière, Vincent Enouf, Maud Vanpeene, Sylvie van der Werf,Alain Delepine |
| EPI_ISL_1526794 | Hospital | National Reference Center for Viruses of Respiratory Infections, Institut Pasteur, Paris | Marion Barbet, Sylvie Behillil, Méline Bizard, Angela Brisebarre, Camille Capel,Frédéric Lemoine,Corinne Maufrais,Christophe Malabat, Louise Lefrançois, Etienne Simon-Lorière, Vincent Enouf, Maud Vanpeene, Sylvie van der Werf,Eleha Guillotel |
| EPI_ISL_1528181 | Hospital | National Reference Center for Viruses of Respiratory Infections, Institut Pasteur, Paris | Marion Barbet, Sylvie Behillil, Frédéric Lemoine, Corinne Maufrais, Christophe Malabat, Méline Bizard, Angela Brisebarre, Camille Capel, Louise Lefrançois, Etienne Simon-Lorière, Vincent Enouf, Maud Vanpeene, Sylvie van der Werf,FrançOise Lunel Fabiani |
| EPI_ISL_1528183, EPI_ISL_1528184 | Armies | National Reference Center for Viruses of Respiratory Infections, Institut Pasteur, Paris | Marion Barbet, Sylvie Behillil, Frédéric Lemoine, Corinne Maufrais, Christophe Malabat, Méline Bizard, Angela Brisebarre, Camille Capel, Louise Lefrançois, Etienne Simon-Lorière, Vincent Enouf, Maud Vanpeene, Sylvie van der Werf,Marine Desroches |
| EPI_ISL_1528185 | Armies | National Reference Center for Viruses of Respiratory Infections, Institut Pasteur, Paris | Marion Barbet, Sylvie Behillil, Frédéric Lemoine, Corinne Maufrais, Christophe Malabat,Amaury Vaysse, Méline Bizard, Angela Brisebarre, Camille Capel, Louise Lefrançois, Etienne Simon-Lorière, Vincent Enouf, Maud Vanpeene, Sylvie van der Werf,Marine Desroches |
| EPI_ISL_1528186, EPI_ISL_1528187 | Hospital | National Reference Center for Viruses of Respiratory Infections, Institut Pasteur, Paris | Marion Barbet, Sylvie Behillil, Frédéric Lemoine, Corinne Maufrais, Christophe Malabat, Méline Bizard, Angela Brisebarre, Camille Capel, Louise Lefrançois, Etienne Simon-Lorière, Vincent Enouf, Maud Vanpeene, Sylvie van der Werf,ValéRie Serazin |
| EPI_ISL_1528188 | Hospital | National Reference Center for Viruses of Respiratory Infections, Institut Pasteur, Paris | Marion Barbet, Sylvie Behillil, Frédéric Lemoine, Corinne Maufrais, Christophe Malabat, Adrien Pain, Méline Bizard, Angela Brisebarre, Camille Capel, Louise Lefrançois, Etienne Simon-Lorière, Vincent Enouf, Maud Vanpeene, Sylvie van der Werf,ValéRie Serazin |
| EPI_ISL_1528189, EPI_ISL_1528190 | Hospital | National Reference Center for Viruses of Respiratory Infections, Institut Pasteur, Paris | Marion Barbet, Sylvie Behillil, Frédéric Lemoine, Corinne Maufrais, Christophe Malabat, Méline Bizard, Angela Brisebarre, Camille Capel, Louise Lefrançois, Etienne Simon-Lorière, Vincent Enouf, Maud Vanpeene, Sylvie van der Werf,ValéRie Serazin |
| EPI_ISL_1528191 | Hospital | National Reference Center for Viruses of Respiratory Infections, Institut Pasteur, Paris | Marion Barbet, Sylvie Behillil, Frédéric Lemoine, Corinne Maufrais, Christophe Malabat,Victoire Baillet, Méline Bizard, Angela Brisebarre, Camille Capel, Louise Lefrançois, Etienne Simon-Lorière, Vincent Enouf, Maud Vanpeene, Sylvie van der Werf,FrançOise Lunel Fabiani |
| EPI_ISL_1528192 | Hospital | National Reference Center for Viruses of Respiratory Infections, Institut Pasteur, Paris | Marion Barbet, Sylvie Behillil, Frédéric Lemoine, Corinne Maufrais, Christophe Malabat,Emmanuelle Permal, Méline Bizard, Angela Brisebarre, Camille Capel, Louise Lefrançois, Etienne Simon-Lorière, Vincent Enouf, Maud Vanpeene, Sylvie van der Werf,FrançOise Lunel Fabiani |
| EPI_ISL_1528193 | Labo Analyses Med | National Reference Center for Viruses of Respiratory Infections, Institut Pasteur, Paris | Marion Barbet, Sylvie Behillil, Frédéric Lemoine, Corinne Maufrais, Christophe Malabat,Victoire Baillet, Méline Bizard, Angela Brisebarre, Camille Capel, Louise Lefrançois, Etienne Simon-Lorière, Vincent Enouf, Maud Vanpeene, Sylvie van der Werf,Pierre-Yves Leonard |
| EPI_ISL_1528194 | Hospital | National Reference Center for Viruses of Respiratory Infections, Institut Pasteur, Paris | Marion Barbet, Sylvie Behillil, Frédéric Lemoine, Corinne Maufrais, Christophe Malabat, Méline Bizard, Angela Brisebarre, Camille Capel, Louise Lefrançois, Etienne Simon-Lorière, Vincent Enouf, Maud Vanpeene, Sylvie van der Werf,Mj Dufour |
| EPI_ISL_1528198 | Hospital | National Reference Center for Viruses of Respiratory | Marion Barbet, Sylvie Behillil, Frédéric Lemoine, Corinne Maufrais, Christophe Malabat, Méline Bizard, Angela Brisebarre, Camille Capel, Louise |

| | | | |
|---|---|---|---|
| | | Infections, Institut Pasteur, Paris | Lefrançois, Etienne Simon-Lorière, Vincent Enouf, Maud Vanpeene, Sylvie van der Werf,H Bisiau |
| EPI_ISL_1528199, EPI_ISL_1528200, EPI_ISL_1528201 | Armies | National Reference Center for Viruses of Respiratory Infections, Institut Pasteur, Paris | Marion Barbet, Sylvie Behillil, Frédéric Lemoine, Corinne Maufrais, Christophe Malabat, Méline Bizard, Angela Brisebarre, Camille Capel, Louise Lefrançois, Etienne Simon-Lorière, Vincent Enouf, Maud Vanpeene, Sylvie van der Werf,Marine Desroches |
| EPI_ISL_1528202 | Labo Analyses Med | National Reference Center for Viruses of Respiratory Infections, Institut Pasteur, Paris | Marion Barbet, Sylvie Behillil, Frédéric Lemoine, Corinne Maufrais, Christophe Malabat, Méline Bizard, Angela Brisebarre, Camille Capel, Louise Lefrançois, Etienne Simon-Lorière, Vincent Enouf, Maud Vanpeene, Sylvie van der Werf,Sarah Schillinger |
| EPI_ISL_1528203, EPI_ISL_1528204, EPI_ISL_1528205, EPI_ISL_1528206, EPI_ISL_1528207 | Hospital | National Reference Center for Viruses of Respiratory Infections, Institut Pasteur, Paris | Marion Barbet, Sylvie Behillil, Frédéric Lemoine, Corinne Maufrais, Christophe Malabat, Méline Bizard, Angela Brisebarre, Camille Capel, Louise Lefrançois, Etienne Simon-Lorière, Vincent Enouf, Maud Vanpeene, Sylvie van der Werf,ValéRie Serazin |
| EPI_ISL_1528208, EPI_ISL_1528209, EPI_ISL_1528210 | Hospital | National Reference Center for Viruses of Respiratory Infections, Institut Pasteur, Paris | Marion Barbet, Sylvie Behillil, Frédéric Lemoine, Corinne Maufrais, Christophe Malabat,Adrien Pain, Méline Bizard, Angela Brisebarre, Camille Capel, Louise Lefrançois, Etienne Simon-Lorière, Vincent Enouf, Maud Vanpeene, Sylvie van der Werf,ValéRie Serazin |
| EPI_ISL_1528211, EPI_ISL_1528212, EPI_ISL_1528213 | Labo Analyses Med | National Reference Center for Viruses of Respiratory Infections, Institut Pasteur, Paris | Marion Barbet, Sylvie Behillil, Frédéric Lemoine, Corinne Maufrais, Christophe Malabat, Méline Bizard, Angela Brisebarre, Camille Capel, Louise Lefrançois, Etienne Simon-Lorière, Vincent Enouf, Maud Vanpeene, Sylvie van der Werf,Pierre-Yves Leonard |
| EPI_ISL_1528215, EPI_ISL_1528216, EPI_ISL_1528217 | Labo Analyses Med | National Reference Center for Viruses of Respiratory Infections, Institut Pasteur, Paris | Marion Barbet, Sylvie Behillil, Frédéric Lemoine, Corinne Maufrais, Christophe Malabat, Méline Bizard, Angela Brisebarre, Camille Capel, Louise Lefrançois, Etienne Simon-Lorière, Vincent Enouf, Maud Vanpeene, Sylvie van der Werf,GréGoire Potiron |
| EPI_ISL_1528218 | Armies | National Reference Center for Viruses of Respiratory Infections, Institut Pasteur, Paris | Marion Barbet, Sylvie Behillil, Frédéric Lemoine, Corinne Maufrais, Christophe Malabat, Méline Bizard, Angela Brisebarre, Camille Capel, Louise Lefrançois, Etienne Simon-Lorière, Vincent Enouf, Maud Vanpeene, Sylvie van der Werf,Marine Desroches |
| EPI_ISL_1528219 | Hospital | National Reference Center for Viruses of Respiratory Infections, Institut Pasteur, Paris | Marion Barbet, Sylvie Behillil, Frédéric Lemoine, Corinne Maufrais, Christophe Malabat, Méline Bizard, Angela Brisebarre, Camille Capel, Louise Lefrançois, Etienne Simon-Lorière, Vincent Enouf, Maud Vanpeene, Sylvie van der Werf,Mj Dufour |
| EPI_ISL_1528220 | Hospital | National Reference Center for Viruses of Respiratory Infections, Institut Pasteur, Paris | Marion Barbet, Sylvie Behillil, Frédéric Lemoine, Corinne Maufrais, Christophe Malabat,Adrien Pain, Méline Bizard, Angela Brisebarre, Camille Capel, Louise Lefrançois, Etienne Simon-Lorière, Vincent Enouf, Maud Vanpeene, Sylvie van der Werf,Isabelle Woercel |
| EPI_ISL_1528221 | Labo Analyses Med | National Reference Center for Viruses of Respiratory Infections, Institut Pasteur, Paris | Marion Barbet, Sylvie Behillil, Frédéric Lemoine, Corinne Maufrais, Christophe Malabat,Victoire Baillet, Méline Bizard, Angela Brisebarre, Camille Capel, Louise Lefrançois, Etienne Simon-Lorière, Vincent Enouf, Maud Vanpeene, Sylvie van der Werf,Jean-FrançOis Comes |
| EPI_ISL_1528230 | Hospital | National Reference Center for Viruses of Respiratory Infections, Institut Pasteur, Paris | Marion Barbet, Sylvie Behillil, Frédéric Lemoine, Corinne Maufrais, Christophe Malabat,Adrien Pain, Méline Bizard, Angela Brisebarre, Camille Capel, Louise Lefrançois, Etienne Simon-Lorière, Vincent Enouf, Maud Vanpeene, Sylvie van der Werf,Isabelle Woercel |
| EPI_ISL_1528231 | Labo Analyses Med | National Reference Center for Viruses of Respiratory Infections, Institut Pasteur, Paris | Marion Barbet, Sylvie Behillil, Frédéric Lemoine, Corinne Maufrais, Christophe Malabat,Amaury Vaysse, Méline Bizard, Angela Brisebarre, Camille Capel, Louise Lefrançois, Etienne Simon-Lorière, Vincent Enouf, Maud Vanpeene, Sylvie van der Werf,Vicky Le |
| EPI_ISL_1528232 | Hospital | National Reference Center for Viruses of Respiratory Infections, Institut Pasteur, Paris | Marion Barbet, Sylvie Behillil, Frédéric Lemoine, Corinne Maufrais, Christophe Malabat,Amaury Vaysse, Méline Bizard, Angela Brisebarre, Camille Capel, Louise Lefrançois, Etienne Simon-Lorière, Vincent Enouf, Maud Vanpeene, Sylvie van der Werf,Rajaonarison |
| EPI_ISL_1528233 | Hospital | National Reference Center for Viruses of Respiratory Infections, Institut Pasteur, Paris | Marion Barbet, Sylvie Behillil, Frédéric Lemoine, Corinne Maufrais, Christophe Malabat,Amaury Vaysse, Méline Bizard, Angela Brisebarre, Camille Capel, Louise Lefrançois, Etienne Simon-Lorière, Vincent Enouf, Maud Vanpeene, Sylvie van der Werf,Pascale Martres |
| EPI_ISL_1528234 | Hospital | National Reference Center for Viruses of Respiratory Infections, Institut Pasteur, Paris | Marion Barbet, Sylvie Behillil, Frédéric Lemoine, Corinne Maufrais, Christophe Malabat, Adrien Pain, Méline Bizard, Angela Brisebarre, Camille Capel, Louise Lefrançois, Etienne Simon-Lorière, Vincent Enouf, Maud Vanpeene, Sylvie van der Werf,ValéRie Serazin |
| EPI_ISL_1528235 | Armies | National Reference Center for Viruses of Respiratory Infections, Institut Pasteur, Paris | Marion Barbet, Sylvie Behillil, Frédéric Lemoine, Corinne Maufrais, Christophe Malabat, Méline Bizard, Angela Brisebarre, Camille Capel, Louise Lefrançois, Etienne Simon-Lorière, Vincent Enouf, Maud Vanpeene, Sylvie van der Werf,Marine Desroches |
| EPI_ISL_1528237, EPI_ISL_1528238 | Labo Analyses Med | National Reference Center for Viruses of Respiratory Infections, Institut Pasteur, Paris | Marion Barbet, Sylvie Behillil, Frédéric Lemoine, Corinne Maufrais, Christophe Malabat, Méline Bizard, Angela Brisebarre, Camille Capel, Louise Lefrançois, Etienne Simon-Lorière, Vincent Enouf, Maud Vanpeene, Sylvie van der Werf,Jean-FrançOis Comes |
| EPI_ISL_1528241 | Hospital | National Reference Center for Viruses of Respiratory Infections, Institut Pasteur, Paris | Marion Barbet, Sylvie Behillil, Frédéric Lemoine, Corinne Maufrais, Christophe Malabat,Victoire Baillet, Méline Bizard, Angela Brisebarre, Camille Capel, Louise Lefrançois, Etienne Simon-Lorière, Vincent Enouf, Maud Vanpeene, Sylvie van der Werf,FrançOise Lunel Fabiani |
| EPI_ISL_1531745, EPI_ISL_1531746, EPI_ISL_1531747, EPI_ISL_1531748, EPI_ISL_1531749, EPI_ISL_1531750, EPI_ISL_1531751, EPI_ISL_1531752, EPI_ISL_1531753, EPI_ISL_1531754, EPI_ISL_1531755, EPI_ISL_1531756, EPI_ISL_1531757, EPI_ISL_1531758, EPI_ISL_1531759, EPI_ISL_1531760, EPI_ISL_1531761, EPI_ISL_1531762, EPI_ISL_1531763, EPI_ISL_1531764, EPI_ISL_1531765, EPI_ISL_1531766, EPI_ISL_1531767, EPI_ISL_1531769 | | | |
| see above | University of Oregon COVID-19 MAP Laboratory | University of Oregon Genomics and Cell Characterization Core Facility (GC3F) | Douglas Turnbull, Ariana White, Jeff Bishop, Jason Carriere |
| EPI_ISL_1532792, EPI_ISL_1532797, EPI_ISL_1532798, EPI_ISL_1532799 | Microbiology Department. Complexo Hospitalario Universitario de Vigo | Microbiology Department. Complexo Hospitalario Universitario de Vigo | Alfaya N, Alonso I, Alvarez M, Cabrera JJ, Carballo R, Cores O, Cortizo S, del-Campo V, Martinez L, Mediero G, Perez S, Potel C, Regueiro B, Rey S, Vasallo FJ |
| EPI_ISL_1534706 | National Institute Health Research and Development | National Institute of Health Research and Development | Subangkit, Hana Apsari Pawestri, Kartika Dewi Puspa, Arie Ardiansyah Nugraha, Hartanti Dian Ikawati, Kristina Nur Andriana Pangesti, Yuni Rukminiati, Ririn Ramadhany, Agustiningsih, Kindi Adam, Holy Arif Wibowo, Triyani Soekarso, Ni Ketut Susilarini, Nurika Hariastuti, Ully Alfi Nikmah, Reni Herman, Nike Susanti, Herna, Tati Febriyanti, Natalie Laurencia Kipuw, Fauzul Muna, Irene Lorinda Indalao, Nelly Puspandari, Vivi Setiawaty. |
| EPI_ISL_1540445 | Hospital | National Reference Center for Viruses of Respiratory Infections, Institut Pasteur, Paris | Marion Barbet, Sylvie Behillil, Frédéric Lemoine, Corinne Maufrais, Christophe Malabat,Amaury Vaysse, Méline Bizard, Angela Brisebarre, Camille Capel, Louise Lefrançois, Etienne Simon-Lorière, Vincent Enouf, Maud Vanpeene, Sylvie van der Werf,Odile Petsaris |
| EPI_ISL_1540446 | Labo Analyses Med | National Reference Center for Viruses of Respiratory Infections, Institut Pasteur, Paris | Marion Barbet, Sylvie Behillil, Frédéric Lemoine, Corinne Maufrais, Christophe Malabat,Amaury Vaysse, Méline Bizard, Angela Brisebarre, Camille Capel, Louise Lefrançois, Etienne Simon-Lorière, Vincent Enouf, Maud Vanpeene, Sylvie van der Werf,Jean-FrançOis Comes |
| EPI_ISL_1540449 | Hospital | National Reference Center for Viruses of Respiratory Infections, Institut Pasteur, Paris | Marion Barbet, Sylvie Behillil, Frédéric Lemoine, Corinne Maufrais, Christophe Malabat,Emmanuelle Permal, Méline Bizard, Angela Brisebarre, Camille Capel, Louise Lefrançois, Etienne Simon-Lorière, Vincent Enouf, Maud Vanpeene, Sylvie van der Werf,Haciba Moudjahed |
| EPI_ISL_1540450 | Hospital | National Reference Center for Viruses of Respiratory Infections, Institut Pasteur, Paris | Marion Barbet, Sylvie Behillil, Frédéric Lemoine, Corinne Maufrais, Christophe Malabat,Emmanuelle Permal, Méline Bizard, Angela Brisebarre, Camille Capel, Louise Lefrançois, Etienne Simon-Lorière, Vincent Enouf, Maud Vanpeene, Sylvie van der Werf,ValéRie Serazin |
| EPI_ISL_1540707 | Universidad Tecnica Particular de Loja | Institute of Microbiology, Universidad San Francisco de Quito | Belén Prado-Vivar, Sully Márquez, Juan José Guadalupe, Monica Becerra-Wong, Bernardo Gutiérrez, Paola Dalgo, Fernando Serrano, Katherine Ojeda, David Zuñiga, Melissa Ortega, Raiza Briceño, Luis Flores, Verónica Barragán, Patricio Rojas-Silva, Gabriel Trueba, Michelle Grunauer, Paúl Cárdenas |
| EPI_ISL_1545633, EPI_ISL_1545635, EPI_ISL_1545636 | CHWAPI - SITE NOTRE DAME | Institut de Pathologie et Genetique (IPG) | Pascale Hilbert, Jérémie Gras |
| EPI_ISL_1545642 | Cabinet Priv | Institut de Pathologie et Genetique (IPG) | Pascale Hilbert, Jérémie Gras |
| EPI_ISL_1545654 | CHWAPI - SITE NOTRE DAME | Institut de Pathologie et Genetique (IPG) | Pascale Hilbert, Jérémie Gras |
| EPI_ISL_1547397, EPI_ISL_1547398 | Centre hospitalier D'AURILLAC | CHU Clermont-Ferrand, service de virologie | Bisseux Maxime, Mirand Audrey, Combes Patricia, Henquell Cécile |
| EPI_ISL_1547399 | Centre Hospitalier Universitaire Clermont-Ferrand | CHU Clermont-Ferrand, service de virologie | Bisseux Maxime, Mirand Audrey, Combes Patricia, Henquell Cécile |
| EPI_ISL_1547400 | HOPITAL DE MONTLUCON | CHU Clermont-Ferrand, service de virologie | Bisseux Maxime, Mirand Audrey, Combes Patricia, Henquell Cécile |
| EPI_ISL_1547401, EPI_ISL_1547402, EPI_ISL_1547403, EPI_ISL_1547404, EPI_ISL_1547405, EPI_ISL_1547406 | Centre Hospitalier Universitaire Clermont-Ferrand | CHU Clermont-Ferrand, service de virologie | Bisseux Maxime, Mirand Audrey, Combes Patricia, Henquell Cécile |
| EPI_ISL_1547407 | Centre hospitalier D'AURILLAC | CHU Clermont-Ferrand, service de virologie | Bisseux Maxime, Mirand Audrey, Combes Patricia, Henquell Cécile |
| EPI_ISL_1547408, EPI_ISL_1547409, EPI_ISL_1547410, EPI_ISL_1547411 | Centre Hospitalier Universitaire Clermont-Ferrand | CHU Clermont-Ferrand, service de virologie | Bisseux Maxime, Mirand Audrey, Combes Patricia, Henquell Cécile |

| | | | |
|---|---|---|---|
| EPI_ISL_1547412 | HOPITAL DU MONT DORE | CHU Clermont-Ferrand, service de virologie | Bisseux Maxime, Mirand Audrey, Combes Patricia, Henquell Cécile |
| EPI_ISL_1547413, EPI_ISL_1547414, EPI_ISL_1547415, EPI_ISL_1547416 | Centre Hospitalier Universitaire Clermont-Ferrand | CHU Clermont-Ferrand, service de virologie | Bisseux Maxime, Mirand Audrey, Combes Patricia, Henquell Cécile |
| EPI_ISL_1547417 | Centre hospitalier D'AURILLAC | CHU Clermont-Ferrand, service de virologie | Bisseux Maxime, Mirand Audrey, Combes Patricia, Henquell Cécile |
| EPI_ISL_1547418, EPI_ISL_1547419, EPI_ISL_1547420 | Centre Hospitalier Universitaire Clermont-Ferrand | CHU Clermont-Ferrand, service de virologie | Bisseux Maxime, Mirand Audrey, Combes Patricia, Henquell Cécile |
| EPI_ISL_1547421 | HOPITAL DE MONTLUCON | CHU Clermont-Ferrand, service de virologie | Bisseux Maxime, Mirand Audrey, Combes Patricia, Henquell Cécile |
| EPI_ISL_1547422, EPI_ISL_1547423, EPI_ISL_1547424, EPI_ISL_1547425 | Centre Hospitalier Universitaire Clermont-Ferrand | CHU Clermont-Ferrand, service de virologie | Bisseux Maxime, Mirand Audrey, Combes Patricia, Henquell Cécile |
| EPI_ISL_1547426 | Centre Hospitalier Paul Ardier | CHU Clermont-Ferrand, service de virologie | Bisseux Maxime, Mirand Audrey, Combes Patricia, Henquell Cécile |
| EPI_ISL_1547427 | HOPITAL DE MONTLUCON | CHU Clermont-Ferrand, service de virologie | Bisseux Maxime, Mirand Audrey, Combes Patricia, Henquell Cécile |
| EPI_ISL_1547428 | Centre Hospitalier Paul Ardier | CHU Clermont-Ferrand, service de virologie | Bisseux Maxime, Mirand Audrey, Combes Patricia, Henquell Cécile |
| EPI_ISL_1547429 | Centre Hospitalier RIOM | CHU Clermont-Ferrand, service de virologie | Bisseux Maxime, Mirand Audrey, Combes Patricia, Henquell Cécile |
| EPI_ISL_1547430, EPI_ISL_1547431, EPI_ISL_1547432, EPI_ISL_1547433 | Centre Hospitalier Universitaire Clermont-Ferrand | CHU Clermont-Ferrand, service de virologie | Bisseux Maxime, Mirand Audrey, Combes Patricia, Henquell Cécile |
| EPI_ISL_1547434 | Centre Hospitalier RIOM | CHU Clermont-Ferrand, service de virologie | Bisseux Maxime, Mirand Audrey, Combes Patricia, Henquell Cécile |
| EPI_ISL_1547435, EPI_ISL_1547436, EPI_ISL_1547437, EPI_ISL_1547438, EPI_ISL_1547439, EPI_ISL_1547440 | Centre Hospitalier Universitaire Clermont-Ferrand | CHU Clermont-Ferrand, service de virologie | Bisseux Maxime, Mirand Audrey, Combes Patricia, Henquell Cécile |
| EPI_ISL_1547441, EPI_ISL_1547442, EPI_ISL_1547443, EPI_ISL_1547444 | CDP COVID MONTLUCON | CHU Clermont-Ferrand, service de virologie | Bisseux Maxime, Mirand Audrey, Combes Patricia, Henquell Cécile |
| EPI_ISL_1547445, EPI_ISL_1547446 | Centre Hospitalier Universitaire Clermont-Ferrand | CHU Clermont-Ferrand, service de virologie | Bisseux Maxime, Mirand Audrey, Combes Patricia, Henquell Cécile |
| EPI_ISL_1547447 | HOPITAL DE VICHY | CHU Clermont-Ferrand, service de virologie | Bisseux Maxime, Mirand Audrey, Combes Patricia, Henquell Cécile |
| EPI_ISL_1547450, EPI_ISL_1547451 | Centre Hospitalier Universitaire Clermont-Ferrand | CHU Clermont-Ferrand, service de virologie | Bisseux Maxime, Mirand Audrey, Combes Patricia, Henquell Cécile |
| EPI_ISL_1547452 | Centre Hospitalier Paul Ardier | CHU Clermont-Ferrand, service de virologie | Bisseux Maxime, Mirand Audrey, Combes Patricia, Henquell Cécile |
| EPI_ISL_1547453 | HOPITAL DU MONT DORE | CHU Clermont-Ferrand, service de virologie | Bisseux Maxime, Mirand Audrey, Combes Patricia, Henquell Cécile |
| EPI_ISL_1547454 | Centre De Prelevement COVID RIOM | CHU Clermont-Ferrand, service de virologie | Bisseux Maxime, Mirand Audrey, Combes Patricia, Henquell Cécile |
| EPI_ISL_1547455 | Centre Hospitalier Universitaire Clermont-Ferrand | CHU Clermont-Ferrand, service de virologie | Bisseux Maxime, Mirand Audrey, Combes Patricia, Henquell Cécile |
| EPI_ISL_1547456 | Centre De Prélévement COVID VICHY | CHU Clermont-Ferrand, service de virologie | Bisseux Maxime, Mirand Audrey, Combes Patricia, Henquell Cécile |
| EPI_ISL_1547457, EPI_ISL_1547458 | Centre Hospitalier Universitaire Clermont-Ferrand | CHU Clermont-Ferrand, service de virologie | Bisseux Maxime, Mirand Audrey, Combes Patricia, Henquell Cécile |
| EPI_ISL_1547459 | HOPITAL DE VICHY | CHU Clermont-Ferrand, service de virologie | Bisseux Maxime, Mirand Audrey, Combes Patricia, Henquell Cécile |
| EPI_ISL_1547460 | Centre Hospitalier RIOM | CHU Clermont-Ferrand, service de virologie | Bisseux Maxime, Mirand Audrey, Combes Patricia, Henquell Cécile |
| EPI_ISL_1547461 | Centre hospitalier D'AURILLAC | CHU Clermont-Ferrand, service de virologie | Bisseux Maxime, Mirand Audrey, Combes Patricia, Henquell Cécile |
| EPI_ISL_1547462 | HOPITAL DE VICHY | CHU Clermont-Ferrand, service de virologie | Bisseux Maxime, Mirand Audrey, Combes Patricia, Henquell Cécile |
| EPI_ISL_1547463, EPI_ISL_1547464 | Centre Hospitalier Universitaire Clermont-Ferrand | CHU Clermont-Ferrand, service de virologie | Bisseux Maxime, Mirand Audrey, Combes Patricia, Henquell Cécile |
| EPI_ISL_1547465, EPI_ISL_1547466, EPI_ISL_1547467 | HOPITAL DE VICHY | CHU Clermont-Ferrand, service de virologie | Bisseux Maxime, Mirand Audrey, Combes Patricia, Henquell Cécile |
| EPI_ISL_1547468 | Centre Hospitalier Universitaire Clermont-Ferrand | CHU Clermont-Ferrand, service de virologie | Bisseux Maxime, Mirand Audrey, Combes Patricia, Henquell Cécile |
| EPI_ISL_1547469 | HOPITAL DE MONTLUCON | CHU Clermont-Ferrand, service de virologie | Bisseux Maxime, Mirand Audrey, Combes Patricia, Henquell Cécile |
| EPI_ISL_1547470 | HOPITAL DE VICHY | CHU Clermont-Ferrand, service de virologie | Bisseux Maxime, Mirand Audrey, Combes Patricia, Henquell Cécile |
| EPI_ISL_1547471 | Centre Hospitalier Universitaire Clermont-Ferrand | CHU Clermont-Ferrand, service de virologie | Bisseux Maxime, Mirand Audrey, Combes Patricia, Henquell Cécile |
| EPI_ISL_1547506 | Cerballiance Pont Audemer | Cerba Lab | Haïm-Boukobza S, Prigent A, Leriche A, Verdurme L, Trombert-Paolantoni S, Lecorche E, Roquebert B |
| EPI_ISL_1547827 | LBM ALPHABIO | LBM ALPHABIO | VINCENT GARCIA, PHILIPPE HALFON |
| EPI_ISL_1549134, EPI_ISL_1549135, EPI_ISL_1549136, EPI_ISL_1549137, EPI_ISL_1549138 | OLVZ Aalst | OLVZ Aalst | Astrid Holderbeke |
| EPI_ISL_1554899 | Hospital | National Reference Center for Viruses of Respiratory Infections, Institut Pasteur, Paris | Marion Barbet, Sylvie Behillil, Méline Bizard, Angela Brisebarre, Camille Capel,Frédéric Lemoine,Corinne Maufrais,Christophe Malabat, Damien Mornico, Louise Lefrançois, Etienne Simon-Lorière, Vincent Enouf, Maud Vanpeene, Sylvie van der Werf,Farfour Eric |
| EPI_ISL_1557219, EPI_ISL_1557220, EPI_ISL_1557221 | Laboratorio Biologia Molecolare Sars Cov2 - UOC Laboratorio Analisi - Servizio Medicina di Laboratorio, Ospedale "San Francesco" - ATS-ASSL Nuoro | Laboratorio specialistico UOC Ematologia - Ospedale "San Francesco" - ATS-ASSL Nuoro | Piras Giovanna, Malune Paolo,Asproni Rosanna,,Fiamma Maura, Monne Maria Itria, Palmas Angelo Domenico, Lo Maglio Iana, Mameli Giuseppe |
| EPI_ISL_1559626 | Diagnostic Genomics Laboratory | Diagnostic Genomics Laboratory | Carolyn Banister, Fadi Abboodi, Ji (Emily) Hao, Diego Altomoare, Michael Shtutman, Phillip Buckhaults, Helmut Albrecht |
| EPI_ISL_1575080, EPI_ISL_1575112 | Cruces University Hospital | Biocruces Bizkaia | Mikel Gallego Rodrigo, Ana Belen de la Hoz Rastrollo |
| EPI_ISL_1577356, EPI_ISL_1577357, EPI_ISL_1577358, EPI_ISL_1577359 | CHU Lille | CHU Lille - Laboratoire de Virologie | AIT YAHYA Emilie, ALIDJINOU Enagnon Kazali, BOCKET Laurence, CREPIN Michel, DEMAY Christophe, ENGELMANN Ilka, GEFFROY Sandrine, GUIGON Aurélie, LAMBERT Valérie, LAZREK Mouna, NOBILLIAUX Florian, PREVOST Brigitte, THUILLIER Caroline, TINEZ Claire ,TCHANTCHOU NJOSSE YANICK |
| EPI_ISL_1577369 | CH Cambrai | CHU Lille - Laboratoire de Virologie | AIT YAHYA Emilie, ALIDJINOU Enagnon Kazali, BOCKET Laurence, CREPIN Michel, DEMAY Christophe, ENGELMANN Ilka, GEFFROY Sandrine, GUIGON Aurélie, LAMBERT Valérie, LAZREK Mouna, NOBILLIAUX Florian, PREVOST Brigitte, THUILLIER Caroline, TINEZ Claire ,TCHANTCHOU NJOSSE YANICK |
| EPI_ISL_1577370, EPI_ISL_1577371, EPI_ISL_1577372 | CHU Lille | CHU Lille - Laboratoire de Virologie | AIT YAHYA Emilie, ALIDJINOU Enagnon Kazali, BOCKET Laurence, CREPIN Michel, DEMAY Christophe, ENGELMANN Ilka, GEFFROY Sandrine, GUIGON Aurélie, LAMBERT Valérie, LAZREK Mouna, NOBILLIAUX Florian, PREVOST Brigitte, THUILLIER Caroline, TINEZ Claire ,TCHANTCHOU NJOSSE YANICK |
| EPI_ISL_1577376 | CH Lens | CHU Lille - Laboratoire de Virologie | AIT YAHYA Emilie, ALIDJINOU Enagnon Kazali, BOCKET Laurence, CREPIN Michel, DEMAY Christophe, ENGELMANN Ilka, GEFFROY Sandrine, GUIGON Aurélie, LAMBERT Valérie, LAZREK Mouna, NOBILLIAUX Florian, PREVOST Brigitte, THUILLIER Caroline, TINEZ Claire ,TCHANTCHOU NJOSSE YANICK |

| | | | |
|---|---|---|---|
| EPI_ISL_1577377, EPI_ISL_1577378, EPI_ISL_1577381, EPI_ISL_1577383, EPI_ISL_1577384, EPI_ISL_1577385, EPI_ISL_1577386, EPI_ISL_1577387, EPI_ISL_1577388 | CHU Lille | CHU Lille - Laboratoire de Virologie | AIT YAHYA Emilie, ALIDJINOU Enagnon Kazali, BOCKET Laurence, CREPIN Michel, DEMAY Christophe, ENGELMANN Ilka, GEFFROY Sandrine, GUIGON Aurélie, LAMBERT Valérie, LAZREK Mouna, NOBILLIAUX Florian, PREVOST Brigitte, THUILLIER Caroline, TINEZ Claire ,TCHANTCHOU NJOSSE YANICK |
| EPI_ISL_1577389 | CH Roubaix | CHU Lille - Laboratoire de Virologie | AIT YAHYA Emilie, ALIDJINOU Enagnon Kazali, BOCKET Laurence, CREPIN Michel, DEMAY Christophe, ENGELMANN Ilka, GEFFROY Sandrine, GUIGON Aurélie, LAMBERT Valérie, LAZREK Mouna, NOBILLIAUX Florian, PREVOST Brigitte, THUILLIER Caroline, TINEZ Claire ,TCHANTCHOU NJOSSE YANICK |
| EPI_ISL_1577906, EPI_ISL_1577907, EPI_ISL_1577911 | Laboratory for Clinical Immunology and Molecular Genetics - University Clinic Golnik Laboratory for Respiratory Microbiology - University Clinic Golnik | Laboratory for Clinical Immunology and Molecular Genetics -University Clinic Golnik | Matija Rijavec, Julij Šelb, Urška Bidovec Stojkovi, Žan Kogovšek, Nina Rupar, Viktorija Tomi, Peter Korošec |
| EPI_ISL_1577915 | PRESIDIO TERRITORIALE CASOLI CASOLI(CHIETI) | Istituto Zooprofilattico Sperimentale dell'Abruzzo e Molise "G. Caporale" | Lorusso A, Marcacci M, Di Domenico M, Ancora M, Curini V, Mangone I, Rinaldi A, Scialabba S, Di Pasquale A, Cammà C, Puglia I, Calistri P, Savini G |
| EPI_ISL_1577916 | SIESP DIPARTIMENTO DI PREVENZIONE CHIETI CHIETI(CHIETI) | Istituto Zooprofilattico Sperimentale dell'Abruzzo e Molise "G. Caporale" | Lorusso A, Marcacci M, Di Domenico M, Ancora M, Curini V, Mangone I, Rinaldi A, Scialabba S, Di Pasquale A, Cammà C, Puglia I, Calistri P, Savini G |
| EPI_ISL_1577917, EPI_ISL_1577918 | SIESP DIPARTIMENTO DI PREVENZIONE TERAMO TERAMO(TERAMO) | Istituto Zooprofilattico Sperimentale dell'Abruzzo e Molise "G. Caporale" | Lorusso A, Marcacci M, Di Domenico M, Ancora M, Curini V, Mangone I, Rinaldi A, Scialabba S, Di Pasquale A, Cammà C, Puglia I, Calistri P, Savini G |
| EPI_ISL_1577919 | SIESP SULMONA SULMONA(L'AQUILA) | Istituto Zooprofilattico Sperimentale dell'Abruzzo e Molise "G. Caporale" | Lorusso A, Marcacci M, Di Domenico M, Ancora M, Curini V, Mangone I, Rinaldi A, Scialabba S, Di Pasquale A, Cammà C, Puglia I, Calistri P, Savini G |
| EPI_ISL_1577920, EPI_ISL_1577921, EPI_ISL_1577922 | DIP. PREV. AVEZZANO SERVIZIO DI IGIENE EPIDEMIOLOGIA E SANITA' PUBBLICA AVEZZANO(L'AQUILA) | Istituto Zooprofilattico Sperimentale dell'Abruzzo e Molise "G. Caporale" | Lorusso A, Marcacci M, Di Domenico M, Ancora M, Curini V, Mangone I, Rinaldi A, Scialabba S, Di Pasquale A, Cammà C, Puglia I, Calistri P, Savini G |
| EPI_ISL_1577923 | SIESP DIPARTIMENTO DI PREVENZIONE TERAMO TERAMO(TERAMO) | Istituto Zooprofilattico Sperimentale dell'Abruzzo e Molise "G. Caporale" | Lorusso A, Marcacci M, Di Domenico M, Ancora M, Curini V, Mangone I, Rinaldi A, Scialabba S, Di Pasquale A, Cammà C, Puglia I, Calistri P, Savini G |
| EPI_ISL_1577924, EPI_ISL_1577925 | USCA Avezzano AVEZZANO(L'AQUILA) | Istituto Zooprofilattico Sperimentale dell'Abruzzo e Molise "G. Caporale" | Lorusso A, Marcacci M, Di Domenico M, Ancora M, Curini V, Mangone I, Rinaldi A, Scialabba S, Di Pasquale A, Cammà C, Puglia I, Calistri P, Savini G |
| EPI_ISL_1577926 | USCA Pescina PESCINA(L'AQUILA) | Istituto Zooprofilattico Sperimentale dell'Abruzzo e Molise "G. Caporale" | Lorusso A, Marcacci M, Di Domenico M, Ancora M, Curini V, Mangone I, Rinaldi A, Scialabba S, Di Pasquale A, Cammà C, Puglia I, Calistri P, Savini G |
| EPI_ISL_1577927 | USCA Tagliacozzo TAGLIACOZZO(L'AQUILA) | Istituto Zooprofilattico Sperimentale dell'Abruzzo e Molise "G. Caporale" | Lorusso A, Marcacci M, Di Domenico M, Ancora M, Curini V, Mangone I, Rinaldi A, Scialabba S, Di Pasquale A, Cammà C, Puglia I, Calistri P, Savini G |
| EPI_ISL_1577928 | SIESP CHIETI - DRIVE IN LANCIANO LANCIANO(CHIETI) | Istituto Zooprofilattico Sperimentale dell'Abruzzo e Molise "G. Caporale" | Lorusso A, Marcacci M, Di Domenico M, Ancora M, Curini V, Mangone I, Rinaldi A, Scialabba S, Di Pasquale A, Cammà C, Puglia I, Calistri P, Savini G |
| EPI_ISL_1577929 | SIESP CHIETI - DRIVE IN VASTO VASTO(CHIETI) | Istituto Zooprofilattico Sperimentale dell'Abruzzo e Molise "G. Caporale" | Lorusso A, Marcacci M, Di Domenico M, Ancora M, Curini V, Mangone I, Rinaldi A, Scialabba S, Di Pasquale A, Cammà C, Puglia I, Calistri P, Savini G |
| EPI_ISL_1577930 | SIESP CHIETI - DRIVE IN CHIETI CHIETI(CHIETI) | Istituto Zooprofilattico Sperimentale dell'Abruzzo e Molise "G. Caporale" | Lorusso A, Marcacci M, Di Domenico M, Ancora M, Curini V, Mangone I, Rinaldi A, Scialabba S, Di Pasquale A, Cammà C, Puglia I, Calistri P, Savini G |
| EPI_ISL_1577931, EPI_ISL_1577932 | SIESP DIPARTIMENTO DI PREVENZIONE CHIETI CHIETI(CHIETI) | Istituto Zooprofilattico Sperimentale dell'Abruzzo e Molise "G. Caporale" | Lorusso A, Marcacci M, Di Domenico M, Ancora M, Curini V, Mangone I, Rinaldi A, Scialabba S, Di Pasquale A, Cammà C, Puglia I, Calistri P, Savini G |
| EPI_ISL_1577933 | SIESP CHIETI - DRIVE IN GISSI GISSI(CHIETI) | Istituto Zooprofilattico Sperimentale dell'Abruzzo e Molise "G. Caporale" | Lorusso A, Marcacci M, Di Domenico M, Ancora M, Curini V, Mangone I, Rinaldi A, Scialabba S, Di Pasquale A, Cammà C, Puglia I, Calistri P, Savini G |
| EPI_ISL_1577934, EPI_ISL_1577935, EPI_ISL_1577936, EPI_ISL_1577937, EPI_ISL_1577938, EPI_ISL_1577939, EPI_ISL_1577940 | SIESP DIPARTIMENTO DI PREVENZIONE TERAMO TERAMO(TERAMO) | Istituto Zooprofilattico Sperimentale dell'Abruzzo e Molise "G. Caporale" | Lorusso A, Marcacci M, Di Domenico M, Ancora M, Curini V, Mangone I, Rinaldi A, Scialabba S, Di Pasquale A, Cammà C, Puglia I, Calistri P, Savini G |
| EPI_ISL_1577941 | Servizio di Prevenzione e Protezione dell'IZSAM TERAMO(TERAMO) | Istituto Zooprofilattico Sperimentale dell'Abruzzo e Molise "G. Caporale" | Lorusso A, Marcacci M, Di Domenico M, Ancora M, Curini V, Mangone I, Rinaldi A, Scialabba S, Di Pasquale A, Cammà C, Puglia I, Calistri P, Savini G |
| EPI_ISL_1577942 | SIESP DIPARTIMENTO DI PREVENZIONE TERAMO TERAMO(TERAMO) | Istituto Zooprofilattico Sperimentale dell'Abruzzo e Molise "G. Caporale" | Lorusso A, Marcacci M, Di Domenico M, Ancora M, Curini V, Mangone I, Rinaldi A, Scialabba S, Di Pasquale A, Cammà C, Puglia I, Calistri P, Savini G |
| EPI_ISL_1577943 | SIESP DIPARTIMENTO DI PREVENZIONE CHIETI CHIETI(CHIETI) | Istituto Zooprofilattico Sperimentale dell'Abruzzo e Molise "G. Caporale" | Lorusso A, Marcacci M, Di Domenico M, Ancora M, Curini V, Mangone I, Rinaldi A, Scialabba S, Di Pasquale A, Cammà C, Puglia I, Calistri P, Savini G |
| EPI_ISL_1577944 | DIP. PREV. AVEZZANO SERVIZIO DI IGIENE EPIDEMIOLOGIA E SANITA' PUBBLICA AVEZZANO(L'AQUILA) | Istituto Zooprofilattico Sperimentale dell'Abruzzo e Molise "G. Caporale" | Lorusso A, Marcacci M, Di Domenico M, Ancora M, Curini V, Mangone I, Rinaldi A, Scialabba S, Di Pasquale A, Cammà C, Puglia I, Calistri P, Savini G |
| EPI_ISL_1577945 | SIESP SULMONA SULMONA(L'AQUILA) | Istituto Zooprofilattico Sperimentale dell'Abruzzo e Molise "G. Caporale" | Lorusso A, Marcacci M, Di Domenico M, Ancora M, Curini V, Mangone I, Rinaldi A, Scialabba S, Di Pasquale A, Cammà C, Puglia I, Calistri P, Savini G |
| EPI_ISL_1577946 | SIESP CHIETI - DRIVE IN ORTONA ORTONA(CHIETI) | Istituto Zooprofilattico Sperimentale dell'Abruzzo e Molise "G. Caporale" | Lorusso A, Marcacci M, Di Domenico M, Ancora M, Curini V, Mangone I, Rinaldi A, Scialabba S, Di Pasquale A, Cammà C, Puglia I, Calistri P, Savini G |
| EPI_ISL_1577947 | SIESP SULMONA SULMONA(L'AQUILA) | Istituto Zooprofilattico Sperimentale dell'Abruzzo e Molise "G. Caporale" | Lorusso A, Marcacci M, Di Domenico M, Ancora M, Curini V, Mangone I, Rinaldi A, Scialabba S, Di Pasquale A, Cammà C, Puglia I, Calistri P, Savini G |
| EPI_ISL_1577948 | DIP. PREV. AVEZZANO SERVIZIO DI IGIENE EPIDEMIOLOGIA E SANITA' PUBBLICA AVEZZANO(L'AQUILA) | Istituto Zooprofilattico Sperimentale dell'Abruzzo e Molise "G. Caporale" | Lorusso A, Marcacci M, Di Domenico M, Ancora M, Curini V, Mangone I, Rinaldi A, Scialabba S, Di Pasquale A, Cammà C, Puglia I, Calistri P, Savini G |
| EPI_ISL_1577949 | SIESP DIPARTIMENTO DI PREVENZIONE TERAMO TERAMO(TERAMO) | Istituto Zooprofilattico Sperimentale dell'Abruzzo e Molise "G. Caporale" | Lorusso A, Marcacci M, Di Domenico M, Ancora M, Curini V, Mangone I, Rinaldi A, Scialabba S, Di Pasquale A, Cammà C, Puglia I, Calistri P, Savini G |
| EPI_ISL_1577950 | SIESP CHIETI - DRIVE IN ORTONA ORTONA(CHIETI) | Istituto Zooprofilattico Sperimentale dell'Abruzzo e Molise "G. Caporale" | Lorusso A, Marcacci M, Di Domenico M, Ancora M, Curini V, Mangone I, Rinaldi A, Scialabba S, Di Pasquale A, Cammà C, Puglia I, Calistri P, Savini G |
| EPI_ISL_1577951 | SIESP DIPARTIMENTO DI PREVENZIONE TERAMO TERAMO(TERAMO) | Istituto Zooprofilattico Sperimentale dell'Abruzzo e Molise "G. Caporale" | Lorusso A, Marcacci M, Di Domenico M, Ancora M, Curini V, Mangone I, Rinaldi A, Scialabba S, Di Pasquale A, Cammà C, Puglia I, Calistri P, Savini G |
| EPI_ISL_1577952 | SIESP SULMONA SULMONA(L'AQUILA) | Istituto Zooprofilattico Sperimentale dell'Abruzzo e Molise "G. Caporale" | Lorusso A, Marcacci M, Di Domenico M, Ancora M, Curini V, Mangone I, Rinaldi A, Scialabba S, Di Pasquale A, Cammà C, Puglia I, Calistri P, Savini G |
| EPI_ISL_1577953 | USCA Avezzano AVEZZANO(L'AQUILA) | Istituto Zooprofilattico Sperimentale dell'Abruzzo e Molise "G. Caporale" | Lorusso A, Marcacci M, Di Domenico M, Ancora M, Curini V, Mangone I, Rinaldi A, Scialabba S, Di Pasquale A, Cammà C, Puglia I, Calistri P, Savini G |

| | | | |
|---|---|---|---|
| EPI_ISL_1577954 | USCA Pescina PESCINA(L'AQUILA) | Istituto Zooprofilattico Sperimentale dell'Abruzzo e Molise "G. Caporale" | Lorusso A, Marcacci M, Di Domenico M, Ancora M, Curini V, Mangone I, Rinaldi A, Scialabba S, Di Pasquale A, Cammà C, Puglia I, Calistri P, Savini G |
| EPI_ISL_1577955 | USCA Sulmona SULMONA(L'AQUILA) | Istituto Zooprofilattico Sperimentale dell'Abruzzo e Molise "G. Caporale" | Lorusso A, Marcacci M, Di Domenico M, Ancora M, Curini V, Mangone I, Rinaldi A, Scialabba S, Di Pasquale A, Cammà C, Puglia I, Calistri P, Savini G |
| EPI_ISL_1577956, EPI_ISL_1577957, EPI_ISL_1577958 | SIESP CHIETI - DRIVE IN ORTONA ORTONA(CHIETI) | Istituto Zooprofilattico Sperimentale dell'Abruzzo e Molise "G. Caporale" | Lorusso A, Marcacci M, Di Domenico M, Ancora M, Curini V, Mangone I, Rinaldi A, Scialabba S, Di Pasquale A, Cammà C, Puglia I, Calistri P, Savini G |
| EPI_ISL_1577959 | USCA Pescina PESCINA(L'AQUILA) | Istituto Zooprofilattico Sperimentale dell'Abruzzo e Molise "G. Caporale" | Lorusso A, Marcacci M, Di Domenico M, Ancora M, Curini V, Mangone I, Rinaldi A, Scialabba S, Di Pasquale A, Cammà C, Puglia I, Calistri P, Savini G |
| EPI_ISL_1577960, EPI_ISL_1577961, EPI_ISL_1577962 | SIESP CHIETI - DRIVE IN LANCIANO LANCIANO(CHIETI) | Istituto Zooprofilattico Sperimentale dell'Abruzzo e Molise "G. Caporale" | Lorusso A, Marcacci M, Di Domenico M, Ancora M, Curini V, Mangone I, Rinaldi A, Scialabba S, Di Pasquale A, Cammà C, Puglia I, Calistri P, Savini G |
| EPI_ISL_1577963 | VILLA NOVECENTO S.R.L LANCIANO(CHIETI) | Istituto Zooprofilattico Sperimentale dell'Abruzzo e Molise "G. Caporale" | Lorusso A, Marcacci M, Di Domenico M, Ancora M, Curini V, Mangone I, Rinaldi A, Scialabba S, Di Pasquale A, Cammà C, Puglia I, Calistri P, Savini G |
| EPI_ISL_1577964 | SIESP DIPARTIMENTO DI PREVENZIONE CHIETI CHIETI(CHIETI) | Istituto Zooprofilattico Sperimentale dell'Abruzzo e Molise "G. Caporale" | Lorusso A, Marcacci M, Di Domenico M, Ancora M, Curini V, Mangone I, Rinaldi A, Scialabba S, Di Pasquale A, Cammà C, Puglia I, Calistri P, Savini G |
| EPI_ISL_1577965 | SIESP CHIETI - DRIVE IN GISSI GISSI(CHIETI) | Istituto Zooprofilattico Sperimentale dell'Abruzzo e Molise "G. Caporale" | Lorusso A, Marcacci M, Di Domenico M, Ancora M, Curini V, Mangone I, Rinaldi A, Scialabba S, Di Pasquale A, Cammà C, Puglia I, Calistri P, Savini G |
| EPI_ISL_1577966 | SIESP CHIETI - DRIVE IN VASTO VASTO(CHIETI) | Istituto Zooprofilattico Sperimentale dell'Abruzzo e Molise "G. Caporale" | Lorusso A, Marcacci M, Di Domenico M, Ancora M, Curini V, Mangone I, Rinaldi A, Scialabba S, Di Pasquale A, Cammà C, Puglia I, Calistri P, Savini G |
| EPI_ISL_1577967 | SIESP CHIETI - DRIVE IN GISSI GISSI(CHIETI) | Istituto Zooprofilattico Sperimentale dell'Abruzzo e Molise "G. Caporale" | Lorusso A, Marcacci M, Di Domenico M, Ancora M, Curini V, Mangone I, Rinaldi A, Scialabba S, Di Pasquale A, Cammà C, Puglia I, Calistri P, Savini G |
| EPI_ISL_1577968 | SIESP CHIETI - DRIVE IN VASTO VASTO(CHIETI) | Istituto Zooprofilattico Sperimentale dell'Abruzzo e Molise "G. Caporale" | Lorusso A, Marcacci M, Di Domenico M, Ancora M, Curini V, Mangone I, Rinaldi A, Scialabba S, Di Pasquale A, Cammà C, Puglia I, Calistri P, Savini G |
| EPI_ISL_1577969, EPI_ISL_1577970, EPI_ISL_1577971 | SIESP CHIETI - DRIVE IN CHIETI CHIETI(CHIETI) | Istituto Zooprofilattico Sperimentale dell'Abruzzo e Molise "G. Caporale" | Lorusso A, Marcacci M, Di Domenico M, Ancora M, Curini V, Mangone I, Rinaldi A, Scialabba S, Di Pasquale A, Cammà C, Puglia I, Calistri P, Savini G |
| EPI_ISL_1577972, EPI_ISL_1577973, EPI_ISL_1577974, EPI_ISL_1577975, EPI_ISL_1577976 | Rsa Residenza PALIZZI VASTO(CHIETI) | Istituto Zooprofilattico Sperimentale dell'Abruzzo e Molise "G. Caporale" | Lorusso A, Marcacci M, Di Domenico M, Ancora M, Curini V, Mangone I, Rinaldi A, Scialabba S, Di Pasquale A, Cammà C, Puglia I, Calistri P, Savini G |
| EPI_ISL_1577977 | RSA GIULIANOVA GIULIANOVA(TERAMO) | Istituto Zooprofilattico Sperimentale dell'Abruzzo e Molise "G. Caporale" | Lorusso A, Marcacci M, Di Domenico M, Ancora M, Curini V, Mangone I, Rinaldi A, Scialabba S, Di Pasquale A, Cammà C, Puglia I, Calistri P, Savini G |
| EPI_ISL_1577978, EPI_ISL_1577979, EPI_ISL_1577980, EPI_ISL_1577981, EPI_ISL_1577982, EPI_ISL_1577983 | SIESP DIPARTIMENTO DI PREVENZIONE TERAMO TERAMO(TERAMO) | Istituto Zooprofilattico Sperimentale dell'Abruzzo e Molise "G. Caporale" | Lorusso A, Marcacci M, Di Domenico M, Ancora M, Curini V, Mangone I, Rinaldi A, Scialabba S, Di Pasquale A, Cammà C, Puglia I, Calistri P, Savini G |
| EPI_ISL_1577984 | SIESP CHIETI - DRIVE IN CHIETI CHIETI(CHIETI) | Istituto Zooprofilattico Sperimentale dell'Abruzzo e Molise "G. Caporale" | Lorusso A, Marcacci M, Di Domenico M, Ancora M, Curini V, Mangone I, Rinaldi A, Scialabba S, Di Pasquale A, Cammà C, Puglia I, Calistri P, Savini G |
| EPI_ISL_1577985 | SIESP CHIETI - DRIVE IN ORTONA ORTONA(CHIETI) | Istituto Zooprofilattico Sperimentale dell'Abruzzo e Molise "G. Caporale" | Lorusso A, Marcacci M, Di Domenico M, Ancora M, Curini V, Mangone I, Rinaldi A, Scialabba S, Di Pasquale A, Cammà C, Puglia I, Calistri P, Savini G |
| EPI_ISL_1577986 | SIESP CHIETI - DRIVE IN CHIETI CHIETI(CHIETI) | Istituto Zooprofilattico Sperimentale dell'Abruzzo e Molise "G. Caporale" | Lorusso A, Marcacci M, Di Domenico M, Ancora M, Curini V, Mangone I, Rinaldi A, Scialabba S, Di Pasquale A, Cammà C, Puglia I, Calistri P, Savini G |
| EPI_ISL_1577987 | SIESP CHIETI - DRIVE IN LANCIANO LANCIANO(CHIETI) | Istituto Zooprofilattico Sperimentale dell'Abruzzo e Molise "G. Caporale" | Lorusso A, Marcacci M, Di Domenico M, Ancora M, Curini V, Mangone I, Rinaldi A, Scialabba S, Di Pasquale A, Cammà C, Puglia I, Calistri P, Savini G |
| EPI_ISL_1577988, EPI_ISL_1577989, EPI_ISL_1577990, EPI_ISL_1577991, EPI_ISL_1577992, EPI_ISL_1577993 | DIP. PREV. AVEZZANO SERVIZIO DI IGIENE EPIDEMIOLOGIA E SANITA' PUBBLICA AVEZZANO(L'AQUILA) | Istituto Zooprofilattico Sperimentale dell'Abruzzo e Molise "G. Caporale" | Lorusso A, Marcacci M, Di Domenico M, Ancora M, Curini V, Mangone I, Rinaldi A, Scialabba S, Di Pasquale A, Cammà C, Puglia I, Calistri P, Savini G |
| EPI_ISL_1577994 | U.O.S.D. Medici Competenti PO Avezzano AVEZZANO(L'AQUILA) | Istituto Zooprofilattico Sperimentale dell'Abruzzo e Molise "G. Caporale" | Lorusso A, Marcacci M, Di Domenico M, Ancora M, Curini V, Mangone I, Rinaldi A, Scialabba S, Di Pasquale A, Cammà C, Puglia I, Calistri P, Savini G |
| EPI_ISL_1577995 | PRESIDIO TERRITORIALE CASOLI CASOLI(CHIETI) | Istituto Zooprofilattico Sperimentale dell'Abruzzo e Molise "G. Caporale" | Lorusso A, Marcacci M, Di Domenico M, Ancora M, Curini V, Mangone I, Rinaldi A, Scialabba S, Di Pasquale A, Cammà C, Puglia I, Calistri P, Savini G |
| EPI_ISL_1577996, EPI_ISL_1577997 | USCA Sulmona SULMONA(L'AQUILA) | Istituto Zooprofilattico Sperimentale dell'Abruzzo e Molise "G. Caporale" | Lorusso A, Marcacci M, Di Domenico M, Ancora M, Curini V, Mangone I, Rinaldi A, Scialabba S, Di Pasquale A, Cammà C, Puglia I, Calistri P, Savini G |
| EPI_ISL_1577998 | SIESP CHIETI - DRIVE IN ORTONA ORTONA(CHIETI) | Istituto Zooprofilattico Sperimentale dell'Abruzzo e Molise "G. Caporale" | Lorusso A, Marcacci M, Di Domenico M, Ancora M, Curini V, Mangone I, Rinaldi A, Scialabba S, Di Pasquale A, Cammà C, Puglia I, Calistri P, Savini G |
| EPI_ISL_1577999, EPI_ISL_1578000, EPI_ISL_1578001 | USCA Castel di Sangro CASTEL DI SANGRO(L'AQUILA) | Istituto Zooprofilattico Sperimentale dell'Abruzzo e Molise "G. Caporale" | Lorusso A, Marcacci M, Di Domenico M, Ancora M, Curini V, Mangone I, Rinaldi A, Scialabba S, Di Pasquale A, Cammà C, Puglia I, Calistri P, Savini G |
| EPI_ISL_1578002, EPI_ISL_1578003 | SIESP CHIETI - DRIVE IN LANCIANO LANCIANO(CHIETI) | Istituto Zooprofilattico Sperimentale dell'Abruzzo e Molise "G. Caporale" | Lorusso A, Marcacci M, Di Domenico M, Ancora M, Curini V, Mangone I, Rinaldi A, Scialabba S, Di Pasquale A, Cammà C, Puglia I, Calistri P, Savini G |
| EPI_ISL_1578004 | USCA Pescina PESCINA(L'AQUILA) | Istituto Zooprofilattico Sperimentale dell'Abruzzo e Molise "G. Caporale" | Lorusso A, Marcacci M, Di Domenico M, Ancora M, Curini V, Mangone I, Rinaldi A, Scialabba S, Di Pasquale A, Cammà C, Puglia I, Calistri P, Savini G |
| EPI_ISL_1578005, EPI_ISL_1578006, EPI_ISL_1578007, EPI_ISL_1578008, EPI_ISL_1578009, EPI_ISL_1578010, EPI_ISL_1578011, EPI_ISL_1578012, EPI_ISL_1578013, EPI_ISL_1578014, EPI_ISL_1578015, EPI_ISL_1578016, EPI_ISL_1578017, EPI_ISL_1578018, EPI_ISL_1578019, EPI_ISL_1578020 | | | |
| see above | OSPEDALE SAN SALVATORE - MEDICINA DI LABORATORIO L'AQUILA(L'AQUILA) | Istituto Zooprofilattico Sperimentale dell'Abruzzo e Molise "G. Caporale" | Lorusso A, Marcacci M, Di Domenico M, Ancora M, Curini V, Mangone I, Rinaldi A, Scialabba S, Di Pasquale A, Cammà C, Puglia I, Calistri P, Savini G |
| EPI_ISL_1578021 | OSPEDALE S.S. ANNUNZIATA CHIETI CHIETI(CHIETI) | Istituto Zooprofilattico Sperimentale dell'Abruzzo e Molise "G. Caporale" | Lorusso A, Marcacci M, Di Domenico M, Ancora M, Curini V, Mangone I, Rinaldi A, Scialabba S, Di Pasquale A, Cammà C, Puglia I, Calistri P, Savini G |
| EPI_ISL_1578022 | SIESP CHIETI - DRIVE IN ORTONA ORTONA(CHIETI) | Istituto Zooprofilattico Sperimentale dell'Abruzzo e Molise "G. Caporale" | Lorusso A, Marcacci M, Di Domenico M, Ancora M, Curini V, Mangone I, Rinaldi A, Scialabba S, Di Pasquale A, Cammà C, Puglia I, Calistri P, Savini G |
| EPI_ISL_1578023 | SIESP CHIETI - DRIVE IN CHIETI CHIETI(CHIETI) | Istituto Zooprofilattico Sperimentale dell'Abruzzo e Molise "G. Caporale" | Lorusso A, Marcacci M, Di Domenico M, Ancora M, Curini V, Mangone I, Rinaldi A, Scialabba S, Di Pasquale A, Cammà C, Puglia I, Calistri P, Savini G |
| EPI_ISL_1578024 | SIESP CHIETI - DRIVE IN ORTONA ORTONA(CHIETI) | Istituto Zooprofilattico Sperimentale dell'Abruzzo e Molise "G. Caporale" | Lorusso A, Marcacci M, Di Domenico M, Ancora M, Curini V, Mangone I, Rinaldi A, Scialabba S, Di Pasquale A, Cammà C, Puglia I, Calistri P, Savini G |
| EPI_ISL_1578025 | SIESP SULMONA SULMONA(L'AQUILA) | Istituto Zooprofilattico Sperimentale dell'Abruzzo e Molise "G. Caporale" | Lorusso A, Marcacci M, Di Domenico M, Ancora M, Curini V, Mangone I, Rinaldi A, Scialabba S, Di Pasquale A, Cammà C, Puglia I, Calistri P, Savini G |
| EPI_ISL_1578026, EPI_ISL_1578027 | USCA Castel di Sangro CASTEL DI SANGRO(L'AQUILA) | Istituto Zooprofilattico Sperimentale dell'Abruzzo e Molise "G. | Lorusso A, Marcacci M, Di Domenico M, Ancora M, Curini V, Mangone I, Rinaldi A, Scialabba S, Di Pasquale A, Cammà C, Puglia I, Calistri P, Savini G |

| | | | |
|---|---|---|---|
| | | Caporale" | |
| EPI_ISL_1578028, EPI_ISL_1578029 | SIESP DIPARTIMENTO DI PREVENZIONE TERAMO TERAMO(TERAMO) | Istituto Zooprofilattico Sperimentale dell'Abruzzo e Molise "G. Caporale" | Lorusso A, Marcacci M, Di Domenico M, Ancora M, Curini V, Mangone I, Rinaldi A, Scialabba S, Di Pasquale A, Cammà C, Puglia I, Calistri P, Savini G |
| EPI_ISL_1578030, EPI_ISL_1578031 | OSPEDALE SAN SALVATORE - MEDICINA DI LABORATORIO L'AQUILA(L'AQUILA) | Istituto Zooprofilattico Sperimentale dell'Abruzzo e Molise "G. Caporale" | Lorusso A, Marcacci M, Di Domenico M, Ancora M, Curini V, Mangone I, Rinaldi A, Scialabba S, Di Pasquale A, Cammà C, Puglia I, Calistri P, Savini G |
| EPI_ISL_1578032, EPI_ISL_1578033 | SIESP CHIETI - DRIVE IN VASTO VASTO(CHIETI) | Istituto Zooprofilattico Sperimentale dell'Abruzzo e Molise "G. Caporale" | Lorusso A, Marcacci M, Di Domenico M, Ancora M, Curini V, Mangone I, Rinaldi A, Scialabba S, Di Pasquale A, Cammà C, Puglia I, Calistri P, Savini G |
| EPI_ISL_1578034, EPI_ISL_1578035 | SIESP CHIETI - DRIVE IN LANCIANO LANCIANO(CHIETI) | Istituto Zooprofilattico Sperimentale dell'Abruzzo e Molise "G. Caporale" | Lorusso A, Marcacci M, Di Domenico M, Ancora M, Curini V, Mangone I, Rinaldi A, Scialabba S, Di Pasquale A, Cammà C, Puglia I, Calistri P, Savini G |
| EPI_ISL_1578036 | VILLA NOVECENTO S.R.L LANCIANO(CHIETI) | Istituto Zooprofilattico Sperimentale dell'Abruzzo e Molise "G. Caporale" | Lorusso A, Marcacci M, Di Domenico M, Ancora M, Curini V, Mangone I, Rinaldi A, Scialabba S, Di Pasquale A, Cammà C, Puglia I, Calistri P, Savini G |
| EPI_ISL_1578037 | PRESIDIO TERRITORIALE CASOLI CASOLI(CHIETI) | Istituto Zooprofilattico Sperimentale dell'Abruzzo e Molise "G. Caporale" | Lorusso A, Marcacci M, Di Domenico M, Ancora M, Curini V, Mangone I, Rinaldi A, Scialabba S, Di Pasquale A, Cammà C, Puglia I, Calistri P, Savini G |
| EPI_ISL_1578038 | SIESP DIPARTIMENTO DI PREVENZIONE CHIETI CHIETI(CHIETI) | Istituto Zooprofilattico Sperimentale dell'Abruzzo e Molise "G. Caporale" | Lorusso A, Marcacci M, Di Domenico M, Ancora M, Curini V, Mangone I, Rinaldi A, Scialabba S, Di Pasquale A, Cammà C, Puglia I, Calistri P, Savini G |
| EPI_ISL_1578114, EPI_ISL_1578115 | Genome Analysis Center, Yamanashi Central Hospital | Genome Analysis Center, Yamanashi Central Hospital | Yosuke Hirotsu |
| EPI_ISL_1579806 | NMVRVI | National Public Health Surveillance Laboratory | Lukas Zemaitis, Migle Gabrielaite, Jelena Razmuk, Svajune Muralyte, Ana Steponkiene, Lukas Vasionis, Danas Baksa |
| EPI_ISL_1580502 | Laboratório de Biologia Molecular do Hospital das Clínicas da Faculdade de Medicina de Botucatu/SP | Laboratórios de Genômica Funcional (FCA/UNESP) e Biologia Molecular (FMB-HC/UNESP) - Rede de Vigilância Genômica (Vigenômica)/UNESP | Patrícia Akemi Assato; Felipe Allan da Silva da Costa; Bianca Cechetto Carlos; Flavia Hebner Barbosa Trovão; Guilherme Targino Valente; Rejane Maria Tommasini Grotto; Jayme A. Souza-Neto. |
| EPI_ISL_1580505, EPI_ISL_1580506 | DIP. PREV. AVEZZANO SERVIZIO DI IGIENE EPIDEMIOLOGIA E SANITA' PUBBLICA AVEZZANO(L'AQUILA) | Istituto Zooprofilattico Sperimentale dell'Abruzzo e Molise "G. Caporale" | Lorusso A, Marcacci M, Di Domenico M, Ancora M, Curini V, Di Lollo Valeria, Mangone I, Rinaldi A, Delli Compagni E, Scialabba S, Caporale M, Di Pasquale A, Cammà C, Puglia I, Calistri P, Savini G |
| EPI_ISL_1580507 | USCA Tagliacozzo TAGLIACOZZO(L'AQUILA) | Istituto Zooprofilattico Sperimentale dell'Abruzzo e Molise "G. Caporale" | Lorusso A, Marcacci M, Di Domenico M, Ancora M, Curini V, Di Lollo Valeria, Mangone I, Rinaldi A, Delli Compagni E, Scialabba S, Caporale M, Di Pasquale A, Cammà C, Puglia I, Calistri P, Savini G |
| EPI_ISL_1580508, EPI_ISL_1580509 | SIESP DIPARTIMENTO DI PREVENZIONE CHIETI CHIETI(CHIETI) | Istituto Zooprofilattico Sperimentale dell'Abruzzo e Molise "G. Caporale" | Lorusso A, Marcacci M, Di Domenico M, Ancora M, Curini V, Di Lollo Valeria, Mangone I, Rinaldi A, Delli Compagni E, Scialabba S, Caporale M, Di Pasquale A, Cammà C, Puglia I, Calistri P, Savini G |
| EPI_ISL_1580510, EPI_ISL_1580511 | SIESP DIPARTIMENTO DI PREVENZIONE TERAMO TERAMO(TERAMO) | Istituto Zooprofilattico Sperimentale dell'Abruzzo e Molise "G. Caporale" | Lorusso A, Marcacci M, Di Domenico M, Ancora M, Curini V, Di Lollo Valeria, Mangone I, Rinaldi A, Delli Compagni E, Scialabba S, Caporale M, Di Pasquale A, Cammà C, Puglia I, Calistri P, Savini G |
| EPI_ISL_1580512 | DIP. PREV. AVEZZANO SERVIZIO DI IGIENE EPIDEMIOLOGIA E SANITA' PUBBLICA AVEZZANO(L'AQUILA) | Istituto Zooprofilattico Sperimentale dell'Abruzzo e Molise "G. Caporale" | Lorusso A, Marcacci M, Di Domenico M, Ancora M, Curini V, Di Lollo Valeria, Mangone I, Rinaldi A, Delli Compagni E, Scialabba S, Caporale M, Di Pasquale A, Cammà C, Puglia I, Calistri P, Savini G |
| EPI_ISL_1580513 | SIESP DIPARTIMENTO DI PREVENZIONE CHIETI CHIETI(CHIETI) | Istituto Zooprofilattico Sperimentale dell'Abruzzo e Molise "G. Caporale" | Lorusso A, Marcacci M, Di Domenico M, Ancora M, Curini V, Di Lollo Valeria, Mangone I, Rinaldi A, Delli Compagni E, Scialabba S, Caporale M, Di Pasquale A, Cammà C, Puglia I, Calistri P, Savini G |
| EPI_ISL_1580514 | DIP. PREV. AVEZZANO SERVIZIO DI IGIENE EPIDEMIOLOGIA E SANITA' PUBBLICA AVEZZANO(L'AQUILA) | Istituto Zooprofilattico Sperimentale dell'Abruzzo e Molise "G. Caporale" | Lorusso A, Marcacci M, Di Domenico M, Ancora M, Curini V, Di Lollo Valeria, Mangone I, Rinaldi A, Delli Compagni E, Scialabba S, Caporale M, Di Pasquale A, Cammà C, Puglia I, Calistri P, Savini G |
| EPI_ISL_1580515 | RESIDENZA PER ANZIANI CASA TUA S.R.L. TORRICELLA PELIGNA(CHIETI) | Istituto Zooprofilattico Sperimentale dell'Abruzzo e Molise "G. Caporale" | Lorusso A, Marcacci M, Di Domenico M, Ancora M, Curini V, Di Lollo Valeria, Mangone I, Rinaldi A, Delli Compagni E, Scialabba S, Caporale M, Di Pasquale A, Cammà C, Puglia I, Calistri P, Savini G |
| EPI_ISL_1580516, EPI_ISL_1580517 | SIESP CHIETI - DRIVE IN CHIETI CHIETI(CHIETI) | Istituto Zooprofilattico Sperimentale dell'Abruzzo e Molise "G. Caporale" | Lorusso A, Marcacci M, Di Domenico M, Ancora M, Curini V, Di Lollo Valeria, Mangone I, Rinaldi A, Delli Compagni E, Scialabba S, Caporale M, Di Pasquale A, Cammà C, Puglia I, Calistri P, Savini G |
| EPI_ISL_1580518, EPI_ISL_1580519 | SIESP DIPARTIMENTO DI PREVENZIONE CHIETI CHIETI(CHIETI) | Istituto Zooprofilattico Sperimentale dell'Abruzzo e Molise "G. Caporale" | Lorusso A, Marcacci M, Di Domenico M, Ancora M, Curini V, Di Lollo Valeria, Mangone I, Rinaldi A, Delli Compagni E, Scialabba S, Caporale M, Di Pasquale A, Cammà C, Puglia I, Calistri P, Savini G |
| EPI_ISL_1580520, EPI_ISL_1580521, EPI_ISL_1580522, EPI_ISL_1580523, EPI_ISL_1580524 | SIESP DIPARTIMENTO DI PREVENZIONE TERAMO TERAMO(TERAMO) | Istituto Zooprofilattico Sperimentale dell'Abruzzo e Molise "G. Caporale" | Lorusso A, Marcacci M, Di Domenico M, Ancora M, Curini V, Di Lollo Valeria, Mangone I, Rinaldi A, Delli Compagni E, Scialabba S, Caporale M, Di Pasquale A, Cammà C, Puglia I, Calistri P, Savini G |
| EPI_ISL_1580525 | RSA GIULIANOVA GIULIANOVA(TERAMO) | Istituto Zooprofilattico Sperimentale dell'Abruzzo e Molise "G. Caporale" | Lorusso A, Marcacci M, Di Domenico M, Ancora M, Curini V, Di Lollo Valeria, Mangone I, Rinaldi A, Delli Compagni E, Scialabba S, Caporale M, Di Pasquale A, Cammà C, Puglia I, Calistri P, Savini G |
| EPI_ISL_1580526 | SIESP DIPARTIMENTO DI PREVENZIONE TERAMO TERAMO(TERAMO) | Istituto Zooprofilattico Sperimentale dell'Abruzzo e Molise "G. Caporale" | Lorusso A, Marcacci M, Di Domenico M, Ancora M, Curini V, Di Lollo Valeria, Mangone I, Rinaldi A, Delli Compagni E, Scialabba S, Caporale M, Di Pasquale A, Cammà C, Puglia I, Calistri P, Savini G |
| EPI_ISL_1580527, EPI_ISL_1580528 | SIESP CHIETI - DRIVE IN VASTO VASTO(CHIETI) | Istituto Zooprofilattico Sperimentale dell'Abruzzo e Molise "G. Caporale" | Lorusso A, Marcacci M, Di Domenico M, Ancora M, Curini V, Di Lollo Valeria, Mangone I, Rinaldi A, Delli Compagni E, Scialabba S, Caporale M, Di Pasquale A, Cammà C, Puglia I, Calistri P, Savini G |
| EPI_ISL_1580529 | SIESP CHIETI - DRIVE IN LANCIANO LANCIANO(CHIETI) | Istituto Zooprofilattico Sperimentale dell'Abruzzo e Molise "G. Caporale" | Lorusso A, Marcacci M, Di Domenico M, Ancora M, Curini V, Di Lollo Valeria, Mangone I, Rinaldi A, Delli Compagni E, Scialabba S, Caporale M, Di Pasquale A, Cammà C, Puglia I, Calistri P, Savini G |
| EPI_ISL_1580530 | SANTA CAMILLA VILLA PINI-RIAB. PSICHIATRICA CHIETI(CHIETI) | Istituto Zooprofilattico Sperimentale dell'Abruzzo e Molise "G. Caporale" | Lorusso A, Marcacci M, Di Domenico M, Ancora M, Curini V, Di Lollo Valeria, Mangone I, Rinaldi A, Delli Compagni E, Scialabba S, Caporale M, Di Pasquale A, Cammà C, Puglia I, Calistri P, Savini G |
| EPI_ISL_1580531 | SIESP SULMONA SULMONA(L'AQUILA) | Istituto Zooprofilattico Sperimentale dell'Abruzzo e Molise "G. Caporale" | Lorusso A, Marcacci M, Di Domenico M, Ancora M, Curini V, Di Lollo Valeria, Mangone I, Rinaldi A, Delli Compagni E, Scialabba S, Caporale M, Di Pasquale A, Cammà C, Puglia I, Calistri P, Savini G |
| EPI_ISL_1580532 | USCA Tagliacozzo TAGLIACOZZO(L'AQUILA) | Istituto Zooprofilattico Sperimentale dell'Abruzzo e Molise "G. Caporale" | Lorusso A, Marcacci M, Di Domenico M, Ancora M, Curini V, Di Lollo Valeria, Mangone I, Rinaldi A, Delli Compagni E, Scialabba S, Caporale M, Di Pasquale A, Cammà C, Puglia I, Calistri P, Savini G |
| EPI_ISL_1580533 | OSPEDALE CIVILE TERAMO - CENTRO TRASFUSIONALE TERAMO(TERAMO) | Istituto Zooprofilattico Sperimentale dell'Abruzzo e Molise "G. Caporale" | Lorusso A, Marcacci M, Di Domenico M, Ancora M, Curini V, Di Lollo Valeria, Mangone I, Rinaldi A, Delli Compagni E, Scialabba S, Caporale M, Di Pasquale A, Cammà C, Puglia I, Calistri P, Savini G |
| EPI_ISL_1580534 | SIESP SULMONA SULMONA(L'AQUILA) | Istituto Zooprofilattico Sperimentale dell'Abruzzo e Molise "G. Caporale" | Lorusso A, Marcacci M, Di Domenico M, Ancora M, Curini V, Di Lollo Valeria, Mangone I, Rinaldi A, Delli Compagni E, Scialabba S, Caporale M, Di Pasquale A, Cammà C, Puglia I, Calistri P, Savini G |
| EPI_ISL_1580535 | SIESP DIPARTIMENTO DI PREVENZIONE TERAMO TERAMO(TERAMO) | Istituto Zooprofilattico Sperimentale dell'Abruzzo e Molise "G. Caporale" | Lorusso A, Marcacci M, Di Domenico M, Ancora M, Curini V, Di Lollo Valeria, Mangone I, Rinaldi A, Delli Compagni E, Scialabba S, Caporale M, Di Pasquale A, Cammà C, Puglia I, Calistri P, Savini G |
| EPI_ISL_1580536 | SIESP CHIETI - DRIVE IN LANCIANO LANCIANO(CHIETI) | Istituto Zooprofilattico Sperimentale dell'Abruzzo e Molise "G. Caporale" | Lorusso A, Marcacci M, Di Domenico M, Ancora M, Curini V, Di Lollo Valeria, Mangone I, Rinaldi A, Delli Compagni E, Scialabba S, Caporale M, Di Pasquale A, Cammà C, Puglia I, Calistri P, Savini G |
| EPI_ISL_1580537 | SIESP DIPARTIMENTO DI PREVENZIONE CHIETI CHIETI(CHIETI) | Istituto Zooprofilattico Sperimentale dell'Abruzzo e Molise "G. Caporale" | Lorusso A, Marcacci M, Di Domenico M, Ancora M, Curini V, Di Lollo Valeria, Mangone I, Rinaldi A, Delli Compagni E, Scialabba S, Caporale M, Di Pasquale A, Cammà C, Puglia I, Calistri P, Savini G |
| EPI_ISL_1580538, EPI_ISL_1580539 | Rsa VILLA SAN PIO LANCIANO(CHIETI) | Istituto Zooprofilattico Sperimentale dell'Abruzzo e Molise "G. | Lorusso A, Marcacci M, Di Domenico M, Ancora M, Curini V, Di Lollo Valeria, Mangone I, Rinaldi A, Delli Compagni E, Scialabba S, Caporale M, Di |

| | | | |
|---|---|---|---|
| | | Caporale" | Pasquale A, Cammà C, Puglia I, Calistri P, Savini G |
| EPI_ISL_1580540 | SIESP DIPARTIMENTO DI PREVENZIONE CHIETI CHIETI(CHIETI) | Istituto Zooprofilattico Sperimentale dell'Abruzzo e Molise "G. Caporale" | Lorusso A, Marcacci M, Di Domenico M, Ancora M, Curini V, Di Lollo Valeria, Mangone I, Rinaldi A, Delli Compagni E, Scialabba S, Caporale M, Di Pasquale A, Cammà C, Puglia I, Calistri P, Savini G |
| EPI_ISL_1580541, EPI_ISL_1580542 | SIESP SULMONA SULMONA(L'AQUILA) | Istituto Zooprofilattico Sperimentale dell'Abruzzo e Molise "G. Caporale" | Lorusso A, Marcacci M, Di Domenico M, Ancora M, Curini V, Di Lollo Valeria, Mangone I, Rinaldi A, Delli Compagni E, Scialabba S, Caporale M, Di Pasquale A, Cammà C, Puglia I, Calistri P, Savini G |
| EPI_ISL_1580543 | PRESIDIO TERRITORIALE CASOLI CASOLI(CHIETI) | Istituto Zooprofilattico Sperimentale dell'Abruzzo e Molise "G. Caporale" | Lorusso A, Marcacci M, Di Domenico M, Ancora M, Curini V, Di Lollo Valeria, Mangone I, Rinaldi A, Delli Compagni E, Scialabba S, Caporale M, Di Pasquale A, Cammà C, Puglia I, Calistri P, Savini G |
| EPI_ISL_1580544, EPI_ISL_1580545, EPI_ISL_1580546, EPI_ISL_1580547 | SIESP CHIETI - DRIVE IN ORTONA ORTONA(CHIETI) | Istituto Zooprofilattico Sperimentale dell'Abruzzo e Molise "G. Caporale" | Lorusso A, Marcacci M, Di Domenico M, Ancora M, Curini V, Di Lollo Valeria, Mangone I, Rinaldi A, Delli Compagni E, Scialabba S, Caporale M, Di Pasquale A, Cammà C, Puglia I, Calistri P, Savini G |
| EPI_ISL_1580548 | SIESP CHIETI - DRIVE IN VASTO VASTO(CHIETI) | Istituto Zooprofilattico Sperimentale dell'Abruzzo e Molise "G. Caporale" | Lorusso A, Marcacci M, Di Domenico M, Ancora M, Curini V, Di Lollo Valeria, Mangone I, Rinaldi A, Delli Compagni E, Scialabba S, Caporale M, Di Pasquale A, Cammà C, Puglia I, Calistri P, Savini G |
| EPI_ISL_1580549 | USCA Avezzano AVEZZANO(L'AQUILA) | Istituto Zooprofilattico Sperimentale dell'Abruzzo e Molise "G. Caporale" | Lorusso A, Marcacci M, Di Domenico M, Ancora M, Curini V, Di Lollo Valeria, Mangone I, Rinaldi A, Delli Compagni E, Scialabba S, Caporale M, Di Pasquale A, Cammà C, Puglia I, Calistri P, Savini G |
| EPI_ISL_1580550 | USCA Pescina PESCINA(L'AQUILA) | Istituto Zooprofilattico Sperimentale dell'Abruzzo e Molise "G. Caporale" | Lorusso A, Marcacci M, Di Domenico M, Ancora M, Curini V, Di Lollo Valeria, Mangone I, Rinaldi A, Delli Compagni E, Scialabba S, Caporale M, Di Pasquale A, Cammà C, Puglia I, Calistri P, Savini G |
| EPI_ISL_1580551 | USCA Tagliacozzo TAGLIACOZZO(L'AQUILA) | Istituto Zooprofilattico Sperimentale dell'Abruzzo e Molise "G. Caporale" | Lorusso A, Marcacci M, Di Domenico M, Ancora M, Curini V, Di Lollo Valeria, Mangone I, Rinaldi A, Delli Compagni E, Scialabba S, Caporale M, Di Pasquale A, Cammà C, Puglia I, Calistri P, Savini G |
| EPI_ISL_1580552 | SIESP DIPARTIMENTO DI PREVENZIONE CHIETI CHIETI(CHIETI) | Istituto Zooprofilattico Sperimentale dell'Abruzzo e Molise "G. Caporale" | Lorusso A, Marcacci M, Di Domenico M, Ancora M, Curini V, Di Lollo Valeria, Mangone I, Rinaldi A, Delli Compagni E, Scialabba S, Caporale M, Di Pasquale A, Cammà C, Puglia I, Calistri P, Savini G |
| EPI_ISL_1580553 | SIESP CHIETI - DRIVE IN CHIETI CHIETI(CHIETI) | Istituto Zooprofilattico Sperimentale dell'Abruzzo e Molise "G. Caporale" | Lorusso A, Marcacci M, Di Domenico M, Ancora M, Curini V, Di Lollo Valeria, Mangone I, Rinaldi A, Delli Compagni E, Scialabba S, Caporale M, Di Pasquale A, Cammà C, Puglia I, Calistri P, Savini G |
| EPI_ISL_1580554, EPI_ISL_1580555, EPI_ISL_1580556 | DIP. PREV. AVEZZANO SERVIZIO DI IGIENE EPIDEMIOLOGIA E SANITA' PUBBLICA AVEZZANO(L'AQUILA) | Istituto Zooprofilattico Sperimentale dell'Abruzzo e Molise "G. Caporale" | Lorusso A, Marcacci M, Di Domenico M, Ancora M, Curini V, Di Lollo Valeria, Mangone I, Rinaldi A, Delli Compagni E, Scialabba S, Caporale M, Di Pasquale A, Cammà C, Puglia I, Calistri P, Savini G |
| EPI_ISL_1580557 | SIESP CHIETI - DRIVE IN GISSI GISSI(CHIETI) | Istituto Zooprofilattico Sperimentale dell'Abruzzo e Molise "G. Caporale" | Lorusso A, Marcacci M, Di Domenico M, Ancora M, Curini V, Di Lollo Valeria, Mangone I, Rinaldi A, Delli Compagni E, Scialabba S, Caporale M, Di Pasquale A, Cammà C, Puglia I, Calistri P, Savini G |
| EPI_ISL_1580558, EPI_ISL_1580559 | SIESP DIPARTIMENTO DI PREVENZIONE TERAMO TERAMO(TERAMO) | Istituto Zooprofilattico Sperimentale dell'Abruzzo e Molise "G. Caporale" | Lorusso A, Marcacci M, Di Domenico M, Ancora M, Curini V, Di Lollo Valeria, Mangone I, Rinaldi A, Delli Compagni E, Scialabba S, Caporale M, Di Pasquale A, Cammà C, Puglia I, Calistri P, Savini G |
| EPI_ISL_1580560, EPI_ISL_1580561 | DIP. PREV. AVEZZANO SERVIZIO DI IGIENE EPIDEMIOLOGIA E SANITA' PUBBLICA AVEZZANO(L'AQUILA) | Istituto Zooprofilattico Sperimentale dell'Abruzzo e Molise "G. Caporale" | Lorusso A, Marcacci M, Di Domenico M, Ancora M, Curini V, Di Lollo Valeria, Mangone I, Rinaldi A, Delli Compagni E, Scialabba S, Caporale M, Di Pasquale A, Cammà C, Puglia I, Calistri P, Savini G |
| EPI_ISL_1580562, EPI_ISL_1580563 | SIESP DIPARTIMENTO DI PREVENZIONE TERAMO TERAMO(TERAMO) | Istituto Zooprofilattico Sperimentale dell'Abruzzo e Molise "G. Caporale" | Lorusso A, Marcacci M, Di Domenico M, Ancora M, Curini V, Di Lollo Valeria, Mangone I, Rinaldi A, Delli Compagni E, Scialabba S, Caporale M, Di Pasquale A, Cammà C, Puglia I, Calistri P, Savini G |
| EPI_ISL_1580564 | RSA GIULIANOVA GIULIANOVA(TERAMO) | Istituto Zooprofilattico Sperimentale dell'Abruzzo e Molise "G. Caporale" | Lorusso A, Marcacci M, Di Domenico M, Ancora M, Curini V, Di Lollo Valeria, Mangone I, Rinaldi A, Delli Compagni E, Scialabba S, Caporale M, Di Pasquale A, Cammà C, Puglia I, Calistri P, Savini G |
| EPI_ISL_1580565, EPI_ISL_1580566, EPI_ISL_1580567, EPI_ISL_1580568, EPI_ISL_1580569, EPI_ISL_1580570, EPI_ISL_1580571 | SIESP DIPARTIMENTO DI PREVENZIONE TERAMO TERAMO(TERAMO) | Istituto Zooprofilattico Sperimentale dell'Abruzzo e Molise "G. Caporale" | Lorusso A, Marcacci M, Di Domenico M, Ancora M, Curini V, Di Lollo Valeria, Mangone I, Rinaldi A, Delli Compagni E, Scialabba S, Caporale M, Di Pasquale A, Cammà C, Puglia I, Calistri P, Savini G |
| EPI_ISL_1580572 | SIESP CHIETI - DRIVE IN VASTO VASTO(CHIETI) | Istituto Zooprofilattico Sperimentale dell'Abruzzo e Molise "G. Caporale" | Lorusso A, Marcacci M, Di Domenico M, Ancora M, Curini V, Di Lollo Valeria, Mangone I, Rinaldi A, Delli Compagni E, Scialabba S, Caporale M, Di Pasquale A, Cammà C, Puglia I, Calistri P, Savini G |
| EPI_ISL_1580573 | RSA GIULIANOVA GIULIANOVA(TERAMO) | Istituto Zooprofilattico Sperimentale dell'Abruzzo e Molise "G. Caporale" | Lorusso A, Marcacci M, Di Domenico M, Ancora M, Curini V, Di Lollo Valeria, Mangone I, Rinaldi A, Delli Compagni E, Scialabba S, Caporale M, Di Pasquale A, Cammà C, Puglia I, Calistri P, Savini G |
| EPI_ISL_1580574, EPI_ISL_1580576, EPI_ISL_1580577 | SIESP DIPARTIMENTO DI PREVENZIONE TERAMO TERAMO(TERAMO) | Istituto Zooprofilattico Sperimentale dell'Abruzzo e Molise "G. Caporale" | Lorusso A, Marcacci M, Di Domenico M, Ancora M, Curini V, Di Lollo Valeria, Mangone I, Rinaldi A, Delli Compagni E, Scialabba S, Caporale M, Di Pasquale A, Cammà C, Puglia I, Calistri P, Savini G |
| EPI_ISL_1580578, EPI_ISL_1580579, EPI_ISL_1580580 | SIESP CHIETI - DRIVE IN VASTO VASTO(CHIETI) | Istituto Zooprofilattico Sperimentale dell'Abruzzo e Molise "G. Caporale" | Lorusso A, Marcacci M, Di Domenico M, Ancora M, Curini V, Di Lollo Valeria, Mangone I, Rinaldi A, Delli Compagni E, Scialabba S, Caporale M, Di Pasquale A, Cammà C, Puglia I, Calistri P, Savini G |
| EPI_ISL_1580581, EPI_ISL_1580582, EPI_ISL_1580583, EPI_ISL_1580584, EPI_ISL_1580585, EPI_ISL_1580586, EPI_ISL_1580587, EPI_ISL_1580588, EPI_ISL_1580589, EPI_ISL_1580590 | SIESP DIPARTIMENTO DI PREVENZIONE TERAMO TERAMO(TERAMO) | Istituto Zooprofilattico Sperimentale dell'Abruzzo e Molise "G. Caporale" | Lorusso A, Marcacci M, Di Domenico M, Ancora M, Curini V, Di Lollo Valeria, Mangone I, Rinaldi A, Delli Compagni E, Scialabba S, Caporale M, Di Pasquale A, Cammà C, Puglia I, Calistri P, Savini G |
| EPI_ISL_1580591, EPI_ISL_1580592 | DIP. PREV. AVEZZANO SERVIZIO DI IGIENE EPIDEMIOLOGIA E SANITA' PUBBLICA AVEZZANO(L'AQUILA) | Istituto Zooprofilattico Sperimentale dell'Abruzzo e Molise "G. Caporale" | Lorusso A, Marcacci M, Di Domenico M, Ancora M, Curini V, Di Lollo Valeria, Mangone I, Rinaldi A, Delli Compagni E, Scialabba S, Caporale M, Di Pasquale A, Cammà C, Puglia I, Calistri P, Savini G |
| EPI_ISL_1580593 | SIESP SULMONA SULMONA(L'AQUILA) | Istituto Zooprofilattico Sperimentale dell'Abruzzo e Molise "G. Caporale" | Lorusso A, Marcacci M, Di Domenico M, Ancora M, Curini V, Di Lollo Valeria, Mangone I, Rinaldi A, Delli Compagni E, Scialabba S, Caporale M, Di Pasquale A, Cammà C, Puglia I, Calistri P, Savini G |
| EPI_ISL_1580594, EPI_ISL_1580595, EPI_ISL_1580596 | SIESP CHIETI - DRIVE IN ORTONA ORTONA(CHIETI) | Istituto Zooprofilattico Sperimentale dell'Abruzzo e Molise "G. Caporale" | Lorusso A, Marcacci M, Di Domenico M, Ancora M, Curini V, Di Lollo Valeria, Mangone I, Rinaldi A, Delli Compagni E, Scialabba S, Caporale M, Di Pasquale A, Cammà C, Puglia I, Calistri P, Savini G |
| EPI_ISL_1580598 | DIP. PREV. AVEZZANO SERVIZIO DI IGIENE EPIDEMIOLOGIA E SANITA' PUBBLICA AVEZZANO(L'AQUILA) | Istituto Zooprofilattico Sperimentale dell'Abruzzo e Molise "G. Caporale" | Lorusso A, Marcacci M, Di Domenico M, Ancora M, Curini V, Di Lollo Valeria, Mangone I, Rinaldi A, Delli Compagni E, Scialabba S, Caporale M, Di Pasquale A, Cammà C, Puglia I, Calistri P, Savini G |
| EPI_ISL_1580599, EPI_ISL_1580600 | SIESP CHIETI - DRIVE IN VASTO VASTO(CHIETI) | Istituto Zooprofilattico Sperimentale dell'Abruzzo e Molise "G. Caporale" | Lorusso A, Marcacci M, Di Domenico M, Ancora M, Curini V, Di Lollo Valeria, Mangone I, Rinaldi A, Delli Compagni E, Scialabba S, Caporale M, Di Pasquale A, Cammà C, Puglia I, Calistri P, Savini G |
| EPI_ISL_1580601 | Servizio di Prevenzione e Protezione dell'IZSAM TERAMO(TERAMO) | Istituto Zooprofilattico Sperimentale dell'Abruzzo e Molise "G. Caporale" | Lorusso A, Marcacci M, Di Domenico M, Ancora M, Curini V, Di Lollo Valeria, Mangone I, Rinaldi A, Delli Compagni E, Scialabba S, Caporale M, Di Pasquale A, Cammà C, Puglia I, Calistri P, Savini G |
| EPI_ISL_1580602 | SIESP CHIETI - DRIVE IN VASTO VASTO(CHIETI) | Istituto Zooprofilattico Sperimentale dell'Abruzzo e Molise "G. Caporale" | Lorusso A, Marcacci M, Di Domenico M, Ancora M, Curini V, Di Lollo Valeria, Mangone I, Rinaldi A, Delli Compagni E, Scialabba S, Caporale M, Di Pasquale A, Cammà C, Puglia I, Calistri P, Savini G |
| EPI_ISL_1580603 | USCA Pescina PESCINA(L'AQUILA) | Istituto Zooprofilattico Sperimentale dell'Abruzzo e Molise "G. Caporale" | Lorusso A, Marcacci M, Di Domenico M, Ancora M, Curini V, Di Lollo Valeria, Mangone I, Rinaldi A, Delli Compagni E, Scialabba S, Caporale M, Di Pasquale A, Cammà C, Puglia I, Calistri P, Savini G |
| EPI_ISL_1580604 | USCA Tagliacozzo TAGLIACOZZO(L'AQUILA) | Istituto Zooprofilattico Sperimentale dell'Abruzzo e Molise "G. Caporale" | Lorusso A, Marcacci M, Di Domenico M, Ancora M, Curini V, Di Lollo Valeria, Mangone I, Rinaldi A, Delli Compagni E, Scialabba S, Caporale M, Di Pasquale A, Cammà C, Puglia I, Calistri P, Savini G |

| | | | |
|---|---|---|---|
| EPI_ISL_1580605, EPI_ISL_1580606, EPI_ISL_1580607 | SIESP CHIETI - DRIVE IN VASTO VASTO(CHIETI) | Istituto Zooprofilattico Sperimentale dell'Abruzzo e Molise "G. Caporale" | Lorusso A, Marcacci M, Di Domenico M, Ancora M, Curini V, Di Lollo Valeria, Mangone I, Rinaldi A, Delli Compagni E, Scialabba S, Caporale M, Di Pasquale A, Cammà C, Puglia I, Calistri P, Savini G |
| EPI_ISL_1580608 | DIP. PREV. AVEZZANO SERVIZIO DI IGIENE EPIDEMIOLOGIA E SANITA' PUBBLICA AVEZZANO(L'AQUILA) | Istituto Zooprofilattico Sperimentale dell'Abruzzo e Molise "G. Caporale" | Lorusso A, Marcacci M, Di Domenico M, Ancora M, Curini V, Di Lollo Valeria, Mangone I, Rinaldi A, Delli Compagni E, Scialabba S, Caporale M, Di Pasquale A, Cammà C, Puglia I, Calistri P, Savini G |
| EPI_ISL_1580609 | SIESP CHIETI - DRIVE IN ATESSA ATESSA(CHIETI) | Istituto Zooprofilattico Sperimentale dell'Abruzzo e Molise "G. Caporale" | Lorusso A, Marcacci M, Di Domenico M, Ancora M, Curini V, Di Lollo Valeria, Mangone I, Rinaldi A, Delli Compagni E, Scialabba S, Caporale M, Di Pasquale A, Cammà C, Puglia I, Calistri P, Savini G |
| EPI_ISL_1580610, EPI_ISL_1580611, EPI_ISL_1580612, EPI_ISL_1580613, EPI_ISL_1580614 | SIESP DIPARTIMENTO DI PREVENZIONE TERAMO TERAMO(TERAMO) | Istituto Zooprofilattico Sperimentale dell'Abruzzo e Molise "G. Caporale" | Lorusso A, Marcacci M, Di Domenico M, Ancora M, Curini V, Di Lollo Valeria, Mangone I, Rinaldi A, Delli Compagni E, Scialabba S, Caporale M, Di Pasquale A, Cammà C, Puglia I, Calistri P, Savini G |
| EPI_ISL_1580615 | USCA Tagliacozzo TAGLIACOZZO(L'AQUILA) | Istituto Zooprofilattico Sperimentale dell'Abruzzo e Molise "G. Caporale" | Lorusso A, Marcacci M, Di Domenico M, Ancora M, Curini V, Di Lollo Valeria, Mangone I, Rinaldi A, Delli Compagni E, Scialabba S, Caporale M, Di Pasquale A, Cammà C, Puglia I, Calistri P, Savini G |
| EPI_ISL_1580616 | SIESP CHIETI - DRIVE IN CHIETI CHIETI(CHIETI) | Istituto Zooprofilattico Sperimentale dell'Abruzzo e Molise "G. Caporale" | Lorusso A, Marcacci M, Di Domenico M, Ancora M, Curini V, Di Lollo Valeria, Mangone I, Rinaldi A, Delli Compagni E, Scialabba S, Caporale M, Di Pasquale A, Cammà C, Puglia I, Calistri P, Savini G |
| EPI_ISL_1580617, EPI_ISL_1580618 | SIESP DIPARTIMENTO DI PREVENZIONE TERAMO TERAMO(TERAMO) | Istituto Zooprofilattico Sperimentale dell'Abruzzo e Molise "G. Caporale" | Lorusso A, Marcacci M, Di Domenico M, Ancora M, Curini V, Di Lollo Valeria, Mangone I, Rinaldi A, Delli Compagni E, Scialabba S, Caporale M, Di Pasquale A, Cammà C, Puglia I, Calistri P, Savini G |
| EPI_ISL_1580619 | PRESIDIO TERRITORIALE CASOLI CASOLI(CHIETI) | Istituto Zooprofilattico Sperimentale dell'Abruzzo e Molise "G. Caporale" | Lorusso A, Marcacci M, Di Domenico M, Ancora M, Curini V, Di Lollo Valeria, Mangone I, Rinaldi A, Delli Compagni E, Scialabba S, Caporale M, Di Pasquale A, Cammà C, Puglia I, Calistri P, Savini G |
| EPI_ISL_1580620 | SIESP DIPARTIMENTO DI PREVENZIONE TERAMO TERAMO(TERAMO) | Istituto Zooprofilattico Sperimentale dell'Abruzzo e Molise "G. Caporale" | Lorusso A, Marcacci M, Di Domenico M, Ancora M, Curini V, Di Lollo Valeria, Mangone I, Rinaldi A, Delli Compagni E, Scialabba S, Caporale M, Di Pasquale A, Cammà C, Puglia I, Calistri P, Savini G |
| EPI_ISL_1580621 | SIESP CHIETI - DRIVE IN GISSI GISSI(CHIETI) | Istituto Zooprofilattico Sperimentale dell'Abruzzo e Molise "G. Caporale" | Lorusso A, Marcacci M, Di Domenico M, Ancora M, Curini V, Di Lollo Valeria, Mangone I, Rinaldi A, Delli Compagni E, Scialabba S, Caporale M, Di Pasquale A, Cammà C, Puglia I, Calistri P, Savini G |
| EPI_ISL_1580622, EPI_ISL_1580623, EPI_ISL_1580624 | SIESP DIPARTIMENTO DI PREVENZIONE TERAMO TERAMO(TERAMO) | Istituto Zooprofilattico Sperimentale dell'Abruzzo e Molise "G. Caporale" | Lorusso A, Marcacci M, Di Domenico M, Ancora M, Curini V, Di Lollo Valeria, Mangone I, Rinaldi A, Delli Compagni E, Scialabba S, Caporale M, Di Pasquale A, Cammà C, Puglia I, Calistri P, Savini G |
| EPI_ISL_1580625, EPI_ISL_1580626 | DIP. PREV. AVEZZANO SERVIZIO DI IGIENE EPIDEMIOLOGIA E SANITA' PUBBLICA AVEZZANO(L'AQUILA) | Istituto Zooprofilattico Sperimentale dell'Abruzzo e Molise "G. Caporale" | Lorusso A, Marcacci M, Di Domenico M, Ancora M, Curini V, Di Lollo Valeria, Mangone I, Rinaldi A, Delli Compagni E, Scialabba S, Caporale M, Di Pasquale A, Cammà C, Puglia I, Calistri P, Savini G |
| EPI_ISL_1580627, EPI_ISL_1580628 | SIESP DIPARTIMENTO DI PREVENZIONE TERAMO TERAMO(TERAMO) | Istituto Zooprofilattico Sperimentale dell'Abruzzo e Molise "G. Caporale" | Lorusso A, Marcacci M, Di Domenico M, Ancora M, Curini V, Di Lollo Valeria, Mangone I, Rinaldi A, Delli Compagni E, Scialabba S, Caporale M, Di Pasquale A, Cammà C, Puglia I, Calistri P, Savini G |
| EPI_ISL_1580629, EPI_ISL_1580630 | SIESP CHIETI - DRIVE IN VASTO VASTO(CHIETI) | Istituto Zooprofilattico Sperimentale dell'Abruzzo e Molise "G. Caporale" | Lorusso A, Marcacci M, Di Domenico M, Ancora M, Curini V, Di Lollo Valeria, Mangone I, Rinaldi A, Delli Compagni E, Scialabba S, Caporale M, Di Pasquale A, Cammà C, Puglia I, Calistri P, Savini G |
| EPI_ISL_1580631 | SIESP CHIETI - DRIVE IN ORTONA ORTONA(CHIETI) | Istituto Zooprofilattico Sperimentale dell'Abruzzo e Molise "G. Caporale" | Lorusso A, Marcacci M, Di Domenico M, Ancora M, Curini V, Di Lollo Valeria, Mangone I, Rinaldi A, Delli Compagni E, Scialabba S, Caporale M, Di Pasquale A, Cammà C, Puglia I, Calistri P, Savini G |
| EPI_ISL_1580632 | SIESP CHIETI - DRIVE IN LANCIANO LANCIANO(CHIETI) | Istituto Zooprofilattico Sperimentale dell'Abruzzo e Molise "G. Caporale" | Lorusso A, Marcacci M, Di Domenico M, Ancora M, Curini V, Di Lollo Valeria, Mangone I, Rinaldi A, Delli Compagni E, Scialabba S, Caporale M, Di Pasquale A, Cammà C, Puglia I, Calistri P, Savini G |
| EPI_ISL_1580633, EPI_ISL_1580634, EPI_ISL_1580635, EPI_ISL_1580636, EPI_ISL_1580637, EPI_ISL_1580638, EPI_ISL_1580639 | SIESP DIPARTIMENTO DI PREVENZIONE TERAMO TERAMO(TERAMO) | Istituto Zooprofilattico Sperimentale dell'Abruzzo e Molise "G. Caporale" | Lorusso A, Marcacci M, Di Domenico M, Ancora M, Curini V, Di Lollo Valeria, Mangone I, Rinaldi A, Delli Compagni E, Scialabba S, Caporale M, Di Pasquale A, Cammà C, Puglia I, Calistri P, Savini G |
| EPI_ISL_1580640 | SIESP CHIETI - DRIVE IN VASTO VASTO(CHIETI) | Istituto Zooprofilattico Sperimentale dell'Abruzzo e Molise "G. Caporale" | Lorusso A, Marcacci M, Di Domenico M, Ancora M, Curini V, Di Lollo Valeria, Mangone I, Rinaldi A, Delli Compagni E, Scialabba S, Caporale M, Di Pasquale A, Cammà C, Puglia I, Calistri P, Savini G |
| EPI_ISL_1580641, EPI_ISL_1580642, EPI_ISL_1580643 | SIESP CHIETI - DRIVE IN CHIETI CHIETI(CHIETI) | Istituto Zooprofilattico Sperimentale dell'Abruzzo e Molise "G. Caporale" | Lorusso A, Marcacci M, Di Domenico M, Ancora M, Curini V, Di Lollo Valeria, Mangone I, Rinaldi A, Delli Compagni E, Scialabba S, Caporale M, Di Pasquale A, Cammà C, Puglia I, Calistri P, Savini G |
| EPI_ISL_1580644 | USCA Pescina PESCINA(L'AQUILA) | Istituto Zooprofilattico Sperimentale dell'Abruzzo e Molise "G. Caporale" | Lorusso A, Marcacci M, Di Domenico M, Ancora M, Curini V, Di Lollo Valeria, Mangone I, Rinaldi A, Delli Compagni E, Scialabba S, Caporale M, Di Pasquale A, Cammà C, Puglia I, Calistri P, Savini G |
| EPI_ISL_1580645 | DIP. PREV. AVEZZANO SERVIZIO DI IGIENE EPIDEMIOLOGIA E SANITA' PUBBLICA AVEZZANO(L'AQUILA) | Istituto Zooprofilattico Sperimentale dell'Abruzzo e Molise "G. Caporale" | Lorusso A, Marcacci M, Di Domenico M, Ancora M, Curini V, Di Lollo Valeria, Mangone I, Rinaldi A, Delli Compagni E, Scialabba S, Caporale M, Di Pasquale A, Cammà C, Puglia I, Calistri P, Savini G |
| EPI_ISL_1580646 | PRESIDIO TERRITORIALE CASOLI CASOLI(CHIETI) | Istituto Zooprofilattico Sperimentale dell'Abruzzo e Molise "G. Caporale" | Lorusso A, Marcacci M, Di Domenico M, Ancora M, Curini V, Di Lollo Valeria, Mangone I, Rinaldi A, Delli Compagni E, Scialabba S, Caporale M, Di Pasquale A, Cammà C, Puglia I, Calistri P, Savini G |
| EPI_ISL_1580647 | SIESP DIPARTIMENTO DI PREVENZIONE CHIETI CHIETI(CHIETI) | Istituto Zooprofilattico Sperimentale dell'Abruzzo e Molise "G. Caporale" | Lorusso A, Marcacci M, Di Domenico M, Ancora M, Curini V, Di Lollo Valeria, Mangone I, Rinaldi A, Delli Compagni E, Scialabba S, Caporale M, Di Pasquale A, Cammà C, Puglia I, Calistri P, Savini G |
| EPI_ISL_1580648 | DIP. PREV. AVEZZANO SERVIZIO DI IGIENE EPIDEMIOLOGIA E SANITA' PUBBLICA AVEZZANO(L'AQUILA) | Istituto Zooprofilattico Sperimentale dell'Abruzzo e Molise "G. Caporale" | Lorusso A, Marcacci M, Di Domenico M, Ancora M, Curini V, Di Lollo Valeria, Mangone I, Rinaldi A, Delli Compagni E, Scialabba S, Caporale M, Di Pasquale A, Cammà C, Puglia I, Calistri P, Savini G |
| EPI_ISL_1580649 | SIESP CHIETI - DRIVE IN ORTONA ORTONA(CHIETI) | Istituto Zooprofilattico Sperimentale dell'Abruzzo e Molise "G. Caporale" | Lorusso A, Marcacci M, Di Domenico M, Ancora M, Curini V, Di Lollo Valeria, Mangone I, Rinaldi A, Delli Compagni E, Scialabba S, Caporale M, Di Pasquale A, Cammà C, Puglia I, Calistri P, Savini G |
| EPI_ISL_1580650, EPI_ISL_1580651 | DIP. PREV. AVEZZANO SERVIZIO DI IGIENE EPIDEMIOLOGIA E SANITA' PUBBLICA AVEZZANO(L'AQUILA) | Istituto Zooprofilattico Sperimentale dell'Abruzzo e Molise "G. Caporale" | Lorusso A, Marcacci M, Di Domenico M, Ancora M, Curini V, Di Lollo Valeria, Mangone I, Rinaldi A, Delli Compagni E, Scialabba S, Caporale M, Di Pasquale A, Cammà C, Puglia I, Calistri P, Savini G |
| EPI_ISL_1580652 | SIESP CHIETI - DRIVE IN GISSI GISSI(CHIETI) | Istituto Zooprofilattico Sperimentale dell'Abruzzo e Molise "G. Caporale" | Lorusso A, Marcacci M, Di Domenico M, Ancora M, Curini V, Di Lollo Valeria, Mangone I, Rinaldi A, Delli Compagni E, Scialabba S, Caporale M, Di Pasquale A, Cammà C, Puglia I, Calistri P, Savini G |
| EPI_ISL_1580653, EPI_ISL_1580654 | SIESP DIPARTIMENTO DI PREVENZIONE TERAMO TERAMO(TERAMO) | Istituto Zooprofilattico Sperimentale dell'Abruzzo e Molise "G. Caporale" | Lorusso A, Marcacci M, Di Domenico M, Ancora M, Curini V, Di Lollo Valeria, Mangone I, Rinaldi A, Delli Compagni E, Scialabba S, Caporale M, Di Pasquale A, Cammà C, Puglia I, Calistri P, Savini G |
| EPI_ISL_1580655, EPI_ISL_1580656 | DIP. PREV. AVEZZANO SERVIZIO DI IGIENE EPIDEMIOLOGIA E SANITA' PUBBLICA AVEZZANO(L'AQUILA) | Istituto Zooprofilattico Sperimentale dell'Abruzzo e Molise "G. Caporale" | Lorusso A, Marcacci M, Di Domenico M, Ancora M, Curini V, Di Lollo Valeria, Mangone I, Rinaldi A, Delli Compagni E, Scialabba S, Caporale M, Di Pasquale A, Cammà C, Puglia I, Calistri P, Savini G |
| EPI_ISL_1580657, EPI_ISL_1580658 | SIESP CHIETI - DRIVE IN LANCIANO LANCIANO(CHIETI) | Istituto Zooprofilattico Sperimentale dell'Abruzzo e Molise "G. Caporale" | Lorusso A, Marcacci M, Di Domenico M, Ancora M, Curini V, Di Lollo Valeria, Mangone I, Rinaldi A, Delli Compagni E, Scialabba S, Caporale M, Di Pasquale A, Cammà C, Puglia I, Calistri P, Savini G |
| EPI_ISL_1580659 | SIESP DIPARTIMENTO DI PREVENZIONE TERAMO TERAMO(TERAMO) | Istituto Zooprofilattico Sperimentale dell'Abruzzo e Molise "G. Caporale" | Lorusso A, Marcacci M, Di Domenico M, Ancora M, Curini V, Di Lollo Valeria, Mangone I, Rinaldi A, Delli Compagni E, Scialabba S, Caporale M, Di Pasquale A, Cammà C, Puglia I, Calistri P, Savini G |
| EPI_ISL_1580660 | SIESP SULMONA SULMONA(L'AQUILA) | Istituto Zooprofilattico Sperimentale dell'Abruzzo e Molise "G. | Lorusso A, Marcacci M, Di Domenico M, Ancora M, Curini V, Di Lollo Valeria, Mangone I, Rinaldi A, Delli Compagni E, Scialabba S, Caporale M, Di |

| | | | |
|---|---|---|---|
| | | Caporale" | Pasquale A, Cammà C, Puglia I, Calistri P, Savini G |
| EPI_ISL_1580661, EPI_ISL_1580662, EPI_ISL_1580663, EPI_ISL_1580664, EPI_ISL_1580665 | SIESP DIPARTIMENTO DI PREVENZIONE TERAMO TERAMO(TERAMO) | Istituto Zooprofilattico Sperimentale dell'Abruzzo e Molise "G. Caporale" | Lorusso A, Marcacci M, Di Domenico M, Ancora M, Curini V, Di Lollo Valeria, Mangone I, Rinaldi A, Delli Compagni E, Scialabba S, Caporale M, Di Pasquale A, Cammà C, Puglia I, Calistri P, Savini G |
| EPI_ISL_1580666 | DIP. PREV. AVEZZANO SERVIZIO DI IGIENE EPIDEMIOLOGIA E SANITA' PUBBLICA AVEZZANO(L'AQUILA) | Istituto Zooprofilattico Sperimentale dell'Abruzzo e Molise "G. Caporale" | Lorusso A, Marcacci M, Di Domenico M, Ancora M, Curini V, Di Lollo Valeria, Mangone I, Rinaldi A, Delli Compagni E, Scialabba S, Caporale M, Di Pasquale A, Cammà C, Puglia I, Calistri P, Savini G |
| EPI_ISL_1582659, EPI_ISL_1582660, EPI_ISL_1582661, EPI_ISL_1582662 | Hospital | National Reference Center for Viruses of Respiratory Infections, Institut Pasteur, Paris | Marion Barbet, Sylvie Behillil, Frédéric Lemoine, Corinne Maufrais, Christophe Malabat, Méline Bizard, Angela Brisebarre, Camille Capel, Louise Lefrançois, Etienne Simon-Lorière, Vincent Enouf, Maud Vanpeene, Sylvie van der Werf,CéCile Farrugia |
| EPI_ISL_1582663 | Hospital | National Reference Center for Viruses of Respiratory Infections, Institut Pasteur, Paris | Marion Barbet, Sylvie Behillil, Frédéric Lemoine, Corinne Maufrais, Christophe Malabat,Gael Millot, Méline Bizard, Angela Brisebarre, Camille Capel, Louise Lefrançois, Etienne Simon-Lorière, Vincent Enouf, Maud Vanpeene, Sylvie van der Werf,CéCile Farrugia |
| EPI_ISL_1582664, EPI_ISL_1582665, EPI_ISL_1582666, EPI_ISL_1582667 | Hospital | National Reference Center for Viruses of Respiratory Infections, Institut Pasteur, Paris | Marion Barbet, Sylvie Behillil, Frédéric Lemoine, Corinne Maufrais, Christophe Malabat, Méline Bizard, Angela Brisebarre, Camille Capel, Louise Lefrançois, Etienne Simon-Lorière, Vincent Enouf, Maud Vanpeene, Sylvie van der Werf,CéCile Farrugia |
| EPI_ISL_1582668 | Hospital | National Reference Center for Viruses of Respiratory Infections, Institut Pasteur, Paris | Marion Barbet, Sylvie Behillil, Frédéric Lemoine, Corinne Maufrais, Christophe Malabat,Gael Millot, Méline Bizard, Angela Brisebarre, Camille Capel, Louise Lefrançois, Etienne Simon-Lorière, Vincent Enouf, Maud Vanpeene, Sylvie van der Werf,CéCile Farrugia |
| EPI_ISL_1582669 | Hospital | National Reference Center for Viruses of Respiratory Infections, Institut Pasteur, Paris | Marion Barbet, Sylvie Behillil, Frédéric Lemoine, Corinne Maufrais, Christophe Malabat,Pierre Lechat, Méline Bizard, Angela Brisebarre, Camille Capel, Louise Lefrançois, Etienne Simon-Lorière, Vincent Enouf, Maud Vanpeene, Sylvie van der Werf,CéCile Farrugia |
| EPI_ISL_1582670, EPI_ISL_1582671, EPI_ISL_1582672, EPI_ISL_1582673, EPI_ISL_1582674, EPI_ISL_1582675, EPI_ISL_1582676, EPI_ISL_1582677, EPI_ISL_1582679, EPI_ISL_1582681, EPI_ISL_1582682 | | | |
| see above | Hospital | National Reference Center for Viruses of Respiratory Infections, Institut Pasteur, Paris | Marion Barbet, Sylvie Behillil, Frédéric Lemoine, Corinne Maufrais, Christophe Malabat, Méline Bizard, Angela Brisebarre, Camille Capel, Louise Lefrançois, Etienne Simon-Lorière, Vincent Enouf, Maud Vanpeene, Sylvie van der Werf,Pascale Martres |
| EPI_ISL_1582683, EPI_ISL_1582684, EPI_ISL_1582685, EPI_ISL_1582686, EPI_ISL_1582687, EPI_ISL_1582688, EPI_ISL_1582689, EPI_ISL_1582690, EPI_ISL_1582691, EPI_ISL_1582692 | Armies | National Reference Center for Viruses of Respiratory Infections, Institut Pasteur, Paris | Marion Barbet, Sylvie Behillil, Frédéric Lemoine, Corinne Maufrais, Christophe Malabat, Méline Bizard, Angela Brisebarre, Camille Capel, Louise Lefrançois, Etienne Simon-Lorière, Vincent Enouf, Maud Vanpeene, Sylvie van der Werf,Marine Desroches |
| EPI_ISL_1582693 | Armies | National Reference Center for Viruses of Respiratory Infections, Institut Pasteur, Paris | Marion Barbet, Sylvie Behillil, Frédéric Lemoine, Corinne Maufrais, Christophe Malabat,Gael Millot, Méline Bizard, Angela Brisebarre, Camille Capel, Louise Lefrançois, Etienne Simon-Lorière, Vincent Enouf, Maud Vanpeene, Sylvie van der Werf,Marine Desroches |
| EPI_ISL_1582695 | Armies | National Reference Center for Viruses of Respiratory Infections, Institut Pasteur, Paris | Marion Barbet, Sylvie Behillil, Frédéric Lemoine, Corinne Maufrais, Christophe Malabat, Méline Bizard, Angela Brisebarre, Camille Capel, Louise Lefrançois, Etienne Simon-Lorière, Vincent Enouf, Maud Vanpeene, Sylvie van der Werf,Marine Desroches |
| EPI_ISL_1582696 | Hospital | National Reference Center for Viruses of Respiratory Infections, Institut Pasteur, Paris | Marion Barbet, Sylvie Behillil, Frédéric Lemoine, Corinne Maufrais, Christophe Malabat,Gael Millot, Méline Bizard, Angela Brisebarre, Camille Capel, Louise Lefrançois, Etienne Simon-Lorière, Vincent Enouf, Maud Vanpeene, Sylvie van der Werf,JéRome Guinard |
| EPI_ISL_1582697, EPI_ISL_1582698, EPI_ISL_1582699, EPI_ISL_1582700, EPI_ISL_1582701, EPI_ISL_1582703, EPI_ISL_1582705 | Hospital | National Reference Center for Viruses of Respiratory Infections, Institut Pasteur, Paris | Marion Barbet, Sylvie Behillil, Frédéric Lemoine, Corinne Maufrais, Christophe Malabat, Méline Bizard, Angela Brisebarre, Camille Capel, Louise Lefrançois, Etienne Simon-Lorière, Vincent Enouf, Maud Vanpeene, Sylvie van der Werf,JéRome Guinard |
| EPI_ISL_1582706 | Hospital | National Reference Center for Viruses of Respiratory Infections, Institut Pasteur, Paris | Marion Barbet, Sylvie Behillil, Frédéric Lemoine, Corinne Maufrais, Christophe Malabat,Gael Millot, Méline Bizard, Angela Brisebarre, Camille Capel, Louise Lefrançois, Etienne Simon-Lorière, Vincent Enouf, Maud Vanpeene, Sylvie van der Werf,JéRome Guinard |
| EPI_ISL_1582711 | Hospital | National Reference Center for Viruses of Respiratory Infections, Institut Pasteur, Paris | Marion Barbet, Sylvie Behillil, Frédéric Lemoine, Corinne Maufrais, Christophe Malabat,Pierre Lechat, Méline Bizard, Angela Brisebarre, Camille Capel, Louise Lefrançois, Etienne Simon-Lorière, Vincent Enouf, Maud Vanpeene, Sylvie van der Werf,Isabelle Gros |
| EPI_ISL_1582714 | Hospital | National Reference Center for Viruses of Respiratory Infections, Institut Pasteur, Paris | Marion Barbet, Sylvie Behillil, Frédéric Lemoine, Corinne Maufrais, Christophe Malabat, Méline Bizard, Angela Brisebarre, Camille Capel, Louise Lefrançois, Etienne Simon-Lorière, Vincent Enouf, Maud Vanpeene, Sylvie van der Werf,Isabelle Gros |
| EPI_ISL_1582715 | Hospital | National Reference Center for Viruses of Respiratory Infections, Institut Pasteur, Paris | Marion Barbet, Sylvie Behillil, Frédéric Lemoine, Corinne Maufrais, Christophe Malabat,Pierre Lechat, Méline Bizard, Angela Brisebarre, Camille Capel, Louise Lefrançois, Etienne Simon-Lorière, Vincent Enouf, Maud Vanpeene, Sylvie van der Werf,Isabelle Gros |
| EPI_ISL_1582717, EPI_ISL_1582718, EPI_ISL_1582719, EPI_ISL_1582721 | Hospital | National Reference Center for Viruses of Respiratory Infections, Institut Pasteur, Paris | Marion Barbet, Sylvie Behillil, Frédéric Lemoine, Corinne Maufrais, Christophe Malabat, Méline Bizard, Angela Brisebarre, Camille Capel, Louise Lefrançois, Etienne Simon-Lorière, Vincent Enouf, Maud Vanpeene, Sylvie van der Werf,Isabelle Gros |
| EPI_ISL_1582723, EPI_ISL_1582725, EPI_ISL_1582726 | Hospital | National Reference Center for Viruses of Respiratory Infections, Institut Pasteur, Paris | Marion Barbet, Sylvie Behillil, Frédéric Lemoine, Corinne Maufrais, Christophe Malabat, Méline Bizard, Angela Brisebarre, Camille Capel, Louise Lefrançois, Etienne Simon-Lorière, Vincent Enouf, Maud Vanpeene, Sylvie van der Werf,StéPhanie Marque Juillet |
| EPI_ISL_1582727 | Hospital | National Reference Center for Viruses of Respiratory Infections, Institut Pasteur, Paris | Marion Barbet, Sylvie Behillil, Frédéric Lemoine, Corinne Maufrais, Christophe Malabat,Gael Millot, Méline Bizard, Angela Brisebarre, Camille Capel, Louise Lefrançois, Etienne Simon-Lorière, Vincent Enouf, Maud Vanpeene, Sylvie van der Werf,StéPhanie Marque Juillet |
| EPI_ISL_1582728 | Hospital | National Reference Center for Viruses of Respiratory Infections, Institut Pasteur, Paris | Marion Barbet, Sylvie Behillil, Frédéric Lemoine, Corinne Maufrais, Christophe Malabat,Pierre Lechat, Méline Bizard, Angela Brisebarre, Camille Capel, Louise Lefrançois, Etienne Simon-Lorière, Vincent Enouf, Maud Vanpeene, Sylvie van der Werf,StéPhanie Marque Juillet |
| EPI_ISL_1582729, EPI_ISL_1582730 | Hospital | National Reference Center for Viruses of Respiratory Infections, Institut Pasteur, Paris | Marion Barbet, Sylvie Behillil, Frédéric Lemoine, Corinne Maufrais, Christophe Malabat, Méline Bizard, Angela Brisebarre, Camille Capel, Louise Lefrançois, Etienne Simon-Lorière, Vincent Enouf, Maud Vanpeene, Sylvie van der Werf,StéPhanie Marque Juillet |
| EPI_ISL_1582731, EPI_ISL_1582732 | Hospital | National Reference Center for Viruses of Respiratory Infections, Institut Pasteur, Paris | Marion Barbet, Sylvie Behillil, Frédéric Lemoine, Corinne Maufrais, Christophe Malabat, Méline Bizard, Angela Brisebarre, Camille Capel, Louise Lefrançois, Etienne Simon-Lorière, Vincent Enouf, Maud Vanpeene, Sylvie van der Werf,Sandrine Castelain |
| EPI_ISL_1582733, EPI_ISL_1582734 | Hospital | National Reference Center for Viruses of Respiratory Infections, Institut Pasteur, Paris | Marion Barbet, Sylvie Behillil, Frédéric Lemoine, Corinne Maufrais, Christophe Malabat,Pierre Lechat, Méline Bizard, Angela Brisebarre, Camille Capel, Louise Lefrançois, Etienne Simon-Lorière, Vincent Enouf, Maud Vanpeene, Sylvie van der Werf,Sandrine Castelain |
| EPI_ISL_1582736, EPI_ISL_1582737 | Hospital | National Reference Center for Viruses of Respiratory Infections, Institut Pasteur, Paris | Marion Barbet, Sylvie Behillil, Frédéric Lemoine, Corinne Maufrais, Christophe Malabat, Méline Bizard, Angela Brisebarre, Camille Capel, Louise Lefrançois, Etienne Simon-Lorière, Vincent Enouf, Maud Vanpeene, Sylvie van der Werf,Sandrine Castelain |
| EPI_ISL_1582738 | Hospital | National Reference Center for Viruses of Respiratory Infections, Institut Pasteur, Paris | Marion Barbet, Sylvie Behillil, Frédéric Lemoine, Corinne Maufrais, Christophe Malabat,Pierre Lechat, Méline Bizard, Angela Brisebarre, Camille Capel, Louise Lefrançois, Etienne Simon-Lorière, Vincent Enouf, Maud Vanpeene, Sylvie van der Werf,Sandrine Castelain |
| EPI_ISL_1582739, EPI_ISL_1582740 | Hospital | National Reference Center for Viruses of Respiratory Infections, Institut Pasteur, Paris | Marion Barbet, Sylvie Behillil, Frédéric Lemoine, Corinne Maufrais, Christophe Malabat, Méline Bizard, Angela Brisebarre, Camille Capel, Louise Lefrançois, Etienne Simon-Lorière, Vincent Enouf, Maud Vanpeene, Sylvie van der Werf,Sandrine Castelain |
| EPI_ISL_1582741, EPI_ISL_1582742, EPI_ISL_1582744, EPI_ISL_1582746 | Hospital | National Reference Center for Viruses of Respiratory Infections, Institut Pasteur, Paris | Marion Barbet, Sylvie Behillil, Frédéric Lemoine, Corinne Maufrais, Christophe Malabat, Méline Bizard, Angela Brisebarre, Camille Capel, Louise Lefrançois, Etienne Simon-Lorière, Vincent Enouf, Maud Vanpeene, Sylvie van der Werf,AuréLie Guigon |
| EPI_ISL_1582747 | Hospital | National Reference Center for Viruses of Respiratory Infections, Institut Pasteur, Paris | Marion Barbet, Sylvie Behillil, Frédéric Lemoine, Corinne Maufrais, Christophe Malabat,Pierre Lechat, Méline Bizard, Angela Brisebarre, Camille Capel, Louise Lefrançois, Etienne Simon-Lorière, Vincent Enouf, Maud Vanpeene, Sylvie van der Werf,Haciba Moudjahed |
| EPI_ISL_1582748, EPI_ISL_1582749, EPI_ISL_1582750 | Hospital | National Reference Center for Viruses of Respiratory Infections, Institut Pasteur, Paris | Marion Barbet, Sylvie Behillil, Frédéric Lemoine, Corinne Maufrais, Christophe Malabat, Méline Bizard, Angela Brisebarre, Camille Capel, Louise Lefrançois, Etienne Simon-Lorière, Vincent Enouf, Maud Vanpeene, Sylvie van der Werf,S. Van Agt |
| EPI_ISL_1582751, EPI_ISL_1582752 | Hospital | National Reference Center for Viruses of Respiratory Infections, Institut Pasteur, Paris | Marion Barbet, Sylvie Behillil, Frédéric Lemoine, Corinne Maufrais, Christophe Malabat, Méline Bizard, Angela Brisebarre, Camille Capel, Louise Lefrançois, Etienne Simon-Lorière, Vincent Enouf, Maud Vanpeene, Sylvie van der Werf,Sylvie Ledru |

| | | | |
|---|---|---|---|
| EPI_ISL_1582753 | Hospital | National Reference Center for Viruses of Respiratory Infections, Institut Pasteur, Paris | Marion Barbet, Sylvie Behillil, Frédéric Lemoine, Corinne Maufrais, Christophe Malabat,Gael Millot, Méline Bizard, Angela Brisebarre, Camille Capel, Louise Lefrançois, Etienne Simon-Lorière, Vincent Enouf, Maud Vanpeene, Sylvie van der Werf,Sylvie Ledru |
| EPI_ISL_1582754 | Hospital | National Reference Center for Viruses of Respiratory Infections, Institut Pasteur, Paris | Marion Barbet, Sylvie Behillil, Frédéric Lemoine, Corinne Maufrais, Christophe Malabat,Pierre Lechat, Méline Bizard, Angela Brisebarre, Camille Capel, Louise Lefrançois, Etienne Simon-Lorière, Vincent Enouf, Maud Vanpeene, Sylvie van der Werf,Franck Griscelli |
| EPI_ISL_1582755 | Hospital | National Reference Center for Viruses of Respiratory Infections, Institut Pasteur, Paris | Marion Barbet, Sylvie Behillil, Frédéric Lemoine, Corinne Maufrais, Christophe Malabat,Gael Millot, Méline Bizard, Angela Brisebarre, Camille Capel, Louise Lefrançois, Etienne Simon-Lorière, Vincent Enouf, Maud Vanpeene, Sylvie van der Werf,Franck Griscelli |
| EPI_ISL_1582757, EPI_ISL_1582758 | Hospital | National Reference Center for Viruses of Respiratory Infections, Institut Pasteur, Paris | Marion Barbet, Sylvie Behillil, Frédéric Lemoine, Corinne Maufrais, Christophe Malabat, Méline Bizard, Angela Brisebarre, Camille Capel, Louise Lefrançois, Etienne Simon-Lorière, Vincent Enouf, Maud Vanpeene, Sylvie van der Werf,Anne Vachee |
| EPI_ISL_1583006, EPI_ISL_1583007 | Institute for Medical Research, Infectious Disease Research Centre, National Institutes of Health, Ministry of Health Malaysia | Institute for Medical Research, Infectious Disease Research Centre, National Institutes of Health, Ministry of Health Malaysia | Suppiah J, Kamel K, Mohd Zawawi Z, Ramly N, Robert F, Thayan R |
| EPI_ISL_1583008, EPI_ISL_1583009, EPI_ISL_1583010, EPI_ISL_1583011, EPI_ISL_1583012, EPI_ISL_1583013, EPI_ISL_1583014, EPI_ISL_1583015 | Hospital | National Reference Center for Viruses of Respiratory Infections, Institut Pasteur, Paris | Marion Barbet, Sylvie Behillil, Frédéric Lemoine, Corinne Maufrais, Christophe Malabat, Méline Bizard, Angela Brisebarre, Camille Capel, Louise Lefrançois, Etienne Simon-Lorière, Vincent Enouf, Maud Vanpeene, Sylvie van der Werf,Emeline Riverain |
| EPI_ISL_1583016, EPI_ISL_1583019, EPI_ISL_1583020, EPI_ISL_1583022, EPI_ISL_1583023, EPI_ISL_1583025, EPI_ISL_1583026 | Hospital | National Reference Center for Viruses of Respiratory Infections, Institut Pasteur, Paris | Marion Barbet, Sylvie Behillil, Frédéric Lemoine, Corinne Maufrais, Christophe Malabat, Méline Bizard, Angela Brisebarre, Camille Capel, Louise Lefrançois, Etienne Simon-Lorière, Vincent Enouf, Maud Vanpeene, Sylvie van der Werf,José Bras Cachinho |
| EPI_ISL_1583027, EPI_ISL_1583028, EPI_ISL_1583029, EPI_ISL_1583030, EPI_ISL_1583031 | Labo Analyses Med | National Reference Center for Viruses of Respiratory Infections, Institut Pasteur, Paris | Marion Barbet, Sylvie Behillil, Frédéric Lemoine, Corinne Maufrais, Christophe Malabat, Méline Bizard, Angela Brisebarre, Camille Capel, Louise Lefrançois, Etienne Simon-Lorière, Vincent Enouf, Maud Vanpeene, Sylvie van der Werf,Lam Alliance Anabio |
| EPI_ISL_1583032, EPI_ISL_1583033, EPI_ISL_1583034, EPI_ISL_1583035, EPI_ISL_1583036, EPI_ISL_1583037, EPI_ISL_1583038, EPI_ISL_1583039, EPI_ISL_1583040, EPI_ISL_1583042 | Hospital | National Reference Center for Viruses of Respiratory Infections, Institut Pasteur, Paris | Marion Barbet, Sylvie Behillil, Frédéric Lemoine, Corinne Maufrais, Christophe Malabat, Méline Bizard, Angela Brisebarre, Camille Capel, Louise Lefrançois, Etienne Simon-Lorière, Vincent Enouf, Maud Vanpeene, Sylvie van der Werf,LéA Pilorge |
| EPI_ISL_1583044, EPI_ISL_1583045, EPI_ISL_1583046, EPI_ISL_1583047, EPI_ISL_1583048 | Labo Analyses Med | National Reference Center for Viruses of Respiratory Infections, Institut Pasteur, Paris | Marion Barbet, Sylvie Behillil, Frédéric Lemoine, Corinne Maufrais, Christophe Malabat, Méline Bizard, Angela Brisebarre, Camille Capel, Louise Lefrançois, Etienne Simon-Lorière, Vincent Enouf, Maud Vanpeene, Sylvie van der Werf,SéVerine Haouisee |
| EPI_ISL_1583049, EPI_ISL_1583050, EPI_ISL_1583051 | Hospital | National Reference Center for Viruses of Respiratory Infections, Institut Pasteur, Paris | Marion Barbet, Sylvie Behillil, Frédéric Lemoine, Corinne Maufrais, Christophe Malabat, Méline Bizard, Angela Brisebarre, Camille Capel, Louise Lefrançois, Etienne Simon-Lorière, Vincent Enouf, Maud Vanpeene, Sylvie van der Werf,Eleha Guillotel |
| EPI_ISL_1583052, EPI_ISL_1583054, EPI_ISL_1583055, EPI_ISL_1583056, EPI_ISL_1583057, EPI_ISL_1583058 | Labo Analyses Med | National Reference Center for Viruses of Respiratory Infections, Institut Pasteur, Paris | Marion Barbet, Sylvie Behillil, Frédéric Lemoine, Corinne Maufrais, Christophe Malabat, Méline Bizard, Angela Brisebarre, Camille Capel, Louise Lefrançois, Etienne Simon-Lorière, Vincent Enouf, Maud Vanpeene, Sylvie van der Werf,GréGoire Potiron |
| EPI_ISL_1583059 | Hospital | National Reference Center for Viruses of Respiratory Infections, Institut Pasteur, Paris | Marion Barbet, Sylvie Behillil, Frédéric Lemoine, Corinne Maufrais, Christophe Malabat, Méline Bizard, Angela Brisebarre, Camille Capel, Louise Lefrançois, Etienne Simon-Lorière, Vincent Enouf, Maud Vanpeene, Sylvie van der Werf,Pauline Morvan |
| EPI_ISL_1583060, EPI_ISL_1583061, EPI_ISL_1583062 | Hospital | National Reference Center for Viruses of Respiratory Infections, Institut Pasteur, Paris | Marion Barbet, Sylvie Behillil, Frédéric Lemoine, Corinne Maufrais, Christophe Malabat, Méline Bizard, Angela Brisebarre, Camille Capel, Louise Lefrançois, Etienne Simon-Lorière, Vincent Enouf, Maud Vanpeene, Sylvie van der Werf,BéAtrice Lesimple |
| EPI_ISL_1583063, EPI_ISL_1583064 | Labo Analyses Med | National Reference Center for Viruses of Respiratory Infections, Institut Pasteur, Paris | Marion Barbet, Sylvie Behillil, Frédéric Lemoine, Corinne Maufrais, Christophe Malabat, Méline Bizard, Angela Brisebarre, Camille Capel, Louise Lefrançois, Etienne Simon-Lorière, Vincent Enouf, Maud Vanpeene, Sylvie van der Werf,Labm Cité Sanitaire |
| EPI_ISL_1583065 | Hospital | National Reference Center for Viruses of Respiratory Infections, Institut Pasteur, Paris | Marion Barbet, Sylvie Behillil, Frédéric Lemoine, Corinne Maufrais, Christophe Malabat, Méline Bizard, Angela Brisebarre, Camille Capel, Louise Lefrançois, Etienne Simon-Lorière, Vincent Enouf, Maud Vanpeene, Sylvie van der Werf,Marc Villemain |
| EPI_ISL_1583066, EPI_ISL_1583067, EPI_ISL_1583068, EPI_ISL_1583069, EPI_ISL_1583070, EPI_ISL_1583071, EPI_ISL_1583072, EPI_ISL_1583073 | Hospital | National Reference Center for Viruses of Respiratory Infections, Institut Pasteur, Paris | Marion Barbet, Sylvie Behillil, Frédéric Lemoine, Corinne Maufrais, Christophe Malabat, Méline Bizard, Angela Brisebarre, Camille Capel, Louise Lefrançois, Etienne Simon-Lorière, Vincent Enouf, Maud Vanpeene, Sylvie van der Werf,Alexandra Ducancelle |
| EPI_ISL_1583075, EPI_ISL_1583076, EPI_ISL_1583077, EPI_ISL_1583079, EPI_ISL_1583080, EPI_ISL_1583081, EPI_ISL_1583082, EPI_ISL_1583083, EPI_ISL_1583084, EPI_ISL_1583085, EPI_ISL_1583086, EPI_ISL_1583087, EPI_ISL_1583088, EPI_ISL_1583089, EPI_ISL_1583090, EPI_ISL_1583091, EPI_ISL_1583092, EPI_ISL_1583093 | | | |
| see above | Labo analyses Med | National Reference Center for Viruses of Respiratory Infections, Institut Pasteur, Paris | Marion Barbet, Sylvie Behillil, Frédéric Lemoine, Corinne Maufrais, Christophe Malabat, Méline Bizard, Angela Brisebarre, Camille Capel, Louise Lefrançois, Etienne Simon-Lorière, Vincent Enouf, Maud Vanpeene, Sylvie van der Werf,Anne Holstein |
| EPI_ISL_1583097, EPI_ISL_1583098, EPI_ISL_1583100, EPI_ISL_1583101 | Hospital | National Reference Center for Viruses of Respiratory Infections, Institut Pasteur, Paris | Marion Barbet, Sylvie Behillil, Frédéric Lemoine, Corinne Maufrais, Christophe Malabat, Méline Bizard, Angela Brisebarre, Camille Capel, Louise Lefrançois, Etienne Simon-Lorière, Vincent Enouf, Maud Vanpeene, Sylvie van der Werf,Sandrine Castelain |
| EPI_ISL_1583102, EPI_ISL_1583103, EPI_ISL_1583104, EPI_ISL_1583105, EPI_ISL_1583106, EPI_ISL_1583107, EPI_ISL_1583108, EPI_ISL_1583109, EPI_ISL_1583110, EPI_ISL_1583111, EPI_ISL_1583112, EPI_ISL_1583113, EPI_ISL_1583114, EPI_ISL_1583115, EPI_ISL_1583116, EPI_ISL_1583117, EPI_ISL_1583118, EPI_ISL_1583119, EPI_ISL_1583120, EPI_ISL_1583121, EPI_ISL_1583122, EPI_ISL_1583123, EPI_ISL_1583124, EPI_ISL_1583127, EPI_ISL_1583128, EPI_ISL_1583129, EPI_ISL_1583130, EPI_ISL_1583131, EPI_ISL_1583132, EPI_ISL_1583133, EPI_ISL_1583134, EPI_ISL_1583136, EPI_ISL_1583137, EPI_ISL_1583138, EPI_ISL_1583139, EPI_ISL_1583140, EPI_ISL_1583141, EPI_ISL_1583142, EPI_ISL_1583143, EPI_ISL_1583144, EPI_ISL_1583146, EPI_ISL_1583147, EPI_ISL_1583148, EPI_ISL_1583150, EPI_ISL_1583151, EPI_ISL_1583152, EPI_ISL_1583153, EPI_ISL_1583154, EPI_ISL_1583156 | | | |
| see above | Hospital | National Reference Center for Viruses of Respiratory Infections, Institut Pasteur, Paris | Marion Barbet, Sylvie Behillil, Frédéric Lemoine, Corinne Maufrais, Christophe Malabat, Méline Bizard, Angela Brisebarre, Camille Capel, Louise Lefrançois, Etienne Simon-Lorière, Vincent Enouf, Maud Vanpeene, Sylvie van der Werf,Nabil Gastli |
| EPI_ISL_1585242 | Institute for Medical Research, Infectious Disease Research Centre, National Institutes of Health, Ministry of Health Malaysia | Institute for Medical Research, Infectious Disease Research Centre, National Institutes of Health, Ministry of Health Malaysia | Suppiah J, Kamel K, Mohd Zawawi Z, Ramly N, Robert F, Thayan R |
| EPI_ISL_1585270, EPI_ISL_1585271 | Labuan Hospital | Institute for Medical Research, Infectious Disease Research Centre, National Institutes of Health, Ministry of Health Malaysia | Suppiah J, Kamel K, Mohd Zawawi Z, Ramly N, Robert F, Thayan R |
| EPI_ISL_1585274, EPI_ISL_1585275 | Virology Unit, AOUP | Virology Unit, AOUP | Marialinda Vatteroni, Susi Frateschi, Mauro Pistello |
| EPI_ISL_1585276 | INT Fondazione Pascale | INT Fondazione Pascale | INT Fondazione Pascale |
| EPI_ISL_1585277 | Virology Unit, AOUP | Virology Unit, AOUP | Marialinda Vatteroni, Susi Frateschi, Mauro Pistello |
| EPI_ISL_1585278 | INT Fondazione Pascale | INT Fondazione Pascale | INT Fondazione Pascale |
| EPI_ISL_1585279, EPI_ISL_1585280 | Virology Unit, AOUP | Virology Unit, AOUP | Marialinda Vatteroni, Susi Frateschi, Mauro Pistello |
| EPI_ISL_1585288 | Virology Unit, AOUP | Virology Unit, AOUP | Marialinda Vatteroni, Susi Frateschi |
| EPI_ISL_1585423 | Centro de Investigación Biomédica del Noreste (CIBIN) | Instituto Nacional de Enfermedades Respiratorias (INER): Centro de Investigación en Enfermedades Infecciosas (CIENI) | Consorcio Mexicano de Vigilancia Genómica (CoViGen-Mex). Authors (in alphabetical order): Julio Elias Alvarado-Yaah, Carlos F. Arias, Santiago Ávila-Ríos, Víctor Hugo Borja-Aburto, Celia Boukadida, Juan Bautista Chale-Dzul, Célida Duque Molina, José Antonio Enciso-Moreno, Gloria Elena Espinosa-Ayala, Fernando Fontove-Herrera, Victor Eduardo Garcia-Arias, Concepción Grajales-Muñiz, Ricardo Grande, Alfredo Herrera-Estrella, Carla |

| | | | |
|---|---|---|---|
| | | | Ivón Herrera-Najera, Pavel Isa, Brenda Irasema Maldonado-Meza, Bernardo Martínez-Miguel, Margarita Matías-Florentino, María Guadalupe de Jesús Mireles-Rivera, Gloria María Molina-Salinas, Hector Montoya-Fuentes, José Esteban Muñoz-Medina, José de Jesús Nuñez-Contreras, Alicia Ocaña-Mondragón, Luis Alberto Ochoa-Carrera, Hector Esteban Paz-Juárez, Francisco Pulido, Helen Haydee Fernanda Ramírez-Plascencia, Angel Gustavo Salas-Lais, Alejandro Sanchez-Flores, Clara Esperanza Santacruz-Tinoco, María Guadalupe Santiago-Mauricio, Nelly Sélem-Mojica, Blanca Taboada, Gloria Vazquez |
| EPI_ISL_1585475, EPI_ISL_1585478, EPI_ISL_1585479 | Unidad de Investigación Biomédica de Zacatecas (UIBZ) | Instituto Nacional de Enfermedades Respiratorias (INER): Centro de Investigación en Enfermedades Infecciosas (CIENI) | Consorcio Mexicano de Vigilancia Genómica (CoViGen-Mex). Authors (in alphabetical order): Julio Elias Alvarado-Yaah, Carlos F. Arias, Santiago Ávila-Ríos, Víctor Hugo Borja-Aburto, Celia Boukadida, Juan Bautista Chale-Dzul, Célida Duque Molina, José Antonio Enciso-Moreno, Gloria Elena Espinosa-Ayala, Fernando Fontove-Herrera, Victor Eduardo Garcia-Arias, Concepción Grajales-Muñiz, Ricardo Grande, Alfredo Herrera-Estrella, Carla Ivón Herrera-Najera, Pavel Isa, Brenda Irasema Maldonado-Meza, Bernardo Martínez-Miguel, Margarita Matías-Florentino, María Guadalupe de Jesús Mireles-Rivera, Gloria María Molina-Salinas, Hector Montoya-Fuentes, José Esteban Muñoz-Medina, José de Jesús Nuñez-Contreras, Alicia Ocaña-Mondragón, Luis Alberto Ochoa-Carrera, Hector Esteban Paz-Juárez, Francisco Pulido, Helen Haydee Fernanda Ramírez-Plascencia, Angel Gustavo Salas-Lais, Alejandro Sanchez-Flores, Clara Esperanza Santacruz-Tinoco, María Guadalupe Santiago-Mauricio, Nelly Sélem-Mojica, Blanca Taboada, Gloria Vazquez |
| EPI_ISL_1585540, EPI_ISL_1585541, EPI_ISL_1585542, EPI_ISL_1585546, EPI_ISL_1585551, EPI_ISL_1585555, EPI_ISL_1585556, EPI_ISL_1585557, EPI_ISL_1585558, EPI_ISL_1585559, EPI_ISL_1585560, EPI_ISL_1585561, EPI_ISL_1585562, EPI_ISL_1585563 | | | |
| see above | Centro de Investigación Biomédica de Occidente (CIBO) | Instituto Nacional de Enfermedades Respiratorias (INER): Centro de Investigación en Enfermedades Infecciosas (CIENI) | Consorcio Mexicano de Vigilancia Genómica (CoViGen-Mex). Authors (in alphabetical order): Julio Elias Alvarado-Yaah, Carlos F. Arias, Santiago Ávila-Ríos, Víctor Hugo Borja-Aburto, Celia Boukadida, Juan Bautista Chale-Dzul, Célida Duque Molina, José Antonio Enciso-Moreno, Gloria Elena Espinosa-Ayala, Fernando Fontove-Herrera, Victor Eduardo Garcia-Arias, Concepción Grajales-Muñiz, Ricardo Grande, Alfredo Herrera-Estrella, Carla Ivón Herrera-Najera, Pavel Isa, Brenda Irasema Maldonado-Meza, Bernardo Martínez-Miguel, Margarita Matías-Florentino, María Guadalupe de Jesús Mireles-Rivera, Gloria María Molina-Salinas, Hector Montoya-Fuentes, José Esteban Muñoz-Medina, José de Jesús Nuñez-Contreras, Alicia Ocaña-Mondragón, Luis Alberto Ochoa-Carrera, Hector Esteban Paz-Juárez, Francisco Pulido, Helen Haydee Fernanda Ramírez-Plascencia, Angel Gustavo Salas-Lais, Alejandro Sanchez-Flores, Clara Esperanza Santacruz-Tinoco, María Guadalupe Santiago-Mauricio, Nelly Sélem-Mojica, Blanca Taboada, Gloria Vazquez |
| EPI_ISL_1585596, EPI_ISL_1585598, EPI_ISL_1585603, EPI_ISL_1585607, EPI_ISL_1585608, EPI_ISL_1585609, EPI_ISL_1585611, EPI_ISL_1585612, EPI_ISL_1585614, EPI_ISL_1585616, EPI_ISL_1585618, EPI_ISL_1585620, EPI_ISL_1585621, EPI_ISL_1585622, EPI_ISL_1585623, EPI_ISL_1585626, EPI_ISL_1585627, EPI_ISL_1585628, EPI_ISL_1585629, EPI_ISL_1585630, EPI_ISL_1585631, EPI_ISL_1585632, EPI_ISL_1585633, EPI_ISL_1585634, EPI_ISL_1585635, EPI_ISL_1585636, EPI_ISL_1585638, EPI_ISL_1585639, EPI_ISL_1585640, EPI_ISL_1585643, EPI_ISL_1585644, EPI_ISL_1585645, EPI_ISL_1585647, EPI_ISL_1585648, EPI_ISL_1585649, EPI_ISL_1585650, EPI_ISL_1585651, EPI_ISL_1585652, EPI_ISL_1585653, EPI_ISL_1585654, EPI_ISL_1585655, EPI_ISL_1585656, EPI_ISL_1585657, EPI_ISL_1585659, EPI_ISL_1585660, EPI_ISL_1585661, EPI_ISL_1585662, EPI_ISL_1585663, EPI_ISL_1585664, EPI_ISL_1585665, EPI_ISL_1585666, EPI_ISL_1585667, EPI_ISL_1585668, EPI_ISL_1585669, EPI_ISL_1585670 | | | |
| see above | Laboratorio Central de Epidemiología (LCE) | Instituto Nacional de Enfermedades Respiratorias (INER): Centro de Investigación en Enfermedades Infecciosas (CIENI) | Consorcio Mexicano de Vigilancia Genómica (CoViGen-Mex). Authors (in alphabetical order): Julio Elias Alvarado-Yaah, Carlos F. Arias, Santiago Ávila-Ríos, Víctor Hugo Borja-Aburto, Celia Boukadida, Juan Bautista Chale-Dzul, Célida Duque Molina, José Antonio Enciso-Moreno, Gloria Elena Espinosa-Ayala, Fernando Fontove-Herrera, Victor Eduardo Garcia-Arias, Concepción Grajales-Muñiz, Ricardo Grande, Alfredo Herrera-Estrella, Carla Ivón Herrera-Najera, Pavel Isa, Brenda Irasema Maldonado-Meza, Bernardo Martínez-Miguel, Margarita Matías-Florentino, María Guadalupe de Jesús Mireles-Rivera, Gloria María Molina-Salinas, Hector Montoya-Fuentes, José Esteban Muñoz-Medina, José de Jesús Nuñez-Contreras, Alicia Ocaña-Mondragón, Luis Alberto Ochoa-Carrera, Hector Esteban Paz-Juárez, Francisco Pulido, Helen Haydee Fernanda Ramírez-Plascencia, Angel Gustavo Salas-Lais, Alejandro Sanchez-Flores, Clara Esperanza Santacruz-Tinoco, María Guadalupe Santiago-Mauricio, Nelly Sélem-Mojica, Blanca Taboada, Gloria Vazquez |
| EPI_ISL_1587582 | Genome Analysis Center, Yamanashi Central Hospital | Genome Analysis Center, Yamanashi Central Hospital | Yosuke Hirotsu |
| EPI_ISL_1588160, EPI_ISL_1588165, EPI_ISL_1588166, EPI_ISL_1588185, EPI_ISL_1588186, EPI_ISL_1588189, EPI_ISL_1588192, EPI_ISL_1588196, EPI_ISL_1588198, EPI_ISL_1588204, EPI_ISL_1588210, EPI_ISL_1588218, EPI_ISL_1588222, EPI_ISL_1588227, EPI_ISL_1588230, EPI_ISL_1588231, EPI_ISL_1588242, EPI_ISL_1588243, EPI_ISL_1588247, EPI_ISL_1588265, EPI_ISL_1588267, EPI_ISL_1588269, EPI_ISL_1588278, EPI_ISL_1588281, EPI_ISL_1588282, EPI_ISL_1588284, EPI_ISL_1588285 | | | |
| see above | NB-Hôpital Georges L. Dumont | National Microbiology Laboratory (NML) | Anna Majer, Shari Tyson, Grace Seo, Philip Mabon, Elsie Grudeski, Rhiannon Huzarewich, Russell Mandes, Anneliese Landgraff, Jennifer Tanner, Natalie Knox, Morag Graham, Gary Van Domselaar, Richard Garceau, Guillaume Desnoyers, Nathalie Bastien, Yan Li, Timothy Booth, Darian Hole, Madison Chapel, Kirsten Biggar, CanCOGeN's metadata curation team, Public Health Agency of Canada CanCOGeN team |
| EPI_ISL_1588522, EPI_ISL_1588524, EPI_ISL_1588584 | National Institute of Public Health | National Institute of Public Health | Helena Jirincova, Jaromira Vecerova, Timotej Suri, Dusan Trnka, Alexander Nagy |
| EPI_ISL_1588900, EPI_ISL_1588901 | Hospital | National Reference Center for Viruses of Respiratory Infections, Institut Pasteur, Paris | Marion Barbet, Sylvie Behillil, Frédéric Lemoine, Corinne Maufrais, Christophe Malabat,Pierre Lechat, Méline Bizard, Angela Brisebarre, Camille Capel, Louise Lefrançois, Etienne Simon-Lorière, Vincent Enouf, Maud Vanpeene, Sylvie van der Werf,StéPhanie Marque Juillet |
| EPI_ISL_1588902, EPI_ISL_1588903 | Hospital | National Reference Center for Viruses of Respiratory Infections, Institut Pasteur, Paris | Marion Barbet, Sylvie Behillil, Frédéric Lemoine, Corinne Maufrais, Christophe Malabat,Pierre Lechat, Méline Bizard, Angela Brisebarre, Camille Capel, Louise Lefrançois, Etienne Simon-Lorière, Vincent Enouf, Maud Vanpeene, Sylvie van der Werf,AuréLie Guigon |
| EPI_ISL_1588998 | Labo Analyses Med | National Reference Center for Viruses of Respiratory Infections, Institut Pasteur, Paris | Marion Barbet, Sylvie Behillil, Frédéric Lemoine, Corinne Maufrais, Christophe Malabat, Méline Bizard, Angela Brisebarre, Camille Capel, Louise Lefrançois, Etienne Simon-Lorière, Vincent Enouf, Maud Vanpeene, Sylvie van der Werf,GréGoire Potiron |
| EPI_ISL_1588999, EPI_ISL_1589000, EPI_ISL_1589001, EPI_ISL_1589002 | Hospital | National Reference Center for Viruses of Respiratory Infections, Institut Pasteur, Paris | Marion Barbet, Sylvie Behillil, Frédéric Lemoine, Corinne Maufrais, Christophe Malabat, Méline Bizard, Angela Brisebarre, Camille Capel, Louise Lefrançois, Etienne Simon-Lorière, Vincent Enouf, Maud Vanpeene, Sylvie van der Werf,Nabil Gastli |
| EPI_ISL_1591253, EPI_ISL_1591254, EPI_ISL_1591255, EPI_ISL_1591256, EPI_ISL_1591258, EPI_ISL_1591259, EPI_ISL_1591261, EPI_ISL_1591262, EPI_ISL_1591263, EPI_ISL_1591264 | Department of Virology, Istituto Zooprofilattico Sperimentale del Lazio e della Toscana (IZSLT) | Department of General Diagnostics; Department of Virology; Istituto Zooprofilattico Sperimentale del Lazio e della Toscana (IZSLT) | Patricia Alba, Giuseppe Manna, Elena L. Diaconu, Fabiola Feltrin, Raffaela Conti, Teresa Scicluna, Virginia Carfora, Antonella Cersini, Alessia Franco, Antonio Battisti. |
| EPI_ISL_1593756, EPI_ISL_1593757, EPI_ISL_1593772 | OCME Office Of Chief Medical Examiner | New York City Public Health Laboratory | Jade Wang, et al. |
| EPI_ISL_1594379 | Hospital | National Reference Center for Viruses of Respiratory Infections, Institut Pasteur, Paris | Marion Barbet, Sylvie Behillil, Frédéric Lemoine, Corinne Maufrais, Christophe Malabat, Méline Bizard, Angela Brisebarre, Camille Capel, Louise Lefrançois, Etienne Simon-Lorière, Vincent Enouf, Maud Vanpeene, Sylvie van der Werf,Anne Cady |
| EPI_ISL_1594380 | Hospital | National Reference Center for Viruses of Respiratory Infections, Institut Pasteur, Paris | Marion Barbet, Sylvie Behillil, Frédéric Lemoine, Corinne Maufrais, Christophe Malabat, Méline Bizard, Angela Brisebarre, Camille Capel, Louise Lefrançois, Etienne Simon-Lorière, Vincent Enouf, Maud Vanpeene, Sylvie van der Werf,Jean-FrançOis Filippi |
| EPI_ISL_1594381, EPI_ISL_1594382, EPI_ISL_1594383, EPI_ISL_1594384, EPI_ISL_1594386, EPI_ISL_1594387, EPI_ISL_1594388, EPI_ISL_1594389, EPI_ISL_1594390, EPI_ISL_1594392, EPI_ISL_1594393 | | | |
| see above | Labo Analyses Med | National Reference Center for Viruses of Respiratory Infections, Institut Pasteur, Paris | Marion Barbet, Sylvie Behillil, Frédéric Lemoine, Corinne Maufrais, Christophe Malabat, Méline Bizard, Angela Brisebarre, Camille Capel, Louise Lefrançois, Etienne Simon-Lorière, Vincent Enouf, Maud Vanpeene, Sylvie van der Werf,Melanie Caron |
| EPI_ISL_1594394, EPI_ISL_1594395 | Labo Analyses Med | National Reference Center for Viruses of Respiratory Infections, Institut Pasteur, Paris | Marion Barbet, Sylvie Behillil, Frédéric Lemoine, Corinne Maufrais, Christophe Malabat, Amaury Vaysse, Méline Bizard, Angela Brisebarre, Camille Capel, Louise Lefrançois, Etienne Simon-Lorière, Vincent Enouf, Maud Vanpeene, Sylvie van der Werf,Melanie Caron |
| EPI_ISL_1594396, EPI_ISL_1594397, EPI_ISL_1594398, EPI_ISL_1594399, EPI_ISL_1594400, EPI_ISL_1594401, EPI_ISL_1594402, EPI_ISL_1594403 | Labo Analyses Med | National Reference Center for Viruses of Respiratory Infections, Institut Pasteur, Paris | Marion Barbet, Sylvie Behillil, Frédéric Lemoine, Corinne Maufrais, Christophe Malabat, Méline Bizard, Angela Brisebarre, Camille Capel, Louise Lefrançois, Etienne Simon-Lorière, Vincent Enouf, Maud Vanpeene, Sylvie van der Werf,Melanie Caron |
| EPI_ISL_1594404 | Labo Analyses Med | National Reference Center for Viruses of Respiratory Infections, Institut Pasteur, Paris | Marion Barbet, Sylvie Behillil, Frédéric Lemoine, Corinne Maufrais, Christophe Malabat, Amaury Vaysse, Méline Bizard, Angela Brisebarre, Camille Capel, Louise Lefrançois, Etienne Simon-Lorière, Vincent Enouf, Maud Vanpeene, Sylvie van der Werf,Melanie Caron |
| EPI_ISL_1594405, EPI_ISL_1594406, EPI_ISL_1594407, EPI_ISL_1594408, EPI_ISL_1594409, EPI_ISL_1594410 | Labo Analyses Med | National Reference Center for Viruses of Respiratory Infections, Institut Pasteur, Paris | Marion Barbet, Sylvie Behillil, Frédéric Lemoine, Corinne Maufrais, Christophe Malabat, Méline Bizard, Angela Brisebarre, Camille Capel, Louise Lefrançois, Etienne Simon-Lorière, Vincent Enouf, Maud Vanpeene, Sylvie van der Werf,Melanie Caron |

| | | | |
|---|---|---|---|
| EPI_ISL_1594411, EPI_ISL_1594412, EPI_ISL_1594413, EPI_ISL_1594414, EPI_ISL_1594415, EPI_ISL_1594416, EPI_ISL_1594417, EPI_ISL_1594418, EPI_ISL_1594419 | Hospital | National Reference Center for Viruses of Respiratory Infections, Institut Pasteur, Paris | Marion Barbet, Sylvie Behillil, Frédéric Lemoine, Corinne Maufrais, Christophe Malabat, Méline Bizard, Angela Brisebarre, Camille Capel, Louise Lefrançois, Etienne Simon-Lorière, Vincent Enouf, Maud Vanpeene, Sylvie van der Werf,CéLine Bressollette |
| EPI_ISL_1594420 | Hospital | National Reference Center for Viruses of Respiratory Infections, Institut Pasteur, Paris | Marion Barbet, Sylvie Behillil, Frédéric Lemoine, Corinne Maufrais, Christophe Malabat, Adrien Pain, Méline Bizard, Angela Brisebarre, Camille Capel, Louise Lefrançois, Etienne Simon-Lorière, Vincent Enouf, Maud Vanpeene, Sylvie van der Werf,CéLine Bressollette |
| EPI_ISL_1594422, EPI_ISL_1594423 | Hospital | National Reference Center for Viruses of Respiratory Infections, Institut Pasteur, Paris | Marion Barbet, Sylvie Behillil, Frédéric Lemoine, Corinne Maufrais, Christophe Malabat, Méline Bizard, Angela Brisebarre, Camille Capel, Louise Lefrançois, Etienne Simon-Lorière, Vincent Enouf, Maud Vanpeene, Sylvie van der Werf,CéLine Bressollette |
| EPI_ISL_1594424 | Hospital | National Reference Center for Viruses of Respiratory Infections, Institut Pasteur, Paris | Marion Barbet, Sylvie Behillil, Frédéric Lemoine, Corinne Maufrais, Christophe Malabat, Adrien Pain, Méline Bizard, Angela Brisebarre, Camille Capel, Louise Lefrançois, Etienne Simon-Lorière, Vincent Enouf, Maud Vanpeene, Sylvie van der Werf,CéLine Bressollette |
| EPI_ISL_1594425 | Hospital | National Reference Center for Viruses of Respiratory Infections, Institut Pasteur, Paris | Marion Barbet, Sylvie Behillil, Frédéric Lemoine, Corinne Maufrais, Christophe Malabat, Méline Bizard, Angela Brisebarre, Camille Capel, Louise Lefrançois, Etienne Simon-Lorière, Vincent Enouf, Maud Vanpeene, Sylvie van der Werf,CéLine Bressollette |
| EPI_ISL_1594426, EPI_ISL_1594427 | Labo Analyses Med | National Reference Center for Viruses of Respiratory Infections, Institut Pasteur, Paris | Marion Barbet, Sylvie Behillil, Frédéric Lemoine, Corinne Maufrais, Christophe Malabat, Adrien Pain, Méline Bizard, Angela Brisebarre, Camille Capel, Louise Lefrançois, Etienne Simon-Lorière, Vincent Enouf, Maud Vanpeene, Sylvie van der Werf,Florent Tomasi |
| EPI_ISL_1594428, EPI_ISL_1594429 | Hospital | National Reference Center for Viruses of Respiratory Infections, Institut Pasteur, Paris | Marion Barbet, Sylvie Behillil, Frédéric Lemoine, Corinne Maufrais, Christophe Malabat, Méline Bizard, Angela Brisebarre, Camille Capel, Louise Lefrançois, Etienne Simon-Lorière, Vincent Enouf, Maud Vanpeene, Sylvie van der Werf,Franck Bernardi |
| EPI_ISL_1594430 | Labo Analyses Med | National Reference Center for Viruses of Respiratory Infections, Institut Pasteur, Paris | Marion Barbet, Sylvie Behillil, Frédéric Lemoine, Corinne Maufrais, Christophe Malabat, Méline Bizard, Angela Brisebarre, Camille Capel, Louise Lefrançois, Etienne Simon-Lorière, Vincent Enouf, Maud Vanpeene, Sylvie van der Werf,J Besson |
| EPI_ISL_1594432, EPI_ISL_1594433 | Labo Analyses Med | National Reference Center for Viruses of Respiratory Infections, Institut Pasteur, Paris | Marion Barbet, Sylvie Behillil, Frédéric Lemoine, Corinne Maufrais, Christophe Malabat, Adrien Pain, Méline Bizard, Angela Brisebarre, Camille Capel, Louise Lefrançois, Etienne Simon-Lorière, Vincent Enouf, Maud Vanpeene, Sylvie van der Werf,Pierre-Yves Leonard |
| EPI_ISL_1594434, EPI_ISL_1594435, EPI_ISL_1594436, EPI_ISL_1594437, EPI_ISL_1594438, EPI_ISL_1594439, EPI_ISL_1594440, EPI_ISL_1594441, EPI_ISL_1594442 | Labo Analyses Med | National Reference Center for Viruses of Respiratory Infections, Institut Pasteur, Paris | Marion Barbet, Sylvie Behillil, Frédéric Lemoine, Corinne Maufrais, Christophe Malabat, Méline Bizard, Angela Brisebarre, Camille Capel, Louise Lefrançois, Etienne Simon-Lorière, Vincent Enouf, Maud Vanpeene, Sylvie van der Werf,Pierre-Yves Leonard |
| EPI_ISL_1594445 | Labo Analyses Med | National Reference Center for Viruses of Respiratory Infections, Institut Pasteur, Paris | Marion Barbet, Sylvie Behillil, Frédéric Lemoine, Corinne Maufrais, Christophe Malabat, Amaury Vaysse, Méline Bizard, Angela Brisebarre, Camille Capel, Louise Lefrançois, Etienne Simon-Lorière, Vincent Enouf, Maud Vanpeene, Sylvie van der Werf,Pierre-Yves Leonard |
| EPI_ISL_1594446, EPI_ISL_1594447, EPI_ISL_1594448, EPI_ISL_1594449 | Labo Analyses Med | National Reference Center for Viruses of Respiratory Infections, Institut Pasteur, Paris | Marion Barbet, Sylvie Behillil, Frédéric Lemoine, Corinne Maufrais, Christophe Malabat, Méline Bizard, Angela Brisebarre, Camille Capel, Louise Lefrançois, Etienne Simon-Lorière, Vincent Enouf, Maud Vanpeene, Sylvie van der Werf,Pierre-Yves Leonard |
| EPI_ISL_1594450 | Labo Analyses Med | National Reference Center for Viruses of Respiratory Infections, Institut Pasteur, Paris | Marion Barbet, Sylvie Behillil, Frédéric Lemoine, Corinne Maufrais, Christophe Malabat, Amaury Vaysse, Méline Bizard, Angela Brisebarre, Camille Capel, Louise Lefrançois, Etienne Simon-Lorière, Vincent Enouf, Maud Vanpeene, Sylvie van der Werf,Pierre-Yves Leonard |
| EPI_ISL_1594451 | Labo Analyses Med | National Reference Center for Viruses of Respiratory Infections, Institut Pasteur, Paris | Marion Barbet, Sylvie Behillil, Frédéric Lemoine, Corinne Maufrais, Christophe Malabat, Méline Bizard, Angela Brisebarre, Camille Capel, Louise Lefrançois, Etienne Simon-Lorière, Vincent Enouf, Maud Vanpeene, Sylvie van der Werf,Pierre-Yves Leonard |
| EPI_ISL_1594452 | Labo Analyses Med | National Reference Center for Viruses of Respiratory Infections, Institut Pasteur, Paris | Marion Barbet, Sylvie Behillil, Frédéric Lemoine, Corinne Maufrais, Christophe Malabat, Amaury Vaysse, Méline Bizard, Angela Brisebarre, Camille Capel, Louise Lefrançois, Etienne Simon-Lorière, Vincent Enouf, Maud Vanpeene, Sylvie van der Werf,Pierre-Yves Leonard |
| EPI_ISL_1595474 | Hospital | National Reference Center for Viruses of Respiratory Infections, Institut Pasteur, Paris | Marion Barbet, Sylvie Behillil, Frédéric Lemoine, Corinne Maufrais, Christophe Malabat, Victoire Baillet, Méline Bizard, Angela Brisebarre, Camille Capel, Louise Lefrançois, Etienne Simon-Lorière, Vincent Enouf, Maud Vanpeene, Sylvie van der Werf,Sandrine Castelain |
| EPI_ISL_1595475 | Hospital | National Reference Center for Viruses of Respiratory Infections, Institut Pasteur, Paris | Marion Barbet, Sylvie Behillil, Frédéric Lemoine, Corinne Maufrais, Christophe Malabat, Victoire Baillet, Méline Bizard, Angela Brisebarre, Camille Capel, Louise Lefrançois, Etienne Simon-Lorière, Vincent Enouf, Maud Vanpeene, Sylvie van der Werf,Pauline Morvan |
| EPI_ISL_1595476, EPI_ISL_1595477 | Hospital | National Reference Center for Viruses of Respiratory Infections, Institut Pasteur, Paris | Marion Barbet, Sylvie Behillil, Frédéric Lemoine, Corinne Maufrais, Christophe Malabat, Victoire Baillet, Méline Bizard, Angela Brisebarre, Camille Capel, Louise Lefrançois, Etienne Simon-Lorière, Vincent Enouf, Maud Vanpeene, Sylvie van der Werf,Dominique Descamps |
| EPI_ISL_1595479 | Labo Analyses Med | National Reference Center for Viruses of Respiratory Infections, Institut Pasteur, Paris | Marion Barbet, Sylvie Behillil, Frédéric Lemoine, Corinne Maufrais, Christophe Malabat, Victoire Baillet, Méline Bizard, Angela Brisebarre, Camille Capel, Louise Lefrançois, Etienne Simon-Lorière, Vincent Enouf, Maud Vanpeene, Sylvie van der Werf,Nelly Luizy |
| EPI_ISL_1595480, EPI_ISL_1595481 | Hospital | National Reference Center for Viruses of Respiratory Infections, Institut Pasteur, Paris | Marion Barbet, Sylvie Behillil, Frédéric Lemoine, Corinne Maufrais, Christophe Malabat, Victoire Baillet, Méline Bizard, Angela Brisebarre, Camille Capel, Louise Lefrançois, Etienne Simon-Lorière, Vincent Enouf, Maud Vanpeene, Sylvie van der Werf,Thibault Guinoiseau |
| EPI_ISL_1595482, EPI_ISL_1595483, EPI_ISL_1595484 | Hospital | National Reference Center for Viruses of Respiratory Infections, Institut Pasteur, Paris | Marion Barbet, Sylvie Behillil, Frédéric Lemoine, Corinne Maufrais, Christophe Malabat, Amaury Vaysse, Méline Bizard, Angela Brisebarre, Camille Capel, Louise Lefrançois, Etienne Simon-Lorière, Vincent Enouf, Maud Vanpeene, Sylvie van der Werf,Eric Farfour |
| EPI_ISL_1595485 | Hospital | National Reference Center for Viruses of Respiratory Infections, Institut Pasteur, Paris | Marion Barbet, Sylvie Behillil, Frédéric Lemoine, Corinne Maufrais, Christophe Malabat, Méline Bizard, Angela Brisebarre, Camille Capel, Louise Lefrançois, Etienne Simon-Lorière, Vincent Enouf, Maud Vanpeene, Sylvie van der Werf,Eric Farfour |
| EPI_ISL_1595486 | Hospital | National Reference Center for Viruses of Respiratory Infections, Institut Pasteur, Paris | Marion Barbet, Sylvie Behillil, Frédéric Lemoine, Corinne Maufrais, Christophe Malabat, Victoire Baillet, Méline Bizard, Angela Brisebarre, Camille Capel, Louise Lefrançois, Etienne Simon-Lorière, Vincent Enouf, Maud Vanpeene, Sylvie van der Werf,Eric Farfour |
| EPI_ISL_1595487, EPI_ISL_1595488, EPI_ISL_1595489, EPI_ISL_1595490, EPI_ISL_1595491, EPI_ISL_1595492, EPI_ISL_1595493, EPI_ISL_1595494, EPI_ISL_1595495, EPI_ISL_1595496 | Hospital | National Reference Center for Viruses of Respiratory Infections, Institut Pasteur, Paris | Marion Barbet, Sylvie Behillil, Frédéric Lemoine, Corinne Maufrais, Christophe Malabat, Méline Bizard, Angela Brisebarre, Camille Capel, Louise Lefrançois, Etienne Simon-Lorière, Vincent Enouf, Maud Vanpeene, Sylvie van der Werf,Anne Cady |
| EPI_ISL_1595497, EPI_ISL_1595498 | Hospital | National Reference Center for Viruses of Respiratory Infections, Institut Pasteur, Paris | Marion Barbet, Sylvie Behillil, Frédéric Lemoine, Corinne Maufrais, Christophe Malabat, Victoire Baillet, Méline Bizard, Angela Brisebarre, Camille Capel, Louise Lefrançois, Etienne Simon-Lorière, Vincent Enouf, Maud Vanpeene, Sylvie van der Werf,Vincent Thibault |
| EPI_ISL_1595499, EPI_ISL_1595500, EPI_ISL_1595501 | Hospital | National Reference Center for Viruses of Respiratory Infections, Institut Pasteur, Paris | Marion Barbet, Sylvie Behillil, Frédéric Lemoine, Corinne Maufrais, Christophe Malabat, Méline Bizard, Angela Brisebarre, Camille Capel, Louise Lefrançois, Etienne Simon-Lorière, Vincent Enouf, Maud Vanpeene, Sylvie van der Werf,Vincent Thibault |
| EPI_ISL_1595502 | Hospital | National Reference Center for Viruses of Respiratory Infections, Institut Pasteur, Paris | Marion Barbet, Sylvie Behillil, Frédéric Lemoine, Corinne Maufrais, Christophe Malabat, Amaury Vaysse, Méline Bizard, Angela Brisebarre, Camille Capel, Louise Lefrançois, Etienne Simon-Lorière, Vincent Enouf, Maud Vanpeene, Sylvie van der Werf,Vincent Thibault |
| EPI_ISL_1595503, EPI_ISL_1595504, EPI_ISL_1595505, EPI_ISL_1595506, EPI_ISL_1595507, EPI_ISL_1595508 | Hospital | National Reference Center for Viruses of Respiratory Infections, Institut Pasteur, Paris | Marion Barbet, Sylvie Behillil, Frédéric Lemoine, Corinne Maufrais, Christophe Malabat, Méline Bizard, Angela Brisebarre, Camille Capel, Louise Lefrançois, Etienne Simon-Lorière, Vincent Enouf, Maud Vanpeene, Sylvie van der Werf,Vincent Thibault |
| EPI_ISL_1595605 | Centro de Investigación Biomédica de Occidente (CIBO) | Instituto Nacional de Enfermedades Respiratorias (INER): Centro de Investigación en Enfermedades Infecciosas (CIENI) | Consorcio Mexicano de Vigilancia Genómica (CoViGen-Mex). Authors (in alphabetical order): Julio Elias Alvarado-Yaah, Carlos F. Arias, Santiago Ávila-Ríos, Víctor Hugo Borja-Aburto, Celia Boukadida, Juan Bautista Chale-Dzul, Célida Duque Molina, José Antonio Enciso-Moreno, Gloria Elena Espinosa-Ayala, Fernando Fontove-Herrera, Victor Eduardo Garcia-Arias, Concepción Grajales-Muñiz, Ricardo Grande, Alfredo Herrera-Estrella, Carla Ivón Herrera-Najera, Pavel Isa, Brenda Irasema Maldonado-Meza, Bernardo Martínez-Miguel, Margarita Matías-Florentino, María Guadalupe de Jesús Mireles-Rivera, Gloria María Molina-Salinas, Hector Montoya-Fuentes, José Esteban Muñoz-Medina, José de Jesús Nuñez-Contreras, Alicia |

| | | | |
|---|---|---|---|
| | | | Ocaña-Mondragón, Luis Alberto Ochoa-Carrera, Hector Esteban Paz-Juárez, Francisco Pulido, Helen Haydee Fernanda Ramírez-Plascencia, Angel Gustavo Salas-Lais, Alejandro Sanchez-Flores, Clara Esperanza Santacruz-Tinoco, María Guadalupe Santiago-Mauricio, Nelly Sélem-Mojica, Blanca Taboada, Gloria Vazquez |
| EPI_ISL_1595722, EPI_ISL_1595723, EPI_ISL_1595724, EPI_ISL_1595725, EPI_ISL_1595726, EPI_ISL_1595727, EPI_ISL_1595728, EPI_ISL_1595729, EPI_ISL_1595730, EPI_ISL_1595732, EPI_ISL_1595733 | | | |
| see above | Microbiology Department. Complexo Hospitalario Universitario de Vigo | Microbiology Department. Complexo Hospitalario Universitario de Vigo | Alfaya N, Alonso I, Alvarez M, Cabrera JJ, Carballo R, Cores O, Cortizo S, del-Campo V, Martinez L, Mediero G, Perez S, Potel C, Regueiro B, Rey S, Vasallo FJ |
| EPI_ISL_1595844 | Compass Laboratory Services, LLC | Compass Laboratory Services | William Budd; PhD |
| EPI_ISL_1595847 | Genome Analysis Center, Yamanashi Central Hospital | Genome Analysis Center, Yamanashi Central Hospital | Yosuke Hirotsu |
| EPI_ISL_1599171, EPI_ISL_1599172, EPI_ISL_1599173, EPI_ISL_1599174, EPI_ISL_1599175, EPI_ISL_1599176, EPI_ISL_1599177, EPI_ISL_1599178 | Cliniques universitaires Saint-Luc | UCLouvain/IREC/MBLG | Jean Ruelle, Lysa Pinsmaye, Benoit Kabamba Mukadi |
| EPI_ISL_1599310 | Public Health Authority of the Slovak Republic | Laboratory of Genomics and Bioinformatics, Comenius University Science Park | Tatiana Sedláková, Diana Rusáková, Miroslav Böhmer, Anna Giová, Jaroslav Budiš, Tomáš Szemes |
| EPI_ISL_1599509, EPI_ISL_1599511, EPI_ISL_1599513, EPI_ISL_1599515, EPI_ISL_1599517, EPI_ISL_1599519, EPI_ISL_1599520, EPI_ISL_1599522, EPI_ISL_1599524, EPI_ISL_1599525, EPI_ISL_1599526, EPI_ISL_1599527 | | | |
| see above | AZ Klina | AZ Klina | Carl Vael - Lynsey Berckmans |
| EPI_ISL_1599528, EPI_ISL_1599529, EPI_ISL_1599530 | Klinisch Laboratorium ZNA | Klinisch Laboratorium ZNA | Verstrepen et al. |
| EPI_ISL_1599531, EPI_ISL_1599532, EPI_ISL_1599533, EPI_ISL_1599534, EPI_ISL_1599536, EPI_ISL_1599537 | Klinisch Laboratorium GZA | Klinisch Laboratorium ZNA | Verstrepen et al. |
| EPI_ISL_1599538, EPI_ISL_1599539, EPI_ISL_1599540 | Klinisch Laboratorium ZNA | Klinisch Laboratorium ZNA | Verstrepen et al. |
| EPI_ISL_1599545, EPI_ISL_1599546, EPI_ISL_1599547, EPI_ISL_1599548 | AZ Klina | AZ Klina | Carl Vael - Lynsey Berckmans |
| EPI_ISL_1599550 | AZ Klina | AZ KLina | Carl Vael - Lynsey Berckmans |
| EPI_ISL_1599551, EPI_ISL_1599856, EPI_ISL_1600921 | AZ Klina | AZ Klina | Carl Vael - Lynsey Berckmans |
| EPI_ISL_1607869 | Department of Virology, Istituto Zooprofilattico Sperimentale del Lazio e della Toscana (IZSLT) | Department of General Diagnostics; Department of Virology; Istituto Zooprofilattico Sperimentale del Lazio e della Toscana (IZSLT) | Patricia Alba, Giuseppe Manna, Elena L. Diaconu, Fabiola Feltrin, Raffaela Conti, Teresa Scicluna, Virginia Carfora, Antonella Cersini, Alessia Franco, Antonio Battisti. |
| EPI_ISL_1607968 | SIESP CHIETI - DRIVE IN LANCIANO LANCIANO(CHIETI) | Istituto Zooprofilattico Sperimentale dell'Abruzzo e Molise "G. Caporale" | Lorusso A, Marcacci M, Di Domenico M, Ancora M, Curini V, Mangone I, Rinaldi A, Scialabba S, Di Pasquale A, Cammà C, Puglia I, Calistri P, Savini G |
| EPI_ISL_1607969 | DIP. PREV. AVEZZANO SERVIZIO DI IGIENE EPIDEMIOLOGIA E SANITA' PUBBLICA AVEZZANO(L'AQUILA) | Istituto Zooprofilattico Sperimentale dell'Abruzzo e Molise "G. Caporale" | Lorusso A, Marcacci M, Di Domenico M, Ancora M, Curini V, Mangone I, Rinaldi A, Scialabba S, Di Pasquale A, Cammà C, Puglia I, Calistri P, Savini G |
| EPI_ISL_1607970, EPI_ISL_1607971 | SIESP CHIETI - DRIVE IN LANCIANO LANCIANO(CHIETI) | Istituto Zooprofilattico Sperimentale dell'Abruzzo e Molise "G. Caporale" | Lorusso A, Marcacci M, Di Domenico M, Ancora M, Curini V, Mangone I, Rinaldi A, Scialabba S, Di Pasquale A, Cammà C, Puglia I, Calistri P, Savini G |
| EPI_ISL_1607972 | SIESP CHIETI - DRIVE IN ATESSA ATESSA(CHIETI) | Istituto Zooprofilattico Sperimentale dell'Abruzzo e Molise "G. Caporale" | Lorusso A, Marcacci M, Di Domenico M, Ancora M, Curini V, Mangone I, Rinaldi A, Scialabba S, Di Pasquale A, Cammà C, Puglia I, Calistri P, Savini G |
| EPI_ISL_1607973 | SIESP DIPARTIMENTO DI PREVENZIONE CHIETI CHIETI(CHIETI) | Istituto Zooprofilattico Sperimentale dell'Abruzzo e Molise "G. Caporale" | Lorusso A, Marcacci M, Di Domenico M, Ancora M, Curini V, Mangone I, Rinaldi A, Scialabba S, Di Pasquale A, Cammà C, Puglia I, Calistri P, Savini G |
| EPI_ISL_1607974 | SIESP CHIETI - DRIVE IN ORTONA ORTONA(CHIETI) | Istituto Zooprofilattico Sperimentale dell'Abruzzo e Molise "G. Caporale" | Lorusso A, Marcacci M, Di Domenico M, Ancora M, Curini V, Mangone I, Rinaldi A, Scialabba S, Di Pasquale A, Cammà C, Puglia I, Calistri P, Savini G |
| EPI_ISL_1607975, EPI_ISL_1607976 | SIESP CHIETI - DRIVE IN CHIETI CHIETI(CHIETI) | Istituto Zooprofilattico Sperimentale dell'Abruzzo e Molise "G. Caporale" | Lorusso A, Marcacci M, Di Domenico M, Ancora M, Curini V, Mangone I, Rinaldi A, Scialabba S, Di Pasquale A, Cammà C, Puglia I, Calistri P, Savini G |
| EPI_ISL_1607977 | SIESP CHIETI - DRIVE IN ORTONA ORTONA(CHIETI) | Istituto Zooprofilattico Sperimentale dell'Abruzzo e Molise "G. Caporale" | Lorusso A, Marcacci M, Di Domenico M, Ancora M, Curini V, Mangone I, Rinaldi A, Scialabba S, Di Pasquale A, Cammà C, Puglia I, Calistri P, Savini G |
| EPI_ISL_1607978 | PRESIDIO TERRITORIALE CASOLI CASOLI(CHIETI) | Istituto Zooprofilattico Sperimentale dell'Abruzzo e Molise "G. Caporale" | Lorusso A, Marcacci M, Di Domenico M, Ancora M, Curini V, Mangone I, Rinaldi A, Scialabba S, Di Pasquale A, Cammà C, Puglia I, Calistri P, Savini G |
| EPI_ISL_1607979 | USCA Pescina PESCINA(L'AQUILA) | Istituto Zooprofilattico Sperimentale dell'Abruzzo e Molise "G. Caporale" | Lorusso A, Marcacci M, Di Domenico M, Ancora M, Curini V, Mangone I, Rinaldi A, Scialabba S, Di Pasquale A, Cammà C, Puglia I, Calistri P, Savini G |
| EPI_ISL_1607980 | SIESP CHIETI - DRIVE IN CHIETI CHIETI(CHIETI) | Istituto Zooprofilattico Sperimentale dell'Abruzzo e Molise "G. Caporale" | Lorusso A, Marcacci M, Di Domenico M, Ancora M, Curini V, Mangone I, Rinaldi A, Scialabba S, Di Pasquale A, Cammà C, Puglia I, Calistri P, Savini G |
| EPI_ISL_1607981 | SIESP CHIETI - DRIVE IN GISSI GISSI(CHIETI) | Istituto Zooprofilattico Sperimentale dell'Abruzzo e Molise "G. Caporale" | Lorusso A, Marcacci M, Di Domenico M, Ancora M, Curini V, Mangone I, Rinaldi A, Scialabba S, Di Pasquale A, Cammà C, Puglia I, Calistri P, Savini G |
| EPI_ISL_1608568 | SIESP DIPARTIMENTO DI PREVENZIONE TERAMO TERAMO(TERAMO) | Istituto Zooprofilattico Sperimentale dell'Abruzzo e Molise "G. Caporale" | Lorusso A, Marcacci M, Di Domenico M, Ancora M, Curini V, Di Lollo Valeria, Mangone I, Rinaldi A, Delli Compagni E, Scialabba S, Caporale M, Di Pasquale A, Cammà C, Puglia I, Calistri P, Savini G |
| EPI_ISL_1608569 | SIESP CHIETI - DRIVE IN LANCIANO LANCIANO(CHIETI) | Istituto Zooprofilattico Sperimentale dell'Abruzzo e Molise "G. Caporale" | Lorusso A, Marcacci M, Di Domenico M, Ancora M, Curini V, Di Lollo Valeria, Mangone I, Rinaldi A, Delli Compagni E, Scialabba S, Caporale M, Di Pasquale A, Cammà C, Puglia I, Calistri P, Savini G |
| EPI_ISL_1608570 | SIESP DIPARTIMENTO DI PREVENZIONE CHIETI CHIETI(CHIETI) | Istituto Zooprofilattico Sperimentale dell'Abruzzo e Molise "G. Caporale" | Lorusso A, Marcacci M, Di Domenico M, Ancora M, Curini V, Di Lollo Valeria, Mangone I, Rinaldi A, Delli Compagni E, Scialabba S, Caporale M, Di Pasquale A, Cammà C, Puglia I, Calistri P, Savini G |
| EPI_ISL_1608571 | SIESP DIPARTIMENTO DI PREVENZIONE TERAMO TERAMO(TERAMO) | Istituto Zooprofilattico Sperimentale dell'Abruzzo e Molise "G. Caporale" | Lorusso A, Marcacci M, Di Domenico M, Ancora M, Curini V, Di Lollo Valeria, Mangone I, Rinaldi A, Delli Compagni E, Scialabba S, Caporale M, Di Pasquale A, Cammà C, Puglia I, Calistri P, Savini G |
| EPI_ISL_1608572 | SIESP DIPARTIMENTO DI PREVENZIONE CHIETI CHIETI(CHIETI) | Istituto Zooprofilattico Sperimentale dell'Abruzzo e Molise "G. Caporale" | Lorusso A, Marcacci M, Di Domenico M, Ancora M, Curini V, Di Lollo Valeria, Mangone I, Rinaldi A, Delli Compagni E, Scialabba S, Caporale M, Di Pasquale A, Cammà C, Puglia I, Calistri P, Savini G |
| EPI_ISL_1608573 | USCA Tagliacozzo TAGLIACOZZO(L'AQUILA) | Istituto Zooprofilattico Sperimentale dell'Abruzzo e Molise "G. Caporale" | Lorusso A, Marcacci M, Di Domenico M, Ancora M, Curini V, Di Lollo Valeria, Mangone I, Rinaldi A, Delli Compagni E, Scialabba S, Caporale M, Di Pasquale A, Cammà C, Puglia I, Calistri P, Savini G |
| EPI_ISL_1608574 | PRESIDIO TERRITORIALE CASOLI CASOLI(CHIETI) | Istituto Zooprofilattico Sperimentale dell'Abruzzo e Molise "G. Caporale" | Lorusso A, Marcacci M, Di Domenico M, Ancora M, Curini V, Di Lollo Valeria, Mangone I, Rinaldi A, Delli Compagni E, Scialabba S, Caporale M, Di Pasquale A, Cammà C, Puglia I, Calistri P, Savini G |

| | | | |
|---|---|---|---|
| EPI_ISL_1608575 | SIESP DIPARTIMENTO DI PREVENZIONE TERAMO TERAMO(TERAMO) | Istituto Zooprofilattico Sperimentale dell'Abruzzo e Molise "G. Caporale" | Lorusso A, Marcacci M, Di Domenico M, Ancora M, Curini V, Di Lollo Valeria, Mangone I, Rinaldi A, Delli Compagni E, Scialabba S, Caporale M, Di Pasquale A, Cammà C, Puglia I, Calistri P, Savini G |
| EPI_ISL_1608576 | USCA Pescina PESCINA(L'AQUILA) | Istituto Zooprofilattico Sperimentale dell'Abruzzo e Molise "G. Caporale" | Lorusso A, Marcacci M, Di Domenico M, Ancora M, Curini V, Di Lollo Valeria, Mangone I, Rinaldi A, Delli Compagni E, Scialabba S, Caporale M, Di Pasquale A, Cammà C, Puglia I, Calistri P, Savini G |
| EPI_ISL_1608577 | USCA Tagliacozzo TAGLIACOZZO(L'AQUILA) | Istituto Zooprofilattico Sperimentale dell'Abruzzo e Molise "G. Caporale" | Lorusso A, Marcacci M, Di Domenico M, Ancora M, Curini V, Di Lollo Valeria, Mangone I, Rinaldi A, Delli Compagni E, Scialabba S, Caporale M, Di Pasquale A, Cammà C, Puglia I, Calistri P, Savini G |
| EPI_ISL_1608578 | SIESP SULMONA SULMONA(L'AQUILA) | Istituto Zooprofilattico Sperimentale dell'Abruzzo e Molise "G. Caporale" | Lorusso A, Marcacci M, Di Domenico M, Ancora M, Curini V, Di Lollo Valeria, Mangone I, Rinaldi A, Delli Compagni E, Scialabba S, Caporale M, Di Pasquale A, Cammà C, Puglia I, Calistri P, Savini G |
| EPI_ISL_1608579 | DIP. PREV. AVEZZANO SERVIZIO DI IGIENE EPIDEMIOLOGIA E SANITA' PUBBLICA AVEZZANO(L'AQUILA) | Istituto Zooprofilattico Sperimentale dell'Abruzzo e Molise "G. Caporale" | Lorusso A, Marcacci M, Di Domenico M, Ancora M, Curini V, Di Lollo Valeria, Mangone I, Rinaldi A, Delli Compagni E, Scialabba S, Caporale M, Di Pasquale A, Cammà C, Puglia I, Calistri P, Savini G |
| EPI_ISL_1608580 | SIESP DIPARTIMENTO DI PREVENZIONE TERAMO TERAMO(TERAMO) | Istituto Zooprofilattico Sperimentale dell'Abruzzo e Molise "G. Caporale" | Lorusso A, Marcacci M, Di Domenico M, Ancora M, Curini V, Di Lollo Valeria, Mangone I, Rinaldi A, Delli Compagni E, Scialabba S, Caporale M, Di Pasquale A, Cammà C, Puglia I, Calistri P, Savini G |
| EPI_ISL_1608581 | SIESP CHIETI - DRIVE IN LANCIANO LANCIANO(CHIETI) | Istituto Zooprofilattico Sperimentale dell'Abruzzo e Molise "G. Caporale" | Lorusso A, Marcacci M, Di Domenico M, Ancora M, Curini V, Di Lollo Valeria, Mangone I, Rinaldi A, Delli Compagni E, Scialabba S, Caporale M, Di Pasquale A, Cammà C, Puglia I, Calistri P, Savini G |
| EPI_ISL_1615665 | National Platform bis UMONS/Jolimont | National Platform bis UMONS/Jolimont | François Dufrasne, Guillaume Bayon-Vicente, Gautier Detry, Ruddy Wattiez |
| EPI_ISL_1619453 | University of Bari Biomedical Sciences and Human Oncology | University of Bari Biomedical Sciences and Human Oncology | Chironna Maria., Sallustio A., Loconsole Daniela., Accogli M. |
| EPI_ISL_1620229, EPI_ISL_1620232, EPI_ISL_1620234, EPI_ISL_1620235 | OLVZ Aalst | OLVZ Aalst | Astrid Holderbeke |
| EPI_ISL_1620238 | University of Bari Biomedical Sciences and Human Oncology | University of Bari Biomedical Sciences and Human Oncology | Chironna M., Sallustio A., loconsole D., Accogli M. |
| EPI_ISL_1620554, EPI_ISL_1620574, EPI_ISL_1620575, EPI_ISL_1620576, EPI_ISL_1620577, EPI_ISL_1620578, EPI_ISL_1620579, EPI_ISL_1620593, EPI_ISL_1620607 | Puerto Rico Department of Health | Centers for Disease Control and Prevention, Dengue Branch | Gilberto A. Santiago, Glenda Gonzalez, Betzabel Flores, Keyla Charriez, Gabriela Paz-Bailey, Jorge L. Munoz-Jordan |
| EPI_ISL_1621624, EPI_ISL_1621626 | Labo analyses Med | National Reference Center for Viruses of Respiratory Infections, Institut Pasteur, Paris | Marion Barbet, Sylvie Behillil, Frédéric Lemoine, Corinne Maufrais, Christophe Malabat, Méline Bizard, Angela Brisebarre, Camille Capel, Louise Lefrançois, Etienne Simon-Lorière, Vincent Enouf, Maud Vanpeene, Sylvie van der Werf,Anne Holstein |
| EPI_ISL_1621627 | Labo analyses Med | National Reference Center for Viruses of Respiratory Infections, Institut Pasteur, Paris | Marion Barbet, Sylvie Behillil, Frédéric Lemoine, Corinne Maufrais, Christophe Malabat, Damien Mornico, Méline Bizard, Angela Brisebarre, Camille Capel, Louise Lefrançois, Etienne Simon-Lorière, Vincent Enouf, Maud Vanpeene, Sylvie van der Werf,Anne Holstein |
| EPI_ISL_1621628, EPI_ISL_1621629, EPI_ISL_1621630 | Labo analyses Med | National Reference Center for Viruses of Respiratory Infections, Institut Pasteur, Paris | Marion Barbet, Sylvie Behillil, Frédéric Lemoine, Corinne Maufrais, Christophe Malabat, Méline Bizard, Angela Brisebarre, Camille Capel, Louise Lefrançois, Etienne Simon-Lorière, Vincent Enouf, Maud Vanpeene, Sylvie van der Werf,Anne Holstein |
| EPI_ISL_1621631 | Labo analyses Med | National Reference Center for Viruses of Respiratory Infections, Institut Pasteur, Paris | Marion Barbet, Sylvie Behillil, Frédéric Lemoine, Corinne Maufrais, Christophe Malabat, Damien Mornico, Méline Bizard, Angela Brisebarre, Camille Capel, Louise Lefrançois, Etienne Simon-Lorière, Vincent Enouf, Maud Vanpeene, Sylvie van der Werf,Anne Holstein |
| EPI_ISL_1621632, EPI_ISL_1621633, EPI_ISL_1621634, EPI_ISL_1621635, EPI_ISL_1621636, EPI_ISL_1621637 | Labo analyses Med | National Reference Center for Viruses of Respiratory Infections, Institut Pasteur, Paris | Marion Barbet, Sylvie Behillil, Frédéric Lemoine, Corinne Maufrais, Christophe Malabat, Méline Bizard, Angela Brisebarre, Camille Capel, Louise Lefrançois, Etienne Simon-Lorière, Vincent Enouf, Maud Vanpeene, Sylvie van der Werf,Anne Holstein |
| EPI_ISL_1621638 | Labo analyses Med | National Reference Center for Viruses of Respiratory Infections, Institut Pasteur, Paris | Marion Barbet, Sylvie Behillil, Frédéric Lemoine, Corinne Maufrais, Christophe Malabat, Damien Mornico, Méline Bizard, Angela Brisebarre, Camille Capel, Louise Lefrançois, Etienne Simon-Lorière, Vincent Enouf, Maud Vanpeene, Sylvie van der Werf,Anne Holstein |
| EPI_ISL_1621639, EPI_ISL_1621640, EPI_ISL_1621641, EPI_ISL_1621642, EPI_ISL_1621643, EPI_ISL_1621644, EPI_ISL_1621645, EPI_ISL_1621646, EPI_ISL_1621647, EPI_ISL_1621648, EPI_ISL_1621649, EPI_ISL_1621650, EPI_ISL_1621651, EPI_ISL_1621652, EPI_ISL_1621653, EPI_ISL_1621654 | | | |
| see above | Labo analyses Med | National Reference Center for Viruses of Respiratory Infections, Institut Pasteur, Paris | Marion Barbet, Sylvie Behillil, Frédéric Lemoine, Corinne Maufrais, Christophe Malabat, Méline Bizard, Angela Brisebarre, Camille Capel, Louise Lefrançois, Etienne Simon-Lorière, Vincent Enouf, Maud Vanpeene, Sylvie van der Werf,Anne Holstein |
| EPI_ISL_1621656 | Labo Analyses Med | National Reference Center for Viruses of Respiratory Infections, Institut Pasteur, Paris | Marion Barbet, Sylvie Behillil, Frédéric Lemoine, Corinne Maufrais, Christophe Malabat, Adrien Pain, Méline Bizard, Angela Brisebarre, Camille Capel, Louise Lefrançois, Etienne Simon-Lorière, Vincent Enouf, Maud Vanpeene, Sylvie van der Werf,Pierre-Yves Leonard |
| EPI_ISL_1621657, EPI_ISL_1621658 | Labo Analyses Med | National Reference Center for Viruses of Respiratory Infections, Institut Pasteur, Paris | Marion Barbet, Sylvie Behillil, Frédéric Lemoine, Corinne Maufrais, Christophe Malabat, Méline Bizard, Angela Brisebarre, Camille Capel, Louise Lefrançois, Etienne Simon-Lorière, Vincent Enouf, Maud Vanpeene, Sylvie van der Werf,Pierre-Yves Leonard |
| EPI_ISL_1621660, EPI_ISL_1621661, EPI_ISL_1621662 | Labo Analyses Med | National Reference Center for Viruses of Respiratory Infections, Institut Pasteur, Paris | Marion Barbet, Sylvie Behillil, Frédéric Lemoine, Corinne Maufrais, Christophe Malabat, Adrien Pain, Méline Bizard, Angela Brisebarre, Camille Capel, Louise Lefrançois, Etienne Simon-Lorière, Vincent Enouf, Maud Vanpeene, Sylvie van der Werf,Pierre-Yves Leonard |
| EPI_ISL_1621663 | Labo Analyses Med | National Reference Center for Viruses of Respiratory Infections, Institut Pasteur, Paris | Marion Barbet, Sylvie Behillil, Frédéric Lemoine, Corinne Maufrais, Christophe Malabat, Méline Bizard, Angela Brisebarre, Camille Capel, Louise Lefrançois, Etienne Simon-Lorière, Vincent Enouf, Maud Vanpeene, Sylvie van der Werf,Pierre-Yves Leonard |
| EPI_ISL_1621664 | Labo Analyses Med | National Reference Center for Viruses of Respiratory Infections, Institut Pasteur, Paris | Marion Barbet, Sylvie Behillil, Frédéric Lemoine, Corinne Maufrais, Christophe Malabat, Adrien Pain, Méline Bizard, Angela Brisebarre, Camille Capel, Louise Lefrançois, Etienne Simon-Lorière, Vincent Enouf, Maud Vanpeene, Sylvie van der Werf,Pierre-Yves Leonard |
| EPI_ISL_1621666, EPI_ISL_1621667 | Labo Analyses Med | National Reference Center for Viruses of Respiratory Infections, Institut Pasteur, Paris | Marion Barbet, Sylvie Behillil, Frédéric Lemoine, Corinne Maufrais, Christophe Malabat, Méline Bizard, Angela Brisebarre, Camille Capel, Louise Lefrançois, Etienne Simon-Lorière, Vincent Enouf, Maud Vanpeene, Sylvie van der Werf,Pierre-Yves Leonard |
| EPI_ISL_1621668 | Labo Analyses Med | National Reference Center for Viruses of Respiratory Infections, Institut Pasteur, Paris | Marion Barbet, Sylvie Behillil, Frédéric Lemoine, Corinne Maufrais, Christophe Malabat, Adrien Pain, Méline Bizard, Angela Brisebarre, Camille Capel, Louise Lefrançois, Etienne Simon-Lorière, Vincent Enouf, Maud Vanpeene, Sylvie van der Werf,Pierre-Yves Leonard |
| EPI_ISL_1621669, EPI_ISL_1621670, EPI_ISL_1621671 | Labo Analyses Med | National Reference Center for Viruses of Respiratory Infections, Institut Pasteur, Paris | Marion Barbet, Sylvie Behillil, Frédéric Lemoine, Corinne Maufrais, Christophe Malabat, Méline Bizard, Angela Brisebarre, Camille Capel, Louise Lefrançois, Etienne Simon-Lorière, Vincent Enouf, Maud Vanpeene, Sylvie van der Werf,Pierre-Yves Leonard |
| EPI_ISL_1621672 | Labo Analyses Med | National Reference Center for Viruses of Respiratory Infections, Institut Pasteur, Paris | Marion Barbet, Sylvie Behillil, Frédéric Lemoine, Corinne Maufrais, Christophe Malabat, Adrien Pain, Méline Bizard, Angela Brisebarre, Camille Capel, Louise Lefrançois, Etienne Simon-Lorière, Vincent Enouf, Maud Vanpeene, Sylvie van der Werf,Pierre-Yves Leonard |
| EPI_ISL_1621673, EPI_ISL_1621675 | Labo Analyses Med | National Reference Center for Viruses of Respiratory Infections, Institut Pasteur, Paris | Marion Barbet, Sylvie Behillil, Frédéric Lemoine, Corinne Maufrais, Christophe Malabat, Méline Bizard, Angela Brisebarre, Camille Capel, Louise Lefrançois, Etienne Simon-Lorière, Vincent Enouf, Maud Vanpeene, Sylvie van der Werf,Pierre-Yves Leonard |
| EPI_ISL_1621676, EPI_ISL_1621677 | Labo Analyses Med | National Reference Center for Viruses of Respiratory Infections, Institut Pasteur, Paris | Marion Barbet, Sylvie Behillil, Frédéric Lemoine, Corinne Maufrais, Christophe Malabat, Adrien Pain, Méline Bizard, Angela Brisebarre, Camille Capel, Louise Lefrançois, Etienne Simon-Lorière, Vincent Enouf, Maud Vanpeene, Sylvie van der Werf,Pierre-Yves Leonard |
| EPI_ISL_1621678 | Labo Analyses Med | National Reference Center for Viruses of Respiratory Infections, Institut Pasteur, Paris | Marion Barbet, Sylvie Behillil, Frédéric Lemoine, Corinne Maufrais, Christophe Malabat, Méline Bizard, Angela Brisebarre, Camille Capel, Louise Lefrançois, Etienne Simon-Lorière, Vincent Enouf, Maud Vanpeene, Sylvie van der Werf,Pierre-Yves Leonard |
| EPI_ISL_1621679, EPI_ISL_1621680, EPI_ISL_1621681, EPI_ISL_1621682 | Labo Analyses Med | National Reference Center for Viruses of Respiratory Infections, Institut Pasteur, Paris | Marion Barbet, Sylvie Behillil, Frédéric Lemoine, Corinne Maufrais, Christophe Malabat, Adrien Pain, Méline Bizard, Angela Brisebarre, Camille Capel, Louise Lefrançois, Etienne Simon-Lorière, Vincent Enouf, Maud Vanpeene, Sylvie van der Werf,Pierre-Yves Leonard |
| EPI_ISL_1621683, EPI_ISL_1621684 | Labo Analyses Med | National Reference Center for Viruses of Respiratory Infections, Institut Pasteur, Paris | Marion Barbet, Sylvie Behillil, Frédéric Lemoine, Corinne Maufrais, Christophe Malabat, Méline Bizard, Angela Brisebarre, Camille Capel, Louise Lefrançois, Etienne Simon-Lorière, Vincent Enouf, Maud Vanpeene, Sylvie van der Werf,Pierre-Yves Leonard |
| EPI_ISL_1621685 | Labo Analyses Med | National Reference Center for Viruses of Respiratory Infections, Institut Pasteur, Paris | Marion Barbet, Sylvie Behillil, Frédéric Lemoine, Corinne Maufrais, Christophe Malabat, Adrien Pain, Méline Bizard, Angela Brisebarre, Camille Capel, Louise Lefrançois, Etienne Simon-Lorière, Vincent Enouf, Maud Vanpeene, Sylvie van der Werf,Pierre-Yves Leonard |

| | | | |
|---|---|---|---|
| EPI_ISL_1621686 | Labo Analyses Med | National Reference Center for Viruses of Respiratory Infections, Institut Pasteur, Paris | Marion Barbet, Sylvie Behillil, Frédéric Lemoine, Corinne Maufrais, Christophe Malabat, Méline Bizard, Angela Brisebarre, Camille Capel, Louise Lefrançois, Etienne Simon-Lorière, Vincent Enouf, Maud Vanpeene, Sylvie van der Werf,Pierre-Yves Leonard |
| EPI_ISL_1621687 | Labo Analyses Med | National Reference Center for Viruses of Respiratory Infections, Institut Pasteur, Paris | Marion Barbet, Sylvie Behillil, Frédéric Lemoine, Corinne Maufrais, Christophe Malabat, Adrien Pain, Méline Bizard, Angela Brisebarre, Camille Capel, Louise Lefrançois, Etienne Simon-Lorière, Vincent Enouf, Maud Vanpeene, Sylvie van der Werf,Pierre-Yves Leonard |
| EPI_ISL_1621688, EPI_ISL_1621689 | Labo Analyses Med | National Reference Center for Viruses of Respiratory Infections, Institut Pasteur, Paris | Marion Barbet, Sylvie Behillil, Frédéric Lemoine, Corinne Maufrais, Christophe Malabat, Méline Bizard, Angela Brisebarre, Camille Capel, Louise Lefrançois, Etienne Simon-Lorière, Vincent Enouf, Maud Vanpeene, Sylvie van der Werf,Pierre-Yves Leonard |
| EPI_ISL_1621690, EPI_ISL_1621691 | Labo Analyses Med | National Reference Center for Viruses of Respiratory Infections, Institut Pasteur, Paris | Marion Barbet, Sylvie Behillil, Frédéric Lemoine, Corinne Maufrais, Christophe Malabat, Adrien Pain, Méline Bizard, Angela Brisebarre, Camille Capel, Louise Lefrançois, Etienne Simon-Lorière, Vincent Enouf, Maud Vanpeene, Sylvie van der Werf,Pierre-Yves Leonard |
| EPI_ISL_1621692, EPI_ISL_1621693, EPI_ISL_1621694 | Labo Analyses Med | National Reference Center for Viruses of Respiratory Infections, Institut Pasteur, Paris | Marion Barbet, Sylvie Behillil, Frédéric Lemoine, Corinne Maufrais, Christophe Malabat, Méline Bizard, Angela Brisebarre, Camille Capel, Louise Lefrançois, Etienne Simon-Lorière, Vincent Enouf, Maud Vanpeene, Sylvie van der Werf,Pierre-Yves Leonard |
| EPI_ISL_1621695, EPI_ISL_1621696 | Labo Analyses Med | National Reference Center for Viruses of Respiratory Infections, Institut Pasteur, Paris | Marion Barbet, Sylvie Behillil, Frédéric Lemoine, Corinne Maufrais, Christophe Malabat, Adrien Pain, Méline Bizard, Angela Brisebarre, Camille Capel, Louise Lefrançois, Etienne Simon-Lorière, Vincent Enouf, Maud Vanpeene, Sylvie van der Werf,Pierre-Yves Leonard |
| EPI_ISL_1621697, EPI_ISL_1621698 | Labo Analyses Med | National Reference Center for Viruses of Respiratory Infections, Institut Pasteur, Paris | Marion Barbet, Sylvie Behillil, Frédéric Lemoine, Corinne Maufrais, Christophe Malabat, Méline Bizard, Angela Brisebarre, Camille Capel, Louise Lefrançois, Etienne Simon-Lorière, Vincent Enouf, Maud Vanpeene, Sylvie van der Werf,Pierre-Yves Leonard |
| EPI_ISL_1621699, EPI_ISL_1621700 | Hospital | National Reference Center for Viruses of Respiratory Infections, Institut Pasteur, Paris | Marion Barbet, Sylvie Behillil, Frédéric Lemoine, Corinne Maufrais, Christophe Malabat, Méline Bizard, Angela Brisebarre, Camille Capel, Louise Lefrançois, Etienne Simon-Lorière, Vincent Enouf, Maud Vanpeene, Sylvie van der Werf,Nabil Gastli |
| EPI_ISL_1621701 | Hospital | National Reference Center for Viruses of Respiratory Infections, Institut Pasteur, Paris | Marion Barbet, Sylvie Behillil, Frédéric Lemoine, Corinne Maufrais, Christophe Malabat, Damien Mornico, Méline Bizard, Angela Brisebarre, Camille Capel, Louise Lefrançois, Etienne Simon-Lorière, Vincent Enouf, Maud Vanpeene, Sylvie van der Werf,Nabil Gastli |
| EPI_ISL_1621702 | Hospital | National Reference Center for Viruses of Respiratory Infections, Institut Pasteur, Paris | Marion Barbet, Sylvie Behillil, Frédéric Lemoine, Corinne Maufrais, Christophe Malabat, Méline Bizard, Angela Brisebarre, Camille Capel, Louise Lefrançois, Etienne Simon-Lorière, Vincent Enouf, Maud Vanpeene, Sylvie van der Werf,Nabil Gastli |
| EPI_ISL_1621703 | Hospital | National Reference Center for Viruses of Respiratory Infections, Institut Pasteur, Paris | Marion Barbet, Sylvie Behillil, Frédéric Lemoine, Corinne Maufrais, Christophe Malabat, Damien Mornico, Méline Bizard, Angela Brisebarre, Camille Capel, Louise Lefrançois, Etienne Simon-Lorière, Vincent Enouf, Maud Vanpeene, Sylvie van der Werf,Nabil Gastli |
| EPI_ISL_1621704, EPI_ISL_1621705, EPI_ISL_1621707 | Hospital | National Reference Center for Viruses of Respiratory Infections, Institut Pasteur, Paris | Marion Barbet, Sylvie Behillil, Frédéric Lemoine, Corinne Maufrais, Christophe Malabat, Méline Bizard, Angela Brisebarre, Camille Capel, Louise Lefrançois, Etienne Simon-Lorière, Vincent Enouf, Maud Vanpeene, Sylvie van der Werf,Nabil Gastli |
| EPI_ISL_1621708 | Hospital | National Reference Center for Viruses of Respiratory Infections, Institut Pasteur, Paris | Marion Barbet, Sylvie Behillil, Frédéric Lemoine, Corinne Maufrais, Christophe Malabat, Damien Mornico, Méline Bizard, Angela Brisebarre, Camille Capel, Louise Lefrançois, Etienne Simon-Lorière, Vincent Enouf, Maud Vanpeene, Sylvie van der Werf,Nabil Gastli |
| EPI_ISL_1621709 | Labo Analyses Med | National Reference Center for Viruses of Respiratory Infections, Institut Pasteur, Paris | Marion Barbet, Sylvie Behillil, Frédéric Lemoine, Corinne Maufrais, Christophe Malabat, Méline Bizard, Angela Brisebarre, Camille Capel, Louise Lefrançois, Etienne Simon-Lorière, Vincent Enouf, Maud Vanpeene, Sylvie van der Werf,Pierre-Yves Leonard |
| EPI_ISL_1621710, EPI_ISL_1621711 | Labo Analyses Med | National Reference Center for Viruses of Respiratory Infections, Institut Pasteur, Paris | Marion Barbet, Sylvie Behillil, Frédéric Lemoine, Corinne Maufrais, Christophe Malabat, Adrien Pain, Méline Bizard, Angela Brisebarre, Camille Capel, Louise Lefrançois, Etienne Simon-Lorière, Vincent Enouf, Maud Vanpeene, Sylvie van der Werf,Pierre-Yves Leonard |
| EPI_ISL_1621712 | Labo Analyses Med | National Reference Center for Viruses of Respiratory Infections, Institut Pasteur, Paris | Marion Barbet, Sylvie Behillil, Frédéric Lemoine, Corinne Maufrais, Christophe Malabat, Méline Bizard, Angela Brisebarre, Camille Capel, Louise Lefrançois, Etienne Simon-Lorière, Vincent Enouf, Maud Vanpeene, Sylvie van der Werf,Pierre-Yves Leonard |
| EPI_ISL_1621713, EPI_ISL_1621714 | Labo Analyses Med | National Reference Center for Viruses of Respiratory Infections, Institut Pasteur, Paris | Marion Barbet, Sylvie Behillil, Frédéric Lemoine, Corinne Maufrais, Christophe Malabat, Adrien Pain, Méline Bizard, Angela Brisebarre, Camille Capel, Louise Lefrançois, Etienne Simon-Lorière, Vincent Enouf, Maud Vanpeene, Sylvie van der Werf,Pierre-Yves Leonard |
| EPI_ISL_1621715 | Labo Analyses Med | National Reference Center for Viruses of Respiratory Infections, Institut Pasteur, Paris | Marion Barbet, Sylvie Behillil, Frédéric Lemoine, Corinne Maufrais, Christophe Malabat, Damien Mornico, Méline Bizard, Angela Brisebarre, Camille Capel, Louise Lefrançois, Etienne Simon-Lorière, Vincent Enouf, Maud Vanpeene, Sylvie van der Werf,Pierre-Yves Leonard |
| EPI_ISL_1621716, EPI_ISL_1621717, EPI_ISL_1621718, EPI_ISL_1621719, EPI_ISL_1621720 | Labo Analyses Med | National Reference Center for Viruses of Respiratory Infections, Institut Pasteur, Paris | Marion Barbet, Sylvie Behillil, Frédéric Lemoine, Corinne Maufrais, Christophe Malabat, Méline Bizard, Angela Brisebarre, Camille Capel, Louise Lefrançois, Etienne Simon-Lorière, Vincent Enouf, Maud Vanpeene, Sylvie van der Werf,Pierre-Yves Leonard |
| EPI_ISL_1621721, EPI_ISL_1621722 | Hospital | National Reference Center for Viruses of Respiratory Infections, Institut Pasteur, Paris | Marion Barbet, Sylvie Behillil, Frédéric Lemoine, Corinne Maufrais, Christophe Malabat, Damien Mornico, Méline Bizard, Angela Brisebarre, Camille Capel, Louise Lefrançois, Etienne Simon-Lorière, Vincent Enouf, Maud Vanpeene, Sylvie van der Werf,Pascale Martres |
| EPI_ISL_1621724, EPI_ISL_1621725, EPI_ISL_1621727, EPI_ISL_1621728, EPI_ISL_1621730, EPI_ISL_1621731, EPI_ISL_1621733 | Hospital | National Reference Center for Viruses of Respiratory Infections, Institut Pasteur, Paris | Marion Barbet, Sylvie Behillil, Frédéric Lemoine, Corinne Maufrais, Christophe Malabat, Méline Bizard, Angela Brisebarre, Camille Capel, Louise Lefrançois, Etienne Simon-Lorière, Vincent Enouf, Maud Vanpeene, Sylvie van der Werf,BenoïT Bergues |
| EPI_ISL_1621734, EPI_ISL_1621736, EPI_ISL_1621737, EPI_ISL_1621739, EPI_ISL_1621740, EPI_ISL_1621742, EPI_ISL_1621743, EPI_ISL_1621746, EPI_ISL_1621748, EPI_ISL_1621750, EPI_ISL_1621753, EPI_ISL_1621754, EPI_ISL_1621756 | | | |
| see above | Labo Analyses Med | National Reference Center for Viruses of Respiratory Infections, Institut Pasteur, Paris | Marion Barbet, Sylvie Behillil, Frédéric Lemoine, Corinne Maufrais, Christophe Malabat, Méline Bizard, Angela Brisebarre, Camille Capel, Louise Lefrançois, Etienne Simon-Lorière, Vincent Enouf, Maud Vanpeene, Sylvie van der Werf |
| EPI_ISL_1621757 | Labo Analyses Med | National Reference Center for Viruses of Respiratory Infections, Institut Pasteur, Paris | Marion Barbet, Sylvie Behillil, Frédéric Lemoine, Corinne Maufrais, Christophe Malabat, Damien Mornico, Méline Bizard, Angela Brisebarre, Camille Capel, Louise Lefrançois, Etienne Simon-Lorière, Vincent Enouf, Maud Vanpeene, Sylvie van der Werf |
| EPI_ISL_1621759 | Labo Analyses Med | National Reference Center for Viruses of Respiratory Infections, Institut Pasteur, Paris | Marion Barbet, Sylvie Behillil, Frédéric Lemoine, Corinne Maufrais, Christophe Malabat, Méline Bizard, Angela Brisebarre, Camille Capel, Louise Lefrançois, Etienne Simon-Lorière, Vincent Enouf, Maud Vanpeene, Sylvie van der Werf |
| EPI_ISL_1621760, EPI_ISL_1621762 | Labo Analyses Med | National Reference Center for Viruses of Respiratory Infections, Institut Pasteur, Paris | Marion Barbet, Sylvie Behillil, Frédéric Lemoine, Corinne Maufrais, Christophe Malabat, Damien Mornico, Méline Bizard, Angela Brisebarre, Camille Capel, Louise Lefrançois, Etienne Simon-Lorière, Vincent Enouf, Maud Vanpeene, Sylvie van der Werf |
| EPI_ISL_1621764, EPI_ISL_1621765 | Labo Analyses med | National Reference Center for Viruses of Respiratory Infections, Institut Pasteur, Paris | Marion Barbet, Sylvie Behillil, Frédéric Lemoine, Corinne Maufrais, Christophe Malabat, Damien Mornico, Méline Bizard, Angela Brisebarre, Camille Capel, Louise Lefrançois, Etienne Simon-Lorière, Vincent Enouf, Maud Vanpeene, Sylvie van der Werf,GréGoire Potiron |
| EPI_ISL_1621766 | Labo Analyses med | National Reference Center for Viruses of Respiratory Infections, Institut Pasteur, Paris | Marion Barbet, Sylvie Behillil, Frédéric Lemoine, Corinne Maufrais, Christophe Malabat, Méline Bizard, Angela Brisebarre, Camille Capel, Louise Lefrançois, Etienne Simon-Lorière, Vincent Enouf, Maud Vanpeene, Sylvie van der Werf,GréGoire Potiron |
| EPI_ISL_1621768, EPI_ISL_1621770, EPI_ISL_1621771 | Labo Analyses med | National Reference Center for Viruses of Respiratory Infections, Institut Pasteur, Paris | Marion Barbet, Sylvie Behillil, Frédéric Lemoine, Corinne Maufrais, Christophe Malabat, Damien Mornico, Méline Bizard, Angela Brisebarre, Camille Capel, Louise Lefrançois, Etienne Simon-Lorière, Vincent Enouf, Maud Vanpeene, Sylvie van der Werf,GréGoire Potiron |
| EPI_ISL_1621773, EPI_ISL_1621774 | Labo Analyses med | National Reference Center for Viruses of Respiratory Infections, Institut Pasteur, Paris | Marion Barbet, Sylvie Behillil, Frédéric Lemoine, Corinne Maufrais, Christophe Malabat, Méline Bizard, Angela Brisebarre, Camille Capel, Louise Lefrançois, Etienne Simon-Lorière, Vincent Enouf, Maud Vanpeene, Sylvie van der Werf,GréGoire Potiron |
| EPI_ISL_1621776 | Labo Analyses med | National Reference Center for Viruses of Respiratory Infections, Institut Pasteur, Paris | Marion Barbet, Sylvie Behillil, Frédéric Lemoine, Corinne Maufrais, Christophe Malabat, Damien Mornico, Méline Bizard, Angela Brisebarre, Camille Capel, Louise Lefrançois, Etienne Simon-Lorière, Vincent Enouf, Maud Vanpeene, Sylvie van der Werf,GréGoire Potiron |
| EPI_ISL_1621778, EPI_ISL_1621779, EPI_ISL_1621781, EPI_ISL_1621782, EPI_ISL_1621784, EPI_ISL_1621785, EPI_ISL_1621787 | Labo Analyses med | National Reference Center for Viruses of Respiratory Infections, Institut Pasteur, Paris | Marion Barbet, Sylvie Behillil, Frédéric Lemoine, Corinne Maufrais, Christophe Malabat, Méline Bizard, Angela Brisebarre, Camille Capel, Louise Lefrançois, Etienne Simon-Lorière, Vincent Enouf, Maud Vanpeene, Sylvie van der Werf,GréGoire Potiron |
| EPI_ISL_1621788 | Labo Analyses med | National Reference Center for Viruses of Respiratory Infections, Institut Pasteur, Paris | Marion Barbet, Sylvie Behillil, Frédéric Lemoine, Corinne Maufrais, Christophe Malabat, Damien Mornico, Méline Bizard, Angela Brisebarre, Camille Capel, Louise Lefrançois, Etienne Simon-Lorière, Vincent Enouf, Maud Vanpeene, Sylvie van der Werf,GréGoire Potiron |
| EPI_ISL_1621790 | Labo Analyses med | National Reference Center for Viruses of Respiratory | Marion Barbet, Sylvie Behillil, Frédéric Lemoine, Corinne Maufrais, Christophe Malabat, Méline Bizard, Angela Brisebarre, Camille Capel, Louise |

| | | | |
|---|---|---|---|
| | | Infections, Institut Pasteur, Paris | Lefrançois, Etienne Simon-Lorière, Vincent Enouf, Maud Vanpeene, Sylvie van der Werf,GréGoire Potiron |
| EPI_ISL_1621793 | Hospital | National Reference Center for Viruses of Respiratory Infections, Institut Pasteur, Paris | Marion Barbet, Sylvie Behillil, Frédéric Lemoine, Corinne Maufrais, Christophe Malabat, Méline Bizard, Angela Brisebarre, Camille Capel, Louise Lefrançois, Etienne Simon-Lorière, Vincent Enouf, Maud Vanpeene, Sylvie van der Werf,Jean-Philippe Emond |
| EPI_ISL_1621794, EPI_ISL_1621796, EPI_ISL_1621797, EPI_ISL_1621799, EPI_ISL_1621800, EPI_ISL_1621802 | Hospital | National Reference Center for Viruses of Respiratory Infections, Institut Pasteur, Paris | Marion Barbet, Sylvie Behillil, Frédéric Lemoine, Corinne Maufrais, Christophe Malabat, Méline Bizard, Angela Brisebarre, Camille Capel, Louise Lefrançois, Etienne Simon-Lorière, Vincent Enouf, Maud Vanpeene, Sylvie van der Werf,CéLine Ramanantsoa |
| EPI_ISL_1621804 | Hospital | National Reference Center for Viruses of Respiratory Infections, Institut Pasteur, Paris | Marion Barbet, Sylvie Behillil, Frédéric Lemoine, Corinne Maufrais, Christophe Malabat, Damien Mornico, Méline Bizard, Angela Brisebarre, Camille Capel, Louise Lefrançois, Etienne Simon-Lorière, Vincent Enouf, Maud Vanpeene, Sylvie van der Werf,Laurence Louvet |
| EPI_ISL_1621805 | Hospital | National Reference Center for Viruses of Respiratory Infections, Institut Pasteur, Paris | Marion Barbet, Sylvie Behillil, Frédéric Lemoine, Corinne Maufrais, Christophe Malabat, Damien Mornico, Méline Bizard, Angela Brisebarre, Camille Capel, Louise Lefrançois, Etienne Simon-Lorière, Vincent Enouf, Maud Vanpeene, Sylvie van der Werf,Pascale Martres |
| EPI_ISL_1621806 | Hospital | National Reference Center for Viruses of Respiratory Infections, Institut Pasteur, Paris | Marion Barbet, Sylvie Behillil, Frédéric Lemoine, Corinne Maufrais, Christophe Malabat, Méline Bizard, Angela Brisebarre, Camille Capel, Louise Lefrançois, Etienne Simon-Lorière, Vincent Enouf, Maud Vanpeene, Sylvie van der Werf,Pascale Martres |
| EPI_ISL_1621808 | Retirement home | National Reference Center for Viruses of Respiratory Infections, Institut Pasteur, Paris | Marion Barbet, Sylvie Behillil, Frédéric Lemoine, Corinne Maufrais, Christophe Malabat, Méline Bizard, Angela Brisebarre, Camille Capel, Louise Lefrançois, Etienne Simon-Lorière, Vincent Enouf, Maud Vanpeene, Sylvie van der Werf,Sylvie Rossignol |
| EPI_ISL_1621810 | Retirement home | National Reference Center for Viruses of Respiratory Infections, Institut Pasteur, Paris | Marion Barbet, Sylvie Behillil, Frédéric Lemoine, Corinne Maufrais, Christophe Malabat, Damien Mornico, Méline Bizard, Angela Brisebarre, Camille Capel, Louise Lefrançois, Etienne Simon-Lorière, Vincent Enouf, Maud Vanpeene, Sylvie van der Werf,Sylvie Rossignol |
| EPI_ISL_1621811, EPI_ISL_1621813, EPI_ISL_1621814, EPI_ISL_1621816, EPI_ISL_1621817, EPI_ISL_1621819 | Retirement home | National Reference Center for Viruses of Respiratory Infections, Institut Pasteur, Paris | Marion Barbet, Sylvie Behillil, Frédéric Lemoine, Corinne Maufrais, Christophe Malabat, Méline Bizard, Angela Brisebarre, Camille Capel, Louise Lefrançois, Etienne Simon-Lorière, Vincent Enouf, Maud Vanpeene, Sylvie van der Werf,Sylvie Rossignol |
| EPI_ISL_1621820 | Hospital | National Reference Center for Viruses of Respiratory Infections, Institut Pasteur, Paris | Marion Barbet, Sylvie Behillil, Frédéric Lemoine, Corinne Maufrais, Christophe Malabat, Méline Bizard, Angela Brisebarre, Camille Capel, Louise Lefrançois, Etienne Simon-Lorière, Vincent Enouf, Maud Vanpeene, Sylvie van der Werf,JéRome Guinard |
| EPI_ISL_1621822 | Hospital | National Reference Center for Viruses of Respiratory Infections, Institut Pasteur, Paris | Marion Barbet, Sylvie Behillil, Frédéric Lemoine, Corinne Maufrais, Christophe Malabat, Adrien Pain, Méline Bizard, Angela Brisebarre, Camille Capel, Louise Lefrançois, Etienne Simon-Lorière, Vincent Enouf, Maud Vanpeene, Sylvie van der Werf,JéRome Guinard |
| EPI_ISL_1621823, EPI_ISL_1621825, EPI_ISL_1621826, EPI_ISL_1621828, EPI_ISL_1621829, EPI_ISL_1621831, EPI_ISL_1621832, EPI_ISL_1621834, EPI_ISL_1621835, EPI_ISL_1621837, EPI_ISL_1621838 | | | |
| see above | Labo Analyses Med | National Reference Center for Viruses of Respiratory Infections, Institut Pasteur, Paris | Marion Barbet, Sylvie Behillil, Frédéric Lemoine, Corinne Maufrais, Christophe Malabat, Méline Bizard, Angela Brisebarre, Camille Capel, Louise Lefrançois, Etienne Simon-Lorière, Vincent Enouf, Maud Vanpeene, Sylvie van der Werf,GréGoire Potiron |
| EPI_ISL_1621840 | Hospital | National Reference Center for Viruses of Respiratory Infections, Institut Pasteur, Paris | Marion Barbet, Sylvie Behillil, Frédéric Lemoine, Corinne Maufrais, Christophe Malabat, Méline Bizard, Angela Brisebarre, Camille Capel, Louise Lefrançois, Etienne Simon-Lorière, Vincent Enouf, Maud Vanpeene, Sylvie van der Werf,Ben Hadj Yahia |
| EPI_ISL_1621841 | Hospital | National Reference Center for Viruses of Respiratory Infections, Institut Pasteur, Paris | Marion Barbet, Sylvie Behillil, Frédéric Lemoine, Corinne Maufrais, Christophe Malabat, Adrien Pain, Méline Bizard, Angela Brisebarre, Camille Capel, Louise Lefrançois, Etienne Simon-Lorière, Vincent Enouf, Maud Vanpeene, Sylvie van der Werf,Ben Hadj Yahia |
| EPI_ISL_1621843 | Hospital | National Reference Center for Viruses of Respiratory Infections, Institut Pasteur, Paris | Marion Barbet, Sylvie Behillil, Frédéric Lemoine, Corinne Maufrais, Christophe Malabat, Méline Bizard, Angela Brisebarre, Camille Capel, Louise Lefrançois, Etienne Simon-Lorière, Vincent Enouf, Maud Vanpeene, Sylvie van der Werf,Ben Hadj Yahia |
| EPI_ISL_1621846, EPI_ISL_1621848, EPI_ISL_1621849 | Retirement home | National Reference Center for Viruses of Respiratory Infections, Institut Pasteur, Paris | Marion Barbet, Sylvie Behillil, Frédéric Lemoine, Corinne Maufrais, Christophe Malabat, Adrien Pain, Méline Bizard, Angela Brisebarre, Camille Capel, Louise Lefrançois, Etienne Simon-Lorière, Vincent Enouf, Maud Vanpeene, Sylvie van der Werf,Bezin |
| EPI_ISL_1622957 | Hospital | National Reference Center for Viruses of Respiratory Infections, Institut Pasteur, Paris | Marion Barbet, Sylvie Behillil, Frédéric Lemoine, Corinne Maufrais, Christophe Malabat, Victoire Baillet, Méline Bizard, Angela Brisebarre, Camille Capel, Louise Lefrançois, Etienne Simon-Lorière, Vincent Enouf, Maud Vanpeene, Sylvie van der Werf,Thibault Guinoiseau |
| EPI_ISL_1622958 | Hospital | National Reference Center for Viruses of Respiratory Infections, Institut Pasteur, Paris | Marion Barbet, Sylvie Behillil, Frédéric Lemoine, Corinne Maufrais, Christophe Malabat, Amaury Vaysse, Méline Bizard, Angela Brisebarre, Camille Capel, Louise Lefrançois, Etienne Simon-Lorière, Vincent Enouf, Maud Vanpeene, Sylvie van der Werf,Eric Farfour |
| EPI_ISL_1623012, EPI_ISL_1623013, EPI_ISL_1623014, EPI_ISL_1623015, EPI_ISL_1623016 | Labo Analyses Med | National Reference Center for Viruses of Respiratory Infections, Institut Pasteur, Paris | Marion Barbet, Sylvie Behillil, Frédéric Lemoine, Corinne Maufrais, Christophe Malabat, Amaury Vaysse, Méline Bizard, Angela Brisebarre, Camille Capel, Louise Lefrançois, Etienne Simon-Lorière, Vincent Enouf, Maud Vanpeene, Sylvie van der Werf,Melanie Caron |
| EPI_ISL_1623017 | Hospital | National Reference Center for Viruses of Respiratory Infections, Institut Pasteur, Paris | Marion Barbet, Sylvie Behillil, Frédéric Lemoine, Corinne Maufrais, Christophe Malabat, Adrien Pain, Méline Bizard, Angela Brisebarre, Camille Capel, Louise Lefrançois, Etienne Simon-Lorière, Vincent Enouf, Maud Vanpeene, Sylvie van der Werf,CéLine Bressollette |
| EPI_ISL_1623018 | Labo Analyses Med | National Reference Center for Viruses of Respiratory Infections, Institut Pasteur, Paris | Marion Barbet, Sylvie Behillil, Frédéric Lemoine, Corinne Maufrais, Christophe Malabat, Adrien Pain, Méline Bizard, Angela Brisebarre, Camille Capel, Louise Lefrançois, Etienne Simon-Lorière, Vincent Enouf, Maud Vanpeene, Sylvie van der Werf,Pierre-Yves Leonard |
| EPI_ISL_1623019 | Labo Analyses Med | National Reference Center for Viruses of Respiratory Infections, Institut Pasteur, Paris | Marion Barbet, Sylvie Behillil, Frédéric Lemoine, Corinne Maufrais, Christophe Malabat, Amaury Vaysse, Méline Bizard, Angela Brisebarre, Camille Capel, Louise Lefrançois, Etienne Simon-Lorière, Vincent Enouf, Maud Vanpeene, Sylvie van der Werf,Pierre-Yves Leonard |
| EPI_ISL_1623029, EPI_ISL_1623030, EPI_ISL_1623031, EPI_ISL_1623032 | Centre Hospitalier Universitaire Clermont-Ferrand | CHU Clermont-Ferrand, service de virologie | Bisseux Maxime, Mirand Audrey, Combes Patricia, Henquell Cécile |
| EPI_ISL_1623033, EPI_ISL_1623034, EPI_ISL_1623035 | Centre hospitalier D'AURILLAC | CHU Clermont-Ferrand, service de virologie | Bisseux Maxime, Mirand Audrey, Combes Patricia, Henquell Cécile |
| EPI_ISL_1623036, EPI_ISL_1623037, EPI_ISL_1623038, EPI_ISL_1623039 | Centre Hospitalier Universitaire Clermont-Ferrand | CHU Clermont-Ferrand, service de virologie | Bisseux Maxime, Mirand Audrey, Combes Patricia, Henquell Cécile |
| EPI_ISL_1623041 | Centre Hospitalier RIOM | CHU Clermont-Ferrand, service de virologie | Bisseux Maxime, Mirand Audrey, Combes Patricia, Henquell Cécile |
| EPI_ISL_1623042, EPI_ISL_1623043 | HOPITAL DU MONT DORE | CHU Clermont-Ferrand, service de virologie | Bisseux Maxime, Mirand Audrey, Combes Patricia, Henquell Cécile |
| EPI_ISL_1623044, EPI_ISL_1623045, EPI_ISL_1623046, EPI_ISL_1623047, EPI_ISL_1623048, EPI_ISL_1623049, EPI_ISL_1623050, EPI_ISL_1623051, EPI_ISL_1623052, EPI_ISL_1623053, EPI_ISL_1623054, EPI_ISL_1623055, EPI_ISL_1623056, EPI_ISL_1623057, EPI_ISL_1623058, EPI_ISL_1623059 | | | |
| see above | Centre Hospitalier Universitaire Clermont-Ferrand | CHU Clermont-Ferrand, service de virologie | Bisseux Maxime, Mirand Audrey, Combes Patricia, Henquell Cécile |
| EPI_ISL_1623067, EPI_ISL_1623068, EPI_ISL_1623069 | Centre Hospitalier Clementel | CHU Clermont-Ferrand, service de virologie | Bisseux Maxime, Mirand Audrey, Combes Patricia, Henquell Cécile |
| EPI_ISL_1623070 | Centre Hospitalier RIOM | CHU Clermont-Ferrand, service de virologie | Bisseux Maxime, Mirand Audrey, Combes Patricia, Henquell Cécile |
| EPI_ISL_1623071, EPI_ISL_1623072, EPI_ISL_1623073, EPI_ISL_1623074, EPI_ISL_1623075 | Centre Hospitalier Clementel | CHU Clermont-Ferrand, service de virologie | Bisseux Maxime, Mirand Audrey, Combes Patricia, Henquell Cécile |
| EPI_ISL_1623076, EPI_ISL_1623077 | Centre Hospitalier Universitaire Clermont-Ferrand | CHU Clermont-Ferrand, service de virologie | Bisseux Maxime, Mirand Audrey, Combes Patricia, Henquell Cécile |
| EPI_ISL_1623078 | HOPITAL DE VICHY | CHU Clermont-Ferrand, service de virologie | Bisseux Maxime, Mirand Audrey, Combes Patricia, Henquell Cécile |
| EPI_ISL_1623079 | HOPITAL DU MONT DORE | CHU Clermont-Ferrand, service de virologie | Bisseux Maxime, Mirand Audrey, Combes Patricia, Henquell Cécile |
| EPI_ISL_1623080, EPI_ISL_1623081, EPI_ISL_1623082, EPI_ISL_1623085 | Centre Hospitalier Universitaire Clermont-Ferrand | CHU Clermont-Ferrand, service de virologie | Bisseux Maxime, Mirand Audrey, Combes Patricia, Henquell Cécile |
| EPI_ISL_1623087 | Centre De Prélévement COVID VICHY | CHU Clermont-Ferrand, service de virologie | Bisseux Maxime, Mirand Audrey, Combes Patricia, Henquell Cécile |

| | | | |
|---|---|---|---|
| EPI_ISL_1623088 | Centre Hospitalier RIOM | CHU Clermont-Ferrand, service de virologie | Bisseux Maxime, Mirand Audrey, Combes Patricia, Henquell Cécile |
| EPI_ISL_1623089, EPI_ISL_1623090, EPI_ISL_1623091, EPI_ISL_1623092 | HOPITAL DE VICHY | CHU Clermont-Ferrand, service de virologie | Bisseux Maxime, Mirand Audrey, Combes Patricia, Henquell Cécile |
| EPI_ISL_1623093 | Centre Hospitalier RIOM | CHU Clermont-Ferrand, service de virologie | Bisseux Maxime, Mirand Audrey, Combes Patricia, Henquell Cécile |
| EPI_ISL_1623094 | Centre Hospitalier Universitaire Clermont-Ferrand | CHU Clermont-Ferrand, service de virologie | Bisseux Maxime, Mirand Audrey, Combes Patricia, Henquell Cécile |
| EPI_ISL_1623095 | HOPITAL DE VICHY | CHU Clermont-Ferrand, service de virologie | Bisseux Maxime, Mirand Audrey, Combes Patricia, Henquell Cécile |
| EPI_ISL_1623658, EPI_ISL_1623659, EPI_ISL_1623660, EPI_ISL_1623661, EPI_ISL_1623662, EPI_ISL_1623663, EPI_ISL_1623664, EPI_ISL_1623667, EPI_ISL_1623668, EPI_ISL_1623669, EPI_ISL_1623670, EPI_ISL_1623671, EPI_ISL_1623672, EPI_ISL_1623673, EPI_ISL_1623674, EPI_ISL_1623675, EPI_ISL_1623676, EPI_ISL_1623677, EPI_ISL_1623678, EPI_ISL_1623679, EPI_ISL_1623681, EPI_ISL_1623682, EPI_ISL_1623683, EPI_ISL_1623684, EPI_ISL_1623685, EPI_ISL_1623686, EPI_ISL_1623688, EPI_ISL_1623691, EPI_ISL_1623692, EPI_ISL_1623693 | | | |
| see above | National Platform bis UMONS/Jolimont | National Platform bis UMONS/Jolimont | François Dufrasne, Guillaume Bayon-Vicente, Gautier Detry, Ruddy Wattiez |
| EPI_ISL_1623816, EPI_ISL_1623817, EPI_ISL_1623818, EPI_ISL_1623819, EPI_ISL_1623820, EPI_ISL_1623821, EPI_ISL_1623822, EPI_ISL_1623823, EPI_ISL_1623824, EPI_ISL_1623825, EPI_ISL_1623826, EPI_ISL_1623827, EPI_ISL_1623828, EPI_ISL_1623829, EPI_ISL_1623830, EPI_ISL_1623831 | | | |
| see above | Baylor Scott & White-Temple | Baylor Scott & White-Temple | Ari Rao, Linden Morales, Kimberly Walker, Marcus Volz, Shelby Hendrickson, Caitlin Maloney |
| EPI_ISL_1623832 | Baylor Scott & White-Grapevine | Baylor Scott & White-Temple | Ari Rao, Linden Morales, Kimberly Walker, Marcus Volz, Shelby Hendrickson, Caitlin Maloney |
| EPI_ISL_1626614 | Instituto Nacional de Medicina Genomica | Instituto Nacional de Medicina Genomica | Hidalgo-Miranda A, Mendoza-Vargas A, Reyes-Grajeda JP, Cedro-Tanda A, Alcaraz N, Gonzalez-Barrera D, Rangel-DeLeon D, Miranda-Ortiz H, Rosas-Escobar P, Canseco Mendez JC, Munguia-Garza P, Ramirez-Vega O, Garcia-Cardenas FJ, Gonzalez-Woge MA, Herrera-Montalvo LA |
| EPI_ISL_1626618, EPI_ISL_1626619 | Laboratorio de Salud Pública Bogota | Gencore - Universidad de los Andes | Luisa Sacristan, David Gonzalez, Silvia Restrepo, Marcela Guevara, Ana Maria Palacio, Johana Hernandez, Gabriela Delgado, Alejandro Gomez |
| EPI_ISL_1626802 | LESP Yucatan | Instituto de Diagnostico y Referencia Epidemiologicos (INDRE) | Claudia Wong-Arambula, Abril Rodriguez-Maldonado, Vanessa Rivero-Arredondo, Ariadna Medina-Benitez, Joaquin Quiroz-Mercado, Sergio Rangel-Guerrero, Natividad Cruz-Ortiz, Tatiana Nunez-Garcia, Gisela Barrera-Badillo, Lucia Hernandez-Rivas, Irma Lopez-Martinez, Ernesto Ramirez-Gonzalez. |
| EPI_ISL_1626803, EPI_ISL_1626804, EPI_ISL_1626805 | LESP Chihuahua | Instituto de Diagnostico y Referencia Epidemiologicos (INDRE) | Claudia Wong-Arambula, Abril Rodriguez-Maldonado, Vanessa Rivero-Arredondo, Ariadna Medina-Benitez, Joaquin Quiroz-Mercado, Sergio Rangel-Guerrero, Natividad Cruz-Ortiz, Tatiana Nunez-Garcia, Gisela Barrera-Badillo, Lucia Hernandez-Rivas, Irma Lopez-Martinez, Ernesto Ramirez-Gonzalez. |
| EPI_ISL_1626809 | LESP Guanajuato | Instituto de Diagnostico y Referencia Epidemiologicos (INDRE) | Claudia Wong-Arambula, Abril Rodriguez-Maldonado, Vanessa Rivero-Arredondo, Ariadna Medina-Benitez, Joaquin Quiroz-Mercado, Sergio Rangel-Guerrero, Natividad Cruz-Ortiz, Tatiana Nunez-Garcia, Gisela Barrera-Badillo, Lucia Hernandez-Rivas, Irma Lopez-Martinez, Ernesto Ramirez-Gonzalez. |
| EPI_ISL_1626811, EPI_ISL_1626812, EPI_ISL_1626813 | LESP Baja California Sur | Instituto de Diagnostico y Referencia Epidemiologicos (INDRE) | Claudia Wong-Arambula, Abril Rodriguez-Maldonado, Vanessa Rivero-Arredondo, Ariadna Medina-Benitez, Joaquin Quiroz-Mercado, Sergio Rangel-Guerrero, Natividad Cruz-Ortiz, Tatiana Nunez-Garcia, Gisela Barrera-Badillo, Lucia Hernandez-Rivas, Irma Lopez-Martinez, Ernesto Ramirez-Gonzalez. |
| EPI_ISL_1626816, EPI_ISL_1626819, EPI_ISL_1626820 | LESP Guanajuato | Instituto de Diagnostico y Referencia Epidemiologicos (INDRE) | Claudia Wong-Arambula, Abril Rodriguez-Maldonado, Vanessa Rivero-Arredondo, Ariadna Medina-Benitez, Joaquin Quiroz-Mercado, Sergio Rangel-Guerrero, Natividad Cruz-Ortiz, Tatiana Nunez-Garcia, Gisela Barrera-Badillo, Lucia Hernandez-Rivas, Irma Lopez-Martinez, Ernesto Ramirez-Gonzalez. |
| EPI_ISL_1626821 | LESP Quintana Roo | Instituto de Diagnostico y Referencia Epidemiologicos (INDRE) | Claudia Wong-Arambula, Abril Rodriguez-Maldonado, Vanessa Rivero-Arredondo, Ariadna Medina-Benitez, Joaquin Quiroz-Mercado, Sergio Rangel-Guerrero, Natividad Cruz-Ortiz, Tatiana Nunez-Garcia, Gisela Barrera-Badillo, Lucia Hernandez-Rivas, Irma Lopez-Martinez, Ernesto Ramirez-Gonzalez. |
| EPI_ISL_1627325, EPI_ISL_1627326, EPI_ISL_1627327, EPI_ISL_1627328, EPI_ISL_1627329, EPI_ISL_1627330, EPI_ISL_1627331, EPI_ISL_1627332, EPI_ISL_1627335, EPI_ISL_1627336, EPI_ISL_1627337, EPI_ISL_1627338, EPI_ISL_1627339, EPI_ISL_1627340, EPI_ISL_1627341, EPI_ISL_1627362 | | | |
| see above | Presidio di Brindisi Di Summa - Perrino | Istituto Zooprofilattico Sperimentale della Puglia e della Basilicata | Parisi A., Bianco A., Capozzi L., Del Sambro L., Simone D., Difato L., Santoro A., Ridolfi D., Giannico A. |
| EPI_ISL_1627363 | Ospedale Santissima Annunziata | Istituto Zooprofilattico Sperimentale della Puglia e della Basilicata | Parisi A., Bianco A., Capozzi L., Del Sambro L., Simone D., Difato L., Ridolfi D., Giannico A, Tampoia M. |
| EPI_ISL_1627405, EPI_ISL_1627406, EPI_ISL_1627407, EPI_ISL_1627408, EPI_ISL_1627409, EPI_ISL_1627410, EPI_ISL_1627411, EPI_ISL_1627412, EPI_ISL_1627414, EPI_ISL_1627415, EPI_ISL_1627416, EPI_ISL_1627417, EPI_ISL_1627418, EPI_ISL_1627419, EPI_ISL_1627420, EPI_ISL_1627421, EPI_ISL_1627422, EPI_ISL_1627423, EPI_ISL_1627424, EPI_ISL_1627425, EPI_ISL_1627426, EPI_ISL_1627427, EPI_ISL_1627428, EPI_ISL_1627429, EPI_ISL_1627430, EPI_ISL_1627431, EPI_ISL_1627432, EPI_ISL_1627433, EPI_ISL_1627434, EPI_ISL_1627435, EPI_ISL_1627436, EPI_ISL_1627437, EPI_ISL_1627438, EPI_ISL_1627439, EPI_ISL_1627440, EPI_ISL_1627441, EPI_ISL_1627442, EPI_ISL_1627443, EPI_ISL_1627444, EPI_ISL_1627445, EPI_ISL_1627446, EPI_ISL_1627447, EPI_ISL_1627451, EPI_ISL_1627452, EPI_ISL_1627453, EPI_ISL_1627454, EPI_ISL_1627455, EPI_ISL_1627456, EPI_ISL_1627457, EPI_ISL_1627458, EPI_ISL_1627459, EPI_ISL_1627460, EPI_ISL_1627461, EPI_ISL_1627462, EPI_ISL_1627463, EPI_ISL_1627464, EPI_ISL_1627465, EPI_ISL_1627466, EPI_ISL_1627467 | | | |
| see above | Baylor Scott & White-Temple | Baylor Scott & White-Temple | Ari Rao, Linden Morales, Kimberly Walker, Marcus Volz, Shelby Hendrickson, Caitlin Maloney |
| EPI_ISL_1628366 | CS III de Patrocinio Paulista | Instituto Adolfo Lutz, Interdiciplinary Procedures Center, Strategic Laboratory | Claudio Tavares Sacchi, Claudia Regina Gonçalves, Erica Valessa Ramos Gomes, Karoline Rodrigues Campos, Caio Vinicius Dias Lopes, Leonardo Jose Tadeu de Araujo, Katia Correa de Oliveira Santos |
| EPI_ISL_1628367 | Santa Casa de Guaira | Instituto Adolfo Lutz, Interdiciplinary Procedures Center, Strategic Laboratory | Claudio Tavares Sacchi, Claudia Regina Gonçalves, Erica Valessa Ramos Gomes, Karoline Rodrigues Campos, Caio Vinicius Dias Lopes, Leonardo Jose Tadeu de Araujo, Katia Correa de Oliveira Santos |
| EPI_ISL_1628373 | UPA De Bebedouro | Instituto Adolfo Lutz, Interdiciplinary Procedures Center, Strategic Laboratory | Claudio Tavares Sacchi, Claudia Regina Gonçalves, Erica Valessa Ramos Gomes, Karoline Rodrigues Campos, Caio Vinicius Dias Lopes, Leonardo Jose Tadeu de Araujo, Katia Correa de Oliveira Santos |
| EPI_ISL_1628375, EPI_ISL_1628376 | Secretaria Municipal de Saude de Guariba | Instituto Adolfo Lutz, Interdiciplinary Procedures Center, Strategic Laboratory | Claudio Tavares Sacchi, Claudia Regina Gonçalves, Erica Valessa Ramos Gomes, Karoline Rodrigues Campos, Caio Vinicius Dias Lopes, Leonardo Jose Tadeu de Araujo, Katia Correa de Oliveira Santos |
| EPI_ISL_1628378 | Vigilancia Epidemiologica Jardinopolis | Instituto Adolfo Lutz, Interdiciplinary Procedures Center, Strategic Laboratory | Claudio Tavares Sacchi, Claudia Regina Gonçalves, Erica Valessa Ramos Gomes, Karoline Rodrigues Campos, Caio Vinicius Dias Lopes, Leonardo Jose Tadeu de Araujo, Katia Correa de Oliveira Santos |
| EPI_ISL_1628380 | Santa Casa De Pitangueiras | Instituto Adolfo Lutz, Interdiciplinary Procedures Center, Strategic Laboratory | Claudio Tavares Sacchi, Claudia Regina Gonçalves, Erica Valessa Ramos Gomes, Karoline Rodrigues Campos, Caio Vinicius Dias Lopes, Leonardo Jose Tadeu de Araujo, Katia Correa de Oliveira Santos |
| EPI_ISL_1628381, EPI_ISL_1628382, EPI_ISL_1628383, EPI_ISL_1628384, EPI_ISL_1628385, EPI_ISL_1628386, EPI_ISL_1628387, EPI_ISL_1628388, EPI_ISL_1628389, EPI_ISL_1628390, EPI_ISL_1628391, EPI_ISL_1628392, EPI_ISL_1628394, EPI_ISL_1628395, EPI_ISL_1628396, EPI_ISL_1628397, EPI_ISL_1628398, EPI_ISL_1628399, EPI_ISL_1628400, EPI_ISL_1628402, EPI_ISL_1628403, EPI_ISL_1628404, EPI_ISL_1628405, EPI_ISL_1628406, EPI_ISL_1628407, EPI_ISL_1628408, EPI_ISL_1628409, EPI_ISL_1628410, EPI_ISL_1628411, EPI_ISL_1628412, EPI_ISL_1628413, EPI_ISL_1628414, EPI_ISL_1628415, EPI_ISL_1628416, EPI_ISL_1628417, EPI_ISL_1628418, EPI_ISL_1628419, EPI_ISL_1628420, EPI_ISL_1628421, EPI_ISL_1628423, EPI_ISL_1628424, EPI_ISL_1628425, EPI_ISL_1628426, EPI_ISL_1628427, EPI_ISL_1628428, EPI_ISL_1628429, EPI_ISL_1628430, EPI_ISL_1628431, EPI_ISL_1628432, EPI_ISL_1628433, EPI_ISL_1628434, EPI_ISL_1628435, EPI_ISL_1628436, EPI_ISL_1628437, EPI_ISL_1628438, EPI_ISL_1628439, EPI_ISL_1628440, EPI_ISL_1628441, EPI_ISL_1628442, EPI_ISL_1628443, EPI_ISL_1628444, EPI_ISL_1628445, EPI_ISL_1628446, EPI_ISL_1628447, EPI_ISL_1628448, EPI_ISL_1628449, EPI_ISL_1628450, EPI_ISL_1628451, EPI_ISL_1628452, EPI_ISL_1628453, EPI_ISL_1628454, EPI_ISL_1628455, EPI_ISL_1628456, EPI_ISL_1628457, EPI_ISL_1628458, EPI_ISL_1628459, EPI_ISL_1628460, EPI_ISL_1628461, EPI_ISL_1628462, EPI_ISL_1628463, EPI_ISL_1628464, EPI_ISL_1628465, EPI_ISL_1628467, EPI_ISL_1628468, EPI_ISL_1628469, EPI_ISL_1628470, EPI_ISL_1628471, EPI_ISL_1628472, EPI_ISL_1628473 | | | |
| see above | NMVRVI | National Public Health Surveillance Laboratory | Lukas Zemaitis, Ingrida Olendraite, Arnoldas Pautienius, Kamile Tamusauskaite, Dovydas Gecys, Laura Pareckaite, Vaiva Lesauskaite, Astra Vitkauskiene |
| EPI_ISL_1629326 | REUNILAB | UMR PIMIT | Dr David A Wilkinson, Dr Patrick Mavingui, Dr Camille Lebarbenchon, Magali Turpin |
| EPI_ISL_1629356, EPI_ISL_1629361, EPI_ISL_1629374, EPI_ISL_1629380 | CERBALLIANCE | UMR PIMIT | Dr David A Wilkinson, Dr Patrick Mavingui, Dr Camille Lebarbenchon, Magali Turpin |
| EPI_ISL_1629430 | CHU | UMR PIMIT | Dr David A Wilkinson, Dr Patrick Mavingui, Dr Camille Lebarbenchon, Magali Turpin |
| EPI_ISL_1629446, EPI_ISL_1629448 | REUNILAB | UMR PIMIT | Dr David A Wilkinson, Dr Patrick Mavingui, Dr Camille Lebarbenchon, Magali Turpin |
| EPI_ISL_1629459 | CHU | UMR PIMIT | Dr David A Wilkinson, Dr Patrick Mavingui, Dr Camille Lebarbenchon, Magali Turpin |
| EPI_ISL_1630166, EPI_ISL_1630253, EPI_ISL_1630254, EPI_ISL_1630273, | National Platform bis UMONS/Jolimont | National Platform bis UMONS/Jolimont | François Dufrasne, Guillaume Bayon-Vicente, Gautier Detry, Ruddy Wattiez |

| | | | |
|---|---|---|---|
| EPI_ISL_1630275, EPI_ISL_1630276 | | | |
| EPI_ISL_1634479, EPI_ISL_1634482, EPI_ISL_1634486, EPI_ISL_1636398 | Institut de Virologie du CHU de Strasbourg | Institut de Virologie du CHU de Strasbourg | Fafi-Kremer Samira, Gallais Floriane, Gantner Pierre, Laugel Elodie, Solis Morgane, Velay Aurélie, Wendling Marie-Josée |
| EPI_ISL_1636500, EPI_ISL_1636501, EPI_ISL_1636503, EPI_ISL_1636504, EPI_ISL_1636505, EPI_ISL_1636506, EPI_ISL_1636507, EPI_ISL_1636508, EPI_ISL_1636509, EPI_ISL_1636510, EPI_ISL_1636512, EPI_ISL_1636514, EPI_ISL_1636516, EPI_ISL_1636517, EPI_ISL_1636518 | | | |
| see above | Althaia. Xarxa Assistencial Universitària de Manresa | IrsiCaixa | Marc Noguera-Julian, Mariona Parera, Maria Casadellà, Pilar Armengol, Francesc Catala-Moll, Roger Paredes, Bonaventura Clotet Gloria Trujillo, Rafel Perez Vidal, Jaume Trape Pujol, Carolina Gonzalez Fernandez, Roger Paredes, Eulalia Grau, Bonaventura Clotet |
| EPI_ISL_1636524 | Microbiology Department, University Hospital Donostia | Microbiology Department, University Hospital Donostia | Marimon JM, Montes M, Piñeiro L, Sorarrain A, Gomez M, Cilla G. |
| EPI_ISL_1647059 | SIESP CHIETI - DRIVE IN ORTONA | Istituto Zooprofilattico Sperimentale dell'Abruzzo e Molise "G. Caporale" | Lorusso A, Marcacci M, Di Domenico M, Ancora M, Curini V, Di Lollo Valeria, Mangone I, Rinaldi A, Delli Compagni E, Scialabba S, Caporale M, Di Pasquale A, Cammà C, Puglia I, Calistri P, Savini G |
| EPI_ISL_1647060 | SIESP CHIETI - DRIVE IN CHIETI | Istituto Zooprofilattico Sperimentale dell'Abruzzo e Molise "G. Caporale" | Lorusso A, Marcacci M, Di Domenico M, Ancora M, Curini V, Di Lollo Valeria, Mangone I, Rinaldi A, Delli Compagni E, Scialabba S, Caporale M, Di Pasquale A, Cammà C, Puglia I, Calistri P, Savini G |
| EPI_ISL_1647061, EPI_ISL_1647062, EPI_ISL_1647063 | SIESP CHIETI - DRIVE IN GISSI | Istituto Zooprofilattico Sperimentale dell'Abruzzo e Molise "G. Caporale" | Lorusso A, Marcacci M, Di Domenico M, Ancora M, Curini V, Di Lollo Valeria, Mangone I, Rinaldi A, Delli Compagni E, Scialabba S, Caporale M, Di Pasquale A, Cammà C, Puglia I, Calistri P, Savini G |
| EPI_ISL_1647064 | SIESP DIPARTIMENTO DI PREVENZIONE CHIETI | Istituto Zooprofilattico Sperimentale dell'Abruzzo e Molise "G. Caporale" | Lorusso A, Marcacci M, Di Domenico M, Ancora M, Curini V, Di Lollo Valeria, Mangone I, Rinaldi A, Delli Compagni E, Scialabba S, Caporale M, Di Pasquale A, Cammà C, Puglia I, Calistri P, Savini G |
| EPI_ISL_1647065 | SIESP CHIETI - DRIVE IN LANCIANO | Istituto Zooprofilattico Sperimentale dell'Abruzzo e Molise "G. Caporale" | Lorusso A, Marcacci M, Di Domenico M, Ancora M, Curini V, Di Lollo Valeria, Mangone I, Rinaldi A, Delli Compagni E, Scialabba S, Caporale M, Di Pasquale A, Cammà C, Puglia I, Calistri P, Savini G |
| EPI_ISL_1647066, EPI_ISL_1647067 | SIESP DIPARTIMENTO DI PREVENZIONE TERAMO | Istituto Zooprofilattico Sperimentale dell'Abruzzo e Molise "G. Caporale" | Lorusso A, Marcacci M, Di Domenico M, Ancora M, Curini V, Di Lollo Valeria, Mangone I, Rinaldi A, Delli Compagni E, Scialabba S, Caporale M, Di Pasquale A, Cammà C, Puglia I, Calistri P, Savini G |
| EPI_ISL_1647068 | SIESP CHIETI - DRIVE IN ORTONA | Istituto Zooprofilattico Sperimentale dell'Abruzzo e Molise "G. Caporale" | Lorusso A, Marcacci M, Di Domenico M, Ancora M, Curini V, Di Lollo Valeria, Mangone I, Rinaldi A, Delli Compagni E, Scialabba S, Caporale M, Di Pasquale A, Cammà C, Puglia I, Calistri P, Savini G |
| EPI_ISL_1647069 | SIESP CHIETI - DRIVE IN CHIETI | Istituto Zooprofilattico Sperimentale dell'Abruzzo e Molise "G. Caporale" | Lorusso A, Marcacci M, Di Domenico M, Ancora M, Curini V, Di Lollo Valeria, Mangone I, Rinaldi A, Delli Compagni E, Scialabba S, Caporale M, Di Pasquale A, Cammà C, Puglia I, Calistri P, Savini G |
| EPI_ISL_1647070 | SIESP CHIETI - DRIVE IN ORTONA | Istituto Zooprofilattico Sperimentale dell'Abruzzo e Molise "G. Caporale" | Lorusso A, Marcacci M, Di Domenico M, Ancora M, Curini V, Di Lollo Valeria, Mangone I, Rinaldi A, Delli Compagni E, Scialabba S, Caporale M, Di Pasquale A, Cammà C, Puglia I, Calistri P, Savini G |
| EPI_ISL_1647071, EPI_ISL_1647072, EPI_ISL_1647073 | SIESP DIPARTIMENTO DI PREVENZIONE TERAMO | Istituto Zooprofilattico Sperimentale dell'Abruzzo e Molise "G. Caporale" | Lorusso A, Marcacci M, Di Domenico M, Ancora M, Curini V, Di Lollo Valeria, Mangone I, Rinaldi A, Delli Compagni E, Scialabba S, Caporale M, Di Pasquale A, Cammà C, Puglia I, Calistri P, Savini G |
| EPI_ISL_1647074 | SIESP CHIETI - DRIVE IN CHIETI | Istituto Zooprofilattico Sperimentale dell'Abruzzo e Molise "G. Caporale" | Lorusso A, Marcacci M, Di Domenico M, Ancora M, Curini V, Di Lollo Valeria, Mangone I, Rinaldi A, Delli Compagni E, Scialabba S, Caporale M, Di Pasquale A, Cammà C, Puglia I, Calistri P, Savini G |
| EPI_ISL_1647075 | SIESP SULMONA | Istituto Zooprofilattico Sperimentale dell'Abruzzo e Molise "G. Caporale" | Lorusso A, Marcacci M, Di Domenico M, Ancora M, Curini V, Di Lollo Valeria, Mangone I, Rinaldi A, Delli Compagni E, Scialabba S, Caporale M, Di Pasquale A, Cammà C, Puglia I, Calistri P, Savini G |
| EPI_ISL_1647076 | SIESP CHIETI - DRIVE IN ORTONA | Istituto Zooprofilattico Sperimentale dell'Abruzzo e Molise "G. Caporale" | Lorusso A, Marcacci M, Di Domenico M, Ancora M, Curini V, Di Lollo Valeria, Mangone I, Rinaldi A, Delli Compagni E, Scialabba S, Caporale M, Di Pasquale A, Cammà C, Puglia I, Calistri P, Savini G |
| EPI_ISL_1647077, EPI_ISL_1647078, EPI_ISL_1647079, EPI_ISL_1647080 | DIP. PREV. AVEZZANO SERVIZIO DI IGIENE EPIDEMIOLOGIA E SANITA' PUBBLICA | Istituto Zooprofilattico Sperimentale dell'Abruzzo e Molise "G. Caporale" | Lorusso A, Marcacci M, Di Domenico M, Ancora M, Curini V, Di Lollo Valeria, Mangone I, Rinaldi A, Delli Compagni E, Scialabba S, Caporale M, Di Pasquale A, Cammà C, Puglia I, Calistri P, Savini G |
| EPI_ISL_1647081, EPI_ISL_1647082 | SIESP SULMONA | Istituto Zooprofilattico Sperimentale dell'Abruzzo e Molise "G. Caporale" | Lorusso A, Marcacci M, Di Domenico M, Ancora M, Curini V, Di Lollo Valeria, Mangone I, Rinaldi A, Delli Compagni E, Scialabba S, Caporale M, Di Pasquale A, Cammà C, Puglia I, Calistri P, Savini G |
| EPI_ISL_1647083 | USCA Castel di Sangro | Istituto Zooprofilattico Sperimentale dell'Abruzzo e Molise "G. Caporale" | Lorusso A, Marcacci M, Di Domenico M, Ancora M, Curini V, Di Lollo Valeria, Mangone I, Rinaldi A, Delli Compagni E, Scialabba S, Caporale M, Di Pasquale A, Cammà C, Puglia I, Calistri P, Savini G |
| EPI_ISL_1647084, EPI_ISL_1647085 | DIP. PREV. AVEZZANO SERVIZIO DI IGIENE EPIDEMIOLOGIA E SANITA' PUBBLICA | Istituto Zooprofilattico Sperimentale dell'Abruzzo e Molise "G. Caporale" | Lorusso A, Marcacci M, Di Domenico M, Ancora M, Curini V, Di Lollo Valeria, Mangone I, Rinaldi A, Delli Compagni E, Scialabba S, Caporale M, Di Pasquale A, Cammà C, Puglia I, Calistri P, Savini G |
| EPI_ISL_1647086 | SIESP CHIETI - DRIVE IN ORTONA | Istituto Zooprofilattico Sperimentale dell'Abruzzo e Molise "G. Caporale" | Lorusso A, Marcacci M, Di Domenico M, Ancora M, Curini V, Di Lollo Valeria, Mangone I, Rinaldi A, Delli Compagni E, Scialabba S, Caporale M, Di Pasquale A, Cammà C, Puglia I, Calistri P, Savini G |
| EPI_ISL_1647087, EPI_ISL_1647088 | SIESP CHIETI - DRIVE IN LANCIANO | Istituto Zooprofilattico Sperimentale dell'Abruzzo e Molise "G. Caporale" | Lorusso A, Marcacci M, Di Domenico M, Ancora M, Curini V, Di Lollo Valeria, Mangone I, Rinaldi A, Delli Compagni E, Scialabba S, Caporale M, Di Pasquale A, Cammà C, Puglia I, Calistri P, Savini G |
| EPI_ISL_1647089, EPI_ISL_1647090 | SIESP CHIETI - DRIVE IN VASTO | Istituto Zooprofilattico Sperimentale dell'Abruzzo e Molise "G. Caporale" | Lorusso A, Marcacci M, Di Domenico M, Ancora M, Curini V, Di Lollo Valeria, Mangone I, Rinaldi A, Delli Compagni E, Scialabba S, Caporale M, Di Pasquale A, Cammà C, Puglia I, Calistri P, Savini G |
| EPI_ISL_1647091, EPI_ISL_1647092 | SIESP CHIETI - DRIVE IN GISSI | Istituto Zooprofilattico Sperimentale dell'Abruzzo e Molise "G. Caporale" | Lorusso A, Marcacci M, Di Domenico M, Ancora M, Curini V, Di Lollo Valeria, Mangone I, Rinaldi A, Delli Compagni E, Scialabba S, Caporale M, Di Pasquale A, Cammà C, Puglia I, Calistri P, Savini G |
| EPI_ISL_1647093, EPI_ISL_1647094, EPI_ISL_1647095, EPI_ISL_1647096, EPI_ISL_1647097, EPI_ISL_1647098, EPI_ISL_1647099 | SIESP DIPARTIMENTO DI PREVENZIONE TERAMO | Istituto Zooprofilattico Sperimentale dell'Abruzzo e Molise "G. Caporale" | Lorusso A, Marcacci M, Di Domenico M, Ancora M, Curini V, Di Lollo Valeria, Mangone I, Rinaldi A, Delli Compagni E, Scialabba S, Caporale M, Di Pasquale A, Cammà C, Puglia I, Calistri P, Savini G |
| EPI_ISL_1647100 | SIESP CHIETI - DRIVE IN CHIETI | Istituto Zooprofilattico Sperimentale dell'Abruzzo e Molise "G. Caporale" | Lorusso A, Marcacci M, Di Domenico M, Ancora M, Curini V, Di Lollo Valeria, Mangone I, Rinaldi A, Delli Compagni E, Scialabba S, Caporale M, Di Pasquale A, Cammà C, Puglia I, Calistri P, Savini G |
| EPI_ISL_1647101 | SIESP CHIETI - DRIVE IN ORTONA | Istituto Zooprofilattico Sperimentale dell'Abruzzo e Molise "G. Caporale" | Lorusso A, Marcacci M, Di Domenico M, Ancora M, Curini V, Di Lollo Valeria, Mangone I, Rinaldi A, Delli Compagni E, Scialabba S, Caporale M, Di Pasquale A, Cammà C, Puglia I, Calistri P, Savini G |
| EPI_ISL_1647102, EPI_ISL_1647103, EPI_ISL_1647104, EPI_ISL_1647105 | SIESP DIPARTIMENTO DI PREVENZIONE TERAMO | Istituto Zooprofilattico Sperimentale dell'Abruzzo e Molise "G. Caporale" | Lorusso A, Marcacci M, Di Domenico M, Ancora M, Curini V, Di Lollo Valeria, Mangone I, Rinaldi A, Delli Compagni E, Scialabba S, Caporale M, Di Pasquale A, Cammà C, Puglia I, Calistri P, Savini G |
| EPI_ISL_1647106 | SIESP CHIETI - DRIVE IN LANCIANO | Istituto Zooprofilattico Sperimentale dell'Abruzzo e Molise "G. Caporale" | Lorusso A, Marcacci M, Di Domenico M, Ancora M, Curini V, Di Lollo Valeria, Mangone I, Rinaldi A, Delli Compagni E, Scialabba S, Caporale M, Di Pasquale A, Cammà C, Puglia I, Calistri P, Savini G |
| EPI_ISL_1647107 | SIESP DIPARTIMENTO DI PREVENZIONE TERAMO | Istituto Zooprofilattico Sperimentale dell'Abruzzo e Molise "G. Caporale" | Lorusso A, Marcacci M, Di Domenico M, Ancora M, Curini V, Di Lollo Valeria, Mangone I, Rinaldi A, Delli Compagni E, Scialabba S, Caporale M, Di Pasquale A, Cammà C, Puglia I, Calistri P, Savini G |
| EPI_ISL_1647108 | SIESP CHIETI - DRIVE IN CHIETI | Istituto Zooprofilattico Sperimentale dell'Abruzzo e Molise "G. Caporale" | Lorusso A, Marcacci M, Di Domenico M, Ancora M, Curini V, Di Lollo Valeria, Mangone I, Rinaldi A, Delli Compagni E, Scialabba S, Caporale M, Di Pasquale A, Cammà C, Puglia I, Calistri P, Savini G |
| EPI_ISL_1647109, EPI_ISL_1647110 | SIESP SULMONA | Istituto Zooprofilattico Sperimentale dell'Abruzzo e Molise "G. Caporale" | Lorusso A, Marcacci M, Di Domenico M, Ancora M, Curini V, Di Lollo Valeria, Mangone I, Rinaldi A, Delli Compagni E, Scialabba S, Caporale M, Di Pasquale A, Cammà C, Puglia I, Calistri P, Savini G |
| EPI_ISL_1647111 | SIESP CHIETI - DRIVE IN LANCIANO | Istituto Zooprofilattico Sperimentale dell'Abruzzo e Molise "G. Caporale" | Lorusso A, Marcacci M, Di Domenico M, Ancora M, Curini V, Di Lollo Valeria, Mangone I, Rinaldi A, Delli Compagni E, Scialabba S, Caporale M, Di Pasquale A, Cammà C, Puglia I, Calistri P, Savini G |
| EPI_ISL_1647112 | DIP. PREV. AVEZZANO SERVIZIO DI IGIENE | Istituto Zooprofilattico Sperimentale dell'Abruzzo e Molise "G. | Lorusso A, Marcacci M, Di Domenico M, Ancora M, Curini V, Di Lollo Valeria, Mangone I, Rinaldi A, Delli Compagni E, Scialabba S, Caporale M, Di |

| | | | |
|---|---|---|---|
| | EPIDEMIOLOGIA E SANITA' PUBBLICA | Caporale" | Pasquale A, Cammà C, Puglia I, Calistri P, Savini G |
| EPI_ISL_1647113, EPI_ISL_1647114 | USCA Sulmona | Istituto Zooprofilattico Sperimentale dell'Abruzzo e Molise "G. Caporale" | Lorusso A, Marcacci M, Di Domenico M, Ancora M, Curini V, Di Lollo Valeria, Mangone I, Rinaldi A, Delli Compagni E, Scialabba S, Caporale M, Di Pasquale A, Cammà C, Puglia I, Calistri P, Savini G |
| EPI_ISL_1647115 | DIP. PREV. AVEZZANO SERVIZIO DI IGIENE EPIDEMIOLOGIA E SANITA' PUBBLICA | Istituto Zooprofilattico Sperimentale dell'Abruzzo e Molise "G. Caporale" | Lorusso A, Marcacci M, Di Domenico M, Ancora M, Curini V, Di Lollo Valeria, Mangone I, Rinaldi A, Delli Compagni E, Scialabba S, Caporale M, Di Pasquale A, Cammà C, Puglia I, Calistri P, Savini G |
| EPI_ISL_1647116 | SIESP DIPARTIMENTO DI PREVENZIONE TERAMO | Istituto Zooprofilattico Sperimentale dell'Abruzzo e Molise "G. Caporale" | Lorusso A, Marcacci M, Di Domenico M, Ancora M, Curini V, Di Lollo Valeria, Mangone I, Rinaldi A, Delli Compagni E, Scialabba S, Caporale M, Di Pasquale A, Cammà C, Puglia I, Calistri P, Savini G |
| EPI_ISL_1647117 | SIESP CHIETI - DRIVE IN LANCIANO | Istituto Zooprofilattico Sperimentale dell'Abruzzo e Molise "G. Caporale" | Lorusso A, Marcacci M, Di Domenico M, Ancora M, Curini V, Di Lollo Valeria, Mangone I, Rinaldi A, Delli Compagni E, Scialabba S, Caporale M, Di Pasquale A, Cammà C, Puglia I, Calistri P, Savini G |
| EPI_ISL_1647118 | SIESP DIPARTIMENTO DI PREVENZIONE CHIETI | Istituto Zooprofilattico Sperimentale dell'Abruzzo e Molise "G. Caporale" | Lorusso A, Marcacci M, Di Domenico M, Ancora M, Curini V, Di Lollo Valeria, Mangone I, Rinaldi A, Delli Compagni E, Scialabba S, Caporale M, Di Pasquale A, Cammà C, Puglia I, Calistri P, Savini G |
| EPI_ISL_1647119, EPI_ISL_1647120 | DIP. PREV. AVEZZANO SERVIZIO DI IGIENE EPIDEMIOLOGIA E SANITA' PUBBLICA | Istituto Zooprofilattico Sperimentale dell'Abruzzo e Molise "G. Caporale" | Lorusso A, Marcacci M, Di Domenico M, Ancora M, Curini V, Di Lollo Valeria, Mangone I, Rinaldi A, Delli Compagni E, Scialabba S, Caporale M, Di Pasquale A, Cammà C, Puglia I, Calistri P, Savini G |
| EPI_ISL_1647121 | SIESP SULMONA | Istituto Zooprofilattico Sperimentale dell'Abruzzo e Molise "G. Caporale" | Lorusso A, Marcacci M, Di Domenico M, Ancora M, Curini V, Di Lollo Valeria, Mangone I, Rinaldi A, Delli Compagni E, Scialabba S, Caporale M, Di Pasquale A, Cammà C, Puglia I, Calistri P, Savini G |
| EPI_ISL_1647122, EPI_ISL_1647123, EPI_ISL_1647124 | DIP. PREV. AVEZZANO SERVIZIO DI IGIENE EPIDEMIOLOGIA E SANITA' PUBBLICA | Istituto Zooprofilattico Sperimentale dell'Abruzzo e Molise "G. Caporale" | Lorusso A, Marcacci M, Di Domenico M, Ancora M, Curini V, Di Lollo Valeria, Mangone I, Rinaldi A, Delli Compagni E, Scialabba S, Caporale M, Di Pasquale A, Cammà C, Puglia I, Calistri P, Savini G |
| EPI_ISL_1647125 | USCA Pescina | Istituto Zooprofilattico Sperimentale dell'Abruzzo e Molise "G. Caporale" | Lorusso A, Marcacci M, Di Domenico M, Ancora M, Curini V, Di Lollo Valeria, Mangone I, Rinaldi A, Delli Compagni E, Scialabba S, Caporale M, Di Pasquale A, Cammà C, Puglia I, Calistri P, Savini G |
| EPI_ISL_1647126 | USCA Avezzano | Istituto Zooprofilattico Sperimentale dell'Abruzzo e Molise "G. Caporale" | Lorusso A, Marcacci M, Di Domenico M, Ancora M, Curini V, Di Lollo Valeria, Mangone I, Rinaldi A, Delli Compagni E, Scialabba S, Caporale M, Di Pasquale A, Cammà C, Puglia I, Calistri P, Savini G |
| EPI_ISL_1647127 | Residenza per Anziani TOLLO | Istituto Zooprofilattico Sperimentale dell'Abruzzo e Molise "G. Caporale" | Lorusso A, Marcacci M, Di Domenico M, Ancora M, Curini V, Di Lollo Valeria, Mangone I, Rinaldi A, Delli Compagni E, Scialabba S, Caporale M, Di Pasquale A, Cammà C, Puglia I, Calistri P, Savini G |
| EPI_ISL_1647128, EPI_ISL_1647129, EPI_ISL_1647131, EPI_ISL_1647133, EPI_ISL_1647134, EPI_ISL_1647135, EPI_ISL_1647136, EPI_ISL_1647137, EPI_ISL_1647138, EPI_ISL_1647139, EPI_ISL_1647140, EPI_ISL_1647141, EPI_ISL_1647142, EPI_ISL_1647143, EPI_ISL_1647144, EPI_ISL_1647145 | | | |
| see above | OSPEDALE SAN SALVATORE - MEDICINA DI LABORATORIO | Istituto Zooprofilattico Sperimentale dell'Abruzzo e Molise "G. Caporale" | Lorusso A, Marcacci M, Di Domenico M, Ancora M, Curini V, Di Lollo Valeria, Mangone I, Rinaldi A, Delli Compagni E, Scialabba S, Caporale M, Di Pasquale A, Cammà C, Puglia I, Calistri P, Savini G |
| EPI_ISL_1647146 | SIESP DIPARTIMENTO DI PREVENZIONE CHIETI | Istituto Zooprofilattico Sperimentale dell'Abruzzo e Molise "G. Caporale" | Lorusso A, Marcacci M, Di Domenico M, Ancora M, Curini V, Di Lollo Valeria, Mangone I, Rinaldi A, Delli Compagni E, Scialabba S, Caporale M, Di Pasquale A, Cammà C, Puglia I, Calistri P, Savini G |
| EPI_ISL_1647147 | SIESP CHIETI - DRIVE IN LANCIANO | Istituto Zooprofilattico Sperimentale dell'Abruzzo e Molise "G. Caporale" | Lorusso A, Marcacci M, Di Domenico M, Ancora M, Curini V, Di Lollo Valeria, Mangone I, Rinaldi A, Delli Compagni E, Scialabba S, Caporale M, Di Pasquale A, Cammà C, Puglia I, Calistri P, Savini G |
| EPI_ISL_1647148, EPI_ISL_1647149, EPI_ISL_1647150, EPI_ISL_1647151 | SIESP DIPARTIMENTO DI PREVENZIONE TERAMO | Istituto Zooprofilattico Sperimentale dell'Abruzzo e Molise "G. Caporale" | Lorusso A, Marcacci M, Di Domenico M, Ancora M, Curini V, Di Lollo Valeria, Mangone I, Rinaldi A, Delli Compagni E, Scialabba S, Caporale M, Di Pasquale A, Cammà C, Puglia I, Calistri P, Savini G |
| EPI_ISL_1647152 | SIESP CHIETI - DRIVE IN CHIETI | Istituto Zooprofilattico Sperimentale dell'Abruzzo e Molise "G. Caporale" | Lorusso A, Marcacci M, Di Domenico M, Ancora M, Curini V, Di Lollo Valeria, Mangone I, Rinaldi A, Delli Compagni E, Scialabba S, Caporale M, Di Pasquale A, Cammà C, Puglia I, Calistri P, Savini G |
| EPI_ISL_1647153 | SIESP SULMONA | Istituto Zooprofilattico Sperimentale dell'Abruzzo e Molise "G. Caporale" | Lorusso A, Marcacci M, Di Domenico M, Ancora M, Curini V, Di Lollo Valeria, Mangone I, Rinaldi A, Delli Compagni E, Scialabba S, Caporale M, Di Pasquale A, Cammà C, Puglia I, Calistri P, Savini G |
| EPI_ISL_1647154 | OSPEDALE CIVILE SANT'OMERO - PRONTO SOCCORSO | Istituto Zooprofilattico Sperimentale dell'Abruzzo e Molise "G. Caporale" | Lorusso A, Marcacci M, Di Domenico M, Ancora M, Curini V, Di Lollo Valeria, Mangone I, Rinaldi A, Delli Compagni E, Scialabba S, Caporale M, Di Pasquale A, Cammà C, Puglia I, Calistri P, Savini G |
| EPI_ISL_1647155 | USCA Avezzano | Istituto Zooprofilattico Sperimentale dell'Abruzzo e Molise "G. Caporale" | Lorusso A, Marcacci M, Di Domenico M, Ancora M, Curini V, Di Lollo Valeria, Mangone I, Rinaldi A, Delli Compagni E, Scialabba S, Caporale M, Di Pasquale A, Cammà C, Puglia I, Calistri P, Savini G |
| EPI_ISL_1647156 | SIESP CHIETI - DRIVE IN VASTO | Istituto Zooprofilattico Sperimentale dell'Abruzzo e Molise "G. Caporale" | Lorusso A, Marcacci M, Di Domenico M, Ancora M, Curini V, Di Lollo Valeria, Mangone I, Rinaldi A, Delli Compagni E, Scialabba S, Caporale M, Di Pasquale A, Cammà C, Puglia I, Calistri P, Savini G |
| EPI_ISL_1647157 | USCA Avezzano | Istituto Zooprofilattico Sperimentale dell'Abruzzo e Molise "G. Caporale" | Lorusso A, Marcacci M, Di Domenico M, Ancora M, Curini V, Di Lollo Valeria, Mangone I, Rinaldi A, Delli Compagni E, Scialabba S, Caporale M, Di Pasquale A, Cammà C, Puglia I, Calistri P, Savini G |
| EPI_ISL_1647158, EPI_ISL_1647159 | SIESP SULMONA | Istituto Zooprofilattico Sperimentale dell'Abruzzo e Molise "G. Caporale" | Lorusso A, Marcacci M, Di Domenico M, Ancora M, Curini V, Di Lollo Valeria, Mangone I, Rinaldi A, Delli Compagni E, Scialabba S, Caporale M, Di Pasquale A, Cammà C, Puglia I, Calistri P, Savini G |
| EPI_ISL_1647160 | SIESP CHIETI - DRIVE IN ORTONA | Istituto Zooprofilattico Sperimentale dell'Abruzzo e Molise "G. Caporale" | Lorusso A, Marcacci M, Di Domenico M, Ancora M, Curini V, Di Lollo Valeria, Mangone I, Rinaldi A, Delli Compagni E, Scialabba S, Caporale M, Di Pasquale A, Cammà C, Puglia I, Calistri P, Savini G |
| EPI_ISL_1647161, EPI_ISL_1647162, EPI_ISL_1647163 | SIESP CHIETI - DRIVE IN VASTO | Istituto Zooprofilattico Sperimentale dell'Abruzzo e Molise "G. Caporale" | Lorusso A, Marcacci M, Di Domenico M, Ancora M, Curini V, Di Lollo Valeria, Mangone I, Rinaldi A, Delli Compagni E, Scialabba S, Caporale M, Di Pasquale A, Cammà C, Puglia I, Calistri P, Savini G |
| EPI_ISL_1647164, EPI_ISL_1647165, EPI_ISL_1647166 | DIP. PREV. AVEZZANO SERVIZIO DI IGIENE EPIDEMIOLOGIA E SANITA' PUBBLICA | Istituto Zooprofilattico Sperimentale dell'Abruzzo e Molise "G. Caporale" | Lorusso A, Marcacci M, Di Domenico M, Ancora M, Curini V, Di Lollo Valeria, Mangone I, Rinaldi A, Delli Compagni E, Scialabba S, Caporale M, Di Pasquale A, Cammà C, Puglia I, Calistri P, Savini G |
| EPI_ISL_1647167 | SIESP CHIETI - DRIVE IN VASTO | Istituto Zooprofilattico Sperimentale dell'Abruzzo e Molise "G. Caporale" | Lorusso A, Marcacci M, Di Domenico M, Ancora M, Curini V, Di Lollo Valeria, Mangone I, Rinaldi A, Delli Compagni E, Scialabba S, Caporale M, Di Pasquale A, Cammà C, Puglia I, Calistri P, Savini G |
| EPI_ISL_1647169, EPI_ISL_1647170, EPI_ISL_1647171 | DIP. PREV. AVEZZANO SERVIZIO DI IGIENE EPIDEMIOLOGIA E SANITA' PUBBLICA | Istituto Zooprofilattico Sperimentale dell'Abruzzo e Molise "G. Caporale" | Lorusso A, Marcacci M, Di Domenico M, Ancora M, Curini V, Di Lollo Valeria, Mangone I, Rinaldi A, Delli Compagni E, Scialabba S, Caporale M, Di Pasquale A, Cammà C, Puglia I, Calistri P, Savini G |
| EPI_ISL_1647172, EPI_ISL_1647173 | SIESP CHIETI - DRIVE IN LANCIANO | Istituto Zooprofilattico Sperimentale dell'Abruzzo e Molise "G. Caporale" | Lorusso A, Marcacci M, Di Domenico M, Ancora M, Curini V, Di Lollo Valeria, Mangone I, Rinaldi A, Delli Compagni E, Scialabba S, Caporale M, Di Pasquale A, Cammà C, Puglia I, Calistri P, Savini G |
| EPI_ISL_1647174 | SIESP CHIETI - DRIVE IN VASTO | Istituto Zooprofilattico Sperimentale dell'Abruzzo e Molise "G. Caporale" | Lorusso A, Marcacci M, Di Domenico M, Ancora M, Curini V, Di Lollo Valeria, Mangone I, Rinaldi A, Delli Compagni E, Scialabba S, Caporale M, Di Pasquale A, Cammà C, Puglia I, Calistri P, Savini G |
| EPI_ISL_1647175, EPI_ISL_1647176 | SIESP DIPARTIMENTO DI PREVENZIONE CHIETI | Istituto Zooprofilattico Sperimentale dell'Abruzzo e Molise "G. Caporale" | Lorusso A, Marcacci M, Di Domenico M, Ancora M, Curini V, Di Lollo Valeria, Mangone I, Rinaldi A, Delli Compagni E, Scialabba S, Caporale M, Di Pasquale A, Cammà C, Puglia I, Calistri P, Savini G |
| EPI_ISL_1647177 | SIESP CHIETI - DRIVE IN VASTO | Istituto Zooprofilattico Sperimentale dell'Abruzzo e Molise "G. Caporale" | Lorusso A, Marcacci M, Di Domenico M, Ancora M, Curini V, Di Lollo Valeria, Mangone I, Rinaldi A, Delli Compagni E, Scialabba S, Caporale M, Di Pasquale A, Cammà C, Puglia I, Calistri P, Savini G |
| EPI_ISL_1647178 | PRESIDIO TERRITORIALE CASOLI | Istituto Zooprofilattico Sperimentale dell'Abruzzo e Molise "G. Caporale" | Lorusso A, Marcacci M, Di Domenico M, Ancora M, Curini V, Di Lollo Valeria, Mangone I, Rinaldi A, Delli Compagni E, Scialabba S, Caporale M, Di Pasquale A, Cammà C, Puglia I, Calistri P, Savini G |
| EPI_ISL_1647179, EPI_ISL_1647180, EPI_ISL_1647181 | SIESP DIPARTIMENTO DI PREVENZIONE TERAMO | Istituto Zooprofilattico Sperimentale dell'Abruzzo e Molise "G. Caporale" | Lorusso A, Marcacci M, Di Domenico M, Ancora M, Curini V, Di Lollo Valeria, Mangone I, Rinaldi A, Delli Compagni E, Scialabba S, Caporale M, Di Pasquale A, Cammà C, Puglia I, Calistri P, Savini G |
| EPI_ISL_1647182 | SIESP CHIETI - DRIVE IN CHIETI | Istituto Zooprofilattico Sperimentale dell'Abruzzo e Molise "G. Caporale" | Lorusso A, Marcacci M, Di Domenico M, Ancora M, Curini V, Di Lollo Valeria, Mangone I, Rinaldi A, Delli Compagni E, Scialabba S, Caporale M, Di Pasquale A, Cammà C, Puglia I, Calistri P, Savini G |

| | | | |
|---|---|---|---|
| EPI_ISL_1647183, EPI_ISL_1647184 | SIESP DIPARTIMENTO DI PREVENZIONE TERAMO | Istituto Zooprofilattico Sperimentale dell'Abruzzo e Molise "G. Caporale" | Lorusso A, Marcacci M, Di Domenico M, Ancora M, Curini V, Di Lollo Valeria, Mangone I, Rinaldi A, Delli Compagni E, Scialabba S, Caporale M, Di Pasquale A, Cammà C, Puglia I, Calistri P, Savini G |
| EPI_ISL_1647185, EPI_ISL_1647186, EPI_ISL_1647187 | SIESP CHIETI - DRIVE IN ORTONA | Istituto Zooprofilattico Sperimentale dell'Abruzzo e Molise "G. Caporale" | Lorusso A, Marcacci M, Di Domenico M, Ancora M, Curini V, Di Lollo Valeria, Mangone I, Rinaldi A, Delli Compagni E, Scialabba S, Caporale M, Di Pasquale A, Cammà C, Puglia I, Calistri P, Savini G |
| EPI_ISL_1651250, EPI_ISL_1651251 | LESP Jalisco | Instituto de Diagnostico y Referencia Epidemiologicos (INDRE) | Claudia Wong-Arambula, Abril Rodriguez-Maldonado, Vanessa Rivero-Arredondo, Ariadna Medina-Benitez, Joaquin Quiroz-Mercado, Sergio Rangel-Guerrero, Natividad Cruz-Ortiz, Tatiana Nunez-Garcia, Gisela Barrera-Badillo, Lucia Hernandez-Rivas, Irma Lopez-Martinez, Ernesto Ramirez-Gonzalez. |
| EPI_ISL_1651262, EPI_ISL_1651263, EPI_ISL_1651266, EPI_ISL_1651267, EPI_ISL_1651268, EPI_ISL_1651269, EPI_ISL_1651270, EPI_ISL_1651271, EPI_ISL_1651272, EPI_ISL_1651273, EPI_ISL_1651274, EPI_ISL_1651276 | | | |
| see above | LESP Morelos | Instituto de Diagnostico y Referencia Epidemiologicos (INDRE) | Claudia Wong-Arambula, Abril Rodriguez-Maldonado, Vanessa Rivero-Arredondo, Ariadna Medina-Benitez, Joaquin Quiroz-Mercado, Sergio Rangel-Guerrero, Natividad Cruz-Ortiz, Tatiana Nunez-Garcia, Gisela Barrera-Badillo, Lucia Hernandez-Rivas, Irma Lopez-Martinez, Ernesto Ramirez-Gonzalez. |
| EPI_ISL_1651279, EPI_ISL_1651281, EPI_ISL_1651282, EPI_ISL_1651283, EPI_ISL_1651284, EPI_ISL_1651285, EPI_ISL_1651286, EPI_ISL_1651287, EPI_ISL_1651876 | LESP Queretaro | Instituto de Diagnostico y Referencia Epidemiologicos (INDRE) | Claudia Wong-Arambula, Abril Rodriguez-Maldonado, Vanessa Rivero-Arredondo, Ariadna Medina-Benitez, Joaquin Quiroz-Mercado, Sergio Rangel-Guerrero, Natividad Cruz-Ortiz, Tatiana Nunez-Garcia, Gisela Barrera-Badillo, Lucia Hernandez-Rivas, Irma Lopez-Martinez, Ernesto Ramirez-Gonzalez. |
| EPI_ISL_1651883, EPI_ISL_1651884, EPI_ISL_1651885, EPI_ISL_1651886 | LESP Yucatan | Instituto de Diagnostico y Referencia Epidemiologicos (INDRE) | Claudia Wong-Arambula, Abril Rodriguez-Maldonado, Vanessa Rivero-Arredondo, Ariadna Medina-Benitez, Joaquin Quiroz-Mercado, Sergio Rangel-Guerrero, Natividad Cruz-Ortiz, Tatiana Nunez-Garcia, Gisela Barrera-Badillo, Lucia Hernandez-Rivas, Irma Lopez-Martinez, Ernesto Ramirez-Gonzalez. |
| EPI_ISL_1651889 | LESP Coahuila | Instituto de Diagnostico y Referencia Epidemiologicos (INDRE) | Claudia Wong-Arambula, Abril Rodriguez-Maldonado, Vanessa Rivero-Arredondo, Ariadna Medina-Benitez, Joaquin Quiroz-Mercado, Sergio Rangel-Guerrero, Natividad Cruz-Ortiz, Tatiana Nunez-Garcia, Gisela Barrera-Badillo, Lucia Hernandez-Rivas, Irma Lopez-Martinez, Ernesto Ramirez-Gonzalez. |
| EPI_ISL_1651890, EPI_ISL_1651891, EPI_ISL_1651892, EPI_ISL_1651893, EPI_ISL_1651894, EPI_ISL_1651895 | LESP Baja California Sur | Instituto de Diagnostico y Referencia Epidemiologicos (INDRE) | Claudia Wong-Arambula, Abril Rodriguez-Maldonado, Vanessa Rivero-Arredondo, Ariadna Medina-Benitez, Joaquin Quiroz-Mercado, Sergio Rangel-Guerrero, Natividad Cruz-Ortiz, Tatiana Nunez-Garcia, Gisela Barrera-Badillo, Lucia Hernandez-Rivas, Irma Lopez-Martinez, Ernesto Ramirez-Gonzalez. |
| EPI_ISL_1651896, EPI_ISL_1651897, EPI_ISL_1651898 | LESP Quintana Roo | Instituto de Diagnostico y Referencia Epidemiologicos (INDRE) | Claudia Wong-Arambula, Abril Rodriguez-Maldonado, Vanessa Rivero-Arredondo, Ariadna Medina-Benitez, Joaquin Quiroz-Mercado, Sergio Rangel-Guerrero, Natividad Cruz-Ortiz, Tatiana Nunez-Garcia, Gisela Barrera-Badillo, Lucia Hernandez-Rivas, Irma Lopez-Martinez, Ernesto Ramirez-Gonzalez. |
| EPI_ISL_1651900 | LESP Baja California Sur | Instituto de Diagnostico y Referencia Epidemiologicos (INDRE) | Claudia Wong-Arambula, Abril Rodriguez-Maldonado, Vanessa Rivero-Arredondo, Ariadna Medina-Benitez, Joaquin Quiroz-Mercado, Sergio Rangel-Guerrero, Natividad Cruz-Ortiz, Tatiana Nunez-Garcia, Gisela Barrera-Badillo, Lucia Hernandez-Rivas, Irma Lopez-Martinez, Ernesto Ramirez-Gonzalez. |
| EPI_ISL_1651904, EPI_ISL_1651906, EPI_ISL_1651908 | LESP Morelos | Instituto de Diagnostico y Referencia Epidemiologicos (INDRE) | Claudia Wong-Arambula, Abril Rodriguez-Maldonado, Vanessa Rivero-Arredondo, Ariadna Medina-Benitez, Joaquin Quiroz-Mercado, Sergio Rangel-Guerrero, Natividad Cruz-Ortiz, Tatiana Nunez-Garcia, Gisela Barrera-Badillo, Lucia Hernandez-Rivas, Irma Lopez-Martinez, Ernesto Ramirez-Gonzalez. |
| EPI_ISL_1653934, EPI_ISL_1653936 | University of Bari Biomedical Sciences and Human Oncology | University of Bari Biomedical Sciences and Human Oncology | Chironna M., Sallustio A., Loconsole D., Accogli M. |
| EPI_ISL_1655918 | Hospital | National Reference Center for Viruses of Respiratory Infections, Institut Pasteur, Paris | Marion Barbet, Sylvie Behillil, Frédéric Lemoine, Corinne Maufrais, Christophe Malabat, Damien Mornico, Méline Bizard, Angela Brisebarre, Camille Capel, Louise Lefrançois, Etienne Simon-Lorière, Vincent Enouf, Maud Vanpeene, Sylvie van der Werf,Odile Petsaris |
| EPI_ISL_1655919 | Hospital | National Reference Center for Viruses of Respiratory Infections, Institut Pasteur, Paris | Marion Barbet, Sylvie Behillil, Frédéric Lemoine, Corinne Maufrais, Christophe Malabat, Damien Mornico, Méline Bizard, Angela Brisebarre, Camille Capel, Louise Lefrançois, Etienne Simon-Lorière, Vincent Enouf, Maud Vanpeene, Sylvie van der Werf,Mj Dufour |
| EPI_ISL_1655920, EPI_ISL_1655921 | Hospital | National Reference Center for Viruses of Respiratory Infections, Institut Pasteur, Paris | Marion Barbet, Sylvie Behillil, Frédéric Lemoine, Corinne Maufrais, Christophe Malabat, Damien Mornico, Méline Bizard, Angela Brisebarre, Camille Capel, Louise Lefrançois, Etienne Simon-Lorière, Vincent Enouf, Maud Vanpeene, Sylvie van der Werf,Odile Petsaris |
| EPI_ISL_1655922 | Labo analyses Med | National Reference Center for Viruses of Respiratory Infections, Institut Pasteur, Paris | Marion Barbet, Sylvie Behillil, Frédéric Lemoine, Corinne Maufrais, Christophe Malabat, Damien Mornico, Méline Bizard, Angela Brisebarre, Camille Capel, Louise Lefrançois, Etienne Simon-Lorière, Vincent Enouf, Maud Vanpeene, Sylvie van der Werf,Anne Holstein |
| EPI_ISL_1655923, EPI_ISL_1655924, EPI_ISL_1655925, EPI_ISL_1655926, EPI_ISL_1655927 | Labo Analyses Med | National Reference Center for Viruses of Respiratory Infections, Institut Pasteur, Paris | Marion Barbet, Sylvie Behillil, Frédéric Lemoine, Corinne Maufrais, Christophe Malabat, Adrien Pain, Méline Bizard, Angela Brisebarre, Camille Capel, Louise Lefrançois, Etienne Simon-Lorière, Vincent Enouf, Maud Vanpeene, Sylvie van der Werf,Pierre-Yves Leonard |
| EPI_ISL_1655928 | Hospital | National Reference Center for Viruses of Respiratory Infections, Institut Pasteur, Paris | Marion Barbet, Sylvie Behillil, Frédéric Lemoine, Corinne Maufrais, Christophe Malabat, Damien Mornico, Méline Bizard, Angela Brisebarre, Camille Capel, Louise Lefrançois, Etienne Simon-Lorière, Vincent Enouf, Maud Vanpeene, Sylvie van der Werf,Nabil Gastli |
| EPI_ISL_1655929 | Labo Analyses Med | National Reference Center for Viruses of Respiratory Infections, Institut Pasteur, Paris | Marion Barbet, Sylvie Behillil, Frédéric Lemoine, Corinne Maufrais, Christophe Malabat, Damien Mornico, Méline Bizard, Angela Brisebarre, Camille Capel, Louise Lefrançois, Etienne Simon-Lorière, Vincent Enouf, Maud Vanpeene, Sylvie van der Werf,Pierre-Yves Leonard |
| EPI_ISL_1655930 | Hospital | National Reference Center for Viruses of Respiratory Infections, Institut Pasteur, Paris | Marion Barbet, Sylvie Behillil, Frédéric Lemoine, Corinne Maufrais, Christophe Malabat, Méline Bizard, Angela Brisebarre, Camille Capel, Louise Lefrançois, Etienne Simon-Lorière, Vincent Enouf, Maud Vanpeene, Sylvie van der Werf,BenoïT Bergues |
| EPI_ISL_1655931 | Labo Analyses Med | National Reference Center for Viruses of Respiratory Infections, Institut Pasteur, Paris | Marion Barbet, Sylvie Behillil, Frédéric Lemoine, Corinne Maufrais, Christophe Malabat, Damien Mornico, Méline Bizard, Angela Brisebarre, Camille Capel, Louise Lefrançois, Etienne Simon-Lorière, Vincent Enouf, Maud Vanpeene, Sylvie van der Werf |
| EPI_ISL_1655932, EPI_ISL_1655933 | Hospital | National Reference Center for Viruses of Respiratory Infections, Institut Pasteur, Paris | Marion Barbet, Sylvie Behillil, Frédéric Lemoine, Corinne Maufrais, Christophe Malabat, Adrien Pain, Méline Bizard, Angela Brisebarre, Camille Capel, Louise Lefrançois, Etienne Simon-Lorière, Vincent Enouf, Maud Vanpeene, Sylvie van der Werf,Ben Hadj Yahia |
| EPI_ISL_1655935 | Department of Virology, Istituto Zooprofilattico Sperimentale del Lazio e della Toscana (IZSLT) | Department of General Diagnostics; Department of Virology; Istituto Zooprofilattico Sperimentale del Lazio e della Toscana (IZSLT) | Patricia Alba, Giuseppe Manna, Elena L. Diaconu, Fabiola Feltrin, Raffaela Conti, Teresa Scicluna, Virginia Carfora, Antonella Cersini, Alessia Franco, Antonio Battisti. |
| EPI_ISL_1656844, EPI_ISL_1656845, EPI_ISL_1656846, EPI_ISL_1656847, EPI_ISL_1656848, EPI_ISL_1656849, EPI_ISL_1656850, EPI_ISL_1656851, EPI_ISL_1656852, EPI_ISL_1656855, EPI_ISL_1656856, EPI_ISL_1656857, EPI_ISL_1656858, EPI_ISL_1656860, EPI_ISL_1656861, EPI_ISL_1656862, EPI_ISL_1656864, EPI_ISL_1656865, EPI_ISL_1656866, EPI_ISL_1656867, EPI_ISL_1656868, EPI_ISL_1656869, EPI_ISL_1656870, EPI_ISL_1656871, EPI_ISL_1656872, EPI_ISL_1656873, EPI_ISL_1656874, EPI_ISL_1656875, EPI_ISL_1656876, EPI_ISL_1656877, EPI_ISL_1656878, EPI_ISL_1656879, EPI_ISL_1656880, EPI_ISL_1656881, EPI_ISL_1656882 | | | |
| see above | NVSPL | National Public Health Surveillance Laboratory | Lukas Zemaitis, Ingrida Olendraite, Arnoldas Pautienius, Kamile Tamusauskaite, Dovydas Gecys, Laura Pareckaite, Vaiva Lesauskaite, Astra Vitkauskiene |
| EPI_ISL_1656883, EPI_ISL_1656884, EPI_ISL_1656885, EPI_ISL_1656886, EPI_ISL_1656887, EPI_ISL_1656888, EPI_ISL_1656889, EPI_ISL_1656890, EPI_ISL_1656891, EPI_ISL_1656893, EPI_ISL_1656894, EPI_ISL_1656895, EPI_ISL_1656896, EPI_ISL_1656897, EPI_ISL_1656898, EPI_ISL_1656899, EPI_ISL_1656900, EPI_ISL_1656901, EPI_ISL_1656902, EPI_ISL_1656903, EPI_ISL_1656904, EPI_ISL_1656905, EPI_ISL_1656906, EPI_ISL_1656907, EPI_ISL_1656908, EPI_ISL_1656909, EPI_ISL_1656910, EPI_ISL_1656911, EPI_ISL_1656912, EPI_ISL_1656913, EPI_ISL_1656914, EPI_ISL_1656915, EPI_ISL_1656916, EPI_ISL_1656917, EPI_ISL_1656918, EPI_ISL_1656919, EPI_ISL_1656920, EPI_ISL_1656921, EPI_ISL_1656922, EPI_ISL_1656923, EPI_ISL_1656924, EPI_ISL_1656925, EPI_ISL_1656926, EPI_ISL_1656927, EPI_ISL_1656928, EPI_ISL_1656929, EPI_ISL_1656930, EPI_ISL_1656931, EPI_ISL_1656932, EPI_ISL_1656933, EPI_ISL_1656934, EPI_ISL_1656935, EPI_ISL_1656936, EPI_ISL_1656939, EPI_ISL_1656940, EPI_ISL_1656941, EPI_ISL_1656942, EPI_ISL_1656943, EPI_ISL_1656944, EPI_ISL_1656945, EPI_ISL_1656946, EPI_ISL_1656947, EPI_ISL_1656948, EPI_ISL_1656949, EPI_ISL_1656950, EPI_ISL_1656951, EPI_ISL_1656952, EPI_ISL_1656953, EPI_ISL_1656954, EPI_ISL_1656955, EPI_ISL_1656956, EPI_ISL_1656957, EPI_ISL_1656958, EPI_ISL_1656959, EPI_ISL_1656960, EPI_ISL_1656961, EPI_ISL_1656962, EPI_ISL_1656963, EPI_ISL_1656964, EPI_ISL_1656965, EPI_ISL_1656966, EPI_ISL_1656967, EPI_ISL_1656968, EPI_ISL_1656969, EPI_ISL_1656970, EPI_ISL_1656971, EPI_ISL_1656972, EPI_ISL_1656974, EPI_ISL_1656975, EPI_ISL_1656976, EPI_ISL_1656977, EPI_ISL_1656978, EPI_ISL_1656979, EPI_ISL_1656980, EPI_ISL_1656981, EPI_ISL_1656982, EPI_ISL_1656983, EPI_ISL_1656984, EPI_ISL_1656985, EPI_ISL_1656986, EPI_ISL_1656987, EPI_ISL_1656988, EPI_ISL_1656989, EPI_ISL_1656990, EPI_ISL_1656991, EPI_ISL_1656992, EPI_ISL_1656993, EPI_ISL_1656994, EPI_ISL_1656995, EPI_ISL_1656996, EPI_ISL_1656997, EPI_ISL_1656998, EPI_ISL_1656999, EPI_ISL_1657000, EPI_ISL_1657001, EPI_ISL_1657002, EPI_ISL_1657003, EPI_ISL_1657004, EPI_ISL_1657005, EPI_ISL_1657007, EPI_ISL_1657008, EPI_ISL_1657009, EPI_ISL_1657010, EPI_ISL_1657011, EPI_ISL_1657012, EPI_ISL_1657013, EPI_ISL_1657014, EPI_ISL_1657015, EPI_ISL_1657016, EPI_ISL_1657017, EPI_ISL_1657018, EPI_ISL_1657019, EPI_ISL_1657020, EPI_ISL_1657021, EPI_ISL_1657022, EPI_ISL_1657023, EPI_ISL_1657024 | | | |
| see above | NMVRVI | National Public Health Surveillance Laboratory | Lukas Zemaitis, Ingrida Olendraite, Arnoldas Pautienius, Kamile Tamusauskaite, Dovydas Gecys, Laura Pareckaite, Vaiva Lesauskaite, Astra Vitkauskiene |

| | | | |
|---|---|---|---|
| EPI_ISL_1657089, EPI_ISL_1657090, EPI_ISL_1657091, EPI_ISL_1657093, EPI_ISL_1657096, EPI_ISL_1657097 | National Institute of Infectious Diseases-Prof. Dr. Matei Bals Molecular Diagnostics Laboratory | National Institute of Infectious Diseases-Prof. Dr. Matei Bals Molecular Diagnostics Laboratory | Corina Casangiu, Leontina Banica, Marius Surleac, Ovidiu Vlaicu, Andreea Tudor, Simona Paraschiv, Dan Otelea |
| EPI_ISL_1658432, EPI_ISL_1658433, EPI_ISL_1658434, EPI_ISL_1658435, EPI_ISL_1658436, EPI_ISL_1658437, EPI_ISL_1658438, EPI_ISL_1658439, EPI_ISL_1658440, EPI_ISL_1658441 | IZSM | IZSM-U.O.C. Virologia | Maurizio Viscardi, Lorena Cardillo,Giovanna Fusco, Esterina De Carlo, Antonio Limone |
| EPI_ISL_1661474, EPI_ISL_1661475, EPI_ISL_1661476, EPI_ISL_1661477, EPI_ISL_1661479, EPI_ISL_1661480 | LESP Morelos | Instituto de Diagnostico y Referencia Epidemiologicos (INDRE) | Claudia Wong-Arambula, Abril Rodriguez-Maldonado, Vanessa Rivero-Arredondo, Ariadna Medina-Benitez, Joaquin Quiroz-Mercado, Sergio Rangel-Guerrero, Natividad Cruz-Ortiz, Tatiana Nunez-Garcia, Gisela Barrera-Badillo, Lucia Hernandez-Rivas, Irma Lopez-Martinez, Ernesto Ramirez-Gonzalez. |
| EPI_ISL_1661481, EPI_ISL_1661482, EPI_ISL_1661483, EPI_ISL_1661484, EPI_ISL_1661485, EPI_ISL_1661486, EPI_ISL_1661487, EPI_ISL_1661488, EPI_ISL_1661489 | LESP Chihuahua | Instituto de Diagnostico y Referencia Epidemiologicos (INDRE) | Claudia Wong-Arambula, Abril Rodriguez-Maldonado, Vanessa Rivero-Arredondo, Ariadna Medina-Benitez, Joaquin Quiroz-Mercado, Sergio Rangel-Guerrero, Natividad Cruz-Ortiz, Tatiana Nunez-Garcia, Gisela Barrera-Badillo, Lucia Hernandez-Rivas, Irma Lopez-Martinez, Ernesto Ramirez-Gonzalez. |
| EPI_ISL_1661491 | LESP Estado de Mexico | Instituto de Diagnostico y Referencia Epidemiologicos (INDRE) | Claudia Wong-Arambula, Abril Rodriguez-Maldonado, Vanessa Rivero-Arredondo, Ariadna Medina-Benitez, Joaquin Quiroz-Mercado, Sergio Rangel-Guerrero, Natividad Cruz-Ortiz, Tatiana Nunez-Garcia, Gisela Barrera-Badillo, Lucia Hernandez-Rivas, Irma Lopez-Martinez, Ernesto Ramirez-Gonzalez. |
| EPI_ISL_1661495, EPI_ISL_1661499, EPI_ISL_1661500, EPI_ISL_1661504, EPI_ISL_1661505, EPI_ISL_1661506, EPI_ISL_1661507, EPI_ISL_1661508, EPI_ISL_1661509, EPI_ISL_1661510, EPI_ISL_1661511, EPI_ISL_1661512, EPI_ISL_1661513, EPI_ISL_1661514, EPI_ISL_1661515, EPI_ISL_1661516, EPI_ISL_1661517, EPI_ISL_1661518, EPI_ISL_1661519, EPI_ISL_1661520 | | | |
| see above | LESP Guanajuato | Instituto de Diagnostico y Referencia Epidemiologicos (INDRE) | Claudia Wong-Arambula, Abril Rodriguez-Maldonado, Vanessa Rivero-Arredondo, Ariadna Medina-Benitez, Joaquin Quiroz-Mercado, Sergio Rangel-Guerrero, Natividad Cruz-Ortiz, Tatiana Nunez-Garcia, Gisela Barrera-Badillo, Lucia Hernandez-Rivas, Irma Lopez-Martinez, Ernesto Ramirez-Gonzalez. |
| EPI_ISL_1661521 | LESP Estado de Mexico | Instituto de Diagnostico y Referencia Epidemiologicos (INDRE) | Claudia Wong-Arambula, Abril Rodriguez-Maldonado, Vanessa Rivero-Arredondo, Ariadna Medina-Benitez, Joaquin Quiroz-Mercado, Sergio Rangel-Guerrero, Natividad Cruz-Ortiz, Tatiana Nunez-Garcia, Gisela Barrera-Badillo, Lucia Hernandez-Rivas, Irma Lopez-Martinez, Ernesto Ramirez-Gonzalez. |
| EPI_ISL_1661522, EPI_ISL_1661523, EPI_ISL_1661524, EPI_ISL_1661525, EPI_ISL_1661526, EPI_ISL_1661527, EPI_ISL_1661528, EPI_ISL_1661529, EPI_ISL_1661530, EPI_ISL_1661531, EPI_ISL_1661532, EPI_ISL_1661533 | | | |
| see above | LESP Queretaro | Instituto de Diagnostico y Referencia Epidemiologicos (INDRE) | Claudia Wong-Arambula, Abril Rodriguez-Maldonado, Vanessa Rivero-Arredondo, Ariadna Medina-Benitez, Joaquin Quiroz-Mercado, Sergio Rangel-Guerrero, Natividad Cruz-Ortiz, Tatiana Nunez-Garcia, Gisela Barrera-Badillo, Lucia Hernandez-Rivas, Irma Lopez-Martinez, Ernesto Ramirez-Gonzalez. |
| EPI_ISL_1661534 | LESP Hidalgo | Instituto de Diagnostico y Referencia Epidemiologicos (INDRE) | Claudia Wong-Arambula, Abril Rodriguez-Maldonado, Vanessa Rivero-Arredondo, Ariadna Medina-Benitez, Joaquin Quiroz-Mercado, Sergio Rangel-Guerrero, Natividad Cruz-Ortiz, Tatiana Nunez-Garcia, Gisela Barrera-Badillo, Lucia Hernandez-Rivas, Irma Lopez-Martinez, Ernesto Ramirez-Gonzalez. |
| EPI_ISL_1661535 | LESP Puebla | Instituto de Diagnostico y Referencia Epidemiologicos (INDRE) | Claudia Wong-Arambula, Abril Rodriguez-Maldonado, Vanessa Rivero-Arredondo, Ariadna Medina-Benitez, Joaquin Quiroz-Mercado, Sergio Rangel-Guerrero, Natividad Cruz-Ortiz, Tatiana Nunez-Garcia, Gisela Barrera-Badillo, Lucia Hernandez-Rivas, Irma Lopez-Martinez, Ernesto Ramirez-Gonzalez. |
| EPI_ISL_1661536, EPI_ISL_1661537, EPI_ISL_1661538 | LESPZacatecas | Instituto de Diagnostico y Referencia Epidemiologicos (INDRE) | Claudia Wong-Arambula, Abril Rodriguez-Maldonado, Vanessa Rivero-Arredondo, Ariadna Medina-Benitez, Joaquin Quiroz-Mercado, Sergio Rangel-Guerrero, Natividad Cruz-Ortiz, Tatiana Nunez-Garcia, Gisela Barrera-Badillo, Lucia Hernandez-Rivas, Irma Lopez-Martinez, Ernesto Ramirez-Gonzalez. |
| EPI_ISL_1661539, EPI_ISL_1661540, EPI_ISL_1661541 | LESP Hidalgo | Instituto de Diagnostico y Referencia Epidemiologicos (INDRE) | Claudia Wong-Arambula, Abril Rodriguez-Maldonado, Vanessa Rivero-Arredondo, Ariadna Medina-Benitez, Joaquin Quiroz-Mercado, Sergio Rangel-Guerrero, Natividad Cruz-Ortiz, Tatiana Nunez-Garcia, Gisela Barrera-Badillo, Lucia Hernandez-Rivas, Irma Lopez-Martinez, Ernesto Ramirez-Gonzalez. |
| EPI_ISL_1661542, EPI_ISL_1661543 | LESP Morelos | Instituto de Diagnostico y Referencia Epidemiologicos (INDRE) | Claudia Wong-Arambula, Abril Rodriguez-Maldonado, Vanessa Rivero-Arredondo, Ariadna Medina-Benitez, Joaquin Quiroz-Mercado, Sergio Rangel-Guerrero, Natividad Cruz-Ortiz, Tatiana Nunez-Garcia, Gisela Barrera-Badillo, Lucia Hernandez-Rivas, Irma Lopez-Martinez, Ernesto Ramirez-Gonzalez. |
| EPI_ISL_1661544, EPI_ISL_1661545, EPI_ISL_1661546, EPI_ISL_1661547 | LESPZacatecas | Instituto de Diagnostico y Referencia Epidemiologicos (INDRE) | Claudia Wong-Arambula, Abril Rodriguez-Maldonado, Vanessa Rivero-Arredondo, Ariadna Medina-Benitez, Joaquin Quiroz-Mercado, Sergio Rangel-Guerrero, Natividad Cruz-Ortiz, Tatiana Nunez-Garcia, Gisela Barrera-Badillo, Lucia Hernandez-Rivas, Irma Lopez-Martinez, Ernesto Ramirez-Gonzalez. |
| EPI_ISL_1661551, EPI_ISL_1661552 | LESP Hidalgo | Instituto de Diagnostico y Referencia Epidemiologicos (INDRE) | Claudia Wong-Arambula, Abril Rodriguez-Maldonado, Vanessa Rivero-Arredondo, Ariadna Medina-Benitez, Joaquin Quiroz-Mercado, Sergio Rangel-Guerrero, Natividad Cruz-Ortiz, Tatiana Nunez-Garcia, Gisela Barrera-Badillo, Lucia Hernandez-Rivas, Irma Lopez-Martinez, Ernesto Ramirez-Gonzalez. |
| EPI_ISL_1661553, EPI_ISL_1661554, EPI_ISL_1661555, EPI_ISL_1661556 | LESP Nuevo Leon | Instituto de Diagnostico y Referencia Epidemiologicos (INDRE) | Claudia Wong-Arambula, Abril Rodriguez-Maldonado, Vanessa Rivero-Arredondo, Ariadna Medina-Benitez, Joaquin Quiroz-Mercado, Sergio Rangel-Guerrero, Natividad Cruz-Ortiz, Tatiana Nunez-Garcia, Gisela Barrera-Badillo, Lucia Hernandez-Rivas, Irma Lopez-Martinez, Ernesto Ramirez-Gonzalez. |
| EPI_ISL_1661557 | LESP San Luis Potosi | Instituto de Diagnostico y Referencia Epidemiologicos (INDRE) | Claudia Wong-Arambula, Abril Rodriguez-Maldonado, Vanessa Rivero-Arredondo, Ariadna Medina-Benitez, Joaquin Quiroz-Mercado, Sergio Rangel-Guerrero, Natividad Cruz-Ortiz, Tatiana Nunez-Garcia, Gisela Barrera-Badillo, Lucia Hernandez-Rivas, Irma Lopez-Martinez, Ernesto Ramirez-Gonzalez. |
| EPI_ISL_1661558, EPI_ISL_1661559, EPI_ISL_1661560, EPI_ISL_1661561, EPI_ISL_1661562, EPI_ISL_1661563, EPI_ISL_1661564 | LESP Nuevo Leon | Instituto de Diagnostico y Referencia Epidemiologicos (INDRE) | Claudia Wong-Arambula, Abril Rodriguez-Maldonado, Vanessa Rivero-Arredondo, Ariadna Medina-Benitez, Joaquin Quiroz-Mercado, Sergio Rangel-Guerrero, Natividad Cruz-Ortiz, Tatiana Nunez-Garcia, Gisela Barrera-Badillo, Lucia Hernandez-Rivas, Irma Lopez-Martinez, Ernesto Ramirez-Gonzalez. |
| EPI_ISL_1661565, EPI_ISL_1661566, EPI_ISL_1661567, EPI_ISL_1661568, EPI_ISL_1661569 | LESP Guanajuato | Instituto de Diagnostico y Referencia Epidemiologicos (INDRE) | Claudia Wong-Arambula, Abril Rodriguez-Maldonado, Vanessa Rivero-Arredondo, Ariadna Medina-Benitez, Joaquin Quiroz-Mercado, Sergio Rangel-Guerrero, Natividad Cruz-Ortiz, Tatiana Nunez-Garcia, Gisela Barrera-Badillo, Lucia Hernandez-Rivas, Irma Lopez-Martinez, Ernesto Ramirez-Gonzalez. |
| EPI_ISL_1661570 | LESP Guerrero | Instituto de Diagnostico y Referencia Epidemiologicos (INDRE) | Claudia Wong-Arambula, Abril Rodriguez-Maldonado, Vanessa Rivero-Arredondo, Ariadna Medina-Benitez, Joaquin Quiroz-Mercado, Sergio Rangel-Guerrero, Natividad Cruz-Ortiz, Tatiana Nunez-Garcia, Gisela Barrera-Badillo, Lucia Hernandez-Rivas, Irma Lopez-Martinez, Ernesto Ramirez-Gonzalez. |
| EPI_ISL_1661919 | IU-Cerrahpasa, Cerrahpasa School of Medicine, COVID-19 Lab | IU-Cerrahpasa, Cerrahpasa School of Medicine, COVID-19 Lab | Mert Kuskucu, Yesim Tuyji Tok, Kenan Midilli |
| EPI_ISL_1661977, EPI_ISL_1661978 | Laboratorio Central de Epidemiologia (LCE) | Unidad de Genomica Avanzada | Consorcio Mexicano de Vigilancia Genomica (CoViGen-Mex). Authors (in alphabetical order): Julio Elias Alvarado-Yaah, Carlos F. Arias, Santiago Ávila-Rios, Victor Hugo Borja-Aburto, Celia Boukadida, Juan Bautista Chale-Dzul, Celida Duque Molina, Jose Antonio Enciso-Moreno, Gloria Elena Espinosa-Ayala, Fernando Fontove-Herrera, Victor Eduardo Garcia-Arias,Concepcion Grajales-Muniz, Ricardo Grande, Alfredo Herrera-Estrella, Carla Ivon Herrera-Najera, Pavel Isa, Brenda Irasema Maldonado-Meza, Bernardo Martinez-Miguel, Margarita Matias-Florentino, Maria Guadalupe de Jesus Mireles-Rivera, Gloria Maria Molina-Salinas, Hector Montoya-Fuentes, Jose Esteban Munoz-Medina, Jose de Jesus Nunez-Contreras, Alicia Ocana-Mondragon, Luis Alberto Ochoa-Carrera, Hector Esteban Paz-Juarez, Francisco Pulido, Helen Haydee Fernanda Ramirez-Plascencia, Angel |

| | | | |
|---|---|---|---|
| | | | Gustavo Salas-Lais, Alejandro Sanchez-Flores, Clara Esperanza Santacruz-Tinoco, Maria Guadalupe Santiago-Mauricio, Nelly Selem-Mojica, Blanca Taboada, Gloria Vazquez |
| EPI_ISL_1661979 | Centro de Investigacion Biomedica de Occidente (CIBO) | Unidad de Genomica Avanzada | Consorcio Mexicano de Vigilancia Genomica (CoViGen-Mex). Authors (in alphabetical order): Julio Elias Alvarado-Yaah, Carlos F. Arias, Santiago Ávila-Rios, Victor Hugo Borja-Aburto, Celia Boukadida, Juan Bautista Chale-Dzul, Celida Duque Molina, Jose Antonio Enciso-Moreno, Gloria Elena Espinosa-Ayala, Fernando Fontove-Herrera, Victor Eduardo Garcia-Arias,Concepcion Grajales-Muniz, Ricardo Grande, Alfredo Herrera-Estrella, Carla Ivon Herrera-Najera, Pavel Isa, Brenda Irasema Maldonado-Meza, Bernardo Martinez-Miguel, Margarita Matias-Florentino, Maria Guadalupe de Jesus Mireles-Rivera, Gloria Maria Molina-Salinas, Hector Montoya-Fuentes, Jose Esteban Munoz-Medina, Jose de Jesus Nunez-Contreras, Alicia Ocana-Mondragon, Luis Alberto Ochoa-Carrera, Hector Esteban Paz-Juarez, Francisco Pulido, Helen Haydee Fernanda Ramirez-Plascencia, Angel Gustavo Salas-Lais, Alejandro Sanchez-Flores, Clara Esperanza Santacruz-Tinoco, Maria Guadalupe Santiago-Mauricio, Nelly Selem-Mojica, Blanca Taboada, Gloria Vazquez |
| EPI_ISL_1661980 | Laboratorio Central de Epidemiologia (LCE) | Unidad de Genomica Avanzada | Consorcio Mexicano de Vigilancia Genomica (CoViGen-Mex). Authors (in alphabetical order): Julio Elias Alvarado-Yaah, Carlos F. Arias, Santiago Ávila-Rios, Victor Hugo Borja-Aburto, Celia Boukadida, Juan Bautista Chale-Dzul, Celida Duque Molina, Jose Antonio Enciso-Moreno, Gloria Elena Espinosa-Ayala, Fernando Fontove-Herrera, Victor Eduardo Garcia-Arias,Concepcion Grajales-Muniz, Ricardo Grande, Alfredo Herrera-Estrella, Carla Ivon Herrera-Najera, Pavel Isa, Brenda Irasema Maldonado-Meza, Bernardo Martinez-Miguel, Margarita Matias-Florentino, Maria Guadalupe de Jesus Mireles-Rivera, Gloria Maria Molina-Salinas, Hector Montoya-Fuentes, Jose Esteban Munoz-Medina, Jose de Jesus Nunez-Contreras, Alicia Ocana-Mondragon, Luis Alberto Ochoa-Carrera, Hector Esteban Paz-Juarez, Francisco Pulido, Helen Haydee Fernanda Ramirez-Plascencia, Angel Gustavo Salas-Lais, Alejandro Sanchez-Flores, Clara Esperanza Santacruz-Tinoco, Maria Guadalupe Santiago-Mauricio, Nelly Selem-Mojica, Blanca Taboada, Gloria Vazquez |
| EPI_ISL_1661982, EPI_ISL_1661983, EPI_ISL_1661984, EPI_ISL_1661985, EPI_ISL_1661986, EPI_ISL_1661988, EPI_ISL_1661990 | Centro de Investigacion Biomedica del Noreste (CIBIN) | Unidad de Genomica Avanzada | Consorcio Mexicano de Vigilancia Genomica (CoViGen-Mex). Authors (in alphabetical order): Julio Elias Alvarado-Yaah, Carlos F. Arias, Santiago Ávila-Rios, Victor Hugo Borja-Aburto, Celia Boukadida, Juan Bautista Chale-Dzul, Celida Duque Molina, Jose Antonio Enciso-Moreno, Gloria Elena Espinosa-Ayala, Fernando Fontove-Herrera, Victor Eduardo Garcia-Arias,Concepcion Grajales-Muniz, Ricardo Grande, Alfredo Herrera-Estrella, Carla Ivon Herrera-Najera, Pavel Isa, Brenda Irasema Maldonado-Meza, Bernardo Martinez-Miguel, Margarita Matias-Florentino, Maria Guadalupe de Jesus Mireles-Rivera, Gloria Maria Molina-Salinas, Hector Montoya-Fuentes, Jose Esteban Munoz-Medina, Jose de Jesus Nunez-Contreras, Alicia Ocana-Mondragon, Luis Alberto Ochoa-Carrera, Hector Esteban Paz-Juarez, Francisco Pulido, Helen Haydee Fernanda Ramirez-Plascencia, Angel Gustavo Salas-Lais, Alejandro Sanchez-Flores, Clara Esperanza Santacruz-Tinoco, Maria Guadalupe Santiago-Mauricio, Nelly Selem-Mojica, Blanca Taboada, Gloria Vazquez |
| EPI_ISL_1661991, EPI_ISL_1661992, EPI_ISL_1661993, EPI_ISL_1661994, EPI_ISL_1661995, EPI_ISL_1661996, EPI_ISL_1661997, EPI_ISL_1661998, EPI_ISL_1661999, EPI_ISL_1662000, EPI_ISL_1662001, EPI_ISL_1662002, EPI_ISL_1662003, EPI_ISL_1662004, EPI_ISL_1662005, EPI_ISL_1662006, EPI_ISL_1662007, EPI_ISL_1662009, EPI_ISL_1662010, EPI_ISL_1662011, EPI_ISL_1662012, EPI_ISL_1662013, EPI_ISL_1662014, EPI_ISL_1662015, EPI_ISL_1662016, EPI_ISL_1662017, EPI_ISL_1662018, EPI_ISL_1662019, EPI_ISL_1662021, EPI_ISL_1662022, EPI_ISL_1662023, EPI_ISL_1662027, EPI_ISL_1662031, EPI_ISL_1662032, EPI_ISL_1662033 | | | |
| see above | Laboratorio Central de Epidemiologia (LCE) | Unidad de Genomica Avanzada | Consorcio Mexicano de Vigilancia Genomica (CoViGen-Mex). Authors (in alphabetical order): Julio Elias Alvarado-Yaah, Carlos F. Arias, Santiago Ávila-Rios, Victor Hugo Borja-Aburto, Celia Boukadida, Juan Bautista Chale-Dzul, Celida Duque Molina, Jose Antonio Enciso-Moreno, Gloria Elena Espinosa-Ayala, Fernando Fontove-Herrera, Victor Eduardo Garcia-Arias,Concepcion Grajales-Muniz, Ricardo Grande, Alfredo Herrera-Estrella, Carla Ivon Herrera-Najera, Pavel Isa, Brenda Irasema Maldonado-Meza, Bernardo Martinez-Miguel, Margarita Matias-Florentino, Maria Guadalupe de Jesus Mireles-Rivera, Gloria Maria Molina-Salinas, Hector Montoya-Fuentes, Jose Esteban Munoz-Medina, Jose de Jesus Nunez-Contreras, Alicia Ocana-Mondragon, Luis Alberto Ochoa-Carrera, Hector Esteban Paz-Juarez, Francisco Pulido, Helen Haydee Fernanda Ramirez-Plascencia, Angel Gustavo Salas-Lais, Alejandro Sanchez-Flores, Clara Esperanza Santacruz-Tinoco, Maria Guadalupe Santiago-Mauricio, Nelly Selem-Mojica, Blanca Taboada, Gloria Vazquez |
| EPI_ISL_1662035, EPI_ISL_1662036, EPI_ISL_1662037, EPI_ISL_1662038, EPI_ISL_1662043, EPI_ISL_1662044, EPI_ISL_1662045, EPI_ISL_1662046, EPI_ISL_1662052, EPI_ISL_1662053, EPI_ISL_1662054, EPI_ISL_1662055, EPI_ISL_1662057 | | | |
| see above | Centro de Investigacion Biomedica del Noreste (CIBIN) | Unidad de Genomica Avanzada | Consorcio Mexicano de Vigilancia Genomica (CoViGen-Mex). Authors (in alphabetical order): Julio Elias Alvarado-Yaah, Carlos F. Arias, Santiago Ávila-Rios, Victor Hugo Borja-Aburto, Celia Boukadida, Juan Bautista Chale-Dzul, Celida Duque Molina, Jose Antonio Enciso-Moreno, Gloria Elena Espinosa-Ayala, Fernando Fontove-Herrera, Victor Eduardo Garcia-Arias,Concepcion Grajales-Muniz, Ricardo Grande, Alfredo Herrera-Estrella, Carla Ivon Herrera-Najera, Pavel Isa, Brenda Irasema Maldonado-Meza, Bernardo Martinez-Miguel, Margarita Matias-Florentino, Maria Guadalupe de Jesus Mireles-Rivera, Gloria Maria Molina-Salinas, Hector Montoya-Fuentes, Jose Esteban Munoz-Medina, Jose de Jesus Nunez-Contreras, Alicia Ocana-Mondragon, Luis Alberto Ochoa-Carrera, Hector Esteban Paz-Juarez, Francisco Pulido, Helen Haydee Fernanda Ramirez-Plascencia, Angel Gustavo Salas-Lais, Alejandro Sanchez-Flores, Clara Esperanza Santacruz-Tinoco, Maria Guadalupe Santiago-Mauricio, Nelly Selem-Mojica, Blanca Taboada, Gloria Vazquez |
| EPI_ISL_1662058, EPI_ISL_1662060, EPI_ISL_1662061 | Laboratorio Central de Epidemiologia (LCE) | Unidad de Genomica Avanzada | Consorcio Mexicano de Vigilancia Genomica (CoViGen-Mex). Authors (in alphabetical order): Julio Elias Alvarado-Yaah, Carlos F. Arias, Santiago Ávila-Rios, Victor Hugo Borja-Aburto, Celia Boukadida, Juan Bautista Chale-Dzul, Celida Duque Molina, Jose Antonio Enciso-Moreno, Gloria Elena Espinosa-Ayala, Fernando Fontove-Herrera, Victor Eduardo Garcia-Arias,Concepcion Grajales-Muniz, Ricardo Grande, Alfredo Herrera-Estrella, Carla Ivon Herrera-Najera, Pavel Isa, Brenda Irasema Maldonado-Meza, Bernardo Martinez-Miguel, Margarita Matias-Florentino, Maria Guadalupe de Jesus Mireles-Rivera, Gloria Maria Molina-Salinas, Hector Montoya-Fuentes, Jose Esteban Munoz-Medina, Jose de Jesus Nunez-Contreras, Alicia Ocana-Mondragon, Luis Alberto Ochoa-Carrera, Hector Esteban Paz-Juarez, Francisco Pulido, Helen Haydee Fernanda Ramirez-Plascencia, Angel Gustavo Salas-Lais, Alejandro Sanchez-Flores, Clara Esperanza Santacruz-Tinoco, Maria Guadalupe Santiago-Mauricio, Nelly Selem-Mojica, Blanca Taboada, Gloria Vazquez |
| EPI_ISL_1662066, EPI_ISL_1662067, EPI_ISL_1662069 | Centro de Investigacion Biomedica de Occidente (CIBO) | Unidad de Genomica Avanzada | Consorcio Mexicano de Vigilancia Genomica (CoViGen-Mex). Authors (in alphabetical order): Julio Elias Alvarado-Yaah, Carlos F. Arias, Santiago Ávila-Rios, Victor Hugo Borja-Aburto, Celia Boukadida, Juan Bautista Chale-Dzul, Celida Duque Molina, Jose Antonio Enciso-Moreno, Gloria Elena Espinosa-Ayala, Fernando Fontove-Herrera, Victor Eduardo Garcia-Arias,Concepcion Grajales-Muniz, Ricardo Grande, Alfredo Herrera-Estrella, Carla Ivon Herrera-Najera, Pavel Isa, Brenda Irasema Maldonado-Meza, Bernardo Martinez-Miguel, Margarita Matias-Florentino, Maria Guadalupe de Jesus Mireles-Rivera, Gloria Maria Molina-Salinas, Hector Montoya-Fuentes, Jose Esteban Munoz-Medina, Jose de Jesus Nunez-Contreras, Alicia Ocana-Mondragon, Luis Alberto Ochoa-Carrera, Hector Esteban Paz-Juarez, Francisco Pulido, Helen Haydee Fernanda Ramirez-Plascencia, Angel Gustavo Salas-Lais, Alejandro Sanchez-Flores, Clara Esperanza Santacruz-Tinoco, Maria Guadalupe Santiago-Mauricio, Nelly Selem-Mojica, Blanca Taboada, Gloria Vazquez |
| EPI_ISL_1662070, EPI_ISL_1662071, EPI_ISL_1662073, EPI_ISL_1662074 | Laboratorio Central de Epidemiologia (LCE) | Unidad de Genomica Avanzada | Consorcio Mexicano de Vigilancia Genomica (CoViGen-Mex). Authors (in alphabetical order): Julio Elias Alvarado-Yaah, Carlos F. Arias, Santiago Ávila-Rios, Victor Hugo Borja-Aburto, Celia Boukadida, Juan Bautista Chale-Dzul, Celida Duque Molina, Jose Antonio Enciso-Moreno, Gloria Elena Espinosa-Ayala, Fernando Fontove-Herrera, Victor Eduardo Garcia-Arias,Concepcion Grajales-Muniz, Ricardo Grande, Alfredo Herrera-Estrella, Carla Ivon Herrera-Najera, Pavel Isa, Brenda Irasema Maldonado-Meza, Bernardo Martinez-Miguel, Margarita Matias-Florentino, Maria Guadalupe de Jesus Mireles-Rivera, Gloria Maria Molina-Salinas, Hector Montoya-Fuentes, Jose Esteban Munoz-Medina, Jose de Jesus Nunez-Contreras, Alicia Ocana-Mondragon, Luis Alberto Ochoa-Carrera, Hector Esteban Paz-Juarez, Francisco Pulido, Helen Haydee Fernanda Ramirez-Plascencia, Angel Gustavo Salas-Lais, Alejandro Sanchez-Flores, Clara Esperanza Santacruz-Tinoco, Maria Guadalupe Santiago-Mauricio, Nelly Selem-Mojica, Blanca Taboada, Gloria Vazquez |
| EPI_ISL_1662080, EPI_ISL_1662081, EPI_ISL_1662084, EPI_ISL_1662085, EPI_ISL_1662087, EPI_ISL_1662089, EPI_ISL_1662090, EPI_ISL_1662093, EPI_ISL_1662095, EPI_ISL_1662097, EPI_ISL_1662100, EPI_ISL_1662102, EPI_ISL_1662103 | | | |
| see above | Centro de Investigacion Biomedica de Occidente (CIBO) | Unidad de Genomica Avanzada | Consorcio Mexicano de Vigilancia Genomica (CoViGen-Mex). Authors (in alphabetical order): Julio Elias Alvarado-Yaah, Carlos F. Arias, Santiago Ávila-Rios, Victor Hugo Borja-Aburto, Celia Boukadida, Juan Bautista Chale-Dzul, Celida Duque Molina, Jose Antonio Enciso-Moreno, Gloria Elena Espinosa-Ayala, Fernando Fontove-Herrera, Victor Eduardo Garcia-Arias,Concepcion Grajales-Muniz, Ricardo Grande, Alfredo Herrera-Estrella, Carla Ivon Herrera-Najera, Pavel Isa, Brenda Irasema Maldonado-Meza, Bernardo Martinez-Miguel, Margarita Matias-Florentino, Maria Guadalupe de Jesus Mireles-Rivera, Gloria Maria Molina-Salinas, Hector Montoya-Fuentes, Jose Esteban Munoz-Medina, Jose de Jesus Nunez-Contreras, Alicia Ocana-Mondragon, Luis Alberto Ochoa-Carrera, Hector Esteban Paz-Juarez, Francisco Pulido, Helen Haydee Fernanda Ramirez-Plascencia, Angel Gustavo Salas-Lais, Alejandro Sanchez-Flores, Clara Esperanza Santacruz-Tinoco, Maria Guadalupe Santiago-Mauricio, Nelly Selem-Mojica, Blanca Taboada, Gloria Vazquez |
| EPI_ISL_1662104, EPI_ISL_1662106, EPI_ISL_1662107, EPI_ISL_1662109, EPI_ISL_1662110, EPI_ISL_1662111, EPI_ISL_1662112, EPI_ISL_1662113, EPI_ISL_1662114, EPI_ISL_1662116, EPI_ISL_1662117, EPI_ISL_1662119, EPI_ISL_1662120 | | | |

| | | | |
|---|---|---|---|
| see above | Laboratorio Central de Epidemiologia (LCE) | Unidad de Genomica Avanzada | Consorcio Mexicano de Vigilancia Genomica (CoViGen-Mex). Authors (in alphabetical order): Julio Elias Alvarado-Yaah, Carlos F. Arias, Santiago Ávila-Rios, Victor Hugo Borja-Aburto, Celia Boukadida, Juan Bautista Chale-Dzul, Celida Duque Molina, Jose Antonio Enciso-Moreno, Gloria Elena Espinosa-Ayala, Fernando Fontove-Herrera, Victor Eduardo Garcia-Arias,Concepcion Grajales-Muniz, Ricardo Grande, Alfredo Herrera-Estrella, Carla Ivon Herrera-Najera, Pavel Isa, Brenda Irasema Maldonado-Meza, Bernardo Martinez-Miguel, Margarita Matias-Florentino, Maria Guadalupe de Jesus Mireles-Rivera, Gloria Maria Molina-Salinas, Hector Montoya-Fuentes, Jose Esteban Munoz-Medina, Jose de Jesus Nunez-Contreras, Alicia Ocana-Mondragon, Luis Alberto Ochoa-Carrera, Hector Esteban Paz-Juarez, Francisco Pulido, Helen Haydee Fernanda Ramirez-Plascencia, Angel Gustavo Salas-Lais, Alejandro Sanchez-Flores, Clara Esperanza Santacruz-Tinoco, Maria Guadalupe Santiago-Mauricio, Nelly Selem-Mojica, Blanca Taboada, Gloria Vazquez |
| EPI_ISL_1662125, EPI_ISL_1662126 | Centro de Investigacion Biomedica de Occidente (CIBO) | Unidad de Genomica Avanzada | Consorcio Mexicano de Vigilancia Genomica (CoViGen-Mex). Authors (in alphabetical order): Julio Elias Alvarado-Yaah, Carlos F. Arias, Santiago Ávila-Rios, Victor Hugo Borja-Aburto, Celia Boukadida, Juan Bautista Chale-Dzul, Celida Duque Molina, Jose Antonio Enciso-Moreno, Gloria Elena Espinosa-Ayala, Fernando Fontove-Herrera, Victor Eduardo Garcia-Arias,Concepcion Grajales-Muniz, Ricardo Grande, Alfredo Herrera-Estrella, Carla Ivon Herrera-Najera, Pavel Isa, Brenda Irasema Maldonado-Meza, Bernardo Martinez-Miguel, Margarita Matias-Florentino, Maria Guadalupe de Jesus Mireles-Rivera, Gloria Maria Molina-Salinas, Hector Montoya-Fuentes, Jose Esteban Munoz-Medina, Jose de Jesus Nunez-Contreras, Alicia Ocana-Mondragon, Luis Alberto Ochoa-Carrera, Hector Esteban Paz-Juarez, Francisco Pulido, Helen Haydee Fernanda Ramirez-Plascencia, Angel Gustavo Salas-Lais, Alejandro Sanchez-Flores, Clara Esperanza Santacruz-Tinoco, Maria Guadalupe Santiago-Mauricio, Nelly Selem-Mojica, Blanca Taboada, Gloria Vazquez |
| EPI_ISL_1662128, EPI_ISL_1662129, EPI_ISL_1662130 | Laboratorio Central de Epidemiologia (LCE) | Unidad de Genomica Avanzada | Consorcio Mexicano de Vigilancia Genomica (CoViGen-Mex). Authors (in alphabetical order): Julio Elias Alvarado-Yaah, Carlos F. Arias, Santiago Ávila-Rios, Victor Hugo Borja-Aburto, Celia Boukadida, Juan Bautista Chale-Dzul, Celida Duque Molina, Jose Antonio Enciso-Moreno, Gloria Elena Espinosa-Ayala, Fernando Fontove-Herrera, Victor Eduardo Garcia-Arias,Concepcion Grajales-Muniz, Ricardo Grande, Alfredo Herrera-Estrella, Carla Ivon Herrera-Najera, Pavel Isa, Brenda Irasema Maldonado-Meza, Bernardo Martinez-Miguel, Margarita Matias-Florentino, Maria Guadalupe de Jesus Mireles-Rivera, Gloria Maria Molina-Salinas, Hector Montoya-Fuentes, Jose Esteban Munoz-Medina, Jose de Jesus Nunez-Contreras, Alicia Ocana-Mondragon, Luis Alberto Ochoa-Carrera, Hector Esteban Paz-Juarez, Francisco Pulido, Helen Haydee Fernanda Ramirez-Plascencia, Angel Gustavo Salas-Lais, Alejandro Sanchez-Flores, Clara Esperanza Santacruz-Tinoco, Maria Guadalupe Santiago-Mauricio, Nelly Selem-Mojica, Blanca Taboada, Gloria Vazquez |
| EPI_ISL_1662133, EPI_ISL_1662136, EPI_ISL_1662138, EPI_ISL_1662139, EPI_ISL_1662140, EPI_ISL_1662141, EPI_ISL_1662142, EPI_ISL_1662143, EPI_ISL_1662144 | Centro de Investigacion Biomedica del Noreste (CIBIN) | Unidad de Genomica Avanzada | Consorcio Mexicano de Vigilancia Genomica (CoViGen-Mex). Authors (in alphabetical order): Julio Elias Alvarado-Yaah, Carlos F. Arias, Santiago Ávila-Rios, Victor Hugo Borja-Aburto, Celia Boukadida, Juan Bautista Chale-Dzul, Celida Duque Molina, Jose Antonio Enciso-Moreno, Gloria Elena Espinosa-Ayala, Fernando Fontove-Herrera, Victor Eduardo Garcia-Arias,Concepcion Grajales-Muniz, Ricardo Grande, Alfredo Herrera-Estrella, Carla Ivon Herrera-Najera, Pavel Isa, Brenda Irasema Maldonado-Meza, Bernardo Martinez-Miguel, Margarita Matias-Florentino, Maria Guadalupe de Jesus Mireles-Rivera, Gloria Maria Molina-Salinas, Hector Montoya-Fuentes, Jose Esteban Munoz-Medina, Jose de Jesus Nunez-Contreras, Alicia Ocana-Mondragon, Luis Alberto Ochoa-Carrera, Hector Esteban Paz-Juarez, Francisco Pulido, Helen Haydee Fernanda Ramirez-Plascencia, Angel Gustavo Salas-Lais, Alejandro Sanchez-Flores, Clara Esperanza Santacruz-Tinoco, Maria Guadalupe Santiago-Mauricio, Nelly Selem-Mojica, Blanca Taboada, Gloria Vazquez |
| EPI_ISL_1662145, EPI_ISL_1662146, EPI_ISL_1662147, EPI_ISL_1662149, EPI_ISL_1662150, EPI_ISL_1662151, EPI_ISL_1662152, EPI_ISL_1662153, EPI_ISL_1662154, EPI_ISL_1662155, EPI_ISL_1662156, EPI_ISL_1662157, EPI_ISL_1662158, EPI_ISL_1662159, EPI_ISL_1662161, EPI_ISL_1662164, EPI_ISL_1662165, EPI_ISL_1662166, EPI_ISL_1662167, EPI_ISL_1662168, EPI_ISL_1662170, EPI_ISL_1662171, EPI_ISL_1662172, EPI_ISL_1662174, EPI_ISL_1662175, EPI_ISL_1662176, EPI_ISL_1662177, EPI_ISL_1662178, EPI_ISL_1662181, EPI_ISL_1662182, EPI_ISL_1662183 | | | |
| see above | Laboratorio Central de Epidemiologia (LCE) | Unidad de Genomica Avanzada | Consorcio Mexicano de Vigilancia Genomica (CoViGen-Mex). Authors (in alphabetical order): Julio Elias Alvarado-Yaah, Carlos F. Arias, Santiago Ávila-Rios, Victor Hugo Borja-Aburto, Celia Boukadida, Juan Bautista Chale-Dzul, Celida Duque Molina, Jose Antonio Enciso-Moreno, Gloria Elena Espinosa-Ayala, Fernando Fontove-Herrera, Victor Eduardo Garcia-Arias,Concepcion Grajales-Muniz, Ricardo Grande, Alfredo Herrera-Estrella, Carla Ivon Herrera-Najera, Pavel Isa, Brenda Irasema Maldonado-Meza, Bernardo Martinez-Miguel, Margarita Matias-Florentino, Maria Guadalupe de Jesus Mireles-Rivera, Gloria Maria Molina-Salinas, Hector Montoya-Fuentes, Jose Esteban Munoz-Medina, Jose de Jesus Nunez-Contreras, Alicia Ocana-Mondragon, Luis Alberto Ochoa-Carrera, Hector Esteban Paz-Juarez, Francisco Pulido, Helen Haydee Fernanda Ramirez-Plascencia, Angel Gustavo Salas-Lais, Alejandro Sanchez-Flores, Clara Esperanza Santacruz-Tinoco, Maria Guadalupe Santiago-Mauricio, Nelly Selem-Mojica, Blanca Taboada, Gloria Vazquez |
| EPI_ISL_1662185 | Unidad de Investigacion Medica de Yucatan (UIMY) | Unidad de Genomica Avanzada | Consorcio Mexicano de Vigilancia Genomica (CoViGen-Mex). Authors (in alphabetical order): Julio Elias Alvarado-Yaah, Carlos F. Arias, Santiago Ávila-Rios, Victor Hugo Borja-Aburto, Celia Boukadida, Juan Bautista Chale-Dzul, Celida Duque Molina, Jose Antonio Enciso-Moreno, Gloria Elena Espinosa-Ayala, Fernando Fontove-Herrera, Victor Eduardo Garcia-Arias,Concepcion Grajales-Muniz, Ricardo Grande, Alfredo Herrera-Estrella, Carla Ivon Herrera-Najera, Pavel Isa, Brenda Irasema Maldonado-Meza, Bernardo Martinez-Miguel, Margarita Matias-Florentino, Maria Guadalupe de Jesus Mireles-Rivera, Gloria Maria Molina-Salinas, Hector Montoya-Fuentes, Jose Esteban Munoz-Medina, Jose de Jesus Nunez-Contreras, Alicia Ocana-Mondragon, Luis Alberto Ochoa-Carrera, Hector Esteban Paz-Juarez, Francisco Pulido, Helen Haydee Fernanda Ramirez-Plascencia, Angel Gustavo Salas-Lais, Alejandro Sanchez-Flores, Clara Esperanza Santacruz-Tinoco, Maria Guadalupe Santiago-Mauricio, Nelly Selem-Mojica, Blanca Taboada, Gloria Vazquez |
| EPI_ISL_1662188, EPI_ISL_1662189, EPI_ISL_1662190, EPI_ISL_1662191, EPI_ISL_1662192, EPI_ISL_1662193 | Centro de Investigacion Biomedica de Occidente (CIBO) | Unidad de Genomica Avanzada | Consorcio Mexicano de Vigilancia Genomica (CoViGen-Mex). Authors (in alphabetical order): Julio Elias Alvarado-Yaah, Carlos F. Arias, Santiago Ávila-Rios, Victor Hugo Borja-Aburto, Celia Boukadida, Juan Bautista Chale-Dzul, Celida Duque Molina, Jose Antonio Enciso-Moreno, Gloria Elena Espinosa-Ayala, Fernando Fontove-Herrera, Victor Eduardo Garcia-Arias,Concepcion Grajales-Muniz, Ricardo Grande, Alfredo Herrera-Estrella, Carla Ivon Herrera-Najera, Pavel Isa, Brenda Irasema Maldonado-Meza, Bernardo Martinez-Miguel, Margarita Matias-Florentino, Maria Guadalupe de Jesus Mireles-Rivera, Gloria Maria Molina-Salinas, Hector Montoya-Fuentes, Jose Esteban Munoz-Medina, Jose de Jesus Nunez-Contreras, Alicia Ocana-Mondragon, Luis Alberto Ochoa-Carrera, Hector Esteban Paz-Juarez, Francisco Pulido, Helen Haydee Fernanda Ramirez-Plascencia, Angel Gustavo Salas-Lais, Alejandro Sanchez-Flores, Clara Esperanza Santacruz-Tinoco, Maria Guadalupe Santiago-Mauricio, Nelly Selem-Mojica, Blanca Taboada, Gloria Vazquez |
| EPI_ISL_1662194, EPI_ISL_1662195, EPI_ISL_1662196 | Laboratorio Central de Epidemiologia (LCE) | Unidad de Genomica Avanzada | Consorcio Mexicano de Vigilancia Genomica (CoViGen-Mex). Authors (in alphabetical order): Julio Elias Alvarado-Yaah, Carlos F. Arias, Santiago Ávila-Rios, Victor Hugo Borja-Aburto, Celia Boukadida, Juan Bautista Chale-Dzul, Celida Duque Molina, Jose Antonio Enciso-Moreno, Gloria Elena Espinosa-Ayala, Fernando Fontove-Herrera, Victor Eduardo Garcia-Arias,Concepcion Grajales-Muniz, Ricardo Grande, Alfredo Herrera-Estrella, Carla Ivon Herrera-Najera, Pavel Isa, Brenda Irasema Maldonado-Meza, Bernardo Martinez-Miguel, Margarita Matias-Florentino, Maria Guadalupe de Jesus Mireles-Rivera, Gloria Maria Molina-Salinas, Hector Montoya-Fuentes, Jose Esteban Munoz-Medina, Jose de Jesus Nunez-Contreras, Alicia Ocana-Mondragon, Luis Alberto Ochoa-Carrera, Hector Esteban Paz-Juarez, Francisco Pulido, Helen Haydee Fernanda Ramirez-Plascencia, Angel Gustavo Salas-Lais, Alejandro Sanchez-Flores, Clara Esperanza Santacruz-Tinoco, Maria Guadalupe Santiago-Mauricio, Nelly Selem-Mojica, Blanca Taboada, Gloria Vazquez |
| EPI_ISL_1662197, EPI_ISL_1662198 | Centro de Investigacion Biomedica de Occidente (CIBO) | Unidad de Genomica Avanzada | Consorcio Mexicano de Vigilancia Genomica (CoViGen-Mex). Authors (in alphabetical order): Julio Elias Alvarado-Yaah, Carlos F. Arias, Santiago Ávila-Rios, Victor Hugo Borja-Aburto, Celia Boukadida, Juan Bautista Chale-Dzul, Celida Duque Molina, Jose Antonio Enciso-Moreno, Gloria Elena Espinosa-Ayala, Fernando Fontove-Herrera, Victor Eduardo Garcia-Arias,Concepcion Grajales-Muniz, Ricardo Grande, Alfredo Herrera-Estrella, Carla Ivon Herrera-Najera, Pavel Isa, Brenda Irasema Maldonado-Meza, Bernardo Martinez-Miguel, Margarita Matias-Florentino, Maria Guadalupe de Jesus Mireles-Rivera, Gloria Maria Molina-Salinas, Hector Montoya-Fuentes, Jose Esteban Munoz-Medina, Jose de Jesus Nunez-Contreras, Alicia Ocana-Mondragon, Luis Alberto Ochoa-Carrera, Hector Esteban Paz-Juarez, Francisco Pulido, Helen Haydee Fernanda Ramirez-Plascencia, Angel Gustavo Salas-Lais, Alejandro Sanchez-Flores, Clara Esperanza Santacruz-Tinoco, Maria Guadalupe Santiago-Mauricio, Nelly Selem-Mojica, Blanca Taboada, Gloria Vazquez |
| EPI_ISL_1662199, EPI_ISL_1662200 | Centro de Investigacion Biomedica del Noreste (CIBIN) | Unidad de Genomica Avanzada | Consorcio Mexicano de Vigilancia Genomica (CoViGen-Mex). Authors (in alphabetical order): Julio Elias Alvarado-Yaah, Carlos F. Arias, Santiago Ávila-Rios, Victor Hugo Borja-Aburto, Celia Boukadida, Juan Bautista Chale-Dzul, Celida Duque Molina, Jose Antonio Enciso-Moreno, Gloria Elena Espinosa-Ayala, Fernando Fontove-Herrera, Victor Eduardo Garcia-Arias,Concepcion Grajales-Muniz, Ricardo Grande, Alfredo Herrera-Estrella, Carla Ivon Herrera-Najera, Pavel Isa, Brenda Irasema Maldonado-Meza, Bernardo Martinez-Miguel, Margarita Matias-Florentino, Maria Guadalupe de Jesus Mireles-Rivera, Gloria Maria Molina-Salinas, Hector Montoya-Fuentes, Jose Esteban Munoz-Medina, Jose de Jesus Nunez-Contreras, Alicia Ocana-Mondragon, Luis Alberto Ochoa-Carrera, Hector Esteban Paz-Juarez, Francisco Pulido, Helen Haydee Fernanda Ramirez-Plascencia, Angel Gustavo Salas-Lais, Alejandro Sanchez-Flores, Clara Esperanza Santacruz-Tinoco, Maria Guadalupe Santiago-Mauricio, Nelly Selem-Mojica, Blanca Taboada, Gloria Vazquez |

| | | | |
|---|---|---|---|
| EPI_ISL_1662201, EPI_ISL_1662202, EPI_ISL_1662203, EPI_ISL_1662204, EPI_ISL_1662206, EPI_ISL_1662207, EPI_ISL_1662208, EPI_ISL_1662209, EPI_ISL_1662210, EPI_ISL_1662211, EPI_ISL_1662212 | | | |
| see above | Laboratorio Central de Epidemiologia (LCE) | Unidad de Genomica Avanzada | Consorcio Mexicano de Vigilancia Genomica (CoViGen-Mex). Authors (in alphabetical order): Julio Elias Alvarado-Yaah, Carlos F. Arias, Santiago Ávila-Rios, Victor Hugo Borja-Aburto, Celia Boukadida, Juan Bautista Chale-Dzul, Celida Duque Molina, Jose Antonio Enciso-Moreno, Gloria Elena Espinosa-Ayala, Fernando Fontove-Herrera, Victor Eduardo Garcia-Arias,Concepcion Grajales-Muniz, Ricardo Grande, Alfredo Herrera-Estrella, Carla Ivon Herrera-Najera, Pavel Isa, Brenda Irasema Maldonado-Meza, Bernardo Martinez-Miguel, Margarita Matias-Florentino, Maria Guadalupe de Jesus Mireles-Rivera, Gloria Maria Molina-Salinas, Hector Montoya-Fuentes, Jose Esteban Munoz-Medina, Jose de Jesus Nunez-Contreras, Alicia Ocana-Mondragon, Luis Alberto Ochoa-Carrera, Hector Esteban Paz-Juarez, Francisco Pulido, Helen Haydee Fernanda Ramirez-Plascencia, Angel Gustavo Salas-Lais, Alejandro Sanchez-Flores, Clara Esperanza Santacruz-Tinoco, Maria Guadalupe Santiago-Mauricio, Nelly Selem-Mojica, Blanca Taboada, Gloria Vazquez |
| EPI_ISL_1662214, EPI_ISL_1662215, EPI_ISL_1662216, EPI_ISL_1662217, EPI_ISL_1662218 | Unidad de Investigacion Medica de Yucatan (UIMY) | Unidad de Genomica Avanzada | Consorcio Mexicano de Vigilancia Genomica (CoViGen-Mex). Authors (in alphabetical order): Julio Elias Alvarado-Yaah, Carlos F. Arias, Santiago Ávila-Rios, Victor Hugo Borja-Aburto, Celia Boukadida, Juan Bautista Chale-Dzul, Celida Duque Molina, Jose Antonio Enciso-Moreno, Gloria Elena Espinosa-Ayala, Fernando Fontove-Herrera, Victor Eduardo Garcia-Arias,Concepcion Grajales-Muniz, Ricardo Grande, Alfredo Herrera-Estrella, Carla Ivon Herrera-Najera, Pavel Isa, Brenda Irasema Maldonado-Meza, Bernardo Martinez-Miguel, Margarita Matias-Florentino, Maria Guadalupe de Jesus Mireles-Rivera, Gloria Maria Molina-Salinas, Hector Montoya-Fuentes, Jose Esteban Munoz-Medina, Jose de Jesus Nunez-Contreras, Alicia Ocana-Mondragon, Luis Alberto Ochoa-Carrera, Hector Esteban Paz-Juarez, Francisco Pulido, Helen Haydee Fernanda Ramirez-Plascencia, Angel Gustavo Salas-Lais, Alejandro Sanchez-Flores, Clara Esperanza Santacruz-Tinoco, Maria Guadalupe Santiago-Mauricio, Nelly Selem-Mojica, Blanca Taboada, Gloria Vazquez |
| EPI_ISL_1662221, EPI_ISL_1662222, EPI_ISL_1662223, EPI_ISL_1662224, EPI_ISL_1662226, EPI_ISL_1662230 | Unidad de Investigacion Biomedica de Zacatecas (UIBZ) | Unidad de Genomica Avanzada | Consorcio Mexicano de Vigilancia Genomica (CoViGen-Mex). Authors (in alphabetical order): Julio Elias Alvarado-Yaah, Carlos F. Arias, Santiago Ávila-Rios, Victor Hugo Borja-Aburto, Celia Boukadida, Juan Bautista Chale-Dzul, Celida Duque Molina, Jose Antonio Enciso-Moreno, Gloria Elena Espinosa-Ayala, Fernando Fontove-Herrera, Victor Eduardo Garcia-Arias,Concepcion Grajales-Muniz, Ricardo Grande, Alfredo Herrera-Estrella, Carla Ivon Herrera-Najera, Pavel Isa, Brenda Irasema Maldonado-Meza, Bernardo Martinez-Miguel, Margarita Matias-Florentino, Maria Guadalupe de Jesus Mireles-Rivera, Gloria Maria Molina-Salinas, Hector Montoya-Fuentes, Jose Esteban Munoz-Medina, Jose de Jesus Nunez-Contreras, Alicia Ocana-Mondragon, Luis Alberto Ochoa-Carrera, Hector Esteban Paz-Juarez, Francisco Pulido, Helen Haydee Fernanda Ramirez-Plascencia, Angel Gustavo Salas-Lais, Alejandro Sanchez-Flores, Clara Esperanza Santacruz-Tinoco, Maria Guadalupe Santiago-Mauricio, Nelly Selem-Mojica, Blanca Taboada, Gloria Vazquez |
| EPI_ISL_1662234 | IU-Cerrahpasa, Cerrahpasa School of Medicine, COVID-19 Lab | IU-Cerrahpasa, Cerrahpasa School of Medicine, COVID-19 Lab | Mert Kuskucu, Yesim Tuyji Tok, Kenan Midilli |
| EPI_ISL_1662240, EPI_ISL_1662468, EPI_ISL_1662529 | Institut de Virologie du CHU de Strasbourg | Institut de Virologie du CHU de Strasbourg | Fafi-Kremer Samira, Gallais Floriane, Gantner Pierre, Laugel Elodie, Solis Morgane, Velay Aurélie, Wendling Marie-Josée |
| EPI_ISL_1662593, EPI_ISL_1662620, EPI_ISL_1663061 | IU-Cerrahpasa, Cerrahpasa School of Medicine, COVID-19 Lab | IU-Cerrahpasa, Cerrahpasa School of Medicine, COVID-19 Lab | Mert Kuskucu, Yesim Tuyji Tok, Kenan Midilli |
| EPI_ISL_1663062 | IU-Cerrahpasa, Cerrahpasa School of Medicine, COVID-19 Lab | Illumina Miseq | Mert Kuskucu, Yesim Tuyji Tok, Kenan Midilli |
| EPI_ISL_1663229 | National Institute of Infectious Diseases-Prof. Dr. Matei Bals Molecular Diagnostics Laboratory | National Institute of Infectious Diseases-Prof. Dr. Matei Bals Molecular Diagnostics Laboratory | Corina Casangiu, Leontina Banica, Marius Surleac, Ovidiu Vlaicu, Andreea Tudor, Simona Paraschiv, Dan Otelea |
| EPI_ISL_1663577, EPI_ISL_1663579 | IU-Cerrahpasa, Cerrahpasa School of Medicine, COVID-19 Lab | IU-Cerrahpasa, Cerrahpasa School of Medicine, COVID-19 Lab | Mert Kuskucu, Yesim Tuyji Tok, Kenan Midilli |
| EPI_ISL_1667474 | Department of Laboratory Medicine, National Taiwan University Hospital | Microbial Genomics Core Lab, National Taiwan University Centers of Genomic and Precision Medicine | Shiou-Hwei Yeh, You-Yu Lin, Ya-Yun Lai, Chiao-Ling Li, Shan-Chwen Chang, Pei-Jer Chen, Sui-Yuan Chang |
| EPI_ISL_1669729, EPI_ISL_1669730, EPI_ISL_1669731, EPI_ISL_1669732, EPI_ISL_1669733, EPI_ISL_1669735, EPI_ISL_1669736, EPI_ISL_1669737, EPI_ISL_1669738, EPI_ISL_1669739, EPI_ISL_1669742, EPI_ISL_1669743, EPI_ISL_1669744, EPI_ISL_1669745, EPI_ISL_1669746, EPI_ISL_1669747, EPI_ISL_1669748, EPI_ISL_1669749, EPI_ISL_1669750, EPI_ISL_1669751, EPI_ISL_1669752, EPI_ISL_1669753 | | | |
| see above | Department of Virology, Istituto Zooprofilattico Sperimentale del Lazio e della Toscana (IZSLT) | Department of General Diagnostics; Department of Virology; Istituto Zooprofilattico Sperimentale del Lazio e della Toscana (IZSLT) | Patricia Alba, Giuseppe Manna, Elena L. Diaconu, Fabiola Feltrin, Raffaela Conti, Teresa Scicluna, Virginia Carfora, Antonella Cersini, Alessia Franco, Antonio Battisti. |
| EPI_ISL_1669826 | Microbiology Department, University Hospital Donostia | Microbiology Department, University Hospital Donostia | Marimon JM, Montes M, Piñeiro L, Sorarrain A, Gomez M, Cilla G. |
| EPI_ISL_1669909, EPI_ISL_1669910, EPI_ISL_1669911, EPI_ISL_1669912, EPI_ISL_1669913, EPI_ISL_1669914, EPI_ISL_1669915, EPI_ISL_1669918, EPI_ISL_1669919, EPI_ISL_1669920 | Azienda Ospedaliera Terni | Istituto Zooprofilattico Sperimentale dell'Abruzzo e Molise "G. Caporale" | Palumbo M, Scaccetti A, Lorusso A, Marcacci M, Di Domenico M, Ancora M, Curini V, Mangone I, Rinaldi A, Scialabba S, Di Pasquale A, Cammà C, Puglia I, Calistri P, Savini G |
| EPI_ISL_1669921, EPI_ISL_1669922, EPI_ISL_1669923, EPI_ISL_1669924, EPI_ISL_1669925 | Istituto Zooprofilattico Sperimentale Umbria e Marche "Togo Rosati" | Istituto Zooprofilattico Sperimentale dell'Abruzzo e Molise "G. Caporale" | Biagetti M, Giammarioli M, Lorusso A, Marcacci M, Di Domenico M, Ancora M, Curini V, Mangone I, Rinaldi A, Scialabba S, Di Pasquale A, Cammà C, Puglia I, Calistri P, Savini G |
| EPI_ISL_1669926, EPI_ISL_1669927, EPI_ISL_1669928 | Azienda USL Umbria 2 | Istituto Zooprofilattico Sperimentale dell'Abruzzo e Molise "G. Caporale" | Proietti A, Pistoni E, Lorusso A, Marcacci M, Di Domenico M, Ancora M, Curini V, Mangone I, Rinaldi A, Scialabba S, Di Pasquale A, Cammà C, Puglia I, Calistri P, Savini G |
| EPI_ISL_1669929 | Azienda Ospedaliera Terni | Istituto Zooprofilattico Sperimentale dell'Abruzzo e Molise "G. Caporale" | Palumbo M, Scaccetti A, Lorusso A, Marcacci M, Di Domenico M, Ancora M, Curini V, Mangone I, Rinaldi A, Scialabba S, Di Pasquale A, Cammà C, Puglia I, Calistri P, Savini G |
| EPI_ISL_1669933, EPI_ISL_1669934, EPI_ISL_1669935, EPI_ISL_1669936, EPI_ISL_1669937, EPI_ISL_1669938, EPI_ISL_1669939 | Università degli Studi di Perugia | Istituto Zooprofilattico Sperimentale dell'Abruzzo e Molise "G. Caporale" | Mencacci A, Camilloni B, Lorusso A, Marcacci M, Di Domenico M, Ancora M, Curini V, Mangone I, Rinaldi A, Scialabba S, Di Pasquale A, Cammà C, Puglia I, Calistri P, Savini G |
| EPI_ISL_1669940 | Azienda Ospedaliera Terni | Istituto Zooprofilattico Sperimentale dell'Abruzzo e Molise "G. Caporale" | Palumbo M, Scaccetti A, Lorusso A, Marcacci M, Di Domenico M, Ancora M, Curini V, Mangone I, Rinaldi A, Scialabba S, Di Pasquale A, Cammà C, Puglia I, Calistri P, Savini G |
| EPI_ISL_1669941, EPI_ISL_1669942, EPI_ISL_1669943, EPI_ISL_1669944, EPI_ISL_1669945, EPI_ISL_1669946, EPI_ISL_1669947 | Università degli Studi di Perugia | Istituto Zooprofilattico Sperimentale dell'Abruzzo e Molise "G. Caporale" | Mencacci A, Camilloni B, Lorusso A, Marcacci M, Di Domenico M, Ancora M, Curini V, Mangone I, Rinaldi A, Scialabba S, Di Pasquale A, Cammà C, Puglia I, Calistri P, Savini G |
| EPI_ISL_1669948, EPI_ISL_1669949, EPI_ISL_1669950, EPI_ISL_1669951, EPI_ISL_1669952, EPI_ISL_1669953 | Azienda Ospedaliera Terni | Istituto Zooprofilattico Sperimentale dell'Abruzzo e Molise "G. Caporale" | Palumbo M, Scaccetti A, Lorusso A, Marcacci M, Di Domenico M, Ancora M, Curini V, Mangone I, Rinaldi A, Scialabba S, Di Pasquale A, Cammà C, Puglia I, Calistri P, Savini G |
| EPI_ISL_1669990 | General Hospital - Ohrid | Laboratory of virology and molecular diagnostics, Institute of Public Health | Kuzmanovska M, Boshevska G, Janchevska E. |
| EPI_ISL_1669999 | General Hospital - Gostivar | Laboratory of virology and molecular diagnostics, Institute of Public Health | Kuzmanovska M, Boshevska G, Janchevska E. |
| EPI_ISL_1670000 | Clinical Center "Mother Theresa" | Laboratory of virology and molecular diagnostics, Institute of | Kuzmanovska M, Boshevska G, Janchevska E. |

| | | | |
|---|---|---|---|
| | | Public Health | |
| EPI_ISL_1670595, EPI_ISL_1670596, EPI_ISL_1670597, EPI_ISL_1670598, EPI_ISL_1670599, EPI_ISL_1670600, EPI_ISL_1670601, EPI_ISL_1670629, EPI_ISL_1670630, EPI_ISL_1670631, EPI_ISL_1670632, EPI_ISL_1670633, EPI_ISL_1670634, EPI_ISL_1670635, EPI_ISL_1670636, EPI_ISL_1670637, EPI_ISL_1670638 | | | |
| see above | Azienda Ospedaliera Terni | Istituto Zooprofilattico Sperimentale dell'Abruzzo e Molise "G. Caporale" | Palumbo M, Scaccetti A, Lorusso A, Marcacci M, Di Domenico M, Ancora M, Curini V, Di Lollo Valeria, Mangone I, Rinaldi A, Delli Compagni E, Scialabba S, Caporale M, Di Pasquale A, Cammà C, Puglia I, Calistri P, Savini G |
| EPI_ISL_1670639, EPI_ISL_1670640, EPI_ISL_1670641, EPI_ISL_1670642, EPI_ISL_1670643, EPI_ISL_1670644 | Laboratorio Analisi Osp. Città di Castello - Azienda USL Umbria1 | Istituto Zooprofilattico Sperimentale dell'Abruzzo e Molise "G. Caporale" | Malagigi V, Tacconi P, Lorusso A, Marcacci M, Di Domenico M, Ancora M, Curini V, Di Lollo Valeria, Mangone I, Rinaldi A, Delli Compagni E, Scialabba S, Caporale M, Di Pasquale A, Cammà C, Puglia I, Calistri P, Savini G |
| EPI_ISL_1670645, EPI_ISL_1670647, EPI_ISL_1670648, EPI_ISL_1670649, EPI_ISL_1670650, EPI_ISL_1670651, EPI_ISL_1670652, EPI_ISL_1670653, EPI_ISL_1670654, EPI_ISL_1670655, EPI_ISL_1670656, EPI_ISL_1670657, EPI_ISL_1670658, EPI_ISL_1670659 | | | |
| see above | Università degli Studi di Perugia | Istituto Zooprofilattico Sperimentale dell'Abruzzo e Molise "G. Caporale" | Mencacci A, Camilloni B, Lorusso A, Marcacci M, Di Domenico M, Ancora M, Curini V, Di Lollo Valeria, Mangone I, Rinaldi A, Delli Compagni E, Scialabba S, Caporale M, Di Pasquale A, Cammà C, Puglia I, Calistri P, Savini G |
| EPI_ISL_1670660, EPI_ISL_1670661, EPI_ISL_1670662 | Istituto Zooprofilattico Sperimentale Umbria e Marche "Togo Rosati" | Istituto Zooprofilattico Sperimentale dell'Abruzzo e Molise "G. Caporale" | Biagetti M, Giammarioli M,Lorusso A, Marcacci M, Di Domenico M, Ancora M, Curini V, Di Lollo Valeria, Mangone I, Rinaldi A, Delli Compagni E, Scialabba S, Caporale M, Di Pasquale A, Cammà C, Puglia I, Calistri P, Savini G |
| EPI_ISL_1670663, EPI_ISL_1670664, EPI_ISL_1670665, EPI_ISL_1670666 | Azienda USL Umbria 2 | Istituto Zooprofilattico Sperimentale dell'Abruzzo e Molise "G. Caporale" | Proietti A, Pistoni E, Lorusso A, Marcacci M, Di Domenico M, Ancora M, Curini V, Di Lollo Valeria, Mangone I, Rinaldi A, Delli Compagni E, Scialabba S, Caporale M, Di Pasquale A, Cammà C, Puglia I, Calistri P, Savini G |
| EPI_ISL_1670709 | Microbiology Department, University Hospital Donostia | Microbiology Department, University Hospital Donostia | Marimon JM, Montes M, Piñeiro L, Sorarrain A, Gomez M, Cilla G. |
| EPI_ISL_1671758 | CH Arras | CHU Lille - Laboratoire de Virologie | AIT YAHYA Emilie, ALIDJINOU Enagnon Kazali, BOCKET Laurence, CREPIN Michel, DEMAY Christophe, ENGELMANN Ilka, GEFFROY Sandrine, GUIGON Aurélie, LAMBERT Valérie, LAZREK Mouna, NOBILLIAUX Florian, PREVOST Brigitte, THUILLIER Caroline, TINEZ Claire |
| EPI_ISL_1671759, EPI_ISL_1671760 | CHU Lille | CHU Lille - Laboratoire de Virologie | AIT YAHYA Emilie, ALIDJINOU Enagnon Kazali, BOCKET Laurence, CREPIN Michel, DEMAY Christophe, ENGELMANN Ilka, GEFFROY Sandrine, GUIGON Aurélie, LAMBERT Valérie, LAZREK Mouna, NOBILLIAUX Florian, PREVOST Brigitte, THUILLIER Caroline, TINEZ Claire |
| EPI_ISL_1671761 | CH Cambrai | CHU Lille - Laboratoire de Virologie | AIT YAHYA Emilie, ALIDJINOU Enagnon Kazali, BOCKET Laurence, CREPIN Michel, DEMAY Christophe, ENGELMANN Ilka, GEFFROY Sandrine, GUIGON Aurélie, LAMBERT Valérie, LAZREK Mouna, NOBILLIAUX Florian, PREVOST Brigitte, THUILLIER Caroline, TINEZ Claire |
| EPI_ISL_1671762 | CH Lens | CHU Lille - Laboratoire de Virologie | AIT YAHYA Emilie, ALIDJINOU Enagnon Kazali, BOCKET Laurence, CREPIN Michel, DEMAY Christophe, ENGELMANN Ilka, GEFFROY Sandrine, GUIGON Aurélie, LAMBERT Valérie, LAZREK Mouna, NOBILLIAUX Florian, PREVOST Brigitte, THUILLIER Caroline, TINEZ Claire |
| EPI_ISL_1671763, EPI_ISL_1671764 | CHU Lille | CHU Lille - Laboratoire de Virologie | AIT YAHYA Emilie, ALIDJINOU Enagnon Kazali, BOCKET Laurence, CREPIN Michel, DEMAY Christophe, ENGELMANN Ilka, GEFFROY Sandrine, GUIGON Aurélie, LAMBERT Valérie, LAZREK Mouna, NOBILLIAUX Florian, PREVOST Brigitte, THUILLIER Caroline, TINEZ Claire |
| EPI_ISL_1671765, EPI_ISL_1671766 | CH Douai | CHU Lille - Laboratoire de Virologie | AIT YAHYA Emilie, ALIDJINOU Enagnon Kazali, BOCKET Laurence, CREPIN Michel, DEMAY Christophe, ENGELMANN Ilka, GEFFROY Sandrine, GUIGON Aurélie, LAMBERT Valérie, LAZREK Mouna, NOBILLIAUX Florian, PREVOST Brigitte, THUILLIER Caroline, TINEZ Claire |
| EPI_ISL_1671767, EPI_ISL_1671768 | CHU Lille | CHU Lille - Laboratoire de Virologie | AIT YAHYA Emilie, ALIDJINOU Enagnon Kazali, BOCKET Laurence, CREPIN Michel, DEMAY Christophe, ENGELMANN Ilka, GEFFROY Sandrine, GUIGON Aurélie, LAMBERT Valérie, LAZREK Mouna, NOBILLIAUX Florian, PREVOST Brigitte, THUILLIER Caroline, TINEZ Claire |
| EPI_ISL_1671769 | CH Roubaix | CHU Lille - Laboratoire de Virologie | AIT YAHYA Emilie, ALIDJINOU Enagnon Kazali, BOCKET Laurence, CREPIN Michel, DEMAY Christophe, ENGELMANN Ilka, GEFFROY Sandrine, GUIGON Aurélie, LAMBERT Valérie, LAZREK Mouna, NOBILLIAUX Florian, PREVOST Brigitte, THUILLIER Caroline, TINEZ Claire |
| EPI_ISL_1671770 | CH Calais | CHU Lille - Laboratoire de Virologie | AIT YAHYA Emilie, ALIDJINOU Enagnon Kazali, BOCKET Laurence, CREPIN Michel, DEMAY Christophe, ENGELMANN Ilka, GEFFROY Sandrine, GUIGON Aurélie, LAMBERT Valérie, LAZREK Mouna, NOBILLIAUX Florian, PREVOST Brigitte, THUILLIER Caroline, TINEZ Claire |
| EPI_ISL_1671771, EPI_ISL_1671772, EPI_ISL_1671773, EPI_ISL_1671774, EPI_ISL_1671775, EPI_ISL_1671776, EPI_ISL_1671777, EPI_ISL_1671778, EPI_ISL_1671779, EPI_ISL_1671780 | CHU Lille | CHU Lille - Laboratoire de Virologie | AIT YAHYA Emilie, ALIDJINOU Enagnon Kazali, BOCKET Laurence, CREPIN Michel, DEMAY Christophe, ENGELMANN Ilka, GEFFROY Sandrine, GUIGON Aurélie, LAMBERT Valérie, LAZREK Mouna, NOBILLIAUX Florian, PREVOST Brigitte, THUILLIER Caroline, TINEZ Claire |
| EPI_ISL_1671781 | CH Dunkerque | CHU Lille - Laboratoire de Virologie | AIT YAHYA Emilie, ALIDJINOU Enagnon Kazali, BOCKET Laurence, CREPIN Michel, DEMAY Christophe, ENGELMANN Ilka, GEFFROY Sandrine, GUIGON Aurélie, LAMBERT Valérie, LAZREK Mouna, NOBILLIAUX Florian, PREVOST Brigitte, THUILLIER Caroline, TINEZ Claire |
| EPI_ISL_1671782 | CHU Lille | CHU Lille - Laboratoire de Virologie | AIT YAHYA Emilie, ALIDJINOU Enagnon Kazali, BOCKET Laurence, CREPIN Michel, DEMAY Christophe, ENGELMANN Ilka, GEFFROY Sandrine, GUIGON Aurélie, LAMBERT Valérie, LAZREK Mouna, NOBILLIAUX Florian, PREVOST Brigitte, THUILLIER Caroline, TINEZ Claire |
| EPI_ISL_1671783 | CH Roubaix | CHU Lille - Laboratoire de Virologie | AIT YAHYA Emilie, ALIDJINOU Enagnon Kazali, BOCKET Laurence, CREPIN Michel, DEMAY Christophe, ENGELMANN Ilka, GEFFROY Sandrine, GUIGON Aurélie, LAMBERT Valérie, LAZREK Mouna, NOBILLIAUX Florian, PREVOST Brigitte, THUILLIER Caroline, TINEZ Claire |
| EPI_ISL_1671784, EPI_ISL_1671785 | CH Boulogne sur mer | CHU Lille - Laboratoire de Virologie | AIT YAHYA Emilie, ALIDJINOU Enagnon Kazali, BOCKET Laurence, CREPIN Michel, DEMAY Christophe, ENGELMANN Ilka, GEFFROY Sandrine, GUIGON Aurélie, LAMBERT Valérie, LAZREK Mouna, NOBILLIAUX Florian, PREVOST Brigitte, THUILLIER Caroline, TINEZ Claire |
| EPI_ISL_1671786, EPI_ISL_1671787 | CH Calais | CHU Lille - Laboratoire de Virologie | AIT YAHYA Emilie, ALIDJINOU Enagnon Kazali, BOCKET Laurence, CREPIN Michel, DEMAY Christophe, ENGELMANN Ilka, GEFFROY Sandrine, GUIGON Aurélie, LAMBERT Valérie, LAZREK Mouna, NOBILLIAUX Florian, PREVOST Brigitte, THUILLIER Caroline, TINEZ Claire |
| EPI_ISL_1671788 | CH Armentières | CHU Lille - Laboratoire de Virologie | AIT YAHYA Emilie, ALIDJINOU Enagnon Kazali, BOCKET Laurence, CREPIN Michel, DEMAY Christophe, ENGELMANN Ilka, GEFFROY Sandrine, GUIGON Aurélie, LAMBERT Valérie, LAZREK Mouna, NOBILLIAUX Florian, PREVOST Brigitte, THUILLIER Caroline, TINEZ Claire |
| EPI_ISL_1671789 | CH Bethune | CHU Lille - Laboratoire de Virologie | AIT YAHYA Emilie, ALIDJINOU Enagnon Kazali, BOCKET Laurence, CREPIN Michel, DEMAY Christophe, ENGELMANN Ilka, GEFFROY Sandrine, GUIGON Aurélie, LAMBERT Valérie, LAZREK Mouna, NOBILLIAUX Florian, PREVOST Brigitte, THUILLIER Caroline, TINEZ Claire |
| EPI_ISL_1671790, EPI_ISL_1671791, EPI_ISL_1671792, EPI_ISL_1671793, EPI_ISL_1671794, EPI_ISL_1671795, EPI_ISL_1671796, EPI_ISL_1671797, EPI_ISL_1671798, EPI_ISL_1671799, EPI_ISL_1671800, EPI_ISL_1671801, EPI_ISL_1671802, EPI_ISL_1671803, EPI_ISL_1671804, EPI_ISL_1671805, EPI_ISL_1671806, EPI_ISL_1671807, EPI_ISL_1671808 | | | |
| see above | CHU Lille | CHU Lille - Laboratoire de Virologie | AIT YAHYA Emilie, ALIDJINOU Enagnon Kazali, BOCKET Laurence, CREPIN Michel, DEMAY Christophe, ENGELMANN Ilka, GEFFROY Sandrine, GUIGON Aurélie, LAMBERT Valérie, LAZREK Mouna, NOBILLIAUX Florian, PREVOST Brigitte, THUILLIER Caroline, TINEZ Claire |
| EPI_ISL_1671818 | CH Bethune | CHU Lille - Laboratoire de Virologie | AIT YAHYA Emilie, ALIDJINOU Enagnon Kazali, BOCKET Laurence, CREPIN Michel, DEMAY Christophe, ENGELMANN Ilka, GEFFROY Sandrine, GUIGON Aurélie, LAMBERT Valérie, LAZREK Mouna, NOBILLIAUX Florian, PREVOST Brigitte, THUILLIER Caroline, TINEZ Claire |
| EPI_ISL_1671819 | CHU Lille | CHU Lille - Laboratoire de Virologie | AIT YAHYA Emilie, ALIDJINOU Enagnon Kazali, BOCKET Laurence, CREPIN Michel, DEMAY Christophe, ENGELMANN Ilka, GEFFROY Sandrine, GUIGON Aurélie, LAMBERT Valérie, LAZREK Mouna, NOBILLIAUX Florian, PREVOST Brigitte, THUILLIER Caroline, TINEZ Claire |
| EPI_ISL_1671820 | CH Roubaix | CHU Lille - Laboratoire de Virologie | AIT YAHYA Emilie, ALIDJINOU Enagnon Kazali, BOCKET Laurence, CREPIN Michel, DEMAY Christophe, ENGELMANN Ilka, GEFFROY Sandrine, GUIGON Aurélie, LAMBERT Valérie, LAZREK Mouna, NOBILLIAUX Florian, PREVOST Brigitte, THUILLIER Caroline, TINEZ Claire |
| EPI_ISL_1671821 | CH Lens | CHU Lille - Laboratoire de Virologie | AIT YAHYA Emilie, ALIDJINOU Enagnon Kazali, BOCKET Laurence, CREPIN Michel, DEMAY Christophe, ENGELMANN Ilka, GEFFROY Sandrine, GUIGON Aurélie, LAMBERT Valérie, LAZREK Mouna, NOBILLIAUX Florian, PREVOST Brigitte, THUILLIER Caroline, TINEZ Claire |
| EPI_ISL_1671822 | CH Roubaix | CHU Lille - Laboratoire de Virologie | AIT YAHYA Emilie, ALIDJINOU Enagnon Kazali, BOCKET Laurence, CREPIN Michel, DEMAY Christophe, ENGELMANN Ilka, GEFFROY Sandrine, GUIGON Aurélie, LAMBERT Valérie, LAZREK Mouna, NOBILLIAUX Florian, PREVOST Brigitte, THUILLIER Caroline, TINEZ Claire |
| EPI_ISL_1671823 | CH Armentières | CHU Lille - Laboratoire de Virologie | AIT YAHYA Emilie, ALIDJINOU Enagnon Kazali, BOCKET Laurence, CREPIN Michel, DEMAY Christophe, ENGELMANN Ilka, GEFFROY Sandrine, GUIGON Aurélie, LAMBERT Valérie, LAZREK Mouna, NOBILLIAUX Florian, PREVOST Brigitte, THUILLIER Caroline, TINEZ Claire |
| EPI_ISL_1671824, EPI_ISL_1671825, EPI_ISL_1671826, EPI_ISL_1671827, EPI_ISL_1671828 | CHU Lille | CHU Lille - Laboratoire de Virologie | AIT YAHYA Emilie, ALIDJINOU Enagnon Kazali, BOCKET Laurence, CREPIN Michel, DEMAY Christophe, ENGELMANN Ilka, GEFFROY Sandrine, GUIGON Aurélie, LAMBERT Valérie, LAZREK Mouna, NOBILLIAUX Florian, PREVOST Brigitte, THUILLIER Caroline, TINEZ Claire |

| | | | |
|---|---|---|---|
| EPI_ISL_1671829 | CH Seclin | CHU Lille - Laboratoire de Virologie | AIT YAHYA Emilie, ALIDJINOU Enagnon Kazali, BOCKET Laurence, CREPIN Michel, DEMAY Christophe, ENGELMANN Ilka, GEFFROY Sandrine, GUIGON Aurélie, LAMBERT Valérie, LAZREK Mouna, NOBILLIAUX Florian, PREVOST Brigitte, THUILLIER Caroline, TINEZ Claire |
| EPI_ISL_1671830 | CHU Lille | CHU Lille - Laboratoire de Virologie | AIT YAHYA Emilie, ALIDJINOU Enagnon Kazali, BOCKET Laurence, CREPIN Michel, DEMAY Christophe, ENGELMANN Ilka, GEFFROY Sandrine, GUIGON Aurélie, LAMBERT Valérie, LAZREK Mouna, NOBILLIAUX Florian, PREVOST Brigitte, THUILLIER Caroline, TINEZ Claire |
| EPI_ISL_1671831 | CH Seclin | CHU Lille - Laboratoire de Virologie | AIT YAHYA Emilie, ALIDJINOU Enagnon Kazali, BOCKET Laurence, CREPIN Michel, DEMAY Christophe, ENGELMANN Ilka, GEFFROY Sandrine, GUIGON Aurélie, LAMBERT Valérie, LAZREK Mouna, NOBILLIAUX Florian, PREVOST Brigitte, THUILLIER Caroline, TINEZ Claire |
| EPI_ISL_1671832, EPI_ISL_1671833, EPI_ISL_1671834 | CH Saint-Omer | CHU Lille - Laboratoire de Virologie | AIT YAHYA Emilie, ALIDJINOU Enagnon Kazali, BOCKET Laurence, CREPIN Michel, DEMAY Christophe, ENGELMANN Ilka, GEFFROY Sandrine, GUIGON Aurélie, LAMBERT Valérie, LAZREK Mouna, NOBILLIAUX Florian, PREVOST Brigitte, THUILLIER Caroline, TINEZ Claire |
| EPI_ISL_1671835, EPI_ISL_1671836 | CH Roubaix | CHU Lille - Laboratoire de Virologie | AIT YAHYA Emilie, ALIDJINOU Enagnon Kazali, BOCKET Laurence, CREPIN Michel, DEMAY Christophe, ENGELMANN Ilka, GEFFROY Sandrine, GUIGON Aurélie, LAMBERT Valérie, LAZREK Mouna, NOBILLIAUX Florian, PREVOST Brigitte, THUILLIER Caroline, TINEZ Claire |
| EPI_ISL_1671837 | CH Tourcoing | CHU Lille - Laboratoire de Virologie | AIT YAHYA Emilie, ALIDJINOU Enagnon Kazali, BOCKET Laurence, CREPIN Michel, DEMAY Christophe, ENGELMANN Ilka, GEFFROY Sandrine, GUIGON Aurélie, LAMBERT Valérie, LAZREK Mouna, NOBILLIAUX Florian, PREVOST Brigitte, THUILLIER Caroline, TINEZ Claire |
| EPI_ISL_1671838, EPI_ISL_1671839 | CHU Lille | CHU Lille - Laboratoire de Virologie | AIT YAHYA Emilie, ALIDJINOU Enagnon Kazali, BOCKET Laurence, CREPIN Michel, DEMAY Christophe, ENGELMANN Ilka, GEFFROY Sandrine, GUIGON Aurélie, LAMBERT Valérie, LAZREK Mouna, NOBILLIAUX Florian, PREVOST Brigitte, THUILLIER Caroline, TINEZ Claire |
| EPI_ISL_1672633 | Hospital Tuanku Jaafar Seremban | Institute for Medical Research, Infectious Disease Research Centre, National Institutes of Health, Ministry of Health Malaysia | Suppiah J, Kamel K, Mohd Zawawi Z, Thayan R |
| EPI_ISL_1672967, EPI_ISL_1672968 | Institut de Virologie du CHU de Strasbourg | Institut de Virologie du CHU de Strasbourg | Fafi-Kremer Samira, Gallais Floriane, Gantner Pierre, Laugel Elodie, Solis Morgane, Velay Aurélie, Wendling Marie-Josée |
| EPI_ISL_1673257 | Hospital Lahad Datu | Institute for Medical Research, Infectious Disease Research Centre, National Institutes of Health, Ministry of Health Malaysia Jalan Setia Murni U13/52, Bandar Setia Alam, 40170 Selangor, Malaysia | Suppiah J, Kamel K, Mohd Zawawi Z, Thayan R |
| EPI_ISL_1673266 | LESP Morelos | Instituto de Diagnostico y Referencia Epidemiologicos (INDRE) | Claudia Wong-Arambula, Abril Rodriguez-Maldonado, Vanessa Rivero-Arredondo, Ariadna Medina-Benitez, Joaquin Quiroz-Mercado, Sergio Rangel-Guerrero, Natividad Cruz-Ortiz, Tatiana Nunez-Garcia, Gisela Barrera-Badillo, Lucia Hernandez-Rivas, Irma Lopez-Martinez, Ernesto Ramirez-Gonzalez. |
| EPI_ISL_1673276 | LESP Jalisco | Instituto de Diagnostico y Referencia Epidemiologicos (INDRE) | Claudia Wong-Arambula, Abril Rodriguez-Maldonado, Vanessa Rivero-Arredondo, Ariadna Medina-Benitez, Joaquin Quiroz-Mercado, Sergio Rangel-Guerrero, Natividad Cruz-Ortiz, Tatiana Nunez-Garcia, Gisela Barrera-Badillo, Lucia Hernandez-Rivas, Irma Lopez-Martinez, Ernesto Ramirez-Gonzalez. |
| EPI_ISL_1673278, EPI_ISL_1673282 | Laboratorio de Virología HUCA | Laboratorio de Virología HUCA | Sandoval M, Castelló C, Gómez de Oña J, Boga JA, Rojo S, Alvarez-Arguelles ME, Abreu F, Costales I, Perez-Martínez Z, Martín-Rodríguez G, Coto E, Melón S |
| EPI_ISL_1673285 | Institut de Virologie du CHU de Strasbourg | Institut de Virologie du CHU de Strasbourg | Fafi-Kremer Samira, Gallais Floriane, Gantner Pierre, Laugel Elodie, Solis Morgane, Velay Aurélie, Wendling Marie-Josée |
| EPI_ISL_1673287 | Institute for Medical Research, Infectious Disease Research Centre, National Institutes of Health, Ministry of Health Malaysia | Institute for Medical Research, Infectious Disease Research Centre, National Institutes of Health, Ministry of Health Malaysia | Suppiah J, Kamel K, Mohd Zawawi Z, Ramly N, Robert F, Thayan R |
| EPI_ISL_1673293, EPI_ISL_1673296, EPI_ISL_1673299 | Institute for Medical Research, Infectious Disease Research Centre, National Institutes of Health, Ministry of Health Malaysia | Institute for Medical Research, Infectious Disease Research Centre, National Institutes of Health, Ministry of Health Malaysia | Suppiah J, Kamel K, Thayan R |
| EPI_ISL_1673301 | Laboratorio de Virología HUCA | Laboratorio de Virología HUCA | Sandoval M, Castelló C, Gómez de Oña J, Boga JA, Rojo S, Alvarez-Arguelles ME, Abreu F, Costales I, Perez-Martínez Z, Martín-Rodríguez G, Coto E, Melón S |
| EPI_ISL_1673304, EPI_ISL_1673336 | Institute for Medical Research, Infectious Disease Research Centre, National Institutes of Health, Ministry of Health Malaysia | Institute for Medical Research, Infectious Disease Research Centre, National Institutes of Health, Ministry of Health Malaysia | Suppiah J, Kamel K, Thayan R |
| EPI_ISL_1673342 | Institute for Medical Research, Infectious Disease Research Centre, National Institutes of Health, Ministry of Health Malaysia | Institute for Medical Research, Infectious Disease Research Centre, National Institutes of Health, Ministry of Health Malaysia | Suppiah J, Kamel K, Mohd Zawawi Z, Thayan R |
| EPI_ISL_1673344 | Institute for Medical Research, Infectious Disease Research Centre, National Institutes of Health, Ministry of Health Malaysia | Institute for Medical Research, Infectious Disease Research Centre, National Institutes of Health, Ministry of Health Malaysia | Suppiah J, Kamel K, Thayan R |
| EPI_ISL_1673345, EPI_ISL_1673420 | Institute for Medical Research, Infectious Disease Research Centre, National Institutes of Health, Ministry of Health Malaysia | Institute for Medical Research, Infectious Disease Research Centre, National Institutes of Health, Ministry of Health Malaysia | Suppiah J, Kamel K, Mohd Zawawi Z, Thayan R |
| EPI_ISL_1673673 | Genomic Research Lab, BCSIR | Genomic Research Lab, BCSIR | Tanjina Akhter Banu, Md. Murshed Hasan Sarkar, Mohammad Samir Uzzaman, Eshrar Osman, Md. Ahasan Habib, Shahina Akter, Abu Sayeed Mohammad Mahmud, Barna Goswami, Iffat Jahan, Md. Saddam Hossain, Mohammad Mohi Uddin, Mohammad Fazle Alam Rabbi, Md Firoz Kabir, Kazi Nadim Hasan, Md. Mizanur Rahman, Md. Abdul Khaleque, Sharif Akhteruzzamani, Md. Salim Khan |
| EPI_ISL_1673680 | Genomic Research Lab, BCSIR | Genomic Research Lab, BCSIR | Tanjina Akhter Banu, Md. Murshed Hasan Sarkar, Mohammad Samir Uzzaman, Eshrar Osman, Md. Ahasan Habib, Shahina Akter, Abu Sayeed Mohammad Mahmud, Barna Goswami, Iffat Jahan, Md. Saddam Hossain, Mohammad Mohi Uddin, Md. Salim Khan |
| EPI_ISL_1675007 | Medicine sentinel network Province | National Reference Center for Viruses of Respiratory Infections, Institut Pasteur, Paris | Marion Barbet, Sylvie Behillil, Méline Bizard, Angela Brisebarre, Camille Capel, Vincent Enouf, Louise Lefrançois, Frédéric Lemoine, Christophe Malabat, Corinne Maufrais, Pierre Lechat, Etienne Simon-Lorière, Maud Vanpeene, Sylvie Van der Werf ,Ludovic Joulaud |
| EPI_ISL_1675008 | Medicine sentinel network IDF | National Reference Center for Viruses of Respiratory Infections, Institut Pasteur, Paris | Marion Barbet, Sylvie Behillil, Méline Bizard, Angela Brisebarre, Camille Capel, Vincent Enouf, Louise Lefrançois, Frédéric Lemoine, Christophe Malabat, Corinne Maufrais, Etienne Simon-Lorière, Maud Vanpeene, Sylvie Van der Werf ,Fatima Afarni |
| EPI_ISL_1675011 | Labo Analyses Med | National Reference Center for Viruses of Respiratory Infections, Institut Pasteur, Paris | Marion Barbet, Sylvie Behillil, Méline Bizard, Angela Brisebarre, Camille Capel, Vincent Enouf, Louise Lefrançois, Frédéric Lemoine, Christophe Malabat, Corinne Maufrais, Etienne Simon-Lorière, Maud Vanpeene, Sylvie Van der Werf ,Antoine Amiot |
| EPI_ISL_1675012 | Hospital | National Reference Center for Viruses of Respiratory Infections, Institut Pasteur, Paris | Marion Barbet, Sylvie Behillil, Méline Bizard, Angela Brisebarre, Camille Capel, Vincent Enouf, Louise Lefrançois, Frédéric Lemoine, Christophe Malabat, Corinne Maufrais, Pierre Lechat, Etienne Simon-Lorière, Maud Vanpeene, Sylvie Van der Werf ,Marianne Burgard |
| EPI_ISL_1675014 | Hospital | National Reference Center for Viruses of Respiratory Infections, Institut Pasteur, Paris | Marion Barbet, Sylvie Behillil, Méline Bizard, Angela Brisebarre, Camille Capel, Vincent Enouf, Louise Lefrançois, Frédéric Lemoine, Christophe Malabat, Corinne Maufrais, Etienne Simon-Lorière, Maud Vanpeene, Sylvie Van der Werf ,Laurence Louvet |
| EPI_ISL_1675015, EPI_ISL_1675017 | Hospital | National Reference Center for Viruses of Respiratory Infections, Institut Pasteur, Paris | Marion Barbet, Sylvie Behillil, Méline Bizard, Angela Brisebarre, Camille Capel, Vincent Enouf, Louise Lefrançois, Frédéric Lemoine, Christophe Malabat, Corinne Maufrais, Pierre Lechat, Etienne Simon-Lorière, Maud Vanpeene, Sylvie Van der Werf ,Pascale Martres |
| EPI_ISL_1675020 | Hospital | National Reference Center for Viruses of Respiratory Infections, Institut Pasteur, Paris | Marion Barbet, Sylvie Behillil, Méline Bizard, Angela Brisebarre, Camille Capel, Vincent Enouf, Louise Lefrançois, Frédéric Lemoine, Christophe Malabat, Corinne Maufrais, Pierre Lechat, Etienne Simon-Lorière, Maud Vanpeene, Sylvie Van der Werf ,Laurence Louvet |
| EPI_ISL_1675026 | Labo Analyses Med | National Reference Center for Viruses of Respiratory Infections, Institut Pasteur, Paris | Marion Barbet, Sylvie Behillil, Méline Bizard, Angela Brisebarre, Camille Capel, Vincent Enouf, Louise Lefrançois, Frédéric Lemoine, Christophe Malabat, Corinne Maufrais, Adrien Pain, Etienne Simon-Lorière, Maud Vanpeene, Sylvie Van der Werf ,Fabienne Artur |

| | | | |
|---|---|---|---|
| EPI_ISL_1675027 | Labo Analyses Med | National Reference Center for Viruses of Respiratory Infections, Institut Pasteur, Paris | Marion Barbet, Sylvie Behillil, Méline Bizard, Angela Brisebarre, Camille Capel, Vincent Enouf, Louise Lefrançois, Frédéric Lemoine, Christophe Malabat, Corinne Maufrais, Adrien Pain, Etienne Simon-Lorière, Maud Vanpeene, Sylvie Van der Werf ,GréGoire Potiron |
| EPI_ISL_1675028, EPI_ISL_1675029 | Labo Analyses Med | National Reference Center for Viruses of Respiratory Infections, Institut Pasteur, Paris | Marion Barbet, Sylvie Behillil, Méline Bizard, Angela Brisebarre, Camille Capel, Vincent Enouf, Louise Lefrançois, Frédéric Lemoine, Christophe Malabat, Corinne Maufrais, Etienne Simon-Lorière, Maud Vanpeene, Sylvie Van der Werf ,GréGoire Potiron |
| EPI_ISL_1675030 | Labo Analyses Med | National Reference Center for Viruses of Respiratory Infections, Institut Pasteur, Paris | Marion Barbet, Sylvie Behillil, Méline Bizard, Angela Brisebarre, Camille Capel, Vincent Enouf, Louise Lefrançois, Frédéric Lemoine, Christophe Malabat, Corinne Maufrais, Adrien Pain, Etienne Simon-Lorière, Maud Vanpeene, Sylvie Van der Werf ,GréGoire Potiron |
| EPI_ISL_1675034 | Hospital | National Reference Center for Viruses of Respiratory Infections, Institut Pasteur, Paris | Marion Barbet, Sylvie Behillil, Méline Bizard, Angela Brisebarre, Camille Capel, Vincent Enouf, Louise Lefrançois, Frédéric Lemoine, Christophe Malabat, Corinne Maufrais, Adrien Pain, Etienne Simon-Lorière, Maud Vanpeene, Sylvie Van der Werf ,CéLine Ramanantsoa |
| EPI_ISL_1675035, EPI_ISL_1675036, EPI_ISL_1675037 | Labo Analyses Med | National Reference Center for Viruses of Respiratory Infections, Institut Pasteur, Paris | Marion Barbet, Sylvie Behillil, Méline Bizard, Angela Brisebarre, Camille Capel, Vincent Enouf, Louise Lefrançois, Frédéric Lemoine, Christophe Malabat, Corinne Maufrais, Etienne Simon-Lorière, Maud Vanpeene, Sylvie Van der Werf ,José Delaval |
| EPI_ISL_1675038 | Hospital | National Reference Center for Viruses of Respiratory Infections, Institut Pasteur, Paris | Marion Barbet, Sylvie Behillil, Méline Bizard, Angela Brisebarre, Camille Capel, Vincent Enouf, Louise Lefrançois, Frédéric Lemoine, Christophe Malabat, Corinne Maufrais, Gael Millot, Etienne Simon-Lorière, Maud Vanpeene, Sylvie Van der Werf ,Nabil Gastli |
| EPI_ISL_1675039, EPI_ISL_1675040, EPI_ISL_1675041, EPI_ISL_1675042, EPI_ISL_1675043 | Hospital | National Reference Center for Viruses of Respiratory Infections, Institut Pasteur, Paris | Marion Barbet, Sylvie Behillil, Méline Bizard, Angela Brisebarre, Camille Capel, Vincent Enouf, Louise Lefrançois, Frédéric Lemoine, Christophe Malabat, Corinne Maufrais, Etienne Simon-Lorière, Maud Vanpeene, Sylvie Van der Werf ,Nabil Gastli |
| EPI_ISL_1675044 | Hospital | National Reference Center for Viruses of Respiratory Infections, Institut Pasteur, Paris | Marion Barbet, Sylvie Behillil, Méline Bizard, Angela Brisebarre, Camille Capel, Vincent Enouf, Louise Lefrançois, Frédéric Lemoine, Christophe Malabat, Corinne Maufrais, Pierre Lechat, Etienne Simon-Lorière, Maud Vanpeene, Sylvie Van der Werf ,Nabil Gastli |
| EPI_ISL_1675045, EPI_ISL_1675046, EPI_ISL_1675047, EPI_ISL_1675048, EPI_ISL_1675049, EPI_ISL_1675050, EPI_ISL_1675051, EPI_ISL_1675052, EPI_ISL_1675053, EPI_ISL_1675054, EPI_ISL_1675055, EPI_ISL_1675056 | | | |
| see above | Hospital | National Reference Center for Viruses of Respiratory Infections, Institut Pasteur, Paris | Marion Barbet, Sylvie Behillil, Méline Bizard, Angela Brisebarre, Camille Capel, Vincent Enouf, Louise Lefrançois, Frédéric Lemoine, Christophe Malabat, Corinne Maufrais, Etienne Simon-Lorière, Maud Vanpeene, Sylvie Van der Werf ,Nabil Gastli |
| EPI_ISL_1675057 | Hospital | National Reference Center for Viruses of Respiratory Infections, Institut Pasteur, Paris | Marion Barbet, Sylvie Behillil, Méline Bizard, Angela Brisebarre, Camille Capel, Vincent Enouf, Louise Lefrançois, Frédéric Lemoine, Christophe Malabat, Corinne Maufrais, Gael Millot, Etienne Simon-Lorière, Maud Vanpeene, Sylvie Van der Werf ,Nabil Gastli |
| EPI_ISL_1675058, EPI_ISL_1675059, EPI_ISL_1675060, EPI_ISL_1675061, EPI_ISL_1675062, EPI_ISL_1675063, EPI_ISL_1675064, EPI_ISL_1675065, EPI_ISL_1675066, EPI_ISL_1675067, EPI_ISL_1675068, EPI_ISL_1675069, EPI_ISL_1675070, EPI_ISL_1675071, EPI_ISL_1675072 | | | |
| see above | Hospital | National Reference Center for Viruses of Respiratory Infections, Institut Pasteur, Paris | Marion Barbet, Sylvie Behillil, Méline Bizard, Angela Brisebarre, Camille Capel, Vincent Enouf, Louise Lefrançois, Frédéric Lemoine, Christophe Malabat, Corinne Maufrais, Etienne Simon-Lorière, Maud Vanpeene, Sylvie Van der Werf ,Nabil Gastli |
| EPI_ISL_1675073 | Armies | National Reference Center for Viruses of Respiratory Infections, Institut Pasteur, Paris | Marion Barbet, Sylvie Behillil, Méline Bizard, Angela Brisebarre, Camille Capel, Vincent Enouf, Louise Lefrançois, Frédéric Lemoine, Christophe Malabat, Corinne Maufrais, Etienne Simon-Lorière, Maud Vanpeene, Sylvie Van der Werf ,Marine Desroches |
| EPI_ISL_1675074 | Hospital | National Reference Center for Viruses of Respiratory Infections, Institut Pasteur, Paris | Marion Barbet, Sylvie Behillil, Méline Bizard, Angela Brisebarre, Camille Capel, Vincent Enouf, Louise Lefrançois, Frédéric Lemoine, Christophe Malabat, Corinne Maufrais, Etienne Simon-Lorière, Maud Vanpeene, Sylvie Van der Werf ,VéRonique Avettand-Fenoel |
| EPI_ISL_1675075 | Hospital | National Reference Center for Viruses of Respiratory Infections, Institut Pasteur, Paris | Marion Barbet, Sylvie Behillil, Méline Bizard, Angela Brisebarre, Camille Capel, Vincent Enouf, Louise Lefrançois, Frédéric Lemoine, Christophe Malabat, Corinne Maufrais, Etienne Simon-Lorière, Maud Vanpeene, Sylvie Van der Werf ,Olivia Raulin |
| EPI_ISL_1675076, EPI_ISL_1675077, EPI_ISL_1675078, EPI_ISL_1675079, EPI_ISL_1675080, EPI_ISL_1675081 | Hospital | National Reference Center for Viruses of Respiratory Infections, Institut Pasteur, Paris | Marion Barbet, Sylvie Behillil, Méline Bizard, Angela Brisebarre, Camille Capel, Vincent Enouf, Louise Lefrançois, Frédéric Lemoine, Christophe Malabat, Corinne Maufrais, Etienne Simon-Lorière, Maud Vanpeene, Sylvie Van der Werf ,LéA Pilorge |
| EPI_ISL_1675083 | Hospital | National Reference Center for Viruses of Respiratory Infections, Institut Pasteur, Paris | Marion Barbet, Sylvie Behillil, Méline Bizard, Angela Brisebarre, Camille Capel, Vincent Enouf, Louise Lefrançois, Frédéric Lemoine, Christophe Malabat, Corinne Maufrais, Etienne Simon-Lorière, Maud Vanpeene, Sylvie Van der Werf ,Pascale Martres |
| EPI_ISL_1675085, EPI_ISL_1675086 | Hospital | National Reference Center for Viruses of Respiratory Infections, Institut Pasteur, Paris | Marion Barbet, Sylvie Behillil, Méline Bizard, Angela Brisebarre, Camille Capel, Vincent Enouf, Louise Lefrançois, Frédéric Lemoine, Christophe Malabat, Corinne Maufrais, Etienne Simon-Lorière, Maud Vanpeene, Sylvie Van der Werf ,Anne Cady |
| EPI_ISL_1675087, EPI_ISL_1675088 | Labo Analyses Med | National Reference Center for Viruses of Respiratory Infections, Institut Pasteur, Paris | Marion Barbet, Sylvie Behillil, Méline Bizard, Angela Brisebarre, Camille Capel, Vincent Enouf, Louise Lefrançois, Frédéric Lemoine, Christophe Malabat, Corinne Maufrais, Etienne Simon-Lorière, Maud Vanpeene, Sylvie Van der Werf ,GréGoire Potiron |
| EPI_ISL_1675089 | Hospital | National Reference Center for Viruses of Respiratory Infections, Institut Pasteur, Paris | Marion Barbet, Sylvie Behillil, Méline Bizard, Angela Brisebarre, Camille Capel, Vincent Enouf, Louise Lefrançois, Frédéric Lemoine, Christophe Malabat, Corinne Maufrais, Etienne Simon-Lorière, Maud Vanpeene, Sylvie Van der Werf ,HéLèNe Fiette |
| EPI_ISL_1675091 | Retirement home | National Reference Center for Viruses of Respiratory Infections, Institut Pasteur, Paris | Marion Barbet, Sylvie Behillil, Méline Bizard, Angela Brisebarre, Camille Capel, Vincent Enouf, Louise Lefrançois, Frédéric Lemoine, Christophe Malabat, Corinne Maufrais, Etienne Simon-Lorière, Maud Vanpeene, Sylvie Van der Werf ,Nouri |
| EPI_ISL_1675092 | Hospital | National Reference Center for Viruses of Respiratory Infections, Institut Pasteur, Paris | Marion Barbet, Sylvie Behillil, Méline Bizard, Angela Brisebarre, Camille Capel, Vincent Enouf, Louise Lefrançois, Frédéric Lemoine, Christophe Malabat, Corinne Maufrais, Etienne Simon-Lorière, Maud Vanpeene, Sylvie Van der Werf ,JéRome Guinard |
| EPI_ISL_1675586, EPI_ISL_1675588, EPI_ISL_1675590, EPI_ISL_1675592, EPI_ISL_1675593, EPI_ISL_1675595, EPI_ISL_1675597, EPI_ISL_1675599, EPI_ISL_1675601, EPI_ISL_1675605, EPI_ISL_1675616, EPI_ISL_1675625, EPI_ISL_1675629, EPI_ISL_1675635, EPI_ISL_1675639, EPI_ISL_1675641, EPI_ISL_1675645, EPI_ISL_1675649, EPI_ISL_1675651, EPI_ISL_1675652 | | | |
| see above | Institute of Molecular and Translational Medicine / Laboratory of Experimental Medicine, Faculty of Medicine and Dentistry, Palacky University and University Hospital Olomouc | Institute of Molecular and Translational Medicine / Laboratory of Experimental Medicine, Faculty of Medicine and Dentistry, Palacky University | Rastislav Slavkovský, Hana Jaworek, Vladimíra Koudeláková, Barbora Blumová, Tomáš Pospíšil, Marián Hajdúch |
| EPI_ISL_1677768 | Gujarat Biotechnology Research Centre (GBRC), Gandhinagar | Gujarat Biotechnology Research Centre | Nitin Savaliya,Dinesh Kumar,Twinkle Soni,Sonal Sharma ,Zuber Saiyed,Ramesh Pandit,Janvi Raval,Zarna Patel,Umang Mishra,Nitesh Shah,Chaitanya Joshi,Madhvi Joshi |
| EPI_ISL_1677769 | Gujarat Biotechnology Research Centre (GBRC), Gandhinagar | Gujarat Biotechnology Research Centre | Dinesh Kumar,Twinkle Soni,Sonal Sharma ,Zuber Saiyed,Ramesh Pandit,Janvi Raval,Zarna Patel,Nitin Savaliya,Umang Mishra,Nitesh Shah,Chaitanya Joshi,Madhvi Joshi |
| EPI_ISL_1678022, EPI_ISL_1678023 | CQRC_QUALITY CONTROL CHEMICAL BIOLOGICAL RISK_AOOR Villa Sofia Cervello Palermo | CQRC_QUALITY CONTROL CHEMICAL BIOLOGICAL RISK_AOOR Villa Sofia Cervello Palermo | Di Gaudio, F., Contino, F., Brunacci, G., |
| EPI_ISL_1678380, EPI_ISL_1678383, EPI_ISL_1678384 | Laboratory for Clinical Immunology and Molecular Genetics - University Clinic Golnik Laboratory for Respiratory Microbiology - University Clinic Golnik | Laboratory for Clinical Immunology and Molecular Genetics -University Clinic Golnik | Matija Rijavec, Julij Šelb, Urška Bidovec Stojkovi, Žan Kogovšek, Nina Rupar, Viktorija Tomi, Peter Korošec |
| EPI_ISL_1678409, EPI_ISL_1678410, EPI_ISL_1678411, EPI_ISL_1678412, EPI_ISL_1678413, EPI_ISL_1678478 | HG Pharma GmbH | Bergthaler laboratory, CeMM Research Center for Molecular Medicine of the Austrian Academy of Sciences | Lukas Endler, Anna Schedl, Fabian Amman, Petr Triska, Thomas Penz, Benedikt Agerer, Maelle Le Moing, Michael Schuster, Bekir Erguner, Jan Laine, Martin Senekowitsch, Christoph Bock, Andreas Bergthaler |
| EPI_ISL_1685361, EPI_ISL_1685369, EPI_ISL_1685370, EPI_ISL_1685371, EPI_ISL_1685372, EPI_ISL_1685374 | Noth Estonia Medical Centre | 1. Laboratory of Communicable Diseases (Estonia); 2. Eurofins Genomics Europe Sequencing GmbH | Liidia Dotsenko et al. |
| EPI_ISL_1685991, EPI_ISL_1685993 | Laboratory of Communicable Diseases | 1. Laboratory of Communicable Diseases (Estonia); 2. Eurofins Genomics Europe Sequencing GmbH | Liidia Dotsenko et al. |
| EPI_ISL_1685995, EPI_ISL_1685997, EPI_ISL_1685998, EPI_ISL_1686000, EPI_ISL_1686001, EPI_ISL_1686004, EPI_ISL_1686007, EPI_ISL_1686010, EPI_ISL_1686014, EPI_ISL_1686016, EPI_ISL_1686018, EPI_ISL_1686019, EPI_ISL_1686021, EPI_ISL_1686022, EPI_ISL_1686024, EPI_ISL_1686025, EPI_ISL_1686027, EPI_ISL_1686028, EPI_ISL_1686030, EPI_ISL_1686031, EPI_ISL_1686033, EPI_ISL_1686034, EPI_ISL_1686036, EPI_ISL_1686037, EPI_ISL_1686039, EPI_ISL_1686040, EPI_ISL_1686042, EPI_ISL_1686043, EPI_ISL_1686045, EPI_ISL_1686046, EPI_ISL_1686048, EPI_ISL_1686049, EPI_ISL_1686050, EPI_ISL_1686052, | | | |

| | | | |
|---|---|---|---|
| EPI_ISL_1686053, EPI_ISL_1686055, EPI_ISL_1686057, EPI_ISL_1686058, EPI_ISL_1686059, EPI_ISL_1686061, EPI_ISL_1686062, EPI_ISL_1686064, EPI_ISL_1686065, EPI_ISL_1686067, EPI_ISL_1686068, EPI_ISL_1686070, EPI_ISL_1686073, EPI_ISL_1686074, EPI_ISL_1686076, EPI_ISL_1686077, EPI_ISL_1686078, EPI_ISL_1686080, EPI_ISL_1686081, EPI_ISL_1686083, EPI_ISL_1686085, EPI_ISL_1686086, EPI_ISL_1686087, EPI_ISL_1686089, EPI_ISL_1686091, EPI_ISL_1686092, EPI_ISL_1686093, EPI_ISL_1686096, EPI_ISL_1686098, EPI_ISL_1686099, EPI_ISL_1686101, EPI_ISL_1686102, EPI_ISL_1686103, EPI_ISL_1686105, EPI_ISL_1686106, EPI_ISL_1686108, EPI_ISL_1686109, EPI_ISL_1686111, EPI_ISL_1686112, EPI_ISL_1686114, EPI_ISL_1686115, EPI_ISL_1686117, EPI_ISL_1686118, EPI_ISL_1686119, EPI_ISL_1686121, EPI_ISL_1686123, EPI_ISL_1686124, EPI_ISL_1686125, EPI_ISL_1686127 | | | |
| see above | Synlab Eesti OÜ | 1. Laboratory of Communicable Diseases (Estonia); 2. Eurofins Genomics Europe Sequencing GmbH | Liidia Dotsenko et al. |
| EPI_ISL_1686129, EPI_ISL_1686130, EPI_ISL_1686131, EPI_ISL_1686133, EPI_ISL_1686134, EPI_ISL_1686136 | Laboratory of Communicable Diseases | 1. Laboratory of Communicable Diseases (Estonia); 2. Eurofins Genomics Europe Sequencing GmbH | Liidia Dotsenko et al. |
| EPI_ISL_1686137, EPI_ISL_1686139, EPI_ISL_1686140, EPI_ISL_1686142, EPI_ISL_1686143, EPI_ISL_1686144, EPI_ISL_1686145, EPI_ISL_1686147, EPI_ISL_1686148, EPI_ISL_1686150, EPI_ISL_1686151, EPI_ISL_1686152, EPI_ISL_1686154, EPI_ISL_1686156, EPI_ISL_1686157, EPI_ISL_1686158, EPI_ISL_1686160, EPI_ISL_1686162, EPI_ISL_1686165, EPI_ISL_1686168, EPI_ISL_1686171, EPI_ISL_1686174, EPI_ISL_1686178, EPI_ISL_1686182, EPI_ISL_1686185, EPI_ISL_1686189, EPI_ISL_1686190, EPI_ISL_1686192, EPI_ISL_1686193, EPI_ISL_1686195, EPI_ISL_1686196, EPI_ISL_1686198, EPI_ISL_1686199, EPI_ISL_1686201, EPI_ISL_1686202, EPI_ISL_1686203, EPI_ISL_1686205, EPI_ISL_1686206, EPI_ISL_1686208, EPI_ISL_1686209, EPI_ISL_1686211, EPI_ISL_1686212, EPI_ISL_1686214, EPI_ISL_1686215, EPI_ISL_1686216, EPI_ISL_1686218, EPI_ISL_1686219, EPI_ISL_1686221, EPI_ISL_1686222, EPI_ISL_1686223, EPI_ISL_1686225, EPI_ISL_1686226, EPI_ISL_1686228, EPI_ISL_1686229, EPI_ISL_1686231, EPI_ISL_1686232, EPI_ISL_1686234, EPI_ISL_1686235, EPI_ISL_1686237, EPI_ISL_1686238, EPI_ISL_1686240, EPI_ISL_1686241, EPI_ISL_1686242, EPI_ISL_1686244, EPI_ISL_1686245, EPI_ISL_1686247, EPI_ISL_1686248, EPI_ISL_1686250, EPI_ISL_1686251, EPI_ISL_1686253, EPI_ISL_1686254, EPI_ISL_1686256, EPI_ISL_1686257, EPI_ISL_1686259, EPI_ISL_1686260, EPI_ISL_1686262, EPI_ISL_1686263, EPI_ISL_1686265, EPI_ISL_1686266, EPI_ISL_1686268, EPI_ISL_1686269, EPI_ISL_1686270, EPI_ISL_1686272, EPI_ISL_1686276 | | | |
| see above | Synlab Eesti OÜ | 1. Laboratory of Communicable Diseases (Estonia); 2. Eurofins Genomics Europe Sequencing GmbH | Liidia Dotsenko et al. |
| EPI_ISL_1694722, EPI_ISL_1694724, EPI_ISL_1694725, EPI_ISL_1694726, EPI_ISL_1694727, EPI_ISL_1694765, EPI_ISL_1694766 | OCME Office Of Chief Medical Examiner | New York City Public Health Laboratory | Jade Wang, et al. |
| EPI_ISL_1696756, EPI_ISL_1696757, EPI_ISL_1696758 | Institute for Medical Research, Infectious Disease Research Centre, National Institutes of Health, Ministry of Health Malaysia | Institute for Medical Research, Infectious Disease Research Centre, National Institutes of Health, Ministry of Health Malaysia | Suppiah J, Kamel K, Mohd Zawawi Z, Thayan R |
| EPI_ISL_1697377 | Institut de Virologie du CHU de Strasbourg | Institut de Virologie du CHU de Strasbourg | Fafi-Kremer Samira, Gallais Floriane, Gantner Pierre, Laugel Elodie, Solis Morgane, Velay Aurélie, Wendling Marie-Josée |
| EPI_ISL_1700564, EPI_ISL_1700566, EPI_ISL_1700567, EPI_ISL_1700568 | Research | National Reference Center for Viruses of Respiratory Infections, Institut Pasteur, Paris | Marion Barbet, Sylvie Behillil, Méline Bizard, Angela Brisebarre, Camille Capel, Vincent Enouf, Louise Lefrançois, Frédéric Lemoine, Christophe Malabat, Corinne Maufrais, Etienne Simon-Lorière, Maud Vanpeene, Sylvie Van der Werf, Marianne Leruez-Ville |
| EPI_ISL_1700569 | Research | National Reference Center for Viruses of Respiratory Infections, Institut Pasteur, Paris | Marion Barbet, Sylvie Behillil, Méline Bizard, Angela Brisebarre, Camille Capel, Vincent Enouf, Louise Lefrançois, Frédéric Lemoine, Christophe Malabat, Corinne Maufrais, Damien Mornico, Etienne Simon-Lorière, Maud Vanpeene, Sylvie Van der Werf, Marianne Leruez-Ville |
| EPI_ISL_1700570, EPI_ISL_1700571, EPI_ISL_1700572, EPI_ISL_1700573, EPI_ISL_1700574, EPI_ISL_1700575, EPI_ISL_1700578, EPI_ISL_1700579, EPI_ISL_1700580 | Research | National Reference Center for Viruses of Respiratory Infections, Institut Pasteur, Paris | Marion Barbet, Sylvie Behillil, Méline Bizard, Angela Brisebarre, Camille Capel, Vincent Enouf, Louise Lefrançois, Frédéric Lemoine, Christophe Malabat, Corinne Maufrais, Etienne Simon-Lorière, Maud Vanpeene, Sylvie Van der Werf, Marianne Leruez-Ville |
| EPI_ISL_1700581 | Research | National Reference Center for Viruses of Respiratory Infections, Institut Pasteur, Paris | Marion Barbet, Sylvie Behillil, Méline Bizard, Angela Brisebarre, Camille Capel, Vincent Enouf, Louise Lefrançois, Frédéric Lemoine, Christophe Malabat, Corinne Maufrais, Damien Mornico, Etienne Simon-Lorière, Maud Vanpeene, Sylvie Van der Werf, Marianne Leruez-Ville |
| EPI_ISL_1700584, EPI_ISL_1700585, EPI_ISL_1700586, EPI_ISL_1700587, EPI_ISL_1700588, EPI_ISL_1700589, EPI_ISL_1700590, EPI_ISL_1700591 | Research | National Reference Center for Viruses of Respiratory Infections, Institut Pasteur, Paris | Marion Barbet, Sylvie Behillil, Méline Bizard, Angela Brisebarre, Camille Capel, Vincent Enouf, Louise Lefrançois, Frédéric Lemoine, Christophe Malabat, Corinne Maufrais, Etienne Simon-Lorière, Maud Vanpeene, Sylvie Van der Werf, Marianne Leruez-Ville |
| EPI_ISL_1700593 | Hospital | National Reference Center for Viruses of Respiratory Infections, Institut Pasteur, Paris | Marion Barbet, Sylvie Behillil, Méline Bizard, Angela Brisebarre, Camille Capel, Vincent Enouf, Louise Lefrançois, Frédéric Lemoine, Christophe Malabat, Corinne Maufrais, Etienne Simon-Lorière, Maud Vanpeene, Sylvie Van der Werf, Haciba Moudjahed |
| EPI_ISL_1700594 | Hospital | National Reference Center for Viruses of Respiratory Infections, Institut Pasteur, Paris | Marion Barbet, Sylvie Behillil, Méline Bizard, Angela Brisebarre, Camille Capel, Vincent Enouf, Louise Lefrançois, Frédéric Lemoine, Christophe Malabat, Corinne Maufrais, Etienne Simon-Lorière, Maud Vanpeene, Sylvie Van der Werf, Sylvaine Bastian |
| EPI_ISL_1700597 | Hospital | National Reference Center for Viruses of Respiratory Infections, Institut Pasteur, Paris | Marion Barbet, Sylvie Behillil, Méline Bizard, Angela Brisebarre, Camille Capel, Vincent Enouf, Louise Lefrançois, Frédéric Lemoine, Christophe Malabat, Corinne Maufrais, Damien Mornico, Etienne Simon-Lorière, Maud Vanpeene, Sylvie Van der Werf, Patricia Stoessel-Thouvenin |
| EPI_ISL_1700599, EPI_ISL_1700600, EPI_ISL_1700601 | Labo Analyses Med | National Reference Center for Viruses of Respiratory Infections, Institut Pasteur, Paris | Marion Barbet, Sylvie Behillil, Méline Bizard, Angela Brisebarre, Camille Capel, Vincent Enouf, Louise Lefrançois, Frédéric Lemoine, Christophe Malabat, Corinne Maufrais, Etienne Simon-Lorière, Maud Vanpeene, Sylvie Van der Werf, BéAtrice Redeker |
| EPI_ISL_1700605, EPI_ISL_1700606, EPI_ISL_1700607, EPI_ISL_1700611, EPI_ISL_1700612, EPI_ISL_1700613, EPI_ISL_1700614 | Labo Analyses Med | National Reference Center for Viruses of Respiratory Infections, Institut Pasteur, Paris | Marion Barbet, Sylvie Behillil, Méline Bizard, Angela Brisebarre, Camille Capel, Vincent Enouf, Louise Lefrançois, Frédéric Lemoine, Christophe Malabat, Corinne Maufrais, Etienne Simon-Lorière, Maud Vanpeene, Sylvie Van der Werf, GréGoire Potiron |
| EPI_ISL_1700615 | Hospital | National Reference Center for Viruses of Respiratory Infections, Institut Pasteur, Paris | Marion Barbet, Sylvie Behillil, Méline Bizard, Angela Brisebarre, Camille Capel, Vincent Enouf, Louise Lefrançois, Frédéric Lemoine, Christophe Malabat, Corinne Maufrais, Damien Mornico, Etienne Simon-Lorière, Maud Vanpeene, Sylvie Van der Werf, Franck Griscelli |
| EPI_ISL_1700617 | Hospital | National Reference Center for Viruses of Respiratory Infections, Institut Pasteur, Paris | Marion Barbet, Sylvie Behillil, Méline Bizard, Angela Brisebarre, Camille Capel, Vincent Enouf, Louise Lefrançois, Frédéric Lemoine, Christophe Malabat, Corinne Maufrais, Damien Mornico, Etienne Simon-Lorière, Maud Vanpeene, Sylvie Van der Werf, Marianne Burgard |
| EPI_ISL_1700625 | Hospital | National Reference Center for Viruses of Respiratory Infections, Institut Pasteur, Paris | Marion Barbet, Sylvie Behillil, Méline Bizard, Angela Brisebarre, Camille Capel, Vincent Enouf, Louise Lefrançois, Frédéric Lemoine, Christophe Malabat, Corinne Maufrais, Etienne Simon-Lorière, Maud Vanpeene, Sylvie Van der Werf, Laurent Bret |
| EPI_ISL_1700627 | Hospital | National Reference Center for Viruses of Respiratory Infections, Institut Pasteur, Paris | Marion Barbet, Sylvie Behillil, Méline Bizard, Angela Brisebarre, Camille Capel, Vincent Enouf, Louise Lefrançois, Frédéric Lemoine, Christophe Malabat, Corinne Maufrais, Etienne Simon-Lorière, Maud Vanpeene, Sylvie Van der Werf, JéRome Guinard |
| EPI_ISL_1700784, EPI_ISL_1700785 | HG Pharma GmbH | Bergthaler laboratory, CeMM Research Center for Molecular Medicine of the Austrian Academy of Sciences | Lukas Endler, Anna Schedl, Fabian Amman, Petr Triska, Thomas Penz, Benedikt Agerer, Maelle Le Moing, Michael Schuster, Bekir Erguner, Jan Laine, Martin Senekowitsch, Christoph Bock, Andreas Bergthaler |
| EPI_ISL_1700806 | LESP Queretaro | Instituto de Diagnostico y Referencia Epidemiologicos (INDRE) | Claudia Wong-Arambula, Abril Rodriguez-Maldonado, Vanessa Rivero-Arredondo, Ariadna Medina-Benitez, Joaquin Quiroz-Mercado, Sergio Rangel-Guerrero, Natividad Cruz-Ortiz, Tatiana Nunez-Garcia, Gisela Barrera-Badillo, Lucia Hernandez-Rivas, Irma Lopez-Martinez, Ernesto Ramirez-Gonzalez. |
| EPI_ISL_1700819 | LESP Baja California Sur | Instituto de Diagnostico y Referencia Epidemiologicos (INDRE) | Claudia Wong-Arambula, Abril Rodriguez-Maldonado, Vanessa Rivero-Arredondo, Ariadna Medina-Benitez, Joaquin Quiroz-Mercado, Sergio Rangel-Guerrero, Natividad Cruz-Ortiz, Tatiana Nunez-Garcia, Gisela Barrera-Badillo, Lucia Hernandez-Rivas, Irma Lopez-Martinez, Ernesto Ramirez-Gonzalez. |
| EPI_ISL_1700821 | LESP Jalisco | Instituto de Diagnostico y Referencia Epidemiologicos (INDRE) | Claudia Wong-Arambula, Abril Rodriguez-Maldonado, Vanessa Rivero-Arredondo, Ariadna Medina-Benitez, Joaquin Quiroz-Mercado, Sergio Rangel-Guerrero, Natividad Cruz-Ortiz, Tatiana Nunez-Garcia, Gisela Barrera-Badillo, Lucia Hernandez-Rivas, Irma Lopez-Martinez, Ernesto Ramirez-Gonzalez. |
| EPI_ISL_1700826 | LESP Morelos | Instituto de Diagnostico y Referencia Epidemiologicos (INDRE) | Claudia Wong-Arambula, Abril Rodriguez-Maldonado, Vanessa Rivero-Arredondo, Ariadna Medina-Benitez, Joaquin Quiroz-Mercado, Sergio Rangel-Guerrero, Natividad Cruz-Ortiz, Tatiana Nunez-Garcia, Gisela Barrera-Badillo, Lucia Hernandez-Rivas, Irma Lopez-Martinez, Ernesto Ramirez-Gonzalez. |
| EPI_ISL_1700828, EPI_ISL_1700830 | LESP Guanajuato | Instituto de Diagnostico y Referencia Epidemiologicos (INDRE) | Claudia Wong-Arambula, Abril Rodriguez-Maldonado, Vanessa Rivero-Arredondo, Ariadna Medina-Benitez, Joaquin Quiroz-Mercado, Sergio Rangel-Guerrero, Natividad Cruz-Ortiz, Tatiana Nunez-Garcia, Gisela Barrera-Badillo, Lucia Hernandez-Rivas, Irma Lopez-Martinez, Ernesto |

| | | | |
|---|---|---|---|
| | | | Ramirez-Gonzalez. |
| EPI_ISL_1700834, EPI_ISL_1700835 | LESP Morelos | Instituto de Diagnostico y Referencia Epidemiologicos (INDRE) | Claudia Wong-Arambula, Abril Rodriguez-Maldonado, Vanessa Rivero-Arredondo, Ariadna Medina-Benitez, Joaquin Quiroz-Mercado, Sergio Rangel-Guerrero, Natividad Cruz-Ortiz, Tatiana Nunez-Garcia, Gisela Barrera-Badillo, Lucia Hernandez-Rivas, Irma Lopez-Martinez, Ernesto Ramirez-Gonzalez. |
| EPI_ISL_1700836 | LESPZacatecas | Instituto de Diagnostico y Referencia Epidemiologicos (INDRE) | Claudia Wong-Arambula, Abril Rodriguez-Maldonado, Vanessa Rivero-Arredondo, Ariadna Medina-Benitez, Joaquin Quiroz-Mercado, Sergio Rangel-Guerrero, Natividad Cruz-Ortiz, Tatiana Nunez-Garcia, Gisela Barrera-Badillo, Lucia Hernandez-Rivas, Irma Lopez-Martinez, Ernesto Ramirez-Gonzalez. |
| EPI_ISL_1700839 | LESP Jalisco | Instituto de Diagnostico y Referencia Epidemiologicos (INDRE) | Claudia Wong-Arambula, Abril Rodriguez-Maldonado, Vanessa Rivero-Arredondo, Ariadna Medina-Benitez, Joaquin Quiroz-Mercado, Sergio Rangel-Guerrero, Natividad Cruz-Ortiz, Tatiana Nunez-Garcia, Gisela Barrera-Badillo, Lucia Hernandez-Rivas, Irma Lopez-Martinez, Ernesto Ramirez-Gonzalez. |
| EPI_ISL_1701615 | Laboratorio de Virología HUCA | Laboratorio de Virología HUCA | Sandoval M, Castelló C, Gómez de Oña J, Boga JA, Rojo S, Alvarez-Arguelles ME, Abreu F, Costales I, Perez-Martínez Z, Martín-Rodríguez G, Coto E, Melón S |
| EPI_ISL_1706277 | Basurto University Hospital Clinical Microbiology Laboratory | Biocruces Bizkaia | Mikel Urrutikoetxea-Gutierrez, Mª Carmen Nieto Toboso, Estibaliz Ugalde Zarraga, Mikel Gallego Rodrigo, Ana de la Hoz, José Luis Díaz de Tuesta del Arco. |
| EPI_ISL_1706360 | Basurto University Hospital Clinical Microbiology Lab | Biocruces Bizkaia | Mikel Urrutikoetxea-Gutierrez, Mª Carmen Nieto Toboso, Estibaliz Ugalde Zarraga, Mikel Gallego Rodrigo, Ana de la Hoz, José Luis Díaz de Tuesta |
| EPI_ISL_1706478 | Centre Hospitalier Universitaire de Rouen Laboratoire de Virologie | Centre Hospitalier Universitaire de Rouen Laboratoire de Virologie | Alice Moisan, Fabienne De Oliveira, Marie Leoz |
| EPI_ISL_1706483 | Centre hospitalier du Belvedère | Centre Hospitalier Universitaire de Rouen Laboratoire de Virologie | Alice Moisan, Fabienne De Oliveira, Marie Leoz |
| EPI_ISL_1706485, EPI_ISL_1706489, EPI_ISL_1706491, EPI_ISL_1706543, EPI_ISL_1706544, EPI_ISL_1706545, EPI_ISL_1706546, EPI_ISL_1706547, EPI_ISL_1706548 | Centre Hospitalier Universitaire de Rouen Laboratoire de Virologie | Centre Hospitalier Universitaire de Rouen Laboratoire de Virologie | Alice Moisan, Fabienne De Oliveira, Marie Leoz |
| EPI_ISL_1706577, EPI_ISL_1706578, EPI_ISL_1706579 | Department of Virology, Istituto Zooprofilattico Sperimentale del Lazio e della Toscana (IZSLT) | Department of General Diagnostics; Department of Virology; Istituto Zooprofilattico Sperimentale del Lazio e della Toscana (IZSLT) | Patricia Alba, Giuseppe Manna, Elena L. Diaconu, Fabiola Feltrin, Raffaela Conti, Teresa Scicluna, Virginia Carfora, Antonella Cersini, Alessia Franco, Antonio Battisti. |
| EPI_ISL_1706598, EPI_ISL_1706599, EPI_ISL_1706600, EPI_ISL_1706602 | Virology Unit, Institut Pasteur du Cambodge | Virology Unit, Institut Pasteur du Cambodge | Jurre Y Siegers, Teyputita Ou, Leakhena Pum, Cecile Troupin, Ly Sovann, Kraing Sidonn, Yi Sengdoeurn, Chin Savuth, Chau Darapheak, Veasna Duong, Erik A Karlsson |
| EPI_ISL_1707072 | Klinisch Laboratorium ZNA | Klinisch Laboratorium ZNA | Verstrepen et al. |
| EPI_ISL_1707540 | SIESP CHIETI - DRIVE IN LANCIANO | Istituto Zooprofilattico Sperimentale dell'Abruzzo e Molise "G. Caporale" | Lorusso A, Marcacci M, Di Domenico M, Ancora M, Curini V, Di Lollo Valeria, Mangone I, Rinaldi A, Delli Compagni E, Scialabba S, Caporale M, Di Pasquale A, Cammà C, Puglia I, Calistri P, Savini G |
| EPI_ISL_1707541, EPI_ISL_1707542, EPI_ISL_1707543 | SIESP CHIETI - DRIVE IN ORTONA | Istituto Zooprofilattico Sperimentale dell'Abruzzo e Molise "G. Caporale" | Lorusso A, Marcacci M, Di Domenico M, Ancora M, Curini V, Di Lollo Valeria, Mangone I, Rinaldi A, Delli Compagni E, Scialabba S, Caporale M, Di Pasquale A, Cammà C, Puglia I, Calistri P, Savini G |
| EPI_ISL_1707544 | SIESP CHIETI - DRIVE IN LANCIANO | Istituto Zooprofilattico Sperimentale dell'Abruzzo e Molise "G. Caporale" | Lorusso A, Marcacci M, Di Domenico M, Ancora M, Curini V, Di Lollo Valeria, Mangone I, Rinaldi A, Delli Compagni E, Scialabba S, Caporale M, Di Pasquale A, Cammà C, Puglia I, Calistri P, Savini G |
| EPI_ISL_1707545 | SIESP DIPARTIMENTO DI PREVENZIONE TERAMO | Istituto Zooprofilattico Sperimentale dell'Abruzzo e Molise "G. Caporale" | Lorusso A, Marcacci M, Di Domenico M, Ancora M, Curini V, Di Lollo Valeria, Mangone I, Rinaldi A, Delli Compagni E, Scialabba S, Caporale M, Di Pasquale A, Cammà C, Puglia I, Calistri P, Savini G |
| EPI_ISL_1707546, EPI_ISL_1707547 | DIP. PREV. AVEZZANO SERVIZIO DI IGIENE EPIDEMIOLOGIA E SANITA' PUBBLICA | Istituto Zooprofilattico Sperimentale dell'Abruzzo e Molise "G. Caporale" | Lorusso A, Marcacci M, Di Domenico M, Ancora M, Curini V, Di Lollo Valeria, Mangone I, Rinaldi A, Delli Compagni E, Scialabba S, Caporale M, Di Pasquale A, Cammà C, Puglia I, Calistri P, Savini G |
| EPI_ISL_1707548 | SIESP DIPARTIMENTO DI PREVENZIONE CHIETI | Istituto Zooprofilattico Sperimentale dell'Abruzzo e Molise "G. Caporale" | Lorusso A, Marcacci M, Di Domenico M, Ancora M, Curini V, Di Lollo Valeria, Mangone I, Rinaldi A, Delli Compagni E, Scialabba S, Caporale M, Di Pasquale A, Cammà C, Puglia I, Calistri P, Savini G |
| EPI_ISL_1707549 | SIESP CHIETI - DRIVE IN VASTO | Istituto Zooprofilattico Sperimentale dell'Abruzzo e Molise "G. Caporale" | Lorusso A, Marcacci M, Di Domenico M, Ancora M, Curini V, Di Lollo Valeria, Mangone I, Rinaldi A, Delli Compagni E, Scialabba S, Caporale M, Di Pasquale A, Cammà C, Puglia I, Calistri P, Savini G |
| EPI_ISL_1707550 | SIESP SULMONA | Istituto Zooprofilattico Sperimentale dell'Abruzzo e Molise "G. Caporale" | Lorusso A, Marcacci M, Di Domenico M, Ancora M, Curini V, Di Lollo Valeria, Mangone I, Rinaldi A, Delli Compagni E, Scialabba S, Caporale M, Di Pasquale A, Cammà C, Puglia I, Calistri P, Savini G |
| EPI_ISL_1707551, EPI_ISL_1707552, EPI_ISL_1707553 | SIESP DIPARTIMENTO DI PREVENZIONE TERAMO | Istituto Zooprofilattico Sperimentale dell'Abruzzo e Molise "G. Caporale" | Lorusso A, Marcacci M, Di Domenico M, Ancora M, Curini V, Di Lollo Valeria, Mangone I, Rinaldi A, Delli Compagni E, Scialabba S, Caporale M, Di Pasquale A, Cammà C, Puglia I, Calistri P, Savini G |
| EPI_ISL_1707554 | USCA Sulmona | Istituto Zooprofilattico Sperimentale dell'Abruzzo e Molise "G. Caporale" | Lorusso A, Marcacci M, Di Domenico M, Ancora M, Curini V, Di Lollo Valeria, Mangone I, Rinaldi A, Delli Compagni E, Scialabba S, Caporale M, Di Pasquale A, Cammà C, Puglia I, Calistri P, Savini G |
| EPI_ISL_1707555, EPI_ISL_1707556 | SIESP DIPARTIMENTO DI PREVENZIONE TERAMO | Istituto Zooprofilattico Sperimentale dell'Abruzzo e Molise "G. Caporale" | Lorusso A, Marcacci M, Di Domenico M, Ancora M, Curini V, Di Lollo Valeria, Mangone I, Rinaldi A, Delli Compagni E, Scialabba S, Caporale M, Di Pasquale A, Cammà C, Puglia I, Calistri P, Savini G |
| EPI_ISL_1707557 | SIESP CHIETI - DRIVE IN CHIETI | Istituto Zooprofilattico Sperimentale dell'Abruzzo e Molise "G. Caporale" | Lorusso A, Marcacci M, Di Domenico M, Ancora M, Curini V, Di Lollo Valeria, Mangone I, Rinaldi A, Delli Compagni E, Scialabba S, Caporale M, Di Pasquale A, Cammà C, Puglia I, Calistri P, Savini G |
| EPI_ISL_1707558 | DIP. PREV. AVEZZANO SERVIZIO DI IGIENE EPIDEMIOLOGIA E SANITA' PUBBLICA | Istituto Zooprofilattico Sperimentale dell'Abruzzo e Molise "G. Caporale" | Lorusso A, Marcacci M, Di Domenico M, Ancora M, Curini V, Di Lollo Valeria, Mangone I, Rinaldi A, Delli Compagni E, Scialabba S, Caporale M, Di Pasquale A, Cammà C, Puglia I, Calistri P, Savini G |
| EPI_ISL_1707559 | USCA Castel di Sangro | Istituto Zooprofilattico Sperimentale dell'Abruzzo e Molise "G. Caporale" | Lorusso A, Marcacci M, Di Domenico M, Ancora M, Curini V, Di Lollo Valeria, Mangone I, Rinaldi A, Delli Compagni E, Scialabba S, Caporale M, Di Pasquale A, Cammà C, Puglia I, Calistri P, Savini G |
| EPI_ISL_1707560 | DIP. PREV. AVEZZANO SERVIZIO DI IGIENE EPIDEMIOLOGIA E SANITA' PUBBLICA | Istituto Zooprofilattico Sperimentale dell'Abruzzo e Molise "G. Caporale" | Lorusso A, Marcacci M, Di Domenico M, Ancora M, Curini V, Di Lollo Valeria, Mangone I, Rinaldi A, Delli Compagni E, Scialabba S, Caporale M, Di Pasquale A, Cammà C, Puglia I, Calistri P, Savini G |
| EPI_ISL_1707561 | USCA Pescina | Istituto Zooprofilattico Sperimentale dell'Abruzzo e Molise "G. Caporale" | Lorusso A, Marcacci M, Di Domenico M, Ancora M, Curini V, Di Lollo Valeria, Mangone I, Rinaldi A, Delli Compagni E, Scialabba S, Caporale M, Di Pasquale A, Cammà C, Puglia I, Calistri P, Savini G |
| EPI_ISL_1707562 | DIP. PREV. AVEZZANO SERVIZIO DI IGIENE EPIDEMIOLOGIA E SANITA' PUBBLICA | Istituto Zooprofilattico Sperimentale dell'Abruzzo e Molise "G. Caporale" | Lorusso A, Marcacci M, Di Domenico M, Ancora M, Curini V, Di Lollo Valeria, Mangone I, Rinaldi A, Delli Compagni E, Scialabba S, Caporale M, Di Pasquale A, Cammà C, Puglia I, Calistri P, Savini G |
| EPI_ISL_1707563 | SIESP DIPARTIMENTO DI PREVENZIONE CHIETI | Istituto Zooprofilattico Sperimentale dell'Abruzzo e Molise "G. Caporale" | Lorusso A, Marcacci M, Di Domenico M, Ancora M, Curini V, Di Lollo Valeria, Mangone I, Rinaldi A, Delli Compagni E, Scialabba S, Caporale M, Di Pasquale A, Cammà C, Puglia I, Calistri P, Savini G |
| EPI_ISL_1707564, EPI_ISL_1707565 | SIESP CHIETI - DRIVE IN VASTO | Istituto Zooprofilattico Sperimentale dell'Abruzzo e Molise "G. Caporale" | Lorusso A, Marcacci M, Di Domenico M, Ancora M, Curini V, Di Lollo Valeria, Mangone I, Rinaldi A, Delli Compagni E, Scialabba S, Caporale M, Di Pasquale A, Cammà C, Puglia I, Calistri P, Savini G |
| EPI_ISL_1707569 | SIESP SULMONA | Istituto Zooprofilattico Sperimentale dell'Abruzzo e Molise "G. Caporale" | Lorusso A, Marcacci M, Di Domenico M, Ancora M, Curini V, Di Lollo Valeria, Mangone I, Rinaldi A, Delli Compagni E, Scialabba S, Caporale M, Di Pasquale A, Cammà C, Puglia I, Calistri P, Savini G |
| EPI_ISL_1707574 | USCA Pescina | Istituto Zooprofilattico Sperimentale dell'Abruzzo e Molise "G. | Lorusso A, Marcacci M, Di Domenico M, Ancora M, Curini V, Di Lollo Valeria, Mangone I, Rinaldi A, Delli Compagni E, Scialabba S, Caporale M, Di |

| | | | |
|---|---|---|---|
| | | Caporale" | Pasquale A, Cammà C, Puglia I, Calistri P, Savini G |
| EPI_ISL_1707578 | SIESP CHIETI - DRIVE IN ORTONA | Istituto Zooprofilattico Sperimentale dell'Abruzzo e Molise "G. Caporale" | Lorusso A, Marcacci M, Di Domenico M, Ancora M, Curini V, Di Lollo Valeria, Mangone I, Rinaldi A, Delli Compagni E, Scialabba S, Caporale M, Di Pasquale A, Cammà C, Puglia I, Calistri P, Savini G |
| EPI_ISL_1707579 | SIESP DIPARTIMENTO DI PREVENZIONE CHIETI | Istituto Zooprofilattico Sperimentale dell'Abruzzo e Molise "G. Caporale" | Lorusso A, Marcacci M, Di Domenico M, Ancora M, Curini V, Di Lollo Valeria, Mangone I, Rinaldi A, Delli Compagni E, Scialabba S, Caporale M, Di Pasquale A, Cammà C, Puglia I, Calistri P, Savini G |
| EPI_ISL_1708640, EPI_ISL_1708641 | Hospital | National Reference Center for Viruses of Respiratory Infections, Institut Pasteur, Paris | Marion Barbet, Sylvie Behillil, Méline Bizard, Angela Brisebarre, Camille Capel, Vincent Enouf, Louise Lefrançois, Frédéric Lemoine, Christophe Malabat, Corinne Maufrais, Etienne Simon-Lorière, Maud Vanpeene, Sylvie Van der Werf ,Haciba Moudjahed |
| EPI_ISL_1710511 | Basurto University Hospital Clinical Microbiology Lab | Biocruces Bizkaia | Mikel Urrutikoetxea-Gutierrez, Mª Carmen Nieto Toboso, Estibaliz Ugalde Zarraga, Mikel Gallego Rodrigo, Ana de la Hoz, José Luis Díaz de Tuesta del Arco. |
| EPI_ISL_1710964, EPI_ISL_1710965 | LESP Ciudad de Mexico | Instituto de Diagnostico y Referencia Epidemiologicos (INDRE) | Claudia Wong-Arambula, Abril Rodriguez-Maldonado, Vanessa Rivero-Arredondo, Ariadna Medina-Benitez, Joaquin Quiroz-Mercado, Sergio Rangel-Guerrero, Natividad Cruz-Ortiz, Tatiana Nunez-Garcia, Gisela Barrera-Badillo, Lucia Hernandez-Rivas, Irma Lopez-Martinez, Ernesto Ramirez-Gonzalez. |
| EPI_ISL_1710967 | LESP Morelos | Instituto de Diagnostico y Referencia Epidemiologicos (INDRE) | Claudia Wong-Arambula, Abril Rodriguez-Maldonado, Vanessa Rivero-Arredondo, Ariadna Medina-Benitez, Joaquin Quiroz-Mercado, Sergio Rangel-Guerrero, Natividad Cruz-Ortiz, Tatiana Nunez-Garcia, Gisela Barrera-Badillo, Lucia Hernandez-Rivas, Irma Lopez-Martinez, Ernesto Ramirez-Gonzalez. |
| EPI_ISL_1710968, EPI_ISL_1710969 | LESP Guanajuato | Instituto de Diagnostico y Referencia Epidemiologicos (INDRE) | Claudia Wong-Arambula, Abril Rodriguez-Maldonado, Vanessa Rivero-Arredondo, Ariadna Medina-Benitez, Joaquin Quiroz-Mercado, Sergio Rangel-Guerrero, Natividad Cruz-Ortiz, Tatiana Nunez-Garcia, Gisela Barrera-Badillo, Lucia Hernandez-Rivas, Irma Lopez-Martinez, Ernesto Ramirez-Gonzalez. |
| EPI_ISL_1710970 | LESPZacatecas | Instituto de Diagnostico y Referencia Epidemiologicos (INDRE) | Claudia Wong-Arambula, Abril Rodriguez-Maldonado, Vanessa Rivero-Arredondo, Ariadna Medina-Benitez, Joaquin Quiroz-Mercado, Sergio Rangel-Guerrero, Natividad Cruz-Ortiz, Tatiana Nunez-Garcia, Gisela Barrera-Badillo, Lucia Hernandez-Rivas, Irma Lopez-Martinez, Ernesto Ramirez-Gonzalez. |
| EPI_ISL_1711976, EPI_ISL_1711977, EPI_ISL_1711978, EPI_ISL_1711979, EPI_ISL_1711980, EPI_ISL_1711981, EPI_ISL_1711982 | Virology Unit, Institut Pasteur du Cambodge | Virology Unit, Institut Pasteur du Cambodge | Jurre Y Siegers, Teyputita Ou, Leakhena Pum, Cecile Troupin, Ly Sovann, Kraing Sidonn, Yi Sengdoeurn, Chin Savuth, Chau Darapheak, Veasna Duong, Erik A Karlsson |
| EPI_ISL_1715140 | UPA Dr Luis Atilio Losi Viana Ribeirao Preto | Instituto Adolfo Lutz, Interdiciplinary Procedures Center, Strategic Laboratory | Claudio Tavares Sacchi, Claudia Regina Gonçalves, Erica Valessa Ramos Gomes, Karoline Rodrigues Campos, Caio Vinicius Dias Lopes, Leonardo Jose Tadeu de Araujo, Katia Correa de Oliveira Santos |
| EPI_ISL_1715142 | CS III de Patrocinio Paulista | Instituto Adolfo Lutz, Interdiciplinary Procedures Center, Strategic Laboratory | Claudio Tavares Sacchi, Claudia Regina Gonçalves, Erica Valessa Ramos Gomes, Karoline Rodrigues Campos, Caio Vinicius Dias Lopes, Leonardo Jose Tadeu de Araujo, Katia Correa de Oliveira Santos |
| EPI_ISL_1715146, EPI_ISL_1715147 | Instituto Adolfo Lutz - Regional de Aracatuba | Instituto Adolfo Lutz, Interdiciplinary Procedures Center, Strategic Laboratory | Claudio Tavares Sacchi, Claudia Regina Gonçalves, Erica Valessa Ramos Gomes, Karoline Rodrigues Campos, Caio Vinicius Dias Lopes, Leonardo Jose Tadeu de Araujo, Katia Correa de Oliveira Santos |
| EPI_ISL_1715148 | Instituto Adolfo Lutz - Regional de Ribeirao Preto | Instituto Adolfo Lutz, Interdiciplinary Procedures Center, Strategic Laboratory | Claudio Tavares Sacchi, Claudia Regina Gonçalves, Erica Valessa Ramos Gomes, Karoline Rodrigues Campos, Caio Vinicius Dias Lopes, Leonardo Jose Tadeu de Araujo, Katia Correa de Oliveira Santos |
| EPI_ISL_1715149, EPI_ISL_1715150, EPI_ISL_1715151, EPI_ISL_1715152, EPI_ISL_1715153, EPI_ISL_1715154, EPI_ISL_1715155, EPI_ISL_1715156, EPI_ISL_1715157, EPI_ISL_1715158 | Instituto Adolfo Lutz Central | Instituto Adolfo Lutz, Interdiciplinary Procedures Center, Strategic Laboratory | Claudio Tavares Sacchi, Claudia Regina Gonçalves, Erica Valessa Ramos Gomes, Karoline Rodrigues Campos, Caio Vinicius Dias Lopes, Leonardo Jose Tadeu de Araujo, Katia Correa de Oliveira Santos |
| EPI_ISL_1715218 | Laboratorio HUB -Azienda Ospedaliero Universitaria - AOU - Cagliari | Laboratorio SPOKE Biologia Molecolare -Azienda Ospedaliero Universitaria - AOU - Cagliari | Germano Orrù, Sara Fais, Valentina Medda, Alessandra Scano, Riccardo Cappai, Ferdinando Coghe |
| EPI_ISL_1715425, EPI_ISL_1715426, EPI_ISL_1715427, EPI_ISL_1715428, EPI_ISL_1715429, EPI_ISL_1715430, EPI_ISL_1715431, EPI_ISL_1715432, EPI_ISL_1715433, EPI_ISL_1715434, EPI_ISL_1715435, EPI_ISL_1715436, EPI_ISL_1715437, EPI_ISL_1715438 | | | |
| see above | University of Rome Tor Vergata: Departm Experim Medicine Chair of Virology | University of Rome Tor Vergata: Departm Experim Medicine Chair of Virology | Maria Concetta Bellocchi, Luca Carioti, Maria Botticelli, Lorenzo Piermatteo, Marco Iannetta, Loredana Sarmati, Massimo Andreoni, Francesca Ceccherini-Silberstein |
| EPI_ISL_1716736 | Laboratory of Immunohematology, Division of Hematology | Greek Genome Center, Biomedical Research Foundation of the Academy of Athens (BRFAA) | Emmanouil Athanasiadis, Ioannis Vatsellas, Theodoros Loupis, Katerina Zoi, Athanasia Mouzaki, Dimitrios Thanos |
| EPI_ISL_1716737 | Central Public Health Lab, National Public Health Organization | Greek Genome Center, Biomedical Research Foundation of the Academy of Athens (BRFAA) | Emmanouil Athanasiadis, Ioannis Vatsellas, Theodoros Loupis, Katerina Zoi, Kyriaki Tryfinopoulou, Dimitrios Thanos |
| EPI_ISL_1718319 | Institut de Virologie du CHU de Strasbourg | Institut de Virologie du CHU de Strasbourg | Fafi-Kremer Samira, Gallais Floriane, Gantner Pierre, Laugel Elodie, Solis Morgane, Velay Aurélie, Wendling Marie-Josée |
| EPI_ISL_1731458, EPI_ISL_1731459, EPI_ISL_1731460 | Centre hospitalier D'AURILLAC | CHU Clermont-Ferrand, service de virologie | Bisseux Maxime, Mirand Audrey, Combes Patricia, Henquell Cécile |
| EPI_ISL_1731575 | Casa de Caridade Sao Vicente de Paulo Cajuru | Instituto Adolfo Lutz, Interdiciplinary Procedures Center, Strategic Laboratory | Claudio Tavares Sacchi, Claudia Regina Gonçalves, Erica Valessa Ramos Gomes, Karoline Rodrigues Campos, Caio Vinicius Dias Lopes, Leonardo Jose Tadeu de Araujo, Katia Correa de Oliveira Santos |
| EPI_ISL_1731577 | Hospital Municipal Dr Mario Gatti Campinas | Instituto Adolfo Lutz, Interdiciplinary Procedures Center, Strategic Laboratory | Claudio Tavares Sacchi, Claudia Regina Gonçalves, Erica Valessa Ramos Gomes, Karoline Rodrigues Campos, Caio Vinicius Dias Lopes, Leonardo Jose Tadeu de Araujo, Katia Correa de Oliveira Santos |
| EPI_ISL_1731578, EPI_ISL_1731579, EPI_ISL_1731580 | Instituto Adolfo Lutz - Regional de Aracatuba | Instituto Adolfo Lutz, Interdiciplinary Procedures Center, Strategic Laboratory | Claudio Tavares Sacchi, Claudia Regina Gonçalves, Erica Valessa Ramos Gomes, Karoline Rodrigues Campos, Caio Vinicius Dias Lopes, Leonardo Jose Tadeu de Araujo, Katia Correa de Oliveira Santos |
| EPI_ISL_1731581, EPI_ISL_1731582 | Instituto Adolfo Lutz - Regional de Ribeirao Preto | Instituto Adolfo Lutz, Interdiciplinary Procedures Center, Strategic Laboratory | Claudio Tavares Sacchi, Claudia Regina Gonçalves, Erica Valessa Ramos Gomes, Karoline Rodrigues Campos, Caio Vinicius Dias Lopes, Leonardo Jose Tadeu de Araujo, Katia Correa de Oliveira Santos |
| EPI_ISL_1731583, EPI_ISL_1731584, EPI_ISL_1731585, EPI_ISL_1731586, EPI_ISL_1731587, EPI_ISL_1731588, EPI_ISL_1731589, EPI_ISL_1731590, EPI_ISL_1731591, EPI_ISL_1731592, EPI_ISL_1731593, EPI_ISL_1731594, EPI_ISL_1731595, EPI_ISL_1731596, EPI_ISL_1731597, EPI_ISL_1731598, EPI_ISL_1731599, EPI_ISL_1731600, EPI_ISL_1731601, EPI_ISL_1731602, EPI_ISL_1731603, EPI_ISL_1731604, EPI_ISL_1731605, EPI_ISL_1731606, EPI_ISL_1731607, EPI_ISL_1731608, EPI_ISL_1731609, EPI_ISL_1731610, EPI_ISL_1731611, EPI_ISL_1731612 | | | |
| see above | Instituto Adolfo Lutz Central | Instituto Adolfo Lutz, Interdiciplinary Procedures Center, Strategic Laboratory | Claudio Tavares Sacchi, Claudia Regina Gonçalves, Erica Valessa Ramos Gomes, Karoline Rodrigues Campos, Caio Vinicius Dias Lopes, Leonardo Jose Tadeu de Araujo, Katia Correa de Oliveira Santos |
| EPI_ISL_1731751, EPI_ISL_1731756 | State Virus Research and Diagnostic Laboratory (VRDL), AIIMS Raipur | State Virus Research and Diagnostic Laboratory (VRDL), AIIMS Raipur | Pushpendra Singh, Kuldeep Sharma, Priyanka Singh, Somya Sharma, Sanjay Singh Negi, Anudita Bhargava |
| EPI_ISL_1731832, EPI_ISL_1731833, EPI_ISL_1731834, EPI_ISL_1731835, EPI_ISL_1731836, EPI_ISL_1731837, EPI_ISL_1731838, EPI_ISL_1731839, EPI_ISL_1731840, EPI_ISL_1731841, EPI_ISL_1731842, EPI_ISL_1731843, EPI_ISL_1731844, EPI_ISL_1731845, EPI_ISL_1731846, EPI_ISL_1731847, EPI_ISL_1731848, EPI_ISL_1731849, EPI_ISL_1731850, EPI_ISL_1731851, EPI_ISL_1731852, EPI_ISL_1731854, EPI_ISL_1731855, EPI_ISL_1731856, EPI_ISL_1731857, EPI_ISL_1731858, EPI_ISL_1731862, EPI_ISL_1731863, EPI_ISL_1731864, EPI_ISL_1731865, EPI_ISL_1731866, EPI_ISL_1731867, EPI_ISL_1731868, EPI_ISL_1731869, EPI_ISL_1731870, EPI_ISL_1731871, EPI_ISL_1731872, EPI_ISL_1731873, EPI_ISL_1731874, EPI_ISL_1731875, EPI_ISL_1731876, EPI_ISL_1731877, EPI_ISL_1731879, EPI_ISL_1731880, EPI_ISL_1731881, EPI_ISL_1731882, EPI_ISL_1731883, EPI_ISL_1731884, EPI_ISL_1731885, EPI_ISL_1731886, EPI_ISL_1731887, EPI_ISL_1731888, EPI_ISL_1731889, EPI_ISL_1731890, EPI_ISL_1731891, EPI_ISL_1731892, EPI_ISL_1731893, EPI_ISL_1731894, EPI_ISL_1731895, EPI_ISL_1731896, EPI_ISL_1731897, EPI_ISL_1731898, EPI_ISL_1731899, EPI_ISL_1731900, EPI_ISL_1731901, EPI_ISL_1731903, EPI_ISL_1731904, EPI_ISL_1731905, EPI_ISL_1731906, EPI_ISL_1731907, EPI_ISL_1731908, EPI_ISL_1731909, EPI_ISL_1731910, EPI_ISL_1731911, EPI_ISL_1731912, EPI_ISL_1731914, EPI_ISL_1731915, EPI_ISL_1731916, EPI_ISL_1731917, EPI_ISL_1731918, EPI_ISL_1731919, EPI_ISL_1731921, EPI_ISL_1731922, EPI_ISL_1731923, EPI_ISL_1731924, EPI_ISL_1731925, EPI_ISL_1731926, EPI_ISL_1731927, EPI_ISL_1731928, EPI_ISL_1731929, EPI_ISL_1731930, EPI_ISL_1731931, EPI_ISL_1731932, EPI_ISL_1731933, EPI_ISL_1731935, EPI_ISL_1731937, EPI_ISL_1731938, EPI_ISL_1731939, EPI_ISL_1731940, EPI_ISL_1731942, EPI_ISL_1731943, EPI_ISL_1731944, EPI_ISL_1731945, EPI_ISL_1731947, EPI_ISL_1731948, EPI_ISL_1731949, EPI_ISL_1731950, EPI_ISL_1731952, EPI_ISL_1731955, EPI_ISL_1731956, EPI_ISL_1731957, EPI_ISL_1731958, EPI_ISL_1731959, EPI_ISL_1731960, EPI_ISL_1731961, EPI_ISL_1731962, EPI_ISL_1731963, EPI_ISL_1731964, EPI_ISL_1731965, EPI_ISL_1731966, EPI_ISL_1731967, EPI_ISL_1731968, EPI_ISL_1731969, EPI_ISL_1731970, EPI_ISL_1731971, EPI_ISL_1731972, EPI_ISL_1731973, EPI_ISL_1731974, EPI_ISL_1731975, EPI_ISL_1731977, EPI_ISL_1731978, EPI_ISL_1731979, EPI_ISL_1731980, EPI_ISL_1731981, EPI_ISL_1731983, EPI_ISL_1731984, EPI_ISL_1731985, EPI_ISL_1731986, EPI_ISL_1731987, EPI_ISL_1731989, EPI_ISL_1731990, EPI_ISL_1731991, EPI_ISL_1731992, EPI_ISL_1731993, EPI_ISL_1731994, EPI_ISL_1731995, EPI_ISL_1731996 | | | |

| | | | |
|---|---|---|---|
| see above | Synlab Eesti OÜ | 1. Laboratory of Communicable Diseases (Estonia); 2. Eurofins Genomics Europe Sequencing GmbH | Liidia Dotsenko et al. |
| EPI_ISL_1731997, EPI_ISL_1731998, EPI_ISL_1731999, EPI_ISL_1732001, EPI_ISL_1732002, EPI_ISL_1732003, EPI_ISL_1732004, EPI_ISL_1732005, EPI_ISL_1732006 | Laboratory of Communicable Diseases | 1. Laboratory of Communicable Diseases (Estonia); 2. Eurofins Genomics Europe Sequencing GmbH | Liidia Dotsenko et al. |
| EPI_ISL_1732007, EPI_ISL_1732008, EPI_ISL_1732010, EPI_ISL_1732011, EPI_ISL_1732012, EPI_ISL_1732013, EPI_ISL_1732014, EPI_ISL_1732015, EPI_ISL_1732016, EPI_ISL_1732017, EPI_ISL_1732018, EPI_ISL_1732019, EPI_ISL_1732021, EPI_ISL_1732022, EPI_ISL_1732024, EPI_ISL_1732025, EPI_ISL_1732026, EPI_ISL_1732027, EPI_ISL_1732028, EPI_ISL_1732030, EPI_ISL_1732031, EPI_ISL_1732032, EPI_ISL_1732033, EPI_ISL_1732034, EPI_ISL_1732035, EPI_ISL_1732036, EPI_ISL_1732037, EPI_ISL_1732038, EPI_ISL_1732039, EPI_ISL_1732040, EPI_ISL_1732041, EPI_ISL_1732042, EPI_ISL_1732043, EPI_ISL_1732044, EPI_ISL_1732045, EPI_ISL_1732046, EPI_ISL_1732047, EPI_ISL_1732048, EPI_ISL_1732049, EPI_ISL_1732050, EPI_ISL_1732051, EPI_ISL_1732052, EPI_ISL_1732053, EPI_ISL_1732054, EPI_ISL_1732055, EPI_ISL_1732056, EPI_ISL_1732057, EPI_ISL_1732058, EPI_ISL_1732059, EPI_ISL_1732060, EPI_ISL_1732061, EPI_ISL_1732062, EPI_ISL_1732063, EPI_ISL_1732064, EPI_ISL_1732065, EPI_ISL_1732066, EPI_ISL_1732067, EPI_ISL_1732068, EPI_ISL_1732069, EPI_ISL_1732070, EPI_ISL_1732071, EPI_ISL_1732072, EPI_ISL_1732073, EPI_ISL_1732074, EPI_ISL_1732075, EPI_ISL_1732076, EPI_ISL_1732077, EPI_ISL_1732078, EPI_ISL_1732079 | | | |
| see above | North Estonia Medical Centre | 1. Laboratory of Communicable Diseases (Estonia); 2. Eurofins Genomics Europe Sequencing GmbH | Liidia Dotsenko et al. |
| EPI_ISL_1732154, EPI_ISL_1732155 | Klinisch Laboratorium ZNA | Klinisch Laboratorium ZNA | Verstrepen et al. |
| EPI_ISL_1732156, EPI_ISL_1732157 | Klinisch Laboratorium GZA | Klinisch Laboratorium ZNA | Verstrepen et al. |
| EPI_ISL_1732158, EPI_ISL_1732159, EPI_ISL_1732160, EPI_ISL_1732161, EPI_ISL_1732162, EPI_ISL_1732163, EPI_ISL_1732164, EPI_ISL_1732165, EPI_ISL_1732166, EPI_ISL_1732167, EPI_ISL_1732168, EPI_ISL_1732169, EPI_ISL_1732170, EPI_ISL_1732171 | | | |
| see above | Klinisch Laboratorium ZNA | Klinisch Laboratorium ZNA | Verstrepen et al. |
| EPI_ISL_1732949, EPI_ISL_1732959, EPI_ISL_1732962, EPI_ISL_1732963, EPI_ISL_1732969, EPI_ISL_1732971, EPI_ISL_1732972, EPI_ISL_1732974, EPI_ISL_1732975, EPI_ISL_1732978, EPI_ISL_1732980, EPI_ISL_1732984, EPI_ISL_1732986, EPI_ISL_1732988, EPI_ISL_1732989, EPI_ISL_1732991, EPI_ISL_1732992, EPI_ISL_1732994, EPI_ISL_1732995, EPI_ISL_1732997, EPI_ISL_1732998, EPI_ISL_1733001, EPI_ISL_1733005, EPI_ISL_1733007, EPI_ISL_1733008, EPI_ISL_1733010, EPI_ISL_1733011, EPI_ISL_1733014, EPI_ISL_1733017, EPI_ISL_1733019, EPI_ISL_1733020, EPI_ISL_1733022, EPI_ISL_1733023, EPI_ISL_1733025, EPI_ISL_1733029, EPI_ISL_1733031, EPI_ISL_1733034, EPI_ISL_1733037, EPI_ISL_1733039, EPI_ISL_1733045, EPI_ISL_1733047, EPI_ISL_1733048, EPI_ISL_1733053, EPI_ISL_1733055, EPI_ISL_1733057, EPI_ISL_1733058, EPI_ISL_1733060, EPI_ISL_1733061, EPI_ISL_1733063, EPI_ISL_1733065 | | | |
| see above | Baylor Scott & White-Temple | Baylor Scott & White-Temple | Ari Rao, Linden Morales, Kimberly Walker, Marcus Volz, Shelby Hendrickson |
| EPI_ISL_1736905 | Basurto University Hospital Microbiology Lab | Biocruces Bizkaia | Mikel Urrutikoetxea-Gutierrez, Mº Carmen Nieto Toboso, Estibaliz Ugalde Zarraga, Mikel Gallego Rodrigo, Ana de la Hoz, Jose Luis Díaz de Tuesta del Arco |
| EPI_ISL_1739306 | Labo Analyses Med | National Reference Center for Viruses of Respiratory Infections, Institut Pasteur, Paris | Marion Barbet, Sylvie Behillil, Méline Bizard, Angela Brisebarre, Camille Capel, Vincent Enouf, Louise Lefrançois, Frédéric Lemoine, Christophe Malabat, Corinne Maufrais, Pierre Lechat, Etienne Simon-Lorière, Maud Vanpeene, Sylvie Van der Werf ,Fabienne Artur |
| EPI_ISL_1739307, EPI_ISL_1739308 | Hospital | National Reference Center for Viruses of Respiratory Infections, Institut Pasteur, Paris | Marion Barbet, Sylvie Behillil, Méline Bizard, Angela Brisebarre, Camille Capel, Vincent Enouf, Louise Lefrançois, Frédéric Lemoine, Christophe Malabat, Corinne Maufrais, Pierre Lechat, Etienne Simon-Lorière, Maud Vanpeene, Sylvie Van der Werf ,Nabil Gastli |
| EPI_ISL_1739309, EPI_ISL_1739310 | Hospital | National Reference Center for Viruses of Respiratory Infections, Institut Pasteur, Paris | Marion Barbet, Sylvie Behillil, Méline Bizard, Angela Brisebarre, Camille Capel, Vincent Enouf, Louise Lefrançois, Frédéric Lemoine, Christophe Malabat, Corinne Maufrais, Gael Millot, Etienne Simon-Lorière, Maud Vanpeene, Sylvie Van der Werf ,Nabil Gastli |
| EPI_ISL_1739311 | Hospital | National Reference Center for Viruses of Respiratory Infections, Institut Pasteur, Paris | Marion Barbet, Sylvie Behillil, Méline Bizard, Angela Brisebarre, Camille Capel, Vincent Enouf, Louise Lefrançois, Frédéric Lemoine, Christophe Malabat, Corinne Maufrais, Etienne Simon-Lorière, Maud Vanpeene, Sylvie Van der Werf ,Nabil Gastli |
| EPI_ISL_1739312 | Hospital | National Reference Center for Viruses of Respiratory Infections, Institut Pasteur, Paris | Marion Barbet, Sylvie Behillil, Méline Bizard, Angela Brisebarre, Camille Capel, Vincent Enouf, Louise Lefrançois, Frédéric Lemoine, Christophe Malabat, Corinne Maufrais, Etienne Simon-Lorière, Maud Vanpeene, Sylvie Van der Werf ,LéA Pilorge |
| EPI_ISL_1739313, EPI_ISL_1739314, EPI_ISL_1739315 | Hospital | National Reference Center for Viruses of Respiratory Infections, Institut Pasteur, Paris | Marion Barbet, Sylvie Behillil, Méline Bizard, Angela Brisebarre, Camille Capel, Vincent Enouf, Louise Lefrançois, Frédéric Lemoine, Christophe Malabat, Corinne Maufrais, Etienne Simon-Lorière, Maud Vanpeene, Sylvie Van der Werf ,Odile Petsaris |
| EPI_ISL_1739316 | Labo Analyses Med | National Reference Center for Viruses of Respiratory Infections, Institut Pasteur, Paris | Marion Barbet, Sylvie Behillil, Méline Bizard, Angela Brisebarre, Camille Capel, Vincent Enouf, Louise Lefrançois, Frédéric Lemoine, Christophe Malabat, Corinne Maufrais, Etienne Simon-Lorière, Maud Vanpeene, Sylvie Van der Werf ,Fabienne Artur |
| EPI_ISL_1739317 | Hospital | National Reference Center for Viruses of Respiratory Infections, Institut Pasteur, Paris | Marion Barbet, Sylvie Behillil, Méline Bizard, Angela Brisebarre, Camille Capel, Vincent Enouf, Louise Lefrançois, Frédéric Lemoine, Christophe Malabat, Corinne Maufrais, Damien Mornico, Etienne Simon-Lorière, Maud Vanpeene, Sylvie Van der Werf ,Odile Petsaris |
| EPI_ISL_1739318, EPI_ISL_1739319, EPI_ISL_1739320, EPI_ISL_1739321 | Research | National Reference Center for Viruses of Respiratory Infections, Institut Pasteur, Paris | Marion Barbet, Sylvie Behillil, Méline Bizard, Angela Brisebarre, Camille Capel, Vincent Enouf, Louise Lefrançois, Frédéric Lemoine, Christophe Malabat, Corinne Maufrais, Damien Mornico, Etienne Simon-Lorière, Maud Vanpeene, Sylvie Van der Werf ,Marianne Leruez-Ville |
| EPI_ISL_1739322 | Hospital | National Reference Center for Viruses of Respiratory Infections, Institut Pasteur, Paris | Marion Barbet, Sylvie Behillil, Méline Bizard, Angela Brisebarre, Camille Capel, Vincent Enouf, Louise Lefrançois, Frédéric Lemoine, Christophe Malabat, Corinne Maufrais, Damien Mornico, Etienne Simon-Lorière, Maud Vanpeene, Sylvie Van der Werf ,Pascale Martres |
| EPI_ISL_1739323 | Hospital | National Reference Center for Viruses of Respiratory Infections, Institut Pasteur, Paris | Marion Barbet, Sylvie Behillil, Méline Bizard, Angela Brisebarre, Camille Capel, Vincent Enouf, Louise Lefrançois, Frédéric Lemoine, Christophe Malabat, Corinne Maufrais, Etienne Simon-Lorière, Maud Vanpeene, Sylvie Van der Werf ,Sylvaine Bastian |
| EPI_ISL_1739324 | Hospital | National Reference Center for Viruses of Respiratory Infections, Institut Pasteur, Paris | Marion Barbet, Sylvie Behillil, Méline Bizard, Angela Brisebarre, Camille Capel, Vincent Enouf, Louise Lefrançois, Frédéric Lemoine, Christophe Malabat, Corinne Maufrais, Damien Mornico, Etienne Simon-Lorière, Maud Vanpeene, Sylvie Van der Werf ,Pascale Martres |
| EPI_ISL_1739325 | Hospital | National Reference Center for Viruses of Respiratory Infections, Institut Pasteur, Paris | Marion Barbet, Sylvie Behillil, Méline Bizard, Angela Brisebarre, Camille Capel, Vincent Enouf, Louise Lefrançois, Frédéric Lemoine, Christophe Malabat, Corinne Maufrais, Damien Mornico, Etienne Simon-Lorière, Maud Vanpeene, Sylvie Van der Werf ,Olivia Raulin |
| EPI_ISL_1739331 | Basurto University Hospital | Biocruces Bizkaia | Mikel Urrutikoetxea-Gutiérrez, Mº Carmen Nieto Toboso, Estibaliz Ugalde Zarraga, Mikel Gallego Rodrigo, Ana de la Hoz, Jose Luis Díaz de Tuesta del Arco |
| EPI_ISL_1743687, EPI_ISL_1743716 | Usansolo-Galdakao University Hospital | Cruces University Hospital | Ana Gual-de-Torrella, Izaskun Alejo-Cancho, Mikel Gallego, Ana Belén de la Hoz |
| EPI_ISL_1749431, EPI_ISL_1749434, EPI_ISL_1749435 | Singapore General Hospital | Department of Microbiology | Nurdyana Abdul Rahman, Kun Lee Lim, Chenhao Li, Sui Sin Goh, Kenneth Xin Long Chan, Kian Sing Chan, Lynette Oon, Kern Rei Chng, Niranjan Nagarajan, Karrie Ko |
| EPI_ISL_1749568 | Public Health Authority of the Slovak Republic | Laboratory of Genomics and Bioinformatics, Comenius University Science Park | Tatiana Sedláková, Diana Rusáková, Miroslav Böhmer, Anna Giová, Jaroslav Budiš, Tomáš Szemes |
| EPI_ISL_1752637 | UBDS Dr Marco Antonio Sahao Vila Virginia | Instituto Adolfo Lutz, Interdiciplinary Procedures Center, Strategic Laboratory | Claudio Tavares Sacchi, Claudia Regina Gonçalves, Erica Valessa Ramos Gomes, Karoline Rodrigues Campos, Caio Vinicius Dias Lopes, Leonardo Jose Tadeu de Araujo, Katia Correa de Oliveira Santos |
| EPI_ISL_1752638 | Casa de Caridade Sao Vicente de Paulo Cajuru | Instituto Adolfo Lutz, Interdiciplinary Procedures Center, Strategic Laboratory | Claudio Tavares Sacchi, Claudia Regina Gonçalves, Erica Valessa Ramos Gomes, Karoline Rodrigues Campos, Caio Vinicius Dias Lopes, Leonardo Jose Tadeu de Araujo, Katia Correa de Oliveira Santos |
| EPI_ISL_1752639 | UPA Dr Luis Atilio Losi Viana Ribeirao Preto | Instituto Adolfo Lutz, Interdiciplinary Procedures Center, Strategic Laboratory | Claudio Tavares Sacchi, Claudia Regina Gonçalves, Erica Valessa Ramos Gomes, Karoline Rodrigues Campos, Caio Vinicius Dias Lopes, Leonardo Jose Tadeu de Araujo, Katia Correa de Oliveira Santos |
| EPI_ISL_1752640 | Hospital Municipal Dr Mario Gatti Campinas | Instituto Adolfo Lutz, Interdiciplinary Procedures Center, Strategic Laboratory | Claudio Tavares Sacchi, Claudia Regina Gonçalves, Erica Valessa Ramos Gomes, Karoline Rodrigues Campos, Caio Vinicius Dias Lopes, Leonardo Jose Tadeu de Araujo, Katia Correa de Oliveira Santos |
| EPI_ISL_1752641, EPI_ISL_1752642, EPI_ISL_1752643 | Instituto Adolfo Lutz - Regional de Aracatuba | Instituto Adolfo Lutz, Interdiciplinary Procedures Center, Strategic Laboratory | Claudio Tavares Sacchi, Claudia Regina Gonçalves, Erica Valessa Ramos Gomes, Karoline Rodrigues Campos, Caio Vinicius Dias Lopes, Leonardo Jose Tadeu de Araujo, Katia Correa de Oliveira Santos |
| EPI_ISL_1752644, EPI_ISL_1752645 | Instituto Adolfo Lutz - Regional de Ribeirao Preto | Instituto Adolfo Lutz, Interdiciplinary Procedures Center, Strategic Laboratory | Claudio Tavares Sacchi, Claudia Regina Gonçalves, Erica Valessa Ramos Gomes, Karoline Rodrigues Campos, Caio Vinicius Dias Lopes, Leonardo Jose Tadeu de Araujo, Katia Correa de Oliveira Santos |
| EPI_ISL_1752646, EPI_ISL_1752647, EPI_ISL_1752648, EPI_ISL_1752655, EPI_ISL_1752656, EPI_ISL_1752658, EPI_ISL_1752659, EPI_ISL_1752660, EPI_ISL_1752661, EPI_ISL_1752663, EPI_ISL_1752664, EPI_ISL_1752665, EPI_ISL_1752666, EPI_ISL_1752667, EPI_ISL_1752668 | | | |

| | | | |
|---|---|---|---|
| see above | Instituto Adolfo Lutz Central | Instituto Adolfo Lutz, Interdiciplinary Procedures Center, Strategic Laboratory | Claudio Tavares Sacchi, Claudia Regina Gonçalves, Erica Valessa Ramos Gomes, Karoline Rodrigues Campos, Caio Vinicius Dias Lopes, Leonardo Jose Tadeu de Araujo, Katia Correa de Oliveira Santos |
| EPI_ISL_1754127, EPI_ISL_1754129, EPI_ISL_1754130, EPI_ISL_1754131, EPI_ISL_1754134, EPI_ISL_1754135 | Servicio Microbiología Hospital La Paz | Servicio Microbiología Hospital La Paz | Fernando Lázaro, Rubén Cáceres, Jesús Mingorance Cruz, Elie Dahdouh |
| EPI_ISL_1754139, EPI_ISL_1754140 | LESP Campeche | Instituto de Diagnostico y Referencia Epidemiologicos (INDRE) | Claudia Wong-Arambula, Abril Rodriguez-Maldonado, Vanessa Rivero-Arredondo, Ariadna Medina-Benitez, Joaquin Quiroz-Mercado, Sergio Rangel-Guerrero, Natividad Cruz-Ortiz, Tatiana Nunez-Garcia, Gisela Barrera-Badillo, Lucia Hernandez-Rivas, Irma Lopez-Martinez, Ernesto Ramirez-Gonzalez. |
| EPI_ISL_1754141 | LESP Baja California | Instituto de Diagnostico y Referencia Epidemiologicos (INDRE) | Claudia Wong-Arambula, Abril Rodriguez-Maldonado, Vanessa Rivero-Arredondo, Ariadna Medina-Benitez, Joaquin Quiroz-Mercado, Sergio Rangel-Guerrero, Natividad Cruz-Ortiz, Tatiana Nunez-Garcia, Gisela Barrera-Badillo, Lucia Hernandez-Rivas, Irma Lopez-Martinez, Ernesto Ramirez-Gonzalez. |
| EPI_ISL_1754233, EPI_ISL_1754236, EPI_ISL_1754249, EPI_ISL_1754341, EPI_ISL_1754343, EPI_ISL_1754345, EPI_ISL_1754349, EPI_ISL_1754351, EPI_ISL_1754356, EPI_ISL_1754359, EPI_ISL_1754360, EPI_ISL_1754363, EPI_ISL_1754364, EPI_ISL_1754370, EPI_ISL_1754371, EPI_ISL_1754382, EPI_ISL_1754384, EPI_ISL_1754387, EPI_ISL_1754395, EPI_ISL_1754396, EPI_ISL_1754586 | | | |
| see above | GH A.CHENEVIER-H.MONDOR | Department of Virology, Henri Mondor University Hospital, Assistance Publique Hôpitaux de Paris, Université Paris-Est Créteil, INSERM U955 | Christophe Rodriguez, Slim Fourati, Vanessa Demontant, Guillaume Gricourt, Melissa N'Debi, Alexandre Soulier, Elisabeth Trawinski, Jean-Michel Pawlotsky |
| EPI_ISL_1754587, EPI_ISL_1754594 | HOPITAL EMILE ROUX | Department of Virology, Henri Mondor University Hospital, Assistance Publique Hôpitaux de Paris, Université Paris-Est Créteil, INSERM U955 | Christophe Rodriguez, Slim Fourati, Vanessa Demontant, Guillaume Gricourt, Melissa N'Debi, Alexandre Soulier, Elisabeth Trawinski, Jean-Michel Pawlotsky |
| EPI_ISL_1754596, EPI_ISL_1754601, EPI_ISL_1754602, EPI_ISL_1754603, EPI_ISL_1754701, EPI_ISL_1754702, EPI_ISL_1754704, EPI_ISL_1754707, EPI_ISL_1754708, EPI_ISL_1754710, EPI_ISL_1754711 | | | |
| see above | GH A.CHENEVIER-H.MONDOR | Department of Virology, Henri Mondor University Hospital, Assistance Publique Hôpitaux de Paris, Université Paris-Est Créteil, INSERM U955 | Christophe Rodriguez, Slim Fourati, Vanessa Demontant, Guillaume Gricourt, Melissa N'Debi, Alexandre Soulier, Elisabeth Trawinski, Jean-Michel Pawlotsky |
| EPI_ISL_1754712 | HOPITAL EMILE ROUX | Department of Virology, Henri Mondor University Hospital, Assistance Publique Hôpitaux de Paris, Université Paris-Est Créteil, INSERM U955 | Christophe Rodriguez, Slim Fourati, Vanessa Demontant, Guillaume Gricourt, Melissa N'Debi, Alexandre Soulier, Elisabeth Trawinski, Jean-Michel Pawlotsky |
| EPI_ISL_1754728, EPI_ISL_1754746, EPI_ISL_1754748, EPI_ISL_1754750 | GH A.CHENEVIER-H.MONDOR | Department of Virology, Henri Mondor University Hospital, Assistance Publique Hôpitaux de Paris, Université Paris-Est Créteil, INSERM U955 | Christophe Rodriguez, Slim Fourati, Vanessa Demontant, Guillaume Gricourt, Melissa N'Debi, Alexandre Soulier, Elisabeth Trawinski, Jean-Michel Pawlotsky |
| EPI_ISL_1754764 | CH.INTERCOMMUNAL DE CRETEIL | Department of Virology, Henri Mondor University Hospital, Assistance Publique Hôpitaux de Paris, Université Paris-Est Créteil, INSERM U955 | Christophe Rodriguez, Slim Fourati, Vanessa Demontant, Guillaume Gricourt, Melissa N'Debi, Alexandre Soulier, Elisabeth Trawinski, Jean-Michel Pawlotsky |
| EPI_ISL_1754793, EPI_ISL_1754807 | GH A.CHENEVIER-H.MONDOR | Department of Virology, Henri Mondor University Hospital, Assistance Publique Hôpitaux de Paris, Université Paris-Est Créteil, INSERM U955 | Christophe Rodriguez, Slim Fourati, Vanessa Demontant, Guillaume Gricourt, Melissa N'Debi, Alexandre Soulier, Elisabeth Trawinski, Jean-Michel Pawlotsky |
| EPI_ISL_1754809 | HOPITAL EMILE ROUX | Department of Virology, Henri Mondor University Hospital, Assistance Publique Hôpitaux de Paris, Université Paris-Est Créteil, INSERM U955 | Christophe Rodriguez, Slim Fourati, Vanessa Demontant, Guillaume Gricourt, Melissa N'Debi, Alexandre Soulier, Elisabeth Trawinski, Jean-Michel Pawlotsky |
| EPI_ISL_1754827, EPI_ISL_1754834, EPI_ISL_1754837, EPI_ISL_1754838, EPI_ISL_1754843, EPI_ISL_1754844, EPI_ISL_1754976, EPI_ISL_1754982, EPI_ISL_1754983, EPI_ISL_1754984, EPI_ISL_1754985, EPI_ISL_1754986, EPI_ISL_1754988, EPI_ISL_1754995 | | | |
| see above | GH A.CHENEVIER-H.MONDOR | Department of Virology, Henri Mondor University Hospital, Assistance Publique Hôpitaux de Paris, Université Paris-Est Créteil, INSERM U955 | Christophe Rodriguez, Slim Fourati, Vanessa Demontant, Guillaume Gricourt, Melissa N'Debi, Alexandre Soulier, Elisabeth Trawinski, Jean-Michel Pawlotsky |
| EPI_ISL_1755031, EPI_ISL_1755032, EPI_ISL_1755033, EPI_ISL_1755036 | GH JOFFRE DUPUYTREN | Department of Virology, Henri Mondor University Hospital, Assistance Publique Hôpitaux de Paris, Université Paris-Est Créteil, INSERM U955 | Christophe Rodriguez, Slim Fourati, Vanessa Demontant, Guillaume Gricourt, Melissa N'Debi, Alexandre Soulier, Elisabeth Trawinski, Jean-Michel Pawlotsky |
| EPI_ISL_1755037, EPI_ISL_1755039, EPI_ISL_1755040, EPI_ISL_1755044, EPI_ISL_1755045, EPI_ISL_1755046, EPI_ISL_1755047, EPI_ISL_1755049, EPI_ISL_1755050, EPI_ISL_1755052, EPI_ISL_1755056, EPI_ISL_1755061, EPI_ISL_1755065, EPI_ISL_1755068, EPI_ISL_1755069, EPI_ISL_1755071, EPI_ISL_1755075, EPI_ISL_1755078 | | | |
| see above | GH A.CHENEVIER-H.MONDOR | Department of Virology, Henri Mondor University Hospital, Assistance Publique Hôpitaux de Paris, Université Paris-Est Créteil, INSERM U955 | Christophe Rodriguez, Slim Fourati, Vanessa Demontant, Guillaume Gricourt, Melissa N'Debi, Alexandre Soulier, Elisabeth Trawinski, Jean-Michel Pawlotsky |
| EPI_ISL_1755080 | GH JOFFRE DUPUYTREN | Department of Virology, Henri Mondor University Hospital, Assistance Publique Hôpitaux de Paris, Université Paris-Est Créteil, INSERM U955 | Christophe Rodriguez, Slim Fourati, Vanessa Demontant, Guillaume Gricourt, Melissa N'Debi, Alexandre Soulier, Elisabeth Trawinski, Jean-Michel Pawlotsky |
| EPI_ISL_1755083, EPI_ISL_1755088 | GH A.CHENEVIER-H.MONDOR | Department of Virology, Henri Mondor University Hospital, Assistance Publique Hôpitaux de Paris, Université Paris-Est Créteil, INSERM U955 | Christophe Rodriguez, Slim Fourati, Vanessa Demontant, Guillaume Gricourt, Melissa N'Debi, Alexandre Soulier, Elisabeth Trawinski, Jean-Michel Pawlotsky |
| EPI_ISL_1755093 | GH JOFFRE DUPUYTREN | Department of Virology, Henri Mondor University Hospital, Assistance Publique Hôpitaux de Paris, Université Paris-Est Créteil, INSERM U955 | Christophe Rodriguez, Slim Fourati, Vanessa Demontant, Guillaume Gricourt, Melissa N'Debi, Alexandre Soulier, Elisabeth Trawinski, Jean-Michel Pawlotsky |
| EPI_ISL_1755095, EPI_ISL_1755096, EPI_ISL_1755309, EPI_ISL_1755319 | GH A.CHENEVIER-H.MONDOR | Department of Virology, Henri Mondor University Hospital, Assistance Publique Hôpitaux de Paris, Université Paris-Est Créteil, INSERM U955 | Christophe Rodriguez, Slim Fourati, Vanessa Demontant, Guillaume Gricourt, Melissa N'Debi, Alexandre Soulier, Elisabeth Trawinski, Jean-Michel Pawlotsky |
| EPI_ISL_1755320 | HOPITAL EMILE ROUX | Department of Virology, Henri Mondor University Hospital, Assistance Publique Hôpitaux de Paris, Université Paris-Est Créteil, INSERM U955 | Christophe Rodriguez, Slim Fourati, Vanessa Demontant, Guillaume Gricourt, Melissa N'Debi, Alexandre Soulier, Elisabeth Trawinski, Jean-Michel Pawlotsky |
| EPI_ISL_1755324, EPI_ISL_1755326, EPI_ISL_1755327, EPI_ISL_1755329, EPI_ISL_1755337, EPI_ISL_1755342 | GH A.CHENEVIER-H.MONDOR | Department of Virology, Henri Mondor University Hospital, Assistance Publique Hôpitaux de Paris, Université Paris-Est Créteil, INSERM U955 | Christophe Rodriguez, Slim Fourati, Vanessa Demontant, Guillaume Gricourt, Melissa N'Debi, Alexandre Soulier, Elisabeth Trawinski, Jean-Michel Pawlotsky |
| EPI_ISL_1755357, EPI_ISL_1755362, EPI_ISL_1755365 | GH JOFFRE DUPUYTREN | Department of Virology, Henri Mondor University Hospital, Assistance Publique Hôpitaux de Paris, Université Paris-Est Créteil, INSERM U955 | Christophe Rodriguez, Slim Fourati, Vanessa Demontant, Guillaume Gricourt, Melissa N'Debi, Alexandre Soulier, Elisabeth Trawinski, Jean-Michel Pawlotsky |
| EPI_ISL_1756111 | Hôpital Pitié-Salpêtrière | Department of Virology, Henri Mondor University Hospital, Assistance Publique Hôpitaux de Paris, Université Paris-Est Créteil, INSERM U955 | Christophe Rodriguez, Slim Fourati, Vanessa Demontant, Guillaume Gricourt, Melissa N'Debi, Alexandre Soulier, Elisabeth Trawinski, Jean-Michel Pawlotsky |

| | | | |
|---|---|---|---|
| EPI_ISL_1756184 | GH A.CHENEVIER-H.MONDOR | Department of Virology, Henri Mondor University Hospital, Assistance Publique Hôpitaux de Paris, Université Paris-Est Créteil, INSERM U955 | Christophe Rodriguez, Slim Fourati, Vanessa Demontant, Guillaume Gricourt, Melissa N'Debi, Alexandre Soulier, Elisabeth Trawinski, Jean-Michel Pawlotsky |
| EPI_ISL_1756197 | HOPITAL EMILE ROUX | Department of Virology, Henri Mondor University Hospital, Assistance Publique Hôpitaux de Paris, Université Paris-Est Créteil, INSERM U955 | Christophe Rodriguez, Slim Fourati, Vanessa Demontant, Guillaume Gricourt, Melissa N'Debi, Alexandre Soulier, Elisabeth Trawinski, Jean-Michel Pawlotsky |
| EPI_ISL_1756198, EPI_ISL_1756199 | GH A.CHENEVIER-H.MONDOR | Department of Virology, Henri Mondor University Hospital, Assistance Publique Hôpitaux de Paris, Université Paris-Est Créteil, INSERM U955 | Christophe Rodriguez, Slim Fourati, Vanessa Demontant, Guillaume Gricourt, Melissa N'Debi, Alexandre Soulier, Elisabeth Trawinski, Jean-Michel Pawlotsky |
| EPI_ISL_1756200, EPI_ISL_1756206, EPI_ISL_1756207, EPI_ISL_1756209 | GH JOFFRE DUPUYTREN | Department of Virology, Henri Mondor University Hospital, Assistance Publique Hôpitaux de Paris, Université Paris-Est Créteil, INSERM U955 | Christophe Rodriguez, Slim Fourati, Vanessa Demontant, Guillaume Gricourt, Melissa N'Debi, Alexandre Soulier, Elisabeth Trawinski, Jean-Michel Pawlotsky |
| EPI_ISL_1756219, EPI_ISL_1756220, EPI_ISL_1756221, EPI_ISL_1756223, EPI_ISL_1756487, EPI_ISL_1756489, EPI_ISL_1756490, EPI_ISL_1756492, EPI_ISL_1756493, EPI_ISL_1756494, EPI_ISL_1756495, EPI_ISL_1756499, EPI_ISL_1756500, EPI_ISL_1756501, EPI_ISL_1756502, EPI_ISL_1756503, EPI_ISL_1756505, EPI_ISL_1756506, EPI_ISL_1756508, EPI_ISL_1756519, EPI_ISL_1756541, EPI_ISL_1756545 | | | |
| see above | GH A.CHENEVIER-H.MONDOR | Department of Virology, Henri Mondor University Hospital, Assistance Publique Hôpitaux de Paris, Université Paris-Est Créteil, INSERM U955 | Christophe Rodriguez, Slim Fourati, Vanessa Demontant, Guillaume Gricourt, Melissa N'Debi, Alexandre Soulier, Elisabeth Trawinski, Jean-Michel Pawlotsky |
| EPI_ISL_1759739, EPI_ISL_1759747 | Institut de Virologie du CHU de Strasbourg | Institut de Virologie du CHU de Strasbourg | Fafi-Kremer Samira, Gallais Floriane, Gantner Pierre, Laugel Elodie, Solis Morgane, Velay Aurélie, Wendling Marie-Josée |
| EPI_ISL_1760251, EPI_ISL_1760253, EPI_ISL_1760254, EPI_ISL_1760255, EPI_ISL_1760261, EPI_ISL_1760262, EPI_ISL_1760263, EPI_ISL_1760264, EPI_ISL_1760265, EPI_ISL_1760266, EPI_ISL_1760267, EPI_ISL_1760268, EPI_ISL_1760269, EPI_ISL_1760270, EPI_ISL_1760271, EPI_ISL_1760272, EPI_ISL_1760273, EPI_ISL_1760274, EPI_ISL_1760275, EPI_ISL_1760276, EPI_ISL_1760277, EPI_ISL_1760278, EPI_ISL_1760279, EPI_ISL_1760280, EPI_ISL_1760281, EPI_ISL_1760282, EPI_ISL_1760283, EPI_ISL_1760284, EPI_ISL_1760285, EPI_ISL_1760286, EPI_ISL_1760287, EPI_ISL_1760288, EPI_ISL_1760289, EPI_ISL_1760290, EPI_ISL_1760291 | | | |
| see above | Centers for Disease Control and Prevention, Dengue Branch | Centers for Disease Control and Prevention, Dengue Branch | Gilberto A. Santiago, Glenda Gonzalez, Betzabel Flores, Keyla Charriez, Gabriela Paz-Bailey, Jorge L. Munoz-Jordan |
| EPI_ISL_1761366, EPI_ISL_1761367, EPI_ISL_1761368, EPI_ISL_1761369 | Tokyo Medical and Dental University | Tokyo Medical and Dental University | Kousuke Tanimoto, Yukie Tanaka, Yoko Nukui, Akinori Kimura, Hiroaki Takeuchi |
| EPI_ISL_1761591, EPI_ISL_1761594, EPI_ISL_1761597, EPI_ISL_1761609 | Althaia. Xarxa Assistencial Universitària de Manresa | IrsiCaixa | Marc Noguera-Julian, Mariona Parera, Maria Casadellà, Pilar Armengol, Francesc Catala-Moll, Roger Paredes, Bonaventura Clotet Gloria Trujillo, Rafel Perez Vidal, Jaume Trape Pujol, Carolina Gonzalez Fernandez, Miquel Micó, Antonia Flor, Eulalia Grau |
| EPI_ISL_1764148 | Klinisch Laboratorium ZNA | Klinisch Laboratorium ZNA | Verstrepen et al. |
| EPI_ISL_1764149 | Klinisch Laboratorium GZA | Klinisch Laboratorium ZNA | Verstrepen et al. |
| EPI_ISL_1764150, EPI_ISL_1764151, EPI_ISL_1764152, EPI_ISL_1764153, EPI_ISL_1764154 | Klinisch Laboratorium ZNA | Klinisch Laboratorium ZNA | Verstrepen et al. |
| EPI_ISL_1786155, EPI_ISL_1786156, EPI_ISL_1786157, EPI_ISL_1786158, EPI_ISL_1786159, EPI_ISL_1786160, EPI_ISL_1786161, EPI_ISL_1786162, EPI_ISL_1786163, EPI_ISL_1786164, EPI_ISL_1786166, EPI_ISL_1786167, EPI_ISL_1786168, EPI_ISL_1786169, EPI_ISL_1786170, EPI_ISL_1786171, EPI_ISL_1786172, EPI_ISL_1786174, EPI_ISL_1786175, EPI_ISL_1786176, EPI_ISL_1786177, EPI_ISL_1786178, EPI_ISL_1786179, EPI_ISL_1786180, EPI_ISL_1786181, EPI_ISL_1786182, EPI_ISL_1786183, EPI_ISL_1786184, EPI_ISL_1786185, EPI_ISL_1786186, EPI_ISL_1786187, EPI_ISL_1786188, EPI_ISL_1786189, EPI_ISL_1786190, EPI_ISL_1786191, EPI_ISL_1786192, EPI_ISL_1786194, EPI_ISL_1786195, EPI_ISL_1786196, EPI_ISL_1786197, EPI_ISL_1786198, EPI_ISL_1786199, EPI_ISL_1786200, EPI_ISL_1786201, EPI_ISL_1786202, EPI_ISL_1786203, EPI_ISL_1786205, EPI_ISL_1786207, EPI_ISL_1786209, EPI_ISL_1786212, EPI_ISL_1786214, EPI_ISL_1786215, EPI_ISL_1786216, EPI_ISL_1786220, EPI_ISL_1786221, EPI_ISL_1786223, EPI_ISL_1786224, EPI_ISL_1786225, EPI_ISL_1786226, EPI_ISL_1786227, EPI_ISL_1786229, EPI_ISL_1786230, EPI_ISL_1786231, EPI_ISL_1786232, EPI_ISL_1786233, EPI_ISL_1786234, EPI_ISL_1786238, EPI_ISL_1786240, EPI_ISL_1786242, EPI_ISL_1786243, EPI_ISL_1786244, EPI_ISL_1786246, EPI_ISL_1786248, EPI_ISL_1786249, EPI_ISL_1786250, EPI_ISL_1786251, EPI_ISL_1786252, EPI_ISL_1786253, EPI_ISL_1786254, EPI_ISL_1786256, EPI_ISL_1786257, EPI_ISL_1786258, EPI_ISL_1786259, EPI_ISL_1786260, EPI_ISL_1786261, EPI_ISL_1786262, EPI_ISL_1786263, EPI_ISL_1786265, EPI_ISL_1786266, EPI_ISL_1786267, EPI_ISL_1786268, EPI_ISL_1786269, EPI_ISL_1786270, EPI_ISL_1786271, EPI_ISL_1786273, EPI_ISL_1786274, EPI_ISL_1786275, EPI_ISL_1786276, EPI_ISL_1786277, EPI_ISL_1786278, EPI_ISL_1786279, EPI_ISL_1786280, EPI_ISL_1786283, EPI_ISL_1786285, EPI_ISL_1786286, EPI_ISL_1786287, EPI_ISL_1786288, EPI_ISL_1786289, EPI_ISL_1786290, EPI_ISL_1786293, EPI_ISL_1786294, EPI_ISL_1786295, EPI_ISL_1786296, EPI_ISL_1786297, EPI_ISL_1786298, EPI_ISL_1786299, EPI_ISL_1786300, EPI_ISL_1786301, EPI_ISL_1786302, EPI_ISL_1786304, EPI_ISL_1786305, EPI_ISL_1786306, EPI_ISL_1786307, EPI_ISL_1786308, EPI_ISL_1786309, EPI_ISL_1786310, EPI_ISL_1786313, EPI_ISL_1786314, EPI_ISL_1786315, EPI_ISL_1786316, EPI_ISL_1786319, EPI_ISL_1786320, EPI_ISL_1786322, EPI_ISL_1786323, EPI_ISL_1786324, EPI_ISL_1786325, EPI_ISL_1786326, EPI_ISL_1786327, EPI_ISL_1786330, EPI_ISL_1786334, EPI_ISL_1786335, EPI_ISL_1786338, EPI_ISL_1786339, EPI_ISL_1786340, EPI_ISL_1786341, EPI_ISL_1786342, EPI_ISL_1786343, EPI_ISL_1786344, EPI_ISL_1786345, EPI_ISL_1786346, EPI_ISL_1786347, EPI_ISL_1786348, EPI_ISL_1786349, EPI_ISL_1786350, EPI_ISL_1786351, EPI_ISL_1786352, EPI_ISL_1786353, EPI_ISL_1786354, EPI_ISL_1786355, EPI_ISL_1786356, EPI_ISL_1786357, EPI_ISL_1786358, EPI_ISL_1786359, EPI_ISL_1786360, EPI_ISL_1786361, EPI_ISL_1786362, EPI_ISL_1786363, EPI_ISL_1786364, EPI_ISL_1786366, EPI_ISL_1786368, EPI_ISL_1786369, EPI_ISL_1786370, EPI_ISL_1786371, EPI_ISL_1786372, EPI_ISL_1786373, EPI_ISL_1786374, EPI_ISL_1786376, EPI_ISL_1786377, EPI_ISL_1786378, EPI_ISL_1786379, EPI_ISL_1786380, EPI_ISL_1786381, EPI_ISL_1786382, EPI_ISL_1786383, EPI_ISL_1786384, EPI_ISL_1786385, EPI_ISL_1786386, EPI_ISL_1786387, EPI_ISL_1786388, EPI_ISL_1786391, EPI_ISL_1786392, EPI_ISL_1786394, EPI_ISL_1786395, EPI_ISL_1786396, EPI_ISL_1786397, EPI_ISL_1786399, EPI_ISL_1786400, EPI_ISL_1786402, EPI_ISL_1786404, EPI_ISL_1786405, EPI_ISL_1786406, EPI_ISL_1786408, EPI_ISL_1786409, EPI_ISL_1786410, EPI_ISL_1786411, EPI_ISL_1786413, EPI_ISL_1786414, EPI_ISL_1786415, EPI_ISL_1786416, EPI_ISL_1786417, EPI_ISL_1786418, EPI_ISL_1786419 | | | |
| see above | National Center of Infectious and Parasitic Diseases | National Center of Infectious and Parasitic Diseases | Alexiev et al |
| EPI_ISL_1787318, EPI_ISL_1788079 | Institute for Medical Research, Infectious Disease Research Centre, National Institutes of Health, Ministry of Health Malaysia | Institute for Medical Research, Infectious Disease Research Centre, National Institutes of Health, Ministry of Health Malaysia | Suppiah J, Kamel K, Mohd Zawawi Z, Azizan MA, Ramly N, Robert F, Thayan R |
| EPI_ISL_1788202, EPI_ISL_1788203 | Cerballiance IDF Lisses | Cerba lab | Bénédicte Roquebert, Emmanuel Lecorche, Laura Verdurme, Sabine Trombert-Paolantoni, Aude Lessenne, Stéphanie Haïm-Boukobza |
| EPI_ISL_1788744 | SIESP DIPARTIMENTO DI PREVENZIONE TERAMO | Istituto Zooprofilattico Sperimentale dell'Abruzzo e Molise "G. Caporale" | Lorusso A, Marcacci M, Di Domenico M, Ancora M, Curini V, Di Lollo Valeria, Mangone I, Rinaldi A, Delli Compagni E, Scialabba S, Caporale M, Di Pasquale A, Cammà C, Puglia I, Calistri P, Savini G |
| EPI_ISL_1788745 | DIP. PREV. AVEZZANO SERVIZIO DI IGIENE EPIDEMIOLOGIA E SANITA' PUBBLICA | Istituto Zooprofilattico Sperimentale dell'Abruzzo e Molise "G. Caporale" | Lorusso A, Marcacci M, Di Domenico M, Ancora M, Curini V, Di Lollo Valeria, Mangone I, Rinaldi A, Delli Compagni E, Scialabba S, Caporale M, Di Pasquale A, Cammà C, Puglia I, Calistri P, Savini G |
| EPI_ISL_1788758 | Research | National Reference Center for Viruses of Respiratory Infections, Institut Pasteur, Paris | Marion Barbet, Sylvie Behillil, Méline Bizard, Angela Brisebarre, Camille Capel, Vincent Enouf, Louise Lefrançois, Frédéric Lemoine, Christophe Malabat, Corinne Maufrais, Amaury Vaysse, Etienne Simon-Lorière, Maud Vanpeene, Sylvie Van der Werf ,Marianne Leruez-Ville |
| EPI_ISL_1788761, EPI_ISL_1788763 | Research | National Reference Center for Viruses of Respiratory Infections, Institut Pasteur, Paris | Marion Barbet, Sylvie Behillil, Méline Bizard, Angela Brisebarre, Camille Capel, Vincent Enouf, Louise Lefrançois, Frédéric Lemoine, Christophe Malabat, Corinne Maufrais, Etienne Simon-Lorière, Maud Vanpeene, Sylvie Van der Werf ,Marianne Leruez-Ville |
| EPI_ISL_1788764, EPI_ISL_1788765, EPI_ISL_1788766 | Research | National Reference Center for Viruses of Respiratory Infections, Institut Pasteur, Paris | Marion Barbet, Sylvie Behillil, Méline Bizard, Angela Brisebarre, Camille Capel, Vincent Enouf, Louise Lefrançois, Frédéric Lemoine, Christophe Malabat, Corinne Maufrais, Amaury Vaysse, Etienne Simon-Lorière, Maud Vanpeene, Sylvie Van der Werf ,Marianne Leruez-Ville |
| EPI_ISL_1788767, EPI_ISL_1788768 | Research | National Reference Center for Viruses of Respiratory Infections, Institut Pasteur, Paris | Marion Barbet, Sylvie Behillil, Méline Bizard, Angela Brisebarre, Camille Capel, Vincent Enouf, Louise Lefrançois, Frédéric Lemoine, Christophe Malabat, Corinne Maufrais, Etienne Simon-Lorière, Maud Vanpeene, Sylvie Van der Werf ,Marianne Leruez-Ville |
| EPI_ISL_1788779 | Hospital | National Reference Center for Viruses of Respiratory Infections, Institut Pasteur, Paris | Marion Barbet, Sylvie Behillil, Méline Bizard, Angela Brisebarre, Camille Capel, Vincent Enouf, Louise Lefrançois, Frédéric Lemoine, Christophe Malabat, Corinne Maufrais, Etienne Simon-Lorière, Maud Vanpeene, Sylvie Van der Werf ,FrançOise Lunel Fabiani |
| EPI_ISL_1788821, EPI_ISL_1788822, EPI_ISL_1788823, EPI_ISL_1788824, EPI_ISL_1788825, EPI_ISL_1788826, EPI_ISL_1788827, EPI_ISL_1788829, EPI_ISL_1788830, EPI_ISL_1788831, EPI_ISL_1788832, EPI_ISL_1788833, EPI_ISL_1788834, EPI_ISL_1788835, EPI_ISL_1788836, EPI_ISL_1788837, EPI_ISL_1788838 | | | |
| see above | Labo Analyses Med | National Reference Center for Viruses of Respiratory Infections, Institut Pasteur, Paris | Marion Barbet, Sylvie Behillil, Méline Bizard, Angela Brisebarre, Camille Capel, Vincent Enouf, Louise Lefrançois, Frédéric Lemoine, Christophe Malabat, Corinne Maufrais, Amaury Vaysse, Etienne Simon-Lorière, Maud Vanpeene, Sylvie Van der Werf ,Catherine Coignard |
| EPI_ISL_1788882, EPI_ISL_1788883, EPI_ISL_1788884 | Hospital | National Reference Center for Viruses of Respiratory Infections, Institut Pasteur, Paris | Marion Barbet, Sylvie Behillil, Méline Bizard, Angela Brisebarre, Camille Capel, Vincent Enouf, Louise Lefrançois, Frédéric Lemoine, Christophe Malabat, Corinne Maufrais, Etienne Simon-Lorière, Maud Vanpeene, Sylvie Van der Werf ,LéA Pilorge |
| EPI_ISL_1788915, EPI_ISL_1788916 | Hospital | National Reference Center for Viruses of Respiratory Infections, Institut Pasteur, Paris | Marion Barbet, Sylvie Behillil, Méline Bizard, Angela Brisebarre, Camille Capel, Vincent Enouf, Louise Lefrançois, Frédéric Lemoine, Christophe Malabat, Corinne Maufrais, Etienne Simon-Lorière, Maud Vanpeene, Sylvie Van der Werf ,Jean Baptiste Bour |
| EPI_ISL_1788971 | Labo Analyses Med | National Reference Center for Viruses of Respiratory Infections, Institut Pasteur, Paris | Marion Barbet, Sylvie Behillil, Méline Bizard, Angela Brisebarre, Camille Capel, Vincent Enouf, Louise Lefrançois, Frédéric Lemoine, Christophe Malabat, Corinne Maufrais, Etienne Simon-Lorière, Maud Vanpeene, Sylvie Van der Werf ,Franck Griscelli |
| EPI_ISL_1789048 | Hospital | National Reference Center for Viruses of Respiratory | Marion Barbet, Sylvie Behillil, Méline Bizard, Angela Brisebarre, Camille Capel, Vincent Enouf, Louise Lefrançois, Frédéric Lemoine, Christophe Malabat, |

| | | | |
|---|---|---|---|
| | | Infections, Institut Pasteur, Paris | Corinne Maufrais, Etienne Simon-Lorière, Maud Vanpeene, Sylvie Van der Werf ,Emilie Przyrowski |
| EPI_ISL_1789049 | Labo Analyses Med | National Reference Center for Viruses of Respiratory Infections, Institut Pasteur, Paris | Marion Barbet, Sylvie Behillil, Méline Bizard, Angela Brisebarre, Camille Capel, Vincent Enouf, Louise Lefrançois, Frédéric Lemoine, Christophe Malabat, Corinne Maufrais, Etienne Simon-Lorière, Maud Vanpeene, Sylvie Van der Werf ,Vicky Le (V) |
| EPI_ISL_1789204 | Institute for Medical Research, Infectious Disease Research Centre, National Institutes of Health, Ministry of Health Malaysia | Institute for Medical Research, Infectious Disease Research Centre, National Institutes of Health, Ministry of Health Malaysia | Suppiah J, Kamel K, Mohd Zawawi Z, Ramly N, Robert F, Thayan R |
| EPI_ISL_1789591, EPI_ISL_1789671, EPI_ISL_1789672, EPI_ISL_1789680, EPI_ISL_1789728, EPI_ISL_1789776, EPI_ISL_1789777, EPI_ISL_1789852, EPI_ISL_1790048, EPI_ISL_1790050, EPI_ISL_1790051, EPI_ISL_1790059 | | | |
| see above | Institut de Virologie du CHU de Strasbourg | Institut de Virologie du CHU de Strasbourg | Fafi-Kremer Samira, Gallais Floriane, Gantner Pierre, Laugel Elodie, Solis Morgane, Velay Aurélie, Wendling Marie-Josée |
| EPI_ISL_1791031 | Virology Unit, AOUP | Virology Unit | Marialinda Vatteroni, Susi Frateschi, Mauro Pistello |
| EPI_ISL_1791475, EPI_ISL_1791476, EPI_ISL_1791477, EPI_ISL_1791478, EPI_ISL_1791479, EPI_ISL_1791480, EPI_ISL_1791481, EPI_ISL_1791482, EPI_ISL_1791483, EPI_ISL_1791484, EPI_ISL_1791485, EPI_ISL_1791486, EPI_ISL_1791487, EPI_ISL_1791493, EPI_ISL_1791495, EPI_ISL_1791496, EPI_ISL_1791497, EPI_ISL_1791498, EPI_ISL_1791499, EPI_ISL_1791500, EPI_ISL_1791501, EPI_ISL_1791502, EPI_ISL_1791503, EPI_ISL_1791504, EPI_ISL_1791505, EPI_ISL_1791506, EPI_ISL_1791507 | | | |
| see above | University of Oregon COVID-19 MAP Laboratory | University of Oregon Genomics and Cell Characterization Core Facility (GC3F) | Douglas Turnbull, Ariana White, Jeff Bishop, Jason Carriere |
| EPI_ISL_1794455 | Laboratorio PGM | Laboratorio de Infectología Molecular, Departamento de Bioquímica y Medicina Molecular,Facultad de Medicina - Universidad Autónoma de Nuevo León | Kame A. Galán-Huerta, María F. Herrera-Saldivar, Natalia Martínez-Acuña, Sonia A. Lozano-Sepúlveda, Daniel Arellanos-Soto, Ana M. Rivas-Estilla, Javier Ramos-Jimenez, Gabriela Elizondo, Eduardo Garza-de-la-Peña |
| EPI_ISL_1797344, EPI_ISL_1797345, EPI_ISL_1797346, EPI_ISL_1797348, EPI_ISL_1797349, EPI_ISL_1797350 | National Center of Infectious and Parasitic Diseases | National Center of Infectious and Parasitic Diseases | Alexiev et al |
| EPI_ISL_1805465 | LESP Queretaro | Instituto de Diagnostico y Referencia Epidemiologicos (INDRE) | Claudia Wong-Arambula, Abril Rodriguez-Maldonado, Vanessa Rivero-Arredondo, Ariadna Medina-Benitez, Joaquin Quiroz-Mercado, Sergio Rangel-Guerrero, Natividad Cruz-Ortiz, Tatiana Nunez-Garcia, Gisela Barrera-Badillo, Lucia Hernandez-Rivas, Irma Lopez-Martinez, Ernesto Ramirez-Gonzalez. |
| EPI_ISL_1805466, EPI_ISL_1805467, EPI_ISL_1805468, EPI_ISL_1805469 | LESP Chihuahua | Instituto de Diagnostico y Referencia Epidemiologicos (INDRE) | Claudia Wong-Arambula, Abril Rodriguez-Maldonado, Vanessa Rivero-Arredondo, Ariadna Medina-Benitez, Joaquin Quiroz-Mercado, Sergio Rangel-Guerrero, Natividad Cruz-Ortiz, Tatiana Nunez-Garcia, Gisela Barrera-Badillo, Lucia Hernandez-Rivas, Irma Lopez-Martinez, Ernesto Ramirez-Gonzalez. |
| EPI_ISL_1805470 | LESP Quintana Roo | Instituto de Diagnostico y Referencia Epidemiologicos (INDRE) | Claudia Wong-Arambula, Abril Rodriguez-Maldonado, Vanessa Rivero-Arredondo, Ariadna Medina-Benitez, Joaquin Quiroz-Mercado, Sergio Rangel-Guerrero, Natividad Cruz-Ortiz, Tatiana Nunez-Garcia, Gisela Barrera-Badillo, Lucia Hernandez-Rivas, Irma Lopez-Martinez, Ernesto Ramirez-Gonzalez. |
| EPI_ISL_1805472 | LESP Sonora | Instituto de Diagnostico y Referencia Epidemiologicos (INDRE) | Claudia Wong-Arambula, Abril Rodriguez-Maldonado, Vanessa Rivero-Arredondo, Ariadna Medina-Benitez, Joaquin Quiroz-Mercado, Sergio Rangel-Guerrero, Natividad Cruz-Ortiz, Tatiana Nunez-Garcia, Gisela Barrera-Badillo, Lucia Hernandez-Rivas, Irma Lopez-Martinez, Ernesto Ramirez-Gonzalez. |
| EPI_ISL_1805473 | LESP Nuevo Leon | Instituto de Diagnostico y Referencia Epidemiologicos (INDRE) | Claudia Wong-Arambula, Abril Rodriguez-Maldonado, Vanessa Rivero-Arredondo, Ariadna Medina-Benitez, Joaquin Quiroz-Mercado, Sergio Rangel-Guerrero, Natividad Cruz-Ortiz, Tatiana Nunez-Garcia, Gisela Barrera-Badillo, Lucia Hernandez-Rivas, Irma Lopez-Martinez, Ernesto Ramirez-Gonzalez. |
| EPI_ISL_1805476, EPI_ISL_1805477, EPI_ISL_1805479 | LESP Chihuahua | Instituto de Diagnostico y Referencia Epidemiologicos (INDRE) | Claudia Wong-Arambula, Abril Rodriguez-Maldonado, Vanessa Rivero-Arredondo, Ariadna Medina-Benitez, Joaquin Quiroz-Mercado, Sergio Rangel-Guerrero, Natividad Cruz-Ortiz, Tatiana Nunez-Garcia, Gisela Barrera-Badillo, Lucia Hernandez-Rivas, Irma Lopez-Martinez, Ernesto Ramirez-Gonzalez. |
| EPI_ISL_1805482 | LESP Oaxaca | Instituto de Diagnostico y Referencia Epidemiologicos (INDRE) | Claudia Wong-Arambula, Abril Rodriguez-Maldonado, Vanessa Rivero-Arredondo, Ariadna Medina-Benitez, Joaquin Quiroz-Mercado, Sergio Rangel-Guerrero, Natividad Cruz-Ortiz, Tatiana Nunez-Garcia, Gisela Barrera-Badillo, Lucia Hernandez-Rivas, Irma Lopez-Martinez, Ernesto Ramirez-Gonzalez. |
| EPI_ISL_1805483 | LESP Hidalgo | Instituto de Diagnostico y Referencia Epidemiologicos (INDRE) | Claudia Wong-Arambula, Abril Rodriguez-Maldonado, Vanessa Rivero-Arredondo, Ariadna Medina-Benitez, Joaquin Quiroz-Mercado, Sergio Rangel-Guerrero, Natividad Cruz-Ortiz, Tatiana Nunez-Garcia, Gisela Barrera-Badillo, Lucia Hernandez-Rivas, Irma Lopez-Martinez, Ernesto Ramirez-Gonzalez. |
| EPI_ISL_1805486 | LESP Ciudad de Mexico | Instituto de Diagnostico y Referencia Epidemiologicos (INDRE) | Claudia Wong-Arambula, Abril Rodriguez-Maldonado, Vanessa Rivero-Arredondo, Ariadna Medina-Benitez, Joaquin Quiroz-Mercado, Sergio Rangel-Guerrero, Natividad Cruz-Ortiz, Tatiana Nunez-Garcia, Gisela Barrera-Badillo, Lucia Hernandez-Rivas, Irma Lopez-Martinez, Ernesto Ramirez-Gonzalez. |
| EPI_ISL_1805490 | LESP Nuevo Leon | Instituto de Diagnostico y Referencia Epidemiologicos (INDRE) | Claudia Wong-Arambula, Abril Rodriguez-Maldonado, Vanessa Rivero-Arredondo, Ariadna Medina-Benitez, Joaquin Quiroz-Mercado, Sergio Rangel-Guerrero, Natividad Cruz-Ortiz, Tatiana Nunez-Garcia, Gisela Barrera-Badillo, Lucia Hernandez-Rivas, Irma Lopez-Martinez, Ernesto Ramirez-Gonzalez. |
| EPI_ISL_1805493 | LESP Baja California | Instituto de Diagnostico y Referencia Epidemiologicos (INDRE) | Claudia Wong-Arambula, Abril Rodriguez-Maldonado, Vanessa Rivero-Arredondo, Ariadna Medina-Benitez, Joaquin Quiroz-Mercado, Sergio Rangel-Guerrero, Natividad Cruz-Ortiz, Tatiana Nunez-Garcia, Gisela Barrera-Badillo, Lucia Hernandez-Rivas, Irma Lopez-Martinez, Ernesto Ramirez-Gonzalez. |
| EPI_ISL_1805498 | LESP Guanajuato | Instituto de Diagnostico y Referencia Epidemiologicos (INDRE) | Claudia Wong-Arambula, Abril Rodriguez-Maldonado, Vanessa Rivero-Arredondo, Ariadna Medina-Benitez, Joaquin Quiroz-Mercado, Sergio Rangel-Guerrero, Natividad Cruz-Ortiz, Tatiana Nunez-Garcia, Gisela Barrera-Badillo, Lucia Hernandez-Rivas, Irma Lopez-Martinez, Ernesto Ramirez-Gonzalez. |
| EPI_ISL_1805499, EPI_ISL_1805500, EPI_ISL_1805503, EPI_ISL_1805504 | LESP Nuevo Leon | Instituto de Diagnostico y Referencia Epidemiologicos (INDRE) | Claudia Wong-Arambula, Abril Rodriguez-Maldonado, Vanessa Rivero-Arredondo, Ariadna Medina-Benitez, Joaquin Quiroz-Mercado, Sergio Rangel-Guerrero, Natividad Cruz-Ortiz, Tatiana Nunez-Garcia, Gisela Barrera-Badillo, Lucia Hernandez-Rivas, Irma Lopez-Martinez, Ernesto Ramirez-Gonzalez. |
| EPI_ISL_1805505, EPI_ISL_1805508 | Institute for Medical Research, Infectious Disease Research Centre, National Institutes of Health, Ministry of Health Malaysia | Institute for Medical Research, Infectious Disease Research Centre, National Institutes of Health, Ministry of Health Malaysia | Suppiah J, Kamel K, Mohd Zawawi Z, Ramly N, Robert F, Thayan R |
| EPI_ISL_1805639 | LDSP FRONTERIZO | Instituto Nacional de Salud- Dirección de Investigación en Salud Pública | Katherine Laiton-Donato, Diego A. Álvarez-Díaz, Carlos Franco-Muñoz, Hector Alejandro Ruiz-Moreno, Paola Rojas, Maria T. Herrera-Sepúlveda, Diego Andrés Prada, Jhonnatan Reales-González, Sheryll Corchuelo, Julian Naizaque, Jorge Rivera, Gerardo Santamaría, Sergio Gomez, Lisseth Pardo, Juan Camilo Martinez, Marta Lopez Blanco, Ángela Alarcon Cruz, Diana Malo, Carmen Osorio, Magdalena Wiesner, Martha Lucia Ospina Martinez, Marcela Mercado-Reyes |
| EPI_ISL_1805747 | National Center of Infectious and Parasitic Diseases | National Center of Infectious and Parasitic Diseases | Alexiev et al |
| EPI_ISL_1807185, EPI_ISL_1807188, EPI_ISL_1807206, EPI_ISL_1807207 | Institute for Medical Research, Infectious Disease Research Centre, National Institutes of Health, Ministry of Health Malaysia | Institute for Medical Research, Infectious Disease Research Centre, National Institutes of Health, Ministry of Health Malaysia | Suppiah J, Kamel K, Mohd Zawawi Z, Ramly N, Robert F, Thayan R |
| EPI_ISL_1807718 | Laboratorio Analisi Praecilia srl | Eurofins Genoma Group | Francesca Spinella, Riccardo Giannico, Valentina Andrioletti, Ettore Cotroneo (Eurofins Genoma Group) |
| EPI_ISL_1811224 | AS Barva | Inciensa, Instituto Costarricense de Investigación y Enseñanza en Nutrición y Salud | Pérez-Corrales C & Kellerman-Montero K |
| EPI_ISL_1811225 | AS Puerto Viejo | Inciensa, Instituto Costarricense de Investigación y | Pérez-Corrales C & Murillo-Bustos H |

| | | | |
|---|---|---|---|
| | | Enseñanza en Nutrición y Salud | |
| EPI_ISL_1811226 | Guapiles | Inciensa, Instituto Costarricense de Investigación y Enseñanza en Nutrición y Salud | Pérez-Corrales C & Cerdas-Quesada C |
| EPI_ISL_1811227 | HNN | Inciensa, Instituto Costarricense de Investigación y Enseñanza en Nutrición y Salud | Pérez-Corrales C & Montero-Chacón L. |
| EPI_ISL_1811228 | Horquetas Rio Frio | Inciensa, Instituto Costarricense de Investigación y Enseñanza en Nutrición y Salud | Pérez-Corrales C & Montiel-Ulloa G |
| EPI_ISL_1811229 | AS Belen Flores | Inciensa, Instituto Costarricense de Investigación y Enseñanza en Nutrición y Salud | Pérez-Corrales C & Centeno-Miranda M |
| EPI_ISL_1811230 | AS Santa Barbara | Inciensa, Instituto Costarricense de Investigación y Enseñanza en Nutrición y Salud | Pérez-Corrales C & Aguilar-Monge R |
| EPI_ISL_1811231 | AS Alajuela Central | Inciensa, Instituto Costarricense de Investigación y Enseñanza en Nutrición y Salud | Pérez-Corrales C |
| EPI_ISL_1811232 | Santo Domingo | Inciensa, Instituto Costarricense de Investigación y Enseñanza en Nutrición y Salud | Pérez-Corrales C & Blanco-Delgado C |
| EPI_ISL_1811233 | San Isidro de Heredia | Inciensa, Instituto Costarricense de Investigación y Enseñanza en Nutrición y Salud | Pérez-Corrales C & Fonseca-Muñoz R |
| EPI_ISL_1811237 | AS Santa Barbara | Inciensa, Instituto Costarricense de Investigación y Enseñanza en Nutrición y Salud | Pérez-Corrales C & Aguilar-Monge R |
| EPI_ISL_1811238 | San Isidro de Heredia | Inciensa, Instituto Costarricense de Investigación y Enseñanza en Nutrición y Salud | Pérez-Corrales C & Fonseca-Muñoz R |
| EPI_ISL_1811239 | AS Alajuela Central | Inciensa, Instituto Costarricense de Investigación y Enseñanza en Nutrición y Salud | Pérez-Corrales C & Mairena-Acuña C |
| EPI_ISL_1811240 | AS Alajuela Central | Inciensa, Instituto Costarricense de Investigación y Enseñanza en Nutrición y Salud | Pérez-Corrales C & Zuñiga-Carvajal P |
| EPI_ISL_1811379, EPI_ISL_1811380, EPI_ISL_1811381, EPI_ISL_1811382, EPI_ISL_1811383, EPI_ISL_1811386, EPI_ISL_1811387, EPI_ISL_1811388, EPI_ISL_1811390, EPI_ISL_1811391, EPI_ISL_1811393, EPI_ISL_1811394, EPI_ISL_1811395, EPI_ISL_1811396, EPI_ISL_1811397, EPI_ISL_1811398, EPI_ISL_1811400, EPI_ISL_1811401, EPI_ISL_1811404, EPI_ISL_1811405, EPI_ISL_1811406, EPI_ISL_1811407, EPI_ISL_1811408, EPI_ISL_1811409, EPI_ISL_1811410, EPI_ISL_1811411, EPI_ISL_1811412, EPI_ISL_1811413, EPI_ISL_1811414, EPI_ISL_1811415, EPI_ISL_1811416, EPI_ISL_1811417, EPI_ISL_1811418, EPI_ISL_1811419, EPI_ISL_1811420, EPI_ISL_1811421, EPI_ISL_1811424, EPI_ISL_1811425, EPI_ISL_1811426, EPI_ISL_1811427, EPI_ISL_1811428, EPI_ISL_1811430, EPI_ISL_1811431, EPI_ISL_1811432, EPI_ISL_1811433, EPI_ISL_1811434, EPI_ISL_1811435, EPI_ISL_1811436, EPI_ISL_1811437, EPI_ISL_1811439, EPI_ISL_1811440, EPI_ISL_1811441, EPI_ISL_1811442, EPI_ISL_1811443, EPI_ISL_1811444, EPI_ISL_1811445, EPI_ISL_1811446, EPI_ISL_1811447, EPI_ISL_1811448, EPI_ISL_1811449, EPI_ISL_1811450, EPI_ISL_1811451, EPI_ISL_1811452, EPI_ISL_1811453, EPI_ISL_1811455, EPI_ISL_1811456, EPI_ISL_1811461, EPI_ISL_1811462, EPI_ISL_1811463, EPI_ISL_1811464, EPI_ISL_1811465, EPI_ISL_1811466, EPI_ISL_1811467, EPI_ISL_1811468, EPI_ISL_1811469, EPI_ISL_1811470, EPI_ISL_1811471, EPI_ISL_1811474, EPI_ISL_1811475, EPI_ISL_1811477, EPI_ISL_1811478, EPI_ISL_1811479, EPI_ISL_1811480, EPI_ISL_1811482, EPI_ISL_1811483, EPI_ISL_1811484, EPI_ISL_1811486, EPI_ISL_1811487, EPI_ISL_1811488, EPI_ISL_1811489, EPI_ISL_1811490, EPI_ISL_1811491, EPI_ISL_1811493, EPI_ISL_1811494, EPI_ISL_1811497, EPI_ISL_1811498, EPI_ISL_1811499, EPI_ISL_1811500, EPI_ISL_1811501, EPI_ISL_1811502, EPI_ISL_1811505, EPI_ISL_1811507, EPI_ISL_1811508, EPI_ISL_1811509, EPI_ISL_1811510, EPI_ISL_1811512, EPI_ISL_1811514, EPI_ISL_1811515, EPI_ISL_1811517, EPI_ISL_1811518, EPI_ISL_1811519, EPI_ISL_1811520, EPI_ISL_1811521, EPI_ISL_1811524, EPI_ISL_1811528, EPI_ISL_1811535, EPI_ISL_1811536, EPI_ISL_1811537, EPI_ISL_1811538, EPI_ISL_1811539, EPI_ISL_1811540, EPI_ISL_1811541, EPI_ISL_1811542, EPI_ISL_1811543, EPI_ISL_1811544 | | | |
| see above | Laboratorio Central de Epidemiología (LCE) | Instituto de Biotecnología de la UNAM | Julio Elias Alvarado-Yaah, Carlos F. Arias, Santiago Ávila-Ríos, Víctor Hugo Borja-Aburto, Celia Boukadida, Juan Bautista Chale-Dzul, Célida Duque-Molina, José Antonio Enciso-Moreno, Gloria Elena Espinosa-Ayala, Fernando Fontove-Herrera, Victor Eduardo Garcia-Arias, Concepción Grajales-Muñiz, Ricardo Grande, Alfredo Herrera-Estrella, Carla Ivón Herrera-Najera, Pavel Isa, Brenda Irasema Maldonado-Meza, Bernardo Martínez-Miguel, Margarita Matías-Florentino, María Guadalupe de Jesús Mireles-Rivera, Gloria María Molina-Salinas, Hector Montoya-Fuentes, José Esteban Muñoz-Medina, José de Jesús Nuñez-Contreras, Alicia Ocaña-Mondragón, Luis Alberto Ochoa-Carrera, Hector Esteban Paz-Juárez, Francisco Pulido, Helen Haydee Fernanda Ramírez-Plascencia, Angel Gustavo Salas-Lais, Alejandro Sanchez-Flores, Clara Esperanza Santacruz-Tinoco, María Guadalupe Santiago-Mauricio, Nelly Sélem-Mojica, Blanca Taboada, Gloria Vazquez |
| EPI_ISL_1817561, EPI_ISL_1817621, EPI_ISL_1817650, EPI_ISL_1817654 | Servicio Microbiología Hospital La Paz | Servicio Microbiología Hospital La Paz | Fernando Lázaro, Rubén Cáceres, Jesús Mingorance Cruz, Elie Dahdouh |
| EPI_ISL_1818780 | CH Saint Philibert | CHU Lille - Laboratoire de Virologie | AIT YAHYA Emilie, ALIDJINOU Enagnon Kazali, BOCKET Laurence, CREPIN Michel, DEMAY Christophe, ENGELMANN Ilka, GEFFROY Sandrine, GUIGON Aurélie, LAMBERT Valérie, LAZREK Mouna, NOBILLIAUX Florian, PREVOST Brigitte, THUILLIER Caroline, TINEZ Claire |
| EPI_ISL_1818781 | CH Seclin | CHU Lille - Laboratoire de Virologie | AIT YAHYA Emilie, ALIDJINOU Enagnon Kazali, BOCKET Laurence, CREPIN Michel, DEMAY Christophe, ENGELMANN Ilka, GEFFROY Sandrine, GUIGON Aurélie, LAMBERT Valérie, LAZREK Mouna, NOBILLIAUX Florian, PREVOST Brigitte, THUILLIER Caroline, TINEZ Claire |
| EPI_ISL_1818783 | CH Boulogne sur mer | CHU Lille - Laboratoire de Virologie | AIT YAHYA Emilie, ALIDJINOU Enagnon Kazali, BOCKET Laurence, CREPIN Michel, DEMAY Christophe, ENGELMANN Ilka, GEFFROY Sandrine, GUIGON Aurélie, LAMBERT Valérie, LAZREK Mouna, NOBILLIAUX Florian, PREVOST Brigitte, THUILLIER Caroline, TINEZ Claire |
| EPI_ISL_1818784 | CHU Lille | CHU Lille - Laboratoire de Virologie | AIT YAHYA Emilie, ALIDJINOU Enagnon Kazali, BOCKET Laurence, CREPIN Michel, DEMAY Christophe, ENGELMANN Ilka, GEFFROY Sandrine, GUIGON Aurélie, LAMBERT Valérie, LAZREK Mouna, NOBILLIAUX Florian, PREVOST Brigitte, THUILLIER Caroline, TINEZ Claire |
| EPI_ISL_1818786 | CH Béthune | CHU Lille - Laboratoire de Virologie | AIT YAHYA Emilie, ALIDJINOU Enagnon Kazali, BOCKET Laurence, CREPIN Michel, DEMAY Christophe, ENGELMANN Ilka, GEFFROY Sandrine, GUIGON Aurélie, LAMBERT Valérie, LAZREK Mouna, NOBILLIAUX Florian, PREVOST Brigitte, THUILLIER Caroline, TINEZ Claire |
| EPI_ISL_1818787 | CHU Lille | CHU Lille - Laboratoire de Virologie | AIT YAHYA Emilie, ALIDJINOU Enagnon Kazali, BOCKET Laurence, CREPIN Michel, DEMAY Christophe, ENGELMANN Ilka, GEFFROY Sandrine, GUIGON Aurélie, LAMBERT Valérie, LAZREK Mouna, NOBILLIAUX Florian, PREVOST Brigitte, THUILLIER Caroline, TINEZ Claire |
| EPI_ISL_1818788 | CH Cambrai | CHU Lille - Laboratoire de Virologie | AIT YAHYA Emilie, ALIDJINOU Enagnon Kazali, BOCKET Laurence, CREPIN Michel, DEMAY Christophe, ENGELMANN Ilka, GEFFROY Sandrine, GUIGON Aurélie, LAMBERT Valérie, LAZREK Mouna, NOBILLIAUX Florian, PREVOST Brigitte, THUILLIER Caroline, TINEZ Claire |
| EPI_ISL_1818791 | CHU Lille | CHU Lille - Laboratoire de Virologie | AIT YAHYA Emilie, ALIDJINOU Enagnon Kazali, BOCKET Laurence, CREPIN Michel, DEMAY Christophe, ENGELMANN Ilka, GEFFROY Sandrine, GUIGON Aurélie, LAMBERT Valérie, LAZREK Mouna, NOBILLIAUX Florian, PREVOST Brigitte, THUILLIER Caroline, TINEZ Claire |
| EPI_ISL_1818792 | CH Douai | CHU Lille - Laboratoire de Virologie | AIT YAHYA Emilie, ALIDJINOU Enagnon Kazali, BOCKET Laurence, CREPIN Michel, DEMAY Christophe, ENGELMANN Ilka, GEFFROY Sandrine, GUIGON Aurélie, LAMBERT Valérie, LAZREK Mouna, NOBILLIAUX Florian, PREVOST Brigitte, THUILLIER Caroline, TINEZ Claire |
| EPI_ISL_1818793 | CH Cambrai | CHU Lille - Laboratoire de Virologie | AIT YAHYA Emilie, ALIDJINOU Enagnon Kazali, BOCKET Laurence, CREPIN Michel, DEMAY Christophe, ENGELMANN Ilka, GEFFROY Sandrine, GUIGON Aurélie, LAMBERT Valérie, LAZREK Mouna, NOBILLIAUX Florian, PREVOST Brigitte, THUILLIER Caroline, TINEZ Claire |
| EPI_ISL_1818795 | CHU Lille | CHU Lille - Laboratoire de Virologie | AIT YAHYA Emilie, ALIDJINOU Enagnon Kazali, BOCKET Laurence, CREPIN Michel, DEMAY Christophe, ENGELMANN Ilka, GEFFROY Sandrine, GUIGON Aurélie, LAMBERT Valérie, LAZREK Mouna, NOBILLIAUX Florian, PREVOST Brigitte, THUILLIER Caroline, TINEZ Claire |
| EPI_ISL_1818796 | CH Armentières | CHU Lille - Laboratoire de Virologie | AIT YAHYA Emilie, ALIDJINOU Enagnon Kazali, BOCKET Laurence, CREPIN Michel, DEMAY Christophe, ENGELMANN Ilka, GEFFROY Sandrine, GUIGON Aurélie, LAMBERT Valérie, LAZREK Mouna, NOBILLIAUX Florian, PREVOST Brigitte, THUILLIER Caroline, TINEZ Claire |
| EPI_ISL_1818797 | CH Dunkerque | CHU Lille - Laboratoire de Virologie | AIT YAHYA Emilie, ALIDJINOU Enagnon Kazali, BOCKET Laurence, CREPIN Michel, DEMAY Christophe, ENGELMANN Ilka, GEFFROY Sandrine, GUIGON Aurélie, LAMBERT Valérie, LAZREK Mouna, NOBILLIAUX Florian, PREVOST Brigitte, THUILLIER Caroline, TINEZ Claire |
| EPI_ISL_1818801 | CH Arras | CHU Lille - Laboratoire de Virologie | AIT YAHYA Emilie, ALIDJINOU Enagnon Kazali, BOCKET Laurence, CREPIN Michel, DEMAY Christophe, ENGELMANN Ilka, GEFFROY Sandrine, GUIGON Aurélie, LAMBERT Valérie, LAZREK Mouna, NOBILLIAUX Florian, PREVOST Brigitte, THUILLIER Caroline, TINEZ Claire |
| EPI_ISL_1818802, EPI_ISL_1818803 | CHU Lille | CHU Lille - Laboratoire de Virologie | AIT YAHYA Emilie, ALIDJINOU Enagnon Kazali, BOCKET Laurence, CREPIN Michel, DEMAY Christophe, ENGELMANN Ilka, GEFFROY Sandrine, GUIGON Aurélie, LAMBERT Valérie, LAZREK Mouna, NOBILLIAUX Florian, PREVOST Brigitte, THUILLIER Caroline, TINEZ Claire |

| | | | |
|---|---|---|---|
| EPI_ISL_1818804 | CH Douai | CHU Lille - Laboratoire de Virologie | AIT YAHYA Emilie, ALIDJINOU Enagnon Kazali, BOCKET Laurence, CREPIN Michel, DEMAY Christophe, ENGELMANN Ilka, GEFFROY Sandrine, GUIGON Aurélie, LAMBERT Valérie, LAZREK Mouna, NOBILLIAUX Florian, PREVOST Brigitte, THUILLIER Caroline, TINEZ Claire |
| EPI_ISL_1818807 | CH Lens | CHU Lille - Laboratoire de Virologie | AIT YAHYA Emilie, ALIDJINOU Enagnon Kazali, BOCKET Laurence, CREPIN Michel, DEMAY Christophe, ENGELMANN Ilka, GEFFROY Sandrine, GUIGON Aurélie, LAMBERT Valérie, LAZREK Mouna, NOBILLIAUX Florian, PREVOST Brigitte, THUILLIER Caroline, TINEZ Claire |
| EPI_ISL_1818808, EPI_ISL_1818809, EPI_ISL_1818810 | CHU Lille | CHU Lille - Laboratoire de Virologie | AIT YAHYA Emilie, ALIDJINOU Enagnon Kazali, BOCKET Laurence, CREPIN Michel, DEMAY Christophe, ENGELMANN Ilka, GEFFROY Sandrine, GUIGON Aurélie, LAMBERT Valérie, LAZREK Mouna, NOBILLIAUX Florian, PREVOST Brigitte, THUILLIER Caroline, TINEZ Claire |
| EPI_ISL_1818811 | CH Seclin | CHU Lille - Laboratoire de Virologie | AIT YAHYA Emilie, ALIDJINOU Enagnon Kazali, BOCKET Laurence, CREPIN Michel, DEMAY Christophe, ENGELMANN Ilka, GEFFROY Sandrine, GUIGON Aurélie, LAMBERT Valérie, LAZREK Mouna, NOBILLIAUX Florian, PREVOST Brigitte, THUILLIER Caroline, TINEZ Claire |
| EPI_ISL_1818812 | CHU Lille | CHU Lille - Laboratoire de Virologie | AIT YAHYA Emilie, ALIDJINOU Enagnon Kazali, BOCKET Laurence, CREPIN Michel, DEMAY Christophe, ENGELMANN Ilka, GEFFROY Sandrine, GUIGON Aurélie, LAMBERT Valérie, LAZREK Mouna, NOBILLIAUX Florian, PREVOST Brigitte, THUILLIER Caroline, TINEZ Claire |
| EPI_ISL_1818813 | CH Armentières | CHU Lille - Laboratoire de Virologie | AIT YAHYA Emilie, ALIDJINOU Enagnon Kazali, BOCKET Laurence, CREPIN Michel, DEMAY Christophe, ENGELMANN Ilka, GEFFROY Sandrine, GUIGON Aurélie, LAMBERT Valérie, LAZREK Mouna, NOBILLIAUX Florian, PREVOST Brigitte, THUILLIER Caroline, TINEZ Claire |
| EPI_ISL_1818814 | CH Tourcoing | CHU Lille - Laboratoire de Virologie | AIT YAHYA Emilie, ALIDJINOU Enagnon Kazali, BOCKET Laurence, CREPIN Michel, DEMAY Christophe, ENGELMANN Ilka, GEFFROY Sandrine, GUIGON Aurélie, LAMBERT Valérie, LAZREK Mouna, NOBILLIAUX Florian, PREVOST Brigitte, THUILLIER Caroline, TINEZ Claire |
| EPI_ISL_1818817, EPI_ISL_1818819 | CHU Lille | CHU Lille - Laboratoire de Virologie | AIT YAHYA Emilie, ALIDJINOU Enagnon Kazali, BOCKET Laurence, CREPIN Michel, DEMAY Christophe, ENGELMANN Ilka, GEFFROY Sandrine, GUIGON Aurélie, LAMBERT Valérie, LAZREK Mouna, NOBILLIAUX Florian, PREVOST Brigitte, THUILLIER Caroline, TINEZ Claire |
| EPI_ISL_1818825 | CH Saint Philibert | CHU Lille - Laboratoire de Virologie | AIT YAHYA Emilie, ALIDJINOU Enagnon Kazali, BOCKET Laurence, CREPIN Michel, DEMAY Christophe, ENGELMANN Ilka, GEFFROY Sandrine, GUIGON Aurélie, LAMBERT Valérie, LAZREK Mouna, NOBILLIAUX Florian, PREVOST Brigitte, THUILLIER Caroline, TINEZ Claire |
| EPI_ISL_1818826, EPI_ISL_1818828, EPI_ISL_1818830 | CHU Lille | CHU Lille - Laboratoire de Virologie | AIT YAHYA Emilie, ALIDJINOU Enagnon Kazali, BOCKET Laurence, CREPIN Michel, DEMAY Christophe, ENGELMANN Ilka, GEFFROY Sandrine, GUIGON Aurélie, LAMBERT Valérie, LAZREK Mouna, NOBILLIAUX Florian, PREVOST Brigitte, THUILLIER Caroline, TINEZ Claire |
| EPI_ISL_1818832 | CH Tourcoing | CHU Lille - Laboratoire de Virologie | AIT YAHYA Emilie, ALIDJINOU Enagnon Kazali, BOCKET Laurence, CREPIN Michel, DEMAY Christophe, ENGELMANN Ilka, GEFFROY Sandrine, GUIGON Aurélie, LAMBERT Valérie, LAZREK Mouna, NOBILLIAUX Florian, PREVOST Brigitte, THUILLIER Caroline, TINEZ Claire |
| EPI_ISL_1818834 | CH Béthune | CHU Lille - Laboratoire de Virologie | AIT YAHYA Emilie, ALIDJINOU Enagnon Kazali, BOCKET Laurence, CREPIN Michel, DEMAY Christophe, ENGELMANN Ilka, GEFFROY Sandrine, GUIGON Aurélie, LAMBERT Valérie, LAZREK Mouna, NOBILLIAUX Florian, PREVOST Brigitte, THUILLIER Caroline, TINEZ Claire |
| EPI_ISL_1818835, EPI_ISL_1818836, EPI_ISL_1818838 | CHU Lille | CHU Lille - Laboratoire de Virologie | AIT YAHYA Emilie, ALIDJINOU Enagnon Kazali, BOCKET Laurence, CREPIN Michel, DEMAY Christophe, ENGELMANN Ilka, GEFFROY Sandrine, GUIGON Aurélie, LAMBERT Valérie, LAZREK Mouna, NOBILLIAUX Florian, PREVOST Brigitte, THUILLIER Caroline, TINEZ Claire |
| EPI_ISL_1818839 | CH Lens | CHU Lille - Laboratoire de Virologie | AIT YAHYA Emilie, ALIDJINOU Enagnon Kazali, BOCKET Laurence, CREPIN Michel, DEMAY Christophe, ENGELMANN Ilka, GEFFROY Sandrine, GUIGON Aurélie, LAMBERT Valérie, LAZREK Mouna, NOBILLIAUX Florian, PREVOST Brigitte, THUILLIER Caroline, TINEZ Claire |
| EPI_ISL_1818840, EPI_ISL_1818842, EPI_ISL_1818843, EPI_ISL_1818844 | CHU Lille | CHU Lille - Laboratoire de Virologie | AIT YAHYA Emilie, ALIDJINOU Enagnon Kazali, BOCKET Laurence, CREPIN Michel, DEMAY Christophe, ENGELMANN Ilka, GEFFROY Sandrine, GUIGON Aurélie, LAMBERT Valérie, LAZREK Mouna, NOBILLIAUX Florian, PREVOST Brigitte, THUILLIER Caroline, TINEZ Claire |
| EPI_ISL_1818846 | CH Calais | CHU Lille - Laboratoire de Virologie | AIT YAHYA Emilie, ALIDJINOU Enagnon Kazali, BOCKET Laurence, CREPIN Michel, DEMAY Christophe, ENGELMANN Ilka, GEFFROY Sandrine, GUIGON Aurélie, LAMBERT Valérie, LAZREK Mouna, NOBILLIAUX Florian, PREVOST Brigitte, THUILLIER Caroline, TINEZ Claire |
| EPI_ISL_1818852 | CHU Lille | CHU Lille - Laboratoire de Virologie | AIT YAHYA Emilie, ALIDJINOU Enagnon Kazali, BOCKET Laurence, CREPIN Michel, DEMAY Christophe, ENGELMANN Ilka, GEFFROY Sandrine, GUIGON Aurélie, LAMBERT Valérie, LAZREK Mouna, NOBILLIAUX Florian, PREVOST Brigitte, THUILLIER Caroline, TINEZ Claire |
| EPI_ISL_1818853 | CH Arras | CHU Lille - Laboratoire de Virologie | AIT YAHYA Emilie, ALIDJINOU Enagnon Kazali, BOCKET Laurence, CREPIN Michel, DEMAY Christophe, ENGELMANN Ilka, GEFFROY Sandrine, GUIGON Aurélie, LAMBERT Valérie, LAZREK Mouna, NOBILLIAUX Florian, PREVOST Brigitte, THUILLIER Caroline, TINEZ Claire |
| EPI_ISL_1818945, EPI_ISL_1818946 | Virology Unit, Institut Pasteur du Cambodge | Virology Unit, Institut Pasteur du Cambodge | Jurre Y Siegers, Cecile Troupin, Ly Sovann, Kraing Sidonn, Yi Sengdoeurn, Chin Savuth, Chau Darapheak, Veasna Duong, Erik A Karlsson |
| EPI_ISL_1821102 | LESP Queretaro | Instituto de Diagnostico y Referencia Epidemiologicos (INDRE) | Claudia Wong-Arambula, Abril Rodriguez-Maldonado, Vanessa Rivero-Arredondo, Ariadna Medina-Benitez, Joaquin Quiroz-Mercado, Sergio Rangel-Guerrero, Natividad Cruz-Ortiz, Tatiana Nunez-Garcia, Gisela Barrera-Badillo, Lucia Hernandez-Rivas, Irma Lopez-Martinez, Ernesto Ramirez-Gonzalez. |
| EPI_ISL_1821103 | LESP Chihuahua | Instituto de Diagnostico y Referencia Epidemiologicos (INDRE) | Claudia Wong-Arambula, Abril Rodriguez-Maldonado, Vanessa Rivero-Arredondo, Ariadna Medina-Benitez, Joaquin Quiroz-Mercado, Sergio Rangel-Guerrero, Natividad Cruz-Ortiz, Tatiana Nunez-Garcia, Gisela Barrera-Badillo, Lucia Hernandez-Rivas, Irma Lopez-Martinez, Ernesto Ramirez-Gonzalez. |
| EPI_ISL_1821104, EPI_ISL_1821105 | LESP Baja California Sur | Instituto de Diagnostico y Referencia Epidemiologicos (INDRE) | Claudia Wong-Arambula, Abril Rodriguez-Maldonado, Vanessa Rivero-Arredondo, Ariadna Medina-Benitez, Joaquin Quiroz-Mercado, Sergio Rangel-Guerrero, Natividad Cruz-Ortiz, Tatiana Nunez-Garcia, Gisela Barrera-Badillo, Lucia Hernandez-Rivas, Irma Lopez-Martinez, Ernesto Ramirez-Gonzalez. |
| EPI_ISL_1821106 | LESP Ciudad de Mexico | Instituto de Diagnostico y Referencia Epidemiologicos (INDRE) | Claudia Wong-Arambula, Abril Rodriguez-Maldonado, Vanessa Rivero-Arredondo, Ariadna Medina-Benitez, Joaquin Quiroz-Mercado, Sergio Rangel-Guerrero, Natividad Cruz-Ortiz, Tatiana Nunez-Garcia, Gisela Barrera-Badillo, Lucia Hernandez-Rivas, Irma Lopez-Martinez, Ernesto Ramirez-Gonzalez. |
| EPI_ISL_1821107 | LESP Colima | Instituto de Diagnostico y Referencia Epidemiologicos (INDRE) | Claudia Wong-Arambula, Abril Rodriguez-Maldonado, Vanessa Rivero-Arredondo, Ariadna Medina-Benitez, Joaquin Quiroz-Mercado, Sergio Rangel-Guerrero, Natividad Cruz-Ortiz, Tatiana Nunez-Garcia, Gisela Barrera-Badillo, Lucia Hernandez-Rivas, Irma Lopez-Martinez, Ernesto Ramirez-Gonzalez. |
| EPI_ISL_1821108, EPI_ISL_1821109, EPI_ISL_1821110, EPI_ISL_1821111 | LESP Guanajuato | Instituto de Diagnostico y Referencia Epidemiologicos (INDRE) | Claudia Wong-Arambula, Abril Rodriguez-Maldonado, Vanessa Rivero-Arredondo, Ariadna Medina-Benitez, Joaquin Quiroz-Mercado, Sergio Rangel-Guerrero, Natividad Cruz-Ortiz, Tatiana Nunez-Garcia, Gisela Barrera-Badillo, Lucia Hernandez-Rivas, Irma Lopez-Martinez, Ernesto Ramirez-Gonzalez. |
| EPI_ISL_1821112, EPI_ISL_1821113, EPI_ISL_1821114, EPI_ISL_1821115, EPI_ISL_1821116, EPI_ISL_1821117, EPI_ISL_1821118 | LESP Nuevo Leon | Instituto de Diagnostico y Referencia Epidemiologicos (INDRE) | Claudia Wong-Arambula, Abril Rodriguez-Maldonado, Vanessa Rivero-Arredondo, Ariadna Medina-Benitez, Joaquin Quiroz-Mercado, Sergio Rangel-Guerrero, Natividad Cruz-Ortiz, Tatiana Nunez-Garcia, Gisela Barrera-Badillo, Lucia Hernandez-Rivas, Irma Lopez-Martinez, Ernesto Ramirez-Gonzalez. |
| EPI_ISL_1821119, EPI_ISL_1821120, EPI_ISL_1821122 | LESP Puebla | Instituto de Diagnostico y Referencia Epidemiologicos (INDRE) | Claudia Wong-Arambula, Abril Rodriguez-Maldonado, Vanessa Rivero-Arredondo, Ariadna Medina-Benitez, Joaquin Quiroz-Mercado, Sergio Rangel-Guerrero, Natividad Cruz-Ortiz, Tatiana Nunez-Garcia, Gisela Barrera-Badillo, Lucia Hernandez-Rivas, Irma Lopez-Martinez, Ernesto Ramirez-Gonzalez. |
| EPI_ISL_1821123 | LESP Guerrero | Instituto de Diagnostico y Referencia Epidemiologicos (INDRE) | Claudia Wong-Arambula, Abril Rodriguez-Maldonado, Vanessa Rivero-Arredondo, Ariadna Medina-Benitez, Joaquin Quiroz-Mercado, Sergio Rangel-Guerrero, Natividad Cruz-Ortiz, Tatiana Nunez-Garcia, Gisela Barrera-Badillo, Lucia Hernandez-Rivas, Irma Lopez-Martinez, Ernesto Ramirez-Gonzalez. |
| EPI_ISL_1821125 | LESP Chihuahua | Instituto de Diagnostico y Referencia Epidemiologicos (INDRE) | Claudia Wong-Arambula, Abril Rodriguez-Maldonado, Vanessa Rivero-Arredondo, Ariadna Medina-Benitez, Joaquin Quiroz-Mercado, Sergio Rangel-Guerrero, Natividad Cruz-Ortiz, Tatiana Nunez-Garcia, Gisela Barrera-Badillo, Lucia Hernandez-Rivas, Irma Lopez-Martinez, Ernesto Ramirez-Gonzalez. |
| EPI_ISL_1821126 | LESP Sonora | Instituto de Diagnostico y Referencia Epidemiologicos (INDRE) | Claudia Wong-Arambula, Abril Rodriguez-Maldonado, Vanessa Rivero-Arredondo, Ariadna Medina-Benitez, Joaquin Quiroz-Mercado, Sergio Rangel-Guerrero, Natividad Cruz-Ortiz, Tatiana Nunez-Garcia, Gisela Barrera-Badillo, Lucia Hernandez-Rivas, Irma Lopez-Martinez, Ernesto Ramirez-Gonzalez. |
| EPI_ISL_1821127 | LESP Chihuahua | Instituto de Diagnostico y Referencia Epidemiologicos (INDRE) | Claudia Wong-Arambula, Abril Rodriguez-Maldonado, Vanessa Rivero-Arredondo, Ariadna Medina-Benitez, Joaquin Quiroz-Mercado, Sergio Rangel-Guerrero, Natividad Cruz-Ortiz, Tatiana Nunez-Garcia, Gisela Barrera-Badillo, Lucia Hernandez-Rivas, Irma Lopez-Martinez, Ernesto |

| | | | |
|---|---|---|---|
| | | | Ramirez-Gonzalez. |
| EPI_ISL_1821128 | LESP Sonora | Instituto de Diagnostico y Referencia Epidemiologicos (INDRE) | Claudia Wong-Arambula, Abril Rodriguez-Maldonado, Vanessa Rivero-Arredondo, Ariadna Medina-Benitez, Joaquin Quiroz-Mercado, Sergio Rangel-Guerrero, Natividad Cruz-Ortiz, Tatiana Nunez-Garcia, Gisela Barrera-Badillo, Lucia Hernandez-Rivas, Irma Lopez-Martinez, Ernesto Ramirez-Gonzalez. |
| EPI_ISL_1821129, EPI_ISL_1821130 | LESP Jalisco | Instituto de Diagnostico y Referencia Epidemiologicos (INDRE) | Claudia Wong-Arambula, Abril Rodriguez-Maldonado, Vanessa Rivero-Arredondo, Ariadna Medina-Benitez, Joaquin Quiroz-Mercado, Sergio Rangel-Guerrero, Natividad Cruz-Ortiz, Tatiana Nunez-Garcia, Gisela Barrera-Badillo, Lucia Hernandez-Rivas, Irma Lopez-Martinez, Ernesto Ramirez-Gonzalez. |
| EPI_ISL_1821133, EPI_ISL_1821134, EPI_ISL_1821136 | LESP Queretaro | Instituto de Diagnostico y Referencia Epidemiologicos (INDRE) | Claudia Wong-Arambula, Abril Rodriguez-Maldonado, Vanessa Rivero-Arredondo, Ariadna Medina-Benitez, Joaquin Quiroz-Mercado, Sergio Rangel-Guerrero, Natividad Cruz-Ortiz, Tatiana Nunez-Garcia, Gisela Barrera-Badillo, Lucia Hernandez-Rivas, Irma Lopez-Martinez, Ernesto Ramirez-Gonzalez. |
| EPI_ISL_1821137, EPI_ISL_1821138, EPI_ISL_1821139, EPI_ISL_1821140, EPI_ISL_1821141, EPI_ISL_1821142, EPI_ISL_1821144, EPI_ISL_1821145, EPI_ISL_1821147, EPI_ISL_1821148 | LESP Morelos | Instituto de Diagnostico y Referencia Epidemiologicos (INDRE) | Claudia Wong-Arambula, Abril Rodriguez-Maldonado, Vanessa Rivero-Arredondo, Ariadna Medina-Benitez, Joaquin Quiroz-Mercado, Sergio Rangel-Guerrero, Natividad Cruz-Ortiz, Tatiana Nunez-Garcia, Gisela Barrera-Badillo, Lucia Hernandez-Rivas, Irma Lopez-Martinez, Ernesto Ramirez-Gonzalez. |
| EPI_ISL_1821149, EPI_ISL_1821150, EPI_ISL_1821151 | LESP Guerrero | Instituto de Diagnostico y Referencia Epidemiologicos (INDRE) | Claudia Wong-Arambula, Abril Rodriguez-Maldonado, Vanessa Rivero-Arredondo, Ariadna Medina-Benitez, Joaquin Quiroz-Mercado, Sergio Rangel-Guerrero, Natividad Cruz-Ortiz, Tatiana Nunez-Garcia, Gisela Barrera-Badillo, Lucia Hernandez-Rivas, Irma Lopez-Martinez, Ernesto Ramirez-Gonzalez. |
| EPI_ISL_1821158, EPI_ISL_1821159, EPI_ISL_1821160 | LESP Tamaulipas | Instituto de Diagnostico y Referencia Epidemiologicos (INDRE) | Claudia Wong-Arambula, Abril Rodriguez-Maldonado, Vanessa Rivero-Arredondo, Ariadna Medina-Benitez, Joaquin Quiroz-Mercado, Sergio Rangel-Guerrero, Natividad Cruz-Ortiz, Tatiana Nunez-Garcia, Gisela Barrera-Badillo, Lucia Hernandez-Rivas, Irma Lopez-Martinez, Ernesto Ramirez-Gonzalez. |
| EPI_ISL_1821162 | LESP Oaxaca | Instituto de Diagnostico y Referencia Epidemiologicos (INDRE) | Claudia Wong-Arambula, Abril Rodriguez-Maldonado, Vanessa Rivero-Arredondo, Ariadna Medina-Benitez, Joaquin Quiroz-Mercado, Sergio Rangel-Guerrero, Natividad Cruz-Ortiz, Tatiana Nunez-Garcia, Gisela Barrera-Badillo, Lucia Hernandez-Rivas, Irma Lopez-Martinez, Ernesto Ramirez-Gonzalez. |
| EPI_ISL_1821163, EPI_ISL_1821164, EPI_ISL_1821165 | LESP Veracruz | Instituto de Diagnostico y Referencia Epidemiologicos (INDRE) | Claudia Wong-Arambula, Abril Rodriguez-Maldonado, Vanessa Rivero-Arredondo, Ariadna Medina-Benitez, Joaquin Quiroz-Mercado, Sergio Rangel-Guerrero, Natividad Cruz-Ortiz, Tatiana Nunez-Garcia, Gisela Barrera-Badillo, Lucia Hernandez-Rivas, Irma Lopez-Martinez, Ernesto Ramirez-Gonzalez. |
| EPI_ISL_1821166 | LESP Tlaxcala | Instituto de Diagnostico y Referencia Epidemiologicos (INDRE) | Claudia Wong-Arambula, Abril Rodriguez-Maldonado, Vanessa Rivero-Arredondo, Ariadna Medina-Benitez, Joaquin Quiroz-Mercado, Sergio Rangel-Guerrero, Natividad Cruz-Ortiz, Tatiana Nunez-Garcia, Gisela Barrera-Badillo, Lucia Hernandez-Rivas, Irma Lopez-Martinez, Ernesto Ramirez-Gonzalez. |
| EPI_ISL_1821169, EPI_ISL_1821170 | LESP Ciudad de Mexico | Instituto de Diagnostico y Referencia Epidemiologicos (INDRE) | Claudia Wong-Arambula, Abril Rodriguez-Maldonado, Vanessa Rivero-Arredondo, Ariadna Medina-Benitez, Joaquin Quiroz-Mercado, Sergio Rangel-Guerrero, Natividad Cruz-Ortiz, Tatiana Nunez-Garcia, Gisela Barrera-Badillo, Lucia Hernandez-Rivas, Irma Lopez-Martinez, Ernesto Ramirez-Gonzalez. |
| EPI_ISL_1821171, EPI_ISL_1821172 | LESP Quintana Roo | Instituto de Diagnostico y Referencia Epidemiologicos (INDRE) | Claudia Wong-Arambula, Abril Rodriguez-Maldonado, Vanessa Rivero-Arredondo, Ariadna Medina-Benitez, Joaquin Quiroz-Mercado, Sergio Rangel-Guerrero, Natividad Cruz-Ortiz, Tatiana Nunez-Garcia, Gisela Barrera-Badillo, Lucia Hernandez-Rivas, Irma Lopez-Martinez, Ernesto Ramirez-Gonzalez. |
| EPI_ISL_1821178, EPI_ISL_1821179 | LESP Zacatecas | Instituto de Diagnostico y Referencia Epidemiologicos (INDRE) | Claudia Wong-Arambula, Abril Rodriguez-Maldonado, Vanessa Rivero-Arredondo, Ariadna Medina-Benitez, Joaquin Quiroz-Mercado, Sergio Rangel-Guerrero, Natividad Cruz-Ortiz, Tatiana Nunez-Garcia, Gisela Barrera-Badillo, Lucia Hernandez-Rivas, Irma Lopez-Martinez, Ernesto Ramirez-Gonzalez. |
| EPI_ISL_1821186 | LESP Jalisco | Instituto de Diagnostico y Referencia Epidemiologicos (INDRE) | Claudia Wong-Arambula, Abril Rodriguez-Maldonado, Vanessa Rivero-Arredondo, Ariadna Medina-Benitez, Joaquin Quiroz-Mercado, Sergio Rangel-Guerrero, Natividad Cruz-Ortiz, Tatiana Nunez-Garcia, Gisela Barrera-Badillo, Lucia Hernandez-Rivas, Irma Lopez-Martinez, Ernesto Ramirez-Gonzalez. |
| EPI_ISL_1821187, EPI_ISL_1821188, EPI_ISL_1821189, EPI_ISL_1821190 | LESP Ciudad de Mexico | Instituto de Diagnostico y Referencia Epidemiologicos (INDRE) | Claudia Wong-Arambula, Abril Rodriguez-Maldonado, Vanessa Rivero-Arredondo, Ariadna Medina-Benitez, Joaquin Quiroz-Mercado, Sergio Rangel-Guerrero, Natividad Cruz-Ortiz, Tatiana Nunez-Garcia, Gisela Barrera-Badillo, Lucia Hernandez-Rivas, Irma Lopez-Martinez, Ernesto Ramirez-Gonzalez. |
| EPI_ISL_1821194, EPI_ISL_1821195, EPI_ISL_1821196 | LESP Hidalgo | Instituto de Diagnostico y Referencia Epidemiologicos (INDRE) | Claudia Wong-Arambula, Abril Rodriguez-Maldonado, Vanessa Rivero-Arredondo, Ariadna Medina-Benitez, Joaquin Quiroz-Mercado, Sergio Rangel-Guerrero, Natividad Cruz-Ortiz, Tatiana Nunez-Garcia, Gisela Barrera-Badillo, Lucia Hernandez-Rivas, Irma Lopez-Martinez, Ernesto Ramirez-Gonzalez. |
| EPI_ISL_1821199 | LESP Sinaloa | Instituto de Diagnostico y Referencia Epidemiologicos (INDRE) | Claudia Wong-Arambula, Abril Rodriguez-Maldonado, Vanessa Rivero-Arredondo, Ariadna Medina-Benitez, Joaquin Quiroz-Mercado, Sergio Rangel-Guerrero, Natividad Cruz-Ortiz, Tatiana Nunez-Garcia, Gisela Barrera-Badillo, Lucia Hernandez-Rivas, Irma Lopez-Martinez, Ernesto Ramirez-Gonzalez. |
| EPI_ISL_1821205 | UPA Dr Luis Atilio Losi Viana Ribeirao Preto | Instituto Adolfo Lutz, Interdiciplinary Procedures Center, Strategic Laboratory | Claudio Tavares Sacchi, Claudia Regina Gonçalves, Erica Valessa Ramos Gomes, Karoline Rodrigues Campos, Caio Vinicius Dias Lopes, Leonardo Jose Tadeu de Araujo |
| EPI_ISL_1821209 | CS de Paulo de Faria | Instituto Adolfo Lutz, Interdiciplinary Procedures Center, Strategic Laboratory | Claudio Tavares Sacchi, Claudia Regina Gonçalves, Erica Valessa Ramos Gomes, Karoline Rodrigues Campos, Caio Vinicius Dias Lopes, Leonardo Jose Tadeu de Araujo |
| EPI_ISL_1821214, EPI_ISL_1821215, EPI_ISL_1821216, EPI_ISL_1821217, EPI_ISL_1821218, EPI_ISL_1821219, EPI_ISL_1821220, EPI_ISL_1821221, EPI_ISL_1821222, EPI_ISL_1821223, EPI_ISL_1821224 | | | |
| see above | Instituto Adolfo Lutz - Regional de Aracatuba | Instituto Adolfo Lutz, Interdiciplinary Procedures Center, Strategic Laboratory | Claudio Tavares Sacchi, Claudia Regina Gonçalves, Erica Valessa Ramos Gomes, Karoline Rodrigues Campos, Caio Vinicius Dias Lopes, Leonardo Jose Tadeu de Araujo |
| EPI_ISL_1821239, EPI_ISL_1821240, EPI_ISL_1821241, EPI_ISL_1821242 | Instituto Adolfo Lutz - Regional de Marilia | Instituto Adolfo Lutz, Interdiciplinary Procedures Center, Strategic Laboratory | Claudio Tavares Sacchi, Claudia Regina Gonçalves, Erica Valessa Ramos Gomes, Karoline Rodrigues Campos, Caio Vinicius Dias Lopes, Leonardo Jose Tadeu de Araujo |
| EPI_ISL_1821246, EPI_ISL_1821247, EPI_ISL_1821248, EPI_ISL_1821249, EPI_ISL_1821250, EPI_ISL_1821251, EPI_ISL_1821252, EPI_ISL_1821253, EPI_ISL_1821254, EPI_ISL_1821255, EPI_ISL_1821256, EPI_ISL_1821257, EPI_ISL_1821258, EPI_ISL_1821259, EPI_ISL_1821260, EPI_ISL_1821261, EPI_ISL_1821262, EPI_ISL_1821263, EPI_ISL_1821264, EPI_ISL_1821265 | | | |
| see above | Instituto Adolfo Lutz - Regional de Ribeirao Preto | Instituto Adolfo Lutz, Interdiciplinary Procedures Center, Strategic Laboratory | Claudio Tavares Sacchi, Claudia Regina Gonçalves, Erica Valessa Ramos Gomes, Karoline Rodrigues Campos, Caio Vinicius Dias Lopes, Leonardo Jose Tadeu de Araujo |
| EPI_ISL_1821266, EPI_ISL_1821267, EPI_ISL_1821268, EPI_ISL_1821269, EPI_ISL_1821270, EPI_ISL_1821271, EPI_ISL_1821272, EPI_ISL_1821273 | Instituto Adolfo Lutz Central | Instituto Adolfo Lutz, Interdiciplinary Procedures Center, Strategic Laboratory | Claudio Tavares Sacchi, Claudia Regina Gonçalves, Erica Valessa Ramos Gomes, Karoline Rodrigues Campos, Caio Vinicius Dias Lopes, Leonardo Jose Tadeu de Araujo |
| EPI_ISL_1821477 | Labo Analyses Med | National Reference Center for Viruses of Respiratory Infections, Institut Pasteur, Paris | Marion Barbet, Sylvie Behillil, Méline Bizard, Angela Brisebarre, Camille Capel, Vincent Enouf, Louise Lefrançois, Frédéric Lemoine, Christophe Malabat, Corinne Maufrais, Etienne Simon-Lorière, Maud Vanpeene, Sylvie Van der Werf ,Pierre Pennacino |
| EPI_ISL_1821480, EPI_ISL_1821481 | Hospital | National Reference Center for Viruses of Respiratory Infections, Institut Pasteur, Paris | Marion Barbet, Sylvie Behillil, Méline Bizard, Angela Brisebarre, Camille Capel, Vincent Enouf, Louise Lefrançois, Frédéric Lemoine, Christophe Malabat, Corinne Maufrais, Etienne Simon-Lorière, Maud Vanpeene, Sylvie Van der Werf ,JéRome Guinard |
| EPI_ISL_1821542 | Labo Analyses Med | National Reference Center for Viruses of Respiratory Infections, Institut Pasteur, Paris | Marion Barbet, Sylvie Behillil, Méline Bizard, Angela Brisebarre, Camille Capel, Vincent Enouf, Louise Lefrançois, Frédéric Lemoine, Christophe Malabat, Corinne Maufrais, Etienne Simon-Lorière, Maud Vanpeene, Sylvie Van der Werf ,GréGoire Potiron |

| | | | |
|---|---|---|---|
| EPI_ISL_1821543 | Hospital | National Reference Center for Viruses of Respiratory Infections, Institut Pasteur, Paris | Marion Barbet, Sylvie Behillil, Méline Bizard, Angela Brisebarre, Camille Capel, Vincent Enouf, Louise Lefrançois, Frédéric Lemoine, Christophe Malabat, Corinne Maufrais, Etienne Simon-Lorière, Maud Vanpeene, Sylvie Van der Werf ,Haciba Moudjahed |
| EPI_ISL_1821546, EPI_ISL_1821547, EPI_ISL_1821548 | Hospital | National Reference Center for Viruses of Respiratory Infections, Institut Pasteur, Paris | Marion Barbet, Sylvie Behillil, Méline Bizard, Angela Brisebarre, Camille Capel, Vincent Enouf, Louise Lefrançois, Frédéric Lemoine, Christophe Malabat, Corinne Maufrais, Damien Mornico, Etienne Simon-Lorière, Maud Vanpeene, Sylvie Van der Werf ,LéA Pilorge |
| EPI_ISL_1821551, EPI_ISL_1821553 | Hospital | National Reference Center for Viruses of Respiratory Infections, Institut Pasteur, Paris | Marion Barbet, Sylvie Behillil, Méline Bizard, Angela Brisebarre, Camille Capel, Vincent Enouf, Louise Lefrançois, Frédéric Lemoine, Christophe Malabat, Corinne Maufrais, Etienne Simon-Lorière, Maud Vanpeene, Sylvie Van der Werf ,JéRome Guinard |
| EPI_ISL_1821562 | Labo Analyses Med | National Reference Center for Viruses of Respiratory Infections, Institut Pasteur, Paris | Marion Barbet, Sylvie Behillil, Méline Bizard, Angela Brisebarre, Camille Capel, Vincent Enouf, Louise Lefrançois, Frédéric Lemoine, Christophe Malabat, Corinne Maufrais, Etienne Simon-Lorière, Maud Vanpeene, Sylvie Van der Werf ,Gilles Desvaux |
| EPI_ISL_1821590, EPI_ISL_1821616, EPI_ISL_1821617, EPI_ISL_1821618 | Hospital | National Reference Center for Viruses of Respiratory Infections, Institut Pasteur, Paris | Marion Barbet, Sylvie Behillil, Méline Bizard, Angela Brisebarre, Camille Capel, Vincent Enouf, Louise Lefrançois, Frédéric Lemoine, Christophe Malabat, Corinne Maufrais, Etienne Simon-Lorière, Maud Vanpeene, Sylvie Van der Werf ,JéRome Guinard |
| EPI_ISL_1821622 | Labo Analyses Med | National Reference Center for Viruses of Respiratory Infections, Institut Pasteur, Paris | Marion Barbet, Sylvie Behillil, Méline Bizard, Angela Brisebarre, Camille Capel, Vincent Enouf, Louise Lefrançois, Frédéric Lemoine, Christophe Malabat, Corinne Maufrais, Etienne Simon-Lorière, Maud Vanpeene, Sylvie Van der Werf ,JéRome Guinard |
| EPI_ISL_1821623 | Labo Analyses Med | National Reference Center for Viruses of Respiratory Infections, Institut Pasteur, Paris | Marion Barbet, Sylvie Behillil, Méline Bizard, Angela Brisebarre, Camille Capel, Vincent Enouf, Louise Lefrançois, Frédéric Lemoine, Christophe Malabat, Corinne Maufrais, Amaury Vaysse, Etienne Simon-Lorière, Maud Vanpeene, Sylvie Van der Werf ,Pauline Garnier |
| EPI_ISL_1821624 | Labo Analyses Med | National Reference Center for Viruses of Respiratory Infections, Institut Pasteur, Paris | Marion Barbet, Sylvie Behillil, Méline Bizard, Angela Brisebarre, Camille Capel, Vincent Enouf, Louise Lefrançois, Frédéric Lemoine, Christophe Malabat, Corinne Maufrais, Etienne Simon-Lorière, Maud Vanpeene, Sylvie Van der Werf ,Pauline Garnier |
| EPI_ISL_1821626 | Labo Analyses Med | National Reference Center for Viruses of Respiratory Infections, Institut Pasteur, Paris | Marion Barbet, Sylvie Behillil, Méline Bizard, Angela Brisebarre, Camille Capel, Vincent Enouf, Louise Lefrançois, Frédéric Lemoine, Christophe Malabat, Corinne Maufrais, Etienne Simon-Lorière, Maud Vanpeene, Sylvie Van der Werf ,Evelyne Bourgerette |
| EPI_ISL_1821627, EPI_ISL_1821628, EPI_ISL_1821629, EPI_ISL_1821630 | Labo Analyses Med | National Reference Center for Viruses of Respiratory Infections, Institut Pasteur, Paris | Marion Barbet, Sylvie Behillil, Méline Bizard, Angela Brisebarre, Camille Capel, Vincent Enouf, Louise Lefrançois, Frédéric Lemoine, Christophe Malabat, Corinne Maufrais, Etienne Simon-Lorière, Maud Vanpeene, Sylvie Van der Werf ,Dominique Rousset |
| EPI_ISL_1821633, EPI_ISL_1821634, EPI_ISL_1821635, EPI_ISL_1821636, EPI_ISL_1821637, EPI_ISL_1821638, EPI_ISL_1821639, EPI_ISL_1821640, EPI_ISL_1821641, EPI_ISL_1821642 | Labo Analyses Med | National Reference Center for Viruses of Respiratory Infections, Institut Pasteur, Paris | Marion Barbet, Sylvie Behillil, Méline Bizard, Angela Brisebarre, Camille Capel, Vincent Enouf, Louise Lefrançois, Frédéric Lemoine, Christophe Malabat, Corinne Maufrais, Emmanuelle Permal, Etienne Simon-Lorière, Maud Vanpeene, Sylvie Van der Werf ,Dominique Rousset |
| EPI_ISL_1821653 | Labo Analyses Med | National Reference Center for Viruses of Respiratory Infections, Institut Pasteur, Paris | Marion Barbet, Sylvie Behillil, Méline Bizard, Angela Brisebarre, Camille Capel, Vincent Enouf, Louise Lefrançois, Frédéric Lemoine, Christophe Malabat, Corinne Maufrais, Amaury Vaysse, Etienne Simon-Lorière, Maud Vanpeene, Sylvie Van der Werf ,Dominique Rousset |
| EPI_ISL_1821657 | Labo Analyses Med | National Reference Center for Viruses of Respiratory Infections, Institut Pasteur, Paris | Marion Barbet, Sylvie Behillil, Méline Bizard, Angela Brisebarre, Camille Capel, Vincent Enouf, Louise Lefrançois, Frédéric Lemoine, Christophe Malabat, Corinne Maufrais, Emmanuelle Permal, Etienne Simon-Lorière, Maud Vanpeene, Sylvie Van der Werf ,Dominique Rousset |
| EPI_ISL_1823181, EPI_ISL_1823183, EPI_ISL_1823184, EPI_ISL_1823185, EPI_ISL_1823186, EPI_ISL_1823187, EPI_ISL_1823188, EPI_ISL_1823189 | Public Institution of Health Care "Diagnostic Center ( Laboratory Testing Center) of Moscow Health Department " | Group of Genomics and Postgenomic Technologies of Central Research Institute of Epidemiology | Samoilov AE, Kaptelova VV, Korneenko EV, Saenko SS, Smirnova YS, Nadtoka MI, Spakova OG, Speranskaya AS, Tivanova EV, Kondrasheva LY, Akimkin VG |
| EPI_ISL_1823190 | Public Institution of Health Care "Diagnostic Center ( Laboratory Testing Center) of Moscow Health Department " | Group of Genomics and Postgenomic Technologies of Central Research Institute of Epidemiology | Samoilov AE, Kaptelova VV, Korneenko EV, Saenko SS, Smirnova YS, Nadtoka MI, Speranskaya AS, Tivanova EV, Kondrasheva LY, Akimkin VG |
| EPI_ISL_1823452, EPI_ISL_1823453, EPI_ISL_1823454, EPI_ISL_1823455, EPI_ISL_1823456, EPI_ISL_1823457, EPI_ISL_1823459, EPI_ISL_1823460, EPI_ISL_1823461, EPI_ISL_1823462, EPI_ISL_1823463, EPI_ISL_1823464, EPI_ISL_1823465, EPI_ISL_1823466, EPI_ISL_1823468, EPI_ISL_1823469, EPI_ISL_1823472, EPI_ISL_1823474, EPI_ISL_1823476, EPI_ISL_1823478, EPI_ISL_1823479, EPI_ISL_1823481, EPI_ISL_1823482, EPI_ISL_1823484, EPI_ISL_1823485, EPI_ISL_1823486, EPI_ISL_1823487, EPI_ISL_1823488, EPI_ISL_1823489, EPI_ISL_1823491, EPI_ISL_1823492, EPI_ISL_1823494, EPI_ISL_1823495, EPI_ISL_1823497, EPI_ISL_1823498, EPI_ISL_1823499, EPI_ISL_1823502, EPI_ISL_1823504, EPI_ISL_1823506, EPI_ISL_1823507, EPI_ISL_1823509, EPI_ISL_1823510, EPI_ISL_1823511, EPI_ISL_1823513, EPI_ISL_1823514, EPI_ISL_1823515, EPI_ISL_1823516, EPI_ISL_1823517, EPI_ISL_1823518, EPI_ISL_1823519, EPI_ISL_1823520, EPI_ISL_1823521, EPI_ISL_1823522, EPI_ISL_1823524, EPI_ISL_1823525, EPI_ISL_1823526, EPI_ISL_1823527, EPI_ISL_1823528, EPI_ISL_1823529, EPI_ISL_1823530, EPI_ISL_1823531, EPI_ISL_1823532, EPI_ISL_1823533, EPI_ISL_1823534, EPI_ISL_1823535, EPI_ISL_1823536, EPI_ISL_1823537, EPI_ISL_1823538, EPI_ISL_1823540, EPI_ISL_1823542, EPI_ISL_1823543, EPI_ISL_1823544, EPI_ISL_1823545, EPI_ISL_1823546, EPI_ISL_1823547, EPI_ISL_1823549, EPI_ISL_1823552, EPI_ISL_1823553, EPI_ISL_1823556, EPI_ISL_1823558, EPI_ISL_1823566 | | | |
| see above | Indiana Animal Disease Diagnostic Laboratory | Carpi Laboratory - Purdue University | Jack Dorman, Ilinca I Ciubotariu, Nicole M Perry, Jobin J Kattoor, Lev Gorenstein, Abebe A Fola, G Kenitra Hendrix, Rebecca P Wilkes, Giovanna Carpi |
| EPI_ISL_1824499, EPI_ISL_1824500 | AZ Klina | AZ Klina | Carl Vael - Lynsey Berckmans |
| EPI_ISL_1824598 | Public Institution of Health Care "Diagnostic Center (Laboratory Testing Center) of Moscow Health Department " | Group of Genomics and Postgenomic Technologies of Central Research Institute of Epidemiology | Samoilov AE, Kaptelova VV, Korneenko EV, Saenko SS, Smirnova YS, Nadtoka MI, Spakova OG, Speranskaya AS, Tivanova EV, Kondrasheva LY, Akimkin VG |
| EPI_ISL_1824606 | RSUP Sanglah | National Institute of Health Research and Development | Vivi Setiawaty, Hana Apsari Pawestri, Subangkit, Kartika Dewi Puspa, Arie Ardiansyah Nugraha, Hartanti Dian Ikawati, Krisna Pangesti, Nelly Puspandari, I Nengah Tony Rustawan, I Gede Ketut Sajinadiyasa |
| EPI_ISL_1824607 | PKC Jagakarsa | National Institute of Health Research and Development | Vivi Setiawaty, Hana Apsari Pawestri, Subangkit, Kartika Dewi Puspa, Arie Ardiansyah Nugraha, Hartanti Dian Ikawati, Krisna Pangesti, Nelly Puspandari |
| EPI_ISL_1824624, EPI_ISL_1824625, EPI_ISL_1824626, EPI_ISL_1824627, EPI_ISL_1824628, EPI_ISL_1824629, EPI_ISL_1824630, EPI_ISL_1824631, EPI_ISL_1824632, EPI_ISL_1824633, EPI_ISL_1824634, EPI_ISL_1824635 | | | |
| see above | COVID Laboratory AOU Messina | Molecular Hepatology Lab AOU Messina | Giuseppina Raffa, Cristina Musolino, Valeria Chines, Daniele Lombardo, Domenico Giosa, Teresa Pollicino. |
| EPI_ISL_1825676, EPI_ISL_1825678, EPI_ISL_1825679, EPI_ISL_1825680, EPI_ISL_1825681, EPI_ISL_1825682, EPI_ISL_1825683, EPI_ISL_1825685, EPI_ISL_1825686 | COVID-19 Detection Lab, Chattogram Veterinary and Animal Sciences University | Genomic Research Lab, Bangladesh Council of Scientific and Industrial Research | Goutam Buddha Das, Tridip Das, Tanvir Ahmad Nizami, Eaftekhar Ahmed Rana, Md. Sirazul Islam, Pronesh Dutta, Sharmin Chowdhury, Md. Morshed Hasan Sarkar, Md. Salim Khan, Paritosh Kumar Biswas |
| EPI_ISL_1827527 | AREA DE SALUD CATEDRAL NORESTE | Inciensa, Instituto Costarricense de Investigación y Enseñanza en Nutrición y Salud | Francisco Duarte, Hebleen Porras, Claudio Soto-Garita, Estela Cordero, Adriana Godínez, Melany Calderón, José Luis Vargas, Mariela Gutiérrez & Joselyn Prado |
| EPI_ISL_1827528, EPI_ISL_1827529 | HOSPITAL DR. WILLIAM ALLEN | Inciensa, Instituto Costarricense de Investigación y Enseñanza en Nutrición y Salud | Francisco Duarte, Hebleen Porras, Claudio Soto-Garita, Estela Cordero, Adriana Godínez, Melany Calderón, José Luis Vargas, Mariela Gutiérrez, Joselyn Prado & Mónica Charpentier-Artavia |
| EPI_ISL_1827532 | AREA DE SALUD MATA REDONDA-HOSPITAL - CLINICA DR. MORENO CAÑAS | Inciensa, Instituto Costarricense de Investigación y Enseñanza en Nutrición y Salud | Francisco Duarte, Hebleen Porras, Claudio Soto-Garita, Estela Cordero, Adriana Godínez, Melany Calderón, José Luis Vargas, Mariela Gutiérrez, Joselyn Prado & María Gabriela Valderrama-Ordoñez |
| EPI_ISL_1827534, EPI_ISL_1827535 | AREA DE SALUD CATEDRAL NORESTE | Inciensa, Instituto Costarricense de Investigación y Enseñanza en Nutrición y Salud | Francisco Duarte, Hebleen Porras, Claudio Soto-Garita, Estela Cordero, Adriana Godínez, Melany Calderón, José Luis Vargas, Mariela Gutiérrez, Joselyn Prado & Mariel López |
| EPI_ISL_1827538 | AREA DE SALUD PEREZ ZELEDON | Inciensa, Instituto Costarricense de Investigación y Enseñanza en Nutrición y Salud | Francisco Duarte, Hebleen Porras, Claudio Soto-Garita, Estela Cordero, Adriana Godínez, Melany Calderón, José Luis Vargas, Mariela Gutiérrez, Joselyn Prado & María Fernanda Matamoros-Madrigal |
| EPI_ISL_1827539, EPI_ISL_1827540 | AREA DE SALUD SIQUIRRES | Inciensa, Instituto Costarricense de Investigación y Enseñanza en Nutrición y Salud | Francisco Duarte, Hebleen Porras, Claudio Soto-Garita, Estela Cordero, Adriana Godínez, Melany Calderón, José Luis Vargas, Mariela Gutiérrez, Joselyn Prado & Ileana Chaves-Peraza |
| EPI_ISL_1827541 | HOSPITAL CIUDAD NEILY | Inciensa, Instituto Costarricense de Investigación y Enseñanza en Nutrición y Salud | Francisco Duarte, Hebleen Porras, Claudio Soto-Garita, Estela Cordero, Adriana Godínez, Melany Calderón, José Luis Vargas, Mariela Gutiérrez, Joselyn Prado & Nelly Castro-Bonilla |
| EPI_ISL_1827543 | HOSPITAL LA ANEXION | Inciensa, Instituto Costarricense de Investigación y Enseñanza en Nutrición y Salud | Francisco Duarte, Hebleen Porras, Claudio Soto-Garita, Estela Cordero, Adriana Godínez, Melany Calderón, José Luis Vargas, Mariela Gutiérrez, Joselyn Prado & Arnoldo Zamora-Villegas |
| EPI_ISL_1827544 | AREA DE SALUD SANTA CRUZ | Inciensa, Instituto Costarricense de Investigación y Enseñanza en Nutrición y Salud | Francisco Duarte, Hebleen Porras, Claudio Soto-Garita, Estela Cordero, Adriana Godínez, Melany Calderón, José Luis Vargas, Mariela Gutiérrez, Joselyn Prado & Arnoldo Zamora-Villegas |

| | | | |
|---|---|---|---|
| EPI_ISL_1831676, EPI_ISL_1831679, EPI_ISL_1831682, EPI_ISL_1831685 | Althaia. Xarxa Assistencial Universitària de Manresa | IrsiCaixa | Marc Noguera-Julian, Mariona Parera, Maria Casadellà, Pilar Armengol, Francesc Catala-Moll, Roger Paredes, Bonaventura Clotet Gloria Trujillo, Rafel Perez Vidal, Jaume Trape Pujol, Carolina Gonzalez Fernandez, Miquel Micó, Roger Paredes, Eulalia Grau, Bonaventura Clotet |
| EPI_ISL_1840869, EPI_ISL_1840870, EPI_ISL_1840871, EPI_ISL_1840872, EPI_ISL_1840873, EPI_ISL_1840874, EPI_ISL_1840875, EPI_ISL_1840877 | Austrian Agency for Health and Food Safety (AGES) | Bergthaler laboratory, CeMM Research Center for Molecular Medicine of the Austrian Academy of Sciences | Lukas Endler, Anna Schedl, Fabian Amman, Petr Triska, Thomas Penz, Benedikt Agerer, Maelle Le Moing, Michael Schuster, Bekir Erguner, Jan Laine, Martin Senekowitsch, Christoph Bock, Andreas Bergthaler |
| EPI_ISL_1840880, EPI_ISL_1840881, EPI_ISL_1840882, EPI_ISL_1840883, EPI_ISL_1840884, EPI_ISL_1840885, EPI_ISL_1840886 | Department of Laboratory Medicine, Division of Clinical Virology, University of Medicine, Vienna | Bergthaler laboratory, CeMM Research Center for Molecular Medicine of the Austrian Academy of Sciences | Lukas Endler, Anna Schedl, Fabian Amman, Petr Triska, Thomas Penz, Benedikt Agerer, Maelle Le Moing, Michael Schuster, Bekir Erguner, Jan Laine, Martin Senekowitsch, Christoph Bock, Andreas Bergthaler |
| EPI_ISL_1840888, EPI_ISL_1840889, EPI_ISL_1840890, EPI_ISL_1840891, EPI_ISL_1840892, EPI_ISL_1840893, EPI_ISL_1840895, EPI_ISL_1840896 | 3. Medizinische Abteilung, Hanusch Krankenhaus | Bergthaler laboratory, CeMM Research Center for Molecular Medicine of the Austrian Academy of Sciences | Lukas Endler, Anna Schedl, Fabian Amman, Petr Triska, Thomas Penz, Benedikt Agerer, Maelle Le Moing, Michael Schuster, Bekir Erguner, Jan Laine, Martin Senekowitsch, Christoph Bock, Andreas Bergthaler |
| EPI_ISL_1840900, EPI_ISL_1840901, EPI_ISL_1840902, EPI_ISL_1840904, EPI_ISL_1840909, EPI_ISL_1840910, EPI_ISL_1840911, EPI_ISL_1840913, EPI_ISL_1840917, EPI_ISL_1840919, EPI_ISL_1840921, EPI_ISL_1840926, EPI_ISL_1840930, EPI_ISL_1840935, EPI_ISL_1840936, EPI_ISL_1840939, EPI_ISL_1840944, EPI_ISL_1840945, EPI_ISL_1840953, EPI_ISL_1840954, EPI_ISL_1840961, EPI_ISL_1840962, EPI_ISL_1840964 | | | |
| see above | Austrian Agency for Health and Food Safety (AGES) | Bergthaler laboratory, CeMM Research Center for Molecular Medicine of the Austrian Academy of Sciences | Lukas Endler, Anna Schedl, Fabian Amman, Petr Triska, Thomas Penz, Benedikt Agerer, Maelle Le Moing, Michael Schuster, Bekir Erguner, Jan Laine, Martin Senekowitsch, Christoph Bock, Andreas Bergthaler |
| EPI_ISL_1843721, EPI_ISL_1843723, EPI_ISL_1843725, EPI_ISL_1843726, EPI_ISL_1843731, EPI_ISL_1843733, EPI_ISL_1843734, EPI_ISL_1843736, EPI_ISL_1843738, EPI_ISL_1843740, EPI_ISL_1843742, EPI_ISL_1843744, EPI_ISL_1843745, EPI_ISL_1843749, EPI_ISL_1843751, EPI_ISL_1843752 | | | |
| see above | Klinisch Laboratorium ZNA | Klinisch Laboratorium ZNA | Verstrepen et al. |
| EPI_ISL_1855854, EPI_ISL_1855855, EPI_ISL_1855856, EPI_ISL_1855858, EPI_ISL_1855860, EPI_ISL_1855861, EPI_ISL_1855870 | Baylor Scott & White-Temple | Baylor Scott & White-Temple | Ari Rao, Linden Morales, Kimberly Walker, Marcus Volz, Shelby Hendrickson |
| EPI_ISL_1856480, EPI_ISL_1856596, EPI_ISL_1856625, EPI_ISL_1856626, EPI_ISL_1856627, EPI_ISL_1856633, EPI_ISL_1856634, EPI_ISL_1856636, EPI_ISL_1856653, EPI_ISL_1856654, EPI_ISL_1856741, EPI_ISL_1856747, EPI_ISL_1856748, EPI_ISL_1856749, EPI_ISL_1856750, EPI_ISL_1856752, EPI_ISL_1856753, EPI_ISL_1856755, EPI_ISL_1856756, EPI_ISL_1856757, EPI_ISL_1856758, EPI_ISL_1856759, EPI_ISL_1856760, EPI_ISL_1856761, EPI_ISL_1856762, EPI_ISL_1856763, EPI_ISL_1856764, EPI_ISL_1856765, EPI_ISL_1856766, EPI_ISL_1856768, EPI_ISL_1856769, EPI_ISL_1856771, EPI_ISL_1856772, EPI_ISL_1856773, EPI_ISL_1856774, EPI_ISL_1856775, EPI_ISL_1856776, EPI_ISL_1856777, EPI_ISL_1856779, EPI_ISL_1856780, EPI_ISL_1856781, EPI_ISL_1856782, EPI_ISL_1856783, EPI_ISL_1856784, EPI_ISL_1856785, EPI_ISL_1856787, EPI_ISL_1856788, EPI_ISL_1856790, EPI_ISL_1856792, EPI_ISL_1856793, EPI_ISL_1856794, EPI_ISL_1856795, EPI_ISL_1856796, EPI_ISL_1856797, EPI_ISL_1856798, EPI_ISL_1856800, EPI_ISL_1856801, EPI_ISL_1856802, EPI_ISL_1856803, EPI_ISL_1856804, EPI_ISL_1856805, EPI_ISL_1856806, EPI_ISL_1856807, EPI_ISL_1856808, EPI_ISL_1856811, EPI_ISL_1856812, EPI_ISL_1856813, EPI_ISL_1856814, EPI_ISL_1856816, EPI_ISL_1856817, EPI_ISL_1856818, EPI_ISL_1856819, EPI_ISL_1856826, EPI_ISL_1856827, EPI_ISL_1856830, EPI_ISL_1856831, EPI_ISL_1856833, EPI_ISL_1856842, EPI_ISL_1856850, EPI_ISL_1856853, EPI_ISL_1856854, EPI_ISL_1856876, EPI_ISL_1856879, EPI_ISL_1856880, EPI_ISL_1856881, EPI_ISL_1856882, EPI_ISL_1856885, EPI_ISL_1856886, EPI_ISL_1856887, EPI_ISL_1856888, EPI_ISL_1856889, EPI_ISL_1856891, EPI_ISL_1856892, EPI_ISL_1856893, EPI_ISL_1856894, EPI_ISL_1856895, EPI_ISL_1856896, EPI_ISL_1856897, EPI_ISL_1856898, EPI_ISL_1856899, EPI_ISL_1856900 | | | |
| see above | UVA Clinical Microbiology Lab | Mathers Lab | Hardik I. Parikh, Katie E. Barry, Amy J. Mathers |
| EPI_ISL_1856993 | Outre mer | National Reference Center for Viruses of Respiratory Infections, Institut Pasteur, Paris | Marion Barbet, Sylvie Behillil, Méline Bizard, Angela Brisebarre, Camille Capel, Vincent Enouf, Louise Lefrançois, Frédéric Lemoine, Christophe Malabat, Corinne Maufrais, Gael Millot, Etienne Simon-Lorière, Maud Vanpeene, Sylvie Van der Werf ,Dominique Rousset |
| EPI_ISL_1857099 | LESP Nuevo Leon | Instituto de Diagnostico y Referencia Epidemiologicos (INDRE) | Claudia Wong-Arambula, Abril Rodriguez-Maldonado, Vanessa Rivero-Arredondo, Ariadna Medina-Benitez, Joaquin Quiroz-Mercado, Sergio Rangel-Guerrero, Natividad Cruz-Ortiz, Tatiana Nunez-Garcia, Gisela Barrera-Badillo, Lucia Hernandez-Rivas, Irma Lopez-Martinez, Ernesto Ramirez-Gonzalez. |
| EPI_ISL_1857100 | LESP Puebla | Instituto de Diagnostico y Referencia Epidemiologicos (INDRE) | Claudia Wong-Arambula, Abril Rodriguez-Maldonado, Vanessa Rivero-Arredondo, Ariadna Medina-Benitez, Joaquin Quiroz-Mercado, Sergio Rangel-Guerrero, Natividad Cruz-Ortiz, Tatiana Nunez-Garcia, Gisela Barrera-Badillo, Lucia Hernandez-Rivas, Irma Lopez-Martinez, Ernesto Ramirez-Gonzalez. |
| EPI_ISL_1857102, EPI_ISL_1857104, EPI_ISL_1857105 | LESP Ciudad de Mexico | Instituto de Diagnostico y Referencia Epidemiologicos (INDRE) | Claudia Wong-Arambula, Abril Rodriguez-Maldonado, Vanessa Rivero-Arredondo, Ariadna Medina-Benitez, Joaquin Quiroz-Mercado, Sergio Rangel-Guerrero, Natividad Cruz-Ortiz, Tatiana Nunez-Garcia, Gisela Barrera-Badillo, Lucia Hernandez-Rivas, Irma Lopez-Martinez, Ernesto Ramirez-Gonzalez. |
| EPI_ISL_1857200 | LESP Tamaulipas | Instituto de Diagnostico y Referencia Epidemiologicos (INDRE) | Claudia Wong-Arambula, Abril Rodriguez-Maldonado, Vanessa Rivero-Arredondo, Ariadna Medina-Benitez, Joaquin Quiroz-Mercado, Sergio Rangel-Guerrero, Natividad Cruz-Ortiz, Tatiana Nunez-Garcia, Gisela Barrera-Badillo, Lucia Hernandez-Rivas, Irma Lopez-Martinez, Ernesto Ramirez-Gonzalez. |
| EPI_ISL_1857201 | LESP Guerrero | Instituto de Diagnostico y Referencia Epidemiologicos (INDRE) | Claudia Wong-Arambula, Abril Rodriguez-Maldonado, Vanessa Rivero-Arredondo, Ariadna Medina-Benitez, Joaquin Quiroz-Mercado, Sergio Rangel-Guerrero, Natividad Cruz-Ortiz, Tatiana Nunez-Garcia, Gisela Barrera-Badillo, Lucia Hernandez-Rivas, Irma Lopez-Martinez, Ernesto Ramirez-Gonzalez. |
| EPI_ISL_1857205, EPI_ISL_1857206 | LESP Yucatan | Instituto de Diagnostico y Referencia Epidemiologicos (INDRE) | Claudia Wong-Arambula, Abril Rodriguez-Maldonado, Vanessa Rivero-Arredondo, Ariadna Medina-Benitez, Joaquin Quiroz-Mercado, Sergio Rangel-Guerrero, Natividad Cruz-Ortiz, Tatiana Nunez-Garcia, Gisela Barrera-Badillo, Lucia Hernandez-Rivas, Irma Lopez-Martinez, Ernesto Ramirez-Gonzalez. |
| EPI_ISL_1857207 | LESP Chiapas | Instituto de Diagnostico y Referencia Epidemiologicos (INDRE) | Claudia Wong-Arambula, Abril Rodriguez-Maldonado, Vanessa Rivero-Arredondo, Ariadna Medina-Benitez, Joaquin Quiroz-Mercado, Sergio Rangel-Guerrero, Natividad Cruz-Ortiz, Tatiana Nunez-Garcia, Gisela Barrera-Badillo, Lucia Hernandez-Rivas, Irma Lopez-Martinez, Ernesto Ramirez-Gonzalez. |
| EPI_ISL_1857209, EPI_ISL_1857210, EPI_ISL_1857211, EPI_ISL_1857215 | LESP Queretaro | Instituto de Diagnostico y Referencia Epidemiologicos (INDRE) | Claudia Wong-Arambula, Abril Rodriguez-Maldonado, Vanessa Rivero-Arredondo, Ariadna Medina-Benitez, Joaquin Quiroz-Mercado, Sergio Rangel-Guerrero, Natividad Cruz-Ortiz, Tatiana Nunez-Garcia, Gisela Barrera-Badillo, Lucia Hernandez-Rivas, Irma Lopez-Martinez, Ernesto Ramirez-Gonzalez. |
| EPI_ISL_1857217 | LESP Ciudad de Mexico | Instituto de Diagnostico y Referencia Epidemiologicos (INDRE) | Claudia Wong-Arambula, Abril Rodriguez-Maldonado, Vanessa Rivero-Arredondo, Ariadna Medina-Benitez, Joaquin Quiroz-Mercado, Sergio Rangel-Guerrero, Natividad Cruz-Ortiz, Tatiana Nunez-Garcia, Gisela Barrera-Badillo, Lucia Hernandez-Rivas, Irma Lopez-Martinez, Ernesto Ramirez-Gonzalez. |
| EPI_ISL_1857219, EPI_ISL_1857221, EPI_ISL_1857222, EPI_ISL_1857223 | LESP Chihuahua | Instituto de Diagnostico y Referencia Epidemiologicos (INDRE) | Claudia Wong-Arambula, Abril Rodriguez-Maldonado, Vanessa Rivero-Arredondo, Ariadna Medina-Benitez, Joaquin Quiroz-Mercado, Sergio Rangel-Guerrero, Natividad Cruz-Ortiz, Tatiana Nunez-Garcia, Gisela Barrera-Badillo, Lucia Hernandez-Rivas, Irma Lopez-Martinez, Ernesto Ramirez-Gonzalez. |
| EPI_ISL_1857224, EPI_ISL_1857225, EPI_ISL_1857226 | LESP Baja California Sur | Instituto de Diagnostico y Referencia Epidemiologicos (INDRE) | Claudia Wong-Arambula, Abril Rodriguez-Maldonado, Vanessa Rivero-Arredondo, Ariadna Medina-Benitez, Joaquin Quiroz-Mercado, Sergio Rangel-Guerrero, Natividad Cruz-Ortiz, Tatiana Nunez-Garcia, Gisela Barrera-Badillo, Lucia Hernandez-Rivas, Irma Lopez-Martinez, Ernesto Ramirez-Gonzalez. |
| EPI_ISL_1857228 | LESP Quintana Roo | Instituto de Diagnostico y Referencia Epidemiologicos (INDRE) | Claudia Wong-Arambula, Abril Rodriguez-Maldonado, Vanessa Rivero-Arredondo, Ariadna Medina-Benitez, Joaquin Quiroz-Mercado, Sergio Rangel-Guerrero, Natividad Cruz-Ortiz, Tatiana Nunez-Garcia, Gisela Barrera-Badillo, Lucia Hernandez-Rivas, Irma Lopez-Martinez, Ernesto Ramirez-Gonzalez. |
| EPI_ISL_1857230 | LESP Jalisco | Instituto de Diagnostico y Referencia Epidemiologicos (INDRE) | Claudia Wong-Arambula, Abril Rodriguez-Maldonado, Vanessa Rivero-Arredondo, Ariadna Medina-Benitez, Joaquin Quiroz-Mercado, Sergio Rangel-Guerrero, Natividad Cruz-Ortiz, Tatiana Nunez-Garcia, Gisela Barrera-Badillo, Lucia Hernandez-Rivas, Irma Lopez-Martinez, Ernesto Ramirez-Gonzalez. |
| EPI_ISL_1857232 | LESP Coahuila | Instituto de Diagnostico y Referencia Epidemiologicos (INDRE) | Claudia Wong-Arambula, Abril Rodriguez-Maldonado, Vanessa Rivero-Arredondo, Ariadna Medina-Benitez, Joaquin Quiroz-Mercado, Sergio Rangel-Guerrero, Natividad Cruz-Ortiz, Tatiana Nunez-Garcia, Gisela Barrera-Badillo, Lucia Hernandez-Rivas, Irma Lopez-Martinez, Ernesto |

| | | | |
|---|---|---|---|
| | | | Ramirez-Gonzalez. |
| EPI_ISL_1857236, EPI_ISL_1857237, EPI_ISL_1857238 | LESP Guanajuato | Instituto de Diagnostico y Referencia Epidemiologicos (INDRE) | Claudia Wong-Arambula, Abril Rodriguez-Maldonado, Vanessa Rivero-Arredondo, Ariadna Medina-Benitez, Joaquin Quiroz-Mercado, Sergio Rangel-Guerrero, Natividad Cruz-Ortiz, Tatiana Nunez-Garcia, Gisela Barrera-Badillo, Lucia Hernandez-Rivas, Irma Lopez-Martinez, Ernesto Ramirez-Gonzalez. |
| EPI_ISL_1857241 | LESP Puebla | Instituto de Diagnostico y Referencia Epidemiologicos (INDRE) | Claudia Wong-Arambula, Abril Rodriguez-Maldonado, Vanessa Rivero-Arredondo, Ariadna Medina-Benitez, Joaquin Quiroz-Mercado, Sergio Rangel-Guerrero, Natividad Cruz-Ortiz, Tatiana Nunez-Garcia, Gisela Barrera-Badillo, Lucia Hernandez-Rivas, Irma Lopez-Martinez, Ernesto Ramirez-Gonzalez. |
| EPI_ISL_1857246 | LESP Queretaro | Instituto de Diagnostico y Referencia Epidemiologicos (INDRE) | Claudia Wong-Arambula, Abril Rodriguez-Maldonado, Vanessa Rivero-Arredondo, Ariadna Medina-Benitez, Joaquin Quiroz-Mercado, Sergio Rangel-Guerrero, Natividad Cruz-Ortiz, Tatiana Nunez-Garcia, Gisela Barrera-Badillo, Lucia Hernandez-Rivas, Irma Lopez-Martinez, Ernesto Ramirez-Gonzalez. |
| EPI_ISL_1857248 | LESP Morelos | Instituto de Diagnostico y Referencia Epidemiologicos (INDRE) | Claudia Wong-Arambula, Abril Rodriguez-Maldonado, Vanessa Rivero-Arredondo, Ariadna Medina-Benitez, Joaquin Quiroz-Mercado, Sergio Rangel-Guerrero, Natividad Cruz-Ortiz, Tatiana Nunez-Garcia, Gisela Barrera-Badillo, Lucia Hernandez-Rivas, Irma Lopez-Martinez, Ernesto Ramirez-Gonzalez. |
| EPI_ISL_1857254 | LESP Tamaulipas | Instituto de Diagnostico y Referencia Epidemiologicos (INDRE) | Claudia Wong-Arambula, Abril Rodriguez-Maldonado, Vanessa Rivero-Arredondo, Ariadna Medina-Benitez, Joaquin Quiroz-Mercado, Sergio Rangel-Guerrero, Natividad Cruz-Ortiz, Tatiana Nunez-Garcia, Gisela Barrera-Badillo, Lucia Hernandez-Rivas, Irma Lopez-Martinez, Ernesto Ramirez-Gonzalez. |
| EPI_ISL_1857265 | LESP Zacatecas | Instituto de Diagnostico y Referencia Epidemiologicos (INDRE) | Claudia Wong-Arambula, Abril Rodriguez-Maldonado, Vanessa Rivero-Arredondo, Ariadna Medina-Benitez, Joaquin Quiroz-Mercado, Sergio Rangel-Guerrero, Natividad Cruz-Ortiz, Tatiana Nunez-Garcia, Gisela Barrera-Badillo, Lucia Hernandez-Rivas, Irma Lopez-Martinez, Ernesto Ramirez-Gonzalez. |
| EPI_ISL_1857269 | LESP Ciudad de Mexico | Instituto de Diagnostico y Referencia Epidemiologicos (INDRE) | Claudia Wong-Arambula, Abril Rodriguez-Maldonado, Vanessa Rivero-Arredondo, Ariadna Medina-Benitez, Joaquin Quiroz-Mercado, Sergio Rangel-Guerrero, Natividad Cruz-Ortiz, Tatiana Nunez-Garcia, Gisela Barrera-Badillo, Lucia Hernandez-Rivas, Irma Lopez-Martinez, Ernesto Ramirez-Gonzalez. |
| EPI_ISL_1904873, EPI_ISL_1904874 | OLVZ Aalst | OLVZ Aalst | Anne Vankeerberghen |
| EPI_ISL_1905079 | Laboratoire Central de Virologie | Laboratoire de Biotechnologie | Mouna Ouadghiri, Tarik Aanniz, Abdelomunim Essabbar, Ghizlane EL Amin, Amal Zouaki, Myriam Seffar, Hakima Kabbaj, Naima El Hafidi, Saaïd Amzazi, Lahcen Belyamani and Azeddine Ibrahimi |
| EPI_ISL_1905210 | AZ St.Jan Brugge-Oostende | AZ SINT-JAN BRUGGE | Marijke Reynders, Patrick Descheemaeker, Laurien Hoornaert, Thomas Van Landschoot, Jorn Hellemans |
| EPI_ISL_1908150 | Provincial Public Health Reference Laboratory (PPHRL) | Provincial Public Health Reference Laboratory (PPHRL) | Andleeb Hanif |
| EPI_ISL_1908788, EPI_ISL_1908789, EPI_ISL_1908790, EPI_ISL_1908791, EPI_ISL_1908792, EPI_ISL_1908793, EPI_ISL_1908794, EPI_ISL_1908795, EPI_ISL_1908796, EPI_ISL_1908797 | Microbiology Department, University Hospital Donostia | Microbiology Department, University Hospital Donostia | Marimon JM, Montes M, Piñeiro L, Sorarrain A, Gomez M, Cilla G. |
| EPI_ISL_1908907 | Provincial Public Health Reference Laboratory | Provincial Public Health Reference Laboratory | ANDLEEB HANIF |
| EPI_ISL_1909829, EPI_ISL_1909830, EPI_ISL_1909832, EPI_ISL_1909834, EPI_ISL_1909835, EPI_ISL_1909836, EPI_ISL_1909837, EPI_ISL_1909838, EPI_ISL_1909839, EPI_ISL_1909840, EPI_ISL_1909841, EPI_ISL_1909842, EPI_ISL_1909843, EPI_ISL_1909844, EPI_ISL_1909845, EPI_ISL_1909846, EPI_ISL_1909847, EPI_ISL_1909850, EPI_ISL_1909851, EPI_ISL_1909852, EPI_ISL_1909853, EPI_ISL_1909854, EPI_ISL_1909855, EPI_ISL_1909856, EPI_ISL_1909857, EPI_ISL_1909858, EPI_ISL_1909859, EPI_ISL_1909860, EPI_ISL_1909861, EPI_ISL_1909862, EPI_ISL_1909863, EPI_ISL_1909864, EPI_ISL_1909865, EPI_ISL_1909866, EPI_ISL_1909867, EPI_ISL_1909868, EPI_ISL_1909869, EPI_ISL_1909870, EPI_ISL_1909871, EPI_ISL_1909872, EPI_ISL_1909873, EPI_ISL_1909874, EPI_ISL_1909875, EPI_ISL_1909876, EPI_ISL_1909877, EPI_ISL_1909878, EPI_ISL_1909879, EPI_ISL_1909880, EPI_ISL_1909881, EPI_ISL_1909882, EPI_ISL_1909883, EPI_ISL_1909884, EPI_ISL_1909885, EPI_ISL_1909886, EPI_ISL_1909887, EPI_ISL_1909888, EPI_ISL_1909889, EPI_ISL_1909890, EPI_ISL_1909891, EPI_ISL_1909892, EPI_ISL_1909893, EPI_ISL_1909894, EPI_ISL_1909895, EPI_ISL_1909896, EPI_ISL_1909897, EPI_ISL_1909898, EPI_ISL_1909899, EPI_ISL_1909900, EPI_ISL_1909901, EPI_ISL_1909902, EPI_ISL_1909903, EPI_ISL_1909904, EPI_ISL_1909905, EPI_ISL_1909906, EPI_ISL_1909907, EPI_ISL_1909908, EPI_ISL_1909909, EPI_ISL_1909911, EPI_ISL_1909912, EPI_ISL_1909913, EPI_ISL_1909914, EPI_ISL_1909915, EPI_ISL_1909916, EPI_ISL_1909917, EPI_ISL_1909918 | | | |
| see above | National Food and Veterinary Risk Assessment Institute (NMVRVI) | National Public Health Surveillance Laboratory | Lukas Zemaitis, Migle Gabrielaite, Jelena Razmuk, Svajune Muralyte, Ana Steponkiene, Lukas Vasionis, Danas Baksa |
| EPI_ISL_1909925, EPI_ISL_1909927, EPI_ISL_1909928, EPI_ISL_1909929, EPI_ISL_1909930, EPI_ISL_1909931, EPI_ISL_1909932, EPI_ISL_1909933, EPI_ISL_1909934, EPI_ISL_1909935, EPI_ISL_1909936, EPI_ISL_1909937, EPI_ISL_1909938, EPI_ISL_1909939, EPI_ISL_1909940, EPI_ISL_1909941, EPI_ISL_1909942, EPI_ISL_1909943, EPI_ISL_1909944, EPI_ISL_1909945, EPI_ISL_1909946, EPI_ISL_1909947, EPI_ISL_1909948, EPI_ISL_1909949, EPI_ISL_1909950, EPI_ISL_1909951, EPI_ISL_1909952, EPI_ISL_1909953, EPI_ISL_1909954, EPI_ISL_1909955, EPI_ISL_1909956, EPI_ISL_1909957, EPI_ISL_1909958, EPI_ISL_1909959, EPI_ISL_1909960, EPI_ISL_1909962, EPI_ISL_1909964, EPI_ISL_1909965, EPI_ISL_1909967, EPI_ISL_1909968, EPI_ISL_1909969, EPI_ISL_1909970, EPI_ISL_1909971, EPI_ISL_1909972, EPI_ISL_1909973, EPI_ISL_1909974, EPI_ISL_1909976, EPI_ISL_1909977, EPI_ISL_1909978, EPI_ISL_1909979, EPI_ISL_1909980, EPI_ISL_1909981, EPI_ISL_1909983, EPI_ISL_1909984, EPI_ISL_1909985, EPI_ISL_1909986, EPI_ISL_1909989, EPI_ISL_1909990, EPI_ISL_1909992, EPI_ISL_1909993, EPI_ISL_1909994, EPI_ISL_1909995, EPI_ISL_1909996, EPI_ISL_1909997, EPI_ISL_1909999, EPI_ISL_1910000, EPI_ISL_1910001, EPI_ISL_1910002, EPI_ISL_1910004, EPI_ISL_1910005, EPI_ISL_1910006, EPI_ISL_1910007, EPI_ISL_1910009, EPI_ISL_1910010, EPI_ISL_1910011, EPI_ISL_1910012, EPI_ISL_1910013, EPI_ISL_1910014, EPI_ISL_1910017, EPI_ISL_1910018, EPI_ISL_1910019, EPI_ISL_1910020, EPI_ISL_1910021, EPI_ISL_1910022, EPI_ISL_1910023, EPI_ISL_1910024, EPI_ISL_1910025, EPI_ISL_1910026, EPI_ISL_1910027, EPI_ISL_1910028, EPI_ISL_1910029, EPI_ISL_1910031, EPI_ISL_1910032, EPI_ISL_1910033, EPI_ISL_1910034, EPI_ISL_1910035, EPI_ISL_1910036, EPI_ISL_1910037, EPI_ISL_1910038, EPI_ISL_1910039, EPI_ISL_1910040, EPI_ISL_1910041, EPI_ISL_1910042, EPI_ISL_1910043, EPI_ISL_1910044, EPI_ISL_1910045, EPI_ISL_1910046, EPI_ISL_1910048, EPI_ISL_1910049, EPI_ISL_1910050, EPI_ISL_1910051, EPI_ISL_1910052, EPI_ISL_1910053, EPI_ISL_1910054, EPI_ISL_1910055, EPI_ISL_1910056, EPI_ISL_1910057, EPI_ISL_1910058, EPI_ISL_1910059, EPI_ISL_1910061, EPI_ISL_1910062, EPI_ISL_1910064, EPI_ISL_1910065, EPI_ISL_1910066, EPI_ISL_1910067, EPI_ISL_1910068, EPI_ISL_1910069, EPI_ISL_1910070, EPI_ISL_1910071, EPI_ISL_1910072, EPI_ISL_1910073, EPI_ISL_1910074, EPI_ISL_1910075, EPI_ISL_1910076, EPI_ISL_1910079, EPI_ISL_1910080, EPI_ISL_1910081, EPI_ISL_1910082, EPI_ISL_1910083, EPI_ISL_1910084, EPI_ISL_1910085, EPI_ISL_1910086, EPI_ISL_1910087, EPI_ISL_1910088, EPI_ISL_1910089, EPI_ISL_1910090, EPI_ISL_1910091, EPI_ISL_1910092, EPI_ISL_1910093, EPI_ISL_1910094, EPI_ISL_1910095, EPI_ISL_1910096, EPI_ISL_1910097, EPI_ISL_1910098, EPI_ISL_1910099, EPI_ISL_1910100, EPI_ISL_1910101, EPI_ISL_1910102, EPI_ISL_1910103, EPI_ISL_1910104, EPI_ISL_1910105, EPI_ISL_1910108, EPI_ISL_1910109, EPI_ISL_1910110, EPI_ISL_1910112, EPI_ISL_1910115, EPI_ISL_1910116, EPI_ISL_1910119, EPI_ISL_1910120, EPI_ISL_1910122, EPI_ISL_1910123, EPI_ISL_1910124, EPI_ISL_1910126, EPI_ISL_1910127, EPI_ISL_1910128, EPI_ISL_1910129, EPI_ISL_1910131, EPI_ISL_1910132, EPI_ISL_1910133, EPI_ISL_1910134, EPI_ISL_1910135, EPI_ISL_1910137, EPI_ISL_1910139, EPI_ISL_1910141, EPI_ISL_1910142, EPI_ISL_1910143, EPI_ISL_1910144, EPI_ISL_1910145, EPI_ISL_1910147, EPI_ISL_1910149, EPI_ISL_1910150, EPI_ISL_1910152, EPI_ISL_1910153, EPI_ISL_1910155, EPI_ISL_1910158, EPI_ISL_1910159, EPI_ISL_1910160, EPI_ISL_1910161, EPI_ISL_1910162, EPI_ISL_1910163, EPI_ISL_1910164, EPI_ISL_1910165, EPI_ISL_1910166, EPI_ISL_1910167, EPI_ISL_1910168, EPI_ISL_1910169, EPI_ISL_1910170, EPI_ISL_1910171, EPI_ISL_1910172, EPI_ISL_1910173, EPI_ISL_1910174, EPI_ISL_1910175, EPI_ISL_1910178, EPI_ISL_1910179, EPI_ISL_1910180, EPI_ISL_1910181, EPI_ISL_1910182, EPI_ISL_1910183, EPI_ISL_1910184, EPI_ISL_1910185, EPI_ISL_1910186, EPI_ISL_1910187, EPI_ISL_1910188, EPI_ISL_1910189, EPI_ISL_1910190, EPI_ISL_1910191, EPI_ISL_1910192, EPI_ISL_1910194, EPI_ISL_1910195, EPI_ISL_1910196, EPI_ISL_1910198, EPI_ISL_1910199, EPI_ISL_1910200, EPI_ISL_1910201, EPI_ISL_1910202, EPI_ISL_1910203, EPI_ISL_1910204, EPI_ISL_1910205, EPI_ISL_1910206, EPI_ISL_1910207, EPI_ISL_1910208, EPI_ISL_1910209, EPI_ISL_1910210, EPI_ISL_1910211, EPI_ISL_1910212, EPI_ISL_1910213, EPI_ISL_1910214, EPI_ISL_1910215, EPI_ISL_1910216, EPI_ISL_1910217, EPI_ISL_1910218, EPI_ISL_1910219, EPI_ISL_1910220, EPI_ISL_1910221, EPI_ISL_1910222, EPI_ISL_1910223, EPI_ISL_1910224, EPI_ISL_1910225, EPI_ISL_1910226, EPI_ISL_1910227, EPI_ISL_1910228, EPI_ISL_1910229, EPI_ISL_1910230, EPI_ISL_1910231, EPI_ISL_1910232, EPI_ISL_1910233, EPI_ISL_1910234, EPI_ISL_1910235, EPI_ISL_1910236, EPI_ISL_1910237, EPI_ISL_1910238, EPI_ISL_1910239, EPI_ISL_1910240, EPI_ISL_1910241, EPI_ISL_1910242, EPI_ISL_1910243, EPI_ISL_1910244, EPI_ISL_1910245, EPI_ISL_1910246, EPI_ISL_1910247, EPI_ISL_1910248, EPI_ISL_1910249, EPI_ISL_1910250, EPI_ISL_1910251, EPI_ISL_1910253, EPI_ISL_1910254, EPI_ISL_1910255, EPI_ISL_1910256, EPI_ISL_1910257, EPI_ISL_1910258, EPI_ISL_1910259, EPI_ISL_1910260, EPI_ISL_1910261, EPI_ISL_1910262, EPI_ISL_1910263, EPI_ISL_1910264, EPI_ISL_1910265, EPI_ISL_1910266, EPI_ISL_1910267, EPI_ISL_1910268, EPI_ISL_1910269, EPI_ISL_1910270, EPI_ISL_1910271, EPI_ISL_1910272, EPI_ISL_1910273, EPI_ISL_1910274, EPI_ISL_1910275, EPI_ISL_1910276, EPI_ISL_1910277, EPI_ISL_1910279, EPI_ISL_1910280, EPI_ISL_1910281, EPI_ISL_1910282, EPI_ISL_1910283, EPI_ISL_1910284, EPI_ISL_1910285, EPI_ISL_1910287, EPI_ISL_1910288, EPI_ISL_1910289, EPI_ISL_1910290, EPI_ISL_1910291, EPI_ISL_1910293, EPI_ISL_1910294, EPI_ISL_1910295, EPI_ISL_1910297, EPI_ISL_1910300, EPI_ISL_1910301, EPI_ISL_1910304, EPI_ISL_1910305, EPI_ISL_1910307, EPI_ISL_1910308, EPI_ISL_1910309, EPI_ISL_1910311, EPI_ISL_1910312, EPI_ISL_1910313, EPI_ISL_1910314, EPI_ISL_1910316, EPI_ISL_1910317, EPI_ISL_1910318, EPI_ISL_1910319, EPI_ISL_1910320, EPI_ISL_1910322, EPI_ISL_1910324, EPI_ISL_1910326, EPI_ISL_1910327, EPI_ISL_1910328, EPI_ISL_1910329, EPI_ISL_1910330, EPI_ISL_1910332, EPI_ISL_1910334, EPI_ISL_1910335, EPI_ISL_1910337, EPI_ISL_1910338, EPI_ISL_1910340, EPI_ISL_1910343, EPI_ISL_1910344, EPI_ISL_1910345, EPI_ISL_1910346, EPI_ISL_1910347, EPI_ISL_1910348, EPI_ISL_1910349, EPI_ISL_1910350, EPI_ISL_1910351, EPI_ISL_1910352, EPI_ISL_1910353, EPI_ISL_1910354, EPI_ISL_1910355, EPI_ISL_1910356, EPI_ISL_1910357, EPI_ISL_1910358, EPI_ISL_1910359, EPI_ISL_1910360, EPI_ISL_1910363, EPI_ISL_1910364, EPI_ISL_1910365, EPI_ISL_1910366, EPI_ISL_1910367, EPI_ISL_1910368, EPI_ISL_1910369, EPI_ISL_1910370, EPI_ISL_1910371, EPI_ISL_1910372, EPI_ISL_1910373, EPI_ISL_1910374, EPI_ISL_1910375, EPI_ISL_1910376, EPI_ISL_1910377, EPI_ISL_1910379, EPI_ISL_1910380, EPI_ISL_1910381, EPI_ISL_1910383, EPI_ISL_1910384 | | | |
| see above | National Institute for Food and Veterinary Risk Assessment (NMVRVI) | National Public Health Surveillance Laboratory | Lukas Zemaitis, Migle Gabrielaite, Jelena Razmuk, Svajune Muralyte, Ana Steponkiene, Lukas Vasionis, Danas Baksa |
| EPI_ISL_1910390, EPI_ISL_1910393 | Iressef Genomics lab | IRESSEF | Souleymane MBOUP, Abdou PADANE, Khadim GUEYE, Papa Alassane DIAW, Birahim Piere NDIAYE, Barada CISSE, Aminata MBOUP, Moustapha MBOW, Ndeye Coumba Toure KANE, Nafisatou LEYE, Gora LO, Ambroise AHOUIDI , Astou Gaye GAYE, Aminata DIA, Yacine DIA |
| EPI_ISL_1910862 | CHU Lille | CHU Lille | AIT YAHYA Emilie, ALIDJINOU Enagnon Kazali, BOCKET Laurence, CREPIN Michel, DEMAY Christophe, ENGELMANN Ilka, GEFFROY Sandrine, GUIGON Aurélie, LAMBERT Valérie, LAZREK Mouna, NOBILLIAUX Florian, PREVOST Brigitte, THUILLIER Caroline, TINEZ Claire |
| EPI_ISL_1910865, EPI_ISL_1910866 | CH Lens | CHU Lille | AIT YAHYA Emilie, ALIDJINOU Enagnon Kazali, BOCKET Laurence, CREPIN Michel, DEMAY Christophe, ENGELMANN Ilka, GEFFROY Sandrine, GUIGON Aurélie, LAMBERT Valérie, LAZREK Mouna, NOBILLIAUX Florian, PREVOST Brigitte, THUILLIER Caroline, TINEZ Claire |
| EPI_ISL_1910868, EPI_ISL_1910869 | CHU Lille | CHU Lille | AIT YAHYA Emilie, ALIDJINOU Enagnon Kazali, BOCKET Laurence, CREPIN Michel, DEMAY Christophe, ENGELMANN Ilka, GEFFROY Sandrine, GUIGON Aurélie, LAMBERT Valérie, LAZREK Mouna, NOBILLIAUX Florian, PREVOST Brigitte, THUILLIER Caroline, TINEZ Claire |
| EPI_ISL_1910870, EPI_ISL_1910871 | CH Boulogne sur mer | CHU Lille | AIT YAHYA Emilie, ALIDJINOU Enagnon Kazali, BOCKET Laurence, CREPIN Michel, DEMAY Christophe, ENGELMANN Ilka, GEFFROY Sandrine, GUIGON Aurélie, LAMBERT Valérie, LAZREK Mouna, NOBILLIAUX Florian, PREVOST Brigitte, THUILLIER Caroline, TINEZ Claire |

| | | | |
|---|---|---|---|
| EPI_ISL_1910873 | CH Calais | CHU Lille | AIT YAHYA Emilie, ALIDJINOU Enagnon Kazali, BOCKET Laurence, CREPIN Michel, DEMAY Christophe, ENGELMANN Ilka, GEFFROY Sandrine, GUIGON Aurélie, LAMBERT Valérie, LAZREK Mouna, NOBILLIAUX Florian, PREVOST Brigitte, THUILLIER Caroline, TINEZ Claire |
| EPI_ISL_1910874 | CH Saint-Philibert | CHU Lille | AIT YAHYA Emilie, ALIDJINOU Enagnon Kazali, BOCKET Laurence, CREPIN Michel, DEMAY Christophe, ENGELMANN Ilka, GEFFROY Sandrine, GUIGON Aurélie, LAMBERT Valérie, LAZREK Mouna, NOBILLIAUX Florian, PREVOST Brigitte, THUILLIER Caroline, TINEZ Claire |
| EPI_ISL_1910875 | CH Roubaix | CHU Lille | AIT YAHYA Emilie, ALIDJINOU Enagnon Kazali, BOCKET Laurence, CREPIN Michel, DEMAY Christophe, ENGELMANN Ilka, GEFFROY Sandrine, GUIGON Aurélie, LAMBERT Valérie, LAZREK Mouna, NOBILLIAUX Florian, PREVOST Brigitte, THUILLIER Caroline, TINEZ Claire |
| EPI_ISL_1910878 | CH Dunkerque | CHU Lille | AIT YAHYA Emilie, ALIDJINOU Enagnon Kazali, BOCKET Laurence, CREPIN Michel, DEMAY Christophe, ENGELMANN Ilka, GEFFROY Sandrine, GUIGON Aurélie, LAMBERT Valérie, LAZREK Mouna, NOBILLIAUX Florian, PREVOST Brigitte, THUILLIER Caroline, TINEZ Claire |
| EPI_ISL_1910880 | CH Cambrai | CHU Lille | AIT YAHYA Emilie, ALIDJINOU Enagnon Kazali, BOCKET Laurence, CREPIN Michel, DEMAY Christophe, ENGELMANN Ilka, GEFFROY Sandrine, GUIGON Aurélie, LAMBERT Valérie, LAZREK Mouna, NOBILLIAUX Florian, PREVOST Brigitte, THUILLIER Caroline, TINEZ Claire |
| EPI_ISL_1910881 | CH Calais | CHU Lille | AIT YAHYA Emilie, ALIDJINOU Enagnon Kazali, BOCKET Laurence, CREPIN Michel, DEMAY Christophe, ENGELMANN Ilka, GEFFROY Sandrine, GUIGON Aurélie, LAMBERT Valérie, LAZREK Mouna, NOBILLIAUX Florian, PREVOST Brigitte, THUILLIER Caroline, TINEZ Claire |
| EPI_ISL_1910882 | CH Bethune | CHU Lille | AIT YAHYA Emilie, ALIDJINOU Enagnon Kazali, BOCKET Laurence, CREPIN Michel, DEMAY Christophe, ENGELMANN Ilka, GEFFROY Sandrine, GUIGON Aurélie, LAMBERT Valérie, LAZREK Mouna, NOBILLIAUX Florian, PREVOST Brigitte, THUILLIER Caroline, TINEZ Claire |
| EPI_ISL_1910884, EPI_ISL_1910885, EPI_ISL_1910886, EPI_ISL_1910887, EPI_ISL_1910889, EPI_ISL_1910890, EPI_ISL_1910891 | CHU Lille | CHU Lille | AIT YAHYA Emilie, ALIDJINOU Enagnon Kazali, BOCKET Laurence, CREPIN Michel, DEMAY Christophe, ENGELMANN Ilka, GEFFROY Sandrine, GUIGON Aurélie, LAMBERT Valérie, LAZREK Mouna, NOBILLIAUX Florian, PREVOST Brigitte, THUILLIER Caroline, TINEZ Claire |
| EPI_ISL_1910892, EPI_ISL_1910893 | CH Tourcoing | CHU Lille | AIT YAHYA Emilie, ALIDJINOU Enagnon Kazali, BOCKET Laurence, CREPIN Michel, DEMAY Christophe, ENGELMANN Ilka, GEFFROY Sandrine, GUIGON Aurélie, LAMBERT Valérie, LAZREK Mouna, NOBILLIAUX Florian, PREVOST Brigitte, THUILLIER Caroline, TINEZ Claire |
| EPI_ISL_1910894 | CH Roubaix | CHU Lille | AIT YAHYA Emilie, ALIDJINOU Enagnon Kazali, BOCKET Laurence, CREPIN Michel, DEMAY Christophe, ENGELMANN Ilka, GEFFROY Sandrine, GUIGON Aurélie, LAMBERT Valérie, LAZREK Mouna, NOBILLIAUX Florian, PREVOST Brigitte, THUILLIER Caroline, TINEZ Claire |
| EPI_ISL_1910895, EPI_ISL_1910896, EPI_ISL_1910897 | CH Arras | CHU Lille | AIT YAHYA Emilie, ALIDJINOU Enagnon Kazali, BOCKET Laurence, CREPIN Michel, DEMAY Christophe, ENGELMANN Ilka, GEFFROY Sandrine, GUIGON Aurélie, LAMBERT Valérie, LAZREK Mouna, NOBILLIAUX Florian, PREVOST Brigitte, THUILLIER Caroline, TINEZ Claire |
| EPI_ISL_1910898, EPI_ISL_1910899 | CH Cambrai | CHU Lille | AIT YAHYA Emilie, ALIDJINOU Enagnon Kazali, BOCKET Laurence, CREPIN Michel, DEMAY Christophe, ENGELMANN Ilka, GEFFROY Sandrine, GUIGON Aurélie, LAMBERT Valérie, LAZREK Mouna, NOBILLIAUX Florian, PREVOST Brigitte, THUILLIER Caroline, TINEZ Claire |
| EPI_ISL_1910900, EPI_ISL_1910903, EPI_ISL_1910905, EPI_ISL_1910906 | CHU Lille | CHU Lille | AIT YAHYA Emilie, ALIDJINOU Enagnon Kazali, BOCKET Laurence, CREPIN Michel, DEMAY Christophe, ENGELMANN Ilka, GEFFROY Sandrine, GUIGON Aurélie, LAMBERT Valérie, LAZREK Mouna, NOBILLIAUX Florian, PREVOST Brigitte, THUILLIER Caroline, TINEZ Claire |
| EPI_ISL_1910907 | CH Tourcoing | CHU Lille | AIT YAHYA Emilie, ALIDJINOU Enagnon Kazali, BOCKET Laurence, CREPIN Michel, DEMAY Christophe, ENGELMANN Ilka, GEFFROY Sandrine, GUIGON Aurélie, LAMBERT Valérie, LAZREK Mouna, NOBILLIAUX Florian, PREVOST Brigitte, THUILLIER Caroline, TINEZ Claire |
| EPI_ISL_1910908 | CH Seclin | CHU Lille | AIT YAHYA Emilie, ALIDJINOU Enagnon Kazali, BOCKET Laurence, CREPIN Michel, DEMAY Christophe, ENGELMANN Ilka, GEFFROY Sandrine, GUIGON Aurélie, LAMBERT Valérie, LAZREK Mouna, NOBILLIAUX Florian, PREVOST Brigitte, THUILLIER Caroline, TINEZ Claire |
| EPI_ISL_1910913, EPI_ISL_1910914 | CHU Lille | CHU Lille | AIT YAHYA Emilie, ALIDJINOU Enagnon Kazali, BOCKET Laurence, CREPIN Michel, DEMAY Christophe, ENGELMANN Ilka, GEFFROY Sandrine, GUIGON Aurélie, LAMBERT Valérie, LAZREK Mouna, NOBILLIAUX Florian, PREVOST Brigitte, THUILLIER Caroline, TINEZ Claire |
| EPI_ISL_1910916 | CH Boulogne sur mer | CHU Lille | AIT YAHYA Emilie, ALIDJINOU Enagnon Kazali, BOCKET Laurence, CREPIN Michel, DEMAY Christophe, ENGELMANN Ilka, GEFFROY Sandrine, GUIGON Aurélie, LAMBERT Valérie, LAZREK Mouna, NOBILLIAUX Florian, PREVOST Brigitte, THUILLIER Caroline, TINEZ Claire |
| EPI_ISL_1910918 | CH Calais | CHU Lille | AIT YAHYA Emilie, ALIDJINOU Enagnon Kazali, BOCKET Laurence, CREPIN Michel, DEMAY Christophe, ENGELMANN Ilka, GEFFROY Sandrine, GUIGON Aurélie, LAMBERT Valérie, LAZREK Mouna, NOBILLIAUX Florian, PREVOST Brigitte, THUILLIER Caroline, TINEZ Claire |
| EPI_ISL_1910920, EPI_ISL_1910921, EPI_ISL_1910922, EPI_ISL_1910923, EPI_ISL_1910924, EPI_ISL_1910925, EPI_ISL_1910926, EPI_ISL_1910927, EPI_ISL_1910928, EPI_ISL_1910929 | CHU Lille | CHU Lille | AIT YAHYA Emilie, ALIDJINOU Enagnon Kazali, BOCKET Laurence, CREPIN Michel, DEMAY Christophe, ENGELMANN Ilka, GEFFROY Sandrine, GUIGON Aurélie, LAMBERT Valérie, LAZREK Mouna, NOBILLIAUX Florian, PREVOST Brigitte, THUILLIER Caroline, TINEZ Claire |
| EPI_ISL_1910930 | CH Seclin | CHU Lille | AIT YAHYA Emilie, ALIDJINOU Enagnon Kazali, BOCKET Laurence, CREPIN Michel, DEMAY Christophe, ENGELMANN Ilka, GEFFROY Sandrine, GUIGON Aurélie, LAMBERT Valérie, LAZREK Mouna, NOBILLIAUX Florian, PREVOST Brigitte, THUILLIER Caroline, TINEZ Claire |
| EPI_ISL_1910931 | CH Roubaix | CHU Lille | AIT YAHYA Emilie, ALIDJINOU Enagnon Kazali, BOCKET Laurence, CREPIN Michel, DEMAY Christophe, ENGELMANN Ilka, GEFFROY Sandrine, GUIGON Aurélie, LAMBERT Valérie, LAZREK Mouna, NOBILLIAUX Florian, PREVOST Brigitte, THUILLIER Caroline, TINEZ Claire |
| EPI_ISL_1910932, EPI_ISL_1910933, EPI_ISL_1910934, EPI_ISL_1910935, EPI_ISL_1910937 | CHU Lille | CHU Lille | AIT YAHYA Emilie, ALIDJINOU Enagnon Kazali, BOCKET Laurence, CREPIN Michel, DEMAY Christophe, ENGELMANN Ilka, GEFFROY Sandrine, GUIGON Aurélie, LAMBERT Valérie, LAZREK Mouna, NOBILLIAUX Florian, PREVOST Brigitte, THUILLIER Caroline, TINEZ Claire |
| EPI_ISL_1910938 | CH Seclin | CHU Lille | AIT YAHYA Emilie, ALIDJINOU Enagnon Kazali, BOCKET Laurence, CREPIN Michel, DEMAY Christophe, ENGELMANN Ilka, GEFFROY Sandrine, GUIGON Aurélie, LAMBERT Valérie, LAZREK Mouna, NOBILLIAUX Florian, PREVOST Brigitte, THUILLIER Caroline, TINEZ Claire |
| EPI_ISL_1910939, EPI_ISL_1910940, EPI_ISL_1910941, EPI_ISL_1910942, EPI_ISL_1910943, EPI_ISL_1910944, EPI_ISL_1910945, EPI_ISL_1910946 | CHU Lille | CHU Lille | AIT YAHYA Emilie, ALIDJINOU Enagnon Kazali, BOCKET Laurence, CREPIN Michel, DEMAY Christophe, ENGELMANN Ilka, GEFFROY Sandrine, GUIGON Aurélie, LAMBERT Valérie, LAZREK Mouna, NOBILLIAUX Florian, PREVOST Brigitte, THUILLIER Caroline, TINEZ Claire |
| EPI_ISL_1910947 | CH Tourcoing | CHU Lille | AIT YAHYA Emilie, ALIDJINOU Enagnon Kazali, BOCKET Laurence, CREPIN Michel, DEMAY Christophe, ENGELMANN Ilka, GEFFROY Sandrine, GUIGON Aurélie, LAMBERT Valérie, LAZREK Mouna, NOBILLIAUX Florian, PREVOST Brigitte, THUILLIER Caroline, TINEZ Claire |
| EPI_ISL_1910948 | CH Cambrai | CHU Lille | AIT YAHYA Emilie, ALIDJINOU Enagnon Kazali, BOCKET Laurence, CREPIN Michel, DEMAY Christophe, ENGELMANN Ilka, GEFFROY Sandrine, GUIGON Aurélie, LAMBERT Valérie, LAZREK Mouna, NOBILLIAUX Florian, PREVOST Brigitte, THUILLIER Caroline, TINEZ Claire |
| EPI_ISL_1911423, EPI_ISL_1911424, EPI_ISL_1911425, EPI_ISL_1911426, EPI_ISL_1911427, EPI_ISL_1911428, EPI_ISL_1911429, EPI_ISL_1911430, EPI_ISL_1911431, EPI_ISL_1911432, EPI_ISL_1911433, EPI_ISL_1911434, EPI_ISL_1911435, EPI_ISL_1911436 | | | |
| see above | NVSPL | National Public Health Surveillance Laboratory | Lukas Zemaitis, Migle Gabrielaite, Jelena Razmuk, Svajune Muralyte, Ana Steponkiene, Lukas Vasionis, Danas Baksa |
| EPI_ISL_1911439, EPI_ISL_1911441, EPI_ISL_1911443, EPI_ISL_1911444, EPI_ISL_1911445, EPI_ISL_1911446, EPI_ISL_1911447, EPI_ISL_1911449, EPI_ISL_1911450, EPI_ISL_1911451, EPI_ISL_1911452, EPI_ISL_1911453, EPI_ISL_1911456, EPI_ISL_1911457, EPI_ISL_1911459, EPI_ISL_1911460, EPI_ISL_1911462, EPI_ISL_1911463, EPI_ISL_1911464, EPI_ISL_1911465, EPI_ISL_1911467, EPI_ISL_1911468, EPI_ISL_1911469, EPI_ISL_1911470, EPI_ISL_1911471, EPI_ISL_1911472, EPI_ISL_1911474, EPI_ISL_1911475, EPI_ISL_1911476, EPI_ISL_1911477, EPI_ISL_1911478, EPI_ISL_1911479, EPI_ISL_1911481, EPI_ISL_1911482, EPI_ISL_1911483, EPI_ISL_1911484, EPI_ISL_1911485, EPI_ISL_1911486, EPI_ISL_1911487, EPI_ISL_1911488, EPI_ISL_1911491, EPI_ISL_1911492, EPI_ISL_1911493, EPI_ISL_1911494, EPI_ISL_1911495, EPI_ISL_1911496, EPI_ISL_1911497, EPI_ISL_1911498, EPI_ISL_1911499, EPI_ISL_1911500, EPI_ISL_1911501, EPI_ISL_1911502, EPI_ISL_1911503, EPI_ISL_1911504, EPI_ISL_1911505, EPI_ISL_1911506, EPI_ISL_1911527, EPI_ISL_1911528, EPI_ISL_1911530, EPI_ISL_1911531, EPI_ISL_1911532, EPI_ISL_1911533, EPI_ISL_1911534, EPI_ISL_1911536, EPI_ISL_1911537, EPI_ISL_1911538, EPI_ISL_1911539, EPI_ISL_1911540, EPI_ISL_1911541, EPI_ISL_1911542, EPI_ISL_1911543, EPI_ISL_1911544, EPI_ISL_1911546, EPI_ISL_1911548, EPI_ISL_1911549, EPI_ISL_1911550, EPI_ISL_1911552, EPI_ISL_1911553, EPI_ISL_1911554, EPI_ISL_1911555, EPI_ISL_1911556, EPI_ISL_1911558, EPI_ISL_1911560, EPI_ISL_1911561, EPI_ISL_1911562, EPI_ISL_1911563, EPI_ISL_1911564, EPI_ISL_1911565, EPI_ISL_1911566, EPI_ISL_1911567, EPI_ISL_1911568, EPI_ISL_1911569, EPI_ISL_1911570, EPI_ISL_1911571, EPI_ISL_1911572, EPI_ISL_1911573, EPI_ISL_1911574, EPI_ISL_1911575, EPI_ISL_1911576, EPI_ISL_1911577, EPI_ISL_1911578, EPI_ISL_1911579, EPI_ISL_1911580, EPI_ISL_1911581, EPI_ISL_1911582, EPI_ISL_1911584, EPI_ISL_1911585, EPI_ISL_1911586, EPI_ISL_1911587, EPI_ISL_1911588, EPI_ISL_1911589, EPI_ISL_1911591, EPI_ISL_1911593, EPI_ISL_1911594, EPI_ISL_1911595, EPI_ISL_1911596, EPI_ISL_1911597, EPI_ISL_1911598, EPI_ISL_1911599, EPI_ISL_1911600, EPI_ISL_1911601, EPI_ISL_1911602, EPI_ISL_1911604, EPI_ISL_1911605, EPI_ISL_1911607, EPI_ISL_1911608, EPI_ISL_1911609, EPI_ISL_1911610, EPI_ISL_1911611, EPI_ISL_1911612, EPI_ISL_1911613, EPI_ISL_1911614, EPI_ISL_1911615, EPI_ISL_1911616 | | | |
| see above | National Institute for Food and Veterinary Risk Assessment (NMVRVI) | National Public Health Surveillance Laboratory | Lukas Zemaitis, Migle Gabrielaite, Jelena Razmuk, Svajune Muralyte, Ana Steponkiene, Lukas Vasionis, Danas Baksa |
| EPI_ISL_1911655, EPI_ISL_1911656, EPI_ISL_1911657, EPI_ISL_1911658, EPI_ISL_1911659, EPI_ISL_1911660, EPI_ISL_1911661, EPI_ISL_1911662, EPI_ISL_1911663, EPI_ISL_1911664, EPI_ISL_1911665, EPI_ISL_1911667, EPI_ISL_1911668, EPI_ISL_1911669, EPI_ISL_1911670, EPI_ISL_1911671, EPI_ISL_1911672, | | | |

| | | | |
|---|---|---|---|
| EPI_ISL_1911673, EPI_ISL_1911674, EPI_ISL_1911675, EPI_ISL_1911676, EPI_ISL_1911677, EPI_ISL_1911678, EPI_ISL_1911679, EPI_ISL_1911680, EPI_ISL_1911681, EPI_ISL_1911682, EPI_ISL_1911683, EPI_ISL_1911684, EPI_ISL_1911685, EPI_ISL_1911686, EPI_ISL_1911687, EPI_ISL_1911689, EPI_ISL_1911690, EPI_ISL_1911691, EPI_ISL_1911692, EPI_ISL_1911693, EPI_ISL_1911694, EPI_ISL_1911695, EPI_ISL_1911696, EPI_ISL_1911698, EPI_ISL_1911700, EPI_ISL_1911701, EPI_ISL_1911702, EPI_ISL_1911704, EPI_ISL_1911705, EPI_ISL_1911706, EPI_ISL_1911707, EPI_ISL_1911708, EPI_ISL_1911709, EPI_ISL_1911710, EPI_ISL_1911712, EPI_ISL_1911713, EPI_ISL_1911714, EPI_ISL_1911715, EPI_ISL_1911716, EPI_ISL_1911717, EPI_ISL_1911719, EPI_ISL_1911720, EPI_ISL_1911721, EPI_ISL_1911722, EPI_ISL_1911723, EPI_ISL_1911724, EPI_ISL_1911725, EPI_ISL_1911727, EPI_ISL_1911728, EPI_ISL_1911729, EPI_ISL_1911730, EPI_ISL_1911732, EPI_ISL_1911733, EPI_ISL_1911734, EPI_ISL_1911735, EPI_ISL_1911736, EPI_ISL_1911738, EPI_ISL_1911739, EPI_ISL_1911740, EPI_ISL_1911741, EPI_ISL_1911742 | | | |
| see above | National Food and Veterinary Risk Assessment Institute (NMVRVI) | National Public Health Surveillance Laboratory | Lukas Zemaitis, Migle Gabrielaite, Jelena Razmuk, Svajune Muralyte, Ana Steponkiene, Lukas Vasionis, Danas Baksa |
| EPI_ISL_1911817, EPI_ISL_1911831, EPI_ISL_1911851, EPI_ISL_1911855, EPI_ISL_1911864, EPI_ISL_1911868, EPI_ISL_1911876, EPI_ISL_1911894, EPI_ISL_1911896 | National Institute for Food and Veterinary Risk Assessment (NMVRVI) | National Public Health Surveillance Laboratory | Lukas Zemaitis, Migle Gabrielaite, Jelena Razmuk, Svajune Muralyte, Ana Steponkiene, Lukas Vasionis, Danas Baksa |
| EPI_ISL_1911917, EPI_ISL_1911947, EPI_ISL_1911948, EPI_ISL_1911949, EPI_ISL_1911950, EPI_ISL_1911951, EPI_ISL_1911952 | Baylor Scott & White-Temple | Baylor Scott & White-Temple | Ari Rao, Linden Morales, Kimberly Walker, Marcus Volz, Shelby Hendrickson |
| EPI_ISL_1914277 | Cliniques universitaires Saint-Luc | UCLouvain/IREC/MBLG | Jean Ruelle, Eleonore Ngyuvula, Benoit Kabamba Mukadi |
| EPI_ISL_1914595 | State Virus Research and Diagnostic Laboratory (VRDL), AIIMS Raipur | State Virus Research and Diagnostic Laboratory (VRDL), AIIMS Raipur | Pushpendra Singh, Kuldeep Sharma, Priyanka Singh, Somya Sharma, Sanjay Singh Negi, Anudita Bhargava |
| EPI_ISL_1915128, EPI_ISL_1915129, EPI_ISL_1915131, EPI_ISL_1915133, EPI_ISL_1915134 | Hospital | National Reference Center for Viruses of Respiratory Infections, Institut Pasteur, Paris | Marion Barbet, Sylvie Behillil, Méline Bizard, Angela Brisebarre, Camille Capel, Vincent Enouf, Louise Lefrançois, Frédéric Lemoine, Christophe Malabat, Corinne Maufrais, Emmanuelle Permal, Etienne Simon-Lorière, Maud Vanpeene, Sylvie Van der Werf ,Eric Farfour |
| EPI_ISL_1915142 | Hospital | National Reference Center for Viruses of Respiratory Infections, Institut Pasteur, Paris | Marion Barbet, Sylvie Behillil, Méline Bizard, Angela Brisebarre, Camille Capel, Vincent Enouf, Louise Lefrançois, Frédéric Lemoine, Christophe Malabat, Corinne Maufrais, Emmanuelle Permal, Etienne Simon-Lorière, Maud Vanpeene, Sylvie Van der Werf ,Farida Moreau Benaoudia |
| EPI_ISL_1915154 | Labo Analyses Med | National Reference Center for Viruses of Respiratory Infections, Institut Pasteur, Paris | Marion Barbet, Sylvie Behillil, Méline Bizard, Angela Brisebarre, Camille Capel, Vincent Enouf, Louise Lefrançois, Frédéric Lemoine, Christophe Malabat, Corinne Maufrais, Emmanuelle Permal, Etienne Simon-Lorière, Maud Vanpeene, Sylvie Van der Werf ,Manuel Zurbaran |
| EPI_ISL_1916411, EPI_ISL_1916414, EPI_ISL_1916421 | CSIR-National Environmental Engineering Research Institute | CSIR-Centre for Cellular and Molecular Biology - INSACOG | Onkar Kulkarni, Lamuk Zaveri, Ara Sreenivas, Sofia Banu, Shreekant Verma, Amareshwar Vodapalli , Viswagithe S L, B Himasri, Sharath Chandra Thota, Karthik Bharadwaj Tallapaka, Krishna Khairnar, Rakesh K Mishra,Divya Tej Sowpati |
| EPI_ISL_1916433, EPI_ISL_1916434, EPI_ISL_1916446, EPI_ISL_1916456, EPI_ISL_1916457, EPI_ISL_1916477, EPI_ISL_1916479, EPI_ISL_1916486, EPI_ISL_1916488, EPI_ISL_1916489 | CSIR-National Environmental Engineering Research Institute | CSIR-Centre for Cellular and Molecular Biology - INSACOG | Ara Sreenivas, Onkar Kulkarni, Lamuk Zaveri, Sofia Banu, Shreekant Verma, Amareshwar Vodapalli , Viswagithe S L, B Himasri, Sharath Chandra Thota, Karthik Bharadwaj Tallapaka, Krishna Khairnar, Rakesh K Mishra,Divya Tej Sowpati |
| EPI_ISL_1919606, EPI_ISL_1919611 | Laboratorio Analisi Osp. Città di Castello - Azienda USL Umbria1 | Istituto Zooprofilattico Sperimentale dell'Abruzzo e Molise "G. Caporale" | Malagigi V, Tacconi P,Lorusso A, Marcacci M, Di Domenico M, Ancora M, Curini V, Di Lollo Valeria, Mangone I, Rinaldi A, Delli Compagni E, Scialabba S, Caporale M, Di Pasquale A, Cammà C, Puglia I, Calistri P, Savini G |
| EPI_ISL_1919612 | Università degli Studi di Perugia | Istituto Zooprofilattico Sperimentale dell'Abruzzo e Molise "G. Caporale" | Mencacci A, Camilloni B, Lorusso A, Marcacci M, Di Domenico M, Ancora M, Curini V, Mangone I, Rinaldi A, Delli Compagni E, Scialabba S, Di Pasquale A, Cammà C, Puglia I, Calistri P, Savini G |
| EPI_ISL_1919613, EPI_ISL_1919614, EPI_ISL_1919615, EPI_ISL_1919616, EPI_ISL_1919617, EPI_ISL_1919618, EPI_ISL_1919619, EPI_ISL_1919620, EPI_ISL_1919621, EPI_ISL_1919622 | Azienda Ospedaliera Terni | Istituto Zooprofilattico Sperimentale dell'Abruzzo e Molise "G. Caporale" | Palumbo M, Scaccetti A, Lorusso A, Marcacci M, Di Domenico M, Ancora M, Curini V, Mangone I, Rinaldi A, Delli Compagni E, Scialabba S, Di Pasquale A, Cammà C, Puglia I, Calistri P, Savini G |
| EPI_ISL_1919623 | Università degli Studi di Perugia | Istituto Zooprofilattico Sperimentale dell'Abruzzo e Molise "G. Caporale" | Mencacci A, Camilloni B, Lorusso A, Marcacci M, Di Domenico M, Ancora M, Curini V, Mangone I, Rinaldi A, Delli Compagni E, Scialabba S, Di Pasquale A, Cammà C, Puglia I, Calistri P, Savini G |
| EPI_ISL_1919624, EPI_ISL_1919625, EPI_ISL_1919626, EPI_ISL_1919627, EPI_ISL_1919628, EPI_ISL_1919629, EPI_ISL_1919630, EPI_ISL_1919632 | Azienda Ospedaliera Terni | Istituto Zooprofilattico Sperimentale dell'Abruzzo e Molise "G. Caporale" | Palumbo M, Scaccetti A, Lorusso A, Marcacci M, Di Domenico M, Ancora M, Curini V, Mangone I, Rinaldi A, Delli Compagni E, Scialabba S, Di Pasquale A, Cammà C, Puglia I, Calistri P, Savini G |
| EPI_ISL_1919639, EPI_ISL_1919640, EPI_ISL_1919643, EPI_ISL_1919644 | Università degli Studi di Perugia | Istituto Zooprofilattico Sperimentale dell'Abruzzo e Molise "G. Caporale" | Mencacci A, Camilloni B, Lorusso A, Marcacci M, Di Domenico M, Ancora M, Curini V, Mangone I, Rinaldi A, Delli Compagni E, Scialabba S, Di Pasquale A, Cammà C, Puglia I, Calistri P, Savini G |
| EPI_ISL_1919651, EPI_ISL_1919652, EPI_ISL_1919653 | Institute for Medical Research, Infectious Disease Research Centre, National Institutes of Health, Ministry of Health Malaysia | Institute for Medical Research, Infectious Disease Research Centre, National Institutes of Health, Ministry of Health Malaysia | Suppiah J, Kamel K, Mohd Zawawi Z, Ramly N, Robert F, Thayan R |
| EPI_ISL_1919745, EPI_ISL_1919746, EPI_ISL_1919747, EPI_ISL_1919748, EPI_ISL_1919749, EPI_ISL_1919751, EPI_ISL_1919752 | Presidio di Brindisi Di Summa - Perrino | Istituto Zooprofilattico Sperimentale della Puglia e della Basilicata | Parisi A., Bianco A., Capozzi L., Del Sambro L., Simone D., Difato L., Santoro A., Ridolfi D., Giannico A. |
| EPI_ISL_1921892, EPI_ISL_1921893 | OLVZ Aalst | OLVZ Aalst | Anne Vankeerberghen |
| EPI_ISL_1936194, EPI_ISL_1936195, EPI_ISL_1936196, EPI_ISL_1936197, EPI_ISL_1936198, EPI_ISL_1936199, EPI_ISL_1936200, EPI_ISL_1936201, EPI_ISL_1936202, EPI_ISL_1936203, EPI_ISL_1936204, EPI_ISL_1936205, EPI_ISL_1936206, EPI_ISL_1936207, EPI_ISL_1936208, EPI_ISL_1936209, EPI_ISL_1936210, EPI_ISL_1936211, EPI_ISL_1936212, EPI_ISL_1936213, EPI_ISL_1936214, EPI_ISL_1936215, EPI_ISL_1936216, EPI_ISL_1936217, EPI_ISL_1936218, EPI_ISL_1936219, EPI_ISL_1936220, EPI_ISL_1936221, EPI_ISL_1936222, EPI_ISL_1936223, EPI_ISL_1936224, EPI_ISL_1936225, EPI_ISL_1936226, EPI_ISL_1936227, EPI_ISL_1936228, EPI_ISL_1936229, EPI_ISL_1936230, EPI_ISL_1936231, EPI_ISL_1936232, EPI_ISL_1936233, EPI_ISL_1936234, EPI_ISL_1936235, EPI_ISL_1936236, EPI_ISL_1936237 | | | |
| see above | Main Chemical Laboratories Egypt Army | Main Chemical Laboratories Egypt Army | Mohamed Seadawy, AbedElrahman Zekri, Mohamed Shamel, Ahmed Gad, Abdullah Salama, Mostfa Elhoseiny |
| EPI_ISL_1936273, EPI_ISL_1936274, EPI_ISL_1936275, EPI_ISL_1936276, EPI_ISL_1936277, EPI_ISL_1936278, EPI_ISL_1936279, EPI_ISL_1936280, EPI_ISL_1936281, EPI_ISL_1936282, EPI_ISL_1936283, EPI_ISL_1936284, EPI_ISL_1936285, EPI_ISL_1936286, EPI_ISL_1936287, EPI_ISL_1936288, EPI_ISL_1936289, EPI_ISL_1936290, EPI_ISL_1936291, EPI_ISL_1936292, EPI_ISL_1936293, EPI_ISL_1936294 | | | |
| see above | Main Chemical Laboratories Egypt Army | Main Chemical Laboratories Egypt Army | Mohamed Seadawy, AbedElrahman Zekri, Mohamed Shamel, Ahmed Gad, Abdullah Salama, Mohamed El-Esawi, Mohamed Abdel-Monem |
| EPI_ISL_1941262 | Basurto University Hospital: Clinical Microbiology Laboratory | Biocruces Bizkaia | Mikel Urrutikoetxea-Gutiérrez, Mª Carmen Nieto Toboso, Estibaliz Ugalde Zarraga, Mikel Gallego Rodrigo, Ana de la Hoz, José Luis Díaz de Tuesta del Arco |
| EPI_ISL_1960074 | ICMT-APARTADO | Universidad Nacional de Colombia - Laboratorio Genómico One Health | Andres F. Cardona-Rios, Maria Stella López, Daniel O. Maldonado-Perez, Simón Villegas Velásquez, Lina M. Hurtado, Laura Silvana Perez, Karl A Ciuoderis, Maria Angélica Maya, Carolina Muñoz-Arango, Celeny Ortiz, Rita Almanza Payares, Idabely Betancur Ortiz, Sandra Ines Cano, Diego A. Álvarez-Díaz, Carlos Franco-Muñoz, Katherine Laiton-Donato, Hector Alejandro Ruiz-Moreno, Marcela Mercado-Reyes, Jorge E. Osorio, Juan P. Hernandez-Ortiz |
| EPI_ISL_1960075 | LDSP | Universidad Nacional de Colombia - Laboratorio Genómico One Health | Simón Villegas Velásquez, Andres F. Cardona-Rios, Maria Stella López, Daniel O. Maldonado-Perez, Lina M. Hurtado, Laura Silvana Perez, Karl A Ciuoderis, Maria Angélica Maya, Carolina Muñoz-Arango, Celeny Ortiz, Rita Almanza Payares, Idabely Betancur Ortiz, Sandra Ines Cano, Diego A. Álvarez-Díaz, Carlos Franco-Muñoz, Katherine Laiton-Donato, Hector Alejandro Ruiz-Moreno, Marcela Mercado-Reyes, Jorge E. Osorio, Juan P. Hernandez-Ortiz |

| | | | |
|---|---|---|---|
| EPI_ISL_1960076 | PECET | Universidad Nacional de Colombia - Laboratorio Genómico One Health | Andres F. Cardona-Rios, Maria Stella López, Daniel O. Maldonado-Perez, Simón Villegas Velásquez, Lina M. Hurtado, Laura Silvana Perez, Karl A Ciuoderis, Maria Angélica Maya, Carolina Muñoz-Arango, Celeny Ortiz, Rita Almanza Payares, Idabely Betancur Ortiz, Sandra Ines Cano, Diego A. Álvarez-Díaz, Carlos Franco-Muñoz, Katherine Laiton-Donato, Hector Alejandro Ruiz-Moreno, Marcela Mercado-Reyes, Jorge E. Osorio, Juan P. Hernandez-Ortiz |
| EPI_ISL_1960078 | LDSP | Universidad Nacional de Colombia - Laboratorio Genómico One Health | Andres F. Cardona-Rios, Maria Stella López, Daniel O. Maldonado-Perez, Simón Villegas Velásquez, Lina M. Hurtado, Laura Silvana Perez, Karl A Ciuoderis, Maria Angélica Maya, Carolina Muñoz-Arango, Celeny Ortiz, Rita Almanza Payares, Idabely Betancur Ortiz, Sandra Ines Cano, Diego A. Álvarez-Díaz, Carlos Franco-Muñoz, Katherine Laiton-Donato, Hector Alejandro Ruiz-Moreno, Marcela Mercado-Reyes, Jorge E. Osorio, Juan P. Hernandez-Ortiz |
| EPI_ISL_1960080 | SYNLAB | Universidad Nacional de Colombia - Laboratorio Genómico One Health | Maria Stella López, Andres F. Cardona-Rios, Daniel O. Maldonado-Perez, Simón Villegas Velásquez, Lina M. Hurtado, Laura Silvana Perez, Karl A Ciuoderis, Maria Angélica Maya, Carolina Muñoz-Arango, Celeny Ortiz, Rita Almanza Payares, Idabely Betancur Ortiz, Sandra Ines Cano, Diego A. Álvarez-Díaz, Carlos Franco-Muñoz, Katherine Laiton-Donato, Hector Alejandro Ruiz-Moreno, Marcela Mercado-Reyes, Jorge E. Osorio, Juan P. Hernandez-Ortiz |
| EPI_ISL_1965217, EPI_ISL_1965218, EPI_ISL_1965219, EPI_ISL_1965220, EPI_ISL_1965221, EPI_ISL_1965222, EPI_ISL_1965223, EPI_ISL_1965224, EPI_ISL_1965225, EPI_ISL_1965226, EPI_ISL_1965227, EPI_ISL_1965228, EPI_ISL_1965229, EPI_ISL_1965230, EPI_ISL_1965231, EPI_ISL_1965232, EPI_ISL_1965233, EPI_ISL_1965234, EPI_ISL_1965235, EPI_ISL_1965237, EPI_ISL_1965238, EPI_ISL_1965239, EPI_ISL_1965240, EPI_ISL_1965241, EPI_ISL_1965242, EPI_ISL_1965243, EPI_ISL_1965244, EPI_ISL_1965245, EPI_ISL_1965246, EPI_ISL_1965247, EPI_ISL_1965248, EPI_ISL_1965249, EPI_ISL_1965250, EPI_ISL_1965252, EPI_ISL_1965253, EPI_ISL_1965254, EPI_ISL_1965255, EPI_ISL_1965256, EPI_ISL_1965257, EPI_ISL_1965258, EPI_ISL_1965259, EPI_ISL_1965260, EPI_ISL_1965261, EPI_ISL_1965263, EPI_ISL_1965264, EPI_ISL_1965265, EPI_ISL_1965266, EPI_ISL_1965267, EPI_ISL_1965268, EPI_ISL_1965269, EPI_ISL_1965270, EPI_ISL_1965271, EPI_ISL_1965272, EPI_ISL_1965273, EPI_ISL_1965274, EPI_ISL_1965275, EPI_ISL_1965276, EPI_ISL_1965277 | | | |
| see above | IPK | Respiratory Infections Laboratory | Lissette Perez , Yahisel Tejero , Yanet Pintos, Dailyn Medero,Odalys Valdes , Mirtha Elisa Aguado , Liena de Regla Ponce |
| EPI_ISL_1965282, EPI_ISL_1965283, EPI_ISL_1965284, EPI_ISL_1965285, EPI_ISL_1965286, EPI_ISL_1965287, EPI_ISL_1965288, EPI_ISL_1965289, EPI_ISL_1965290, EPI_ISL_1965291, EPI_ISL_1965292, EPI_ISL_1965293, EPI_ISL_1965294, EPI_ISL_1965295, EPI_ISL_1965296, EPI_ISL_1965297, EPI_ISL_1965298, EPI_ISL_1965299, EPI_ISL_1965300, EPI_ISL_1965301, EPI_ISL_1965302, EPI_ISL_1965303, EPI_ISL_1965304, EPI_ISL_1965305, EPI_ISL_1965306, EPI_ISL_1965307, EPI_ISL_1965308, EPI_ISL_1965309, EPI_ISL_1965310, EPI_ISL_1965311, EPI_ISL_1965312, EPI_ISL_1965313, EPI_ISL_1965315, EPI_ISL_1965316, EPI_ISL_1965317, EPI_ISL_1965318, EPI_ISL_1965319, EPI_ISL_1965320, EPI_ISL_1965321, EPI_ISL_1965322, EPI_ISL_1965323, EPI_ISL_1965324, EPI_ISL_1965325, EPI_ISL_1965326, EPI_ISL_1965327, EPI_ISL_1965330, EPI_ISL_1965331, EPI_ISL_1965332, EPI_ISL_1965333, EPI_ISL_1965334, EPI_ISL_1965335, EPI_ISL_1965336, EPI_ISL_1965337, EPI_ISL_1965338 | | | |
| see above | IPK | Respiratory Infections Laboratory | Lissette Perez, Yahisel Tejero, Yanet Pintos, Daylin Mederos, Odalys Valdes, Mirtha Elisa Aguado, Liena Ponce |
| EPI_ISL_1965496, EPI_ISL_1965497, EPI_ISL_1965499, EPI_ISL_1965501, EPI_ISL_1965502, EPI_ISL_1965503, EPI_ISL_1965507, EPI_ISL_1965508, EPI_ISL_1965511, EPI_ISL_1965512, EPI_ISL_1965513, EPI_ISL_1965515, EPI_ISL_1965517, EPI_ISL_1965519, EPI_ISL_1965520, EPI_ISL_1965523, EPI_ISL_1965524, EPI_ISL_1965525, EPI_ISL_1965527, EPI_ISL_1965528, EPI_ISL_1965529, EPI_ISL_1965530, EPI_ISL_1965531, EPI_ISL_1965533, EPI_ISL_1965534, EPI_ISL_1965551, EPI_ISL_1965552, EPI_ISL_1965553, EPI_ISL_1965554, EPI_ISL_1965555, EPI_ISL_1965556, EPI_ISL_1965560, EPI_ISL_1965561, EPI_ISL_1965564, EPI_ISL_1965569, EPI_ISL_1965572, EPI_ISL_1965576, EPI_ISL_1965581, EPI_ISL_1965582, EPI_ISL_1965583, EPI_ISL_1965584, EPI_ISL_1965585, EPI_ISL_1965586, EPI_ISL_1965589, EPI_ISL_1965590, EPI_ISL_1965591, EPI_ISL_1965594, EPI_ISL_1965595, EPI_ISL_1965607, EPI_ISL_1965611, EPI_ISL_1965612, EPI_ISL_1965614, EPI_ISL_1965621, EPI_ISL_1965628, EPI_ISL_1965630, EPI_ISL_1965633, EPI_ISL_1965634, EPI_ISL_1965636, EPI_ISL_1965637, EPI_ISL_1965641 | | | |
| see above | HG Pharma GmbH | Bergthaler laboratory, CeMM Research Center for Molecular Medicine of the Austrian Academy of Sciences | Lukas Endler, Anna Schedl, Fabian Amman, Petr Triska, Thomas Penz, Benedikt Agerer, Maelle Le Moing, Michael Schuster, Bekir Erguner, Jan Laine, Martin Senekowitsch, Christoph Bock, Andreas Bergthaler |
| EPI_ISL_1965643, EPI_ISL_1965645, EPI_ISL_1965647, EPI_ISL_1965648, EPI_ISL_1965649, EPI_ISL_1965650, EPI_ISL_1965651, EPI_ISL_1965652, EPI_ISL_1965660, EPI_ISL_1965661, EPI_ISL_1965662, EPI_ISL_1965663, EPI_ISL_1965664 | | | |
| see above | Elling group, Institute of Molecular Biotechnology (IMBA) | Bergthaler laboratory, CeMM Research Center for Molecular Medicine of the Austrian Academy of Sciences | Lukas Endler, Anna Schedl, Fabian Amman, Petr Triska, Thomas Penz, Benedikt Agerer, Maelle Le Moing, Michael Schuster, Bekir Erguner, Jan Laine, Martin Senekowitsch, Christoph Bock, Andreas Bergthaler |
| EPI_ISL_1965667 | Institute for Laboratory Diagnostics and Microbiology, Klinikum Klagenfurt am Wörthersee | Bergthaler laboratory, CeMM Research Center for Molecular Medicine of the Austrian Academy of Sciences | Lukas Endler, Anna Schedl, Fabian Amman, Petr Triska, Thomas Penz, Benedikt Agerer, Maelle Le Moing, Michael Schuster, Bekir Erguner, Jan Laine, Martin Senekowitsch, Christoph Bock, Andreas Bergthaler |
| EPI_ISL_1968629, EPI_ISL_1968632 | Provincial Public Health Reference Laboratory | Provincial Public Health Reference Laboratory | ANDLEEB HANIF |
| EPI_ISL_1969078, EPI_ISL_1969079 | Main Chemical Laboratories Egypt Army | Main Chemical Laboratories Egypt Army | Mohamed Seadawy, AbedElrahman Zekri, Mohamed Shamel, Ahmed Gad, Abdullah Salama, Mostfa Elhoseiny |
| EPI_ISL_1972107, EPI_ISL_1972109, EPI_ISL_1972111, EPI_ISL_1972113, EPI_ISL_1972114, EPI_ISL_1972115, EPI_ISL_1972116 | AZ St.Jan Brugge-Oostende | AZ SINT-JAN BRUGGE | Marijke Reynders, Patrick Descheemaeker, Laurien Hoornaert, Thomas Van Landschoot, Jorn Hellemans |
| EPI_ISL_1972899, EPI_ISL_1972943, EPI_ISL_1972947 | Usansolo-Galdakao University Hospital | Cruces University Hospital | Ana Gual-de-Torrella, Izaskun Alejo-Cancho, Mikel Gallego, Ana Belén de la Hoz |
| EPI_ISL_1973326, EPI_ISL_1973327, EPI_ISL_1973328 | Bintulu Hospital | Institute for Medical Research, Infectious Disease Research Centre, National Institutes of Health, Ministry of Health Malaysia | Suppiah J, Kamel K, Mohd Zawawi Z, Azizan MA, Ramly N, Robert F, Thayan R |
| EPI_ISL_1993548 | Molecular Diagnostic Laboratory (Hormozghan University of Medical Sciences) | National Influenza Center | NZ Shafiei Jandaghi, V Salimi, A Nejati, K Sadeghi, J Yavarian, N Ghavvami,F Ajaminejad and T Mokhtari Azad |
| EPI_ISL_1993551, EPI_ISL_1993556 | National Influenza Center, Virology Department | National Influenza Center | K Sadeghi, A Nejati, J Yavarian, NZ Shafiei Jandaghi, V Salimi, F Ajaminejad,N Ghavvami and T Mokhtari Azad |
| EPI_ISL_1993557 | National Influenza Center, Virology Department | National Influenza Center | V Salimi,NZ Shafiei Jandaghi, J Yavarian, A Nejati, K Sadeghi, N Ghavvami,F Ajaminejad and T Mokhtari Azad |
| EPI_ISL_2001127 | GMERS, Government Medical College, Gandhinagar | Gujarat Biotechnology Research Centre | Zuber Saiyed,Ramesh Pandit,Janvi Raval,Zarna Patel,Nitin Savaliya,Dinesh Kumar,Twinkle Soni,Sonal Sharma ,Gaurishankar Shrimali,Umang Mishra,Nitesh Shah,Chaitanya Joshi,Madhvi Joshi |
| EPI_ISL_2001128 | GMERS, Government Medical College, Gandhinagar | Gujarat Biotechnology Research Centre | Ramesh Pandit,Janvi Raval,Zarna Patel,Nitin Savaliya,Dinesh Kumar,Twinkle Soni,Sonal Sharma ,Zuber Saiyed,Gaurishankar Shrimali,Umang Mishra,Nitesh Shah,Chaitanya Joshi,Madhvi Joshi |
| EPI_ISL_2001129 | GMERS, Government Medical College, Gandhinagar | Gujarat Biotechnology Research Centre | Janvi Raval,Zarna Patel,Nitin Savaliya,Dinesh Kumar,Twinkle Soni,Sonal Sharma ,Zuber Saiyed,Ramesh Pandit,Gaurishankar Shrimali,Umang Mishra,Nitesh Shah,Chaitanya Joshi,Madhvi Joshi |
| EPI_ISL_2001130 | GMERS, Government Medical College, Gandhinagar | Gujarat Biotechnology Research Centre | Zarna Patel,Nitin Savaliya,Dinesh Kumar,Twinkle Soni,Sonal Sharma ,Zuber Saiyed,Ramesh Pandit,Janvi Raval,Gaurishankar Shrimali,Umang Mishra,Nitesh Shah,Chaitanya Joshi,Madhvi Joshi |
| EPI_ISL_2001131 | GMERS, Government Medical College, Gandhinagar | Gujarat Biotechnology Research Centre | Nitin Savaliya,Dinesh Kumar,Twinkle Soni,Sonal Sharma ,Zuber Saiyed,Ramesh Pandit,Janvi Raval,Zarna Patel,Gaurishankar Shrimali,Umang Mishra,Nitesh Shah,Chaitanya Joshi,Madhvi Joshi |
| EPI_ISL_2001132 | GMERS, Government Medical College, Gandhinagar | Gujarat Biotechnology Research Centre | Dinesh Kumar,Twinkle Soni,Sonal Sharma ,Zuber Saiyed,Ramesh Pandit,Janvi Raval,Zarna Patel,Nitin Savaliya,Gaurishankar Shrimali,Umang Mishra,Nitesh Shah,Chaitanya Joshi,Madhvi Joshi |
| EPI_ISL_2001133 | GMERS, Government Medical College, Gandhinagar | Gujarat Biotechnology Research Centre | Twinkle Soni,Sonal Sharma ,Zuber Saiyed,Ramesh Pandit,Janvi Raval,Zarna Patel,Nitin Savaliya,Dinesh Kumar,Gaurishankar Shrimali,Umang Mishra,Nitesh Shah,Chaitanya Joshi,Madhvi Joshi |
| EPI_ISL_2001134 | GMERS, Government Medical College, Gandhinagar | Gujarat Biotechnology Research Centre | Sonal Sharma ,Zuber Saiyed,Ramesh Pandit,Janvi Raval,Zarna Patel,Nitin Savaliya,Dinesh Kumar,Twinkle Soni,Gaurishankar Shrimali,Umang Mishra,Nitesh Shah,Chaitanya Joshi,Madhvi Joshi |
| EPI_ISL_2001135 | GMERS, Government Medical College, Gandhinagar | Gujarat Biotechnology Research Centre | Zuber Saiyed,Ramesh Pandit,Janvi Raval,Zarna Patel,Nitin Savaliya,Dinesh Kumar,Twinkle Soni,Sonal Sharma ,Gaurishankar Shrimali,Umang Mishra,Nitesh Shah,Chaitanya Joshi,Madhvi Joshi |
| EPI_ISL_2001136 | GMERS, Government Medical College, Gandhinagar | Gujarat Biotechnology Research Centre | Ramesh Pandit,Janvi Raval,Zarna Patel,Nitin Savaliya,Dinesh Kumar,Twinkle Soni,Sonal Sharma ,Zuber Saiyed,Gaurishankar Shrimali,Umang Mishra,Nitesh Shah,Chaitanya Joshi,Madhvi Joshi |
| EPI_ISL_2001137 | GMERS, Government Medical College, Gandhinagar | Gujarat Biotechnology Research Centre | Janvi Raval,Zarna Patel,Nitin Savaliya,Dinesh Kumar,Twinkle Soni,Sonal Sharma ,Zuber Saiyed,Ramesh Pandit,Gaurishankar Shrimali,Umang Mishra,Nitesh Shah,Chaitanya Joshi,Madhvi Joshi |
| EPI_ISL_2001138 | GMERS, Government Medical College, Gandhinagar | Gujarat Biotechnology Research Centre | Zarna Patel,Nitin Savaliya,Dinesh Kumar,Twinkle Soni,Sonal Sharma ,Zuber Saiyed,Ramesh Pandit,Janvi Raval,Gaurishankar Shrimali,Umang Mishra,Nitesh Shah,Chaitanya Joshi,Madhvi Joshi |

| | | | |
|---|---|---|---|
| EPI_ISL_2001139 | GMERS, Government Medical College, Gandhinagar | Gujarat Biotechnology Research Centre | Nitin Savaliya,Dinesh Kumar,Twinkle Soni,Sonal Sharma ,Zuber Saiyed,Ramesh Pandit,Janvi Raval,Zarna Patel,Gaurishankar Shrimali,Umang Mishra,Nitesh Shah,Chaitanya Joshi,Madhvi Joshi |
| EPI_ISL_2001140 | GMERS, Government Medical College, Gandhinagar | Gujarat Biotechnology Research Centre | Dinesh Kumar,Twinkle Soni,Sonal Sharma ,Zuber Saiyed,Ramesh Pandit,Janvi Raval,Zarna Patel,Nitin Savaliya,Gaurishankar Shrimali,Umang Mishra,Nitesh Shah,Chaitanya Joshi,Madhvi Joshi |
| EPI_ISL_2001141 | GMERS, Government Medical College, Gandhinagar | Gujarat Biotechnology Research Centre | Twinkle Soni,Sonal Sharma ,Zuber Saiyed,Ramesh Pandit,Janvi Raval,Zarna Patel,Nitin Savaliya,Dinesh Kumar,Gaurishankar Shrimali,Umang Mishra,Nitesh Shah,Chaitanya Joshi,Madhvi Joshi |
| EPI_ISL_2001142 | GMERS, Government Medical College, Gandhinagar | Gujarat Biotechnology Research Centre | Sonal Sharma ,Zuber Saiyed,Ramesh Pandit,Janvi Raval,Zarna Patel,Nitin Savaliya,Dinesh Kumar,Twinkle Soni,Gaurishankar Shrimali,Umang Mishra,Nitesh Shah,Chaitanya Joshi,Madhvi Joshi |
| EPI_ISL_2001143 | GMERS, Government Medical College, Gandhinagar | Gujarat Biotechnology Research Centre | Zuber Saiyed,Ramesh Pandit,Janvi Raval,Zarna Patel,Nitin Savaliya,Dinesh Kumar,Twinkle Soni,Sonal Sharma ,Gaurishankar Shrimali,Umang Mishra,Nitesh Shah,Chaitanya Joshi,Madhvi Joshi |
| EPI_ISL_2001144 | GMERS, Government Medical College, Gandhinagar | Gujarat Biotechnology Research Centre | Ramesh Pandit,Janvi Raval,Zarna Patel,Nitin Savaliya,Dinesh Kumar,Twinkle Soni,Sonal Sharma ,Zuber Saiyed,Gaurishankar Shrimali,Umang Mishra,Nitesh Shah,Chaitanya Joshi,Madhvi Joshi |
| EPI_ISL_2001145 | GMERS, Government Medical College, Gandhinagar | Gujarat Biotechnology Research Centre | Janvi Raval,Zarna Patel,Nitin Savaliya,Dinesh Kumar,Twinkle Soni,Sonal Sharma ,Zuber Saiyed,Ramesh Pandit,Gaurishankar Shrimali,Umang Mishra,Nitesh Shah,Chaitanya Joshi,Madhvi Joshi |
| EPI_ISL_2001146 | GMERS, Government Medical College, Gandhinagar | Gujarat Biotechnology Research Centre | Zarna Patel,Nitin Savaliya,Dinesh Kumar,Twinkle Soni,Sonal Sharma ,Zuber Saiyed,Ramesh Pandit,Janvi Raval,Gaurishankar Shrimali,Umang Mishra,Nitesh Shah,Chaitanya Joshi,Madhvi Joshi |
| EPI_ISL_2001147 | GMERS, Government Medical College, Gandhinagar | Gujarat Biotechnology Research Centre | Nitin Savaliya,Dinesh Kumar,Twinkle Soni,Sonal Sharma ,Zuber Saiyed,Ramesh Pandit,Janvi Raval,Zarna Patel,Gaurishankar Shrimali,Umang Mishra,Nitesh Shah,Chaitanya Joshi,Madhvi Joshi |
| EPI_ISL_2001148 | GMERS, Government Medical College, Gandhinagar | Gujarat Biotechnology Research Centre | Dinesh Kumar,Twinkle Soni,Sonal Sharma ,Zuber Saiyed,Ramesh Pandit,Janvi Raval,Zarna Patel,Nitin Savaliya,Gaurishankar Shrimali,Umang Mishra,Nitesh Shah,Chaitanya Joshi,Madhvi Joshi |
| EPI_ISL_2001178 | GMERS, Government Medical College (GMC), Gotri | Gujarat Biotechnology Research Centre | Ramesh Pandit,Janvi Raval,Zarna Patel,Nitin Savaliya,Dinesh Kumar,Twinkle Soni,Sonal Sharma ,Zuber Saiyed,Bithika Duttaroy,Umang Mishra,Nitesh Shah,Chaitanya Joshi,Madhvi Joshi |
| EPI_ISL_2001179 | GMERS, Government Medical College (GMC), Gotri | Gujarat Biotechnology Research Centre | Janvi Raval,Zarna Patel,Nitin Savaliya,Dinesh Kumar,Twinkle Soni,Sonal Sharma ,Zuber Saiyed,Ramesh Pandit,Bithika Duttaroy,Umang Mishra,Nitesh Shah,Chaitanya Joshi,Madhvi Joshi |
| EPI_ISL_2001180 | GMERS, Government Medical College (GMC), Gotri | Gujarat Biotechnology Research Centre | Zarna Patel,Nitin Savaliya,Dinesh Kumar,Twinkle Soni,Sonal Sharma ,Zuber Saiyed,Ramesh Pandit,Janvi Raval,Bithika Duttaroy,Umang Mishra,Nitesh Shah,Chaitanya Joshi,Madhvi Joshi |
| EPI_ISL_2001183 | GMERS, Government Medical College (GMC), Gotri | Gujarat Biotechnology Research Centre | Twinkle Soni,Sonal Sharma ,Zuber Saiyed,Ramesh Pandit,Janvi Raval,Zarna Patel,Nitin Savaliya,Dinesh Kumar,Bithika Duttaroy,Umang Mishra,Nitesh Shah,Chaitanya Joshi,Madhvi Joshi |
| EPI_ISL_2001184 | GMERS, Government Medical College (GMC), Gotri | Gujarat Biotechnology Research Centre | Sonal Sharma ,Zuber Saiyed,Ramesh Pandit,Janvi Raval,Zarna Patel,Nitin Savaliya,Dinesh Kumar,Twinkle Soni,Bithika Duttaroy,Umang Mishra,Nitesh Shah,Chaitanya Joshi,Madhvi Joshi |
| EPI_ISL_2001185 | GMERS, Government Medical College (GMC), Gotri | Gujarat Biotechnology Research Centre | Zuber Saiyed,Ramesh Pandit,Janvi Raval,Zarna Patel,Nitin Savaliya,Dinesh Kumar,Twinkle Soni,Sonal Sharma ,Bithika Duttaroy,Umang Mishra,Nitesh Shah,Chaitanya Joshi,Madhvi Joshi |
| EPI_ISL_2001186 | GMERS, Government Medical College (GMC), Gotri | Gujarat Biotechnology Research Centre | Ramesh Pandit,Janvi Raval,Zarna Patel,Nitin Savaliya,Dinesh Kumar,Twinkle Soni,Sonal Sharma ,Zuber Saiyed,Bithika Duttaroy,Umang Mishra,Nitesh Shah,Chaitanya Joshi,Madhvi Joshi |
| EPI_ISL_2001187 | VRDL, Goverment Medical College (GMC), Surat | Gujarat Biotechnology Research Centre | Janvi Raval,Zarna Patel,Nitin Savaliya,Dinesh Kumar,Twinkle Soni,Sonal Sharma ,Zuber Saiyed,Ramesh Pandit,Neeta Khandelwal,Umang Mishra,Nitesh Shah,Chaitanya Joshi,Madhvi Joshi |
| EPI_ISL_2001188 | VRDL, Goverment Medical College (GMC), Surat | Gujarat Biotechnology Research Centre | Zarna Patel,Nitin Savaliya,Dinesh Kumar,Twinkle Soni,Sonal Sharma ,Zuber Saiyed,Ramesh Pandit,Janvi Raval,Neeta Khandelwal,Umang Mishra,Nitesh Shah,Chaitanya Joshi,Madhvi Joshi |
| EPI_ISL_2001199 | VRDL, Goverment Medical College (GMC), Surat | Gujarat Biotechnology Research Centre | Twinkle Soni,Sonal Sharma ,Zuber Saiyed,Ramesh Pandit,Janvi Raval,Zarna Patel,Nitin Savaliya,Dinesh Kumar,Neeta Khandelwal,Umang Mishra,Nitesh Shah,Chaitanya Joshi,Madhvi Joshi |
| EPI_ISL_2001200 | VRDL, Goverment Medical College (GMC), Surat | Gujarat Biotechnology Research Centre | Sonal Sharma ,Zuber Saiyed,Ramesh Pandit,Janvi Raval,Zarna Patel,Nitin Savaliya,Dinesh Kumar,Twinkle Soni,Neeta Khandelwal,Umang Mishra,Nitesh Shah,Chaitanya Joshi,Madhvi Joshi |
| EPI_ISL_2001205 | VRDL, Goverment Medical College (GMC), Surat | Gujarat Biotechnology Research Centre | Nitin Savaliya,Dinesh Kumar,Twinkle Soni,Sonal Sharma ,Zuber Saiyed,Ramesh Pandit,Janvi Raval,Zarna Patel,Neeta Khandelwal,Umang Mishra,Nitesh Shah,Chaitanya Joshi,Madhvi Joshi |
| EPI_ISL_2001206 | VRDL, Goverment Medical College (GMC), Surat | Gujarat Biotechnology Research Centre | Dinesh Kumar,Twinkle Soni,Sonal Sharma ,Zuber Saiyed,Ramesh Pandit,Janvi Raval,Zarna Patel,Nitin Savaliya,Neeta Khandelwal,Umang Mishra,Nitesh Shah,Chaitanya Joshi,Madhvi Joshi |
| EPI_ISL_2001207 | VRDL, Goverment Medical College (GMC), Surat | Gujarat Biotechnology Research Centre | Twinkle Soni,Sonal Sharma ,Zuber Saiyed,Ramesh Pandit,Janvi Raval,Zarna Patel,Nitin Savaliya,Dinesh Kumar,Neeta Khandelwal,Umang Mishra,Nitesh Shah,Chaitanya Joshi,Madhvi Joshi |
| EPI_ISL_2001208 | VRDL, Goverment Medical College (GMC), Surat | Gujarat Biotechnology Research Centre | Sonal Sharma ,Zuber Saiyed,Ramesh Pandit,Janvi Raval,Zarna Patel,Nitin Savaliya,Dinesh Kumar,Twinkle Soni,Neeta Khandelwal,Umang Mishra,Nitesh Shah,Chaitanya Joshi,Madhvi Joshi |
| EPI_ISL_2001209 | VRDL, Goverment Medical College (GMC), Surat | Gujarat Biotechnology Research Centre | Zuber Saiyed,Ramesh Pandit,Janvi Raval,Zarna Patel,Nitin Savaliya,Dinesh Kumar,Twinkle Soni,Sonal Sharma ,Neeta Khandelwal,Umang Mishra,Nitesh Shah,Chaitanya Joshi,Madhvi Joshi |
| EPI_ISL_2001210 | VRDL, Goverment Medical College (GMC), Surat | Gujarat Biotechnology Research Centre | Ramesh Pandit,Janvi Raval,Zarna Patel,Nitin Savaliya,Dinesh Kumar,Twinkle Soni,Sonal Sharma ,Zuber Saiyed,Neeta Khandelwal,Umang Mishra,Nitesh Shah,Chaitanya Joshi,Madhvi Joshi |
| EPI_ISL_2001211 | VRDL, Goverment Medical College (GMC), Surat | Gujarat Biotechnology Research Centre | Janvi Raval,Zarna Patel,Nitin Savaliya,Dinesh Kumar,Twinkle Soni,Sonal Sharma ,Zuber Saiyed,Ramesh Pandit,Neeta Khandelwal,Umang Mishra,Nitesh Shah,Chaitanya Joshi,Madhvi Joshi |
| EPI_ISL_2001212 | VRDL, Goverment Medical College (GMC), Surat | Gujarat Biotechnology Research Centre | Zarna Patel,Nitin Savaliya,Dinesh Kumar,Twinkle Soni,Sonal Sharma ,Zuber Saiyed,Ramesh Pandit,Janvi Raval,Neeta Khandelwal,Umang Mishra,Nitesh Shah,Chaitanya Joshi,Madhvi Joshi |
| EPI_ISL_2001297 | USCA Pescina | Istituto Zooprofilattico Sperimentale dell'Abruzzo e Molise "G. Caporale" | Lorusso A, Marcacci M, Di Domenico M, Ancora M, Curini V, Di Lollo Valeria, Mangone I, Rinaldi A, Delli Compagni E, Scialabba S, Caporale M, Di Pasquale A, Cammà C, Puglia I, Calistri P, Savini G |
| EPI_ISL_2003114, EPI_ISL_2003115, EPI_ISL_2003116, EPI_ISL_2003117, EPI_ISL_2003118, EPI_ISL_2003119, EPI_ISL_2003120, EPI_ISL_2003121, EPI_ISL_2003122, EPI_ISL_2003123, EPI_ISL_2003124, EPI_ISL_2003125, EPI_ISL_2003126, EPI_ISL_2003127, EPI_ISL_2003128, EPI_ISL_2003129, EPI_ISL_2003130, EPI_ISL_2003131 | | | |
| see above | Instituto Adolfo Lutz Central | Instituto Adolfo Lutz, Interdiciplinary Procedures Center, Strategic Laboratory | Claudio Tavares Sacchi, Claudia Regina Gonçalves, Erica Valessa Ramos Gomes, Karoline Rodrigues Campos, Caio Vinicius Dias Lopes, Leonardo Jose Tadeu de Araujo |
| EPI_ISL_2003139, EPI_ISL_2003140, EPI_ISL_2003141, EPI_ISL_2003142, EPI_ISL_2003143, EPI_ISL_2003144, EPI_ISL_2003145, EPI_ISL_2003146, EPI_ISL_2003147, EPI_ISL_2003148, EPI_ISL_2003149 | | | |
| see above | Instituto Adolfo Lutz - Regional de Santos | Instituto Adolfo Lutz, Interdiciplinary Procedures Center, Strategic Laboratory | Claudio Tavares Sacchi, Claudia Regina Gonçalves, Erica Valessa Ramos Gomes, Karoline Rodrigues Campos, Caio Vinicius Dias Lopes, Leonardo Jose Tadeu de Araujo |
| EPI_ISL_2003151 | Instituto Adolfo Lutz - Regional de Bauru | Instituto Adolfo Lutz, Interdiciplinary Procedures Center, Strategic Laboratory | Claudio Tavares Sacchi, Claudia Regina Gonçalves, Erica Valessa Ramos Gomes, Karoline Rodrigues Campos, Caio Vinicius Dias Lopes, Leonardo Jose Tadeu de Araujo |
| EPI_ISL_2003152 | Instituto Adolfo Lutz - Regional de Marilia | Instituto Adolfo Lutz, Interdiciplinary Procedures Center, | Claudio Tavares Sacchi, Claudia Regina Gonçalves, Erica Valessa Ramos Gomes, Karoline Rodrigues Campos, Caio Vinicius Dias Lopes, Leonardo Jose |

| | | | |
|---|---|---|---|
| | | Strategic Laboratory | Tadeu de Araujo |
| EPI_ISL_2003162, EPI_ISL_2003163, EPI_ISL_2003164, EPI_ISL_2003165, EPI_ISL_2003166, EPI_ISL_2003167, EPI_ISL_2003168, EPI_ISL_2003169, EPI_ISL_2003170 | Instituto Adolfo Lutz - Regional de Campinas | Instituto Adolfo Lutz, Interdiciplinary Procedures Center, Strategic Laboratory | Claudio Tavares Sacchi, Claudia Regina Gonçalves, Erica Valessa Ramos Gomes, Karoline Rodrigues Campos, Caio Vinicius Dias Lopes, Leonardo Jose Tadeu de Araujo |
| EPI_ISL_2006966, EPI_ISL_2006968, EPI_ISL_2006969, EPI_ISL_2006970, EPI_ISL_2006973, EPI_ISL_2006975, EPI_ISL_2006976, EPI_ISL_2006984 | The Ohio State University Applied Microbiology Services Laboratory | The Ohio State University Applied Microbiology Services Laboratory | Seth A. Faith PhD |
| EPI_ISL_2009137, EPI_ISL_2009138, EPI_ISL_2009139, EPI_ISL_2009140, EPI_ISL_2009141, EPI_ISL_2009142, EPI_ISL_2009143, EPI_ISL_2009166, EPI_ISL_2009167, EPI_ISL_2009168, EPI_ISL_2009169, EPI_ISL_2009176, EPI_ISL_2009177 | | | |
| see above | MSK Microbiology Lab | MSK Microbiology Lab | Esther Babady, Tracy McMillen, Krupa Jani |
| EPI_ISL_2017728 | VRDL, Goverment Medical College (GMC), Surat | Gujarat Biotechnology Research Centre | Nitin Savaliya,Dinesh Kumar,Twinkle Soni,Sonal Sharma ,Zuber Saiyed,Ramesh Pandit,Janvi Raval,Zarna Patel,Neeta Khandelwal,Umang Mishra,Nitesh Shah,Chaitanya Joshi,Madhvi Joshi |
| EPI_ISL_2017729 | VRDL, Goverment Medical College (GMC), Surat | Gujarat Biotechnology Research Centre | Dinesh Kumar,Twinkle Soni,Sonal Sharma ,Zuber Saiyed,Ramesh Pandit,Janvi Raval,Zarna Patel,Nitin Savaliya,Neeta Khandelwal,Umang Mishra,Nitesh Shah,Chaitanya Joshi,Madhvi Joshi |
| EPI_ISL_2017730 | VRDL, Goverment Medical College (GMC), Surat | Gujarat Biotechnology Research Centre | Twinkle Soni,Sonal Sharma ,Zuber Saiyed,Ramesh Pandit,Janvi Raval,Zarna Patel,Nitin Savaliya,Dinesh Kumar,Neeta Khandelwal,Umang Mishra,Nitesh Shah,Chaitanya Joshi,Madhvi Joshi |
| EPI_ISL_2017731 | VRDL, Goverment Medical College (GMC), Surat | Gujarat Biotechnology Research Centre | Sonal Sharma ,Zuber Saiyed,Ramesh Pandit,Janvi Raval,Zarna Patel,Nitin Savaliya,Dinesh Kumar,Twinkle Soni,Neeta Khandelwal,Umang Mishra,Nitesh Shah,Chaitanya Joshi,Madhvi Joshi |
| EPI_ISL_2017732 | VRDL, Goverment Medical College (GMC), Surat | Gujarat Biotechnology Research Centre | Zuber Saiyed,Ramesh Pandit,Janvi Raval,Zarna Patel,Nitin Savaliya,Dinesh Kumar,Twinkle Soni,Sonal Sharma ,Neeta Khandelwal,Umang Mishra,Nitesh Shah,Chaitanya Joshi,Madhvi Joshi |
| EPI_ISL_2017733 | VRDL, Goverment Medical College (GMC), Surat | Gujarat Biotechnology Research Centre | Ramesh Pandit,Janvi Raval,Zarna Patel,Nitin Savaliya,Dinesh Kumar,Twinkle Soni,Sonal Sharma ,Zuber Saiyed,Neeta Khandelwal,Umang Mishra,Nitesh Shah,Chaitanya Joshi,Madhvi Joshi |
| EPI_ISL_2017734 | VRDL, Goverment Medical College (GMC), Surat | Gujarat Biotechnology Research Centre | Janvi Raval,Zarna Patel,Nitin Savaliya,Dinesh Kumar,Twinkle Soni,Sonal Sharma ,Zuber Saiyed,Ramesh Pandit,Neeta Khandelwal,Umang Mishra,Nitesh Shah,Chaitanya Joshi,Madhvi Joshi |
| EPI_ISL_2017735 | VRDL, Goverment Medical College (GMC), Surat | Gujarat Biotechnology Research Centre | Zarna Patel,Nitin Savaliya,Dinesh Kumar,Twinkle Soni,Sonal Sharma ,Zuber Saiyed,Ramesh Pandit,Janvi Raval,Neeta Khandelwal,Umang Mishra,Nitesh Shah,Chaitanya Joshi,Madhvi Joshi |
| EPI_ISL_2017736 | VRDL, Goverment Medical College (GMC), Surat | Gujarat Biotechnology Research Centre | Nitin Savaliya,Dinesh Kumar,Twinkle Soni,Sonal Sharma ,Zuber Saiyed,Ramesh Pandit,Janvi Raval,Zarna Patel,Neeta Khandelwal,Umang Mishra,Nitesh Shah,Chaitanya Joshi,Madhvi Joshi |
| EPI_ISL_2017737 | VRDL, Goverment Medical College (GMC), Surat | Gujarat Biotechnology Research Centre | Dinesh Kumar,Twinkle Soni,Sonal Sharma ,Zuber Saiyed,Ramesh Pandit,Janvi Raval,Zarna Patel,Nitin Savaliya,Neeta Khandelwal,Umang Mishra,Nitesh Shah,Chaitanya Joshi,Madhvi Joshi |
| EPI_ISL_2017738 | VRDL, Goverment Medical College (GMC), Surat | Gujarat Biotechnology Research Centre | Twinkle Soni,Sonal Sharma ,Zuber Saiyed,Ramesh Pandit,Janvi Raval,Zarna Patel,Nitin Savaliya,Dinesh Kumar,Neeta Khandelwal,Umang Mishra,Nitesh Shah,Chaitanya Joshi,Madhvi Joshi |
| EPI_ISL_2017739 | VRDL, Goverment Medical College (GMC), Surat | Gujarat Biotechnology Research Centre | Sonal Sharma ,Zuber Saiyed,Ramesh Pandit,Janvi Raval,Zarna Patel,Nitin Savaliya,Dinesh Kumar,Twinkle Soni,Neeta Khandelwal,Umang Mishra,Nitesh Shah,Chaitanya Joshi,Madhvi Joshi |
| EPI_ISL_2017742 | VRDL, Goverment Medical College (GMC), Surat | Gujarat Biotechnology Research Centre | Janvi Raval,Zarna Patel,Nitin Savaliya,Dinesh Kumar,Twinkle Soni,Sonal Sharma ,Zuber Saiyed,Ramesh Pandit,Neeta Khandelwal,Umang Mishra,Nitesh Shah,Chaitanya Joshi,Madhvi Joshi |
| EPI_ISL_2017743 | VRDL, Goverment Medical College (GMC), Surat | Gujarat Biotechnology Research Centre | Zarna Patel,Nitin Savaliya,Dinesh Kumar,Twinkle Soni,Sonal Sharma ,Zuber Saiyed,Ramesh Pandit,Janvi Raval,Neeta Khandelwal,Umang Mishra,Nitesh Shah,Chaitanya Joshi,Madhvi Joshi |
| EPI_ISL_2017744 | VRDL, Goverment Medical College (GMC), Surat | Gujarat Biotechnology Research Centre | Nitin Savaliya,Dinesh Kumar,Twinkle Soni,Sonal Sharma ,Zuber Saiyed,Ramesh Pandit,Janvi Raval,Zarna Patel,Neeta Khandelwal,Umang Mishra,Nitesh Shah,Chaitanya Joshi,Madhvi Joshi |
| EPI_ISL_2017745 | VRDL, Goverment Medical College (GMC), Surat | Gujarat Biotechnology Research Centre | Dinesh Kumar,Twinkle Soni,Sonal Sharma ,Zuber Saiyed,Ramesh Pandit,Janvi Raval,Zarna Patel,Nitin Savaliya,Neeta Khandelwal,Umang Mishra,Nitesh Shah,Chaitanya Joshi,Madhvi Joshi |
| EPI_ISL_2017746 | VRDL, Goverment Medical College (GMC) and Hospital, Valsad | Gujarat Biotechnology Research Centre | Twinkle Soni,Sonal Sharma ,Zuber Saiyed,Ramesh Pandit,Janvi Raval,Zarna Patel,Nitin Savaliya,Dinesh Kumar,Vicky Gandhi,Umang Mishra,Nitesh Shah,Chaitanya Joshi,Madhvi Joshi |
| EPI_ISL_2017747 | VRDL, Goverment Medical College (GMC) and Hospital, Valsad | Gujarat Biotechnology Research Centre | Sonal Sharma ,Zuber Saiyed,Ramesh Pandit,Janvi Raval,Zarna Patel,Nitin Savaliya,Dinesh Kumar,Twinkle Soni,Vicky Gandhi,Umang Mishra,Nitesh Shah,Chaitanya Joshi,Madhvi Joshi |
| EPI_ISL_2017750 | VRDL, Goverment Medical College (GMC) and Hospital, Valsad | Gujarat Biotechnology Research Centre | Janvi Raval,Zarna Patel,Nitin Savaliya,Dinesh Kumar,Twinkle Soni,Sonal Sharma ,Zuber Saiyed,Ramesh Pandit,Vicky Gandhi,Umang Mishra,Nitesh Shah,Chaitanya Joshi,Madhvi Joshi |
| EPI_ISL_2017751 | VRDL, Goverment Medical College (GMC) and Hospital, Valsad | Gujarat Biotechnology Research Centre | Zarna Patel,Nitin Savaliya,Dinesh Kumar,Twinkle Soni,Sonal Sharma ,Zuber Saiyed,Ramesh Pandit,Janvi Raval,Vicky Gandhi,Umang Mishra,Nitesh Shah,Chaitanya Joshi,Madhvi Joshi |
| EPI_ISL_2017754 | GMERS, Government Medical College (GMC), Gotri | Gujarat Biotechnology Research Centre | Twinkle Soni,Sonal Sharma ,Zuber Saiyed,Ramesh Pandit,Janvi Raval,Zarna Patel,Nitin Savaliya,Dinesh Kumar,Bithika Duttaroy,Umang Mishra,Nitesh Shah,Chaitanya Joshi,Madhvi Joshi |
| EPI_ISL_2017755 | GMERS, Government Medical College (GMC), Gotri | Gujarat Biotechnology Research Centre | Sonal Sharma ,Zuber Saiyed,Ramesh Pandit,Janvi Raval,Zarna Patel,Nitin Savaliya,Dinesh Kumar,Twinkle Soni,Bithika Duttaroy,Umang Mishra,Nitesh Shah,Chaitanya Joshi,Madhvi Joshi |
| EPI_ISL_2017756 | GMERS, Government Medical College (GMC), Gotri | Gujarat Biotechnology Research Centre | Zuber Saiyed,Ramesh Pandit,Janvi Raval,Zarna Patel,Nitin Savaliya,Dinesh Kumar,Twinkle Soni,Sonal Sharma ,Bithika Duttaroy,Umang Mishra,Nitesh Shah,Chaitanya Joshi,Madhvi Joshi |
| EPI_ISL_2017757 | GMERS, Government Medical College (GMC), Gotri | Gujarat Biotechnology Research Centre | Ramesh Pandit,Janvi Raval,Zarna Patel,Nitin Savaliya,Dinesh Kumar,Twinkle Soni,Sonal Sharma ,Zuber Saiyed,Bithika Duttaroy,Umang Mishra,Nitesh Shah,Chaitanya Joshi,Madhvi Joshi |
| EPI_ISL_2017758 | GMERS, Government Medical College (GMC), Gotri | Gujarat Biotechnology Research Centre | Janvi Raval,Zarna Patel,Nitin Savaliya,Dinesh Kumar,Twinkle Soni,Sonal Sharma ,Zuber Saiyed,Ramesh Pandit,Bithika Duttaroy,Umang Mishra,Nitesh Shah,Chaitanya Joshi,Madhvi Joshi |
| EPI_ISL_2017790, EPI_ISL_2017791, EPI_ISL_2017792, EPI_ISL_2017793, EPI_ISL_2017794, EPI_ISL_2017795, EPI_ISL_2017796, EPI_ISL_2017797, EPI_ISL_2017798, EPI_ISL_2017799, EPI_ISL_2017800, EPI_ISL_2017801, EPI_ISL_2017802, EPI_ISL_2017803, EPI_ISL_2017804, EPI_ISL_2017805, EPI_ISL_2017807, EPI_ISL_2017808, EPI_ISL_2017809, EPI_ISL_2017810, EPI_ISL_2017811, EPI_ISL_2017812, EPI_ISL_2017813, EPI_ISL_2017814, EPI_ISL_2017815, EPI_ISL_2017816, EPI_ISL_2017817, EPI_ISL_2017818, EPI_ISL_2017819, EPI_ISL_2017820, EPI_ISL_2017821, EPI_ISL_2017822, EPI_ISL_2017823, EPI_ISL_2017824, EPI_ISL_2017825, EPI_ISL_2017826, EPI_ISL_2017827, EPI_ISL_2017828, EPI_ISL_2017829, EPI_ISL_2017830, EPI_ISL_2017831, EPI_ISL_2017832, EPI_ISL_2017833, EPI_ISL_2017834, EPI_ISL_2017835, EPI_ISL_2017836, EPI_ISL_2017837, EPI_ISL_2017838, EPI_ISL_2017839, EPI_ISL_2017840, EPI_ISL_2017841, EPI_ISL_2017842, EPI_ISL_2017843, EPI_ISL_2017844, EPI_ISL_2017845, EPI_ISL_2017846, EPI_ISL_2017847, EPI_ISL_2017848, EPI_ISL_2017849, EPI_ISL_2017850, EPI_ISL_2017851 | | | |
| see above | Viral Respiratory Infections Laboratory, Cantacuzino National Military-Medical Institute | Cantacuzino Institute Virology | Luiza Ustea, Nicoleta Paraschiv, Catalina Pascu, Sorin Dinu, Mihaela Oprea, Mihaela Lazar |
| EPI_ISL_2017852 | "St. Parascheva" Infectious Diseases Clinical Hospital | Cantacuzino Institute Virology | Luiza Ustea, Nicoleta Paraschiv, Catalina Pascu, Sorin Dinu, Mihaela Oprea, Mihaela Lazar |
| EPI_ISL_2017853 | Viral Respiratory Infections Laboratory, Cantacuzino National Military-Medical Institute | Cantacuzino Institute Virology | Luiza Ustea, Nicoleta Paraschiv, Catalina Pascu, Sorin Dinu, Mihaela Oprea, Mihaela Lazar |

| | | | |
|---|---|---|---|
| EPI_ISL_2017854 | "St. Parascheva" Infectious Diseases Clinical Hospital | Cantacuzino Institute Virology | Luiza Ustea, Nicoleta Paraschiv, Catalina Pascu, Sorin Dinu, Mihaela Oprea, Mihaela Lazar |
| EPI_ISL_2017855, EPI_ISL_2017856, EPI_ISL_2017857, EPI_ISL_2017858, EPI_ISL_2017861, EPI_ISL_2017862 | Viral Respiratory Infections Laboratory, Cantacuzino National Military-Medical Institute | Cantacuzino Institute Virology | Luiza Ustea, Nicoleta Paraschiv, Catalina Pascu, Sorin Dinu, Mihaela Oprea, Mihaela Lazar |
| EPI_ISL_2017863, EPI_ISL_2017864, EPI_ISL_2017865, EPI_ISL_2017866, EPI_ISL_2017867 | "St. Parascheva" Infectious Diseases Clinical Hospital | Cantacuzino Institute Virology | Luiza Ustea, Nicoleta Paraschiv, Catalina Pascu, Sorin Dinu, Mihaela Oprea, Mihaela Lazar |
| EPI_ISL_2017868, EPI_ISL_2017869, EPI_ISL_2017871 | Viral Respiratory Infections Laboratory, Cantacuzino National Military-Medical Institute | Cantacuzino Institute Virology | Luiza Ustea, Nicoleta Paraschiv, Catalina Pascu, Sorin Dinu, Mihaela Oprea, Mihaela Lazar |
| EPI_ISL_2018922, EPI_ISL_2018923, EPI_ISL_2018924, EPI_ISL_2018925, EPI_ISL_2018926, EPI_ISL_2018927, EPI_ISL_2018928, EPI_ISL_2018929, EPI_ISL_2018930, EPI_ISL_2018931, EPI_ISL_2018932, EPI_ISL_2018933, EPI_ISL_2018934, EPI_ISL_2018935, EPI_ISL_2018936, EPI_ISL_2018937, EPI_ISL_2018938, EPI_ISL_2018943 | | | |
| see above | University of Rome Tor Vergata: Departm Experim Medicine Chair of Virology | University of Rome Tor Vergata: Departm Experim Medicine Chair of Virology | Maria Concetta Bellocchi, Luca Carioti, Maria Botticelli, Lorenzo Piermatteo, Rossana Scutari, Marco Iannetta, Loredana Sarmati, Massimo Andreoni, Francesca Ceccherini-Silberstein |
| EPI_ISL_2020238, EPI_ISL_2020239, EPI_ISL_2020240 | Bintulu Hospital | Institute for Medical Research, Infectious Disease Research Centre, National Institutes of Health, Ministry of Health Malaysia | Suppiah J, Kamel K, Mohd Zawawi Z, Azizan MA, Ramly N, Robert F, Thayan R |
| EPI_ISL_2020863 | Institut de Virologie du CHU de Strasbourg | Institut de Virologie du CHU de Strasbourg | Fafi-Kremer Samira, Gallais Floriane, Gantner Pierre, Laugel Elodie, Solis Morgane, Velay Aurélie, Wendling Marie-Josée |
| EPI_ISL_2023429, EPI_ISL_2023430, EPI_ISL_2023431, EPI_ISL_2023432, EPI_ISL_2023433, EPI_ISL_2023434, EPI_ISL_2023435, EPI_ISL_2023438, EPI_ISL_2023439, EPI_ISL_2023441, EPI_ISL_2023442, EPI_ISL_2023443, EPI_ISL_2023444, EPI_ISL_2023446, EPI_ISL_2023447, EPI_ISL_2023449 | | | |
| see above | Microbiology Department, University Hospital Donostia | Microbiology Department, University Hospital Donostia | Marimon JM, Montes M, Piñeiro L, Sorarrain A, Gomez M, Cilla G. |
| EPI_ISL_2029634 | Outre mer | National Reference Center for Viruses of Respiratory Infections, Institut Pasteur, Paris | Marion Barbet, Sylvie Behillil, Méline Bizard, Angela Brisebarre, Camille Capel, Vincent Enouf, Louise Lefrançois, Frédéric Lemoine, Christophe Malabat, Corinne Maufrais, Etienne Simon-Lorière, Maud Vanpeene, Sylvie Van der Werf ,Patrice Combe |
| EPI_ISL_2030016 | Hospital | National Reference Center for Viruses of Respiratory Infections, Institut Pasteur, Paris | Marion Barbet, Sylvie Behillil, Méline Bizard, Angela Brisebarre, Camille Capel, Vincent Enouf, Louise Lefrançois, Frédéric Lemoine, Christophe Malabat, Corinne Maufrais, Etienne Simon-Lorière, Maud Vanpeene, Sylvie Van der Werf ,CéLine Bressollette |
| EPI_ISL_2030084, EPI_ISL_2030085 | Hospital | National Reference Center for Viruses of Respiratory Infections, Institut Pasteur, Paris | Marion Barbet, Sylvie Behillil, Méline Bizard, Angela Brisebarre, Camille Capel, Vincent Enouf, Louise Lefrançois, Frédéric Lemoine, Christophe Malabat, Corinne Maufrais, Etienne Simon-Lorière, Maud Vanpeene, Sylvie Van der Werf ,EugèNe Castelain |
| EPI_ISL_2030088 | Hospital | National Reference Center for Viruses of Respiratory Infections, Institut Pasteur, Paris | Marion Barbet, Sylvie Behillil, Méline Bizard, Angela Brisebarre, Camille Capel, Vincent Enouf, Louise Lefrançois, Frédéric Lemoine, Christophe Malabat, Corinne Maufrais, Amaury Vaysse, Etienne Simon-Lorière, Maud Vanpeene, Sylvie Van der Werf ,CéLine Ramanantsoa |
| EPI_ISL_2030122 | Hospital | National Reference Center for Viruses of Respiratory Infections, Institut Pasteur, Paris | Marion Barbet, Sylvie Behillil, Méline Bizard, Angela Brisebarre, Camille Capel, Vincent Enouf, Louise Lefrançois, Frédéric Lemoine, Christophe Malabat, Corinne Maufrais, Etienne Simon-Lorière, Maud Vanpeene, Sylvie Van der Werf ,CéLine Ramanantsoa |
| EPI_ISL_2030166 | Hospital | National Reference Center for Viruses of Respiratory Infections, Institut Pasteur, Paris | Marion Barbet, Sylvie Behillil, Méline Bizard, Angela Brisebarre, Camille Capel, Vincent Enouf, Louise Lefrançois, Frédéric Lemoine, Christophe Malabat, Corinne Maufrais, Etienne Simon-Lorière, Maud Vanpeene, Sylvie Van der Werf ,EugèNe Castelain |
| EPI_ISL_2033747 | Translational Health Science and Technology Institute -ESIC medical college and hospital, Faridabad | THSTI Bioassay laboratory | Naseem Ahmed Khan, Heena Shaman, Jigme Wangchuk, Anil Pandey, Akshay Kanakan, Ranjeet Maurya, Janani Srinivasa Vasudevan, Neha Jha, Priyanka Mehta, Rajesh Pandey, Guruprasad R Medigeshi |
| EPI_ISL_2035722, EPI_ISL_2035723, EPI_ISL_2035725, EPI_ISL_2035726, EPI_ISL_2035727, EPI_ISL_2035728 | Public Institution of Health Care "Diagnostic Center ( Laboratory Testing Center) of Moscow Health Department " | Group of Genomics and Postgenomic Technologies of Central Research Institute of Epidemiology | Samoilov AE, Kaptelova VV, Korneenko EV, Saenko SS, Smirnova YS, Nadtoka MI, Spakova OG, Speranskaya AS, Tivanova EV, Kondrasheva LY, Akimkin VG |
| EPI_ISL_2040582, EPI_ISL_2040584 | CHU Nantes Virology | CHU Nantes Virology | Celine Bressollette-Bodin, Thomas Drumel, Audrey Rodallec, Berthe-Marie Imbert-Marcille |
| EPI_ISL_2047860 | Azienda USL Umbria 2 | Istituto Zooprofilattico Sperimentale dell'Abruzzo e Molise "G. Caporale" | Proietti A, Pistoni E, Lorusso A, Marcacci M, Di Domenico M, Ancora M, Curini V, Di Lollo Valeria, Mangone I, Rinaldi A, Delli Compagni E, Scialabba S, Caporale M, Di Pasquale A, Cammà C, Puglia I, Calistri P, Savini G |
| EPI_ISL_2047861 | Università degli Studi di Perugia | Istituto Zooprofilattico Sperimentale dell'Abruzzo e Molise "G. Caporale" | Mencacci A, Camilloni B, Lorusso A, Marcacci M, Di Domenico M, Ancora M, Curini V, Di Lollo Valeria, Mangone I, Rinaldi A, Delli Compagni E, Scialabba S, Caporale M, Di Pasquale A, Cammà C, Puglia I, Calistri P, Savini G |
| EPI_ISL_2047862, EPI_ISL_2047863 | Laboratorio Analisi Osp. Città di Castello - Azienda USL Umbria1 | Istituto Zooprofilattico Sperimentale dell'Abruzzo e Molise "G. Caporale" | Malagigi V, Tacconi P, Lorusso A, Marcacci M, Di Domenico M, Ancora M, Curini V, Di Lollo Valeria, Mangone I, Rinaldi A, Delli Compagni E, Scialabba S, Caporale M, Di Pasquale A, Cammà C, Puglia I, Calistri P, Savini G |
| EPI_ISL_2047865 | Università degli Studi di Perugia | Istituto Zooprofilattico Sperimentale dell'Abruzzo e Molise "G. Caporale" | Mencacci A, Camilloni B, Lorusso A, Marcacci M, Di Domenico M, Ancora M, Curini V, Mangone I, Rinaldi A, Delli Compagni E, Scialabba S, Di Pasquale A, Cammà C, Puglia I, Calistri P, Savini G |
| EPI_ISL_2047866 | Università degli Studi di Perugia | Istituto Zooprofilattico Sperimentale dell'Abruzzo e Molise "G. Caporale" | Mencacci A, Camilloni B, Lorusso A, Marcacci M, Di Domenico M, Ancora M, Curini V, Di Lollo Valeria, Mangone I, Rinaldi A, Delli Compagni E, Scialabba S, Caporale M, Di Pasquale A, Cammà C, Puglia I, Calistri P, Savini G |
| EPI_ISL_2047867 | Istituto Zooprofilattico Sperimentale Umbria e Marche "Togo Rosati" | Istituto Zooprofilattico Sperimentale dell'Abruzzo e Molise "G. Caporale" | Biagetti M, Giammarioli M,Lorusso A, Marcacci M, Di Domenico M, Ancora M, Curini V, Di Lollo Valeria, Mangone I, Rinaldi A, Delli Compagni E, Scialabba S, Caporale M, Di Pasquale A, Cammà C, Puglia I, Calistri P, Savini G |
| EPI_ISL_2047868 | Azienda USL Umbria 2 | Istituto Zooprofilattico Sperimentale dell'Abruzzo e Molise "G. Caporale" | Proietti A, Pistoni E, Lorusso A, Marcacci M, Di Domenico M, Ancora M, Curini V, Di Lollo Valeria, Mangone I, Rinaldi A, Delli Compagni E, Scialabba S, Caporale M, Di Pasquale A, Cammà C, Puglia I, Calistri P, Savini G |
| EPI_ISL_2047869 | Azienda Ospedaliera Terni | Istituto Zooprofilattico Sperimentale dell'Abruzzo e Molise "G. Caporale" | Palumbo M, Scaccetti A, Lorusso A, Marcacci M, Di Domenico M, Ancora M, Curini V, Mangone I, Rinaldi A, Delli Compagni E, Scialabba S, Di Pasquale A, Cammà C, Puglia I, Calistri P, Savini G |
| EPI_ISL_2047870 | Istituto Zooprofilattico Sperimentale Umbria e Marche "Togo Rosati" | Istituto Zooprofilattico Sperimentale dell'Abruzzo e Molise "G. Caporale" | Biagetti M, Giammarioli M, Lorusso A, Marcacci M, Di Domenico M, Ancora M, Curini V, Mangone I, Rinaldi A, Delli Compagni E, Scialabba S, Di Pasquale A, Cammà C, Puglia I, Calistri P, Savini G |
| EPI_ISL_2081192, EPI_ISL_2081193 | Premier Laboratory Services | BioInfoExperts LLC | Cheryl Metzler, Valerie Murray, Rebecca Rose, David Nolan, Susanna Lamers |
| EPI_ISL_2083045 | Baylor Scott & White-Temple | Baylor Scott & White-Temple | Ari Rao, Linden Morales, Kimberly Walker, Marcus Volz, Shelby Hendrickson |
| EPI_ISL_2086122, EPI_ISL_2086123, EPI_ISL_2086124, EPI_ISL_2086126, EPI_ISL_2086127, EPI_ISL_2086128, EPI_ISL_2086129, EPI_ISL_2086130, EPI_ISL_2086131, EPI_ISL_2086132, EPI_ISL_2086133, EPI_ISL_2086134, EPI_ISL_2086135, EPI_ISL_2086136, EPI_ISL_2086137, EPI_ISL_2086167 | | | |
| see above | Microbiology Department. Complexo Hospitalario Universitario de Vigo | Microbiology Department. Complexo Hospitalario Universitario de Vigo | Alfaya N, Alonso I, Alvarez M, Cabrera JJ, Carballo R, Cores O, Cortizo S, del-Campo V, Martinez L, Mediero G, Perez S, Potel C, Regueiro B, Rey S, Vasallo FJ |
| EPI_ISL_2086610, EPI_ISL_2086611, EPI_ISL_2086612, EPI_ISL_2086613, EPI_ISL_2086614, EPI_ISL_2086615, EPI_ISL_2086616, EPI_ISL_2086617, EPI_ISL_2086618, EPI_ISL_2086619, EPI_ISL_2086620, EPI_ISL_2086621, EPI_ISL_2086622, EPI_ISL_2086765, EPI_ISL_2086766, EPI_ISL_2086767 | | | |
| see above | LABCOVID_HCPA | LABRESIS_HCPA | Wink PL, Martins AF, Volpato F, Monteiro F, Zavascki AP, Barth AL |
| EPI_ISL_2090728 | Institut de Virologie du CHU de Strasbourg | Institut de Virologie du CHU de Strasbourg | Fafi-Kremer Samira, Gallais Floriane, Gantner Pierre, Laugel Elodie, Solis Morgane, Velay Aurélie, Wendling Marie-Josée |
| EPI_ISL_2090759 | CHU NORD | UMR PIMIT | Dr David A Wilkinson, Dr Patrick Mavingui, Dr Camille Lebarbenchon, Magali Turpin |
| EPI_ISL_2090785 | SAINT BENOIT | UMR PIMIT | Dr David A Wilkinson, Dr Patrick Mavingui, Dr Camille Lebarbenchon, Magali Turpin |
| EPI_ISL_2090821, EPI_ISL_2090828, EPI_ISL_2090829, EPI_ISL_2090832, EPI_ISL_2090837, EPI_ISL_2090841, EPI_ISL_2090842, EPI_ISL_2090844, EPI_ISL_2090850, EPI_ISL_2090861, EPI_ISL_2090862 | | | |
| see above | REUNILAB | UMR PIMIT | Dr David A Wilkinson, Dr Patrick Mavingui, Dr Camille Lebarbenchon, Magali Turpin |
| EPI_ISL_2090864 | CHU NORD | UMR PIMIT | Dr David A Wilkinson, Dr Patrick Mavingui, Dr Camille Lebarbenchon, Magali Turpin |

| | | | |
|---|---|---|---|
| EPI_ISL_2090866, EPI_ISL_2090867, EPI_ISL_2090869, EPI_ISL_2090879, EPI_ISL_2090880, EPI_ISL_2090882 | REUNILAB | UMR PIMIT | Dr David A Wilkinson, Dr Patrick Mavingui, Dr Camille Lebarbenchon, Magali Turpin |
| EPI_ISL_2091023, EPI_ISL_2091025, EPI_ISL_2091120 | Institut de Virologie du CHU de Strasbourg | Institut de Virologie du CHU de Strasbourg | Fafi-Kremer Samira, Gallais Floriane, Gantner Pierre, Laugel Elodie, Solis Morgane, Velay Aurélie, Wendling Marie-Josée |
| EPI_ISL_2091144, EPI_ISL_2091146, EPI_ISL_2091147, EPI_ISL_2091148, EPI_ISL_2091149, EPI_ISL_2091150, EPI_ISL_2091151, EPI_ISL_2091152, EPI_ISL_2091153, EPI_ISL_2091154, EPI_ISL_2091155, EPI_ISL_2091156, EPI_ISL_2091157, EPI_ISL_2091158, EPI_ISL_2091159 | | | |
| see above | Laboratorio Central de Epidemiología (LCE) | Centro de Investigación en Enfermedades Infecciosas (CIENI), Instituto Nacional de Enfermedades Respiratorias (INER) | Consorcio Mexicano de Vigilancia Genómica (CoViGen-Mex). Authors (in alphabetical order): Julio Elias Alvarado-Yaah, Carlos F. Arias, Santiago Ávila-Ríos, Víctor Hugo Borja-Aburto, Celia Boukadida, Juan Bautista Chale-Dzul, Célida Duque Molina, José Antonio Enciso-Moreno, Gloria Elena Espinosa-Ayala, Fernando Fontove-Herrera, Victor Eduardo Garcia-Arias, Concepción Grajales-Muñiz, Ricardo Grande, Alfredo Herrera-Estrella, Carla Ivón Herrera-Najera, Pavel Isa, Brenda Irasema Maldonado-Meza, Bernardo Martínez-Miguel, Margarita Matías-Florentino, María Guadalupe de Jesús Mireles-Rivera, Gloria María Molina-Salinas, Hector Montoya-Fuentes, José Esteban Muñoz-Medina, José de Jesús Nuñez-Contreras, Alicia Ocaña-Mondragón, Luis Alberto Ochoa-Carrera, Hector Esteban Paz-Juárez, Francisco Pulido, Helen Haydee Fernanda Ramírez-Plascencia, Angel Gustavo Salas-Lais, Alejandro Sanchez-Flores, Clara Esperanza Santacruz-Tinoco, María Guadalupe Santiago-Mauricio, Nelly Sélem-Mojica, Blanca Taboada, Gloria Vazquez |
| EPI_ISL_2091160, EPI_ISL_2091161, EPI_ISL_2091162, EPI_ISL_2091163, EPI_ISL_2091164, EPI_ISL_2091165, EPI_ISL_2091166, EPI_ISL_2091167, EPI_ISL_2091168, EPI_ISL_2091169, EPI_ISL_2091170, EPI_ISL_2091171, EPI_ISL_2091172, EPI_ISL_2091173, EPI_ISL_2091174, EPI_ISL_2091175, EPI_ISL_2091176, EPI_ISL_2091177, EPI_ISL_2091178, EPI_ISL_2091179, EPI_ISL_2091180, EPI_ISL_2091182, EPI_ISL_2091183, EPI_ISL_2091184, EPI_ISL_2091185, EPI_ISL_2091186, EPI_ISL_2091187, EPI_ISL_2091188, EPI_ISL_2091190, EPI_ISL_2091191, EPI_ISL_2091192, EPI_ISL_2091193, EPI_ISL_2091194, EPI_ISL_2091195, EPI_ISL_2091196, EPI_ISL_2091197, EPI_ISL_2091198, EPI_ISL_2091199, EPI_ISL_2091213 | | | |
| see above | Centro de Investigación Biomédica del Noreste (CIBIN) | Centro de Investigación en Enfermedades Infecciosas (CIENI), Instituto Nacional de Enfermedades Respiratorias (INER) | Consorcio Mexicano de Vigilancia Genómica (CoViGen-Mex). Authors (in alphabetical order): Julio Elias Alvarado-Yaah, Carlos F. Arias, Santiago Ávila-Ríos, Víctor Hugo Borja-Aburto, Celia Boukadida, Juan Bautista Chale-Dzul, Célida Duque Molina, José Antonio Enciso-Moreno, Gloria Elena Espinosa-Ayala, Fernando Fontove-Herrera, Victor Eduardo Garcia-Arias, Concepción Grajales-Muñiz, Ricardo Grande, Alfredo Herrera-Estrella, Carla Ivón Herrera-Najera, Pavel Isa, Brenda Irasema Maldonado-Meza, Bernardo Martínez-Miguel, Margarita Matías-Florentino, María Guadalupe de Jesús Mireles-Rivera, Gloria María Molina-Salinas, Hector Montoya-Fuentes, José Esteban Muñoz-Medina, José de Jesús Nuñez-Contreras, Alicia Ocaña-Mondragón, Luis Alberto Ochoa-Carrera, Hector Esteban Paz-Juárez, Francisco Pulido, Helen Haydee Fernanda Ramírez-Plascencia, Angel Gustavo Salas-Lais, Alejandro Sanchez-Flores, Clara Esperanza Santacruz-Tinoco, María Guadalupe Santiago-Mauricio, Nelly Sélem-Mojica, Blanca Taboada, Gloria Vazquez |
| EPI_ISL_2091225, EPI_ISL_2091226, EPI_ISL_2091227, EPI_ISL_2091228, EPI_ISL_2091229, EPI_ISL_2091230, EPI_ISL_2091231, EPI_ISL_2091232, EPI_ISL_2091233, EPI_ISL_2091234, EPI_ISL_2091235, EPI_ISL_2091236, EPI_ISL_2091237, EPI_ISL_2091238, EPI_ISL_2091239, EPI_ISL_2091240, EPI_ISL_2091241, EPI_ISL_2091242, EPI_ISL_2091243, EPI_ISL_2091244, EPI_ISL_2091245, EPI_ISL_2091246, EPI_ISL_2091248, EPI_ISL_2091249, EPI_ISL_2091250 | | | |
| see above | Unidad de Investigación Médica de Yucatán (UIMY) | Centro de Investigación en Enfermedades Infecciosas (CIENI), Instituto Nacional de Enfermedades Respiratorias (INER) | Consorcio Mexicano de Vigilancia Genómica (CoViGen-Mex). Authors (in alphabetical order): Julio Elias Alvarado-Yaah, Carlos F. Arias, Santiago Ávila-Ríos, Víctor Hugo Borja-Aburto, Celia Boukadida, Juan Bautista Chale-Dzul, Célida Duque Molina, José Antonio Enciso-Moreno, Gloria Elena Espinosa-Ayala, Fernando Fontove-Herrera, Victor Eduardo Garcia-Arias, Concepción Grajales-Muñiz, Ricardo Grande, Alfredo Herrera-Estrella, Carla Ivón Herrera-Najera, Pavel Isa, Brenda Irasema Maldonado-Meza, Bernardo Martínez-Miguel, Margarita Matías-Florentino, María Guadalupe de Jesús Mireles-Rivera, Gloria María Molina-Salinas, Hector Montoya-Fuentes, José Esteban Muñoz-Medina, José de Jesús Nuñez-Contreras, Alicia Ocaña-Mondragón, Luis Alberto Ochoa-Carrera, Hector Esteban Paz-Juárez, Francisco Pulido, Helen Haydee Fernanda Ramírez-Plascencia, Angel Gustavo Salas-Lais, Alejandro Sanchez-Flores, Clara Esperanza Santacruz-Tinoco, María Guadalupe Santiago-Mauricio, Nelly Sélem-Mojica, Blanca Taboada, Gloria Vazquez |
| EPI_ISL_2091258, EPI_ISL_2091259, EPI_ISL_2091260, EPI_ISL_2091261, EPI_ISL_2091262, EPI_ISL_2091263, EPI_ISL_2091264, EPI_ISL_2091265, EPI_ISL_2091266, EPI_ISL_2091267, EPI_ISL_2091268, EPI_ISL_2091269, EPI_ISL_2091270, EPI_ISL_2091271, EPI_ISL_2091272, EPI_ISL_2091273, EPI_ISL_2091274, EPI_ISL_2091275, EPI_ISL_2091276, EPI_ISL_2091278, EPI_ISL_2091279, EPI_ISL_2091280, EPI_ISL_2091283, EPI_ISL_2091287, EPI_ISL_2091288, EPI_ISL_2091298, EPI_ISL_2091300, EPI_ISL_2091306 | | | |
| see above | Centro de Investigación Biomédica de Occidente (CIBO) | Centro de Investigación en Enfermedades Infecciosas (CIENI), Instituto Nacional de Enfermedades Respiratorias (INER) | Consorcio Mexicano de Vigilancia Genómica (CoViGen-Mex). Authors (in alphabetical order): Julio Elias Alvarado-Yaah, Carlos F. Arias, Santiago Ávila-Ríos, Víctor Hugo Borja-Aburto, Celia Boukadida, Juan Bautista Chale-Dzul, Célida Duque Molina, José Antonio Enciso-Moreno, Gloria Elena Espinosa-Ayala, Fernando Fontove-Herrera, Victor Eduardo Garcia-Arias, Concepción Grajales-Muñiz, Ricardo Grande, Alfredo Herrera-Estrella, Carla Ivón Herrera-Najera, Pavel Isa, Brenda Irasema Maldonado-Meza, Bernardo Martínez-Miguel, Margarita Matías-Florentino, María Guadalupe de Jesús Mireles-Rivera, Gloria María Molina-Salinas, Hector Montoya-Fuentes, José Esteban Muñoz-Medina, José de Jesús Nuñez-Contreras, Alicia Ocaña-Mondragón, Luis Alberto Ochoa-Carrera, Hector Esteban Paz-Juárez, Francisco Pulido, Helen Haydee Fernanda Ramírez-Plascencia, Angel Gustavo Salas-Lais, Alejandro Sanchez-Flores, Clara Esperanza Santacruz-Tinoco, María Guadalupe Santiago-Mauricio, Nelly Sélem-Mojica, Blanca Taboada, Gloria Vazquez |
| EPI_ISL_2091320, EPI_ISL_2091321, EPI_ISL_2091322, EPI_ISL_2091323, EPI_ISL_2091324, EPI_ISL_2091325, EPI_ISL_2091326, EPI_ISL_2091327, EPI_ISL_2091328, EPI_ISL_2091329, EPI_ISL_2091330, EPI_ISL_2091331, EPI_ISL_2091332, EPI_ISL_2091333, EPI_ISL_2091334, EPI_ISL_2091335, EPI_ISL_2091336, EPI_ISL_2091337, EPI_ISL_2091338, EPI_ISL_2091339, EPI_ISL_2091340, EPI_ISL_2091341, EPI_ISL_2091342, EPI_ISL_2091343, EPI_ISL_2091348, EPI_ISL_2091354, EPI_ISL_2091355, EPI_ISL_2091356, EPI_ISL_2091357, EPI_ISL_2091358, EPI_ISL_2091359, EPI_ISL_2091360, EPI_ISL_2091413, EPI_ISL_2091441 | | | |
| see above | Laboratorio Central de Epidemiología (LCE) | Centro de Investigación en Enfermedades Infecciosas (CIENI), Instituto Nacional de Enfermedades Respiratorias (INER) | Consorcio Mexicano de Vigilancia Genómica (CoViGen-Mex). Authors (in alphabetical order): Julio Elias Alvarado-Yaah, Carlos F. Arias, Santiago Ávila-Ríos, Víctor Hugo Borja-Aburto, Celia Boukadida, Juan Bautista Chale-Dzul, Célida Duque Molina, José Antonio Enciso-Moreno, Gloria Elena Espinosa-Ayala, Fernando Fontove-Herrera, Victor Eduardo Garcia-Arias, Concepción Grajales-Muñiz, Ricardo Grande, Alfredo Herrera-Estrella, Carla Ivón Herrera-Najera, Pavel Isa, Brenda Irasema Maldonado-Meza, Bernardo Martínez-Miguel, Margarita Matías-Florentino, María Guadalupe de Jesús Mireles-Rivera, Gloria María Molina-Salinas, Hector Montoya-Fuentes, José Esteban Muñoz-Medina, José de Jesús Nuñez-Contreras, Alicia Ocaña-Mondragón, Luis Alberto Ochoa-Carrera, Hector Esteban Paz-Juárez, Francisco Pulido, Helen Haydee Fernanda Ramírez-Plascencia, Angel Gustavo Salas-Lais, Alejandro Sanchez-Flores, Clara Esperanza Santacruz-Tinoco, María Guadalupe Santiago-Mauricio, Nelly Sélem-Mojica, Blanca Taboada, Gloria Vazquez |
| EPI_ISL_2092872 | CQRC_QUALITY CONTROL CHEMICAL BIOLOGICAL RISK_AOOR Villa Sofia Cervello Palermo | CQRC_QUALITY CONTROL CHEMICAL BIOLOGICAL RISK_AOOR Villa Sofia Cervello Palermo | Di Gaudio, F., Brunacci, G., Contino, F., |
| EPI_ISL_2094417, EPI_ISL_2094418, EPI_ISL_2094419, EPI_ISL_2094420, EPI_ISL_2094421 | cerballiance-HDF | Cerba lab | Emmanuel Lecorche, Sabine Trombert-Paolantoni, Claire Felloni, Anne Mainardi, Laura Verdurme, Jean-Michel Damien, Stéphanie Haïm-Boukobza, Bénédicte Roquebert |
| EPI_ISL_2094422, EPI_ISL_2094423, EPI_ISL_2094424, EPI_ISL_2094425, EPI_ISL_2094426, EPI_ISL_2094427, EPI_ISL_2094428, EPI_ISL_2094429, EPI_ISL_2094430, EPI_ISL_2094431, EPI_ISL_2094432, EPI_ISL_2094433, EPI_ISL_2094434, EPI_ISL_2094435, EPI_ISL_2094436 | | | |
| see above | cerballiance-IDF | Cerba lab | Emmanuel Lecorche, Sabine Trombert-Paolantoni, Aude Lessenne, Kader Merah, Thierry Collin, Laura Verdurme, Patrice Herisson, Stéphanie Haïm-Boukobza, Bénédicte Roquebert |
| EPI_ISL_2094437, EPI_ISL_2094438 | cerballiance-PACA | Cerba lab | Emmanuel Lecorche, Sabine Trombert-Paolantoni, Laurence Prots, Jean-Christophe Roig, Edouard Delaunay, Isabelle Butin, Laura Verdurme, Bruno Roure, Stéphanie Haïm-Boukobza, Bénédicte Roquebert |
| EPI_ISL_2095823, EPI_ISL_2095824, EPI_ISL_2095825, EPI_ISL_2095826, EPI_ISL_2095827 | NMVRVI | National Public Health Surveillance Laboratory | Lukas Zemaitis, Migle Gabrielaite, Jelena Razmuk, Svajune Muralyte, Ana Steponkiene, Lukas Vasionis, Danas Baksa |
| EPI_ISL_2095843 | RezusLT | National Public Health Surveillance Laboratory | Lukas Zemaitis, Migle Gabrielaite, Jelena Razmuk, Svajune Muralyte, Ana Steponkiene, Lukas Vasionis, Danas Baksa |
| EPI_ISL_2095845 | Vilniaus universitetas | National Public Health Surveillance Laboratory | Lukas Zemaitis, Migle Gabrielaite, Jelena Razmuk, Svajune Muralyte, Ana Steponkiene, Lukas Vasionis, Danas Baksa |
| EPI_ISL_2095846, EPI_ISL_2095847 | Klaipedos universitetine ligonine | National Public Health Surveillance Laboratory | Lukas Zemaitis, Migle Gabrielaite, Jelena Razmuk, Svajune Muralyte, Ana Steponkiene, Lukas Vasionis, Danas Baksa |
| EPI_ISL_2095851, EPI_ISL_2095853, EPI_ISL_2095854, EPI_ISL_2095855, EPI_ISL_2095856, EPI_ISL_2095857, EPI_ISL_2095859 | RezusLT | National Public Health Surveillance Laboratory | Lukas Zemaitis, Migle Gabrielaite, Jelena Razmuk, Svajune Muralyte, Ana Steponkiene, Lukas Vasionis, Danas Baksa |
| EPI_ISL_2095861, EPI_ISL_2095862, EPI_ISL_2095864, EPI_ISL_2095865, EPI_ISL_2095866, EPI_ISL_2095867, EPI_ISL_2095868, EPI_ISL_2095869, EPI_ISL_2095870, EPI_ISL_2095871, EPI_ISL_2095872, EPI_ISL_2095873, EPI_ISL_2095874, EPI_ISL_2095875, EPI_ISL_2095876 | | | |

| | | | |
|---|---|---|---|
| see above | Vilniaus universitetas | National Public Health Surveillance Laboratory | Lukas Zemaitis, Migle Gabrielaite, Jelena Razmuk, Svajune Muralyte, Ana Steponkiene, Lukas Vasionis, Danas Baksa |
| EPI_ISL_2095887 | Klaipedos universitetine ligonine | National Public Health Surveillance Laboratory | Lukas Zemaitis, Migle Gabrielaite, Jelena Razmuk, Svajune Muralyte, Ana Steponkiene, Lukas Vasionis, Danas Baksa |
| EPI_ISL_2095890 | Vilniaus universitetas | National Public Health Surveillance Laboratory | Lukas Zemaitis, Migle Gabrielaite, Jelena Razmuk, Svajune Muralyte, Ana Steponkiene, Lukas Vasionis, Danas Baksa |
| EPI_ISL_2095891 | RezusLT | National Public Health Surveillance Laboratory | Lukas Zemaitis, Migle Gabrielaite, Jelena Razmuk, Svajune Muralyte, Ana Steponkiene, Lukas Vasionis, Danas Baksa |
| EPI_ISL_2095893, EPI_ISL_2095894, EPI_ISL_2095898, EPI_ISL_2095899, EPI_ISL_2095900, EPI_ISL_2095903, EPI_ISL_2095904 | Klaipedos universitetine ligonine | National Public Health Surveillance Laboratory | Lukas Zemaitis, Migle Gabrielaite, Jelena Razmuk, Svajune Muralyte, Ana Steponkiene, Lukas Vasionis, Danas Baksa |
| EPI_ISL_2095907, EPI_ISL_2095908 | RezusLT | National Public Health Surveillance Laboratory | Lukas Zemaitis, Migle Gabrielaite, Jelena Razmuk, Svajune Muralyte, Ana Steponkiene, Lukas Vasionis, Danas Baksa |
| EPI_ISL_2095909, EPI_ISL_2095910 | Vilniaus universitetas | National Public Health Surveillance Laboratory | Lukas Zemaitis, Migle Gabrielaite, Jelena Razmuk, Svajune Muralyte, Ana Steponkiene, Lukas Vasionis, Danas Baksa |
| EPI_ISL_2095911 | Klaipedos universitetine ligonine | National Public Health Surveillance Laboratory | Lukas Zemaitis, Migle Gabrielaite, Jelena Razmuk, Svajune Muralyte, Ana Steponkiene, Lukas Vasionis, Danas Baksa |
| EPI_ISL_2095917, EPI_ISL_2095919, EPI_ISL_2095920, EPI_ISL_2095921 | National Food and Veterinary Risk Assessment Institute | National Public Health Surveillance Laboratory | Lukas Zemaitis, Migle Gabrielaite, Jelena Razmuk, Svajune Muralyte, Ana Steponkiene, Lukas Vasionis, Danas Baksa |
| EPI_ISL_2095922, EPI_ISL_2095923, EPI_ISL_2095924, EPI_ISL_2095925, EPI_ISL_2095926, EPI_ISL_2095927, EPI_ISL_2095928, EPI_ISL_2095929, EPI_ISL_2095930, EPI_ISL_2095931, EPI_ISL_2095932, EPI_ISL_2095933, EPI_ISL_2095934, EPI_ISL_2095935, EPI_ISL_2095936, EPI_ISL_2095937, EPI_ISL_2095938, EPI_ISL_2095939, EPI_ISL_2095940, EPI_ISL_2095941, EPI_ISL_2095942, EPI_ISL_2095943, EPI_ISL_2095944, EPI_ISL_2095945 | | | |
| see above | National Public Health Surveillance Laboratory | National Public Health Surveillance Laboratory | Lukas Zemaitis, Migle Gabrielaite, Jelena Razmuk, Svajune Muralyte, Ana Steponkiene, Lukas Vasionis, Danas Baksa |
| EPI_ISL_2095946, EPI_ISL_2095947, EPI_ISL_2095948, EPI_ISL_2095949, EPI_ISL_2095950, EPI_ISL_2095951, EPI_ISL_2095952, EPI_ISL_2095953, EPI_ISL_2095954, EPI_ISL_2095955, EPI_ISL_2095956, EPI_ISL_2095957, EPI_ISL_2095958, EPI_ISL_2095959, EPI_ISL_2095960, EPI_ISL_2095961, EPI_ISL_2095962, EPI_ISL_2095963, EPI_ISL_2095964, EPI_ISL_2095965, EPI_ISL_2095966, EPI_ISL_2095967, EPI_ISL_2095968, EPI_ISL_2095969 | | | |
| see above | National Food and Veterinary Risk Assessment Institute | National Public Health Surveillance Laboratory | Lukas Zemaitis, Migle Gabrielaite, Jelena Razmuk, Svajune Muralyte, Ana Steponkiene, Lukas Vasionis, Danas Baksa |
| EPI_ISL_2095970, EPI_ISL_2095971, EPI_ISL_2095972, EPI_ISL_2095973, EPI_ISL_2095974 | National Public Health Surveillance Laboratory | National Public Health Surveillance Laboratory | Lukas Zemaitis, Migle Gabrielaite, Jelena Razmuk, Svajune Muralyte, Ana Steponkiene, Lukas Vasionis, Danas Baksa |
| EPI_ISL_2095975, EPI_ISL_2095976, EPI_ISL_2095977, EPI_ISL_2095978, EPI_ISL_2095979 | National Food and Veterinary Risk Assessment Institute | National Public Health Surveillance Laboratory | Lukas Zemaitis, Migle Gabrielaite, Jelena Razmuk, Svajune Muralyte, Ana Steponkiene, Lukas Vasionis, Danas Baksa |
| EPI_ISL_2095982, EPI_ISL_2095983, EPI_ISL_2095984 | National Public Health Surveillance Laboratory | National Public Health Surveillance Laboratory | Lukas Zemaitis, Migle Gabrielaite, Jelena Razmuk, Svajune Muralyte, Ana Steponkiene, Lukas Vasionis, Danas Baksa |
| EPI_ISL_2095985 | Nacionaline visuomenes sveikatos prieziuros laboratorija | National Public Health Surveillance Laboratory | Lukas Zemaitis, Migle Gabrielaite, Jelena Razmuk, Svajune Muralyte, Ana Steponkiene, Lukas Vasionis, Danas Baksa |
| EPI_ISL_2095986, EPI_ISL_2095987, EPI_ISL_2095988, EPI_ISL_2095989 | National Public Health Surveillance Laboratory | National Public Health Surveillance Laboratory | Lukas Zemaitis, Migle Gabrielaite, Jelena Razmuk, Svajune Muralyte, Ana Steponkiene, Lukas Vasionis, Danas Baksa |
| EPI_ISL_2095990 | Nacionaline visuomenes sveikatos prieziuros laboratorija | National Public Health Surveillance Laboratory | Lukas Zemaitis, Migle Gabrielaite, Jelena Razmuk, Svajune Muralyte, Ana Steponkiene, Lukas Vasionis, Danas Baksa |
| EPI_ISL_2095991, EPI_ISL_2095992, EPI_ISL_2095993, EPI_ISL_2095994, EPI_ISL_2095995, EPI_ISL_2095996, EPI_ISL_2095997, EPI_ISL_2095998, EPI_ISL_2095999 | National Public Health Surveillance Laboratory | National Public Health Surveillance Laboratory | Lukas Zemaitis, Migle Gabrielaite, Jelena Razmuk, Svajune Muralyte, Ana Steponkiene, Lukas Vasionis, Danas Baksa |
| EPI_ISL_2096000, EPI_ISL_2096001, EPI_ISL_2096002 | National Food and Veterinary Risk Assessment Institute | National Public Health Surveillance Laboratory | Lukas Zemaitis, Migle Gabrielaite, Jelena Razmuk, Svajune Muralyte, Ana Steponkiene, Lukas Vasionis, Danas Baksa |
| EPI_ISL_2096992, EPI_ISL_2096993, EPI_ISL_2096994, EPI_ISL_2096995, EPI_ISL_2096996 | Viral Respiratory Infections Laboratory, Cantacuzino National Military-Medical Institute | Cantacuzino Institute Virology | Luiza Ustea, Nicoleta Paraschiv, Catalina Pascu, Sorin Dinu, Mihaela Oprea, Mihaela Lazar |
| EPI_ISL_2096997, EPI_ISL_2096998 | "St. Parascheva" Infectious Diseases Clinical Hospital | Cantacuzino Institute Virology | Luiza Ustea, Nicoleta Paraschiv, Catalina Pascu, Sorin Dinu, Mihaela Oprea, Mihaela Lazar |
| EPI_ISL_2097000, EPI_ISL_2097001, EPI_ISL_2097002, EPI_ISL_2097010, EPI_ISL_2097011, EPI_ISL_2097012, EPI_ISL_2097015, EPI_ISL_2097016, EPI_ISL_2097018, EPI_ISL_2097019, EPI_ISL_2097020, EPI_ISL_2097021, EPI_ISL_2097024, EPI_ISL_2097025, EPI_ISL_2097026, EPI_ISL_2097027, EPI_ISL_2097028, EPI_ISL_2097029, EPI_ISL_2097030, EPI_ISL_2097031, EPI_ISL_2097032, EPI_ISL_2097033, EPI_ISL_2097035, EPI_ISL_2097037, EPI_ISL_2097041, EPI_ISL_2097042, EPI_ISL_2097043, EPI_ISL_2097044, EPI_ISL_2097045, EPI_ISL_2097046, EPI_ISL_2097047, EPI_ISL_2097048, EPI_ISL_2097052, EPI_ISL_2097053, EPI_ISL_2097054, EPI_ISL_2097055, EPI_ISL_2097056, EPI_ISL_2097057, EPI_ISL_2097058, EPI_ISL_2097060, EPI_ISL_2097061, EPI_ISL_2097063, EPI_ISL_2097064, EPI_ISL_2097065, EPI_ISL_2097066, EPI_ISL_2097067, EPI_ISL_2097068, EPI_ISL_2097069, EPI_ISL_2097070, EPI_ISL_2097071, EPI_ISL_2097072, EPI_ISL_2097073, EPI_ISL_2097074, EPI_ISL_2097075, EPI_ISL_2097076, EPI_ISL_2097078, EPI_ISL_2097079, EPI_ISL_2097082 | | | |
| see above | Synlab Eesti OÜ | 1. Laboratory of Communicable Diseases (Estonia); 2. Eurofins Genomics Europe Sequencing GmbH | Liidia Dotsenko et al. |
| EPI_ISL_2098714, EPI_ISL_2098715, EPI_ISL_2098721, EPI_ISL_2098722 | State Virus Research and Diagnostic Laboratory (VRDL), AIIMS Raipur | State Virus Research and Diagnostic Laboratory (VRDL), AIIMS Raipur | Pushpendra Singh, Kuldeep Sharma, Priyanka Singh, Somya Sharma, MD Rafiiqullah Khan, Jiyan Chandrawanshi, Sanjay Singh Negi, Anudita Bhargava |
| EPI_ISL_2099345, EPI_ISL_2099346, EPI_ISL_2099358, EPI_ISL_2099359, EPI_ISL_2099360, EPI_ISL_2099361, EPI_ISL_2099362, EPI_ISL_2099363, EPI_ISL_2099364, EPI_ISL_2099365, EPI_ISL_2099366, EPI_ISL_2099367, EPI_ISL_2099368, EPI_ISL_2099369, EPI_ISL_2099370, EPI_ISL_2099371, EPI_ISL_2099372, EPI_ISL_2099373, EPI_ISL_2099374, EPI_ISL_2099375, EPI_ISL_2099376, EPI_ISL_2099377, EPI_ISL_2099378, EPI_ISL_2099379, EPI_ISL_2099380, EPI_ISL_2099381, EPI_ISL_2099382, EPI_ISL_2099383, EPI_ISL_2099384, EPI_ISL_2099385, EPI_ISL_2099386, EPI_ISL_2099387, EPI_ISL_2099388, EPI_ISL_2099389, EPI_ISL_2099390, EPI_ISL_2099391, EPI_ISL_2099392, EPI_ISL_2099393, EPI_ISL_2099394, EPI_ISL_2099395, EPI_ISL_2099396, EPI_ISL_2099397, EPI_ISL_2099398, EPI_ISL_2099399, EPI_ISL_2099400, EPI_ISL_2099401, EPI_ISL_2099402, EPI_ISL_2099403, EPI_ISL_2099404, EPI_ISL_2099405, EPI_ISL_2099406, EPI_ISL_2099407, EPI_ISL_2099408, EPI_ISL_2099409, EPI_ISL_2099410, EPI_ISL_2099411, EPI_ISL_2099412, EPI_ISL_2099413, EPI_ISL_2099414, EPI_ISL_2099415, EPI_ISL_2099416, EPI_ISL_2099417, EPI_ISL_2099418, EPI_ISL_2099419, EPI_ISL_2099420, EPI_ISL_2099421, EPI_ISL_2099423, EPI_ISL_2099424, EPI_ISL_2099425, EPI_ISL_2099426, EPI_ISL_2099429, EPI_ISL_2099453, EPI_ISL_2099454, EPI_ISL_2099458, EPI_ISL_2099459, EPI_ISL_2099460, EPI_ISL_2099461, EPI_ISL_2099462, EPI_ISL_2099463, EPI_ISL_2099464, EPI_ISL_2099465, EPI_ISL_2099466, EPI_ISL_2099467, EPI_ISL_2099468, EPI_ISL_2099469, EPI_ISL_2099470, EPI_ISL_2099471, EPI_ISL_2099472, EPI_ISL_2099473, EPI_ISL_2099474, EPI_ISL_2099475, EPI_ISL_2099476, EPI_ISL_2099477, EPI_ISL_2099478, EPI_ISL_2099479, EPI_ISL_2099480, EPI_ISL_2099481, EPI_ISL_2099483, EPI_ISL_2099485, EPI_ISL_2099486, EPI_ISL_2099487 | | | |
| see above | National Center of Infectious and Parasitic Diseases | National Center of Infectious and Parasitic Diseases | Alexiev, Ivanov, Korsun, Stoitsova, Philipova, Dimitrova, Grigorova L., Hristova, Donchev, Stoykov, Trifonova, Dobrinov, Grigorova I., Kantardjiev |
| EPI_ISL_2100241, EPI_ISL_2100242 | Viral Respiratory Infections Laboratory, Cantacuzino National Military-Medical Institute | Cantacuzino Institute Virology | Sorin Dinu,Carmen Cherciu, Mihaela Oprea, Mihaela Lazar |
| EPI_ISL_2100243, EPI_ISL_2100244, EPI_ISL_2100245, EPI_ISL_2100246, EPI_ISL_2100247 | Clinical Hospital of Infectious and Tropical Diseases "Dr. Victor Babes" | Cantacuzino Institute Virology | Sorin Dinu,Carmen Cherciu, Mihaela Oprea, Mihaela Lazar |
| EPI_ISL_2100447, EPI_ISL_2100450, EPI_ISL_2100452, EPI_ISL_2100453, EPI_ISL_2100454, EPI_ISL_2100456, EPI_ISL_2100457, EPI_ISL_2100458, EPI_ISL_2100459, EPI_ISL_2100461, EPI_ISL_2100462, EPI_ISL_2100463, EPI_ISL_2100464, EPI_ISL_2100465, EPI_ISL_2100466, EPI_ISL_2100467, EPI_ISL_2100468, EPI_ISL_2100469, EPI_ISL_2100470, EPI_ISL_2100471, EPI_ISL_2100472, EPI_ISL_2100473 | | | |
| see above | University of Health Sciences | Quadram Institute Bioscience | Muhammad Bilal Sarwar, Sidra-tul-muntaha, Muhammad Roman, Alam Khan, Almina Shafiq, Nadeem Afzal, Shah Jahan, Javed Akram, Dave J. Baker, Gemma L. Kay, Alp Aydin, Thanh Le-Viet, Steven Rudder, Ana P. Tedim, Anastasia Kolyva, Maria Diaz, Leonardo de Oliveira Martins, Nabil-Fareed Alikhan, Lizzie Meadows, Rachael Stanley, Ngozi Elumogo, Muhammed Yasir, Nicholas M. Thomson, Alexander J Trotter, Rachel Gilroy, Samuel Bloomfield, Claire Stuart, Andrew Bell, Reenesh Prakash, Samir Dervisevic, Alison E. Mather, John Wain, Mark Webber, Andrew J. Page, Justin O'Grady |
| EPI_ISL_2101147 | Molecular Biology Laboratory, Faculty Medicine and Health Sciences, Warmadewa University | Eijkman Institute for Molecular Biology, National Agency for Research and Innovation; Molecular Biology Laboratory, Faculty Medicine and Health Sciences, Warmadewa University | Sukma Oktavianthi, Lidwina Priliani, Edison Johar, Frilasita A Yudhaputri, Muhammad Rezki Rasyak, Willy Agustine, Hidayat Trimarsanto, Lydia V. Panggalo, Iskandar Adnan, Sri Masyeni, Erly Sintya, Safarina G Malik, Khin Saw Myint, Amin Soebandrio |

| | | | |
|---|---|---|---|
| EPI_ISL_2101148 | Molecular Biology Laboratory, Faculty Medicine and Health Sciences, Warmadewa University | Eijkman Institute for Molecular Biology, National Agency for Research and Innovation; Molecular Biology Laboratory, Faculty Medicine and Health Sciences, Warmadewa University | Lidwina Priliani, Edison Johar, Frilasita A Yudhaputri, Muhammad Rezki Rasyak, Willy Agustine, Hidayat Trimarsanto, Lydia V. Panggalo, Iskandar Adnan, Sukma Oktavianthi, Sri Masyeni, Erly Sintya, Safarina G Malik, Khin Saw Myint, Amin Soebandrio |
| EPI_ISL_2101149 | Molecular Biology Laboratory, Faculty Medicine and Health Sciences, Warmadewa University | Eijkman Institute for Molecular Biology, National Agency for Research and Innovation; Molecular Biology Laboratory, Faculty Medicine and Health Sciences, Warmadewa University | Edison Johar, Frilasita A Yudhaputri, Muhammad Rezki Rasyak, Willy Agustine, Hidayat Trimarsanto, Lydia V. Panggalo, Iskandar Adnan, Sukma Oktavianthi, Lidwina Priliani, Sri Masyeni, Erly Sintya, Safarina G Malik, Khin Saw Myint, Amin Soebandrio |
| EPI_ISL_2101150 | Molecular Biology Laboratory, Faculty Medicine and Health Sciences, Warmadewa University | Eijkman Institute for Molecular Biology, National Agency for Research and Innovation; Molecular Biology Laboratory, Faculty Medicine and Health Sciences, Warmadewa University | Frilasita A Yudhaputri, Muhammad Rezki Rasyak, Willy Agustine, Hidayat Trimarsanto, Lydia V. Panggalo, Iskandar Adnan, Sukma Oktavianthi, Lidwina Priliani, Edison Johar, Sri Masyeni, Erly Sintya, Safarina G Malik, Khin Saw Myint, Amin Soebandrio |
| EPI_ISL_2101151 | Molecular Biology Laboratory, Faculty Medicine and Health Sciences, Warmadewa University | Eijkman Institute for Molecular Biology, National Agency for Research and Innovation; Molecular Biology Laboratory, Faculty Medicine and Health Sciences, Warmadewa University | Willy Agustine, Hidayat Trimarsanto, Lydia V. Panggalo, Iskandar Adnan, Sukma Oktavianthi, Lidwina Priliani, Edison Johar, Frilasita A Yudhaputri, Muhammad Rezki Rasyak, Sri Masyeni, Erly Sintya, Safarina G Malik, Khin Saw Myint, Amin Soebandrio |
| EPI_ISL_2101152 | Molecular Biology Laboratory, Faculty Medicine and Health Sciences, Warmadewa University | Eijkman Institute for Molecular Biology, National Agency for Research and Innovation; Molecular Biology Laboratory, Faculty Medicine and Health Sciences, Warmadewa University | Hidayat Trimarsanto, Lydia V. Panggalo, Iskandar Adnan, Sukma Oktavianthi, Lidwina Priliani, Edison Johar, Frilasita A Yudhaputri, Muhammad Rezki Rasyak, Willy Agustine, Sri Masyeni, Erly Sintya, Safarina G Malik, Khin Saw Myint, Amin Soebandrio |
| EPI_ISL_2101153 | Molecular Biology Laboratory, Faculty Medicine and Health Sciences, Warmadewa University | Eijkman Institute for Molecular Biology, National Agency for Research and Innovation; Molecular Biology Laboratory, Faculty Medicine and Health Sciences, Warmadewa University | Frilasita A Yudhaputri, Muhammad Rezki Rasyak, Willy Agustine, Hidayat Trimarsanto, Lydia V. Panggalo, Iskandar Adnan, Sukma Oktavianthi, Lidwina Priliani, Edison Johar, Sri Masyeni, Erly Sintya, Safarina G Malik, Khin Saw Myint, Amin Soebandrio |
| EPI_ISL_2101154 | Molecular Biology Laboratory, Faculty Medicine and Health Sciences, Warmadewa University | Eijkman Institute for Molecular Biology, National Agency for Research and Innovation; Molecular Biology Laboratory, Faculty Medicine and Health Sciences, Warmadewa University | Sukma Oktavianthi, Lidwina Priliani, Edison Johar, Frilasita A Yudhaputri, Muhammad Rezki Rasyak, Willy Agustine, Hidayat Trimarsanto, Lydia V. Panggalo, Iskandar Adnan, Sri Masyeni, Erly Sintya, Safarina G Malik, Khin Saw Myint, Amin Soebandrio |
| EPI_ISL_2101155 | Molecular Biology Laboratory, Faculty Medicine and Health Sciences, Warmadewa University | Eijkman Institute for Molecular Biology, National Agency for Research and Innovation; Molecular Biology Laboratory, Faculty Medicine and Health Sciences, Warmadewa University | Lidwina Priliani, Edison Johar, Frilasita A Yudhaputri, Muhammad Rezki Rasyak, Willy Agustine, Hidayat Trimarsanto, Lydia V. Panggalo, Iskandar Adnan, Sukma Oktavianthi, Sri Masyeni, Erly Sintya, Safarina G Malik, Khin Saw Myint, Amin Soebandrio |
| EPI_ISL_2101156 | Molecular Biology Laboratory, Faculty Medicine and Health Sciences, Warmadewa University | Eijkman Institute for Molecular Biology, National Agency for Research and Innovation; Molecular Biology Laboratory, Faculty Medicine and Health Sciences, Warmadewa University | Edison Johar, Frilasita A Yudhaputri, Muhammad Rezki Rasyak, Willy Agustine, Hidayat Trimarsanto, Lydia V. Panggalo, Iskandar Adnan, Sukma Oktavianthi, Lidwina Priliani, Sri Masyeni, Erly Sintya, Safarina G Malik, Khin Saw Myint, Amin Soebandrio |
| EPI_ISL_2101157 | Molecular Biology Laboratory, Faculty Medicine and Health Sciences, Warmadewa University | Eijkman Institute for Molecular Biology, National Agency for Research and Innovation; Molecular Biology Laboratory, Faculty Medicine and Health Sciences, Warmadewa University | Frilasita A Yudhaputri, Muhammad Rezki Rasyak, Willy Agustine, Hidayat Trimarsanto, Lydia V. Panggalo, Iskandar Adnan, Sukma Oktavianthi, Lidwina Priliani, Edison Johar, Sri Masyeni, Erly Sintya, Safarina G Malik, Khin Saw Myint, Amin Soebandrio |
| EPI_ISL_2101158 | Molecular Biology Laboratory, Faculty Medicine and Health Sciences, Warmadewa University | Eijkman Institute for Molecular Biology, National Agency for Research and Innovation; Molecular Biology Laboratory, Faculty Medicine and Health Sciences, Warmadewa University | Willy Agustine, Hidayat Trimarsanto, Lydia V. Panggalo, Iskandar Adnan, Sukma Oktavianthi, Lidwina Priliani, Edison Johar, Frilasita A Yudhaputri, Muhammad Rezki Rasyak, Sri Masyeni, Erly Sintya, Safarina G Malik, Khin Saw Myint, Amin Soebandrio |
| EPI_ISL_2101159 | Molecular Biology Laboratory, Faculty Medicine and Health Sciences, Warmadewa University | Eijkman Institute for Molecular Biology, National Agency for Research and Innovation; Molecular Biology Laboratory, Faculty Medicine and Health Sciences, Warmadewa University | Hidayat Trimarsanto, Lydia V. Panggalo, Iskandar Adnan, Sukma Oktavianthi, Lidwina Priliani, Edison Johar, Frilasita A Yudhaputri, Muhammad Rezki Rasyak, Willy Agustine, Sri Masyeni, Erly Sintya, Safarina G Malik, Khin Saw Myint, Amin Soebandrio |
| EPI_ISL_2101160 | Molecular Biology Laboratory, Faculty Medicine and Health Sciences, Warmadewa University | Eijkman Institute for Molecular Biology, National Agency for Research and Innovation; Molecular Biology Laboratory, Faculty Medicine and Health Sciences, Warmadewa University | Lydia V. Panggalo, Iskandar Adnan, Sukma Oktavianthi, Lidwina Priliani, Edison Johar, Frilasita A Yudhaputri, Muhammad Rezki Rasyak, Willy Agustine, Hidayat Trimarsanto, Sri Masyeni, Erly Sintya, Safarina G Malik, Khin Saw Myint, Amin Soebandrio |
| EPI_ISL_2101161 | Molecular Biology Laboratory, Faculty Medicine and Health Sciences, Warmadewa University | Eijkman Institute for Molecular Biology, National Agency for Research and Innovation; Molecular Biology Laboratory, Faculty Medicine and Health Sciences, Warmadewa University | Iskandar Adnan, Sukma Oktavianthi, Lidwina Priliani, Edison Johar, Frilasita A Yudhaputri, Muhammad Rezki Rasyak, Willy Agustine, Hidayat Trimarsanto, Lydia V. Panggalo, Sri Masyeni, Erly Sintya, Safarina G Malik, Khin Saw Myint, Amin Soebandrio |
| EPI_ISL_2101162 | Molecular Biology Laboratory, Faculty Medicine and Health Sciences, Warmadewa University | Eijkman Institute for Molecular Biology, National Agency for Research and Innovation; Molecular Biology Laboratory, Faculty Medicine and Health Sciences, Warmadewa University | Sukma Oktavianthi, Lidwina Priliani, Edison Johar, Frilasita A Yudhaputri, Muhammad Rezki Rasyak, Willy Agustine, Hidayat Trimarsanto, Lydia V. Panggalo, Iskandar Adnan, Sri Masyeni, Erly Sintya, Safarina G Malik, Khin Saw Myint, Amin Soebandrio |
| EPI_ISL_2101163 | Molecular Biology Laboratory, Faculty Medicine and Health Sciences, Warmadewa University | Eijkman Institute for Molecular Biology, National Agency for Research and Innovation; Molecular Biology Laboratory, Faculty Medicine and Health Sciences, Warmadewa University | Iskandar Adnan, Sukma Oktavianthi, Lidwina Priliani, Edison Johar, Frilasita A Yudhaputri, Muhammad Rezki Rasyak, Willy Agustine, Hidayat Trimarsanto, Lydia V. Panggalo, Sri Masyeni, Erly Sintya, Safarina G Malik, Khin Saw Myint, Amin Soebandrio |
| EPI_ISL_2101164 | Molecular Biology Laboratory, Faculty Medicine and Health Sciences, Warmadewa University | Eijkman Institute for Molecular Biology, National Agency for Research and Innovation; Molecular Biology Laboratory, Faculty Medicine and Health Sciences, Warmadewa University | Sukma Oktavianthi, Lidwina Priliani, Edison Johar, Frilasita A Yudhaputri, Muhammad Rezki Rasyak, Willy Agustine, Hidayat Trimarsanto, Lydia V. Panggalo, Iskandar Adnan, Sri Masyeni, Erly Sintya, Safarina G Malik, Khin Saw Myint, Amin Soebandrio |
| EPI_ISL_2101165 | Molecular Biology Laboratory, Faculty Medicine and Health Sciences, Warmadewa University | Eijkman Institute for Molecular Biology, National Agency for Research and Innovation; Molecular Biology Laboratory, Faculty Medicine and Health Sciences, Warmadewa University | Lidwina Priliani, Edison Johar, Frilasita A Yudhaputri, Muhammad Rezki Rasyak, Willy Agustine, Hidayat Trimarsanto, Lydia V. Panggalo, Iskandar Adnan, Sukma Oktavianthi, Sri Masyeni, Erly Sintya, Safarina G Malik, Khin Saw Myint, Amin Soebandrio |
| EPI_ISL_2101166 | Molecular Biology Laboratory, Faculty Medicine and Health Sciences, Warmadewa University | Eijkman Institute for Molecular Biology, National Agency for Research and Innovation; Molecular Biology Laboratory, Faculty Medicine and Health Sciences, Warmadewa University | Edison Johar, Frilasita A Yudhaputri, Muhammad Rezki Rasyak, Willy Agustine, Hidayat Trimarsanto, Lydia V. Panggalo, Iskandar Adnan, Sukma Oktavianthi, Lidwina Priliani, Sri Masyeni, Erly Sintya, Safarina G Malik, Khin Saw Myint, Amin Soebandrio |
| EPI_ISL_2101167 | Molecular Biology Laboratory, Faculty Medicine and Health | Eijkman Institute for Molecular Biology, National Agency for | Frilasita A Yudhaputri, Muhammad Rezki Rasyak, Willy Agustine, Hidayat Trimarsanto, Lydia V. Panggalo, Iskandar Adnan, Sukma Oktavianthi, Lidwina |

| | | | |
|---|---|---|---|
| | Sciences, Warmadewa University | Research and Innovation; Molecular Biology Laboratory, Faculty Medicine and Health Sciences, Warmadewa University | Priliani, Edison Johar, Sri Masyeni, Erly Sintya, Safarina G Malik, Khin Saw Myint, Amin Soebandrio |
| EPI_ISL_2101168 | Molecular Biology Laboratory, Faculty Medicine and Health Sciences, Warmadewa University | Eijkman Institute for Molecular Biology, National Agency for Research and Innovation; Molecular Biology Laboratory, Faculty Medicine and Health Sciences, Warmadewa University | Willy Agustine, Hidayat Trimarsanto, Lydia V. Panggalo, Iskandar Adnan, Sukma Oktavianthi, Lidwina Priliani, Edison Johar, Frilasita A Yudhaputri, Muhammad Rezki Rasyak, Sri Masyeni, Erly Sintya, Safarina G Malik, Khin Saw Myint, Amin Soebandrio |
| EPI_ISL_2101169 | Molecular Biology Laboratory, Faculty Medicine and Health Sciences, Warmadewa University | Eijkman Institute for Molecular Biology, National Agency for Research and Innovation; Molecular Biology Laboratory, Faculty Medicine and Health Sciences, Warmadewa University | Hidayat Trimarsanto, Lydia V. Panggalo, Iskandar Adnan, Sukma Oktavianthi, Lidwina Priliani, Edison Johar, Frilasita A Yudhaputri, Muhammad Rezki Rasyak, Willy Agustine, Sri Masyeni, Erly Sintya, Safarina G Malik, Khin Saw Myint, Amin Soebandrio |
| EPI_ISL_2101877 | LESP Chihuahua | Instituto de Diagnostico y Referencia Epidemiologicos (INDRE) | Claudia Wong-Arambula, Abril Rodriguez-Maldonado, Vanessa Rivero-Arredondo, Ariadna Medina-Benitez, Joaquin Quiroz-Mercado, Sergio Rangel-Guerrero, Natividad Cruz-Ortiz, Tatiana Nunez-Garcia, Gisela Barrera-Badillo, Lucia Hernandez-Rivas, Irma Lopez-Martinez, Ernesto Ramirez-Gonzalez. |
| EPI_ISL_2103374 | AREA DE SALUD NANDAYURE | Inciensa, Instituto Costarricense de Investigación y Enseñanza en Nutrición y Salud | Francisco Duarte, Hebleen Porras, Claudio Soto-Garita, Estela Cordero, Adriana Godínez, Melany Calderón, José Luis Vargas, Mariela Gutiérrez, Joselyn Prado, Caterina Guzmán, Nazareth Ruiz & Arnoldo Zamora-Villegas |
| EPI_ISL_2103375 | AREA DE SALUD GUATUSO | Inciensa, Instituto Costarricense de Investigación y Enseñanza en Nutrición y Salud | Francisco Duarte, Hebleen Porras, Claudio Soto-Garita, Estela Cordero, Adriana Godínez, Melany Calderón, José Luis Vargas, Mariela Gutiérrez, Joselyn Prado, Caterina Guzmán, Nazareth Ruiz & Francisco Chacón-Valverde |
| EPI_ISL_2103376 | HOSPITAL DR. WILLIAM ALLEN | Inciensa, Instituto Costarricense de Investigación y Enseñanza en Nutrición y Salud | Francisco Duarte, Hebleen Porras, Claudio Soto-Garita, Estela Cordero, Adriana Godínez, Melany Calderón, José Luis Vargas, Mariela Gutiérrez, Joselyn Prado, Caterina Guzmán, Nazareth Ruiz & Mónica Charpentier-Artavia |
| EPI_ISL_2103377 | HOSPITAL METROPOLITANO | Inciensa, Instituto Costarricense de Investigación y Enseñanza en Nutrición y Salud | Francisco Duarte, Hebleen Porras, Claudio Soto-Garita, Estela Cordero, Adriana Godínez, Melany Calderón, José Luis Vargas, Mariela Gutiérrez, Joselyn Prado, Caterina Guzmán, Nazareth Ruiz & Margarita Lee-Lui |
| EPI_ISL_2103380 | HOSPITAL DR. FERNANDO ESCALANTE PRADILLA | Inciensa, Instituto Costarricense de Investigación y Enseñanza en Nutrición y Salud | Francisco Duarte, Hebleen Porras, Claudio Soto-Garita, Estela Cordero, Adriana Godínez, Melany Calderón, José Luis Vargas, Mariela Gutiérrez, Joselyn Prado, Caterina Guzmán, Nazareth Ruiz & Lizzeth Blanco |
| EPI_ISL_2103381 | HOSPITAL DR. RAFAEL A. CALDERON GUARDIA | Inciensa, Instituto Costarricense de Investigación y Enseñanza en Nutrición y Salud | Francisco Duarte, Hebleen Porras, Claudio Soto-Garita, Estela Cordero, Adriana Godínez, Melany Calderón, José Luis Vargas, Mariela Gutiérrez, Joselyn Prado, Caterina Guzmán, Nazareth Ruiz & Fabian Salas-Flores |
| EPI_ISL_2103382 | HOSPITAL METROPOLITANO | Inciensa, Instituto Costarricense de Investigación y Enseñanza en Nutrición y Salud | Francisco Duarte, Hebleen Porras, Claudio Soto-Garita, Estela Cordero, Adriana Godínez, Melany Calderón, José Luis Vargas, Mariela Gutiérrez, Joselyn Prado, Caterina Guzmán, Nazareth Ruiz & Margarita Lee-Lui |
| EPI_ISL_2103383 | AREA DE SALUD MORAVIA | Inciensa, Instituto Costarricense de Investigación y Enseñanza en Nutrición y Salud | Francisco Duarte, Hebleen Porras, Claudio Soto-Garita, Estela Cordero, Adriana Godínez, Melany Calderón, José Luis Vargas, Mariela Gutiérrez, Caterina Guzmán, Nazareth Ruiz & Joselyn Prado |
| EPI_ISL_2103384 | AREA DE SALUD SIQUIRRES | Inciensa, Instituto Costarricense de Investigación y Enseñanza en Nutrición y Salud | Francisco Duarte, Hebleen Porras, Claudio Soto-Garita, Estela Cordero, Adriana Godínez, Melany Calderón, José Luis Vargas, Mariela Gutiérrez, Joselyn Prado, Caterina Guzmán, Nazareth Ruiz & Ileana Chaves-Peraza |
| EPI_ISL_2103385 | AREA DE SALUD OREAMUNO-PACAYAS-TIERRA BLANCA | Inciensa, Instituto Costarricense de Investigación y Enseñanza en Nutrición y Salud | Francisco Duarte, Hebleen Porras, Claudio Soto-Garita, Estela Cordero, Adriana Godínez, Melany Calderón, José Luis Vargas, Mariela Gutiérrez, Joselyn Prado, Caterina Guzmán, Nazareth Ruiz & Carolina Loría Acosta |
| EPI_ISL_2103386 | HOSPITAL METROPOLITANO | Inciensa, Instituto Costarricense de Investigación y Enseñanza en Nutrición y Salud | Francisco Duarte, Hebleen Porras, Claudio Soto-Garita, Estela Cordero, Adriana Godínez, Melany Calderón, José Luis Vargas, Mariela Gutiérrez, Joselyn Prado, Caterina Guzmán, Nazareth Ruiz & Margarita Lee-Lui |
| EPI_ISL_2103387 | HOSPITAL DR. MAX TERAN VALLS | Inciensa, Instituto Costarricense de Investigación y Enseñanza en Nutrición y Salud | Francisco Duarte, Hebleen Porras, Claudio Soto-Garita, Estela Cordero, Adriana Godínez, Melany Calderón, José Luis Vargas, Mariela Gutiérrez, Joselyn Prado, Caterina Guzmán, Nazareth Ruiz & María José Gómez-Umaña |
| EPI_ISL_2103388 | HOSPITAL MONSEÑOR SANABRIA | Inciensa, Instituto Costarricense de Investigación y Enseñanza en Nutrición y Salud | Francisco Duarte, Hebleen Porras, Claudio Soto-Garita, Estela Cordero, Adriana Godínez, Melany Calderón, José Luis Vargas, Mariela Gutiérrez, Joselyn Prado, Caterina Guzmán, Nazareth Ruiz & María José Gómez-Umaña |
| EPI_ISL_2103389, EPI_ISL_2103390 | AREA DE SALUD OROTINA-SAN MATEO | Inciensa, Instituto Costarricense de Investigación y Enseñanza en Nutrición y Salud | Francisco Duarte, Hebleen Porras, Claudio Soto-Garita, Estela Cordero, Adriana Godínez, Melany Calderón, José Luis Vargas, Mariela Gutiérrez, Joselyn Prado, Caterina Guzmán, Nazareth Ruiz & María José Gómez-Umaña |
| EPI_ISL_2103391 | HOSPITAL DR. MAX TERAN VALLS | Inciensa, Instituto Costarricense de Investigación y Enseñanza en Nutrición y Salud | Francisco Duarte, Hebleen Porras, Claudio Soto-Garita, Estela Cordero, Adriana Godínez, Melany Calderón, José Luis Vargas, Mariela Gutiérrez, Joselyn Prado, Caterina Guzmán, Nazareth Ruiz & María José Gómez-Umaña |
| EPI_ISL_2103392 | AREA DE SALUD MATA REDONDA-HOSPITAL - CLINICA DR. MORENO CAÑAS | Inciensa, Instituto Costarricense de Investigación y Enseñanza en Nutrición y Salud | Francisco Duarte, Hebleen Porras, Claudio Soto-Garita, Estela Cordero, Adriana Godínez, Melany Calderón, José Luis Vargas, Mariela Gutiérrez, Joselyn Prado, Caterina Guzmán, Nazareth Ruiz & María Gabriela Valderrama-Ordoñez |
| EPI_ISL_2103393, EPI_ISL_2103394, EPI_ISL_2103395 | AREA DE SALUD PAVAS (COOPESALUD) | Inciensa, Instituto Costarricense de Investigación y Enseñanza en Nutrición y Salud | Francisco Duarte, Hebleen Porras, Claudio Soto-Garita, Estela Cordero, Adriana Godínez, Melany Calderón, José Luis Vargas, Mariela Gutiérrez, Joselyn Prado, Caterina Guzmán, Nazareth Ruiz & Mariel López |
| EPI_ISL_2103396 | AREA DE SALUD CORONADO | Inciensa, Instituto Costarricense de Investigación y Enseñanza en Nutrición y Salud | Francisco Duarte, Hebleen Porras, Claudio Soto-Garita, Estela Cordero, Adriana Godínez, Melany Calderón, José Luis Vargas, Mariela Gutiérrez, Caterina Guzmán, Nazareth Ruiz & Joselyn Prado |
| EPI_ISL_2103397 | AREA DE SALUD ALAJUELITA | Inciensa, Instituto Costarricense de Investigación y Enseñanza en Nutrición y Salud | Francisco Duarte, Hebleen Porras, Claudio Soto-Garita, Estela Cordero, Adriana Godínez, Melany Calderón, José Luis Vargas, Mariela Gutiérrez, Joselyn Prado, Caterina Guzmán, Nazareth Ruiz & Jonathan Rojas |
| EPI_ISL_2103398 | AREA DE SALUD GOICOECHEA 2 - CLINICA DR. JIMENEZ NUÑEZ | Inciensa, Instituto Costarricense de Investigación y Enseñanza en Nutrición y Salud | Francisco Duarte, Hebleen Porras, Claudio Soto-Garita, Estela Cordero, Adriana Godínez, Melany Calderón, José Luis Vargas, Mariela Gutiérrez, Joselyn Prado, Caterina Guzmán, Nazareth Ruiz & Mariel López |
| EPI_ISL_2103399, EPI_ISL_2103400 | HOSPITAL METROPOLITANO | Inciensa, Instituto Costarricense de Investigación y Enseñanza en Nutrición y Salud | Francisco Duarte, Hebleen Porras, Claudio Soto-Garita, Estela Cordero, Adriana Godínez, Melany Calderón, José Luis Vargas, Mariela Gutiérrez, Joselyn Prado, Caterina Guzmán, Nazareth Ruiz & Margarita Lee-Lui |
| EPI_ISL_2103401 | HOSPITAL SAN VICENTE DE PAUL | Inciensa, Instituto Costarricense de Investigación y Enseñanza en Nutrición y Salud | Francisco Duarte, Hebleen Porras, Claudio Soto-Garita, Estela Cordero, Adriana Godínez, Melany Calderón, José Luis Vargas, Mariela Gutiérrez, Joselyn Prado, Caterina Guzmán, Nazareth Ruiz & Silvia Sáenz-García |
| EPI_ISL_2103402 | HOSPITAL DR. MAX TERAN VALLS | Inciensa, Instituto Costarricense de Investigación y Enseñanza en Nutrición y Salud | Francisco Duarte, Hebleen Porras, Claudio Soto-Garita, Estela Cordero, Adriana Godínez, Melany Calderón, José Luis Vargas, Mariela Gutiérrez, Joselyn Prado, Caterina Guzmán, Nazareth Ruiz & María José Gómez-Umaña |
| EPI_ISL_2103403 | AREA DE SALUD CHACARITA - CLINICA DR. FRANCISCO QUINTANA | Inciensa, Instituto Costarricense de Investigación y Enseñanza en Nutrición y Salud | Francisco Duarte, Hebleen Porras, Claudio Soto-Garita, Estela Cordero, Adriana Godínez, Melany Calderón, José Luis Vargas, Mariela Gutiérrez, Joselyn Prado, Caterina Guzmán, Nazareth Ruiz & María José Gómez-Umaña |
| EPI_ISL_2103404 | HOSPITAL DR. MAX TERAN VALLS | Inciensa, Instituto Costarricense de Investigación y Enseñanza en Nutrición y Salud | Francisco Duarte, Hebleen Porras, Claudio Soto-Garita, Estela Cordero, Adriana Godínez, Melany Calderón, José Luis Vargas, Mariela Gutiérrez, Joselyn Prado, Caterina Guzmán, Nazareth Ruiz & María José Gómez-Umaña |
| EPI_ISL_2103405 | AREA DE SALUD PAVAS (COOPESALUD) | Inciensa, Instituto Costarricense de Investigación y Enseñanza en Nutrición y Salud | Francisco Duarte, Hebleen Porras, Claudio Soto-Garita, Estela Cordero, Adriana Godínez, Melany Calderón, José Luis Vargas, Mariela Gutiérrez, Joselyn Prado, Caterina Guzmán, Nazareth Ruiz & Nicole Vargas-Víquez |
| EPI_ISL_2103406 | HOSPITAL SAN FRANCISCO DE ASIS | Inciensa, Instituto Costarricense de Investigación y Enseñanza en Nutrición y Salud | Francisco Duarte, Hebleen Porras, Claudio Soto-Garita, Estela Cordero, Adriana Godínez, Melany Calderón, José Luis Vargas, Mariela Gutiérrez, Caterina Guzmán, Nazareth Ruiz & Joselyn Prado |
| EPI_ISL_2103407 | HOSPITAL SAN VICENTE DE PAUL | Inciensa, Instituto Costarricense de Investigación y Enseñanza en Nutrición y Salud | Francisco Duarte, Hebleen Porras, Claudio Soto-Garita, Estela Cordero, Adriana Godínez, Melany Calderón, José Luis Vargas, Mariela Gutiérrez, Joselyn Prado, Caterina Guzmán, Nazareth Ruiz & Silvia Sáenz-García |
| EPI_ISL_2103408 | AREA DE SALUD SAN ISIDRO | Inciensa, Instituto Costarricense de Investigación y Enseñanza en Nutrición y Salud | Francisco Duarte, Hebleen Porras, Claudio Soto-Garita, Estela Cordero, Adriana Godínez, Melany Calderón, José Luis Vargas, Mariela Gutiérrez, Joselyn Prado, Caterina Guzmán, Nazareth Ruiz & Mariel López |

| | | | |
|---|---|---|---|
| EPI_ISL_2103409 | AREA DE SALUD FORTUNA | Inciensa, Instituto Costarricense de Investigación y Enseñanza en Nutrición y Salud | Francisco Duarte, Hebleen Porras, Claudio Soto-Garita, Estela Cordero, Adriana Godínez, Melany Calderón, José Luis Vargas, Mariela Gutiérrez, Joselyn Prado, Caterina Guzmán, Nazareth Ruiz & Carolina Arrieta Rojas |
| EPI_ISL_2103411 | HOSPITAL GOLFITO MANUEL MORA VALVERDE | Inciensa, Instituto Costarricense de Investigación y Enseñanza en Nutrición y Salud | Francisco Duarte, Hebleen Porras, Claudio Soto-Garita, Estela Cordero, Adriana Godínez, Melany Calderón, José Luis Vargas, Mariela Gutiérrez, Joselyn Prado, Caterina Guzmán, Nazareth Ruiz & Lizzeth Blanco |
| EPI_ISL_2103412 | HOSPITAL DR. FERNANDO ESCALANTE PRADILLA | Inciensa, Instituto Costarricense de Investigación y Enseñanza en Nutrición y Salud | Francisco Duarte, Hebleen Porras, Claudio Soto-Garita, Estela Cordero, Adriana Godínez, Melany Calderón, José Luis Vargas, Mariela Gutiérrez, Caterina Guzmán, Nazareth Ruiz & Joselyn Prado |
| EPI_ISL_2106191 | Dr. Gernot Walder GmbH | Dr. Gernot Walder GmbH | Sissy T. Lamprecht-Sonnleitner, Stefanie Sonnleitner |
| EPI_ISL_2106229, EPI_ISL_2106230, EPI_ISL_2106231 | National Center of Infectious and Parasitic Diseases | National Center of Infectious and Parasitic Diseases | Alexiev, Ivanov, Korsun, Stoitsova, Philipova, Dimitrova, Grigorova L., Hristova, Donchev, Stoykov, Trifonova, Dobrinov, Grigorova I., Kantardjiev |
| EPI_ISL_2107117, EPI_ISL_2107118, EPI_ISL_2107119, EPI_ISL_2107120, EPI_ISL_2107122, EPI_ISL_2107123, EPI_ISL_2107124, EPI_ISL_2107125, EPI_ISL_2107126, EPI_ISL_2107127, EPI_ISL_2107128, EPI_ISL_2107129, EPI_ISL_2107130, EPI_ISL_2107131, EPI_ISL_2107132, EPI_ISL_2107133, EPI_ISL_2107134, EPI_ISL_2107135, EPI_ISL_2107136, EPI_ISL_2107137, EPI_ISL_2107138, EPI_ISL_2107139, EPI_ISL_2107141, EPI_ISL_2107142, EPI_ISL_2107143, EPI_ISL_2107144, EPI_ISL_2107145, EPI_ISL_2107146, EPI_ISL_2107147, EPI_ISL_2107148, EPI_ISL_2107149, EPI_ISL_2107150, EPI_ISL_2107151, EPI_ISL_2107152, EPI_ISL_2107153, EPI_ISL_2107154, EPI_ISL_2107155, EPI_ISL_2107156, EPI_ISL_2107157, EPI_ISL_2107158, EPI_ISL_2107159, EPI_ISL_2107160, EPI_ISL_2107161, EPI_ISL_2107162, EPI_ISL_2107163, EPI_ISL_2107164, EPI_ISL_2107165, EPI_ISL_2107166, EPI_ISL_2107167, EPI_ISL_2107169, EPI_ISL_2107170, EPI_ISL_2107171, EPI_ISL_2107172, EPI_ISL_2107173, EPI_ISL_2107174, EPI_ISL_2107175, EPI_ISL_2107176, EPI_ISL_2107177, EPI_ISL_2107178, EPI_ISL_2107179, EPI_ISL_2107181, EPI_ISL_2107182, EPI_ISL_2107183, EPI_ISL_2107185, EPI_ISL_2107186, EPI_ISL_2107187, EPI_ISL_2107188, EPI_ISL_2107189, EPI_ISL_2107190, EPI_ISL_2107192, EPI_ISL_2107194, EPI_ISL_2107195, EPI_ISL_2107196, EPI_ISL_2107197, EPI_ISL_2107198, EPI_ISL_2107199, EPI_ISL_2107200, EPI_ISL_2107201, EPI_ISL_2107202, EPI_ISL_2107204, EPI_ISL_2107205, EPI_ISL_2107206, EPI_ISL_2107207, EPI_ISL_2107208, EPI_ISL_2107209, EPI_ISL_2107211, EPI_ISL_2107212, EPI_ISL_2107214, EPI_ISL_2107215, EPI_ISL_2107216, EPI_ISL_2107217, EPI_ISL_2107218, EPI_ISL_2107219, EPI_ISL_2107220, EPI_ISL_2107221, EPI_ISL_2107222, EPI_ISL_2107223, EPI_ISL_2107224, EPI_ISL_2107225, EPI_ISL_2107226, EPI_ISL_2107227, EPI_ISL_2107228, EPI_ISL_2107230, EPI_ISL_2107231, EPI_ISL_2107232, EPI_ISL_2107234, EPI_ISL_2107235, EPI_ISL_2107236, EPI_ISL_2107237, EPI_ISL_2107238, EPI_ISL_2107239, EPI_ISL_2107241, EPI_ISL_2107242, EPI_ISL_2107243, EPI_ISL_2107244, EPI_ISL_2107245, EPI_ISL_2107246, EPI_ISL_2107247, EPI_ISL_2107248, EPI_ISL_2107249, EPI_ISL_2107250, EPI_ISL_2107251, EPI_ISL_2107252, EPI_ISL_2107253, EPI_ISL_2107254, EPI_ISL_2107256, EPI_ISL_2107257, EPI_ISL_2107258, EPI_ISL_2107259, EPI_ISL_2107260, EPI_ISL_2107261, EPI_ISL_2107262, EPI_ISL_2107263, EPI_ISL_2107264, EPI_ISL_2107265, EPI_ISL_2107266, EPI_ISL_2107267, EPI_ISL_2107268, EPI_ISL_2107269, EPI_ISL_2107270, EPI_ISL_2107271, EPI_ISL_2107272, EPI_ISL_2107273, EPI_ISL_2107274, EPI_ISL_2107275, EPI_ISL_2107276, EPI_ISL_2107277, EPI_ISL_2107278, EPI_ISL_2107279, EPI_ISL_2107280, EPI_ISL_2107281, EPI_ISL_2107282, EPI_ISL_2107283, EPI_ISL_2107284, EPI_ISL_2107285, EPI_ISL_2107286, EPI_ISL_2107288 | | | |
| see above | Synlab Eesti OÜ | 1. Laboratory of Communicable Diseases (Estonia); 2. Eurofins Genomics Europe Sequencing GmbH | Liidia Dotsenko et al. |
| EPI_ISL_2107432, EPI_ISL_2107433 | Department of Intensive Care Medicine, and clinical lab., Osaka Women's and Children's Hospital | Department of Developmental Medicine, Research Institute, Osaka Women's and Children's Hospital | Itaru Yanagihara, Yukiko Nakura, Muneyuki Takeuchi, Shinobu Ida |
| EPI_ISL_2107437 | Dr. Yoshitaka Tamura Department of Clinical Laboratory, Osaka Habikino Medical Center | Department of Developmental Medicine, Research Institute, Osaka Women's and Children's Hospital | Itaru Yanagihara, Yukiko Nakura |
| EPI_ISL_2107438 | Dr. Yoshitaka Tamura Department of Clinical Laboratory, Osaka Habikino Medical | Department of Developmental Medicine, Research Institute, Osaka Women's and Children's Hospital | Itaru Yanagihara, Yukiko Nakura |
| EPI_ISL_2107439, EPI_ISL_2107440, EPI_ISL_2107441, EPI_ISL_2107460, EPI_ISL_2107461, EPI_ISL_2107462, EPI_ISL_2107463 | Dr. Yoshitaka Tamura Department of Clinical Laboratory, Osaka Habikino Medical Center | Department of Developmental Medicine, Research Institute, Osaka Women's and Children's Hospital | Itaru Yanagihara, Yukiko Nakura |
| EPI_ISL_2107464 | Dr. Yoshitaka Tamura Department of Clinical Laboratory, Osaka Habikino Medical Center Dr. Yoshitaka Tamura Department of Clinical Laboratory, Osaka Habikino Medical Center | Department of Developmental Medicine, Research Institute, Osaka Women's and Children's Hospital | Itaru Yanagihara, Yukiko Nakura |
| EPI_ISL_2107465, EPI_ISL_2107466, EPI_ISL_2107467, EPI_ISL_2107468 | Dr. Yoshitaka Tamura Department of Clinical Laboratory, Osaka Habikino Medical Center | Department of Developmental Medicine, Research Institute, Osaka Women's and Children's Hospital | Itaru Yanagihara, Yukiko Nakura |
| EPI_ISL_2107508, EPI_ISL_2107509, EPI_ISL_2107510, EPI_ISL_2107511, EPI_ISL_2107512 | Medical Laboratory Sciences, Arab American University | Medical Laboratory Sciences, Arab American University | Dumaidi,K., Al-Jawabreh,A., Al-Jawabreh,H., Ereqat,S., Nasereddin,A. |
| EPI_ISL_2107513, EPI_ISL_2107514, EPI_ISL_2107515, EPI_ISL_2107516, EPI_ISL_2107517, EPI_ISL_2107518, EPI_ISL_2107519 | Biochemistry and Molecular Biology Department-Faculty of Medicine, Al-Quds University | Biochemistry and Molecular Biology Department-Faculty of Medicine, Al-Quds University | Ereqat,S., Naserddin,A., Al-Jawabreh,A. |
| EPI_ISL_2107520, EPI_ISL_2107521, EPI_ISL_2107522, EPI_ISL_2107523, EPI_ISL_2107524, EPI_ISL_2107525, EPI_ISL_2107526 | Medical Laboratory Sciences, Arab American University | Medical Laboratory Sciences, Arab American University | Al-Jawabreh,A., Ereqat,S., Dumaidi,K., Al-Jawabreh,H., Nasereddin,A. |
| EPI_ISL_2107984, EPI_ISL_2113123 | Institut de Virologie du CHU de Strasbourg | Institut de Virologie du CHU de Strasbourg | Fafi-Kremer Samira, Gallais Floriane, Gantner Pierre, Laugel Elodie, Solis Morgane, Velay Aurélie, Wendling Marie-Josée |
| EPI_ISL_2131179, EPI_ISL_2131365, EPI_ISL_2131366, EPI_ISL_2131444, EPI_ISL_2131445, EPI_ISL_2131452 | AZ Klina | AZ Klina | Carl Vael - Lynsey Berckmans |
| EPI_ISL_2133385 | CHU Nantes Virology | CH La Roche sur Yon | Celine Bressollette-Bodin, Thomas Drumel, Audrey Rodallec, Berthe-Marie Imbert-Marcille, Cécile Le Boterff |
| EPI_ISL_2133386 | CHU Nantes Virology | CHU Nantes Virology | Celine Bressollette-Bodin, Thomas Drumel, Audrey Rodallec, Berthe-Marie Imbert-Marcille |
| EPI_ISL_2137037 | Institut de Virologie du CHU de Strasbourg | Institut de Virologie du CHU de Strasbourg | Fafi-Kremer Samira, Gallais Floriane, Gantner Pierre, Laugel Elodie, Solis Morgane, Velay Aurélie, Wendling Marie-Josée |
| EPI_ISL_2139910 | LESP Nuevo Leon | Instituto de Diagnostico y Referencia Epidemiologicos (INDRE) | Claudia Wong-Arambula, Abril Rodriguez-Maldonado, Vanessa Rivero-Arredondo, Ariadna Medina-Benitez, Joaquin Quiroz-Mercado, Sergio Rangel-Guerrero, Natividad Cruz-Ortiz, Tatiana Nunez-Garcia, Gisela Barrera-Badillo, Lucia Hernandez-Rivas, Irma Lopez-Martinez, Ernesto Ramirez-Gonzalez. |
| EPI_ISL_2142747 | MRCG at LSHTM Genomics lab | MRCG at LSHTM Genomics lab | Abdul Karim sesay, Abdoulie Kanteh, Jarra Manneh, Mariama Kujabi, Bakary Sanyang, Damiri Damilari, Sainabou laye Ndure |
| EPI_ISL_2151120 | Università di Parma, Laboratorio di Igiene e Sanità Pubblica | Istituto Zooprofilattico Sperimentale della Lombardia e dell'Emilia Romagna (IZSLER), Risk Analysis and Genomic Epidemiology Unit | Maria Eugenia Colucci, Licia Veronesi, Paola Affanni, Marina Morganti, Ilaria Menozzi, Erika Scaltriti, Stefano Pongolini |
| EPI_ISL_2153725, EPI_ISL_2153727, EPI_ISL_2153732, EPI_ISL_2153733, EPI_ISL_2153741, EPI_ISL_2153743, EPI_ISL_2153744, EPI_ISL_2153745, EPI_ISL_2153747, EPI_ISL_2153749, EPI_ISL_2153752, EPI_ISL_2153754, EPI_ISL_2153758, EPI_ISL_2153759, EPI_ISL_2153761, EPI_ISL_2153765, EPI_ISL_2153767, EPI_ISL_2153769, EPI_ISL_2153771, EPI_ISL_2153772, EPI_ISL_2153773, EPI_ISL_2153774, EPI_ISL_2153775, EPI_ISL_2153776, EPI_ISL_2153777, EPI_ISL_2153778, EPI_ISL_2153779, EPI_ISL_2153781, EPI_ISL_2153782, EPI_ISL_2153783, EPI_ISL_2153784, EPI_ISL_2153788, EPI_ISL_2153789, EPI_ISL_2153790, EPI_ISL_2153791, EPI_ISL_2153793, EPI_ISL_2153794, EPI_ISL_2153795, EPI_ISL_2153797, EPI_ISL_2153798, EPI_ISL_2153800, EPI_ISL_2153801, EPI_ISL_2153803, EPI_ISL_2153804, EPI_ISL_2153805 | | | |
| see above | The Ohio State University Applied Microbiology Services Laboratory | The Ohio State University Applied Microbiology Services Laboratory | Seth A. Faith PhD |
| EPI_ISL_2155037 | SYNLAB -ANGEL DIAGNOSTICA | Instituto Nacional de Salud- Dirección de Investigación en Salud Pública | Katherine Laiton-Donato, Diego A. Álvarez-Díaz, Carlos Franco-Muñoz, Hector Alejandro Ruiz-Moreno, Paola Rojas, Maria T. Herrera-Sepúlveda, Diego Andrés Prada, Jhonnatan Reales-González, Sheryll Corchuelo, Julian Naizaque, Jorge Rivera, Gerardo Santamaría, Sergio Gomez, Lisseth Pardo, Juan Camilo Martinez, Marta Lopez Blanco, Ángela Alarcon Cruz, Diana Malo, Carmen Osorio, Magdalena Wiesner, Martha Lucia Ospina Martinez, Marcela Mercado-Reyes |
| EPI_ISL_2157318 | LESP Chihuahua | Instituto de Diagnostico y Referencia Epidemiologicos (INDRE) | Claudia Wong-Arambula, Abril Rodriguez-Maldonado, Vanessa Rivero-Arredondo, Ariadna Medina-Benitez, Joaquin Quiroz-Mercado, Sergio Rangel-Guerrero, Natividad Cruz-Ortiz, Tatiana Nunez-Garcia, Gisela Barrera-Badillo, Lucia Hernandez-Rivas, Irma Lopez-Martinez, Ernesto Ramirez-Gonzalez. |

| | | | |
|---|---|---|---|
| EPI_ISL_2157516, EPI_ISL_2157517 | Laboratório Central de Saude Publica do Estado do Paraná (LACEN/PR) | Laboratory of Respiratory Viruses and Measles, Oswaldo Cruz Institute, FIOCRUZ | Paola Resende, Luciana Appolinario, Fernando Motta, Anna Carolina Paixao, Ana Carolina Mendonca, Alice Sampaio Rocha, Renata Serrano Lopes, Maria do Carmo Debur, Taina Venas, Elisa Cavalcante Pereira, Irina Nastassja Riediger, Marilda Siqueira on behalf of the Fiocruz COVID-19 Genomic Surveillance Network |
| EPI_ISL_2157527 | Lboratorio Central de Saude Publica do Estado do Parana (LACEN/PR) | Laboratory of Respiratory Viruses and Measles, Oswaldo Cruz Institute, FIOCRUZ | Paola Resende, Luciana Appolinario, Fernando Motta, Anna Carolina Paixao, Ana Carolina Mendonca, Alice Sampaio Rocha, Taina Venas, Elisa Cavalcante Pereira, Renata Serrano Lopes, Irina Riediger, Marilda Siqueira on behalf of the Fiocruz COVID-19 Genomic Surveillance Network |
| EPI_ISL_2158136 | LESP Chihuahua | Instituto de Diagnostico y Referencia Epidemiologicos (INDRE) | Claudia Wong-Arambula, Abril Rodriguez-Maldonado, Vanessa Rivero-Arredondo, Ariadna Medina-Benitez, Joaquin Quiroz-Mercado, Sergio Rangel-Guerrero, Natividad Cruz-Ortiz, Tatiana Nunez-Garcia, Gisela Barrera-Badillo, Lucia Hernandez-Rivas, Irma Lopez-Martinez, Ernesto Ramirez-Gonzalez. |
| EPI_ISL_2158242 | LESP Queretaro /BIMODI | Instituto de Diagnostico y Referencia Epidemiologicos (INDRE) | Claudia Wong-Arambula, Abril Rodriguez-Maldonado, Vanessa Rivero-Arredondo, Ariadna Medina-Benitez, Joaquin Quiroz-Mercado, Sergio Rangel-Guerrero, Natividad Cruz-Ortiz, Tatiana Nunez-Garcia, Gisela Barrera-Badillo, Lucia Hernandez-Rivas, Irma Lopez-Martinez, Ernesto Ramirez-Gonzalez. |
| EPI_ISL_2161783, EPI_ISL_2161784, EPI_ISL_2161785, EPI_ISL_2161786, EPI_ISL_2161787 | NL-Dr. Leonard A. Miller Centre for Health Services | National Microbiology Laboratory (NML) | Anna Majer, Shari Tyson, Grace Seo, Philip Mabon, Elsie Grudeski, Rhiannon Huzarewich, Russell Mandes, Anneliese Landgraff, Jennifer Tanner, Natalie Knox, Morag Graham, Gary Van Domselaar, Robert Needle, Yang Yu, Adel Malek, Laura Gilbert, George Zahariadis, Nathalie Bastien, Yan Li, Timothy Booth, Darian Hole, Madison Chapel, Kirsten Biggar, Kerri Smith, CanCOGeN's metadata curation team, Public Health Agency of Canada CanCOGeN team |
| EPI_ISL_2162810 | Klinisch Laboratorium ZNA | Klinisch Laboratorium ZNA | Verstrepen et al. |
| EPI_ISL_2171268, EPI_ISL_2171607 | Institut de Virologie du CHU de Strasbourg | Institut de Virologie du CHU de Strasbourg | Fafi-Kremer Samira, Gallais Floriane, Gantner Pierre, Laugel Elodie, Solis Morgane, Velay Aurélie, Wendling Marie-Josée |
| EPI_ISL_2178413 | Hospital | National Reference Center for Viruses of Respiratory Infections, Institut Pasteur, Paris | Marion Barbet, Sylvie Behillil, Méline Bizard, Angela Brisebarre, Camille Capel, Vincent Enouf, Louise Lefrançois, Frédéric Lemoine, Christophe Malabat, Corinne Maufrais, Amaury Vaysse, Etienne Simon-Lorière, Maud Vanpeene, Sylvie Van der Werf ,LéA Pilorge |
| EPI_ISL_2178418 | Hospital | National Reference Center for Viruses of Respiratory Infections, Institut Pasteur, Paris | Marion Barbet, Sylvie Behillil, Méline Bizard, Angela Brisebarre, Camille Capel, Vincent Enouf, Louise Lefrançois, Frédéric Lemoine, Christophe Malabat, Corinne Maufrais, Etienne Simon-Lorière, Maud Vanpeene, Sylvie Van der Werf ,Odile Petsaris |
| EPI_ISL_2178420 | Hospital | National Reference Center for Viruses of Respiratory Infections, Institut Pasteur, Paris | Marion Barbet, Sylvie Behillil, Méline Bizard, Angela Brisebarre, Camille Capel, Vincent Enouf, Louise Lefrançois, Frédéric Lemoine, Christophe Malabat, Corinne Maufrais, Amaury Vaysse, Etienne Simon-Lorière, Maud Vanpeene, Sylvie Van der Werf ,LéA Pilorge |
| EPI_ISL_2178425, EPI_ISL_2178429 | Research | National Reference Center for Viruses of Respiratory Infections, Institut Pasteur, Paris | Marion Barbet, Sylvie Behillil, Méline Bizard, Angela Brisebarre, Camille Capel, Vincent Enouf, Louise Lefrançois, Frédéric Lemoine, Christophe Malabat, Corinne Maufrais, Amaury Vaysse, Etienne Simon-Lorière, Maud Vanpeene, Sylvie Van der Werf ,Marianne Leruez-Ville |
| EPI_ISL_2178432 | Labo Analyses Med | National Reference Center for Viruses of Respiratory Infections, Institut Pasteur, Paris | Marion Barbet, Sylvie Behillil, Méline Bizard, Angela Brisebarre, Camille Capel, Vincent Enouf, Louise Lefrançois, Frédéric Lemoine, Christophe Malabat, Corinne Maufrais, Amaury Vaysse, Etienne Simon-Lorière, Maud Vanpeene, Sylvie Van der Werf ,Odile Petsaris |
| EPI_ISL_2178439 | Hospital | National Reference Center for Viruses of Respiratory Infections, Institut Pasteur, Paris | Marion Barbet, Sylvie Behillil, Méline Bizard, Angela Brisebarre, Camille Capel, Vincent Enouf, Louise Lefrançois, Frédéric Lemoine, Christophe Malabat, Corinne Maufrais, Damien Mornico, Etienne Simon-Lorière, Maud Vanpeene, Sylvie Van der Werf ,Odile Petsaris |
| EPI_ISL_2178440, EPI_ISL_2178441 | Hospital | National Reference Center for Viruses of Respiratory Infections, Institut Pasteur, Paris | Marion Barbet, Sylvie Behillil, Méline Bizard, Angela Brisebarre, Camille Capel, Vincent Enouf, Louise Lefrançois, Frédéric Lemoine, Christophe Malabat, Corinne Maufrais, Adrien Pain, Etienne Simon-Lorière, Maud Vanpeene, Sylvie Van der Werf ,JéRome Guinard |
| EPI_ISL_2178445 | Hospital | National Reference Center for Viruses of Respiratory Infections, Institut Pasteur, Paris | Marion Barbet, Sylvie Behillil, Méline Bizard, Angela Brisebarre, Camille Capel, Vincent Enouf, Louise Lefrançois, Frédéric Lemoine, Christophe Malabat, Corinne Maufrais, Emmanuelle Permal, Etienne Simon-Lorière, Maud Vanpeene, Sylvie Van der Werf ,Raphael Guiheneuf |
| EPI_ISL_2178476 | Hospital | National Reference Center for Viruses of Respiratory Infections, Institut Pasteur, Paris | Marion Barbet, Sylvie Behillil, Méline Bizard, Angela Brisebarre, Camille Capel, Vincent Enouf, Louise Lefrançois, Frédéric Lemoine, Christophe Malabat, Corinne Maufrais, Victoire Baillet, Etienne Simon-Lorière, Maud Vanpeene, Sylvie Van der Werf ,Hery-Zo Rakotomalala |
| EPI_ISL_2178725 | Centro de Investigación Biomédica del Noreste (CIBIN) | Centro de Investigación en Enfermedades Infecciosas (CIENI), Instituto Nacional de Enfermedades Respiratorias (INER) | Consorcio Mexicano de Vigilancia Genómica (CoViGen-Mex). Authors (in alphabetical order): Julio Elias Alvarado-Yaah, Carlos F. Arias, Santiago Ávila-Ríos, Víctor Hugo Borja-Aburto, Celia Boukadida, Juan Bautista Chale-Dzul, Célida Duque Molina, José Antonio Enciso-Moreno, Gloria Elena Espinosa-Ayala, Fernando Fontove-Herrera, Victor Eduardo Garcia-Arias, Concepción Grajales-Muñiz, Ricardo Grande, Alfredo Herrera-Estrella, Carla Ivón Herrera-Najera, Pavel Isa, Brenda Irasema Maldonado-Meza, Bernardo Martínez-Miguel, Margarita Matías-Florentino, María Guadalupe de Jesús Mireles-Rivera, Gloria María Molina-Salinas, Hector Montoya-Fuentes, José Esteban Muñoz-Medina, José de Jesús Nuñez-Contreras, Alicia Ocaña-Mondragón, Luis Alberto Ochoa-Carrera, Hector Esteban Paz-Juárez, Francisco Pulido, Helen Haydee Fernanda Ramírez-Plascencia, Angel Gustavo Salas-Lais, Alejandro Sanchez-Flores, Clara Esperanza Santacruz-Tinoco, María Guadalupe Santiago-Mauricio, Nelly Sélem-Mojica, Blanca Taboada, Gloria Vazquez |
| EPI_ISL_2178727 | Unidad de Investigación Médica de Yucatán (UIMY) | Centro de Investigación en Enfermedades Infecciosas (CIENI), Instituto Nacional de Enfermedades Respiratorias (INER) | Consorcio Mexicano de Vigilancia Genómica (CoViGen-Mex). Authors (in alphabetical order): Julio Elias Alvarado-Yaah, Carlos F. Arias, Santiago Ávila-Ríos, Víctor Hugo Borja-Aburto, Celia Boukadida, Juan Bautista Chale-Dzul, Célida Duque Molina, José Antonio Enciso-Moreno, Gloria Elena Espinosa-Ayala, Fernando Fontove-Herrera, Victor Eduardo Garcia-Arias, Concepción Grajales-Muñiz, Ricardo Grande, Alfredo Herrera-Estrella, Carla Ivón Herrera-Najera, Pavel Isa, Brenda Irasema Maldonado-Meza, Bernardo Martínez-Miguel, Margarita Matías-Florentino, María Guadalupe de Jesús Mireles-Rivera, Gloria María Molina-Salinas, Hector Montoya-Fuentes, José Esteban Muñoz-Medina, José de Jesús Nuñez-Contreras, Alicia Ocaña-Mondragón, Luis Alberto Ochoa-Carrera, Hector Esteban Paz-Juárez, Francisco Pulido, Helen Haydee Fernanda Ramírez-Plascencia, Angel Gustavo Salas-Lais, Alejandro Sanchez-Flores, Clara Esperanza Santacruz-Tinoco, María Guadalupe Santiago-Mauricio, Nelly Sélem-Mojica, Blanca Taboada, Gloria Vazquez |
| EPI_ISL_2178729 | Centro de Investigación Biomédica de Occidente (CIBO) | Centro de Investigación en Enfermedades Infecciosas (CIENI), Instituto Nacional de Enfermedades Respiratorias (INER) | Consorcio Mexicano de Vigilancia Genómica (CoViGen-Mex). Authors (in alphabetical order): Julio Elias Alvarado-Yaah, Carlos F. Arias, Santiago Ávila-Ríos, Víctor Hugo Borja-Aburto, Celia Boukadida, Juan Bautista Chale-Dzul, Célida Duque Molina, José Antonio Enciso-Moreno, Gloria Elena Espinosa-Ayala, Fernando Fontove-Herrera, Victor Eduardo Garcia-Arias, Concepción Grajales-Muñiz, Ricardo Grande, Alfredo Herrera-Estrella, Carla Ivón Herrera-Najera, Pavel Isa, Brenda Irasema Maldonado-Meza, Bernardo Martínez-Miguel, Margarita Matías-Florentino, María Guadalupe de Jesús Mireles-Rivera, Gloria María Molina-Salinas, Hector Montoya-Fuentes, José Esteban Muñoz-Medina, José de Jesús Nuñez-Contreras, Alicia Ocaña-Mondragón, Luis Alberto Ochoa-Carrera, Hector Esteban Paz-Juárez, Francisco Pulido, Helen Haydee Fernanda Ramírez-Plascencia, Angel Gustavo Salas-Lais, Alejandro Sanchez-Flores, Clara Esperanza Santacruz-Tinoco, María Guadalupe Santiago-Mauricio, Nelly Sélem-Mojica, Blanca Taboada, Gloria Vazquez |
| EPI_ISL_2178730, EPI_ISL_2178731, EPI_ISL_2178732 | Laboratorio Central de Epidemiología (LCE) | Centro de Investigación en Enfermedades Infecciosas (CIENI), Instituto Nacional de Enfermedades Respiratorias (INER) | Consorcio Mexicano de Vigilancia Genómica (CoViGen-Mex). Authors (in alphabetical order): Julio Elias Alvarado-Yaah, Carlos F. Arias, Santiago Ávila-Ríos, Víctor Hugo Borja-Aburto, Celia Boukadida, Juan Bautista Chale-Dzul, Célida Duque Molina, José Antonio Enciso-Moreno, Gloria Elena Espinosa-Ayala, Fernando Fontove-Herrera, Victor Eduardo Garcia-Arias, Concepción Grajales-Muñiz, Ricardo Grande, Alfredo Herrera-Estrella, Carla Ivón Herrera-Najera, Pavel Isa, Brenda Irasema Maldonado-Meza, Bernardo Martínez-Miguel, Margarita Matías-Florentino, María Guadalupe de Jesús Mireles-Rivera, Gloria María Molina-Salinas, Hector Montoya-Fuentes, José Esteban Muñoz-Medina, José de Jesús Nuñez-Contreras, Alicia Ocaña-Mondragón, Luis Alberto Ochoa-Carrera, Hector Esteban Paz-Juárez, Francisco Pulido, Helen Haydee Fernanda Ramírez-Plascencia, Angel Gustavo Salas-Lais, Alejandro Sanchez-Flores, Clara Esperanza Santacruz-Tinoco, María Guadalupe Santiago-Mauricio, Nelly Sélem-Mojica, Blanca Taboada, Gloria Vazquez |
| EPI_ISL_2179375, EPI_ISL_2179381, EPI_ISL_2179383, EPI_ISL_2179386, EPI_ISL_2179387, EPI_ISL_2179388 | GH A.CHENEVIER-H.MONDOR | Department of Virology, Henri Mondor University Hospital, Assistance Publique Hôpitaux de Paris, Université Paris-Est Créteil, INSERM U955 | Christophe Rodriguez, Slim Fourati, Vanessa Demontant, Guillaume Gricourt, Melissa N'Debi, Alexandre Soulier, Elisabeth Trawinski, Jean-Michel Pawlotsky |
| EPI_ISL_2179389 | GH JOFFRE DUPUYTREN | Department of Virology, Henri Mondor University Hospital, Assistance Publique Hôpitaux de Paris, Université Paris-Est Créteil, INSERM U955 | Christophe Rodriguez, Slim Fourati, Vanessa Demontant, Guillaume Gricourt, Melissa N'Debi, Alexandre Soulier, Elisabeth Trawinski, Jean-Michel Pawlotsky |
| EPI_ISL_2179396, EPI_ISL_2179401, EPI_ISL_2179402, EPI_ISL_2179414, EPI_ISL_2179449, EPI_ISL_2179450, | GH A.CHENEVIER-H.MONDOR | Department of Virology, Henri Mondor University Hospital, Assistance Publique Hôpitaux de Paris, Université Paris-Est Créteil, INSERM U955 | Christophe Rodriguez, Slim Fourati, Vanessa Demontant, Guillaume Gricourt, Melissa N'Debi, Alexandre Soulier, Elisabeth Trawinski, Jean-Michel Pawlotsky |

| | | | |
|---|---|---|---|
| EPI_ISL_2179452 | | | |
| EPI_ISL_2179456 | GH JOFFRE DUPUYTREN | Department of Virology, Henri Mondor University Hospital, Assistance Publique Hôpitaux de Paris, Université Paris-Est Créteil, INSERM U955 | Christophe Rodriguez, Slim Fourati, Vanessa Demontant, Guillaume Gricourt, Melissa N'Debi, Alexandre Soulier, Elisabeth Trawinski, Jean-Michel Pawlotsky |
| EPI_ISL_2179463 | HOPITAL EMILE ROUX | Department of Virology, Henri Mondor University Hospital, Assistance Publique Hôpitaux de Paris, Université Paris-Est Créteil, INSERM U955 | Christophe Rodriguez, Slim Fourati, Vanessa Demontant, Guillaume Gricourt, Melissa N'Debi, Alexandre Soulier, Elisabeth Trawinski, Jean-Michel Pawlotsky |
| EPI_ISL_2179467 | GH A.CHENEVIER-H.MONDOR | Department of Virology, Henri Mondor University Hospital, Assistance Publique Hôpitaux de Paris, Université Paris-Est Créteil, INSERM U955 | Christophe Rodriguez, Slim Fourati, Vanessa Demontant, Guillaume Gricourt, Melissa N'Debi, Alexandre Soulier, Elisabeth Trawinski, Jean-Michel Pawlotsky |
| EPI_ISL_2179468 | HOPITAL EMILE ROUX | Department of Virology, Henri Mondor University Hospital, Assistance Publique Hôpitaux de Paris, Université Paris-Est Créteil, INSERM U955 | Christophe Rodriguez, Slim Fourati, Vanessa Demontant, Guillaume Gricourt, Melissa N'Debi, Alexandre Soulier, Elisabeth Trawinski, Jean-Michel Pawlotsky |
| EPI_ISL_2179473, EPI_ISL_2179475, EPI_ISL_2179484, EPI_ISL_2179485, EPI_ISL_2179486, EPI_ISL_2179487, EPI_ISL_2179615 | GH A.CHENEVIER-H.MONDOR | Department of Virology, Henri Mondor University Hospital, Assistance Publique Hôpitaux de Paris, Université Paris-Est Créteil, INSERM U955 | Christophe Rodriguez, Slim Fourati, Vanessa Demontant, Guillaume Gricourt, Melissa N'Debi, Alexandre Soulier, Elisabeth Trawinski, Jean-Michel Pawlotsky |
| EPI_ISL_2179616 | GH JOFFRE DUPUYTREN | Department of Virology, Henri Mondor University Hospital, Assistance Publique Hôpitaux de Paris, Université Paris-Est Créteil, INSERM U955 | Christophe Rodriguez, Slim Fourati, Vanessa Demontant, Guillaume Gricourt, Melissa N'Debi, Alexandre Soulier, Elisabeth Trawinski, Jean-Michel Pawlotsky |
| EPI_ISL_2179620 | GH A.CHENEVIER-H.MONDOR | Department of Virology, Henri Mondor University Hospital, Assistance Publique Hôpitaux de Paris, Université Paris-Est Créteil, INSERM U955 | Christophe Rodriguez, Slim Fourati, Vanessa Demontant, Guillaume Gricourt, Melissa N'Debi, Alexandre Soulier, Elisabeth Trawinski, Jean-Michel Pawlotsky |
| EPI_ISL_2179622 | HOPITAL EMILE ROUX | Department of Virology, Henri Mondor University Hospital, Assistance Publique Hôpitaux de Paris, Université Paris-Est Créteil, INSERM U955 | Christophe Rodriguez, Slim Fourati, Vanessa Demontant, Guillaume Gricourt, Melissa N'Debi, Alexandre Soulier, Elisabeth Trawinski, Jean-Michel Pawlotsky |
| EPI_ISL_2179625, EPI_ISL_2179626 | GH A.CHENEVIER-H.MONDOR | Department of Virology, Henri Mondor University Hospital, Assistance Publique Hôpitaux de Paris, Université Paris-Est Créteil, INSERM U955 | Christophe Rodriguez, Slim Fourati, Vanessa Demontant, Guillaume Gricourt, Melissa N'Debi, Alexandre Soulier, Elisabeth Trawinski, Jean-Michel Pawlotsky |
| EPI_ISL_2180057, EPI_ISL_2180058, EPI_ISL_2180059, EPI_ISL_2180060, EPI_ISL_2180061, EPI_ISL_2180062, EPI_ISL_2180064, EPI_ISL_2180065, EPI_ISL_2180066, EPI_ISL_2180067, EPI_ISL_2180068, EPI_ISL_2180069, EPI_ISL_2180070, EPI_ISL_2180071, EPI_ISL_2180072, EPI_ISL_2180073, EPI_ISL_2180074, EPI_ISL_2180075, EPI_ISL_2180076, EPI_ISL_2180078, EPI_ISL_2180079, EPI_ISL_2180080, EPI_ISL_2180081, EPI_ISL_2180082, EPI_ISL_2180083, EPI_ISL_2180084, EPI_ISL_2180085, EPI_ISL_2180086, EPI_ISL_2180087, EPI_ISL_2180088, EPI_ISL_2180089, EPI_ISL_2180090, EPI_ISL_2180091, EPI_ISL_2180092, EPI_ISL_2180093, EPI_ISL_2180094, EPI_ISL_2180096, EPI_ISL_2180097, EPI_ISL_2180098, EPI_ISL_2180099, EPI_ISL_2180100, EPI_ISL_2180101, EPI_ISL_2180102, EPI_ISL_2180103, EPI_ISL_2180104, EPI_ISL_2180106, EPI_ISL_2180107, EPI_ISL_2180108, EPI_ISL_2180109, EPI_ISL_2180110, EPI_ISL_2180111, EPI_ISL_2180112, EPI_ISL_2180113, EPI_ISL_2180114, EPI_ISL_2180115, EPI_ISL_2180116, EPI_ISL_2180117, EPI_ISL_2180118, EPI_ISL_2180119, EPI_ISL_2180120, EPI_ISL_2180121, EPI_ISL_2180122, EPI_ISL_2180123, EPI_ISL_2180124, EPI_ISL_2180126, EPI_ISL_2180127, EPI_ISL_2180128, EPI_ISL_2180129, EPI_ISL_2180130, EPI_ISL_2180131, EPI_ISL_2180132, EPI_ISL_2180133, EPI_ISL_2180135, EPI_ISL_2180136, EPI_ISL_2180137, EPI_ISL_2180139, EPI_ISL_2180140, EPI_ISL_2180141, EPI_ISL_2180142, EPI_ISL_2180143, EPI_ISL_2180145, EPI_ISL_2180146, EPI_ISL_2180147, EPI_ISL_2180148, EPI_ISL_2180149, EPI_ISL_2180153, EPI_ISL_2180155, EPI_ISL_2180156, EPI_ISL_2180158, EPI_ISL_2180159, EPI_ISL_2180161, EPI_ISL_2180162, EPI_ISL_2180164, EPI_ISL_2180165, EPI_ISL_2180166, EPI_ISL_2180167, EPI_ISL_2180168, EPI_ISL_2180169, EPI_ISL_2180170, EPI_ISL_2180171, EPI_ISL_2180172, EPI_ISL_2180173, EPI_ISL_2180174, EPI_ISL_2180176, EPI_ISL_2180179, EPI_ISL_2180180, EPI_ISL_2180182, EPI_ISL_2180183, EPI_ISL_2180184, EPI_ISL_2180185, EPI_ISL_2180186, EPI_ISL_2180187, EPI_ISL_2180189, EPI_ISL_2180190, EPI_ISL_2180191, EPI_ISL_2180192, EPI_ISL_2180193, EPI_ISL_2180194, EPI_ISL_2180196, EPI_ISL_2180197, EPI_ISL_2180198, EPI_ISL_2180199, EPI_ISL_2180200, EPI_ISL_2180201, EPI_ISL_2180202, EPI_ISL_2180203, EPI_ISL_2180205, EPI_ISL_2180206, EPI_ISL_2180207, EPI_ISL_2180208, EPI_ISL_2180209, EPI_ISL_2180212, EPI_ISL_2180213, EPI_ISL_2180214, EPI_ISL_2180215, EPI_ISL_2180216, EPI_ISL_2180217, EPI_ISL_2180218, EPI_ISL_2180219, EPI_ISL_2180220, EPI_ISL_2180221, EPI_ISL_2180222, EPI_ISL_2180223, EPI_ISL_2180224, EPI_ISL_2180225, EPI_ISL_2180226, EPI_ISL_2180227, EPI_ISL_2180228, EPI_ISL_2180229, EPI_ISL_2180230, EPI_ISL_2180231, EPI_ISL_2180232, EPI_ISL_2180235, EPI_ISL_2180236, EPI_ISL_2180240, EPI_ISL_2180241, EPI_ISL_2180242, EPI_ISL_2180243, EPI_ISL_2180244, EPI_ISL_2180247, EPI_ISL_2180248, EPI_ISL_2180249, EPI_ISL_2180250, EPI_ISL_2180251, EPI_ISL_2180253, EPI_ISL_2180254, EPI_ISL_2180257, EPI_ISL_2180259, EPI_ISL_2180260, EPI_ISL_2180262, EPI_ISL_2180263, EPI_ISL_2180264, EPI_ISL_2180265, EPI_ISL_2180266, EPI_ISL_2180267, EPI_ISL_2180268, EPI_ISL_2180269, EPI_ISL_2180270, EPI_ISL_2180271, EPI_ISL_2180272, EPI_ISL_2180273, EPI_ISL_2180274, EPI_ISL_2180275, EPI_ISL_2180277, EPI_ISL_2180278, EPI_ISL_2180280, EPI_ISL_2180282, EPI_ISL_2180283, EPI_ISL_2180284, EPI_ISL_2180285, EPI_ISL_2180286, EPI_ISL_2180287, EPI_ISL_2180291, EPI_ISL_2180292, EPI_ISL_2180293, EPI_ISL_2180297, EPI_ISL_2180298, EPI_ISL_2180299, EPI_ISL_2180301, EPI_ISL_2180304, EPI_ISL_2180309, EPI_ISL_2180310, EPI_ISL_2180338, EPI_ISL_2180339, EPI_ISL_2180340, EPI_ISL_2180341, EPI_ISL_2180342, EPI_ISL_2180343, EPI_ISL_2180344, EPI_ISL_2180345, EPI_ISL_2180347, EPI_ISL_2180348, EPI_ISL_2180349, EPI_ISL_2180351, EPI_ISL_2180352, EPI_ISL_2180353, EPI_ISL_2180354, EPI_ISL_2180355, EPI_ISL_2180356, EPI_ISL_2180357, EPI_ISL_2180358, EPI_ISL_2180359, EPI_ISL_2180360, EPI_ISL_2180361, EPI_ISL_2180362, EPI_ISL_2180363, EPI_ISL_2180364, EPI_ISL_2180365, EPI_ISL_2180366, EPI_ISL_2180367, EPI_ISL_2180369, EPI_ISL_2180370 | | | |
| see above | National Center of Infectious and Parasitic Diseases | National Center of Infectious and Parasitic Diseases | Alexiev et al |
| EPI_ISL_2188485 | Labo Analyses Med | National Reference Center for Viruses of Respiratory Infections, Institut Pasteur, Paris | Marion Barbet, Sylvie Behillil, Méline Bizard, Angela Brisebarre, Camille Capel, Vincent Enouf, Louise Lefrançois, Frédéric Lemoine, Christophe Malabat, Corinne Maufrais, Gael Millot, Etienne Simon-Lorière, Maud Vanpeene, Sylvie Van der Werf ,Adamou Lagare |
| EPI_ISL_2188869 | The Lord's Grace Medical and Industrial Clinic | Philippine Genome Center | Francis A. Tablizo, Kenneth M. Kim, Maria Sofia L. Yangzon, Elcid Aaron R. Pangilinan, Renato Jacinto Q. Mantaring, Benedict A. Maralit, Marc Edsel C. Ayes, Eva Maria Cutiongco-de la Paz, Alethea R. de Guzman, Jan Michael C. Yap, Jo-Hannah S. Llames, Sheila Mae M. Araiza, Kris P. Punayan, Irish Coleen A. Asin, Candice Francheska B. Tambaoan, Asia Louisa U. Chong, Karol Sophia Agape R. Padilla, Rianna Patricia S. Cruz, Carlo M. Lapid, El King D. Morado, Joshua Gregor A. Dizon, Razel Nikka M. Hao, Arianne A. Zamora, Devon Ray Pacial, Juan Antonio R. Magalang, Marissa Alejandria, Celia Carlos, Anna Ong-Lim, Edsel Maurice Salvaña, John Q. Wong, Jaime C. Montoya, Maria Rosario Singh-Vergeire and Cynthia P. Saloma |
| EPI_ISL_2188874 | Pasig City Children's Hospital - Child's Hope | Philippine Genome Center | Francis A. Tablizo, Kenneth M. Kim, Maria Sofia L. Yangzon, Elcid Aaron R. Pangilinan, Renato Jacinto Q. Mantaring, Benedict A. Maralit, Marc Edsel C. Ayes, Eva Maria Cutiongco-de la Paz, Alethea R. de Guzman, Jan Michael C. Yap, Jo-Hannah S. Llames, Sheila Mae M. Araiza, Kris P. Punayan, Irish Coleen A. Asin, Candice Francheska B. Tambaoan, Asia Louisa U. Chong, Karol Sophia Agape R. Padilla, Rianna Patricia S. Cruz, Carlo M. Lapid, El King D. Morado, Joshua Gregor A. Dizon, Razel Nikka M. Hao, Arianne A. Zamora, Devon Ray Pacial, Juan Antonio R. Magalang, Marissa Alejandria, Celia Carlos, Anna Ong-Lim, Edsel Maurice Salvaña, John Q. Wong, Jaime C. Montoya, Maria Rosario Singh-Vergeire and Cynthia P. Saloma |
| EPI_ISL_2188875 | Sta. Rosa CHO- GENEXPERT LABORATORY | Philippine Genome Center | Francis A. Tablizo, Kenneth M. Kim, Maria Sofia L. Yangzon, Elcid Aaron R. Pangilinan, Renato Jacinto Q. Mantaring, Benedict A. Maralit, Marc Edsel C. Ayes, Eva Maria Cutiongco-de la Paz, Alethea R. de Guzman, Jan Michael C. Yap, Jo-Hannah S. Llames, Sheila Mae M. Araiza, Kris P. Punayan, Irish Coleen A. Asin, Candice Francheska B. Tambaoan, Asia Louisa U. Chong, Karol Sophia Agape R. Padilla, Rianna Patricia S. Cruz, Carlo M. Lapid, El King D. Morado, Joshua Gregor A. Dizon, Razel Nikka M. Hao, Arianne A. Zamora, Devon Ray Pacial, Juan Antonio R. Magalang, Marissa Alejandria, Celia Carlos, Anna Ong-Lim, Edsel Maurice Salvaña, John Q. Wong, Jaime C. Montoya, Maria Rosario Singh-Vergeire and Cynthia P. Saloma |
| EPI_ISL_2188876 | Qualimed Hospital Sta. Rosa | Philippine Genome Center | Francis A. Tablizo, Kenneth M. Kim, Maria Sofia L. Yangzon, Elcid Aaron R. Pangilinan, Renato Jacinto Q. Mantaring, Benedict A. Maralit, Marc Edsel C. Ayes, Eva Maria Cutiongco-de la Paz, Alethea R. de Guzman, Jan Michael C. Yap, Jo-Hannah S. Llames, Sheila Mae M. Araiza, Kris P. Punayan, Irish Coleen A. Asin, Candice Francheska B. Tambaoan, Asia Louisa U. Chong, Karol Sophia Agape R. Padilla, Rianna Patricia S. Cruz, Carlo M. Lapid, El King D. Morado, Joshua Gregor A. Dizon, Razel Nikka M. Hao, Arianne A. Zamora, Devon Ray Pacial, Juan Antonio R. Magalang, Marissa Alejandria, Celia Carlos, Anna Ong-Lim, Edsel Maurice Salvaña, John Q. Wong, Jaime C. Montoya, Maria Rosario Singh-Vergeire and Cynthia P. Saloma |
| EPI_ISL_2188877 | Pasig City Children's Hospital - Child's Hope | Philippine Genome Center | Francis A. Tablizo, Kenneth M. Kim, Maria Sofia L. Yangzon, Elcid Aaron R. Pangilinan, Renato Jacinto Q. Mantaring, Benedict A. Maralit, Marc Edsel C. Ayes, Eva Maria Cutiongco-de la Paz, Alethea R. de Guzman, Jan Michael C. Yap, Jo-Hannah S. Llames, Sheila Mae M. Araiza, Kris P. Punayan, Irish Coleen A. Asin, Candice Francheska B. Tambaoan, Asia Louisa U. Chong, Karol Sophia Agape R. Padilla, Rianna Patricia S. Cruz, Carlo M. Lapid, El King D. Morado, Joshua Gregor A. Dizon, Razel Nikka M. Hao, Arianne A. Zamora, Devon Ray Pacial, Juan Antonio R. Magalang, Marissa Alejandria, Celia Carlos, Anna Ong-Lim, Edsel Maurice Salvaña, John Q. Wong, Jaime C. Montoya, Maria Rosario Singh-Vergeire and Cynthia P. Saloma |
| EPI_ISL_2188878 | Las PiÃ±as General Hospital and Salellite Trauma Center | Philippine Genome Center | Francis A. Tablizo, Kenneth M. Kim, Maria Sofia L. Yangzon, Elcid Aaron R. Pangilinan, Renato Jacinto Q. Mantaring, Benedict A. Maralit, Marc Edsel C. Ayes, Eva Maria Cutiongco-de la Paz, Alethea R. de Guzman, Jan Michael C. Yap, Jo-Hannah S. Llames, Sheila Mae M. Araiza, Kris P. Punayan, Irish Coleen A. Asin, Candice Francheska B. Tambaoan, Asia Louisa U. Chong, Karol Sophia Agape R. Padilla, Rianna Patricia S. Cruz, Carlo M. Lapid, El King D. Morado, Joshua Gregor A. Dizon, Razel Nikka M. Hao, Arianne A. Zamora, Devon Ray Pacial, Juan Antonio R. Magalang, Marissa Alejandria, Celia Carlos, Anna Ong-Lim, Edsel Maurice Salvaña, John Q. Wong, Jaime C. Montoya, Maria Rosario Singh-Vergeire and Cynthia P. Saloma |
| EPI_ISL_2188881, EPI_ISL_2188883 | The Lord's Grace Medical and Industrial Clinic | Philippine Genome Center | Francis A. Tablizo, Kenneth M. Kim, Maria Sofia L. Yangzon, Elcid Aaron R. Pangilinan, Renato Jacinto Q. Mantaring, Benedict A. Maralit, Marc Edsel C. Ayes, Eva Maria Cutiongco-de la Paz, Alethea R. de Guzman, Jan Michael C. Yap, Jo-Hannah S. Llames, Sheila Mae M. Araiza, Kris P. Punayan, Irish |

| | | | |
|---|---|---|---|
| | | | Coleen A. Asin, Candice Francheska B. Tambaoan, Asia Louisa U. Chong, Karol Sophia Agape R. Padilla, Rianna Patricia S. Cruz, Carlo M. Lapid, El King D. Morado, Joshua Gregor A. Dizon, Razel Nikka M. Hao, Arianne A. Zamora, Devon Ray Pacial, Juan Antonio R. Magalang, Marissa Alejandria, Celia Carlos, Anna Ong-Lim, Edsel Maurice Salvaña, John Q. Wong, Jaime C. Montoya, Maria Rosario Singh-Vergeire and Cynthia P. Saloma |
| EPI_ISL_2188884 | The Medical City | Philippine Genome Center | Francis A. Tablizo, Kenneth M. Kim, Maria Sofia L. Yangzon, Elcid Aaron R. Pangilinan, Renato Jacinto Q. Mantaring, Benedict A. Maralit, Marc Edsel C. Ayes, Eva Maria Cutiongco-de la Paz, Alethea R. de Guzman, Jan Michael C. Yap, Jo-Hannah S. Llames, Sheila Mae M. Araiza, Kris P. Punayan, Irish Coleen A. Asin, Candice Francheska B. Tambaoan, Asia Louisa U. Chong, Karol Sophia Agape R. Padilla, Rianna Patricia S. Cruz, Carlo M. Lapid, El King D. Morado, Joshua Gregor A. Dizon, Razel Nikka M. Hao, Arianne A. Zamora, Devon Ray Pacial, Juan Antonio R. Magalang, Marissa Alejandria, Celia Carlos, Anna Ong-Lim, Edsel Maurice Salvaña, John Q. Wong, Jaime C. Montoya, Maria Rosario Singh-Vergeire and Cynthia P. Saloma |
| EPI_ISL_2188885 | Qualimed Hospital Sta. Rosa | Philippine Genome Center | Francis A. Tablizo, Kenneth M. Kim, Maria Sofia L. Yangzon, Elcid Aaron R. Pangilinan, Renato Jacinto Q. Mantaring, Benedict A. Maralit, Marc Edsel C. Ayes, Eva Maria Cutiongco-de la Paz, Alethea R. de Guzman, Jan Michael C. Yap, Jo-Hannah S. Llames, Sheila Mae M. Araiza, Kris P. Punayan, Irish Coleen A. Asin, Candice Francheska B. Tambaoan, Asia Louisa U. Chong, Karol Sophia Agape R. Padilla, Rianna Patricia S. Cruz, Carlo M. Lapid, El King D. Morado, Joshua Gregor A. Dizon, Razel Nikka M. Hao, Arianne A. Zamora, Devon Ray Pacial, Juan Antonio R. Magalang, Marissa Alejandria, Celia Carlos, Anna Ong-Lim, Edsel Maurice Salvaña, John Q. Wong, Jaime C. Montoya, Maria Rosario Singh-Vergeire and Cynthia P. Saloma |
| EPI_ISL_2188887 | Philippine General Hospital (PGH) | Philippine Genome Center | Francis A. Tablizo, Kenneth M. Kim, Maria Sofia L. Yangzon, Elcid Aaron R. Pangilinan, Renato Jacinto Q. Mantaring, Benedict A. Maralit, Marc Edsel C. Ayes, Eva Maria Cutiongco-de la Paz, Alethea R. de Guzman, Jan Michael C. Yap, Jo-Hannah S. Llames, Sheila Mae M. Araiza, Kris P. Punayan, Irish Coleen A. Asin, Candice Francheska B. Tambaoan, Asia Louisa U. Chong, Karol Sophia Agape R. Padilla, Rianna Patricia S. Cruz, Carlo M. Lapid, El King D. Morado, Joshua Gregor A. Dizon, Razel Nikka M. Hao, Arianne A. Zamora, Devon Ray Pacial, Juan Antonio R. Magalang, Marissa Alejandria, Celia Carlos, Anna Ong-Lim, Edsel Maurice Salvaña, John Q. Wong, Jaime C. Montoya, Maria Rosario Singh-Vergeire and Cynthia P. Saloma |
| EPI_ISL_2188889 | The Lord's Grace Medical and Industrial Clinic | Philippine Genome Center | Francis A. Tablizo, Kenneth M. Kim, Maria Sofia L. Yangzon, Elcid Aaron R. Pangilinan, Renato Jacinto Q. Mantaring, Benedict A. Maralit, Marc Edsel C. Ayes, Eva Maria Cutiongco-de la Paz, Alethea R. de Guzman, Jan Michael C. Yap, Jo-Hannah S. Llames, Sheila Mae M. Araiza, Kris P. Punayan, Irish Coleen A. Asin, Candice Francheska B. Tambaoan, Asia Louisa U. Chong, Karol Sophia Agape R. Padilla, Rianna Patricia S. Cruz, Carlo M. Lapid, El King D. Morado, Joshua Gregor A. Dizon, Razel Nikka M. Hao, Arianne A. Zamora, Devon Ray Pacial, Juan Antonio R. Magalang, Marissa Alejandria, Celia Carlos, Anna Ong-Lim, Edsel Maurice Salvaña, John Q. Wong, Jaime C. Montoya, Maria Rosario Singh-Vergeire and Cynthia P. Saloma |
| EPI_ISL_2188891 | University of the Philippines National Institutes of Health (UP NIH) | Philippine Genome Center | Francis A. Tablizo, Kenneth M. Kim, Maria Sofia L. Yangzon, Elcid Aaron R. Pangilinan, Renato Jacinto Q. Mantaring, Benedict A. Maralit, Marc Edsel C. Ayes, Eva Maria Cutiongco-de la Paz, Alethea R. de Guzman, Jan Michael C. Yap, Jo-Hannah S. Llames, Sheila Mae M. Araiza, Kris P. Punayan, Irish Coleen A. Asin, Candice Francheska B. Tambaoan, Asia Louisa U. Chong, Karol Sophia Agape R. Padilla, Rianna Patricia S. Cruz, Carlo M. Lapid, El King D. Morado, Joshua Gregor A. Dizon, Razel Nikka M. Hao, Arianne A. Zamora, Devon Ray Pacial, Juan Antonio R. Magalang, Marissa Alejandria, Celia Carlos, Anna Ong-Lim, Edsel Maurice Salvaña, John Q. Wong, Jaime C. Montoya, Maria Rosario Singh-Vergeire and Cynthia P. Saloma |
| EPI_ISL_2188892 | Philippine Red Cross - Port Area | Philippine Genome Center | Francis A. Tablizo, Kenneth M. Kim, Maria Sofia L. Yangzon, Elcid Aaron R. Pangilinan, Renato Jacinto Q. Mantaring, Benedict A. Maralit, Marc Edsel C. Ayes, Eva Maria Cutiongco-de la Paz, Alethea R. de Guzman, Jan Michael C. Yap, Jo-Hannah S. Llames, Sheila Mae M. Araiza, Kris P. Punayan, Irish Coleen A. Asin, Candice Francheska B. Tambaoan, Asia Louisa U. Chong, Karol Sophia Agape R. Padilla, Rianna Patricia S. Cruz, Carlo M. Lapid, El King D. Morado, Joshua Gregor A. Dizon, Razel Nikka M. Hao, Arianne A. Zamora, Devon Ray Pacial, Juan Antonio R. Magalang, Marissa Alejandria, Celia Carlos, Anna Ong-Lim, Edsel Maurice Salvaña, John Q. Wong, Jaime C. Montoya, Maria Rosario Singh-Vergeire and Cynthia P. Saloma |
| EPI_ISL_2188893 | The Lord's Grace Medical and Industrial Clinic | Philippine Genome Center | Francis A. Tablizo, Kenneth M. Kim, Maria Sofia L. Yangzon, Elcid Aaron R. Pangilinan, Renato Jacinto Q. Mantaring, Benedict A. Maralit, Marc Edsel C. Ayes, Eva Maria Cutiongco-de la Paz, Alethea R. de Guzman, Jan Michael C. Yap, Jo-Hannah S. Llames, Sheila Mae M. Araiza, Kris P. Punayan, Irish Coleen A. Asin, Candice Francheska B. Tambaoan, Asia Louisa U. Chong, Karol Sophia Agape R. Padilla, Rianna Patricia S. Cruz, Carlo M. Lapid, El King D. Morado, Joshua Gregor A. Dizon, Razel Nikka M. Hao, Arianne A. Zamora, Devon Ray Pacial, Juan Antonio R. Magalang, Marissa Alejandria, Celia Carlos, Anna Ong-Lim, Edsel Maurice Salvaña, John Q. Wong, Jaime C. Montoya, Maria Rosario Singh-Vergeire and Cynthia P. Saloma |
| EPI_ISL_2188895 | Sta. Rosa CHO- GENEXPERT LABORATORY | Philippine Genome Center | Francis A. Tablizo, Kenneth M. Kim, Maria Sofia L. Yangzon, Elcid Aaron R. Pangilinan, Renato Jacinto Q. Mantaring, Benedict A. Maralit, Marc Edsel C. Ayes, Eva Maria Cutiongco-de la Paz, Alethea R. de Guzman, Jan Michael C. Yap, Jo-Hannah S. Llames, Sheila Mae M. Araiza, Kris P. Punayan, Irish Coleen A. Asin, Candice Francheska B. Tambaoan, Asia Louisa U. Chong, Karol Sophia Agape R. Padilla, Rianna Patricia S. Cruz, Carlo M. Lapid, El King D. Morado, Joshua Gregor A. Dizon, Razel Nikka M. Hao, Arianne A. Zamora, Devon Ray Pacial, Juan Antonio R. Magalang, Marissa Alejandria, Celia Carlos, Anna Ong-Lim, Edsel Maurice Salvaña, John Q. Wong, Jaime C. Montoya, Maria Rosario Singh-Vergeire and Cynthia P. Saloma |
| EPI_ISL_2188896 | Philippine Red Cross - National Blood Center | Philippine Genome Center | Francis A. Tablizo, Kenneth M. Kim, Maria Sofia L. Yangzon, Elcid Aaron R. Pangilinan, Renato Jacinto Q. Mantaring, Benedict A. Maralit, Marc Edsel C. Ayes, Eva Maria Cutiongco-de la Paz, Alethea R. de Guzman, Jan Michael C. Yap, Jo-Hannah S. Llames, Sheila Mae M. Araiza, Kris P. Punayan, Irish Coleen A. Asin, Candice Francheska B. Tambaoan, Asia Louisa U. Chong, Karol Sophia Agape R. Padilla, Rianna Patricia S. Cruz, Carlo M. Lapid, El King D. Morado, Joshua Gregor A. Dizon, Razel Nikka M. Hao, Arianne A. Zamora, Devon Ray Pacial, Juan Antonio R. Magalang, Marissa Alejandria, Celia Carlos, Anna Ong-Lim, Edsel Maurice Salvaña, John Q. Wong, Jaime C. Montoya, Maria Rosario Singh-Vergeire and Cynthia P. Saloma |
| EPI_ISL_2188897 | The Lord's Grace Medical and Industrial Clinic | Philippine Genome Center | Francis A. Tablizo, Kenneth M. Kim, Maria Sofia L. Yangzon, Elcid Aaron R. Pangilinan, Renato Jacinto Q. Mantaring, Benedict A. Maralit, Marc Edsel C. Ayes, Eva Maria Cutiongco-de la Paz, Alethea R. de Guzman, Jan Michael C. Yap, Jo-Hannah S. Llames, Sheila Mae M. Araiza, Kris P. Punayan, Irish Coleen A. Asin, Candice Francheska B. Tambaoan, Asia Louisa U. Chong, Karol Sophia Agape R. Padilla, Rianna Patricia S. Cruz, Carlo M. Lapid, El King D. Morado, Joshua Gregor A. Dizon, Razel Nikka M. Hao, Arianne A. Zamora, Devon Ray Pacial, Juan Antonio R. Magalang, Marissa Alejandria, Celia Carlos, Anna Ong-Lim, Edsel Maurice Salvaña, John Q. Wong, Jaime C. Montoya, Maria Rosario Singh-Vergeire and Cynthia P. Saloma |
| EPI_ISL_2188898 | Philippine Red Cross - Port Area | Philippine Genome Center | Francis A. Tablizo, Kenneth M. Kim, Maria Sofia L. Yangzon, Elcid Aaron R. Pangilinan, Renato Jacinto Q. Mantaring, Benedict A. Maralit, Marc Edsel C. Ayes, Eva Maria Cutiongco-de la Paz, Alethea R. de Guzman, Jan Michael C. Yap, Jo-Hannah S. Llames, Sheila Mae M. Araiza, Kris P. Punayan, Irish Coleen A. Asin, Candice Francheska B. Tambaoan, Asia Louisa U. Chong, Karol Sophia Agape R. Padilla, Rianna Patricia S. Cruz, Carlo M. Lapid, El King D. Morado, Joshua Gregor A. Dizon, Razel Nikka M. Hao, Arianne A. Zamora, Devon Ray Pacial, Juan Antonio R. Magalang, Marissa Alejandria, Celia Carlos, Anna Ong-Lim, Edsel Maurice Salvaña, John Q. Wong, Jaime C. Montoya, Maria Rosario Singh-Vergeire and Cynthia P. Saloma |
| EPI_ISL_2188899 | The Medical City | Philippine Genome Center | Francis A. Tablizo, Kenneth M. Kim, Maria Sofia L. Yangzon, Elcid Aaron R. Pangilinan, Renato Jacinto Q. Mantaring, Benedict A. Maralit, Marc Edsel C. Ayes, Eva Maria Cutiongco-de la Paz, Alethea R. de Guzman, Jan Michael C. Yap, Jo-Hannah S. Llames, Sheila Mae M. Araiza, Kris P. Punayan, Irish Coleen A. Asin, Candice Francheska B. Tambaoan, Asia Louisa U. Chong, Karol Sophia Agape R. Padilla, Rianna Patricia S. Cruz, Carlo M. Lapid, El King D. Morado, Joshua Gregor A. Dizon, Razel Nikka M. Hao, Arianne A. Zamora, Devon Ray Pacial, Juan Antonio R. Magalang, Marissa Alejandria, Celia Carlos, Anna Ong-Lim, Edsel Maurice Salvaña, John Q. Wong, Jaime C. Montoya, Maria Rosario Singh-Vergeire and Cynthia P. Saloma |
| EPI_ISL_2188901 | The Lord's Grace Medical and Industrial Clinic | Philippine Genome Center | Francis A. Tablizo, Kenneth M. Kim, Maria Sofia L. Yangzon, Elcid Aaron R. Pangilinan, Renato Jacinto Q. Mantaring, Benedict A. Maralit, Marc Edsel C. Ayes, Eva Maria Cutiongco-de la Paz, Alethea R. de Guzman, Jan Michael C. Yap, Jo-Hannah S. Llames, Sheila Mae M. Araiza, Kris P. Punayan, Irish Coleen A. Asin, Candice Francheska B. Tambaoan, Asia Louisa U. Chong, Karol Sophia Agape R. Padilla, Rianna Patricia S. Cruz, Carlo M. Lapid, El King D. Morado, Joshua Gregor A. Dizon, Razel Nikka M. Hao, Arianne A. Zamora, Devon Ray Pacial, Juan Antonio R. Magalang, Marissa Alejandria, Celia Carlos, Anna Ong-Lim, Edsel Maurice Salvaña, John Q. Wong, Jaime C. Montoya, Maria Rosario Singh-Vergeire and Cynthia P. Saloma |
| EPI_ISL_2188902 | Philippine Red Cross - Clark Molecular Laboratory | Philippine Genome Center | Francis A. Tablizo, Kenneth M. Kim, Maria Sofia L. Yangzon, Elcid Aaron R. Pangilinan, Renato Jacinto Q. Mantaring, Benedict A. Maralit, Marc Edsel C. Ayes, Eva Maria Cutiongco-de la Paz, Alethea R. de Guzman, Jan Michael C. Yap, Jo-Hannah S. Llames, Sheila Mae M. Araiza, Kris P. Punayan, Irish Coleen A. Asin, Candice Francheska B. Tambaoan, Asia Louisa U. Chong, Karol Sophia Agape R. Padilla, Rianna Patricia S. Cruz, Carlo M. Lapid, El King D. Morado, Joshua Gregor A. Dizon, Razel Nikka M. Hao, Arianne A. Zamora, Devon Ray Pacial, Juan Antonio R. Magalang, Marissa Alejandria, Celia Carlos, Anna Ong-Lim, Edsel Maurice Salvaña, John Q. Wong, Jaime C. Montoya, Maria Rosario Singh-Vergeire and Cynthia P. Saloma |
| EPI_ISL_2188903 | Sta. Rosa CHO- GENEXPERT LABORATORY | Philippine Genome Center | Francis A. Tablizo, Kenneth M. Kim, Maria Sofia L. Yangzon, Elcid Aaron R. Pangilinan, Renato Jacinto Q. Mantaring, Benedict A. Maralit, Marc Edsel C. Ayes, Eva Maria Cutiongco-de la Paz, Alethea R. de Guzman, Jan Michael C. Yap, Jo-Hannah S. Llames, Sheila Mae M. Araiza, Kris P. Punayan, Irish Coleen A. Asin, Candice Francheska B. Tambaoan, Asia Louisa U. Chong, Karol Sophia Agape R. Padilla, Rianna Patricia S. Cruz, Carlo M. Lapid, El King D. Morado, Joshua Gregor A. Dizon, Razel Nikka M. Hao, Arianne A. Zamora, Devon Ray Pacial, Juan Antonio R. Magalang, Marissa Alejandria, Celia Carlos, Anna Ong-Lim, Edsel Maurice Salvaña, John Q. Wong, Jaime C. Montoya, Maria Rosario Singh-Vergeire and Cynthia P. Saloma |
| EPI_ISL_2188904 | The Medical City | Philippine Genome Center | Francis A. Tablizo, Kenneth M. Kim, Maria Sofia L. Yangzon, Elcid Aaron R. Pangilinan, Renato Jacinto Q. Mantaring, Benedict A. Maralit, Marc Edsel C. |

| | | | |
|---|---|---|---|
| | | | Ayes, Eva Maria Cutiongco-de la Paz, Alethea R. de Guzman, Jan Michael C. Yap, Jo-Hannah S. Llames, Sheila Mae M. Araiza, Kris P. Punayan, Irish Coleen A. Asin, Candice Francheska B. Tambaoan, Asia Louisa U. Chong, Karol Sophia Agape R. Padilla, Rianna Patricia S. Cruz, Carlo M. Lapid, El King D. Morado, Joshua Gregor A. Dizon, Razel Nikka M. Hao, Arianne A. Zamora, Devon Ray Pacial, Juan Antonio R. Magalang, Marissa Alejandria, Celia Carlos, Anna Ong-Lim, Edsel Maurice Salvaña, John Q. Wong, Jaime C. Montoya, Maria Rosario Singh-Vergeire and Cynthia P. Saloma |
| EPI_ISL_2188905 | University of the Philippines National Institutes of Health (UP NIH) | Philippine Genome Center | Francis A. Tablizo, Kenneth M. Kim, Maria Sofia L. Yangzon, Elcid Aaron R. Pangilinan, Renato Jacinto Q. Mantaring, Benedict A. Maralit, Marc Edsel C. Ayes, Eva Maria Cutiongco-de la Paz, Alethea R. de Guzman, Jan Michael C. Yap, Jo-Hannah S. Llames, Sheila Mae M. Araiza, Kris P. Punayan, Irish Coleen A. Asin, Candice Francheska B. Tambaoan, Asia Louisa U. Chong, Karol Sophia Agape R. Padilla, Rianna Patricia S. Cruz, Carlo M. Lapid, El King D. Morado, Joshua Gregor A. Dizon, Razel Nikka M. Hao, Arianne A. Zamora, Devon Ray Pacial, Juan Antonio R. Magalang, Marissa Alejandria, Celia Carlos, Anna Ong-Lim, Edsel Maurice Salvaña, John Q. Wong, Jaime C. Montoya, Maria Rosario Singh-Vergeire and Cynthia P. Saloma |
| EPI_ISL_2188907 | Pasig City Children's Hospital - Child's Hope | Philippine Genome Center | Francis A. Tablizo, Kenneth M. Kim, Maria Sofia L. Yangzon, Elcid Aaron R. Pangilinan, Renato Jacinto Q. Mantaring, Benedict A. Maralit, Marc Edsel C. Ayes, Eva Maria Cutiongco-de la Paz, Alethea R. de Guzman, Jan Michael C. Yap, Jo-Hannah S. Llames, Sheila Mae M. Araiza, Kris P. Punayan, Irish Coleen A. Asin, Candice Francheska B. Tambaoan, Asia Louisa U. Chong, Karol Sophia Agape R. Padilla, Rianna Patricia S. Cruz, Carlo M. Lapid, El King D. Morado, Joshua Gregor A. Dizon, Razel Nikka M. Hao, Arianne A. Zamora, Devon Ray Pacial, Juan Antonio R. Magalang, Marissa Alejandria, Celia Carlos, Anna Ong-Lim, Edsel Maurice Salvaña, John Q. Wong, Jaime C. Montoya, Maria Rosario Singh-Vergeire and Cynthia P. Saloma |
| EPI_ISL_2188911 | Sta. Rosa CHO- GENEXPERT LABORATORY | Philippine Genome Center | Francis A. Tablizo, Kenneth M. Kim, Maria Sofia L. Yangzon, Elcid Aaron R. Pangilinan, Renato Jacinto Q. Mantaring, Benedict A. Maralit, Marc Edsel C. Ayes, Eva Maria Cutiongco-de la Paz, Alethea R. de Guzman, Jan Michael C. Yap, Jo-Hannah S. Llames, Sheila Mae M. Araiza, Kris P. Punayan, Irish Coleen A. Asin, Candice Francheska B. Tambaoan, Asia Louisa U. Chong, Karol Sophia Agape R. Padilla, Rianna Patricia S. Cruz, Carlo M. Lapid, El King D. Morado, Joshua Gregor A. Dizon, Razel Nikka M. Hao, Arianne A. Zamora, Devon Ray Pacial, Juan Antonio R. Magalang, Marissa Alejandria, Celia Carlos, Anna Ong-Lim, Edsel Maurice Salvaña, John Q. Wong, Jaime C. Montoya, Maria Rosario Singh-Vergeire and Cynthia P. Saloma |
| EPI_ISL_2188912 | The Medical City | Philippine Genome Center | Francis A. Tablizo, Kenneth M. Kim, Maria Sofia L. Yangzon, Elcid Aaron R. Pangilinan, Renato Jacinto Q. Mantaring, Benedict A. Maralit, Marc Edsel C. Ayes, Eva Maria Cutiongco-de la Paz, Alethea R. de Guzman, Jan Michael C. Yap, Jo-Hannah S. Llames, Sheila Mae M. Araiza, Kris P. Punayan, Irish Coleen A. Asin, Candice Francheska B. Tambaoan, Asia Louisa U. Chong, Karol Sophia Agape R. Padilla, Rianna Patricia S. Cruz, Carlo M. Lapid, El King D. Morado, Joshua Gregor A. Dizon, Razel Nikka M. Hao, Arianne A. Zamora, Devon Ray Pacial, Juan Antonio R. Magalang, Marissa Alejandria, Celia Carlos, Anna Ong-Lim, Edsel Maurice Salvaña, John Q. Wong, Jaime C. Montoya, Maria Rosario Singh-Vergeire and Cynthia P. Saloma |
| EPI_ISL_2188914 | Philippine Red Cross - Port Area | Philippine Genome Center | Francis A. Tablizo, Kenneth M. Kim, Maria Sofia L. Yangzon, Elcid Aaron R. Pangilinan, Renato Jacinto Q. Mantaring, Benedict A. Maralit, Marc Edsel C. Ayes, Eva Maria Cutiongco-de la Paz, Alethea R. de Guzman, Jan Michael C. Yap, Jo-Hannah S. Llames, Sheila Mae M. Araiza, Kris P. Punayan, Irish Coleen A. Asin, Candice Francheska B. Tambaoan, Asia Louisa U. Chong, Karol Sophia Agape R. Padilla, Rianna Patricia S. Cruz, Carlo M. Lapid, El King D. Morado, Joshua Gregor A. Dizon, Razel Nikka M. Hao, Arianne A. Zamora, Devon Ray Pacial, Juan Antonio R. Magalang, Marissa Alejandria, Celia Carlos, Anna Ong-Lim, Edsel Maurice Salvaña, John Q. Wong, Jaime C. Montoya, Maria Rosario Singh-Vergeire and Cynthia P. Saloma |
| EPI_ISL_2188916 | Philippine Red Cross - National Blood Center | Philippine Genome Center | Francis A. Tablizo, Kenneth M. Kim, Maria Sofia L. Yangzon, Elcid Aaron R. Pangilinan, Renato Jacinto Q. Mantaring, Benedict A. Maralit, Marc Edsel C. Ayes, Eva Maria Cutiongco-de la Paz, Alethea R. de Guzman, Jan Michael C. Yap, Jo-Hannah S. Llames, Sheila Mae M. Araiza, Kris P. Punayan, Irish Coleen A. Asin, Candice Francheska B. Tambaoan, Asia Louisa U. Chong, Karol Sophia Agape R. Padilla, Rianna Patricia S. Cruz, Carlo M. Lapid, El King D. Morado, Joshua Gregor A. Dizon, Razel Nikka M. Hao, Arianne A. Zamora, Devon Ray Pacial, Juan Antonio R. Magalang, Marissa Alejandria, Celia Carlos, Anna Ong-Lim, Edsel Maurice Salvaña, John Q. Wong, Jaime C. Montoya, Maria Rosario Singh-Vergeire and Cynthia P. Saloma |
| EPI_ISL_2188917 | The Lord's Grace Medical and Industrial Clinic | Philippine Genome Center | Francis A. Tablizo, Kenneth M. Kim, Maria Sofia L. Yangzon, Elcid Aaron R. Pangilinan, Renato Jacinto Q. Mantaring, Benedict A. Maralit, Marc Edsel C. Ayes, Eva Maria Cutiongco-de la Paz, Alethea R. de Guzman, Jan Michael C. Yap, Jo-Hannah S. Llames, Sheila Mae M. Araiza, Kris P. Punayan, Irish Coleen A. Asin, Candice Francheska B. Tambaoan, Asia Louisa U. Chong, Karol Sophia Agape R. Padilla, Rianna Patricia S. Cruz, Carlo M. Lapid, El King D. Morado, Joshua Gregor A. Dizon, Razel Nikka M. Hao, Arianne A. Zamora, Devon Ray Pacial, Juan Antonio R. Magalang, Marissa Alejandria, Celia Carlos, Anna Ong-Lim, Edsel Maurice Salvaña, John Q. Wong, Jaime C. Montoya, Maria Rosario Singh-Vergeire and Cynthia P. Saloma |
| EPI_ISL_2188918 | Mary Mediatrix Medical Center | Philippine Genome Center | Francis A. Tablizo, Kenneth M. Kim, Maria Sofia L. Yangzon, Elcid Aaron R. Pangilinan, Renato Jacinto Q. Mantaring, Benedict A. Maralit, Marc Edsel C. Ayes, Eva Maria Cutiongco-de la Paz, Alethea R. de Guzman, Jan Michael C. Yap, Jo-Hannah S. Llames, Sheila Mae M. Araiza, Kris P. Punayan, Irish Coleen A. Asin, Candice Francheska B. Tambaoan, Asia Louisa U. Chong, Karol Sophia Agape R. Padilla, Rianna Patricia S. Cruz, Carlo M. Lapid, El King D. Morado, Joshua Gregor A. Dizon, Razel Nikka M. Hao, Arianne A. Zamora, Devon Ray Pacial, Juan Antonio R. Magalang, Marissa Alejandria, Celia Carlos, Anna Ong-Lim, Edsel Maurice Salvaña, John Q. Wong, Jaime C. Montoya, Maria Rosario Singh-Vergeire and Cynthia P. Saloma |
| EPI_ISL_2188920 | Sta. Ana Hospital | Philippine Genome Center | Francis A. Tablizo, Kenneth M. Kim, Maria Sofia L. Yangzon, Elcid Aaron R. Pangilinan, Renato Jacinto Q. Mantaring, Benedict A. Maralit, Marc Edsel C. Ayes, Eva Maria Cutiongco-de la Paz, Alethea R. de Guzman, Jan Michael C. Yap, Jo-Hannah S. Llames, Sheila Mae M. Araiza, Kris P. Punayan, Irish Coleen A. Asin, Candice Francheska B. Tambaoan, Asia Louisa U. Chong, Karol Sophia Agape R. Padilla, Rianna Patricia S. Cruz, Carlo M. Lapid, El King D. Morado, Joshua Gregor A. Dizon, Razel Nikka M. Hao, Arianne A. Zamora, Devon Ray Pacial, Juan Antonio R. Magalang, Marissa Alejandria, Celia Carlos, Anna Ong-Lim, Edsel Maurice Salvaña, John Q. Wong, Jaime C. Montoya, Maria Rosario Singh-Vergeire and Cynthia P. Saloma |
| EPI_ISL_2188924 | The Lord's Grace Medical and Industrial Clinic | Philippine Genome Center | Francis A. Tablizo, Kenneth M. Kim, Maria Sofia L. Yangzon, Elcid Aaron R. Pangilinan, Renato Jacinto Q. Mantaring, Benedict A. Maralit, Marc Edsel C. Ayes, Eva Maria Cutiongco-de la Paz, Alethea R. de Guzman, Jan Michael C. Yap, Jo-Hannah S. Llames, Sheila Mae M. Araiza, Kris P. Punayan, Irish Coleen A. Asin, Candice Francheska B. Tambaoan, Asia Louisa U. Chong, Karol Sophia Agape R. Padilla, Rianna Patricia S. Cruz, Carlo M. Lapid, El King D. Morado, Joshua Gregor A. Dizon, Razel Nikka M. Hao, Arianne A. Zamora, Devon Ray Pacial, Juan Antonio R. Magalang, Marissa Alejandria, Celia Carlos, Anna Ong-Lim, Edsel Maurice Salvaña, John Q. Wong, Jaime C. Montoya, Maria Rosario Singh-Vergeire and Cynthia P. Saloma |
| EPI_ISL_2188925 | Pasig City Children's Hospital - Child's Hope | Philippine Genome Center | Francis A. Tablizo, Kenneth M. Kim, Maria Sofia L. Yangzon, Elcid Aaron R. Pangilinan, Renato Jacinto Q. Mantaring, Benedict A. Maralit, Marc Edsel C. Ayes, Eva Maria Cutiongco-de la Paz, Alethea R. de Guzman, Jan Michael C. Yap, Jo-Hannah S. Llames, Sheila Mae M. Araiza, Kris P. Punayan, Irish Coleen A. Asin, Candice Francheska B. Tambaoan, Asia Louisa U. Chong, Karol Sophia Agape R. Padilla, Rianna Patricia S. Cruz, Carlo M. Lapid, El King D. Morado, Joshua Gregor A. Dizon, Razel Nikka M. Hao, Arianne A. Zamora, Devon Ray Pacial, Juan Antonio R. Magalang, Marissa Alejandria, Celia Carlos, Anna Ong-Lim, Edsel Maurice Salvaña, John Q. Wong, Jaime C. Montoya, Maria Rosario Singh-Vergeire and Cynthia P. Saloma |
| EPI_ISL_2188927 | Philippine Red Cross - National Blood Center | Philippine Genome Center | Francis A. Tablizo, Kenneth M. Kim, Maria Sofia L. Yangzon, Elcid Aaron R. Pangilinan, Renato Jacinto Q. Mantaring, Benedict A. Maralit, Marc Edsel C. Ayes, Eva Maria Cutiongco-de la Paz, Alethea R. de Guzman, Jan Michael C. Yap, Jo-Hannah S. Llames, Sheila Mae M. Araiza, Kris P. Punayan, Irish Coleen A. Asin, Candice Francheska B. Tambaoan, Asia Louisa U. Chong, Karol Sophia Agape R. Padilla, Rianna Patricia S. Cruz, Carlo M. Lapid, El King D. Morado, Joshua Gregor A. Dizon, Razel Nikka M. Hao, Arianne A. Zamora, Devon Ray Pacial, Juan Antonio R. Magalang, Marissa Alejandria, Celia Carlos, Anna Ong-Lim, Edsel Maurice Salvaña, John Q. Wong, Jaime C. Montoya, Maria Rosario Singh-Vergeire and Cynthia P. Saloma |
| EPI_ISL_2188928, EPI_ISL_2188931 | Pasig City Children's Hospital - Child's Hope | Philippine Genome Center | Francis A. Tablizo, Kenneth M. Kim, Maria Sofia L. Yangzon, Elcid Aaron R. Pangilinan, Renato Jacinto Q. Mantaring, Benedict A. Maralit, Marc Edsel C. Ayes, Eva Maria Cutiongco-de la Paz, Alethea R. de Guzman, Jan Michael C. Yap, Jo-Hannah S. Llames, Sheila Mae M. Araiza, Kris P. Punayan, Irish Coleen A. Asin, Candice Francheska B. Tambaoan, Asia Louisa U. Chong, Karol Sophia Agape R. Padilla, Rianna Patricia S. Cruz, Carlo M. Lapid, El King D. Morado, Joshua Gregor A. Dizon, Razel Nikka M. Hao, Arianne A. Zamora, Devon Ray Pacial, Juan Antonio R. Magalang, Marissa Alejandria, Celia Carlos, Anna Ong-Lim, Edsel Maurice Salvaña, John Q. Wong, Jaime C. Montoya, Maria Rosario Singh-Vergeire and Cynthia P. Saloma |
| EPI_ISL_2188932 | The Lord's Grace Medical and Industrial Clinic | Philippine Genome Center | Francis A. Tablizo, Kenneth M. Kim, Maria Sofia L. Yangzon, Elcid Aaron R. Pangilinan, Renato Jacinto Q. Mantaring, Benedict A. Maralit, Marc Edsel C. Ayes, Eva Maria Cutiongco-de la Paz, Alethea R. de Guzman, Jan Michael C. Yap, Jo-Hannah S. Llames, Sheila Mae M. Araiza, Kris P. Punayan, Irish Coleen A. Asin, Candice Francheska B. Tambaoan, Asia Louisa U. Chong, Karol Sophia Agape R. Padilla, Rianna Patricia S. Cruz, Carlo M. Lapid, El King D. Morado, Joshua Gregor A. Dizon, Razel Nikka M. Hao, Arianne A. Zamora, Devon Ray Pacial, Juan Antonio R. Magalang, Marissa Alejandria, Celia Carlos, Anna Ong-Lim, Edsel Maurice Salvaña, John Q. Wong, Jaime C. Montoya, Maria Rosario Singh-Vergeire and Cynthia P. Saloma |
| EPI_ISL_2188933 | Philippine Red Cross - Port Area | Philippine Genome Center | Francis A. Tablizo, Kenneth M. Kim, Maria Sofia L. Yangzon, Elcid Aaron R. Pangilinan, Renato Jacinto Q. Mantaring, Benedict A. Maralit, Marc Edsel C. Ayes, Eva Maria Cutiongco-de la Paz, Alethea R. de Guzman, Jan Michael C. Yap, Jo-Hannah S. Llames, Sheila Mae M. Araiza, Kris P. Punayan, Irish Coleen A. Asin, Candice Francheska B. Tambaoan, Asia Louisa U. Chong, Karol Sophia Agape R. Padilla, Rianna Patricia S. Cruz, Carlo M. Lapid, El King D. Morado, Joshua Gregor A. Dizon, Razel Nikka M. Hao, Arianne A. Zamora, Devon Ray Pacial, Juan Antonio R. Magalang, Marissa Alejandria, Celia Carlos, Anna Ong-Lim, Edsel Maurice Salvaña, John Q. Wong, Jaime C. Montoya, Maria Rosario Singh-Vergeire and Cynthia P. Saloma |

| | | | |
|---|---|---|---|
| EPI_ISL_2188937 | The Medical City | Philippine Genome Center | Francis A. Tablizo, Kenneth M. Kim, Maria Sofia L. Yangzon, Elcid Aaron R. Pangilinan, Renato Jacinto Q. Mantaring, Benedict A. Maralit, Marc Edsel C. Ayes, Eva Maria Cutiongco-de la Paz, Alethea R. de Guzman, Jan Michael C. Yap, Jo-Hannah S. Llames, Sheila Mae M. Araiza, Kris P. Punayan, Irish Coleen A. Asin, Candice Francheska B. Tambaoan, Asia Louisa U. Chong, Karol Sophia Agape R. Padilla, Rianna Patricia S. Cruz, Carlo M. Lapid, El King D. Morado, Joshua Gregor A. Dizon, Razel Nikka M. Hao, Arianne A. Zamora, Devon Ray Pacial, Juan Antonio R. Magalang, Marissa Alejandria, Celia Carlos, Anna Ong-Lim, Edsel Maurice Salvaña, John Q. Wong, Jaime C. Montoya, Maria Rosario Singh-Vergeire and Cynthia P. Saloma |
| EPI_ISL_2188938 | Philippine Red Cross - Port Area | Philippine Genome Center | Francis A. Tablizo, Kenneth M. Kim, Maria Sofia L. Yangzon, Elcid Aaron R. Pangilinan, Renato Jacinto Q. Mantaring, Benedict A. Maralit, Marc Edsel C. Ayes, Eva Maria Cutiongco-de la Paz, Alethea R. de Guzman, Jan Michael C. Yap, Jo-Hannah S. Llames, Sheila Mae M. Araiza, Kris P. Punayan, Irish Coleen A. Asin, Candice Francheska B. Tambaoan, Asia Louisa U. Chong, Karol Sophia Agape R. Padilla, Rianna Patricia S. Cruz, Carlo M. Lapid, El King D. Morado, Joshua Gregor A. Dizon, Razel Nikka M. Hao, Arianne A. Zamora, Devon Ray Pacial, Juan Antonio R. Magalang, Marissa Alejandria, Celia Carlos, Anna Ong-Lim, Edsel Maurice Salvaña, John Q. Wong, Jaime C. Montoya, Maria Rosario Singh-Vergeire and Cynthia P. Saloma |
| EPI_ISL_2188939 | Philippine General Hospital (PGH) | Philippine Genome Center | Francis A. Tablizo, Kenneth M. Kim, Maria Sofia L. Yangzon, Elcid Aaron R. Pangilinan, Renato Jacinto Q. Mantaring, Benedict A. Maralit, Marc Edsel C. Ayes, Eva Maria Cutiongco-de la Paz, Alethea R. de Guzman, Jan Michael C. Yap, Jo-Hannah S. Llames, Sheila Mae M. Araiza, Kris P. Punayan, Irish Coleen A. Asin, Candice Francheska B. Tambaoan, Asia Louisa U. Chong, Karol Sophia Agape R. Padilla, Rianna Patricia S. Cruz, Carlo M. Lapid, El King D. Morado, Joshua Gregor A. Dizon, Razel Nikka M. Hao, Arianne A. Zamora, Devon Ray Pacial, Juan Antonio R. Magalang, Marissa Alejandria, Celia Carlos, Anna Ong-Lim, Edsel Maurice Salvaña, John Q. Wong, Jaime C. Montoya, Maria Rosario Singh-Vergeire and Cynthia P. Saloma |
| EPI_ISL_2188940 | South Super Highway Molecular Diagnostic Laboratory | Philippine Genome Center | Francis A. Tablizo, Kenneth M. Kim, Maria Sofia L. Yangzon, Elcid Aaron R. Pangilinan, Renato Jacinto Q. Mantaring, Benedict A. Maralit, Marc Edsel C. Ayes, Eva Maria Cutiongco-de la Paz, Alethea R. de Guzman, Jan Michael C. Yap, Jo-Hannah S. Llames, Sheila Mae M. Araiza, Kris P. Punayan, Irish Coleen A. Asin, Candice Francheska B. Tambaoan, Asia Louisa U. Chong, Karol Sophia Agape R. Padilla, Rianna Patricia S. Cruz, Carlo M. Lapid, El King D. Morado, Joshua Gregor A. Dizon, Razel Nikka M. Hao, Arianne A. Zamora, Devon Ray Pacial, Juan Antonio R. Magalang, Marissa Alejandria, Celia Carlos, Anna Ong-Lim, Edsel Maurice Salvaña, John Q. Wong, Jaime C. Montoya, Maria Rosario Singh-Vergeire and Cynthia P. Saloma |
| EPI_ISL_2188948 | The Medical City | Philippine Genome Center | Francis A. Tablizo, Kenneth M. Kim, Maria Sofia L. Yangzon, Elcid Aaron R. Pangilinan, Renato Jacinto Q. Mantaring, Benedict A. Maralit, Marc Edsel C. Ayes, Eva Maria Cutiongco-de la Paz, Alethea R. de Guzman, Jan Michael C. Yap, Jo-Hannah S. Llames, Sheila Mae M. Araiza, Kris P. Punayan, Irish Coleen A. Asin, Candice Francheska B. Tambaoan, Asia Louisa U. Chong, Karol Sophia Agape R. Padilla, Rianna Patricia S. Cruz, Carlo M. Lapid, El King D. Morado, Joshua Gregor A. Dizon, Razel Nikka M. Hao, Arianne A. Zamora, Devon Ray Pacial, Juan Antonio R. Magalang, Marissa Alejandria, Celia Carlos, Anna Ong-Lim, Edsel Maurice Salvaña, John Q. Wong, Jaime C. Montoya, Maria Rosario Singh-Vergeire and Cynthia P. Saloma |
| EPI_ISL_2188949 | Philippine Red Cross - Port Area | Philippine Genome Center | Francis A. Tablizo, Kenneth M. Kim, Maria Sofia L. Yangzon, Elcid Aaron R. Pangilinan, Renato Jacinto Q. Mantaring, Benedict A. Maralit, Marc Edsel C. Ayes, Eva Maria Cutiongco-de la Paz, Alethea R. de Guzman, Jan Michael C. Yap, Jo-Hannah S. Llames, Sheila Mae M. Araiza, Kris P. Punayan, Irish Coleen A. Asin, Candice Francheska B. Tambaoan, Asia Louisa U. Chong, Karol Sophia Agape R. Padilla, Rianna Patricia S. Cruz, Carlo M. Lapid, El King D. Morado, Joshua Gregor A. Dizon, Razel Nikka M. Hao, Arianne A. Zamora, Devon Ray Pacial, Juan Antonio R. Magalang, Marissa Alejandria, Celia Carlos, Anna Ong-Lim, Edsel Maurice Salvaña, John Q. Wong, Jaime C. Montoya, Maria Rosario Singh-Vergeire and Cynthia P. Saloma |
| EPI_ISL_2188950 | Philippine General Hospital (PGH) | Philippine Genome Center | Francis A. Tablizo, Kenneth M. Kim, Maria Sofia L. Yangzon, Elcid Aaron R. Pangilinan, Renato Jacinto Q. Mantaring, Benedict A. Maralit, Marc Edsel C. Ayes, Eva Maria Cutiongco-de la Paz, Alethea R. de Guzman, Jan Michael C. Yap, Jo-Hannah S. Llames, Sheila Mae M. Araiza, Kris P. Punayan, Irish Coleen A. Asin, Candice Francheska B. Tambaoan, Asia Louisa U. Chong, Karol Sophia Agape R. Padilla, Rianna Patricia S. Cruz, Carlo M. Lapid, El King D. Morado, Joshua Gregor A. Dizon, Razel Nikka M. Hao, Arianne A. Zamora, Devon Ray Pacial, Juan Antonio R. Magalang, Marissa Alejandria, Celia Carlos, Anna Ong-Lim, Edsel Maurice Salvaña, John Q. Wong, Jaime C. Montoya, Maria Rosario Singh-Vergeire and Cynthia P. Saloma |
| EPI_ISL_2188951 | Prime Care Alpha Covid-19 Testing Laboratory | Philippine Genome Center | Francis A. Tablizo, Kenneth M. Kim, Maria Sofia L. Yangzon, Elcid Aaron R. Pangilinan, Renato Jacinto Q. Mantaring, Benedict A. Maralit, Marc Edsel C. Ayes, Eva Maria Cutiongco-de la Paz, Alethea R. de Guzman, Jan Michael C. Yap, Jo-Hannah S. Llames, Sheila Mae M. Araiza, Kris P. Punayan, Irish Coleen A. Asin, Candice Francheska B. Tambaoan, Asia Louisa U. Chong, Karol Sophia Agape R. Padilla, Rianna Patricia S. Cruz, Carlo M. Lapid, El King D. Morado, Joshua Gregor A. Dizon, Razel Nikka M. Hao, Arianne A. Zamora, Devon Ray Pacial, Juan Antonio R. Magalang, Marissa Alejandria, Celia Carlos, Anna Ong-Lim, Edsel Maurice Salvaña, John Q. Wong, Jaime C. Montoya, Maria Rosario Singh-Vergeire and Cynthia P. Saloma |
| EPI_ISL_2188952 | Philippine Red Cross - Port Area | Philippine Genome Center | Francis A. Tablizo, Kenneth M. Kim, Maria Sofia L. Yangzon, Elcid Aaron R. Pangilinan, Renato Jacinto Q. Mantaring, Benedict A. Maralit, Marc Edsel C. Ayes, Eva Maria Cutiongco-de la Paz, Alethea R. de Guzman, Jan Michael C. Yap, Jo-Hannah S. Llames, Sheila Mae M. Araiza, Kris P. Punayan, Irish Coleen A. Asin, Candice Francheska B. Tambaoan, Asia Louisa U. Chong, Karol Sophia Agape R. Padilla, Rianna Patricia S. Cruz, Carlo M. Lapid, El King D. Morado, Joshua Gregor A. Dizon, Razel Nikka M. Hao, Arianne A. Zamora, Devon Ray Pacial, Juan Antonio R. Magalang, Marissa Alejandria, Celia Carlos, Anna Ong-Lim, Edsel Maurice Salvaña, John Q. Wong, Jaime C. Montoya, Maria Rosario Singh-Vergeire and Cynthia P. Saloma |
| EPI_ISL_2188953 | The Lord's Grace Medical and Industrial Clinic | Philippine Genome Center | Francis A. Tablizo, Kenneth M. Kim, Maria Sofia L. Yangzon, Elcid Aaron R. Pangilinan, Renato Jacinto Q. Mantaring, Benedict A. Maralit, Marc Edsel C. Ayes, Eva Maria Cutiongco-de la Paz, Alethea R. de Guzman, Jan Michael C. Yap, Jo-Hannah S. Llames, Sheila Mae M. Araiza, Kris P. Punayan, Irish Coleen A. Asin, Candice Francheska B. Tambaoan, Asia Louisa U. Chong, Karol Sophia Agape R. Padilla, Rianna Patricia S. Cruz, Carlo M. Lapid, El King D. Morado, Joshua Gregor A. Dizon, Razel Nikka M. Hao, Arianne A. Zamora, Devon Ray Pacial, Juan Antonio R. Magalang, Marissa Alejandria, Celia Carlos, Anna Ong-Lim, Edsel Maurice Salvaña, John Q. Wong, Jaime C. Montoya, Maria Rosario Singh-Vergeire and Cynthia P. Saloma |
| EPI_ISL_2188954 | The Medical City | Philippine Genome Center | Francis A. Tablizo, Kenneth M. Kim, Maria Sofia L. Yangzon, Elcid Aaron R. Pangilinan, Renato Jacinto Q. Mantaring, Benedict A. Maralit, Marc Edsel C. Ayes, Eva Maria Cutiongco-de la Paz, Alethea R. de Guzman, Jan Michael C. Yap, Jo-Hannah S. Llames, Sheila Mae M. Araiza, Kris P. Punayan, Irish Coleen A. Asin, Candice Francheska B. Tambaoan, Asia Louisa U. Chong, Karol Sophia Agape R. Padilla, Rianna Patricia S. Cruz, Carlo M. Lapid, El King D. Morado, Joshua Gregor A. Dizon, Razel Nikka M. Hao, Arianne A. Zamora, Devon Ray Pacial, Juan Antonio R. Magalang, Marissa Alejandria, Celia Carlos, Anna Ong-Lim, Edsel Maurice Salvaña, John Q. Wong, Jaime C. Montoya, Maria Rosario Singh-Vergeire and Cynthia P. Saloma |
| EPI_ISL_2188955 | The Lord's Grace Medical and Industrial Clinic | Philippine Genome Center | Francis A. Tablizo, Kenneth M. Kim, Maria Sofia L. Yangzon, Elcid Aaron R. Pangilinan, Renato Jacinto Q. Mantaring, Benedict A. Maralit, Marc Edsel C. Ayes, Eva Maria Cutiongco-de la Paz, Alethea R. de Guzman, Jan Michael C. Yap, Jo-Hannah S. Llames, Sheila Mae M. Araiza, Kris P. Punayan, Irish Coleen A. Asin, Candice Francheska B. Tambaoan, Asia Louisa U. Chong, Karol Sophia Agape R. Padilla, Rianna Patricia S. Cruz, Carlo M. Lapid, El King D. Morado, Joshua Gregor A. Dizon, Razel Nikka M. Hao, Arianne A. Zamora, Devon Ray Pacial, Juan Antonio R. Magalang, Marissa Alejandria, Celia Carlos, Anna Ong-Lim, Edsel Maurice Salvaña, John Q. Wong, Jaime C. Montoya, Maria Rosario Singh-Vergeire and Cynthia P. Saloma |
| EPI_ISL_2188956 | The Medical City | Philippine Genome Center | Francis A. Tablizo, Kenneth M. Kim, Maria Sofia L. Yangzon, Elcid Aaron R. Pangilinan, Renato Jacinto Q. Mantaring, Benedict A. Maralit, Marc Edsel C. Ayes, Eva Maria Cutiongco-de la Paz, Alethea R. de Guzman, Jan Michael C. Yap, Jo-Hannah S. Llames, Sheila Mae M. Araiza, Kris P. Punayan, Irish Coleen A. Asin, Candice Francheska B. Tambaoan, Asia Louisa U. Chong, Karol Sophia Agape R. Padilla, Rianna Patricia S. Cruz, Carlo M. Lapid, El King D. Morado, Joshua Gregor A. Dizon, Razel Nikka M. Hao, Arianne A. Zamora, Devon Ray Pacial, Juan Antonio R. Magalang, Marissa Alejandria, Celia Carlos, Anna Ong-Lim, Edsel Maurice Salvaña, John Q. Wong, Jaime C. Montoya, Maria Rosario Singh-Vergeire and Cynthia P. Saloma |
| EPI_ISL_2188957 | The Lord's Grace Medical and Industrial Clinic | Philippine Genome Center | Francis A. Tablizo, Kenneth M. Kim, Maria Sofia L. Yangzon, Elcid Aaron R. Pangilinan, Renato Jacinto Q. Mantaring, Benedict A. Maralit, Marc Edsel C. Ayes, Eva Maria Cutiongco-de la Paz, Alethea R. de Guzman, Jan Michael C. Yap, Jo-Hannah S. Llames, Sheila Mae M. Araiza, Kris P. Punayan, Irish Coleen A. Asin, Candice Francheska B. Tambaoan, Asia Louisa U. Chong, Karol Sophia Agape R. Padilla, Rianna Patricia S. Cruz, Carlo M. Lapid, El King D. Morado, Joshua Gregor A. Dizon, Razel Nikka M. Hao, Arianne A. Zamora, Devon Ray Pacial, Juan Antonio R. Magalang, Marissa Alejandria, Celia Carlos, Anna Ong-Lim, Edsel Maurice Salvaña, John Q. Wong, Jaime C. Montoya, Maria Rosario Singh-Vergeire and Cynthia P. Saloma |
| EPI_ISL_2188961 | Philippine Red Cross - Clark Molecular Laboratory | Philippine Genome Center | Francis A. Tablizo, Kenneth M. Kim, Maria Sofia L. Yangzon, Elcid Aaron R. Pangilinan, Renato Jacinto Q. Mantaring, Benedict A. Maralit, Marc Edsel C. Ayes, Eva Maria Cutiongco-de la Paz, Alethea R. de Guzman, Jan Michael C. Yap, Jo-Hannah S. Llames, Sheila Mae M. Araiza, Kris P. Punayan, Irish Coleen A. Asin, Candice Francheska B. Tambaoan, Asia Louisa U. Chong, Karol Sophia Agape R. Padilla, Rianna Patricia S. Cruz, Carlo M. Lapid, El King D. Morado, Joshua Gregor A. Dizon, Razel Nikka M. Hao, Arianne A. Zamora, Devon Ray Pacial, Juan Antonio R. Magalang, Marissa Alejandria, Celia Carlos, Anna Ong-Lim, Edsel Maurice Salvaña, John Q. Wong, Jaime C. Montoya, Maria Rosario Singh-Vergeire and Cynthia P. Saloma |
| EPI_ISL_2188962 | Parkway Medical and Diagnostic Center | Philippine Genome Center | Francis A. Tablizo, Kenneth M. Kim, Maria Sofia L. Yangzon, Elcid Aaron R. Pangilinan, Renato Jacinto Q. Mantaring, Benedict A. Maralit, Marc Edsel C. Ayes, Eva Maria Cutiongco-de la Paz, Alethea R. de Guzman, Jan Michael C. Yap, Jo-Hannah S. Llames, Sheila Mae M. Araiza, Kris P. Punayan, Irish Coleen A. Asin, Candice Francheska B. Tambaoan, Asia Louisa U. Chong, Karol Sophia Agape R. Padilla, Rianna Patricia S. Cruz, Carlo M. Lapid, El King D. Morado, Joshua Gregor A. Dizon, Razel Nikka M. Hao, Arianne A. Zamora, Devon Ray Pacial, Juan Antonio R. Magalang, Marissa Alejandria, Celia Carlos, Anna Ong-Lim, Edsel Maurice Salvaña, John Q. Wong, Jaime C. Montoya, Maria Rosario Singh-Vergeire and Cynthia P. Saloma |

| | | | |
|---|---|---|---|
| EPI_ISL_2188963 | The Medical City | Philippine Genome Center | Francis A. Tablizo, Kenneth M. Kim, Maria Sofia L. Yangzon, Elcid Aaron R. Pangilinan, Renato Jacinto Q. Mantaring, Benedict A. Maralit, Marc Edsel C. Ayes, Eva Maria Cutiongco-de la Paz, Alethea R. de Guzman, Jan Michael C. Yap, Jo-Hannah S. Llames, Sheila Mae M. Araiza, Kris P. Punayan, Irish Coleen A. Asin, Candice Francheska B. Tambaoan, Asia Louisa U. Chong, Karol Sophia Agape R. Padilla, Rianna Patricia S. Cruz, Carlo M. Lapid, El King D. Morado, Joshua Gregor A. Dizon, Razel Nikka M. Hao, Arianne A. Zamora, Devon Ray Pacial, Juan Antonio R. Magalang, Marissa Alejandria, Celia Carlos, Anna Ong-Lim, Edsel Maurice Salvaña, John Q. Wong, Jaime C. Montoya, Maria Rosario Singh-Vergeire and Cynthia P. Saloma |
| EPI_ISL_2188964 | University of the Philippines National Institutes of Health (UP NIH) | Philippine Genome Center | Francis A. Tablizo, Kenneth M. Kim, Maria Sofia L. Yangzon, Elcid Aaron R. Pangilinan, Renato Jacinto Q. Mantaring, Benedict A. Maralit, Marc Edsel C. Ayes, Eva Maria Cutiongco-de la Paz, Alethea R. de Guzman, Jan Michael C. Yap, Jo-Hannah S. Llames, Sheila Mae M. Araiza, Kris P. Punayan, Irish Coleen A. Asin, Candice Francheska B. Tambaoan, Asia Louisa U. Chong, Karol Sophia Agape R. Padilla, Rianna Patricia S. Cruz, Carlo M. Lapid, El King D. Morado, Joshua Gregor A. Dizon, Razel Nikka M. Hao, Arianne A. Zamora, Devon Ray Pacial, Juan Antonio R. Magalang, Marissa Alejandria, Celia Carlos, Anna Ong-Lim, Edsel Maurice Salvaña, John Q. Wong, Jaime C. Montoya, Maria Rosario Singh-Vergeire and Cynthia P. Saloma |
| EPI_ISL_2188965, EPI_ISL_2188966 | The Lord's Grace Medical and Industrial Clinic | Philippine Genome Center | Francis A. Tablizo, Kenneth M. Kim, Maria Sofia L. Yangzon, Elcid Aaron R. Pangilinan, Renato Jacinto Q. Mantaring, Benedict A. Maralit, Marc Edsel C. Ayes, Eva Maria Cutiongco-de la Paz, Alethea R. de Guzman, Jan Michael C. Yap, Jo-Hannah S. Llames, Sheila Mae M. Araiza, Kris P. Punayan, Irish Coleen A. Asin, Candice Francheska B. Tambaoan, Asia Louisa U. Chong, Karol Sophia Agape R. Padilla, Rianna Patricia S. Cruz, Carlo M. Lapid, El King D. Morado, Joshua Gregor A. Dizon, Razel Nikka M. Hao, Arianne A. Zamora, Devon Ray Pacial, Juan Antonio R. Magalang, Marissa Alejandria, Celia Carlos, Anna Ong-Lim, Edsel Maurice Salvaña, John Q. Wong, Jaime C. Montoya, Maria Rosario Singh-Vergeire and Cynthia P. Saloma |
| EPI_ISL_2188969 | Philippine Red Cross - National Blood Center | Philippine Genome Center | Francis A. Tablizo, Kenneth M. Kim, Maria Sofia L. Yangzon, Elcid Aaron R. Pangilinan, Renato Jacinto Q. Mantaring, Benedict A. Maralit, Marc Edsel C. Ayes, Eva Maria Cutiongco-de la Paz, Alethea R. de Guzman, Jan Michael C. Yap, Jo-Hannah S. Llames, Sheila Mae M. Araiza, Kris P. Punayan, Irish Coleen A. Asin, Candice Francheska B. Tambaoan, Asia Louisa U. Chong, Karol Sophia Agape R. Padilla, Rianna Patricia S. Cruz, Carlo M. Lapid, El King D. Morado, Joshua Gregor A. Dizon, Razel Nikka M. Hao, Arianne A. Zamora, Devon Ray Pacial, Juan Antonio R. Magalang, Marissa Alejandria, Celia Carlos, Anna Ong-Lim, Edsel Maurice Salvaña, John Q. Wong, Jaime C. Montoya, Maria Rosario Singh-Vergeire and Cynthia P. Saloma |
| EPI_ISL_2188970 | Sta. Rosa CHO- GENEXPERT LABORATORY | Philippine Genome Center | Francis A. Tablizo, Kenneth M. Kim, Maria Sofia L. Yangzon, Elcid Aaron R. Pangilinan, Renato Jacinto Q. Mantaring, Benedict A. Maralit, Marc Edsel C. Ayes, Eva Maria Cutiongco-de la Paz, Alethea R. de Guzman, Jan Michael C. Yap, Jo-Hannah S. Llames, Sheila Mae M. Araiza, Kris P. Punayan, Irish Coleen A. Asin, Candice Francheska B. Tambaoan, Asia Louisa U. Chong, Karol Sophia Agape R. Padilla, Rianna Patricia S. Cruz, Carlo M. Lapid, El King D. Morado, Joshua Gregor A. Dizon, Razel Nikka M. Hao, Arianne A. Zamora, Devon Ray Pacial, Juan Antonio R. Magalang, Marissa Alejandria, Celia Carlos, Anna Ong-Lim, Edsel Maurice Salvaña, John Q. Wong, Jaime C. Montoya, Maria Rosario Singh-Vergeire and Cynthia P. Saloma |
| EPI_ISL_2188971 | The Lord's Grace Medical and Industrial Clinic | Philippine Genome Center | Francis A. Tablizo, Kenneth M. Kim, Maria Sofia L. Yangzon, Elcid Aaron R. Pangilinan, Renato Jacinto Q. Mantaring, Benedict A. Maralit, Marc Edsel C. Ayes, Eva Maria Cutiongco-de la Paz, Alethea R. de Guzman, Jan Michael C. Yap, Jo-Hannah S. Llames, Sheila Mae M. Araiza, Kris P. Punayan, Irish Coleen A. Asin, Candice Francheska B. Tambaoan, Asia Louisa U. Chong, Karol Sophia Agape R. Padilla, Rianna Patricia S. Cruz, Carlo M. Lapid, El King D. Morado, Joshua Gregor A. Dizon, Razel Nikka M. Hao, Arianne A. Zamora, Devon Ray Pacial, Juan Antonio R. Magalang, Marissa Alejandria, Celia Carlos, Anna Ong-Lim, Edsel Maurice Salvaña, John Q. Wong, Jaime C. Montoya, Maria Rosario Singh-Vergeire and Cynthia P. Saloma |
| EPI_ISL_2188973 | Philippine Red Cross - National Blood Center | Philippine Genome Center | Francis A. Tablizo, Kenneth M. Kim, Maria Sofia L. Yangzon, Elcid Aaron R. Pangilinan, Renato Jacinto Q. Mantaring, Benedict A. Maralit, Marc Edsel C. Ayes, Eva Maria Cutiongco-de la Paz, Alethea R. de Guzman, Jan Michael C. Yap, Jo-Hannah S. Llames, Sheila Mae M. Araiza, Kris P. Punayan, Irish Coleen A. Asin, Candice Francheska B. Tambaoan, Asia Louisa U. Chong, Karol Sophia Agape R. Padilla, Rianna Patricia S. Cruz, Carlo M. Lapid, El King D. Morado, Joshua Gregor A. Dizon, Razel Nikka M. Hao, Arianne A. Zamora, Devon Ray Pacial, Juan Antonio R. Magalang, Marissa Alejandria, Celia Carlos, Anna Ong-Lim, Edsel Maurice Salvaña, John Q. Wong, Jaime C. Montoya, Maria Rosario Singh-Vergeire and Cynthia P. Saloma |
| EPI_ISL_2188975 | The Medical City | Philippine Genome Center | Francis A. Tablizo, Kenneth M. Kim, Maria Sofia L. Yangzon, Elcid Aaron R. Pangilinan, Renato Jacinto Q. Mantaring, Benedict A. Maralit, Marc Edsel C. Ayes, Eva Maria Cutiongco-de la Paz, Alethea R. de Guzman, Jan Michael C. Yap, Jo-Hannah S. Llames, Sheila Mae M. Araiza, Kris P. Punayan, Irish Coleen A. Asin, Candice Francheska B. Tambaoan, Asia Louisa U. Chong, Karol Sophia Agape R. Padilla, Rianna Patricia S. Cruz, Carlo M. Lapid, El King D. Morado, Joshua Gregor A. Dizon, Razel Nikka M. Hao, Arianne A. Zamora, Devon Ray Pacial, Juan Antonio R. Magalang, Marissa Alejandria, Celia Carlos, Anna Ong-Lim, Edsel Maurice Salvaña, John Q. Wong, Jaime C. Montoya, Maria Rosario Singh-Vergeire and Cynthia P. Saloma |
| EPI_ISL_2188977 | Philippine Red Cross - Port Area | Philippine Genome Center | Francis A. Tablizo, Kenneth M. Kim, Maria Sofia L. Yangzon, Elcid Aaron R. Pangilinan, Renato Jacinto Q. Mantaring, Benedict A. Maralit, Marc Edsel C. Ayes, Eva Maria Cutiongco-de la Paz, Alethea R. de Guzman, Jan Michael C. Yap, Jo-Hannah S. Llames, Sheila Mae M. Araiza, Kris P. Punayan, Irish Coleen A. Asin, Candice Francheska B. Tambaoan, Asia Louisa U. Chong, Karol Sophia Agape R. Padilla, Rianna Patricia S. Cruz, Carlo M. Lapid, El King D. Morado, Joshua Gregor A. Dizon, Razel Nikka M. Hao, Arianne A. Zamora, Devon Ray Pacial, Juan Antonio R. Magalang, Marissa Alejandria, Celia Carlos, Anna Ong-Lim, Edsel Maurice Salvaña, John Q. Wong, Jaime C. Montoya, Maria Rosario Singh-Vergeire and Cynthia P. Saloma |
| EPI_ISL_2188978 | The Lord's Grace Medical and Industrial Clinic | Philippine Genome Center | Francis A. Tablizo, Kenneth M. Kim, Maria Sofia L. Yangzon, Elcid Aaron R. Pangilinan, Renato Jacinto Q. Mantaring, Benedict A. Maralit, Marc Edsel C. Ayes, Eva Maria Cutiongco-de la Paz, Alethea R. de Guzman, Jan Michael C. Yap, Jo-Hannah S. Llames, Sheila Mae M. Araiza, Kris P. Punayan, Irish Coleen A. Asin, Candice Francheska B. Tambaoan, Asia Louisa U. Chong, Karol Sophia Agape R. Padilla, Rianna Patricia S. Cruz, Carlo M. Lapid, El King D. Morado, Joshua Gregor A. Dizon, Razel Nikka M. Hao, Arianne A. Zamora, Devon Ray Pacial, Juan Antonio R. Magalang, Marissa Alejandria, Celia Carlos, Anna Ong-Lim, Edsel Maurice Salvaña, John Q. Wong, Jaime C. Montoya, Maria Rosario Singh-Vergeire and Cynthia P. Saloma |
| EPI_ISL_2188979 | Mary Mediatrix Medical Center | Philippine Genome Center | Francis A. Tablizo, Kenneth M. Kim, Maria Sofia L. Yangzon, Elcid Aaron R. Pangilinan, Renato Jacinto Q. Mantaring, Benedict A. Maralit, Marc Edsel C. Ayes, Eva Maria Cutiongco-de la Paz, Alethea R. de Guzman, Jan Michael C. Yap, Jo-Hannah S. Llames, Sheila Mae M. Araiza, Kris P. Punayan, Irish Coleen A. Asin, Candice Francheska B. Tambaoan, Asia Louisa U. Chong, Karol Sophia Agape R. Padilla, Rianna Patricia S. Cruz, Carlo M. Lapid, El King D. Morado, Joshua Gregor A. Dizon, Razel Nikka M. Hao, Arianne A. Zamora, Devon Ray Pacial, Juan Antonio R. Magalang, Marissa Alejandria, Celia Carlos, Anna Ong-Lim, Edsel Maurice Salvaña, John Q. Wong, Jaime C. Montoya, Maria Rosario Singh-Vergeire and Cynthia P. Saloma |
| EPI_ISL_2188980 | The Lord's Grace Medical and Industrial Clinic | Philippine Genome Center | Francis A. Tablizo, Kenneth M. Kim, Maria Sofia L. Yangzon, Elcid Aaron R. Pangilinan, Renato Jacinto Q. Mantaring, Benedict A. Maralit, Marc Edsel C. Ayes, Eva Maria Cutiongco-de la Paz, Alethea R. de Guzman, Jan Michael C. Yap, Jo-Hannah S. Llames, Sheila Mae M. Araiza, Kris P. Punayan, Irish Coleen A. Asin, Candice Francheska B. Tambaoan, Asia Louisa U. Chong, Karol Sophia Agape R. Padilla, Rianna Patricia S. Cruz, Carlo M. Lapid, El King D. Morado, Joshua Gregor A. Dizon, Razel Nikka M. Hao, Arianne A. Zamora, Devon Ray Pacial, Juan Antonio R. Magalang, Marissa Alejandria, Celia Carlos, Anna Ong-Lim, Edsel Maurice Salvaña, John Q. Wong, Jaime C. Montoya, Maria Rosario Singh-Vergeire and Cynthia P. Saloma |
| EPI_ISL_2188983 | Jose B. Lingad Memorial General Hospital | Philippine Genome Center | Francis A. Tablizo, Kenneth M. Kim, Maria Sofia L. Yangzon, Elcid Aaron R. Pangilinan, Renato Jacinto Q. Mantaring, Benedict A. Maralit, Marc Edsel C. Ayes, Eva Maria Cutiongco-de la Paz, Alethea R. de Guzman, Jan Michael C. Yap, Jo-Hannah S. Llames, Sheila Mae M. Araiza, Kris P. Punayan, Irish Coleen A. Asin, Candice Francheska B. Tambaoan, Asia Louisa U. Chong, Karol Sophia Agape R. Padilla, Rianna Patricia S. Cruz, Carlo M. Lapid, El King D. Morado, Joshua Gregor A. Dizon, Razel Nikka M. Hao, Arianne A. Zamora, Devon Ray Pacial, Juan Antonio R. Magalang, Marissa Alejandria, Celia Carlos, Anna Ong-Lim, Edsel Maurice Salvaña, John Q. Wong, Jaime C. Montoya, Maria Rosario Singh-Vergeire and Cynthia P. Saloma |
| EPI_ISL_2188984 | Philippine General Hospital (PGH) | Philippine Genome Center | Francis A. Tablizo, Kenneth M. Kim, Maria Sofia L. Yangzon, Elcid Aaron R. Pangilinan, Renato Jacinto Q. Mantaring, Benedict A. Maralit, Marc Edsel C. Ayes, Eva Maria Cutiongco-de la Paz, Alethea R. de Guzman, Jan Michael C. Yap, Jo-Hannah S. Llames, Sheila Mae M. Araiza, Kris P. Punayan, Irish Coleen A. Asin, Candice Francheska B. Tambaoan, Asia Louisa U. Chong, Karol Sophia Agape R. Padilla, Rianna Patricia S. Cruz, Carlo M. Lapid, El King D. Morado, Joshua Gregor A. Dizon, Razel Nikka M. Hao, Arianne A. Zamora, Devon Ray Pacial, Juan Antonio R. Magalang, Marissa Alejandria, Celia Carlos, Anna Ong-Lim, Edsel Maurice Salvaña, John Q. Wong, Jaime C. Montoya, Maria Rosario Singh-Vergeire and Cynthia P. Saloma |
| EPI_ISL_2188985 | The Lord's Grace Medical and Industrial Clinic | Philippine Genome Center | Francis A. Tablizo, Kenneth M. Kim, Maria Sofia L. Yangzon, Elcid Aaron R. Pangilinan, Renato Jacinto Q. Mantaring, Benedict A. Maralit, Marc Edsel C. Ayes, Eva Maria Cutiongco-de la Paz, Alethea R. de Guzman, Jan Michael C. Yap, Jo-Hannah S. Llames, Sheila Mae M. Araiza, Kris P. Punayan, Irish Coleen A. Asin, Candice Francheska B. Tambaoan, Asia Louisa U. Chong, Karol Sophia Agape R. Padilla, Rianna Patricia S. Cruz, Carlo M. Lapid, El King D. Morado, Joshua Gregor A. Dizon, Razel Nikka M. Hao, Arianne A. Zamora, Devon Ray Pacial, Juan Antonio R. Magalang, Marissa Alejandria, Celia Carlos, Anna Ong-Lim, Edsel Maurice Salvaña, John Q. Wong, Jaime C. Montoya, Maria Rosario Singh-Vergeire and Cynthia P. Saloma |
| EPI_ISL_2188986 | The Medical City | Philippine Genome Center | Francis A. Tablizo, Kenneth M. Kim, Maria Sofia L. Yangzon, Elcid Aaron R. Pangilinan, Renato Jacinto Q. Mantaring, Benedict A. Maralit, Marc Edsel C. Ayes, Eva Maria Cutiongco-de la Paz, Alethea R. de Guzman, Jan Michael C. Yap, Jo-Hannah S. Llames, Sheila Mae M. Araiza, Kris P. Punayan, Irish Coleen A. Asin, Candice Francheska B. Tambaoan, Asia Louisa U. Chong, Karol Sophia Agape R. Padilla, Rianna Patricia S. Cruz, Carlo M. Lapid, El King D. Morado, Joshua Gregor A. Dizon, Razel Nikka M. Hao, Arianne A. Zamora, Devon Ray Pacial, Juan Antonio R. Magalang, Marissa Alejandria, Celia Carlos, Anna Ong-Lim, Edsel Maurice Salvaña, John Q. Wong, Jaime C. Montoya, Maria Rosario Singh-Vergeire and Cynthia P. Saloma |

| | | | |
|---|---|---|---|
| EPI_ISL_2188987 | The Lord's Grace Medical and Industrial Clinic | Philippine Genome Center | Francis A. Tablizo, Kenneth M. Kim, Maria Sofia L. Yangzon, Elcid Aaron R. Pangilinan, Renato Jacinto Q. Mantaring, Benedict A. Maralit, Marc Edsel C. Ayes, Eva Maria Cutiongco-de la Paz, Alethea R. de Guzman, Jan Michael C. Yap, Jo-Hannah S. Llames, Sheila Mae M. Araiza, Kris P. Punayan, Irish Coleen A. Asin, Candice Francheska B. Tambaoan, Asia Louisa U. Chong, Karol Sophia Agape R. Padilla, Rianna Patricia S. Cruz, Carlo M. Lapid, El King D. Morado, Joshua Gregor A. Dizon, Razel Nikka M. Hao, Arianne A. Zamora, Devon Ray Pacial, Juan Antonio R. Magalang, Marissa Alejandria, Celia Carlos, Anna Ong-Lim, Edsel Maurice Salvaña, John Q. Wong, Jaime C. Montoya, Maria Rosario Singh-Vergeire and Cynthia P. Saloma |
| EPI_ISL_2188988, EPI_ISL_2188989 | The Medical City | Philippine Genome Center | Francis A. Tablizo, Kenneth M. Kim, Maria Sofia L. Yangzon, Elcid Aaron R. Pangilinan, Renato Jacinto Q. Mantaring, Benedict A. Maralit, Marc Edsel C. Ayes, Eva Maria Cutiongco-de la Paz, Alethea R. de Guzman, Jan Michael C. Yap, Jo-Hannah S. Llames, Sheila Mae M. Araiza, Kris P. Punayan, Irish Coleen A. Asin, Candice Francheska B. Tambaoan, Asia Louisa U. Chong, Karol Sophia Agape R. Padilla, Rianna Patricia S. Cruz, Carlo M. Lapid, El King D. Morado, Joshua Gregor A. Dizon, Razel Nikka M. Hao, Arianne A. Zamora, Devon Ray Pacial, Juan Antonio R. Magalang, Marissa Alejandria, Celia Carlos, Anna Ong-Lim, Edsel Maurice Salvaña, John Q. Wong, Jaime C. Montoya, Maria Rosario Singh-Vergeire and Cynthia P. Saloma |
| EPI_ISL_2188990 | Sta. Ana Hospital | Philippine Genome Center | Francis A. Tablizo, Kenneth M. Kim, Maria Sofia L. Yangzon, Elcid Aaron R. Pangilinan, Renato Jacinto Q. Mantaring, Benedict A. Maralit, Marc Edsel C. Ayes, Eva Maria Cutiongco-de la Paz, Alethea R. de Guzman, Jan Michael C. Yap, Jo-Hannah S. Llames, Sheila Mae M. Araiza, Kris P. Punayan, Irish Coleen A. Asin, Candice Francheska B. Tambaoan, Asia Louisa U. Chong, Karol Sophia Agape R. Padilla, Rianna Patricia S. Cruz, Carlo M. Lapid, El King D. Morado, Joshua Gregor A. Dizon, Razel Nikka M. Hao, Arianne A. Zamora, Devon Ray Pacial, Juan Antonio R. Magalang, Marissa Alejandria, Celia Carlos, Anna Ong-Lim, Edsel Maurice Salvaña, John Q. Wong, Jaime C. Montoya, Maria Rosario Singh-Vergeire and Cynthia P. Saloma |
| EPI_ISL_2188991 | The Medical City | Philippine Genome Center | Francis A. Tablizo, Kenneth M. Kim, Maria Sofia L. Yangzon, Elcid Aaron R. Pangilinan, Renato Jacinto Q. Mantaring, Benedict A. Maralit, Marc Edsel C. Ayes, Eva Maria Cutiongco-de la Paz, Alethea R. de Guzman, Jan Michael C. Yap, Jo-Hannah S. Llames, Sheila Mae M. Araiza, Kris P. Punayan, Irish Coleen A. Asin, Candice Francheska B. Tambaoan, Asia Louisa U. Chong, Karol Sophia Agape R. Padilla, Rianna Patricia S. Cruz, Carlo M. Lapid, El King D. Morado, Joshua Gregor A. Dizon, Razel Nikka M. Hao, Arianne A. Zamora, Devon Ray Pacial, Juan Antonio R. Magalang, Marissa Alejandria, Celia Carlos, Anna Ong-Lim, Edsel Maurice Salvaña, John Q. Wong, Jaime C. Montoya, Maria Rosario Singh-Vergeire and Cynthia P. Saloma |
| EPI_ISL_2188994 | Butuan Medical Center | Philippine Genome Center | Francis A. Tablizo, Kenneth M. Kim, Maria Sofia L. Yangzon, Elcid Aaron R. Pangilinan, Renato Jacinto Q. Mantaring, Benedict A. Maralit, Marc Edsel C. Ayes, Eva Maria Cutiongco-de la Paz, Alethea R. de Guzman, Jan Michael C. Yap, Jo-Hannah S. Llames, Sheila Mae M. Araiza, Kris P. Punayan, Irish Coleen A. Asin, Candice Francheska B. Tambaoan, Asia Louisa U. Chong, Karol Sophia Agape R. Padilla, Rianna Patricia S. Cruz, Carlo M. Lapid, El King D. Morado, Joshua Gregor A. Dizon, Razel Nikka M. Hao, Arianne A. Zamora, Devon Ray Pacial, Juan Antonio R. Magalang, Marissa Alejandria, Celia Carlos, Anna Ong-Lim, Edsel Maurice Salvaña, John Q. Wong, Jaime C. Montoya, Maria Rosario Singh-Vergeire and Cynthia P. Saloma |
| EPI_ISL_2188995 | The Medical City | Philippine Genome Center | Francis A. Tablizo, Kenneth M. Kim, Maria Sofia L. Yangzon, Elcid Aaron R. Pangilinan, Renato Jacinto Q. Mantaring, Benedict A. Maralit, Marc Edsel C. Ayes, Eva Maria Cutiongco-de la Paz, Alethea R. de Guzman, Jan Michael C. Yap, Jo-Hannah S. Llames, Sheila Mae M. Araiza, Kris P. Punayan, Irish Coleen A. Asin, Candice Francheska B. Tambaoan, Asia Louisa U. Chong, Karol Sophia Agape R. Padilla, Rianna Patricia S. Cruz, Carlo M. Lapid, El King D. Morado, Joshua Gregor A. Dizon, Razel Nikka M. Hao, Arianne A. Zamora, Devon Ray Pacial, Juan Antonio R. Magalang, Marissa Alejandria, Celia Carlos, Anna Ong-Lim, Edsel Maurice Salvaña, John Q. Wong, Jaime C. Montoya, Maria Rosario Singh-Vergeire and Cynthia P. Saloma |
| EPI_ISL_2188996 | Mary Mediatrix Medical Center | Philippine Genome Center | Francis A. Tablizo, Kenneth M. Kim, Maria Sofia L. Yangzon, Elcid Aaron R. Pangilinan, Renato Jacinto Q. Mantaring, Benedict A. Maralit, Marc Edsel C. Ayes, Eva Maria Cutiongco-de la Paz, Alethea R. de Guzman, Jan Michael C. Yap, Jo-Hannah S. Llames, Sheila Mae M. Araiza, Kris P. Punayan, Irish Coleen A. Asin, Candice Francheska B. Tambaoan, Asia Louisa U. Chong, Karol Sophia Agape R. Padilla, Rianna Patricia S. Cruz, Carlo M. Lapid, El King D. Morado, Joshua Gregor A. Dizon, Razel Nikka M. Hao, Arianne A. Zamora, Devon Ray Pacial, Juan Antonio R. Magalang, Marissa Alejandria, Celia Carlos, Anna Ong-Lim, Edsel Maurice Salvaña, John Q. Wong, Jaime C. Montoya, Maria Rosario Singh-Vergeire and Cynthia P. Saloma |
| EPI_ISL_2188998 | The Medical City | Philippine Genome Center | Francis A. Tablizo, Kenneth M. Kim, Maria Sofia L. Yangzon, Elcid Aaron R. Pangilinan, Renato Jacinto Q. Mantaring, Benedict A. Maralit, Marc Edsel C. Ayes, Eva Maria Cutiongco-de la Paz, Alethea R. de Guzman, Jan Michael C. Yap, Jo-Hannah S. Llames, Sheila Mae M. Araiza, Kris P. Punayan, Irish Coleen A. Asin, Candice Francheska B. Tambaoan, Asia Louisa U. Chong, Karol Sophia Agape R. Padilla, Rianna Patricia S. Cruz, Carlo M. Lapid, El King D. Morado, Joshua Gregor A. Dizon, Razel Nikka M. Hao, Arianne A. Zamora, Devon Ray Pacial, Juan Antonio R. Magalang, Marissa Alejandria, Celia Carlos, Anna Ong-Lim, Edsel Maurice Salvaña, John Q. Wong, Jaime C. Montoya, Maria Rosario Singh-Vergeire and Cynthia P. Saloma |
| EPI_ISL_2188999 | Laguna Provincial Hospital - San Pablo District Hospital | Philippine Genome Center | Francis A. Tablizo, Kenneth M. Kim, Maria Sofia L. Yangzon, Elcid Aaron R. Pangilinan, Renato Jacinto Q. Mantaring, Benedict A. Maralit, Marc Edsel C. Ayes, Eva Maria Cutiongco-de la Paz, Alethea R. de Guzman, Jan Michael C. Yap, Jo-Hannah S. Llames, Sheila Mae M. Araiza, Kris P. Punayan, Irish Coleen A. Asin, Candice Francheska B. Tambaoan, Asia Louisa U. Chong, Karol Sophia Agape R. Padilla, Rianna Patricia S. Cruz, Carlo M. Lapid, El King D. Morado, Joshua Gregor A. Dizon, Razel Nikka M. Hao, Arianne A. Zamora, Devon Ray Pacial, Juan Antonio R. Magalang, Marissa Alejandria, Celia Carlos, Anna Ong-Lim, Edsel Maurice Salvaña, John Q. Wong, Jaime C. Montoya, Maria Rosario Singh-Vergeire and Cynthia P. Saloma |
| EPI_ISL_2189000 | Prime Care Alpha Covid-19 Testing Laboratory | Philippine Genome Center | Francis A. Tablizo, Kenneth M. Kim, Maria Sofia L. Yangzon, Elcid Aaron R. Pangilinan, Renato Jacinto Q. Mantaring, Benedict A. Maralit, Marc Edsel C. Ayes, Eva Maria Cutiongco-de la Paz, Alethea R. de Guzman, Jan Michael C. Yap, Jo-Hannah S. Llames, Sheila Mae M. Araiza, Kris P. Punayan, Irish Coleen A. Asin, Candice Francheska B. Tambaoan, Asia Louisa U. Chong, Karol Sophia Agape R. Padilla, Rianna Patricia S. Cruz, Carlo M. Lapid, El King D. Morado, Joshua Gregor A. Dizon, Razel Nikka M. Hao, Arianne A. Zamora, Devon Ray Pacial, Juan Antonio R. Magalang, Marissa Alejandria, Celia Carlos, Anna Ong-Lim, Edsel Maurice Salvaña, John Q. Wong, Jaime C. Montoya, Maria Rosario Singh-Vergeire and Cynthia P. Saloma |
| EPI_ISL_2189001 | The Lord's Grace Medical and Industrial Clinic | Philippine Genome Center | Francis A. Tablizo, Kenneth M. Kim, Maria Sofia L. Yangzon, Elcid Aaron R. Pangilinan, Renato Jacinto Q. Mantaring, Benedict A. Maralit, Marc Edsel C. Ayes, Eva Maria Cutiongco-de la Paz, Alethea R. de Guzman, Jan Michael C. Yap, Jo-Hannah S. Llames, Sheila Mae M. Araiza, Kris P. Punayan, Irish Coleen A. Asin, Candice Francheska B. Tambaoan, Asia Louisa U. Chong, Karol Sophia Agape R. Padilla, Rianna Patricia S. Cruz, Carlo M. Lapid, El King D. Morado, Joshua Gregor A. Dizon, Razel Nikka M. Hao, Arianne A. Zamora, Devon Ray Pacial, Juan Antonio R. Magalang, Marissa Alejandria, Celia Carlos, Anna Ong-Lim, Edsel Maurice Salvaña, John Q. Wong, Jaime C. Montoya, Maria Rosario Singh-Vergeire and Cynthia P. Saloma |
| EPI_ISL_2189002 | Mary Mediatrix Medical Center | Philippine Genome Center | Francis A. Tablizo, Kenneth M. Kim, Maria Sofia L. Yangzon, Elcid Aaron R. Pangilinan, Renato Jacinto Q. Mantaring, Benedict A. Maralit, Marc Edsel C. Ayes, Eva Maria Cutiongco-de la Paz, Alethea R. de Guzman, Jan Michael C. Yap, Jo-Hannah S. Llames, Sheila Mae M. Araiza, Kris P. Punayan, Irish Coleen A. Asin, Candice Francheska B. Tambaoan, Asia Louisa U. Chong, Karol Sophia Agape R. Padilla, Rianna Patricia S. Cruz, Carlo M. Lapid, El King D. Morado, Joshua Gregor A. Dizon, Razel Nikka M. Hao, Arianne A. Zamora, Devon Ray Pacial, Juan Antonio R. Magalang, Marissa Alejandria, Celia Carlos, Anna Ong-Lim, Edsel Maurice Salvaña, John Q. Wong, Jaime C. Montoya, Maria Rosario Singh-Vergeire and Cynthia P. Saloma |
| EPI_ISL_2189003 | Philippine Red Cross - Port Area | Philippine Genome Center | Francis A. Tablizo, Kenneth M. Kim, Maria Sofia L. Yangzon, Elcid Aaron R. Pangilinan, Renato Jacinto Q. Mantaring, Benedict A. Maralit, Marc Edsel C. Ayes, Eva Maria Cutiongco-de la Paz, Alethea R. de Guzman, Jan Michael C. Yap, Jo-Hannah S. Llames, Sheila Mae M. Araiza, Kris P. Punayan, Irish Coleen A. Asin, Candice Francheska B. Tambaoan, Asia Louisa U. Chong, Karol Sophia Agape R. Padilla, Rianna Patricia S. Cruz, Carlo M. Lapid, El King D. Morado, Joshua Gregor A. Dizon, Razel Nikka M. Hao, Arianne A. Zamora, Devon Ray Pacial, Juan Antonio R. Magalang, Marissa Alejandria, Celia Carlos, Anna Ong-Lim, Edsel Maurice Salvaña, John Q. Wong, Jaime C. Montoya, Maria Rosario Singh-Vergeire and Cynthia P. Saloma |
| EPI_ISL_2189004, EPI_ISL_2189006 | The Lord's Grace Medical and Industrial Clinic | Philippine Genome Center | Francis A. Tablizo, Kenneth M. Kim, Maria Sofia L. Yangzon, Elcid Aaron R. Pangilinan, Renato Jacinto Q. Mantaring, Benedict A. Maralit, Marc Edsel C. Ayes, Eva Maria Cutiongco-de la Paz, Alethea R. de Guzman, Jan Michael C. Yap, Jo-Hannah S. Llames, Sheila Mae M. Araiza, Kris P. Punayan, Irish Coleen A. Asin, Candice Francheska B. Tambaoan, Asia Louisa U. Chong, Karol Sophia Agape R. Padilla, Rianna Patricia S. Cruz, Carlo M. Lapid, El King D. Morado, Joshua Gregor A. Dizon, Razel Nikka M. Hao, Arianne A. Zamora, Devon Ray Pacial, Juan Antonio R. Magalang, Marissa Alejandria, Celia Carlos, Anna Ong-Lim, Edsel Maurice Salvaña, John Q. Wong, Jaime C. Montoya, Maria Rosario Singh-Vergeire and Cynthia P. Saloma |
| EPI_ISL_2189007 | The Medical City | Philippine Genome Center | Francis A. Tablizo, Kenneth M. Kim, Maria Sofia L. Yangzon, Elcid Aaron R. Pangilinan, Renato Jacinto Q. Mantaring, Benedict A. Maralit, Marc Edsel C. Ayes, Eva Maria Cutiongco-de la Paz, Alethea R. de Guzman, Jan Michael C. Yap, Jo-Hannah S. Llames, Sheila Mae M. Araiza, Kris P. Punayan, Irish Coleen A. Asin, Candice Francheska B. Tambaoan, Asia Louisa U. Chong, Karol Sophia Agape R. Padilla, Rianna Patricia S. Cruz, Carlo M. Lapid, El King D. Morado, Joshua Gregor A. Dizon, Razel Nikka M. Hao, Arianne A. Zamora, Devon Ray Pacial, Juan Antonio R. Magalang, Marissa Alejandria, Celia Carlos, Anna Ong-Lim, Edsel Maurice Salvaña, John Q. Wong, Jaime C. Montoya, Maria Rosario Singh-Vergeire and Cynthia P. Saloma |
| EPI_ISL_2189014 | Philippine Red Cross - Port Area | Philippine Genome Center | Francis A. Tablizo, Kenneth M. Kim, Maria Sofia L. Yangzon, Elcid Aaron R. Pangilinan, Renato Jacinto Q. Mantaring, Benedict A. Maralit, Marc Edsel C. Ayes, Eva Maria Cutiongco-de la Paz, Alethea R. de Guzman, Jan Michael C. Yap, Jo-Hannah S. Llames, Sheila Mae M. Araiza, Kris P. Punayan, Irish Coleen A. Asin, Candice Francheska B. Tambaoan, Asia Louisa U. Chong, Karol Sophia Agape R. Padilla, Rianna Patricia S. Cruz, Carlo M. Lapid, El King D. Morado, Joshua Gregor A. Dizon, Razel Nikka M. Hao, Arianne A. Zamora, Devon Ray Pacial, Juan Antonio R. Magalang, Marissa Alejandria, Celia Carlos, Anna Ong-Lim, Edsel Maurice Salvaña, John Q. Wong, Jaime C. Montoya, Maria Rosario Singh-Vergeire and Cynthia P. Saloma |

| | | | |
|---|---|---|---|
| EPI_ISL_2189015 | Pasig City Children's Hospital - Child's Hope | Philippine Genome Center | Francis A. Tablizo, Kenneth M. Kim, Maria Sofia L. Yangzon, Elcid Aaron R. Pangilinan, Renato Jacinto Q. Mantaring, Benedict A. Maralit, Marc Edsel C. Ayes, Eva Maria Cutiongco-de la Paz, Alethea R. de Guzman, Jan Michael C. Yap, Jo-Hannah S. Llames, Sheila Mae M. Araiza, Kris P. Punayan, Irish Coleen A. Asin, Candice Francheska B. Tambaoan, Asia Louisa U. Chong, Karol Sophia Agape R. Padilla, Rianna Patricia S. Cruz, Carlo M. Lapid, El King D. Morado, Joshua Gregor A. Dizon, Razel Nikka M. Hao, Arianne A. Zamora, Devon Ray Pacial, Juan Antonio R. Magalang, Marissa Alejandria, Celia Carlos, Anna Ong-Lim, Edsel Maurice Salvaña, John Q. Wong, Jaime C. Montoya, Maria Rosario Singh-Vergeire and Cynthia P. Saloma |
| EPI_ISL_2189016 | Philippine Red Cross - Port Area | Philippine Genome Center | Francis A. Tablizo, Kenneth M. Kim, Maria Sofia L. Yangzon, Elcid Aaron R. Pangilinan, Renato Jacinto Q. Mantaring, Benedict A. Maralit, Marc Edsel C. Ayes, Eva Maria Cutiongco-de la Paz, Alethea R. de Guzman, Jan Michael C. Yap, Jo-Hannah S. Llames, Sheila Mae M. Araiza, Kris P. Punayan, Irish Coleen A. Asin, Candice Francheska B. Tambaoan, Asia Louisa U. Chong, Karol Sophia Agape R. Padilla, Rianna Patricia S. Cruz, Carlo M. Lapid, El King D. Morado, Joshua Gregor A. Dizon, Razel Nikka M. Hao, Arianne A. Zamora, Devon Ray Pacial, Juan Antonio R. Magalang, Marissa Alejandria, Celia Carlos, Anna Ong-Lim, Edsel Maurice Salvaña, John Q. Wong, Jaime C. Montoya, Maria Rosario Singh-Vergeire and Cynthia P. Saloma |
| EPI_ISL_2189019 | The Lord's Grace Medical and Industrial Clinic | Philippine Genome Center | Francis A. Tablizo, Kenneth M. Kim, Maria Sofia L. Yangzon, Elcid Aaron R. Pangilinan, Renato Jacinto Q. Mantaring, Benedict A. Maralit, Marc Edsel C. Ayes, Eva Maria Cutiongco-de la Paz, Alethea R. de Guzman, Jan Michael C. Yap, Jo-Hannah S. Llames, Sheila Mae M. Araiza, Kris P. Punayan, Irish Coleen A. Asin, Candice Francheska B. Tambaoan, Asia Louisa U. Chong, Karol Sophia Agape R. Padilla, Rianna Patricia S. Cruz, Carlo M. Lapid, El King D. Morado, Joshua Gregor A. Dizon, Razel Nikka M. Hao, Arianne A. Zamora, Devon Ray Pacial, Juan Antonio R. Magalang, Marissa Alejandria, Celia Carlos, Anna Ong-Lim, Edsel Maurice Salvaña, John Q. Wong, Jaime C. Montoya, Maria Rosario Singh-Vergeire and Cynthia P. Saloma |
| EPI_ISL_2189021 | Parkway Medical and Diagnostic Center | Philippine Genome Center | Francis A. Tablizo, Kenneth M. Kim, Maria Sofia L. Yangzon, Elcid Aaron R. Pangilinan, Renato Jacinto Q. Mantaring, Benedict A. Maralit, Marc Edsel C. Ayes, Eva Maria Cutiongco-de la Paz, Alethea R. de Guzman, Jan Michael C. Yap, Jo-Hannah S. Llames, Sheila Mae M. Araiza, Kris P. Punayan, Irish Coleen A. Asin, Candice Francheska B. Tambaoan, Asia Louisa U. Chong, Karol Sophia Agape R. Padilla, Rianna Patricia S. Cruz, Carlo M. Lapid, El King D. Morado, Joshua Gregor A. Dizon, Razel Nikka M. Hao, Arianne A. Zamora, Devon Ray Pacial, Juan Antonio R. Magalang, Marissa Alejandria, Celia Carlos, Anna Ong-Lim, Edsel Maurice Salvaña, John Q. Wong, Jaime C. Montoya, Maria Rosario Singh-Vergeire and Cynthia P. Saloma |
| EPI_ISL_2189022 | Philippine General Hospital (PGH) | Philippine Genome Center | Francis A. Tablizo, Kenneth M. Kim, Maria Sofia L. Yangzon, Elcid Aaron R. Pangilinan, Renato Jacinto Q. Mantaring, Benedict A. Maralit, Marc Edsel C. Ayes, Eva Maria Cutiongco-de la Paz, Alethea R. de Guzman, Jan Michael C. Yap, Jo-Hannah S. Llames, Sheila Mae M. Araiza, Kris P. Punayan, Irish Coleen A. Asin, Candice Francheska B. Tambaoan, Asia Louisa U. Chong, Karol Sophia Agape R. Padilla, Rianna Patricia S. Cruz, Carlo M. Lapid, El King D. Morado, Joshua Gregor A. Dizon, Razel Nikka M. Hao, Arianne A. Zamora, Devon Ray Pacial, Juan Antonio R. Magalang, Marissa Alejandria, Celia Carlos, Anna Ong-Lim, Edsel Maurice Salvaña, John Q. Wong, Jaime C. Montoya, Maria Rosario Singh-Vergeire and Cynthia P. Saloma |
| EPI_ISL_2189024 | South Super Highway Molecular Diagnostic Laboratory | Philippine Genome Center | Francis A. Tablizo, Kenneth M. Kim, Maria Sofia L. Yangzon, Elcid Aaron R. Pangilinan, Renato Jacinto Q. Mantaring, Benedict A. Maralit, Marc Edsel C. Ayes, Eva Maria Cutiongco-de la Paz, Alethea R. de Guzman, Jan Michael C. Yap, Jo-Hannah S. Llames, Sheila Mae M. Araiza, Kris P. Punayan, Irish Coleen A. Asin, Candice Francheska B. Tambaoan, Asia Louisa U. Chong, Karol Sophia Agape R. Padilla, Rianna Patricia S. Cruz, Carlo M. Lapid, El King D. Morado, Joshua Gregor A. Dizon, Razel Nikka M. Hao, Arianne A. Zamora, Devon Ray Pacial, Juan Antonio R. Magalang, Marissa Alejandria, Celia Carlos, Anna Ong-Lim, Edsel Maurice Salvaña, John Q. Wong, Jaime C. Montoya, Maria Rosario Singh-Vergeire and Cynthia P. Saloma |
| EPI_ISL_2189025 | Mary Mediatrix Medical Center | Philippine Genome Center | Francis A. Tablizo, Kenneth M. Kim, Maria Sofia L. Yangzon, Elcid Aaron R. Pangilinan, Renato Jacinto Q. Mantaring, Benedict A. Maralit, Marc Edsel C. Ayes, Eva Maria Cutiongco-de la Paz, Alethea R. de Guzman, Jan Michael C. Yap, Jo-Hannah S. Llames, Sheila Mae M. Araiza, Kris P. Punayan, Irish Coleen A. Asin, Candice Francheska B. Tambaoan, Asia Louisa U. Chong, Karol Sophia Agape R. Padilla, Rianna Patricia S. Cruz, Carlo M. Lapid, El King D. Morado, Joshua Gregor A. Dizon, Razel Nikka M. Hao, Arianne A. Zamora, Devon Ray Pacial, Juan Antonio R. Magalang, Marissa Alejandria, Celia Carlos, Anna Ong-Lim, Edsel Maurice Salvaña, John Q. Wong, Jaime C. Montoya, Maria Rosario Singh-Vergeire and Cynthia P. Saloma |
| EPI_ISL_2189028 | Qualimed Hospital Sta. Rosa | Philippine Genome Center | Francis A. Tablizo, Kenneth M. Kim, Maria Sofia L. Yangzon, Elcid Aaron R. Pangilinan, Renato Jacinto Q. Mantaring, Benedict A. Maralit, Marc Edsel C. Ayes, Eva Maria Cutiongco-de la Paz, Alethea R. de Guzman, Jan Michael C. Yap, Jo-Hannah S. Llames, Sheila Mae M. Araiza, Kris P. Punayan, Irish Coleen A. Asin, Candice Francheska B. Tambaoan, Asia Louisa U. Chong, Karol Sophia Agape R. Padilla, Rianna Patricia S. Cruz, Carlo M. Lapid, El King D. Morado, Joshua Gregor A. Dizon, Razel Nikka M. Hao, Arianne A. Zamora, Devon Ray Pacial, Juan Antonio R. Magalang, Marissa Alejandria, Celia Carlos, Anna Ong-Lim, Edsel Maurice Salvaña, John Q. Wong, Jaime C. Montoya, Maria Rosario Singh-Vergeire and Cynthia P. Saloma |
| EPI_ISL_2189032 | The Lord's Grace Medical and Industrial Clinic | Philippine Genome Center | Francis A. Tablizo, Kenneth M. Kim, Maria Sofia L. Yangzon, Elcid Aaron R. Pangilinan, Renato Jacinto Q. Mantaring, Benedict A. Maralit, Marc Edsel C. Ayes, Eva Maria Cutiongco-de la Paz, Alethea R. de Guzman, Jan Michael C. Yap, Jo-Hannah S. Llames, Sheila Mae M. Araiza, Kris P. Punayan, Irish Coleen A. Asin, Candice Francheska B. Tambaoan, Asia Louisa U. Chong, Karol Sophia Agape R. Padilla, Rianna Patricia S. Cruz, Carlo M. Lapid, El King D. Morado, Joshua Gregor A. Dizon, Razel Nikka M. Hao, Arianne A. Zamora, Devon Ray Pacial, Juan Antonio R. Magalang, Marissa Alejandria, Celia Carlos, Anna Ong-Lim, Edsel Maurice Salvaña, John Q. Wong, Jaime C. Montoya, Maria Rosario Singh-Vergeire and Cynthia P. Saloma |
| EPI_ISL_2189034 | Pasig City Children's Hospital - Child's Hope | Philippine Genome Center | Francis A. Tablizo, Kenneth M. Kim, Maria Sofia L. Yangzon, Elcid Aaron R. Pangilinan, Renato Jacinto Q. Mantaring, Benedict A. Maralit, Marc Edsel C. Ayes, Eva Maria Cutiongco-de la Paz, Alethea R. de Guzman, Jan Michael C. Yap, Jo-Hannah S. Llames, Sheila Mae M. Araiza, Kris P. Punayan, Irish Coleen A. Asin, Candice Francheska B. Tambaoan, Asia Louisa U. Chong, Karol Sophia Agape R. Padilla, Rianna Patricia S. Cruz, Carlo M. Lapid, El King D. Morado, Joshua Gregor A. Dizon, Razel Nikka M. Hao, Arianne A. Zamora, Devon Ray Pacial, Juan Antonio R. Magalang, Marissa Alejandria, Celia Carlos, Anna Ong-Lim, Edsel Maurice Salvaña, John Q. Wong, Jaime C. Montoya, Maria Rosario Singh-Vergeire and Cynthia P. Saloma |
| EPI_ISL_2189035 | Philippine Red Cross - Port Area | Philippine Genome Center | Francis A. Tablizo, Kenneth M. Kim, Maria Sofia L. Yangzon, Elcid Aaron R. Pangilinan, Renato Jacinto Q. Mantaring, Benedict A. Maralit, Marc Edsel C. Ayes, Eva Maria Cutiongco-de la Paz, Alethea R. de Guzman, Jan Michael C. Yap, Jo-Hannah S. Llames, Sheila Mae M. Araiza, Kris P. Punayan, Irish Coleen A. Asin, Candice Francheska B. Tambaoan, Asia Louisa U. Chong, Karol Sophia Agape R. Padilla, Rianna Patricia S. Cruz, Carlo M. Lapid, El King D. Morado, Joshua Gregor A. Dizon, Razel Nikka M. Hao, Arianne A. Zamora, Devon Ray Pacial, Juan Antonio R. Magalang, Marissa Alejandria, Celia Carlos, Anna Ong-Lim, Edsel Maurice Salvaña, John Q. Wong, Jaime C. Montoya, Maria Rosario Singh-Vergeire and Cynthia P. Saloma |
| EPI_ISL_2189039 | The Medical City | Philippine Genome Center | Francis A. Tablizo, Kenneth M. Kim, Maria Sofia L. Yangzon, Elcid Aaron R. Pangilinan, Renato Jacinto Q. Mantaring, Benedict A. Maralit, Marc Edsel C. Ayes, Eva Maria Cutiongco-de la Paz, Alethea R. de Guzman, Jan Michael C. Yap, Jo-Hannah S. Llames, Sheila Mae M. Araiza, Kris P. Punayan, Irish Coleen A. Asin, Candice Francheska B. Tambaoan, Asia Louisa U. Chong, Karol Sophia Agape R. Padilla, Rianna Patricia S. Cruz, Carlo M. Lapid, El King D. Morado, Joshua Gregor A. Dizon, Razel Nikka M. Hao, Arianne A. Zamora, Devon Ray Pacial, Juan Antonio R. Magalang, Marissa Alejandria, Celia Carlos, Anna Ong-Lim, Edsel Maurice Salvaña, John Q. Wong, Jaime C. Montoya, Maria Rosario Singh-Vergeire and Cynthia P. Saloma |
| EPI_ISL_2189040 | Qualimed Hospital Sta. Rosa | Philippine Genome Center | Francis A. Tablizo, Kenneth M. Kim, Maria Sofia L. Yangzon, Elcid Aaron R. Pangilinan, Renato Jacinto Q. Mantaring, Benedict A. Maralit, Marc Edsel C. Ayes, Eva Maria Cutiongco-de la Paz, Alethea R. de Guzman, Jan Michael C. Yap, Jo-Hannah S. Llames, Sheila Mae M. Araiza, Kris P. Punayan, Irish Coleen A. Asin, Candice Francheska B. Tambaoan, Asia Louisa U. Chong, Karol Sophia Agape R. Padilla, Rianna Patricia S. Cruz, Carlo M. Lapid, El King D. Morado, Joshua Gregor A. Dizon, Razel Nikka M. Hao, Arianne A. Zamora, Devon Ray Pacial, Juan Antonio R. Magalang, Marissa Alejandria, Celia Carlos, Anna Ong-Lim, Edsel Maurice Salvaña, John Q. Wong, Jaime C. Montoya, Maria Rosario Singh-Vergeire and Cynthia P. Saloma |
| EPI_ISL_2189041 | The Medical City | Philippine Genome Center | Francis A. Tablizo, Kenneth M. Kim, Maria Sofia L. Yangzon, Elcid Aaron R. Pangilinan, Renato Jacinto Q. Mantaring, Benedict A. Maralit, Marc Edsel C. Ayes, Eva Maria Cutiongco-de la Paz, Alethea R. de Guzman, Jan Michael C. Yap, Jo-Hannah S. Llames, Sheila Mae M. Araiza, Kris P. Punayan, Irish Coleen A. Asin, Candice Francheska B. Tambaoan, Asia Louisa U. Chong, Karol Sophia Agape R. Padilla, Rianna Patricia S. Cruz, Carlo M. Lapid, El King D. Morado, Joshua Gregor A. Dizon, Razel Nikka M. Hao, Arianne A. Zamora, Devon Ray Pacial, Juan Antonio R. Magalang, Marissa Alejandria, Celia Carlos, Anna Ong-Lim, Edsel Maurice Salvaña, John Q. Wong, Jaime C. Montoya, Maria Rosario Singh-Vergeire and Cynthia P. Saloma |
| EPI_ISL_2189043, EPI_ISL_2189044, EPI_ISL_2189045 | The Lord's Grace Medical and Industrial Clinic | Philippine Genome Center | Francis A. Tablizo, Kenneth M. Kim, Maria Sofia L. Yangzon, Elcid Aaron R. Pangilinan, Renato Jacinto Q. Mantaring, Benedict A. Maralit, Marc Edsel C. Ayes, Eva Maria Cutiongco-de la Paz, Alethea R. de Guzman, Jan Michael C. Yap, Jo-Hannah S. Llames, Sheila Mae M. Araiza, Kris P. Punayan, Irish Coleen A. Asin, Candice Francheska B. Tambaoan, Asia Louisa U. Chong, Karol Sophia Agape R. Padilla, Rianna Patricia S. Cruz, Carlo M. Lapid, El King D. Morado, Joshua Gregor A. Dizon, Razel Nikka M. Hao, Arianne A. Zamora, Devon Ray Pacial, Juan Antonio R. Magalang, Marissa Alejandria, Celia Carlos, Anna Ong-Lim, Edsel Maurice Salvaña, John Q. Wong, Jaime C. Montoya, Maria Rosario Singh-Vergeire and Cynthia P. Saloma |
| EPI_ISL_2189047 | Philippine Red Cross - Port Area | Philippine Genome Center | Francis A. Tablizo, Kenneth M. Kim, Maria Sofia L. Yangzon, Elcid Aaron R. Pangilinan, Renato Jacinto Q. Mantaring, Benedict A. Maralit, Marc Edsel C. Ayes, Eva Maria Cutiongco-de la Paz, Alethea R. de Guzman, Jan Michael C. Yap, Jo-Hannah S. Llames, Sheila Mae M. Araiza, Kris P. Punayan, Irish Coleen A. Asin, Candice Francheska B. Tambaoan, Asia Louisa U. Chong, Karol Sophia Agape R. Padilla, Rianna Patricia S. Cruz, Carlo M. Lapid, El King D. Morado, Joshua Gregor A. Dizon, Razel Nikka M. Hao, Arianne A. Zamora, Devon Ray Pacial, Juan Antonio R. Magalang, Marissa Alejandria, Celia Carlos, Anna Ong-Lim, Edsel Maurice Salvaña, John Q. Wong, Jaime C. Montoya, Maria Rosario Singh-Vergeire and Cynthia P. Saloma |

| | | | |
|---|---|---|---|
| EPI_ISL_2189049 | The Lord's Grace Medical and Industrial Clinic | Philippine Genome Center | Francis A. Tablizo, Kenneth M. Kim, Maria Sofia L. Yangzon, Elcid Aaron R. Pangilinan, Renato Jacinto Q. Mantaring, Benedict A. Maralit, Marc Edsel C. Ayes, Eva Maria Cutiongco-de la Paz, Alethea R. de Guzman, Jan Michael C. Yap, Jo-Hannah S. Llames, Sheila Mae M. Araiza, Kris P. Punayan, Irish Coleen A. Asin, Candice Francheska B. Tambaoan, Asia Louisa U. Chong, Karol Sophia Agape R. Padilla, Rianna Patricia S. Cruz, Carlo M. Lapid, El King D. Morado, Joshua Gregor A. Dizon, Razel Nikka M. Hao, Arianne A. Zamora, Devon Ray Pacial, Juan Antonio R. Magalang, Marissa Alejandria, Celia Carlos, Anna Ong-Lim, Edsel Maurice Salvaña, John Q. Wong, Jaime C. Montoya, Maria Rosario Singh-Vergeire and Cynthia P. Saloma |
| EPI_ISL_2189050 | Pasig City Children's Hospital - Child's Hope | Philippine Genome Center | Francis A. Tablizo, Kenneth M. Kim, Maria Sofia L. Yangzon, Elcid Aaron R. Pangilinan, Renato Jacinto Q. Mantaring, Benedict A. Maralit, Marc Edsel C. Ayes, Eva Maria Cutiongco-de la Paz, Alethea R. de Guzman, Jan Michael C. Yap, Jo-Hannah S. Llames, Sheila Mae M. Araiza, Kris P. Punayan, Irish Coleen A. Asin, Candice Francheska B. Tambaoan, Asia Louisa U. Chong, Karol Sophia Agape R. Padilla, Rianna Patricia S. Cruz, Carlo M. Lapid, El King D. Morado, Joshua Gregor A. Dizon, Razel Nikka M. Hao, Arianne A. Zamora, Devon Ray Pacial, Juan Antonio R. Magalang, Marissa Alejandria, Celia Carlos, Anna Ong-Lim, Edsel Maurice Salvaña, John Q. Wong, Jaime C. Montoya, Maria Rosario Singh-Vergeire and Cynthia P. Saloma |
| EPI_ISL_2189051, EPI_ISL_2189053 | The Medical City | Philippine Genome Center | Francis A. Tablizo, Kenneth M. Kim, Maria Sofia L. Yangzon, Elcid Aaron R. Pangilinan, Renato Jacinto Q. Mantaring, Benedict A. Maralit, Marc Edsel C. Ayes, Eva Maria Cutiongco-de la Paz, Alethea R. de Guzman, Jan Michael C. Yap, Jo-Hannah S. Llames, Sheila Mae M. Araiza, Kris P. Punayan, Irish Coleen A. Asin, Candice Francheska B. Tambaoan, Asia Louisa U. Chong, Karol Sophia Agape R. Padilla, Rianna Patricia S. Cruz, Carlo M. Lapid, El King D. Morado, Joshua Gregor A. Dizon, Razel Nikka M. Hao, Arianne A. Zamora, Devon Ray Pacial, Juan Antonio R. Magalang, Marissa Alejandria, Celia Carlos, Anna Ong-Lim, Edsel Maurice Salvaña, John Q. Wong, Jaime C. Montoya, Maria Rosario Singh-Vergeire and Cynthia P. Saloma |
| EPI_ISL_2189054 | The Lord's Grace Medical and Industrial Clinic | Philippine Genome Center | Francis A. Tablizo, Kenneth M. Kim, Maria Sofia L. Yangzon, Elcid Aaron R. Pangilinan, Renato Jacinto Q. Mantaring, Benedict A. Maralit, Marc Edsel C. Ayes, Eva Maria Cutiongco-de la Paz, Alethea R. de Guzman, Jan Michael C. Yap, Jo-Hannah S. Llames, Sheila Mae M. Araiza, Kris P. Punayan, Irish Coleen A. Asin, Candice Francheska B. Tambaoan, Asia Louisa U. Chong, Karol Sophia Agape R. Padilla, Rianna Patricia S. Cruz, Carlo M. Lapid, El King D. Morado, Joshua Gregor A. Dizon, Razel Nikka M. Hao, Arianne A. Zamora, Devon Ray Pacial, Juan Antonio R. Magalang, Marissa Alejandria, Celia Carlos, Anna Ong-Lim, Edsel Maurice Salvaña, John Q. Wong, Jaime C. Montoya, Maria Rosario Singh-Vergeire and Cynthia P. Saloma |
| EPI_ISL_2189055, EPI_ISL_2189056 | Philippine Red Cross - Port Area | Philippine Genome Center | Francis A. Tablizo, Kenneth M. Kim, Maria Sofia L. Yangzon, Elcid Aaron R. Pangilinan, Renato Jacinto Q. Mantaring, Benedict A. Maralit, Marc Edsel C. Ayes, Eva Maria Cutiongco-de la Paz, Alethea R. de Guzman, Jan Michael C. Yap, Jo-Hannah S. Llames, Sheila Mae M. Araiza, Kris P. Punayan, Irish Coleen A. Asin, Candice Francheska B. Tambaoan, Asia Louisa U. Chong, Karol Sophia Agape R. Padilla, Rianna Patricia S. Cruz, Carlo M. Lapid, El King D. Morado, Joshua Gregor A. Dizon, Razel Nikka M. Hao, Arianne A. Zamora, Devon Ray Pacial, Juan Antonio R. Magalang, Marissa Alejandria, Celia Carlos, Anna Ong-Lim, Edsel Maurice Salvaña, John Q. Wong, Jaime C. Montoya, Maria Rosario Singh-Vergeire and Cynthia P. Saloma |
| EPI_ISL_2189058 | Ospital ng ParaÃ±aque II | Philippine Genome Center | Francis A. Tablizo, Kenneth M. Kim, Maria Sofia L. Yangzon, Elcid Aaron R. Pangilinan, Renato Jacinto Q. Mantaring, Benedict A. Maralit, Marc Edsel C. Ayes, Eva Maria Cutiongco-de la Paz, Alethea R. de Guzman, Jan Michael C. Yap, Jo-Hannah S. Llames, Sheila Mae M. Araiza, Kris P. Punayan, Irish Coleen A. Asin, Candice Francheska B. Tambaoan, Asia Louisa U. Chong, Karol Sophia Agape R. Padilla, Rianna Patricia S. Cruz, Carlo M. Lapid, El King D. Morado, Joshua Gregor A. Dizon, Razel Nikka M. Hao, Arianne A. Zamora, Devon Ray Pacial, Juan Antonio R. Magalang, Marissa Alejandria, Celia Carlos, Anna Ong-Lim, Edsel Maurice Salvaña, John Q. Wong, Jaime C. Montoya, Maria Rosario Singh-Vergeire and Cynthia P. Saloma |
| EPI_ISL_2189059 | Philippine Red Cross - Clark Molecular Laboratory | Philippine Genome Center | Francis A. Tablizo, Kenneth M. Kim, Maria Sofia L. Yangzon, Elcid Aaron R. Pangilinan, Renato Jacinto Q. Mantaring, Benedict A. Maralit, Marc Edsel C. Ayes, Eva Maria Cutiongco-de la Paz, Alethea R. de Guzman, Jan Michael C. Yap, Jo-Hannah S. Llames, Sheila Mae M. Araiza, Kris P. Punayan, Irish Coleen A. Asin, Candice Francheska B. Tambaoan, Asia Louisa U. Chong, Karol Sophia Agape R. Padilla, Rianna Patricia S. Cruz, Carlo M. Lapid, El King D. Morado, Joshua Gregor A. Dizon, Razel Nikka M. Hao, Arianne A. Zamora, Devon Ray Pacial, Juan Antonio R. Magalang, Marissa Alejandria, Celia Carlos, Anna Ong-Lim, Edsel Maurice Salvaña, John Q. Wong, Jaime C. Montoya, Maria Rosario Singh-Vergeire and Cynthia P. Saloma |
| EPI_ISL_2189061 | The Lord's Grace Medical and Industrial Clinic | Philippine Genome Center | Francis A. Tablizo, Kenneth M. Kim, Maria Sofia L. Yangzon, Elcid Aaron R. Pangilinan, Renato Jacinto Q. Mantaring, Benedict A. Maralit, Marc Edsel C. Ayes, Eva Maria Cutiongco-de la Paz, Alethea R. de Guzman, Jan Michael C. Yap, Jo-Hannah S. Llames, Sheila Mae M. Araiza, Kris P. Punayan, Irish Coleen A. Asin, Candice Francheska B. Tambaoan, Asia Louisa U. Chong, Karol Sophia Agape R. Padilla, Rianna Patricia S. Cruz, Carlo M. Lapid, El King D. Morado, Joshua Gregor A. Dizon, Razel Nikka M. Hao, Arianne A. Zamora, Devon Ray Pacial, Juan Antonio R. Magalang, Marissa Alejandria, Celia Carlos, Anna Ong-Lim, Edsel Maurice Salvaña, John Q. Wong, Jaime C. Montoya, Maria Rosario Singh-Vergeire and Cynthia P. Saloma |
| EPI_ISL_2189062 | The Medical City | Philippine Genome Center | Francis A. Tablizo, Kenneth M. Kim, Maria Sofia L. Yangzon, Elcid Aaron R. Pangilinan, Renato Jacinto Q. Mantaring, Benedict A. Maralit, Marc Edsel C. Ayes, Eva Maria Cutiongco-de la Paz, Alethea R. de Guzman, Jan Michael C. Yap, Jo-Hannah S. Llames, Sheila Mae M. Araiza, Kris P. Punayan, Irish Coleen A. Asin, Candice Francheska B. Tambaoan, Asia Louisa U. Chong, Karol Sophia Agape R. Padilla, Rianna Patricia S. Cruz, Carlo M. Lapid, El King D. Morado, Joshua Gregor A. Dizon, Razel Nikka M. Hao, Arianne A. Zamora, Devon Ray Pacial, Juan Antonio R. Magalang, Marissa Alejandria, Celia Carlos, Anna Ong-Lim, Edsel Maurice Salvaña, John Q. Wong, Jaime C. Montoya, Maria Rosario Singh-Vergeire and Cynthia P. Saloma |
| EPI_ISL_2189063 | Mary Mediatrix Medical Center | Philippine Genome Center | Francis A. Tablizo, Kenneth M. Kim, Maria Sofia L. Yangzon, Elcid Aaron R. Pangilinan, Renato Jacinto Q. Mantaring, Benedict A. Maralit, Marc Edsel C. Ayes, Eva Maria Cutiongco-de la Paz, Alethea R. de Guzman, Jan Michael C. Yap, Jo-Hannah S. Llames, Sheila Mae M. Araiza, Kris P. Punayan, Irish Coleen A. Asin, Candice Francheska B. Tambaoan, Asia Louisa U. Chong, Karol Sophia Agape R. Padilla, Rianna Patricia S. Cruz, Carlo M. Lapid, El King D. Morado, Joshua Gregor A. Dizon, Razel Nikka M. Hao, Arianne A. Zamora, Devon Ray Pacial, Juan Antonio R. Magalang, Marissa Alejandria, Celia Carlos, Anna Ong-Lim, Edsel Maurice Salvaña, John Q. Wong, Jaime C. Montoya, Maria Rosario Singh-Vergeire and Cynthia P. Saloma |
| EPI_ISL_2189065 | First Aide Diagnostic Center | Philippine Genome Center | Francis A. Tablizo, Kenneth M. Kim, Maria Sofia L. Yangzon, Elcid Aaron R. Pangilinan, Renato Jacinto Q. Mantaring, Benedict A. Maralit, Marc Edsel C. Ayes, Eva Maria Cutiongco-de la Paz, Alethea R. de Guzman, Jan Michael C. Yap, Jo-Hannah S. Llames, Sheila Mae M. Araiza, Kris P. Punayan, Irish Coleen A. Asin, Candice Francheska B. Tambaoan, Asia Louisa U. Chong, Karol Sophia Agape R. Padilla, Rianna Patricia S. Cruz, Carlo M. Lapid, El King D. Morado, Joshua Gregor A. Dizon, Razel Nikka M. Hao, Arianne A. Zamora, Devon Ray Pacial, Juan Antonio R. Magalang, Marissa Alejandria, Celia Carlos, Anna Ong-Lim, Edsel Maurice Salvaña, John Q. Wong, Jaime C. Montoya, Maria Rosario Singh-Vergeire and Cynthia P. Saloma |
| EPI_ISL_2189067, EPI_ISL_2189069 | The Medical City | Philippine Genome Center | Francis A. Tablizo, Kenneth M. Kim, Maria Sofia L. Yangzon, Elcid Aaron R. Pangilinan, Renato Jacinto Q. Mantaring, Benedict A. Maralit, Marc Edsel C. Ayes, Eva Maria Cutiongco-de la Paz, Alethea R. de Guzman, Jan Michael C. Yap, Jo-Hannah S. Llames, Sheila Mae M. Araiza, Kris P. Punayan, Irish Coleen A. Asin, Candice Francheska B. Tambaoan, Asia Louisa U. Chong, Karol Sophia Agape R. Padilla, Rianna Patricia S. Cruz, Carlo M. Lapid, El King D. Morado, Joshua Gregor A. Dizon, Razel Nikka M. Hao, Arianne A. Zamora, Devon Ray Pacial, Juan Antonio R. Magalang, Marissa Alejandria, Celia Carlos, Anna Ong-Lim, Edsel Maurice Salvaña, John Q. Wong, Jaime C. Montoya, Maria Rosario Singh-Vergeire and Cynthia P. Saloma |
| EPI_ISL_2189070 | Philippine Red Cross - National Blood Center | Philippine Genome Center | Francis A. Tablizo, Kenneth M. Kim, Maria Sofia L. Yangzon, Elcid Aaron R. Pangilinan, Renato Jacinto Q. Mantaring, Benedict A. Maralit, Marc Edsel C. Ayes, Eva Maria Cutiongco-de la Paz, Alethea R. de Guzman, Jan Michael C. Yap, Jo-Hannah S. Llames, Sheila Mae M. Araiza, Kris P. Punayan, Irish Coleen A. Asin, Candice Francheska B. Tambaoan, Asia Louisa U. Chong, Karol Sophia Agape R. Padilla, Rianna Patricia S. Cruz, Carlo M. Lapid, El King D. Morado, Joshua Gregor A. Dizon, Razel Nikka M. Hao, Arianne A. Zamora, Devon Ray Pacial, Juan Antonio R. Magalang, Marissa Alejandria, Celia Carlos, Anna Ong-Lim, Edsel Maurice Salvaña, John Q. Wong, Jaime C. Montoya, Maria Rosario Singh-Vergeire and Cynthia P. Saloma |
| EPI_ISL_2189073 | The Medical City | Philippine Genome Center | Francis A. Tablizo, Kenneth M. Kim, Maria Sofia L. Yangzon, Elcid Aaron R. Pangilinan, Renato Jacinto Q. Mantaring, Benedict A. Maralit, Marc Edsel C. Ayes, Eva Maria Cutiongco-de la Paz, Alethea R. de Guzman, Jan Michael C. Yap, Jo-Hannah S. Llames, Sheila Mae M. Araiza, Kris P. Punayan, Irish Coleen A. Asin, Candice Francheska B. Tambaoan, Asia Louisa U. Chong, Karol Sophia Agape R. Padilla, Rianna Patricia S. Cruz, Carlo M. Lapid, El King D. Morado, Joshua Gregor A. Dizon, Razel Nikka M. Hao, Arianne A. Zamora, Devon Ray Pacial, Juan Antonio R. Magalang, Marissa Alejandria, Celia Carlos, Anna Ong-Lim, Edsel Maurice Salvaña, John Q. Wong, Jaime C. Montoya, Maria Rosario Singh-Vergeire and Cynthia P. Saloma |
| EPI_ISL_2189078 | The Lord's Grace Medical and Industrial Clinic | Philippine Genome Center | Francis A. Tablizo, Kenneth M. Kim, Maria Sofia L. Yangzon, Elcid Aaron R. Pangilinan, Renato Jacinto Q. Mantaring, Benedict A. Maralit, Marc Edsel C. Ayes, Eva Maria Cutiongco-de la Paz, Alethea R. de Guzman, Jan Michael C. Yap, Jo-Hannah S. Llames, Sheila Mae M. Araiza, Kris P. Punayan, Irish Coleen A. Asin, Candice Francheska B. Tambaoan, Asia Louisa U. Chong, Karol Sophia Agape R. Padilla, Rianna Patricia S. Cruz, Carlo M. Lapid, El King D. Morado, Joshua Gregor A. Dizon, Razel Nikka M. Hao, Arianne A. Zamora, Devon Ray Pacial, Juan Antonio R. Magalang, Marissa Alejandria, Celia Carlos, Anna Ong-Lim, Edsel Maurice Salvaña, John Q. Wong, Jaime C. Montoya, Maria Rosario Singh-Vergeire and Cynthia P. Saloma |
| EPI_ISL_2189080, EPI_ISL_2189084 | Pasig City Children's Hospital - Child's Hope | Philippine Genome Center | Francis A. Tablizo, Kenneth M. Kim, Maria Sofia L. Yangzon, Elcid Aaron R. Pangilinan, Renato Jacinto Q. Mantaring, Benedict A. Maralit, Marc Edsel C. Ayes, Eva Maria Cutiongco-de la Paz, Alethea R. de Guzman, Jan Michael C. Yap, Jo-Hannah S. Llames, Sheila Mae M. Araiza, Kris P. Punayan, Irish Coleen A. Asin, Candice Francheska B. Tambaoan, Asia Louisa U. Chong, Karol Sophia Agape R. Padilla, Rianna Patricia S. Cruz, Carlo M. Lapid, El King D. Morado, Joshua Gregor A. Dizon, Razel Nikka M. Hao, Arianne A. Zamora, Devon Ray Pacial, Juan Antonio R. Magalang, Marissa Alejandria, Celia Carlos, Anna Ong-Lim, Edsel Maurice Salvaña, John Q. Wong, Jaime C. Montoya, Maria Rosario Singh-Vergeire and Cynthia P. Saloma |

| | | | |
|---|---|---|---|
| EPI_ISL_2189086, EPI_ISL_2189087 | The Lord's Grace Medical and Industrial Clinic | Philippine Genome Center | Francis A. Tablizo, Kenneth M. Kim, Maria Sofia L. Yangzon, Elcid Aaron R. Pangilinan, Renato Jacinto Q. Mantaring, Benedict A. Maralit, Marc Edsel C. Ayes, Eva Maria Cutiongco-de la Paz, Alethea R. de Guzman, Jan Michael C. Yap, Jo-Hannah S. Llames, Sheila Mae M. Araiza, Kris P. Punayan, Irish Coleen A. Asin, Candice Francheska B. Tambaoan, Asia Louisa U. Chong, Karol Sophia Agape R. Padilla, Rianna Patricia S. Cruz, Carlo M. Lapid, El King D. Morado, Joshua Gregor A. Dizon, Razel Nikka M. Hao, Arianne A. Zamora, Devon Ray Pacial, Juan Antonio R. Magalang, Marissa Alejandria, Celia Carlos, Anna Ong-Lim, Edsel Maurice Salvaña, John Q. Wong, Jaime C. Montoya, Maria Rosario Singh-Vergeire and Cynthia P. Saloma |
| EPI_ISL_2189088 | Safeguard DNA Diagnostic, Inc. | Philippine Genome Center | Francis A. Tablizo, Kenneth M. Kim, Maria Sofia L. Yangzon, Elcid Aaron R. Pangilinan, Renato Jacinto Q. Mantaring, Benedict A. Maralit, Marc Edsel C. Ayes, Eva Maria Cutiongco-de la Paz, Alethea R. de Guzman, Jan Michael C. Yap, Jo-Hannah S. Llames, Sheila Mae M. Araiza, Kris P. Punayan, Irish Coleen A. Asin, Candice Francheska B. Tambaoan, Asia Louisa U. Chong, Karol Sophia Agape R. Padilla, Rianna Patricia S. Cruz, Carlo M. Lapid, El King D. Morado, Joshua Gregor A. Dizon, Razel Nikka M. Hao, Arianne A. Zamora, Devon Ray Pacial, Juan Antonio R. Magalang, Marissa Alejandria, Celia Carlos, Anna Ong-Lim, Edsel Maurice Salvaña, John Q. Wong, Jaime C. Montoya, Maria Rosario Singh-Vergeire and Cynthia P. Saloma |
| EPI_ISL_2189089 | Mary Mediatrix Medical Center | Philippine Genome Center | Francis A. Tablizo, Kenneth M. Kim, Maria Sofia L. Yangzon, Elcid Aaron R. Pangilinan, Renato Jacinto Q. Mantaring, Benedict A. Maralit, Marc Edsel C. Ayes, Eva Maria Cutiongco-de la Paz, Alethea R. de Guzman, Jan Michael C. Yap, Jo-Hannah S. Llames, Sheila Mae M. Araiza, Kris P. Punayan, Irish Coleen A. Asin, Candice Francheska B. Tambaoan, Asia Louisa U. Chong, Karol Sophia Agape R. Padilla, Rianna Patricia S. Cruz, Carlo M. Lapid, El King D. Morado, Joshua Gregor A. Dizon, Razel Nikka M. Hao, Arianne A. Zamora, Devon Ray Pacial, Juan Antonio R. Magalang, Marissa Alejandria, Celia Carlos, Anna Ong-Lim, Edsel Maurice Salvaña, John Q. Wong, Jaime C. Montoya, Maria Rosario Singh-Vergeire and Cynthia P. Saloma |
| EPI_ISL_2189090 | The Lord's Grace Medical and Industrial Clinic | Philippine Genome Center | Francis A. Tablizo, Kenneth M. Kim, Maria Sofia L. Yangzon, Elcid Aaron R. Pangilinan, Renato Jacinto Q. Mantaring, Benedict A. Maralit, Marc Edsel C. Ayes, Eva Maria Cutiongco-de la Paz, Alethea R. de Guzman, Jan Michael C. Yap, Jo-Hannah S. Llames, Sheila Mae M. Araiza, Kris P. Punayan, Irish Coleen A. Asin, Candice Francheska B. Tambaoan, Asia Louisa U. Chong, Karol Sophia Agape R. Padilla, Rianna Patricia S. Cruz, Carlo M. Lapid, El King D. Morado, Joshua Gregor A. Dizon, Razel Nikka M. Hao, Arianne A. Zamora, Devon Ray Pacial, Juan Antonio R. Magalang, Marissa Alejandria, Celia Carlos, Anna Ong-Lim, Edsel Maurice Salvaña, John Q. Wong, Jaime C. Montoya, Maria Rosario Singh-Vergeire and Cynthia P. Saloma |
| EPI_ISL_2189091 | Philippine Red Cross - Clark Molecular Laboratory | Philippine Genome Center | Francis A. Tablizo, Kenneth M. Kim, Maria Sofia L. Yangzon, Elcid Aaron R. Pangilinan, Renato Jacinto Q. Mantaring, Benedict A. Maralit, Marc Edsel C. Ayes, Eva Maria Cutiongco-de la Paz, Alethea R. de Guzman, Jan Michael C. Yap, Jo-Hannah S. Llames, Sheila Mae M. Araiza, Kris P. Punayan, Irish Coleen A. Asin, Candice Francheska B. Tambaoan, Asia Louisa U. Chong, Karol Sophia Agape R. Padilla, Rianna Patricia S. Cruz, Carlo M. Lapid, El King D. Morado, Joshua Gregor A. Dizon, Razel Nikka M. Hao, Arianne A. Zamora, Devon Ray Pacial, Juan Antonio R. Magalang, Marissa Alejandria, Celia Carlos, Anna Ong-Lim, Edsel Maurice Salvaña, John Q. Wong, Jaime C. Montoya, Maria Rosario Singh-Vergeire and Cynthia P. Saloma |
| EPI_ISL_2189093, EPI_ISL_2189094 | The Lord's Grace Medical and Industrial Clinic | Philippine Genome Center | Francis A. Tablizo, Kenneth M. Kim, Maria Sofia L. Yangzon, Elcid Aaron R. Pangilinan, Renato Jacinto Q. Mantaring, Benedict A. Maralit, Marc Edsel C. Ayes, Eva Maria Cutiongco-de la Paz, Alethea R. de Guzman, Jan Michael C. Yap, Jo-Hannah S. Llames, Sheila Mae M. Araiza, Kris P. Punayan, Irish Coleen A. Asin, Candice Francheska B. Tambaoan, Asia Louisa U. Chong, Karol Sophia Agape R. Padilla, Rianna Patricia S. Cruz, Carlo M. Lapid, El King D. Morado, Joshua Gregor A. Dizon, Razel Nikka M. Hao, Arianne A. Zamora, Devon Ray Pacial, Juan Antonio R. Magalang, Marissa Alejandria, Celia Carlos, Anna Ong-Lim, Edsel Maurice Salvaña, John Q. Wong, Jaime C. Montoya, Maria Rosario Singh-Vergeire and Cynthia P. Saloma |
| EPI_ISL_2189095 | Philippine Red Cross - National Blood Center | Philippine Genome Center | Francis A. Tablizo, Kenneth M. Kim, Maria Sofia L. Yangzon, Elcid Aaron R. Pangilinan, Renato Jacinto Q. Mantaring, Benedict A. Maralit, Marc Edsel C. Ayes, Eva Maria Cutiongco-de la Paz, Alethea R. de Guzman, Jan Michael C. Yap, Jo-Hannah S. Llames, Sheila Mae M. Araiza, Kris P. Punayan, Irish Coleen A. Asin, Candice Francheska B. Tambaoan, Asia Louisa U. Chong, Karol Sophia Agape R. Padilla, Rianna Patricia S. Cruz, Carlo M. Lapid, El King D. Morado, Joshua Gregor A. Dizon, Razel Nikka M. Hao, Arianne A. Zamora, Devon Ray Pacial, Juan Antonio R. Magalang, Marissa Alejandria, Celia Carlos, Anna Ong-Lim, Edsel Maurice Salvaña, John Q. Wong, Jaime C. Montoya, Maria Rosario Singh-Vergeire and Cynthia P. Saloma |
| EPI_ISL_2189097 | Prime Care Alpha Covid-19 Testing Laboratory | Philippine Genome Center | Francis A. Tablizo, Kenneth M. Kim, Maria Sofia L. Yangzon, Elcid Aaron R. Pangilinan, Renato Jacinto Q. Mantaring, Benedict A. Maralit, Marc Edsel C. Ayes, Eva Maria Cutiongco-de la Paz, Alethea R. de Guzman, Jan Michael C. Yap, Jo-Hannah S. Llames, Sheila Mae M. Araiza, Kris P. Punayan, Irish Coleen A. Asin, Candice Francheska B. Tambaoan, Asia Louisa U. Chong, Karol Sophia Agape R. Padilla, Rianna Patricia S. Cruz, Carlo M. Lapid, El King D. Morado, Joshua Gregor A. Dizon, Razel Nikka M. Hao, Arianne A. Zamora, Devon Ray Pacial, Juan Antonio R. Magalang, Marissa Alejandria, Celia Carlos, Anna Ong-Lim, Edsel Maurice Salvaña, John Q. Wong, Jaime C. Montoya, Maria Rosario Singh-Vergeire and Cynthia P. Saloma |
| EPI_ISL_2189098, EPI_ISL_2189100 | The Lord's Grace Medical and Industrial Clinic | Philippine Genome Center | Francis A. Tablizo, Kenneth M. Kim, Maria Sofia L. Yangzon, Elcid Aaron R. Pangilinan, Renato Jacinto Q. Mantaring, Benedict A. Maralit, Marc Edsel C. Ayes, Eva Maria Cutiongco-de la Paz, Alethea R. de Guzman, Jan Michael C. Yap, Jo-Hannah S. Llames, Sheila Mae M. Araiza, Kris P. Punayan, Irish Coleen A. Asin, Candice Francheska B. Tambaoan, Asia Louisa U. Chong, Karol Sophia Agape R. Padilla, Rianna Patricia S. Cruz, Carlo M. Lapid, El King D. Morado, Joshua Gregor A. Dizon, Razel Nikka M. Hao, Arianne A. Zamora, Devon Ray Pacial, Juan Antonio R. Magalang, Marissa Alejandria, Celia Carlos, Anna Ong-Lim, Edsel Maurice Salvaña, John Q. Wong, Jaime C. Montoya, Maria Rosario Singh-Vergeire and Cynthia P. Saloma |
| EPI_ISL_2189101, EPI_ISL_2189102 | The Medical City | Philippine Genome Center | Francis A. Tablizo, Kenneth M. Kim, Maria Sofia L. Yangzon, Elcid Aaron R. Pangilinan, Renato Jacinto Q. Mantaring, Benedict A. Maralit, Marc Edsel C. Ayes, Eva Maria Cutiongco-de la Paz, Alethea R. de Guzman, Jan Michael C. Yap, Jo-Hannah S. Llames, Sheila Mae M. Araiza, Kris P. Punayan, Irish Coleen A. Asin, Candice Francheska B. Tambaoan, Asia Louisa U. Chong, Karol Sophia Agape R. Padilla, Rianna Patricia S. Cruz, Carlo M. Lapid, El King D. Morado, Joshua Gregor A. Dizon, Razel Nikka M. Hao, Arianne A. Zamora, Devon Ray Pacial, Juan Antonio R. Magalang, Marissa Alejandria, Celia Carlos, Anna Ong-Lim, Edsel Maurice Salvaña, John Q. Wong, Jaime C. Montoya, Maria Rosario Singh-Vergeire and Cynthia P. Saloma |
| EPI_ISL_2189104 | Pasig City Children's Hospital - Child's Hope | Philippine Genome Center | Francis A. Tablizo, Kenneth M. Kim, Maria Sofia L. Yangzon, Elcid Aaron R. Pangilinan, Renato Jacinto Q. Mantaring, Benedict A. Maralit, Marc Edsel C. Ayes, Eva Maria Cutiongco-de la Paz, Alethea R. de Guzman, Jan Michael C. Yap, Jo-Hannah S. Llames, Sheila Mae M. Araiza, Kris P. Punayan, Irish Coleen A. Asin, Candice Francheska B. Tambaoan, Asia Louisa U. Chong, Karol Sophia Agape R. Padilla, Rianna Patricia S. Cruz, Carlo M. Lapid, El King D. Morado, Joshua Gregor A. Dizon, Razel Nikka M. Hao, Arianne A. Zamora, Devon Ray Pacial, Juan Antonio R. Magalang, Marissa Alejandria, Celia Carlos, Anna Ong-Lim, Edsel Maurice Salvaña, John Q. Wong, Jaime C. Montoya, Maria Rosario Singh-Vergeire and Cynthia P. Saloma |
| EPI_ISL_2189105 | Philippine Red Cross - National Blood Center | Philippine Genome Center | Francis A. Tablizo, Kenneth M. Kim, Maria Sofia L. Yangzon, Elcid Aaron R. Pangilinan, Renato Jacinto Q. Mantaring, Benedict A. Maralit, Marc Edsel C. Ayes, Eva Maria Cutiongco-de la Paz, Alethea R. de Guzman, Jan Michael C. Yap, Jo-Hannah S. Llames, Sheila Mae M. Araiza, Kris P. Punayan, Irish Coleen A. Asin, Candice Francheska B. Tambaoan, Asia Louisa U. Chong, Karol Sophia Agape R. Padilla, Rianna Patricia S. Cruz, Carlo M. Lapid, El King D. Morado, Joshua Gregor A. Dizon, Razel Nikka M. Hao, Arianne A. Zamora, Devon Ray Pacial, Juan Antonio R. Magalang, Marissa Alejandria, Celia Carlos, Anna Ong-Lim, Edsel Maurice Salvaña, John Q. Wong, Jaime C. Montoya, Maria Rosario Singh-Vergeire and Cynthia P. Saloma |
| EPI_ISL_2189110 | The Medical City | Philippine Genome Center | Francis A. Tablizo, Kenneth M. Kim, Maria Sofia L. Yangzon, Elcid Aaron R. Pangilinan, Renato Jacinto Q. Mantaring, Benedict A. Maralit, Marc Edsel C. Ayes, Eva Maria Cutiongco-de la Paz, Alethea R. de Guzman, Jan Michael C. Yap, Jo-Hannah S. Llames, Sheila Mae M. Araiza, Kris P. Punayan, Irish Coleen A. Asin, Candice Francheska B. Tambaoan, Asia Louisa U. Chong, Karol Sophia Agape R. Padilla, Rianna Patricia S. Cruz, Carlo M. Lapid, El King D. Morado, Joshua Gregor A. Dizon, Razel Nikka M. Hao, Arianne A. Zamora, Devon Ray Pacial, Juan Antonio R. Magalang, Marissa Alejandria, Celia Carlos, Anna Ong-Lim, Edsel Maurice Salvaña, John Q. Wong, Jaime C. Montoya, Maria Rosario Singh-Vergeire and Cynthia P. Saloma |
| EPI_ISL_2189112 | South Super Highway Molecular Diagnostic Laboratory | Philippine Genome Center | Francis A. Tablizo, Kenneth M. Kim, Maria Sofia L. Yangzon, Elcid Aaron R. Pangilinan, Renato Jacinto Q. Mantaring, Benedict A. Maralit, Marc Edsel C. Ayes, Eva Maria Cutiongco-de la Paz, Alethea R. de Guzman, Jan Michael C. Yap, Jo-Hannah S. Llames, Sheila Mae M. Araiza, Kris P. Punayan, Irish Coleen A. Asin, Candice Francheska B. Tambaoan, Asia Louisa U. Chong, Karol Sophia Agape R. Padilla, Rianna Patricia S. Cruz, Carlo M. Lapid, El King D. Morado, Joshua Gregor A. Dizon, Razel Nikka M. Hao, Arianne A. Zamora, Devon Ray Pacial, Juan Antonio R. Magalang, Marissa Alejandria, Celia Carlos, Anna Ong-Lim, Edsel Maurice Salvaña, John Q. Wong, Jaime C. Montoya, Maria Rosario Singh-Vergeire and Cynthia P. Saloma |
| EPI_ISL_2189114 | The Medical City | Philippine Genome Center | Francis A. Tablizo, Kenneth M. Kim, Maria Sofia L. Yangzon, Elcid Aaron R. Pangilinan, Renato Jacinto Q. Mantaring, Benedict A. Maralit, Marc Edsel C. Ayes, Eva Maria Cutiongco-de la Paz, Alethea R. de Guzman, Jan Michael C. Yap, Jo-Hannah S. Llames, Sheila Mae M. Araiza, Kris P. Punayan, Irish Coleen A. Asin, Candice Francheska B. Tambaoan, Asia Louisa U. Chong, Karol Sophia Agape R. Padilla, Rianna Patricia S. Cruz, Carlo M. Lapid, El King D. Morado, Joshua Gregor A. Dizon, Razel Nikka M. Hao, Arianne A. Zamora, Devon Ray Pacial, Juan Antonio R. Magalang, Marissa Alejandria, Celia Carlos, Anna Ong-Lim, Edsel Maurice Salvaña, John Q. Wong, Jaime C. Montoya, Maria Rosario Singh-Vergeire and Cynthia P. Saloma |
| EPI_ISL_2189115 | Butuan Medical Center | Philippine Genome Center | Francis A. Tablizo, Kenneth M. Kim, Maria Sofia L. Yangzon, Elcid Aaron R. Pangilinan, Renato Jacinto Q. Mantaring, Benedict A. Maralit, Marc Edsel C. Ayes, Eva Maria Cutiongco-de la Paz, Alethea R. de Guzman, Jan Michael C. Yap, Jo-Hannah S. Llames, Sheila Mae M. Araiza, Kris P. Punayan, Irish Coleen A. Asin, Candice Francheska B. Tambaoan, Asia Louisa U. Chong, Karol Sophia Agape R. Padilla, Rianna Patricia S. Cruz, Carlo M. Lapid, El King D. Morado, Joshua Gregor A. Dizon, Razel Nikka M. Hao, Arianne A. Zamora, Devon Ray Pacial, Juan Antonio R. Magalang, Marissa Alejandria, Celia Carlos, Anna Ong-Lim, Edsel Maurice Salvaña, John Q. Wong, Jaime C. Montoya, Maria Rosario Singh-Vergeire and Cynthia P. Saloma |

| | | | |
|---|---|---|---|
| EPI_ISL_2189116 | Philippine General Hospital (PGH) | Philippine Genome Center | Francis A. Tablizo, Kenneth M. Kim, Maria Sofia L. Yangzon, Elcid Aaron R. Pangilinan, Renato Jacinto Q. Mantaring, Benedict A. Maralit, Marc Edsel C. Ayes, Eva Maria Cutiongco-de la Paz, Alethea R. de Guzman, Jan Michael C. Yap, Jo-Hannah S. Llames, Sheila Mae M. Araiza, Kris P. Punayan, Irish Coleen A. Asin, Candice Francheska B. Tambaoan, Asia Louisa U. Chong, Karol Sophia Agape R. Padilla, Rianna Patricia S. Cruz, Carlo M. Lapid, El King D. Morado, Joshua Gregor A. Dizon, Razel Nikka M. Hao, Arianne A. Zamora, Devon Ray Pacial, Juan Antonio R. Magalang, Marissa Alejandria, Celia Carlos, Anna Ong-Lim, Edsel Maurice Salvaña, John Q. Wong, Jaime C. Montoya, Maria Rosario Singh-Vergeire and Cynthia P. Saloma |
| EPI_ISL_2189118 | South Super Highway Molecular Diagnostic Laboratory | Philippine Genome Center | Francis A. Tablizo, Kenneth M. Kim, Maria Sofia L. Yangzon, Elcid Aaron R. Pangilinan, Renato Jacinto Q. Mantaring, Benedict A. Maralit, Marc Edsel C. Ayes, Eva Maria Cutiongco-de la Paz, Alethea R. de Guzman, Jan Michael C. Yap, Jo-Hannah S. Llames, Sheila Mae M. Araiza, Kris P. Punayan, Irish Coleen A. Asin, Candice Francheska B. Tambaoan, Asia Louisa U. Chong, Karol Sophia Agape R. Padilla, Rianna Patricia S. Cruz, Carlo M. Lapid, El King D. Morado, Joshua Gregor A. Dizon, Razel Nikka M. Hao, Arianne A. Zamora, Devon Ray Pacial, Juan Antonio R. Magalang, Marissa Alejandria, Celia Carlos, Anna Ong-Lim, Edsel Maurice Salvaña, John Q. Wong, Jaime C. Montoya, Maria Rosario Singh-Vergeire and Cynthia P. Saloma |
| EPI_ISL_2189119 | The Medical City | Philippine Genome Center | Francis A. Tablizo, Kenneth M. Kim, Maria Sofia L. Yangzon, Elcid Aaron R. Pangilinan, Renato Jacinto Q. Mantaring, Benedict A. Maralit, Marc Edsel C. Ayes, Eva Maria Cutiongco-de la Paz, Alethea R. de Guzman, Jan Michael C. Yap, Jo-Hannah S. Llames, Sheila Mae M. Araiza, Kris P. Punayan, Irish Coleen A. Asin, Candice Francheska B. Tambaoan, Asia Louisa U. Chong, Karol Sophia Agape R. Padilla, Rianna Patricia S. Cruz, Carlo M. Lapid, El King D. Morado, Joshua Gregor A. Dizon, Razel Nikka M. Hao, Arianne A. Zamora, Devon Ray Pacial, Juan Antonio R. Magalang, Marissa Alejandria, Celia Carlos, Anna Ong-Lim, Edsel Maurice Salvaña, John Q. Wong, Jaime C. Montoya, Maria Rosario Singh-Vergeire and Cynthia P. Saloma |
| EPI_ISL_2189120 | Prime Care Alpha Covid-19 Testing Laboratory | Philippine Genome Center | Francis A. Tablizo, Kenneth M. Kim, Maria Sofia L. Yangzon, Elcid Aaron R. Pangilinan, Renato Jacinto Q. Mantaring, Benedict A. Maralit, Marc Edsel C. Ayes, Eva Maria Cutiongco-de la Paz, Alethea R. de Guzman, Jan Michael C. Yap, Jo-Hannah S. Llames, Sheila Mae M. Araiza, Kris P. Punayan, Irish Coleen A. Asin, Candice Francheska B. Tambaoan, Asia Louisa U. Chong, Karol Sophia Agape R. Padilla, Rianna Patricia S. Cruz, Carlo M. Lapid, El King D. Morado, Joshua Gregor A. Dizon, Razel Nikka M. Hao, Arianne A. Zamora, Devon Ray Pacial, Juan Antonio R. Magalang, Marissa Alejandria, Celia Carlos, Anna Ong-Lim, Edsel Maurice Salvaña, John Q. Wong, Jaime C. Montoya, Maria Rosario Singh-Vergeire and Cynthia P. Saloma |
| EPI_ISL_2189121, EPI_ISL_2189122 | The Lord's Grace Medical and Industrial Clinic | Philippine Genome Center | Francis A. Tablizo, Kenneth M. Kim, Maria Sofia L. Yangzon, Elcid Aaron R. Pangilinan, Renato Jacinto Q. Mantaring, Benedict A. Maralit, Marc Edsel C. Ayes, Eva Maria Cutiongco-de la Paz, Alethea R. de Guzman, Jan Michael C. Yap, Jo-Hannah S. Llames, Sheila Mae M. Araiza, Kris P. Punayan, Irish Coleen A. Asin, Candice Francheska B. Tambaoan, Asia Louisa U. Chong, Karol Sophia Agape R. Padilla, Rianna Patricia S. Cruz, Carlo M. Lapid, El King D. Morado, Joshua Gregor A. Dizon, Razel Nikka M. Hao, Arianne A. Zamora, Devon Ray Pacial, Juan Antonio R. Magalang, Marissa Alejandria, Celia Carlos, Anna Ong-Lim, Edsel Maurice Salvaña, John Q. Wong, Jaime C. Montoya, Maria Rosario Singh-Vergeire and Cynthia P. Saloma |
| EPI_ISL_2189123 | Sta. Rosa CHO- GENEXPERT LABORATORY | Philippine Genome Center | Francis A. Tablizo, Kenneth M. Kim, Maria Sofia L. Yangzon, Elcid Aaron R. Pangilinan, Renato Jacinto Q. Mantaring, Benedict A. Maralit, Marc Edsel C. Ayes, Eva Maria Cutiongco-de la Paz, Alethea R. de Guzman, Jan Michael C. Yap, Jo-Hannah S. Llames, Sheila Mae M. Araiza, Kris P. Punayan, Irish Coleen A. Asin, Candice Francheska B. Tambaoan, Asia Louisa U. Chong, Karol Sophia Agape R. Padilla, Rianna Patricia S. Cruz, Carlo M. Lapid, El King D. Morado, Joshua Gregor A. Dizon, Razel Nikka M. Hao, Arianne A. Zamora, Devon Ray Pacial, Juan Antonio R. Magalang, Marissa Alejandria, Celia Carlos, Anna Ong-Lim, Edsel Maurice Salvaña, John Q. Wong, Jaime C. Montoya, Maria Rosario Singh-Vergeire and Cynthia P. Saloma |
| EPI_ISL_2189124 | Philippine Red Cross - Port Area | Philippine Genome Center | Francis A. Tablizo, Kenneth M. Kim, Maria Sofia L. Yangzon, Elcid Aaron R. Pangilinan, Renato Jacinto Q. Mantaring, Benedict A. Maralit, Marc Edsel C. Ayes, Eva Maria Cutiongco-de la Paz, Alethea R. de Guzman, Jan Michael C. Yap, Jo-Hannah S. Llames, Sheila Mae M. Araiza, Kris P. Punayan, Irish Coleen A. Asin, Candice Francheska B. Tambaoan, Asia Louisa U. Chong, Karol Sophia Agape R. Padilla, Rianna Patricia S. Cruz, Carlo M. Lapid, El King D. Morado, Joshua Gregor A. Dizon, Razel Nikka M. Hao, Arianne A. Zamora, Devon Ray Pacial, Juan Antonio R. Magalang, Marissa Alejandria, Celia Carlos, Anna Ong-Lim, Edsel Maurice Salvaña, John Q. Wong, Jaime C. Montoya, Maria Rosario Singh-Vergeire and Cynthia P. Saloma |
| EPI_ISL_2189125 | The Lord's Grace Medical and Industrial Clinic | Philippine Genome Center | Francis A. Tablizo, Kenneth M. Kim, Maria Sofia L. Yangzon, Elcid Aaron R. Pangilinan, Renato Jacinto Q. Mantaring, Benedict A. Maralit, Marc Edsel C. Ayes, Eva Maria Cutiongco-de la Paz, Alethea R. de Guzman, Jan Michael C. Yap, Jo-Hannah S. Llames, Sheila Mae M. Araiza, Kris P. Punayan, Irish Coleen A. Asin, Candice Francheska B. Tambaoan, Asia Louisa U. Chong, Karol Sophia Agape R. Padilla, Rianna Patricia S. Cruz, Carlo M. Lapid, El King D. Morado, Joshua Gregor A. Dizon, Razel Nikka M. Hao, Arianne A. Zamora, Devon Ray Pacial, Juan Antonio R. Magalang, Marissa Alejandria, Celia Carlos, Anna Ong-Lim, Edsel Maurice Salvaña, John Q. Wong, Jaime C. Montoya, Maria Rosario Singh-Vergeire and Cynthia P. Saloma |
| EPI_ISL_2189126 | Philippine Red Cross - National Blood Center | Philippine Genome Center | Francis A. Tablizo, Kenneth M. Kim, Maria Sofia L. Yangzon, Elcid Aaron R. Pangilinan, Renato Jacinto Q. Mantaring, Benedict A. Maralit, Marc Edsel C. Ayes, Eva Maria Cutiongco-de la Paz, Alethea R. de Guzman, Jan Michael C. Yap, Jo-Hannah S. Llames, Sheila Mae M. Araiza, Kris P. Punayan, Irish Coleen A. Asin, Candice Francheska B. Tambaoan, Asia Louisa U. Chong, Karol Sophia Agape R. Padilla, Rianna Patricia S. Cruz, Carlo M. Lapid, El King D. Morado, Joshua Gregor A. Dizon, Razel Nikka M. Hao, Arianne A. Zamora, Devon Ray Pacial, Juan Antonio R. Magalang, Marissa Alejandria, Celia Carlos, Anna Ong-Lim, Edsel Maurice Salvaña, John Q. Wong, Jaime C. Montoya, Maria Rosario Singh-Vergeire and Cynthia P. Saloma |
| EPI_ISL_2189128 | The Medical City | Philippine Genome Center | Francis A. Tablizo, Kenneth M. Kim, Maria Sofia L. Yangzon, Elcid Aaron R. Pangilinan, Renato Jacinto Q. Mantaring, Benedict A. Maralit, Marc Edsel C. Ayes, Eva Maria Cutiongco-de la Paz, Alethea R. de Guzman, Jan Michael C. Yap, Jo-Hannah S. Llames, Sheila Mae M. Araiza, Kris P. Punayan, Irish Coleen A. Asin, Candice Francheska B. Tambaoan, Asia Louisa U. Chong, Karol Sophia Agape R. Padilla, Rianna Patricia S. Cruz, Carlo M. Lapid, El King D. Morado, Joshua Gregor A. Dizon, Razel Nikka M. Hao, Arianne A. Zamora, Devon Ray Pacial, Juan Antonio R. Magalang, Marissa Alejandria, Celia Carlos, Anna Ong-Lim, Edsel Maurice Salvaña, John Q. Wong, Jaime C. Montoya, Maria Rosario Singh-Vergeire and Cynthia P. Saloma |
| EPI_ISL_2189129 | Qualimed Hospital Sta. Rosa | Philippine Genome Center | Francis A. Tablizo, Kenneth M. Kim, Maria Sofia L. Yangzon, Elcid Aaron R. Pangilinan, Renato Jacinto Q. Mantaring, Benedict A. Maralit, Marc Edsel C. Ayes, Eva Maria Cutiongco-de la Paz, Alethea R. de Guzman, Jan Michael C. Yap, Jo-Hannah S. Llames, Sheila Mae M. Araiza, Kris P. Punayan, Irish Coleen A. Asin, Candice Francheska B. Tambaoan, Asia Louisa U. Chong, Karol Sophia Agape R. Padilla, Rianna Patricia S. Cruz, Carlo M. Lapid, El King D. Morado, Joshua Gregor A. Dizon, Razel Nikka M. Hao, Arianne A. Zamora, Devon Ray Pacial, Juan Antonio R. Magalang, Marissa Alejandria, Celia Carlos, Anna Ong-Lim, Edsel Maurice Salvaña, John Q. Wong, Jaime C. Montoya, Maria Rosario Singh-Vergeire and Cynthia P. Saloma |
| EPI_ISL_2189131 | National Kidney and Transplant Institute (NKTI) | Philippine Genome Center | Francis A. Tablizo, Kenneth M. Kim, Maria Sofia L. Yangzon, Elcid Aaron R. Pangilinan, Renato Jacinto Q. Mantaring, Benedict A. Maralit, Marc Edsel C. Ayes, Eva Maria Cutiongco-de la Paz, Alethea R. de Guzman, Jan Michael C. Yap, Jo-Hannah S. Llames, Sheila Mae M. Araiza, Kris P. Punayan, Irish Coleen A. Asin, Candice Francheska B. Tambaoan, Asia Louisa U. Chong, Karol Sophia Agape R. Padilla, Rianna Patricia S. Cruz, Carlo M. Lapid, El King D. Morado, Joshua Gregor A. Dizon, Razel Nikka M. Hao, Arianne A. Zamora, Devon Ray Pacial, Juan Antonio R. Magalang, Marissa Alejandria, Celia Carlos, Anna Ong-Lim, Edsel Maurice Salvaña, John Q. Wong, Jaime C. Montoya, Maria Rosario Singh-Vergeire and Cynthia P. Saloma |
| EPI_ISL_2189132 | The Medical City | Philippine Genome Center | Francis A. Tablizo, Kenneth M. Kim, Maria Sofia L. Yangzon, Elcid Aaron R. Pangilinan, Renato Jacinto Q. Mantaring, Benedict A. Maralit, Marc Edsel C. Ayes, Eva Maria Cutiongco-de la Paz, Alethea R. de Guzman, Jan Michael C. Yap, Jo-Hannah S. Llames, Sheila Mae M. Araiza, Kris P. Punayan, Irish Coleen A. Asin, Candice Francheska B. Tambaoan, Asia Louisa U. Chong, Karol Sophia Agape R. Padilla, Rianna Patricia S. Cruz, Carlo M. Lapid, El King D. Morado, Joshua Gregor A. Dizon, Razel Nikka M. Hao, Arianne A. Zamora, Devon Ray Pacial, Juan Antonio R. Magalang, Marissa Alejandria, Celia Carlos, Anna Ong-Lim, Edsel Maurice Salvaña, John Q. Wong, Jaime C. Montoya, Maria Rosario Singh-Vergeire and Cynthia P. Saloma |
| EPI_ISL_2189133 | University of the Philippines National Institutes of Health (UP NIH) | Philippine Genome Center | Francis A. Tablizo, Kenneth M. Kim, Maria Sofia L. Yangzon, Elcid Aaron R. Pangilinan, Renato Jacinto Q. Mantaring, Benedict A. Maralit, Marc Edsel C. Ayes, Eva Maria Cutiongco-de la Paz, Alethea R. de Guzman, Jan Michael C. Yap, Jo-Hannah S. Llames, Sheila Mae M. Araiza, Kris P. Punayan, Irish Coleen A. Asin, Candice Francheska B. Tambaoan, Asia Louisa U. Chong, Karol Sophia Agape R. Padilla, Rianna Patricia S. Cruz, Carlo M. Lapid, El King D. Morado, Joshua Gregor A. Dizon, Razel Nikka M. Hao, Arianne A. Zamora, Devon Ray Pacial, Juan Antonio R. Magalang, Marissa Alejandria, Celia Carlos, Anna Ong-Lim, Edsel Maurice Salvaña, John Q. Wong, Jaime C. Montoya, Maria Rosario Singh-Vergeire and Cynthia P. Saloma |
| EPI_ISL_2189134 | The Medical City | Philippine Genome Center | Francis A. Tablizo, Kenneth M. Kim, Maria Sofia L. Yangzon, Elcid Aaron R. Pangilinan, Renato Jacinto Q. Mantaring, Benedict A. Maralit, Marc Edsel C. Ayes, Eva Maria Cutiongco-de la Paz, Alethea R. de Guzman, Jan Michael C. Yap, Jo-Hannah S. Llames, Sheila Mae M. Araiza, Kris P. Punayan, Irish Coleen A. Asin, Candice Francheska B. Tambaoan, Asia Louisa U. Chong, Karol Sophia Agape R. Padilla, Rianna Patricia S. Cruz, Carlo M. Lapid, El King D. Morado, Joshua Gregor A. Dizon, Razel Nikka M. Hao, Arianne A. Zamora, Devon Ray Pacial, Juan Antonio R. Magalang, Marissa Alejandria, Celia Carlos, Anna Ong-Lim, Edsel Maurice Salvaña, John Q. Wong, Jaime C. Montoya, Maria Rosario Singh-Vergeire and Cynthia P. Saloma |
| EPI_ISL_2189135 | Philippine Red Cross - Clark Molecular Laboratory | Philippine Genome Center | Francis A. Tablizo, Kenneth M. Kim, Maria Sofia L. Yangzon, Elcid Aaron R. Pangilinan, Renato Jacinto Q. Mantaring, Benedict A. Maralit, Marc Edsel C. Ayes, Eva Maria Cutiongco-de la Paz, Alethea R. de Guzman, Jan Michael C. Yap, Jo-Hannah S. Llames, Sheila Mae M. Araiza, Kris P. Punayan, Irish |

| | | | |
|---|---|---|---|
| | | | Coleen A. Asin, Candice Francheska B. Tambaoan, Asia Louisa U. Chong, Karol Sophia Agape R. Padilla, Rianna Patricia S. Cruz, Carlo M. Lapid, El King D. Morado, Joshua Gregor A. Dizon, Razel Nikka M. Hao, Arianne A. Zamora, Devon Ray Pacial, Juan Antonio R. Magalang, Marissa Alejandria, Celia Carlos, Anna Ong-Lim, Edsel Maurice Salvaña, John Q. Wong, Jaime C. Montoya, Maria Rosario Singh-Vergeire and Cynthia P. Saloma |
| EPI_ISL_2189136 | Philippine Red Cross - Port Area | Philippine Genome Center | Francis A. Tablizo, Kenneth M. Kim, Maria Sofia L. Yangzon, Elcid Aaron R. Pangilinan, Renato Jacinto Q. Mantaring, Benedict A. Maralit, Marc Edsel C. Ayes, Eva Maria Cutiongco-de la Paz, Alethea R. de Guzman, Jan Michael C. Yap, Jo-Hannah S. Llames, Sheila Mae M. Araiza, Kris P. Punayan, Irish Coleen A. Asin, Candice Francheska B. Tambaoan, Asia Louisa U. Chong, Karol Sophia Agape R. Padilla, Rianna Patricia S. Cruz, Carlo M. Lapid, El King D. Morado, Joshua Gregor A. Dizon, Razel Nikka M. Hao, Arianne A. Zamora, Devon Ray Pacial, Juan Antonio R. Magalang, Marissa Alejandria, Celia Carlos, Anna Ong-Lim, Edsel Maurice Salvaña, John Q. Wong, Jaime C. Montoya, Maria Rosario Singh-Vergeire and Cynthia P. Saloma |
| EPI_ISL_2189137 | The Medical City | Philippine Genome Center | Francis A. Tablizo, Kenneth M. Kim, Maria Sofia L. Yangzon, Elcid Aaron R. Pangilinan, Renato Jacinto Q. Mantaring, Benedict A. Maralit, Marc Edsel C. Ayes, Eva Maria Cutiongco-de la Paz, Alethea R. de Guzman, Jan Michael C. Yap, Jo-Hannah S. Llames, Sheila Mae M. Araiza, Kris P. Punayan, Irish Coleen A. Asin, Candice Francheska B. Tambaoan, Asia Louisa U. Chong, Karol Sophia Agape R. Padilla, Rianna Patricia S. Cruz, Carlo M. Lapid, El King D. Morado, Joshua Gregor A. Dizon, Razel Nikka M. Hao, Arianne A. Zamora, Devon Ray Pacial, Juan Antonio R. Magalang, Marissa Alejandria, Celia Carlos, Anna Ong-Lim, Edsel Maurice Salvaña, John Q. Wong, Jaime C. Montoya, Maria Rosario Singh-Vergeire and Cynthia P. Saloma |
| EPI_ISL_2189138 | Sta. Ana Hospital | Philippine Genome Center | Francis A. Tablizo, Kenneth M. Kim, Maria Sofia L. Yangzon, Elcid Aaron R. Pangilinan, Renato Jacinto Q. Mantaring, Benedict A. Maralit, Marc Edsel C. Ayes, Eva Maria Cutiongco-de la Paz, Alethea R. de Guzman, Jan Michael C. Yap, Jo-Hannah S. Llames, Sheila Mae M. Araiza, Kris P. Punayan, Irish Coleen A. Asin, Candice Francheska B. Tambaoan, Asia Louisa U. Chong, Karol Sophia Agape R. Padilla, Rianna Patricia S. Cruz, Carlo M. Lapid, El King D. Morado, Joshua Gregor A. Dizon, Razel Nikka M. Hao, Arianne A. Zamora, Devon Ray Pacial, Juan Antonio R. Magalang, Marissa Alejandria, Celia Carlos, Anna Ong-Lim, Edsel Maurice Salvaña, John Q. Wong, Jaime C. Montoya, Maria Rosario Singh-Vergeire and Cynthia P. Saloma |
| EPI_ISL_2189142 | The Medical City | Philippine Genome Center | Francis A. Tablizo, Kenneth M. Kim, Maria Sofia L. Yangzon, Elcid Aaron R. Pangilinan, Renato Jacinto Q. Mantaring, Benedict A. Maralit, Marc Edsel C. Ayes, Eva Maria Cutiongco-de la Paz, Alethea R. de Guzman, Jan Michael C. Yap, Jo-Hannah S. Llames, Sheila Mae M. Araiza, Kris P. Punayan, Irish Coleen A. Asin, Candice Francheska B. Tambaoan, Asia Louisa U. Chong, Karol Sophia Agape R. Padilla, Rianna Patricia S. Cruz, Carlo M. Lapid, El King D. Morado, Joshua Gregor A. Dizon, Razel Nikka M. Hao, Arianne A. Zamora, Devon Ray Pacial, Juan Antonio R. Magalang, Marissa Alejandria, Celia Carlos, Anna Ong-Lim, Edsel Maurice Salvaña, John Q. Wong, Jaime C. Montoya, Maria Rosario Singh-Vergeire and Cynthia P. Saloma |
| EPI_ISL_2189144 | Sta. Rosa CHO- GENEXPERT LABORATORY | Philippine Genome Center | Francis A. Tablizo, Kenneth M. Kim, Maria Sofia L. Yangzon, Elcid Aaron R. Pangilinan, Renato Jacinto Q. Mantaring, Benedict A. Maralit, Marc Edsel C. Ayes, Eva Maria Cutiongco-de la Paz, Alethea R. de Guzman, Jan Michael C. Yap, Jo-Hannah S. Llames, Sheila Mae M. Araiza, Kris P. Punayan, Irish Coleen A. Asin, Candice Francheska B. Tambaoan, Asia Louisa U. Chong, Karol Sophia Agape R. Padilla, Rianna Patricia S. Cruz, Carlo M. Lapid, El King D. Morado, Joshua Gregor A. Dizon, Razel Nikka M. Hao, Arianne A. Zamora, Devon Ray Pacial, Juan Antonio R. Magalang, Marissa Alejandria, Celia Carlos, Anna Ong-Lim, Edsel Maurice Salvaña, John Q. Wong, Jaime C. Montoya, Maria Rosario Singh-Vergeire and Cynthia P. Saloma |
| EPI_ISL_2189145 | Baguio General Hospital Medical Center (BGHMC) | Philippine Genome Center | Francis A. Tablizo, Kenneth M. Kim, Maria Sofia L. Yangzon, Elcid Aaron R. Pangilinan, Renato Jacinto Q. Mantaring, Benedict A. Maralit, Marc Edsel C. Ayes, Eva Maria Cutiongco-de la Paz, Alethea R. de Guzman, Jan Michael C. Yap, Jo-Hannah S. Llames, Sheila Mae M. Araiza, Kris P. Punayan, Irish Coleen A. Asin, Candice Francheska B. Tambaoan, Asia Louisa U. Chong, Karol Sophia Agape R. Padilla, Rianna Patricia S. Cruz, Carlo M. Lapid, El King D. Morado, Joshua Gregor A. Dizon, Razel Nikka M. Hao, Arianne A. Zamora, Devon Ray Pacial, Juan Antonio R. Magalang, Marissa Alejandria, Celia Carlos, Anna Ong-Lim, Edsel Maurice Salvaña, John Q. Wong, Jaime C. Montoya, Maria Rosario Singh-Vergeire and Cynthia P. Saloma |
| EPI_ISL_2189146 | Philippine Red Cross - National Blood Center | Philippine Genome Center | Francis A. Tablizo, Kenneth M. Kim, Maria Sofia L. Yangzon, Elcid Aaron R. Pangilinan, Renato Jacinto Q. Mantaring, Benedict A. Maralit, Marc Edsel C. Ayes, Eva Maria Cutiongco-de la Paz, Alethea R. de Guzman, Jan Michael C. Yap, Jo-Hannah S. Llames, Sheila Mae M. Araiza, Kris P. Punayan, Irish Coleen A. Asin, Candice Francheska B. Tambaoan, Asia Louisa U. Chong, Karol Sophia Agape R. Padilla, Rianna Patricia S. Cruz, Carlo M. Lapid, El King D. Morado, Joshua Gregor A. Dizon, Razel Nikka M. Hao, Arianne A. Zamora, Devon Ray Pacial, Juan Antonio R. Magalang, Marissa Alejandria, Celia Carlos, Anna Ong-Lim, Edsel Maurice Salvaña, John Q. Wong, Jaime C. Montoya, Maria Rosario Singh-Vergeire and Cynthia P. Saloma |
| EPI_ISL_2189147 | Safeguard DNA Diagnostic, Inc. | Philippine Genome Center | Francis A. Tablizo, Kenneth M. Kim, Maria Sofia L. Yangzon, Elcid Aaron R. Pangilinan, Renato Jacinto Q. Mantaring, Benedict A. Maralit, Marc Edsel C. Ayes, Eva Maria Cutiongco-de la Paz, Alethea R. de Guzman, Jan Michael C. Yap, Jo-Hannah S. Llames, Sheila Mae M. Araiza, Kris P. Punayan, Irish Coleen A. Asin, Candice Francheska B. Tambaoan, Asia Louisa U. Chong, Karol Sophia Agape R. Padilla, Rianna Patricia S. Cruz, Carlo M. Lapid, El King D. Morado, Joshua Gregor A. Dizon, Razel Nikka M. Hao, Arianne A. Zamora, Devon Ray Pacial, Juan Antonio R. Magalang, Marissa Alejandria, Celia Carlos, Anna Ong-Lim, Edsel Maurice Salvaña, John Q. Wong, Jaime C. Montoya, Maria Rosario Singh-Vergeire and Cynthia P. Saloma |
| EPI_ISL_2189148 | Pasig City Children's Hospital - Child's Hope | Philippine Genome Center | Francis A. Tablizo, Kenneth M. Kim, Maria Sofia L. Yangzon, Elcid Aaron R. Pangilinan, Renato Jacinto Q. Mantaring, Benedict A. Maralit, Marc Edsel C. Ayes, Eva Maria Cutiongco-de la Paz, Alethea R. de Guzman, Jan Michael C. Yap, Jo-Hannah S. Llames, Sheila Mae M. Araiza, Kris P. Punayan, Irish Coleen A. Asin, Candice Francheska B. Tambaoan, Asia Louisa U. Chong, Karol Sophia Agape R. Padilla, Rianna Patricia S. Cruz, Carlo M. Lapid, El King D. Morado, Joshua Gregor A. Dizon, Razel Nikka M. Hao, Arianne A. Zamora, Devon Ray Pacial, Juan Antonio R. Magalang, Marissa Alejandria, Celia Carlos, Anna Ong-Lim, Edsel Maurice Salvaña, John Q. Wong, Jaime C. Montoya, Maria Rosario Singh-Vergeire and Cynthia P. Saloma |
| EPI_ISL_2189149 | Philippine Red Cross - Port Area | Philippine Genome Center | Francis A. Tablizo, Kenneth M. Kim, Maria Sofia L. Yangzon, Elcid Aaron R. Pangilinan, Renato Jacinto Q. Mantaring, Benedict A. Maralit, Marc Edsel C. Ayes, Eva Maria Cutiongco-de la Paz, Alethea R. de Guzman, Jan Michael C. Yap, Jo-Hannah S. Llames, Sheila Mae M. Araiza, Kris P. Punayan, Irish Coleen A. Asin, Candice Francheska B. Tambaoan, Asia Louisa U. Chong, Karol Sophia Agape R. Padilla, Rianna Patricia S. Cruz, Carlo M. Lapid, El King D. Morado, Joshua Gregor A. Dizon, Razel Nikka M. Hao, Arianne A. Zamora, Devon Ray Pacial, Juan Antonio R. Magalang, Marissa Alejandria, Celia Carlos, Anna Ong-Lim, Edsel Maurice Salvaña, John Q. Wong, Jaime C. Montoya, Maria Rosario Singh-Vergeire and Cynthia P. Saloma |
| EPI_ISL_2189150 | The Medical City | Philippine Genome Center | Francis A. Tablizo, Kenneth M. Kim, Maria Sofia L. Yangzon, Elcid Aaron R. Pangilinan, Renato Jacinto Q. Mantaring, Benedict A. Maralit, Marc Edsel C. Ayes, Eva Maria Cutiongco-de la Paz, Alethea R. de Guzman, Jan Michael C. Yap, Jo-Hannah S. Llames, Sheila Mae M. Araiza, Kris P. Punayan, Irish Coleen A. Asin, Candice Francheska B. Tambaoan, Asia Louisa U. Chong, Karol Sophia Agape R. Padilla, Rianna Patricia S. Cruz, Carlo M. Lapid, El King D. Morado, Joshua Gregor A. Dizon, Razel Nikka M. Hao, Arianne A. Zamora, Devon Ray Pacial, Juan Antonio R. Magalang, Marissa Alejandria, Celia Carlos, Anna Ong-Lim, Edsel Maurice Salvaña, John Q. Wong, Jaime C. Montoya, Maria Rosario Singh-Vergeire and Cynthia P. Saloma |
| EPI_ISL_2189153 | Sta. Rosa CHO- GENEXPERT LABORATORY | Philippine Genome Center | Francis A. Tablizo, Kenneth M. Kim, Maria Sofia L. Yangzon, Elcid Aaron R. Pangilinan, Renato Jacinto Q. Mantaring, Benedict A. Maralit, Marc Edsel C. Ayes, Eva Maria Cutiongco-de la Paz, Alethea R. de Guzman, Jan Michael C. Yap, Jo-Hannah S. Llames, Sheila Mae M. Araiza, Kris P. Punayan, Irish Coleen A. Asin, Candice Francheska B. Tambaoan, Asia Louisa U. Chong, Karol Sophia Agape R. Padilla, Rianna Patricia S. Cruz, Carlo M. Lapid, El King D. Morado, Joshua Gregor A. Dizon, Razel Nikka M. Hao, Arianne A. Zamora, Devon Ray Pacial, Juan Antonio R. Magalang, Marissa Alejandria, Celia Carlos, Anna Ong-Lim, Edsel Maurice Salvaña, John Q. Wong, Jaime C. Montoya, Maria Rosario Singh-Vergeire and Cynthia P. Saloma |
| EPI_ISL_2189154 | Philippine Red Cross - National Blood Center | Philippine Genome Center | Francis A. Tablizo, Kenneth M. Kim, Maria Sofia L. Yangzon, Elcid Aaron R. Pangilinan, Renato Jacinto Q. Mantaring, Benedict A. Maralit, Marc Edsel C. Ayes, Eva Maria Cutiongco-de la Paz, Alethea R. de Guzman, Jan Michael C. Yap, Jo-Hannah S. Llames, Sheila Mae M. Araiza, Kris P. Punayan, Irish Coleen A. Asin, Candice Francheska B. Tambaoan, Asia Louisa U. Chong, Karol Sophia Agape R. Padilla, Rianna Patricia S. Cruz, Carlo M. Lapid, El King D. Morado, Joshua Gregor A. Dizon, Razel Nikka M. Hao, Arianne A. Zamora, Devon Ray Pacial, Juan Antonio R. Magalang, Marissa Alejandria, Celia Carlos, Anna Ong-Lim, Edsel Maurice Salvaña, John Q. Wong, Jaime C. Montoya, Maria Rosario Singh-Vergeire and Cynthia P. Saloma |
| EPI_ISL_2189155 | Sta. Rosa CHO- GENEXPERT LABORATORY | Philippine Genome Center | Francis A. Tablizo, Kenneth M. Kim, Maria Sofia L. Yangzon, Elcid Aaron R. Pangilinan, Renato Jacinto Q. Mantaring, Benedict A. Maralit, Marc Edsel C. Ayes, Eva Maria Cutiongco-de la Paz, Alethea R. de Guzman, Jan Michael C. Yap, Jo-Hannah S. Llames, Sheila Mae M. Araiza, Kris P. Punayan, Irish Coleen A. Asin, Candice Francheska B. Tambaoan, Asia Louisa U. Chong, Karol Sophia Agape R. Padilla, Rianna Patricia S. Cruz, Carlo M. Lapid, El King D. Morado, Joshua Gregor A. Dizon, Razel Nikka M. Hao, Arianne A. Zamora, Devon Ray Pacial, Juan Antonio R. Magalang, Marissa Alejandria, Celia Carlos, Anna Ong-Lim, Edsel Maurice Salvaña, John Q. Wong, Jaime C. Montoya, Maria Rosario Singh-Vergeire and Cynthia P. Saloma |
| EPI_ISL_2189156 | Sta. Ana Hospital | Philippine Genome Center | Francis A. Tablizo, Kenneth M. Kim, Maria Sofia L. Yangzon, Elcid Aaron R. Pangilinan, Renato Jacinto Q. Mantaring, Benedict A. Maralit, Marc Edsel C. Ayes, Eva Maria Cutiongco-de la Paz, Alethea R. de Guzman, Jan Michael C. Yap, Jo-Hannah S. Llames, Sheila Mae M. Araiza, Kris P. Punayan, Irish Coleen A. Asin, Candice Francheska B. Tambaoan, Asia Louisa U. Chong, Karol Sophia Agape R. Padilla, Rianna Patricia S. Cruz, Carlo M. Lapid, El King D. Morado, Joshua Gregor A. Dizon, Razel Nikka M. Hao, Arianne A. Zamora, Devon Ray Pacial, Juan Antonio R. Magalang, Marissa Alejandria, Celia Carlos, Anna Ong-Lim, Edsel Maurice Salvaña, John Q. Wong, Jaime C. Montoya, Maria Rosario Singh-Vergeire and Cynthia P. Saloma |
| EPI_ISL_2189159 | Philippine Red Cross - National Blood Center | Philippine Genome Center | Francis A. Tablizo, Kenneth M. Kim, Maria Sofia L. Yangzon, Elcid Aaron R. Pangilinan, Renato Jacinto Q. Mantaring, Benedict A. Maralit, Marc Edsel C. |

| | | | |
|---|---|---|---|
| | | | Ayes, Eva Maria Cutiongco-de la Paz, Alethea R. de Guzman, Jan Michael C. Yap, Jo-Hannah S. Llames, Sheila Mae M. Araiza, Kris P. Punayan, Irish Coleen A. Asin, Candice Francheska B. Tambaoan, Asia Louisa U. Chong, Karol Sophia Agape R. Padilla, Rianna Patricia S. Cruz, Carlo M. Lapid, El King D. Morado, Joshua Gregor A. Dizon, Razel Nikka M. Hao, Arianne A. Zamora, Devon Ray Pacial, Juan Antonio R. Magalang, Marissa Alejandria, Celia Carlos, Anna Ong-Lim, Edsel Maurice Salvaña, John Q. Wong, Jaime C. Montoya, Maria Rosario Singh-Vergeire and Cynthia P. Saloma |
| EPI_ISL_2189161 | The Medical City | Philippine Genome Center | Francis A. Tablizo, Kenneth M. Kim, Maria Sofia L. Yangzon, Elcid Aaron R. Pangilinan, Renato Jacinto Q. Mantaring, Benedict A. Maralit, Marc Edsel C. Ayes, Eva Maria Cutiongco-de la Paz, Alethea R. de Guzman, Jan Michael C. Yap, Jo-Hannah S. Llames, Sheila Mae M. Araiza, Kris P. Punayan, Irish Coleen A. Asin, Candice Francheska B. Tambaoan, Asia Louisa U. Chong, Karol Sophia Agape R. Padilla, Rianna Patricia S. Cruz, Carlo M. Lapid, El King D. Morado, Joshua Gregor A. Dizon, Razel Nikka M. Hao, Arianne A. Zamora, Devon Ray Pacial, Juan Antonio R. Magalang, Marissa Alejandria, Celia Carlos, Anna Ong-Lim, Edsel Maurice Salvaña, John Q. Wong, Jaime C. Montoya, Maria Rosario Singh-Vergeire and Cynthia P. Saloma |
| EPI_ISL_2189162 | Sta. Ana Hospital | Philippine Genome Center | Francis A. Tablizo, Kenneth M. Kim, Maria Sofia L. Yangzon, Elcid Aaron R. Pangilinan, Renato Jacinto Q. Mantaring, Benedict A. Maralit, Marc Edsel C. Ayes, Eva Maria Cutiongco-de la Paz, Alethea R. de Guzman, Jan Michael C. Yap, Jo-Hannah S. Llames, Sheila Mae M. Araiza, Kris P. Punayan, Irish Coleen A. Asin, Candice Francheska B. Tambaoan, Asia Louisa U. Chong, Karol Sophia Agape R. Padilla, Rianna Patricia S. Cruz, Carlo M. Lapid, El King D. Morado, Joshua Gregor A. Dizon, Razel Nikka M. Hao, Arianne A. Zamora, Devon Ray Pacial, Juan Antonio R. Magalang, Marissa Alejandria, Celia Carlos, Anna Ong-Lim, Edsel Maurice Salvaña, John Q. Wong, Jaime C. Montoya, Maria Rosario Singh-Vergeire and Cynthia P. Saloma |
| EPI_ISL_2189163 | Philippine Red Cross - Port Area | Philippine Genome Center | Francis A. Tablizo, Kenneth M. Kim, Maria Sofia L. Yangzon, Elcid Aaron R. Pangilinan, Renato Jacinto Q. Mantaring, Benedict A. Maralit, Marc Edsel C. Ayes, Eva Maria Cutiongco-de la Paz, Alethea R. de Guzman, Jan Michael C. Yap, Jo-Hannah S. Llames, Sheila Mae M. Araiza, Kris P. Punayan, Irish Coleen A. Asin, Candice Francheska B. Tambaoan, Asia Louisa U. Chong, Karol Sophia Agape R. Padilla, Rianna Patricia S. Cruz, Carlo M. Lapid, El King D. Morado, Joshua Gregor A. Dizon, Razel Nikka M. Hao, Arianne A. Zamora, Devon Ray Pacial, Juan Antonio R. Magalang, Marissa Alejandria, Celia Carlos, Anna Ong-Lim, Edsel Maurice Salvaña, John Q. Wong, Jaime C. Montoya, Maria Rosario Singh-Vergeire and Cynthia P. Saloma |
| EPI_ISL_2189166 | Sta. Rosa CHO- GENEXPERT LABORATORY | Philippine Genome Center | Francis A. Tablizo, Kenneth M. Kim, Maria Sofia L. Yangzon, Elcid Aaron R. Pangilinan, Renato Jacinto Q. Mantaring, Benedict A. Maralit, Marc Edsel C. Ayes, Eva Maria Cutiongco-de la Paz, Alethea R. de Guzman, Jan Michael C. Yap, Jo-Hannah S. Llames, Sheila Mae M. Araiza, Kris P. Punayan, Irish Coleen A. Asin, Candice Francheska B. Tambaoan, Asia Louisa U. Chong, Karol Sophia Agape R. Padilla, Rianna Patricia S. Cruz, Carlo M. Lapid, El King D. Morado, Joshua Gregor A. Dizon, Razel Nikka M. Hao, Arianne A. Zamora, Devon Ray Pacial, Juan Antonio R. Magalang, Marissa Alejandria, Celia Carlos, Anna Ong-Lim, Edsel Maurice Salvaña, John Q. Wong, Jaime C. Montoya, Maria Rosario Singh-Vergeire and Cynthia P. Saloma |
| EPI_ISL_2189167 | The Medical City | Philippine Genome Center | Francis A. Tablizo, Kenneth M. Kim, Maria Sofia L. Yangzon, Elcid Aaron R. Pangilinan, Renato Jacinto Q. Mantaring, Benedict A. Maralit, Marc Edsel C. Ayes, Eva Maria Cutiongco-de la Paz, Alethea R. de Guzman, Jan Michael C. Yap, Jo-Hannah S. Llames, Sheila Mae M. Araiza, Kris P. Punayan, Irish Coleen A. Asin, Candice Francheska B. Tambaoan, Asia Louisa U. Chong, Karol Sophia Agape R. Padilla, Rianna Patricia S. Cruz, Carlo M. Lapid, El King D. Morado, Joshua Gregor A. Dizon, Razel Nikka M. Hao, Arianne A. Zamora, Devon Ray Pacial, Juan Antonio R. Magalang, Marissa Alejandria, Celia Carlos, Anna Ong-Lim, Edsel Maurice Salvaña, John Q. Wong, Jaime C. Montoya, Maria Rosario Singh-Vergeire and Cynthia P. Saloma |
| EPI_ISL_2189168 | The Lord's Grace Medical and Industrial Clinic | Philippine Genome Center | Francis A. Tablizo, Kenneth M. Kim, Maria Sofia L. Yangzon, Elcid Aaron R. Pangilinan, Renato Jacinto Q. Mantaring, Benedict A. Maralit, Marc Edsel C. Ayes, Eva Maria Cutiongco-de la Paz, Alethea R. de Guzman, Jan Michael C. Yap, Jo-Hannah S. Llames, Sheila Mae M. Araiza, Kris P. Punayan, Irish Coleen A. Asin, Candice Francheska B. Tambaoan, Asia Louisa U. Chong, Karol Sophia Agape R. Padilla, Rianna Patricia S. Cruz, Carlo M. Lapid, El King D. Morado, Joshua Gregor A. Dizon, Razel Nikka M. Hao, Arianne A. Zamora, Devon Ray Pacial, Juan Antonio R. Magalang, Marissa Alejandria, Celia Carlos, Anna Ong-Lim, Edsel Maurice Salvaña, John Q. Wong, Jaime C. Montoya, Maria Rosario Singh-Vergeire and Cynthia P. Saloma |
| EPI_ISL_2189169 | The Medical City | Philippine Genome Center | Francis A. Tablizo, Kenneth M. Kim, Maria Sofia L. Yangzon, Elcid Aaron R. Pangilinan, Renato Jacinto Q. Mantaring, Benedict A. Maralit, Marc Edsel C. Ayes, Eva Maria Cutiongco-de la Paz, Alethea R. de Guzman, Jan Michael C. Yap, Jo-Hannah S. Llames, Sheila Mae M. Araiza, Kris P. Punayan, Irish Coleen A. Asin, Candice Francheska B. Tambaoan, Asia Louisa U. Chong, Karol Sophia Agape R. Padilla, Rianna Patricia S. Cruz, Carlo M. Lapid, El King D. Morado, Joshua Gregor A. Dizon, Razel Nikka M. Hao, Arianne A. Zamora, Devon Ray Pacial, Juan Antonio R. Magalang, Marissa Alejandria, Celia Carlos, Anna Ong-Lim, Edsel Maurice Salvaña, John Q. Wong, Jaime C. Montoya, Maria Rosario Singh-Vergeire and Cynthia P. Saloma |
| EPI_ISL_2189170 | Mary Mediatrix Medical Center | Philippine Genome Center | Francis A. Tablizo, Kenneth M. Kim, Maria Sofia L. Yangzon, Elcid Aaron R. Pangilinan, Renato Jacinto Q. Mantaring, Benedict A. Maralit, Marc Edsel C. Ayes, Eva Maria Cutiongco-de la Paz, Alethea R. de Guzman, Jan Michael C. Yap, Jo-Hannah S. Llames, Sheila Mae M. Araiza, Kris P. Punayan, Irish Coleen A. Asin, Candice Francheska B. Tambaoan, Asia Louisa U. Chong, Karol Sophia Agape R. Padilla, Rianna Patricia S. Cruz, Carlo M. Lapid, El King D. Morado, Joshua Gregor A. Dizon, Razel Nikka M. Hao, Arianne A. Zamora, Devon Ray Pacial, Juan Antonio R. Magalang, Marissa Alejandria, Celia Carlos, Anna Ong-Lim, Edsel Maurice Salvaña, John Q. Wong, Jaime C. Montoya, Maria Rosario Singh-Vergeire and Cynthia P. Saloma |
| EPI_ISL_2189172 | Philippine Red Cross - National Blood Center | Philippine Genome Center | Francis A. Tablizo, Kenneth M. Kim, Maria Sofia L. Yangzon, Elcid Aaron R. Pangilinan, Renato Jacinto Q. Mantaring, Benedict A. Maralit, Marc Edsel C. Ayes, Eva Maria Cutiongco-de la Paz, Alethea R. de Guzman, Jan Michael C. Yap, Jo-Hannah S. Llames, Sheila Mae M. Araiza, Kris P. Punayan, Irish Coleen A. Asin, Candice Francheska B. Tambaoan, Asia Louisa U. Chong, Karol Sophia Agape R. Padilla, Rianna Patricia S. Cruz, Carlo M. Lapid, El King D. Morado, Joshua Gregor A. Dizon, Razel Nikka M. Hao, Arianne A. Zamora, Devon Ray Pacial, Juan Antonio R. Magalang, Marissa Alejandria, Celia Carlos, Anna Ong-Lim, Edsel Maurice Salvaña, John Q. Wong, Jaime C. Montoya, Maria Rosario Singh-Vergeire and Cynthia P. Saloma |
| EPI_ISL_2189173 | Philippine Red Cross - Port Area | Philippine Genome Center | Francis A. Tablizo, Kenneth M. Kim, Maria Sofia L. Yangzon, Elcid Aaron R. Pangilinan, Renato Jacinto Q. Mantaring, Benedict A. Maralit, Marc Edsel C. Ayes, Eva Maria Cutiongco-de la Paz, Alethea R. de Guzman, Jan Michael C. Yap, Jo-Hannah S. Llames, Sheila Mae M. Araiza, Kris P. Punayan, Irish Coleen A. Asin, Candice Francheska B. Tambaoan, Asia Louisa U. Chong, Karol Sophia Agape R. Padilla, Rianna Patricia S. Cruz, Carlo M. Lapid, El King D. Morado, Joshua Gregor A. Dizon, Razel Nikka M. Hao, Arianne A. Zamora, Devon Ray Pacial, Juan Antonio R. Magalang, Marissa Alejandria, Celia Carlos, Anna Ong-Lim, Edsel Maurice Salvaña, John Q. Wong, Jaime C. Montoya, Maria Rosario Singh-Vergeire and Cynthia P. Saloma |
| EPI_ISL_2189174 | First Aide Diagnostic Center | Philippine Genome Center | Francis A. Tablizo, Kenneth M. Kim, Maria Sofia L. Yangzon, Elcid Aaron R. Pangilinan, Renato Jacinto Q. Mantaring, Benedict A. Maralit, Marc Edsel C. Ayes, Eva Maria Cutiongco-de la Paz, Alethea R. de Guzman, Jan Michael C. Yap, Jo-Hannah S. Llames, Sheila Mae M. Araiza, Kris P. Punayan, Irish Coleen A. Asin, Candice Francheska B. Tambaoan, Asia Louisa U. Chong, Karol Sophia Agape R. Padilla, Rianna Patricia S. Cruz, Carlo M. Lapid, El King D. Morado, Joshua Gregor A. Dizon, Razel Nikka M. Hao, Arianne A. Zamora, Devon Ray Pacial, Juan Antonio R. Magalang, Marissa Alejandria, Celia Carlos, Anna Ong-Lim, Edsel Maurice Salvaña, John Q. Wong, Jaime C. Montoya, Maria Rosario Singh-Vergeire and Cynthia P. Saloma |
| EPI_ISL_2189175 | Philippine Red Cross - National Blood Center | Philippine Genome Center | Francis A. Tablizo, Kenneth M. Kim, Maria Sofia L. Yangzon, Elcid Aaron R. Pangilinan, Renato Jacinto Q. Mantaring, Benedict A. Maralit, Marc Edsel C. Ayes, Eva Maria Cutiongco-de la Paz, Alethea R. de Guzman, Jan Michael C. Yap, Jo-Hannah S. Llames, Sheila Mae M. Araiza, Kris P. Punayan, Irish Coleen A. Asin, Candice Francheska B. Tambaoan, Asia Louisa U. Chong, Karol Sophia Agape R. Padilla, Rianna Patricia S. Cruz, Carlo M. Lapid, El King D. Morado, Joshua Gregor A. Dizon, Razel Nikka M. Hao, Arianne A. Zamora, Devon Ray Pacial, Juan Antonio R. Magalang, Marissa Alejandria, Celia Carlos, Anna Ong-Lim, Edsel Maurice Salvaña, John Q. Wong, Jaime C. Montoya, Maria Rosario Singh-Vergeire and Cynthia P. Saloma |
| EPI_ISL_2189178 | Philippine Red Cross - Port Area | Philippine Genome Center | Francis A. Tablizo, Kenneth M. Kim, Maria Sofia L. Yangzon, Elcid Aaron R. Pangilinan, Renato Jacinto Q. Mantaring, Benedict A. Maralit, Marc Edsel C. Ayes, Eva Maria Cutiongco-de la Paz, Alethea R. de Guzman, Jan Michael C. Yap, Jo-Hannah S. Llames, Sheila Mae M. Araiza, Kris P. Punayan, Irish Coleen A. Asin, Candice Francheska B. Tambaoan, Asia Louisa U. Chong, Karol Sophia Agape R. Padilla, Rianna Patricia S. Cruz, Carlo M. Lapid, El King D. Morado, Joshua Gregor A. Dizon, Razel Nikka M. Hao, Arianne A. Zamora, Devon Ray Pacial, Juan Antonio R. Magalang, Marissa Alejandria, Celia Carlos, Anna Ong-Lim, Edsel Maurice Salvaña, John Q. Wong, Jaime C. Montoya, Maria Rosario Singh-Vergeire and Cynthia P. Saloma |
| EPI_ISL_2189179 | Sta. Rosa CHO- GENEXPERT LABORATORY | Philippine Genome Center | Francis A. Tablizo, Kenneth M. Kim, Maria Sofia L. Yangzon, Elcid Aaron R. Pangilinan, Renato Jacinto Q. Mantaring, Benedict A. Maralit, Marc Edsel C. Ayes, Eva Maria Cutiongco-de la Paz, Alethea R. de Guzman, Jan Michael C. Yap, Jo-Hannah S. Llames, Sheila Mae M. Araiza, Kris P. Punayan, Irish Coleen A. Asin, Candice Francheska B. Tambaoan, Asia Louisa U. Chong, Karol Sophia Agape R. Padilla, Rianna Patricia S. Cruz, Carlo M. Lapid, El King D. Morado, Joshua Gregor A. Dizon, Razel Nikka M. Hao, Arianne A. Zamora, Devon Ray Pacial, Juan Antonio R. Magalang, Marissa Alejandria, Celia Carlos, Anna Ong-Lim, Edsel Maurice Salvaña, John Q. Wong, Jaime C. Montoya, Maria Rosario Singh-Vergeire and Cynthia P. Saloma |
| EPI_ISL_2189180 | Pasig City Children's Hospital - Child's Hope | Philippine Genome Center | Francis A. Tablizo, Kenneth M. Kim, Maria Sofia L. Yangzon, Elcid Aaron R. Pangilinan, Renato Jacinto Q. Mantaring, Benedict A. Maralit, Marc Edsel C. Ayes, Eva Maria Cutiongco-de la Paz, Alethea R. de Guzman, Jan Michael C. Yap, Jo-Hannah S. Llames, Sheila Mae M. Araiza, Kris P. Punayan, Irish Coleen A. Asin, Candice Francheska B. Tambaoan, Asia Louisa U. Chong, Karol Sophia Agape R. Padilla, Rianna Patricia S. Cruz, Carlo M. Lapid, El King D. Morado, Joshua Gregor A. Dizon, Razel Nikka M. Hao, Arianne A. Zamora, Devon Ray Pacial, Juan Antonio R. Magalang, Marissa Alejandria, Celia Carlos, Anna Ong-Lim, Edsel Maurice Salvaña, John Q. Wong, Jaime C. Montoya, Maria Rosario Singh-Vergeire and Cynthia P. Saloma |

| | | | |
|---|---|---|---|
| EPI_ISL_2189181 | Qualimed Hospital Sta. Rosa | Philippine Genome Center | Francis A. Tablizo, Kenneth M. Kim, Maria Sofia L. Yangzon, Elcid Aaron R. Pangilinan, Renato Jacinto Q. Mantaring, Benedict A. Maralit, Marc Edsel C. Ayes, Eva Maria Cutiongco-de la Paz, Alethea R. de Guzman, Jan Michael C. Yap, Jo-Hannah S. Llames, Sheila Mae M. Araiza, Kris P. Punayan, Irish Coleen A. Asin, Candice Francheska B. Tambaoan, Asia Louisa U. Chong, Karol Sophia Agape R. Padilla, Rianna Patricia S. Cruz, Carlo M. Lapid, El King D. Morado, Joshua Gregor A. Dizon, Razel Nikka M. Hao, Arianne A. Zamora, Devon Ray Pacial, Juan Antonio R. Magalang, Marissa Alejandria, Celia Carlos, Anna Ong-Lim, Edsel Maurice Salvaña, John Q. Wong, Jaime C. Montoya, Maria Rosario Singh-Vergeire and Cynthia P. Saloma |
| EPI_ISL_2189183 | Philippine Red Cross - Port Area | Philippine Genome Center | Francis A. Tablizo, Kenneth M. Kim, Maria Sofia L. Yangzon, Elcid Aaron R. Pangilinan, Renato Jacinto Q. Mantaring, Benedict A. Maralit, Marc Edsel C. Ayes, Eva Maria Cutiongco-de la Paz, Alethea R. de Guzman, Jan Michael C. Yap, Jo-Hannah S. Llames, Sheila Mae M. Araiza, Kris P. Punayan, Irish Coleen A. Asin, Candice Francheska B. Tambaoan, Asia Louisa U. Chong, Karol Sophia Agape R. Padilla, Rianna Patricia S. Cruz, Carlo M. Lapid, El King D. Morado, Joshua Gregor A. Dizon, Razel Nikka M. Hao, Arianne A. Zamora, Devon Ray Pacial, Juan Antonio R. Magalang, Marissa Alejandria, Celia Carlos, Anna Ong-Lim, Edsel Maurice Salvaña, John Q. Wong, Jaime C. Montoya, Maria Rosario Singh-Vergeire and Cynthia P. Saloma |
| EPI_ISL_2189184 | Pasig City Children's Hospital - Child's Hope | Philippine Genome Center | Francis A. Tablizo, Kenneth M. Kim, Maria Sofia L. Yangzon, Elcid Aaron R. Pangilinan, Renato Jacinto Q. Mantaring, Benedict A. Maralit, Marc Edsel C. Ayes, Eva Maria Cutiongco-de la Paz, Alethea R. de Guzman, Jan Michael C. Yap, Jo-Hannah S. Llames, Sheila Mae M. Araiza, Kris P. Punayan, Irish Coleen A. Asin, Candice Francheska B. Tambaoan, Asia Louisa U. Chong, Karol Sophia Agape R. Padilla, Rianna Patricia S. Cruz, Carlo M. Lapid, El King D. Morado, Joshua Gregor A. Dizon, Razel Nikka M. Hao, Arianne A. Zamora, Devon Ray Pacial, Juan Antonio R. Magalang, Marissa Alejandria, Celia Carlos, Anna Ong-Lim, Edsel Maurice Salvaña, John Q. Wong, Jaime C. Montoya, Maria Rosario Singh-Vergeire and Cynthia P. Saloma |
| EPI_ISL_2189185 | Philippine Red Cross - National Blood Center | Philippine Genome Center | Francis A. Tablizo, Kenneth M. Kim, Maria Sofia L. Yangzon, Elcid Aaron R. Pangilinan, Renato Jacinto Q. Mantaring, Benedict A. Maralit, Marc Edsel C. Ayes, Eva Maria Cutiongco-de la Paz, Alethea R. de Guzman, Jan Michael C. Yap, Jo-Hannah S. Llames, Sheila Mae M. Araiza, Kris P. Punayan, Irish Coleen A. Asin, Candice Francheska B. Tambaoan, Asia Louisa U. Chong, Karol Sophia Agape R. Padilla, Rianna Patricia S. Cruz, Carlo M. Lapid, El King D. Morado, Joshua Gregor A. Dizon, Razel Nikka M. Hao, Arianne A. Zamora, Devon Ray Pacial, Juan Antonio R. Magalang, Marissa Alejandria, Celia Carlos, Anna Ong-Lim, Edsel Maurice Salvaña, John Q. Wong, Jaime C. Montoya, Maria Rosario Singh-Vergeire and Cynthia P. Saloma |
| EPI_ISL_2189186 | The Medical City | Philippine Genome Center | Francis A. Tablizo, Kenneth M. Kim, Maria Sofia L. Yangzon, Elcid Aaron R. Pangilinan, Renato Jacinto Q. Mantaring, Benedict A. Maralit, Marc Edsel C. Ayes, Eva Maria Cutiongco-de la Paz, Alethea R. de Guzman, Jan Michael C. Yap, Jo-Hannah S. Llames, Sheila Mae M. Araiza, Kris P. Punayan, Irish Coleen A. Asin, Candice Francheska B. Tambaoan, Asia Louisa U. Chong, Karol Sophia Agape R. Padilla, Rianna Patricia S. Cruz, Carlo M. Lapid, El King D. Morado, Joshua Gregor A. Dizon, Razel Nikka M. Hao, Arianne A. Zamora, Devon Ray Pacial, Juan Antonio R. Magalang, Marissa Alejandria, Celia Carlos, Anna Ong-Lim, Edsel Maurice Salvaña, John Q. Wong, Jaime C. Montoya, Maria Rosario Singh-Vergeire and Cynthia P. Saloma |
| EPI_ISL_2189189 | The Lord's Grace Medical and Industrial Clinic | Philippine Genome Center | Francis A. Tablizo, Kenneth M. Kim, Maria Sofia L. Yangzon, Elcid Aaron R. Pangilinan, Renato Jacinto Q. Mantaring, Benedict A. Maralit, Marc Edsel C. Ayes, Eva Maria Cutiongco-de la Paz, Alethea R. de Guzman, Jan Michael C. Yap, Jo-Hannah S. Llames, Sheila Mae M. Araiza, Kris P. Punayan, Irish Coleen A. Asin, Candice Francheska B. Tambaoan, Asia Louisa U. Chong, Karol Sophia Agape R. Padilla, Rianna Patricia S. Cruz, Carlo M. Lapid, El King D. Morado, Joshua Gregor A. Dizon, Razel Nikka M. Hao, Arianne A. Zamora, Devon Ray Pacial, Juan Antonio R. Magalang, Marissa Alejandria, Celia Carlos, Anna Ong-Lim, Edsel Maurice Salvaña, John Q. Wong, Jaime C. Montoya, Maria Rosario Singh-Vergeire and Cynthia P. Saloma |
| EPI_ISL_2189190 | Sta. Rosa CHO- GENEXPERT LABORATORY | Philippine Genome Center | Francis A. Tablizo, Kenneth M. Kim, Maria Sofia L. Yangzon, Elcid Aaron R. Pangilinan, Renato Jacinto Q. Mantaring, Benedict A. Maralit, Marc Edsel C. Ayes, Eva Maria Cutiongco-de la Paz, Alethea R. de Guzman, Jan Michael C. Yap, Jo-Hannah S. Llames, Sheila Mae M. Araiza, Kris P. Punayan, Irish Coleen A. Asin, Candice Francheska B. Tambaoan, Asia Louisa U. Chong, Karol Sophia Agape R. Padilla, Rianna Patricia S. Cruz, Carlo M. Lapid, El King D. Morado, Joshua Gregor A. Dizon, Razel Nikka M. Hao, Arianne A. Zamora, Devon Ray Pacial, Juan Antonio R. Magalang, Marissa Alejandria, Celia Carlos, Anna Ong-Lim, Edsel Maurice Salvaña, John Q. Wong, Jaime C. Montoya, Maria Rosario Singh-Vergeire and Cynthia P. Saloma |
| EPI_ISL_2189191 | Qualimed Hospital Sta. Rosa | Philippine Genome Center | Francis A. Tablizo, Kenneth M. Kim, Maria Sofia L. Yangzon, Elcid Aaron R. Pangilinan, Renato Jacinto Q. Mantaring, Benedict A. Maralit, Marc Edsel C. Ayes, Eva Maria Cutiongco-de la Paz, Alethea R. de Guzman, Jan Michael C. Yap, Jo-Hannah S. Llames, Sheila Mae M. Araiza, Kris P. Punayan, Irish Coleen A. Asin, Candice Francheska B. Tambaoan, Asia Louisa U. Chong, Karol Sophia Agape R. Padilla, Rianna Patricia S. Cruz, Carlo M. Lapid, El King D. Morado, Joshua Gregor A. Dizon, Razel Nikka M. Hao, Arianne A. Zamora, Devon Ray Pacial, Juan Antonio R. Magalang, Marissa Alejandria, Celia Carlos, Anna Ong-Lim, Edsel Maurice Salvaña, John Q. Wong, Jaime C. Montoya, Maria Rosario Singh-Vergeire and Cynthia P. Saloma |
| EPI_ISL_2189192 | Mary Mediatrix Medical Center | Philippine Genome Center | Francis A. Tablizo, Kenneth M. Kim, Maria Sofia L. Yangzon, Elcid Aaron R. Pangilinan, Renato Jacinto Q. Mantaring, Benedict A. Maralit, Marc Edsel C. Ayes, Eva Maria Cutiongco-de la Paz, Alethea R. de Guzman, Jan Michael C. Yap, Jo-Hannah S. Llames, Sheila Mae M. Araiza, Kris P. Punayan, Irish Coleen A. Asin, Candice Francheska B. Tambaoan, Asia Louisa U. Chong, Karol Sophia Agape R. Padilla, Rianna Patricia S. Cruz, Carlo M. Lapid, El King D. Morado, Joshua Gregor A. Dizon, Razel Nikka M. Hao, Arianne A. Zamora, Devon Ray Pacial, Juan Antonio R. Magalang, Marissa Alejandria, Celia Carlos, Anna Ong-Lim, Edsel Maurice Salvaña, John Q. Wong, Jaime C. Montoya, Maria Rosario Singh-Vergeire and Cynthia P. Saloma |
| EPI_ISL_2189193 | University of the Philippines National Institutes of Health (UP NIH) | Philippine Genome Center | Francis A. Tablizo, Kenneth M. Kim, Maria Sofia L. Yangzon, Elcid Aaron R. Pangilinan, Renato Jacinto Q. Mantaring, Benedict A. Maralit, Marc Edsel C. Ayes, Eva Maria Cutiongco-de la Paz, Alethea R. de Guzman, Jan Michael C. Yap, Jo-Hannah S. Llames, Sheila Mae M. Araiza, Kris P. Punayan, Irish Coleen A. Asin, Candice Francheska B. Tambaoan, Asia Louisa U. Chong, Karol Sophia Agape R. Padilla, Rianna Patricia S. Cruz, Carlo M. Lapid, El King D. Morado, Joshua Gregor A. Dizon, Razel Nikka M. Hao, Arianne A. Zamora, Devon Ray Pacial, Juan Antonio R. Magalang, Marissa Alejandria, Celia Carlos, Anna Ong-Lim, Edsel Maurice Salvaña, John Q. Wong, Jaime C. Montoya, Maria Rosario Singh-Vergeire and Cynthia P. Saloma |
| EPI_ISL_2189197 | Pasig City Children's Hospital - Child's Hope | Philippine Genome Center | Francis A. Tablizo, Kenneth M. Kim, Maria Sofia L. Yangzon, Elcid Aaron R. Pangilinan, Renato Jacinto Q. Mantaring, Benedict A. Maralit, Marc Edsel C. Ayes, Eva Maria Cutiongco-de la Paz, Alethea R. de Guzman, Jan Michael C. Yap, Jo-Hannah S. Llames, Sheila Mae M. Araiza, Kris P. Punayan, Irish Coleen A. Asin, Candice Francheska B. Tambaoan, Asia Louisa U. Chong, Karol Sophia Agape R. Padilla, Rianna Patricia S. Cruz, Carlo M. Lapid, El King D. Morado, Joshua Gregor A. Dizon, Razel Nikka M. Hao, Arianne A. Zamora, Devon Ray Pacial, Juan Antonio R. Magalang, Marissa Alejandria, Celia Carlos, Anna Ong-Lim, Edsel Maurice Salvaña, John Q. Wong, Jaime C. Montoya, Maria Rosario Singh-Vergeire and Cynthia P. Saloma |
| EPI_ISL_2189199 | Philippine Red Cross - National Blood Center | Philippine Genome Center | Francis A. Tablizo, Kenneth M. Kim, Maria Sofia L. Yangzon, Elcid Aaron R. Pangilinan, Renato Jacinto Q. Mantaring, Benedict A. Maralit, Marc Edsel C. Ayes, Eva Maria Cutiongco-de la Paz, Alethea R. de Guzman, Jan Michael C. Yap, Jo-Hannah S. Llames, Sheila Mae M. Araiza, Kris P. Punayan, Irish Coleen A. Asin, Candice Francheska B. Tambaoan, Asia Louisa U. Chong, Karol Sophia Agape R. Padilla, Rianna Patricia S. Cruz, Carlo M. Lapid, El King D. Morado, Joshua Gregor A. Dizon, Razel Nikka M. Hao, Arianne A. Zamora, Devon Ray Pacial, Juan Antonio R. Magalang, Marissa Alejandria, Celia Carlos, Anna Ong-Lim, Edsel Maurice Salvaña, John Q. Wong, Jaime C. Montoya, Maria Rosario Singh-Vergeire and Cynthia P. Saloma |
| EPI_ISL_2189201 | Sta. Ana Hospital | Philippine Genome Center | Francis A. Tablizo, Kenneth M. Kim, Maria Sofia L. Yangzon, Elcid Aaron R. Pangilinan, Renato Jacinto Q. Mantaring, Benedict A. Maralit, Marc Edsel C. Ayes, Eva Maria Cutiongco-de la Paz, Alethea R. de Guzman, Jan Michael C. Yap, Jo-Hannah S. Llames, Sheila Mae M. Araiza, Kris P. Punayan, Irish Coleen A. Asin, Candice Francheska B. Tambaoan, Asia Louisa U. Chong, Karol Sophia Agape R. Padilla, Rianna Patricia S. Cruz, Carlo M. Lapid, El King D. Morado, Joshua Gregor A. Dizon, Razel Nikka M. Hao, Arianne A. Zamora, Devon Ray Pacial, Juan Antonio R. Magalang, Marissa Alejandria, Celia Carlos, Anna Ong-Lim, Edsel Maurice Salvaña, John Q. Wong, Jaime C. Montoya, Maria Rosario Singh-Vergeire and Cynthia P. Saloma |
| EPI_ISL_2189206 | First Aide Diagnostic Center | Philippine Genome Center | Francis A. Tablizo, Kenneth M. Kim, Maria Sofia L. Yangzon, Elcid Aaron R. Pangilinan, Renato Jacinto Q. Mantaring, Benedict A. Maralit, Marc Edsel C. Ayes, Eva Maria Cutiongco-de la Paz, Alethea R. de Guzman, Jan Michael C. Yap, Jo-Hannah S. Llames, Sheila Mae M. Araiza, Kris P. Punayan, Irish Coleen A. Asin, Candice Francheska B. Tambaoan, Asia Louisa U. Chong, Karol Sophia Agape R. Padilla, Rianna Patricia S. Cruz, Carlo M. Lapid, El King D. Morado, Joshua Gregor A. Dizon, Razel Nikka M. Hao, Arianne A. Zamora, Devon Ray Pacial, Juan Antonio R. Magalang, Marissa Alejandria, Celia Carlos, Anna Ong-Lim, Edsel Maurice Salvaña, John Q. Wong, Jaime C. Montoya, Maria Rosario Singh-Vergeire and Cynthia P. Saloma |
| EPI_ISL_2189209, EPI_ISL_2189212 | The Lord's Grace Medical and Industrial Clinic | Philippine Genome Center | Francis A. Tablizo, Kenneth M. Kim, Maria Sofia L. Yangzon, Elcid Aaron R. Pangilinan, Renato Jacinto Q. Mantaring, Benedict A. Maralit, Marc Edsel C. Ayes, Eva Maria Cutiongco-de la Paz, Alethea R. de Guzman, Jan Michael C. Yap, Jo-Hannah S. Llames, Sheila Mae M. Araiza, Kris P. Punayan, Irish Coleen A. Asin, Candice Francheska B. Tambaoan, Asia Louisa U. Chong, Karol Sophia Agape R. Padilla, Rianna Patricia S. Cruz, Carlo M. Lapid, El King D. Morado, Joshua Gregor A. Dizon, Razel Nikka M. Hao, Arianne A. Zamora, Devon Ray Pacial, Juan Antonio R. Magalang, Marissa Alejandria, Celia Carlos, Anna Ong-Lim, Edsel Maurice Salvaña, John Q. Wong, Jaime C. Montoya, Maria Rosario Singh-Vergeire and Cynthia P. Saloma |
| EPI_ISL_2189213 | Philippine Red Cross - Port Area | Philippine Genome Center | Francis A. Tablizo, Kenneth M. Kim, Maria Sofia L. Yangzon, Elcid Aaron R. Pangilinan, Renato Jacinto Q. Mantaring, Benedict A. Maralit, Marc Edsel C. Ayes, Eva Maria Cutiongco-de la Paz, Alethea R. de Guzman, Jan Michael C. Yap, Jo-Hannah S. Llames, Sheila Mae M. Araiza, Kris P. Punayan, Irish Coleen A. Asin, Candice Francheska B. Tambaoan, Asia Louisa U. Chong, Karol Sophia Agape R. Padilla, Rianna Patricia S. Cruz, Carlo M. Lapid, El King D. Morado, Joshua Gregor A. Dizon, Razel Nikka M. Hao, Arianne A. Zamora, Devon Ray Pacial, Juan Antonio R. Magalang, Marissa Alejandria, Celia Carlos, Anna Ong-Lim, Edsel Maurice Salvaña, John Q. Wong, Jaime C. Montoya, Maria Rosario Singh-Vergeire and Cynthia P. Saloma |

| | | | |
|---|---|---|---|
| EPI_ISL_2189214 | The Lord's Grace Medical and Industrial Clinic | Philippine Genome Center | Francis A. Tablizo, Kenneth M. Kim, Maria Sofia L. Yangzon, Elcid Aaron R. Pangilinan, Renato Jacinto Q. Mantaring, Benedict A. Maralit, Marc Edsel C. Ayes, Eva Maria Cutiongco-de la Paz, Alethea R. de Guzman, Jan Michael C. Yap, Jo-Hannah S. Llames, Sheila Mae M. Araiza, Kris P. Punayan, Irish Coleen A. Asin, Candice Francheska B. Tambaoan, Asia Louisa U. Chong, Karol Sophia Agape R. Padilla, Rianna Patricia S. Cruz, Carlo M. Lapid, El King D. Morado, Joshua Gregor A. Dizon, Razel Nikka M. Hao, Arianne A. Zamora, Devon Ray Pacial, Juan Antonio R. Magalang, Marissa Alejandria, Celia Carlos, Anna Ong-Lim, Edsel Maurice Salvaña, John Q. Wong, Jaime C. Montoya, Maria Rosario Singh-Vergeire and Cynthia P. Saloma |
| EPI_ISL_2189215, EPI_ISL_2189217 | The Medical City | Philippine Genome Center | Francis A. Tablizo, Kenneth M. Kim, Maria Sofia L. Yangzon, Elcid Aaron R. Pangilinan, Renato Jacinto Q. Mantaring, Benedict A. Maralit, Marc Edsel C. Ayes, Eva Maria Cutiongco-de la Paz, Alethea R. de Guzman, Jan Michael C. Yap, Jo-Hannah S. Llames, Sheila Mae M. Araiza, Kris P. Punayan, Irish Coleen A. Asin, Candice Francheska B. Tambaoan, Asia Louisa U. Chong, Karol Sophia Agape R. Padilla, Rianna Patricia S. Cruz, Carlo M. Lapid, El King D. Morado, Joshua Gregor A. Dizon, Razel Nikka M. Hao, Arianne A. Zamora, Devon Ray Pacial, Juan Antonio R. Magalang, Marissa Alejandria, Celia Carlos, Anna Ong-Lim, Edsel Maurice Salvaña, John Q. Wong, Jaime C. Montoya, Maria Rosario Singh-Vergeire and Cynthia P. Saloma |
| EPI_ISL_2189218 | Philippine Red Cross - Port Area | Philippine Genome Center | Francis A. Tablizo, Kenneth M. Kim, Maria Sofia L. Yangzon, Elcid Aaron R. Pangilinan, Renato Jacinto Q. Mantaring, Benedict A. Maralit, Marc Edsel C. Ayes, Eva Maria Cutiongco-de la Paz, Alethea R. de Guzman, Jan Michael C. Yap, Jo-Hannah S. Llames, Sheila Mae M. Araiza, Kris P. Punayan, Irish Coleen A. Asin, Candice Francheska B. Tambaoan, Asia Louisa U. Chong, Karol Sophia Agape R. Padilla, Rianna Patricia S. Cruz, Carlo M. Lapid, El King D. Morado, Joshua Gregor A. Dizon, Razel Nikka M. Hao, Arianne A. Zamora, Devon Ray Pacial, Juan Antonio R. Magalang, Marissa Alejandria, Celia Carlos, Anna Ong-Lim, Edsel Maurice Salvaña, John Q. Wong, Jaime C. Montoya, Maria Rosario Singh-Vergeire and Cynthia P. Saloma |
| EPI_ISL_2189219 | Qualimed Hospital Sta. Rosa | Philippine Genome Center | Francis A. Tablizo, Kenneth M. Kim, Maria Sofia L. Yangzon, Elcid Aaron R. Pangilinan, Renato Jacinto Q. Mantaring, Benedict A. Maralit, Marc Edsel C. Ayes, Eva Maria Cutiongco-de la Paz, Alethea R. de Guzman, Jan Michael C. Yap, Jo-Hannah S. Llames, Sheila Mae M. Araiza, Kris P. Punayan, Irish Coleen A. Asin, Candice Francheska B. Tambaoan, Asia Louisa U. Chong, Karol Sophia Agape R. Padilla, Rianna Patricia S. Cruz, Carlo M. Lapid, El King D. Morado, Joshua Gregor A. Dizon, Razel Nikka M. Hao, Arianne A. Zamora, Devon Ray Pacial, Juan Antonio R. Magalang, Marissa Alejandria, Celia Carlos, Anna Ong-Lim, Edsel Maurice Salvaña, John Q. Wong, Jaime C. Montoya, Maria Rosario Singh-Vergeire and Cynthia P. Saloma |
| EPI_ISL_2189220 | Pasig City Children's Hospital - Child's Hope | Philippine Genome Center | Francis A. Tablizo, Kenneth M. Kim, Maria Sofia L. Yangzon, Elcid Aaron R. Pangilinan, Renato Jacinto Q. Mantaring, Benedict A. Maralit, Marc Edsel C. Ayes, Eva Maria Cutiongco-de la Paz, Alethea R. de Guzman, Jan Michael C. Yap, Jo-Hannah S. Llames, Sheila Mae M. Araiza, Kris P. Punayan, Irish Coleen A. Asin, Candice Francheska B. Tambaoan, Asia Louisa U. Chong, Karol Sophia Agape R. Padilla, Rianna Patricia S. Cruz, Carlo M. Lapid, El King D. Morado, Joshua Gregor A. Dizon, Razel Nikka M. Hao, Arianne A. Zamora, Devon Ray Pacial, Juan Antonio R. Magalang, Marissa Alejandria, Celia Carlos, Anna Ong-Lim, Edsel Maurice Salvaña, John Q. Wong, Jaime C. Montoya, Maria Rosario Singh-Vergeire and Cynthia P. Saloma |
| EPI_ISL_2189221 | The Lord's Grace Medical and Industrial Clinic | Philippine Genome Center | Francis A. Tablizo, Kenneth M. Kim, Maria Sofia L. Yangzon, Elcid Aaron R. Pangilinan, Renato Jacinto Q. Mantaring, Benedict A. Maralit, Marc Edsel C. Ayes, Eva Maria Cutiongco-de la Paz, Alethea R. de Guzman, Jan Michael C. Yap, Jo-Hannah S. Llames, Sheila Mae M. Araiza, Kris P. Punayan, Irish Coleen A. Asin, Candice Francheska B. Tambaoan, Asia Louisa U. Chong, Karol Sophia Agape R. Padilla, Rianna Patricia S. Cruz, Carlo M. Lapid, El King D. Morado, Joshua Gregor A. Dizon, Razel Nikka M. Hao, Arianne A. Zamora, Devon Ray Pacial, Juan Antonio R. Magalang, Marissa Alejandria, Celia Carlos, Anna Ong-Lim, Edsel Maurice Salvaña, John Q. Wong, Jaime C. Montoya, Maria Rosario Singh-Vergeire and Cynthia P. Saloma |
| EPI_ISL_2189222 | South Super Highway Molecular Diagnostic Laboratory | Philippine Genome Center | Francis A. Tablizo, Kenneth M. Kim, Maria Sofia L. Yangzon, Elcid Aaron R. Pangilinan, Renato Jacinto Q. Mantaring, Benedict A. Maralit, Marc Edsel C. Ayes, Eva Maria Cutiongco-de la Paz, Alethea R. de Guzman, Jan Michael C. Yap, Jo-Hannah S. Llames, Sheila Mae M. Araiza, Kris P. Punayan, Irish Coleen A. Asin, Candice Francheska B. Tambaoan, Asia Louisa U. Chong, Karol Sophia Agape R. Padilla, Rianna Patricia S. Cruz, Carlo M. Lapid, El King D. Morado, Joshua Gregor A. Dizon, Razel Nikka M. Hao, Arianne A. Zamora, Devon Ray Pacial, Juan Antonio R. Magalang, Marissa Alejandria, Celia Carlos, Anna Ong-Lim, Edsel Maurice Salvaña, John Q. Wong, Jaime C. Montoya, Maria Rosario Singh-Vergeire and Cynthia P. Saloma |
| EPI_ISL_2189223 | Mary Mediatrix Medical Center | Philippine Genome Center | Francis A. Tablizo, Kenneth M. Kim, Maria Sofia L. Yangzon, Elcid Aaron R. Pangilinan, Renato Jacinto Q. Mantaring, Benedict A. Maralit, Marc Edsel C. Ayes, Eva Maria Cutiongco-de la Paz, Alethea R. de Guzman, Jan Michael C. Yap, Jo-Hannah S. Llames, Sheila Mae M. Araiza, Kris P. Punayan, Irish Coleen A. Asin, Candice Francheska B. Tambaoan, Asia Louisa U. Chong, Karol Sophia Agape R. Padilla, Rianna Patricia S. Cruz, Carlo M. Lapid, El King D. Morado, Joshua Gregor A. Dizon, Razel Nikka M. Hao, Arianne A. Zamora, Devon Ray Pacial, Juan Antonio R. Magalang, Marissa Alejandria, Celia Carlos, Anna Ong-Lim, Edsel Maurice Salvaña, John Q. Wong, Jaime C. Montoya, Maria Rosario Singh-Vergeire and Cynthia P. Saloma |
| EPI_ISL_2189224 | Pasig City Children's Hospital - Child's Hope | Philippine Genome Center | Francis A. Tablizo, Kenneth M. Kim, Maria Sofia L. Yangzon, Elcid Aaron R. Pangilinan, Renato Jacinto Q. Mantaring, Benedict A. Maralit, Marc Edsel C. Ayes, Eva Maria Cutiongco-de la Paz, Alethea R. de Guzman, Jan Michael C. Yap, Jo-Hannah S. Llames, Sheila Mae M. Araiza, Kris P. Punayan, Irish Coleen A. Asin, Candice Francheska B. Tambaoan, Asia Louisa U. Chong, Karol Sophia Agape R. Padilla, Rianna Patricia S. Cruz, Carlo M. Lapid, El King D. Morado, Joshua Gregor A. Dizon, Razel Nikka M. Hao, Arianne A. Zamora, Devon Ray Pacial, Juan Antonio R. Magalang, Marissa Alejandria, Celia Carlos, Anna Ong-Lim, Edsel Maurice Salvaña, John Q. Wong, Jaime C. Montoya, Maria Rosario Singh-Vergeire and Cynthia P. Saloma |
| EPI_ISL_2189226, EPI_ISL_2189227 | The Medical City | Philippine Genome Center | Francis A. Tablizo, Kenneth M. Kim, Maria Sofia L. Yangzon, Elcid Aaron R. Pangilinan, Renato Jacinto Q. Mantaring, Benedict A. Maralit, Marc Edsel C. Ayes, Eva Maria Cutiongco-de la Paz, Alethea R. de Guzman, Jan Michael C. Yap, Jo-Hannah S. Llames, Sheila Mae M. Araiza, Kris P. Punayan, Irish Coleen A. Asin, Candice Francheska B. Tambaoan, Asia Louisa U. Chong, Karol Sophia Agape R. Padilla, Rianna Patricia S. Cruz, Carlo M. Lapid, El King D. Morado, Joshua Gregor A. Dizon, Razel Nikka M. Hao, Arianne A. Zamora, Devon Ray Pacial, Juan Antonio R. Magalang, Marissa Alejandria, Celia Carlos, Anna Ong-Lim, Edsel Maurice Salvaña, John Q. Wong, Jaime C. Montoya, Maria Rosario Singh-Vergeire and Cynthia P. Saloma |
| EPI_ISL_2189228 | South Super Highway Molecular Diagnostic Laboratory | Philippine Genome Center | Francis A. Tablizo, Kenneth M. Kim, Maria Sofia L. Yangzon, Elcid Aaron R. Pangilinan, Renato Jacinto Q. Mantaring, Benedict A. Maralit, Marc Edsel C. Ayes, Eva Maria Cutiongco-de la Paz, Alethea R. de Guzman, Jan Michael C. Yap, Jo-Hannah S. Llames, Sheila Mae M. Araiza, Kris P. Punayan, Irish Coleen A. Asin, Candice Francheska B. Tambaoan, Asia Louisa U. Chong, Karol Sophia Agape R. Padilla, Rianna Patricia S. Cruz, Carlo M. Lapid, El King D. Morado, Joshua Gregor A. Dizon, Razel Nikka M. Hao, Arianne A. Zamora, Devon Ray Pacial, Juan Antonio R. Magalang, Marissa Alejandria, Celia Carlos, Anna Ong-Lim, Edsel Maurice Salvaña, John Q. Wong, Jaime C. Montoya, Maria Rosario Singh-Vergeire and Cynthia P. Saloma |
| EPI_ISL_2189229 | The Lord's Grace Medical and Industrial Clinic | Philippine Genome Center | Francis A. Tablizo, Kenneth M. Kim, Maria Sofia L. Yangzon, Elcid Aaron R. Pangilinan, Renato Jacinto Q. Mantaring, Benedict A. Maralit, Marc Edsel C. Ayes, Eva Maria Cutiongco-de la Paz, Alethea R. de Guzman, Jan Michael C. Yap, Jo-Hannah S. Llames, Sheila Mae M. Araiza, Kris P. Punayan, Irish Coleen A. Asin, Candice Francheska B. Tambaoan, Asia Louisa U. Chong, Karol Sophia Agape R. Padilla, Rianna Patricia S. Cruz, Carlo M. Lapid, El King D. Morado, Joshua Gregor A. Dizon, Razel Nikka M. Hao, Arianne A. Zamora, Devon Ray Pacial, Juan Antonio R. Magalang, Marissa Alejandria, Celia Carlos, Anna Ong-Lim, Edsel Maurice Salvaña, John Q. Wong, Jaime C. Montoya, Maria Rosario Singh-Vergeire and Cynthia P. Saloma |
| EPI_ISL_2189230 | Sta. Rosa CHO- GENEXPERT LABORATORY | Philippine Genome Center | Francis A. Tablizo, Kenneth M. Kim, Maria Sofia L. Yangzon, Elcid Aaron R. Pangilinan, Renato Jacinto Q. Mantaring, Benedict A. Maralit, Marc Edsel C. Ayes, Eva Maria Cutiongco-de la Paz, Alethea R. de Guzman, Jan Michael C. Yap, Jo-Hannah S. Llames, Sheila Mae M. Araiza, Kris P. Punayan, Irish Coleen A. Asin, Candice Francheska B. Tambaoan, Asia Louisa U. Chong, Karol Sophia Agape R. Padilla, Rianna Patricia S. Cruz, Carlo M. Lapid, El King D. Morado, Joshua Gregor A. Dizon, Razel Nikka M. Hao, Arianne A. Zamora, Devon Ray Pacial, Juan Antonio R. Magalang, Marissa Alejandria, Celia Carlos, Anna Ong-Lim, Edsel Maurice Salvaña, John Q. Wong, Jaime C. Montoya, Maria Rosario Singh-Vergeire and Cynthia P. Saloma |
| EPI_ISL_2189231 | The Medical City | Philippine Genome Center | Francis A. Tablizo, Kenneth M. Kim, Maria Sofia L. Yangzon, Elcid Aaron R. Pangilinan, Renato Jacinto Q. Mantaring, Benedict A. Maralit, Marc Edsel C. Ayes, Eva Maria Cutiongco-de la Paz, Alethea R. de Guzman, Jan Michael C. Yap, Jo-Hannah S. Llames, Sheila Mae M. Araiza, Kris P. Punayan, Irish Coleen A. Asin, Candice Francheska B. Tambaoan, Asia Louisa U. Chong, Karol Sophia Agape R. Padilla, Rianna Patricia S. Cruz, Carlo M. Lapid, El King D. Morado, Joshua Gregor A. Dizon, Razel Nikka M. Hao, Arianne A. Zamora, Devon Ray Pacial, Juan Antonio R. Magalang, Marissa Alejandria, Celia Carlos, Anna Ong-Lim, Edsel Maurice Salvaña, John Q. Wong, Jaime C. Montoya, Maria Rosario Singh-Vergeire and Cynthia P. Saloma |
| EPI_ISL_2189232, EPI_ISL_2189234 | Pasig City Children's Hospital - Child's Hope | Philippine Genome Center | Francis A. Tablizo, Kenneth M. Kim, Maria Sofia L. Yangzon, Elcid Aaron R. Pangilinan, Renato Jacinto Q. Mantaring, Benedict A. Maralit, Marc Edsel C. Ayes, Eva Maria Cutiongco-de la Paz, Alethea R. de Guzman, Jan Michael C. Yap, Jo-Hannah S. Llames, Sheila Mae M. Araiza, Kris P. Punayan, Irish Coleen A. Asin, Candice Francheska B. Tambaoan, Asia Louisa U. Chong, Karol Sophia Agape R. Padilla, Rianna Patricia S. Cruz, Carlo M. Lapid, El King D. Morado, Joshua Gregor A. Dizon, Razel Nikka M. Hao, Arianne A. Zamora, Devon Ray Pacial, Juan Antonio R. Magalang, Marissa Alejandria, Celia Carlos, Anna Ong-Lim, Edsel Maurice Salvaña, John Q. Wong, Jaime C. Montoya, Maria Rosario Singh-Vergeire and Cynthia P. Saloma |
| EPI_ISL_2189235, EPI_ISL_2189236, EPI_ISL_2189237, EPI_ISL_2189239, EPI_ISL_2189240 | The Lord's Grace Medical and Industrial Clinic | Philippine Genome Center | Francis A. Tablizo, Kenneth M. Kim, Maria Sofia L. Yangzon, Elcid Aaron R. Pangilinan, Renato Jacinto Q. Mantaring, Benedict A. Maralit, Marc Edsel C. Ayes, Eva Maria Cutiongco-de la Paz, Alethea R. de Guzman, Jan Michael C. Yap, Jo-Hannah S. Llames, Sheila Mae M. Araiza, Kris P. Punayan, Irish Coleen A. Asin, Candice Francheska B. Tambaoan, Asia Louisa U. Chong, Karol Sophia Agape R. Padilla, Rianna Patricia S. Cruz, Carlo M. Lapid, El King D. Morado, Joshua Gregor A. Dizon, Razel Nikka M. Hao, Arianne A. Zamora, Devon Ray Pacial, Juan Antonio R. Magalang, Marissa Alejandria, Celia Carlos, Anna Ong-Lim, Edsel Maurice Salvaña, John Q. Wong, Jaime C. Montoya, Maria Rosario Singh-Vergeire and Cynthia P. Saloma |

| | | | |
|---|---|---|---|
| EPI_ISL_2189242 | Philippine Red Cross - National Blood Center | Philippine Genome Center | Francis A. Tablizo, Kenneth M. Kim, Maria Sofia L. Yangzon, Elcid Aaron R. Pangilinan, Renato Jacinto Q. Mantaring, Benedict A. Maralit, Marc Edsel C. Ayes, Eva Maria Cutiongco-de la Paz, Alethea R. de Guzman, Jan Michael C. Yap, Jo-Hannah S. Llames, Sheila Mae M. Araiza, Kris P. Punayan, Irish Coleen A. Asin, Candice Francheska B. Tambaoan, Asia Louisa U. Chong, Karol Sophia Agape R. Padilla, Rianna Patricia S. Cruz, Carlo M. Lapid, El King D. Morado, Joshua Gregor A. Dizon, Razel Nikka M. Hao, Arianne A. Zamora, Devon Ray Pacial, Juan Antonio R. Magalang, Marissa Alejandria, Celia Carlos, Anna Ong-Lim, Edsel Maurice Salvaña, John Q. Wong, Jaime C. Montoya, Maria Rosario Singh-Vergeire and Cynthia P. Saloma |
| EPI_ISL_2189243 | Philippine Red Cross - Port Area | Philippine Genome Center | Francis A. Tablizo, Kenneth M. Kim, Maria Sofia L. Yangzon, Elcid Aaron R. Pangilinan, Renato Jacinto Q. Mantaring, Benedict A. Maralit, Marc Edsel C. Ayes, Eva Maria Cutiongco-de la Paz, Alethea R. de Guzman, Jan Michael C. Yap, Jo-Hannah S. Llames, Sheila Mae M. Araiza, Kris P. Punayan, Irish Coleen A. Asin, Candice Francheska B. Tambaoan, Asia Louisa U. Chong, Karol Sophia Agape R. Padilla, Rianna Patricia S. Cruz, Carlo M. Lapid, El King D. Morado, Joshua Gregor A. Dizon, Razel Nikka M. Hao, Arianne A. Zamora, Devon Ray Pacial, Juan Antonio R. Magalang, Marissa Alejandria, Celia Carlos, Anna Ong-Lim, Edsel Maurice Salvaña, John Q. Wong, Jaime C. Montoya, Maria Rosario Singh-Vergeire and Cynthia P. Saloma |
| EPI_ISL_2189244 | The Lord's Grace Medical and Industrial Clinic | Philippine Genome Center | Francis A. Tablizo, Kenneth M. Kim, Maria Sofia L. Yangzon, Elcid Aaron R. Pangilinan, Renato Jacinto Q. Mantaring, Benedict A. Maralit, Marc Edsel C. Ayes, Eva Maria Cutiongco-de la Paz, Alethea R. de Guzman, Jan Michael C. Yap, Jo-Hannah S. Llames, Sheila Mae M. Araiza, Kris P. Punayan, Irish Coleen A. Asin, Candice Francheska B. Tambaoan, Asia Louisa U. Chong, Karol Sophia Agape R. Padilla, Rianna Patricia S. Cruz, Carlo M. Lapid, El King D. Morado, Joshua Gregor A. Dizon, Razel Nikka M. Hao, Arianne A. Zamora, Devon Ray Pacial, Juan Antonio R. Magalang, Marissa Alejandria, Celia Carlos, Anna Ong-Lim, Edsel Maurice Salvaña, John Q. Wong, Jaime C. Montoya, Maria Rosario Singh-Vergeire and Cynthia P. Saloma |
| EPI_ISL_2189245 | Pasig City Children's Hospital - Child's Hope | Philippine Genome Center | Francis A. Tablizo, Kenneth M. Kim, Maria Sofia L. Yangzon, Elcid Aaron R. Pangilinan, Renato Jacinto Q. Mantaring, Benedict A. Maralit, Marc Edsel C. Ayes, Eva Maria Cutiongco-de la Paz, Alethea R. de Guzman, Jan Michael C. Yap, Jo-Hannah S. Llames, Sheila Mae M. Araiza, Kris P. Punayan, Irish Coleen A. Asin, Candice Francheska B. Tambaoan, Asia Louisa U. Chong, Karol Sophia Agape R. Padilla, Rianna Patricia S. Cruz, Carlo M. Lapid, El King D. Morado, Joshua Gregor A. Dizon, Razel Nikka M. Hao, Arianne A. Zamora, Devon Ray Pacial, Juan Antonio R. Magalang, Marissa Alejandria, Celia Carlos, Anna Ong-Lim, Edsel Maurice Salvaña, John Q. Wong, Jaime C. Montoya, Maria Rosario Singh-Vergeire and Cynthia P. Saloma |
| EPI_ISL_2189247 | Sta. Ana Hospital | Philippine Genome Center | Francis A. Tablizo, Kenneth M. Kim, Maria Sofia L. Yangzon, Elcid Aaron R. Pangilinan, Renato Jacinto Q. Mantaring, Benedict A. Maralit, Marc Edsel C. Ayes, Eva Maria Cutiongco-de la Paz, Alethea R. de Guzman, Jan Michael C. Yap, Jo-Hannah S. Llames, Sheila Mae M. Araiza, Kris P. Punayan, Irish Coleen A. Asin, Candice Francheska B. Tambaoan, Asia Louisa U. Chong, Karol Sophia Agape R. Padilla, Rianna Patricia S. Cruz, Carlo M. Lapid, El King D. Morado, Joshua Gregor A. Dizon, Razel Nikka M. Hao, Arianne A. Zamora, Devon Ray Pacial, Juan Antonio R. Magalang, Marissa Alejandria, Celia Carlos, Anna Ong-Lim, Edsel Maurice Salvaña, John Q. Wong, Jaime C. Montoya, Maria Rosario Singh-Vergeire and Cynthia P. Saloma |
| EPI_ISL_2189248 | Philippine Red Cross - National Blood Center | Philippine Genome Center | Francis A. Tablizo, Kenneth M. Kim, Maria Sofia L. Yangzon, Elcid Aaron R. Pangilinan, Renato Jacinto Q. Mantaring, Benedict A. Maralit, Marc Edsel C. Ayes, Eva Maria Cutiongco-de la Paz, Alethea R. de Guzman, Jan Michael C. Yap, Jo-Hannah S. Llames, Sheila Mae M. Araiza, Kris P. Punayan, Irish Coleen A. Asin, Candice Francheska B. Tambaoan, Asia Louisa U. Chong, Karol Sophia Agape R. Padilla, Rianna Patricia S. Cruz, Carlo M. Lapid, El King D. Morado, Joshua Gregor A. Dizon, Razel Nikka M. Hao, Arianne A. Zamora, Devon Ray Pacial, Juan Antonio R. Magalang, Marissa Alejandria, Celia Carlos, Anna Ong-Lim, Edsel Maurice Salvaña, John Q. Wong, Jaime C. Montoya, Maria Rosario Singh-Vergeire and Cynthia P. Saloma |
| EPI_ISL_2189249 | University of the Philippines National Institutes of Health (UP NIH) | Philippine Genome Center | Francis A. Tablizo, Kenneth M. Kim, Maria Sofia L. Yangzon, Elcid Aaron R. Pangilinan, Renato Jacinto Q. Mantaring, Benedict A. Maralit, Marc Edsel C. Ayes, Eva Maria Cutiongco-de la Paz, Alethea R. de Guzman, Jan Michael C. Yap, Jo-Hannah S. Llames, Sheila Mae M. Araiza, Kris P. Punayan, Irish Coleen A. Asin, Candice Francheska B. Tambaoan, Asia Louisa U. Chong, Karol Sophia Agape R. Padilla, Rianna Patricia S. Cruz, Carlo M. Lapid, El King D. Morado, Joshua Gregor A. Dizon, Razel Nikka M. Hao, Arianne A. Zamora, Devon Ray Pacial, Juan Antonio R. Magalang, Marissa Alejandria, Celia Carlos, Anna Ong-Lim, Edsel Maurice Salvaña, John Q. Wong, Jaime C. Montoya, Maria Rosario Singh-Vergeire and Cynthia P. Saloma |
| EPI_ISL_2189250 | Philippine Red Cross - Clark Molecular Laboratory | Philippine Genome Center | Francis A. Tablizo, Kenneth M. Kim, Maria Sofia L. Yangzon, Elcid Aaron R. Pangilinan, Renato Jacinto Q. Mantaring, Benedict A. Maralit, Marc Edsel C. Ayes, Eva Maria Cutiongco-de la Paz, Alethea R. de Guzman, Jan Michael C. Yap, Jo-Hannah S. Llames, Sheila Mae M. Araiza, Kris P. Punayan, Irish Coleen A. Asin, Candice Francheska B. Tambaoan, Asia Louisa U. Chong, Karol Sophia Agape R. Padilla, Rianna Patricia S. Cruz, Carlo M. Lapid, El King D. Morado, Joshua Gregor A. Dizon, Razel Nikka M. Hao, Arianne A. Zamora, Devon Ray Pacial, Juan Antonio R. Magalang, Marissa Alejandria, Celia Carlos, Anna Ong-Lim, Edsel Maurice Salvaña, John Q. Wong, Jaime C. Montoya, Maria Rosario Singh-Vergeire and Cynthia P. Saloma |
| EPI_ISL_2189251 | Pasig City Children's Hospital - Child's Hope | Philippine Genome Center | Francis A. Tablizo, Kenneth M. Kim, Maria Sofia L. Yangzon, Elcid Aaron R. Pangilinan, Renato Jacinto Q. Mantaring, Benedict A. Maralit, Marc Edsel C. Ayes, Eva Maria Cutiongco-de la Paz, Alethea R. de Guzman, Jan Michael C. Yap, Jo-Hannah S. Llames, Sheila Mae M. Araiza, Kris P. Punayan, Irish Coleen A. Asin, Candice Francheska B. Tambaoan, Asia Louisa U. Chong, Karol Sophia Agape R. Padilla, Rianna Patricia S. Cruz, Carlo M. Lapid, El King D. Morado, Joshua Gregor A. Dizon, Razel Nikka M. Hao, Arianne A. Zamora, Devon Ray Pacial, Juan Antonio R. Magalang, Marissa Alejandria, Celia Carlos, Anna Ong-Lim, Edsel Maurice Salvaña, John Q. Wong, Jaime C. Montoya, Maria Rosario Singh-Vergeire and Cynthia P. Saloma |
| EPI_ISL_2189254 | The Lord's Grace Medical and Industrial Clinic | Philippine Genome Center | Francis A. Tablizo, Kenneth M. Kim, Maria Sofia L. Yangzon, Elcid Aaron R. Pangilinan, Renato Jacinto Q. Mantaring, Benedict A. Maralit, Marc Edsel C. Ayes, Eva Maria Cutiongco-de la Paz, Alethea R. de Guzman, Jan Michael C. Yap, Jo-Hannah S. Llames, Sheila Mae M. Araiza, Kris P. Punayan, Irish Coleen A. Asin, Candice Francheska B. Tambaoan, Asia Louisa U. Chong, Karol Sophia Agape R. Padilla, Rianna Patricia S. Cruz, Carlo M. Lapid, El King D. Morado, Joshua Gregor A. Dizon, Razel Nikka M. Hao, Arianne A. Zamora, Devon Ray Pacial, Juan Antonio R. Magalang, Marissa Alejandria, Celia Carlos, Anna Ong-Lim, Edsel Maurice Salvaña, John Q. Wong, Jaime C. Montoya, Maria Rosario Singh-Vergeire and Cynthia P. Saloma |
| EPI_ISL_2189256 | Qualimed Hospital Sta. Rosa | Philippine Genome Center | Francis A. Tablizo, Kenneth M. Kim, Maria Sofia L. Yangzon, Elcid Aaron R. Pangilinan, Renato Jacinto Q. Mantaring, Benedict A. Maralit, Marc Edsel C. Ayes, Eva Maria Cutiongco-de la Paz, Alethea R. de Guzman, Jan Michael C. Yap, Jo-Hannah S. Llames, Sheila Mae M. Araiza, Kris P. Punayan, Irish Coleen A. Asin, Candice Francheska B. Tambaoan, Asia Louisa U. Chong, Karol Sophia Agape R. Padilla, Rianna Patricia S. Cruz, Carlo M. Lapid, El King D. Morado, Joshua Gregor A. Dizon, Razel Nikka M. Hao, Arianne A. Zamora, Devon Ray Pacial, Juan Antonio R. Magalang, Marissa Alejandria, Celia Carlos, Anna Ong-Lim, Edsel Maurice Salvaña, John Q. Wong, Jaime C. Montoya, Maria Rosario Singh-Vergeire and Cynthia P. Saloma |
| EPI_ISL_2189257 | Sta. Ana Hospital | Philippine Genome Center | Francis A. Tablizo, Kenneth M. Kim, Maria Sofia L. Yangzon, Elcid Aaron R. Pangilinan, Renato Jacinto Q. Mantaring, Benedict A. Maralit, Marc Edsel C. Ayes, Eva Maria Cutiongco-de la Paz, Alethea R. de Guzman, Jan Michael C. Yap, Jo-Hannah S. Llames, Sheila Mae M. Araiza, Kris P. Punayan, Irish Coleen A. Asin, Candice Francheska B. Tambaoan, Asia Louisa U. Chong, Karol Sophia Agape R. Padilla, Rianna Patricia S. Cruz, Carlo M. Lapid, El King D. Morado, Joshua Gregor A. Dizon, Razel Nikka M. Hao, Arianne A. Zamora, Devon Ray Pacial, Juan Antonio R. Magalang, Marissa Alejandria, Celia Carlos, Anna Ong-Lim, Edsel Maurice Salvaña, John Q. Wong, Jaime C. Montoya, Maria Rosario Singh-Vergeire and Cynthia P. Saloma |
| EPI_ISL_2189258 | Qualimed Hospital Sta. Rosa | Philippine Genome Center | Francis A. Tablizo, Kenneth M. Kim, Maria Sofia L. Yangzon, Elcid Aaron R. Pangilinan, Renato Jacinto Q. Mantaring, Benedict A. Maralit, Marc Edsel C. Ayes, Eva Maria Cutiongco-de la Paz, Alethea R. de Guzman, Jan Michael C. Yap, Jo-Hannah S. Llames, Sheila Mae M. Araiza, Kris P. Punayan, Irish Coleen A. Asin, Candice Francheska B. Tambaoan, Asia Louisa U. Chong, Karol Sophia Agape R. Padilla, Rianna Patricia S. Cruz, Carlo M. Lapid, El King D. Morado, Joshua Gregor A. Dizon, Razel Nikka M. Hao, Arianne A. Zamora, Devon Ray Pacial, Juan Antonio R. Magalang, Marissa Alejandria, Celia Carlos, Anna Ong-Lim, Edsel Maurice Salvaña, John Q. Wong, Jaime C. Montoya, Maria Rosario Singh-Vergeire and Cynthia P. Saloma |
| EPI_ISL_2189259 | Philippine Red Cross - National Blood Center | Philippine Genome Center | Francis A. Tablizo, Kenneth M. Kim, Maria Sofia L. Yangzon, Elcid Aaron R. Pangilinan, Renato Jacinto Q. Mantaring, Benedict A. Maralit, Marc Edsel C. Ayes, Eva Maria Cutiongco-de la Paz, Alethea R. de Guzman, Jan Michael C. Yap, Jo-Hannah S. Llames, Sheila Mae M. Araiza, Kris P. Punayan, Irish Coleen A. Asin, Candice Francheska B. Tambaoan, Asia Louisa U. Chong, Karol Sophia Agape R. Padilla, Rianna Patricia S. Cruz, Carlo M. Lapid, El King D. Morado, Joshua Gregor A. Dizon, Razel Nikka M. Hao, Arianne A. Zamora, Devon Ray Pacial, Juan Antonio R. Magalang, Marissa Alejandria, Celia Carlos, Anna Ong-Lim, Edsel Maurice Salvaña, John Q. Wong, Jaime C. Montoya, Maria Rosario Singh-Vergeire and Cynthia P. Saloma |
| EPI_ISL_2189260 | The Lord's Grace Medical and Industrial Clinic | Philippine Genome Center | Francis A. Tablizo, Kenneth M. Kim, Maria Sofia L. Yangzon, Elcid Aaron R. Pangilinan, Renato Jacinto Q. Mantaring, Benedict A. Maralit, Marc Edsel C. Ayes, Eva Maria Cutiongco-de la Paz, Alethea R. de Guzman, Jan Michael C. Yap, Jo-Hannah S. Llames, Sheila Mae M. Araiza, Kris P. Punayan, Irish Coleen A. Asin, Candice Francheska B. Tambaoan, Asia Louisa U. Chong, Karol Sophia Agape R. Padilla, Rianna Patricia S. Cruz, Carlo M. Lapid, El King D. Morado, Joshua Gregor A. Dizon, Razel Nikka M. Hao, Arianne A. Zamora, Devon Ray Pacial, Juan Antonio R. Magalang, Marissa Alejandria, Celia Carlos, Anna Ong-Lim, Edsel Maurice Salvaña, John Q. Wong, Jaime C. Montoya, Maria Rosario Singh-Vergeire and Cynthia P. Saloma |
| EPI_ISL_2189261 | The Medical City | Philippine Genome Center | Francis A. Tablizo, Kenneth M. Kim, Maria Sofia L. Yangzon, Elcid Aaron R. Pangilinan, Renato Jacinto Q. Mantaring, Benedict A. Maralit, Marc Edsel C. Ayes, Eva Maria Cutiongco-de la Paz, Alethea R. de Guzman, Jan Michael C. Yap, Jo-Hannah S. Llames, Sheila Mae M. Araiza, Kris P. Punayan, Irish Coleen A. Asin, Candice Francheska B. Tambaoan, Asia Louisa U. Chong, Karol Sophia Agape R. Padilla, Rianna Patricia S. Cruz, Carlo M. Lapid, El King D. Morado, Joshua Gregor A. Dizon, Razel Nikka M. Hao, Arianne A. Zamora, Devon Ray Pacial, Juan Antonio R. Magalang, Marissa Alejandria, Celia Carlos, Anna Ong-Lim, Edsel Maurice Salvaña, John Q. Wong, Jaime C. Montoya, Maria Rosario Singh-Vergeire and Cynthia P. Saloma |

| | | | |
|---|---|---|---|
| EPI_ISL_2189262 | Philippine Red Cross - National Blood Center | Philippine Genome Center | Francis A. Tablizo, Kenneth M. Kim, Maria Sofia L. Yangzon, Elcid Aaron R. Pangilinan, Renato Jacinto Q. Mantaring, Benedict A. Maralit, Marc Edsel C. Ayes, Eva Maria Cutiongco-de la Paz, Alethea R. de Guzman, Jan Michael C. Yap, Jo-Hannah S. Llames, Sheila Mae M. Araiza, Kris P. Punayan, Irish Coleen A. Asin, Candice Francheska B. Tambaoan, Asia Louisa U. Chong, Karol Sophia Agape R. Padilla, Rianna Patricia S. Cruz, Carlo M. Lapid, El King D. Morado, Joshua Gregor A. Dizon, Razel Nikka M. Hao, Arianne A. Zamora, Devon Ray Pacial, Juan Antonio R. Magalang, Marissa Alejandria, Celia Carlos, Anna Ong-Lim, Edsel Maurice Salvaña, John Q. Wong, Jaime C. Montoya, Maria Rosario Singh-Vergeire and Cynthia P. Saloma |
| EPI_ISL_2189263, EPI_ISL_2189264 | Pasig City Children's Hospital - Child's Hope | Philippine Genome Center | Francis A. Tablizo, Kenneth M. Kim, Maria Sofia L. Yangzon, Elcid Aaron R. Pangilinan, Renato Jacinto Q. Mantaring, Benedict A. Maralit, Marc Edsel C. Ayes, Eva Maria Cutiongco-de la Paz, Alethea R. de Guzman, Jan Michael C. Yap, Jo-Hannah S. Llames, Sheila Mae M. Araiza, Kris P. Punayan, Irish Coleen A. Asin, Candice Francheska B. Tambaoan, Asia Louisa U. Chong, Karol Sophia Agape R. Padilla, Rianna Patricia S. Cruz, Carlo M. Lapid, El King D. Morado, Joshua Gregor A. Dizon, Razel Nikka M. Hao, Arianne A. Zamora, Devon Ray Pacial, Juan Antonio R. Magalang, Marissa Alejandria, Celia Carlos, Anna Ong-Lim, Edsel Maurice Salvaña, John Q. Wong, Jaime C. Montoya, Maria Rosario Singh-Vergeire and Cynthia P. Saloma |
| EPI_ISL_2189265 | The Medical City | Philippine Genome Center | Francis A. Tablizo, Kenneth M. Kim, Maria Sofia L. Yangzon, Elcid Aaron R. Pangilinan, Renato Jacinto Q. Mantaring, Benedict A. Maralit, Marc Edsel C. Ayes, Eva Maria Cutiongco-de la Paz, Alethea R. de Guzman, Jan Michael C. Yap, Jo-Hannah S. Llames, Sheila Mae M. Araiza, Kris P. Punayan, Irish Coleen A. Asin, Candice Francheska B. Tambaoan, Asia Louisa U. Chong, Karol Sophia Agape R. Padilla, Rianna Patricia S. Cruz, Carlo M. Lapid, El King D. Morado, Joshua Gregor A. Dizon, Razel Nikka M. Hao, Arianne A. Zamora, Devon Ray Pacial, Juan Antonio R. Magalang, Marissa Alejandria, Celia Carlos, Anna Ong-Lim, Edsel Maurice Salvaña, John Q. Wong, Jaime C. Montoya, Maria Rosario Singh-Vergeire and Cynthia P. Saloma |
| EPI_ISL_2189268 | Philippine General Hospital (PGH) | Philippine Genome Center | Francis A. Tablizo, Kenneth M. Kim, Maria Sofia L. Yangzon, Elcid Aaron R. Pangilinan, Renato Jacinto Q. Mantaring, Benedict A. Maralit, Marc Edsel C. Ayes, Eva Maria Cutiongco-de la Paz, Alethea R. de Guzman, Jan Michael C. Yap, Jo-Hannah S. Llames, Sheila Mae M. Araiza, Kris P. Punayan, Irish Coleen A. Asin, Candice Francheska B. Tambaoan, Asia Louisa U. Chong, Karol Sophia Agape R. Padilla, Rianna Patricia S. Cruz, Carlo M. Lapid, El King D. Morado, Joshua Gregor A. Dizon, Razel Nikka M. Hao, Arianne A. Zamora, Devon Ray Pacial, Juan Antonio R. Magalang, Marissa Alejandria, Celia Carlos, Anna Ong-Lim, Edsel Maurice Salvaña, John Q. Wong, Jaime C. Montoya, Maria Rosario Singh-Vergeire and Cynthia P. Saloma |
| EPI_ISL_2189269, EPI_ISL_2189270 | The Medical City | Philippine Genome Center | Francis A. Tablizo, Kenneth M. Kim, Maria Sofia L. Yangzon, Elcid Aaron R. Pangilinan, Renato Jacinto Q. Mantaring, Benedict A. Maralit, Marc Edsel C. Ayes, Eva Maria Cutiongco-de la Paz, Alethea R. de Guzman, Jan Michael C. Yap, Jo-Hannah S. Llames, Sheila Mae M. Araiza, Kris P. Punayan, Irish Coleen A. Asin, Candice Francheska B. Tambaoan, Asia Louisa U. Chong, Karol Sophia Agape R. Padilla, Rianna Patricia S. Cruz, Carlo M. Lapid, El King D. Morado, Joshua Gregor A. Dizon, Razel Nikka M. Hao, Arianne A. Zamora, Devon Ray Pacial, Juan Antonio R. Magalang, Marissa Alejandria, Celia Carlos, Anna Ong-Lim, Edsel Maurice Salvaña, John Q. Wong, Jaime C. Montoya, Maria Rosario Singh-Vergeire and Cynthia P. Saloma |
| EPI_ISL_2189271 | Qualimed Hospital Sta. Rosa | Philippine Genome Center | Francis A. Tablizo, Kenneth M. Kim, Maria Sofia L. Yangzon, Elcid Aaron R. Pangilinan, Renato Jacinto Q. Mantaring, Benedict A. Maralit, Marc Edsel C. Ayes, Eva Maria Cutiongco-de la Paz, Alethea R. de Guzman, Jan Michael C. Yap, Jo-Hannah S. Llames, Sheila Mae M. Araiza, Kris P. Punayan, Irish Coleen A. Asin, Candice Francheska B. Tambaoan, Asia Louisa U. Chong, Karol Sophia Agape R. Padilla, Rianna Patricia S. Cruz, Carlo M. Lapid, El King D. Morado, Joshua Gregor A. Dizon, Razel Nikka M. Hao, Arianne A. Zamora, Devon Ray Pacial, Juan Antonio R. Magalang, Marissa Alejandria, Celia Carlos, Anna Ong-Lim, Edsel Maurice Salvaña, John Q. Wong, Jaime C. Montoya, Maria Rosario Singh-Vergeire and Cynthia P. Saloma |
| EPI_ISL_2189275 | The Medical City | Philippine Genome Center | Francis A. Tablizo, Kenneth M. Kim, Maria Sofia L. Yangzon, Elcid Aaron R. Pangilinan, Renato Jacinto Q. Mantaring, Benedict A. Maralit, Marc Edsel C. Ayes, Eva Maria Cutiongco-de la Paz, Alethea R. de Guzman, Jan Michael C. Yap, Jo-Hannah S. Llames, Sheila Mae M. Araiza, Kris P. Punayan, Irish Coleen A. Asin, Candice Francheska B. Tambaoan, Asia Louisa U. Chong, Karol Sophia Agape R. Padilla, Rianna Patricia S. Cruz, Carlo M. Lapid, El King D. Morado, Joshua Gregor A. Dizon, Razel Nikka M. Hao, Arianne A. Zamora, Devon Ray Pacial, Juan Antonio R. Magalang, Marissa Alejandria, Celia Carlos, Anna Ong-Lim, Edsel Maurice Salvaña, John Q. Wong, Jaime C. Montoya, Maria Rosario Singh-Vergeire and Cynthia P. Saloma |
| EPI_ISL_2189276 | The Lord's Grace Medical and Industrial Clinic | Philippine Genome Center | Francis A. Tablizo, Kenneth M. Kim, Maria Sofia L. Yangzon, Elcid Aaron R. Pangilinan, Renato Jacinto Q. Mantaring, Benedict A. Maralit, Marc Edsel C. Ayes, Eva Maria Cutiongco-de la Paz, Alethea R. de Guzman, Jan Michael C. Yap, Jo-Hannah S. Llames, Sheila Mae M. Araiza, Kris P. Punayan, Irish Coleen A. Asin, Candice Francheska B. Tambaoan, Asia Louisa U. Chong, Karol Sophia Agape R. Padilla, Rianna Patricia S. Cruz, Carlo M. Lapid, El King D. Morado, Joshua Gregor A. Dizon, Razel Nikka M. Hao, Arianne A. Zamora, Devon Ray Pacial, Juan Antonio R. Magalang, Marissa Alejandria, Celia Carlos, Anna Ong-Lim, Edsel Maurice Salvaña, John Q. Wong, Jaime C. Montoya, Maria Rosario Singh-Vergeire and Cynthia P. Saloma |
| EPI_ISL_2189277 | Philippine Red Cross - Clark Molecular Laboratory | Philippine Genome Center | Francis A. Tablizo, Kenneth M. Kim, Maria Sofia L. Yangzon, Elcid Aaron R. Pangilinan, Renato Jacinto Q. Mantaring, Benedict A. Maralit, Marc Edsel C. Ayes, Eva Maria Cutiongco-de la Paz, Alethea R. de Guzman, Jan Michael C. Yap, Jo-Hannah S. Llames, Sheila Mae M. Araiza, Kris P. Punayan, Irish Coleen A. Asin, Candice Francheska B. Tambaoan, Asia Louisa U. Chong, Karol Sophia Agape R. Padilla, Rianna Patricia S. Cruz, Carlo M. Lapid, El King D. Morado, Joshua Gregor A. Dizon, Razel Nikka M. Hao, Arianne A. Zamora, Devon Ray Pacial, Juan Antonio R. Magalang, Marissa Alejandria, Celia Carlos, Anna Ong-Lim, Edsel Maurice Salvaña, John Q. Wong, Jaime C. Montoya, Maria Rosario Singh-Vergeire and Cynthia P. Saloma |
| EPI_ISL_2189279 | Northern Mindanao TB Regional Center | Philippine Genome Center | Francis A. Tablizo, Kenneth M. Kim, Maria Sofia L. Yangzon, Elcid Aaron R. Pangilinan, Renato Jacinto Q. Mantaring, Benedict A. Maralit, Marc Edsel C. Ayes, Eva Maria Cutiongco-de la Paz, Alethea R. de Guzman, Jan Michael C. Yap, Jo-Hannah S. Llames, Sheila Mae M. Araiza, Kris P. Punayan, Irish Coleen A. Asin, Candice Francheska B. Tambaoan, Asia Louisa U. Chong, Karol Sophia Agape R. Padilla, Rianna Patricia S. Cruz, Carlo M. Lapid, El King D. Morado, Joshua Gregor A. Dizon, Razel Nikka M. Hao, Arianne A. Zamora, Devon Ray Pacial, Juan Antonio R. Magalang, Marissa Alejandria, Celia Carlos, Anna Ong-Lim, Edsel Maurice Salvaña, John Q. Wong, Jaime C. Montoya, Maria Rosario Singh-Vergeire and Cynthia P. Saloma |
| EPI_ISL_2189283 | The Lord's Grace Medical and Industrial Clinic | Philippine Genome Center | Francis A. Tablizo, Kenneth M. Kim, Maria Sofia L. Yangzon, Elcid Aaron R. Pangilinan, Renato Jacinto Q. Mantaring, Benedict A. Maralit, Marc Edsel C. Ayes, Eva Maria Cutiongco-de la Paz, Alethea R. de Guzman, Jan Michael C. Yap, Jo-Hannah S. Llames, Sheila Mae M. Araiza, Kris P. Punayan, Irish Coleen A. Asin, Candice Francheska B. Tambaoan, Asia Louisa U. Chong, Karol Sophia Agape R. Padilla, Rianna Patricia S. Cruz, Carlo M. Lapid, El King D. Morado, Joshua Gregor A. Dizon, Razel Nikka M. Hao, Arianne A. Zamora, Devon Ray Pacial, Juan Antonio R. Magalang, Marissa Alejandria, Celia Carlos, Anna Ong-Lim, Edsel Maurice Salvaña, John Q. Wong, Jaime C. Montoya, Maria Rosario Singh-Vergeire and Cynthia P. Saloma |
| EPI_ISL_2189284 | First Aide Diagnostic Center | Philippine Genome Center | Francis A. Tablizo, Kenneth M. Kim, Maria Sofia L. Yangzon, Elcid Aaron R. Pangilinan, Renato Jacinto Q. Mantaring, Benedict A. Maralit, Marc Edsel C. Ayes, Eva Maria Cutiongco-de la Paz, Alethea R. de Guzman, Jan Michael C. Yap, Jo-Hannah S. Llames, Sheila Mae M. Araiza, Kris P. Punayan, Irish Coleen A. Asin, Candice Francheska B. Tambaoan, Asia Louisa U. Chong, Karol Sophia Agape R. Padilla, Rianna Patricia S. Cruz, Carlo M. Lapid, El King D. Morado, Joshua Gregor A. Dizon, Razel Nikka M. Hao, Arianne A. Zamora, Devon Ray Pacial, Juan Antonio R. Magalang, Marissa Alejandria, Celia Carlos, Anna Ong-Lim, Edsel Maurice Salvaña, John Q. Wong, Jaime C. Montoya, Maria Rosario Singh-Vergeire and Cynthia P. Saloma |
| EPI_ISL_2189285 | The Lord's Grace Medical and Industrial Clinic | Philippine Genome Center | Francis A. Tablizo, Kenneth M. Kim, Maria Sofia L. Yangzon, Elcid Aaron R. Pangilinan, Renato Jacinto Q. Mantaring, Benedict A. Maralit, Marc Edsel C. Ayes, Eva Maria Cutiongco-de la Paz, Alethea R. de Guzman, Jan Michael C. Yap, Jo-Hannah S. Llames, Sheila Mae M. Araiza, Kris P. Punayan, Irish Coleen A. Asin, Candice Francheska B. Tambaoan, Asia Louisa U. Chong, Karol Sophia Agape R. Padilla, Rianna Patricia S. Cruz, Carlo M. Lapid, El King D. Morado, Joshua Gregor A. Dizon, Razel Nikka M. Hao, Arianne A. Zamora, Devon Ray Pacial, Juan Antonio R. Magalang, Marissa Alejandria, Celia Carlos, Anna Ong-Lim, Edsel Maurice Salvaña, John Q. Wong, Jaime C. Montoya, Maria Rosario Singh-Vergeire and Cynthia P. Saloma |
| EPI_ISL_2189286 | Pasig City Children's Hospital - Child's Hope | Philippine Genome Center | Francis A. Tablizo, Kenneth M. Kim, Maria Sofia L. Yangzon, Elcid Aaron R. Pangilinan, Renato Jacinto Q. Mantaring, Benedict A. Maralit, Marc Edsel C. Ayes, Eva Maria Cutiongco-de la Paz, Alethea R. de Guzman, Jan Michael C. Yap, Jo-Hannah S. Llames, Sheila Mae M. Araiza, Kris P. Punayan, Irish Coleen A. Asin, Candice Francheska B. Tambaoan, Asia Louisa U. Chong, Karol Sophia Agape R. Padilla, Rianna Patricia S. Cruz, Carlo M. Lapid, El King D. Morado, Joshua Gregor A. Dizon, Razel Nikka M. Hao, Arianne A. Zamora, Devon Ray Pacial, Juan Antonio R. Magalang, Marissa Alejandria, Celia Carlos, Anna Ong-Lim, Edsel Maurice Salvaña, John Q. Wong, Jaime C. Montoya, Maria Rosario Singh-Vergeire and Cynthia P. Saloma |
| EPI_ISL_2189289 | The Lord's Grace Medical and Industrial Clinic | Philippine Genome Center | Francis A. Tablizo, Kenneth M. Kim, Maria Sofia L. Yangzon, Elcid Aaron R. Pangilinan, Renato Jacinto Q. Mantaring, Benedict A. Maralit, Marc Edsel C. Ayes, Eva Maria Cutiongco-de la Paz, Alethea R. de Guzman, Jan Michael C. Yap, Jo-Hannah S. Llames, Sheila Mae M. Araiza, Kris P. Punayan, Irish Coleen A. Asin, Candice Francheska B. Tambaoan, Asia Louisa U. Chong, Karol Sophia Agape R. Padilla, Rianna Patricia S. Cruz, Carlo M. Lapid, El King D. Morado, Joshua Gregor A. Dizon, Razel Nikka M. Hao, Arianne A. Zamora, Devon Ray Pacial, Juan Antonio R. Magalang, Marissa Alejandria, Celia Carlos, Anna Ong-Lim, Edsel Maurice Salvaña, John Q. Wong, Jaime C. Montoya, Maria Rosario Singh-Vergeire and Cynthia P. Saloma |
| EPI_ISL_2189290 | Pasig City Children's Hospital - Child's Hope | Philippine Genome Center | Francis A. Tablizo, Kenneth M. Kim, Maria Sofia L. Yangzon, Elcid Aaron R. Pangilinan, Renato Jacinto Q. Mantaring, Benedict A. Maralit, Marc Edsel C. Ayes, Eva Maria Cutiongco-de la Paz, Alethea R. de Guzman, Jan Michael C. Yap, Jo-Hannah S. Llames, Sheila Mae M. Araiza, Kris P. Punayan, Irish Coleen A. Asin, Candice Francheska B. Tambaoan, Asia Louisa U. Chong, Karol Sophia Agape R. Padilla, Rianna Patricia S. Cruz, Carlo M. Lapid, El King D. Morado, Joshua Gregor A. Dizon, Razel Nikka M. Hao, Arianne A. Zamora, Devon Ray Pacial, Juan Antonio R. Magalang, Marissa Alejandria, Celia Carlos, Anna Ong-Lim, Edsel Maurice Salvaña, John Q. Wong, Jaime C. Montoya, Maria Rosario Singh-Vergeire and Cynthia P. Saloma |

| | | | |
|---|---|---|---|
| EPI_ISL_2189292 | The Lord's Grace Medical and Industrial Clinic | Philippine Genome Center | Francis A. Tablizo, Kenneth M. Kim, Maria Sofia L. Yangzon, Elcid Aaron R. Pangilinan, Renato Jacinto Q. Mantaring, Benedict A. Maralit, Marc Edsel C. Ayes, Eva Maria Cutiongco-de la Paz, Alethea R. de Guzman, Jan Michael C. Yap, Jo-Hannah S. Llames, Sheila Mae M. Araiza, Kris P. Punayan, Irish Coleen A. Asin, Candice Francheska B. Tambaoan, Asia Louisa U. Chong, Karol Sophia Agape R. Padilla, Rianna Patricia S. Cruz, Carlo M. Lapid, El King D. Morado, Joshua Gregor A. Dizon, Razel Nikka M. Hao, Arianne A. Zamora, Devon Ray Pacial, Juan Antonio R. Magalang, Marissa Alejandria, Celia Carlos, Anna Ong-Lim, Edsel Maurice Salvaña, John Q. Wong, Jaime C. Montoya, Maria Rosario Singh-Vergeire and Cynthia P. Saloma |
| EPI_ISL_2189293 | South Super Highway Molecular Diagnostic Laboratory | Philippine Genome Center | Francis A. Tablizo, Kenneth M. Kim, Maria Sofia L. Yangzon, Elcid Aaron R. Pangilinan, Renato Jacinto Q. Mantaring, Benedict A. Maralit, Marc Edsel C. Ayes, Eva Maria Cutiongco-de la Paz, Alethea R. de Guzman, Jan Michael C. Yap, Jo-Hannah S. Llames, Sheila Mae M. Araiza, Kris P. Punayan, Irish Coleen A. Asin, Candice Francheska B. Tambaoan, Asia Louisa U. Chong, Karol Sophia Agape R. Padilla, Rianna Patricia S. Cruz, Carlo M. Lapid, El King D. Morado, Joshua Gregor A. Dizon, Razel Nikka M. Hao, Arianne A. Zamora, Devon Ray Pacial, Juan Antonio R. Magalang, Marissa Alejandria, Celia Carlos, Anna Ong-Lim, Edsel Maurice Salvaña, John Q. Wong, Jaime C. Montoya, Maria Rosario Singh-Vergeire and Cynthia P. Saloma |
| EPI_ISL_2189294 | The Lord's Grace Medical and Industrial Clinic | Philippine Genome Center | Francis A. Tablizo, Kenneth M. Kim, Maria Sofia L. Yangzon, Elcid Aaron R. Pangilinan, Renato Jacinto Q. Mantaring, Benedict A. Maralit, Marc Edsel C. Ayes, Eva Maria Cutiongco-de la Paz, Alethea R. de Guzman, Jan Michael C. Yap, Jo-Hannah S. Llames, Sheila Mae M. Araiza, Kris P. Punayan, Irish Coleen A. Asin, Candice Francheska B. Tambaoan, Asia Louisa U. Chong, Karol Sophia Agape R. Padilla, Rianna Patricia S. Cruz, Carlo M. Lapid, El King D. Morado, Joshua Gregor A. Dizon, Razel Nikka M. Hao, Arianne A. Zamora, Devon Ray Pacial, Juan Antonio R. Magalang, Marissa Alejandria, Celia Carlos, Anna Ong-Lim, Edsel Maurice Salvaña, John Q. Wong, Jaime C. Montoya, Maria Rosario Singh-Vergeire and Cynthia P. Saloma |
| EPI_ISL_2189297 | National Kidney and Transplant Institute (NKTI) | Philippine Genome Center | Francis A. Tablizo, Kenneth M. Kim, Maria Sofia L. Yangzon, Elcid Aaron R. Pangilinan, Renato Jacinto Q. Mantaring, Benedict A. Maralit, Marc Edsel C. Ayes, Eva Maria Cutiongco-de la Paz, Alethea R. de Guzman, Jan Michael C. Yap, Jo-Hannah S. Llames, Sheila Mae M. Araiza, Kris P. Punayan, Irish Coleen A. Asin, Candice Francheska B. Tambaoan, Asia Louisa U. Chong, Karol Sophia Agape R. Padilla, Rianna Patricia S. Cruz, Carlo M. Lapid, El King D. Morado, Joshua Gregor A. Dizon, Razel Nikka M. Hao, Arianne A. Zamora, Devon Ray Pacial, Juan Antonio R. Magalang, Marissa Alejandria, Celia Carlos, Anna Ong-Lim, Edsel Maurice Salvaña, John Q. Wong, Jaime C. Montoya, Maria Rosario Singh-Vergeire and Cynthia P. Saloma |
| EPI_ISL_2189301 | Philippine Red Cross - Port Area | Philippine Genome Center | Francis A. Tablizo, Kenneth M. Kim, Maria Sofia L. Yangzon, Elcid Aaron R. Pangilinan, Renato Jacinto Q. Mantaring, Benedict A. Maralit, Marc Edsel C. Ayes, Eva Maria Cutiongco-de la Paz, Alethea R. de Guzman, Jan Michael C. Yap, Jo-Hannah S. Llames, Sheila Mae M. Araiza, Kris P. Punayan, Irish Coleen A. Asin, Candice Francheska B. Tambaoan, Asia Louisa U. Chong, Karol Sophia Agape R. Padilla, Rianna Patricia S. Cruz, Carlo M. Lapid, El King D. Morado, Joshua Gregor A. Dizon, Razel Nikka M. Hao, Arianne A. Zamora, Devon Ray Pacial, Juan Antonio R. Magalang, Marissa Alejandria, Celia Carlos, Anna Ong-Lim, Edsel Maurice Salvaña, John Q. Wong, Jaime C. Montoya, Maria Rosario Singh-Vergeire and Cynthia P. Saloma |
| EPI_ISL_2189302 | Mariano Marcos Memorial Hospital Medical Center | Philippine Genome Center | Francis A. Tablizo, Kenneth M. Kim, Maria Sofia L. Yangzon, Elcid Aaron R. Pangilinan, Renato Jacinto Q. Mantaring, Benedict A. Maralit, Marc Edsel C. Ayes, Eva Maria Cutiongco-de la Paz, Alethea R. de Guzman, Jan Michael C. Yap, Jo-Hannah S. Llames, Sheila Mae M. Araiza, Kris P. Punayan, Irish Coleen A. Asin, Candice Francheska B. Tambaoan, Asia Louisa U. Chong, Karol Sophia Agape R. Padilla, Rianna Patricia S. Cruz, Carlo M. Lapid, El King D. Morado, Joshua Gregor A. Dizon, Razel Nikka M. Hao, Arianne A. Zamora, Devon Ray Pacial, Juan Antonio R. Magalang, Marissa Alejandria, Celia Carlos, Anna Ong-Lim, Edsel Maurice Salvaña, John Q. Wong, Jaime C. Montoya, Maria Rosario Singh-Vergeire and Cynthia P. Saloma |
| EPI_ISL_2189303 | First Aide Diagnostic Center | Philippine Genome Center | Francis A. Tablizo, Kenneth M. Kim, Maria Sofia L. Yangzon, Elcid Aaron R. Pangilinan, Renato Jacinto Q. Mantaring, Benedict A. Maralit, Marc Edsel C. Ayes, Eva Maria Cutiongco-de la Paz, Alethea R. de Guzman, Jan Michael C. Yap, Jo-Hannah S. Llames, Sheila Mae M. Araiza, Kris P. Punayan, Irish Coleen A. Asin, Candice Francheska B. Tambaoan, Asia Louisa U. Chong, Karol Sophia Agape R. Padilla, Rianna Patricia S. Cruz, Carlo M. Lapid, El King D. Morado, Joshua Gregor A. Dizon, Razel Nikka M. Hao, Arianne A. Zamora, Devon Ray Pacial, Juan Antonio R. Magalang, Marissa Alejandria, Celia Carlos, Anna Ong-Lim, Edsel Maurice Salvaña, John Q. Wong, Jaime C. Montoya, Maria Rosario Singh-Vergeire and Cynthia P. Saloma |
| EPI_ISL_2189304 | Philippine Red Cross - National Blood Center | Philippine Genome Center | Francis A. Tablizo, Kenneth M. Kim, Maria Sofia L. Yangzon, Elcid Aaron R. Pangilinan, Renato Jacinto Q. Mantaring, Benedict A. Maralit, Marc Edsel C. Ayes, Eva Maria Cutiongco-de la Paz, Alethea R. de Guzman, Jan Michael C. Yap, Jo-Hannah S. Llames, Sheila Mae M. Araiza, Kris P. Punayan, Irish Coleen A. Asin, Candice Francheska B. Tambaoan, Asia Louisa U. Chong, Karol Sophia Agape R. Padilla, Rianna Patricia S. Cruz, Carlo M. Lapid, El King D. Morado, Joshua Gregor A. Dizon, Razel Nikka M. Hao, Arianne A. Zamora, Devon Ray Pacial, Juan Antonio R. Magalang, Marissa Alejandria, Celia Carlos, Anna Ong-Lim, Edsel Maurice Salvaña, John Q. Wong, Jaime C. Montoya, Maria Rosario Singh-Vergeire and Cynthia P. Saloma |
| EPI_ISL_2189305, EPI_ISL_2189307 | The Lord's Grace Medical and Industrial Clinic | Philippine Genome Center | Francis A. Tablizo, Kenneth M. Kim, Maria Sofia L. Yangzon, Elcid Aaron R. Pangilinan, Renato Jacinto Q. Mantaring, Benedict A. Maralit, Marc Edsel C. Ayes, Eva Maria Cutiongco-de la Paz, Alethea R. de Guzman, Jan Michael C. Yap, Jo-Hannah S. Llames, Sheila Mae M. Araiza, Kris P. Punayan, Irish Coleen A. Asin, Candice Francheska B. Tambaoan, Asia Louisa U. Chong, Karol Sophia Agape R. Padilla, Rianna Patricia S. Cruz, Carlo M. Lapid, El King D. Morado, Joshua Gregor A. Dizon, Razel Nikka M. Hao, Arianne A. Zamora, Devon Ray Pacial, Juan Antonio R. Magalang, Marissa Alejandria, Celia Carlos, Anna Ong-Lim, Edsel Maurice Salvaña, John Q. Wong, Jaime C. Montoya, Maria Rosario Singh-Vergeire and Cynthia P. Saloma |
| EPI_ISL_2189308 | Sta. Rosa CHO- GENEXPERT LABORATORY | Philippine Genome Center | Francis A. Tablizo, Kenneth M. Kim, Maria Sofia L. Yangzon, Elcid Aaron R. Pangilinan, Renato Jacinto Q. Mantaring, Benedict A. Maralit, Marc Edsel C. Ayes, Eva Maria Cutiongco-de la Paz, Alethea R. de Guzman, Jan Michael C. Yap, Jo-Hannah S. Llames, Sheila Mae M. Araiza, Kris P. Punayan, Irish Coleen A. Asin, Candice Francheska B. Tambaoan, Asia Louisa U. Chong, Karol Sophia Agape R. Padilla, Rianna Patricia S. Cruz, Carlo M. Lapid, El King D. Morado, Joshua Gregor A. Dizon, Razel Nikka M. Hao, Arianne A. Zamora, Devon Ray Pacial, Juan Antonio R. Magalang, Marissa Alejandria, Celia Carlos, Anna Ong-Lim, Edsel Maurice Salvaña, John Q. Wong, Jaime C. Montoya, Maria Rosario Singh-Vergeire and Cynthia P. Saloma |
| EPI_ISL_2189309, EPI_ISL_2189310 | Mary Mediatrix Medical Center | Philippine Genome Center | Francis A. Tablizo, Kenneth M. Kim, Maria Sofia L. Yangzon, Elcid Aaron R. Pangilinan, Renato Jacinto Q. Mantaring, Benedict A. Maralit, Marc Edsel C. Ayes, Eva Maria Cutiongco-de la Paz, Alethea R. de Guzman, Jan Michael C. Yap, Jo-Hannah S. Llames, Sheila Mae M. Araiza, Kris P. Punayan, Irish Coleen A. Asin, Candice Francheska B. Tambaoan, Asia Louisa U. Chong, Karol Sophia Agape R. Padilla, Rianna Patricia S. Cruz, Carlo M. Lapid, El King D. Morado, Joshua Gregor A. Dizon, Razel Nikka M. Hao, Arianne A. Zamora, Devon Ray Pacial, Juan Antonio R. Magalang, Marissa Alejandria, Celia Carlos, Anna Ong-Lim, Edsel Maurice Salvaña, John Q. Wong, Jaime C. Montoya, Maria Rosario Singh-Vergeire and Cynthia P. Saloma |
| EPI_ISL_2189311, EPI_ISL_2189312 | Tarlac Provincial Hospital - Molecular Pathology Laboratory | Philippine Genome Center | Francis A. Tablizo, Kenneth M. Kim, Maria Sofia L. Yangzon, Elcid Aaron R. Pangilinan, Renato Jacinto Q. Mantaring, Benedict A. Maralit, Marc Edsel C. Ayes, Eva Maria Cutiongco-de la Paz, Alethea R. de Guzman, Jan Michael C. Yap, Jo-Hannah S. Llames, Sheila Mae M. Araiza, Kris P. Punayan, Irish Coleen A. Asin, Candice Francheska B. Tambaoan, Asia Louisa U. Chong, Karol Sophia Agape R. Padilla, Rianna Patricia S. Cruz, Carlo M. Lapid, El King D. Morado, Joshua Gregor A. Dizon, Razel Nikka M. Hao, Arianne A. Zamora, Devon Ray Pacial, Juan Antonio R. Magalang, Marissa Alejandria, Celia Carlos, Anna Ong-Lim, Edsel Maurice Salvaña, John Q. Wong, Jaime C. Montoya, Maria Rosario Singh-Vergeire and Cynthia P. Saloma |
| EPI_ISL_2189313 | University of the Philippines National Institutes of Health (UP NIH) | Philippine Genome Center | Francis A. Tablizo, Kenneth M. Kim, Maria Sofia L. Yangzon, Elcid Aaron R. Pangilinan, Renato Jacinto Q. Mantaring, Benedict A. Maralit, Marc Edsel C. Ayes, Eva Maria Cutiongco-de la Paz, Alethea R. de Guzman, Jan Michael C. Yap, Jo-Hannah S. Llames, Sheila Mae M. Araiza, Kris P. Punayan, Irish Coleen A. Asin, Candice Francheska B. Tambaoan, Asia Louisa U. Chong, Karol Sophia Agape R. Padilla, Rianna Patricia S. Cruz, Carlo M. Lapid, El King D. Morado, Joshua Gregor A. Dizon, Razel Nikka M. Hao, Arianne A. Zamora, Devon Ray Pacial, Juan Antonio R. Magalang, Marissa Alejandria, Celia Carlos, Anna Ong-Lim, Edsel Maurice Salvaña, John Q. Wong, Jaime C. Montoya, Maria Rosario Singh-Vergeire and Cynthia P. Saloma |
| EPI_ISL_2189315, EPI_ISL_2189316, EPI_ISL_2189317 | The Lord's Grace Medical and Industrial Clinic | Philippine Genome Center | Francis A. Tablizo, Kenneth M. Kim, Maria Sofia L. Yangzon, Elcid Aaron R. Pangilinan, Renato Jacinto Q. Mantaring, Benedict A. Maralit, Marc Edsel C. Ayes, Eva Maria Cutiongco-de la Paz, Alethea R. de Guzman, Jan Michael C. Yap, Jo-Hannah S. Llames, Sheila Mae M. Araiza, Kris P. Punayan, Irish Coleen A. Asin, Candice Francheska B. Tambaoan, Asia Louisa U. Chong, Karol Sophia Agape R. Padilla, Rianna Patricia S. Cruz, Carlo M. Lapid, El King D. Morado, Joshua Gregor A. Dizon, Razel Nikka M. Hao, Arianne A. Zamora, Devon Ray Pacial, Juan Antonio R. Magalang, Marissa Alejandria, Celia Carlos, Anna Ong-Lim, Edsel Maurice Salvaña, John Q. Wong, Jaime C. Montoya, Maria Rosario Singh-Vergeire and Cynthia P. Saloma |
| EPI_ISL_2189319 | The Medical City | Philippine Genome Center | Francis A. Tablizo, Kenneth M. Kim, Maria Sofia L. Yangzon, Elcid Aaron R. Pangilinan, Renato Jacinto Q. Mantaring, Benedict A. Maralit, Marc Edsel C. Ayes, Eva Maria Cutiongco-de la Paz, Alethea R. de Guzman, Jan Michael C. Yap, Jo-Hannah S. Llames, Sheila Mae M. Araiza, Kris P. Punayan, Irish Coleen A. Asin, Candice Francheska B. Tambaoan, Asia Louisa U. Chong, Karol Sophia Agape R. Padilla, Rianna Patricia S. Cruz, Carlo M. Lapid, El King D. Morado, Joshua Gregor A. Dizon, Razel Nikka M. Hao, Arianne A. Zamora, Devon Ray Pacial, Juan Antonio R. Magalang, Marissa Alejandria, Celia Carlos, Anna Ong-Lim, Edsel Maurice Salvaña, John Q. Wong, Jaime C. Montoya, Maria Rosario Singh-Vergeire and Cynthia P. Saloma |
| EPI_ISL_2189321 | Laguna Provincial Hospital - San Pablo District Hospital | Philippine Genome Center | Francis A. Tablizo, Kenneth M. Kim, Maria Sofia L. Yangzon, Elcid Aaron R. Pangilinan, Renato Jacinto Q. Mantaring, Benedict A. Maralit, Marc Edsel C. Ayes, Eva Maria Cutiongco-de la Paz, Alethea R. de Guzman, Jan Michael C. Yap, Jo-Hannah S. Llames, Sheila Mae M. Araiza, Kris P. Punayan, Irish Coleen A. Asin, Candice Francheska B. Tambaoan, Asia Louisa U. Chong, Karol Sophia Agape R. Padilla, Rianna Patricia S. Cruz, Carlo M. Lapid, El King D. Morado, Joshua Gregor A. Dizon, Razel Nikka M. Hao, Arianne A. Zamora, Devon Ray Pacial, Juan Antonio R. Magalang, Marissa Alejandria, Celia Carlos, Anna Ong-Lim, Edsel Maurice Salvaña, John Q. Wong, Jaime C. Montoya, Maria Rosario Singh-Vergeire and Cynthia P. Saloma |

| | | | |
|---|---|---|---|
| EPI_ISL_2189322 | The Medical City | Philippine Genome Center | Francis A. Tablizo, Kenneth M. Kim, Maria Sofia L. Yangzon, Elcid Aaron R. Pangilinan, Renato Jacinto Q. Mantaring, Benedict A. Maralit, Marc Edsel C. Ayes, Eva Maria Cutiongco-de la Paz, Alethea R. de Guzman, Jan Michael C. Yap, Jo-Hannah S. Llames, Sheila Mae M. Araiza, Kris P. Punayan, Irish Coleen A. Asin, Candice Francheska B. Tambaoan, Asia Louisa U. Chong, Karol Sophia Agape R. Padilla, Rianna Patricia S. Cruz, Carlo M. Lapid, El King D. Morado, Joshua Gregor A. Dizon, Razel Nikka M. Hao, Arianne A. Zamora, Devon Ray Pacial, Juan Antonio R. Magalang, Marissa Alejandria, Celia Carlos, Anna Ong-Lim, Edsel Maurice Salvaña, John Q. Wong, Jaime C. Montoya, Maria Rosario Singh-Vergeire and Cynthia P. Saloma |
| EPI_ISL_2189323, EPI_ISL_2189324 | The Lord's Grace Medical and Industrial Clinic | Philippine Genome Center | Francis A. Tablizo, Kenneth M. Kim, Maria Sofia L. Yangzon, Elcid Aaron R. Pangilinan, Renato Jacinto Q. Mantaring, Benedict A. Maralit, Marc Edsel C. Ayes, Eva Maria Cutiongco-de la Paz, Alethea R. de Guzman, Jan Michael C. Yap, Jo-Hannah S. Llames, Sheila Mae M. Araiza, Kris P. Punayan, Irish Coleen A. Asin, Candice Francheska B. Tambaoan, Asia Louisa U. Chong, Karol Sophia Agape R. Padilla, Rianna Patricia S. Cruz, Carlo M. Lapid, El King D. Morado, Joshua Gregor A. Dizon, Razel Nikka M. Hao, Arianne A. Zamora, Devon Ray Pacial, Juan Antonio R. Magalang, Marissa Alejandria, Celia Carlos, Anna Ong-Lim, Edsel Maurice Salvaña, John Q. Wong, Jaime C. Montoya, Maria Rosario Singh-Vergeire and Cynthia P. Saloma |
| EPI_ISL_2189325 | Sta. Rosa CHO- GENEXPERT LABORATORY | Philippine Genome Center | Francis A. Tablizo, Kenneth M. Kim, Maria Sofia L. Yangzon, Elcid Aaron R. Pangilinan, Renato Jacinto Q. Mantaring, Benedict A. Maralit, Marc Edsel C. Ayes, Eva Maria Cutiongco-de la Paz, Alethea R. de Guzman, Jan Michael C. Yap, Jo-Hannah S. Llames, Sheila Mae M. Araiza, Kris P. Punayan, Irish Coleen A. Asin, Candice Francheska B. Tambaoan, Asia Louisa U. Chong, Karol Sophia Agape R. Padilla, Rianna Patricia S. Cruz, Carlo M. Lapid, El King D. Morado, Joshua Gregor A. Dizon, Razel Nikka M. Hao, Arianne A. Zamora, Devon Ray Pacial, Juan Antonio R. Magalang, Marissa Alejandria, Celia Carlos, Anna Ong-Lim, Edsel Maurice Salvaña, John Q. Wong, Jaime C. Montoya, Maria Rosario Singh-Vergeire and Cynthia P. Saloma |
| EPI_ISL_2189327 | The Medical City | Philippine Genome Center | Francis A. Tablizo, Kenneth M. Kim, Maria Sofia L. Yangzon, Elcid Aaron R. Pangilinan, Renato Jacinto Q. Mantaring, Benedict A. Maralit, Marc Edsel C. Ayes, Eva Maria Cutiongco-de la Paz, Alethea R. de Guzman, Jan Michael C. Yap, Jo-Hannah S. Llames, Sheila Mae M. Araiza, Kris P. Punayan, Irish Coleen A. Asin, Candice Francheska B. Tambaoan, Asia Louisa U. Chong, Karol Sophia Agape R. Padilla, Rianna Patricia S. Cruz, Carlo M. Lapid, El King D. Morado, Joshua Gregor A. Dizon, Razel Nikka M. Hao, Arianne A. Zamora, Devon Ray Pacial, Juan Antonio R. Magalang, Marissa Alejandria, Celia Carlos, Anna Ong-Lim, Edsel Maurice Salvaña, John Q. Wong, Jaime C. Montoya, Maria Rosario Singh-Vergeire and Cynthia P. Saloma |
| EPI_ISL_2189330 | Philippine Red Cross - National Blood Center | Philippine Genome Center | Francis A. Tablizo, Kenneth M. Kim, Maria Sofia L. Yangzon, Elcid Aaron R. Pangilinan, Renato Jacinto Q. Mantaring, Benedict A. Maralit, Marc Edsel C. Ayes, Eva Maria Cutiongco-de la Paz, Alethea R. de Guzman, Jan Michael C. Yap, Jo-Hannah S. Llames, Sheila Mae M. Araiza, Kris P. Punayan, Irish Coleen A. Asin, Candice Francheska B. Tambaoan, Asia Louisa U. Chong, Karol Sophia Agape R. Padilla, Rianna Patricia S. Cruz, Carlo M. Lapid, El King D. Morado, Joshua Gregor A. Dizon, Razel Nikka M. Hao, Arianne A. Zamora, Devon Ray Pacial, Juan Antonio R. Magalang, Marissa Alejandria, Celia Carlos, Anna Ong-Lim, Edsel Maurice Salvaña, John Q. Wong, Jaime C. Montoya, Maria Rosario Singh-Vergeire and Cynthia P. Saloma |
| EPI_ISL_2189331 | South Super Highway Molecular Diagnostic Laboratory | Philippine Genome Center | Francis A. Tablizo, Kenneth M. Kim, Maria Sofia L. Yangzon, Elcid Aaron R. Pangilinan, Renato Jacinto Q. Mantaring, Benedict A. Maralit, Marc Edsel C. Ayes, Eva Maria Cutiongco-de la Paz, Alethea R. de Guzman, Jan Michael C. Yap, Jo-Hannah S. Llames, Sheila Mae M. Araiza, Kris P. Punayan, Irish Coleen A. Asin, Candice Francheska B. Tambaoan, Asia Louisa U. Chong, Karol Sophia Agape R. Padilla, Rianna Patricia S. Cruz, Carlo M. Lapid, El King D. Morado, Joshua Gregor A. Dizon, Razel Nikka M. Hao, Arianne A. Zamora, Devon Ray Pacial, Juan Antonio R. Magalang, Marissa Alejandria, Celia Carlos, Anna Ong-Lim, Edsel Maurice Salvaña, John Q. Wong, Jaime C. Montoya, Maria Rosario Singh-Vergeire and Cynthia P. Saloma |
| EPI_ISL_2189332 | The Lord's Grace Medical and Industrial Clinic | Philippine Genome Center | Francis A. Tablizo, Kenneth M. Kim, Maria Sofia L. Yangzon, Elcid Aaron R. Pangilinan, Renato Jacinto Q. Mantaring, Benedict A. Maralit, Marc Edsel C. Ayes, Eva Maria Cutiongco-de la Paz, Alethea R. de Guzman, Jan Michael C. Yap, Jo-Hannah S. Llames, Sheila Mae M. Araiza, Kris P. Punayan, Irish Coleen A. Asin, Candice Francheska B. Tambaoan, Asia Louisa U. Chong, Karol Sophia Agape R. Padilla, Rianna Patricia S. Cruz, Carlo M. Lapid, El King D. Morado, Joshua Gregor A. Dizon, Razel Nikka M. Hao, Arianne A. Zamora, Devon Ray Pacial, Juan Antonio R. Magalang, Marissa Alejandria, Celia Carlos, Anna Ong-Lim, Edsel Maurice Salvaña, John Q. Wong, Jaime C. Montoya, Maria Rosario Singh-Vergeire and Cynthia P. Saloma |
| EPI_ISL_2189333 | Laguna Provincial Hospital - San Pablo District Hospital | Philippine Genome Center | Francis A. Tablizo, Kenneth M. Kim, Maria Sofia L. Yangzon, Elcid Aaron R. Pangilinan, Renato Jacinto Q. Mantaring, Benedict A. Maralit, Marc Edsel C. Ayes, Eva Maria Cutiongco-de la Paz, Alethea R. de Guzman, Jan Michael C. Yap, Jo-Hannah S. Llames, Sheila Mae M. Araiza, Kris P. Punayan, Irish Coleen A. Asin, Candice Francheska B. Tambaoan, Asia Louisa U. Chong, Karol Sophia Agape R. Padilla, Rianna Patricia S. Cruz, Carlo M. Lapid, El King D. Morado, Joshua Gregor A. Dizon, Razel Nikka M. Hao, Arianne A. Zamora, Devon Ray Pacial, Juan Antonio R. Magalang, Marissa Alejandria, Celia Carlos, Anna Ong-Lim, Edsel Maurice Salvaña, John Q. Wong, Jaime C. Montoya, Maria Rosario Singh-Vergeire and Cynthia P. Saloma |
| EPI_ISL_2189334 | Philippine Red Cross - Port Area | Philippine Genome Center | Francis A. Tablizo, Kenneth M. Kim, Maria Sofia L. Yangzon, Elcid Aaron R. Pangilinan, Renato Jacinto Q. Mantaring, Benedict A. Maralit, Marc Edsel C. Ayes, Eva Maria Cutiongco-de la Paz, Alethea R. de Guzman, Jan Michael C. Yap, Jo-Hannah S. Llames, Sheila Mae M. Araiza, Kris P. Punayan, Irish Coleen A. Asin, Candice Francheska B. Tambaoan, Asia Louisa U. Chong, Karol Sophia Agape R. Padilla, Rianna Patricia S. Cruz, Carlo M. Lapid, El King D. Morado, Joshua Gregor A. Dizon, Razel Nikka M. Hao, Arianne A. Zamora, Devon Ray Pacial, Juan Antonio R. Magalang, Marissa Alejandria, Celia Carlos, Anna Ong-Lim, Edsel Maurice Salvaña, John Q. Wong, Jaime C. Montoya, Maria Rosario Singh-Vergeire and Cynthia P. Saloma |
| EPI_ISL_2189335 | The Lord's Grace Medical and Industrial Clinic | Philippine Genome Center | Francis A. Tablizo, Kenneth M. Kim, Maria Sofia L. Yangzon, Elcid Aaron R. Pangilinan, Renato Jacinto Q. Mantaring, Benedict A. Maralit, Marc Edsel C. Ayes, Eva Maria Cutiongco-de la Paz, Alethea R. de Guzman, Jan Michael C. Yap, Jo-Hannah S. Llames, Sheila Mae M. Araiza, Kris P. Punayan, Irish Coleen A. Asin, Candice Francheska B. Tambaoan, Asia Louisa U. Chong, Karol Sophia Agape R. Padilla, Rianna Patricia S. Cruz, Carlo M. Lapid, El King D. Morado, Joshua Gregor A. Dizon, Razel Nikka M. Hao, Arianne A. Zamora, Devon Ray Pacial, Juan Antonio R. Magalang, Marissa Alejandria, Celia Carlos, Anna Ong-Lim, Edsel Maurice Salvaña, John Q. Wong, Jaime C. Montoya, Maria Rosario Singh-Vergeire and Cynthia P. Saloma |
| EPI_ISL_2189336, EPI_ISL_2189338 | The Medical City | Philippine Genome Center | Francis A. Tablizo, Kenneth M. Kim, Maria Sofia L. Yangzon, Elcid Aaron R. Pangilinan, Renato Jacinto Q. Mantaring, Benedict A. Maralit, Marc Edsel C. Ayes, Eva Maria Cutiongco-de la Paz, Alethea R. de Guzman, Jan Michael C. Yap, Jo-Hannah S. Llames, Sheila Mae M. Araiza, Kris P. Punayan, Irish Coleen A. Asin, Candice Francheska B. Tambaoan, Asia Louisa U. Chong, Karol Sophia Agape R. Padilla, Rianna Patricia S. Cruz, Carlo M. Lapid, El King D. Morado, Joshua Gregor A. Dizon, Razel Nikka M. Hao, Arianne A. Zamora, Devon Ray Pacial, Juan Antonio R. Magalang, Marissa Alejandria, Celia Carlos, Anna Ong-Lim, Edsel Maurice Salvaña, John Q. Wong, Jaime C. Montoya, Maria Rosario Singh-Vergeire and Cynthia P. Saloma |
| EPI_ISL_2189339 | Laguna Provincial Hospital - San Pablo District Hospital | Philippine Genome Center | Francis A. Tablizo, Kenneth M. Kim, Maria Sofia L. Yangzon, Elcid Aaron R. Pangilinan, Renato Jacinto Q. Mantaring, Benedict A. Maralit, Marc Edsel C. Ayes, Eva Maria Cutiongco-de la Paz, Alethea R. de Guzman, Jan Michael C. Yap, Jo-Hannah S. Llames, Sheila Mae M. Araiza, Kris P. Punayan, Irish Coleen A. Asin, Candice Francheska B. Tambaoan, Asia Louisa U. Chong, Karol Sophia Agape R. Padilla, Rianna Patricia S. Cruz, Carlo M. Lapid, El King D. Morado, Joshua Gregor A. Dizon, Razel Nikka M. Hao, Arianne A. Zamora, Devon Ray Pacial, Juan Antonio R. Magalang, Marissa Alejandria, Celia Carlos, Anna Ong-Lim, Edsel Maurice Salvaña, John Q. Wong, Jaime C. Montoya, Maria Rosario Singh-Vergeire and Cynthia P. Saloma |
| EPI_ISL_2189340 | The Lord's Grace Medical and Industrial Clinic | Philippine Genome Center | Francis A. Tablizo, Kenneth M. Kim, Maria Sofia L. Yangzon, Elcid Aaron R. Pangilinan, Renato Jacinto Q. Mantaring, Benedict A. Maralit, Marc Edsel C. Ayes, Eva Maria Cutiongco-de la Paz, Alethea R. de Guzman, Jan Michael C. Yap, Jo-Hannah S. Llames, Sheila Mae M. Araiza, Kris P. Punayan, Irish Coleen A. Asin, Candice Francheska B. Tambaoan, Asia Louisa U. Chong, Karol Sophia Agape R. Padilla, Rianna Patricia S. Cruz, Carlo M. Lapid, El King D. Morado, Joshua Gregor A. Dizon, Razel Nikka M. Hao, Arianne A. Zamora, Devon Ray Pacial, Juan Antonio R. Magalang, Marissa Alejandria, Celia Carlos, Anna Ong-Lim, Edsel Maurice Salvaña, John Q. Wong, Jaime C. Montoya, Maria Rosario Singh-Vergeire and Cynthia P. Saloma |
| EPI_ISL_2189344 | Pasig City Children's Hospital - Child's Hope | Philippine Genome Center | Francis A. Tablizo, Kenneth M. Kim, Maria Sofia L. Yangzon, Elcid Aaron R. Pangilinan, Renato Jacinto Q. Mantaring, Benedict A. Maralit, Marc Edsel C. Ayes, Eva Maria Cutiongco-de la Paz, Alethea R. de Guzman, Jan Michael C. Yap, Jo-Hannah S. Llames, Sheila Mae M. Araiza, Kris P. Punayan, Irish Coleen A. Asin, Candice Francheska B. Tambaoan, Asia Louisa U. Chong, Karol Sophia Agape R. Padilla, Rianna Patricia S. Cruz, Carlo M. Lapid, El King D. Morado, Joshua Gregor A. Dizon, Razel Nikka M. Hao, Arianne A. Zamora, Devon Ray Pacial, Juan Antonio R. Magalang, Marissa Alejandria, Celia Carlos, Anna Ong-Lim, Edsel Maurice Salvaña, John Q. Wong, Jaime C. Montoya, Maria Rosario Singh-Vergeire and Cynthia P. Saloma |
| EPI_ISL_2189345 | Philippine Red Cross - National Blood Center | Philippine Genome Center | Francis A. Tablizo, Kenneth M. Kim, Maria Sofia L. Yangzon, Elcid Aaron R. Pangilinan, Renato Jacinto Q. Mantaring, Benedict A. Maralit, Marc Edsel C. Ayes, Eva Maria Cutiongco-de la Paz, Alethea R. de Guzman, Jan Michael C. Yap, Jo-Hannah S. Llames, Sheila Mae M. Araiza, Kris P. Punayan, Irish Coleen A. Asin, Candice Francheska B. Tambaoan, Asia Louisa U. Chong, Karol Sophia Agape R. Padilla, Rianna Patricia S. Cruz, Carlo M. Lapid, El King D. Morado, Joshua Gregor A. Dizon, Razel Nikka M. Hao, Arianne A. Zamora, Devon Ray Pacial, Juan Antonio R. Magalang, Marissa Alejandria, Celia Carlos, Anna Ong-Lim, Edsel Maurice Salvaña, John Q. Wong, Jaime C. Montoya, Maria Rosario Singh-Vergeire and Cynthia P. Saloma |
| EPI_ISL_2189346 | Philippine Red Cross - Port Area | Philippine Genome Center | Francis A. Tablizo, Kenneth M. Kim, Maria Sofia L. Yangzon, Elcid Aaron R. Pangilinan, Renato Jacinto Q. Mantaring, Benedict A. Maralit, Marc Edsel C. Ayes, Eva Maria Cutiongco-de la Paz, Alethea R. de Guzman, Jan Michael C. Yap, Jo-Hannah S. Llames, Sheila Mae M. Araiza, Kris P. Punayan, Irish Coleen A. Asin, Candice Francheska B. Tambaoan, Asia Louisa U. Chong, Karol Sophia Agape R. Padilla, Rianna Patricia S. Cruz, Carlo M. Lapid, El King D. Morado, Joshua Gregor A. Dizon, Razel Nikka M. Hao, Arianne A. Zamora, Devon Ray Pacial, Juan Antonio R. Magalang, Marissa Alejandria, Celia Carlos, Anna Ong-Lim, Edsel Maurice Salvaña, John Q. Wong, Jaime C. Montoya, Maria Rosario Singh-Vergeire and Cynthia P. Saloma |

| | | | |
|---|---|---|---|
| EPI_ISL_2189347 | Philippine General Hospital (PGH) | Philippine Genome Center | Francis A. Tablizo, Kenneth M. Kim, Maria Sofia L. Yangzon, Elcid Aaron R. Pangilinan, Renato Jacinto Q. Mantaring, Benedict A. Maralit, Marc Edsel C. Ayes, Eva Maria Cutiongco-de la Paz, Alethea R. de Guzman, Jan Michael C. Yap, Jo-Hannah S. Llames, Sheila Mae M. Araiza, Kris P. Punayan, Irish Coleen A. Asin, Candice Francheska B. Tambaoan, Asia Louisa U. Chong, Karol Sophia Agape R. Padilla, Rianna Patricia S. Cruz, Carlo M. Lapid, El King D. Morado, Joshua Gregor A. Dizon, Razel Nikka M. Hao, Arianne A. Zamora, Devon Ray Pacial, Juan Antonio R. Magalang, Marissa Alejandria, Celia Carlos, Anna Ong-Lim, Edsel Maurice Salvaña, John Q. Wong, Jaime C. Montoya, Maria Rosario Singh-Vergeire and Cynthia P. Saloma |
| EPI_ISL_2189349 | Philippine Red Cross - Port Area | Philippine Genome Center | Francis A. Tablizo, Kenneth M. Kim, Maria Sofia L. Yangzon, Elcid Aaron R. Pangilinan, Renato Jacinto Q. Mantaring, Benedict A. Maralit, Marc Edsel C. Ayes, Eva Maria Cutiongco-de la Paz, Alethea R. de Guzman, Jan Michael C. Yap, Jo-Hannah S. Llames, Sheila Mae M. Araiza, Kris P. Punayan, Irish Coleen A. Asin, Candice Francheska B. Tambaoan, Asia Louisa U. Chong, Karol Sophia Agape R. Padilla, Rianna Patricia S. Cruz, Carlo M. Lapid, El King D. Morado, Joshua Gregor A. Dizon, Razel Nikka M. Hao, Arianne A. Zamora, Devon Ray Pacial, Juan Antonio R. Magalang, Marissa Alejandria, Celia Carlos, Anna Ong-Lim, Edsel Maurice Salvaña, John Q. Wong, Jaime C. Montoya, Maria Rosario Singh-Vergeire and Cynthia P. Saloma |
| EPI_ISL_2189352 | Philippine Red Cross - National Blood Center | Philippine Genome Center | Francis A. Tablizo, Kenneth M. Kim, Maria Sofia L. Yangzon, Elcid Aaron R. Pangilinan, Renato Jacinto Q. Mantaring, Benedict A. Maralit, Marc Edsel C. Ayes, Eva Maria Cutiongco-de la Paz, Alethea R. de Guzman, Jan Michael C. Yap, Jo-Hannah S. Llames, Sheila Mae M. Araiza, Kris P. Punayan, Irish Coleen A. Asin, Candice Francheska B. Tambaoan, Asia Louisa U. Chong, Karol Sophia Agape R. Padilla, Rianna Patricia S. Cruz, Carlo M. Lapid, El King D. Morado, Joshua Gregor A. Dizon, Razel Nikka M. Hao, Arianne A. Zamora, Devon Ray Pacial, Juan Antonio R. Magalang, Marissa Alejandria, Celia Carlos, Anna Ong-Lim, Edsel Maurice Salvaña, John Q. Wong, Jaime C. Montoya, Maria Rosario Singh-Vergeire and Cynthia P. Saloma |
| EPI_ISL_2189353 | Laguna Provincial Hospital - San Pablo District Hospital | Philippine Genome Center | Francis A. Tablizo, Kenneth M. Kim, Maria Sofia L. Yangzon, Elcid Aaron R. Pangilinan, Renato Jacinto Q. Mantaring, Benedict A. Maralit, Marc Edsel C. Ayes, Eva Maria Cutiongco-de la Paz, Alethea R. de Guzman, Jan Michael C. Yap, Jo-Hannah S. Llames, Sheila Mae M. Araiza, Kris P. Punayan, Irish Coleen A. Asin, Candice Francheska B. Tambaoan, Asia Louisa U. Chong, Karol Sophia Agape R. Padilla, Rianna Patricia S. Cruz, Carlo M. Lapid, El King D. Morado, Joshua Gregor A. Dizon, Razel Nikka M. Hao, Arianne A. Zamora, Devon Ray Pacial, Juan Antonio R. Magalang, Marissa Alejandria, Celia Carlos, Anna Ong-Lim, Edsel Maurice Salvaña, John Q. Wong, Jaime C. Montoya, Maria Rosario Singh-Vergeire and Cynthia P. Saloma |
| EPI_ISL_2189355 | University of the Philippines National Institutes of Health (UP NIH) | Philippine Genome Center | Francis A. Tablizo, Kenneth M. Kim, Maria Sofia L. Yangzon, Elcid Aaron R. Pangilinan, Renato Jacinto Q. Mantaring, Benedict A. Maralit, Marc Edsel C. Ayes, Eva Maria Cutiongco-de la Paz, Alethea R. de Guzman, Jan Michael C. Yap, Jo-Hannah S. Llames, Sheila Mae M. Araiza, Kris P. Punayan, Irish Coleen A. Asin, Candice Francheska B. Tambaoan, Asia Louisa U. Chong, Karol Sophia Agape R. Padilla, Rianna Patricia S. Cruz, Carlo M. Lapid, El King D. Morado, Joshua Gregor A. Dizon, Razel Nikka M. Hao, Arianne A. Zamora, Devon Ray Pacial, Juan Antonio R. Magalang, Marissa Alejandria, Celia Carlos, Anna Ong-Lim, Edsel Maurice Salvaña, John Q. Wong, Jaime C. Montoya, Maria Rosario Singh-Vergeire and Cynthia P. Saloma |
| EPI_ISL_2189356 | The Medical City | Philippine Genome Center | Francis A. Tablizo, Kenneth M. Kim, Maria Sofia L. Yangzon, Elcid Aaron R. Pangilinan, Renato Jacinto Q. Mantaring, Benedict A. Maralit, Marc Edsel C. Ayes, Eva Maria Cutiongco-de la Paz, Alethea R. de Guzman, Jan Michael C. Yap, Jo-Hannah S. Llames, Sheila Mae M. Araiza, Kris P. Punayan, Irish Coleen A. Asin, Candice Francheska B. Tambaoan, Asia Louisa U. Chong, Karol Sophia Agape R. Padilla, Rianna Patricia S. Cruz, Carlo M. Lapid, El King D. Morado, Joshua Gregor A. Dizon, Razel Nikka M. Hao, Arianne A. Zamora, Devon Ray Pacial, Juan Antonio R. Magalang, Marissa Alejandria, Celia Carlos, Anna Ong-Lim, Edsel Maurice Salvaña, John Q. Wong, Jaime C. Montoya, Maria Rosario Singh-Vergeire and Cynthia P. Saloma |
| EPI_ISL_2189357 | Philippine General Hospital (PGH) | Philippine Genome Center | Francis A. Tablizo, Kenneth M. Kim, Maria Sofia L. Yangzon, Elcid Aaron R. Pangilinan, Renato Jacinto Q. Mantaring, Benedict A. Maralit, Marc Edsel C. Ayes, Eva Maria Cutiongco-de la Paz, Alethea R. de Guzman, Jan Michael C. Yap, Jo-Hannah S. Llames, Sheila Mae M. Araiza, Kris P. Punayan, Irish Coleen A. Asin, Candice Francheska B. Tambaoan, Asia Louisa U. Chong, Karol Sophia Agape R. Padilla, Rianna Patricia S. Cruz, Carlo M. Lapid, El King D. Morado, Joshua Gregor A. Dizon, Razel Nikka M. Hao, Arianne A. Zamora, Devon Ray Pacial, Juan Antonio R. Magalang, Marissa Alejandria, Celia Carlos, Anna Ong-Lim, Edsel Maurice Salvaña, John Q. Wong, Jaime C. Montoya, Maria Rosario Singh-Vergeire and Cynthia P. Saloma |
| EPI_ISL_2189359 | Laguna Provincial Hospital - San Pablo District Hospital | Philippine Genome Center | Francis A. Tablizo, Kenneth M. Kim, Maria Sofia L. Yangzon, Elcid Aaron R. Pangilinan, Renato Jacinto Q. Mantaring, Benedict A. Maralit, Marc Edsel C. Ayes, Eva Maria Cutiongco-de la Paz, Alethea R. de Guzman, Jan Michael C. Yap, Jo-Hannah S. Llames, Sheila Mae M. Araiza, Kris P. Punayan, Irish Coleen A. Asin, Candice Francheska B. Tambaoan, Asia Louisa U. Chong, Karol Sophia Agape R. Padilla, Rianna Patricia S. Cruz, Carlo M. Lapid, El King D. Morado, Joshua Gregor A. Dizon, Razel Nikka M. Hao, Arianne A. Zamora, Devon Ray Pacial, Juan Antonio R. Magalang, Marissa Alejandria, Celia Carlos, Anna Ong-Lim, Edsel Maurice Salvaña, John Q. Wong, Jaime C. Montoya, Maria Rosario Singh-Vergeire and Cynthia P. Saloma |
| EPI_ISL_2189361, EPI_ISL_2189363 | Philippine General Hospital (PGH) | Philippine Genome Center | Francis A. Tablizo, Kenneth M. Kim, Maria Sofia L. Yangzon, Elcid Aaron R. Pangilinan, Renato Jacinto Q. Mantaring, Benedict A. Maralit, Marc Edsel C. Ayes, Eva Maria Cutiongco-de la Paz, Alethea R. de Guzman, Jan Michael C. Yap, Jo-Hannah S. Llames, Sheila Mae M. Araiza, Kris P. Punayan, Irish Coleen A. Asin, Candice Francheska B. Tambaoan, Asia Louisa U. Chong, Karol Sophia Agape R. Padilla, Rianna Patricia S. Cruz, Carlo M. Lapid, El King D. Morado, Joshua Gregor A. Dizon, Razel Nikka M. Hao, Arianne A. Zamora, Devon Ray Pacial, Juan Antonio R. Magalang, Marissa Alejandria, Celia Carlos, Anna Ong-Lim, Edsel Maurice Salvaña, John Q. Wong, Jaime C. Montoya, Maria Rosario Singh-Vergeire and Cynthia P. Saloma |
| EPI_ISL_2189365 | Philippine Red Cross - Port Area | Philippine Genome Center | Francis A. Tablizo, Kenneth M. Kim, Maria Sofia L. Yangzon, Elcid Aaron R. Pangilinan, Renato Jacinto Q. Mantaring, Benedict A. Maralit, Marc Edsel C. Ayes, Eva Maria Cutiongco-de la Paz, Alethea R. de Guzman, Jan Michael C. Yap, Jo-Hannah S. Llames, Sheila Mae M. Araiza, Kris P. Punayan, Irish Coleen A. Asin, Candice Francheska B. Tambaoan, Asia Louisa U. Chong, Karol Sophia Agape R. Padilla, Rianna Patricia S. Cruz, Carlo M. Lapid, El King D. Morado, Joshua Gregor A. Dizon, Razel Nikka M. Hao, Arianne A. Zamora, Devon Ray Pacial, Juan Antonio R. Magalang, Marissa Alejandria, Celia Carlos, Anna Ong-Lim, Edsel Maurice Salvaña, John Q. Wong, Jaime C. Montoya, Maria Rosario Singh-Vergeire and Cynthia P. Saloma |
| EPI_ISL_2189366 | Philippine Red Cross - Clark Molecular Laboratory | Philippine Genome Center | Francis A. Tablizo, Kenneth M. Kim, Maria Sofia L. Yangzon, Elcid Aaron R. Pangilinan, Renato Jacinto Q. Mantaring, Benedict A. Maralit, Marc Edsel C. Ayes, Eva Maria Cutiongco-de la Paz, Alethea R. de Guzman, Jan Michael C. Yap, Jo-Hannah S. Llames, Sheila Mae M. Araiza, Kris P. Punayan, Irish Coleen A. Asin, Candice Francheska B. Tambaoan, Asia Louisa U. Chong, Karol Sophia Agape R. Padilla, Rianna Patricia S. Cruz, Carlo M. Lapid, El King D. Morado, Joshua Gregor A. Dizon, Razel Nikka M. Hao, Arianne A. Zamora, Devon Ray Pacial, Juan Antonio R. Magalang, Marissa Alejandria, Celia Carlos, Anna Ong-Lim, Edsel Maurice Salvaña, John Q. Wong, Jaime C. Montoya, Maria Rosario Singh-Vergeire and Cynthia P. Saloma |
| EPI_ISL_2189367, EPI_ISL_2189368 | The Medical City | Philippine Genome Center | Francis A. Tablizo, Kenneth M. Kim, Maria Sofia L. Yangzon, Elcid Aaron R. Pangilinan, Renato Jacinto Q. Mantaring, Benedict A. Maralit, Marc Edsel C. Ayes, Eva Maria Cutiongco-de la Paz, Alethea R. de Guzman, Jan Michael C. Yap, Jo-Hannah S. Llames, Sheila Mae M. Araiza, Kris P. Punayan, Irish Coleen A. Asin, Candice Francheska B. Tambaoan, Asia Louisa U. Chong, Karol Sophia Agape R. Padilla, Rianna Patricia S. Cruz, Carlo M. Lapid, El King D. Morado, Joshua Gregor A. Dizon, Razel Nikka M. Hao, Arianne A. Zamora, Devon Ray Pacial, Juan Antonio R. Magalang, Marissa Alejandria, Celia Carlos, Anna Ong-Lim, Edsel Maurice Salvaña, John Q. Wong, Jaime C. Montoya, Maria Rosario Singh-Vergeire and Cynthia P. Saloma |
| EPI_ISL_2189369 | First Aide Diagnostic Center | Philippine Genome Center | Francis A. Tablizo, Kenneth M. Kim, Maria Sofia L. Yangzon, Elcid Aaron R. Pangilinan, Renato Jacinto Q. Mantaring, Benedict A. Maralit, Marc Edsel C. Ayes, Eva Maria Cutiongco-de la Paz, Alethea R. de Guzman, Jan Michael C. Yap, Jo-Hannah S. Llames, Sheila Mae M. Araiza, Kris P. Punayan, Irish Coleen A. Asin, Candice Francheska B. Tambaoan, Asia Louisa U. Chong, Karol Sophia Agape R. Padilla, Rianna Patricia S. Cruz, Carlo M. Lapid, El King D. Morado, Joshua Gregor A. Dizon, Razel Nikka M. Hao, Arianne A. Zamora, Devon Ray Pacial, Juan Antonio R. Magalang, Marissa Alejandria, Celia Carlos, Anna Ong-Lim, Edsel Maurice Salvaña, John Q. Wong, Jaime C. Montoya, Maria Rosario Singh-Vergeire and Cynthia P. Saloma |
| EPI_ISL_2189370 | Pasig City Children's Hospital - Child's Hope | Philippine Genome Center | Francis A. Tablizo, Kenneth M. Kim, Maria Sofia L. Yangzon, Elcid Aaron R. Pangilinan, Renato Jacinto Q. Mantaring, Benedict A. Maralit, Marc Edsel C. Ayes, Eva Maria Cutiongco-de la Paz, Alethea R. de Guzman, Jan Michael C. Yap, Jo-Hannah S. Llames, Sheila Mae M. Araiza, Kris P. Punayan, Irish Coleen A. Asin, Candice Francheska B. Tambaoan, Asia Louisa U. Chong, Karol Sophia Agape R. Padilla, Rianna Patricia S. Cruz, Carlo M. Lapid, El King D. Morado, Joshua Gregor A. Dizon, Razel Nikka M. Hao, Arianne A. Zamora, Devon Ray Pacial, Juan Antonio R. Magalang, Marissa Alejandria, Celia Carlos, Anna Ong-Lim, Edsel Maurice Salvaña, John Q. Wong, Jaime C. Montoya, Maria Rosario Singh-Vergeire and Cynthia P. Saloma |
| EPI_ISL_2189371 | The Lord's Grace Medical and Industrial Clinic | Philippine Genome Center | Francis A. Tablizo, Kenneth M. Kim, Maria Sofia L. Yangzon, Elcid Aaron R. Pangilinan, Renato Jacinto Q. Mantaring, Benedict A. Maralit, Marc Edsel C. Ayes, Eva Maria Cutiongco-de la Paz, Alethea R. de Guzman, Jan Michael C. Yap, Jo-Hannah S. Llames, Sheila Mae M. Araiza, Kris P. Punayan, Irish Coleen A. Asin, Candice Francheska B. Tambaoan, Asia Louisa U. Chong, Karol Sophia Agape R. Padilla, Rianna Patricia S. Cruz, Carlo M. Lapid, El King D. Morado, Joshua Gregor A. Dizon, Razel Nikka M. Hao, Arianne A. Zamora, Devon Ray Pacial, Juan Antonio R. Magalang, Marissa Alejandria, Celia Carlos, Anna Ong-Lim, Edsel Maurice Salvaña, John Q. Wong, Jaime C. Montoya, Maria Rosario Singh-Vergeire and Cynthia P. Saloma |
| EPI_ISL_2189372, EPI_ISL_2189375 | The Medical City | Philippine Genome Center | Francis A. Tablizo, Kenneth M. Kim, Maria Sofia L. Yangzon, Elcid Aaron R. Pangilinan, Renato Jacinto Q. Mantaring, Benedict A. Maralit, Marc Edsel C. Ayes, Eva Maria Cutiongco-de la Paz, Alethea R. de Guzman, Jan Michael C. Yap, Jo-Hannah S. Llames, Sheila Mae M. Araiza, Kris P. Punayan, Irish |

| | | | |
|---|---|---|---|
| | | | Coleen A. Asin, Candice Francheska B. Tambaoan, Asia Louisa U. Chong, Karol Sophia Agape R. Padilla, Rianna Patricia S. Cruz, Carlo M. Lapid, El King D. Morado, Joshua Gregor A. Dizon, Razel Nikka M. Hao, Arianne A. Zamora, Devon Ray Pacial, Juan Antonio R. Magalang, Marissa Alejandria, Celia Carlos, Anna Ong-Lim, Edsel Maurice Salvaña, John Q. Wong, Jaime C. Montoya, Maria Rosario Singh-Vergeire and Cynthia P. Saloma |
| EPI_ISL_2189376 | Philippine General Hospital (PGH) | Philippine Genome Center | Francis A. Tablizo, Kenneth M. Kim, Maria Sofia L. Yangzon, Elcid Aaron R. Pangilinan, Renato Jacinto Q. Mantaring, Benedict A. Maralit, Marc Edsel C. Ayes, Eva Maria Cutiongco-de la Paz, Alethea R. de Guzman, Jan Michael C. Yap, Jo-Hannah S. Llames, Sheila Mae M. Araiza, Kris P. Punayan, Irish Coleen A. Asin, Candice Francheska B. Tambaoan, Asia Louisa U. Chong, Karol Sophia Agape R. Padilla, Rianna Patricia S. Cruz, Carlo M. Lapid, El King D. Morado, Joshua Gregor A. Dizon, Razel Nikka M. Hao, Arianne A. Zamora, Devon Ray Pacial, Juan Antonio R. Magalang, Marissa Alejandria, Celia Carlos, Anna Ong-Lim, Edsel Maurice Salvaña, John Q. Wong, Jaime C. Montoya, Maria Rosario Singh-Vergeire and Cynthia P. Saloma |
| EPI_ISL_2189378 | University of the Philippines National Institutes of Health (UP NIH) | Philippine Genome Center | Francis A. Tablizo, Kenneth M. Kim, Maria Sofia L. Yangzon, Elcid Aaron R. Pangilinan, Renato Jacinto Q. Mantaring, Benedict A. Maralit, Marc Edsel C. Ayes, Eva Maria Cutiongco-de la Paz, Alethea R. de Guzman, Jan Michael C. Yap, Jo-Hannah S. Llames, Sheila Mae M. Araiza, Kris P. Punayan, Irish Coleen A. Asin, Candice Francheska B. Tambaoan, Asia Louisa U. Chong, Karol Sophia Agape R. Padilla, Rianna Patricia S. Cruz, Carlo M. Lapid, El King D. Morado, Joshua Gregor A. Dizon, Razel Nikka M. Hao, Arianne A. Zamora, Devon Ray Pacial, Juan Antonio R. Magalang, Marissa Alejandria, Celia Carlos, Anna Ong-Lim, Edsel Maurice Salvaña, John Q. Wong, Jaime C. Montoya, Maria Rosario Singh-Vergeire and Cynthia P. Saloma |
| EPI_ISL_2189379 | The Medical City | Philippine Genome Center | Francis A. Tablizo, Kenneth M. Kim, Maria Sofia L. Yangzon, Elcid Aaron R. Pangilinan, Renato Jacinto Q. Mantaring, Benedict A. Maralit, Marc Edsel C. Ayes, Eva Maria Cutiongco-de la Paz, Alethea R. de Guzman, Jan Michael C. Yap, Jo-Hannah S. Llames, Sheila Mae M. Araiza, Kris P. Punayan, Irish Coleen A. Asin, Candice Francheska B. Tambaoan, Asia Louisa U. Chong, Karol Sophia Agape R. Padilla, Rianna Patricia S. Cruz, Carlo M. Lapid, El King D. Morado, Joshua Gregor A. Dizon, Razel Nikka M. Hao, Arianne A. Zamora, Devon Ray Pacial, Juan Antonio R. Magalang, Marissa Alejandria, Celia Carlos, Anna Ong-Lim, Edsel Maurice Salvaña, John Q. Wong, Jaime C. Montoya, Maria Rosario Singh-Vergeire and Cynthia P. Saloma |
| EPI_ISL_2189381, EPI_ISL_2189383 | Pasig City Children's Hospital - Child's Hope | Philippine Genome Center | Francis A. Tablizo, Kenneth M. Kim, Maria Sofia L. Yangzon, Elcid Aaron R. Pangilinan, Renato Jacinto Q. Mantaring, Benedict A. Maralit, Marc Edsel C. Ayes, Eva Maria Cutiongco-de la Paz, Alethea R. de Guzman, Jan Michael C. Yap, Jo-Hannah S. Llames, Sheila Mae M. Araiza, Kris P. Punayan, Irish Coleen A. Asin, Candice Francheska B. Tambaoan, Asia Louisa U. Chong, Karol Sophia Agape R. Padilla, Rianna Patricia S. Cruz, Carlo M. Lapid, El King D. Morado, Joshua Gregor A. Dizon, Razel Nikka M. Hao, Arianne A. Zamora, Devon Ray Pacial, Juan Antonio R. Magalang, Marissa Alejandria, Celia Carlos, Anna Ong-Lim, Edsel Maurice Salvaña, John Q. Wong, Jaime C. Montoya, Maria Rosario Singh-Vergeire and Cynthia P. Saloma |
| EPI_ISL_2189384 | The Medical City | Philippine Genome Center | Francis A. Tablizo, Kenneth M. Kim, Maria Sofia L. Yangzon, Elcid Aaron R. Pangilinan, Renato Jacinto Q. Mantaring, Benedict A. Maralit, Marc Edsel C. Ayes, Eva Maria Cutiongco-de la Paz, Alethea R. de Guzman, Jan Michael C. Yap, Jo-Hannah S. Llames, Sheila Mae M. Araiza, Kris P. Punayan, Irish Coleen A. Asin, Candice Francheska B. Tambaoan, Asia Louisa U. Chong, Karol Sophia Agape R. Padilla, Rianna Patricia S. Cruz, Carlo M. Lapid, El King D. Morado, Joshua Gregor A. Dizon, Razel Nikka M. Hao, Arianne A. Zamora, Devon Ray Pacial, Juan Antonio R. Magalang, Marissa Alejandria, Celia Carlos, Anna Ong-Lim, Edsel Maurice Salvaña, John Q. Wong, Jaime C. Montoya, Maria Rosario Singh-Vergeire and Cynthia P. Saloma |
| EPI_ISL_2189385 | Pasig City Children's Hospital - Child's Hope | Philippine Genome Center | Francis A. Tablizo, Kenneth M. Kim, Maria Sofia L. Yangzon, Elcid Aaron R. Pangilinan, Renato Jacinto Q. Mantaring, Benedict A. Maralit, Marc Edsel C. Ayes, Eva Maria Cutiongco-de la Paz, Alethea R. de Guzman, Jan Michael C. Yap, Jo-Hannah S. Llames, Sheila Mae M. Araiza, Kris P. Punayan, Irish Coleen A. Asin, Candice Francheska B. Tambaoan, Asia Louisa U. Chong, Karol Sophia Agape R. Padilla, Rianna Patricia S. Cruz, Carlo M. Lapid, El King D. Morado, Joshua Gregor A. Dizon, Razel Nikka M. Hao, Arianne A. Zamora, Devon Ray Pacial, Juan Antonio R. Magalang, Marissa Alejandria, Celia Carlos, Anna Ong-Lim, Edsel Maurice Salvaña, John Q. Wong, Jaime C. Montoya, Maria Rosario Singh-Vergeire and Cynthia P. Saloma |
| EPI_ISL_2189386 | University of the Philippines National Institutes of Health (UP NIH) | Philippine Genome Center | Francis A. Tablizo, Kenneth M. Kim, Maria Sofia L. Yangzon, Elcid Aaron R. Pangilinan, Renato Jacinto Q. Mantaring, Benedict A. Maralit, Marc Edsel C. Ayes, Eva Maria Cutiongco-de la Paz, Alethea R. de Guzman, Jan Michael C. Yap, Jo-Hannah S. Llames, Sheila Mae M. Araiza, Kris P. Punayan, Irish Coleen A. Asin, Candice Francheska B. Tambaoan, Asia Louisa U. Chong, Karol Sophia Agape R. Padilla, Rianna Patricia S. Cruz, Carlo M. Lapid, El King D. Morado, Joshua Gregor A. Dizon, Razel Nikka M. Hao, Arianne A. Zamora, Devon Ray Pacial, Juan Antonio R. Magalang, Marissa Alejandria, Celia Carlos, Anna Ong-Lim, Edsel Maurice Salvaña, John Q. Wong, Jaime C. Montoya, Maria Rosario Singh-Vergeire and Cynthia P. Saloma |
| EPI_ISL_2189391, EPI_ISL_2189392 | The Lord's Grace Medical and Industrial Clinic | Philippine Genome Center | Francis A. Tablizo, Kenneth M. Kim, Maria Sofia L. Yangzon, Elcid Aaron R. Pangilinan, Renato Jacinto Q. Mantaring, Benedict A. Maralit, Marc Edsel C. Ayes, Eva Maria Cutiongco-de la Paz, Alethea R. de Guzman, Jan Michael C. Yap, Jo-Hannah S. Llames, Sheila Mae M. Araiza, Kris P. Punayan, Irish Coleen A. Asin, Candice Francheska B. Tambaoan, Asia Louisa U. Chong, Karol Sophia Agape R. Padilla, Rianna Patricia S. Cruz, Carlo M. Lapid, El King D. Morado, Joshua Gregor A. Dizon, Razel Nikka M. Hao, Arianne A. Zamora, Devon Ray Pacial, Juan Antonio R. Magalang, Marissa Alejandria, Celia Carlos, Anna Ong-Lim, Edsel Maurice Salvaña, John Q. Wong, Jaime C. Montoya, Maria Rosario Singh-Vergeire and Cynthia P. Saloma |
| EPI_ISL_2189393 | Philippine Red Cross - National Blood Center | Philippine Genome Center | Francis A. Tablizo, Kenneth M. Kim, Maria Sofia L. Yangzon, Elcid Aaron R. Pangilinan, Renato Jacinto Q. Mantaring, Benedict A. Maralit, Marc Edsel C. Ayes, Eva Maria Cutiongco-de la Paz, Alethea R. de Guzman, Jan Michael C. Yap, Jo-Hannah S. Llames, Sheila Mae M. Araiza, Kris P. Punayan, Irish Coleen A. Asin, Candice Francheska B. Tambaoan, Asia Louisa U. Chong, Karol Sophia Agape R. Padilla, Rianna Patricia S. Cruz, Carlo M. Lapid, El King D. Morado, Joshua Gregor A. Dizon, Razel Nikka M. Hao, Arianne A. Zamora, Devon Ray Pacial, Juan Antonio R. Magalang, Marissa Alejandria, Celia Carlos, Anna Ong-Lim, Edsel Maurice Salvaña, John Q. Wong, Jaime C. Montoya, Maria Rosario Singh-Vergeire and Cynthia P. Saloma |
| EPI_ISL_2189394 | The Medical City | Philippine Genome Center | Francis A. Tablizo, Kenneth M. Kim, Maria Sofia L. Yangzon, Elcid Aaron R. Pangilinan, Renato Jacinto Q. Mantaring, Benedict A. Maralit, Marc Edsel C. Ayes, Eva Maria Cutiongco-de la Paz, Alethea R. de Guzman, Jan Michael C. Yap, Jo-Hannah S. Llames, Sheila Mae M. Araiza, Kris P. Punayan, Irish Coleen A. Asin, Candice Francheska B. Tambaoan, Asia Louisa U. Chong, Karol Sophia Agape R. Padilla, Rianna Patricia S. Cruz, Carlo M. Lapid, El King D. Morado, Joshua Gregor A. Dizon, Razel Nikka M. Hao, Arianne A. Zamora, Devon Ray Pacial, Juan Antonio R. Magalang, Marissa Alejandria, Celia Carlos, Anna Ong-Lim, Edsel Maurice Salvaña, John Q. Wong, Jaime C. Montoya, Maria Rosario Singh-Vergeire and Cynthia P. Saloma |
| EPI_ISL_2189395 | Philippine General Hospital (PGH) | Philippine Genome Center | Francis A. Tablizo, Kenneth M. Kim, Maria Sofia L. Yangzon, Elcid Aaron R. Pangilinan, Renato Jacinto Q. Mantaring, Benedict A. Maralit, Marc Edsel C. Ayes, Eva Maria Cutiongco-de la Paz, Alethea R. de Guzman, Jan Michael C. Yap, Jo-Hannah S. Llames, Sheila Mae M. Araiza, Kris P. Punayan, Irish Coleen A. Asin, Candice Francheska B. Tambaoan, Asia Louisa U. Chong, Karol Sophia Agape R. Padilla, Rianna Patricia S. Cruz, Carlo M. Lapid, El King D. Morado, Joshua Gregor A. Dizon, Razel Nikka M. Hao, Arianne A. Zamora, Devon Ray Pacial, Juan Antonio R. Magalang, Marissa Alejandria, Celia Carlos, Anna Ong-Lim, Edsel Maurice Salvaña, John Q. Wong, Jaime C. Montoya, Maria Rosario Singh-Vergeire and Cynthia P. Saloma |
| EPI_ISL_2189396 | South Super Highway Molecular Diagnostic Laboratory | Philippine Genome Center | Francis A. Tablizo, Kenneth M. Kim, Maria Sofia L. Yangzon, Elcid Aaron R. Pangilinan, Renato Jacinto Q. Mantaring, Benedict A. Maralit, Marc Edsel C. Ayes, Eva Maria Cutiongco-de la Paz, Alethea R. de Guzman, Jan Michael C. Yap, Jo-Hannah S. Llames, Sheila Mae M. Araiza, Kris P. Punayan, Irish Coleen A. Asin, Candice Francheska B. Tambaoan, Asia Louisa U. Chong, Karol Sophia Agape R. Padilla, Rianna Patricia S. Cruz, Carlo M. Lapid, El King D. Morado, Joshua Gregor A. Dizon, Razel Nikka M. Hao, Arianne A. Zamora, Devon Ray Pacial, Juan Antonio R. Magalang, Marissa Alejandria, Celia Carlos, Anna Ong-Lim, Edsel Maurice Salvaña, John Q. Wong, Jaime C. Montoya, Maria Rosario Singh-Vergeire and Cynthia P. Saloma |
| EPI_ISL_2189397 | Safeguard DNA Diagnostic, Inc. | Philippine Genome Center | Francis A. Tablizo, Kenneth M. Kim, Maria Sofia L. Yangzon, Elcid Aaron R. Pangilinan, Renato Jacinto Q. Mantaring, Benedict A. Maralit, Marc Edsel C. Ayes, Eva Maria Cutiongco-de la Paz, Alethea R. de Guzman, Jan Michael C. Yap, Jo-Hannah S. Llames, Sheila Mae M. Araiza, Kris P. Punayan, Irish Coleen A. Asin, Candice Francheska B. Tambaoan, Asia Louisa U. Chong, Karol Sophia Agape R. Padilla, Rianna Patricia S. Cruz, Carlo M. Lapid, El King D. Morado, Joshua Gregor A. Dizon, Razel Nikka M. Hao, Arianne A. Zamora, Devon Ray Pacial, Juan Antonio R. Magalang, Marissa Alejandria, Celia Carlos, Anna Ong-Lim, Edsel Maurice Salvaña, John Q. Wong, Jaime C. Montoya, Maria Rosario Singh-Vergeire and Cynthia P. Saloma |
| EPI_ISL_2189398 | Laguna Provincial Hospital - San Pablo District Hospital | Philippine Genome Center | Francis A. Tablizo, Kenneth M. Kim, Maria Sofia L. Yangzon, Elcid Aaron R. Pangilinan, Renato Jacinto Q. Mantaring, Benedict A. Maralit, Marc Edsel C. Ayes, Eva Maria Cutiongco-de la Paz, Alethea R. de Guzman, Jan Michael C. Yap, Jo-Hannah S. Llames, Sheila Mae M. Araiza, Kris P. Punayan, Irish Coleen A. Asin, Candice Francheska B. Tambaoan, Asia Louisa U. Chong, Karol Sophia Agape R. Padilla, Rianna Patricia S. Cruz, Carlo M. Lapid, El King D. Morado, Joshua Gregor A. Dizon, Razel Nikka M. Hao, Arianne A. Zamora, Devon Ray Pacial, Juan Antonio R. Magalang, Marissa Alejandria, Celia Carlos, Anna Ong-Lim, Edsel Maurice Salvaña, John Q. Wong, Jaime C. Montoya, Maria Rosario Singh-Vergeire and Cynthia P. Saloma |
| EPI_ISL_2189399 | First Aide Diagnostic Center | Philippine Genome Center | Francis A. Tablizo, Kenneth M. Kim, Maria Sofia L. Yangzon, Elcid Aaron R. Pangilinan, Renato Jacinto Q. Mantaring, Benedict A. Maralit, Marc Edsel C. Ayes, Eva Maria Cutiongco-de la Paz, Alethea R. de Guzman, Jan Michael C. Yap, Jo-Hannah S. Llames, Sheila Mae M. Araiza, Kris P. Punayan, Irish Coleen A. Asin, Candice Francheska B. Tambaoan, Asia Louisa U. Chong, Karol Sophia Agape R. Padilla, Rianna Patricia S. Cruz, Carlo M. Lapid, El King D. Morado, Joshua Gregor A. Dizon, Razel Nikka M. Hao, Arianne A. Zamora, Devon Ray Pacial, Juan Antonio R. Magalang, Marissa Alejandria, Celia Carlos, Anna Ong-Lim, Edsel Maurice Salvaña, John Q. Wong, Jaime C. Montoya, Maria Rosario Singh-Vergeire and Cynthia P. Saloma |
| EPI_ISL_2189400 | The Lord's Grace Medical and Industrial Clinic | Philippine Genome Center | Francis A. Tablizo, Kenneth M. Kim, Maria Sofia L. Yangzon, Elcid Aaron R. Pangilinan, Renato Jacinto Q. Mantaring, Benedict A. Maralit, Marc Edsel C. |

| | | | |
|---|---|---|---|
| | | | Ayes, Eva Maria Cutiongco-de la Paz, Alethea R. de Guzman, Jan Michael C. Yap, Jo-Hannah S. Llames, Sheila Mae M. Araiza, Kris P. Punayan, Irish Coleen A. Asin, Candice Francheska B. Tambaoan, Asia Louisa U. Chong, Karol Sophia Agape R. Padilla, Rianna Patricia S. Cruz, Carlo M. Lapid, El King D. Morado, Joshua Gregor A. Dizon, Razel Nikka M. Hao, Arianne A. Zamora, Devon Ray Pacial, Juan Antonio R. Magalang, Marissa Alejandria, Celia Carlos, Anna Ong-Lim, Edsel Maurice Salvaña, John Q. Wong, Jaime C. Montoya, Maria Rosario Singh-Vergeire and Cynthia P. Saloma |
| EPI_ISL_2189402 | Sta. Rosa CHO- GENEXPERT LABORATORY | Philippine Genome Center | Francis A. Tablizo, Kenneth M. Kim, Maria Sofia L. Yangzon, Elcid Aaron R. Pangilinan, Renato Jacinto Q. Mantaring, Benedict A. Maralit, Marc Edsel C. Ayes, Eva Maria Cutiongco-de la Paz, Alethea R. de Guzman, Jan Michael C. Yap, Jo-Hannah S. Llames, Sheila Mae M. Araiza, Kris P. Punayan, Irish Coleen A. Asin, Candice Francheska B. Tambaoan, Asia Louisa U. Chong, Karol Sophia Agape R. Padilla, Rianna Patricia S. Cruz, Carlo M. Lapid, El King D. Morado, Joshua Gregor A. Dizon, Razel Nikka M. Hao, Arianne A. Zamora, Devon Ray Pacial, Juan Antonio R. Magalang, Marissa Alejandria, Celia Carlos, Anna Ong-Lim, Edsel Maurice Salvaña, John Q. Wong, Jaime C. Montoya, Maria Rosario Singh-Vergeire and Cynthia P. Saloma |
| EPI_ISL_2189404 | The Medical City | Philippine Genome Center | Francis A. Tablizo, Kenneth M. Kim, Maria Sofia L. Yangzon, Elcid Aaron R. Pangilinan, Renato Jacinto Q. Mantaring, Benedict A. Maralit, Marc Edsel C. Ayes, Eva Maria Cutiongco-de la Paz, Alethea R. de Guzman, Jan Michael C. Yap, Jo-Hannah S. Llames, Sheila Mae M. Araiza, Kris P. Punayan, Irish Coleen A. Asin, Candice Francheska B. Tambaoan, Asia Louisa U. Chong, Karol Sophia Agape R. Padilla, Rianna Patricia S. Cruz, Carlo M. Lapid, El King D. Morado, Joshua Gregor A. Dizon, Razel Nikka M. Hao, Arianne A. Zamora, Devon Ray Pacial, Juan Antonio R. Magalang, Marissa Alejandria, Celia Carlos, Anna Ong-Lim, Edsel Maurice Salvaña, John Q. Wong, Jaime C. Montoya, Maria Rosario Singh-Vergeire and Cynthia P. Saloma |
| EPI_ISL_2189406 | The Lord's Grace Medical and Industrial Clinic | Philippine Genome Center | Francis A. Tablizo, Kenneth M. Kim, Maria Sofia L. Yangzon, Elcid Aaron R. Pangilinan, Renato Jacinto Q. Mantaring, Benedict A. Maralit, Marc Edsel C. Ayes, Eva Maria Cutiongco-de la Paz, Alethea R. de Guzman, Jan Michael C. Yap, Jo-Hannah S. Llames, Sheila Mae M. Araiza, Kris P. Punayan, Irish Coleen A. Asin, Candice Francheska B. Tambaoan, Asia Louisa U. Chong, Karol Sophia Agape R. Padilla, Rianna Patricia S. Cruz, Carlo M. Lapid, El King D. Morado, Joshua Gregor A. Dizon, Razel Nikka M. Hao, Arianne A. Zamora, Devon Ray Pacial, Juan Antonio R. Magalang, Marissa Alejandria, Celia Carlos, Anna Ong-Lim, Edsel Maurice Salvaña, John Q. Wong, Jaime C. Montoya, Maria Rosario Singh-Vergeire and Cynthia P. Saloma |
| EPI_ISL_2189407 | Philippine Red Cross - National Blood Center | Philippine Genome Center | Francis A. Tablizo, Kenneth M. Kim, Maria Sofia L. Yangzon, Elcid Aaron R. Pangilinan, Renato Jacinto Q. Mantaring, Benedict A. Maralit, Marc Edsel C. Ayes, Eva Maria Cutiongco-de la Paz, Alethea R. de Guzman, Jan Michael C. Yap, Jo-Hannah S. Llames, Sheila Mae M. Araiza, Kris P. Punayan, Irish Coleen A. Asin, Candice Francheska B. Tambaoan, Asia Louisa U. Chong, Karol Sophia Agape R. Padilla, Rianna Patricia S. Cruz, Carlo M. Lapid, El King D. Morado, Joshua Gregor A. Dizon, Razel Nikka M. Hao, Arianne A. Zamora, Devon Ray Pacial, Juan Antonio R. Magalang, Marissa Alejandria, Celia Carlos, Anna Ong-Lim, Edsel Maurice Salvaña, John Q. Wong, Jaime C. Montoya, Maria Rosario Singh-Vergeire and Cynthia P. Saloma |
| EPI_ISL_2189408 | The Medical City | Philippine Genome Center | Francis A. Tablizo, Kenneth M. Kim, Maria Sofia L. Yangzon, Elcid Aaron R. Pangilinan, Renato Jacinto Q. Mantaring, Benedict A. Maralit, Marc Edsel C. Ayes, Eva Maria Cutiongco-de la Paz, Alethea R. de Guzman, Jan Michael C. Yap, Jo-Hannah S. Llames, Sheila Mae M. Araiza, Kris P. Punayan, Irish Coleen A. Asin, Candice Francheska B. Tambaoan, Asia Louisa U. Chong, Karol Sophia Agape R. Padilla, Rianna Patricia S. Cruz, Carlo M. Lapid, El King D. Morado, Joshua Gregor A. Dizon, Razel Nikka M. Hao, Arianne A. Zamora, Devon Ray Pacial, Juan Antonio R. Magalang, Marissa Alejandria, Celia Carlos, Anna Ong-Lim, Edsel Maurice Salvaña, John Q. Wong, Jaime C. Montoya, Maria Rosario Singh-Vergeire and Cynthia P. Saloma |
| EPI_ISL_2189410 | IOM- Manila Health Centre Laboratory | Philippine Genome Center | Francis A. Tablizo, Kenneth M. Kim, Maria Sofia L. Yangzon, Elcid Aaron R. Pangilinan, Renato Jacinto Q. Mantaring, Benedict A. Maralit, Marc Edsel C. Ayes, Eva Maria Cutiongco-de la Paz, Alethea R. de Guzman, Jan Michael C. Yap, Jo-Hannah S. Llames, Sheila Mae M. Araiza, Kris P. Punayan, Irish Coleen A. Asin, Candice Francheska B. Tambaoan, Asia Louisa U. Chong, Karol Sophia Agape R. Padilla, Rianna Patricia S. Cruz, Carlo M. Lapid, El King D. Morado, Joshua Gregor A. Dizon, Razel Nikka M. Hao, Arianne A. Zamora, Devon Ray Pacial, Juan Antonio R. Magalang, Marissa Alejandria, Celia Carlos, Anna Ong-Lim, Edsel Maurice Salvaña, John Q. Wong, Jaime C. Montoya, Maria Rosario Singh-Vergeire and Cynthia P. Saloma |
| EPI_ISL_2189411 | Qualimed Hospital Sta. Rosa | Philippine Genome Center | Francis A. Tablizo, Kenneth M. Kim, Maria Sofia L. Yangzon, Elcid Aaron R. Pangilinan, Renato Jacinto Q. Mantaring, Benedict A. Maralit, Marc Edsel C. Ayes, Eva Maria Cutiongco-de la Paz, Alethea R. de Guzman, Jan Michael C. Yap, Jo-Hannah S. Llames, Sheila Mae M. Araiza, Kris P. Punayan, Irish Coleen A. Asin, Candice Francheska B. Tambaoan, Asia Louisa U. Chong, Karol Sophia Agape R. Padilla, Rianna Patricia S. Cruz, Carlo M. Lapid, El King D. Morado, Joshua Gregor A. Dizon, Razel Nikka M. Hao, Arianne A. Zamora, Devon Ray Pacial, Juan Antonio R. Magalang, Marissa Alejandria, Celia Carlos, Anna Ong-Lim, Edsel Maurice Salvaña, John Q. Wong, Jaime C. Montoya, Maria Rosario Singh-Vergeire and Cynthia P. Saloma |
| EPI_ISL_2189412 | The Lord's Grace Medical and Industrial Clinic | Philippine Genome Center | Francis A. Tablizo, Kenneth M. Kim, Maria Sofia L. Yangzon, Elcid Aaron R. Pangilinan, Renato Jacinto Q. Mantaring, Benedict A. Maralit, Marc Edsel C. Ayes, Eva Maria Cutiongco-de la Paz, Alethea R. de Guzman, Jan Michael C. Yap, Jo-Hannah S. Llames, Sheila Mae M. Araiza, Kris P. Punayan, Irish Coleen A. Asin, Candice Francheska B. Tambaoan, Asia Louisa U. Chong, Karol Sophia Agape R. Padilla, Rianna Patricia S. Cruz, Carlo M. Lapid, El King D. Morado, Joshua Gregor A. Dizon, Razel Nikka M. Hao, Arianne A. Zamora, Devon Ray Pacial, Juan Antonio R. Magalang, Marissa Alejandria, Celia Carlos, Anna Ong-Lim, Edsel Maurice Salvaña, John Q. Wong, Jaime C. Montoya, Maria Rosario Singh-Vergeire and Cynthia P. Saloma |
| EPI_ISL_2189413 | Laguna Provincial Hospital - San Pablo District Hospital | Philippine Genome Center | Francis A. Tablizo, Kenneth M. Kim, Maria Sofia L. Yangzon, Elcid Aaron R. Pangilinan, Renato Jacinto Q. Mantaring, Benedict A. Maralit, Marc Edsel C. Ayes, Eva Maria Cutiongco-de la Paz, Alethea R. de Guzman, Jan Michael C. Yap, Jo-Hannah S. Llames, Sheila Mae M. Araiza, Kris P. Punayan, Irish Coleen A. Asin, Candice Francheska B. Tambaoan, Asia Louisa U. Chong, Karol Sophia Agape R. Padilla, Rianna Patricia S. Cruz, Carlo M. Lapid, El King D. Morado, Joshua Gregor A. Dizon, Razel Nikka M. Hao, Arianne A. Zamora, Devon Ray Pacial, Juan Antonio R. Magalang, Marissa Alejandria, Celia Carlos, Anna Ong-Lim, Edsel Maurice Salvaña, John Q. Wong, Jaime C. Montoya, Maria Rosario Singh-Vergeire and Cynthia P. Saloma |
| EPI_ISL_2189415 | The Lord's Grace Medical and Industrial Clinic | Philippine Genome Center | Francis A. Tablizo, Kenneth M. Kim, Maria Sofia L. Yangzon, Elcid Aaron R. Pangilinan, Renato Jacinto Q. Mantaring, Benedict A. Maralit, Marc Edsel C. Ayes, Eva Maria Cutiongco-de la Paz, Alethea R. de Guzman, Jan Michael C. Yap, Jo-Hannah S. Llames, Sheila Mae M. Araiza, Kris P. Punayan, Irish Coleen A. Asin, Candice Francheska B. Tambaoan, Asia Louisa U. Chong, Karol Sophia Agape R. Padilla, Rianna Patricia S. Cruz, Carlo M. Lapid, El King D. Morado, Joshua Gregor A. Dizon, Razel Nikka M. Hao, Arianne A. Zamora, Devon Ray Pacial, Juan Antonio R. Magalang, Marissa Alejandria, Celia Carlos, Anna Ong-Lim, Edsel Maurice Salvaña, John Q. Wong, Jaime C. Montoya, Maria Rosario Singh-Vergeire and Cynthia P. Saloma |
| EPI_ISL_2189416 | Pasig City Children's Hospital - Child's Hope | Philippine Genome Center | Francis A. Tablizo, Kenneth M. Kim, Maria Sofia L. Yangzon, Elcid Aaron R. Pangilinan, Renato Jacinto Q. Mantaring, Benedict A. Maralit, Marc Edsel C. Ayes, Eva Maria Cutiongco-de la Paz, Alethea R. de Guzman, Jan Michael C. Yap, Jo-Hannah S. Llames, Sheila Mae M. Araiza, Kris P. Punayan, Irish Coleen A. Asin, Candice Francheska B. Tambaoan, Asia Louisa U. Chong, Karol Sophia Agape R. Padilla, Rianna Patricia S. Cruz, Carlo M. Lapid, El King D. Morado, Joshua Gregor A. Dizon, Razel Nikka M. Hao, Arianne A. Zamora, Devon Ray Pacial, Juan Antonio R. Magalang, Marissa Alejandria, Celia Carlos, Anna Ong-Lim, Edsel Maurice Salvaña, John Q. Wong, Jaime C. Montoya, Maria Rosario Singh-Vergeire and Cynthia P. Saloma |
| EPI_ISL_2189418 | The Medical City | Philippine Genome Center | Francis A. Tablizo, Kenneth M. Kim, Maria Sofia L. Yangzon, Elcid Aaron R. Pangilinan, Renato Jacinto Q. Mantaring, Benedict A. Maralit, Marc Edsel C. Ayes, Eva Maria Cutiongco-de la Paz, Alethea R. de Guzman, Jan Michael C. Yap, Jo-Hannah S. Llames, Sheila Mae M. Araiza, Kris P. Punayan, Irish Coleen A. Asin, Candice Francheska B. Tambaoan, Asia Louisa U. Chong, Karol Sophia Agape R. Padilla, Rianna Patricia S. Cruz, Carlo M. Lapid, El King D. Morado, Joshua Gregor A. Dizon, Razel Nikka M. Hao, Arianne A. Zamora, Devon Ray Pacial, Juan Antonio R. Magalang, Marissa Alejandria, Celia Carlos, Anna Ong-Lim, Edsel Maurice Salvaña, John Q. Wong, Jaime C. Montoya, Maria Rosario Singh-Vergeire and Cynthia P. Saloma |
| EPI_ISL_2189422 | Sta. Ana Hospital | Philippine Genome Center | Francis A. Tablizo, Kenneth M. Kim, Maria Sofia L. Yangzon, Elcid Aaron R. Pangilinan, Renato Jacinto Q. Mantaring, Benedict A. Maralit, Marc Edsel C. Ayes, Eva Maria Cutiongco-de la Paz, Alethea R. de Guzman, Jan Michael C. Yap, Jo-Hannah S. Llames, Sheila Mae M. Araiza, Kris P. Punayan, Irish Coleen A. Asin, Candice Francheska B. Tambaoan, Asia Louisa U. Chong, Karol Sophia Agape R. Padilla, Rianna Patricia S. Cruz, Carlo M. Lapid, El King D. Morado, Joshua Gregor A. Dizon, Razel Nikka M. Hao, Arianne A. Zamora, Devon Ray Pacial, Juan Antonio R. Magalang, Marissa Alejandria, Celia Carlos, Anna Ong-Lim, Edsel Maurice Salvaña, John Q. Wong, Jaime C. Montoya, Maria Rosario Singh-Vergeire and Cynthia P. Saloma |
| EPI_ISL_2189423 | University of the Philippines National Institutes of Health (UP NIH) | Philippine Genome Center | Francis A. Tablizo, Kenneth M. Kim, Maria Sofia L. Yangzon, Elcid Aaron R. Pangilinan, Renato Jacinto Q. Mantaring, Benedict A. Maralit, Marc Edsel C. Ayes, Eva Maria Cutiongco-de la Paz, Alethea R. de Guzman, Jan Michael C. Yap, Jo-Hannah S. Llames, Sheila Mae M. Araiza, Kris P. Punayan, Irish Coleen A. Asin, Candice Francheska B. Tambaoan, Asia Louisa U. Chong, Karol Sophia Agape R. Padilla, Rianna Patricia S. Cruz, Carlo M. Lapid, El King D. Morado, Joshua Gregor A. Dizon, Razel Nikka M. Hao, Arianne A. Zamora, Devon Ray Pacial, Juan Antonio R. Magalang, Marissa Alejandria, Celia Carlos, Anna Ong-Lim, Edsel Maurice Salvaña, John Q. Wong, Jaime C. Montoya, Maria Rosario Singh-Vergeire and Cynthia P. Saloma |
| EPI_ISL_2189425 | The Medical City | Philippine Genome Center | Francis A. Tablizo, Kenneth M. Kim, Maria Sofia L. Yangzon, Elcid Aaron R. Pangilinan, Renato Jacinto Q. Mantaring, Benedict A. Maralit, Marc Edsel C. Ayes, Eva Maria Cutiongco-de la Paz, Alethea R. de Guzman, Jan Michael C. Yap, Jo-Hannah S. Llames, Sheila Mae M. Araiza, Kris P. Punayan, Irish Coleen A. Asin, Candice Francheska B. Tambaoan, Asia Louisa U. Chong, Karol Sophia Agape R. Padilla, Rianna Patricia S. Cruz, Carlo M. Lapid, El King D. Morado, Joshua Gregor A. Dizon, Razel Nikka M. Hao, Arianne A. Zamora, Devon Ray Pacial, Juan Antonio R. Magalang, Marissa Alejandria, Celia Carlos, Anna Ong-Lim, Edsel Maurice Salvaña, John Q. Wong, Jaime C. Montoya, Maria Rosario Singh-Vergeire and Cynthia P. Saloma |

| | | | |
|---|---|---|---|
| EPI_ISL_2189427, EPI_ISL_2189428, EPI_ISL_2189429 | The Lord's Grace Medical and Industrial Clinic | Philippine Genome Center | Francis A. Tablizo, Kenneth M. Kim, Maria Sofia L. Yangzon, Elcid Aaron R. Pangilinan, Renato Jacinto Q. Mantaring, Benedict A. Maralit, Marc Edsel C. Ayes, Eva Maria Cutiongco-de la Paz, Alethea R. de Guzman, Jan Michael C. Yap, Jo-Hannah S. Llames, Sheila Mae M. Araiza, Kris P. Punayan, Irish Coleen A. Asin, Candice Francheska B. Tambaoan, Asia Louisa U. Chong, Karol Sophia Agape R. Padilla, Rianna Patricia S. Cruz, Carlo M. Lapid, El King D. Morado, Joshua Gregor A. Dizon, Razel Nikka M. Hao, Arianne A. Zamora, Devon Ray Pacial, Juan Antonio R. Magalang, Marissa Alejandria, Celia Carlos, Anna Ong-Lim, Edsel Maurice Salvaña, John Q. Wong, Jaime C. Montoya, Maria Rosario Singh-Vergeire and Cynthia P. Saloma |
| EPI_ISL_2189431 | The Medical City | Philippine Genome Center | Francis A. Tablizo, Kenneth M. Kim, Maria Sofia L. Yangzon, Elcid Aaron R. Pangilinan, Renato Jacinto Q. Mantaring, Benedict A. Maralit, Marc Edsel C. Ayes, Eva Maria Cutiongco-de la Paz, Alethea R. de Guzman, Jan Michael C. Yap, Jo-Hannah S. Llames, Sheila Mae M. Araiza, Kris P. Punayan, Irish Coleen A. Asin, Candice Francheska B. Tambaoan, Asia Louisa U. Chong, Karol Sophia Agape R. Padilla, Rianna Patricia S. Cruz, Carlo M. Lapid, El King D. Morado, Joshua Gregor A. Dizon, Razel Nikka M. Hao, Arianne A. Zamora, Devon Ray Pacial, Juan Antonio R. Magalang, Marissa Alejandria, Celia Carlos, Anna Ong-Lim, Edsel Maurice Salvaña, John Q. Wong, Jaime C. Montoya, Maria Rosario Singh-Vergeire and Cynthia P. Saloma |
| EPI_ISL_2189432 | Pasig City Children's Hospital - Child's Hope | Philippine Genome Center | Francis A. Tablizo, Kenneth M. Kim, Maria Sofia L. Yangzon, Elcid Aaron R. Pangilinan, Renato Jacinto Q. Mantaring, Benedict A. Maralit, Marc Edsel C. Ayes, Eva Maria Cutiongco-de la Paz, Alethea R. de Guzman, Jan Michael C. Yap, Jo-Hannah S. Llames, Sheila Mae M. Araiza, Kris P. Punayan, Irish Coleen A. Asin, Candice Francheska B. Tambaoan, Asia Louisa U. Chong, Karol Sophia Agape R. Padilla, Rianna Patricia S. Cruz, Carlo M. Lapid, El King D. Morado, Joshua Gregor A. Dizon, Razel Nikka M. Hao, Arianne A. Zamora, Devon Ray Pacial, Juan Antonio R. Magalang, Marissa Alejandria, Celia Carlos, Anna Ong-Lim, Edsel Maurice Salvaña, John Q. Wong, Jaime C. Montoya, Maria Rosario Singh-Vergeire and Cynthia P. Saloma |
| EPI_ISL_2189433 | Sta. Rosa CHO- GENEXPERT LABORATORY | Philippine Genome Center | Francis A. Tablizo, Kenneth M. Kim, Maria Sofia L. Yangzon, Elcid Aaron R. Pangilinan, Renato Jacinto Q. Mantaring, Benedict A. Maralit, Marc Edsel C. Ayes, Eva Maria Cutiongco-de la Paz, Alethea R. de Guzman, Jan Michael C. Yap, Jo-Hannah S. Llames, Sheila Mae M. Araiza, Kris P. Punayan, Irish Coleen A. Asin, Candice Francheska B. Tambaoan, Asia Louisa U. Chong, Karol Sophia Agape R. Padilla, Rianna Patricia S. Cruz, Carlo M. Lapid, El King D. Morado, Joshua Gregor A. Dizon, Razel Nikka M. Hao, Arianne A. Zamora, Devon Ray Pacial, Juan Antonio R. Magalang, Marissa Alejandria, Celia Carlos, Anna Ong-Lim, Edsel Maurice Salvaña, John Q. Wong, Jaime C. Montoya, Maria Rosario Singh-Vergeire and Cynthia P. Saloma |
| EPI_ISL_2189435 | The Medical City | Philippine Genome Center | Francis A. Tablizo, Kenneth M. Kim, Maria Sofia L. Yangzon, Elcid Aaron R. Pangilinan, Renato Jacinto Q. Mantaring, Benedict A. Maralit, Marc Edsel C. Ayes, Eva Maria Cutiongco-de la Paz, Alethea R. de Guzman, Jan Michael C. Yap, Jo-Hannah S. Llames, Sheila Mae M. Araiza, Kris P. Punayan, Irish Coleen A. Asin, Candice Francheska B. Tambaoan, Asia Louisa U. Chong, Karol Sophia Agape R. Padilla, Rianna Patricia S. Cruz, Carlo M. Lapid, El King D. Morado, Joshua Gregor A. Dizon, Razel Nikka M. Hao, Arianne A. Zamora, Devon Ray Pacial, Juan Antonio R. Magalang, Marissa Alejandria, Celia Carlos, Anna Ong-Lim, Edsel Maurice Salvaña, John Q. Wong, Jaime C. Montoya, Maria Rosario Singh-Vergeire and Cynthia P. Saloma |
| EPI_ISL_2189437 | Pasig City Children's Hospital - Child's Hope | Philippine Genome Center | Francis A. Tablizo, Kenneth M. Kim, Maria Sofia L. Yangzon, Elcid Aaron R. Pangilinan, Renato Jacinto Q. Mantaring, Benedict A. Maralit, Marc Edsel C. Ayes, Eva Maria Cutiongco-de la Paz, Alethea R. de Guzman, Jan Michael C. Yap, Jo-Hannah S. Llames, Sheila Mae M. Araiza, Kris P. Punayan, Irish Coleen A. Asin, Candice Francheska B. Tambaoan, Asia Louisa U. Chong, Karol Sophia Agape R. Padilla, Rianna Patricia S. Cruz, Carlo M. Lapid, El King D. Morado, Joshua Gregor A. Dizon, Razel Nikka M. Hao, Arianne A. Zamora, Devon Ray Pacial, Juan Antonio R. Magalang, Marissa Alejandria, Celia Carlos, Anna Ong-Lim, Edsel Maurice Salvaña, John Q. Wong, Jaime C. Montoya, Maria Rosario Singh-Vergeire and Cynthia P. Saloma |
| EPI_ISL_2189439 | The Medical City | Philippine Genome Center | Francis A. Tablizo, Kenneth M. Kim, Maria Sofia L. Yangzon, Elcid Aaron R. Pangilinan, Renato Jacinto Q. Mantaring, Benedict A. Maralit, Marc Edsel C. Ayes, Eva Maria Cutiongco-de la Paz, Alethea R. de Guzman, Jan Michael C. Yap, Jo-Hannah S. Llames, Sheila Mae M. Araiza, Kris P. Punayan, Irish Coleen A. Asin, Candice Francheska B. Tambaoan, Asia Louisa U. Chong, Karol Sophia Agape R. Padilla, Rianna Patricia S. Cruz, Carlo M. Lapid, El King D. Morado, Joshua Gregor A. Dizon, Razel Nikka M. Hao, Arianne A. Zamora, Devon Ray Pacial, Juan Antonio R. Magalang, Marissa Alejandria, Celia Carlos, Anna Ong-Lim, Edsel Maurice Salvaña, John Q. Wong, Jaime C. Montoya, Maria Rosario Singh-Vergeire and Cynthia P. Saloma |
| EPI_ISL_2189441 | Mary Mediatrix Medical Center | Philippine Genome Center | Francis A. Tablizo, Kenneth M. Kim, Maria Sofia L. Yangzon, Elcid Aaron R. Pangilinan, Renato Jacinto Q. Mantaring, Benedict A. Maralit, Marc Edsel C. Ayes, Eva Maria Cutiongco-de la Paz, Alethea R. de Guzman, Jan Michael C. Yap, Jo-Hannah S. Llames, Sheila Mae M. Araiza, Kris P. Punayan, Irish Coleen A. Asin, Candice Francheska B. Tambaoan, Asia Louisa U. Chong, Karol Sophia Agape R. Padilla, Rianna Patricia S. Cruz, Carlo M. Lapid, El King D. Morado, Joshua Gregor A. Dizon, Razel Nikka M. Hao, Arianne A. Zamora, Devon Ray Pacial, Juan Antonio R. Magalang, Marissa Alejandria, Celia Carlos, Anna Ong-Lim, Edsel Maurice Salvaña, John Q. Wong, Jaime C. Montoya, Maria Rosario Singh-Vergeire and Cynthia P. Saloma |
| EPI_ISL_2189444 | The Lord's Grace Medical and Industrial Clinic | Philippine Genome Center | Francis A. Tablizo, Kenneth M. Kim, Maria Sofia L. Yangzon, Elcid Aaron R. Pangilinan, Renato Jacinto Q. Mantaring, Benedict A. Maralit, Marc Edsel C. Ayes, Eva Maria Cutiongco-de la Paz, Alethea R. de Guzman, Jan Michael C. Yap, Jo-Hannah S. Llames, Sheila Mae M. Araiza, Kris P. Punayan, Irish Coleen A. Asin, Candice Francheska B. Tambaoan, Asia Louisa U. Chong, Karol Sophia Agape R. Padilla, Rianna Patricia S. Cruz, Carlo M. Lapid, El King D. Morado, Joshua Gregor A. Dizon, Razel Nikka M. Hao, Arianne A. Zamora, Devon Ray Pacial, Juan Antonio R. Magalang, Marissa Alejandria, Celia Carlos, Anna Ong-Lim, Edsel Maurice Salvaña, John Q. Wong, Jaime C. Montoya, Maria Rosario Singh-Vergeire and Cynthia P. Saloma |
| EPI_ISL_2189448 | Qualimed Hospital Sta. Rosa | Philippine Genome Center | Francis A. Tablizo, Kenneth M. Kim, Maria Sofia L. Yangzon, Elcid Aaron R. Pangilinan, Renato Jacinto Q. Mantaring, Benedict A. Maralit, Marc Edsel C. Ayes, Eva Maria Cutiongco-de la Paz, Alethea R. de Guzman, Jan Michael C. Yap, Jo-Hannah S. Llames, Sheila Mae M. Araiza, Kris P. Punayan, Irish Coleen A. Asin, Candice Francheska B. Tambaoan, Asia Louisa U. Chong, Karol Sophia Agape R. Padilla, Rianna Patricia S. Cruz, Carlo M. Lapid, El King D. Morado, Joshua Gregor A. Dizon, Razel Nikka M. Hao, Arianne A. Zamora, Devon Ray Pacial, Juan Antonio R. Magalang, Marissa Alejandria, Celia Carlos, Anna Ong-Lim, Edsel Maurice Salvaña, John Q. Wong, Jaime C. Montoya, Maria Rosario Singh-Vergeire and Cynthia P. Saloma |
| EPI_ISL_2189449 | Sta. Ana Hospital | Philippine Genome Center | Francis A. Tablizo, Kenneth M. Kim, Maria Sofia L. Yangzon, Elcid Aaron R. Pangilinan, Renato Jacinto Q. Mantaring, Benedict A. Maralit, Marc Edsel C. Ayes, Eva Maria Cutiongco-de la Paz, Alethea R. de Guzman, Jan Michael C. Yap, Jo-Hannah S. Llames, Sheila Mae M. Araiza, Kris P. Punayan, Irish Coleen A. Asin, Candice Francheska B. Tambaoan, Asia Louisa U. Chong, Karol Sophia Agape R. Padilla, Rianna Patricia S. Cruz, Carlo M. Lapid, El King D. Morado, Joshua Gregor A. Dizon, Razel Nikka M. Hao, Arianne A. Zamora, Devon Ray Pacial, Juan Antonio R. Magalang, Marissa Alejandria, Celia Carlos, Anna Ong-Lim, Edsel Maurice Salvaña, John Q. Wong, Jaime C. Montoya, Maria Rosario Singh-Vergeire and Cynthia P. Saloma |
| EPI_ISL_2189451 | Pasig City Children's Hospital - Child's Hope | Philippine Genome Center | Francis A. Tablizo, Kenneth M. Kim, Maria Sofia L. Yangzon, Elcid Aaron R. Pangilinan, Renato Jacinto Q. Mantaring, Benedict A. Maralit, Marc Edsel C. Ayes, Eva Maria Cutiongco-de la Paz, Alethea R. de Guzman, Jan Michael C. Yap, Jo-Hannah S. Llames, Sheila Mae M. Araiza, Kris P. Punayan, Irish Coleen A. Asin, Candice Francheska B. Tambaoan, Asia Louisa U. Chong, Karol Sophia Agape R. Padilla, Rianna Patricia S. Cruz, Carlo M. Lapid, El King D. Morado, Joshua Gregor A. Dizon, Razel Nikka M. Hao, Arianne A. Zamora, Devon Ray Pacial, Juan Antonio R. Magalang, Marissa Alejandria, Celia Carlos, Anna Ong-Lim, Edsel Maurice Salvaña, John Q. Wong, Jaime C. Montoya, Maria Rosario Singh-Vergeire and Cynthia P. Saloma |
| EPI_ISL_2189453 | South Super Highway Molecular Diagnostic Laboratory | Philippine Genome Center | Francis A. Tablizo, Kenneth M. Kim, Maria Sofia L. Yangzon, Elcid Aaron R. Pangilinan, Renato Jacinto Q. Mantaring, Benedict A. Maralit, Marc Edsel C. Ayes, Eva Maria Cutiongco-de la Paz, Alethea R. de Guzman, Jan Michael C. Yap, Jo-Hannah S. Llames, Sheila Mae M. Araiza, Kris P. Punayan, Irish Coleen A. Asin, Candice Francheska B. Tambaoan, Asia Louisa U. Chong, Karol Sophia Agape R. Padilla, Rianna Patricia S. Cruz, Carlo M. Lapid, El King D. Morado, Joshua Gregor A. Dizon, Razel Nikka M. Hao, Arianne A. Zamora, Devon Ray Pacial, Juan Antonio R. Magalang, Marissa Alejandria, Celia Carlos, Anna Ong-Lim, Edsel Maurice Salvaña, John Q. Wong, Jaime C. Montoya, Maria Rosario Singh-Vergeire and Cynthia P. Saloma |
| EPI_ISL_2189455, EPI_ISL_2189459 | Philippine Red Cross - Port Area | Philippine Genome Center | Francis A. Tablizo, Kenneth M. Kim, Maria Sofia L. Yangzon, Elcid Aaron R. Pangilinan, Renato Jacinto Q. Mantaring, Benedict A. Maralit, Marc Edsel C. Ayes, Eva Maria Cutiongco-de la Paz, Alethea R. de Guzman, Jan Michael C. Yap, Jo-Hannah S. Llames, Sheila Mae M. Araiza, Kris P. Punayan, Irish Coleen A. Asin, Candice Francheska B. Tambaoan, Asia Louisa U. Chong, Karol Sophia Agape R. Padilla, Rianna Patricia S. Cruz, Carlo M. Lapid, El King D. Morado, Joshua Gregor A. Dizon, Razel Nikka M. Hao, Arianne A. Zamora, Devon Ray Pacial, Juan Antonio R. Magalang, Marissa Alejandria, Celia Carlos, Anna Ong-Lim, Edsel Maurice Salvaña, John Q. Wong, Jaime C. Montoya, Maria Rosario Singh-Vergeire and Cynthia P. Saloma |
| EPI_ISL_2189460 | First Aide Diagnostic Center | Philippine Genome Center | Francis A. Tablizo, Kenneth M. Kim, Maria Sofia L. Yangzon, Elcid Aaron R. Pangilinan, Renato Jacinto Q. Mantaring, Benedict A. Maralit, Marc Edsel C. Ayes, Eva Maria Cutiongco-de la Paz, Alethea R. de Guzman, Jan Michael C. Yap, Jo-Hannah S. Llames, Sheila Mae M. Araiza, Kris P. Punayan, Irish Coleen A. Asin, Candice Francheska B. Tambaoan, Asia Louisa U. Chong, Karol Sophia Agape R. Padilla, Rianna Patricia S. Cruz, Carlo M. Lapid, El King D. Morado, Joshua Gregor A. Dizon, Razel Nikka M. Hao, Arianne A. Zamora, Devon Ray Pacial, Juan Antonio R. Magalang, Marissa Alejandria, Celia Carlos, Anna Ong-Lim, Edsel Maurice Salvaña, John Q. Wong, Jaime C. Montoya, Maria Rosario Singh-Vergeire and Cynthia P. Saloma |
| EPI_ISL_2189462 | South Super Highway Molecular Diagnostic Laboratory | Philippine Genome Center | Francis A. Tablizo, Kenneth M. Kim, Maria Sofia L. Yangzon, Elcid Aaron R. Pangilinan, Renato Jacinto Q. Mantaring, Benedict A. Maralit, Marc Edsel C. Ayes, Eva Maria Cutiongco-de la Paz, Alethea R. de Guzman, Jan Michael C. Yap, Jo-Hannah S. Llames, Sheila Mae M. Araiza, Kris P. Punayan, Irish Coleen A. Asin, Candice Francheska B. Tambaoan, Asia Louisa U. Chong, Karol Sophia Agape R. Padilla, Rianna Patricia S. Cruz, Carlo M. Lapid, El King D. Morado, Joshua Gregor A. Dizon, Razel Nikka M. Hao, Arianne A. Zamora, Devon Ray Pacial, Juan Antonio R. Magalang, Marissa Alejandria, Celia Carlos, Anna Ong-Lim, Edsel Maurice Salvaña, John Q. Wong, Jaime C. Montoya, Maria Rosario Singh-Vergeire and Cynthia P. Saloma |

| | | | |
|---|---|---|---|
| EPI_ISL_2189463 | The Lord's Grace Medical and Industrial Clinic | Philippine Genome Center | Francis A. Tablizo, Kenneth M. Kim, Maria Sofia L. Yangzon, Elcid Aaron R. Pangilinan, Renato Jacinto Q. Mantaring, Benedict A. Maralit, Marc Edsel C. Ayes, Eva Maria Cutiongco-de la Paz, Alethea R. de Guzman, Jan Michael C. Yap, Jo-Hannah S. Llames, Sheila Mae M. Araiza, Kris P. Punayan, Irish Coleen A. Asin, Candice Francheska B. Tambaoan, Asia Louisa U. Chong, Karol Sophia Agape R. Padilla, Rianna Patricia S. Cruz, Carlo M. Lapid, El King D. Morado, Joshua Gregor A. Dizon, Razel Nikka M. Hao, Arianne A. Zamora, Devon Ray Pacial, Juan Antonio R. Magalang, Marissa Alejandria, Celia Carlos, Anna Ong-Lim, Edsel Maurice Salvaña, John Q. Wong, Jaime C. Montoya, Maria Rosario Singh-Vergeire and Cynthia P. Saloma |
| EPI_ISL_2189464 | Philippine Red Cross - Clark Molecular Laboratory | Philippine Genome Center | Francis A. Tablizo, Kenneth M. Kim, Maria Sofia L. Yangzon, Elcid Aaron R. Pangilinan, Renato Jacinto Q. Mantaring, Benedict A. Maralit, Marc Edsel C. Ayes, Eva Maria Cutiongco-de la Paz, Alethea R. de Guzman, Jan Michael C. Yap, Jo-Hannah S. Llames, Sheila Mae M. Araiza, Kris P. Punayan, Irish Coleen A. Asin, Candice Francheska B. Tambaoan, Asia Louisa U. Chong, Karol Sophia Agape R. Padilla, Rianna Patricia S. Cruz, Carlo M. Lapid, El King D. Morado, Joshua Gregor A. Dizon, Razel Nikka M. Hao, Arianne A. Zamora, Devon Ray Pacial, Juan Antonio R. Magalang, Marissa Alejandria, Celia Carlos, Anna Ong-Lim, Edsel Maurice Salvaña, John Q. Wong, Jaime C. Montoya, Maria Rosario Singh-Vergeire and Cynthia P. Saloma |
| EPI_ISL_2189465 | Jose B. Lingad Memorial General Hospital | Philippine Genome Center | Francis A. Tablizo, Kenneth M. Kim, Maria Sofia L. Yangzon, Elcid Aaron R. Pangilinan, Renato Jacinto Q. Mantaring, Benedict A. Maralit, Marc Edsel C. Ayes, Eva Maria Cutiongco-de la Paz, Alethea R. de Guzman, Jan Michael C. Yap, Jo-Hannah S. Llames, Sheila Mae M. Araiza, Kris P. Punayan, Irish Coleen A. Asin, Candice Francheska B. Tambaoan, Asia Louisa U. Chong, Karol Sophia Agape R. Padilla, Rianna Patricia S. Cruz, Carlo M. Lapid, El King D. Morado, Joshua Gregor A. Dizon, Razel Nikka M. Hao, Arianne A. Zamora, Devon Ray Pacial, Juan Antonio R. Magalang, Marissa Alejandria, Celia Carlos, Anna Ong-Lim, Edsel Maurice Salvaña, John Q. Wong, Jaime C. Montoya, Maria Rosario Singh-Vergeire and Cynthia P. Saloma |
| EPI_ISL_2189466 | Pasig City Children's Hospital - Child's Hope | Philippine Genome Center | Francis A. Tablizo, Kenneth M. Kim, Maria Sofia L. Yangzon, Elcid Aaron R. Pangilinan, Renato Jacinto Q. Mantaring, Benedict A. Maralit, Marc Edsel C. Ayes, Eva Maria Cutiongco-de la Paz, Alethea R. de Guzman, Jan Michael C. Yap, Jo-Hannah S. Llames, Sheila Mae M. Araiza, Kris P. Punayan, Irish Coleen A. Asin, Candice Francheska B. Tambaoan, Asia Louisa U. Chong, Karol Sophia Agape R. Padilla, Rianna Patricia S. Cruz, Carlo M. Lapid, El King D. Morado, Joshua Gregor A. Dizon, Razel Nikka M. Hao, Arianne A. Zamora, Devon Ray Pacial, Juan Antonio R. Magalang, Marissa Alejandria, Celia Carlos, Anna Ong-Lim, Edsel Maurice Salvaña, John Q. Wong, Jaime C. Montoya, Maria Rosario Singh-Vergeire and Cynthia P. Saloma |
| EPI_ISL_2189467 | The Medical City | Philippine Genome Center | Francis A. Tablizo, Kenneth M. Kim, Maria Sofia L. Yangzon, Elcid Aaron R. Pangilinan, Renato Jacinto Q. Mantaring, Benedict A. Maralit, Marc Edsel C. Ayes, Eva Maria Cutiongco-de la Paz, Alethea R. de Guzman, Jan Michael C. Yap, Jo-Hannah S. Llames, Sheila Mae M. Araiza, Kris P. Punayan, Irish Coleen A. Asin, Candice Francheska B. Tambaoan, Asia Louisa U. Chong, Karol Sophia Agape R. Padilla, Rianna Patricia S. Cruz, Carlo M. Lapid, El King D. Morado, Joshua Gregor A. Dizon, Razel Nikka M. Hao, Arianne A. Zamora, Devon Ray Pacial, Juan Antonio R. Magalang, Marissa Alejandria, Celia Carlos, Anna Ong-Lim, Edsel Maurice Salvaña, John Q. Wong, Jaime C. Montoya, Maria Rosario Singh-Vergeire and Cynthia P. Saloma |
| EPI_ISL_2189468 | Pasig City Children's Hospital - Child's Hope | Philippine Genome Center | Francis A. Tablizo, Kenneth M. Kim, Maria Sofia L. Yangzon, Elcid Aaron R. Pangilinan, Renato Jacinto Q. Mantaring, Benedict A. Maralit, Marc Edsel C. Ayes, Eva Maria Cutiongco-de la Paz, Alethea R. de Guzman, Jan Michael C. Yap, Jo-Hannah S. Llames, Sheila Mae M. Araiza, Kris P. Punayan, Irish Coleen A. Asin, Candice Francheska B. Tambaoan, Asia Louisa U. Chong, Karol Sophia Agape R. Padilla, Rianna Patricia S. Cruz, Carlo M. Lapid, El King D. Morado, Joshua Gregor A. Dizon, Razel Nikka M. Hao, Arianne A. Zamora, Devon Ray Pacial, Juan Antonio R. Magalang, Marissa Alejandria, Celia Carlos, Anna Ong-Lim, Edsel Maurice Salvaña, John Q. Wong, Jaime C. Montoya, Maria Rosario Singh-Vergeire and Cynthia P. Saloma |
| EPI_ISL_2189469 | The Lord's Grace Medical and Industrial Clinic | Philippine Genome Center | Francis A. Tablizo, Kenneth M. Kim, Maria Sofia L. Yangzon, Elcid Aaron R. Pangilinan, Renato Jacinto Q. Mantaring, Benedict A. Maralit, Marc Edsel C. Ayes, Eva Maria Cutiongco-de la Paz, Alethea R. de Guzman, Jan Michael C. Yap, Jo-Hannah S. Llames, Sheila Mae M. Araiza, Kris P. Punayan, Irish Coleen A. Asin, Candice Francheska B. Tambaoan, Asia Louisa U. Chong, Karol Sophia Agape R. Padilla, Rianna Patricia S. Cruz, Carlo M. Lapid, El King D. Morado, Joshua Gregor A. Dizon, Razel Nikka M. Hao, Arianne A. Zamora, Devon Ray Pacial, Juan Antonio R. Magalang, Marissa Alejandria, Celia Carlos, Anna Ong-Lim, Edsel Maurice Salvaña, John Q. Wong, Jaime C. Montoya, Maria Rosario Singh-Vergeire and Cynthia P. Saloma |
| EPI_ISL_2189471 | University of the Philippines National Institutes of Health (UP NIH) | Philippine Genome Center | Francis A. Tablizo, Kenneth M. Kim, Maria Sofia L. Yangzon, Elcid Aaron R. Pangilinan, Renato Jacinto Q. Mantaring, Benedict A. Maralit, Marc Edsel C. Ayes, Eva Maria Cutiongco-de la Paz, Alethea R. de Guzman, Jan Michael C. Yap, Jo-Hannah S. Llames, Sheila Mae M. Araiza, Kris P. Punayan, Irish Coleen A. Asin, Candice Francheska B. Tambaoan, Asia Louisa U. Chong, Karol Sophia Agape R. Padilla, Rianna Patricia S. Cruz, Carlo M. Lapid, El King D. Morado, Joshua Gregor A. Dizon, Razel Nikka M. Hao, Arianne A. Zamora, Devon Ray Pacial, Juan Antonio R. Magalang, Marissa Alejandria, Celia Carlos, Anna Ong-Lim, Edsel Maurice Salvaña, John Q. Wong, Jaime C. Montoya, Maria Rosario Singh-Vergeire and Cynthia P. Saloma |
| EPI_ISL_2189473 | The Medical City | Philippine Genome Center | Francis A. Tablizo, Kenneth M. Kim, Maria Sofia L. Yangzon, Elcid Aaron R. Pangilinan, Renato Jacinto Q. Mantaring, Benedict A. Maralit, Marc Edsel C. Ayes, Eva Maria Cutiongco-de la Paz, Alethea R. de Guzman, Jan Michael C. Yap, Jo-Hannah S. Llames, Sheila Mae M. Araiza, Kris P. Punayan, Irish Coleen A. Asin, Candice Francheska B. Tambaoan, Asia Louisa U. Chong, Karol Sophia Agape R. Padilla, Rianna Patricia S. Cruz, Carlo M. Lapid, El King D. Morado, Joshua Gregor A. Dizon, Razel Nikka M. Hao, Arianne A. Zamora, Devon Ray Pacial, Juan Antonio R. Magalang, Marissa Alejandria, Celia Carlos, Anna Ong-Lim, Edsel Maurice Salvaña, John Q. Wong, Jaime C. Montoya, Maria Rosario Singh-Vergeire and Cynthia P. Saloma |
| EPI_ISL_2189474 | The Lord's Grace Medical and Industrial Clinic | Philippine Genome Center | Francis A. Tablizo, Kenneth M. Kim, Maria Sofia L. Yangzon, Elcid Aaron R. Pangilinan, Renato Jacinto Q. Mantaring, Benedict A. Maralit, Marc Edsel C. Ayes, Eva Maria Cutiongco-de la Paz, Alethea R. de Guzman, Jan Michael C. Yap, Jo-Hannah S. Llames, Sheila Mae M. Araiza, Kris P. Punayan, Irish Coleen A. Asin, Candice Francheska B. Tambaoan, Asia Louisa U. Chong, Karol Sophia Agape R. Padilla, Rianna Patricia S. Cruz, Carlo M. Lapid, El King D. Morado, Joshua Gregor A. Dizon, Razel Nikka M. Hao, Arianne A. Zamora, Devon Ray Pacial, Juan Antonio R. Magalang, Marissa Alejandria, Celia Carlos, Anna Ong-Lim, Edsel Maurice Salvaña, John Q. Wong, Jaime C. Montoya, Maria Rosario Singh-Vergeire and Cynthia P. Saloma |
| EPI_ISL_2189475 | Mariano Marcos Memorial Hospital Medical Center | Philippine Genome Center | Francis A. Tablizo, Kenneth M. Kim, Maria Sofia L. Yangzon, Elcid Aaron R. Pangilinan, Renato Jacinto Q. Mantaring, Benedict A. Maralit, Marc Edsel C. Ayes, Eva Maria Cutiongco-de la Paz, Alethea R. de Guzman, Jan Michael C. Yap, Jo-Hannah S. Llames, Sheila Mae M. Araiza, Kris P. Punayan, Irish Coleen A. Asin, Candice Francheska B. Tambaoan, Asia Louisa U. Chong, Karol Sophia Agape R. Padilla, Rianna Patricia S. Cruz, Carlo M. Lapid, El King D. Morado, Joshua Gregor A. Dizon, Razel Nikka M. Hao, Arianne A. Zamora, Devon Ray Pacial, Juan Antonio R. Magalang, Marissa Alejandria, Celia Carlos, Anna Ong-Lim, Edsel Maurice Salvaña, John Q. Wong, Jaime C. Montoya, Maria Rosario Singh-Vergeire and Cynthia P. Saloma |
| EPI_ISL_2189476 | Philippine Red Cross - Clark Molecular Laboratory | Philippine Genome Center | Francis A. Tablizo, Kenneth M. Kim, Maria Sofia L. Yangzon, Elcid Aaron R. Pangilinan, Renato Jacinto Q. Mantaring, Benedict A. Maralit, Marc Edsel C. Ayes, Eva Maria Cutiongco-de la Paz, Alethea R. de Guzman, Jan Michael C. Yap, Jo-Hannah S. Llames, Sheila Mae M. Araiza, Kris P. Punayan, Irish Coleen A. Asin, Candice Francheska B. Tambaoan, Asia Louisa U. Chong, Karol Sophia Agape R. Padilla, Rianna Patricia S. Cruz, Carlo M. Lapid, El King D. Morado, Joshua Gregor A. Dizon, Razel Nikka M. Hao, Arianne A. Zamora, Devon Ray Pacial, Juan Antonio R. Magalang, Marissa Alejandria, Celia Carlos, Anna Ong-Lim, Edsel Maurice Salvaña, John Q. Wong, Jaime C. Montoya, Maria Rosario Singh-Vergeire and Cynthia P. Saloma |
| EPI_ISL_2189477 | Mary Mediatrix Medical Center | Philippine Genome Center | Francis A. Tablizo, Kenneth M. Kim, Maria Sofia L. Yangzon, Elcid Aaron R. Pangilinan, Renato Jacinto Q. Mantaring, Benedict A. Maralit, Marc Edsel C. Ayes, Eva Maria Cutiongco-de la Paz, Alethea R. de Guzman, Jan Michael C. Yap, Jo-Hannah S. Llames, Sheila Mae M. Araiza, Kris P. Punayan, Irish Coleen A. Asin, Candice Francheska B. Tambaoan, Asia Louisa U. Chong, Karol Sophia Agape R. Padilla, Rianna Patricia S. Cruz, Carlo M. Lapid, El King D. Morado, Joshua Gregor A. Dizon, Razel Nikka M. Hao, Arianne A. Zamora, Devon Ray Pacial, Juan Antonio R. Magalang, Marissa Alejandria, Celia Carlos, Anna Ong-Lim, Edsel Maurice Salvaña, John Q. Wong, Jaime C. Montoya, Maria Rosario Singh-Vergeire and Cynthia P. Saloma |
| EPI_ISL_2189478 | Philippine Red Cross - National Blood Center | Philippine Genome Center | Francis A. Tablizo, Kenneth M. Kim, Maria Sofia L. Yangzon, Elcid Aaron R. Pangilinan, Renato Jacinto Q. Mantaring, Benedict A. Maralit, Marc Edsel C. Ayes, Eva Maria Cutiongco-de la Paz, Alethea R. de Guzman, Jan Michael C. Yap, Jo-Hannah S. Llames, Sheila Mae M. Araiza, Kris P. Punayan, Irish Coleen A. Asin, Candice Francheska B. Tambaoan, Asia Louisa U. Chong, Karol Sophia Agape R. Padilla, Rianna Patricia S. Cruz, Carlo M. Lapid, El King D. Morado, Joshua Gregor A. Dizon, Razel Nikka M. Hao, Arianne A. Zamora, Devon Ray Pacial, Juan Antonio R. Magalang, Marissa Alejandria, Celia Carlos, Anna Ong-Lim, Edsel Maurice Salvaña, John Q. Wong, Jaime C. Montoya, Maria Rosario Singh-Vergeire and Cynthia P. Saloma |
| EPI_ISL_2189479 | The Lord's Grace Medical and Industrial Clinic | Philippine Genome Center | Francis A. Tablizo, Kenneth M. Kim, Maria Sofia L. Yangzon, Elcid Aaron R. Pangilinan, Renato Jacinto Q. Mantaring, Benedict A. Maralit, Marc Edsel C. Ayes, Eva Maria Cutiongco-de la Paz, Alethea R. de Guzman, Jan Michael C. Yap, Jo-Hannah S. Llames, Sheila Mae M. Araiza, Kris P. Punayan, Irish Coleen A. Asin, Candice Francheska B. Tambaoan, Asia Louisa U. Chong, Karol Sophia Agape R. Padilla, Rianna Patricia S. Cruz, Carlo M. Lapid, El King D. Morado, Joshua Gregor A. Dizon, Razel Nikka M. Hao, Arianne A. Zamora, Devon Ray Pacial, Juan Antonio R. Magalang, Marissa Alejandria, Celia Carlos, Anna Ong-Lim, Edsel Maurice Salvaña, John Q. Wong, Jaime C. Montoya, Maria Rosario Singh-Vergeire and Cynthia P. Saloma |
| EPI_ISL_2189482 | Sta. Rosa CHO- GENEXPERT LABORATORY | Philippine Genome Center | Francis A. Tablizo, Kenneth M. Kim, Maria Sofia L. Yangzon, Elcid Aaron R. Pangilinan, Renato Jacinto Q. Mantaring, Benedict A. Maralit, Marc Edsel C. Ayes, Eva Maria Cutiongco-de la Paz, Alethea R. de Guzman, Jan Michael C. Yap, Jo-Hannah S. Llames, Sheila Mae M. Araiza, Kris P. Punayan, Irish Coleen A. Asin, Candice Francheska B. Tambaoan, Asia Louisa U. Chong, Karol Sophia Agape R. Padilla, Rianna Patricia S. Cruz, Carlo M. Lapid, El King D. Morado, Joshua Gregor A. Dizon, Razel Nikka M. Hao, Arianne A. Zamora, Devon Ray Pacial, Juan Antonio R. Magalang, Marissa Alejandria, Celia Carlos, Anna Ong-Lim, Edsel Maurice Salvaña, John Q. Wong, Jaime C. Montoya, Maria Rosario Singh-Vergeire and Cynthia P. Saloma |

| | | | |
|---|---|---|---|
| EPI_ISL_2189483 | The Lord's Grace Medical and Industrial Clinic | Philippine Genome Center | Francis A. Tablizo, Kenneth M. Kim, Maria Sofia L. Yangzon, Elcid Aaron R. Pangilinan, Renato Jacinto Q. Mantaring, Benedict A. Maralit, Marc Edsel C. Ayes, Eva Maria Cutiongco-de la Paz, Alethea R. de Guzman, Jan Michael C. Yap, Jo-Hannah S. Llames, Sheila Mae M. Araiza, Kris P. Punayan, Irish Coleen A. Asin, Candice Francheska B. Tambaoan, Asia Louisa U. Chong, Karol Sophia Agape R. Padilla, Rianna Patricia S. Cruz, Carlo M. Lapid, El King D. Morado, Joshua Gregor A. Dizon, Razel Nikka M. Hao, Arianne A. Zamora, Devon Ray Pacial, Juan Antonio R. Magalang, Marissa Alejandria, Celia Carlos, Anna Ong-Lim, Edsel Maurice Salvaña, John Q. Wong, Jaime C. Montoya, Maria Rosario Singh-Vergeire and Cynthia P. Saloma |
| EPI_ISL_2189484 | Parkway Medical and Diagnostic Center | Philippine Genome Center | Francis A. Tablizo, Kenneth M. Kim, Maria Sofia L. Yangzon, Elcid Aaron R. Pangilinan, Renato Jacinto Q. Mantaring, Benedict A. Maralit, Marc Edsel C. Ayes, Eva Maria Cutiongco-de la Paz, Alethea R. de Guzman, Jan Michael C. Yap, Jo-Hannah S. Llames, Sheila Mae M. Araiza, Kris P. Punayan, Irish Coleen A. Asin, Candice Francheska B. Tambaoan, Asia Louisa U. Chong, Karol Sophia Agape R. Padilla, Rianna Patricia S. Cruz, Carlo M. Lapid, El King D. Morado, Joshua Gregor A. Dizon, Razel Nikka M. Hao, Arianne A. Zamora, Devon Ray Pacial, Juan Antonio R. Magalang, Marissa Alejandria, Celia Carlos, Anna Ong-Lim, Edsel Maurice Salvaña, John Q. Wong, Jaime C. Montoya, Maria Rosario Singh-Vergeire and Cynthia P. Saloma |
| EPI_ISL_2189485, EPI_ISL_2189489 | The Lord's Grace Medical and Industrial Clinic | Philippine Genome Center | Francis A. Tablizo, Kenneth M. Kim, Maria Sofia L. Yangzon, Elcid Aaron R. Pangilinan, Renato Jacinto Q. Mantaring, Benedict A. Maralit, Marc Edsel C. Ayes, Eva Maria Cutiongco-de la Paz, Alethea R. de Guzman, Jan Michael C. Yap, Jo-Hannah S. Llames, Sheila Mae M. Araiza, Kris P. Punayan, Irish Coleen A. Asin, Candice Francheska B. Tambaoan, Asia Louisa U. Chong, Karol Sophia Agape R. Padilla, Rianna Patricia S. Cruz, Carlo M. Lapid, El King D. Morado, Joshua Gregor A. Dizon, Razel Nikka M. Hao, Arianne A. Zamora, Devon Ray Pacial, Juan Antonio R. Magalang, Marissa Alejandria, Celia Carlos, Anna Ong-Lim, Edsel Maurice Salvaña, John Q. Wong, Jaime C. Montoya, Maria Rosario Singh-Vergeire and Cynthia P. Saloma |
| EPI_ISL_2189490 | Philippine Red Cross - Clark Molecular Laboratory | Philippine Genome Center | Francis A. Tablizo, Kenneth M. Kim, Maria Sofia L. Yangzon, Elcid Aaron R. Pangilinan, Renato Jacinto Q. Mantaring, Benedict A. Maralit, Marc Edsel C. Ayes, Eva Maria Cutiongco-de la Paz, Alethea R. de Guzman, Jan Michael C. Yap, Jo-Hannah S. Llames, Sheila Mae M. Araiza, Kris P. Punayan, Irish Coleen A. Asin, Candice Francheska B. Tambaoan, Asia Louisa U. Chong, Karol Sophia Agape R. Padilla, Rianna Patricia S. Cruz, Carlo M. Lapid, El King D. Morado, Joshua Gregor A. Dizon, Razel Nikka M. Hao, Arianne A. Zamora, Devon Ray Pacial, Juan Antonio R. Magalang, Marissa Alejandria, Celia Carlos, Anna Ong-Lim, Edsel Maurice Salvaña, John Q. Wong, Jaime C. Montoya, Maria Rosario Singh-Vergeire and Cynthia P. Saloma |
| EPI_ISL_2189491, EPI_ISL_2189494 | The Lord's Grace Medical and Industrial Clinic | Philippine Genome Center | Francis A. Tablizo, Kenneth M. Kim, Maria Sofia L. Yangzon, Elcid Aaron R. Pangilinan, Renato Jacinto Q. Mantaring, Benedict A. Maralit, Marc Edsel C. Ayes, Eva Maria Cutiongco-de la Paz, Alethea R. de Guzman, Jan Michael C. Yap, Jo-Hannah S. Llames, Sheila Mae M. Araiza, Kris P. Punayan, Irish Coleen A. Asin, Candice Francheska B. Tambaoan, Asia Louisa U. Chong, Karol Sophia Agape R. Padilla, Rianna Patricia S. Cruz, Carlo M. Lapid, El King D. Morado, Joshua Gregor A. Dizon, Razel Nikka M. Hao, Arianne A. Zamora, Devon Ray Pacial, Juan Antonio R. Magalang, Marissa Alejandria, Celia Carlos, Anna Ong-Lim, Edsel Maurice Salvaña, John Q. Wong, Jaime C. Montoya, Maria Rosario Singh-Vergeire and Cynthia P. Saloma |
| EPI_ISL_2189497 | The Medical City | Philippine Genome Center | Francis A. Tablizo, Kenneth M. Kim, Maria Sofia L. Yangzon, Elcid Aaron R. Pangilinan, Renato Jacinto Q. Mantaring, Benedict A. Maralit, Marc Edsel C. Ayes, Eva Maria Cutiongco-de la Paz, Alethea R. de Guzman, Jan Michael C. Yap, Jo-Hannah S. Llames, Sheila Mae M. Araiza, Kris P. Punayan, Irish Coleen A. Asin, Candice Francheska B. Tambaoan, Asia Louisa U. Chong, Karol Sophia Agape R. Padilla, Rianna Patricia S. Cruz, Carlo M. Lapid, El King D. Morado, Joshua Gregor A. Dizon, Razel Nikka M. Hao, Arianne A. Zamora, Devon Ray Pacial, Juan Antonio R. Magalang, Marissa Alejandria, Celia Carlos, Anna Ong-Lim, Edsel Maurice Salvaña, John Q. Wong, Jaime C. Montoya, Maria Rosario Singh-Vergeire and Cynthia P. Saloma |
| EPI_ISL_2189499 | Pasig City Children's Hospital - Child's Hope | Philippine Genome Center | Francis A. Tablizo, Kenneth M. Kim, Maria Sofia L. Yangzon, Elcid Aaron R. Pangilinan, Renato Jacinto Q. Mantaring, Benedict A. Maralit, Marc Edsel C. Ayes, Eva Maria Cutiongco-de la Paz, Alethea R. de Guzman, Jan Michael C. Yap, Jo-Hannah S. Llames, Sheila Mae M. Araiza, Kris P. Punayan, Irish Coleen A. Asin, Candice Francheska B. Tambaoan, Asia Louisa U. Chong, Karol Sophia Agape R. Padilla, Rianna Patricia S. Cruz, Carlo M. Lapid, El King D. Morado, Joshua Gregor A. Dizon, Razel Nikka M. Hao, Arianne A. Zamora, Devon Ray Pacial, Juan Antonio R. Magalang, Marissa Alejandria, Celia Carlos, Anna Ong-Lim, Edsel Maurice Salvaña, John Q. Wong, Jaime C. Montoya, Maria Rosario Singh-Vergeire and Cynthia P. Saloma |
| EPI_ISL_2189500, EPI_ISL_2189501 | The Medical City | Philippine Genome Center | Francis A. Tablizo, Kenneth M. Kim, Maria Sofia L. Yangzon, Elcid Aaron R. Pangilinan, Renato Jacinto Q. Mantaring, Benedict A. Maralit, Marc Edsel C. Ayes, Eva Maria Cutiongco-de la Paz, Alethea R. de Guzman, Jan Michael C. Yap, Jo-Hannah S. Llames, Sheila Mae M. Araiza, Kris P. Punayan, Irish Coleen A. Asin, Candice Francheska B. Tambaoan, Asia Louisa U. Chong, Karol Sophia Agape R. Padilla, Rianna Patricia S. Cruz, Carlo M. Lapid, El King D. Morado, Joshua Gregor A. Dizon, Razel Nikka M. Hao, Arianne A. Zamora, Devon Ray Pacial, Juan Antonio R. Magalang, Marissa Alejandria, Celia Carlos, Anna Ong-Lim, Edsel Maurice Salvaña, John Q. Wong, Jaime C. Montoya, Maria Rosario Singh-Vergeire and Cynthia P. Saloma |
| EPI_ISL_2189503, EPI_ISL_2189504, EPI_ISL_2189506, EPI_ISL_2189507, EPI_ISL_2189508 | The Lord's Grace Medical and Industrial Clinic | Philippine Genome Center | Francis A. Tablizo, Kenneth M. Kim, Maria Sofia L. Yangzon, Elcid Aaron R. Pangilinan, Renato Jacinto Q. Mantaring, Benedict A. Maralit, Marc Edsel C. Ayes, Eva Maria Cutiongco-de la Paz, Alethea R. de Guzman, Jan Michael C. Yap, Jo-Hannah S. Llames, Sheila Mae M. Araiza, Kris P. Punayan, Irish Coleen A. Asin, Candice Francheska B. Tambaoan, Asia Louisa U. Chong, Karol Sophia Agape R. Padilla, Rianna Patricia S. Cruz, Carlo M. Lapid, El King D. Morado, Joshua Gregor A. Dizon, Razel Nikka M. Hao, Arianne A. Zamora, Devon Ray Pacial, Juan Antonio R. Magalang, Marissa Alejandria, Celia Carlos, Anna Ong-Lim, Edsel Maurice Salvaña, John Q. Wong, Jaime C. Montoya, Maria Rosario Singh-Vergeire and Cynthia P. Saloma |
| EPI_ISL_2189510, EPI_ISL_2189511 | Philippine Red Cross - Port Area | Philippine Genome Center | Francis A. Tablizo, Kenneth M. Kim, Maria Sofia L. Yangzon, Elcid Aaron R. Pangilinan, Renato Jacinto Q. Mantaring, Benedict A. Maralit, Marc Edsel C. Ayes, Eva Maria Cutiongco-de la Paz, Alethea R. de Guzman, Jan Michael C. Yap, Jo-Hannah S. Llames, Sheila Mae M. Araiza, Kris P. Punayan, Irish Coleen A. Asin, Candice Francheska B. Tambaoan, Asia Louisa U. Chong, Karol Sophia Agape R. Padilla, Rianna Patricia S. Cruz, Carlo M. Lapid, El King D. Morado, Joshua Gregor A. Dizon, Razel Nikka M. Hao, Arianne A. Zamora, Devon Ray Pacial, Juan Antonio R. Magalang, Marissa Alejandria, Celia Carlos, Anna Ong-Lim, Edsel Maurice Salvaña, John Q. Wong, Jaime C. Montoya, Maria Rosario Singh-Vergeire and Cynthia P. Saloma |
| EPI_ISL_2189512 | Pasig City Children's Hospital - Child's Hope | Philippine Genome Center | Francis A. Tablizo, Kenneth M. Kim, Maria Sofia L. Yangzon, Elcid Aaron R. Pangilinan, Renato Jacinto Q. Mantaring, Benedict A. Maralit, Marc Edsel C. Ayes, Eva Maria Cutiongco-de la Paz, Alethea R. de Guzman, Jan Michael C. Yap, Jo-Hannah S. Llames, Sheila Mae M. Araiza, Kris P. Punayan, Irish Coleen A. Asin, Candice Francheska B. Tambaoan, Asia Louisa U. Chong, Karol Sophia Agape R. Padilla, Rianna Patricia S. Cruz, Carlo M. Lapid, El King D. Morado, Joshua Gregor A. Dizon, Razel Nikka M. Hao, Arianne A. Zamora, Devon Ray Pacial, Juan Antonio R. Magalang, Marissa Alejandria, Celia Carlos, Anna Ong-Lim, Edsel Maurice Salvaña, John Q. Wong, Jaime C. Montoya, Maria Rosario Singh-Vergeire and Cynthia P. Saloma |
| EPI_ISL_2189513 | Philippine Red Cross - Port Area | Philippine Genome Center | Francis A. Tablizo, Kenneth M. Kim, Maria Sofia L. Yangzon, Elcid Aaron R. Pangilinan, Renato Jacinto Q. Mantaring, Benedict A. Maralit, Marc Edsel C. Ayes, Eva Maria Cutiongco-de la Paz, Alethea R. de Guzman, Jan Michael C. Yap, Jo-Hannah S. Llames, Sheila Mae M. Araiza, Kris P. Punayan, Irish Coleen A. Asin, Candice Francheska B. Tambaoan, Asia Louisa U. Chong, Karol Sophia Agape R. Padilla, Rianna Patricia S. Cruz, Carlo M. Lapid, El King D. Morado, Joshua Gregor A. Dizon, Razel Nikka M. Hao, Arianne A. Zamora, Devon Ray Pacial, Juan Antonio R. Magalang, Marissa Alejandria, Celia Carlos, Anna Ong-Lim, Edsel Maurice Salvaña, John Q. Wong, Jaime C. Montoya, Maria Rosario Singh-Vergeire and Cynthia P. Saloma |
| EPI_ISL_2189514 | The Lord's Grace Medical and Industrial Clinic | Philippine Genome Center | Francis A. Tablizo, Kenneth M. Kim, Maria Sofia L. Yangzon, Elcid Aaron R. Pangilinan, Renato Jacinto Q. Mantaring, Benedict A. Maralit, Marc Edsel C. Ayes, Eva Maria Cutiongco-de la Paz, Alethea R. de Guzman, Jan Michael C. Yap, Jo-Hannah S. Llames, Sheila Mae M. Araiza, Kris P. Punayan, Irish Coleen A. Asin, Candice Francheska B. Tambaoan, Asia Louisa U. Chong, Karol Sophia Agape R. Padilla, Rianna Patricia S. Cruz, Carlo M. Lapid, El King D. Morado, Joshua Gregor A. Dizon, Razel Nikka M. Hao, Arianne A. Zamora, Devon Ray Pacial, Juan Antonio R. Magalang, Marissa Alejandria, Celia Carlos, Anna Ong-Lim, Edsel Maurice Salvaña, John Q. Wong, Jaime C. Montoya, Maria Rosario Singh-Vergeire and Cynthia P. Saloma |
| EPI_ISL_2189515, EPI_ISL_2189517 | Pasig City Children's Hospital - Child's Hope | Philippine Genome Center | Francis A. Tablizo, Kenneth M. Kim, Maria Sofia L. Yangzon, Elcid Aaron R. Pangilinan, Renato Jacinto Q. Mantaring, Benedict A. Maralit, Marc Edsel C. Ayes, Eva Maria Cutiongco-de la Paz, Alethea R. de Guzman, Jan Michael C. Yap, Jo-Hannah S. Llames, Sheila Mae M. Araiza, Kris P. Punayan, Irish Coleen A. Asin, Candice Francheska B. Tambaoan, Asia Louisa U. Chong, Karol Sophia Agape R. Padilla, Rianna Patricia S. Cruz, Carlo M. Lapid, El King D. Morado, Joshua Gregor A. Dizon, Razel Nikka M. Hao, Arianne A. Zamora, Devon Ray Pacial, Juan Antonio R. Magalang, Marissa Alejandria, Celia Carlos, Anna Ong-Lim, Edsel Maurice Salvaña, John Q. Wong, Jaime C. Montoya, Maria Rosario Singh-Vergeire and Cynthia P. Saloma |
| EPI_ISL_2189519 | Philippine General Hospital (PGH) | Philippine Genome Center | Francis A. Tablizo, Kenneth M. Kim, Maria Sofia L. Yangzon, Elcid Aaron R. Pangilinan, Renato Jacinto Q. Mantaring, Benedict A. Maralit, Marc Edsel C. Ayes, Eva Maria Cutiongco-de la Paz, Alethea R. de Guzman, Jan Michael C. Yap, Jo-Hannah S. Llames, Sheila Mae M. Araiza, Kris P. Punayan, Irish Coleen A. Asin, Candice Francheska B. Tambaoan, Asia Louisa U. Chong, Karol Sophia Agape R. Padilla, Rianna Patricia S. Cruz, Carlo M. Lapid, El King D. Morado, Joshua Gregor A. Dizon, Razel Nikka M. Hao, Arianne A. Zamora, Devon Ray Pacial, Juan Antonio R. Magalang, Marissa Alejandria, Celia Carlos, Anna Ong-Lim, Edsel Maurice Salvaña, John Q. Wong, Jaime C. Montoya, Maria Rosario Singh-Vergeire and Cynthia P. Saloma |
| EPI_ISL_2189520 | Philippine Red Cross - Port Area | Philippine Genome Center | Francis A. Tablizo, Kenneth M. Kim, Maria Sofia L. Yangzon, Elcid Aaron R. Pangilinan, Renato Jacinto Q. Mantaring, Benedict A. Maralit, Marc Edsel C. Ayes, Eva Maria Cutiongco-de la Paz, Alethea R. de Guzman, Jan Michael C. Yap, Jo-Hannah S. Llames, Sheila Mae M. Araiza, Kris P. Punayan, Irish Coleen A. Asin, Candice Francheska B. Tambaoan, Asia Louisa U. Chong, Karol Sophia Agape R. Padilla, Rianna Patricia S. Cruz, Carlo M. Lapid, El King D. Morado, Joshua Gregor A. Dizon, Razel Nikka M. Hao, Arianne A. Zamora, Devon Ray Pacial, Juan Antonio R. Magalang, Marissa Alejandria, Celia Carlos, Anna Ong-Lim, Edsel Maurice Salvaña, John Q. Wong, Jaime C. Montoya, Maria Rosario Singh-Vergeire and Cynthia P. Saloma |

| | | | |
|---|---|---|---|
| EPI_ISL_2189521 | South Super Highway Molecular Diagnostic Laboratory | Philippine Genome Center | Francis A. Tablizo, Kenneth M. Kim, Maria Sofia L. Yangzon, Elcid Aaron R. Pangilinan, Renato Jacinto Q. Mantaring, Benedict A. Maralit, Marc Edsel C. Ayes, Eva Maria Cutiongco-de la Paz, Alethea R. de Guzman, Jan Michael C. Yap, Jo-Hannah S. Llames, Sheila Mae M. Araiza, Kris P. Punayan, Irish Coleen A. Asin, Candice Francheska B. Tambaoan, Asia Louisa U. Chong, Karol Sophia Agape R. Padilla, Rianna Patricia S. Cruz, Carlo M. Lapid, El King D. Morado, Joshua Gregor A. Dizon, Razel Nikka M. Hao, Arianne A. Zamora, Devon Ray Pacial, Juan Antonio R. Magalang, Marissa Alejandria, Celia Carlos, Anna Ong-Lim, Edsel Maurice Salvaña, John Q. Wong, Jaime C. Montoya, Maria Rosario Singh-Vergeire and Cynthia P. Saloma |
| EPI_ISL_2189522 | Mariano Marcos Memorial Hospital Medical Center | Philippine Genome Center | Francis A. Tablizo, Kenneth M. Kim, Maria Sofia L. Yangzon, Elcid Aaron R. Pangilinan, Renato Jacinto Q. Mantaring, Benedict A. Maralit, Marc Edsel C. Ayes, Eva Maria Cutiongco-de la Paz, Alethea R. de Guzman, Jan Michael C. Yap, Jo-Hannah S. Llames, Sheila Mae M. Araiza, Kris P. Punayan, Irish Coleen A. Asin, Candice Francheska B. Tambaoan, Asia Louisa U. Chong, Karol Sophia Agape R. Padilla, Rianna Patricia S. Cruz, Carlo M. Lapid, El King D. Morado, Joshua Gregor A. Dizon, Razel Nikka M. Hao, Arianne A. Zamora, Devon Ray Pacial, Juan Antonio R. Magalang, Marissa Alejandria, Celia Carlos, Anna Ong-Lim, Edsel Maurice Salvaña, John Q. Wong, Jaime C. Montoya, Maria Rosario Singh-Vergeire and Cynthia P. Saloma |
| EPI_ISL_2189526 | Pasig City Children's Hospital - Child's Hope | Philippine Genome Center | Francis A. Tablizo, Kenneth M. Kim, Maria Sofia L. Yangzon, Elcid Aaron R. Pangilinan, Renato Jacinto Q. Mantaring, Benedict A. Maralit, Marc Edsel C. Ayes, Eva Maria Cutiongco-de la Paz, Alethea R. de Guzman, Jan Michael C. Yap, Jo-Hannah S. Llames, Sheila Mae M. Araiza, Kris P. Punayan, Irish Coleen A. Asin, Candice Francheska B. Tambaoan, Asia Louisa U. Chong, Karol Sophia Agape R. Padilla, Rianna Patricia S. Cruz, Carlo M. Lapid, El King D. Morado, Joshua Gregor A. Dizon, Razel Nikka M. Hao, Arianne A. Zamora, Devon Ray Pacial, Juan Antonio R. Magalang, Marissa Alejandria, Celia Carlos, Anna Ong-Lim, Edsel Maurice Salvaña, John Q. Wong, Jaime C. Montoya, Maria Rosario Singh-Vergeire and Cynthia P. Saloma |
| EPI_ISL_2189527 | Qualimed Hospital Sta. Rosa | Philippine Genome Center | Francis A. Tablizo, Kenneth M. Kim, Maria Sofia L. Yangzon, Elcid Aaron R. Pangilinan, Renato Jacinto Q. Mantaring, Benedict A. Maralit, Marc Edsel C. Ayes, Eva Maria Cutiongco-de la Paz, Alethea R. de Guzman, Jan Michael C. Yap, Jo-Hannah S. Llames, Sheila Mae M. Araiza, Kris P. Punayan, Irish Coleen A. Asin, Candice Francheska B. Tambaoan, Asia Louisa U. Chong, Karol Sophia Agape R. Padilla, Rianna Patricia S. Cruz, Carlo M. Lapid, El King D. Morado, Joshua Gregor A. Dizon, Razel Nikka M. Hao, Arianne A. Zamora, Devon Ray Pacial, Juan Antonio R. Magalang, Marissa Alejandria, Celia Carlos, Anna Ong-Lim, Edsel Maurice Salvaña, John Q. Wong, Jaime C. Montoya, Maria Rosario Singh-Vergeire and Cynthia P. Saloma |
| EPI_ISL_2191524, EPI_ISL_2191530, EPI_ISL_2191533, EPI_ISL_2191536, EPI_ISL_2191541 | Microbiology Department, University Hospital Donostia | Microbiology Department, University Hospital Donostia | Marimon JM, Montes M, Piñeiro L, Grandioso D, Martin-Peñaranda T, Gomez M, Cilla G. |
| EPI_ISL_2191878 | Arcispedale Santa Maria Nuova, Autoimmunità, Allergologia e Biotecnologie Innovative | Istituto Zooprofilattico Sperimentale della Lombardia e dell'Emilia Romagna (IZSLER), Risk Analysis and Genomic Epidemiology Unit | Alessandro Zerbini, Lucia Belloni, Stefania Croci, Marina Morganti, Ilaria Menozzi, Erika Scaltriti, Stefano Pongolini |
| EPI_ISL_2193349, EPI_ISL_2193350, EPI_ISL_2193362 | Institut de Virologie du CHU de Strasbourg | Institut de Virologie du CHU de Strasbourg | Fafi-Kremer Samira, Gallais Floriane, Gantner Pierre, Laugel Elodie, Solis Morgane, Velay Aurélie, Wendling Marie-Josée |
| EPI_ISL_2193956, EPI_ISL_2193957, EPI_ISL_2193958, EPI_ISL_2193960, EPI_ISL_2193990, EPI_ISL_2193993, EPI_ISL_2193995, EPI_ISL_2194017, EPI_ISL_2194018, EPI_ISL_2194020 | CHU Nantes Virology | CHU Nantes Virology | Celine Bressollette-Bodin, Thomas Drumel, Audrey Rodallec, Berthe-Marie Imbert-Marcille |
| EPI_ISL_2196257 | Laboratorio Central de Saude Publica do Estado de Minas Gerais (LACEN/MG) | Laboratory of Respiratory Viruses and Measles, Oswaldo Cruz Institute, FIOCRUZ | Paola Resende, Luciana Appolinario, Fernando Motta, Anna Carolina Paixao, Ana Carolina Mendonca, Alice Sampaio Rocha, Taina Venas, Elisa Cavalcante Pereira, Renata Serrano Lopes, Andre Felipe Leal Bernardes, Marilda Siqueira on behalf of the Fiocruz COVID-19 Genomic Surveillance Network |
| EPI_ISL_2196259 | Lboratorio Central de Saude Publica do Estado do Parana (LACEN/PR) | Laboratory of Respiratory Viruses and Measles, Oswaldo Cruz Institute, FIOCRUZ | Paola Resende, Luciana Appolinario, Fernando Motta, Anna Carolina Paixao, Ana Carolina Mendonca, Alice Sampaio Rocha, Taina Venas, Elisa Cavalcante Pereira, Renata Serrano Lopes, Irina Riediger, Marilda Siqueira on behalf of the Fiocruz COVID-19 Genomic Surveillance Network |
| EPI_ISL_2196322 | Labortorio Central de Saude Publica do Estado do Rio de Janeiro (LACEN/RJ) | Laboratory of Respiratory Viruses and Measles, Oswaldo Cruz Institute, FIOCRUZ | Paola Resende, Luciana Appolinario, Fernando Motta, Anna Carolina Paixao, Ana Carolina Mendonca, Alice Sampaio Rocha, Taina Venas, Elisa Cavalcante Pereira, Renata Serrano Lopes, Andrea Cony Cavalcanti, Marilda Siqueira on behalf of the Fiocruz COVID-19 Genomic Surveillance Network |
| EPI_ISL_2196366 | Microbiology Department - University Hospital Brussel | Microbiology Department - University Hospital Brussel | Oriane Soetens, Thomas Demuyser, Lucie Seyler |
| EPI_ISL_2200079, EPI_ISL_2200082, EPI_ISL_2200084, EPI_ISL_2200085, EPI_ISL_2200087, EPI_ISL_2200088, EPI_ISL_2200089, EPI_ISL_2200094, EPI_ISL_2200096, EPI_ISL_2200099, EPI_ISL_2200100, EPI_ISL_2200101, EPI_ISL_2200103, EPI_ISL_2200107, EPI_ISL_2200109, EPI_ISL_2200110, EPI_ISL_2200112, EPI_ISL_2200115, EPI_ISL_2200120, EPI_ISL_2200123, EPI_ISL_2200124, EPI_ISL_2200128, EPI_ISL_2200131, EPI_ISL_2200133, EPI_ISL_2200134, EPI_ISL_2200135, EPI_ISL_2200138, EPI_ISL_2200141, EPI_ISL_2200142, EPI_ISL_2200143, EPI_ISL_2200144, EPI_ISL_2200145, EPI_ISL_2200146, EPI_ISL_2200147, EPI_ISL_2200149, EPI_ISL_2200150, EPI_ISL_2200152 | | | |
| see above | The Ohio State University Applied Microbiology Services Laboratory | The Ohio State University Applied Microbiology Services Laboratory | Seth A. Faith PhD |
| EPI_ISL_2214702, EPI_ISL_2214883, EPI_ISL_2214888, EPI_ISL_2214972 | Centre Hospitalier Universitaire Clermont-Ferrand | CHU Clermont-Ferrand, service de virologie | Bisseux Maxime, Mirand Audrey, Combes Patricia, Henquell Cécile |
| EPI_ISL_2225607 | CHU Pontchaillou | CHU Pontchaillou | GROLHIER Claire, DENOUAL Florent, ETCHEVERRY Amandine, SASSI Mohamed, JAGLINE Steven, FEBREAU Christine, PRONIER Charlotte, GALIBERT Marie Dominique, DE TAYRAC Marie, THIBAULT Vincent |
| EPI_ISL_2226213, EPI_ISL_2226214 | Università di Parma, Laboratorio di Igiene e Sanità Pubblica | Istituto Zooprofilattico Sperimentale della Lombardia e dell'Emilia Romagna (IZSLER), Risk Analysis and Genomic Epidemiology Unit | Maria Eugenia Colucci, Licia Veronesi, Paola Affanni, Marina Morganti, Ilaria Menozzi, Erika Scaltriti, Stefano Pongolini |
| EPI_ISL_2226265, EPI_ISL_2226266 | BIO86 CHAUMONT | CHU Poitiers | Caroline MICHAUD, Maxime PICHON, Manon PRAT, Valentin BON-BARET, Birama N'DIAYE, Magali GARCIA, Agnes BEBY-DEFAUX, Nicolas LEVEQUE |
| EPI_ISL_2226267 | CH NIORT | CHU Poitiers | Caroline MICHAUD, Maxime PICHON, Manon PRAT, Valentin BON-BARET, Birama N'DIAYE, Magali GARCIA, Agnes BEBY-DEFAUX, Nicolas LEVEQUE |
| EPI_ISL_2226268 | CHU POITIERS | CHU Poitiers | Caroline MICHAUD, Maxime PICHON, Manon PRAT, Valentin BON-BARET, Birama N'DIAYE, Magali GARCIA, Agnes BEBY-DEFAUX, Nicolas LEVEQUE |
| EPI_ISL_2226269 | MEDILAB | CHU Poitiers | Caroline MICHAUD, Maxime PICHON, Manon PRAT, Valentin BON-BARET, Birama N'DIAYE, Magali GARCIA, Agnes BEBY-DEFAUX, Nicolas LEVEQUE |
| EPI_ISL_2226270, EPI_ISL_2226271 | BIO86 CHAUMONT | CHU Poitiers | Caroline MICHAUD, Maxime PICHON, Manon PRAT, Valentin BON-BARET, Birama N'DIAYE, Magali GARCIA, Agnes BEBY-DEFAUX, Nicolas LEVEQUE |
| EPI_ISL_2226272 | LABOFFICE | CHU Poitiers | Caroline MICHAUD, Maxime PICHON, Manon PRAT, Valentin BON-BARET, Birama N'DIAYE, Magali GARCIA, Agnes BEBY-DEFAUX, Nicolas LEVEQUE |
| EPI_ISL_2226273, EPI_ISL_2226274 | BIO86 CHAUMONT | CHU Poitiers | Caroline MICHAUD, Maxime PICHON, Manon PRAT, Valentin BON-BARET, Birama N'DIAYE, Magali GARCIA, Agnes BEBY-DEFAUX, Nicolas LEVEQUE |
| EPI_ISL_2226276 | MEDILAB | CHU Poitiers | Caroline MICHAUD, Maxime PICHON, Manon PRAT, Valentin BON-BARET, Birama N'DIAYE, Magali GARCIA, Agnes BEBY-DEFAUX, Nicolas LEVEQUE |
| EPI_ISL_2226277, EPI_ISL_2226278 | BIO86 CHAUMONT | CHU Poitiers | Caroline MICHAUD, Maxime PICHON, Manon PRAT, Valentin BON-BARET, Birama N'DIAYE, Magali GARCIA, Agnes BEBY-DEFAUX, Nicolas LEVEQUE |
| EPI_ISL_2226280 | CH NIORT | CHU Poitiers | Caroline MICHAUD, Maxime PICHON, Manon PRAT, Valentin BON-BARET, Birama N'DIAYE, Magali GARCIA, Agnes BEBY-DEFAUX, Nicolas LEVEQUE |
| EPI_ISL_2226282 | BIO86 CHAUMONT | CHU Poitiers | Caroline MICHAUD, Maxime PICHON, Manon PRAT, Valentin BON-BARET, Birama N'DIAYE, Magali GARCIA, Agnes BEBY-DEFAUX, Nicolas |

| | | | |
|---|---|---|---|
| | | | LEVEQUE |
| EPI_ISL_2226284 | MEDILAB | CHU Poitiers | Caroline MICHAUD, Maxime PICHON, Manon PRAT, Valentin BON-BARET, Birama N'DIAYE, Magali GARCIA, Agnes BEBY-DEFAUX, Nicolas LEVEQUE |
| EPI_ISL_2226285 | CH LA ROCHELLE | CHU Poitiers | Caroline MICHAUD, Maxime PICHON, Manon PRAT, Valentin BON-BARET, Birama N'DIAYE, Magali GARCIA, Agnes BEBY-DEFAUX, Nicolas LEVEQUE |
| EPI_ISL_2226286 | MEDILAB | CHU Poitiers | Caroline MICHAUD, Maxime PICHON, Manon PRAT, Valentin BON-BARET, Birama N'DIAYE, Magali GARCIA, Agnes BEBY-DEFAUX, Nicolas LEVEQUE |
| EPI_ISL_2226287 | CHU POITIERS | CHU Poitiers | Caroline MICHAUD, Maxime PICHON, Manon PRAT, Valentin BON-BARET, Birama N'DIAYE, Magali GARCIA, Agnes BEBY-DEFAUX, Nicolas LEVEQUE |
| EPI_ISL_2226288 | CH ANGOULEME | CHU Poitiers | Caroline MICHAUD, Maxime PICHON, Manon PRAT, Valentin BON-BARET, Birama N'DIAYE, Magali GARCIA, Agnes BEBY-DEFAUX, Nicolas LEVEQUE |
| EPI_ISL_2226289, EPI_ISL_2226290 | MEDILAB | CHU Poitiers | Caroline MICHAUD, Maxime PICHON, Manon PRAT, Valentin BON-BARET, Birama N'DIAYE, Magali GARCIA, Agnes BEBY-DEFAUX, Nicolas LEVEQUE |
| EPI_ISL_2226291 | CH ANGOULEME | CHU Poitiers | Caroline MICHAUD, Maxime PICHON, Manon PRAT, Valentin BON-BARET, Birama N'DIAYE, Magali GARCIA, Agnes BEBY-DEFAUX, Nicolas LEVEQUE |
| EPI_ISL_2226292 | LABOFFICE | CHU Poitiers | Caroline MICHAUD, Maxime PICHON, Manon PRAT, Valentin BON-BARET, Birama N'DIAYE, Magali GARCIA, Agnes BEBY-DEFAUX, Nicolas LEVEQUE |
| EPI_ISL_2226293 | MEDILAB | CHU Poitiers | Caroline MICHAUD, Maxime PICHON, Manon PRAT, Valentin BON-BARET, Birama N'DIAYE, Magali GARCIA, Agnes BEBY-DEFAUX, Nicolas LEVEQUE |
| EPI_ISL_2226294 | LABOFFICE | CHU Poitiers | Caroline MICHAUD, Maxime PICHON, Manon PRAT, Valentin BON-BARET, Birama N'DIAYE, Magali GARCIA, Agnes BEBY-DEFAUX, Nicolas LEVEQUE |
| EPI_ISL_2226295, EPI_ISL_2226296 | MEDILAB | CHU Poitiers | Caroline MICHAUD, Maxime PICHON, Manon PRAT, Valentin BON-BARET, Birama N'DIAYE, Magali GARCIA, Agnes BEBY-DEFAUX, Nicolas LEVEQUE |
| EPI_ISL_2226298 | CHU POITIERS | CHU Poitiers | Caroline MICHAUD, Maxime PICHON, Manon PRAT, Valentin BON-BARET, Birama N'DIAYE, Magali GARCIA, Agnes BEBY-DEFAUX, Nicolas LEVEQUE |
| EPI_ISL_2226300 | MEDILAB | CHU Poitiers | Caroline MICHAUD, Maxime PICHON, Manon PRAT, Valentin BON-BARET, Birama N'DIAYE, Magali GARCIA, Agnes BEBY-DEFAUX, Nicolas LEVEQUE |
| EPI_ISL_2226301 | BIO86 CHAUMONT | CHU Poitiers | Caroline MICHAUD, Maxime PICHON, Manon PRAT, Valentin BON-BARET, Birama N'DIAYE, Magali GARCIA, Agnes BEBY-DEFAUX, Nicolas LEVEQUE |
| EPI_ISL_2226303 | CH SAINTONGE | CHU Poitiers | Caroline MICHAUD, Maxime PICHON, Manon PRAT, Valentin BON-BARET, Birama N'DIAYE, Magali GARCIA, Agnes BEBY-DEFAUX, Nicolas LEVEQUE |
| EPI_ISL_2226304 | BIO86 CHAUMONT | CHU Poitiers | Caroline MICHAUD, Maxime PICHON, Manon PRAT, Valentin BON-BARET, Birama N'DIAYE, Magali GARCIA, Agnes BEBY-DEFAUX, Nicolas LEVEQUE |
| EPI_ISL_2226307 | CH HENRI LABORIT | CHU Poitiers | Caroline MICHAUD, Maxime PICHON, Manon PRAT, Valentin BON-BARET, Birama N'DIAYE, Magali GARCIA, Agnes BEBY-DEFAUX, Nicolas LEVEQUE |
| EPI_ISL_2226309 | CHU POITIERS | CHU Poitiers | Caroline MICHAUD, Maxime PICHON, Manon PRAT, Valentin BON-BARET, Birama N'DIAYE, Magali GARCIA, Agnes BEBY-DEFAUX, Nicolas LEVEQUE |
| EPI_ISL_2226311 | CH HENRI LABORIT | CHU Poitiers | Caroline MICHAUD, Maxime PICHON, Manon PRAT, Valentin BON-BARET, Birama N'DIAYE, Magali GARCIA, Agnes BEBY-DEFAUX, Nicolas LEVEQUE |
| EPI_ISL_2226315 | CH LA ROCHELLE | CHU Poitiers | Caroline MICHAUD, Maxime PICHON, Manon PRAT, Valentin BON-BARET, Birama N'DIAYE, Magali GARCIA, Agnes BEBY-DEFAUX, Nicolas LEVEQUE |
| EPI_ISL_2226323 | CHU POITIERS | CHU Poitiers | Caroline MICHAUD, Maxime PICHON, Manon PRAT, Valentin BON-BARET, Birama N'DIAYE, Magali GARCIA, Agnes BEBY-DEFAUX, Nicolas LEVEQUE |
| EPI_ISL_2226324 | CH NIORT | CHU Poitiers | Caroline MICHAUD, Maxime PICHON, Manon PRAT, Valentin BON-BARET, Birama N'DIAYE, Magali GARCIA, Agnes BEBY-DEFAUX, Nicolas LEVEQUE |
| EPI_ISL_2226342 | CH ANGOULEME | CHU Poitiers | Caroline MICHAUD, Maxime PICHON, Manon PRAT, Valentin BON-BARET, Birama N'DIAYE, Magali GARCIA, Agnes BEBY-DEFAUX, Nicolas LEVEQUE |
| EPI_ISL_2226351 | CHU POITIERS | CHU Poitiers | Caroline MICHAUD, Maxime PICHON, Manon PRAT, Valentin BON-BARET, Birama N'DIAYE, Magali GARCIA, Agnes BEBY-DEFAUX, Nicolas LEVEQUE |
| EPI_ISL_2226353, EPI_ISL_2226366 | MEDILAB | CHU Poitiers | Caroline MICHAUD, Maxime PICHON, Manon PRAT, Valentin BON-BARET, Birama N'DIAYE, Magali GARCIA, Agnes BEBY-DEFAUX, Nicolas LEVEQUE |
| EPI_ISL_2226368 | CHU POITIERS | CHU Poitiers | Caroline MICHAUD, Maxime PICHON, Manon PRAT, Valentin BON-BARET, Birama N'DIAYE, Magali GARCIA, Agnes BEBY-DEFAUX, Nicolas LEVEQUE |
| EPI_ISL_2226369 | CH LA ROCHELLE | CHU Poitiers | Caroline MICHAUD, Maxime PICHON, Manon PRAT, Valentin BON-BARET, Birama N'DIAYE, Magali GARCIA, Agnes BEBY-DEFAUX, Nicolas LEVEQUE |
| EPI_ISL_2226371 | LABOFFICE | CHU Poitiers | Caroline MICHAUD, Maxime PICHON, Manon PRAT, Valentin BON-BARET, Birama N'DIAYE, Magali GARCIA, Agnes BEBY-DEFAUX, Nicolas LEVEQUE |
| EPI_ISL_2226372 | MEDILAB | CHU Poitiers | Caroline MICHAUD, Maxime PICHON, Manon PRAT, Valentin BON-BARET, Birama N'DIAYE, Magali GARCIA, Agnes BEBY-DEFAUX, Nicolas LEVEQUE |
| EPI_ISL_2226676, EPI_ISL_2226677, EPI_ISL_2226678, EPI_ISL_2226679, EPI_ISL_2226680, EPI_ISL_2226681, EPI_ISL_2226683, EPI_ISL_2226684, EPI_ISL_2226685, EPI_ISL_2226686, EPI_ISL_2226687, EPI_ISL_2226688, EPI_ISL_2226689, EPI_ISL_2226691, EPI_ISL_2226692, EPI_ISL_2226694, EPI_ISL_2226696, EPI_ISL_2226697, EPI_ISL_2226699, EPI_ISL_2226702 | | | |
| see above | ADMED Microbiologie | Genomics and Transcriptomics, Philip Morris International | Reto Lienhard, Marie-Lise Tritten, Emmanuel Guedj, Nicolas Sierro, Rémi Dulize, David Bornand, Mehdi Auberson, Maxime Berthouzoz, Nikolai Ivanov, Manuel Peitsch |
| EPI_ISL_2227205, EPI_ISL_2227206, EPI_ISL_2227207 | CH ARMENTIERES | CHU Lille - Laboratoire de Virologie | AIT YAHYA Emilie, ALIDJINOU Enagnon Kazali, BOCKET Laurence, CREPIN Michel, DEMAY Christophe, ENGELMANN Ilka, GEFFROY Sandrine, GUIGON Aurélie, LAMBERT Valérie, LAZREK Mouna, NOBILLIAUX Florian, PREVOST Brigitte, THUILLIER Caroline, TINEZ Claire ,TCHANTCHOU NJOSSE YANICK |
| EPI_ISL_2227210 | CH BETHUNE | CHU Lille - Laboratoire de Virologie | AIT YAHYA Emilie, ALIDJINOU Enagnon Kazali, BOCKET Laurence, CREPIN Michel, DEMAY Christophe, ENGELMANN Ilka, GEFFROY Sandrine, GUIGON Aurélie, LAMBERT Valérie, LAZREK Mouna, NOBILLIAUX Florian, PREVOST Brigitte, THUILLIER Caroline, TINEZ Claire ,TCHANTCHOU NJOSSE YANICK |
| EPI_ISL_2227233, EPI_ISL_2227238 | CH ARRAS | CHU Lille - Laboratoire de Virologie | AIT YAHYA Emilie, ALIDJINOU Enagnon Kazali, BOCKET Laurence, CREPIN Michel, DEMAY Christophe, ENGELMANN Ilka, GEFFROY Sandrine, GUIGON Aurélie, LAMBERT Valérie, LAZREK Mouna, NOBILLIAUX Florian, PREVOST Brigitte, THUILLIER Caroline, TINEZ Claire ,TCHANTCHOU |

| | | | |
|---|---|---|---|
| | | | NJOSSE YANICK |
| EPI_ISL_2227745, EPI_ISL_2227746 | Microbiology Department - University Hospital Brussel | Microbiology Department - University Hospital Brussel | Oriane Soetens, Thomas Demuyser, Lucie Seyler |
| EPI_ISL_2228076, EPI_ISL_2228081, EPI_ISL_2228083, EPI_ISL_2228085, EPI_ISL_2228086 | Microbiology Department, University Hospital Donostia | Microbiology Department, University Hospital Donostia | Marimon JM, Montes M, Piñeiro L, Grandioso D, Martin-Peñaranda T, Gomez M, Cilla G. |
| EPI_ISL_2232728, EPI_ISL_2232729, EPI_ISL_2232730, EPI_ISL_2232731, EPI_ISL_2232732, EPI_ISL_2232733, EPI_ISL_2232734, EPI_ISL_2232735, EPI_ISL_2232736, EPI_ISL_2232737, EPI_ISL_2232738, EPI_ISL_2232739, EPI_ISL_2232740, EPI_ISL_2232741, EPI_ISL_2232743, EPI_ISL_2232744, EPI_ISL_2232745, EPI_ISL_2232746, EPI_ISL_2232747, EPI_ISL_2232748, EPI_ISL_2232749, EPI_ISL_2232750, EPI_ISL_2232751, EPI_ISL_2232752 | | | |
| see above | Department of Laboratory Medicine, Division of Clinical Virology, University of Medicine, Vienna | Bergthaler laboratory, CeMM Research Center for Molecular Medicine of the Austrian Academy of Sciences | Lukas Endler, Anna Schedl, Fabian Amman, Petr Triska, Thomas Penz, Benedikt Agerer, Maelle Le Moing, Michael Schuster, Bekir Erguner, Jan Laine, Martin Senekowitsch, Christoph Bock, Andreas Bergthaler |
| EPI_ISL_2232772, EPI_ISL_2232775, EPI_ISL_2232776, EPI_ISL_2232777, EPI_ISL_2232787, EPI_ISL_2232793, EPI_ISL_2232794, EPI_ISL_2232797, EPI_ISL_2232804, EPI_ISL_2232807, EPI_ISL_2232808, EPI_ISL_2232814, EPI_ISL_2232818, EPI_ISL_2232819, EPI_ISL_2232826, EPI_ISL_2232830, EPI_ISL_2232831, EPI_ISL_2232841, EPI_ISL_2232842, EPI_ISL_2232849, EPI_ISL_2232852, EPI_ISL_2232853, EPI_ISL_2232854, EPI_ISL_2232855, EPI_ISL_2232860, EPI_ISL_2232862, EPI_ISL_2232863, EPI_ISL_2232873, EPI_ISL_2232875, EPI_ISL_2232876, EPI_ISL_2232880, EPI_ISL_2232884, EPI_ISL_2232885, EPI_ISL_2232892, EPI_ISL_2232893, EPI_ISL_2232898, EPI_ISL_2232904, EPI_ISL_2232905, EPI_ISL_2232906, EPI_ISL_2232909, EPI_ISL_2232914, EPI_ISL_2232915, EPI_ISL_2232916, EPI_ISL_2232920, EPI_ISL_2232922, EPI_ISL_2232951, EPI_ISL_2232952 | | | |
| see above | Austrian Agency for Health and Food Safety (AGES) | Bergthaler laboratory, CeMM Research Center for Molecular Medicine of the Austrian Academy of Sciences | Lukas Endler, Anna Schedl, Fabian Amman, Petr Triska, Thomas Penz, Benedikt Agerer, Maelle Le Moing, Michael Schuster, Bekir Erguner, Jan Laine, Martin Senekowitsch, Christoph Bock, Andreas Bergthaler |
| EPI_ISL_2235301, EPI_ISL_2235302, EPI_ISL_2235304 | AZ St.Jan Brugge-Oostende AV | AZ SINT-JAN BRUGGE | Marijke Reynders, Patrick Descheemaeker, Laurien Hoornaert, Thomas Van Landschoot, Jorn Hellemans, Marie Madeleine Chabert-Consen |
| EPI_ISL_2259256, EPI_ISL_2259258, EPI_ISL_2259259, EPI_ISL_2259260, EPI_ISL_2259261, EPI_ISL_2259262, EPI_ISL_2259266, EPI_ISL_2259267 | National Center of Infectious and Parasitic Diseases | National Center of Infectious and Parasitic Diseases | Alexiev et al |
| EPI_ISL_2259288 | GH JOFFRE DUPUYTREN | Department of Virology, Henri Mondor University Hospital, Assistance Publique Hôpitaux de Paris, Université Paris-Est Créteil, INSERM U955 | Christophe Rodriguez, Slim Fourati, Vanessa Demontant, Guillaume Gricourt, Melissa N'Debi, Alexandre Soulier, Elisabeth Trawinski, Jean-Michel Pawlotsky |
| EPI_ISL_2259290 | HOPITAL EMILE ROUX | Department of Virology, Henri Mondor University Hospital, Assistance Publique Hôpitaux de Paris, Université Paris-Est Créteil, INSERM U955 | Christophe Rodriguez, Slim Fourati, Vanessa Demontant, Guillaume Gricourt, Melissa N'Debi, Alexandre Soulier, Elisabeth Trawinski, Jean-Michel Pawlotsky |
| EPI_ISL_2259292, EPI_ISL_2259293 | GH JOFFRE DUPUYTREN | Department of Virology, Henri Mondor University Hospital, Assistance Publique Hôpitaux de Paris, Université Paris-Est Créteil, INSERM U955 | Christophe Rodriguez, Slim Fourati, Vanessa Demontant, Guillaume Gricourt, Melissa N'Debi, Alexandre Soulier, Elisabeth Trawinski, Jean-Michel Pawlotsky |
| EPI_ISL_2259294, EPI_ISL_2259295 | GH A.CHENEVIER-H.MONDOR | Department of Virology, Henri Mondor University Hospital, Assistance Publique Hôpitaux de Paris, Université Paris-Est Créteil, INSERM U955 | Christophe Rodriguez, Slim Fourati, Vanessa Demontant, Guillaume Gricourt, Melissa N'Debi, Alexandre Soulier, Elisabeth Trawinski, Jean-Michel Pawlotsky |
| EPI_ISL_2259296 | HOPITAL EMILE ROUX | Department of Virology, Henri Mondor University Hospital, Assistance Publique Hôpitaux de Paris, Université Paris-Est Créteil, INSERM U955 | Christophe Rodriguez, Slim Fourati, Vanessa Demontant, Guillaume Gricourt, Melissa N'Debi, Alexandre Soulier, Elisabeth Trawinski, Jean-Michel Pawlotsky |
| EPI_ISL_2259302, EPI_ISL_2259304 | GH A.CHENEVIER-H.MONDOR | Department of Virology, Henri Mondor University Hospital, Assistance Publique Hôpitaux de Paris, Université Paris-Est Créteil, INSERM U955 | Christophe Rodriguez, Slim Fourati, Vanessa Demontant, Guillaume Gricourt, Melissa N'Debi, Alexandre Soulier, Elisabeth Trawinski, Jean-Michel Pawlotsky |
| EPI_ISL_2262546, EPI_ISL_2262548, EPI_ISL_2262550, EPI_ISL_2262552, EPI_ISL_2262554, EPI_ISL_2262564, EPI_ISL_2262575 | OCME Office Of Chief Medical Examiner | New York City Public Health Laboratory | Jade Wang, et al. |
| EPI_ISL_2272958 | AREA DE SALUD SIQUIRRES | Inciensa, Instituto Costarricense de Investigación y Enseñanza en Nutrición y Salud | Francisco Duarte, Hebleen Porras, Claudio Soto-Garita, Estela Cordero, Adriana Godínez, Melany Calderón, José Luis Vargas, Mariela Gutiérrez, Joselyn Prado & Ileana Chaves-Peraza |
| EPI_ISL_2272961, EPI_ISL_2272962 | HOSPITAL MONSEÑOR SANABRIA | Inciensa, Instituto Costarricense de Investigación y Enseñanza en Nutrición y Salud | Francisco Duarte, Hebleen Porras, Claudio Soto-Garita, Estela Cordero, Adriana Godínez, Melany Calderón, José Luis Vargas, Mariela Gutiérrez, Joselyn Prado & María José Gómez-Umaña |
| EPI_ISL_2272963 | AREA DE SALUD COBANO | Inciensa, Instituto Costarricense de Investigación y Enseñanza en Nutrición y Salud | Francisco Duarte, Hebleen Porras, Claudio Soto-Garita, Estela Cordero, Adriana Godínez, Melany Calderón, José Luis Vargas, Mariela Gutiérrez, Joselyn Prado & María José Gómez-Umaña |
| EPI_ISL_2272964 | HOSPITAL MONSEÑOR SANABRIA | Inciensa, Instituto Costarricense de Investigación y Enseñanza en Nutrición y Salud | Francisco Duarte, Hebleen Porras, Claudio Soto-Garita, Estela Cordero, Adriana Godínez, Melany Calderón, José Luis Vargas, Mariela Gutiérrez, Joselyn Prado & María José Gómez-Umaña |
| EPI_ISL_2272965 | AREA DE SALUD HEREDIA-CUBUJUQUI - CLINICA DR. FRANCISCO BOLAÑOS | Inciensa, Instituto Costarricense de Investigación y Enseñanza en Nutrición y Salud | Francisco Duarte, Hebleen Porras, Claudio Soto-Garita, Estela Cordero, Adriana Godínez, Melany Calderón, José Luis Vargas, Mariela Gutiérrez, Joselyn Prado & Juan Carlos Villalobos-Ugalde |
| EPI_ISL_2274115, EPI_ISL_2274117, EPI_ISL_2274128, EPI_ISL_2274130 | Laboratorio Central de Saude Publica do Estado do Rio Grande do Sul (LACEN-RS) | Laboratory of Respiratory Viruses and Measles, Oswaldo Cruz Institute, FIOCRUZ | Paola Resende, Luciana Appolinario, Fernando Motta, Anna Carolina Paixao, Ana Carolina Mendonca, Alice Sampaio Rocha, Taina Venas, Elisa Cavalcante Pereira, Renata Serrano Lopes, Tatiana Schaffer Gregianini, Richard Salvato, Marilda Siqueira on behalf of the Fiocruz COVID-19 Genomic Surveillance Network |
| EPI_ISL_2278010 | Institut National d'Hygiène (INH) | Unité Mixte Internationale TransVIHMI (UMI 233 IRD - U1175 INSERM - Université de Montpellier)IRD (Institut de recherche pour le développement) | Mounerou SALOU, Christelle BUTEL, Wembo A. HALATOKO,Issaka Maman, Abla A. KONOU, Amivi EHLAN, Adodo SADJI, Kokou TEGUENI,Sidonie A.M.KAGNISSODE, Akoélé SILIADIN, Alassane OURO-MEDELI, Messanh DOUFFAN,Déléma MABA,Sika DOSSIM, Améyo DORKENOO, Mireille PRINCE-DAVID,Anoumou DAGNRA,Laetitia SERRANO,Ahidjo AYOUBA,Eric DELAPORTE, Martine PEETERS |
| EPI_ISL_2279610, EPI_ISL_2279611 | Hospital | National Reference Center for Viruses of Respiratory Infections, Institut Pasteur, Paris | Marion Barbet, Sylvie Behillil, Méline Bizard, Angela Brisebarre, Camille Capel, Vincent Enouf, Louise Lefrançois, Frédéric Lemoine, Christophe Malabat, Corinne Maufrais, Amaury Vaysse, Etienne Simon-Lorière, Maud Vanpeene, Sylvie Van der Werf ,StéPhanie Marque Juillet |
| EPI_ISL_2279612, EPI_ISL_2279613 | Hospital | National Reference Center for Viruses of Respiratory Infections, Institut Pasteur, Paris | Marion Barbet, Sylvie Behillil, Méline Bizard, Angela Brisebarre, Camille Capel, Vincent Enouf, Louise Lefrançois, Frédéric Lemoine, Christophe Malabat, Corinne Maufrais, Etienne Simon-Lorière, Maud Vanpeene, Sylvie Van der Werf ,StéPhanie Marque Juillet |
| EPI_ISL_2279615, EPI_ISL_2279616 | Hospital | National Reference Center for Viruses of Respiratory Infections, Institut Pasteur, Paris | Marion Barbet, Sylvie Behillil, Méline Bizard, Angela Brisebarre, Camille Capel, Vincent Enouf, Louise Lefrançois, Frédéric Lemoine, Christophe Malabat, Corinne Maufrais, Amaury Vaysse, Etienne Simon-Lorière, Maud Vanpeene, Sylvie Van der Werf ,StéPhanie Marque Juillet |
| EPI_ISL_2279617, EPI_ISL_2279618, EPI_ISL_2279619 | Hospital | National Reference Center for Viruses of Respiratory Infections, Institut Pasteur, Paris | Marion Barbet, Sylvie Behillil, Méline Bizard, Angela Brisebarre, Camille Capel, Vincent Enouf, Louise Lefrançois, Frédéric Lemoine, Christophe Malabat, Corinne Maufrais, Etienne Simon-Lorière, Maud Vanpeene, Sylvie Van der Werf ,StéPhanie Marque Juillet |
| EPI_ISL_2279620 | Hospital | National Reference Center for Viruses of Respiratory Infections, Institut Pasteur, Paris | Marion Barbet, Sylvie Behillil, Méline Bizard, Angela Brisebarre, Camille Capel, Vincent Enouf, Louise Lefrançois, Frédéric Lemoine, Christophe Malabat, Corinne Maufrais, Ammanuelle Permal, Etienne Simon-Lorière, Maud Vanpeene, Sylvie Van der Werf ,StéPhanie Marque Juillet |
| EPI_ISL_2279621, EPI_ISL_2279624, EPI_ISL_2279626 | Hospital | National Reference Center for Viruses of Respiratory Infections, Institut Pasteur, Paris | Marion Barbet, Sylvie Behillil, Méline Bizard, Angela Brisebarre, Camille Capel, Vincent Enouf, Louise Lefrançois, Frédéric Lemoine, Christophe Malabat, Corinne Maufrais, Etienne Simon-Lorière, Maud Vanpeene, Sylvie Van der Werf ,StéPhanie Marque Juillet |
| EPI_ISL_2279628 | Hospital | National Reference Center for Viruses of Respiratory Infections, Institut Pasteur, Paris | Marion Barbet, Sylvie Behillil, Méline Bizard, Angela Brisebarre, Camille Capel, Vincent Enouf, Louise Lefrançois, Frédéric Lemoine, Christophe Malabat, Corinne Maufrais, Amaury Vaysse, Etienne Simon-Lorière, Maud Vanpeene, Sylvie Van der Werf ,StéPhanie Marque Juillet |
| EPI_ISL_2279640 | Hospital | National Reference Center for Viruses of Respiratory | Marion Barbet, Sylvie Behillil, Méline Bizard, Angela Brisebarre, Camille Capel, Vincent Enouf, Louise Lefrançois, Frédéric Lemoine, Christophe Malabat, |

| | | | |
|---|---|---|---|
| | | Infections, Institut Pasteur, Paris | Corinne Maufrais, Etienne Simon-Lorière, Maud Vanpeene, Sylvie Van der Werf ,StéPhanie Marque Juillet |
| EPI_ISL_2283681, EPI_ISL_2283682 | LESP Oaxaca | Instituto de Diagnostico y Referencia Epidemiologicos (INDRE) | Claudia Wong-Arambula, Abril Rodriguez-Maldonado, Vanessa Rivero-Arredondo, Ariadna Medina-Benitez, Joaquin Quiroz-Mercado, Sergio Rangel-Guerrero, Natividad Cruz-Ortiz, Tatiana Nunez-Garcia, Gisela Barrera-Badillo, Lucia Hernandez-Rivas, Irma Lopez-Martinez, Ernesto Ramirez-Gonzalez. |
| EPI_ISL_2283695 | LESP Michoacan | Instituto de Diagnostico y Referencia Epidemiologicos (INDRE) | Claudia Wong-Arambula, Abril Rodriguez-Maldonado, Vanessa Rivero-Arredondo, Ariadna Medina-Benitez, Joaquin Quiroz-Mercado, Sergio Rangel-Guerrero, Natividad Cruz-Ortiz, Tatiana Nunez-Garcia, Gisela Barrera-Badillo, Lucia Hernandez-Rivas, Irma Lopez-Martinez, Ernesto Ramirez-Gonzalez. |
| EPI_ISL_2283698, EPI_ISL_2283699, EPI_ISL_2283719 | LESP Yucatan | Instituto de Diagnostico y Referencia Epidemiologicos (INDRE) | Claudia Wong-Arambula, Abril Rodriguez-Maldonado, Vanessa Rivero-Arredondo, Ariadna Medina-Benitez, Joaquin Quiroz-Mercado, Sergio Rangel-Guerrero, Natividad Cruz-Ortiz, Tatiana Nunez-Garcia, Gisela Barrera-Badillo, Lucia Hernandez-Rivas, Irma Lopez-Martinez, Ernesto Ramirez-Gonzalez. |
| EPI_ISL_2283723, EPI_ISL_2283724 | LESP Nuevo Leon | Instituto de Diagnostico y Referencia Epidemiologicos (INDRE) | Claudia Wong-Arambula, Abril Rodriguez-Maldonado, Vanessa Rivero-Arredondo, Ariadna Medina-Benitez, Joaquin Quiroz-Mercado, Sergio Rangel-Guerrero, Natividad Cruz-Ortiz, Tatiana Nunez-Garcia, Gisela Barrera-Badillo, Lucia Hernandez-Rivas, Irma Lopez-Martinez, Ernesto Ramirez-Gonzalez. |
| EPI_ISL_2283725 | LESP Chihuahua | Instituto de Diagnostico y Referencia Epidemiologicos (INDRE) | Claudia Wong-Arambula, Abril Rodriguez-Maldonado, Vanessa Rivero-Arredondo, Ariadna Medina-Benitez, Joaquin Quiroz-Mercado, Sergio Rangel-Guerrero, Natividad Cruz-Ortiz, Tatiana Nunez-Garcia, Gisela Barrera-Badillo, Lucia Hernandez-Rivas, Irma Lopez-Martinez, Ernesto Ramirez-Gonzalez. |
| EPI_ISL_2285108 | Microbiology Department. Complexo Hospitalario Universitario de Vigo | Microbiology Department. Complexo Hospitalario Universitario de Vigo | Alfaya N, Alonso I, Alvarez M, Cabrera JJ, Carballo R, Cores O, Cortizo S, del-Campo V, Martinez L, Mediero G, Perez S, Potel C, Regueiro B, Rey S, Vasallo FJ |
| EPI_ISL_2285843, EPI_ISL_2285844, EPI_ISL_2285845, EPI_ISL_2285846, EPI_ISL_2285848 | National Influenza Centre | National Influenza Centre | William K. Ampofo, Michael Marks, Ivy A. Asante, Sharon Hsu, Benjamiin B. Lindsey, Benjamin H. Foulkes, Mildred Adusei-Poku, Linda Boatemaa, Lorreta Kwasah, Joseph Oliver-Commey, Ernest Asiedu, Franklin Asiedu-Bekoe, Gordon Awandare, Joyce Ngoi, Dennis Laryea, Mathew D. Parker, Thushan I de Silva, |
| EPI_ISL_2289131, EPI_ISL_2289132, EPI_ISL_2289133, EPI_ISL_2289134, EPI_ISL_2289135, EPI_ISL_2289136, EPI_ISL_2289138, EPI_ISL_2289139, EPI_ISL_2289140, EPI_ISL_2289141, EPI_ISL_2289142, EPI_ISL_2289143, EPI_ISL_2289144 | | | |
| see above | Laboratori de Referencia de Catalunya | Laboratori de Referencia de Catalunya | Ramirez A., Bellosillo B., Padilla E., Vilas A., Hernandez JJ., Canal M. |
| EPI_ISL_2293415, EPI_ISL_2293416, EPI_ISL_2293417, EPI_ISL_2293418, EPI_ISL_2293419, EPI_ISL_2293420, EPI_ISL_2293424, EPI_ISL_2293426, EPI_ISL_2293427, EPI_ISL_2293428, EPI_ISL_2293429, EPI_ISL_2293430, EPI_ISL_2293431 | | | |
| see above | Research | National Reference Center for Viruses of Respiratory Infections, Institut Pasteur, Paris | Marion Barbet, Sylvie Behillil, Méline Bizard, Angela Brisebarre, Camille Capel, Vincent Enouf, Louise Lefrançois, Frédéric Lemoine, Christophe Malabat, Corinne Maufrais, Etienne Simon-Lorière, Maud Vanpeene, Sylvie Van der Werf ,Marianne Leruez-Ville |
| EPI_ISL_2308479, EPI_ISL_2308480, EPI_ISL_2308481, EPI_ISL_2308483, EPI_ISL_2308485, EPI_ISL_2308486, EPI_ISL_2308487, EPI_ISL_2308488, EPI_ISL_2308489, EPI_ISL_2308490, EPI_ISL_2308491, EPI_ISL_2308492, EPI_ISL_2308493, EPI_ISL_2308495 | | | |
| see above | Laboratori de Referencia de Catalunya | Laboratori de Referencia de Catalunya | Ramirez A., Bellosillo B., Padilla E., Vilas A., Hernandez JJ., Canal M. |
| EPI_ISL_2312658 | Centre Hospitalier Universitaire Clermont-Ferrand | CHU Clermont-Ferrand, service de virologie | Bisseux Maxime, Mirand Audrey, Combes Patricia, Henquell Cécile |
| EPI_ISL_2312659, EPI_ISL_2312661, EPI_ISL_2312662 | HOPITAL DE VICHY | CHU Clermont-Ferrand, service de virologie | Bisseux Maxime, Mirand Audrey, Combes Patricia, Henquell Cécile |
| EPI_ISL_2321586 | BIOREFERENCIA | Universidad Nacional de Colombia - Laboratorio Genómico One Health | Simón Villegas Velásquez, Andres F. Cardona-Rios, Maria Stella López, Daniel O. Maldonado-Perez, Lina M. Hurtado, Laura Silvana Perez, Karl A Ciuoderis, Maria Angélica Maya, Carolina Muñoz-Arango, Celeny Ortiz, Rita Almanza Payares, Idabely Betancur Ortiz, Sandra Ines Cano, Diego A. Álvarez-Díaz, Carlos Franco-Muñoz, Katherine Laiton-Donato, Hector Alejandro Ruiz-Moreno, Marcela Mercado-Reyes, Jorge E. Osorio, Juan P. Hernandez-Ortiz |
| EPI_ISL_2322757 | Collection of WHO National Influenza Center, N.F. Gamaleya National Research Center for Epidemiology and Microbiology of the Ministry of Health of the Russian Federation | Laboratory of genome analysis, N.F. Gamaleya National Research Center for Epidemiology and Microbiology of the Ministry of Health of the Russian Federation | Olga Voronina, Ekaterina Aksenova, Natalia Ryzhova, Marina Kunda, Kirill Krasnoslobodtsev, Elena Burtseva, Denis Logunov |
| EPI_ISL_2323006 | cerballiance-HDF | Cerba lab | Emmanuel Lecorche, Sabine Trombert-Paolantoni, Claire Felloni, Anne Mainardi, Laura Verdurme, Jean-Michel Damien, Stéphanie Haïm-Boukobza, Bénédicte Roquebert |
| EPI_ISL_2323007, EPI_ISL_2323008 | cerballiance-IDF | Cerba lab | Emmanuel Lecorche, Sabine Trombert-Paolantoni, Aude Lessenne, Kader Merah, Thierry Collin, Laura Verdurme, Patrice Herisson, Stéphanie Haïm-Boukobza, Bénédicte Roquebert |
| EPI_ISL_2323009 | cerballiance-HDF | Cerba lab | Emmanuel Lecorche, Sabine Trombert-Paolantoni, Claire Felloni, Anne Mainardi, Laura Verdurme, Jean-Michel Damien, Stéphanie Haïm-Boukobza, Bénédicte Roquebert |
| EPI_ISL_2323010 | cerballiance-IDF | Cerba lab | Emmanuel Lecorche, Sabine Trombert-Paolantoni, Aude Lessenne, Kader Merah, Thierry Collin, Laura Verdurme, Patrice Herisson, Stéphanie Haïm-Boukobza, Bénédicte Roquebert |
| EPI_ISL_2323011 | cerballiance-HDF | Cerba lab | Emmanuel Lecorche, Sabine Trombert-Paolantoni, Claire Felloni, Anne Mainardi, Laura Verdurme, Jean-Michel Damien, Stéphanie Haïm-Boukobza, Bénédicte Roquebert |
| EPI_ISL_2323012, EPI_ISL_2323013, EPI_ISL_2323015 | cerballiance-IDF | Cerba lab | Emmanuel Lecorche, Sabine Trombert-Paolantoni, Aude Lessenne, Kader Merah, Thierry Collin, Laura Verdurme, Patrice Herisson, Stéphanie Haïm-Boukobza, Bénédicte Roquebert |
| EPI_ISL_2323016 | cerballiance-ARA | Cerba lab | Emmanuel Lecorche, Sabine Trombert-Paolantoni, Emmanuel Chanard, Samuel Granjon, Laura Verdurme, Charles Leclerc, Stéphanie Haïm-Boukobza, Bénédicte Roquebert |
| EPI_ISL_2323467 | CH ANGOULEME | CHU Poitiers | Caroline MICHAUD, Maxime PICHON, Manon PRAT, Valentin BON-BARET, Birama N'DIAYE, Magali GARCIA, Agnes BEBY-DEFAUX, Nicolas LEVEQUE |
| EPI_ISL_2324262, EPI_ISL_2324263 | ILV Kärnten | Bergthaler laboratory, CeMM Research Center for Molecular Medicine of the Austrian Academy of Sciences | Lukas Endler, Anna Schedl, Fabian Amman, Petr Triska, Thomas Penz, Benedikt Agerer, Maelle Le Moing, Michael Schuster, Bekir Erguner, Jan Laine, Martin Senekowitsch, Christoph Bock, Andreas Bergthaler |
| EPI_ISL_2324347 | Austrian Agency for Health and Food Safety (AGES) | Bergthaler laboratory, CeMM Research Center for Molecular Medicine of the Austrian Academy of Sciences | Lukas Endler, Anna Schedl, Fabian Amman, Petr Triska, Thomas Penz, Benedikt Agerer, Maelle Le Moing, Michael Schuster, Bekir Erguner, Jan Laine, Martin Senekowitsch, Christoph Bock, Andreas Bergthaler |
| EPI_ISL_2325180 | Microbiology Department, University Hospital Donostia | Microbiology Department, University Hospital Donostia | Marimon JM, Montes M, Piñeiro L, Grandioso D, Martin-Peñaranda T, Gomez M, Cilla G. |
| EPI_ISL_2340872 | LESP Michoacan | Instituto de Diagnostico y Referencia Epidemiologicos (INDRE) | Claudia Wong-Arambula, Abril Rodriguez-Maldonado, Vanessa Rivero-Arredondo, Ariadna Medina-Benitez, Joaquin Quiroz-Mercado, Sergio Rangel-Guerrero, Natividad Cruz-Ortiz, Tatiana Nunez-Garcia, Gisela Barrera-Badillo, Lucia Hernandez-Rivas, Irma Lopez-Martinez, Ernesto Ramirez-Gonzalez. |
| EPI_ISL_2340874 | LESP Yucatan | Instituto de Diagnostico y Referencia Epidemiologicos (INDRE) | Claudia Wong-Arambula, Abril Rodriguez-Maldonado, Vanessa Rivero-Arredondo, Ariadna Medina-Benitez, Joaquin Quiroz-Mercado, Sergio Rangel-Guerrero, Natividad Cruz-Ortiz, Tatiana Nunez-Garcia, Gisela Barrera-Badillo, Lucia Hernandez-Rivas, Irma Lopez-Martinez, Ernesto Ramirez-Gonzalez. |
| EPI_ISL_2340890, EPI_ISL_2340892 | LESP Michoacan | Instituto de Diagnostico y Referencia Epidemiologicos (INDRE) | Claudia Wong-Arambula, Abril Rodriguez-Maldonado, Vanessa Rivero-Arredondo, Ariadna Medina-Benitez, Joaquin Quiroz-Mercado, Sergio Rangel-Guerrero, Natividad Cruz-Ortiz, Tatiana Nunez-Garcia, Gisela Barrera-Badillo, Lucia Hernandez-Rivas, Irma Lopez-Martinez, Ernesto Ramirez-Gonzalez. |
| EPI_ISL_2340915, EPI_ISL_2340916, EPI_ISL_2340917, EPI_ISL_2340918, EPI_ISL_2340919, EPI_ISL_2340920, EPI_ISL_2340921, EPI_ISL_2340922, EPI_ISL_2340923, EPI_ISL_2340924, EPI_ISL_2340925, EPI_ISL_2340926, EPI_ISL_2340927, EPI_ISL_2340928, EPI_ISL_2340929, EPI_ISL_2340930, EPI_ISL_2340931, EPI_ISL_2340932, EPI_ISL_2340933, EPI_ISL_2340934, EPI_ISL_2340935, EPI_ISL_2340936, EPI_ISL_2340937, EPI_ISL_2340938, EPI_ISL_2340939, EPI_ISL_2340940, EPI_ISL_2340941, EPI_ISL_2340998, EPI_ISL_2340999, EPI_ISL_2341000, EPI_ISL_2341001, EPI_ISL_2341024, EPI_ISL_2341025, EPI_ISL_2341026, EPI_ISL_2341027 | | | |

| | | | |
|---|---|---|---|
| see above | LESP Yucatan | Instituto de Diagnostico y Referencia Epidemiologicos (INDRE) | Claudia Wong-Arambula, Abril Rodriguez-Maldonado, Vanessa Rivero-Arredondo, Ariadna Medina-Benitez, Joaquin Quiroz-Mercado, Sergio Rangel-Guerrero, Natividad Cruz-Ortiz, Tatiana Nunez-Garcia, Gisela Barrera-Badillo, Lucia Hernandez-Rivas, Irma Lopez-Martinez, Ernesto Ramirez-Gonzalez. |
| EPI_ISL_2341347, EPI_ISL_2341348, EPI_ISL_2341350, EPI_ISL_2341353, EPI_ISL_2341355, EPI_ISL_2341356, EPI_ISL_2341358, EPI_ISL_2341359, EPI_ISL_2341361, EPI_ISL_2341362, EPI_ISL_2341363, EPI_ISL_2341364, EPI_ISL_2341365, EPI_ISL_2341366, EPI_ISL_2341367, EPI_ISL_2341368, EPI_ISL_2341369, EPI_ISL_2341370, EPI_ISL_2341371, EPI_ISL_2341372, EPI_ISL_2341373, EPI_ISL_2341374, EPI_ISL_2341375, EPI_ISL_2341376, EPI_ISL_2341377, EPI_ISL_2341378, EPI_ISL_2341379, EPI_ISL_2341380, EPI_ISL_2341381, EPI_ISL_2341382, EPI_ISL_2341383, EPI_ISL_2341384, EPI_ISL_2341385, EPI_ISL_2341386, EPI_ISL_2341387, EPI_ISL_2341388, EPI_ISL_2341389, EPI_ISL_2341390, EPI_ISL_2341391, EPI_ISL_2341392, EPI_ISL_2341393, EPI_ISL_2341394, EPI_ISL_2341395, EPI_ISL_2341396, EPI_ISL_2341397, EPI_ISL_2341398, EPI_ISL_2341399, EPI_ISL_2341400, EPI_ISL_2341401, EPI_ISL_2341402, EPI_ISL_2341403, EPI_ISL_2341404, EPI_ISL_2341405, EPI_ISL_2341406, EPI_ISL_2341407, EPI_ISL_2341408, EPI_ISL_2341409, EPI_ISL_2341410, EPI_ISL_2341411 | | | |
| see above | The Ohio State University Applied Microbiology Services Laboratory | The Ohio State University Applied Microbiology Services Laboratory | Seth A. Faith PhD |
| EPI_ISL_2343001, EPI_ISL_2343002, EPI_ISL_2343004 | LESP Yucatan | Instituto de Diagnostico y Referencia Epidemiologicos (INDRE) | Claudia Wong-Arambula, Abril Rodriguez-Maldonado, Vanessa Rivero-Arredondo, Ariadna Medina-Benitez, Joaquin Quiroz-Mercado, Sergio Rangel-Guerrero, Natividad Cruz-Ortiz, Tatiana Nunez-Garcia, Gisela Barrera-Badillo, Lucia Hernandez-Rivas, Irma Lopez-Martinez, Ernesto Ramirez-Gonzalez. |
| EPI_ISL_2343007 | LESP Ciudad de Mexico | Instituto de Diagnostico y Referencia Epidemiologicos (INDRE) | Claudia Wong-Arambula, Abril Rodriguez-Maldonado, Vanessa Rivero-Arredondo, Ariadna Medina-Benitez, Joaquin Quiroz-Mercado, Sergio Rangel-Guerrero, Natividad Cruz-Ortiz, Tatiana Nunez-Garcia, Gisela Barrera-Badillo, Lucia Hernandez-Rivas, Irma Lopez-Martinez, Ernesto Ramirez-Gonzalez. |
| EPI_ISL_2343018 | LESP Yucatan | Instituto de Diagnostico y Referencia Epidemiologicos (INDRE) | Claudia Wong-Arambula, Abril Rodriguez-Maldonado, Vanessa Rivero-Arredondo, Ariadna Medina-Benitez, Joaquin Quiroz-Mercado, Sergio Rangel-Guerrero, Natividad Cruz-Ortiz, Tatiana Nunez-Garcia, Gisela Barrera-Badillo, Lucia Hernandez-Rivas, Irma Lopez-Martinez, Ernesto Ramirez-Gonzalez. |
| EPI_ISL_2348705, EPI_ISL_2348729 | Gyncentrum | Gyncentrum | Emilia Morawiec, Pawel Czerwinski, Adam Pudelko, Aleksandra Skubis-Sikora, Magdalena Samul, Agnieszka Polak, Celina Kruszniewska-Rajs, Tomasz Wasik, Robert Wojtyczka, Jolanta Bartosiewicz- Wasik, Maria Maklasinska-Majdanik |
| EPI_ISL_2352604, EPI_ISL_2352688, EPI_ISL_2354075, EPI_ISL_2354478 | Dr. Gernot Walder GmbH | Dr. Gernot Walder GmbH | Sissy T. Sonnleitner |
| EPI_ISL_2360255 | Molecular Diagnosis Laboratory | National Influenza Center | J Yavarian, NZ Shafiei Jandaghi, V Salimi, A Nejati, K Sadeghi, Hamed Gouklani, Masoumeh Moosazadeh, F Ajaminejad,N Ghavvami and T Mokhtari Azad |
| EPI_ISL_2362673, EPI_ISL_2362676 | Microbiology and Virology Unit, Florence Careggi University Hospital | Microbiology and Virology Unit, Florence Careggi University Hospital | Vincenzo Di Pilato, Marco Coppi, Fabio Morecchiato, Noemi Aiezza, Ilaria Baccani, Nicla Giovacchini, Alberto Antonelli, Emanuele Gori, Gian Maria Rossolini |
| EPI_ISL_2375798 | Laboratório de Microbiologia Molecular - Universidade FEEVALE | Molecular Microbiology Laboratory | Alana Witt Hansen, Fágner Henrique Heldt, Fernando Rosado Spilki, Flávio Silveira, Juliana Schons Gularte, Juliane Deise Fleck, Mariana Soares da Silva, Meriane Demoliner, Matheus Nunes Weber, Paula Rodrigues de Almeida, Micheli Filippi. |
| EPI_ISL_2375899 | Programa de Oncovirologia, Instituto Nacional de Câncer | Programa de Oncovirologia, Instituto Nacional de Câncer | Livia R. Goes, Juliana D. Siqueira, Marianne M. Garrido, Brunna M. Alves, Ana Cristina P. M. Pereira, Claudia Cicala, James Arthos, João P.B. Viola, Marcelo A. Soares |
| EPI_ISL_2379012, EPI_ISL_2379013, EPI_ISL_2379025, EPI_ISL_2379029, EPI_ISL_2379031, EPI_ISL_2379034, EPI_ISL_2379035, EPI_ISL_2379044, EPI_ISL_2379045, EPI_ISL_2379046, EPI_ISL_2379050, EPI_ISL_2379052, EPI_ISL_2379053, EPI_ISL_2379054, EPI_ISL_2379055, EPI_ISL_2379056, EPI_ISL_2379058, EPI_ISL_2379061, EPI_ISL_2379062, EPI_ISL_2379064, EPI_ISL_2379065, EPI_ISL_2379066, EPI_ISL_2379072, EPI_ISL_2379073, EPI_ISL_2379074, EPI_ISL_2379075, EPI_ISL_2379076, EPI_ISL_2379077, EPI_ISL_2379079, EPI_ISL_2379080, EPI_ISL_2379081, EPI_ISL_2379082, EPI_ISL_2379084, EPI_ISL_2379086, EPI_ISL_2379087, EPI_ISL_2379089, EPI_ISL_2379090, EPI_ISL_2379091, EPI_ISL_2379092, EPI_ISL_2379093, EPI_ISL_2379094, EPI_ISL_2379095, EPI_ISL_2379096, EPI_ISL_2379097, EPI_ISL_2379098, EPI_ISL_2379101, EPI_ISL_2379102, EPI_ISL_2379104, EPI_ISL_2379105, EPI_ISL_2379106, EPI_ISL_2379107, EPI_ISL_2379108, EPI_ISL_2379110, EPI_ISL_2379111, EPI_ISL_2379113, EPI_ISL_2379114, EPI_ISL_2379115, EPI_ISL_2379116, EPI_ISL_2379117, EPI_ISL_2379118, EPI_ISL_2379119, EPI_ISL_2379121, EPI_ISL_2379122, EPI_ISL_2379123, EPI_ISL_2379124, EPI_ISL_2379125, EPI_ISL_2379126, EPI_ISL_2379145, EPI_ISL_2379146, EPI_ISL_2379147, EPI_ISL_2379148, EPI_ISL_2379149, EPI_ISL_2379150, EPI_ISL_2379151, EPI_ISL_2379152, EPI_ISL_2379153, EPI_ISL_2379154, EPI_ISL_2379155, EPI_ISL_2379156, EPI_ISL_2379182, EPI_ISL_2379183, EPI_ISL_2379184, EPI_ISL_2379185, EPI_ISL_2379186, EPI_ISL_2379187, EPI_ISL_2379188, EPI_ISL_2379189, EPI_ISL_2379190, EPI_ISL_2379191, EPI_ISL_2379197, EPI_ISL_2379199, EPI_ISL_2379200, EPI_ISL_2379201, EPI_ISL_2379202, EPI_ISL_2379203, EPI_ISL_2379204, EPI_ISL_2379205, EPI_ISL_2379206, EPI_ISL_2379207, EPI_ISL_2379208, EPI_ISL_2379209, EPI_ISL_2379210 | | | |
| see above | National Center of Infectious and Parasitic Diseases | National Center of Infectious and Parasitic Diseases | Alexiev et al |
| EPI_ISL_2381995, EPI_ISL_2381996, EPI_ISL_2381997 | AZ Zeno Knokke-Blankenberge | AZ Sint Jan | Marijke Reynders, Patrick Descheemaeker, Laurien Hoornaert, Thomas Van Landschoot, Marie Madeleine Chabert-Consen, Jorn Hellemans |
| EPI_ISL_2382753, EPI_ISL_2382754, EPI_ISL_2382755, EPI_ISL_2382757, EPI_ISL_2382758, EPI_ISL_2382759, EPI_ISL_2382760, EPI_ISL_2382761, EPI_ISL_2382762, EPI_ISL_2382763, EPI_ISL_2382764, EPI_ISL_2382765, EPI_ISL_2382767, EPI_ISL_2382769, EPI_ISL_2382771, EPI_ISL_2382772, EPI_ISL_2382773, EPI_ISL_2382774, EPI_ISL_2382775, EPI_ISL_2382776, EPI_ISL_2382777, EPI_ISL_2382778, EPI_ISL_2382779, EPI_ISL_2382780, EPI_ISL_2382781, EPI_ISL_2382782, EPI_ISL_2382783, EPI_ISL_2382784, EPI_ISL_2382785, EPI_ISL_2382786, EPI_ISL_2382787, EPI_ISL_2382788, EPI_ISL_2382789, EPI_ISL_2382790, EPI_ISL_2382791, EPI_ISL_2382792, EPI_ISL_2382794, EPI_ISL_2382795, EPI_ISL_2382796, EPI_ISL_2382797, EPI_ISL_2382798, EPI_ISL_2382799, EPI_ISL_2382800, EPI_ISL_2382801, EPI_ISL_2382802, EPI_ISL_2382803 | | | |
| see above | The Ohio State University Applied Microbiology Services Laboratory | The Ohio State University Applied Microbiology Services Laboratory | Seth A. Faith PhD |
| EPI_ISL_2386155, EPI_ISL_2386159 | Botswana Harvard HIV Reference Laboratory | Botswana Harvard HIV Reference Laboratory | Sikhulile Moyo, Wonderful T. Choga, Pamela Smith-Lawrence, Kgomotso Moruisi, Dorcas Maruapula, Keoratile Ntshambiwa, Thongbotho Mphoyakgosi, Boitumelo Zuze, Botshelo Radibe, Legodile Kooepile, Ontlametse T. Bareng, Roger Shapiro, Shahin Lockman, Joseph Makhema, Madisa Mine, Mosepele Mosepele, Simani Gaseitsiwe |
| EPI_ISL_2391479 | N.H.L Municipal Medical College, Ahmedabad | Gujarat Biotechnology Research Centre | Zuber Saiyed,Ramesh Pandit,Janvi Raval,Zarna Patel,Nitin Savaliya,Dinesh Kumar,Twinkle Soni,Sonal Sharma ,Jayshri Pethani,Umang Mishra,Nitesh Shah,Chaitanya Joshi,Madhvi Joshi |
| EPI_ISL_2391480 | N.H.L Municipal Medical College, Ahmedabad | Gujarat Biotechnology Research Centre | Ramesh Pandit,Janvi Raval,Zarna Patel,Nitin Savaliya,Dinesh Kumar,Twinkle Soni,Sonal Sharma ,Zuber Saiyed,Jayshri Pethani,Umang Mishra,Nitesh Shah,Chaitanya Joshi,Madhvi Joshi |
| EPI_ISL_2391485 | N.H.L Municipal Medical College, Ahmedabad | Gujarat Biotechnology Research Centre | Twinkle Soni,Sonal Sharma ,Zuber Saiyed,Ramesh Pandit,Janvi Raval,Zarna Patel,Nitin Savaliya,Dinesh Kumar,Jayshri Pethani,Umang Mishra,Nitesh Shah,Chaitanya Joshi,Madhvi Joshi |
| EPI_ISL_2391486 | N.H.L Municipal Medical College, Ahmedabad | Gujarat Biotechnology Research Centre | Sonal Sharma ,Zuber Saiyed,Ramesh Pandit,Janvi Raval,Zarna Patel,Nitin Savaliya,Dinesh Kumar,Twinkle Soni,Jayshri Pethani,Umang Mishra,Nitesh Shah,Chaitanya Joshi,Madhvi Joshi |
| EPI_ISL_2391487 | N.H.L Municipal Medical College, Ahmedabad | Gujarat Biotechnology Research Centre | Zuber Saiyed,Ramesh Pandit,Janvi Raval,Zarna Patel,Nitin Savaliya,Dinesh Kumar,Twinkle Soni,Sonal Sharma ,Jayshri Pethani,Umang Mishra,Nitesh Shah,Chaitanya Joshi,Madhvi Joshi |
| EPI_ISL_2391488 | BJ Medical College and Civil Hospital, Ahmedabad | Gujarat Biotechnology Research Centre | Janvi Raval,Zarna Patel,Nitin Savaliya,Dinesh Kumar,Twinkle Soni,Sonal Sharma ,Zuber Saiyed,Ramesh Pandit,Pranay Shah,Umang Mishra,Nitesh Shah,Chaitanya Joshi,Madhvi Joshi |
| EPI_ISL_2391507, EPI_ISL_2391523 | Centro de Investigación Biomédica del Noreste (CIBIN) | Instituto de Biotecnología de la UNAM | Consorcio Mexicano de Vigilancia Genómica (CoViGen-Mex). Authors (in alphabetical order): Julio Elias Alvarado-Yaah, Carlos F. Arias, Santiago Ávila-Ríos, Eduardo Becerril-Vargas, Víctor Hugo Borja-Aburto, Celia Boukadida, Cristóbal Cháidez-Quiróz, Juan Bautista Chale-Dzul, Ricardo Ciria Merce, Andreu Comas-García, Célida Duque Molina, Julissa Enciso-Ibarra, José Antonio Enciso-Moreno, Gloria Elena Espinosa-Ayala, Fernando Fontove-Herrera, Daniel Fregoso-Rueda, Victor Eduardo Garcia-Arias, Alejandra Garcia-Gasca, Bruno Gomez-Gil, Jean Pierre González, Irvin González-López, Concepción Grajales-Muñiz, Ricardo Grande, Rosa María Gutierrez Rios, Alejandra Hernández-Terán, Alfredo Herrera-Estrella, Carla Ivón Herrera-Najera, Pavel Isa, Daniel Lira Morales, Susana Lopez, Antonio Loza Román, Brenda Irasema Maldonado-Meza, Bernardo Martínez-Miguel, José Arturo Martínez-Orozco, Célida Martínez- Rodríguez, Margarita Matías-Florentino, Fidencio Mejía-Nepomuceno, María Guadalupe de Jesús Mireles-Rivera, Gloria María Molina-Salinas, Hector Montoya-Fuentes, , Mario Mújica-Sánchez, José Esteban Muñoz-Medina, José de Jesús Nuñez-Contreras, Alicia Ocaña-Mondragón, Luis Alberto Ochoa-Carrera, Hector Esteban Paz-Juárez, Francisco Pulido, Helen Haydee Fernanda Ramírez-Plascencia, Jorge Salas-Hernández, Angel Gustavo Salas-Lais, Alejandro Sanchez-Flores, Clara Esperanza Santacruz-Tinoco, María Guadalupe Santiago-Mauricio, Selene Zárate, Nelly Sélem-Mojica, Blanca Taboada, Gloria Vazquez, Joel Armando Vázquez-Pérez. |
| EPI_ISL_2391537 | Unidad de Investigación Médica de Yucatán (UIMY) | Instituto de Biotecnología de la UNAM | Consorcio Mexicano de Vigilancia Genómica (CoViGen-Mex). Authors (in alphabetical order): Julio Elias Alvarado-Yaah, Carlos F. Arias, Santiago Ávila-Ríos, Eduardo Becerril-Vargas, Víctor Hugo Borja-Aburto, Celia Boukadida, Cristóbal Cháidez-Quiróz, Juan Bautista Chale-Dzul, Ricardo Ciria Merce, Andreu Comas-García, Célida Duque Molina, Julissa Enciso-Ibarra, José Antonio Enciso-Moreno, Gloria Elena Espinosa-Ayala, Fernando Fontove-Herrera, Daniel Fregoso-Rueda, Victor Eduardo Garcia-Arias, Alejandra Garcia-Gasca, Bruno Gomez-Gil, Jean Pierre González, Irvin González-López, Concepción Grajales-Muñiz, Ricardo Grande, Rosa María Gutierrez Rios, Alejandra Hernández-Terán, Alfredo Herrera-Estrella, Carla |

| | | | |
|---|---|---|---|
| | | | Ivón Herrera-Najera, Pavel Isa, Daniel Lira Morales, Susana Lopez, Antonio Loza Román, Brenda Irasema Maldonado-Meza, Bernardo Martínez-Miguel, José Arturo Martínez-Orozco, Célida Martínez- Rodríguez, Margarita Matías-Florentino, Fidencio Mejía-Nepomuceno, María Guadalupe de Jesús Mireles-Rivera, Gloria María Molina-Salinas, Hector Montoya-Fuentes, , Mario Mújica-Sánchez, José Esteban Muñoz-Medina, José de Jesús Nuñez-Contreras, Alicia Ocaña-Mondragón, Luis Alberto Ochoa-Carrera, Hector Esteban Paz-Juárez, Francisco Pulido, Helen Haydee Fernanda Ramírez-Plascencia, Jorge Salas-Hernández, Angel Gustavo Salas-Lais, Alejandro Sanchez-Flores, Clara Esperanza Santacruz-Tinoco, María Guadalupe Santiago-Mauricio, Selene Zárate, Nelly Sélem-Mojica, Blanca Taboada, Gloria Vazquez, Joel Armando Vázquez-Pérez. |
| EPI_ISL_2391544, EPI_ISL_2391568 | Centro de Investigación Biomédica de Occidente (CIBO) | Instituto de Biotecnología de la UNAM | Consorcio Mexicano de Vigilancia Genómica (CoViGen-Mex). Authors (in alphabetical order): Julio Elias Alvarado-Yaah, Carlos F. Arias, Santiago Ávila-Ríos, Eduardo Becerril-Vargas, Víctor Hugo Borja-Aburto, Celia Boukadida, Cristóbal Cháidez-Quiróz, Juan Bautista Chale-Dzul, Ricardo Ciria Merce, Andreu Comas-García, Célida Duque Molina, Julissa Enciso-Ibarra, José Antonio Enciso-Moreno, Gloria Elena Espinosa-Ayala, Fernando Fontove-Herrera, Daniel Fregoso-Rueda, Victor Eduardo Garcia-Arias, Alejandra Garcia-Gasca, Bruno Gomez-Gil, Jean Pierre González, Irvin González-López, Concepción Grajales-Muñiz, Ricardo Grande, Rosa María Gutierrez Rios, Alejandra Hernández-Terán, Alfredo Herrera-Estrella, Carla Ivón Herrera-Najera, Pavel Isa, Daniel Lira Morales, Susana Lopez, Antonio Loza Román, Brenda Irasema Maldonado-Meza, Bernardo Martínez-Miguel, José Arturo Martínez-Orozco, Célida Martínez- Rodríguez, Margarita Matías-Florentino, Fidencio Mejía-Nepomuceno, María Guadalupe de Jesús Mireles-Rivera, Gloria María Molina-Salinas, Hector Montoya-Fuentes, , Mario Mújica-Sánchez, José Esteban Muñoz-Medina, José de Jesús Nuñez-Contreras, Alicia Ocaña-Mondragón, Luis Alberto Ochoa-Carrera, Hector Esteban Paz-Juárez, Francisco Pulido, Helen Haydee Fernanda Ramírez-Plascencia, Jorge Salas-Hernández, Angel Gustavo Salas-Lais, Alejandro Sanchez-Flores, Clara Esperanza Santacruz-Tinoco, María Guadalupe Santiago-Mauricio, Selene Zárate, Nelly Sélem-Mojica, Blanca Taboada, Gloria Vazquez, Joel Armando Vázquez-Pérez. |
| EPI_ISL_2391578, EPI_ISL_2391581 | Unidad de Investigación Médica de Yucatán (UIMY) | Instituto de Biotecnología de la UNAM | Consorcio Mexicano de Vigilancia Genómica (CoViGen-Mex). Authors (in alphabetical order): Julio Elias Alvarado-Yaah, Carlos F. Arias, Santiago Ávila-Ríos, Eduardo Becerril-Vargas, Víctor Hugo Borja-Aburto, Celia Boukadida, Cristóbal Cháidez-Quiróz, Juan Bautista Chale-Dzul, Ricardo Ciria Merce, Andreu Comas-García, Célida Duque Molina, Julissa Enciso-Ibarra, José Antonio Enciso-Moreno, Gloria Elena Espinosa-Ayala, Fernando Fontove-Herrera, Daniel Fregoso-Rueda, Victor Eduardo Garcia-Arias, Alejandra Garcia-Gasca, Bruno Gomez-Gil, Jean Pierre González, Irvin González-López, Concepción Grajales-Muñiz, Ricardo Grande, Rosa María Gutierrez Rios, Alejandra Hernández-Terán, Alfredo Herrera-Estrella, Carla Ivón Herrera-Najera, Pavel Isa, Daniel Lira Morales, Susana Lopez, Antonio Loza Román, Brenda Irasema Maldonado-Meza, Bernardo Martínez-Miguel, José Arturo Martínez-Orozco, Célida Martínez- Rodríguez, Margarita Matías-Florentino, Fidencio Mejía-Nepomuceno, María Guadalupe de Jesús Mireles-Rivera, Gloria María Molina-Salinas, Hector Montoya-Fuentes, , Mario Mújica-Sánchez, José Esteban Muñoz-Medina, José de Jesús Nuñez-Contreras, Alicia Ocaña-Mondragón, Luis Alberto Ochoa-Carrera, Hector Esteban Paz-Juárez, Francisco Pulido, Helen Haydee Fernanda Ramírez-Plascencia, Jorge Salas-Hernández, Angel Gustavo Salas-Lais, Alejandro Sanchez-Flores, Clara Esperanza Santacruz-Tinoco, María Guadalupe Santiago-Mauricio, Selene Zárate, Nelly Sélem-Mojica, Blanca Taboada, Gloria Vazquez, Joel Armando Vázquez-Pérez. |
| EPI_ISL_2391619 | Laboratorio Central de Epidemiología (LCE) | Instituto de Biotecnología de la UNAM | Consorcio Mexicano de Vigilancia Genómica (CoViGen-Mex). Authors (in alphabetical order): Julio Elias Alvarado-Yaah, Carlos F. Arias, Santiago Ávila-Ríos, Eduardo Becerril-Vargas, Víctor Hugo Borja-Aburto, Celia Boukadida, Cristóbal Cháidez-Quiróz, Juan Bautista Chale-Dzul, Ricardo Ciria Merce, Andreu Comas-García, Célida Duque Molina, Julissa Enciso-Ibarra, José Antonio Enciso-Moreno, Gloria Elena Espinosa-Ayala, Fernando Fontove-Herrera, Daniel Fregoso-Rueda, Victor Eduardo Garcia-Arias, Alejandra Garcia-Gasca, Bruno Gomez-Gil, Jean Pierre González, Irvin González-López, Concepción Grajales-Muñiz, Ricardo Grande, Rosa María Gutierrez Rios, Alejandra Hernández-Terán, Alfredo Herrera-Estrella, Carla Ivón Herrera-Najera, Pavel Isa, Daniel Lira Morales, Susana Lopez, Antonio Loza Román, Brenda Irasema Maldonado-Meza, Bernardo Martínez-Miguel, José Arturo Martínez-Orozco, Célida Martínez- Rodríguez, Margarita Matías-Florentino, Fidencio Mejía-Nepomuceno, María Guadalupe de Jesús Mireles-Rivera, Gloria María Molina-Salinas, Hector Montoya-Fuentes, , Mario Mújica-Sánchez, José Esteban Muñoz-Medina, José de Jesús Nuñez-Contreras, Alicia Ocaña-Mondragón, Luis Alberto Ochoa-Carrera, Hector Esteban Paz-Juárez, Francisco Pulido, Helen Haydee Fernanda Ramírez-Plascencia, Jorge Salas-Hernández, Angel Gustavo Salas-Lais, Alejandro Sanchez-Flores, Clara Esperanza Santacruz-Tinoco, María Guadalupe Santiago-Mauricio, Selene Zárate, Nelly Sélem-Mojica, Blanca Taboada, Gloria Vazquez, Joel Armando Vázquez-Pérez. |
| EPI_ISL_2400383 | BJ Medical College and Civil Hospital, Ahmedabad | Gujarat Biotechnology Research Centre | Ramesh Pandit,Janvi Raval,Zarna Patel,Nitin Savaliya,Dinesh Kumar,Twinkle Soni,Sonal Sharma ,Zuber Saiyed,Pranay Shah,Umang Mishra,Nitesh Shah,Chaitanya Joshi,Madhvi Joshi |
| EPI_ISL_2402633, EPI_ISL_2402634, EPI_ISL_2402638, EPI_ISL_2402639, EPI_ISL_2402640, EPI_ISL_2402641, EPI_ISL_2402642, EPI_ISL_2402644, EPI_ISL_2402645, EPI_ISL_2402646, EPI_ISL_2402647, EPI_ISL_2402648, EPI_ISL_2402649, EPI_ISL_2402650, EPI_ISL_2402651, EPI_ISL_2402652, EPI_ISL_2402653, EPI_ISL_2402654, EPI_ISL_2402655, EPI_ISL_2402656, EPI_ISL_2402657, EPI_ISL_2402658, EPI_ISL_2402659, EPI_ISL_2402660, EPI_ISL_2402661, EPI_ISL_2402662, EPI_ISL_2402663, EPI_ISL_2402664, EPI_ISL_2402665, EPI_ISL_2402675, EPI_ISL_2402676, EPI_ISL_2402677, EPI_ISL_2402678, EPI_ISL_2402679, EPI_ISL_2402680, EPI_ISL_2402681, EPI_ISL_2402682, EPI_ISL_2402683, EPI_ISL_2402684, EPI_ISL_2402686, EPI_ISL_2402687, EPI_ISL_2402688, EPI_ISL_2402689, EPI_ISL_2402690, EPI_ISL_2402691, EPI_ISL_2402692, EPI_ISL_2402693, EPI_ISL_2402694, EPI_ISL_2402695, EPI_ISL_2402696, EPI_ISL_2402697, EPI_ISL_2402698, EPI_ISL_2402699, EPI_ISL_2402700, EPI_ISL_2402701, EPI_ISL_2402702, EPI_ISL_2402703, EPI_ISL_2402704, EPI_ISL_2402705, EPI_ISL_2402706, EPI_ISL_2402707, EPI_ISL_2402708, EPI_ISL_2402709, EPI_ISL_2402710, EPI_ISL_2402711, EPI_ISL_2402712, EPI_ISL_2402713, EPI_ISL_2402714, EPI_ISL_2402715, EPI_ISL_2402716, EPI_ISL_2402717, EPI_ISL_2402718, EPI_ISL_2402719, EPI_ISL_2402720, EPI_ISL_2402721, EPI_ISL_2402722, EPI_ISL_2402723, EPI_ISL_2402724, EPI_ISL_2402725, EPI_ISL_2402726, EPI_ISL_2402727, EPI_ISL_2402728, EPI_ISL_2402729, EPI_ISL_2402730, EPI_ISL_2402731, EPI_ISL_2402732, EPI_ISL_2402733, EPI_ISL_2402734, EPI_ISL_2402735, EPI_ISL_2402736, EPI_ISL_2402737, EPI_ISL_2402739, EPI_ISL_2402740, EPI_ISL_2402741, EPI_ISL_2402742, EPI_ISL_2402743, EPI_ISL_2402744, EPI_ISL_2402745, EPI_ISL_2402746, EPI_ISL_2402747, EPI_ISL_2402748, EPI_ISL_2402750, EPI_ISL_2402751, EPI_ISL_2402752, EPI_ISL_2402753, EPI_ISL_2402754, EPI_ISL_2402755, EPI_ISL_2402756, EPI_ISL_2402758, EPI_ISL_2402759, EPI_ISL_2402760, EPI_ISL_2402761, EPI_ISL_2402762, EPI_ISL_2402763, EPI_ISL_2402764, EPI_ISL_2402765, EPI_ISL_2402766, EPI_ISL_2402767, EPI_ISL_2402768, EPI_ISL_2402771 | | | |
| see above | Gyncentrum | Gyncentrum | Emilia Morawiec, Pawe Czerwiski, Adam Pudeko, Aleksandra Skubis-Sikora, Magdalena Samul, Agnieszka Polak, Celina Kruszniewska-Rajs, Tomasz Wsik, Robert Wojtyczka, Jolanta Bartosiewicz- Wsik, Maria Maklasiska-Majdanik |
| EPI_ISL_2405120 | N.H.L Municipal Medical College, Ahmedabad | Gujarat Biotechnology Research Centre | Nitin Savaliya,Dinesh Kumar,Twinkle Soni,Sonal Sharma ,Zuber Saiyed,Ramesh Pandit,Janvi Raval,Zarna Patel,Jayshri Pethani,Umang Mishra,Nitesh Shah,Chaitanya Joshi,Madhvi Joshi |
| EPI_ISL_2405121 | N.H.L Municipal Medical College, Ahmedabad | Gujarat Biotechnology Research Centre | Dinesh Kumar,Twinkle Soni,Sonal Sharma ,Zuber Saiyed,Ramesh Pandit,Janvi Raval,Zarna Patel,Nitin Savaliya,Jayshri Pethani,Umang Mishra,Nitesh Shah,Chaitanya Joshi,Madhvi Joshi |
| EPI_ISL_2405123 | N.H.L Municipal Medical College, Ahmedabad | Gujarat Biotechnology Research Centre | Sonal Sharma ,Zuber Saiyed,Ramesh Pandit,Janvi Raval,Zarna Patel,Nitin Savaliya,Dinesh Kumar,Twinkle Soni,Jayshri Pethani,Umang Mishra,Nitesh Shah,Chaitanya Joshi,Madhvi Joshi |
| EPI_ISL_2405124 | N.H.L Municipal Medical College, Ahmedabad | Gujarat Biotechnology Research Centre | Zuber Saiyed,Ramesh Pandit,Janvi Raval,Zarna Patel,Nitin Savaliya,Dinesh Kumar,Twinkle Soni,Sonal Sharma ,Jayshri Pethani,Umang Mishra,Nitesh Shah,Chaitanya Joshi,Madhvi Joshi |
| EPI_ISL_2422416 | cerballiance-NOR | Cerba lab | Emmanuel Lecorche, Sabine Trombert-Paolantoni, Gwénolé Prigent, Antoine Prigent, Francois Xavier Le Guillou, Alexandre Leriche, Laura Verdurme, Sylvain Metge, Stéphanie Haïm-Boukobza, Bénédicte Roquebert |
| EPI_ISL_2422417 | cerballiance-HDF | Cerba lab | Emmanuel Lecorche, Sabine Trombert-Paolantoni, Claire Felloni, Anne Mainardi, Laura Verdurme, Jean-Michel Damien, Stéphanie Haïm-Boukobza, Bénédicte Roquebert |
| EPI_ISL_2422419 | cerballiance-NOR | Cerba lab | Emmanuel Lecorche, Sabine Trombert-Paolantoni, Gwénolé Prigent, Antoine Prigent, Francois Xavier Le Guillou, Alexandre Leriche, Laura Verdurme, Sylvain Metge, Stéphanie Haïm-Boukobza, Bénédicte Roquebert |
| EPI_ISL_2422421 | cerballiance-HDF | Cerba lab | Emmanuel Lecorche, Sabine Trombert-Paolantoni, Claire Felloni, Anne Mainardi, Laura Verdurme, Jean-Michel Damien, Stéphanie Haïm-Boukobza, Bénédicte Roquebert |
| EPI_ISL_2422422, EPI_ISL_2422424 | cerballiance-NOR | Cerba lab | Emmanuel Lecorche, Sabine Trombert-Paolantoni, Gwénolé Prigent, Antoine Prigent, Francois Xavier Le Guillou, Alexandre Leriche, Laura Verdurme, Sylvain Metge, Stéphanie Haïm-Boukobza, Bénédicte Roquebert |
| EPI_ISL_2422425, EPI_ISL_2422426 | cerballiance-ARA | Cerba lab | Emmanuel Lecorche, Sabine Trombert-Paolantoni, Emmanuel Chanard, Samuel Granjon, Laura Verdurme, Charles Leclerc, Stéphanie Haïm-Boukobza, Bénédicte Roquebert |
| EPI_ISL_2422427 | cerballiance-NOR | Cerba lab | Emmanuel Lecorche, Sabine Trombert-Paolantoni, Gwénolé Prigent, Antoine Prigent, Francois Xavier Le Guillou, Alexandre Leriche, Laura Verdurme, Sylvain Metge, Stéphanie Haïm-Boukobza, Bénédicte Roquebert |
| EPI_ISL_2422429 | cerballiance-ARA | Cerba lab | Emmanuel Lecorche, Sabine Trombert-Paolantoni, Emmanuel Chanard, Samuel Granjon, Laura Verdurme, Charles Leclerc, Stéphanie Haïm-Boukobza, Bénédicte Roquebert |
| EPI_ISL_2423010 | San Diego County Public Health Laboratory | Andersen lab at Scripps Research | SEARCH Alliance San Diego with Ashleigh Murphy, Jovan Shephard, Brett Austin |
| EPI_ISL_2425544 | cerballiance-ARA | Cerba lab | Emmanuel Lecorche, Sabine Trombert-Paolantoni, Emmanuel Chanard, Samuel Granjon, Laura Verdurme, Charles Leclerc, Stéphanie Haïm-Boukobza, |

| | | | |
|---|---|---|---|
| | | | Bénédicte Roquebert |
| EPI_ISL_2425545, EPI_ISL_2425546 | cerballiance-HDF | Cerba lab | Emmanuel Lecorche, Sabine Trombert-Paolantoni, Claire Felloni, Anne Mainardi, Laura Verdurme, Jean-Michel Damien, Stéphanie Haïm-Boukobza, Bénédicte Roquebert |
| EPI_ISL_2425547, EPI_ISL_2425548 | cerballiance-ARA | Cerba lab | Emmanuel Lecorche, Sabine Trombert-Paolantoni, Emmanuel Chanard, Samuel Granjon, Laura Verdurme, Charles Leclerc, Stéphanie Haïm-Boukobza, Bénédicte Roquebert |
| EPI_ISL_2425549 | cerballiance-HDF | Cerba lab | Emmanuel Lecorche, Sabine Trombert-Paolantoni, Claire Felloni, Anne Mainardi, Laura Verdurme, Jean-Michel Damien, Stéphanie Haïm-Boukobza, Bénédicte Roquebert |
| EPI_ISL_2425550 | cerballiance-ARA | Cerba lab | Emmanuel Lecorche, Sabine Trombert-Paolantoni, Emmanuel Chanard, Samuel Granjon, Laura Verdurme, Charles Leclerc, Stéphanie Haïm-Boukobza, Bénédicte Roquebert |
| EPI_ISL_2425551, EPI_ISL_2425552, EPI_ISL_2425553 | cerballiance-NOR | Cerba lab | Emmanuel Lecorche, Sabine Trombert-Paolantoni, Gwénolé Prigent, Antoine Prigent, Francois Xavier Le Guillou, Alexandre Leriche, Laura Verdurme, Sylvain Metge, Stéphanie Haïm-Boukobza, Bénédicte Roquebert |
| EPI_ISL_2425554 | cerballiance-OCC | Cerba lab | Emmanuel Lecorche, Sabine Trombert-Paolantoni, Anne-Claire Strzelecki, Laura Verdurme, Laurent Escudie, Stéphanie Haïm-Boukobza, Bénédicte Roquebert |
| EPI_ISL_2425555 | cerballiance-HDF | Cerba lab | Emmanuel Lecorche, Sabine Trombert-Paolantoni, Claire Felloni, Anne Mainardi, Laura Verdurme, Jean-Michel Damien, Stéphanie Haïm-Boukobza, Bénédicte Roquebert |
| EPI_ISL_2425556 | cerballiance-IDF | Cerba lab | Emmanuel Lecorche, Sabine Trombert-Paolantoni, Aude Lessenne, Kader Merah, Thierry Collin, Laura Verdurme, Patrice Herisson, Stéphanie Haïm-Boukobza, Bénédicte Roquebert |
| EPI_ISL_2425557 | cerballiance-OCC | Cerba lab | Emmanuel Lecorche, Sabine Trombert-Paolantoni, Anne-Claire Strzelecki, Laura Verdurme, Laurent Escudie, Stéphanie Haïm-Boukobza, Bénédicte Roquebert |
| EPI_ISL_2425558 | cerballiance-ARA | Cerba lab | Emmanuel Lecorche, Sabine Trombert-Paolantoni, Emmanuel Chanard, Samuel Granjon, Laura Verdurme, Charles Leclerc, Stéphanie Haïm-Boukobza, Bénédicte Roquebert |
| EPI_ISL_2425559, EPI_ISL_2425560 | cerballiance-HDF | Cerba lab | Emmanuel Lecorche, Sabine Trombert-Paolantoni, Claire Felloni, Anne Mainardi, Laura Verdurme, Jean-Michel Damien, Stéphanie Haïm-Boukobza, Bénédicte Roquebert |
| EPI_ISL_2425561 | cerballiance-ARA | Cerba lab | Emmanuel Lecorche, Sabine Trombert-Paolantoni, Emmanuel Chanard, Samuel Granjon, Laura Verdurme, Charles Leclerc, Stéphanie Haïm-Boukobza, Bénédicte Roquebert |
| EPI_ISL_2426147, EPI_ISL_2426148, EPI_ISL_2426149 | Clinical Virology Laboratory, Institute of Liver and Biliary Sciences | ILBS - IGIB | Ekta Gupta, Sheetalnath Rooge, Pratibha Kale, Amit Pandey, Chhagan Bihari Sharma, Partha Chattopadhyay, Priti Devi, Ranjeet Maurya, Neha Jha, Priyanka Mehta, Manish Kumar, Pooja Sharma, Sheeba Saifi, Aparna S, Sarfaraz Alam, Bharathram Uppili, Mohammed Faruq, Anurag Agrawal, Rajesh Pandey, Shiv Kumar Sarin |
| EPI_ISL_2426206, EPI_ISL_2426212, EPI_ISL_2426213 | All India Institute of Medical Sciences, Delhi | CSIR Institute of Genomics and Integrative Biology | Urvashi B Singh, Mercy Rophina, Rama Chaudhry, Vigneshwar Senthivel, Kiran Bala, Rahul C. Bhoyar, Bani Jolly, Nayer Jamshed, Mohamed Imran,Ritu Gupta, Praveen Aggarwal, Mohit Kumar Divakar, Subrata Sinha , Arvinden VR, Anjali Bajaj, Afra Shamnath, Abhinav Jain, COVID CBNAAT CORE GROUP , Vinod Scaria, Sridhar Sivasubbu, Randeep Guleria |
| EPI_ISL_2426858 | Centers for Disease Control and Prevention, Dengue Branch | Centers for Disease Control and Prevention, Dengue Branch | Gilberto A. Santiago, Glenda Gonzalez, Betzabel Flores, Keyla Charriez, Gabriela Paz-Bailey, Jorge L. Munoz-Jordan |
| EPI_ISL_2426861, EPI_ISL_2426862, EPI_ISL_2426863, EPI_ISL_2426864 | Puerto Rico Department of Health | Centers for Disease Control and Prevention, Dengue Branch | Gilberto A. Santiago, Glenda Gonzalez, Betzabel Flores, Keyla Charriez, Gabriela Paz-Bailey, Jorge L. Munoz-Jordan |
| EPI_ISL_2426873, EPI_ISL_2426874 | Centers for Disease Control and Prevention, Dengue Branch | Centers for Disease Control and Prevention, Dengue Branch | Gilberto A. Santiago, Glenda Gonzalez, Betzabel Flores, Keyla Charriez, Gabriela Paz-Bailey, Jorge L. Munoz-Jordan |
| EPI_ISL_2426880, EPI_ISL_2426881 | Puerto Rico Department of Health | Centers for Disease Control and Prevention, Dengue Branch | Gilberto A. Santiago, Glenda Gonzalez, Betzabel Flores, Keyla Charriez, Gabriela Paz-Bailey, Jorge L. Munoz-Jordan |
| EPI_ISL_2426887, EPI_ISL_2426888, EPI_ISL_2426889, EPI_ISL_2426890 | Centers for Disease Control and Prevention, Dengue Branch | Centers for Disease Control and Prevention, Dengue Branch | Gilberto A. Santiago, Glenda Gonzalez, Betzabel Flores, Keyla Charriez, Gabriela Paz-Bailey, Jorge L. Munoz-Jordan |
| EPI_ISL_2426892, EPI_ISL_2426896, EPI_ISL_2426897 | Puerto Rico Department of Health | Centers for Disease Control and Prevention, Dengue Branch | Gilberto A. Santiago, Glenda Gonzalez, Betzabel Flores, Keyla Charriez, Gabriela Paz-Bailey, Jorge L. Munoz-Jordan |
| EPI_ISL_2439136 | 65Biocev | Charles University, Faculty of Science, BIOCEV, OMICS Genomics | Vladimír Hampl, Štpánka Hrdá, Ruth Tachezy, Blanka Hamplová, Sebastian Cristian Treitli, Zoltán Füssy, Magdalena Janáová, Ingrid Poláková, Jana Šmahelová, Jí Novák |
| EPI_ISL_2441780 | CHU de la Réunion - Nord | Laboratoire de virologie, CNR arbovirus Associé, Chu de la Réunion | Etienne Frumence, Marie-Christine Jaffar Bandjee, Nicolas Traversier, Sabrina Petit Genet |
| EPI_ISL_2443502 | Baylor Scott & White-Temple | Baylor Scott & White-Temple | Ari Rao, Linden Morales, Kimberly Walker, Marcus Volz, Shelby Johnson |
| EPI_ISL_2443530, EPI_ISL_2443531, EPI_ISL_2443532, EPI_ISL_2443533, EPI_ISL_2443534, EPI_ISL_2443535, EPI_ISL_2443536, EPI_ISL_2443537, EPI_ISL_2443538 | Lboratorio Central de Saude Publica do Estado do Parana (LACEN/PR) | Laboratory of Respiratory Viruses and Measles, Oswaldo Cruz Institute, FIOCRUZ | Paola Resende, Luciana Appolinario, Fernando Motta, Anna Carolina Paixao, Ana Carolina Mendonca, Alice Sampaio Rocha, Taina Venas, Elisa Cavalcante Pereira, Renata Serrano Lopes, Irina Riediger, Marilda Siqueira on behalf of the Fiocruz COVID-19 Genomic Surveillance Network |
| EPI_ISL_2443551, EPI_ISL_2443552 | Labortorio Central de Saude Publica do Estado de Santa Catarina (LACEN/SC) | Laboratory of Respiratory Viruses and Measles, Oswaldo Cruz Institute, FIOCRUZ | Paola Resende, Luciana Appolinario, Fernando Motta, Anna Carolina Paixao, Ana Carolina Mendonca, Alice Sampaio Rocha, Taina Venas, Elisa Cavalcante Pereira, Renata Serrano Lopes, Darcita Buerger Rovaris, Sandra Bianchini Fernandes, Marilda Siqueira on behalf of the Fiocruz COVID-19 Genomic Surveillance Network |
| EPI_ISL_2443778, EPI_ISL_2443779, EPI_ISL_2443780 | The Ohio State University Applied Microbiology Services Laboratory | The Ohio State University Applied Microbiology Services Laboratory | Seth A. Faith PhD |
| EPI_ISL_2462862, EPI_ISL_2462863, EPI_ISL_2462864, EPI_ISL_2462865 | AZ Sint-Lucas Gent | AZ Sint-Lucas Gent | Jos Van Acker, Charlotte Verfaillie, Elke Vanlaere, Klara De Rauw |
| EPI_ISL_2484769, EPI_ISL_2484771 | Temporary Specimen Collection Centre at the AsiaWorld-Expo | Hong Kong Department of Health | Alan K.L. Tsang, Gannon C.K. Mak, Peter K.C. Cheng, Peter C.W. Yip, Ken H.L. Ng, Edman T.K. Lam, Rickjason C.W. Chan, Dominic N.C. Tsang |
| EPI_ISL_2484772 | HOME QUARANTINE TASKFORCE | Hong Kong Department of Health | Alan K.L. Tsang, Gannon C.K. Mak, Peter K.C. Cheng, Peter C.W. Yip, Ken H.L. Ng, Edman T.K. Lam, Rickjason C.W. Chan, Dominic N.C. Tsang |
| EPI_ISL_2484773 | North Lantau Hospital | Hong Kong Department of Health | Alan K.L. Tsang, Gannon C.K. Mak, Peter K.C. Cheng, Peter C.W. Yip, Ken H.L. Ng, Edman T.K. Lam, Rickjason C.W. Chan, Dominic N.C. Tsang |
| EPI_ISL_2484774 | Pamela Youde Nethersole Eastern Hospital | Hong Kong Department of Health | Alan K.L. Tsang, Gannon C.K. Mak, Peter K.C. Cheng, Peter C.W. Yip, Ken H.L. Ng, Edman T.K. Lam, Rickjason C.W. Chan, Dominic N.C. Tsang |
| EPI_ISL_2484775 | HOME QUARANTINE TASKFORCE | Hong Kong Department of Health | Alan K.L. Tsang, Gannon C.K. Mak, Peter K.C. Cheng, Peter C.W. Yip, Ken H.L. Ng, Edman T.K. Lam, Rickjason C.W. Chan, Dominic N.C. Tsang |
| EPI_ISL_2484776 | Temporary Specimen Collection Centre at the AsiaWorld-Expo | Hong Kong Department of Health | Alan K.L. Tsang, Gannon C.K. Mak, Peter K.C. Cheng, Peter C.W. Yip, Ken H.L. Ng, Edman T.K. Lam, Rickjason C.W. Chan, Dominic N.C. Tsang |
| EPI_ISL_2484777, EPI_ISL_2484778, EPI_ISL_2484780, EPI_ISL_2484781, EPI_ISL_2484782, EPI_ISL_2484783, EPI_ISL_2484785 | HOME QUARANTINE TASKFORCE | Hong Kong Department of Health | Alan K.L. Tsang, Gannon C.K. Mak, Peter K.C. Cheng, Peter C.W. Yip, Ken H.L. Ng, Edman T.K. Lam, Rickjason C.W. Chan, Dominic N.C. Tsang |
| EPI_ISL_2488812 | LACEN - Laboratório Central de Saúde Pública do Rio Grande do Norte | Evandro Chagas Institute | Santos, M.C.; Silva, A.M.; Junior, W.D.C.; Barbagelata, L.S.; Ferreira, J.A.; Sousa, E.M.A.; da Silva, P.S.; Pinheiro, K.C.; L.C.; Sousa Junior, E.C. |
| EPI_ISL_2491631 | Centre Hospitalier Universitaire Clermont-Ferrand | CHU Clermont-Ferrand, service de virologie | Bisseux Maxime, Mirand Audrey, Combes Patricia, Henquell Cécile |

| | | | |
|---|---|---|---|
| EPI_ISL_2495448, EPI_ISL_2495449, EPI_ISL_2495450, EPI_ISL_2495451, EPI_ISL_2495452, EPI_ISL_2495453, EPI_ISL_2495454, EPI_ISL_2495455, EPI_ISL_2495456, EPI_ISL_2495457, EPI_ISL_2495458, EPI_ISL_2495459, EPI_ISL_2495460, EPI_ISL_2495461, EPI_ISL_2495462, EPI_ISL_2495463, EPI_ISL_2495464, EPI_ISL_2495465, EPI_ISL_2495466, EPI_ISL_2495467, EPI_ISL_2495468, EPI_ISL_2495469, EPI_ISL_2495470, EPI_ISL_2495471, EPI_ISL_2495472, EPI_ISL_2495473, EPI_ISL_2495475, EPI_ISL_2495476, EPI_ISL_2495477, EPI_ISL_2495478, EPI_ISL_2495479, EPI_ISL_2495480, EPI_ISL_2495481, EPI_ISL_2495482, EPI_ISL_2495511, EPI_ISL_2495512, EPI_ISL_2495513, EPI_ISL_2495514, EPI_ISL_2495515, EPI_ISL_2495516, EPI_ISL_2495517, EPI_ISL_2495518, EPI_ISL_2495519, EPI_ISL_2495520, EPI_ISL_2495521, EPI_ISL_2495522, EPI_ISL_2495523, EPI_ISL_2495524, EPI_ISL_2495525, EPI_ISL_2495526, EPI_ISL_2495527, EPI_ISL_2495528, EPI_ISL_2495529, EPI_ISL_2495530, EPI_ISL_2495531, EPI_ISL_2495532, EPI_ISL_2495533, EPI_ISL_2495534, EPI_ISL_2495535, EPI_ISL_2495536, EPI_ISL_2495537, EPI_ISL_2495538, EPI_ISL_2495539, EPI_ISL_2495540, EPI_ISL_2495541, EPI_ISL_2495542, EPI_ISL_2495543, EPI_ISL_2495544, EPI_ISL_2495545, EPI_ISL_2495546, EPI_ISL_2495547, EPI_ISL_2495548, EPI_ISL_2495549, EPI_ISL_2495550, EPI_ISL_2495551, EPI_ISL_2495552, EPI_ISL_2495553, EPI_ISL_2495556, EPI_ISL_2495557, EPI_ISL_2495558, EPI_ISL_2495559, EPI_ISL_2495560, EPI_ISL_2495561, EPI_ISL_2495562, EPI_ISL_2495563, EPI_ISL_2495564, EPI_ISL_2495565, EPI_ISL_2495593, EPI_ISL_2495594, EPI_ISL_2495595, EPI_ISL_2495616, EPI_ISL_2495620, EPI_ISL_2495722, EPI_ISL_2495723, EPI_ISL_2495724, EPI_ISL_2495725, EPI_ISL_2495726, EPI_ISL_2495727, EPI_ISL_2495728, EPI_ISL_2495729, EPI_ISL_2495730, EPI_ISL_2495731, EPI_ISL_2495732, EPI_ISL_2495733, EPI_ISL_2495734 | | | |
| see above | MB-Cadham Provincial laboratory | National Microbiology Laboratory (NML) | Anna Majer, Shari Tyson, Grace Seo, Philip Mabon, Elsie Grudeski, Rhiannon Huzarewich, Russell Mandes, Anneliese Landgraff, Jennifer Tanner, Natalie Knox, Morag Graham, Gary Van Domselaar, Paul Van Caeseele, Jared Bullard, David Alexander, Kerry Dust, Nathalie Bastien, Yan Li, Timothy Booth, Darian Hole, Madison Chapel, Kirsten Biggar, CanCOGeN's metadata curation team, Public Health Agency of Canada CanCOGeN team |
| EPI_ISL_2500929, EPI_ISL_2500931 | Instituto Nacional de Salud | Instituto Nacional de Salud- Dirección de Investigación en Salud Pública | Katherine Laiton-Donato, Diego A. Álvarez-Díaz, Carlos Franco-Muñoz, Hector Alejandro Ruiz-Moreno, Paola Rojas, Maria T. Herrera-Sepúlveda, Diego Andrés Prada, Jhonnatan Reales-González, Sheryll Corchuelo, Julian Naizaque, Jorge Rivera, Gerardo Santamaría, Sergio Gomez, Lisseth Pardo, Juan Camilo Martinez, Marta Lopez Blanco, Ángela Alarcon Cruz, Diana Malo, Carmen Osorio, Magdalena Wiesner, Martha Lucia Ospina Martinez, Marcela Mercado-Reyes |
| EPI_ISL_2500969 | HOSPITAL DEPARTAMENTAL MARIA INMACULADA | Instituto Nacional de Salud- Dirección de Investigación en Salud Pública | Katherine Laiton-Donato, Diego A. Álvarez-Díaz, Carlos Franco-Muñoz, Hector Alejandro Ruiz-Moreno, Paola Rojas, Maria T. Herrera-Sepúlveda, Diego Andrés Prada, Jhonnatan Reales-González, Sheryll Corchuelo, Julian Naizaque, Jorge Rivera, Gerardo Santamaría, Sergio Gomez, Lisseth Pardo, Juan Camilo Martinez, Marta Lopez Blanco, Ángela Alarcon Cruz, Diana Malo, Carmen Osorio, Magdalena Wiesner, Martha Lucia Ospina Martinez, Marcela Mercado-Reyes |
| EPI_ISL_2501731, EPI_ISL_2501732, EPI_ISL_2501733, EPI_ISL_2501734, EPI_ISL_2501735 | Premier Laboratory Services | BioInfoExperts LLC | Cheryl Metzler, Valerie Murray, Rebecca Rose, David Nolan, Susanna Lamers |
| EPI_ISL_2502617 | U.O. Igiene, Ospedale Policlinico San Martino | U.O. Igiene, Ospedale Policlinico San Martino | Bruzzone Bianca, Caligiuri Patrizia, De Pace Vanessa, Domnich Alexander, Orsi Andrea, Ricucci Valentina, Icardi Giancarlo |
| EPI_ISL_2502618 | S.C. Laboratorio di analisi, Ente ospedaliero "Ospedali Galliera" Genova | U.O. Igiene, Ospedale Policlinico San Martino | Bruzzone Bianca, Caligiuri Patrizia, De Pace Vanessa, Domnich Alexander, Innocenti Lionello, Orsi Andrea, Ricucci Valentina, Icardi Giancarlo |
| EPI_ISL_2502619, EPI_ISL_2502621 | U.O. Igiene, Ospedale Policlinico San Martino | U.O. Igiene, Ospedale Policlinico San Martino | Bruzzone Bianca, Caligiuri Patrizia, De Pace Vanessa, Domnich Alexander, Orsi Andrea, Ricucci Valentina, Icardi Giancarlo |
| EPI_ISL_2509935 | N.H.L Municipal Medical College, Ahmedabad | Gujarat Biotechnology Research Centre | Twinkle Soni,Sonal Sharma ,Zuber Saiyed,Ramesh Pandit,Janvi Raval,Zarna Patel,Nitin Savaliya,Dinesh Kumar,Jayshri Pethani,Umang Mishra,Nitesh Shah,Chaitanya Joshi,Madhvi Joshi |
| EPI_ISL_2509936 | N.H.L Municipal Medical College, Ahmedabad | Gujarat Biotechnology Research Centre | Sonal Sharma ,Zuber Saiyed,Ramesh Pandit,Janvi Raval,Zarna Patel,Nitin Savaliya,Dinesh Kumar,Twinkle Soni,Jayshri Pethani,Umang Mishra,Nitesh Shah,Chaitanya Joshi,Madhvi Joshi |
| EPI_ISL_2509937 | N.H.L Municipal Medical College, Ahmedabad | Gujarat Biotechnology Research Centre | Zuber Saiyed,Ramesh Pandit,Janvi Raval,Zarna Patel,Nitin Savaliya,Dinesh Kumar,Twinkle Soni,Sonal Sharma ,Jayshri Pethani,Umang Mishra,Nitesh Shah,Chaitanya Joshi,Madhvi Joshi |
| EPI_ISL_2516963, EPI_ISL_2516968, EPI_ISL_2516969, EPI_ISL_2516970, EPI_ISL_2516971, EPI_ISL_2516972, EPI_ISL_2516973 | Microbiology Department. Complexo Hospitalario Universitario de Vigo | Microbiology Department. Complexo Hospitalario Universitario de Vigo | Alfaya N, Alonso I, Alvarez M, Cabrera JJ, Carballo R, Cores O, Cortizo S, del-Campo V, Martinez L, Mediero G, Perez S, Potel C, Regueiro B, Rey S, Vasallo FJ |
| EPI_ISL_2530111, EPI_ISL_2530120, EPI_ISL_2530122, EPI_ISL_2530124, EPI_ISL_2530126, EPI_ISL_2530128, EPI_ISL_2530129, EPI_ISL_2530131, EPI_ISL_2530132, EPI_ISL_2530137, EPI_ISL_2530139, EPI_ISL_2530141, EPI_ISL_2530146, EPI_ISL_2530148, EPI_ISL_2530149, EPI_ISL_2530151, EPI_ISL_2530154, EPI_ISL_2530155, EPI_ISL_2530157, EPI_ISL_2530163, EPI_ISL_2530165, EPI_ISL_2530167, EPI_ISL_2530169, EPI_ISL_2530170, EPI_ISL_2530172, EPI_ISL_2530176, EPI_ISL_2530265, EPI_ISL_2530272, EPI_ISL_2530273, EPI_ISL_2530275, EPI_ISL_2530276, EPI_ISL_2530295, EPI_ISL_2530317, EPI_ISL_2530319, EPI_ISL_2530321, EPI_ISL_2530322, EPI_ISL_2530324, EPI_ISL_2530326, EPI_ISL_2530330, EPI_ISL_2530334, EPI_ISL_2530336, EPI_ISL_2530337, EPI_ISL_2530339, EPI_ISL_2530340, EPI_ISL_2530344, EPI_ISL_2530349, EPI_ISL_2530353, EPI_ISL_2530357, EPI_ISL_2530360, EPI_ISL_2530364, EPI_ISL_2530366, EPI_ISL_2530367, EPI_ISL_2530369, EPI_ISL_2530371 | | | |
| see above | Hematopathology Laboratory, ACTREC, TMC | Hematopathology Laboratory, ACTREC, TMC | Hematopathology Laboratory, ACTREC |
| EPI_ISL_2534023, EPI_ISL_2534024, EPI_ISL_2534025, EPI_ISL_2534026, EPI_ISL_2534027, EPI_ISL_2534028, EPI_ISL_2534029, EPI_ISL_2534030, EPI_ISL_2534031, EPI_ISL_2534032, EPI_ISL_2534033, EPI_ISL_2534034 | | | |
| see above | S.C. Patologia Clinica, Ospedale Sant'Andrea, ASL 5 | U.O. Igiene, Ospedale Policlinico San Martino | Bruzzone Bianca, Battolla Enrico, Caligiuri Patrizia, De Pace Vanessa, Domnich Alexander, Orsi Andrea, Ricucci Valentina, Icardi Giancarlo |
| EPI_ISL_2534035, EPI_ISL_2534036 | U.O.C. Laboratorio Analisi, Ospedale G. Gaslini IRCCS Genova | U.O. Igiene, Ospedale Policlinico San Martino | Bruzzone Bianca, Caligiuri Patrizia, De Pace Vanessa, Domnich Alexander, Orsi Andrea, Ricucci Valentina, Tripodi Gino, Icardi Giancarlo |
| EPI_ISL_2534039, EPI_ISL_2534040 | S.S.D. Microbiologia, Stabilimento ospedaliero di Sanremo, ASL 1 Liguria | U.O. Igiene, Ospedale Policlinico San Martino | Bruzzone Bianca, Caligiuri Patrizia, De Pace Vanessa, Domnich Alexander, Dusi Pier Andrea, Orsi Andrea, Ricucci Valentina, Icardi Giancarlo |
| EPI_ISL_2534041, EPI_ISL_2534042, EPI_ISL_2534043 | Laboratorio di Patologia Clinica, Ospedale San Paolo in Valloria, ASL 2 Liguria | U.O. Igiene, Ospedale Policlinico San Martino | Bruzzone Bianca, Caligiuri Patrizia, De Pace Vanessa, Domnich Alexander, Lillo Flavia, Orsi Andrea, Ricucci Valentina, Icardi Giancarlo |
| EPI_ISL_2534044, EPI_ISL_2534045, EPI_ISL_2534046, EPI_ISL_2534047, EPI_ISL_2534048 | S.S.D. Microbiologia, Ospedale Santa Corona di Pietra Ligure, ASL 2 Liguria | U.O. Igiene, Ospedale Policlinico San Martino | Bruzzone Bianca, Caligiuri Patrizia, De Pace Vanessa, Domnich Alexander, Orsi Andrea, Ricucci Valentina, Valle Caterina, Icardi Giancarlo |
| EPI_ISL_2534049 | S.S.D. Microbiologia, Stabilimento ospedaliero di Sanremo, ASL 1 Liguria | U.O. Igiene, Ospedale Policlinico San Martino | Bruzzone Bianca, Caligiuri Patrizia, De Pace Vanessa, Domnich Alexander, Dusi Pier Andrea, Orsi Andrea, Ricucci Valentina, Icardi Giancarlo |
| EPI_ISL_2534050 | Laboratorio di Patologia Clinica, Ospedale San Paolo in Valloria, ASL 2 Liguria | U.O. Igiene, Ospedale Policlinico San Martino | Bruzzone Bianca, Caligiuri Patrizia, De Pace Vanessa, Domnich Alexander, Lillo Flavia, Orsi Andrea, Ricucci Valentina, Icardi Giancarlo |
| EPI_ISL_2534051 | U.O.C. Laboratorio Analisi, Ospedale G. Gaslini IRCCS Genova | U.O. Igiene, Ospedale Policlinico San Martino | Bruzzone Bianca, Caligiuri Patrizia, De Pace Vanessa, Domnich Alexander, Orsi Andrea, Ricucci Valentina, Tripodi Gino, Icardi Giancarlo |
| EPI_ISL_2534052, EPI_ISL_2534053 | U.O. Igiene, Ospedale Policlinico San Martino | U.O. Igiene, Ospedale Policlinico San Martino | Bruzzone Bianca, Caligiuri Patrizia, De Pace Vanessa, Domnich Alexander, Orsi Andrea, Ricucci Valentina, Icardi Giancarlo |
| EPI_ISL_2534054, EPI_ISL_2534055, EPI_ISL_2534056 | S.C. Laboratorio Analisi, ASL 3 Liguria | U.O. Igiene, Ospedale Policlinico San Martino | Bruzzone Bianca, Caligiuri Patrizia, De Pace Vanessa, Domnich Alexander, Orsi Andrea, Ricucci Valentina, Spitaleri Antonino, Icardi Giancarlo |
| EPI_ISL_2534077 | S.C. Patologia Clinica, Ospedale Sant'Andrea, ASL 5 | U.O. Igiene, Ospedale Policlinico San Martino | Bruzzone Bianca, Battolla Enrico, Caligiuri Patrizia, De Pace Vanessa, Domnich Alexander, Orsi Andrea, Ricucci Valentina, Icardi Giancarlo |
| EPI_ISL_2535792 | National Influenza Centre | National Influenza Centre | William K. Ampofo, Michael Marks, Ivy A. Asante, Sharon Hsu, Benjamiin B. Lindsey, Benjamin H. Foulkes, Mildred Adusei-Poku, Linda Boatemaa, Lorreta Kwasah, Joseph Oliver-Commey, Ernest Asiedu, Franklin Asiedu-Bekoe, Gordon Awandare, Joyce Ngoi, Dennis Laryea, Mathew D. Parker, Thushan I de Silva, |
| EPI_ISL_2536262, EPI_ISL_2536264 | Labortorio Central de Saude Publica do Estado de Santa Catarina (LACEN/SC) | Laboratory of Respiratory Viruses and Measles, Oswaldo Cruz Institute, FIOCRUZ | Paola Resende, Luciana Appolinario, Fernando Motta, Anna Carolina Paixao, Ana Carolina Mendonca, Alice Sampaio Rocha, Taina Venas, Elisa Cavalcante Pereira, Renata Serrano Lopes, Darcita Buerger Rovaris, Sandra Bianchini Fernandes, Marilda Siqueira on behalf of the Fiocruz COVID-19 Genomic Surveillance Network |
| EPI_ISL_2536311, EPI_ISL_2536313, EPI_ISL_2536321, EPI_ISL_2536322 | Laboratorio Central de Saude Publica do Estado da Paraiba (LACEN-PB) | Laboratory of Respiratory Viruses and Measles, Oswaldo Cruz Institute, FIOCRUZ | Paola Resende, Luciana Appolinario, Fernando Motta, Anna Carolina Paixao, Ana Carolina Mendonca, Alice Sampaio Rocha, Taina Venas, Elisa Cavalcante Pereira, Renata Serrano Lopes, Joao Felipe Bezerra, Dalane Loudal Florentino Teixeira, Marilda Siqueira on behalf of the Fiocruz COVID-19 Genomic Surveillance Network |
| EPI_ISL_2544193 | Provincial Public Health Reference Laboratory (PPHRL) P& SHD | Provincial Public Health Reference Laboratory (PPHRL) P& SHD | ANDLEEB HANIF |

| | | | |
|---|---|---|---|
| EPI_ISL_2544286, EPI_ISL_2544494, EPI_ISL_2544496 | Baker lab at Loyola Medical Center | Baker lab at Loyola Medical Center | Xufang Deng, PhD, Monika Evdokimova, BS, Amornrat O'Brien, PhD, Cynthia Rowe, PhD, Nina M. Clark, MD, Amanda Harrington, PhD, Gail E. Reid, MD, Susan Uprichard, PhD, and Susan C. Baker, PhD |
| EPI_ISL_2550530, EPI_ISL_2550538, EPI_ISL_2550540, EPI_ISL_2550541, EPI_ISL_2550542, EPI_ISL_2550543, EPI_ISL_2550546, EPI_ISL_2550551, EPI_ISL_2550557 | NB-Hôpital Georges L. Dumont | National Microbiology Laboratory (NML) | Anna Majer, Shari Tyson, Grace Seo, Philip Mabon, Elsie Grudeski, Rhiannon Huzarewich, Russell Mandes, Anneliese Landgraff, Jennifer Tanner, Natalie Knox, Morag Graham, Gary Van Domselaar, Richard Garceau, Guillaume Desnoyers, Nathalie Bastien, Yan Li, Timothy Booth, Darian Hole, Madison Chapel, Kirsten Biggar, CanCOGeN's metadata curation team, Public Health Agency of Canada CanCOGeN team |
| EPI_ISL_2562035, EPI_ISL_2562036 | DPM Diagnostics | Cerba lab | Emmanuel Lecorche, Sabine Trombert-Paolantoni, Aude Lessenne, Kader Merah, Thierry Collin, Laura Verdurme, Patrice Herisson, Stéphanie Haïm-Boukobza, Bénédicte Roquebert |
| EPI_ISL_2562039, EPI_ISL_2562040, EPI_ISL_2562041, EPI_ISL_2562042, EPI_ISL_2562043, EPI_ISL_2562044 | cerballiance-IDF | Cerba lab | Emmanuel Lecorche, Sabine Trombert-Paolantoni, Aude Lessenne, Kader Merah, Thierry Collin, Laura Verdurme, Patrice Herisson, Stéphanie Haïm-Boukobza, Bénédicte Roquebert |
| EPI_ISL_2565859, EPI_ISL_2565860 | S.C. Laboratorio Analisi, Ospedale di Lavagna, ASL 4 Liguria | U.O. Igiene, Ospedale Policlinico San Martino | Bruzzone Bianca, Bandettini Roberto, Caligiuri Patrizia, De Pace Vanessa, Domnich Alexander, Orsi Andrea, Ricucci Valentina, Icardi Giancarlo |
| EPI_ISL_2566237 | SC (UCO) Igiene e Sanità Pubblica, ASUGI, Trieste and SC Microbiologia e Virologia, ASFO, Pordenone | ARGO Laboratorio Genomica ed Epigenomica | Licastro D, Dal Monego S, Degasperi M, Marcello A, Segat L, De Rosa R, Braida C, Barbone F, D'Agaro P |
| EPI_ISL_2566238 | SC (UCO) Igiene e Sanità Pubblica, ASUGI, Trieste | ARGO Laboratorio Genomica ed Epigenomica | Licastro D, Dal Monego S, Degasperi M, Marcello A, Maggiore A, Busetti M, Segat L, Piscianz E, Braida C, Barbone F, D'Agaro P |
| EPI_ISL_2600353 | Temporary Specimen Collection Centre at the AsiaWorld-Expo | Hong Kong Department of Health | Alan K.L. Tsang, Gannon C.K. Mak, Peter K.C. Cheng, Peter C.W. Yip, Ken H.L. Ng, Edman T.K. Lam, Rickjason C.W. Chan, Dominic N.C. Tsang |
| EPI_ISL_2600362 | Princess Margaret Hospital | Hong Kong Department of Health | Alan K.L. Tsang, Gannon C.K. Mak, Peter K.C. Cheng, Peter C.W. Yip, Ken H.L. Ng, Edman T.K. Lam, Rickjason C.W. Chan, Dominic N.C. Tsang |
| EPI_ISL_2600363, EPI_ISL_2600364 | HOME QUARANTINE TASKFORCE | Hong Kong Department of Health | Alan K.L. Tsang, Gannon C.K. Mak, Peter K.C. Cheng, Peter C.W. Yip, Ken H.L. Ng, Edman T.K. Lam, Rickjason C.W. Chan, Dominic N.C. Tsang |
| EPI_ISL_2600953, EPI_ISL_2600954, EPI_ISL_2600955, EPI_ISL_2600956, EPI_ISL_2600957, EPI_ISL_2600958, EPI_ISL_2600959, EPI_ISL_2600962, EPI_ISL_2600963 | Hematopathology Laboratory, ACTREC, TMC | Hematopathology Laboratory, ACTREC, TMC | Hematopathology Laboratory, ACTREC |
| EPI_ISL_2601589, EPI_ISL_2601590, EPI_ISL_2601592, EPI_ISL_2601593, EPI_ISL_2601594, EPI_ISL_2601595, EPI_ISL_2601596, EPI_ISL_2601597, EPI_ISL_2601598, EPI_ISL_2601600, EPI_ISL_2601601, EPI_ISL_2601602, EPI_ISL_2601603, EPI_ISL_2601604, EPI_ISL_2601605, EPI_ISL_2601606, EPI_ISL_2601607, EPI_ISL_2601608, EPI_ISL_2601609 | | | |
| see above | Laboratory of Communicable Diseases | 1. Laboratory of Communicable Diseases (Estonia); 2. Eurofins Genomics Europe Sequencing GmbH | Liidia Dotsenko et al. |
| EPI_ISL_2601613, EPI_ISL_2601614, EPI_ISL_2601615, EPI_ISL_2601616, EPI_ISL_2601617, EPI_ISL_2601618, EPI_ISL_2601620, EPI_ISL_2601621, EPI_ISL_2601624, EPI_ISL_2601625, EPI_ISL_2601627, EPI_ISL_2601629, EPI_ISL_2601630, EPI_ISL_2601631, EPI_ISL_2601632, EPI_ISL_2601633, EPI_ISL_2601634, EPI_ISL_2601635, EPI_ISL_2601637 | | | |
| see above | North Estonia Medical Centre | 1. Laboratory of Communicable Diseases (Estonia); 2. Eurofins Genomics Europe Sequencing GmbH | Liidia Dotsenko et al. |
| EPI_ISL_2601652, EPI_ISL_2601653, EPI_ISL_2601654 | Pärnu Hospital | 1. Laboratory of Communicable Diseases (Estonia); 2. Eurofins Genomics Europe Sequencing GmbH | Liidia Dotsenko et al. |
| EPI_ISL_2603132, EPI_ISL_2603133, EPI_ISL_2603137, EPI_ISL_2603138, EPI_ISL_2603139, EPI_ISL_2603141, EPI_ISL_2603143, EPI_ISL_2603144, EPI_ISL_2603146, EPI_ISL_2603147, EPI_ISL_2603148, EPI_ISL_2603153, EPI_ISL_2603154, EPI_ISL_2603156, EPI_ISL_2603160, EPI_ISL_2603162, EPI_ISL_2603163, EPI_ISL_2603177, EPI_ISL_2603179, EPI_ISL_2603180, EPI_ISL_2603181 | | | |
| see above | Grupo de Investigación en Enfermedades Tropicales del Ejército (GINETEJ), Laboratorio de Referencia e Investigación, Dirección de Sanidad Ejército, Bogotá, Colombia | Centro de Investigaciones en Microbiología y Biotecnología-UR (CIMBIUR), Facultad de Ciencias Naturales, Universidad del Rosario, Bogotá, Colombia | Sergio Castañeda, Nathalia Ballesteros, Marina Muñoz, Luz H. Patiño, Claudia Méndez, Carolina Oliveros, Julie Pérez, Lorena Albarracín, Elizabeth K. Márquez, María Teresa Alvarado, Frank de los Santos Ortiz, Yanira Romero, Camilo A. Correa-Cárdenas, Maria Clara Duque, Sergio Gutiérrez-Riveros, Zulma Cucunubá, Juan David Ramírez |
| EPI_ISL_2603427, EPI_ISL_2603428, EPI_ISL_2603429, EPI_ISL_2603431, EPI_ISL_2603438, EPI_ISL_2603440 | Lboratorio Central de Saude Publica do Estado do Parana (LACEN/PR) | Laboratory of Respiratory Viruses and Measles, Oswaldo Cruz Institute, FIOCRUZ | Paola Resende, Luciana Appolinario, Fernando Motta, Anna Carolina Paixao, Ana Carolina Mendonca, Alice Sampaio Rocha, Taina Venas, Elisa Cavalcante Pereira, Renata Serrano Lopes, Irina Riediger, Marilda Siqueira on behalf of the Fiocruz COVID-19 Genomic Surveillance Network |
| EPI_ISL_2612626, EPI_ISL_2612628 | Hospital | National Reference Center for Viruses of Respiratory Infections, Institut Pasteur, Paris | Marion Barbet, Sylvie Behillil, Méline Bizard, Angela Brisebarre, Camille Capel, Vincent Enouf, Louise Lefrançois, Frédéric Lemoine, Christophe Malabat, Corinne Maufrais, Amaury Vaysse, Etienne Simon-Lorière, Maud Vanpeene, Sylvie Van der Werf ,Alexandra Ducancelle |
| EPI_ISL_2614520, EPI_ISL_2614521, EPI_ISL_2614522, EPI_ISL_2614523, EPI_ISL_2614524, EPI_ISL_2614525, EPI_ISL_2614526, EPI_ISL_2614527, EPI_ISL_2614528, EPI_ISL_2614529, EPI_ISL_2614530, EPI_ISL_2614531, EPI_ISL_2614532, EPI_ISL_2614533, EPI_ISL_2614534, EPI_ISL_2614535, EPI_ISL_2614536, EPI_ISL_2614537, EPI_ISL_2614538 | | | |
| see above | Instituto Adolfo Lutz - Regional de Campinas | Instituto Adolfo Lutz, Interdiciplinary Procedures Center, Strategic Laboratory | Claudio Tavares Sacchi, Claudia Regina Gonçalves, Erica Valessa Ramos Gomes, Karoline Rodrigues Campos, Caio Vinicius Dias Lopes, Leonardo Jose Tadeu de Araujo |
| EPI_ISL_2614539 | Centro de Diagnostico SECEDI | Instituto Adolfo Lutz, Interdiciplinary Procedures Center, Strategic Laboratory | Claudio Tavares Sacchi, Claudia Regina Gonçalves, Erica Valessa Ramos Gomes, Karoline Rodrigues Campos, Caio Vinicius Dias Lopes, Leonardo Jose Tadeu de Araujo |
| EPI_ISL_2614544, EPI_ISL_2614545, EPI_ISL_2614546, EPI_ISL_2614547, EPI_ISL_2614548, EPI_ISL_2614549, EPI_ISL_2614550, EPI_ISL_2614551, EPI_ISL_2614552, EPI_ISL_2614553, EPI_ISL_2614554, EPI_ISL_2614555, EPI_ISL_2614556, EPI_ISL_2614562, EPI_ISL_2614563, EPI_ISL_2614564 | | | |
| see above | IAL Presidente Prudente | Instituto Adolfo Lutz, Interdiciplinary Procedures Center, Strategic Laboratory | Claudio Tavares Sacchi, Claudia Regina Gonçalves, Erica Valessa Ramos Gomes, Karoline Rodrigues Campos, Caio Vinicius Dias Lopes, Leonardo Jose Tadeu de Araujo |
| EPI_ISL_2614566 | AMA Dr. Cesar Antunes da Rocha | Instituto Adolfo Lutz, Interdiciplinary Procedures Center, Strategic Laboratory | Claudio Tavares Sacchi, Claudia Regina Gonçalves, Erica Valessa Ramos Gomes, Karoline Rodrigues Campos, Caio Vinicius Dias Lopes, Leonardo Jose Tadeu de Araujo |
| EPI_ISL_2614567 | UPA de Jandira | Instituto Adolfo Lutz, Interdiciplinary Procedures Center, Strategic Laboratory | Claudio Tavares Sacchi, Claudia Regina Gonçalves, Erica Valessa Ramos Gomes, Karoline Rodrigues Campos, Caio Vinicius Dias Lopes, Leonardo Jose Tadeu de Araujo |
| EPI_ISL_2614568 | PS Jose Ibrahin | Instituto Adolfo Lutz, Interdiciplinary Procedures Center, Strategic Laboratory | Claudio Tavares Sacchi, Claudia Regina Gonçalves, Erica Valessa Ramos Gomes, Karoline Rodrigues Campos, Caio Vinicius Dias Lopes, Leonardo Jose Tadeu de Araujo |
| EPI_ISL_2614569 | Hospital Universitário da USP | Instituto Adolfo Lutz, Interdiciplinary Procedures Center, Strategic Laboratory | Claudio Tavares Sacchi, Claudia Regina Gonçalves, Erica Valessa Ramos Gomes, Karoline Rodrigues Campos, Caio Vinicius Dias Lopes, Leonardo Jose Tadeu de Araujo |
| EPI_ISL_2614570 | Hospital Sagrada Família- Unidade Maua | Instituto Adolfo Lutz, Interdiciplinary Procedures Center, Strategic Laboratory | Claudio Tavares Sacchi, Claudia Regina Gonçalves, Erica Valessa Ramos Gomes, Karoline Rodrigues Campos, Caio Vinicius Dias Lopes, Leonardo Jose Tadeu de Araujo |
| EPI_ISL_2614571 | Pronto-Socorro Municipal Dr. Lauro Ribas Braga | Instituto Adolfo Lutz, Interdiciplinary Procedures Center, Strategic Laboratory | Claudio Tavares Sacchi, Claudia Regina Gonçalves, Erica Valessa Ramos Gomes, Karoline Rodrigues Campos, Caio Vinicius Dias Lopes, Leonardo Jose Tadeu de Araujo |
| EPI_ISL_2614584 | UBS Jardim Jovaia | Instituto Adolfo Lutz, Interdiciplinary Procedures Center, Strategic Laboratory | Claudio Tavares Sacchi, Claudia Regina Gonçalves, Erica Valessa Ramos Gomes, Karoline Rodrigues Campos, Caio Vinicius Dias Lopes, Leonardo Jose Tadeu de Araujo |
| EPI_ISL_2614585 | Laboratorio Municipal De Suzano | Instituto Adolfo Lutz, Interdiciplinary Procedures Center, Strategic Laboratory | Claudio Tavares Sacchi, Claudia Regina Gonçalves, Erica Valessa Ramos Gomes, Karoline Rodrigues Campos, Caio Vinicius Dias Lopes, Leonardo Jose Tadeu de Araujo |
| EPI_ISL_2614602 | Instituto Biologico | Instituto Adolfo Lutz, Interdiciplinary Procedures Center, Strategic Laboratory | Claudio Tavares Sacchi, Claudia Regina Gonçalves, Erica Valessa Ramos Gomes, Karoline Rodrigues Campos, Caio Vinicius Dias Lopes, Leonardo Jose Tadeu de Araujo |

| | | | |
|---|---|---|---|
| EPI_ISL_2614614, EPI_ISL_2614615, EPI_ISL_2614616 | Instituto Adolfo Lutz - Regional de Presidente Prudente | Instituto Adolfo Lutz, Interdiciplinary Procedures Center, Strategic Laboratory | Claudio Tavares Sacchi, Claudia Regina Gonçalves, Erica Valessa Ramos Gomes, Karoline Rodrigues Campos, Caio Vinicius Dias Lopes, Leonardo Jose Tadeu de Araujo |
| EPI_ISL_2616368 | Elling group, Institute of Molecular Biotechnology (IMBA) | Bergthaler laboratory, CeMM Research Center for Molecular Medicine of the Austrian Academy of Sciences | Lukas Endler, Anna Schedl, Fabian Amman, Petr Triska, Matthew Thornton, Thomas Penz, Benedikt Agerer, Maelle Le Moing, Michael Schuster, Bekir Erguner, Jan Laine, Martin Senekowitsch, Christoph Bock, Andreas Bergthaler |
| EPI_ISL_2622008, EPI_ISL_2622009, EPI_ISL_2622010, EPI_ISL_2622011, EPI_ISL_2622012, EPI_ISL_2622013, EPI_ISL_2622015, EPI_ISL_2622016 | cerballiance-IDF | Cerba lab | Emmanuel Lecorche, Sabine Trombert-Paolantoni, Aude Lessenne, Kader Merah, Thierry Collin, Laura Verdurme, Patrice Herisson, Stéphanie Haïm-Boukobza, Bénédicte Roquebert |
| EPI_ISL_2622017 | cerballiance-Oise | Cerba lab | Emmanuel Lecorche, Sabine Trombert-Paolantoni, Aude Lessenne, Kader Merah, Thierry Collin, Laura Verdurme, Patrice Herisson, Stéphanie Haïm-Boukobza, Bénédicte Roquebert |
| EPI_ISL_2622018, EPI_ISL_2622019, EPI_ISL_2622020, EPI_ISL_2622021, EPI_ISL_2622022, EPI_ISL_2622023, EPI_ISL_2622024, EPI_ISL_2622025, EPI_ISL_2622026, EPI_ISL_2622027, EPI_ISL_2622028, EPI_ISL_2622029, EPI_ISL_2622030, EPI_ISL_2622031, EPI_ISL_2622032, EPI_ISL_2622033 | | | |
| see above | cerballiance-IDF | Cerba lab | Emmanuel Lecorche, Sabine Trombert-Paolantoni, Aude Lessenne, Kader Merah, Thierry Collin, Laura Verdurme, Patrice Herisson, Stéphanie Haïm-Boukobza, Bénédicte Roquebert |
| EPI_ISL_2622034 | cerballiance-Oise | Cerba lab | Emmanuel Lecorche, Sabine Trombert-Paolantoni, Aude Lessenne, Kader Merah, Thierry Collin, Laura Verdurme, Patrice Herisson, Stéphanie Haïm-Boukobza, Bénédicte Roquebert |
| EPI_ISL_2622035, EPI_ISL_2622036, EPI_ISL_2622038, EPI_ISL_2622039, EPI_ISL_2622040, EPI_ISL_2622041, EPI_ISL_2622042 | cerballiance-IDF | Cerba lab | Emmanuel Lecorche, Sabine Trombert-Paolantoni, Aude Lessenne, Kader Merah, Thierry Collin, Laura Verdurme, Patrice Herisson, Stéphanie Haïm-Boukobza, Bénédicte Roquebert |
| EPI_ISL_2622044 | cerballiance-Oise | Cerba lab | Emmanuel Lecorche, Sabine Trombert-Paolantoni, Aude Lessenne, Kader Merah, Thierry Collin, Laura Verdurme, Patrice Herisson, Stéphanie Haïm-Boukobza, Bénédicte Roquebert |
| EPI_ISL_2626192, EPI_ISL_2626193 | Molecular diagnostic laboratory of Federal Budget Institution of Science "Central Research Institute of Epidemiology" of The Federal Service on Customers' Rights Protection and Human Well-being Surveillance | Group of Genomics and Postgenomic Technologies of Central Research Institute of Epidemiology | Samoilov A.E., Kaptelova V.V., Saenko S.S.,Sinitsyn S.O., Korneenko E.V., Smirnova Y.S., Nadtoka M.I., Speranskaya A.S., Tivanova E.V., Shipulina O.Y., Kondrasheva L.Y., Akimkin V.G. |
| EPI_ISL_2626443, EPI_ISL_2626444, EPI_ISL_2626445, EPI_ISL_2626446, EPI_ISL_2626447, EPI_ISL_2626448 | cerballiance-IDF | Cerba lab | Emmanuel Lecorche, Sabine Trombert-Paolantoni, Aude Lessenne, Kader Merah, Thierry Collin, Laura Verdurme, Patrice Herisson, Stéphanie Haïm-Boukobza, Bénédicte Roquebert |
| EPI_ISL_2627957 | Hospital | National Reference Center for Viruses of Respiratory Infections, Institut Pasteur, Paris | Marion Barbet, Sylvie Behillil, Méline Bizard, Angela Brisebarre, Camille Capel, Vincent Enouf, Louise Lefrançois, Frédéric Lemoine, Christophe Malabat, Corinne Maufrais, Etienne Simon-Lorière, Maud Vanpeene, Sylvie Van der Werf ,Franck Griscelli |
| EPI_ISL_2629002, EPI_ISL_2629003, EPI_ISL_2629004, EPI_ISL_2629005, EPI_ISL_2629006, EPI_ISL_2629007, EPI_ISL_2629008, EPI_ISL_2629009, EPI_ISL_2629011, EPI_ISL_2629012, EPI_ISL_2629013, EPI_ISL_2629014, EPI_ISL_2629015, EPI_ISL_2629016, EPI_ISL_2629017, EPI_ISL_2629018, EPI_ISL_2629019, EPI_ISL_2629020, EPI_ISL_2629021, EPI_ISL_2629023 | | | |
| see above | HEGP - Laboratoire de Virologie | HEGP - Laboratoire de Virologie | David Veyer, Sandrine Imbeaud, Julien Puech, Hélène Péré, Maxime Wack, Nicolas Robillard |
| EPI_ISL_2629024, EPI_ISL_2629025, EPI_ISL_2629027, EPI_ISL_2629028, EPI_ISL_2629029, EPI_ISL_2629030, EPI_ISL_2629031 | Plateforme Covid IDF, site Broussais | HEGP - Laboratoire de Virologie | David Veyer, Sandrine Imbeaud, Julien Puech, Nabil Gastli, Hélène Péré, Maxime Wack, Nicolas Robillard |
| EPI_ISL_2629032, EPI_ISL_2629034, EPI_ISL_2629035, EPI_ISL_2629036, EPI_ISL_2629037, EPI_ISL_2629038, EPI_ISL_2629039, EPI_ISL_2629040, EPI_ISL_2629041, EPI_ISL_2629042, EPI_ISL_2629043, EPI_ISL_2629044 | | | |
| see above | HEGP - Laboratoire de Virologie | HEGP - Laboratoire de Virologie | David Veyer, Sandrine Imbeaud, Julien Puech, Hélène Péré, Maxime Wack, Nicolas Robillard |
| EPI_ISL_2629045, EPI_ISL_2629046, EPI_ISL_2629047, EPI_ISL_2629048, EPI_ISL_2629049, EPI_ISL_2629050, EPI_ISL_2629051 | Plateforme Covid IDF, site Broussais | HEGP - Laboratoire de Virologie | David Veyer, Sandrine Imbeaud, Julien Puech, Nabil Gastli, Hélène Péré, Maxime Wack, Nicolas Robillard |
| EPI_ISL_2629052 | Plateforme Covid IDF, site Broussais | HEGP - Laboratoire de Virologie | David Veyer, Sandrine Imbeaud, Julien Puech, Hélène Péré, Maxime Wack, Nicolas Robillard |
| EPI_ISL_2629053, EPI_ISL_2629055, EPI_ISL_2629056, EPI_ISL_2629057, EPI_ISL_2629058, EPI_ISL_2629059, EPI_ISL_2629060, EPI_ISL_2629061, EPI_ISL_2629062, EPI_ISL_2629063, EPI_ISL_2629064, EPI_ISL_2629065, EPI_ISL_2629066, EPI_ISL_2629067, EPI_ISL_2629068, EPI_ISL_2629069, EPI_ISL_2629075, EPI_ISL_2629076, EPI_ISL_2629077, EPI_ISL_2629081, EPI_ISL_2629082, EPI_ISL_2629083, EPI_ISL_2629084, EPI_ISL_2629085, EPI_ISL_2629086, EPI_ISL_2629087, EPI_ISL_2629088, EPI_ISL_2629089, EPI_ISL_2629090, EPI_ISL_2629091, EPI_ISL_2629092, EPI_ISL_2629093 | | | |
| see above | HEGP - Laboratoire de Virologie | HEGP - Laboratoire de Virologie | David Veyer, Sandrine Imbeaud, Julien Puech, Hélène Péré, Maxime Wack, Nicolas Robillard |
| EPI_ISL_2629164, EPI_ISL_2629166, EPI_ISL_2629167, EPI_ISL_2629168, EPI_ISL_2629169, EPI_ISL_2629170, EPI_ISL_2629172, EPI_ISL_2629173, EPI_ISL_2629174, EPI_ISL_2629175, EPI_ISL_2629176, EPI_ISL_2629177, EPI_ISL_2629178, EPI_ISL_2629179, EPI_ISL_2629180, EPI_ISL_2629181, EPI_ISL_2629183, EPI_ISL_2629184, EPI_ISL_2629186 | | | |
| see above | Turkish Society of Internal Medicine | Gen Era Diagnostics Corporation Life Science Research and Molecular Diagnostics | Serhat Ünal |
| EPI_ISL_2648110, EPI_ISL_2648111, EPI_ISL_2648112, EPI_ISL_2648113 | Instituto Nacional de Enfermedades Respiratorias (INER) | Centro de Investigación en Enfermedades Infecciosas (CIENI), Instituto Nacional de Enfermedades Respiratorias (INER) | Consorcio Mexicano de Vigilancia Genómica (CoViGen-Mex). Authors (in alphabetical order): Julio Elias Alvarado-Yaah, Carlos F. Arias, Santiago Ávila-Ríos, Eduardo Becerril-Vargas, Víctor Hugo Borja-Aburto, Celia Boukadida, Cristóbal Cháidez-Quiróz, Juan Bautista Chale-Dzul, Ricardo Ciria Merce, Andreu Comas-García, Célida Duque Molina, Julissa Enciso-Ibarra, José Antonio Enciso-Moreno, Gloria Elena Espinosa-Ayala, Fernando Fontove-Herrera, Daniel Fregoso-Rueda, Victor Eduardo Garcia-Arias, Alejandra Garcia-Gasca, Bruno Gomez-Gil, Jean Pierre González, Irvin González-López, Concepción Grajales-Muñiz, Ricardo Grande, Rosa María Gutierrez Rios, Alejandra Hernández-Terán, Alfredo Herrera-Estrella, Carla Ivón Herrera-Najera, Pavel Isa, Daniel Lira Morales, Susana Lopez, Antonio Loza Román, Brenda Irasema Maldonado-Meza, Bernardo Martínez-Miguel, José Arturo Martínez-Orozco, Célida Martínez- Rodríguez, Margarita Matías-Florentino, Fidencio Mejía-Nepomuceno, María Guadalupe de Jesús Mireles-Rivera, Gloria María Molina-Salinas, Hector Montoya-Fuentes, , Mario Mújica-Sánchez, José Esteban Muñoz-Medina, José de Jesús Nuñez-Contreras, Alicia Ocaña-Mondragón, Luis Alberto Ochoa-Carrera, Hector Esteban Paz-Juárez, Marissa Pérez-García, Francisco Pulido, Helen Haydee Fernanda Ramírez-Plascencia, Jorge Salas-Hernández, Angel Gustavo Salas-Lais, Alejandro Sanchez-Flores, Clara Esperanza Santacruz-Tinoco, María Guadalupe Santiago-Mauricio, Selene Zárate, Nelly Sélem-Mojica, Blanca Taboada, Gloria Vazquez, Joel Armando Vázquez-Pérez. |
| EPI_ISL_2657513, EPI_ISL_2657514, EPI_ISL_2657515, EPI_ISL_2657516 | Dr. Gernot Walder GmbH | Bergthaler laboratory, CeMM Research Center for Molecular Medicine of the Austrian Academy of Sciences | Lukas Endler, Anna Schedl, Fabian Amman, Petr Triska, Matthew Thornton, Thomas Penz, Benedikt Agerer, Maelle Le Moing, Michael Schuster, Bekir Erguner, Jan Laine, Martin Senekowitsch, Christoph Bock, Andreas Bergthaler |
| EPI_ISL_2657681 | AREA DE SALUD CHACARITA - CLINICA DR. FRANCISCO QUINTANA | Inciensa, Instituto Costarricense de Investigación y Enseñanza en Nutrición y Salud | Francisco Duarte, Hebleen Porras, Claudio Soto-Garita, Estela Cordero, Adriana Godínez, Melany Calderón, José Luis Vargas, Mariela Gutiérrez, Joselyn Prado, Caterina Guzmán, Nazareth Ruiz & María José Gómez-Umaña |
| EPI_ISL_2657682, EPI_ISL_2657683 | HOSPITAL METROPOLITANO | Inciensa, Instituto Costarricense de Investigación y Enseñanza en Nutrición y Salud | Francisco Duarte, Hebleen Porras, Claudio Soto-Garita, Estela Cordero, Adriana Godínez, Melany Calderón, José Luis Vargas, Mariela Gutiérrez, Joselyn Prado, Caterina Guzmán, Nazareth Ruiz & Margarita Lee-Lui |
| EPI_ISL_2657684 | LABORATORIO CLINICO LABIN | Inciensa, Instituto Costarricense de Investigación y Enseñanza en Nutrición y Salud | Francisco Duarte, Hebleen Porras, Claudio Soto-Garita, Estela Cordero, Adriana Godínez, Melany Calderón, José Luis Vargas, Mariela Gutiérrez, Joselyn Prado, Caterina Guzmán, Nazareth Ruiz & Pei Ling Chan Ma |
| EPI_ISL_2661778 | Laboratorio Central de Saude Publica do Estado do Rio Grande do Sul (LACEN-RS) | Laboratory of Respiratory Viruses and Measles, Oswaldo Cruz Institute, FIOCRUZ | Paola Resende, Luciana Appolinario, Fernando Motta, Anna Carolina Paixao, Ana Carolina Mendonca, Alice Sampaio Rocha, Taina Venas, Elisa Cavalcante Pereira, Renata Serrano Lopes, Tatiana Schaffer Gregianini, Richard Salvato, Marilda Siqueira on behalf of the Fiocruz COVID-19 Genomic |

| | | | |
|---|---|---|---|
| | | | Surveillance Network |
| EPI_ISL_2671891, EPI_ISL_2671898 | Juntendo University Hospital | Juntendo University Hospital | Koji Tuchiya, Yoshie Hosaka, Yoko Tabe |
| EPI_ISL_2672210 | INSP, Bamako | Malaria Research and Training Center-Parasito | Antoine Dara, Ibrehima Guindo, Amadou Daou, Demba Koita, Abdoulaye Djimde |
| EPI_ISL_2681394, EPI_ISL_2681395, EPI_ISL_2681396, EPI_ISL_2681397, EPI_ISL_2681398, EPI_ISL_2681399 | Clinic of infective diseases | Laboratory of virology and molecular diagnostics, Institute of Public Health | Kuzmanovska M, Boshevska G, Janchevska E. |
| EPI_ISL_2688278, EPI_ISL_2688279, EPI_ISL_2688280, EPI_ISL_2688281, EPI_ISL_2688282, EPI_ISL_2688283, EPI_ISL_2688288 | Puerto Rico Department of Health | Centers for Disease Control and Prevention, Dengue Branch | Gilberto A. Santiago, Glenda Gonzalez, Betzabel Flores, Keyla Charriez, Gabriela Paz-Bailey, Jorge L. Munoz-Jordan |
| EPI_ISL_2688626, EPI_ISL_2688632 | Centre Hospitalier Universitaire de Rouen Laboratoire de Virologie | Centre Hospitalier Universitaire de Rouen Laboratoire de Virologie | Alice Moisan, Fabienne De Oliveira, Marie Leoz |
| EPI_ISL_2690312, EPI_ISL_2690423, EPI_ISL_2690668, EPI_ISL_2690670, EPI_ISL_2690671, EPI_ISL_2690673, EPI_ISL_2690675, EPI_ISL_2690677, EPI_ISL_2690683, EPI_ISL_2691074, EPI_ISL_2691077, EPI_ISL_2691079, EPI_ISL_2691080, EPI_ISL_2691082, EPI_ISL_2691088 | | | |
| see above | MSK Microbiology Lab | MSK Microbiology Lab | Esther Babady, Tracy McMillen, Krupa Jani |
| EPI_ISL_2691102, EPI_ISL_2691103, EPI_ISL_2691104, EPI_ISL_2691105, EPI_ISL_2691106 | Instituto Adolfo Lutz - Regional de Marilia | Instituto Adolfo Lutz, Interdiciplinary Procedures Center, Strategic Laboratory | Claudio Tavares Sacchi, Claudia Regina Gonçalves, Erica Valessa Ramos Gomes, Karoline Rodrigues Campos, Caio Vinicius Dias Lopes, Leonardo Jose Tadeu de Araujo |
| EPI_ISL_2691107, EPI_ISL_2691121 | Instituto Adolfo Lutz - Regional de Bauru | Instituto Adolfo Lutz, Interdiciplinary Procedures Center, Strategic Laboratory | Claudio Tavares Sacchi, Claudia Regina Gonçalves, Erica Valessa Ramos Gomes, Karoline Rodrigues Campos, Caio Vinicius Dias Lopes, Leonardo Jose Tadeu de Araujo |
| EPI_ISL_2691149, EPI_ISL_2691150 | Intituto Adolfo Lutz - Regional de Sorocaba | Instituto Adolfo Lutz, Interdiciplinary Procedures Center, Strategic Laboratory | Claudio Tavares Sacchi, Claudia Regina Gonçalves, Erica Valessa Ramos Gomes, Karoline Rodrigues Campos, Caio Vinicius Dias Lopes, Leonardo Jose Tadeu de Araujo |
| EPI_ISL_2691215 | LACEN do Estado de Goias | Instituto Adolfo Lutz, Interdiciplinary Procedures Center, Strategic Laboratory | Claudio Tavares Sacchi, Claudia Regina Gonçalves, Erica Valessa Ramos Gomes, Karoline Rodrigues Campos, Caio Vinicius Dias Lopes, Leonardo Jose Tadeu de Araujo |
| EPI_ISL_2695785 | National Influenza Center (NIC). Virology department. Institut National d'Hygiène. | National Influenza Center (NIC). Virology department. Institut National d'Hygiène. | Hicham OUMZIL, Hassan IHAZMAD, Abderrahmane BIMOUHEN, Zakia REGRAGUI, Samira BENKERROUM, fatima EL FALAKI and Mohamed RHAJAOUI |
| EPI_ISL_2713213, EPI_ISL_2713214, EPI_ISL_2713215, EPI_ISL_2713216, EPI_ISL_2713219, EPI_ISL_2713227, EPI_ISL_2713228, EPI_ISL_2713229, EPI_ISL_2713230, EPI_ISL_2713231, EPI_ISL_2713232, EPI_ISL_2713233, EPI_ISL_2713252, EPI_ISL_2713253, EPI_ISL_2713254 | | | |
| see above | Department of Health Technology and Informatics, The Hong Kong Polytechnic University | Department of Health Technology and Informatics, The Hong Kong Polytechnic University | Gilman Kit-Hang Siu, Lam-Kwong Lee, Kenneth Siu-Sing Leung, Jake Siu-Lun Leung, Wing-Hei Lo, Timothy Ting-Leung Ng, Chloe Toi-Mei Chan, Hoi-Ching Jim, Ivan Tak-Fai Wong, Kingsley King-Gee Tam, Hiu-Yin Lao, Denise Sze-Hang Wong, Alan Ka-Lun Wu, Miranda Chong-Yee Yau, Yvette Wai-Man Lai, Kitty Sau-Chun Fung, Sandy Ka-Yee Chau, Barry Kin-Chung Wong, Wing-Kin To, Kristine Luk, Alex Yat-Man Ho, Tak-Lun Que, Kam-Tong Yip, Wing Cheong Yam, David Ho-Keung Shum, Shea Ping Yip |
| EPI_ISL_2728535, EPI_ISL_2728541 | Laboratorio de Infectologia y Virologia Molecular | Laboratory of Molecular Virology, School of Medicine, Pontificia Universidad Catolica de Chile | Catalina Pardo-Roa, Constanza Martinez-Valdevenito, Jennifer Angulo, Tamara Garcia-Salum, Estefany Poblete, Maria Jose Avendano, Leonardo I. Almonacid, Ana Maria Contreras, Carlos Palma, Jorge Levican, Eileen Serrano, Constanza Maldonado, M. Belen Leyton, Erick Salinas, Andres E. Munoz-Marcos, Francisco Melo, Marcela Ferres, Alejandro Bhrun, Magdalena Vera, Rafael A. Medina |
| EPI_ISL_2756433, EPI_ISL_2756434, EPI_ISL_2756444, EPI_ISL_2756445, EPI_ISL_2756446, EPI_ISL_2756447, EPI_ISL_2756457, EPI_ISL_2756459, EPI_ISL_2756460, EPI_ISL_2756461, EPI_ISL_2756462, EPI_ISL_2756463, EPI_ISL_2756464, EPI_ISL_2756465, EPI_ISL_2756466, EPI_ISL_2756469, EPI_ISL_2756470, EPI_ISL_2756471, EPI_ISL_2756472, EPI_ISL_2756473, EPI_ISL_2756476, EPI_ISL_2756488, EPI_ISL_2756489, EPI_ISL_2756490, EPI_ISL_2756491, EPI_ISL_2756492, EPI_ISL_2756493 | | | |
| see above | Instituto Adolfo Lutz Central | Instituto Adolfo Lutz, Interdiciplinary Procedures Center, Strategic Laboratory | Claudio Tavares Sacchi, Claudia Regina Gonçalves, Erica Valessa Ramos Gomes, Karoline Rodrigues Campos, Caio Vinicius Dias Lopes, Leonardo Jose Tadeu de Araujo |
| EPI_ISL_2757943, EPI_ISL_2757945, EPI_ISL_2757946, EPI_ISL_2757947 | Department of Virology, Istituto Zooprofilattico Sperimentale del Lazio e della Toscana (IZSLT) | Department of General Diagnostics; Department of Virology; Istituto Zooprofilattico Sperimentale del Lazio e della Toscana (IZSLT) | Patricia Alba, Giuseppe Manna, Elena L. Diaconu, Fabiola Feltrin, Raffaela Conti, Teresa Scicluna, Virginia Carfora, Antonella Cersini, Alessia Franco, Antonio Battisti. |
| EPI_ISL_2779310, EPI_ISL_2779311, EPI_ISL_2779312 | Lanet hospital | USAMRD-A, Basic Science Laboratory | Gathii Kimita, Josphat Nyataya, Esther Omuseni, Faith Sigei, Alan Lemtudo, Eric Muthanje, Brian Andika, Rehema Liyai, Rachel Githii, Clement Masakwe, Stephen Ochola, George Awinda, Carol Kifude, Beth Mutai, John Waitumbi |
| EPI_ISL_2779319, EPI_ISL_2779320 | Jaramogi Oginga Odinga Teaching and Referral Hospital | USAMRD-A, Basic Science Laboratory | Gathii Kimita, Josphat Nyataya, Esther Omuseni, Faith Sigei, Alan Lemtudo, Eric Muthanje, Brian Andika, Rehema Liyai, Rachel Githii, Clement Masakwe, Stephen Ochola, George Awinda, Carol Kifude, Beth Mutai, John Waitumbi |
| EPI_ISL_2779321 | Bungoma County Referral Hospital | USAMRD-A, Basic Science Laboratory | Gathii Kimita, Josphat Nyataya, Esther Omuseni, Faith Sigei, Alan Lemtudo, Eric Muthanje, Brian Andika, Rehema Liyai, Rachel Githii, Clement Masakwe, Stephen Ochola, George Awinda, Carol Kifude, Beth Mutai, John Waitumbi |
| EPI_ISL_2779323 | KEMRI/DEID Surveillance Site (Isebani Sub-County Hospital) | USAMRD-A, Basic Science Laboratory | Gathii Kimita, Josphat Nyataya, Esther Omuseni, Faith Sigei, Alan Lemtudo, Eric Muthanje, Brian Andika, Rehema Liyai, Rachel Githii, Clement Masakwe, Stephen Ochola, George Awinda, Carol Kifude, Beth Mutai, John Waitumbi |
| EPI_ISL_2779324 | KCRH KEMRI/DEID Surveillance Site | USAMRD-A, Basic Science Laboratory | Gathii Kimita, Josphat Nyataya, Esther Omuseni, Faith Sigei, Alan Lemtudo, Eric Muthanje, Brian Andika, Rehema Liyai, Rachel Githii, Clement Masakwe, Stephen Ochola, George Awinda, Carol Kifude, Beth Mutai, John Waitumbi |
| EPI_ISL_2779325, EPI_ISL_2779326 | KEMRI/DEID Surveillance Site (Lamu County Hospital) | USAMRD-A, Basic Science Laboratory | Gathii Kimita, Josphat Nyataya, Esther Omuseni, Faith Sigei, Alan Lemtudo, Eric Muthanje, Brian Andika, Rehema Liyai, Rachel Githii, Clement Masakwe, Stephen Ochola, George Awinda, Carol Kifude, Beth Mutai, John Waitumbi |
| EPI_ISL_2779327 | Bungoma County Referral Hospital | USAMRD-A, Basic Science Laboratory | Gathii Kimita, Josphat Nyataya, Esther Omuseni, Faith Sigei, Alan Lemtudo, Eric Muthanje, Brian Andika, Rehema Liyai, Rachel Githii, Clement Masakwe, Stephen Ochola, George Awinda, Carol Kifude, Beth Mutai, John Waitumbi |
| EPI_ISL_2779328, EPI_ISL_2779329 | Life Care Hospital (Bungoma) | USAMRD-A, Basic Science Laboratory | Gathii Kimita, Josphat Nyataya, Esther Omuseni, Faith Sigei, Alan Lemtudo, Eric Muthanje, Brian Andika, Rehema Liyai, Rachel Githii, Clement Masakwe, Stephen Ochola, George Awinda, Carol Kifude, Beth Mutai, John Waitumbi |
| EPI_ISL_2779330 | Homabay County Referral Hospital | USAMRD-A, Basic Science Laboratory | Gathii Kimita, Josphat Nyataya, Esther Omuseni, Faith Sigei, Alan Lemtudo, Eric Muthanje, Brian Andika, Rehema Liyai, Rachel Githii, Clement Masakwe, Stephen Ochola, George Awinda, Carol Kifude, Beth Mutai, John Waitumbi |
| EPI_ISL_2779331, EPI_ISL_2779332, EPI_ISL_2779333 | KCRH KEMRI/DEID Surveillance Site | USAMRD-A, Basic Science Laboratory | Gathii Kimita, Josphat Nyataya, Esther Omuseni, Faith Sigei, Alan Lemtudo, Eric Muthanje, Brian Andika, Rehema Liyai, Rachel Githii, Clement Masakwe, Stephen Ochola, George Awinda, Carol Kifude, Beth Mutai, John Waitumbi |
| EPI_ISL_2779334, EPI_ISL_2779335, EPI_ISL_2779336 | KCRH KEMRI-DEID Surveillance Site | USAMRD-A, Basic Science Laboratory | Gathii Kimita, Josphat Nyataya, Esther Omuseni, Faith Sigei, Alan Lemtudo, Eric Muthanje, Brian Andika, Rehema Liyai, Rachel Githii, Clement Masakwe, Stephen Ochola, George Awinda, Carol Kifude, Beth Mutai, John Waitumbi |
| EPI_ISL_2779337 | KCRH KEMRI/DEID Surveillance Site | USAMRD-A, Basic Science Laboratory | Gathii Kimita, Josphat Nyataya, Esther Omuseni, Faith Sigei, Alan Lemtudo, Eric Muthanje, Brian Andika, Rehema Liyai, Rachel Githii, Clement Masakwe, Stephen Ochola, George Awinda, Carol Kifude, Beth Mutai, John Waitumbi |
| EPI_ISL_2779338, EPI_ISL_2779339, EPI_ISL_2779340, EPI_ISL_2779341, EPI_ISL_2779342 | Nyamira County Referral Hopital | USAMRD-A, Basic Science Laboratory | Gathii Kimita, Josphat Nyataya, Esther Omuseni, Faith Sigei, Alan Lemtudo, Eric Muthanje, Brian Andika, Rehema Liyai, Rachel Githii, Clement Masakwe, Stephen Ochola, George Awinda, Carol Kifude, Beth Mutai, John Waitumbi |
| EPI_ISL_2779343 | Vihiga County referral hospital | USAMRD-A, Basic Science Laboratory | Gathii Kimita, Josphat Nyataya, Esther Omuseni, Faith Sigei, Alan Lemtudo, Eric Muthanje, Brian Andika, Rehema Liyai, Rachel Githii, Clement Masakwe, Stephen Ochola, George Awinda, Carol Kifude, Beth Mutai, John Waitumbi |
| EPI_ISL_2779344 | KEMRI/DEID Surveillance Site (Lamu County Hospital) | USAMRD-A, Basic Science Laboratory | Gathii Kimita, Josphat Nyataya, Esther Omuseni, Faith Sigei, Alan Lemtudo, Eric Muthanje, Brian Andika, Rehema Liyai, Rachel Githii, Clement Masakwe, Stephen Ochola, George Awinda, Carol Kifude, Beth Mutai, John Waitumbi |

| | | | |
|---|---|---|---|
| EPI_ISL_2779345 | Bungoma County Referral Hospital | USAMRD-A, Basic Science Laboratory | Gathii Kimita, Josphat Nyataya, Esther Omuseni, Faith Sigei, Alan Lemtudo, Eric Muthanje, Brian Andika, Rehema Liyai, Rachel Githii, Clement Masakwe, Stephen Ochola, George Awinda, Carol Kifude, Beth Mutai, John Waitumbi |
| EPI_ISL_2779346 | Homabay County Referral Hospital | USAMRD-A, Basic Science Laboratory | Gathii Kimita, Josphat Nyataya, Esther Omuseni, Faith Sigei, Alan Lemtudo, Eric Muthanje, Brian Andika, Rehema Liyai, Rachel Githii, Clement Masakwe, Stephen Ochola, George Awinda, Carol Kifude, Beth Mutai, John Waitumbi |
| EPI_ISL_2779347 | KCRH KEMRI-DEID Surveillance Site | USAMRD-A, Basic Science Laboratory | Gathii Kimita, Josphat Nyataya, Esther Omuseni, Faith Sigei, Alan Lemtudo, Eric Muthanje, Brian Andika, Rehema Liyai, Rachel Githii, Clement Masakwe, Stephen Ochola, George Awinda, Carol Kifude, Beth Mutai, John Waitumbi |
| EPI_ISL_2790232, EPI_ISL_2790233, EPI_ISL_2790234, EPI_ISL_2790235 | CHWAPI - SITE NOTRE DAME | Institut de Pathologie et Genetique (IPG) | Pascale Hilbert, Jérémie Gras |
| EPI_ISL_2828022, EPI_ISL_2828023 | Unidad de Investigación Molecular (UNIMOL) | Instituto Nacional de Salud- Dirección de Investigación en Salud Pública | Katherine Laiton-Donato, Diego A. Álvarez-Díaz, Carlos Franco-Muñoz, Hector Alejandro Ruiz-Moreno, Paola Rojas, Maria T. Herrera-Sepúlveda, Diego Andrés Prada, Jhonnatan Reales-González, Sheryll Corchuelo, Julian Naizaque, Jorge Rivera, Gerardo Santamaría, Sergio Gomez, Lisseth Pardo, Juan Camilo Martinez, Marta Lopez Blanco, Ángela Alarcon Cruz, Diana Malo, Carmen Osorio, Magdalena Wiesner, Martha Lucia Ospina Martinez, Marcela Mercado-Reyes |
| EPI_ISL_2828687, EPI_ISL_2828688, EPI_ISL_2828689 | Instituto Adolfo Lutz - Regional de Campinas | Instituto Adolfo Lutz, Interdiciplinary Procedures Center, Strategic Laboratory | Claudio Tavares Sacchi, Claudia Regina Gonçalves, Erica Valessa Ramos Gomes, Karoline Rodrigues Campos, Caio Vinicius Dias Lopes |
| EPI_ISL_2860796, EPI_ISL_2860802, EPI_ISL_2860803, EPI_ISL_2860804, EPI_ISL_2860805, EPI_ISL_2860806, EPI_ISL_2860807, EPI_ISL_2860808, EPI_ISL_2860809, EPI_ISL_2860810, EPI_ISL_2860819 | | | |
| see above | Department of Health Technology and Informatics, The Hong Kong Polytechnic University | Department of Health Technology and Informatics, The Hong Kong Polytechnic University | Gilman Kit-Hang Siu, Lam-Kwong Lee, Kenneth Siu-Sing Leung, Jake Siu-Lun Leung, Wing-Hei Lo, Timothy Ting-Leung Ng, Chloe Toi-Mei Chan, Hoi-Ching Jim, Ivan Tak-Fai Wong, Kingsley King-Gee Tam, Hiu-Yin Lao, Denise Sze-Hang Wong, Alan Ka-Lun Wu, Miranda Chong-Yee Yau, Yvette Wai-Man Lai, Kitty Sau-Chun Fung, Sandy Ka-Yee Chau, Barry Kin-Chung Wong, Wing-Kin To, Kristine Luk, Alex Yat-Man Ho, Tak-Lun Que, Kam-Tong Yip, Wing Cheong Yam, David Ho-Keung Shum, Shea Ping Yip |
| EPI_ISL_2873844, EPI_ISL_2873852 | National Influenza Centre | National Influenza Centre | William K. Ampofo, Michael Marks, Ivy A. Asante, Sharon Hsu, Benjamiin B. Lindsey, Benjamin H. Foulkes, Mildred Adusei-Poku, Linda Boatemaa, Lorreta Kwasah, Joseph Oliver-Commey, Ernest Asiedu, Franklin Asiedu-Bekoe, Gordon Awandare, Joyce Ngoi, Dennis Laryea, Mathew D. Parker, Thushan I de Silva, |

We gratefully acknowledge the following Authors from the Originating laboratories responsible for obtaining the specimens, as well as the Submitting laboratories where the genome data were generated and shared via GISAID, on which this research is based.

All Submitters of data may be contacted directly via www.gisaid.org

Authors are sorted alphabetically.

| Accession ID | Originating Laboratory | Submitting Laboratory | Authors |
|---|---|---|---|
| EPI_ISL_1595848 | Genome Analysis Center, Yamanashi Central Hospital | Genome Analysis Center, Yamanashi Central Hospital | Yosuke Hirotsu |
| EPI_ISL_1601578, EPI_ISL_1601807 | AZ Klina | AZ Klina | Carl Vael - Lynsey Berckmans |
| EPI_ISL_1623040, EPI_ISL_1623060, EPI_ISL_1623061, EPI_ISL_1623062 | Centre Hospitalier Universitaire Clermont-Ferrand | CHU Clermont-Ferrand, service de virologie | Bisseux Maxime, Mirand Audrey, Combes Patricia, Henquell Cécile |
| EPI_ISL_1623063 | Centre Hospitalier Clementel | CHU Clermont-Ferrand, service de virologie | Bisseux Maxime, Mirand Audrey, Combes Patricia, Henquell Cécile |
| EPI_ISL_1623064 | HOPITAL DE VICHY | CHU Clermont-Ferrand, service de virologie | Bisseux Maxime, Mirand Audrey, Combes Patricia, Henquell Cécile |
| EPI_ISL_1623065, EPI_ISL_1623066, EPI_ISL_1623086 | Centre Hospitalier Universitaire Clermont-Ferrand | CHU Clermont-Ferrand, service de virologie | Bisseux Maxime, Mirand Audrey, Combes Patricia, Henquell Cécile |
| EPI_ISL_1628343 | CHC Andrée Rosemon | Institut Pasteur de la Guyane | Anne Lavergne, Dominique Rousset, A. Salmier |
| EPI_ISL_1657100 | National Institute of Infectious Diseases-Prof. Dr. Matei Bals Molecular Diagnostics Laboratory | National Institute of Infectious Diseases-Prof. Dr. Matei Bals Molecular Diagnostics Laboratory | Corina Casangiu, Leontina Banica, Marius Surleac, Ovidiu Vlaicu, Andreea Tudor, Simona Paraschiv, Dan Otelea |
| EPI_ISL_1660337, EPI_ISL_1660338 | Pro-Vitam Diagnostics and Research Laboratory | Pro-Vitam Diagnostics and Research Laboratory | Szilard N. Fejer, Istvan Horvath, Monika Korodi, Kinga Rakosi, Zsuzsanna Jenei |
| EPI_ISL_1667475 | Department of Laboratory Medicine, National Taiwan University Hospital | Microbial Genomics Core Lab, National Taiwan University Centers of Genomic and Precision Medicine | Shiou-Hwei Yeh, You-Yu Lin, Ya-Yun Lai, Chiao-Ling Li, Shan-Chwen Chang, Pei-Jer Chen, Sui-Yuan Chang |
| EPI_ISL_1669760 | Genome Analysis Center, Yamanashi Central Hospital | Genome Analysis Center, Yamanashi Central Hospital | Yosuke Hirotsu |
| EPI_ISL_1694769, EPI_ISL_1694770, EPI_ISL_1694772 | OCME Office Of Chief Medical Examiner | New York City Public Health Laboratory | Jade Wang, et al. |
| EPI_ISL_1697433, EPI_ISL_1697434, EPI_ISL_1697435, EPI_ISL_1697436, EPI_ISL_1697437, EPI_ISL_1697438 | Bioptická laborato, s.r.o. | Bioptická laborato, s.r.o. | Petr Šteiner, Tomáš Vanek, Nikola Bílá, Martina Putzová, Michaela íhová, Silva Vondráková |
| EPI_ISL_1700598 | Hospital | National Reference Center for Viruses of Respiratory Infections, Institut Pasteur, Paris | Marion Barbet, Sylvie Behillil, Méline Bizard, Angela Brisebarre, Camille Capel, Vincent Enouf, Louise Lefrançois, Frédéric Lemoine, Christophe Malabat, Corinne Maufrais, Damien Mornico, Etienne Simon-Lorière, Maud Vanpeene, Sylvie Van der Werf, Marianne Burgard |
| EPI_ISL_1700602 | Hospital | National Reference Center for Viruses of Respiratory Infections, Institut Pasteur, Paris | Marion Barbet, Sylvie Behillil, Méline Bizard, Angela Brisebarre, Camille Capel, Vincent Enouf, Louise Lefrançois, Frédéric Lemoine, Christophe Malabat, Corinne Maufrais, Damien Mornico, Etienne Simon-Lorière, Maud Vanpeene, Sylvie Van der Werf, Pascale Martres |
| EPI_ISL_1700609, EPI_ISL_1700610 | Labo Analyses Med | National Reference Center for Viruses of Respiratory Infections, Institut Pasteur, Paris | Marion Barbet, Sylvie Behillil, Méline Bizard, Angela Brisebarre, Camille Capel, Vincent Enouf, Louise Lefrançois, Frédéric Lemoine, Christophe Malabat, Corinne Maufrais, Etienne Simon-Lorière, Maud Vanpeene, Sylvie Van der Werf, GréGoire Potiron |
| EPI_ISL_1700616 | Hospital | National Reference Center for Viruses of Respiratory Infections, Institut Pasteur, Paris | Marion Barbet, Sylvie Behillil, Méline Bizard, Angela Brisebarre, Camille Capel, Vincent Enouf, Louise Lefrançois, Frédéric Lemoine, Christophe Malabat, Corinne Maufrais, Etienne Simon-Lorière, Maud Vanpeene, Sylvie Van der Werf, Maxime Paluch |
| EPI_ISL_1700618, EPI_ISL_1700619, EPI_ISL_1700620, EPI_ISL_1700621, EPI_ISL_1700622 | Hospital | National Reference Center for Viruses of Respiratory Infections, Institut Pasteur, Paris | Marion Barbet, Sylvie Behillil, Méline Bizard, Angela Brisebarre, Camille Capel, Vincent Enouf, Louise Lefrançois, Frédéric Lemoine, Christophe Malabat, Corinne Maufrais, Etienne Simon-Lorière, Maud Vanpeene, Sylvie Van der Werf, ValéRie Serazin |
| EPI_ISL_1700628, EPI_ISL_1700629 | Hospital | National Reference Center for Viruses of Respiratory Infections, Institut Pasteur, Paris | Marion Barbet, Sylvie Behillil, Méline Bizard, Angela Brisebarre, Camille Capel, Vincent Enouf, Louise Lefrançois, Frédéric Lemoine, Christophe Malabat, Corinne Maufrais, Etienne Simon-Lorière, Maud Vanpeene, Sylvie Van der Werf, JéRome Guinard |
| EPI_ISL_1700631 | Hospital | National Reference Center for Viruses of Respiratory Infections, Institut Pasteur, Paris | Marion Barbet, Sylvie Behillil, Méline Bizard, Angela Brisebarre, Camille Capel, Vincent Enouf, Louise Lefrançois, Frédéric Lemoine, Christophe Malabat, Corinne Maufrais, Damien Mornico, Etienne Simon-Lorière, Maud Vanpeene, Sylvie Van der Werf, JéRome Guinard |
| EPI_ISL_1700632, EPI_ISL_1700633, EPI_ISL_1700634, EPI_ISL_1700635 | Hospital | National Reference Center for Viruses of Respiratory Infections, Institut Pasteur, Paris | Marion Barbet, Sylvie Behillil, Méline Bizard, Angela Brisebarre, Camille Capel, Vincent Enouf, Louise Lefrançois, Frédéric Lemoine, Christophe Malabat, Corinne Maufrais, Etienne Simon-Lorière, Maud Vanpeene, Sylvie Van der Werf, JéRome Guinard |
| EPI_ISL_1700636 | Armies | National Reference Center for Viruses of Respiratory Infections, Institut Pasteur, Paris | Marion Barbet, Sylvie Behillil, Méline Bizard, Angela Brisebarre, Camille Capel, Vincent Enouf, Louise Lefrançois, Frédéric Lemoine, Christophe Malabat, Corinne Maufrais, Damien Mornico, Etienne Simon-Lorière, Maud Vanpeene, Sylvie Van der Werf, Marine Desroches |
| EPI_ISL_1700637, EPI_ISL_1700638, EPI_ISL_1700639, EPI_ISL_1700640 | Armies | National Reference Center for Viruses of Respiratory Infections, Institut Pasteur, Paris | Marion Barbet, Sylvie Behillil, Méline Bizard, Angela Brisebarre, Camille Capel, Vincent Enouf, Louise Lefrançois, Frédéric Lemoine, Christophe Malabat, Corinne Maufrais, Etienne Simon-Lorière, Maud Vanpeene, Sylvie Van der Werf, Marine Desroches |
| EPI_ISL_1700641 | Armies | National Reference Center for Viruses of Respiratory Infections, Institut Pasteur, Paris | Marion Barbet, Sylvie Behillil, Méline Bizard, Angela Brisebarre, Camille Capel, Vincent Enouf, Louise Lefrançois, Frédéric Lemoine, Christophe Malabat, Corinne Maufrais, Damien Mornico, Etienne Simon-Lorière, Maud Vanpeene, Sylvie Van der Werf, Marine Desroches |
| EPI_ISL_1700642, EPI_ISL_1700643 | Armies | National Reference Center for Viruses of Respiratory Infections, Institut Pasteur, Paris | Marion Barbet, Sylvie Behillil, Méline Bizard, Angela Brisebarre, Camille Capel, Vincent Enouf, Louise Lefrançois, Frédéric Lemoine, Christophe Malabat, Corinne Maufrais, Etienne Simon-Lorière, Maud Vanpeene, Sylvie Van der Werf, Marine Desroches |
| EPI_ISL_1700644 | Hospital | National Reference Center for Viruses of Respiratory Infections, Institut Pasteur, Paris | Marion Barbet, Sylvie Behillil, Méline Bizard, Angela Brisebarre, Camille Capel, Vincent Enouf, Louise Lefrançois, Frédéric Lemoine, Christophe Malabat, Corinne Maufrais, Adrien Pain, Etienne Simon-Lorière, Maud Vanpeene, Sylvie Van der Werf, CéCile Farrugia |
| EPI_ISL_1700645 | Hospital | National Reference Center for Viruses of Respiratory Infections, Institut Pasteur, Paris | Marion Barbet, Sylvie Behillil, Méline Bizard, Angela Brisebarre, Camille Capel, Vincent Enouf, Louise Lefrançois, Frédéric Lemoine, Christophe Malabat, Corinne Maufrais, Etienne Simon-Lorière, Maud Vanpeene, Sylvie Van der Werf, CéCile Farrugia |
| EPI_ISL_1700646 | Hospital | National Reference Center for Viruses of Respiratory Infections, Institut Pasteur, Paris | Marion Barbet, Sylvie Behillil, Méline Bizard, Angela Brisebarre, Camille Capel, Vincent Enouf, Louise Lefrançois, Frédéric Lemoine, Christophe Malabat, Corinne Maufrais, Damien Mornico, Etienne Simon-Lorière, Maud Vanpeene, Sylvie Van der Werf, Pascale Martres |
| EPI_ISL_1700647, EPI_ISL_1700648 | Hospital | National Reference Center for Viruses of Respiratory Infections, Institut Pasteur, Paris | Marion Barbet, Sylvie Behillil, Méline Bizard, Angela Brisebarre, Camille Capel, Vincent Enouf, Louise Lefrançois, Frédéric Lemoine, Christophe Malabat, Corinne Maufrais, Adrien Pain, Etienne Simon-Lorière, Maud Vanpeene, Sylvie Van der Werf, Pascale Martres |
| EPI_ISL_1700649 | Hospital | National Reference Center for Viruses of Respiratory Infections, Institut Pasteur, Paris | Marion Barbet, Sylvie Behillil, Méline Bizard, Angela Brisebarre, Camille Capel, Vincent Enouf, Louise Lefrançois, Frédéric Lemoine, Christophe Malabat, Corinne Maufrais, Damien Mornico, Etienne Simon-Lorière, Maud Vanpeene, Sylvie Van der Werf, Pascale Martres |
| EPI_ISL_1700650 | Hospital | National Reference Center for Viruses of Respiratory Infections, Institut Pasteur, Paris | Marion Barbet, Sylvie Behillil, Méline Bizard, Angela Brisebarre, Camille Capel, Vincent Enouf, Louise Lefrançois, Frédéric Lemoine, Christophe Malabat, Corinne Maufrais, Adrien Pain, Etienne Simon-Lorière, Maud Vanpeene, Sylvie Van der Werf, Pascale Martres |
| EPI_ISL_1700652, EPI_ISL_1700654, EPI_ISL_1700655, EPI_ISL_1700656, EPI_ISL_1700657, EPI_ISL_1700658, | Hospital | National Reference Center for Viruses of Respiratory Infections, Institut Pasteur, Paris | Marion Barbet, Sylvie Behillil, Méline Bizard, Angela Brisebarre, Camille Capel, Vincent Enouf, Louise Lefrançois, Frédéric Lemoine, Christophe Malabat, Corinne Maufrais, Etienne Simon-Lorière, Maud Vanpeene, Sylvie Van der Werf, Pascale Martres |

| | | | |
|---|---|---|---|
| EPI_ISL_1700659 | | | |
| EPI_ISL_1700660 | Hospital | National Reference Center for Viruses of Respiratory Infections, Institut Pasteur, Paris | Marion Barbet, Sylvie Behillil, Méline Bizard, Angela Brisebarre, Camille Capel, Vincent Enouf, Louise Lefrançois, Frédéric Lemoine, Christophe Malabat, Corinne Maufrais, Damien Mornico, Etienne Simon-Lorière, Maud Vanpeene, Sylvie Van der Werf, CéCile Farrugia |
| EPI_ISL_1700661, EPI_ISL_1700662, EPI_ISL_1700663, EPI_ISL_1700664, EPI_ISL_1700665, EPI_ISL_1700666 | Hospital | National Reference Center for Viruses of Respiratory Infections, Institut Pasteur, Paris | Marion Barbet, Sylvie Behillil, Méline Bizard, Angela Brisebarre, Camille Capel, Vincent Enouf, Louise Lefrançois, Frédéric Lemoine, Christophe Malabat, Corinne Maufrais, Etienne Simon-Lorière, Maud Vanpeene, Sylvie Van der Werf, CéCile Farrugia |
| EPI_ISL_1700667 | Hospital | National Reference Center for Viruses of Respiratory Infections, Institut Pasteur, Paris | Marion Barbet, Sylvie Behillil, Méline Bizard, Angela Brisebarre, Camille Capel, Vincent Enouf, Louise Lefrançois, Frédéric Lemoine, Christophe Malabat, Corinne Maufrais, Adrien Pain, Etienne Simon-Lorière, Maud Vanpeene, Sylvie Van der Werf, CéCile Farrugia |
| EPI_ISL_1701391, EPI_ISL_1701394 | INT Fondazione Pascale | INT Fondazione Pascale | INT Fondazione Pascale |
| EPI_ISL_1706601 | Virology Unit, Institut Pasteur du Cambodge | Virology Unit, Institut Pasteur du Cambodge | Jurre Y Siegers, Teyputita Ou, Leakhena Pum, Cecile Troupin, Ly Sovann, Kraing Sidonn, Yi Sengdoeurn, Chin Savuth, Chau Darapheak, Veasna Duong, Erik A Karlsson |
| EPI_ISL_1707566, EPI_ISL_1707567, EPI_ISL_1707568, EPI_ISL_1707570, EPI_ISL_1707571, EPI_ISL_1707572, EPI_ISL_1707573 | SIESP DIPARTIMENTO DI PREVENZIONE TERAMO | Istituto Zooprofilattico Sperimentale dell'Abruzzo e Molise "G. Caporale" | Lorusso A, Marcacci M, Di Domenico M, Ancora M, Curini V, Di Lollo Valeria, Mangone I, Rinaldi A, Delli Compagni E, Scialabba S, Caporale M, Di Pasquale A, Cammà C, Puglia I, Calistri P, Savini G |
| EPI_ISL_1707575 | SIESP CHIETI - DRIVE IN LANCIANO | Istituto Zooprofilattico Sperimentale dell'Abruzzo e Molise "G. Caporale" | Lorusso A, Marcacci M, Di Domenico M, Ancora M, Curini V, Di Lollo Valeria, Mangone I, Rinaldi A, Delli Compagni E, Scialabba S, Caporale M, Di Pasquale A, Cammà C, Puglia I, Calistri P, Savini G |
| EPI_ISL_1707576 | SIESP CHIETI - DRIVE IN CHIETI | Istituto Zooprofilattico Sperimentale dell'Abruzzo e Molise "G. Caporale" | Lorusso A, Marcacci M, Di Domenico M, Ancora M, Curini V, Di Lollo Valeria, Mangone I, Rinaldi A, Delli Compagni E, Scialabba S, Caporale M, Di Pasquale A, Cammà C, Puglia I, Calistri P, Savini G |
| EPI_ISL_1707577 | SIESP CHIETI - DRIVE IN ORTONA | Istituto Zooprofilattico Sperimentale dell'Abruzzo e Molise "G. Caporale" | Lorusso A, Marcacci M, Di Domenico M, Ancora M, Curini V, Di Lollo Valeria, Mangone I, Rinaldi A, Delli Compagni E, Scialabba S, Caporale M, Di Pasquale A, Cammà C, Puglia I, Calistri P, Savini G |
| EPI_ISL_1707580, EPI_ISL_1707581 | USCA Tagliacozzo | Istituto Zooprofilattico Sperimentale dell'Abruzzo e Molise "G. Caporale" | Lorusso A, Marcacci M, Di Domenico M, Ancora M, Curini V, Di Lollo Valeria, Mangone I, Rinaldi A, Delli Compagni E, Scialabba S, Caporale M, Di Pasquale A, Cammà C, Puglia I, Calistri P, Savini G |
| EPI_ISL_1707582, EPI_ISL_1707583 | SIESP CHIETI - DRIVE IN VASTO | Istituto Zooprofilattico Sperimentale dell'Abruzzo e Molise "G. Caporale" | Lorusso A, Marcacci M, Di Domenico M, Ancora M, Curini V, Di Lollo Valeria, Mangone I, Rinaldi A, Delli Compagni E, Scialabba S, Caporale M, Di Pasquale A, Cammà C, Puglia I, Calistri P, Savini G |
| EPI_ISL_1707584 | SIESP CHIETI - DRIVE IN GISSI | Istituto Zooprofilattico Sperimentale dell'Abruzzo e Molise "G. Caporale" | Lorusso A, Marcacci M, Di Domenico M, Ancora M, Curini V, Di Lollo Valeria, Mangone I, Rinaldi A, Delli Compagni E, Scialabba S, Caporale M, Di Pasquale A, Cammà C, Puglia I, Calistri P, Savini G |
| EPI_ISL_1707585 | SIESP CHIETI - DRIVE IN LANCIANO | Istituto Zooprofilattico Sperimentale dell'Abruzzo e Molise "G. Caporale" | Lorusso A, Marcacci M, Di Domenico M, Ancora M, Curini V, Di Lollo Valeria, Mangone I, Rinaldi A, Delli Compagni E, Scialabba S, Caporale M, Di Pasquale A, Cammà C, Puglia I, Calistri P, Savini G |
| EPI_ISL_1707586 | SIESP CHIETI - DRIVE IN ORTONA | Istituto Zooprofilattico Sperimentale dell'Abruzzo e Molise "G. Caporale" | Lorusso A, Marcacci M, Di Domenico M, Ancora M, Curini V, Di Lollo Valeria, Mangone I, Rinaldi A, Delli Compagni E, Scialabba S, Caporale M, Di Pasquale A, Cammà C, Puglia I, Calistri P, Savini G |
| EPI_ISL_1707587 | SIESP CHIETI - DRIVE IN LANCIANO | Istituto Zooprofilattico Sperimentale dell'Abruzzo e Molise "G. Caporale" | Lorusso A, Marcacci M, Di Domenico M, Ancora M, Curini V, Di Lollo Valeria, Mangone I, Rinaldi A, Delli Compagni E, Scialabba S, Caporale M, Di Pasquale A, Cammà C, Puglia I, Calistri P, Savini G |
| EPI_ISL_1707588, EPI_ISL_1707589, EPI_ISL_1707590 | SIESP DIPARTIMENTO DI PREVENZIONE CHIETI | Istituto Zooprofilattico Sperimentale dell'Abruzzo e Molise "G. Caporale" | Lorusso A, Marcacci M, Di Domenico M, Ancora M, Curini V, Di Lollo Valeria, Mangone I, Rinaldi A, Delli Compagni E, Scialabba S, Caporale M, Di Pasquale A, Cammà C, Puglia I, Calistri P, Savini G |
| EPI_ISL_1707591, EPI_ISL_1707592, EPI_ISL_1707593, EPI_ISL_1707594, EPI_ISL_1707595, EPI_ISL_1707596, EPI_ISL_1707597, EPI_ISL_1707598, EPI_ISL_1707599, EPI_ISL_1707600 | SIESP DIPARTIMENTO DI PREVENZIONE TERAMO | Istituto Zooprofilattico Sperimentale dell'Abruzzo e Molise "G. Caporale" | Lorusso A, Marcacci M, Di Domenico M, Ancora M, Curini V, Di Lollo Valeria, Mangone I, Rinaldi A, Delli Compagni E, Scialabba S, Caporale M, Di Pasquale A, Cammà C, Puglia I, Calistri P, Savini G |
| EPI_ISL_1707601, EPI_ISL_1707602, EPI_ISL_1707603, EPI_ISL_1707604, EPI_ISL_1707605, EPI_ISL_1707606, EPI_ISL_1707607 | DIP. PREV. AVEZZANO SERVIZIO DI IGIENE EPIDEMIOLOGIA E SANITA' PUBBLICA | Istituto Zooprofilattico Sperimentale dell'Abruzzo e Molise "G. Caporale" | Lorusso A, Marcacci M, Di Domenico M, Ancora M, Curini V, Di Lollo Valeria, Mangone I, Rinaldi A, Delli Compagni E, Scialabba S, Caporale M, Di Pasquale A, Cammà C, Puglia I, Calistri P, Savini G |
| EPI_ISL_1707608, EPI_ISL_1707609 | USCA Avezzano | Istituto Zooprofilattico Sperimentale dell'Abruzzo e Molise "G. Caporale" | Lorusso A, Marcacci M, Di Domenico M, Ancora M, Curini V, Di Lollo Valeria, Mangone I, Rinaldi A, Delli Compagni E, Scialabba S, Caporale M, Di Pasquale A, Cammà C, Puglia I, Calistri P, Savini G |
| EPI_ISL_1707610, EPI_ISL_1707611 | USCA Castel di Sangro | Istituto Zooprofilattico Sperimentale dell'Abruzzo e Molise "G. Caporale" | Lorusso A, Marcacci M, Di Domenico M, Ancora M, Curini V, Di Lollo Valeria, Mangone I, Rinaldi A, Delli Compagni E, Scialabba S, Caporale M, Di Pasquale A, Cammà C, Puglia I, Calistri P, Savini G |
| EPI_ISL_1707612, EPI_ISL_1707613 | SIESP CHIETI - DRIVE IN ORTONA | Istituto Zooprofilattico Sperimentale dell'Abruzzo e Molise "G. Caporale" | Lorusso A, Marcacci M, Di Domenico M, Ancora M, Curini V, Di Lollo Valeria, Mangone I, Rinaldi A, Delli Compagni E, Scialabba S, Caporale M, Di Pasquale A, Cammà C, Puglia I, Calistri P, Savini G |
| EPI_ISL_1707614 | SIESP CHIETI - DRIVE IN VASTO | Istituto Zooprofilattico Sperimentale dell'Abruzzo e Molise "G. Caporale" | Lorusso A, Marcacci M, Di Domenico M, Ancora M, Curini V, Di Lollo Valeria, Mangone I, Rinaldi A, Delli Compagni E, Scialabba S, Caporale M, Di Pasquale A, Cammà C, Puglia I, Calistri P, Savini G |
| EPI_ISL_1707615, EPI_ISL_1707616 | SIESP CHIETI - DRIVE IN GISSI | Istituto Zooprofilattico Sperimentale dell'Abruzzo e Molise "G. Caporale" | Lorusso A, Marcacci M, Di Domenico M, Ancora M, Curini V, Di Lollo Valeria, Mangone I, Rinaldi A, Delli Compagni E, Scialabba S, Caporale M, Di Pasquale A, Cammà C, Puglia I, Calistri P, Savini G |
| EPI_ISL_1707617 | SIESP DIPARTIMENTO DI PREVENZIONE CHIETI | Istituto Zooprofilattico Sperimentale dell'Abruzzo e Molise "G. Caporale" | Lorusso A, Marcacci M, Di Domenico M, Ancora M, Curini V, Di Lollo Valeria, Mangone I, Rinaldi A, Delli Compagni E, Scialabba S, Caporale M, Di Pasquale A, Cammà C, Puglia I, Calistri P, Savini G |
| EPI_ISL_1707618 | SIESP DIPARTIMENTO DI PREVENZIONE TERAMO | Istituto Zooprofilattico Sperimentale dell'Abruzzo e Molise "G. Caporale" | Lorusso A, Marcacci M, Di Domenico M, Ancora M, Curini V, Di Lollo Valeria, Mangone I, Rinaldi A, Delli Compagni E, Scialabba S, Caporale M, Di Pasquale A, Cammà C, Puglia I, Calistri P, Savini G |
| EPI_ISL_1707619, EPI_ISL_1707620, EPI_ISL_1707621, EPI_ISL_1707622, EPI_ISL_1707623, EPI_ISL_1707624, EPI_ISL_1707625, EPI_ISL_1707626, EPI_ISL_1707627, EPI_ISL_1707628, EPI_ISL_1707629, EPI_ISL_1707630, EPI_ISL_1707631, EPI_ISL_1707632, EPI_ISL_1707633, EPI_ISL_1707634, EPI_ISL_1707635, EPI_ISL_1707636, EPI_ISL_1707637, EPI_ISL_1707638, EPI_ISL_1707639, EPI_ISL_1707640 | | | |
| see above | OSPEDALE SAN SALVATORE - MEDICINA DI LABORATORIO | Istituto Zooprofilattico Sperimentale dell'Abruzzo e Molise "G. Caporale" | Lorusso A, Marcacci M, Di Domenico M, Ancora M, Curini V, Di Lollo Valeria, Mangone I, Rinaldi A, Delli Compagni E, Scialabba S, Caporale M, Di Pasquale A, Cammà C, Puglia I, Calistri P, Savini G |
| EPI_ISL_1707641, EPI_ISL_1707642, EPI_ISL_1707643, EPI_ISL_1707644, EPI_ISL_1707645, EPI_ISL_1707646, EPI_ISL_1707647 | SIESP DIPARTIMENTO DI PREVENZIONE TERAMO | Istituto Zooprofilattico Sperimentale dell'Abruzzo e Molise "G. Caporale" | Lorusso A, Marcacci M, Di Domenico M, Ancora M, Curini V, Di Lollo Valeria, Mangone I, Rinaldi A, Delli Compagni E, Scialabba S, Caporale M, Di Pasquale A, Cammà C, Puglia I, Calistri P, Savini G |
| EPI_ISL_1707648 | SIESP CHIETI - DRIVE IN ORTONA | Istituto Zooprofilattico Sperimentale dell'Abruzzo e Molise "G. Caporale" | Lorusso A, Marcacci M, Di Domenico M, Ancora M, Curini V, Di Lollo Valeria, Mangone I, Rinaldi A, Delli Compagni E, Scialabba S, Caporale M, Di Pasquale A, Cammà C, Puglia I, Calistri P, Savini G |
| EPI_ISL_1707649 | SIESP DIPARTIMENTO DI PREVENZIONE TERAMO | Istituto Zooprofilattico Sperimentale dell'Abruzzo e Molise "G. Caporale" | Lorusso A, Marcacci M, Di Domenico M, Ancora M, Curini V, Di Lollo Valeria, Mangone I, Rinaldi A, Delli Compagni E, Scialabba S, Caporale M, Di Pasquale A, Cammà C, Puglia I, Calistri P, Savini G |
| EPI_ISL_1707650 | SIESP CHIETI - DRIVE IN ORTONA | Istituto Zooprofilattico Sperimentale dell'Abruzzo e Molise "G. Caporale" | Lorusso A, Marcacci M, Di Domenico M, Ancora M, Curini V, Di Lollo Valeria, Mangone I, Rinaldi A, Delli Compagni E, Scialabba S, Caporale M, Di Pasquale A, Cammà C, Puglia I, Calistri P, Savini G |

| | | | |
|---|---|---|---|
| EPI_ISL_1707651 | SIESP CHIETI - DRIVE IN LANCIANO | Istituto Zooprofilattico Sperimentale dell'Abruzzo e Molise "G. Caporale" | Lorusso A, Marcacci M, Di Domenico M, Ancora M, Curini V, Di Lollo Valeria, Mangone I, Rinaldi A, Delli Compagni E, Scialabba S, Caporale M, Di Pasquale A, Cammà C, Puglia I, Calistri P, Savini G |
| EPI_ISL_1707652, EPI_ISL_1707653, EPI_ISL_1707654 | DIP. PREV. AVEZZANO SERVIZIO DI IGIENE EPIDEMIOLOGIA E SANITA' PUBBLICA | Istituto Zooprofilattico Sperimentale dell'Abruzzo e Molise "G. Caporale" | Lorusso A, Marcacci M, Di Domenico M, Ancora M, Curini V, Di Lollo Valeria, Mangone I, Rinaldi A, Delli Compagni E, Scialabba S, Caporale M, Di Pasquale A, Cammà C, Puglia I, Calistri P, Savini G |
| EPI_ISL_1707655 | SIESP DIPARTIMENTO DI PREVENZIONE TERAMO | Istituto Zooprofilattico Sperimentale dell'Abruzzo e Molise "G. Caporale" | Lorusso A, Marcacci M, Di Domenico M, Ancora M, Curini V, Di Lollo Valeria, Mangone I, Rinaldi A, Delli Compagni E, Scialabba S, Caporale M, Di Pasquale A, Cammà C, Puglia I, Calistri P, Savini G |
| EPI_ISL_1707656, EPI_ISL_1707657 | DIP. PREV. AVEZZANO SERVIZIO DI IGIENE EPIDEMIOLOGIA E SANITA' PUBBLICA | Istituto Zooprofilattico Sperimentale dell'Abruzzo e Molise "G. Caporale" | Lorusso A, Marcacci M, Di Domenico M, Ancora M, Curini V, Di Lollo Valeria, Mangone I, Rinaldi A, Delli Compagni E, Scialabba S, Caporale M, Di Pasquale A, Cammà C, Puglia I, Calistri P, Savini G |
| EPI_ISL_1707658, EPI_ISL_1707659 | USCA Tagliacozzo | Istituto Zooprofilattico Sperimentale dell'Abruzzo e Molise "G. Caporale" | Lorusso A, Marcacci M, Di Domenico M, Ancora M, Curini V, Di Lollo Valeria, Mangone I, Rinaldi A, Delli Compagni E, Scialabba S, Caporale M, Di Pasquale A, Cammà C, Puglia I, Calistri P, Savini G |
| EPI_ISL_1707660, EPI_ISL_1707661 | DIP. PREV. AVEZZANO SERVIZIO DI IGIENE EPIDEMIOLOGIA E SANITA' PUBBLICA | Istituto Zooprofilattico Sperimentale dell'Abruzzo e Molise "G. Caporale" | Lorusso A, Marcacci M, Di Domenico M, Ancora M, Curini V, Di Lollo Valeria, Mangone I, Rinaldi A, Delli Compagni E, Scialabba S, Caporale M, Di Pasquale A, Cammà C, Puglia I, Calistri P, Savini G |
| EPI_ISL_1707662, EPI_ISL_1707663 | USCA Pescina | Istituto Zooprofilattico Sperimentale dell'Abruzzo e Molise "G. Caporale" | Lorusso A, Marcacci M, Di Domenico M, Ancora M, Curini V, Di Lollo Valeria, Mangone I, Rinaldi A, Delli Compagni E, Scialabba S, Caporale M, Di Pasquale A, Cammà C, Puglia I, Calistri P, Savini G |
| EPI_ISL_1707664, EPI_ISL_1707665 | DIP. PREV. AVEZZANO SERVIZIO DI IGIENE EPIDEMIOLOGIA E SANITA' PUBBLICA | Istituto Zooprofilattico Sperimentale dell'Abruzzo e Molise "G. Caporale" | Lorusso A, Marcacci M, Di Domenico M, Ancora M, Curini V, Di Lollo Valeria, Mangone I, Rinaldi A, Delli Compagni E, Scialabba S, Caporale M, Di Pasquale A, Cammà C, Puglia I, Calistri P, Savini G |
| EPI_ISL_1707666 | USCA Avezzano | Istituto Zooprofilattico Sperimentale dell'Abruzzo e Molise "G. Caporale" | Lorusso A, Marcacci M, Di Domenico M, Ancora M, Curini V, Di Lollo Valeria, Mangone I, Rinaldi A, Delli Compagni E, Scialabba S, Caporale M, Di Pasquale A, Cammà C, Puglia I, Calistri P, Savini G |
| EPI_ISL_1707667 | SIESP DIPARTIMENTO DI PREVENZIONE TERAMO | Istituto Zooprofilattico Sperimentale dell'Abruzzo e Molise "G. Caporale" | Lorusso A, Marcacci M, Di Domenico M, Ancora M, Curini V, Di Lollo Valeria, Mangone I, Rinaldi A, Delli Compagni E, Scialabba S, Caporale M, Di Pasquale A, Cammà C, Puglia I, Calistri P, Savini G |
| EPI_ISL_1707668 | SIESP CHIETI - DRIVE IN ORTONA | Istituto Zooprofilattico Sperimentale dell'Abruzzo e Molise "G. Caporale" | Lorusso A, Marcacci M, Di Domenico M, Ancora M, Curini V, Di Lollo Valeria, Mangone I, Rinaldi A, Delli Compagni E, Scialabba S, Caporale M, Di Pasquale A, Cammà C, Puglia I, Calistri P, Savini G |
| EPI_ISL_1707669, EPI_ISL_1707670, EPI_ISL_1707671 | SIESP CHIETI - DRIVE IN VASTO | Istituto Zooprofilattico Sperimentale dell'Abruzzo e Molise "G. Caporale" | Lorusso A, Marcacci M, Di Domenico M, Ancora M, Curini V, Di Lollo Valeria, Mangone I, Rinaldi A, Delli Compagni E, Scialabba S, Caporale M, Di Pasquale A, Cammà C, Puglia I, Calistri P, Savini G |
| EPI_ISL_1707672 | SIESP DIPARTIMENTO DI PREVENZIONE TERAMO | Istituto Zooprofilattico Sperimentale dell'Abruzzo e Molise "G. Caporale" | Lorusso A, Marcacci M, Di Domenico M, Ancora M, Curini V, Di Lollo Valeria, Mangone I, Rinaldi A, Delli Compagni E, Scialabba S, Caporale M, Di Pasquale A, Cammà C, Puglia I, Calistri P, Savini G |
| EPI_ISL_1707673 | SIESP CHIETI - DRIVE IN ORTONA | Istituto Zooprofilattico Sperimentale dell'Abruzzo e Molise "G. Caporale" | Lorusso A, Marcacci M, Di Domenico M, Ancora M, Curini V, Di Lollo Valeria, Mangone I, Rinaldi A, Delli Compagni E, Scialabba S, Caporale M, Di Pasquale A, Cammà C, Puglia I, Calistri P, Savini G |
| EPI_ISL_1707674 | OSPEDALE SAN SALVATORE - MEDICINA DI LABORATORIO | Istituto Zooprofilattico Sperimentale dell'Abruzzo e Molise "G. Caporale" | Lorusso A, Marcacci M, Di Domenico M, Ancora M, Curini V, Di Lollo Valeria, Mangone I, Rinaldi A, Delli Compagni E, Scialabba S, Caporale M, Di Pasquale A, Cammà C, Puglia I, Calistri P, Savini G |
| EPI_ISL_1707689, EPI_ISL_1707690, EPI_ISL_1707691, EPI_ISL_1707692 | Instituto Adolfo Lutz - Regional de Taubate | Instituto Adolfo Lutz, Interdiciplinary Procedures Center, Strategic Laboratory | Claudio Tavares Sacchi, Claudia Regina Gonçalves, Erica Valessa Ramos Gomes, Karoline Rodrigues Campos, Caio Vinicius Dias Lopes, Leonardo Jose Tadeu de Araujo, Katia Correa de Oliveira Santos |
| EPI_ISL_1708622 | Hospital | National Reference Center for Viruses of Respiratory Infections, Institut Pasteur, Paris | Marion Barbet, Sylvie Behillil, Méline Bizard, Angela Brisebarre, Camille Capel, Vincent Enouf, Louise Lefrançois, Frédéric Lemoine, Christophe Malabat, Corinne Maufrais, Etienne Simon-Lorière, Maud Vanpeene, Sylvie Van der Werf ,CéCile Farrugia |
| EPI_ISL_1708623, EPI_ISL_1708624, EPI_ISL_1708625, EPI_ISL_1708626, EPI_ISL_1708627 | Hospital | National Reference Center for Viruses of Respiratory Infections, Institut Pasteur, Paris | Marion Barbet, Sylvie Behillil, Méline Bizard, Angela Brisebarre, Camille Capel, Vincent Enouf, Louise Lefrançois, Frédéric Lemoine, Christophe Malabat, Corinne Maufrais, Etienne Simon-Lorière, Maud Vanpeene, Sylvie Van der Werf ,Pierre Alain Billy |
| EPI_ISL_1708629 | Hospital | National Reference Center for Viruses of Respiratory Infections, Institut Pasteur, Paris | Marion Barbet, Sylvie Behillil, Méline Bizard, Angela Brisebarre, Camille Capel, Vincent Enouf, Louise Lefrançois, Frédéric Lemoine, Christophe Malabat, Corinne Maufrais, Amaury Vaysse, Etienne Simon-Lorière, Maud Vanpeene, Sylvie Van der Werf ,Pierre Alain Billy |
| EPI_ISL_1708630 | Hospital | National Reference Center for Viruses of Respiratory Infections, Institut Pasteur, Paris | Marion Barbet, Sylvie Behillil, Méline Bizard, Angela Brisebarre, Camille Capel, Vincent Enouf, Louise Lefrançois, Frédéric Lemoine, Christophe Malabat, Corinne Maufrais, Etienne Simon-Lorière, Maud Vanpeene, Sylvie Van der Werf ,Pierre Alain Billy |
| EPI_ISL_1708632, EPI_ISL_1708633 | Hospital | National Reference Center for Viruses of Respiratory Infections, Institut Pasteur, Paris | Marion Barbet, Sylvie Behillil, Méline Bizard, Angela Brisebarre, Camille Capel, Vincent Enouf, Louise Lefrançois, Frédéric Lemoine, Christophe Malabat, Corinne Maufrais, Etienne Simon-Lorière, Maud Vanpeene, Sylvie Van der Werf ,Eric Farfour |
| EPI_ISL_1708634, EPI_ISL_1708636 | Hospital | National Reference Center for Viruses of Respiratory Infections, Institut Pasteur, Paris | Marion Barbet, Sylvie Behillil, Méline Bizard, Angela Brisebarre, Camille Capel, Vincent Enouf, Louise Lefrançois, Frédéric Lemoine, Christophe Malabat, Corinne Maufrais, Emmanuelle Permal, Etienne Simon-Lorière, Maud Vanpeene, Sylvie Van der Werf ,Eric Farfour |
| EPI_ISL_1708637 | Hospital | National Reference Center for Viruses of Respiratory Infections, Institut Pasteur, Paris | Marion Barbet, Sylvie Behillil, Méline Bizard, Angela Brisebarre, Camille Capel, Vincent Enouf, Louise Lefrançois, Frédéric Lemoine, Christophe Malabat, Corinne Maufrais, Amaury Vaysse, Etienne Simon-Lorière, Maud Vanpeene, Sylvie Van der Werf ,StéPhanie Marque Juillet |
| EPI_ISL_1708639 | Hospital | National Reference Center for Viruses of Respiratory Infections, Institut Pasteur, Paris | Marion Barbet, Sylvie Behillil, Méline Bizard, Angela Brisebarre, Camille Capel, Vincent Enouf, Louise Lefrançois, Frédéric Lemoine, Christophe Malabat, Corinne Maufrais, Etienne Simon-Lorière, Maud Vanpeene, Sylvie Van der Werf ,StéPhanie Marque Juillet |
| EPI_ISL_1708642 | Hospital | National Reference Center for Viruses of Respiratory Infections, Institut Pasteur, Paris | Marion Barbet, Sylvie Behillil, Méline Bizard, Angela Brisebarre, Camille Capel, Vincent Enouf, Louise Lefrançois, Frédéric Lemoine, Christophe Malabat, Corinne Maufrais, Amaury Vaysse, Etienne Simon-Lorière, Maud Vanpeene, Sylvie Van der Werf ,Sylvie Ledru |
| EPI_ISL_1708643 | Hospital | National Reference Center for Viruses of Respiratory Infections, Institut Pasteur, Paris | Marion Barbet, Sylvie Behillil, Méline Bizard, Angela Brisebarre, Camille Capel, Vincent Enouf, Louise Lefrançois, Frédéric Lemoine, Christophe Malabat, Corinne Maufrais, Etienne Simon-Lorière, Maud Vanpeene, Sylvie Van der Werf ,BenoïT Bergues |
| EPI_ISL_1708644 | Hospital | National Reference Center for Viruses of Respiratory Infections, Institut Pasteur, Paris | Marion Barbet, Sylvie Behillil, Méline Bizard, Angela Brisebarre, Camille Capel, Vincent Enouf, Louise Lefrançois, Frédéric Lemoine, Christophe Malabat, Corinne Maufrais, Amaury Vaysse, Etienne Simon-Lorière, Maud Vanpeene, Sylvie Van der Werf ,BenoïT Bergues |
| EPI_ISL_1708646, EPI_ISL_1708647, EPI_ISL_1708648 | Hospital | National Reference Center for Viruses of Respiratory Infections, Institut Pasteur, Paris | Marion Barbet, Sylvie Behillil, Méline Bizard, Angela Brisebarre, Camille Capel, Vincent Enouf, Louise Lefrançois, Frédéric Lemoine, Christophe Malabat, Corinne Maufrais, Etienne Simon-Lorière, Maud Vanpeene, Sylvie Van der Werf ,Sandrine Castelain |
| EPI_ISL_1708651 | Hospital | National Reference Center for Viruses of Respiratory Infections, Institut Pasteur, Paris | Marion Barbet, Sylvie Behillil, Méline Bizard, Angela Brisebarre, Camille Capel, Vincent Enouf, Louise Lefrançois, Frédéric Lemoine, Christophe Malabat, Corinne Maufrais, Amaury Vaysse, Etienne Simon-Lorière, Maud Vanpeene, Sylvie Van der Werf ,Sandrine Castelain |
| EPI_ISL_1708654, EPI_ISL_1708655 | Hospital | National Reference Center for Viruses of Respiratory Infections, Institut Pasteur, Paris | Marion Barbet, Sylvie Behillil, Méline Bizard, Angela Brisebarre, Camille Capel, Vincent Enouf, Louise Lefrançois, Frédéric Lemoine, Christophe Malabat, Corinne Maufrais, Etienne Simon-Lorière, Maud Vanpeene, Sylvie Van der Werf ,Ilka Engelmann |
| EPI_ISL_1708656, EPI_ISL_1708657, EPI_ISL_1708658 | Hospital | National Reference Center for Viruses of Respiratory Infections, Institut Pasteur, Paris | Marion Barbet, Sylvie Behillil, Méline Bizard, Angela Brisebarre, Camille Capel, Vincent Enouf, Louise Lefrançois, Frédéric Lemoine, Christophe Malabat, Corinne Maufrais, Etienne Simon-Lorière, Maud Vanpeene, Sylvie Van der Werf ,Anne Vachee |
| EPI_ISL_1708742 | Imeda Hospital | Imeda Hospital | Johan Frans, Dagmar Obbels, Hanne Valgaeren |
| EPI_ISL_1711983, EPI_ISL_1711984, EPI_ISL_1711985, EPI_ISL_1711986, EPI_ISL_1711987, EPI_ISL_1711988, EPI_ISL_1711989, EPI_ISL_1711990, EPI_ISL_1711991, EPI_ISL_1711992, EPI_ISL_1711993, EPI_ISL_1711994, EPI_ISL_1711995, EPI_ISL_1711996, EPI_ISL_1711997, EPI_ISL_1711998 | | | |
| see above | Virology Unit, Institut Pasteur du Cambodge | Virology Unit, Institut Pasteur du Cambodge | Jurre Y Siegers, Teyputita Ou, Leakhena Pum, Cecile Troupin, Ly Sovann, Kraing Sidonn, Yi Sengdoeurn, Chin Savuth, Chau Darapheak, Veasna Duong, Erik A Karlsson |
| EPI_ISL_1731455 | Centre Hospitalier Universitaire Clermont-Ferrand | CHU Clermont-Ferrand, service de virologie | Bisseux Maxime, Mirand Audrey, Combes Patricia, Henquell Cécile |
| EPI_ISL_1731456, EPI_ISL_1731457 | HOPITAL DE VICHY | CHU Clermont-Ferrand, service de virologie | Bisseux Maxime, Mirand Audrey, Combes Patricia, Henquell Cécile |
| EPI_ISL_1731461 | Centre hospitalier D'AURILLAC | CHU Clermont-Ferrand, service de virologie | Bisseux Maxime, Mirand Audrey, Combes Patricia, Henquell Cécile |

| | | | |
|---|---|---|---|
| EPI_ISL_1731462 | Centre Hospitalier Universitaire Clermont-Ferrand | CHU Clermont-Ferrand, service de virologie | Bisseux Maxime, Mirand Audrey, Combes Patricia, Henquell Cécile |
| EPI_ISL_1731463, EPI_ISL_1731464, EPI_ISL_1731465 | HOPITAL DE VICHY | CHU Clermont-Ferrand, service de virologie | Bisseux Maxime, Mirand Audrey, Combes Patricia, Henquell Cécile |
| EPI_ISL_1731466 | Centre hospitalier D'AURILLAC | CHU Clermont-Ferrand, service de virologie | Bisseux Maxime, Mirand Audrey, Combes Patricia, Henquell Cécile |
| EPI_ISL_1731467 | Centre Hospitalier Paul Ardier | CHU Clermont-Ferrand, service de virologie | Bisseux Maxime, Mirand Audrey, Combes Patricia, Henquell Cécile |
| EPI_ISL_1731468, EPI_ISL_1731469 | Centre Hospitalier Universitaire Clermont-Ferrand | CHU Clermont-Ferrand, service de virologie | Bisseux Maxime, Mirand Audrey, Combes Patricia, Henquell Cécile |
| EPI_ISL_1731470 | HOPITAL DE VICHY | CHU Clermont-Ferrand, service de virologie | Bisseux Maxime, Mirand Audrey, Combes Patricia, Henquell Cécile |
| EPI_ISL_1731471 | Centre hospitalier D'AURILLAC | CHU Clermont-Ferrand, service de virologie | Bisseux Maxime, Mirand Audrey, Combes Patricia, Henquell Cécile |
| EPI_ISL_1731472, EPI_ISL_1731473 | Centre Hospitalier Universitaire Clermont-Ferrand | CHU Clermont-Ferrand, service de virologie | Bisseux Maxime, Mirand Audrey, Combes Patricia, Henquell Cécile |
| EPI_ISL_1731474 | HOPITAL DE MONTLUCON | CHU Clermont-Ferrand, service de virologie | Bisseux Maxime, Mirand Audrey, Combes Patricia, Henquell Cécile |
| EPI_ISL_1731475, EPI_ISL_1731476 | Centre Hospitalier Universitaire Clermont-Ferrand | CHU Clermont-Ferrand, service de virologie | Bisseux Maxime, Mirand Audrey, Combes Patricia, Henquell Cécile |
| EPI_ISL_1731477 | HOPITAL DE VICHY | CHU Clermont-Ferrand, service de virologie | Bisseux Maxime, Mirand Audrey, Combes Patricia, Henquell Cécile |
| EPI_ISL_1731478, EPI_ISL_1731479, EPI_ISL_1731480 | Centre Hospitalier Universitaire Clermont-Ferrand | CHU Clermont-Ferrand, service de virologie | Bisseux Maxime, Mirand Audrey, Combes Patricia, Henquell Cécile |
| EPI_ISL_1731481, EPI_ISL_1731482 | Centre hospitalier D'AURILLAC | CHU Clermont-Ferrand, service de virologie | Bisseux Maxime, Mirand Audrey, Combes Patricia, Henquell Cécile |
| EPI_ISL_1731483, EPI_ISL_1731485, EPI_ISL_1731486, EPI_ISL_1731487, EPI_ISL_1731488, EPI_ISL_1731489, EPI_ISL_1731490 | Centre Hospitalier Universitaire Clermont-Ferrand | CHU Clermont-Ferrand, service de virologie | Bisseux Maxime, Mirand Audrey, Combes Patricia, Henquell Cécile |
| EPI_ISL_1731491, EPI_ISL_1731492 | HOPITAL DE VICHY | CHU Clermont-Ferrand, service de virologie | Bisseux Maxime, Mirand Audrey, Combes Patricia, Henquell Cécile |
| EPI_ISL_1731493, EPI_ISL_1731494, EPI_ISL_1731495, EPI_ISL_1731496, EPI_ISL_1731497 | Centre Hospitalier Universitaire Clermont-Ferrand | CHU Clermont-Ferrand, service de virologie | Bisseux Maxime, Mirand Audrey, Combes Patricia, Henquell Cécile |
| EPI_ISL_1731498, EPI_ISL_1731499, EPI_ISL_1731500, EPI_ISL_1731501, EPI_ISL_1731502 | Centre Hospitalier RIOM | CHU Clermont-Ferrand, service de virologie | Bisseux Maxime, Mirand Audrey, Combes Patricia, Henquell Cécile |
| EPI_ISL_1731503, EPI_ISL_1731504 | HOPITAL DU MONT DORE | CHU Clermont-Ferrand, service de virologie | Bisseux Maxime, Mirand Audrey, Combes Patricia, Henquell Cécile |
| EPI_ISL_1731507, EPI_ISL_1731508, EPI_ISL_1731509 | Centre Hospitalier Universitaire Clermont-Ferrand | CHU Clermont-Ferrand, service de virologie | Bisseux Maxime, Mirand Audrey, Combes Patricia, Henquell Cécile |
| EPI_ISL_1731510, EPI_ISL_1731511, EPI_ISL_1731512 | Centre De Prelevement COVID RIOM | CHU Clermont-Ferrand, service de virologie | Bisseux Maxime, Mirand Audrey, Combes Patricia, Henquell Cécile |
| EPI_ISL_1731514, EPI_ISL_1731515, EPI_ISL_1731516 | Centre Hospitalier Universitaire Clermont-Ferrand | CHU Clermont-Ferrand, service de virologie | Bisseux Maxime, Mirand Audrey, Combes Patricia, Henquell Cécile |
| EPI_ISL_1731517 | Centre De Prelevement COVID RIOM | CHU Clermont-Ferrand, service de virologie | Bisseux Maxime, Mirand Audrey, Combes Patricia, Henquell Cécile |
| EPI_ISL_1731518 | Centre Jean Perrin | CHU Clermont-Ferrand, service de virologie | Bisseux Maxime, Mirand Audrey, Combes Patricia, Henquell Cécile |
| EPI_ISL_1731519, EPI_ISL_1731520, EPI_ISL_1731521, EPI_ISL_1731522 | Centre De Prelevement COVID RIOM | CHU Clermont-Ferrand, service de virologie | Bisseux Maxime, Mirand Audrey, Combes Patricia, Henquell Cécile |
| EPI_ISL_1731523 | Centre Hospitalier Universitaire Clermont-Ferrand | CHU Clermont-Ferrand, service de virologie | Bisseux Maxime, Mirand Audrey, Combes Patricia, Henquell Cécile |
| EPI_ISL_1738807 | Institute of Microbiology, Universidad San Francisco de Quito | Institute of Microbiology, Universidad San Francisco de Quito | Sully Márquez, Belén Prado-Vivar, Juan José Guadalupe, Monica Becerra-Wong, Bernardo Gutiérrez, Manuel Jaramillo, Verónica Barragán, Patricio Rojas-Silva, Gabriel Trueba, Michelle Grunauer, Paúl Cárdenas |
| EPI_ISL_1739244 | Institute of Microbiology, Universidad San Francisco de Quito | Institute of Microbiology, Universidad San Francisco de Quito | Sully Márquez, Belén Prado-Vivar, Juan José Guadalupe, Monica Becerra-Wong, Bernardo Gutiérrez, Gabriela Zambrano, Verónica Barragán, Patricio Rojas-Silva, Gabriel Trueba, Michelle Grunauer, Paúl Cárdenas |
| EPI_ISL_1739326 | Hospital | National Reference Center for Viruses of Respiratory Infections, Institut Pasteur, Paris | Marion Barbet, Sylvie Behillil, Méline Bizard, Angela Brisebarre, Camille Capel, Vincent Enouf, Louise Lefrançois, Frédéric Lemoine, Christophe Malabat, Corinne Maufrais, Damien Mornico, Etienne Simon-Lorière, Maud Vanpeene, Sylvie Van der Werf ,ValéRie Serazin |
| EPI_ISL_1739327 | Hospital | National Reference Center for Viruses of Respiratory Infections, Institut Pasteur, Paris | Marion Barbet, Sylvie Behillil, Méline Bizard, Angela Brisebarre, Camille Capel, Vincent Enouf, Louise Lefrançois, Frédéric Lemoine, Christophe Malabat, Corinne Maufrais, Adrien Pain, Etienne Simon-Lorière, Maud Vanpeene, Sylvie Van der Werf ,Pascale Martres |
| EPI_ISL_1739328, EPI_ISL_1739329 | Hospital | National Reference Center for Viruses of Respiratory Infections, Institut Pasteur, Paris | Marion Barbet, Sylvie Behillil, Méline Bizard, Angela Brisebarre, Camille Capel, Vincent Enouf, Louise Lefrançois, Frédéric Lemoine, Christophe Malabat, Corinne Maufrais, Adrien Pain, Etienne Simon-Lorière, Maud Vanpeene, Sylvie Van der Werf ,CéCile Farrugia |
| EPI_ISL_1739330 | Basurto University Hospital Microbiology Lab | Biocruces Bizkaia | Mikel Urrutikoetxea-Gutierrez, Mº Carmen Nieto Toboso, Estibaliz Ugalde Zarraga, Mikel Gallego Rodrigo, Ana de la Hoz, Jose Luís Díaz de Tuesta del Arco |
| EPI_ISL_1745681, EPI_ISL_1745682, EPI_ISL_1745683, EPI_ISL_1745684, EPI_ISL_1745685, EPI_ISL_1745686, EPI_ISL_1745687, EPI_ISL_1745688, EPI_ISL_1745690, EPI_ISL_1745691, EPI_ISL_1745692, EPI_ISL_1745693, EPI_ISL_1745694, EPI_ISL_1745695 | | | |
| see above | Institute of Molecular and Translational Medicine / Laboratory of Experimental Medicine, Faculty of Medicine and Dentistry, Palacky University and University Hospital Olomouc | Institute of Molecular and Translational Medicine / Laboratory of Experimental Medicine, Faculty of Medicine and Dentistry, Palacky University | Rastislav Slavkovský, Hana Jaworek, Vladimíra Koudeláková, Barbora Blumová, Tomáš Pospíšíl, Marián Hajdúch |
| EPI_ISL_1745696, EPI_ISL_1745697, EPI_ISL_1745698, EPI_ISL_1745699, EPI_ISL_1745700 | Department of Microbiology, Faculty of Medicine and Dentistry, Palacky University and University Hospital Olomouc | Institute of Molecular and Translational Medicine / Laboratory of Experimental Medicine, Faculty of Medicine and Dentistry, Palacky University | Rastislav Slavkovský, Hana Jaworek, Vladimíra Koudeláková, Barbora Blumová, Tomáš Pospíšíl, Marián Hajdúch, Pavel Sauer |
| EPI_ISL_1745701, EPI_ISL_1745702, EPI_ISL_1745703, EPI_ISL_1745704, EPI_ISL_1745705, EPI_ISL_1745706, EPI_ISL_1745707, EPI_ISL_1745708, EPI_ISL_1745709, EPI_ISL_1745710, EPI_ISL_1745711, EPI_ISL_1745712, EPI_ISL_1745713, EPI_ISL_1745714 | | | |
| see above | Institute of Molecular and Translational Medicine / Laboratory of Experimental Medicine, Faculty of Medicine and Dentistry, Palacky University and University Hospital Olomouc | Institute of Molecular and Translational Medicine / Laboratory of Experimental Medicine, Faculty of Medicine and Dentistry, Palacky University | Rastislav Slavkovský, Hana Jaworek, Vladimíra Koudeláková, Barbora Blumová, Tomáš Pospíšíl, Marián Hajdúch |
| EPI_ISL_1749430, EPI_ISL_1749436, EPI_ISL_1749437 | Singapore General Hospital | Department of Microbiology | Nurdyana Abdul Rahman, Kun Lee Lim, Chenhao Li, Sui Sin Goh, Kenneth Xin Long Chan, Kian Sing Chan, Lynette Oon, Kern Rei Chng, Niranjan Nagarajan, Karrie Ko |
| EPI_ISL_1752678, EPI_ISL_1752679, EPI_ISL_1752680, EPI_ISL_1752681, EPI_ISL_1752683, EPI_ISL_1752684, EPI_ISL_1752685, EPI_ISL_1752686, EPI_ISL_1752687, EPI_ISL_1752688 | Bioptická laborato, s.r.o. | Bioptická laborato, s.r.o. | Petr Šteiner, Tomáš Vanek, Nikola Bílá, Martina Putzová, Michaela íhová, Silva Vondráková |

| | | | |
|---|---|---|---|
| EPI_ISL_1756222, EPI_ISL_1756225, EPI_ISL_1756229, EPI_ISL_1756231, EPI_ISL_1756232, EPI_ISL_1756235, EPI_ISL_1756237, EPI_ISL_1756239, EPI_ISL_1756240 | GH A.CHENEVIER-H.MONDOR | Department of Virology, Henri Mondor University Hospital, Assistance Publique Hôpitaux de Paris, Université Paris-Est Créteil, INSERM U955 | Christophe Rodriguez, Slim Fourati, Vanessa Demontant, Guillaume Gricourt, Melissa N'Debi, Alexandre Soulier, Elisabeth Trawinski, Jean-Michel Pawlotsky |
| EPI_ISL_1756866 | UAB InMedica | Vilnius University Hospital Santaros Klinikos, Center of Laboratory Medicine | Ingrida Olendraite, Gytis Dudas, Daniel Naumovas, Dovile Ezerskyte, Rimvydas Norvilas, Ligita Raugaite, Monika Katenaite, Mindaugas Stoskus, Laimonas Griskevicius |
| EPI_ISL_1757326, EPI_ISL_1757327, EPI_ISL_1757329, EPI_ISL_1757337, EPI_ISL_1757344, EPI_ISL_1757349, EPI_ISL_1757356, EPI_ISL_1757360, EPI_ISL_1757383, EPI_ISL_1757385 | GH A.CHENEVIER-H.MONDOR | Department of Virology, Henri Mondor University Hospital, Assistance Publique Hôpitaux de Paris, Université Paris-Est Créteil, INSERM U955 | Christophe Rodriguez, Slim Fourati, Vanessa Demontant, Guillaume Gricourt, Melissa N'Debi, Alexandre Soulier, Elisabeth Trawinski, Jean-Michel Pawlotsky |
| EPI_ISL_1757395 | GH JOFFRE DUPUYTREN | Department of Virology, Henri Mondor University Hospital, Assistance Publique Hôpitaux de Paris, Université Paris-Est Créteil, INSERM U955 | Christophe Rodriguez, Slim Fourati, Vanessa Demontant, Guillaume Gricourt, Melissa N'Debi, Alexandre Soulier, Elisabeth Trawinski, Jean-Michel Pawlotsky |
| EPI_ISL_1757417 | HOPITAL EMILE ROUX | Department of Virology, Henri Mondor University Hospital, Assistance Publique Hôpitaux de Paris, Université Paris-Est Créteil, INSERM U955 | Christophe Rodriguez, Slim Fourati, Vanessa Demontant, Guillaume Gricourt, Melissa N'Debi, Alexandre Soulier, Elisabeth Trawinski, Jean-Michel Pawlotsky |
| EPI_ISL_1757599 | Hôpital Pitié-Salpêtrière | Department of Virology, Henri Mondor University Hospital, Assistance Publique Hôpitaux de Paris, Université Paris-Est Créteil, INSERM U955 | Christophe Rodriguez, Slim Fourati, Vanessa Demontant, Guillaume Gricourt, Melissa N'Debi, Alexandre Soulier, Elisabeth Trawinski, Jean-Michel Pawlotsky |
| EPI_ISL_1760209 | COVID Laboratory AOU Messina | Molecular Hepatology Lab AOU Messina | Giuseppina Raffa, Cristina Musolino, Valeria Chines, Daniele Lombardo, Teresa Pollicino |
| EPI_ISL_1761554, EPI_ISL_1761555, EPI_ISL_1761560, EPI_ISL_1761561, EPI_ISL_1761562, EPI_ISL_1761563, EPI_ISL_1761564, EPI_ISL_1761565, EPI_ISL_1761566, EPI_ISL_1761567, EPI_ISL_1761568, EPI_ISL_1761569, EPI_ISL_1761570, EPI_ISL_1761571, EPI_ISL_1761572, EPI_ISL_1761573, EPI_ISL_1761574, EPI_ISL_1761575, EPI_ISL_1761576, EPI_ISL_1761577, EPI_ISL_1761578, EPI_ISL_1761579, EPI_ISL_1761580 | | | |
| see above | P.O.CARDARELLI | P.O.CARDARELLI | Scutellà M, Felice V, Niro G |
| EPI_ISL_1785081, EPI_ISL_1785082 | SIESP DIPARTIMENTO DI PREVENZIONE TERAMO | Istituto Zooprofilattico Sperimentale dell'Abruzzo e Molise "G. Caporale" | Lorusso A, Marcacci M, Di Domenico M, Ancora M, Curini V, Di Lollo Valeria, Mangone I, Rinaldi A, Delli Compagni E, Scialabba S, Caporale M, Di Pasquale A, Cammà C, Puglia I, Calistri P, Savini G |
| EPI_ISL_1785083, EPI_ISL_1785084 | SIESP CHIETI - DRIVE IN CHIETI | Istituto Zooprofilattico Sperimentale dell'Abruzzo e Molise "G. Caporale" | Lorusso A, Marcacci M, Di Domenico M, Ancora M, Curini V, Di Lollo Valeria, Mangone I, Rinaldi A, Delli Compagni E, Scialabba S, Caporale M, Di Pasquale A, Cammà C, Puglia I, Calistri P, Savini G |
| EPI_ISL_1785085, EPI_ISL_1785086, EPI_ISL_1785087, EPI_ISL_1785088, EPI_ISL_1785089, EPI_ISL_1785090, EPI_ISL_1785091, EPI_ISL_1785092, EPI_ISL_1785093, EPI_ISL_1785094, EPI_ISL_1785095, EPI_ISL_1785096, EPI_ISL_1785097, EPI_ISL_1785098, EPI_ISL_1785099, EPI_ISL_1785100, EPI_ISL_1785101, EPI_ISL_1785102, EPI_ISL_1785103, EPI_ISL_1785104, EPI_ISL_1785105, EPI_ISL_1785106, EPI_ISL_1785107, EPI_ISL_1785108, EPI_ISL_1785109, EPI_ISL_1785110, EPI_ISL_1785111, EPI_ISL_1785112, EPI_ISL_1785113, EPI_ISL_1785114 | | | |
| see above | SIESP DIPARTIMENTO DI PREVENZIONE TERAMO | Istituto Zooprofilattico Sperimentale dell'Abruzzo e Molise "G. Caporale" | Lorusso A, Marcacci M, Di Domenico M, Ancora M, Curini V, Di Lollo Valeria, Mangone I, Rinaldi A, Delli Compagni E, Scialabba S, Caporale M, Di Pasquale A, Cammà C, Puglia I, Calistri P, Savini G |
| EPI_ISL_1785115 | Casa di Cura Di Lorenzo | Istituto Zooprofilattico Sperimentale dell'Abruzzo e Molise "G. Caporale" | Lorusso A, Marcacci M, Di Domenico M, Ancora M, Curini V, Di Lollo Valeria, Mangone I, Rinaldi A, Delli Compagni E, Scialabba S, Caporale M, Di Pasquale A, Cammà C, Puglia I, Calistri P, Savini G |
| EPI_ISL_1785116, EPI_ISL_1785117, EPI_ISL_1785118 | SIESP CHIETI - DRIVE IN LANCIANO | Istituto Zooprofilattico Sperimentale dell'Abruzzo e Molise "G. Caporale" | Lorusso A, Marcacci M, Di Domenico M, Ancora M, Curini V, Di Lollo Valeria, Mangone I, Rinaldi A, Delli Compagni E, Scialabba S, Caporale M, Di Pasquale A, Cammà C, Puglia I, Calistri P, Savini G |
| EPI_ISL_1785119 | SIESP SULMONA | Istituto Zooprofilattico Sperimentale dell'Abruzzo e Molise "G. Caporale" | Lorusso A, Marcacci M, Di Domenico M, Ancora M, Curini V, Di Lollo Valeria, Mangone I, Rinaldi A, Delli Compagni E, Scialabba S, Caporale M, Di Pasquale A, Cammà C, Puglia I, Calistri P, Savini G |
| EPI_ISL_1785120, EPI_ISL_1785121 | SIESP CHIETI - DRIVE IN LANCIANO | Istituto Zooprofilattico Sperimentale dell'Abruzzo e Molise "G. Caporale" | Lorusso A, Marcacci M, Di Domenico M, Ancora M, Curini V, Di Lollo Valeria, Mangone I, Rinaldi A, Delli Compagni E, Scialabba S, Caporale M, Di Pasquale A, Cammà C, Puglia I, Calistri P, Savini G |
| EPI_ISL_1785122 | SIESP DIPARTIMENTO DI PREVENZIONE CHIETI | Istituto Zooprofilattico Sperimentale dell'Abruzzo e Molise "G. Caporale" | Lorusso A, Marcacci M, Di Domenico M, Ancora M, Curini V, Di Lollo Valeria, Mangone I, Rinaldi A, Delli Compagni E, Scialabba S, Caporale M, Di Pasquale A, Cammà C, Puglia I, Calistri P, Savini G |
| EPI_ISL_1785124 | SIESP SULMONA | Istituto Zooprofilattico Sperimentale dell'Abruzzo e Molise "G. Caporale" | Lorusso A, Marcacci M, Di Domenico M, Ancora M, Curini V, Di Lollo Valeria, Mangone I, Rinaldi A, Delli Compagni E, Scialabba S, Caporale M, Di Pasquale A, Cammà C, Puglia I, Calistri P, Savini G |
| EPI_ISL_1785125, EPI_ISL_1785126, EPI_ISL_1785127, EPI_ISL_1785128, EPI_ISL_1785129, EPI_ISL_1785130 | DIP. PREV. AVEZZANO SERVIZIO DI IGIENE EPIDEMIOLOGIA E SANITA' PUBBLICA | Istituto Zooprofilattico Sperimentale dell'Abruzzo e Molise "G. Caporale" | Lorusso A, Marcacci M, Di Domenico M, Ancora M, Curini V, Di Lollo Valeria, Mangone I, Rinaldi A, Delli Compagni E, Scialabba S, Caporale M, Di Pasquale A, Cammà C, Puglia I, Calistri P, Savini G |
| EPI_ISL_1785131 | USCA Pescina | Istituto Zooprofilattico Sperimentale dell'Abruzzo e Molise "G. Caporale" | Lorusso A, Marcacci M, Di Domenico M, Ancora M, Curini V, Di Lollo Valeria, Mangone I, Rinaldi A, Delli Compagni E, Scialabba S, Caporale M, Di Pasquale A, Cammà C, Puglia I, Calistri P, Savini G |
| EPI_ISL_1785132, EPI_ISL_1785133 | OSPEDALE "F. RENZETTI" LANCIANO - MEDICINA INTERNA | Istituto Zooprofilattico Sperimentale dell'Abruzzo e Molise "G. Caporale" | Lorusso A, Marcacci M, Di Domenico M, Ancora M, Curini V, Di Lollo Valeria, Mangone I, Rinaldi A, Delli Compagni E, Scialabba S, Caporale M, Di Pasquale A, Cammà C, Puglia I, Calistri P, Savini G |
| EPI_ISL_1785134, EPI_ISL_1785135, EPI_ISL_1785136, EPI_ISL_1785137 | SIESP CHIETI - DRIVE IN LANCIANO | Istituto Zooprofilattico Sperimentale dell'Abruzzo e Molise "G. Caporale" | Lorusso A, Marcacci M, Di Domenico M, Ancora M, Curini V, Di Lollo Valeria, Mangone I, Rinaldi A, Delli Compagni E, Scialabba S, Caporale M, Di Pasquale A, Cammà C, Puglia I, Calistri P, Savini G |
| EPI_ISL_1785138 | SIESP DIPARTIMENTO DI PREVENZIONE CHIETI | Istituto Zooprofilattico Sperimentale dell'Abruzzo e Molise "G. Caporale" | Lorusso A, Marcacci M, Di Domenico M, Ancora M, Curini V, Di Lollo Valeria, Mangone I, Rinaldi A, Delli Compagni E, Scialabba S, Caporale M, Di Pasquale A, Cammà C, Puglia I, Calistri P, Savini G |
| EPI_ISL_1785139, EPI_ISL_1785140, EPI_ISL_1785141, EPI_ISL_1785142, EPI_ISL_1785143 | SIESP CHIETI - DRIVE IN VASTO | Istituto Zooprofilattico Sperimentale dell'Abruzzo e Molise "G. Caporale" | Lorusso A, Marcacci M, Di Domenico M, Ancora M, Curini V, Di Lollo Valeria, Mangone I, Rinaldi A, Delli Compagni E, Scialabba S, Caporale M, Di Pasquale A, Cammà C, Puglia I, Calistri P, Savini G |
| EPI_ISL_1785144 | SIESP CHIETI - DRIVE IN LANCIANO | Istituto Zooprofilattico Sperimentale dell'Abruzzo e Molise "G. Caporale" | Lorusso A, Marcacci M, Di Domenico M, Ancora M, Curini V, Di Lollo Valeria, Mangone I, Rinaldi A, Delli Compagni E, Scialabba S, Caporale M, Di Pasquale A, Cammà C, Puglia I, Calistri P, Savini G |
| EPI_ISL_1785145 | SIESP CHIETI - DRIVE IN VASTO | Istituto Zooprofilattico Sperimentale dell'Abruzzo e Molise "G. Caporale" | Lorusso A, Marcacci M, Di Domenico M, Ancora M, Curini V, Di Lollo Valeria, Mangone I, Rinaldi A, Delli Compagni E, Scialabba S, Caporale M, Di Pasquale A, Cammà C, Puglia I, Calistri P, Savini G |
| EPI_ISL_1785146, EPI_ISL_1785147 | SIESP CHIETI - DRIVE IN GISSI | Istituto Zooprofilattico Sperimentale dell'Abruzzo e Molise "G. Caporale" | Lorusso A, Marcacci M, Di Domenico M, Ancora M, Curini V, Di Lollo Valeria, Mangone I, Rinaldi A, Delli Compagni E, Scialabba S, Caporale M, Di Pasquale A, Cammà C, Puglia I, Calistri P, Savini G |
| EPI_ISL_1785148, EPI_ISL_1785149 | SIESP DIPARTIMENTO DI PREVENZIONE TERAMO | Istituto Zooprofilattico Sperimentale dell'Abruzzo e Molise "G. Caporale" | Lorusso A, Marcacci M, Di Domenico M, Ancora M, Curini V, Di Lollo Valeria, Mangone I, Rinaldi A, Delli Compagni E, Scialabba S, Caporale M, Di Pasquale A, Cammà C, Puglia I, Calistri P, Savini G |
| EPI_ISL_1785150 | SIESP CHIETI - DRIVE IN ORTONA | Istituto Zooprofilattico Sperimentale dell'Abruzzo e Molise "G. Caporale" | Lorusso A, Marcacci M, Di Domenico M, Ancora M, Curini V, Di Lollo Valeria, Mangone I, Rinaldi A, Delli Compagni E, Scialabba S, Caporale M, Di Pasquale A, Cammà C, Puglia I, Calistri P, Savini G |
| EPI_ISL_1785151 | SIESP DIPARTIMENTO DI PREVENZIONE TERAMO | Istituto Zooprofilattico Sperimentale dell'Abruzzo e Molise "G. Caporale" | Lorusso A, Marcacci M, Di Domenico M, Ancora M, Curini V, Di Lollo Valeria, Mangone I, Rinaldi A, Delli Compagni E, Scialabba S, Caporale M, Di Pasquale A, Cammà C, Puglia I, Calistri P, Savini G |
| EPI_ISL_1785152 | SIESP CHIETI - DRIVE IN VASTO | Istituto Zooprofilattico Sperimentale dell'Abruzzo e Molise "G. | Lorusso A, Marcacci M, Di Domenico M, Ancora M, Curini V, Di Lollo Valeria, Mangone I, Rinaldi A, Delli Compagni E, Scialabba S, Caporale M, Di |

| | | | |
|---|---|---|---|
| | | Caporale" | Pasquale A, Cammà C, Puglia I, Calistri P, Savini G |
| EPI_ISL_1785153, EPI_ISL_1785154 | SIESP CHIETI - DRIVE IN CHIETI | Istituto Zooprofilattico Sperimentale dell'Abruzzo e Molise "G. Caporale" | Lorusso A, Marcacci M, Di Domenico M, Ancora M, Curini V, Di Lollo Valeria, Mangone I, Rinaldi A, Delli Compagni E, Scialabba S, Caporale M, Di Pasquale A, Cammà C, Puglia I, Calistri P, Savini G |
| EPI_ISL_1785155 | DIP. PREV. AVEZZANO SERVIZIO DI IGIENE EPIDEMIOLOGIA E SANITA' PUBBLICA | Istituto Zooprofilattico Sperimentale dell'Abruzzo e Molise "G. Caporale" | Lorusso A, Marcacci M, Di Domenico M, Ancora M, Curini V, Di Lollo Valeria, Mangone I, Rinaldi A, Delli Compagni E, Scialabba S, Caporale M, Di Pasquale A, Cammà C, Puglia I, Calistri P, Savini G |
| EPI_ISL_1785156 | USCA Castel di Sangro | Istituto Zooprofilattico Sperimentale dell'Abruzzo e Molise "G. Caporale" | Lorusso A, Marcacci M, Di Domenico M, Ancora M, Curini V, Di Lollo Valeria, Mangone I, Rinaldi A, Delli Compagni E, Scialabba S, Caporale M, Di Pasquale A, Cammà C, Puglia I, Calistri P, Savini G |
| EPI_ISL_1785157 | SIESP CHIETI - DRIVE IN LANCIANO | Istituto Zooprofilattico Sperimentale dell'Abruzzo e Molise "G. Caporale" | Lorusso A, Marcacci M, Di Domenico M, Ancora M, Curini V, Di Lollo Valeria, Mangone I, Rinaldi A, Delli Compagni E, Scialabba S, Caporale M, Di Pasquale A, Cammà C, Puglia I, Calistri P, Savini G |
| EPI_ISL_1785158 | SIESP DIPARTIMENTO DI PREVENZIONE TERAMO | Istituto Zooprofilattico Sperimentale dell'Abruzzo e Molise "G. Caporale" | Lorusso A, Marcacci M, Di Domenico M, Ancora M, Curini V, Di Lollo Valeria, Mangone I, Rinaldi A, Delli Compagni E, Scialabba S, Caporale M, Di Pasquale A, Cammà C, Puglia I, Calistri P, Savini G |
| EPI_ISL_1785159, EPI_ISL_1785160 | OSPEDALE S.S. ANNUNZIATA CHIETI | Istituto Zooprofilattico Sperimentale dell'Abruzzo e Molise "G. Caporale" | Lorusso A, Marcacci M, Di Domenico M, Ancora M, Curini V, Di Lollo Valeria, Mangone I, Rinaldi A, Delli Compagni E, Scialabba S, Caporale M, Di Pasquale A, Cammà C, Puglia I, Calistri P, Savini G |
| EPI_ISL_1785161, EPI_ISL_1785162, EPI_ISL_1785163 | SIESP DIPARTIMENTO DI PREVENZIONE TERAMO | Istituto Zooprofilattico Sperimentale dell'Abruzzo e Molise "G. Caporale" | Lorusso A, Marcacci M, Di Domenico M, Ancora M, Curini V, Di Lollo Valeria, Mangone I, Rinaldi A, Delli Compagni E, Scialabba S, Caporale M, Di Pasquale A, Cammà C, Puglia I, Calistri P, Savini G |
| EPI_ISL_1785164 | SIESP CHIETI - DRIVE IN ORTONA | Istituto Zooprofilattico Sperimentale dell'Abruzzo e Molise "G. Caporale" | Lorusso A, Marcacci M, Di Domenico M, Ancora M, Curini V, Di Lollo Valeria, Mangone I, Rinaldi A, Delli Compagni E, Scialabba S, Caporale M, Di Pasquale A, Cammà C, Puglia I, Calistri P, Savini G |
| EPI_ISL_1785165, EPI_ISL_1785166, EPI_ISL_1785167 | SIESP DIPARTIMENTO DI PREVENZIONE TERAMO | Istituto Zooprofilattico Sperimentale dell'Abruzzo e Molise "G. Caporale" | Lorusso A, Marcacci M, Di Domenico M, Ancora M, Curini V, Di Lollo Valeria, Mangone I, Rinaldi A, Delli Compagni E, Scialabba S, Caporale M, Di Pasquale A, Cammà C, Puglia I, Calistri P, Savini G |
| EPI_ISL_1785168 | DIP. PREV. AVEZZANO SERVIZIO DI IGIENE EPIDEMIOLOGIA E SANITA' PUBBLICA | Istituto Zooprofilattico Sperimentale dell'Abruzzo e Molise "G. Caporale" | Lorusso A, Marcacci M, Di Domenico M, Ancora M, Curini V, Di Lollo Valeria, Mangone I, Rinaldi A, Delli Compagni E, Scialabba S, Caporale M, Di Pasquale A, Cammà C, Puglia I, Calistri P, Savini G |
| EPI_ISL_1785169 | SIESP SULMONA | Istituto Zooprofilattico Sperimentale dell'Abruzzo e Molise "G. Caporale" | Lorusso A, Marcacci M, Di Domenico M, Ancora M, Curini V, Di Lollo Valeria, Mangone I, Rinaldi A, Delli Compagni E, Scialabba S, Caporale M, Di Pasquale A, Cammà C, Puglia I, Calistri P, Savini G |
| EPI_ISL_1785170 | SIESP DIPARTIMENTO DI PREVENZIONE CHIETI | Istituto Zooprofilattico Sperimentale dell'Abruzzo e Molise "G. Caporale" | Lorusso A, Marcacci M, Di Domenico M, Ancora M, Curini V, Di Lollo Valeria, Mangone I, Rinaldi A, Delli Compagni E, Scialabba S, Caporale M, Di Pasquale A, Cammà C, Puglia I, Calistri P, Savini G |
| EPI_ISL_1785171, EPI_ISL_1785172 | DIP. PREV. AVEZZANO SERVIZIO DI IGIENE EPIDEMIOLOGIA E SANITA' PUBBLICA | Istituto Zooprofilattico Sperimentale dell'Abruzzo e Molise "G. Caporale" | Lorusso A, Marcacci M, Di Domenico M, Ancora M, Curini V, Di Lollo Valeria, Mangone I, Rinaldi A, Delli Compagni E, Scialabba S, Caporale M, Di Pasquale A, Cammà C, Puglia I, Calistri P, Savini G |
| EPI_ISL_1785173 | SIESP SULMONA | Istituto Zooprofilattico Sperimentale dell'Abruzzo e Molise "G. Caporale" | Lorusso A, Marcacci M, Di Domenico M, Ancora M, Curini V, Di Lollo Valeria, Mangone I, Rinaldi A, Delli Compagni E, Scialabba S, Caporale M, Di Pasquale A, Cammà C, Puglia I, Calistri P, Savini G |
| EPI_ISL_1785174 | USCA Castel di Sangro | Istituto Zooprofilattico Sperimentale dell'Abruzzo e Molise "G. Caporale" | Lorusso A, Marcacci M, Di Domenico M, Ancora M, Curini V, Di Lollo Valeria, Mangone I, Rinaldi A, Delli Compagni E, Scialabba S, Caporale M, Di Pasquale A, Cammà C, Puglia I, Calistri P, Savini G |
| EPI_ISL_1785175 | SIESP CHIETI - DRIVE IN LANCIANO | Istituto Zooprofilattico Sperimentale dell'Abruzzo e Molise "G. Caporale" | Lorusso A, Marcacci M, Di Domenico M, Ancora M, Curini V, Di Lollo Valeria, Mangone I, Rinaldi A, Delli Compagni E, Scialabba S, Caporale M, Di Pasquale A, Cammà C, Puglia I, Calistri P, Savini G |
| EPI_ISL_1785176 | PRESIDIO TERRITORIALE CASOLI | Istituto Zooprofilattico Sperimentale dell'Abruzzo e Molise "G. Caporale" | Lorusso A, Marcacci M, Di Domenico M, Ancora M, Curini V, Di Lollo Valeria, Mangone I, Rinaldi A, Delli Compagni E, Scialabba S, Caporale M, Di Pasquale A, Cammà C, Puglia I, Calistri P, Savini G |
| EPI_ISL_1785177, EPI_ISL_1785178 | SIESP CHIETI - DRIVE IN GISSI | Istituto Zooprofilattico Sperimentale dell'Abruzzo e Molise "G. Caporale" | Lorusso A, Marcacci M, Di Domenico M, Ancora M, Curini V, Di Lollo Valeria, Mangone I, Rinaldi A, Delli Compagni E, Scialabba S, Caporale M, Di Pasquale A, Cammà C, Puglia I, Calistri P, Savini G |
| EPI_ISL_1785179, EPI_ISL_1785180, EPI_ISL_1785181, EPI_ISL_1785182, EPI_ISL_1785183, EPI_ISL_1785184, EPI_ISL_1785185, EPI_ISL_1785186, EPI_ISL_1785187, EPI_ISL_1785188, EPI_ISL_1785189, EPI_ISL_1785190, EPI_ISL_1785191 | | | |
| see above | SIESP DIPARTIMENTO DI PREVENZIONE TERAMO | Istituto Zooprofilattico Sperimentale dell'Abruzzo e Molise "G. Caporale" | Lorusso A, Marcacci M, Di Domenico M, Ancora M, Curini V, Di Lollo Valeria, Mangone I, Rinaldi A, Delli Compagni E, Scialabba S, Caporale M, Di Pasquale A, Cammà C, Puglia I, Calistri P, Savini G |
| EPI_ISL_1785192 | SIESP CHIETI - DRIVE IN CHIETI | Istituto Zooprofilattico Sperimentale dell'Abruzzo e Molise "G. Caporale" | Lorusso A, Marcacci M, Di Domenico M, Ancora M, Curini V, Di Lollo Valeria, Mangone I, Rinaldi A, Delli Compagni E, Scialabba S, Caporale M, Di Pasquale A, Cammà C, Puglia I, Calistri P, Savini G |
| EPI_ISL_1785193 | SIESP SULMONA | Istituto Zooprofilattico Sperimentale dell'Abruzzo e Molise "G. Caporale" | Lorusso A, Marcacci M, Di Domenico M, Ancora M, Curini V, Di Lollo Valeria, Mangone I, Rinaldi A, Delli Compagni E, Scialabba S, Caporale M, Di Pasquale A, Cammà C, Puglia I, Calistri P, Savini G |
| EPI_ISL_1785194, EPI_ISL_1785195 | SIESP CHIETI - DRIVE IN ORTONA | Istituto Zooprofilattico Sperimentale dell'Abruzzo e Molise "G. Caporale" | Lorusso A, Marcacci M, Di Domenico M, Ancora M, Curini V, Di Lollo Valeria, Mangone I, Rinaldi A, Delli Compagni E, Scialabba S, Caporale M, Di Pasquale A, Cammà C, Puglia I, Calistri P, Savini G |
| EPI_ISL_1785196, EPI_ISL_1785197 | USCA Tagliacozzo | Istituto Zooprofilattico Sperimentale dell'Abruzzo e Molise "G. Caporale" | Lorusso A, Marcacci M, Di Domenico M, Ancora M, Curini V, Di Lollo Valeria, Mangone I, Rinaldi A, Delli Compagni E, Scialabba S, Caporale M, Di Pasquale A, Cammà C, Puglia I, Calistri P, Savini G |
| EPI_ISL_1785198 | SIESP DIPARTIMENTO DI PREVENZIONE CHIETI | Istituto Zooprofilattico Sperimentale dell'Abruzzo e Molise "G. Caporale" | Lorusso A, Marcacci M, Di Domenico M, Ancora M, Curini V, Di Lollo Valeria, Mangone I, Rinaldi A, Delli Compagni E, Scialabba S, Caporale M, Di Pasquale A, Cammà C, Puglia I, Calistri P, Savini G |
| EPI_ISL_1785199 | SIESP CHIETI - DRIVE IN CHIETI | Istituto Zooprofilattico Sperimentale dell'Abruzzo e Molise "G. Caporale" | Lorusso A, Marcacci M, Di Domenico M, Ancora M, Curini V, Di Lollo Valeria, Mangone I, Rinaldi A, Delli Compagni E, Scialabba S, Caporale M, Di Pasquale A, Cammà C, Puglia I, Calistri P, Savini G |
| EPI_ISL_1785200, EPI_ISL_1785201 | SIESP CHIETI - DRIVE IN GISSI | Istituto Zooprofilattico Sperimentale dell'Abruzzo e Molise "G. Caporale" | Lorusso A, Marcacci M, Di Domenico M, Ancora M, Curini V, Di Lollo Valeria, Mangone I, Rinaldi A, Delli Compagni E, Scialabba S, Caporale M, Di Pasquale A, Cammà C, Puglia I, Calistri P, Savini G |
| EPI_ISL_1785202, EPI_ISL_1785203 | SIESP CHIETI - DRIVE IN LANCIANO | Istituto Zooprofilattico Sperimentale dell'Abruzzo e Molise "G. Caporale" | Lorusso A, Marcacci M, Di Domenico M, Ancora M, Curini V, Di Lollo Valeria, Mangone I, Rinaldi A, Delli Compagni E, Scialabba S, Caporale M, Di Pasquale A, Cammà C, Puglia I, Calistri P, Savini G |
| EPI_ISL_1785204, EPI_ISL_1785205, EPI_ISL_1785206, EPI_ISL_1785207, EPI_ISL_1785208, EPI_ISL_1785209, EPI_ISL_1785210, EPI_ISL_1785211, EPI_ISL_1785212, EPI_ISL_1785213 | Università degli Studi di Perugia | Istituto Zooprofilattico Sperimentale dell'Abruzzo e Molise "G. Caporale" | Mencacci A, Camilloni B,Lorusso A, Marcacci M, Di Domenico M, Ancora M, Curini V, Di Lollo Valeria, Mangone I, Rinaldi A, Delli Compagni E, Scialabba S, Caporale M, Di Pasquale A, Cammà C, Puglia I, Calistri P, Savini G |
| EPI_ISL_1785214 | Laboratorio Analisi Osp. Città di Castello - Azienda USL Umbria1 | Istituto Zooprofilattico Sperimentale dell'Abruzzo e Molise "G. Caporale" | Malagigi V, Tacconi P, Lorusso A, Marcacci M, Di Domenico M, Ancora M, Curini V, Di Lollo Valeria, Mangone I, Rinaldi A, Delli Compagni E, Scialabba S, Caporale M, Di Pasquale A, Cammà C, Puglia I, Calistri P, Savini G |
| EPI_ISL_1785215, EPI_ISL_1785216, EPI_ISL_1785217, EPI_ISL_1785218, EPI_ISL_1785219, EPI_ISL_1785220, EPI_ISL_1785221, EPI_ISL_1785222, EPI_ISL_1785223 | Università degli Studi di Perugia | Istituto Zooprofilattico Sperimentale dell'Abruzzo e Molise "G. Caporale" | Mencacci A, Camilloni B,Lorusso A, Marcacci M, Di Domenico M, Ancora M, Curini V, Di Lollo Valeria, Mangone I, Rinaldi A, Delli Compagni E, Scialabba S, Caporale M, Di Pasquale A, Cammà C, Puglia I, Calistri P, Savini G |
| EPI_ISL_1785224 | Laboratorio Analisi Osp. Città di Castello - Azienda USL Umbria1 | Istituto Zooprofilattico Sperimentale dell'Abruzzo e Molise "G. Caporale" | Malagigi V, Tacconi P, Lorusso A, Marcacci M, Di Domenico M, Ancora M, Curini V, Di Lollo Valeria, Mangone I, Rinaldi A, Delli Compagni E, Scialabba S, Caporale M, Di Pasquale A, Cammà C, Puglia I, Calistri P, Savini G |
| EPI_ISL_1785225, EPI_ISL_1785226, | Università degli Studi di Perugia | Istituto Zooprofilattico Sperimentale dell'Abruzzo e Molise "G. | Mencacci A, Camilloni B,Lorusso A, Marcacci M, Di Domenico M, Ancora M, Curini V, Di Lollo Valeria, Mangone I, Rinaldi A, Delli Compagni E, Scialabba |

| | | | |
|---|---|---|---|
| EPI_ISL_1785227 | | Caporale" | S, Caporale M, Di Pasquale A, Cammà C, Puglia I, Calistri P, Savini G |
| EPI_ISL_1785228, EPI_ISL_1785229, EPI_ISL_1785230 | Laboratorio Analisi Osp. Città di Castello - Azienda USL Umbria1 | Istituto Zooprofilattico Sperimentale dell'Abruzzo e Molise "G. Caporale" | Malagigi V, Tacconi P, Lorusso A, Marcacci M, Di Domenico M, Ancora M, Curini V, Di Lollo Valeria, Mangone I, Rinaldi A, Delli Compagni E, Scialabba S, Caporale M, Di Pasquale A, Cammà C, Puglia I, Calistri P, Savini G |
| EPI_ISL_1785231, EPI_ISL_1785232, EPI_ISL_1785233 | Azienda Ospedaliera Terni | Istituto Zooprofilattico Sperimentale dell'Abruzzo e Molise "G. Caporale" | Palumbo M, Scaccetti A, Lorusso A, Marcacci M, Di Domenico M, Ancora M, Curini V, Di Lollo Valeria, Mangone I, Rinaldi A, Delli Compagni E, Scialabba S, Caporale M, Di Pasquale A, Cammà C, Puglia I, Calistri P, Savini G |
| EPI_ISL_1785234 | Laboratorio Analisi Osp. Città di Castello - Azienda USL Umbria1 | Istituto Zooprofilattico Sperimentale dell'Abruzzo e Molise "G. Caporale" | Malagigi V, Tacconi P, Lorusso A, Marcacci M, Di Domenico M, Ancora M, Curini V, Di Lollo Valeria, Mangone I, Rinaldi A, Delli Compagni E, Scialabba S, Caporale M, Di Pasquale A, Cammà C, Puglia I, Calistri P, Savini G |
| EPI_ISL_1785235, EPI_ISL_1785236, EPI_ISL_1785237, EPI_ISL_1785238, EPI_ISL_1785239, EPI_ISL_1785240, EPI_ISL_1785241, EPI_ISL_1785242, EPI_ISL_1785243, EPI_ISL_1785244 | Azienda Ospedaliera Terni | Istituto Zooprofilattico Sperimentale dell'Abruzzo e Molise "G. Caporale" | Palumbo M, Scaccetti A, Lorusso A, Marcacci M, Di Domenico M, Ancora M, Curini V, Di Lollo Valeria, Mangone I, Rinaldi A, Delli Compagni E, Scialabba S, Caporale M, Di Pasquale A, Cammà C, Puglia I, Calistri P, Savini G |
| EPI_ISL_1785245 | Laboratorio Analisi Osp. Città di Castello - Azienda USL Umbria1 | Istituto Zooprofilattico Sperimentale dell'Abruzzo e Molise "G. Caporale" | Malagigi V, Tacconi P, Lorusso A, Marcacci M, Di Domenico M, Ancora M, Curini V, Di Lollo Valeria, Mangone I, Rinaldi A, Delli Compagni E, Scialabba S, Caporale M, Di Pasquale A, Cammà C, Puglia I, Calistri P, Savini G |
| EPI_ISL_1785246 | Azienda Ospedaliera Terni | Istituto Zooprofilattico Sperimentale dell'Abruzzo e Molise "G. Caporale" | Palumbo M, Scaccetti A, Lorusso A, Marcacci M, Di Domenico M, Ancora M, Curini V, Di Lollo Valeria, Mangone I, Rinaldi A, Delli Compagni E, Scialabba S, Caporale M, Di Pasquale A, Cammà C, Puglia I, Calistri P, Savini G |
| EPI_ISL_1785247, EPI_ISL_1785248, EPI_ISL_1785249, EPI_ISL_1785250 | Università degli Studi di Perugia | Istituto Zooprofilattico Sperimentale dell'Abruzzo e Molise "G. Caporale" | Mencacci A, Camilloni B,Lorusso A, Marcacci M, Di Domenico M, Ancora M, Curini V, Di Lollo Valeria, Mangone I, Rinaldi A, Delli Compagni E, Scialabba S, Caporale M, Di Pasquale A, Cammà C, Puglia I, Calistri P, Savini G |
| EPI_ISL_1785251 | Laboratorio Analisi Osp. Città di Castello - Azienda USL Umbria1 | Istituto Zooprofilattico Sperimentale dell'Abruzzo e Molise "G. Caporale" | Malagigi V, Tacconi P, Lorusso A, Marcacci M, Di Domenico M, Ancora M, Curini V, Di Lollo Valeria, Mangone I, Rinaldi A, Delli Compagni E, Scialabba S, Caporale M, Di Pasquale A, Cammà C, Puglia I, Calistri P, Savini G |
| EPI_ISL_1787254, EPI_ISL_1787317, EPI_ISL_1787320, EPI_ISL_1787321, EPI_ISL_1787323, EPI_ISL_1787324, EPI_ISL_1787689, EPI_ISL_1787983 | Institute for Medical Research, Infectious Disease Research Centre, National Institutes of Health, Ministry of Health Malaysia | Institute for Medical Research, Infectious Disease Research Centre, National Institutes of Health, Ministry of Health Malaysia | Suppiah J, Kamel K, Mohd Zawawi Z, Azizan MA, Ramly N, Robert F, Thayan R |
| EPI_ISL_1788746 | DIP. PREV. AVEZZANO SERVIZIO DI IGIENE EPIDEMIOLOGIA E SANITA' PUBBLICA | Istituto Zooprofilattico Sperimentale dell'Abruzzo e Molise "G. Caporale" | Lorusso A, Marcacci M, Di Domenico M, Ancora M, Curini V, Di Lollo Valeria, Mangone I, Rinaldi A, Delli Compagni E, Scialabba S, Caporale M, Di Pasquale A, Cammà C, Puglia I, Calistri P, Savini G |
| EPI_ISL_1788747 | SIESP DIPARTIMENTO DI PREVENZIONE TERAMO | Istituto Zooprofilattico Sperimentale dell'Abruzzo e Molise "G. Caporale" | Lorusso A, Marcacci M, Di Domenico M, Ancora M, Curini V, Di Lollo Valeria, Mangone I, Rinaldi A, Delli Compagni E, Scialabba S, Caporale M, Di Pasquale A, Cammà C, Puglia I, Calistri P, Savini G |
| EPI_ISL_1788748, EPI_ISL_1788749, EPI_ISL_1788750, EPI_ISL_1788751 | DIP. PREV. AVEZZANO SERVIZIO DI IGIENE EPIDEMIOLOGIA E SANITA' PUBBLICA | Istituto Zooprofilattico Sperimentale dell'Abruzzo e Molise "G. Caporale" | Lorusso A, Marcacci M, Di Domenico M, Ancora M, Curini V, Di Lollo Valeria, Mangone I, Rinaldi A, Delli Compagni E, Scialabba S, Caporale M, Di Pasquale A, Cammà C, Puglia I, Calistri P, Savini G |
| EPI_ISL_1788752 | USCA Castel di Sangro | Istituto Zooprofilattico Sperimentale dell'Abruzzo e Molise "G. Caporale" | Lorusso A, Marcacci M, Di Domenico M, Ancora M, Curini V, Di Lollo Valeria, Mangone I, Rinaldi A, Delli Compagni E, Scialabba S, Caporale M, Di Pasquale A, Cammà C, Puglia I, Calistri P, Savini G |
| EPI_ISL_1788753, EPI_ISL_1788754 | SIESP CHIETI - DRIVE IN ORTONA | Istituto Zooprofilattico Sperimentale dell'Abruzzo e Molise "G. Caporale" | Lorusso A, Marcacci M, Di Domenico M, Ancora M, Curini V, Di Lollo Valeria, Mangone I, Rinaldi A, Delli Compagni E, Scialabba S, Caporale M, Di Pasquale A, Cammà C, Puglia I, Calistri P, Savini G |
| EPI_ISL_1788755 | OSPEDALE SAN SALVATORE - MEDICINA DI LABORATORIO | Istituto Zooprofilattico Sperimentale dell'Abruzzo e Molise "G. Caporale" | Lorusso A, Marcacci M, Di Domenico M, Ancora M, Curini V, Di Lollo Valeria, Mangone I, Rinaldi A, Delli Compagni E, Scialabba S, Caporale M, Di Pasquale A, Cammà C, Puglia I, Calistri P, Savini G |
| EPI_ISL_1788756 | SIESP CHIETI - DRIVE IN CHIETI | Istituto Zooprofilattico Sperimentale dell'Abruzzo e Molise "G. Caporale" | Lorusso A, Marcacci M, Di Domenico M, Ancora M, Curini V, Di Lollo Valeria, Mangone I, Rinaldi A, Delli Compagni E, Scialabba S, Caporale M, Di Pasquale A, Cammà C, Puglia I, Calistri P, Savini G |
| EPI_ISL_1788769, EPI_ISL_1788770, EPI_ISL_1788772, EPI_ISL_1788773, EPI_ISL_1788774, EPI_ISL_1788775, EPI_ISL_1788776, EPI_ISL_1788777, EPI_ISL_1788778 | Labo Analyses Med | National Reference Center for Viruses of Respiratory Infections, Institut Pasteur, Paris | Marion Barbet, Sylvie Behillil, Méline Bizard, Angela Brisebarre, Camille Capel, Vincent Enouf, Louise Lefrançois, Frédéric Lemoine, Christophe Malabat, Corinne Maufrais, Etienne Simon-Lorière, Maud Vanpeene, Sylvie Van der Werf ,Bruno Berdin |
| EPI_ISL_1788791, EPI_ISL_1788792, EPI_ISL_1788794, EPI_ISL_1788796, EPI_ISL_1788797, EPI_ISL_1788798 | Hospital | National Reference Center for Viruses of Respiratory Infections, Institut Pasteur, Paris | Marion Barbet, Sylvie Behillil, Méline Bizard, Angela Brisebarre, Camille Capel, Vincent Enouf, Louise Lefrançois, Frédéric Lemoine, Christophe Malabat, Corinne Maufrais, Etienne Simon-Lorière, Maud Vanpeene, Sylvie Van der Werf ,Alexandra Ducancelle |
| EPI_ISL_1788799 | Hospital | National Reference Center for Viruses of Respiratory Infections, Institut Pasteur, Paris | Marion Barbet, Sylvie Behillil, Méline Bizard, Angela Brisebarre, Camille Capel, Vincent Enouf, Louise Lefrançois, Frédéric Lemoine, Christophe Malabat, Corinne Maufrais, Amaury Vaysse, Etienne Simon-Lorière, Maud Vanpeene, Sylvie Van der Werf ,Alexandra Ducancelle |
| EPI_ISL_1788800, EPI_ISL_1788801, EPI_ISL_1788802, EPI_ISL_1788803 | Hospital | National Reference Center for Viruses of Respiratory Infections, Institut Pasteur, Paris | Marion Barbet, Sylvie Behillil, Méline Bizard, Angela Brisebarre, Camille Capel, Vincent Enouf, Louise Lefrançois, Frédéric Lemoine, Christophe Malabat, Corinne Maufrais, Etienne Simon-Lorière, Maud Vanpeene, Sylvie Van der Werf ,Alexandra Ducancelle |
| EPI_ISL_1788804, EPI_ISL_1788805, EPI_ISL_1788806, EPI_ISL_1788807, EPI_ISL_1788808, EPI_ISL_1788809, EPI_ISL_1788810 | Labo Analyses Med | National Reference Center for Viruses of Respiratory Infections, Institut Pasteur, Paris | Marion Barbet, Sylvie Behillil, Méline Bizard, Angela Brisebarre, Camille Capel, Vincent Enouf, Louise Lefrançois, Frédéric Lemoine, Christophe Malabat, Corinne Maufrais, Etienne Simon-Lorière, Maud Vanpeene, Sylvie Van der Werf ,Serge Ney |
| EPI_ISL_1788811 | Hospital | National Reference Center for Viruses of Respiratory Infections, Institut Pasteur, Paris | Marion Barbet, Sylvie Behillil, Méline Bizard, Angela Brisebarre, Camille Capel, Vincent Enouf, Louise Lefrançois, Frédéric Lemoine, Christophe Malabat, Corinne Maufrais, Amaury Vaysse, Etienne Simon-Lorière, Maud Vanpeene, Sylvie Van der Werf ,Franck Bernardi |
| EPI_ISL_1788817 | Labo Analyses Med | National Reference Center for Viruses of Respiratory Infections, Institut Pasteur, Paris | Marion Barbet, Sylvie Behillil, Méline Bizard, Angela Brisebarre, Camille Capel, Vincent Enouf, Louise Lefrançois, Frédéric Lemoine, Christophe Malabat, Corinne Maufrais, Amaury Vaysse, Etienne Simon-Lorière, Maud Vanpeene, Sylvie Van der Werf ,SéVerine Haouisee |
| EPI_ISL_1788818, EPI_ISL_1788819, EPI_ISL_1788820 | Labo Analyses Med | National Reference Center for Viruses of Respiratory Infections, Institut Pasteur, Paris | Marion Barbet, Sylvie Behillil, Méline Bizard, Angela Brisebarre, Camille Capel, Vincent Enouf, Louise Lefrançois, Frédéric Lemoine, Christophe Malabat, Corinne Maufrais, Etienne Simon-Lorière, Maud Vanpeene, Sylvie Van der Werf ,SéVerine Haouisee |
| EPI_ISL_1788839 | Labo Analyses med | National Reference Center for Viruses of Respiratory Infections, Institut Pasteur, Paris | Marion Barbet, Sylvie Behillil, Méline Bizard, Angela Brisebarre, Camille Capel, Vincent Enouf, Louise Lefrançois, Frédéric Lemoine, Christophe Malabat, Corinne Maufrais, Amaury Vaysse, Etienne Simon-Lorière, Maud Vanpeene, Sylvie Van der Werf ,Anne Holstein |
| EPI_ISL_1788840, EPI_ISL_1788841, EPI_ISL_1788844, EPI_ISL_1788845, EPI_ISL_1788847, EPI_ISL_1788849, EPI_ISL_1788850, EPI_ISL_1788851, EPI_ISL_1788852, EPI_ISL_1788853, EPI_ISL_1788854 | | | |
| see above | Labo Analyses med | National Reference Center for Viruses of Respiratory Infections, Institut Pasteur, Paris | Marion Barbet, Sylvie Behillil, Méline Bizard, Angela Brisebarre, Camille Capel, Vincent Enouf, Louise Lefrançois, Frédéric Lemoine, Christophe Malabat, Corinne Maufrais, Etienne Simon-Lorière, Maud Vanpeene, Sylvie Van der Werf ,Anne Holstein |
| EPI_ISL_1788855 | Labo Analyses med | National Reference Center for Viruses of Respiratory Infections, Institut Pasteur, Paris | Marion Barbet, Sylvie Behillil, Méline Bizard, Angela Brisebarre, Camille Capel, Vincent Enouf, Louise Lefrançois, Frédéric Lemoine, Christophe Malabat, Corinne Maufrais, Amaury Vaysse, Etienne Simon-Lorière, Maud Vanpeene, Sylvie Van der Werf ,Anne Holstein |
| EPI_ISL_1788856, EPI_ISL_1788857, EPI_ISL_1788858, EPI_ISL_1788859 | Labo Analyses med | National Reference Center for Viruses of Respiratory Infections, Institut Pasteur, Paris | Marion Barbet, Sylvie Behillil, Méline Bizard, Angela Brisebarre, Camille Capel, Vincent Enouf, Louise Lefrançois, Frédéric Lemoine, Christophe Malabat, Corinne Maufrais, Etienne Simon-Lorière, Maud Vanpeene, Sylvie Van der Werf ,Anne Holstein |
| EPI_ISL_1788860, EPI_ISL_1788861 | Labo Analyses med | National Reference Center for Viruses of Respiratory Infections, Institut Pasteur, Paris | Marion Barbet, Sylvie Behillil, Méline Bizard, Angela Brisebarre, Camille Capel, Vincent Enouf, Louise Lefrançois, Frédéric Lemoine, Christophe Malabat, Corinne Maufrais, Amaury Vaysse, Etienne Simon-Lorière, Maud Vanpeene, Sylvie Van der Werf ,Anne Holstein |
| EPI_ISL_1788862, EPI_ISL_1788863, | Labo Analyses med | National Reference Center for Viruses of Respiratory | Marion Barbet, Sylvie Behillil, Méline Bizard, Angela Brisebarre, Camille Capel, Vincent Enouf, Louise Lefrançois, Frédéric Lemoine, Christophe Malabat, |

| | | | |
|---|---|---|---|
| EPI_ISL_1788864, EPI_ISL_1788865 | | Infections, Institut Pasteur, Paris | Corinne Maufrais, Etienne Simon-Lorière, Maud Vanpeene, Sylvie Van der Werf ,Anne Holstein |
| EPI_ISL_1788866 | Labo Analyses med | National Reference Center for Viruses of Respiratory Infections, Institut Pasteur, Paris | Marion Barbet, Sylvie Behillil, Méline Bizard, Angela Brisebarre, Camille Capel, Vincent Enouf, Louise Lefrançois, Frédéric Lemoine, Christophe Malabat, Corinne Maufrais, Amaury Vaysse, Etienne Simon-Lorière, Maud Vanpeene, Sylvie Van der Werf ,Anne Holstein |
| EPI_ISL_1788867 | Labo Analyses med | National Reference Center for Viruses of Respiratory Infections, Institut Pasteur, Paris | Marion Barbet, Sylvie Behillil, Méline Bizard, Angela Brisebarre, Camille Capel, Vincent Enouf, Louise Lefrançois, Frédéric Lemoine, Christophe Malabat, Corinne Maufrais, Etienne Simon-Lorière, Maud Vanpeene, Sylvie Van der Werf ,Anne Holstein |
| EPI_ISL_1788868 | Labo Analyses med | National Reference Center for Viruses of Respiratory Infections, Institut Pasteur, Paris | Marion Barbet, Sylvie Behillil, Méline Bizard, Angela Brisebarre, Camille Capel, Vincent Enouf, Louise Lefrançois, Frédéric Lemoine, Christophe Malabat, Corinne Maufrais, Amaury Vaysse, Etienne Simon-Lorière, Maud Vanpeene, Sylvie Van der Werf ,Anne Holstein |
| EPI_ISL_1788870, EPI_ISL_1788871, EPI_ISL_1788872, EPI_ISL_1788873, EPI_ISL_1788874, EPI_ISL_1788875, EPI_ISL_1788876, EPI_ISL_1788877 | Labo Analyses med | National Reference Center for Viruses of Respiratory Infections, Institut Pasteur, Paris | Marion Barbet, Sylvie Behillil, Méline Bizard, Angela Brisebarre, Camille Capel, Vincent Enouf, Louise Lefrançois, Frédéric Lemoine, Christophe Malabat, Corinne Maufrais, Etienne Simon-Lorière, Maud Vanpeene, Sylvie Van der Werf ,Anne Holstein |
| EPI_ISL_1788878 | Labo Analyses med | National Reference Center for Viruses of Respiratory Infections, Institut Pasteur, Paris | Marion Barbet, Sylvie Behillil, Méline Bizard, Angela Brisebarre, Camille Capel, Vincent Enouf, Louise Lefrançois, Frédéric Lemoine, Christophe Malabat, Corinne Maufrais, Amaury Vaysse, Etienne Simon-Lorière, Maud Vanpeene, Sylvie Van der Werf ,Anne Holstein |
| EPI_ISL_1788879, EPI_ISL_1788880, EPI_ISL_1788881 | Labo Analyses med | National Reference Center for Viruses of Respiratory Infections, Institut Pasteur, Paris | Marion Barbet, Sylvie Behillil, Méline Bizard, Angela Brisebarre, Camille Capel, Vincent Enouf, Louise Lefrançois, Frédéric Lemoine, Christophe Malabat, Corinne Maufrais, Etienne Simon-Lorière, Maud Vanpeene, Sylvie Van der Werf ,Anne Holstein |
| EPI_ISL_1788885, EPI_ISL_1788886, EPI_ISL_1788889, EPI_ISL_1788890, EPI_ISL_1788891 | Hospital | National Reference Center for Viruses of Respiratory Infections, Institut Pasteur, Paris | Marion Barbet, Sylvie Behillil, Méline Bizard, Angela Brisebarre, Camille Capel, Vincent Enouf, Louise Lefrançois, Frédéric Lemoine, Christophe Malabat, Corinne Maufrais, Etienne Simon-Lorière, Maud Vanpeene, Sylvie Van der Werf ,LéA Pilorge |
| EPI_ISL_1788892, EPI_ISL_1788893, EPI_ISL_1788894, EPI_ISL_1788895, EPI_ISL_1788896, EPI_ISL_1788897, EPI_ISL_1788898, EPI_ISL_1788900, EPI_ISL_1788901 | Hospital | National Reference Center for Viruses of Respiratory Infections, Institut Pasteur, Paris | Marion Barbet, Sylvie Behillil, Méline Bizard, Angela Brisebarre, Camille Capel, Vincent Enouf, Louise Lefrançois, Frédéric Lemoine, Christophe Malabat, Corinne Maufrais, Etienne Simon-Lorière, Maud Vanpeene, Sylvie Van der Werf ,Vincent Thibault |
| EPI_ISL_1788902, EPI_ISL_1788903, EPI_ISL_1788904, EPI_ISL_1788905, EPI_ISL_1788906, EPI_ISL_1788907, EPI_ISL_1788908 | Hospital | National Reference Center for Viruses of Respiratory Infections, Institut Pasteur, Paris | Marion Barbet, Sylvie Behillil, Méline Bizard, Angela Brisebarre, Camille Capel, Vincent Enouf, Louise Lefrançois, Frédéric Lemoine, Christophe Malabat, Corinne Maufrais, Etienne Simon-Lorière, Maud Vanpeene, Sylvie Van der Werf ,Anne Cady |
| EPI_ISL_1788909, EPI_ISL_1788910 | Hospital | National Reference Center for Viruses of Respiratory Infections, Institut Pasteur, Paris | Marion Barbet, Sylvie Behillil, Méline Bizard, Angela Brisebarre, Camille Capel, Vincent Enouf, Louise Lefrançois, Frédéric Lemoine, Christophe Malabat, Corinne Maufrais, Etienne Simon-Lorière, Maud Vanpeene, Sylvie Van der Werf ,BéAtrice Lesimple |
| EPI_ISL_1788911, EPI_ISL_1788912 | Hospital | National Reference Center for Viruses of Respiratory Infections, Institut Pasteur, Paris | Marion Barbet, Sylvie Behillil, Méline Bizard, Angela Brisebarre, Camille Capel, Vincent Enouf, Louise Lefrançois, Frédéric Lemoine, Christophe Malabat, Corinne Maufrais, Etienne Simon-Lorière, Maud Vanpeene, Sylvie Van der Werf ,Alix Irimia |
| EPI_ISL_1788913 | Labo Analyses Med | National Reference Center for Viruses of Respiratory Infections, Institut Pasteur, Paris | Marion Barbet, Sylvie Behillil, Méline Bizard, Angela Brisebarre, Camille Capel, Vincent Enouf, Louise Lefrançois, Frédéric Lemoine, Christophe Malabat, Corinne Maufrais, Etienne Simon-Lorière, Maud Vanpeene, Sylvie Van der Werf ,Claude Jorion |
| EPI_ISL_1788917, EPI_ISL_1788918, EPI_ISL_1788919, EPI_ISL_1788920, EPI_ISL_1788921, EPI_ISL_1788922, EPI_ISL_1788923, EPI_ISL_1788924, EPI_ISL_1788925 | Hospital | National Reference Center for Viruses of Respiratory Infections, Institut Pasteur, Paris | Marion Barbet, Sylvie Behillil, Méline Bizard, Angela Brisebarre, Camille Capel, Vincent Enouf, Louise Lefrançois, Frédéric Lemoine, Christophe Malabat, Corinne Maufrais, Etienne Simon-Lorière, Maud Vanpeene, Sylvie Van der Werf ,Jean Baptiste Bour |
| EPI_ISL_1788926, EPI_ISL_1788927, EPI_ISL_1788928, EPI_ISL_1788929, EPI_ISL_1788930, EPI_ISL_1788931, EPI_ISL_1788932, EPI_ISL_1788933, EPI_ISL_1788934, EPI_ISL_1788935, EPI_ISL_1788936, EPI_ISL_1788937, EPI_ISL_1788938, EPI_ISL_1788939, EPI_ISL_1788940, EPI_ISL_1788941, EPI_ISL_1788942, EPI_ISL_1788943, EPI_ISL_1788944, EPI_ISL_1788945, EPI_ISL_1788946, EPI_ISL_1788947, EPI_ISL_1788948, EPI_ISL_1788949 | | | |
| see above | Labo Analyses Med | National Reference Center for Viruses of Respiratory Infections, Institut Pasteur, Paris | Marion Barbet, Sylvie Behillil, Méline Bizard, Angela Brisebarre, Camille Capel, Vincent Enouf, Louise Lefrançois, Frédéric Lemoine, Christophe Malabat, Corinne Maufrais, Etienne Simon-Lorière, Maud Vanpeene, Sylvie Van der Werf ,GréGoire Potiron |
| EPI_ISL_1788950, EPI_ISL_1788951, EPI_ISL_1788952, EPI_ISL_1788953, EPI_ISL_1788954, EPI_ISL_1788955, EPI_ISL_1788956, EPI_ISL_1788957, EPI_ISL_1788958 | Hospital | National Reference Center for Viruses of Respiratory Infections, Institut Pasteur, Paris | Marion Barbet, Sylvie Behillil, Méline Bizard, Angela Brisebarre, Camille Capel, Vincent Enouf, Louise Lefrançois, Frédéric Lemoine, Christophe Malabat, Corinne Maufrais, Etienne Simon-Lorière, Maud Vanpeene, Sylvie Van der Werf ,Eleha Guillotel |
| EPI_ISL_1788959, EPI_ISL_1788961, EPI_ISL_1788962, EPI_ISL_1788963, EPI_ISL_1788964, EPI_ISL_1788965, EPI_ISL_1788966, EPI_ISL_1788967, EPI_ISL_1788968, EPI_ISL_1788969 | Hospital | National Reference Center for Viruses of Respiratory Infections, Institut Pasteur, Paris | Marion Barbet, Sylvie Behillil, Méline Bizard, Angela Brisebarre, Camille Capel, Vincent Enouf, Louise Lefrançois, Frédéric Lemoine, Christophe Malabat, Corinne Maufrais, Etienne Simon-Lorière, Maud Vanpeene, Sylvie Van der Werf ,Esther Gyde |
| EPI_ISL_1788970 | Hospital | National Reference Center for Viruses of Respiratory Infections, Institut Pasteur, Paris | Marion Barbet, Sylvie Behillil, Méline Bizard, Angela Brisebarre, Camille Capel, Vincent Enouf, Louise Lefrançois, Frédéric Lemoine, Christophe Malabat, Corinne Maufrais, Etienne Simon-Lorière, Maud Vanpeene, Sylvie Van der Werf ,Pascale Martres |
| EPI_ISL_1788972, EPI_ISL_1788973 | Hospital | National Reference Center for Viruses of Respiratory Infections, Institut Pasteur, Paris | Marion Barbet, Sylvie Behillil, Méline Bizard, Angela Brisebarre, Camille Capel, Vincent Enouf, Louise Lefrançois, Frédéric Lemoine, Christophe Malabat, Corinne Maufrais, Etienne Simon-Lorière, Maud Vanpeene, Sylvie Van der Werf ,Thibault Guinoiseau |
| EPI_ISL_1788974, EPI_ISL_1788975, EPI_ISL_1788976, EPI_ISL_1788977, EPI_ISL_1788978, EPI_ISL_1788979, EPI_ISL_1788980, EPI_ISL_1788981, EPI_ISL_1788982, EPI_ISL_1788983, EPI_ISL_1788984, EPI_ISL_1788985, EPI_ISL_1788986, EPI_ISL_1788987, EPI_ISL_1788988, EPI_ISL_1788989, EPI_ISL_1788990, EPI_ISL_1788991, EPI_ISL_1788992, EPI_ISL_1788994, EPI_ISL_1788995, EPI_ISL_1788996, EPI_ISL_1788997, EPI_ISL_1788998, EPI_ISL_1788999, EPI_ISL_1789000, EPI_ISL_1789001, EPI_ISL_1789002, EPI_ISL_1789003, EPI_ISL_1789004, EPI_ISL_1789005, EPI_ISL_1789006, EPI_ISL_1789007, EPI_ISL_1789008 | | | |
| see above | Labo Analyses Med | National Reference Center for Viruses of Respiratory Infections, Institut Pasteur, Paris | Marion Barbet, Sylvie Behillil, Méline Bizard, Angela Brisebarre, Camille Capel, Vincent Enouf, Louise Lefrançois, Frédéric Lemoine, Christophe Malabat, Corinne Maufrais, Etienne Simon-Lorière, Maud Vanpeene, Sylvie Van der Werf ,Nabil Gastli |
| EPI_ISL_1789009, EPI_ISL_1789010, EPI_ISL_1789011, EPI_ISL_1789012, EPI_ISL_1789013, EPI_ISL_1789014, EPI_ISL_1789015, EPI_ISL_1789016, EPI_ISL_1789017, EPI_ISL_1789018, EPI_ISL_1789019 | | | |
| see above | Labo Analyses Med | National Reference Center for Viruses of Respiratory Infections, Institut Pasteur, Paris | Marion Barbet, Sylvie Behillil, Méline Bizard, Angela Brisebarre, Camille Capel, Vincent Enouf, Louise Lefrançois, Frédéric Lemoine, Christophe Malabat, Corinne Maufrais, Etienne Simon-Lorière, Maud Vanpeene, Sylvie Van der Werf ,Jean-FrançOis Comes |
| EPI_ISL_1789020 | Labo Analyses Med | National Reference Center for Viruses of Respiratory Infections, Institut Pasteur, Paris | Marion Barbet, Sylvie Behillil, Méline Bizard, Angela Brisebarre, Camille Capel, Vincent Enouf, Louise Lefrançois, Frédéric Lemoine, Christophe Malabat, Corinne Maufrais, Etienne Simon-Lorière, Maud Vanpeene, Sylvie Van der Werf ,Anne Cady |
| EPI_ISL_1789021, EPI_ISL_1789022, EPI_ISL_1789023, EPI_ISL_1789024, EPI_ISL_1789025, EPI_ISL_1789026, EPI_ISL_1789027, EPI_ISL_1789028, EPI_ISL_1789029, EPI_ISL_1789030, EPI_ISL_1789031, EPI_ISL_1789032, EPI_ISL_1789034, EPI_ISL_1789036 | | | |
| see above | Hospital | National Reference Center for Viruses of Respiratory Infections, Institut Pasteur, Paris | Marion Barbet, Sylvie Behillil, Méline Bizard, Angela Brisebarre, Camille Capel, Vincent Enouf, Louise Lefrançois, Frédéric Lemoine, Christophe Malabat, Corinne Maufrais, Etienne Simon-Lorière, Maud Vanpeene, Sylvie Van der Werf ,CéLine Bressollette |
| EPI_ISL_1789038 | Hospital | National Reference Center for Viruses of Respiratory Infections, Institut Pasteur, Paris | Marion Barbet, Sylvie Behillil, Méline Bizard, Angela Brisebarre, Camille Capel, Vincent Enouf, Louise Lefrançois, Frédéric Lemoine, Christophe Malabat, Corinne Maufrais, Etienne Simon-Lorière, Maud Vanpeene, Sylvie Van der Werf ,Ben Hadj Yahia |
| EPI_ISL_1789039 | Hospital | National Reference Center for Viruses of Respiratory Infections, Institut Pasteur, Paris | Marion Barbet, Sylvie Behillil, Méline Bizard, Angela Brisebarre, Camille Capel, Vincent Enouf, Louise Lefrançois, Frédéric Lemoine, Christophe Malabat, Corinne Maufrais, Etienne Simon-Lorière, Maud Vanpeene, Sylvie Van der Werf ,Patricia Stoessel-Thouvenin |
| EPI_ISL_1789040, EPI_ISL_1789041, | Hospital | National Reference Center for Viruses of Respiratory | Marion Barbet, Sylvie Behillil, Méline Bizard, Angela Brisebarre, Camille Capel, Vincent Enouf, Louise Lefrançois, Frédéric Lemoine, Christophe Malabat, |

| | | | |
|---|---|---|---|
| EPI_ISL_1789042, EPI_ISL_1789043, EPI_ISL_1789044, EPI_ISL_1789045, EPI_ISL_1789046, EPI_ISL_1789047 | | Infections, Institut Pasteur, Paris | Corinne Maufrais, Etienne Simon-Lorière, Maud Vanpeene, Sylvie Van der Werf ,Dominique Descamps |
| EPI_ISL_1789050, EPI_ISL_1789051, EPI_ISL_1789052 | Hospital | National Reference Center for Viruses of Respiratory Infections, Institut Pasteur, Paris | Marion Barbet, Sylvie Behillil, Méline Bizard, Angela Brisebarre, Camille Capel, Vincent Enouf, Louise Lefrançois, Frédéric Lemoine, Christophe Malabat, Corinne Maufrais, Etienne Simon-Lorière, Maud Vanpeene, Sylvie Van der Werf ,André-Guy Combremont |
| EPI_ISL_1789053, EPI_ISL_1789054, EPI_ISL_1789055, EPI_ISL_1789056 | Hospital | National Reference Center for Viruses of Respiratory Infections, Institut Pasteur, Paris | Marion Barbet, Sylvie Behillil, Méline Bizard, Angela Brisebarre, Camille Capel, Vincent Enouf, Louise Lefrançois, Frédéric Lemoine, Christophe Malabat, Corinne Maufrais, Etienne Simon-Lorière, Maud Vanpeene, Sylvie Van der Werf ,CéLine Ramanantsoa |
| EPI_ISL_1789057, EPI_ISL_1789058, EPI_ISL_1789059, EPI_ISL_1789060, EPI_ISL_1789061, EPI_ISL_1789062, EPI_ISL_1789063 | Labo Analyses Med | National Reference Center for Viruses of Respiratory Infections, Institut Pasteur, Paris | Marion Barbet, Sylvie Behillil, Méline Bizard, Angela Brisebarre, Camille Capel, Vincent Enouf, Louise Lefrançois, Frédéric Lemoine, Christophe Malabat, Corinne Maufrais, Etienne Simon-Lorière, Maud Vanpeene, Sylvie Van der Werf ,GréGoire Potiron |
| EPI_ISL_1789064, EPI_ISL_1789065, EPI_ISL_1789066, EPI_ISL_1789067, EPI_ISL_1789068, EPI_ISL_1789069, EPI_ISL_1789071, EPI_ISL_1789072, EPI_ISL_1789073, EPI_ISL_1789074, EPI_ISL_1789075, EPI_ISL_1789076, EPI_ISL_1789077, EPI_ISL_1789078, EPI_ISL_1789079, EPI_ISL_1789080 | | | |
| see above | Labo Analyses Med | National Reference Center for Viruses of Respiratory Infections, Institut Pasteur, Paris | Marion Barbet, Sylvie Behillil, Méline Bizard, Angela Brisebarre, Camille Capel, Vincent Enouf, Louise Lefrançois, Frédéric Lemoine, Christophe Malabat, Corinne Maufrais, Etienne Simon-Lorière, Maud Vanpeene, Sylvie Van der Werf ,Jean-FrançOis Comes |
| EPI_ISL_1789081, EPI_ISL_1789082, EPI_ISL_1789083 | Hospital | National Reference Center for Viruses of Respiratory Infections, Institut Pasteur, Paris | Marion Barbet, Sylvie Behillil, Méline Bizard, Angela Brisebarre, Camille Capel, Vincent Enouf, Louise Lefrançois, Frédéric Lemoine, Christophe Malabat, Corinne Maufrais, Etienne Simon-Lorière, Maud Vanpeene, Sylvie Van der Werf ,Pascale Martres |
| EPI_ISL_1789084, EPI_ISL_1789086, EPI_ISL_1789087, EPI_ISL_1789088 | Hospital | National Reference Center for Viruses of Respiratory Infections, Institut Pasteur, Paris | Marion Barbet, Sylvie Behillil, Méline Bizard, Angela Brisebarre, Camille Capel, Vincent Enouf, Louise Lefrançois, Frédéric Lemoine, Christophe Malabat, Corinne Maufrais, Etienne Simon-Lorière, Maud Vanpeene, Sylvie Van der Werf ,Nelly Luizy |
| EPI_ISL_1789101 | Hospital | National Reference Center for Viruses of Respiratory Infections, Institut Pasteur, Paris | Marion Barbet, Sylvie Behillil, Méline Bizard, Angela Brisebarre, Camille Capel, Vincent Enouf, Louise Lefrançois, Frédéric Lemoine, Christophe Malabat, Corinne Maufrais, Etienne Simon-Lorière, Maud Vanpeene, Sylvie Van der Werf ,JéRome Guinard |
| EPI_ISL_1789104 | Labo Analyses Med | National Reference Center for Viruses of Respiratory Infections, Institut Pasteur, Paris | Marion Barbet, Sylvie Behillil, Méline Bizard, Angela Brisebarre, Camille Capel, Vincent Enouf, Louise Lefrançois, Frédéric Lemoine, Christophe Malabat, Corinne Maufrais, Etienne Simon-Lorière, Maud Vanpeene, Sylvie Van der Werf ,Adrien Fabre |
| EPI_ISL_1789109, EPI_ISL_1789110 | Labo Analyses Med | National Reference Center for Viruses of Respiratory Infections, Institut Pasteur, Paris | Marion Barbet, Sylvie Behillil, Méline Bizard, Angela Brisebarre, Camille Capel, Vincent Enouf, Louise Lefrançois, Frédéric Lemoine, Christophe Malabat, Corinne Maufrais, Etienne Simon-Lorière, Maud Vanpeene, Sylvie Van der Werf ,GréGoire Potiron |
| EPI_ISL_1789111 | Labo Analyses Med | National Reference Center for Viruses of Respiratory Infections, Institut Pasteur, Paris | Marion Barbet, Sylvie Behillil, Méline Bizard, Angela Brisebarre, Camille Capel, Vincent Enouf, Louise Lefrançois, Frédéric Lemoine, Christophe Malabat, Corinne Maufrais, Etienne Simon-Lorière, Maud Vanpeene, Sylvie Van der Werf ,Selas Medilys |
| EPI_ISL_1789203 | Institute for Medical Research, Infectious Disease Research Centre, National Institutes of Health, Ministry of Health Malaysia | Institute for Medical Research, Infectious Disease Research Centre, National Institutes of Health, Ministry of Health Malaysia | Suppiah J, Kamel K, Mohd Zawawi Z, Ramly N, Robert F, Thayan R |
| EPI_ISL_1789488, EPI_ISL_1789489, EPI_ISL_1789490, EPI_ISL_1789491, EPI_ISL_1789492, EPI_ISL_1789494, EPI_ISL_1789495, EPI_ISL_1789496, EPI_ISL_1789497 | Bioptická laborato, s.r.o. | Bioptická laborato, s.r.o. | Petr Šteiner, Tomáš Vanek, Nikola Bílá, Martina Putzová, Michaela íhová, Silva Vondráková |
| EPI_ISL_1789527, EPI_ISL_1789529, EPI_ISL_1789531, EPI_ISL_1789532, EPI_ISL_1789533, EPI_ISL_1789536, EPI_ISL_1789537, EPI_ISL_1789538 | Department of Virology, Istituto Zooprofilattico Sperimentale del Lazio e della Toscana (IZSLT) | Department of General Diagnostics; Department of Virology; Istituto Zooprofilattico Sperimentale del Lazio e della Toscana (IZSLT) | Patricia Alba, Giuseppe Manna, Elena L. Diaconu, Fabiola Feltrin, Raffaela Conti, Teresa Scicluna, Virginia Carfora, Antonella Cersini, Alessia Franco, Antonio Battisti. |
| EPI_ISL_1789541, EPI_ISL_1789542 | WHO National Influenza Centre Russian Federation | WHO National Influenza Centre Russian Federation | Andrey Komissarov, Artem Fadeev, Kseniya Komissarova, Oula Mansour, Mikhail Bakaev, Tamila Musaeva, Maria Timofeeva, Veronika Eder, Maria Pisareva, Daria Danilenko, Ksenia Safina, Elena Nabieva, Georgii Bazykin, Dmitry Lioznov |
| EPI_ISL_1789673 | Pro-Vitam Diagnostics and Research Laboratory | Pro-Vitam Diagnostics and Research Laboratory | Szilard N. Fejer, Istvan Horvath, Monika Korodi, Kinga Rakosi, Zsuzsanna Jenei |
| EPI_ISL_1789696, EPI_ISL_1789697, EPI_ISL_1789698, EPI_ISL_1789699 | Laboratorio PGM | Laboratorio de Infectología Molecular, Departamento de Bioquímica y Medicina Molecular,Facultad de Medicina - Universidad Autónoma de Nuevo León | Kame A. Galán-Huerta, María F. Herrera-Saldivar, Natalia Martínez-Acuña, Sonia A. Lozano-Sepúlveda, Daniel Arellanos-Soto, Ana M. Rivas-Estilla, Javier Ramos-Jimenez, Gabriela Elizondo, Eduardo Garza-de-la-Peña |
| EPI_ISL_1790054, EPI_ISL_1790056 | Virology Unit, AOUP | Virology Unit, AOUP | Marialinda Vatteroni, Susi Frateschi, Mauro Pistello |
| EPI_ISL_1790060 | Virology Unit, AOUP | Virology Unit | Marialinda Vatteroni, Susi Frateschi, Mauro Pistello |
| EPI_ISL_1790212, EPI_ISL_1791023 | Virology Unit, AOUP | Virology Unit, AOUP | Marialinda Vatteroni, Susi Frateschi, Mauro Pistello |
| EPI_ISL_1791027 | COVID Laboratory AOU Messina | Molecular Hepatology Lab AOU Messina | Giuseppina Raffa, Cristina Musolino, Valeria Chines, Daniele Lombardo, Teresa Pollicino |
| EPI_ISL_1791029 | Virology Unit, AOUP | Virology Unit, AOUP | Marialinda Vatteroni, Susi Frateschi, Mauro Pistello |
| EPI_ISL_1791030 | COVID Laboratory AOU Messina | Molecular Hepatology Lab AOU Messina | Giuseppina Raffa, Cristina Musolino, Valeria Chines, Daniele Lombardo, Teresa Pollicino |
| EPI_ISL_1791033, EPI_ISL_1791035 | Virology Unit, AOUP | Virology Unit, AOUP | Marialinda Vatteroni, Susi Frateschi, Mauro Pistello |
| EPI_ISL_1791037, EPI_ISL_1791039, EPI_ISL_1791040, EPI_ISL_1791041, EPI_ISL_1791043, EPI_ISL_1791044, EPI_ISL_1791049, EPI_ISL_1791051, EPI_ISL_1791054, EPI_ISL_1791056, EPI_ISL_1791057, EPI_ISL_1791058, EPI_ISL_1791060, EPI_ISL_1791061, EPI_ISL_1791062, EPI_ISL_1791063 | | | |
| see above | COVID Laboratory AOU Messina | Molecular Hepatology Lab AOU Messina | Giuseppina Raffa, Cristina Musolino, Valeria Chines, Daniele Lombardo, Teresa Pollicino |
| EPI_ISL_1791488, EPI_ISL_1791489, EPI_ISL_1791491, EPI_ISL_1791509, EPI_ISL_1791511, EPI_ISL_1791512, EPI_ISL_1791513, EPI_ISL_1791514, EPI_ISL_1791515, EPI_ISL_1791517, EPI_ISL_1791518, EPI_ISL_1791519, EPI_ISL_1791520, EPI_ISL_1791521, EPI_ISL_1791522, EPI_ISL_1791523, EPI_ISL_1791524, EPI_ISL_1791526, EPI_ISL_1791527, EPI_ISL_1791528, EPI_ISL_1791529, EPI_ISL_1791530, EPI_ISL_1791531, EPI_ISL_1791532, EPI_ISL_1791533, EPI_ISL_1791534, EPI_ISL_1791535, EPI_ISL_1791536, EPI_ISL_1791537, EPI_ISL_1791539, EPI_ISL_1791540, EPI_ISL_1791541, EPI_ISL_1791542, EPI_ISL_1791543, EPI_ISL_1791544, EPI_ISL_1791546 | | | |
| see above | University of Oregon COVID-19 MAP Laboratory | University of Oregon Genomics and Cell Characterization Core Facility (GC3F) | Douglas Turnbull, Ariana White, Jeff Bishop, Jason Carriere |
| EPI_ISL_1792926 | Microbiologia e Virologia Cotugno | Microbiologia e Virologia Cotugno | Antonio Grimaldi Patrizia Annunziata Francesco Panariello Biancamaria Pierri Claudia Tiberio Teresa Giuliano Anna Perfetti Valentina Bouche Chiara Colantuono Maria Concetta Cuomo Denise Di Concilio Lucio Di Filippo Anna Manfredi Marcello Salvi Antonio Limone Luigi Atripaldi Pellegrino Cerino Andrea Ballabio Davide Cacchiarelli |
| EPI_ISL_1792927 | Texas Children's Hospital | Texas Children's Microbiome Center | Ruth Ann Luna, Jennifer K. Spinler, Jessica Runge, Sabeen Raza, James Dunn, Ila Singh, James Versalovic, Sridevi Devaraj |
| EPI_ISL_1793646, EPI_ISL_1793647, EPI_ISL_1793649, EPI_ISL_1793650, EPI_ISL_1793652, EPI_ISL_1793653 | Genome Analysis Center, Yamanashi Central Hospital | Genome Analysis Center, Yamanashi Central Hospital | Yosuke Hirotsu |
| EPI_ISL_1797436, EPI_ISL_1797437 | WHO National Influenza Centre Russian Federation | WHO National Influenza Centre Russian Federation | Andrey Komissarov, Artem Fadeev, Kseniya Komissarova, Oula Mansour, Mikhail Bakaev, Tamila Musaeva, Maria Timofeeva, Veronika Eder, Maria Pisareva, Daria Danilenko, Ksenia Safina, Elena Nabieva, Georgii Bazykin, Dmitry Lioznov |
| EPI_ISL_1797438 | CLINIQUE N-D DE GRACE | Institut de Pathologie et Genetique (IPG) | Pascale Hilbert, Jérémie Gras |
| EPI_ISL_1797619, EPI_ISL_1797620, EPI_ISL_1797622, EPI_ISL_1797623, EPI_ISL_1797624, EPI_ISL_1797625 | Department of Health Technology and Informatics, The Hong Kong Polytechnic University | Department of Health Technology and Informatics, The Hong Kong Polytechnic University | Gilman Kit-Hang Siu, Lam-Kwong Lee, Kenneth Siu-Sing Leung, Jake Siu-Lun Leung, Wing-Hei Lo, Timothy Ting-Leung Ng, Chloe Toi-Mei Chan, Hoi-Ching Jim, Kingsley King-Gee Tam, Hiu-Yin Lao, Denise Sze-Hang Wong, Alan Ka-Lun Wu, Miranda Chong-Yee Yau, Yvette Wai-Man Lai, Kitty Sau-Chun Fung, Sandy Ka-Yee Chau, Barry Kin-Chung Wong, Wing-Kin To, Kristine Luk, Alex Yat-Man Ho, Tak-Lun Que, Kam-Tong Yip, Wing Cheong |

| | | | |
|---|---|---|---|
| | | | Yam, David Ho-Keung Shum, Shea Ping Yip |
| EPI_ISL_1803231, EPI_ISL_1803232 | CQRC_QUALITY CONTROL CHEMICAL BIOLOGICAL RISK_AOOR Villa Sofia Cervello Palermo | CQRC_QUALITY CONTROL CHEMICAL BIOLOGICAL RISK_AOOR Villa Sofia Cervello Palermo | Di Gaudio, F., Brunacci, G., Contino, F., |
| EPI_ISL_1803233, EPI_ISL_1803236 | CQRC_QUALITY CONTROL CHEMICAL BIOLOGICAL RISK_AOOR Villa Sofia Cervello Palermo | CQRC_QUALITY CONTROL CHEMICAL BIOLOGICAL RISK_AOOR Villa Sofia Cervello Palermo | Di Gaudio, F., Contino, F., Brunacci, G., |
| EPI_ISL_1803239 | CQRC_QUALITY CONTROL CHEMICAL BIOLOGICAL RISK_AOOR Villa Sofia Cervello Palermo | CQRC_QUALITY CONTROL CHEMICAL BIOLOGICAL RISK_AOOR Villa Sofia Cervello Palermo | Di Gaudio, F., Brunacci, G., Contino, F., |
| EPI_ISL_1803244 | CQRC_QUALITY CONTROL CHEMICAL BIOLOGICAL RISK_AOOR Villa Sofia Cervello Palermo | CQRC_QUALITY CONTROL CHEMICAL BIOLOGICAL RISK_AOOR Villa Sofia Cervello Palermo | Di Gaudio, F., Contino, F., Brunacci, G., |
| EPI_ISL_1803245 | CQRC_QUALITY CONTROL CHEMICAL BIOLOGICAL RISK_AOOR Villa Sofia Cervello Palermo | CQRC_QUALITY CONTROL CHEMICAL BIOLOGICAL RISK_AOOR Villa Sofia Cervello Palermo | Di Gaudio, F., Brunacci, G., Contino, F., |
| EPI_ISL_1803246, EPI_ISL_1803249, EPI_ISL_1803251, EPI_ISL_1803254 | CQRC_QUALITY CONTROL CHEMICAL BIOLOGICAL RISK_AOOR Villa Sofia Cervello Palermo | CQRC_QUALITY CONTROL CHEMICAL BIOLOGICAL RISK_AOOR Villa Sofia Cervello Palermo | Di Gaudio, F., Contino, F., Brunacci, G., |
| EPI_ISL_1803255 | CQRC_QUALITY CONTROL CHEMICAL BIOLOGICAL RISK_AOOR Villa Sofia Cervello Palermo | CQRC_QUALITY CONTROL CHEMICAL BIOLOGICAL RISK_AOOR Villa Sofia Cervello Palermo | Di Gaudio, F., Brunacci, G., Contino, F., |
| EPI_ISL_1803258 | CQRC_QUALITY CONTROL CHEMICAL BIOLOGICAL RISK_AOOR Villa Sofia Cervello Palermo | CQRC_QUALITY CONTROL CHEMICAL BIOLOGICAL RISK_AOOR Villa Sofia Cervello Palermo | Di Gaudio, F., Contino, F., Brunacci, G. |
| EPI_ISL_1803261, EPI_ISL_1803262 | CQRC_QUALITY CONTROL CHEMICAL BIOLOGICAL RISK_AOOR Villa Sofia Cervello Palermo | CQRC_QUALITY CONTROL CHEMICAL BIOLOGICAL RISK_AOOR Villa Sofia Cervello Palermo | Di Gaudio, F., Contino, F., Brunacci, G., |
| EPI_ISL_1803264, EPI_ISL_1803269, EPI_ISL_1803272, EPI_ISL_1803274, EPI_ISL_1803275, EPI_ISL_1803278, EPI_ISL_1803282, EPI_ISL_1803284 | CQRC_QUALITY CONTROL CHEMICAL BIOLOGICAL RISK_AOOR Villa Sofia Cervello Palermo | CQRC_QUALITY CONTROL CHEMICAL BIOLOGICAL RISK_AOOR Villa Sofia Cervello Palermo | Di Gaudio, F., Brunacci, G., Contino, F., |
| EPI_ISL_1803286 | CQRC_QUALITY CONTROL CHEMICAL BIOLOGICAL RISK_AOOR Villa Sofia Cervello Palermo | CQRC_QUALITY CONTROL CHEMICAL BIOLOGICAL RISK_AOOR Villa Sofia Cervello Palermo | Di Gaudio, F., Contino, F., Orlando, V., Buffa, V., Bonvissuto, M., Di Chiara, V., Todaro, I., Seidita, G., Di Sclafani, A., Lungari, A., Brunacci, G. |
| EPI_ISL_1803288 | CQRC_QUALITY CONTROL CHEMICAL BIOLOGICAL RISK_AOOR Villa Sofia Cervello Palermo | CQRC_QUALITY CONTROL CHEMICAL BIOLOGICAL RISK_AOOR Villa Sofia Cervello Palermo | Di Gaudio, F., Brunacci, G., Contino, F., |
| EPI_ISL_1803291, EPI_ISL_1803294 | CQRC_QUALITY CONTROL CHEMICAL BIOLOGICAL RISK_AOOR Villa Sofia Cervello Palermo | CQRC_QUALITY CONTROL CHEMICAL BIOLOGICAL RISK_AOOR Villa Sofia Cervello Palermo | Di Gaudio, F., Contino, F., Brunacci, G., |
| EPI_ISL_1804821, EPI_ISL_1804822, EPI_ISL_1804823, EPI_ISL_1804824, EPI_ISL_1804825, EPI_ISL_1804826, EPI_ISL_1804827, EPI_ISL_1804828, EPI_ISL_1804829, EPI_ISL_1804830, EPI_ISL_1804831, EPI_ISL_1804832, EPI_ISL_1804833, EPI_ISL_1804834, EPI_ISL_1804835, EPI_ISL_1804836, EPI_ISL_1804837, EPI_ISL_1804838, EPI_ISL_1804839, EPI_ISL_1804840, EPI_ISL_1804841, EPI_ISL_1804842 | | | |
| see above | Bioptická laborato, s.r.o. | Bioptická laborato, s.r.o. | Petr Šteiner, Tomáš Vanek, Nikola Bílá, Martina Putzová, Michaela íhová, Silva Vondráková |
| EPI_ISL_1804970 | CQRC_QUALITY CONTROL CHEMICAL BIOLOGICAL RISK_AOOR Villa Sofia Cervello Palermo | CQRC_QUALITY CONTROL CHEMICAL BIOLOGICAL RISK_AOOR Villa Sofia Cervello Palermo | Di Gaudio, F., Contino, F., Brunacci, G., |
| EPI_ISL_1804972, EPI_ISL_1804973 | CQRC_QUALITY CONTROL CHEMICAL BIOLOGICAL RISK_AOOR Villa Sofia Cervello Palermo | CQRC_QUALITY CONTROL CHEMICAL BIOLOGICAL RISK_AOOR Villa Sofia Cervello Palermo | Di Gaudio, F., Brunacci, G., Contino, F., |
| EPI_ISL_1804978 | CQRC_QUALITY CONTROL CHEMICAL BIOLOGICAL RISK_AOOR Villa Sofia Cervello Palermo | CQRC_QUALITY CONTROL CHEMICAL BIOLOGICAL RISK_AOOR Villa Sofia Cervello Palermo | Di Gaudio, F., Contino, F., Brunacci, G., |
| EPI_ISL_1804979 | CQRC_QUALITY CONTROL CHEMICAL BIOLOGICAL RISK_AOOR Villa Sofia Cervello Palermo | CQRC_QUALITY CONTROL CHEMICAL BIOLOGICAL RISK_AOOR Villa Sofia Cervello Palermo | Di Gaudio, F., Brunacci, G., Contino, F., |
| EPI_ISL_1804985 | CQRC_QUALITY CONTROL CHEMICAL BIOLOGICAL RISK_AOOR Villa Sofia Cervello Palermo | CQRC_QUALITY CONTROL CHEMICAL BIOLOGICAL RISK_AOOR Villa Sofia Cervello Palermo | Di Gaudio, F., Contino, F., Brunacci, G., |
| EPI_ISL_1805480 | LESP Ciudad de Mexico | Instituto de Diagnostico y Referencia Epidemiologicos (INDRE) | Claudia Wong-Arambula, Abril Rodriguez-Maldonado, Vanessa Rivero-Arredondo, Ariadna Medina-Benitez, Joaquin Quiroz-Mercado, Sergio Rangel-Guerrero, Natividad Cruz-Ortiz, Tatiana Nunez-Garcia, Gisela Barrera-Badillo, Lucia Hernandez-Rivas, Irma Lopez-Martinez, Ernesto Ramirez-Gonzalez. |
| EPI_ISL_1805491 | LESP Jalisco | Instituto de Diagnostico y Referencia Epidemiologicos (INDRE) | Claudia Wong-Arambula, Abril Rodriguez-Maldonado, Vanessa Rivero-Arredondo, Ariadna Medina-Benitez, Joaquin Quiroz-Mercado, Sergio Rangel-Guerrero, Natividad Cruz-Ortiz, Tatiana Nunez-Garcia, Gisela Barrera-Badillo, Lucia Hernandez-Rivas, Irma Lopez-Martinez, Ernesto Ramirez-Gonzalez. |
| EPI_ISL_1805494 | LESPZacatecas | Instituto de Diagnostico y Referencia Epidemiologicos (INDRE) | Claudia Wong-Arambula, Abril Rodriguez-Maldonado, Vanessa Rivero-Arredondo, Ariadna Medina-Benitez, Joaquin Quiroz-Mercado, Sergio Rangel-Guerrero, Natividad Cruz-Ortiz, Tatiana Nunez-Garcia, Gisela Barrera-Badillo, Lucia Hernandez-Rivas, Irma Lopez-Martinez, Ernesto Ramirez-Gonzalez. |
| EPI_ISL_1805501, EPI_ISL_1805502 | LESP Yucatan | Instituto de Diagnostico y Referencia Epidemiologicos (INDRE) | Claudia Wong-Arambula, Abril Rodriguez-Maldonado, Vanessa Rivero-Arredondo, Ariadna Medina-Benitez, Joaquin Quiroz-Mercado, Sergio Rangel-Guerrero, Natividad Cruz-Ortiz, Tatiana Nunez-Garcia, Gisela Barrera-Badillo, Lucia Hernandez-Rivas, Irma Lopez-Martinez, Ernesto Ramirez-Gonzalez. |
| EPI_ISL_1807186 | Institute for Medical Research, Infectious Disease Research Centre, National Institutes of Health, Ministry of Health Malaysia | Institute for Medical Research, Infectious Disease Research Centre, National Institutes of Health, Ministry of Health Malaysia | Suppiah J, Kamel K, Mohd Zawawi Z, Ramly N, Robert F, Thayan R |
| EPI_ISL_1810831, EPI_ISL_1810832, EPI_ISL_1810833, EPI_ISL_1810834, EPI_ISL_1810835, EPI_ISL_1810841, EPI_ISL_1810842, EPI_ISL_1810843, EPI_ISL_1810844, EPI_ISL_1810845, EPI_ISL_1810846, EPI_ISL_1810847, EPI_ISL_1810849, EPI_ISL_1810850, EPI_ISL_1810853, EPI_ISL_1810856, EPI_ISL_1810859, EPI_ISL_1810863, EPI_ISL_1810864, EPI_ISL_1810865, EPI_ISL_1810866, EPI_ISL_1810867, EPI_ISL_1810868, EPI_ISL_1810869, EPI_ISL_1810872, EPI_ISL_1810873, EPI_ISL_1810874, EPI_ISL_1810875, EPI_ISL_1810879, EPI_ISL_1810881, EPI_ISL_1810882, EPI_ISL_1810884, EPI_ISL_1810885, EPI_ISL_1810886, EPI_ISL_1810887, EPI_ISL_1810888, EPI_ISL_1810891, EPI_ISL_1810892, EPI_ISL_1810893, EPI_ISL_1810894, EPI_ISL_1810895, EPI_ISL_1810896, EPI_ISL_1810898, EPI_ISL_1810899, EPI_ISL_1810900, EPI_ISL_1810905, EPI_ISL_1810907, EPI_ISL_1810908, EPI_ISL_1810910, EPI_ISL_1810911, EPI_ISL_1810912, EPI_ISL_1810917, EPI_ISL_1810918, EPI_ISL_1810919, EPI_ISL_1810921, EPI_ISL_1810923 | | | |
| see above | InDRE | Instituto Nacional de Medicina Genomica | Hidalgo-Miranda A, Mendoza-Vargas A, Reyes-Grajeda JP, Cedro-Tanda A, Gisela Barrera-Badillo, Irma Lopez-Martinez, Jose Ernesto Ramirez González, Gonzalez-Barrera D, Rangel-DeLeon D, Munguia-Garza P, Garcia-Cardenas FJ, Gonzalez-Woge MA, Herrera-Montalvo LA. |
| EPI_ISL_1810925 | SISP - Servizio Igiene e Sanità Pubblica-ASL AT | Genomics facility CAAD, Università del Piemonte Orientale (UPO) | Marta Mellai, Marta Allesina, Nadia Barizzone, Maurizio Penna, Nadia Ardovini, Stefano Mosca, Sandra D'Alfonso |
| EPI_ISL_1810930, EPI_ISL_1810931, EPI_ISL_1810932 | SISP - Servizio Igiene e Sanità Pubblica-ASL AT | Genomics Facility CAAD - Università del Piemonte Orientale (UPO) | Marta Mellai, Marta Allesina, Nadia Barizzone, Maurizio Penna, Nadia Ardovini, Stefano Mosca, Sandra D'Alfonso |
| EPI_ISL_1811223 | COVID Laboratory AOU Messina | Molecular Hepatology Lab AOU Messina | Giuseppina Raffa, Cristina Musolino, Valeria Chines, Daniele Lombardo, Teresa Pollicino |
| EPI_ISL_1811234 | AS Santa Barbara | Inciensa, Instituto Costarricense de Investigación y Enseñanza en Nutrición y Salud | Pérez-Corrales C & Aguilar-Monge R |
| EPI_ISL_1811235 | HNN | Inciensa, Instituto Costarricense de Investigación y Enseñanza en Nutrición y Salud | Pérez-Corrales C & Morazán-Fernandez D |

| | | | |
|---|---|---|---|
| EPI_ISL_1811236 | HNN | Inciensa, Instituto Costarricense de Investigación y Enseñanza en Nutrición y Salud | Pérez-Corrales C & Catalán-Marín F |
| EPI_ISL_1811525, EPI_ISL_1811526, EPI_ISL_1811527, EPI_ISL_1811529, EPI_ISL_1811530, EPI_ISL_1811531, EPI_ISL_1811532, EPI_ISL_1811533, EPI_ISL_1811534 | Laboratorio Central de Epidemiología (LCE) | Instituto de Biotecnología de la UNAM | Julio Elias Alvarado-Yaah, Carlos F. Arias, Santiago Ávila-Ríos, Víctor Hugo Borja-Aburto, Celia Boukadida, Juan Bautista Chale-Dzul, Célida Duque-Molina, José Antonio Enciso-Moreno, Gloria Elena Espinosa-Ayala, Fernando Fontove-Herrera, Victor Eduardo Garcia-Arias, Concepción Grajales-Muñiz, Ricardo Grande, Alfredo Herrera-Estrella, Carla Ivón Herrera-Najera, Pavel Isa, Brenda Irasema Maldonado-Meza, Bernardo Martínez-Miguel, Margarita Matías-Florentino, María Guadalupe de Jesús Mireles-Rivera, Gloria María Molina-Salinas, Hector Montoya-Fuentes, José Esteban Muñoz-Medina, José de Jesús Nuñez-Contreras, Alicia Ocaña-Mondragón, Luis Alberto Ochoa-Carrera, Hector Esteban Paz-Juárez, Francisco Pulido, Helen Haydee Fernanda Ramírez-Plascencia, Angel Gustavo Salas-Lais, Alejandro Sanchez-Flores, Clara Esperanza Santacruz-Tinoco, María Guadalupe Santiago-Mauricio, Nelly Sélem-Mojica, Blanca Taboada, Gloria Vazquez |
| EPI_ISL_1818785 | CH Roubaix | CHU Lille - Laboratoire de Virologie | AIT YAHYA Emilie, ALIDJINOU Enagnon Kazali, BOCKET Laurence, CREPIN Michel, DEMAY Christophe, ENGELMANN Ilka, GEFFROY Sandrine, GUIGON Aurélie, LAMBERT Valérie, LAZREK Mouna, NOBILLIAUX Florian, PREVOST Brigitte, THUILLIER Caroline, TINEZ Claire |
| EPI_ISL_1818794 | CH Saint Philibert | CHU Lille - Laboratoire de Virologie | AIT YAHYA Emilie, ALIDJINOU Enagnon Kazali, BOCKET Laurence, CREPIN Michel, DEMAY Christophe, ENGELMANN Ilka, GEFFROY Sandrine, GUIGON Aurélie, LAMBERT Valérie, LAZREK Mouna, NOBILLIAUX Florian, PREVOST Brigitte, THUILLIER Caroline, TINEZ Claire |
| EPI_ISL_1818798 | CHU Lille | CHU Lille - Laboratoire de Virologie | AIT YAHYA Emilie, ALIDJINOU Enagnon Kazali, BOCKET Laurence, CREPIN Michel, DEMAY Christophe, ENGELMANN Ilka, GEFFROY Sandrine, GUIGON Aurélie, LAMBERT Valérie, LAZREK Mouna, NOBILLIAUX Florian, PREVOST Brigitte, THUILLIER Caroline, TINEZ Claire |
| EPI_ISL_1818799 | CH Lens | CHU Lille - Laboratoire de Virologie | AIT YAHYA Emilie, ALIDJINOU Enagnon Kazali, BOCKET Laurence, CREPIN Michel, DEMAY Christophe, ENGELMANN Ilka, GEFFROY Sandrine, GUIGON Aurélie, LAMBERT Valérie, LAZREK Mouna, NOBILLIAUX Florian, PREVOST Brigitte, THUILLIER Caroline, TINEZ Claire |
| EPI_ISL_1818800 | CHU Lille | CHU Lille - Laboratoire de Virologie | AIT YAHYA Emilie, ALIDJINOU Enagnon Kazali, BOCKET Laurence, CREPIN Michel, DEMAY Christophe, ENGELMANN Ilka, GEFFROY Sandrine, GUIGON Aurélie, LAMBERT Valérie, LAZREK Mouna, NOBILLIAUX Florian, PREVOST Brigitte, THUILLIER Caroline, TINEZ Claire |
| EPI_ISL_1818806 | CH Dunkerque | CHU Lille - Laboratoire de Virologie | AIT YAHYA Emilie, ALIDJINOU Enagnon Kazali, BOCKET Laurence, CREPIN Michel, DEMAY Christophe, ENGELMANN Ilka, GEFFROY Sandrine, GUIGON Aurélie, LAMBERT Valérie, LAZREK Mouna, NOBILLIAUX Florian, PREVOST Brigitte, THUILLIER Caroline, TINEZ Claire |
| EPI_ISL_1818815, EPI_ISL_1818816, EPI_ISL_1818820, EPI_ISL_1818821 | CHU Lille | CHU Lille - Laboratoire de Virologie | AIT YAHYA Emilie, ALIDJINOU Enagnon Kazali, BOCKET Laurence, CREPIN Michel, DEMAY Christophe, ENGELMANN Ilka, GEFFROY Sandrine, GUIGON Aurélie, LAMBERT Valérie, LAZREK Mouna, NOBILLIAUX Florian, PREVOST Brigitte, THUILLIER Caroline, TINEZ Claire |
| EPI_ISL_1818822 | CH Arras | CHU Lille - Laboratoire de Virologie | AIT YAHYA Emilie, ALIDJINOU Enagnon Kazali, BOCKET Laurence, CREPIN Michel, DEMAY Christophe, ENGELMANN Ilka, GEFFROY Sandrine, GUIGON Aurélie, LAMBERT Valérie, LAZREK Mouna, NOBILLIAUX Florian, PREVOST Brigitte, THUILLIER Caroline, TINEZ Claire |
| EPI_ISL_1818824, EPI_ISL_1818827, EPI_ISL_1818829, EPI_ISL_1818831, EPI_ISL_1818833, EPI_ISL_1818837, EPI_ISL_1818841, EPI_ISL_1818847 | CHU Lille | CHU Lille - Laboratoire de Virologie | AIT YAHYA Emilie, ALIDJINOU Enagnon Kazali, BOCKET Laurence, CREPIN Michel, DEMAY Christophe, ENGELMANN Ilka, GEFFROY Sandrine, GUIGON Aurélie, LAMBERT Valérie, LAZREK Mouna, NOBILLIAUX Florian, PREVOST Brigitte, THUILLIER Caroline, TINEZ Claire |
| EPI_ISL_1818848 | CH Armentières | CHU Lille - Laboratoire de Virologie | AIT YAHYA Emilie, ALIDJINOU Enagnon Kazali, BOCKET Laurence, CREPIN Michel, DEMAY Christophe, ENGELMANN Ilka, GEFFROY Sandrine, GUIGON Aurélie, LAMBERT Valérie, LAZREK Mouna, NOBILLIAUX Florian, PREVOST Brigitte, THUILLIER Caroline, TINEZ Claire |
| EPI_ISL_1818850 | CH Roubaix | CHU Lille - Laboratoire de Virologie | AIT YAHYA Emilie, ALIDJINOU Enagnon Kazali, BOCKET Laurence, CREPIN Michel, DEMAY Christophe, ENGELMANN Ilka, GEFFROY Sandrine, GUIGON Aurélie, LAMBERT Valérie, LAZREK Mouna, NOBILLIAUX Florian, PREVOST Brigitte, THUILLIER Caroline, TINEZ Claire |
| EPI_ISL_1818947, EPI_ISL_1818948, EPI_ISL_1818949, EPI_ISL_1818950, EPI_ISL_1818951, EPI_ISL_1818952, EPI_ISL_1818953, EPI_ISL_1818954, EPI_ISL_1818955, EPI_ISL_1818956, EPI_ISL_1818957, EPI_ISL_1818958, EPI_ISL_1818959, EPI_ISL_1818960, EPI_ISL_1818961, EPI_ISL_1818962, EPI_ISL_1818963, EPI_ISL_1818964, EPI_ISL_1818965, EPI_ISL_1818966 | | | |
| see above | Virology Unit, Institut Pasteur du Cambodge | Virology Unit, Institut Pasteur du Cambodge | Jurre Y Siegers, Cecile Troupin, Ly Sovann, Kraing Sidonn, Yi Sengdoeurn, Chin Savuth, Chau Darapheak, Veasna Duong, Erik A Karlsson |
| EPI_ISL_1819188, EPI_ISL_1819189, EPI_ISL_1819192, EPI_ISL_1819193, EPI_ISL_1819194, EPI_ISL_1819196, EPI_ISL_1819197, EPI_ISL_1819198 | Ospedale di Venere | Istituto Zooprofilattico Sperimentale della Puglia e della Basilicata | Parisi A., Bianco A., Capozzi L., Del Sambro L., Simone D., Iacobellis M., Ridolfi D., Giannico A. |
| EPI_ISL_1819199, EPI_ISL_1819200, EPI_ISL_1819201, EPI_ISL_1819202, EPI_ISL_1819203, EPI_ISL_1819204 | Presidio di Brindisi Di Summa - Perrino | Istituto Zooprofilattico Sperimentale della Puglia e della Basilicata | Parisi A., Bianco A., Capozzi L., Del Sambro L., Simone D., Difato L., Santoro A., Ridolfi D., Giannico A. |
| EPI_ISL_1819205, EPI_ISL_1819206, EPI_ISL_1819207, EPI_ISL_1819208, EPI_ISL_1819209, EPI_ISL_1819211, EPI_ISL_1819212 | Ospedale Santa Caterina Novella | Istituto Zooprofilattico Sperimentale della Puglia e della Basilicata | Parisi A., Bianco A., Capozzi L., Del Sambro L., Simone D., Difato L., Bruno A. R. |
| EPI_ISL_1819213, EPI_ISL_1819215, EPI_ISL_1819216, EPI_ISL_1819217, EPI_ISL_1819218, EPI_ISL_1819219, EPI_ISL_1819221 | Ospedale Vito Fazzi | Istituto Zooprofilattico Sperimentale della Puglia e della Basilicata | Parisi A., Bianco A., Capozzi L., Del Sambro L., Simone D., Difato L., Lobreglio G. |
| EPI_ISL_1819222, EPI_ISL_1819223, EPI_ISL_1819224 | Ospedale Generale Regionale F. Miulli | Istituto Zooprofilattico Sperimentale della Puglia e della Basilicata | Parisi A., Bianco A., Capozzi L., Del Sambro L., Simone D., Ceci E. |
| EPI_ISL_1819225, EPI_ISL_1819227, EPI_ISL_1819229, EPI_ISL_1819230, EPI_ISL_1819231, EPI_ISL_1819232, EPI_ISL_1819235 | Ospedale Santissima Annunziata | Istituto Zooprofilattico Sperimentale della Puglia e della Basilicata | Parisi A., Bianco A., Capozzi L., Del Sambro L., Simone D., Difato L., Ridolfi D., Giannico A, Tampoia M. |
| EPI_ISL_1819237, EPI_ISL_1819238, EPI_ISL_1819239, EPI_ISL_1819240, EPI_ISL_1819241, EPI_ISL_1819242, EPI_ISL_1819244 | I.R.C.C.S. "S. De Bellis" - Ente Ospedaliero | Istituto Zooprofilattico Sperimentale della Puglia e della Basilicata | Parisi A., Bianco A., Capozzi L., Del Sambro L., Simone D., Lippolis A., Notarnicola M. |
| EPI_ISL_1819245, EPI_ISL_1819246, EPI_ISL_1819247 | Ospedale Casa Sollievo della Sofferenza | Istituto Zooprofilattico Sperimentale della Puglia e della Basilicata | Parisi A., Bianco A., Capozzi L., Del Sambro L., Simone D., Difato L., Ridolfi D., Giannico A, Labonia M. |
| EPI_ISL_1819248 | Ospedale Santissima Annunziata | Istituto Zooprofilattico Sperimentale della Puglia e della Basilicata | Parisi A., Bianco A., Capozzi L., Del Sambro L., Simone D., Difato L., Ridolfi D., Giannico A, Tampoia M. |
| EPI_ISL_1819250, EPI_ISL_1819251, EPI_ISL_1819252, EPI_ISL_1819253, EPI_ISL_1819254, EPI_ISL_1819255, EPI_ISL_1819256 | Presidio di Brindisi Di Summa - Perrino | Istituto Zooprofilattico Sperimentale della Puglia e della Basilicata | Parisi A., Bianco A., Capozzi L., Del Sambro L., Simone D., Difato L., Santoro A., Ridolfi D., Giannico A. |
| EPI_ISL_1819257 | Ospedale Generale Regionale F. Miulli | Istituto Zooprofilattico Sperimentale della Puglia e della Basilicata | Parisi A., Bianco A., Capozzi L., Del Sambro L., Simone D., Ceci E. |
| EPI_ISL_1820961 | IMAT | Instituto Nacional de Salud- Dirección de Investigación en Salud Pública | Katherine Laiton-Donato, Diego A. Álvarez-Díaz, Carlos Franco-Muñoz, Hector Alejandro Ruiz-Moreno, Paola Rojas, Maria T. Herrera-Sepúlveda, Diego Andrés Prada, Jhonnatan Reales-González, Sheryll Corchuelo, Julian Naizaque, Jorge Rivera, Gerardo Santamaría, Sergio Gomez, Lisseth Pardo, Juan Camilo Martinez, Marta Lopez Blanco, Ángela Alarcon Cruz, Diana Malo, Carmen Osorio, Magdalena Wiesner, Martha Lucia Ospina Martinez, Marcela |

| | | | |
|---|---|---|---|
| | | | Mercado-Reyes |
| EPI_ISL_1820962 | Clinica los Nogales | Instituto Nacional de Salud- Dirección de Investigación en Salud Pública | Katherine Laiton-Donato, Diego A. Álvarez-Díaz, Carlos Franco-Muñoz, Hector Alejandro Ruiz-Moreno, Paola Rojas, Maria T. Herrera-Sepúlveda, Diego Andrés Prada, Jhonnatan Reales-González, Sheryll Corchuelo, Julian Naizaque, Jorge Rivera, Gerardo Santamaría, Sergio Gomez, Lisseth Pardo, Juan Camilo Martinez, Marta Lopez Blanco, Ángela Alarcon Cruz, Diana Malo, Carmen Osorio, Magdalena Wiesner, Martha Lucia Ospina Martinez, Marcela Mercado-Reyes |
| EPI_ISL_1821124 | LESP Puebla | Instituto de Diagnostico y Referencia Epidemiologicos (INDRE) | Claudia Wong-Arambula, Abril Rodriguez-Maldonado, Vanessa Rivero-Arredondo, Ariadna Medina-Benitez, Joaquin Quiroz-Mercado, Sergio Rangel-Guerrero, Natividad Cruz-Ortiz, Tatiana Nunez-Garcia, Gisela Barrera-Badillo, Lucia Hernandez-Rivas, Irma Lopez-Martinez, Ernesto Ramirez-Gonzalez. |
| EPI_ISL_1821131 | LESP Jalisco | Instituto de Diagnostico y Referencia Epidemiologicos (INDRE) | Claudia Wong-Arambula, Abril Rodriguez-Maldonado, Vanessa Rivero-Arredondo, Ariadna Medina-Benitez, Joaquin Quiroz-Mercado, Sergio Rangel-Guerrero, Natividad Cruz-Ortiz, Tatiana Nunez-Garcia, Gisela Barrera-Badillo, Lucia Hernandez-Rivas, Irma Lopez-Martinez, Ernesto Ramirez-Gonzalez. |
| EPI_ISL_1821135 | LESP Queretaro | Instituto de Diagnostico y Referencia Epidemiologicos (INDRE) | Claudia Wong-Arambula, Abril Rodriguez-Maldonado, Vanessa Rivero-Arredondo, Ariadna Medina-Benitez, Joaquin Quiroz-Mercado, Sergio Rangel-Guerrero, Natividad Cruz-Ortiz, Tatiana Nunez-Garcia, Gisela Barrera-Badillo, Lucia Hernandez-Rivas, Irma Lopez-Martinez, Ernesto Ramirez-Gonzalez. |
| EPI_ISL_1821152, EPI_ISL_1821153, EPI_ISL_1821154, EPI_ISL_1821155 | LESP Yucatan | Instituto de Diagnostico y Referencia Epidemiologicos (INDRE) | Claudia Wong-Arambula, Abril Rodriguez-Maldonado, Vanessa Rivero-Arredondo, Ariadna Medina-Benitez, Joaquin Quiroz-Mercado, Sergio Rangel-Guerrero, Natividad Cruz-Ortiz, Tatiana Nunez-Garcia, Gisela Barrera-Badillo, Lucia Hernandez-Rivas, Irma Lopez-Martinez, Ernesto Ramirez-Gonzalez. |
| EPI_ISL_1821156 | LESP Sinaloa | Instituto de Diagnostico y Referencia Epidemiologicos (INDRE) | Claudia Wong-Arambula, Abril Rodriguez-Maldonado, Vanessa Rivero-Arredondo, Ariadna Medina-Benitez, Joaquin Quiroz-Mercado, Sergio Rangel-Guerrero, Natividad Cruz-Ortiz, Tatiana Nunez-Garcia, Gisela Barrera-Badillo, Lucia Hernandez-Rivas, Irma Lopez-Martinez, Ernesto Ramirez-Gonzalez. |
| EPI_ISL_1821161 | LESP Nayarit | Instituto de Diagnostico y Referencia Epidemiologicos (INDRE) | Claudia Wong-Arambula, Abril Rodriguez-Maldonado, Vanessa Rivero-Arredondo, Ariadna Medina-Benitez, Joaquin Quiroz-Mercado, Sergio Rangel-Guerrero, Natividad Cruz-Ortiz, Tatiana Nunez-Garcia, Gisela Barrera-Badillo, Lucia Hernandez-Rivas, Irma Lopez-Martinez, Ernesto Ramirez-Gonzalez. |
| EPI_ISL_1821168 | LESP Chihuahua | Instituto de Diagnostico y Referencia Epidemiologicos (INDRE) | Claudia Wong-Arambula, Abril Rodriguez-Maldonado, Vanessa Rivero-Arredondo, Ariadna Medina-Benitez, Joaquin Quiroz-Mercado, Sergio Rangel-Guerrero, Natividad Cruz-Ortiz, Tatiana Nunez-Garcia, Gisela Barrera-Badillo, Lucia Hernandez-Rivas, Irma Lopez-Martinez, Ernesto Ramirez-Gonzalez. |
| EPI_ISL_1821173, EPI_ISL_1821174, EPI_ISL_1821175, EPI_ISL_1821176, EPI_ISL_1821177 | LESP Zacatecas | Instituto de Diagnostico y Referencia Epidemiologicos (INDRE) | Claudia Wong-Arambula, Abril Rodriguez-Maldonado, Vanessa Rivero-Arredondo, Ariadna Medina-Benitez, Joaquin Quiroz-Mercado, Sergio Rangel-Guerrero, Natividad Cruz-Ortiz, Tatiana Nunez-Garcia, Gisela Barrera-Badillo, Lucia Hernandez-Rivas, Irma Lopez-Martinez, Ernesto Ramirez-Gonzalez. |
| EPI_ISL_1821180, EPI_ISL_1821181, EPI_ISL_1821182, EPI_ISL_1821183, EPI_ISL_1821184 | LESP Baja California Sur | Instituto de Diagnostico y Referencia Epidemiologicos (INDRE) | Claudia Wong-Arambula, Abril Rodriguez-Maldonado, Vanessa Rivero-Arredondo, Ariadna Medina-Benitez, Joaquin Quiroz-Mercado, Sergio Rangel-Guerrero, Natividad Cruz-Ortiz, Tatiana Nunez-Garcia, Gisela Barrera-Badillo, Lucia Hernandez-Rivas, Irma Lopez-Martinez, Ernesto Ramirez-Gonzalez. |
| EPI_ISL_1821185 | LESP Jalisco | Instituto de Diagnostico y Referencia Epidemiologicos (INDRE) | Claudia Wong-Arambula, Abril Rodriguez-Maldonado, Vanessa Rivero-Arredondo, Ariadna Medina-Benitez, Joaquin Quiroz-Mercado, Sergio Rangel-Guerrero, Natividad Cruz-Ortiz, Tatiana Nunez-Garcia, Gisela Barrera-Badillo, Lucia Hernandez-Rivas, Irma Lopez-Martinez, Ernesto Ramirez-Gonzalez. |
| EPI_ISL_1821201, EPI_ISL_1821202, EPI_ISL_1821203 | LESP Baja California | Instituto de Diagnostico y Referencia Epidemiologicos (INDRE) | Claudia Wong-Arambula, Abril Rodriguez-Maldonado, Vanessa Rivero-Arredondo, Ariadna Medina-Benitez, Joaquin Quiroz-Mercado, Sergio Rangel-Guerrero, Natividad Cruz-Ortiz, Tatiana Nunez-Garcia, Gisela Barrera-Badillo, Lucia Hernandez-Rivas, Irma Lopez-Martinez, Ernesto Ramirez-Gonzalez. |
| EPI_ISL_1821475, EPI_ISL_1821476 | Labo Analyses Med | National Reference Center for Viruses of Respiratory Infections, Institut Pasteur, Paris | Marion Barbet, Sylvie Behillil, Méline Bizard, Angela Brisebarre, Camille Capel, Vincent Enouf, Louise Lefrançois, Frédéric Lemoine, Christophe Malabat, Corinne Maufrais, Etienne Simon-Lorière, Maud Vanpeene, Sylvie Van der Werf ,Sophie Chalmin |
| EPI_ISL_1821478 | Labo Analyses Med | National Reference Center for Viruses of Respiratory Infections, Institut Pasteur, Paris | Marion Barbet, Sylvie Behillil, Méline Bizard, Angela Brisebarre, Camille Capel, Vincent Enouf, Louise Lefrançois, Frédéric Lemoine, Christophe Malabat, Corinne Maufrais, Etienne Simon-Lorière, Maud Vanpeene, Sylvie Van der Werf ,Pierre Pennacino |
| EPI_ISL_1821479 | Hospital | National Reference Center for Viruses of Respiratory Infections, Institut Pasteur, Paris | Marion Barbet, Sylvie Behillil, Méline Bizard, Angela Brisebarre, Camille Capel, Vincent Enouf, Louise Lefrançois, Frédéric Lemoine, Christophe Malabat, Corinne Maufrais, Etienne Simon-Lorière, Maud Vanpeene, Sylvie Van der Werf ,LéA Pilorge |
| EPI_ISL_1821482, EPI_ISL_1821483 | Hospital | National Reference Center for Viruses of Respiratory Infections, Institut Pasteur, Paris | Marion Barbet, Sylvie Behillil, Méline Bizard, Angela Brisebarre, Camille Capel, Vincent Enouf, Louise Lefrançois, Frédéric Lemoine, Christophe Malabat, Corinne Maufrais, Etienne Simon-Lorière, Maud Vanpeene, Sylvie Van der Werf ,JéRome Guinard |
| EPI_ISL_1821484, EPI_ISL_1821485, EPI_ISL_1821486, EPI_ISL_1821487, EPI_ISL_1821488 | Hospital | National Reference Center for Viruses of Respiratory Infections, Institut Pasteur, Paris | Marion Barbet, Sylvie Behillil, Méline Bizard, Angela Brisebarre, Camille Capel, Vincent Enouf, Louise Lefrançois, Frédéric Lemoine, Christophe Malabat, Corinne Maufrais, Etienne Simon-Lorière, Maud Vanpeene, Sylvie Van der Werf ,FréDéRique Canis |
| EPI_ISL_1821489 | Hospital | National Reference Center for Viruses of Respiratory Infections, Institut Pasteur, Paris | Marion Barbet, Sylvie Behillil, Méline Bizard, Angela Brisebarre, Camille Capel, Vincent Enouf, Louise Lefrançois, Frédéric Lemoine, Christophe Malabat, Corinne Maufrais, Etienne Simon-Lorière, Maud Vanpeene, Sylvie Van der Werf ,Haciba Moudjahed |
| EPI_ISL_1821490, EPI_ISL_1821491, EPI_ISL_1821492, EPI_ISL_1821493 | Labo Analyses Med | National Reference Center for Viruses of Respiratory Infections, Institut Pasteur, Paris | Marion Barbet, Sylvie Behillil, Méline Bizard, Angela Brisebarre, Camille Capel, Vincent Enouf, Louise Lefrançois, Frédéric Lemoine, Christophe Malabat, Corinne Maufrais, Etienne Simon-Lorière, Maud Vanpeene, Sylvie Van der Werf ,GréGoire Potiron |
| EPI_ISL_1821494, EPI_ISL_1821495 | Labo Analyses Med | National Reference Center for Viruses of Respiratory Infections, Institut Pasteur, Paris | Marion Barbet, Sylvie Behillil, Méline Bizard, Angela Brisebarre, Camille Capel, Vincent Enouf, Louise Lefrançois, Frédéric Lemoine, Christophe Malabat, Corinne Maufrais, Etienne Simon-Lorière, Maud Vanpeene, Sylvie Van der Werf ,Sophie Chalmin |
| EPI_ISL_1821496, EPI_ISL_1821497 | Labo Analyses Med | National Reference Center for Viruses of Respiratory Infections, Institut Pasteur, Paris | Marion Barbet, Sylvie Behillil, Méline Bizard, Angela Brisebarre, Camille Capel, Vincent Enouf, Louise Lefrançois, Frédéric Lemoine, Christophe Malabat, Corinne Maufrais, Damien Mornico, Etienne Simon-Lorière, Maud Vanpeene, Sylvie Van der Werf ,Sophie Chalmin |
| EPI_ISL_1821498, EPI_ISL_1821499 | Labo Analyses Med | National Reference Center for Viruses of Respiratory Infections, Institut Pasteur, Paris | Marion Barbet, Sylvie Behillil, Méline Bizard, Angela Brisebarre, Camille Capel, Vincent Enouf, Louise Lefrançois, Frédéric Lemoine, Christophe Malabat, Corinne Maufrais, Etienne Simon-Lorière, Maud Vanpeene, Sylvie Van der Werf ,Sophie Chalmin |
| EPI_ISL_1821500 | Labo Analyses Med | National Reference Center for Viruses of Respiratory Infections, Institut Pasteur, Paris | Marion Barbet, Sylvie Behillil, Méline Bizard, Angela Brisebarre, Camille Capel, Vincent Enouf, Louise Lefrançois, Frédéric Lemoine, Christophe Malabat, Corinne Maufrais, Damien Mornico, Etienne Simon-Lorière, Maud Vanpeene, Sylvie Van der Werf ,Sophie Chalmin |
| EPI_ISL_1821501, EPI_ISL_1821502, EPI_ISL_1821503, EPI_ISL_1821504, EPI_ISL_1821505, EPI_ISL_1821506, EPI_ISL_1821507, EPI_ISL_1821508, EPI_ISL_1821509 | Labo Analyses Med | National Reference Center for Viruses of Respiratory Infections, Institut Pasteur, Paris | Marion Barbet, Sylvie Behillil, Méline Bizard, Angela Brisebarre, Camille Capel, Vincent Enouf, Louise Lefrançois, Frédéric Lemoine, Christophe Malabat, Corinne Maufrais, Etienne Simon-Lorière, Maud Vanpeene, Sylvie Van der Werf ,Sophie Chalmin |
| EPI_ISL_1821510 | Labo Analyses Med | National Reference Center for Viruses of Respiratory Infections, Institut Pasteur, Paris | Marion Barbet, Sylvie Behillil, Méline Bizard, Angela Brisebarre, Camille Capel, Vincent Enouf, Louise Lefrançois, Frédéric Lemoine, Christophe Malabat, Corinne Maufrais, Damien Mornico, Etienne Simon-Lorière, Maud Vanpeene, Sylvie Van der Werf ,Sophie Chalmin |
| EPI_ISL_1821511, EPI_ISL_1821512, EPI_ISL_1821513, EPI_ISL_1821514, EPI_ISL_1821515, EPI_ISL_1821516, EPI_ISL_1821517 | Labo Analyses Med | National Reference Center for Viruses of Respiratory Infections, Institut Pasteur, Paris | Marion Barbet, Sylvie Behillil, Méline Bizard, Angela Brisebarre, Camille Capel, Vincent Enouf, Louise Lefrançois, Frédéric Lemoine, Christophe Malabat, Corinne Maufrais, Etienne Simon-Lorière, Maud Vanpeene, Sylvie Van der Werf ,Sophie Chalmin |
| EPI_ISL_1821518 | Labo Analyses Med | National Reference Center for Viruses of Respiratory Infections, Institut Pasteur, Paris | Marion Barbet, Sylvie Behillil, Méline Bizard, Angela Brisebarre, Camille Capel, Vincent Enouf, Louise Lefrançois, Frédéric Lemoine, Christophe Malabat, Corinne Maufrais, Damien Mornico, Etienne Simon-Lorière, Maud Vanpeene, Sylvie Van der Werf ,Sophie Chalmin |

| | | | |
|---|---|---|---|
| EPI_ISL_1821519, EPI_ISL_1821520, EPI_ISL_1821521, EPI_ISL_1821522, EPI_ISL_1821523, EPI_ISL_1821524, EPI_ISL_1821525, EPI_ISL_1821526 | Labo Analyses Med | National Reference Center for Viruses of Respiratory Infections, Institut Pasteur, Paris | Marion Barbet, Sylvie Behillil, Méline Bizard, Angela Brisebarre, Camille Capel, Vincent Enouf, Louise Lefrançois, Frédéric Lemoine, Christophe Malabat, Corinne Maufrais, Etienne Simon-Lorière, Maud Vanpeene, Sylvie Van der Werf ,Sophie Chalmin |
| EPI_ISL_1821527 | Labo Analyses Med | National Reference Center for Viruses of Respiratory Infections, Institut Pasteur, Paris | Marion Barbet, Sylvie Behillil, Méline Bizard, Angela Brisebarre, Camille Capel, Vincent Enouf, Louise Lefrançois, Frédéric Lemoine, Christophe Malabat, Corinne Maufrais, Damien Mornico, Etienne Simon-Lorière, Maud Vanpeene, Sylvie Van der Werf ,Sophie Chalmin |
| EPI_ISL_1821528, EPI_ISL_1821529 | Labo Analyses Med | National Reference Center for Viruses of Respiratory Infections, Institut Pasteur, Paris | Marion Barbet, Sylvie Behillil, Méline Bizard, Angela Brisebarre, Camille Capel, Vincent Enouf, Louise Lefrançois, Frédéric Lemoine, Christophe Malabat, Corinne Maufrais, Etienne Simon-Lorière, Maud Vanpeene, Sylvie Van der Werf ,Sophie Chalmin |
| EPI_ISL_1821530 | Labo Analyses Med | National Reference Center for Viruses of Respiratory Infections, Institut Pasteur, Paris | Marion Barbet, Sylvie Behillil, Méline Bizard, Angela Brisebarre, Camille Capel, Vincent Enouf, Louise Lefrançois, Frédéric Lemoine, Christophe Malabat, Corinne Maufrais, Damien Mornico, Etienne Simon-Lorière, Maud Vanpeene, Sylvie Van der Werf ,Sophie Chalmin |
| EPI_ISL_1821531, EPI_ISL_1821532 | Labo Analyses Med | National Reference Center for Viruses of Respiratory Infections, Institut Pasteur, Paris | Marion Barbet, Sylvie Behillil, Méline Bizard, Angela Brisebarre, Camille Capel, Vincent Enouf, Louise Lefrançois, Frédéric Lemoine, Christophe Malabat, Corinne Maufrais, Etienne Simon-Lorière, Maud Vanpeene, Sylvie Van der Werf ,Sophie Chalmin |
| EPI_ISL_1821533 | Labo Analyses Med | National Reference Center for Viruses of Respiratory Infections, Institut Pasteur, Paris | Marion Barbet, Sylvie Behillil, Méline Bizard, Angela Brisebarre, Camille Capel, Vincent Enouf, Louise Lefrançois, Frédéric Lemoine, Christophe Malabat, Corinne Maufrais, Damien Mornico, Etienne Simon-Lorière, Maud Vanpeene, Sylvie Van der Werf ,Sophie Chalmin |
| EPI_ISL_1821534, EPI_ISL_1821535 | Labo Analyses Med | National Reference Center for Viruses of Respiratory Infections, Institut Pasteur, Paris | Marion Barbet, Sylvie Behillil, Méline Bizard, Angela Brisebarre, Camille Capel, Vincent Enouf, Louise Lefrançois, Frédéric Lemoine, Christophe Malabat, Corinne Maufrais, Etienne Simon-Lorière, Maud Vanpeene, Sylvie Van der Werf ,Sophie Chalmin |
| EPI_ISL_1821536, EPI_ISL_1821537 | Labo Analyses Med | National Reference Center for Viruses of Respiratory Infections, Institut Pasteur, Paris | Marion Barbet, Sylvie Behillil, Méline Bizard, Angela Brisebarre, Camille Capel, Vincent Enouf, Louise Lefrançois, Frédéric Lemoine, Christophe Malabat, Corinne Maufrais, Etienne Simon-Lorière, Maud Vanpeene, Sylvie Van der Werf ,Jean-Philippe Rault |
| EPI_ISL_1821538, EPI_ISL_1821544 | Labo Analyses Med | National Reference Center for Viruses of Respiratory Infections, Institut Pasteur, Paris | Marion Barbet, Sylvie Behillil, Méline Bizard, Angela Brisebarre, Camille Capel, Vincent Enouf, Louise Lefrançois, Frédéric Lemoine, Christophe Malabat, Corinne Maufrais, Etienne Simon-Lorière, Maud Vanpeene, Sylvie Van der Werf ,Sophie Chalmin |
| EPI_ISL_1821545 | Labo Analyses Med | National Reference Center for Viruses of Respiratory Infections, Institut Pasteur, Paris | Marion Barbet, Sylvie Behillil, Méline Bizard, Angela Brisebarre, Camille Capel, Vincent Enouf, Louise Lefrançois, Frédéric Lemoine, Christophe Malabat, Corinne Maufrais, Etienne Simon-Lorière, Maud Vanpeene, Sylvie Van der Werf ,Pierre Pennacino |
| EPI_ISL_1821549 | Hospital | National Reference Center for Viruses of Respiratory Infections, Institut Pasteur, Paris | Marion Barbet, Sylvie Behillil, Méline Bizard, Angela Brisebarre, Camille Capel, Vincent Enouf, Louise Lefrançois, Frédéric Lemoine, Christophe Malabat, Corinne Maufrais, Damien Mornico, Etienne Simon-Lorière, Maud Vanpeene, Sylvie Van der Werf ,LéA Pilorge |
| EPI_ISL_1821550 | Hospital | National Reference Center for Viruses of Respiratory Infections, Institut Pasteur, Paris | Marion Barbet, Sylvie Behillil, Méline Bizard, Angela Brisebarre, Camille Capel, Vincent Enouf, Louise Lefrançois, Frédéric Lemoine, Christophe Malabat, Corinne Maufrais, Etienne Simon-Lorière, Maud Vanpeene, Sylvie Van der Werf ,JéRéMy Jaffre |
| EPI_ISL_1821552, EPI_ISL_1821554 | Hospital | National Reference Center for Viruses of Respiratory Infections, Institut Pasteur, Paris | Marion Barbet, Sylvie Behillil, Méline Bizard, Angela Brisebarre, Camille Capel, Vincent Enouf, Louise Lefrançois, Frédéric Lemoine, Christophe Malabat, Corinne Maufrais, Etienne Simon-Lorière, Maud Vanpeene, Sylvie Van der Werf ,JéRome Guinard |
| EPI_ISL_1821555, EPI_ISL_1821556, EPI_ISL_1821557, EPI_ISL_1821558 | Hospital | National Reference Center for Viruses of Respiratory Infections, Institut Pasteur, Paris | Marion Barbet, Sylvie Behillil, Méline Bizard, Angela Brisebarre, Camille Capel, Vincent Enouf, Louise Lefrançois, Frédéric Lemoine, Christophe Malabat, Corinne Maufrais, Etienne Simon-Lorière, Maud Vanpeene, Sylvie Van der Werf ,FréDéRique Canis |
| EPI_ISL_1821559 | Labo Analyses Med | National Reference Center for Viruses of Respiratory Infections, Institut Pasteur, Paris | Marion Barbet, Sylvie Behillil, Méline Bizard, Angela Brisebarre, Camille Capel, Vincent Enouf, Louise Lefrançois, Frédéric Lemoine, Christophe Malabat, Corinne Maufrais, Etienne Simon-Lorière, Maud Vanpeene, Sylvie Van der Werf ,OphéLie Said-Delattre |
| EPI_ISL_1821560 | Hospital | National Reference Center for Viruses of Respiratory Infections, Institut Pasteur, Paris | Marion Barbet, Sylvie Behillil, Méline Bizard, Angela Brisebarre, Camille Capel, Vincent Enouf, Louise Lefrançois, Frédéric Lemoine, Christophe Malabat, Corinne Maufrais, Etienne Simon-Lorière, Maud Vanpeene, Sylvie Van der Werf ,Nelly Luizy |
| EPI_ISL_1821561 | Labo Analyses Med | National Reference Center for Viruses of Respiratory Infections, Institut Pasteur, Paris | Marion Barbet, Sylvie Behillil, Méline Bizard, Angela Brisebarre, Camille Capel, Vincent Enouf, Louise Lefrançois, Frédéric Lemoine, Christophe Malabat, Corinne Maufrais, Damien Mornico, Etienne Simon-Lorière, Maud Vanpeene, Sylvie Van der Werf ,StéPhane Romand |
| EPI_ISL_1821563 | Labo Analyses Med | National Reference Center for Viruses of Respiratory Infections, Institut Pasteur, Paris | Marion Barbet, Sylvie Behillil, Méline Bizard, Angela Brisebarre, Camille Capel, Vincent Enouf, Louise Lefrançois, Frédéric Lemoine, Christophe Malabat, Corinne Maufrais, Etienne Simon-Lorière, Maud Vanpeene, Sylvie Van der Werf ,Gilles Desvaux |
| EPI_ISL_1821564, EPI_ISL_1821565 | Labo Analyses Med | National Reference Center for Viruses of Respiratory Infections, Institut Pasteur, Paris | Marion Barbet, Sylvie Behillil, Méline Bizard, Angela Brisebarre, Camille Capel, Vincent Enouf, Louise Lefrançois, Frédéric Lemoine, Christophe Malabat, Corinne Maufrais, Etienne Simon-Lorière, Maud Vanpeene, Sylvie Van der Werf ,GréGoire Potiron |
| EPI_ISL_1821566, EPI_ISL_1821567, EPI_ISL_1821568, EPI_ISL_1821569, EPI_ISL_1821570, EPI_ISL_1821571, EPI_ISL_1821572, EPI_ISL_1821573, EPI_ISL_1821574, EPI_ISL_1821575, EPI_ISL_1821576, EPI_ISL_1821577, EPI_ISL_1821578, EPI_ISL_1821579, EPI_ISL_1821580, EPI_ISL_1821581, EPI_ISL_1821582, EPI_ISL_1821583, EPI_ISL_1821584, EPI_ISL_1821585, EPI_ISL_1821586 | | | |
| see above | Labo Analyses Med | National Reference Center for Viruses of Respiratory Infections, Institut Pasteur, Paris | Marion Barbet, Sylvie Behillil, Méline Bizard, Angela Brisebarre, Camille Capel, Vincent Enouf, Louise Lefrançois, Frédéric Lemoine, Christophe Malabat, Corinne Maufrais, Etienne Simon-Lorière, Maud Vanpeene, Sylvie Van der Werf ,Sophie Chalmin |
| EPI_ISL_1821587 | Labo Analyses Med | National Reference Center for Viruses of Respiratory Infections, Institut Pasteur, Paris | Marion Barbet, Sylvie Behillil, Méline Bizard, Angela Brisebarre, Camille Capel, Vincent Enouf, Louise Lefrançois, Frédéric Lemoine, Christophe Malabat, Corinne Maufrais, Etienne Simon-Lorière, Maud Vanpeene, Sylvie Van der Werf ,Sylvaine Bastian |
| EPI_ISL_1821588 | Labo Analyses Med | National Reference Center for Viruses of Respiratory Infections, Institut Pasteur, Paris | Marion Barbet, Sylvie Behillil, Méline Bizard, Angela Brisebarre, Camille Capel, Vincent Enouf, Louise Lefrançois, Frédéric Lemoine, Christophe Malabat, Corinne Maufrais, Emmanuelle Permal, Etienne Simon-Lorière, Maud Vanpeene, Sylvie Van der Werf ,GréGoire Potiron |
| EPI_ISL_1821589 | Labo Analyses Med | National Reference Center for Viruses of Respiratory Infections, Institut Pasteur, Paris | Marion Barbet, Sylvie Behillil, Méline Bizard, Angela Brisebarre, Camille Capel, Vincent Enouf, Louise Lefrançois, Frédéric Lemoine, Christophe Malabat, Corinne Maufrais, Etienne Simon-Lorière, Maud Vanpeene, Sylvie Van der Werf ,Sophie Girard (S) |
| EPI_ISL_1821591 | Hospital | National Reference Center for Viruses of Respiratory Infections, Institut Pasteur, Paris | Marion Barbet, Sylvie Behillil, Méline Bizard, Angela Brisebarre, Camille Capel, Vincent Enouf, Louise Lefrançois, Frédéric Lemoine, Christophe Malabat, Corinne Maufrais, Damien Mornico, Etienne Simon-Lorière, Maud Vanpeene, Sylvie Van der Werf ,Laurence Louvet |
| EPI_ISL_1821592 | Hospital | National Reference Center for Viruses of Respiratory Infections, Institut Pasteur, Paris | Marion Barbet, Sylvie Behillil, Méline Bizard, Angela Brisebarre, Camille Capel, Vincent Enouf, Louise Lefrançois, Frédéric Lemoine, Christophe Malabat, Corinne Maufrais, Emmanuelle Permal, Etienne Simon-Lorière, Maud Vanpeene, Sylvie Van der Werf ,Guillaume Gautier |
| EPI_ISL_1821606, EPI_ISL_1821607 | Hospital | National Reference Center for Viruses of Respiratory Infections, Institut Pasteur, Paris | Marion Barbet, Sylvie Behillil, Méline Bizard, Angela Brisebarre, Camille Capel, Vincent Enouf, Louise Lefrançois, Frédéric Lemoine, Christophe Malabat, Corinne Maufrais, Damien Mornico, Etienne Simon-Lorière, Maud Vanpeene, Sylvie Van der Werf ,LéA Pilorge |
| EPI_ISL_1821608 | Hospital | National Reference Center for Viruses of Respiratory Infections, Institut Pasteur, Paris | Marion Barbet, Sylvie Behillil, Méline Bizard, Angela Brisebarre, Camille Capel, Vincent Enouf, Louise Lefrançois, Frédéric Lemoine, Christophe Malabat, Corinne Maufrais, Damien Mornico, Etienne Simon-Lorière, Maud Vanpeene, Sylvie Van der Werf ,Haciba Moudjahed |
| EPI_ISL_1821609 | Hospital | National Reference Center for Viruses of Respiratory Infections, Institut Pasteur, Paris | Marion Barbet, Sylvie Behillil, Méline Bizard, Angela Brisebarre, Camille Capel, Vincent Enouf, Louise Lefrançois, Frédéric Lemoine, Christophe Malabat, Corinne Maufrais, Damien Mornico, Etienne Simon-Lorière, Maud Vanpeene, Sylvie Van der Werf ,Pascale Martres |
| EPI_ISL_1821610, EPI_ISL_1821611 | Labo Analyses Med | National Reference Center for Viruses of Respiratory Infections, Institut Pasteur, Paris | Marion Barbet, Sylvie Behillil, Méline Bizard, Angela Brisebarre, Camille Capel, Vincent Enouf, Louise Lefrançois, Frédéric Lemoine, Christophe Malabat, Corinne Maufrais, Damien Mornico, Etienne Simon-Lorière, Maud Vanpeene, Sylvie Van der Werf ,Sophie Chalmin |
| EPI_ISL_1821612 | Labo Analyses Med | National Reference Center for Viruses of Respiratory Infections, Institut Pasteur, Paris | Marion Barbet, Sylvie Behillil, Méline Bizard, Angela Brisebarre, Camille Capel, Vincent Enouf, Louise Lefrançois, Frédéric Lemoine, Christophe Malabat, Corinne Maufrais, Etienne Simon-Lorière, Maud Vanpeene, Sylvie Van der Werf ,Sophie Chalmin |
| EPI_ISL_1821613, EPI_ISL_1821614, EPI_ISL_1821615 | Labo Analyses Med | National Reference Center for Viruses of Respiratory Infections, Institut Pasteur, Paris | Marion Barbet, Sylvie Behillil, Méline Bizard, Angela Brisebarre, Camille Capel, Vincent Enouf, Louise Lefrançois, Frédéric Lemoine, Christophe Malabat, Corinne Maufrais, Etienne Simon-Lorière, Maud Vanpeene, Sylvie Van der Werf ,Franck Ennouchi |
| EPI_ISL_1821619, EPI_ISL_1821620 | Hospital | National Reference Center for Viruses of Respiratory Infections, Institut Pasteur, Paris | Marion Barbet, Sylvie Behillil, Méline Bizard, Angela Brisebarre, Camille Capel, Vincent Enouf, Louise Lefrançois, Frédéric Lemoine, Christophe Malabat, Corinne Maufrais, Etienne Simon-Lorière, Maud Vanpeene, Sylvie Van der Werf ,FréDéRique Canis |
| EPI_ISL_1821621 | Hospital | National Reference Center for Viruses of Respiratory Infections, Institut Pasteur, Paris | Marion Barbet, Sylvie Behillil, Méline Bizard, Angela Brisebarre, Camille Capel, Vincent Enouf, Louise Lefrançois, Frédéric Lemoine, Christophe Malabat, Corinne Maufrais, Etienne Simon-Lorière, Maud Vanpeene, Sylvie Van der Werf ,Haciba Moudjahed |
| EPI_ISL_1821631, EPI_ISL_1821632 | Labo Analyses Med | National Reference Center for Viruses of Respiratory | Marion Barbet, Sylvie Behillil, Méline Bizard, Angela Brisebarre, Camille Capel, Vincent Enouf, Louise Lefrançois, Frédéric Lemoine, Christophe Malabat, |

| | | | |
|---|---|---|---|
| | | Infections, Institut Pasteur, Paris | Corinne Maufrais, Etienne Simon-Lorière, Maud Vanpeene, Sylvie Van der Werf ,Dominique Rousset |
| EPI_ISL_1821643, EPI_ISL_1821644, EPI_ISL_1821645, EPI_ISL_1821646, EPI_ISL_1821647, EPI_ISL_1821648, EPI_ISL_1821649, EPI_ISL_1821650, EPI_ISL_1821651 | Labo Analyses Med | National Reference Center for Viruses of Respiratory Infections, Institut Pasteur, Paris | Marion Barbet, Sylvie Behillil, Méline Bizard, Angela Brisebarre, Camille Capel, Vincent Enouf, Louise Lefrançois, Frédéric Lemoine, Christophe Malabat, Corinne Maufrais, Emmanuelle Permal, Etienne Simon-Lorière, Maud Vanpeene, Sylvie Van der Werf ,Dominique Rousset |
| EPI_ISL_1821652, EPI_ISL_1821654 | Labo Analyses Med | National Reference Center for Viruses of Respiratory Infections, Institut Pasteur, Paris | Marion Barbet, Sylvie Behillil, Méline Bizard, Angela Brisebarre, Camille Capel, Vincent Enouf, Louise Lefrançois, Frédéric Lemoine, Christophe Malabat, Corinne Maufrais, Amaury Vaysse, Etienne Simon-Lorière, Maud Vanpeene, Sylvie Van der Werf ,Dominique Rousset |
| EPI_ISL_1821659 | Labo Analyses Med | National Reference Center for Viruses of Respiratory Infections, Institut Pasteur, Paris | Marion Barbet, Sylvie Behillil, Méline Bizard, Angela Brisebarre, Camille Capel, Vincent Enouf, Louise Lefrançois, Frédéric Lemoine, Christophe Malabat, Corinne Maufrais, Amaury Vaysse, Etienne Simon-Lorière, Maud Vanpeene, Sylvie Van der Werf ,Marianne Burgard |
| EPI_ISL_1821752, EPI_ISL_1821753, EPI_ISL_1821754, EPI_ISL_1821755, EPI_ISL_1821756, EPI_ISL_1821757, EPI_ISL_1821758, EPI_ISL_1821759, EPI_ISL_1821760 | Laboratorio di Riferimento Regionale della Sicilia Occidentale per l'Emergenza COVID-19 | Laboratorio di Riferimento Regionale della Sicilia Occidentale per l'Emergenza COVID-19 | Fabio Tramuto, Stefano Reale, Carmelo Massimo Maida, Silvia Scibetta, Daniela Di Naro, Giulia Randazzo, Walter Mazzucco, Giorgio Graziano, Vincenzo Restivo, Claudio Costantino, Francesco Vitale |
| EPI_ISL_1821795, EPI_ISL_1821803, EPI_ISL_1821811, EPI_ISL_1821816, EPI_ISL_1821818, EPI_ISL_1821824, EPI_ISL_1821825, EPI_ISL_1821826, EPI_ISL_1821827 | GH A.CHENEVIER-H.MONDOR | Department of Virology, Henri Mondor University Hospital, Assistance Publique Hôpitaux de Paris, Université Paris-Est Créteil, INSERM U955 | Christophe Rodriguez, Slim Fourati, Vanessa Demontant, Guillaume Gricourt, Melissa N'Debi, Alexandre Soulier, Elisabeth Trawinski, Jean-Michel Pawlotsky |
| EPI_ISL_1821834 | GH JOFFRE DUPUYTREN | Department of Virology, Henri Mondor University Hospital, Assistance Publique Hôpitaux de Paris, Université Paris-Est Créteil, INSERM U955 | Christophe Rodriguez, Slim Fourati, Vanessa Demontant, Guillaume Gricourt, Melissa N'Debi, Alexandre Soulier, Elisabeth Trawinski, Jean-Michel Pawlotsky |
| EPI_ISL_1821842, EPI_ISL_1821845, EPI_ISL_1821847, EPI_ISL_1821848, EPI_ISL_1821852, EPI_ISL_1821854, EPI_ISL_1821855, EPI_ISL_1821856, EPI_ISL_1821857, EPI_ISL_1821860, EPI_ISL_1821863, EPI_ISL_1821866, EPI_ISL_1821871 | | | |
| see above | GH A.CHENEVIER-H.MONDOR | Department of Virology, Henri Mondor University Hospital, Assistance Publique Hôpitaux de Paris, Université Paris-Est Créteil, INSERM U955 | Christophe Rodriguez, Slim Fourati, Vanessa Demontant, Guillaume Gricourt, Melissa N'Debi, Alexandre Soulier, Elisabeth Trawinski, Jean-Michel Pawlotsky |
| EPI_ISL_1821883, EPI_ISL_1821884 | Ipoh Public Health Laboratory (MKAI), Ministry of Health Malaysia | Institute for Medical Research, Infectious Disease Research Centre, National Institutes of Health, Ministry of Health Malaysia | Suppiah J, Kamel K, Mohd Zawawi Z, Thayan R |
| EPI_ISL_1822600 | Baylor Scott & White-Irving | Baylor Scott & White - Temple | Ari Rao, Linden Morales, Kimberly Walker, Marcus Volz, Shelby Hendrickson |
| EPI_ISL_1824148, EPI_ISL_1824150 | The Ohio State University Applied Microbiology Services Laboratory | The Ohio State University Applied Microbiology Services Laboratory | Seth A. Faith PhD |
| EPI_ISL_1824179, EPI_ISL_1824180, EPI_ISL_1824181, EPI_ISL_1824182, EPI_ISL_1824183, EPI_ISL_1824184, EPI_ISL_1824185, EPI_ISL_1824186, EPI_ISL_1824187, EPI_ISL_1824188 | Grupo de Investigación en Enfermedades Tropicales del Ejército (GINETEJ), Laboratorio de Referencia e Investigación, Dirección de Sanidad Ejército, Bogotá, Colombia | Centro de Investigaciones en Microbiología y Biotecnología-UR (CIMBIUR), Facultad de Ciencias Naturales, Universidad del Rosario, Bogotá, Colombia | Sergio Castañeda, Nathalia Ballesteros, Marina Muñoz, Luz H. Patiño, Claudia Méndez, Carolina Oliveros, Julie Pérez, Lorena Albarracín, Elizabeth K. Márquez, María Teresa Alvarado, Frank de los Santos Ortiz, Yanira Romero, Camilo A. Correa-Cárdenas, Maria Clara Duque, Sergio Gutíerrez-Riveros, Juan David Ramírez |
| EPI_ISL_1824596, EPI_ISL_1824597, EPI_ISL_1824599, EPI_ISL_1824600 | AZ Klina | AZ Klina | Carl Vael - Lynsey Berckmans |
| EPI_ISL_1824604 | PRVKP FK UI | National Institute of Health Research and Development | Vivi Setiawaty, Hana Apsari Pawestri, Subangkit, Kartika Dewi Puspa, Arie Ardiansyah Nugraha, Hartanti Dian Ikawati, Krisna Pangesti, Nelly Puspandari |
| EPI_ISL_1824615 | RSUD dr. Hasri Ainun Habibie Gorontalo | National Institute of Health Research and Development | Hana Apsari Pawestri, Subangkit, Kartika Dewi Puspa, Arie Ardiansyah Nugraha, Hartanti Dian Ikawati, Krisna Pangesti, Nelly Puspandari, Vivi Setiawaty |
| EPI_ISL_1824619, EPI_ISL_1824623, EPI_ISL_1824637, EPI_ISL_1824638, EPI_ISL_1824639, EPI_ISL_1824679, EPI_ISL_1824681 | AZ Klina | AZ Klina | Carl Vael - Lynsey Berckmans |
| EPI_ISL_1824682, EPI_ISL_1824683, EPI_ISL_1824684 | VITHAS Lab | Microbiology Department. Complexo Hospitalario Universitario de Vigo | Camacho AT, Perez S, Regueiro B |
| EPI_ISL_1824685, EPI_ISL_1824698, EPI_ISL_1824699 | AZ Klina | AZ Klina | Carl Vael - Lynsey Berckmans |
| EPI_ISL_1831675, EPI_ISL_1831677, EPI_ISL_1831678, EPI_ISL_1831680, EPI_ISL_1831681, EPI_ISL_1831683, EPI_ISL_1831684, EPI_ISL_1831687, EPI_ISL_1831691, EPI_ISL_1831692, EPI_ISL_1831693 | | | |
| see above | Althaia. Xarxa Assistencial Universitària de Manresa | IrsiCaixa | Marc Noguera-Julian, Mariona Parera, Maria Casadellà, Pilar Armengol, Francesc Catala-Moll, Roger Paredes, Bonaventura Clotet Gloria Trujillo, Rafel Perez Vidal, Jaume Trape Pujol, Carolina Gonzalez Fernandez, Miquel Micó, Roger Paredes, Eulalia Grau, Bonaventura Clotet |
| EPI_ISL_1840243, EPI_ISL_1840245, EPI_ISL_1840247 | Pro-Vitam Diagnostics and Research Laboratory | Pro-Vitam Diagnostics and Research Laboratory | Szilard N. Fejer, Istvan Horvath, Monika Korodi, Kinga Rakosi, Zsuzsanna Jenei |
| EPI_ISL_1840693 | Baylor Scott & White-Temple | Baylor Scott & White-Temple | Ari Rao, Linden Morales, Kimberly Walker, Marcus Volz, Shelby Hendrickson |
| EPI_ISL_1840903, EPI_ISL_1840905, EPI_ISL_1840907, EPI_ISL_1840908, EPI_ISL_1840912, EPI_ISL_1840914, EPI_ISL_1840915, EPI_ISL_1840916, EPI_ISL_1840918, EPI_ISL_1840920, EPI_ISL_1840922, EPI_ISL_1840923, EPI_ISL_1840924, EPI_ISL_1840925, EPI_ISL_1840927, EPI_ISL_1840928, EPI_ISL_1840929, EPI_ISL_1840931, EPI_ISL_1840932, EPI_ISL_1840933, EPI_ISL_1840934, EPI_ISL_1840937, EPI_ISL_1840938, EPI_ISL_1840940, EPI_ISL_1840941, EPI_ISL_1840942, EPI_ISL_1840946, EPI_ISL_1840947, EPI_ISL_1840948, EPI_ISL_1840949, EPI_ISL_1840950, EPI_ISL_1840951, EPI_ISL_1840952, EPI_ISL_1840955, EPI_ISL_1840956, EPI_ISL_1840957, EPI_ISL_1840959, EPI_ISL_1840960, EPI_ISL_1840963, EPI_ISL_1840965, EPI_ISL_1840966, EPI_ISL_1840967, EPI_ISL_1840968 | | | |
| see above | Austrian Agency for Health and Food Safety (AGES) | Bergthaler laboratory, CeMM Research Center for Molecular Medicine of the Austrian Academy of Sciences | Lukas Endler, Anna Schedl, Fabian Amman, Petr Triska, Thomas Penz, Benedikt Agerer, Maelle Le Moing, Michael Schuster, Bekir Erguner, Jan Laine, Martin Senekowitsch, Christoph Bock, Andreas Bergthaler |
| EPI_ISL_1841083 | SIESP CHIETI - DRIVE IN LANCIANO | Istituto Zooprofilattico Sperimentale dell'Abruzzo e Molise "G. Caporale" | Lorusso A, Marcacci M, Di Domenico M, Ancora M, Curini V, Di Lollo Valeria, Mangone I, Rinaldi A, Delli Compagni E, Scialabba S, Caporale M, Di Pasquale A, Cammà C, Puglia I, Calistri P, Savini G |
| EPI_ISL_1841084, EPI_ISL_1841085, EPI_ISL_1841086 | SIESP DIPARTIMENTO DI PREVENZIONE TERAMO | Istituto Zooprofilattico Sperimentale dell'Abruzzo e Molise "G. Caporale" | Lorusso A, Marcacci M, Di Domenico M, Ancora M, Curini V, Di Lollo Valeria, Mangone I, Rinaldi A, Delli Compagni E, Scialabba S, Caporale M, Di Pasquale A, Cammà C, Puglia I, Calistri P, Savini G |
| EPI_ISL_1841087 | SIESP CHIETI - DRIVE IN LANCIANO | Istituto Zooprofilattico Sperimentale dell'Abruzzo e Molise "G. Caporale" | Lorusso A, Marcacci M, Di Domenico M, Ancora M, Curini V, Di Lollo Valeria, Mangone I, Rinaldi A, Delli Compagni E, Scialabba S, Caporale M, Di Pasquale A, Cammà C, Puglia I, Calistri P, Savini G |
| EPI_ISL_1841088 | PRESIDIO TERRITORIALE CASOLI | Istituto Zooprofilattico Sperimentale dell'Abruzzo e Molise "G. Caporale" | Lorusso A, Marcacci M, Di Domenico M, Ancora M, Curini V, Di Lollo Valeria, Mangone I, Rinaldi A, Delli Compagni E, Scialabba S, Caporale M, Di Pasquale A, Cammà C, Puglia I, Calistri P, Savini G |
| EPI_ISL_1841089 | USCA Tagliacozzo | Istituto Zooprofilattico Sperimentale dell'Abruzzo e Molise "G. Caporale" | Lorusso A, Marcacci M, Di Domenico M, Ancora M, Curini V, Di Lollo Valeria, Mangone I, Rinaldi A, Delli Compagni E, Scialabba S, Caporale M, Di Pasquale A, Cammà C, Puglia I, Calistri P, Savini G |
| EPI_ISL_1841090 | SIESP DIPARTIMENTO DI PREVENZIONE TERAMO | Istituto Zooprofilattico Sperimentale dell'Abruzzo e Molise "G. Caporale" | Lorusso A, Marcacci M, Di Domenico M, Ancora M, Curini V, Di Lollo Valeria, Mangone I, Rinaldi A, Delli Compagni E, Scialabba S, Caporale M, Di Pasquale A, Cammà C, Puglia I, Calistri P, Savini G |

| | | | |
|---|---|---|---|
| EPI_ISL_1841091 | DIP. PREV. AVEZZANO SERVIZIO DI IGIENE EPIDEMIOLOGIA E SANITA' PUBBLICA | Istituto Zooprofilattico Sperimentale dell'Abruzzo e Molise "G. Caporale" | Lorusso A, Marcacci M, Di Domenico M, Ancora M, Curini V, Di Lollo Valeria, Mangone I, Rinaldi A, Delli Compagni E, Scialabba S, Caporale M, Di Pasquale A, Cammà C, Puglia I, Calistri P, Savini G |
| EPI_ISL_1841092 | U.O.S.D. Medici Competenti PO Avezzano | Istituto Zooprofilattico Sperimentale dell'Abruzzo e Molise "G. Caporale" | Lorusso A, Marcacci M, Di Domenico M, Ancora M, Curini V, Di Lollo Valeria, Mangone I, Rinaldi A, Delli Compagni E, Scialabba S, Caporale M, Di Pasquale A, Cammà C, Puglia I, Calistri P, Savini G |
| EPI_ISL_1841093 | SIESP CHIETI - DRIVE IN LANCIANO | Istituto Zooprofilattico Sperimentale dell'Abruzzo e Molise "G. Caporale" | Lorusso A, Marcacci M, Di Domenico M, Ancora M, Curini V, Di Lollo Valeria, Mangone I, Rinaldi A, Delli Compagni E, Scialabba S, Caporale M, Di Pasquale A, Cammà C, Puglia I, Calistri P, Savini G |
| EPI_ISL_1841094, EPI_ISL_1841095 | SIESP CHIETI - DRIVE IN ORTONA | Istituto Zooprofilattico Sperimentale dell'Abruzzo e Molise "G. Caporale" | Lorusso A, Marcacci M, Di Domenico M, Ancora M, Curini V, Di Lollo Valeria, Mangone I, Rinaldi A, Delli Compagni E, Scialabba S, Caporale M, Di Pasquale A, Cammà C, Puglia I, Calistri P, Savini G |
| EPI_ISL_1841096, EPI_ISL_1841097, EPI_ISL_1841098, EPI_ISL_1841099 | DIP. PREV. AVEZZANO SERVIZIO DI IGIENE EPIDEMIOLOGIA E SANITA' PUBBLICA | Istituto Zooprofilattico Sperimentale dell'Abruzzo e Molise "G. Caporale" | Lorusso A, Marcacci M, Di Domenico M, Ancora M, Curini V, Di Lollo Valeria, Mangone I, Rinaldi A, Delli Compagni E, Scialabba S, Caporale M, Di Pasquale A, Cammà C, Puglia I, Calistri P, Savini G |
| EPI_ISL_1841100 | OSPEDALE SAN SALVATORE - MEDICINA DI LABORATORIO | Istituto Zooprofilattico Sperimentale dell'Abruzzo e Molise "G. Caporale" | Lorusso A, Marcacci M, Di Domenico M, Ancora M, Curini V, Di Lollo Valeria, Mangone I, Rinaldi A, Delli Compagni E, Scialabba S, Caporale M, Di Pasquale A, Cammà C, Puglia I, Calistri P, Savini G |
| EPI_ISL_1841101, EPI_ISL_1841102, EPI_ISL_1841103, EPI_ISL_1841104, EPI_ISL_1841105, EPI_ISL_1841106, EPI_ISL_1841107, EPI_ISL_1841108, EPI_ISL_1841109, EPI_ISL_1841110, EPI_ISL_1841111, EPI_ISL_1841112, EPI_ISL_1841113, EPI_ISL_1841114, EPI_ISL_1841115, EPI_ISL_1841116, EPI_ISL_1841117, EPI_ISL_1841118, EPI_ISL_1841119, EPI_ISL_1841120, EPI_ISL_1841121 | | | |
| see above | SIESP DIPARTIMENTO DI PREVENZIONE TERAMO | Istituto Zooprofilattico Sperimentale dell'Abruzzo e Molise "G. Caporale" | Lorusso A, Marcacci M, Di Domenico M, Ancora M, Curini V, Di Lollo Valeria, Mangone I, Rinaldi A, Delli Compagni E, Scialabba S, Caporale M, Di Pasquale A, Cammà C, Puglia I, Calistri P, Savini G |
| EPI_ISL_1841122 | SIESP CHIETI - DRIVE IN CHIETI | Istituto Zooprofilattico Sperimentale dell'Abruzzo e Molise "G. Caporale" | Lorusso A, Marcacci M, Di Domenico M, Ancora M, Curini V, Di Lollo Valeria, Mangone I, Rinaldi A, Delli Compagni E, Scialabba S, Caporale M, Di Pasquale A, Cammà C, Puglia I, Calistri P, Savini G |
| EPI_ISL_1841123 | SIESP CHIETI - DRIVE IN ORTONA | Istituto Zooprofilattico Sperimentale dell'Abruzzo e Molise "G. Caporale" | Lorusso A, Marcacci M, Di Domenico M, Ancora M, Curini V, Di Lollo Valeria, Mangone I, Rinaldi A, Delli Compagni E, Scialabba S, Caporale M, Di Pasquale A, Cammà C, Puglia I, Calistri P, Savini G |
| EPI_ISL_1841127, EPI_ISL_1841128 | SIESP SULMONA | Istituto Zooprofilattico Sperimentale dell'Abruzzo e Molise "G. Caporale" | Lorusso A, Marcacci M, Di Domenico M, Ancora M, Curini V, Di Lollo Valeria, Mangone I, Rinaldi A, Delli Compagni E, Scialabba S, Caporale M, Di Pasquale A, Cammà C, Puglia I, Calistri P, Savini G |
| EPI_ISL_1841129 | SIESP DIPARTIMENTO DI PREVENZIONE CHIETI | Istituto Zooprofilattico Sperimentale dell'Abruzzo e Molise "G. Caporale" | Lorusso A, Marcacci M, Di Domenico M, Ancora M, Curini V, Di Lollo Valeria, Mangone I, Rinaldi A, Delli Compagni E, Scialabba S, Caporale M, Di Pasquale A, Cammà C, Puglia I, Calistri P, Savini G |
| EPI_ISL_1841130, EPI_ISL_1841131 | SIESP CHIETI - DRIVE IN LANCIANO | Istituto Zooprofilattico Sperimentale dell'Abruzzo e Molise "G. Caporale" | Lorusso A, Marcacci M, Di Domenico M, Ancora M, Curini V, Di Lollo Valeria, Mangone I, Rinaldi A, Delli Compagni E, Scialabba S, Caporale M, Di Pasquale A, Cammà C, Puglia I, Calistri P, Savini G |
| EPI_ISL_1841132, EPI_ISL_1841133, EPI_ISL_1841134 | SIESP DIPARTIMENTO DI PREVENZIONE TERAMO | Istituto Zooprofilattico Sperimentale dell'Abruzzo e Molise "G. Caporale" | Lorusso A, Marcacci M, Di Domenico M, Ancora M, Curini V, Di Lollo Valeria, Mangone I, Rinaldi A, Delli Compagni E, Scialabba S, Caporale M, Di Pasquale A, Cammà C, Puglia I, Calistri P, Savini G |
| EPI_ISL_1841135 | SIESP CHIETI - DRIVE IN VASTO | Istituto Zooprofilattico Sperimentale dell'Abruzzo e Molise "G. Caporale" | Lorusso A, Marcacci M, Di Domenico M, Ancora M, Curini V, Di Lollo Valeria, Mangone I, Rinaldi A, Delli Compagni E, Scialabba S, Caporale M, Di Pasquale A, Cammà C, Puglia I, Calistri P, Savini G |
| EPI_ISL_1841136 | USCA Castel di Sangro | Istituto Zooprofilattico Sperimentale dell'Abruzzo e Molise "G. Caporale" | Lorusso A, Marcacci M, Di Domenico M, Ancora M, Curini V, Di Lollo Valeria, Mangone I, Rinaldi A, Delli Compagni E, Scialabba S, Caporale M, Di Pasquale A, Cammà C, Puglia I, Calistri P, Savini G |
| EPI_ISL_1841137, EPI_ISL_1841138, EPI_ISL_1841139 | SIESP CHIETI - DRIVE IN CHIETI | Istituto Zooprofilattico Sperimentale dell'Abruzzo e Molise "G. Caporale" | Lorusso A, Marcacci M, Di Domenico M, Ancora M, Curini V, Di Lollo Valeria, Mangone I, Rinaldi A, Delli Compagni E, Scialabba S, Caporale M, Di Pasquale A, Cammà C, Puglia I, Calistri P, Savini G |
| EPI_ISL_1841140 | USCA Castel di Sangro | Istituto Zooprofilattico Sperimentale dell'Abruzzo e Molise "G. Caporale" | Lorusso A, Marcacci M, Di Domenico M, Ancora M, Curini V, Di Lollo Valeria, Mangone I, Rinaldi A, Delli Compagni E, Scialabba S, Caporale M, Di Pasquale A, Cammà C, Puglia I, Calistri P, Savini G |
| EPI_ISL_1841141 | SIESP CHIETI - DRIVE IN LANCIANO | Istituto Zooprofilattico Sperimentale dell'Abruzzo e Molise "G. Caporale" | Lorusso A, Marcacci M, Di Domenico M, Ancora M, Curini V, Di Lollo Valeria, Mangone I, Rinaldi A, Delli Compagni E, Scialabba S, Caporale M, Di Pasquale A, Cammà C, Puglia I, Calistri P, Savini G |
| EPI_ISL_1841142, EPI_ISL_1841143, EPI_ISL_1841144, EPI_ISL_1841145, EPI_ISL_1841146, EPI_ISL_1841147, EPI_ISL_1841148, EPI_ISL_1841149, EPI_ISL_1841150, EPI_ISL_1841151, EPI_ISL_1841153 | | | |
| see above | SIESP DIPARTIMENTO DI PREVENZIONE TERAMO | Istituto Zooprofilattico Sperimentale dell'Abruzzo e Molise "G. Caporale" | Lorusso A, Marcacci M, Di Domenico M, Ancora M, Curini V, Di Lollo Valeria, Mangone I, Rinaldi A, Delli Compagni E, Scialabba S, Caporale M, Di Pasquale A, Cammà C, Puglia I, Calistri P, Savini G |
| EPI_ISL_1841154 | SIESP CHIETI - DRIVE IN VASTO | Istituto Zooprofilattico Sperimentale dell'Abruzzo e Molise "G. Caporale" | Lorusso A, Marcacci M, Di Domenico M, Ancora M, Curini V, Di Lollo Valeria, Mangone I, Rinaldi A, Delli Compagni E, Scialabba S, Caporale M, Di Pasquale A, Cammà C, Puglia I, Calistri P, Savini G |
| EPI_ISL_1841155, EPI_ISL_1841156, EPI_ISL_1841157 | SIESP DIPARTIMENTO DI PREVENZIONE TERAMO | Istituto Zooprofilattico Sperimentale dell'Abruzzo e Molise "G. Caporale" | Lorusso A, Marcacci M, Di Domenico M, Ancora M, Curini V, Di Lollo Valeria, Mangone I, Rinaldi A, Delli Compagni E, Scialabba S, Caporale M, Di Pasquale A, Cammà C, Puglia I, Calistri P, Savini G |
| EPI_ISL_1841158 | DIP. PREV. AVEZZANO SERVIZIO DI IGIENE EPIDEMIOLOGIA E SANITA' PUBBLICA | Istituto Zooprofilattico Sperimentale dell'Abruzzo e Molise "G. Caporale" | Lorusso A, Marcacci M, Di Domenico M, Ancora M, Curini V, Di Lollo Valeria, Mangone I, Rinaldi A, Delli Compagni E, Scialabba S, Caporale M, Di Pasquale A, Cammà C, Puglia I, Calistri P, Savini G |
| EPI_ISL_1841159 | USCA Tagliacozzo | Istituto Zooprofilattico Sperimentale dell'Abruzzo e Molise "G. Caporale" | Lorusso A, Marcacci M, Di Domenico M, Ancora M, Curini V, Di Lollo Valeria, Mangone I, Rinaldi A, Delli Compagni E, Scialabba S, Caporale M, Di Pasquale A, Cammà C, Puglia I, Calistri P, Savini G |
| EPI_ISL_1841160 | SIESP CHIETI - DRIVE IN GISSI | Istituto Zooprofilattico Sperimentale dell'Abruzzo e Molise "G. Caporale" | Lorusso A, Marcacci M, Di Domenico M, Ancora M, Curini V, Di Lollo Valeria, Mangone I, Rinaldi A, Delli Compagni E, Scialabba S, Caporale M, Di Pasquale A, Cammà C, Puglia I, Calistri P, Savini G |
| EPI_ISL_1841161, EPI_ISL_1841162 | SIESP DIPARTIMENTO DI PREVENZIONE CHIETI | Istituto Zooprofilattico Sperimentale dell'Abruzzo e Molise "G. Caporale" | Lorusso A, Marcacci M, Di Domenico M, Ancora M, Curini V, Di Lollo Valeria, Mangone I, Rinaldi A, Delli Compagni E, Scialabba S, Caporale M, Di Pasquale A, Cammà C, Puglia I, Calistri P, Savini G |
| EPI_ISL_1841163 | SIESP CHIETI - DRIVE IN ORTONA | Istituto Zooprofilattico Sperimentale dell'Abruzzo e Molise "G. Caporale" | Lorusso A, Marcacci M, Di Domenico M, Ancora M, Curini V, Di Lollo Valeria, Mangone I, Rinaldi A, Delli Compagni E, Scialabba S, Caporale M, Di Pasquale A, Cammà C, Puglia I, Calistri P, Savini G |
| EPI_ISL_1841164, EPI_ISL_1841165, EPI_ISL_1841166, EPI_ISL_1841167, EPI_ISL_1841168, EPI_ISL_1841169 | SIESP DIPARTIMENTO DI PREVENZIONE TERAMO | Istituto Zooprofilattico Sperimentale dell'Abruzzo e Molise "G. Caporale" | Lorusso A, Marcacci M, Di Domenico M, Ancora M, Curini V, Di Lollo Valeria, Mangone I, Rinaldi A, Delli Compagni E, Scialabba S, Caporale M, Di Pasquale A, Cammà C, Puglia I, Calistri P, Savini G |
| EPI_ISL_1841170 | SIESP CHIETI - DRIVE IN CHIETI | Istituto Zooprofilattico Sperimentale dell'Abruzzo e Molise "G. Caporale" | Lorusso A, Marcacci M, Di Domenico M, Ancora M, Curini V, Di Lollo Valeria, Mangone I, Rinaldi A, Delli Compagni E, Scialabba S, Caporale M, Di Pasquale A, Cammà C, Puglia I, Calistri P, Savini G |
| EPI_ISL_1841171, EPI_ISL_1841172 | USCA Tagliacozzo | Istituto Zooprofilattico Sperimentale dell'Abruzzo e Molise "G. Caporale" | Lorusso A, Marcacci M, Di Domenico M, Ancora M, Curini V, Di Lollo Valeria, Mangone I, Rinaldi A, Delli Compagni E, Scialabba S, Caporale M, Di Pasquale A, Cammà C, Puglia I, Calistri P, Savini G |
| EPI_ISL_1841173 | DIP. PREV. AVEZZANO SERVIZIO DI IGIENE EPIDEMIOLOGIA E SANITA' PUBBLICA | Istituto Zooprofilattico Sperimentale dell'Abruzzo e Molise "G. Caporale" | Lorusso A, Marcacci M, Di Domenico M, Ancora M, Curini V, Di Lollo Valeria, Mangone I, Rinaldi A, Delli Compagni E, Scialabba S, Caporale M, Di Pasquale A, Cammà C, Puglia I, Calistri P, Savini G |
| EPI_ISL_1841174 | SIESP DIPARTIMENTO DI PREVENZIONE TERAMO | Istituto Zooprofilattico Sperimentale dell'Abruzzo e Molise "G. Caporale" | Lorusso A, Marcacci M, Di Domenico M, Ancora M, Curini V, Di Lollo Valeria, Mangone I, Rinaldi A, Delli Compagni E, Scialabba S, Caporale M, Di Pasquale A, Cammà C, Puglia I, Calistri P, Savini G |
| EPI_ISL_1841175 | SIESP CHIETI - DRIVE IN LANCIANO | Istituto Zooprofilattico Sperimentale dell'Abruzzo e Molise "G. Caporale" | Lorusso A, Marcacci M, Di Domenico M, Ancora M, Curini V, Di Lollo Valeria, Mangone I, Rinaldi A, Delli Compagni E, Scialabba S, Caporale M, Di Pasquale A, Cammà C, Puglia I, Calistri P, Savini G |
| EPI_ISL_1841176, EPI_ISL_1841177, EPI_ISL_1841178, EPI_ISL_1841179, | SIESP DIPARTIMENTO DI PREVENZIONE TERAMO | Istituto Zooprofilattico Sperimentale dell'Abruzzo e Molise "G. Caporale" | Lorusso A, Marcacci M, Di Domenico M, Ancora M, Curini V, Di Lollo Valeria, Mangone I, Rinaldi A, Delli Compagni E, Scialabba S, Caporale M, Di Pasquale A, Cammà C, Puglia I, Calistri P, Savini G |

| | | | |
|---|---|---|---|
| EPI_ISL_1841180, EPI_ISL_1841181 | | | |
| EPI_ISL_1841182 | SIESP SULMONA | Istituto Zooprofilattico Sperimentale dell'Abruzzo e Molise "G. Caporale" | Lorusso A, Marcacci M, Di Domenico M, Ancora M, Curini V, Di Lollo Valeria, Mangone I, Rinaldi A, Delli Compagni E, Scialabba S, Caporale M, Di Pasquale A, Cammà C, Puglia I, Calistri P, Savini G |
| EPI_ISL_1841183 | SIESP DIPARTIMENTO DI PREVENZIONE TERAMO | Istituto Zooprofilattico Sperimentale dell'Abruzzo e Molise "G. Caporale" | Lorusso A, Marcacci M, Di Domenico M, Ancora M, Curini V, Di Lollo Valeria, Mangone I, Rinaldi A, Delli Compagni E, Scialabba S, Caporale M, Di Pasquale A, Cammà C, Puglia I, Calistri P, Savini G |
| EPI_ISL_1841184 | DIP. PREV. AVEZZANO SERVIZIO DI IGIENE EPIDEMIOLOGIA E SANITA' PUBBLICA | Istituto Zooprofilattico Sperimentale dell'Abruzzo e Molise "G. Caporale" | Lorusso A, Marcacci M, Di Domenico M, Ancora M, Curini V, Di Lollo Valeria, Mangone I, Rinaldi A, Delli Compagni E, Scialabba S, Caporale M, Di Pasquale A, Cammà C, Puglia I, Calistri P, Savini G |
| EPI_ISL_1841185, EPI_ISL_1841186, EPI_ISL_1841187 | SIESP DIPARTIMENTO DI PREVENZIONE TERAMO | Istituto Zooprofilattico Sperimentale dell'Abruzzo e Molise "G. Caporale" | Lorusso A, Marcacci M, Di Domenico M, Ancora M, Curini V, Di Lollo Valeria, Mangone I, Rinaldi A, Delli Compagni E, Scialabba S, Caporale M, Di Pasquale A, Cammà C, Puglia I, Calistri P, Savini G |
| EPI_ISL_1841188 | SIESP CHIETI - DRIVE IN LANCIANO | Istituto Zooprofilattico Sperimentale dell'Abruzzo e Molise "G. Caporale" | Lorusso A, Marcacci M, Di Domenico M, Ancora M, Curini V, Di Lollo Valeria, Mangone I, Rinaldi A, Delli Compagni E, Scialabba S, Caporale M, Di Pasquale A, Cammà C, Puglia I, Calistri P, Savini G |
| EPI_ISL_1841189, EPI_ISL_1841190 | SIESP DIPARTIMENTO DI PREVENZIONE TERAMO | Istituto Zooprofilattico Sperimentale dell'Abruzzo e Molise "G. Caporale" | Lorusso A, Marcacci M, Di Domenico M, Ancora M, Curini V, Di Lollo Valeria, Mangone I, Rinaldi A, Delli Compagni E, Scialabba S, Caporale M, Di Pasquale A, Cammà C, Puglia I, Calistri P, Savini G |
| EPI_ISL_1841191 | SIESP CHIETI - DRIVE IN ORTONA | Istituto Zooprofilattico Sperimentale dell'Abruzzo e Molise "G. Caporale" | Lorusso A, Marcacci M, Di Domenico M, Ancora M, Curini V, Di Lollo Valeria, Mangone I, Rinaldi A, Delli Compagni E, Scialabba S, Caporale M, Di Pasquale A, Cammà C, Puglia I, Calistri P, Savini G |
| EPI_ISL_1841192, EPI_ISL_1841193 | SIESP CHIETI - DRIVE IN CHIETI | Istituto Zooprofilattico Sperimentale dell'Abruzzo e Molise "G. Caporale" | Lorusso A, Marcacci M, Di Domenico M, Ancora M, Curini V, Di Lollo Valeria, Mangone I, Rinaldi A, Delli Compagni E, Scialabba S, Caporale M, Di Pasquale A, Cammà C, Puglia I, Calistri P, Savini G |
| EPI_ISL_1841194 | SIESP CHIETI - DRIVE IN LANCIANO | Istituto Zooprofilattico Sperimentale dell'Abruzzo e Molise "G. Caporale" | Lorusso A, Marcacci M, Di Domenico M, Ancora M, Curini V, Di Lollo Valeria, Mangone I, Rinaldi A, Delli Compagni E, Scialabba S, Caporale M, Di Pasquale A, Cammà C, Puglia I, Calistri P, Savini G |
| EPI_ISL_1841195 | SIESP CHIETI - DRIVE IN VASTO | Istituto Zooprofilattico Sperimentale dell'Abruzzo e Molise "G. Caporale" | Lorusso A, Marcacci M, Di Domenico M, Ancora M, Curini V, Di Lollo Valeria, Mangone I, Rinaldi A, Delli Compagni E, Scialabba S, Caporale M, Di Pasquale A, Cammà C, Puglia I, Calistri P, Savini G |
| EPI_ISL_1841196 | DIP. PREV. AVEZZANO SERVIZIO DI IGIENE EPIDEMIOLOGIA E SANITA' PUBBLICA | Istituto Zooprofilattico Sperimentale dell'Abruzzo e Molise "G. Caporale" | Lorusso A, Marcacci M, Di Domenico M, Ancora M, Curini V, Di Lollo Valeria, Mangone I, Rinaldi A, Delli Compagni E, Scialabba S, Caporale M, Di Pasquale A, Cammà C, Puglia I, Calistri P, Savini G |
| EPI_ISL_1841197 | USCA Tagliacozzo | Istituto Zooprofilattico Sperimentale dell'Abruzzo e Molise "G. Caporale" | Lorusso A, Marcacci M, Di Domenico M, Ancora M, Curini V, Di Lollo Valeria, Mangone I, Rinaldi A, Delli Compagni E, Scialabba S, Caporale M, Di Pasquale A, Cammà C, Puglia I, Calistri P, Savini G |
| EPI_ISL_1841198, EPI_ISL_1841199 | DIP. PREV. AVEZZANO SERVIZIO DI IGIENE EPIDEMIOLOGIA E SANITA' PUBBLICA | Istituto Zooprofilattico Sperimentale dell'Abruzzo e Molise "G. Caporale" | Lorusso A, Marcacci M, Di Domenico M, Ancora M, Curini V, Di Lollo Valeria, Mangone I, Rinaldi A, Delli Compagni E, Scialabba S, Caporale M, Di Pasquale A, Cammà C, Puglia I, Calistri P, Savini G |
| EPI_ISL_1841200 | USCA Tagliacozzo | Istituto Zooprofilattico Sperimentale dell'Abruzzo e Molise "G. Caporale" | Lorusso A, Marcacci M, Di Domenico M, Ancora M, Curini V, Di Lollo Valeria, Mangone I, Rinaldi A, Delli Compagni E, Scialabba S, Caporale M, Di Pasquale A, Cammà C, Puglia I, Calistri P, Savini G |
| EPI_ISL_1841201 | USCA Pescina | Istituto Zooprofilattico Sperimentale dell'Abruzzo e Molise "G. Caporale" | Lorusso A, Marcacci M, Di Domenico M, Ancora M, Curini V, Di Lollo Valeria, Mangone I, Rinaldi A, Delli Compagni E, Scialabba S, Caporale M, Di Pasquale A, Cammà C, Puglia I, Calistri P, Savini G |
| EPI_ISL_1841202 | PRESIDIO OSPEDALIERO TAGLIACOZZO | Istituto Zooprofilattico Sperimentale dell'Abruzzo e Molise "G. Caporale" | Lorusso A, Marcacci M, Di Domenico M, Ancora M, Curini V, Di Lollo Valeria, Mangone I, Rinaldi A, Delli Compagni E, Scialabba S, Caporale M, Di Pasquale A, Cammà C, Puglia I, Calistri P, Savini G |
| EPI_ISL_1841203, EPI_ISL_1841204 | SIESP DIPARTIMENTO DI PREVENZIONE TERAMO | Istituto Zooprofilattico Sperimentale dell'Abruzzo e Molise "G. Caporale" | Lorusso A, Marcacci M, Di Domenico M, Ancora M, Curini V, Di Lollo Valeria, Mangone I, Rinaldi A, Delli Compagni E, Scialabba S, Caporale M, Di Pasquale A, Cammà C, Puglia I, Calistri P, Savini G |
| EPI_ISL_1841205, EPI_ISL_1841206, EPI_ISL_1841207, EPI_ISL_1841208, EPI_ISL_1841209 | SIESP CHIETI - DRIVE IN LANCIANO | Istituto Zooprofilattico Sperimentale dell'Abruzzo e Molise "G. Caporale" | Lorusso A, Marcacci M, Di Domenico M, Ancora M, Curini V, Di Lollo Valeria, Mangone I, Rinaldi A, Delli Compagni E, Scialabba S, Caporale M, Di Pasquale A, Cammà C, Puglia I, Calistri P, Savini G |
| EPI_ISL_1841210, EPI_ISL_1841211 | SIESP DIPARTIMENTO DI PREVENZIONE TERAMO | Istituto Zooprofilattico Sperimentale dell'Abruzzo e Molise "G. Caporale" | Lorusso A, Marcacci M, Di Domenico M, Ancora M, Curini V, Di Lollo Valeria, Mangone I, Rinaldi A, Delli Compagni E, Scialabba S, Caporale M, Di Pasquale A, Cammà C, Puglia I, Calistri P, Savini G |
| EPI_ISL_1841212 | USCA Castel di Sangro | Istituto Zooprofilattico Sperimentale dell'Abruzzo e Molise "G. Caporale" | Lorusso A, Marcacci M, Di Domenico M, Ancora M, Curini V, Di Lollo Valeria, Mangone I, Rinaldi A, Delli Compagni E, Scialabba S, Caporale M, Di Pasquale A, Cammà C, Puglia I, Calistri P, Savini G |
| EPI_ISL_1841213 | SIESP DIPARTIMENTO DI PREVENZIONE TERAMO | Istituto Zooprofilattico Sperimentale dell'Abruzzo e Molise "G. Caporale" | Lorusso A, Marcacci M, Di Domenico M, Ancora M, Curini V, Di Lollo Valeria, Mangone I, Rinaldi A, Delli Compagni E, Scialabba S, Caporale M, Di Pasquale A, Cammà C, Puglia I, Calistri P, Savini G |
| EPI_ISL_1841214 | DIP. PREV. AVEZZANO SERVIZIO DI IGIENE EPIDEMIOLOGIA E SANITA' PUBBLICA | Istituto Zooprofilattico Sperimentale dell'Abruzzo e Molise "G. Caporale" | Lorusso A, Marcacci M, Di Domenico M, Ancora M, Curini V, Di Lollo Valeria, Mangone I, Rinaldi A, Delli Compagni E, Scialabba S, Caporale M, Di Pasquale A, Cammà C, Puglia I, Calistri P, Savini G |
| EPI_ISL_1841215 | USCA Pescina | Istituto Zooprofilattico Sperimentale dell'Abruzzo e Molise "G. Caporale" | Lorusso A, Marcacci M, Di Domenico M, Ancora M, Curini V, Di Lollo Valeria, Mangone I, Rinaldi A, Delli Compagni E, Scialabba S, Caporale M, Di Pasquale A, Cammà C, Puglia I, Calistri P, Savini G |
| EPI_ISL_1841216 | DIP. PREV. AVEZZANO SERVIZIO DI IGIENE EPIDEMIOLOGIA E SANITA' PUBBLICA | Istituto Zooprofilattico Sperimentale dell'Abruzzo e Molise "G. Caporale" | Lorusso A, Marcacci M, Di Domenico M, Ancora M, Curini V, Di Lollo Valeria, Mangone I, Rinaldi A, Delli Compagni E, Scialabba S, Caporale M, Di Pasquale A, Cammà C, Puglia I, Calistri P, Savini G |
| EPI_ISL_1841217 | SIESP DIPARTIMENTO DI PREVENZIONE TERAMO | Istituto Zooprofilattico Sperimentale dell'Abruzzo e Molise "G. Caporale" | Lorusso A, Marcacci M, Di Domenico M, Ancora M, Curini V, Di Lollo Valeria, Mangone I, Rinaldi A, Delli Compagni E, Scialabba S, Caporale M, Di Pasquale A, Cammà C, Puglia I, Calistri P, Savini G |
| EPI_ISL_1841218, EPI_ISL_1841219 | USCA Avezzano | Istituto Zooprofilattico Sperimentale dell'Abruzzo e Molise "G. Caporale" | Lorusso A, Marcacci M, Di Domenico M, Ancora M, Curini V, Di Lollo Valeria, Mangone I, Rinaldi A, Delli Compagni E, Scialabba S, Caporale M, Di Pasquale A, Cammà C, Puglia I, Calistri P, Savini G |
| EPI_ISL_1841220 | DIP. PREV. AVEZZANO SERVIZIO DI IGIENE EPIDEMIOLOGIA E SANITA' PUBBLICA | Istituto Zooprofilattico Sperimentale dell'Abruzzo e Molise "G. Caporale" | Lorusso A, Marcacci M, Di Domenico M, Ancora M, Curini V, Di Lollo Valeria, Mangone I, Rinaldi A, Delli Compagni E, Scialabba S, Caporale M, Di Pasquale A, Cammà C, Puglia I, Calistri P, Savini G |
| EPI_ISL_1841221 | OSPEDALE CIVILE SANT'OMERO - PRONTO SOCCORSO | Istituto Zooprofilattico Sperimentale dell'Abruzzo e Molise "G. Caporale" | Lorusso A, Marcacci M, Di Domenico M, Ancora M, Curini V, Di Lollo Valeria, Mangone I, Rinaldi A, Delli Compagni E, Scialabba S, Caporale M, Di Pasquale A, Cammà C, Puglia I, Calistri P, Savini G |
| EPI_ISL_1841222, EPI_ISL_1841223, EPI_ISL_1841224, EPI_ISL_1841225 | SIESP CHIETI - DRIVE IN LANCIANO | Istituto Zooprofilattico Sperimentale dell'Abruzzo e Molise "G. Caporale" | Lorusso A, Marcacci M, Di Domenico M, Ancora M, Curini V, Di Lollo Valeria, Mangone I, Rinaldi A, Delli Compagni E, Scialabba S, Caporale M, Di Pasquale A, Cammà C, Puglia I, Calistri P, Savini G |
| EPI_ISL_1841226 | SIESP DIPARTIMENTO DI PREVENZIONE TERAMO | Istituto Zooprofilattico Sperimentale dell'Abruzzo e Molise "G. Caporale" | Lorusso A, Marcacci M, Di Domenico M, Ancora M, Curini V, Di Lollo Valeria, Mangone I, Rinaldi A, Delli Compagni E, Scialabba S, Caporale M, Di Pasquale A, Cammà C, Puglia I, Calistri P, Savini G |
| EPI_ISL_1841227 | SIESP CHIETI - DRIVE IN ORTONA | Istituto Zooprofilattico Sperimentale dell'Abruzzo e Molise "G. Caporale" | Lorusso A, Marcacci M, Di Domenico M, Ancora M, Curini V, Di Lollo Valeria, Mangone I, Rinaldi A, Delli Compagni E, Scialabba S, Caporale M, Di Pasquale A, Cammà C, Puglia I, Calistri P, Savini G |
| EPI_ISL_1841228 | Baylor Scott & White-Temple | Baylor Scott & White-Temple | Ari Rao, Linden Morales, Kimberly Walker, Marcus Volz, Shelby Hendrickson |
| EPI_ISL_1843754, EPI_ISL_1843756, EPI_ISL_1843758, EPI_ISL_1843759, EPI_ISL_1843761, EPI_ISL_1843763, EPI_ISL_1843765, EPI_ISL_1843767, EPI_ISL_1843769, EPI_ISL_1843771, EPI_ISL_1843774, EPI_ISL_1843776, EPI_ISL_1843778, EPI_ISL_1843779 | | | |
| see above | Klinisch Laboratorium ZNA | Klinisch Laboratorium ZNA | Verstrepen et al. |
| EPI_ISL_1851394, EPI_ISL_1851399 | SIESP DIPARTIMENTO DI PREVENZIONE TERAMO | Istituto Zooprofilattico Sperimentale dell'Abruzzo e Molise "G. Caporale" | Lorusso A, Marcacci M, Di Domenico M, Ancora M, Curini V, Di Lollo Valeria, Mangone I, Rinaldi A, Delli Compagni E, Scialabba S, Caporale M, Di Pasquale A, Cammà C, Puglia I, Calistri P, Savini G |
| EPI_ISL_1851405 | DIP. PREV. AVEZZANO SERVIZIO DI IGIENE | Istituto Zooprofilattico Sperimentale dell'Abruzzo e Molise "G. | Lorusso A, Marcacci M, Di Domenico M, Ancora M, Curini V, Di Lollo Valeria, Mangone I, Rinaldi A, Delli Compagni E, Scialabba S, Caporale M, Di |

| | | | |
|---|---|---|---|
| | EPIDEMIOLOGIA E SANITA' PUBBLICA | Caporale" | Pasquale A, Cammà C, Puglia I, Calistri P, Savini G |
| EPI_ISL_1851410 | OSPEDALE "F. RENZETTI" LANCIANO - MEDICINA INTERNA | Istituto Zooprofilattico Sperimentale dell'Abruzzo e Molise "G. Caporale" | Lorusso A, Marcacci M, Di Domenico M, Ancora M, Curini V, Di Lollo Valeria, Mangone I, Rinaldi A, Delli Compagni E, Scialabba S, Caporale M, Di Pasquale A, Cammà C, Puglia I, Calistri P, Savini G |
| EPI_ISL_1851416 | SIESP CHIETI - DRIVE IN LANCIANO | Istituto Zooprofilattico Sperimentale dell'Abruzzo e Molise "G. Caporale" | Lorusso A, Marcacci M, Di Domenico M, Ancora M, Curini V, Di Lollo Valeria, Mangone I, Rinaldi A, Delli Compagni E, Scialabba S, Caporale M, Di Pasquale A, Cammà C, Puglia I, Calistri P, Savini G |
| EPI_ISL_1851421 | SIESP DIPARTIMENTO DI PREVENZIONE CHIETI | Istituto Zooprofilattico Sperimentale dell'Abruzzo e Molise "G. Caporale" | Lorusso A, Marcacci M, Di Domenico M, Ancora M, Curini V, Di Lollo Valeria, Mangone I, Rinaldi A, Delli Compagni E, Scialabba S, Caporale M, Di Pasquale A, Cammà C, Puglia I, Calistri P, Savini G |
| EPI_ISL_1851815 | SIESP DIPARTIMENTO DI PREVENZIONE TERAMO | Istituto Zooprofilattico Sperimentale dell'Abruzzo e Molise "G. Caporale" | Lorusso A, Marcacci M, Di Domenico M, Ancora M, Curini V, Di Lollo Valeria, Mangone I, Rinaldi A, Delli Compagni E, Scialabba S, Caporale M, Di Pasquale A, Cammà C, Puglia I, Calistri P, Savini G |
| EPI_ISL_1851820 | SIESP CHIETI - DRIVE IN ORTONA | Istituto Zooprofilattico Sperimentale dell'Abruzzo e Molise "G. Caporale" | Lorusso A, Marcacci M, Di Domenico M, Ancora M, Curini V, Di Lollo Valeria, Mangone I, Rinaldi A, Delli Compagni E, Scialabba S, Caporale M, Di Pasquale A, Cammà C, Puglia I, Calistri P, Savini G |
| EPI_ISL_1851826 | SIESP DIPARTIMENTO DI PREVENZIONE TERAMO | Istituto Zooprofilattico Sperimentale dell'Abruzzo e Molise "G. Caporale" | Lorusso A, Marcacci M, Di Domenico M, Ancora M, Curini V, Di Lollo Valeria, Mangone I, Rinaldi A, Delli Compagni E, Scialabba S, Caporale M, Di Pasquale A, Cammà C, Puglia I, Calistri P, Savini G |
| EPI_ISL_1851831 | DIP. PREV. AVEZZANO SERVIZIO DI IGIENE EPIDEMIOLOGIA E SANITA' PUBBLICA | Istituto Zooprofilattico Sperimentale dell'Abruzzo e Molise "G. Caporale" | Lorusso A, Marcacci M, Di Domenico M, Ancora M, Curini V, Di Lollo Valeria, Mangone I, Rinaldi A, Delli Compagni E, Scialabba S, Caporale M, Di Pasquale A, Cammà C, Puglia I, Calistri P, Savini G |
| EPI_ISL_1851836 | PRESIDIO TERRITORIALE CASOLI | Istituto Zooprofilattico Sperimentale dell'Abruzzo e Molise "G. Caporale" | Lorusso A, Marcacci M, Di Domenico M, Ancora M, Curini V, Di Lollo Valeria, Mangone I, Rinaldi A, Delli Compagni E, Scialabba S, Caporale M, Di Pasquale A, Cammà C, Puglia I, Calistri P, Savini G |
| EPI_ISL_1851842 | SIESP CHIETI - DRIVE IN CHIETI | Istituto Zooprofilattico Sperimentale dell'Abruzzo e Molise "G. Caporale" | Lorusso A, Marcacci M, Di Domenico M, Ancora M, Curini V, Di Lollo Valeria, Mangone I, Rinaldi A, Delli Compagni E, Scialabba S, Caporale M, Di Pasquale A, Cammà C, Puglia I, Calistri P, Savini G |
| EPI_ISL_1851847 | DIP. PREV. AVEZZANO SERVIZIO DI IGIENE EPIDEMIOLOGIA E SANITA' PUBBLICA | Istituto Zooprofilattico Sperimentale dell'Abruzzo e Molise "G. Caporale" | Lorusso A, Marcacci M, Di Domenico M, Ancora M, Curini V, Di Lollo Valeria, Mangone I, Rinaldi A, Delli Compagni E, Scialabba S, Caporale M, Di Pasquale A, Cammà C, Puglia I, Calistri P, Savini G |
| EPI_ISL_1851852 | SIESP CHIETI - DRIVE IN ORTONA | Istituto Zooprofilattico Sperimentale dell'Abruzzo e Molise "G. Caporale" | Lorusso A, Marcacci M, Di Domenico M, Ancora M, Curini V, Di Lollo Valeria, Mangone I, Rinaldi A, Delli Compagni E, Scialabba S, Caporale M, Di Pasquale A, Cammà C, Puglia I, Calistri P, Savini G |
| EPI_ISL_1851857 | SIESP CHIETI - DRIVE IN GISSI | Istituto Zooprofilattico Sperimentale dell'Abruzzo e Molise "G. Caporale" | Lorusso A, Marcacci M, Di Domenico M, Ancora M, Curini V, Di Lollo Valeria, Mangone I, Rinaldi A, Delli Compagni E, Scialabba S, Caporale M, Di Pasquale A, Cammà C, Puglia I, Calistri P, Savini G |
| EPI_ISL_1852192 | Università degli Studi di Perugia | Istituto Zooprofilattico Sperimentale dell'Abruzzo e Molise "G. Caporale" | Mencacci A, Camilloni B,Lorusso A, Marcacci M, Di Domenico M, Ancora M, Curini V, Di Lollo Valeria, Mangone I, Rinaldi A, Delli Compagni E, Scialabba S, Caporale M, Di Pasquale A, Cammà C, Puglia I, Calistri P, Savini G |
| EPI_ISL_1852197 | Laboratorio Analisi Osp. Città di Castello - Azienda USL Umbria1 | Istituto Zooprofilattico Sperimentale dell'Abruzzo e Molise "G. Caporale" | Malagigi V, Tacconi P, Lorusso A, Marcacci M, Di Domenico M, Ancora M, Curini V, Di Lollo Valeria, Mangone I, Rinaldi A, Delli Compagni E, Scialabba S, Caporale M, Di Pasquale A, Cammà C, Puglia I, Calistri P, Savini G |
| EPI_ISL_1852563 | Ospedale Santissima Annunziata | Istituto Zooprofilattico Sperimentale della Puglia e della Basilicata | Parisi A., Bianco A., Capozzi L., Del Sambro L., Simone D., Difato L., Ridolfi D., Giannico A, Tampoia M. |
| EPI_ISL_1852568 | Presidio di Brindisi Di Summa - Perrino | Istituto Zooprofilattico Sperimentale della Puglia e della Basilicata | Parisi A., Bianco A., Capozzi L., Del Sambro L., Simone D., Difato L., Santoro A., Ridolfi D., Giannico A. |
| EPI_ISL_1853279, EPI_ISL_1853280, EPI_ISL_1853281, EPI_ISL_1853282, EPI_ISL_1853283 | Centre Hospitalier Universitaire Clermont-Ferrand | CHU Clermont-Ferrand, service de virologie | Bisseux Maxime, Mirand Audrey, Combes Patricia, Henquell Cécile |
| EPI_ISL_1853285 | Centre Hospitalier Paul Ardier | CHU Clermont-Ferrand, service de virologie | Bisseux Maxime, Mirand Audrey, Combes Patricia, Henquell Cécile |
| EPI_ISL_1853286 | Centre Hospitalier Universitaire Clermont-Ferrand | CHU Clermont-Ferrand, service de virologie | Bisseux Maxime, Mirand Audrey, Combes Patricia, Henquell Cécile |
| EPI_ISL_1853287 | HOPITAL DE VICHY | CHU Clermont-Ferrand, service de virologie | Bisseux Maxime, Mirand Audrey, Combes Patricia, Henquell Cécile |
| EPI_ISL_1853288, EPI_ISL_1853289, EPI_ISL_1853290 | Centre hospitalier D'AURILLAC | CHU Clermont-Ferrand, service de virologie | Bisseux Maxime, Mirand Audrey, Combes Patricia, Henquell Cécile |
| EPI_ISL_1853291, EPI_ISL_1853292, EPI_ISL_1853293 | Centre Hospitalier Universitaire Clermont-Ferrand | CHU Clermont-Ferrand, service de virologie | Bisseux Maxime, Mirand Audrey, Combes Patricia, Henquell Cécile |
| EPI_ISL_1853294 | HOPITAL DE VICHY | CHU Clermont-Ferrand, service de virologie | Bisseux Maxime, Mirand Audrey, Combes Patricia, Henquell Cécile |
| EPI_ISL_1853295 | Centre Hospitalier Paul Ardier | CHU Clermont-Ferrand, service de virologie | Bisseux Maxime, Mirand Audrey, Combes Patricia, Henquell Cécile |
| EPI_ISL_1853296 | Centre Hospitalier Universitaire Clermont-Ferrand | CHU Clermont-Ferrand, service de virologie | Bisseux Maxime, Mirand Audrey, Combes Patricia, Henquell Cécile |
| EPI_ISL_1853297 | Centre hospitalier D'AURILLAC | CHU Clermont-Ferrand, service de virologie | Bisseux Maxime, Mirand Audrey, Combes Patricia, Henquell Cécile |
| EPI_ISL_1853298, EPI_ISL_1853299, EPI_ISL_1853301 | Centre Hospitalier Universitaire Clermont-Ferrand | CHU Clermont-Ferrand, service de virologie | Bisseux Maxime, Mirand Audrey, Combes Patricia, Henquell Cécile |
| EPI_ISL_1853304 | HOPITAL DE VICHY | CHU Clermont-Ferrand, service de virologie | Bisseux Maxime, Mirand Audrey, Combes Patricia, Henquell Cécile |
| EPI_ISL_1853306 | HOPITAL DE MONTLUCON | CHU Clermont-Ferrand, service de virologie | Bisseux Maxime, Mirand Audrey, Combes Patricia, Henquell Cécile |
| EPI_ISL_1853308, EPI_ISL_1853310, EPI_ISL_1853313, EPI_ISL_1853315 | Centre Hospitalier Universitaire Clermont-Ferrand | CHU Clermont-Ferrand, service de virologie | Bisseux Maxime, Mirand Audrey, Combes Patricia, Henquell Cécile |
| EPI_ISL_1853317 | Centre Hospitalier Billom | CHU Clermont-Ferrand, service de virologie | Bisseux Maxime, Mirand Audrey, Combes Patricia, Henquell Cécile |
| EPI_ISL_1853319, EPI_ISL_1853322, EPI_ISL_1853324 | Centre Hospitalier Universitaire Clermont-Ferrand | CHU Clermont-Ferrand, service de virologie | Bisseux Maxime, Mirand Audrey, Combes Patricia, Henquell Cécile |
| EPI_ISL_1853326 | Centre hospitalier D'AURILLAC | CHU Clermont-Ferrand, service de virologie | Bisseux Maxime, Mirand Audrey, Combes Patricia, Henquell Cécile |
| EPI_ISL_1853328, EPI_ISL_1853330 | Centre Hospitalier Universitaire Clermont-Ferrand | CHU Clermont-Ferrand, service de virologie | Bisseux Maxime, Mirand Audrey, Combes Patricia, Henquell Cécile |
| EPI_ISL_1853333 | HOPITAL DE MONTLUCON | CHU Clermont-Ferrand, service de virologie | Bisseux Maxime, Mirand Audrey, Combes Patricia, Henquell Cécile |
| EPI_ISL_1853335 | Centre Hospitalier Universitaire Clermont-Ferrand | CHU Clermont-Ferrand, service de virologie | Bisseux Maxime, Mirand Audrey, Combes Patricia, Henquell Cécile |
| EPI_ISL_1853337, EPI_ISL_1853339 | HOPITAL DE VICHY | CHU Clermont-Ferrand, service de virologie | Bisseux Maxime, Mirand Audrey, Combes Patricia, Henquell Cécile |
| EPI_ISL_1853341, EPI_ISL_1853343, EPI_ISL_1853346 | Centre Hospitalier Universitaire Clermont-Ferrand | CHU Clermont-Ferrand, service de virologie | Bisseux Maxime, Mirand Audrey, Combes Patricia, Henquell Cécile |
| EPI_ISL_1853350 | HOPITAL DE VICHY | CHU Clermont-Ferrand, service de virologie | Bisseux Maxime, Mirand Audrey, Combes Patricia, Henquell Cécile |
| EPI_ISL_1853352, EPI_ISL_1853355, EPI_ISL_1853357 | HOPITAL DE MONTLUCON | CHU Clermont-Ferrand, service de virologie | Bisseux Maxime, Mirand Audrey, Combes Patricia, Henquell Cécile |
| EPI_ISL_1853359, EPI_ISL_1853362 | Centre Hospitalier Universitaire Clermont-Ferrand | CHU Clermont-Ferrand, service de virologie | Bisseux Maxime, Mirand Audrey, Combes Patricia, Henquell Cécile |

| | | | |
|---|---|---|---|
| EPI_ISL_1853366, EPI_ISL_1853368 | Centre Hospitalier RIOM | CHU Clermont-Ferrand, service de virologie | Bisseux Maxime, Mirand Audrey, Combes Patricia, Henquell Cécile |
| EPI_ISL_1853370, EPI_ISL_1853372, EPI_ISL_1853374, EPI_ISL_1853377 | Centre Hospitalier Universitaire Clermont-Ferrand | CHU Clermont-Ferrand, service de virologie | Bisseux Maxime, Mirand Audrey, Combes Patricia, Henquell Cécile |
| EPI_ISL_1853379 | Centre De Prelevement COVID RIOM | CHU Clermont-Ferrand, service de virologie | Bisseux Maxime, Mirand Audrey, Combes Patricia, Henquell Cécile |
| EPI_ISL_1853381 | Centre De Prélévement COVID VICHY | CHU Clermont-Ferrand, service de virologie | Bisseux Maxime, Mirand Audrey, Combes Patricia, Henquell Cécile |
| EPI_ISL_1853383 | HOPITAL DE VICHY | CHU Clermont-Ferrand, service de virologie | Bisseux Maxime, Mirand Audrey, Combes Patricia, Henquell Cécile |
| EPI_ISL_1853386 | Centre De Prélévement COVID VICHY | CHU Clermont-Ferrand, service de virologie | Bisseux Maxime, Mirand Audrey, Combes Patricia, Henquell Cécile |
| EPI_ISL_1853388 | center de prélévement COVID ISSOIRE | CHU Clermont-Ferrand, service de virologie | Bisseux Maxime, Mirand Audrey, Combes Patricia, Henquell Cécile |
| EPI_ISL_1853390 | Centre De Prélévement COVID VICHY | CHU Clermont-Ferrand, service de virologie | Bisseux Maxime, Mirand Audrey, Combes Patricia, Henquell Cécile |
| EPI_ISL_1853393, EPI_ISL_1853395 | Centre Hospitalier Universitaire Clermont-Ferrand | CHU Clermont-Ferrand, service de virologie | Bisseux Maxime, Mirand Audrey, Combes Patricia, Henquell Cécile |
| EPI_ISL_1853397, EPI_ISL_1853400 | Centre Jean Perrin | CHU Clermont-Ferrand, service de virologie | Bisseux Maxime, Mirand Audrey, Combes Patricia, Henquell Cécile |
| EPI_ISL_1853402, EPI_ISL_1853404 | center de prélévement COVID ISSOIRE | CHU Clermont-Ferrand, service de virologie | Bisseux Maxime, Mirand Audrey, Combes Patricia, Henquell Cécile |
| EPI_ISL_1853407, EPI_ISL_1853409, EPI_ISL_1853411, EPI_ISL_1853413, EPI_ISL_1853416 | Centre Hospitalier Universitaire Clermont-Ferrand | CHU Clermont-Ferrand, service de virologie | Bisseux Maxime, Mirand Audrey, Combes Patricia, Henquell Cécile |
| EPI_ISL_1853418 | Centre De Prélévement COVID VICHY | CHU Clermont-Ferrand, service de virologie | Bisseux Maxime, Mirand Audrey, Combes Patricia, Henquell Cécile |
| EPI_ISL_1853419 | CDP COVID MONTLUCON | CHU Clermont-Ferrand, service de virologie | Bisseux Maxime, Mirand Audrey, Combes Patricia, Henquell Cécile |
| EPI_ISL_1853420 | Centre Hospitalier Universitaire Clermont-Ferrand | CHU Clermont-Ferrand, service de virologie | Bisseux Maxime, Mirand Audrey, Combes Patricia, Henquell Cécile |
| EPI_ISL_1855850, EPI_ISL_1855852, EPI_ISL_1855864, EPI_ISL_1855866, EPI_ISL_1855868, EPI_ISL_1855869, EPI_ISL_1855872, EPI_ISL_1855874, EPI_ISL_1855875, EPI_ISL_1855879, EPI_ISL_1855880, EPI_ISL_1855883, EPI_ISL_1855886, EPI_ISL_1855887, EPI_ISL_1855889, EPI_ISL_1855890, EPI_ISL_1855891, EPI_ISL_1855892, EPI_ISL_1855893, EPI_ISL_1855895, EPI_ISL_1855896, EPI_ISL_1855897, EPI_ISL_1855898, EPI_ISL_1855900, EPI_ISL_1855905, EPI_ISL_1855907, EPI_ISL_1855908, EPI_ISL_1855909, EPI_ISL_1855910 | | | |
| see above | Baylor Scott & White-Temple | Baylor Scott & White-Temple | Ari Rao, Linden Morales, Kimberly Walker, Marcus Volz, Shelby Hendrickson |
| EPI_ISL_1856983 | Outre mer | National Reference Center for Viruses of Respiratory Infections, Institut Pasteur, Paris | Marion Barbet, Sylvie Behillil, Méline Bizard, Angela Brisebarre, Camille Capel, Vincent Enouf, Louise Lefrançois, Frédéric Lemoine, Christophe Malabat, Corinne Maufrais, Etienne Simon-Lorière, Maud Vanpeene, Sylvie Van der Werf ,Dominique Rousset |
| EPI_ISL_1856984, EPI_ISL_1856985, EPI_ISL_1856986, EPI_ISL_1856987 | Outre mer | National Reference Center for Viruses of Respiratory Infections, Institut Pasteur, Paris | Marion Barbet, Sylvie Behillil, Méline Bizard, Angela Brisebarre, Camille Capel, Vincent Enouf, Louise Lefrançois, Frédéric Lemoine, Christophe Malabat, Corinne Maufrais, Pierre Lechat, Etienne Simon-Lorière, Maud Vanpeene, Sylvie Van der Werf ,Dominique Rousset |
| EPI_ISL_1856988 | Outre mer | National Reference Center for Viruses of Respiratory Infections, Institut Pasteur, Paris | Marion Barbet, Sylvie Behillil, Méline Bizard, Angela Brisebarre, Camille Capel, Vincent Enouf, Louise Lefrançois, Frédéric Lemoine, Christophe Malabat, Corinne Maufrais, Etienne Simon-Lorière, Maud Vanpeene, Sylvie Van der Werf ,Dominique Rousset |
| EPI_ISL_1856989, EPI_ISL_1856990 | Outre mer | National Reference Center for Viruses of Respiratory Infections, Institut Pasteur, Paris | Marion Barbet, Sylvie Behillil, Méline Bizard, Angela Brisebarre, Camille Capel, Vincent Enouf, Louise Lefrançois, Frédéric Lemoine, Christophe Malabat, Corinne Maufrais, Pierre Lechat, Etienne Simon-Lorière, Maud Vanpeene, Sylvie Van der Werf ,Dominique Rousset |
| EPI_ISL_1856991, EPI_ISL_1856992 | Outre mer | National Reference Center for Viruses of Respiratory Infections, Institut Pasteur, Paris | Marion Barbet, Sylvie Behillil, Méline Bizard, Angela Brisebarre, Camille Capel, Vincent Enouf, Louise Lefrançois, Frédéric Lemoine, Christophe Malabat, Corinne Maufrais, Etienne Simon-Lorière, Maud Vanpeene, Sylvie Van der Werf ,Dominique Rousset |
| EPI_ISL_1856994, EPI_ISL_1856995 | Labo Analyses Med | National Reference Center for Viruses of Respiratory Infections, Institut Pasteur, Paris | Marion Barbet, Sylvie Behillil, Méline Bizard, Angela Brisebarre, Camille Capel, Vincent Enouf, Louise Lefrançois, Frédéric Lemoine, Christophe Malabat, Corinne Maufrais, Etienne Simon-Lorière, Maud Vanpeene, Sylvie Van der Werf ,Sophie Girard |
| EPI_ISL_1856996 | Labo Analyses Med | National Reference Center for Viruses of Respiratory Infections, Institut Pasteur, Paris | Marion Barbet, Sylvie Behillil, Méline Bizard, Angela Brisebarre, Camille Capel, Vincent Enouf, Louise Lefrançois, Frédéric Lemoine, Christophe Malabat, Corinne Maufrais, Gael Millot, Etienne Simon-Lorière, Maud Vanpeene, Sylvie Van der Werf ,Fabienne Artur |
| EPI_ISL_1856997 | Labo Analyses Med | National Reference Center for Viruses of Respiratory Infections, Institut Pasteur, Paris | Marion Barbet, Sylvie Behillil, Méline Bizard, Angela Brisebarre, Camille Capel, Vincent Enouf, Louise Lefrançois, Frédéric Lemoine, Christophe Malabat, Corinne Maufrais, Gael Millot, Etienne Simon-Lorière, Maud Vanpeene, Sylvie Van der Werf ,Christine Laroche |
| EPI_ISL_1856999, EPI_ISL_1857001 | Labo Analyses Med | National Reference Center for Viruses of Respiratory Infections, Institut Pasteur, Paris | Marion Barbet, Sylvie Behillil, Méline Bizard, Angela Brisebarre, Camille Capel, Vincent Enouf, Louise Lefrançois, Frédéric Lemoine, Christophe Malabat, Corinne Maufrais, Etienne Simon-Lorière, Maud Vanpeene, Sylvie Van der Werf ,Nabil Gastli |
| EPI_ISL_1857002 | Labo Analyses Med | National Reference Center for Viruses of Respiratory Infections, Institut Pasteur, Paris | Marion Barbet, Sylvie Behillil, Méline Bizard, Angela Brisebarre, Camille Capel, Vincent Enouf, Louise Lefrançois, Frédéric Lemoine, Christophe Malabat, Corinne Maufrais, Pierre Lechat, Etienne Simon-Lorière, Maud Vanpeene, Sylvie Van der Werf ,Nabil Gastli |
| EPI_ISL_1857003 | Labo Analyses Med | National Reference Center for Viruses of Respiratory Infections, Institut Pasteur, Paris | Marion Barbet, Sylvie Behillil, Méline Bizard, Angela Brisebarre, Camille Capel, Vincent Enouf, Louise Lefrançois, Frédéric Lemoine, Christophe Malabat, Corinne Maufrais, Gael Millot, Etienne Simon-Lorière, Maud Vanpeene, Sylvie Van der Werf ,Nabil Gastli |
| EPI_ISL_1857004, EPI_ISL_1857005 | Labo Analyses Med | National Reference Center for Viruses of Respiratory Infections, Institut Pasteur, Paris | Marion Barbet, Sylvie Behillil, Méline Bizard, Angela Brisebarre, Camille Capel, Vincent Enouf, Louise Lefrançois, Frédéric Lemoine, Christophe Malabat, Corinne Maufrais, Etienne Simon-Lorière, Maud Vanpeene, Sylvie Van der Werf ,Nabil Gastli |
| EPI_ISL_1857006 | Labo Analyses Med | National Reference Center for Viruses of Respiratory Infections, Institut Pasteur, Paris | Marion Barbet, Sylvie Behillil, Méline Bizard, Angela Brisebarre, Camille Capel, Vincent Enouf, Louise Lefrançois, Frédéric Lemoine, Christophe Malabat, Corinne Maufrais, Gael Millot, Etienne Simon-Lorière, Maud Vanpeene, Sylvie Van der Werf ,Nabil Gastli |
| EPI_ISL_1857007, EPI_ISL_1857008, EPI_ISL_1857009, EPI_ISL_1857010, EPI_ISL_1857011, EPI_ISL_1857012, EPI_ISL_1857013, EPI_ISL_1857014, EPI_ISL_1857015 | Labo Analyses Med | National Reference Center for Viruses of Respiratory Infections, Institut Pasteur, Paris | Marion Barbet, Sylvie Behillil, Méline Bizard, Angela Brisebarre, Camille Capel, Vincent Enouf, Louise Lefrançois, Frédéric Lemoine, Christophe Malabat, Corinne Maufrais, Etienne Simon-Lorière, Maud Vanpeene, Sylvie Van der Werf ,Nabil Gastli |
| EPI_ISL_1857016 | Labo Analyses Med | National Reference Center for Viruses of Respiratory Infections, Institut Pasteur, Paris | Marion Barbet, Sylvie Behillil, Méline Bizard, Angela Brisebarre, Camille Capel, Vincent Enouf, Louise Lefrançois, Frédéric Lemoine, Christophe Malabat, Corinne Maufrais, Pierre Lechat, Etienne Simon-Lorière, Maud Vanpeene, Sylvie Van der Werf ,Nabil Gastli |
| EPI_ISL_1857017, EPI_ISL_1857018, EPI_ISL_1857019, EPI_ISL_1857020, EPI_ISL_1857021 | Labo Analyses Med | National Reference Center for Viruses of Respiratory Infections, Institut Pasteur, Paris | Marion Barbet, Sylvie Behillil, Méline Bizard, Angela Brisebarre, Camille Capel, Vincent Enouf, Louise Lefrançois, Frédéric Lemoine, Christophe Malabat, Corinne Maufrais, Etienne Simon-Lorière, Maud Vanpeene, Sylvie Van der Werf ,Nabil Gastli |
| EPI_ISL_1857022 | Labo Analyses Med | National Reference Center for Viruses of Respiratory Infections, Institut Pasteur, Paris | Marion Barbet, Sylvie Behillil, Méline Bizard, Angela Brisebarre, Camille Capel, Vincent Enouf, Louise Lefrançois, Frédéric Lemoine, Christophe Malabat, Corinne Maufrais, Pierre Lechat, Etienne Simon-Lorière, Maud Vanpeene, Sylvie Van der Werf ,Nabil Gastli |
| EPI_ISL_1857024, EPI_ISL_1857025 | Labo Analyses Med | National Reference Center for Viruses of Respiratory Infections, Institut Pasteur, Paris | Marion Barbet, Sylvie Behillil, Méline Bizard, Angela Brisebarre, Camille Capel, Vincent Enouf, Louise Lefrançois, Frédéric Lemoine, Christophe Malabat, Corinne Maufrais, Etienne Simon-Lorière, Maud Vanpeene, Sylvie Van der Werf ,Nabil Gastli |
| EPI_ISL_1857026 | Labo Analyses Med | National Reference Center for Viruses of Respiratory Infections, Institut Pasteur, Paris | Marion Barbet, Sylvie Behillil, Méline Bizard, Angela Brisebarre, Camille Capel, Vincent Enouf, Louise Lefrançois, Frédéric Lemoine, Christophe Malabat, Corinne Maufrais, Pierre Lechat, Etienne Simon-Lorière, Maud Vanpeene, Sylvie Van der Werf ,Nabil Gastli |
| EPI_ISL_1857027 | Labo Analyses Med | National Reference Center for Viruses of Respiratory Infections, Institut Pasteur, Paris | Marion Barbet, Sylvie Behillil, Méline Bizard, Angela Brisebarre, Camille Capel, Vincent Enouf, Louise Lefrançois, Frédéric Lemoine, Christophe Malabat, Corinne Maufrais, Etienne Simon-Lorière, Maud Vanpeene, Sylvie Van der Werf ,Nabil Gastli |
| EPI_ISL_1857028 | Labo Analyses Med | National Reference Center for Viruses of Respiratory Infections, Institut Pasteur, Paris | Marion Barbet, Sylvie Behillil, Méline Bizard, Angela Brisebarre, Camille Capel, Vincent Enouf, Louise Lefrançois, Frédéric Lemoine, Christophe Malabat, Corinne Maufrais, Pierre Lechat, Etienne Simon-Lorière, Maud Vanpeene, Sylvie Van der Werf ,Nabil Gastli |
| EPI_ISL_1857029, EPI_ISL_1857030, EPI_ISL_1857031, EPI_ISL_1857032, | Labo Analyses Med | National Reference Center for Viruses of Respiratory Infections, Institut Pasteur, Paris | Marion Barbet, Sylvie Behillil, Méline Bizard, Angela Brisebarre, Camille Capel, Vincent Enouf, Louise Lefrançois, Frédéric Lemoine, Christophe Malabat, Corinne Maufrais, Etienne Simon-Lorière, Maud Vanpeene, Sylvie Van der Werf ,Nabil Gastli |

| | | | |
|---|---|---|---|
| EPI_ISL_1857033, EPI_ISL_1857034 | | | |
| EPI_ISL_1857103 | LESP Ciudad de Mexico | Instituto de Diagnostico y Referencia Epidemiologicos (INDRE) | Claudia Wong-Arambula, Abril Rodriguez-Maldonado, Vanessa Rivero-Arredondo, Ariadna Medina-Benitez, Joaquin Quiroz-Mercado, Sergio Rangel-Guerrero, Natividad Cruz-Ortiz, Tatiana Nunez-Garcia, Gisela Barrera-Badillo, Lucia Hernandez-Rivas, Irma Lopez-Martinez, Ernesto Ramirez-Gonzalez. |
| EPI_ISL_1857243 | LESP Puebla | Instituto de Diagnostico y Referencia Epidemiologicos (INDRE) | Claudia Wong-Arambula, Abril Rodriguez-Maldonado, Vanessa Rivero-Arredondo, Ariadna Medina-Benitez, Joaquin Quiroz-Mercado, Sergio Rangel-Guerrero, Natividad Cruz-Ortiz, Tatiana Nunez-Garcia, Gisela Barrera-Badillo, Lucia Hernandez-Rivas, Irma Lopez-Martinez, Ernesto Ramirez-Gonzalez. |
| EPI_ISL_1857244 | LESP Jalisco | Instituto de Diagnostico y Referencia Epidemiologicos (INDRE) | Claudia Wong-Arambula, Abril Rodriguez-Maldonado, Vanessa Rivero-Arredondo, Ariadna Medina-Benitez, Joaquin Quiroz-Mercado, Sergio Rangel-Guerrero, Natividad Cruz-Ortiz, Tatiana Nunez-Garcia, Gisela Barrera-Badillo, Lucia Hernandez-Rivas, Irma Lopez-Martinez, Ernesto Ramirez-Gonzalez. |
| EPI_ISL_1857257 | LESP Tlaxcala | Instituto de Diagnostico y Referencia Epidemiologicos (INDRE) | Claudia Wong-Arambula, Abril Rodriguez-Maldonado, Vanessa Rivero-Arredondo, Ariadna Medina-Benitez, Joaquin Quiroz-Mercado, Sergio Rangel-Guerrero, Natividad Cruz-Ortiz, Tatiana Nunez-Garcia, Gisela Barrera-Badillo, Lucia Hernandez-Rivas, Irma Lopez-Martinez, Ernesto Ramirez-Gonzalez. |
| EPI_ISL_1857262 | LESP Chihuahua | Instituto de Diagnostico y Referencia Epidemiologicos (INDRE) | Claudia Wong-Arambula, Abril Rodriguez-Maldonado, Vanessa Rivero-Arredondo, Ariadna Medina-Benitez, Joaquin Quiroz-Mercado, Sergio Rangel-Guerrero, Natividad Cruz-Ortiz, Tatiana Nunez-Garcia, Gisela Barrera-Badillo, Lucia Hernandez-Rivas, Irma Lopez-Martinez, Ernesto Ramirez-Gonzalez. |
| EPI_ISL_1857272, EPI_ISL_1857273, EPI_ISL_1857274 | LESP Ciudad de Mexico | Instituto de Diagnostico y Referencia Epidemiologicos (INDRE) | Claudia Wong-Arambula, Abril Rodriguez-Maldonado, Vanessa Rivero-Arredondo, Ariadna Medina-Benitez, Joaquin Quiroz-Mercado, Sergio Rangel-Guerrero, Natividad Cruz-Ortiz, Tatiana Nunez-Garcia, Gisela Barrera-Badillo, Lucia Hernandez-Rivas, Irma Lopez-Martinez, Ernesto Ramirez-Gonzalez. |
| EPI_ISL_1857488 | ALPHABIO Laboratory | ALPHABIO Laboratory | Vincent GARCIA |
| EPI_ISL_1904862, EPI_ISL_1904863, EPI_ISL_1904864, EPI_ISL_1904865, EPI_ISL_1904866, EPI_ISL_1904867, EPI_ISL_1904868, EPI_ISL_1904869, EPI_ISL_1904870, EPI_ISL_1904871, EPI_ISL_1904872 | | | |
| see above | OLVZ Aalst | OLVZ Aalst | Anne Vankeerberghen |
| EPI_ISL_1905190, EPI_ISL_1905192, EPI_ISL_1905195, EPI_ISL_1905197, EPI_ISL_1905198, EPI_ISL_1905200, EPI_ISL_1905201, EPI_ISL_1905212, EPI_ISL_1905214, EPI_ISL_1905216 | AZ St.Jan Brugge-Oostende | AZ SINT-JAN BRUGGE | Marijke Reynders, Patrick Descheemaeker, Laurien Hoornaert, Thomas Van Landschoot, Jorn Hellemans |
| EPI_ISL_1909803, EPI_ISL_1909804, EPI_ISL_1909810, EPI_ISL_1909811, EPI_ISL_1909815, EPI_ISL_1909818 | Institute of Molecular and Translational Medicine / Laboratory of Experimental Medicine, Faculty of Medicine and Dentistry, Palacky University and University Hospital Olomouc | Institute of Molecular and Translational Medicine / Laboratory of Experimental Medicine, Faculty of Medicine and Dentistry, Palacky University | Rastislav Slavkovský, Hana Jaworek, Vladimíra Koudeláková, Barbora Blumová, Tomáš Pospíšíl, Marián Hajdúch |
| EPI_ISL_1909822 | Imeda Hospital | Imeda Hospital | Johan Frans, Dagmar Obbels, Hanne Valgaeren |
| EPI_ISL_1911084, EPI_ISL_1911085, EPI_ISL_1911086, EPI_ISL_1911087, EPI_ISL_1911088, EPI_ISL_1911089, EPI_ISL_1911090, EPI_ISL_1911092, EPI_ISL_1911093, EPI_ISL_1911094, EPI_ISL_1911095, EPI_ISL_1911096, EPI_ISL_1911097, EPI_ISL_1911098, EPI_ISL_1911099, EPI_ISL_1911100, EPI_ISL_1911101 | | | |
| see above | The Ohio State University Applied Microbiology Services Laboratory | The Ohio State University Applied Microbiology Services Laboratory | Seth A. Faith PhD |
| EPI_ISL_1911195, EPI_ISL_1911196, EPI_ISL_1911197, EPI_ISL_1911250 | Zhejiang Provincial Center for Disease Control and Prevention Zhoushan Center for Disease Prevention and Contol | Zhejiang Province Center of Disease Control and prevention | Yanjun Zhang, Bing WU, Hongling Wang |
| EPI_ISL_1911437, EPI_ISL_1911438 | NVSPL | National Public Health Surveillance Laboratory | Lukas Zemaitis, Migle Gabrielaite, Jelena Razmuk, Svajune Muralyte, Ana Steponkiene, Lukas Vasionis, Danas Baksa |
| EPI_ISL_1911440, EPI_ISL_1911442, EPI_ISL_1911448, EPI_ISL_1911454, EPI_ISL_1911455, EPI_ISL_1911458, EPI_ISL_1911461, EPI_ISL_1911466, EPI_ISL_1911473, EPI_ISL_1911480, EPI_ISL_1911489, EPI_ISL_1911490, EPI_ISL_1911507, EPI_ISL_1911529, EPI_ISL_1911547, EPI_ISL_1911557, EPI_ISL_1911559, EPI_ISL_1911590, EPI_ISL_1911603, EPI_ISL_1911606, EPI_ISL_1911617, EPI_ISL_1911814, EPI_ISL_1911815, EPI_ISL_1911816, EPI_ISL_1911818, EPI_ISL_1911819, EPI_ISL_1911821, EPI_ISL_1911822, EPI_ISL_1911823, EPI_ISL_1911824, EPI_ISL_1911825, EPI_ISL_1911826, EPI_ISL_1911827, EPI_ISL_1911828, EPI_ISL_1911829, EPI_ISL_1911830, EPI_ISL_1911832, EPI_ISL_1911833, EPI_ISL_1911834, EPI_ISL_1911835, EPI_ISL_1911836, EPI_ISL_1911837, EPI_ISL_1911838, EPI_ISL_1911839, EPI_ISL_1911840, EPI_ISL_1911841, EPI_ISL_1911842, EPI_ISL_1911843, EPI_ISL_1911845, EPI_ISL_1911846, EPI_ISL_1911847, EPI_ISL_1911848, EPI_ISL_1911849, EPI_ISL_1911850, EPI_ISL_1911852, EPI_ISL_1911853, EPI_ISL_1911854, EPI_ISL_1911856, EPI_ISL_1911857, EPI_ISL_1911858, EPI_ISL_1911859, EPI_ISL_1911860, EPI_ISL_1911861, EPI_ISL_1911862, EPI_ISL_1911863, EPI_ISL_1911865, EPI_ISL_1911866, EPI_ISL_1911869, EPI_ISL_1911870, EPI_ISL_1911871, EPI_ISL_1911872, EPI_ISL_1911874, EPI_ISL_1911875, EPI_ISL_1911877, EPI_ISL_1911878, EPI_ISL_1911879, EPI_ISL_1911880, EPI_ISL_1911881, EPI_ISL_1911882, EPI_ISL_1911883, EPI_ISL_1911884, EPI_ISL_1911885, EPI_ISL_1911886, EPI_ISL_1911887, EPI_ISL_1911888, EPI_ISL_1911889, EPI_ISL_1911890, EPI_ISL_1911891, EPI_ISL_1911892, EPI_ISL_1911893, EPI_ISL_1911895, EPI_ISL_1911897, EPI_ISL_1911898, EPI_ISL_1911899, EPI_ISL_1911900 | | | |
| see above | National Institute for Food and Veterinary Risk Assessment (NMVRVI) | National Public Health Surveillance Laboratory | Lukas Zemaitis, Migle Gabrielaite, Jelena Razmuk, Svajune Muralyte, Ana Steponkiene, Lukas Vasionis, Danas Baksa |
| EPI_ISL_1911902, EPI_ISL_1911903, EPI_ISL_1911904, EPI_ISL_1911905, EPI_ISL_1911906, EPI_ISL_1911908, EPI_ISL_1911909, EPI_ISL_1911911, EPI_ISL_1911912, EPI_ISL_1911913, EPI_ISL_1911914, EPI_ISL_1911915, EPI_ISL_1911916, EPI_ISL_1911919, EPI_ISL_1911921, EPI_ISL_1911923, EPI_ISL_1911924, EPI_ISL_1911925, EPI_ISL_1911926, EPI_ISL_1911927, EPI_ISL_1911928, EPI_ISL_1911929, EPI_ISL_1911930, EPI_ISL_1911935, EPI_ISL_1911936, EPI_ISL_1911938, EPI_ISL_1911939, EPI_ISL_1911940, EPI_ISL_1911942, EPI_ISL_1911943, EPI_ISL_1911945, EPI_ISL_1911946, EPI_ISL_1911954, EPI_ISL_1911955 | | | |
| see above | Baylor Scott & White-Temple | Baylor Scott & White-Temple | Ari Rao, Linden Morales, Kimberly Walker, Marcus Volz, Shelby Hendrickson |
| EPI_ISL_1913255 | Laboratorio de Virología HUCA | Laboratorio de Virología HUCA | Sandoval M, Castelló C, Gómez de Oña J, Boga JA, Rojo S, Alvarez-Arguelles ME, Abreu F, Costales I, Perez-Martínez Z, Martín-Rodríguez G, Coto E, Melón S |
| EPI_ISL_1914278, EPI_ISL_1914279, EPI_ISL_1914280 | Cliniques universitaires Saint-Luc | UCLouvain/IREC/MBLG | Jean Ruelle, Eleonore Ngyuvula, Benoit Kabamba Mukadi |
| EPI_ISL_1914380 | Laboratorio de Virología HUCA | Laboratorio de Virología HUCA | Sandoval M, Castelló C, Gómez de Oña J, Boga JA, Rojo S, Alvarez-Arguelles ME, Abreu F, Costales I, Perez-Martínez Z, Martín-Rodríguez G, Coto E, Melón S |
| EPI_ISL_1915122 | Hospital | National Reference Center for Viruses of Respiratory Infections, Institut Pasteur, Paris | Marion Barbet, Sylvie Behillil, Méline Bizard, Angela Brisebarre, Camille Capel, Vincent Enouf, Louise Lefrançois, Frédéric Lemoine, Christophe Malabat, Corinne Maufrais, Emmanuelle Permal, Etienne Simon-Lorière, Maud Vanpeene, Sylvie Van der Werf ,Geoffroy Lorin De La Grandmaison |
| EPI_ISL_1915124, EPI_ISL_1915125, EPI_ISL_1915126 | Labo Analyses Med | National Reference Center for Viruses of Respiratory Infections, Institut Pasteur, Paris | Marion Barbet, Sylvie Behillil, Méline Bizard, Angela Brisebarre, Camille Capel, Vincent Enouf, Louise Lefrançois, Frédéric Lemoine, Christophe Malabat, Corinne Maufrais, Emmanuelle Permal, Etienne Simon-Lorière, Maud Vanpeene, Sylvie Van der Werf ,Nabil Gastli |
| EPI_ISL_1915127, EPI_ISL_1915130, EPI_ISL_1915132, EPI_ISL_1915136 | Hospital | National Reference Center for Viruses of Respiratory Infections, Institut Pasteur, Paris | Marion Barbet, Sylvie Behillil, Méline Bizard, Angela Brisebarre, Camille Capel, Vincent Enouf, Louise Lefrançois, Frédéric Lemoine, Christophe Malabat, Corinne Maufrais, Emmanuelle Permal, Etienne Simon-Lorière, Maud Vanpeene, Sylvie Van der Werf ,Eric Farfour |
| EPI_ISL_1915137, EPI_ISL_1915138, EPI_ISL_1915139 | Hospital | National Reference Center for Viruses of Respiratory Infections, Institut Pasteur, Paris | Marion Barbet, Sylvie Behillil, Méline Bizard, Angela Brisebarre, Camille Capel, Vincent Enouf, Louise Lefrançois, Frédéric Lemoine, Christophe Malabat, Corinne Maufrais, Emmanuelle Permal, Etienne Simon-Lorière, Maud Vanpeene, Sylvie Van der Werf ,Haciba Moudjahed |
| EPI_ISL_1915143 | Hospital | National Reference Center for Viruses of Respiratory Infections, Institut Pasteur, Paris | Marion Barbet, Sylvie Behillil, Méline Bizard, Angela Brisebarre, Camille Capel, Vincent Enouf, Louise Lefrançois, Frédéric Lemoine, Christophe Malabat, Corinne Maufrais, Emmanuelle Permal, Etienne Simon-Lorière, Maud Vanpeene, Sylvie Van der Werf ,Fribourg |
| EPI_ISL_1915144 | Labo Analyses Med | National Reference Center for Viruses of Respiratory Infections, Institut Pasteur, Paris | Marion Barbet, Sylvie Behillil, Méline Bizard, Angela Brisebarre, Camille Capel, Vincent Enouf, Louise Lefrançois, Frédéric Lemoine, Christophe Malabat, Corinne Maufrais, Emmanuelle Permal, Etienne Simon-Lorière, Maud Vanpeene, Sylvie Van der Werf ,Selas Medilys |
| EPI_ISL_1915145 | Hospital | National Reference Center for Viruses of Respiratory Infections, Institut Pasteur, Paris | Marion Barbet, Sylvie Behillil, Méline Bizard, Angela Brisebarre, Camille Capel, Vincent Enouf, Louise Lefrançois, Frédéric Lemoine, Christophe Malabat, Corinne Maufrais, Emmanuelle Permal, Etienne Simon-Lorière, Maud Vanpeene, Sylvie Van der Werf ,Pauline Garnier |
| EPI_ISL_1915146 | Labo Analyses Med | National Reference Center for Viruses of Respiratory Infections, Institut Pasteur, Paris | Marion Barbet, Sylvie Behillil, Méline Bizard, Angela Brisebarre, Camille Capel, Vincent Enouf, Louise Lefrançois, Frédéric Lemoine, Christophe Malabat, Corinne Maufrais, Emmanuelle Permal, Etienne Simon-Lorière, Maud Vanpeene, Sylvie Van der Werf ,T Flao |
| EPI_ISL_1915147, EPI_ISL_1915149 | Hospital | National Reference Center for Viruses of Respiratory Infections, Institut Pasteur, Paris | Marion Barbet, Sylvie Behillil, Méline Bizard, Angela Brisebarre, Camille Capel, Vincent Enouf, Louise Lefrançois, Frédéric Lemoine, Christophe Malabat, Corinne Maufrais, Emmanuelle Permal, Etienne Simon-Lorière, Maud Vanpeene, Sylvie Van der Werf ,Marianne Burgard |

| | | | |
|---|---|---|---|
| EPI_ISL_1915150 | Hospital | National Reference Center for Viruses of Respiratory Infections, Institut Pasteur, Paris | Marion Barbet, Sylvie Behillil, Méline Bizard, Angela Brisebarre, Camille Capel, Vincent Enouf, Louise Lefrançois, Frédéric Lemoine, Christophe Malabat, Corinne Maufrais, Emmanuelle Permal, Etienne Simon-Lorière, Maud Vanpeene, Sylvie Van der Werf ,CéLine Ramanantsoa |
| EPI_ISL_1915151, EPI_ISL_1915152, EPI_ISL_1915153 | Labo Analyses Med | National Reference Center for Viruses of Respiratory Infections, Institut Pasteur, Paris | Marion Barbet, Sylvie Behillil, Méline Bizard, Angela Brisebarre, Camille Capel, Vincent Enouf, Louise Lefrançois, Frédéric Lemoine, Christophe Malabat, Corinne Maufrais, Emmanuelle Permal, Etienne Simon-Lorière, Maud Vanpeene, Sylvie Van der Werf ,Anne Grattard |
| EPI_ISL_1915156 | Labo Analyses Med | National Reference Center for Viruses of Respiratory Infections, Institut Pasteur, Paris | Marion Barbet, Sylvie Behillil, Méline Bizard, Angela Brisebarre, Camille Capel, Vincent Enouf, Louise Lefrançois, Frédéric Lemoine, Christophe Malabat, Corinne Maufrais, Emmanuelle Permal, Etienne Simon-Lorière, Maud Vanpeene, Sylvie Van der Werf ,Fabienne Artur |
| EPI_ISL_1915157 | Hospital | National Reference Center for Viruses of Respiratory Infections, Institut Pasteur, Paris | Marion Barbet, Sylvie Behillil, Méline Bizard, Angela Brisebarre, Camille Capel, Vincent Enouf, Louise Lefrançois, Frédéric Lemoine, Christophe Malabat, Corinne Maufrais, Emmanuelle Permal, Etienne Simon-Lorière, Maud Vanpeene, Sylvie Van der Werf ,Marianne Burgard |
| EPI_ISL_1915158, EPI_ISL_1915159, EPI_ISL_1915160, EPI_ISL_1915161, EPI_ISL_1915162, EPI_ISL_1915163, EPI_ISL_1915165, EPI_ISL_1915166, EPI_ISL_1915167, EPI_ISL_1915173, EPI_ISL_1915175 | | | |
| see above | Labo Analyses Med | National Reference Center for Viruses of Respiratory Infections, Institut Pasteur, Paris | Marion Barbet, Sylvie Behillil, Méline Bizard, Angela Brisebarre, Camille Capel, Vincent Enouf, Louise Lefrançois, Frédéric Lemoine, Christophe Malabat, Corinne Maufrais, Emmanuelle Permal, Etienne Simon-Lorière, Maud Vanpeene, Sylvie Van der Werf ,Florent Tomasi |
| EPI_ISL_1915179 | Hospital | National Reference Center for Viruses of Respiratory Infections, Institut Pasteur, Paris | Marion Barbet, Sylvie Behillil, Méline Bizard, Angela Brisebarre, Camille Capel, Vincent Enouf, Louise Lefrançois, Frédéric Lemoine, Christophe Malabat, Corinne Maufrais, Emmanuelle Permal, Etienne Simon-Lorière, Maud Vanpeene, Sylvie Van der Werf ,Haciba Moudjahed |
| EPI_ISL_1915180, EPI_ISL_1915181, EPI_ISL_1915182 | Hospital | National Reference Center for Viruses of Respiratory Infections, Institut Pasteur, Paris | Marion Barbet, Sylvie Behillil, Méline Bizard, Angela Brisebarre, Camille Capel, Vincent Enouf, Louise Lefrançois, Frédéric Lemoine, Christophe Malabat, Corinne Maufrais, Emmanuelle Permal, Etienne Simon-Lorière, Maud Vanpeene, Sylvie Van der Werf ,Pascale Martres |
| EPI_ISL_1915185 | Hospital | National Reference Center for Viruses of Respiratory Infections, Institut Pasteur, Paris | Marion Barbet, Sylvie Behillil, Méline Bizard, Angela Brisebarre, Camille Capel, Vincent Enouf, Louise Lefrançois, Frédéric Lemoine, Christophe Malabat, Corinne Maufrais, Emmanuelle Permal, Etienne Simon-Lorière, Maud Vanpeene, Sylvie Van der Werf ,Franck Griscelli |
| EPI_ISL_1915187 | Labo Analyses Med | National Reference Center for Viruses of Respiratory Infections, Institut Pasteur, Paris | Marion Barbet, Sylvie Behillil, Méline Bizard, Angela Brisebarre, Camille Capel, Vincent Enouf, Louise Lefrançois, Frédéric Lemoine, Christophe Malabat, Corinne Maufrais, Emmanuelle Permal, Etienne Simon-Lorière, Maud Vanpeene, Sylvie Van der Werf ,StéPhane Romand |
| EPI_ISL_1915190 | Labo Analyses Med | National Reference Center for Viruses of Respiratory Infections, Institut Pasteur, Paris | Marion Barbet, Sylvie Behillil, Méline Bizard, Angela Brisebarre, Camille Capel, Vincent Enouf, Louise Lefrançois, Frédéric Lemoine, Christophe Malabat, Corinne Maufrais, Emmanuelle Permal, Etienne Simon-Lorière, Maud Vanpeene, Sylvie Van der Werf ,Vicky Le |
| EPI_ISL_1915191 | Labo Analyses Med | National Reference Center for Viruses of Respiratory Infections, Institut Pasteur, Paris | Marion Barbet, Sylvie Behillil, Méline Bizard, Angela Brisebarre, Camille Capel, Vincent Enouf, Louise Lefrançois, Frédéric Lemoine, Christophe Malabat, Corinne Maufrais, Emmanuelle Permal, Etienne Simon-Lorière, Maud Vanpeene, Sylvie Van der Werf ,Gilles Desvaux |
| EPI_ISL_1917493 | SIESP CHIETI - DRIVE IN VASTO | Istituto Zooprofilattico Sperimentale dell'Abruzzo e Molise "G. Caporale" | Lorusso A, Marcacci M, Di Domenico M, Ancora M, Curini V, Di Lollo Valeria, Mangone I, Rinaldi A, Delli Compagni E, Scialabba S, Caporale M, Di Pasquale A, Cammà C, Puglia I, Calistri P, Savini G |
| EPI_ISL_1917494 | RSA GIULIANOVA | Istituto Zooprofilattico Sperimentale dell'Abruzzo e Molise "G. Caporale" | Lorusso A, Marcacci M, Di Domenico M, Ancora M, Curini V, Di Lollo Valeria, Mangone I, Rinaldi A, Delli Compagni E, Scialabba S, Caporale M, Di Pasquale A, Cammà C, Puglia I, Calistri P, Savini G |
| EPI_ISL_1917495 | DIP. PREV. AVEZZANO SERVIZIO DI IGIENE EPIDEMIOLOGIA E SANITA' PUBBLICA | Istituto Zooprofilattico Sperimentale dell'Abruzzo e Molise "G. Caporale" | Lorusso A, Marcacci M, Di Domenico M, Ancora M, Curini V, Di Lollo Valeria, Mangone I, Rinaldi A, Delli Compagni E, Scialabba S, Caporale M, Di Pasquale A, Cammà C, Puglia I, Calistri P, Savini G |
| EPI_ISL_1917496 | USCA Avezzano | Istituto Zooprofilattico Sperimentale dell'Abruzzo e Molise "G. Caporale" | Lorusso A, Marcacci M, Di Domenico M, Ancora M, Curini V, Di Lollo Valeria, Mangone I, Rinaldi A, Delli Compagni E, Scialabba S, Caporale M, Di Pasquale A, Cammà C, Puglia I, Calistri P, Savini G |
| EPI_ISL_1917497 | USCA Tagliacozzo | Istituto Zooprofilattico Sperimentale dell'Abruzzo e Molise "G. Caporale" | Lorusso A, Marcacci M, Di Domenico M, Ancora M, Curini V, Di Lollo Valeria, Mangone I, Rinaldi A, Delli Compagni E, Scialabba S, Caporale M, Di Pasquale A, Cammà C, Puglia I, Calistri P, Savini G |
| EPI_ISL_1917498, EPI_ISL_1917499, EPI_ISL_1917500, EPI_ISL_1917501 | SIESP CHIETI - DRIVE IN LANCIANO | Istituto Zooprofilattico Sperimentale dell'Abruzzo e Molise "G. Caporale" | Lorusso A, Marcacci M, Di Domenico M, Ancora M, Curini V, Di Lollo Valeria, Mangone I, Rinaldi A, Delli Compagni E, Scialabba S, Caporale M, Di Pasquale A, Cammà C, Puglia I, Calistri P, Savini G |
| EPI_ISL_1917502 | SIESP DIPARTIMENTO DI PREVENZIONE CHIETI | Istituto Zooprofilattico Sperimentale dell'Abruzzo e Molise "G. Caporale" | Lorusso A, Marcacci M, Di Domenico M, Ancora M, Curini V, Di Lollo Valeria, Mangone I, Rinaldi A, Delli Compagni E, Scialabba S, Caporale M, Di Pasquale A, Cammà C, Puglia I, Calistri P, Savini G |
| EPI_ISL_1917503 | SIESP CHIETI - DRIVE IN ORTONA | Istituto Zooprofilattico Sperimentale dell'Abruzzo e Molise "G. Caporale" | Lorusso A, Marcacci M, Di Domenico M, Ancora M, Curini V, Di Lollo Valeria, Mangone I, Rinaldi A, Delli Compagni E, Scialabba S, Caporale M, Di Pasquale A, Cammà C, Puglia I, Calistri P, Savini G |
| EPI_ISL_1917504 | SIESP DIPARTIMENTO DI PREVENZIONE CHIETI | Istituto Zooprofilattico Sperimentale dell'Abruzzo e Molise "G. Caporale" | Lorusso A, Marcacci M, Di Domenico M, Ancora M, Curini V, Di Lollo Valeria, Mangone I, Rinaldi A, Delli Compagni E, Scialabba S, Caporale M, Di Pasquale A, Cammà C, Puglia I, Calistri P, Savini G |
| EPI_ISL_1917505, EPI_ISL_1917506 | DIP. PREV. AVEZZANO SERVIZIO DI IGIENE EPIDEMIOLOGIA E SANITA' PUBBLICA | Istituto Zooprofilattico Sperimentale dell'Abruzzo e Molise "G. Caporale" | Lorusso A, Marcacci M, Di Domenico M, Ancora M, Curini V, Di Lollo Valeria, Mangone I, Rinaldi A, Delli Compagni E, Scialabba S, Caporale M, Di Pasquale A, Cammà C, Puglia I, Calistri P, Savini G |
| EPI_ISL_1917507 | SIESP SULMONA | Istituto Zooprofilattico Sperimentale dell'Abruzzo e Molise "G. Caporale" | Lorusso A, Marcacci M, Di Domenico M, Ancora M, Curini V, Di Lollo Valeria, Mangone I, Rinaldi A, Delli Compagni E, Scialabba S, Caporale M, Di Pasquale A, Cammà C, Puglia I, Calistri P, Savini G |
| EPI_ISL_1917508, EPI_ISL_1917509 | SIESP DIPARTIMENTO DI PREVENZIONE TERAMO | Istituto Zooprofilattico Sperimentale dell'Abruzzo e Molise "G. Caporale" | Lorusso A, Marcacci M, Di Domenico M, Ancora M, Curini V, Di Lollo Valeria, Mangone I, Rinaldi A, Delli Compagni E, Scialabba S, Caporale M, Di Pasquale A, Cammà C, Puglia I, Calistri P, Savini G |
| EPI_ISL_1917510 | SIESP CHIETI - DRIVE IN VASTO | Istituto Zooprofilattico Sperimentale dell'Abruzzo e Molise "G. Caporale" | Lorusso A, Marcacci M, Di Domenico M, Ancora M, Curini V, Di Lollo Valeria, Mangone I, Rinaldi A, Delli Compagni E, Scialabba S, Caporale M, Di Pasquale A, Cammà C, Puglia I, Calistri P, Savini G |
| EPI_ISL_1917511 | SIESP CHIETI - DRIVE IN CHIETI | Istituto Zooprofilattico Sperimentale dell'Abruzzo e Molise "G. Caporale" | Lorusso A, Marcacci M, Di Domenico M, Ancora M, Curini V, Di Lollo Valeria, Mangone I, Rinaldi A, Delli Compagni E, Scialabba S, Caporale M, Di Pasquale A, Cammà C, Puglia I, Calistri P, Savini G |
| EPI_ISL_1917512 | SIESP CHIETI - DRIVE IN LANCIANO | Istituto Zooprofilattico Sperimentale dell'Abruzzo e Molise "G. Caporale" | Lorusso A, Marcacci M, Di Domenico M, Ancora M, Curini V, Di Lollo Valeria, Mangone I, Rinaldi A, Delli Compagni E, Scialabba S, Caporale M, Di Pasquale A, Cammà C, Puglia I, Calistri P, Savini G |
| EPI_ISL_1917513 | PRESIDIO TERRITORIALE CASOLI | Istituto Zooprofilattico Sperimentale dell'Abruzzo e Molise "G. Caporale" | Lorusso A, Marcacci M, Di Domenico M, Ancora M, Curini V, Di Lollo Valeria, Mangone I, Rinaldi A, Delli Compagni E, Scialabba S, Caporale M, Di Pasquale A, Cammà C, Puglia I, Calistri P, Savini G |
| EPI_ISL_1917514 | SIESP DIPARTIMENTO DI PREVENZIONE CHIETI | Istituto Zooprofilattico Sperimentale dell'Abruzzo e Molise "G. Caporale" | Lorusso A, Marcacci M, Di Domenico M, Ancora M, Curini V, Di Lollo Valeria, Mangone I, Rinaldi A, Delli Compagni E, Scialabba S, Caporale M, Di Pasquale A, Cammà C, Puglia I, Calistri P, Savini G |
| EPI_ISL_1917515 | SIESP DIPARTIMENTO DI PREVENZIONE TERAMO | Istituto Zooprofilattico Sperimentale dell'Abruzzo e Molise "G. Caporale" | Lorusso A, Marcacci M, Di Domenico M, Ancora M, Curini V, Di Lollo Valeria, Mangone I, Rinaldi A, Delli Compagni E, Scialabba S, Caporale M, Di Pasquale A, Cammà C, Puglia I, Calistri P, Savini G |
| EPI_ISL_1917516 | USCA Tagliacozzo | Istituto Zooprofilattico Sperimentale dell'Abruzzo e Molise "G. Caporale" | Lorusso A, Marcacci M, Di Domenico M, Ancora M, Curini V, Di Lollo Valeria, Mangone I, Rinaldi A, Delli Compagni E, Scialabba S, Caporale M, Di Pasquale A, Cammà C, Puglia I, Calistri P, Savini G |
| EPI_ISL_1917517, EPI_ISL_1917518, EPI_ISL_1917519 | DIP. PREV. AVEZZANO SERVIZIO DI IGIENE EPIDEMIOLOGIA E SANITA' PUBBLICA | Istituto Zooprofilattico Sperimentale dell'Abruzzo e Molise "G. Caporale" | Lorusso A, Marcacci M, Di Domenico M, Ancora M, Curini V, Di Lollo Valeria, Mangone I, Rinaldi A, Delli Compagni E, Scialabba S, Caporale M, Di Pasquale A, Cammà C, Puglia I, Calistri P, Savini G |
| EPI_ISL_1917520 | SIESP DIPARTIMENTO DI PREVENZIONE TERAMO | Istituto Zooprofilattico Sperimentale dell'Abruzzo e Molise "G. Caporale" | Lorusso A, Marcacci M, Di Domenico M, Ancora M, Curini V, Di Lollo Valeria, Mangone I, Rinaldi A, Delli Compagni E, Scialabba S, Caporale M, Di Pasquale A, Cammà C, Puglia I, Calistri P, Savini G |
| EPI_ISL_1917521, EPI_ISL_1917522 | SIESP CHIETI - DRIVE IN VASTO | Istituto Zooprofilattico Sperimentale dell'Abruzzo e Molise "G. Caporale" | Lorusso A, Marcacci M, Di Domenico M, Ancora M, Curini V, Di Lollo Valeria, Mangone I, Rinaldi A, Delli Compagni E, Scialabba S, Caporale M, Di Pasquale A, Cammà C, Puglia I, Calistri P, Savini G |
| EPI_ISL_1917523 | SIESP DIPARTIMENTO DI PREVENZIONE CHIETI | Istituto Zooprofilattico Sperimentale dell'Abruzzo e Molise "G. Caporale" | Lorusso A, Marcacci M, Di Domenico M, Ancora M, Curini V, Di Lollo Valeria, Mangone I, Rinaldi A, Delli Compagni E, Scialabba S, Caporale M, Di Pasquale A, Cammà C, Puglia I, Calistri P, Savini G |
| EPI_ISL_1917524 | SIESP CHIETI - DRIVE IN GISSI | Istituto Zooprofilattico Sperimentale dell'Abruzzo e Molise "G. | Lorusso A, Marcacci M, Di Domenico M, Ancora M, Curini V, Di Lollo Valeria, Mangone I, Rinaldi A, Delli Compagni E, Scialabba S, Caporale M, Di |

| | | | |
|---|---|---|---|
| | | Caporale" | Pasquale A, Cammà C, Puglia I, Calistri P, Savini G |
| EPI_ISL_1917525, EPI_ISL_1917526 | SIESP DIPARTIMENTO DI PREVENZIONE TERAMO | Istituto Zooprofilattico Sperimentale dell'Abruzzo e Molise "G. Caporale" | Lorusso A, Marcacci M, Di Domenico M, Ancora M, Curini V, Di Lollo Valeria, Mangone I, Rinaldi A, Delli Compagni E, Scialabba S, Caporale M, Di Pasquale A, Cammà C, Puglia I, Calistri P, Savini G |
| EPI_ISL_1917527 | SIESP CHIETI - DRIVE IN LANCIANO | Istituto Zooprofilattico Sperimentale dell'Abruzzo e Molise "G. Caporale" | Lorusso A, Marcacci M, Di Domenico M, Ancora M, Curini V, Di Lollo Valeria, Mangone I, Rinaldi A, Delli Compagni E, Scialabba S, Caporale M, Di Pasquale A, Cammà C, Puglia I, Calistri P, Savini G |
| EPI_ISL_1917528 | SIESP DIPARTIMENTO DI PREVENZIONE TERAMO | Istituto Zooprofilattico Sperimentale dell'Abruzzo e Molise "G. Caporale" | Lorusso A, Marcacci M, Di Domenico M, Ancora M, Curini V, Di Lollo Valeria, Mangone I, Rinaldi A, Delli Compagni E, Scialabba S, Caporale M, Di Pasquale A, Cammà C, Puglia I, Calistri P, Savini G |
| EPI_ISL_1917530 | USCA Avezzano | Istituto Zooprofilattico Sperimentale dell'Abruzzo e Molise "G. Caporale" | Lorusso A, Marcacci M, Di Domenico M, Ancora M, Curini V, Di Lollo Valeria, Mangone I, Rinaldi A, Delli Compagni E, Scialabba S, Caporale M, Di Pasquale A, Cammà C, Puglia I, Calistri P, Savini G |
| EPI_ISL_1917531 | SIESP SULMONA | Istituto Zooprofilattico Sperimentale dell'Abruzzo e Molise "G. Caporale" | Lorusso A, Marcacci M, Di Domenico M, Ancora M, Curini V, Di Lollo Valeria, Mangone I, Rinaldi A, Delli Compagni E, Scialabba S, Caporale M, Di Pasquale A, Cammà C, Puglia I, Calistri P, Savini G |
| EPI_ISL_1919272, EPI_ISL_1919273 | Ipoh Public Health Laboratory (MKAI), Ministry of Health Malaysia | Institute for Medical Research, Infectious Disease Research Centre, National Institutes of Health, Ministry of Health Malaysia | Suppiah J, Kamel K, Mohd Zawawi Z, Thayan R |
| EPI_ISL_1919422 | Ipoh Public Health Laboratory (MKAI), Ministry of Health Malaysia | Institute for Medical Research, Infectious Disease Research Centre, National Institutes of Health, Ministry | Suppiah J, Kamel K, Mohd Zawawi Z, Thayan R |
| EPI_ISL_1919602, EPI_ISL_1919603, EPI_ISL_1919604 | Università degli Studi di Perugia | Istituto Zooprofilattico Sperimentale dell'Abruzzo e Molise "G. Caporale" | Mencacci A, Camilloni B, Lorusso A, Marcacci M, Di Domenico M, Ancora M, Curini V, Mangone I, Rinaldi A, Delli Compagni E, Scialabba S, Di Pasquale A, Cammà C, Puglia I, Calistri P, Savini G |
| EPI_ISL_1919605, EPI_ISL_1919607 | Laboratorio Analisi Osp. Città di Castello - Azienda USL Umbria1 | Istituto Zooprofilattico Sperimentale dell'Abruzzo e Molise "G. Caporale" | Malagigi V, Tacconi P,Lorusso A, Marcacci M, Di Domenico M, Ancora M, Curini V, Di Lollo Valeria, Mangone I, Rinaldi A, Delli Compagni E, Scialabba S, Caporale M, Di Pasquale A, Cammà C, Puglia I, Calistri P, Savini G |
| EPI_ISL_1919631 | Azienda Ospedaliera Terni | Istituto Zooprofilattico Sperimentale dell'Abruzzo e Molise "G. Caporale" | Palumbo M, Scaccetti A, Lorusso A, Marcacci M, Di Domenico M, Ancora M, Curini V, Mangone I, Rinaldi A, Delli Compagni E, Scialabba S, Di Pasquale A, Cammà C, Puglia I, Calistri P, Savini G |
| EPI_ISL_1919633 | Università degli Studi di Perugia | Istituto Zooprofilattico Sperimentale dell'Abruzzo e Molise "G. Caporale" | Mencacci A, Camilloni B, Lorusso A, Marcacci M, Di Domenico M, Ancora M, Curini V, Mangone I, Rinaldi A, Delli Compagni E, Scialabba S, Di Pasquale A, Cammà C, Puglia I, Calistri P, Savini G |
| EPI_ISL_1919634, EPI_ISL_1919635, EPI_ISL_1919636, EPI_ISL_1919637, EPI_ISL_1919638 | Istituto Zooprofilattico Sperimentale Umbria e Marche "Togo Rosati" | Istituto Zooprofilattico Sperimentale dell'Abruzzo e Molise "G. Caporale" | Biagetti M, Giammarioli M, Lorusso A, Marcacci M, Di Domenico M, Ancora M, Curini V, Mangone I, Rinaldi A, Delli Compagni E, Scialabba S, Di Pasquale A, Cammà C, Puglia I, Calistri P, Savini G |
| EPI_ISL_1919641, EPI_ISL_1919642 | Università degli Studi di Perugia | Istituto Zooprofilattico Sperimentale dell'Abruzzo e Molise "G. Caporale" | Mencacci A, Camilloni B, Lorusso A, Marcacci M, Di Domenico M, Ancora M, Curini V, Mangone I, Rinaldi A, Delli Compagni E, Scialabba S, Di Pasquale A, Cammà C, Puglia I, Calistri P, Savini G |
| EPI_ISL_1919645, EPI_ISL_1919646 | Ipoh Public Health Laboratory (MKAI), Ministry of Health Malaysia | Institute for Medical Research, Infectious Disease Research Centre, National Institutes of Health, Ministry | Suppiah J, Kamel K, Mohd Zawawi Z, Thayan R |
| EPI_ISL_1919647, EPI_ISL_1919648, EPI_ISL_1919649 | Ipoh Public Health Laboratory (MKAI), Ministry of Health Malaysia | Institute for Medical Research, Infectious Disease Research Centre, National Institutes of Health, Ministry of Health Malaysia | Suppiah J, Kamel K, Mohd Zawawi Z, Thayan R |
| EPI_ISL_1919650, EPI_ISL_1919654 | Institute for Medical Research, Infectious Disease Research Centre, National Institutes of Health, Ministry of Health Malaysia | Institute for Medical Research, Infectious Disease Research Centre, National Institutes of Health, Ministry of Health Malaysia | Suppiah J, Kamel K, Mohd Zawawi Z, Ramly N, Robert F, Thayan R |
| EPI_ISL_1919754, EPI_ISL_1919755, EPI_ISL_1919757 | Ospedale Santa Caterina Novella | Istituto Zooprofilattico Sperimentale della Puglia e della Basilicata | Parisi A., Bianco A., Capozzi L., Del Sambro L., Simone D., Difato L., Bruno A. |
| EPI_ISL_1919858, EPI_ISL_1919859 | Ipoh Public Health Laboratory (MKAI), Ministry of Health Malaysia | Institute for Medical Research, Infectious Disease Research Centre, National Institutes of Health, Ministry of Health Malaysia | Suppiah J, Kamel K, Mohd Zawawi Z, Thayan R |
| EPI_ISL_1919860 | Institute for Medical Research, Infectious Disease Research Centre, National Institutes of Health, Ministry of Health Malaysia | Institute for Medical Research, Infectious Disease Research Centre, National Institutes of Health, Ministry of Health Malaysia | Suppiah J, Kamel K, Mohd Zawawi Z, Azizan MA, Ramly N, Robert F, Thayan R |
| EPI_ISL_1919861 | Duchess of Kent Hospital, Sandakan | Institute for Medical Research, Infectious Disease Research Centre, National Institutes of Health, Ministry of Health Malaysia | Suppiah J, Kamel K, Mohd Zawawi Z, Thayan R |
| EPI_ISL_1919862, EPI_ISL_1919863 | Tawau Hospital | Institute for Medical Research, Infectious Disease Research Centre, National Institutes of Health, Ministry of Health Malaysia | Suppiah J, Kamel K, Mohd Zawawi Z, Thayan R |
| EPI_ISL_1919944, EPI_ISL_1919945, EPI_ISL_1919946, EPI_ISL_1919947, EPI_ISL_1919949, EPI_ISL_1919950, EPI_ISL_1919986, EPI_ISL_1919987 | Ipoh Public Health Laboratory (MKAI), Ministry of Health Malaysia | Institute for Medical Research, Infectious Disease Research Centre, National Institutes of Health, Ministry of Health Malaysia | Suppiah J, Kamel K, Mohd Zawawi Z, Thayan R |
| EPI_ISL_1920176 | CH SAINT-PHILIBERT | CHU Lille | AIT YAHYA Emilie, ALIDJINOU Enagnon Kazali, BOCKET Laurence, CREPIN Michel, DEMAY Christophe, ENGELMANN Ilka, GEFFROY Sandrine, GUIGON Aurélie, LAMBERT Valérie, LAZREK Mouna, NOBILLIAUX Florian, PREVOST Brigitte, THUILLIER Caroline, TINEZ Claire |
| EPI_ISL_1920177 | CH ARRAS | CHU Lille | AIT YAHYA Emilie, ALIDJINOU Enagnon Kazali, BOCKET Laurence, CREPIN Michel, DEMAY Christophe, ENGELMANN Ilka, GEFFROY Sandrine, GUIGON Aurélie, LAMBERT Valérie, LAZREK Mouna, NOBILLIAUX Florian, PREVOST Brigitte, THUILLIER Caroline, TINEZ Claire |
| EPI_ISL_1920179, EPI_ISL_1920180 | CHU Lille | CHU Lille | AIT YAHYA Emilie, ALIDJINOU Enagnon Kazali, BOCKET Laurence, CREPIN Michel, DEMAY Christophe, ENGELMANN Ilka, GEFFROY Sandrine, GUIGON Aurélie, LAMBERT Valérie, LAZREK Mouna, NOBILLIAUX Florian, PREVOST Brigitte, THUILLIER Caroline, TINEZ Claire |
| EPI_ISL_1920182 | CH ARMENTIERES | CHU Lille | AIT YAHYA Emilie, ALIDJINOU Enagnon Kazali, BOCKET Laurence, CREPIN Michel, DEMAY Christophe, ENGELMANN Ilka, GEFFROY Sandrine, GUIGON Aurélie, LAMBERT Valérie, LAZREK Mouna, NOBILLIAUX Florian, PREVOST Brigitte, THUILLIER Caroline, TINEZ Claire |
| EPI_ISL_1920183, EPI_ISL_1920184 | CH CAMBRAI | CHU Lille | AIT YAHYA Emilie, ALIDJINOU Enagnon Kazali, BOCKET Laurence, CREPIN Michel, DEMAY Christophe, ENGELMANN Ilka, GEFFROY Sandrine, GUIGON Aurélie, LAMBERT Valérie, LAZREK Mouna, NOBILLIAUX Florian, PREVOST Brigitte, THUILLIER Caroline, TINEZ Claire |
| EPI_ISL_1920185, EPI_ISL_1920186, EPI_ISL_1920187, EPI_ISL_1920188, EPI_ISL_1920189, EPI_ISL_1920190, EPI_ISL_1920192, EPI_ISL_1920193, EPI_ISL_1920194 | CHU Lille | CHU Lille | AIT YAHYA Emilie, ALIDJINOU Enagnon Kazali, BOCKET Laurence, CREPIN Michel, DEMAY Christophe, ENGELMANN Ilka, GEFFROY Sandrine, GUIGON Aurélie, LAMBERT Valérie, LAZREK Mouna, NOBILLIAUX Florian, PREVOST Brigitte, THUILLIER Caroline, TINEZ Claire |
| EPI_ISL_1920195 | CH ARRAS | CHU Lille | AIT YAHYA Emilie, ALIDJINOU Enagnon Kazali, BOCKET Laurence, CREPIN Michel, DEMAY Christophe, ENGELMANN Ilka, GEFFROY Sandrine, GUIGON Aurélie, LAMBERT Valérie, LAZREK Mouna, NOBILLIAUX Florian, PREVOST Brigitte, THUILLIER Caroline, TINEZ Claire |
| EPI_ISL_1920196 | CH SAINT-PHILIBERT | CHU Lille | AIT YAHYA Emilie, ALIDJINOU Enagnon Kazali, BOCKET Laurence, CREPIN Michel, DEMAY Christophe, ENGELMANN Ilka, GEFFROY Sandrine, |

| | | | |
|---|---|---|---|
| | | | GUIGON Aurélie, LAMBERT Valérie, LAZREK Mouna, NOBILLIAUX Florian, PREVOST Brigitte, THUILLIER Caroline, TINEZ Claire |
| EPI_ISL_1920197 | CH DOUAI | CHU Lille | AIT YAHYA Emilie, ALIDJINOU Enagnon Kazali, BOCKET Laurence, CREPIN Michel, DEMAY Christophe, ENGELMANN Ilka, GEFFROY Sandrine, GUIGON Aurélie, LAMBERT Valérie, LAZREK Mouna, NOBILLIAUX Florian, PREVOST Brigitte, THUILLIER Caroline, TINEZ Claire |
| EPI_ISL_1920200, EPI_ISL_1920201, EPI_ISL_1920202 | CHU Lille | CHU Lille | AIT YAHYA Emilie, ALIDJINOU Enagnon Kazali, BOCKET Laurence, CREPIN Michel, DEMAY Christophe, ENGELMANN Ilka, GEFFROY Sandrine, GUIGON Aurélie, LAMBERT Valérie, LAZREK Mouna, NOBILLIAUX Florian, PREVOST Brigitte, THUILLIER Caroline, TINEZ Claire |
| EPI_ISL_1920203, EPI_ISL_1920204 | CH ROUBAIX | CHU Lille | AIT YAHYA Emilie, ALIDJINOU Enagnon Kazali, BOCKET Laurence, CREPIN Michel, DEMAY Christophe, ENGELMANN Ilka, GEFFROY Sandrine, GUIGON Aurélie, LAMBERT Valérie, LAZREK Mouna, NOBILLIAUX Florian, PREVOST Brigitte, THUILLIER Caroline, TINEZ Claire |
| EPI_ISL_1920205, EPI_ISL_1920207, EPI_ISL_1920208, EPI_ISL_1920209, EPI_ISL_1920210, EPI_ISL_1920211, EPI_ISL_1920212, EPI_ISL_1920214 | CHU Lille | CHU Lille | AIT YAHYA Emilie, ALIDJINOU Enagnon Kazali, BOCKET Laurence, CREPIN Michel, DEMAY Christophe, ENGELMANN Ilka, GEFFROY Sandrine, GUIGON Aurélie, LAMBERT Valérie, LAZREK Mouna, NOBILLIAUX Florian, PREVOST Brigitte, THUILLIER Caroline, TINEZ Claire |
| EPI_ISL_1920215, EPI_ISL_1920216 | CH DUNKERQUE | CHU Lille | AIT YAHYA Emilie, ALIDJINOU Enagnon Kazali, BOCKET Laurence, CREPIN Michel, DEMAY Christophe, ENGELMANN Ilka, GEFFROY Sandrine, GUIGON Aurélie, LAMBERT Valérie, LAZREK Mouna, NOBILLIAUX Florian, PREVOST Brigitte, THUILLIER Caroline, TINEZ Claire |
| EPI_ISL_1920217 | CH TOURCOING | CHU Lille | AIT YAHYA Emilie, ALIDJINOU Enagnon Kazali, BOCKET Laurence, CREPIN Michel, DEMAY Christophe, ENGELMANN Ilka, GEFFROY Sandrine, GUIGON Aurélie, LAMBERT Valérie, LAZREK Mouna, NOBILLIAUX Florian, PREVOST Brigitte, THUILLIER Caroline, TINEZ Claire |
| EPI_ISL_1920218 | CH ARRAS | CHU Lille | AIT YAHYA Emilie, ALIDJINOU Enagnon Kazali, BOCKET Laurence, CREPIN Michel, DEMAY Christophe, ENGELMANN Ilka, GEFFROY Sandrine, GUIGON Aurélie, LAMBERT Valérie, LAZREK Mouna, NOBILLIAUX Florian, PREVOST Brigitte, THUILLIER Caroline, TINEZ Claire |
| EPI_ISL_1920219, EPI_ISL_1920220 | CH ROUBAIX | CHU Lille | AIT YAHYA Emilie, ALIDJINOU Enagnon Kazali, BOCKET Laurence, CREPIN Michel, DEMAY Christophe, ENGELMANN Ilka, GEFFROY Sandrine, GUIGON Aurélie, LAMBERT Valérie, LAZREK Mouna, NOBILLIAUX Florian, PREVOST Brigitte, THUILLIER Caroline, TINEZ Claire |
| EPI_ISL_1920221, EPI_ISL_1920222 | CH BETHUNE | CHU Lille | AIT YAHYA Emilie, ALIDJINOU Enagnon Kazali, BOCKET Laurence, CREPIN Michel, DEMAY Christophe, ENGELMANN Ilka, GEFFROY Sandrine, GUIGON Aurélie, LAMBERT Valérie, LAZREK Mouna, NOBILLIAUX Florian, PREVOST Brigitte, THUILLIER Caroline, TINEZ Claire |
| EPI_ISL_1920223, EPI_ISL_1920224, EPI_ISL_1920225, EPI_ISL_1920226 | CH ARMENTIERES | CHU Lille | AIT YAHYA Emilie, ALIDJINOU Enagnon Kazali, BOCKET Laurence, CREPIN Michel, DEMAY Christophe, ENGELMANN Ilka, GEFFROY Sandrine, GUIGON Aurélie, LAMBERT Valérie, LAZREK Mouna, NOBILLIAUX Florian, PREVOST Brigitte, THUILLIER Caroline, TINEZ Claire |
| EPI_ISL_1920227 | CH TOURCOING | CHU Lille | AIT YAHYA Emilie, ALIDJINOU Enagnon Kazali, BOCKET Laurence, CREPIN Michel, DEMAY Christophe, ENGELMANN Ilka, GEFFROY Sandrine, GUIGON Aurélie, LAMBERT Valérie, LAZREK Mouna, NOBILLIAUX Florian, PREVOST Brigitte, THUILLIER Caroline, TINEZ Claire |
| EPI_ISL_1920228, EPI_ISL_1920229, EPI_ISL_1920230 | CH LENS | CHU Lille | AIT YAHYA Emilie, ALIDJINOU Enagnon Kazali, BOCKET Laurence, CREPIN Michel, DEMAY Christophe, ENGELMANN Ilka, GEFFROY Sandrine, GUIGON Aurélie, LAMBERT Valérie, LAZREK Mouna, NOBILLIAUX Florian, PREVOST Brigitte, THUILLIER Caroline, TINEZ Claire |
| EPI_ISL_1920231 | CH BOULOGNE-SUR-MER | CHU Lille | AIT YAHYA Emilie, ALIDJINOU Enagnon Kazali, BOCKET Laurence, CREPIN Michel, DEMAY Christophe, ENGELMANN Ilka, GEFFROY Sandrine, GUIGON Aurélie, LAMBERT Valérie, LAZREK Mouna, NOBILLIAUX Florian, PREVOST Brigitte, THUILLIER Caroline, TINEZ Claire |
| EPI_ISL_1920233, EPI_ISL_1920234, EPI_ISL_1920235, EPI_ISL_1920237, EPI_ISL_1920238, EPI_ISL_1920239, EPI_ISL_1920240, EPI_ISL_1920241 | CHU Lille | CHU Lille | AIT YAHYA Emilie, ALIDJINOU Enagnon Kazali, BOCKET Laurence, CREPIN Michel, DEMAY Christophe, ENGELMANN Ilka, GEFFROY Sandrine, GUIGON Aurélie, LAMBERT Valérie, LAZREK Mouna, NOBILLIAUX Florian, PREVOST Brigitte, THUILLIER Caroline, TINEZ Claire |
| EPI_ISL_1920503, EPI_ISL_1920504, EPI_ISL_1920506, EPI_ISL_1920507, EPI_ISL_1920508, EPI_ISL_1920509 | Laboratori de Referencia de Catalunya | Laboratori de Referencia de Catalunya | Ramirez A., Bellosillo B., Padilla E., Vilas A., Hernandez JJ., Canal M. |
| EPI_ISL_1921767, EPI_ISL_1921769 | Ipoh Public Health Laboratory (MKAI), Ministry of Health Malaysia | Institute for Medical Research, Infectious Disease Research Centre, National Institutes of Health, Ministry of Health Malaysia | Suppiah J, Kamel K, Mohd Zawawi Z, Thayan R |
| EPI_ISL_1921894, EPI_ISL_1921896 | OLVZ Aalst | OLVZ Aalst | Anne Vankeerberghen |
| EPI_ISL_1922125 | University of Bari Biomedical Sciences and Human Oncology | University of Bari Biomedical Sciences and Human Oncology | Chironna M., Sallustio A., Loconsole D., Accogli M. |
| EPI_ISL_1935600, EPI_ISL_1935601, EPI_ISL_1935602, EPI_ISL_1935603, EPI_ISL_1935604, EPI_ISL_1935605 | Cliniques universitaires Saint-Luc | UCLouvain/IREC/MBLG | Jean Ruelle, Eleonore Ngyuvula, Benoit Kabamba Mukadi |
| EPI_ISL_1936102, EPI_ISL_1936103, EPI_ISL_1936104, EPI_ISL_1936105, EPI_ISL_1936106, EPI_ISL_1936107, EPI_ISL_1936108, EPI_ISL_1936109, EPI_ISL_1936110, EPI_ISL_1936111, EPI_ISL_1936112, EPI_ISL_1936113, EPI_ISL_1936114, EPI_ISL_1936115, EPI_ISL_1936116, EPI_ISL_1936117, EPI_ISL_1936118, EPI_ISL_1936119, EPI_ISL_1936120, EPI_ISL_1936121, EPI_ISL_1936122, EPI_ISL_1936123, EPI_ISL_1936124, EPI_ISL_1936125, EPI_ISL_1936126, EPI_ISL_1936127, EPI_ISL_1936128, EPI_ISL_1936129, EPI_ISL_1936130, EPI_ISL_1936131, EPI_ISL_1936132, EPI_ISL_1936133, EPI_ISL_1936134, EPI_ISL_1936135, EPI_ISL_1936136, EPI_ISL_1936137, EPI_ISL_1936138, EPI_ISL_1936139, EPI_ISL_1936140, EPI_ISL_1936141, EPI_ISL_1936142, EPI_ISL_1936143, EPI_ISL_1936144, EPI_ISL_1936145 | | | |
| see above | Main Chemical Laboratories Egypt Army | Main Chemical Laboratories Egypt Army | Mohamed Seadawy, AbedElrahman Zekri, Mohamed Shamel, Ahmed Gad, Abdullah Salama, Mervat Hassan, Sabah Ahmed |
| EPI_ISL_1936240, EPI_ISL_1936241, EPI_ISL_1936242, EPI_ISL_1936243, EPI_ISL_1936244, EPI_ISL_1936245, EPI_ISL_1936246, EPI_ISL_1936247, EPI_ISL_1936248, EPI_ISL_1936249, EPI_ISL_1936250, EPI_ISL_1936251, EPI_ISL_1936252, EPI_ISL_1936253, EPI_ISL_1936254, EPI_ISL_1936255, EPI_ISL_1936256, EPI_ISL_1936257, EPI_ISL_1936258, EPI_ISL_1936259, EPI_ISL_1936260, EPI_ISL_1936261, EPI_ISL_1936262, EPI_ISL_1936263, EPI_ISL_1936264, EPI_ISL_1936265, EPI_ISL_1936266, EPI_ISL_1936267, EPI_ISL_1936268, EPI_ISL_1936269, EPI_ISL_1936270, EPI_ISL_1936271, EPI_ISL_1936272 | | | |
| see above | Main Chemical Laboratories Egypt Army | Main Chemical Laboratories Egypt Army | Mohamed Seadawy, AbedElrahman Zekri, Mohamed Shamel, Ahmed Gad, Bassem Elharty, Mostfa Elhoseiny |
| EPI_ISL_1939827, EPI_ISL_1939830, EPI_ISL_1939831, EPI_ISL_1939833, EPI_ISL_1939834, EPI_ISL_1939835, EPI_ISL_1939837, EPI_ISL_1939838 | Laboratory for Clinical Immunology and Molecular Genetics - University Clinic Golnik Laboratory for Respiratory Microbiology - University Clinic Golnik | Laboratory for Clinical Immunology and Molecular Genetics -University Clinic Golnik | Matija Rijavec, Julij Šelb, Urška Bidovec Stojkovi, Žan Kogovšek, Nina Rupar, Viktorija Tomi, Peter Korošec |
| EPI_ISL_1941875 | Hospital Universitario Cruces | Biocruces Bizkaia | Ana Belen de la Hoz, Mikel Gallego |
| EPI_ISL_1942243 | CQRC_QUALITY CONTROL CHEMICAL BIOLOGICAL RISK_AOOR Villa Sofia Cervello Palermo | CQRC_QUALITY CONTROL CHEMICAL BIOLOGICAL RISK_AOOR Villa Sofia Cervello Palermo | Di Gaudio, F., Contino, F., Orlando, V., Buffa, V., Bonvissuto, M., Di Chiara, V., Seidita, G.,Todaro, I., Di Sclafani, A., Lungari, A., Brunacci,G. |
| EPI_ISL_1942249 | Genome Centre | Genome Centre | Md. Shazid Hasan, Shovon Lal Sarkar, Ali Ahsan Setu, Tanay Chakrabarty, A. S. M. Rubayet Ul Alam, M. Shaminur Rahman, Ovinu Kibria Islam, M. Tanvir Islam, Hassan M. Al-Emran, Iqbal Kabir Jahid, M. Anwar Hossain |
| EPI_ISL_1960077 | LDSP | Universidad Nacional de Colombia - Laboratorio Genómico One Health | Simón Villegas Velásquez, Andres F. Cardona-Rios, Maria Stella López, Daniel O. Maldonado-Perez, Lina M. Hurtado, Laura Silvana Perez, Karl A Ciuoderis, Maria Angélica Maya, Carolina Muñoz-Arango, Celeny Ortiz, Rita Almanza Payares, Idabely Betancur Ortiz, Sandra Ines Cano, Diego A. Álvarez-Díaz, Carlos Franco-Muñoz, Katherine Laiton-Donato, Hector Alejandro Ruiz-Moreno, Marcela Mercado-Reyes, Jorge E. Osorio, Juan P. Hernandez-Ortiz |
| EPI_ISL_1960079 | ICMT-APARTADO | Universidad Nacional de Colombia - Laboratorio Genómico One Health | Andres F. Cardona-Rios, Maria Stella López, Daniel O. Maldonado-Perez, Simón Villegas Velásquez, Lina M. Hurtado, Laura Silvana Perez, Karl A Ciuoderis, Maria Angélica Maya, Carolina Muñoz-Arango, Celeny Ortiz, Rita Almanza Payares, Idabely Betancur Ortiz, Sandra Ines Cano, Diego A. Álvarez-Díaz, Carlos Franco-Muñoz, Katherine Laiton-Donato, Hector Alejandro Ruiz-Moreno, Marcela Mercado-Reyes, Jorge E. Osorio, Juan P. Hernandez-Ortiz |
| EPI_ISL_1960081 | LDSP | Universidad Nacional de Colombia - Laboratorio Genómico One Health | Simón Villegas Velásquez, Andres F. Cardona-Rios, Maria Stella López, Daniel O. Maldonado-Perez, Lina M. Hurtado, Laura Silvana Perez, Karl A Ciuoderis, Maria Angélica Maya, Carolina Muñoz-Arango, Celeny Ortiz, Rita Almanza Payares, Idabely Betancur Ortiz, Sandra Ines Cano, Diego A. Álvarez-Díaz, Carlos Franco-Muñoz, Katherine Laiton-Donato, Hector Alejandro Ruiz-Moreno, Marcela Mercado-Reyes, Jorge E. Osorio, Juan P. Hernandez-Ortiz |

| | | | |
|---|---|---|---|
| EPI_ISL_1960082 | UDEA | Universidad Nacional de Colombia - Laboratorio Genómico One Health | Andres F. Cardona-Rios, Maria Stella López, Daniel O. Maldonado-Perez, Simón Villegas Velásquez, Lina M. Hurtado, Laura Silvana Perez, Karl A Ciuoderis, Maria Angélica Maya, Carolina Muñoz-Arango, Celeny Ortiz, Rita Almanza Payares, Idabely Betancur Ortiz, Sandra Ines Cano, Diego A. Álvarez-Díaz, Carlos Franco-Muñoz, Katherine Laiton-Donato, Hector Alejandro Ruiz-Moreno, Marcela Mercado-Reyes, Jorge E. Osorio, Juan P. Hernandez-Ortiz |
| EPI_ISL_1960213, EPI_ISL_1960214 | UAB Diagnostikos laboratorija | National Public Health Surveillance Laboratory | Lukas Zemaitis, Migle Gabrielaite, Jelena Razmuk, Svajune Muralyte, Ana Steponkiene, Lukas Vasionis, Danas Baksa |
| EPI_ISL_1960215 | UAB Medicina practica laboratorija | National Public Health Surveillance Laboratory | Lukas Zemaitis, Migle Gabrielaite, Jelena Razmuk, Svajune Muralyte, Ana Steponkiene, Lukas Vasionis, Danas Baksa |
| EPI_ISL_1960217 | UAB Diagnostikos laboratorija | National Public Health Surveillance Laboratory | Lukas Zemaitis, Migle Gabrielaite, Jelena Razmuk, Svajune Muralyte, Ana Steponkiene, Lukas Vasionis, Danas Baksa |
| EPI_ISL_1960220, EPI_ISL_1960221, EPI_ISL_1960222, EPI_ISL_1960223 | Nacionalinis maisto ir veterinarijos rizikos vertinimo institutas | National Public Health Surveillance Laboratory | Lukas Zemaitis, Migle Gabrielaite, Jelena Razmuk, Svajune Muralyte, Ana Steponkiene, Lukas Vasionis, Danas Baksa |
| EPI_ISL_1960224 | UAB Diagnostikos laboratorija | National Public Health Surveillance Laboratory | Lukas Zemaitis, Migle Gabrielaite, Jelena Razmuk, Svajune Muralyte, Ana Steponkiene, Lukas Vasionis, Danas Baksa |
| EPI_ISL_1960225, EPI_ISL_1960226 | Nacionalinis maisto ir veterinarijos rizikos vertinimo institutas | National Public Health Surveillance Laboratory | Lukas Zemaitis, Migle Gabrielaite, Jelena Razmuk, Svajune Muralyte, Ana Steponkiene, Lukas Vasionis, Danas Baksa |
| EPI_ISL_1960227, EPI_ISL_1960228, EPI_ISL_1960229, EPI_ISL_1960230, EPI_ISL_1960231, EPI_ISL_1960232, EPI_ISL_1960233, EPI_ISL_1960234, EPI_ISL_1960235, EPI_ISL_1960236, EPI_ISL_1960237, EPI_ISL_1960238, EPI_ISL_1960239 | | | |
| see above | Nacionaline visuomenes sveikatos prieziuros laboratorija | National Public Health Surveillance Laboratory | Lukas Zemaitis, Migle Gabrielaite, Jelena Razmuk, Svajune Muralyte, Ana Steponkiene, Lukas Vasionis, Danas Baksa |
| EPI_ISL_1960240, EPI_ISL_1960241, EPI_ISL_1960242, EPI_ISL_1960243, EPI_ISL_1960244, EPI_ISL_1960245, EPI_ISL_1960246, EPI_ISL_1960247, EPI_ISL_1960248, EPI_ISL_1960249, EPI_ISL_1960250, EPI_ISL_1960251, EPI_ISL_1960252, EPI_ISL_1960253, EPI_ISL_1960254, EPI_ISL_1960255, EPI_ISL_1960256, EPI_ISL_1960257, EPI_ISL_1960258, EPI_ISL_1960259, EPI_ISL_1960260, EPI_ISL_1960261, EPI_ISL_1960262, EPI_ISL_1960263, EPI_ISL_1960264, EPI_ISL_1960265, EPI_ISL_1960266, EPI_ISL_1960267, EPI_ISL_1960268, EPI_ISL_1960269, EPI_ISL_1960270, EPI_ISL_1960271, EPI_ISL_1960272, EPI_ISL_1960273, EPI_ISL_1960274, EPI_ISL_1960275, EPI_ISL_1960276, EPI_ISL_1960277, EPI_ISL_1960278, EPI_ISL_1960279, EPI_ISL_1960280, EPI_ISL_1960281, EPI_ISL_1960282, EPI_ISL_1960283, EPI_ISL_1960284, EPI_ISL_1960285, EPI_ISL_1960286, EPI_ISL_1960287, EPI_ISL_1960288, EPI_ISL_1960289, EPI_ISL_1960290, EPI_ISL_1960291, EPI_ISL_1960292, EPI_ISL_1960293, EPI_ISL_1960294, EPI_ISL_1960295, EPI_ISL_1960296, EPI_ISL_1960297, EPI_ISL_1960298, EPI_ISL_1960299, EPI_ISL_1960300, EPI_ISL_1960301, EPI_ISL_1960302, EPI_ISL_1960303, EPI_ISL_1960304, EPI_ISL_1960305, EPI_ISL_1960306, EPI_ISL_1960307, EPI_ISL_1960308, EPI_ISL_1960309, EPI_ISL_1960310, EPI_ISL_1960311, EPI_ISL_1960312, EPI_ISL_1960313, EPI_ISL_1960314, EPI_ISL_1960315, EPI_ISL_1960316, EPI_ISL_1960317, EPI_ISL_1960318, EPI_ISL_1960319, EPI_ISL_1960320, EPI_ISL_1960321, EPI_ISL_1960322, EPI_ISL_1960323, EPI_ISL_1960324, EPI_ISL_1960325, EPI_ISL_1960326, EPI_ISL_1960327, EPI_ISL_1960328, EPI_ISL_1960329, EPI_ISL_1960330, EPI_ISL_1960331, EPI_ISL_1960332, EPI_ISL_1960333, EPI_ISL_1960334, EPI_ISL_1960335, EPI_ISL_1960336, EPI_ISL_1960337, EPI_ISL_1960338, EPI_ISL_1960339, EPI_ISL_1960340, EPI_ISL_1960341, EPI_ISL_1960342, EPI_ISL_1960343, EPI_ISL_1960344, EPI_ISL_1960345, EPI_ISL_1960346, EPI_ISL_1960347, EPI_ISL_1960348, EPI_ISL_1960349, EPI_ISL_1960350, EPI_ISL_1960351, EPI_ISL_1960352, EPI_ISL_1960353, EPI_ISL_1960354, EPI_ISL_1960355, EPI_ISL_1960356, EPI_ISL_1960357, EPI_ISL_1960358, EPI_ISL_1960359, EPI_ISL_1960360, EPI_ISL_1960361, EPI_ISL_1960362, EPI_ISL_1960363, EPI_ISL_1960364, EPI_ISL_1960365, EPI_ISL_1960366, EPI_ISL_1960367, EPI_ISL_1960368, EPI_ISL_1960369, EPI_ISL_1960370, EPI_ISL_1960371, EPI_ISL_1960372, EPI_ISL_1960373, EPI_ISL_1960374, EPI_ISL_1960375, EPI_ISL_1960376, EPI_ISL_1960377, EPI_ISL_1960378, EPI_ISL_1960379, EPI_ISL_1960380, EPI_ISL_1960381, EPI_ISL_1960382, EPI_ISL_1960383, EPI_ISL_1960384, EPI_ISL_1960385, EPI_ISL_1960386, EPI_ISL_1960387, EPI_ISL_1960388, EPI_ISL_1960389, EPI_ISL_1960390, EPI_ISL_1960391, EPI_ISL_1960392, EPI_ISL_1960393, EPI_ISL_1960394, EPI_ISL_1960395, EPI_ISL_1960396, EPI_ISL_1960397, EPI_ISL_1960398, EPI_ISL_1960399, EPI_ISL_1960400, EPI_ISL_1960401, EPI_ISL_1960402, EPI_ISL_1960403, EPI_ISL_1960404, EPI_ISL_1960405, EPI_ISL_1960406, EPI_ISL_1960407, EPI_ISL_1960408, EPI_ISL_1960409, EPI_ISL_1960410, EPI_ISL_1960411, EPI_ISL_1960412, EPI_ISL_1960413, EPI_ISL_1960414, EPI_ISL_1960415, EPI_ISL_1960416, EPI_ISL_1960417 | | | |
| see above | Nacionalinis maisto ir veterinarijos rizikos vertinimo institutas | National Public Health Surveillance Laboratory | Lukas Zemaitis, Migle Gabrielaite, Jelena Razmuk, Svajune Muralyte, Ana Steponkiene, Lukas Vasionis, Danas Baksa |
| EPI_ISL_1960418, EPI_ISL_1960419, EPI_ISL_1960420, EPI_ISL_1960421, EPI_ISL_1960422, EPI_ISL_1960423, EPI_ISL_1960424, EPI_ISL_1960425, EPI_ISL_1960426, EPI_ISL_1960427 | UAB "Rezus.lt" | National Public Health Surveillance Laboratory | Lukas Zemaitis, Migle Gabrielaite, Jelena Razmuk, Svajune Muralyte, Ana Steponkiene, Lukas Vasionis, Danas Baksa |
| EPI_ISL_1960428, EPI_ISL_1960429, EPI_ISL_1960430, EPI_ISL_1960431, EPI_ISL_1960432, EPI_ISL_1960433, EPI_ISL_1960434, EPI_ISL_1960435, EPI_ISL_1960436, EPI_ISL_1960437, EPI_ISL_1960438, EPI_ISL_1960439, EPI_ISL_1960440, EPI_ISL_1960441, EPI_ISL_1960442, EPI_ISL_1960443, EPI_ISL_1960444, EPI_ISL_1960445, EPI_ISL_1960446, EPI_ISL_1960447, EPI_ISL_1960448, EPI_ISL_1960449, EPI_ISL_1960450, EPI_ISL_1960451, EPI_ISL_1960452, EPI_ISL_1960453, EPI_ISL_1960454, EPI_ISL_1960455, EPI_ISL_1960456, EPI_ISL_1960457, EPI_ISL_1960458, EPI_ISL_1960459, EPI_ISL_1960460, EPI_ISL_1960461, EPI_ISL_1960462, EPI_ISL_1960463, EPI_ISL_1960464, EPI_ISL_1960465, EPI_ISL_1960466, EPI_ISL_1960467, EPI_ISL_1960468, EPI_ISL_1960469, EPI_ISL_1960470, EPI_ISL_1960471, EPI_ISL_1960472, EPI_ISL_1960473, EPI_ISL_1960474, EPI_ISL_1960475, EPI_ISL_1960476, EPI_ISL_1960477, EPI_ISL_1960478, EPI_ISL_1960479, EPI_ISL_1960480, EPI_ISL_1960481, EPI_ISL_1960482, EPI_ISL_1960483, EPI_ISL_1960484, EPI_ISL_1960485, EPI_ISL_1960486, EPI_ISL_1960487, EPI_ISL_1960488, EPI_ISL_1960489, EPI_ISL_1960490, EPI_ISL_1960491, EPI_ISL_1960492, EPI_ISL_1960493, EPI_ISL_1960494, EPI_ISL_1960495, EPI_ISL_1960496 | | | |
| see above | UAB Diagnostikos laboratorija | National Public Health Surveillance Laboratory | Lukas Zemaitis, Migle Gabrielaite, Jelena Razmuk, Svajune Muralyte, Ana Steponkiene, Lukas Vasionis, Danas Baksa |
| EPI_ISL_1960497, EPI_ISL_1960498, EPI_ISL_1960499, EPI_ISL_1960500, EPI_ISL_1960501, EPI_ISL_1960502, EPI_ISL_1960503, EPI_ISL_1960504, EPI_ISL_1960505, EPI_ISL_1960506, EPI_ISL_1960507, EPI_ISL_1960508, EPI_ISL_1960509, EPI_ISL_1960510, EPI_ISL_1960511, EPI_ISL_1960512, EPI_ISL_1960513, EPI_ISL_1960514, EPI_ISL_1960515, EPI_ISL_1960516, EPI_ISL_1960517, EPI_ISL_1960518, EPI_ISL_1960519, EPI_ISL_1960520 | | | |
| see above | UAB Medicina practica laboratorija | National Public Health Surveillance Laboratory | Lukas Zemaitis, Migle Gabrielaite, Jelena Razmuk, Svajune Muralyte, Ana Steponkiene, Lukas Vasionis, Danas Baksa |
| EPI_ISL_1960521, EPI_ISL_1960522, EPI_ISL_1960523, EPI_ISL_1960524, EPI_ISL_1960525 | Viesoji istaiga Klaipedos universitetine ligonine | National Public Health Surveillance Laboratory | Lukas Zemaitis, Migle Gabrielaite, Jelena Razmuk, Svajune Muralyte, Ana Steponkiene, Lukas Vasionis, Danas Baksa |
| EPI_ISL_1960526, EPI_ISL_1960527, EPI_ISL_1960528, EPI_ISL_1960529, EPI_ISL_1960530, EPI_ISL_1960531 | Nacionaline visuomenes sveikatos prieziuros laboratorija | National Public Health Surveillance Laboratory | Lukas Zemaitis, Migle Gabrielaite, Jelena Razmuk, Svajune Muralyte, Ana Steponkiene, Lukas Vasionis, Danas Baksa |
| EPI_ISL_1960532, EPI_ISL_1960533, EPI_ISL_1960534, EPI_ISL_1960535, EPI_ISL_1960536, EPI_ISL_1960537, EPI_ISL_1960538, EPI_ISL_1960539, EPI_ISL_1960540, EPI_ISL_1960541, EPI_ISL_1960542, EPI_ISL_1960543, EPI_ISL_1960544, EPI_ISL_1960545, EPI_ISL_1960546, EPI_ISL_1960547, EPI_ISL_1960548, EPI_ISL_1960549, EPI_ISL_1960550, EPI_ISL_1960551, EPI_ISL_1960552, EPI_ISL_1960553, EPI_ISL_1960554, EPI_ISL_1960555, EPI_ISL_1960556, EPI_ISL_1960557, EPI_ISL_1960558 | | | |
| see above | Nacionalinis maisto ir veterinarijos rizikos vertinimo institutas | National Public Health Surveillance Laboratory | Lukas Zemaitis, Migle Gabrielaite, Jelena Razmuk, Svajune Muralyte, Ana Steponkiene, Lukas Vasionis, Danas Baksa |
| EPI_ISL_1960559, EPI_ISL_1960560 | UAB "Rezus.lt" | National Public Health Surveillance Laboratory | Lukas Zemaitis, Migle Gabrielaite, Jelena Razmuk, Svajune Muralyte, Ana Steponkiene, Lukas Vasionis, Danas Baksa |
| EPI_ISL_1960561, EPI_ISL_1960562, EPI_ISL_1960563, EPI_ISL_1960564, EPI_ISL_1960565, EPI_ISL_1960566, EPI_ISL_1960567, EPI_ISL_1960568 | UAB Diagnostikos laboratorija | National Public Health Surveillance Laboratory | Lukas Zemaitis, Migle Gabrielaite, Jelena Razmuk, Svajune Muralyte, Ana Steponkiene, Lukas Vasionis, Danas Baksa |
| EPI_ISL_1960569, EPI_ISL_1960570, EPI_ISL_1960571, EPI_ISL_1960572, EPI_ISL_1960573, EPI_ISL_1960574, EPI_ISL_1960575, EPI_ISL_1960576, EPI_ISL_1960577, EPI_ISL_1960578 | UAB Medicina practica laboratorija | National Public Health Surveillance Laboratory | Lukas Zemaitis, Migle Gabrielaite, Jelena Razmuk, Svajune Muralyte, Ana Steponkiene, Lukas Vasionis, Danas Baksa |
| EPI_ISL_1960579 | Viesoji istaiga Klaipedos universitetine ligonine | National Public Health Surveillance Laboratory | Lukas Zemaitis, Migle Gabrielaite, Jelena Razmuk, Svajune Muralyte, Ana Steponkiene, Lukas Vasionis, Danas Baksa |
| EPI_ISL_1960580 | UAB Medicina practica laboratorija | National Public Health Surveillance Laboratory | Lukas Zemaitis, Migle Gabrielaite, Jelena Razmuk, Svajune Muralyte, Ana Steponkiene, Lukas Vasionis, Danas Baksa |
| EPI_ISL_1960581 | Nacionalinis maisto ir veterinarijos rizikos vertinimo institutas | National Public Health Surveillance Laboratory | Lukas Zemaitis, Migle Gabrielaite, Jelena Razmuk, Svajune Muralyte, Ana Steponkiene, Lukas Vasionis, Danas Baksa |
| EPI_ISL_1960603 | Viesoji istaiga Klaipedos universitetine ligonine | National Public Health Surveillance Laboratory | Lukas Zemaitis, Migle Gabrielaite, Jelena Razmuk, Svajune Muralyte, Ana Steponkiene, Lukas Vasionis, Danas Baksa |
| EPI_ISL_1960604 | UAB Diagnostikos laboratorija | National Public Health Surveillance Laboratory | Lukas Zemaitis, Migle Gabrielaite, Jelena Razmuk, Svajune Muralyte, Ana Steponkiene, Lukas Vasionis, Danas Baksa |
| EPI_ISL_1960606, EPI_ISL_1960607, EPI_ISL_1960608, EPI_ISL_1960609, EPI_ISL_1960610, EPI_ISL_1960611, EPI_ISL_1960612, EPI_ISL_1960613, EPI_ISL_1960614, EPI_ISL_1960615, EPI_ISL_1960616 | | | |
| see above | Nacionaline visuomenes sveikatos prieziuros laboratorija | National Public Health Surveillance Laboratory | Lukas Zemaitis, Migle Gabrielaite, Jelena Razmuk, Svajune Muralyte, Ana Steponkiene, Lukas Vasionis, Danas Baksa |
| EPI_ISL_1960617 | Viesoji istaiga Klaipedos universitetine ligonine | National Public Health Surveillance Laboratory | Lukas Zemaitis, Migle Gabrielaite, Jelena Razmuk, Svajune Muralyte, Ana Steponkiene, Lukas Vasionis, Danas Baksa |
| EPI_ISL_1960618, EPI_ISL_1960619, EPI_ISL_1960620 | Nacionalinis maisto ir veterinarijos rizikos vertinimo institutas | National Public Health Surveillance Laboratory | Lukas Zemaitis, Migle Gabrielaite, Jelena Razmuk, Svajune Muralyte, Ana Steponkiene, Lukas Vasionis, Danas Baksa |
| EPI_ISL_1960621, EPI_ISL_1960622, EPI_ISL_1960623, EPI_ISL_1960624, EPI_ISL_1960625, EPI_ISL_1960626 | UAB Diagnostikos laboratorija | National Public Health Surveillance Laboratory | Lukas Zemaitis, Migle Gabrielaite, Jelena Razmuk, Svajune Muralyte, Ana Steponkiene, Lukas Vasionis, Danas Baksa |

| | | | |
|---|---|---|---|
| EPI_ISL_1960627, EPI_ISL_1960628, EPI_ISL_1960629, EPI_ISL_1960630, EPI_ISL_1960631 | Viesoji istaiga Klaipedos universitetine ligonine | National Public Health Surveillance Laboratory | Lukas Zemaitis, Migle Gabrielaite, Jelena Razmuk, Svajune Muralyte, Ana Steponkiene, Lukas Vasionis, Danas Baksa |
| EPI_ISL_1965328, EPI_ISL_1965329, EPI_ISL_1965339, EPI_ISL_1965340, EPI_ISL_1965341, EPI_ISL_1965342, EPI_ISL_1965343, EPI_ISL_1965344, EPI_ISL_1965345, EPI_ISL_1965346, EPI_ISL_1965347, EPI_ISL_1965348, EPI_ISL_1965349, EPI_ISL_1965350, EPI_ISL_1965352, EPI_ISL_1965353, EPI_ISL_1965354, EPI_ISL_1965355, EPI_ISL_1965356, EPI_ISL_1965357, EPI_ISL_1965358, EPI_ISL_1965359, EPI_ISL_1965360, EPI_ISL_1965361, EPI_ISL_1965362, EPI_ISL_1965363, EPI_ISL_1965364, EPI_ISL_1965366, EPI_ISL_1965367, EPI_ISL_1965368, EPI_ISL_1965369, EPI_ISL_1965370, EPI_ISL_1965371, EPI_ISL_1965372, EPI_ISL_1965373, EPI_ISL_1965374, EPI_ISL_1965375, EPI_ISL_1965376, EPI_ISL_1965377, EPI_ISL_1965378, EPI_ISL_1965379, EPI_ISL_1965380, EPI_ISL_1965381, EPI_ISL_1965382, EPI_ISL_1965383, EPI_ISL_1965384, EPI_ISL_1965385, EPI_ISL_1965386, EPI_ISL_1965387, EPI_ISL_1965388, EPI_ISL_1965389, EPI_ISL_1965390, EPI_ISL_1965391, EPI_ISL_1965392, EPI_ISL_1965393, EPI_ISL_1965394, EPI_ISL_1965395, EPI_ISL_1965396, EPI_ISL_1965397, EPI_ISL_1965398, EPI_ISL_1965399, EPI_ISL_1965400, EPI_ISL_1965401, EPI_ISL_1965402, EPI_ISL_1965403, EPI_ISL_1965404, EPI_ISL_1965405, EPI_ISL_1965406, EPI_ISL_1965407, EPI_ISL_1965408, EPI_ISL_1965409, EPI_ISL_1965410, EPI_ISL_1965411, EPI_ISL_1965412, EPI_ISL_1965413, EPI_ISL_1965414, EPI_ISL_1965415, EPI_ISL_1965417, EPI_ISL_1965418, EPI_ISL_1965419, EPI_ISL_1965420, EPI_ISL_1965421, EPI_ISL_1965422, EPI_ISL_1965423, EPI_ISL_1965424, EPI_ISL_1965425, EPI_ISL_1965426, EPI_ISL_1965427, EPI_ISL_1965429, EPI_ISL_1965430, EPI_ISL_1965431, EPI_ISL_1965432, EPI_ISL_1965433, EPI_ISL_1965434, EPI_ISL_1965435, EPI_ISL_1965436, EPI_ISL_1965437 | | | |
| see above | IPK | Respiratory Infections Laboratory | Lissette Perez, Yahisel Tejero, Yanet Pintos, Daylin Mederos, Odalys Valdes, Mirtha Elisa Aguado, Liena Ponce |
| EPI_ISL_1965438, EPI_ISL_1965439, EPI_ISL_1965442, EPI_ISL_1965443, EPI_ISL_1965444, EPI_ISL_1965445, EPI_ISL_1965446, EPI_ISL_1965447, EPI_ISL_1965448, EPI_ISL_1965449, EPI_ISL_1965450, EPI_ISL_1965451, EPI_ISL_1965452, EPI_ISL_1965453, EPI_ISL_1965454, EPI_ISL_1965455, EPI_ISL_1965456, EPI_ISL_1965457, EPI_ISL_1965458, EPI_ISL_1965459, EPI_ISL_1965460, EPI_ISL_1965461, EPI_ISL_1965462, EPI_ISL_1965463, EPI_ISL_1965464, EPI_ISL_1965465, EPI_ISL_1965466, EPI_ISL_1965467, EPI_ISL_1965468, EPI_ISL_1965470, EPI_ISL_1965471, EPI_ISL_1965472, EPI_ISL_1965473, EPI_ISL_1965474, EPI_ISL_1965475, EPI_ISL_1965477, EPI_ISL_1965478, EPI_ISL_1965479, EPI_ISL_1965480, EPI_ISL_1965481, EPI_ISL_1965483, EPI_ISL_1965484, EPI_ISL_1965485, EPI_ISL_1965486 | | | |
| see above | HG Pharma GmbH | Bergthaler laboratory, CeMM Research Center for Molecular Medicine of the Austrian Academy of Sciences | Lukas Endler, Anna Schedl, Fabian Amman, Petr Triska, Thomas Penz, Benedikt Agerer, Maelle Le Moing, Michael Schuster, Bekir Erguner, Jan Laine, Martin Senekowitsch, Christoph Bock, Andreas Bergthaler |
| EPI_ISL_1965488 | Department of Laboratory Medicine, Division of Clinical Virology, University of Medicine, Vienna | Bergthaler laboratory, CeMM Research Center for Molecular Medicine of the Austrian Academy of Sciences | Lukas Endler, Anna Schedl, Fabian Amman, Petr Triska, Thomas Penz, Benedikt Agerer, Maelle Le Moing, Michael Schuster, Bekir Erguner, Jan Laine, Martin Senekowitsch, Christoph Bock, Andreas Bergthaler |
| EPI_ISL_1965665, EPI_ISL_1965666 | Institute for Laboratory Diagnostics and Microbiology, Klinikum Klagenfurt am Wörthersee | Bergthaler laboratory, CeMM Research Center for Molecular Medicine of the Austrian Academy of Sciences | Lukas Endler, Anna Schedl, Fabian Amman, Petr Triska, Thomas Penz, Benedikt Agerer, Maelle Le Moing, Michael Schuster, Bekir Erguner, Jan Laine, Martin Senekowitsch, Christoph Bock, Andreas Bergthaler |
| EPI_ISL_1965714, EPI_ISL_1965716, EPI_ISL_1965717, EPI_ISL_1965719, EPI_ISL_1965720 | 3. Medizinische Abteilung, Hanusch Krankenhaus | Bergthaler laboratory, CeMM Research Center for Molecular Medicine of the Austrian Academy of Sciences | Lukas Endler, Anna Schedl, Fabian Amman, Petr Triska, Thomas Penz, Benedikt Agerer, Maelle Le Moing, Michael Schuster, Bekir Erguner, Jan Laine, Martin Senekowitsch, Christoph Bock, Andreas Bergthaler |
| EPI_ISL_1968980 | Omics Sciences Laboratory | Omics Sciences Laboratory | Derly Andrade Molina, Rubén Armas González, Gabriel Morey León, Darlyn Amaya, Katheryn Sacheri Viteri, Juan Carlos Fernández Cadena |
| EPI_ISL_1969080, EPI_ISL_1969081 | Main Chemical Laboratories Egypt Army | Main Chemical Laboratories Egypt Army | Mohamed Seadawy, AbedElrahman Zekri, Mohamed Shamel, Ahmed Gad, Abdullah Salama, Mostfa Elhoseiny |
| EPI_ISL_1969669, EPI_ISL_1969670, EPI_ISL_1969671, EPI_ISL_1969672, EPI_ISL_1969673, EPI_ISL_1969674, EPI_ISL_1969675, EPI_ISL_1969676, EPI_ISL_1969677, EPI_ISL_1969678, EPI_ISL_1969679, EPI_ISL_1969680, EPI_ISL_1969681, EPI_ISL_1969682, EPI_ISL_1969683, EPI_ISL_1969684, EPI_ISL_1969685, EPI_ISL_1969686, EPI_ISL_1969687, EPI_ISL_1969688, EPI_ISL_1969689 | | | |
| see above | Virology Unit, Institut Pasteur du Cambodge | Virology Unit, Institut Pasteur du Cambodge | Jurre Y Siegers, Cecile Troupin, Ly Sovann, Kraing Sidonn, Yi Sengdoeurn, Chin Savuth, Chau Darapheak, Veasna Duong, Erik A Karlsson |
| EPI_ISL_1969690 | Singapore General Hospital | Department of Microbiology | Nurdyana Abdul Rahman, Kenneth Xin Long Chan, Suphavilai Chayaporn, Kun Lee Lim, Sui Sin Goh, Kian Sing Chan, Lynette Oon, Kern Rei Chng, James Sim Heng Chiak, Niranjan Nagarajan, Karrie Ko |
| EPI_ISL_1969933 | National Institute of Infectious Diseases-Prof. Dr. Matei Bals Molecular Diagnostics Laboratory | National Institute of Infectious Diseases-Prof. Dr. Matei Bals Molecular Diagnostics Laboratory | Corina Casangiu, Leontina Banica, Marius Surleac, Ovidiu Vlaicu, Andreea Tudor, Simona Paraschiv, Dan Otelea |
| EPI_ISL_1970692 | CHWAPI - SITE NOTRE DAME | Institut de Pathologie et Genetique (IPG) | Pascale Hilbert, Jérémie Gras |
| EPI_ISL_1970693 | CLINIQUE N-D DE GRACE | Institut de Pathologie et Genetique (IPG) | Pascale Hilbert, Jérémie Gras |
| EPI_ISL_1970694, EPI_ISL_1970702 | CHWAPI - SITE NOTRE DAME | Institut de Pathologie et Genetique (IPG) | Pascale Hilbert, Jérémie Gras |
| EPI_ISL_1970836, EPI_ISL_1970837 | Centro de Investigación en Enfermedades Infecciosas (CIENI), Instituto Nacional de Enfermedades Respiratorias (INER) | Centro de Investigación en Enfermedades Infecciosas (CIENI), Instituto Nacional de Enfermedades Respiratorias (INER) | Celia Boukadida, Carlos F. Arias, Santiago Ávila-Ríos, Eduardo Becerril-Vargas, Ricardo Grande, Alejandra Hernández-Terán, Pavel Isa, José Arturo Martínez-Orozco, Margarita Matías-Florentino, Fidencio Mejía-Nepomuceno, Mario Mújica-Sánchez, Hector Esteban Paz-Juárez, Francisco Pulido, Jorge Salas-Hernández, Alejandro Sanchez-Flores, Blanca Taboada, Joel Armando Vázquez-Pérez and Consorcio Mexicano de Vigilancia Genómica (CoViGen-Mex) |
| EPI_ISL_1972117 | AZ St.Jan Brugge-Oostende | AZ SINT-JAN BRUGGE | Marijke Reynders, Patrick Descheemaeker, Laurien Hoornaert, Thomas Van Landschoot, Jorn Hellemans |
| EPI_ISL_1972118 | AZ Zeno | AZ SINT-JAN BRUGGE | Marijke Reynders, Patrick Descheemaeker, Laurien Hoornaert, Thomas Van Landschoot, Jorn Hellemans |
| EPI_ISL_1972119, EPI_ISL_1972120, EPI_ISL_1972121, EPI_ISL_1972122, EPI_ISL_1972123, EPI_ISL_1972124 | AZ St.Jan Brugge-Oostende | AZ SINT-JAN BRUGGE | Marijke Reynders, Patrick Descheemaeker, Laurien Hoornaert, Thomas Van Landschoot, Jorn Hellemans |
| EPI_ISL_1972125, EPI_ISL_1972126, EPI_ISL_1972127, EPI_ISL_1972128, EPI_ISL_1972129 | AZ Zeno | AZ SINT-JAN BRUGGE | Marijke Reynders, Patrick Descheemaeker, Laurien Hoornaert, Thomas Van Landschoot, Jorn Hellemans |
| EPI_ISL_1972238, EPI_ISL_1972239, EPI_ISL_1972240, EPI_ISL_1972241, EPI_ISL_1972242, EPI_ISL_1972243, EPI_ISL_1972245, EPI_ISL_1972248, EPI_ISL_1972249, EPI_ISL_1972254, EPI_ISL_1972255, EPI_ISL_1972256, EPI_ISL_1972257, EPI_ISL_1972260, EPI_ISL_1972261, EPI_ISL_1972264, EPI_ISL_1972266, EPI_ISL_1972267, EPI_ISL_1972268, EPI_ISL_1972269, EPI_ISL_1972270, EPI_ISL_1972271, EPI_ISL_1972273, EPI_ISL_1972274, EPI_ISL_1972275, EPI_ISL_1972278, EPI_ISL_1972279, EPI_ISL_1972280, EPI_ISL_1972281, EPI_ISL_1972287, EPI_ISL_1972289, EPI_ISL_1972290, EPI_ISL_1972291, EPI_ISL_1972293, EPI_ISL_1972294, EPI_ISL_1972295, EPI_ISL_1972296, EPI_ISL_1972297, EPI_ISL_1972298, EPI_ISL_1972299, EPI_ISL_1972300, EPI_ISL_1972304 | | | |
| see above | Baylor Scott & White-Temple | Baylor Scott & White-Temple | Ari Rao, Linden Morales, Kimberly Walker, Marcus Volz, Shelby Hendrickson |
| EPI_ISL_1972356 | Sungai Buloh Hospital | Institute for Medical Research, Infectious Disease Research Centre, National Institutes of Health, Ministry of Health Malaysia | Suppiah J, Kamel K, Mohd Zawawi Z, Azizan MA, Ramly N, Robert F, Thayan R |
| EPI_ISL_1972365, EPI_ISL_1972533 | Sultanah Bahiyah Hospital, Alor Setar | Institute for Medical Research, Infectious Disease Research Centre, National Institutes of Health, Ministry of Health Malaysia | Suppiah J, Kamel K, Mohd Zawawi Z, Azizan MA, Ramly N, Robert F, Thayan R |
| EPI_ISL_1972534, EPI_ISL_1972535, EPI_ISL_1972536, EPI_ISL_1972537 | Institute for Medical Research, Infectious Disease Research Centre, National Institutes of Health, Ministry of Health Malaysia | Institute for Medical Research, Infectious Disease Research Centre, National Institutes of Health, Ministry of Health Malaysia | Suppiah J, Kamel K, Mohd Zawawi Z, Azizan MA, Ramly N, Robert F, Thayan R |
| EPI_ISL_1972539, EPI_ISL_1972540, EPI_ISL_1972547 | Tampin Hospital | Institute for Medical Research, Infectious Disease Research Centre, National Institutes of Health, Ministry of Health Malaysia | Suppiah J, Kamel K, Mohd Zawawi Z, Azizan MA, Ramly N, Robert F, Thayan R |
| EPI_ISL_1972548 | Klinik Kesihatan Lurah Bilut (Felda) | Institute for Medical Research, Infectious Disease Research Centre, National Institutes of Health, Ministry of Health Malaysia | Suppiah J, Kamel K, Mohd Zawawi Z, Azizan MA, Ramly N, Robert F, Thayan R |
| EPI_ISL_1972549, EPI_ISL_1972550 | Klinik Kesihatan Bentong | Institute for Medical Research, Infectious Disease Research Centre, National Institutes of Health, Ministry of Health Malaysia | Suppiah J, Kamel K, Mohd Zawawi Z, Azizan MA, Ramly N, Robert F, Thayan R |
| EPI_ISL_1972551, EPI_ISL_1972552 | Klinik Kesihatan Lurah Bilut (Felda) | Institute for Medical Research, Infectious Disease Research Centre, National Institutes of Health, Ministry of Health Malaysia | Suppiah J, Kamel K, Mohd Zawawi Z, Azizan MA, Ramly N, Robert F, Thayan R |
| EPI_ISL_1972553 | Klinik Kesihatan Karak | Institute for Medical Research, Infectious Disease Research | Suppiah J, Kamel K, Mohd Zawawi Z, Azizan MA, Ramly N, Robert F, Thayan R |

| | | | |
|---|---|---|---|
| | | Centre, National Institutes of Health, Ministry of Health Malaysia | |
| EPI_ISL_1972906, EPI_ISL_1972907, EPI_ISL_1972913, EPI_ISL_1972914, EPI_ISL_1972915, EPI_ISL_1972948 | Usansolo-Galdakao University Hospital | Cruces University Hospital | Ana Gual-de-Torrella, Izaskun Alejo-Cancho, Mikel Gallego, Ana Belén de la Hoz |
| EPI_ISL_1973324 | Kota Bharu Public Health Laboratory | Institute for Medical Research, Infectious Disease Research Centre, National Institutes of Health, Ministry of Health Malaysia | uppiah J, Kamel K, Mohd Zawawi Z, Azizan MA, Ramly N, Robert F, Thayan R |
| EPI_ISL_1973325 | Bintulu Hospital | Institute for Medical Research, Infectious Disease Research Centre, National Institutes of Health, Ministry of Health Malaysia | Suppiah J, Kamel K, Mohd Zawawi Z, Azizan MA, Ramly N, Robert F, Thayan R |
| EPI_ISL_1993547 | National Influenza Center, Virology Department | National Influenza Center | V Salimi,NZ Shafiei Jandaghi, J Yavarian, A Nejati, K Sadeghi, N Ghavvami,F Ajaminejad and T Mokhtari Azad |
| EPI_ISL_1993549 | National Influenza Center, Virology Department | National Influenza Center | J Yavarian,K Sadeghi, NZ Shafiei Jandaghi, V Salimi, A Nejati, N Ghavvami,F Ajaminejad and T Mokhtari Azad |
| EPI_ISL_1993550 | National Influenza Center, Virology Department | National Influenza Center | A Nejati, J Yavarian,K Sadeghi, NZ Shafiei Jandaghi, V Salimi, N Ghavvami,F Ajaminejad and T Mokhtari Azad |
| EPI_ISL_1993552 | National Influenza Center, Virology Department | National Influenza Center | J Yavarian, NZ Shafiei Jandaghi, V Salimi, A Nejati, K Sadeghi, F Ajaminejad,N Ghavvami and T Mokhtari Azad |
| EPI_ISL_1993553 | National Influenza Center, Virology Department | National Influenza Center | NZ Shafiei Jandaghi, V Salimi, A Nejati, K Sadeghi, J Yavarian, F Ajaminejad,N Ghavvami and T Mokhtari Azad |
| EPI_ISL_1993554 | National Influenza Center, Virology Department | National Influenza Center | V Salimi,NZ Shafiei Jandaghi, J Yavarian, A Nejati, K Sadeghi, N Ghavvami,F Ajaminejad and T Mokhtari Azad |
| EPI_ISL_1993555 | National Influenza Center, Virology Department | National Influenza Center | A Nejati, J Yavarian,K Sadeghi, NZ Shafiei Jandaghi, V Salimi, N Ghavvami,F Ajaminejad and T Mokhtari Azad |
| EPI_ISL_1993955, EPI_ISL_1993956, EPI_ISL_1993957, EPI_ISL_1993958, EPI_ISL_1993959, EPI_ISL_1993960, EPI_ISL_1993961, EPI_ISL_1993962, EPI_ISL_1993963, EPI_ISL_1993964, EPI_ISL_1993965, EPI_ISL_1993966, EPI_ISL_1993967, EPI_ISL_1993968, EPI_ISL_1993969, EPI_ISL_1993970, EPI_ISL_1993971, EPI_ISL_1993972, EPI_ISL_1993973, EPI_ISL_1993974, EPI_ISL_1993977, EPI_ISL_1993979, EPI_ISL_1993980, EPI_ISL_1993981, EPI_ISL_1993982, EPI_ISL_1993983, EPI_ISL_1993984, EPI_ISL_1993985, EPI_ISL_1993986, EPI_ISL_1993987, EPI_ISL_1993988, EPI_ISL_1993989 | | | |
| see above | CHU Lille | CHU Lille | AIT YAHYA Emilie, ALIDJINOU Enagnon Kazali, BOCKET Laurence, CREPIN Michel, DEMAY Christophe, ENGELMANN Ilka, GEFFROY Sandrine, GUIGON Aurélie, LAMBERT Valérie, LAZREK Mouna, NOBILLIAUX Florian, PREVOST Brigitte, THUILLIER Caroline, TINEZ Claire |
| EPI_ISL_1993990, EPI_ISL_1993991, EPI_ISL_1993992, EPI_ISL_1993993, EPI_ISL_1993994, EPI_ISL_1993995 | CH Armentieres | CHU Lille | AIT YAHYA Emilie, ALIDJINOU Enagnon Kazali, BOCKET Laurence, CREPIN Michel, DEMAY Christophe, ENGELMANN Ilka, GEFFROY Sandrine, GUIGON Aurélie, LAMBERT Valérie, LAZREK Mouna, NOBILLIAUX Florian, PREVOST Brigitte, THUILLIER Caroline, TINEZ Claire |
| EPI_ISL_1993996, EPI_ISL_1993997, EPI_ISL_1993998, EPI_ISL_1993999 | CH Bethunes | CHU Lille | AIT YAHYA Emilie, ALIDJINOU Enagnon Kazali, BOCKET Laurence, CREPIN Michel, DEMAY Christophe, ENGELMANN Ilka, GEFFROY Sandrine, GUIGON Aurélie, LAMBERT Valérie, LAZREK Mouna, NOBILLIAUX Florian, PREVOST Brigitte, THUILLIER Caroline, TINEZ Claire |
| EPI_ISL_1994000, EPI_ISL_1994001, EPI_ISL_1994002 | CH Boulogne Sur Mer | CHU Lille | AIT YAHYA Emilie, ALIDJINOU Enagnon Kazali, BOCKET Laurence, CREPIN Michel, DEMAY Christophe, ENGELMANN Ilka, GEFFROY Sandrine, GUIGON Aurélie, LAMBERT Valérie, LAZREK Mouna, NOBILLIAUX Florian, PREVOST Brigitte, THUILLIER Caroline, TINEZ Claire |
| EPI_ISL_1994003, EPI_ISL_1994004 | CH Valenciennes | CHU Lille | AIT YAHYA Emilie, ALIDJINOU Enagnon Kazali, BOCKET Laurence, CREPIN Michel, DEMAY Christophe, ENGELMANN Ilka, GEFFROY Sandrine, GUIGON Aurélie, LAMBERT Valérie, LAZREK Mouna, NOBILLIAUX Florian, PREVOST Brigitte, THUILLIER Caroline, TINEZ Claire |
| EPI_ISL_1994005, EPI_ISL_1994006 | CH St Omer | CHU Lille | AIT YAHYA Emilie, ALIDJINOU Enagnon Kazali, BOCKET Laurence, CREPIN Michel, DEMAY Christophe, ENGELMANN Ilka, GEFFROY Sandrine, GUIGON Aurélie, LAMBERT Valérie, LAZREK Mouna, NOBILLIAUX Florian, PREVOST Brigitte, THUILLIER Caroline, TINEZ Claire |
| EPI_ISL_1994007, EPI_ISL_1994008 | CH Calais | CHU Lille | AIT YAHYA Emilie, ALIDJINOU Enagnon Kazali, BOCKET Laurence, CREPIN Michel, DEMAY Christophe, ENGELMANN Ilka, GEFFROY Sandrine, GUIGON Aurélie, LAMBERT Valérie, LAZREK Mouna, NOBILLIAUX Florian, PREVOST Brigitte, THUILLIER Caroline, TINEZ Claire |
| EPI_ISL_1994009, EPI_ISL_1994010 | CH Cambrai | CHU Lille | AIT YAHYA Emilie, ALIDJINOU Enagnon Kazali, BOCKET Laurence, CREPIN Michel, DEMAY Christophe, ENGELMANN Ilka, GEFFROY Sandrine, GUIGON Aurélie, LAMBERT Valérie, LAZREK Mouna, NOBILLIAUX Florian, PREVOST Brigitte, THUILLIER Caroline, TINEZ Claire |
| EPI_ISL_1994011 | CH Douai | CHU Lille | AIT YAHYA Emilie, ALIDJINOU Enagnon Kazali, BOCKET Laurence, CREPIN Michel, DEMAY Christophe, ENGELMANN Ilka, GEFFROY Sandrine, GUIGON Aurélie, LAMBERT Valérie, LAZREK Mouna, NOBILLIAUX Florian, PREVOST Brigitte, THUILLIER Caroline, TINEZ Claire |
| EPI_ISL_1994013, EPI_ISL_1994014 | CH Dunkerque | CHU Lille | AIT YAHYA Emilie, ALIDJINOU Enagnon Kazali, BOCKET Laurence, CREPIN Michel, DEMAY Christophe, ENGELMANN Ilka, GEFFROY Sandrine, GUIGON Aurélie, LAMBERT Valérie, LAZREK Mouna, NOBILLIAUX Florian, PREVOST Brigitte, THUILLIER Caroline, TINEZ Claire |
| EPI_ISL_1994015, EPI_ISL_1994016 | CH Lens | CHU Lille | AIT YAHYA Emilie, ALIDJINOU Enagnon Kazali, BOCKET Laurence, CREPIN Michel, DEMAY Christophe, ENGELMANN Ilka, GEFFROY Sandrine, GUIGON Aurélie, LAMBERT Valérie, LAZREK Mouna, NOBILLIAUX Florian, PREVOST Brigitte, THUILLIER Caroline, TINEZ Claire |
| EPI_ISL_1994017, EPI_ISL_1994018, EPI_ISL_1994019, EPI_ISL_1994020 | CH Roubaix | CHU Lille | AIT YAHYA Emilie, ALIDJINOU Enagnon Kazali, BOCKET Laurence, CREPIN Michel, DEMAY Christophe, ENGELMANN Ilka, GEFFROY Sandrine, GUIGON Aurélie, LAMBERT Valérie, LAZREK Mouna, NOBILLIAUX Florian, PREVOST Brigitte, THUILLIER Caroline, TINEZ Claire |
| EPI_ISL_1994021 | CH Seclin | CHU Lille | AIT YAHYA Emilie, ALIDJINOU Enagnon Kazali, BOCKET Laurence, CREPIN Michel, DEMAY Christophe, ENGELMANN Ilka, GEFFROY Sandrine, GUIGON Aurélie, LAMBERT Valérie, LAZREK Mouna, NOBILLIAUX Florian, PREVOST Brigitte, THUILLIER Caroline, TINEZ Claire |
| EPI_ISL_1994023 | CHU Lille | CHU Lille | AIT YAHYA Emilie, ALIDJINOU Enagnon Kazali, BOCKET Laurence, CREPIN Michel, DEMAY Christophe, ENGELMANN Ilka, GEFFROY Sandrine, GUIGON Aurélie, LAMBERT Valérie, LAZREK Mouna, NOBILLIAUX Florian, PREVOST Brigitte, THUILLIER Caroline, TINEZ Claire |
| EPI_ISL_1994024, EPI_ISL_1994025, EPI_ISL_1994027 | CH St Philibert | CHU Lille | AIT YAHYA Emilie, ALIDJINOU Enagnon Kazali, BOCKET Laurence, CREPIN Michel, DEMAY Christophe, ENGELMANN Ilka, GEFFROY Sandrine, GUIGON Aurélie, LAMBERT Valérie, LAZREK Mouna, NOBILLIAUX Florian, PREVOST Brigitte, THUILLIER Caroline, TINEZ Claire |
| EPI_ISL_1994029 | CH Tourcoing | CHU Lille | AIT YAHYA Emilie, ALIDJINOU Enagnon Kazali, BOCKET Laurence, CREPIN Michel, DEMAY Christophe, ENGELMANN Ilka, GEFFROY Sandrine, GUIGON Aurélie, LAMBERT Valérie, LAZREK Mouna, NOBILLIAUX Florian, PREVOST Brigitte, THUILLIER Caroline, TINEZ Claire |
| EPI_ISL_2001298 | USCA Avezzano | Istituto Zooprofilattico Sperimentale dell'Abruzzo e Molise "G. Caporale" | Lorusso A, Marcacci M, Di Domenico M, Ancora M, Curini V, Di Lollo Valeria, Mangone I, Rinaldi A, Delli Compagni E, Scialabba S, Caporale M, Di Pasquale A, Cammà C, Puglia I, Calistri P, Savini G |
| EPI_ISL_2001299 | SIESP CHIETI - DRIVE IN ORTONA | Istituto Zooprofilattico Sperimentale dell'Abruzzo e Molise "G. Caporale" | Lorusso A, Marcacci M, Di Domenico M, Ancora M, Curini V, Di Lollo Valeria, Mangone I, Rinaldi A, Delli Compagni E, Scialabba S, Caporale M, Di Pasquale A, Cammà C, Puglia I, Calistri P, Savini G |
| EPI_ISL_2001300, EPI_ISL_2001301, EPI_ISL_2001302 | USCA Castel Di Sangro | Istituto Zooprofilattico Sperimentale dell'Abruzzo e Molise "G. Caporale" | Lorusso A, Marcacci M, Di Domenico M, Ancora M, Curini V, Di Lollo Valeria, Mangone I, Rinaldi A, Delli Compagni E, Scialabba S, Caporale M, Di Pasquale A, Cammà C, Puglia I, Calistri P, Savini G |
| EPI_ISL_2001303, EPI_ISL_2001304 | SIESP CHIETI - DRIVE IN GISSI | Istituto Zooprofilattico Sperimentale dell'Abruzzo e Molise "G. Caporale" | Lorusso A, Marcacci M, Di Domenico M, Ancora M, Curini V, Di Lollo Valeria, Mangone I, Rinaldi A, Delli Compagni E, Scialabba S, Caporale M, Di Pasquale A, Cammà C, Puglia I, Calistri P, Savini G |
| EPI_ISL_2001305 | SIESP CHIETI - DRIVE IN VASTO | Istituto Zooprofilattico Sperimentale dell'Abruzzo e Molise "G. Caporale" | Lorusso A, Marcacci M, Di Domenico M, Ancora M, Curini V, Di Lollo Valeria, Mangone I, Rinaldi A, Delli Compagni E, Scialabba S, Caporale M, Di Pasquale A, Cammà C, Puglia I, Calistri P, Savini G |
| EPI_ISL_2001306, EPI_ISL_2001307, EPI_ISL_2001308, EPI_ISL_2001309, EPI_ISL_2001311 | SIESP CHIETI - DRIVE IN LANCIANO | Istituto Zooprofilattico Sperimentale dell'Abruzzo e Molise "G. Caporale" | Lorusso A, Marcacci M, Di Domenico M, Ancora M, Curini V, Di Lollo Valeria, Mangone I, Rinaldi A, Delli Compagni E, Scialabba S, Caporale M, Di Pasquale A, Cammà C, Puglia I, Calistri P, Savini G |
| EPI_ISL_2001312 | SIESP CHIETI - DRIVE IN VASTO | Istituto Zooprofilattico Sperimentale dell'Abruzzo e Molise "G. Caporale" | Lorusso A, Marcacci M, Di Domenico M, Ancora M, Curini V, Di Lollo Valeria, Mangone I, Rinaldi A, Delli Compagni E, Scialabba S, Caporale M, Di Pasquale A, Cammà C, Puglia I, Calistri P, Savini G |
| EPI_ISL_2001313 | SIESP CHIETI - DRIVE IN LANCIANO | Istituto Zooprofilattico Sperimentale dell'Abruzzo e Molise "G. Caporale" | Lorusso A, Marcacci M, Di Domenico M, Ancora M, Curini V, Di Lollo Valeria, Mangone I, Rinaldi A, Delli Compagni E, Scialabba S, Caporale M, Di Pasquale A, Cammà C, Puglia I, Calistri P, Savini G |

| | | | |
|---|---|---|---|
| EPI_ISL_2001314 | PRESIDIO TERRITORIALE CASOLI | Istituto Zooprofilattico Sperimentale dell'Abruzzo e Molise "G. Caporale" | Lorusso A, Marcacci M, Di Domenico M, Ancora M, Curini V, Di Lollo Valeria, Mangone I, Rinaldi A, Delli Compagni E, Scialabba S, Caporale M, Di Pasquale A, Cammà C, Puglia I, Calistri P, Savini G |
| EPI_ISL_2001315 | SIESP CHIETI - DRIVE IN VASTO | Istituto Zooprofilattico Sperimentale dell'Abruzzo e Molise "G. Caporale" | Lorusso A, Marcacci M, Di Domenico M, Ancora M, Curini V, Di Lollo Valeria, Mangone I, Rinaldi A, Delli Compagni E, Scialabba S, Caporale M, Di Pasquale A, Cammà C, Puglia I, Calistri P, Savini G |
| EPI_ISL_2001316 | SIESP CHIETI - DRIVE IN LANCIANO | Istituto Zooprofilattico Sperimentale dell'Abruzzo e Molise "G. Caporale" | Lorusso A, Marcacci M, Di Domenico M, Ancora M, Curini V, Di Lollo Valeria, Mangone I, Rinaldi A, Delli Compagni E, Scialabba S, Caporale M, Di Pasquale A, Cammà C, Puglia I, Calistri P, Savini G |
| EPI_ISL_2001317 | SIESP CHIETI - DRIVE IN ORTONA | Istituto Zooprofilattico Sperimentale dell'Abruzzo e Molise "G. Caporale" | Lorusso A, Marcacci M, Di Domenico M, Ancora M, Curini V, Di Lollo Valeria, Mangone I, Rinaldi A, Delli Compagni E, Scialabba S, Caporale M, Di Pasquale A, Cammà C, Puglia I, Calistri P, Savini G |
| EPI_ISL_2001319, EPI_ISL_2001320, EPI_ISL_2001321, EPI_ISL_2001322, EPI_ISL_2001323, EPI_ISL_2001324, EPI_ISL_2001325 | SIESP DIPARTIMENTO DI PREVENZIONE TERAMO | Istituto Zooprofilattico Sperimentale dell'Abruzzo e Molise "G. Caporale" | Lorusso A, Marcacci M, Di Domenico M, Ancora M, Curini V, Di Lollo Valeria, Mangone I, Rinaldi A, Delli Compagni E, Scialabba S, Caporale M, Di Pasquale A, Cammà C, Puglia I, Calistri P, Savini G |
| EPI_ISL_2001326 | SIESP SULMONA | Istituto Zooprofilattico Sperimentale dell'Abruzzo e Molise "G. Caporale" | Lorusso A, Marcacci M, Di Domenico M, Ancora M, Curini V, Di Lollo Valeria, Mangone I, Rinaldi A, Delli Compagni E, Scialabba S, Caporale M, Di Pasquale A, Cammà C, Puglia I, Calistri P, Savini G |
| EPI_ISL_2001327, EPI_ISL_2001328 | SIESP DIPARTIMENTO DI PREVENZIONE TERAMO | Istituto Zooprofilattico Sperimentale dell'Abruzzo e Molise "G. Caporale" | Lorusso A, Marcacci M, Di Domenico M, Ancora M, Curini V, Di Lollo Valeria, Mangone I, Rinaldi A, Delli Compagni E, Scialabba S, Caporale M, Di Pasquale A, Cammà C, Puglia I, Calistri P, Savini G |
| EPI_ISL_2001329, EPI_ISL_2001331 | SIESP CHIETI - DRIVE IN LANCIANO | Istituto Zooprofilattico Sperimentale dell'Abruzzo e Molise "G. Caporale" | Lorusso A, Marcacci M, Di Domenico M, Ancora M, Curini V, Di Lollo Valeria, Mangone I, Rinaldi A, Delli Compagni E, Scialabba S, Caporale M, Di Pasquale A, Cammà C, Puglia I, Calistri P, Savini G |
| EPI_ISL_2001332, EPI_ISL_2001333 | SIESP DIPARTIMENTO DI PREVENZIONE TERAMO | Istituto Zooprofilattico Sperimentale dell'Abruzzo e Molise "G. Caporale" | Lorusso A, Marcacci M, Di Domenico M, Ancora M, Curini V, Di Lollo Valeria, Mangone I, Rinaldi A, Delli Compagni E, Scialabba S, Caporale M, Di Pasquale A, Cammà C, Puglia I, Calistri P, Savini G |
| EPI_ISL_2001334, EPI_ISL_2001335, EPI_ISL_2001336, EPI_ISL_2001337 | SIESP CHIETI - DRIVE IN ORTONA | Istituto Zooprofilattico Sperimentale dell'Abruzzo e Molise "G. Caporale" | Lorusso A, Marcacci M, Di Domenico M, Ancora M, Curini V, Di Lollo Valeria, Mangone I, Rinaldi A, Delli Compagni E, Scialabba S, Caporale M, Di Pasquale A, Cammà C, Puglia I, Calistri P, Savini G |
| EPI_ISL_2001338 | SIESP DIPARTIMENTO DI PREVENZIONE CHIETI | Istituto Zooprofilattico Sperimentale dell'Abruzzo e Molise "G. Caporale" | Lorusso A, Marcacci M, Di Domenico M, Ancora M, Curini V, Di Lollo Valeria, Mangone I, Rinaldi A, Delli Compagni E, Scialabba S, Caporale M, Di Pasquale A, Cammà C, Puglia I, Calistri P, Savini G |
| EPI_ISL_2001339 | SIESP CHIETI - DRIVE IN VASTO | Istituto Zooprofilattico Sperimentale dell'Abruzzo e Molise "G. Caporale" | Lorusso A, Marcacci M, Di Domenico M, Ancora M, Curini V, Di Lollo Valeria, Mangone I, Rinaldi A, Delli Compagni E, Scialabba S, Caporale M, Di Pasquale A, Cammà C, Puglia I, Calistri P, Savini G |
| EPI_ISL_2001340 | SIESP CHIETI - DRIVE IN GISSI | Istituto Zooprofilattico Sperimentale dell'Abruzzo e Molise "G. Caporale" | Lorusso A, Marcacci M, Di Domenico M, Ancora M, Curini V, Di Lollo Valeria, Mangone I, Rinaldi A, Delli Compagni E, Scialabba S, Caporale M, Di Pasquale A, Cammà C, Puglia I, Calistri P, Savini G |
| EPI_ISL_2001341 | SIESP CHIETI - DRIVE IN VASTO | Istituto Zooprofilattico Sperimentale dell'Abruzzo e Molise "G. Caporale" | Lorusso A, Marcacci M, Di Domenico M, Ancora M, Curini V, Di Lollo Valeria, Mangone I, Rinaldi A, Delli Compagni E, Scialabba S, Caporale M, Di Pasquale A, Cammà C, Puglia I, Calistri P, Savini G |
| EPI_ISL_2001342, EPI_ISL_2001343, EPI_ISL_2001344, EPI_ISL_2001345 | SIESP SULMONA | Istituto Zooprofilattico Sperimentale dell'Abruzzo e Molise "G. Caporale" | Lorusso A, Marcacci M, Di Domenico M, Ancora M, Curini V, Di Lollo Valeria, Mangone I, Rinaldi A, Delli Compagni E, Scialabba S, Caporale M, Di Pasquale A, Cammà C, Puglia I, Calistri P, Savini G |
| EPI_ISL_2001346, EPI_ISL_2001347 | SIESP DIPARTIMENTO DI PREVENZIONE TERAMO | Istituto Zooprofilattico Sperimentale dell'Abruzzo e Molise "G. Caporale" | Lorusso A, Marcacci M, Di Domenico M, Ancora M, Curini V, Di Lollo Valeria, Mangone I, Rinaldi A, Delli Compagni E, Scialabba S, Caporale M, Di Pasquale A, Cammà C, Puglia I, Calistri P, Savini G |
| EPI_ISL_2001348 | DIP. PREV. AVEZZANO SERVIZIO DI IGIENE EPIDEMIOLOGIA E SANITA' PUBBLICA | Istituto Zooprofilattico Sperimentale dell'Abruzzo e Molise "G. Caporale" | Lorusso A, Marcacci M, Di Domenico M, Ancora M, Curini V, Di Lollo Valeria, Mangone I, Rinaldi A, Delli Compagni E, Scialabba S, Caporale M, Di Pasquale A, Cammà C, Puglia I, Calistri P, Savini G |
| EPI_ISL_2001349, EPI_ISL_2001350 | SIESP DIPARTIMENTO DI PREVENZIONE TERAMO | Istituto Zooprofilattico Sperimentale dell'Abruzzo e Molise "G. Caporale" | Lorusso A, Marcacci M, Di Domenico M, Ancora M, Curini V, Di Lollo Valeria, Mangone I, Rinaldi A, Delli Compagni E, Scialabba S, Caporale M, Di Pasquale A, Cammà C, Puglia I, Calistri P, Savini G |
| EPI_ISL_2001351, EPI_ISL_2001352, EPI_ISL_2001353 | DIP. PREV. AVEZZANO SERVIZIO DI IGIENE EPIDEMIOLOGIA E SANITA' PUBBLICA | Istituto Zooprofilattico Sperimentale dell'Abruzzo e Molise "G. Caporale" | Lorusso A, Marcacci M, Di Domenico M, Ancora M, Curini V, Di Lollo Valeria, Mangone I, Rinaldi A, Delli Compagni E, Scialabba S, Caporale M, Di Pasquale A, Cammà C, Puglia I, Calistri P, Savini G |
| EPI_ISL_2001354 | SIESP DIPARTIMENTO DI PREVENZIONE TERAMO | Istituto Zooprofilattico Sperimentale dell'Abruzzo e Molise "G. Caporale" | Lorusso A, Marcacci M, Di Domenico M, Ancora M, Curini V, Di Lollo Valeria, Mangone I, Rinaldi A, Delli Compagni E, Scialabba S, Caporale M, Di Pasquale A, Cammà C, Puglia I, Calistri P, Savini G |
| EPI_ISL_2001355 | DIP. PREV. AVEZZANO SERVIZIO DI IGIENE EPIDEMIOLOGIA E SANITA' PUBBLICA | Istituto Zooprofilattico Sperimentale dell'Abruzzo e Molise "G. Caporale" | Lorusso A, Marcacci M, Di Domenico M, Ancora M, Curini V, Di Lollo Valeria, Mangone I, Rinaldi A, Delli Compagni E, Scialabba S, Caporale M, Di Pasquale A, Cammà C, Puglia I, Calistri P, Savini G |
| EPI_ISL_2001356 | SIESP DIPARTIMENTO DI PREVENZIONE TERAMO | Istituto Zooprofilattico Sperimentale dell'Abruzzo e Molise "G. Caporale" | Lorusso A, Marcacci M, Di Domenico M, Ancora M, Curini V, Di Lollo Valeria, Mangone I, Rinaldi A, Delli Compagni E, Scialabba S, Caporale M, Di Pasquale A, Cammà C, Puglia I, Calistri P, Savini G |
| EPI_ISL_2001357 | DIP. PREV. AVEZZANO SERVIZIO DI IGIENE EPIDEMIOLOGIA E SANITA' PUBBLICA | Istituto Zooprofilattico Sperimentale dell'Abruzzo e Molise "G. Caporale" | Lorusso A, Marcacci M, Di Domenico M, Ancora M, Curini V, Di Lollo Valeria, Mangone I, Rinaldi A, Delli Compagni E, Scialabba S, Caporale M, Di Pasquale A, Cammà C, Puglia I, Calistri P, Savini G |
| EPI_ISL_2001358, EPI_ISL_2001359, EPI_ISL_2001360 | SIESP DIPARTIMENTO DI PREVENZIONE TERAMO | Istituto Zooprofilattico Sperimentale dell'Abruzzo e Molise "G. Caporale" | Lorusso A, Marcacci M, Di Domenico M, Ancora M, Curini V, Di Lollo Valeria, Mangone I, Rinaldi A, Delli Compagni E, Scialabba S, Caporale M, Di Pasquale A, Cammà C, Puglia I, Calistri P, Savini G |
| EPI_ISL_2001361 | DIP. PREV. AVEZZANO SERVIZIO DI IGIENE EPIDEMIOLOGIA E SANITA' PUBBLICA | Istituto Zooprofilattico Sperimentale dell'Abruzzo e Molise "G. Caporale" | Lorusso A, Marcacci M, Di Domenico M, Ancora M, Curini V, Di Lollo Valeria, Mangone I, Rinaldi A, Delli Compagni E, Scialabba S, Caporale M, Di Pasquale A, Cammà C, Puglia I, Calistri P, Savini G |
| EPI_ISL_2001362, EPI_ISL_2001363, EPI_ISL_2001364 | SIESP CHIETI - DRIVE IN ORTONA | Istituto Zooprofilattico Sperimentale dell'Abruzzo e Molise "G. Caporale" | Lorusso A, Marcacci M, Di Domenico M, Ancora M, Curini V, Di Lollo Valeria, Mangone I, Rinaldi A, Delli Compagni E, Scialabba S, Caporale M, Di Pasquale A, Cammà C, Puglia I, Calistri P, Savini G |
| EPI_ISL_2001365 | SIESP DIPARTIMENTO DI PREVENZIONE TERAMO | Istituto Zooprofilattico Sperimentale dell'Abruzzo e Molise "G. Caporale" | Lorusso A, Marcacci M, Di Domenico M, Ancora M, Curini V, Di Lollo Valeria, Mangone I, Rinaldi A, Delli Compagni E, Scialabba S, Caporale M, Di Pasquale A, Cammà C, Puglia I, Calistri P, Savini G |
| EPI_ISL_2001366 | SIESP CHIETI - DRIVE IN ORTONA | Istituto Zooprofilattico Sperimentale dell'Abruzzo e Molise "G. Caporale" | Lorusso A, Marcacci M, Di Domenico M, Ancora M, Curini V, Di Lollo Valeria, Mangone I, Rinaldi A, Delli Compagni E, Scialabba S, Caporale M, Di Pasquale A, Cammà C, Puglia I, Calistri P, Savini G |
| EPI_ISL_2001367 | SIESP DIPARTIMENTO DI PREVENZIONE CHIETI | Istituto Zooprofilattico Sperimentale dell'Abruzzo e Molise "G. Caporale" | Lorusso A, Marcacci M, Di Domenico M, Ancora M, Curini V, Di Lollo Valeria, Mangone I, Rinaldi A, Delli Compagni E, Scialabba S, Caporale M, Di Pasquale A, Cammà C, Puglia I, Calistri P, Savini G |
| EPI_ISL_2001368 | SIESP DIPARTIMENTO DI PREVENZIONE TERAMO | Istituto Zooprofilattico Sperimentale dell'Abruzzo e Molise "G. Caporale" | Lorusso A, Marcacci M, Di Domenico M, Ancora M, Curini V, Di Lollo Valeria, Mangone I, Rinaldi A, Delli Compagni E, Scialabba S, Caporale M, Di Pasquale A, Cammà C, Puglia I, Calistri P, Savini G |
| EPI_ISL_2001369 | SIESP DIPARTIMENTO DI PREVENZIONE CHIETI | Istituto Zooprofilattico Sperimentale dell'Abruzzo e Molise "G. Caporale" | Lorusso A, Marcacci M, Di Domenico M, Ancora M, Curini V, Di Lollo Valeria, Mangone I, Rinaldi A, Delli Compagni E, Scialabba S, Caporale M, Di Pasquale A, Cammà C, Puglia I, Calistri P, Savini G |
| EPI_ISL_2001370 | SIESP CHIETI - DRIVE IN LANCIANO | Istituto Zooprofilattico Sperimentale dell'Abruzzo e Molise "G. Caporale" | Lorusso A, Marcacci M, Di Domenico M, Ancora M, Curini V, Di Lollo Valeria, Mangone I, Rinaldi A, Delli Compagni E, Scialabba S, Caporale M, Di Pasquale A, Cammà C, Puglia I, Calistri P, Savini G |
| EPI_ISL_2001371 | SIESP DIPARTIMENTO DI PREVENZIONE TERAMO | Istituto Zooprofilattico Sperimentale dell'Abruzzo e Molise "G. Caporale" | Lorusso A, Marcacci M, Di Domenico M, Ancora M, Curini V, Di Lollo Valeria, Mangone I, Rinaldi A, Delli Compagni E, Scialabba S, Caporale M, Di Pasquale A, Cammà C, Puglia I, Calistri P, Savini G |
| EPI_ISL_2001372 | SIESP CHIETI - DRIVE IN VASTO | Istituto Zooprofilattico Sperimentale dell'Abruzzo e Molise "G. Caporale" | Lorusso A, Marcacci M, Di Domenico M, Ancora M, Curini V, Di Lollo Valeria, Mangone I, Rinaldi A, Delli Compagni E, Scialabba S, Caporale M, Di Pasquale A, Cammà C, Puglia I, Calistri P, Savini G |
| EPI_ISL_2001373, EPI_ISL_2001374 | SIESP CHIETI - DRIVE IN LANCIANO | Istituto Zooprofilattico Sperimentale dell'Abruzzo e Molise "G. | Lorusso A, Marcacci M, Di Domenico M, Ancora M, Curini V, Di Lollo Valeria, Mangone I, Rinaldi A, Delli Compagni E, Scialabba S, Caporale M, Di |

| | | | |
|---|---|---|---|
| | | Caporale" | Pasquale A, Cammà C, Puglia I, Calistri P, Savini G |
| EPI_ISL_2001375 | SIESP DIPARTIMENTO DI PREVENZIONE CHIETI | Istituto Zooprofilattico Sperimentale dell'Abruzzo e Molise "G. Caporale" | Lorusso A, Marcacci M, Di Domenico M, Ancora M, Curini V, Di Lollo Valeria, Mangone I, Rinaldi A, Delli Compagni E, Scialabba S, Caporale M, Di Pasquale A, Cammà C, Puglia I, Calistri P, Savini G |
| EPI_ISL_2001376, EPI_ISL_2001377 | SIESP CHIETI - DRIVE IN LANCIANO | Istituto Zooprofilattico Sperimentale dell'Abruzzo e Molise "G. Caporale" | Lorusso A, Marcacci M, Di Domenico M, Ancora M, Curini V, Di Lollo Valeria, Mangone I, Rinaldi A, Delli Compagni E, Scialabba S, Caporale M, Di Pasquale A, Cammà C, Puglia I, Calistri P, Savini G |
| EPI_ISL_2001378 | SIESP DIPARTIMENTO DI PREVENZIONE TERAMO | Istituto Zooprofilattico Sperimentale dell'Abruzzo e Molise "G. Caporale" | Lorusso A, Marcacci M, Di Domenico M, Ancora M, Curini V, Di Lollo Valeria, Mangone I, Rinaldi A, Delli Compagni E, Scialabba S, Caporale M, Di Pasquale A, Cammà C, Puglia I, Calistri P, Savini G |
| EPI_ISL_2001379 | RSA GIULIANOVA | Istituto Zooprofilattico Sperimentale dell'Abruzzo e Molise "G. Caporale" | Lorusso A, Marcacci M, Di Domenico M, Ancora M, Curini V, Di Lollo Valeria, Mangone I, Rinaldi A, Delli Compagni E, Scialabba S, Caporale M, Di Pasquale A, Cammà C, Puglia I, Calistri P, Savini G |
| EPI_ISL_2001380, EPI_ISL_2001381, EPI_ISL_2001382 | SIESP DIPARTIMENTO DI PREVENZIONE TERAMO | Istituto Zooprofilattico Sperimentale dell'Abruzzo e Molise "G. Caporale" | Lorusso A, Marcacci M, Di Domenico M, Ancora M, Curini V, Di Lollo Valeria, Mangone I, Rinaldi A, Delli Compagni E, Scialabba S, Caporale M, Di Pasquale A, Cammà C, Puglia I, Calistri P, Savini G |
| EPI_ISL_2001383 | SIESP CHIETI - DRIVE IN LANCIANO | Istituto Zooprofilattico Sperimentale dell'Abruzzo e Molise "G. Caporale" | Lorusso A, Marcacci M, Di Domenico M, Ancora M, Curini V, Di Lollo Valeria, Mangone I, Rinaldi A, Delli Compagni E, Scialabba S, Caporale M, Di Pasquale A, Cammà C, Puglia I, Calistri P, Savini G |
| EPI_ISL_2001384 | SIESP CHIETI - DRIVE IN GISSI | Istituto Zooprofilattico Sperimentale dell'Abruzzo e Molise "G. Caporale" | Lorusso A, Marcacci M, Di Domenico M, Ancora M, Curini V, Di Lollo Valeria, Mangone I, Rinaldi A, Delli Compagni E, Scialabba S, Caporale M, Di Pasquale A, Cammà C, Puglia I, Calistri P, Savini G |
| EPI_ISL_2001385, EPI_ISL_2001386 | SIESP CHIETI - DRIVE IN ORTONA | Istituto Zooprofilattico Sperimentale dell'Abruzzo e Molise "G. Caporale" | Lorusso A, Marcacci M, Di Domenico M, Ancora M, Curini V, Di Lollo Valeria, Mangone I, Rinaldi A, Delli Compagni E, Scialabba S, Caporale M, Di Pasquale A, Cammà C, Puglia I, Calistri P, Savini G |
| EPI_ISL_2001387 | OSPEDALE SAN SALVATORE - MEDICINA INTERNA | Istituto Zooprofilattico Sperimentale dell'Abruzzo e Molise "G. Caporale" | Lorusso A, Marcacci M, Di Domenico M, Ancora M, Curini V, Di Lollo Valeria, Mangone I, Rinaldi A, Delli Compagni E, Scialabba S, Caporale M, Di Pasquale A, Cammà C, Puglia I, Calistri P, Savini G |
| EPI_ISL_2001388, EPI_ISL_2001389 | SIESP CHIETI - DRIVE IN LANCIANO | Istituto Zooprofilattico Sperimentale dell'Abruzzo e Molise "G. Caporale" | Lorusso A, Marcacci M, Di Domenico M, Ancora M, Curini V, Di Lollo Valeria, Mangone I, Rinaldi A, Delli Compagni E, Scialabba S, Caporale M, Di Pasquale A, Cammà C, Puglia I, Calistri P, Savini G |
| EPI_ISL_2001390 | USCA Avezzano | Istituto Zooprofilattico Sperimentale dell'Abruzzo e Molise "G. Caporale" | Lorusso A, Marcacci M, Di Domenico M, Ancora M, Curini V, Di Lollo Valeria, Mangone I, Rinaldi A, Delli Compagni E, Scialabba S, Caporale M, Di Pasquale A, Cammà C, Puglia I, Calistri P, Savini G |
| EPI_ISL_2001391 | PRESIDIO TERRITORIALE CASOLI | Istituto Zooprofilattico Sperimentale dell'Abruzzo e Molise "G. Caporale" | Lorusso A, Marcacci M, Di Domenico M, Ancora M, Curini V, Di Lollo Valeria, Mangone I, Rinaldi A, Delli Compagni E, Scialabba S, Caporale M, Di Pasquale A, Cammà C, Puglia I, Calistri P, Savini G |
| EPI_ISL_2001392, EPI_ISL_2001393, EPI_ISL_2001394, EPI_ISL_2001395, EPI_ISL_2001396, EPI_ISL_2001397, EPI_ISL_2001398 | SIESP DIPARTIMENTO DI PREVENZIONE TERAMO | Istituto Zooprofilattico Sperimentale dell'Abruzzo e Molise "G. Caporale" | Lorusso A, Marcacci M, Di Domenico M, Ancora M, Curini V, Di Lollo Valeria, Mangone I, Rinaldi A, Delli Compagni E, Scialabba S, Caporale M, Di Pasquale A, Cammà C, Puglia I, Calistri P, Savini G |
| EPI_ISL_2001399 | SIESP DIPARTIMENTO DI PREVENZIONE CHIETI | Istituto Zooprofilattico Sperimentale dell'Abruzzo e Molise "G. Caporale" | Lorusso A, Marcacci M, Di Domenico M, Ancora M, Curini V, Di Lollo Valeria, Mangone I, Rinaldi A, Delli Compagni E, Scialabba S, Caporale M, Di Pasquale A, Cammà C, Puglia I, Calistri P, Savini G |
| EPI_ISL_2001400, EPI_ISL_2001401 | SIESP DIPARTIMENTO DI PREVENZIONE TERAMO | Istituto Zooprofilattico Sperimentale dell'Abruzzo e Molise "G. Caporale" | Lorusso A, Marcacci M, Di Domenico M, Ancora M, Curini V, Di Lollo Valeria, Mangone I, Rinaldi A, Delli Compagni E, Scialabba S, Caporale M, Di Pasquale A, Cammà C, Puglia I, Calistri P, Savini G |
| EPI_ISL_2001402, EPI_ISL_2001403, EPI_ISL_2001404 | SIESP CHIETI - DRIVE IN LANCIANO | Istituto Zooprofilattico Sperimentale dell'Abruzzo e Molise "G. Caporale" | Lorusso A, Marcacci M, Di Domenico M, Ancora M, Curini V, Di Lollo Valeria, Mangone I, Rinaldi A, Delli Compagni E, Scialabba S, Caporale M, Di Pasquale A, Cammà C, Puglia I, Calistri P, Savini G |
| EPI_ISL_2001405 | DIP. PREV. AVEZZANO SERVIZIO DI IGIENE EPIDEMIOLOGIA E SANITA' PUBBLICA | Istituto Zooprofilattico Sperimentale dell'Abruzzo e Molise "G. Caporale" | Lorusso A, Marcacci M, Di Domenico M, Ancora M, Curini V, Di Lollo Valeria, Mangone I, Rinaldi A, Delli Compagni E, Scialabba S, Caporale M, Di Pasquale A, Cammà C, Puglia I, Calistri P, Savini G |
| EPI_ISL_2001406 | SIESP SULMONA | Istituto Zooprofilattico Sperimentale dell'Abruzzo e Molise "G. Caporale" | Lorusso A, Marcacci M, Di Domenico M, Ancora M, Curini V, Di Lollo Valeria, Mangone I, Rinaldi A, Delli Compagni E, Scialabba S, Caporale M, Di Pasquale A, Cammà C, Puglia I, Calistri P, Savini G |
| EPI_ISL_2001407 | SIESP CHIETI - DRIVE IN ORTONA | Istituto Zooprofilattico Sperimentale dell'Abruzzo e Molise "G. Caporale" | Lorusso A, Marcacci M, Di Domenico M, Ancora M, Curini V, Di Lollo Valeria, Mangone I, Rinaldi A, Delli Compagni E, Scialabba S, Caporale M, Di Pasquale A, Cammà C, Puglia I, Calistri P, Savini G |
| EPI_ISL_2001408 | SIESP SULMONA | Istituto Zooprofilattico Sperimentale dell'Abruzzo e Molise "G. Caporale" | Lorusso A, Marcacci M, Di Domenico M, Ancora M, Curini V, Di Lollo Valeria, Mangone I, Rinaldi A, Delli Compagni E, Scialabba S, Caporale M, Di Pasquale A, Cammà C, Puglia I, Calistri P, Savini G |
| EPI_ISL_2001409 | SIESP CHIETI - DRIVE IN ORTONA | Istituto Zooprofilattico Sperimentale dell'Abruzzo e Molise "G. Caporale" | Lorusso A, Marcacci M, Di Domenico M, Ancora M, Curini V, Di Lollo Valeria, Mangone I, Rinaldi A, Delli Compagni E, Scialabba S, Caporale M, Di Pasquale A, Cammà C, Puglia I, Calistri P, Savini G |
| EPI_ISL_2001410, EPI_ISL_2001411 | DIP. PREV. AVEZZANO SERVIZIO DI IGIENE EPIDEMIOLOGIA E SANITA' PUBBLICA | Istituto Zooprofilattico Sperimentale dell'Abruzzo e Molise "G. Caporale" | Lorusso A, Marcacci M, Di Domenico M, Ancora M, Curini V, Di Lollo Valeria, Mangone I, Rinaldi A, Delli Compagni E, Scialabba S, Caporale M, Di Pasquale A, Cammà C, Puglia I, Calistri P, Savini G |
| EPI_ISL_2001412 | SIESP CHIETI - DRIVE IN CHIETI | Istituto Zooprofilattico Sperimentale dell'Abruzzo e Molise "G. Caporale" | Lorusso A, Marcacci M, Di Domenico M, Ancora M, Curini V, Di Lollo Valeria, Mangone I, Rinaldi A, Delli Compagni E, Scialabba S, Caporale M, Di Pasquale A, Cammà C, Puglia I, Calistri P, Savini G |
| EPI_ISL_2001413 | DIP. PREV. AVEZZANO SERVIZIO DI IGIENE EPIDEMIOLOGIA E SANITA' PUBBLICA | Istituto Zooprofilattico Sperimentale dell'Abruzzo e Molise "G. Caporale" | Lorusso A, Marcacci M, Di Domenico M, Ancora M, Curini V, Di Lollo Valeria, Mangone I, Rinaldi A, Delli Compagni E, Scialabba S, Caporale M, Di Pasquale A, Cammà C, Puglia I, Calistri P, Savini G |
| EPI_ISL_2001414, EPI_ISL_2001415 | SIESP DIPARTIMENTO DI PREVENZIONE CHIETI | Istituto Zooprofilattico Sperimentale dell'Abruzzo e Molise "G. Caporale" | Lorusso A, Marcacci M, Di Domenico M, Ancora M, Curini V, Di Lollo Valeria, Mangone I, Rinaldi A, Delli Compagni E, Scialabba S, Caporale M, Di Pasquale A, Cammà C, Puglia I, Calistri P, Savini G |
| EPI_ISL_2001416 | SIESP DIPARTIMENTO DI PREVENZIONE TERAMO | Istituto Zooprofilattico Sperimentale dell'Abruzzo e Molise "G. Caporale" | Lorusso A, Marcacci M, Di Domenico M, Ancora M, Curini V, Di Lollo Valeria, Mangone I, Rinaldi A, Delli Compagni E, Scialabba S, Caporale M, Di Pasquale A, Cammà C, Puglia I, Calistri P, Savini G |
| EPI_ISL_2001417 | PRESIDIO TERRITORIALE CASOLI | Istituto Zooprofilattico Sperimentale dell'Abruzzo e Molise "G. Caporale" | Lorusso A, Marcacci M, Di Domenico M, Ancora M, Curini V, Di Lollo Valeria, Mangone I, Rinaldi A, Delli Compagni E, Scialabba S, Caporale M, Di Pasquale A, Cammà C, Puglia I, Calistri P, Savini G |
| EPI_ISL_2001418 | SIESP CHIETI - DRIVE IN VASTO | Istituto Zooprofilattico Sperimentale dell'Abruzzo e Molise "G. Caporale" | Lorusso A, Marcacci M, Di Domenico M, Ancora M, Curini V, Di Lollo Valeria, Mangone I, Rinaldi A, Delli Compagni E, Scialabba S, Caporale M, Di Pasquale A, Cammà C, Puglia I, Calistri P, Savini G |
| EPI_ISL_2001419 | SIESP DIPARTIMENTO DI PREVENZIONE CHIETI | Istituto Zooprofilattico Sperimentale dell'Abruzzo e Molise "G. Caporale" | Lorusso A, Marcacci M, Di Domenico M, Ancora M, Curini V, Di Lollo Valeria, Mangone I, Rinaldi A, Delli Compagni E, Scialabba S, Caporale M, Di Pasquale A, Cammà C, Puglia I, Calistri P, Savini G |
| EPI_ISL_2001420 | SIESP DIPARTIMENTO DI PREVENZIONE TERAMO | Istituto Zooprofilattico Sperimentale dell'Abruzzo e Molise "G. Caporale" | Lorusso A, Marcacci M, Di Domenico M, Ancora M, Curini V, Di Lollo Valeria, Mangone I, Rinaldi A, Delli Compagni E, Scialabba S, Caporale M, Di Pasquale A, Cammà C, Puglia I, Calistri P, Savini G |
| EPI_ISL_2001421 | SIESP CHIETI - DRIVE IN LANCIANO | Istituto Zooprofilattico Sperimentale dell'Abruzzo e Molise "G. Caporale" | Lorusso A, Marcacci M, Di Domenico M, Ancora M, Curini V, Di Lollo Valeria, Mangone I, Rinaldi A, Delli Compagni E, Scialabba S, Caporale M, Di Pasquale A, Cammà C, Puglia I, Calistri P, Savini G |
| EPI_ISL_2001422, EPI_ISL_2001423 | SIESP CHIETI - DRIVE IN ORTONA | Istituto Zooprofilattico Sperimentale dell'Abruzzo e Molise "G. Caporale" | Lorusso A, Marcacci M, Di Domenico M, Ancora M, Curini V, Di Lollo Valeria, Mangone I, Rinaldi A, Delli Compagni E, Scialabba S, Caporale M, Di Pasquale A, Cammà C, Puglia I, Calistri P, Savini G |
| EPI_ISL_2001424, EPI_ISL_2001425 | SIESP CHIETI - DRIVE IN GISSI | Istituto Zooprofilattico Sperimentale dell'Abruzzo e Molise "G. Caporale" | Lorusso A, Marcacci M, Di Domenico M, Ancora M, Curini V, Di Lollo Valeria, Mangone I, Rinaldi A, Delli Compagni E, Scialabba S, Caporale M, Di Pasquale A, Cammà C, Puglia I, Calistri P, Savini G |
| EPI_ISL_2001426 | SIESP DIPARTIMENTO DI PREVENZIONE TERAMO | Istituto Zooprofilattico Sperimentale dell'Abruzzo e Molise "G. Caporale" | Lorusso A, Marcacci M, Di Domenico M, Ancora M, Curini V, Di Lollo Valeria, Mangone I, Rinaldi A, Delli Compagni E, Scialabba S, Caporale M, Di Pasquale A, Cammà C, Puglia I, Calistri P, Savini G |

| | | | |
|---|---|---|---|
| EPI_ISL_2001427 | SIESP DIPARTIMENTO DI PREVENZIONE CHIETI | Istituto Zooprofilattico Sperimentale dell'Abruzzo e Molise "G. Caporale" | Lorusso A, Marcacci M, Di Domenico M, Ancora M, Curini V, Di Lollo Valeria, Mangone I, Rinaldi A, Delli Compagni E, Scialabba S, Caporale M, Di Pasquale A, Cammà C, Puglia I, Calistri P, Savini G |
| EPI_ISL_2001428 | SIESP DIPARTIMENTO DI PREVENZIONE TERAMO | Istituto Zooprofilattico Sperimentale dell'Abruzzo e Molise "G. Caporale" | Lorusso A, Marcacci M, Di Domenico M, Ancora M, Curini V, Di Lollo Valeria, Mangone I, Rinaldi A, Delli Compagni E, Scialabba S, Caporale M, Di Pasquale A, Cammà C, Puglia I, Calistri P, Savini G |
| EPI_ISL_2001431, EPI_ISL_2001432, EPI_ISL_2001433, EPI_ISL_2001434, EPI_ISL_2001435, EPI_ISL_2001436, EPI_ISL_2001437, EPI_ISL_2001438, EPI_ISL_2001439, EPI_ISL_2001440, EPI_ISL_2001441, EPI_ISL_2001442, EPI_ISL_2001443, EPI_ISL_2001444, EPI_ISL_2001445, EPI_ISL_2001446, EPI_ISL_2001447 | | | |
| see above | Università degli Studi di Perugia | Istituto Zooprofilattico Sperimentale dell'Abruzzo e Molise "G. Caporale" | Mencacci A, Camilloni B, Lorusso A, Marcacci M, Di Domenico M, Ancora M, Curini V, Mangone I, Rinaldi A, Scialabba S, Di Pasquale A, Cammà C, Puglia I, Calistri P, Savini G |
| EPI_ISL_2001448, EPI_ISL_2001449, EPI_ISL_2001450, EPI_ISL_2001451, EPI_ISL_2001452 | Laboratorio Analisi Osp. Città di Castello - Azienda USL Umbria1 | Istituto Zooprofilattico Sperimentale dell'Abruzzo e Molise "G. Caporale" | Malagigi V, Tacconi P, Lorusso A, Marcacci M, Di Domenico M, Ancora M, Curini V, Mangone I, Rinaldi A, Scialabba S, Di Pasquale A, Cammà C, Puglia I, Calistri P, Savini G |
| EPI_ISL_2001453, EPI_ISL_2001454, EPI_ISL_2001455, EPI_ISL_2001456, EPI_ISL_2001457, EPI_ISL_2001458, EPI_ISL_2001459, EPI_ISL_2001460, EPI_ISL_2001461, EPI_ISL_2001462, EPI_ISL_2001463, EPI_ISL_2001464, EPI_ISL_2001465, EPI_ISL_2001466, EPI_ISL_2001467, EPI_ISL_2001468, EPI_ISL_2001469, EPI_ISL_2001470, EPI_ISL_2001471, EPI_ISL_2001472, EPI_ISL_2001473 | | | |
| see above | Azienda Ospedaliera Terni | Istituto Zooprofilattico Sperimentale dell'Abruzzo e Molise "G. Caporale" | Palumbo M, Scaccetti A, Lorusso A, Marcacci M, Di Domenico M, Ancora M, Curini V, Mangone I, Rinaldi A, Scialabba S, Di Pasquale A, Cammà C, Puglia I, Calistri P, Savini G |
| EPI_ISL_2001474, EPI_ISL_2001475, EPI_ISL_2001476, EPI_ISL_2001477, EPI_ISL_2001478, EPI_ISL_2001479, EPI_ISL_2001480 | Azienda USL Umbria 2 | Istituto Zooprofilattico Sperimentale dell'Abruzzo e Molise "G. Caporale" | Proietti A, Pistoni E,Lorusso A, Marcacci M, Di Domenico M, Ancora M, Curini V, Mangone I, Rinaldi A, Scialabba S, Di Pasquale A, Cammà C, Puglia I, Calistri P, Savini G |
| EPI_ISL_2002598 | Centre Hospitalier Universitaire Clermont-Ferrand | CHU Clermont-Ferrand, service de virologie | Bisseux Maxime, Mirand Audrey, Combes Patricia, Henquell Cécile |
| EPI_ISL_2002599 | HOPITAL DE VICHY | CHU Clermont-Ferrand, service de virologie | Bisseux Maxime, Mirand Audrey, Combes Patricia, Henquell Cécile |
| EPI_ISL_2002600, EPI_ISL_2002601, EPI_ISL_2002602, EPI_ISL_2002603, EPI_ISL_2002604, EPI_ISL_2002605, EPI_ISL_2002606, EPI_ISL_2002607 | Centre Hospitalier Universitaire Clermont-Ferrand | CHU Clermont-Ferrand, service de virologie | Bisseux Maxime, Mirand Audrey, Combes Patricia, Henquell Cécile |
| EPI_ISL_2002608 | HOPITAL DE MONTLUCON | CHU Clermont-Ferrand, service de virologie | Bisseux Maxime, Mirand Audrey, Combes Patricia, Henquell Cécile |
| EPI_ISL_2002610, EPI_ISL_2002611 | Centre Hospitalier Universitaire Clermont-Ferrand | CHU Clermont-Ferrand, service de virologie | Bisseux Maxime, Mirand Audrey, Combes Patricia, Henquell Cécile |
| EPI_ISL_2002613 | Centre hospitalier D'AURILLAC | CHU Clermont-Ferrand, service de virologie | Bisseux Maxime, Mirand Audrey, Combes Patricia, Henquell Cécile |
| EPI_ISL_2002614, EPI_ISL_2002615, EPI_ISL_2002616 | Centre Hospitalier Universitaire Clermont-Ferrand | CHU Clermont-Ferrand, service de virologie | Bisseux Maxime, Mirand Audrey, Combes Patricia, Henquell Cécile |
| EPI_ISL_2002617 | Centre Hospitalier RIOM | CHU Clermont-Ferrand, service de virologie | Bisseux Maxime, Mirand Audrey, Combes Patricia, Henquell Cécile |
| EPI_ISL_2002618 | Centre Hospitalier Universitaire Clermont-Ferrand | CHU Clermont-Ferrand, service de virologie | Bisseux Maxime, Mirand Audrey, Combes Patricia, Henquell Cécile |
| EPI_ISL_2002619 | Centre Hospitalier Paul Ardier | CHU Clermont-Ferrand, service de virologie | Bisseux Maxime, Mirand Audrey, Combes Patricia, Henquell Cécile |
| EPI_ISL_2002620 | Centre Hospitalier Billom | CHU Clermont-Ferrand, service de virologie | Bisseux Maxime, Mirand Audrey, Combes Patricia, Henquell Cécile |
| EPI_ISL_2002621, EPI_ISL_2002622, EPI_ISL_2002623, EPI_ISL_2002624, EPI_ISL_2002625 | Centre Hospitalier Universitaire Clermont-Ferrand | CHU Clermont-Ferrand, service de virologie | Bisseux Maxime, Mirand Audrey, Combes Patricia, Henquell Cécile |
| EPI_ISL_2002626 | Centre hospitalier D'AURILLAC | CHU Clermont-Ferrand, service de virologie | Bisseux Maxime, Mirand Audrey, Combes Patricia, Henquell Cécile |
| EPI_ISL_2002627 | Centre Hospitalier Universitaire Clermont-Ferrand | CHU Clermont-Ferrand, service de virologie | Bisseux Maxime, Mirand Audrey, Combes Patricia, Henquell Cécile |
| EPI_ISL_2002628 | Centre Hospitalier Paul Ardier | CHU Clermont-Ferrand, service de virologie | Bisseux Maxime, Mirand Audrey, Combes Patricia, Henquell Cécile |
| EPI_ISL_2002629, EPI_ISL_2002630, EPI_ISL_2002631 | Centre Hospitalier Universitaire Clermont-Ferrand | CHU Clermont-Ferrand, service de virologie | Bisseux Maxime, Mirand Audrey, Combes Patricia, Henquell Cécile |
| EPI_ISL_2002632 | HOPITAL DE VICHY | CHU Clermont-Ferrand, service de virologie | Bisseux Maxime, Mirand Audrey, Combes Patricia, Henquell Cécile |
| EPI_ISL_2002633 | HOPITAL DE MONTLUCON | CHU Clermont-Ferrand, service de virologie | Bisseux Maxime, Mirand Audrey, Combes Patricia, Henquell Cécile |
| EPI_ISL_2002634, EPI_ISL_2002635 | Centre Hospitalier Universitaire Clermont-Ferrand | CHU Clermont-Ferrand, service de virologie | Bisseux Maxime, Mirand Audrey, Combes Patricia, Henquell Cécile |
| EPI_ISL_2002636 | Centre hospitalier D'AURILLAC | CHU Clermont-Ferrand, service de virologie | Bisseux Maxime, Mirand Audrey, Combes Patricia, Henquell Cécile |
| EPI_ISL_2002637 | HOPITAL DE VICHY | CHU Clermont-Ferrand, service de virologie | Bisseux Maxime, Mirand Audrey, Combes Patricia, Henquell Cécile |
| EPI_ISL_2002638 | HOPITAL DE MONTLUCON | CHU Clermont-Ferrand, service de virologie | Bisseux Maxime, Mirand Audrey, Combes Patricia, Henquell Cécile |
| EPI_ISL_2002639 | Centre Hospitalier RIOM | CHU Clermont-Ferrand, service de virologie | Bisseux Maxime, Mirand Audrey, Combes Patricia, Henquell Cécile |
| EPI_ISL_2002640, EPI_ISL_2002641, EPI_ISL_2002642, EPI_ISL_2002643, EPI_ISL_2002644, EPI_ISL_2002645, EPI_ISL_2002646, EPI_ISL_2002647, EPI_ISL_2002648, EPI_ISL_2002649, EPI_ISL_2002650, EPI_ISL_2002651 | | | |
| see above | Centre Hospitalier Universitaire Clermont-Ferrand | CHU Clermont-Ferrand, service de virologie | Bisseux Maxime, Mirand Audrey, Combes Patricia, Henquell Cécile |
| EPI_ISL_2002652 | Centre De Prelevement COVID RIOM | CHU Clermont-Ferrand, service de virologie | Bisseux Maxime, Mirand Audrey, Combes Patricia, Henquell Cécile |
| EPI_ISL_2002653 | Centre De Prélévement COVID VICHY | CHU Clermont-Ferrand, service de virologie | Bisseux Maxime, Mirand Audrey, Combes Patricia, Henquell Cécile |
| EPI_ISL_2002654 | Centre De Prelevement COVID RIOM | CHU Clermont-Ferrand, service de virologie | Bisseux Maxime, Mirand Audrey, Combes Patricia, Henquell Cécile |
| EPI_ISL_2002655 | Centre Hospitalier Universitaire Clermont-Ferrand | CHU Clermont-Ferrand, service de virologie | Bisseux Maxime, Mirand Audrey, Combes Patricia, Henquell Cécile |
| EPI_ISL_2002656 | HOPITAL DU MONT DORE | CHU Clermont-Ferrand, service de virologie | Bisseux Maxime, Mirand Audrey, Combes Patricia, Henquell Cécile |
| EPI_ISL_2002657 | center de prélévement COVID ISSOIRE | CHU Clermont-Ferrand, service de virologie | Bisseux Maxime, Mirand Audrey, Combes Patricia, Henquell Cécile |
| EPI_ISL_2002658 | Centre Hospitalier Universitaire Clermont-Ferrand | CHU Clermont-Ferrand, service de virologie | Bisseux Maxime, Mirand Audrey, Combes Patricia, Henquell Cécile |
| EPI_ISL_2002659 | CDP COVID MONTLUCON | CHU Clermont-Ferrand, service de virologie | Bisseux Maxime, Mirand Audrey, Combes Patricia, Henquell Cécile |
| EPI_ISL_2002660, EPI_ISL_2002662, EPI_ISL_2002663, EPI_ISL_2002664, EPI_ISL_2002665, EPI_ISL_2002666 | Centre Hospitalier Universitaire Clermont-Ferrand | CHU Clermont-Ferrand, service de virologie | Bisseux Maxime, Mirand Audrey, Combes Patricia, Henquell Cécile |
| EPI_ISL_2003132, EPI_ISL_2003133 | Instituto Adolfo Lutz Central | Instituto Adolfo Lutz, Interdiciplinary Procedures Center, Strategic Laboratory | Claudio Tavares Sacchi, Claudia Regina Gonçalves, Erica Valessa Ramos Gomes, Karoline Rodrigues Campos, Caio Vinicius Dias Lopes, Leonardo Jose Tadeu de Araujo |
| EPI_ISL_2003981 | Basurto University Hospital: Clinical Microbiology Laboratory | Biocruces Bizkaia | Mikel Urrutikoetxea-Gutiérrez, Mª Carmen Nieto Toboso, Estibaliz Ugalde Zarraga, Mikel Gallego Rodrigo, Ana de la Hoz, José Luis Díaz de Tuesta del Arco |

| | | | |
|---|---|---|---|
| EPI_ISL_2004066 | INT Fondazione Pascale | INT Fondazione Pascale | INT Fondazione Pascale |
| EPI_ISL_2006928, EPI_ISL_2006929, EPI_ISL_2006930, EPI_ISL_2006931, EPI_ISL_2006932, EPI_ISL_2006933, EPI_ISL_2006934, EPI_ISL_2006935, EPI_ISL_2006937, EPI_ISL_2006938, EPI_ISL_2006939, EPI_ISL_2006940, EPI_ISL_2006941, EPI_ISL_2006943, EPI_ISL_2006944, EPI_ISL_2006946, EPI_ISL_2006947, EPI_ISL_2006948, EPI_ISL_2006950, EPI_ISL_2006951, EPI_ISL_2006953, EPI_ISL_2006954, EPI_ISL_2006955, EPI_ISL_2006956, EPI_ISL_2006957, EPI_ISL_2006958, EPI_ISL_2006959, EPI_ISL_2006960, EPI_ISL_2006961, EPI_ISL_2006965, EPI_ISL_2006990, EPI_ISL_2006991, EPI_ISL_2006994, EPI_ISL_2006995, EPI_ISL_2006998, EPI_ISL_2006999, EPI_ISL_2007000, EPI_ISL_2007001, EPI_ISL_2007002, EPI_ISL_2007004, EPI_ISL_2007005, EPI_ISL_2007008, EPI_ISL_2007009 | | | |
| see above | The Ohio State University Applied Microbiology Services Laboratory | The Ohio State University Applied Microbiology Services Laboratory | Seth A. Faith PhD |
| EPI_ISL_2007802, EPI_ISL_2007803, EPI_ISL_2007804, EPI_ISL_2007805, EPI_ISL_2007806, EPI_ISL_2007807, EPI_ISL_2007808, EPI_ISL_2007809, EPI_ISL_2007810, EPI_ISL_2007811, EPI_ISL_2007812, EPI_ISL_2007813, EPI_ISL_2007814, EPI_ISL_2007815, EPI_ISL_2007816, EPI_ISL_2007817, EPI_ISL_2007818, EPI_ISL_2007819, EPI_ISL_2007820, EPI_ISL_2007821, EPI_ISL_2007822, EPI_ISL_2007823, EPI_ISL_2007824, EPI_ISL_2007829, EPI_ISL_2007830, EPI_ISL_2007831, EPI_ISL_2007832, EPI_ISL_2007833, EPI_ISL_2007834 | | | |
| see above | University of Oregon COVID-19 MAP Laboratory | University of Oregon Genomics and Cell Characterization Core Facility (GC3F) | Douglas Turnbull, Ariana White, Jeff Bishop, Jason Carriere, Jason Sydes, Peter Batzel |
| EPI_ISL_2007879 | Laboratoire Hôpital Sainte Musse | Laboratoire Hôpital Sainte Musse | Lionel Chollet |
| EPI_ISL_2011683 | CHU Angoulême | CHU Poitiers | Maxime PICHON, Caroline MICHAUD, Manon PRAT, Birama N'DIAYE, Agnes BEBY-DEFAUX, Nicolas LEVEQUE |
| EPI_ISL_2013035 | Plateforme Covid IDF, site Broussais | HEGP - Laboratoire de Virologie | David Veyer, Sandrine Imbeaud, Julien Puech, Nabil Gastli, Hélène Péré, Maxime Wack, Nicolas Robillard |
| EPI_ISL_2017485, EPI_ISL_2017486, EPI_ISL_2017487, EPI_ISL_2017488, EPI_ISL_2017489, EPI_ISL_2017490, EPI_ISL_2017491, EPI_ISL_2017492, EPI_ISL_2017493, EPI_ISL_2017494, EPI_ISL_2017495, EPI_ISL_2017496, EPI_ISL_2017497, EPI_ISL_2017498, EPI_ISL_2017499, EPI_ISL_2017500, EPI_ISL_2017501, EPI_ISL_2017502 | | | |
| see above | OLVZ Aalst | OLVZ Aalst | Anne Vankeerberghen |
| EPI_ISL_2017759 | N.H.L Municipal Medical College, Ahmedabad | Gujarat Biotechnology Research Centre | Zarna Patel,Nitin Savaliya,Dinesh Kumar,Twinkle Soni,Sonal Sharma ,Zuber Saiyed,Ramesh Pandit,Janvi Raval,Jayshri Pethani,Umang Mishra,Nitesh Shah,Chaitanya Joshi,Madhvi Joshi |
| EPI_ISL_2017760 | N.H.L Municipal Medical College, Ahmedabad | Gujarat Biotechnology Research Centre | Nitin Savaliya,Dinesh Kumar,Twinkle Soni,Sonal Sharma ,Zuber Saiyed,Ramesh Pandit,Janvi Raval,Zarna Patel,Jayshri Pethani,Umang Mishra,Nitesh Shah,Chaitanya Joshi,Madhvi Joshi |
| EPI_ISL_2017761 | N.H.L Municipal Medical College, Ahmedabad | Gujarat Biotechnology Research Centre | Dinesh Kumar,Twinkle Soni,Sonal Sharma ,Zuber Saiyed,Ramesh Pandit,Janvi Raval,Zarna Patel,Nitin Savaliya,Jayshri Pethani,Umang Mishra,Nitesh Shah,Chaitanya Joshi,Madhvi Joshi |
| EPI_ISL_2017762 | N.H.L Municipal Medical College, Ahmedabad | Gujarat Biotechnology Research Centre | Twinkle Soni,Sonal Sharma ,Zuber Saiyed,Ramesh Pandit,Janvi Raval,Zarna Patel,Nitin Savaliya,Dinesh Kumar,Jayshri Pethani,Umang Mishra,Nitesh Shah,Chaitanya Joshi,Madhvi Joshi |
| EPI_ISL_2017765 | BJ Medical College and Civil Hospital, Ahmedabad | Gujarat Biotechnology Research Centre | Ramesh Pandit,Janvi Raval,Zarna Patel,Nitin Savaliya,Dinesh Kumar,Twinkle Soni,Sonal Sharma ,Zuber Saiyed,Pranay Shah,Umang Mishra,Nitesh Shah,Chaitanya Joshi,Madhvi Joshi |
| EPI_ISL_2017766 | BJ Medical College and Civil Hospital, Ahmedabad | Gujarat Biotechnology Research Centre | Janvi Raval,Zarna Patel,Nitin Savaliya,Dinesh Kumar,Twinkle Soni,Sonal Sharma ,Zuber Saiyed,Ramesh Pandit,Pranay Shah,Umang Mishra,Nitesh Shah,Chaitanya Joshi,Madhvi Joshi |
| EPI_ISL_2017767 | BJ Medical College and Civil Hospital, Ahmedabad | Gujarat Biotechnology Research Centre | Zarna Patel,Nitin Savaliya,Dinesh Kumar,Twinkle Soni,Sonal Sharma ,Zuber Saiyed,Ramesh Pandit,Janvi Raval,Pranay Shah,Umang Mishra,Nitesh Shah,Chaitanya Joshi,Madhvi Joshi |
| EPI_ISL_2017768 | BJ Medical College and Civil Hospital, Ahmedabad | Gujarat Biotechnology Research Centre | Nitin Savaliya,Dinesh Kumar,Twinkle Soni,Sonal Sharma ,Zuber Saiyed,Ramesh Pandit,Janvi Raval,Zarna Patel,Pranay Shah,Umang Mishra,Nitesh Shah,Chaitanya Joshi,Madhvi Joshi |
| EPI_ISL_2017769 | BJ Medical College and Civil Hospital, Ahmedabad | Gujarat Biotechnology Research Centre | Dinesh Kumar,Twinkle Soni,Sonal Sharma ,Zuber Saiyed,Ramesh Pandit,Janvi Raval,Zarna Patel,Nitin Savaliya,Pranay Shah,Umang Mishra,Nitesh Shah,Chaitanya Joshi,Madhvi Joshi |
| EPI_ISL_2017770 | BJ Medical College and Civil Hospital, Ahmedabad | Gujarat Biotechnology Research Centre | Twinkle Soni,Sonal Sharma ,Zuber Saiyed,Ramesh Pandit,Janvi Raval,Zarna Patel,Nitin Savaliya,Dinesh Kumar,Pranay Shah,Umang Mishra,Nitesh Shah,Chaitanya Joshi,Madhvi Joshi |
| EPI_ISL_2017771 | BJ Medical College and Civil Hospital, Ahmedabad | Gujarat Biotechnology Research Centre | Sonal Sharma ,Zuber Saiyed,Ramesh Pandit,Janvi Raval,Zarna Patel,Nitin Savaliya,Dinesh Kumar,Twinkle Soni,Pranay Shah,Umang Mishra,Nitesh Shah,Chaitanya Joshi,Madhvi Joshi |
| EPI_ISL_2017772 | BJ Medical College and Civil Hospital, Ahmedabad | Gujarat Biotechnology Research Centre | Zuber Saiyed,Ramesh Pandit,Janvi Raval,Zarna Patel,Nitin Savaliya,Dinesh Kumar,Twinkle Soni,Sonal Sharma ,Pranay Shah,Umang Mishra,Nitesh Shah,Chaitanya Joshi,Madhvi Joshi |
| EPI_ISL_2017774 | BJ Medical College and Civil Hospital, Ahmedabad | Gujarat Biotechnology Research Centre | Janvi Raval,Zarna Patel,Nitin Savaliya,Dinesh Kumar,Twinkle Soni,Sonal Sharma ,Zuber Saiyed,Ramesh Pandit,Pranay Shah,Umang Mishra,Nitesh Shah,Chaitanya Joshi,Madhvi Joshi |
| EPI_ISL_2017775 | BJ Medical College and Civil Hospital, Ahmedabad | Gujarat Biotechnology Research Centre | Zarna Patel,Nitin Savaliya,Dinesh Kumar,Twinkle Soni,Sonal Sharma ,Zuber Saiyed,Ramesh Pandit,Janvi Raval,Pranay Shah,Umang Mishra,Nitesh Shah,Chaitanya Joshi,Madhvi Joshi |
| EPI_ISL_2017776 | BJ Medical College and Civil Hospital, Ahmedabad | Gujarat Biotechnology Research Centre | Nitin Savaliya,Dinesh Kumar,Twinkle Soni,Sonal Sharma ,Zuber Saiyed,Ramesh Pandit,Janvi Raval,Zarna Patel,Pranay Shah,Umang Mishra,Nitesh Shah,Chaitanya Joshi,Madhvi Joshi |
| EPI_ISL_2017777 | BJ Medical College and Civil Hospital, Ahmedabad | Gujarat Biotechnology Research Centre | Dinesh Kumar,Twinkle Soni,Sonal Sharma ,Zuber Saiyed,Ramesh Pandit,Janvi Raval,Zarna Patel,Nitin Savaliya,Pranay Shah,Umang Mishra,Nitesh Shah,Chaitanya Joshi,Madhvi Joshi |
| EPI_ISL_2017778 | BJ Medical College and Civil Hospital, Ahmedabad | Gujarat Biotechnology Research Centre | Twinkle Soni,Sonal Sharma ,Zuber Saiyed,Ramesh Pandit,Janvi Raval,Zarna Patel,Nitin Savaliya,Dinesh Kumar,Pranay Shah,Umang Mishra,Nitesh Shah,Chaitanya Joshi,Madhvi Joshi |
| EPI_ISL_2017779 | BJ Medical College and Civil Hospital, Ahmedabad | Gujarat Biotechnology Research Centre | Sonal Sharma ,Zuber Saiyed,Ramesh Pandit,Janvi Raval,Zarna Patel,Nitin Savaliya,Dinesh Kumar,Twinkle Soni,Pranay Shah,Umang Mishra,Nitesh Shah,Chaitanya Joshi,Madhvi Joshi |
| EPI_ISL_2017780 | BJ Medical College and Civil Hospital, Ahmedabad | Gujarat Biotechnology Research Centre | Zuber Saiyed,Ramesh Pandit,Janvi Raval,Zarna Patel,Nitin Savaliya,Dinesh Kumar,Twinkle Soni,Sonal Sharma ,Pranay Shah,Umang Mishra,Nitesh Shah,Chaitanya Joshi,Madhvi Joshi |
| EPI_ISL_2017781 | BJ Medical College and Civil Hospital, Ahmedabad | Gujarat Biotechnology Research Centre | Ramesh Pandit,Janvi Raval,Zarna Patel,Nitin Savaliya,Dinesh Kumar,Twinkle Soni,Sonal Sharma ,Zuber Saiyed,Pranay Shah,Umang Mishra,Nitesh Shah,Chaitanya Joshi,Madhvi Joshi |
| EPI_ISL_2017782 | BJ Medical College and Civil Hospital, Ahmedabad | Gujarat Biotechnology Research Centre | Janvi Raval,Zarna Patel,Nitin Savaliya,Dinesh Kumar,Twinkle Soni,Sonal Sharma ,Zuber Saiyed,Ramesh Pandit,Pranay Shah,Umang Mishra,Nitesh Shah,Chaitanya Joshi,Madhvi Joshi |
| EPI_ISL_2017783 | BJ Medical College and Civil Hospital, Ahmedabad | Gujarat Biotechnology Research Centre | Zarna Patel,Nitin Savaliya,Dinesh Kumar,Twinkle Soni,Sonal Sharma ,Zuber Saiyed,Ramesh Pandit,Janvi Raval,Pranay Shah,Umang Mishra,Nitesh Shah,Chaitanya Joshi,Madhvi Joshi |
| EPI_ISL_2017784 | BJ Medical College and Civil Hospital, Ahmedabad | Gujarat Biotechnology Research Centre | Nitin Savaliya,Dinesh Kumar,Twinkle Soni,Sonal Sharma ,Zuber Saiyed,Ramesh Pandit,Janvi Raval,Zarna Patel,Pranay Shah,Umang Mishra,Nitesh Shah,Chaitanya Joshi,Madhvi Joshi |
| EPI_ISL_2017785 | BJ Medical College and Civil Hospital, Ahmedabad | Gujarat Biotechnology Research Centre | Dinesh Kumar,Twinkle Soni,Sonal Sharma ,Zuber Saiyed,Ramesh Pandit,Janvi Raval,Zarna Patel,Nitin Savaliya,Pranay Shah,Umang Mishra,Nitesh Shah,Chaitanya Joshi,Madhvi Joshi |
| EPI_ISL_2017786 | BJ Medical College and Civil Hospital, Ahmedabad | Gujarat Biotechnology Research Centre | Twinkle Soni,Sonal Sharma ,Zuber Saiyed,Ramesh Pandit,Janvi Raval,Zarna Patel,Nitin Savaliya,Dinesh Kumar,Pranay Shah,Umang Mishra,Nitesh Shah,Chaitanya Joshi,Madhvi Joshi |
| EPI_ISL_2017787 | BJ Medical College and Civil Hospital, Ahmedabad | Gujarat Biotechnology Research Centre | Sonal Sharma ,Zuber Saiyed,Ramesh Pandit,Janvi Raval,Zarna Patel,Nitin Savaliya,Dinesh Kumar,Twinkle Soni,Pranay Shah,Umang Mishra,Nitesh Shah,Chaitanya Joshi,Madhvi Joshi |
| EPI_ISL_2017788 | BJ Medical College and Civil Hospital, Ahmedabad | Gujarat Biotechnology Research Centre | Zuber Saiyed,Ramesh Pandit,Janvi Raval,Zarna Patel,Nitin Savaliya,Dinesh Kumar,Twinkle Soni,Sonal Sharma ,Pranay Shah,Umang Mishra,Nitesh Shah,Chaitanya Joshi,Madhvi Joshi |

| | | | |
|---|---|---|---|
| EPI_ISL_2017789 | Gujarat Biotechnology Research Centre | Gujarat Biotechnology Research Centre | Ramesh Pandit,Janvi Raval,Zarna Patel,Nitin Savaliya,Dinesh Kumar,Twinkle Soni,Sonal Sharma ,Zuber Saiyed,Umang Mishra,Nitesh Shah,Chaitanya Joshi,Madhvi Joshi |
| EPI_ISL_2018939, EPI_ISL_2018940, EPI_ISL_2018941, EPI_ISL_2018942, EPI_ISL_2018944, EPI_ISL_2018945, EPI_ISL_2018946, EPI_ISL_2018947, EPI_ISL_2018948, EPI_ISL_2018949, EPI_ISL_2018950, EPI_ISL_2018951, EPI_ISL_2018952, EPI_ISL_2018953, EPI_ISL_2018954, EPI_ISL_2018955, EPI_ISL_2018956, EPI_ISL_2018957, EPI_ISL_2018958, EPI_ISL_2018959, EPI_ISL_2018960, EPI_ISL_2018961, EPI_ISL_2018962 | | | |
| see above | University of Rome Tor Vergata: Departm Experim Medicine Chair of Virology | University of Rome Tor Vergata: Departm Experim Medicine Chair of Virology | Maria Concetta Bellocchi, Luca Carioti, Maria Botticelli, Lorenzo Piermatteo, Rossana Scutari, Marco Iannetta, Loredana Sarmati, Massimo Andreoni, Francesca Ceccherini-Silberstein |
| EPI_ISL_2020075, EPI_ISL_2020076, EPI_ISL_2020078, EPI_ISL_2020165, EPI_ISL_2020166, EPI_ISL_2020167 | Kota Bharu Public Health Laboratory | Institute for Medical Research, Infectious Disease Research Centre, National Institutes of Health, Ministry of Health Malaysia | Suppiah J, Kamel K, Mohd Zawawi Z, Azizan MA, Ramly N, Robert F, Thayan R |
| EPI_ISL_2020930 | Urban Public Health Centre (UPHC), SECTOR-2, Gandhinagar | Gujarat Biotechnology Research Centre | Amit Kanani,Twinkle Soni,Sonal Sharma ,Zuber Saiyed,Ramesh Pandit,Janvi Raval,Zarna Patel,Nitin Savaliya,Dinesh Kumar,Bhargav Waghela,Bhumika Prajapati,Tejendra Purohit ,Nilen Patel ,Kalpesh Goswami,Amrutlal Patel,Umang Mishra,Nitesh Shah,Chaitanya Joshi,Madhvi Joshi |
| EPI_ISL_2020931 | Urban Public Health Centre (UPHC), SECTOR-2, Gandhinagar | Gujarat Biotechnology Research Centre | Amit Kanani,Sonal Sharma ,Zuber Saiyed,Ramesh Pandit,Janvi Raval,Zarna Patel,Nitin Savaliya,Dinesh Kumar,Twinkle Soni,Bhargav Waghela,Bhumika Prajapati,Tejendra Purohit ,Nilen Patel ,Kalpesh Goswami,Amrutlal Patel,Umang Mishra,Nitesh Shah,Chaitanya Joshi,Madhvi Joshi |
| EPI_ISL_2020932 | Urban Public Health Centre (UPHC), SECTOR-2, Gandhinagar | Gujarat Biotechnology Research Centre | Zuber Saiyed,Ramesh Pandit,Janvi Raval,Zarna Patel,Nitin Savaliya,Dinesh Kumar,Twinkle Soni,Sonal Sharma ,Bhargav Waghela,Bhumika Prajapati,Tejendra Purohit ,Nilen Patel ,Amit Kanani,Kalpesh Goswami,Amrutlal Patel,Umang Mishra,Nitesh Shah,Chaitanya Joshi,Madhvi Joshi |
| EPI_ISL_2020933 | Urban Public Health Centre (UPHC), SECTOR-2, Gandhinagar | Gujarat Biotechnology Research Centre | Ramesh Pandit,Janvi Raval,Zarna Patel,Nitin Savaliya,Dinesh Kumar,Twinkle Soni,Sonal Sharma ,Zuber Saiyed,Bhargav Waghela,Bhumika Prajapati,Tejendra Purohit ,Nilen Patel ,Amit Kanani,Kalpesh Goswami,Amrutlal Patel,Umang Mishra,Nitesh Shah,Chaitanya Joshi,Madhvi Joshi |
| EPI_ISL_2020934 | Urban Public Health Centre (UPHC), SECTOR-2, Gandhinagar | Gujarat Biotechnology Research Centre | Janvi Raval,Zarna Patel,Nitin Savaliya,Dinesh Kumar,Twinkle Soni,Sonal Sharma ,Zuber Saiyed,Ramesh Pandit,Bhargav Waghela,Bhumika Prajapati,Tejendra Purohit ,Nilen Patel ,Amit Kanani,Kalpesh Goswami,Amrutlal Patel,Umang Mishra,Nitesh Shah,Chaitanya Joshi,Madhvi Joshi |
| EPI_ISL_2020935 | Urban Public Health Centre (UPHC), SECTOR-2, Gandhinagar | Gujarat Biotechnology Research Centre | Zarna Patel,Nitin Savaliya,Dinesh Kumar,Twinkle Soni,Sonal Sharma ,Zuber Saiyed,Ramesh Pandit,Janvi Raval,Bhargav Waghela,Bhumika Prajapati,Tejendra Purohit ,Nilen Patel ,Amit Kanani,Kalpesh Goswami,Amrutlal Patel,Umang Mishra,Nitesh Shah,Chaitanya Joshi,Madhvi Joshi |
| EPI_ISL_2020936 | Urban Public Health Centre (UPHC), SECTOR-2, Gandhinagar | Gujarat Biotechnology Research Centre | Nitin Savaliya,Dinesh Kumar,Twinkle Soni,Sonal Sharma ,Zuber Saiyed,Ramesh Pandit,Janvi Raval,Zarna Patel,Bhargav Waghela,Bhumika Prajapati,Tejendra Purohit ,Nilen Patel ,Amit Kanani,Kalpesh Goswami,Amrutlal Patel,Umang Mishra,Nitesh Shah,Chaitanya Joshi,Madhvi Joshi |
| EPI_ISL_2020941 | Urban Public Health Centre (UPHC), SECTOR-2, Gandhinagar | Gujarat Biotechnology Research Centre | Ramesh Pandit,Janvi Raval,Zarna Patel,Nitin Savaliya,Dinesh Kumar,Twinkle Soni,Sonal Sharma ,Zuber Saiyed,Bhargav Waghela,Bhumika Prajapati,Tejendra Purohit ,Nilen Patel ,Amit Kanani,Kalpesh Goswami,Amrutlal Patel,Umang Mishra,Nitesh Shah,Chaitanya Joshi,Madhvi Joshi |
| EPI_ISL_2020942 | Urban Public Health Centre (UPHC), SECTOR-2, Gandhinagar | Gujarat Biotechnology Research Centre | Janvi Raval,Zarna Patel,Nitin Savaliya,Dinesh Kumar,Twinkle Soni,Sonal Sharma ,Zuber Saiyed,Ramesh Pandit,Bhargav Waghela,Bhumika Prajapati,Tejendra Purohit ,Nilen Patel ,Amit Kanani,Kalpesh Goswami,Amrutlal Patel,Umang Mishra,Nitesh Shah,Chaitanya Joshi,Madhvi Joshi |
| EPI_ISL_2020943 | Urban Public Health Centre (UPHC), SECTOR-2, Gandhinagar | Gujarat Biotechnology Research Centre | Zarna Patel,Nitin Savaliya,Dinesh Kumar,Twinkle Soni,Sonal Sharma ,Zuber Saiyed,Ramesh Pandit,Janvi Raval,Bhargav Waghela,Bhumika Prajapati,Tejendra Purohit ,Nilen Patel ,Amit Kanani,Kalpesh Goswami,Amrutlal Patel,Umang Mishra,Nitesh Shah,Chaitanya Joshi,Madhvi Joshi |
| EPI_ISL_2020945 | Urban Public Health Centre (UPHC), SECTOR-2, Gandhinagar | Gujarat Biotechnology Research Centre | Bhumika Prajapati,Dinesh Kumar,Twinkle Soni,Sonal Sharma ,Zuber Saiyed,Ramesh Pandit,Janvi Raval,Zarna Patel,Nitin Savaliya,Bhargav Waghela,Tejendra Purohit ,Nilen Patel ,Amit Kanani,Kalpesh Goswami,Amrutlal Patel,Umang Mishra,Nitesh Shah,Chaitanya Joshi,Madhvi Joshi |
| EPI_ISL_2020946 | Urban Public Health Centre (UPHC), SECTOR-2, Gandhinagar | Gujarat Biotechnology Research Centre | Bhumika Prajapati,Twinkle Soni,Sonal Sharma ,Zuber Saiyed,Ramesh Pandit,Janvi Raval,Zarna Patel,Nitin Savaliya,Dinesh Kumar,Bhargav Waghela,Tejendra Purohit ,Nilen Patel ,Amit Kanani,Kalpesh Goswami,Amrutlal Patel,Umang Mishra,Nitesh Shah,Chaitanya Joshi,Madhvi Joshi |
| EPI_ISL_2020947 | Urban Public Health Centre (UPHC), SECTOR-2, Gandhinagar | Gujarat Biotechnology Research Centre | Sonal Sharma ,Zuber Saiyed,Ramesh Pandit,Janvi Raval,Zarna Patel,Nitin Savaliya,Dinesh Kumar,Twinkle Soni,Bhargav Waghela,Bhumika Prajapati,Tejendra Purohit ,Nilen Patel ,Amit Kanani,Kalpesh Goswami,Amrutlal Patel,Umang Mishra,Nitesh Shah,Chaitanya Joshi,Madhvi Joshi |
| EPI_ISL_2020950 | Urban Public Health Centre (UPHC), SECTOR-2, Gandhinagar | Gujarat Biotechnology Research Centre | Janvi Raval,Zarna Patel,Nitin Savaliya,Dinesh Kumar,Twinkle Soni,Sonal Sharma ,Zuber Saiyed,Ramesh Pandit,Bhargav Waghela,Bhumika Prajapati,Tejendra Purohit ,Nilen Patel ,Amit Kanani,Kalpesh Goswami,Amrutlal Patel,Umang Mishra,Nitesh Shah,Chaitanya Joshi,Madhvi Joshi |
| EPI_ISL_2020951 | Urban Public Health Centre (UPHC), SECTOR-2, Gandhinagar | Gujarat Biotechnology Research Centre | Zarna Patel,Nitin Savaliya,Dinesh Kumar,Twinkle Soni,Sonal Sharma ,Zuber Saiyed,Ramesh Pandit,Janvi Raval,Bhargav Waghela,Bhumika Prajapati,Tejendra Purohit ,Nilen Patel ,Amit Kanani,Kalpesh Goswami,Amrutlal Patel,Umang Mishra,Nitesh Shah,Chaitanya Joshi,Madhvi Joshi |
| EPI_ISL_2020952 | Urban Public Health Centre (UPHC), SECTOR-2, Gandhinagar | Gujarat Biotechnology Research Centre | Nitin Savaliya,Dinesh Kumar,Twinkle Soni,Sonal Sharma ,Zuber Saiyed,Ramesh Pandit,Janvi Raval,Zarna Patel,Bhargav Waghela,Bhumika Prajapati,Tejendra Purohit ,Nilen Patel ,Amit Kanani,Kalpesh Goswami,Amrutlal Patel,Umang Mishra,Nitesh Shah,Chaitanya Joshi,Madhvi Joshi |
| EPI_ISL_2020953 | Urban Public Health Centre (UPHC), SECTOR-2, Gandhinagar | Gujarat Biotechnology Research Centre | Dinesh Kumar,Twinkle Soni,Sonal Sharma ,Zuber Saiyed,Ramesh Pandit,Janvi Raval,Zarna Patel,Nitin Savaliya,Bhargav Waghela,Bhumika Prajapati,Tejendra Purohit ,Nilen Patel ,Amit Kanani,Kalpesh Goswami,Amrutlal Patel,Umang Mishra,Nitesh Shah,Chaitanya Joshi,Madhvi Joshi |
| EPI_ISL_2020954 | Urban Public Health Centre (UPHC), SECTOR-2, Gandhinagar | Gujarat Biotechnology Research Centre | Twinkle Soni,Sonal Sharma ,Zuber Saiyed,Ramesh Pandit,Janvi Raval,Zarna Patel,Nitin Savaliya,Dinesh Kumar,Bhargav Waghela,Bhumika Prajapati,Tejendra Purohit ,Nilen Patel ,Amit Kanani,Kalpesh Goswami,Amrutlal Patel,Umang Mishra,Nitesh Shah,Chaitanya Joshi,Madhvi Joshi |
| EPI_ISL_2020955 | Urban Public Health Centre (UPHC), SECTOR-2, Gandhinagar | Gujarat Biotechnology Research Centre | Nilen Patel ,Sonal Sharma ,Zuber Saiyed,Ramesh Pandit,Janvi Raval,Zarna Patel,Nitin Savaliya,Dinesh Kumar,Twinkle Soni,Bhargav Waghela,Bhumika Prajapati,Tejendra Purohit ,Amit Kanani,Kalpesh Goswami,Amrutlal Patel,Umang Mishra,Nitesh Shah,Chaitanya Joshi,Madhvi Joshi |
| EPI_ISL_2020956 | Urban Public Health Centre (UPHC), SECTOR-2, Gandhinagar | Gujarat Biotechnology Research Centre | Nilen Patel ,Zuber Saiyed,Ramesh Pandit,Janvi Raval,Zarna Patel,Nitin Savaliya,Dinesh Kumar,Twinkle Soni,Sonal Sharma ,Bhargav Waghela,Bhumika Prajapati,Tejendra Purohit ,Amit Kanani,Kalpesh Goswami,Amrutlal Patel,Umang Mishra,Nitesh Shah,Chaitanya Joshi,Madhvi Joshi |
| EPI_ISL_2020957 | Urban Public Health Centre (UPHC), SECTOR-2, Gandhinagar | Gujarat Biotechnology Research Centre | Kalpesh Goswami,Nitin Savaliya,Dinesh Kumar,Twinkle Soni,Sonal Sharma ,Zuber Saiyed,Ramesh Pandit,Janvi Raval,Zarna Patel,Bhargav Waghela,Bhumika Prajapati,Tejendra Purohit ,Nilen Patel ,Amit Kanani,Amrutlal Patel,Umang Mishra,Nitesh Shah,Chaitanya Joshi,Madhvi Joshi |
| EPI_ISL_2020958 | Urban Public Health Centre (UPHC), SECTOR-2, Gandhinagar | Gujarat Biotechnology Research Centre | Kalpesh Goswami,Dinesh Kumar,Twinkle Soni,Sonal Sharma ,Zuber Saiyed,Ramesh Pandit,Janvi Raval,Zarna Patel,Nitin Savaliya,Bhargav Waghela,Bhumika Prajapati,Tejendra Purohit ,Nilen Patel ,Amit Kanani,Amrutlal Patel,Umang Mishra,Nitesh Shah,Chaitanya Joshi,Madhvi Joshi |
| EPI_ISL_2023034, EPI_ISL_2023037, EPI_ISL_2023039, EPI_ISL_2023042, EPI_ISL_2023047 | Department of Virology, Istituto Zooprofilattico Sperimentale del Lazio e della Toscana (IZSLT) | Department of General Diagnostics; Department of Virology; Istituto Zooprofilattico Sperimentale del Lazio e della Toscana (IZSLT) | Patricia Alba, Giuseppe Manna, Elena L. Diaconu, Fabiola Feltrin, Raffaela Conti, Teresa Scicluna, Virginia Carfora, Antonella Cersini, Alessia Franco, Antonio Battisti. |
| EPI_ISL_2023049 | Laboratorio di Microbiologia e Virologia, Università Vita-Salute San Raffaele, Milano | Laboratorio di Microbiologia e Virologia, Università Vita-Salute San Raffaele, Milano | Michela Sampaolo, Elena Criscuolo, Matteo Castelli, Roberto Ferrarese, Roberta Antonia Diotti, Enzo Boeri, Massimo Locatelli, Massimo Clementi, Nicasio Mancini, Nicola Clementi |
| EPI_ISL_2023437 | Microbiology Department, University Hospital Donostia | Microbiology Department, University Hospital Donostia | Marimon JM, Montes M, Piñeiro L, Sorarrain A, Gomez M, Cilla G. |
| EPI_ISL_2029113, EPI_ISL_2029114 | HEGP - Laboratoire de Virologie | HEGP - Laboratoire de Virologie | David Veyer, Sandrine Imbeaud, Julien Puech, Hélène Péré, Maxime Wack, Nicolas Robillard |
| EPI_ISL_2029115 | USCA Tagliacozzo | Istituto Zooprofilattico Sperimentale dell'Abruzzo e Molise "G. Caporale" | Lorusso A, Marcacci M, Di Domenico M, Ancora M, Curini V, Di Lollo Valeria, Mangone I, Rinaldi A, Delli Compagni E, Scialabba S, Caporale M, Di Pasquale A, Cammà C, Puglia I, Calistri P, Savini G |
| EPI_ISL_2029116 | SIESP CHIETI - DRIVE IN VASTO | Istituto Zooprofilattico Sperimentale dell'Abruzzo e Molise "G. Caporale" | Lorusso A, Marcacci M, Di Domenico M, Ancora M, Curini V, Di Lollo Valeria, Mangone I, Rinaldi A, Delli Compagni E, Scialabba S, Caporale M, Di Pasquale A, Cammà C, Puglia I, Calistri P, Savini G |
| EPI_ISL_2029604 | Hospital | National Reference Center for Viruses of Respiratory Infections, Institut Pasteur, Paris | Marion Barbet, Sylvie Behillil, Méline Bizard, Angela Brisebarre, Camille Capel, Vincent Enouf, Louise Lefrançois, Frédéric Lemoine, Christophe Malabat, Corinne Maufrais, Etienne Simon-Lorière, Maud Vanpeene, Sylvie Van der Werf ,Isabelle Gros |
| EPI_ISL_2029605 | Hospital | National Reference Center for Viruses of Respiratory Infections, Institut Pasteur, Paris | Marion Barbet, Sylvie Behillil, Méline Bizard, Angela Brisebarre, Camille Capel, Vincent Enouf, Louise Lefrançois, Frédéric Lemoine, Christophe Malabat, Corinne Maufrais, Etienne Simon-Lorière, Maud Vanpeene, Sylvie Van der Werf ,Sylvie Ledru |
| EPI_ISL_2029606 | Hospital | National Reference Center for Viruses of Respiratory Infections, Institut Pasteur, Paris | Marion Barbet, Sylvie Behillil, Méline Bizard, Angela Brisebarre, Camille Capel, Vincent Enouf, Louise Lefrançois, Frédéric Lemoine, Christophe Malabat, Corinne Maufrais, Amaury Vaysse, Etienne Simon-Lorière, Maud Vanpeene, Sylvie Van der Werf ,André-Guy Combremont |

| | | | |
|---|---|---|---|
| EPI_ISL_2029608 | Hospital | National Reference Center for Viruses of Respiratory Infections, Institut Pasteur, Paris | Marion Barbet, Sylvie Behillil, Méline Bizard, Angela Brisebarre, Camille Capel, Vincent Enouf, Louise Lefrançois, Frédéric Lemoine, Christophe Malabat, Corinne Maufrais, Etienne Simon-Lorière, Maud Vanpeene, Sylvie Van der Werf ,Alexandra Ducancelle |
| EPI_ISL_2029609 | Labo Analyses Med | National Reference Center for Viruses of Respiratory Infections, Institut Pasteur, Paris | Marion Barbet, Sylvie Behillil, Méline Bizard, Angela Brisebarre, Camille Capel, Vincent Enouf, Louise Lefrançois, Frédéric Lemoine, Christophe Malabat, Corinne Maufrais, Etienne Simon-Lorière, Maud Vanpeene, Sylvie Van der Werf ,Pierre-Yves Leonard |
| EPI_ISL_2029610 | Hospital | National Reference Center for Viruses of Respiratory Infections, Institut Pasteur, Paris | Marion Barbet, Sylvie Behillil, Méline Bizard, Angela Brisebarre, Camille Capel, Vincent Enouf, Louise Lefrançois, Frédéric Lemoine, Christophe Malabat, Corinne Maufrais, Etienne Simon-Lorière, Maud Vanpeene, Sylvie Van der Werf ,Pascale Martres |
| EPI_ISL_2029611 | Hospital | National Reference Center for Viruses of Respiratory Infections, Institut Pasteur, Paris | Marion Barbet, Sylvie Behillil, Méline Bizard, Angela Brisebarre, Camille Capel, Vincent Enouf, Louise Lefrançois, Frédéric Lemoine, Christophe Malabat, Corinne Maufrais, Etienne Simon-Lorière, Maud Vanpeene, Sylvie Van der Werf ,Isabelle Gros |
| EPI_ISL_2029612 | Hospital | National Reference Center for Viruses of Respiratory Infections, Institut Pasteur, Paris | Marion Barbet, Sylvie Behillil, Méline Bizard, Angela Brisebarre, Camille Capel, Vincent Enouf, Louise Lefrançois, Frédéric Lemoine, Christophe Malabat, Corinne Maufrais, Etienne Simon-Lorière, Maud Vanpeene, Sylvie Van der Werf ,CéLine Ramanantsoa |
| EPI_ISL_2029613 | Labo Analyses Med | National Reference Center for Viruses of Respiratory Infections, Institut Pasteur, Paris | Marion Barbet, Sylvie Behillil, Méline Bizard, Angela Brisebarre, Camille Capel, Vincent Enouf, Louise Lefrançois, Frédéric Lemoine, Christophe Malabat, Corinne Maufrais, Etienne Simon-Lorière, Maud Vanpeene, Sylvie Van der Werf ,Pierre-Yves Leonard |
| EPI_ISL_2029614 | Labo Analyses Med | National Reference Center for Viruses of Respiratory Infections, Institut Pasteur, Paris | Marion Barbet, Sylvie Behillil, Méline Bizard, Angela Brisebarre, Camille Capel, Vincent Enouf, Louise Lefrançois, Frédéric Lemoine, Christophe Malabat, Corinne Maufrais, Etienne Simon-Lorière, Maud Vanpeene, Sylvie Van der Werf ,Brieuc Gestin |
| EPI_ISL_2029616 | Hospital | National Reference Center for Viruses of Respiratory Infections, Institut Pasteur, Paris | Marion Barbet, Sylvie Behillil, Méline Bizard, Angela Brisebarre, Camille Capel, Vincent Enouf, Louise Lefrançois, Frédéric Lemoine, Christophe Malabat, Corinne Maufrais, Etienne Simon-Lorière, Maud Vanpeene, Sylvie Van der Werf ,LéA Pilorge |
| EPI_ISL_2029617 | Labo Analyses Med | National Reference Center for Viruses of Respiratory Infections, Institut Pasteur, Paris | Marion Barbet, Sylvie Behillil, Méline Bizard, Angela Brisebarre, Camille Capel, Vincent Enouf, Louise Lefrançois, Frédéric Lemoine, Christophe Malabat, Corinne Maufrais, Etienne Simon-Lorière, Maud Vanpeene, Sylvie Van der Werf ,SéVerine Haouisee |
| EPI_ISL_2029619 | Labo Analyses Med | National Reference Center for Viruses of Respiratory Infections, Institut Pasteur, Paris | Marion Barbet, Sylvie Behillil, Méline Bizard, Angela Brisebarre, Camille Capel, Vincent Enouf, Louise Lefrançois, Frédéric Lemoine, Christophe Malabat, Corinne Maufrais, Etienne Simon-Lorière, Maud Vanpeene, Sylvie Van der Werf ,Pierre-Yves Leonard |
| EPI_ISL_2029620 | Hospital | National Reference Center for Viruses of Respiratory Infections, Institut Pasteur, Paris | Marion Barbet, Sylvie Behillil, Méline Bizard, Angela Brisebarre, Camille Capel, Vincent Enouf, Louise Lefrançois, Frédéric Lemoine, Christophe Malabat, Corinne Maufrais, Amaury Vaysse, Etienne Simon-Lorière, Maud Vanpeene, Sylvie Van der Werf ,CéLine Bressollette |
| EPI_ISL_2029622 | Labo Analyses Med | National Reference Center for Viruses of Respiratory Infections, Institut Pasteur, Paris | Marion Barbet, Sylvie Behillil, Méline Bizard, Angela Brisebarre, Camille Capel, Vincent Enouf, Louise Lefrançois, Frédéric Lemoine, Christophe Malabat, Corinne Maufrais, Emmanuelle Permal, Etienne Simon-Lorière, Maud Vanpeene, Sylvie Van der Werf ,Anne Holstein |
| EPI_ISL_2029623 | Labo Analyses Med | National Reference Center for Viruses of Respiratory Infections, Institut Pasteur, Paris | Marion Barbet, Sylvie Behillil, Méline Bizard, Angela Brisebarre, Camille Capel, Vincent Enouf, Louise Lefrançois, Frédéric Lemoine, Christophe Malabat, Corinne Maufrais, Etienne Simon-Lorière, Maud Vanpeene, Sylvie Van der Werf ,Delphine MammÈS |
| EPI_ISL_2029625 | Labo Analyses Med | National Reference Center for Viruses of Respiratory Infections, Institut Pasteur, Paris | Marion Barbet, Sylvie Behillil, Méline Bizard, Angela Brisebarre, Camille Capel, Vincent Enouf, Louise Lefrançois, Frédéric Lemoine, Christophe Malabat, Corinne Maufrais, Etienne Simon-Lorière, Maud Vanpeene, Sylvie Van der Werf ,Pierre-Yves Leonard |
| EPI_ISL_2029626 | Labo Analyses Med | National Reference Center for Viruses of Respiratory Infections, Institut Pasteur, Paris | Marion Barbet, Sylvie Behillil, Méline Bizard, Angela Brisebarre, Camille Capel, Vincent Enouf, Louise Lefrançois, Frédéric Lemoine, Christophe Malabat, Corinne Maufrais, Etienne Simon-Lorière, Maud Vanpeene, Sylvie Van der Werf ,Brieuc Gestin |
| EPI_ISL_2029627 | Labo Analyses Med | National Reference Center for Viruses of Respiratory Infections, Institut Pasteur, Paris | Marion Barbet, Sylvie Behillil, Méline Bizard, Angela Brisebarre, Camille Capel, Vincent Enouf, Louise Lefrançois, Frédéric Lemoine, Christophe Malabat, Corinne Maufrais, Etienne Simon-Lorière, Maud Vanpeene, Sylvie Van der Werf ,GréGoire Potiron |
| EPI_ISL_2029628 | Labo Analyses Med | National Reference Center for Viruses of Respiratory Infections, Institut Pasteur, Paris | Marion Barbet, Sylvie Behillil, Méline Bizard, Angela Brisebarre, Camille Capel, Vincent Enouf, Louise Lefrançois, Frédéric Lemoine, Christophe Malabat, Corinne Maufrais, Etienne Simon-Lorière, Maud Vanpeene, Sylvie Van der Werf ,Pierre-Yves Leonard |
| EPI_ISL_2029630 | Hospital | National Reference Center for Viruses of Respiratory Infections, Institut Pasteur, Paris | Marion Barbet, Sylvie Behillil, Méline Bizard, Angela Brisebarre, Camille Capel, Vincent Enouf, Louise Lefrançois, Frédéric Lemoine, Christophe Malabat, Corinne Maufrais, Etienne Simon-Lorière, Maud Vanpeene, Sylvie Van der Werf ,L. Courdavault |
| EPI_ISL_2029631 | Labo Analyses Med | National Reference Center for Viruses of Respiratory Infections, Institut Pasteur, Paris | Marion Barbet, Sylvie Behillil, Méline Bizard, Angela Brisebarre, Camille Capel, Vincent Enouf, Louise Lefrançois, Frédéric Lemoine, Christophe Malabat, Corinne Maufrais, Emmanuelle Permal, Etienne Simon-Lorière, Maud Vanpeene, Sylvie Van der Werf ,Anne Holstein |
| EPI_ISL_2029632 | Hospital | National Reference Center for Viruses of Respiratory Infections, Institut Pasteur, Paris | Marion Barbet, Sylvie Behillil, Méline Bizard, Angela Brisebarre, Camille Capel, Vincent Enouf, Louise Lefrançois, Frédéric Lemoine, Christophe Malabat, Corinne Maufrais, Etienne Simon-Lorière, Maud Vanpeene, Sylvie Van der Werf ,Thibault Guinoiseau |
| EPI_ISL_2029637 | Labo Analyses Med | National Reference Center for Viruses of Respiratory Infections, Institut Pasteur, Paris | Marion Barbet, Sylvie Behillil, Méline Bizard, Angela Brisebarre, Camille Capel, Vincent Enouf, Louise Lefrançois, Frédéric Lemoine, Christophe Malabat, Corinne Maufrais, Etienne Simon-Lorière, Maud Vanpeene, Sylvie Van der Werf ,Anne Holstein |
| EPI_ISL_2029638 | Labo Analyses Med | National Reference Center for Viruses of Respiratory Infections, Institut Pasteur, Paris | Marion Barbet, Sylvie Behillil, Méline Bizard, Angela Brisebarre, Camille Capel, Vincent Enouf, Louise Lefrançois, Frédéric Lemoine, Christophe Malabat, Corinne Maufrais, Etienne Simon-Lorière, Maud Vanpeene, Sylvie Van der Werf ,Pierre-Yves Leonard |
| EPI_ISL_2029639 | Labo Analyses Med | National Reference Center for Viruses of Respiratory Infections, Institut Pasteur, Paris | Marion Barbet, Sylvie Behillil, Méline Bizard, Angela Brisebarre, Camille Capel, Vincent Enouf, Louise Lefrançois, Frédéric Lemoine, Christophe Malabat, Corinne Maufrais, Etienne Simon-Lorière, Maud Vanpeene, Sylvie Van der Werf ,Bruno Berdin |
| EPI_ISL_2029640, EPI_ISL_2029641 | Labo Analyses Med | National Reference Center for Viruses of Respiratory Infections, Institut Pasteur, Paris | Marion Barbet, Sylvie Behillil, Méline Bizard, Angela Brisebarre, Camille Capel, Vincent Enouf, Louise Lefrançois, Frédéric Lemoine, Christophe Malabat, Corinne Maufrais, Etienne Simon-Lorière, Maud Vanpeene, Sylvie Van der Werf ,GréGoire Potiron |
| EPI_ISL_2029642 | Labo Analyses Med | National Reference Center for Viruses of Respiratory Infections, Institut Pasteur, Paris | Marion Barbet, Sylvie Behillil, Méline Bizard, Angela Brisebarre, Camille Capel, Vincent Enouf, Louise Lefrançois, Frédéric Lemoine, Christophe Malabat, Corinne Maufrais, Etienne Simon-Lorière, Maud Vanpeene, Sylvie Van der Werf ,Arditi Cocco |
| EPI_ISL_2029643 | Hospital | National Reference Center for Viruses of Respiratory Infections, Institut Pasteur, Paris | Marion Barbet, Sylvie Behillil, Méline Bizard, Angela Brisebarre, Camille Capel, Vincent Enouf, Louise Lefrançois, Frédéric Lemoine, Christophe Malabat, Corinne Maufrais, Etienne Simon-Lorière, Maud Vanpeene, Sylvie Van der Werf ,Mj Dufour |
| EPI_ISL_2029644 | Hospital | National Reference Center for Viruses of Respiratory Infections, Institut Pasteur, Paris | Marion Barbet, Sylvie Behillil, Méline Bizard, Angela Brisebarre, Camille Capel, Vincent Enouf, Louise Lefrançois, Frédéric Lemoine, Christophe Malabat, Corinne Maufrais, Etienne Simon-Lorière, Maud Vanpeene, Sylvie Van der Werf ,Nabil Gastli |
| EPI_ISL_2029645 | Labo Analyses Med | National Reference Center for Viruses of Respiratory Infections, Institut Pasteur, Paris | Marion Barbet, Sylvie Behillil, Méline Bizard, Angela Brisebarre, Camille Capel, Vincent Enouf, Louise Lefrançois, Frédéric Lemoine, Christophe Malabat, Corinne Maufrais, Etienne Simon-Lorière, Maud Vanpeene, Sylvie Van der Werf ,Pierre-Yves Leonard |
| EPI_ISL_2029646 | Hospital | National Reference Center for Viruses of Respiratory Infections, Institut Pasteur, Paris | Marion Barbet, Sylvie Behillil, Méline Bizard, Angela Brisebarre, Camille Capel, Vincent Enouf, Louise Lefrançois, Frédéric Lemoine, Christophe Malabat, Corinne Maufrais, Etienne Simon-Lorière, Maud Vanpeene, Sylvie Van der Werf ,CéLine Bressollette |
| EPI_ISL_2029647 | Hospital | National Reference Center for Viruses of Respiratory Infections, Institut Pasteur, Paris | Marion Barbet, Sylvie Behillil, Méline Bizard, Angela Brisebarre, Camille Capel, Vincent Enouf, Louise Lefrançois, Frédéric Lemoine, Christophe Malabat, Corinne Maufrais, Etienne Simon-Lorière, Maud Vanpeene, Sylvie Van der Werf ,Benjamin Maneglie |
| EPI_ISL_2029648 | Hospital | National Reference Center for Viruses of Respiratory Infections, Institut Pasteur, Paris | Marion Barbet, Sylvie Behillil, Méline Bizard, Angela Brisebarre, Camille Capel, Vincent Enouf, Louise Lefrançois, Frédéric Lemoine, Christophe Malabat, Corinne Maufrais, Etienne Simon-Lorière, Maud Vanpeene, Sylvie Van der Werf ,CéCile Farrugia |
| EPI_ISL_2029650 | Labo Analyses Med | National Reference Center for Viruses of Respiratory Infections, Institut Pasteur, Paris | Marion Barbet, Sylvie Behillil, Méline Bizard, Angela Brisebarre, Camille Capel, Vincent Enouf, Louise Lefrançois, Frédéric Lemoine, Christophe Malabat, Corinne Maufrais, Pierre Lechat, Etienne Simon-Lorière, Maud Vanpeene, Sylvie Van der Werf ,Ronan Pichard |
| EPI_ISL_2029651, EPI_ISL_2029652 | Labo Analyses Med | National Reference Center for Viruses of Respiratory Infections, Institut Pasteur, Paris | Marion Barbet, Sylvie Behillil, Méline Bizard, Angela Brisebarre, Camille Capel, Vincent Enouf, Louise Lefrançois, Frédéric Lemoine, Christophe Malabat, Corinne Maufrais, Etienne Simon-Lorière, Maud Vanpeene, Sylvie Van der Werf ,Anne Holstein |
| EPI_ISL_2029653 | Hospital | National Reference Center for Viruses of Respiratory Infections, Institut Pasteur, Paris | Marion Barbet, Sylvie Behillil, Méline Bizard, Angela Brisebarre, Camille Capel, Vincent Enouf, Louise Lefrançois, Frédéric Lemoine, Christophe Malabat, Corinne Maufrais, Etienne Simon-Lorière, Maud Vanpeene, Sylvie Van der Werf ,CéLine Bressollette |
| EPI_ISL_2029654 | Labo Analyses Med | National Reference Center for Viruses of Respiratory Infections, Institut Pasteur, Paris | Marion Barbet, Sylvie Behillil, Méline Bizard, Angela Brisebarre, Camille Capel, Vincent Enouf, Louise Lefrançois, Frédéric Lemoine, Christophe Malabat, Corinne Maufrais, Etienne Simon-Lorière, Maud Vanpeene, Sylvie Van der Werf |

| | | | |
|---|---|---|---|
| EPI_ISL_2029655 | Labo Analyses Med | National Reference Center for Viruses of Respiratory Infections, Institut Pasteur, Paris | Marion Barbet, Sylvie Behillil, Méline Bizard, Angela Brisebarre, Camille Capel, Vincent Enouf, Louise Lefrançois, Frédéric Lemoine, Christophe Malabat, Corinne Maufrais, Etienne Simon-Lorière, Maud Vanpeene, Sylvie Van der Werf ,Brieuc Gestin |
| EPI_ISL_2029656 | Hospital | National Reference Center for Viruses of Respiratory Infections, Institut Pasteur, Paris | Marion Barbet, Sylvie Behillil, Méline Bizard, Angela Brisebarre, Camille Capel, Vincent Enouf, Louise Lefrançois, Frédéric Lemoine, Christophe Malabat, Corinne Maufrais, Etienne Simon-Lorière, Maud Vanpeene, Sylvie Van der Werf ,LéA Pilorge |
| EPI_ISL_2029657 | Labo Analyses Med | National Reference Center for Viruses of Respiratory Infections, Institut Pasteur, Paris | Marion Barbet, Sylvie Behillil, Méline Bizard, Angela Brisebarre, Camille Capel, Vincent Enouf, Louise Lefrançois, Frédéric Lemoine, Christophe Malabat, Corinne Maufrais, Etienne Simon-Lorière, Maud Vanpeene, Sylvie Van der Werf ,Ronan Pichard |
| EPI_ISL_2029658, EPI_ISL_2029659, EPI_ISL_2029660 | Labo Analyses Med | National Reference Center for Viruses of Respiratory Infections, Institut Pasteur, Paris | Marion Barbet, Sylvie Behillil, Méline Bizard, Angela Brisebarre, Camille Capel, Vincent Enouf, Louise Lefrançois, Frédéric Lemoine, Christophe Malabat, Corinne Maufrais, Etienne Simon-Lorière, Maud Vanpeene, Sylvie Van der Werf ,GréGoire Potiron |
| EPI_ISL_2029661 | Labo Analyses Med | National Reference Center for Viruses of Respiratory Infections, Institut Pasteur, Paris | Marion Barbet, Sylvie Behillil, Méline Bizard, Angela Brisebarre, Camille Capel, Vincent Enouf, Louise Lefrançois, Frédéric Lemoine, Christophe Malabat, Corinne Maufrais, Etienne Simon-Lorière, Maud Vanpeene, Sylvie Van der Werf ,Arditi Cocco |
| EPI_ISL_2029662 | Labo Analyses Med | National Reference Center for Viruses of Respiratory Infections, Institut Pasteur, Paris | Marion Barbet, Sylvie Behillil, Méline Bizard, Angela Brisebarre, Camille Capel, Vincent Enouf, Louise Lefrançois, Frédéric Lemoine, Christophe Malabat, Corinne Maufrais, Etienne Simon-Lorière, Maud Vanpeene, Sylvie Van der Werf ,Franck Ennouchi |
| EPI_ISL_2029663 | Labo Analyses Med | National Reference Center for Viruses of Respiratory Infections, Institut Pasteur, Paris | Marion Barbet, Sylvie Behillil, Méline Bizard, Angela Brisebarre, Camille Capel, Vincent Enouf, Louise Lefrançois, Frédéric Lemoine, Christophe Malabat, Corinne Maufrais, Etienne Simon-Lorière, Maud Vanpeene, Sylvie Van der Werf ,Anne Holstein |
| EPI_ISL_2029664 | Hospital | National Reference Center for Viruses of Respiratory Infections, Institut Pasteur, Paris | Marion Barbet, Sylvie Behillil, Méline Bizard, Angela Brisebarre, Camille Capel, Vincent Enouf, Louise Lefrançois, Frédéric Lemoine, Christophe Malabat, Corinne Maufrais, Etienne Simon-Lorière, Maud Vanpeene, Sylvie Van der Werf ,Nabil Gastli |
| EPI_ISL_2029665 | Labo analyses med | National Reference Center for Viruses of Respiratory Infections, Institut Pasteur, Paris | Marion Barbet, Sylvie Behillil, Méline Bizard, Angela Brisebarre, Camille Capel, Vincent Enouf, Louise Lefrançois, Frédéric Lemoine, Christophe Malabat, Corinne Maufrais, Etienne Simon-Lorière, Maud Vanpeene, Sylvie Van der Werf ,Jean-FrançOis Comes |
| EPI_ISL_2029669 | Labo Analyses Med | National Reference Center for Viruses of Respiratory Infections, Institut Pasteur, Paris | Marion Barbet, Sylvie Behillil, Méline Bizard, Angela Brisebarre, Camille Capel, Vincent Enouf, Louise Lefrançois, Frédéric Lemoine, Christophe Malabat, Corinne Maufrais, Etienne Simon-Lorière, Maud Vanpeene, Sylvie Van der Werf ,Brieuc Gestin |
| EPI_ISL_2029670 | Hospital | National Reference Center for Viruses of Respiratory Infections, Institut Pasteur, Paris | Marion Barbet, Sylvie Behillil, Méline Bizard, Angela Brisebarre, Camille Capel, Vincent Enouf, Louise Lefrançois, Frédéric Lemoine, Christophe Malabat, Corinne Maufrais, Etienne Simon-Lorière, Maud Vanpeene, Sylvie Van der Werf ,LéA Pilorge |
| EPI_ISL_2029671, EPI_ISL_2029672, EPI_ISL_2029673 | Labo Analyses Med | National Reference Center for Viruses of Respiratory Infections, Institut Pasteur, Paris | Marion Barbet, Sylvie Behillil, Méline Bizard, Angela Brisebarre, Camille Capel, Vincent Enouf, Louise Lefrançois, Frédéric Lemoine, Christophe Malabat, Corinne Maufrais, Etienne Simon-Lorière, Maud Vanpeene, Sylvie Van der Werf ,Ronan Pichard |
| EPI_ISL_2029674 | Labo Analyses Med | National Reference Center for Viruses of Respiratory Infections, Institut Pasteur, Paris | Marion Barbet, Sylvie Behillil, Méline Bizard, Angela Brisebarre, Camille Capel, Vincent Enouf, Louise Lefrançois, Frédéric Lemoine, Christophe Malabat, Corinne Maufrais, Etienne Simon-Lorière, Maud Vanpeene, Sylvie Van der Werf ,GréGoire Potiron |
| EPI_ISL_2029676 | Labo Analyses Med | National Reference Center for Viruses of Respiratory Infections, Institut Pasteur, Paris | Marion Barbet, Sylvie Behillil, Méline Bizard, Angela Brisebarre, Camille Capel, Vincent Enouf, Louise Lefrançois, Frédéric Lemoine, Christophe Malabat, Corinne Maufrais, Pierre Lechat, Etienne Simon-Lorière, Maud Vanpeene, Sylvie Van der Werf ,GréGoire Potiron |
| EPI_ISL_2029677 | Labo Analyses Med | National Reference Center for Viruses of Respiratory Infections, Institut Pasteur, Paris | Marion Barbet, Sylvie Behillil, Méline Bizard, Angela Brisebarre, Camille Capel, Vincent Enouf, Louise Lefrançois, Frédéric Lemoine, Christophe Malabat, Corinne Maufrais, Etienne Simon-Lorière, Maud Vanpeene, Sylvie Van der Werf ,Arditi Cocco |
| EPI_ISL_2029679 | Hospital | National Reference Center for Viruses of Respiratory Infections, Institut Pasteur, Paris | Marion Barbet, Sylvie Behillil, Méline Bizard, Angela Brisebarre, Camille Capel, Vincent Enouf, Louise Lefrançois, Frédéric Lemoine, Christophe Malabat, Corinne Maufrais, Etienne Simon-Lorière, Maud Vanpeene, Sylvie Van der Werf ,EugèNe Castelain |
| EPI_ISL_2029680 | Labo Analyses Med | National Reference Center for Viruses of Respiratory Infections, Institut Pasteur, Paris | Marion Barbet, Sylvie Behillil, Méline Bizard, Angela Brisebarre, Camille Capel, Vincent Enouf, Louise Lefrançois, Frédéric Lemoine, Christophe Malabat, Corinne Maufrais, Etienne Simon-Lorière, Maud Vanpeene, Sylvie Van der Werf |
| EPI_ISL_2029681 | Labo Analyses Med | National Reference Center for Viruses of Respiratory Infections, Institut Pasteur, Paris | Marion Barbet, Sylvie Behillil, Méline Bizard, Angela Brisebarre, Camille Capel, Vincent Enouf, Louise Lefrançois, Frédéric Lemoine, Christophe Malabat, Corinne Maufrais, Etienne Simon-Lorière, Maud Vanpeene, Sylvie Van der Werf ,Brieuc Gestin |
| EPI_ISL_2029683 | Hospital | National Reference Center for Viruses of Respiratory Infections, Institut Pasteur, Paris | Marion Barbet, Sylvie Behillil, Méline Bizard, Angela Brisebarre, Camille Capel, Vincent Enouf, Louise Lefrançois, Frédéric Lemoine, Christophe Malabat, Corinne Maufrais, Etienne Simon-Lorière, Maud Vanpeene, Sylvie Van der Werf ,LéA Pilorge |
| EPI_ISL_2029684 | Labo Analyses Med | National Reference Center for Viruses of Respiratory Infections, Institut Pasteur, Paris | Marion Barbet, Sylvie Behillil, Méline Bizard, Angela Brisebarre, Camille Capel, Vincent Enouf, Louise Lefrançois, Frédéric Lemoine, Christophe Malabat, Corinne Maufrais, Etienne Simon-Lorière, Maud Vanpeene, Sylvie Van der Werf ,Arditi Cocco |
| EPI_ISL_2029685 | Labo analyses med | National Reference Center for Viruses of Respiratory Infections, Institut Pasteur, Paris | Marion Barbet, Sylvie Behillil, Méline Bizard, Angela Brisebarre, Camille Capel, Vincent Enouf, Louise Lefrançois, Frédéric Lemoine, Christophe Malabat, Corinne Maufrais, Etienne Simon-Lorière, Maud Vanpeene, Sylvie Van der Werf ,Jean-FrançOis Comes |
| EPI_ISL_2029686 | Labo Analyses Med | National Reference Center for Viruses of Respiratory Infections, Institut Pasteur, Paris | Marion Barbet, Sylvie Behillil, Méline Bizard, Angela Brisebarre, Camille Capel, Vincent Enouf, Louise Lefrançois, Frédéric Lemoine, Christophe Malabat, Corinne Maufrais, Etienne Simon-Lorière, Maud Vanpeene, Sylvie Van der Werf ,Pierre-Yves Leonard |
| EPI_ISL_2029687 | Labo Analyses Med | National Reference Center for Viruses of Respiratory Infections, Institut Pasteur, Paris | Marion Barbet, Sylvie Behillil, Méline Bizard, Angela Brisebarre, Camille Capel, Vincent Enouf, Louise Lefrançois, Frédéric Lemoine, Christophe Malabat, Corinne Maufrais, Etienne Simon-Lorière, Maud Vanpeene, Sylvie Van der Werf ,Pierre Netzer |
| EPI_ISL_2029688, EPI_ISL_2029689 | Armies | National Reference Center for Viruses of Respiratory Infections, Institut Pasteur, Paris | Marion Barbet, Sylvie Behillil, Méline Bizard, Angela Brisebarre, Camille Capel, Vincent Enouf, Louise Lefrançois, Frédéric Lemoine, Christophe Malabat, Corinne Maufrais, Etienne Simon-Lorière, Maud Vanpeene, Sylvie Van der Werf ,Marine Desroches |
| EPI_ISL_2029691 | Labo Analyses Med | National Reference Center for Viruses of Respiratory Infections, Institut Pasteur, Paris | Marion Barbet, Sylvie Behillil, Méline Bizard, Angela Brisebarre, Camille Capel, Vincent Enouf, Louise Lefrançois, Frédéric Lemoine, Christophe Malabat, Corinne Maufrais, Etienne Simon-Lorière, Maud Vanpeene, Sylvie Van der Werf ,Brieuc Gestin |
| EPI_ISL_2029692 | Hospital | National Reference Center for Viruses of Respiratory Infections, Institut Pasteur, Paris | Marion Barbet, Sylvie Behillil, Méline Bizard, Angela Brisebarre, Camille Capel, Vincent Enouf, Louise Lefrançois, Frédéric Lemoine, Christophe Malabat, Corinne Maufrais, Etienne Simon-Lorière, Maud Vanpeene, Sylvie Van der Werf ,Anne Cady |
| EPI_ISL_2029693 | Labo Analyses Med | National Reference Center for Viruses of Respiratory Infections, Institut Pasteur, Paris | Marion Barbet, Sylvie Behillil, Méline Bizard, Angela Brisebarre, Camille Capel, Vincent Enouf, Louise Lefrançois, Frédéric Lemoine, Christophe Malabat, Corinne Maufrais, Etienne Simon-Lorière, Maud Vanpeene, Sylvie Van der Werf ,Franck Ennouchi |
| EPI_ISL_2029694 | Labo Analyses Med | National Reference Center for Viruses of Respiratory Infections, Institut Pasteur, Paris | Marion Barbet, Sylvie Behillil, Méline Bizard, Angela Brisebarre, Camille Capel, Vincent Enouf, Louise Lefrançois, Frédéric Lemoine, Christophe Malabat, Corinne Maufrais, Etienne Simon-Lorière, Maud Vanpeene, Sylvie Van der Werf ,Anne Holstein |
| EPI_ISL_2029695 | Hospital | National Reference Center for Viruses of Respiratory Infections, Institut Pasteur, Paris | Marion Barbet, Sylvie Behillil, Méline Bizard, Angela Brisebarre, Camille Capel, Vincent Enouf, Louise Lefrançois, Frédéric Lemoine, Christophe Malabat, Corinne Maufrais, Etienne Simon-Lorière, Maud Vanpeene, Sylvie Van der Werf ,Nabil Gastli |
| EPI_ISL_2029696 | Hospital | National Reference Center for Viruses of Respiratory Infections, Institut Pasteur, Paris | Marion Barbet, Sylvie Behillil, Méline Bizard, Angela Brisebarre, Camille Capel, Vincent Enouf, Louise Lefrançois, Frédéric Lemoine, Christophe Malabat, Corinne Maufrais, Etienne Simon-Lorière, Maud Vanpeene, Sylvie Van der Werf ,Ilka Engelmann |
| EPI_ISL_2029697 | Armies | National Reference Center for Viruses of Respiratory Infections, Institut Pasteur, Paris | Marion Barbet, Sylvie Behillil, Méline Bizard, Angela Brisebarre, Camille Capel, Vincent Enouf, Louise Lefrançois, Frédéric Lemoine, Christophe Malabat, Corinne Maufrais, Victoire Baillet, Etienne Simon-Lorière, Maud Vanpeene, Sylvie Van der Werf ,Marine Desroches |
| EPI_ISL_2029698 | Hospital | National Reference Center for Viruses of Respiratory Infections, Institut Pasteur, Paris | Marion Barbet, Sylvie Behillil, Méline Bizard, Angela Brisebarre, Camille Capel, Vincent Enouf, Louise Lefrançois, Frédéric Lemoine, Christophe Malabat, Corinne Maufrais, Etienne Simon-Lorière, Maud Vanpeene, Sylvie Van der Werf ,ValéRie Serazin |
| EPI_ISL_2029699 | Labo Analyses Med | National Reference Center for Viruses of Respiratory Infections, Institut Pasteur, Paris | Marion Barbet, Sylvie Behillil, Méline Bizard, Angela Brisebarre, Camille Capel, Vincent Enouf, Louise Lefrançois, Frédéric Lemoine, Christophe Malabat, Corinne Maufrais, Pierre Lechat, Etienne Simon-Lorière, Maud Vanpeene, Sylvie Van der Werf ,Franck Ennouchi |
| EPI_ISL_2029701, EPI_ISL_2029702 | Labo Analyses Med | National Reference Center for Viruses of Respiratory Infections, Institut Pasteur, Paris | Marion Barbet, Sylvie Behillil, Méline Bizard, Angela Brisebarre, Camille Capel, Vincent Enouf, Louise Lefrançois, Frédéric Lemoine, Christophe Malabat, Corinne Maufrais, Etienne Simon-Lorière, Maud Vanpeene, Sylvie Van der Werf ,Brieuc Gestin |
| EPI_ISL_2029703 | Labo Analyses Med | National Reference Center for Viruses of Respiratory Infections, Institut Pasteur, Paris | Marion Barbet, Sylvie Behillil, Méline Bizard, Angela Brisebarre, Camille Capel, Vincent Enouf, Louise Lefrançois, Frédéric Lemoine, Christophe Malabat, Corinne Maufrais, Etienne Simon-Lorière, Maud Vanpeene, Sylvie Van der Werf ,Franck Ennouchi |

| | | | |
|---|---|---|---|
| EPI_ISL_2029704 | Hospital | National Reference Center for Viruses of Respiratory Infections, Institut Pasteur, Paris | Marion Barbet, Sylvie Behillil, Méline Bizard, Angela Brisebarre, Camille Capel, Vincent Enouf, Louise Lefrançois, Frédéric Lemoine, Christophe Malabat, Corinne Maufrais, Etienne Simon-Lorière, Maud Vanpeene, Sylvie Van der Werf ,Alexandra Ducancelle |
| EPI_ISL_2029705, EPI_ISL_2029706, EPI_ISL_2029707 | Labo Analyses Med | National Reference Center for Viruses of Respiratory Infections, Institut Pasteur, Paris | Marion Barbet, Sylvie Behillil, Méline Bizard, Angela Brisebarre, Camille Capel, Vincent Enouf, Louise Lefrançois, Frédéric Lemoine, Christophe Malabat, Corinne Maufrais, Etienne Simon-Lorière, Maud Vanpeene, Sylvie Van der Werf ,Anne Holstein |
| EPI_ISL_2029708 | Labo Analyses Med | National Reference Center for Viruses of Respiratory Infections, Institut Pasteur, Paris | Marion Barbet, Sylvie Behillil, Méline Bizard, Angela Brisebarre, Camille Capel, Vincent Enouf, Louise Lefrançois, Frédéric Lemoine, Christophe Malabat, Corinne Maufrais, Etienne Simon-Lorière, Maud Vanpeene, Sylvie Van der Werf ,Delphine MammÈS |
| EPI_ISL_2029710 | Labo analyses med | National Reference Center for Viruses of Respiratory Infections, Institut Pasteur, Paris | Marion Barbet, Sylvie Behillil, Méline Bizard, Angela Brisebarre, Camille Capel, Vincent Enouf, Louise Lefrançois, Frédéric Lemoine, Christophe Malabat, Corinne Maufrais, Etienne Simon-Lorière, Maud Vanpeene, Sylvie Van der Werf ,Jean-FrançOis Comes |
| EPI_ISL_2029711 | Outre mer | National Reference Center for Viruses of Respiratory Infections, Institut Pasteur, Paris | Marion Barbet, Sylvie Behillil, Méline Bizard, Angela Brisebarre, Camille Capel, Vincent Enouf, Louise Lefrançois, Frédéric Lemoine, Christophe Malabat, Corinne Maufrais, Amaury Vaysse, Etienne Simon-Lorière, Maud Vanpeene, Sylvie Van der Werf ,Patrice Combe |
| EPI_ISL_2029713 | Labo Analyses Med | National Reference Center for Viruses of Respiratory Infections, Institut Pasteur, Paris | Marion Barbet, Sylvie Behillil, Méline Bizard, Angela Brisebarre, Camille Capel, Vincent Enouf, Louise Lefrançois, Frédéric Lemoine, Christophe Malabat, Corinne Maufrais, Etienne Simon-Lorière, Maud Vanpeene, Sylvie Van der Werf ,Pierre Netzer |
| EPI_ISL_2029715, EPI_ISL_2029716 | Labo Analyses Med | National Reference Center for Viruses of Respiratory Infections, Institut Pasteur, Paris | Marion Barbet, Sylvie Behillil, Méline Bizard, Angela Brisebarre, Camille Capel, Vincent Enouf, Louise Lefrançois, Frédéric Lemoine, Christophe Malabat, Corinne Maufrais, Etienne Simon-Lorière, Maud Vanpeene, Sylvie Van der Werf ,Brieuc Gestin |
| EPI_ISL_2029718, EPI_ISL_2029719 | Labo Analyses Med | National Reference Center for Viruses of Respiratory Infections, Institut Pasteur, Paris | Marion Barbet, Sylvie Behillil, Méline Bizard, Angela Brisebarre, Camille Capel, Vincent Enouf, Louise Lefrançois, Frédéric Lemoine, Christophe Malabat, Corinne Maufrais, Etienne Simon-Lorière, Maud Vanpeene, Sylvie Van der Werf ,Anne Holstein |
| EPI_ISL_2029720 | Hospital | National Reference Center for Viruses of Respiratory Infections, Institut Pasteur, Paris | Marion Barbet, Sylvie Behillil, Méline Bizard, Angela Brisebarre, Camille Capel, Vincent Enouf, Louise Lefrançois, Frédéric Lemoine, Christophe Malabat, Corinne Maufrais, Etienne Simon-Lorière, Maud Vanpeene, Sylvie Van der Werf ,Dominique Descamps |
| EPI_ISL_2029721 | Labo analyses med | National Reference Center for Viruses of Respiratory Infections, Institut Pasteur, Paris | Marion Barbet, Sylvie Behillil, Méline Bizard, Angela Brisebarre, Camille Capel, Vincent Enouf, Louise Lefrançois, Frédéric Lemoine, Christophe Malabat, Corinne Maufrais, Etienne Simon-Lorière, Maud Vanpeene, Sylvie Van der Werf ,Jean-FrançOis Comes |
| EPI_ISL_2029722 | Labo Analyses Med | National Reference Center for Viruses of Respiratory Infections, Institut Pasteur, Paris | Marion Barbet, Sylvie Behillil, Méline Bizard, Angela Brisebarre, Camille Capel, Vincent Enouf, Louise Lefrançois, Frédéric Lemoine, Christophe Malabat, Corinne Maufrais, Amaury Vaysse, Etienne Simon-Lorière, Maud Vanpeene, Sylvie Van der Werf ,Pierre-Yves Leonard |
| EPI_ISL_2029724 | Armies | National Reference Center for Viruses of Respiratory Infections, Institut Pasteur, Paris | Marion Barbet, Sylvie Behillil, Méline Bizard, Angela Brisebarre, Camille Capel, Vincent Enouf, Louise Lefrançois, Frédéric Lemoine, Christophe Malabat, Corinne Maufrais, Etienne Simon-Lorière, Maud Vanpeene, Sylvie Van der Werf ,Marine Desroches |
| EPI_ISL_2029725 | Hospital | National Reference Center for Viruses of Respiratory Infections, Institut Pasteur, Paris | Marion Barbet, Sylvie Behillil, Méline Bizard, Angela Brisebarre, Camille Capel, Vincent Enouf, Louise Lefrançois, Frédéric Lemoine, Christophe Malabat, Corinne Maufrais, Etienne Simon-Lorière, Maud Vanpeene, Sylvie Van der Werf ,Isabelle Gros |
| EPI_ISL_2029726 | Labo Analyses Med | National Reference Center for Viruses of Respiratory Infections, Institut Pasteur, Paris | Marion Barbet, Sylvie Behillil, Méline Bizard, Angela Brisebarre, Camille Capel, Vincent Enouf, Louise Lefrançois, Frédéric Lemoine, Christophe Malabat, Corinne Maufrais, Etienne Simon-Lorière, Maud Vanpeene, Sylvie Van der Werf ,Arditi Cocco |
| EPI_ISL_2029727 | Labo Analyses Med | National Reference Center for Viruses of Respiratory Infections, Institut Pasteur, Paris | Marion Barbet, Sylvie Behillil, Méline Bizard, Angela Brisebarre, Camille Capel, Vincent Enouf, Louise Lefrançois, Frédéric Lemoine, Christophe Malabat, Corinne Maufrais, Pierre Lechat, Etienne Simon-Lorière, Maud Vanpeene, Sylvie Van der Werf ,Jean Boyer |
| EPI_ISL_2029728 | Labo Analyses Med | National Reference Center for Viruses of Respiratory Infections, Institut Pasteur, Paris | Marion Barbet, Sylvie Behillil, Méline Bizard, Angela Brisebarre, Camille Capel, Vincent Enouf, Louise Lefrançois, Frédéric Lemoine, Christophe Malabat, Corinne Maufrais, Etienne Simon-Lorière, Maud Vanpeene, Sylvie Van der Werf ,Pierre-Yves Leonard |
| EPI_ISL_2029729, EPI_ISL_2029730 | Labo Analyses Med | National Reference Center for Viruses of Respiratory Infections, Institut Pasteur, Paris | Marion Barbet, Sylvie Behillil, Méline Bizard, Angela Brisebarre, Camille Capel, Vincent Enouf, Louise Lefrançois, Frédéric Lemoine, Christophe Malabat, Corinne Maufrais, Etienne Simon-Lorière, Maud Vanpeene, Sylvie Van der Werf ,Bruno Berdin |
| EPI_ISL_2029732 | Labo Analyses Med | National Reference Center for Viruses of Respiratory Infections, Institut Pasteur, Paris | Marion Barbet, Sylvie Behillil, Méline Bizard, Angela Brisebarre, Camille Capel, Vincent Enouf, Louise Lefrançois, Frédéric Lemoine, Christophe Malabat, Corinne Maufrais, Etienne Simon-Lorière, Maud Vanpeene, Sylvie Van der Werf ,GréGoire Potiron |
| EPI_ISL_2029733 | Outre mer | National Reference Center for Viruses of Respiratory Infections, Institut Pasteur, Paris | Marion Barbet, Sylvie Behillil, Méline Bizard, Angela Brisebarre, Camille Capel, Vincent Enouf, Louise Lefrançois, Frédéric Lemoine, Christophe Malabat, Corinne Maufrais, Etienne Simon-Lorière, Maud Vanpeene, Sylvie Van der Werf ,Patrice Combe |
| EPI_ISL_2029734 | Labo Analyses Med | National Reference Center for Viruses of Respiratory Infections, Institut Pasteur, Paris | Marion Barbet, Sylvie Behillil, Méline Bizard, Angela Brisebarre, Camille Capel, Vincent Enouf, Louise Lefrançois, Frédéric Lemoine, Christophe Malabat, Corinne Maufrais, Amaury Vaysse, Etienne Simon-Lorière, Maud Vanpeene, Sylvie Van der Werf ,Pierre Netzer |
| EPI_ISL_2029735 | Hospital | National Reference Center for Viruses of Respiratory Infections, Institut Pasteur, Paris | Marion Barbet, Sylvie Behillil, Méline Bizard, Angela Brisebarre, Camille Capel, Vincent Enouf, Louise Lefrançois, Frédéric Lemoine, Christophe Malabat, Corinne Maufrais, Etienne Simon-Lorière, Maud Vanpeene, Sylvie Van der Werf ,Mignard |
| EPI_ISL_2029736, EPI_ISL_2029737 | Hospital | National Reference Center for Viruses of Respiratory Infections, Institut Pasteur, Paris | Marion Barbet, Sylvie Behillil, Méline Bizard, Angela Brisebarre, Camille Capel, Vincent Enouf, Louise Lefrançois, Frédéric Lemoine, Christophe Malabat, Corinne Maufrais, Etienne Simon-Lorière, Maud Vanpeene, Sylvie Van der Werf ,L. Courdavault |
| EPI_ISL_2029738 | Labo Analyses Med | National Reference Center for Viruses of Respiratory Infections, Institut Pasteur, Paris | Marion Barbet, Sylvie Behillil, Méline Bizard, Angela Brisebarre, Camille Capel, Vincent Enouf, Louise Lefrançois, Frédéric Lemoine, Christophe Malabat, Corinne Maufrais, Etienne Simon-Lorière, Maud Vanpeene, Sylvie Van der Werf ,Arditi Cocco |
| EPI_ISL_2029739 | Labo Analyses Med | National Reference Center for Viruses of Respiratory Infections, Institut Pasteur, Paris | Marion Barbet, Sylvie Behillil, Méline Bizard, Angela Brisebarre, Camille Capel, Vincent Enouf, Louise Lefrançois, Frédéric Lemoine, Christophe Malabat, Corinne Maufrais, Pierre Lechat, Etienne Simon-Lorière, Maud Vanpeene, Sylvie Van der Werf ,Arditi Cocco |
| EPI_ISL_2029740 | Labo Analyses Med | National Reference Center for Viruses of Respiratory Infections, Institut Pasteur, Paris | Marion Barbet, Sylvie Behillil, Méline Bizard, Angela Brisebarre, Camille Capel, Vincent Enouf, Louise Lefrançois, Frédéric Lemoine, Christophe Malabat, Corinne Maufrais, Pierre Lechat, Etienne Simon-Lorière, Maud Vanpeene, Sylvie Van der Werf ,Jean Boyer |
| EPI_ISL_2029741 | Labo Analyses Med | National Reference Center for Viruses of Respiratory Infections, Institut Pasteur, Paris | Marion Barbet, Sylvie Behillil, Méline Bizard, Angela Brisebarre, Camille Capel, Vincent Enouf, Louise Lefrançois, Frédéric Lemoine, Christophe Malabat, Corinne Maufrais, Etienne Simon-Lorière, Maud Vanpeene, Sylvie Van der Werf ,Anne Holstein |
| EPI_ISL_2029743 | Hospital | National Reference Center for Viruses of Respiratory Infections, Institut Pasteur, Paris | Marion Barbet, Sylvie Behillil, Méline Bizard, Angela Brisebarre, Camille Capel, Vincent Enouf, Louise Lefrançois, Frédéric Lemoine, Christophe Malabat, Corinne Maufrais, Etienne Simon-Lorière, Maud Vanpeene, Sylvie Van der Werf ,FréDéRique Canis |
| EPI_ISL_2029744 | Labo Analyses Med | National Reference Center for Viruses of Respiratory Infections, Institut Pasteur, Paris | Marion Barbet, Sylvie Behillil, Méline Bizard, Angela Brisebarre, Camille Capel, Vincent Enouf, Louise Lefrançois, Frédéric Lemoine, Christophe Malabat, Corinne Maufrais, Etienne Simon-Lorière, Maud Vanpeene, Sylvie Van der Werf ,Pierre-Yves Leonard |
| EPI_ISL_2029745 | Labo Analyses Med | National Reference Center for Viruses of Respiratory Infections, Institut Pasteur, Paris | Marion Barbet, Sylvie Behillil, Méline Bizard, Angela Brisebarre, Camille Capel, Vincent Enouf, Louise Lefrançois, Frédéric Lemoine, Christophe Malabat, Corinne Maufrais, Etienne Simon-Lorière, Maud Vanpeene, Sylvie Van der Werf ,Anne Holstein |
| EPI_ISL_2029746, EPI_ISL_2029747, EPI_ISL_2029748 | Hospital | National Reference Center for Viruses of Respiratory Infections, Institut Pasteur, Paris | Marion Barbet, Sylvie Behillil, Méline Bizard, Angela Brisebarre, Camille Capel, Vincent Enouf, Louise Lefrançois, Frédéric Lemoine, Christophe Malabat, Corinne Maufrais, Etienne Simon-Lorière, Maud Vanpeene, Sylvie Van der Werf ,Nabil Gastli |
| EPI_ISL_2029749 | Labo Analyses Med | National Reference Center for Viruses of Respiratory Infections, Institut Pasteur, Paris | Marion Barbet, Sylvie Behillil, Méline Bizard, Angela Brisebarre, Camille Capel, Vincent Enouf, Louise Lefrançois, Frédéric Lemoine, Christophe Malabat, Corinne Maufrais, Amaury Vaysse, Etienne Simon-Lorière, Maud Vanpeene, Sylvie Van der Werf ,Pierre-Yves Leonard |
| EPI_ISL_2029750 | Labo Analyses Med | National Reference Center for Viruses of Respiratory Infections, Institut Pasteur, Paris | Marion Barbet, Sylvie Behillil, Méline Bizard, Angela Brisebarre, Camille Capel, Vincent Enouf, Louise Lefrançois, Frédéric Lemoine, Christophe Malabat, Corinne Maufrais, Etienne Simon-Lorière, Maud Vanpeene, Sylvie Van der Werf ,Pierre-Yves Leonard |
| EPI_ISL_2029751 | Labo Analyses Med | National Reference Center for Viruses of Respiratory Infections, Institut Pasteur, Paris | Marion Barbet, Sylvie Behillil, Méline Bizard, Angela Brisebarre, Camille Capel, Vincent Enouf, Louise Lefrançois, Frédéric Lemoine, Christophe Malabat, Corinne Maufrais, Etienne Simon-Lorière, Maud Vanpeene, Sylvie Van der Werf ,Judith Zerah |
| EPI_ISL_2029752 | Labo Analyses Med | National Reference Center for Viruses of Respiratory Infections, Institut Pasteur, Paris | Marion Barbet, Sylvie Behillil, Méline Bizard, Angela Brisebarre, Camille Capel, Vincent Enouf, Louise Lefrançois, Frédéric Lemoine, Christophe Malabat, Corinne Maufrais, Etienne Simon-Lorière, Maud Vanpeene, Sylvie Van der Werf ,Anne Holstein |
| EPI_ISL_2029753 | Hospital | National Reference Center for Viruses of Respiratory Infections, Institut Pasteur, Paris | Marion Barbet, Sylvie Behillil, Méline Bizard, Angela Brisebarre, Camille Capel, Vincent Enouf, Louise Lefrançois, Frédéric Lemoine, Christophe Malabat, Corinne Maufrais, Etienne Simon-Lorière, Maud Vanpeene, Sylvie Van der Werf ,Nabil Gastli |

| | | | |
|---|---|---|---|
| EPI_ISL_2029754 | Hospital | National Reference Center for Viruses of Respiratory Infections, Institut Pasteur, Paris | Marion Barbet, Sylvie Behillil, Méline Bizard, Angela Brisebarre, Camille Capel, Vincent Enouf, Louise Lefrançois, Frédéric Lemoine, Christophe Malabat, Corinne Maufrais, Etienne Simon-Lorière, Maud Vanpeene, Sylvie Van der Werf ,Ilka Engelmann |
| EPI_ISL_2029755, EPI_ISL_2029756 | Labo Analyses Med | National Reference Center for Viruses of Respiratory Infections, Institut Pasteur, Paris | Marion Barbet, Sylvie Behillil, Méline Bizard, Angela Brisebarre, Camille Capel, Vincent Enouf, Louise Lefrançois, Frédéric Lemoine, Christophe Malabat, Corinne Maufrais, Etienne Simon-Lorière, Maud Vanpeene, Sylvie Van der Werf ,Pierre-Yves Leonard |
| EPI_ISL_2029758 | Labo Analyses Med | National Reference Center for Viruses of Respiratory Infections, Institut Pasteur, Paris | Marion Barbet, Sylvie Behillil, Méline Bizard, Angela Brisebarre, Camille Capel, Vincent Enouf, Louise Lefrançois, Frédéric Lemoine, Christophe Malabat, Corinne Maufrais, Etienne Simon-Lorière, Maud Vanpeene, Sylvie Van der Werf ,Brieuc Gestin |
| EPI_ISL_2029759 | Hospital | National Reference Center for Viruses of Respiratory Infections, Institut Pasteur, Paris | Marion Barbet, Sylvie Behillil, Méline Bizard, Angela Brisebarre, Camille Capel, Vincent Enouf, Louise Lefrançois, Frédéric Lemoine, Christophe Malabat, Corinne Maufrais, Amaury Vaysse, Etienne Simon-Lorière, Maud Vanpeene, Sylvie Van der Werf ,Dominique Descamps |
| EPI_ISL_2029761 | Labo Analyses Med | National Reference Center for Viruses of Respiratory Infections, Institut Pasteur, Paris | Marion Barbet, Sylvie Behillil, Méline Bizard, Angela Brisebarre, Camille Capel, Vincent Enouf, Louise Lefrançois, Frédéric Lemoine, Christophe Malabat, Corinne Maufrais, Victoire Baillet, Etienne Simon-Lorière, Maud Vanpeene, Sylvie Van der Werf ,Judith Zerah |
| EPI_ISL_2029763 | Hospital | National Reference Center for Viruses of Respiratory Infections, Institut Pasteur, Paris | Marion Barbet, Sylvie Behillil, Méline Bizard, Angela Brisebarre, Camille Capel, Vincent Enouf, Louise Lefrançois, Frédéric Lemoine, Christophe Malabat, Corinne Maufrais, Amaury Vaysse, Etienne Simon-Lorière, Maud Vanpeene, Sylvie Van der Werf ,EugèNe Castelain |
| EPI_ISL_2029764 | Labo Analyses Med | National Reference Center for Viruses of Respiratory Infections, Institut Pasteur, Paris | Marion Barbet, Sylvie Behillil, Méline Bizard, Angela Brisebarre, Camille Capel, Vincent Enouf, Louise Lefrançois, Frédéric Lemoine, Christophe Malabat, Corinne Maufrais, Etienne Simon-Lorière, Maud Vanpeene, Sylvie Van der Werf ,Brieuc Gestin |
| EPI_ISL_2029765 | Hospital | National Reference Center for Viruses of Respiratory Infections, Institut Pasteur, Paris | Marion Barbet, Sylvie Behillil, Méline Bizard, Angela Brisebarre, Camille Capel, Vincent Enouf, Louise Lefrançois, Frédéric Lemoine, Christophe Malabat, Corinne Maufrais, Etienne Simon-Lorière, Maud Vanpeene, Sylvie Van der Werf ,LéA Pilorge |
| EPI_ISL_2029766 | Labo Analyses Med | National Reference Center for Viruses of Respiratory Infections, Institut Pasteur, Paris | Marion Barbet, Sylvie Behillil, Méline Bizard, Angela Brisebarre, Camille Capel, Vincent Enouf, Louise Lefrançois, Frédéric Lemoine, Christophe Malabat, Corinne Maufrais, Gael Millot, Etienne Simon-Lorière, Maud Vanpeene, Sylvie Van der Werf ,GréGoire Potiron |
| EPI_ISL_2029767, EPI_ISL_2029768 | Labo Analyses Med | National Reference Center for Viruses of Respiratory Infections, Institut Pasteur, Paris | Marion Barbet, Sylvie Behillil, Méline Bizard, Angela Brisebarre, Camille Capel, Vincent Enouf, Louise Lefrançois, Frédéric Lemoine, Christophe Malabat, Corinne Maufrais, Pierre Lechat, Etienne Simon-Lorière, Maud Vanpeene, Sylvie Van der Werf ,GréGoire Potiron |
| EPI_ISL_2029769 | Hospital | National Reference Center for Viruses of Respiratory Infections, Institut Pasteur, Paris | Marion Barbet, Sylvie Behillil, Méline Bizard, Angela Brisebarre, Camille Capel, Vincent Enouf, Louise Lefrançois, Frédéric Lemoine, Christophe Malabat, Corinne Maufrais, Etienne Simon-Lorière, Maud Vanpeene, Sylvie Van der Werf ,Anne Cady |
| EPI_ISL_2029770 | Hospital | National Reference Center for Viruses of Respiratory Infections, Institut Pasteur, Paris | Marion Barbet, Sylvie Behillil, Méline Bizard, Angela Brisebarre, Camille Capel, Vincent Enouf, Louise Lefrançois, Frédéric Lemoine, Christophe Malabat, Corinne Maufrais, Etienne Simon-Lorière, Maud Vanpeene, Sylvie Van der Werf ,L. Courdavault |
| EPI_ISL_2029771 | Labo Analyses Med | National Reference Center for Viruses of Respiratory Infections, Institut Pasteur, Paris | Marion Barbet, Sylvie Behillil, Méline Bizard, Angela Brisebarre, Camille Capel, Vincent Enouf, Louise Lefrançois, Frédéric Lemoine, Christophe Malabat, Corinne Maufrais, Gael Millot, Etienne Simon-Lorière, Maud Vanpeene, Sylvie Van der Werf ,Arditi Cocco |
| EPI_ISL_2029773, EPI_ISL_2029774 | Hospital | National Reference Center for Viruses of Respiratory Infections, Institut Pasteur, Paris | Marion Barbet, Sylvie Behillil, Méline Bizard, Angela Brisebarre, Camille Capel, Vincent Enouf, Louise Lefrançois, Frédéric Lemoine, Christophe Malabat, Corinne Maufrais, Etienne Simon-Lorière, Maud Vanpeene, Sylvie Van der Werf ,Nabil Gastli |
| EPI_ISL_2029775 | Labo Analyses Med | National Reference Center for Viruses of Respiratory Infections, Institut Pasteur, Paris | Marion Barbet, Sylvie Behillil, Méline Bizard, Angela Brisebarre, Camille Capel, Vincent Enouf, Louise Lefrançois, Frédéric Lemoine, Christophe Malabat, Corinne Maufrais, Etienne Simon-Lorière, Maud Vanpeene, Sylvie Van der Werf |
| EPI_ISL_2029776 | Labo Analyses Med | National Reference Center for Viruses of Respiratory Infections, Institut Pasteur, Paris | Marion Barbet, Sylvie Behillil, Méline Bizard, Angela Brisebarre, Camille Capel, Vincent Enouf, Louise Lefrançois, Frédéric Lemoine, Christophe Malabat, Corinne Maufrais, Etienne Simon-Lorière, Maud Vanpeene, Sylvie Van der Werf ,Ronan Pichard |
| EPI_ISL_2029777 | Labo Analyses Med | National Reference Center for Viruses of Respiratory Infections, Institut Pasteur, Paris | Marion Barbet, Sylvie Behillil, Méline Bizard, Angela Brisebarre, Camille Capel, Vincent Enouf, Louise Lefrançois, Frédéric Lemoine, Christophe Malabat, Corinne Maufrais, Etienne Simon-Lorière, Maud Vanpeene, Sylvie Van der Werf ,GréGoire Potiron |
| EPI_ISL_2029778 | Hospital | National Reference Center for Viruses of Respiratory Infections, Institut Pasteur, Paris | Marion Barbet, Sylvie Behillil, Méline Bizard, Angela Brisebarre, Camille Capel, Vincent Enouf, Louise Lefrançois, Frédéric Lemoine, Christophe Malabat, Corinne Maufrais, Etienne Simon-Lorière, Maud Vanpeene, Sylvie Van der Werf ,L. Courdavault |
| EPI_ISL_2029779 | Hospital | National Reference Center for Viruses of Respiratory Infections, Institut Pasteur, Paris | Marion Barbet, Sylvie Behillil, Méline Bizard, Angela Brisebarre, Camille Capel, Vincent Enouf, Louise Lefrançois, Frédéric Lemoine, Christophe Malabat, Corinne Maufrais, Etienne Simon-Lorière, Maud Vanpeene, Sylvie Van der Werf ,Nabil Gastli |
| EPI_ISL_2029780 | Labo Analyses Med | National Reference Center for Viruses of Respiratory Infections, Institut Pasteur, Paris | Marion Barbet, Sylvie Behillil, Méline Bizard, Angela Brisebarre, Camille Capel, Vincent Enouf, Louise Lefrançois, Frédéric Lemoine, Christophe Malabat, Corinne Maufrais, Etienne Simon-Lorière, Maud Vanpeene, Sylvie Van der Werf ,Pierre-Yves Leonard |
| EPI_ISL_2029781 | Outre mer | National Reference Center for Viruses of Respiratory Infections, Institut Pasteur, Paris | Marion Barbet, Sylvie Behillil, Méline Bizard, Angela Brisebarre, Camille Capel, Vincent Enouf, Louise Lefrançois, Frédéric Lemoine, Christophe Malabat, Corinne Maufrais, Etienne Simon-Lorière, Maud Vanpeene, Sylvie Van der Werf ,Patrice Combe |
| EPI_ISL_2029782 | Armies | National Reference Center for Viruses of Respiratory Infections, Institut Pasteur, Paris | Marion Barbet, Sylvie Behillil, Méline Bizard, Angela Brisebarre, Camille Capel, Vincent Enouf, Louise Lefrançois, Frédéric Lemoine, Christophe Malabat, Corinne Maufrais, Etienne Simon-Lorière, Maud Vanpeene, Sylvie Van der Werf ,Marine Desroches |
| EPI_ISL_2029784 | Hospital | National Reference Center for Viruses of Respiratory Infections, Institut Pasteur, Paris | Marion Barbet, Sylvie Behillil, Méline Bizard, Angela Brisebarre, Camille Capel, Vincent Enouf, Louise Lefrançois, Frédéric Lemoine, Christophe Malabat, Corinne Maufrais, Etienne Simon-Lorière, Maud Vanpeene, Sylvie Van der Werf ,L. Courdavault |
| EPI_ISL_2029785 | Hospital | National Reference Center for Viruses of Respiratory Infections, Institut Pasteur, Paris | Marion Barbet, Sylvie Behillil, Méline Bizard, Angela Brisebarre, Camille Capel, Vincent Enouf, Louise Lefrançois, Frédéric Lemoine, Christophe Malabat, Corinne Maufrais, Etienne Simon-Lorière, Maud Vanpeene, Sylvie Van der Werf ,Christine Febreau |
| EPI_ISL_2029786 | Labo Analyses Med | National Reference Center for Viruses of Respiratory Infections, Institut Pasteur, Paris | Marion Barbet, Sylvie Behillil, Méline Bizard, Angela Brisebarre, Camille Capel, Vincent Enouf, Louise Lefrançois, Frédéric Lemoine, Christophe Malabat, Corinne Maufrais, Etienne Simon-Lorière, Maud Vanpeene, Sylvie Van der Werf ,Anne Holstein |
| EPI_ISL_2029788 | Labo Analyses Med | National Reference Center for Viruses of Respiratory Infections, Institut Pasteur, Paris | Marion Barbet, Sylvie Behillil, Méline Bizard, Angela Brisebarre, Camille Capel, Vincent Enouf, Louise Lefrançois, Frédéric Lemoine, Christophe Malabat, Corinne Maufrais, Etienne Simon-Lorière, Maud Vanpeene, Sylvie Van der Werf ,Ronan Pichard |
| EPI_ISL_2029789 | Labo Analyses Med | National Reference Center for Viruses of Respiratory Infections, Institut Pasteur, Paris | Marion Barbet, Sylvie Behillil, Méline Bizard, Angela Brisebarre, Camille Capel, Vincent Enouf, Louise Lefrançois, Frédéric Lemoine, Christophe Malabat, Corinne Maufrais, Etienne Simon-Lorière, Maud Vanpeene, Sylvie Van der Werf ,GréGoire Potiron |
| EPI_ISL_2029790 | Labo Analyses Med | National Reference Center for Viruses of Respiratory Infections, Institut Pasteur, Paris | Marion Barbet, Sylvie Behillil, Méline Bizard, Angela Brisebarre, Camille Capel, Vincent Enouf, Louise Lefrançois, Frédéric Lemoine, Christophe Malabat, Corinne Maufrais, Pierre Lechat, Etienne Simon-Lorière, Maud Vanpeene, Sylvie Van der Werf ,GréGoire Potiron |
| EPI_ISL_2029791 | Hospital | National Reference Center for Viruses of Respiratory Infections, Institut Pasteur, Paris | Marion Barbet, Sylvie Behillil, Méline Bizard, Angela Brisebarre, Camille Capel, Vincent Enouf, Louise Lefrançois, Frédéric Lemoine, Christophe Malabat, Corinne Maufrais, Etienne Simon-Lorière, Maud Vanpeene, Sylvie Van der Werf ,L. Courdavault |
| EPI_ISL_2029792 | Labo Analyses Med | National Reference Center for Viruses of Respiratory Infections, Institut Pasteur, Paris | Marion Barbet, Sylvie Behillil, Méline Bizard, Angela Brisebarre, Camille Capel, Vincent Enouf, Louise Lefrançois, Frédéric Lemoine, Christophe Malabat, Corinne Maufrais, Pierre Lechat, Etienne Simon-Lorière, Maud Vanpeene, Sylvie Van der Werf ,Anne Holstein |
| EPI_ISL_2029794 | Labo Analyses Med | National Reference Center for Viruses of Respiratory Infections, Institut Pasteur, Paris | Marion Barbet, Sylvie Behillil, Méline Bizard, Angela Brisebarre, Camille Capel, Vincent Enouf, Louise Lefrançois, Frédéric Lemoine, Christophe Malabat, Corinne Maufrais, Etienne Simon-Lorière, Maud Vanpeene, Sylvie Van der Werf ,Pierre-Yves Leonard |
| EPI_ISL_2029797 | Labo Analyses Med | National Reference Center for Viruses of Respiratory Infections, Institut Pasteur, Paris | Marion Barbet, Sylvie Behillil, Méline Bizard, Angela Brisebarre, Camille Capel, Vincent Enouf, Louise Lefrançois, Frédéric Lemoine, Christophe Malabat, Corinne Maufrais, Etienne Simon-Lorière, Maud Vanpeene, Sylvie Van der Werf ,Brieuc Gestin |
| EPI_ISL_2029798, EPI_ISL_2029799 | Labo Analyses Med | National Reference Center for Viruses of Respiratory Infections, Institut Pasteur, Paris | Marion Barbet, Sylvie Behillil, Méline Bizard, Angela Brisebarre, Camille Capel, Vincent Enouf, Louise Lefrançois, Frédéric Lemoine, Christophe Malabat, Corinne Maufrais, Etienne Simon-Lorière, Maud Vanpeene, Sylvie Van der Werf ,GréGoire Potiron |
| EPI_ISL_2029801 | Labo Analyses Med | National Reference Center for Viruses of Respiratory Infections, Institut Pasteur, Paris | Marion Barbet, Sylvie Behillil, Méline Bizard, Angela Brisebarre, Camille Capel, Vincent Enouf, Louise Lefrançois, Frédéric Lemoine, Christophe Malabat, Corinne Maufrais, Etienne Simon-Lorière, Maud Vanpeene, Sylvie Van der Werf ,Pierre-Yves Leonard |
| EPI_ISL_2029802 | Labo Analyses Med | National Reference Center for Viruses of Respiratory Infections, Institut Pasteur, Paris | Marion Barbet, Sylvie Behillil, Méline Bizard, Angela Brisebarre, Camille Capel, Vincent Enouf, Louise Lefrançois, Frédéric Lemoine, Christophe Malabat, Corinne Maufrais, Etienne Simon-Lorière, Maud Vanpeene, Sylvie Van der Werf |

| | | | |
|---|---|---|---|
| EPI_ISL_2029803 | Armies | National Reference Center for Viruses of Respiratory Infections, Institut Pasteur, Paris | Marion Barbet, Sylvie Behillil, Méline Bizard, Angela Brisebarre, Camille Capel, Vincent Enouf, Louise Lefrançois, Frédéric Lemoine, Christophe Malabat, Corinne Maufrais, Etienne Simon-Lorière, Maud Vanpeene, Sylvie Van der Werf ,Marine Desroches |
| EPI_ISL_2029804 | Hospital | National Reference Center for Viruses of Respiratory Infections, Institut Pasteur, Paris | Marion Barbet, Sylvie Behillil, Méline Bizard, Angela Brisebarre, Camille Capel, Vincent Enouf, Louise Lefrançois, Frédéric Lemoine, Christophe Malabat, Corinne Maufrais, Etienne Simon-Lorière, Maud Vanpeene, Sylvie Van der Werf ,Gandon |
| EPI_ISL_2029805 | Labo Analyses Med | National Reference Center for Viruses of Respiratory Infections, Institut Pasteur, Paris | Marion Barbet, Sylvie Behillil, Méline Bizard, Angela Brisebarre, Camille Capel, Vincent Enouf, Louise Lefrançois, Frédéric Lemoine, Christophe Malabat, Corinne Maufrais, Etienne Simon-Lorière, Maud Vanpeene, Sylvie Van der Werf ,Arditi Cocco |
| EPI_ISL_2029807 | Hospital | National Reference Center for Viruses of Respiratory Infections, Institut Pasteur, Paris | Marion Barbet, Sylvie Behillil, Méline Bizard, Angela Brisebarre, Camille Capel, Vincent Enouf, Louise Lefrançois, Frédéric Lemoine, Christophe Malabat, Corinne Maufrais, Etienne Simon-Lorière, Maud Vanpeene, Sylvie Van der Werf ,Pascale Martres |
| EPI_ISL_2029808, EPI_ISL_2029809 | Hospital | National Reference Center for Viruses of Respiratory Infections, Institut Pasteur, Paris | Marion Barbet, Sylvie Behillil, Méline Bizard, Angela Brisebarre, Camille Capel, Vincent Enouf, Louise Lefrançois, Frédéric Lemoine, Christophe Malabat, Corinne Maufrais, Etienne Simon-Lorière, Maud Vanpeene, Sylvie Van der Werf ,Nabil Gastli |
| EPI_ISL_2029810 | Hospital | National Reference Center for Viruses of Respiratory Infections, Institut Pasteur, Paris | Marion Barbet, Sylvie Behillil, Méline Bizard, Angela Brisebarre, Camille Capel, Vincent Enouf, Louise Lefrançois, Frédéric Lemoine, Christophe Malabat, Corinne Maufrais, Etienne Simon-Lorière, Maud Vanpeene, Sylvie Van der Werf ,FréDéRique Canis |
| EPI_ISL_2029812 | Hospital | National Reference Center for Viruses of Respiratory Infections, Institut Pasteur, Paris | Marion Barbet, Sylvie Behillil, Méline Bizard, Angela Brisebarre, Camille Capel, Vincent Enouf, Louise Lefrançois, Frédéric Lemoine, Christophe Malabat, Corinne Maufrais, Etienne Simon-Lorière, Maud Vanpeene, Sylvie Van der Werf ,CéLine Ramanantsoa |
| EPI_ISL_2029813 | Outre mer | National Reference Center for Viruses of Respiratory Infections, Institut Pasteur, Paris | Marion Barbet, Sylvie Behillil, Méline Bizard, Angela Brisebarre, Camille Capel, Vincent Enouf, Louise Lefrançois, Frédéric Lemoine, Christophe Malabat, Corinne Maufrais, Etienne Simon-Lorière, Maud Vanpeene, Sylvie Van der Werf ,Patrice Combe |
| EPI_ISL_2029815 | Armies | National Reference Center for Viruses of Respiratory Infections, Institut Pasteur, Paris | Marion Barbet, Sylvie Behillil, Méline Bizard, Angela Brisebarre, Camille Capel, Vincent Enouf, Louise Lefrançois, Frédéric Lemoine, Christophe Malabat, Corinne Maufrais, Etienne Simon-Lorière, Maud Vanpeene, Sylvie Van der Werf ,Marine Desroches |
| EPI_ISL_2029816 | Labo Analyses Med | National Reference Center for Viruses of Respiratory Infections, Institut Pasteur, Paris | Marion Barbet, Sylvie Behillil, Méline Bizard, Angela Brisebarre, Camille Capel, Vincent Enouf, Louise Lefrançois, Frédéric Lemoine, Christophe Malabat, Corinne Maufrais, Etienne Simon-Lorière, Maud Vanpeene, Sylvie Van der Werf ,Brieuc Gestin |
| EPI_ISL_2029817 | Hospital | National Reference Center for Viruses of Respiratory Infections, Institut Pasteur, Paris | Marion Barbet, Sylvie Behillil, Méline Bizard, Angela Brisebarre, Camille Capel, Vincent Enouf, Louise Lefrançois, Frédéric Lemoine, Christophe Malabat, Corinne Maufrais, Etienne Simon-Lorière, Maud Vanpeene, Sylvie Van der Werf ,Pascale Martres |
| EPI_ISL_2029818 | Hospital | National Reference Center for Viruses of Respiratory Infections, Institut Pasteur, Paris | Marion Barbet, Sylvie Behillil, Méline Bizard, Angela Brisebarre, Camille Capel, Vincent Enouf, Louise Lefrançois, Frédéric Lemoine, Christophe Malabat, Corinne Maufrais, Etienne Simon-Lorière, Maud Vanpeene, Sylvie Van der Werf ,Nabil Gastli |
| EPI_ISL_2029819 | Labo analyses med | National Reference Center for Viruses of Respiratory Infections, Institut Pasteur, Paris | Marion Barbet, Sylvie Behillil, Méline Bizard, Angela Brisebarre, Camille Capel, Vincent Enouf, Louise Lefrançois, Frédéric Lemoine, Christophe Malabat, Corinne Maufrais, Etienne Simon-Lorière, Maud Vanpeene, Sylvie Van der Werf ,Jean-FrançOis Comes |
| EPI_ISL_2029820 | Labo Analyses Med | National Reference Center for Viruses of Respiratory Infections, Institut Pasteur, Paris | Marion Barbet, Sylvie Behillil, Méline Bizard, Angela Brisebarre, Camille Capel, Vincent Enouf, Louise Lefrançois, Frédéric Lemoine, Christophe Malabat, Corinne Maufrais, Etienne Simon-Lorière, Maud Vanpeene, Sylvie Van der Werf ,Pierre-Yves Leonard |
| EPI_ISL_2029821 | Hospital | National Reference Center for Viruses of Respiratory Infections, Institut Pasteur, Paris | Marion Barbet, Sylvie Behillil, Méline Bizard, Angela Brisebarre, Camille Capel, Vincent Enouf, Louise Lefrançois, Frédéric Lemoine, Christophe Malabat, Corinne Maufrais, Etienne Simon-Lorière, Maud Vanpeene, Sylvie Van der Werf ,Benjamin Maneglie |
| EPI_ISL_2029822 | Labo Analyses Med | National Reference Center for Viruses of Respiratory Infections, Institut Pasteur, Paris | Marion Barbet, Sylvie Behillil, Méline Bizard, Angela Brisebarre, Camille Capel, Vincent Enouf, Louise Lefrançois, Frédéric Lemoine, Christophe Malabat, Corinne Maufrais, Etienne Simon-Lorière, Maud Vanpeene, Sylvie Van der Werf ,Ronan Pichard |
| EPI_ISL_2029823 | Labo Analyses Med | National Reference Center for Viruses of Respiratory Infections, Institut Pasteur, Paris | Marion Barbet, Sylvie Behillil, Méline Bizard, Angela Brisebarre, Camille Capel, Vincent Enouf, Louise Lefrançois, Frédéric Lemoine, Christophe Malabat, Corinne Maufrais, Etienne Simon-Lorière, Maud Vanpeene, Sylvie Van der Werf ,GréGoire Potiron |
| EPI_ISL_2029824 | Labo Analyses Med | National Reference Center for Viruses of Respiratory Infections, Institut Pasteur, Paris | Marion Barbet, Sylvie Behillil, Méline Bizard, Angela Brisebarre, Camille Capel, Vincent Enouf, Louise Lefrançois, Frédéric Lemoine, Christophe Malabat, Corinne Maufrais, Etienne Simon-Lorière, Maud Vanpeene, Sylvie Van der Werf ,Anne Holstein |
| EPI_ISL_2029825 | Hospital | National Reference Center for Viruses of Respiratory Infections, Institut Pasteur, Paris | Marion Barbet, Sylvie Behillil, Méline Bizard, Angela Brisebarre, Camille Capel, Vincent Enouf, Louise Lefrançois, Frédéric Lemoine, Christophe Malabat, Corinne Maufrais, Etienne Simon-Lorière, Maud Vanpeene, Sylvie Van der Werf ,Nabil Gastli |
| EPI_ISL_2029826 | Labo Analyses Med | National Reference Center for Viruses of Respiratory Infections, Institut Pasteur, Paris | Marion Barbet, Sylvie Behillil, Méline Bizard, Angela Brisebarre, Camille Capel, Vincent Enouf, Louise Lefrançois, Frédéric Lemoine, Christophe Malabat, Corinne Maufrais, Amaury Vaysse, Etienne Simon-Lorière, Maud Vanpeene, Sylvie Van der Werf ,Pierre-Yves Leonard |
| EPI_ISL_2029827 | Labo Analyses Med | National Reference Center for Viruses of Respiratory Infections, Institut Pasteur, Paris | Marion Barbet, Sylvie Behillil, Méline Bizard, Angela Brisebarre, Camille Capel, Vincent Enouf, Louise Lefrançois, Frédéric Lemoine, Christophe Malabat, Corinne Maufrais, Etienne Simon-Lorière, Maud Vanpeene, Sylvie Van der Werf ,Pierre-Yves Leonard |
| EPI_ISL_2029829 | Hospital | National Reference Center for Viruses of Respiratory Infections, Institut Pasteur, Paris | Marion Barbet, Sylvie Behillil, Méline Bizard, Angela Brisebarre, Camille Capel, Vincent Enouf, Louise Lefrançois, Frédéric Lemoine, Christophe Malabat, Corinne Maufrais, Etienne Simon-Lorière, Maud Vanpeene, Sylvie Van der Werf ,Isabelle Gros |
| EPI_ISL_2029830 | Hospital | National Reference Center for Viruses of Respiratory Infections, Institut Pasteur, Paris | Marion Barbet, Sylvie Behillil, Méline Bizard, Angela Brisebarre, Camille Capel, Vincent Enouf, Louise Lefrançois, Frédéric Lemoine, Christophe Malabat, Corinne Maufrais, Etienne Simon-Lorière, Maud Vanpeene, Sylvie Van der Werf ,CéCile Farrugia |
| EPI_ISL_2029831 | Labo Analyses Med | National Reference Center for Viruses of Respiratory Infections, Institut Pasteur, Paris | Marion Barbet, Sylvie Behillil, Méline Bizard, Angela Brisebarre, Camille Capel, Vincent Enouf, Louise Lefrançois, Frédéric Lemoine, Christophe Malabat, Corinne Maufrais, Etienne Simon-Lorière, Maud Vanpeene, Sylvie Van der Werf |
| EPI_ISL_2029832 | Hospital | National Reference Center for Viruses of Respiratory Infections, Institut Pasteur, Paris | Marion Barbet, Sylvie Behillil, Méline Bizard, Angela Brisebarre, Camille Capel, Vincent Enouf, Louise Lefrançois, Frédéric Lemoine, Christophe Malabat, Corinne Maufrais, Etienne Simon-Lorière, Maud Vanpeene, Sylvie Van der Werf ,L. Courdavault |
| EPI_ISL_2029833 | Labo Analyses Med | National Reference Center for Viruses of Respiratory Infections, Institut Pasteur, Paris | Marion Barbet, Sylvie Behillil, Méline Bizard, Angela Brisebarre, Camille Capel, Vincent Enouf, Louise Lefrançois, Frédéric Lemoine, Christophe Malabat, Corinne Maufrais, Amaury Vaysse, Etienne Simon-Lorière, Maud Vanpeene, Sylvie Van der Werf ,Pierre-Yves Leonard |
| EPI_ISL_2029834, EPI_ISL_2029835 | Labo Analyses Med | National Reference Center for Viruses of Respiratory Infections, Institut Pasteur, Paris | Marion Barbet, Sylvie Behillil, Méline Bizard, Angela Brisebarre, Camille Capel, Vincent Enouf, Louise Lefrançois, Frédéric Lemoine, Christophe Malabat, Corinne Maufrais, Etienne Simon-Lorière, Maud Vanpeene, Sylvie Van der Werf ,Pierre-Yves Leonard |
| EPI_ISL_2029836 | Hospital | National Reference Center for Viruses of Respiratory Infections, Institut Pasteur, Paris | Marion Barbet, Sylvie Behillil, Méline Bizard, Angela Brisebarre, Camille Capel, Vincent Enouf, Louise Lefrançois, Frédéric Lemoine, Christophe Malabat, Corinne Maufrais, Etienne Simon-Lorière, Maud Vanpeene, Sylvie Van der Werf ,CéLine Bressollette |
| EPI_ISL_2029838 | Hospital | National Reference Center for Viruses of Respiratory Infections, Institut Pasteur, Paris | Marion Barbet, Sylvie Behillil, Méline Bizard, Angela Brisebarre, Camille Capel, Vincent Enouf, Louise Lefrançois, Frédéric Lemoine, Christophe Malabat, Corinne Maufrais, Etienne Simon-Lorière, Maud Vanpeene, Sylvie Van der Werf ,Gandon |
| EPI_ISL_2029839 | Labo Analyses Med | National Reference Center for Viruses of Respiratory Infections, Institut Pasteur, Paris | Marion Barbet, Sylvie Behillil, Méline Bizard, Angela Brisebarre, Camille Capel, Vincent Enouf, Louise Lefrançois, Frédéric Lemoine, Christophe Malabat, Corinne Maufrais, Etienne Simon-Lorière, Maud Vanpeene, Sylvie Van der Werf ,Ronan Pichard |
| EPI_ISL_2029840 | Labo Analyses Med | National Reference Center for Viruses of Respiratory Infections, Institut Pasteur, Paris | Marion Barbet, Sylvie Behillil, Méline Bizard, Angela Brisebarre, Camille Capel, Vincent Enouf, Louise Lefrançois, Frédéric Lemoine, Christophe Malabat, Corinne Maufrais, Etienne Simon-Lorière, Maud Vanpeene, Sylvie Van der Werf ,GréGoire Potiron |
| EPI_ISL_2029841 | Hospital | National Reference Center for Viruses of Respiratory Infections, Institut Pasteur, Paris | Marion Barbet, Sylvie Behillil, Méline Bizard, Angela Brisebarre, Camille Capel, Vincent Enouf, Louise Lefrançois, Frédéric Lemoine, Christophe Malabat, Corinne Maufrais, Etienne Simon-Lorière, Maud Vanpeene, Sylvie Van der Werf ,Anne Cady |
| EPI_ISL_2029843 | Labo Analyses Med | National Reference Center for Viruses of Respiratory Infections, Institut Pasteur, Paris | Marion Barbet, Sylvie Behillil, Méline Bizard, Angela Brisebarre, Camille Capel, Vincent Enouf, Louise Lefrançois, Frédéric Lemoine, Christophe Malabat, Corinne Maufrais, Etienne Simon-Lorière, Maud Vanpeene, Sylvie Van der Werf ,Anne Holstein |
| EPI_ISL_2029844 | Hospital | National Reference Center for Viruses of Respiratory Infections, Institut Pasteur, Paris | Marion Barbet, Sylvie Behillil, Méline Bizard, Angela Brisebarre, Camille Capel, Vincent Enouf, Louise Lefrançois, Frédéric Lemoine, Christophe Malabat, Corinne Maufrais, Etienne Simon-Lorière, Maud Vanpeene, Sylvie Van der Werf ,CéLine Bressollette |
| EPI_ISL_2029845 | Labo Analyses Med | National Reference Center for Viruses of Respiratory Infections, Institut Pasteur, Paris | Marion Barbet, Sylvie Behillil, Méline Bizard, Angela Brisebarre, Camille Capel, Vincent Enouf, Louise Lefrançois, Frédéric Lemoine, Christophe Malabat, Corinne Maufrais, Etienne Simon-Lorière, Maud Vanpeene, Sylvie Van der Werf ,Arditi Cocco |

| | | | |
|---|---|---|---|
| EPI_ISL_2029846 | Hospital | National Reference Center for Viruses of Respiratory Infections, Institut Pasteur, Paris | Marion Barbet, Sylvie Behillil, Méline Bizard, Angela Brisebarre, Camille Capel, Vincent Enouf, Louise Lefrançois, Frédéric Lemoine, Christophe Malabat, Corinne Maufrais, Etienne Simon-Lorière, Maud Vanpeene, Sylvie Van der Werf ,Nabil Gastli |
| EPI_ISL_2029847 | Labo Analyses Med | National Reference Center for Viruses of Respiratory Infections, Institut Pasteur, Paris | Marion Barbet, Sylvie Behillil, Méline Bizard, Angela Brisebarre, Camille Capel, Vincent Enouf, Louise Lefrançois, Frédéric Lemoine, Christophe Malabat, Corinne Maufrais, Etienne Simon-Lorière, Maud Vanpeene, Sylvie Van der Werf ,Pierre-Yves Leonard |
| EPI_ISL_2029848 | Labo Analyses Med | National Reference Center for Viruses of Respiratory Infections, Institut Pasteur, Paris | Marion Barbet, Sylvie Behillil, Méline Bizard, Angela Brisebarre, Camille Capel, Vincent Enouf, Louise Lefrançois, Frédéric Lemoine, Christophe Malabat, Corinne Maufrais, Etienne Simon-Lorière, Maud Vanpeene, Sylvie Van der Werf |
| EPI_ISL_2029849 | Labo Analyses Med | National Reference Center for Viruses of Respiratory Infections, Institut Pasteur, Paris | Marion Barbet, Sylvie Behillil, Méline Bizard, Angela Brisebarre, Camille Capel, Vincent Enouf, Louise Lefrançois, Frédéric Lemoine, Christophe Malabat, Corinne Maufrais, Etienne Simon-Lorière, Maud Vanpeene, Sylvie Van der Werf ,Ronan Pichard |
| EPI_ISL_2029851 | Hospital | National Reference Center for Viruses of Respiratory Infections, Institut Pasteur, Paris | Marion Barbet, Sylvie Behillil, Méline Bizard, Angela Brisebarre, Camille Capel, Vincent Enouf, Louise Lefrançois, Frédéric Lemoine, Christophe Malabat, Corinne Maufrais, Etienne Simon-Lorière, Maud Vanpeene, Sylvie Van der Werf ,Ilka Engelmann |
| EPI_ISL_2029852 | Hospital | National Reference Center for Viruses of Respiratory Infections, Institut Pasteur, Paris | Marion Barbet, Sylvie Behillil, Méline Bizard, Angela Brisebarre, Camille Capel, Vincent Enouf, Louise Lefrançois, Frédéric Lemoine, Christophe Malabat, Corinne Maufrais, Pierre Lechat, Etienne Simon-Lorière, Maud Vanpeene, Sylvie Van der Werf ,LéA Pilorge |
| EPI_ISL_2029853, EPI_ISL_2029854 | Labo Analyses Med | National Reference Center for Viruses of Respiratory Infections, Institut Pasteur, Paris | Marion Barbet, Sylvie Behillil, Méline Bizard, Angela Brisebarre, Camille Capel, Vincent Enouf, Louise Lefrançois, Frédéric Lemoine, Christophe Malabat, Corinne Maufrais, Etienne Simon-Lorière, Maud Vanpeene, Sylvie Van der Werf ,Anne Holstein |
| EPI_ISL_2029855 | Labo Analyses Med | National Reference Center for Viruses of Respiratory Infections, Institut Pasteur, Paris | Marion Barbet, Sylvie Behillil, Méline Bizard, Angela Brisebarre, Camille Capel, Vincent Enouf, Louise Lefrançois, Frédéric Lemoine, Christophe Malabat, Corinne Maufrais, Etienne Simon-Lorière, Maud Vanpeene, Sylvie Van der Werf ,Pierre-Yves Leonard |
| EPI_ISL_2029856 | Hospital | National Reference Center for Viruses of Respiratory Infections, Institut Pasteur, Paris | Marion Barbet, Sylvie Behillil, Méline Bizard, Angela Brisebarre, Camille Capel, Vincent Enouf, Louise Lefrançois, Frédéric Lemoine, Christophe Malabat, Corinne Maufrais, Amaury Vaysse, Etienne Simon-Lorière, Maud Vanpeene, Sylvie Van der Werf ,CéLine Bressollette |
| EPI_ISL_2029858 | Labo Analyses Med | National Reference Center for Viruses of Respiratory Infections, Institut Pasteur, Paris | Marion Barbet, Sylvie Behillil, Méline Bizard, Angela Brisebarre, Camille Capel, Vincent Enouf, Louise Lefrançois, Frédéric Lemoine, Christophe Malabat, Corinne Maufrais, Etienne Simon-Lorière, Maud Vanpeene, Sylvie Van der Werf ,Brieuc Gestin |
| EPI_ISL_2029859 | Labo Analyses Med | National Reference Center for Viruses of Respiratory Infections, Institut Pasteur, Paris | Marion Barbet, Sylvie Behillil, Méline Bizard, Angela Brisebarre, Camille Capel, Vincent Enouf, Louise Lefrançois, Frédéric Lemoine, Christophe Malabat, Corinne Maufrais, Etienne Simon-Lorière, Maud Vanpeene, Sylvie Van der Werf ,Franck Ennouchi |
| EPI_ISL_2029860 | Labo Analyses Med | National Reference Center for Viruses of Respiratory Infections, Institut Pasteur, Paris | Marion Barbet, Sylvie Behillil, Méline Bizard, Angela Brisebarre, Camille Capel, Vincent Enouf, Louise Lefrançois, Frédéric Lemoine, Christophe Malabat, Corinne Maufrais, Etienne Simon-Lorière, Maud Vanpeene, Sylvie Van der Werf |
| EPI_ISL_2029861 | Labo Analyses Med | National Reference Center for Viruses of Respiratory Infections, Institut Pasteur, Paris | Marion Barbet, Sylvie Behillil, Méline Bizard, Angela Brisebarre, Camille Capel, Vincent Enouf, Louise Lefrançois, Frédéric Lemoine, Christophe Malabat, Corinne Maufrais, Etienne Simon-Lorière, Maud Vanpeene, Sylvie Van der Werf ,Anne Holstein |
| EPI_ISL_2029863 | Armies | National Reference Center for Viruses of Respiratory Infections, Institut Pasteur, Paris | Marion Barbet, Sylvie Behillil, Méline Bizard, Angela Brisebarre, Camille Capel, Vincent Enouf, Louise Lefrançois, Frédéric Lemoine, Christophe Malabat, Corinne Maufrais, Etienne Simon-Lorière, Maud Vanpeene, Sylvie Van der Werf ,Marine Desroches |
| EPI_ISL_2029864 | Labo Analyses Med | National Reference Center for Viruses of Respiratory Infections, Institut Pasteur, Paris | Marion Barbet, Sylvie Behillil, Méline Bizard, Angela Brisebarre, Camille Capel, Vincent Enouf, Louise Lefrançois, Frédéric Lemoine, Christophe Malabat, Corinne Maufrais, Etienne Simon-Lorière, Maud Vanpeene, Sylvie Van der Werf ,GréGoire Potiron |
| EPI_ISL_2029865 | Hospital | National Reference Center for Viruses of Respiratory Infections, Institut Pasteur, Paris | Marion Barbet, Sylvie Behillil, Méline Bizard, Angela Brisebarre, Camille Capel, Vincent Enouf, Louise Lefrançois, Frédéric Lemoine, Christophe Malabat, Corinne Maufrais, Pierre Lechat, Etienne Simon-Lorière, Maud Vanpeene, Sylvie Van der Werf ,L. Courdavault |
| EPI_ISL_2029866 | Labo analyses med | National Reference Center for Viruses of Respiratory Infections, Institut Pasteur, Paris | Marion Barbet, Sylvie Behillil, Méline Bizard, Angela Brisebarre, Camille Capel, Vincent Enouf, Louise Lefrançois, Frédéric Lemoine, Christophe Malabat, Corinne Maufrais, Amaury Vaysse, Etienne Simon-Lorière, Maud Vanpeene, Sylvie Van der Werf ,Jean-FrançOis Comes |
| EPI_ISL_2029867 | Labo Analyses Med | National Reference Center for Viruses of Respiratory Infections, Institut Pasteur, Paris | Marion Barbet, Sylvie Behillil, Méline Bizard, Angela Brisebarre, Camille Capel, Vincent Enouf, Louise Lefrançois, Frédéric Lemoine, Christophe Malabat, Corinne Maufrais, Etienne Simon-Lorière, Maud Vanpeene, Sylvie Van der Werf ,SéVerine Haouisee |
| EPI_ISL_2029868 | Hospital | National Reference Center for Viruses of Respiratory Infections, Institut Pasteur, Paris | Marion Barbet, Sylvie Behillil, Méline Bizard, Angela Brisebarre, Camille Capel, Vincent Enouf, Louise Lefrançois, Frédéric Lemoine, Christophe Malabat, Corinne Maufrais, Etienne Simon-Lorière, Maud Vanpeene, Sylvie Van der Werf ,Nabil Gastli |
| EPI_ISL_2029869 | Hospital | National Reference Center for Viruses of Respiratory Infections, Institut Pasteur, Paris | Marion Barbet, Sylvie Behillil, Méline Bizard, Angela Brisebarre, Camille Capel, Vincent Enouf, Louise Lefrançois, Frédéric Lemoine, Christophe Malabat, Corinne Maufrais, Etienne Simon-Lorière, Maud Vanpeene, Sylvie Van der Werf ,Farida Benouadi |
| EPI_ISL_2029870 | Labo Analyses Med | National Reference Center for Viruses of Respiratory Infections, Institut Pasteur, Paris | Marion Barbet, Sylvie Behillil, Méline Bizard, Angela Brisebarre, Camille Capel, Vincent Enouf, Louise Lefrançois, Frédéric Lemoine, Christophe Malabat, Corinne Maufrais, Emmanuelle Permal, Etienne Simon-Lorière, Maud Vanpeene, Sylvie Van der Werf ,Brieuc Gestin |
| EPI_ISL_2029871 | Labo Analyses Med | National Reference Center for Viruses of Respiratory Infections, Institut Pasteur, Paris | Marion Barbet, Sylvie Behillil, Méline Bizard, Angela Brisebarre, Camille Capel, Vincent Enouf, Louise Lefrançois, Frédéric Lemoine, Christophe Malabat, Corinne Maufrais, Etienne Simon-Lorière, Maud Vanpeene, Sylvie Van der Werf ,Anne Holstein |
| EPI_ISL_2029872 | Labo analyses med | National Reference Center for Viruses of Respiratory Infections, Institut Pasteur, Paris | Marion Barbet, Sylvie Behillil, Méline Bizard, Angela Brisebarre, Camille Capel, Vincent Enouf, Louise Lefrançois, Frédéric Lemoine, Christophe Malabat, Corinne Maufrais, Etienne Simon-Lorière, Maud Vanpeene, Sylvie Van der Werf ,Jean-FrançOis Comes |
| EPI_ISL_2029875 | Labo Analyses Med | National Reference Center for Viruses of Respiratory Infections, Institut Pasteur, Paris | Marion Barbet, Sylvie Behillil, Méline Bizard, Angela Brisebarre, Camille Capel, Vincent Enouf, Louise Lefrançois, Frédéric Lemoine, Christophe Malabat, Corinne Maufrais, Etienne Simon-Lorière, Maud Vanpeene, Sylvie Van der Werf ,Pierre-Yves Leonard |
| EPI_ISL_2029876 | Labo Analyses Med | National Reference Center for Viruses of Respiratory Infections, Institut Pasteur, Paris | Marion Barbet, Sylvie Behillil, Méline Bizard, Angela Brisebarre, Camille Capel, Vincent Enouf, Louise Lefrançois, Frédéric Lemoine, Christophe Malabat, Corinne Maufrais, Etienne Simon-Lorière, Maud Vanpeene, Sylvie Van der Werf ,Valentin Robbe |
| EPI_ISL_2029877 | Labo Analyses Med | National Reference Center for Viruses of Respiratory Infections, Institut Pasteur, Paris | Marion Barbet, Sylvie Behillil, Méline Bizard, Angela Brisebarre, Camille Capel, Vincent Enouf, Louise Lefrançois, Frédéric Lemoine, Christophe Malabat, Corinne Maufrais, Victoire Baillet, Etienne Simon-Lorière, Maud Vanpeene, Sylvie Van der Werf ,Judith Zerah |
| EPI_ISL_2029878, EPI_ISL_2029879 | Hospital | National Reference Center for Viruses of Respiratory Infections, Institut Pasteur, Paris | Marion Barbet, Sylvie Behillil, Méline Bizard, Angela Brisebarre, Camille Capel, Vincent Enouf, Louise Lefrançois, Frédéric Lemoine, Christophe Malabat, Corinne Maufrais, Etienne Simon-Lorière, Maud Vanpeene, Sylvie Van der Werf ,L. Courdavault |
| EPI_ISL_2029880 | Labo Analyses Med | National Reference Center for Viruses of Respiratory Infections, Institut Pasteur, Paris | Marion Barbet, Sylvie Behillil, Méline Bizard, Angela Brisebarre, Camille Capel, Vincent Enouf, Louise Lefrançois, Frédéric Lemoine, Christophe Malabat, Corinne Maufrais, Etienne Simon-Lorière, Maud Vanpeene, Sylvie Van der Werf ,Arditi Cocco |
| EPI_ISL_2029881 | Labo analyses med | National Reference Center for Viruses of Respiratory Infections, Institut Pasteur, Paris | Marion Barbet, Sylvie Behillil, Méline Bizard, Angela Brisebarre, Camille Capel, Vincent Enouf, Louise Lefrançois, Frédéric Lemoine, Christophe Malabat, Corinne Maufrais, Etienne Simon-Lorière, Maud Vanpeene, Sylvie Van der Werf ,Jean-FrançOis Comes |
| EPI_ISL_2029882, EPI_ISL_2029883 | Labo Analyses Med | National Reference Center for Viruses of Respiratory Infections, Institut Pasteur, Paris | Marion Barbet, Sylvie Behillil, Méline Bizard, Angela Brisebarre, Camille Capel, Vincent Enouf, Louise Lefrançois, Frédéric Lemoine, Christophe Malabat, Corinne Maufrais, Etienne Simon-Lorière, Maud Vanpeene, Sylvie Van der Werf ,Pierre-Yves Leonard |
| EPI_ISL_2029884 | Hospital | National Reference Center for Viruses of Respiratory Infections, Institut Pasteur, Paris | Marion Barbet, Sylvie Behillil, Méline Bizard, Angela Brisebarre, Camille Capel, Vincent Enouf, Louise Lefrançois, Frédéric Lemoine, Christophe Malabat, Corinne Maufrais, Amaury Vaysse, Etienne Simon-Lorière, Maud Vanpeene, Sylvie Van der Werf ,CéLine Bressollette |
| EPI_ISL_2029885 | Hospital | National Reference Center for Viruses of Respiratory Infections, Institut Pasteur, Paris | Marion Barbet, Sylvie Behillil, Méline Bizard, Angela Brisebarre, Camille Capel, Vincent Enouf, Louise Lefrançois, Frédéric Lemoine, Christophe Malabat, Corinne Maufrais, Etienne Simon-Lorière, Maud Vanpeene, Sylvie Van der Werf ,CéLine Bressollette |
| EPI_ISL_2029886, EPI_ISL_2029887 | Labo Analyses Med | National Reference Center for Viruses of Respiratory Infections, Institut Pasteur, Paris | Marion Barbet, Sylvie Behillil, Méline Bizard, Angela Brisebarre, Camille Capel, Vincent Enouf, Louise Lefrançois, Frédéric Lemoine, Christophe Malabat, Corinne Maufrais, Etienne Simon-Lorière, Maud Vanpeene, Sylvie Van der Werf ,Brieuc Gestin |
| EPI_ISL_2029889 | Labo Analyses Med | National Reference Center for Viruses of Respiratory Infections, Institut Pasteur, Paris | Marion Barbet, Sylvie Behillil, Méline Bizard, Angela Brisebarre, Camille Capel, Vincent Enouf, Louise Lefrançois, Frédéric Lemoine, Christophe Malabat, Corinne Maufrais, Etienne Simon-Lorière, Maud Vanpeene, Sylvie Van der Werf ,GréGoire Potiron |
| EPI_ISL_2029890, EPI_ISL_2029891 | Labo Analyses Med | National Reference Center for Viruses of Respiratory Infections, Institut Pasteur, Paris | Marion Barbet, Sylvie Behillil, Méline Bizard, Angela Brisebarre, Camille Capel, Vincent Enouf, Louise Lefrançois, Frédéric Lemoine, Christophe Malabat, Corinne Maufrais, Etienne Simon-Lorière, Maud Vanpeene, Sylvie Van der Werf ,Arditi Cocco |

| | | | |
|---|---|---|---|
| EPI_ISL_2029892 | Labo Analyses Med | National Reference Center for Viruses of Respiratory Infections, Institut Pasteur, Paris | Marion Barbet, Sylvie Behillil, Méline Bizard, Angela Brisebarre, Camille Capel, Vincent Enouf, Louise Lefrançois, Frédéric Lemoine, Christophe Malabat, Corinne Maufrais, Pierre Lechat, Etienne Simon-Lorière, Maud Vanpeene, Sylvie Van der Werf ,Franck Ennouchi |
| EPI_ISL_2029893, EPI_ISL_2029894 | Labo analyses med | National Reference Center for Viruses of Respiratory Infections, Institut Pasteur, Paris | Marion Barbet, Sylvie Behillil, Méline Bizard, Angela Brisebarre, Camille Capel, Vincent Enouf, Louise Lefrançois, Frédéric Lemoine, Christophe Malabat, Corinne Maufrais, Etienne Simon-Lorière, Maud Vanpeene, Sylvie Van der Werf ,Jean-FrançOis Comes |
| EPI_ISL_2029896 | Hospital | National Reference Center for Viruses of Respiratory Infections, Institut Pasteur, Paris | Marion Barbet, Sylvie Behillil, Méline Bizard, Angela Brisebarre, Camille Capel, Vincent Enouf, Louise Lefrançois, Frédéric Lemoine, Christophe Malabat, Corinne Maufrais, Etienne Simon-Lorière, Maud Vanpeene, Sylvie Van der Werf ,JéRome Guinard |
| EPI_ISL_2029897 | Hospital | National Reference Center for Viruses of Respiratory Infections, Institut Pasteur, Paris | Marion Barbet, Sylvie Behillil, Méline Bizard, Angela Brisebarre, Camille Capel, Vincent Enouf, Louise Lefrançois, Frédéric Lemoine, Christophe Malabat, Corinne Maufrais, Emmanuelle Permal, Etienne Simon-Lorière, Maud Vanpeene, Sylvie Van der Werf ,Isabelle Gros |
| EPI_ISL_2029898 | Labo Analyses Med | National Reference Center for Viruses of Respiratory Infections, Institut Pasteur, Paris | Marion Barbet, Sylvie Behillil, Méline Bizard, Angela Brisebarre, Camille Capel, Vincent Enouf, Louise Lefrançois, Frédéric Lemoine, Christophe Malabat, Corinne Maufrais, Etienne Simon-Lorière, Maud Vanpeene, Sylvie Van der Werf ,GréGoire Potiron |
| EPI_ISL_2029899 | Labo Analyses Med | National Reference Center for Viruses of Respiratory Infections, Institut Pasteur, Paris | Marion Barbet, Sylvie Behillil, Méline Bizard, Angela Brisebarre, Camille Capel, Vincent Enouf, Louise Lefrançois, Frédéric Lemoine, Christophe Malabat, Corinne Maufrais, Etienne Simon-Lorière, Maud Vanpeene, Sylvie Van der Werf ,Anne Holstein |
| EPI_ISL_2029900 | Labo Analyses Med | National Reference Center for Viruses of Respiratory Infections, Institut Pasteur, Paris | Marion Barbet, Sylvie Behillil, Méline Bizard, Angela Brisebarre, Camille Capel, Vincent Enouf, Louise Lefrançois, Frédéric Lemoine, Christophe Malabat, Corinne Maufrais, Emmanuelle Permal, Etienne Simon-Lorière, Maud Vanpeene, Sylvie Van der Werf ,Anne Holstein |
| EPI_ISL_2029901 | Labo Analyses Med | National Reference Center for Viruses of Respiratory Infections, Institut Pasteur, Paris | Marion Barbet, Sylvie Behillil, Méline Bizard, Angela Brisebarre, Camille Capel, Vincent Enouf, Louise Lefrançois, Frédéric Lemoine, Christophe Malabat, Corinne Maufrais, Etienne Simon-Lorière, Maud Vanpeene, Sylvie Van der Werf ,Anne Holstein |
| EPI_ISL_2029904 | Hospital | National Reference Center for Viruses of Respiratory Infections, Institut Pasteur, Paris | Marion Barbet, Sylvie Behillil, Méline Bizard, Angela Brisebarre, Camille Capel, Vincent Enouf, Louise Lefrançois, Frédéric Lemoine, Christophe Malabat, Corinne Maufrais, Etienne Simon-Lorière, Maud Vanpeene, Sylvie Van der Werf ,JéRome Guinard |
| EPI_ISL_2029905 | Labo Analyses Med | National Reference Center for Viruses of Respiratory Infections, Institut Pasteur, Paris | Marion Barbet, Sylvie Behillil, Méline Bizard, Angela Brisebarre, Camille Capel, Vincent Enouf, Louise Lefrançois, Frédéric Lemoine, Christophe Malabat, Corinne Maufrais, Etienne Simon-Lorière, Maud Vanpeene, Sylvie Van der Werf |
| EPI_ISL_2029906 | Labo Analyses Med | National Reference Center for Viruses of Respiratory Infections, Institut Pasteur, Paris | Marion Barbet, Sylvie Behillil, Méline Bizard, Angela Brisebarre, Camille Capel, Vincent Enouf, Louise Lefrançois, Frédéric Lemoine, Christophe Malabat, Corinne Maufrais, Etienne Simon-Lorière, Maud Vanpeene, Sylvie Van der Werf ,Brieuc Gestin |
| EPI_ISL_2029908, EPI_ISL_2029909, EPI_ISL_2029910 | Labo Analyses Med | National Reference Center for Viruses of Respiratory Infections, Institut Pasteur, Paris | Marion Barbet, Sylvie Behillil, Méline Bizard, Angela Brisebarre, Camille Capel, Vincent Enouf, Louise Lefrançois, Frédéric Lemoine, Christophe Malabat, Corinne Maufrais, Etienne Simon-Lorière, Maud Vanpeene, Sylvie Van der Werf ,Anne Holstein |
| EPI_ISL_2029911 | Labo analyses med | National Reference Center for Viruses of Respiratory Infections, Institut Pasteur, Paris | Marion Barbet, Sylvie Behillil, Méline Bizard, Angela Brisebarre, Camille Capel, Vincent Enouf, Louise Lefrançois, Frédéric Lemoine, Christophe Malabat, Corinne Maufrais, Amaury Vaysse, Etienne Simon-Lorière, Maud Vanpeene, Sylvie Van der Werf ,Jean-FrançOis Comes |
| EPI_ISL_2029912 | Labo Analyses Med | National Reference Center for Viruses of Respiratory Infections, Institut Pasteur, Paris | Marion Barbet, Sylvie Behillil, Méline Bizard, Angela Brisebarre, Camille Capel, Vincent Enouf, Louise Lefrançois, Frédéric Lemoine, Christophe Malabat, Corinne Maufrais, Etienne Simon-Lorière, Maud Vanpeene, Sylvie Van der Werf ,Pierre-Yves Leonard |
| EPI_ISL_2029915 | Labo Analyses Med | National Reference Center for Viruses of Respiratory Infections, Institut Pasteur, Paris | Marion Barbet, Sylvie Behillil, Méline Bizard, Angela Brisebarre, Camille Capel, Vincent Enouf, Louise Lefrançois, Frédéric Lemoine, Christophe Malabat, Corinne Maufrais, Etienne Simon-Lorière, Maud Vanpeene, Sylvie Van der Werf ,Brieuc Gestin |
| EPI_ISL_2029916 | Labo Analyses Med | National Reference Center for Viruses of Respiratory Infections, Institut Pasteur, Paris | Marion Barbet, Sylvie Behillil, Méline Bizard, Angela Brisebarre, Camille Capel, Vincent Enouf, Louise Lefrançois, Frédéric Lemoine, Christophe Malabat, Corinne Maufrais, Etienne Simon-Lorière, Maud Vanpeene, Sylvie Van der Werf ,Elodie Etienne |
| EPI_ISL_2029918 | Labo Analyses Med | National Reference Center for Viruses of Respiratory Infections, Institut Pasteur, Paris | Marion Barbet, Sylvie Behillil, Méline Bizard, Angela Brisebarre, Camille Capel, Vincent Enouf, Louise Lefrançois, Frédéric Lemoine, Christophe Malabat, Corinne Maufrais, Etienne Simon-Lorière, Maud Vanpeene, Sylvie Van der Werf ,Brieuc Gestin |
| EPI_ISL_2029919 | Labo Analyses Med | National Reference Center for Viruses of Respiratory Infections, Institut Pasteur, Paris | Marion Barbet, Sylvie Behillil, Méline Bizard, Angela Brisebarre, Camille Capel, Vincent Enouf, Louise Lefrançois, Frédéric Lemoine, Christophe Malabat, Corinne Maufrais, Etienne Simon-Lorière, Maud Vanpeene, Sylvie Van der Werf ,GréGoire Potiron |
| EPI_ISL_2029920 | Hospital | National Reference Center for Viruses of Respiratory Infections, Institut Pasteur, Paris | Marion Barbet, Sylvie Behillil, Méline Bizard, Angela Brisebarre, Camille Capel, Vincent Enouf, Louise Lefrançois, Frédéric Lemoine, Christophe Malabat, Corinne Maufrais, Etienne Simon-Lorière, Maud Vanpeene, Sylvie Van der Werf ,L. Courdavault |
| EPI_ISL_2029921 | Labo Analyses Med | National Reference Center for Viruses of Respiratory Infections, Institut Pasteur, Paris | Marion Barbet, Sylvie Behillil, Méline Bizard, Angela Brisebarre, Camille Capel, Vincent Enouf, Louise Lefrançois, Frédéric Lemoine, Christophe Malabat, Corinne Maufrais, Pierre Lechat, Etienne Simon-Lorière, Maud Vanpeene, Sylvie Van der Werf ,Arditi Cocco |
| EPI_ISL_2029923 | Labo Analyses Med | National Reference Center for Viruses of Respiratory Infections, Institut Pasteur, Paris | Marion Barbet, Sylvie Behillil, Méline Bizard, Angela Brisebarre, Camille Capel, Vincent Enouf, Louise Lefrançois, Frédéric Lemoine, Christophe Malabat, Corinne Maufrais, Pierre Lechat, Etienne Simon-Lorière, Maud Vanpeene, Sylvie Van der Werf ,Anne Holstein |
| EPI_ISL_2029924 | Hospital | National Reference Center for Viruses of Respiratory Infections, Institut Pasteur, Paris | Marion Barbet, Sylvie Behillil, Méline Bizard, Angela Brisebarre, Camille Capel, Vincent Enouf, Louise Lefrançois, Frédéric Lemoine, Christophe Malabat, Corinne Maufrais, Amaury Vaysse, Etienne Simon-Lorière, Maud Vanpeene, Sylvie Van der Werf ,CéLine Ramanantsoa |
| EPI_ISL_2029926 | Labo Analyses Med | National Reference Center for Viruses of Respiratory Infections, Institut Pasteur, Paris | Marion Barbet, Sylvie Behillil, Méline Bizard, Angela Brisebarre, Camille Capel, Vincent Enouf, Louise Lefrançois, Frédéric Lemoine, Christophe Malabat, Corinne Maufrais, Etienne Simon-Lorière, Maud Vanpeene, Sylvie Van der Werf ,Ronan Pichard |
| EPI_ISL_2029927 | Labo Analyses Med | National Reference Center for Viruses of Respiratory Infections, Institut Pasteur, Paris | Marion Barbet, Sylvie Behillil, Méline Bizard, Angela Brisebarre, Camille Capel, Vincent Enouf, Louise Lefrançois, Frédéric Lemoine, Christophe Malabat, Corinne Maufrais, Pierre Lechat, Etienne Simon-Lorière, Maud Vanpeene, Sylvie Van der Werf ,Ronan Pichard |
| EPI_ISL_2029928, EPI_ISL_2029929 | Hospital | National Reference Center for Viruses of Respiratory Infections, Institut Pasteur, Paris | Marion Barbet, Sylvie Behillil, Méline Bizard, Angela Brisebarre, Camille Capel, Vincent Enouf, Louise Lefrançois, Frédéric Lemoine, Christophe Malabat, Corinne Maufrais, Etienne Simon-Lorière, Maud Vanpeene, Sylvie Van der Werf ,Pascale Martres |
| EPI_ISL_2029930, EPI_ISL_2029931, EPI_ISL_2029932 | Hospital | National Reference Center for Viruses of Respiratory Infections, Institut Pasteur, Paris | Marion Barbet, Sylvie Behillil, Méline Bizard, Angela Brisebarre, Camille Capel, Vincent Enouf, Louise Lefrançois, Frédéric Lemoine, Christophe Malabat, Corinne Maufrais, Etienne Simon-Lorière, Maud Vanpeene, Sylvie Van der Werf ,Nabil Gastli |
| EPI_ISL_2029933 | Hospital | National Reference Center for Viruses of Respiratory Infections, Institut Pasteur, Paris | Marion Barbet, Sylvie Behillil, Méline Bizard, Angela Brisebarre, Camille Capel, Vincent Enouf, Louise Lefrançois, Frédéric Lemoine, Christophe Malabat, Corinne Maufrais, Etienne Simon-Lorière, Maud Vanpeene, Sylvie Van der Werf ,Sylvie Ledru |
| EPI_ISL_2029934 | Hospital | National Reference Center for Viruses of Respiratory Infections, Institut Pasteur, Paris | Marion Barbet, Sylvie Behillil, Méline Bizard, Angela Brisebarre, Camille Capel, Vincent Enouf, Louise Lefrançois, Frédéric Lemoine, Christophe Malabat, Corinne Maufrais, Etienne Simon-Lorière, Maud Vanpeene, Sylvie Van der Werf ,CéLine Bressollette |
| EPI_ISL_2029935, EPI_ISL_2029937 | Labo Analyses Med | National Reference Center for Viruses of Respiratory Infections, Institut Pasteur, Paris | Marion Barbet, Sylvie Behillil, Méline Bizard, Angela Brisebarre, Camille Capel, Vincent Enouf, Louise Lefrançois, Frédéric Lemoine, Christophe Malabat, Corinne Maufrais, Etienne Simon-Lorière, Maud Vanpeene, Sylvie Van der Werf ,Brieuc Gestin |
| EPI_ISL_2029938 | Labo Analyses Med | National Reference Center for Viruses of Respiratory Infections, Institut Pasteur, Paris | Marion Barbet, Sylvie Behillil, Méline Bizard, Angela Brisebarre, Camille Capel, Vincent Enouf, Louise Lefrançois, Frédéric Lemoine, Christophe Malabat, Corinne Maufrais, Etienne Simon-Lorière, Maud Vanpeene, Sylvie Van der Werf ,Anne Holstein |
| EPI_ISL_2029939 | Hospital | National Reference Center for Viruses of Respiratory Infections, Institut Pasteur, Paris | Marion Barbet, Sylvie Behillil, Méline Bizard, Angela Brisebarre, Camille Capel, Vincent Enouf, Louise Lefrançois, Frédéric Lemoine, Christophe Malabat, Corinne Maufrais, Etienne Simon-Lorière, Maud Vanpeene, Sylvie Van der Werf ,Odile Morvan |
| EPI_ISL_2029940 | Hospital | National Reference Center for Viruses of Respiratory Infections, Institut Pasteur, Paris | Marion Barbet, Sylvie Behillil, Méline Bizard, Angela Brisebarre, Camille Capel, Vincent Enouf, Louise Lefrançois, Frédéric Lemoine, Christophe Malabat, Corinne Maufrais, Etienne Simon-Lorière, Maud Vanpeene, Sylvie Van der Werf ,Pascale Martres |
| EPI_ISL_2029941 | Hospital | National Reference Center for Viruses of Respiratory Infections, Institut Pasteur, Paris | Marion Barbet, Sylvie Behillil, Méline Bizard, Angela Brisebarre, Camille Capel, Vincent Enouf, Louise Lefrançois, Frédéric Lemoine, Christophe Malabat, Corinne Maufrais, Etienne Simon-Lorière, Maud Vanpeene, Sylvie Van der Werf ,Nabil Gastli |
| EPI_ISL_2029942 | Labo Analyses Med | National Reference Center for Viruses of Respiratory Infections, Institut Pasteur, Paris | Marion Barbet, Sylvie Behillil, Méline Bizard, Angela Brisebarre, Camille Capel, Vincent Enouf, Louise Lefrançois, Frédéric Lemoine, Christophe Malabat, Corinne Maufrais, Etienne Simon-Lorière, Maud Vanpeene, Sylvie Van der Werf ,Pierre-Yves Leonard |
| EPI_ISL_2029943 | Outre mer | National Reference Center for Viruses of Respiratory Infections, Institut Pasteur, Paris | Marion Barbet, Sylvie Behillil, Méline Bizard, Angela Brisebarre, Camille Capel, Vincent Enouf, Louise Lefrançois, Frédéric Lemoine, Christophe Malabat, Corinne Maufrais, Etienne Simon-Lorière, Maud Vanpeene, Sylvie Van der Werf ,Patrice Combe |

| | | | |
|---|---|---|---|
| EPI_ISL_2029944 | Labo Analyses Med | National Reference Center for Viruses of Respiratory Infections, Institut Pasteur, Paris | Marion Barbet, Sylvie Behillil, Méline Bizard, Angela Brisebarre, Camille Capel, Vincent Enouf, Louise Lefrançois, Frédéric Lemoine, Christophe Malabat, Corinne Maufrais, Pierre Lechat, Etienne Simon-Lorière, Maud Vanpeene, Sylvie Van der Werf ,GréGoire Potiron |
| EPI_ISL_2029945 | Labo Analyses Med | National Reference Center for Viruses of Respiratory Infections, Institut Pasteur, Paris | Marion Barbet, Sylvie Behillil, Méline Bizard, Angela Brisebarre, Camille Capel, Vincent Enouf, Louise Lefrançois, Frédéric Lemoine, Christophe Malabat, Corinne Maufrais, Etienne Simon-Lorière, Maud Vanpeene, Sylvie Van der Werf ,Anne Holstein |
| EPI_ISL_2029946 | Labo Analyses Med | National Reference Center for Viruses of Respiratory Infections, Institut Pasteur, Paris | Marion Barbet, Sylvie Behillil, Méline Bizard, Angela Brisebarre, Camille Capel, Vincent Enouf, Louise Lefrançois, Frédéric Lemoine, Christophe Malabat, Corinne Maufrais, Etienne Simon-Lorière, Maud Vanpeene, Sylvie Van der Werf ,Pierre-Yves Leonard |
| EPI_ISL_2029947 | Outre mer | National Reference Center for Viruses of Respiratory Infections, Institut Pasteur, Paris | Marion Barbet, Sylvie Behillil, Méline Bizard, Angela Brisebarre, Camille Capel, Vincent Enouf, Louise Lefrançois, Frédéric Lemoine, Christophe Malabat, Corinne Maufrais, Amaury Vaysse, Etienne Simon-Lorière, Maud Vanpeene, Sylvie Van der Werf ,Patrice Combe |
| EPI_ISL_2029948 | Hospital | National Reference Center for Viruses of Respiratory Infections, Institut Pasteur, Paris | Marion Barbet, Sylvie Behillil, Méline Bizard, Angela Brisebarre, Camille Capel, Vincent Enouf, Louise Lefrançois, Frédéric Lemoine, Christophe Malabat, Corinne Maufrais, Etienne Simon-Lorière, Maud Vanpeene, Sylvie Van der Werf ,Isabelle Gros |
| EPI_ISL_2029949 | Labo Analyses Med | National Reference Center for Viruses of Respiratory Infections, Institut Pasteur, Paris | Marion Barbet, Sylvie Behillil, Méline Bizard, Angela Brisebarre, Camille Capel, Vincent Enouf, Louise Lefrançois, Frédéric Lemoine, Christophe Malabat, Corinne Maufrais, Etienne Simon-Lorière, Maud Vanpeene, Sylvie Van der Werf |
| EPI_ISL_2029950, EPI_ISL_2029951 | Labo Analyses Med | National Reference Center for Viruses of Respiratory Infections, Institut Pasteur, Paris | Marion Barbet, Sylvie Behillil, Méline Bizard, Angela Brisebarre, Camille Capel, Vincent Enouf, Louise Lefrançois, Frédéric Lemoine, Christophe Malabat, Corinne Maufrais, Etienne Simon-Lorière, Maud Vanpeene, Sylvie Van der Werf ,Brieuc Gestin |
| EPI_ISL_2029952 | Hospital | National Reference Center for Viruses of Respiratory Infections, Institut Pasteur, Paris | Marion Barbet, Sylvie Behillil, Méline Bizard, Angela Brisebarre, Camille Capel, Vincent Enouf, Louise Lefrançois, Frédéric Lemoine, Christophe Malabat, Corinne Maufrais, Etienne Simon-Lorière, Maud Vanpeene, Sylvie Van der Werf ,LéA Pilorge |
| EPI_ISL_2029953 | Labo Analyses Med | National Reference Center for Viruses of Respiratory Infections, Institut Pasteur, Paris | Marion Barbet, Sylvie Behillil, Méline Bizard, Angela Brisebarre, Camille Capel, Vincent Enouf, Louise Lefrançois, Frédéric Lemoine, Christophe Malabat, Corinne Maufrais, Etienne Simon-Lorière, Maud Vanpeene, Sylvie Van der Werf ,Ronan Pichard |
| EPI_ISL_2029954 | Labo Analyses Med | National Reference Center for Viruses of Respiratory Infections, Institut Pasteur, Paris | Marion Barbet, Sylvie Behillil, Méline Bizard, Angela Brisebarre, Camille Capel, Vincent Enouf, Louise Lefrançois, Frédéric Lemoine, Christophe Malabat, Corinne Maufrais, Pierre Lechat, Etienne Simon-Lorière, Maud Vanpeene, Sylvie Van der Werf ,GréGoire Potiron |
| EPI_ISL_2029955 | Labo Analyses Med | National Reference Center for Viruses of Respiratory Infections, Institut Pasteur, Paris | Marion Barbet, Sylvie Behillil, Méline Bizard, Angela Brisebarre, Camille Capel, Vincent Enouf, Louise Lefrançois, Frédéric Lemoine, Christophe Malabat, Corinne Maufrais, Etienne Simon-Lorière, Maud Vanpeene, Sylvie Van der Werf ,GréGoire Potiron |
| EPI_ISL_2029956 | Hospital | National Reference Center for Viruses of Respiratory Infections, Institut Pasteur, Paris | Marion Barbet, Sylvie Behillil, Méline Bizard, Angela Brisebarre, Camille Capel, Vincent Enouf, Louise Lefrançois, Frédéric Lemoine, Christophe Malabat, Corinne Maufrais, Etienne Simon-Lorière, Maud Vanpeene, Sylvie Van der Werf ,Anne Cady |
| EPI_ISL_2029957, EPI_ISL_2029958, EPI_ISL_2029959 | Labo Analyses Med | National Reference Center for Viruses of Respiratory Infections, Institut Pasteur, Paris | Marion Barbet, Sylvie Behillil, Méline Bizard, Angela Brisebarre, Camille Capel, Vincent Enouf, Louise Lefrançois, Frédéric Lemoine, Christophe Malabat, Corinne Maufrais, Etienne Simon-Lorière, Maud Vanpeene, Sylvie Van der Werf ,Arditi Cocco |
| EPI_ISL_2029960 | Labo Analyses Med | National Reference Center for Viruses of Respiratory Infections, Institut Pasteur, Paris | Marion Barbet, Sylvie Behillil, Méline Bizard, Angela Brisebarre, Camille Capel, Vincent Enouf, Louise Lefrançois, Frédéric Lemoine, Christophe Malabat, Corinne Maufrais, Etienne Simon-Lorière, Maud Vanpeene, Sylvie Van der Werf ,Anne Holstein |
| EPI_ISL_2029961 | Labo Analyses Med | National Reference Center for Viruses of Respiratory Infections, Institut Pasteur, Paris | Marion Barbet, Sylvie Behillil, Méline Bizard, Angela Brisebarre, Camille Capel, Vincent Enouf, Louise Lefrançois, Frédéric Lemoine, Christophe Malabat, Corinne Maufrais, Etienne Simon-Lorière, Maud Vanpeene, Sylvie Van der Werf ,Delphine MammÈS |
| EPI_ISL_2029962 | Hospital | National Reference Center for Viruses of Respiratory Infections, Institut Pasteur, Paris | Marion Barbet, Sylvie Behillil, Méline Bizard, Angela Brisebarre, Camille Capel, Vincent Enouf, Louise Lefrançois, Frédéric Lemoine, Christophe Malabat, Corinne Maufrais, Etienne Simon-Lorière, Maud Vanpeene, Sylvie Van der Werf ,Pascale Martres |
| EPI_ISL_2029963, EPI_ISL_2029964 | Labo Analyses Med | National Reference Center for Viruses of Respiratory Infections, Institut Pasteur, Paris | Marion Barbet, Sylvie Behillil, Méline Bizard, Angela Brisebarre, Camille Capel, Vincent Enouf, Louise Lefrançois, Frédéric Lemoine, Christophe Malabat, Corinne Maufrais, Etienne Simon-Lorière, Maud Vanpeene, Sylvie Van der Werf ,Pierre-Yves Leonard |
| EPI_ISL_2029965 | Hospital | National Reference Center for Viruses of Respiratory Infections, Institut Pasteur, Paris | Marion Barbet, Sylvie Behillil, Méline Bizard, Angela Brisebarre, Camille Capel, Vincent Enouf, Louise Lefrançois, Frédéric Lemoine, Christophe Malabat, Corinne Maufrais, Etienne Simon-Lorière, Maud Vanpeene, Sylvie Van der Werf ,CéLine Bressollette |
| EPI_ISL_2029966 | Labo Analyses Med | National Reference Center for Viruses of Respiratory Infections, Institut Pasteur, Paris | Marion Barbet, Sylvie Behillil, Méline Bizard, Angela Brisebarre, Camille Capel, Vincent Enouf, Louise Lefrançois, Frédéric Lemoine, Christophe Malabat, Corinne Maufrais, Etienne Simon-Lorière, Maud Vanpeene, Sylvie Van der Werf |
| EPI_ISL_2029967, EPI_ISL_2029968, EPI_ISL_2029969 | Labo Analyses Med | National Reference Center for Viruses of Respiratory Infections, Institut Pasteur, Paris | Marion Barbet, Sylvie Behillil, Méline Bizard, Angela Brisebarre, Camille Capel, Vincent Enouf, Louise Lefrançois, Frédéric Lemoine, Christophe Malabat, Corinne Maufrais, Etienne Simon-Lorière, Maud Vanpeene, Sylvie Van der Werf ,Brieuc Gestin |
| EPI_ISL_2029971 | Hospital | National Reference Center for Viruses of Respiratory Infections, Institut Pasteur, Paris | Marion Barbet, Sylvie Behillil, Méline Bizard, Angela Brisebarre, Camille Capel, Vincent Enouf, Louise Lefrançois, Frédéric Lemoine, Christophe Malabat, Corinne Maufrais, Etienne Simon-Lorière, Maud Vanpeene, Sylvie Van der Werf ,L. Courdavault |
| EPI_ISL_2029972 | Hospital | National Reference Center for Viruses of Respiratory Infections, Institut Pasteur, Paris | Marion Barbet, Sylvie Behillil, Méline Bizard, Angela Brisebarre, Camille Capel, Vincent Enouf, Louise Lefrançois, Frédéric Lemoine, Christophe Malabat, Corinne Maufrais, Etienne Simon-Lorière, Maud Vanpeene, Sylvie Van der Werf ,Pascale Martres |
| EPI_ISL_2029973 | Labo Analyses Med | National Reference Center for Viruses of Respiratory Infections, Institut Pasteur, Paris | Marion Barbet, Sylvie Behillil, Méline Bizard, Angela Brisebarre, Camille Capel, Vincent Enouf, Louise Lefrançois, Frédéric Lemoine, Christophe Malabat, Corinne Maufrais, Etienne Simon-Lorière, Maud Vanpeene, Sylvie Van der Werf ,Anne Holstein |
| EPI_ISL_2029975, EPI_ISL_2029976 | Labo Analyses Med | National Reference Center for Viruses of Respiratory Infections, Institut Pasteur, Paris | Marion Barbet, Sylvie Behillil, Méline Bizard, Angela Brisebarre, Camille Capel, Vincent Enouf, Louise Lefrançois, Frédéric Lemoine, Christophe Malabat, Corinne Maufrais, Etienne Simon-Lorière, Maud Vanpeene, Sylvie Van der Werf ,Pierre-Yves Leonard |
| EPI_ISL_2029978, EPI_ISL_2029979 | Labo Analyses Med | National Reference Center for Viruses of Respiratory Infections, Institut Pasteur, Paris | Marion Barbet, Sylvie Behillil, Méline Bizard, Angela Brisebarre, Camille Capel, Vincent Enouf, Louise Lefrançois, Frédéric Lemoine, Christophe Malabat, Corinne Maufrais, Etienne Simon-Lorière, Maud Vanpeene, Sylvie Van der Werf ,Anne Holstein |
| EPI_ISL_2029980 | Hospital | National Reference Center for Viruses of Respiratory Infections, Institut Pasteur, Paris | Marion Barbet, Sylvie Behillil, Méline Bizard, Angela Brisebarre, Camille Capel, Vincent Enouf, Louise Lefrançois, Frédéric Lemoine, Christophe Malabat, Corinne Maufrais, Etienne Simon-Lorière, Maud Vanpeene, Sylvie Van der Werf ,Thibault Guinoiseau |
| EPI_ISL_2029981 | Labo Analyses Med | National Reference Center for Viruses of Respiratory Infections, Institut Pasteur, Paris | Marion Barbet, Sylvie Behillil, Méline Bizard, Angela Brisebarre, Camille Capel, Vincent Enouf, Louise Lefrançois, Frédéric Lemoine, Christophe Malabat, Corinne Maufrais, Etienne Simon-Lorière, Maud Vanpeene, Sylvie Van der Werf ,Valentin Robbe |
| EPI_ISL_2029982 | Labo Analyses Med | National Reference Center for Viruses of Respiratory Infections, Institut Pasteur, Paris | Marion Barbet, Sylvie Behillil, Méline Bizard, Angela Brisebarre, Camille Capel, Vincent Enouf, Louise Lefrançois, Frédéric Lemoine, Christophe Malabat, Corinne Maufrais, Etienne Simon-Lorière, Maud Vanpeene, Sylvie Van der Werf ,Bruno Berdin |
| EPI_ISL_2029984 | Hospital | National Reference Center for Viruses of Respiratory Infections, Institut Pasteur, Paris | Marion Barbet, Sylvie Behillil, Méline Bizard, Angela Brisebarre, Camille Capel, Vincent Enouf, Louise Lefrançois, Frédéric Lemoine, Christophe Malabat, Corinne Maufrais, Etienne Simon-Lorière, Maud Vanpeene, Sylvie Van der Werf ,Anne Cady |
| EPI_ISL_2029985 | Hospital | National Reference Center for Viruses of Respiratory Infections, Institut Pasteur, Paris | Marion Barbet, Sylvie Behillil, Méline Bizard, Angela Brisebarre, Camille Capel, Vincent Enouf, Louise Lefrançois, Frédéric Lemoine, Christophe Malabat, Corinne Maufrais, Etienne Simon-Lorière, Maud Vanpeene, Sylvie Van der Werf ,Nabil Gastli |
| EPI_ISL_2029986 | Hospital | National Reference Center for Viruses of Respiratory Infections, Institut Pasteur, Paris | Marion Barbet, Sylvie Behillil, Méline Bizard, Angela Brisebarre, Camille Capel, Vincent Enouf, Louise Lefrançois, Frédéric Lemoine, Christophe Malabat, Corinne Maufrais, Etienne Simon-Lorière, Maud Vanpeene, Sylvie Van der Werf ,Anne Vachee |
| EPI_ISL_2029987, EPI_ISL_2029988 | Labo Analyses Med | National Reference Center for Viruses of Respiratory Infections, Institut Pasteur, Paris | Marion Barbet, Sylvie Behillil, Méline Bizard, Angela Brisebarre, Camille Capel, Vincent Enouf, Louise Lefrançois, Frédéric Lemoine, Christophe Malabat, Corinne Maufrais, Etienne Simon-Lorière, Maud Vanpeene, Sylvie Van der Werf ,Pierre-Yves Leonard |
| EPI_ISL_2029989 | Labo Analyses Med | National Reference Center for Viruses of Respiratory Infections, Institut Pasteur, Paris | Marion Barbet, Sylvie Behillil, Méline Bizard, Angela Brisebarre, Camille Capel, Vincent Enouf, Louise Lefrançois, Frédéric Lemoine, Christophe Malabat, Corinne Maufrais, Amaury Vaysse, Etienne Simon-Lorière, Maud Vanpeene, Sylvie Van der Werf ,Pierre-Yves Leonard |
| EPI_ISL_2029990 | Hospital | National Reference Center for Viruses of Respiratory Infections, Institut Pasteur, Paris | Marion Barbet, Sylvie Behillil, Méline Bizard, Angela Brisebarre, Camille Capel, Vincent Enouf, Louise Lefrançois, Frédéric Lemoine, Christophe Malabat, Corinne Maufrais, Amaury Vaysse, Etienne Simon-Lorière, Maud Vanpeene, Sylvie Van der Werf ,CéLine Ramanantsoa |
| EPI_ISL_2029992 | Armies | National Reference Center for Viruses of Respiratory Infections, Institut Pasteur, Paris | Marion Barbet, Sylvie Behillil, Méline Bizard, Angela Brisebarre, Camille Capel, Vincent Enouf, Louise Lefrançois, Frédéric Lemoine, Christophe Malabat, Corinne Maufrais, Etienne Simon-Lorière, Maud Vanpeene, Sylvie Van der Werf ,Marine Desroches |

| | | | |
|---|---|---|---|
| EPI_ISL_2029993, EPI_ISL_2029994 | Labo Analyses Med | National Reference Center for Viruses of Respiratory Infections, Institut Pasteur, Paris | Marion Barbet, Sylvie Behillil, Méline Bizard, Angela Brisebarre, Camille Capel, Vincent Enouf, Louise Lefrançois, Frédéric Lemoine, Christophe Malabat, Corinne Maufrais, Etienne Simon-Lorière, Maud Vanpeene, Sylvie Van der Werf ,Ronan Pichard |
| EPI_ISL_2029995 | Labo Analyses Med | National Reference Center for Viruses of Respiratory Infections, Institut Pasteur, Paris | Marion Barbet, Sylvie Behillil, Méline Bizard, Angela Brisebarre, Camille Capel, Vincent Enouf, Louise Lefrançois, Frédéric Lemoine, Christophe Malabat, Corinne Maufrais, Etienne Simon-Lorière, Maud Vanpeene, Sylvie Van der Werf ,Arditi Cocco |
| EPI_ISL_2029996 | Labo analyses med | National Reference Center for Viruses of Respiratory Infections, Institut Pasteur, Paris | Marion Barbet, Sylvie Behillil, Méline Bizard, Angela Brisebarre, Camille Capel, Vincent Enouf, Louise Lefrançois, Frédéric Lemoine, Christophe Malabat, Corinne Maufrais, Etienne Simon-Lorière, Maud Vanpeene, Sylvie Van der Werf ,Jean-FrançOis Comes |
| EPI_ISL_2029998 | Hospital | National Reference Center for Viruses of Respiratory Infections, Institut Pasteur, Paris | Marion Barbet, Sylvie Behillil, Méline Bizard, Angela Brisebarre, Camille Capel, Vincent Enouf, Louise Lefrançois, Frédéric Lemoine, Christophe Malabat, Corinne Maufrais, Etienne Simon-Lorière, Maud Vanpeene, Sylvie Van der Werf ,Mignard |
| EPI_ISL_2029999 | Labo Analyses Med | National Reference Center for Viruses of Respiratory Infections, Institut Pasteur, Paris | Marion Barbet, Sylvie Behillil, Méline Bizard, Angela Brisebarre, Camille Capel, Vincent Enouf, Louise Lefrançois, Frédéric Lemoine, Christophe Malabat, Corinne Maufrais, Amaury Vaysse, Etienne Simon-Lorière, Maud Vanpeene, Sylvie Van der Werf ,Brieuc Gestin |
| EPI_ISL_2030000, EPI_ISL_2030001 | Labo Analyses Med | National Reference Center for Viruses of Respiratory Infections, Institut Pasteur, Paris | Marion Barbet, Sylvie Behillil, Méline Bizard, Angela Brisebarre, Camille Capel, Vincent Enouf, Louise Lefrançois, Frédéric Lemoine, Christophe Malabat, Corinne Maufrais, Etienne Simon-Lorière, Maud Vanpeene, Sylvie Van der Werf ,Brieuc Gestin |
| EPI_ISL_2030002 | Hospital | National Reference Center for Viruses of Respiratory Infections, Institut Pasteur, Paris | Marion Barbet, Sylvie Behillil, Méline Bizard, Angela Brisebarre, Camille Capel, Vincent Enouf, Louise Lefrançois, Frédéric Lemoine, Christophe Malabat, Corinne Maufrais, Etienne Simon-Lorière, Maud Vanpeene, Sylvie Van der Werf ,Anne Cady |
| EPI_ISL_2030003 | Hospital | National Reference Center for Viruses of Respiratory Infections, Institut Pasteur, Paris | Marion Barbet, Sylvie Behillil, Méline Bizard, Angela Brisebarre, Camille Capel, Vincent Enouf, Louise Lefrançois, Frédéric Lemoine, Christophe Malabat, Corinne Maufrais, Etienne Simon-Lorière, Maud Vanpeene, Sylvie Van der Werf ,Nabil Gastli |
| EPI_ISL_2030005 | Labo Analyses Med | National Reference Center for Viruses of Respiratory Infections, Institut Pasteur, Paris | Marion Barbet, Sylvie Behillil, Méline Bizard, Angela Brisebarre, Camille Capel, Vincent Enouf, Louise Lefrançois, Frédéric Lemoine, Christophe Malabat, Corinne Maufrais, Etienne Simon-Lorière, Maud Vanpeene, Sylvie Van der Werf ,Ronan Pichard |
| EPI_ISL_2030006 | Hospital | National Reference Center for Viruses of Respiratory Infections, Institut Pasteur, Paris | Marion Barbet, Sylvie Behillil, Méline Bizard, Angela Brisebarre, Camille Capel, Vincent Enouf, Louise Lefrançois, Frédéric Lemoine, Christophe Malabat, Corinne Maufrais, Amaury Vaysse, Etienne Simon-Lorière, Maud Vanpeene, Sylvie Van der Werf ,CéLine Bressollette |
| EPI_ISL_2030007 | Hospital | National Reference Center for Viruses of Respiratory Infections, Institut Pasteur, Paris | Marion Barbet, Sylvie Behillil, Méline Bizard, Angela Brisebarre, Camille Capel, Vincent Enouf, Louise Lefrançois, Frédéric Lemoine, Christophe Malabat, Corinne Maufrais, Etienne Simon-Lorière, Maud Vanpeene, Sylvie Van der Werf ,EugèNe Castelain |
| EPI_ISL_2030008 | Labo Analyses Med | National Reference Center for Viruses of Respiratory Infections, Institut Pasteur, Paris | Marion Barbet, Sylvie Behillil, Méline Bizard, Angela Brisebarre, Camille Capel, Vincent Enouf, Louise Lefrançois, Frédéric Lemoine, Christophe Malabat, Corinne Maufrais, Etienne Simon-Lorière, Maud Vanpeene, Sylvie Van der Werf ,GréGoire Potiron |
| EPI_ISL_2030009 | Hospital | National Reference Center for Viruses of Respiratory Infections, Institut Pasteur, Paris | Marion Barbet, Sylvie Behillil, Méline Bizard, Angela Brisebarre, Camille Capel, Vincent Enouf, Louise Lefrançois, Frédéric Lemoine, Christophe Malabat, Corinne Maufrais, Etienne Simon-Lorière, Maud Vanpeene, Sylvie Van der Werf ,Marianne Burgard |
| EPI_ISL_2030010 | Hospital | National Reference Center for Viruses of Respiratory Infections, Institut Pasteur, Paris | Marion Barbet, Sylvie Behillil, Méline Bizard, Angela Brisebarre, Camille Capel, Vincent Enouf, Louise Lefrançois, Frédéric Lemoine, Christophe Malabat, Corinne Maufrais, Pierre Lechat, Etienne Simon-Lorière, Maud Vanpeene, Sylvie Van der Werf ,Christine Febreau |
| EPI_ISL_2030011 | Labo Analyses Med | National Reference Center for Viruses of Respiratory Infections, Institut Pasteur, Paris | Marion Barbet, Sylvie Behillil, Méline Bizard, Angela Brisebarre, Camille Capel, Vincent Enouf, Louise Lefrançois, Frédéric Lemoine, Christophe Malabat, Corinne Maufrais, Etienne Simon-Lorière, Maud Vanpeene, Sylvie Van der Werf ,Anne Holstein |
| EPI_ISL_2030012 | Hospital | National Reference Center for Viruses of Respiratory Infections, Institut Pasteur, Paris | Marion Barbet, Sylvie Behillil, Méline Bizard, Angela Brisebarre, Camille Capel, Vincent Enouf, Louise Lefrançois, Frédéric Lemoine, Christophe Malabat, Corinne Maufrais, Etienne Simon-Lorière, Maud Vanpeene, Sylvie Van der Werf ,Odile Morvan |
| EPI_ISL_2030013 | Labo Analyses Med | National Reference Center for Viruses of Respiratory Infections, Institut Pasteur, Paris | Marion Barbet, Sylvie Behillil, Méline Bizard, Angela Brisebarre, Camille Capel, Vincent Enouf, Louise Lefrançois, Frédéric Lemoine, Christophe Malabat, Corinne Maufrais, Etienne Simon-Lorière, Maud Vanpeene, Sylvie Van der Werf ,Delphine MammÈS |
| EPI_ISL_2030014 | Hospital | National Reference Center for Viruses of Respiratory Infections, Institut Pasteur, Paris | Marion Barbet, Sylvie Behillil, Méline Bizard, Angela Brisebarre, Camille Capel, Vincent Enouf, Louise Lefrançois, Frédéric Lemoine, Christophe Malabat, Corinne Maufrais, Etienne Simon-Lorière, Maud Vanpeene, Sylvie Van der Werf ,Pascale Martres |
| EPI_ISL_2030015 | Hospital | National Reference Center for Viruses of Respiratory Infections, Institut Pasteur, Paris | Marion Barbet, Sylvie Behillil, Méline Bizard, Angela Brisebarre, Camille Capel, Vincent Enouf, Louise Lefrançois, Frédéric Lemoine, Christophe Malabat, Corinne Maufrais, Amaury Vaysse, Etienne Simon-Lorière, Maud Vanpeene, Sylvie Van der Werf ,CéLine Bressollette |
| EPI_ISL_2030017 | Labo Analyses Med | National Reference Center for Viruses of Respiratory Infections, Institut Pasteur, Paris | Marion Barbet, Sylvie Behillil, Méline Bizard, Angela Brisebarre, Camille Capel, Vincent Enouf, Louise Lefrançois, Frédéric Lemoine, Christophe Malabat, Corinne Maufrais, Etienne Simon-Lorière, Maud Vanpeene, Sylvie Van der Werf ,Bruno Berdin |
| EPI_ISL_2030018 | Labo Analyses Med | National Reference Center for Viruses of Respiratory Infections, Institut Pasteur, Paris | Marion Barbet, Sylvie Behillil, Méline Bizard, Angela Brisebarre, Camille Capel, Vincent Enouf, Louise Lefrançois, Frédéric Lemoine, Christophe Malabat, Corinne Maufrais, Pierre Lechat, Etienne Simon-Lorière, Maud Vanpeene, Sylvie Van der Werf ,Arditi Cocco |
| EPI_ISL_2030019 | Labo Analyses Med | National Reference Center for Viruses of Respiratory Infections, Institut Pasteur, Paris | Marion Barbet, Sylvie Behillil, Méline Bizard, Angela Brisebarre, Camille Capel, Vincent Enouf, Louise Lefrançois, Frédéric Lemoine, Christophe Malabat, Corinne Maufrais, Etienne Simon-Lorière, Maud Vanpeene, Sylvie Van der Werf ,Anne Holstein |
| EPI_ISL_2030020 | Hospital | National Reference Center for Viruses of Respiratory Infections, Institut Pasteur, Paris | Marion Barbet, Sylvie Behillil, Méline Bizard, Angela Brisebarre, Camille Capel, Vincent Enouf, Louise Lefrançois, Frédéric Lemoine, Christophe Malabat, Corinne Maufrais, Etienne Simon-Lorière, Maud Vanpeene, Sylvie Van der Werf ,Nabil Gastli |
| EPI_ISL_2030021 | Labo analyses med | National Reference Center for Viruses of Respiratory Infections, Institut Pasteur, Paris | Marion Barbet, Sylvie Behillil, Méline Bizard, Angela Brisebarre, Camille Capel, Vincent Enouf, Louise Lefrançois, Frédéric Lemoine, Christophe Malabat, Corinne Maufrais, Etienne Simon-Lorière, Maud Vanpeene, Sylvie Van der Werf ,Jean-FrançOis Comes |
| EPI_ISL_2030022 | Labo analyses med | National Reference Center for Viruses of Respiratory Infections, Institut Pasteur, Paris | Marion Barbet, Sylvie Behillil, Méline Bizard, Angela Brisebarre, Camille Capel, Vincent Enouf, Louise Lefrançois, Frédéric Lemoine, Christophe Malabat, Corinne Maufrais, Amaury Vaysse, Etienne Simon-Lorière, Maud Vanpeene, Sylvie Van der Werf ,Jean-FrançOis Comes |
| EPI_ISL_2030023 | Hospital | National Reference Center for Viruses of Respiratory Infections, Institut Pasteur, Paris | Marion Barbet, Sylvie Behillil, Méline Bizard, Angela Brisebarre, Camille Capel, Vincent Enouf, Louise Lefrançois, Frédéric Lemoine, Christophe Malabat, Corinne Maufrais, Etienne Simon-Lorière, Maud Vanpeene, Sylvie Van der Werf ,ValéRie Serazin |
| EPI_ISL_2030024 | Hospital | National Reference Center for Viruses of Respiratory Infections, Institut Pasteur, Paris | Marion Barbet, Sylvie Behillil, Méline Bizard, Angela Brisebarre, Camille Capel, Vincent Enouf, Louise Lefrançois, Frédéric Lemoine, Christophe Malabat, Corinne Maufrais, Emmanuelle Permal, Etienne Simon-Lorière, Maud Vanpeene, Sylvie Van der Werf ,Isabelle Gros |
| EPI_ISL_2030025, EPI_ISL_2030026 | Labo Analyses Med | National Reference Center for Viruses of Respiratory Infections, Institut Pasteur, Paris | Marion Barbet, Sylvie Behillil, Méline Bizard, Angela Brisebarre, Camille Capel, Vincent Enouf, Louise Lefrançois, Frédéric Lemoine, Christophe Malabat, Corinne Maufrais, Etienne Simon-Lorière, Maud Vanpeene, Sylvie Van der Werf ,GréGoire Potiron |
| EPI_ISL_2030028 | Hospital | National Reference Center for Viruses of Respiratory Infections, Institut Pasteur, Paris | Marion Barbet, Sylvie Behillil, Méline Bizard, Angela Brisebarre, Camille Capel, Vincent Enouf, Louise Lefrançois, Frédéric Lemoine, Christophe Malabat, Corinne Maufrais, Etienne Simon-Lorière, Maud Vanpeene, Sylvie Van der Werf ,Jean Baptiste Bour |
| EPI_ISL_2030030 | Labo Analyses Med | National Reference Center for Viruses of Respiratory Infections, Institut Pasteur, Paris | Marion Barbet, Sylvie Behillil, Méline Bizard, Angela Brisebarre, Camille Capel, Vincent Enouf, Louise Lefrançois, Frédéric Lemoine, Christophe Malabat, Corinne Maufrais, Pierre Lechat, Etienne Simon-Lorière, Maud Vanpeene, Sylvie Van der Werf ,Anne Holstein |
| EPI_ISL_2030031 | Hospital | National Reference Center for Viruses of Respiratory Infections, Institut Pasteur, Paris | Marion Barbet, Sylvie Behillil, Méline Bizard, Angela Brisebarre, Camille Capel, Vincent Enouf, Louise Lefrançois, Frédéric Lemoine, Christophe Malabat, Corinne Maufrais, Etienne Simon-Lorière, Maud Vanpeene, Sylvie Van der Werf ,CéLine Ramanantsoa |
| EPI_ISL_2030032 | Labo Analyses Med | National Reference Center for Viruses of Respiratory Infections, Institut Pasteur, Paris | Marion Barbet, Sylvie Behillil, Méline Bizard, Angela Brisebarre, Camille Capel, Vincent Enouf, Louise Lefrançois, Frédéric Lemoine, Christophe Malabat, Corinne Maufrais, Etienne Simon-Lorière, Maud Vanpeene, Sylvie Van der Werf ,Bruno Berdin |
| EPI_ISL_2030033 | Labo Analyses Med | National Reference Center for Viruses of Respiratory Infections, Institut Pasteur, Paris | Marion Barbet, Sylvie Behillil, Méline Bizard, Angela Brisebarre, Camille Capel, Vincent Enouf, Louise Lefrançois, Frédéric Lemoine, Christophe Malabat, Corinne Maufrais, Etienne Simon-Lorière, Maud Vanpeene, Sylvie Van der Werf ,GréGoire Potiron |
| EPI_ISL_2030034 | Labo Analyses Med | National Reference Center for Viruses of Respiratory Infections, Institut Pasteur, Paris | Marion Barbet, Sylvie Behillil, Méline Bizard, Angela Brisebarre, Camille Capel, Vincent Enouf, Louise Lefrançois, Frédéric Lemoine, Christophe Malabat, Corinne Maufrais, Gael Millot, Etienne Simon-Lorière, Maud Vanpeene, Sylvie Van der Werf ,GréGoire Potiron |
| EPI_ISL_2030035 | Hospital | National Reference Center for Viruses of Respiratory Infections, Institut Pasteur, Paris | Marion Barbet, Sylvie Behillil, Méline Bizard, Angela Brisebarre, Camille Capel, Vincent Enouf, Louise Lefrançois, Frédéric Lemoine, Christophe Malabat, Corinne Maufrais, Etienne Simon-Lorière, Maud Vanpeene, Sylvie Van der Werf ,Laura Djamdjian |

| | | | |
|---|---|---|---|
| EPI_ISL_2030037 | Labo analyses med | National Reference Center for Viruses of Respiratory Infections, Institut Pasteur, Paris | Marion Barbet, Sylvie Behillil, Méline Bizard, Angela Brisebarre, Camille Capel, Vincent Enouf, Louise Lefrançois, Frédéric Lemoine, Christophe Malabat, Corinne Maufrais, Etienne Simon-Lorière, Maud Vanpeene, Sylvie Van der Werf ,Jean-FrançOis Comes |
| EPI_ISL_2030038 | Hospital | National Reference Center for Viruses of Respiratory Infections, Institut Pasteur, Paris | Marion Barbet, Sylvie Behillil, Méline Bizard, Angela Brisebarre, Camille Capel, Vincent Enouf, Louise Lefrançois, Frédéric Lemoine, Christophe Malabat, Corinne Maufrais, Etienne Simon-Lorière, Maud Vanpeene, Sylvie Van der Werf ,EugèNe Castelain |
| EPI_ISL_2030039 | Hospital | National Reference Center for Viruses of Respiratory Infections, Institut Pasteur, Paris | Marion Barbet, Sylvie Behillil, Méline Bizard, Angela Brisebarre, Camille Capel, Vincent Enouf, Louise Lefrançois, Frédéric Lemoine, Christophe Malabat, Corinne Maufrais, Etienne Simon-Lorière, Maud Vanpeene, Sylvie Van der Werf ,Laura Djamdjian |
| EPI_ISL_2030040 | Hospital | National Reference Center for Viruses of Respiratory Infections, Institut Pasteur, Paris | Marion Barbet, Sylvie Behillil, Méline Bizard, Angela Brisebarre, Camille Capel, Vincent Enouf, Louise Lefrançois, Frédéric Lemoine, Christophe Malabat, Corinne Maufrais, Etienne Simon-Lorière, Maud Vanpeene, Sylvie Van der Werf ,Nabil Gastli |
| EPI_ISL_2030041 | Hospital | National Reference Center for Viruses of Respiratory Infections, Institut Pasteur, Paris | Marion Barbet, Sylvie Behillil, Méline Bizard, Angela Brisebarre, Camille Capel, Vincent Enouf, Louise Lefrançois, Frédéric Lemoine, Christophe Malabat, Corinne Maufrais, Etienne Simon-Lorière, Maud Vanpeene, Sylvie Van der Werf ,André-Guy Combremont |
| EPI_ISL_2030042 | Labo Analyses Med | National Reference Center for Viruses of Respiratory Infections, Institut Pasteur, Paris | Marion Barbet, Sylvie Behillil, Méline Bizard, Angela Brisebarre, Camille Capel, Vincent Enouf, Louise Lefrançois, Frédéric Lemoine, Christophe Malabat, Corinne Maufrais, Etienne Simon-Lorière, Maud Vanpeene, Sylvie Van der Werf ,Pierre-Yves Leonard |
| EPI_ISL_2030045 | Hospital | National Reference Center for Viruses of Respiratory Infections, Institut Pasteur, Paris | Marion Barbet, Sylvie Behillil, Méline Bizard, Angela Brisebarre, Camille Capel, Vincent Enouf, Louise Lefrançois, Frédéric Lemoine, Christophe Malabat, Corinne Maufrais, Etienne Simon-Lorière, Maud Vanpeene, Sylvie Van der Werf ,FréDéRique Canis |
| EPI_ISL_2030048 | Labo Analyses Med | National Reference Center for Viruses of Respiratory Infections, Institut Pasteur, Paris | Marion Barbet, Sylvie Behillil, Méline Bizard, Angela Brisebarre, Camille Capel, Vincent Enouf, Louise Lefrançois, Frédéric Lemoine, Christophe Malabat, Corinne Maufrais, Etienne Simon-Lorière, Maud Vanpeene, Sylvie Van der Werf ,Arditi Cocco |
| EPI_ISL_2030049 | Labo Analyses Med | National Reference Center for Viruses of Respiratory Infections, Institut Pasteur, Paris | Marion Barbet, Sylvie Behillil, Méline Bizard, Angela Brisebarre, Camille Capel, Vincent Enouf, Louise Lefrançois, Frédéric Lemoine, Christophe Malabat, Corinne Maufrais, Etienne Simon-Lorière, Maud Vanpeene, Sylvie Van der Werf ,Pierre-Yves Leonard |
| EPI_ISL_2030050 | Hospital | National Reference Center for Viruses of Respiratory Infections, Institut Pasteur, Paris | Marion Barbet, Sylvie Behillil, Méline Bizard, Angela Brisebarre, Camille Capel, Vincent Enouf, Louise Lefrançois, Frédéric Lemoine, Christophe Malabat, Corinne Maufrais, Damien Mornico, Etienne Simon-Lorière, Maud Vanpeene, Sylvie Van der Werf ,Pascale Martres |
| EPI_ISL_2030052 | Labo Analyses Med | National Reference Center for Viruses of Respiratory Infections, Institut Pasteur, Paris | Marion Barbet, Sylvie Behillil, Méline Bizard, Angela Brisebarre, Camille Capel, Vincent Enouf, Louise Lefrançois, Frédéric Lemoine, Christophe Malabat, Corinne Maufrais, Etienne Simon-Lorière, Maud Vanpeene, Sylvie Van der Werf ,Arditi Cocco |
| EPI_ISL_2030053, EPI_ISL_2030054 | Labo Analyses Med | National Reference Center for Viruses of Respiratory Infections, Institut Pasteur, Paris | Marion Barbet, Sylvie Behillil, Méline Bizard, Angela Brisebarre, Camille Capel, Vincent Enouf, Louise Lefrançois, Frédéric Lemoine, Christophe Malabat, Corinne Maufrais, Etienne Simon-Lorière, Maud Vanpeene, Sylvie Van der Werf ,Anne Holstein |
| EPI_ISL_2030055 | Labo analyses med | National Reference Center for Viruses of Respiratory Infections, Institut Pasteur, Paris | Marion Barbet, Sylvie Behillil, Méline Bizard, Angela Brisebarre, Camille Capel, Vincent Enouf, Louise Lefrançois, Frédéric Lemoine, Christophe Malabat, Corinne Maufrais, Amaury Vaysse, Etienne Simon-Lorière, Maud Vanpeene, Sylvie Van der Werf ,Jean-FrançOis Comes |
| EPI_ISL_2030056 | Labo Analyses Med | National Reference Center for Viruses of Respiratory Infections, Institut Pasteur, Paris | Marion Barbet, Sylvie Behillil, Méline Bizard, Angela Brisebarre, Camille Capel, Vincent Enouf, Louise Lefrançois, Frédéric Lemoine, Christophe Malabat, Corinne Maufrais, Etienne Simon-Lorière, Maud Vanpeene, Sylvie Van der Werf ,Pierre-Yves Leonard |
| EPI_ISL_2030058 | Hospital | National Reference Center for Viruses of Respiratory Infections, Institut Pasteur, Paris | Marion Barbet, Sylvie Behillil, Méline Bizard, Angela Brisebarre, Camille Capel, Vincent Enouf, Louise Lefrançois, Frédéric Lemoine, Christophe Malabat, Corinne Maufrais, Emmanuelle Permal, Etienne Simon-Lorière, Maud Vanpeene, Sylvie Van der Werf ,Benjamin Maneglie |
| EPI_ISL_2030059 | Labo Analyses Med | National Reference Center for Viruses of Respiratory Infections, Institut Pasteur, Paris | Marion Barbet, Sylvie Behillil, Méline Bizard, Angela Brisebarre, Camille Capel, Vincent Enouf, Louise Lefrançois, Frédéric Lemoine, Christophe Malabat, Corinne Maufrais, Etienne Simon-Lorière, Maud Vanpeene, Sylvie Van der Werf ,Bruno Berdin |
| EPI_ISL_2030061, EPI_ISL_2030062 | Labo Analyses Med | National Reference Center for Viruses of Respiratory Infections, Institut Pasteur, Paris | Marion Barbet, Sylvie Behillil, Méline Bizard, Angela Brisebarre, Camille Capel, Vincent Enouf, Louise Lefrançois, Frédéric Lemoine, Christophe Malabat, Corinne Maufrais, Etienne Simon-Lorière, Maud Vanpeene, Sylvie Van der Werf ,Ronan Pichard |
| EPI_ISL_2030063 | Labo Analyses Med | National Reference Center for Viruses of Respiratory Infections, Institut Pasteur, Paris | Marion Barbet, Sylvie Behillil, Méline Bizard, Angela Brisebarre, Camille Capel, Vincent Enouf, Louise Lefrançois, Frédéric Lemoine, Christophe Malabat, Corinne Maufrais, Amaury Vaysse, Etienne Simon-Lorière, Maud Vanpeene, Sylvie Van der Werf ,Pierre-Yves Leonard |
| EPI_ISL_2030064 | Hospital | National Reference Center for Viruses of Respiratory Infections, Institut Pasteur, Paris | Marion Barbet, Sylvie Behillil, Méline Bizard, Angela Brisebarre, Camille Capel, Vincent Enouf, Louise Lefrançois, Frédéric Lemoine, Christophe Malabat, Corinne Maufrais, Etienne Simon-Lorière, Maud Vanpeene, Sylvie Van der Werf ,EugèNe Castelain |
| EPI_ISL_2030065 | Labo Analyses Med | National Reference Center for Viruses of Respiratory Infections, Institut Pasteur, Paris | Marion Barbet, Sylvie Behillil, Méline Bizard, Angela Brisebarre, Camille Capel, Vincent Enouf, Louise Lefrançois, Frédéric Lemoine, Christophe Malabat, Corinne Maufrais, Etienne Simon-Lorière, Maud Vanpeene, Sylvie Van der Werf ,Ronan Pichard |
| EPI_ISL_2030066 | Hospital | National Reference Center for Viruses of Respiratory Infections, Institut Pasteur, Paris | Marion Barbet, Sylvie Behillil, Méline Bizard, Angela Brisebarre, Camille Capel, Vincent Enouf, Louise Lefrançois, Frédéric Lemoine, Christophe Malabat, Corinne Maufrais, Pierre Lechat, Etienne Simon-Lorière, Maud Vanpeene, Sylvie Van der Werf ,Laura Djamdjian |
| EPI_ISL_2030067 | Hospital | National Reference Center for Viruses of Respiratory Infections, Institut Pasteur, Paris | Marion Barbet, Sylvie Behillil, Méline Bizard, Angela Brisebarre, Camille Capel, Vincent Enouf, Louise Lefrançois, Frédéric Lemoine, Christophe Malabat, Corinne Maufrais, Etienne Simon-Lorière, Maud Vanpeene, Sylvie Van der Werf ,L. Courdavault |
| EPI_ISL_2030068 | Labo Analyses Med | National Reference Center for Viruses of Respiratory Infections, Institut Pasteur, Paris | Marion Barbet, Sylvie Behillil, Méline Bizard, Angela Brisebarre, Camille Capel, Vincent Enouf, Louise Lefrançois, Frédéric Lemoine, Christophe Malabat, Corinne Maufrais, Etienne Simon-Lorière, Maud Vanpeene, Sylvie Van der Werf ,Delphine MammÈS |
| EPI_ISL_2030069 | Hospital | National Reference Center for Viruses of Respiratory Infections, Institut Pasteur, Paris | Marion Barbet, Sylvie Behillil, Méline Bizard, Angela Brisebarre, Camille Capel, Vincent Enouf, Louise Lefrançois, Frédéric Lemoine, Christophe Malabat, Corinne Maufrais, Etienne Simon-Lorière, Maud Vanpeene, Sylvie Van der Werf ,FréDéRique Canis |
| EPI_ISL_2030070 | Hospital | National Reference Center for Viruses of Respiratory Infections, Institut Pasteur, Paris | Marion Barbet, Sylvie Behillil, Méline Bizard, Angela Brisebarre, Camille Capel, Vincent Enouf, Louise Lefrançois, Frédéric Lemoine, Christophe Malabat, Corinne Maufrais, Amaury Vaysse, Etienne Simon-Lorière, Maud Vanpeene, Sylvie Van der Werf ,Pascale Martres |
| EPI_ISL_2030071 | Labo analyses med | National Reference Center for Viruses of Respiratory Infections, Institut Pasteur, Paris | Marion Barbet, Sylvie Behillil, Méline Bizard, Angela Brisebarre, Camille Capel, Vincent Enouf, Louise Lefrançois, Frédéric Lemoine, Christophe Malabat, Corinne Maufrais, Etienne Simon-Lorière, Maud Vanpeene, Sylvie Van der Werf ,Jean-FrançOis Comes |
| EPI_ISL_2030072 | Hospital | National Reference Center for Viruses of Respiratory Infections, Institut Pasteur, Paris | Marion Barbet, Sylvie Behillil, Méline Bizard, Angela Brisebarre, Camille Capel, Vincent Enouf, Louise Lefrançois, Frédéric Lemoine, Christophe Malabat, Corinne Maufrais, Etienne Simon-Lorière, Maud Vanpeene, Sylvie Van der Werf ,CéLine Bressollette |
| EPI_ISL_2030073 | Labo Analyses Med | National Reference Center for Viruses of Respiratory Infections, Institut Pasteur, Paris | Marion Barbet, Sylvie Behillil, Méline Bizard, Angela Brisebarre, Camille Capel, Vincent Enouf, Louise Lefrançois, Frédéric Lemoine, Christophe Malabat, Corinne Maufrais, Pierre Lechat, Etienne Simon-Lorière, Maud Vanpeene, Sylvie Van der Werf ,GréGoire Potiron |
| EPI_ISL_2030074 | Hospital | National Reference Center for Viruses of Respiratory Infections, Institut Pasteur, Paris | Marion Barbet, Sylvie Behillil, Méline Bizard, Angela Brisebarre, Camille Capel, Vincent Enouf, Louise Lefrançois, Frédéric Lemoine, Christophe Malabat, Corinne Maufrais, Etienne Simon-Lorière, Maud Vanpeene, Sylvie Van der Werf ,Isabelle Gros |
| EPI_ISL_2030075 | Labo Analyses Med | National Reference Center for Viruses of Respiratory Infections, Institut Pasteur, Paris | Marion Barbet, Sylvie Behillil, Méline Bizard, Angela Brisebarre, Camille Capel, Vincent Enouf, Louise Lefrançois, Frédéric Lemoine, Christophe Malabat, Corinne Maufrais, Pierre Lechat, Etienne Simon-Lorière, Maud Vanpeene, Sylvie Van der Werf ,Anne Holstein |
| EPI_ISL_2030076 | Hospital | National Reference Center for Viruses of Respiratory Infections, Institut Pasteur, Paris | Marion Barbet, Sylvie Behillil, Méline Bizard, Angela Brisebarre, Camille Capel, Vincent Enouf, Louise Lefrançois, Frédéric Lemoine, Christophe Malabat, Corinne Maufrais, Etienne Simon-Lorière, Maud Vanpeene, Sylvie Van der Werf ,Anne Cady |
| EPI_ISL_2030078 | Hospital | National Reference Center for Viruses of Respiratory Infections, Institut Pasteur, Paris | Marion Barbet, Sylvie Behillil, Méline Bizard, Angela Brisebarre, Camille Capel, Vincent Enouf, Louise Lefrançois, Frédéric Lemoine, Christophe Malabat, Corinne Maufrais, Etienne Simon-Lorière, Maud Vanpeene, Sylvie Van der Werf ,CéLine Bressollette |
| EPI_ISL_2030079, EPI_ISL_2030080 | Hospital | National Reference Center for Viruses of Respiratory Infections, Institut Pasteur, Paris | Marion Barbet, Sylvie Behillil, Méline Bizard, Angela Brisebarre, Camille Capel, Vincent Enouf, Louise Lefrançois, Frédéric Lemoine, Christophe Malabat, Corinne Maufrais, Etienne Simon-Lorière, Maud Vanpeene, Sylvie Van der Werf ,Pauline Garnier |
| EPI_ISL_2030081 | Hospital | National Reference Center for Viruses of Respiratory Infections, Institut Pasteur, Paris | Marion Barbet, Sylvie Behillil, Méline Bizard, Angela Brisebarre, Camille Capel, Vincent Enouf, Louise Lefrançois, Frédéric Lemoine, Christophe Malabat, Corinne Maufrais, Etienne Simon-Lorière, Maud Vanpeene, Sylvie Van der Werf ,ValéRie Serazin |
| EPI_ISL_2030083 | Hospital | National Reference Center for Viruses of Respiratory Infections, Institut Pasteur, Paris | Marion Barbet, Sylvie Behillil, Méline Bizard, Angela Brisebarre, Camille Capel, Vincent Enouf, Louise Lefrançois, Frédéric Lemoine, Christophe Malabat, Corinne Maufrais, Etienne Simon-Lorière, Maud Vanpeene, Sylvie Van der Werf ,EugèNe Castelain |

| | | | |
|---|---|---|---|
| EPI_ISL_2030086 | Labo analyses med | National Reference Center for Viruses of Respiratory Infections, Institut Pasteur, Paris | Marion Barbet, Sylvie Behillil, Méline Bizard, Angela Brisebarre, Camille Capel, Vincent Enouf, Louise Lefrançois, Frédéric Lemoine, Christophe Malabat, Corinne Maufrais, Amaury Vaysse, Etienne Simon-Lorière, Maud Vanpeene, Sylvie Van der Werf ,Jean-FrançOis Comes |
| EPI_ISL_2030087 | Hospital | National Reference Center for Viruses of Respiratory Infections, Institut Pasteur, Paris | Marion Barbet, Sylvie Behillil, Méline Bizard, Angela Brisebarre, Camille Capel, Vincent Enouf, Louise Lefrançois, Frédéric Lemoine, Christophe Malabat, Corinne Maufrais, Etienne Simon-Lorière, Maud Vanpeene, Sylvie Van der Werf ,CéLine Ramanantsoa |
| EPI_ISL_2030089 | Hospital | National Reference Center for Viruses of Respiratory Infections, Institut Pasteur, Paris | Marion Barbet, Sylvie Behillil, Méline Bizard, Angela Brisebarre, Camille Capel, Vincent Enouf, Louise Lefrançois, Frédéric Lemoine, Christophe Malabat, Corinne Maufrais, Etienne Simon-Lorière, Maud Vanpeene, Sylvie Van der Werf ,ValéRie Serazin |
| EPI_ISL_2030090 | Labo Analyses Med | National Reference Center for Viruses of Respiratory Infections, Institut Pasteur, Paris | Marion Barbet, Sylvie Behillil, Méline Bizard, Angela Brisebarre, Camille Capel, Vincent Enouf, Louise Lefrançois, Frédéric Lemoine, Christophe Malabat, Corinne Maufrais, Etienne Simon-Lorière, Maud Vanpeene, Sylvie Van der Werf ,GréGoire Potiron |
| EPI_ISL_2030091 | Hospital | National Reference Center for Viruses of Respiratory Infections, Institut Pasteur, Paris | Marion Barbet, Sylvie Behillil, Méline Bizard, Angela Brisebarre, Camille Capel, Vincent Enouf, Louise Lefrançois, Frédéric Lemoine, Christophe Malabat, Corinne Maufrais, Etienne Simon-Lorière, Maud Vanpeene, Sylvie Van der Werf ,Christine Febreau |
| EPI_ISL_2030093 | Hospital | National Reference Center for Viruses of Respiratory Infections, Institut Pasteur, Paris | Marion Barbet, Sylvie Behillil, Méline Bizard, Angela Brisebarre, Camille Capel, Vincent Enouf, Louise Lefrançois, Frédéric Lemoine, Christophe Malabat, Corinne Maufrais, Etienne Simon-Lorière, Maud Vanpeene, Sylvie Van der Werf ,Pauline Garnier |
| EPI_ISL_2030094 | Hospital | National Reference Center for Viruses of Respiratory Infections, Institut Pasteur, Paris | Marion Barbet, Sylvie Behillil, Méline Bizard, Angela Brisebarre, Camille Capel, Vincent Enouf, Louise Lefrançois, Frédéric Lemoine, Christophe Malabat, Corinne Maufrais, Amaury Vaysse, Etienne Simon-Lorière, Maud Vanpeene, Sylvie Van der Werf ,Pauline Garnier |
| EPI_ISL_2030095 | Labo analyses med | National Reference Center for Viruses of Respiratory Infections, Institut Pasteur, Paris | Marion Barbet, Sylvie Behillil, Méline Bizard, Angela Brisebarre, Camille Capel, Vincent Enouf, Louise Lefrançois, Frédéric Lemoine, Christophe Malabat, Corinne Maufrais, Etienne Simon-Lorière, Maud Vanpeene, Sylvie Van der Werf ,Jean-FrançOis Comes |
| EPI_ISL_2030098 | Hospital | National Reference Center for Viruses of Respiratory Infections, Institut Pasteur, Paris | Marion Barbet, Sylvie Behillil, Méline Bizard, Angela Brisebarre, Camille Capel, Vincent Enouf, Louise Lefrançois, Frédéric Lemoine, Christophe Malabat, Corinne Maufrais, Etienne Simon-Lorière, Maud Vanpeene, Sylvie Van der Werf ,Kahla |
| EPI_ISL_2030101 | Labo Analyses Med | National Reference Center for Viruses of Respiratory Infections, Institut Pasteur, Paris | Marion Barbet, Sylvie Behillil, Méline Bizard, Angela Brisebarre, Camille Capel, Vincent Enouf, Louise Lefrançois, Frédéric Lemoine, Christophe Malabat, Corinne Maufrais, Etienne Simon-Lorière, Maud Vanpeene, Sylvie Van der Werf ,Brieuc Gestin |
| EPI_ISL_2030102 | Hospital | National Reference Center for Viruses of Respiratory Infections, Institut Pasteur, Paris | Marion Barbet, Sylvie Behillil, Méline Bizard, Angela Brisebarre, Camille Capel, Vincent Enouf, Louise Lefrançois, Frédéric Lemoine, Christophe Malabat, Corinne Maufrais, Etienne Simon-Lorière, Maud Vanpeene, Sylvie Van der Werf ,Pascale Martres |
| EPI_ISL_2030103, EPI_ISL_2030104 | Hospital | National Reference Center for Viruses of Respiratory Infections, Institut Pasteur, Paris | Marion Barbet, Sylvie Behillil, Méline Bizard, Angela Brisebarre, Camille Capel, Vincent Enouf, Louise Lefrançois, Frédéric Lemoine, Christophe Malabat, Corinne Maufrais, Etienne Simon-Lorière, Maud Vanpeene, Sylvie Van der Werf ,FréDéRique Canis |
| EPI_ISL_2030105 | Labo Analyses Med | National Reference Center for Viruses of Respiratory Infections, Institut Pasteur, Paris | Marion Barbet, Sylvie Behillil, Méline Bizard, Angela Brisebarre, Camille Capel, Vincent Enouf, Louise Lefrançois, Frédéric Lemoine, Christophe Malabat, Corinne Maufrais, Amaury Vaysse, Etienne Simon-Lorière, Maud Vanpeene, Sylvie Van der Werf ,Pierre-Yves Leonard |
| EPI_ISL_2030106 | Labo Analyses Med | National Reference Center for Viruses of Respiratory Infections, Institut Pasteur, Paris | Marion Barbet, Sylvie Behillil, Méline Bizard, Angela Brisebarre, Camille Capel, Vincent Enouf, Louise Lefrançois, Frédéric Lemoine, Christophe Malabat, Corinne Maufrais, Etienne Simon-Lorière, Maud Vanpeene, Sylvie Van der Werf ,Anne Holstein |
| EPI_ISL_2030107 | Hospital | National Reference Center for Viruses of Respiratory Infections, Institut Pasteur, Paris | Marion Barbet, Sylvie Behillil, Méline Bizard, Angela Brisebarre, Camille Capel, Vincent Enouf, Louise Lefrançois, Frédéric Lemoine, Christophe Malabat, Corinne Maufrais, Etienne Simon-Lorière, Maud Vanpeene, Sylvie Van der Werf ,FréDéRique Canis |
| EPI_ISL_2030108 | Hospital | National Reference Center for Viruses of Respiratory Infections, Institut Pasteur, Paris | Marion Barbet, Sylvie Behillil, Méline Bizard, Angela Brisebarre, Camille Capel, Vincent Enouf, Louise Lefrançois, Frédéric Lemoine, Christophe Malabat, Corinne Maufrais, Etienne Simon-Lorière, Maud Vanpeene, Sylvie Van der Werf ,CéLine Ramanantsoa |
| EPI_ISL_2030110 | Hospital | National Reference Center for Viruses of Respiratory Infections, Institut Pasteur, Paris | Marion Barbet, Sylvie Behillil, Méline Bizard, Angela Brisebarre, Camille Capel, Vincent Enouf, Louise Lefrançois, Frédéric Lemoine, Christophe Malabat, Corinne Maufrais, Etienne Simon-Lorière, Maud Vanpeene, Sylvie Van der Werf ,EugèNe Castelain |
| EPI_ISL_2030111 | Labo Analyses Med | National Reference Center for Viruses of Respiratory Infections, Institut Pasteur, Paris | Marion Barbet, Sylvie Behillil, Méline Bizard, Angela Brisebarre, Camille Capel, Vincent Enouf, Louise Lefrançois, Frédéric Lemoine, Christophe Malabat, Corinne Maufrais, Etienne Simon-Lorière, Maud Vanpeene, Sylvie Van der Werf ,Bruno Berdin |
| EPI_ISL_2030112 | Hospital | National Reference Center for Viruses of Respiratory Infections, Institut Pasteur, Paris | Marion Barbet, Sylvie Behillil, Méline Bizard, Angela Brisebarre, Camille Capel, Vincent Enouf, Louise Lefrançois, Frédéric Lemoine, Christophe Malabat, Corinne Maufrais, Etienne Simon-Lorière, Maud Vanpeene, Sylvie Van der Werf ,LéA Pilorge |
| EPI_ISL_2030115 | Hospital | National Reference Center for Viruses of Respiratory Infections, Institut Pasteur, Paris | Marion Barbet, Sylvie Behillil, Méline Bizard, Angela Brisebarre, Camille Capel, Vincent Enouf, Louise Lefrançois, Frédéric Lemoine, Christophe Malabat, Corinne Maufrais, Etienne Simon-Lorière, Maud Vanpeene, Sylvie Van der Werf ,Anne Cady |
| EPI_ISL_2030116 | Hospital | National Reference Center for Viruses of Respiratory Infections, Institut Pasteur, Paris | Marion Barbet, Sylvie Behillil, Méline Bizard, Angela Brisebarre, Camille Capel, Vincent Enouf, Louise Lefrançois, Frédéric Lemoine, Christophe Malabat, Corinne Maufrais, Etienne Simon-Lorière, Maud Vanpeene, Sylvie Van der Werf ,Sylvie Ledru |
| EPI_ISL_2030118 | Hospital | National Reference Center for Viruses of Respiratory Infections, Institut Pasteur, Paris | Marion Barbet, Sylvie Behillil, Méline Bizard, Angela Brisebarre, Camille Capel, Vincent Enouf, Louise Lefrançois, Frédéric Lemoine, Christophe Malabat, Corinne Maufrais, Etienne Simon-Lorière, Maud Vanpeene, Sylvie Van der Werf ,JéRome Guinard |
| EPI_ISL_2030119 | Hospital | National Reference Center for Viruses of Respiratory Infections, Institut Pasteur, Paris | Marion Barbet, Sylvie Behillil, Méline Bizard, Angela Brisebarre, Camille Capel, Vincent Enouf, Louise Lefrançois, Frédéric Lemoine, Christophe Malabat, Corinne Maufrais, Etienne Simon-Lorière, Maud Vanpeene, Sylvie Van der Werf ,Laura Djamdjian |
| EPI_ISL_2030120 | Hospital | National Reference Center for Viruses of Respiratory Infections, Institut Pasteur, Paris | Marion Barbet, Sylvie Behillil, Méline Bizard, Angela Brisebarre, Camille Capel, Vincent Enouf, Louise Lefrançois, Frédéric Lemoine, Christophe Malabat, Corinne Maufrais, Pierre Lechat, Etienne Simon-Lorière, Maud Vanpeene, Sylvie Van der Werf ,L. Courdavault |
| EPI_ISL_2030121 | Hospital | National Reference Center for Viruses of Respiratory Infections, Institut Pasteur, Paris | Marion Barbet, Sylvie Behillil, Méline Bizard, Angela Brisebarre, Camille Capel, Vincent Enouf, Louise Lefrançois, Frédéric Lemoine, Christophe Malabat, Corinne Maufrais, Etienne Simon-Lorière, Maud Vanpeene, Sylvie Van der Werf ,L. Courdavault |
| EPI_ISL_2030125 | Hospital | National Reference Center for Viruses of Respiratory Infections, Institut Pasteur, Paris | Marion Barbet, Sylvie Behillil, Méline Bizard, Angela Brisebarre, Camille Capel, Vincent Enouf, Louise Lefrançois, Frédéric Lemoine, Christophe Malabat, Corinne Maufrais, Etienne Simon-Lorière, Maud Vanpeene, Sylvie Van der Werf ,Pascale Martres |
| EPI_ISL_2030126 | Hospital | National Reference Center for Viruses of Respiratory Infections, Institut Pasteur, Paris | Marion Barbet, Sylvie Behillil, Méline Bizard, Angela Brisebarre, Camille Capel, Vincent Enouf, Louise Lefrançois, Frédéric Lemoine, Christophe Malabat, Corinne Maufrais, Etienne Simon-Lorière, Maud Vanpeene, Sylvie Van der Werf ,CéLine Bressollette |
| EPI_ISL_2030128 | Hospital | National Reference Center for Viruses of Respiratory Infections, Institut Pasteur, Paris | Marion Barbet, Sylvie Behillil, Méline Bizard, Angela Brisebarre, Camille Capel, Vincent Enouf, Louise Lefrançois, Frédéric Lemoine, Christophe Malabat, Corinne Maufrais, Etienne Simon-Lorière, Maud Vanpeene, Sylvie Van der Werf ,Kahla |
| EPI_ISL_2030129, EPI_ISL_2030130 | Labo Analyses Med | National Reference Center for Viruses of Respiratory Infections, Institut Pasteur, Paris | Marion Barbet, Sylvie Behillil, Méline Bizard, Angela Brisebarre, Camille Capel, Vincent Enouf, Louise Lefrançois, Frédéric Lemoine, Christophe Malabat, Corinne Maufrais, Etienne Simon-Lorière, Maud Vanpeene, Sylvie Van der Werf ,Brieuc Gestin |
| EPI_ISL_2030132 | Labo Analyses Med | National Reference Center for Viruses of Respiratory Infections, Institut Pasteur, Paris | Marion Barbet, Sylvie Behillil, Méline Bizard, Angela Brisebarre, Camille Capel, Vincent Enouf, Louise Lefrançois, Frédéric Lemoine, Christophe Malabat, Corinne Maufrais, Etienne Simon-Lorière, Maud Vanpeene, Sylvie Van der Werf ,Anne Holstein |
| EPI_ISL_2030133 | Hospital | National Reference Center for Viruses of Respiratory Infections, Institut Pasteur, Paris | Marion Barbet, Sylvie Behillil, Méline Bizard, Angela Brisebarre, Camille Capel, Vincent Enouf, Louise Lefrançois, Frédéric Lemoine, Christophe Malabat, Corinne Maufrais, Etienne Simon-Lorière, Maud Vanpeene, Sylvie Van der Werf ,Benjamin Maneglie |
| EPI_ISL_2030134 | Hospital | National Reference Center for Viruses of Respiratory Infections, Institut Pasteur, Paris | Marion Barbet, Sylvie Behillil, Méline Bizard, Angela Brisebarre, Camille Capel, Vincent Enouf, Louise Lefrançois, Frédéric Lemoine, Christophe Malabat, Corinne Maufrais, Etienne Simon-Lorière, Maud Vanpeene, Sylvie Van der Werf ,BéAtrice Lesimple |
| EPI_ISL_2030135 | Hospital | National Reference Center for Viruses of Respiratory Infections, Institut Pasteur, Paris | Marion Barbet, Sylvie Behillil, Méline Bizard, Angela Brisebarre, Camille Capel, Vincent Enouf, Louise Lefrançois, Frédéric Lemoine, Christophe Malabat, Corinne Maufrais, Etienne Simon-Lorière, Maud Vanpeene, Sylvie Van der Werf ,FréDéRique Canis |
| EPI_ISL_2030136 | Hospital | National Reference Center for Viruses of Respiratory Infections, Institut Pasteur, Paris | Marion Barbet, Sylvie Behillil, Méline Bizard, Angela Brisebarre, Camille Capel, Vincent Enouf, Louise Lefrançois, Frédéric Lemoine, Christophe Malabat, Corinne Maufrais, Etienne Simon-Lorière, Maud Vanpeene, Sylvie Van der Werf ,ValéRie Serazin |
| EPI_ISL_2030137 | Hospital | National Reference Center for Viruses of Respiratory Infections, Institut Pasteur, Paris | Marion Barbet, Sylvie Behillil, Méline Bizard, Angela Brisebarre, Camille Capel, Vincent Enouf, Louise Lefrançois, Frédéric Lemoine, Christophe Malabat, Corinne Maufrais, Gael Millot, Etienne Simon-Lorière, Maud Vanpeene, Sylvie Van der Werf ,Christine Febreau |

| | | | |
|---|---|---|---|
| EPI_ISL_2030138 | Hospital | National Reference Center for Viruses of Respiratory Infections, Institut Pasteur, Paris | Marion Barbet, Sylvie Behillil, Méline Bizard, Angela Brisebarre, Camille Capel, Vincent Enouf, Louise Lefrançois, Frédéric Lemoine, Christophe Malabat, Corinne Maufrais, Etienne Simon-Lorière, Maud Vanpeene, Sylvie Van der Werf ,Odile Morvan |
| EPI_ISL_2030139, EPI_ISL_2030140 | Hospital | National Reference Center for Viruses of Respiratory Infections, Institut Pasteur, Paris | Marion Barbet, Sylvie Behillil, Méline Bizard, Angela Brisebarre, Camille Capel, Vincent Enouf, Louise Lefrançois, Frédéric Lemoine, Christophe Malabat, Corinne Maufrais, Etienne Simon-Lorière, Maud Vanpeene, Sylvie Van der Werf ,L. Courdavault |
| EPI_ISL_2030141 | Hospital | National Reference Center for Viruses of Respiratory Infections, Institut Pasteur, Paris | Marion Barbet, Sylvie Behillil, Méline Bizard, Angela Brisebarre, Camille Capel, Vincent Enouf, Louise Lefrançois, Frédéric Lemoine, Christophe Malabat, Corinne Maufrais, Etienne Simon-Lorière, Maud Vanpeene, Sylvie Van der Werf ,FréDéRique Canis |
| EPI_ISL_2030142 | Hospital | National Reference Center for Viruses of Respiratory Infections, Institut Pasteur, Paris | Marion Barbet, Sylvie Behillil, Méline Bizard, Angela Brisebarre, Camille Capel, Vincent Enouf, Louise Lefrançois, Frédéric Lemoine, Christophe Malabat, Corinne Maufrais, Amaury Vaysse, Etienne Simon-Lorière, Maud Vanpeene, Sylvie Van der Werf ,Patricia Stoessel-Thouvenin |
| EPI_ISL_2030143 | Labo Analyses Med | National Reference Center for Viruses of Respiratory Infections, Institut Pasteur, Paris | Marion Barbet, Sylvie Behillil, Méline Bizard, Angela Brisebarre, Camille Capel, Vincent Enouf, Louise Lefrançois, Frédéric Lemoine, Christophe Malabat, Corinne Maufrais, Etienne Simon-Lorière, Maud Vanpeene, Sylvie Van der Werf ,Pierre-Yves Leonard |
| EPI_ISL_2030144 | Hospital | National Reference Center for Viruses of Respiratory Infections, Institut Pasteur, Paris | Marion Barbet, Sylvie Behillil, Méline Bizard, Angela Brisebarre, Camille Capel, Vincent Enouf, Louise Lefrançois, Frédéric Lemoine, Christophe Malabat, Corinne Maufrais, Etienne Simon-Lorière, Maud Vanpeene, Sylvie Van der Werf ,Odile Morvan |
| EPI_ISL_2030145 | Hospital | National Reference Center for Viruses of Respiratory Infections, Institut Pasteur, Paris | Marion Barbet, Sylvie Behillil, Méline Bizard, Angela Brisebarre, Camille Capel, Vincent Enouf, Louise Lefrançois, Frédéric Lemoine, Christophe Malabat, Corinne Maufrais, Etienne Simon-Lorière, Maud Vanpeene, Sylvie Van der Werf ,Nabil Gastli |
| EPI_ISL_2030146 | Hospital | National Reference Center for Viruses of Respiratory Infections, Institut Pasteur, Paris | Marion Barbet, Sylvie Behillil, Méline Bizard, Angela Brisebarre, Camille Capel, Vincent Enouf, Louise Lefrançois, Frédéric Lemoine, Christophe Malabat, Corinne Maufrais, Etienne Simon-Lorière, Maud Vanpeene, Sylvie Van der Werf ,JéRome Guinard |
| EPI_ISL_2030147 | Hospital | National Reference Center for Viruses of Respiratory Infections, Institut Pasteur, Paris | Marion Barbet, Sylvie Behillil, Méline Bizard, Angela Brisebarre, Camille Capel, Vincent Enouf, Louise Lefrançois, Frédéric Lemoine, Christophe Malabat, Corinne Maufrais, Pierre Lechat, Etienne Simon-Lorière, Maud Vanpeene, Sylvie Van der Werf ,Laura Djamdjian |
| EPI_ISL_2030148 | Hospital | National Reference Center for Viruses of Respiratory Infections, Institut Pasteur, Paris | Marion Barbet, Sylvie Behillil, Méline Bizard, Angela Brisebarre, Camille Capel, Vincent Enouf, Louise Lefrançois, Frédéric Lemoine, Christophe Malabat, Corinne Maufrais, Etienne Simon-Lorière, Maud Vanpeene, Sylvie Van der Werf ,Benjamin Maneglie |
| EPI_ISL_2030149 | Labo Analyses Med | National Reference Center for Viruses of Respiratory Infections, Institut Pasteur, Paris | Marion Barbet, Sylvie Behillil, Méline Bizard, Angela Brisebarre, Camille Capel, Vincent Enouf, Louise Lefrançois, Frédéric Lemoine, Christophe Malabat, Corinne Maufrais, Etienne Simon-Lorière, Maud Vanpeene, Sylvie Van der Werf ,Pierre-Yves Leonard |
| EPI_ISL_2030150 | Hospital | National Reference Center for Viruses of Respiratory Infections, Institut Pasteur, Paris | Marion Barbet, Sylvie Behillil, Méline Bizard, Angela Brisebarre, Camille Capel, Vincent Enouf, Louise Lefrançois, Frédéric Lemoine, Christophe Malabat, Corinne Maufrais, Etienne Simon-Lorière, Maud Vanpeene, Sylvie Van der Werf ,CéLine Bressollette |
| EPI_ISL_2030151 | Hospital | National Reference Center for Viruses of Respiratory Infections, Institut Pasteur, Paris | Marion Barbet, Sylvie Behillil, Méline Bizard, Angela Brisebarre, Camille Capel, Vincent Enouf, Louise Lefrançois, Frédéric Lemoine, Christophe Malabat, Corinne Maufrais, Etienne Simon-Lorière, Maud Vanpeene, Sylvie Van der Werf ,EugèNe Castelain |
| EPI_ISL_2030152 | Labo Analyses Med | National Reference Center for Viruses of Respiratory Infections, Institut Pasteur, Paris | Marion Barbet, Sylvie Behillil, Méline Bizard, Angela Brisebarre, Camille Capel, Vincent Enouf, Louise Lefrançois, Frédéric Lemoine, Christophe Malabat, Corinne Maufrais, Etienne Simon-Lorière, Maud Vanpeene, Sylvie Van der Werf ,Brieuc Gestin |
| EPI_ISL_2030153 | Hospital | National Reference Center for Viruses of Respiratory Infections, Institut Pasteur, Paris | Marion Barbet, Sylvie Behillil, Méline Bizard, Angela Brisebarre, Camille Capel, Vincent Enouf, Louise Lefrançois, Frédéric Lemoine, Christophe Malabat, Corinne Maufrais, Etienne Simon-Lorière, Maud Vanpeene, Sylvie Van der Werf ,Laura Djamdjian |
| EPI_ISL_2030154 | Hospital | National Reference Center for Viruses of Respiratory Infections, Institut Pasteur, Paris | Marion Barbet, Sylvie Behillil, Méline Bizard, Angela Brisebarre, Camille Capel, Vincent Enouf, Louise Lefrançois, Frédéric Lemoine, Christophe Malabat, Corinne Maufrais, Etienne Simon-Lorière, Maud Vanpeene, Sylvie Van der Werf ,CéLine Ramanantsoa |
| EPI_ISL_2030155 | Labo Analyses Med | National Reference Center for Viruses of Respiratory Infections, Institut Pasteur, Paris | Marion Barbet, Sylvie Behillil, Méline Bizard, Angela Brisebarre, Camille Capel, Vincent Enouf, Louise Lefrançois, Frédéric Lemoine, Christophe Malabat, Corinne Maufrais, Etienne Simon-Lorière, Maud Vanpeene, Sylvie Van der Werf ,Cyrile Bonnet |
| EPI_ISL_2030157 | Labo Analyses Med | National Reference Center for Viruses of Respiratory Infections, Institut Pasteur, Paris | Marion Barbet, Sylvie Behillil, Méline Bizard, Angela Brisebarre, Camille Capel, Vincent Enouf, Louise Lefrançois, Frédéric Lemoine, Christophe Malabat, Corinne Maufrais, Etienne Simon-Lorière, Maud Vanpeene, Sylvie Van der Werf ,GréGoire Potiron |
| EPI_ISL_2030158 | Labo Analyses Med | National Reference Center for Viruses of Respiratory Infections, Institut Pasteur, Paris | Marion Barbet, Sylvie Behillil, Méline Bizard, Angela Brisebarre, Camille Capel, Vincent Enouf, Louise Lefrançois, Frédéric Lemoine, Christophe Malabat, Corinne Maufrais, Etienne Simon-Lorière, Maud Vanpeene, Sylvie Van der Werf ,Pierre-Yves Leonard |
| EPI_ISL_2030159 | Hospital | National Reference Center for Viruses of Respiratory Infections, Institut Pasteur, Paris | Marion Barbet, Sylvie Behillil, Méline Bizard, Angela Brisebarre, Camille Capel, Vincent Enouf, Louise Lefrançois, Frédéric Lemoine, Christophe Malabat, Corinne Maufrais, Etienne Simon-Lorière, Maud Vanpeene, Sylvie Van der Werf ,ValéRie Serazin |
| EPI_ISL_2030160 | Hospital | National Reference Center for Viruses of Respiratory Infections, Institut Pasteur, Paris | Marion Barbet, Sylvie Behillil, Méline Bizard, Angela Brisebarre, Camille Capel, Vincent Enouf, Louise Lefrançois, Frédéric Lemoine, Christophe Malabat, Corinne Maufrais, Etienne Simon-Lorière, Maud Vanpeene, Sylvie Van der Werf ,Mignard |
| EPI_ISL_2030161 | Labo Analyses Med | National Reference Center for Viruses of Respiratory Infections, Institut Pasteur, Paris | Marion Barbet, Sylvie Behillil, Méline Bizard, Angela Brisebarre, Camille Capel, Vincent Enouf, Louise Lefrançois, Frédéric Lemoine, Christophe Malabat, Corinne Maufrais, Etienne Simon-Lorière, Maud Vanpeene, Sylvie Van der Werf ,Arditi Cocco |
| EPI_ISL_2030162 | Hospital | National Reference Center for Viruses of Respiratory Infections, Institut Pasteur, Paris | Marion Barbet, Sylvie Behillil, Méline Bizard, Angela Brisebarre, Camille Capel, Vincent Enouf, Louise Lefrançois, Frédéric Lemoine, Christophe Malabat, Corinne Maufrais, Etienne Simon-Lorière, Maud Vanpeene, Sylvie Van der Werf ,E. Bichier |
| EPI_ISL_2030163 | Hospital | National Reference Center for Viruses of Respiratory Infections, Institut Pasteur, Paris | Marion Barbet, Sylvie Behillil, Méline Bizard, Angela Brisebarre, Camille Capel, Vincent Enouf, Louise Lefrançois, Frédéric Lemoine, Christophe Malabat, Corinne Maufrais, Etienne Simon-Lorière, Maud Vanpeene, Sylvie Van der Werf ,EugèNe Castelain |
| EPI_ISL_2030164 | Labo Analyses Med | National Reference Center for Viruses of Respiratory Infections, Institut Pasteur, Paris | Marion Barbet, Sylvie Behillil, Méline Bizard, Angela Brisebarre, Camille Capel, Vincent Enouf, Louise Lefrançois, Frédéric Lemoine, Christophe Malabat, Corinne Maufrais, Etienne Simon-Lorière, Maud Vanpeene, Sylvie Van der Werf ,Brieuc Gestin |
| EPI_ISL_2030165, EPI_ISL_2030167 | Labo Analyses Med | National Reference Center for Viruses of Respiratory Infections, Institut Pasteur, Paris | Marion Barbet, Sylvie Behillil, Méline Bizard, Angela Brisebarre, Camille Capel, Vincent Enouf, Louise Lefrançois, Frédéric Lemoine, Christophe Malabat, Corinne Maufrais, Etienne Simon-Lorière, Maud Vanpeene, Sylvie Van der Werf ,GréGoire Potiron |
| EPI_ISL_2030168 | Hospital | National Reference Center for Viruses of Respiratory Infections, Institut Pasteur, Paris | Marion Barbet, Sylvie Behillil, Méline Bizard, Angela Brisebarre, Camille Capel, Vincent Enouf, Louise Lefrançois, Frédéric Lemoine, Christophe Malabat, Corinne Maufrais, Etienne Simon-Lorière, Maud Vanpeene, Sylvie Van der Werf ,Gandon |
| EPI_ISL_2030169 | Labo Analyses Med | National Reference Center for Viruses of Respiratory Infections, Institut Pasteur, Paris | Marion Barbet, Sylvie Behillil, Méline Bizard, Angela Brisebarre, Camille Capel, Vincent Enouf, Louise Lefrançois, Frédéric Lemoine, Christophe Malabat, Corinne Maufrais, Etienne Simon-Lorière, Maud Vanpeene, Sylvie Van der Werf ,GréGoire Potiron |
| EPI_ISL_2030170 | Labo Analyses Med | National Reference Center for Viruses of Respiratory Infections, Institut Pasteur, Paris | Marion Barbet, Sylvie Behillil, Méline Bizard, Angela Brisebarre, Camille Capel, Vincent Enouf, Louise Lefrançois, Frédéric Lemoine, Christophe Malabat, Corinne Maufrais, Etienne Simon-Lorière, Maud Vanpeene, Sylvie Van der Werf ,Arditi Cocco |
| EPI_ISL_2030172 | Hospital | National Reference Center for Viruses of Respiratory Infections, Institut Pasteur, Paris | Marion Barbet, Sylvie Behillil, Méline Bizard, Angela Brisebarre, Camille Capel, Vincent Enouf, Louise Lefrançois, Frédéric Lemoine, Christophe Malabat, Corinne Maufrais, Etienne Simon-Lorière, Maud Vanpeene, Sylvie Van der Werf ,Dominique Descamps |
| EPI_ISL_2030173 | Hospital | National Reference Center for Viruses of Respiratory Infections, Institut Pasteur, Paris | Marion Barbet, Sylvie Behillil, Méline Bizard, Angela Brisebarre, Camille Capel, Vincent Enouf, Louise Lefrançois, Frédéric Lemoine, Christophe Malabat, Corinne Maufrais, Etienne Simon-Lorière, Maud Vanpeene, Sylvie Van der Werf ,André-Guy Combremont |
| EPI_ISL_2030174 | Hospital | National Reference Center for Viruses of Respiratory Infections, Institut Pasteur, Paris | Marion Barbet, Sylvie Behillil, Méline Bizard, Angela Brisebarre, Camille Capel, Vincent Enouf, Louise Lefrançois, Frédéric Lemoine, Christophe Malabat, Corinne Maufrais, Etienne Simon-Lorière, Maud Vanpeene, Sylvie Van der Werf ,EugèNe Castelain |
| EPI_ISL_2030175 | Labo Analyses Med | National Reference Center for Viruses of Respiratory Infections, Institut Pasteur, Paris | Marion Barbet, Sylvie Behillil, Méline Bizard, Angela Brisebarre, Camille Capel, Vincent Enouf, Louise Lefrançois, Frédéric Lemoine, Christophe Malabat, Corinne Maufrais, Etienne Simon-Lorière, Maud Vanpeene, Sylvie Van der Werf ,GréGoire Potiron |
| EPI_ISL_2030176 | Hospital | National Reference Center for Viruses of Respiratory Infections, Institut Pasteur, Paris | Marion Barbet, Sylvie Behillil, Méline Bizard, Angela Brisebarre, Camille Capel, Vincent Enouf, Louise Lefrançois, Frédéric Lemoine, Christophe Malabat, Corinne Maufrais, Etienne Simon-Lorière, Maud Vanpeene, Sylvie Van der Werf ,Kahla |
| EPI_ISL_2030177 | Hospital | National Reference Center for Viruses of Respiratory Infections, Institut Pasteur, Paris | Marion Barbet, Sylvie Behillil, Méline Bizard, Angela Brisebarre, Camille Capel, Vincent Enouf, Louise Lefrançois, Frédéric Lemoine, Christophe Malabat, Corinne Maufrais, Etienne Simon-Lorière, Maud Vanpeene, Sylvie Van der Werf ,Jean Baptiste Bour |

| | | | |
|---|---|---|---|
| EPI_ISL_2030178 | Labo Analyses Med | National Reference Center for Viruses of Respiratory Infections, Institut Pasteur, Paris | Marion Barbet, Sylvie Behillil, Méline Bizard, Angela Brisebarre, Camille Capel, Vincent Enouf, Louise Lefrançois, Frédéric Lemoine, Christophe Malabat, Corinne Maufrais, Etienne Simon-Lorière, Maud Vanpeene, Sylvie Van der Werf ,Cyrile Bonnet |
| EPI_ISL_2030179 | Hospital | National Reference Center for Viruses of Respiratory Infections, Institut Pasteur, Paris | Marion Barbet, Sylvie Behillil, Méline Bizard, Angela Brisebarre, Camille Capel, Vincent Enouf, Louise Lefrançois, Frédéric Lemoine, Christophe Malabat, Corinne Maufrais, Etienne Simon-Lorière, Maud Vanpeene, Sylvie Van der Werf ,Benjamin Maneglie |
| EPI_ISL_2033158, EPI_ISL_2033160, EPI_ISL_2036092 | Microbiology Department, University Hospital Donostia | Microbiology Department, University Hospital Donostia | Marimon JM, Montes M, Piñeiro L, Sorarrain A, Gomez M, Cilla G. |
| EPI_ISL_2036263, EPI_ISL_2036266, EPI_ISL_2036268 | Laboratori de Referencia de Catalunya | Laboratori de Referencia de Catalunya | Ramirez A., Bellosillo B., Padilla E., Vilas A., Hernandez JJ., Canal M. |
| EPI_ISL_2037414, EPI_ISL_2037443 | Basurto University Hospital: Clinical Microbiology Laboratory | Biocruces Bizkaia | Mikel Urrutikoetxea-Gutiérrez, Mª Carmen Nieto Toboso, Estibaliz Ugalde Zarraga, Mikel Gallego Rodrigo, Ana de la Hoz, José Luis Díaz de Tuesta del Arco |
| EPI_ISL_2038898 | Federal Budget Health Care Institution "Center of Hygiene and Epidemiology in Ryazan region" | Group of Genomics and Postgenomic Technologies of Central Research Institute of Epidemiology | Samoilov AE, Kaptelova VV, Korneenko EV, Saenko SS, Smirnova YS, Nadtoka MI, Baranova SV, Speranskaya AS, Tivanova EV, Kondrasheva LY, Akimkin VG |
| EPI_ISL_2040227, EPI_ISL_2040229 | Cliniques universitaires Saint-Luc | UCLouvain/IREC/MBLG | Jean Ruelle, Eleonore Ngyuvula, Benoit Kabamba Mukadi |
| EPI_ISL_2047871 | Università degli Studi di Perugia | Istituto Zooprofilattico Sperimentale dell'Abruzzo e Molise "G. Caporale" | Mencacci A, Camilloni B,Lorusso A, Marcacci M, Di Domenico M, Ancora M, Curini V, Di Lollo Valeria, Mangone I, Rinaldi A, Delli Compagni E, Scialabba S, Caporale M, Di Pasquale A, Cammà C, Puglia I, Calistri P, Savini G |
| EPI_ISL_2047872 | Laboratorio Analisi Osp. Città di Castello - Azienda USL Umbria1 | Istituto Zooprofilattico Sperimentale dell'Abruzzo e Molise "G. Caporale" | Malagigi V, Tacconi P, Lorusso A, Marcacci M, Di Domenico M, Ancora M, Curini V, Di Lollo Valeria, Mangone I, Rinaldi A, Delli Compagni E, Scialabba S, Caporale M, Di Pasquale A, Cammà C, Puglia I, Calistri P, Savini G |
| EPI_ISL_2080934 | Department of Virology, Istituto Zooprofilattico Sperimentale del Lazio e della Toscana (IZSLT) | Department of General Diagnostics; Department of Virology; Istituto Zooprofilattico Sperimentale del Lazio e della Toscana (IZSLT) | Patricia Alba, Giuseppe Manna, Elena L. Diaconu, Fabiola Feltrin, Raffaela Conti, Teresa Scicluna, Virginia Carfora, Antonella Cersini, Alessia Franco, Antonio Battisti. |
| EPI_ISL_2083016, EPI_ISL_2083017, EPI_ISL_2083018, EPI_ISL_2083019, EPI_ISL_2083020, EPI_ISL_2083022, EPI_ISL_2083025, EPI_ISL_2083026, EPI_ISL_2083027, EPI_ISL_2083028, EPI_ISL_2083029, EPI_ISL_2083030, EPI_ISL_2083031, EPI_ISL_2083032 | | | |
| see above | Baylor Scott & White-Temple | Baylor Scott & White-Temple | Ari Rao, Linden Morales, Kimberly Walker, Marcus Volz, Shelby Hendrickson |
| EPI_ISL_2083033 | Baylor Scott & White-Grapevine | Baylor Scott & White-Temple | Ari Rao, Linden Morales, Kimberly Walker, Marcus Volz, Shelby Hendrickson |
| EPI_ISL_2083034, EPI_ISL_2083035, EPI_ISL_2083036, EPI_ISL_2083037, EPI_ISL_2083038, EPI_ISL_2083039, EPI_ISL_2083040, EPI_ISL_2083041, EPI_ISL_2083042, EPI_ISL_2083043, EPI_ISL_2083046, EPI_ISL_2083047, EPI_ISL_2083048, EPI_ISL_2083049, EPI_ISL_2083052, EPI_ISL_2083055, EPI_ISL_2083056, EPI_ISL_2083057, EPI_ISL_2083058, EPI_ISL_2083059, EPI_ISL_2083060, EPI_ISL_2083061, EPI_ISL_2083062, EPI_ISL_2083063, EPI_ISL_2083064, EPI_ISL_2083065, EPI_ISL_2083067, EPI_ISL_2083068, EPI_ISL_2083069, EPI_ISL_2083070, EPI_ISL_2083071, EPI_ISL_2083072, EPI_ISL_2083073, EPI_ISL_2083074, EPI_ISL_2083076, EPI_ISL_2083077, EPI_ISL_2083079, EPI_ISL_2083080, EPI_ISL_2083081, EPI_ISL_2083082, EPI_ISL_2083083, EPI_ISL_2083084, EPI_ISL_2083085 | | | |
| see above | Baylor Scott & White-Temple | Baylor Scott & White-Temple | Ari Rao, Linden Morales, Kimberly Walker, Marcus Volz, Shelby Hendrickson |
| EPI_ISL_2086138, EPI_ISL_2086139, EPI_ISL_2086140, EPI_ISL_2086141, EPI_ISL_2086142, EPI_ISL_2086143, EPI_ISL_2086144, EPI_ISL_2086145, EPI_ISL_2086146, EPI_ISL_2086147, EPI_ISL_2086148, EPI_ISL_2086149, EPI_ISL_2086151, EPI_ISL_2086152, EPI_ISL_2086153, EPI_ISL_2086154, EPI_ISL_2086155, EPI_ISL_2086156, EPI_ISL_2086157, EPI_ISL_2086158, EPI_ISL_2086159, EPI_ISL_2086160, EPI_ISL_2086162, EPI_ISL_2086163, EPI_ISL_2086164, EPI_ISL_2086165 | | | |
| see above | Microbiology Department. Complexo Hospitalario Universitario de Vigo | Microbiology Department. Complexo Hospitalario Universitario de Vigo | Alfaya N, Alonso I, Alvarez M, Cabrera JJ, Carballo R, Cores O, Cortizo S, del-Campo V, Martinez L, Mediero G, Perez S, Potel C, Regueiro B, Rey S, Vasallo FJ |
| EPI_ISL_2086166 | VITHAS Lab | Microbiology Department. Complexo Hospitalario Universitario de Vigo | Camacho AT, Perez S, Regueiro B |
| EPI_ISL_2086168, EPI_ISL_2086169, EPI_ISL_2086170, EPI_ISL_2086171, EPI_ISL_2086172, EPI_ISL_2086173, EPI_ISL_2086174 | Microbiology Department. Complexo Hospitalario Universitario de Vigo | Microbiology Department. Complexo Hospitalario Universitario de Vigo | Alfaya N, Alonso I, Alvarez M, Cabrera JJ, Carballo R, Cores O, Cortizo S, del-Campo V, Martinez L, Mediero G, Perez S, Potel C, Regueiro B, Rey S, Vasallo FJ |
| EPI_ISL_2086292, EPI_ISL_2086293, EPI_ISL_2086294 | Laboratori de Referencia de Catalunya | Laboratori de Referencia de Catalunya | Ramirez A., Bellosillo B., Padilla E., Vilas A., Hernandez JJ., Canal M. |
| EPI_ISL_2090772 | CERBALLIANCE | UMR PIMIT | Dr David A Wilkinson, Dr Patrick Mavingui, Dr Camille Lebarbenchon, Magali Turpin |
| EPI_ISL_2090779 | REUNILAB | UMR PIMIT | Dr David A Wilkinson, Dr Patrick Mavingui, Dr Camille Lebarbenchon, Magali Turpin |
| EPI_ISL_2090781 | CERBALLIANCE | UMR PIMIT | Dr David A Wilkinson, Dr Patrick Mavingui, Dr Camille Lebarbenchon, Magali Turpin |
| EPI_ISL_2090784, EPI_ISL_2090789, EPI_ISL_2090792, EPI_ISL_2090796, EPI_ISL_2090797 | REUNILAB | UMR PIMIT | Dr David A Wilkinson, Dr Patrick Mavingui, Dr Camille Lebarbenchon, Magali Turpin |
| EPI_ISL_2090801 | CERBALLIANCE | UMR PIMIT | Dr David A Wilkinson, Dr Patrick Mavingui, Dr Camille Lebarbenchon, Magali Turpin |
| EPI_ISL_2090802, EPI_ISL_2090808, EPI_ISL_2090809, EPI_ISL_2090872 | REUNILAB | UMR PIMIT | Dr David A Wilkinson, Dr Patrick Mavingui, Dr Camille Lebarbenchon, Magali Turpin |
| EPI_ISL_2090886, EPI_ISL_2090887, EPI_ISL_2090889, EPI_ISL_2090890, EPI_ISL_2091000 | Institute for Medical Research, Infectious Disease Research Centre, National Institutes of Health, Ministry of Health Malaysia | Institute for Medical Research, Infectious Disease Research Centre, National Institutes of Health, Ministry of Health Malaysia | Suppiah J, Kamel K, Mohd Zawawi Z, Azizan MA, Ramly N, Robert F, Thayan R |
| EPI_ISL_2091014 | Hospital Universitario Cruces | Biocruces Bizkaia | Mikel Gallego, Ana Belen de la Hoz |
| EPI_ISL_2091016 | Laboratorio Biologia Molecolare Sars Cov2 - UOC Laboratorio Analisi - Servizio Medicina di Laboratorio, Ospedale "San Francesco" - ATS-ASSL Nuoro | Laboratorio specialistico UOC Ematologia - Ospedale "San Francesco" - ATS-ASSL Nuoro | Giovanna Piras, Rosanna Asproni,Paolo Malune,Franco Carta,Pira Giovanna,Fancello Patrizia, Rosu Valentina, Maura Fiamma, Maria Itria Monne, Maria Cristina Garau, Giuseppe Mameli |
| EPI_ISL_2091020 | Sungai Buloh Hospital | Institute for Medical Research, Infectious Disease Research Centre, National Institutes of Health, Ministry of Health Malaysia | Suppiah J, Kamel K, Mohd Zawawi Z, Azizan MA, Ramly N, Robert F, Thayan R |
| EPI_ISL_2091022 | Johor Bahru Public Health Laboratory | Institute for Medical Research, Infectious Disease Research Centre, National Institutes of Health, Ministry of Health Malaysia | Suppiah J, Kamel K, Mohd Zawawi Z, Azizan MA, Ramly N, Robert F, Thayan R |
| EPI_ISL_2091024 | Sultanah Aminah Hospital, Johor Bahru | Institute for Medical Research, Infectious Disease Research Centre, National Institutes of Health, Ministry of Health Malaysia | Suppiah J, Kamel K, Mohd Zawawi Z, Azizan MA, Ramly N, Robert F, Thayan R |
| EPI_ISL_2091121, EPI_ISL_2091122, EPI_ISL_2091123, EPI_ISL_2091124, EPI_ISL_2091125, EPI_ISL_2091126, EPI_ISL_2091127, EPI_ISL_2091129, EPI_ISL_2091130 | Laboratorio di Genetica Medica, Matera, Italy | Laboratorio di Genetica Medica, Matera, Italy | Domenico Dell'Edera, Arianna Allegretti |
| EPI_ISL_2091200, EPI_ISL_2091201, EPI_ISL_2091202, EPI_ISL_2091203, EPI_ISL_2091204, EPI_ISL_2091205, EPI_ISL_2091206, EPI_ISL_2091207, EPI_ISL_2091208, EPI_ISL_2091209, EPI_ISL_2091210, EPI_ISL_2091211, EPI_ISL_2091212, EPI_ISL_2091214, EPI_ISL_2091215, EPI_ISL_2091218, EPI_ISL_2091219, EPI_ISL_2091220, EPI_ISL_2091221, EPI_ISL_2091222, EPI_ISL_2091223, EPI_ISL_2091224 | | | |

| | | | |
|---|---|---|---|
| see above | Centro de Investigación Biomédica del Noreste (CIBIN) | Centro de Investigación en Enfermedades Infecciosas (CIENI), Instituto Nacional de Enfermedades Respiratorias (INER) | Consorcio Mexicano de Vigilancia Genómica (CoViGen-Mex). Authors (in alphabetical order): Julio Elias Alvarado-Yaah, Carlos F. Arias, Santiago Ávila-Ríos, Víctor Hugo Borja-Aburto, Celia Boukadida, Juan Bautista Chale-Dzul, Célida Duque Molina, José Antonio Enciso-Moreno, Gloria Elena Espinosa-Ayala, Fernando Fontove-Herrera, Victor Eduardo Garcia-Arias, Concepción Grajales-Muñiz, Ricardo Grande, Alfredo Herrera-Estrella, Carla Ivón Herrera-Najera, Pavel Isa, Brenda Irasema Maldonado-Meza, Bernardo Martínez-Miguel, Margarita Matías-Florentino, María Guadalupe de Jesús Mireles-Rivera, Gloria María Molina-Salinas, Hector Montoya-Fuentes, José Esteban Muñoz-Medina, José de Jesús Nuñez-Contreras, Alicia Ocaña-Mondragón, Luis Alberto Ochoa-Carrera, Hector Esteban Paz-Juárez, Francisco Pulido, Helen Haydee Fernanda Ramírez-Plascencia, Angel Gustavo Salas-Lais, Alejandro Sanchez-Flores, Clara Esperanza Santacruz-Tinoco, María Guadalupe Santiago-Mauricio, Nelly Sélem-Mojica, Blanca Taboada, Gloria Vazquez |
| EPI_ISL_2091247, EPI_ISL_2091251, EPI_ISL_2091252, EPI_ISL_2091253, EPI_ISL_2091254, EPI_ISL_2091255, EPI_ISL_2091256, EPI_ISL_2091257 | Unidad de Investigación Médica de Yucatán (UIMY) | Centro de Investigación en Enfermedades Infecciosas (CIENI), Instituto Nacional de Enfermedades Respiratorias (INER) | Consorcio Mexicano de Vigilancia Genómica (CoViGen-Mex). Authors (in alphabetical order): Julio Elias Alvarado-Yaah, Carlos F. Arias, Santiago Ávila-Ríos, Víctor Hugo Borja-Aburto, Celia Boukadida, Juan Bautista Chale-Dzul, Célida Duque Molina, José Antonio Enciso-Moreno, Gloria Elena Espinosa-Ayala, Fernando Fontove-Herrera, Victor Eduardo Garcia-Arias, Concepción Grajales-Muñiz, Ricardo Grande, Alfredo Herrera-Estrella, Carla Ivón Herrera-Najera, Pavel Isa, Brenda Irasema Maldonado-Meza, Bernardo Martínez-Miguel, Margarita Matías-Florentino, María Guadalupe de Jesús Mireles-Rivera, Gloria María Molina-Salinas, Hector Montoya-Fuentes, José Esteban Muñoz-Medina, José de Jesús Nuñez-Contreras, Alicia Ocaña-Mondragón, Luis Alberto Ochoa-Carrera, Hector Esteban Paz-Juárez, Francisco Pulido, Helen Haydee Fernanda Ramírez-Plascencia, Angel Gustavo Salas-Lais, Alejandro Sanchez-Flores, Clara Esperanza Santacruz-Tinoco, María Guadalupe Santiago-Mauricio, Nelly Sélem-Mojica, Blanca Taboada, Gloria Vazquez |
| EPI_ISL_2091281, EPI_ISL_2091282, EPI_ISL_2091284, EPI_ISL_2091285, EPI_ISL_2091286, EPI_ISL_2091289, EPI_ISL_2091292, EPI_ISL_2091293, EPI_ISL_2091294, EPI_ISL_2091295, EPI_ISL_2091296, EPI_ISL_2091297, EPI_ISL_2091299, EPI_ISL_2091301, EPI_ISL_2091302, EPI_ISL_2091303, EPI_ISL_2091304, EPI_ISL_2091305, EPI_ISL_2091307, EPI_ISL_2091308, EPI_ISL_2091309, EPI_ISL_2091310, EPI_ISL_2091311, EPI_ISL_2091312, EPI_ISL_2091313, EPI_ISL_2091314, EPI_ISL_2091315, EPI_ISL_2091316, EPI_ISL_2091317, EPI_ISL_2091318, EPI_ISL_2091319 | | | |
| see above | Centro de Investigación Biomédica de Occidente (CIBO) | Centro de Investigación en Enfermedades Infecciosas (CIENI), Instituto Nacional de Enfermedades Respiratorias (INER) | Consorcio Mexicano de Vigilancia Genómica (CoViGen-Mex). Authors (in alphabetical order): Julio Elias Alvarado-Yaah, Carlos F. Arias, Santiago Ávila-Ríos, Víctor Hugo Borja-Aburto, Celia Boukadida, Juan Bautista Chale-Dzul, Célida Duque Molina, José Antonio Enciso-Moreno, Gloria Elena Espinosa-Ayala, Fernando Fontove-Herrera, Victor Eduardo Garcia-Arias, Concepción Grajales-Muñiz, Ricardo Grande, Alfredo Herrera-Estrella, Carla Ivón Herrera-Najera, Pavel Isa, Brenda Irasema Maldonado-Meza, Bernardo Martínez-Miguel, Margarita Matías-Florentino, María Guadalupe de Jesús Mireles-Rivera, Gloria María Molina-Salinas, Hector Montoya-Fuentes, José Esteban Muñoz-Medina, José de Jesús Nuñez-Contreras, Alicia Ocaña-Mondragón, Luis Alberto Ochoa-Carrera, Hector Esteban Paz-Juárez, Francisco Pulido, Helen Haydee Fernanda Ramírez-Plascencia, Angel Gustavo Salas-Lais, Alejandro Sanchez-Flores, Clara Esperanza Santacruz-Tinoco, María Guadalupe Santiago-Mauricio, Nelly Sélem-Mojica, Blanca Taboada, Gloria Vazquez |
| EPI_ISL_2091344, EPI_ISL_2091345, EPI_ISL_2091346, EPI_ISL_2091347, EPI_ISL_2091349, EPI_ISL_2091350, EPI_ISL_2091351, EPI_ISL_2091352, EPI_ISL_2091353, EPI_ISL_2091361, EPI_ISL_2091362, EPI_ISL_2091363, EPI_ISL_2091364, EPI_ISL_2091365, EPI_ISL_2091366, EPI_ISL_2091367, EPI_ISL_2091368, EPI_ISL_2091369, EPI_ISL_2091370, EPI_ISL_2091371, EPI_ISL_2091372, EPI_ISL_2091373, EPI_ISL_2091374, EPI_ISL_2091375, EPI_ISL_2091376, EPI_ISL_2091377, EPI_ISL_2091378, EPI_ISL_2091379, EPI_ISL_2091381, EPI_ISL_2091382, EPI_ISL_2091383, EPI_ISL_2091384, EPI_ISL_2091385, EPI_ISL_2091386, EPI_ISL_2091387, EPI_ISL_2091388, EPI_ISL_2091389, EPI_ISL_2091390, EPI_ISL_2091391, EPI_ISL_2091392, EPI_ISL_2091393, EPI_ISL_2091394, EPI_ISL_2091395, EPI_ISL_2091396, EPI_ISL_2091397, EPI_ISL_2091398, EPI_ISL_2091399, EPI_ISL_2091400, EPI_ISL_2091401, EPI_ISL_2091402, EPI_ISL_2091403, EPI_ISL_2091404, EPI_ISL_2091405, EPI_ISL_2091406, EPI_ISL_2091407, EPI_ISL_2091408, EPI_ISL_2091409, EPI_ISL_2091410, EPI_ISL_2091411, EPI_ISL_2091412, EPI_ISL_2091414, EPI_ISL_2091415, EPI_ISL_2091416, EPI_ISL_2091417, EPI_ISL_2091418, EPI_ISL_2091419, EPI_ISL_2091420, EPI_ISL_2091421, EPI_ISL_2091422, EPI_ISL_2091423, EPI_ISL_2091424, EPI_ISL_2091425, EPI_ISL_2091426, EPI_ISL_2091427, EPI_ISL_2091428, EPI_ISL_2091429, EPI_ISL_2091430, EPI_ISL_2091431, EPI_ISL_2091432, EPI_ISL_2091433, EPI_ISL_2091434, EPI_ISL_2091435, EPI_ISL_2091436, EPI_ISL_2091437, EPI_ISL_2091438, EPI_ISL_2091439, EPI_ISL_2091442, EPI_ISL_2091443, EPI_ISL_2091444, EPI_ISL_2091445, EPI_ISL_2091446, EPI_ISL_2091447, EPI_ISL_2091448, EPI_ISL_2091449, EPI_ISL_2091450, EPI_ISL_2091451, EPI_ISL_2091452, EPI_ISL_2091453, EPI_ISL_2091454, EPI_ISL_2091455, EPI_ISL_2091456 | | | |
| see above | Laboratorio Central de Epidemiología (LCE) | Centro de Investigación en Enfermedades Infecciosas (CIENI), Instituto Nacional de Enfermedades Respiratorias (INER) | Consorcio Mexicano de Vigilancia Genómica (CoViGen-Mex). Authors (in alphabetical order): Julio Elias Alvarado-Yaah, Carlos F. Arias, Santiago Ávila-Ríos, Víctor Hugo Borja-Aburto, Celia Boukadida, Juan Bautista Chale-Dzul, Célida Duque Molina, José Antonio Enciso-Moreno, Gloria Elena Espinosa-Ayala, Fernando Fontove-Herrera, Victor Eduardo Garcia-Arias, Concepción Grajales-Muñiz, Ricardo Grande, Alfredo Herrera-Estrella, Carla Ivón Herrera-Najera, Pavel Isa, Brenda Irasema Maldonado-Meza, Bernardo Martínez-Miguel, Margarita Matías-Florentino, María Guadalupe de Jesús Mireles-Rivera, Gloria María Molina-Salinas, Hector Montoya-Fuentes, José Esteban Muñoz-Medina, José de Jesús Nuñez-Contreras, Alicia Ocaña-Mondragón, Luis Alberto Ochoa-Carrera, Hector Esteban Paz-Juárez, Francisco Pulido, Helen Haydee Fernanda Ramírez-Plascencia, Angel Gustavo Salas-Lais, Alejandro Sanchez-Flores, Clara Esperanza Santacruz-Tinoco, María Guadalupe Santiago-Mauricio, Nelly Sélem-Mojica, Blanca Taboada, Gloria Vazquez |
| EPI_ISL_2094505 | LESP Jalisco | Instituto de Diagnostico y Referencia Epidemiologicos (INDRE) | Claudia Wong-Arambula, Abril Rodriguez-Maldonado, Vanessa Rivero-Arredondo, Ariadna Medina-Benitez, Joaquin Quiroz-Mercado, Sergio Rangel-Guerrero, Natividad Cruz-Ortiz, Tatiana Nunez-Garcia, Gisela Barrera-Badillo, Lucia Hernandez-Rivas, Irma Lopez-Martinez, Ernesto Ramirez-Gonzalez. |
| EPI_ISL_2095848, EPI_ISL_2095849, EPI_ISL_2095850 | Klaipedos universitetine ligonine | National Public Health Surveillance Laboratory | Lukas Zemaitis, Migle Gabrielaite, Jelena Razmuk, Svajune Muralyte, Ana Steponkiene, Lukas Vasionis, Danas Baksa |
| EPI_ISL_2095852, EPI_ISL_2095858, EPI_ISL_2095860 | RezusLT | National Public Health Surveillance Laboratory | Lukas Zemaitis, Migle Gabrielaite, Jelena Razmuk, Svajune Muralyte, Ana Steponkiene, Lukas Vasionis, Danas Baksa |
| EPI_ISL_2095877, EPI_ISL_2095878, EPI_ISL_2095879, EPI_ISL_2095880, EPI_ISL_2095881, EPI_ISL_2095882, EPI_ISL_2095883 | Vilniaus universitetas | National Public Health Surveillance Laboratory | Lukas Zemaitis, Migle Gabrielaite, Jelena Razmuk, Svajune Muralyte, Ana Steponkiene, Lukas Vasionis, Danas Baksa |
| EPI_ISL_2095885, EPI_ISL_2095886 | Klaipedos universitetine ligonine | National Public Health Surveillance Laboratory | Lukas Zemaitis, Migle Gabrielaite, Jelena Razmuk, Svajune Muralyte, Ana Steponkiene, Lukas Vasionis, Danas Baksa |
| EPI_ISL_2095888, EPI_ISL_2095889, EPI_ISL_2095895 | RezusLT | National Public Health Surveillance Laboratory | Lukas Zemaitis, Migle Gabrielaite, Jelena Razmuk, Svajune Muralyte, Ana Steponkiene, Lukas Vasionis, Danas Baksa |
| EPI_ISL_2095896, EPI_ISL_2095897 | Vilniaus universitetas | National Public Health Surveillance Laboratory | Lukas Zemaitis, Migle Gabrielaite, Jelena Razmuk, Svajune Muralyte, Ana Steponkiene, Lukas Vasionis, Danas Baksa |
| EPI_ISL_2095901, EPI_ISL_2095902 | Klaipedos universitetine ligonine | National Public Health Surveillance Laboratory | Lukas Zemaitis, Migle Gabrielaite, Jelena Razmuk, Svajune Muralyte, Ana Steponkiene, Lukas Vasionis, Danas Baksa |
| EPI_ISL_2095905, EPI_ISL_2095906 | RezusLT | National Public Health Surveillance Laboratory | Lukas Zemaitis, Migle Gabrielaite, Jelena Razmuk, Svajune Muralyte, Ana Steponkiene, Lukas Vasionis, Danas Baksa |
| EPI_ISL_2097003, EPI_ISL_2097004, EPI_ISL_2097005, EPI_ISL_2097006, EPI_ISL_2097007, EPI_ISL_2097008, EPI_ISL_2097009, EPI_ISL_2097036, EPI_ISL_2097038, EPI_ISL_2097039, EPI_ISL_2097040, EPI_ISL_2097080, EPI_ISL_2097083, EPI_ISL_2097084, EPI_ISL_2097085, EPI_ISL_2097086, EPI_ISL_2097087, EPI_ISL_2097088, EPI_ISL_2097089, EPI_ISL_2097090, EPI_ISL_2097091, EPI_ISL_2097092, EPI_ISL_2097093, EPI_ISL_2097094, EPI_ISL_2097095, EPI_ISL_2097097, EPI_ISL_2097098, EPI_ISL_2097099, EPI_ISL_2097101, EPI_ISL_2097102, EPI_ISL_2097103, EPI_ISL_2097104, EPI_ISL_2097105, EPI_ISL_2097106, EPI_ISL_2097107, EPI_ISL_2097108, EPI_ISL_2097109, EPI_ISL_2097110, EPI_ISL_2097111, EPI_ISL_2097112, EPI_ISL_2097113, EPI_ISL_2097114, EPI_ISL_2097116, EPI_ISL_2097117, EPI_ISL_2097118, EPI_ISL_2097119, EPI_ISL_2097120, EPI_ISL_2097122, EPI_ISL_2097123, EPI_ISL_2097124, EPI_ISL_2097126, EPI_ISL_2097127, EPI_ISL_2097128, EPI_ISL_2097129, EPI_ISL_2097130, EPI_ISL_2097131, EPI_ISL_2097132, EPI_ISL_2097133, EPI_ISL_2097134, EPI_ISL_2097135, EPI_ISL_2097136, EPI_ISL_2097137, EPI_ISL_2097138, EPI_ISL_2097139, EPI_ISL_2097140, EPI_ISL_2097141, EPI_ISL_2097142, EPI_ISL_2097143, EPI_ISL_2097144, EPI_ISL_2097145, EPI_ISL_2097146, EPI_ISL_2097147, EPI_ISL_2097149, EPI_ISL_2097151, EPI_ISL_2097152, EPI_ISL_2097153, EPI_ISL_2097154, EPI_ISL_2097155, EPI_ISL_2097157, EPI_ISL_2097158, EPI_ISL_2097159, EPI_ISL_2097161, EPI_ISL_2097163, EPI_ISL_2097164, EPI_ISL_2097165, EPI_ISL_2097166, EPI_ISL_2097167 | | | |
| see above | Synlab Eesti OÜ | 1. Laboratory of Communicable Diseases (Estonia); 2. Eurofins Genomics Europe Sequencing GmbH | Liidia Dotsenko et al. |
| EPI_ISL_2097315, EPI_ISL_2097316 | Cliniques universitaires Saint-Luc | UCLouvain/IREC/MBLG | Jean Ruelle, Eleonore Ngyuvula, Benoit Kabamba Mukadi |
| EPI_ISL_2098710 | State Virus Research and Diagnostic Laboratory (VRDL), AIIMS Raipur | State Virus Research and Diagnostic Laboratory (VRDL), AIIMS Raipur | Pushpendra Singh, Kuldeep Sharma, Priyanka Singh, Somya Sharma, MD Rafiiqullah Khan, Jiyan Chandrawanshi, Sanjay Singh Negi, Anudita Bhargava |
| EPI_ISL_2098949 | Laboratorio Biologia Molecolare Sars Cov2 - UOC Laboratorio Analisi - Servizio Medicina di Laboratorio, Ospedale "San Francesco" - ATS-ASSL Nuoro | Laboratorio specialistico UOC Ematologia - Ospedale "San Francesco" - ATS-ASSL Nuoro | Piras Giovanna, Asproni Rosanna,Paolo Malune, , Monne Maria Itria, Palmas Angelo Domenico, Fiamma Maura, Garau Maria Cristina, Mameli Giuseppe |
| EPI_ISL_2099337, EPI_ISL_2099422, EPI_ISL_2099427, EPI_ISL_2099428, EPI_ISL_2099430, EPI_ISL_2099431, EPI_ISL_2099432, EPI_ISL_2099433, EPI_ISL_2099434, EPI_ISL_2099435, EPI_ISL_2099436, EPI_ISL_2099437, EPI_ISL_2099438, EPI_ISL_2099439, EPI_ISL_2099440, EPI_ISL_2099441, EPI_ISL_2099442, EPI_ISL_2099443, EPI_ISL_2099444, EPI_ISL_2099445, EPI_ISL_2099446, EPI_ISL_2099447, EPI_ISL_2099448, EPI_ISL_2099449, EPI_ISL_2099450, EPI_ISL_2099451, EPI_ISL_2099452, EPI_ISL_2099482, EPI_ISL_2099484, EPI_ISL_2099488, EPI_ISL_2099489, EPI_ISL_2099490, EPI_ISL_2099491, EPI_ISL_2099492, EPI_ISL_2099493, EPI_ISL_2099494, EPI_ISL_2099495, EPI_ISL_2099496, EPI_ISL_2099497 | | | |
| see above | National Center of Infectious and Parasitic Diseases | National Center of Infectious and Parasitic Diseases | Alexiev, Ivanov, Korsun, Stoitsova, Philipova, Dimitrova, Grigorova L., Hristova, Donchev, Stoykov, Trifonova, Dobrinov, Grigorova I., Kantardjiev |
| EPI_ISL_2099853 | National Institute of Infectious Diseases-Prof. Dr. Matei Bals | National Institute of Infectious Diseases-Prof. Dr. Matei Bals | Corina Casangiu, Leontina Banica, Marius Surleac, Ovidiu Vlaicu, Andreea Tudor, Simona Paraschiv, Dan Otelea |

| | | | |
|---|---|---|---|
| | Molecular Diagnostics Laboratory | Molecular Diagnostics Laboratory | |
| EPI_ISL_2100220, EPI_ISL_2100221, EPI_ISL_2100222, EPI_ISL_2100223 | Laboratori de Referencia de Catalunya | Laboratori de Referencia de Catalunya | Ramirez A., Bellosillo B., Padilla E., Vilas A., Hernandez JJ., Canal M. |
| EPI_ISL_2100252, EPI_ISL_2100253 | Imelda Ziekenhuis | Imelda Ziekenhuis | Johan Frans, Hanne Valgaeren, Dagmar Obbels |
| EPI_ISL_2100475, EPI_ISL_2100476, EPI_ISL_2100477, EPI_ISL_2100478, EPI_ISL_2100479, EPI_ISL_2100480, EPI_ISL_2100481, EPI_ISL_2100482, EPI_ISL_2100483, EPI_ISL_2100484, EPI_ISL_2100485, EPI_ISL_2100486, EPI_ISL_2100487, EPI_ISL_2100488, EPI_ISL_2100489, EPI_ISL_2100490, EPI_ISL_2100491, EPI_ISL_2100492, EPI_ISL_2100493, EPI_ISL_2100494, EPI_ISL_2100496, EPI_ISL_2100497, EPI_ISL_2100498, EPI_ISL_2100499, EPI_ISL_2100500, EPI_ISL_2100501, EPI_ISL_2100502, EPI_ISL_2100503, EPI_ISL_2100504, EPI_ISL_2100505, EPI_ISL_2100506, EPI_ISL_2100507, EPI_ISL_2100508, EPI_ISL_2100509 | | | |
| see above | University of Health Sciences | Quadram Institute Bioscience | Muhammad Bilal Sarwar, Sidra-tul-muntaha, Muhammad Roman, Alam Khan, Almina Shafiq, Nadeem Afzal, Shah Jahan, Javed Akram, Dave J. Baker, Gemma L. Kay, Alp Aydin, Thanh Le-Viet, Steven Rudder, Ana P. Tedim, Anastasia Kolyva, Maria Diaz, Leonardo de Oliveira Martins, Nabil-Fareed Alikhan, Lizzie Meadows, Rachael Stanley, Ngozi Elumogo, Muhammed Yasir, Nicholas M. Thomson, Alexander J Trotter, Rachel Gilroy, Samuel Bloomfield, Claire Stuart, Andrew Bell, Reenesh Prakash, Samir Dervisevic, Alison E. Mather, John Wain, Mark Webber, Andrew J. Page, Justin O'Grady |
| EPI_ISL_2100679, EPI_ISL_2100681, EPI_ISL_2100682, EPI_ISL_2100687, EPI_ISL_2100693, EPI_ISL_2100700 | All India Institute of Medical Sciences, Ansari Nagar Delhi | CSIR-Institute of Genomics and Integrative Biology | Pooja Sharma*, Bharathram Uppilli*, Animesh Ray, Sushma Rajpoot, Saruchi Wadhwa, Asangala Kamai, Devasish Desai, Manish Kumar, Satish Swain, Sarafaraz Alam, Umang Arora, Sheeba Saifi, Mohammed Ahmed, Ayush Goel, Ashwin Varadarajan, Naveet Wig, Rajesh Pandey, Mohammed Faruq |
| EPI_ISL_2100871, EPI_ISL_2100872, EPI_ISL_2100873, EPI_ISL_2100874, EPI_ISL_2100875, EPI_ISL_2100876, EPI_ISL_2100877, EPI_ISL_2100878, EPI_ISL_2100879, EPI_ISL_2100966, EPI_ISL_2100968 | | | |
| see above | The Ohio State University Applied Microbiology Services Laboratory | The Ohio State University Applied Microbiology Services Laboratory | Seth A. Faith PhD |
| EPI_ISL_2101870 | LESP Morelos | Instituto de Diagnostico y Referencia Epidemiologicos (INDRE) | Claudia Wong-Arambula, Abril Rodriguez-Maldonado, Vanessa Rivero-Arredondo, Ariadna Medina-Benitez, Joaquin Quiroz-Mercado, Sergio Rangel-Guerrero, Natividad Cruz-Ortiz, Tatiana Nunez-Garcia, Gisela Barrera-Badillo, Lucia Hernandez-Rivas, Irma Lopez-Martinez, Ernesto Ramirez-Gonzalez. |
| EPI_ISL_2101871, EPI_ISL_2101872, EPI_ISL_2101873, EPI_ISL_2101874, EPI_ISL_2101875, EPI_ISL_2101876, EPI_ISL_2101878 | LESP Chihuahua | Instituto de Diagnostico y Referencia Epidemiologicos (INDRE) | Claudia Wong-Arambula, Abril Rodriguez-Maldonado, Vanessa Rivero-Arredondo, Ariadna Medina-Benitez, Joaquin Quiroz-Mercado, Sergio Rangel-Guerrero, Natividad Cruz-Ortiz, Tatiana Nunez-Garcia, Gisela Barrera-Badillo, Lucia Hernandez-Rivas, Irma Lopez-Martinez, Ernesto Ramirez-Gonzalez. |
| EPI_ISL_2101879 | LESP Quintana Roo | Instituto de Diagnostico y Referencia Epidemiologicos (INDRE) | Claudia Wong-Arambula, Abril Rodriguez-Maldonado, Vanessa Rivero-Arredondo, Ariadna Medina-Benitez, Joaquin Quiroz-Mercado, Sergio Rangel-Guerrero, Natividad Cruz-Ortiz, Tatiana Nunez-Garcia, Gisela Barrera-Badillo, Lucia Hernandez-Rivas, Irma Lopez-Martinez, Ernesto Ramirez-Gonzalez. |
| EPI_ISL_2101880 | LESP Baja California | Instituto de Diagnostico y Referencia Epidemiologicos (INDRE) | Claudia Wong-Arambula, Abril Rodriguez-Maldonado, Vanessa Rivero-Arredondo, Ariadna Medina-Benitez, Joaquin Quiroz-Mercado, Sergio Rangel-Guerrero, Natividad Cruz-Ortiz, Tatiana Nunez-Garcia, Gisela Barrera-Badillo, Lucia Hernandez-Rivas, Irma Lopez-Martinez, Ernesto Ramirez-Gonzalez. |
| EPI_ISL_2101881, EPI_ISL_2101882, EPI_ISL_2101883, EPI_ISL_2101884, EPI_ISL_2101887, EPI_ISL_2101888, EPI_ISL_2101890 | LESP Chihuahua | Instituto de Diagnostico y Referencia Epidemiologicos (INDRE) | Claudia Wong-Arambula, Abril Rodriguez-Maldonado, Vanessa Rivero-Arredondo, Ariadna Medina-Benitez, Joaquin Quiroz-Mercado, Sergio Rangel-Guerrero, Natividad Cruz-Ortiz, Tatiana Nunez-Garcia, Gisela Barrera-Badillo, Lucia Hernandez-Rivas, Irma Lopez-Martinez, Ernesto Ramirez-Gonzalez. |
| EPI_ISL_2101891 | LESP Jalisco | Instituto de Diagnostico y Referencia Epidemiologicos (INDRE) | Claudia Wong-Arambula, Abril Rodriguez-Maldonado, Vanessa Rivero-Arredondo, Ariadna Medina-Benitez, Joaquin Quiroz-Mercado, Sergio Rangel-Guerrero, Natividad Cruz-Ortiz, Tatiana Nunez-Garcia, Gisela Barrera-Badillo, Lucia Hernandez-Rivas, Irma Lopez-Martinez, Ernesto Ramirez-Gonzalez. |
| EPI_ISL_2101892 | LESP Veracruz | Instituto de Diagnostico y Referencia Epidemiologicos (INDRE) | Claudia Wong-Arambula, Abril Rodriguez-Maldonado, Vanessa Rivero-Arredondo, Ariadna Medina-Benitez, Joaquin Quiroz-Mercado, Sergio Rangel-Guerrero, Natividad Cruz-Ortiz, Tatiana Nunez-Garcia, Gisela Barrera-Badillo, Lucia Hernandez-Rivas, Irma Lopez-Martinez, Ernesto Ramirez-Gonzalez. |
| EPI_ISL_2101893, EPI_ISL_2101894 | LESP Guanajuato | Instituto de Diagnostico y Referencia Epidemiologicos (INDRE) | Claudia Wong-Arambula, Abril Rodriguez-Maldonado, Vanessa Rivero-Arredondo, Ariadna Medina-Benitez, Joaquin Quiroz-Mercado, Sergio Rangel-Guerrero, Natividad Cruz-Ortiz, Tatiana Nunez-Garcia, Gisela Barrera-Badillo, Lucia Hernandez-Rivas, Irma Lopez-Martinez, Ernesto Ramirez-Gonzalez. |
| EPI_ISL_2101895 | LESP Jalisco | Instituto de Diagnostico y Referencia Epidemiologicos (INDRE) | Claudia Wong-Arambula, Abril Rodriguez-Maldonado, Vanessa Rivero-Arredondo, Ariadna Medina-Benitez, Joaquin Quiroz-Mercado, Sergio Rangel-Guerrero, Natividad Cruz-Ortiz, Tatiana Nunez-Garcia, Gisela Barrera-Badillo, Lucia Hernandez-Rivas, Irma Lopez-Martinez, Ernesto Ramirez-Gonzalez. |
| EPI_ISL_2101896 | LESP Sinaloa | Instituto de Diagnostico y Referencia Epidemiologicos (INDRE) | Claudia Wong-Arambula, Abril Rodriguez-Maldonado, Vanessa Rivero-Arredondo, Ariadna Medina-Benitez, Joaquin Quiroz-Mercado, Sergio Rangel-Guerrero, Natividad Cruz-Ortiz, Tatiana Nunez-Garcia, Gisela Barrera-Badillo, Lucia Hernandez-Rivas, Irma Lopez-Martinez, Ernesto Ramirez-Gonzalez. |
| EPI_ISL_2101897 | LESP Sonora | Instituto de Diagnostico y Referencia Epidemiologicos (INDRE) | Claudia Wong-Arambula, Abril Rodriguez-Maldonado, Vanessa Rivero-Arredondo, Ariadna Medina-Benitez, Joaquin Quiroz-Mercado, Sergio Rangel-Guerrero, Natividad Cruz-Ortiz, Tatiana Nunez-Garcia, Gisela Barrera-Badillo, Lucia Hernandez-Rivas, Irma Lopez-Martinez, Ernesto Ramirez-Gonzalez. |
| EPI_ISL_2101898, EPI_ISL_2101899 | LESP Quintana Roo | Instituto de Diagnostico y Referencia Epidemiologicos (INDRE) | Claudia Wong-Arambula, Abril Rodriguez-Maldonado, Vanessa Rivero-Arredondo, Ariadna Medina-Benitez, Joaquin Quiroz-Mercado, Sergio Rangel-Guerrero, Natividad Cruz-Ortiz, Tatiana Nunez-Garcia, Gisela Barrera-Badillo, Lucia Hernandez-Rivas, Irma Lopez-Martinez, Ernesto Ramirez-Gonzalez. |
| EPI_ISL_2101900 | LESP Jalisco | Instituto de Diagnostico y Referencia Epidemiologicos (INDRE) | Claudia Wong-Arambula, Abril Rodriguez-Maldonado, Vanessa Rivero-Arredondo, Ariadna Medina-Benitez, Joaquin Quiroz-Mercado, Sergio Rangel-Guerrero, Natividad Cruz-Ortiz, Tatiana Nunez-Garcia, Gisela Barrera-Badillo, Lucia Hernandez-Rivas, Irma Lopez-Martinez, Ernesto Ramirez-Gonzalez. |
| EPI_ISL_2101901 | LESP Coahuila | Instituto de Diagnostico y Referencia Epidemiologicos (INDRE) | Claudia Wong-Arambula, Abril Rodriguez-Maldonado, Vanessa Rivero-Arredondo, Ariadna Medina-Benitez, Joaquin Quiroz-Mercado, Sergio Rangel-Guerrero, Natividad Cruz-Ortiz, Tatiana Nunez-Garcia, Gisela Barrera-Badillo, Lucia Hernandez-Rivas, Irma Lopez-Martinez, Ernesto Ramirez-Gonzalez. |
| EPI_ISL_2101906, EPI_ISL_2101907, EPI_ISL_2101908, EPI_ISL_2101909, EPI_ISL_2101911, EPI_ISL_2101912 | LESP Ciudad de Mexico | Instituto de Diagnostico y Referencia Epidemiologicos (INDRE) | Claudia Wong-Arambula, Abril Rodriguez-Maldonado, Vanessa Rivero-Arredondo, Ariadna Medina-Benitez, Joaquin Quiroz-Mercado, Sergio Rangel-Guerrero, Natividad Cruz-Ortiz, Tatiana Nunez-Garcia, Gisela Barrera-Badillo, Lucia Hernandez-Rivas, Irma Lopez-Martinez, Ernesto Ramirez-Gonzalez. |
| EPI_ISL_2103413 | AREA DE SALUD SAN FRANCISCO-SAN ANTONIO (COOPESANA) | Inciensa, Instituto Costarricense de Investigación y Enseñanza en Nutrición y Salud | Francisco Duarte, Hebleen Porras, Claudio Soto-Garita, Estela Cordero, Adriana Godínez, Melany Calderón, José Luis Vargas, Mariela Gutiérrez, Joselyn Prado, Caterina Guzmán, Nazareth Ruiz & Mariel López |
| EPI_ISL_2103415 | CLINICA CATOLICA | Inciensa, Instituto Costarricense de Investigación y Enseñanza en Nutrición y Salud | Francisco Duarte, Hebleen Porras, Claudio Soto-Garita, Estela Cordero, Adriana Godínez, Melany Calderón, José Luis Vargas, Mariela Gutiérrez, Joselyn Prado, Caterina Guzmán, Nazareth Ruiz & Mariel López |
| EPI_ISL_2105681, EPI_ISL_2105682, EPI_ISL_2105683, EPI_ISL_2105684, EPI_ISL_2105685, EPI_ISL_2105686, EPI_ISL_2105687 | LESP Guanajuato | Instituto de Diagnostico y Referencia Epidemiologicos (INDRE) | Claudia Wong-Arambula, Abril Rodriguez-Maldonado, Vanessa Rivero-Arredondo, Ariadna Medina-Benitez, Joaquin Quiroz-Mercado, Sergio Rangel-Guerrero, Natividad Cruz-Ortiz, Tatiana Nunez-Garcia, Gisela Barrera-Badillo, Lucia Hernandez-Rivas, Irma Lopez-Martinez, Ernesto Ramirez-Gonzalez. |
| EPI_ISL_2105688 | LESP Baja California | Instituto de Diagnostico y Referencia Epidemiologicos (INDRE) | Claudia Wong-Arambula, Abril Rodriguez-Maldonado, Vanessa Rivero-Arredondo, Ariadna Medina-Benitez, Joaquin Quiroz-Mercado, Sergio Rangel-Guerrero, Natividad Cruz-Ortiz, Tatiana Nunez-Garcia, Gisela Barrera-Badillo, Lucia Hernandez-Rivas, Irma Lopez-Martinez, Ernesto Ramirez-Gonzalez. |

| | | | |
|---|---|---|---|
| EPI_ISL_2105689, EPI_ISL_2105690, EPI_ISL_2105691, EPI_ISL_2105692 | LESP Quintana Roo | Instituto de Diagnostico y Referencia Epidemiologicos (INDRE) | Claudia Wong-Arambula, Abril Rodriguez-Maldonado, Vanessa Rivero-Arredondo, Ariadna Medina-Benitez, Joaquin Quiroz-Mercado, Sergio Rangel-Guerrero, Natividad Cruz-Ortiz, Tatiana Nunez-Garcia, Gisela Barrera-Badillo, Lucia Hernandez-Rivas, Irma Lopez-Martinez, Ernesto Ramirez-Gonzalez. |
| EPI_ISL_2105716, EPI_ISL_2105717, EPI_ISL_2105718 | LESP Ciudad de Mexico | Instituto de Diagnostico y Referencia Epidemiologicos (INDRE) | Claudia Wong-Arambula, Abril Rodriguez-Maldonado, Vanessa Rivero-Arredondo, Ariadna Medina-Benitez, Joaquin Quiroz-Mercado, Sergio Rangel-Guerrero, Natividad Cruz-Ortiz, Tatiana Nunez-Garcia, Gisela Barrera-Badillo, Lucia Hernandez-Rivas, Irma Lopez-Martinez, Ernesto Ramirez-Gonzalez. |
| EPI_ISL_2105944, EPI_ISL_2105950, EPI_ISL_2105951 | Molecular Diagnostics Pathology Department Mater Dei Hospital Malta | Molecular Diagnostics Pathology Department Mater Dei Hospital Malta | G Zahra, R Borg, C Cilia, L Grech |
| EPI_ISL_2106232, EPI_ISL_2106233 | National Center of Infectious and Parasitic Diseases | National Center of Infectious and Parasitic Diseases | Alexiev, Ivanov, Korsun, Stoitsova, Philipova, Dimitrova, Grigorova L., Hristova, Donchev, Stoykov, Trifonova, Dobrinov, Grigorova I., Kantardjiev |
| EPI_ISL_2106234, EPI_ISL_2106235 | Siem Reap Provincial Laboratory | Virology Unit, Institut Pasteur du Cambodge | Jurre Y Siegers, Cecile Troupin, Leakhena Pum, Lamleav Leak, Ly Sovann, Kraing Sidonn, Yi Sengdoeurn, Chin Savuth, Chau Darapheak, Veasna Duong, Erik A Karlsson |
| EPI_ISL_2106236, EPI_ISL_2106237 | Virology Unit, Institut Pasteur du Cambodge | Virology Unit, Institut Pasteur du Cambodge | Jurre Y Siegers, Cecile Troupin, Leakhena Pum, Ly Sovann, Kraing Sidonn, Yi Sengdoeurn, Chin Savuth, Chau Darapheak, Veasna Duong, Erik A Karlsson |
| EPI_ISL_2106238, EPI_ISL_2106239, EPI_ISL_2106240, EPI_ISL_2106241, EPI_ISL_2106242 | Siem Reap Provincial Laboratory | Virology Unit, Institut Pasteur du Cambodge | Jurre Y Siegers, Cecile Troupin, Leakhena Pum, Lamleav Leak, Ly Sovann, Kraing Sidonn, Yi Sengdoeurn, Chin Savuth, Chau Darapheak, Veasna Duong, Erik A Karlsson |
| EPI_ISL_2106243, EPI_ISL_2106244, EPI_ISL_2106245, EPI_ISL_2106246, EPI_ISL_2106247, EPI_ISL_2106248, EPI_ISL_2106249, EPI_ISL_2106250, EPI_ISL_2106251, EPI_ISL_2106253, EPI_ISL_2106261, EPI_ISL_2106262, EPI_ISL_2106263 | | | |
| see above | Virology Unit, Institut Pasteur du Cambodge | Virology Unit, Institut Pasteur du Cambodge | Jurre Y Siegers, Cecile Troupin, Leakhena Pum, Ly Sovann, Kraing Sidonn, Yi Sengdoeurn, Chin Savuth, Chau Darapheak, Veasna Duong, Erik A Karlsson |
| EPI_ISL_2106264, EPI_ISL_2106265, EPI_ISL_2106266, EPI_ISL_2106267, EPI_ISL_2106268, EPI_ISL_2106269, EPI_ISL_2106270, EPI_ISL_2106271 | Cambodian National Public Health Laboratory, National Institute of Public Health | Virology Unit, Institut Pasteur du Cambodge | Jurre Y Siegers, Cecile Troupin, Leakhena Pum, Ly Sovann, Kraing Sidonn, Yi Sengdoeurn, Chin Savuth, Chau Darapheak, Veasna Duong, Erik A Karlsson |
| EPI_ISL_2106272 | Siem Reap Provincial Laboratory | Virology Unit, Institut Pasteur du Cambodge | Jurre Y Siegers, Cecile Troupin, Leakhena Pum, Lamleav Leak, Ly Sovann, Kraing Sidonn, Yi Sengdoeurn, Chin Savuth, Chau Darapheak, Veasna Duong, Erik A Karlsson |
| EPI_ISL_2106273, EPI_ISL_2106274, EPI_ISL_2106275 | Virology Unit, Institut Pasteur du Cambodge | Virology Unit, Institut Pasteur du Cambodge | Jurre Y Siegers, Cecile Troupin, Leakhena Pum, Ly Sovann, Kraing Sidonn, Yi Sengdoeurn, Chin Savuth, Chau Darapheak, Veasna Duong, Erik A Karlsson |
| EPI_ISL_2110643 | Institut Pasteur du Maroc | Functional Genomics platform/CNRST | Melloul Marouane, Chouati Taha, Hemlali Mouhsine, Alaoui Sanaa-amine, Rfaki Abderrazak Touil Nadia, Elannaz Hicham, Ennibi Khalid, Youbi Mohammed, MERABET MOUAD, NOURLIL Jalal, Maaroufi Abderrahmane, EL FAHIME Elmostafa |
| EPI_ISL_2116499 | Basurto University Hospital: Clinical Microbiology Laboratory | Biocruces Bizkaia | Mikel Urrutikoetxea-Gutiérrez, Mª Carmen Nieto Toboso, Estibaliz Ugalde Zarraga, Mikel Gallego Rodrigo, Ana de la Hoz, José Luis Díaz de Tuesta del Arco |
| EPI_ISL_2131175, EPI_ISL_2131176, EPI_ISL_2131177 | AZ Klina | AZ Klina | Carl Vael - Lynsey Berckmans |
| EPI_ISL_2131269, EPI_ISL_2131272, EPI_ISL_2131273 | OLVZ Aalst | OLVZ Aalst | Astrid Holderbeke |
| EPI_ISL_2131367 | SIESP CHIETI - DRIVE IN LANCIANO | Istituto Zooprofilattico Sperimentale dell'Abruzzo e Molise "G. Caporale" | Lorusso A, Marcacci M, Di Domenico M, Ancora M, Curini V, Di Lollo Valeria, Mangone I, Rinaldi A, Delli Compagni E, Scialabba S, Caporale M, Di Pasquale A, Cammà C, Puglia I, Calistri P, Savini G |
| EPI_ISL_2131368, EPI_ISL_2131369, EPI_ISL_2131370 | SIESP DIPARTIMENTO DI PREVENZIONE TERAMO | Istituto Zooprofilattico Sperimentale dell'Abruzzo e Molise "G. Caporale" | Lorusso A, Marcacci M, Di Domenico M, Ancora M, Curini V, Di Lollo Valeria, Mangone I, Rinaldi A, Delli Compagni E, Scialabba S, Caporale M, Di Pasquale A, Cammà C, Puglia I, Calistri P, Savini G |
| EPI_ISL_2131371 | USCA Pescina | Istituto Zooprofilattico Sperimentale dell'Abruzzo e Molise "G. Caporale" | Lorusso A, Marcacci M, Di Domenico M, Ancora M, Curini V, Di Lollo Valeria, Mangone I, Rinaldi A, Delli Compagni E, Scialabba S, Caporale M, Di Pasquale A, Cammà C, Puglia I, Calistri P, Savini G |
| EPI_ISL_2131372 | USCA Avezzano | Istituto Zooprofilattico Sperimentale dell'Abruzzo e Molise "G. Caporale" | Lorusso A, Marcacci M, Di Domenico M, Ancora M, Curini V, Di Lollo Valeria, Mangone I, Rinaldi A, Delli Compagni E, Scialabba S, Caporale M, Di Pasquale A, Cammà C, Puglia I, Calistri P, Savini G |
| EPI_ISL_2131373 | SIESP CHIETI - DRIVE IN GISSI | Istituto Zooprofilattico Sperimentale dell'Abruzzo e Molise "G. Caporale" | Lorusso A, Marcacci M, Di Domenico M, Ancora M, Curini V, Di Lollo Valeria, Mangone I, Rinaldi A, Delli Compagni E, Scialabba S, Caporale M, Di Pasquale A, Cammà C, Puglia I, Calistri P, Savini G |
| EPI_ISL_2131374 | SIESP DIPARTIMENTO DI PREVENZIONE CHIETI | Istituto Zooprofilattico Sperimentale dell'Abruzzo e Molise "G. Caporale" | Lorusso A, Marcacci M, Di Domenico M, Ancora M, Curini V, Di Lollo Valeria, Mangone I, Rinaldi A, Delli Compagni E, Scialabba S, Caporale M, Di Pasquale A, Cammà C, Puglia I, Calistri P, Savini G |
| EPI_ISL_2131375 | SIESP CHIETI - DRIVE IN ORTONA | Istituto Zooprofilattico Sperimentale dell'Abruzzo e Molise "G. Caporale" | Lorusso A, Marcacci M, Di Domenico M, Ancora M, Curini V, Di Lollo Valeria, Mangone I, Rinaldi A, Delli Compagni E, Scialabba S, Caporale M, Di Pasquale A, Cammà C, Puglia I, Calistri P, Savini G |
| EPI_ISL_2131376, EPI_ISL_2131377 | SIESP CHIETI - DRIVE IN LANCIANO | Istituto Zooprofilattico Sperimentale dell'Abruzzo e Molise "G. Caporale" | Lorusso A, Marcacci M, Di Domenico M, Ancora M, Curini V, Di Lollo Valeria, Mangone I, Rinaldi A, Delli Compagni E, Scialabba S, Caporale M, Di Pasquale A, Cammà C, Puglia I, Calistri P, Savini G |
| EPI_ISL_2131378 | USCA Tagliacozzo | Istituto Zooprofilattico Sperimentale dell'Abruzzo e Molise "G. Caporale" | Lorusso A, Marcacci M, Di Domenico M, Ancora M, Curini V, Di Lollo Valeria, Mangone I, Rinaldi A, Delli Compagni E, Scialabba S, Caporale M, Di Pasquale A, Cammà C, Puglia I, Calistri P, Savini G |
| EPI_ISL_2131379, EPI_ISL_2131380, EPI_ISL_2131381 | SIESP CHIETI - DRIVE IN LANCIANO | Istituto Zooprofilattico Sperimentale dell'Abruzzo e Molise "G. Caporale" | Lorusso A, Marcacci M, Di Domenico M, Ancora M, Curini V, Di Lollo Valeria, Mangone I, Rinaldi A, Delli Compagni E, Scialabba S, Caporale M, Di Pasquale A, Cammà C, Puglia I, Calistri P, Savini G |
| EPI_ISL_2131382, EPI_ISL_2131383, EPI_ISL_2131384, EPI_ISL_2131385, EPI_ISL_2131386, EPI_ISL_2131387, EPI_ISL_2131388, EPI_ISL_2131389, EPI_ISL_2131390, EPI_ISL_2131391, EPI_ISL_2131392, EPI_ISL_2131393 | | | |
| see above | SIESP DIPARTIMENTO DI PREVENZIONE TERAMO | Istituto Zooprofilattico Sperimentale dell'Abruzzo e Molise "G. Caporale" | Lorusso A, Marcacci M, Di Domenico M, Ancora M, Curini V, Di Lollo Valeria, Mangone I, Rinaldi A, Delli Compagni E, Scialabba S, Caporale M, Di Pasquale A, Cammà C, Puglia I, Calistri P, Savini G |
| EPI_ISL_2131394 | USCA Sulmona | Istituto Zooprofilattico Sperimentale dell'Abruzzo e Molise "G. Caporale" | Lorusso A, Marcacci M, Di Domenico M, Ancora M, Curini V, Di Lollo Valeria, Mangone I, Rinaldi A, Delli Compagni E, Scialabba S, Caporale M, Di Pasquale A, Cammà C, Puglia I, Calistri P, Savini G |
| EPI_ISL_2131396, EPI_ISL_2131397 | SIESP CHIETI - DRIVE IN LANCIANO | Istituto Zooprofilattico Sperimentale dell'Abruzzo e Molise "G. Caporale" | Lorusso A, Marcacci M, Di Domenico M, Ancora M, Curini V, Di Lollo Valeria, Mangone I, Rinaldi A, Delli Compagni E, Scialabba S, Caporale M, Di Pasquale A, Cammà C, Puglia I, Calistri P, Savini G |
| EPI_ISL_2131398, EPI_ISL_2131399, EPI_ISL_2131400 | SIESP CHIETI - DRIVE IN VASTO | Istituto Zooprofilattico Sperimentale dell'Abruzzo e Molise "G. Caporale" | Lorusso A, Marcacci M, Di Domenico M, Ancora M, Curini V, Di Lollo Valeria, Mangone I, Rinaldi A, Delli Compagni E, Scialabba S, Caporale M, Di Pasquale A, Cammà C, Puglia I, Calistri P, Savini G |
| EPI_ISL_2131401 | SIESP DIPARTIMENTO DI PREVENZIONE CHIETI | Istituto Zooprofilattico Sperimentale dell'Abruzzo e Molise "G. Caporale" | Lorusso A, Marcacci M, Di Domenico M, Ancora M, Curini V, Di Lollo Valeria, Mangone I, Rinaldi A, Delli Compagni E, Scialabba S, Caporale M, Di Pasquale A, Cammà C, Puglia I, Calistri P, Savini G |
| EPI_ISL_2131406 | SIESP CHIETI - DRIVE IN VASTO | Istituto Zooprofilattico Sperimentale dell'Abruzzo e Molise "G. Caporale" | Lorusso A, Marcacci M, Di Domenico M, Ancora M, Curini V, Di Lollo Valeria, Mangone I, Rinaldi A, Delli Compagni E, Scialabba S, Caporale M, Di Pasquale A, Cammà C, Puglia I, Calistri P, Savini G |
| EPI_ISL_2131417 | SIESP CHIETI - DRIVE IN GISSI | Istituto Zooprofilattico Sperimentale dell'Abruzzo e Molise "G. Caporale" | Lorusso A, Marcacci M, Di Domenico M, Ancora M, Curini V, Di Lollo Valeria, Mangone I, Rinaldi A, Delli Compagni E, Scialabba S, Caporale M, Di Pasquale A, Cammà C, Puglia I, Calistri P, Savini G |
| EPI_ISL_2131428, EPI_ISL_2131438 | SIESP DIPARTIMENTO DI PREVENZIONE CHIETI | Istituto Zooprofilattico Sperimentale dell'Abruzzo e Molise "G. | Lorusso A, Marcacci M, Di Domenico M, Ancora M, Curini V, Di Lollo Valeria, Mangone I, Rinaldi A, Delli Compagni E, Scialabba S, Caporale M, Di |

| | | | |
|---|---|---|---|
| | | Caporale" | Pasquale A, Cammà C, Puglia I, Calistri P, Savini G |
| EPI_ISL_2131439, EPI_ISL_2131440 | DIP. PREV. AVEZZANO SERVIZIO DI IGIENE EPIDEMIOLOGIA E SANITA' PUBBLICA | Istituto Zooprofilattico Sperimentale dell'Abruzzo e Molise "G. Caporale" | Lorusso A, Marcacci M, Di Domenico M, Ancora M, Curini V, Di Lollo Valeria, Mangone I, Rinaldi A, Delli Compagni E, Scialabba S, Caporale M, Di Pasquale A, Cammà C, Puglia I, Calistri P, Savini G |
| EPI_ISL_2131441, EPI_ISL_2131442 | SIESP CHIETI - DRIVE IN LANCIANO | Istituto Zooprofilattico Sperimentale dell'Abruzzo e Molise "G. Caporale" | Lorusso A, Marcacci M, Di Domenico M, Ancora M, Curini V, Di Lollo Valeria, Mangone I, Rinaldi A, Delli Compagni E, Scialabba S, Caporale M, Di Pasquale A, Cammà C, Puglia I, Calistri P, Savini G |
| EPI_ISL_2131449, EPI_ISL_2131450 | CHU NIMES | CHU NIMES | Stephan Robin, Marie-josée Carles, Sophie Bravo, Agathe Boudet |
| EPI_ISL_2131455, EPI_ISL_2131457 | Laboratorio Analisi Osp. Città di Castello - Azienda USL Umbria1 | Istituto Zooprofilattico Sperimentale dell'Abruzzo e Molise "G. Caporale" | Malagigi V, Tacconi P, Lorusso A, Marcacci M, Di Domenico M, Ancora M, Curini V, Mangone I, Rinaldi A, Delli Compagni E, Scialabba S, Di Pasquale A, Cammà C, Puglia I, Calistri P, Savini G |
| EPI_ISL_2131459, EPI_ISL_2131460, EPI_ISL_2131461, EPI_ISL_2131462, EPI_ISL_2131463, EPI_ISL_2131464, EPI_ISL_2131465, EPI_ISL_2131466, EPI_ISL_2131467, EPI_ISL_2131468, EPI_ISL_2131469, EPI_ISL_2131470, EPI_ISL_2131471, EPI_ISL_2131472, EPI_ISL_2131473, EPI_ISL_2131474, EPI_ISL_2131475, EPI_ISL_2131476, EPI_ISL_2131477, EPI_ISL_2131478, EPI_ISL_2131479, EPI_ISL_2131480, EPI_ISL_2131481, EPI_ISL_2131482, EPI_ISL_2131483, EPI_ISL_2131484, EPI_ISL_2131485 | | | |
| see above | Università degli Studi di Perugia | Istituto Zooprofilattico Sperimentale dell'Abruzzo e Molise "G. Caporale" | Mencacci A, Camilloni B, Lorusso A, Marcacci M, Di Domenico M, Ancora M, Curini V, Mangone I, Rinaldi A, Delli Compagni E, Scialabba S, Di Pasquale A, Cammà C, Puglia I, Calistri P, Savini G |
| EPI_ISL_2131486, EPI_ISL_2131487, EPI_ISL_2131488, EPI_ISL_2131489, EPI_ISL_2131490, EPI_ISL_2131491, EPI_ISL_2131492, EPI_ISL_2131493, EPI_ISL_2131494, EPI_ISL_2131495, EPI_ISL_2131496, EPI_ISL_2131497, EPI_ISL_2131498, EPI_ISL_2131499, EPI_ISL_2131500, EPI_ISL_2131501, EPI_ISL_2131502, EPI_ISL_2131503, EPI_ISL_2131504, EPI_ISL_2131505, EPI_ISL_2131506, EPI_ISL_2131507, EPI_ISL_2131508 | | | |
| see above | Azienda Ospedaliera Terni | Istituto Zooprofilattico Sperimentale dell'Abruzzo e Molise "G. Caporale" | Palumbo M, Scaccetti A, Lorusso A, Marcacci M, Di Domenico M, Ancora M, Curini V, Mangone I, Rinaldi A, Delli Compagni E, Scialabba S, Di Pasquale A, Cammà C, Puglia I, Calistri P, Savini G |
| EPI_ISL_2131806, EPI_ISL_2131833 | AZ Klina | AZ Klina | Carl Vael - Lynsey Berckmans |
| EPI_ISL_2134719, EPI_ISL_2134720, EPI_ISL_2134721, EPI_ISL_2134722 | Centre Hospitalier Universitaire Clermont-Ferrand | CHU Clermont-Ferrand, service de virologie | Bisseux Maxime, Mirand Audrey, Combes Patricia, Henquell Cécile |
| EPI_ISL_2134723 | EHPAD | CHU Clermont-Ferrand, service de virologie | Bisseux Maxime, Mirand Audrey, Combes Patricia, Henquell Cécile |
| EPI_ISL_2134725 | Centre Hospitalier Paul Ardier | CHU Clermont-Ferrand, service de virologie | Bisseux Maxime, Mirand Audrey, Combes Patricia, Henquell Cécile |
| EPI_ISL_2134727 | Centre Hospitalier Universitaire Clermont-Ferrand | CHU Clermont-Ferrand, service de virologie | Bisseux Maxime, Mirand Audrey, Combes Patricia, Henquell Cécile |
| EPI_ISL_2134728 | HOPITAL DE VICHY | CHU Clermont-Ferrand, service de virologie | Bisseux Maxime, Mirand Audrey, Combes Patricia, Henquell Cécile |
| EPI_ISL_2134729 | Centre hospitalier D'AURILLAC | CHU Clermont-Ferrand, service de virologie | Bisseux Maxime, Mirand Audrey, Combes Patricia, Henquell Cécile |
| EPI_ISL_2134730 | Centre Hospitalier Universitaire Clermont-Ferrand | CHU Clermont-Ferrand, service de virologie | Bisseux Maxime, Mirand Audrey, Combes Patricia, Henquell Cécile |
| EPI_ISL_2134731 | Centre Hospitalier RIOM | CHU Clermont-Ferrand, service de virologie | Bisseux Maxime, Mirand Audrey, Combes Patricia, Henquell Cécile |
| EPI_ISL_2134732 | HOPITAL DE VICHY | CHU Clermont-Ferrand, service de virologie | Bisseux Maxime, Mirand Audrey, Combes Patricia, Henquell Cécile |
| EPI_ISL_2134733 | Centre Hospitalier Universitaire Clermont-Ferrand | CHU Clermont-Ferrand, service de virologie | Bisseux Maxime, Mirand Audrey, Combes Patricia, Henquell Cécile |
| EPI_ISL_2134738 | HOPITAL DE VICHY | CHU Clermont-Ferrand, service de virologie | Bisseux Maxime, Mirand Audrey, Combes Patricia, Henquell Cécile |
| EPI_ISL_2134742 | Centre hospitalier D'AURILLAC | CHU Clermont-Ferrand, service de virologie | Bisseux Maxime, Mirand Audrey, Combes Patricia, Henquell Cécile |
| EPI_ISL_2134751, EPI_ISL_2134752 | Centre Hospitalier RIOM | CHU Clermont-Ferrand, service de virologie | Bisseux Maxime, Mirand Audrey, Combes Patricia, Henquell Cécile |
| EPI_ISL_2134754, EPI_ISL_2134755, EPI_ISL_2134756, EPI_ISL_2134757, EPI_ISL_2134758 | Centre Hospitalier Universitaire Clermont-Ferrand | CHU Clermont-Ferrand, service de virologie | Bisseux Maxime, Mirand Audrey, Combes Patricia, Henquell Cécile |
| EPI_ISL_2134761, EPI_ISL_2134762 | HOPITAL DU MONT DORE | CHU Clermont-Ferrand, service de virologie | Bisseux Maxime, Mirand Audrey, Combes Patricia, Henquell Cécile |
| EPI_ISL_2134764, EPI_ISL_2134765 | CDP COVID MONTLUCON | CHU Clermont-Ferrand, service de virologie | Bisseux Maxime, Mirand Audrey, Combes Patricia, Henquell Cécile |
| EPI_ISL_2134767, EPI_ISL_2134768, EPI_ISL_2134769 | Centre Hospitalier Universitaire Clermont-Ferrand | CHU Clermont-Ferrand, service de virologie | Bisseux Maxime, Mirand Audrey, Combes Patricia, Henquell Cécile |
| EPI_ISL_2134771 | Centre Hospitalier Billom | CHU Clermont-Ferrand, service de virologie | Bisseux Maxime, Mirand Audrey, Combes Patricia, Henquell Cécile |
| EPI_ISL_2134772, EPI_ISL_2134773, EPI_ISL_2134780 | Centre Hospitalier Universitaire Clermont-Ferrand | CHU Clermont-Ferrand, service de virologie | Bisseux Maxime, Mirand Audrey, Combes Patricia, Henquell Cécile |
| EPI_ISL_2134781 | EHPAD | CHU Clermont-Ferrand, service de virologie | Bisseux Maxime, Mirand Audrey, Combes Patricia, Henquell Cécile |
| EPI_ISL_2134782 | Centre Hospitalier Universitaire Clermont-Ferrand | CHU Clermont-Ferrand, service de virologie | Bisseux Maxime, Mirand Audrey, Combes Patricia, Henquell Cécile |
| EPI_ISL_2134783 | HOPITAL DU MONT DORE | CHU Clermont-Ferrand, service de virologie | Bisseux Maxime, Mirand Audrey, Combes Patricia, Henquell Cécile |
| EPI_ISL_2136324 | UNIGEM | Universidad Nacional de Colombia - Laboratorio Genómico One Health | Maria Stella López, Andres F. Cardona-Rios, Daniel O. Maldonado-Perez, Simón Villegas Velásquez, Lina M. Hurtado, Laura Silvana Perez, Karl A Ciuoderis, Maria Angélica Maya, Carolina Muñoz-Arango, Celeny Ortiz, Rita Almanza Payares, Idabely Betancur Ortiz, Sandra Ines Cano, Diego A. Álvarez-Díaz, Carlos Franco-Muñoz, Katherine Laiton-Donato, Hector Alejandro Ruiz-Moreno, Marcela Mercado-Reyes, Jorge E. Osorio, Juan P. Hernandez-Ortiz |
| EPI_ISL_2136329 | UNIGEM | Universidad Nacional de Colombia - Laboratorio Genómico One Health | Simón Villegas Velásquez, Andres F. Cardona-Rios, Maria Stella López, Daniel O. Maldonado-Perez, Lina M. Hurtado, Laura Silvana Perez, Karl A Ciuoderis, Maria Angélica Maya, Carolina Muñoz-Arango, Celeny Ortiz, Rita Almanza Payares, Idabely Betancur Ortiz, Sandra Ines Cano, Diego A. Álvarez-Díaz, Carlos Franco-Muñoz, Katherine Laiton-Donato, Hector Alejandro Ruiz-Moreno, Marcela Mercado-Reyes, Jorge E. Osorio, Juan P. Hernandez-Ortiz |
| EPI_ISL_2136334 | UNIGEM | Universidad Nacional de Colombia - Laboratorio Genómico One Health | Andres F. Cardona-Rios, Maria Stella López, Daniel O. Maldonado-Perez, Simón Villegas Velásquez, Lina M. Hurtado, Laura Silvana Perez, Karl A Ciuoderis, Maria Angélica Maya, Carolina Muñoz-Arango, Celeny Ortiz, Rita Almanza Payares, Idabely Betancur Ortiz, Sandra Ines Cano, Diego A. Álvarez-Díaz, Carlos Franco-Muñoz, Katherine Laiton-Donato, Hector Alejandro Ruiz-Moreno, Marcela Mercado-Reyes, Jorge E. Osorio, Juan P. Hernandez-Ortiz |
| EPI_ISL_2137020, EPI_ISL_2137021, EPI_ISL_2137023, EPI_ISL_2137024, EPI_ISL_2137030, EPI_ISL_2137031 | Althaia. Xarxa Assistencial Universitària de Manresa | IrsiCaixa | Marc Noguera-Julian, Mariona Parera, Maria Casadellà, Pilar Armengol, Francesc Catala-Moll, Roger Paredes, Bonaventura Clotet Gloria Trujillo, Rafel Perez Vidal, Jaume Trape Pujol, Carolina Gonzalez Fernandez, Miquel Micó, Roger Paredes, Eulalia Grau, Bonaventura Clotet |
| EPI_ISL_2137045, EPI_ISL_2137047, EPI_ISL_2137378, EPI_ISL_2137379, EPI_ISL_2137380, EPI_ISL_2137724, EPI_ISL_2137725, EPI_ISL_2137733 | AZ Klina | AZ Klina | Carl Vael - Lynsey Berckmans |
| EPI_ISL_2137768 | Università degli Studi di Perugia | Istituto Zooprofilattico Sperimentale dell'Abruzzo e Molise "G. Caporale" | Mencacci A, Camilloni B, Lorusso A, Marcacci M, Di Domenico M, Ancora M, Curini V, Mangone I, Rinaldi A, Delli Compagni E, Scialabba S, Di Pasquale A, Cammà C, Puglia I, Calistri P, Savini G |
| EPI_ISL_2139908, EPI_ISL_2139909 | LESP Baja California | Instituto de Diagnostico y Referencia Epidemiologicos (INDRE) | Claudia Wong-Arambula, Abril Rodriguez-Maldonado, Vanessa Rivero-Arredondo, Ariadna Medina-Benitez, Joaquin Quiroz-Mercado, Sergio Rangel-Guerrero, Natividad Cruz-Ortiz, Tatiana Nunez-Garcia, Gisela Barrera-Badillo, Lucia Hernandez-Rivas, Irma Lopez-Martinez, Ernesto Ramirez-Gonzalez. |

| | | | |
|---|---|---|---|
| EPI_ISL_2139911 | LESP Guanajuato | Instituto de Diagnostico y Referencia Epidemiologicos (INDRE) | Claudia Wong-Arambula, Abril Rodriguez-Maldonado, Vanessa Rivero-Arredondo, Ariadna Medina-Benitez, Joaquin Quiroz-Mercado, Sergio Rangel-Guerrero, Natividad Cruz-Ortiz, Tatiana Nunez-Garcia, Gisela Barrera-Badillo, Lucia Hernandez-Rivas, Irma Lopez-Martinez, Ernesto Ramirez-Gonzalez. |
| EPI_ISL_2139912 | LESP Colima | Instituto de Diagnostico y Referencia Epidemiologicos (INDRE) | Claudia Wong-Arambula, Abril Rodriguez-Maldonado, Vanessa Rivero-Arredondo, Ariadna Medina-Benitez, Joaquin Quiroz-Mercado, Sergio Rangel-Guerrero, Natividad Cruz-Ortiz, Tatiana Nunez-Garcia, Gisela Barrera-Badillo, Lucia Hernandez-Rivas, Irma Lopez-Martinez, Ernesto Ramirez-Gonzalez. |
| EPI_ISL_2139913 | LESP Baja California Sur | Instituto de Diagnostico y Referencia Epidemiologicos (INDRE) | Claudia Wong-Arambula, Abril Rodriguez-Maldonado, Vanessa Rivero-Arredondo, Ariadna Medina-Benitez, Joaquin Quiroz-Mercado, Sergio Rangel-Guerrero, Natividad Cruz-Ortiz, Tatiana Nunez-Garcia, Gisela Barrera-Badillo, Lucia Hernandez-Rivas, Irma Lopez-Martinez, Ernesto Ramirez-Gonzalez. |
| EPI_ISL_2139915 | LESP Jalisco | Instituto de Diagnostico y Referencia Epidemiologicos (INDRE) | Claudia Wong-Arambula, Abril Rodriguez-Maldonado, Vanessa Rivero-Arredondo, Ariadna Medina-Benitez, Joaquin Quiroz-Mercado, Sergio Rangel-Guerrero, Natividad Cruz-Ortiz, Tatiana Nunez-Garcia, Gisela Barrera-Badillo, Lucia Hernandez-Rivas, Irma Lopez-Martinez, Ernesto Ramirez-Gonzalez. |
| EPI_ISL_2139916 | LESP Chihuahua | Instituto de Diagnostico y Referencia Epidemiologicos (INDRE) | Claudia Wong-Arambula, Abril Rodriguez-Maldonado, Vanessa Rivero-Arredondo, Ariadna Medina-Benitez, Joaquin Quiroz-Mercado, Sergio Rangel-Guerrero, Natividad Cruz-Ortiz, Tatiana Nunez-Garcia, Gisela Barrera-Badillo, Lucia Hernandez-Rivas, Irma Lopez-Martinez, Ernesto Ramirez-Gonzalez. |
| EPI_ISL_2139918 | LESP Sinaloa | Instituto de Diagnostico y Referencia Epidemiologicos (INDRE) | Claudia Wong-Arambula, Abril Rodriguez-Maldonado, Vanessa Rivero-Arredondo, Ariadna Medina-Benitez, Joaquin Quiroz-Mercado, Sergio Rangel-Guerrero, Natividad Cruz-Ortiz, Tatiana Nunez-Garcia, Gisela Barrera-Badillo, Lucia Hernandez-Rivas, Irma Lopez-Martinez, Ernesto Ramirez-Gonzalez. |
| EPI_ISL_2139919 | LESP Jalisco | Instituto de Diagnostico y Referencia Epidemiologicos (INDRE) | Claudia Wong-Arambula, Abril Rodriguez-Maldonado, Vanessa Rivero-Arredondo, Ariadna Medina-Benitez, Joaquin Quiroz-Mercado, Sergio Rangel-Guerrero, Natividad Cruz-Ortiz, Tatiana Nunez-Garcia, Gisela Barrera-Badillo, Lucia Hernandez-Rivas, Irma Lopez-Martinez, Ernesto Ramirez-Gonzalez. |
| EPI_ISL_2139922 | LESP Ciudad de Mexico | Instituto de Diagnostico y Referencia Epidemiologicos (INDRE) | Claudia Wong-Arambula, Abril Rodriguez-Maldonado, Vanessa Rivero-Arredondo, Ariadna Medina-Benitez, Joaquin Quiroz-Mercado, Sergio Rangel-Guerrero, Natividad Cruz-Ortiz, Tatiana Nunez-Garcia, Gisela Barrera-Badillo, Lucia Hernandez-Rivas, Irma Lopez-Martinez, Ernesto Ramirez-Gonzalez. |
| EPI_ISL_2140730 | OLVZ Aalst | OLVZ Aalst | Astrid Holderbeke |
| EPI_ISL_2140877, EPI_ISL_2140879, EPI_ISL_2140880 | The Ohio State University Applied Microbiology Services Laboratory | The Ohio State University Applied Microbiology Services Laboratory | Seth A. Faith PhD |
| EPI_ISL_2143385, EPI_ISL_2143386, EPI_ISL_2143388 | CHU LILLE | CHU Lille - Laboratoire de Virologie | AIT YAHYA Emilie, ALIDJINOU Enagnon Kazali, BOCKET Laurence, CREPIN Michel, DEMAY Christophe, ENGELMANN Ilka, GEFFROY Sandrine, GUIGON Aurélie, LAMBERT Valérie, LAZREK Mouna, NOBILLIAUX Florian, PREVOST Brigitte, THUILLIER Caroline, TINEZ Claire |
| EPI_ISL_2143390 | CH ARRAS | CHU Lille - Laboratoire de Virologie | AIT YAHYA Emilie, ALIDJINOU Enagnon Kazali, BOCKET Laurence, CREPIN Michel, DEMAY Christophe, ENGELMANN Ilka, GEFFROY Sandrine, GUIGON Aurélie, LAMBERT Valérie, LAZREK Mouna, NOBILLIAUX Florian, PREVOST Brigitte, THUILLIER Caroline, TINEZ Claire |
| EPI_ISL_2143391 | CHU LILLE | CHU Lille - Laboratoire de Virologie | AIT YAHYA Emilie, ALIDJINOU Enagnon Kazali, BOCKET Laurence, CREPIN Michel, DEMAY Christophe, ENGELMANN Ilka, GEFFROY Sandrine, GUIGON Aurélie, LAMBERT Valérie, LAZREK Mouna, NOBILLIAUX Florian, PREVOST Brigitte, THUILLIER Caroline, TINEZ Claire |
| EPI_ISL_2143393 | CH CAMBRAI | CHU Lille - Laboratoire de Virologie | AIT YAHYA Emilie, ALIDJINOU Enagnon Kazali, BOCKET Laurence, CREPIN Michel, DEMAY Christophe, ENGELMANN Ilka, GEFFROY Sandrine, GUIGON Aurélie, LAMBERT Valérie, LAZREK Mouna, NOBILLIAUX Florian, PREVOST Brigitte, THUILLIER Caroline, TINEZ Claire |
| EPI_ISL_2143395 | CH DOUAI | CHU Lille - Laboratoire de Virologie | AIT YAHYA Emilie, ALIDJINOU Enagnon Kazali, BOCKET Laurence, CREPIN Michel, DEMAY Christophe, ENGELMANN Ilka, GEFFROY Sandrine, GUIGON Aurélie, LAMBERT Valérie, LAZREK Mouna, NOBILLIAUX Florian, PREVOST Brigitte, THUILLIER Caroline, TINEZ Claire |
| EPI_ISL_2143396, EPI_ISL_2143398 | CHU LILLE | CHU Lille - Laboratoire de Virologie | AIT YAHYA Emilie, ALIDJINOU Enagnon Kazali, BOCKET Laurence, CREPIN Michel, DEMAY Christophe, ENGELMANN Ilka, GEFFROY Sandrine, GUIGON Aurélie, LAMBERT Valérie, LAZREK Mouna, NOBILLIAUX Florian, PREVOST Brigitte, THUILLIER Caroline, TINEZ Claire |
| EPI_ISL_2143400 | CH TOURCOING | CHU Lille - Laboratoire de Virologie | AIT YAHYA Emilie, ALIDJINOU Enagnon Kazali, BOCKET Laurence, CREPIN Michel, DEMAY Christophe, ENGELMANN Ilka, GEFFROY Sandrine, GUIGON Aurélie, LAMBERT Valérie, LAZREK Mouna, NOBILLIAUX Florian, PREVOST Brigitte, THUILLIER Caroline, TINEZ Claire |
| EPI_ISL_2143403, EPI_ISL_2143404 | CHU LILLE | CHU Lille - Laboratoire de Virologie | AIT YAHYA Emilie, ALIDJINOU Enagnon Kazali, BOCKET Laurence, CREPIN Michel, DEMAY Christophe, ENGELMANN Ilka, GEFFROY Sandrine, GUIGON Aurélie, LAMBERT Valérie, LAZREK Mouna, NOBILLIAUX Florian, PREVOST Brigitte, THUILLIER Caroline, TINEZ Claire |
| EPI_ISL_2143406 | CH CALAIS | CHU Lille - Laboratoire de Virologie | AIT YAHYA Emilie, ALIDJINOU Enagnon Kazali, BOCKET Laurence, CREPIN Michel, DEMAY Christophe, ENGELMANN Ilka, GEFFROY Sandrine, GUIGON Aurélie, LAMBERT Valérie, LAZREK Mouna, NOBILLIAUX Florian, PREVOST Brigitte, THUILLIER Caroline, TINEZ Claire |
| EPI_ISL_2143408 | CH ARMENTIERES | CHU Lille - Laboratoire de Virologie | AIT YAHYA Emilie, ALIDJINOU Enagnon Kazali, BOCKET Laurence, CREPIN Michel, DEMAY Christophe, ENGELMANN Ilka, GEFFROY Sandrine, GUIGON Aurélie, LAMBERT Valérie, LAZREK Mouna, NOBILLIAUX Florian, PREVOST Brigitte, THUILLIER Caroline, TINEZ Claire |
| EPI_ISL_2143409, EPI_ISL_2143411 | CHU LILLE | CHU Lille - Laboratoire de Virologie | AIT YAHYA Emilie, ALIDJINOU Enagnon Kazali, BOCKET Laurence, CREPIN Michel, DEMAY Christophe, ENGELMANN Ilka, GEFFROY Sandrine, GUIGON Aurélie, LAMBERT Valérie, LAZREK Mouna, NOBILLIAUX Florian, PREVOST Brigitte, THUILLIER Caroline, TINEZ Claire |
| EPI_ISL_2143413 | CH ROUBAIX | CHU Lille - Laboratoire de Virologie | AIT YAHYA Emilie, ALIDJINOU Enagnon Kazali, BOCKET Laurence, CREPIN Michel, DEMAY Christophe, ENGELMANN Ilka, GEFFROY Sandrine, GUIGON Aurélie, LAMBERT Valérie, LAZREK Mouna, NOBILLIAUX Florian, PREVOST Brigitte, THUILLIER Caroline, TINEZ Claire |
| EPI_ISL_2143416 | CHU LILLE | CHU Lille - Laboratoire de Virologie | AIT YAHYA Emilie, ALIDJINOU Enagnon Kazali, BOCKET Laurence, CREPIN Michel, DEMAY Christophe, ENGELMANN Ilka, GEFFROY Sandrine, GUIGON Aurélie, LAMBERT Valérie, LAZREK Mouna, NOBILLIAUX Florian, PREVOST Brigitte, THUILLIER Caroline, TINEZ Claire |
| EPI_ISL_2143417 | CH SAINT-PHILIBERT | CHU Lille - Laboratoire de Virologie | AIT YAHYA Emilie, ALIDJINOU Enagnon Kazali, BOCKET Laurence, CREPIN Michel, DEMAY Christophe, ENGELMANN Ilka, GEFFROY Sandrine, GUIGON Aurélie, LAMBERT Valérie, LAZREK Mouna, NOBILLIAUX Florian, PREVOST Brigitte, THUILLIER Caroline, TINEZ Claire |
| EPI_ISL_2143419 | CH ARRAS | CHU Lille - Laboratoire de Virologie | AIT YAHYA Emilie, ALIDJINOU Enagnon Kazali, BOCKET Laurence, CREPIN Michel, DEMAY Christophe, ENGELMANN Ilka, GEFFROY Sandrine, GUIGON Aurélie, LAMBERT Valérie, LAZREK Mouna, NOBILLIAUX Florian, PREVOST Brigitte, THUILLIER Caroline, TINEZ Claire |
| EPI_ISL_2143421 | CH CAMBRAI | CHU Lille - Laboratoire de Virologie | AIT YAHYA Emilie, ALIDJINOU Enagnon Kazali, BOCKET Laurence, CREPIN Michel, DEMAY Christophe, ENGELMANN Ilka, GEFFROY Sandrine, GUIGON Aurélie, LAMBERT Valérie, LAZREK Mouna, NOBILLIAUX Florian, PREVOST Brigitte, THUILLIER Caroline, TINEZ Claire |
| EPI_ISL_2143422 | CH ROUBAIX | CHU Lille - Laboratoire de Virologie | AIT YAHYA Emilie, ALIDJINOU Enagnon Kazali, BOCKET Laurence, CREPIN Michel, DEMAY Christophe, ENGELMANN Ilka, GEFFROY Sandrine, GUIGON Aurélie, LAMBERT Valérie, LAZREK Mouna, NOBILLIAUX Florian, PREVOST Brigitte, THUILLIER Caroline, TINEZ Claire |
| EPI_ISL_2143424 | CH ARMENTIERES | CHU Lille - Laboratoire de Virologie | AIT YAHYA Emilie, ALIDJINOU Enagnon Kazali, BOCKET Laurence, CREPIN Michel, DEMAY Christophe, ENGELMANN Ilka, GEFFROY Sandrine, GUIGON Aurélie, LAMBERT Valérie, LAZREK Mouna, NOBILLIAUX Florian, PREVOST Brigitte, THUILLIER Caroline, TINEZ Claire |
| EPI_ISL_2143426 | CH TOURCOING | CHU Lille - Laboratoire de Virologie | AIT YAHYA Emilie, ALIDJINOU Enagnon Kazali, BOCKET Laurence, CREPIN Michel, DEMAY Christophe, ENGELMANN Ilka, GEFFROY Sandrine, GUIGON Aurélie, LAMBERT Valérie, LAZREK Mouna, NOBILLIAUX Florian, PREVOST Brigitte, THUILLIER Caroline, TINEZ Claire |
| EPI_ISL_2143427 | CH DUNKERQUE | CHU Lille - Laboratoire de Virologie | AIT YAHYA Emilie, ALIDJINOU Enagnon Kazali, BOCKET Laurence, CREPIN Michel, DEMAY Christophe, ENGELMANN Ilka, GEFFROY Sandrine, GUIGON Aurélie, LAMBERT Valérie, LAZREK Mouna, NOBILLIAUX Florian, PREVOST Brigitte, THUILLIER Caroline, TINEZ Claire |
| EPI_ISL_2143429, EPI_ISL_2143433, EPI_ISL_2143435, EPI_ISL_2143437, EPI_ISL_2143440, EPI_ISL_2143442 | CHU LILLE | CHU Lille - Laboratoire de Virologie | AIT YAHYA Emilie, ALIDJINOU Enagnon Kazali, BOCKET Laurence, CREPIN Michel, DEMAY Christophe, ENGELMANN Ilka, GEFFROY Sandrine, GUIGON Aurélie, LAMBERT Valérie, LAZREK Mouna, NOBILLIAUX Florian, PREVOST Brigitte, THUILLIER Caroline, TINEZ Claire |
| EPI_ISL_2143443, EPI_ISL_2143445 | CH ROUBAIX | CHU Lille - Laboratoire de Virologie | AIT YAHYA Emilie, ALIDJINOU Enagnon Kazali, BOCKET Laurence, CREPIN Michel, DEMAY Christophe, ENGELMANN Ilka, GEFFROY Sandrine, GUIGON Aurélie, LAMBERT Valérie, LAZREK Mouna, NOBILLIAUX Florian, PREVOST Brigitte, THUILLIER Caroline, TINEZ Claire |

| | | | |
|---|---|---|---|
| EPI_ISL_2143446 | CH ARMENTIERES | CHU Lille - Laboratoire de Virologie | AIT YAHYA Emilie, ALIDJINOU Enagnon Kazali, BOCKET Laurence, CREPIN Michel, DEMAY Christophe, ENGELMANN Ilka, GEFFROY Sandrine, GUIGON Aurélie, LAMBERT Valérie, LAZREK Mouna, NOBILLIAUX Florian, PREVOST Brigitte, THUILLIER Caroline, TINEZ Claire |
| EPI_ISL_2143448, EPI_ISL_2143450, EPI_ISL_2143451, EPI_ISL_2143453, EPI_ISL_2143454, EPI_ISL_2143456 | CHU LILLE | CHU Lille - Laboratoire de Virologie | AIT YAHYA Emilie, ALIDJINOU Enagnon Kazali, BOCKET Laurence, CREPIN Michel, DEMAY Christophe, ENGELMANN Ilka, GEFFROY Sandrine, GUIGON Aurélie, LAMBERT Valérie, LAZREK Mouna, NOBILLIAUX Florian, PREVOST Brigitte, THUILLIER Caroline, TINEZ Claire |
| EPI_ISL_2143459 | CH CAMBRAI | CHU Lille - Laboratoire de Virologie | AIT YAHYA Emilie, ALIDJINOU Enagnon Kazali, BOCKET Laurence, CREPIN Michel, DEMAY Christophe, ENGELMANN Ilka, GEFFROY Sandrine, GUIGON Aurélie, LAMBERT Valérie, LAZREK Mouna, NOBILLIAUX Florian, PREVOST Brigitte, THUILLIER Caroline, TINEZ Claire |
| EPI_ISL_2143460, EPI_ISL_2143462 | CH ARRAS | CHU Lille - Laboratoire de Virologie | AIT YAHYA Emilie, ALIDJINOU Enagnon Kazali, BOCKET Laurence, CREPIN Michel, DEMAY Christophe, ENGELMANN Ilka, GEFFROY Sandrine, GUIGON Aurélie, LAMBERT Valérie, LAZREK Mouna, NOBILLIAUX Florian, PREVOST Brigitte, THUILLIER Caroline, TINEZ Claire |
| EPI_ISL_2143464 | CH CALAIS | CHU Lille - Laboratoire de Virologie | AIT YAHYA Emilie, ALIDJINOU Enagnon Kazali, BOCKET Laurence, CREPIN Michel, DEMAY Christophe, ENGELMANN Ilka, GEFFROY Sandrine, GUIGON Aurélie, LAMBERT Valérie, LAZREK Mouna, NOBILLIAUX Florian, PREVOST Brigitte, THUILLIER Caroline, TINEZ Claire |
| EPI_ISL_2143466 | CH DUNKERQUE | CHU Lille - Laboratoire de Virologie | AIT YAHYA Emilie, ALIDJINOU Enagnon Kazali, BOCKET Laurence, CREPIN Michel, DEMAY Christophe, ENGELMANN Ilka, GEFFROY Sandrine, GUIGON Aurélie, LAMBERT Valérie, LAZREK Mouna, NOBILLIAUX Florian, PREVOST Brigitte, THUILLIER Caroline, TINEZ Claire |
| EPI_ISL_2143467 | CHU LILLE | CHU Lille - Laboratoire de Virologie | AIT YAHYA Emilie, ALIDJINOU Enagnon Kazali, BOCKET Laurence, CREPIN Michel, DEMAY Christophe, ENGELMANN Ilka, GEFFROY Sandrine, GUIGON Aurélie, LAMBERT Valérie, LAZREK Mouna, NOBILLIAUX Florian, PREVOST Brigitte, THUILLIER Caroline, TINEZ Claire |
| EPI_ISL_2143469 | CH SAINT-PHILIBERT | CHU Lille - Laboratoire de Virologie | AIT YAHYA Emilie, ALIDJINOU Enagnon Kazali, BOCKET Laurence, CREPIN Michel, DEMAY Christophe, ENGELMANN Ilka, GEFFROY Sandrine, GUIGON Aurélie, LAMBERT Valérie, LAZREK Mouna, NOBILLIAUX Florian, PREVOST Brigitte, THUILLIER Caroline, TINEZ Claire |
| EPI_ISL_2143471 | CH ARMENTIERES | CHU Lille - Laboratoire de Virologie | AIT YAHYA Emilie, ALIDJINOU Enagnon Kazali, BOCKET Laurence, CREPIN Michel, DEMAY Christophe, ENGELMANN Ilka, GEFFROY Sandrine, GUIGON Aurélie, LAMBERT Valérie, LAZREK Mouna, NOBILLIAUX Florian, PREVOST Brigitte, THUILLIER Caroline, TINEZ Claire |
| EPI_ISL_2143472 | CHU LILLE | CHU Lille - Laboratoire de Virologie | AIT YAHYA Emilie, ALIDJINOU Enagnon Kazali, BOCKET Laurence, CREPIN Michel, DEMAY Christophe, ENGELMANN Ilka, GEFFROY Sandrine, GUIGON Aurélie, LAMBERT Valérie, LAZREK Mouna, NOBILLIAUX Florian, PREVOST Brigitte, THUILLIER Caroline, TINEZ Claire |
| EPI_ISL_2143474 | CH ARMENTIERES | CHU Lille - Laboratoire de Virologie | AIT YAHYA Emilie, ALIDJINOU Enagnon Kazali, BOCKET Laurence, CREPIN Michel, DEMAY Christophe, ENGELMANN Ilka, GEFFROY Sandrine, GUIGON Aurélie, LAMBERT Valérie, LAZREK Mouna, NOBILLIAUX Florian, PREVOST Brigitte, THUILLIER Caroline, TINEZ Claire |
| EPI_ISL_2143475 | CH ARRAS | CHU Lille - Laboratoire de Virologie | AIT YAHYA Emilie, ALIDJINOU Enagnon Kazali, BOCKET Laurence, CREPIN Michel, DEMAY Christophe, ENGELMANN Ilka, GEFFROY Sandrine, GUIGON Aurélie, LAMBERT Valérie, LAZREK Mouna, NOBILLIAUX Florian, PREVOST Brigitte, THUILLIER Caroline, TINEZ Claire |
| EPI_ISL_2143477 | CH VALENCIENNES | CHU Lille - Laboratoire de Virologie | AIT YAHYA Emilie, ALIDJINOU Enagnon Kazali, BOCKET Laurence, CREPIN Michel, DEMAY Christophe, ENGELMANN Ilka, GEFFROY Sandrine, GUIGON Aurélie, LAMBERT Valérie, LAZREK Mouna, NOBILLIAUX Florian, PREVOST Brigitte, THUILLIER Caroline, TINEZ Claire |
| EPI_ISL_2143479, EPI_ISL_2143480 | CHU LILLE | CHU Lille - Laboratoire de Virologie | AIT YAHYA Emilie, ALIDJINOU Enagnon Kazali, BOCKET Laurence, CREPIN Michel, DEMAY Christophe, ENGELMANN Ilka, GEFFROY Sandrine, GUIGON Aurélie, LAMBERT Valérie, LAZREK Mouna, NOBILLIAUX Florian, PREVOST Brigitte, THUILLIER Caroline, TINEZ Claire |
| EPI_ISL_2143482, EPI_ISL_2143483 | CH TOURCOING | CHU Lille - Laboratoire de Virologie | AIT YAHYA Emilie, ALIDJINOU Enagnon Kazali, BOCKET Laurence, CREPIN Michel, DEMAY Christophe, ENGELMANN Ilka, GEFFROY Sandrine, GUIGON Aurélie, LAMBERT Valérie, LAZREK Mouna, NOBILLIAUX Florian, PREVOST Brigitte, THUILLIER Caroline, TINEZ Claire |
| EPI_ISL_2143485 | CH ARMENTIERES | CHU Lille - Laboratoire de Virologie | AIT YAHYA Emilie, ALIDJINOU Enagnon Kazali, BOCKET Laurence, CREPIN Michel, DEMAY Christophe, ENGELMANN Ilka, GEFFROY Sandrine, GUIGON Aurélie, LAMBERT Valérie, LAZREK Mouna, NOBILLIAUX Florian, PREVOST Brigitte, THUILLIER Caroline, TINEZ Claire |
| EPI_ISL_2143486, EPI_ISL_2143490 | CHU LILLE | CHU Lille - Laboratoire de Virologie | AIT YAHYA Emilie, ALIDJINOU Enagnon Kazali, BOCKET Laurence, CREPIN Michel, DEMAY Christophe, ENGELMANN Ilka, GEFFROY Sandrine, GUIGON Aurélie, LAMBERT Valérie, LAZREK Mouna, NOBILLIAUX Florian, PREVOST Brigitte, THUILLIER Caroline, TINEZ Claire |
| EPI_ISL_2143497 | CH BETHUNE | CHU Lille - Laboratoire de Virologie | AIT YAHYA Emilie, ALIDJINOU Enagnon Kazali, BOCKET Laurence, CREPIN Michel, DEMAY Christophe, ENGELMANN Ilka, GEFFROY Sandrine, GUIGON Aurélie, LAMBERT Valérie, LAZREK Mouna, NOBILLIAUX Florian, PREVOST Brigitte, THUILLIER Caroline, TINEZ Claire |
| EPI_ISL_2143499, EPI_ISL_2143502 | CHU LILLE | CHU Lille - Laboratoire de Virologie | AIT YAHYA Emilie, ALIDJINOU Enagnon Kazali, BOCKET Laurence, CREPIN Michel, DEMAY Christophe, ENGELMANN Ilka, GEFFROY Sandrine, GUIGON Aurélie, LAMBERT Valérie, LAZREK Mouna, NOBILLIAUX Florian, PREVOST Brigitte, THUILLIER Caroline, TINEZ Claire |
| EPI_ISL_2143503 | CH ARRAS | CHU Lille - Laboratoire de Virologie | AIT YAHYA Emilie, ALIDJINOU Enagnon Kazali, BOCKET Laurence, CREPIN Michel, DEMAY Christophe, ENGELMANN Ilka, GEFFROY Sandrine, GUIGON Aurélie, LAMBERT Valérie, LAZREK Mouna, NOBILLIAUX Florian, PREVOST Brigitte, THUILLIER Caroline, TINEZ Claire |
| EPI_ISL_2143506 | CHU LILLE | CHU Lille - Laboratoire de Virologie | AIT YAHYA Emilie, ALIDJINOU Enagnon Kazali, BOCKET Laurence, CREPIN Michel, DEMAY Christophe, ENGELMANN Ilka, GEFFROY Sandrine, GUIGON Aurélie, LAMBERT Valérie, LAZREK Mouna, NOBILLIAUX Florian, PREVOST Brigitte, THUILLIER Caroline, TINEZ Claire |
| EPI_ISL_2143508 | CH CAMBRAI | CHU Lille - Laboratoire de Virologie | AIT YAHYA Emilie, ALIDJINOU Enagnon Kazali, BOCKET Laurence, CREPIN Michel, DEMAY Christophe, ENGELMANN Ilka, GEFFROY Sandrine, GUIGON Aurélie, LAMBERT Valérie, LAZREK Mouna, NOBILLIAUX Florian, PREVOST Brigitte, THUILLIER Caroline, TINEZ Claire |
| EPI_ISL_2143511 | CH DOUAI | CHU Lille - Laboratoire de Virologie | AIT YAHYA Emilie, ALIDJINOU Enagnon Kazali, BOCKET Laurence, CREPIN Michel, DEMAY Christophe, ENGELMANN Ilka, GEFFROY Sandrine, GUIGON Aurélie, LAMBERT Valérie, LAZREK Mouna, NOBILLIAUX Florian, PREVOST Brigitte, THUILLIER Caroline, TINEZ Claire |
| EPI_ISL_2143513, EPI_ISL_2143516 | CHU LILLE | CHU Lille - Laboratoire de Virologie | AIT YAHYA Emilie, ALIDJINOU Enagnon Kazali, BOCKET Laurence, CREPIN Michel, DEMAY Christophe, ENGELMANN Ilka, GEFFROY Sandrine, GUIGON Aurélie, LAMBERT Valérie, LAZREK Mouna, NOBILLIAUX Florian, PREVOST Brigitte, THUILLIER Caroline, TINEZ Claire |
| EPI_ISL_2143519 | CH TOURCOING | CHU Lille - Laboratoire de Virologie | AIT YAHYA Emilie, ALIDJINOU Enagnon Kazali, BOCKET Laurence, CREPIN Michel, DEMAY Christophe, ENGELMANN Ilka, GEFFROY Sandrine, GUIGON Aurélie, LAMBERT Valérie, LAZREK Mouna, NOBILLIAUX Florian, PREVOST Brigitte, THUILLIER Caroline, TINEZ Claire |
| EPI_ISL_2143524, EPI_ISL_2143526 | CHU LILLE | CHU Lille - Laboratoire de Virologie | AIT YAHYA Emilie, ALIDJINOU Enagnon Kazali, BOCKET Laurence, CREPIN Michel, DEMAY Christophe, ENGELMANN Ilka, GEFFROY Sandrine, GUIGON Aurélie, LAMBERT Valérie, LAZREK Mouna, NOBILLIAUX Florian, PREVOST Brigitte, THUILLIER Caroline, TINEZ Claire |
| EPI_ISL_2143528 | CH CALAIS | CHU Lille - Laboratoire de Virologie | AIT YAHYA Emilie, ALIDJINOU Enagnon Kazali, BOCKET Laurence, CREPIN Michel, DEMAY Christophe, ENGELMANN Ilka, GEFFROY Sandrine, GUIGON Aurélie, LAMBERT Valérie, LAZREK Mouna, NOBILLIAUX Florian, PREVOST Brigitte, THUILLIER Caroline, TINEZ Claire |
| EPI_ISL_2143531 | CH ARMENTIERES | CHU Lille - Laboratoire de Virologie | AIT YAHYA Emilie, ALIDJINOU Enagnon Kazali, BOCKET Laurence, CREPIN Michel, DEMAY Christophe, ENGELMANN Ilka, GEFFROY Sandrine, GUIGON Aurélie, LAMBERT Valérie, LAZREK Mouna, NOBILLIAUX Florian, PREVOST Brigitte, THUILLIER Caroline, TINEZ Claire |
| EPI_ISL_2143533, EPI_ISL_2143536 | CHU LILLE | CHU Lille - Laboratoire de Virologie | AIT YAHYA Emilie, ALIDJINOU Enagnon Kazali, BOCKET Laurence, CREPIN Michel, DEMAY Christophe, ENGELMANN Ilka, GEFFROY Sandrine, GUIGON Aurélie, LAMBERT Valérie, LAZREK Mouna, NOBILLIAUX Florian, PREVOST Brigitte, THUILLIER Caroline, TINEZ Claire |
| EPI_ISL_2143539 | CH ROUBAIX | CHU Lille - Laboratoire de Virologie | AIT YAHYA Emilie, ALIDJINOU Enagnon Kazali, BOCKET Laurence, CREPIN Michel, DEMAY Christophe, ENGELMANN Ilka, GEFFROY Sandrine, GUIGON Aurélie, LAMBERT Valérie, LAZREK Mouna, NOBILLIAUX Florian, PREVOST Brigitte, THUILLIER Caroline, TINEZ Claire |
| EPI_ISL_2143541 | CHU LILLE | CHU Lille - Laboratoire de Virologie | AIT YAHYA Emilie, ALIDJINOU Enagnon Kazali, BOCKET Laurence, CREPIN Michel, DEMAY Christophe, ENGELMANN Ilka, GEFFROY Sandrine, GUIGON Aurélie, LAMBERT Valérie, LAZREK Mouna, NOBILLIAUX Florian, PREVOST Brigitte, THUILLIER Caroline, TINEZ Claire |
| EPI_ISL_2143544 | CH SAINT-PHILIBERT | CHU Lille - Laboratoire de Virologie | AIT YAHYA Emilie, ALIDJINOU Enagnon Kazali, BOCKET Laurence, CREPIN Michel, DEMAY Christophe, ENGELMANN Ilka, GEFFROY Sandrine, GUIGON Aurélie, LAMBERT Valérie, LAZREK Mouna, NOBILLIAUX Florian, PREVOST Brigitte, THUILLIER Caroline, TINEZ Claire |
| EPI_ISL_2143546 | CH ARRAS | CHU Lille - Laboratoire de Virologie | AIT YAHYA Emilie, ALIDJINOU Enagnon Kazali, BOCKET Laurence, CREPIN Michel, DEMAY Christophe, ENGELMANN Ilka, GEFFROY Sandrine, GUIGON Aurélie, LAMBERT Valérie, LAZREK Mouna, NOBILLIAUX Florian, PREVOST Brigitte, THUILLIER Caroline, TINEZ Claire |
| EPI_ISL_2143548 | CH CAMBRAI | CHU Lille - Laboratoire de Virologie | AIT YAHYA Emilie, ALIDJINOU Enagnon Kazali, BOCKET Laurence, CREPIN Michel, DEMAY Christophe, ENGELMANN Ilka, GEFFROY Sandrine, GUIGON Aurélie, LAMBERT Valérie, LAZREK Mouna, NOBILLIAUX Florian, PREVOST Brigitte, THUILLIER Caroline, TINEZ Claire |
| EPI_ISL_2143551 | CH ROUBAIX | CHU Lille - Laboratoire de Virologie | AIT YAHYA Emilie, ALIDJINOU Enagnon Kazali, BOCKET Laurence, CREPIN Michel, DEMAY Christophe, ENGELMANN Ilka, GEFFROY Sandrine, GUIGON Aurélie, LAMBERT Valérie, LAZREK Mouna, NOBILLIAUX Florian, PREVOST Brigitte, THUILLIER Caroline, TINEZ Claire |

| | | | |
|---|---|---|---|
| EPI_ISL_2143553 | CH ARMENTIERES | CHU Lille - Laboratoire de Virologie | AIT YAHYA Emilie, ALIDJINOU Enagnon Kazali, BOCKET Laurence, CREPIN Michel, DEMAY Christophe, ENGELMANN Ilka, GEFFROY Sandrine, GUIGON Aurélie, LAMBERT Valérie, LAZREK Mouna, NOBILLIAUX Florian, PREVOST Brigitte, THUILLIER Caroline, TINEZ Claire |
| EPI_ISL_2143556 | CH TOURCOING | CHU Lille - Laboratoire de Virologie | AIT YAHYA Emilie, ALIDJINOU Enagnon Kazali, BOCKET Laurence, CREPIN Michel, DEMAY Christophe, ENGELMANN Ilka, GEFFROY Sandrine, GUIGON Aurélie, LAMBERT Valérie, LAZREK Mouna, NOBILLIAUX Florian, PREVOST Brigitte, THUILLIER Caroline, TINEZ Claire |
| EPI_ISL_2143559 | CH DUNKERQUE | CHU Lille - Laboratoire de Virologie | AIT YAHYA Emilie, ALIDJINOU Enagnon Kazali, BOCKET Laurence, CREPIN Michel, DEMAY Christophe, ENGELMANN Ilka, GEFFROY Sandrine, GUIGON Aurélie, LAMBERT Valérie, LAZREK Mouna, NOBILLIAUX Florian, PREVOST Brigitte, THUILLIER Caroline, TINEZ Claire |
| EPI_ISL_2143561, EPI_ISL_2143570, EPI_ISL_2143572, EPI_ISL_2143575, EPI_ISL_2143577, EPI_ISL_2143579 | CHU LILLE | CHU Lille - Laboratoire de Virologie | AIT YAHYA Emilie, ALIDJINOU Enagnon Kazali, BOCKET Laurence, CREPIN Michel, DEMAY Christophe, ENGELMANN Ilka, GEFFROY Sandrine, GUIGON Aurélie, LAMBERT Valérie, LAZREK Mouna, NOBILLIAUX Florian, PREVOST Brigitte, THUILLIER Caroline, TINEZ Claire |
| EPI_ISL_2143581, EPI_ISL_2143583 | CH ROUBAIX | CHU Lille - Laboratoire de Virologie | AIT YAHYA Emilie, ALIDJINOU Enagnon Kazali, BOCKET Laurence, CREPIN Michel, DEMAY Christophe, ENGELMANN Ilka, GEFFROY Sandrine, GUIGON Aurélie, LAMBERT Valérie, LAZREK Mouna, NOBILLIAUX Florian, PREVOST Brigitte, THUILLIER Caroline, TINEZ Claire |
| EPI_ISL_2143585 | CH ARMENTIERES | CHU Lille - Laboratoire de Virologie | AIT YAHYA Emilie, ALIDJINOU Enagnon Kazali, BOCKET Laurence, CREPIN Michel, DEMAY Christophe, ENGELMANN Ilka, GEFFROY Sandrine, GUIGON Aurélie, LAMBERT Valérie, LAZREK Mouna, NOBILLIAUX Florian, PREVOST Brigitte, THUILLIER Caroline, TINEZ Claire |
| EPI_ISL_2143587, EPI_ISL_2143589, EPI_ISL_2143590, EPI_ISL_2143592, EPI_ISL_2143594, EPI_ISL_2143596 | CHU LILLE | CHU Lille - Laboratoire de Virologie | AIT YAHYA Emilie, ALIDJINOU Enagnon Kazali, BOCKET Laurence, CREPIN Michel, DEMAY Christophe, ENGELMANN Ilka, GEFFROY Sandrine, GUIGON Aurélie, LAMBERT Valérie, LAZREK Mouna, NOBILLIAUX Florian, PREVOST Brigitte, THUILLIER Caroline, TINEZ Claire |
| EPI_ISL_2143599 | CH CAMBRAI | CHU Lille - Laboratoire de Virologie | AIT YAHYA Emilie, ALIDJINOU Enagnon Kazali, BOCKET Laurence, CREPIN Michel, DEMAY Christophe, ENGELMANN Ilka, GEFFROY Sandrine, GUIGON Aurélie, LAMBERT Valérie, LAZREK Mouna, NOBILLIAUX Florian, PREVOST Brigitte, THUILLIER Caroline, TINEZ Claire |
| EPI_ISL_2143601, EPI_ISL_2143603 | CH ARRAS | CHU Lille - Laboratoire de Virologie | AIT YAHYA Emilie, ALIDJINOU Enagnon Kazali, BOCKET Laurence, CREPIN Michel, DEMAY Christophe, ENGELMANN Ilka, GEFFROY Sandrine, GUIGON Aurélie, LAMBERT Valérie, LAZREK Mouna, NOBILLIAUX Florian, PREVOST Brigitte, THUILLIER Caroline, TINEZ Claire |
| EPI_ISL_2143605 | CH CALAIS | CHU Lille - Laboratoire de Virologie | AIT YAHYA Emilie, ALIDJINOU Enagnon Kazali, BOCKET Laurence, CREPIN Michel, DEMAY Christophe, ENGELMANN Ilka, GEFFROY Sandrine, GUIGON Aurélie, LAMBERT Valérie, LAZREK Mouna, NOBILLIAUX Florian, PREVOST Brigitte, THUILLIER Caroline, TINEZ Claire |
| EPI_ISL_2143607 | CH DUNKERQUE | CHU Lille - Laboratoire de Virologie | AIT YAHYA Emilie, ALIDJINOU Enagnon Kazali, BOCKET Laurence, CREPIN Michel, DEMAY Christophe, ENGELMANN Ilka, GEFFROY Sandrine, GUIGON Aurélie, LAMBERT Valérie, LAZREK Mouna, NOBILLIAUX Florian, PREVOST Brigitte, THUILLIER Caroline, TINEZ Claire |
| EPI_ISL_2143608 | CHU LILLE | CHU Lille - Laboratoire de Virologie | AIT YAHYA Emilie, ALIDJINOU Enagnon Kazali, BOCKET Laurence, CREPIN Michel, DEMAY Christophe, ENGELMANN Ilka, GEFFROY Sandrine, GUIGON Aurélie, LAMBERT Valérie, LAZREK Mouna, NOBILLIAUX Florian, PREVOST Brigitte, THUILLIER Caroline, TINEZ Claire |
| EPI_ISL_2143610 | CH SAINT-PHILIBERT | CHU Lille - Laboratoire de Virologie | AIT YAHYA Emilie, ALIDJINOU Enagnon Kazali, BOCKET Laurence, CREPIN Michel, DEMAY Christophe, ENGELMANN Ilka, GEFFROY Sandrine, GUIGON Aurélie, LAMBERT Valérie, LAZREK Mouna, NOBILLIAUX Florian, PREVOST Brigitte, THUILLIER Caroline, TINEZ Claire |
| EPI_ISL_2143613 | CH ARMENTIERES | CHU Lille - Laboratoire de Virologie | AIT YAHYA Emilie, ALIDJINOU Enagnon Kazali, BOCKET Laurence, CREPIN Michel, DEMAY Christophe, ENGELMANN Ilka, GEFFROY Sandrine, GUIGON Aurélie, LAMBERT Valérie, LAZREK Mouna, NOBILLIAUX Florian, PREVOST Brigitte, THUILLIER Caroline, TINEZ Claire |
| EPI_ISL_2143614 | CHU LILLE | CHU Lille - Laboratoire de Virologie | AIT YAHYA Emilie, ALIDJINOU Enagnon Kazali, BOCKET Laurence, CREPIN Michel, DEMAY Christophe, ENGELMANN Ilka, GEFFROY Sandrine, GUIGON Aurélie, LAMBERT Valérie, LAZREK Mouna, NOBILLIAUX Florian, PREVOST Brigitte, THUILLIER Caroline, TINEZ Claire |
| EPI_ISL_2143616 | CH ARMENTIERES | CHU Lille - Laboratoire de Virologie | AIT YAHYA Emilie, ALIDJINOU Enagnon Kazali, BOCKET Laurence, CREPIN Michel, DEMAY Christophe, ENGELMANN Ilka, GEFFROY Sandrine, GUIGON Aurélie, LAMBERT Valérie, LAZREK Mouna, NOBILLIAUX Florian, PREVOST Brigitte, THUILLIER Caroline, TINEZ Claire |
| EPI_ISL_2143618 | CH ARRAS | CHU Lille - Laboratoire de Virologie | AIT YAHYA Emilie, ALIDJINOU Enagnon Kazali, BOCKET Laurence, CREPIN Michel, DEMAY Christophe, ENGELMANN Ilka, GEFFROY Sandrine, GUIGON Aurélie, LAMBERT Valérie, LAZREK Mouna, NOBILLIAUX Florian, PREVOST Brigitte, THUILLIER Caroline, TINEZ Claire |
| EPI_ISL_2143620 | CH VALENCIENNES | CHU Lille - Laboratoire de Virologie | AIT YAHYA Emilie, ALIDJINOU Enagnon Kazali, BOCKET Laurence, CREPIN Michel, DEMAY Christophe, ENGELMANN Ilka, GEFFROY Sandrine, GUIGON Aurélie, LAMBERT Valérie, LAZREK Mouna, NOBILLIAUX Florian, PREVOST Brigitte, THUILLIER Caroline, TINEZ Claire |
| EPI_ISL_2143622, EPI_ISL_2143624 | CHU LILLE | CHU Lille - Laboratoire de Virologie | AIT YAHYA Emilie, ALIDJINOU Enagnon Kazali, BOCKET Laurence, CREPIN Michel, DEMAY Christophe, ENGELMANN Ilka, GEFFROY Sandrine, GUIGON Aurélie, LAMBERT Valérie, LAZREK Mouna, NOBILLIAUX Florian, PREVOST Brigitte, THUILLIER Caroline, TINEZ Claire |
| EPI_ISL_2143626, EPI_ISL_2143629 | CH TOURCOING | CHU Lille - Laboratoire de Virologie | AIT YAHYA Emilie, ALIDJINOU Enagnon Kazali, BOCKET Laurence, CREPIN Michel, DEMAY Christophe, ENGELMANN Ilka, GEFFROY Sandrine, GUIGON Aurélie, LAMBERT Valérie, LAZREK Mouna, NOBILLIAUX Florian, PREVOST Brigitte, THUILLIER Caroline, TINEZ Claire |
| EPI_ISL_2143630 | CH ARMENTIERES | CHU Lille - Laboratoire de Virologie | AIT YAHYA Emilie, ALIDJINOU Enagnon Kazali, BOCKET Laurence, CREPIN Michel, DEMAY Christophe, ENGELMANN Ilka, GEFFROY Sandrine, GUIGON Aurélie, LAMBERT Valérie, LAZREK Mouna, NOBILLIAUX Florian, PREVOST Brigitte, THUILLIER Caroline, TINEZ Claire |
| EPI_ISL_2143633, EPI_ISL_2143636 | CHU LILLE | CHU Lille - Laboratoire de Virologie | AIT YAHYA Emilie, ALIDJINOU Enagnon Kazali, BOCKET Laurence, CREPIN Michel, DEMAY Christophe, ENGELMANN Ilka, GEFFROY Sandrine, GUIGON Aurélie, LAMBERT Valérie, LAZREK Mouna, NOBILLIAUX Florian, PREVOST Brigitte, THUILLIER Caroline, TINEZ Claire |
| EPI_ISL_2143643 | CH BETHUNE | CHU Lille - Laboratoire de Virologie | AIT YAHYA Emilie, ALIDJINOU Enagnon Kazali, BOCKET Laurence, CREPIN Michel, DEMAY Christophe, ENGELMANN Ilka, GEFFROY Sandrine, GUIGON Aurélie, LAMBERT Valérie, LAZREK Mouna, NOBILLIAUX Florian, PREVOST Brigitte, THUILLIER Caroline, TINEZ Claire |
| EPI_ISL_2143645 | CHU LILLE | CHU Lille - Laboratoire de Virologie | AIT YAHYA Emilie, ALIDJINOU Enagnon Kazali, BOCKET Laurence, CREPIN Michel, DEMAY Christophe, ENGELMANN Ilka, GEFFROY Sandrine, GUIGON Aurélie, LAMBERT Valérie, LAZREK Mouna, NOBILLIAUX Florian, PREVOST Brigitte, THUILLIER Caroline, TINEZ Claire |
| EPI_ISL_2151335 | Basurto University Hospital Clinical Microbiology Laboratory | Biocruces Bizkaia | Mikel Urrutikoetxea-Gutierrez, Mª Carmen Nieto Toboso, Estibaliz Ugalde Zarraga, Mikel Gallego Rodrigo, Ana de la Hoz, José Luis Díaz de Tuesta del Arco |
| EPI_ISL_2151405, EPI_ISL_2151406, EPI_ISL_2151407, EPI_ISL_2151408, EPI_ISL_2151409, EPI_ISL_2151411 | OCME Office Of Chief Medical Examiner | New York City Public Health Laboratory | Jade Wang, et al. |
| EPI_ISL_2152365, EPI_ISL_2152373, EPI_ISL_2152376, EPI_ISL_2152377, EPI_ISL_2152378 | GH A.CHENEVIER-H.MONDOR | Department of Virology, Henri Mondor University Hospital, Assistance Publique Hôpitaux de Paris, Université Paris-Est Créteil, INSERM U955 | Christophe Rodriguez, Slim Fourati, Vanessa Demontant, Guillaume Gricourt, Melissa N'Debi, Alexandre Soulier, Elisabeth Trawinski, Jean-Michel Pawlotsky |
| EPI_ISL_2152435 | Usansolo-Galdakao University Hospital | Cruces University Hospital | Ana Gual-de-Torrella, Izaskun Alejo-Cancho, Mikel Gallego, Ana Belén de la Hoz |
| EPI_ISL_2152539, EPI_ISL_2152541, EPI_ISL_2152542 | GH A.CHENEVIER-H.MONDOR | Department of Virology, Henri Mondor University Hospital, Assistance Publique Hôpitaux de Paris, Université Paris-Est Créteil, INSERM U955 | Christophe Rodriguez, Slim Fourati, Vanessa Demontant, Guillaume Gricourt, Melissa N'Debi, Alexandre Soulier, Elisabeth Trawinski, Jean-Michel Pawlotsky |
| EPI_ISL_2152588 | HOPITAL G. CLEMENCEAU | Department of Virology, Henri Mondor University Hospital, Assistance Publique Hôpitaux de Paris, Université Paris-Est Créteil, INSERM U955 | Christophe Rodriguez, Slim Fourati, Vanessa Demontant, Guillaume Gricourt, Melissa N'Debi, Alexandre Soulier, Elisabeth Trawinski, Jean-Michel Pawlotsky |
| EPI_ISL_2152591 | GH A.CHENEVIER-H.MONDOR | Department of Virology, Henri Mondor University Hospital, Assistance Publique Hôpitaux de Paris, Université Paris-Est Créteil, INSERM U955 | Christophe Rodriguez, Slim Fourati, Vanessa Demontant, Guillaume Gricourt, Melissa N'Debi, Alexandre Soulier, Elisabeth Trawinski, Jean-Michel Pawlotsky |
| EPI_ISL_2153433 | Laboratory of Microbiology, National Reference Lab, Charles Nicolle Hospital; 2-University of Tunis ElManar, Faculty of Medicine of Tunis, LR99ES09, Tunis, Tunisia | Clinical and Experimental Pharmacology Lab, LR16SP02, National Center of Pharmacovigilance, University of Tunis El Manar, Tunis, Tunisia. 2-Neurodegenerative diseases and psychiatric troubles, LR18SP03, Razi Hospital, University of | Sameh Trabelsi, Nissaf Ben Alaya, Mouna Ben Sassi, Sana Ferjani, Salma Abid, Mouna Safer, Roua Ben Othman, Sarra Chamman, Imen Mdini, Manel Ben Sassi, Imen Kacem, Maher Kharrat, Alia BenKahla, Jalila Ben Khelil, Riadh Daghfous, Riadh Gouider, Raja Mahfoudh, Ilhem Boutiba-Ben Boubaker. |

| | | | |
|---|---|---|---|
| | | Tunis El Manar, Tunis, Tunisia. 3- Ministry of Health, National Observatory of New and Emerging Diseases, 1006, Tunis, Tunisia | |
| EPI_ISL_2154331 | Laboratory of Microbiology, National Reference Lab, Charles Nicolle Hospital; 2-University of Tunis ElManar, Faculty of Medicine of Tunis, LR99ES09, Tunis, Tunisia | Clinical and Experimental Pharmacology Lab, LR16SP02, National Center of Pharmacovigilance, University of Tunis El Manar, Tunis, Tunisia. 2-Neurodegenerative diseases and psychiatric troubles, LR18SP03, Razi Hospital, University of Tunis El Manar, Tunis, Tunisia. 3- Ministry of Health, National Observatory of New and Emerging Diseases, 1006, Tunis, Tunisia | Sameh Trabelsi, Nissaf Ben Alaya, Mouna Ben Sassi, Sana Ferjani, Salma Abid, Mouna Safer, Roua Ben Othman, Sarra Chamman, Imen Mdini, Manel Ben Sassi, Imen Kacem, Maher Kharrat, Alia BenKahla, Jalila Ben Khelil, Riadh Daghfous, Riadh Gouider, Taha Maatoug, Ilhem Boutiba-Ben Boubaker. |
| EPI_ISL_2157310 | LESP Nuevo Leon | Instituto de Diagnostico y Referencia Epidemiologicos (INDRE) | Claudia Wong-Arambula, Abril Rodriguez-Maldonado, Vanessa Rivero-Arredondo, Ariadna Medina-Benitez, Joaquin Quiroz-Mercado, Sergio Rangel-Guerrero, Natividad Cruz-Ortiz, Tatiana Nunez-Garcia, Gisela Barrera-Badillo, Lucia Hernandez-Rivas, Irma Lopez-Martinez, Ernesto Ramirez-Gonzalez. |
| EPI_ISL_2157313 | LESP Queretaro | Instituto de Diagnostico y Referencia Epidemiologicos (INDRE) | Claudia Wong-Arambula, Abril Rodriguez-Maldonado, Vanessa Rivero-Arredondo, Ariadna Medina-Benitez, Joaquin Quiroz-Mercado, Sergio Rangel-Guerrero, Natividad Cruz-Ortiz, Tatiana Nunez-Garcia, Gisela Barrera-Badillo, Lucia Hernandez-Rivas, Irma Lopez-Martinez, Ernesto Ramirez-Gonzalez. |
| EPI_ISL_2157315 | LESP Morelos | Instituto de Diagnostico y Referencia Epidemiologicos (INDRE) | Claudia Wong-Arambula, Abril Rodriguez-Maldonado, Vanessa Rivero-Arredondo, Ariadna Medina-Benitez, Joaquin Quiroz-Mercado, Sergio Rangel-Guerrero, Natividad Cruz-Ortiz, Tatiana Nunez-Garcia, Gisela Barrera-Badillo, Lucia Hernandez-Rivas, Irma Lopez-Martinez, Ernesto Ramirez-Gonzalez. |
| EPI_ISL_2157317 | LESP Chihuahua | Instituto de Diagnostico y Referencia Epidemiologicos (INDRE) | Claudia Wong-Arambula, Abril Rodriguez-Maldonado, Vanessa Rivero-Arredondo, Ariadna Medina-Benitez, Joaquin Quiroz-Mercado, Sergio Rangel-Guerrero, Natividad Cruz-Ortiz, Tatiana Nunez-Garcia, Gisela Barrera-Badillo, Lucia Hernandez-Rivas, Irma Lopez-Martinez, Ernesto Ramirez-Gonzalez. |
| EPI_ISL_2157319 | LESP Coahuila | Instituto de Diagnostico y Referencia Epidemiologicos (INDRE) | Claudia Wong-Arambula, Abril Rodriguez-Maldonado, Vanessa Rivero-Arredondo, Ariadna Medina-Benitez, Joaquin Quiroz-Mercado, Sergio Rangel-Guerrero, Natividad Cruz-Ortiz, Tatiana Nunez-Garcia, Gisela Barrera-Badillo, Lucia Hernandez-Rivas, Irma Lopez-Martinez, Ernesto Ramirez-Gonzalez. |
| EPI_ISL_2157321 | LESP Sonora | Instituto de Diagnostico y Referencia Epidemiologicos (INDRE) | Claudia Wong-Arambula, Abril Rodriguez-Maldonado, Vanessa Rivero-Arredondo, Ariadna Medina-Benitez, Joaquin Quiroz-Mercado, Sergio Rangel-Guerrero, Natividad Cruz-Ortiz, Tatiana Nunez-Garcia, Gisela Barrera-Badillo, Lucia Hernandez-Rivas, Irma Lopez-Martinez, Ernesto Ramirez-Gonzalez. |
| EPI_ISL_2157322 | LESP Chihuahua | Instituto de Diagnostico y Referencia Epidemiologicos (INDRE) | Claudia Wong-Arambula, Abril Rodriguez-Maldonado, Vanessa Rivero-Arredondo, Ariadna Medina-Benitez, Joaquin Quiroz-Mercado, Sergio Rangel-Guerrero, Natividad Cruz-Ortiz, Tatiana Nunez-Garcia, Gisela Barrera-Badillo, Lucia Hernandez-Rivas, Irma Lopez-Martinez, Ernesto Ramirez-Gonzalez. |
| EPI_ISL_2157324, EPI_ISL_2157325, EPI_ISL_2157326 | LESP Guanajuato | Instituto de Diagnostico y Referencia Epidemiologicos (INDRE) | Claudia Wong-Arambula, Abril Rodriguez-Maldonado, Vanessa Rivero-Arredondo, Ariadna Medina-Benitez, Joaquin Quiroz-Mercado, Sergio Rangel-Guerrero, Natividad Cruz-Ortiz, Tatiana Nunez-Garcia, Gisela Barrera-Badillo, Lucia Hernandez-Rivas, Irma Lopez-Martinez, Ernesto Ramirez-Gonzalez. |
| EPI_ISL_2157328 | LESP Baja California | Instituto de Diagnostico y Referencia Epidemiologicos (INDRE) | Claudia Wong-Arambula, Abril Rodriguez-Maldonado, Vanessa Rivero-Arredondo, Ariadna Medina-Benitez, Joaquin Quiroz-Mercado, Sergio Rangel-Guerrero, Natividad Cruz-Ortiz, Tatiana Nunez-Garcia, Gisela Barrera-Badillo, Lucia Hernandez-Rivas, Irma Lopez-Martinez, Ernesto Ramirez-Gonzalez. |
| EPI_ISL_2157331 | LESP Oaxaca | Instituto de Diagnostico y Referencia Epidemiologicos (INDRE) | Claudia Wong-Arambula, Abril Rodriguez-Maldonado, Vanessa Rivero-Arredondo, Ariadna Medina-Benitez, Joaquin Quiroz-Mercado, Sergio Rangel-Guerrero, Natividad Cruz-Ortiz, Tatiana Nunez-Garcia, Gisela Barrera-Badillo, Lucia Hernandez-Rivas, Irma Lopez-Martinez, Ernesto Ramirez-Gonzalez. |
| EPI_ISL_2157333 | LESP Ciudad de Mexico | Instituto de Diagnostico y Referencia Epidemiologicos (INDRE) | Claudia Wong-Arambula, Abril Rodriguez-Maldonado, Vanessa Rivero-Arredondo, Ariadna Medina-Benitez, Joaquin Quiroz-Mercado, Sergio Rangel-Guerrero, Natividad Cruz-Ortiz, Tatiana Nunez-Garcia, Gisela Barrera-Badillo, Lucia Hernandez-Rivas, Irma Lopez-Martinez, Ernesto Ramirez-Gonzalez. |
| EPI_ISL_2158108 | LESP Baja California | Instituto de Diagnostico y Referencia Epidemiologicos (INDRE) | Claudia Wong-Arambula, Abril Rodriguez-Maldonado, Vanessa Rivero-Arredondo, Ariadna Medina-Benitez, Joaquin Quiroz-Mercado, Sergio Rangel-Guerrero, Natividad Cruz-Ortiz, Tatiana Nunez-Garcia, Gisela Barrera-Badillo, Lucia Hernandez-Rivas, Irma Lopez-Martinez, Ernesto Ramirez-Gonzalez. |
| EPI_ISL_2158109, EPI_ISL_2158110, EPI_ISL_2158111, EPI_ISL_2158112, EPI_ISL_2158113 | LESP Nuevo Leon | Instituto de Diagnostico y Referencia Epidemiologicos (INDRE) | Claudia Wong-Arambula, Abril Rodriguez-Maldonado, Vanessa Rivero-Arredondo, Ariadna Medina-Benitez, Joaquin Quiroz-Mercado, Sergio Rangel-Guerrero, Natividad Cruz-Ortiz, Tatiana Nunez-Garcia, Gisela Barrera-Badillo, Lucia Hernandez-Rivas, Irma Lopez-Martinez, Ernesto Ramirez-Gonzalez. |
| EPI_ISL_2158114 | LESP Tamaulipas | Instituto de Diagnostico y Referencia Epidemiologicos (INDRE) | Claudia Wong-Arambula, Abril Rodriguez-Maldonado, Vanessa Rivero-Arredondo, Ariadna Medina-Benitez, Joaquin Quiroz-Mercado, Sergio Rangel-Guerrero, Natividad Cruz-Ortiz, Tatiana Nunez-Garcia, Gisela Barrera-Badillo, Lucia Hernandez-Rivas, Irma Lopez-Martinez, Ernesto Ramirez-Gonzalez. |
| EPI_ISL_2158115, EPI_ISL_2158116, EPI_ISL_2158117, EPI_ISL_2158118, EPI_ISL_2158119, EPI_ISL_2158120 | LESP Nuevo Leon | Instituto de Diagnostico y Referencia Epidemiologicos (INDRE) | Claudia Wong-Arambula, Abril Rodriguez-Maldonado, Vanessa Rivero-Arredondo, Ariadna Medina-Benitez, Joaquin Quiroz-Mercado, Sergio Rangel-Guerrero, Natividad Cruz-Ortiz, Tatiana Nunez-Garcia, Gisela Barrera-Badillo, Lucia Hernandez-Rivas, Irma Lopez-Martinez, Ernesto Ramirez-Gonzalez. |
| EPI_ISL_2158122, EPI_ISL_2158123, EPI_ISL_2158124, EPI_ISL_2158125, EPI_ISL_2158126, EPI_ISL_2158127 | LESP Hidalgo | Instituto de Diagnostico y Referencia Epidemiologicos (INDRE) | Claudia Wong-Arambula, Abril Rodriguez-Maldonado, Vanessa Rivero-Arredondo, Ariadna Medina-Benitez, Joaquin Quiroz-Mercado, Sergio Rangel-Guerrero, Natividad Cruz-Ortiz, Tatiana Nunez-Garcia, Gisela Barrera-Badillo, Lucia Hernandez-Rivas, Irma Lopez-Martinez, Ernesto Ramirez-Gonzalez. |
| EPI_ISL_2158128, EPI_ISL_2158129, EPI_ISL_2158130 | LESP Puebla | Instituto de Diagnostico y Referencia Epidemiologicos (INDRE) | Claudia Wong-Arambula, Abril Rodriguez-Maldonado, Vanessa Rivero-Arredondo, Ariadna Medina-Benitez, Joaquin Quiroz-Mercado, Sergio Rangel-Guerrero, Natividad Cruz-Ortiz, Tatiana Nunez-Garcia, Gisela Barrera-Badillo, Lucia Hernandez-Rivas, Irma Lopez-Martinez, Ernesto Ramirez-Gonzalez. |
| EPI_ISL_2158131 | LESP Tlaxcala | Instituto de Diagnostico y Referencia Epidemiologicos (INDRE) | Claudia Wong-Arambula, Abril Rodriguez-Maldonado, Vanessa Rivero-Arredondo, Ariadna Medina-Benitez, Joaquin Quiroz-Mercado, Sergio Rangel-Guerrero, Natividad Cruz-Ortiz, Tatiana Nunez-Garcia, Gisela Barrera-Badillo, Lucia Hernandez-Rivas, Irma Lopez-Martinez, Ernesto Ramirez-Gonzalez. |
| EPI_ISL_2158132, EPI_ISL_2158133, EPI_ISL_2158134 | LESP Guanajuato | Instituto de Diagnostico y Referencia Epidemiologicos (INDRE) | Claudia Wong-Arambula, Abril Rodriguez-Maldonado, Vanessa Rivero-Arredondo, Ariadna Medina-Benitez, Joaquin Quiroz-Mercado, Sergio Rangel-Guerrero, Natividad Cruz-Ortiz, Tatiana Nunez-Garcia, Gisela Barrera-Badillo, Lucia Hernandez-Rivas, Irma Lopez-Martinez, Ernesto Ramirez-Gonzalez. |
| EPI_ISL_2158135 | LESP Jalisco | Instituto de Diagnostico y Referencia Epidemiologicos (INDRE) | Claudia Wong-Arambula, Abril Rodriguez-Maldonado, Vanessa Rivero-Arredondo, Ariadna Medina-Benitez, Joaquin Quiroz-Mercado, Sergio Rangel-Guerrero, Natividad Cruz-Ortiz, Tatiana Nunez-Garcia, Gisela Barrera-Badillo, Lucia Hernandez-Rivas, Irma Lopez-Martinez, Ernesto Ramirez-Gonzalez. |
| EPI_ISL_2158137 | LESP Nuevo Leon | Instituto de Diagnostico y Referencia Epidemiologicos (INDRE) | Claudia Wong-Arambula, Abril Rodriguez-Maldonado, Vanessa Rivero-Arredondo, Ariadna Medina-Benitez, Joaquin Quiroz-Mercado, Sergio Rangel-Guerrero, Natividad Cruz-Ortiz, Tatiana Nunez-Garcia, Gisela Barrera-Badillo, Lucia Hernandez-Rivas, Irma Lopez-Martinez, Ernesto Ramirez-Gonzalez. |
| EPI_ISL_2158138, EPI_ISL_2158139, EPI_ISL_2158140 | LESP Colima | Instituto de Diagnostico y Referencia Epidemiologicos (INDRE) | Claudia Wong-Arambula, Abril Rodriguez-Maldonado, Vanessa Rivero-Arredondo, Ariadna Medina-Benitez, Joaquin Quiroz-Mercado, Sergio Rangel-Guerrero, Natividad Cruz-Ortiz, Tatiana Nunez-Garcia, Gisela Barrera-Badillo, Lucia Hernandez-Rivas, Irma Lopez-Martinez, Ernesto Ramirez-Gonzalez. |

| | | | |
|---|---|---|---|
| EPI_ISL_2158141 | LESP Queretaro | Instituto de Diagnostico y Referencia Epidemiologicos (INDRE) | Claudia Wong-Arambula, Abril Rodriguez-Maldonado, Vanessa Rivero-Arredondo, Ariadna Medina-Benitez, Joaquin Quiroz-Mercado, Sergio Rangel-Guerrero, Natividad Cruz-Ortiz, Tatiana Nunez-Garcia, Gisela Barrera-Badillo, Lucia Hernandez-Rivas, Irma Lopez-Martinez, Ernesto Ramirez-Gonzalez. |
| EPI_ISL_2158142 | LESP Veracruz | Instituto de Diagnostico y Referencia Epidemiologicos (INDRE) | Claudia Wong-Arambula, Abril Rodriguez-Maldonado, Vanessa Rivero-Arredondo, Ariadna Medina-Benitez, Joaquin Quiroz-Mercado, Sergio Rangel-Guerrero, Natividad Cruz-Ortiz, Tatiana Nunez-Garcia, Gisela Barrera-Badillo, Lucia Hernandez-Rivas, Irma Lopez-Martinez, Ernesto Ramirez-Gonzalez. |
| EPI_ISL_2158143, EPI_ISL_2158144, EPI_ISL_2158145, EPI_ISL_2158146, EPI_ISL_2158147, EPI_ISL_2158148, EPI_ISL_2158149, EPI_ISL_2158150 | LESP Queretaro | Instituto de Diagnostico y Referencia Epidemiologicos (INDRE) | Claudia Wong-Arambula, Abril Rodriguez-Maldonado, Vanessa Rivero-Arredondo, Ariadna Medina-Benitez, Joaquin Quiroz-Mercado, Sergio Rangel-Guerrero, Natividad Cruz-Ortiz, Tatiana Nunez-Garcia, Gisela Barrera-Badillo, Lucia Hernandez-Rivas, Irma Lopez-Martinez, Ernesto Ramirez-Gonzalez. |
| EPI_ISL_2158151 | LESP Puebla | Instituto de Diagnostico y Referencia Epidemiologicos (INDRE) | Claudia Wong-Arambula, Abril Rodriguez-Maldonado, Vanessa Rivero-Arredondo, Ariadna Medina-Benitez, Joaquin Quiroz-Mercado, Sergio Rangel-Guerrero, Natividad Cruz-Ortiz, Tatiana Nunez-Garcia, Gisela Barrera-Badillo, Lucia Hernandez-Rivas, Irma Lopez-Martinez, Ernesto Ramirez-Gonzalez. |
| EPI_ISL_2158152, EPI_ISL_2158153, EPI_ISL_2158155, EPI_ISL_2158156, EPI_ISL_2158157, EPI_ISL_2158158, EPI_ISL_2158159, EPI_ISL_2158161 | LESP Morelos | Instituto de Diagnostico y Referencia Epidemiologicos (INDRE) | Claudia Wong-Arambula, Abril Rodriguez-Maldonado, Vanessa Rivero-Arredondo, Ariadna Medina-Benitez, Joaquin Quiroz-Mercado, Sergio Rangel-Guerrero, Natividad Cruz-Ortiz, Tatiana Nunez-Garcia, Gisela Barrera-Badillo, Lucia Hernandez-Rivas, Irma Lopez-Martinez, Ernesto Ramirez-Gonzalez. |
| EPI_ISL_2158162, EPI_ISL_2158163, EPI_ISL_2158164, EPI_ISL_2158165 | LESP Chihuahua | Instituto de Diagnostico y Referencia Epidemiologicos (INDRE) | Claudia Wong-Arambula, Abril Rodriguez-Maldonado, Vanessa Rivero-Arredondo, Ariadna Medina-Benitez, Joaquin Quiroz-Mercado, Sergio Rangel-Guerrero, Natividad Cruz-Ortiz, Tatiana Nunez-Garcia, Gisela Barrera-Badillo, Lucia Hernandez-Rivas, Irma Lopez-Martinez, Ernesto Ramirez-Gonzalez. |
| EPI_ISL_2158166 | LESP Ciudad de Mexico | Instituto de Diagnostico y Referencia Epidemiologicos (INDRE) | Claudia Wong-Arambula, Abril Rodriguez-Maldonado, Vanessa Rivero-Arredondo, Ariadna Medina-Benitez, Joaquin Quiroz-Mercado, Sergio Rangel-Guerrero, Natividad Cruz-Ortiz, Tatiana Nunez-Garcia, Gisela Barrera-Badillo, Lucia Hernandez-Rivas, Irma Lopez-Martinez, Ernesto Ramirez-Gonzalez. |
| EPI_ISL_2158167, EPI_ISL_2158168 | LESP Chihuahua | Instituto de Diagnostico y Referencia Epidemiologicos (INDRE) | Claudia Wong-Arambula, Abril Rodriguez-Maldonado, Vanessa Rivero-Arredondo, Ariadna Medina-Benitez, Joaquin Quiroz-Mercado, Sergio Rangel-Guerrero, Natividad Cruz-Ortiz, Tatiana Nunez-Garcia, Gisela Barrera-Badillo, Lucia Hernandez-Rivas, Irma Lopez-Martinez, Ernesto Ramirez-Gonzalez. |
| EPI_ISL_2158171, EPI_ISL_2158172 | LESP Jalisco | Instituto de Diagnostico y Referencia Epidemiologicos (INDRE) | Claudia Wong-Arambula, Abril Rodriguez-Maldonado, Vanessa Rivero-Arredondo, Ariadna Medina-Benitez, Joaquin Quiroz-Mercado, Sergio Rangel-Guerrero, Natividad Cruz-Ortiz, Tatiana Nunez-Garcia, Gisela Barrera-Badillo, Lucia Hernandez-Rivas, Irma Lopez-Martinez, Ernesto Ramirez-Gonzalez. |
| EPI_ISL_2158173 | LESP Sinaloa | Instituto de Diagnostico y Referencia Epidemiologicos (INDRE) | Claudia Wong-Arambula, Abril Rodriguez-Maldonado, Vanessa Rivero-Arredondo, Ariadna Medina-Benitez, Joaquin Quiroz-Mercado, Sergio Rangel-Guerrero, Natividad Cruz-Ortiz, Tatiana Nunez-Garcia, Gisela Barrera-Badillo, Lucia Hernandez-Rivas, Irma Lopez-Martinez, Ernesto Ramirez-Gonzalez. |
| EPI_ISL_2158174, EPI_ISL_2158175, EPI_ISL_2158176, EPI_ISL_2158177, EPI_ISL_2158178, EPI_ISL_2158179, EPI_ISL_2158181, EPI_ISL_2158183, EPI_ISL_2158184 | LESP Morelos | Instituto de Diagnostico y Referencia Epidemiologicos (INDRE) | Claudia Wong-Arambula, Abril Rodriguez-Maldonado, Vanessa Rivero-Arredondo, Ariadna Medina-Benitez, Joaquin Quiroz-Mercado, Sergio Rangel-Guerrero, Natividad Cruz-Ortiz, Tatiana Nunez-Garcia, Gisela Barrera-Badillo, Lucia Hernandez-Rivas, Irma Lopez-Martinez, Ernesto Ramirez-Gonzalez. |
| EPI_ISL_2158186, EPI_ISL_2158187, EPI_ISL_2158188, EPI_ISL_2158189, EPI_ISL_2158190, EPI_ISL_2158191, EPI_ISL_2158192, EPI_ISL_2158193, EPI_ISL_2158194 | LESP Chihuahua | Instituto de Diagnostico y Referencia Epidemiologicos (INDRE) | Claudia Wong-Arambula, Abril Rodriguez-Maldonado, Vanessa Rivero-Arredondo, Ariadna Medina-Benitez, Joaquin Quiroz-Mercado, Sergio Rangel-Guerrero, Natividad Cruz-Ortiz, Tatiana Nunez-Garcia, Gisela Barrera-Badillo, Lucia Hernandez-Rivas, Irma Lopez-Martinez, Ernesto Ramirez-Gonzalez. |
| EPI_ISL_2158197 | LESP Nuevo Leon | Instituto de Diagnostico y Referencia Epidemiologicos (INDRE) | Claudia Wong-Arambula, Abril Rodriguez-Maldonado, Vanessa Rivero-Arredondo, Ariadna Medina-Benitez, Joaquin Quiroz-Mercado, Sergio Rangel-Guerrero, Natividad Cruz-Ortiz, Tatiana Nunez-Garcia, Gisela Barrera-Badillo, Lucia Hernandez-Rivas, Irma Lopez-Martinez, Ernesto Ramirez-Gonzalez. |
| EPI_ISL_2158198 | LESP Chihuahua | Instituto de Diagnostico y Referencia Epidemiologicos (INDRE) | Claudia Wong-Arambula, Abril Rodriguez-Maldonado, Vanessa Rivero-Arredondo, Ariadna Medina-Benitez, Joaquin Quiroz-Mercado, Sergio Rangel-Guerrero, Natividad Cruz-Ortiz, Tatiana Nunez-Garcia, Gisela Barrera-Badillo, Lucia Hernandez-Rivas, Irma Lopez-Martinez, Ernesto Ramirez-Gonzalez. |
| EPI_ISL_2158199, EPI_ISL_2158200 | LESP Puebla | Instituto de Diagnostico y Referencia Epidemiologicos (INDRE) | Claudia Wong-Arambula, Abril Rodriguez-Maldonado, Vanessa Rivero-Arredondo, Ariadna Medina-Benitez, Joaquin Quiroz-Mercado, Sergio Rangel-Guerrero, Natividad Cruz-Ortiz, Tatiana Nunez-Garcia, Gisela Barrera-Badillo, Lucia Hernandez-Rivas, Irma Lopez-Martinez, Ernesto Ramirez-Gonzalez. |
| EPI_ISL_2158201, EPI_ISL_2158202 | LESP Jalisco | Instituto de Diagnostico y Referencia Epidemiologicos (INDRE) | Claudia Wong-Arambula, Abril Rodriguez-Maldonado, Vanessa Rivero-Arredondo, Ariadna Medina-Benitez, Joaquin Quiroz-Mercado, Sergio Rangel-Guerrero, Natividad Cruz-Ortiz, Tatiana Nunez-Garcia, Gisela Barrera-Badillo, Lucia Hernandez-Rivas, Irma Lopez-Martinez, Ernesto Ramirez-Gonzalez. |
| EPI_ISL_2158203 | LESP Hidalgo | Instituto de Diagnostico y Referencia Epidemiologicos (INDRE) | Claudia Wong-Arambula, Abril Rodriguez-Maldonado, Vanessa Rivero-Arredondo, Ariadna Medina-Benitez, Joaquin Quiroz-Mercado, Sergio Rangel-Guerrero, Natividad Cruz-Ortiz, Tatiana Nunez-Garcia, Gisela Barrera-Badillo, Lucia Hernandez-Rivas, Irma Lopez-Martinez, Ernesto Ramirez-Gonzalez. |
| EPI_ISL_2158204 | LESP Nayarit | Instituto de Diagnostico y Referencia Epidemiologicos (INDRE) | Claudia Wong-Arambula, Abril Rodriguez-Maldonado, Vanessa Rivero-Arredondo, Ariadna Medina-Benitez, Joaquin Quiroz-Mercado, Sergio Rangel-Guerrero, Natividad Cruz-Ortiz, Tatiana Nunez-Garcia, Gisela Barrera-Badillo, Lucia Hernandez-Rivas, Irma Lopez-Martinez, Ernesto Ramirez-Gonzalez. |
| EPI_ISL_2158205 | LESP Veracruz | Instituto de Diagnostico y Referencia Epidemiologicos (INDRE) | Claudia Wong-Arambula, Abril Rodriguez-Maldonado, Vanessa Rivero-Arredondo, Ariadna Medina-Benitez, Joaquin Quiroz-Mercado, Sergio Rangel-Guerrero, Natividad Cruz-Ortiz, Tatiana Nunez-Garcia, Gisela Barrera-Badillo, Lucia Hernandez-Rivas, Irma Lopez-Martinez, Ernesto Ramirez-Gonzalez. |
| EPI_ISL_2158206 | LESP Tamaulipas | Instituto de Diagnostico y Referencia Epidemiologicos (INDRE) | Claudia Wong-Arambula, Abril Rodriguez-Maldonado, Vanessa Rivero-Arredondo, Ariadna Medina-Benitez, Joaquin Quiroz-Mercado, Sergio Rangel-Guerrero, Natividad Cruz-Ortiz, Tatiana Nunez-Garcia, Gisela Barrera-Badillo, Lucia Hernandez-Rivas, Irma Lopez-Martinez, Ernesto Ramirez-Gonzalez. |
| EPI_ISL_2158207, EPI_ISL_2158208, EPI_ISL_2158209, EPI_ISL_2158210, EPI_ISL_2158211, EPI_ISL_2158212, EPI_ISL_2158213, EPI_ISL_2158214, EPI_ISL_2158215, EPI_ISL_2158216, EPI_ISL_2158217, EPI_ISL_2158218, EPI_ISL_2158219, EPI_ISL_2158220, EPI_ISL_2158221, EPI_ISL_2158222, EPI_ISL_2158223, EPI_ISL_2158224, EPI_ISL_2158225 | | | |
| see above | LESP Guanajuato | Instituto de Diagnostico y Referencia Epidemiologicos (INDRE) | Claudia Wong-Arambula, Abril Rodriguez-Maldonado, Vanessa Rivero-Arredondo, Ariadna Medina-Benitez, Joaquin Quiroz-Mercado, Sergio Rangel-Guerrero, Natividad Cruz-Ortiz, Tatiana Nunez-Garcia, Gisela Barrera-Badillo, Lucia Hernandez-Rivas, Irma Lopez-Martinez, Ernesto Ramirez-Gonzalez. |
| EPI_ISL_2158226 | LESP Puebla | Instituto de Diagnostico y Referencia Epidemiologicos (INDRE) | Claudia Wong-Arambula, Abril Rodriguez-Maldonado, Vanessa Rivero-Arredondo, Ariadna Medina-Benitez, Joaquin Quiroz-Mercado, Sergio Rangel-Guerrero, Natividad Cruz-Ortiz, Tatiana Nunez-Garcia, Gisela Barrera-Badillo, Lucia Hernandez-Rivas, Irma Lopez-Martinez, Ernesto Ramirez-Gonzalez. |
| EPI_ISL_2158227 | LESP Hidalgo | Instituto de Diagnostico y Referencia Epidemiologicos (INDRE) | Claudia Wong-Arambula, Abril Rodriguez-Maldonado, Vanessa Rivero-Arredondo, Ariadna Medina-Benitez, Joaquin Quiroz-Mercado, Sergio Rangel-Guerrero, Natividad Cruz-Ortiz, Tatiana Nunez-Garcia, Gisela Barrera-Badillo, Lucia Hernandez-Rivas, Irma Lopez-Martinez, Ernesto |

| | | | |
|---|---|---|---|
| | | | Ramirez-Gonzalez. |
| EPI_ISL_2158228 | LESP Veracruz | Instituto de Diagnostico y Referencia Epidemiologicos (INDRE) | Claudia Wong-Arambula, Abril Rodriguez-Maldonado, Vanessa Rivero-Arredondo, Ariadna Medina-Benitez, Joaquin Quiroz-Mercado, Sergio Rangel-Guerrero, Natividad Cruz-Ortiz, Tatiana Nunez-Garcia, Gisela Barrera-Badillo, Lucia Hernandez-Rivas, Irma Lopez-Martinez, Ernesto Ramirez-Gonzalez. |
| EPI_ISL_2158229 | LESP Tamaulipas | Instituto de Diagnostico y Referencia Epidemiologicos (INDRE) | Claudia Wong-Arambula, Abril Rodriguez-Maldonado, Vanessa Rivero-Arredondo, Ariadna Medina-Benitez, Joaquin Quiroz-Mercado, Sergio Rangel-Guerrero, Natividad Cruz-Ortiz, Tatiana Nunez-Garcia, Gisela Barrera-Badillo, Lucia Hernandez-Rivas, Irma Lopez-Martinez, Ernesto Ramirez-Gonzalez. |
| EPI_ISL_2158230 | LESP Chiapas | Instituto de Diagnostico y Referencia Epidemiologicos (INDRE) | Claudia Wong-Arambula, Abril Rodriguez-Maldonado, Vanessa Rivero-Arredondo, Ariadna Medina-Benitez, Joaquin Quiroz-Mercado, Sergio Rangel-Guerrero, Natividad Cruz-Ortiz, Tatiana Nunez-Garcia, Gisela Barrera-Badillo, Lucia Hernandez-Rivas, Irma Lopez-Martinez, Ernesto Ramirez-Gonzalez. |
| EPI_ISL_2158232 | LESP Guerrero | Instituto de Diagnostico y Referencia Epidemiologicos (INDRE) | Claudia Wong-Arambula, Abril Rodriguez-Maldonado, Vanessa Rivero-Arredondo, Ariadna Medina-Benitez, Joaquin Quiroz-Mercado, Sergio Rangel-Guerrero, Natividad Cruz-Ortiz, Tatiana Nunez-Garcia, Gisela Barrera-Badillo, Lucia Hernandez-Rivas, Irma Lopez-Martinez, Ernesto Ramirez-Gonzalez. |
| EPI_ISL_2158233 | LESP Jalisco | Instituto de Diagnostico y Referencia Epidemiologicos (INDRE) | Claudia Wong-Arambula, Abril Rodriguez-Maldonado, Vanessa Rivero-Arredondo, Ariadna Medina-Benitez, Joaquin Quiroz-Mercado, Sergio Rangel-Guerrero, Natividad Cruz-Ortiz, Tatiana Nunez-Garcia, Gisela Barrera-Badillo, Lucia Hernandez-Rivas, Irma Lopez-Martinez, Ernesto Ramirez-Gonzalez. |
| EPI_ISL_2158234 | LESP Coahuila | Instituto de Diagnostico y Referencia Epidemiologicos (INDRE) | Claudia Wong-Arambula, Abril Rodriguez-Maldonado, Vanessa Rivero-Arredondo, Ariadna Medina-Benitez, Joaquin Quiroz-Mercado, Sergio Rangel-Guerrero, Natividad Cruz-Ortiz, Tatiana Nunez-Garcia, Gisela Barrera-Badillo, Lucia Hernandez-Rivas, Irma Lopez-Martinez, Ernesto Ramirez-Gonzalez. |
| EPI_ISL_2158249 | LESP Guanajuato | Instituto de Diagnostico y Referencia Epidemiologicos (INDRE) | Claudia Wong-Arambula, Abril Rodriguez-Maldonado, Vanessa Rivero-Arredondo, Ariadna Medina-Benitez, Joaquin Quiroz-Mercado, Sergio Rangel-Guerrero, Natividad Cruz-Ortiz, Tatiana Nunez-Garcia, Gisela Barrera-Badillo, Lucia Hernandez-Rivas, Irma Lopez-Martinez, Ernesto Ramirez-Gonzalez. |
| EPI_ISL_2158257, EPI_ISL_2158258, EPI_ISL_2158259, EPI_ISL_2158260, EPI_ISL_2158261, EPI_ISL_2158262 | LESP Ciudad de Mexico | Instituto de Diagnostico y Referencia Epidemiologicos (INDRE) | Claudia Wong-Arambula, Abril Rodriguez-Maldonado, Vanessa Rivero-Arredondo, Ariadna Medina-Benitez, Joaquin Quiroz-Mercado, Sergio Rangel-Guerrero, Natividad Cruz-Ortiz, Tatiana Nunez-Garcia, Gisela Barrera-Badillo, Lucia Hernandez-Rivas, Irma Lopez-Martinez, Ernesto Ramirez-Gonzalez. |
| EPI_ISL_2158310 | LESP Jalisco | Instituto de Diagnostico y Referencia Epidemiologicos (INDRE) | Claudia Wong-Arambula, Abril Rodriguez-Maldonado, Vanessa Rivero-Arredondo, Ariadna Medina-Benitez, Joaquin Quiroz-Mercado, Sergio Rangel-Guerrero, Natividad Cruz-Ortiz, Tatiana Nunez-Garcia, Gisela Barrera-Badillo, Lucia Hernandez-Rivas, Irma Lopez-Martinez, Ernesto Ramirez-Gonzalez. |
| EPI_ISL_2158311 | LESP Chihuahua | Instituto de Diagnostico y Referencia Epidemiologicos (INDRE) | Claudia Wong-Arambula, Abril Rodriguez-Maldonado, Vanessa Rivero-Arredondo, Ariadna Medina-Benitez, Joaquin Quiroz-Mercado, Sergio Rangel-Guerrero, Natividad Cruz-Ortiz, Tatiana Nunez-Garcia, Gisela Barrera-Badillo, Lucia Hernandez-Rivas, Irma Lopez-Martinez, Ernesto Ramirez-Gonzalez. |
| EPI_ISL_2158312, EPI_ISL_2158313 | LESP Nuevo Leon | Instituto de Diagnostico y Referencia Epidemiologicos (INDRE) | Claudia Wong-Arambula, Abril Rodriguez-Maldonado, Vanessa Rivero-Arredondo, Ariadna Medina-Benitez, Joaquin Quiroz-Mercado, Sergio Rangel-Guerrero, Natividad Cruz-Ortiz, Tatiana Nunez-Garcia, Gisela Barrera-Badillo, Lucia Hernandez-Rivas, Irma Lopez-Martinez, Ernesto Ramirez-Gonzalez. |
| EPI_ISL_2158314 | LESP Morelos | Instituto de Diagnostico y Referencia Epidemiologicos (INDRE) | Claudia Wong-Arambula, Abril Rodriguez-Maldonado, Vanessa Rivero-Arredondo, Ariadna Medina-Benitez, Joaquin Quiroz-Mercado, Sergio Rangel-Guerrero, Natividad Cruz-Ortiz, Tatiana Nunez-Garcia, Gisela Barrera-Badillo, Lucia Hernandez-Rivas, Irma Lopez-Martinez, Ernesto Ramirez-Gonzalez. |
| EPI_ISL_2158315, EPI_ISL_2158316, EPI_ISL_2158317, EPI_ISL_2158318 | LESP Ciudad de Mexico | Instituto de Diagnostico y Referencia Epidemiologicos (INDRE) | Claudia Wong-Arambula, Abril Rodriguez-Maldonado, Vanessa Rivero-Arredondo, Ariadna Medina-Benitez, Joaquin Quiroz-Mercado, Sergio Rangel-Guerrero, Natividad Cruz-Ortiz, Tatiana Nunez-Garcia, Gisela Barrera-Badillo, Lucia Hernandez-Rivas, Irma Lopez-Martinez, Ernesto Ramirez-Gonzalez. |
| EPI_ISL_2158319 | LESP Nuevo Leon | Instituto de Diagnostico y Referencia Epidemiologicos (INDRE) | Claudia Wong-Arambula, Abril Rodriguez-Maldonado, Vanessa Rivero-Arredondo, Ariadna Medina-Benitez, Joaquin Quiroz-Mercado, Sergio Rangel-Guerrero, Natividad Cruz-Ortiz, Tatiana Nunez-Garcia, Gisela Barrera-Badillo, Lucia Hernandez-Rivas, Irma Lopez-Martinez, Ernesto Ramirez-Gonzalez. |
| EPI_ISL_2158320 | LESP Chihuahua | Instituto de Diagnostico y Referencia Epidemiologicos (INDRE) | Claudia Wong-Arambula, Abril Rodriguez-Maldonado, Vanessa Rivero-Arredondo, Ariadna Medina-Benitez, Joaquin Quiroz-Mercado, Sergio Rangel-Guerrero, Natividad Cruz-Ortiz, Tatiana Nunez-Garcia, Gisela Barrera-Badillo, Lucia Hernandez-Rivas, Irma Lopez-Martinez, Ernesto Ramirez-Gonzalez. |
| EPI_ISL_2158321 | LESP Queretaro | Instituto de Diagnostico y Referencia Epidemiologicos (INDRE) | Claudia Wong-Arambula, Abril Rodriguez-Maldonado, Vanessa Rivero-Arredondo, Ariadna Medina-Benitez, Joaquin Quiroz-Mercado, Sergio Rangel-Guerrero, Natividad Cruz-Ortiz, Tatiana Nunez-Garcia, Gisela Barrera-Badillo, Lucia Hernandez-Rivas, Irma Lopez-Martinez, Ernesto Ramirez-Gonzalez. |
| EPI_ISL_2158353 | LDSP CUNDINAMARCA | Instituto Nacional de Salud- Dirección de Investigación en Salud Pública | Katherine Laiton-Donato, Diego A. Álvarez-Díaz, Carlos Franco-Muñoz, Hector Alejandro Ruiz-Moreno, Paola Rojas, Maria T. Herrera-Sepúlveda, Diego Andrés Prada, Jhonnatan Reales-González, Sheryll Corchuelo, Julian Naizaque, Jorge Rivera, Gerardo Santamaría, Sergio Gomez, Lisseth Pardo, Juan Camilo Martinez, Marta Lopez Blanco, Ángela Alarcon Cruz, Diana Malo, Carmen Osorio, Magdalena Wiesner, Martha Lucia Ospina Martinez, Marcela Mercado-Reyes |
| EPI_ISL_2158354, EPI_ISL_2158355 | Lab. Virología y Genética Universidad Simón Bolívar | Instituto Nacional de Salud- Dirección de Investigación en Salud Pública | Katherine Laiton-Donato, Diego A. Álvarez-Díaz, Carlos Franco-Muñoz, Hector Alejandro Ruiz-Moreno, Paola Rojas, Maria T. Herrera-Sepúlveda, Diego Andrés Prada, Jhonnatan Reales-González, Sheryll Corchuelo, Julian Naizaque, Jorge Rivera, Gerardo Santamaría, Sergio Gomez, Lisseth Pardo, Juan Camilo Martinez, Marta Lopez Blanco, Ángela Alarcon Cruz, Diana Malo, Carmen Osorio, Magdalena Wiesner, Martha Lucia Ospina Martinez, Marcela Mercado-Reyes |
| EPI_ISL_2161788, EPI_ISL_2161789, EPI_ISL_2161790, EPI_ISL_2161793, EPI_ISL_2161794, EPI_ISL_2161795, EPI_ISL_2161797, EPI_ISL_2161798, EPI_ISL_2161800, EPI_ISL_2161801, EPI_ISL_2161803, EPI_ISL_2161804, EPI_ISL_2161806, EPI_ISL_2161809, EPI_ISL_2161811 | | | |
| see above | NL-Dr. Leonard A. Miller Centre for Health Services | National Microbiology Laboratory (NML) | Anna Majer, Shari Tyson, Grace Seo, Philip Mabon, Elsie Grudeski, Rhiannon Huzarewich, Russell Mandes, Anneliese Landgraff, Jennifer Tanner, Natalie Knox, Morag Graham, Gary Van Domselaar, Robert Needle, Yang Yu, Adel Malek, Laura Gilbert, George Zahariadis, Nathalie Bastien, Yan Li, Timothy Booth, Darian Hole, Madison Chapel, Kirsten Biggar, Kerri Smith, CanCOGeN's metadata curation team, Public Health Agency of Canada CanCOGeN team |
| EPI_ISL_2162811, EPI_ISL_2162812, EPI_ISL_2162813, EPI_ISL_2162814, EPI_ISL_2162815, EPI_ISL_2162816, EPI_ISL_2162817, EPI_ISL_2162818, EPI_ISL_2162819, EPI_ISL_2162820, EPI_ISL_2162821, EPI_ISL_2162822, EPI_ISL_2162823, EPI_ISL_2162824, EPI_ISL_2162825, EPI_ISL_2162826, EPI_ISL_2162827, EPI_ISL_2162828, EPI_ISL_2162829, EPI_ISL_2162830 | | | |
| see above | Klinisch Laboratorium ZNA | Klinisch Laboratorium ZNA | Verstrepen et al. |
| EPI_ISL_2164238 | Basurto University Hospital: Clinical Microbiology Laboratory | Biocruces Bizkaia | Mikel Urrutikoetxea-Gutiérrez, Mª Carmen Nieto Toboso, Estibaliz Ugalde Zarraga, Mikel Gallego Rodrigo, Ana de la Hoz, José Luis Díaz de Tuesta del Arco |
| EPI_ISL_2164421 | Basurto University Hospital: Clinical Microbiology Laboratory | Biocruces Bizkaia | Basurto University Hospital: Clinical Microbiology Laboratory |
| EPI_ISL_2178411 | Labo Analyses Med | National Reference Center for Viruses of Respiratory Infections, Institut Pasteur, Paris | Marion Barbet, Sylvie Behillil, Méline Bizard, Angela Brisebarre, Camille Capel, Vincent Enouf, Louise Lefrançois, Frédéric Lemoine, Christophe Malabat, Corinne Maufrais, Etienne Simon-Lorière, Maud Vanpeene, Sylvie Van der Werf ,Bruno Berdin |
| EPI_ISL_2178412 | Labo Analyses Med | National Reference Center for Viruses of Respiratory Infections, Institut Pasteur, Paris | Marion Barbet, Sylvie Behillil, Méline Bizard, Angela Brisebarre, Camille Capel, Vincent Enouf, Louise Lefrançois, Frédéric Lemoine, Christophe Malabat, Corinne Maufrais, Etienne Simon-Lorière, Maud Vanpeene, Sylvie Van der Werf ,Nabil Gastli |
| EPI_ISL_2178414 | Labo Analyses med | National Reference Center for Viruses of Respiratory Infections, Institut Pasteur, Paris | Marion Barbet, Sylvie Behillil, Méline Bizard, Angela Brisebarre, Camille Capel, Vincent Enouf, Louise Lefrançois, Frédéric Lemoine, Christophe Malabat, Corinne Maufrais, Amaury Vaysse, Etienne Simon-Lorière, Maud Vanpeene, Sylvie Van der Werf ,Anne Holstein |

| | | | |
|---|---|---|---|
| EPI_ISL_2178415 | Hospital | National Reference Center for Viruses of Respiratory Infections, Institut Pasteur, Paris | Marion Barbet, Sylvie Behillil, Méline Bizard, Angela Brisebarre, Camille Capel, Vincent Enouf, Louise Lefrançois, Frédéric Lemoine, Christophe Malabat, Corinne Maufrais, Amaury Vaysse, Etienne Simon-Lorière, Maud Vanpeene, Sylvie Van der Werf ,LéA Pilorge |
| EPI_ISL_2178416 | Labo Analyses med | National Reference Center for Viruses of Respiratory Infections, Institut Pasteur, Paris | Marion Barbet, Sylvie Behillil, Méline Bizard, Angela Brisebarre, Camille Capel, Vincent Enouf, Louise Lefrançois, Frédéric Lemoine, Christophe Malabat, Corinne Maufrais, Amaury Vaysse, Etienne Simon-Lorière, Maud Vanpeene, Sylvie Van der Werf ,Anne Holstein |
| EPI_ISL_2178417 | Hospital | National Reference Center for Viruses of Respiratory Infections, Institut Pasteur, Paris | Marion Barbet, Sylvie Behillil, Méline Bizard, Angela Brisebarre, Camille Capel, Vincent Enouf, Louise Lefrançois, Frédéric Lemoine, Christophe Malabat, Corinne Maufrais, Etienne Simon-Lorière, Maud Vanpeene, Sylvie Van der Werf ,Eleha Guillotel |
| EPI_ISL_2178419 | Hospital | National Reference Center for Viruses of Respiratory Infections, Institut Pasteur, Paris | Marion Barbet, Sylvie Behillil, Méline Bizard, Angela Brisebarre, Camille Capel, Vincent Enouf, Louise Lefrançois, Frédéric Lemoine, Christophe Malabat, Corinne Maufrais, Etienne Simon-Lorière, Maud Vanpeene, Sylvie Van der Werf ,CéLine Bressollette |
| EPI_ISL_2178421, EPI_ISL_2178422 | Hospital | National Reference Center for Viruses of Respiratory Infections, Institut Pasteur, Paris | Marion Barbet, Sylvie Behillil, Méline Bizard, Angela Brisebarre, Camille Capel, Vincent Enouf, Louise Lefrançois, Frédéric Lemoine, Christophe Malabat, Corinne Maufrais, Amaury Vaysse, Etienne Simon-Lorière, Maud Vanpeene, Sylvie Van der Werf ,Alexandra Ducancelle |
| EPI_ISL_2178423 | Hospital | National Reference Center for Viruses of Respiratory Infections, Institut Pasteur, Paris | Marion Barbet, Sylvie Behillil, Méline Bizard, Angela Brisebarre, Camille Capel, Vincent Enouf, Louise Lefrançois, Frédéric Lemoine, Christophe Malabat, Corinne Maufrais, Etienne Simon-Lorière, Maud Vanpeene, Sylvie Van der Werf ,CéLine Bressollette |
| EPI_ISL_2178424 | Labo Analyses Med | National Reference Center for Viruses of Respiratory Infections, Institut Pasteur, Paris | Marion Barbet, Sylvie Behillil, Méline Bizard, Angela Brisebarre, Camille Capel, Vincent Enouf, Louise Lefrançois, Frédéric Lemoine, Christophe Malabat, Corinne Maufrais, Etienne Simon-Lorière, Maud Vanpeene, Sylvie Van der Werf ,GréGoire Potiron |
| EPI_ISL_2178426, EPI_ISL_2178427 | Labo Analyses med | National Reference Center for Viruses of Respiratory Infections, Institut Pasteur, Paris | Marion Barbet, Sylvie Behillil, Méline Bizard, Angela Brisebarre, Camille Capel, Vincent Enouf, Louise Lefrançois, Frédéric Lemoine, Christophe Malabat, Corinne Maufrais, Amaury Vaysse, Etienne Simon-Lorière, Maud Vanpeene, Sylvie Van der Werf ,Anne Holstein |
| EPI_ISL_2178428 | Labo Analyses Med | National Reference Center for Viruses of Respiratory Infections, Institut Pasteur, Paris | Marion Barbet, Sylvie Behillil, Méline Bizard, Angela Brisebarre, Camille Capel, Vincent Enouf, Louise Lefrançois, Frédéric Lemoine, Christophe Malabat, Corinne Maufrais, Etienne Simon-Lorière, Maud Vanpeene, Sylvie Van der Werf ,Jean-FrançOis Comes |
| EPI_ISL_2178430, EPI_ISL_2178431 | Labo Analyses med | National Reference Center for Viruses of Respiratory Infections, Institut Pasteur, Paris | Marion Barbet, Sylvie Behillil, Méline Bizard, Angela Brisebarre, Camille Capel, Vincent Enouf, Louise Lefrançois, Frédéric Lemoine, Christophe Malabat, Corinne Maufrais, Amaury Vaysse, Etienne Simon-Lorière, Maud Vanpeene, Sylvie Van der Werf ,Anne Holstein |
| EPI_ISL_2178433 | Labo Analyses Med | National Reference Center for Viruses of Respiratory Infections, Institut Pasteur, Paris | Marion Barbet, Sylvie Behillil, Méline Bizard, Angela Brisebarre, Camille Capel, Vincent Enouf, Louise Lefrançois, Frédéric Lemoine, Christophe Malabat, Corinne Maufrais, Amaury Vaysse, Etienne Simon-Lorière, Maud Vanpeene, Sylvie Van der Werf ,Anne Cady |
| EPI_ISL_2178434 | Labo Analyses Med | National Reference Center for Viruses of Respiratory Infections, Institut Pasteur, Paris | Marion Barbet, Sylvie Behillil, Méline Bizard, Angela Brisebarre, Camille Capel, Vincent Enouf, Louise Lefrançois, Frédéric Lemoine, Christophe Malabat, Corinne Maufrais, Etienne Simon-Lorière, Maud Vanpeene, Sylvie Van der Werf ,Sophie Chalmin |
| EPI_ISL_2178435, EPI_ISL_2178436, EPI_ISL_2178437 | Labo Analyses Med | National Reference Center for Viruses of Respiratory Infections, Institut Pasteur, Paris | Marion Barbet, Sylvie Behillil, Méline Bizard, Angela Brisebarre, Camille Capel, Vincent Enouf, Louise Lefrançois, Frédéric Lemoine, Christophe Malabat, Corinne Maufrais, Damien Mornico, Etienne Simon-Lorière, Maud Vanpeene, Sylvie Van der Werf ,Sophie Chalmin |
| EPI_ISL_2178438 | Hospital | National Reference Center for Viruses of Respiratory Infections, Institut Pasteur, Paris | Marion Barbet, Sylvie Behillil, Méline Bizard, Angela Brisebarre, Camille Capel, Vincent Enouf, Louise Lefrançois, Frédéric Lemoine, Christophe Malabat, Corinne Maufrais, Damien Mornico, Etienne Simon-Lorière, Maud Vanpeene, Sylvie Van der Werf ,JéRome Guinard |
| EPI_ISL_2178442, EPI_ISL_2178443 | Labo Analyses Med | National Reference Center for Viruses of Respiratory Infections, Institut Pasteur, Paris | Marion Barbet, Sylvie Behillil, Méline Bizard, Angela Brisebarre, Camille Capel, Vincent Enouf, Louise Lefrançois, Frédéric Lemoine, Christophe Malabat, Corinne Maufrais, Pierre Lechat, Etienne Simon-Lorière, Maud Vanpeene, Sylvie Van der Werf ,Nabil Gastli |
| EPI_ISL_2178446 | Medicine Sentinel Network Province | National Reference Center for Viruses of Respiratory Infections, Institut Pasteur, Paris | Marion Barbet, Sylvie Behillil, Méline Bizard, Angela Brisebarre, Camille Capel, Vincent Enouf, Louise Lefrançois, Frédéric Lemoine, Christophe Malabat, Corinne Maufrais, Emmanuelle Permal, Etienne Simon-Lorière, Maud Vanpeene, Sylvie Van der Werf ,Jean-Christian Bientz |
| EPI_ISL_2178447, EPI_ISL_2178448 | Hospital | National Reference Center for Viruses of Respiratory Infections, Institut Pasteur, Paris | Marion Barbet, Sylvie Behillil, Méline Bizard, Angela Brisebarre, Camille Capel, Vincent Enouf, Louise Lefrançois, Frédéric Lemoine, Christophe Malabat, Corinne Maufrais, Emmanuelle Permal, Etienne Simon-Lorière, Maud Vanpeene, Sylvie Van der Werf ,Odile Petsaris |
| EPI_ISL_2178449 | Labo Analyses Med | National Reference Center for Viruses of Respiratory Infections, Institut Pasteur, Paris | Marion Barbet, Sylvie Behillil, Méline Bizard, Angela Brisebarre, Camille Capel, Vincent Enouf, Louise Lefrançois, Frédéric Lemoine, Christophe Malabat, Corinne Maufrais, Emmanuelle Permal, Etienne Simon-Lorière, Maud Vanpeene, Sylvie Van der Werf ,J. Simart |
| EPI_ISL_2178450 | Hospital | National Reference Center for Viruses of Respiratory Infections, Institut Pasteur, Paris | Marion Barbet, Sylvie Behillil, Méline Bizard, Angela Brisebarre, Camille Capel, Vincent Enouf, Louise Lefrançois, Frédéric Lemoine, Christophe Malabat, Corinne Maufrais, Emmanuelle Permal, Etienne Simon-Lorière, Maud Vanpeene, Sylvie Van der Werf ,Pascale Martres |
| EPI_ISL_2178451 | Labo Analyses Med | National Reference Center for Viruses of Respiratory Infections, Institut Pasteur, Paris | Marion Barbet, Sylvie Behillil, Méline Bizard, Angela Brisebarre, Camille Capel, Vincent Enouf, Louise Lefrançois, Frédéric Lemoine, Christophe Malabat, Corinne Maufrais, Emmanuelle Permal, Etienne Simon-Lorière, Maud Vanpeene, Sylvie Van der Werf ,Jean-FrançOis Comes |
| EPI_ISL_2178466 | Labo Analyses Med | National Reference Center for Viruses of Respiratory Infections, Institut Pasteur, Paris | Marion Barbet, Sylvie Behillil, Méline Bizard, Angela Brisebarre, Camille Capel, Vincent Enouf, Louise Lefrançois, Frédéric Lemoine, Christophe Malabat, Corinne Maufrais, Emmanuelle Permal, Etienne Simon-Lorière, Maud Vanpeene, Sylvie Van der Werf ,Bruno Berdin |
| EPI_ISL_2178467 | Labo Analyses Med | National Reference Center for Viruses of Respiratory Infections, Institut Pasteur, Paris | Marion Barbet, Sylvie Behillil, Méline Bizard, Angela Brisebarre, Camille Capel, Vincent Enouf, Louise Lefrançois, Frédéric Lemoine, Christophe Malabat, Corinne Maufrais, Amaury Vaysse, Etienne Simon-Lorière, Maud Vanpeene, Sylvie Van der Werf ,Pierre-Yves Leonard |
| EPI_ISL_2178469 | Hospital | National Reference Center for Viruses of Respiratory Infections, Institut Pasteur, Paris | Marion Barbet, Sylvie Behillil, Méline Bizard, Angela Brisebarre, Camille Capel, Vincent Enouf, Louise Lefrançois, Frédéric Lemoine, Christophe Malabat, Corinne Maufrais, Etienne Simon-Lorière, Maud Vanpeene, Sylvie Van der Werf ,FréDéRique Canis |
| EPI_ISL_2178471 | Labo Analyses Med | National Reference Center for Viruses of Respiratory Infections, Institut Pasteur, Paris | Marion Barbet, Sylvie Behillil, Méline Bizard, Angela Brisebarre, Camille Capel, Vincent Enouf, Louise Lefrançois, Frédéric Lemoine, Christophe Malabat, Corinne Maufrais, Pierre Lechat, Etienne Simon-Lorière, Maud Vanpeene, Sylvie Van der Werf ,Arditi Cocco |
| EPI_ISL_2178472 | Hospital | National Reference Center for Viruses of Respiratory Infections, Institut Pasteur, Paris | Marion Barbet, Sylvie Behillil, Méline Bizard, Angela Brisebarre, Camille Capel, Vincent Enouf, Louise Lefrançois, Frédéric Lemoine, Christophe Malabat, Corinne Maufrais, Etienne Simon-Lorière, Maud Vanpeene, Sylvie Van der Werf ,Nabil Gastli |
| EPI_ISL_2178473 | Hospital | National Reference Center for Viruses of Respiratory Infections, Institut Pasteur, Paris | Marion Barbet, Sylvie Behillil, Méline Bizard, Angela Brisebarre, Camille Capel, Vincent Enouf, Louise Lefrançois, Frédéric Lemoine, Christophe Malabat, Corinne Maufrais, Etienne Simon-Lorière, Maud Vanpeene, Sylvie Van der Werf ,Thibault Guinoiseau |
| EPI_ISL_2178474 | Hospital | National Reference Center for Viruses of Respiratory Infections, Institut Pasteur, Paris | Marion Barbet, Sylvie Behillil, Méline Bizard, Angela Brisebarre, Camille Capel, Vincent Enouf, Louise Lefrançois, Frédéric Lemoine, Christophe Malabat, Corinne Maufrais, Etienne Simon-Lorière, Maud Vanpeene, Sylvie Van der Werf ,Nabil Gastli |
| EPI_ISL_2178475 | Labo Analyses Med | National Reference Center for Viruses of Respiratory Infections, Institut Pasteur, Paris | Marion Barbet, Sylvie Behillil, Méline Bizard, Angela Brisebarre, Camille Capel, Vincent Enouf, Louise Lefrançois, Frédéric Lemoine, Christophe Malabat, Corinne Maufrais, Amaury Vaysse, Etienne Simon-Lorière, Maud Vanpeene, Sylvie Van der Werf ,Brieuc Gestin |
| EPI_ISL_2178477 | Labo Analyses Med | National Reference Center for Viruses of Respiratory Infections, Institut Pasteur, Paris | Marion Barbet, Sylvie Behillil, Méline Bizard, Angela Brisebarre, Camille Capel, Vincent Enouf, Louise Lefrançois, Frédéric Lemoine, Christophe Malabat, Corinne Maufrais, Pierre Lechat, Etienne Simon-Lorière, Maud Vanpeene, Sylvie Van der Werf ,Franck Ennouchi |
| EPI_ISL_2178478, EPI_ISL_2178479 | Labo Analyses Med | National Reference Center for Viruses of Respiratory Infections, Institut Pasteur, Paris | Marion Barbet, Sylvie Behillil, Méline Bizard, Angela Brisebarre, Camille Capel, Vincent Enouf, Louise Lefrançois, Frédéric Lemoine, Christophe Malabat, Corinne Maufrais, Amaury Vaysse, Etienne Simon-Lorière, Maud Vanpeene, Sylvie Van der Werf ,Pierre-Yves Leonard |
| EPI_ISL_2178480 | Labo analyses med | National Reference Center for Viruses of Respiratory Infections, Institut Pasteur, Paris | Marion Barbet, Sylvie Behillil, Méline Bizard, Angela Brisebarre, Camille Capel, Vincent Enouf, Louise Lefrançois, Frédéric Lemoine, Christophe Malabat, Corinne Maufrais, Amaury Vaysse, Etienne Simon-Lorière, Maud Vanpeene, Sylvie Van der Werf ,Jean-FrançOis Comes |
| EPI_ISL_2178481 | Hospital | National Reference Center for Viruses of Respiratory Infections, Institut Pasteur, Paris | Marion Barbet, Sylvie Behillil, Méline Bizard, Angela Brisebarre, Camille Capel, Vincent Enouf, Louise Lefrançois, Frédéric Lemoine, Christophe Malabat, Corinne Maufrais, Etienne Simon-Lorière, Maud Vanpeene, Sylvie Van der Werf ,EugèNe Castelain |
| EPI_ISL_2178482 | Hospital | National Reference Center for Viruses of Respiratory Infections, Institut Pasteur, Paris | Marion Barbet, Sylvie Behillil, Méline Bizard, Angela Brisebarre, Camille Capel, Vincent Enouf, Louise Lefrançois, Frédéric Lemoine, Christophe Malabat, Corinne Maufrais, Victoire Baillet, Etienne Simon-Lorière, Maud Vanpeene, Sylvie Van der Werf ,ValéRie Serazin |
| EPI_ISL_2178483 | Labo analyses med | National Reference Center for Viruses of Respiratory Infections, Institut Pasteur, Paris | Marion Barbet, Sylvie Behillil, Méline Bizard, Angela Brisebarre, Camille Capel, Vincent Enouf, Louise Lefrançois, Frédéric Lemoine, Christophe Malabat, Corinne Maufrais, Amaury Vaysse, Etienne Simon-Lorière, Maud Vanpeene, Sylvie Van der Werf ,Jean-FrançOis Comes |
| EPI_ISL_2178484, EPI_ISL_2178485 | Labo Analyses Med | National Reference Center for Viruses of Respiratory Infections, Institut Pasteur, Paris | Marion Barbet, Sylvie Behillil, Méline Bizard, Angela Brisebarre, Camille Capel, Vincent Enouf, Louise Lefrançois, Frédéric Lemoine, Christophe Malabat, Corinne Maufrais, Pierre Lechat, Etienne Simon-Lorière, Maud Vanpeene, Sylvie Van der Werf ,GréGoire Potiron |

| | | | |
|---|---|---|---|
| EPI_ISL_2178486 | Labo Analyses Med | National Reference Center for Viruses of Respiratory Infections, Institut Pasteur, Paris | Marion Barbet, Sylvie Behillil, Méline Bizard, Angela Brisebarre, Camille Capel, Vincent Enouf, Louise Lefrançois, Frédéric Lemoine, Christophe Malabat, Corinne Maufrais, Pierre Lechat, Etienne Simon-Lorière, Maud Vanpeene, Sylvie Van der Werf ,Arditi Cocco |
| EPI_ISL_2178487 | Labo Analyses Med | National Reference Center for Viruses of Respiratory Infections, Institut Pasteur, Paris | Marion Barbet, Sylvie Behillil, Méline Bizard, Angela Brisebarre, Camille Capel, Vincent Enouf, Louise Lefrançois, Frédéric Lemoine, Christophe Malabat, Corinne Maufrais, Amaury Vaysse, Etienne Simon-Lorière, Maud Vanpeene, Sylvie Van der Werf ,Valentin Robbe |
| EPI_ISL_2178488 | Hospital | National Reference Center for Viruses of Respiratory Infections, Institut Pasteur, Paris | Marion Barbet, Sylvie Behillil, Méline Bizard, Angela Brisebarre, Camille Capel, Vincent Enouf, Louise Lefrançois, Frédéric Lemoine, Christophe Malabat, Corinne Maufrais, Amaury Vaysse, Etienne Simon-Lorière, Maud Vanpeene, Sylvie Van der Werf ,Ilka Engelmann |
| EPI_ISL_2178489, EPI_ISL_2178490 | Hospital | National Reference Center for Viruses of Respiratory Infections, Institut Pasteur, Paris | Marion Barbet, Sylvie Behillil, Méline Bizard, Angela Brisebarre, Camille Capel, Vincent Enouf, Louise Lefrançois, Frédéric Lemoine, Christophe Malabat, Corinne Maufrais, Amaury Vaysse, Etienne Simon-Lorière, Maud Vanpeene, Sylvie Van der Werf ,CéLine Bressollette |
| EPI_ISL_2178491 | Labo Analyses Med | National Reference Center for Viruses of Respiratory Infections, Institut Pasteur, Paris | Marion Barbet, Sylvie Behillil, Méline Bizard, Angela Brisebarre, Camille Capel, Vincent Enouf, Louise Lefrançois, Frédéric Lemoine, Christophe Malabat, Corinne Maufrais, Amaury Vaysse, Etienne Simon-Lorière, Maud Vanpeene, Sylvie Van der Werf |
| EPI_ISL_2178492 | Labo Analyses Med | National Reference Center for Viruses of Respiratory Infections, Institut Pasteur, Paris | Marion Barbet, Sylvie Behillil, Méline Bizard, Angela Brisebarre, Camille Capel, Vincent Enouf, Louise Lefrançois, Frédéric Lemoine, Christophe Malabat, Corinne Maufrais, Emmanuelle Permal, Etienne Simon-Lorière, Maud Vanpeene, Sylvie Van der Werf ,Brieuc Gestin |
| EPI_ISL_2178493 | Labo Analyses Med | National Reference Center for Viruses of Respiratory Infections, Institut Pasteur, Paris | Marion Barbet, Sylvie Behillil, Méline Bizard, Angela Brisebarre, Camille Capel, Vincent Enouf, Louise Lefrançois, Frédéric Lemoine, Christophe Malabat, Corinne Maufrais, Amaury Vaysse, Etienne Simon-Lorière, Maud Vanpeene, Sylvie Van der Werf ,Brieuc Gestin |
| EPI_ISL_2178494 | Labo Analyses Med | National Reference Center for Viruses of Respiratory Infections, Institut Pasteur, Paris | Marion Barbet, Sylvie Behillil, Méline Bizard, Angela Brisebarre, Camille Capel, Vincent Enouf, Louise Lefrançois, Frédéric Lemoine, Christophe Malabat, Corinne Maufrais, Pierre Lechat, Etienne Simon-Lorière, Maud Vanpeene, Sylvie Van der Werf ,Anne Holstein |
| EPI_ISL_2178495, EPI_ISL_2178496 | Labo Analyses Med | National Reference Center for Viruses of Respiratory Infections, Institut Pasteur, Paris | Marion Barbet, Sylvie Behillil, Méline Bizard, Angela Brisebarre, Camille Capel, Vincent Enouf, Louise Lefrançois, Frédéric Lemoine, Christophe Malabat, Corinne Maufrais, Emmanuelle Permal, Etienne Simon-Lorière, Maud Vanpeene, Sylvie Van der Werf ,Anne Holstein |
| EPI_ISL_2178497 | Labo Analyses Med | National Reference Center for Viruses of Respiratory Infections, Institut Pasteur, Paris | Marion Barbet, Sylvie Behillil, Méline Bizard, Angela Brisebarre, Camille Capel, Vincent Enouf, Louise Lefrançois, Frédéric Lemoine, Christophe Malabat, Corinne Maufrais, Amaury Vaysse, Etienne Simon-Lorière, Maud Vanpeene, Sylvie Van der Werf ,Pierre-Yves Leonard |
| EPI_ISL_2178498 | Hospital | National Reference Center for Viruses of Respiratory Infections, Institut Pasteur, Paris | Marion Barbet, Sylvie Behillil, Méline Bizard, Angela Brisebarre, Camille Capel, Vincent Enouf, Louise Lefrançois, Frédéric Lemoine, Christophe Malabat, Corinne Maufrais, Amaury Vaysse, Etienne Simon-Lorière, Maud Vanpeene, Sylvie Van der Werf ,CéLine Bressollette |
| EPI_ISL_2178499 | Hospital | National Reference Center for Viruses of Respiratory Infections, Institut Pasteur, Paris | Marion Barbet, Sylvie Behillil, Méline Bizard, Angela Brisebarre, Camille Capel, Vincent Enouf, Louise Lefrançois, Frédéric Lemoine, Christophe Malabat, Corinne Maufrais, Emmanuelle Permal, Etienne Simon-Lorière, Maud Vanpeene, Sylvie Van der Werf ,Ali Si-Mohammed |
| EPI_ISL_2178500 | Hospital | National Reference Center for Viruses of Respiratory Infections, Institut Pasteur, Paris | Marion Barbet, Sylvie Behillil, Méline Bizard, Angela Brisebarre, Camille Capel, Vincent Enouf, Louise Lefrançois, Frédéric Lemoine, Christophe Malabat, Corinne Maufrais, Pierre Lechat, Etienne Simon-Lorière, Maud Vanpeene, Sylvie Van der Werf ,Jean Baptiste Bour |
| EPI_ISL_2178501 | Hospital | National Reference Center for Viruses of Respiratory Infections, Institut Pasteur, Paris | Marion Barbet, Sylvie Behillil, Méline Bizard, Angela Brisebarre, Camille Capel, Vincent Enouf, Louise Lefrançois, Frédéric Lemoine, Christophe Malabat, Corinne Maufrais, Amaury Vaysse, Etienne Simon-Lorière, Maud Vanpeene, Sylvie Van der Werf ,CéLine Bressollette |
| EPI_ISL_2178502 | Hospital | National Reference Center for Viruses of Respiratory Infections, Institut Pasteur, Paris | Marion Barbet, Sylvie Behillil, Méline Bizard, Angela Brisebarre, Camille Capel, Vincent Enouf, Louise Lefrançois, Frédéric Lemoine, Christophe Malabat, Corinne Maufrais, Etienne Simon-Lorière, Maud Vanpeene, Sylvie Van der Werf ,Nabil Gastli |
| EPI_ISL_2178503 | Labo Analyses Med | National Reference Center for Viruses of Respiratory Infections, Institut Pasteur, Paris | Marion Barbet, Sylvie Behillil, Méline Bizard, Angela Brisebarre, Camille Capel, Vincent Enouf, Louise Lefrançois, Frédéric Lemoine, Christophe Malabat, Corinne Maufrais, Etienne Simon-Lorière, Maud Vanpeene, Sylvie Van der Werf ,Ronan Pichard |
| EPI_ISL_2178504 | Hospital | National Reference Center for Viruses of Respiratory Infections, Institut Pasteur, Paris | Marion Barbet, Sylvie Behillil, Méline Bizard, Angela Brisebarre, Camille Capel, Vincent Enouf, Louise Lefrançois, Frédéric Lemoine, Christophe Malabat, Corinne Maufrais, Victoire Baillet, Etienne Simon-Lorière, Maud Vanpeene, Sylvie Van der Werf ,A Hochart |
| EPI_ISL_2178505, EPI_ISL_2178506 | Hospital | National Reference Center for Viruses of Respiratory Infections, Institut Pasteur, Paris | Marion Barbet, Sylvie Behillil, Méline Bizard, Angela Brisebarre, Camille Capel, Vincent Enouf, Louise Lefrançois, Frédéric Lemoine, Christophe Malabat, Corinne Maufrais, Amaury Vaysse, Etienne Simon-Lorière, Maud Vanpeene, Sylvie Van der Werf ,Ilka Engelmann |
| EPI_ISL_2178507 | Hospital | National Reference Center for Viruses of Respiratory Infections, Institut Pasteur, Paris | Marion Barbet, Sylvie Behillil, Méline Bizard, Angela Brisebarre, Camille Capel, Vincent Enouf, Louise Lefrançois, Frédéric Lemoine, Christophe Malabat, Corinne Maufrais, Amaury Vaysse, Etienne Simon-Lorière, Maud Vanpeene, Sylvie Van der Werf ,Dominique Descamps |
| EPI_ISL_2178667, EPI_ISL_2178668, EPI_ISL_2178669, EPI_ISL_2178670, EPI_ISL_2178671, EPI_ISL_2178672, EPI_ISL_2178673, EPI_ISL_2178674, EPI_ISL_2178675, EPI_ISL_2178676, EPI_ISL_2178677, EPI_ISL_2178678, EPI_ISL_2178679, EPI_ISL_2178680, EPI_ISL_2178681, EPI_ISL_2178682, EPI_ISL_2178683, EPI_ISL_2178684, EPI_ISL_2178685, EPI_ISL_2178686, EPI_ISL_2178687, EPI_ISL_2178688, EPI_ISL_2178689, EPI_ISL_2178690, EPI_ISL_2178691, EPI_ISL_2178692, EPI_ISL_2178693, EPI_ISL_2178694, EPI_ISL_2178695, EPI_ISL_2178696, EPI_ISL_2178697, EPI_ISL_2178698, EPI_ISL_2178699, EPI_ISL_2178700, EPI_ISL_2178701, EPI_ISL_2178702, EPI_ISL_2178703, EPI_ISL_2178704, EPI_ISL_2178705, EPI_ISL_2178706, EPI_ISL_2178707, EPI_ISL_2178708, EPI_ISL_2178709, EPI_ISL_2178710, EPI_ISL_2178711, EPI_ISL_2178712, EPI_ISL_2178713, EPI_ISL_2178714, EPI_ISL_2178715, EPI_ISL_2178716, EPI_ISL_2178717, EPI_ISL_2178718, EPI_ISL_2178719, EPI_ISL_2178720, EPI_ISL_2178721, EPI_ISL_2178722, EPI_ISL_2178723, EPI_ISL_2178724 | | | |
| see above | Tropical Medicine Institute Pedro Kouri | Respiratory Infections Laboratory | Yahisel Tejero Suarez, Lissette Perez Santos, Yanet Pintos Saavedra, Dailyn Medero Díaz, Odalys Valdes, Mirtha Elisa Aguado, Liena Ponce |
| EPI_ISL_2178726 | Centro de Investigación Biomédica del Noreste (CIBIN) | Centro de Investigación en Enfermedades Infecciosas (CIENI), Instituto Nacional de Enfermedades Respiratorias (INER) | Consorcio Mexicano de Vigilancia Genómica (CoViGen-Mex). Authors (in alphabetical order): Julio Elias Alvarado-Yaah, Carlos F. Arias, Santiago Ávila-Ríos, Víctor Hugo Borja-Aburto, Celia Boukadida, Juan Bautista Chale-Dzul, Célida Duque Molina, José Antonio Enciso-Moreno, Gloria Elena Espinosa-Ayala, Fernando Fontove-Herrera, Victor Eduardo Garcia-Arias, Concepción Grajales-Muñiz, Ricardo Grande, Alfredo Herrera-Estrella, Carla Ivón Herrera-Najera, Pavel Isa, Brenda Irasema Maldonado-Meza, Bernardo Martínez-Miguel, Margarita Matías-Florentino, María Guadalupe de Jesús Mireles-Rivera, Gloria María Molina-Salinas, Hector Montoya-Fuentes, José Esteban Muñoz-Medina, José de Jesús Nuñez-Contreras, Alicia Ocaña-Mondragón, Luis Alberto Ochoa-Carrera, Hector Esteban Paz-Juárez, Francisco Pulido, Helen Haydee Fernanda Ramírez-Plascencia, Angel Gustavo Salas-Lais, Alejandro Sanchez-Flores, Clara Esperanza Santacruz-Tinoco, María Guadalupe Santiago-Mauricio, Nelly Sélem-Mojica, Blanca Taboada, Gloria Vazquez |
| EPI_ISL_2178728 | Unidad de Investigación Médica de Yucatán (UIMY) | Centro de Investigación en Enfermedades Infecciosas (CIENI), Instituto Nacional de Enfermedades Respiratorias (INER) | Consorcio Mexicano de Vigilancia Genómica (CoViGen-Mex). Authors (in alphabetical order): Julio Elias Alvarado-Yaah, Carlos F. Arias, Santiago Ávila-Ríos, Víctor Hugo Borja-Aburto, Celia Boukadida, Juan Bautista Chale-Dzul, Célida Duque Molina, José Antonio Enciso-Moreno, Gloria Elena Espinosa-Ayala, Fernando Fontove-Herrera, Victor Eduardo Garcia-Arias, Concepción Grajales-Muñiz, Ricardo Grande, Alfredo Herrera-Estrella, Carla Ivón Herrera-Najera, Pavel Isa, Brenda Irasema Maldonado-Meza, Bernardo Martínez-Miguel, Margarita Matías-Florentino, María Guadalupe de Jesús Mireles-Rivera, Gloria María Molina-Salinas, Hector Montoya-Fuentes, José Esteban Muñoz-Medina, José de Jesús Nuñez-Contreras, Alicia Ocaña-Mondragón, Luis Alberto Ochoa-Carrera, Hector Esteban Paz-Juárez, Francisco Pulido, Helen Haydee Fernanda Ramírez-Plascencia, Angel Gustavo Salas-Lais, Alejandro Sanchez-Flores, Clara Esperanza Santacruz-Tinoco, María Guadalupe Santiago-Mauricio, Nelly Sélem-Mojica, Blanca Taboada, Gloria Vazquez |
| EPI_ISL_2178733, EPI_ISL_2178734, EPI_ISL_2178735, EPI_ISL_2178736, EPI_ISL_2178737, EPI_ISL_2178738, EPI_ISL_2178739, EPI_ISL_2178740, EPI_ISL_2178741 | Laboratorio Central de Epidemiología (LCE) | Centro de Investigación en Enfermedades Infecciosas (CIENI), Instituto Nacional de Enfermedades Respiratorias (INER) | Consorcio Mexicano de Vigilancia Genómica (CoViGen-Mex). Authors (in alphabetical order): Julio Elias Alvarado-Yaah, Carlos F. Arias, Santiago Ávila-Ríos, Víctor Hugo Borja-Aburto, Celia Boukadida, Juan Bautista Chale-Dzul, Célida Duque Molina, José Antonio Enciso-Moreno, Gloria Elena Espinosa-Ayala, Fernando Fontove-Herrera, Victor Eduardo Garcia-Arias, Concepción Grajales-Muñiz, Ricardo Grande, Alfredo Herrera-Estrella, Carla Ivón Herrera-Najera, Pavel Isa, Brenda Irasema Maldonado-Meza, Bernardo Martínez-Miguel, Margarita Matías-Florentino, María Guadalupe de Jesús Mireles-Rivera, Gloria María Molina-Salinas, Hector Montoya-Fuentes, José Esteban Muñoz-Medina, José de Jesús Nuñez-Contreras, Alicia Ocaña-Mondragón, Luis Alberto Ochoa-Carrera, Hector Esteban Paz-Juárez, Francisco Pulido, Helen Haydee Fernanda Ramírez-Plascencia, Angel Gustavo Salas-Lais, Alejandro Sanchez-Flores, Clara Esperanza Santacruz-Tinoco, María Guadalupe Santiago-Mauricio, Nelly Sélem-Mojica, Blanca Taboada, Gloria Vazquez |
| EPI_ISL_2178941, EPI_ISL_2178942, EPI_ISL_2178943, EPI_ISL_2178944, EPI_ISL_2178945 | Klinisch Laboratorium ZNA | Klinisch Laboratorium ZNA | Verstrepen et al. |
| EPI_ISL_2179606, EPI_ISL_2179630 | GH A.CHENEVIER-H.MONDOR | Department of Virology, Henri Mondor University Hospital, Assistance Publique Hôpitaux de Paris, Université Paris-Est Créteil, INSERM U955 | Christophe Rodriguez, Slim Fourati, Vanessa Demontant, Guillaume Gricourt, Melissa N'Debi, Alexandre Soulier, Elisabeth Trawinski, Jean-Michel Pawlotsky |
| EPI_ISL_2179727, EPI_ISL_2179759, | Servicio Microbiología Hospital La Paz | Servicio Microbiología Hospital La Paz | Fernando Lázaro, Rubén Cáceres, Jesús Mingorance Cruz, Elie Dahdouh |

| | | | |
|---|---|---|---|
| EPI_ISL_2179763 | | | |
| EPI_ISL_2180312, EPI_ISL_2180313, EPI_ISL_2180314, EPI_ISL_2180316, EPI_ISL_2180317, EPI_ISL_2180318, EPI_ISL_2180319, EPI_ISL_2180320, EPI_ISL_2180321, EPI_ISL_2180322, EPI_ISL_2180323, EPI_ISL_2180324, EPI_ISL_2180325, EPI_ISL_2180327, EPI_ISL_2180328, EPI_ISL_2180371, EPI_ISL_2180372, EPI_ISL_2180375, EPI_ISL_2180377, EPI_ISL_2180379, EPI_ISL_2180380, EPI_ISL_2180381, EPI_ISL_2180382, EPI_ISL_2180383, EPI_ISL_2180384, EPI_ISL_2180386, EPI_ISL_2180387, EPI_ISL_2180388, EPI_ISL_2180389, EPI_ISL_2180391, EPI_ISL_2180392, EPI_ISL_2180393, EPI_ISL_2180394, EPI_ISL_2180395, EPI_ISL_2180396, EPI_ISL_2180397, EPI_ISL_2180398, EPI_ISL_2180399, EPI_ISL_2180400, EPI_ISL_2180401, EPI_ISL_2180402, EPI_ISL_2180403, EPI_ISL_2180404, EPI_ISL_2180405, EPI_ISL_2180406, EPI_ISL_2180407, EPI_ISL_2180408, EPI_ISL_2180409, EPI_ISL_2180410, EPI_ISL_2180411, EPI_ISL_2180412, EPI_ISL_2180413, EPI_ISL_2180414, EPI_ISL_2180415, EPI_ISL_2180416, EPI_ISL_2180417, EPI_ISL_2180418, EPI_ISL_2180419, EPI_ISL_2180420, EPI_ISL_2180421, EPI_ISL_2180422, EPI_ISL_2180423, EPI_ISL_2180424, EPI_ISL_2180425, EPI_ISL_2180426, EPI_ISL_2180427, EPI_ISL_2180429, EPI_ISL_2180430, EPI_ISL_2180431, EPI_ISL_2180432, EPI_ISL_2180433, EPI_ISL_2180434, EPI_ISL_2180435, EPI_ISL_2180436, EPI_ISL_2180437, EPI_ISL_2180438, EPI_ISL_2180439, EPI_ISL_2180441, EPI_ISL_2180443, EPI_ISL_2180444, EPI_ISL_2180445, EPI_ISL_2180446, EPI_ISL_2180449, EPI_ISL_2180450, EPI_ISL_2180451, EPI_ISL_2180452 | | | |
| see above | National Center of Infectious and Parasitic Diseases | National Center of Infectious and Parasitic Diseases | Alexiev et al |
| EPI_ISL_2187013 | Servicio de Microbiología, Hospital General Universitario de Ciudad Real | Servicio de Microbiología, Hospital General Universitario de Ciudad Real | Jose Martinez Alarcon, Soledad Illescas |
| EPI_ISL_2187984 | HLAGYN - Laboratorio de Imunologia de Transplantes de Goias | HLAGYN - Laboratorio de Imunologia de Transplantes de Goias | Fernando Antonio Vinhal dos Santos, Erika Lopes Rocha Batista, Alessandro Leonardo Alvares Magalhaes, Frederico Rodrigues Vinhal, Sabrina Sara Moreira Duarte, Lucas Carlos Gomes Pereira, Daniel Ferreira de Sousa |
| EPI_ISL_2188316 | Medicine Sentinel NetworkIdf | National Reference Center for Viruses of Respiratory Infections, Institut Pasteur, Paris | Marion Barbet, Sylvie Behillil, Méline Bizard, Angela Brisebarre, Camille Capel, Vincent Enouf, Louise Lefrançois, Frédéric Lemoine, Christophe Malabat, Corinne Maufrais, Etienne Simon-Lorière, Maud Vanpeene, Sylvie Van der Werf ,Patricia Lefebure |
| EPI_ISL_2188317, EPI_ISL_2188318 | Hospital | National Reference Center for Viruses of Respiratory Infections, Institut Pasteur, Paris | Marion Barbet, Sylvie Behillil, Méline Bizard, Angela Brisebarre, Camille Capel, Vincent Enouf, Louise Lefrançois, Frédéric Lemoine, Christophe Malabat, Corinne Maufrais, Etienne Simon-Lorière, Maud Vanpeene, Sylvie Van der Werf ,Pascale Martres |
| EPI_ISL_2188319, EPI_ISL_2188320, EPI_ISL_2188321 | Hospital | National Reference Center for Viruses of Respiratory Infections, Institut Pasteur, Paris | Marion Barbet, Sylvie Behillil, Méline Bizard, Angela Brisebarre, Camille Capel, Vincent Enouf, Louise Lefrançois, Frédéric Lemoine, Christophe Malabat, Corinne Maufrais, Etienne Simon-Lorière, Maud Vanpeene, Sylvie Van der Werf ,Sylvaine Bastian |
| EPI_ISL_2188322 | Hospital | National Reference Center for Viruses of Respiratory Infections, Institut Pasteur, Paris | Marion Barbet, Sylvie Behillil, Méline Bizard, Angela Brisebarre, Camille Capel, Vincent Enouf, Louise Lefrançois, Frédéric Lemoine, Christophe Malabat, Corinne Maufrais, Etienne Simon-Lorière, Maud Vanpeene, Sylvie Van der Werf ,Anne Cady |
| EPI_ISL_2188324 | Hospital | National Reference Center for Viruses of Respiratory Infections, Institut Pasteur, Paris | Marion Barbet, Sylvie Behillil, Méline Bizard, Angela Brisebarre, Camille Capel, Vincent Enouf, Louise Lefrançois, Frédéric Lemoine, Christophe Malabat, Corinne Maufrais, Etienne Simon-Lorière, Maud Vanpeene, Sylvie Van der Werf ,Alexandra Ducancelle |
| EPI_ISL_2188327 | Hospital | National Reference Center for Viruses of Respiratory Infections, Institut Pasteur, Paris | Marion Barbet, Sylvie Behillil, Méline Bizard, Angela Brisebarre, Camille Capel, Vincent Enouf, Louise Lefrançois, Frédéric Lemoine, Christophe Malabat, Corinne Maufrais, Etienne Simon-Lorière, Maud Vanpeene, Sylvie Van der Werf ,Schoenauer |
| EPI_ISL_2188328, EPI_ISL_2188329 | Labo Analyses Med | National Reference Center for Viruses of Respiratory Infections, Institut Pasteur, Paris | Marion Barbet, Sylvie Behillil, Méline Bizard, Angela Brisebarre, Camille Capel, Vincent Enouf, Louise Lefrançois, Frédéric Lemoine, Christophe Malabat, Corinne Maufrais, Damien Mornico, Etienne Simon-Lorière, Maud Vanpeene, Sylvie Van der Werf ,Sophie Chalmin |
| EPI_ISL_2188330, EPI_ISL_2188331, EPI_ISL_2188332, EPI_ISL_2188333, EPI_ISL_2188334, EPI_ISL_2188335, EPI_ISL_2188336, EPI_ISL_2188337 | Labo Analyses Med | National Reference Center for Viruses of Respiratory Infections, Institut Pasteur, Paris | Marion Barbet, Sylvie Behillil, Méline Bizard, Angela Brisebarre, Camille Capel, Vincent Enouf, Louise Lefrançois, Frédéric Lemoine, Christophe Malabat, Corinne Maufrais, Etienne Simon-Lorière, Maud Vanpeene, Sylvie Van der Werf |
| EPI_ISL_2188338 | Hospital | National Reference Center for Viruses of Respiratory Infections, Institut Pasteur, Paris | Marion Barbet, Sylvie Behillil, Méline Bizard, Angela Brisebarre, Camille Capel, Vincent Enouf, Louise Lefrançois, Frédéric Lemoine, Christophe Malabat, Corinne Maufrais, Etienne Simon-Lorière, Maud Vanpeene, Sylvie Van der Werf ,Pascale Martres |
| EPI_ISL_2188339 | Hospital | National Reference Center for Viruses of Respiratory Infections, Institut Pasteur, Paris | Marion Barbet, Sylvie Behillil, Méline Bizard, Angela Brisebarre, Camille Capel, Vincent Enouf, Louise Lefrançois, Frédéric Lemoine, Christophe Malabat, Corinne Maufrais, Damien Mornico, Etienne Simon-Lorière, Maud Vanpeene, Sylvie Van der Werf ,Pascale Martres |
| EPI_ISL_2188345, EPI_ISL_2188346, EPI_ISL_2188347, EPI_ISL_2188348, EPI_ISL_2188349, EPI_ISL_2188351 | Hospital | National Reference Center for Viruses of Respiratory Infections, Institut Pasteur, Paris | Marion Barbet, Sylvie Behillil, Méline Bizard, Angela Brisebarre, Camille Capel, Vincent Enouf, Louise Lefrançois, Frédéric Lemoine, Christophe Malabat, Corinne Maufrais, Etienne Simon-Lorière, Maud Vanpeene, Sylvie Van der Werf ,Eric Farfour |
| EPI_ISL_2188352 | Hospital | National Reference Center for Viruses of Respiratory Infections, Institut Pasteur, Paris | Marion Barbet, Sylvie Behillil, Méline Bizard, Angela Brisebarre, Camille Capel, Vincent Enouf, Louise Lefrançois, Frédéric Lemoine, Christophe Malabat, Corinne Maufrais, Etienne Simon-Lorière, Maud Vanpeene, Sylvie Van der Werf ,Laura Djamdjian |
| EPI_ISL_2188354 | Labo Analyses Med | National Reference Center for Viruses of Respiratory Infections, Institut Pasteur, Paris | Marion Barbet, Sylvie Behillil, Méline Bizard, Angela Brisebarre, Camille Capel, Vincent Enouf, Louise Lefrançois, Frédéric Lemoine, Christophe Malabat, Corinne Maufrais, Damien Mornico, Etienne Simon-Lorière, Maud Vanpeene, Sylvie Van der Werf ,Franck Ennouchi |
| EPI_ISL_2188355 | Labo Analyses Med | National Reference Center for Viruses of Respiratory Infections, Institut Pasteur, Paris | Marion Barbet, Sylvie Behillil, Méline Bizard, Angela Brisebarre, Camille Capel, Vincent Enouf, Louise Lefrançois, Frédéric Lemoine, Christophe Malabat, Corinne Maufrais, Etienne Simon-Lorière, Maud Vanpeene, Sylvie Van der Werf ,Franck Ennouchi |
| EPI_ISL_2188356, EPI_ISL_2188357 | Labo Analyses Med | National Reference Center for Viruses of Respiratory Infections, Institut Pasteur, Paris | Marion Barbet, Sylvie Behillil, Méline Bizard, Angela Brisebarre, Camille Capel, Vincent Enouf, Louise Lefrançois, Frédéric Lemoine, Christophe Malabat, Corinne Maufrais, Damien Mornico, Etienne Simon-Lorière, Maud Vanpeene, Sylvie Van der Werf ,Franck Ennouchi |
| EPI_ISL_2188360, EPI_ISL_2188362, EPI_ISL_2188363 | Hospital | National Reference Center for Viruses of Respiratory Infections, Institut Pasteur, Paris | Marion Barbet, Sylvie Behillil, Méline Bizard, Angela Brisebarre, Camille Capel, Vincent Enouf, Louise Lefrançois, Frédéric Lemoine, Christophe Malabat, Corinne Maufrais, Etienne Simon-Lorière, Maud Vanpeene, Sylvie Van der Werf ,Marianne Burgard |
| EPI_ISL_2188364, EPI_ISL_2188365, EPI_ISL_2188366, EPI_ISL_2188367 | Labo Analyses Med | National Reference Center for Viruses of Respiratory Infections, Institut Pasteur, Paris | Marion Barbet, Sylvie Behillil, Méline Bizard, Angela Brisebarre, Camille Capel, Vincent Enouf, Louise Lefrançois, Frédéric Lemoine, Christophe Malabat, Corinne Maufrais, Etienne Simon-Lorière, Maud Vanpeene, Sylvie Van der Werf ,Claude Jorion |
| EPI_ISL_2188368, EPI_ISL_2188369, EPI_ISL_2188370 | Hospital | National Reference Center for Viruses of Respiratory Infections, Institut Pasteur, Paris | Marion Barbet, Sylvie Behillil, Méline Bizard, Angela Brisebarre, Camille Capel, Vincent Enouf, Louise Lefrançois, Frédéric Lemoine, Christophe Malabat, Corinne Maufrais, Etienne Simon-Lorière, Maud Vanpeene, Sylvie Van der Werf ,Laurence Louvet |
| EPI_ISL_2188371 | Labo Analyses Med | National Reference Center for Viruses of Respiratory Infections, Institut Pasteur, Paris | Marion Barbet, Sylvie Behillil, Méline Bizard, Angela Brisebarre, Camille Capel, Vincent Enouf, Louise Lefrançois, Frédéric Lemoine, Christophe Malabat, Corinne Maufrais, Etienne Simon-Lorière, Maud Vanpeene, Sylvie Van der Werf ,J Besson |
| EPI_ISL_2188372 | Labo Analyses Med | National Reference Center for Viruses of Respiratory Infections, Institut Pasteur, Paris | Marion Barbet, Sylvie Behillil, Méline Bizard, Angela Brisebarre, Camille Capel, Vincent Enouf, Louise Lefrançois, Frédéric Lemoine, Christophe Malabat, Corinne Maufrais, Etienne Simon-Lorière, Maud Vanpeene, Sylvie Van der Werf ,GréGoire Potiron |
| EPI_ISL_2188373, EPI_ISL_2188374, EPI_ISL_2188375, EPI_ISL_2188376 | Hospital | National Reference Center for Viruses of Respiratory Infections, Institut Pasteur, Paris | Marion Barbet, Sylvie Behillil, Méline Bizard, Angela Brisebarre, Camille Capel, Vincent Enouf, Louise Lefrançois, Frédéric Lemoine, Christophe Malabat, Corinne Maufrais, Etienne Simon-Lorière, Maud Vanpeene, Sylvie Van der Werf ,Nabil Gastli |
| EPI_ISL_2188377, EPI_ISL_2188378 | Hospital | National Reference Center for Viruses of Respiratory Infections, Institut Pasteur, Paris | Marion Barbet, Sylvie Behillil, Méline Bizard, Angela Brisebarre, Camille Capel, Vincent Enouf, Louise Lefrançois, Frédéric Lemoine, Christophe Malabat, Corinne Maufrais, Damien Mornico, Etienne Simon-Lorière, Maud Vanpeene, Sylvie Van der Werf ,Nabil Gastli |
| EPI_ISL_2188379 | Hospital | National Reference Center for Viruses of Respiratory Infections, Institut Pasteur, Paris | Marion Barbet, Sylvie Behillil, Méline Bizard, Angela Brisebarre, Camille Capel, Vincent Enouf, Louise Lefrançois, Frédéric Lemoine, Christophe Malabat, Corinne Maufrais, Etienne Simon-Lorière, Maud Vanpeene, Sylvie Van der Werf ,Nabil Gastli |
| EPI_ISL_2188381 | Hospital | National Reference Center for Viruses of Respiratory Infections, Institut Pasteur, Paris | Marion Barbet, Sylvie Behillil, Méline Bizard, Angela Brisebarre, Camille Capel, Vincent Enouf, Louise Lefrançois, Frédéric Lemoine, Christophe Malabat, Corinne Maufrais, Victoire Baillet, Etienne Simon-Lorière, Maud Vanpeene, Sylvie Van der Werf ,Nabil Gastli |
| EPI_ISL_2188382, EPI_ISL_2188383, EPI_ISL_2188384, EPI_ISL_2188385 | Hospital | National Reference Center for Viruses of Respiratory Infections, Institut Pasteur, Paris | Marion Barbet, Sylvie Behillil, Méline Bizard, Angela Brisebarre, Camille Capel, Vincent Enouf, Louise Lefrançois, Frédéric Lemoine, Christophe Malabat, Corinne Maufrais, Etienne Simon-Lorière, Maud Vanpeene, Sylvie Van der Werf ,Nabil Gastli |
| EPI_ISL_2188386 | Hospital | National Reference Center for Viruses of Respiratory Infections, Institut Pasteur, Paris | Marion Barbet, Sylvie Behillil, Méline Bizard, Angela Brisebarre, Camille Capel, Vincent Enouf, Louise Lefrançois, Frédéric Lemoine, Christophe Malabat, Corinne Maufrais, Etienne Simon-Lorière, Maud Vanpeene, Sylvie Van der Werf ,Laurence Louvet |
| EPI_ISL_2188387 | Labo Analyses Med | National Reference Center for Viruses of Respiratory Infections, Institut Pasteur, Paris | Marion Barbet, Sylvie Behillil, Méline Bizard, Angela Brisebarre, Camille Capel, Vincent Enouf, Louise Lefrançois, Frédéric Lemoine, Christophe Malabat, Corinne Maufrais, Victoire Baillet, Etienne Simon-Lorière, Maud Vanpeene, Sylvie Van der Werf ,Jean-FrançOis Comes |
| EPI_ISL_2188388 | Labo Analyses Med | National Reference Center for Viruses of Respiratory Infections, Institut Pasteur, Paris | Marion Barbet, Sylvie Behillil, Méline Bizard, Angela Brisebarre, Camille Capel, Vincent Enouf, Louise Lefrançois, Frédéric Lemoine, Christophe Malabat, Corinne Maufrais, Etienne Simon-Lorière, Maud Vanpeene, Sylvie Van der Werf ,StéPhane Romand |

| | | | |
|---|---|---|---|
| EPI_ISL_2188389, EPI_ISL_2188390, EPI_ISL_2188391 | Labo Analyses Med | National Reference Center for Viruses of Respiratory Infections, Institut Pasteur, Paris | Marion Barbet, Sylvie Behillil, Méline Bizard, Angela Brisebarre, Camille Capel, Vincent Enouf, Louise Lefrançois, Frédéric Lemoine, Christophe Malabat, Corinne Maufrais, Etienne Simon-Lorière, Maud Vanpeene, Sylvie Van der Werf ,Jean-Baptiste Foulquier |
| EPI_ISL_2188392 | Labo Analyses Med | National Reference Center for Viruses of Respiratory Infections, Institut Pasteur, Paris | Marion Barbet, Sylvie Behillil, Méline Bizard, Angela Brisebarre, Camille Capel, Vincent Enouf, Louise Lefrançois, Frédéric Lemoine, Christophe Malabat, Corinne Maufrais, Victoire Baillet, Etienne Simon-Lorière, Maud Vanpeene, Sylvie Van der Werf ,Jean-Baptiste Foulquier |
| EPI_ISL_2188393, EPI_ISL_2188394, EPI_ISL_2188395, EPI_ISL_2188396, EPI_ISL_2188397, EPI_ISL_2188398, EPI_ISL_2188399 | Labo Analyses Med | National Reference Center for Viruses of Respiratory Infections, Institut Pasteur, Paris | Marion Barbet, Sylvie Behillil, Méline Bizard, Angela Brisebarre, Camille Capel, Vincent Enouf, Louise Lefrançois, Frédéric Lemoine, Christophe Malabat, Corinne Maufrais, Etienne Simon-Lorière, Maud Vanpeene, Sylvie Van der Werf ,Jean-Baptiste Foulquier |
| EPI_ISL_2188400, EPI_ISL_2188402, EPI_ISL_2188403 | Labo Analyses Med | National Reference Center for Viruses of Respiratory Infections, Institut Pasteur, Paris | Marion Barbet, Sylvie Behillil, Méline Bizard, Angela Brisebarre, Camille Capel, Vincent Enouf, Louise Lefrançois, Frédéric Lemoine, Christophe Malabat, Corinne Maufrais, Victoire Baillet, Etienne Simon-Lorière, Maud Vanpeene, Sylvie Van der Werf ,Jean-Baptiste Foulquier |
| EPI_ISL_2188404, EPI_ISL_2188405, EPI_ISL_2188406, EPI_ISL_2188407, EPI_ISL_2188408 | Labo Analyses Med | National Reference Center for Viruses of Respiratory Infections, Institut Pasteur, Paris | Marion Barbet, Sylvie Behillil, Méline Bizard, Angela Brisebarre, Camille Capel, Vincent Enouf, Louise Lefrançois, Frédéric Lemoine, Christophe Malabat, Corinne Maufrais, Etienne Simon-Lorière, Maud Vanpeene, Sylvie Van der Werf ,Jean-Baptiste Foulquier |
| EPI_ISL_2188409 | Labo Analyses Med | National Reference Center for Viruses of Respiratory Infections, Institut Pasteur, Paris | Marion Barbet, Sylvie Behillil, Méline Bizard, Angela Brisebarre, Camille Capel, Vincent Enouf, Louise Lefrançois, Frédéric Lemoine, Christophe Malabat, Corinne Maufrais, Amaury Vaysse, Etienne Simon-Lorière, Maud Vanpeene, Sylvie Van der Werf ,Jean-Baptiste Foulquier |
| EPI_ISL_2188410 | Labo Analyses Med | National Reference Center for Viruses of Respiratory Infections, Institut Pasteur, Paris | Marion Barbet, Sylvie Behillil, Méline Bizard, Angela Brisebarre, Camille Capel, Vincent Enouf, Louise Lefrançois, Frédéric Lemoine, Christophe Malabat, Corinne Maufrais, Etienne Simon-Lorière, Maud Vanpeene, Sylvie Van der Werf ,Jean-Baptiste Foulquier |
| EPI_ISL_2188412, EPI_ISL_2188413 | Hospital | National Reference Center for Viruses of Respiratory Infections, Institut Pasteur, Paris | Marion Barbet, Sylvie Behillil, Méline Bizard, Angela Brisebarre, Camille Capel, Vincent Enouf, Louise Lefrançois, Frédéric Lemoine, Christophe Malabat, Corinne Maufrais, Etienne Simon-Lorière, Maud Vanpeene, Sylvie Van der Werf ,Marianne Burgard |
| EPI_ISL_2188414 | Hospital | National Reference Center for Viruses of Respiratory Infections, Institut Pasteur, Paris | Marion Barbet, Sylvie Behillil, Méline Bizard, Angela Brisebarre, Camille Capel, Vincent Enouf, Louise Lefrançois, Frédéric Lemoine, Christophe Malabat, Corinne Maufrais, Etienne Simon-Lorière, Maud Vanpeene, Sylvie Van der Werf ,Jacques Fourgeaud |
| EPI_ISL_2188417 | Hospital | National Reference Center for Viruses of Respiratory Infections, Institut Pasteur, Paris | Marion Barbet, Sylvie Behillil, Méline Bizard, Angela Brisebarre, Camille Capel, Vincent Enouf, Louise Lefrançois, Frédéric Lemoine, Christophe Malabat, Corinne Maufrais, Etienne Simon-Lorière, Maud Vanpeene, Sylvie Van der Werf ,JéRome Guinard |
| EPI_ISL_2188418 | Hospital | National Reference Center for Viruses of Respiratory Infections, Institut Pasteur, Paris | Marion Barbet, Sylvie Behillil, Méline Bizard, Angela Brisebarre, Camille Capel, Vincent Enouf, Louise Lefrançois, Frédéric Lemoine, Christophe Malabat, Corinne Maufrais, Amaury Vaysse, Etienne Simon-Lorière, Maud Vanpeene, Sylvie Van der Werf ,Raphael Guiheneuf |
| EPI_ISL_2188419 | Hospital | National Reference Center for Viruses of Respiratory Infections, Institut Pasteur, Paris | Marion Barbet, Sylvie Behillil, Méline Bizard, Angela Brisebarre, Camille Capel, Vincent Enouf, Louise Lefrançois, Frédéric Lemoine, Christophe Malabat, Corinne Maufrais, Etienne Simon-Lorière, Maud Vanpeene, Sylvie Van der Werf ,Raphael Guiheneuf |
| EPI_ISL_2188420 | Hospital | National Reference Center for Viruses of Respiratory Infections, Institut Pasteur, Paris | Marion Barbet, Sylvie Behillil, Méline Bizard, Angela Brisebarre, Camille Capel, Vincent Enouf, Louise Lefrançois, Frédéric Lemoine, Christophe Malabat, Corinne Maufrais, Etienne Simon-Lorière, Maud Vanpeene, Sylvie Van der Werf ,Anne Cady |
| EPI_ISL_2188422 | Hospital | National Reference Center for Viruses of Respiratory Infections, Institut Pasteur, Paris | Marion Barbet, Sylvie Behillil, Méline Bizard, Angela Brisebarre, Camille Capel, Vincent Enouf, Louise Lefrançois, Frédéric Lemoine, Christophe Malabat, Corinne Maufrais, Etienne Simon-Lorière, Maud Vanpeene, Sylvie Van der Werf ,Franck Griscelli |
| EPI_ISL_2188423, EPI_ISL_2188424 | Hospital | National Reference Center for Viruses of Respiratory Infections, Institut Pasteur, Paris | Marion Barbet, Sylvie Behillil, Méline Bizard, Angela Brisebarre, Camille Capel, Vincent Enouf, Louise Lefrançois, Frédéric Lemoine, Christophe Malabat, Corinne Maufrais, Etienne Simon-Lorière, Maud Vanpeene, Sylvie Van der Werf ,VéRonique Avettand-Fenoel |
| EPI_ISL_2188425 | Labo Analyses Med | National Reference Center for Viruses of Respiratory Infections, Institut Pasteur, Paris | Marion Barbet, Sylvie Behillil, Méline Bizard, Angela Brisebarre, Camille Capel, Vincent Enouf, Louise Lefrançois, Frédéric Lemoine, Christophe Malabat, Corinne Maufrais, Etienne Simon-Lorière, Maud Vanpeene, Sylvie Van der Werf ,Gilles Desvaux |
| EPI_ISL_2188427 | Hospital | National Reference Center for Viruses of Respiratory Infections, Institut Pasteur, Paris | Marion Barbet, Sylvie Behillil, Méline Bizard, Angela Brisebarre, Camille Capel, Vincent Enouf, Louise Lefrançois, Frédéric Lemoine, Christophe Malabat, Corinne Maufrais, Amaury Vaysse, Etienne Simon-Lorière, Maud Vanpeene, Sylvie Van der Werf ,Alain Delepine |
| EPI_ISL_2188428 | Labo Analyses Med | National Reference Center for Viruses of Respiratory Infections, Institut Pasteur, Paris | Marion Barbet, Sylvie Behillil, Méline Bizard, Angela Brisebarre, Camille Capel, Vincent Enouf, Louise Lefrançois, Frédéric Lemoine, Christophe Malabat, Corinne Maufrais, Amaury Vaysse, Etienne Simon-Lorière, Maud Vanpeene, Sylvie Van der Werf ,GéRaldine Bonnaudet |
| EPI_ISL_2188434 | Labo Analyses Med | National Reference Center for Viruses of Respiratory Infections, Institut Pasteur, Paris | Marion Barbet, Sylvie Behillil, Méline Bizard, Angela Brisebarre, Camille Capel, Vincent Enouf, Louise Lefrançois, Frédéric Lemoine, Christophe Malabat, Corinne Maufrais, Amaury Vaysse, Etienne Simon-Lorière, Maud Vanpeene, Sylvie Van der Werf ,F Bondu |
| EPI_ISL_2188440, EPI_ISL_2188441, EPI_ISL_2188443, EPI_ISL_2188444, EPI_ISL_2188445, EPI_ISL_2188447, EPI_ISL_2188448, EPI_ISL_2188449, EPI_ISL_2188450, EPI_ISL_2188451, EPI_ISL_2188452, EPI_ISL_2188453, EPI_ISL_2188454, EPI_ISL_2188455, EPI_ISL_2188456, EPI_ISL_2188457, EPI_ISL_2188458, EPI_ISL_2188459, EPI_ISL_2188460, EPI_ISL_2188461, EPI_ISL_2188462, EPI_ISL_2188463, EPI_ISL_2188465, EPI_ISL_2188466, EPI_ISL_2188470, EPI_ISL_2188472, EPI_ISL_2188473, EPI_ISL_2188474, EPI_ISL_2188475, EPI_ISL_2188476, EPI_ISL_2188477 | | | |
| see above | Hospital | National Reference Center for Viruses of Respiratory Infections, Institut Pasteur, Paris | Marion Barbet, Sylvie Behillil, Méline Bizard, Angela Brisebarre, Camille Capel, Vincent Enouf, Louise Lefrançois, Frédéric Lemoine, Christophe Malabat, Corinne Maufrais, Etienne Simon-Lorière, Maud Vanpeene, Sylvie Van der Werf ,Nabil Gastli |
| EPI_ISL_2188478, EPI_ISL_2188480 | Hospital | National Reference Center for Viruses of Respiratory Infections, Institut Pasteur, Paris | Marion Barbet, Sylvie Behillil, Méline Bizard, Angela Brisebarre, Camille Capel, Vincent Enouf, Louise Lefrançois, Frédéric Lemoine, Christophe Malabat, Corinne Maufrais, Etienne Simon-Lorière, Maud Vanpeene, Sylvie Van der Werf ,StéPhanie Marque Juillet |
| EPI_ISL_2189530, EPI_ISL_2189532, EPI_ISL_2189533, EPI_ISL_2189534, EPI_ISL_2189535, EPI_ISL_2189536, EPI_ISL_2189537, EPI_ISL_2189538, EPI_ISL_2189540, EPI_ISL_2189542 | Institute of Molecular and Translational Medicine / Laboratory of Experimental Medicine, Faculty of Medicine and Dentistry, Palacky University and University Hospital Olomouc | Institute of Molecular and Translational Medicine / Laboratory of Experimental Medicine, Faculty of Medicine and Dentistry, Palacky University | Rastislav Slavkovský, Hana Jaworek, Vladimíra Koudeláková, Barbora Blumová, Tomáš Pospíšil, Marián Hajdúch |
| EPI_ISL_2189546, EPI_ISL_2189547, EPI_ISL_2189549, EPI_ISL_2189550 | Institute of Molecular and Translational Medicine / Laboratory of Experimental Medicine, Faculty of Medicine and Dentistry, Palacky University and University Hospital Olomouc | Institute of Molecular and Translational Medicine / Laboratory of Experimental Medicine, Faculty of Medicine and Dentistry, Palacky University | Rastislav Slavkovský, Hana Jaworek, Vladimíra Koudeláková, Barbora Blumová, Tomáš Pospíšil, Marián Hajdúch, Pavel Sauer |
| EPI_ISL_2189551, EPI_ISL_2189552, EPI_ISL_2189553, EPI_ISL_2189554, EPI_ISL_2189555, EPI_ISL_2189556, EPI_ISL_2189557, EPI_ISL_2189558, EPI_ISL_2189559, EPI_ISL_2189560, EPI_ISL_2189561, EPI_ISL_2189562, EPI_ISL_2189564, EPI_ISL_2189565, EPI_ISL_2189566, EPI_ISL_2189567 | | | |
| see above | Institute of Molecular and Translational Medicine / Laboratory of Experimental Medicine, Faculty of Medicine and Dentistry, Palacky University and University Hospital Olomouc | Institute of Molecular and Translational Medicine / Laboratory of Experimental Medicine, Faculty of Medicine and Dentistry, Palacky University | Rastislav Slavkovský, Hana Jaworek, Vladimíra Koudeláková, Barbora Blumová, Tomáš Pospíšil, Marián Hajdúch |
| EPI_ISL_2190098 | National Institute of Infectious Diseases-Prof. Dr. Matei Bals Molecular Diagnostics Laboratory | National Institute of Infectious Diseases-Prof. Dr. Matei Bals Molecular Diagnostics Laboratory | Corina Casangiu, Leontina Banica, Marius Surleac, Ovidiu Vlaicu, Andreea Tudor, Simona Paraschiv, Dan Otelea |
| EPI_ISL_2191526, EPI_ISL_2191528, EPI_ISL_2191532, EPI_ISL_2191535, EPI_ISL_2191538, EPI_ISL_2191539 | Microbiology Department, University Hospital Donostia | Microbiology Department, University Hospital Donostia | Marimon JM, Montes M, Piñeiro L, Grandioso D, Martin-Peñaranda T, Gomez M, Cilla G. |
| EPI_ISL_2193232, EPI_ISL_2193233, EPI_ISL_2193234, EPI_ISL_2193235, EPI_ISL_2193236, EPI_ISL_2193237, EPI_ISL_2193238, EPI_ISL_2193239, EPI_ISL_2193240, EPI_ISL_2193241 | Baylor Scott & White-Temple | Baylor Scott & White-Temple | Ari Rao, Linden Morales, Kimberly Walker, Marcus Volz, Shelby Hendrickson |
| EPI_ISL_2193352 | SC (UCO) Igiene e Sanità Pubblica, ASUGI, Trieste | ARGO Laboratorio Genomica ed Epigenomica | Licastro D, Dal Monego S, Degasperi M, Marcello A, Segat L, Piscianz E, D'Agaro P |
| EPI_ISL_2193353, EPI_ISL_2193382, EPI_ISL_2193383, EPI_ISL_2193768 | Institut de Virologie du CHU de Strasbourg | Institut de Virologie du CHU de Strasbourg | Fafi-Kremer Samira, Gallais Floriane, Gantner Pierre, Laugel Elodie, Solis Morgane, Velay Aurélie, Wendling Marie-Josée |

| | | | |
|---|---|---|---|
| EPI_ISL_2193780 | SC (UCO) Igiene e Sanità Pubblica, ASUGI, Trieste | ARGO Laboratorio Genomica ed Epigenomica | Licastro D, Dal Monego S, Degasperi M, Marcello A, Segat L, Piscianz E, D'Agaro P |
| EPI_ISL_2193812 | Institut de Virologie du CHU de Strasbourg | Institut de Virologie du CHU de Strasbourg | Fafi-Kremer Samira, Gallais Floriane, Gantner Pierre, Laugel Elodie, Solis Morgane, Velay Aurélie, Wendling Marie-Josée |
| EPI_ISL_2193963 | SC (UCO) Igiene e Sanità Pubblica, ASUGI, Trieste | ARGO Laboratorio Genomica ed Epigenomica | Licastro D, Dal Monego S, Degasperi M, Marcello A, Segat L, Piscianz E, D'Agaro P |
| EPI_ISL_2193965 | Institut de Virologie du CHU de Strasbourg | Institut de Virologie du CHU de Strasbourg | Fafi-Kremer Samira, Gallais Floriane, Gantner Pierre, Laugel Elodie, Solis Morgane, Velay Aurélie, Wendling Marie-Josée |
| EPI_ISL_2193986, EPI_ISL_2193988 | SC (UCO) Igiene e Sanità Pubblica, ASUGI, Trieste | ARGO Laboratorio Genomica ed Epigenomica | Licastro D, Dal Monego S, Degasperi M, Marcello A, Segat L, Piscianz E, D'Agaro P |
| EPI_ISL_2193997 | Institut de Virologie du CHU de Strasbourg | Institut de Virologie du CHU de Strasbourg | Fafi-Kremer Samira, Gallais Floriane, Gantner Pierre, Laugel Elodie, Solis Morgane, Velay Aurélie, Wendling Marie-Josée |
| EPI_ISL_2194006, EPI_ISL_2194012 | CHU Nantes Virology | CHU Nantes Virology | Celine Bressollette-Bodin, Thomas Drumel, Audrey Rodallec, Berthe-Marie Imbert-Marcille |
| EPI_ISL_2194015 | Institut de Virologie du CHU de Strasbourg | Institut de Virologie du CHU de Strasbourg | Fafi-Kremer Samira, Gallais Floriane, Gantner Pierre, Laugel Elodie, Solis Morgane, Velay Aurélie, Wendling Marie-Josée |
| EPI_ISL_2194022, EPI_ISL_2194032, EPI_ISL_2194034 | CHU Nantes Virology | CHU Nantes Virology | Celine Bressollette-Bodin, Thomas Drumel, Audrey Rodallec, Berthe-Marie Imbert-Marcille |
| EPI_ISL_2194100, EPI_ISL_2194217 | Institut de Virologie du CHU de Strasbourg | Institut de Virologie du CHU de Strasbourg | Fafi-Kremer Samira, Gallais Floriane, Gantner Pierre, Laugel Elodie, Solis Morgane, Velay Aurélie, Wendling Marie-Josée |
| EPI_ISL_2194220, EPI_ISL_2194539 | SC (UCO) Igiene e Sanità Pubblica, ASUGI, Trieste | ARGO Laboratorio Genomica ed Epigenomica | Licastro D, Dal Monego S, Degasperi M, Marcello A, Segat L, Piscianz E, D'Agaro P |
| EPI_ISL_2194540 | CHU Lille - Laboratoire de Virologie | CHU Lille - Laboratoire de Virologie | AIT YAHYA Emilie, ALIDJINOU Enagnon Kazali, BOCKET Laurence, CREPIN Michel, DEMAY Christophe, ENGELMANN Ilka, GEFFROY Sandrine, GUIGON Aurélie, LAMBERT Valérie, LAZREK Mouna, NOBILLIAUX Florian, PREVOST Brigitte, THUILLIER Caroline, TINEZ Claire ,TCHANTCHOU NJOSSE YANICK |
| EPI_ISL_2194541, EPI_ISL_2200868, EPI_ISL_2201320 | SC (UCO) Igiene e Sanità Pubblica, ASUGI, Trieste | ARGO Laboratorio Genomica ed Epigenomica | Licastro D, Dal Monego S, Degasperi M, Marcello A, Segat L, Piscianz E, D'Agaro P |
| EPI_ISL_2212318 | SC (UCO) Igiene e Sanità Pubblica, ASUGI, Trieste and SC Microbiologia e Virologia, ASFO, Pordenone | ARGO Laboratorio Genomica ed Epigenomica | Licastro D, Dal Monego S, Degasperi M, Marcello A, Segat L, Piscianz E, De Rosa R, D'Agaro P |
| EPI_ISL_2212980 | CHU Lille | CHU Lille | AIT YAHYA Emilie, ALIDJINOU Enagnon Kazali, BOCKET Laurence, CREPIN Michel, DEMAY Christophe, ENGELMANN Ilka, GEFFROY Sandrine, GUIGON Aurélie, LAMBERT Valérie, LAZREK Mouna, NOBILLIAUX Florian, PREVOST Brigitte, THUILLIER Caroline, TINEZ Claire |
| EPI_ISL_2212983 | CH St Philibert | CHU Lille | AIT YAHYA Emilie, ALIDJINOU Enagnon Kazali, BOCKET Laurence, CREPIN Michel, DEMAY Christophe, ENGELMANN Ilka, GEFFROY Sandrine, GUIGON Aurélie, LAMBERT Valérie, LAZREK Mouna, NOBILLIAUX Florian, PREVOST Brigitte, THUILLIER Caroline, TINEZ Claire |
| EPI_ISL_2212985 | CHU Lille | CHU Lille | AIT YAHYA Emilie, ALIDJINOU Enagnon Kazali, BOCKET Laurence, CREPIN Michel, DEMAY Christophe, ENGELMANN Ilka, GEFFROY Sandrine, GUIGON Aurélie, LAMBERT Valérie, LAZREK Mouna, NOBILLIAUX Florian, PREVOST Brigitte, THUILLIER Caroline, TINEZ Claire |
| EPI_ISL_2212988, EPI_ISL_2212990 | CH Calais | CHU Lille | AIT YAHYA Emilie, ALIDJINOU Enagnon Kazali, BOCKET Laurence, CREPIN Michel, DEMAY Christophe, ENGELMANN Ilka, GEFFROY Sandrine, GUIGON Aurélie, LAMBERT Valérie, LAZREK Mouna, NOBILLIAUX Florian, PREVOST Brigitte, THUILLIER Caroline, TINEZ Claire |
| EPI_ISL_2212993, EPI_ISL_2212995, EPI_ISL_2212997, EPI_ISL_2213000, EPI_ISL_2213002, EPI_ISL_2213005, EPI_ISL_2213006 | CHU Lille | CHU Lille | AIT YAHYA Emilie, ALIDJINOU Enagnon Kazali, BOCKET Laurence, CREPIN Michel, DEMAY Christophe, ENGELMANN Ilka, GEFFROY Sandrine, GUIGON Aurélie, LAMBERT Valérie, LAZREK Mouna, NOBILLIAUX Florian, PREVOST Brigitte, THUILLIER Caroline, TINEZ Claire |
| EPI_ISL_2213008 | CH Roubaix | CHU Lille | AIT YAHYA Emilie, ALIDJINOU Enagnon Kazali, BOCKET Laurence, CREPIN Michel, DEMAY Christophe, ENGELMANN Ilka, GEFFROY Sandrine, GUIGON Aurélie, LAMBERT Valérie, LAZREK Mouna, NOBILLIAUX Florian, PREVOST Brigitte, THUILLIER Caroline, TINEZ Claire |
| EPI_ISL_2213011 | CH Seclin | CHU Lille | AIT YAHYA Emilie, ALIDJINOU Enagnon Kazali, BOCKET Laurence, CREPIN Michel, DEMAY Christophe, ENGELMANN Ilka, GEFFROY Sandrine, GUIGON Aurélie, LAMBERT Valérie, LAZREK Mouna, NOBILLIAUX Florian, PREVOST Brigitte, THUILLIER Caroline, TINEZ Claire |
| EPI_ISL_2213013 | CH Armentieres | CHU Lille | AIT YAHYA Emilie, ALIDJINOU Enagnon Kazali, BOCKET Laurence, CREPIN Michel, DEMAY Christophe, ENGELMANN Ilka, GEFFROY Sandrine, GUIGON Aurélie, LAMBERT Valérie, LAZREK Mouna, NOBILLIAUX Florian, PREVOST Brigitte, THUILLIER Caroline, TINEZ Claire |
| EPI_ISL_2213016 | CH Lens | CHU Lille | AIT YAHYA Emilie, ALIDJINOU Enagnon Kazali, BOCKET Laurence, CREPIN Michel, DEMAY Christophe, ENGELMANN Ilka, GEFFROY Sandrine, GUIGON Aurélie, LAMBERT Valérie, LAZREK Mouna, NOBILLIAUX Florian, PREVOST Brigitte, THUILLIER Caroline, TINEZ Claire |
| EPI_ISL_2213305, EPI_ISL_2214131 | SC (UCO) Igiene e Sanità Pubblica, ASUGI, Trieste and SC Microbiologia e Virologia, ASFO, Pordenone | ARGO Laboratorio Genomica ed Epigenomica | Licastro D, Dal Monego S, Degasperi M, Marcello A, Segat L, Piscianz E, De Rosa R, D'Agaro P |
| EPI_ISL_2214696, EPI_ISL_2214701, EPI_ISL_2214704, EPI_ISL_2214706, EPI_ISL_2214713, EPI_ISL_2214717 | Centre Hospitalier Universitaire Clermont-Ferrand | CHU Clermont-Ferrand, service de virologie | Bisseux Maxime, Mirand Audrey, Combes Patricia, Henquell Cécile |
| EPI_ISL_2214865 | SC (UCO) Igiene e Sanità Pubblica, ASUGI, Trieste and SC Microbiologia e Virologia, ASFO, Pordenone | ARGO Laboratorio Genomica ed Epigenomica | Licastro D, Dal Monego S, Degasperi M, Marcello A, Segat L, Piscianz E, De Rosa R, D'Agaro P |
| EPI_ISL_2214900 | Centre Hospitalier Universitaire Clermont-Ferrand | CHU Clermont-Ferrand, service de virologie | Bisseux Maxime, Mirand Audrey, Combes Patricia, Henquell Cécile |
| EPI_ISL_2214978 | HOPITAL DE VICHY | CHU Clermont-Ferrand, service de virologie | Bisseux Maxime, Mirand Audrey, Combes Patricia, Henquell Cécile |
| EPI_ISL_2217042 | SC (UCO) Igiene e Sanità Pubblica, ASUGI, Trieste and SC Microbiologia e Virologia, ASFO, Pordenone | ARGO Laboratorio Genomica ed Epigenomica | Licastro D, Dal Monego S, Degasperi M, Marcello A, Segat L, Piscianz E, De Rosa R, D'Agaro P |
| EPI_ISL_2225602, EPI_ISL_2225603, EPI_ISL_2225604, EPI_ISL_2225605 | CHU Pontchaillou | CHU Pontchaillou | GROLHIER Claire, DENOUAL Florent, ETCHEVERRY Amandine, SASSI Mohamed, JAGLINE Steven, FEBREAU Christine, PRONIER Charlotte, GALIBERT Marie Dominique, DE TAYRAC Marie, THIBAULT Vincent |
| EPI_ISL_2225610 | Basurto University Hospital: Clinical Microbiology Laboratory | Biocruces Bizkaia | Mikel Urrutikoetxea-Gutiérrez, Mª Carmen Nieto Toboso, Estibaliz Ugalde Zarraga, Mikel Gallego Rodrigo, Ana de la Hoz, José Luis Díaz de Tuesta del Arco |
| EPI_ISL_2226308 | BIO86 CHAUMONT | CHU Poitiers | Caroline MICHAUD, Maxime PICHON, Manon PRAT, Valentin BON-BARET, Birama N'DIAYE, Magali GARCIA, Agnes BEBY-DEFAUX, Nicolas LEVEQUE |
| EPI_ISL_2226310 | CH ANGOULEME | CHU Poitiers | Caroline MICHAUD, Maxime PICHON, Manon PRAT, Valentin BON-BARET, Birama N'DIAYE, Magali GARCIA, Agnes BEBY-DEFAUX, Nicolas LEVEQUE |
| EPI_ISL_2226312 | CH SAINTONGE | CHU Poitiers | Caroline MICHAUD, Maxime PICHON, Manon PRAT, Valentin BON-BARET, Birama N'DIAYE, Magali GARCIA, Agnes BEBY-DEFAUX, Nicolas LEVEQUE |
| EPI_ISL_2226314, EPI_ISL_2226316 | BIO86 CHAUMONT | CHU Poitiers | Caroline MICHAUD, Maxime PICHON, Manon PRAT, Valentin BON-BARET, Birama N'DIAYE, Magali GARCIA, Agnes BEBY-DEFAUX, Nicolas LEVEQUE |
| EPI_ISL_2226317, EPI_ISL_2226318, EPI_ISL_2226320 | MEDILAB | CHU Poitiers | Caroline MICHAUD, Maxime PICHON, Manon PRAT, Valentin BON-BARET, Birama N'DIAYE, Magali GARCIA, Agnes BEBY-DEFAUX, Nicolas LEVEQUE |
| EPI_ISL_2226321, EPI_ISL_2226322 | BIO86 CHAUMONT | CHU Poitiers | Caroline MICHAUD, Maxime PICHON, Manon PRAT, Valentin BON-BARET, Birama N'DIAYE, Magali GARCIA, Agnes BEBY-DEFAUX, Nicolas LEVEQUE |
| EPI_ISL_2226327, EPI_ISL_2226328, EPI_ISL_2226330, EPI_ISL_2226332 | MEDILAB | CHU Poitiers | Caroline MICHAUD, Maxime PICHON, Manon PRAT, Valentin BON-BARET, Birama N'DIAYE, Magali GARCIA, Agnes BEBY-DEFAUX, Nicolas LEVEQUE |
| EPI_ISL_2226335 | CH SAINTONGE | CHU Poitiers | Caroline MICHAUD, Maxime PICHON, Manon PRAT, Valentin BON-BARET, Birama N'DIAYE, Magali GARCIA, Agnes BEBY-DEFAUX, Nicolas |

| | | | |
|---|---|---|---|
| | | | LEVEQUE |
| EPI_ISL_2226337, EPI_ISL_2226341 | CHU POITIERS | CHU Poitiers | Caroline MICHAUD, Maxime PICHON, Manon PRAT, Valentin BON-BARET, Birama N'DIAYE, Magali GARCIA, Agnes BEBY-DEFAUX, Nicolas LEVEQUE |
| EPI_ISL_2226343 | CH SAINTONGE | CHU Poitiers | Caroline MICHAUD, Maxime PICHON, Manon PRAT, Valentin BON-BARET, Birama N'DIAYE, Magali GARCIA, Agnes BEBY-DEFAUX, Nicolas LEVEQUE |
| EPI_ISL_2226344, EPI_ISL_2226345 | CH NIORT | CHU Poitiers | Caroline MICHAUD, Maxime PICHON, Manon PRAT, Valentin BON-BARET, Birama N'DIAYE, Magali GARCIA, Agnes BEBY-DEFAUX, Nicolas LEVEQUE |
| EPI_ISL_2226346 | CHU POITIERS | CHU Poitiers | Caroline MICHAUD, Maxime PICHON, Manon PRAT, Valentin BON-BARET, Birama N'DIAYE, Magali GARCIA, Agnes BEBY-DEFAUX, Nicolas LEVEQUE |
| EPI_ISL_2226347 | CH SAINTONGE | CHU Poitiers | Caroline MICHAUD, Maxime PICHON, Manon PRAT, Valentin BON-BARET, Birama N'DIAYE, Magali GARCIA, Agnes BEBY-DEFAUX, Nicolas LEVEQUE |
| EPI_ISL_2226348 | CHU POITIERS | CHU Poitiers | Caroline MICHAUD, Maxime PICHON, Manon PRAT, Valentin BON-BARET, Birama N'DIAYE, Magali GARCIA, Agnes BEBY-DEFAUX, Nicolas LEVEQUE |
| EPI_ISL_2226349, EPI_ISL_2226350 | MEDILAB | CHU Poitiers | Caroline MICHAUD, Maxime PICHON, Manon PRAT, Valentin BON-BARET, Birama N'DIAYE, Magali GARCIA, Agnes BEBY-DEFAUX, Nicolas LEVEQUE |
| EPI_ISL_2226352 | BIO86 CHAUMONT | CHU Poitiers | Caroline MICHAUD, Maxime PICHON, Manon PRAT, Valentin BON-BARET, Birama N'DIAYE, Magali GARCIA, Agnes BEBY-DEFAUX, Nicolas LEVEQUE |
| EPI_ISL_2226354 | CH SAINTONGE | CHU Poitiers | Caroline MICHAUD, Maxime PICHON, Manon PRAT, Valentin BON-BARET, Birama N'DIAYE, Magali GARCIA, Agnes BEBY-DEFAUX, Nicolas LEVEQUE |
| EPI_ISL_2226355, EPI_ISL_2226356 | BIO86 CHAUMONT | CHU Poitiers | Caroline MICHAUD, Maxime PICHON, Manon PRAT, Valentin BON-BARET, Birama N'DIAYE, Magali GARCIA, Agnes BEBY-DEFAUX, Nicolas LEVEQUE |
| EPI_ISL_2226357 | CH LA ROCHELLE | CHU Poitiers | Caroline MICHAUD, Maxime PICHON, Manon PRAT, Valentin BON-BARET, Birama N'DIAYE, Magali GARCIA, Agnes BEBY-DEFAUX, Nicolas LEVEQUE |
| EPI_ISL_2226358 | CH SAINTONGE | CHU Poitiers | Caroline MICHAUD, Maxime PICHON, Manon PRAT, Valentin BON-BARET, Birama N'DIAYE, Magali GARCIA, Agnes BEBY-DEFAUX, Nicolas LEVEQUE |
| EPI_ISL_2226359 | CH NIORT | CHU Poitiers | Caroline MICHAUD, Maxime PICHON, Manon PRAT, Valentin BON-BARET, Birama N'DIAYE, Magali GARCIA, Agnes BEBY-DEFAUX, Nicolas LEVEQUE |
| EPI_ISL_2226360 | MEDILAB | CHU Poitiers | Caroline MICHAUD, Maxime PICHON, Manon PRAT, Valentin BON-BARET, Birama N'DIAYE, Magali GARCIA, Agnes BEBY-DEFAUX, Nicolas LEVEQUE |
| EPI_ISL_2226361 | CH SAINTONGE | CHU Poitiers | Caroline MICHAUD, Maxime PICHON, Manon PRAT, Valentin BON-BARET, Birama N'DIAYE, Magali GARCIA, Agnes BEBY-DEFAUX, Nicolas LEVEQUE |
| EPI_ISL_2226362, EPI_ISL_2226363, EPI_ISL_2226364 | CHU POITIERS | CHU Poitiers | Caroline MICHAUD, Maxime PICHON, Manon PRAT, Valentin BON-BARET, Birama N'DIAYE, Magali GARCIA, Agnes BEBY-DEFAUX, Nicolas LEVEQUE |
| EPI_ISL_2226365 | LABOFFICE | CHU Poitiers | Caroline MICHAUD, Maxime PICHON, Manon PRAT, Valentin BON-BARET, Birama N'DIAYE, Magali GARCIA, Agnes BEBY-DEFAUX, Nicolas LEVEQUE |
| EPI_ISL_2226367 | MEDILAB | CHU Poitiers | Caroline MICHAUD, Maxime PICHON, Manon PRAT, Valentin BON-BARET, Birama N'DIAYE, Magali GARCIA, Agnes BEBY-DEFAUX, Nicolas LEVEQUE |
| EPI_ISL_2226370 | CHU POITIERS | CHU Poitiers | Caroline MICHAUD, Maxime PICHON, Manon PRAT, Valentin BON-BARET, Birama N'DIAYE, Magali GARCIA, Agnes BEBY-DEFAUX, Nicolas LEVEQUE |
| EPI_ISL_2226373 | BIO86 CHAUMONT | CHU Poitiers | Caroline MICHAUD, Maxime PICHON, Manon PRAT, Valentin BON-BARET, Birama N'DIAYE, Magali GARCIA, Agnes BEBY-DEFAUX, Nicolas LEVEQUE |
| EPI_ISL_2226374 | MEDILAB | CHU Poitiers | Caroline MICHAUD, Maxime PICHON, Manon PRAT, Valentin BON-BARET, Birama N'DIAYE, Magali GARCIA, Agnes BEBY-DEFAUX, Nicolas LEVEQUE |
| EPI_ISL_2226375 | BIO86 CHAUMONT | CHU Poitiers | Caroline MICHAUD, Maxime PICHON, Manon PRAT, Valentin BON-BARET, Birama N'DIAYE, Magali GARCIA, Agnes BEBY-DEFAUX, Nicolas LEVEQUE |
| EPI_ISL_2226376, EPI_ISL_2226377 | CH ANGOULEME | CHU Poitiers | Caroline MICHAUD, Maxime PICHON, Manon PRAT, Valentin BON-BARET, Birama N'DIAYE, Magali GARCIA, Agnes BEBY-DEFAUX, Nicolas LEVEQUE |
| EPI_ISL_2226378 | CH NIORT | CHU Poitiers | Caroline MICHAUD, Maxime PICHON, Manon PRAT, Valentin BON-BARET, Birama N'DIAYE, Magali GARCIA, Agnes BEBY-DEFAUX, Nicolas LEVEQUE |
| EPI_ISL_2226590 | Hospital | National Reference Center for Viruses of Respiratory Infections, Institut Pasteur, Paris | Marion Barbet, Sylvie Behillil, Méline Bizard, Angela Brisebarre, Camille Capel, Vincent Enouf, Louise Lefrançois, Frédéric Lemoine, Christophe Malabat, Corinne Maufrais, Emmanuelle Permal, Etienne Simon-Lorière, Maud Vanpeene, Sylvie Van der Werf ,Sabine Trombert |
| EPI_ISL_2226648 | BBTKLPP Surabaya | Institute of Tropical Disease, Universitas Airlangga | Aldise M Nastri, Krisnoadi Rahardjo, Jezzy R Dewantari, Rima R Prasetya, Acub Zaenal, Rosidi Roslan, Zahrotunnisa, Gatot Soegiarto, Laksmi Wulandari, Resti Yudhawati, Yasuko Mori, Soetjipto, Kazufumi Shimizu, Maria I Lusida |
| EPI_ISL_2226706, EPI_ISL_2226708, EPI_ISL_2226715, EPI_ISL_2226718, EPI_ISL_2226719, EPI_ISL_2226721, EPI_ISL_2226722, EPI_ISL_2226723, EPI_ISL_2226724, EPI_ISL_2226725, EPI_ISL_2226727, EPI_ISL_2226728, EPI_ISL_2226731, EPI_ISL_2226732, EPI_ISL_2226733, EPI_ISL_2226734, EPI_ISL_2226736, EPI_ISL_2226737, EPI_ISL_2226739, EPI_ISL_2226740, EPI_ISL_2226742, EPI_ISL_2226745, EPI_ISL_2226748, EPI_ISL_2226750, EPI_ISL_2226751, EPI_ISL_2226752, EPI_ISL_2226754, EPI_ISL_2226759, EPI_ISL_2226761, EPI_ISL_2226764, EPI_ISL_2226765 | | | |
| see above | ADMED Microbiologie | Genomics and Transcriptomics, Philip Morris International | Reto Lienhard, Marie-Lise Tritten, Emmanuel Guedj, Nicolas Sierro, Rémi Dulize, David Bornand, Mehdi Auberson, Maxime Berthouzoz, Nikolai Ivanov, Manuel Peitsch |
| EPI_ISL_2227208 | CH SECLIN | CHU Lille - Laboratoire de Virologie | AIT YAHYA Emilie, ALIDJINOU Enagnon Kazali, BOCKET Laurence, CREPIN Michel, DEMAY Christophe, ENGELMANN Ilka, GEFFROY Sandrine, GUIGON Aurélie, LAMBERT Valérie, LAZREK Mouna, NOBILLIAUX Florian, PREVOST Brigitte, THUILLIER Caroline, TINEZ Claire ,TCHANTCHOU NJOSSE YANICK |
| EPI_ISL_2227209 | CH BETHUNE | CHU Lille - Laboratoire de Virologie | AIT YAHYA Emilie, ALIDJINOU Enagnon Kazali, BOCKET Laurence, CREPIN Michel, DEMAY Christophe, ENGELMANN Ilka, GEFFROY Sandrine, GUIGON Aurélie, LAMBERT Valérie, LAZREK Mouna, NOBILLIAUX Florian, PREVOST Brigitte, THUILLIER Caroline, TINEZ Claire ,TCHANTCHOU NJOSSE YANICK |
| EPI_ISL_2227211 | CH LENS | CHU Lille - Laboratoire de Virologie | AIT YAHYA Emilie, ALIDJINOU Enagnon Kazali, BOCKET Laurence, CREPIN Michel, DEMAY Christophe, ENGELMANN Ilka, GEFFROY Sandrine, GUIGON Aurélie, LAMBERT Valérie, LAZREK Mouna, NOBILLIAUX Florian, PREVOST Brigitte, THUILLIER Caroline, TINEZ Claire ,TCHANTCHOU NJOSSE YANICK |
| EPI_ISL_2227212, EPI_ISL_2227213, EPI_ISL_2227214 | CHU LILLE | CHU Lille - Laboratoire de Virologie | AIT YAHYA Emilie, ALIDJINOU Enagnon Kazali, BOCKET Laurence, CREPIN Michel, DEMAY Christophe, ENGELMANN Ilka, GEFFROY Sandrine, GUIGON Aurélie, LAMBERT Valérie, LAZREK Mouna, NOBILLIAUX Florian, PREVOST Brigitte, THUILLIER Caroline, TINEZ Claire ,TCHANTCHOU NJOSSE YANICK |
| EPI_ISL_2227215 | CH BOULOGNE-SUR-MER | CHU Lille - Laboratoire de Virologie | AIT YAHYA Emilie, ALIDJINOU Enagnon Kazali, BOCKET Laurence, CREPIN Michel, DEMAY Christophe, ENGELMANN Ilka, GEFFROY Sandrine, GUIGON Aurélie, LAMBERT Valérie, LAZREK Mouna, NOBILLIAUX Florian, PREVOST Brigitte, THUILLIER Caroline, TINEZ Claire ,TCHANTCHOU |

| | | | |
|---|---|---|---|
| | | | NJOSSE YANICK |
| EPI_ISL_2227216 | CH SECLIN | CHU Lille - Laboratoire de Virologie | AIT YAHYA Emilie, ALIDJINOU Enagnon Kazali, BOCKET Laurence, CREPIN Michel, DEMAY Christophe, ENGELMANN Ilka, GEFFROY Sandrine, GUIGON Aurélie, LAMBERT Valérie, LAZREK Mouna, NOBILLIAUX Florian, PREVOST Brigitte, THUILLIER Caroline, TINEZ Claire ,TCHANTCHOU NJOSSE YANICK |
| EPI_ISL_2227217 | CH DUNKERQUE | CHU Lille - Laboratoire de Virologie | AIT YAHYA Emilie, ALIDJINOU Enagnon Kazali, BOCKET Laurence, CREPIN Michel, DEMAY Christophe, ENGELMANN Ilka, GEFFROY Sandrine, GUIGON Aurélie, LAMBERT Valérie, LAZREK Mouna, NOBILLIAUX Florian, PREVOST Brigitte, THUILLIER Caroline, TINEZ Claire ,TCHANTCHOU NJOSSE YANICK |
| EPI_ISL_2227219, EPI_ISL_2227220, EPI_ISL_2227221, EPI_ISL_2227222, EPI_ISL_2227223 | CHU LILLE | CHU Lille - Laboratoire de Virologie | AIT YAHYA Emilie, ALIDJINOU Enagnon Kazali, BOCKET Laurence, CREPIN Michel, DEMAY Christophe, ENGELMANN Ilka, GEFFROY Sandrine, GUIGON Aurélie, LAMBERT Valérie, LAZREK Mouna, NOBILLIAUX Florian, PREVOST Brigitte, THUILLIER Caroline, TINEZ Claire ,TCHANTCHOU NJOSSE YANICK |
| EPI_ISL_2227224 | CH DUNKERQUE | CHU Lille - Laboratoire de Virologie | AIT YAHYA Emilie, ALIDJINOU Enagnon Kazali, BOCKET Laurence, CREPIN Michel, DEMAY Christophe, ENGELMANN Ilka, GEFFROY Sandrine, GUIGON Aurélie, LAMBERT Valérie, LAZREK Mouna, NOBILLIAUX Florian, PREVOST Brigitte, THUILLIER Caroline, TINEZ Claire ,TCHANTCHOU NJOSSE YANICK |
| EPI_ISL_2227225, EPI_ISL_2227226 | CH TOURCOING | CHU Lille - Laboratoire de Virologie | AIT YAHYA Emilie, ALIDJINOU Enagnon Kazali, BOCKET Laurence, CREPIN Michel, DEMAY Christophe, ENGELMANN Ilka, GEFFROY Sandrine, GUIGON Aurélie, LAMBERT Valérie, LAZREK Mouna, NOBILLIAUX Florian, PREVOST Brigitte, THUILLIER Caroline, TINEZ Claire ,TCHANTCHOU NJOSSE YANICK |
| EPI_ISL_2227227 | CH ARRAS | CHU Lille - Laboratoire de Virologie | AIT YAHYA Emilie, ALIDJINOU Enagnon Kazali, BOCKET Laurence, CREPIN Michel, DEMAY Christophe, ENGELMANN Ilka, GEFFROY Sandrine, GUIGON Aurélie, LAMBERT Valérie, LAZREK Mouna, NOBILLIAUX Florian, PREVOST Brigitte, THUILLIER Caroline, TINEZ Claire ,TCHANTCHOU NJOSSE YANICK |
| EPI_ISL_2227228, EPI_ISL_2227229, EPI_ISL_2227230, EPI_ISL_2227231, EPI_ISL_2227232, EPI_ISL_2227234, EPI_ISL_2227235 | CHU LILLE | CHU Lille - Laboratoire de Virologie | AIT YAHYA Emilie, ALIDJINOU Enagnon Kazali, BOCKET Laurence, CREPIN Michel, DEMAY Christophe, ENGELMANN Ilka, GEFFROY Sandrine, GUIGON Aurélie, LAMBERT Valérie, LAZREK Mouna, NOBILLIAUX Florian, PREVOST Brigitte, THUILLIER Caroline, TINEZ Claire ,TCHANTCHOU NJOSSE YANICK |
| EPI_ISL_2227236 | CH CALAIS | CHU Lille - Laboratoire de Virologie | AIT YAHYA Emilie, ALIDJINOU Enagnon Kazali, BOCKET Laurence, CREPIN Michel, DEMAY Christophe, ENGELMANN Ilka, GEFFROY Sandrine, GUIGON Aurélie, LAMBERT Valérie, LAZREK Mouna, NOBILLIAUX Florian, PREVOST Brigitte, THUILLIER Caroline, TINEZ Claire ,TCHANTCHOU NJOSSE YANICK |
| EPI_ISL_2227237 | CHU LILLE | CHU Lille - Laboratoire de Virologie | AIT YAHYA Emilie, ALIDJINOU Enagnon Kazali, BOCKET Laurence, CREPIN Michel, DEMAY Christophe, ENGELMANN Ilka, GEFFROY Sandrine, GUIGON Aurélie, LAMBERT Valérie, LAZREK Mouna, NOBILLIAUX Florian, PREVOST Brigitte, THUILLIER Caroline, TINEZ Claire ,TCHANTCHOU NJOSSE YANICK |
| EPI_ISL_2227239 | CH ROUBAIX | CHU Lille - Laboratoire de Virologie | AIT YAHYA Emilie, ALIDJINOU Enagnon Kazali, BOCKET Laurence, CREPIN Michel, DEMAY Christophe, ENGELMANN Ilka, GEFFROY Sandrine, GUIGON Aurélie, LAMBERT Valérie, LAZREK Mouna, NOBILLIAUX Florian, PREVOST Brigitte, THUILLIER Caroline, TINEZ Claire ,TCHANTCHOU NJOSSE YANICK |
| EPI_ISL_2227240 | CH CALAIS | CHU Lille - Laboratoire de Virologie | AIT YAHYA Emilie, ALIDJINOU Enagnon Kazali, BOCKET Laurence, CREPIN Michel, DEMAY Christophe, ENGELMANN Ilka, GEFFROY Sandrine, GUIGON Aurélie, LAMBERT Valérie, LAZREK Mouna, NOBILLIAUX Florian, PREVOST Brigitte, THUILLIER Caroline, TINEZ Claire ,TCHANTCHOU NJOSSE YANICK |
| EPI_ISL_2227241, EPI_ISL_2227242, EPI_ISL_2227243, EPI_ISL_2227244, EPI_ISL_2227245 | CHU LILLE | CHU Lille - Laboratoire de Virologie | AIT YAHYA Emilie, ALIDJINOU Enagnon Kazali, BOCKET Laurence, CREPIN Michel, DEMAY Christophe, ENGELMANN Ilka, GEFFROY Sandrine, GUIGON Aurélie, LAMBERT Valérie, LAZREK Mouna, NOBILLIAUX Florian, PREVOST Brigitte, THUILLIER Caroline, TINEZ Claire ,TCHANTCHOU NJOSSE YANICK |
| EPI_ISL_2227246 | CH ROUBAIX | CHU Lille - Laboratoire de Virologie | AIT YAHYA Emilie, ALIDJINOU Enagnon Kazali, BOCKET Laurence, CREPIN Michel, DEMAY Christophe, ENGELMANN Ilka, GEFFROY Sandrine, GUIGON Aurélie, LAMBERT Valérie, LAZREK Mouna, NOBILLIAUX Florian, PREVOST Brigitte, THUILLIER Caroline, TINEZ Claire ,TCHANTCHOU NJOSSE YANICK |
| EPI_ISL_2227247, EPI_ISL_2227248, EPI_ISL_2227249, EPI_ISL_2227250, EPI_ISL_2227251, EPI_ISL_2227252, EPI_ISL_2227253 | CHU LILLE | CHU Lille - Laboratoire de Virologie | AIT YAHYA Emilie, ALIDJINOU Enagnon Kazali, BOCKET Laurence, CREPIN Michel, DEMAY Christophe, ENGELMANN Ilka, GEFFROY Sandrine, GUIGON Aurélie, LAMBERT Valérie, LAZREK Mouna, NOBILLIAUX Florian, PREVOST Brigitte, THUILLIER Caroline, TINEZ Claire ,TCHANTCHOU NJOSSE YANICK |
| EPI_ISL_2227254 | CH ROUBAIX | CHU Lille - Laboratoire de Virologie | AIT YAHYA Emilie, ALIDJINOU Enagnon Kazali, BOCKET Laurence, CREPIN Michel, DEMAY Christophe, ENGELMANN Ilka, GEFFROY Sandrine, GUIGON Aurélie, LAMBERT Valérie, LAZREK Mouna, NOBILLIAUX Florian, PREVOST Brigitte, THUILLIER Caroline, TINEZ Claire ,TCHANTCHOU NJOSSE YANICK |
| EPI_ISL_2227255, EPI_ISL_2227256, EPI_ISL_2227257, EPI_ISL_2227258, EPI_ISL_2227259, EPI_ISL_2227260, EPI_ISL_2227261 | CHU LILLE | CHU Lille - Laboratoire de Virologie | AIT YAHYA Emilie, ALIDJINOU Enagnon Kazali, BOCKET Laurence, CREPIN Michel, DEMAY Christophe, ENGELMANN Ilka, GEFFROY Sandrine, GUIGON Aurélie, LAMBERT Valérie, LAZREK Mouna, NOBILLIAUX Florian, PREVOST Brigitte, THUILLIER Caroline, TINEZ Claire ,TCHANTCHOU NJOSSE YANICK |
| EPI_ISL_2227262 | CH TOURCOING | CHU Lille - Laboratoire de Virologie | AIT YAHYA Emilie, ALIDJINOU Enagnon Kazali, BOCKET Laurence, CREPIN Michel, DEMAY Christophe, ENGELMANN Ilka, GEFFROY Sandrine, GUIGON Aurélie, LAMBERT Valérie, LAZREK Mouna, NOBILLIAUX Florian, PREVOST Brigitte, THUILLIER Caroline, TINEZ Claire ,TCHANTCHOU NJOSSE YANICK |
| EPI_ISL_2227263 | CH ROUBAIX | CHU Lille - Laboratoire de Virologie | AIT YAHYA Emilie, ALIDJINOU Enagnon Kazali, BOCKET Laurence, CREPIN Michel, DEMAY Christophe, ENGELMANN Ilka, GEFFROY Sandrine, GUIGON Aurélie, LAMBERT Valérie, LAZREK Mouna, NOBILLIAUX Florian, PREVOST Brigitte, THUILLIER Caroline, TINEZ Claire ,TCHANTCHOU NJOSSE YANICK |
| EPI_ISL_2227264 | CH TOURCOING | CHU Lille - Laboratoire de Virologie | AIT YAHYA Emilie, ALIDJINOU Enagnon Kazali, BOCKET Laurence, CREPIN Michel, DEMAY Christophe, ENGELMANN Ilka, GEFFROY Sandrine, GUIGON Aurélie, LAMBERT Valérie, LAZREK Mouna, NOBILLIAUX Florian, PREVOST Brigitte, THUILLIER Caroline, TINEZ Claire ,TCHANTCHOU NJOSSE YANICK |
| EPI_ISL_2227265 | CH ROUBAIX | CHU Lille - Laboratoire de Virologie | AIT YAHYA Emilie, ALIDJINOU Enagnon Kazali, BOCKET Laurence, CREPIN Michel, DEMAY Christophe, ENGELMANN Ilka, GEFFROY Sandrine, GUIGON Aurélie, LAMBERT Valérie, LAZREK Mouna, NOBILLIAUX Florian, PREVOST Brigitte, THUILLIER Caroline, TINEZ Claire ,TCHANTCHOU NJOSSE YANICK |
| EPI_ISL_2227266 | CH SAINT-OMER | CHU Lille - Laboratoire de Virologie | AIT YAHYA Emilie, ALIDJINOU Enagnon Kazali, BOCKET Laurence, CREPIN Michel, DEMAY Christophe, ENGELMANN Ilka, GEFFROY Sandrine, GUIGON Aurélie, LAMBERT Valérie, LAZREK Mouna, NOBILLIAUX Florian, PREVOST Brigitte, THUILLIER Caroline, TINEZ Claire ,TCHANTCHOU NJOSSE YANICK |
| EPI_ISL_2227267 | CH LENS | CHU Lille - Laboratoire de Virologie | AIT YAHYA Emilie, ALIDJINOU Enagnon Kazali, BOCKET Laurence, CREPIN Michel, DEMAY Christophe, ENGELMANN Ilka, GEFFROY Sandrine, GUIGON Aurélie, LAMBERT Valérie, LAZREK Mouna, NOBILLIAUX Florian, PREVOST Brigitte, THUILLIER Caroline, TINEZ Claire ,TCHANTCHOU NJOSSE YANICK |
| EPI_ISL_2227460, EPI_ISL_2227461 | Institute of Molecular and Translational Medicine / Laboratory of Experimental Medicine, Faculty of Medicine and Dentistry, Palacky University and University Hospital Olomouc | Institute of Molecular and Translational Medicine / Laboratory of Experimental Medicine, Faculty of Medicine and Dentistry, Palacky University | Rastislav Slavkovský, Hana Jaworek, Vladimíra Koudeláková, Barbora Blumová, Tomáš Pospíšíl, Marián Hajdúch |
| EPI_ISL_2227748 | Microbiology Department - University Hospital Brussel | Microbiology Department - University Hospital Brussel | Oriane Soetens, Thomas Demuyser, Lucie Seyler |
| EPI_ISL_2228045, EPI_ISL_2228046, EPI_ISL_2228047, EPI_ISL_2228048, EPI_ISL_2228049, EPI_ISL_2228050, EPI_ISL_2228051, EPI_ISL_2228052, EPI_ISL_2228053, EPI_ISL_2228054, EPI_ISL_2228055, EPI_ISL_2228056, EPI_ISL_2228057, EPI_ISL_2228058, EPI_ISL_2228059, EPI_ISL_2228060, EPI_ISL_2228061 | | | |

| | | | |
|---|---|---|---|
| see above | Klinisch Laboratorium ZNA | Klinisch Laboratorium ZNA | Verstrepen et al. |
| EPI_ISL_2228077, EPI_ISL_2228078, EPI_ISL_2228079, EPI_ISL_2228080, EPI_ISL_2228082, EPI_ISL_2228084, EPI_ISL_2228087 | Microbiology Department, University Hospital Donostia | Microbiology Department, University Hospital Donostia | Marimon JM, Montes M, Piñeiro L, Grandioso D, Martin-Peñaranda T, Gomez M, Cilla G. |
| EPI_ISL_2228429 | Basurto University Hospital: Clinical Microbiology Laboratory | Biocruces Bizkaia | Mikel Urrutikoetxea-Gutiérrez, Mª Carmen Nieto Toboso, Estibaliz Ugalde Zarraga, Mikel Gallego Rodrigo, Ana de la Hoz, José Luis Díaz de Tuesta del Arco |
| EPI_ISL_2228683, EPI_ISL_2228925 | GH A.CHENEVIER-H.MONDOR | Department of Virology, Henri Mondor University Hospital, Assistance Publique Hôpitaux de Paris, Université Paris-Est Créteil, INSERM U955 | Christophe Rodriguez, Slim Fourati, Vanessa Demontant, Guillaume Gricourt, Melissa N'Debi, Alexandre Soulier, Elisabeth Trawinski, Jean-Michel Pawlotsky |
| EPI_ISL_2231091, EPI_ISL_2231092, EPI_ISL_2231093, EPI_ISL_2231094, EPI_ISL_2231095, EPI_ISL_2231096, EPI_ISL_2231097, EPI_ISL_2231098, EPI_ISL_2231099, EPI_ISL_2231100, EPI_ISL_2231101, EPI_ISL_2231102, EPI_ISL_2231103 | | | |
| see above | Sharp HealthCare Laboratory | Andersen lab at Scripps Research | SEARCH Alliance San Diego with Aaron Harding, Jacquelyn Berumen, Cathy Woerle, Liam McGinnis, Art Mendoza, Omid Bakhtar |
| EPI_ISL_2231104, EPI_ISL_2231105, EPI_ISL_2231106, EPI_ISL_2231107, EPI_ISL_2231108, EPI_ISL_2231109, EPI_ISL_2231110, EPI_ISL_2231111, EPI_ISL_2231112, EPI_ISL_2231113, EPI_ISL_2231114, EPI_ISL_2231115, EPI_ISL_2231116, EPI_ISL_2231117, EPI_ISL_2231118, EPI_ISL_2231119, EPI_ISL_2231120, EPI_ISL_2231121, EPI_ISL_2231122, EPI_ISL_2231123, EPI_ISL_2231124, EPI_ISL_2231125, EPI_ISL_2231126, EPI_ISL_2231127, EPI_ISL_2231128 | | | |
| see above | San Diego County Public Health Laboratory | Andersen lab at Scripps Research | SEARCH Alliance San Diego with Ashleigh Murphy, Jovan Shephard, Brett Austin |
| EPI_ISL_2231129, EPI_ISL_2231130, EPI_ISL_2231131, EPI_ISL_2231132, EPI_ISL_2231133, EPI_ISL_2231134, EPI_ISL_2231135, EPI_ISL_2231136, EPI_ISL_2231137, EPI_ISL_2231138, EPI_ISL_2231139, EPI_ISL_2231140, EPI_ISL_2231141, EPI_ISL_2231142, EPI_ISL_2231143, EPI_ISL_2231144, EPI_ISL_2231145, EPI_ISL_2231146, EPI_ISL_2231147, EPI_ISL_2231148, EPI_ISL_2231149, EPI_ISL_2231150, EPI_ISL_2231151, EPI_ISL_2231155, EPI_ISL_2231156, EPI_ISL_2231158, EPI_ISL_2231159, EPI_ISL_2231160 | | | |
| see above | Sharp HealthCare Laboratory | Andersen lab at Scripps Research | SEARCH Alliance San Diego with Aaron Harding, Jacquelyn Berumen, Cathy Woerle, Liam McGinnis, Art Mendoza, Omid Bakhtar |
| EPI_ISL_2231161 | San Diego County Public Health Laboratory | Andersen lab at Scripps Research | SEARCH Alliance San Diego with Ashleigh Murphy, Jovan Shephard, Brett Austin |
| EPI_ISL_2231162, EPI_ISL_2231163, EPI_ISL_2231164, EPI_ISL_2231165, EPI_ISL_2231166, EPI_ISL_2231167, EPI_ISL_2231168, EPI_ISL_2231169, EPI_ISL_2231170, EPI_ISL_2231176, EPI_ISL_2231177 | | | |
| see above | Sharp HealthCare Laboratory | Andersen lab at Scripps Research | SEARCH Alliance San Diego with Aaron Harding, Jacquelyn Berumen, Cathy Woerle, Liam McGinnis, Art Mendoza, Omid Bakhtar |
| EPI_ISL_2231178 | San Diego County Public Health Laboratory | Andersen lab at Scripps Research | SEARCH Alliance San Diego with Ashleigh Murphy, Jovan Shephard, Brett Austin |
| EPI_ISL_2231179, EPI_ISL_2231180, EPI_ISL_2231181, EPI_ISL_2231182, EPI_ISL_2231183, EPI_ISL_2231184, EPI_ISL_2231185, EPI_ISL_2231195, EPI_ISL_2231198, EPI_ISL_2231205, EPI_ISL_2231206, EPI_ISL_2231318, EPI_ISL_2231319, EPI_ISL_2231320 | | | |
| see above | Sharp HealthCare Laboratory | Andersen lab at Scripps Research | SEARCH Alliance San Diego with Aaron Harding, Jacquelyn Berumen, Cathy Woerle, Liam McGinnis, Art Mendoza, Omid Bakhtar |
| EPI_ISL_2231435, EPI_ISL_2231442 | San Diego County Public Health Laboratory | Andersen lab at Scripps Research | SEARCH Alliance San Diego with Ashleigh Murphy, Jovan Shephard, Brett Austin |
| EPI_ISL_2231526, EPI_ISL_2231527, EPI_ISL_2231530 | Sharp HealthCare Laboratory | Andersen lab at Scripps Research | SEARCH Alliance San Diego with Aaron Harding, Jacquelyn Berumen, Cathy Woerle, Liam McGinnis, Art Mendoza, Omid Bakhtar |
| EPI_ISL_2232757, EPI_ISL_2232758, EPI_ISL_2232760, EPI_ISL_2232761, EPI_ISL_2232762, EPI_ISL_2232763, EPI_ISL_2232764 | Department of Laboratory Medicine, Division of Clinical Virology, University of Medicine, Vienna | Bergthaler laboratory, CeMM Research Center for Molecular Medicine of the Austrian Academy of Sciences | Lukas Endler, Anna Schedl, Fabian Amman, Petr Triska, Thomas Penz, Benedikt Agerer, Maelle Le Moing, Michael Schuster, Bekir Erguner, Jan Laine, Martin Senekowitsch, Christoph Bock, Andreas Bergthaler |
| EPI_ISL_2232765, EPI_ISL_2232766, EPI_ISL_2232767, EPI_ISL_2232768, EPI_ISL_2232769, EPI_ISL_2232770, EPI_ISL_2232771, EPI_ISL_2232773, EPI_ISL_2232774, EPI_ISL_2232778, EPI_ISL_2232780, EPI_ISL_2232781, EPI_ISL_2232782, EPI_ISL_2232783, EPI_ISL_2232785, EPI_ISL_2232786, EPI_ISL_2232788, EPI_ISL_2232789, EPI_ISL_2232790, EPI_ISL_2232791, EPI_ISL_2232792, EPI_ISL_2232795, EPI_ISL_2232796, EPI_ISL_2232798, EPI_ISL_2232799, EPI_ISL_2232800, EPI_ISL_2232801, EPI_ISL_2232802, EPI_ISL_2232803, EPI_ISL_2232806, EPI_ISL_2232809, EPI_ISL_2232810, EPI_ISL_2232811, EPI_ISL_2232812, EPI_ISL_2232813, EPI_ISL_2232815, EPI_ISL_2232816, EPI_ISL_2232817, EPI_ISL_2232820, EPI_ISL_2232821, EPI_ISL_2232822, EPI_ISL_2232823, EPI_ISL_2232824, EPI_ISL_2232825, EPI_ISL_2232827, EPI_ISL_2232828, EPI_ISL_2232829, EPI_ISL_2232833, EPI_ISL_2232834, EPI_ISL_2232835, EPI_ISL_2232836, EPI_ISL_2232838, EPI_ISL_2232839, EPI_ISL_2232840, EPI_ISL_2232843, EPI_ISL_2232844, EPI_ISL_2232845, EPI_ISL_2232846, EPI_ISL_2232847, EPI_ISL_2232848, EPI_ISL_2232850, EPI_ISL_2232851, EPI_ISL_2232856, EPI_ISL_2232858, EPI_ISL_2232859, EPI_ISL_2232861, EPI_ISL_2232865, EPI_ISL_2232869, EPI_ISL_2232870, EPI_ISL_2232871, EPI_ISL_2232872, EPI_ISL_2232879, EPI_ISL_2232881, EPI_ISL_2232882, EPI_ISL_2232887, EPI_ISL_2232889, EPI_ISL_2232890, EPI_ISL_2232895, EPI_ISL_2232896, EPI_ISL_2232897, EPI_ISL_2232899, EPI_ISL_2232901, EPI_ISL_2232902, EPI_ISL_2232903, EPI_ISL_2232907, EPI_ISL_2232908, EPI_ISL_2232910, EPI_ISL_2232911, EPI_ISL_2232912, EPI_ISL_2232919, EPI_ISL_2232953 | | | |
| see above | Austrian Agency for Health and Food Safety (AGES) | Bergthaler laboratory, CeMM Research Center for Molecular Medicine of the Austrian Academy of Sciences | Lukas Endler, Anna Schedl, Fabian Amman, Petr Triska, Thomas Penz, Benedikt Agerer, Maelle Le Moing, Michael Schuster, Bekir Erguner, Jan Laine, Martin Senekowitsch, Christoph Bock, Andreas Bergthaler |
| EPI_ISL_2232982, EPI_ISL_2232985, EPI_ISL_2232986, EPI_ISL_2232987, EPI_ISL_2232988 | 3. Medizinische Abteilung, Hanusch Krankenhaus | Bergthaler laboratory, CeMM Research Center for Molecular Medicine of the Austrian Academy of Sciences | Lukas Endler, Anna Schedl, Fabian Amman, Petr Triska, Thomas Penz, Benedikt Agerer, Maelle Le Moing, Michael Schuster, Bekir Erguner, Jan Laine, Martin Senekowitsch, Christoph Bock, Andreas Bergthaler |
| EPI_ISL_2233089 | RSUD Abdul Moeis | National Institute of Health Research and Development | Vivi Setiawaty, Hartanti Dian Ikawati, Hana Apsari Pawestri, Subangkit, Kartika Dewi Puspa, Arie Ardiansyah Nugraha, Triyani Soekarso, Krisna Pangesti, Nelly Puspandari |
| EPI_ISL_2233094 | Labkesda Prov Kal Tim | National Institute of Health Research and Development | Kartika Dewi Puspa, Subangkit, Arie Ardiansyah Nugraha, Hartanti Dian Ikawati, Hana Apsari Pawestri, Triyani Soekarso, Krisna Pangesti, Nelly Puspandari, Vivi Setiawaty |
| EPI_ISL_2233145, EPI_ISL_2233146 | National Centre For Cell Science | National Centre For Cell Science - INSACOG | Dhiraj Paul, Shivang P. Bhanushali, Mitali Inamdar, Mohak P Gujare, Manoj Kumar Bhat, Ajay Pillai, INSACOG Consortium team, Yogesh Shouche |
| EPI_ISL_2233382 | Hospital Labuan | Institute for Medical Research, Infectious Disease Research Centre, National Institutes of Health, Ministry of Health Malaysia | Suppiah J, Kamel K, Mohd Zawawi Z, Azizan MA, Ramly N, Robert F, Thayan R |
| EPI_ISL_2233384 | Johor Bahru Public Health Laboratory | Institute for Medical Research, Infectious Disease Research Centre, National Institutes of Health, Ministry of Health Malaysia | Suppiah J, Kamel K, Mohd Zawawi Z, Azizan MA, Ramly N, Robert F, Thayan R |
| EPI_ISL_2235297, EPI_ISL_2235298, EPI_ISL_2235303 | AZ St.Jan Brugge-Oostende AV | AZ SINT-JAN BRUGGE | Marijke Reynders, Patrick Descheemaeker, Laurien Hoornaert, Thomas Van Landschoot, Jorn Hellemans, Marie Madeleine Chabert-Consen |
| EPI_ISL_2235305 | AZ Zeno Knokke-Blankenberge | AZ SINT-JAN BRUGGE | Marijke Reynders, Patrick Descheemaeker, Laurien Hoornaert, Thomas Van Landschoot, Jorn Hellemans, Marie Madeleine Chabert-Consen |
| EPI_ISL_2235307 | AZ St.Jan Brugge-Oostende AV | AZ SINT-JAN BRUGGE | Marijke Reynders, Patrick Descheemaeker, Laurien Hoornaert, Thomas Van Landschoot, Jorn Hellemans, Marie Madeleine Chabert-Consen |
| EPI_ISL_2235315 | SIESP CHIETI - DRIVE IN ORTONA | Istituto Zooprofilattico Sperimentale dell'Abruzzo e Molise "G. Caporale" | Lorusso A, Marcacci M, Di Domenico M, Ancora M, Curini V, Di Lollo Valeria, Mangone I, Rinaldi A, Delli Compagni E, Scialabba S, Caporale M, Di Pasquale A, Cammà C, Puglia I, Calistri P, Savini G |
| EPI_ISL_2235316, EPI_ISL_2235317, EPI_ISL_2235318, EPI_ISL_2235319, EPI_ISL_2235320 | SIESP DIPARTIMENTO DI PREVENZIONE TERAMO | Istituto Zooprofilattico Sperimentale dell'Abruzzo e Molise "G. Caporale" | Lorusso A, Marcacci M, Di Domenico M, Ancora M, Curini V, Di Lollo Valeria, Mangone I, Rinaldi A, Delli Compagni E, Scialabba S, Caporale M, Di Pasquale A, Cammà C, Puglia I, Calistri P, Savini G |
| EPI_ISL_2235321, EPI_ISL_2235322 | RSA GIULIANOVA | Istituto Zooprofilattico Sperimentale dell'Abruzzo e Molise "G. Caporale" | Lorusso A, Marcacci M, Di Domenico M, Ancora M, Curini V, Di Lollo Valeria, Mangone I, Rinaldi A, Delli Compagni E, Scialabba S, Caporale M, Di Pasquale A, Cammà C, Puglia I, Calistri P, Savini G |
| EPI_ISL_2235323, EPI_ISL_2235324 | SIESP CHIETI - DRIVE IN LANCIANO | Istituto Zooprofilattico Sperimentale dell'Abruzzo e Molise "G. Caporale" | Lorusso A, Marcacci M, Di Domenico M, Ancora M, Curini V, Di Lollo Valeria, Mangone I, Rinaldi A, Delli Compagni E, Scialabba S, Caporale M, Di Pasquale A, Cammà C, Puglia I, Calistri P, Savini G |
| EPI_ISL_2235325 | USCA Avezzano | Istituto Zooprofilattico Sperimentale dell'Abruzzo e Molise "G. Caporale" | Lorusso A, Marcacci M, Di Domenico M, Ancora M, Curini V, Di Lollo Valeria, Mangone I, Rinaldi A, Delli Compagni E, Scialabba S, Caporale M, Di Pasquale A, Cammà C, Puglia I, Calistri P, Savini G |
| EPI_ISL_2235326, EPI_ISL_2235327 | SIESP CHIETI - DRIVE IN LANCIANO | Istituto Zooprofilattico Sperimentale dell'Abruzzo e Molise "G. Caporale" | Lorusso A, Marcacci M, Di Domenico M, Ancora M, Curini V, Di Lollo Valeria, Mangone I, Rinaldi A, Delli Compagni E, Scialabba S, Caporale M, Di Pasquale A, Cammà C, Puglia I, Calistri P, Savini G |

| | | | |
|---|---|---|---|
| EPI_ISL_2235328, EPI_ISL_2235329 | SIESP CHIETI - DRIVE IN CHIETI | Istituto Zooprofilattico Sperimentale dell'Abruzzo e Molise "G. Caporale" | Lorusso A, Marcacci M, Di Domenico M, Ancora M, Curini V, Di Lollo Valeria, Mangone I, Rinaldi A, Delli Compagni E, Scialabba S, Caporale M, Di Pasquale A, Cammà C, Puglia I, Calistri P, Savini G |
| EPI_ISL_2235330 | SIESP DIPARTIMENTO DI PREVENZIONE TERAMO | Istituto Zooprofilattico Sperimentale dell'Abruzzo e Molise "G. Caporale" | Lorusso A, Marcacci M, Di Domenico M, Ancora M, Curini V, Di Lollo Valeria, Mangone I, Rinaldi A, Delli Compagni E, Scialabba S, Caporale M, Di Pasquale A, Cammà C, Puglia I, Calistri P, Savini G |
| EPI_ISL_2235331 | USCA Avezzano | Istituto Zooprofilattico Sperimentale dell'Abruzzo e Molise "G. Caporale" | Lorusso A, Marcacci M, Di Domenico M, Ancora M, Curini V, Di Lollo Valeria, Mangone I, Rinaldi A, Delli Compagni E, Scialabba S, Caporale M, Di Pasquale A, Cammà C, Puglia I, Calistri P, Savini G |
| EPI_ISL_2235332 | USCA Pescina | Istituto Zooprofilattico Sperimentale dell'Abruzzo e Molise "G. Caporale" | Lorusso A, Marcacci M, Di Domenico M, Ancora M, Curini V, Di Lollo Valeria, Mangone I, Rinaldi A, Delli Compagni E, Scialabba S, Caporale M, Di Pasquale A, Cammà C, Puglia I, Calistri P, Savini G |
| EPI_ISL_2235333 | SIESP CHIETI - DRIVE IN LANCIANO | Istituto Zooprofilattico Sperimentale dell'Abruzzo e Molise "G. Caporale" | Lorusso A, Marcacci M, Di Domenico M, Ancora M, Curini V, Di Lollo Valeria, Mangone I, Rinaldi A, Delli Compagni E, Scialabba S, Caporale M, Di Pasquale A, Cammà C, Puglia I, Calistri P, Savini G |
| EPI_ISL_2235334 | SIESP CHIETI - DRIVE IN VASTO | Istituto Zooprofilattico Sperimentale dell'Abruzzo e Molise "G. Caporale" | Lorusso A, Marcacci M, Di Domenico M, Ancora M, Curini V, Di Lollo Valeria, Mangone I, Rinaldi A, Delli Compagni E, Scialabba S, Caporale M, Di Pasquale A, Cammà C, Puglia I, Calistri P, Savini G |
| EPI_ISL_2235335 | SIESP CHIETI - DRIVE IN GISSI | Istituto Zooprofilattico Sperimentale dell'Abruzzo e Molise "G. Caporale" | Lorusso A, Marcacci M, Di Domenico M, Ancora M, Curini V, Di Lollo Valeria, Mangone I, Rinaldi A, Delli Compagni E, Scialabba S, Caporale M, Di Pasquale A, Cammà C, Puglia I, Calistri P, Savini G |
| EPI_ISL_2235336 | USCA Avezzano | Istituto Zooprofilattico Sperimentale dell'Abruzzo e Molise "G. Caporale" | Lorusso A, Marcacci M, Di Domenico M, Ancora M, Curini V, Di Lollo Valeria, Mangone I, Rinaldi A, Delli Compagni E, Scialabba S, Caporale M, Di Pasquale A, Cammà C, Puglia I, Calistri P, Savini G |
| EPI_ISL_2235337, EPI_ISL_2235338, EPI_ISL_2235339, EPI_ISL_2235340 | SIESP CHIETI - DRIVE IN LANCIANO | Istituto Zooprofilattico Sperimentale dell'Abruzzo e Molise "G. Caporale" | Lorusso A, Marcacci M, Di Domenico M, Ancora M, Curini V, Di Lollo Valeria, Mangone I, Rinaldi A, Delli Compagni E, Scialabba S, Caporale M, Di Pasquale A, Cammà C, Puglia I, Calistri P, Savini G |
| EPI_ISL_2235341 | SIESP DIPARTIMENTO DI PREVENZIONE CHIETI | Istituto Zooprofilattico Sperimentale dell'Abruzzo e Molise "G. Caporale" | Lorusso A, Marcacci M, Di Domenico M, Ancora M, Curini V, Di Lollo Valeria, Mangone I, Rinaldi A, Delli Compagni E, Scialabba S, Caporale M, Di Pasquale A, Cammà C, Puglia I, Calistri P, Savini G |
| EPI_ISL_2235342 | DIP. PREV. AVEZZANO SERVIZIO DI IGIENE EPIDEMIOLOGIA E SANITA' PUBBLICA | Istituto Zooprofilattico Sperimentale dell'Abruzzo e Molise "G. Caporale" | Lorusso A, Marcacci M, Di Domenico M, Ancora M, Curini V, Di Lollo Valeria, Mangone I, Rinaldi A, Delli Compagni E, Scialabba S, Caporale M, Di Pasquale A, Cammà C, Puglia I, Calistri P, Savini G |
| EPI_ISL_2235343 | SIESP CHIETI - DRIVE IN GISSI | Istituto Zooprofilattico Sperimentale dell'Abruzzo e Molise "G. Caporale" | Lorusso A, Marcacci M, Di Domenico M, Ancora M, Curini V, Di Lollo Valeria, Mangone I, Rinaldi A, Delli Compagni E, Scialabba S, Caporale M, Di Pasquale A, Cammà C, Puglia I, Calistri P, Savini G |
| EPI_ISL_2235344, EPI_ISL_2235345, EPI_ISL_2235346 | SIESP DIPARTIMENTO DI PREVENZIONE TERAMO | Istituto Zooprofilattico Sperimentale dell'Abruzzo e Molise "G. Caporale" | Lorusso A, Marcacci M, Di Domenico M, Ancora M, Curini V, Di Lollo Valeria, Mangone I, Rinaldi A, Delli Compagni E, Scialabba S, Caporale M, Di Pasquale A, Cammà C, Puglia I, Calistri P, Savini G |
| EPI_ISL_2235347 | USCA Tagliacozzo | Istituto Zooprofilattico Sperimentale dell'Abruzzo e Molise "G. Caporale" | Lorusso A, Marcacci M, Di Domenico M, Ancora M, Curini V, Di Lollo Valeria, Mangone I, Rinaldi A, Delli Compagni E, Scialabba S, Caporale M, Di Pasquale A, Cammà C, Puglia I, Calistri P, Savini G |
| EPI_ISL_2235348, EPI_ISL_2235349 | SIESP DIPARTIMENTO DI PREVENZIONE TERAMO | Istituto Zooprofilattico Sperimentale dell'Abruzzo e Molise "G. Caporale" | Lorusso A, Marcacci M, Di Domenico M, Ancora M, Curini V, Di Lollo Valeria, Mangone I, Rinaldi A, Delli Compagni E, Scialabba S, Caporale M, Di Pasquale A, Cammà C, Puglia I, Calistri P, Savini G |
| EPI_ISL_2235350, EPI_ISL_2235351, EPI_ISL_2235352, EPI_ISL_2235353 | SIESP CHIETI - DRIVE IN LANCIANO | Istituto Zooprofilattico Sperimentale dell'Abruzzo e Molise "G. Caporale" | Lorusso A, Marcacci M, Di Domenico M, Ancora M, Curini V, Di Lollo Valeria, Mangone I, Rinaldi A, Delli Compagni E, Scialabba S, Caporale M, Di Pasquale A, Cammà C, Puglia I, Calistri P, Savini G |
| EPI_ISL_2235354, EPI_ISL_2235355, EPI_ISL_2235356, EPI_ISL_2235357 | SIESP DIPARTIMENTO DI PREVENZIONE TERAMO | Istituto Zooprofilattico Sperimentale dell'Abruzzo e Molise "G. Caporale" | Lorusso A, Marcacci M, Di Domenico M, Ancora M, Curini V, Di Lollo Valeria, Mangone I, Rinaldi A, Delli Compagni E, Scialabba S, Caporale M, Di Pasquale A, Cammà C, Puglia I, Calistri P, Savini G |
| EPI_ISL_2235358 | OSPEDALE CIVILE SANT'OMERO - PRONTO SOCCORSO | Istituto Zooprofilattico Sperimentale dell'Abruzzo e Molise "G. Caporale" | Lorusso A, Marcacci M, Di Domenico M, Ancora M, Curini V, Di Lollo Valeria, Mangone I, Rinaldi A, Delli Compagni E, Scialabba S, Caporale M, Di Pasquale A, Cammà C, Puglia I, Calistri P, Savini G |
| EPI_ISL_2235359, EPI_ISL_2235360 | SIESP CHIETI - DRIVE IN CHIETI | Istituto Zooprofilattico Sperimentale dell'Abruzzo e Molise "G. Caporale" | Lorusso A, Marcacci M, Di Domenico M, Ancora M, Curini V, Di Lollo Valeria, Mangone I, Rinaldi A, Delli Compagni E, Scialabba S, Caporale M, Di Pasquale A, Cammà C, Puglia I, Calistri P, Savini G |
| EPI_ISL_2235361 | SIESP DIPARTIMENTO DI PREVENZIONE TERAMO | Istituto Zooprofilattico Sperimentale dell'Abruzzo e Molise "G. Caporale" | Lorusso A, Marcacci M, Di Domenico M, Ancora M, Curini V, Di Lollo Valeria, Mangone I, Rinaldi A, Delli Compagni E, Scialabba S, Caporale M, Di Pasquale A, Cammà C, Puglia I, Calistri P, Savini G |
| EPI_ISL_2235362, EPI_ISL_2235363, EPI_ISL_2235364, EPI_ISL_2235365, EPI_ISL_2235366 | SIESP CHIETI - DRIVE IN ORTONA | Istituto Zooprofilattico Sperimentale dell'Abruzzo e Molise "G. Caporale" | Lorusso A, Marcacci M, Di Domenico M, Ancora M, Curini V, Di Lollo Valeria, Mangone I, Rinaldi A, Delli Compagni E, Scialabba S, Caporale M, Di Pasquale A, Cammà C, Puglia I, Calistri P, Savini G |
| EPI_ISL_2235367 | USCA Avezzano | Istituto Zooprofilattico Sperimentale dell'Abruzzo e Molise "G. Caporale" | Lorusso A, Marcacci M, Di Domenico M, Ancora M, Curini V, Di Lollo Valeria, Mangone I, Rinaldi A, Delli Compagni E, Scialabba S, Caporale M, Di Pasquale A, Cammà C, Puglia I, Calistri P, Savini G |
| EPI_ISL_2235368 | DIP. PREV. AVEZZANO SERVIZIO DI IGIENE EPIDEMIOLOGIA E SANITA' PUBBLICA | Istituto Zooprofilattico Sperimentale dell'Abruzzo e Molise "G. Caporale" | Lorusso A, Marcacci M, Di Domenico M, Ancora M, Curini V, Di Lollo Valeria, Mangone I, Rinaldi A, Delli Compagni E, Scialabba S, Caporale M, Di Pasquale A, Cammà C, Puglia I, Calistri P, Savini G |
| EPI_ISL_2235369 | SIESP CHIETI - DRIVE IN LANCIANO | Istituto Zooprofilattico Sperimentale dell'Abruzzo e Molise "G. Caporale" | Lorusso A, Marcacci M, Di Domenico M, Ancora M, Curini V, Di Lollo Valeria, Mangone I, Rinaldi A, Delli Compagni E, Scialabba S, Caporale M, Di Pasquale A, Cammà C, Puglia I, Calistri P, Savini G |
| EPI_ISL_2235370 | SIESP DIPARTIMENTO DI PREVENZIONE CHIETI | Istituto Zooprofilattico Sperimentale dell'Abruzzo e Molise "G. Caporale" | Lorusso A, Marcacci M, Di Domenico M, Ancora M, Curini V, Di Lollo Valeria, Mangone I, Rinaldi A, Delli Compagni E, Scialabba S, Caporale M, Di Pasquale A, Cammà C, Puglia I, Calistri P, Savini G |
| EPI_ISL_2235371 | DIP. PREV. AVEZZANO SERVIZIO DI IGIENE EPIDEMIOLOGIA E SANITA' PUBBLICA | Istituto Zooprofilattico Sperimentale dell'Abruzzo e Molise "G. Caporale" | Lorusso A, Marcacci M, Di Domenico M, Ancora M, Curini V, Di Lollo Valeria, Mangone I, Rinaldi A, Delli Compagni E, Scialabba S, Caporale M, Di Pasquale A, Cammà C, Puglia I, Calistri P, Savini G |
| EPI_ISL_2235372, EPI_ISL_2235373, EPI_ISL_2235374, EPI_ISL_2235375, EPI_ISL_2235376 | SIESP DIPARTIMENTO DI PREVENZIONE TERAMO | Istituto Zooprofilattico Sperimentale dell'Abruzzo e Molise "G. Caporale" | Lorusso A, Marcacci M, Di Domenico M, Ancora M, Curini V, Di Lollo Valeria, Mangone I, Rinaldi A, Delli Compagni E, Scialabba S, Caporale M, Di Pasquale A, Cammà C, Puglia I, Calistri P, Savini G |
| EPI_ISL_2235378 | DIP. PREV. AVEZZANO SERVIZIO DI IGIENE EPIDEMIOLOGIA E SANITA' PUBBLICA | Istituto Zooprofilattico Sperimentale dell'Abruzzo e Molise "G. Caporale" | Lorusso A, Marcacci M, Di Domenico M, Ancora M, Curini V, Di Lollo Valeria, Mangone I, Rinaldi A, Delli Compagni E, Scialabba S, Caporale M, Di Pasquale A, Cammà C, Puglia I, Calistri P, Savini G |
| EPI_ISL_2235380 | SIESP DIPARTIMENTO DI PREVENZIONE TERAMO | Istituto Zooprofilattico Sperimentale dell'Abruzzo e Molise "G. Caporale" | Lorusso A, Marcacci M, Di Domenico M, Ancora M, Curini V, Di Lollo Valeria, Mangone I, Rinaldi A, Delli Compagni E, Scialabba S, Caporale M, Di Pasquale A, Cammà C, Puglia I, Calistri P, Savini G |
| EPI_ISL_2235381 | USCA Castel di Sangro | Istituto Zooprofilattico Sperimentale dell'Abruzzo e Molise "G. Caporale" | Lorusso A, Marcacci M, Di Domenico M, Ancora M, Curini V, Di Lollo Valeria, Mangone I, Rinaldi A, Delli Compagni E, Scialabba S, Caporale M, Di Pasquale A, Cammà C, Puglia I, Calistri P, Savini G |
| EPI_ISL_2235382 | SIESP CHIETI - DRIVE IN LANCIANO | Istituto Zooprofilattico Sperimentale dell'Abruzzo e Molise "G. Caporale" | Lorusso A, Marcacci M, Di Domenico M, Ancora M, Curini V, Di Lollo Valeria, Mangone I, Rinaldi A, Delli Compagni E, Scialabba S, Caporale M, Di Pasquale A, Cammà C, Puglia I, Calistri P, Savini G |
| EPI_ISL_2235383 | SIESP CHIETI - DRIVE IN VASTO | Istituto Zooprofilattico Sperimentale dell'Abruzzo e Molise "G. Caporale" | Lorusso A, Marcacci M, Di Domenico M, Ancora M, Curini V, Di Lollo Valeria, Mangone I, Rinaldi A, Delli Compagni E, Scialabba S, Caporale M, Di Pasquale A, Cammà C, Puglia I, Calistri P, Savini G |
| EPI_ISL_2235384, EPI_ISL_2235385 | SIESP CHIETI - DRIVE IN LANCIANO | Istituto Zooprofilattico Sperimentale dell'Abruzzo e Molise "G. Caporale" | Lorusso A, Marcacci M, Di Domenico M, Ancora M, Curini V, Di Lollo Valeria, Mangone I, Rinaldi A, Delli Compagni E, Scialabba S, Caporale M, Di Pasquale A, Cammà C, Puglia I, Calistri P, Savini G |
| EPI_ISL_2235386 | SIESP CHIETI - DRIVE IN CHIETI | Istituto Zooprofilattico Sperimentale dell'Abruzzo e Molise "G. Caporale" | Lorusso A, Marcacci M, Di Domenico M, Ancora M, Curini V, Di Lollo Valeria, Mangone I, Rinaldi A, Delli Compagni E, Scialabba S, Caporale M, Di Pasquale A, Cammà C, Puglia I, Calistri P, Savini G |
| EPI_ISL_2235387, EPI_ISL_2235388, | SIESP DIPARTIMENTO DI PREVENZIONE CHIETI | Istituto Zooprofilattico Sperimentale dell'Abruzzo e Molise "G. | Lorusso A, Marcacci M, Di Domenico M, Ancora M, Curini V, Di Lollo Valeria, Mangone I, Rinaldi A, Delli Compagni E, Scialabba S, Caporale M, Di |

| | | | |
|---|---|---|---|
| EPI_ISL_2235389, EPI_ISL_2235390 | | Caporale" | Pasquale A, Cammà C, Puglia I, Calistri P, Savini G |
| EPI_ISL_2235391 | SIESP CHIETI - DRIVE IN ORTONA | Istituto Zooprofilattico Sperimentale dell'Abruzzo e Molise "G. Caporale" | Lorusso A, Marcacci M, Di Domenico M, Ancora M, Curini V, Di Lollo Valeria, Mangone I, Rinaldi A, Delli Compagni E, Scialabba S, Caporale M, Di Pasquale A, Cammà C, Puglia I, Calistri P, Savini G |
| EPI_ISL_2235392 | SIESP DIPARTIMENTO DI PREVENZIONE CHIETI | Istituto Zooprofilattico Sperimentale dell'Abruzzo e Molise "G. Caporale" | Lorusso A, Marcacci M, Di Domenico M, Ancora M, Curini V, Di Lollo Valeria, Mangone I, Rinaldi A, Delli Compagni E, Scialabba S, Caporale M, Di Pasquale A, Cammà C, Puglia I, Calistri P, Savini G |
| EPI_ISL_2235393, EPI_ISL_2235394, EPI_ISL_2235395, EPI_ISL_2235396 | SIESP DIPARTIMENTO DI PREVENZIONE TERAMO | Istituto Zooprofilattico Sperimentale dell'Abruzzo e Molise "G. Caporale" | Lorusso A, Marcacci M, Di Domenico M, Ancora M, Curini V, Di Lollo Valeria, Mangone I, Rinaldi A, Delli Compagni E, Scialabba S, Caporale M, Di Pasquale A, Cammà C, Puglia I, Calistri P, Savini G |
| EPI_ISL_2235397 | SIESP CHIETI - DRIVE IN ORTONA | Istituto Zooprofilattico Sperimentale dell'Abruzzo e Molise "G. Caporale" | Lorusso A, Marcacci M, Di Domenico M, Ancora M, Curini V, Di Lollo Valeria, Mangone I, Rinaldi A, Delli Compagni E, Scialabba S, Caporale M, Di Pasquale A, Cammà C, Puglia I, Calistri P, Savini G |
| EPI_ISL_2235398, EPI_ISL_2235399, EPI_ISL_2235400 | SIESP DIPARTIMENTO DI PREVENZIONE TERAMO | Istituto Zooprofilattico Sperimentale dell'Abruzzo e Molise "G. Caporale" | Lorusso A, Marcacci M, Di Domenico M, Ancora M, Curini V, Di Lollo Valeria, Mangone I, Rinaldi A, Delli Compagni E, Scialabba S, Caporale M, Di Pasquale A, Cammà C, Puglia I, Calistri P, Savini G |
| EPI_ISL_2235401 | SIESP CHIETI - DRIVE IN VASTO | Istituto Zooprofilattico Sperimentale dell'Abruzzo e Molise "G. Caporale" | Lorusso A, Marcacci M, Di Domenico M, Ancora M, Curini V, Di Lollo Valeria, Mangone I, Rinaldi A, Delli Compagni E, Scialabba S, Caporale M, Di Pasquale A, Cammà C, Puglia I, Calistri P, Savini G |
| EPI_ISL_2235402 | SIESP CHIETI - DRIVE IN ORTONA | Istituto Zooprofilattico Sperimentale dell'Abruzzo e Molise "G. Caporale" | Lorusso A, Marcacci M, Di Domenico M, Ancora M, Curini V, Di Lollo Valeria, Mangone I, Rinaldi A, Delli Compagni E, Scialabba S, Caporale M, Di Pasquale A, Cammà C, Puglia I, Calistri P, Savini G |
| EPI_ISL_2235403 | SIESP CHIETI - DRIVE IN CHIETI | Istituto Zooprofilattico Sperimentale dell'Abruzzo e Molise "G. Caporale" | Lorusso A, Marcacci M, Di Domenico M, Ancora M, Curini V, Di Lollo Valeria, Mangone I, Rinaldi A, Delli Compagni E, Scialabba S, Caporale M, Di Pasquale A, Cammà C, Puglia I, Calistri P, Savini G |
| EPI_ISL_2235404 | SIESP SULMONA | Istituto Zooprofilattico Sperimentale dell'Abruzzo e Molise "G. Caporale" | Lorusso A, Marcacci M, Di Domenico M, Ancora M, Curini V, Di Lollo Valeria, Mangone I, Rinaldi A, Delli Compagni E, Scialabba S, Caporale M, Di Pasquale A, Cammà C, Puglia I, Calistri P, Savini G |
| EPI_ISL_2235405 | DIP. PREV. AVEZZANO SERVIZIO DI IGIENE EPIDEMIOLOGIA E SANITA' PUBBLICA | Istituto Zooprofilattico Sperimentale dell'Abruzzo e Molise "G. Caporale" | Lorusso A, Marcacci M, Di Domenico M, Ancora M, Curini V, Di Lollo Valeria, Mangone I, Rinaldi A, Delli Compagni E, Scialabba S, Caporale M, Di Pasquale A, Cammà C, Puglia I, Calistri P, Savini G |
| EPI_ISL_2235406 | USCA Sulmona | Istituto Zooprofilattico Sperimentale dell'Abruzzo e Molise "G. Caporale" | Lorusso A, Marcacci M, Di Domenico M, Ancora M, Curini V, Di Lollo Valeria, Mangone I, Rinaldi A, Delli Compagni E, Scialabba S, Caporale M, Di Pasquale A, Cammà C, Puglia I, Calistri P, Savini G |
| EPI_ISL_2235438 | Università degli Studi di Perugia | Istituto Zooprofilattico Sperimentale dell'Abruzzo e Molise "G. Caporale" | Mencacci A, Camilloni B, Lorusso A, Marcacci M, Di Domenico M, Ancora M, Curini V, Di Lollo Valeria, Mangone I, Rinaldi A, Delli Compagni E, Scialabba S, Caporale M, Di Pasquale A, Cammà C, Puglia I, Calistri P, Savini G |
| EPI_ISL_2235464, EPI_ISL_2235465, EPI_ISL_2235466, EPI_ISL_2235467, EPI_ISL_2235468, EPI_ISL_2235469, EPI_ISL_2235470, EPI_ISL_2235471, EPI_ISL_2235472, EPI_ISL_2235475, EPI_ISL_2235484 | | | |
| see above | Azienda Ospedaliera Terni | Istituto Zooprofilattico Sperimentale dell'Abruzzo e Molise "G. Caporale" | Palumbo M, Scaccetti A, Lorusso A, Marcacci M, Di Domenico M, Ancora M, Curini V, Di Lollo Valeria, Mangone I, Rinaldi A, Delli Compagni E, Scialabba S, Caporale M, Di Pasquale A, Cammà C, Puglia I, Calistri P, Savini G |
| EPI_ISL_2235486 | Azienda USL Umbria 3 | Istituto Zooprofilattico Sperimentale dell'Abruzzo e Molise "G. Caporale" | Proietti A, Pistoni E, Lorusso A, Marcacci M, Di Domenico M, Ancora M, Curini V, Di Lollo Valeria, Mangone I, Rinaldi A, Delli Compagni E, Scialabba S, Caporale M, Di Pasquale A, Cammà C, Puglia I, Calistri P, Savini G |
| EPI_ISL_2235487 | Azienda USL Umbria 4 | Istituto Zooprofilattico Sperimentale dell'Abruzzo e Molise "G. Caporale" | Proietti A, Pistoni E, Lorusso A, Marcacci M, Di Domenico M, Ancora M, Curini V, Di Lollo Valeria, Mangone I, Rinaldi A, Delli Compagni E, Scialabba S, Caporale M, Di Pasquale A, Cammà C, Puglia I, Calistri P, Savini G |
| EPI_ISL_2235511, EPI_ISL_2235512, EPI_ISL_2235513 | Genome Analysis Center, Yamanashi Central Hospital | Genome Analysis Center, Yamanashi Central Hospital | Yosuke Hirotsu |
| EPI_ISL_2235527 | Laboratorio Biologia Molecolare Sars Cov2 - UOC Laboratorio Analisi - Servizio Medicina di Laboratorio, Ospedale "San Francesco" - ATS-ASSL Nuoro | Laboratorio specialistico UOC Ematologia - Ospedale "San Francesco" - ATS-ASSL Nuoro | Piras Giovanna, Asproni Rosanna, Fiamma Maura, Monne Maria Itria, Palmas Angelo Domenico, Garau Cristina, Mameli Giuseppe |
| EPI_ISL_2253126 | National Center of Infectious and Parasitic Diseases | National Center of Infectious and Parasitic Diseases | Alexiev et al |
| EPI_ISL_2253197, EPI_ISL_2253212, EPI_ISL_2253213, EPI_ISL_2253216, EPI_ISL_2253218, EPI_ISL_2253219, EPI_ISL_2253221, EPI_ISL_2254123, EPI_ISL_2254124, EPI_ISL_2254128, EPI_ISL_2254129, EPI_ISL_2254132, EPI_ISL_2254495, EPI_ISL_2254499 | | | |
| see above | Institut de Virologie du CHU de Strasbourg | Institut de Virologie du CHU de Strasbourg | Fafi-Kremer Samira, Gallais Floriane, Gantner Pierre, Laugel Elodie, Solis Morgane, Velay Aurélie, Wendling Marie-Josée |
| EPI_ISL_2258207 | RSUD dr Soetomo Surabaya | National Institute of Health Research and Development | Hana Apsari Pawestri, Kartika Dewi Puspa, Arie Ardiansyah Nugraha, Subangkit, Hartanti Dian Ikawati, Triyani Soekarso, Krisna Pangesti, Nelly Puspandari, Vivi Setiawaty |
| EPI_ISL_2258213, EPI_ISL_2258218 | RSUD Arifin Ahmad Riau | National Institute of Health Research and Development | Hana Apsari Pawestri, Hartanti Dian Ikawati, Subangkit, Kartika Dewi Puspa, Arie Ardiansyah Nugraha, Triyani Soekarso, Krisna Pangesti, Nelly Puspandari, Fajri Marinda, Vivi Setiawaty |
| EPI_ISL_2258367 | Laboratorio Biologia Molecolare Sars Cov2 - UOC Laboratorio Analisi - Servizio Medicina di Laboratorio, Ospedale "San Francesco" - ATS-ASSL Nuoro | Laboratorio specialistico UOC Ematologia - Ospedale "San Francesco" - ATS-ASSL Nuoro | Piras Giovanna, Monne Maria Itria, Asproni Rosanna,,Fiamma Maura, Palmas Angelo Domenico, Mameli Giuseppe |
| EPI_ISL_2258368 | Laboratorio Biologia Molecolare Sars Cov2 - UOC Laboratorio Analisi - Servizio Medicina di Laboratorio, Ospedale "San Francesco" - ATS-ASSL Nuoro | Laboratorio specialistico UOC Ematologia - Ospedale "San Francesco" - ATS-ASSL Nuoro | Asproni Rosanna, Carta Franco, Pira Giovanna, Fancello Patrizia, Rosu Valentina, Fiamma Maura, Garau Maria Cristina,Mameli Giuseppe |
| EPI_ISL_2259087 | Hospital | National Reference Center for Viruses of Respiratory Infections, Institut Pasteur, Paris | Marion Barbet, Sylvie Behillil, Méline Bizard, Angela Brisebarre, Camille Capel, Vincent Enouf, Louise Lefrançois, Frédéric Lemoine, Christophe Malabat, Corinne Maufrais, Damien Mornico, Etienne Simon-Lorière, Maud Vanpeene, Sylvie Van der Werf ,Nabil Gastli |
| EPI_ISL_2259088, EPI_ISL_2259089 | Hospital | National Reference Center for Viruses of Respiratory Infections, Institut Pasteur, Paris | Marion Barbet, Sylvie Behillil, Méline Bizard, Angela Brisebarre, Camille Capel, Vincent Enouf, Louise Lefrançois, Frédéric Lemoine, Christophe Malabat, Corinne Maufrais, Amaury Vaysse, Etienne Simon-Lorière, Maud Vanpeene, Sylvie Van der Werf ,Maxime Paluch |
| EPI_ISL_2259090 | Hospital | National Reference Center for Viruses of Respiratory Infections, Institut Pasteur, Paris | Marion Barbet, Sylvie Behillil, Méline Bizard, Angela Brisebarre, Camille Capel, Vincent Enouf, Louise Lefrançois, Frédéric Lemoine, Christophe Malabat, Corinne Maufrais, Etienne Simon-Lorière, Maud Vanpeene, Sylvie Van der Werf ,Nabil Gastli |
| EPI_ISL_2259091 | Labo Analyses Med | National Reference Center for Viruses of Respiratory Infections, Institut Pasteur, Paris | Marion Barbet, Sylvie Behillil, Méline Bizard, Angela Brisebarre, Camille Capel, Vincent Enouf, Louise Lefrançois, Frédéric Lemoine, Christophe Malabat, Corinne Maufrais, Damien Mornico, Etienne Simon-Lorière, Maud Vanpeene, Sylvie Van der Werf ,Fabienne Artur |
| EPI_ISL_2259092 | Labo Analyses Med | National Reference Center for Viruses of Respiratory Infections, Institut Pasteur, Paris | Marion Barbet, Sylvie Behillil, Méline Bizard, Angela Brisebarre, Camille Capel, Vincent Enouf, Louise Lefrançois, Frédéric Lemoine, Christophe Malabat, Corinne Maufrais, Etienne Simon-Lorière, Maud Vanpeene, Sylvie Van der Werf ,Fabienne Artur |
| EPI_ISL_2259093 | Labo Analyses Med | National Reference Center for Viruses of Respiratory Infections, Institut Pasteur, Paris | Marion Barbet, Sylvie Behillil, Méline Bizard, Angela Brisebarre, Camille Capel, Vincent Enouf, Louise Lefrançois, Frédéric Lemoine, Christophe Malabat, Corinne Maufrais, Victoire Baillet, Etienne Simon-Lorière, Maud Vanpeene, Sylvie Van der Werf ,Jean-Baptiste Foulquier |
| EPI_ISL_2259094 | Hospital | National Reference Center for Viruses of Respiratory Infections, Institut Pasteur, Paris | Marion Barbet, Sylvie Behillil, Méline Bizard, Angela Brisebarre, Camille Capel, Vincent Enouf, Louise Lefrançois, Frédéric Lemoine, Christophe Malabat, Corinne Maufrais, Damien Mornico, Etienne Simon-Lorière, Maud Vanpeene, Sylvie Van der Werf ,Pierre Alain Billy |
| EPI_ISL_2259095 | Hospital | National Reference Center for Viruses of Respiratory Infections, Institut Pasteur, Paris | Marion Barbet, Sylvie Behillil, Méline Bizard, Angela Brisebarre, Camille Capel, Vincent Enouf, Louise Lefrançois, Frédéric Lemoine, Christophe Malabat, Corinne Maufrais, Etienne Simon-Lorière, Maud Vanpeene, Sylvie Van der Werf ,Nabil Gastli |
| EPI_ISL_2259255, EPI_ISL_2259257, EPI_ISL_2259263, EPI_ISL_2259264, EPI_ISL_2259265 | National Center of Infectious and Parasitic Diseases | National Center of Infectious and Parasitic Diseases | Alexiev et al |
| EPI_ISL_2259324, EPI_ISL_2259330, | GH A.CHENEVIER-H.MONDOR | Department of Virology, Henri Mondor University Hospital, | Christophe Rodriguez, Slim Fourati, Vanessa Demontant, Guillaume Gricourt, Melissa N'Debi, Alexandre Soulier, Elisabeth Trawinski, Jean-Michel |

| | | | |
|---|---|---|---|
| EPI_ISL_2259339, EPI_ISL_2259362, EPI_ISL_2259363, EPI_ISL_2259364 | | Assistance Publique Hôpitaux de Paris, Université Paris-Est Créteil, INSERM U955 | Pawlotsky |
| EPI_ISL_2259413, EPI_ISL_2259414, EPI_ISL_2259415, EPI_ISL_2259416, EPI_ISL_2259417, EPI_ISL_2259418, EPI_ISL_2259419, EPI_ISL_2259420, EPI_ISL_2259421, EPI_ISL_2259422, EPI_ISL_2259423 | | | |
| see above | Klinisch Laboratorium ZNA | Klinisch Laboratorium ZNA | Verstrepen et al. |
| EPI_ISL_2262294 | RSUD Brebes | National Institute of Health Research and Development | Hana Apsari Pawestri, Kartika Dewi Puspa, Arie Ardiansyah Nugraha, Subangkit, Hartanti Dian Ikawati, Triyani Soekarso, Krisna Pangesti, Nelly Puspandari, Vivi Setiawaty |
| EPI_ISL_2262665, EPI_ISL_2262686 | OCME Office Of Chief Medical Examiner | New York City Public Health Laboratory | Jade Wang, et al. |
| EPI_ISL_2264766 | Ospedale Santa Caterina Novella | Istituto Zooprofilattico Sperimentale della Puglia e della Basilicata | Parisi A., Bianco A., Capozzi L., Del Sambro L., Simone D., Difato L., Bruno A. R. |
| EPI_ISL_2264767, EPI_ISL_2264769, EPI_ISL_2264771, EPI_ISL_2264775 | Istituto Zooprofilattico Sperimentale della Puglia e della Basilicata | Istituto Zooprofilattico Sperimentale della Puglia e della Basilicata | Parisi A., Bianco A., Capozzi L., Del Sambro L., Simone D., Difato L., Pace L., Galante D. |
| EPI_ISL_2272789, EPI_ISL_2272790, EPI_ISL_2272792, EPI_ISL_2272793, EPI_ISL_2272794, EPI_ISL_2272795, EPI_ISL_2272796, EPI_ISL_2272798, EPI_ISL_2272799, EPI_ISL_2272800, EPI_ISL_2272802, EPI_ISL_2272803, EPI_ISL_2272804, EPI_ISL_2272805, EPI_ISL_2272808, EPI_ISL_2272809, EPI_ISL_2272810, EPI_ISL_2272813, EPI_ISL_2272814, EPI_ISL_2272815, EPI_ISL_2272816, EPI_ISL_2272817, EPI_ISL_2272818, EPI_ISL_2272819, EPI_ISL_2272820, EPI_ISL_2272821, EPI_ISL_2272824, EPI_ISL_2272825, EPI_ISL_2272826, EPI_ISL_2272827, EPI_ISL_2272828, EPI_ISL_2272829, EPI_ISL_2272831, EPI_ISL_2272832, EPI_ISL_2272833, EPI_ISL_2272834, EPI_ISL_2272835, EPI_ISL_2272836, EPI_ISL_2272838, EPI_ISL_2272839, EPI_ISL_2272840, EPI_ISL_2272841, EPI_ISL_2272842, EPI_ISL_2272843, EPI_ISL_2272844, EPI_ISL_2272845, EPI_ISL_2272846, EPI_ISL_2272849, EPI_ISL_2272850, EPI_ISL_2272851, EPI_ISL_2272852, EPI_ISL_2272853, EPI_ISL_2272854, EPI_ISL_2272855, EPI_ISL_2272856, EPI_ISL_2272857, EPI_ISL_2272858, EPI_ISL_2272859, EPI_ISL_2272860, EPI_ISL_2272861, EPI_ISL_2272862, EPI_ISL_2272863, EPI_ISL_2272864, EPI_ISL_2272865, EPI_ISL_2272866, EPI_ISL_2272868, EPI_ISL_2272869, EPI_ISL_2272870, EPI_ISL_2272871, EPI_ISL_2272872, EPI_ISL_2272873, EPI_ISL_2272875, EPI_ISL_2272876, EPI_ISL_2272877, EPI_ISL_2272878, EPI_ISL_2272879, EPI_ISL_2272880, EPI_ISL_2272882, EPI_ISL_2272883, EPI_ISL_2272884, EPI_ISL_2272885, EPI_ISL_2272886, EPI_ISL_2272887, EPI_ISL_2272888, EPI_ISL_2272889, EPI_ISL_2272891, EPI_ISL_2272892, EPI_ISL_2272893, EPI_ISL_2272894, EPI_ISL_2272895, EPI_ISL_2272896, EPI_ISL_2272897, EPI_ISL_2272899, EPI_ISL_2272900, EPI_ISL_2272901, EPI_ISL_2272902, EPI_ISL_2272903, EPI_ISL_2272904, EPI_ISL_2272905, EPI_ISL_2272907, EPI_ISL_2272908, EPI_ISL_2272909, EPI_ISL_2272910, EPI_ISL_2272911, EPI_ISL_2272912, EPI_ISL_2272913, EPI_ISL_2272914, EPI_ISL_2272915, EPI_ISL_2272917, EPI_ISL_2272918, EPI_ISL_2272919, EPI_ISL_2272920, EPI_ISL_2272921, EPI_ISL_2272923, EPI_ISL_2272924, EPI_ISL_2272925, EPI_ISL_2272926, EPI_ISL_2272927, EPI_ISL_2272930, EPI_ISL_2272931, EPI_ISL_2272932, EPI_ISL_2272933, EPI_ISL_2272936, EPI_ISL_2272937 | | | |
| see above | JIPMER Puducherry | inStem NCBS - INSACOG | Uma Ramakrishnan Dasaradhi Palakodeti Aswin SaiNarain |
| EPI_ISL_2272959 | HOSPITAL DR. ENRIQUE BALTODANO BRICEÑO | Inciensa, Instituto Costarricense de Investigación y Enseñanza en Nutrición y Salud | Francisco Duarte, Hebleen Porras, Claudio Soto-Garita, Estela Cordero, Adriana Godínez, Melany Calderón, José Luis Vargas, Mariela Gutiérrez, Joselyn Prado & Arnoldo Zamora-Villegas |
| EPI_ISL_2272960 | AREA DE SALUD LA CRUZ | Inciensa, Instituto Costarricense de Investigación y Enseñanza en Nutrición y Salud | Francisco Duarte, Hebleen Porras, Claudio Soto-Garita, Estela Cordero, Adriana Godínez, Melany Calderón, José Luis Vargas, Mariela Gutiérrez, Joselyn Prado & Arnoldo Zamora-Villegas |
| EPI_ISL_2272966 | AREA DE SALUD HATILLO - CLINICA DR. SOLON NUÑEZ | Inciensa, Instituto Costarricense de Investigación y Enseñanza en Nutrición y Salud | Francisco Duarte, Hebleen Porras, Claudio Soto-Garita, Estela Cordero, Adriana Godínez, Melany Calderón, José Luis Vargas, Mariela Gutiérrez, Joselyn Prado & Roberto Brilla-Ferrer |
| EPI_ISL_2272967, EPI_ISL_2272968 | AREA DE SALUD MATA REDONDA-HOSPITAL - CLINICA DR. MORENO CAÑAS | Inciensa, Instituto Costarricense de Investigación y Enseñanza en Nutrición y Salud | Francisco Duarte, Hebleen Porras, Claudio Soto-Garita, Estela Cordero, Adriana Godínez, Melany Calderón, José Luis Vargas, Mariela Gutiérrez, Joselyn Prado & María Gabriela Valderrama-Ordoñez |
| EPI_ISL_2272969 | HOSPITAL SAN VICENTE DE PAUL | Inciensa, Instituto Costarricense de Investigación y Enseñanza en Nutrición y Salud | Francisco Duarte, Hebleen Porras, Claudio Soto-Garita, Estela Cordero, Adriana Godínez, Melany Calderón, José Luis Vargas, Mariela Gutiérrez, Joselyn Prado & Silvia Sáenz-García |
| EPI_ISL_2272970 | HOSPITAL DR. CARLOS LUIS VALVERDE VEGA | Inciensa, Instituto Costarricense de Investigación y Enseñanza en Nutrición y Salud | Francisco Duarte, Hebleen Porras, Claudio Soto-Garita, Estela Cordero, Adriana Godínez, Melany Calderón, José Luis Vargas, Mariela Gutiérrez, Joselyn Prado & Juan Carlos Villalobos-Ugalde |
| EPI_ISL_2272971 | HOSPITAL DR. WILLIAM ALLEN | Inciensa, Instituto Costarricense de Investigación y Enseñanza en Nutrición y Salud | Francisco Duarte, Hebleen Porras, Claudio Soto-Garita, Estela Cordero, Adriana Godínez, Melany Calderón, José Luis Vargas, Mariela Gutiérrez & Joselyn Prado |
| EPI_ISL_2272972 | HOSPITAL DR. FERNANDO ESCALANTE PRADILLA | Inciensa, Instituto Costarricense de Investigación y Enseñanza en Nutrición y Salud | Francisco Duarte, Hebleen Porras, Claudio Soto-Garita, Estela Cordero, Adriana Godínez, Melany Calderón, José Luis Vargas, Mariela Gutiérrez, Joselyn Prado & María Fernanda Matamoros Ceciliano |
| EPI_ISL_2272973, EPI_ISL_2272974 | HOSPITAL SAN JUAN DE DIOS | Inciensa, Instituto Costarricense de Investigación y Enseñanza en Nutrición y Salud | Francisco Duarte, Hebleen Porras, Claudio Soto-Garita, Estela Cordero, Adriana Godínez, Melany Calderón, José Luis Vargas, Mariela Gutiérrez, Joselyn Prado & Andony Cordero-Jiménez |
| EPI_ISL_2272975 | AREA DE SALUD PAVAS (COOPESALUD) | Inciensa, Instituto Costarricense de Investigación y Enseñanza en Nutrición y Salud | Francisco Duarte, Hebleen Porras, Claudio Soto-Garita, Estela Cordero, Adriana Godínez, Melany Calderón, José Luis Vargas, Mariela Gutiérrez, Joselyn Prado & Andony Cordero-Jiménez |
| EPI_ISL_2272976 | AREA DE SALUD SAN JUAN-SAN DIEGO-CONCEPCION 2 | Inciensa, Instituto Costarricense de Investigación y Enseñanza en Nutrición y Salud | Francisco Duarte, Hebleen Porras, Claudio Soto-Garita, Estela Cordero, Adriana Godínez, Melany Calderón, José Luis Vargas, Mariela Gutiérrez, Joselyn Prado & Mariel López |
| EPI_ISL_2272977 | AREA DE SALUD SIQUIRRES | Inciensa, Instituto Costarricense de Investigación y Enseñanza en Nutrición y Salud | Francisco Duarte, Hebleen Porras, Claudio Soto-Garita, Estela Cordero, Adriana Godínez, Melany Calderón, José Luis Vargas, Mariela Gutiérrez, Joselyn Prado & Ileana Chaves-Peraza |
| EPI_ISL_2272978 | AREA DE SALUD OREAMUNO-PACAYAS-TIERRA BLANCA | Inciensa, Instituto Costarricense de Investigación y Enseñanza en Nutrición y Salud | Francisco Duarte, Hebleen Porras, Claudio Soto-Garita, Estela Cordero, Adriana Godínez, Melany Calderón, José Luis Vargas, Mariela Gutiérrez, Joselyn Prado & Carolina Loría Acosta |
| EPI_ISL_2272979 | LABORATORIO CLINICO LABIN | Inciensa, Instituto Costarricense de Investigación y Enseñanza en Nutrición y Salud | Francisco Duarte, Hebleen Porras, Claudio Soto-Garita, Estela Cordero, Adriana Godínez, Melany Calderón, José Luis Vargas, Mariela Gutiérrez, Joselyn Prado & Pei Ling Chan Ma |
| EPI_ISL_2272980, EPI_ISL_2272981 | AREA DE SALUD CARRILLO | Inciensa, Instituto Costarricense de Investigación y Enseñanza en Nutrición y Salud | Francisco Duarte, Hebleen Porras, Claudio Soto-Garita, Estela Cordero, Adriana Godínez, Melany Calderón, José Luis Vargas, Mariela Gutiérrez, Joselyn Prado & Adriana Bemúdez Espinoza |
| EPI_ISL_2272982 | HOSPITAL DR. ENRIQUE BALTODANO BRICEÑO | Inciensa, Instituto Costarricense de Investigación y Enseñanza en Nutrición y Salud | Francisco Duarte, Hebleen Porras, Claudio Soto-Garita, Estela Cordero, Adriana Godínez, Melany Calderón, José Luis Vargas, Mariela Gutiérrez, Joselyn Prado & Adriana Bemúdez Espinoza |
| EPI_ISL_2272983, EPI_ISL_2272984 | AREA DE SALUD GOICOECHEA 2 - CLINICA DR. JIMENEZ NUÑEZ | Inciensa, Instituto Costarricense de Investigación y Enseñanza en Nutrición y Salud | Francisco Duarte, Hebleen Porras, Claudio Soto-Garita, Estela Cordero, Adriana Godínez, Melany Calderón, José Luis Vargas, Mariela Gutiérrez, Joselyn Prado & Juan Carlos Cartes Parra |
| EPI_ISL_2272985 | HOSPITAL CIUDAD NEILY | Inciensa, Instituto Costarricense de Investigación y Enseñanza en Nutrición y Salud | Francisco Duarte, Hebleen Porras, Claudio Soto-Garita, Estela Cordero, Adriana Godínez, Melany Calderón, José Luis Vargas, Mariela Gutiérrez, Joselyn Prado & Raúl Zeledón-Mayorga |
| EPI_ISL_2272986 | HOSPITAL DR. MAX PERALTA JIMENEZ | Inciensa, Instituto Costarricense de Investigación y Enseñanza en Nutrición y Salud | Francisco Duarte, Hebleen Porras, Claudio Soto-Garita, Estela Cordero, Adriana Godínez, Melany Calderón, José Luis Vargas, Mariela Gutiérrez, Joselyn Prado & Mariana Piedra Calderón |
| EPI_ISL_2272987 | AREA DE SALUD COBANO | Inciensa, Instituto Costarricense de Investigación y Enseñanza en Nutrición y Salud | Francisco Duarte, Hebleen Porras, Claudio Soto-Garita, Estela Cordero, Adriana Godínez, Melany Calderón, José Luis Vargas, Mariela Gutiérrez, Joselyn Prado & María José Gómez-Umaña |
| EPI_ISL_2272988 | AREA DE SALUD MORAVIA | Inciensa, Instituto Costarricense de Investigación y Enseñanza en Nutrición y Salud | Francisco Duarte, Hebleen Porras, Claudio Soto-Garita, Estela Cordero, Adriana Godínez, Melany Calderón, José Luis Vargas, Mariela Gutiérrez, Joselyn Prado & Mariel López |
| EPI_ISL_2272989 | AREA DE SALUD CATEDRAL NORESTE | Inciensa, Instituto Costarricense de Investigación y Enseñanza en Nutrición y Salud | Francisco Duarte, Hebleen Porras, Claudio Soto-Garita, Estela Cordero, Adriana Godínez, Melany Calderón, José Luis Vargas, Mariela Gutiérrez, Joselyn Prado & Mariel López |
| EPI_ISL_2272990 | HOSPITAL DR. RAFAEL A. CALDERON GUARDIA | Inciensa, Instituto Costarricense de Investigación y Enseñanza en Nutrición y Salud | Francisco Duarte, Hebleen Porras, Claudio Soto-Garita, Estela Cordero, Adriana Godínez, Melany Calderón, José Luis Vargas, Mariela Gutiérrez, Joselyn Prado & Fabian Salas-Flores |
| EPI_ISL_2272991 | AREA DE SALUD FORTUNA | Inciensa, Instituto Costarricense de Investigación y Enseñanza en Nutrición y Salud | Francisco Duarte, Hebleen Porras, Claudio Soto-Garita, Estela Cordero, Adriana Godínez, Melany Calderón, José Luis Vargas, Mariela Gutiérrez, Joselyn Prado & Carolina Arrieta Rojas |
| EPI_ISL_2272992 | AREA DE SALUD PURISCAL-TURRUBARES | Inciensa, Instituto Costarricense de Investigación y Enseñanza en Nutrición y Salud | Francisco Duarte, Hebleen Porras, Claudio Soto-Garita, Estela Cordero, Adriana Godínez, Melany Calderón, José Luis Vargas, Mariela Gutiérrez & Joselyn Prado |
| EPI_ISL_2272993 | AREA DE SALUD SANTO DOMINGO | Inciensa, Instituto Costarricense de Investigación y | Francisco Duarte, Hebleen Porras, Claudio Soto-Garita, Estela Cordero, Adriana Godínez, Melany Calderón, José Luis Vargas, Mariela Gutiérrez, Joselyn |

| | | | |
|---|---|---|---|
| | | Enseñanza en Nutrición y Salud | Prado & Margarita Lee-Lui |
| EPI_ISL_2272994 | HOSPITAL SAN JUAN DE DIOS | Inciensa, Instituto Costarricense de Investigación y Enseñanza en Nutrición y Salud | Francisco Duarte, Hebleen Porras, Claudio Soto-Garita, Estela Cordero, Adriana Godínez, Melany Calderón, José Luis Vargas, Mariela Gutiérrez, Joselyn Prado & Andony Cordero-Jiménez |
| EPI_ISL_2272995 | AREA DE SALUD PAVAS (COOPESALUD) | Inciensa, Instituto Costarricense de Investigación y Enseñanza en Nutrición y Salud | Francisco Duarte, Hebleen Porras, Claudio Soto-Garita, Estela Cordero, Adriana Godínez, Melany Calderón, José Luis Vargas, Mariela Gutiérrez, Joselyn Prado & Andony Cordero-Jiménez |
| EPI_ISL_2272996, EPI_ISL_2272997 | AREA DE SALUD ZAPOTE-CATEDRAL - CLINICA DR. CARLOS DURAN | Inciensa, Instituto Costarricense de Investigación y Enseñanza en Nutrición y Salud | Francisco Duarte, Hebleen Porras, Claudio Soto-Garita, Estela Cordero, Adriana Godínez, Melany Calderón, José Luis Vargas, Mariela Gutiérrez, Joselyn Prado & Andony Cordero-Jiménez |
| EPI_ISL_2272998 | AREA DE SALUD DESAMPARADOS 2 (COOPESALUD) | Inciensa, Instituto Costarricense de Investigación y Enseñanza en Nutrición y Salud | Francisco Duarte, Hebleen Porras, Claudio Soto-Garita, Estela Cordero, Adriana Godínez, Melany Calderón, José Luis Vargas, Mariela Gutiérrez, Joselyn Prado & Andony Cordero-Jiménez |
| EPI_ISL_2272999 | AREA DE SALUD DESAMPARADOS 1 - CLINICA DR. MARCIAL FALLAS | Inciensa, Instituto Costarricense de Investigación y Enseñanza en Nutrición y Salud | Francisco Duarte, Hebleen Porras, Claudio Soto-Garita, Estela Cordero, Adriana Godínez, Melany Calderón, José Luis Vargas, Mariela Gutiérrez, Joselyn Prado & Andony Cordero-Jiménez |
| EPI_ISL_2273000 | AREA DE SALUD PAVAS (COOPESALUD) | Inciensa, Instituto Costarricense de Investigación y Enseñanza en Nutrición y Salud | Francisco Duarte, Hebleen Porras, Claudio Soto-Garita, Estela Cordero, Adriana Godínez, Melany Calderón, José Luis Vargas, Mariela Gutiérrez, Joselyn Prado & Andony Cordero-Jiménez |
| EPI_ISL_2273001 | AREA DE SALUD CORONADO | Inciensa, Instituto Costarricense de Investigación y Enseñanza en Nutrición y Salud | Francisco Duarte, Hebleen Porras, Claudio Soto-Garita, Estela Cordero, Adriana Godínez, Melany Calderón, José Luis Vargas, Mariela Gutiérrez, Joselyn Prado & Mariel López |
| EPI_ISL_2273002, EPI_ISL_2273003 | AREA DE SALUD CATEDRAL NORESTE | Inciensa, Instituto Costarricense de Investigación y Enseñanza en Nutrición y Salud | Francisco Duarte, Hebleen Porras, Claudio Soto-Garita, Estela Cordero, Adriana Godínez, Melany Calderón, José Luis Vargas, Mariela Gutiérrez, Joselyn Prado & Mariel López |
| EPI_ISL_2273004 | AREA DE SALUD SAN JUAN-SAN DIEGO-CONCEPCION 2 | Inciensa, Instituto Costarricense de Investigación y Enseñanza en Nutrición y Salud | Francisco Duarte, Hebleen Porras, Claudio Soto-Garita, Estela Cordero, Adriana Godínez, Melany Calderón, José Luis Vargas, Mariela Gutiérrez, Joselyn Prado & Mariel López |
| EPI_ISL_2274482, EPI_ISL_2274488, EPI_ISL_2274490, EPI_ISL_2274573, EPI_ISL_2274655, EPI_ISL_2274715, EPI_ISL_2274972, EPI_ISL_2274974, EPI_ISL_2274975, EPI_ISL_2274976 | Institut de Virologie du CHU de Strasbourg | Institut de Virologie du CHU de Strasbourg | Fafi-Kremer Samira, Gallais Floriane, Gantner Pierre, Laugel Elodie, Solis Morgane, Velay Aurélie, Wendling Marie-Josée |
| EPI_ISL_2279629, EPI_ISL_2279630, EPI_ISL_2279633, EPI_ISL_2279636, EPI_ISL_2279637, EPI_ISL_2279638, EPI_ISL_2279639, EPI_ISL_2279641 | Hospital | National Reference Center for Viruses of Respiratory Infections, Institut Pasteur, Paris | Marion Barbet, Sylvie Behillil, Méline Bizard, Angela Brisebarre, Camille Capel, Vincent Enouf, Louise Lefrançois, Frédéric Lemoine, Christophe Malabat, Corinne Maufrais, Etienne Simon-Lorière, Maud Vanpeene, Sylvie Van der Werf ,StéPhanie Marque Juillet |
| EPI_ISL_2279642 | Hospital | National Reference Center for Viruses of Respiratory Infections, Institut Pasteur, Paris | Marion Barbet, Sylvie Behillil, Méline Bizard, Angela Brisebarre, Camille Capel, Vincent Enouf, Louise Lefrançois, Frédéric Lemoine, Christophe Malabat, Corinne Maufrais, Amaury Vaysse, Etienne Simon-Lorière, Maud Vanpeene, Sylvie Van der Werf ,StéPhanie Marque Juillet |
| EPI_ISL_2279643, EPI_ISL_2279644 | Hospital | National Reference Center for Viruses of Respiratory Infections, Institut Pasteur, Paris | Marion Barbet, Sylvie Behillil, Méline Bizard, Angela Brisebarre, Camille Capel, Vincent Enouf, Louise Lefrançois, Frédéric Lemoine, Christophe Malabat, Corinne Maufrais, Etienne Simon-Lorière, Maud Vanpeene, Sylvie Van der Werf ,StéPhanie Marque Juillet |
| EPI_ISL_2279645 | Hospital | National Reference Center for Viruses of Respiratory Infections, Institut Pasteur, Paris | Marion Barbet, Sylvie Behillil, Méline Bizard, Angela Brisebarre, Camille Capel, Vincent Enouf, Louise Lefrançois, Frédéric Lemoine, Christophe Malabat, Corinne Maufrais, Amaury Vaysse, Etienne Simon-Lorière, Maud Vanpeene, Sylvie Van der Werf ,StéPhanie Marque Juillet |
| EPI_ISL_2279646 | Hospital | National Reference Center for Viruses of Respiratory Infections, Institut Pasteur, Paris | Marion Barbet, Sylvie Behillil, Méline Bizard, Angela Brisebarre, Camille Capel, Vincent Enouf, Louise Lefrançois, Frédéric Lemoine, Christophe Malabat, Corinne Maufrais, Etienne Simon-Lorière, Maud Vanpeene, Sylvie Van der Werf ,StéPhanie Marque Juillet |
| EPI_ISL_2279648, EPI_ISL_2279649, EPI_ISL_2279650 | Labo Analyses Med | National Reference Center for Viruses of Respiratory Infections, Institut Pasteur, Paris | Marion Barbet, Sylvie Behillil, Méline Bizard, Angela Brisebarre, Camille Capel, Vincent Enouf, Louise Lefrançois, Frédéric Lemoine, Christophe Malabat, Corinne Maufrais, Etienne Simon-Lorière, Maud Vanpeene, Sylvie Van der Werf ,Anne-Laure Garand |
| EPI_ISL_2279651, EPI_ISL_2279652, EPI_ISL_2279653, EPI_ISL_2279654 | Hospital | National Reference Center for Viruses of Respiratory Infections, Institut Pasteur, Paris | Marion Barbet, Sylvie Behillil, Méline Bizard, Angela Brisebarre, Camille Capel, Vincent Enouf, Louise Lefrançois, Frédéric Lemoine, Christophe Malabat, Corinne Maufrais, Etienne Simon-Lorière, Maud Vanpeene, Sylvie Van der Werf ,Fatiha Najioullah |
| EPI_ISL_2279655, EPI_ISL_2279656, EPI_ISL_2279657, EPI_ISL_2279658, EPI_ISL_2279659, EPI_ISL_2279660, EPI_ISL_2279662, EPI_ISL_2279663, EPI_ISL_2279664, EPI_ISL_2279665, EPI_ISL_2279666, EPI_ISL_2279667 | | | |
| see above | Labo analyses med | National Reference Center for Viruses of Respiratory Infections, Institut Pasteur, Paris | Marion Barbet, Sylvie Behillil, Méline Bizard, Angela Brisebarre, Camille Capel, Vincent Enouf, Louise Lefrançois, Frédéric Lemoine, Christophe Malabat, Corinne Maufrais, Etienne Simon-Lorière, Maud Vanpeene, Sylvie Van der Werf ,OphéLie Said-Delattre |
| EPI_ISL_2279668, EPI_ISL_2279669 | Labo analyses med | National Reference Center for Viruses of Respiratory Infections, Institut Pasteur, Paris | Marion Barbet, Sylvie Behillil, Méline Bizard, Angela Brisebarre, Camille Capel, Vincent Enouf, Louise Lefrançois, Frédéric Lemoine, Christophe Malabat, Corinne Maufrais, Victoire Baillet, Etienne Simon-Lorière, Maud Vanpeene, Sylvie Van der Werf ,OphéLie Said-Delattre |
| EPI_ISL_2279670, EPI_ISL_2279671, EPI_ISL_2279672, EPI_ISL_2279673, EPI_ISL_2279674, EPI_ISL_2279675, EPI_ISL_2279676, EPI_ISL_2279677, EPI_ISL_2279678, EPI_ISL_2279679, EPI_ISL_2279681, EPI_ISL_2279682, EPI_ISL_2279683, EPI_ISL_2279684, EPI_ISL_2279686, EPI_ISL_2279687 | | | |
| see above | Labo analyses med | National Reference Center for Viruses of Respiratory Infections, Institut Pasteur, Paris | Marion Barbet, Sylvie Behillil, Méline Bizard, Angela Brisebarre, Camille Capel, Vincent Enouf, Louise Lefrançois, Frédéric Lemoine, Christophe Malabat, Corinne Maufrais, Etienne Simon-Lorière, Maud Vanpeene, Sylvie Van der Werf ,OphéLie Said-Delattre |
| EPI_ISL_2279688 | Labo analyses med | National Reference Center for Viruses of Respiratory Infections, Institut Pasteur, Paris | Marion Barbet, Sylvie Behillil, Méline Bizard, Angela Brisebarre, Camille Capel, Vincent Enouf, Louise Lefrançois, Frédéric Lemoine, Christophe Malabat, Corinne Maufrais, Victoire Baillet, Etienne Simon-Lorière, Maud Vanpeene, Sylvie Van der Werf ,OphéLie Said-Delattre |
| EPI_ISL_2279690, EPI_ISL_2279691, EPI_ISL_2279692, EPI_ISL_2279693 | Labo analyses med | National Reference Center for Viruses of Respiratory Infections, Institut Pasteur, Paris | Marion Barbet, Sylvie Behillil, Méline Bizard, Angela Brisebarre, Camille Capel, Vincent Enouf, Louise Lefrançois, Frédéric Lemoine, Christophe Malabat, Corinne Maufrais, Etienne Simon-Lorière, Maud Vanpeene, Sylvie Van der Werf ,OphéLie Said-Delattre |
| EPI_ISL_2279695 | Labo analyses med | National Reference Center for Viruses of Respiratory Infections, Institut Pasteur, Paris | Marion Barbet, Sylvie Behillil, Méline Bizard, Angela Brisebarre, Camille Capel, Vincent Enouf, Louise Lefrançois, Frédéric Lemoine, Christophe Malabat, Corinne Maufrais, Victoire Baillet, Etienne Simon-Lorière, Maud Vanpeene, Sylvie Van der Werf ,OphéLie Said-Delattre |
| EPI_ISL_2279697 | Labo analyses med | National Reference Center for Viruses of Respiratory Infections, Institut Pasteur, Paris | Marion Barbet, Sylvie Behillil, Méline Bizard, Angela Brisebarre, Camille Capel, Vincent Enouf, Louise Lefrançois, Frédéric Lemoine, Christophe Malabat, Corinne Maufrais, Etienne Simon-Lorière, Maud Vanpeene, Sylvie Van der Werf ,OphéLie Said-Delattre |
| EPI_ISL_2280079 | Laboratorio Biologia Molecolare Sars Cov2 - UOC Laboratorio Analisi - Servizio Medicina di Laboratorio, Ospedale "San Francesco" - ATS-ASSL Nuoro | Laboratorio Specialistico UOC Ematologia, Ospedale "San Francesco" - ATS-ASSL Nuoro | Piras Giovanna, Asproni Rosanna, Monne Maria Itria, Fiamma Maura,, Palmas Angelo Domenico, Mameli Giuseppe |
| EPI_ISL_2281281, EPI_ISL_2281443, EPI_ISL_2281674, EPI_ISL_2281890, EPI_ISL_2282208, EPI_ISL_2282228, EPI_ISL_2282229, EPI_ISL_2282230, EPI_ISL_2282231, EPI_ISL_2282232, EPI_ISL_2282233, EPI_ISL_2282234, EPI_ISL_2282247, EPI_ISL_2282248 | | | |
| see above | Azienda Ospedaliera Pugliese Ciaccio di Catanzaro SOC Microbiologia e Virologia | Azienda Ospedaliera Pugliese Ciaccio di Catanzaro SOC Microbiologia e Virologia | Rossana Tallerico Federica Pasceri Marco De Fazio Ilenia Talotta Giuseppina Panduri Pasquale Minchella |
| EPI_ISL_2283683, EPI_ISL_2283684, EPI_ISL_2283685 | LESP Oaxaca | Instituto de Diagnostico y Referencia Epidemiologicos (INDRE) | Claudia Wong-Arambula, Abril Rodriguez-Maldonado, Vanessa Rivero-Arredondo, Ariadna Medina-Benitez, Joaquin Quiroz-Mercado, Sergio Rangel-Guerrero, Natividad Cruz-Ortiz, Tatiana Nunez-Garcia, Gisela Barrera-Badillo, Lucia Hernandez-Rivas, Irma Lopez-Martinez, Ernesto Ramirez-Gonzalez. |
| EPI_ISL_2283687 | LESP Tamaulipas | Instituto de Diagnostico y Referencia Epidemiologicos (INDRE) | Claudia Wong-Arambula, Abril Rodriguez-Maldonado, Vanessa Rivero-Arredondo, Ariadna Medina-Benitez, Joaquin Quiroz-Mercado, Sergio Rangel-Guerrero, Natividad Cruz-Ortiz, Tatiana Nunez-Garcia, Gisela Barrera-Badillo, Lucia Hernandez-Rivas, Irma Lopez-Martinez, Ernesto Ramirez-Gonzalez. |
| EPI_ISL_2283688 | LESP Hidalgo | Instituto de Diagnostico y Referencia Epidemiologicos (INDRE) | Claudia Wong-Arambula, Abril Rodriguez-Maldonado, Vanessa Rivero-Arredondo, Ariadna Medina-Benitez, Joaquin Quiroz-Mercado, Sergio Rangel-Guerrero, Natividad Cruz-Ortiz, Tatiana Nunez-Garcia, Gisela Barrera-Badillo, Lucia Hernandez-Rivas, Irma Lopez-Martinez, Ernesto Ramirez-Gonzalez. |

| | | | |
|---|---|---|---|
| EPI_ISL_2283689 | LESP Campeche | Instituto de Diagnostico y Referencia Epidemiologicos (INDRE) | Claudia Wong-Arambula, Abril Rodriguez-Maldonado, Vanessa Rivero-Arredondo, Ariadna Medina-Benitez, Joaquin Quiroz-Mercado, Sergio Rangel-Guerrero, Natividad Cruz-Ortiz, Tatiana Nunez-Garcia, Gisela Barrera-Badillo, Lucia Hernandez-Rivas, Irma Lopez-Martinez, Ernesto Ramirez-Gonzalez. |
| EPI_ISL_2283690 | LESP Hidalgo | Instituto de Diagnostico y Referencia Epidemiologicos (INDRE) | Claudia Wong-Arambula, Abril Rodriguez-Maldonado, Vanessa Rivero-Arredondo, Ariadna Medina-Benitez, Joaquin Quiroz-Mercado, Sergio Rangel-Guerrero, Natividad Cruz-Ortiz, Tatiana Nunez-Garcia, Gisela Barrera-Badillo, Lucia Hernandez-Rivas, Irma Lopez-Martinez, Ernesto Ramirez-Gonzalez. |
| EPI_ISL_2283691, EPI_ISL_2283692 | LESP Chihuahua | Instituto de Diagnostico y Referencia Epidemiologicos (INDRE) | Claudia Wong-Arambula, Abril Rodriguez-Maldonado, Vanessa Rivero-Arredondo, Ariadna Medina-Benitez, Joaquin Quiroz-Mercado, Sergio Rangel-Guerrero, Natividad Cruz-Ortiz, Tatiana Nunez-Garcia, Gisela Barrera-Badillo, Lucia Hernandez-Rivas, Irma Lopez-Martinez, Ernesto Ramirez-Gonzalez. |
| EPI_ISL_2283693 | LESP Tamaulipas | Instituto de Diagnostico y Referencia Epidemiologicos (INDRE) | Claudia Wong-Arambula, Abril Rodriguez-Maldonado, Vanessa Rivero-Arredondo, Ariadna Medina-Benitez, Joaquin Quiroz-Mercado, Sergio Rangel-Guerrero, Natividad Cruz-Ortiz, Tatiana Nunez-Garcia, Gisela Barrera-Badillo, Lucia Hernandez-Rivas, Irma Lopez-Martinez, Ernesto Ramirez-Gonzalez. |
| EPI_ISL_2283700, EPI_ISL_2283701, EPI_ISL_2283702, EPI_ISL_2283703 | LESP Sinaloa | Instituto de Diagnostico y Referencia Epidemiologicos (INDRE) | Claudia Wong-Arambula, Abril Rodriguez-Maldonado, Vanessa Rivero-Arredondo, Ariadna Medina-Benitez, Joaquin Quiroz-Mercado, Sergio Rangel-Guerrero, Natividad Cruz-Ortiz, Tatiana Nunez-Garcia, Gisela Barrera-Badillo, Lucia Hernandez-Rivas, Irma Lopez-Martinez, Ernesto Ramirez-Gonzalez. |
| EPI_ISL_2283705 | LESP Ciudad de Mexico | Instituto de Diagnostico y Referencia Epidemiologicos (INDRE) | Claudia Wong-Arambula, Abril Rodriguez-Maldonado, Vanessa Rivero-Arredondo, Ariadna Medina-Benitez, Joaquin Quiroz-Mercado, Sergio Rangel-Guerrero, Natividad Cruz-Ortiz, Tatiana Nunez-Garcia, Gisela Barrera-Badillo, Lucia Hernandez-Rivas, Irma Lopez-Martinez, Ernesto Ramirez-Gonzalez. |
| EPI_ISL_2283709, EPI_ISL_2283710 | LESP Sinaloa | Instituto de Diagnostico y Referencia Epidemiologicos (INDRE) | Claudia Wong-Arambula, Abril Rodriguez-Maldonado, Vanessa Rivero-Arredondo, Ariadna Medina-Benitez, Joaquin Quiroz-Mercado, Sergio Rangel-Guerrero, Natividad Cruz-Ortiz, Tatiana Nunez-Garcia, Gisela Barrera-Badillo, Lucia Hernandez-Rivas, Irma Lopez-Martinez, Ernesto Ramirez-Gonzalez. |
| EPI_ISL_2283711, EPI_ISL_2283713, EPI_ISL_2283715, EPI_ISL_2283716 | LESP Tamaulipas | Instituto de Diagnostico y Referencia Epidemiologicos (INDRE) | Claudia Wong-Arambula, Abril Rodriguez-Maldonado, Vanessa Rivero-Arredondo, Ariadna Medina-Benitez, Joaquin Quiroz-Mercado, Sergio Rangel-Guerrero, Natividad Cruz-Ortiz, Tatiana Nunez-Garcia, Gisela Barrera-Badillo, Lucia Hernandez-Rivas, Irma Lopez-Martinez, Ernesto Ramirez-Gonzalez. |
| EPI_ISL_2283717 | LESP Guanajuato | Instituto de Diagnostico y Referencia Epidemiologicos (INDRE) | Claudia Wong-Arambula, Abril Rodriguez-Maldonado, Vanessa Rivero-Arredondo, Ariadna Medina-Benitez, Joaquin Quiroz-Mercado, Sergio Rangel-Guerrero, Natividad Cruz-Ortiz, Tatiana Nunez-Garcia, Gisela Barrera-Badillo, Lucia Hernandez-Rivas, Irma Lopez-Martinez, Ernesto Ramirez-Gonzalez. |
| EPI_ISL_2283720 | LESP Queretaro | Instituto de Diagnostico y Referencia Epidemiologicos (INDRE) | Claudia Wong-Arambula, Abril Rodriguez-Maldonado, Vanessa Rivero-Arredondo, Ariadna Medina-Benitez, Joaquin Quiroz-Mercado, Sergio Rangel-Guerrero, Natividad Cruz-Ortiz, Tatiana Nunez-Garcia, Gisela Barrera-Badillo, Lucia Hernandez-Rivas, Irma Lopez-Martinez, Ernesto Ramirez-Gonzalez. |
| EPI_ISL_2283721 | LESP Guanajuato | Instituto de Diagnostico y Referencia Epidemiologicos (INDRE) | Claudia Wong-Arambula, Abril Rodriguez-Maldonado, Vanessa Rivero-Arredondo, Ariadna Medina-Benitez, Joaquin Quiroz-Mercado, Sergio Rangel-Guerrero, Natividad Cruz-Ortiz, Tatiana Nunez-Garcia, Gisela Barrera-Badillo, Lucia Hernandez-Rivas, Irma Lopez-Martinez, Ernesto Ramirez-Gonzalez. |
| EPI_ISL_2285110, EPI_ISL_2285111, EPI_ISL_2285127, EPI_ISL_2285128, EPI_ISL_2285129, EPI_ISL_2285130, EPI_ISL_2285131, EPI_ISL_2285132, EPI_ISL_2285133, EPI_ISL_2285134, EPI_ISL_2285156 | | | |
| see above | Microbiology Department. Complexo Hospitalario Universitario de Vigo | Microbiology Department. Complexo Hospitalario Universitario de Vigo | Alfaya N, Alonso I, Alvarez M, Cabrera JJ, Carballo R, Cores O, Cortizo S, del-Campo V, Martinez L, Mediero G, Perez S, Potel C, Regueiro B, Rey S, Vasallo FJ |
| EPI_ISL_2285339, EPI_ISL_2285340, EPI_ISL_2285341, EPI_ISL_2285342, EPI_ISL_2285343, EPI_ISL_2285344, EPI_ISL_2285345, EPI_ISL_2285346, EPI_ISL_2285347, EPI_ISL_2285348, EPI_ISL_2285349, EPI_ISL_2285350, EPI_ISL_2285351, EPI_ISL_2285352, EPI_ISL_2285353, EPI_ISL_2285354, EPI_ISL_2285355, EPI_ISL_2285356, EPI_ISL_2285357, EPI_ISL_2285358, EPI_ISL_2285359, EPI_ISL_2285360, EPI_ISL_2285361, EPI_ISL_2285362, EPI_ISL_2285366, EPI_ISL_2285367, EPI_ISL_2285368, EPI_ISL_2285369 | | | |
| see above | University of Rome Tor Vergata: Departm Experim Medicine Chair of Virology | University of Rome Tor Vergata: Departm Experim Medicine Chair of Virology | Maria Concetta Bellocchi, Luca Carioti, Maria Botticelli, Lorenzo Piermatteo, Rossana Scutari, Marco Iannetta, Loredana Sarmati, Massimo Andreoni, Francesca Ceccherini-Silberstein |
| EPI_ISL_2285756, EPI_ISL_2285757, EPI_ISL_2285758, EPI_ISL_2285759, EPI_ISL_2285760, EPI_ISL_2285761 | Laboratori de Referencia de Catalunya | Laboratori de Referencia de Catalunya | Ramirez A., Bellosillo B., Padilla E., Vilas A., Hernandez JJ., Canal M. |
| EPI_ISL_2285852, EPI_ISL_2285853, EPI_ISL_2285862, EPI_ISL_2285866, EPI_ISL_2285869 | National Influenza Centre | National Influenza Centre | William K. Ampofo, Michael Marks, Ivy A. Asante, Sharon Hsu, Benjamiin B. Lindsey, Benjamin H. Foulkes, Mildred Adusei-Poku, Linda Boatemaa, Lorreta Kwasah, Joseph Oliver-Commey, Ernest Asiedu, Franklin Asiedu-Bekoe, Gordon Awandare, Joyce Ngoi, Dennis Laryea, Mathew D. Parker, Thushan I de Silva, |
| EPI_ISL_2293257 | AZ Sint Maarten | Imelda Ziekenhuis | Johan Frans, Hanne Valgaeren, Dagmar Obbels |
| EPI_ISL_2293266, EPI_ISL_2293267, EPI_ISL_2293268, EPI_ISL_2293269, EPI_ISL_2293270 | Labo analyses med | National Reference Center for Viruses of Respiratory Infections, Institut Pasteur, Paris | Marion Barbet, Sylvie Behillil, Méline Bizard, Angela Brisebarre, Camille Capel, Vincent Enouf, Louise Lefrançois, Frédéric Lemoine, Christophe Malabat, Corinne Maufrais, Etienne Simon-Lorière, Maud Vanpeene, Sylvie Van der Werf ,OphéLie Said-Delattre |
| EPI_ISL_2293271 | Hospital | National Reference Center for Viruses of Respiratory Infections, Institut Pasteur, Paris | Marion Barbet, Sylvie Behillil, Méline Bizard, Angela Brisebarre, Camille Capel, Vincent Enouf, Louise Lefrançois, Frédéric Lemoine, Christophe Malabat, Corinne Maufrais, Etienne Simon-Lorière, Maud Vanpeene, Sylvie Van der Werf ,José-Manuel Ramirez |
| EPI_ISL_2293272 | Labo Analyses Med | National Reference Center for Viruses of Respiratory Infections, Institut Pasteur, Paris | Marion Barbet, Sylvie Behillil, Méline Bizard, Angela Brisebarre, Camille Capel, Vincent Enouf, Louise Lefrançois, Frédéric Lemoine, Christophe Malabat, Corinne Maufrais, Etienne Simon-Lorière, Maud Vanpeene, Sylvie Van der Werf ,Fabrice Thomas |
| EPI_ISL_2293275 | Hospital | National Reference Center for Viruses of Respiratory Infections, Institut Pasteur, Paris | Marion Barbet, Sylvie Behillil, Méline Bizard, Angela Brisebarre, Camille Capel, Vincent Enouf, Louise Lefrançois, Frédéric Lemoine, Christophe Malabat, Corinne Maufrais, Pierre Lechat, Etienne Simon-Lorière, Maud Vanpeene, Sylvie Van der Werf ,Jacques Fourgeaud |
| EPI_ISL_2293277, EPI_ISL_2293278, EPI_ISL_2293279, EPI_ISL_2293280 | Labo analyses med | National Reference Center for Viruses of Respiratory Infections, Institut Pasteur, Paris | Marion Barbet, Sylvie Behillil, Méline Bizard, Angela Brisebarre, Camille Capel, Vincent Enouf, Louise Lefrançois, Frédéric Lemoine, Christophe Malabat, Corinne Maufrais, Etienne Simon-Lorière, Maud Vanpeene, Sylvie Van der Werf ,Hervé Dupont |
| EPI_ISL_2293281 | Hospital | National Reference Center for Viruses of Respiratory Infections, Institut Pasteur, Paris | Marion Barbet, Sylvie Behillil, Méline Bizard, Angela Brisebarre, Camille Capel, Vincent Enouf, Louise Lefrançois, Frédéric Lemoine, Christophe Malabat, Corinne Maufrais, Pierre Lechat, Etienne Simon-Lorière, Maud Vanpeene, Sylvie Van der Werf ,HéLèNe Roger |
| EPI_ISL_2293283 | Hospital | National Reference Center for Viruses of Respiratory Infections, Institut Pasteur, Paris | Marion Barbet, Sylvie Behillil, Méline Bizard, Angela Brisebarre, Camille Capel, Vincent Enouf, Louise Lefrançois, Frédéric Lemoine, Christophe Malabat, Corinne Maufrais, Etienne Simon-Lorière, Maud Vanpeene, Sylvie Van der Werf ,Salomon |
| EPI_ISL_2293292, EPI_ISL_2293293, EPI_ISL_2293295, EPI_ISL_2293296, EPI_ISL_2293297 | Hospital | National Reference Center for Viruses of Respiratory Infections, Institut Pasteur, Paris | Marion Barbet, Sylvie Behillil, Méline Bizard, Angela Brisebarre, Camille Capel, Vincent Enouf, Louise Lefrançois, Frédéric Lemoine, Christophe Malabat, Corinne Maufrais, Etienne Simon-Lorière, Maud Vanpeene, Sylvie Van der Werf ,Odile Petsaris |
| EPI_ISL_2293298, EPI_ISL_2293299 | Hospital | National Reference Center for Viruses of Respiratory Infections, Institut Pasteur, Paris | Marion Barbet, Sylvie Behillil, Méline Bizard, Angela Brisebarre, Camille Capel, Vincent Enouf, Louise Lefrançois, Frédéric Lemoine, Christophe Malabat, Corinne Maufrais, Pierre Lechat, Etienne Simon-Lorière, Maud Vanpeene, Sylvie Van der Werf ,Odile Petsaris |
| EPI_ISL_2293300 | Hospital | National Reference Center for Viruses of Respiratory Infections, Institut Pasteur, Paris | Marion Barbet, Sylvie Behillil, Méline Bizard, Angela Brisebarre, Camille Capel, Vincent Enouf, Louise Lefrançois, Frédéric Lemoine, Christophe Malabat, Corinne Maufrais, Etienne Simon-Lorière, Maud Vanpeene, Sylvie Van der Werf ,Odile Petsaris |
| EPI_ISL_2293302 | Hospital | National Reference Center for Viruses of Respiratory Infections, Institut Pasteur, Paris | Marion Barbet, Sylvie Behillil, Méline Bizard, Angela Brisebarre, Camille Capel, Vincent Enouf, Louise Lefrançois, Frédéric Lemoine, Christophe Malabat, Corinne Maufrais, Pierre Lechat, Etienne Simon-Lorière, Maud Vanpeene, Sylvie Van der Werf ,Samuel Kouyoumdjian |
| EPI_ISL_2293310, EPI_ISL_2293311, EPI_ISL_2293312 | Hospital | National Reference Center for Viruses of Respiratory Infections, Institut Pasteur, Paris | Marion Barbet, Sylvie Behillil, Méline Bizard, Angela Brisebarre, Camille Capel, Vincent Enouf, Louise Lefrançois, Frédéric Lemoine, Christophe Malabat, Corinne Maufrais, Etienne Simon-Lorière, Maud Vanpeene, Sylvie Van der Werf ,Franck Griscelli |

| | | | |
|---|---|---|---|
| EPI_ISL_2293328, EPI_ISL_2293329, EPI_ISL_2293330, EPI_ISL_2293332, EPI_ISL_2293334, EPI_ISL_2293335, EPI_ISL_2293336, EPI_ISL_2293337, EPI_ISL_2293338, EPI_ISL_2293339, EPI_ISL_2293340, EPI_ISL_2293341, EPI_ISL_2293342, EPI_ISL_2293343 | | | |
| see above | Outre mer | National Reference Center for Viruses of Respiratory Infections, Institut Pasteur, Paris | Marion Barbet, Sylvie Behillil, Méline Bizard, Angela Brisebarre, Camille Capel, Vincent Enouf, Louise Lefrançois, Frédéric Lemoine, Christophe Malabat, Corinne Maufrais, Pierre Lechat, Etienne Simon-Lorière, Maud Vanpeene, Sylvie Van der Werf ,Dominique Rousset |
| EPI_ISL_2293344 | Outre mer | National Reference Center for Viruses of Respiratory Infections, Institut Pasteur, Paris | Marion Barbet, Sylvie Behillil, Méline Bizard, Angela Brisebarre, Camille Capel, Vincent Enouf, Louise Lefrançois, Frédéric Lemoine, Christophe Malabat, Corinne Maufrais, Pierre Lechat, Etienne Simon-Lorière, Maud Vanpeene, Sylvie Van der Werf ,Dominique Rousset |
| EPI_ISL_2293345, EPI_ISL_2293346, EPI_ISL_2293347, EPI_ISL_2293348, EPI_ISL_2293349, EPI_ISL_2293351, EPI_ISL_2293352, EPI_ISL_2293354, EPI_ISL_2293355, EPI_ISL_2293356, EPI_ISL_2293357, EPI_ISL_2293358, EPI_ISL_2293359, EPI_ISL_2293360, EPI_ISL_2293361, EPI_ISL_2293362, EPI_ISL_2293363, EPI_ISL_2293364, EPI_ISL_2293365, EPI_ISL_2293367, EPI_ISL_2293368, EPI_ISL_2293369, EPI_ISL_2293370, EPI_ISL_2293371, EPI_ISL_2293373, EPI_ISL_2293374, EPI_ISL_2293375, EPI_ISL_2293376, EPI_ISL_2293377, EPI_ISL_2293378, EPI_ISL_2293379, EPI_ISL_2293380, EPI_ISL_2293381, EPI_ISL_2293382 | | | |
| see above | Outre mer | National Reference Center for Viruses of Respiratory Infections, Institut Pasteur, Paris | Marion Barbet, Sylvie Behillil, Méline Bizard, Angela Brisebarre, Camille Capel, Vincent Enouf, Louise Lefrançois, Frédéric Lemoine, Christophe Malabat, Corinne Maufrais, Pierre Lechat, Etienne Simon-Lorière, Maud Vanpeene, Sylvie Van der Werf ,Dominique Rousset |
| EPI_ISL_2293425, EPI_ISL_2293432, EPI_ISL_2293433 | Research | National Reference Center for Viruses of Respiratory Infections, Institut Pasteur, Paris | Marion Barbet, Sylvie Behillil, Méline Bizard, Angela Brisebarre, Camille Capel, Vincent Enouf, Louise Lefrançois, Frédéric Lemoine, Christophe Malabat, Corinne Maufrais, Etienne Simon-Lorière, Maud Vanpeene, Sylvie Van der Werf ,Marianne Leruez-Ville |
| EPI_ISL_2296059 | LESP Oaxaca | Instituto de Diagnostico y Referencia Epidemiologicos (INDRE) | Claudia Wong-Arambula, Abril Rodriguez-Maldonado, Vanessa Rivero-Arredondo, Ariadna Medina-Benitez, Joaquin Quiroz-Mercado, Sergio Rangel-Guerrero, Natividad Cruz-Ortiz, Tatiana Nunez-Garcia, Gisela Barrera-Badillo, Lucia Hernandez-Rivas, Irma Lopez-Martinez, Ernesto Ramirez-Gonzalez. |
| EPI_ISL_2296060 | LESP Ciudad de Mexico | Instituto de Diagnostico y Referencia Epidemiologicos (INDRE) | Claudia Wong-Arambula, Abril Rodriguez-Maldonado, Vanessa Rivero-Arredondo, Ariadna Medina-Benitez, Joaquin Quiroz-Mercado, Sergio Rangel-Guerrero, Natividad Cruz-Ortiz, Tatiana Nunez-Garcia, Gisela Barrera-Badillo, Lucia Hernandez-Rivas, Irma Lopez-Martinez, Ernesto Ramirez-Gonzalez. |
| EPI_ISL_2296061, EPI_ISL_2296062 | LESP Quintana Roo | Instituto de Diagnostico y Referencia Epidemiologicos (INDRE) | Claudia Wong-Arambula, Abril Rodriguez-Maldonado, Vanessa Rivero-Arredondo, Ariadna Medina-Benitez, Joaquin Quiroz-Mercado, Sergio Rangel-Guerrero, Natividad Cruz-Ortiz, Tatiana Nunez-Garcia, Gisela Barrera-Badillo, Lucia Hernandez-Rivas, Irma Lopez-Martinez, Ernesto Ramirez-Gonzalez. |
| EPI_ISL_2296065 | LESP Tamaulipas | Instituto de Diagnostico y Referencia Epidemiologicos (INDRE) | Claudia Wong-Arambula, Abril Rodriguez-Maldonado, Vanessa Rivero-Arredondo, Ariadna Medina-Benitez, Joaquin Quiroz-Mercado, Sergio Rangel-Guerrero, Natividad Cruz-Ortiz, Tatiana Nunez-Garcia, Gisela Barrera-Badillo, Lucia Hernandez-Rivas, Irma Lopez-Martinez, Ernesto Ramirez-Gonzalez. |
| EPI_ISL_2296066 | LESP Michoacan | Instituto de Diagnostico y Referencia Epidemiologicos (INDRE) | Claudia Wong-Arambula, Abril Rodriguez-Maldonado, Vanessa Rivero-Arredondo, Ariadna Medina-Benitez, Joaquin Quiroz-Mercado, Sergio Rangel-Guerrero, Natividad Cruz-Ortiz, Tatiana Nunez-Garcia, Gisela Barrera-Badillo, Lucia Hernandez-Rivas, Irma Lopez-Martinez, Ernesto Ramirez-Gonzalez. |
| EPI_ISL_2296068 | LESP Nuevo Leon | Instituto de Diagnostico y Referencia Epidemiologicos (INDRE) | Claudia Wong-Arambula, Abril Rodriguez-Maldonado, Vanessa Rivero-Arredondo, Ariadna Medina-Benitez, Joaquin Quiroz-Mercado, Sergio Rangel-Guerrero, Natividad Cruz-Ortiz, Tatiana Nunez-Garcia, Gisela Barrera-Badillo, Lucia Hernandez-Rivas, Irma Lopez-Martinez, Ernesto Ramirez-Gonzalez. |
| EPI_ISL_2296069 | LESP Guanajuato | Instituto de Diagnostico y Referencia Epidemiologicos (INDRE) | Claudia Wong-Arambula, Abril Rodriguez-Maldonado, Vanessa Rivero-Arredondo, Ariadna Medina-Benitez, Joaquin Quiroz-Mercado, Sergio Rangel-Guerrero, Natividad Cruz-Ortiz, Tatiana Nunez-Garcia, Gisela Barrera-Badillo, Lucia Hernandez-Rivas, Irma Lopez-Martinez, Ernesto Ramirez-Gonzalez. |
| EPI_ISL_2296070 | LESP Jalisco | Instituto de Diagnostico y Referencia Epidemiologicos (INDRE) | Claudia Wong-Arambula, Abril Rodriguez-Maldonado, Vanessa Rivero-Arredondo, Ariadna Medina-Benitez, Joaquin Quiroz-Mercado, Sergio Rangel-Guerrero, Natividad Cruz-Ortiz, Tatiana Nunez-Garcia, Gisela Barrera-Badillo, Lucia Hernandez-Rivas, Irma Lopez-Martinez, Ernesto Ramirez-Gonzalez. |
| EPI_ISL_2296072, EPI_ISL_2296075, EPI_ISL_2296079, EPI_ISL_2296080, EPI_ISL_2296081 | LESP Ciudad de Mexico | Instituto de Diagnostico y Referencia Epidemiologicos (INDRE) | Claudia Wong-Arambula, Abril Rodriguez-Maldonado, Vanessa Rivero-Arredondo, Ariadna Medina-Benitez, Joaquin Quiroz-Mercado, Sergio Rangel-Guerrero, Natividad Cruz-Ortiz, Tatiana Nunez-Garcia, Gisela Barrera-Badillo, Lucia Hernandez-Rivas, Irma Lopez-Martinez, Ernesto Ramirez-Gonzalez. |
| EPI_ISL_2296082 | LESP Estado de Mexico | Instituto de Diagnostico y Referencia Epidemiologicos (INDRE) | Claudia Wong-Arambula, Abril Rodriguez-Maldonado, Vanessa Rivero-Arredondo, Ariadna Medina-Benitez, Joaquin Quiroz-Mercado, Sergio Rangel-Guerrero, Natividad Cruz-Ortiz, Tatiana Nunez-Garcia, Gisela Barrera-Badillo, Lucia Hernandez-Rivas, Irma Lopez-Martinez, Ernesto Ramirez-Gonzalez. |
| EPI_ISL_2296083 | LESP Ciudad de Mexico | Instituto de Diagnostico y Referencia Epidemiologicos (INDRE) | Claudia Wong-Arambula, Abril Rodriguez-Maldonado, Vanessa Rivero-Arredondo, Ariadna Medina-Benitez, Joaquin Quiroz-Mercado, Sergio Rangel-Guerrero, Natividad Cruz-Ortiz, Tatiana Nunez-Garcia, Gisela Barrera-Badillo, Lucia Hernandez-Rivas, Irma Lopez-Martinez, Ernesto Ramirez-Gonzalez. |
| EPI_ISL_2296084 | LESP Tamaulipas | Instituto de Diagnostico y Referencia Epidemiologicos (INDRE) | Claudia Wong-Arambula, Abril Rodriguez-Maldonado, Vanessa Rivero-Arredondo, Ariadna Medina-Benitez, Joaquin Quiroz-Mercado, Sergio Rangel-Guerrero, Natividad Cruz-Ortiz, Tatiana Nunez-Garcia, Gisela Barrera-Badillo, Lucia Hernandez-Rivas, Irma Lopez-Martinez, Ernesto Ramirez-Gonzalez. |
| EPI_ISL_2296085 | LESP Chiapas | Instituto de Diagnostico y Referencia Epidemiologicos (INDRE) | Claudia Wong-Arambula, Abril Rodriguez-Maldonado, Vanessa Rivero-Arredondo, Ariadna Medina-Benitez, Joaquin Quiroz-Mercado, Sergio Rangel-Guerrero, Natividad Cruz-Ortiz, Tatiana Nunez-Garcia, Gisela Barrera-Badillo, Lucia Hernandez-Rivas, Irma Lopez-Martinez, Ernesto Ramirez-Gonzalez. |
| EPI_ISL_2296086 | LESP Baja California | Instituto de Diagnostico y Referencia Epidemiologicos (INDRE) | Claudia Wong-Arambula, Abril Rodriguez-Maldonado, Vanessa Rivero-Arredondo, Ariadna Medina-Benitez, Joaquin Quiroz-Mercado, Sergio Rangel-Guerrero, Natividad Cruz-Ortiz, Tatiana Nunez-Garcia, Gisela Barrera-Badillo, Lucia Hernandez-Rivas, Irma Lopez-Martinez, Ernesto Ramirez-Gonzalez. |
| EPI_ISL_2296087 | LESP Quintana Roo | Instituto de Diagnostico y Referencia Epidemiologicos (INDRE) | Claudia Wong-Arambula, Abril Rodriguez-Maldonado, Vanessa Rivero-Arredondo, Ariadna Medina-Benitez, Joaquin Quiroz-Mercado, Sergio Rangel-Guerrero, Natividad Cruz-Ortiz, Tatiana Nunez-Garcia, Gisela Barrera-Badillo, Lucia Hernandez-Rivas, Irma Lopez-Martinez, Ernesto Ramirez-Gonzalez. |
| EPI_ISL_2296089, EPI_ISL_2296090, EPI_ISL_2296091 | LESP Yucatan | Instituto de Diagnostico y Referencia Epidemiologicos (INDRE) | Claudia Wong-Arambula, Abril Rodriguez-Maldonado, Vanessa Rivero-Arredondo, Ariadna Medina-Benitez, Joaquin Quiroz-Mercado, Sergio Rangel-Guerrero, Natividad Cruz-Ortiz, Tatiana Nunez-Garcia, Gisela Barrera-Badillo, Lucia Hernandez-Rivas, Irma Lopez-Martinez, Ernesto Ramirez-Gonzalez. |
| EPI_ISL_2296093 | LESP Guanajuato | Instituto de Diagnostico y Referencia Epidemiologicos (INDRE) | Claudia Wong-Arambula, Abril Rodriguez-Maldonado, Vanessa Rivero-Arredondo, Ariadna Medina-Benitez, Joaquin Quiroz-Mercado, Sergio Rangel-Guerrero, Natividad Cruz-Ortiz, Tatiana Nunez-Garcia, Gisela Barrera-Badillo, Lucia Hernandez-Rivas, Irma Lopez-Martinez, Ernesto Ramirez-Gonzalez. |
| EPI_ISL_2296094 | LESP Morelos | Instituto de Diagnostico y Referencia Epidemiologicos (INDRE) | Claudia Wong-Arambula, Abril Rodriguez-Maldonado, Vanessa Rivero-Arredondo, Ariadna Medina-Benitez, Joaquin Quiroz-Mercado, Sergio Rangel-Guerrero, Natividad Cruz-Ortiz, Tatiana Nunez-Garcia, Gisela Barrera-Badillo, Lucia Hernandez-Rivas, Irma Lopez-Martinez, Ernesto Ramirez-Gonzalez. |
| EPI_ISL_2296095 | LESP Guanajuato | Instituto de Diagnostico y Referencia Epidemiologicos (INDRE) | Claudia Wong-Arambula, Abril Rodriguez-Maldonado, Vanessa Rivero-Arredondo, Ariadna Medina-Benitez, Joaquin Quiroz-Mercado, Sergio Rangel-Guerrero, Natividad Cruz-Ortiz, Tatiana Nunez-Garcia, Gisela Barrera-Badillo, Lucia Hernandez-Rivas, Irma Lopez-Martinez, Ernesto Ramirez-Gonzalez. |
| EPI_ISL_2296097 | LESP Hidalgo | Instituto de Diagnostico y Referencia Epidemiologicos (INDRE) | Claudia Wong-Arambula, Abril Rodriguez-Maldonado, Vanessa Rivero-Arredondo, Ariadna Medina-Benitez, Joaquin Quiroz-Mercado, Sergio Rangel-Guerrero, Natividad Cruz-Ortiz, Tatiana Nunez-Garcia, Gisela Barrera-Badillo, Lucia Hernandez-Rivas, Irma Lopez-Martinez, Ernesto Ramirez-Gonzalez. |
| EPI_ISL_2296099 | LESP Michoacan | Instituto de Diagnostico y Referencia Epidemiologicos (INDRE) | Claudia Wong-Arambula, Abril Rodriguez-Maldonado, Vanessa Rivero-Arredondo, Ariadna Medina-Benitez, Joaquin Quiroz-Mercado, Sergio Rangel-Guerrero, Natividad Cruz-Ortiz, Tatiana Nunez-Garcia, Gisela Barrera-Badillo, Lucia Hernandez-Rivas, Irma Lopez-Martinez, Ernesto Ramirez-Gonzalez. |

| | | | |
|---|---|---|---|
| EPI_ISL_2296100 | LESP Colima | Instituto de Diagnostico y Referencia Epidemiologicos (INDRE) | Claudia Wong-Arambula, Abril Rodriguez-Maldonado, Vanessa Rivero-Arredondo, Ariadna Medina-Benitez, Joaquin Quiroz-Mercado, Sergio Rangel-Guerrero, Natividad Cruz-Ortiz, Tatiana Nunez-Garcia, Gisela Barrera-Badillo, Lucia Hernandez-Rivas, Irma Lopez-Martinez, Ernesto Ramirez-Gonzalez. |
| EPI_ISL_2296102, EPI_ISL_2296105 | LESP Sinaloa | Instituto de Diagnostico y Referencia Epidemiologicos (INDRE) | Claudia Wong-Arambula, Abril Rodriguez-Maldonado, Vanessa Rivero-Arredondo, Ariadna Medina-Benitez, Joaquin Quiroz-Mercado, Sergio Rangel-Guerrero, Natividad Cruz-Ortiz, Tatiana Nunez-Garcia, Gisela Barrera-Badillo, Lucia Hernandez-Rivas, Irma Lopez-Martinez, Ernesto Ramirez-Gonzalez. |
| EPI_ISL_2296106, EPI_ISL_2296107 | LESP Guanajuato | Instituto de Diagnostico y Referencia Epidemiologicos (INDRE) | Claudia Wong-Arambula, Abril Rodriguez-Maldonado, Vanessa Rivero-Arredondo, Ariadna Medina-Benitez, Joaquin Quiroz-Mercado, Sergio Rangel-Guerrero, Natividad Cruz-Ortiz, Tatiana Nunez-Garcia, Gisela Barrera-Badillo, Lucia Hernandez-Rivas, Irma Lopez-Martinez, Ernesto Ramirez-Gonzalez. |
| EPI_ISL_2296108 | LESP Yucatan | Instituto de Diagnostico y Referencia Epidemiologicos (INDRE) | Claudia Wong-Arambula, Abril Rodriguez-Maldonado, Vanessa Rivero-Arredondo, Ariadna Medina-Benitez, Joaquin Quiroz-Mercado, Sergio Rangel-Guerrero, Natividad Cruz-Ortiz, Tatiana Nunez-Garcia, Gisela Barrera-Badillo, Lucia Hernandez-Rivas, Irma Lopez-Martinez, Ernesto Ramirez-Gonzalez. |
| EPI_ISL_2296109 | LESP Jalisco | Instituto de Diagnostico y Referencia Epidemiologicos (INDRE) | Claudia Wong-Arambula, Abril Rodriguez-Maldonado, Vanessa Rivero-Arredondo, Ariadna Medina-Benitez, Joaquin Quiroz-Mercado, Sergio Rangel-Guerrero, Natividad Cruz-Ortiz, Tatiana Nunez-Garcia, Gisela Barrera-Badillo, Lucia Hernandez-Rivas, Irma Lopez-Martinez, Ernesto Ramirez-Gonzalez. |
| EPI_ISL_2304003, EPI_ISL_2304004, EPI_ISL_2304005, EPI_ISL_2304006, EPI_ISL_2304007, EPI_ISL_2304011, EPI_ISL_2304012, EPI_ISL_2304023, EPI_ISL_2304025 | University of Oregon COVID-19 MAP Laboratory | University of Oregon Genomics and Cell Charactericzation Core Facility (GC3F) | Douglas Turnbull, Ariana White, Jeff Bishop, Jason Carriere, Jason Sydes, Peter Batzel |
| EPI_ISL_2307122, EPI_ISL_2307127, EPI_ISL_2307129, EPI_ISL_2307131 | State Virus Research and Diagnostic Laboratory (VRDL), AIIMS Raipur | State Virus Research and Diagnostic Laboratory (VRDL), AIIMS Raipur | Pushpendra Singh, Kuldeep Sharma, Priyanka Singh, Somya Sharma, MD Rafiiqullah Khan, Jiyan Chandrawanshi, Sanjay Singh Negi, Anudita Bhargava |
| EPI_ISL_2308277, EPI_ISL_2308278, EPI_ISL_2308279, EPI_ISL_2308280, EPI_ISL_2308281 | Laboratori de Referencia de Catalunya | Laboratori de Referencia de Catalunya | Ramirez A., Bellosillo B., Padilla E., Vilas A., Hernandez JJ., Canal M. |
| EPI_ISL_2312663 | HOPITAL DE VICHY | CHU Clermont-Ferrand, service de virologie | Bisseux Maxime, Mirand Audrey, Combes Patricia, Henquell Cécile |
| EPI_ISL_2312664 | Centre Hospitalier Universitaire Clermont-Ferrand | CHU Clermont-Ferrand, service de virologie | Bisseux Maxime, Mirand Audrey, Combes Patricia, Henquell Cécile |
| EPI_ISL_2321598 | UNIGEM | Universidad Nacional de Colombia - Laboratorio Genómico One Health | Simón Villegas Velásquez, Andres F. Cardona-Rios, Maria Stella López, Daniel O. Maldonado-Perez, Lina M. Hurtado, Laura Silvana Perez, Karl A Ciuoderis, Maria Angélica Maya, Carolina Muñoz-Arango, Celeny Ortiz, Rita Almanza Payares, Idabely Betancur Ortiz, Sandra Ines Cano, Diego A. Álvarez-Díaz, Carlos Franco-Muñoz, Katherine Laiton-Donato, Hector Alejandro Ruiz-Moreno, Marcela Mercado-Reyes, Jorge E. Osorio, Juan P. Hernandez-Ortiz |
| EPI_ISL_2321600 | UNIGEM | Universidad Nacional de Colombia - Laboratorio Genómico One Health | Andres F. Cardona-Rios, Maria Stella López, Daniel O. Maldonado-Perez, Simón Villegas Velásquez, Lina M. Hurtado, Laura Silvana Perez, Karl A Ciuoderis, Maria Angélica Maya, Carolina Muñoz-Arango, Celeny Ortiz, Rita Almanza Payares, Idabely Betancur Ortiz, Sandra Ines Cano, Diego A. Álvarez-Díaz, Carlos Franco-Muñoz, Katherine Laiton-Donato, Hector Alejandro Ruiz-Moreno, Marcela Mercado-Reyes, Jorge E. Osorio, Juan P. Hernandez-Ortiz |
| EPI_ISL_2321623, EPI_ISL_2321624 | UNIGEM | Universidad Nacional de Colombia - Laboratorio Genómico One Health | Maria Stella López, Andres F. Cardona-Rios, Daniel O. Maldonado-Perez, Simón Villegas Velásquez, Lina M. Hurtado, Laura Silvana Perez, Karl A Ciuoderis, Maria Angélica Maya, Carolina Muñoz-Arango, Celeny Ortiz, Rita Almanza Payares, Idabely Betancur Ortiz, Sandra Ines Cano, Diego A. Álvarez-Díaz, Carlos Franco-Muñoz, Katherine Laiton-Donato, Hector Alejandro Ruiz-Moreno, Marcela Mercado-Reyes, Jorge E. Osorio, Juan P. Hernandez-Ortiz |
| EPI_ISL_2321627 | SOMER | Universidad Nacional de Colombia - Laboratorio Genómico One Health | Simón Villegas Velásquez, Andres F. Cardona-Rios, Maria Stella López, Daniel O. Maldonado-Perez, Lina M. Hurtado, Laura Silvana Perez, Karl A Ciuoderis, Maria Angélica Maya, Carolina Muñoz-Arango, Celeny Ortiz, Rita Almanza Payares, Idabely Betancur Ortiz, Sandra Ines Cano, Diego A. Álvarez-Díaz, Carlos Franco-Muñoz, Katherine Laiton-Donato, Hector Alejandro Ruiz-Moreno, Marcela Mercado-Reyes, Jorge E. Osorio, Juan P. Hernandez-Ortiz |
| EPI_ISL_2322760 | Collection of WHO National Influenza Center, N.F. Gamaleya National Research Center for Epidemiology and Microbiology of the Ministry of Health of the Russian Federation | Laboratory of genome analysis, N.F. Gamaleya National Research Center for Epidemiology and Microbiology of the Ministry of Health of the Russian Federation | Olga Voronina, Ekaterina Aksenova, Natalia Ryzhova, Marina Kunda, Kirill Krasnoslobodtsev, Elena Burtseva, Denis Logunov |
| EPI_ISL_2323464, EPI_ISL_2323465, EPI_ISL_2323469 | MEDILAB | CHU Poitiers | Caroline MICHAUD, Maxime PICHON, Manon PRAT, Valentin BON-BARET, Birama N'DIAYE, Magali GARCIA, Agnes BEBY-DEFAUX, Nicolas LEVEQUE |
| EPI_ISL_2323470 | CHU POITIERS | CHU Poitiers | Caroline MICHAUD, Maxime PICHON, Manon PRAT, Valentin BON-BARET, Birama N'DIAYE, Magali GARCIA, Agnes BEBY-DEFAUX, Nicolas LEVEQUE |
| EPI_ISL_2323472 | CH NIORT | CHU Poitiers | Caroline MICHAUD, Maxime PICHON, Manon PRAT, Valentin BON-BARET, Birama N'DIAYE, Magali GARCIA, Agnes BEBY-DEFAUX, Nicolas LEVEQUE |
| EPI_ISL_2323474, EPI_ISL_2323477 | CH SAINTONGE | CHU Poitiers | Caroline MICHAUD, Maxime PICHON, Manon PRAT, Valentin BON-BARET, Birama N'DIAYE, Magali GARCIA, Agnes BEBY-DEFAUX, Nicolas LEVEQUE |
| EPI_ISL_2323478 | CH ROCHEFORT | CHU Poitiers | Caroline MICHAUD, Maxime PICHON, Manon PRAT, Valentin BON-BARET, Birama N'DIAYE, Magali GARCIA, Agnes BEBY-DEFAUX, Nicolas LEVEQUE |
| EPI_ISL_2323480, EPI_ISL_2323482 | BIO86 CHAUMONT | CHU Poitiers | Caroline MICHAUD, Maxime PICHON, Manon PRAT, Valentin BON-BARET, Birama N'DIAYE, Magali GARCIA, Agnes BEBY-DEFAUX, Nicolas LEVEQUE |
| EPI_ISL_2324265 | ILV Kärnten | Bergthaler laboratory, CeMM Research Center for Molecular Medicine of the Austrian Academy of Sciences | Lukas Endler, Anna Schedl, Fabian Amman, Petr Triska, Thomas Penz, Benedikt Agerer, Maelle Le Moing, Michael Schuster, Bekir Erguner, Jan Laine, Martin Senekowitsch, Christoph Bock, Andreas Bergthaler |
| EPI_ISL_2324270, EPI_ISL_2324271, EPI_ISL_2324273 | Institute for Medical and Chemical Laboratory Diagnostics, Kepler Universitätsklinikum | Bergthaler laboratory, CeMM Research Center for Molecular Medicine of the Austrian Academy of Sciences | Lukas Endler, Anna Schedl, Fabian Amman, Petr Triska, Thomas Penz, Benedikt Agerer, Maelle Le Moing, Michael Schuster, Bekir Erguner, Jan Laine, Martin Senekowitsch, Christoph Bock, Andreas Bergthaler |
| EPI_ISL_2324297, EPI_ISL_2324298, EPI_ISL_2324299, EPI_ISL_2324300, EPI_ISL_2324313, EPI_ISL_2324314, EPI_ISL_2324315, EPI_ISL_2324316, EPI_ISL_2324328, EPI_ISL_2324329, EPI_ISL_2324331, EPI_ISL_2324333, EPI_ISL_2324338, EPI_ISL_2324339, EPI_ISL_2324340, EPI_ISL_2324343, EPI_ISL_2324344, EPI_ISL_2324345, EPI_ISL_2324346, EPI_ISL_2324348, EPI_ISL_2324349, EPI_ISL_2324350, EPI_ISL_2324351, EPI_ISL_2324352, EPI_ISL_2324353, EPI_ISL_2324360, EPI_ISL_2324373, EPI_ISL_2324374, EPI_ISL_2324375, EPI_ISL_2324376, EPI_ISL_2324377, EPI_ISL_2324378, EPI_ISL_2324379, EPI_ISL_2324380, EPI_ISL_2324381, EPI_ISL_2324382 | | | |
| see above | Austrian Agency for Health and Food Safety (AGES) | Bergthaler laboratory, CeMM Research Center for Molecular Medicine of the Austrian Academy of Sciences | Lukas Endler, Anna Schedl, Fabian Amman, Petr Triska, Thomas Penz, Benedikt Agerer, Maelle Le Moing, Michael Schuster, Bekir Erguner, Jan Laine, Martin Senekowitsch, Christoph Bock, Andreas Bergthaler |
| EPI_ISL_2325177, EPI_ISL_2325178, EPI_ISL_2325179, EPI_ISL_2325181, EPI_ISL_2325182, EPI_ISL_2325183, EPI_ISL_2325184, EPI_ISL_2332580, EPI_ISL_2332581, EPI_ISL_2332582, EPI_ISL_2332583, EPI_ISL_2332584, EPI_ISL_2332585, EPI_ISL_2332586, EPI_ISL_2332587, EPI_ISL_2332588, EPI_ISL_2332589, EPI_ISL_2332590, EPI_ISL_2332591, EPI_ISL_2332592, EPI_ISL_2332594, EPI_ISL_2332595, EPI_ISL_2332596, EPI_ISL_2332597, EPI_ISL_2332598, EPI_ISL_2332599, EPI_ISL_2332600, EPI_ISL_2332601, EPI_ISL_2332602, EPI_ISL_2332603, EPI_ISL_2332604 | | | |
| see above | Microbiology Department, University Hospital Donostia | Microbiology Department, University Hospital Donostia | Marimon JM, Montes M, Piñeiro L, Grandioso D, Martin-Peñaranda T, Gomez M, Cilla G. |
| EPI_ISL_2332704, EPI_ISL_2332705, EPI_ISL_2332706, EPI_ISL_2332707, EPI_ISL_2332708, EPI_ISL_2332709, EPI_ISL_2332710, EPI_ISL_2332711, EPI_ISL_2332712, EPI_ISL_2332713, EPI_ISL_2332714, EPI_ISL_2332715, EPI_ISL_2332716, EPI_ISL_2332717, EPI_ISL_2332718, EPI_ISL_2332719, EPI_ISL_2332720, EPI_ISL_2332721, EPI_ISL_2332722, EPI_ISL_2332723, EPI_ISL_2332724, EPI_ISL_2332725, EPI_ISL_2332726, EPI_ISL_2332727, EPI_ISL_2332728, EPI_ISL_2332729, EPI_ISL_2332730, EPI_ISL_2332731, EPI_ISL_2332732, EPI_ISL_2332733, EPI_ISL_2332734, EPI_ISL_2332735, EPI_ISL_2332736, EPI_ISL_2332737, EPI_ISL_2332738, EPI_ISL_2332739, EPI_ISL_2332740, EPI_ISL_2332741, EPI_ISL_2332742, EPI_ISL_2332743, EPI_ISL_2332744, EPI_ISL_2332745, EPI_ISL_2332746, EPI_ISL_2332747, EPI_ISL_2332748, EPI_ISL_2332749, EPI_ISL_2332750, EPI_ISL_2332751, EPI_ISL_2332752, EPI_ISL_2332753, EPI_ISL_2332754, EPI_ISL_2332755, EPI_ISL_2332756, EPI_ISL_2332757, EPI_ISL_2332758, EPI_ISL_2332759, EPI_ISL_2332760, EPI_ISL_2332761, EPI_ISL_2332762, EPI_ISL_2332763, EPI_ISL_2332764, EPI_ISL_2332765 | | | |

| | | | |
|---|---|---|---|
| see above | National Centre For Cell Science | National Centre For Cell Science - INSACOG | Dhiraj Paul, Shivang P. Bhanushali, Mitali Inamdar, Mohak P Gujare, Manoj Kumar Bhat, Ajay Pillai, INSACOG Consortium team, Yogesh Shouche |
| EPI_ISL_2333665, EPI_ISL_2333666 | Istituto Zooprofilattico Sperimentale della Lombardia e dell'Emilia Romagna (IZSLER), Sede Territoriale di Modena | Istituto Zooprofilattico Sperimentale della Lombardia e dell'Emilia Romagna (IZSLER), Risk Analysis and Genomic Epidemiology Unit | Carra Elena, Gianluca Rugna, Marina Morganti, Ilaria Menozzi, Erika Scaltriti, Stefano Pongolini |
| EPI_ISL_2333783 | GH A.CHENEVIER-H.MONDOR | Department of Virology, Henri Mondor University Hospital, Assistance Publique Hôpitaux de Paris, Université Paris-Est Créteil, INSERM U955 | Christophe Rodriguez, Slim Fourati, Vanessa Demontant, Guillaume Gricourt, Melissa N'Debi, Alexandre Soulier, Elisabeth Trawinski, Jean-Michel Pawlotsky |
| EPI_ISL_2334499, EPI_ISL_2334500, EPI_ISL_2334501, EPI_ISL_2334502, EPI_ISL_2334503, EPI_ISL_2334504, EPI_ISL_2334505, EPI_ISL_2334506, EPI_ISL_2334507, EPI_ISL_2334508, EPI_ISL_2334509, EPI_ISL_2334510, EPI_ISL_2334511, EPI_ISL_2334512, EPI_ISL_2334513, EPI_ISL_2334514, EPI_ISL_2334515, EPI_ISL_2334516, EPI_ISL_2334517, EPI_ISL_2335018, EPI_ISL_2335019, EPI_ISL_2335020 | | | |
| see above | Nacionalinis maisto ir veterinarijos rizikos vertinimo institutas | National Public Health Surveillance Laboratory | Lukas Zemaitis, Migle Gabrielaite, Jelena Razmuk, Svajune Muralyte, Ana Steponkiene, Lukas Vasionis, Danas Baksa |
| EPI_ISL_2339258 | CH SAINT-PHILIBERT | CHU Lille - Laboratoire de Virologie | AIT YAHYA Emilie, ALIDJINOU Enagnon Kazali, BOCKET Laurence, CREPIN Michel, DEMAY Christophe, ENGELMANN Ilka, GEFFROY Sandrine, GUIGON Aurélie, LAMBERT Valérie, LAZREK Mouna, NOBILLIAUX Florian, PREVOST Brigitte, THUILLIER Caroline, TINEZ Claire |
| EPI_ISL_2339259, EPI_ISL_2339260, EPI_ISL_2339261 | CHU LILLE | CHU Lille - Laboratoire de Virologie | AIT YAHYA Emilie, ALIDJINOU Enagnon Kazali, BOCKET Laurence, CREPIN Michel, DEMAY Christophe, ENGELMANN Ilka, GEFFROY Sandrine, GUIGON Aurélie, LAMBERT Valérie, LAZREK Mouna, NOBILLIAUX Florian, PREVOST Brigitte, THUILLIER Caroline, TINEZ Claire |
| EPI_ISL_2339262 | CH DOUAI | CHU Lille - Laboratoire de Virologie | AIT YAHYA Emilie, ALIDJINOU Enagnon Kazali, BOCKET Laurence, CREPIN Michel, DEMAY Christophe, ENGELMANN Ilka, GEFFROY Sandrine, GUIGON Aurélie, LAMBERT Valérie, LAZREK Mouna, NOBILLIAUX Florian, PREVOST Brigitte, THUILLIER Caroline, TINEZ Claire |
| EPI_ISL_2339263 | CH TOURCOING | CHU Lille - Laboratoire de Virologie | AIT YAHYA Emilie, ALIDJINOU Enagnon Kazali, BOCKET Laurence, CREPIN Michel, DEMAY Christophe, ENGELMANN Ilka, GEFFROY Sandrine, GUIGON Aurélie, LAMBERT Valérie, LAZREK Mouna, NOBILLIAUX Florian, PREVOST Brigitte, THUILLIER Caroline, TINEZ Claire |
| EPI_ISL_2339264, EPI_ISL_2339265 | CHU LILLE | CHU Lille - Laboratoire de Virologie | AIT YAHYA Emilie, ALIDJINOU Enagnon Kazali, BOCKET Laurence, CREPIN Michel, DEMAY Christophe, ENGELMANN Ilka, GEFFROY Sandrine, GUIGON Aurélie, LAMBERT Valérie, LAZREK Mouna, NOBILLIAUX Florian, PREVOST Brigitte, THUILLIER Caroline, TINEZ Claire |
| EPI_ISL_2339266 | CH SAINT-PHILIBERT | CHU Lille - Laboratoire de Virologie | AIT YAHYA Emilie, ALIDJINOU Enagnon Kazali, BOCKET Laurence, CREPIN Michel, DEMAY Christophe, ENGELMANN Ilka, GEFFROY Sandrine, GUIGON Aurélie, LAMBERT Valérie, LAZREK Mouna, NOBILLIAUX Florian, PREVOST Brigitte, THUILLIER Caroline, TINEZ Claire |
| EPI_ISL_2339267 | CH CAMBRAI | CHU Lille - Laboratoire de Virologie | AIT YAHYA Emilie, ALIDJINOU Enagnon Kazali, BOCKET Laurence, CREPIN Michel, DEMAY Christophe, ENGELMANN Ilka, GEFFROY Sandrine, GUIGON Aurélie, LAMBERT Valérie, LAZREK Mouna, NOBILLIAUX Florian, PREVOST Brigitte, THUILLIER Caroline, TINEZ Claire |
| EPI_ISL_2339268, EPI_ISL_2339269 | CHU LILLE | CHU Lille - Laboratoire de Virologie | AIT YAHYA Emilie, ALIDJINOU Enagnon Kazali, BOCKET Laurence, CREPIN Michel, DEMAY Christophe, ENGELMANN Ilka, GEFFROY Sandrine, GUIGON Aurélie, LAMBERT Valérie, LAZREK Mouna, NOBILLIAUX Florian, PREVOST Brigitte, THUILLIER Caroline, TINEZ Claire |
| EPI_ISL_2339270 | CH LENS | CHU Lille - Laboratoire de Virologie | AIT YAHYA Emilie, ALIDJINOU Enagnon Kazali, BOCKET Laurence, CREPIN Michel, DEMAY Christophe, ENGELMANN Ilka, GEFFROY Sandrine, GUIGON Aurélie, LAMBERT Valérie, LAZREK Mouna, NOBILLIAUX Florian, PREVOST Brigitte, THUILLIER Caroline, TINEZ Claire |
| EPI_ISL_2339271 | CHU LILLE | CHU Lille - Laboratoire de Virologie | AIT YAHYA Emilie, ALIDJINOU Enagnon Kazali, BOCKET Laurence, CREPIN Michel, DEMAY Christophe, ENGELMANN Ilka, GEFFROY Sandrine, GUIGON Aurélie, LAMBERT Valérie, LAZREK Mouna, NOBILLIAUX Florian, PREVOST Brigitte, THUILLIER Caroline, TINEZ Claire |
| EPI_ISL_2339272 | CH LENS | CHU Lille - Laboratoire de Virologie | AIT YAHYA Emilie, ALIDJINOU Enagnon Kazali, BOCKET Laurence, CREPIN Michel, DEMAY Christophe, ENGELMANN Ilka, GEFFROY Sandrine, GUIGON Aurélie, LAMBERT Valérie, LAZREK Mouna, NOBILLIAUX Florian, PREVOST Brigitte, THUILLIER Caroline, TINEZ Claire |
| EPI_ISL_2339273 | CH DOUAI | CHU Lille - Laboratoire de Virologie | AIT YAHYA Emilie, ALIDJINOU Enagnon Kazali, BOCKET Laurence, CREPIN Michel, DEMAY Christophe, ENGELMANN Ilka, GEFFROY Sandrine, GUIGON Aurélie, LAMBERT Valérie, LAZREK Mouna, NOBILLIAUX Florian, PREVOST Brigitte, THUILLIER Caroline, TINEZ Claire |
| EPI_ISL_2339274 | CHU LILLE | CHU Lille - Laboratoire de Virologie | AIT YAHYA Emilie, ALIDJINOU Enagnon Kazali, BOCKET Laurence, CREPIN Michel, DEMAY Christophe, ENGELMANN Ilka, GEFFROY Sandrine, GUIGON Aurélie, LAMBERT Valérie, LAZREK Mouna, NOBILLIAUX Florian, PREVOST Brigitte, THUILLIER Caroline, TINEZ Claire |
| EPI_ISL_2339776, EPI_ISL_2339777, EPI_ISL_2339778, EPI_ISL_2339780, EPI_ISL_2339782, EPI_ISL_2339783, EPI_ISL_2339784, EPI_ISL_2339786, EPI_ISL_2339787, EPI_ISL_2339788, EPI_ISL_2339789, EPI_ISL_2339790, EPI_ISL_2339791, EPI_ISL_2339792, EPI_ISL_2339793, EPI_ISL_2339794 | | | |
| see above | Grupo de Investigación en Enfermedades Tropicales del Ejército (GINETEJ), Laboratorio de Referencia e Investigación, Dirección de Sanidad Ejército, Bogotá, Colombia | Centro de Investigaciones en Microbiología y Biotecnología-UR (CIMBIUR), Facultad de Ciencias Naturales, Universidad del Rosario, Bogotá, Colombia | Nathalia Ballesteros, Sergio Castañeda, Marina Muñoz, Luz H. Patiño, Claudia Méndez, Carolina Oliveros, Julie Pérez, Lorena Albarracín, Elizabeth K. Márquez, María Teresa Alvarado, Frank de los Santos Ortiz, Yanira Romero, Camilo A. Correa-Cárdenas, Maria Clara Duque, Sergio Gutíerrez-Riveros, Zulma Cucunubá, Juan David Ramírez |
| EPI_ISL_2340864, EPI_ISL_2340866 | LESP Guanajuato | Instituto de Diagnostico y Referencia Epidemiologicos (INDRE) | Claudia Wong-Arambula, Abril Rodriguez-Maldonado, Vanessa Rivero-Arredondo, Ariadna Medina-Benitez, Joaquin Quiroz-Mercado, Sergio Rangel-Guerrero, Natividad Cruz-Ortiz, Tatiana Nunez-Garcia, Gisela Barrera-Badillo, Lucia Hernandez-Rivas, Irma Lopez-Martinez, Ernesto Ramirez-Gonzalez. |
| EPI_ISL_2340867 | LESP Nuevo Leon | Instituto de Diagnostico y Referencia Epidemiologicos (INDRE) | Claudia Wong-Arambula, Abril Rodriguez-Maldonado, Vanessa Rivero-Arredondo, Ariadna Medina-Benitez, Joaquin Quiroz-Mercado, Sergio Rangel-Guerrero, Natividad Cruz-Ortiz, Tatiana Nunez-Garcia, Gisela Barrera-Badillo, Lucia Hernandez-Rivas, Irma Lopez-Martinez, Ernesto Ramirez-Gonzalez. |
| EPI_ISL_2340868, EPI_ISL_2340869, EPI_ISL_2340870 | LESP Michoacan | Instituto de Diagnostico y Referencia Epidemiologicos (INDRE) | Claudia Wong-Arambula, Abril Rodriguez-Maldonado, Vanessa Rivero-Arredondo, Ariadna Medina-Benitez, Joaquin Quiroz-Mercado, Sergio Rangel-Guerrero, Natividad Cruz-Ortiz, Tatiana Nunez-Garcia, Gisela Barrera-Badillo, Lucia Hernandez-Rivas, Irma Lopez-Martinez, Ernesto Ramirez-Gonzalez. |
| EPI_ISL_2340871 | LESP Oaxaca | Instituto de Diagnostico y Referencia Epidemiologicos (INDRE) | Claudia Wong-Arambula, Abril Rodriguez-Maldonado, Vanessa Rivero-Arredondo, Ariadna Medina-Benitez, Joaquin Quiroz-Mercado, Sergio Rangel-Guerrero, Natividad Cruz-Ortiz, Tatiana Nunez-Garcia, Gisela Barrera-Badillo, Lucia Hernandez-Rivas, Irma Lopez-Martinez, Ernesto Ramirez-Gonzalez. |
| EPI_ISL_2340873 | LESP Michoacan | Instituto de Diagnostico y Referencia Epidemiologicos (INDRE) | Claudia Wong-Arambula, Abril Rodriguez-Maldonado, Vanessa Rivero-Arredondo, Ariadna Medina-Benitez, Joaquin Quiroz-Mercado, Sergio Rangel-Guerrero, Natividad Cruz-Ortiz, Tatiana Nunez-Garcia, Gisela Barrera-Badillo, Lucia Hernandez-Rivas, Irma Lopez-Martinez, Ernesto Ramirez-Gonzalez. |
| EPI_ISL_2340875 | LESP Sinaloa | Instituto de Diagnostico y Referencia Epidemiologicos (INDRE) | Claudia Wong-Arambula, Abril Rodriguez-Maldonado, Vanessa Rivero-Arredondo, Ariadna Medina-Benitez, Joaquin Quiroz-Mercado, Sergio Rangel-Guerrero, Natividad Cruz-Ortiz, Tatiana Nunez-Garcia, Gisela Barrera-Badillo, Lucia Hernandez-Rivas, Irma Lopez-Martinez, Ernesto Ramirez-Gonzalez. |
| EPI_ISL_2340876 | LESP Queretaro | Instituto de Diagnostico y Referencia Epidemiologicos (INDRE) | Claudia Wong-Arambula, Abril Rodriguez-Maldonado, Vanessa Rivero-Arredondo, Ariadna Medina-Benitez, Joaquin Quiroz-Mercado, Sergio Rangel-Guerrero, Natividad Cruz-Ortiz, Tatiana Nunez-Garcia, Gisela Barrera-Badillo, Lucia Hernandez-Rivas, Irma Lopez-Martinez, Ernesto Ramirez-Gonzalez. |
| EPI_ISL_2340877, EPI_ISL_2340878 | LESP Tamaulipas | Instituto de Diagnostico y Referencia Epidemiologicos (INDRE) | Claudia Wong-Arambula, Abril Rodriguez-Maldonado, Vanessa Rivero-Arredondo, Ariadna Medina-Benitez, Joaquin Quiroz-Mercado, Sergio Rangel-Guerrero, Natividad Cruz-Ortiz, Tatiana Nunez-Garcia, Gisela Barrera-Badillo, Lucia Hernandez-Rivas, Irma Lopez-Martinez, Ernesto Ramirez-Gonzalez. |
| EPI_ISL_2340879, EPI_ISL_2340880 | LESP Guanajuato | Instituto de Diagnostico y Referencia Epidemiologicos (INDRE) | Claudia Wong-Arambula, Abril Rodriguez-Maldonado, Vanessa Rivero-Arredondo, Ariadna Medina-Benitez, Joaquin Quiroz-Mercado, Sergio Rangel-Guerrero, Natividad Cruz-Ortiz, Tatiana Nunez-Garcia, Gisela Barrera-Badillo, Lucia Hernandez-Rivas, Irma Lopez-Martinez, Ernesto Ramirez-Gonzalez. |
| EPI_ISL_2340881 | LESP Queretaro | Instituto de Diagnostico y Referencia Epidemiologicos (INDRE) | Claudia Wong-Arambula, Abril Rodriguez-Maldonado, Vanessa Rivero-Arredondo, Ariadna Medina-Benitez, Joaquin Quiroz-Mercado, Sergio Rangel-Guerrero, Natividad Cruz-Ortiz, Tatiana Nunez-Garcia, Gisela Barrera-Badillo, Lucia Hernandez-Rivas, Irma Lopez-Martinez, Ernesto Ramirez-Gonzalez. |
| EPI_ISL_2340882 | LESP Ciudad de Mexico | Instituto de Diagnostico y Referencia Epidemiologicos (INDRE) | Claudia Wong-Arambula, Abril Rodriguez-Maldonado, Vanessa Rivero-Arredondo, Ariadna Medina-Benitez, Joaquin Quiroz-Mercado, Sergio Rangel-Guerrero, Natividad Cruz-Ortiz, Tatiana Nunez-Garcia, Gisela Barrera-Badillo, Lucia Hernandez-Rivas, Irma Lopez-Martinez, Ernesto |

| | | | |
|---|---|---|---|
| | | | Ramirez-Gonzalez. |
| EPI_ISL_2340883 | LESP Queretaro | Instituto de Diagnostico y Referencia Epidemiologicos (INDRE) | Claudia Wong-Arambula, Abril Rodriguez-Maldonado, Vanessa Rivero-Arredondo, Ariadna Medina-Benitez, Joaquin Quiroz-Mercado, Sergio Rangel-Guerrero, Natividad Cruz-Ortiz, Tatiana Nunez-Garcia, Gisela Barrera-Badillo, Lucia Hernandez-Rivas, Irma Lopez-Martinez, Ernesto Ramirez-Gonzalez. |
| EPI_ISL_2340886 | LESP Tamaulipas | Instituto de Diagnostico y Referencia Epidemiologicos (INDRE) | Claudia Wong-Arambula, Abril Rodriguez-Maldonado, Vanessa Rivero-Arredondo, Ariadna Medina-Benitez, Joaquin Quiroz-Mercado, Sergio Rangel-Guerrero, Natividad Cruz-Ortiz, Tatiana Nunez-Garcia, Gisela Barrera-Badillo, Lucia Hernandez-Rivas, Irma Lopez-Martinez, Ernesto Ramirez-Gonzalez. |
| EPI_ISL_2340888, EPI_ISL_2340889 | LESP Chihuahua | Instituto de Diagnostico y Referencia Epidemiologicos (INDRE) | Claudia Wong-Arambula, Abril Rodriguez-Maldonado, Vanessa Rivero-Arredondo, Ariadna Medina-Benitez, Joaquin Quiroz-Mercado, Sergio Rangel-Guerrero, Natividad Cruz-Ortiz, Tatiana Nunez-Garcia, Gisela Barrera-Badillo, Lucia Hernandez-Rivas, Irma Lopez-Martinez, Ernesto Ramirez-Gonzalez. |
| EPI_ISL_2340891, EPI_ISL_2340893, EPI_ISL_2340894, EPI_ISL_2340895, EPI_ISL_2340896, EPI_ISL_2340897, EPI_ISL_2340898, EPI_ISL_2340899, EPI_ISL_2340900, EPI_ISL_2340901, EPI_ISL_2340902 | | | |
| see above | LESP Michoacan | Instituto de Diagnostico y Referencia Epidemiologicos (INDRE) | Claudia Wong-Arambula, Abril Rodriguez-Maldonado, Vanessa Rivero-Arredondo, Ariadna Medina-Benitez, Joaquin Quiroz-Mercado, Sergio Rangel-Guerrero, Natividad Cruz-Ortiz, Tatiana Nunez-Garcia, Gisela Barrera-Badillo, Lucia Hernandez-Rivas, Irma Lopez-Martinez, Ernesto Ramirez-Gonzalez. |
| EPI_ISL_2340942, EPI_ISL_2340943, EPI_ISL_2340944, EPI_ISL_2340945, EPI_ISL_2340946, EPI_ISL_2340947, EPI_ISL_2340948, EPI_ISL_2340949 | LESP Guanajuato | Instituto de Diagnostico y Referencia Epidemiologicos (INDRE) | Claudia Wong-Arambula, Abril Rodriguez-Maldonado, Vanessa Rivero-Arredondo, Ariadna Medina-Benitez, Joaquin Quiroz-Mercado, Sergio Rangel-Guerrero, Natividad Cruz-Ortiz, Tatiana Nunez-Garcia, Gisela Barrera-Badillo, Lucia Hernandez-Rivas, Irma Lopez-Martinez, Ernesto Ramirez-Gonzalez. |
| EPI_ISL_2340950, EPI_ISL_2340951, EPI_ISL_2340952, EPI_ISL_2340953, EPI_ISL_2340954 | LESP Puebla | Instituto de Diagnostico y Referencia Epidemiologicos (INDRE) | Claudia Wong-Arambula, Abril Rodriguez-Maldonado, Vanessa Rivero-Arredondo, Ariadna Medina-Benitez, Joaquin Quiroz-Mercado, Sergio Rangel-Guerrero, Natividad Cruz-Ortiz, Tatiana Nunez-Garcia, Gisela Barrera-Badillo, Lucia Hernandez-Rivas, Irma Lopez-Martinez, Ernesto Ramirez-Gonzalez. |
| EPI_ISL_2340955, EPI_ISL_2340956, EPI_ISL_2340957, EPI_ISL_2340958 | LESP Hidalgo | Instituto de Diagnostico y Referencia Epidemiologicos (INDRE) | Claudia Wong-Arambula, Abril Rodriguez-Maldonado, Vanessa Rivero-Arredondo, Ariadna Medina-Benitez, Joaquin Quiroz-Mercado, Sergio Rangel-Guerrero, Natividad Cruz-Ortiz, Tatiana Nunez-Garcia, Gisela Barrera-Badillo, Lucia Hernandez-Rivas, Irma Lopez-Martinez, Ernesto Ramirez-Gonzalez. |
| EPI_ISL_2340959 | LESP Queretaro | Instituto de Diagnostico y Referencia Epidemiologicos (INDRE) | Claudia Wong-Arambula, Abril Rodriguez-Maldonado, Vanessa Rivero-Arredondo, Ariadna Medina-Benitez, Joaquin Quiroz-Mercado, Sergio Rangel-Guerrero, Natividad Cruz-Ortiz, Tatiana Nunez-Garcia, Gisela Barrera-Badillo, Lucia Hernandez-Rivas, Irma Lopez-Martinez, Ernesto Ramirez-Gonzalez. |
| EPI_ISL_2340960, EPI_ISL_2340961 | LESP Sinaloa | Instituto de Diagnostico y Referencia Epidemiologicos (INDRE) | Claudia Wong-Arambula, Abril Rodriguez-Maldonado, Vanessa Rivero-Arredondo, Ariadna Medina-Benitez, Joaquin Quiroz-Mercado, Sergio Rangel-Guerrero, Natividad Cruz-Ortiz, Tatiana Nunez-Garcia, Gisela Barrera-Badillo, Lucia Hernandez-Rivas, Irma Lopez-Martinez, Ernesto Ramirez-Gonzalez. |
| EPI_ISL_2340962, EPI_ISL_2340963, EPI_ISL_2340964, EPI_ISL_2340965, EPI_ISL_2340966, EPI_ISL_2340967, EPI_ISL_2340968 | LESP Queretaro | Instituto de Diagnostico y Referencia Epidemiologicos (INDRE) | Claudia Wong-Arambula, Abril Rodriguez-Maldonado, Vanessa Rivero-Arredondo, Ariadna Medina-Benitez, Joaquin Quiroz-Mercado, Sergio Rangel-Guerrero, Natividad Cruz-Ortiz, Tatiana Nunez-Garcia, Gisela Barrera-Badillo, Lucia Hernandez-Rivas, Irma Lopez-Martinez, Ernesto Ramirez-Gonzalez. |
| EPI_ISL_2340969, EPI_ISL_2340970, EPI_ISL_2340971 | LESP Veracruz | Instituto de Diagnostico y Referencia Epidemiologicos (INDRE) | Claudia Wong-Arambula, Abril Rodriguez-Maldonado, Vanessa Rivero-Arredondo, Ariadna Medina-Benitez, Joaquin Quiroz-Mercado, Sergio Rangel-Guerrero, Natividad Cruz-Ortiz, Tatiana Nunez-Garcia, Gisela Barrera-Badillo, Lucia Hernandez-Rivas, Irma Lopez-Martinez, Ernesto Ramirez-Gonzalez. |
| EPI_ISL_2340972, EPI_ISL_2340973 | LESP Sinaloa | Instituto de Diagnostico y Referencia Epidemiologicos (INDRE) | Claudia Wong-Arambula, Abril Rodriguez-Maldonado, Vanessa Rivero-Arredondo, Ariadna Medina-Benitez, Joaquin Quiroz-Mercado, Sergio Rangel-Guerrero, Natividad Cruz-Ortiz, Tatiana Nunez-Garcia, Gisela Barrera-Badillo, Lucia Hernandez-Rivas, Irma Lopez-Martinez, Ernesto Ramirez-Gonzalez. |
| EPI_ISL_2340974, EPI_ISL_2340975 | LESP Tamaulipas | Instituto de Diagnostico y Referencia Epidemiologicos (INDRE) | Claudia Wong-Arambula, Abril Rodriguez-Maldonado, Vanessa Rivero-Arredondo, Ariadna Medina-Benitez, Joaquin Quiroz-Mercado, Sergio Rangel-Guerrero, Natividad Cruz-Ortiz, Tatiana Nunez-Garcia, Gisela Barrera-Badillo, Lucia Hernandez-Rivas, Irma Lopez-Martinez, Ernesto Ramirez-Gonzalez. |
| EPI_ISL_2340976, EPI_ISL_2340977 | LESP Ciudad de Mexico | Instituto de Diagnostico y Referencia Epidemiologicos (INDRE) | Claudia Wong-Arambula, Abril Rodriguez-Maldonado, Vanessa Rivero-Arredondo, Ariadna Medina-Benitez, Joaquin Quiroz-Mercado, Sergio Rangel-Guerrero, Natividad Cruz-Ortiz, Tatiana Nunez-Garcia, Gisela Barrera-Badillo, Lucia Hernandez-Rivas, Irma Lopez-Martinez, Ernesto Ramirez-Gonzalez. |
| EPI_ISL_2340978 | LESP Sonora | Instituto de Diagnostico y Referencia Epidemiologicos (INDRE) | Claudia Wong-Arambula, Abril Rodriguez-Maldonado, Vanessa Rivero-Arredondo, Ariadna Medina-Benitez, Joaquin Quiroz-Mercado, Sergio Rangel-Guerrero, Natividad Cruz-Ortiz, Tatiana Nunez-Garcia, Gisela Barrera-Badillo, Lucia Hernandez-Rivas, Irma Lopez-Martinez, Ernesto Ramirez-Gonzalez. |
| EPI_ISL_2340979 | LESP Baja California | Instituto de Diagnostico y Referencia Epidemiologicos (INDRE) | Claudia Wong-Arambula, Abril Rodriguez-Maldonado, Vanessa Rivero-Arredondo, Ariadna Medina-Benitez, Joaquin Quiroz-Mercado, Sergio Rangel-Guerrero, Natividad Cruz-Ortiz, Tatiana Nunez-Garcia, Gisela Barrera-Badillo, Lucia Hernandez-Rivas, Irma Lopez-Martinez, Ernesto Ramirez-Gonzalez. |
| EPI_ISL_2340980, EPI_ISL_2340981, EPI_ISL_2340982, EPI_ISL_2340983, EPI_ISL_2340984, EPI_ISL_2340985, EPI_ISL_2340986, EPI_ISL_2340987, EPI_ISL_2340988, EPI_ISL_2340989, EPI_ISL_2340990, EPI_ISL_2340991, EPI_ISL_2340992, EPI_ISL_2340993, EPI_ISL_2340994, EPI_ISL_2340995, EPI_ISL_2340996 | | | |
| see above | LESP Guanajuato | Instituto de Diagnostico y Referencia Epidemiologicos (INDRE) | Claudia Wong-Arambula, Abril Rodriguez-Maldonado, Vanessa Rivero-Arredondo, Ariadna Medina-Benitez, Joaquin Quiroz-Mercado, Sergio Rangel-Guerrero, Natividad Cruz-Ortiz, Tatiana Nunez-Garcia, Gisela Barrera-Badillo, Lucia Hernandez-Rivas, Irma Lopez-Martinez, Ernesto Ramirez-Gonzalez. |
| EPI_ISL_2340997 | LESP Tamaulipas | Instituto de Diagnostico y Referencia Epidemiologicos (INDRE) | Claudia Wong-Arambula, Abril Rodriguez-Maldonado, Vanessa Rivero-Arredondo, Ariadna Medina-Benitez, Joaquin Quiroz-Mercado, Sergio Rangel-Guerrero, Natividad Cruz-Ortiz, Tatiana Nunez-Garcia, Gisela Barrera-Badillo, Lucia Hernandez-Rivas, Irma Lopez-Martinez, Ernesto Ramirez-Gonzalez. |
| EPI_ISL_2341002, EPI_ISL_2341003, EPI_ISL_2341004, EPI_ISL_2341005 | LESP Yucatan | Instituto de Diagnostico y Referencia Epidemiologicos (INDRE) | Claudia Wong-Arambula, Abril Rodriguez-Maldonado, Vanessa Rivero-Arredondo, Ariadna Medina-Benitez, Joaquin Quiroz-Mercado, Sergio Rangel-Guerrero, Natividad Cruz-Ortiz, Tatiana Nunez-Garcia, Gisela Barrera-Badillo, Lucia Hernandez-Rivas, Irma Lopez-Martinez, Ernesto Ramirez-Gonzalez. |
| EPI_ISL_2341006 | LESP Ciudad de Mexico | Instituto de Diagnostico y Referencia Epidemiologicos (INDRE) | Claudia Wong-Arambula, Abril Rodriguez-Maldonado, Vanessa Rivero-Arredondo, Ariadna Medina-Benitez, Joaquin Quiroz-Mercado, Sergio Rangel-Guerrero, Natividad Cruz-Ortiz, Tatiana Nunez-Garcia, Gisela Barrera-Badillo, Lucia Hernandez-Rivas, Irma Lopez-Martinez, Ernesto Ramirez-Gonzalez. |
| EPI_ISL_2341007 | LESP Jalisco | Instituto de Diagnostico y Referencia Epidemiologicos (INDRE) | Claudia Wong-Arambula, Abril Rodriguez-Maldonado, Vanessa Rivero-Arredondo, Ariadna Medina-Benitez, Joaquin Quiroz-Mercado, Sergio Rangel-Guerrero, Natividad Cruz-Ortiz, Tatiana Nunez-Garcia, Gisela Barrera-Badillo, Lucia Hernandez-Rivas, Irma Lopez-Martinez, Ernesto Ramirez-Gonzalez. |
| EPI_ISL_2341008 | LESP Queretaro | Instituto de Diagnostico y Referencia Epidemiologicos (INDRE) | Claudia Wong-Arambula, Abril Rodriguez-Maldonado, Vanessa Rivero-Arredondo, Ariadna Medina-Benitez, Joaquin Quiroz-Mercado, Sergio Rangel-Guerrero, Natividad Cruz-Ortiz, Tatiana Nunez-Garcia, Gisela Barrera-Badillo, Lucia Hernandez-Rivas, Irma Lopez-Martinez, Ernesto Ramirez-Gonzalez. |
| EPI_ISL_2341009, EPI_ISL_2341010, EPI_ISL_2341011, EPI_ISL_2341012 | LESP Guanajuato | Instituto de Diagnostico y Referencia Epidemiologicos (INDRE) | Claudia Wong-Arambula, Abril Rodriguez-Maldonado, Vanessa Rivero-Arredondo, Ariadna Medina-Benitez, Joaquin Quiroz-Mercado, Sergio Rangel-Guerrero, Natividad Cruz-Ortiz, Tatiana Nunez-Garcia, Gisela Barrera-Badillo, Lucia Hernandez-Rivas, Irma Lopez-Martinez, Ernesto Ramirez-Gonzalez. |

| | | | |
|---|---|---|---|
| EPI_ISL_2341013 | LESP Baja California Sur | Instituto de Diagnostico y Referencia Epidemiologicos (INDRE) | Claudia Wong-Arambula, Abril Rodriguez-Maldonado, Vanessa Rivero-Arredondo, Ariadna Medina-Benitez, Joaquin Quiroz-Mercado, Sergio Rangel-Guerrero, Natividad Cruz-Ortiz, Tatiana Nunez-Garcia, Gisela Barrera-Badillo, Lucia Hernandez-Rivas, Irma Lopez-Martinez, Ernesto Ramirez-Gonzalez. |
| EPI_ISL_2341016, EPI_ISL_2341017, EPI_ISL_2341018, EPI_ISL_2341019 | LESP Guanajuato | Instituto de Diagnostico y Referencia Epidemiologicos (INDRE) | Claudia Wong-Arambula, Abril Rodriguez-Maldonado, Vanessa Rivero-Arredondo, Ariadna Medina-Benitez, Joaquin Quiroz-Mercado, Sergio Rangel-Guerrero, Natividad Cruz-Ortiz, Tatiana Nunez-Garcia, Gisela Barrera-Badillo, Lucia Hernandez-Rivas, Irma Lopez-Martinez, Ernesto Ramirez-Gonzalez. |
| EPI_ISL_2341021 | LESP Nuevo Leon | Instituto de Diagnostico y Referencia Epidemiologicos (INDRE) | Claudia Wong-Arambula, Abril Rodriguez-Maldonado, Vanessa Rivero-Arredondo, Ariadna Medina-Benitez, Joaquin Quiroz-Mercado, Sergio Rangel-Guerrero, Natividad Cruz-Ortiz, Tatiana Nunez-Garcia, Gisela Barrera-Badillo, Lucia Hernandez-Rivas, Irma Lopez-Martinez, Ernesto Ramirez-Gonzalez. |
| EPI_ISL_2341023 | LESP Campeche | Instituto de Diagnostico y Referencia Epidemiologicos (INDRE) | Claudia Wong-Arambula, Abril Rodriguez-Maldonado, Vanessa Rivero-Arredondo, Ariadna Medina-Benitez, Joaquin Quiroz-Mercado, Sergio Rangel-Guerrero, Natividad Cruz-Ortiz, Tatiana Nunez-Garcia, Gisela Barrera-Badillo, Lucia Hernandez-Rivas, Irma Lopez-Martinez, Ernesto Ramirez-Gonzalez. |
| EPI_ISL_2342996, EPI_ISL_2342997 | LESP Ciudad de Mexico | Instituto de Diagnostico y Referencia Epidemiologicos (INDRE) | Claudia Wong-Arambula, Abril Rodriguez-Maldonado, Vanessa Rivero-Arredondo, Ariadna Medina-Benitez, Joaquin Quiroz-Mercado, Sergio Rangel-Guerrero, Natividad Cruz-Ortiz, Tatiana Nunez-Garcia, Gisela Barrera-Badillo, Lucia Hernandez-Rivas, Irma Lopez-Martinez, Ernesto Ramirez-Gonzalez. |
| EPI_ISL_2342998 | LESP Queretaro | Instituto de Diagnostico y Referencia Epidemiologicos (INDRE) | Claudia Wong-Arambula, Abril Rodriguez-Maldonado, Vanessa Rivero-Arredondo, Ariadna Medina-Benitez, Joaquin Quiroz-Mercado, Sergio Rangel-Guerrero, Natividad Cruz-Ortiz, Tatiana Nunez-Garcia, Gisela Barrera-Badillo, Lucia Hernandez-Rivas, Irma Lopez-Martinez, Ernesto Ramirez-Gonzalez. |
| EPI_ISL_2343006 | LESP Ciudad de Mexico | Instituto de Diagnostico y Referencia Epidemiologicos (INDRE) | Claudia Wong-Arambula, Abril Rodriguez-Maldonado, Vanessa Rivero-Arredondo, Ariadna Medina-Benitez, Joaquin Quiroz-Mercado, Sergio Rangel-Guerrero, Natividad Cruz-Ortiz, Tatiana Nunez-Garcia, Gisela Barrera-Badillo, Lucia Hernandez-Rivas, Irma Lopez-Martinez, Ernesto Ramirez-Gonzalez. |
| EPI_ISL_2343008 | LESP Tamaulipas | Instituto de Diagnostico y Referencia Epidemiologicos (INDRE) | Claudia Wong-Arambula, Abril Rodriguez-Maldonado, Vanessa Rivero-Arredondo, Ariadna Medina-Benitez, Joaquin Quiroz-Mercado, Sergio Rangel-Guerrero, Natividad Cruz-Ortiz, Tatiana Nunez-Garcia, Gisela Barrera-Badillo, Lucia Hernandez-Rivas, Irma Lopez-Martinez, Ernesto Ramirez-Gonzalez. |
| EPI_ISL_2343009 | LESP Ciudad de Mexico | Instituto de Diagnostico y Referencia Epidemiologicos (INDRE) | Claudia Wong-Arambula, Abril Rodriguez-Maldonado, Vanessa Rivero-Arredondo, Ariadna Medina-Benitez, Joaquin Quiroz-Mercado, Sergio Rangel-Guerrero, Natividad Cruz-Ortiz, Tatiana Nunez-Garcia, Gisela Barrera-Badillo, Lucia Hernandez-Rivas, Irma Lopez-Martinez, Ernesto Ramirez-Gonzalez. |
| EPI_ISL_2343011, EPI_ISL_2343012, EPI_ISL_2343013, EPI_ISL_2343014, EPI_ISL_2343015, EPI_ISL_2343016 | LESP Michoacan | Instituto de Diagnostico y Referencia Epidemiologicos (INDRE) | Claudia Wong-Arambula, Abril Rodriguez-Maldonado, Vanessa Rivero-Arredondo, Ariadna Medina-Benitez, Joaquin Quiroz-Mercado, Sergio Rangel-Guerrero, Natividad Cruz-Ortiz, Tatiana Nunez-Garcia, Gisela Barrera-Badillo, Lucia Hernandez-Rivas, Irma Lopez-Martinez, Ernesto Ramirez-Gonzalez. |
| EPI_ISL_2343019 | LESP Queretaro | Instituto de Diagnostico y Referencia Epidemiologicos (INDRE) | Claudia Wong-Arambula, Abril Rodriguez-Maldonado, Vanessa Rivero-Arredondo, Ariadna Medina-Benitez, Joaquin Quiroz-Mercado, Sergio Rangel-Guerrero, Natividad Cruz-Ortiz, Tatiana Nunez-Garcia, Gisela Barrera-Badillo, Lucia Hernandez-Rivas, Irma Lopez-Martinez, Ernesto Ramirez-Gonzalez. |
| EPI_ISL_2343021 | LESP Veracruz | Instituto de Diagnostico y Referencia Epidemiologicos (INDRE) | Claudia Wong-Arambula, Abril Rodriguez-Maldonado, Vanessa Rivero-Arredondo, Ariadna Medina-Benitez, Joaquin Quiroz-Mercado, Sergio Rangel-Guerrero, Natividad Cruz-Ortiz, Tatiana Nunez-Garcia, Gisela Barrera-Badillo, Lucia Hernandez-Rivas, Irma Lopez-Martinez, Ernesto Ramirez-Gonzalez. |
| EPI_ISL_2343022 | LESP Sinaloa | Instituto de Diagnostico y Referencia Epidemiologicos (INDRE) | Claudia Wong-Arambula, Abril Rodriguez-Maldonado, Vanessa Rivero-Arredondo, Ariadna Medina-Benitez, Joaquin Quiroz-Mercado, Sergio Rangel-Guerrero, Natividad Cruz-Ortiz, Tatiana Nunez-Garcia, Gisela Barrera-Badillo, Lucia Hernandez-Rivas, Irma Lopez-Martinez, Ernesto Ramirez-Gonzalez. |
| EPI_ISL_2343023 | LESP Tamaulipas | Instituto de Diagnostico y Referencia Epidemiologicos (INDRE) | Claudia Wong-Arambula, Abril Rodriguez-Maldonado, Vanessa Rivero-Arredondo, Ariadna Medina-Benitez, Joaquin Quiroz-Mercado, Sergio Rangel-Guerrero, Natividad Cruz-Ortiz, Tatiana Nunez-Garcia, Gisela Barrera-Badillo, Lucia Hernandez-Rivas, Irma Lopez-Martinez, Ernesto Ramirez-Gonzalez. |
| EPI_ISL_2343024 | LESP Chiapas | Instituto de Diagnostico y Referencia Epidemiologicos (INDRE) | Claudia Wong-Arambula, Abril Rodriguez-Maldonado, Vanessa Rivero-Arredondo, Ariadna Medina-Benitez, Joaquin Quiroz-Mercado, Sergio Rangel-Guerrero, Natividad Cruz-Ortiz, Tatiana Nunez-Garcia, Gisela Barrera-Badillo, Lucia Hernandez-Rivas, Irma Lopez-Martinez, Ernesto Ramirez-Gonzalez. |
| EPI_ISL_2343025 | LESP Sonora | Instituto de Diagnostico y Referencia Epidemiologicos (INDRE) | Claudia Wong-Arambula, Abril Rodriguez-Maldonado, Vanessa Rivero-Arredondo, Ariadna Medina-Benitez, Joaquin Quiroz-Mercado, Sergio Rangel-Guerrero, Natividad Cruz-Ortiz, Tatiana Nunez-Garcia, Gisela Barrera-Badillo, Lucia Hernandez-Rivas, Irma Lopez-Martinez, Ernesto Ramirez-Gonzalez. |
| EPI_ISL_2343026, EPI_ISL_2343027, EPI_ISL_2343028 | LESP Guanajuato | Instituto de Diagnostico y Referencia Epidemiologicos (INDRE) | Claudia Wong-Arambula, Abril Rodriguez-Maldonado, Vanessa Rivero-Arredondo, Ariadna Medina-Benitez, Joaquin Quiroz-Mercado, Sergio Rangel-Guerrero, Natividad Cruz-Ortiz, Tatiana Nunez-Garcia, Gisela Barrera-Badillo, Lucia Hernandez-Rivas, Irma Lopez-Martinez, Ernesto Ramirez-Gonzalez. |
| EPI_ISL_2343029 | LESP Yucatan | Instituto de Diagnostico y Referencia Epidemiologicos (INDRE) | Claudia Wong-Arambula, Abril Rodriguez-Maldonado, Vanessa Rivero-Arredondo, Ariadna Medina-Benitez, Joaquin Quiroz-Mercado, Sergio Rangel-Guerrero, Natividad Cruz-Ortiz, Tatiana Nunez-Garcia, Gisela Barrera-Badillo, Lucia Hernandez-Rivas, Irma Lopez-Martinez, Ernesto Ramirez-Gonzalez. |
| EPI_ISL_2346436, EPI_ISL_2346437, EPI_ISL_2346438, EPI_ISL_2346439, EPI_ISL_2346440, EPI_ISL_2346441, EPI_ISL_2346442, EPI_ISL_2346443, EPI_ISL_2346444, EPI_ISL_2346445, EPI_ISL_2346446, EPI_ISL_2346447, EPI_ISL_2346448, EPI_ISL_2346449, EPI_ISL_2346450, EPI_ISL_2346451, EPI_ISL_2346452, EPI_ISL_2346453, EPI_ISL_2346454, EPI_ISL_2346455, EPI_ISL_2346456, EPI_ISL_2346457, EPI_ISL_2346458, EPI_ISL_2346459, EPI_ISL_2346460, EPI_ISL_2346461, EPI_ISL_2346462, EPI_ISL_2346463, EPI_ISL_2346464, EPI_ISL_2346466, EPI_ISL_2346467, EPI_ISL_2346468, EPI_ISL_2346469, EPI_ISL_2346470 | | | |
| see above | Instituto Nacional de Enfermedades Respiratorias (INER) | Centro de Investigación en Enfermedades Infecciosas (CIENI), Instituto Nacional de Enfermedades Respiratorias (INER) | Consorcio Mexicano de Vigilancia Genómica (CoViGen-Mex). Authors (in alphabetical order): Julio Elias Alvarado-Yaah, Carlos F. Arias, Santiago Ávila-Ríos, Eduardo Becerril-Vargas, Víctor Hugo Borja-Aburto, Celia Boukadida, Juan Bautista Chale-Dzul, Célida Duque Molina, José Antonio Enciso-Moreno, Gloria Elena Espinosa-Ayala, Fernando Fontove-Herrera, Victor Eduardo Garcia-Arias, Concepción Grajales-Muñiz, Ricardo Grande, Alejandra Hernández-Terán, Alfredo Herrera-Estrella, Carla Ivón Herrera-Najera, Pavel Isa, Brenda Irasema Maldonado-Meza, Bernardo Martínez-Miguel, José Arturo Martínez-Orozco, Margarita Matías-Florentino, Fidencio Mejía-Nepomuceno, María Guadalupe de Jesús Mireles-Rivera, Gloria María Molina-Salinas, Hector Montoya-Fuentes, Mario Mújica-Sánchez, José Esteban Muñoz-Medina, José de Jesús Nuñez-Contreras, Alicia Ocaña-Mondragón, Luis Alberto Ochoa-Carrera, Hector Esteban Paz-Juárez, Francisco Pulido, Helen Haydee Fernanda Ramírez-Plascencia, Angel Gustavo Salas-Lais, Alejandro Sanchez-Flores, Clara Esperanza Santacruz-Tinoco, María Guadalupe Santiago-Mauricio, Jorge Salas-Hernández, Nelly Sélem-Mojica, Blanca Taboada, Gloria Vazquez, Joel Armando Vázquez-Pérez |
| EPI_ISL_2348730 | Hospital Dr. Horacio Heller | Laboratorio Central Mg. Luis Alfredo Pianciola on behalf of 'Proyecto Argentino Interinstitucional de genomica de SARS-CoV-2' (PAIS Consortium) | L Pianciola, M Mazzeo, C Ziehm, C Pintos, M Fernandez, J Ousset, C Rastellini |
| EPI_ISL_2348731 | Hospital de Area Villa la Angostura Dr. Oscar Arraiz | Laboratorio Central Mg. Luis Alfredo Pianciola on behalf of 'Proyecto Argentino Interinstitucional de genomica de SARS-CoV-2' (PAIS Consortium) | L Pianciola, M Mazzeo, C Ziehm, C Pintos, M Fernandez, J Ousset, C Rastellini |
| EPI_ISL_2348732 | Hospital Provincial Neuquen Dr. Eduardo Castro Rendon | Laboratorio Central Mg. Luis Alfredo Pianciola on behalf of 'Proyecto Argentino Interinstitucional de genomica de SARS-CoV-2' (PAIS Consortium) | L Pianciola, M Mazzeo, C Ziehm, C Pintos, M Fernandez, J Ousset, C Rastellini |
| EPI_ISL_2348733 | Clinica CMIC | Laboratorio Central Mg. Luis Alfredo Pianciola on behalf of 'Proyecto Argentino Interinstitucional de genomica de | L Pianciola, M Mazzeo, C Ziehm, C Pintos, M Fernandez, J Ousset, C Rastellini |

| | | | |
|---|---|---|---|
| | | SARS-CoV-2' (PAIS Consortium) | |
| EPI_ISL_2348734 | Hospital Regional Ramon Carillo | Laboratorio Central Mg. Luis Alfredo Pianciola on behalf of 'Proyecto Argentino Interinstitucional de genomica de SARS-CoV-2' (PAIS Consortium) | L Pianciola, M Mazzeo, C Ziehm, C Pintos, M Fernandez, J Ousset, C Rastellini |
| EPI_ISL_2348735 | Hospital de Area Junin de los Andes | Laboratorio Central Mg. Luis Alfredo Pianciola on behalf of 'Proyecto Argentino Interinstitucional de genomica de SARS-CoV-2' (PAIS Consortium) | L Pianciola, M Mazzeo, C Ziehm, C Pintos, M Fernandez, J Ousset, C Rastellini |
| EPI_ISL_2348736 | Hospital Regional Ramon Carillo | Laboratorio Central Mg. Luis Alfredo Pianciola on behalf of 'Proyecto Argentino Interinstitucional de genomica de SARS-CoV-2' (PAIS Consortium) | L Pianciola, M Mazzeo, C Ziehm, C Pintos, M Fernandez, J Ousset, C Rastellini |
| EPI_ISL_2348737 | Clinica Chapelco | Laboratorio Central Mg. Luis Alfredo Pianciola on behalf of 'Proyecto Argentino Interinstitucional de genomica de SARS-CoV-2' (PAIS Consortium) | L Pianciola, M Mazzeo, C Ziehm, C Pintos, M Fernandez, J Ousset, C Rastellini |
| EPI_ISL_2348738 | Trailer Operativo Detectar | Laboratorio Central Mg. Luis Alfredo Pianciola on behalf of 'Proyecto Argentino Interinstitucional de genomica de SARS-CoV-2' (PAIS Consortium) | L Pianciola, M Mazzeo, C Ziehm, C Pintos, M Fernandez, J Ousset, C Rastellini |
| EPI_ISL_2348739 | Hospital de Area Plottier | Laboratorio Central Mg. Luis Alfredo Pianciola on behalf of 'Proyecto Argentino Interinstitucional de genomica de SARS-CoV-2' (PAIS Consortium) | L Pianciola, M Mazzeo, C Ziehm, C Pintos, M Fernandez, J Ousset, C Rastellini |
| EPI_ISL_2348740 | Hospital de Area Alumine | Laboratorio Central Mg. Luis Alfredo Pianciola on behalf of 'Proyecto Argentino Interinstitucional de genomica de SARS-CoV-2' (PAIS Consortium) | L Pianciola, M Mazzeo, C Ziehm, C Pintos, M Fernandez, J Ousset, C Rastellini |
| EPI_ISL_2348741 | Hospital de Area Plottier | Laboratorio Central Mg. Luis Alfredo Pianciola on behalf of 'Proyecto Argentino Interinstitucional de genomica de SARS-CoV-2' (PAIS Consortium) | L Pianciola, M Mazzeo, C Ziehm, C Pintos, M Fernandez, J Ousset, C Rastellini |
| EPI_ISL_2349552 | National Scientific Center for Especially Dangerous Infections (NSCEDI) | Kazakh National Agrarian University (KazNAU); TreeGene LLP Genetic Laboratory; National Scientific Center for Especially Dangerous Infections (NSCEDI) | Tabynov Kaissar, Strochkov Vitaliy, Tabynov Kairat, Beloussov Vyacheslav, Sandybayev Nurlan, Turebekov Nurkeldy, Granica Joanna, Yerubayev Toktassyn, Yespolov Tlektes |
| EPI_ISL_2349693, EPI_ISL_2349694 | Genome Analysis Center, Yamanashi Central Hospital | Genome Analysis Center, Yamanashi Central Hospital | Yosuke Hirotsu |
| EPI_ISL_2350787, EPI_ISL_2350788, EPI_ISL_2350789, EPI_ISL_2350790 | Translational Health Science and Technology Institute -ESIC medical college and hospital, Faridabad | THSTI Bioassay laboratory | Naseem Ahmed Khan, Heena Shaman, Janmejay Singh, Kamal Pargai, Jigme Wangchuk, Anil Pandey, Akshay Kanakan, Ranjeet Maurya, Janani Srinivasa Vasudevan, Neha Jha, Priyanka Mehta, Rajesh Pandey, Guruprasad R Medigeshi |
| EPI_ISL_2350818, EPI_ISL_2350821, EPI_ISL_2350822, EPI_ISL_2351216, EPI_ISL_2351217, EPI_ISL_2351223, EPI_ISL_2351281, EPI_ISL_2351392, EPI_ISL_2351398, EPI_ISL_2351857, EPI_ISL_2352024, EPI_ISL_2352687, EPI_ISL_2354480, EPI_ISL_2354500, EPI_ISL_2356914, EPI_ISL_2357229, EPI_ISL_2357231, EPI_ISL_2357235, EPI_ISL_2357715, EPI_ISL_2357732 | | | |
| see above | Institut de Virologie du CHU de Strasbourg | Institut de Virologie du CHU de Strasbourg | Fafi-Kremer Samira, Gallais Floriane, Gantner Pierre, Laugel Elodie, Solis Morgane, Velay Aurélie, Wendling Marie-Josée |
| EPI_ISL_2358033 | Laboratoire Hôpital Sainte Musse | Laboratoire Hôpital Sainte Musse | Lionel Chollet |
| EPI_ISL_2360252 | Shahid Ghodossi Laboratory | National Influenza Center | J Yavarian,K Sadeghi, NZ Shafiei Jandaghi, V Salimi, A Nejati, N Ghavvami,F Ajaminejad and T Mokhtari Azad |
| EPI_ISL_2360253 | Maku Molecular Laboratory | National Influenza Center | A Nejati, J Yavarian,K Sadeghi, NZ Shafiei Jandaghi, V Salimi, N Ghavvami,F Ajaminejad and T Mokhtari Azad |
| EPI_ISL_2360254 | Shahid Ghodossi Laboratory | National Influenza Center | K Sadeghi, A Nejati, J Yavarian, NZ Shafiei Jandaghi, V Salimi,F Ajaminejad,N Ghavvami and T Mokhtari Azad |
| EPI_ISL_2360256 | Boshehr University of Medical Sciences | National Influenza Center | NZ Shafiei Jandaghi, V Salimi, A Nejati, K Sadeghi, J Yavarian, Mostafa Salehi-Vaziri, F Ajaminejad,N Ghavvami and T Mokhtari Azad |
| EPI_ISL_2360257 | Boshehr University of Medical Sciences | National Influenza Center | V Salimi,NZ Shafiei Jandaghi, J Yavarian, A Nejati, K Sadeghi,Mostafa Salehi-Vaziri , N Ghavvami,F Ajaminejad and T Mokhtari Azad |
| EPI_ISL_2361358 | Institute of Molecular and Translational Medicine / Laboratory of Experimental Medicine, Faculty of Medicine and Dentistry, Palacky University and University Hospital Olomouc | Institute of Molecular and Translational Medicine / Laboratory of Experimental Medicine, Faculty of Medicine and Dentistry, Palacky University | Rastislav Slavkovský, Hana Jaworek, Vladimíra Koudeláková, Barbora Blumová, Tomáš Pospíšil, Marián Hajdúch |
| EPI_ISL_2361359 | Institute of Molecular and Translational Medicine / Laboratory of Experimental Medicine, Faculty of Medicine and Dentistry, Palacky University and University Hospital Olomouc | Institute of Molecular and Translational Medicine / Laboratory of Experimental Medicine, Faculty of Medicine and Dentistry, Palacky University | Rastislav Slavkovský, Hana Jaworek, Vladimíra Koudeláková, Barbora Blumová, Tomáš Pospíšil, Marián Hajdúch, Pavel Sauer |
| EPI_ISL_2361360 | Institute of Molecular and Translational Medicine / Laboratory of Experimental Medicine, Faculty of Medicine and Dentistry, Palacky University and University Hospital Olomouc | Institute of Molecular and Translational Medicine / Laboratory of Experimental Medicine, Faculty of Medicine and Dentistry, Palacky University | Rastislav Slavkovský, Hana Jaworek, Vladimíra Koudeláková, Barbora Blumová, Tomáš Pospíšil, Marián Hajdúch |
| EPI_ISL_2361502, EPI_ISL_2361521, EPI_ISL_2361522, EPI_ISL_2361618, EPI_ISL_2361692, EPI_ISL_2361712, EPI_ISL_2361744, EPI_ISL_2362041, EPI_ISL_2362042, EPI_ISL_2362043, EPI_ISL_2362044 | | | |
| see above | Gyncentrum | Gyncentrum | Emilia Morawiec, Pawel Czerwinski, Adam Pudelko, Aleksandra Skubis-Sikora, Magdalena Samul, Agnieszka Polak, Celina Kruszniewska-Rajs, Tomasz Wasik, Robert Wojtyczka, Jolanta Bartosiewicz- Wasik, Maria Maklasinska-Majdanik |
| EPI_ISL_2362674, EPI_ISL_2362677, EPI_ISL_2362680, EPI_ISL_2362684 | Microbiology and Virology Unit, Florence Careggi University Hospital | Microbiology and Virology Unit, Florence Careggi University Hospital | Vincenzo Di Pilato, Marco Coppi, Fabio Morecchiato, Noemi Aiezza, Ilaria Baccani, Nicla Giovacchini, Alberto Antonelli, Emanuele Gori, Gian Maria Rossolini |
| EPI_ISL_2363778 | Microbiology and Virology Laboratory, "Policlinico Riuniti, Azienda Ospedaliero Universitaria, Foggia" | Microbiology and Virology Laboratory, "Policlinico Riuniti, Azienda Ospedaliero Universitaria, Foggia" | Fabio Arena, Settimia Altamura, Donatella Cedola, Valeria Delli Carri, Annamaria D'Aprile, Maria Rosaria Lipsi, Loris Micelli, Maria Pia Patrizio, Daniela Pisanelli, Tiziana Rollo, Maurizio Margaglione and Rosella De Nittis. |
| EPI_ISL_2365882, EPI_ISL_2365886, EPI_ISL_2365887 | Microbiology Department. Complexo Hospitalario Universitario de Vigo | Microbiology Department. Complexo Hospitalario Universitario de Vigo | Alfaya N, Alonso I, Alvarez M, Cabrera JJ, Carballo R, Cores O, Cortizo S, del-Campo V, Martinez L, Mediero G, Perez S, Potel C, Regueiro B, Rey S, Vasallo FJ |
| EPI_ISL_2367261, EPI_ISL_2367262, EPI_ISL_2367467, EPI_ISL_2367468, EPI_ISL_2367563 | Institut de Virologie du CHU de Strasbourg | Institut de Virologie du CHU de Strasbourg | Fafi-Kremer Samira, Gallais Floriane, Gantner Pierre, Laugel Elodie, Solis Morgane, Velay Aurélie, Wendling Marie-Josée |
| EPI_ISL_2374154, EPI_ISL_2374155, EPI_ISL_2374156, EPI_ISL_2374157, EPI_ISL_2374158 | Research | National Reference Center for Viruses of Respiratory Infections, Institut Pasteur, Paris | Marion Barbet, Sylvie Behillil, Méline Bizard, Angela Brisebarre, Camille Capel, Vincent Enouf, Louise Lefrançois, Frédéric Lemoine, Christophe Malabat, Corinne Maufrais, Etienne Simon-Lorière, Maud Vanpeene, Sylvie Van der Werf ,Marianne Leruez-Ville |
| EPI_ISL_2374162, EPI_ISL_2374163, EPI_ISL_2374164, EPI_ISL_2374165, EPI_ISL_2374166, EPI_ISL_2374167, EPI_ISL_2374168 | Labo Analyses Med | National Reference Center for Viruses of Respiratory Infections, Institut Pasteur, Paris | Marion Barbet, Sylvie Behillil, Méline Bizard, Angela Brisebarre, Camille Capel, Vincent Enouf, Louise Lefrançois, Frédéric Lemoine, Christophe Malabat, Corinne Maufrais, Etienne Simon-Lorière, Maud Vanpeene, Sylvie Van der Werf ,Anne Lise Bravetti |
| EPI_ISL_2374169 | Hospital | National Reference Center for Viruses of Respiratory Infections, Institut Pasteur, Paris | Marion Barbet, Sylvie Behillil, Méline Bizard, Angela Brisebarre, Camille Capel, Vincent Enouf, Louise Lefrançois, Frédéric Lemoine, Christophe Malabat, Corinne Maufrais, Etienne Simon-Lorière, Maud Vanpeene, Sylvie Van der Werf ,Laurence Louvet |
| EPI_ISL_2374180, EPI_ISL_2374181 | Hospital | National Reference Center for Viruses of Respiratory Infections, Institut Pasteur, Paris | Marion Barbet, Sylvie Behillil, Méline Bizard, Angela Brisebarre, Camille Capel, Vincent Enouf, Louise Lefrançois, Frédéric Lemoine, Christophe Malabat, Corinne Maufrais, Etienne Simon-Lorière, Maud Vanpeene, Sylvie Van der Werf ,Sandrine Castelain |

| | | | |
|---|---|---|---|
| EPI_ISL_2374390, EPI_ISL_2374393 | Hospital | National Reference Center for Viruses of Respiratory Infections, Institut Pasteur, Paris | Marion Barbet, Sylvie Behillil, Méline Bizard, Angela Brisebarre, Camille Capel, Vincent Enouf, Louise Lefrançois, Frédéric Lemoine, Christophe Malabat, Corinne Maufrais, Emmanuelle Permal, Etienne Simon-Lorière, Maud Vanpeene, Sylvie Van der Werf ,Alexandra Ducancelle |
| EPI_ISL_2374394 | Hospital | National Reference Center for Viruses of Respiratory Infections, Institut Pasteur, Paris | Marion Barbet, Sylvie Behillil, Méline Bizard, Angela Brisebarre, Camille Capel, Vincent Enouf, Louise Lefrançois, Frédéric Lemoine, Christophe Malabat, Corinne Maufrais, Emmanuelle Permal, Etienne Simon-Lorière, Maud Vanpeene, Sylvie Van der Werf ,CéLine Ramanantsoa |
| EPI_ISL_2374453 | Hospital | National Reference Center for Viruses of Respiratory Infections, Institut Pasteur, Paris | Marion Barbet, Sylvie Behillil, Méline Bizard, Angela Brisebarre, Camille Capel, Vincent Enouf, Louise Lefrançois, Frédéric Lemoine, Christophe Malabat, Corinne Maufrais, Emmanuelle Permal, Etienne Simon-Lorière, Maud Vanpeene, Sylvie Van der Werf ,Axelle Paquin |
| EPI_ISL_2374472, EPI_ISL_2374473 | Labo Analyses Med | National Reference Center for Viruses of Respiratory Infections, Institut Pasteur, Paris | Marion Barbet, Sylvie Behillil, Méline Bizard, Angela Brisebarre, Camille Capel, Vincent Enouf, Louise Lefrançois, Frédéric Lemoine, Christophe Malabat, Corinne Maufrais, Emmanuelle Permal, Etienne Simon-Lorière, Maud Vanpeene, Sylvie Van der Werf ,Odile Petsaris |
| EPI_ISL_2374478 | Hospital | National Reference Center for Viruses of Respiratory Infections, Institut Pasteur, Paris | Marion Barbet, Sylvie Behillil, Méline Bizard, Angela Brisebarre, Camille Capel, Vincent Enouf, Louise Lefrançois, Frédéric Lemoine, Christophe Malabat, Corinne Maufrais, Emmanuelle Permal, Etienne Simon-Lorière, Maud Vanpeene, Sylvie Van der Werf ,Haciba Moudjahed |
| EPI_ISL_2374479 | Labo Analyses Med | National Reference Center for Viruses of Respiratory Infections, Institut Pasteur, Paris | Marion Barbet, Sylvie Behillil, Méline Bizard, Angela Brisebarre, Camille Capel, Vincent Enouf, Louise Lefrançois, Frédéric Lemoine, Christophe Malabat, Corinne Maufrais, Emmanuelle Permal, Etienne Simon-Lorière, Maud Vanpeene, Sylvie Van der Werf ,Corgiat |
| EPI_ISL_2376227, EPI_ISL_2376229 | Imeda Hospital | Imeda Hospital | Johan Frans, Dagmar Obbels, Hanne Valgaeren |
| EPI_ISL_2379157, EPI_ISL_2379158, EPI_ISL_2379159, EPI_ISL_2379161, EPI_ISL_2379162, EPI_ISL_2379163, EPI_ISL_2379164, EPI_ISL_2379165, EPI_ISL_2379166, EPI_ISL_2379167, EPI_ISL_2379168, EPI_ISL_2379169, EPI_ISL_2379170, EPI_ISL_2379171, EPI_ISL_2379172, EPI_ISL_2379173, EPI_ISL_2379174, EPI_ISL_2379175, EPI_ISL_2379176, EPI_ISL_2379178, EPI_ISL_2379180, EPI_ISL_2379181, EPI_ISL_2379192, EPI_ISL_2379193, EPI_ISL_2379194, EPI_ISL_2379195, EPI_ISL_2379196, EPI_ISL_2379198, EPI_ISL_2379211, EPI_ISL_2379212, EPI_ISL_2379213, EPI_ISL_2379214, EPI_ISL_2379215, EPI_ISL_2379216, EPI_ISL_2379217, EPI_ISL_2379218, EPI_ISL_2379219, EPI_ISL_2379221, EPI_ISL_2379222, EPI_ISL_2379223, EPI_ISL_2379224, EPI_ISL_2379225, EPI_ISL_2379226, EPI_ISL_2379227, EPI_ISL_2379228, EPI_ISL_2379229, EPI_ISL_2379230, EPI_ISL_2379231, EPI_ISL_2379232, EPI_ISL_2379233, EPI_ISL_2379234, EPI_ISL_2379235, EPI_ISL_2379236, EPI_ISL_2379237 | | | |
| see above | National Center of Infectious and Parasitic Diseases | National Center of Infectious and Parasitic Diseases | Alexiev et al |
| EPI_ISL_2379433, EPI_ISL_2379443, EPI_ISL_2379650 | Institute for Medical Research, Infectious Disease Research Centre, National Institutes of Health, Ministry of Health Malaysia | Institute for Medical Research, Infectious Disease Research Centre, National Institutes of Health, Ministry of Health Malaysia | Suppiah J, Kamel K, Mohd Zawawi Z, Azizan MA, Ramly N, Robert F, Thayan R |
| EPI_ISL_2380500, EPI_ISL_2380544, EPI_ISL_2380626, EPI_ISL_2380680, EPI_ISL_2380725, EPI_ISL_2380858 | Institut de Virologie du CHU de Strasbourg | Institut de Virologie du CHU de Strasbourg | Fafi-Kremer Samira, Gallais Floriane, Gantner Pierre, Laugel Elodie, Solis Morgane, Velay Aurélie, Wendling Marie-Josée |
| EPI_ISL_2380902 | Institut de Virologie du CHU de StrasbourgInstitut de Virologie du CHU de Strasbourg | Institut de Virologie du CHU de Strasbourg | Fafi-Kremer Samira, Gallais Floriane, Gantner Pierre, Laugel Elodie, Solis Morgane, Velay Aurélie, Wendling Marie-Josée |
| EPI_ISL_2381414, EPI_ISL_2381415 | Institut de Virologie du CHU de Strasbourg | Institut de Virologie du CHU de Strasbourg | Fafi-Kremer Samira, Gallais Floriane, Gantner Pierre, Laugel Elodie, Solis Morgane, Velay Aurélie, Wendling Marie-Josée |
| EPI_ISL_2381998, EPI_ISL_2381999 | AZ Zeno Knokke-Blankenberge | AZ Sint Jan | Marijke Reynders, Patrick Descheemaeker, Laurien Hoornaert, Thomas Van Landschoot, Marie Madeleine Chabert-Consen, Jorn Hellemans |
| EPI_ISL_2382005, EPI_ISL_2382006 | AZ St.Jan campus Brugge | AZ Sint Jan | Marijke Reynders, Patrick Descheemaeker, Laurien Hoornaert, Thomas Van Landschoot, Marie Madeleine Chabert-Consen, Jorn Hellemans |
| EPI_ISL_2382043, EPI_ISL_2382045, EPI_ISL_2382046, EPI_ISL_2382047, EPI_ISL_2382048, EPI_ISL_2382049, EPI_ISL_2382050, EPI_ISL_2382051, EPI_ISL_2382052, EPI_ISL_2382053, EPI_ISL_2382055, EPI_ISL_2382057 | | | |
| see above | Klinikum Wels-Grieskirchen | Bergthaler laboratory, CeMM Research Center for Molecular Medicine of the Austrian Academy of Sciences | Lukas Endler, Anna Schedl, Fabian Amman, Petr Triska, Thomas Penz, Benedikt Agerer, Maelle Le Moing, Michael Schuster, Bekir Erguner, Jan Laine, Martin Senekowitsch, Christoph Bock, Andreas Bergthaler |
| EPI_ISL_2382295, EPI_ISL_2382296, EPI_ISL_2382297, EPI_ISL_2382299, EPI_ISL_2382300, EPI_ISL_2382301, EPI_ISL_2382302, EPI_ISL_2382303, EPI_ISL_2382304, EPI_ISL_2382305 | Uconn Health Clinical Microbiology | Brenton Graveley Lab, UConn Health | Lori Avery, Christina Caron, Stacey Ferraro, Kimberly Hayes, Michael Senh, Sara Olson, Xintao Wei, Brenton Graveley |
| EPI_ISL_2389632 | B.J. Medical College and Civil Hospital, Ahmedabad | Gujarat Biotechnology Research Centre | Zuber Saiyed,Ramesh Pandit,Janvi Raval,Zarna Patel,Nitin Savaliya,Dinesh Kumar,Twinkle Soni,Sonal Sharma ,Pranay Shah,Umang Mishra,Nitesh Shah,Chaitanya Joshi,Madhvi Joshi |
| EPI_ISL_2389634 | B.J. Medical College and Civil Hospital, Ahmedabad | Gujarat Biotechnology Research Centre | Ramesh Pandit,Janvi Raval,Zarna Patel,Nitin Savaliya,Dinesh Kumar,Twinkle Soni,Sonal Sharma ,Zuber Saiyed,Pranay Shah,Umang Mishra,Nitesh Shah,Chaitanya Joshi,Madhvi Joshi |
| EPI_ISL_2389636 | B.J. Medical College and Civil Hospital, Ahmedabad | Gujarat Biotechnology Research Centre | Janvi Raval,Zarna Patel,Nitin Savaliya,Dinesh Kumar,Twinkle Soni,Sonal Sharma ,Zuber Saiyed,Ramesh Pandit,Pranay Shah,Umang Mishra,Nitesh Shah,Chaitanya Joshi,Madhvi Joshi |
| EPI_ISL_2389638 | B.J. Medical College and Civil Hospital, Ahmedabad | Gujarat Biotechnology Research Centre | Zarna Patel,Nitin Savaliya,Dinesh Kumar,Twinkle Soni,Sonal Sharma ,Zuber Saiyed,Ramesh Pandit,Janvi Raval,Pranay Shah,Umang Mishra,Nitesh Shah,Chaitanya Joshi,Madhvi Joshi |
| EPI_ISL_2389640 | B.J. Medical College and Civil Hospital, Ahmedabad | Gujarat Biotechnology Research Centre | Nitin Savaliya,Dinesh Kumar,Twinkle Soni,Sonal Sharma ,Zuber Saiyed,Ramesh Pandit,Janvi Raval,Zarna Patel,Pranay Shah,Umang Mishra,Nitesh Shah,Chaitanya Joshi,Madhvi Joshi |
| EPI_ISL_2389641 | B.J. Medical College and Civil Hospital, Ahmedabad | Gujarat Biotechnology Research Centre | Dinesh Kumar,Twinkle Soni,Sonal Sharma ,Zuber Saiyed,Ramesh Pandit,Janvi Raval,Zarna Patel,Nitin Savaliya,Pranay Shah,Umang Mishra,Nitesh Shah,Chaitanya Joshi,Madhvi Joshi |
| EPI_ISL_2389643 | B.J. Medical College and Civil Hospital, Ahmedabad | Gujarat Biotechnology Research Centre | Twinkle Soni,Sonal Sharma ,Zuber Saiyed,Ramesh Pandit,Janvi Raval,Zarna Patel,Nitin Savaliya,Dinesh Kumar,Pranay Shah,Umang Mishra,Nitesh Shah,Chaitanya Joshi,Madhvi Joshi |
| EPI_ISL_2389645 | B.J. Medical College and Civil Hospital, Ahmedabad | Gujarat Biotechnology Research Centre | Sonal Sharma ,Zuber Saiyed,Ramesh Pandit,Janvi Raval,Zarna Patel,Nitin Savaliya,Dinesh Kumar,Twinkle Soni,Pranay Shah,Umang Mishra,Nitesh Shah,Chaitanya Joshi,Madhvi Joshi |
| EPI_ISL_2389647 | B.J. Medical College and Civil Hospital, Ahmedabad | Gujarat Biotechnology Research Centre | Zuber Saiyed,Ramesh Pandit,Janvi Raval,Zarna Patel,Nitin Savaliya,Dinesh Kumar,Twinkle Soni,Sonal Sharma ,Pranay Shah,Umang Mishra,Nitesh Shah,Chaitanya Joshi,Madhvi Joshi |
| EPI_ISL_2389649 | B.J. Medical College and Civil Hospital, Ahmedabad | Gujarat Biotechnology Research Centre | Ramesh Pandit,Janvi Raval,Zarna Patel,Nitin Savaliya,Dinesh Kumar,Twinkle Soni,Sonal Sharma ,Zuber Saiyed,Pranay Shah,Umang Mishra,Nitesh Shah,Chaitanya Joshi,Madhvi Joshi |
| EPI_ISL_2389651 | B.J. Medical College and Civil Hospital, Ahmedabad | Gujarat Biotechnology Research Centre | Janvi Raval,Zarna Patel,Nitin Savaliya,Dinesh Kumar,Twinkle Soni,Sonal Sharma ,Zuber Saiyed,Ramesh Pandit,Pranay Shah,Umang Mishra,Nitesh Shah,Chaitanya Joshi,Madhvi Joshi |
| EPI_ISL_2389653 | B.J. Medical College and Civil Hospital, Ahmedabad | Gujarat Biotechnology Research Centre | Zarna Patel,Nitin Savaliya,Dinesh Kumar,Twinkle Soni,Sonal Sharma ,Zuber Saiyed,Ramesh Pandit,Janvi Raval,Pranay Shah,Umang Mishra,Nitesh Shah,Chaitanya Joshi,Madhvi Joshi |
| EPI_ISL_2389655 | B.J. Medical College and Civil Hospital, Ahmedabad | Gujarat Biotechnology Research Centre | Nitin Savaliya,Dinesh Kumar,Twinkle Soni,Sonal Sharma ,Zuber Saiyed,Ramesh Pandit,Janvi Raval,Zarna Patel,Pranay Shah,Umang Mishra,Nitesh Shah,Chaitanya Joshi,Madhvi Joshi |
| EPI_ISL_2389657 | B.J. Medical College and Civil Hospital, Ahmedabad | Gujarat Biotechnology Research Centre | Dinesh Kumar,Twinkle Soni,Sonal Sharma ,Zuber Saiyed,Ramesh Pandit,Janvi Raval,Zarna Patel,Nitin Savaliya,Pranay Shah,Umang Mishra,Nitesh Shah,Chaitanya Joshi,Madhvi Joshi |
| EPI_ISL_2391478 | N.H.L Municipal Medical College, Ahmedabad | Gujarat Biotechnology Research Centre | Sonal Sharma ,Zuber Saiyed,Ramesh Pandit,Janvi Raval,Zarna Patel,Nitin Savaliya,Dinesh Kumar,Twinkle Soni,Jayshri Pethani,Umang Mishra,Nitesh Shah,Chaitanya Joshi,Madhvi Joshi |
| EPI_ISL_2391481 | N.H.L Municipal Medical College, Ahmedabad | Gujarat Biotechnology Research Centre | Janvi Raval,Zarna Patel,Nitin Savaliya,Dinesh Kumar,Twinkle Soni,Sonal Sharma ,Zuber Saiyed,Ramesh Pandit,Jayshri Pethani,Umang Mishra,Nitesh Shah,Chaitanya Joshi,Madhvi Joshi |
| EPI_ISL_2391482 | N.H.L Municipal Medical College, Ahmedabad | Gujarat Biotechnology Research Centre | Zarna Patel,Nitin Savaliya,Dinesh Kumar,Twinkle Soni,Sonal Sharma ,Zuber Saiyed,Ramesh Pandit,Janvi Raval,Jayshri Pethani,Umang Mishra,Nitesh |

| | | | |
|---|---|---|---|
| | | | Shah,Chaitanya Joshi,Madhvi Joshi |
| EPI_ISL_2391483 | N.H.L Municipal Medical College, Ahmedabad | Gujarat Biotechnology Research Centre | Nitin Savaliya,Dinesh Kumar,Twinkle Soni,Sonal Sharma ,Zuber Saiyed,Ramesh Pandit,Janvi Raval,Zarna Patel,Jayshri Pethani,Umang Mishra,Nitesh Shah,Chaitanya Joshi,Madhvi Joshi |
| EPI_ISL_2391484 | N.H.L Municipal Medical College, Ahmedabad | Gujarat Biotechnology Research Centre | Dinesh Kumar,Twinkle Soni,Sonal Sharma ,Zuber Saiyed,Ramesh Pandit,Janvi Raval,Zarna Patel,Nitin Savaliya,Jayshri Pethani,Umang Mishra,Nitesh Shah,Chaitanya Joshi,Madhvi Joshi |
| EPI_ISL_2391489 | GMERS, Government Medical College (GMC), Gotri | Gujarat Biotechnology Research Centre | Zarna Patel,Nitin Savaliya,Dinesh Kumar,Twinkle Soni,Sonal Sharma ,Zuber Saiyed,Ramesh Pandit,Janvi Raval,Bithika Duttaroy,Umang Mishra,Nitesh Shah,Chaitanya Joshi,Madhvi Joshi |
| EPI_ISL_2391490 | GMERS, Government Medical College (GMC), Gotri | Gujarat Biotechnology Research Centre | Nitin Savaliya,Dinesh Kumar,Twinkle Soni,Sonal Sharma ,Zuber Saiyed,Ramesh Pandit,Janvi Raval,Zarna Patel,Bithika Duttaroy,Umang Mishra,Nitesh Shah,Chaitanya Joshi,Madhvi Joshi |
| EPI_ISL_2391491 | GMERS, Government Medical College (GMC), Gotri | Gujarat Biotechnology Research Centre | Dinesh Kumar,Twinkle Soni,Sonal Sharma ,Zuber Saiyed,Ramesh Pandit,Janvi Raval,Zarna Patel,Nitin Savaliya,Bithika Duttaroy,Umang Mishra,Nitesh Shah,Chaitanya Joshi,Madhvi Joshi |
| EPI_ISL_2391494 | GMERS, Government Medical College (GMC), Gotri | Gujarat Biotechnology Research Centre | Zuber Saiyed,Ramesh Pandit,Janvi Raval,Zarna Patel,Nitin Savaliya,Dinesh Kumar,Twinkle Soni,Sonal Sharma ,Bithika Duttaroy,Umang Mishra,Nitesh Shah,Chaitanya Joshi,Madhvi Joshi |
| EPI_ISL_2391495 | GMERS, Government Medical College (GMC), Gotri | Gujarat Biotechnology Research Centre | Ramesh Pandit,Janvi Raval,Zarna Patel,Nitin Savaliya,Dinesh Kumar,Twinkle Soni,Sonal Sharma ,Zuber Saiyed,Bithika Duttaroy,Umang Mishra,Nitesh Shah,Chaitanya Joshi,Madhvi Joshi |
| EPI_ISL_2391496 | GMERS, Government Medical College (GMC), Gotri | Gujarat Biotechnology Research Centre | Janvi Raval,Zarna Patel,Nitin Savaliya,Dinesh Kumar,Twinkle Soni,Sonal Sharma ,Zuber Saiyed,Ramesh Pandit,Bithika Duttaroy,Umang Mishra,Nitesh Shah,Chaitanya Joshi,Madhvi Joshi |
| EPI_ISL_2391502 | GMERS, Government Medical College (GMC), Gotri | Gujarat Biotechnology Research Centre | Zuber Saiyed,Ramesh Pandit,Janvi Raval,Zarna Patel,Nitin Savaliya,Dinesh Kumar,Twinkle Soni,Sonal Sharma ,Bithika Duttaroy,Umang Mishra,Nitesh Shah,Chaitanya Joshi,Madhvi Joshi |
| EPI_ISL_2391503 | GMERS, Government Medical College (GMC), Gotri | Gujarat Biotechnology Research Centre | Ramesh Pandit,Janvi Raval,Zarna Patel,Nitin Savaliya,Dinesh Kumar,Twinkle Soni,Sonal Sharma ,Zuber Saiyed,Bithika Duttaroy,Umang Mishra,Nitesh Shah,Chaitanya Joshi,Madhvi Joshi |
| EPI_ISL_2391508, EPI_ISL_2391509, EPI_ISL_2391510, EPI_ISL_2391511 | Laboratorio Central de Epidemiología (LCE) | Instituto de Biotecnología de la UNAM | Consorcio Mexicano de Vigilancia Genómica (CoViGen-Mex). Authors (in alphabetical order): Julio Elias Alvarado-Yaah, Carlos F. Arias, Santiago Ávila-Ríos, Eduardo Becerril-Vargas, Víctor Hugo Borja-Aburto, Celia Boukadida, Cristóbal Cháidez-Quiróz, Juan Bautista Chale-Dzul, Ricardo Ciria Merce, Andreu Comas-García, Célida Duque Molina, Julissa Enciso-Ibarra, José Antonio Enciso-Moreno, Gloria Elena Espinosa-Ayala, Fernando Fontove-Herrera, Daniel Fregoso-Rueda, Victor Eduardo Garcia-Arias, Alejandra Garcia-Gasca, Bruno Gomez-Gil, Jean Pierre González, Irvin González-López, Concepción Grajales-Muñiz, Ricardo Grande, Rosa María Gutierrez Rios, Alejandra Hernández-Terán, Alfredo Herrera-Estrella, Carla Ivón Herrera-Najera, Pavel Isa, Daniel Lira Morales, Susana Lopez, Antonio Loza Román, Brenda Irasema Maldonado-Meza, Bernardo Martínez-Miguel, José Arturo Martínez-Orozco, Célida Martínez- Rodríguez, Margarita Matías-Florentino, Fidencio Mejía-Nepomuceno, María Guadalupe de Jesús Mireles-Rivera, Gloria María Molina-Salinas, Hector Montoya-Fuentes, , Mario Mújica-Sánchez, José Esteban Muñoz-Medina, José de Jesús Nuñez-Contreras, Alicia Ocaña-Mondragón, Luis Alberto Ochoa-Carrera, Hector Esteban Paz-Juárez, Francisco Pulido, Helen Haydee Fernanda Ramírez-Plascencia, Jorge Salas-Hernández, Angel Gustavo Salas-Lais, Alejandro Sanchez-Flores, Clara Esperanza Santacruz-Tinoco, María Guadalupe Santiago-Mauricio, Selene Zárate, Nelly Sélem-Mojica, Blanca Taboada, Gloria Vazquez, Joel Armando Vázquez-Pérez. |
| EPI_ISL_2391512 | Unidad de Investigación Médica de Yucatán (UIMY) | Instituto de Biotecnología de la UNAM | Consorcio Mexicano de Vigilancia Genómica (CoViGen-Mex). Authors (in alphabetical order): Julio Elias Alvarado-Yaah, Carlos F. Arias, Santiago Ávila-Ríos, Eduardo Becerril-Vargas, Víctor Hugo Borja-Aburto, Celia Boukadida, Cristóbal Cháidez-Quiróz, Juan Bautista Chale-Dzul, Ricardo Ciria Merce, Andreu Comas-García, Célida Duque Molina, Julissa Enciso-Ibarra, José Antonio Enciso-Moreno, Gloria Elena Espinosa-Ayala, Fernando Fontove-Herrera, Daniel Fregoso-Rueda, Victor Eduardo Garcia-Arias, Alejandra Garcia-Gasca, Bruno Gomez-Gil, Jean Pierre González, Irvin González-López, Concepción Grajales-Muñiz, Ricardo Grande, Rosa María Gutierrez Rios, Alejandra Hernández-Terán, Alfredo Herrera-Estrella, Carla Ivón Herrera-Najera, Pavel Isa, Daniel Lira Morales, Susana Lopez, Antonio Loza Román, Brenda Irasema Maldonado-Meza, Bernardo Martínez-Miguel, José Arturo Martínez-Orozco, Célida Martínez- Rodríguez, Margarita Matías-Florentino, Fidencio Mejía-Nepomuceno, María Guadalupe de Jesús Mireles-Rivera, Gloria María Molina-Salinas, Hector Montoya-Fuentes, , Mario Mújica-Sánchez, José Esteban Muñoz-Medina, José de Jesús Nuñez-Contreras, Alicia Ocaña-Mondragón, Luis Alberto Ochoa-Carrera, Hector Esteban Paz-Juárez, Francisco Pulido, Helen Haydee Fernanda Ramírez-Plascencia, Jorge Salas-Hernández, Angel Gustavo Salas-Lais, Alejandro Sanchez-Flores, Clara Esperanza Santacruz-Tinoco, María Guadalupe Santiago-Mauricio, Selene Zárate, Nelly Sélem-Mojica, Blanca Taboada, Gloria Vazquez, Joel Armando Vázquez-Pérez. |
| EPI_ISL_2391513, EPI_ISL_2391514, EPI_ISL_2391515, EPI_ISL_2391516, EPI_ISL_2391517, EPI_ISL_2391518, EPI_ISL_2391519, EPI_ISL_2391520, EPI_ISL_2391521, EPI_ISL_2391522, EPI_ISL_2391524, EPI_ISL_2391525, EPI_ISL_2391526, EPI_ISL_2391527, EPI_ISL_2391528, EPI_ISL_2391529, EPI_ISL_2391530, EPI_ISL_2391531, EPI_ISL_2391534, EPI_ISL_2391535, EPI_ISL_2391538, EPI_ISL_2391539, EPI_ISL_2391541, EPI_ISL_2391542, EPI_ISL_2391543, EPI_ISL_2391545, EPI_ISL_2391546, EPI_ISL_2391547, EPI_ISL_2391548, EPI_ISL_2391549, EPI_ISL_2391550, EPI_ISL_2391551, EPI_ISL_2391552, EPI_ISL_2391553, EPI_ISL_2391554, EPI_ISL_2391555, EPI_ISL_2391556, EPI_ISL_2391558, EPI_ISL_2391559, EPI_ISL_2391560, EPI_ISL_2391561, EPI_ISL_2391562, EPI_ISL_2391563, EPI_ISL_2391564 | | | |
| see above | Laboratorio Central de Epidemiología (LCE) | Instituto de Biotecnología de la UNAM | Consorcio Mexicano de Vigilancia Genómica (CoViGen-Mex). Authors (in alphabetical order): Julio Elias Alvarado-Yaah, Carlos F. Arias, Santiago Ávila-Ríos, Eduardo Becerril-Vargas, Víctor Hugo Borja-Aburto, Celia Boukadida, Cristóbal Cháidez-Quiróz, Juan Bautista Chale-Dzul, Ricardo Ciria Merce, Andreu Comas-García, Célida Duque Molina, Julissa Enciso-Ibarra, José Antonio Enciso-Moreno, Gloria Elena Espinosa-Ayala, Fernando Fontove-Herrera, Daniel Fregoso-Rueda, Victor Eduardo Garcia-Arias, Alejandra Garcia-Gasca, Bruno Gomez-Gil, Jean Pierre González, Irvin González-López, Concepción Grajales-Muñiz, Ricardo Grande, Rosa María Gutierrez Rios, Alejandra Hernández-Terán, Alfredo Herrera-Estrella, Carla Ivón Herrera-Najera, Pavel Isa, Daniel Lira Morales, Susana Lopez, Antonio Loza Román, Brenda Irasema Maldonado-Meza, Bernardo Martínez-Miguel, José Arturo Martínez-Orozco, Célida Martínez- Rodríguez, Margarita Matías-Florentino, Fidencio Mejía-Nepomuceno, María Guadalupe de Jesús Mireles-Rivera, Gloria María Molina-Salinas, Hector Montoya-Fuentes, , Mario Mújica-Sánchez, José Esteban Muñoz-Medina, José de Jesús Nuñez-Contreras, Alicia Ocaña-Mondragón, Luis Alberto Ochoa-Carrera, Hector Esteban Paz-Juárez, Francisco Pulido, Helen Haydee Fernanda Ramírez-Plascencia, Jorge Salas-Hernández, Angel Gustavo Salas-Lais, Alejandro Sanchez-Flores, Clara Esperanza Santacruz-Tinoco, María Guadalupe Santiago-Mauricio, Selene Zárate, Nelly Sélem-Mojica, Blanca Taboada, Gloria Vazquez, Joel Armando Vázquez-Pérez. |
| EPI_ISL_2391565 | Centro de Investigación Biomédica de Occidente (CIBO) | Instituto de Biotecnología de la UNAM | Consorcio Mexicano de Vigilancia Genómica (CoViGen-Mex). Authors (in alphabetical order): Julio Elias Alvarado-Yaah, Carlos F. Arias, Santiago Ávila-Ríos, Eduardo Becerril-Vargas, Víctor Hugo Borja-Aburto, Celia Boukadida, Cristóbal Cháidez-Quiróz, Juan Bautista Chale-Dzul, Ricardo Ciria Merce, Andreu Comas-García, Célida Duque Molina, Julissa Enciso-Ibarra, José Antonio Enciso-Moreno, Gloria Elena Espinosa-Ayala, Fernando Fontove-Herrera, Daniel Fregoso-Rueda, Victor Eduardo Garcia-Arias, Alejandra Garcia-Gasca, Bruno Gomez-Gil, Jean Pierre González, Irvin González-López, Concepción Grajales-Muñiz, Ricardo Grande, Rosa María Gutierrez Rios, Alejandra Hernández-Terán, Alfredo Herrera-Estrella, Carla Ivón Herrera-Najera, Pavel Isa, Daniel Lira Morales, Susana Lopez, Antonio Loza Román, Brenda Irasema Maldonado-Meza, Bernardo Martínez-Miguel, José Arturo Martínez-Orozco, Célida Martínez- Rodríguez, Margarita Matías-Florentino, Fidencio Mejía-Nepomuceno, María Guadalupe de Jesús Mireles-Rivera, Gloria María Molina-Salinas, Hector Montoya-Fuentes, , Mario Mújica-Sánchez, José Esteban Muñoz-Medina, José de Jesús Nuñez-Contreras, Alicia Ocaña-Mondragón, Luis Alberto Ochoa-Carrera, Hector Esteban Paz-Juárez, Francisco Pulido, Helen Haydee Fernanda Ramírez-Plascencia, Jorge Salas-Hernández, Angel Gustavo Salas-Lais, Alejandro Sanchez-Flores, Clara Esperanza Santacruz-Tinoco, María Guadalupe Santiago-Mauricio, Selene Zárate, Nelly Sélem-Mojica, Blanca Taboada, Gloria Vazquez, Joel Armando Vázquez-Pérez. |
| EPI_ISL_2391566, EPI_ISL_2391567, EPI_ISL_2391569, EPI_ISL_2391570, EPI_ISL_2391571, EPI_ISL_2391572, EPI_ISL_2391573, EPI_ISL_2391574, EPI_ISL_2391576, EPI_ISL_2391577, EPI_ISL_2391579, EPI_ISL_2391582, EPI_ISL_2391583, EPI_ISL_2391584, EPI_ISL_2391585 | | | |
| see above | Laboratorio Central de Epidemiología (LCE) | Instituto de Biotecnología de la UNAM | Consorcio Mexicano de Vigilancia Genómica (CoViGen-Mex). Authors (in alphabetical order): Julio Elias Alvarado-Yaah, Carlos F. Arias, Santiago Ávila-Ríos, Eduardo Becerril-Vargas, Víctor Hugo Borja-Aburto, Celia Boukadida, Cristóbal Cháidez-Quiróz, Juan Bautista Chale-Dzul, Ricardo Ciria Merce, Andreu Comas-García, Célida Duque Molina, Julissa Enciso-Ibarra, José Antonio Enciso-Moreno, Gloria Elena Espinosa-Ayala, Fernando Fontove-Herrera, Daniel Fregoso-Rueda, Victor Eduardo Garcia-Arias, Alejandra Garcia-Gasca, Bruno Gomez-Gil, Jean Pierre González, Irvin González-López, Concepción Grajales-Muñiz, Ricardo Grande, Rosa María Gutierrez Rios, Alejandra Hernández-Terán, Alfredo Herrera-Estrella, Carla Ivón Herrera-Najera, Pavel Isa, Daniel Lira Morales, Susana Lopez, Antonio Loza Román, Brenda Irasema Maldonado-Meza, Bernardo Martínez-Miguel, José Arturo Martínez-Orozco, Célida Martínez- Rodríguez, Margarita Matías-Florentino, Fidencio Mejía-Nepomuceno, María Guadalupe de Jesús Mireles-Rivera, Gloria María Molina-Salinas, Hector Montoya-Fuentes, , Mario Mújica-Sánchez, José Esteban Muñoz-Medina, José de Jesús |

| | | | |
|---|---|---|---|
| | | | Nuñez-Contreras, Alicia Ocaña-Mondragón, Luis Alberto Ochoa-Carrera, Hector Esteban Paz-Juárez, Francisco Pulido, Helen Haydee Fernanda Ramírez-Plascencia, Jorge Salas-Hernández, Angel Gustavo Salas-Lais, Alejandro Sanchez-Flores, Clara Esperanza Santacruz-Tinoco, María Guadalupe Santiago-Mauricio, Selene Zárate, Nelly Sélem-Mojica, Blanca Taboada, Gloria Vazquez, Joel Armando Vázquez-Pérez. |
| EPI_ISL_2391588 | Centro de Investigación Biomédica del Noreste (CIBIN) | Instituto de Biotecnología de la UNAM | Consorcio Mexicano de Vigilancia Genómica (CoViGen-Mex). Authors (in alphabetical order): Julio Elias Alvarado-Yaah, Carlos F. Arias, Santiago Ávila-Ríos, Eduardo Becerril-Vargas, Víctor Hugo Borja-Aburto, Celia Boukadida, Cristóbal Cháidez-Quiróz, Juan Bautista Chale-Dzul, Ricardo Ciria Merce, Andreu Comas-García, Célida Duque Molina, Julissa Enciso-Ibarra, José Antonio Enciso-Moreno, Gloria Elena Espinosa-Ayala, Fernando Fontove-Herrera, Daniel Fregoso-Rueda, Victor Eduardo Garcia-Arias, Alejandra Garcia-Gasca, Bruno Gomez-Gil, Jean Pierre González, Irvin González-López, Concepción Grajales-Muñiz, Ricardo Grande, Rosa María Gutierrez Rios, Alejandra Hernández-Terán, Alfredo Herrera-Estrella, Carla Ivón Herrera-Najera, Pavel Isa, Daniel Lira Morales, Susana Lopez, Antonio Loza Román, Brenda Irasema Maldonado-Meza, Bernardo Martínez-Miguel, José Arturo Martínez-Orozco, Célida Martínez- Rodríguez, Margarita Matías-Florentino, Fidencio Mejía-Nepomuceno, María Guadalupe de Jesús Mireles-Rivera, Gloria María Molina-Salinas, Hector Montoya-Fuentes, , Mario Mújica-Sánchez, José Esteban Muñoz-Medina, José de Jesús Nuñez-Contreras, Alicia Ocaña-Mondragón, Luis Alberto Ochoa-Carrera, Hector Esteban Paz-Juárez, Francisco Pulido, Helen Haydee Fernanda Ramírez-Plascencia, Jorge Salas-Hernández, Angel Gustavo Salas-Lais, Alejandro Sanchez-Flores, Clara Esperanza Santacruz-Tinoco, María Guadalupe Santiago-Mauricio, Selene Zárate, Nelly Sélem-Mojica, Blanca Taboada, Gloria Vazquez, Joel Armando Vázquez-Pérez. |
| EPI_ISL_2391590, EPI_ISL_2391591, EPI_ISL_2391592, EPI_ISL_2391593, EPI_ISL_2391595, EPI_ISL_2391596, EPI_ISL_2391597, EPI_ISL_2391599, EPI_ISL_2391600, EPI_ISL_2391601, EPI_ISL_2391602, EPI_ISL_2391603, EPI_ISL_2391604, EPI_ISL_2391605, EPI_ISL_2391606, EPI_ISL_2391608, EPI_ISL_2391609, EPI_ISL_2391610, EPI_ISL_2391611, EPI_ISL_2391612, EPI_ISL_2391613, EPI_ISL_2391614, EPI_ISL_2391615, EPI_ISL_2391616, EPI_ISL_2391618, EPI_ISL_2391620, EPI_ISL_2391621, EPI_ISL_2391622, EPI_ISL_2391624, EPI_ISL_2391627, EPI_ISL_2391629, EPI_ISL_2391630, EPI_ISL_2391631, EPI_ISL_2391633, EPI_ISL_2391634, EPI_ISL_2391635, EPI_ISL_2391636, EPI_ISL_2391637, EPI_ISL_2391638, EPI_ISL_2391640, EPI_ISL_2391641, EPI_ISL_2391642, EPI_ISL_2391644, EPI_ISL_2391645, EPI_ISL_2391646, EPI_ISL_2391648, EPI_ISL_2391649, EPI_ISL_2391652 | | | |
| see above | Laboratorio Central de Epidemiología (LCE) | Instituto de Biotecnología de la UNAM | Consorcio Mexicano de Vigilancia Genómica (CoViGen-Mex). Authors (in alphabetical order): Julio Elias Alvarado-Yaah, Carlos F. Arias, Santiago Ávila-Ríos, Eduardo Becerril-Vargas, Víctor Hugo Borja-Aburto, Celia Boukadida, Cristóbal Cháidez-Quiróz, Juan Bautista Chale-Dzul, Ricardo Ciria Merce, Andreu Comas-García, Célida Duque Molina, Julissa Enciso-Ibarra, José Antonio Enciso-Moreno, Gloria Elena Espinosa-Ayala, Fernando Fontove-Herrera, Daniel Fregoso-Rueda, Victor Eduardo Garcia-Arias, Alejandra Garcia-Gasca, Bruno Gomez-Gil, Jean Pierre González, Irvin González-López, Concepción Grajales-Muñiz, Ricardo Grande, Rosa María Gutierrez Rios, Alejandra Hernández-Terán, Alfredo Herrera-Estrella, Carla Ivón Herrera-Najera, Pavel Isa, Daniel Lira Morales, Susana Lopez, Antonio Loza Román, Brenda Irasema Maldonado-Meza, Bernardo Martínez-Miguel, José Arturo Martínez-Orozco, Célida Martínez- Rodríguez, Margarita Matías-Florentino, Fidencio Mejía-Nepomuceno, María Guadalupe de Jesús Mireles-Rivera, Gloria María Molina-Salinas, Hector Montoya-Fuentes, , Mario Mújica-Sánchez, José Esteban Muñoz-Medina, José de Jesús Nuñez-Contreras, Alicia Ocaña-Mondragón, Luis Alberto Ochoa-Carrera, Hector Esteban Paz-Juárez, Francisco Pulido, Helen Haydee Fernanda Ramírez-Plascencia, Jorge Salas-Hernández, Angel Gustavo Salas-Lais, Alejandro Sanchez-Flores, Clara Esperanza Santacruz-Tinoco, María Guadalupe Santiago-Mauricio, Selene Zárate, Nelly Sélem-Mojica, Blanca Taboada, Gloria Vazquez, Joel Armando Vázquez-Pérez. |
| EPI_ISL_2397159, EPI_ISL_2397160, EPI_ISL_2397161, EPI_ISL_2397162, EPI_ISL_2397163, EPI_ISL_2397164, EPI_ISL_2397165 | Klinisch Laboratorium ZNA | Klinisch Laboratorium ZNA | Verstrepen et al. |
| EPI_ISL_2400428 | OCME Office Of Chief Medical Examiner | New York City Public Health Laboratory | Jade Wang, et al. |
| EPI_ISL_2400443, EPI_ISL_2400444, EPI_ISL_2400449, EPI_ISL_2400450, EPI_ISL_2400451, EPI_ISL_2400452, EPI_ISL_2400455, EPI_ISL_2400457, EPI_ISL_2400459 | Molecular Biology Laboratory, Westchester Medical Center | Genomics Core Laboratory, New York Medical College | Weihua Huang, Utsav Pandey, Jian Zhuge, Changhong Yin |
| EPI_ISL_2400493, EPI_ISL_2400497 | Azarian Lab at Burnett School of Biomedical Sciences | Azarian Lab at Burnett School of Biomedical Sciences | Catherine Johnston, Eleonora Cella, Taj Azarian |
| EPI_ISL_2401933, EPI_ISL_2401934, EPI_ISL_2401935, EPI_ISL_2401936, EPI_ISL_2401937, EPI_ISL_2401938, EPI_ISL_2401939, EPI_ISL_2401940, EPI_ISL_2401942, EPI_ISL_2401943, EPI_ISL_2401944, EPI_ISL_2401945, EPI_ISL_2401946, EPI_ISL_2401947, EPI_ISL_2401948, EPI_ISL_2401949, EPI_ISL_2401951, EPI_ISL_2401952, EPI_ISL_2401953, EPI_ISL_2401954, EPI_ISL_2401955, EPI_ISL_2401956, EPI_ISL_2401958, EPI_ISL_2401959, EPI_ISL_2401960, EPI_ISL_2401963, EPI_ISL_2401964, EPI_ISL_2401965, EPI_ISL_2401966, EPI_ISL_2401967, EPI_ISL_2401968, EPI_ISL_2401971, EPI_ISL_2401973, EPI_ISL_2401974, EPI_ISL_2401977, EPI_ISL_2401978, EPI_ISL_2401979, EPI_ISL_2401980, EPI_ISL_2401981, EPI_ISL_2401982, EPI_ISL_2401984, EPI_ISL_2401987, EPI_ISL_2401988, EPI_ISL_2401989, EPI_ISL_2401990, EPI_ISL_2401991, EPI_ISL_2401992, EPI_ISL_2401993, EPI_ISL_2401994, EPI_ISL_2401995, EPI_ISL_2401996, EPI_ISL_2401997, EPI_ISL_2401998, EPI_ISL_2401999, EPI_ISL_2402002, EPI_ISL_2402003, EPI_ISL_2402004, EPI_ISL_2402005, EPI_ISL_2402006, EPI_ISL_2402008, EPI_ISL_2402009, EPI_ISL_2402011, EPI_ISL_2402012, EPI_ISL_2402014 | | | |
| see above | Centro de Investigación Biomédica del Noreste (CIBIN) | Unidad de Genomica Avanzada | Consorcio Mexicano de Vigilancia Genomica (CoViGen-Mex). Authors (in alphabetical order): Julio Elias Alvarado-Yaah, Carlos F. Arias, Santiago Avila-Rios, Victor Hugo Borja-Aburto, Celia Boukadida, Juan Bautista Chale-Dzul , Jose Antonio Enciso-Moreno, Gloria Elena Espinoza-Ayala, Fernando Fontove-Herrera, Concepcion Grajales-Muniz, Ricardo Grande, Alfredo Herrera-Estrella, Carla Ivon Herrera-Najera, Pavel Isa, Brenda Irasema Maldonado-Meza, Bernardo Martinez-Miguel, Margarita Matias-Florentino, Maria Guadalupe de Jesus Mireles-Rivera, Gloria Maria Molina-Salinas, Hector Montoya-Fuentes, Jose Esteban Munoz-Medina, Jose de Jesus Nunez-Contreras, Alicia Ocana-Mondragon, Luis Alberto Ochoa-Carrera, Hector Esteban Paz-Juarez, Francisco Pulido, Helen Haydee Fernanda Ramirez-Plascencia, Angel Gustavo Salas-Lais, Jorge Ivan Salinal-Nevarez, Alejandro Sanchez-Flores, Clara Esperanza Santacruz-Tinoco, Maria Guadalupe Santiago-Mauricio, Nelly Selem-Mojica, Blanca Taboada, Gloria Vazquez |
| EPI_ISL_2402018, EPI_ISL_2402019, EPI_ISL_2402021, EPI_ISL_2402023, EPI_ISL_2402024, EPI_ISL_2402025, EPI_ISL_2402026, EPI_ISL_2402027, EPI_ISL_2402028, EPI_ISL_2402033, EPI_ISL_2402036, EPI_ISL_2402037, EPI_ISL_2402039, EPI_ISL_2402040, EPI_ISL_2402041, EPI_ISL_2402042, EPI_ISL_2402043, EPI_ISL_2402044, EPI_ISL_2402045, EPI_ISL_2402046, EPI_ISL_2402048, EPI_ISL_2402049, EPI_ISL_2402051, EPI_ISL_2402054, EPI_ISL_2402055, EPI_ISL_2402056, EPI_ISL_2402057, EPI_ISL_2402058, EPI_ISL_2402060, EPI_ISL_2402061, EPI_ISL_2402062, EPI_ISL_2402063, EPI_ISL_2402064, EPI_ISL_2402065 | | | |
| see above | Unidad de Investigación Médica de Yucatán (UIMY) | Unidad de Genomica Avanzada | Consorcio Mexicano de Vigilancia Genomica (CoViGen-Mex). Authors (in alphabetical order): Julio Elias Alvarado-Yaah, Carlos F. Arias, Santiago Avila-Rios, Victor Hugo Borja-Aburto, Celia Boukadida, Juan Bautista Chale-Dzul , Jose Antonio Enciso-Moreno, Gloria Elena Espinoza-Ayala, Fernando Fontove-Herrera, Concepcion Grajales-Muniz, Ricardo Grande, Alfredo Herrera-Estrella, Carla Ivon Herrera-Najera, Pavel Isa, Brenda Irasema Maldonado-Meza, Bernardo Martinez-Miguel, Margarita Matias-Florentino, Maria Guadalupe de Jesus Mireles-Rivera, Gloria Maria Molina-Salinas, Hector Montoya-Fuentes, Jose Esteban Munoz-Medina, Jose de Jesus Nunez-Contreras, Alicia Ocana-Mondragon, Luis Alberto Ochoa-Carrera, Hector Esteban Paz-Juarez, Francisco Pulido, Helen Haydee Fernanda Ramirez-Plascencia, Angel Gustavo Salas-Lais, Jorge Ivan Salinal-Nevarez, Alejandro Sanchez-Flores, Clara Esperanza Santacruz-Tinoco, Maria Guadalupe Santiago-Mauricio, Nelly Selem-Mojica, Blanca Taboada, Gloria Vazquez |
| EPI_ISL_2402066, EPI_ISL_2402069, EPI_ISL_2402070, EPI_ISL_2402072, EPI_ISL_2402074, EPI_ISL_2402075, EPI_ISL_2402077, EPI_ISL_2402079, EPI_ISL_2402080, EPI_ISL_2402082, EPI_ISL_2402083, EPI_ISL_2402084, EPI_ISL_2402085, EPI_ISL_2402086, EPI_ISL_2402087, EPI_ISL_2402089, EPI_ISL_2402090, EPI_ISL_2402091, EPI_ISL_2402093, EPI_ISL_2402094, EPI_ISL_2402097, EPI_ISL_2402098, EPI_ISL_2402099, EPI_ISL_2402100, EPI_ISL_2402101, EPI_ISL_2402102, EPI_ISL_2402103, EPI_ISL_2402104, EPI_ISL_2402105, EPI_ISL_2402107, EPI_ISL_2402108, EPI_ISL_2402109, EPI_ISL_2402111, EPI_ISL_2402112, EPI_ISL_2402113, EPI_ISL_2402116, EPI_ISL_2402118, EPI_ISL_2402121, EPI_ISL_2402122, EPI_ISL_2402123, EPI_ISL_2402124, EPI_ISL_2402127, EPI_ISL_2402128, EPI_ISL_2402129, EPI_ISL_2402130, EPI_ISL_2402131 | | | |
| see above | Unidad de Investigación Biomédica de Zacatecas (UIBZ) | Unidad de Genomica Avanzada | Consorcio Mexicano de Vigilancia Genomica (CoViGen-Mex). Authors (in alphabetical order): Julio Elias Alvarado-Yaah, Carlos F. Arias, Santiago Avila-Rios, Victor Hugo Borja-Aburto, Celia Boukadida, Juan Bautista Chale-Dzul , Jose Antonio Enciso-Moreno, Gloria Elena Espinoza-Ayala, Fernando Fontove-Herrera, Concepcion Grajales-Muniz, Ricardo Grande, Alfredo Herrera-Estrella, Carla Ivon Herrera-Najera, Pavel Isa, Brenda Irasema Maldonado-Meza, Bernardo Martinez-Miguel, Margarita Matias-Florentino, Maria Guadalupe de Jesus Mireles-Rivera, Gloria Maria Molina-Salinas, Hector Montoya-Fuentes, Jose Esteban Munoz-Medina, Jose de Jesus Nunez-Contreras, Alicia Ocana-Mondragon, Luis Alberto Ochoa-Carrera, Hector Esteban Paz-Juarez, Francisco Pulido, Helen Haydee Fernanda Ramirez-Plascencia, Angel Gustavo Salas-Lais, Jorge Ivan Salinal-Nevarez, Alejandro Sanchez-Flores, Clara Esperanza Santacruz-Tinoco, Maria Guadalupe Santiago-Mauricio, Nelly Selem-Mojica, Blanca Taboada, Gloria Vazquez |
| EPI_ISL_2402132, EPI_ISL_2402133, EPI_ISL_2402138, EPI_ISL_2402139, EPI_ISL_2402140, EPI_ISL_2402142, EPI_ISL_2402144, EPI_ISL_2402145, EPI_ISL_2402147, EPI_ISL_2402148, EPI_ISL_2402149, EPI_ISL_2402150, EPI_ISL_2402151, EPI_ISL_2402152, EPI_ISL_2402153, EPI_ISL_2402154, EPI_ISL_2402155, EPI_ISL_2402156, EPI_ISL_2402158, EPI_ISL_2402159, EPI_ISL_2402160, EPI_ISL_2402161, EPI_ISL_2402162, EPI_ISL_2402163, EPI_ISL_2402164, EPI_ISL_2402165, EPI_ISL_2402166, EPI_ISL_2402167, EPI_ISL_2402168, EPI_ISL_2402169, EPI_ISL_2402171, EPI_ISL_2402172, EPI_ISL_2402174, EPI_ISL_2402175, EPI_ISL_2402178 | | | |
| see above | Centro de Investigación Biomédica de Occidente (CIBO) | Unidad de Genomica Avanzada | Consorcio Mexicano de Vigilancia Genomica (CoViGen-Mex). Authors (in alphabetical order): Julio Elias Alvarado-Yaah, Carlos F. Arias, Santiago Avila-Rios, Victor Hugo Borja-Aburto, Celia Boukadida, Juan Bautista Chale-Dzul , Jose Antonio Enciso-Moreno, Gloria Elena Espinoza-Ayala, Fernando Fontove-Herrera, Concepcion Grajales-Muniz, Ricardo Grande, Alfredo Herrera-Estrella, Carla Ivon Herrera-Najera, Pavel Isa, Brenda Irasema Maldonado-Meza, Bernardo Martinez-Miguel, Margarita Matias-Florentino, Maria Guadalupe de Jesus Mireles-Rivera, Gloria Maria Molina-Salinas, Hector Montoya-Fuentes, Jose Esteban Munoz-Medina, Jose de Jesus Nunez-Contreras, Alicia Ocana-Mondragon, Luis Alberto Ochoa-Carrera, Hector Esteban Paz-Juarez, Francisco Pulido, Helen Haydee Fernanda Ramirez-Plascencia, Angel Gustavo Salas-Lais, Jorge Ivan Salinal-Nevarez, Alejandro Sanchez-Flores, Clara Esperanza Santacruz-Tinoco, Maria Guadalupe Santiago-Mauricio, Nelly Selem-Mojica, Blanca Taboada, Gloria Vazquez |

| | | | |
|---|---|---|---|
| EPI_ISL_2402179, EPI_ISL_2402180, EPI_ISL_2402182, EPI_ISL_2402184, EPI_ISL_2402185, EPI_ISL_2402188, EPI_ISL_2402190, EPI_ISL_2402193, EPI_ISL_2402194, EPI_ISL_2402195, EPI_ISL_2402196, EPI_ISL_2402197, EPI_ISL_2402198, EPI_ISL_2402199, EPI_ISL_2402200, EPI_ISL_2402201, EPI_ISL_2402202, EPI_ISL_2402203, EPI_ISL_2402204, EPI_ISL_2402206, EPI_ISL_2402207, EPI_ISL_2402208, EPI_ISL_2402209, EPI_ISL_2402211, EPI_ISL_2402214, EPI_ISL_2402216, EPI_ISL_2402217, EPI_ISL_2402218, EPI_ISL_2402219, EPI_ISL_2402220, EPI_ISL_2402224, EPI_ISL_2402225, EPI_ISL_2402226, EPI_ISL_2402227 | | | |
| see above | Laboratorio Central de Epidemiologí-a (LCE) | Unidad de Genomica Avanzada | Consorcio Mexicano de Vigilancia Genomica (CoViGen-Mex). Authors (in alphabetical order): Julio Elias Alvarado-Yaah, Carlos F. Arias, Santiago Avila-Rios, Victor Hugo Borja-Aburto, Celia Boukadida, Juan Bautista Chale-Dzul , Jose Antonio Enciso-Moreno, Gloria Elena Espinoza-Ayala, Fernando Fontove-Herrera, Concepcion Grajales-Muniz, Ricardo Grande, Alfredo Herrera-Estrella, Carla Ivon Herrera-Najera, Pavel Isa, Brenda Irasema Maldonado-Meza, Bernardo Martinez-Miguel, Margarita Matias-Florentino, Maria Guadalupe de Jesus Mireles-Rivera, Gloria Maria Molina-Salinas, Hector Montoya-Fuentes, Jose Esteban Munoz-Medina, Jose de Jesus Nunez-Contreras, Alicia Ocana-Mondragon, Luis Alberto Ochoa-Carrera, Hector Esteban Paz-Juarez, Francisco Pulido, Helen Haydee Fernanda Ramirez-Plascencia, Angel Gustavo Salas-Lais, Jorge Ivan Salinal-Nevarez, Alejandro Sanchez-Flores, Clara Esperanza Santacruz-Tinoco, Maria Guadalupe Santiago-Mauricio, Nelly Selem-Mojica, Blanca Taboada, Gloria Vazquez |
| EPI_ISL_2402667, EPI_ISL_2402668, EPI_ISL_2402669, EPI_ISL_2402670, EPI_ISL_2402769, EPI_ISL_2402770, EPI_ISL_2402772, EPI_ISL_2402773, EPI_ISL_2402774, EPI_ISL_2402775, EPI_ISL_2402776, EPI_ISL_2402777, EPI_ISL_2402778, EPI_ISL_2402779, EPI_ISL_2402780, EPI_ISL_2402781, EPI_ISL_2402782, EPI_ISL_2402783, EPI_ISL_2402784, EPI_ISL_2402786, EPI_ISL_2402787, EPI_ISL_2402788, EPI_ISL_2402789, EPI_ISL_2402790, EPI_ISL_2402791, EPI_ISL_2402792, EPI_ISL_2402793, EPI_ISL_2402794, EPI_ISL_2402795, EPI_ISL_2402796, EPI_ISL_2402797, EPI_ISL_2402798, EPI_ISL_2402799, EPI_ISL_2402800, EPI_ISL_2402801, EPI_ISL_2402802, EPI_ISL_2402803, EPI_ISL_2402804, EPI_ISL_2402805, EPI_ISL_2402806, EPI_ISL_2402807, EPI_ISL_2402808, EPI_ISL_2402809 | | | |
| see above | Gyncentrum | Gyncentrum | Emilia Morawiec, Pawe Czerwiski, Adam Pudeko, Aleksandra Skubis-Sikora, Magdalena Samul, Agnieszka Polak, Celina Kruszniewska-Rajs, Tomasz Wsik, Robert Wojtyczka, Jolanta Bartosiewicz- Wsik, Maria Maklasiska-Majdanik |
| EPI_ISL_2405118 | N.H.L Municipal Medical College, Ahmedabad | Gujarat Biotechnology Research Centre | Janvi Raval,Zarna Patel,Nitin Savaliya,Dinesh Kumar,Twinkle Soni,Sonal Sharma ,Zuber Saiyed,Ramesh Pandit,Jayshri Pethani,Umang Mishra,Nitesh Shah,Chaitanya Joshi,Madhvi Joshi |
| EPI_ISL_2405125 | GMERS Government Medical College (GMC), Gotri | Gujarat Biotechnology Research Centre | Ramesh Pandit,Janvi Raval,Zarna Patel,Nitin Savaliya,Dinesh Kumar,Twinkle Soni,Sonal Sharma ,Zuber Saiyed,Bithika Duttaroy,Umang Mishra,Nitesh Shah,Chaitanya Joshi,Madhvi Joshi |
| EPI_ISL_2405126 | GMERS Government Medical College (GMC), Gotri | Gujarat Biotechnology Research Centre | Janvi Raval,Zarna Patel,Nitin Savaliya,Dinesh Kumar,Twinkle Soni,Sonal Sharma ,Zuber Saiyed,Ramesh Pandit,Bithika Duttaroy,Umang Mishra,Nitesh Shah,Chaitanya Joshi,Madhvi Joshi |
| EPI_ISL_2405127 | BJ Medical College and Civil Hospital, Ahmedabad | Gujarat Biotechnology Research Centre | Zarna Patel,Nitin Savaliya,Dinesh Kumar,Twinkle Soni,Sonal Sharma ,Zuber Saiyed,Ramesh Pandit,Janvi Raval,Pranay Shah,Umang Mishra,Nitesh Shah,Chaitanya Joshi,Madhvi Joshi |
| EPI_ISL_2405128 | BJ Medical College and Civil Hospital, Ahmedabad | Gujarat Biotechnology Research Centre | Nitin Savaliya,Dinesh Kumar,Twinkle Soni,Sonal Sharma ,Zuber Saiyed,Ramesh Pandit,Janvi Raval,Zarna Patel,Pranay Shah,Umang Mishra,Nitesh Shah,Chaitanya Joshi,Madhvi Joshi |
| EPI_ISL_2405129 | BJ Medical College and Civil Hospital, Ahmedabad | Gujarat Biotechnology Research Centre | Dinesh Kumar,Twinkle Soni,Sonal Sharma ,Zuber Saiyed,Ramesh Pandit,Janvi Raval,Zarna Patel,Nitin Savaliya,Pranay Shah,Umang Mishra,Nitesh Shah,Chaitanya Joshi,Madhvi Joshi |
| EPI_ISL_2405130 | BJ Medical College and Civil Hospital, Ahmedabad | Gujarat Biotechnology Research Centre | Twinkle Soni,Sonal Sharma ,Zuber Saiyed,Ramesh Pandit,Janvi Raval,Zarna Patel,Nitin Savaliya,Dinesh Kumar,Pranay Shah,Umang Mishra,Nitesh Shah,Chaitanya Joshi,Madhvi Joshi |
| EPI_ISL_2405131 | BJ Medical College and Civil Hospital, Ahmedabad | Gujarat Biotechnology Research Centre | Sonal Sharma ,Zuber Saiyed,Ramesh Pandit,Janvi Raval,Zarna Patel,Nitin Savaliya,Dinesh Kumar,Twinkle Soni,Pranay Shah,Umang Mishra,Nitesh Shah,Chaitanya Joshi,Madhvi Joshi |
| EPI_ISL_2405132 | BJ Medical College and Civil Hospital, Ahmedabad | Gujarat Biotechnology Research Centre | Zuber Saiyed,Ramesh Pandit,Janvi Raval,Zarna Patel,Nitin Savaliya,Dinesh Kumar,Twinkle Soni,Sonal Sharma ,Pranay Shah,Umang Mishra,Nitesh Shah,Chaitanya Joshi,Madhvi Joshi |
| EPI_ISL_2405133 | BJ Medical College and Civil Hospital, Ahmedabad | Gujarat Biotechnology Research Centre | Ramesh Pandit,Janvi Raval,Zarna Patel,Nitin Savaliya,Dinesh Kumar,Twinkle Soni,Sonal Sharma ,Zuber Saiyed,Pranay Shah,Umang Mishra,Nitesh Shah,Chaitanya Joshi,Madhvi Joshi |
| EPI_ISL_2405134 | BJ Medical College and Civil Hospital, Ahmedabad | Gujarat Biotechnology Research Centre | Janvi Raval,Zarna Patel,Nitin Savaliya,Dinesh Kumar,Twinkle Soni,Sonal Sharma ,Zuber Saiyed,Ramesh Pandit,Pranay Shah,Umang Mishra,Nitesh Shah,Chaitanya Joshi,Madhvi Joshi |
| EPI_ISL_2405135 | BJ Medical College and Civil Hospital, Ahmedabad | Gujarat Biotechnology Research Centre | Zarna Patel,Nitin Savaliya,Dinesh Kumar,Twinkle Soni,Sonal Sharma ,Zuber Saiyed,Ramesh Pandit,Janvi Raval,Pranay Shah,Umang Mishra,Nitesh Shah,Chaitanya Joshi,Madhvi Joshi |
| EPI_ISL_2405136 | BJ Medical College and Civil Hospital, Ahmedabad | Gujarat Biotechnology Research Centre | Nitin Savaliya,Dinesh Kumar,Twinkle Soni,Sonal Sharma ,Zuber Saiyed,Ramesh Pandit,Janvi Raval,Zarna Patel,Pranay Shah,Umang Mishra,Nitesh Shah,Chaitanya Joshi,Madhvi Joshi |
| EPI_ISL_2405137 | BJ Medical College and Civil Hospital, Ahmedabad | Gujarat Biotechnology Research Centre | Dinesh Kumar,Twinkle Soni,Sonal Sharma ,Zuber Saiyed,Ramesh Pandit,Janvi Raval,Zarna Patel,Nitin Savaliya,Pranay Shah,Umang Mishra,Nitesh Shah,Chaitanya Joshi,Madhvi Joshi |
| EPI_ISL_2405138 | BJ Medical College and Civil Hospital, Ahmedabad | Gujarat Biotechnology Research Centre | Twinkle Soni,Sonal Sharma ,Zuber Saiyed,Ramesh Pandit,Janvi Raval,Zarna Patel,Nitin Savaliya,Dinesh Kumar,Pranay Shah,Umang Mishra,Nitesh Shah,Chaitanya Joshi,Madhvi Joshi |
| EPI_ISL_2405139 | BJ Medical College and Civil Hospital, Ahmedabad | Gujarat Biotechnology Research Centre | Sonal Sharma ,Zuber Saiyed,Ramesh Pandit,Janvi Raval,Zarna Patel,Nitin Savaliya,Dinesh Kumar,Twinkle Soni,Pranay Shah,Umang Mishra,Nitesh Shah,Chaitanya Joshi,Madhvi Joshi |
| EPI_ISL_2405140 | BJ Medical College and Civil Hospital, Ahmedabad | Gujarat Biotechnology Research Centre | Zuber Saiyed,Ramesh Pandit,Janvi Raval,Zarna Patel,Nitin Savaliya,Dinesh Kumar,Twinkle Soni,Sonal Sharma ,Pranay Shah,Umang Mishra,Nitesh Shah,Chaitanya Joshi,Madhvi Joshi |
| EPI_ISL_2405141 | BJ Medical College and Civil Hospital, Ahmedabad | Gujarat Biotechnology Research Centre | Ramesh Pandit,Janvi Raval,Zarna Patel,Nitin Savaliya,Dinesh Kumar,Twinkle Soni,Sonal Sharma ,Zuber Saiyed,Pranay Shah,Umang Mishra,Nitesh Shah,Chaitanya Joshi,Madhvi Joshi |
| EPI_ISL_2405142 | BJ Medical College and Civil Hospital, Ahmedabad | Gujarat Biotechnology Research Centre | Janvi Raval,Zarna Patel,Nitin Savaliya,Dinesh Kumar,Twinkle Soni,Sonal Sharma ,Zuber Saiyed,Ramesh Pandit,Pranay Shah,Umang Mishra,Nitesh Shah,Chaitanya Joshi,Madhvi Joshi |
| EPI_ISL_2405143 | BJ Medical College and Civil Hospital, Ahmedabad | Gujarat Biotechnology Research Centre | Zarna Patel,Nitin Savaliya,Dinesh Kumar,Twinkle Soni,Sonal Sharma ,Zuber Saiyed,Ramesh Pandit,Janvi Raval,Pranay Shah,Umang Mishra,Nitesh Shah,Chaitanya Joshi,Madhvi Joshi |
| EPI_ISL_2405144 | BJ Medical College and Civil Hospital, Ahmedabad | Gujarat Biotechnology Research Centre | Nitin Savaliya,Dinesh Kumar,Twinkle Soni,Sonal Sharma ,Zuber Saiyed,Ramesh Pandit,Janvi Raval,Zarna Patel,Pranay Shah,Umang Mishra,Nitesh Shah,Chaitanya Joshi,Madhvi Joshi |
| EPI_ISL_2408062 | Usansolo-Galdakao University Hospital | Cruces University Hospital | Ana Gual-de-Torrella, Izaskun Alejo-Cancho, Mikel Gallego, Ana Belén de la Hoz |
| EPI_ISL_2408064, EPI_ISL_2408065, EPI_ISL_2408066, EPI_ISL_2408067 | University of Bari Biomedical Sciences and Human Oncology | University of Bari Biomedical Sciences and Human Oncology | Chironna M., Sallustio A., Loconsole D., Accogli M. |
| EPI_ISL_2421108, EPI_ISL_2421111 | cerballiance-IDF | Cerba lab | Emmanuel Lecorche, Sabine Trombert-Paolantoni, Aude Lessenne, Kader Merah, Thierry Collin, Laura Verdurme, Patrice Herisson, Stéphanie Haïm-Boukobza, Bénédicte Roquebert |
| EPI_ISL_2421113 | cerballiance-PACA | Cerba lab | Emmanuel Lecorche, Sabine Trombert-Paolantoni, Laurence Prots, Jean-Christophe Roig, Edouard Delaunay, Isabelle Butin, Laura Verdurme, Bruno Roure, Stéphanie Haïm-Boukobza, Bénédicte Roquebert |
| EPI_ISL_2421114, EPI_ISL_2421115, EPI_ISL_2421118 | cerballiance-IDF | Cerba lab | Emmanuel Lecorche, Sabine Trombert-Paolantoni, Aude Lessenne, Kader Merah, Thierry Collin, Laura Verdurme, Patrice Herisson, Stéphanie Haïm-Boukobza, Bénédicte Roquebert |
| EPI_ISL_2421120 | cerballiance-Normandie | Cerba lab | Emmanuel Lecorche, Sabine Trombert-Paolantoni, Gwénolé Prigent, Antoine Prigent, Francois Xavier Le Guillou, Alexandre Leriche, Laura Verdurme, Sylvain Metge, Stéphanie Haïm-Boukobza, Bénédicte Roquebert |
| EPI_ISL_2421121 | cerballiance-IDF | Cerba lab | Emmanuel Lecorche, Sabine Trombert-Paolantoni, Aude Lessenne, Kader Merah, Thierry Collin, Laura Verdurme, Patrice Herisson, Stéphanie Haïm-Boukobza, Bénédicte Roquebert |
| EPI_ISL_2421123 | cerballiance-PACA | Cerba lab | Emmanuel Lecorche, Sabine Trombert-Paolantoni, Laurence Prots, Jean-Christophe Roig, Edouard Delaunay, Isabelle Butin, Laura Verdurme, Bruno Roure, Stéphanie Haïm-Boukobza, Bénédicte Roquebert |

| | | | |
|---|---|---|---|
| EPI_ISL_2421124, EPI_ISL_2421125, EPI_ISL_2421127, EPI_ISL_2421129, EPI_ISL_2421130, EPI_ISL_2421131, EPI_ISL_2421134, EPI_ISL_2421135, EPI_ISL_2421136, EPI_ISL_2421138 | cerballiance-IDF | Cerba lab | Emmanuel Lecorche, Sabine Trombert-Paolantoni, Aude Lessenne, Kader Merah, Thierry Collin, Laura Verdurme, Patrice Herisson, Stéphanie Haïm-Boukobza, Bénédicte Roquebert |
| EPI_ISL_2421278, EPI_ISL_2421283, EPI_ISL_2421285, EPI_ISL_2421286, EPI_ISL_2421288 | Laboratori de Referencia de Catalunya | Laboratori de Referencia de Catalunya | Ramirez A., Bellosillo B., Padilla E., Vilas A., Hernandez JJ., Canal M. |
| EPI_ISL_2422827, EPI_ISL_2422833 | Sharp HealthCare Laboratory | Andersen lab at Scripps Research | SEARCH Alliance San Diego with Aaron Harding, Jacquelyn Berumen, Cathy Woerle, Liam McGinnis, Art Mendoza, Omid Bakhtar |
| EPI_ISL_2422976, EPI_ISL_2422977, EPI_ISL_2422978, EPI_ISL_2422979, EPI_ISL_2422980, EPI_ISL_2422981, EPI_ISL_2422982, EPI_ISL_2422983, EPI_ISL_2422984, EPI_ISL_2422985, EPI_ISL_2422986, EPI_ISL_2422987, EPI_ISL_2422988, EPI_ISL_2422989, EPI_ISL_2422991, EPI_ISL_2422992, EPI_ISL_2422993, EPI_ISL_2422995, EPI_ISL_2422997, EPI_ISL_2422998, EPI_ISL_2423003, EPI_ISL_2423004, EPI_ISL_2423005, EPI_ISL_2423006, EPI_ISL_2423007, EPI_ISL_2423011, EPI_ISL_2423013, EPI_ISL_2423014, EPI_ISL_2423015, EPI_ISL_2423016, EPI_ISL_2423017, EPI_ISL_2423018, EPI_ISL_2423019, EPI_ISL_2423020, EPI_ISL_2423021, EPI_ISL_2423022, EPI_ISL_2423023, EPI_ISL_2423025, EPI_ISL_2423026, EPI_ISL_2423027, EPI_ISL_2423029, EPI_ISL_2423031, EPI_ISL_2423032 | | | |
| see above | San Diego County Public Health Laboratory | Andersen lab at Scripps Research | SEARCH Alliance San Diego with Ashleigh Murphy, Jovan Shephard, Brett Austin |
| EPI_ISL_2424135 | Clinical Virology Laboratory, Institute of Liver and Biliary Sciences | ILBS - IGIB | Ekta Gupta, Sheetalnath Rooge, Pratibha Kale, Amit Pandey, Chhagan Bihari Sharma, Partha Chattopadhyay, Priti Devi, Ranjeet Maurya, Neha Jha, Priyanka Mehta, Manish Kumar, Pooja Sharma, Sheeba Saifi, Aparna S, Sarfaraz Alam, Bharathram Uppili, Mohammed Faruq, Anurag Agrawal, Rajesh Pandey, Shiv Kumar Sarin |
| EPI_ISL_2425562, EPI_ISL_2425563, EPI_ISL_2425564 | cerballiance-IDF | Cerba lab | Emmanuel Lecorche, Sabine Trombert-Paolantoni, Aude Lessenne, Kader Merah, Thierry Collin, Laura Verdurme, Patrice Herisson, Stéphanie Haïm-Boukobza, Bénédicte Roquebert |
| EPI_ISL_2425566 | cerballiance-PACA | Cerba lab | Emmanuel Lecorche, Sabine Trombert-Paolantoni, Laurence Prots, Jean-Christophe Roig, Edouard Delaunay, Isabelle Butin, Laura Verdurme, Bruno Roure, Stéphanie Haïm-Boukobza, Bénédicte Roquebert |
| EPI_ISL_2425567, EPI_ISL_2425571, EPI_ISL_2425572, EPI_ISL_2425575, EPI_ISL_2425576, EPI_ISL_2425578, EPI_ISL_2425580, EPI_ISL_2425583, EPI_ISL_2425586, EPI_ISL_2425587, EPI_ISL_2425590, EPI_ISL_2425593, EPI_ISL_2425594 | | | |
| see above | cerballiance-IDF | Cerba lab | Emmanuel Lecorche, Sabine Trombert-Paolantoni, Aude Lessenne, Kader Merah, Thierry Collin, Laura Verdurme, Patrice Herisson, Stéphanie Haïm-Boukobza, Bénédicte Roquebert |
| EPI_ISL_2425596 | cerballiance-Normandie | Cerba lab | Emmanuel Lecorche, Sabine Trombert-Paolantoni, Gwénolé Prigent, Antoine Prigent, Francois Xavier Le Guillou, Alexandre Leriche, Laura Verdurme, Sylvain Metge, Stéphanie Haïm-Boukobza, Bénédicte Roquebert |
| EPI_ISL_2425598, EPI_ISL_2425599 | cerballiance-PACA | Cerba lab | Emmanuel Lecorche, Sabine Trombert-Paolantoni, Laurence Prots, Jean-Christophe Roig, Edouard Delaunay, Isabelle Butin, Laura Verdurme, Bruno Roure, Stéphanie Haïm-Boukobza, Bénédicte Roquebert |
| EPI_ISL_2425604, EPI_ISL_2425605, EPI_ISL_2425606, EPI_ISL_2425607, EPI_ISL_2425609 | cerballiance-IDF | Cerba lab | Emmanuel Lecorche, Sabine Trombert-Paolantoni, Aude Lessenne, Kader Merah, Thierry Collin, Laura Verdurme, Patrice Herisson, Stéphanie Haïm-Boukobza, Bénédicte Roquebert |
| EPI_ISL_2425610 | cerballiance-PACA | Cerba lab | Emmanuel Lecorche, Sabine Trombert-Paolantoni, Laurence Prots, Jean-Christophe Roig, Edouard Delaunay, Isabelle Butin, Laura Verdurme, Bruno Roure, Stéphanie Haïm-Boukobza, Bénédicte Roquebert |
| EPI_ISL_2425611 | cerballiance-IDF | Cerba lab | Emmanuel Lecorche, Sabine Trombert-Paolantoni, Aude Lessenne, Kader Merah, Thierry Collin, Laura Verdurme, Patrice Herisson, Stéphanie Haïm-Boukobza, Bénédicte Roquebert |
| EPI_ISL_2425612 | cerballiance-HDF | Cerba lab | Emmanuel Lecorche, Sabine Trombert-Paolantoni, Claire Felloni, Anne Mainardi, Laura Verdurme, Jean-Michel Damien, Stéphanie Haïm-Boukobza, Bénédicte Roquebert |
| EPI_ISL_2426150, EPI_ISL_2426151, EPI_ISL_2426152, EPI_ISL_2426153, EPI_ISL_2426154, EPI_ISL_2426155, EPI_ISL_2426156, EPI_ISL_2426157, EPI_ISL_2426158, EPI_ISL_2426159, EPI_ISL_2426160, EPI_ISL_2426161, EPI_ISL_2426162, EPI_ISL_2426163, EPI_ISL_2426165, EPI_ISL_2426166, EPI_ISL_2426167, EPI_ISL_2426168, EPI_ISL_2426170, EPI_ISL_2426171, EPI_ISL_2426174, EPI_ISL_2426175, EPI_ISL_2426176, EPI_ISL_2426177, EPI_ISL_2426178, EPI_ISL_2426179, EPI_ISL_2426182, EPI_ISL_2426183, EPI_ISL_2426186, EPI_ISL_2426189 | | | |
| see above | Clinical Virology Laboratory, Institute of Liver and Biliary Sciences | ILBS - IGIB | Ekta Gupta, Sheetalnath Rooge, Pratibha Kale, Amit Pandey, Chhagan Bihari Sharma, Partha Chattopadhyay, Priti Devi, Ranjeet Maurya, Neha Jha, Priyanka Mehta, Manish Kumar, Pooja Sharma, Sheeba Saifi, Aparna S, Sarfaraz Alam, Bharathram Uppili, Mohammed Faruq, Anurag Agrawal, Rajesh Pandey, Shiv Kumar Sarin |
| EPI_ISL_2426222, EPI_ISL_2426226, EPI_ISL_2426227, EPI_ISL_2426230, EPI_ISL_2426231, EPI_ISL_2426246 | All India Institute of Medical Sciences, Delhi | CSIR Institute of Genomics and Integrative Biology | Urvashi B Singh, Mercy Rophina, Rama Chaudhry, Vigneshwar Senthivel, Kiran Bala, Rahul C. Bhoyar, Bani Jolly, Nayer Jamshed, Mohamed Imran,Ritu Gupta, Praveen Aggarwal, Mohit Kumar Divakar, Subrata Sinha , Arvinden VR, Anjali Bajaj, Afra Shamnath, Abhinav Jain, COVID CBNAAT CORE GROUP , Vinod Scaria, Sridhar Sivasubbu, Randeep Guleria |
| EPI_ISL_2426855, EPI_ISL_2426856, EPI_ISL_2426857, EPI_ISL_2426859, EPI_ISL_2426860 | Centers for Disease Control and Prevention, Dengue Branch | Centers for Disease Control and Prevention, Dengue Branch | Gilberto A. Santiago, Glenda Gonzalez, Betzabel Flores, Keyla Charriez, Gabriela Paz-Bailey, Jorge L. Munoz-Jordan |
| EPI_ISL_2426865 | Puerto Rico Department of Health | Centers for Disease Control and Prevention, Dengue Branch | Gilberto A. Santiago, Glenda Gonzalez, Betzabel Flores, Keyla Charriez, Gabriela Paz-Bailey, Jorge L. Munoz-Jordan |
| EPI_ISL_2426866, EPI_ISL_2426867, EPI_ISL_2426868, EPI_ISL_2426869, EPI_ISL_2426870, EPI_ISL_2426871, EPI_ISL_2426872, EPI_ISL_2426875, EPI_ISL_2426876, EPI_ISL_2426877, EPI_ISL_2426878, EPI_ISL_2426879, EPI_ISL_2426882, EPI_ISL_2426883, EPI_ISL_2426884, EPI_ISL_2426885, EPI_ISL_2426886, EPI_ISL_2426891, EPI_ISL_2426893, EPI_ISL_2426894, EPI_ISL_2426895 | | | |
| see above | Centers for Disease Control and Prevention, Dengue Branch | Centers for Disease Control and Prevention, Dengue Branch | Gilberto A. Santiago, Glenda Gonzalez, Betzabel Flores, Keyla Charriez, Gabriela Paz-Bailey, Jorge L. Munoz-Jordan |
| EPI_ISL_2429133, EPI_ISL_2429139, EPI_ISL_2429140 | Basurto University Hospital: Clinical Microbiology Laboratory | Biocruces Bizkaia | Mikel Urrutikoetxea-Gutiérrez, Mª Carmen Nieto Toboso, Estibaliz Ugalde Zarraga, Mikel Gallego Rodrigo, Ana de la Hoz, José Luis Díaz de Tuesta del Arco |
| EPI_ISL_2429142 | Institute for Medical Research, Infectious Disease Research Centre, National Institutes of Health, Ministry of Health Malaysia | Institute for Medical Research, Infectious Disease Research Centre, National Institutes of Health, Ministry of Health Malaysia | Suppiah J, Kamel K, Mohd Zawawi Z, Azizan MA, Ramly N, Robert F, Thayan R |
| EPI_ISL_2429146 | FarmaLab Indoutama Halim | National Institute of Health Research and Development | Vivi Setiawaty, Hana Apsari Pawestri, Hartanti Dian Ikawati, Subangkit, Kartika Dewi Puspa, Arie Ardiansyah Nugraha, Triyani Soekarso, Krisna Pangesti, Nelly Puspandari |
| EPI_ISL_2432937, EPI_ISL_2432938, EPI_ISL_2432939, EPI_ISL_2432940, EPI_ISL_2432941, EPI_ISL_2432942, EPI_ISL_2432943, EPI_ISL_2432944, EPI_ISL_2432945, EPI_ISL_2432946, EPI_ISL_2432947, EPI_ISL_2432948, EPI_ISL_2432949, EPI_ISL_2432950, EPI_ISL_2432952 | | | |
| see above | Microbiology Department, University Hospital Donostia | Microbiology Department, University Hospital Donostia | Marimon JM, Montes M, Piñeiro L, Grandioso D, Martin-Peñaranda T, Gomez M, Cilla G. |
| EPI_ISL_2441782 | CHU de la Réunion - Sud | Laboratoire de virologie, CNR arbovirus Associé, Chu de la Réunion | Etienne Frumence, Marie-Christine Jaffar Bandjee, Nicolas Traversier, Sabrina Petit Genet |
| EPI_ISL_2441784 | CHU de la Réunion - Nord | Laboratoire de virologie, CNR arbovirus Associé, Chu de la Réunion | Etienne Frumence, Marie-Christine Jaffar Bandjee, Nicolas Traversier, Sabrina Petit Genet |
| EPI_ISL_2441787, EPI_ISL_2441788, EPI_ISL_2441789 | CHU de la Réunion - Sud | Laboratoire de virologie, CNR arbovirus Associé, Chu de la Réunion | Etienne Frumence, Marie-Christine Jaffar Bandjee, Nicolas Traversier, Sabrina Petit Genet |
| EPI_ISL_2441793, EPI_ISL_2441794 | CHU de la Réunion - Nord | Laboratoire de virologie, CNR arbovirus Associé, Chu de la Réunion | Etienne Frumence, Marie-Christine Jaffar Bandjee, Nicolas Traversier, Sabrina Petit Genet |
| EPI_ISL_2441795 | CHU de la Réunion - Sud | Laboratoire de virologie, CNR arbovirus Associé, Chu de la Réunion | Etienne Frumence, Marie-Christine Jaffar Bandjee, Nicolas Traversier, Sabrina Petit Genet |
| EPI_ISL_2441796, EPI_ISL_2441797, | CHU de la Réunion - Nord | Laboratoire de virologie, CNR arbovirus Associé, Chu de la | Etienne Frumence, Marie-Christine Jaffar Bandjee, Nicolas Traversier, Sabrina Petit Genet |

| | | | |
|---|---|---|---|
| EPI_ISL_2441798 | | Réunion | |
| EPI_ISL_2441799, EPI_ISL_2441800 | CHU de la Réunion - Sud | Laboratoire de virologie, CNR arbovirus Associé, Chu de la Réunion | Etienne Frumence, Marie-Christine Jaffar Bandjee, Nicolas Traversier, Sabrina Petit Genet |
| EPI_ISL_2441801 | CHU de la Réunion - Nord | Laboratoire de virologie, CNR arbovirus Associé, Chu de la Réunion | Etienne Frumence, Marie-Christine Jaffar Bandjee, Nicolas Traversier, Sabrina Petit Genet |
| EPI_ISL_2441803 | CHU de la Réunion - Sud | Laboratoire de virologie, CNR arbovirus Associé, Chu de la Réunion | Etienne Frumence, Marie-Christine Jaffar Bandjee, Nicolas Traversier, Sabrina Petit Genet |
| EPI_ISL_2443528 | Lboratorio Central de Saude Publica do Estado do Parana (LACEN/PR) | Laboratory of Respiratory Viruses and Measles, Oswaldo Cruz Institute, FIOCRUZ | Paola Resende, Luciana Appolinario, Fernando Motta, Anna Carolina Paixao, Ana Carolina Mendonca, Alice Sampaio Rocha, Taina Venas, Elisa Cavalcante Pereira, Renata Serrano Lopes, Irina Riediger, Marilda Siqueira on behalf of the Fiocruz COVID-19 Genomic Surveillance Network |
| EPI_ISL_2443529 | Labortorio Central de Saude Publica do Estado de Santa Catarina (LACEN/SC) | Laboratory of Respiratory Viruses and Measles, Oswaldo Cruz Institute, FIOCRUZ | Paola Resende, Luciana Appolinario, Fernando Motta, Anna Carolina Paixao, Ana Carolina Mendonca, Alice Sampaio Rocha, Taina Venas, Elisa Cavalcante Pereira, Renata Serrano Lopes, Darcita Buerger Rovaris, Sandra Bianchini Fernandes, Marilda Siqueira on behalf of the Fiocruz COVID-19 Genomic Surveillance Network |
| EPI_ISL_2443539, EPI_ISL_2443540, EPI_ISL_2443541, EPI_ISL_2443542, EPI_ISL_2443543, EPI_ISL_2443544, EPI_ISL_2443545, EPI_ISL_2443547, EPI_ISL_2443548, EPI_ISL_2443549, EPI_ISL_2443550 | | | |
| see above | Lboratorio Central de Saude Publica do Estado do Parana (LACEN/PR) | Laboratory of Respiratory Viruses and Measles, Oswaldo Cruz Institute, FIOCRUZ | Paola Resende, Luciana Appolinario, Fernando Motta, Anna Carolina Paixao, Ana Carolina Mendonca, Alice Sampaio Rocha, Taina Venas, Elisa Cavalcante Pereira, Renata Serrano Lopes, Irina Riediger, Marilda Siqueira on behalf of the Fiocruz COVID-19 Genomic Surveillance Network |
| EPI_ISL_2448624 | CHU Poitiers | CHU Poitiers | Caroline MICHAUD, Maxime PICHON, Manon PRAT, Valentin BON-BARET, Birama N'DIAYE, Magali GARCIA, Agnes BEBY-DEFAUX, Nicolas LEVEQUE |
| EPI_ISL_2448626, EPI_ISL_2448632 | CH Angoulême | CHU Poitiers | Caroline MICHAUD, Maxime PICHON, Manon PRAT, Valentin BON-BARET, Birama N'DIAYE, Magali GARCIA, Agnes BEBY-DEFAUX, Nicolas LEVEQUE |
| EPI_ISL_2450815, EPI_ISL_2450816, EPI_ISL_2450817, EPI_ISL_2450818, EPI_ISL_2450819, EPI_ISL_2450820, EPI_ISL_2450821, EPI_ISL_2450822 | S.C. Patologia Clinica, Ospedale Sant'Andrea, ASL 5 | U.O. Igiene, Ospedale Policlinico San Martino | Bruzzone Bianca, Battolla Enrico, Caligiuri Patrizia, De Pace Vanessa, Domnich Alexander, Orsi Andrea, Ricucci Valentina, Icardi Giancarlo on behalf of SARS-CoV-2 ITALIAN RESEARCH ENTERPRISE-(SCIRE) Collaborative Group |
| EPI_ISL_2450823 | S.C. Laboratorio Analisi, Ospedale di Lavagna, ASL 4 Liguria | U.O. Igiene, Ospedale Policlinico San Martino | Bruzzone Bianca, Bandettini Roberto, Caligiuri Patrizia, De Pace Vanessa, Domnich Alexander, Orsi Andrea, Ricucci Valentina, Icardi Giancarlo on behalf of SARS-CoV-2 ITALIAN RESEARCH ENTERPRISE-(SCIRE) Collaborative Group |
| EPI_ISL_2450824 | S.S.D. Microbiologia, Stabilimento ospedaliero di Sanremo, ASL 1 Liguria | U.O. Igiene, Ospedale Policlinico San Martino | Bruzzone Bianca, Caligiuri Patrizia, De Pace Vanessa, Domnich Alexander, Dusi Pier Andrea, Orsi Andrea, Ricucci Valentina, Icardi Giancarlo on behalf of SARS-CoV-2 ITALIAN RESEARCH ENTERPRISE-(SCIRE) Collaborative Group |
| EPI_ISL_2450825 | S.C. Laboratorio Analisi, Ospedale di Lavagna, ASL 4 Liguria | U.O. Igiene, Ospedale Policlinico San Martino | Bruzzone Bianca, Bandettini Roberto, Caligiuri Patrizia, De Pace Vanessa, Domnich Alexander, Orsi Andrea, Ricucci Valentina, Icardi Giancarlo on behalf of SARS-CoV-2 ITALIAN RESEARCH ENTERPRISE-(SCIRE) Collaborative Group |
| EPI_ISL_2450826 | U.O. Igiene, Ospedale Policlinico San Martino | U.O. Igiene, Ospedale Policlinico San Martino | Bruzzone Bianca, Caligiuri Patrizia, De Pace Vanessa, Domnich Alexander, Orsi Andrea, Ricucci Valentina, Spitaleri Antonino, Icardi Giancarlo on behalf of SARS-CoV-2 ITALIAN RESEARCH ENTERPRISE-(SCIRE) Collaborative Group |
| EPI_ISL_2450827, EPI_ISL_2450828, EPI_ISL_2450829, EPI_ISL_2450830, EPI_ISL_2450831, EPI_ISL_2450832, EPI_ISL_2450833, EPI_ISL_2450834, EPI_ISL_2450835, EPI_ISL_2450836, EPI_ISL_2450837, EPI_ISL_2450838, EPI_ISL_2450839, EPI_ISL_2450840 | | | |
| see above | S.C. Laboratorio Analisi, ASL 3 Liguria | U.O. Igiene, Ospedale Policlinico San Martino | Bruzzone Bianca, Caligiuri Patrizia, De Pace Vanessa, Domnich Alexander, Orsi Andrea, Ricucci Valentina, Spitaleri Antonino, Icardi Giancarlo on behalf of SARS-CoV-2 ITALIAN RESEARCH ENTERPRISE-(SCIRE) Collaborative Group |
| EPI_ISL_2450841, EPI_ISL_2450842 | S.S.D. Microbiologia, Stabilimento ospedaliero di Sanremo, ASL 1 Liguria | U.O. Igiene, Ospedale Policlinico San Martino | Bruzzone Bianca, Caligiuri Patrizia, De Pace Vanessa, Domnich Alexander, Dusi Pier Andrea, Orsi Andrea, Ricucci Valentina, Icardi Giancarlo on behalf of SARS-CoV-2 ITALIAN RESEARCH ENTERPRISE-(SCIRE) Collaborative Group |
| EPI_ISL_2450843, EPI_ISL_2450844 | S.S.D. Microbiologia, Ospedale Santa Corona di Pietra Ligure, ASL 2 Liguria | U.O. Igiene, Ospedale Policlinico San Martino | Bruzzone Bianca, Caligiuri Patrizia, De Pace Vanessa, Domnich Alexander, Orsi Andrea, Ricucci Valentina, Valle Caterina, Icardi Giancarlo on behalf of SARS-CoV-2 ITALIAN RESEARCH ENTERPRISE-(SCIRE) Collaborative Group |
| EPI_ISL_2450845 | Laboratorio di Patologia Clinica, Ospedale San Paolo in Valloria, ASL 2 Liguria | U.O. Igiene, Ospedale Policlinico San Martino | Bruzzone Bianca, Caligiuri Patrizia, De Pace Vanessa, Domnich Alexander, Lillo Flavia, Orsi Andrea, Ricucci Valentina, Icardi Giancarlo on behalf of SARS-CoV-2 ITALIAN RESEARCH ENTERPRISE-(SCIRE) Collaborative Group |
| EPI_ISL_2450846 | U.O.C. Laboratorio Analisi, Ospedale G. Gaslini IRCCS Genova | U.O. Igiene, Ospedale Policlinico San Martino | Bruzzone Bianca, Caligiuri Patrizia, De Pace Vanessa, Domnich Alexander, Orsi Andrea, Ricucci Valentina, Tripodi Gino, Icardi Giancarlo on behalf of SARS-CoV-2 ITALIAN RESEARCH ENTERPRISE-(SCIRE) Collaborative Group |
| EPI_ISL_2450847 | S.C. Laboratorio Analisi, ASL 3 Liguria | U.O. Igiene, Ospedale Policlinico San Martino | Bruzzone Bianca, Caligiuri Patrizia, De Pace Vanessa, Domnich Alexander, Orsi Andrea, Ricucci Valentina, Spitaleri Antonino, Icardi Giancarlo on behalf of SARS-CoV-2 ITALIAN RESEARCH ENTERPRISE-(SCIRE) Collaborative Group |
| EPI_ISL_2450848, EPI_ISL_2450849, EPI_ISL_2450850, EPI_ISL_2450851 | Laboratorio di Patologia Clinica, Ospedale San Paolo in Valloria, ASL 2 Liguria | U.O. Igiene, Ospedale Policlinico San Martino | Bruzzone Bianca, Caligiuri Patrizia, De Pace Vanessa, Domnich Alexander, Lillo Flavia, Orsi Andrea, Ricucci Valentina, Icardi Giancarlo on behalf of SARS-CoV-2 ITALIAN RESEARCH ENTERPRISE-(SCIRE) Collaborative Group |
| EPI_ISL_2455241 | CICESE-BSL3 | CICESE-BSL3 | Cervantes Karla, Gonzalez Ricardo, Martinez Miguel, Galindo Clara and Licea Alexei |
| EPI_ISL_2455242 | CICESE-BSL3 | CICESE-BSL3 | Karla Cervantes, Ricardo Gonzalez, Miguel Martinez, Clara Galindo and Alexei Licea |
| EPI_ISL_2455264, EPI_ISL_2455327 | Centers for Disease Control, R.O.C. (Taiwan) | Centers for Disease Control, R.O.C. (Taiwan) | Ji-Rong Yang, Yu-Chi-Lin, Jung-Jung Mu, Ming-Tsan-Liu |
| EPI_ISL_2455896, EPI_ISL_2455897, EPI_ISL_2455898, EPI_ISL_2455899, EPI_ISL_2455900, EPI_ISL_2455901, EPI_ISL_2455902, EPI_ISL_2455905, EPI_ISL_2455907, EPI_ISL_2455908 | LESP Queretaro | Instituto de Diagnostico y Referencia Epidemiologicos (INDRE) | Claudia Wong-Arambula, Abril Rodriguez-Maldonado, Vanessa Rivero-Arredondo, Ariadna Medina-Benitez, Joaquin Quiroz-Mercado, Sergio Rangel-Guerrero, Natividad Cruz-Ortiz, Tatiana Nunez-Garcia, Gisela Barrera-Badillo, Lucia Hernandez-Rivas, Irma Lopez-Martinez, Ernesto Ramirez-Gonzalez. |
| EPI_ISL_2455914, EPI_ISL_2455915 | LESP Chihuahua | Instituto de Diagnostico y Referencia Epidemiologicos (INDRE) | Claudia Wong-Arambula, Abril Rodriguez-Maldonado, Vanessa Rivero-Arredondo, Ariadna Medina-Benitez, Joaquin Quiroz-Mercado, Sergio Rangel-Guerrero, Natividad Cruz-Ortiz, Tatiana Nunez-Garcia, Gisela Barrera-Badillo, Lucia Hernandez-Rivas, Irma Lopez-Martinez, Ernesto Ramirez-Gonzalez. |
| EPI_ISL_2455916, EPI_ISL_2455918 | LESP Tamaulipas | Instituto de Diagnostico y Referencia Epidemiologicos (INDRE) | Claudia Wong-Arambula, Abril Rodriguez-Maldonado, Vanessa Rivero-Arredondo, Ariadna Medina-Benitez, Joaquin Quiroz-Mercado, Sergio Rangel-Guerrero, Natividad Cruz-Ortiz, Tatiana Nunez-Garcia, Gisela Barrera-Badillo, Lucia Hernandez-Rivas, Irma Lopez-Martinez, Ernesto Ramirez-Gonzalez. |
| EPI_ISL_2455927, EPI_ISL_2455928, EPI_ISL_2455929 | LESP Chihuahua | Instituto de Diagnostico y Referencia Epidemiologicos (INDRE) | Claudia Wong-Arambula, Abril Rodriguez-Maldonado, Vanessa Rivero-Arredondo, Ariadna Medina-Benitez, Joaquin Quiroz-Mercado, Sergio Rangel-Guerrero, Natividad Cruz-Ortiz, Tatiana Nunez-Garcia, Gisela Barrera-Badillo, Lucia Hernandez-Rivas, Irma Lopez-Martinez, Ernesto Ramirez-Gonzalez. |
| EPI_ISL_2455964, EPI_ISL_2455965, EPI_ISL_2455969, EPI_ISL_2455970, EPI_ISL_2455971, EPI_ISL_2455972, EPI_ISL_2455973, EPI_ISL_2455975 | LESP Hidalgo | Instituto de Diagnostico y Referencia Epidemiologicos (INDRE) | Claudia Wong-Arambula, Abril Rodriguez-Maldonado, Vanessa Rivero-Arredondo, Ariadna Medina-Benitez, Joaquin Quiroz-Mercado, Sergio Rangel-Guerrero, Natividad Cruz-Ortiz, Tatiana Nunez-Garcia, Gisela Barrera-Badillo, Lucia Hernandez-Rivas, Irma Lopez-Martinez, Ernesto Ramirez-Gonzalez. |
| EPI_ISL_2455977, EPI_ISL_2455978 | LESP Sinaloa | Instituto de Diagnostico y Referencia Epidemiologicos (INDRE) | Claudia Wong-Arambula, Abril Rodriguez-Maldonado, Vanessa Rivero-Arredondo, Ariadna Medina-Benitez, Joaquin Quiroz-Mercado, Sergio Rangel-Guerrero, Natividad Cruz-Ortiz, Tatiana Nunez-Garcia, Gisela Barrera-Badillo, Lucia Hernandez-Rivas, Irma Lopez-Martinez, Ernesto Ramirez-Gonzalez. |
| EPI_ISL_2455979 | LESP Morelos | Instituto de Diagnostico y Referencia Epidemiologicos (INDRE) | Claudia Wong-Arambula, Abril Rodriguez-Maldonado, Vanessa Rivero-Arredondo, Ariadna Medina-Benitez, Joaquin Quiroz-Mercado, Sergio Rangel-Guerrero, Natividad Cruz-Ortiz, Tatiana Nunez-Garcia, Gisela Barrera-Badillo, Lucia Hernandez-Rivas, Irma Lopez-Martinez, Ernesto |

| | | | |
|---|---|---|---|
| | | | Ramirez-Gonzalez. |
| EPI_ISL_2455989 | LESP Queretaro | Instituto de Diagnostico y Referencia Epidemiologicos (INDRE) | Claudia Wong-Arambula, Abril Rodriguez-Maldonado, Vanessa Rivero-Arredondo, Ariadna Medina-Benitez, Joaquin Quiroz-Mercado, Sergio Rangel-Guerrero, Natividad Cruz-Ortiz, Tatiana Nunez-Garcia, Gisela Barrera-Badillo, Lucia Hernandez-Rivas, Irma Lopez-Martinez, Ernesto Ramirez-Gonzalez. |
| EPI_ISL_2455990, EPI_ISL_2455991 | LESP Guanajuato | Instituto de Diagnostico y Referencia Epidemiologicos (INDRE) | Claudia Wong-Arambula, Abril Rodriguez-Maldonado, Vanessa Rivero-Arredondo, Ariadna Medina-Benitez, Joaquin Quiroz-Mercado, Sergio Rangel-Guerrero, Natividad Cruz-Ortiz, Tatiana Nunez-Garcia, Gisela Barrera-Badillo, Lucia Hernandez-Rivas, Irma Lopez-Martinez, Ernesto Ramirez-Gonzalez. |
| EPI_ISL_2462866, EPI_ISL_2462867 | AZ Sint-Lucas Gent | AZ Sint-Lucas Gent | Jos Van Acker, Charlotte Verfaillie, Elke Vanlaere, Klara De Rauw |
| EPI_ISL_2462903 | Labo Analyses Med | National Reference Center for Viruses of Respiratory Infections, Institut Pasteur, Paris | Marion Barbet, Sylvie Behillil, Méline Bizard, Angela Brisebarre, Camille Capel, Vincent Enouf, Louise Lefrançois, Frédéric Lemoine, Christophe Malabat, Corinne Maufrais, Etienne Simon-Lorière, Maud Vanpeene, Sylvie Van der Werf ,Anne Lise Bravetti |
| EPI_ISL_2462924 | Hospital | National Reference Center for Viruses of Respiratory Infections, Institut Pasteur, Paris | Marion Barbet, Sylvie Behillil, Méline Bizard, Angela Brisebarre, Camille Capel, Vincent Enouf, Louise Lefrançois, Frédéric Lemoine, Christophe Malabat, Corinne Maufrais, Etienne Simon-Lorière, Maud Vanpeene, Sylvie Van der Werf ,Marianne Burgard |
| EPI_ISL_2462961 | Hospital | National Reference Center for Viruses of Respiratory Infections, Institut Pasteur, Paris | Marion Barbet, Sylvie Behillil, Méline Bizard, Angela Brisebarre, Camille Capel, Vincent Enouf, Louise Lefrançois, Frédéric Lemoine, Christophe Malabat, Corinne Maufrais, Etienne Simon-Lorière, Maud Vanpeene, Sylvie Van der Werf ,Axelle Paquin |
| EPI_ISL_2464255 | Hospital | National Reference Center for Viruses of Respiratory Infections, Institut Pasteur, Paris | Marion Barbet, Sylvie Behillil, Méline Bizard, Angela Brisebarre, Camille Capel, Vincent Enouf, Louise Lefrançois, Frédéric Lemoine, Christophe Malabat, Corinne Maufrais, Etienne Simon-Lorière, Maud Vanpeene, Sylvie Van der Werf ,Sandrine Castelain |
| EPI_ISL_2466132, EPI_ISL_2466133, EPI_ISL_2466135 | Laboratorio Central de Saude Publica do Estado do Rio Grande do Sul (LACEN-RS) | Laboratory of Respiratory Viruses and Measles, Oswaldo Cruz Institute, FIOCRUZ | Paola Resende, Luciana Appolinario, Fernando Motta, Anna Carolina Paixao, Ana Carolina Mendonca, Alice Sampaio Rocha, Taina Venas, Elisa Cavalcante Pereira, Renata Serrano Lopes, Tatiana Schaffer Gregianini, Richard Salvato, Marilda Siqueira on behalf of the Fiocruz COVID-19 Genomic Surveillance Network |
| EPI_ISL_2466343, EPI_ISL_2466344, EPI_ISL_2466345 | AZ Zeno Knokke-Blankenberge | AZ Sint Jan | Marijke Reynders, Patrick Descheemaeker, Laurien Hoornaert, Thomas Van Landschoot, Marie Madeleine Chabert-Consen, Jorn Hellemans |
| EPI_ISL_2476413, EPI_ISL_2476415 | LESP Guanajuato | Instituto de Diagnostico y Referencia Epidemiologicos (INDRE) | Claudia Wong-Arambula, Abril Rodriguez-Maldonado, Vanessa Rivero-Arredondo, Ariadna Medina-Benitez, Joaquin Quiroz-Mercado, Sergio Rangel-Guerrero, Natividad Cruz-Ortiz, Tatiana Nunez-Garcia, Gisela Barrera-Badillo, Lucia Hernandez-Rivas, Irma Lopez-Martinez, Ernesto Ramirez-Gonzalez. |
| EPI_ISL_2476416 | LESP Tamaulipas | Instituto de Diagnostico y Referencia Epidemiologicos (INDRE) | Claudia Wong-Arambula, Abril Rodriguez-Maldonado, Vanessa Rivero-Arredondo, Ariadna Medina-Benitez, Joaquin Quiroz-Mercado, Sergio Rangel-Guerrero, Natividad Cruz-Ortiz, Tatiana Nunez-Garcia, Gisela Barrera-Badillo, Lucia Hernandez-Rivas, Irma Lopez-Martinez, Ernesto Ramirez-Gonzalez. |
| EPI_ISL_2476424 | LESP Guanajuato | Instituto de Diagnostico y Referencia Epidemiologicos (INDRE) | Claudia Wong-Arambula, Abril Rodriguez-Maldonado, Vanessa Rivero-Arredondo, Ariadna Medina-Benitez, Joaquin Quiroz-Mercado, Sergio Rangel-Guerrero, Natividad Cruz-Ortiz, Tatiana Nunez-Garcia, Gisela Barrera-Badillo, Lucia Hernandez-Rivas, Irma Lopez-Martinez, Ernesto Ramirez-Gonzalez. |
| EPI_ISL_2476425, EPI_ISL_2476433 | LESP Tamaulipas | Instituto de Diagnostico y Referencia Epidemiologicos (INDRE) | Claudia Wong-Arambula, Abril Rodriguez-Maldonado, Vanessa Rivero-Arredondo, Ariadna Medina-Benitez, Joaquin Quiroz-Mercado, Sergio Rangel-Guerrero, Natividad Cruz-Ortiz, Tatiana Nunez-Garcia, Gisela Barrera-Badillo, Lucia Hernandez-Rivas, Irma Lopez-Martinez, Ernesto Ramirez-Gonzalez. |
| EPI_ISL_2479877, EPI_ISL_2479878 | LESP Queretaro | Instituto de Diagnostico y Referencia Epidemiologicos (INDRE) | Claudia Wong-Arambula, Abril Rodriguez-Maldonado, Vanessa Rivero-Arredondo, Ariadna Medina-Benitez, Joaquin Quiroz-Mercado, Sergio Rangel-Guerrero, Natividad Cruz-Ortiz, Tatiana Nunez-Garcia, Gisela Barrera-Badillo, Lucia Hernandez-Rivas, Irma Lopez-Martinez, Ernesto Ramirez-Gonzalez. |
| EPI_ISL_2479892 | LESP Colima | Instituto de Diagnostico y Referencia Epidemiologicos (INDRE) | Claudia Wong-Arambula, Abril Rodriguez-Maldonado, Vanessa Rivero-Arredondo, Ariadna Medina-Benitez, Joaquin Quiroz-Mercado, Sergio Rangel-Guerrero, Natividad Cruz-Ortiz, Tatiana Nunez-Garcia, Gisela Barrera-Badillo, Lucia Hernandez-Rivas, Irma Lopez-Martinez, Ernesto Ramirez-Gonzalez. |
| EPI_ISL_2479901 | LESP Hidalgo | Instituto de Diagnostico y Referencia Epidemiologicos (INDRE) | Claudia Wong-Arambula, Abril Rodriguez-Maldonado, Vanessa Rivero-Arredondo, Ariadna Medina-Benitez, Joaquin Quiroz-Mercado, Sergio Rangel-Guerrero, Natividad Cruz-Ortiz, Tatiana Nunez-Garcia, Gisela Barrera-Badillo, Lucia Hernandez-Rivas, Irma Lopez-Martinez, Ernesto Ramirez-Gonzalez. |
| EPI_ISL_2479904 | LESP Puebla | Instituto de Diagnostico y Referencia Epidemiologicos (INDRE) | Claudia Wong-Arambula, Abril Rodriguez-Maldonado, Vanessa Rivero-Arredondo, Ariadna Medina-Benitez, Joaquin Quiroz-Mercado, Sergio Rangel-Guerrero, Natividad Cruz-Ortiz, Tatiana Nunez-Garcia, Gisela Barrera-Badillo, Lucia Hernandez-Rivas, Irma Lopez-Martinez, Ernesto Ramirez-Gonzalez. |
| EPI_ISL_2484164 | LESP Chihuahua | Instituto de Diagnostico y Referencia Epidemiologicos (INDRE) | Claudia Wong-Arambula, Abril Rodriguez-Maldonado, Vanessa Rivero-Arredondo, Ariadna Medina-Benitez, Joaquin Quiroz-Mercado, Sergio Rangel-Guerrero, Natividad Cruz-Ortiz, Tatiana Nunez-Garcia, Gisela Barrera-Badillo, Lucia Hernandez-Rivas, Irma Lopez-Martinez, Ernesto Ramirez-Gonzalez. |
| EPI_ISL_2484165 | LESP Yucatan | Instituto de Diagnostico y Referencia Epidemiologicos (INDRE) | Claudia Wong-Arambula, Abril Rodriguez-Maldonado, Vanessa Rivero-Arredondo, Ariadna Medina-Benitez, Joaquin Quiroz-Mercado, Sergio Rangel-Guerrero, Natividad Cruz-Ortiz, Tatiana Nunez-Garcia, Gisela Barrera-Badillo, Lucia Hernandez-Rivas, Irma Lopez-Martinez, Ernesto Ramirez-Gonzalez. |
| EPI_ISL_2484596, EPI_ISL_2484598, EPI_ISL_2484604, EPI_ISL_2484606, EPI_ISL_2484608, EPI_ISL_2484612, EPI_ISL_2484613, EPI_ISL_2484620 | NL-Dr. Leonard A. Miller Centre for Health Services | National Microbiology Laboratory (NML) | Anna Majer, Shari Tyson, Grace Seo, Philip Mabon, Elsie Grudeski, Rhiannon Huzarewich, Russell Mandes, Anneliese Landgraff, Jennifer Tanner, Natalie Knox, Morag Graham, Gary Van Domselaar, Robert Needle, Yang Yu, Adel Malek, Laura Gilbert, George Zahariadis, Nathalie Bastien, Yan Li, Timothy Booth, Darian Hole, Madison Chapel, Kirsten Biggar, Kerri Smith, CanCOGeN's metadata curation team, Public Health Agency of Canada CanCOGeN team |
| EPI_ISL_2484787 | Temporary Specimen Collection Centre at the AsiaWorld-Expo | Hong Kong Department of Health | Alan K.L. Tsang, Gannon C.K. Mak, Peter K.C. Cheng, Peter C.W. Yip, Ken H.L. Ng, Edman T.K. Lam, Rickjason C.W. Chan, Dominic N.C. Tsang |
| EPI_ISL_2484788 | COMMUNITY TESTING CENTRE | Hong Kong Department of Health | Alan K.L. Tsang, Gannon C.K. Mak, Peter K.C. Cheng, Peter C.W. Yip, Ken H.L. Ng, Edman T.K. Lam, Rickjason C.W. Chan, Dominic N.C. Tsang |
| EPI_ISL_2484789 | Temporary Specimen Collection Centre at the AsiaWorld-Expo | Hong Kong Department of Health | Alan K.L. Tsang, Gannon C.K. Mak, Peter K.C. Cheng, Peter C.W. Yip, Ken H.L. Ng, Edman T.K. Lam, Rickjason C.W. Chan, Dominic N.C. Tsang |
| EPI_ISL_2484790 | HOME QUARANTINE TASKFORCE | Hong Kong Department of Health | Alan K.L. Tsang, Gannon C.K. Mak, Peter K.C. Cheng, Peter C.W. Yip, Ken H.L. Ng, Edman T.K. Lam, Rickjason C.W. Chan, Dominic N.C. Tsang |
| EPI_ISL_2484791 | Penny's Bay Quarantine Centre | Hong Kong Department of Health | Alan K.L. Tsang, Gannon C.K. Mak, Peter K.C. Cheng, Peter C.W. Yip, Ken H.L. Ng, Edman T.K. Lam, Rickjason C.W. Chan, Dominic N.C. Tsang |
| EPI_ISL_2484793, EPI_ISL_2484794, EPI_ISL_2484795 | HOME QUARANTINE TASKFORCE | Hong Kong Department of Health | Alan K.L. Tsang, Gannon C.K. Mak, Peter K.C. Cheng, Peter C.W. Yip, Ken H.L. Ng, Edman T.K. Lam, Rickjason C.W. Chan, Dominic N.C. Tsang |
| EPI_ISL_2484796 | COMMUNITY TESTING CENTRE | Hong Kong Department of Health | Alan K.L. Tsang, Gannon C.K. Mak, Peter K.C. Cheng, Peter C.W. Yip, Ken H.L. Ng, Edman T.K. Lam, Rickjason C.W. Chan, Dominic N.C. Tsang |
| EPI_ISL_2484797 | Penny's Bay Quarantine Centre | Hong Kong Department of Health | Alan K.L. Tsang, Gannon C.K. Mak, Peter K.C. Cheng, Peter C.W. Yip, Ken H.L. Ng, Edman T.K. Lam, Rickjason C.W. Chan, Dominic N.C. Tsang |
| EPI_ISL_2484798 | COMMUNITY TESTING CENTRE | Hong Kong Department of Health | Alan K.L. Tsang, Gannon C.K. Mak, Peter K.C. Cheng, Peter C.W. Yip, Ken H.L. Ng, Edman T.K. Lam, Rickjason C.W. Chan, Dominic N.C. Tsang |
| EPI_ISL_2488760 | Policlinico Universitario Campus Bio-Medico, Rome, Italy | Policlinico Universitario Campus Bio-Medico, Rome, Italy | Silvia Angeletti, Marta Giovanetti, Marta Fogolori, Lucia de Florio, Maria Francesconi, Roberta Veralli, Francesca Antonelli, Elisabetta Riva, Massimo Ciccozzi |
| EPI_ISL_2490312, EPI_ISL_2490314, EPI_ISL_2490317, EPI_ISL_2490318, EPI_ISL_2490319, EPI_ISL_2490323, EPI_ISL_2490324 | Centro de Investigación Biomédica del Noreste (CIBIN) | Centro de Investigación en Enfermedades Infecciosas (CIENI), Instituto Nacional de Enfermedades Respiratorias (INER) | Consorcio Mexicano de Vigilancia Genómica (CoViGen-Mex). Authors (in alphabetical order): Julio Elias Alvarado-Yaah, Carlos F. Arias, Santiago Ávila-Ríos, Eduardo Becerril-Vargas, Víctor Hugo Borja-Aburto, Celia Boukadida, Cristóbal Cháidez-Quiróz, Juan Bautista Chale-Dzul, Ricardo Ciria Merce, Andreu Comas-García, Célida Duque Molina, Julissa Enciso-Ibarra, José Antonio Enciso-Moreno, Gloria Elena Espinosa-Ayala, Fernando Fontove-Herrera, Daniel Fregoso-Rueda, Victor Eduardo Garcia-Arias, Alejandra Garcia-Gasca, Bruno Gomez-Gil, Jean Pierre González, Irvin González-López, Concepción Grajales-Muñiz, Ricardo Grande, Rosa María Gutierrez Rios, Alejandra Hernández-Terán, Alfredo Herrera-Estrella, Carla |

| | | | |
|---|---|---|---|
| | | | Ivón Herrera-Najera, Pavel Isa, Daniel Lira Morales, Susana Lopez, Antonio Loza Román, Brenda Irasema Maldonado-Meza, Bernardo Martínez-Miguel, José Arturo Martínez-Orozco, Célida Martínez- Rodríguez, Margarita Matías-Florentino, Fidencio Mejía-Nepomuceno, María Guadalupe de Jesús Mireles-Rivera, Gloria María Molina-Salinas, Hector Montoya-Fuentes, , Mario Mújica-Sánchez, José Esteban Muñoz-Medina, José de Jesús Nuñez-Contreras, Alicia Ocaña-Mondragón, Luis Alberto Ochoa-Carrera, Hector Esteban Paz-Juárez, Marissa Perez-Garcia, Francisco Pulido, Helen Haydee Fernanda Ramírez-Plascencia, Jorge Salas-Hernández, Angel Gustavo Salas-Lais, Alejandro Sanchez-Flores, Clara Esperanza Santacruz-Tinoco, María Guadalupe Santiago-Mauricio, Selene Zárate, Nelly Sélem-Mojica, Blanca Taboada, Gloria Vazquez, Joel Armando Vázquez-Pérez. |
| EPI_ISL_2490327 | Unidad de Investigación Médica de Yucatán (UIMY) | Centro de Investigación en Enfermedades Infecciosas (CIENI), Instituto Nacional de Enfermedades Respiratorias (INER) | Consorcio Mexicano de Vigilancia Genómica (CoViGen-Mex). Authors (in alphabetical order): Julio Elias Alvarado-Yaah, Carlos F. Arias, Santiago Ávila-Ríos, Eduardo Becerril-Vargas, Víctor Hugo Borja-Aburto, Celia Boukadida, Cristóbal Cháidez-Quiróz, Juan Bautista Chale-Dzul, Ricardo Ciria Merce, Andreu Comas-García, Célida Duque Molina, Julissa Enciso-Ibarra, José Antonio Enciso-Moreno, Gloria Elena Espinosa-Ayala, Fernando Fontove-Herrera, Daniel Fregoso-Rueda, Victor Eduardo Garcia-Arias, Alejandra Garcia-Gasca, Bruno Gomez-Gil, Jean Pierre González, Irvin González-López, Concepción Grajales-Muñiz, Ricardo Grande, Rosa María Gutierrez Rios, Alejandra Hernández-Terán, Alfredo Herrera-Estrella, Carla Ivón Herrera-Najera, Pavel Isa, Daniel Lira Morales, Susana Lopez, Antonio Loza Román, Brenda Irasema Maldonado-Meza, Bernardo Martínez-Miguel, José Arturo Martínez-Orozco, Célida Martínez- Rodríguez, Margarita Matías-Florentino, Fidencio Mejía-Nepomuceno, María Guadalupe de Jesús Mireles-Rivera, Gloria María Molina-Salinas, Hector Montoya-Fuentes, , Mario Mújica-Sánchez, José Esteban Muñoz-Medina, José de Jesús Nuñez-Contreras, Alicia Ocaña-Mondragón, Luis Alberto Ochoa-Carrera, Hector Esteban Paz-Juárez, Marissa Perez-Garcia, Francisco Pulido, Helen Haydee Fernanda Ramírez-Plascencia, Jorge Salas-Hernández, Angel Gustavo Salas-Lais, Alejandro Sanchez-Flores, Clara Esperanza Santacruz-Tinoco, María Guadalupe Santiago-Mauricio, Selene Zárate, Nelly Sélem-Mojica, Blanca Taboada, Gloria Vazquez, Joel Armando Vázquez-Pérez. |
| EPI_ISL_2490334, EPI_ISL_2490337, EPI_ISL_2490338 | Centro de Investigación Biomédica del Noreste (CIBIN) | Centro de Investigación en Enfermedades Infecciosas (CIENI), Instituto Nacional de Enfermedades Respiratorias (INER) | Consorcio Mexicano de Vigilancia Genómica (CoViGen-Mex). Authors (in alphabetical order): Julio Elias Alvarado-Yaah, Carlos F. Arias, Santiago Ávila-Ríos, Eduardo Becerril-Vargas, Víctor Hugo Borja-Aburto, Celia Boukadida, Cristóbal Cháidez-Quiróz, Juan Bautista Chale-Dzul, Ricardo Ciria Merce, Andreu Comas-García, Célida Duque Molina, Julissa Enciso-Ibarra, José Antonio Enciso-Moreno, Gloria Elena Espinosa-Ayala, Fernando Fontove-Herrera, Daniel Fregoso-Rueda, Victor Eduardo Garcia-Arias, Alejandra Garcia-Gasca, Bruno Gomez-Gil, Jean Pierre González, Irvin González-López, Concepción Grajales-Muñiz, Ricardo Grande, Rosa María Gutierrez Rios, Alejandra Hernández-Terán, Alfredo Herrera-Estrella, Carla Ivón Herrera-Najera, Pavel Isa, Daniel Lira Morales, Susana Lopez, Antonio Loza Román, Brenda Irasema Maldonado-Meza, Bernardo Martínez-Miguel, José Arturo Martínez-Orozco, Célida Martínez- Rodríguez, Margarita Matías-Florentino, Fidencio Mejía-Nepomuceno, María Guadalupe de Jesús Mireles-Rivera, Gloria María Molina-Salinas, Hector Montoya-Fuentes, , Mario Mújica-Sánchez, José Esteban Muñoz-Medina, José de Jesús Nuñez-Contreras, Alicia Ocaña-Mondragón, Luis Alberto Ochoa-Carrera, Hector Esteban Paz-Juárez, Marissa Perez-Garcia, Francisco Pulido, Helen Haydee Fernanda Ramírez-Plascencia, Jorge Salas-Hernández, Angel Gustavo Salas-Lais, Alejandro Sanchez-Flores, Clara Esperanza Santacruz-Tinoco, María Guadalupe Santiago-Mauricio, Selene Zárate, Nelly Sélem-Mojica, Blanca Taboada, Gloria Vazquez, Joel Armando Vázquez-Pérez. |
| EPI_ISL_2490343, EPI_ISL_2490344, EPI_ISL_2490345, EPI_ISL_2490346, EPI_ISL_2490357 | Unidad de Investigación Médica de Yucatán (UIMY) | Centro de Investigación en Enfermedades Infecciosas (CIENI), Instituto Nacional de Enfermedades Respiratorias (INER) | Consorcio Mexicano de Vigilancia Genómica (CoViGen-Mex). Authors (in alphabetical order): Julio Elias Alvarado-Yaah, Carlos F. Arias, Santiago Ávila-Ríos, Eduardo Becerril-Vargas, Víctor Hugo Borja-Aburto, Celia Boukadida, Cristóbal Cháidez-Quiróz, Juan Bautista Chale-Dzul, Ricardo Ciria Merce, Andreu Comas-García, Célida Duque Molina, Julissa Enciso-Ibarra, José Antonio Enciso-Moreno, Gloria Elena Espinosa-Ayala, Fernando Fontove-Herrera, Daniel Fregoso-Rueda, Victor Eduardo Garcia-Arias, Alejandra Garcia-Gasca, Bruno Gomez-Gil, Jean Pierre González, Irvin González-López, Concepción Grajales-Muñiz, Ricardo Grande, Rosa María Gutierrez Rios, Alejandra Hernández-Terán, Alfredo Herrera-Estrella, Carla Ivón Herrera-Najera, Pavel Isa, Daniel Lira Morales, Susana Lopez, Antonio Loza Román, Brenda Irasema Maldonado-Meza, Bernardo Martínez-Miguel, José Arturo Martínez-Orozco, Célida Martínez- Rodríguez, Margarita Matías-Florentino, Fidencio Mejía-Nepomuceno, María Guadalupe de Jesús Mireles-Rivera, Gloria María Molina-Salinas, Hector Montoya-Fuentes, , Mario Mújica-Sánchez, José Esteban Muñoz-Medina, José de Jesús Nuñez-Contreras, Alicia Ocaña-Mondragón, Luis Alberto Ochoa-Carrera, Hector Esteban Paz-Juárez, Marissa Perez-Garcia, Francisco Pulido, Helen Haydee Fernanda Ramírez-Plascencia, Jorge Salas-Hernández, Angel Gustavo Salas-Lais, Alejandro Sanchez-Flores, Clara Esperanza Santacruz-Tinoco, María Guadalupe Santiago-Mauricio, Selene Zárate, Nelly Sélem-Mojica, Blanca Taboada, Gloria Vazquez, Joel Armando Vázquez-Pérez. |
| EPI_ISL_2490358 | Unidad de Investigación Biomédica de Zacatecas (UIBZ) | Centro de Investigación en Enfermedades Infecciosas (CIENI), Instituto Nacional de Enfermedades Respiratorias (INER) | Consorcio Mexicano de Vigilancia Genómica (CoViGen-Mex). Authors (in alphabetical order): Julio Elias Alvarado-Yaah, Carlos F. Arias, Santiago Ávila-Ríos, Eduardo Becerril-Vargas, Víctor Hugo Borja-Aburto, Celia Boukadida, Cristóbal Cháidez-Quiróz, Juan Bautista Chale-Dzul, Ricardo Ciria Merce, Andreu Comas-García, Célida Duque Molina, Julissa Enciso-Ibarra, José Antonio Enciso-Moreno, Gloria Elena Espinosa-Ayala, Fernando Fontove-Herrera, Daniel Fregoso-Rueda, Victor Eduardo Garcia-Arias, Alejandra Garcia-Gasca, Bruno Gomez-Gil, Jean Pierre González, Irvin González-López, Concepción Grajales-Muñiz, Ricardo Grande, Rosa María Gutierrez Rios, Alejandra Hernández-Terán, Alfredo Herrera-Estrella, Carla Ivón Herrera-Najera, Pavel Isa, Daniel Lira Morales, Susana Lopez, Antonio Loza Román, Brenda Irasema Maldonado-Meza, Bernardo Martínez-Miguel, José Arturo Martínez-Orozco, Célida Martínez- Rodríguez, Margarita Matías-Florentino, Fidencio Mejía-Nepomuceno, María Guadalupe de Jesús Mireles-Rivera, Gloria María Molina-Salinas, Hector Montoya-Fuentes, , Mario Mújica-Sánchez, José Esteban Muñoz-Medina, José de Jesús Nuñez-Contreras, Alicia Ocaña-Mondragón, Luis Alberto Ochoa-Carrera, Hector Esteban Paz-Juárez, Marissa Perez-Garcia, Francisco Pulido, Helen Haydee Fernanda Ramírez-Plascencia, Jorge Salas-Hernández, Angel Gustavo Salas-Lais, Alejandro Sanchez-Flores, Clara Esperanza Santacruz-Tinoco, María Guadalupe Santiago-Mauricio, Selene Zárate, Nelly Sélem-Mojica, Blanca Taboada, Gloria Vazquez, Joel Armando Vázquez-Pérez. |
| EPI_ISL_2490359, EPI_ISL_2490360, EPI_ISL_2490361, EPI_ISL_2490364, EPI_ISL_2490365 | Centro de Investigación Biomédica del Noreste (CIBIN) | Centro de Investigación en Enfermedades Infecciosas (CIENI), Instituto Nacional de Enfermedades Respiratorias (INER) | Consorcio Mexicano de Vigilancia Genómica (CoViGen-Mex). Authors (in alphabetical order): Julio Elias Alvarado-Yaah, Carlos F. Arias, Santiago Ávila-Ríos, Eduardo Becerril-Vargas, Víctor Hugo Borja-Aburto, Celia Boukadida, Cristóbal Cháidez-Quiróz, Juan Bautista Chale-Dzul, Ricardo Ciria Merce, Andreu Comas-García, Célida Duque Molina, Julissa Enciso-Ibarra, José Antonio Enciso-Moreno, Gloria Elena Espinosa-Ayala, Fernando Fontove-Herrera, Daniel Fregoso-Rueda, Victor Eduardo Garcia-Arias, Alejandra Garcia-Gasca, Bruno Gomez-Gil, Jean Pierre González, Irvin González-López, Concepción Grajales-Muñiz, Ricardo Grande, Rosa María Gutierrez Rios, Alejandra Hernández-Terán, Alfredo Herrera-Estrella, Carla Ivón Herrera-Najera, Pavel Isa, Daniel Lira Morales, Susana Lopez, Antonio Loza Román, Brenda Irasema Maldonado-Meza, Bernardo Martínez-Miguel, José Arturo Martínez-Orozco, Célida Martínez- Rodríguez, Margarita Matías-Florentino, Fidencio Mejía-Nepomuceno, María Guadalupe de Jesús Mireles-Rivera, Gloria María Molina-Salinas, Hector Montoya-Fuentes, , Mario Mújica-Sánchez, José Esteban Muñoz-Medina, José de Jesús Nuñez-Contreras, Alicia Ocaña-Mondragón, Luis Alberto Ochoa-Carrera, Hector Esteban Paz-Juárez, Marissa Perez-Garcia, Francisco Pulido, Helen Haydee Fernanda Ramírez-Plascencia, Jorge Salas-Hernández, Angel Gustavo Salas-Lais, Alejandro Sanchez-Flores, Clara Esperanza Santacruz-Tinoco, María Guadalupe Santiago-Mauricio, Selene Zárate, Nelly Sélem-Mojica, Blanca Taboada, Gloria Vazquez, Joel Armando Vázquez-Pérez. |
| EPI_ISL_2490367, EPI_ISL_2490368, EPI_ISL_2490369, EPI_ISL_2490370, EPI_ISL_2490371, EPI_ISL_2490372 | Unidad de Investigación Médica de Yucatán (UIMY) | Centro de Investigación en Enfermedades Infecciosas (CIENI), Instituto Nacional de Enfermedades Respiratorias (INER) | Consorcio Mexicano de Vigilancia Genómica (CoViGen-Mex). Authors (in alphabetical order): Julio Elias Alvarado-Yaah, Carlos F. Arias, Santiago Ávila-Ríos, Eduardo Becerril-Vargas, Víctor Hugo Borja-Aburto, Celia Boukadida, Cristóbal Cháidez-Quiróz, Juan Bautista Chale-Dzul, Ricardo Ciria Merce, Andreu Comas-García, Célida Duque Molina, Julissa Enciso-Ibarra, José Antonio Enciso-Moreno, Gloria Elena Espinosa-Ayala, Fernando Fontove-Herrera, Daniel Fregoso-Rueda, Victor Eduardo Garcia-Arias, Alejandra Garcia-Gasca, Bruno Gomez-Gil, Jean Pierre González, Irvin González-López, Concepción Grajales-Muñiz, Ricardo Grande, Rosa María Gutierrez Rios, Alejandra Hernández-Terán, Alfredo Herrera-Estrella, Carla Ivón Herrera-Najera, Pavel Isa, Daniel Lira Morales, Susana Lopez, Antonio Loza Román, Brenda Irasema Maldonado-Meza, Bernardo Martínez-Miguel, José Arturo Martínez-Orozco, Célida Martínez- Rodríguez, Margarita Matías-Florentino, Fidencio Mejía-Nepomuceno, María Guadalupe de Jesús Mireles-Rivera, Gloria María Molina-Salinas, Hector Montoya-Fuentes, , Mario Mújica-Sánchez, José Esteban Muñoz-Medina, José de Jesús Nuñez-Contreras, Alicia Ocaña-Mondragón, Luis Alberto Ochoa-Carrera, Hector Esteban Paz-Juárez, Marissa Perez-Garcia, Francisco Pulido, Helen Haydee Fernanda Ramírez-Plascencia, Jorge Salas-Hernández, Angel Gustavo Salas-Lais, Alejandro Sanchez-Flores, Clara Esperanza Santacruz-Tinoco, María Guadalupe Santiago-Mauricio, Selene Zárate, Nelly Sélem-Mojica, Blanca Taboada, Gloria Vazquez, Joel Armando Vázquez-Pérez. |
| EPI_ISL_2490385 | Unidad de Investigación Biomédica de Zacatecas (UIBZ) | Centro de Investigación en Enfermedades Infecciosas (CIENI), Instituto Nacional de Enfermedades Respiratorias (INER) | Consorcio Mexicano de Vigilancia Genómica (CoViGen-Mex). Authors (in alphabetical order): Julio Elias Alvarado-Yaah, Carlos F. Arias, Santiago Ávila-Ríos, Eduardo Becerril-Vargas, Víctor Hugo Borja-Aburto, Celia Boukadida, Cristóbal Cháidez-Quiróz, Juan Bautista Chale-Dzul, Ricardo Ciria Merce, Andreu Comas-García, Célida Duque Molina, Julissa Enciso-Ibarra, José Antonio Enciso-Moreno, Gloria Elena Espinosa-Ayala, Fernando Fontove-Herrera, Daniel Fregoso-Rueda, Victor Eduardo Garcia-Arias, Alejandra Garcia-Gasca, Bruno Gomez-Gil, Jean Pierre González, Irvin González-López, Concepción Grajales-Muñiz, Ricardo Grande, Rosa María Gutierrez Rios, Alejandra Hernández-Terán, Alfredo Herrera-Estrella, Carla |

| | | | |
|---|---|---|---|
| | | | Ivón Herrera-Najera, Pavel Isa, Daniel Lira Morales, Susana Lopez, Antonio Loza Román, Brenda Irasema Maldonado-Meza, Bernardo Martínez-Miguel, José Arturo Martínez-Orozco, Célida Martínez- Rodríguez, Margarita Matías-Florentino, Fidencio Mejía-Nepomuceno, María Guadalupe de Jesús Mireles-Rivera, Gloria María Molina-Salinas, Hector Montoya-Fuentes, , Mario Mújica-Sánchez, José Esteban Muñoz-Medina, José de Jesús Nuñez-Contreras, Alicia Ocaña-Mondragón, Luis Alberto Ochoa-Carrera, Hector Esteban Paz-Juárez, Marissa Perez-Garcia, Francisco Pulido, Helen Haydee Fernanda Ramírez-Plascencia, Jorge Salas-Hernández, Angel Gustavo Salas-Lais, Alejandro Sanchez-Flores, Clara Esperanza Santacruz-Tinoco, María Guadalupe Santiago-Mauricio, Selene Zárate, Nelly Sélem-Mojica, Blanca Taboada, Gloria Vazquez, Joel Armando Vázquez-Pérez. |
| EPI_ISL_2490394 | Centro de Investigación Biomédica del Noreste (CIBIN) | Centro de Investigación en Enfermedades Infecciosas (CIENI), Instituto Nacional de Enfermedades Respiratorias (INER) | Consorcio Mexicano de Vigilancia Genómica (CoViGen-Mex). Authors (in alphabetical order): Julio Elias Alvarado-Yaah, Carlos F. Arias, Santiago Ávila-Ríos, Eduardo Becerril-Vargas, Víctor Hugo Borja-Aburto, Celia Boukadida, Cristóbal Cháidez-Quiróz, Juan Bautista Chale-Dzul, Ricardo Ciria Merce, Andreu Comas-García, Célida Duque Molina, Julissa Enciso-Ibarra, José Antonio Enciso-Moreno, Gloria Elena Espinosa-Ayala, Fernando Fontove-Herrera, Daniel Fregoso-Rueda, Victor Eduardo Garcia-Arias, Alejandra Garcia-Gasca, Bruno Gomez-Gil, Jean Pierre González, Irvin González-López, Concepción Grajales-Muñiz, Ricardo Grande, Rosa María Gutierrez Rios, Alejandra Hernández-Terán, Alfredo Herrera-Estrella, Carla Ivón Herrera-Najera, Pavel Isa, Daniel Lira Morales, Susana Lopez, Antonio Loza Román, Brenda Irasema Maldonado-Meza, Bernardo Martínez-Miguel, José Arturo Martínez-Orozco, Célida Martínez- Rodríguez, Margarita Matías-Florentino, Fidencio Mejía-Nepomuceno, María Guadalupe de Jesús Mireles-Rivera, Gloria María Molina-Salinas, Hector Montoya-Fuentes, , Mario Mújica-Sánchez, José Esteban Muñoz-Medina, José de Jesús Nuñez-Contreras, Alicia Ocaña-Mondragón, Luis Alberto Ochoa-Carrera, Hector Esteban Paz-Juárez, Marissa Perez-Garcia, Francisco Pulido, Helen Haydee Fernanda Ramírez-Plascencia, Jorge Salas-Hernández, Angel Gustavo Salas-Lais, Alejandro Sanchez-Flores, Clara Esperanza Santacruz-Tinoco, María Guadalupe Santiago-Mauricio, Selene Zárate, Nelly Sélem-Mojica, Blanca Taboada, Gloria Vazquez, Joel Armando Vázquez-Pérez. |
| EPI_ISL_2490395 | Centro de Investigación Biomédica de Occidente (CIBO) | Centro de Investigación en Enfermedades Infecciosas (CIENI), Instituto Nacional de Enfermedades Respiratorias (INER) | Consorcio Mexicano de Vigilancia Genómica (CoViGen-Mex). Authors (in alphabetical order): Julio Elias Alvarado-Yaah, Carlos F. Arias, Santiago Ávila-Ríos, Eduardo Becerril-Vargas, Víctor Hugo Borja-Aburto, Celia Boukadida, Cristóbal Cháidez-Quiróz, Juan Bautista Chale-Dzul, Ricardo Ciria Merce, Andreu Comas-García, Célida Duque Molina, Julissa Enciso-Ibarra, José Antonio Enciso-Moreno, Gloria Elena Espinosa-Ayala, Fernando Fontove-Herrera, Daniel Fregoso-Rueda, Victor Eduardo Garcia-Arias, Alejandra Garcia-Gasca, Bruno Gomez-Gil, Jean Pierre González, Irvin González-López, Concepción Grajales-Muñiz, Ricardo Grande, Rosa María Gutierrez Rios, Alejandra Hernández-Terán, Alfredo Herrera-Estrella, Carla Ivón Herrera-Najera, Pavel Isa, Daniel Lira Morales, Susana Lopez, Antonio Loza Román, Brenda Irasema Maldonado-Meza, Bernardo Martínez-Miguel, José Arturo Martínez-Orozco, Célida Martínez- Rodríguez, Margarita Matías-Florentino, Fidencio Mejía-Nepomuceno, María Guadalupe de Jesús Mireles-Rivera, Gloria María Molina-Salinas, Hector Montoya-Fuentes, , Mario Mújica-Sánchez, José Esteban Muñoz-Medina, José de Jesús Nuñez-Contreras, Alicia Ocaña-Mondragón, Luis Alberto Ochoa-Carrera, Hector Esteban Paz-Juárez, Marissa Perez-Garcia, Francisco Pulido, Helen Haydee Fernanda Ramírez-Plascencia, Jorge Salas-Hernández, Angel Gustavo Salas-Lais, Alejandro Sanchez-Flores, Clara Esperanza Santacruz-Tinoco, María Guadalupe Santiago-Mauricio, Selene Zárate, Nelly Sélem-Mojica, Blanca Taboada, Gloria Vazquez, Joel Armando Vázquez-Pérez. |
| EPI_ISL_2490409 | Centro de Investigación Biomédica del Noreste (CIBIN) | Centro de Investigación en Enfermedades Infecciosas (CIENI), Instituto Nacional de Enfermedades Respiratorias (INER) | Consorcio Mexicano de Vigilancia Genómica (CoViGen-Mex). Authors (in alphabetical order): Julio Elias Alvarado-Yaah, Carlos F. Arias, Santiago Ávila-Ríos, Eduardo Becerril-Vargas, Víctor Hugo Borja-Aburto, Celia Boukadida, Cristóbal Cháidez-Quiróz, Juan Bautista Chale-Dzul, Ricardo Ciria Merce, Andreu Comas-García, Célida Duque Molina, Julissa Enciso-Ibarra, José Antonio Enciso-Moreno, Gloria Elena Espinosa-Ayala, Fernando Fontove-Herrera, Daniel Fregoso-Rueda, Victor Eduardo Garcia-Arias, Alejandra Garcia-Gasca, Bruno Gomez-Gil, Jean Pierre González, Irvin González-López, Concepción Grajales-Muñiz, Ricardo Grande, Rosa María Gutierrez Rios, Alejandra Hernández-Terán, Alfredo Herrera-Estrella, Carla Ivón Herrera-Najera, Pavel Isa, Daniel Lira Morales, Susana Lopez, Antonio Loza Román, Brenda Irasema Maldonado-Meza, Bernardo Martínez-Miguel, José Arturo Martínez-Orozco, Célida Martínez- Rodríguez, Margarita Matías-Florentino, Fidencio Mejía-Nepomuceno, María Guadalupe de Jesús Mireles-Rivera, Gloria María Molina-Salinas, Hector Montoya-Fuentes, , Mario Mújica-Sánchez, José Esteban Muñoz-Medina, José de Jesús Nuñez-Contreras, Alicia Ocaña-Mondragón, Luis Alberto Ochoa-Carrera, Hector Esteban Paz-Juárez, Marissa Perez-Garcia, Francisco Pulido, Helen Haydee Fernanda Ramírez-Plascencia, Jorge Salas-Hernández, Angel Gustavo Salas-Lais, Alejandro Sanchez-Flores, Clara Esperanza Santacruz-Tinoco, María Guadalupe Santiago-Mauricio, Selene Zárate, Nelly Sélem-Mojica, Blanca Taboada, Gloria Vazquez, Joel Armando Vázquez-Pérez. |
| EPI_ISL_2490417 | Laboratorio Central de Epidemiología (LCE) | Centro de Investigación en Enfermedades Infecciosas (CIENI), Instituto Nacional de Enfermedades Respiratorias (INER) | Consorcio Mexicano de Vigilancia Genómica (CoViGen-Mex). Authors (in alphabetical order): Julio Elias Alvarado-Yaah, Carlos F. Arias, Santiago Ávila-Ríos, Eduardo Becerril-Vargas, Víctor Hugo Borja-Aburto, Celia Boukadida, Cristóbal Cháidez-Quiróz, Juan Bautista Chale-Dzul, Ricardo Ciria Merce, Andreu Comas-García, Célida Duque Molina, Julissa Enciso-Ibarra, José Antonio Enciso-Moreno, Gloria Elena Espinosa-Ayala, Fernando Fontove-Herrera, Daniel Fregoso-Rueda, Victor Eduardo Garcia-Arias, Alejandra Garcia-Gasca, Bruno Gomez-Gil, Jean Pierre González, Irvin González-López, Concepción Grajales-Muñiz, Ricardo Grande, Rosa María Gutierrez Rios, Alejandra Hernández-Terán, Alfredo Herrera-Estrella, Carla Ivón Herrera-Najera, Pavel Isa, Daniel Lira Morales, Susana Lopez, Antonio Loza Román, Brenda Irasema Maldonado-Meza, Bernardo Martínez-Miguel, José Arturo Martínez-Orozco, Célida Martínez- Rodríguez, Margarita Matías-Florentino, Fidencio Mejía-Nepomuceno, María Guadalupe de Jesús Mireles-Rivera, Gloria María Molina-Salinas, Hector Montoya-Fuentes, , Mario Mújica-Sánchez, José Esteban Muñoz-Medina, José de Jesús Nuñez-Contreras, Alicia Ocaña-Mondragón, Luis Alberto Ochoa-Carrera, Hector Esteban Paz-Juárez, Marissa Perez-Garcia, Francisco Pulido, Helen Haydee Fernanda Ramírez-Plascencia, Jorge Salas-Hernández, Angel Gustavo Salas-Lais, Alejandro Sanchez-Flores, Clara Esperanza Santacruz-Tinoco, María Guadalupe Santiago-Mauricio, Selene Zárate, Nelly Sélem-Mojica, Blanca Taboada, Gloria Vazquez, Joel Armando Vázquez-Pérez. |
| EPI_ISL_2490418, EPI_ISL_2490419, EPI_ISL_2490420 | Unidad de Investigación Médica de Yucatán (UIMY) | Centro de Investigación en Enfermedades Infecciosas (CIENI), Instituto Nacional de Enfermedades Respiratorias (INER) | Consorcio Mexicano de Vigilancia Genómica (CoViGen-Mex). Authors (in alphabetical order): Julio Elias Alvarado-Yaah, Carlos F. Arias, Santiago Ávila-Ríos, Eduardo Becerril-Vargas, Víctor Hugo Borja-Aburto, Celia Boukadida, Cristóbal Cháidez-Quiróz, Juan Bautista Chale-Dzul, Ricardo Ciria Merce, Andreu Comas-García, Célida Duque Molina, Julissa Enciso-Ibarra, José Antonio Enciso-Moreno, Gloria Elena Espinosa-Ayala, Fernando Fontove-Herrera, Daniel Fregoso-Rueda, Victor Eduardo Garcia-Arias, Alejandra Garcia-Gasca, Bruno Gomez-Gil, Jean Pierre González, Irvin González-López, Concepción Grajales-Muñiz, Ricardo Grande, Rosa María Gutierrez Rios, Alejandra Hernández-Terán, Alfredo Herrera-Estrella, Carla Ivón Herrera-Najera, Pavel Isa, Daniel Lira Morales, Susana Lopez, Antonio Loza Román, Brenda Irasema Maldonado-Meza, Bernardo Martínez-Miguel, José Arturo Martínez-Orozco, Célida Martínez- Rodríguez, Margarita Matías-Florentino, Fidencio Mejía-Nepomuceno, María Guadalupe de Jesús Mireles-Rivera, Gloria María Molina-Salinas, Hector Montoya-Fuentes, , Mario Mújica-Sánchez, José Esteban Muñoz-Medina, José de Jesús Nuñez-Contreras, Alicia Ocaña-Mondragón, Luis Alberto Ochoa-Carrera, Hector Esteban Paz-Juárez, Marissa Perez-Garcia, Francisco Pulido, Helen Haydee Fernanda Ramírez-Plascencia, Jorge Salas-Hernández, Angel Gustavo Salas-Lais, Alejandro Sanchez-Flores, Clara Esperanza Santacruz-Tinoco, María Guadalupe Santiago-Mauricio, Selene Zárate, Nelly Sélem-Mojica, Blanca Taboada, Gloria Vazquez, Joel Armando Vázquez-Pérez. |
| EPI_ISL_2490427, EPI_ISL_2490428, EPI_ISL_2490429, EPI_ISL_2490430, EPI_ISL_2490431, EPI_ISL_2490432 | Centro de Investigación Biomédica del Noreste (CIBIN) | Centro de Investigación en Enfermedades Infecciosas (CIENI), Instituto Nacional de Enfermedades Respiratorias (INER) | Consorcio Mexicano de Vigilancia Genómica (CoViGen-Mex). Authors (in alphabetical order): Julio Elias Alvarado-Yaah, Carlos F. Arias, Santiago Ávila-Ríos, Eduardo Becerril-Vargas, Víctor Hugo Borja-Aburto, Celia Boukadida, Cristóbal Cháidez-Quiróz, Juan Bautista Chale-Dzul, Ricardo Ciria Merce, Andreu Comas-García, Célida Duque Molina, Julissa Enciso-Ibarra, José Antonio Enciso-Moreno, Gloria Elena Espinosa-Ayala, Fernando Fontove-Herrera, Daniel Fregoso-Rueda, Victor Eduardo Garcia-Arias, Alejandra Garcia-Gasca, Bruno Gomez-Gil, Jean Pierre González, Irvin González-López, Concepción Grajales-Muñiz, Ricardo Grande, Rosa María Gutierrez Rios, Alejandra Hernández-Terán, Alfredo Herrera-Estrella, Carla Ivón Herrera-Najera, Pavel Isa, Daniel Lira Morales, Susana Lopez, Antonio Loza Román, Brenda Irasema Maldonado-Meza, Bernardo Martínez-Miguel, José Arturo Martínez-Orozco, Célida Martínez- Rodríguez, Margarita Matías-Florentino, Fidencio Mejía-Nepomuceno, María Guadalupe de Jesús Mireles-Rivera, Gloria María Molina-Salinas, Hector Montoya-Fuentes, , Mario Mújica-Sánchez, José Esteban Muñoz-Medina, José de Jesús Nuñez-Contreras, Alicia Ocaña-Mondragón, Luis Alberto Ochoa-Carrera, Hector Esteban Paz-Juárez, Marissa Perez-Garcia, Francisco Pulido, Helen Haydee Fernanda Ramírez-Plascencia, Jorge Salas-Hernández, Angel Gustavo Salas-Lais, Alejandro Sanchez-Flores, Clara Esperanza Santacruz-Tinoco, María Guadalupe Santiago-Mauricio, Selene Zárate, Nelly Sélem-Mojica, Blanca Taboada, Gloria Vazquez, Joel Armando Vázquez-Pérez. |
| EPI_ISL_2490436 | Laboratorio Central de Epidemiología (LCE) | Centro de Investigación en Enfermedades Infecciosas (CIENI), Instituto Nacional de Enfermedades Respiratorias (INER) | Consorcio Mexicano de Vigilancia Genómica (CoViGen-Mex). Authors (in alphabetical order): Julio Elias Alvarado-Yaah, Carlos F. Arias, Santiago Ávila-Ríos, Eduardo Becerril-Vargas, Víctor Hugo Borja-Aburto, Celia Boukadida, Cristóbal Cháidez-Quiróz, Juan Bautista Chale-Dzul, Ricardo Ciria Merce, Andreu Comas-García, Célida Duque Molina, Julissa Enciso-Ibarra, José Antonio Enciso-Moreno, Gloria Elena Espinosa-Ayala, Fernando Fontove-Herrera, Daniel Fregoso-Rueda, Victor Eduardo Garcia-Arias, Alejandra Garcia-Gasca, Bruno Gomez-Gil, Jean Pierre González, Irvin González-López, Concepción Grajales-Muñiz, Ricardo Grande, Rosa María Gutierrez Rios, Alejandra Hernández-Terán, Alfredo Herrera-Estrella, Carla |

| | | | |
|---|---|---|---|
| | | | Ivón Herrera-Najera, Pavel Isa, Daniel Lira Morales, Susana Lopez, Antonio Loza Román, Brenda Irasema Maldonado-Meza, Bernardo Martínez-Miguel, José Arturo Martínez-Orozco, Célida Martínez- Rodríguez, Margarita Matías-Florentino, Fidencio Mejía-Nepomuceno, María Guadalupe de Jesús Mireles-Rivera, Gloria María Molina-Salinas, Hector Montoya-Fuentes, , Mario Mújica-Sánchez, José Esteban Muñoz-Medina, José de Jesús Nuñez-Contreras, Alicia Ocaña-Mondragón, Luis Alberto Ochoa-Carrera, Hector Esteban Paz-Juárez, Marissa Perez-Garcia, Francisco Pulido, Helen Haydee Fernanda Ramírez-Plascencia, Jorge Salas-Hernández, Angel Gustavo Salas-Lais, Alejandro Sanchez-Flores, Clara Esperanza Santacruz-Tinoco, María Guadalupe Santiago-Mauricio, Selene Zárate, Nelly Sélem-Mojica, Blanca Taboada, Gloria Vazquez, Joel Armando Vázquez-Pérez. |
| EPI_ISL_2490439, EPI_ISL_2490440, EPI_ISL_2490441, EPI_ISL_2490442, EPI_ISL_2490443, EPI_ISL_2490444, EPI_ISL_2490445, EPI_ISL_2490446, EPI_ISL_2490447, EPI_ISL_2490448, EPI_ISL_2490449, EPI_ISL_2490450, EPI_ISL_2490451, EPI_ISL_2490453 | | | |
| see above | Centro de Investigación Biomédica del Noreste (CIBIN) | Centro de Investigación en Enfermedades Infecciosas (CIENI), Instituto Nacional de Enfermedades Respiratorias (INER) | Consorcio Mexicano de Vigilancia Genómica (CoViGen-Mex). Authors (in alphabetical order): Julio Elias Alvarado-Yaah, Carlos F. Arias, Santiago Ávila-Ríos, Eduardo Becerril-Vargas, Víctor Hugo Borja-Aburto, Celia Boukadida, Cristóbal Cháidez-Quiróz, Juan Bautista Chale-Dzul, Ricardo Ciria Merce, Andreu Comas-García, Célida Duque Molina, Julissa Enciso-Ibarra, José Antonio Enciso-Moreno, Gloria Elena Espinosa-Ayala, Fernando Fontove-Herrera, Daniel Fregoso-Rueda, Victor Eduardo Garcia-Arias, Alejandra Garcia-Gasca, Bruno Gomez-Gil, Jean Pierre González, Irvin González-López, Concepción Grajales-Muñiz, Ricardo Grande, Rosa María Gutierrez Rios, Alejandra Hernández-Terán, Alfredo Herrera-Estrella, Carla Ivón Herrera-Najera, Pavel Isa, Daniel Lira Morales, Susana Lopez, Antonio Loza Román, Brenda Irasema Maldonado-Meza, Bernardo Martínez-Miguel, José Arturo Martínez-Orozco, Célida Martínez- Rodríguez, Margarita Matías-Florentino, Fidencio Mejía-Nepomuceno, María Guadalupe de Jesús Mireles-Rivera, Gloria María Molina-Salinas, Hector Montoya-Fuentes, , Mario Mújica-Sánchez, José Esteban Muñoz-Medina, José de Jesús Nuñez-Contreras, Alicia Ocaña-Mondragón, Luis Alberto Ochoa-Carrera, Hector Esteban Paz-Juárez, Marissa Perez-Garcia, Francisco Pulido, Helen Haydee Fernanda Ramírez-Plascencia, Jorge Salas-Hernández, Angel Gustavo Salas-Lais, Alejandro Sanchez-Flores, Clara Esperanza Santacruz-Tinoco, María Guadalupe Santiago-Mauricio, Selene Zárate, Nelly Sélem-Mojica, Blanca Taboada, Gloria Vazquez, Joel Armando Vázquez-Pérez. |
| EPI_ISL_2490466, EPI_ISL_2490467, EPI_ISL_2490468, EPI_ISL_2490469, EPI_ISL_2490470, EPI_ISL_2490471, EPI_ISL_2490472, EPI_ISL_2490473, EPI_ISL_2490474, EPI_ISL_2490475, EPI_ISL_2490476, EPI_ISL_2490477, EPI_ISL_2490478 | | | |
| see above | Unidad de Investigación Médica de Yucatán (UIMY) | Centro de Investigación en Enfermedades Infecciosas (CIENI), Instituto Nacional de Enfermedades Respiratorias (INER) | Consorcio Mexicano de Vigilancia Genómica (CoViGen-Mex). Authors (in alphabetical order): Julio Elias Alvarado-Yaah, Carlos F. Arias, Santiago Ávila-Ríos, Eduardo Becerril-Vargas, Víctor Hugo Borja-Aburto, Celia Boukadida, Cristóbal Cháidez-Quiróz, Juan Bautista Chale-Dzul, Ricardo Ciria Merce, Andreu Comas-García, Célida Duque Molina, Julissa Enciso-Ibarra, José Antonio Enciso-Moreno, Gloria Elena Espinosa-Ayala, Fernando Fontove-Herrera, Daniel Fregoso-Rueda, Victor Eduardo Garcia-Arias, Alejandra Garcia-Gasca, Bruno Gomez-Gil, Jean Pierre González, Irvin González-López, Concepción Grajales-Muñiz, Ricardo Grande, Rosa María Gutierrez Rios, Alejandra Hernández-Terán, Alfredo Herrera-Estrella, Carla Ivón Herrera-Najera, Pavel Isa, Daniel Lira Morales, Susana Lopez, Antonio Loza Román, Brenda Irasema Maldonado-Meza, Bernardo Martínez-Miguel, José Arturo Martínez-Orozco, Célida Martínez- Rodríguez, Margarita Matías-Florentino, Fidencio Mejía-Nepomuceno, María Guadalupe de Jesús Mireles-Rivera, Gloria María Molina-Salinas, Hector Montoya-Fuentes, , Mario Mújica-Sánchez, José Esteban Muñoz-Medina, José de Jesús Nuñez-Contreras, Alicia Ocaña-Mondragón, Luis Alberto Ochoa-Carrera, Hector Esteban Paz-Juárez, Marissa Perez-Garcia, Francisco Pulido, Helen Haydee Fernanda Ramírez-Plascencia, Jorge Salas-Hernández, Angel Gustavo Salas-Lais, Alejandro Sanchez-Flores, Clara Esperanza Santacruz-Tinoco, María Guadalupe Santiago-Mauricio, Selene Zárate, Nelly Sélem-Mojica, Blanca Taboada, Gloria Vazquez, Joel Armando Vázquez-Pérez. |
| EPI_ISL_2490493, EPI_ISL_2490494, EPI_ISL_2490495, EPI_ISL_2490497, EPI_ISL_2490499 | Centro de Investigación Biomédica de Occidente (CIBO) | Centro de Investigación en Enfermedades Infecciosas (CIENI), Instituto Nacional de Enfermedades Respiratorias (INER) | Consorcio Mexicano de Vigilancia Genómica (CoViGen-Mex). Authors (in alphabetical order): Julio Elias Alvarado-Yaah, Carlos F. Arias, Santiago Ávila-Ríos, Eduardo Becerril-Vargas, Víctor Hugo Borja-Aburto, Celia Boukadida, Cristóbal Cháidez-Quiróz, Juan Bautista Chale-Dzul, Ricardo Ciria Merce, Andreu Comas-García, Célida Duque Molina, Julissa Enciso-Ibarra, José Antonio Enciso-Moreno, Gloria Elena Espinosa-Ayala, Fernando Fontove-Herrera, Daniel Fregoso-Rueda, Victor Eduardo Garcia-Arias, Alejandra Garcia-Gasca, Bruno Gomez-Gil, Jean Pierre González, Irvin González-López, Concepción Grajales-Muñiz, Ricardo Grande, Rosa María Gutierrez Rios, Alejandra Hernández-Terán, Alfredo Herrera-Estrella, Carla Ivón Herrera-Najera, Pavel Isa, Daniel Lira Morales, Susana Lopez, Antonio Loza Román, Brenda Irasema Maldonado-Meza, Bernardo Martínez-Miguel, José Arturo Martínez-Orozco, Célida Martínez- Rodríguez, Margarita Matías-Florentino, Fidencio Mejía-Nepomuceno, María Guadalupe de Jesús Mireles-Rivera, Gloria María Molina-Salinas, Hector Montoya-Fuentes, , Mario Mújica-Sánchez, José Esteban Muñoz-Medina, José de Jesús Nuñez-Contreras, Alicia Ocaña-Mondragón, Luis Alberto Ochoa-Carrera, Hector Esteban Paz-Juárez, Marissa Perez-Garcia, Francisco Pulido, Helen Haydee Fernanda Ramírez-Plascencia, Jorge Salas-Hernández, Angel Gustavo Salas-Lais, Alejandro Sanchez-Flores, Clara Esperanza Santacruz-Tinoco, María Guadalupe Santiago-Mauricio, Selene Zárate, Nelly Sélem-Mojica, Blanca Taboada, Gloria Vazquez, Joel Armando Vázquez-Pérez. |
| EPI_ISL_2490500, EPI_ISL_2490501, EPI_ISL_2490502, EPI_ISL_2490503, EPI_ISL_2490504, EPI_ISL_2490505, EPI_ISL_2490506, EPI_ISL_2490507, EPI_ISL_2490508, EPI_ISL_2490509, EPI_ISL_2490510 | | | |
| see above | Unidad de Investigación Biomédica de Zacatecas (UIBZ) | Centro de Investigación en Enfermedades Infecciosas (CIENI), Instituto Nacional de Enfermedades Respiratorias (INER) | Consorcio Mexicano de Vigilancia Genómica (CoViGen-Mex). Authors (in alphabetical order): Julio Elias Alvarado-Yaah, Carlos F. Arias, Santiago Ávila-Ríos, Eduardo Becerril-Vargas, Víctor Hugo Borja-Aburto, Celia Boukadida, Cristóbal Cháidez-Quiróz, Juan Bautista Chale-Dzul, Ricardo Ciria Merce, Andreu Comas-García, Célida Duque Molina, Julissa Enciso-Ibarra, José Antonio Enciso-Moreno, Gloria Elena Espinosa-Ayala, Fernando Fontove-Herrera, Daniel Fregoso-Rueda, Victor Eduardo Garcia-Arias, Alejandra Garcia-Gasca, Bruno Gomez-Gil, Jean Pierre González, Irvin González-López, Concepción Grajales-Muñiz, Ricardo Grande, Rosa María Gutierrez Rios, Alejandra Hernández-Terán, Alfredo Herrera-Estrella, Carla Ivón Herrera-Najera, Pavel Isa, Daniel Lira Morales, Susana Lopez, Antonio Loza Román, Brenda Irasema Maldonado-Meza, Bernardo Martínez-Miguel, José Arturo Martínez-Orozco, Célida Martínez- Rodríguez, Margarita Matías-Florentino, Fidencio Mejía-Nepomuceno, María Guadalupe de Jesús Mireles-Rivera, Gloria María Molina-Salinas, Hector Montoya-Fuentes, , Mario Mújica-Sánchez, José Esteban Muñoz-Medina, José de Jesús Nuñez-Contreras, Alicia Ocaña-Mondragón, Luis Alberto Ochoa-Carrera, Hector Esteban Paz-Juárez, Marissa Perez-Garcia, Francisco Pulido, Helen Haydee Fernanda Ramírez-Plascencia, Jorge Salas-Hernández, Angel Gustavo Salas-Lais, Alejandro Sanchez-Flores, Clara Esperanza Santacruz-Tinoco, María Guadalupe Santiago-Mauricio, Selene Zárate, Nelly Sélem-Mojica, Blanca Taboada, Gloria Vazquez, Joel Armando Vázquez-Pérez. |
| EPI_ISL_2490558, EPI_ISL_2490559, EPI_ISL_2490560, EPI_ISL_2490561, EPI_ISL_2490563, EPI_ISL_2490583 | Laboratorio Central de Epidemiología (LCE) | Centro de Investigación en Enfermedades Infecciosas (CIENI), Instituto Nacional de Enfermedades Respiratorias (INER) | Consorcio Mexicano de Vigilancia Genómica (CoViGen-Mex). Authors (in alphabetical order): Julio Elias Alvarado-Yaah, Carlos F. Arias, Santiago Ávila-Ríos, Eduardo Becerril-Vargas, Víctor Hugo Borja-Aburto, Celia Boukadida, Cristóbal Cháidez-Quiróz, Juan Bautista Chale-Dzul, Ricardo Ciria Merce, Andreu Comas-García, Célida Duque Molina, Julissa Enciso-Ibarra, José Antonio Enciso-Moreno, Gloria Elena Espinosa-Ayala, Fernando Fontove-Herrera, Daniel Fregoso-Rueda, Victor Eduardo Garcia-Arias, Alejandra Garcia-Gasca, Bruno Gomez-Gil, Jean Pierre González, Irvin González-López, Concepción Grajales-Muñiz, Ricardo Grande, Rosa María Gutierrez Rios, Alejandra Hernández-Terán, Alfredo Herrera-Estrella, Carla Ivón Herrera-Najera, Pavel Isa, Daniel Lira Morales, Susana Lopez, Antonio Loza Román, Brenda Irasema Maldonado-Meza, Bernardo Martínez-Miguel, José Arturo Martínez-Orozco, Célida Martínez- Rodríguez, Margarita Matías-Florentino, Fidencio Mejía-Nepomuceno, María Guadalupe de Jesús Mireles-Rivera, Gloria María Molina-Salinas, Hector Montoya-Fuentes, , Mario Mújica-Sánchez, José Esteban Muñoz-Medina, José de Jesús Nuñez-Contreras, Alicia Ocaña-Mondragón, Luis Alberto Ochoa-Carrera, Hector Esteban Paz-Juárez, Marissa Perez-Garcia, Francisco Pulido, Helen Haydee Fernanda Ramírez-Plascencia, Jorge Salas-Hernández, Angel Gustavo Salas-Lais, Alejandro Sanchez-Flores, Clara Esperanza Santacruz-Tinoco, María Guadalupe Santiago-Mauricio, Selene Zárate, Nelly Sélem-Mojica, Blanca Taboada, Gloria Vazquez, Joel Armando Vázquez-Pérez. |
| EPI_ISL_2490592, EPI_ISL_2490593, EPI_ISL_2490594 | Instituto Nacional de Enfermedades Respiratorias (INER) | Centro de Investigación en Enfermedades Infecciosas (CIENI), Instituto Nacional de Enfermedades Respiratorias (INER) | Consorcio Mexicano de Vigilancia Genómica (CoViGen-Mex). Authors (in alphabetical order): Julio Elias Alvarado-Yaah, Carlos F. Arias, Santiago Ávila-Ríos, Eduardo Becerril-Vargas, Víctor Hugo Borja-Aburto, Celia Boukadida, Cristóbal Cháidez-Quiróz, Juan Bautista Chale-Dzul, Ricardo Ciria Merce, Andreu Comas-García, Célida Duque Molina, Julissa Enciso-Ibarra, José Antonio Enciso-Moreno, Gloria Elena Espinosa-Ayala, Fernando Fontove-Herrera, Daniel Fregoso-Rueda, Victor Eduardo Garcia-Arias, Alejandra Garcia-Gasca, Bruno Gomez-Gil, Jean Pierre González, Irvin González-López, Concepción Grajales-Muñiz, Ricardo Grande, Rosa María Gutierrez Rios, Alejandra Hernández-Terán, Alfredo Herrera-Estrella, Carla Ivón Herrera-Najera, Pavel Isa, Daniel Lira Morales, Susana Lopez, Antonio Loza Román, Brenda Irasema Maldonado-Meza, Bernardo Martínez-Miguel, José Arturo Martínez-Orozco, Célida Martínez- Rodríguez, Margarita Matías-Florentino, Fidencio Mejía-Nepomuceno, María Guadalupe de Jesús Mireles-Rivera, Gloria María Molina-Salinas, Hector Montoya-Fuentes, , Mario Mújica-Sánchez, José Esteban Muñoz-Medina, José de Jesús Nuñez-Contreras, Alicia Ocaña-Mondragón, Luis Alberto Ochoa-Carrera, Hector Esteban Paz-Juárez, Marissa Perez-Garcia, Francisco Pulido, Helen Haydee Fernanda Ramírez-Plascencia, Jorge Salas-Hernández, Angel Gustavo Salas-Lais, Alejandro Sanchez-Flores, Clara Esperanza Santacruz-Tinoco, María Guadalupe Santiago-Mauricio, Selene Zárate, Nelly Sélem-Mojica, Blanca Taboada, Gloria Vazquez, Joel Armando Vázquez-Pérez. |

| | | | |
|---|---|---|---|
| EPI_ISL_2491632 | Centre Hospitalier Universitaire Clermont-Ferrand | CHU Clermont-Ferrand, service de virologie | Bisseux Maxime, Mirand Audrey, Combes Patricia, Henquell Cécile |
| EPI_ISL_2492441, EPI_ISL_2492442, EPI_ISL_2492443, EPI_ISL_2492444, EPI_ISL_2492445, EPI_ISL_2492446, EPI_ISL_2492447, EPI_ISL_2492448, EPI_ISL_2492449, EPI_ISL_2492450, EPI_ISL_2492451, EPI_ISL_2492452, EPI_ISL_2492453, EPI_ISL_2492454, EPI_ISL_2492455, EPI_ISL_2492456, EPI_ISL_2492457, EPI_ISL_2492458, EPI_ISL_2492459, EPI_ISL_2492460, EPI_ISL_2492461 | | | |
| see above | LESP Queretaro | Instituto de Diagnostico y Referencia Epidemiologicos (INDRE) | Claudia Wong-Arambula, Abril Rodriguez-Maldonado, Vanessa Rivero-Arredondo, Ariadna Medina-Benitez, Joaquin Quiroz-Mercado, Sergio Rangel-Guerrero, Natividad Cruz-Ortiz, Tatiana Nunez-Garcia, Gisela Barrera-Badillo, Lucia Hernandez-Rivas, Irma Lopez-Martinez, Ernesto Ramirez-Gonzalez. |
| EPI_ISL_2492464 | LESP Nuevo Leon | Instituto de Diagnostico y Referencia Epidemiologicos (INDRE) | Claudia Wong-Arambula, Abril Rodriguez-Maldonado, Vanessa Rivero-Arredondo, Ariadna Medina-Benitez, Joaquin Quiroz-Mercado, Sergio Rangel-Guerrero, Natividad Cruz-Ortiz, Tatiana Nunez-Garcia, Gisela Barrera-Badillo, Lucia Hernandez-Rivas, Irma Lopez-Martinez, Ernesto Ramirez-Gonzalez. |
| EPI_ISL_2492465 | LESP Guanajuato | Instituto de Diagnostico y Referencia Epidemiologicos (INDRE) | Claudia Wong-Arambula, Abril Rodriguez-Maldonado, Vanessa Rivero-Arredondo, Ariadna Medina-Benitez, Joaquin Quiroz-Mercado, Sergio Rangel-Guerrero, Natividad Cruz-Ortiz, Tatiana Nunez-Garcia, Gisela Barrera-Badillo, Lucia Hernandez-Rivas, Irma Lopez-Martinez, Ernesto Ramirez-Gonzalez. |
| EPI_ISL_2492467 | LESP Nuevo Leon | Instituto de Diagnostico y Referencia Epidemiologicos (INDRE) | Claudia Wong-Arambula, Abril Rodriguez-Maldonado, Vanessa Rivero-Arredondo, Ariadna Medina-Benitez, Joaquin Quiroz-Mercado, Sergio Rangel-Guerrero, Natividad Cruz-Ortiz, Tatiana Nunez-Garcia, Gisela Barrera-Badillo, Lucia Hernandez-Rivas, Irma Lopez-Martinez, Ernesto Ramirez-Gonzalez. |
| EPI_ISL_2492472 | LESP Oaxaca | Instituto de Diagnostico y Referencia Epidemiologicos (INDRE) | Claudia Wong-Arambula, Abril Rodriguez-Maldonado, Vanessa Rivero-Arredondo, Ariadna Medina-Benitez, Joaquin Quiroz-Mercado, Sergio Rangel-Guerrero, Natividad Cruz-Ortiz, Tatiana Nunez-Garcia, Gisela Barrera-Badillo, Lucia Hernandez-Rivas, Irma Lopez-Martinez, Ernesto Ramirez-Gonzalez. |
| EPI_ISL_2492473 | LESP San Luis Potosi | Instituto de Diagnostico y Referencia Epidemiologicos (INDRE) | Claudia Wong-Arambula, Abril Rodriguez-Maldonado, Vanessa Rivero-Arredondo, Ariadna Medina-Benitez, Joaquin Quiroz-Mercado, Sergio Rangel-Guerrero, Natividad Cruz-Ortiz, Tatiana Nunez-Garcia, Gisela Barrera-Badillo, Lucia Hernandez-Rivas, Irma Lopez-Martinez, Ernesto Ramirez-Gonzalez. |
| EPI_ISL_2492485 | LESP Guanajuato | Instituto de Diagnostico y Referencia Epidemiologicos (INDRE) | Claudia Wong-Arambula, Abril Rodriguez-Maldonado, Vanessa Rivero-Arredondo, Ariadna Medina-Benitez, Joaquin Quiroz-Mercado, Sergio Rangel-Guerrero, Natividad Cruz-Ortiz, Tatiana Nunez-Garcia, Gisela Barrera-Badillo, Lucia Hernandez-Rivas, Irma Lopez-Martinez, Ernesto Ramirez-Gonzalez. |
| EPI_ISL_2492486 | LESP Nuevo Leon | Instituto de Diagnostico y Referencia Epidemiologicos (INDRE) | Claudia Wong-Arambula, Abril Rodriguez-Maldonado, Vanessa Rivero-Arredondo, Ariadna Medina-Benitez, Joaquin Quiroz-Mercado, Sergio Rangel-Guerrero, Natividad Cruz-Ortiz, Tatiana Nunez-Garcia, Gisela Barrera-Badillo, Lucia Hernandez-Rivas, Irma Lopez-Martinez, Ernesto Ramirez-Gonzalez. |
| EPI_ISL_2492488 | LESP Puebla | Instituto de Diagnostico y Referencia Epidemiologicos (INDRE) | Claudia Wong-Arambula, Abril Rodriguez-Maldonado, Vanessa Rivero-Arredondo, Ariadna Medina-Benitez, Joaquin Quiroz-Mercado, Sergio Rangel-Guerrero, Natividad Cruz-Ortiz, Tatiana Nunez-Garcia, Gisela Barrera-Badillo, Lucia Hernandez-Rivas, Irma Lopez-Martinez, Ernesto Ramirez-Gonzalez. |
| EPI_ISL_2495483, EPI_ISL_2495484, EPI_ISL_2495485, EPI_ISL_2495486, EPI_ISL_2495487, EPI_ISL_2495488, EPI_ISL_2495489, EPI_ISL_2495490, EPI_ISL_2495491, EPI_ISL_2495555, EPI_ISL_2495566, EPI_ISL_2495567, EPI_ISL_2495568, EPI_ISL_2495569, EPI_ISL_2495570, EPI_ISL_2495571, EPI_ISL_2495572, EPI_ISL_2495573, EPI_ISL_2495574, EPI_ISL_2495575, EPI_ISL_2495576, EPI_ISL_2495577, EPI_ISL_2495578, EPI_ISL_2495579, EPI_ISL_2495580, EPI_ISL_2495581, EPI_ISL_2495582, EPI_ISL_2495583, EPI_ISL_2495584, EPI_ISL_2495585, EPI_ISL_2495596, EPI_ISL_2495601, EPI_ISL_2495602, EPI_ISL_2495604, EPI_ISL_2495621, EPI_ISL_2495627, EPI_ISL_2495667, EPI_ISL_2495668, EPI_ISL_2495669, EPI_ISL_2495670, EPI_ISL_2495671, EPI_ISL_2495672, EPI_ISL_2495673, EPI_ISL_2495674, EPI_ISL_2495675, EPI_ISL_2495676, EPI_ISL_2495677, EPI_ISL_2495678, EPI_ISL_2495679, EPI_ISL_2495680, EPI_ISL_2495681, EPI_ISL_2495682, EPI_ISL_2495735, EPI_ISL_2495736, EPI_ISL_2495737, EPI_ISL_2495738, EPI_ISL_2495739, EPI_ISL_2495740, EPI_ISL_2495741, EPI_ISL_2495742 | | | |
| see above | MB-Cadham Provincial laboratory | National Microbiology Laboratory (NML) | Anna Majer, Shari Tyson, Grace Seo, Philip Mabon, Elsie Grudeski, Rhiannon Huzarewich, Russell Mandes, Anneliese Landgraff, Jennifer Tanner, Natalie Knox, Morag Graham, Gary Van Domselaar, Paul Van Caeseele, Jared Bullard, David Alexander, Kerry Dust, Nathalie Bastien, Yan Li, Timothy Booth, Darian Hole, Madison Chapel, Kirsten Biggar, CanCOGeN's metadata curation team, Public Health Agency of Canada CanCOGeN team |
| EPI_ISL_2495931, EPI_ISL_2495932 | LESP Queretaro | Instituto de Diagnostico y Referencia Epidemiologicos (INDRE) | Claudia Wong-Arambula, Abril Rodriguez-Maldonado, Vanessa Rivero-Arredondo, Ariadna Medina-Benitez, Joaquin Quiroz-Mercado, Sergio Rangel-Guerrero, Natividad Cruz-Ortiz, Tatiana Nunez-Garcia, Gisela Barrera-Badillo, Lucia Hernandez-Rivas, Irma Lopez-Martinez, Ernesto Ramirez-Gonzalez. |
| EPI_ISL_2495933, EPI_ISL_2495935 | LESP Morelos | Instituto de Diagnostico y Referencia Epidemiologicos (INDRE) | Claudia Wong-Arambula, Abril Rodriguez-Maldonado, Vanessa Rivero-Arredondo, Ariadna Medina-Benitez, Joaquin Quiroz-Mercado, Sergio Rangel-Guerrero, Natividad Cruz-Ortiz, Tatiana Nunez-Garcia, Gisela Barrera-Badillo, Lucia Hernandez-Rivas, Irma Lopez-Martinez, Ernesto Ramirez-Gonzalez. |
| EPI_ISL_2495936 | LESP Michoacan | Instituto de Diagnostico y Referencia Epidemiologicos (INDRE) | Claudia Wong-Arambula, Abril Rodriguez-Maldonado, Vanessa Rivero-Arredondo, Ariadna Medina-Benitez, Joaquin Quiroz-Mercado, Sergio Rangel-Guerrero, Natividad Cruz-Ortiz, Tatiana Nunez-Garcia, Gisela Barrera-Badillo, Lucia Hernandez-Rivas, Irma Lopez-Martinez, Ernesto Ramirez-Gonzalez. |
| EPI_ISL_2495937, EPI_ISL_2495940, EPI_ISL_2495941, EPI_ISL_2495942, EPI_ISL_2495943, EPI_ISL_2495944, EPI_ISL_2495945, EPI_ISL_2495946, EPI_ISL_2495947 | LESP Queretaro | Instituto de Diagnostico y Referencia Epidemiologicos (INDRE) | Claudia Wong-Arambula, Abril Rodriguez-Maldonado, Vanessa Rivero-Arredondo, Ariadna Medina-Benitez, Joaquin Quiroz-Mercado, Sergio Rangel-Guerrero, Natividad Cruz-Ortiz, Tatiana Nunez-Garcia, Gisela Barrera-Badillo, Lucia Hernandez-Rivas, Irma Lopez-Martinez, Ernesto Ramirez-Gonzalez. |
| EPI_ISL_2495963 | LESP Nuevo Leon | Instituto de Diagnostico y Referencia Epidemiologicos (INDRE) | Claudia Wong-Arambula, Abril Rodriguez-Maldonado, Vanessa Rivero-Arredondo, Ariadna Medina-Benitez, Joaquin Quiroz-Mercado, Sergio Rangel-Guerrero, Natividad Cruz-Ortiz, Tatiana Nunez-Garcia, Gisela Barrera-Badillo, Lucia Hernandez-Rivas, Irma Lopez-Martinez, Ernesto Ramirez-Gonzalez. |
| EPI_ISL_2495964 | LESP Guanajuato | Instituto de Diagnostico y Referencia Epidemiologicos (INDRE) | Claudia Wong-Arambula, Abril Rodriguez-Maldonado, Vanessa Rivero-Arredondo, Ariadna Medina-Benitez, Joaquin Quiroz-Mercado, Sergio Rangel-Guerrero, Natividad Cruz-Ortiz, Tatiana Nunez-Garcia, Gisela Barrera-Badillo, Lucia Hernandez-Rivas, Irma Lopez-Martinez, Ernesto Ramirez-Gonzalez. |
| EPI_ISL_2495969 | LESP Queretaro | Instituto de Diagnostico y Referencia Epidemiologicos (INDRE) | Claudia Wong-Arambula, Abril Rodriguez-Maldonado, Vanessa Rivero-Arredondo, Ariadna Medina-Benitez, Joaquin Quiroz-Mercado, Sergio Rangel-Guerrero, Natividad Cruz-Ortiz, Tatiana Nunez-Garcia, Gisela Barrera-Badillo, Lucia Hernandez-Rivas, Irma Lopez-Martinez, Ernesto Ramirez-Gonzalez. |
| EPI_ISL_2495970 | LESP Michoacan | Instituto de Diagnostico y Referencia Epidemiologicos (INDRE) | Claudia Wong-Arambula, Abril Rodriguez-Maldonado, Vanessa Rivero-Arredondo, Ariadna Medina-Benitez, Joaquin Quiroz-Mercado, Sergio Rangel-Guerrero, Natividad Cruz-Ortiz, Tatiana Nunez-Garcia, Gisela Barrera-Badillo, Lucia Hernandez-Rivas, Irma Lopez-Martinez, Ernesto Ramirez-Gonzalez. |
| EPI_ISL_2495993 | LESP Campeche | Instituto de Diagnostico y Referencia Epidemiologicos (INDRE) | Claudia Wong-Arambula, Abril Rodriguez-Maldonado, Vanessa Rivero-Arredondo, Ariadna Medina-Benitez, Joaquin Quiroz-Mercado, Sergio Rangel-Guerrero, Natividad Cruz-Ortiz, Tatiana Nunez-Garcia, Gisela Barrera-Badillo, Lucia Hernandez-Rivas, Irma Lopez-Martinez, Ernesto Ramirez-Gonzalez. |
| EPI_ISL_2495995, EPI_ISL_2495996 | LESP Morelos | Instituto de Diagnostico y Referencia Epidemiologicos (INDRE) | Claudia Wong-Arambula, Abril Rodriguez-Maldonado, Vanessa Rivero-Arredondo, Ariadna Medina-Benitez, Joaquin Quiroz-Mercado, Sergio Rangel-Guerrero, Natividad Cruz-Ortiz, Tatiana Nunez-Garcia, Gisela Barrera-Badillo, Lucia Hernandez-Rivas, Irma Lopez-Martinez, Ernesto Ramirez-Gonzalez. |
| EPI_ISL_2495997 | LESP Guanajuato | Instituto de Diagnostico y Referencia Epidemiologicos (INDRE) | Claudia Wong-Arambula, Abril Rodriguez-Maldonado, Vanessa Rivero-Arredondo, Ariadna Medina-Benitez, Joaquin Quiroz-Mercado, Sergio Rangel-Guerrero, Natividad Cruz-Ortiz, Tatiana Nunez-Garcia, Gisela Barrera-Badillo, Lucia Hernandez-Rivas, Irma Lopez-Martinez, Ernesto Ramirez-Gonzalez. |
| EPI_ISL_2495998 | LESP Nuevo Leon | Instituto de Diagnostico y Referencia Epidemiologicos (INDRE) | Claudia Wong-Arambula, Abril Rodriguez-Maldonado, Vanessa Rivero-Arredondo, Ariadna Medina-Benitez, Joaquin Quiroz-Mercado, Sergio Rangel-Guerrero, Natividad Cruz-Ortiz, Tatiana Nunez-Garcia, Gisela Barrera-Badillo, Lucia Hernandez-Rivas, Irma Lopez-Martinez, Ernesto Ramirez-Gonzalez. |

| | | | |
|---|---|---|---|
| EPI_ISL_2496000 | LESP Guanajuato | Instituto de Diagnostico y Referencia Epidemiologicos (INDRE) | Claudia Wong-Arambula, Abril Rodriguez-Maldonado, Vanessa Rivero-Arredondo, Ariadna Medina-Benitez, Joaquin Quiroz-Mercado, Sergio Rangel-Guerrero, Natividad Cruz-Ortiz, Tatiana Nunez-Garcia, Gisela Barrera-Badillo, Lucia Hernandez-Rivas, Irma Lopez-Martinez, Ernesto Ramirez-Gonzalez. |
| EPI_ISL_2496001 | LESP Puebla | Instituto de Diagnostico y Referencia Epidemiologicos (INDRE) | Claudia Wong-Arambula, Abril Rodriguez-Maldonado, Vanessa Rivero-Arredondo, Ariadna Medina-Benitez, Joaquin Quiroz-Mercado, Sergio Rangel-Guerrero, Natividad Cruz-Ortiz, Tatiana Nunez-Garcia, Gisela Barrera-Badillo, Lucia Hernandez-Rivas, Irma Lopez-Martinez, Ernesto Ramirez-Gonzalez. |
| EPI_ISL_2496002 | LESP Guanajuato | Instituto de Diagnostico y Referencia Epidemiologicos (INDRE) | Claudia Wong-Arambula, Abril Rodriguez-Maldonado, Vanessa Rivero-Arredondo, Ariadna Medina-Benitez, Joaquin Quiroz-Mercado, Sergio Rangel-Guerrero, Natividad Cruz-Ortiz, Tatiana Nunez-Garcia, Gisela Barrera-Badillo, Lucia Hernandez-Rivas, Irma Lopez-Martinez, Ernesto Ramirez-Gonzalez. |
| EPI_ISL_2496006, EPI_ISL_2496007, EPI_ISL_2496010, EPI_ISL_2496011 | LESP Michoacan | Instituto de Diagnostico y Referencia Epidemiologicos (INDRE) | Claudia Wong-Arambula, Abril Rodriguez-Maldonado, Vanessa Rivero-Arredondo, Ariadna Medina-Benitez, Joaquin Quiroz-Mercado, Sergio Rangel-Guerrero, Natividad Cruz-Ortiz, Tatiana Nunez-Garcia, Gisela Barrera-Badillo, Lucia Hernandez-Rivas, Irma Lopez-Martinez, Ernesto Ramirez-Gonzalez. |
| EPI_ISL_2500928, EPI_ISL_2500933, EPI_ISL_2500934, EPI_ISL_2500935, EPI_ISL_2500936, EPI_ISL_2500937, EPI_ISL_2500939, EPI_ISL_2500940, EPI_ISL_2500941, EPI_ISL_2500942, EPI_ISL_2500943, EPI_ISL_2500945, EPI_ISL_2500947, EPI_ISL_2500948, EPI_ISL_2500951, EPI_ISL_2500952, EPI_ISL_2500953, EPI_ISL_2500954, EPI_ISL_2500955, EPI_ISL_2500956, EPI_ISL_2500960, EPI_ISL_2500961, EPI_ISL_2500962, EPI_ISL_2500963, EPI_ISL_2500964, EPI_ISL_2500965 | | | |
| see above | Instituto Nacional de Salud | Instituto Nacional de Salud- Dirección de Investigación en Salud Pública | Katherine Laiton-Donato, Diego A. Álvarez-Díaz, Carlos Franco-Muñoz, Hector Alejandro Ruiz-Moreno, Paola Rojas, Maria T. Herrera-Sepúlveda, Diego Andrés Prada, Jhonnatan Reales-González, Sheryll Corchuelo, Julian Naizaque, Jorge Rivera, Gerardo Santamaría, Sergio Gomez, Lisseth Pardo, Juan Camilo Martinez, Marta Lopez Blanco, Ángela Alarcon Cruz, Diana Malo, Carmen Osorio, Magdalena Wiesner, Martha Lucia Ospina Martinez, Marcela Mercado-Reyes |
| EPI_ISL_2501736, EPI_ISL_2501737, EPI_ISL_2501738 | Premier Laboratory Services | BioInfoExperts LLC | Cheryl Metzler, Valerie Murray, Rebecca Rose, David Nolan, Susanna Lamers |
| EPI_ISL_2502622, EPI_ISL_2502623, EPI_ISL_2502624, EPI_ISL_2502625 | S.C. Patologia Clinica, Ospedale Sant'Andrea, ASL 5 | U.O. Igiene, Ospedale Policlinico San Martino | Bruzzone Bianca, Battolla Enrico, Caligiuri Patrizia, De Pace Vanessa, Domnich Alexander, Orsi Andrea, Ricucci Valentina, Icardi Giancarlo |
| EPI_ISL_2502626 | U.O. Igiene, Ospedale Policlinico San Martino | U.O. Igiene, Ospedale Policlinico San Martino | Bruzzone Bianca, Caligiuri Patrizia, De Pace Vanessa, Domnich Alexander, Orsi Andrea, Ricucci Valentina, Icardi Giancarlo |
| EPI_ISL_2502669, EPI_ISL_2502672, EPI_ISL_2502673, EPI_ISL_2502674 | CQRC_QUALITY CONTROL CHEMICAL BIOLOGICAL RISK_AOOR Villa Sofia Cervello Palermo | CQRC_QUALITY CONTROL CHEMICAL BIOLOGICAL RISK_AOOR Villa Sofia Cervello Palermo | Di Gaudio, F., Contino, F., Brunacci, G. |
| EPI_ISL_2502675, EPI_ISL_2502676, EPI_ISL_2502677, EPI_ISL_2502678, EPI_ISL_2502679, EPI_ISL_2502680, EPI_ISL_2502681, EPI_ISL_2502682, EPI_ISL_2502683, EPI_ISL_2502684, EPI_ISL_2502685, EPI_ISL_2502686 | | | |
| see above | CQRC_QUALITY CONTROL CHEMICAL BIOLOGICAL RISK_AOOR Villa Sofia Cervello Palermo | CQRC_QUALITY CONTROL CHEMICAL BIOLOGICAL RISK_AOOR Villa Sofia Cervello Palermo | Di Gaudio, F., Brunacci, G., Contino, F. |
| EPI_ISL_2502688, EPI_ISL_2502689, EPI_ISL_2502690, EPI_ISL_2502691, EPI_ISL_2502692, EPI_ISL_2502693, EPI_ISL_2502694, EPI_ISL_2502695, EPI_ISL_2502696, EPI_ISL_2502697, EPI_ISL_2502698, EPI_ISL_2502699, EPI_ISL_2502700, EPI_ISL_2502701, EPI_ISL_2502702, EPI_ISL_2502703, EPI_ISL_2502704, EPI_ISL_2502705, EPI_ISL_2502707, EPI_ISL_2502709, EPI_ISL_2502710, EPI_ISL_2502711, EPI_ISL_2502712, EPI_ISL_2502713, EPI_ISL_2502714, EPI_ISL_2502715, EPI_ISL_2502716 | | | |
| see above | Microbiology Department, University Hospital Donostia | Microbiology Department, University Hospital Donostia | Marimon JM, Montes M, Piñeiro L, Grandioso D, Martin-Peñaranda T, Gomez M, Cilla G. |
| EPI_ISL_2502718, EPI_ISL_2502719 | HOSPITAL SAN VICENTE DE PAUL | Inciensa, Instituto Costarricense de InvestigaciOn y Enseñanza en NutriciOn y Salud | Francisco Duarte, Hebleen Porras, Claudio Soto-Garita, Estela Cordero, Adriana Godínez, Melany CalderOn, Jose Luis Vargas, Mariela Gutierrez, Joselyn Prado & Silvia Saenz-Garcia |
| EPI_ISL_2502720 | AREA DE SALUD MORAVIA | Inciensa, Instituto Costarricense de InvestigaciOn y Enseñanza en NutriciOn y Salud | Francisco Duarte, Hebleen Porras, Claudio Soto-Garita, Estela Cordero, Adriana Godínez, Melany CalderOn, Jose Luis Vargas, Mariela Gutierrez, Joselyn Prado & Mariel LOpez |
| EPI_ISL_2502721 | HOSPITAL DR. MAX PERALTA JIMENEZ | Inciensa, Instituto Costarricense de InvestigaciOn y Enseñanza en NutriciOn y Salud | Francisco Duarte, Hebleen Porras, Claudio Soto-Garita, Estela Cordero, Adriana Godínez, Melany CalderOn, Jose Luis Vargas, Mariela Gutierrez, Joselyn Prado & Pamela Serrano-Valerín |
| EPI_ISL_2502722 | AREA DE SALUD CATEDRAL NORESTE | Inciensa, Instituto Costarricense de InvestigaciOn y Enseñanza en NutriciOn y Salud | Francisco Duarte, Hebleen Porras, Claudio Soto-Garita, Estela Cordero, Adriana Godínez, Melany CalderOn, Jose Luis Vargas, Mariela Gutierrez, Joselyn Prado & Mariel LOpez |
| EPI_ISL_2502723 | HOSPITAL DR. FERNANDO ESCALANTE PRADILLA | Inciensa, Instituto Costarricense de InvestigaciOn y Enseñanza en NutriciOn y Salud | Francisco Duarte, Hebleen Porras, Claudio Soto-Garita, Estela Cordero, Adriana Godínez, Melany CalderOn, Jose Luis Vargas, Mariela Gutierrez, Joselyn Prado & Lizeth Blanco-Porras |
| EPI_ISL_2502724 | HOSPITAL SAN JUAN DE DIOS | Inciensa, Instituto Costarricense de InvestigaciOn y Enseñanza en NutriciOn y Salud | Francisco Duarte, Hebleen Porras, Claudio Soto-Garita, Estela Cordero, Adriana Godínez, Melany CalderOn, Jose Luis Vargas, Mariela Gutierrez, Joselyn Prado & Andony Cordero-Jimenez |
| EPI_ISL_2502725 | AREA DE SALUD CURRIDABAT 2 | Inciensa, Instituto Costarricense de InvestigaciOn y Enseñanza en NutriciOn y Salud | Francisco Duarte, Hebleen Porras, Claudio Soto-Garita, Estela Cordero, Adriana Godínez, Melany CalderOn, Jose Luis Vargas, Mariela Gutierrez, Joselyn Prado & Mariel LOpez |
| EPI_ISL_2502727 | AREA DE SALUD SAN JUAN-SAN DIEGO-CONCEPCION 2 | Inciensa, Instituto Costarricense de InvestigaciOn y Enseñanza en NutriciOn y Salud | Francisco Duarte, Hebleen Porras, Claudio Soto-Garita, Estela Cordero, Adriana Godínez, Melany CalderOn, Jose Luis Vargas, Mariela Gutierrez, Joselyn Prado & Mariel LOpez |
| EPI_ISL_2502728 | AREA DE SALUD CARRILLO | Inciensa, Instituto Costarricense de InvestigaciOn y Enseñanza en NutriciOn y Salud | Francisco Duarte, Hebleen Porras, Claudio Soto-Garita, Estela Cordero, Adriana Godínez, Melany CalderOn, Jose Luis Vargas, Mariela Gutierrez, Joselyn Prado & Ivanna Krize-Morún |
| EPI_ISL_2502729 | HOSPITAL LA ANEXION | Inciensa, Instituto Costarricense de InvestigaciOn y Enseñanza en NutriciOn y Salud | Francisco Duarte, Hebleen Porras, Claudio Soto-Garita, Estela Cordero, Adriana Godínez, Melany CalderOn, Jose Luis Vargas, Mariela Gutierrez, Joselyn Prado & Ivanna Krize-Morún |
| EPI_ISL_2502730 | O.I.J. MORGUE JUDICIAL | Inciensa, Instituto Costarricense de InvestigaciOn y Enseñanza en NutriciOn y Salud | Francisco Duarte, Hebleen Porras, Claudio Soto-Garita, Estela Cordero, Adriana Godínez, Melany CalderOn, Jose Luis Vargas, Mariela Gutierrez, Joselyn Prado & Grettel Chanto-ChacOn |
| EPI_ISL_2502731 | CLINICA BIBLICA | Inciensa, Instituto Costarricense de InvestigaciOn y Enseñanza en NutriciOn y Salud | Francisco Duarte, Hebleen Porras, Claudio Soto-Garita, Estela Cordero, Adriana Godínez, Melany CalderOn, Jose Luis Vargas, Mariela Gutierrez, Joselyn Prado & Karla Gutierrez-González |
| EPI_ISL_2502732 | AREA DE SALUD OREAMUNO-PACAYAS-TIERRA BLANCA | Inciensa, Instituto Costarricense de InvestigaciOn y Enseñanza en NutriciOn y Salud | Francisco Duarte, Hebleen Porras, Claudio Soto-Garita, Estela Cordero, Adriana Godínez, Melany CalderOn, Jose Luis Vargas, Mariela Gutierrez, Joselyn Prado & Carolina Loría-Acosta |
| EPI_ISL_2502733 | HOSPITAL GUAPILES | Inciensa, Instituto Costarricense de InvestigaciOn y Enseñanza en NutriciOn y Salud | Francisco Duarte, Hebleen Porras, Claudio Soto-Garita, Estela Cordero, Adriana Godínez, Melany CalderOn, Jose Luis Vargas, Mariela Gutierrez, Joselyn Prado & Cesar Cerdas-Quesada |
| EPI_ISL_2502734 | AREA DE SALUD HATILLO - CLINICA DR. SOLON NUÑEZ | Inciensa, Instituto Costarricense de InvestigaciOn y Enseñanza en NutriciOn y Salud | Francisco Duarte, Hebleen Porras, Claudio Soto-Garita, Estela Cordero, Adriana Godínez, Melany CalderOn, Jose Luis Vargas, Mariela Gutierrez, Joselyn Prado & Jessica Mora-Sanchez |
| EPI_ISL_2502736 | AREA DE SALUD GUATUSO | Inciensa, Instituto Costarricense de InvestigaciOn y Enseñanza en NutriciOn y Salud | Francisco Duarte, Hebleen Porras, Claudio Soto-Garita, Estela Cordero, Adriana Godínez, Melany CalderOn, Jose Luis Vargas, Mariela Gutierrez, Joselyn Prado & Francisco ChacOn-Valverde |
| EPI_ISL_2502737 | AREA DE SALUD SIQUIRRES | Inciensa, Instituto Costarricense de InvestigaciOn y Enseñanza en NutriciOn y Salud | Francisco Duarte, Hebleen Porras, Claudio Soto-Garita, Estela Cordero, Adriana Godínez, Melany CalderOn, Jose Luis Vargas, Mariela Gutierrez, Joselyn Prado & Ileana Cháves-Peraza |
| EPI_ISL_2509329 | Dr. Yoshitaka Tamura Department of Clinical Laboratory, Osaka Habikino Medical Center | Department of Developmental Medicine, Research Institute, Osaka Women's and Children's Hospital | Itaru Yanagihara, Yukiko Nakura |
| EPI_ISL_2509337 | Dr. Yoshitaka Tamura Department of Clinical Laboratory, Osaka Habikino Medical Center | Department of Developmental Medicine, Research Institute, Osaka Women's and Children's Hospital Department of Developmental Medicine, Research Institute, Osaka Women's and Children's Hospital | Itaru Yanagihara, Yukiko Nakura |

| | | | |
|---|---|---|---|
| EPI_ISL_2509372 | Dr. Yoshitaka Tamura Department of Clinical Laboratory, Osaka Habikino Medical Center | Department of Developmental Medicine, Research Institute, Osaka Women's and Children's Hospital | Itaru Yanagihara, Yukiko Nakura |
| EPI_ISL_2509934 | N.H.L Municipal Medical College, Ahmedabad | Gujarat Biotechnology Research Centre | Dinesh Kumar,Twinkle Soni,Sonal Sharma ,Zuber Saiyed,Ramesh Pandit,Janvi Raval,Zarna Patel,Nitin Savaliya,Jayshri Pethani,Umang Mishra,Nitesh Shah,Chaitanya Joshi,Madhvi Joshi |
| EPI_ISL_2509938 | BJ Medical College and Civil Hospital, Ahmedabad | Gujarat Biotechnology Research Centre | Ramesh Pandit,Janvi Raval,Zarna Patel,Nitin Savaliya,Dinesh Kumar,Twinkle Soni,Sonal Sharma ,Zuber Saiyed,Pranay Shah,Umang Mishra,Nitesh Shah,Chaitanya Joshi,Madhvi Joshi |
| EPI_ISL_2509939 | N.H.L Municipal Medical College, Ahmedabad | Gujarat Biotechnology Research Centre | Janvi Raval,Zarna Patel,Nitin Savaliya,Dinesh Kumar,Twinkle Soni,Sonal Sharma ,Zuber Saiyed,Ramesh Pandit,Jayshri Pethani,Umang Mishra,Nitesh Shah,Chaitanya Joshi,Madhvi Joshi |
| EPI_ISL_2509940 | N.H.L Municipal Medical College, Ahmedabad | Gujarat Biotechnology Research Centre | Zarna Patel,Nitin Savaliya,Dinesh Kumar,Twinkle Soni,Sonal Sharma ,Zuber Saiyed,Ramesh Pandit,Janvi Raval,Jayshri Pethani,Umang Mishra,Nitesh Shah,Chaitanya Joshi,Madhvi Joshi |
| EPI_ISL_2509941 | N.H.L Municipal Medical College, Ahmedabad | Gujarat Biotechnology Research Centre | Nitin Savaliya,Dinesh Kumar,Twinkle Soni,Sonal Sharma ,Zuber Saiyed,Ramesh Pandit,Janvi Raval,Zarna Patel,Jayshri Pethani,Umang Mishra,Nitesh Shah,Chaitanya Joshi,Madhvi Joshi |
| EPI_ISL_2509942 | N.H.L Municipal Medical College, Ahmedabad | Gujarat Biotechnology Research Centre | Dinesh Kumar,Twinkle Soni,Sonal Sharma ,Zuber Saiyed,Ramesh Pandit,Janvi Raval,Zarna Patel,Nitin Savaliya,Jayshri Pethani,Umang Mishra,Nitesh Shah,Chaitanya Joshi,Madhvi Joshi |
| EPI_ISL_2509944 | N.H.L Municipal Medical College, Ahmedabad | Gujarat Biotechnology Research Centre | Sonal Sharma ,Zuber Saiyed,Ramesh Pandit,Janvi Raval,Zarna Patel,Nitin Savaliya,Dinesh Kumar,Twinkle Soni,Jayshri Pethani,Umang Mishra,Nitesh Shah,Chaitanya Joshi,Madhvi Joshi |
| EPI_ISL_2509945 | N.H.L Municipal Medical College, Ahmedabad | Gujarat Biotechnology Research Centre | Zuber Saiyed,Ramesh Pandit,Janvi Raval,Zarna Patel,Nitin Savaliya,Dinesh Kumar,Twinkle Soni,Sonal Sharma ,Jayshri Pethani,Umang Mishra,Nitesh Shah,Chaitanya Joshi,Madhvi Joshi |
| EPI_ISL_2509946 | N.H.L Municipal Medical College, Ahmedabad | Gujarat Biotechnology Research Centre | Ramesh Pandit,Janvi Raval,Zarna Patel,Nitin Savaliya,Dinesh Kumar,Twinkle Soni,Sonal Sharma ,Zuber Saiyed,Jayshri Pethani,Umang Mishra,Nitesh Shah,Chaitanya Joshi,Madhvi Joshi |
| EPI_ISL_2509947 | N.H.L Municipal Medical College, Ahmedabad | Gujarat Biotechnology Research Centre | Janvi Raval,Zarna Patel,Nitin Savaliya,Dinesh Kumar,Twinkle Soni,Sonal Sharma ,Zuber Saiyed,Ramesh Pandit,Jayshri Pethani,Umang Mishra,Nitesh Shah,Chaitanya Joshi,Madhvi Joshi |
| EPI_ISL_2509948 | N.H.L Municipal Medical College, Ahmedabad | Gujarat Biotechnology Research Centre | Zarna Patel,Nitin Savaliya,Dinesh Kumar,Twinkle Soni,Sonal Sharma ,Zuber Saiyed,Ramesh Pandit,Janvi Raval,Jayshri Pethani,Umang Mishra,Nitesh Shah,Chaitanya Joshi,Madhvi Joshi |
| EPI_ISL_2509951 | N.H.L Municipal Medical College, Ahmedabad | Gujarat Biotechnology Research Centre | Twinkle Soni,Sonal Sharma ,Zuber Saiyed,Ramesh Pandit,Janvi Raval,Zarna Patel,Nitin Savaliya,Dinesh Kumar,Jayshri Pethani,Umang Mishra,Nitesh Shah,Chaitanya Joshi,Madhvi Joshi |
| EPI_ISL_2509952 | N.H.L Municipal Medical College, Ahmedabad | Gujarat Biotechnology Research Centre | Sonal Sharma ,Zuber Saiyed,Ramesh Pandit,Janvi Raval,Zarna Patel,Nitin Savaliya,Dinesh Kumar,Twinkle Soni,Jayshri Pethani,Umang Mishra,Nitesh Shah,Chaitanya Joshi,Madhvi Joshi |
| EPI_ISL_2509953 | N.H.L Municipal Medical College, Ahmedabad | Gujarat Biotechnology Research Centre | Zuber Saiyed,Ramesh Pandit,Janvi Raval,Zarna Patel,Nitin Savaliya,Dinesh Kumar,Twinkle Soni,Sonal Sharma ,Jayshri Pethani,Umang Mishra,Nitesh Shah,Chaitanya Joshi,Madhvi Joshi |
| EPI_ISL_2509954 | N.H.L Municipal Medical College, Ahmedabad | Gujarat Biotechnology Research Centre | Ramesh Pandit,Janvi Raval,Zarna Patel,Nitin Savaliya,Dinesh Kumar,Twinkle Soni,Sonal Sharma ,Zuber Saiyed,Jayshri Pethani,Umang Mishra,Nitesh Shah,Chaitanya Joshi,Madhvi Joshi |
| EPI_ISL_2509955 | N.H.L Municipal Medical College, Ahmedabad | Gujarat Biotechnology Research Centre | Janvi Raval,Zarna Patel,Nitin Savaliya,Dinesh Kumar,Twinkle Soni,Sonal Sharma ,Zuber Saiyed,Ramesh Pandit,Jayshri Pethani,Umang Mishra,Nitesh Shah,Chaitanya Joshi,Madhvi Joshi |
| EPI_ISL_2509956 | N.H.L Municipal Medical College, Ahmedabad | Gujarat Biotechnology Research Centre | Zarna Patel,Nitin Savaliya,Dinesh Kumar,Twinkle Soni,Sonal Sharma ,Zuber Saiyed,Ramesh Pandit,Janvi Raval,Jayshri Pethani,Umang Mishra,Nitesh Shah,Chaitanya Joshi,Madhvi Joshi |
| EPI_ISL_2509957 | N.H.L Municipal Medical College, Ahmedabad | Gujarat Biotechnology Research Centre | Nitin Savaliya,Dinesh Kumar,Twinkle Soni,Sonal Sharma ,Zuber Saiyed,Ramesh Pandit,Janvi Raval,Zarna Patel,Jayshri Pethani,Umang Mishra,Nitesh Shah,Chaitanya Joshi,Madhvi Joshi |
| EPI_ISL_2509958 | N.H.L Municipal Medical College, Ahmedabad | Gujarat Biotechnology Research Centre | Dinesh Kumar,Twinkle Soni,Sonal Sharma ,Zuber Saiyed,Ramesh Pandit,Janvi Raval,Zarna Patel,Nitin Savaliya,Jayshri Pethani,Umang Mishra,Nitesh Shah,Chaitanya Joshi,Madhvi Joshi |
| EPI_ISL_2516976 | Microbiology Department. Complexo Hospitalario Universitario de Vigo | Microbiology Department. Complexo Hospitalario Universitario de Vigo | Alfaya N, Alonso I, Alvarez M, Cabrera JJ, Carballo R, Cores O, Cortizo S, del-Campo V, Martinez L, Mediero G, Perez S, Potel C, Regueiro B, Rey S, Vasallo FJ |
| EPI_ISL_2529850, EPI_ISL_2529852, EPI_ISL_2529854, EPI_ISL_2529856, EPI_ISL_2529858, EPI_ISL_2529861 | Baylor College of Medicine/ GCID | Baylor College of Medicine/ GCID | Timothy Farinholt, Qing Meng, Qinchang Meng, Vipin Menon, Xiang Qin, Harsha Doddapaneni, Paige Farinholt, Charu Agrawal, Donna Muzny, Pedro A. Piedra, Richard A. Gibbs, Joseph Pertosino |
| EPI_ISL_2530152, EPI_ISL_2530177, EPI_ISL_2530179, EPI_ISL_2530180, EPI_ISL_2530182, EPI_ISL_2530183, EPI_ISL_2530185, EPI_ISL_2530186, EPI_ISL_2530187, EPI_ISL_2530188, EPI_ISL_2530190, EPI_ISL_2530191, EPI_ISL_2530193, EPI_ISL_2530195, EPI_ISL_2530196, EPI_ISL_2530199, EPI_ISL_2530201, EPI_ISL_2530202, EPI_ISL_2530204, EPI_ISL_2530206, EPI_ISL_2530207, EPI_ISL_2530209, EPI_ISL_2530210, EPI_ISL_2530212, EPI_ISL_2530214, EPI_ISL_2530216, EPI_ISL_2530217, EPI_ISL_2530219, EPI_ISL_2530221, EPI_ISL_2530222, EPI_ISL_2530224, EPI_ISL_2530226, EPI_ISL_2530227, EPI_ISL_2530229, EPI_ISL_2530230, EPI_ISL_2530232, EPI_ISL_2530233, EPI_ISL_2530235, EPI_ISL_2530236, EPI_ISL_2530238, EPI_ISL_2530239, EPI_ISL_2530241, EPI_ISL_2530243, EPI_ISL_2530245, EPI_ISL_2530246, EPI_ISL_2530248, EPI_ISL_2530250, EPI_ISL_2530252, EPI_ISL_2530253, EPI_ISL_2530255, EPI_ISL_2530267, EPI_ISL_2530268, EPI_ISL_2530270, EPI_ISL_2530278, EPI_ISL_2530279, EPI_ISL_2530281, EPI_ISL_2530283, EPI_ISL_2530285, EPI_ISL_2530286, EPI_ISL_2530299, EPI_ISL_2530304, EPI_ISL_2530308, EPI_ISL_2530373, EPI_ISL_2530374, EPI_ISL_2530376, EPI_ISL_2530378, EPI_ISL_2530379 | | | |
| see above | Hematopathology Laboratory, ACTREC, TMC | Hematopathology Laboratory, ACTREC, TMC | Hematopathology Laboratory, ACTREC |
| EPI_ISL_2531855 | Hospital | National Reference Center for Viruses of Respiratory Infections, Institut Pasteur, Paris | Marion Barbet, Sylvie Behillil, Méline Bizard, Angela Brisebarre, Camille Capel, Vincent Enouf, Louise Lefrançois, Frédéric Lemoine, Christophe Malabat, Corinne Maufrais, Victoire Baillet, Etienne Simon-Lorière, Maud Vanpeene, Sylvie Van der Werf ,Alice Moisan |
| EPI_ISL_2531859, EPI_ISL_2531864, EPI_ISL_2531868 | Hospital | National Reference Center for Viruses of Respiratory Infections, Institut Pasteur, Paris | Marion Barbet, Sylvie Behillil, Méline Bizard, Angela Brisebarre, Camille Capel, Vincent Enouf, Louise Lefrançois, Frédéric Lemoine, Christophe Malabat, Corinne Maufrais, Etienne Simon-Lorière, Maud Vanpeene, Sylvie Van der Werf ,Alice Moisan |
| EPI_ISL_2532379, EPI_ISL_2532380, EPI_ISL_2532381, EPI_ISL_2532382, EPI_ISL_2532383, EPI_ISL_2532384, EPI_ISL_2532385, EPI_ISL_2532386, EPI_ISL_2532387 | Labo Analyses Med | National Reference Center for Viruses of Respiratory Infections, Institut Pasteur, Paris | Marion Barbet, Sylvie Behillil, Méline Bizard, Angela Brisebarre, Camille Capel, Vincent Enouf, Louise Lefrançois, Frédéric Lemoine, Christophe Malabat, Corinne Maufrais, Etienne Simon-Lorière, Maud Vanpeene, Sylvie Van der Werf ,Jean-Christophe Denis |
| EPI_ISL_2532394, EPI_ISL_2532395, EPI_ISL_2532396 | Hospital | National Reference Center for Viruses of Respiratory Infections, Institut Pasteur, Paris | Marion Barbet, Sylvie Behillil, Méline Bizard, Angela Brisebarre, Camille Capel, Vincent Enouf, Louise Lefrançois, Frédéric Lemoine, Christophe Malabat, Corinne Maufrais, Etienne Simon-Lorière, Maud Vanpeene, Sylvie Van der Werf ,Patrice Combe |
| EPI_ISL_2533930, EPI_ISL_2533931 | CIAD-Hermosillo | Immunology Laboratory | Consorcio Mexicano de Vigilancia Genómica (CoViGen-Mex). Authors (in alphabetical order): Julio Elias Alvarado-Yaah, Carlos F. Arias, Santiago Ávila-Ríos, Eduardo Becerril-Vargas, Víctor Hugo Borja-Aburto, Celia Boukadida, Cristóbal Cháidez-Quiróz, Juan Bautista Chale-Dzul, Ricardo Ciria Merce, Andreu Comas-García, Célida Duque Molina, Julissa Enciso-Ibarra, José Antonio Enciso-Moreno, Gloria Elena Espinosa-Ayala, Fernando Fontove-Herrera, Daniel Fregoso-Rueda, Victor Eduardo García-Arias, Alejandra García-Gasca, Bruno Gómez-Gil, Jean Pierre González, Irvin González-López, Concepción Grajales-Muñiz, Ricardo Grande, Rosa María Gutiérrez Rios, Jesús Hernández, Alejandra Hernández-Terán, Alfredo Herrera-Estrella, Carla Ivón Herrera-Najera, Pavel Isa, Verónica Mata-Haro, Daniel Lira Morales, Susana Lopez, Antonio Loza Román, Brenda Irasema Maldonado-Meza, Bernardo Martínez-Miguel, José Arturo Martínez-Orozco, Célida Martínez- Rodríguez, Margarita Matías-Florentino, Fidencio Mejía-Nepomuceno, María Guadalupe de Jesús Mireles-Rivera, Gloria María Molina-Salinas, Hector Montoya-Fuentes, , Mario Mújica-Sánchez, José Esteban Muñoz-Medina, José de Jesús Nuñez-Contreras, Alicia Ocaña-Mondragón, Luis Alberto Ochoa-Carrera, Hector Esteban Paz-Juárez, Marissa Perez-Garcia, Francisco Pulido, Helen Haydee Fernanda Ramírez-Plascencia, Jorge Salas-Hernández, Angel Gustavo Salas-Lais, Alejandro Sánchez-Flores, Clara Esperanza Santacruz-Tinoco, María Guadalupe Santiago-Mauricio, Selene Zárate, Nelly Sélem-Mojica, Blanca Taboada, Gloria Vazquez, Joel Armando Vázquez-Pérez. |

| | | | |
|---|---|---|---|
| EPI_ISL_2535471, EPI_ISL_2535472, EPI_ISL_2535473, EPI_ISL_2535474, EPI_ISL_2535475 | Hospital | National Reference Center for Viruses of Respiratory Infections, Institut Pasteur, Paris | Marion Barbet, Sylvie Behillil, Méline Bizard, Angela Brisebarre, Camille Capel, Vincent Enouf, Louise Lefrançois, Frédéric Lemoine, Christophe Malabat, Corinne Maufrais, Etienne Simon-Lorière, Maud Vanpeene, Sylvie Van der Werf ,LéA Pilorge |
| EPI_ISL_2536230, EPI_ISL_2536263 | Labortorio Central de Saude Publica do Estado de Santa Catarina (LACEN/SC) | Laboratory of Respiratory Viruses and Measles, Oswaldo Cruz Institute, FIOCRUZ | Paola Resende, Luciana Appolinario, Fernando Motta, Anna Carolina Paixao, Ana Carolina Mendonca, Alice Sampaio Rocha, Taina Venas, Elisa Cavalcante Pereira, Renata Serrano Lopes, Darcita Buerger Rovaris, Sandra Bianchini Fernandes, Marilda Siqueira on behalf of the Fiocruz COVID-19 Genomic Surveillance Network |
| EPI_ISL_2536323 | Laboratorio Central de Saude Publica do Estado da Paraiba (LACEN-PB) | Laboratory of Respiratory Viruses and Measles, Oswaldo Cruz Institute, FIOCRUZ | Paola Resende, Luciana Appolinario, Fernando Motta, Anna Carolina Paixao, Ana Carolina Mendonca, Alice Sampaio Rocha, Taina Venas, Elisa Cavalcante Pereira, Renata Serrano Lopes, Joao Felipe Bezerra, Dalane Loudal Florentino Teixeira, Marilda Siqueira on behalf of the Fiocruz COVID-19 Genomic Surveillance Network |
| EPI_ISL_2537481, EPI_ISL_2537482, EPI_ISL_2537483 | Department of Intensive Care Medicine, and clinical lab., Osaka Women's and Children's Hospital Drs. Muneyuki Takeuchi, Shinobu Ida. | Department of Developmental Medicine, Research Institute, Osaka Women's and Children's Hospital | Itaru Yanagihara, Yukiko Nakura |
| EPI_ISL_2543563 | Provincial Public Health Reference Laboratory (PPHRL) P& SHD | Provincial Public Health Reference Laboratory (PPHRL) P& SHD | ANDLEEB HANIF |
| EPI_ISL_2543891 | N.H.L Municipal Medical College, Ahmedabad | Gujarat Biotechnology Research Centre | Sonal Sharma ,Zuber Saiyed,Ramesh Pandit,Janvi Raval,Zarna Patel,Nitin Savaliya,Dinesh Kumar,Twinkle Soni,Jayshri Pethani,Umang Mishra,Nitesh Shah,Chaitanya Joshi,Madhvi Joshi |
| EPI_ISL_2543892 | N.H.L Municipal Medical College, Ahmedabad | Gujarat Biotechnology Research Centre | Zuber Saiyed,Ramesh Pandit,Janvi Raval,Zarna Patel,Nitin Savaliya,Dinesh Kumar,Twinkle Soni,Sonal Sharma ,Jayshri Pethani,Umang Mishra,Nitesh Shah,Chaitanya Joshi,Madhvi Joshi |
| EPI_ISL_2543893 | N.H.L Municipal Medical College, Ahmedabad | Gujarat Biotechnology Research Centre | Ramesh Pandit,Janvi Raval,Zarna Patel,Nitin Savaliya,Dinesh Kumar,Twinkle Soni,Sonal Sharma ,Zuber Saiyed,Jayshri Pethani,Umang Mishra,Nitesh Shah,Chaitanya Joshi,Madhvi Joshi |
| EPI_ISL_2543895 | N.H.L Municipal Medical College, Ahmedabad | Gujarat Biotechnology Research Centre | Zarna Patel,Nitin Savaliya,Dinesh Kumar,Twinkle Soni,Sonal Sharma ,Zuber Saiyed,Ramesh Pandit,Janvi Raval,Jayshri Pethani,Umang Mishra,Nitesh Shah,Chaitanya Joshi,Madhvi Joshi |
| EPI_ISL_2543898 | N.H.L Municipal Medical College, Ahmedabad | Gujarat Biotechnology Research Centre | Twinkle Soni,Sonal Sharma ,Zuber Saiyed,Ramesh Pandit,Janvi Raval,Zarna Patel,Nitin Savaliya,Dinesh Kumar,Jayshri Pethani,Umang Mishra,Nitesh Shah,Chaitanya Joshi,Madhvi Joshi |
| EPI_ISL_2544198, EPI_ISL_2544200 | Provincial Public Health Reference Laboratory (PPHRL) P& SHD | Provincial Public Health Reference Laboratory (PPHRL) P& SHD | ANDLEEB HANIF |
| EPI_ISL_2544498, EPI_ISL_2544499, EPI_ISL_2544501, EPI_ISL_2544507, EPI_ISL_2544510, EPI_ISL_2544512, EPI_ISL_2544513, EPI_ISL_2544524, EPI_ISL_2544525 | Baker lab at Loyola Medical Center | Baker lab at Loyola Medical Center | Xufang Deng, PhD, Monika Evdokimova, BS, Amornrat O'Brien, PhD, Cynthia Rowe, PhD, Nina M. Clark, MD, Amanda Harrington, PhD, Gail E. Reid, MD, Susan Uprichard, PhD, and Susan C. Baker, PhD |
| EPI_ISL_2544814 | GMERS Government Medical College (GMC), Gandhinagar | Gujarat Biotechnology Research Centre | Janvi Raval,Zarna Patel,Nitin Savaliya,Dinesh Kumar,Twinkle Soni,Sonal Sharma ,Zuber Saiyed,Ramesh Pandit,Gaurishankar Shrimali,Umang Mishra,Nitesh Shah,Chaitanya Joshi,Madhvi Joshi |
| EPI_ISL_2544815 | GMERS Government Medical College (GMC), Gandhinagar | Gujarat Biotechnology Research Centre | Zarna Patel,Nitin Savaliya,Dinesh Kumar,Twinkle Soni,Sonal Sharma ,Zuber Saiyed,Ramesh Pandit,Janvi Raval,Gaurishankar Shrimali,Umang Mishra,Nitesh Shah,Chaitanya Joshi,Madhvi Joshi |
| EPI_ISL_2544816 | GMERS Government Medical College (GMC), Gandhinagar | Gujarat Biotechnology Research Centre | Nitin Savaliya,Dinesh Kumar,Twinkle Soni,Sonal Sharma ,Zuber Saiyed,Ramesh Pandit,Janvi Raval,Zarna Patel,Gaurishankar Shrimali,Umang Mishra,Nitesh Shah,Chaitanya Joshi,Madhvi Joshi |
| EPI_ISL_2544817 | GMERS Government Medical College (GMC), Gandhinagar | Gujarat Biotechnology Research Centre | Dinesh Kumar,Twinkle Soni,Sonal Sharma ,Zuber Saiyed,Ramesh Pandit,Janvi Raval,Zarna Patel,Nitin Savaliya,Gaurishankar Shrimali,Umang Mishra,Nitesh Shah,Chaitanya Joshi,Madhvi Joshi |
| EPI_ISL_2544818 | GMERS Government Medical College (GMC), Gandhinagar | Gujarat Biotechnology Research Centre | Twinkle Soni,Sonal Sharma ,Zuber Saiyed,Ramesh Pandit,Janvi Raval,Zarna Patel,Nitin Savaliya,Dinesh Kumar,Gaurishankar Shrimali,Umang Mishra,Nitesh Shah,Chaitanya Joshi,Madhvi Joshi |
| EPI_ISL_2544819 | GMERS Government Medical College (GMC), Gandhinagar | Gujarat Biotechnology Research Centre | Sonal Sharma ,Zuber Saiyed,Ramesh Pandit,Janvi Raval,Zarna Patel,Nitin Savaliya,Dinesh Kumar,Twinkle Soni,Gaurishankar Shrimali,Umang Mishra,Nitesh Shah,Chaitanya Joshi,Madhvi Joshi |
| EPI_ISL_2544836 | GMERS Government Medical College (GMC), Gotri | Gujarat Biotechnology Research Centre | Zuber Saiyed,Ramesh Pandit,Janvi Raval,Zarna Patel,Nitin Savaliya,Dinesh Kumar,Twinkle Soni,Sonal Sharma ,Bithika Duttaroy,Umang Mishra,Nitesh Shah,Chaitanya Joshi,Madhvi Joshi |
| EPI_ISL_2545680, EPI_ISL_2545681 | LESP Tamaulipas | Instituto de Diagnostico y Referencia Epidemiologicos (INDRE) | Claudia Wong-Arambula, Abril Rodriguez-Maldonado, Vanessa Rivero-Arredondo, Ariadna Medina-Benitez, Joaquin Quiroz-Mercado, Sergio Rangel-Guerrero, Natividad Cruz-Ortiz, Tatiana Nunez-Garcia, Gisela Barrera-Badillo, Lucia Hernandez-Rivas, Irma Lopez-Martinez, Ernesto Ramirez-Gonzalez. |
| EPI_ISL_2545682, EPI_ISL_2545700, EPI_ISL_2545701 | LESP Queretaro | Instituto de Diagnostico y Referencia Epidemiologicos (INDRE) | Claudia Wong-Arambula, Abril Rodriguez-Maldonado, Vanessa Rivero-Arredondo, Ariadna Medina-Benitez, Joaquin Quiroz-Mercado, Sergio Rangel-Guerrero, Natividad Cruz-Ortiz, Tatiana Nunez-Garcia, Gisela Barrera-Badillo, Lucia Hernandez-Rivas, Irma Lopez-Martinez, Ernesto Ramirez-Gonzalez. |
| EPI_ISL_2545707 | LESP Chihuahua | Instituto de Diagnostico y Referencia Epidemiologicos (INDRE) | Claudia Wong-Arambula, Abril Rodriguez-Maldonado, Vanessa Rivero-Arredondo, Ariadna Medina-Benitez, Joaquin Quiroz-Mercado, Sergio Rangel-Guerrero, Natividad Cruz-Ortiz, Tatiana Nunez-Garcia, Gisela Barrera-Badillo, Lucia Hernandez-Rivas, Irma Lopez-Martinez, Ernesto Ramirez-Gonzalez. |
| EPI_ISL_2545712 | LESP Tamaulipas | Instituto de Diagnostico y Referencia Epidemiologicos (INDRE) | Claudia Wong-Arambula, Abril Rodriguez-Maldonado, Vanessa Rivero-Arredondo, Ariadna Medina-Benitez, Joaquin Quiroz-Mercado, Sergio Rangel-Guerrero, Natividad Cruz-Ortiz, Tatiana Nunez-Garcia, Gisela Barrera-Badillo, Lucia Hernandez-Rivas, Irma Lopez-Martinez, Ernesto Ramirez-Gonzalez. |
| EPI_ISL_2545713, EPI_ISL_2545714, EPI_ISL_2545716, EPI_ISL_2545717, EPI_ISL_2545718, EPI_ISL_2545719, EPI_ISL_2545720 | LESP Queretaro | Instituto de Diagnostico y Referencia Epidemiologicos (INDRE) | Claudia Wong-Arambula, Abril Rodriguez-Maldonado, Vanessa Rivero-Arredondo, Ariadna Medina-Benitez, Joaquin Quiroz-Mercado, Sergio Rangel-Guerrero, Natividad Cruz-Ortiz, Tatiana Nunez-Garcia, Gisela Barrera-Badillo, Lucia Hernandez-Rivas, Irma Lopez-Martinez, Ernesto Ramirez-Gonzalez. |
| EPI_ISL_2545820 | Laboratory of Communicable Diseases | 1. Laboratory of Communicable Diseases (Estonia); 2. Eurofins Genomics Europe Sequencing GmbH | Liidia Dotsenko et al. |

EPI_ISL_2545822, EPI_ISL_2545823, EPI_ISL_2545824, EPI_ISL_2545825, EPI_ISL_2545826, EPI_ISL_2545827, EPI_ISL_2545828, EPI_ISL_2545829, EPI_ISL_2545830, EPI_ISL_2545832, EPI_ISL_2545833, EPI_ISL_2545834, EPI_ISL_2545835, EPI_ISL_2545836, EPI_ISL_2545838, EPI_ISL_2545839, EPI_ISL_2545840, EPI_ISL_2545841, EPI_ISL_2545842, EPI_ISL_2545843, EPI_ISL_2545844, EPI_ISL_2545845, EPI_ISL_2545846, EPI_ISL_2545847, EPI_ISL_2545848, EPI_ISL_2545849, EPI_ISL_2545850, EPI_ISL_2545851, EPI_ISL_2545854, EPI_ISL_2545855, EPI_ISL_2545856, EPI_ISL_2545857, EPI_ISL_2545858, EPI_ISL_2545859, EPI_ISL_2545860, EPI_ISL_2545861, EPI_ISL_2545862, EPI_ISL_2545863, EPI_ISL_2545864, EPI_ISL_2545865, EPI_ISL_2545867, EPI_ISL_2545868, EPI_ISL_2545869, EPI_ISL_2545870, EPI_ISL_2545871, EPI_ISL_2545872, EPI_ISL_2545873, EPI_ISL_2545874, EPI_ISL_2545875, EPI_ISL_2545877, EPI_ISL_2545878, EPI_ISL_2545879, EPI_ISL_2545880, EPI_ISL_2545881, EPI_ISL_2545882, EPI_ISL_2545883, EPI_ISL_2545884, EPI_ISL_2545885, EPI_ISL_2545886, EPI_ISL_2545887, EPI_ISL_2545888, EPI_ISL_2545889, EPI_ISL_2545890, EPI_ISL_2545891, EPI_ISL_2545892, EPI_ISL_2545893, EPI_ISL_2545894, EPI_ISL_2545895, EPI_ISL_2545896, EPI_ISL_2545897, EPI_ISL_2545898, EPI_ISL_2545899, EPI_ISL_2545900, EPI_ISL_2545901, EPI_ISL_2545902, EPI_ISL_2545903, EPI_ISL_2545904, EPI_ISL_2545905, EPI_ISL_2545906, EPI_ISL_2545907, EPI_ISL_2545908, EPI_ISL_2545909, EPI_ISL_2545910, EPI_ISL_2545911, EPI_ISL_2545913, EPI_ISL_2545914, EPI_ISL_2545915, EPI_ISL_2545916, EPI_ISL_2545918, EPI_ISL_2545919, EPI_ISL_2545920, EPI_ISL_2545921, EPI_ISL_2545922, EPI_ISL_2545925, EPI_ISL_2545926, EPI_ISL_2545927, EPI_ISL_2545928, EPI_ISL_2545930, EPI_ISL_2545931, EPI_ISL_2545932, EPI_ISL_2545933, EPI_ISL_2545934, EPI_ISL_2545935, EPI_ISL_2545936, EPI_ISL_2545937, EPI_ISL_2545938, EPI_ISL_2545939, EPI_ISL_2545941, EPI_ISL_2545942, EPI_ISL_2545943, EPI_ISL_2545944, EPI_ISL_2545945, EPI_ISL_2545946, EPI_ISL_2545947, EPI_ISL_2545949, EPI_ISL_2545950, EPI_ISL_2545951, EPI_ISL_2545952, EPI_ISL_2545953, EPI_ISL_2545954, EPI_ISL_2545956, EPI_ISL_2545957, EPI_ISL_2545958, EPI_ISL_2545959, EPI_ISL_2545960, EPI_ISL_2545961, EPI_ISL_2545962, EPI_ISL_2545965, EPI_ISL_2545967, EPI_ISL_2545968, EPI_ISL_2545969, EPI_ISL_2545970, EPI_ISL_2545971, EPI_ISL_2545972, EPI_ISL_2545973, EPI_ISL_2545974, EPI_ISL_2545975, EPI_ISL_2545976, EPI_ISL_2545977, EPI_ISL_2545978, EPI_ISL_2545979, EPI_ISL_2545981, EPI_ISL_2545982, EPI_ISL_2545983, EPI_ISL_2545984, EPI_ISL_2545985, EPI_ISL_2545986, EPI_ISL_2545987, EPI_ISL_2545989, EPI_ISL_2546902, EPI_ISL_2546903, EPI_ISL_2546904, EPI_ISL_2546905, EPI_ISL_2546906, EPI_ISL_2546907, EPI_ISL_2546908, EPI_ISL_2546909, EPI_ISL_2546910, EPI_ISL_2546911, EPI_ISL_2546912, EPI_ISL_2546913, EPI_ISL_2546914, EPI_ISL_2546915, EPI_ISL_2546916, EPI_ISL_2546917, EPI_ISL_2546918, EPI_ISL_2546919, EPI_ISL_2546920, EPI_ISL_2546921, EPI_ISL_2546922, EPI_ISL_2546923, EPI_ISL_2546924, EPI_ISL_2546925, EPI_ISL_2546926, EPI_ISL_2546927, EPI_ISL_2546928, EPI_ISL_2546929, EPI_ISL_2546930, EPI_ISL_2546931, EPI_ISL_2546932, EPI_ISL_2546933, EPI_ISL_2546934, EPI_ISL_2546935, EPI_ISL_2546936, EPI_ISL_2546937, EPI_ISL_2546938, EPI_ISL_2546939, EPI_ISL_2546940, EPI_ISL_2546941, EPI_ISL_2546942, EPI_ISL_2546943, EPI_ISL_2546944, EPI_ISL_2546945, EPI_ISL_2546946, EPI_ISL_2546947, EPI_ISL_2546948, EPI_ISL_2546949, EPI_ISL_2546950, EPI_ISL_2546951, EPI_ISL_2546952, EPI_ISL_2546953, EPI_ISL_2546954, EPI_ISL_2546955, EPI_ISL_2546956,

| | | | |
|---|---|---|---|
| EPI_ISL_2546957, EPI_ISL_2546958, EPI_ISL_2546959, EPI_ISL_2546960, EPI_ISL_2546961, EPI_ISL_2546962, EPI_ISL_2546963, EPI_ISL_2546964, EPI_ISL_2546965, EPI_ISL_2546966, EPI_ISL_2546967, EPI_ISL_2546968, EPI_ISL_2546969, EPI_ISL_2546970, EPI_ISL_2546971, EPI_ISL_2546972, EPI_ISL_2546973, EPI_ISL_2546974, EPI_ISL_2546975, EPI_ISL_2546976, EPI_ISL_2546977, EPI_ISL_2546978, EPI_ISL_2546979, EPI_ISL_2546980, EPI_ISL_2546981, EPI_ISL_2546982, EPI_ISL_2546983, EPI_ISL_2546985, EPI_ISL_2546986, EPI_ISL_2546988, EPI_ISL_2546989, EPI_ISL_2546990, EPI_ISL_2546991, EPI_ISL_2546992, EPI_ISL_2546993, EPI_ISL_2546994, EPI_ISL_2546996, EPI_ISL_2546998, EPI_ISL_2546999, EPI_ISL_2547001, EPI_ISL_2547002, EPI_ISL_2547003, EPI_ISL_2547004, EPI_ISL_2547005, EPI_ISL_2547007, EPI_ISL_2547008, EPI_ISL_2547009, EPI_ISL_2547010, EPI_ISL_2547011, EPI_ISL_2547012, EPI_ISL_2547013, EPI_ISL_2547014, EPI_ISL_2547015, EPI_ISL_2547016, EPI_ISL_2547018, EPI_ISL_2547019, EPI_ISL_2547020, EPI_ISL_2547021, EPI_ISL_2547023, EPI_ISL_2547024, EPI_ISL_2547025, EPI_ISL_2547026, EPI_ISL_2547027, EPI_ISL_2547028, EPI_ISL_2547030, EPI_ISL_2547031, EPI_ISL_2547032, EPI_ISL_2547033, EPI_ISL_2547034, EPI_ISL_2547035, EPI_ISL_2547036, EPI_ISL_2547037, EPI_ISL_2547038, EPI_ISL_2547039, EPI_ISL_2547040, EPI_ISL_2547041, EPI_ISL_2547042, EPI_ISL_2547043, EPI_ISL_2547044, EPI_ISL_2547045, EPI_ISL_2547046, EPI_ISL_2547047, EPI_ISL_2547048, EPI_ISL_2547049, EPI_ISL_2547051, EPI_ISL_2547052, EPI_ISL_2547053, EPI_ISL_2547054, EPI_ISL_2547055, EPI_ISL_2547056, EPI_ISL_2547057, EPI_ISL_2547060, EPI_ISL_2547061, EPI_ISL_2547062, EPI_ISL_2547063, EPI_ISL_2547064, EPI_ISL_2547065, EPI_ISL_2547066, EPI_ISL_2547067, EPI_ISL_2547068 | | | |
| see above | Synlab Eesti OÜ | 1. Laboratory of Communicable Diseases (Estonia); 2. Eurofins Genomics Europe Sequencing GmbH | Liidia Dotsenko et al. |
| EPI_ISL_2550572, EPI_ISL_2550573, EPI_ISL_2550574, EPI_ISL_2550575, EPI_ISL_2550580, EPI_ISL_2550581, EPI_ISL_2550582, EPI_ISL_2550586, EPI_ISL_2550588, EPI_ISL_2550589, EPI_ISL_2550590, EPI_ISL_2550591, EPI_ISL_2550592 | | | |
| see above | NB-Hôpital Georges L. Dumont | National Microbiology Laboratory (NML) | Anna Majer, Shari Tyson, Grace Seo, Philip Mabon, Elsie Grudeski, Rhiannon Huzarewich, Russell Mandes, Anneliese Landgraff, Jennifer Tanner, Natalie Knox, Morag Graham, Gary Van Domselaar, Richard Garceau, Guillaume Desnoyers, Nathalie Bastien, Yan Li, Timothy Booth, Darian Hole, Madison Chapel, Kirsten Biggar, CanCOGeN's metadata curation team, Public Health Agency of Canada CanCOGeN team |
| EPI_ISL_2557359, EPI_ISL_2557360, EPI_ISL_2557367, EPI_ISL_2557368, EPI_ISL_2557372, EPI_ISL_2557374 | Lboratorio Central de Saude Publica do Estado do Parana (LACEN/PR) | Laboratory of Respiratory Viruses and Measles, Oswaldo Cruz Institute, FIOCRUZ | Paola Resende, Luciana Appolinario, Fernando Motta, Anna Carolina Paixao, Ana Carolina Mendonca, Alice Sampaio Rocha, Taina Venas, Elisa Cavalcante Pereira, Renata Serrano Lopes, Irina Riediger, Marilda Siqueira on behalf of the Fiocruz COVID-19 Genomic Surveillance Network |
| EPI_ISL_2559261 | LESP Jalisco | Instituto de Diagnostico y Referencia Epidemiologicos (INDRE) | Claudia Wong-Arambula, Abril Rodriguez-Maldonado, Vanessa Rivero-Arredondo, Ariadna Medina-Benitez, Joaquin Quiroz-Mercado, Sergio Rangel-Guerrero, Natividad Cruz-Ortiz, Tatiana Nunez-Garcia, Gisela Barrera-Badillo, Lucia Hernandez-Rivas, Irma Lopez-Martinez, Ernesto Ramirez-Gonzalez. |
| EPI_ISL_2559263, EPI_ISL_2559265, EPI_ISL_2559267, EPI_ISL_2559269 | LESP Chihuahua | Instituto de Diagnostico y Referencia Epidemiologicos (INDRE) | Claudia Wong-Arambula, Abril Rodriguez-Maldonado, Vanessa Rivero-Arredondo, Ariadna Medina-Benitez, Joaquin Quiroz-Mercado, Sergio Rangel-Guerrero, Natividad Cruz-Ortiz, Tatiana Nunez-Garcia, Gisela Barrera-Badillo, Lucia Hernandez-Rivas, Irma Lopez-Martinez, Ernesto Ramirez-Gonzalez. |
| EPI_ISL_2559279 | LESP Veracruz | Instituto de Diagnostico y Referencia Epidemiologicos (INDRE) | Claudia Wong-Arambula, Abril Rodriguez-Maldonado, Vanessa Rivero-Arredondo, Ariadna Medina-Benitez, Joaquin Quiroz-Mercado, Sergio Rangel-Guerrero, Natividad Cruz-Ortiz, Tatiana Nunez-Garcia, Gisela Barrera-Badillo, Lucia Hernandez-Rivas, Irma Lopez-Martinez, Ernesto Ramirez-Gonzalez. |
| EPI_ISL_2559281 | LESP Tamaulipas | Instituto de Diagnostico y Referencia Epidemiologicos (INDRE) | Claudia Wong-Arambula, Abril Rodriguez-Maldonado, Vanessa Rivero-Arredondo, Ariadna Medina-Benitez, Joaquin Quiroz-Mercado, Sergio Rangel-Guerrero, Natividad Cruz-Ortiz, Tatiana Nunez-Garcia, Gisela Barrera-Badillo, Lucia Hernandez-Rivas, Irma Lopez-Martinez, Ernesto Ramirez-Gonzalez. |
| EPI_ISL_2559283, EPI_ISL_2559285, EPI_ISL_2559287, EPI_ISL_2559289, EPI_ISL_2559291, EPI_ISL_2559293, EPI_ISL_2559295, EPI_ISL_2559297, EPI_ISL_2559299, EPI_ISL_2559302, EPI_ISL_2559303, EPI_ISL_2559305, EPI_ISL_2559307, EPI_ISL_2559309, EPI_ISL_2559311, EPI_ISL_2559314, EPI_ISL_2559316, EPI_ISL_2559317, EPI_ISL_2559319, EPI_ISL_2559321, EPI_ISL_2559323, EPI_ISL_2559325, EPI_ISL_2559327, EPI_ISL_2559329, EPI_ISL_2559331, EPI_ISL_2559333, EPI_ISL_2559336, EPI_ISL_2559338, EPI_ISL_2559342, EPI_ISL_2559343, EPI_ISL_2559346, EPI_ISL_2559347, EPI_ISL_2559350, EPI_ISL_2559352 | | | |
| see above | LESP Queretaro | Instituto de Diagnostico y Referencia Epidemiologicos (INDRE) | Claudia Wong-Arambula, Abril Rodriguez-Maldonado, Vanessa Rivero-Arredondo, Ariadna Medina-Benitez, Joaquin Quiroz-Mercado, Sergio Rangel-Guerrero, Natividad Cruz-Ortiz, Tatiana Nunez-Garcia, Gisela Barrera-Badillo, Lucia Hernandez-Rivas, Irma Lopez-Martinez, Ernesto Ramirez-Gonzalez. |
| EPI_ISL_2559431, EPI_ISL_2559433 | LESP Michoacan | Instituto de Diagnostico y Referencia Epidemiologicos (INDRE) | Claudia Wong-Arambula, Abril Rodriguez-Maldonado, Vanessa Rivero-Arredondo, Ariadna Medina-Benitez, Joaquin Quiroz-Mercado, Sergio Rangel-Guerrero, Natividad Cruz-Ortiz, Tatiana Nunez-Garcia, Gisela Barrera-Badillo, Lucia Hernandez-Rivas, Irma Lopez-Martinez, Ernesto Ramirez-Gonzalez. |
| EPI_ISL_2561981, EPI_ISL_2561982, EPI_ISL_2561983 | cerballiance-IDF | Cerba lab | Emmanuel Lecorche, Sabine Trombert-Paolantoni, Aude Lessenne, Kader Merah, Thierry Collin, Laura Verdurme, Patrice Herisson, Stéphanie Haïm-Boukobza, Bénédicte Roquebert |
| EPI_ISL_2561984, EPI_ISL_2561985 | cerballiance-cote d'azur | Cerba lab | Emmanuel Lecorche, Sabine Trombert-Paolantoni, Laura Verdurme, Stéphanie Haïm-Boukobza, Bénédicte Roquebert |
| EPI_ISL_2561986 | cerballiance-IDF | Cerba lab | Emmanuel Lecorche, Sabine Trombert-Paolantoni, Aude Lessenne, Kader Merah, Thierry Collin, Laura Verdurme, Patrice Herisson, Stéphanie Haïm-Boukobza, Bénédicte Roquebert |
| EPI_ISL_2561987 | LABM Lescaroux | Cerba lab | Emmanuel Lecorche, Sabine Trombert-Paolantoni, Laura Verdurme, Stéphanie Haïm-Boukobza, Bénédicte Roquebert |
| EPI_ISL_2561988 | CH Marie Lannelongue | Cerba lab | Emmanuel Lecorche, Sabine Trombert-Paolantoni, Laura Verdurme, Stéphanie Haïm-Boukobza, Bénédicte Roquebert |
| EPI_ISL_2561989 | cerballiance-IDF | Cerba lab | Emmanuel Lecorche, Sabine Trombert-Paolantoni, Laura Verdurme, Stéphanie Haïm-Boukobza, Bénédicte Roquebert |
| EPI_ISL_2561990 | LABM Lescaroux | Cerba lab | Emmanuel Lecorche, Sabine Trombert-Paolantoni, Laura Verdurme, Stéphanie Haïm-Boukobza, Bénédicte Roquebert |
| EPI_ISL_2561991 | cerballiance-IDF | Cerba lab | Emmanuel Lecorche, Sabine Trombert-Paolantoni, Aude Lessenne, Kader Merah, Thierry Collin, Laura Verdurme, Patrice Herisson, Stéphanie Haïm-Boukobza, Bénédicte Roquebert |
| EPI_ISL_2561992 | cerballiance-centre val de loire | Cerba lab | Emmanuel Lecorche, Sabine Trombert-Paolantoni, Aude Lessenne, Kader Merah, Thierry Collin, Laura Verdurme, Patrice Herisson, Stéphanie Haïm-Boukobza, Bénédicte Roquebert |
| EPI_ISL_2561993, EPI_ISL_2561994, EPI_ISL_2561995, EPI_ISL_2561996, EPI_ISL_2561997, EPI_ISL_2561998, EPI_ISL_2561999, EPI_ISL_2562000, EPI_ISL_2562001, EPI_ISL_2562002, EPI_ISL_2562003 | | | |
| see above | cerballiance-IDF | Cerba lab | Emmanuel Lecorche, Sabine Trombert-Paolantoni, Aude Lessenne, Kader Merah, Thierry Collin, Laura Verdurme, Patrice Herisson, Stéphanie Haïm-Boukobza, Bénédicte Roquebert |
| EPI_ISL_2562004 | cerballiance-cote d'azur | Cerba lab | Emmanuel Lecorche, Sabine Trombert-Paolantoni, Laura Verdurme, Stéphanie Haïm-Boukobza, Bénédicte Roquebert |
| EPI_ISL_2562005 | cerballiance-centre val de loire | Cerba lab | Emmanuel Lecorche, Sabine Trombert-Paolantoni, Aude Lessenne, Kader Merah, Thierry Collin, Laura Verdurme, Patrice Herisson, Stéphanie Haïm-Boukobza, Bénédicte Roquebert |
| EPI_ISL_2562006 | cerballiance-IDF | Cerba lab | Emmanuel Lecorche, Sabine Trombert-Paolantoni, Aude Lessenne, Kader Merah, Thierry Collin, Laura Verdurme, Patrice Herisson, Stéphanie Haïm-Boukobza, Bénédicte Roquebert |
| EPI_ISL_2562007 | cerballiance-cote d'azur | Cerba lab | Emmanuel Lecorche, Sabine Trombert-Paolantoni, Laura Verdurme, Stéphanie Haïm-Boukobza, Bénédicte Roquebert |
| EPI_ISL_2562008 | cerballiance-IDF | Cerba lab | Emmanuel Lecorche, Sabine Trombert-Paolantoni, Aude Lessenne, Kader Merah, Thierry Collin, Laura Verdurme, Patrice Herisson, Stéphanie Haïm-Boukobza, Bénédicte Roquebert |
| EPI_ISL_2562009 | cerballiance-centre val de loire | Cerba lab | Emmanuel Lecorche, Sabine Trombert-Paolantoni, Aude Lessenne, Kader Merah, Thierry Collin, Laura Verdurme, Patrice Herisson, Stéphanie Haïm-Boukobza, Bénédicte Roquebert |
| EPI_ISL_2562010 | cerballiance-IDF | Cerba lab | Emmanuel Lecorche, Sabine Trombert-Paolantoni, Aude Lessenne, Kader Merah, Thierry Collin, Laura Verdurme, Patrice Herisson, Stéphanie Haïm-Boukobza, Bénédicte Roquebert |
| EPI_ISL_2562011 | cerballiance-centre val de loire | Cerba lab | Emmanuel Lecorche, Sabine Trombert-Paolantoni, Aude Lessenne, Kader Merah, Thierry Collin, Laura Verdurme, Patrice Herisson, Stéphanie Haïm-Boukobza, Bénédicte Roquebert |
| EPI_ISL_2562012, EPI_ISL_2562013, EPI_ISL_2562014 | cerballiance-IDF | Cerba lab | Emmanuel Lecorche, Sabine Trombert-Paolantoni, Aude Lessenne, Kader Merah, Thierry Collin, Laura Verdurme, Patrice Herisson, Stéphanie Haïm-Boukobza, Bénédicte Roquebert |
| EPI_ISL_2562015 | cerballiance-HDF | Cerba lab | Emmanuel Lecorche, Sabine Trombert-Paolantoni, Aude Lessenne, Kader Merah, Thierry Collin, Laura Verdurme, Patrice Herisson, Stéphanie Haïm-Boukobza, Bénédicte Roquebert |
| EPI_ISL_2562016 | cerballiance-centre val de loire | Cerba lab | Emmanuel Lecorche, Sabine Trombert-Paolantoni, Aude Lessenne, Kader Merah, Thierry Collin, Laura Verdurme, Patrice Herisson, Stéphanie Haïm-Boukobza, Bénédicte Roquebert |

| | | | |
|---|---|---|---|
| EPI_ISL_2562017 | cerballiance-HDF | Cerba lab | Emmanuel Lecorche, Sabine Trombert-Paolantoni, Aude Lessenne, Kader Merah, Thierry Collin, Laura Verdurme, Patrice Herisson, Stéphanie Haïm-Boukobza, Bénédicte Roquebert |
| EPI_ISL_2562018 | cerballiance-IDF | Cerba lab | Emmanuel Lecorche, Sabine Trombert-Paolantoni, Aude Lessenne, Kader Merah, Thierry Collin, Laura Verdurme, Patrice Herisson, Stéphanie Haïm-Boukobza, Bénédicte Roquebert |
| EPI_ISL_2562019, EPI_ISL_2562020, EPI_ISL_2562023, EPI_ISL_2562024, EPI_ISL_2562025, EPI_ISL_2562026 | cerballiance-HDF | Cerba lab | Emmanuel Lecorche, Sabine Trombert-Paolantoni, Aude Lessenne, Kader Merah, Thierry Collin, Laura Verdurme, Patrice Herisson, Stéphanie Haïm-Boukobza, Bénédicte Roquebert |
| EPI_ISL_2562027 | cerballiance-IDF | Cerba lab | Emmanuel Lecorche, Sabine Trombert-Paolantoni, Aude Lessenne, Kader Merah, Thierry Collin, Laura Verdurme, Patrice Herisson, Stéphanie Haïm-Boukobza, Bénédicte Roquebert |
| EPI_ISL_2562028 | DPM Diagnostics | Cerba lab | Emmanuel Lecorche, Sabine Trombert-Paolantoni, Aude Lessenne, Kader Merah, Thierry Collin, Laura Verdurme, Patrice Herisson, Stéphanie Haïm-Boukobza, Bénédicte Roquebert |
| EPI_ISL_2562029, EPI_ISL_2562030 | CH Franck Joly | Cerba lab | Emmanuel Lecorche, Sabine Trombert-Paolantoni, Aude Lessenne, Kader Merah, Thierry Collin, Laura Verdurme, Patrice Herisson, Stéphanie Haïm-Boukobza, Bénédicte Roquebert |
| EPI_ISL_2562031 | CH Princesse Grace | Cerba lab | Emmanuel Lecorche, Sabine Trombert-Paolantoni, Aude Lessenne, Kader Merah, Thierry Collin, Laura Verdurme, Patrice Herisson, Stéphanie Haïm-Boukobza, Bénédicte Roquebert |
| EPI_ISL_2562038 | cerballiance-Artois | Cerba lab | Emmanuel Lecorche, Sabine Trombert-Paolantoni, Aude Lessenne, Kader Merah, Thierry Collin, Laura Verdurme, Patrice Herisson, Stéphanie Haïm-Boukobza, Bénédicte Roquebert |
| EPI_ISL_2562045 | CH Princesse Grace | Cerba lab | Emmanuel Lecorche, Sabine Trombert-Paolantoni, Aude Lessenne, Kader Merah, Thierry Collin, Laura Verdurme, Patrice Herisson, Stéphanie Haïm-Boukobza, Bénédicte Roquebert |
| EPI_ISL_2562087 | GMERS Government Medical College (GMC), Gotri | Gujarat Biotechnology Research Centre | Zarna Patel,Nitin Savaliya,Dinesh Kumar,Twinkle Soni,Sonal Sharma ,Zuber Saiyed,Ramesh Pandit,Janvi Raval,Bithika Duttaroy,Umang Mishra,Nitesh Shah,Chaitanya Joshi,Madhvi Joshi |
| EPI_ISL_2562088 | GMERS Government Medical College (GMC), Gotri | Gujarat Biotechnology Research Centre | Nitin Savaliya,Dinesh Kumar,Twinkle Soni,Sonal Sharma ,Zuber Saiyed,Ramesh Pandit,Janvi Raval,Zarna Patel,Bithika Duttaroy,Umang Mishra,Nitesh Shah,Chaitanya Joshi,Madhvi Joshi |
| EPI_ISL_2562089 | GMERS Government Medical College (GMC), Gotri | Gujarat Biotechnology Research Centre | Dinesh Kumar,Twinkle Soni,Sonal Sharma ,Zuber Saiyed,Ramesh Pandit,Janvi Raval,Zarna Patel,Nitin Savaliya,Bithika Duttaroy,Umang Mishra,Nitesh Shah,Chaitanya Joshi,Madhvi Joshi |
| EPI_ISL_2562091 | GMERS Government Medical College (GMC), Gotri | Gujarat Biotechnology Research Centre | Sonal Sharma ,Zuber Saiyed,Ramesh Pandit,Janvi Raval,Zarna Patel,Nitin Savaliya,Dinesh Kumar,Twinkle Soni,Bithika Duttaroy,Umang Mishra,Nitesh Shah,Chaitanya Joshi,Madhvi Joshi |
| EPI_ISL_2562092 | GMERS Government Medical College (GMC), Gotri | Gujarat Biotechnology Research Centre | Zuber Saiyed,Ramesh Pandit,Janvi Raval,Zarna Patel,Nitin Savaliya,Dinesh Kumar,Twinkle Soni,Sonal Sharma ,Bithika Duttaroy,Umang Mishra,Nitesh Shah,Chaitanya Joshi,Madhvi Joshi |
| EPI_ISL_2562093 | GMERS Government Medical College (GMC), Gotri | Gujarat Biotechnology Research Centre | Ramesh Pandit,Janvi Raval,Zarna Patel,Nitin Savaliya,Dinesh Kumar,Twinkle Soni,Sonal Sharma ,Zuber Saiyed,Bithika Duttaroy,Umang Mishra,Nitesh Shah,Chaitanya Joshi,Madhvi Joshi |
| EPI_ISL_2562098 | GMERS Government Medical College (GMC), Gotri | Gujarat Biotechnology Research Centre | Twinkle Soni,Sonal Sharma ,Zuber Saiyed,Ramesh Pandit,Janvi Raval,Zarna Patel,Nitin Savaliya,Dinesh Kumar,Bithika Duttaroy,Umang Mishra,Nitesh Shah,Chaitanya Joshi,Madhvi Joshi |
| EPI_ISL_2562099 | GMERS Government Medical College (GMC), Gotri | Gujarat Biotechnology Research Centre | Sonal Sharma ,Zuber Saiyed,Ramesh Pandit,Janvi Raval,Zarna Patel,Nitin Savaliya,Dinesh Kumar,Twinkle Soni,Bithika Duttaroy,Umang Mishra,Nitesh Shah,Chaitanya Joshi,Madhvi Joshi |
| EPI_ISL_2562100 | GMERS Government Medical College (GMC), Gotri | Gujarat Biotechnology Research Centre | Zuber Saiyed,Ramesh Pandit,Janvi Raval,Zarna Patel,Nitin Savaliya,Dinesh Kumar,Twinkle Soni,Sonal Sharma ,Bithika Duttaroy,Umang Mishra,Nitesh Shah,Chaitanya Joshi,Madhvi Joshi |
| EPI_ISL_2562103 | GMERS Government Medical College (GMC), Gotri | Gujarat Biotechnology Research Centre | Zarna Patel,Nitin Savaliya,Dinesh Kumar,Twinkle Soni,Sonal Sharma ,Zuber Saiyed,Ramesh Pandit,Janvi Raval,Bithika Duttaroy,Umang Mishra,Nitesh Shah,Chaitanya Joshi,Madhvi Joshi |
| EPI_ISL_2566210, EPI_ISL_2566211, EPI_ISL_2566212, EPI_ISL_2566213, EPI_ISL_2566214 | SC (UCO) Igiene e Sanità Pubblica, ASUGI, Trieste | ARGO Laboratorio Genomica ed Epigenomica | Licastro D, Dal Monego S, Degasperi M, Marcello A, Breda A, Fontana F, Segat L, Piscianz E, Braida C, Barbone F, D'Agaro P |
| EPI_ISL_2566215, EPI_ISL_2566216, EPI_ISL_2566217, EPI_ISL_2566218, EPI_ISL_2566219, EPI_ISL_2566220, EPI_ISL_2566221, EPI_ISL_2566222 | SC (UCO) Igiene e Sanità Pubblica, ASUGI, Trieste and SC Microbiologia e Virologia, ASFO, Pordenone | ARGO Laboratorio Genomica ed Epigenomica | Licastro D, Dal Monego S, Degasperi M, Marcello A, Segat L, De Rosa R, Braida C, Barbone F, D'Agaro P |
| EPI_ISL_2566223, EPI_ISL_2566225, EPI_ISL_2566226, EPI_ISL_2566227, EPI_ISL_2566228, EPI_ISL_2566229 | SC (UCO) Igiene e Sanità Pubblica, ASUGI, Trieste | ARGO Laboratorio Genomica ed Epigenomica | Licastro D, Dal Monego S, Degasperi M, Marcello A, Maggiore A, Busetti M, Segat L, Piscianz E, Braida C, Barbone F, D'Agaro P |
| EPI_ISL_2566231, EPI_ISL_2566232, EPI_ISL_2566233, EPI_ISL_2566234, EPI_ISL_2566235 | SC (UCO) Igiene e Sanità Pubblica, ASUGI, Trieste and SOC Igiene ed Epidemiologia Clinica, laboratorio di Virologia ,ASUFC, Udine | ARGO Laboratorio Genomica ed Epigenomica | Licastro D, Dal Monego S, Degasperi M, Marcello A, Segat L, Pipan C, Braida C, Barbone F, D'Agaro P |
| EPI_ISL_2566236 | SC (UCO) Igiene e Sanità Pubblica, ASUGI, Trieste and SC Microbiologia e Virologia, ASFO, Pordenone | ARGO Laboratorio Genomica ed Epigenomica | Licastro D, Dal Monego S, Degasperi M, Marcello A, Segat L, De Rosa R, Braida C, Barbone F, D'Agaro P |
| EPI_ISL_2566239, EPI_ISL_2566240, EPI_ISL_2566241 | SC (UCO) Igiene e Sanità Pubblica, ASUGI, Trieste | ARGO Laboratorio Genomica ed Epigenomica | Licastro D, Dal Monego S, Degasperi M, Marcello A, Maggiore A, Busetti M, Segat L, Piscianz E, Braida C, Barbone F, D'Agaro P |
| EPI_ISL_2566242 | SC (UCO) Igiene e Sanità Pubblica, ASUGI, Trieste | ARGO Laboratorio Genomica ed Epigenomica | Licastro D, Dal Monego S, Degasperi M, Marcello A, Breda A, Fontana F, Segat L, Piscianz E, Braida C, Barbone F, D'Agaro P |
| EPI_ISL_2567592, EPI_ISL_2567593, EPI_ISL_2567594, EPI_ISL_2567595, EPI_ISL_2567596, EPI_ISL_2567597, EPI_ISL_2567598, EPI_ISL_2567599, EPI_ISL_2567600, EPI_ISL_2567601 | SC (UCO) Igiene e Sanità Pubblica, ASUGI, Trieste | ARGO Laboratorio Genomica ed Epigenomica | Licastro D, Dal Monego S, Degasperi M, Marcello A, Segat L, Piscianz E, Braida C, Barbone F, D'Agaro P |
| EPI_ISL_2567602, EPI_ISL_2567603, EPI_ISL_2567604, EPI_ISL_2567605 | SC (UCO) Igiene e Sanità Pubblica, ASUGI, Trieste and SC Microbiologia e Virologia, ASFO, Pordenone | ARGO Laboratorio Genomica ed Epigenomica | Licastro D, Dal Monego S, Degasperi M, Marcello A, Segat L, De Rosa R, Braida C, Barbone F, D'Agaro P |
| EPI_ISL_2567606, EPI_ISL_2567607, EPI_ISL_2567608, EPI_ISL_2567609 | SC (UCO) Igiene e Sanità Pubblica, ASUGI, Trieste | ARGO Laboratorio Genomica ed Epigenomica | Licastro D, Dal Monego S, Degasperi M, Marcello A, Segat L, Piscianz E, Braida C, Barbone F, D'Agaro P |
| EPI_ISL_2567610, EPI_ISL_2567611, EPI_ISL_2567612 | SC (UCO) Igiene e Sanità Pubblica, ASUGI, Trieste and SC Microbiologia e Virologia, ASFO, Pordenone | ARGO Laboratorio Genomica ed Epigenomica | Licastro D, Dal Monego S, Degasperi M, Marcello A, Segat L, De Rosa R, Braida C, Barbone F, D'Agaro P |
| EPI_ISL_2600375 | cerballiance-IDF | Cerba lab | Emmanuel Lecorche, Sabine Trombert-Paolantoni, Aude Lessenne, Kader Merah, Thierry Collin, Laura Verdurme, Patrice Herisson, Stéphanie Haïm-Boukobza, Bénédicte Roquebert |
| EPI_ISL_2600376 | cerballiance-centre val de loire | Cerba lab | Emmanuel Lecorche, Sabine Trombert-Paolantoni, Aude Lessenne, Kader Merah, Thierry Collin, Laura Verdurme, Patrice Herisson, Stéphanie Haïm-Boukobza, Bénédicte Roquebert |

| | | | |
|---|---|---|---|
| EPI_ISL_2600377 | cerballiance-HDF | Cerba lab | Emmanuel Lecorche, Sabine Trombert-Paolantoni, Aude Lessenne, Kader Merah, Thierry Collin, Laura Verdurme, Patrice Herisson, Stéphanie Haïm-Boukobza, Bénédicte Roquebert |
| EPI_ISL_2600960, EPI_ISL_2600961 | Hematopathology Laboratory, ACTREC, TMC | Hematopathology Laboratory, ACTREC, TMC | Hematopathology Laboratory, ACTREC |
| EPI_ISL_2601414, EPI_ISL_2601415, EPI_ISL_2601416, EPI_ISL_2601418, EPI_ISL_2601420, EPI_ISL_2601421, EPI_ISL_2601422, EPI_ISL_2601423, EPI_ISL_2601429, EPI_ISL_2601430, EPI_ISL_2601431, EPI_ISL_2601432, EPI_ISL_2601433, EPI_ISL_2601434, EPI_ISL_2601435, EPI_ISL_2601436, EPI_ISL_2601438, EPI_ISL_2601439, EPI_ISL_2601440, EPI_ISL_2601441, EPI_ISL_2601443, EPI_ISL_2601444, EPI_ISL_2601445, EPI_ISL_2601446, EPI_ISL_2601448, EPI_ISL_2601449, EPI_ISL_2601450, EPI_ISL_2601452, EPI_ISL_2601454, EPI_ISL_2601455, EPI_ISL_2601456, EPI_ISL_2601457, EPI_ISL_2601458, EPI_ISL_2601459, EPI_ISL_2601460, EPI_ISL_2601462, EPI_ISL_2601463, EPI_ISL_2601464, EPI_ISL_2601465, EPI_ISL_2601466, EPI_ISL_2601468, EPI_ISL_2601469, EPI_ISL_2601470, EPI_ISL_2601471, EPI_ISL_2601472, EPI_ISL_2601474, EPI_ISL_2601476, EPI_ISL_2601478, EPI_ISL_2601479, EPI_ISL_2601481, EPI_ISL_2601482, EPI_ISL_2601483, EPI_ISL_2601486, EPI_ISL_2601488, EPI_ISL_2601489, EPI_ISL_2601491, EPI_ISL_2601492, EPI_ISL_2601494, EPI_ISL_2601495, EPI_ISL_2601496, EPI_ISL_2601497, EPI_ISL_2601498, EPI_ISL_2601499, EPI_ISL_2601500, EPI_ISL_2601501, EPI_ISL_2601502, EPI_ISL_2601503, EPI_ISL_2601504, EPI_ISL_2601505, EPI_ISL_2601506, EPI_ISL_2601507, EPI_ISL_2601508, EPI_ISL_2601509, EPI_ISL_2601510, EPI_ISL_2601511, EPI_ISL_2601512, EPI_ISL_2601513, EPI_ISL_2601515, EPI_ISL_2601516, EPI_ISL_2601517, EPI_ISL_2601518, EPI_ISL_2601519, EPI_ISL_2601520, EPI_ISL_2601521, EPI_ISL_2601522, EPI_ISL_2601523, EPI_ISL_2601524, EPI_ISL_2601525, EPI_ISL_2601526, EPI_ISL_2601527, EPI_ISL_2601528, EPI_ISL_2601529, EPI_ISL_2601530, EPI_ISL_2601531, EPI_ISL_2601532, EPI_ISL_2601533, EPI_ISL_2601534, EPI_ISL_2601535, EPI_ISL_2601536, EPI_ISL_2601537, EPI_ISL_2601538, EPI_ISL_2601539, EPI_ISL_2601540, EPI_ISL_2601541, EPI_ISL_2601542, EPI_ISL_2601543, EPI_ISL_2601546, EPI_ISL_2601547, EPI_ISL_2601548, EPI_ISL_2601550, EPI_ISL_2601551, EPI_ISL_2601552, EPI_ISL_2601553, EPI_ISL_2601554, EPI_ISL_2601555, EPI_ISL_2601556, EPI_ISL_2601557, EPI_ISL_2601558, EPI_ISL_2601559, EPI_ISL_2601560, EPI_ISL_2601561, EPI_ISL_2601562, EPI_ISL_2601563, EPI_ISL_2601564, EPI_ISL_2601565, EPI_ISL_2601566, EPI_ISL_2601567, EPI_ISL_2601569, EPI_ISL_2601570, EPI_ISL_2601571, EPI_ISL_2601572, EPI_ISL_2601573, EPI_ISL_2601574, EPI_ISL_2601575, EPI_ISL_2601576, EPI_ISL_2601577 | | | |
| see above | Synlab Eesti OÜ | 1. Laboratory of Communicable Diseases (Estonia); 2. Eurofins Genomics Europe Sequencing GmbH | Liidia Dotsenko et al. |
| EPI_ISL_2601610, EPI_ISL_2601611, EPI_ISL_2601612 | Laboratory of Communicable Diseases | 1. Laboratory of Communicable Diseases (Estonia); 2. Eurofins Genomics Europe Sequencing GmbH | Liidia Dotsenko et al. |
| EPI_ISL_2601619, EPI_ISL_2601636, EPI_ISL_2601638, EPI_ISL_2601639, EPI_ISL_2601640, EPI_ISL_2601641, EPI_ISL_2601642, EPI_ISL_2601643, EPI_ISL_2601644, EPI_ISL_2601645, EPI_ISL_2601646, EPI_ISL_2601647, EPI_ISL_2601648, EPI_ISL_2601649, EPI_ISL_2601650, EPI_ISL_2601651 | | | |
| see above | North Estonia Medical Centre | 1. Laboratory of Communicable Diseases (Estonia); 2. Eurofins Genomics Europe Sequencing GmbH | Liidia Dotsenko et al. |
| EPI_ISL_2601655, EPI_ISL_2601656, EPI_ISL_2601658, EPI_ISL_2601659, EPI_ISL_2601660 | Pärnu Hospital | 1. Laboratory of Communicable Diseases (Estonia); 2. Eurofins Genomics Europe Sequencing GmbH | Liidia Dotsenko et al. |
| EPI_ISL_2603131 | Grupo de Investigación en Enfermedades Tropicales del Ejército (GINETEJ), Laboratorio de Referencia e Investigación, Dirección de Sanidad Ejército, Bogotá, Colombia | Centro de Investigaciones en Microbiología y Biotecnología-UR (CIMBIUR), Facultad de Ciencias Naturales, Universidad del Rosario, Bogotá, Colombia | Sergio Castañeda, Nathalia Ballesteros, Marina Muñoz, Luz H. Patiño, Claudia Méndez, Carolina Oliveros, Julie Pérez, Lorena Albarracín, Elizabeth K. Márquez, María Teresa Alvarado, Frank de los Santos Ortiz, Yanira Romero, Camilo A. Correa-Cárdenas, Maria Clara Duque, Sergio Gutíerrez-Riveros, Zulma Cucunubá, Juan David Ramírez |
| EPI_ISL_2603425, EPI_ISL_2603426, EPI_ISL_2603430 | Lboratorio Central de Saude Publica do Estado do Parana (LACEN/PR) | Laboratory of Respiratory Viruses and Measles, Oswaldo Cruz Institute, FIOCRUZ | Paola Resende, Luciana Appolinario, Fernando Motta, Anna Carolina Paixao, Ana Carolina Mendonca, Alice Sampaio Rocha, Taina Venas, Elisa Cavalcante Pereira, Renata Serrano Lopes, Irina Riediger, Marilda Siqueira on behalf of the Fiocruz COVID-19 Genomic Surveillance Network |
| EPI_ISL_2603434, EPI_ISL_2603435, EPI_ISL_2603436 | Laboratorio Central de Saude Publica do Estado do Rio Grande do Sul (LACEN-RS) | Laboratory of Respiratory Viruses and Measles, Oswaldo Cruz Institute, FIOCRUZ | Paola Resende, Luciana Appolinario, Fernando Motta, Anna Carolina Paixao, Ana Carolina Mendonca, Alice Sampaio Rocha, Taina Venas, Elisa Cavalcante Pereira, Renata Serrano Lopes, Anderson Brandao Leite, Marilda Siqueira on behalf of the Fiocruz COVID-19 Genomic Surveillance Network |
| EPI_ISL_2603437, EPI_ISL_2603439 | Lboratorio Central de Saude Publica do Estado do Parana (LACEN/PR) | Laboratory of Respiratory Viruses and Measles, Oswaldo Cruz Institute, FIOCRUZ | Paola Resende, Luciana Appolinario, Fernando Motta, Anna Carolina Paixao, Ana Carolina Mendonca, Alice Sampaio Rocha, Taina Venas, Elisa Cavalcante Pereira, Renata Serrano Lopes, Irina Riediger, Marilda Siqueira on behalf of the Fiocruz COVID-19 Genomic Surveillance Network |
| EPI_ISL_2603445, EPI_ISL_2603449, EPI_ISL_2603450 | Laboratorio Central de Saude Publica do Estado do Rio Grande do Sul (LACEN-RS) | Laboratory of Respiratory Viruses and Measles, Oswaldo Cruz Institute, FIOCRUZ | Paola Resende, Luciana Appolinario, Fernando Motta, Anna Carolina Paixao, Ana Carolina Mendonca, Alice Sampaio Rocha, Taina Venas, Elisa Cavalcante Pereira, Renata Serrano Lopes, Anderson Brandao Leite, Marilda Siqueira on behalf of the Fiocruz COVID-19 Genomic Surveillance Network |
| EPI_ISL_2611665, EPI_ISL_2611667, EPI_ISL_2611668, EPI_ISL_2611671 | HOME QUARANTINE TASKFORCE | Hong Kong Department of Health | Alan K.L. Tsang, Peter C.W. Yip, Ken H.L. Ng, Edman T.K. Lam, Rickjason C.W. Chan, Dominic N.C. Tsang |
| EPI_ISL_2612627 | Hospital | National Reference Center for Viruses of Respiratory Infections, Institut Pasteur, Paris | Marion Barbet, Sylvie Behillil, Méline Bizard, Angela Brisebarre, Camille Capel, Vincent Enouf, Louise Lefrançois, Frédéric Lemoine, Christophe Malabat, Corinne Maufrais, Etienne Simon-Lorière, Maud Vanpeene, Sylvie Van der Werf ,Alexandra Ducancelle |
| EPI_ISL_2613602, EPI_ISL_2613603, EPI_ISL_2613604, EPI_ISL_2613605, EPI_ISL_2613606, EPI_ISL_2613607, EPI_ISL_2613608 | Dipartimento di Medicina di Laboratorio, Azienda sanitaria universitaria Friuli Centrale (ASU FC) | Dipartimento di Medicina di Laboratorio, Azienda sanitaria universitaria Friuli Centrale (ASU FC) | Corrado Pipan, Catia Mio, Chiara Dal Secco, Stefania Marzinotto, Francesco Curcio |
| EPI_ISL_2613816, EPI_ISL_2613817, EPI_ISL_2613822 | Lboratorio Central de Saude Publica do Estado do Parana (LACEN/PR) | Laboratory of Respiratory Viruses and Measles, Oswaldo Cruz Institute, FIOCRUZ | Paola Resende, Luciana Appolinario, Fernando Motta, Anna Carolina Paixao, Ana Carolina Mendonca, Alice Sampaio Rocha, Taina Venas, Elisa Cavalcante Pereira, Renata Serrano Lopes, Irina Riediger, Marilda Siqueira on behalf of the Fiocruz COVID-19 Genomic Surveillance Network |
| EPI_ISL_2614565 | Hospital de Base de Sao Jose do Rio Preto | Instituto Adolfo Lutz, Interdiciplinary Procedures Center, Strategic Laboratory | Claudio Tavares Sacchi, Claudia Regina Gonçalves, Erica Valessa Ramos Gomes, Karoline Rodrigues Campos, Caio Vinicius Dias Lopes, Leonardo Jose Tadeu de Araujo |
| EPI_ISL_2614572 | Hospital Municipal Ermelino Matarazzo | Instituto Adolfo Lutz, Interdiciplinary Procedures Center, Strategic Laboratory | Claudio Tavares Sacchi, Claudia Regina Gonçalves, Erica Valessa Ramos Gomes, Karoline Rodrigues Campos, Caio Vinicius Dias Lopes, Leonardo Jose Tadeu de Araujo |
| EPI_ISL_2614573 | Pronto Socorro Andre Sacco - Pestana | Instituto Adolfo Lutz, Interdiciplinary Procedures Center, Strategic Laboratory | Claudio Tavares Sacchi, Claudia Regina Gonçalves, Erica Valessa Ramos Gomes, Karoline Rodrigues Campos, Caio Vinicius Dias Lopes, Leonardo Jose Tadeu de Araujo |
| EPI_ISL_2614574 | UPA Akira Tada | Instituto Adolfo Lutz, Interdiciplinary Procedures Center, Strategic Laboratory | Claudio Tavares Sacchi, Claudia Regina Gonçalves, Erica Valessa Ramos Gomes, Karoline Rodrigues Campos, Caio Vinicius Dias Lopes, Leonardo Jose Tadeu de Araujo |
| EPI_ISL_2614575 | Hospital das Clinicas Luzia de Pinho Melo | Instituto Adolfo Lutz, Interdiciplinary Procedures Center, Strategic Laboratory | Claudio Tavares Sacchi, Claudia Regina Gonçalves, Erica Valessa Ramos Gomes, Karoline Rodrigues Campos, Caio Vinicius Dias Lopes, Leonardo Jose Tadeu de Araujo |
| EPI_ISL_2614576 | PS Municipal Dr Alceu Lot | Instituto Adolfo Lutz, Interdiciplinary Procedures Center, Strategic Laboratory | Claudio Tavares Sacchi, Claudia Regina Gonçalves, Erica Valessa Ramos Gomes, Karoline Rodrigues Campos, Caio Vinicius Dias Lopes, Leonardo Jose Tadeu de Araujo |
| EPI_ISL_2614577 | Hospital Municipal Antonio Giglio | Instituto Adolfo Lutz, Interdiciplinary Procedures Center, Strategic Laboratory | Claudio Tavares Sacchi, Claudia Regina Gonçalves, Erica Valessa Ramos Gomes, Karoline Rodrigues Campos, Caio Vinicius Dias Lopes, Leonardo Jose Tadeu de Araujo |
| EPI_ISL_2614578 | UBS São Lourenco da Serra | Instituto Adolfo Lutz, Interdiciplinary Procedures Center, Strategic Laboratory | Claudio Tavares Sacchi, Claudia Regina Gonçalves, Erica Valessa Ramos Gomes, Karoline Rodrigues Campos, Caio Vinicius Dias Lopes, Leonardo Jose Tadeu de Araujo |
| EPI_ISL_2614579 | PS Municipal Dr Alceu Lot | Instituto Adolfo Lutz, Interdiciplinary Procedures Center, Strategic Laboratory | Claudio Tavares Sacchi, Claudia Regina Gonçalves, Erica Valessa Ramos Gomes, Karoline Rodrigues Campos, Caio Vinicius Dias Lopes, Leonardo Jose Tadeu de Araujo |
| EPI_ISL_2614580 | Hospital Geral de Pedreira | Instituto Adolfo Lutz, Interdiciplinary Procedures Center, Strategic Laboratory | Claudio Tavares Sacchi, Claudia Regina Gonçalves, Erica Valessa Ramos Gomes, Karoline Rodrigues Campos, Caio Vinicius Dias Lopes, Leonardo Jose Tadeu de Araujo |
| EPI_ISL_2614581 | Hospital Geral do Grajau | Instituto Adolfo Lutz, Interdiciplinary Procedures Center, Strategic Laboratory | Claudio Tavares Sacchi, Claudia Regina Gonçalves, Erica Valessa Ramos Gomes, Karoline Rodrigues Campos, Caio Vinicius Dias Lopes, Leonardo Jose Tadeu de Araujo |
| EPI_ISL_2614582 | Unidade Mista de Saude | Instituto Adolfo Lutz, Interdiciplinary Procedures Center, Strategic Laboratory | Claudio Tavares Sacchi, Claudia Regina Gonçalves, Erica Valessa Ramos Gomes, Karoline Rodrigues Campos, Caio Vinicius Dias Lopes, Leonardo Jose Tadeu de Araujo |
| EPI_ISL_2614583 | Laboratorio Municipal De Suzano | Instituto Adolfo Lutz, Interdiciplinary Procedures Center, Strategic Laboratory | Claudio Tavares Sacchi, Claudia Regina Gonçalves, Erica Valessa Ramos Gomes, Karoline Rodrigues Campos, Caio Vinicius Dias Lopes, Leonardo Jose Tadeu de Araujo |
| EPI_ISL_2614586 | Vigilância Sanitaria E Epidemiologica | Instituto Adolfo Lutz, Interdiciplinary Procedures Center, Strategic Laboratory | Claudio Tavares Sacchi, Claudia Regina Gonçalves, Erica Valessa Ramos Gomes, Karoline Rodrigues Campos, Caio Vinicius Dias Lopes, Leonardo Jose Tadeu de Araujo |
| EPI_ISL_2614587, EPI_ISL_2614588 | UPA Vila Santa Catarina | Instituto Adolfo Lutz, Interdiciplinary Procedures Center, Strategic Laboratory | Claudio Tavares Sacchi, Claudia Regina Gonçalves, Erica Valessa Ramos Gomes, Karoline Rodrigues Campos, Caio Vinicius Dias Lopes, Leonardo Jose Tadeu de Araujo |

| | | | |
|---|---|---|---|
| EPI_ISL_2614589 | Laboratorio Municipal De Suzano | Instituto Adolfo Lutz, Interdiciplinary Procedures Center, Strategic Laboratory | Claudio Tavares Sacchi, Claudia Regina Gonçalves, Erica Valessa Ramos Gomes, Karoline Rodrigues Campos, Caio Vinicius Dias Lopes, Leonardo Jose Tadeu de Araujo |
| EPI_ISL_2614590 | Hospital Municipal Dr. Guido Guida | Instituto Adolfo Lutz, Interdiciplinary Procedures Center, Strategic Laboratory | Claudio Tavares Sacchi, Claudia Regina Gonçalves, Erica Valessa Ramos Gomes, Karoline Rodrigues Campos, Caio Vinicius Dias Lopes, Leonardo Jose Tadeu de Araujo |
| EPI_ISL_2614591 | Hospital Municipal Enf Antonio Policarpo de Oliveira | Instituto Adolfo Lutz, Interdiciplinary Procedures Center, Strategic Laboratory | Claudio Tavares Sacchi, Claudia Regina Gonçalves, Erica Valessa Ramos Gomes, Karoline Rodrigues Campos, Caio Vinicius Dias Lopes, Leonardo Jose Tadeu de Araujo |
| EPI_ISL_2614592 | UBS Silvio Joao Luiz de Lucia | Instituto Adolfo Lutz, Interdiciplinary Procedures Center, Strategic Laboratory | Claudio Tavares Sacchi, Claudia Regina Gonçalves, Erica Valessa Ramos Gomes, Karoline Rodrigues Campos, Caio Vinicius Dias Lopes, Leonardo Jose Tadeu de Araujo |
| EPI_ISL_2614593 | UBS Maria Pia de Oliveira | Instituto Adolfo Lutz, Interdiciplinary Procedures Center, Strategic Laboratory | Claudio Tavares Sacchi, Claudia Regina Gonçalves, Erica Valessa Ramos Gomes, Karoline Rodrigues Campos, Caio Vinicius Dias Lopes, Leonardo Jose Tadeu de Araujo |
| EPI_ISL_2614594 | Hospital Regional de Cotia | Instituto Adolfo Lutz, Interdiciplinary Procedures Center, Strategic Laboratory | Claudio Tavares Sacchi, Claudia Regina Gonçalves, Erica Valessa Ramos Gomes, Karoline Rodrigues Campos, Caio Vinicius Dias Lopes, Leonardo Jose Tadeu de Araujo |
| EPI_ISL_2614595 | UPA Vila Santa Catarina | Instituto Adolfo Lutz, Interdiciplinary Procedures Center, Strategic Laboratory | Claudio Tavares Sacchi, Claudia Regina Gonçalves, Erica Valessa Ramos Gomes, Karoline Rodrigues Campos, Caio Vinicius Dias Lopes, Leonardo Jose Tadeu de Araujo |
| EPI_ISL_2614596 | UBS Atalaia | Instituto Adolfo Lutz, Interdiciplinary Procedures Center, Strategic Laboratory | Claudio Tavares Sacchi, Claudia Regina Gonçalves, Erica Valessa Ramos Gomes, Karoline Rodrigues Campos, Caio Vinicius Dias Lopes, Leonardo Jose Tadeu de Araujo |
| EPI_ISL_2614597 | Hospital Municipal de Mogi das Cruzes | Instituto Adolfo Lutz, Interdiciplinary Procedures Center, Strategic Laboratory | Claudio Tavares Sacchi, Claudia Regina Gonçalves, Erica Valessa Ramos Gomes, Karoline Rodrigues Campos, Caio Vinicius Dias Lopes, Leonardo Jose Tadeu de Araujo |
| EPI_ISL_2614598, EPI_ISL_2614599 | Hospital de Campanha Covid-19 Mairipora | Instituto Adolfo Lutz, Interdiciplinary Procedures Center, Strategic Laboratory | Claudio Tavares Sacchi, Claudia Regina Gonçalves, Erica Valessa Ramos Gomes, Karoline Rodrigues Campos, Caio Vinicius Dias Lopes, Leonardo Jose Tadeu de Araujo |
| EPI_ISL_2614600 | Hospital Geral de Guaianazes | Instituto Adolfo Lutz, Interdiciplinary Procedures Center, Strategic Laboratory | Claudio Tavares Sacchi, Claudia Regina Gonçalves, Erica Valessa Ramos Gomes, Karoline Rodrigues Campos, Caio Vinicius Dias Lopes, Leonardo Jose Tadeu de Araujo |
| EPI_ISL_2614601 | UPA Akira Tada | Instituto Adolfo Lutz, Interdiciplinary Procedures Center, Strategic Laboratory | Claudio Tavares Sacchi, Claudia Regina Gonçalves, Erica Valessa Ramos Gomes, Karoline Rodrigues Campos, Caio Vinicius Dias Lopes, Leonardo Jose Tadeu de Araujo |
| EPI_ISL_2614618, EPI_ISL_2614620, EPI_ISL_2614622 | Instituto Adolfo Lutz - Regional de Presidente Prudente | Instituto Adolfo Lutz, Interdiciplinary Procedures Center, Strategic Laboratory | Claudio Tavares Sacchi, Claudia Regina Gonçalves, Erica Valessa Ramos Gomes, Karoline Rodrigues Campos, Caio Vinicius Dias Lopes, Leonardo Jose Tadeu de Araujo |
| EPI_ISL_2614626, EPI_ISL_2614627, EPI_ISL_2614628 | Instituto Adolfo Lutz - Regional de Marilia | Instituto Adolfo Lutz, Interdiciplinary Procedures Center, Strategic Laboratory | Claudio Tavares Sacchi, Claudia Regina Gonçalves, Erica Valessa Ramos Gomes, Karoline Rodrigues Campos, Caio Vinicius Dias Lopes, Leonardo Jose Tadeu de Araujo |
| EPI_ISL_2614629, EPI_ISL_2614630, EPI_ISL_2614631, EPI_ISL_2614632, EPI_ISL_2614633 | Dr. Yoshitaka Tamura Department of Clinical Laboratory, Osaka Habikino Medical Center | Department of Developmental Medicine, Research Institute, Osaka Women's and Children's Hospital | Itaru Yanagihara, Yukiko Nakura |
| EPI_ISL_2616337, EPI_ISL_2616347, EPI_ISL_2616363, EPI_ISL_2616366, EPI_ISL_2616367 | Elling group, Institute of Molecular Biotechnology (IMBA) | Bergthaler laboratory, CeMM Research Center for Molecular Medicine of the Austrian Academy of Sciences | Lukas Endler, Anna Schedl, Fabian Amman, Petr Triska, Matthew Thornton, Thomas Penz, Benedikt Agerer, Maelle Le Moing, Michael Schuster, Bekir Erguner, Jan Laine, Martin Senekowitsch, Christoph Bock, Andreas Bergthaler |
| EPI_ISL_2616369, EPI_ISL_2616370, EPI_ISL_2616373, EPI_ISL_2616379, EPI_ISL_2616380, EPI_ISL_2616383, EPI_ISL_2616389, EPI_ISL_2616396, EPI_ISL_2616397, EPI_ISL_2616398, EPI_ISL_2616399, EPI_ISL_2616400, EPI_ISL_2616401, EPI_ISL_2616402, EPI_ISL_2616403, EPI_ISL_2616404, EPI_ISL_2616410, EPI_ISL_2616411, EPI_ISL_2616412, EPI_ISL_2616413, EPI_ISL_2616414, EPI_ISL_2616415, EPI_ISL_2616416, EPI_ISL_2616417, EPI_ISL_2616418, EPI_ISL_2616419, EPI_ISL_2616420, EPI_ISL_2616422, EPI_ISL_2616423, EPI_ISL_2616424, EPI_ISL_2616425, EPI_ISL_2616426, EPI_ISL_2616427, EPI_ISL_2616428, EPI_ISL_2616429, EPI_ISL_2616430, EPI_ISL_2616432, EPI_ISL_2616433, EPI_ISL_2616434, EPI_ISL_2616435, EPI_ISL_2616448, EPI_ISL_2616451, EPI_ISL_2616452, EPI_ISL_2616453, EPI_ISL_2616454, EPI_ISL_2616455, EPI_ISL_2616457, EPI_ISL_2616458, EPI_ISL_2616459, EPI_ISL_2616460, EPI_ISL_2616461, EPI_ISL_2616462, EPI_ISL_2616463, EPI_ISL_2616465, EPI_ISL_2616466, EPI_ISL_2616467, EPI_ISL_2616468, EPI_ISL_2616470, EPI_ISL_2616471, EPI_ISL_2616473, EPI_ISL_2616474, EPI_ISL_2616475, EPI_ISL_2616476, EPI_ISL_2616477, EPI_ISL_2616478, EPI_ISL_2616479, EPI_ISL_2616480, EPI_ISL_2616481, EPI_ISL_2616482, EPI_ISL_2616483, EPI_ISL_2616484, EPI_ISL_2616487, EPI_ISL_2616488, EPI_ISL_2616490, EPI_ISL_2616491, EPI_ISL_2616492, EPI_ISL_2616493, EPI_ISL_2616495, EPI_ISL_2616496, EPI_ISL_2616498, EPI_ISL_2616499, EPI_ISL_2616501, EPI_ISL_2616503, EPI_ISL_2616504, EPI_ISL_2616505, EPI_ISL_2616506, EPI_ISL_2616507, EPI_ISL_2616508, EPI_ISL_2616509, EPI_ISL_2616511, EPI_ISL_2616512, EPI_ISL_2616513, EPI_ISL_2616514, EPI_ISL_2616515, EPI_ISL_2616516, EPI_ISL_2616517, EPI_ISL_2616518, EPI_ISL_2616519, EPI_ISL_2616520, EPI_ISL_2616521 | | | |
| see above | Gyncentrum | Gyncentrum | Emilia Morawiec, Pawe Czerwiski, Adam Pudeko, Aleksandra Skubis-Sikora, Magdalena Samul, Agnieszka Polak, Celina Kruszniewska-Rajs, Tomasz Wsik, Robert Wojtyczka, Jolanta Bartosiewicz- Wsik, Maria Maklasiska-Majdanik |
| EPI_ISL_2617090 | Africa_CDC - Malawi P6 S2 | CERI, Centre for Epidemic Response and Innvoation, Stellenbosch University and KRISP, KZN Research Innovation and Sequencing Platform, UKZN. | Mvula B, Chilima B, Chiwaula M, Mwangomba W, Panja L, Kasambara W, Auld A, Kim L, Kampira E, Kaba M, Wadonda N, Maida A, Giandhari J, Pillay S, Naidoo Y, Lessells R, Emmanuel SJ, Tegally H, Wilkinson E, de Oliveira T |
| EPI_ISL_2617882, EPI_ISL_2617883, EPI_ISL_2617884, EPI_ISL_2617886, EPI_ISL_2617887, EPI_ISL_2617892, EPI_ISL_2617893, EPI_ISL_2617895, EPI_ISL_2617896, EPI_ISL_2617897, EPI_ISL_2617898, EPI_ISL_2617899, EPI_ISL_2617900, EPI_ISL_2617901, EPI_ISL_2617906, EPI_ISL_2617908, EPI_ISL_2617909, EPI_ISL_2617912, EPI_ISL_2617914, EPI_ISL_2617915, EPI_ISL_2617918, EPI_ISL_2617919 | | | |
| see above | LABCOVID_HCPA | LABRESIS_HCPA | Wink PL, Martins AF, Volpato F, Monteiro F, Zavascki AP, Barth AL |
| EPI_ISL_2620718, EPI_ISL_2620728, EPI_ISL_2620744, EPI_ISL_2620745, EPI_ISL_2620901 | Ipoh Public Health Laboratory (MKAI), Ministry of Health Malaysia | Institute for Medical Research, Infectious Disease Research Centre, National Institutes of Health, Ministry of Health Malaysia | Suppiah J, Kamel K, Mohd Zawawi Z, Azizan MA, Ramly N, Robert F, Thayan R |
| EPI_ISL_2621164, EPI_ISL_2621165 | ICMT-APARTADO | Universidad Nacional de Colombia - Laboratorio Genómico One Health | Andres F. Cardona-Rios, Maria Stella López, Daniel O. Maldonado-Perez, Simón Villegas Velásquez, Lina M. Hurtado, Laura Silvana Perez, Karl A Ciuoderis, Maria Angélica Maya, Carolina Muñoz-Arango, Celeny Ortiz, Rita Almanza Payares, Idabely Betancur Ortiz, Sandra Ines Cano, Diego A. Álvarez-Díaz, Carlos Franco-Muñoz, Katherine Laiton-Donato, Hector Alejandro Ruiz-Moreno, Marcela Mercado-Reyes, Jorge E. Osorio, Juan P. Hernandez-Ortiz |
| EPI_ISL_2621222, EPI_ISL_2621223, EPI_ISL_2621224, EPI_ISL_2621225, EPI_ISL_2621226, EPI_ISL_2621227, EPI_ISL_2621228, EPI_ISL_2621230, EPI_ISL_2621233, EPI_ISL_2621234, EPI_ISL_2621235, EPI_ISL_2621236, EPI_ISL_2621237, EPI_ISL_2621238, EPI_ISL_2621239, EPI_ISL_2621240, EPI_ISL_2621241, EPI_ISL_2621242, EPI_ISL_2621243, EPI_ISL_2621244, EPI_ISL_2621245, EPI_ISL_2621246, EPI_ISL_2621247, EPI_ISL_2621249, EPI_ISL_2621250, EPI_ISL_2621251, EPI_ISL_2621252, EPI_ISL_2621253, EPI_ISL_2621256, EPI_ISL_2621257, EPI_ISL_2621259, EPI_ISL_2621262, EPI_ISL_2621263, EPI_ISL_2621264, EPI_ISL_2621265, EPI_ISL_2621267, EPI_ISL_2621269, EPI_ISL_2621270, EPI_ISL_2621272, EPI_ISL_2621274, EPI_ISL_2621275, EPI_ISL_2621277, EPI_ISL_2621278, EPI_ISL_2621280, EPI_ISL_2621281, EPI_ISL_2621282, EPI_ISL_2621283, EPI_ISL_2621284, EPI_ISL_2621285, EPI_ISL_2621286, EPI_ISL_2621287, EPI_ISL_2621288, EPI_ISL_2621289, EPI_ISL_2621290, EPI_ISL_2621291, EPI_ISL_2621292, EPI_ISL_2621293, EPI_ISL_2621294, EPI_ISL_2621296, EPI_ISL_2621297, EPI_ISL_2621298, EPI_ISL_2621299, EPI_ISL_2621300, EPI_ISL_2621301, EPI_ISL_2621302, EPI_ISL_2621303, EPI_ISL_2621304, EPI_ISL_2621305, EPI_ISL_2621306, EPI_ISL_2621307, EPI_ISL_2621308, EPI_ISL_2621309, EPI_ISL_2621310, EPI_ISL_2621311, EPI_ISL_2621312, EPI_ISL_2621314, EPI_ISL_2621316, EPI_ISL_2621318, EPI_ISL_2621319, EPI_ISL_2621320, EPI_ISL_2621321, EPI_ISL_2621322, EPI_ISL_2621323, EPI_ISL_2621324, EPI_ISL_2621326, EPI_ISL_2621327, EPI_ISL_2621328, EPI_ISL_2621329, EPI_ISL_2621330, EPI_ISL_2621331, EPI_ISL_2621332, EPI_ISL_2621333, EPI_ISL_2621334, EPI_ISL_2621336, EPI_ISL_2621338, EPI_ISL_2621339, EPI_ISL_2621340, EPI_ISL_2621342, EPI_ISL_2621343, EPI_ISL_2621344, EPI_ISL_2621345, EPI_ISL_2621346, EPI_ISL_2621347, EPI_ISL_2621348, EPI_ISL_2621349, EPI_ISL_2621350, EPI_ISL_2621351, EPI_ISL_2621352, EPI_ISL_2621353, EPI_ISL_2621354, EPI_ISL_2621355, EPI_ISL_2621356, EPI_ISL_2621357, EPI_ISL_2621358, EPI_ISL_2621359, EPI_ISL_2621360, EPI_ISL_2621361, EPI_ISL_2621362, EPI_ISL_2621363, EPI_ISL_2621364, EPI_ISL_2621365, EPI_ISL_2621366, EPI_ISL_2621368, EPI_ISL_2621369, EPI_ISL_2621370, EPI_ISL_2621371, EPI_ISL_2621372, EPI_ISL_2621373, EPI_ISL_2621375, EPI_ISL_2621376, EPI_ISL_2621377, EPI_ISL_2621378, EPI_ISL_2621379, EPI_ISL_2621380, EPI_ISL_2621382, EPI_ISL_2621383, EPI_ISL_2621384, EPI_ISL_2621385, EPI_ISL_2621386, EPI_ISL_2621387, EPI_ISL_2621388, EPI_ISL_2621389, EPI_ISL_2621390, EPI_ISL_2621391, EPI_ISL_2621392, EPI_ISL_2621394, EPI_ISL_2621395, EPI_ISL_2621396, EPI_ISL_2621397, EPI_ISL_2621398, EPI_ISL_2621399, EPI_ISL_2621400, EPI_ISL_2621401, EPI_ISL_2621402, EPI_ISL_2621403, EPI_ISL_2621404, EPI_ISL_2621405, EPI_ISL_2621406, EPI_ISL_2621407, EPI_ISL_2621408, EPI_ISL_2621409, EPI_ISL_2621410, EPI_ISL_2621411, EPI_ISL_2621413, EPI_ISL_2621414, EPI_ISL_2621415, EPI_ISL_2621416, EPI_ISL_2621417, EPI_ISL_2621418, EPI_ISL_2621419, EPI_ISL_2621420, EPI_ISL_2621421, EPI_ISL_2621422, EPI_ISL_2621423, EPI_ISL_2621424, EPI_ISL_2621425, EPI_ISL_2621426, EPI_ISL_2621427, EPI_ISL_2621428, EPI_ISL_2621430, EPI_ISL_2621431, EPI_ISL_2621432, EPI_ISL_2621433, EPI_ISL_2621434, EPI_ISL_2621435, EPI_ISL_2621436, EPI_ISL_2621438, EPI_ISL_2621439, EPI_ISL_2621440, EPI_ISL_2621441, EPI_ISL_2621442, EPI_ISL_2621443, EPI_ISL_2621444, EPI_ISL_2621445, EPI_ISL_2621446, EPI_ISL_2621448, EPI_ISL_2621449, EPI_ISL_2621452, EPI_ISL_2621453, EPI_ISL_2621455, EPI_ISL_2621456, EPI_ISL_2621458, EPI_ISL_2621462, EPI_ISL_2621463, EPI_ISL_2621468 | | | |
| see above | National Center of Infectious and Parasitic Diseases | National Center of Infectious and Parasitic Diseases | Alexiev et al |
| EPI_ISL_2621621, EPI_ISL_2621622, EPI_ISL_2621623, EPI_ISL_2621624, EPI_ISL_2621626, EPI_ISL_2621627, EPI_ISL_2621628, EPI_ISL_2621629, EPI_ISL_2621630, EPI_ISL_2621631 | Ipoh Public Health Laboratory (MKAI), Ministry of Health Malaysia | Institute for Medical Research, Infectious Disease Research Centre, National Institutes of Health, Ministry of Health Malaysia | Suppiah J, Kamel K, Mohd Zawawi Z, Azizan MA, Ramly N, Robert F, Thayan R |

| | | | |
|---|---|---|---|
| EPI_ISL_2621664 | Kulim Hospital | Institute for Medical Research, Infectious Disease Research Centre, National Institutes of Health, Ministry of Health Malaysia | Suppiah J, Kamel K, Mohd Zawawi Z, Azizan MA, Ramly N, Robert F, Thayan R |
| EPI_ISL_2622163 | Covid-19 laboratory, Institute of Human Genetics, Polish Academy of Sciences | Covid-19 laboratory, Institute of Human Genetics, Polish Academy of Sciences | Szymon Hryhorowicz, Adam Ustaszewski, Emilia Lis, Marta Kaczmarek-Ry, Ewa Zitkiewicz, Micha Witt, Andrzej Pawski |
| EPI_ISL_2625621 | Johor Bahru Public Health Laboratory | Institute for Medical Research, Infectious Disease Research Centre, National Institutes of Health, Ministry of Health Malaysia | Suppiah J, Kamel K, Mohd Zawawi Z, Azizan MA, Ramly N, Robert F, Thayan R |
| EPI_ISL_2625623 | Sungai Buloh Hospital | Institute for Medical Research, Infectious Disease Research Centre, National Institutes of Health, Ministry of Health Malaysia | Suppiah J, Kamel K, Mohd Zawawi Z, Azizan MA, Ramly N, Robert F, Thayan R |
| EPI_ISL_2625624, EPI_ISL_2625625, EPI_ISL_2625626, EPI_ISL_2625627, EPI_ISL_2625628, EPI_ISL_2625629, EPI_ISL_2625630, EPI_ISL_2625631 | Johor Bahru Public Health Laboratory | Institute for Medical Research, Infectious Disease Research Centre, National Institutes of Health, Ministry of Health Malaysia | Suppiah J, Kamel K, Mohd Zawawi Z, Azizan MA, Ramly N, Robert F, Thayan R |
| EPI_ISL_2625633 | Institute for Medical Research, Infectious Disease Research Centre, National Institutes of Health, Ministry of Health Malaysia | Institute for Medical Research, Infectious Disease Research Centre, National Institutes of Health, Ministry of Health Malaysia | Suppiah J, Kamel K, Mohd Zawawi Z, Azizan MA, Ramly N, Robert F, Thayan R |
| EPI_ISL_2625637 | Sultanah Aminah Hospital, Johor Bahru | Institute for Medical Research, Infectious Disease Research Centre, National Institutes of Health, Ministry of Health Malaysia | Suppiah J, Kamel K, Mohd Zawawi Z, Azizan MA, Ramly N, Robert F, Thayan R |
| EPI_ISL_2626223, EPI_ISL_2626225, EPI_ISL_2626226, EPI_ISL_2626229, EPI_ISL_2626230, EPI_ISL_2626232, EPI_ISL_2626233, EPI_ISL_2626234, EPI_ISL_2626236, EPI_ISL_2626237, EPI_ISL_2626239, EPI_ISL_2626240, EPI_ISL_2626241, EPI_ISL_2626242, EPI_ISL_2626243, EPI_ISL_2626245 | | | |
| see above | Molecular diagnostic laboratory of Federal Budget Institution of Science "Central Research Institute of Epidemiology" of The Federal Service on Customers' Rights Protection and Human Well-being Surveillance | Group of Genomics and Postgenomic Technologies of Central Research Institute of Epidemiology | Samoilov A.E., Kaptelova V.V., Korneenko E.V., Sinitsyn S.O., Saenko S.S., Nadtoka M.I., Speranskaya A.S., Tivanova E.V., Kondrasheva L.Y., Smirnova Y.S., Cherkashina A.S., Akimkin V.G. |
| EPI_ISL_2626267, EPI_ISL_2626268, EPI_ISL_2626270, EPI_ISL_2626271, EPI_ISL_2626272, EPI_ISL_2626273, EPI_ISL_2626274, EPI_ISL_2626275, EPI_ISL_2626276, EPI_ISL_2626277, EPI_ISL_2626278, EPI_ISL_2626280, EPI_ISL_2626281, EPI_ISL_2626283, EPI_ISL_2626285, EPI_ISL_2626286, EPI_ISL_2626287, EPI_ISL_2626288, EPI_ISL_2626289, EPI_ISL_2626292, EPI_ISL_2626293, EPI_ISL_2626294, EPI_ISL_2626300 | | | |
| see above | Molecular diagnostic laboratory of Federal Budget Institution of Science "Central Research Institute of Epidemiology" of The Federal Service on Customers' Rights Protection and Human Well-being Surveillance | Group of Genomics and Postgenomic Technologies of Central Research Institute of Epidemiology | Samoilov A.E., Kaptelova V.V., Korneenko E.V., Sinitsyn S.O., Saenko S.S., Nadtoka M.I., Speranskaya A.S., Tivanova E.V., Kondrasheva L.Y., Smirnova Y.S., Akimkin V.G. |
| EPI_ISL_2626432, EPI_ISL_2626434 | Laboratori de Referencia de Catalunya | Laboratori de Referencia de Catalunya | Ramirez A., Bellosillo B., Padilla E., Vilas A., Hernandez JJ., Canal M. |
| EPI_ISL_2626451 | AZ Zeno Knokke-Blankenberge | AZ Sint Jan | Marijke Reynders, Patrick Descheemaeker, Laurien Hoornaert, Thomas Van Landschoot, Marie Madeleine Chabert-Consen, Jorn Hellemans |
| EPI_ISL_2628278, EPI_ISL_2628279, EPI_ISL_2628280 | Hospital | National Reference Center for Viruses of Respiratory Infections, Institut Pasteur, Paris | Marion Barbet, Sylvie Behillil, Méline Bizard, Angela Brisebarre, Camille Capel, Vincent Enouf, Louise Lefrançois, Frédéric Lemoine, Christophe Malabat, Corinne Maufrais, Etienne Simon-Lorière, Maud Vanpeene, Sylvie Van der Werf ,Marianne Coste-Burel |
| EPI_ISL_2628281 | Hospital | National Reference Center for Viruses of Respiratory Infections, Institut Pasteur, Paris | Marion Barbet, Sylvie Behillil, Méline Bizard, Angela Brisebarre, Camille Capel, Vincent Enouf, Louise Lefrançois, Frédéric Lemoine, Christophe Malabat, Corinne Maufrais, Adrien Pain, Etienne Simon-Lorière, Maud Vanpeene, Sylvie Van der Werf ,Marianne Coste-Burel |
| EPI_ISL_2628282 | Hospital | National Reference Center for Viruses of Respiratory Infections, Institut Pasteur, Paris | Marion Barbet, Sylvie Behillil, Méline Bizard, Angela Brisebarre, Camille Capel, Vincent Enouf, Louise Lefrançois, Frédéric Lemoine, Christophe Malabat, Corinne Maufrais, Etienne Simon-Lorière, Maud Vanpeene, Sylvie Van der Werf ,Marianne Coste-Burel |
| EPI_ISL_2629094, EPI_ISL_2629095, EPI_ISL_2629096, EPI_ISL_2629098, EPI_ISL_2629099, EPI_ISL_2629100, EPI_ISL_2629102, EPI_ISL_2629103, EPI_ISL_2629104, EPI_ISL_2629105, EPI_ISL_2629106, EPI_ISL_2629107, EPI_ISL_2629108, EPI_ISL_2629109, EPI_ISL_2629110, EPI_ISL_2629111, EPI_ISL_2629112, EPI_ISL_2629113, EPI_ISL_2629114, EPI_ISL_2629115, EPI_ISL_2629116, EPI_ISL_2629117, EPI_ISL_2629118, EPI_ISL_2629119, EPI_ISL_2629120, EPI_ISL_2629121, EPI_ISL_2629123, EPI_ISL_2629124, EPI_ISL_2629125, EPI_ISL_2629126, EPI_ISL_2629127, EPI_ISL_2629128, EPI_ISL_2629129, EPI_ISL_2629130, EPI_ISL_2629131, EPI_ISL_2629132, EPI_ISL_2629133, EPI_ISL_2629134, EPI_ISL_2629135, EPI_ISL_2629136, EPI_ISL_2629137, EPI_ISL_2629138, EPI_ISL_2629139, EPI_ISL_2629140, EPI_ISL_2629142, EPI_ISL_2629143, EPI_ISL_2629144, EPI_ISL_2629145, EPI_ISL_2629146, EPI_ISL_2629147, EPI_ISL_2629148, EPI_ISL_2629149, EPI_ISL_2629150, EPI_ISL_2629151, EPI_ISL_2629152, EPI_ISL_2629153, EPI_ISL_2629155 | | | |
| see above | HEGP - Laboratoire de Virologie | HEGP - Laboratoire de Virologie | David Veyer, Sandrine Imbeaud, Julien Puech, Hélène Péré, Maxime Wack, Nicolas Robillard |
| EPI_ISL_2629156, EPI_ISL_2629157, EPI_ISL_2629160, EPI_ISL_2629161, EPI_ISL_2629162, EPI_ISL_2629163 | Plateforme Covid IDF, site Broussais | HEGP - Laboratoire de Virologie | David Veyer, Sandrine Imbeaud, Julien Puech, Nabil Gastli, Hélène Péré, Maxime Wack, Nicolas Robillard |
| EPI_ISL_2642550, EPI_ISL_2642551 | AOP Policlinico " G. Rodolico- S. Marco" PO Rodolico Catania | AOP Policlinico " G. Rodolico- S. Marco" PO Rodolico Catania and BIOMETEC UNICT | Stefani and Scalia |
| EPI_ISL_2645440, EPI_ISL_2645441, EPI_ISL_2645442, EPI_ISL_2645443, EPI_ISL_2645488, EPI_ISL_2645489, EPI_ISL_2645490 | Lboratorio Central de Saude Publica do Estado do Parana (LACEN/PR) | Laboratory of Respiratory Viruses and Measles, Oswaldo Cruz Institute, FIOCRUZ | Paola Resende, Luciana Appolinario, Fernando Motta, Anna Carolina Paixao, Ana Carolina Mendonca, Alice Sampaio Rocha, Taina Venas, Elisa Cavalcante Pereira, Renata Serrano Lopes, Irina Riediger, Marilda Siqueira on behalf of the Fiocruz COVID-19 Genomic Surveillance Network |
| EPI_ISL_2648058, EPI_ISL_2648059, EPI_ISL_2648060, EPI_ISL_2648061, EPI_ISL_2648062, EPI_ISL_2648063, EPI_ISL_2648064, EPI_ISL_2648065, EPI_ISL_2648068, EPI_ISL_2648069, EPI_ISL_2648070, EPI_ISL_2648071, EPI_ISL_2648072, EPI_ISL_2648073, EPI_ISL_2648074, EPI_ISL_2648075, EPI_ISL_2648076, EPI_ISL_2648077, EPI_ISL_2648078, EPI_ISL_2648079, EPI_ISL_2648080, EPI_ISL_2648081, EPI_ISL_2648082, EPI_ISL_2648083 | | | |
| see above | National Center of Infectious and Parasitic Diseases | National Center of Infectious and Parasitic Diseases | Alexiev et al |
| EPI_ISL_2648217 | Debswana Orapa Mine Hospital Laboratory | Botswana Harvard HIV Reference Laboratory | Sikhulile Moyo, Wonderful T. Choga, Dorcas Maruapula, Keoratile Ntshambiwa, Lesedi Magama, Mpo Molapisi, Koketso Maotwe, Thongbotho Mphoyakgosi, Boitumelo Zuze, Botshelo Radibe, Legodile Kooepile, Ontlametse T. Bareng, Roger Shapiro, Shahin Lockman, Joseph Makhema, Madisa Mine, Mosepele Mosepele, Simani Gaseitsiwe |
| EPI_ISL_2651363 | CHU de la Réunion - Nord | Laboratoire de virologie, CNR arbovirus Associé, Chu de la Réunion | Etienne Frumence, Marie-Christine Jaffar Bandjee, Nicolas Traversier, Sabrina Petit Genet |
| EPI_ISL_2652164 | Clinical Emergency County Hospital, Craiova | Cantacuzino Institute Virology | Carmen Cherciu, Luiza Ustea, Nicoleta Paraschiv, Sorin Dinu, Mihaela Oprea, Mihaela Lazar |
| EPI_ISL_2657566 | Medizinische Universität Graz | Bergthaler laboratory, CeMM Research Center for Molecular Medicine of the Austrian Academy of Sciences | Lukas Endler, Anna Schedl, Fabian Amman, Petr Triska, Matthew Thornton, Thomas Penz, Benedikt Agerer, Maelle Le Moing, Michael Schuster, Bekir Erguner, Jan Laine, Martin Senekowitsch, Christoph Bock, Andreas Bergthaler |
| EPI_ISL_2657685 | HOSPITAL METROPOLITANO | Inciensa, Instituto Costarricense de Investigación y Enseñanza en Nutrición y Salud | Francisco Duarte, Hebleen Porras, Claudio Soto-Garita, Estela Cordero, Adriana Godínez, Melany Calderón, José Luis Vargas, Mariela Gutiérrez, Joselyn Prado & Margarita Lee-Lui |
| EPI_ISL_2657686 | INCIENSA | Inciensa, Instituto Costarricense de Investigación y Enseñanza en Nutrición y Salud | Francisco Duarte, Hebleen Porras, Claudio Soto-Garita, Estela Cordero, Adriana Godínez, Melany Calderón, José Luis Vargas, Mariela Gutiérrez, Joselyn Prado & Grettel Chanto-Chacón |
| EPI_ISL_2657866 | CARVAJAL LABORATORIO IPS | Instituto Nacional de Salud- Dirección de Investigación en Salud Pública | Katherine Laiton-Donato, Diego A. Álvarez-Díaz, Carlos Franco-Muñoz, Hector Alejandro Ruiz-Moreno, Paola Rojas, Maria T. Herrera-Sepúlveda, Diego Andrés Prada, Jhonnatan Reales-González, Sheryll Corchuelo, Julian Naizaque, Jorge Rivera, Gerardo Santamaría, Sergio Gomez, Lisseth Pardo, Juan Camilo Martinez, Marta Lopez Blanco, Ángela Alarcon Cruz, Diana Malo, Carmen Osorio, Magdalena Wiesner, Martha Lucia Ospina Martinez, Marcela Mercado-Reyes |

| | | | |
|---|---|---|---|
| EPI_ISL_2657872 | LABORATORIO CLINICA IMBANACO | Instituto Nacional de Salud- Dirección de Investigación en Salud Pública | Katherine Laiton-Donato, Diego A. Álvarez-Díaz, Carlos Franco-Muñoz, Hector Alejandro Ruiz-Moreno, Paola Rojas, Maria T. Herrera-Sepúlveda, Diego Andrés Prada, Jhonnatan Reales-González, Sheryll Corchuelo, Julian Naizaque, Jorge Rivera, Gerardo Santamaría, Sergio Gomez, Lisseth Pardo, Juan Camilo Martinez, Marta Lopez Blanco, Ángela Alarcon Cruz, Diana Malo, Carmen Osorio, Magdalena Wiesner, Martha Lucia Ospina Martinez, Marcela Mercado-Reyes |
| EPI_ISL_2657877 | Instituto de Diagnóstico Médico | Instituto Nacional de Salud- Dirección de Investigación en Salud Pública | Katherine Laiton-Donato, Diego A. Álvarez-Díaz, Carlos Franco-Muñoz, Hector Alejandro Ruiz-Moreno, Paola Rojas, Maria T. Herrera-Sepúlveda, Diego Andrés Prada, Jhonnatan Reales-González, Sheryll Corchuelo, Julian Naizaque, Jorge Rivera, Gerardo Santamaría, Sergio Gomez, Lisseth Pardo, Juan Camilo Martinez, Marta Lopez Blanco, Ángela Alarcon Cruz, Diana Malo, Carmen Osorio, Magdalena Wiesner, Martha Lucia Ospina Martinez, Marcela Mercado-Reyes |
| EPI_ISL_2658264 | AREA DE SALUD TURRIALBA-JIMENEZ | Inciensa, Instituto Costarricense de Investigación y Enseñanza en Nutrición y Salud | Francisco Duarte, Hebleen Porras, Claudio Soto-Garita, Estela Cordero, Adriana Godínez, Melany Calderón, José Luis Vargas, Mariela Gutiérrez & Joselyn Prado |
| EPI_ISL_2658265 | AREA DE SALUD CAÑAS | Inciensa, Instituto Costarricense de Investigación y Enseñanza en Nutrición y Salud | Francisco Duarte, Hebleen Porras, Claudio Soto-Garita, Estela Cordero, Adriana Godínez, Melany Calderón, José Luis Vargas, Mariela Gutiérrez, Joselyn Prado & Adriana Bermúdez Espinoza |
| EPI_ISL_2658266 | HOSPITAL DR. RAFAEL A. CALDERON GUARDIA | Inciensa, Instituto Costarricense de Investigación y Enseñanza en Nutrición y Salud | Francisco Duarte, Hebleen Porras, Claudio Soto-Garita, Estela Cordero, Adriana Godínez, Melany Calderón, José Luis Vargas, Mariela Gutiérrez, Joselyn Prado & Fabian Salas Flores |
| EPI_ISL_2658267 | AREA DE SALUD PAVAS (COOPESALUD) | Inciensa, Instituto Costarricense de Investigación y Enseñanza en Nutrición y Salud | Francisco Duarte, Hebleen Porras, Claudio Soto-Garita, Estela Cordero, Adriana Godínez, Melany Calderón, José Luis Vargas, Mariela Gutiérrez, Joselyn Prado & Andony Cordero-Jiménez |
| EPI_ISL_2658268 | AREA DE SALUD CARTAGO | Inciensa, Instituto Costarricense de Investigación y Enseñanza en Nutrición y Salud | Francisco Duarte, Hebleen Porras, Claudio Soto-Garita, Estela Cordero, Adriana Godínez, Melany Calderón, José Luis Vargas, Mariela Gutiérrez, Joselyn Prado & Tania Garro Vargas |
| EPI_ISL_2658269 | AREA DE SALUD ZAPOTE-CATEDRAL - CLINICA DR. CARLOS DURAN | Inciensa, Instituto Costarricense de Investigación y Enseñanza en Nutrición y Salud | Francisco Duarte, Hebleen Porras, Claudio Soto-Garita, Estela Cordero, Adriana Godínez, Melany Calderón, José Luis Vargas, Mariela Gutiérrez, Joselyn Prado & Andony Cordero-Jiménez |
| EPI_ISL_2658271 | AREA DE SALUD CHOMES - MONTEVERDE | Inciensa, Instituto Costarricense de Investigación y Enseñanza en Nutrición y Salud | Francisco Duarte, Hebleen Porras, Claudio Soto-Garita, Estela Cordero, Adriana Godínez, Melany Calderón, José Luis Vargas, Mariela Gutiérrez, Joselyn Prado & Javier Zárate-León |
| EPI_ISL_2658272 | AREA DE SALUD SAN RAFAEL | Inciensa, Instituto Costarricense de Investigación y Enseñanza en Nutrición y Salud | Francisco Duarte, Hebleen Porras, Claudio Soto-Garita, Estela Cordero, Adriana Godínez, Melany Calderón, José Luis Vargas, Mariela Gutiérrez, Joselyn Prado & Javier Zárate-León |
| EPI_ISL_2658273, EPI_ISL_2658274 | HOSPITAL MONSEÑOR SANABRIA | Inciensa, Instituto Costarricense de Investigación y Enseñanza en Nutrición y Salud | Francisco Duarte, Hebleen Porras, Claudio Soto-Garita, Estela Cordero, Adriana Godínez, Melany Calderón, José Luis Vargas, Mariela Gutiérrez, Joselyn Prado & Javier Zárate-León |
| EPI_ISL_2658275 | AREA DE SALUD SAN RAFAEL | Inciensa, Instituto Costarricense de Investigación y Enseñanza en Nutrición y Salud | Francisco Duarte, Hebleen Porras, Claudio Soto-Garita, Estela Cordero, Adriana Godínez, Melany Calderón, José Luis Vargas, Mariela Gutiérrez, Joselyn Prado & Javier Zárate-León |
| EPI_ISL_2658276 | AREA DE SALUD EL GUARCO | Inciensa, Instituto Costarricense de Investigación y Enseñanza en Nutrición y Salud | Francisco Duarte, Hebleen Porras, Claudio Soto-Garita, Estela Cordero, Adriana Godínez, Melany Calderón, José Luis Vargas, Mariela Gutiérrez, Joselyn Prado & Pamela Serrano Valerín |
| EPI_ISL_2658277 | HOSPITAL SAN VICENTE DE PAUL | Inciensa, Instituto Costarricense de Investigación y Enseñanza en Nutrición y Salud | Francisco Duarte, Hebleen Porras, Claudio Soto-Garita, Estela Cordero, Adriana Godínez, Melany Calderón, José Luis Vargas, Mariela Gutiérrez, Joselyn Prado & Laura Brizuela Cruz |
| EPI_ISL_2658278 | HOSPITAL SAN FRANCISCO DE ASIS | Inciensa, Instituto Costarricense de Investigación y Enseñanza en Nutrición y Salud | Francisco Duarte, Hebleen Porras, Claudio Soto-Garita, Estela Cordero, Adriana Godínez, Melany Calderón, José Luis Vargas, Mariela Gutiérrez, Joselyn Prado & Juan Carlos Villalobos-Ugalde |
| EPI_ISL_2658279 | AREA DE SALUD CORONADO | Inciensa, Instituto Costarricense de Investigación y Enseñanza en Nutrición y Salud | Francisco Duarte, Hebleen Porras, Claudio Soto-Garita, Estela Cordero, Adriana Godínez, Melany Calderón, José Luis Vargas, Mariela Gutiérrez, Joselyn Prado & Mariel López |
| EPI_ISL_2658280 | AREA DE SALUD SAN JUAN-SAN DIEGO-CONCEPCION 2 | Inciensa, Instituto Costarricense de Investigación y Enseñanza en Nutrición y Salud | Francisco Duarte, Hebleen Porras, Claudio Soto-Garita, Estela Cordero, Adriana Godínez, Melany Calderón, José Luis Vargas, Mariela Gutiérrez, Joselyn Prado & Mariel López |
| EPI_ISL_2658281 | HOSPITAL DR. MAX PERALTA JIMENEZ | Inciensa, Instituto Costarricense de Investigación y Enseñanza en Nutrición y Salud | Francisco Duarte, Hebleen Porras, Claudio Soto-Garita, Estela Cordero, Adriana Godínez, Melany Calderón, José Luis Vargas, Mariela Gutiérrez, Joselyn Prado & Pamela Serrano Valerín |
| EPI_ISL_2663126, EPI_ISL_2663129, EPI_ISL_2663145 | Ochsner Health | Bioinfoexperts, LLC | Judy Crabtree, Amy Feehan, Rebecca Rose, David J. Nolan, Tessa LaFleur, Julia-Garcia-Diaz, Lucio Miele, Susanna L. Lamers |
| EPI_ISL_2665396 | Baylor University Medical Center | Baylor Scott & White Medical Center - Temple | Ari Rao, Linden Morales, Kimberly Walker, Marcus Volz, Shelby Johnson |
| EPI_ISL_2673521 | MOH - Jaber Al-Ahmad Hospital (Innovation Research Laboratory) | MOH - Jaber Al-Ahmad Hospital (Innovation Research Laboratory) | Salman Al-Sabah , Mohammad Alghounaim |
| EPI_ISL_2674282, EPI_ISL_2674283 | LABORATORIO CONTINENTAL | Instituto Nacional de Salud- Dirección de Investigación en Salud Pública | Katherine Laiton-Donato, Diego A. Álvarez-Díaz, Carlos Franco-Muñoz, Hector Alejandro Ruiz-Moreno, Paola Rojas, Maria T. Herrera-Sepúlveda, Diego Andrés Prada, Jhonnatan Reales-González, Sheryll Corchuelo, Julian Naizaque, Jorge Rivera, Gerardo Santamaría, Sergio Gomez, Lisseth Pardo, Juan Camilo Martinez, Marta Lopez Blanco, Ángela Alarcon Cruz, Diana Malo, Carmen Osorio, Magdalena Wiesner, Martha Lucia Ospina Martinez, Marcela Mercado-Reyes |
| EPI_ISL_2674284 | LDSP CESAR | Instituto Nacional de Salud- Dirección de Investigación en Salud Pública | Katherine Laiton-Donato, Diego A. Álvarez-Díaz, Carlos Franco-Muñoz, Hector Alejandro Ruiz-Moreno, Paola Rojas, Maria T. Herrera-Sepúlveda, Diego Andrés Prada, Jhonnatan Reales-González, Sheryll Corchuelo, Julian Naizaque, Jorge Rivera, Gerardo Santamaría, Sergio Gomez, Lisseth Pardo, Juan Camilo Martinez, Marta Lopez Blanco, Ángela Alarcon Cruz, Diana Malo, Carmen Osorio, Magdalena Wiesner, Martha Lucia Ospina Martinez, Marcela Mercado-Reyes |
| EPI_ISL_2674286, EPI_ISL_2674287, EPI_ISL_2674288 | LABORATORIO BIO-TEST | Instituto Nacional de Salud- Dirección de Investigación en Salud Pública | Katherine Laiton-Donato, Diego A. Álvarez-Díaz, Carlos Franco-Muñoz, Hector Alejandro Ruiz-Moreno, Paola Rojas, Maria T. Herrera-Sepúlveda, Diego Andrés Prada, Jhonnatan Reales-González, Sheryll Corchuelo, Julian Naizaque, Jorge Rivera, Gerardo Santamaría, Sergio Gomez, Lisseth Pardo, Juan Camilo Martinez, Marta Lopez Blanco, Ángela Alarcon Cruz, Diana Malo, Carmen Osorio, Magdalena Wiesner, Martha Lucia Ospina Martinez, Marcela Mercado-Reyes |
| EPI_ISL_2674290, EPI_ISL_2674291, EPI_ISL_2674292, EPI_ISL_2674293 | LABORATORIO CLINICO MASVIDA DE LA COSTA | Instituto Nacional de Salud- Dirección de Investigación en Salud Pública | Katherine Laiton-Donato, Diego A. Álvarez-Díaz, Carlos Franco-Muñoz, Hector Alejandro Ruiz-Moreno, Paola Rojas, Maria T. Herrera-Sepúlveda, Diego Andrés Prada, Jhonnatan Reales-González, Sheryll Corchuelo, Julian Naizaque, Jorge Rivera, Gerardo Santamaría, Sergio Gomez, Lisseth Pardo, Juan Camilo Martinez, Marta Lopez Blanco, Ángela Alarcon Cruz, Diana Malo, Carmen Osorio, Magdalena Wiesner, Martha Lucia Ospina Martinez, Marcela Mercado-Reyes |
| EPI_ISL_2674296 | LDSP CALDAS | Instituto Nacional de Salud- Dirección de Investigación en Salud Pública | Katherine Laiton-Donato, Diego A. Álvarez-Díaz, Carlos Franco-Muñoz, Hector Alejandro Ruiz-Moreno, Paola Rojas, Maria T. Herrera-Sepúlveda, Diego Andrés Prada, Jhonnatan Reales-González, Sheryll Corchuelo, Julian Naizaque, Jorge Rivera, Gerardo Santamaría, Sergio Gomez, Lisseth Pardo, Juan Camilo Martinez, Marta Lopez Blanco, Ángela Alarcon Cruz, Diana Malo, Carmen Osorio, Magdalena Wiesner, Martha Lucia Ospina Martinez, Marcela Mercado-Reyes |
| EPI_ISL_2674313 | COMFANDI - SAN NICOLAS | Instituto Nacional de Salud- Dirección de Investigación en Salud Pública | Katherine Laiton-Donato, Diego A. Álvarez-Díaz, Carlos Franco-Muñoz, Hector Alejandro Ruiz-Moreno, Paola Rojas, Maria T. Herrera-Sepúlveda, Diego Andrés Prada, Jhonnatan Reales-González, Sheryll Corchuelo, Julian Naizaque, Jorge Rivera, Gerardo Santamaría, Sergio Gomez, Lisseth Pardo, Juan Camilo Martinez, Marta Lopez Blanco, Ángela Alarcon Cruz, Diana Malo, Carmen Osorio, Magdalena Wiesner, Martha Lucia Ospina Martinez, Marcela Mercado-Reyes |
| EPI_ISL_2674317 | LDSP TOLIMA | Instituto Nacional de Salud- Dirección de Investigación en Salud Pública | Katherine Laiton-Donato, Diego A. Álvarez-Díaz, Carlos Franco-Muñoz, Hector Alejandro Ruiz-Moreno, Paola Rojas, Maria T. Herrera-Sepúlveda, Diego Andrés Prada, Jhonnatan Reales-González, Sheryll Corchuelo, Julian Naizaque, Jorge Rivera, Gerardo Santamaría, Sergio Gomez, Lisseth Pardo, Juan Camilo Martinez, Marta Lopez Blanco, Ángela Alarcon Cruz, Diana Malo, Carmen Osorio, Magdalena Wiesner, Martha Lucia Ospina Martinez, Marcela Mercado-Reyes |

| | | | |
|---|---|---|---|
| EPI_ISL_2674325 | LABORATORIO DE INFECCION E INMUNIDAD | Instituto Nacional de Salud- Dirección de Investigación en Salud Pública | Katherine Laiton-Donato, Diego A. Álvarez-Díaz, Carlos Franco-Muñoz, Hector Alejandro Ruiz-Moreno, Paola Rojas, Maria T. Herrera-Sepúlveda, Diego Andrés Prada, Jhonnatan Reales-González, Sheryll Corchuelo, Julian Naizaque, Jorge Rivera, Gerardo Santamaría, Sergio Gomez, Lisseth Pardo, Juan Camilo Martinez, Marta Lopez Blanco, Ángela Alarcon Cruz, Diana Malo, Carmen Osorio, Magdalena Wiesner, Martha Lucia Ospina Martinez, Marcela Mercado-Reyes |
| EPI_ISL_2674339 | HOSPITAL COMUNAL MALVINAS | Instituto Nacional de Salud- Dirección de Investigación en Salud Pública | Katherine Laiton-Donato, Diego A. Álvarez-Díaz, Carlos Franco-Muñoz, Hector Alejandro Ruiz-Moreno, Paola Rojas, Maria T. Herrera-Sepúlveda, Diego Andrés Prada, Jhonnatan Reales-González, Sheryll Corchuelo, Julian Naizaque, Jorge Rivera, Gerardo Santamaría, Sergio Gomez, Lisseth Pardo, Juan Camilo Martinez, Marta Lopez Blanco, Ángela Alarcon Cruz, Diana Malo, Carmen Osorio, Magdalena Wiesner, Martha Lucia Ospina Martinez, Marcela Mercado-Reyes |
| EPI_ISL_2674342, EPI_ISL_2674343 | IPS YENNY ZORAYA SALAZAR | Instituto Nacional de Salud- Dirección de Investigación en Salud Pública | Katherine Laiton-Donato, Diego A. Álvarez-Díaz, Carlos Franco-Muñoz, Hector Alejandro Ruiz-Moreno, Paola Rojas, Maria T. Herrera-Sepúlveda, Diego Andrés Prada, Jhonnatan Reales-González, Sheryll Corchuelo, Julian Naizaque, Jorge Rivera, Gerardo Santamaría, Sergio Gomez, Lisseth Pardo, Juan Camilo Martinez, Marta Lopez Blanco, Ángela Alarcon Cruz, Diana Malo, Carmen Osorio, Magdalena Wiesner, Martha Lucia Ospina Martinez, Marcela Mercado-Reyes |
| EPI_ISL_2674656, EPI_ISL_2674657 | LESP Guanajuato | Instituto de Diagnostico y Referencia Epidemiologicos (INDRE) | Claudia Wong-Arambula, Abril Rodriguez-Maldonado, Vanessa Rivero-Arredondo, Ariadna Medina-Benitez, Joaquin Quiroz-Mercado, Sergio Rangel-Guerrero, Natividad Cruz-Ortiz, Tatiana Nunez-Garcia, Gisela Barrera-Badillo, Lucia Hernandez-Rivas, Irma Lopez-Martinez, Ernesto Ramirez-Gonzalez. |
| EPI_ISL_2674702 | LESP Yucatan | Instituto de Diagnostico y Referencia Epidemiologicos (INDRE) | Claudia Wong-Arambula, Abril Rodriguez-Maldonado, Vanessa Rivero-Arredondo, Ariadna Medina-Benitez, Joaquin Quiroz-Mercado, Sergio Rangel-Guerrero, Natividad Cruz-Ortiz, Tatiana Nunez-Garcia, Gisela Barrera-Badillo, Lucia Hernandez-Rivas, Irma Lopez-Martinez, Ernesto Ramirez-Gonzalez. |
| EPI_ISL_2674717 | LESP Sonora | Instituto de Diagnostico y Referencia Epidemiologicos (INDRE) | Claudia Wong-Arambula, Abril Rodriguez-Maldonado, Vanessa Rivero-Arredondo, Ariadna Medina-Benitez, Joaquin Quiroz-Mercado, Sergio Rangel-Guerrero, Natividad Cruz-Ortiz, Tatiana Nunez-Garcia, Gisela Barrera-Badillo, Lucia Hernandez-Rivas, Irma Lopez-Martinez, Ernesto Ramirez-Gonzalez. |
| EPI_ISL_2681053 | Centro de Investigación Biomédica del Noreste (CIBIN) | Instituto de Biotecnología de la UNAM | Consorcio Mexicano de Vigilancia Genómica (CoViGen-Mex). Authors (in alphabetical order): Julio Elias Alvarado-Yaah, Carlos F. Arias, Santiago Ávila-Ríos, Eduardo Becerril-Vargas, Víctor Hugo Borja-Aburto, Celia Boukadida, Cristóbal Cháidez-Quiróz, Juan Bautista Chale-Dzul, Ricardo Ciria Merce, Andreu Comas-García, Célida Duque Molina, Julissa Enciso-Ibarra, José Antonio Enciso-Moreno, Gloria Elena Espinosa-Ayala, Fernando Fontove-Herrera, Daniel Fregoso-Rueda, Victor Eduardo García-Arias, Alejandra García-Gasca, Bruno Gómez-Gil, Jean Pierre González, Irvin González-López, Concepción Grajales-Muñiz, Ricardo Grande, Rosa María Gutiérrez Rios, Jesús Hernández, Alejandra Hernández-Terán, Alfredo Herrera-Estrella, Carla Ivón Herrera-Najera, Pavel Isa, Verónica Mata-Haro, Daniel Lira Morales, Susana Lopez, Antonio Loza Román, Brenda Irasema Maldonado-Meza, Bernardo Martínez-Miguel, José Arturo Martínez-Orozco, Célida Martínez- Rodríguez, Margarita Matías-Florentino, Fidencio Mejía-Nepomuceno, María Guadalupe de Jesús Mireles-Rivera, Gloria María Molina-Salinas, Hector Montoya-Fuentes, , Mario Mújica-Sánchez, José Esteban Muñoz-Medina, José de Jesús Nuñez-Contreras, Alicia Ocaña-Mondragón, Luis Alberto Ochoa-Carrera, Hector Esteban Paz-Juárez, Marissa Perez-Garcia, Francisco Pulido, Helen Haydee Fernanda Ramírez-Plascencia, Jorge Salas-Hernández, Angel Gustavo Salas-Lais, Alejandro Sánchez-Flores, Clara Esperanza Santacruz-Tinoco, María Guadalupe Santiago-Mauricio, Selene Zárate, Nelly Sélem-Mojica, Blanca Taboada, Gloria Vazquez, Joel Armando Vázquez-Pérez. |
| EPI_ISL_2681409, EPI_ISL_2681410, EPI_ISL_2681412, EPI_ISL_2681413 | General Hospital "8th of September" | Laboratory of virology and molecular diagnostics, Institute of Public Health | Kuzmanovska M, Boshevska G, Janchevska E. |
| EPI_ISL_2681440 | General Hospital Kochani | Laboratory of virology and molecular diagnostics, Institute of Public Health | Kuzmanovska M, Boshevska G, Janchevska E. |
| EPI_ISL_2683956 | Sungai Buloh Hospital | Institute for Medical Research, Infectious Disease Research Centre, National Institutes of Health, Ministry of Health Malaysia | Suppiah J, Kamel K, Mohd Zawawi Z, Azizan MA, Ramly N, Robert F, Thayan R |
| EPI_ISL_2687731 | Public Health Authority of the Slovak Republic | Public Health Authority of the Slovak Republic | Barbora Kotvasová, Lucia Ševíková, Terézia Vrabová, Anna Giová, Elena Tichá, Miroslav Böhmer, Tomáš Szemes, Pavol Mišenko |
| EPI_ISL_2688261, EPI_ISL_2688262, EPI_ISL_2688263, EPI_ISL_2688264, EPI_ISL_2688265, EPI_ISL_2688266, EPI_ISL_2688268, EPI_ISL_2688269, EPI_ISL_2688270, EPI_ISL_2688272, EPI_ISL_2688273, EPI_ISL_2688275, EPI_ISL_2688276 | | | |
| see above | Centers for Disease Control and Prevention, Dengue Branch | Centers for Disease Control and Prevention, Dengue Branch | Gilberto A. Santiago, Glenda Gonzalez, Betzabel Flores, Keyla Charriez, Gabriela Paz-Bailey, Jorge L. Munoz-Jordan |
| EPI_ISL_2688284, EPI_ISL_2688285, EPI_ISL_2688286, EPI_ISL_2688287, EPI_ISL_2688289, EPI_ISL_2688290, EPI_ISL_2688291, EPI_ISL_2688292, EPI_ISL_2688293, EPI_ISL_2688294, EPI_ISL_2688295, EPI_ISL_2688296, EPI_ISL_2688297, EPI_ISL_2688298, EPI_ISL_2688299, EPI_ISL_2688300, EPI_ISL_2688301, EPI_ISL_2688302, EPI_ISL_2688303 | | | |
| see above | Puerto Rico Department of Health | Centers for Disease Control and Prevention, Dengue Branch | Gilberto A. Santiago, Glenda Gonzalez, Betzabel Flores, Keyla Charriez, Gabriela Paz-Bailey, Jorge L. Munoz-Jordan |
| EPI_ISL_2688656, EPI_ISL_2688661, EPI_ISL_2688687 | Centre Hospitalier Universitaire de Rouen Laboratoire de Virologie | Centre Hospitalier Universitaire de Rouen Laboratoire de Virologie | Alice Moisan, Fabienne De Oliveira, Marie Leoz |
| EPI_ISL_2688957 | LESP Chihuahua | Instituto de Diagnostico y Referencia Epidemiologicos (INDRE) | Claudia Wong-Arambula, Abril Rodriguez-Maldonado, Vanessa Rivero-Arredondo, Ariadna Medina-Benitez, Joaquin Quiroz-Mercado, Sergio Rangel-Guerrero, Natividad Cruz-Ortiz, Tatiana Nunez-Garcia, Gisela Barrera-Badillo, Lucia Hernandez-Rivas, Irma Lopez-Martinez, Ernesto Ramirez-Gonzalez. |
| EPI_ISL_2690137 | Ochsner Health | Bioinfoexperts, LLC | Judy Crabtree, Amy Feehan, Rebecca Rose, David J. Nolan, Tessa LaFleur, Julia-Garcia-Diaz, Lucio Miele, Susanna L. Lamers |
| EPI_ISL_2690321, EPI_ISL_2690326, EPI_ISL_2690329, EPI_ISL_2690684, EPI_ISL_2690685, EPI_ISL_2690691, EPI_ISL_2690694 | MSK Microbiology Lab | MSK Microbiology Lab | Esther Babady, Tracy McMillen, Krupa Jani |
| EPI_ISL_2691089, EPI_ISL_2691090, EPI_ISL_2691091, EPI_ISL_2691092, EPI_ISL_2691093, EPI_ISL_2691094, EPI_ISL_2691095 | Instituto Adolfo Lutz - Regional de Marilia | Instituto Adolfo Lutz, Interdiciplinary Procedures Center, Strategic Laboratory | Claudio Tavares Sacchi, Claudia Regina Gonçalves, Erica Valessa Ramos Gomes, Karoline Rodrigues Campos, Caio Vinicius Dias Lopes, Leonardo Jose Tadeu de Araujo |
| EPI_ISL_2691108, EPI_ISL_2691109, EPI_ISL_2691113, EPI_ISL_2691114, EPI_ISL_2691115, EPI_ISL_2691116, EPI_ISL_2691117, EPI_ISL_2691118, EPI_ISL_2691119, EPI_ISL_2691120, EPI_ISL_2691122 | | | |
| see above | Instituto Adolfo Lutz - Regional de Bauru | Instituto Adolfo Lutz, Interdiciplinary Procedures Center, Strategic Laboratory | Claudio Tavares Sacchi, Claudia Regina Gonçalves, Erica Valessa Ramos Gomes, Karoline Rodrigues Campos, Caio Vinicius Dias Lopes, Leonardo Jose Tadeu de Araujo |
| EPI_ISL_2691123, EPI_ISL_2691124, EPI_ISL_2691125 | Instituto Adolfo Lutz - Regional de Ribeirao Preto | Instituto Adolfo Lutz, Interdiciplinary Procedures Center, Strategic Laboratory | Claudio Tavares Sacchi, Claudia Regina Gonçalves, Erica Valessa Ramos Gomes, Karoline Rodrigues Campos, Caio Vinicius Dias Lopes, Leonardo Jose Tadeu de Araujo |
| EPI_ISL_2691126, EPI_ISL_2691127, EPI_ISL_2691128, EPI_ISL_2691129, EPI_ISL_2691130, EPI_ISL_2691131, EPI_ISL_2691132, EPI_ISL_2691133, EPI_ISL_2691134, EPI_ISL_2691135, EPI_ISL_2691136, EPI_ISL_2691137, EPI_ISL_2691138, EPI_ISL_2691139, EPI_ISL_2691140, EPI_ISL_2691141, EPI_ISL_2691142, EPI_ISL_2691143, EPI_ISL_2691144 | | | |
| see above | Instituto Adolfo Lutz Central | Instituto Adolfo Lutz, Interdiciplinary Procedures Center, Strategic Laboratory | Claudio Tavares Sacchi, Claudia Regina Gonçalves, Erica Valessa Ramos Gomes, Karoline Rodrigues Campos, Caio Vinicius Dias Lopes, Leonardo Jose Tadeu de Araujo |
| EPI_ISL_2691146, EPI_ISL_2691147, EPI_ISL_2691148 | Intituto Adolfo Lutz - Regional de Sorocaba | Instituto Adolfo Lutz, Interdiciplinary Procedures Center, Strategic Laboratory | Claudio Tavares Sacchi, Claudia Regina Gonçalves, Erica Valessa Ramos Gomes, Karoline Rodrigues Campos, Caio Vinicius Dias Lopes, Leonardo Jose Tadeu de Araujo |
| EPI_ISL_2691157, EPI_ISL_2691158 | Instituo Adolfo Lutz Central | Instituto Adolfo Lutz, Interdiciplinary Procedures Center, Strategic Laboratory | Claudio Tavares Sacchi, Claudia Regina Gonçalves, Erica Valessa Ramos Gomes, Karoline Rodrigues Campos, Caio Vinicius Dias Lopes, Leonardo Jose Tadeu de Araujo |
| EPI_ISL_2691206, EPI_ISL_2691207, | LACEN do Estado de Goias | Instituto Adolfo Lutz, Interdiciplinary Procedures Center, | Claudio Tavares Sacchi, Claudia Regina Gonçalves, Erica Valessa Ramos Gomes, Karoline Rodrigues Campos, Caio Vinicius Dias Lopes, Leonardo Jose |

| | | | |
|---|---|---|---|
| EPI_ISL_2691208, EPI_ISL_2691212, EPI_ISL_2691213, EPI_ISL_2691214, EPI_ISL_2691216, EPI_ISL_2691217, EPI_ISL_2691218 | | Strategic Laboratory | Tadeu de Araujo |
| EPI_ISL_2692981 | Clinical Microbiology Laboratory, Faculty of Medicine, Universitas Indonesia | Clinical Microbiology Laboratory, Faculty of Medicine, Universitas Indonesia | Ibrahim F, Natasha A, Yasmon A, Fitrhiyah, Saharman YR, Karuniawati A, Sudarmono P. |
| EPI_ISL_2693000, EPI_ISL_2693004 | TSGH-CP molecular lab | TSGH-CP molecular lab | Cherng-Lih Perng, Ming-Jr JIAN, Chih-Kai Chang, Jung-Chung Lin, Kuo-Ming Yeh, Chien-Wen Chen, Sheng-Kang Chiu, Hsing-Yi Chung, Shih-Hung Tsai, Kuo-Sheng Hung, Tien-Yao Chang, Feng-Yee Chang, Hung-Sheng Shang |
| EPI_ISL_2695877 | Dr. Yoshitaka Tamura Department of Clinical Laboratory, Osaka Habikino Medical Center | Department of Developmental Medicine, Research Institute, Osaka Women's and Children's Hospital | Itaru Yanagihara, Yukiko Nakur |
| EPI_ISL_2695884 | Department of Developmental Medicine, Research Institute, Osaka Women's and Children's Hospital | Department of Developmental Medicine, Research Institute, Osaka Women's and Children's Hospital | Itaru Yanagihara, Yukiko Nakura |
| EPI_ISL_2709689 | LESP Nuevo Leon | Instituto de Diagnostico y Referencia Epidemiologicos (INDRE) | Claudia Wong-Arambula, Abril Rodriguez-Maldonado, Vanessa Rivero-Arredondo, Ariadna Medina-Benitez, Joaquin Quiroz-Mercado, Sergio Rangel-Guerrero, Natividad Cruz-Ortiz, Tatiana Nunez-Garcia, Gisela Barrera-Badillo, Lucia Hernandez-Rivas, Irma Lopez-Martinez, Ernesto Ramirez-Gonzalez. |
| EPI_ISL_2713234, EPI_ISL_2713235, EPI_ISL_2713236, EPI_ISL_2713237, EPI_ISL_2713238, EPI_ISL_2713239, EPI_ISL_2713240, EPI_ISL_2713241, EPI_ISL_2713242, EPI_ISL_2713243, EPI_ISL_2713244, EPI_ISL_2713245, EPI_ISL_2713246, EPI_ISL_2713247, EPI_ISL_2713248, EPI_ISL_2713249, EPI_ISL_2713250, EPI_ISL_2713251, EPI_ISL_2713255, EPI_ISL_2713256 | | | |
| see above | Department of Health Technology and Informatics, The Hong Kong Polytechnic University | Department of Health Technology and Informatics, The Hong Kong Polytechnic University | Gilman Kit-Hang Siu, Lam-Kwong Lee, Kenneth Siu-Sing Leung, Jake Siu-Lun Leung, Wing-Hei Lo, Timothy Ting-Leung Ng, Chloe Toi-Mei Chan, Hoi-Ching Jim, Ivan Tak-Fai Wong, Kingsley King-Gee Tam, Hiu-Yin Lao, Denise Sze-Hang Wong, Alan Ka-Lun Wu, Miranda Chong-Yee Yau, Yvette Wai-Man Lai, Kitty Sau-Chun Fung, Sandy Ka-Yee Chau, Barry Kin-Chung Wong, Wing-Kin To, Kristine Luk, Alex Yat-Man Ho, Tak-Lun Que, Kam-Tong Yip, Wing Cheong Yam, David Ho-Keung Shum, Shea Ping Yip |
| EPI_ISL_2726970, EPI_ISL_2726972 | National Institute of Public Health | National Institute of Public Health | Helena Jirincova, Jaromira Vecerova, Timotej Suri, Dusan Trnka, Alexander Nagy |
| EPI_ISL_2728536, EPI_ISL_2728538, EPI_ISL_2728543, EPI_ISL_2728544, EPI_ISL_2728548, EPI_ISL_2728552, EPI_ISL_2728553, EPI_ISL_2728555, EPI_ISL_2728556, EPI_ISL_2728564, EPI_ISL_2728565, EPI_ISL_2728566 | | | |
| see above | Laboratorio de Infectologia y Virologia Molecular | Laboratory of Molecular Virology, School of Medicine, Pontificia Universidad Catolica de Chile | Catalina Pardo-Roa, Constanza Martinez-Valdevenito, Jennifer Angulo, Tamara Garcia-Salum, Estefany Poblete, Maria Jose Avendano, Leonardo I. Almonacid, Ana Maria Contreras, Carlos Palma, Jorge Levican, Eileen Serrano, Constanza Maldonado, M. Belen Leyton, Erick Salinas, Andres E. Munoz-Marcos, Francisco Melo, Marcela Ferres, Alejandro Bhrun, Magdalena Vera, Rafael A. Medina |
| EPI_ISL_2731270, EPI_ISL_2731272, EPI_ISL_2731273 | SC (UCO) Igiene e Sanità Pubblica, ASUGI, Trieste | ARGO Laboratorio Genomica ed Epigenomica | Licastro D, Dal Monego S, Degasperi M, Marcello A, Segat L, Piscianz E, Busetti M, Fontana F, Braida C, Barbone F, D'Agaro P |
| EPI_ISL_2756426, EPI_ISL_2756427, EPI_ISL_2756428, EPI_ISL_2756429, EPI_ISL_2756430, EPI_ISL_2756435 | Instituto Adolfo Lutz Central | Instituto Adolfo Lutz, Interdiciplinary Procedures Center, Strategic Laboratory | Claudio Tavares Sacchi, Claudia Regina Gonçalves, Erica Valessa Ramos Gomes, Karoline Rodrigues Campos, Caio Vinicius Dias Lopes, Leonardo Jose Tadeu de Araujo |
| EPI_ISL_2756521 | Instituto de Investigaciones Biomédicas en Retrovirus y SIDA (INBIRS), Universidad de Buenos Aires (UBA)-CONICET, Buenos Aires, Argentina. | INEI - ANLIS. Instituto Nacional de Enfermedades Infecciosas, Administración Nacional de. Laboratorios e institutos de Salud Dr. Carlos G. Malbran | Augusto Varese, Ignacio Mazzitelli, Valeria Ochoa, Federico Remes Lenicov, Ana Ceballos, Jorge Geffner |
| EPI_ISL_2756556 | Instituto de Investigaciones Biomédicas en Retrovirus y SIDA (INBIRS), Universidad de Buenos Aires (UBA)-CONICET, Buenos Aires, Argentina. | INEI - ANLIS. Instituto Nacional de Enfermedades Infecciosas, Administración Nacional de. Laboratorios e institutos de Salud "Dr. Carlos G. Malbran". | Augusto Varese, Ignacio Mazzitelli, Valeria Ochoa, Federico Remes Lenicov, Ana Ceballos, Jorge Geffner. |
| EPI_ISL_2756604 | Instituto de Investigaciones Biomédicas en Retrovirus y SIDA (INBIRS), Universidad de Buenos Aires (UBA)-CONICET, Buenos Aires, Argentina. | INEI - ANLIS. Instituto Nacional de Enfermedades Infecciosas, Administración Nacional de. Laboratorios e institutos de Salud "Dr. Carlos G. Malbran"./INBIRS. | Augusto Varese, Ignacio Mazzitelli, Valeria Ochoa, Federico Remes Lenicov, Ana Ceballos, Jorge Geffner. |
| EPI_ISL_2769378, EPI_ISL_2769380 | Centre Hospitalier Universitaire de Rouen Laboratoire de Virologie | Centre Hospitalier Universitaire de Rouen Laboratoire de Virologie | Alice Moisan, Fabienne De Oliveira, Marie Leoz |
| EPI_ISL_2779639 | State Key Laboratory of Emerging Infectious Diseases, The University of Hong Kong | The University of Hong Kong | Lifeng Li, Yiu Man Cheung, Huachen Zhu, Yi Guan |
| EPI_ISL_2788720, EPI_ISL_2788721, EPI_ISL_2788722, EPI_ISL_2788727, EPI_ISL_2788728, EPI_ISL_2788729, EPI_ISL_2788730, EPI_ISL_2788734, EPI_ISL_2788735, EPI_ISL_2788747, EPI_ISL_2788748, EPI_ISL_2788749, EPI_ISL_2788750, EPI_ISL_2788758, EPI_ISL_2788759, EPI_ISL_2788760, EPI_ISL_2788765, EPI_ISL_2788767, EPI_ISL_2788768, EPI_ISL_2788769, EPI_ISL_2788770, EPI_ISL_2788771, EPI_ISL_2788772, EPI_ISL_2788775, EPI_ISL_2788776, EPI_ISL_2788802, EPI_ISL_2788813, EPI_ISL_2788814, EPI_ISL_2788820, EPI_ISL_2788823, EPI_ISL_2788838, EPI_ISL_2788839, EPI_ISL_2788840, EPI_ISL_2788841, EPI_ISL_2788852, EPI_ISL_2788853, EPI_ISL_2788854, EPI_ISL_2788861, EPI_ISL_2788862, EPI_ISL_2788864 | | | |
| see above | Synlab Eesti OÜ | 1. Laboratory of Communicable Diseases (Estonia); 2. Eurofins Genomics Europe Sequencing GmbH | Liidia Dotsenko et al. |
| EPI_ISL_2811865, EPI_ISL_2811866, EPI_ISL_2811867, EPI_ISL_2811868, EPI_ISL_2811869 | Kuala Lumpur International Healthcare Centre (KLIHC) | Institute for Medical Research, Infectious Disease Research Centre, National Institutes of Health, Ministry of Health Malaysia | Suppiah J, Kamel K, Mohd Zawawi Z, Azizan MA, Ramly N, Robert F, Thayan R |
| EPI_ISL_2811870 | Kuala Lumpur International Healthcare Centre (KLIHC) | Institute for Medical Research, Infectious Disease Research Centre, National Institutes of Health, Ministry of Health Malaysia | Suppiah J, Kamel K, Mohd Zawawi Z, Azizan MA, Ramly N, Robert F, Thaya |
| EPI_ISL_2811871 | Pasir Mas Town Health Clinic | Institute for Medical Research, Infectious Disease Research Centre, National Institutes of Health, Ministry of Health Malaysia | Suppiah J, Kamel K, Mohd Zawawi Z, Azizan MA, Ramly N, Robert F, Thayan R |
| EPI_ISL_2811872 | Tumpat Hospital | Institute for Medical Research, Infectious Disease Research Centre, National Institutes of Health, Ministry of Health Malaysia | Suppiah J, Kamel K, Mohd Zawawi Z, Azizan MA, Ramly N, Robert F, Thayan R |
| EPI_ISL_2811873 | Pasir Mas Town Health Clinic | Institute for Medical Research, Infectious Disease Research Centre, National Institutes of Health, Ministry of Health Malaysia | Suppiah J, Kamel K, Mohd Zawawi Z, Azizan MA, Ramly N, Robert F, Thayan R |
| EPI_ISL_2811874 | Klinik Kesihatan Pasir Akar | Institute for Medical Research, Infectious Disease Research Centre, National Institutes of Health, Ministry of Health Malaysia | Suppiah J, Kamel K, Mohd Zawawi Z, Azizan MA, Ramly N, Robert F, Thayan R |
| EPI_ISL_2811875 | Universiti Sains Malaysia Hospital | Institute for Medical Research, Infectious Disease Research Centre, National Institutes of Health, Ministry of Health Malaysia | Suppiah J, Kamel K, Mohd Zawawi Z, Azizan MA, Ramly N, Robert F, Thayan R |
| EPI_ISL_2811876, EPI_ISL_2811877 | Melaka Hospital | Institute for Medical Research, Infectious Disease Research Centre, National Institutes of Health, Ministry of Health Malaysia | Suppiah J, Kamel K, Mohd Zawawi Z, Azizan MA, Ramly N, Robert F, Thayan R |

| | | | |
|---|---|---|---|
| EPI_ISL_2811878, EPI_ISL_2811879, EPI_ISL_2811880, EPI_ISL_2811881, EPI_ISL_2811882 | Sultanah Nora Ismail Hospital, Batu Pahat | Institute for Medical Research, Infectious Disease Research Centre, National Institutes of Health, Ministry of Health Malaysia | Suppiah J, Kamel K, Mohd Zawawi Z, Azizan MA, Ramly N, Robert F, Thayan R |
| EPI_ISL_2811884 | Institute for Medical Research, Infectious Disease Research Centre, National Institutes of Health, Ministry of Health Malaysia | Institute for Medical Research, Infectious Disease Research Centre, National Institutes of Health, Ministry of Health Malaysia | Suppiah J, Kamel K, Mohd Zawawi Z, Azizan MA, Ramly N, Robert F, Thayan R |
| EPI_ISL_2811885 | Sultanah Bahiyah Hospital, Alor Setar | Institute for Medical Research, Infectious Disease Research Centre, National Institutes of Health, Ministry of Health Malaysia | Suppiah J, Kamel K, Mohd Zawawi Z, Azizan MA, Ramly N, Robert F, Thayan R |
| EPI_ISL_2815377 | Ipoh Public Health Laboratory (MKAI), Ministry of Health Malaysia | Institute for Medical Research, Infectious Disease Research Centre, National Institutes of Health, Ministry of Health Malaysia | Suppiah J, Kamel K, Mohd Zawawi Z, Azizan MA, Ramly N, Robert F, Thayan R |
| EPI_ISL_2828040 | CARVAJAL LABORATORIO IPS | Instituto Nacional de Salud- Dirección de Investigación en Salud Pública | Katherine Laiton-Donato, Diego A. Álvarez-Díaz, Carlos Franco-Muñoz, Hector Alejandro Ruiz-Moreno, Paola Rojas, Maria T. Herrera-Sepúlveda, Diego Andrés Prada, Jhonnatan Reales-González, Sheryll Corchuelo, Julian Naizaque, Jorge Rivera, Gerardo Santamaría, Sergio Gomez, Lisseth Pardo, Juan Camilo Martinez, Marta Lopez Blanco, Ángela Alarcon Cruz, Diana Malo, Carmen Osorio, Magdalena Wiesner, Martha Lucia Ospina Martinez, Marcela Mercado-Reyes |
| EPI_ISL_2828661, EPI_ISL_2828662, EPI_ISL_2828666, EPI_ISL_2828667 | Instituo Adolfo Lutz - Reginal de Ribeirao Preto | Instituto Adolfo Lutz, Interdiciplinary Procedures Center, Strategic Laboratory | Claudio Tavares Sacchi, Claudia Regina Gonçalves, Erica Valessa Ramos Gomes, Karoline Rodrigues Campos, Caio Vinicius Dias Lopes |
| EPI_ISL_2828686, EPI_ISL_2828690, EPI_ISL_2828691, EPI_ISL_2828692, EPI_ISL_2828693, EPI_ISL_2828694, EPI_ISL_2828695 | Instituto Adolfo Lutz - Regional de Campinas | Instituto Adolfo Lutz, Interdiciplinary Procedures Center, Strategic Laboratory | Claudio Tavares Sacchi, Claudia Regina Gonçalves, Erica Valessa Ramos Gomes, Karoline Rodrigues Campos, Caio Vinicius Dias Lopes |
| EPI_ISL_2828696, EPI_ISL_2828697, EPI_ISL_2828699, EPI_ISL_2828700, EPI_ISL_2828701, EPI_ISL_2828702, EPI_ISL_2828703 | Instituto Adolfo Lutz - Regional de Santos | Instituto Adolfo Lutz, Interdiciplinary Procedures Center, Strategic Laboratory | Claudio Tavares Sacchi, Claudia Regina Gonçalves, Erica Valessa Ramos Gomes, Karoline Rodrigues Campos, Caio Vinicius Dias Lopes |
| EPI_ISL_2832344, EPI_ISL_2832346, EPI_ISL_2832350, EPI_ISL_2832352 | Department of Virology, Istituto Zooprofilattico Sperimentale del Lazio e della Toscana (IZSLT) | Department of General Diagnostics; Department of Virology; Istituto Zooprofilattico Sperimentale del Lazio e della Toscana (IZSLT) | Patricia Alba, Giuseppe Manna, Elena L. Diaconu, Fabiola Feltrin, Raffaela Conti, Teresa Scicluna, Virginia Carfora, Antonella Cersini, Alessia Franco, Antonio Battisti. |
| EPI_ISL_2840614, EPI_ISL_2840615 | S.C. Laboratorio Analisi, Ospedale di Lavagna, ASL 4 Liguria | U.O. Igiene, Ospedale Policlinico San Martino | Bruzzone Bianca, Bandettini Roberto, Caligiuri Patrizia, De Pace Vanessa, Domnich Alexander, Orsi Andrea, Ricucci Valentina, Icardi Giancarlo on behalf of SARS-CoV-2 ITALIAN RESEARCH ENTERPRISE-(SCIRE) Collaborative Group |
| EPI_ISL_2840616 | S.C. Laboratorio Analisi, ASL 3 Liguria | U.O. Igiene, Ospedale Policlinico San Martino | Bruzzone Bianca, Caligiuri Patrizia, De Pace Vanessa, Domnich Alexander, Orsi Andrea, Ricucci Valentina, Spitaleri Antonino, Icardi Giancarlo on behalf of SARS-CoV-2 ITALIAN RESEARCH ENTERPRISE-(SCIRE) Collaborative Group |
| EPI_ISL_2841553, EPI_ISL_2841554, EPI_ISL_2841555 | Department of Virology, Istituto Zooprofilattico Sperimentale del Lazio e della Toscana (IZSLT) | Department of General Diagnostics; Department of Virology; Istituto Zooprofilattico Sperimentale del Lazio e della Toscana (IZSLT) | Patricia Alba, Giuseppe Manna, Elena L. Diaconu, Fabiola Feltrin, Raffaela Conti, Teresa Scicluna, Virginia Carfora, Antonella Cersini, Alessia Franco, Antonio Battisti. |
| EPI_ISL_2841605 | UBS Silvio Joao Luiz de Lucia | Instituto Adolfo Lutz, Interdiciplinary Procedures Center, Strategic Laboratory | Claudio Tavares Sacchi, Claudia Regina Gonçalves, Erica Valessa Ramos Gomes, Karoline Rodrigues Campos, Caio Vinicius Dias Lopes |
| EPI_ISL_2841606 | Pronto Atendimento de Vargem Grande Paulista | Instituto Adolfo Lutz, Interdiciplinary Procedures Center, Strategic Laboratory | Claudio Tavares Sacchi, Claudia Regina Gonçalves, Erica Valessa Ramos Gomes, Karoline Rodrigues Campos, Caio Vinicius Dias Lopes |
| EPI_ISL_2841951, EPI_ISL_2841952, EPI_ISL_2841971, EPI_ISL_2841972 | National Center of Infectious and Parasitic Diseases | National Center of Infectious and Parasitic Diseases | Alexiev et al |
| EPI_ISL_2854049, EPI_ISL_2854050, EPI_ISL_2854051 | Melaka Hospital | Institute for Medical Research, Infectious Disease Research Centre, National Institutes of Health, Ministry of Health Malaysia | Suppiah J, Kamel K, Mohd Zawawi Z, Azizan MA, Ramly N, Robert F, Anasir MI, Thayan R |
| EPI_ISL_2854695 | Dinkes Mojokerto | National Institute of Health Research and Development | Vivi Setiawaty, Hana Apsari Pawestri, Subangkit, Kartika Dewi Puspa, Arie Ardiansyah Nugraha, Hartanti Dian Ikawati, Krisna Pangesti, Triyani Soekarso, Nelly Puspandari |
| EPI_ISL_2854703 | RS Umum Kartika Pulomas | National Institute of Health Research and Development | Vivi Setiawaty, Hana Apsari Pawestri, Subangkit, Kartika Dewi Puspa, Arie Ardiansyah Nugraha, Hartanti Dian Ikawati, Krisna Pangesti, Triyani Soekarso, Nelly Puspandari |
| EPI_ISL_2854712 | RS Hermina Depok | National Institute of Health Research and Development | Subangkit, Hana Apsari Pawestri, , Kartika Dewi Puspa, Arie Ardiansyah Nugraha, Hartanti Dian Ikawati, Krisna Pangesti, Triyani Soekarso, Nelly Puspandari, Vivi Setiawaty |
| EPI_ISL_2854734 | Labkesda Prov DKI Jakarta | National Institute of Health Research and Development | Subangkit, Hana Apsari Pawestri, , Kartika Dewi Puspa, Arie Ardiansyah Nugraha, Hartanti Dian Ikawati, Krisna Pangesti, Triyani Soekarso, Nelly Puspandari, Vivi Setiawaty |
| EPI_ISL_2854754 | BPPK Aceh | National Institute of Health Research and Development | Hana Apsari Pawestri, Subangkit, Kartika Dewi Puspa, Arie Ardiansyah Nugraha, Hartanti Dian Ikawati, Krisna Pangesti, Triyani Soekarso, Nelly Puspandari, Vivi Setiawaty |
| EPI_ISL_2859305, EPI_ISL_2859317, EPI_ISL_2859324, EPI_ISL_2859330, EPI_ISL_2859334, EPI_ISL_2859349, EPI_ISL_2859352, EPI_ISL_2859356, EPI_ISL_2859359, EPI_ISL_2859367 | Zamboanga City Medical Center | Philippine Genome Center | Francis A. Tablizo, Kenneth M. Kim, Maria Sofia L. Yangzon, Elcid Aaron R. Pangilinan, Renato Jacinto Q. Mantaring, Benedict A. Maralit, Marc Edsel C. Ayes, Eva Maria Cutiongco-de la Paz, Alethea R. de Guzman, Jan Michael C. Yap, Jo-Hannah S. Llames, Sheila Mae M. Araiza, Kris P. Punayan, Karol Sophia Agape R. Padilla, Lindsay Claire D.L. Carandang, Jarvin E. Nipales, Zipporah Mariebelle R. Enriquez, Diomedes A. Carino, Yvonne Valerie Austria, Nina Francesca Bustamante, Marielle M Gamboa, Zyrel V. Mollejon, Krisitna Patriz Dela Cruz, Ma. Exanil Plantig, Rianna Patricia S. Cruz, Carlo M. Lapid, El King D. Morado, Joshua Gregor A. Dizon, Razel Nikka M. Hao, Arianne A. Zamora, Devon Ray Pacial, Juan Antonio R. Magalang, Henrietta Marie Rodriguez, Marissa Alejandria, Celia Carlos, Anna Ong-Lim, Edsel Maurice Salvaña, John Q. Wong, Jaime C. Montoya, Maria Rosario Singh-Vergeire and Cynthia P. Saloma |
| EPI_ISL_2860812, EPI_ISL_2860820, EPI_ISL_2860821 | Department of Health Technology and Informatics, The Hong Kong Polytechnic University | Department of Health Technology and Informatics, The Hong Kong Polytechnic University | Gilman Kit-Hang Siu, Lam-Kwong Lee, Kenneth Siu-Sing Leung, Jake Siu-Lun Leung, Wing-Hei Lo, Timothy Ting-Leung Ng, Chloe Toi-Mei Chan, Hoi-Ching Jim, Ivan Tak-Fai Wong, Kingsley King-Gee Tam, Hiu-Yin Lao, Denise Sze-Hang Wong, Alan Ka-Lun Wu, Miranda Chong-Yee Yau, Yvette Wai-Man Lai, Kitty Sau-Chun Fung, Sandy Ka-Yee Chau, Barry Kin-Chung Wong, Wing-Kin To, Kristine Luk, Alex Yat-Man Ho, Tak-Lun Que, Kam-Tong Yip, Wing Cheong Yam, David Ho-Keung Shum, Shea Ping Yip |
| EPI_ISL_2863795, EPI_ISL_2863855, EPI_ISL_2863856, EPI_ISL_2863857, EPI_ISL_2863858, EPI_ISL_2863859, EPI_ISL_2863860, EPI_ISL_2863861, EPI_ISL_2863862, EPI_ISL_2863863, EPI_ISL_2863864, EPI_ISL_2863865, EPI_ISL_2863866, EPI_ISL_2863869, EPI_ISL_2863870, EPI_ISL_2863872, EPI_ISL_2863873, EPI_ISL_2863874, EPI_ISL_2863875, EPI_ISL_2863876, EPI_ISL_2863877, EPI_ISL_2863878, EPI_ISL_2863879, EPI_ISL_2863880, EPI_ISL_2863881, EPI_ISL_2863882, EPI_ISL_2863883, EPI_ISL_2863884, EPI_ISL_2863885, EPI_ISL_2863886, EPI_ISL_2863887, EPI_ISL_2863888, EPI_ISL_2863890, EPI_ISL_2863893, EPI_ISL_2863895, EPI_ISL_2863896, EPI_ISL_2863898, EPI_ISL_2863899 | | | |
| see above | Lboratorio Central de Saude Publica do Estado do Parana (LACEN/PR) | Laboratory of Respiratory Viruses and Measles, Oswaldo Cruz Institute, FIOCRUZ | Paola Resende, Luciana Appolinario, Fernando Motta, Anna Carolina Paixao, Ana Carolina Mendonca, Alice Sampaio Rocha, Taina Venas, Elisa Cavalcante Pereira, Renata Serrano Lopes, Agatha Cristinne Prudencio Soares, Ighor Leonardo Arantes Gomes, Irina Riediger, Marilda Siqueira on behalf of the Fiocruz COVID-19 Genomic Surveillance Network |
| EPI_ISL_2868925 | BTKLPP Kelas I Manado | National Institute of Health Research and Development | Vivi Setiawaty, Hana Apsari Pawestri, Subangkit, Kartika Dewi Puspa, Arie Ardiansyah Nugraha, Hartanti Dian Ikawati, Krisna Pangesti, Triyani Soekarso, Nelly Puspandari |

| | | | |
|---|---|---|---|
| EPI_ISL_2868941 | DINKES Prov Kalimantan Timur | National Institute of Health Research and Development | Vivi Setiawaty, Hana Apsari Pawestri, Subangkit, Kartika Dewi Puspa, Arie Ardiansyah Nugraha, Hartanti Dian Ikawati, Krisna Pangesti, Triyani Soekarso, Nelly Puspandari |
| EPI_ISL_2873865, EPI_ISL_2873866 | National Influenza Centre | National Influenza Centre | William K. Ampofo, Michael Marks, Ivy A. Asante, Sharon Hsu, Benjamiin B. Lindsey, Benjamin H. Foulkes, Mildred Adusei-Poku, Linda Boatemaa, Lorreta Kwasah, Joseph Oliver-Commey, Ernest Asiedu, Franklin Asiedu-Bekoe, Gordon Awandare, Joyce Ngoi, Dennis Laryea, Mathew D. Parker, Thushan I de Silva, |

We gratefully acknowledge the following Authors from the Originating laboratories responsible for obtaining the specimens, as well as the Submitting laboratories where the genome data were generated and shared via GISAID, on which this research is based.

All Submitters of data may be contacted directly via www.gisaid.org

Authors are sorted alphabetically.

| Accession ID | Originating Laboratory | Submitting Laboratory | Authors |
|---|---|---|---|
| EPI_ISL_1972570, EPI_ISL_2001050 | Hospital Sharp | Microbial Genomics Laboratory | Bruno Gomez-Gil, Julissa Enciso-Ibarra, Alejandra Garcia-Gasca, Daniel Fregoso-Rueda |
| EPI_ISL_2036272 | Genome Centre | Genome Centre | Shovon Lal Sarkar, Md. Shazid Hasan, Ali Ahsan Setu, Prosanto Kumar Das, A. S. M. Rubayet Ul Alam, Ovinu Kibria Islam, Md. Tanvir Islam, Hassan Md. Al-Emran, Iqbal Kabir Jahid, M. Anwar Hossain |
| EPI_ISL_2106252, EPI_ISL_2106254, EPI_ISL_2106255, EPI_ISL_2106256, EPI_ISL_2106257, EPI_ISL_2106258, EPI_ISL_2106259, EPI_ISL_2106260, EPI_ISL_2106276, EPI_ISL_2106277, EPI_ISL_2106278, EPI_ISL_2106279, EPI_ISL_2106280, EPI_ISL_2106281, EPI_ISL_2106282 | | | |
| see above | Virology Unit, Institut Pasteur du Cambodge | Virology Unit, Institut Pasteur du Cambodge | Jurre Y Siegers, Cecile Troupin, Leakhena Pum, Ly Sovann, Kraing Sidonn, Yi Sengdoeurn, Chin Savuth, Chau Darapheak, Veasna Duong, Erik A Karlsson |
| EPI_ISL_2114672, EPI_ISL_2115909, EPI_ISL_2116339 | Molecular Diagnostics Pathology Department Mater Dei Hospital Malta | Molecular Diagnostics Pathology Department Mater Dei Hospital Malta | G Zahra, R Borg, C Cilia, L Grech |
| EPI_ISL_2131274, EPI_ISL_2131277, EPI_ISL_2131278, EPI_ISL_2131279, EPI_ISL_2131280, EPI_ISL_2131281, EPI_ISL_2131282 | OLVZ Aalst | OLVZ Aalst | Astrid Holderbeke |
| EPI_ISL_2131395 | SIESP SULMONA | Istituto Zooprofilattico Sperimentale dell'Abruzzo e Molise "G. Caporale" | Lorusso A, Marcacci M, Di Domenico M, Ancora M, Curini V, Di Lollo Valeria, Mangone I, Rinaldi A, Delli Compagni E, Scialabba S, Caporale M, Di Pasquale A, Cammà C, Puglia I, Calistri P, Savini G |
| EPI_ISL_2131402, EPI_ISL_2131403 | SIESP DIPARTIMENTO DI PREVENZIONE TERAMO | Istituto Zooprofilattico Sperimentale dell'Abruzzo e Molise "G. Caporale" | Lorusso A, Marcacci M, Di Domenico M, Ancora M, Curini V, Di Lollo Valeria, Mangone I, Rinaldi A, Delli Compagni E, Scialabba S, Caporale M, Di Pasquale A, Cammà C, Puglia I, Calistri P, Savini G |
| EPI_ISL_2131404 | SIESP CHIETI - DRIVE IN CHIETI | Istituto Zooprofilattico Sperimentale dell'Abruzzo e Molise "G. Caporale" | Lorusso A, Marcacci M, Di Domenico M, Ancora M, Curini V, Di Lollo Valeria, Mangone I, Rinaldi A, Delli Compagni E, Scialabba S, Caporale M, Di Pasquale A, Cammà C, Puglia I, Calistri P, Savini G |
| EPI_ISL_2131405 | SIESP CHIETI - DRIVE IN ORTONA | Istituto Zooprofilattico Sperimentale dell'Abruzzo e Molise "G. Caporale" | Lorusso A, Marcacci M, Di Domenico M, Ancora M, Curini V, Di Lollo Valeria, Mangone I, Rinaldi A, Delli Compagni E, Scialabba S, Caporale M, Di Pasquale A, Cammà C, Puglia I, Calistri P, Savini G |
| EPI_ISL_2131407, EPI_ISL_2131408 | SIESP DIPARTIMENTO DI PREVENZIONE TERAMO | Istituto Zooprofilattico Sperimentale dell'Abruzzo e Molise "G. Caporale" | Lorusso A, Marcacci M, Di Domenico M, Ancora M, Curini V, Di Lollo Valeria, Mangone I, Rinaldi A, Delli Compagni E, Scialabba S, Caporale M, Di Pasquale A, Cammà C, Puglia I, Calistri P, Savini G |
| EPI_ISL_2131409, EPI_ISL_2131410 | SIESP SULMONA | Istituto Zooprofilattico Sperimentale dell'Abruzzo e Molise "G. Caporale" | Lorusso A, Marcacci M, Di Domenico M, Ancora M, Curini V, Di Lollo Valeria, Mangone I, Rinaldi A, Delli Compagni E, Scialabba S, Caporale M, Di Pasquale A, Cammà C, Puglia I, Calistri P, Savini G |
| EPI_ISL_2131411, EPI_ISL_2131412 | SIESP DIPARTIMENTO DI PREVENZIONE TERAMO | Istituto Zooprofilattico Sperimentale dell'Abruzzo e Molise "G. Caporale" | Lorusso A, Marcacci M, Di Domenico M, Ancora M, Curini V, Di Lollo Valeria, Mangone I, Rinaldi A, Delli Compagni E, Scialabba S, Caporale M, Di Pasquale A, Cammà C, Puglia I, Calistri P, Savini G |
| EPI_ISL_2131413 | SIESP CHIETI - DRIVE IN LANCIANO | Istituto Zooprofilattico Sperimentale dell'Abruzzo e Molise "G. Caporale" | Lorusso A, Marcacci M, Di Domenico M, Ancora M, Curini V, Di Lollo Valeria, Mangone I, Rinaldi A, Delli Compagni E, Scialabba S, Caporale M, Di Pasquale A, Cammà C, Puglia I, Calistri P, Savini G |
| EPI_ISL_2131414 | DIP. PREV. AVEZZANO SERVIZIO DI IGIENE EPIDEMIOLOGIA E SANITA' PUBBLICA | Istituto Zooprofilattico Sperimentale dell'Abruzzo e Molise "G. Caporale" | Lorusso A, Marcacci M, Di Domenico M, Ancora M, Curini V, Di Lollo Valeria, Mangone I, Rinaldi A, Delli Compagni E, Scialabba S, Caporale M, Di Pasquale A, Cammà C, Puglia I, Calistri P, Savini G |
| EPI_ISL_2131415 | SIESP CHIETI - DRIVE IN LANCIANO | Istituto Zooprofilattico Sperimentale dell'Abruzzo e Molise "G. Caporale" | Lorusso A, Marcacci M, Di Domenico M, Ancora M, Curini V, Di Lollo Valeria, Mangone I, Rinaldi A, Delli Compagni E, Scialabba S, Caporale M, Di Pasquale A, Cammà C, Puglia I, Calistri P, Savini G |
| EPI_ISL_2131416 | USCA Avezzano | Istituto Zooprofilattico Sperimentale dell'Abruzzo e Molise "G. Caporale" | Lorusso A, Marcacci M, Di Domenico M, Ancora M, Curini V, Di Lollo Valeria, Mangone I, Rinaldi A, Delli Compagni E, Scialabba S, Caporale M, Di Pasquale A, Cammà C, Puglia I, Calistri P, Savini G |
| EPI_ISL_2131418 | USCA Pescina | Istituto Zooprofilattico Sperimentale dell'Abruzzo e Molise "G. Caporale" | Lorusso A, Marcacci M, Di Domenico M, Ancora M, Curini V, Di Lollo Valeria, Mangone I, Rinaldi A, Delli Compagni E, Scialabba S, Caporale M, Di Pasquale A, Cammà C, Puglia I, Calistri P, Savini G |
| EPI_ISL_2131419 | USCA Tagliacozzo | Istituto Zooprofilattico Sperimentale dell'Abruzzo e Molise "G. Caporale" | Lorusso A, Marcacci M, Di Domenico M, Ancora M, Curini V, Di Lollo Valeria, Mangone I, Rinaldi A, Delli Compagni E, Scialabba S, Caporale M, Di Pasquale A, Cammà C, Puglia I, Calistri P, Savini G |
| EPI_ISL_2131420 | SIESP CHIETI - DRIVE IN VASTO | Istituto Zooprofilattico Sperimentale dell'Abruzzo e Molise "G. Caporale" | Lorusso A, Marcacci M, Di Domenico M, Ancora M, Curini V, Di Lollo Valeria, Mangone I, Rinaldi A, Delli Compagni E, Scialabba S, Caporale M, Di Pasquale A, Cammà C, Puglia I, Calistri P, Savini G |
| EPI_ISL_2131421, EPI_ISL_2131422, EPI_ISL_2131423, EPI_ISL_2131424, EPI_ISL_2131425, EPI_ISL_2131426 | SIESP DIPARTIMENTO DI PREVENZIONE TERAMO | Istituto Zooprofilattico Sperimentale dell'Abruzzo e Molise "G. Caporale" | Lorusso A, Marcacci M, Di Domenico M, Ancora M, Curini V, Di Lollo Valeria, Mangone I, Rinaldi A, Delli Compagni E, Scialabba S, Caporale M, Di Pasquale A, Cammà C, Puglia I, Calistri P, Savini G |
| EPI_ISL_2131427 | SIESP CHIETI - DRIVE IN CHIETI | Istituto Zooprofilattico Sperimentale dell'Abruzzo e Molise "G. Caporale" | Lorusso A, Marcacci M, Di Domenico M, Ancora M, Curini V, Di Lollo Valeria, Mangone I, Rinaldi A, Delli Compagni E, Scialabba S, Caporale M, Di Pasquale A, Cammà C, Puglia I, Calistri P, Savini G |
| EPI_ISL_2131429, EPI_ISL_2131430 | SIESP DIPARTIMENTO DI PREVENZIONE TERAMO | Istituto Zooprofilattico Sperimentale dell'Abruzzo e Molise "G. Caporale" | Lorusso A, Marcacci M, Di Domenico M, Ancora M, Curini V, Di Lollo Valeria, Mangone I, Rinaldi A, Delli Compagni E, Scialabba S, Caporale M, Di Pasquale A, Cammà C, Puglia I, Calistri P, Savini G |
| EPI_ISL_2131431, EPI_ISL_2131432, EPI_ISL_2131433, EPI_ISL_2131434 | SIESP CHIETI - DRIVE IN LANCIANO | Istituto Zooprofilattico Sperimentale dell'Abruzzo e Molise "G. Caporale" | Lorusso A, Marcacci M, Di Domenico M, Ancora M, Curini V, Di Lollo Valeria, Mangone I, Rinaldi A, Delli Compagni E, Scialabba S, Caporale M, Di Pasquale A, Cammà C, Puglia I, Calistri P, Savini G |
| EPI_ISL_2131435, EPI_ISL_2131436, EPI_ISL_2131437 | DIP. PREV. AVEZZANO SERVIZIO DI IGIENE EPIDEMIOLOGIA E SANITA' PUBBLICA | Istituto Zooprofilattico Sperimentale dell'Abruzzo e Molise "G. Caporale" | Lorusso A, Marcacci M, Di Domenico M, Ancora M, Curini V, Di Lollo Valeria, Mangone I, Rinaldi A, Delli Compagni E, Scialabba S, Caporale M, Di Pasquale A, Cammà C, Puglia I, Calistri P, Savini G |
| EPI_ISL_2131805 | Microbiology Department, Lu`an Center for Disease Control and Prevention | Microbiology Department, Lu`an Center for Disease Control and Prevention | Chen Zhichao, Gao Dawei, Yang Wei, Chong Hongwei, Zhang Feng, Zhang Limei, Zhang Qin, Zhu Rui, Li Zhaoyang, Fan Yuzhen |
| EPI_ISL_2131843, EPI_ISL_2132221 | Microbiology Department, Lu`an Center for Disease Control and Prevention | Microbiology Department, Lu`an Center for Disease Control and Prevention | Chen Zhichao, Gao Dawei, Yang Wei, Chen Beilei, Chang Hongwei, Zhang Feng, Zhang Limei, Zhang Qin, Zhu Rui, Li Zhaoyang, Fan Yuzhen. |
| EPI_ISL_2134734, EPI_ISL_2134735, EPI_ISL_2134736, EPI_ISL_2134737 | Centre Hospitalier Universitaire Clermont-Ferrand | CHU Clermont-Ferrand, service de virologie | Bisseux Maxime, Mirand Audrey, Combes Patricia, Henquell Cécile |
| EPI_ISL_2134739 | HOPITAL DE VICHY | CHU Clermont-Ferrand, service de virologie | Bisseux Maxime, Mirand Audrey, Combes Patricia, Henquell Cécile |
| EPI_ISL_2134743 | EHPAD | CHU Clermont-Ferrand, service de virologie | Bisseux Maxime, Mirand Audrey, Combes Patricia, Henquell Cécile |
| EPI_ISL_2134744, EPI_ISL_2134745, | Centre Hospitalier Universitaire Clermont-Ferrand | CHU Clermont-Ferrand, service de virologie | Bisseux Maxime, Mirand Audrey, Combes Patricia, Henquell Cécile |

| | | | |
|---|---|---|---|
| EPI_ISL_2134746, EPI_ISL_2134747, EPI_ISL_2134748, EPI_ISL_2134749 | | | |
| EPI_ISL_2134750 | Centre Hospitalier RIOM | CHU Clermont-Ferrand, service de virologie | Bisseux Maxime, Mirand Audrey, Combes Patricia, Henquell Cécile |
| EPI_ISL_2134753 | Centre Hospitalier Universitaire Clermont-Ferrand | CHU Clermont-Ferrand, service de virologie | Bisseux Maxime, Mirand Audrey, Combes Patricia, Henquell Cécile |
| EPI_ISL_2134759, EPI_ISL_2134760 | Centre De Prelevement COVID RIOM | CHU Clermont-Ferrand, service de virologie | Bisseux Maxime, Mirand Audrey, Combes Patricia, Henquell Cécile |
| EPI_ISL_2134763 | HOPITAL DU MONT DORE | CHU Clermont-Ferrand, service de virologie | Bisseux Maxime, Mirand Audrey, Combes Patricia, Henquell Cécile |
| EPI_ISL_2134766, EPI_ISL_2134775, EPI_ISL_2134776 | Centre Hospitalier Universitaire Clermont-Ferrand | CHU Clermont-Ferrand, service de virologie | Bisseux Maxime, Mirand Audrey, Combes Patricia, Henquell Cécile |
| EPI_ISL_2134777 | Centre Hospitalier Paul Ardier | CHU Clermont-Ferrand, service de virologie | Bisseux Maxime, Mirand Audrey, Combes Patricia, Henquell Cécile |
| EPI_ISL_2134778, EPI_ISL_2134779 | Centre Hospitalier Universitaire Clermont-Ferrand | CHU Clermont-Ferrand, service de virologie | Bisseux Maxime, Mirand Audrey, Combes Patricia, Henquell Cécile |
| EPI_ISL_2137734, EPI_ISL_2137735, EPI_ISL_2137736 | AZ Klina | AZ Klina | Carl Vael - Lynsey Berckmans |
| EPI_ISL_2140882, EPI_ISL_2140884 | The Ohio State University Applied Microbiology Services Laboratory | The Ohio State University Applied Microbiology Services Laboratory | Seth A. Faith PhD |
| EPI_ISL_2146023, EPI_ISL_2146025, EPI_ISL_2146027, EPI_ISL_2146030, EPI_ISL_2146031 | Department of Virology, Istituto Zooprofilattico Sperimentale del Lazio e della Toscana (IZSLT) | Department of General Diagnostics; Department of Virology; Istituto Zooprofilattico Sperimentale del Lazio e della Toscana (IZSLT) | Patricia Alba, Giuseppe Manna, Elena L. Diaconu, Fabiola Feltrin, Raffaela Conti, Teresa Scicluna, Virginia Carfora, Antonella Cersini, Alessia Franco, Antonio Battisti. |
| EPI_ISL_2171267 | Dhulikhel Hospital, Kathmandu University Hospital | Molecular and Genomics Research Lab, Dhulikhel Hospital, Kathmandu University Hospital | Rajeev Shrestha, Nishan Katuwal, Navin Adhikari, Dipesh Tamrakar, Meghnath Dhimal, Pradip Gyanwali, Saroj Bhattarai, Surendra Kumar Madhup |
| EPI_ISL_2178978, EPI_ISL_2178979, EPI_ISL_2178980, EPI_ISL_2178981, EPI_ISL_2178982 | OLVZ Aalst | OLVZ Aalst | Astrid Holderbeke |
| EPI_ISL_2179190 | OCME Office Of Chief Medical Examiner | New York City Public Health Laboratory | Jade Wang, et al. |
| EPI_ISL_2188481, EPI_ISL_2188482, EPI_ISL_2188483 | Hospital | National Reference Center for Viruses of Respiratory Infections, Institut Pasteur, Paris | Marion Barbet, Sylvie Behillil, Méline Bizard, Angela Brisebarre, Camille Capel, Vincent Enouf, Louise Lefrançois, Frédéric Lemoine, Christophe Malabat, Corinne Maufrais, Etienne Simon-Lorière, Maud Vanpeene, Sylvie Van der Werf ,StéPhanie Marque Juillet |
| EPI_ISL_2190110, EPI_ISL_2190111, EPI_ISL_2190121, EPI_ISL_2190122 | National Institute of Infectious Diseases-Prof. Dr. Matei Bals Molecular Diagnostics Laboratory | National Institute of Infectious Diseases-Prof. Dr. Matei Bals Molecular Diagnostics Laboratory | Corina Casangiu, Leontina Banica, Marius Surleac, Ovidiu Vlaicu, Andreea Tudor, Simona Paraschiv, Dan Otelea |
| EPI_ISL_2193243, EPI_ISL_2193247, EPI_ISL_2193248, EPI_ISL_2193249, EPI_ISL_2193250, EPI_ISL_2193251, EPI_ISL_2193253, EPI_ISL_2193254, EPI_ISL_2193255, EPI_ISL_2193256, EPI_ISL_2193258, EPI_ISL_2193259, EPI_ISL_2193260, EPI_ISL_2193261, EPI_ISL_2193262, EPI_ISL_2193263, EPI_ISL_2193264, EPI_ISL_2193265, EPI_ISL_2193266, EPI_ISL_2193273, EPI_ISL_2193274, EPI_ISL_2193275, EPI_ISL_2193276, EPI_ISL_2193277, EPI_ISL_2193282, EPI_ISL_2193284, EPI_ISL_2193285 | | | |
| see above | Baylor Scott & White-Temple | Baylor Scott & White-Temple | Ari Rao, Linden Morales, Kimberly Walker, Marcus Volz, Shelby Hendrickson |
| EPI_ISL_2193288, EPI_ISL_2193289, EPI_ISL_2193291, EPI_ISL_2193294, EPI_ISL_2193295, EPI_ISL_2193296, EPI_ISL_2193297, EPI_ISL_2193298, EPI_ISL_2193299, EPI_ISL_2193300 | Laboratorio Microbiologia PO Cardarelli | Laboratorio Microbiologia PO Cardarelli | Felice V., Niro G. Scutella' M. |
| EPI_ISL_2200064, EPI_ISL_2200156, EPI_ISL_2200157, EPI_ISL_2202178, EPI_ISL_2203011, EPI_ISL_2203964, EPI_ISL_2205561, EPI_ISL_2208894 | SC (UCO) Igiene e Sanità Pubblica, ASUGI, Trieste | ARGO Laboratorio Genomica ed Epigenomica | Licastro D, Dal Monego S, Degasperi M, Marcello A, Segat L, Piscianz E, D'Agaro P |
| EPI_ISL_2211115 | SC (UCO) Igiene e Sanità Pubblica, ASUGI, Trieste and SC Microbiologia e Virologia, ASFO, Pordenone | ARGO Laboratorio Genomica ed Epigenomica | Licastro D, Dal Monego S, Degasperi M, Marcello A, Segat L, Piscianz E, De Rosa R, D'Agaro P |
| EPI_ISL_2214657, EPI_ISL_2214662, EPI_ISL_2214666, EPI_ISL_2214672 | Centre Hospitalier Universitaire Clermont-Ferrand | CHU Clermont-Ferrand, service de virologie | Bisseux Maxime, Mirand Audrey, Combes Patricia, Henquell Cécile |
| EPI_ISL_2214676 | HOPITAL DE VICHY | CHU Clermont-Ferrand, service de virologie | Bisseux Maxime, Mirand Audrey, Combes Patricia, Henquell Cécile |
| EPI_ISL_2214680 | Centre hospitalier D'AURILLAC | CHU Clermont-Ferrand, service de virologie | Bisseux Maxime, Mirand Audrey, Combes Patricia, Henquell Cécile |
| EPI_ISL_2214685 | Centre Hospitalier Universitaire Clermont-Ferrand | CHU Clermont-Ferrand, service de virologie | Bisseux Maxime, Mirand Audrey, Combes Patricia, Henquell Cécile |
| EPI_ISL_2214690 | center de prélévement COVID ISSOIRE | CHU Clermont-Ferrand, service de virologie | Bisseux Maxime, Mirand Audrey, Combes Patricia, Henquell Cécile |
| EPI_ISL_2214692, EPI_ISL_2214694, EPI_ISL_2214699 | Centre Hospitalier Universitaire Clermont-Ferrand | CHU Clermont-Ferrand, service de virologie | Bisseux Maxime, Mirand Audrey, Combes Patricia, Henquell Cécile |
| EPI_ISL_2214721 | Centre Hospitalier Billom | CHU Clermont-Ferrand, service de virologie | Bisseux Maxime, Mirand Audrey, Combes Patricia, Henquell Cécile |
| EPI_ISL_2214727 | Centre Hospitalier Universitaire Clermont-Ferrand | CHU Clermont-Ferrand, service de virologie | Bisseux Maxime, Mirand Audrey, Combes Patricia, Henquell Cécile |
| EPI_ISL_2214731 | HOPITAL DE VICHY | CHU Clermont-Ferrand, service de virologie | Bisseux Maxime, Mirand Audrey, Combes Patricia, Henquell Cécile |
| EPI_ISL_2214736 | Centre Hospitalier Universitaire Clermont-Ferrand | CHU Clermont-Ferrand, service de virologie | Bisseux Maxime, Mirand Audrey, Combes Patricia, Henquell Cécile |
| EPI_ISL_2214739 | EHPAD | CHU Clermont-Ferrand, service de virologie | Bisseux Maxime, Mirand Audrey, Combes Patricia, Henquell Cécile |
| EPI_ISL_2214742 | Centre Hospitalier Universitaire Clermont-Ferrand | CHU Clermont-Ferrand, service de virologie | Bisseux Maxime, Mirand Audrey, Combes Patricia, Henquell Cécile |
| EPI_ISL_2214747, EPI_ISL_2214753, EPI_ISL_2214758 | HOPITAL DE VICHY | CHU Clermont-Ferrand, service de virologie | Bisseux Maxime, Mirand Audrey, Combes Patricia, Henquell Cécile |
| EPI_ISL_2214763, EPI_ISL_2214769, EPI_ISL_2214774 | Centre Hospitalier Universitaire Clermont-Ferrand | CHU Clermont-Ferrand, service de virologie | Bisseux Maxime, Mirand Audrey, Combes Patricia, Henquell Cécile |
| EPI_ISL_2214779 | HOPITAL DE VICHY | CHU Clermont-Ferrand, service de virologie | Bisseux Maxime, Mirand Audrey, Combes Patricia, Henquell Cécile |
| EPI_ISL_2214783, EPI_ISL_2214788 | Centre Hospitalier Universitaire Clermont-Ferrand | CHU Clermont-Ferrand, service de virologie | Bisseux Maxime, Mirand Audrey, Combes Patricia, Henquell Cécile |
| EPI_ISL_2214794 | Centre Hospitalier Billom | CHU Clermont-Ferrand, service de virologie | Bisseux Maxime, Mirand Audrey, Combes Patricia, Henquell Cécile |
| EPI_ISL_2214798 | HOPITAL DE VICHY | CHU Clermont-Ferrand, service de virologie | Bisseux Maxime, Mirand Audrey, Combes Patricia, Henquell Cécile |
| EPI_ISL_2214804 | Centre Hospitalier RIOM | CHU Clermont-Ferrand, service de virologie | Bisseux Maxime, Mirand Audrey, Combes Patricia, Henquell Cécile |
| EPI_ISL_2214808 | Centre Hospitalier Paul Ardier | CHU Clermont-Ferrand, service de virologie | Bisseux Maxime, Mirand Audrey, Combes Patricia, Henquell Cécile |
| EPI_ISL_2214814 | Centre Hospitalier RIOM | CHU Clermont-Ferrand, service de virologie | Bisseux Maxime, Mirand Audrey, Combes Patricia, Henquell Cécile |
| EPI_ISL_2214819, EPI_ISL_2214825, EPI_ISL_2214830, EPI_ISL_2214836, EPI_ISL_2214841, EPI_ISL_2214845, EPI_ISL_2214853, EPI_ISL_2214856, EPI_ISL_2214861, EPI_ISL_2214868, EPI_ISL_2214873, EPI_ISL_2214879 | | | |

| | | | |
|---|---|---|---|
| see above | Centre Hospitalier Universitaire Clermont-Ferrand | CHU Clermont-Ferrand, service de virologie | Bisseux Maxime, Mirand Audrey, Combes Patricia, Henquell Cécile |
| EPI_ISL_2214895 | Centre hospitalier D'AURILLAC | CHU Clermont-Ferrand, service de virologie | Bisseux Maxime, Mirand Audrey, Combes Patricia, Henquell Cécile |
| EPI_ISL_2215527, EPI_ISL_2216284 | SC (UCO) Igiene e Sanità Pubblica, ASUGI, Trieste and SC Microbiologia e Virologia, ASFO, Pordenone | ARGO Laboratorio Genomica ed Epigenomica | Licastro D, Dal Monego S, Degasperi M, Marcello A, Segat L, Piscianz E, De Rosa R, D'Agaro P |
| EPI_ISL_2225611 | Basurto University Hospital: Clinical Microbiology Laboratory | Biocruces Bizkaia | Mikel Urrutikoetxea-Gutiérrez, Mª Carmen Nieto Toboso, Estibaliz Ugalde Zarraga, Mikel Gallego Rodrigo, Ana de la Hoz, José Luis Díaz de Tuesta del Arco |
| EPI_ISL_2228062, EPI_ISL_2228063, EPI_ISL_2228064, EPI_ISL_2228065, EPI_ISL_2228066, EPI_ISL_2228067, EPI_ISL_2228068, EPI_ISL_2228069, EPI_ISL_2228070, EPI_ISL_2228071, EPI_ISL_2228072, EPI_ISL_2228073, EPI_ISL_2228074 | | | |
| see above | OLVZ Aalst | OLVZ Aalst | Anne Vankeerberghen |
| EPI_ISL_2228430 | Basurto University Hospital: Clinical Microbiology Laboratory | Biocruces Bizkaia | Mikel Urrutikoetxea-Gutiérrez, Mª Carmen Nieto Toboso, Estibaliz Ugalde Zarraga, Mikel Gallego Rodrigo, Ana de la Hoz, José Luis Díaz de Tuesta del Arco |
| EPI_ISL_2228654, EPI_ISL_2228655, EPI_ISL_2228656 | GH A.CHENEVIER-H.MONDOR | Department of Virology, Henri Mondor University Hospital, Assistance Publique Hôpitaux de Paris, Université Paris-Est Créteil, INSERM U955 | Christophe Rodriguez, Slim Fourati, Vanessa Demontant, Guillaume Gricourt, Melissa N'Debi, Alexandre Soulier, Elisabeth Trawinski, Jean-Michel Pawlotsky |
| EPI_ISL_2228657 | HOPITAL EMILE ROUX | Department of Virology, Henri Mondor University Hospital, Assistance Publique Hôpitaux de Paris, Université Paris-Est Créteil, INSERM U955 | Christophe Rodriguez, Slim Fourati, Vanessa Demontant, Guillaume Gricourt, Melissa N'Debi, Alexandre Soulier, Elisabeth Trawinski, Jean-Michel Pawlotsky |
| EPI_ISL_2228666 | GH A.CHENEVIER-H.MONDOR | Department of Virology, Henri Mondor University Hospital, Assistance Publique Hôpitaux de Paris, Université Paris-Est Créteil, INSERM U955 | Christophe Rodriguez, Slim Fourati, Vanessa Demontant, Guillaume Gricourt, Melissa N'Debi, Alexandre Soulier, Elisabeth Trawinski, Jean-Michel Pawlotsky |
| EPI_ISL_2228669 | HOPITAL EMILE ROUX | Department of Virology, Henri Mondor University Hospital, Assistance Publique Hôpitaux de Paris, Université Paris-Est Créteil, INSERM U955 | Christophe Rodriguez, Slim Fourati, Vanessa Demontant, Guillaume Gricourt, Melissa N'Debi, Alexandre Soulier, Elisabeth Trawinski, Jean-Michel Pawlotsky |
| EPI_ISL_2228728, EPI_ISL_2228862, EPI_ISL_2228873, EPI_ISL_2228937 | GH A.CHENEVIER-H.MONDOR | Department of Virology, Henri Mondor University Hospital, Assistance Publique Hôpitaux de Paris, Université Paris-Est Créteil, INSERM U955 | Christophe Rodriguez, Slim Fourati, Vanessa Demontant, Guillaume Gricourt, Melissa N'Debi, Alexandre Soulier, Elisabeth Trawinski, Jean-Michel Pawlotsky |
| EPI_ISL_2230773, EPI_ISL_2230774, EPI_ISL_2230775, EPI_ISL_2230776, EPI_ISL_2230777, EPI_ISL_2230778, EPI_ISL_2230779, EPI_ISL_2230780, EPI_ISL_2230781, EPI_ISL_2230782, EPI_ISL_2230783, EPI_ISL_2230784, EPI_ISL_2230785, EPI_ISL_2230786, EPI_ISL_2230787, EPI_ISL_2230788, EPI_ISL_2230789, EPI_ISL_2230790, EPI_ISL_2230791, EPI_ISL_2230792 | | | |
| see above | Cliniques universitaires Saint-Luc | UCLouvain/IREC/MBLG | Jean Ruelle, Ophélie Simon, Nicolas Pinte, Benoit Kabamba Mukadi |
| EPI_ISL_2231157 | Sharp HealthCare Laboratory | Andersen lab at Scripps Research | SEARCH Alliance San Diego with Aaron Harding, Jacquelyn Berumen, Cathy Woerle, Liam McGinnis, Art Mendoza, Omid Bakhtar |
| EPI_ISL_2231565, EPI_ISL_2231566, EPI_ISL_2231567, EPI_ISL_2231568, EPI_ISL_2231569 | Battambang Provincial Laboratory | Virology Unit, Institut Pasteur du Cambodge | Jurre Y Siegers, Cecile Troupin, Leakhena Pum, Chiek Sivhour, Ly Sovann, Kraing Sidonn, Yi Sengdoeurn, Chin Savuth, Chau Darapheak, Veasna Duong, Erik A Karlsson |
| EPI_ISL_2231570, EPI_ISL_2231571, EPI_ISL_2231572 | Kampong Cham Rapid Response Team | Virology Unit, Institut Pasteur du Cambodge | Jurre Y Siegers, Cecile Troupin, Leakhena Pum, Ly Sovann, Kraing Sidonn, Yi Sengdoeurn, Chin Savuth, Chau Darapheak, Veasna Duong, Erik A Karlsson |
| EPI_ISL_2231573, EPI_ISL_2231574, EPI_ISL_2231575, EPI_ISL_2231576, EPI_ISL_2231577, EPI_ISL_2231578, EPI_ISL_2231579, EPI_ISL_2231580, EPI_ISL_2231581, EPI_ISL_2231582, EPI_ISL_2231583, EPI_ISL_2231584, EPI_ISL_2231585, EPI_ISL_2231586, EPI_ISL_2231587, EPI_ISL_2231588 | | | |
| see above | Virology Unit, Institut Pasteur du Cambodge | Virology Unit, Institut Pasteur du Cambodge | Jurre Y Siegers, Cecile Troupin, Leakhena Pum, Ly Sovann, Kraing Sidonn, Yi Sengdoeurn, Chin Savuth, Chau Darapheak, Veasna Duong, Erik A Karlsson |
| EPI_ISL_2232923 | Institute for Laboratory Diagnostics and Microbiology, Klinikum Klagenfurt am Wörthersee | Bergthaler laboratory, CeMM Research Center for Molecular Medicine of the Austrian Academy of Sciences | Lukas Endler, Anna Schedl, Fabian Amman, Petr Triska, Thomas Penz, Benedikt Agerer, Maelle Le Moing, Michael Schuster, Bekir Erguner, Jan Laine, Martin Senekowitsch, Christoph Bock, Andreas Bergthaler |
| EPI_ISL_2233087 | Hotel Horison | National Institute of Health Research and Development | Hartanti Dian Ikawati, Hana Apsari Pawestri, Subangkit, Kartika Dewi Puspa, Arie Ardiansyah Nugraha, Triyani Soekarso, Krisna Pangesti, Nelly Puspandari, Vivi Setiawaty |
| EPI_ISL_2233383 | Gribbles Pathology Malaysia | Institute for Medical Research, Infectious Disease Research Centre, National Institutes of Health, Ministry of Health Malaysia | Suppiah J, Kamel K, Mohd Zawawi Z, Azizan MA, Ramly N, Robert F, Thayan R |
| EPI_ISL_2235294, EPI_ISL_2235308, EPI_ISL_2235309 | AZ Zeno Knokke-Blankenberge | AZ SINT-JAN BRUGGE | Marijke Reynders, Patrick Descheemaeker, Laurien Hoornaert, Thomas Van Landschoot, Jorn Hellemans, Marie Madeleine Chabert-Consen |
| EPI_ISL_2235407 | DIP. PREV. AVEZZANO SERVIZIO DI IGIENE EPIDEMIOLOGIA E SANITA' PUBBLICA | Istituto Zooprofilattico Sperimentale dell'Abruzzo e Molise "G. Caporale" | Lorusso A, Marcacci M, Di Domenico M, Ancora M, Curini V, Di Lollo Valeria, Mangone I, Rinaldi A, Delli Compagni E, Scialabba S, Caporale M, Di Pasquale A, Cammà C, Puglia I, Calistri P, Savini G |
| EPI_ISL_2235408 | USCA Pescina | Istituto Zooprofilattico Sperimentale dell'Abruzzo e Molise "G. Caporale" | Lorusso A, Marcacci M, Di Domenico M, Ancora M, Curini V, Di Lollo Valeria, Mangone I, Rinaldi A, Delli Compagni E, Scialabba S, Caporale M, Di Pasquale A, Cammà C, Puglia I, Calistri P, Savini G |
| EPI_ISL_2235409 | SIESP CHIETI - DRIVE IN CHIETI | Istituto Zooprofilattico Sperimentale dell'Abruzzo e Molise "G. Caporale" | Lorusso A, Marcacci M, Di Domenico M, Ancora M, Curini V, Di Lollo Valeria, Mangone I, Rinaldi A, Delli Compagni E, Scialabba S, Caporale M, Di Pasquale A, Cammà C, Puglia I, Calistri P, Savini G |
| EPI_ISL_2235410 | SIESP CHIETI - DRIVE IN GISSI | Istituto Zooprofilattico Sperimentale dell'Abruzzo e Molise "G. Caporale" | Lorusso A, Marcacci M, Di Domenico M, Ancora M, Curini V, Di Lollo Valeria, Mangone I, Rinaldi A, Delli Compagni E, Scialabba S, Caporale M, Di Pasquale A, Cammà C, Puglia I, Calistri P, Savini G |
| EPI_ISL_2235411 | DIP. PREV. AVEZZANO SERVIZIO DI IGIENE EPIDEMIOLOGIA E SANITA' PUBBLICA | Istituto Zooprofilattico Sperimentale dell'Abruzzo e Molise "G. Caporale" | Lorusso A, Marcacci M, Di Domenico M, Ancora M, Curini V, Di Lollo Valeria, Mangone I, Rinaldi A, Delli Compagni E, Scialabba S, Caporale M, Di Pasquale A, Cammà C, Puglia I, Calistri P, Savini G |
| EPI_ISL_2235412, EPI_ISL_2235413 | SIESP CHIETI - DRIVE IN VASTO | Istituto Zooprofilattico Sperimentale dell'Abruzzo e Molise "G. Caporale" | Lorusso A, Marcacci M, Di Domenico M, Ancora M, Curini V, Di Lollo Valeria, Mangone I, Rinaldi A, Delli Compagni E, Scialabba S, Caporale M, Di Pasquale A, Cammà C, Puglia I, Calistri P, Savini G |
| EPI_ISL_2235414, EPI_ISL_2235415, EPI_ISL_2235416 | SIESP DIPARTIMENTO DI PREVENZIONE TERAMO | Istituto Zooprofilattico Sperimentale dell'Abruzzo e Molise "G. Caporale" | Lorusso A, Marcacci M, Di Domenico M, Ancora M, Curini V, Di Lollo Valeria, Mangone I, Rinaldi A, Delli Compagni E, Scialabba S, Caporale M, Di Pasquale A, Cammà C, Puglia I, Calistri P, Savini G |
| EPI_ISL_2235417 | DIP. PREV. AVEZZANO SERVIZIO DI IGIENE EPIDEMIOLOGIA E SANITA' PUBBLICA | Istituto Zooprofilattico Sperimentale dell'Abruzzo e Molise "G. Caporale" | Lorusso A, Marcacci M, Di Domenico M, Ancora M, Curini V, Di Lollo Valeria, Mangone I, Rinaldi A, Delli Compagni E, Scialabba S, Caporale M, Di Pasquale A, Cammà C, Puglia I, Calistri P, Savini G |
| EPI_ISL_2235418, EPI_ISL_2235419 | SIESP DIPARTIMENTO DI PREVENZIONE TERAMO | Istituto Zooprofilattico Sperimentale dell'Abruzzo e Molise "G. Caporale" | Lorusso A, Marcacci M, Di Domenico M, Ancora M, Curini V, Di Lollo Valeria, Mangone I, Rinaldi A, Delli Compagni E, Scialabba S, Caporale M, Di Pasquale A, Cammà C, Puglia I, Calistri P, Savini G |
| EPI_ISL_2235420, EPI_ISL_2235421, EPI_ISL_2235422 | SIESP DIPARTIMENTO DI PREVENZIONE CHIETI | Istituto Zooprofilattico Sperimentale dell'Abruzzo e Molise "G. Caporale" | Lorusso A, Marcacci M, Di Domenico M, Ancora M, Curini V, Di Lollo Valeria, Mangone I, Rinaldi A, Delli Compagni E, Scialabba S, Caporale M, Di Pasquale A, Cammà C, Puglia I, Calistri P, Savini G |
| EPI_ISL_2235423 | SIESP CHIETI - DRIVE IN VASTO | Istituto Zooprofilattico Sperimentale dell'Abruzzo e Molise "G. Caporale" | Lorusso A, Marcacci M, Di Domenico M, Ancora M, Curini V, Di Lollo Valeria, Mangone I, Rinaldi A, Delli Compagni E, Scialabba S, Caporale M, Di Pasquale A, Cammà C, Puglia I, Calistri P, Savini G |
| EPI_ISL_2235424 | SIESP CHIETI - DRIVE IN LANCIANO | Istituto Zooprofilattico Sperimentale dell'Abruzzo e Molise "G. Caporale" | Lorusso A, Marcacci M, Di Domenico M, Ancora M, Curini V, Di Lollo Valeria, Mangone I, Rinaldi A, Delli Compagni E, Scialabba S, Caporale M, Di Pasquale A, Cammà C, Puglia I, Calistri P, Savini G |

| | | | |
|---|---|---|---|
| EPI_ISL_2235425 | SIESP CHIETI - DRIVE IN GISSI | Istituto Zooprofilattico Sperimentale dell'Abruzzo e Molise "G. Caporale" | Lorusso A, Marcacci M, Di Domenico M, Ancora M, Curini V, Di Lollo Valeria, Mangone I, Rinaldi A, Delli Compagni E, Scialabba S, Caporale M, Di Pasquale A, Cammà C, Puglia I, Calistri P, Savini G |
| EPI_ISL_2235426, EPI_ISL_2235427 | SIESP DIPARTIMENTO DI PREVENZIONE CHIETI | Istituto Zooprofilattico Sperimentale dell'Abruzzo e Molise "G. Caporale" | Lorusso A, Marcacci M, Di Domenico M, Ancora M, Curini V, Di Lollo Valeria, Mangone I, Rinaldi A, Delli Compagni E, Scialabba S, Caporale M, Di Pasquale A, Cammà C, Puglia I, Calistri P, Savini G |
| EPI_ISL_2235428 | SIESP SULMONA | Istituto Zooprofilattico Sperimentale dell'Abruzzo e Molise "G. Caporale" | Lorusso A, Marcacci M, Di Domenico M, Ancora M, Curini V, Di Lollo Valeria, Mangone I, Rinaldi A, Delli Compagni E, Scialabba S, Caporale M, Di Pasquale A, Cammà C, Puglia I, Calistri P, Savini G |
| EPI_ISL_2235429, EPI_ISL_2235430 | USCA Avezzano | Istituto Zooprofilattico Sperimentale dell'Abruzzo e Molise "G. Caporale" | Lorusso A, Marcacci M, Di Domenico M, Ancora M, Curini V, Di Lollo Valeria, Mangone I, Rinaldi A, Delli Compagni E, Scialabba S, Caporale M, Di Pasquale A, Cammà C, Puglia I, Calistri P, Savini G |
| EPI_ISL_2235431 | USCA Sulmona | Istituto Zooprofilattico Sperimentale dell'Abruzzo e Molise "G. Caporale" | Lorusso A, Marcacci M, Di Domenico M, Ancora M, Curini V, Di Lollo Valeria, Mangone I, Rinaldi A, Delli Compagni E, Scialabba S, Caporale M, Di Pasquale A, Cammà C, Puglia I, Calistri P, Savini G |
| EPI_ISL_2235432 | SIESP CHIETI - DRIVE IN LANCIANO | Istituto Zooprofilattico Sperimentale dell'Abruzzo e Molise "G. Caporale" | Lorusso A, Marcacci M, Di Domenico M, Ancora M, Curini V, Di Lollo Valeria, Mangone I, Rinaldi A, Delli Compagni E, Scialabba S, Caporale M, Di Pasquale A, Cammà C, Puglia I, Calistri P, Savini G |
| EPI_ISL_2235433, EPI_ISL_2235434, EPI_ISL_2235435 | SIESP CHIETI - DRIVE IN CHIETI | Istituto Zooprofilattico Sperimentale dell'Abruzzo e Molise "G. Caporale" | Lorusso A, Marcacci M, Di Domenico M, Ancora M, Curini V, Di Lollo Valeria, Mangone I, Rinaldi A, Delli Compagni E, Scialabba S, Caporale M, Di Pasquale A, Cammà C, Puglia I, Calistri P, Savini G |
| EPI_ISL_2235436 | SIESP CHIETI - DRIVE IN LANCIANO | Istituto Zooprofilattico Sperimentale dell'Abruzzo e Molise "G. Caporale" | Lorusso A, Marcacci M, Di Domenico M, Ancora M, Curini V, Di Lollo Valeria, Mangone I, Rinaldi A, Delli Compagni E, Scialabba S, Caporale M, Di Pasquale A, Cammà C, Puglia I, Calistri P, Savini G |
| EPI_ISL_2235439, EPI_ISL_2235440, EPI_ISL_2235441, EPI_ISL_2235442, EPI_ISL_2235443, EPI_ISL_2235444, EPI_ISL_2235446, EPI_ISL_2235447, EPI_ISL_2235448, EPI_ISL_2235449, EPI_ISL_2235450, EPI_ISL_2235451, EPI_ISL_2235452, EPI_ISL_2235453, EPI_ISL_2235454, EPI_ISL_2235455, EPI_ISL_2235456, EPI_ISL_2235457, EPI_ISL_2235458, EPI_ISL_2235459, EPI_ISL_2235460, EPI_ISL_2235461, EPI_ISL_2235462, EPI_ISL_2235463 | | | |
| see above | Università degli Studi di Perugia | Istituto Zooprofilattico Sperimentale dell'Abruzzo e Molise "G. Caporale" | Mencacci A, Camilloni B, Lorusso A, Marcacci M, Di Domenico M, Ancora M, Curini V, Di Lollo Valeria, Mangone I, Rinaldi A, Delli Compagni E, Scialabba S, Caporale M, Di Pasquale A, Cammà C, Puglia I, Calistri P, Savini G |
| EPI_ISL_2235473, EPI_ISL_2235474, EPI_ISL_2235476, EPI_ISL_2235477, EPI_ISL_2235478, EPI_ISL_2235479, EPI_ISL_2235480, EPI_ISL_2235481, EPI_ISL_2235482, EPI_ISL_2235483, EPI_ISL_2235485 | | | |
| see above | Azienda Ospedaliera Terni | Istituto Zooprofilattico Sperimentale dell'Abruzzo e Molise "G. Caporale" | Palumbo M, Scaccetti A, Lorusso A, Marcacci M, Di Domenico M, Ancora M, Curini V, Di Lollo Valeria, Mangone I, Rinaldi A, Delli Compagni E, Scialabba S, Caporale M, Di Pasquale A, Cammà C, Puglia I, Calistri P, Savini G |
| EPI_ISL_2235488 | Azienda USL Umbria 5 | Istituto Zooprofilattico Sperimentale dell'Abruzzo e Molise "G. Caporale" | Proietti A, Pistoni E, Lorusso A, Marcacci M, Di Domenico M, Ancora M, Curini V, Di Lollo Valeria, Mangone I, Rinaldi A, Delli Compagni E, Scialabba S, Caporale M, Di Pasquale A, Cammà C, Puglia I, Calistri P, Savini G |
| EPI_ISL_2235489 | Azienda USL Umbria 6 | Istituto Zooprofilattico Sperimentale dell'Abruzzo e Molise "G. Caporale" | Proietti A, Pistoni E, Lorusso A, Marcacci M, Di Domenico M, Ancora M, Curini V, Di Lollo Valeria, Mangone I, Rinaldi A, Delli Compagni E, Scialabba S, Caporale M, Di Pasquale A, Cammà C, Puglia I, Calistri P, Savini G |
| EPI_ISL_2235490 | Azienda USL Umbria 7 | Istituto Zooprofilattico Sperimentale dell'Abruzzo e Molise "G. Caporale" | Proietti A, Pistoni E, Lorusso A, Marcacci M, Di Domenico M, Ancora M, Curini V, Di Lollo Valeria, Mangone I, Rinaldi A, Delli Compagni E, Scialabba S, Caporale M, Di Pasquale A, Cammà C, Puglia I, Calistri P, Savini G |
| EPI_ISL_2235491 | Azienda USL Umbria 8 | Istituto Zooprofilattico Sperimentale dell'Abruzzo e Molise "G. Caporale" | Proietti A, Pistoni E, Lorusso A, Marcacci M, Di Domenico M, Ancora M, Curini V, Di Lollo Valeria, Mangone I, Rinaldi A, Delli Compagni E, Scialabba S, Caporale M, Di Pasquale A, Cammà C, Puglia I, Calistri P, Savini G |
| EPI_ISL_2235492 | Azienda USL Umbria 9 | Istituto Zooprofilattico Sperimentale dell'Abruzzo e Molise "G. Caporale" | Proietti A, Pistoni E, Lorusso A, Marcacci M, Di Domenico M, Ancora M, Curini V, Di Lollo Valeria, Mangone I, Rinaldi A, Delli Compagni E, Scialabba S, Caporale M, Di Pasquale A, Cammà C, Puglia I, Calistri P, Savini G |
| EPI_ISL_2235493 | Azienda USL Umbria 10 | Istituto Zooprofilattico Sperimentale dell'Abruzzo e Molise "G. Caporale" | Proietti A, Pistoni E, Lorusso A, Marcacci M, Di Domenico M, Ancora M, Curini V, Di Lollo Valeria, Mangone I, Rinaldi A, Delli Compagni E, Scialabba S, Caporale M, Di Pasquale A, Cammà C, Puglia I, Calistri P, Savini G |
| EPI_ISL_2245014 | Università degli Studi di Perugia | Istituto Zooprofilattico Sperimentale dell'Abruzzo e Molise "G. Caporale" | Mencacci A, Camilloni B, Lorusso A, Marcacci M, Di Domenico M, Ancora M, Curini V, Di Lollo Valeria, Mangone I, Rinaldi A, Delli Compagni E, Scialabba S, Caporale M, Di Pasquale A, Cammà C, Puglia I, Calistri P, Savini G |
| EPI_ISL_2245017 | Basurto University Hospital: Clinical Microbiology Laboratory | Biocruces Bizkaia | Mikel Urrutikoetxea-Gutiérrez, Mª Carmen Nieto Toboso, Estibaliz Ugalde Zarraga, Mikel Gallego Rodrigo, Ana de la Hoz, José Luis Díaz de Tuesta del Arco |
| EPI_ISL_2246272, EPI_ISL_2246280, EPI_ISL_2246281, EPI_ISL_2246283 | Department of Virology, Istituto Zooprofilattico Sperimentale del Lazio e della Toscana (IZSLT) | Department of General Diagnostics; Department of Virology; Istituto Zooprofilattico Sperimentale del Lazio e della Toscana (IZSLT) | Patricia Alba, Giuseppe Manna, Elena L. Diaconu, Fabiola Feltrin, Raffaela Conti, Teresa Scicluna, Virginia Carfora, Antonella Cersini, Alessia Franco, Antonio Battisti. |
| EPI_ISL_2246829, EPI_ISL_2246830, EPI_ISL_2246831, EPI_ISL_2246832, EPI_ISL_2246833 | LESP Michoacan | Instituto de Diagnostico y Referencia Epidemiologicos (INDRE) | Claudia Wong-Arambula, Abril Rodriguez-Maldonado, Vanessa Rivero-Arredondo, Ariadna Medina-Benitez, Joaquin Quiroz-Mercado, Sergio Rangel-Guerrero, Natividad Cruz-Ortiz, Tatiana Nunez-Garcia, Gisela Barrera-Badillo, Lucia Hernandez-Rivas, Irma Lopez-Martinez, Ernesto Ramirez-Gonzalez. |
| EPI_ISL_2246834, EPI_ISL_2246835, EPI_ISL_2246836, EPI_ISL_2246837 | LESP Sinaloa | Instituto de Diagnostico y Referencia Epidemiologicos (INDRE) | Claudia Wong-Arambula, Abril Rodriguez-Maldonado, Vanessa Rivero-Arredondo, Ariadna Medina-Benitez, Joaquin Quiroz-Mercado, Sergio Rangel-Guerrero, Natividad Cruz-Ortiz, Tatiana Nunez-Garcia, Gisela Barrera-Badillo, Lucia Hernandez-Rivas, Irma Lopez-Martinez, Ernesto Ramirez-Gonzalez. |
| EPI_ISL_2246838 | LESP Ciudad de Mexico | Instituto de Diagnostico y Referencia Epidemiologicos (INDRE) | Claudia Wong-Arambula, Abril Rodriguez-Maldonado, Vanessa Rivero-Arredondo, Ariadna Medina-Benitez, Joaquin Quiroz-Mercado, Sergio Rangel-Guerrero, Natividad Cruz-Ortiz, Tatiana Nunez-Garcia, Gisela Barrera-Badillo, Lucia Hernandez-Rivas, Irma Lopez-Martinez, Ernesto Ramirez-Gonzalez. |
| EPI_ISL_2246839 | LESP Nuevo Leon | Instituto de Diagnostico y Referencia Epidemiologicos (INDRE) | Claudia Wong-Arambula, Abril Rodriguez-Maldonado, Vanessa Rivero-Arredondo, Ariadna Medina-Benitez, Joaquin Quiroz-Mercado, Sergio Rangel-Guerrero, Natividad Cruz-Ortiz, Tatiana Nunez-Garcia, Gisela Barrera-Badillo, Lucia Hernandez-Rivas, Irma Lopez-Martinez, Ernesto Ramirez-Gonzalez. |
| EPI_ISL_2249246, EPI_ISL_2249247, EPI_ISL_2249249, EPI_ISL_2249252, EPI_ISL_2249253 | Hospital Sharp | Microbial Genomics Laboratory | Bruno Gomez-Gil, Julissa Enciso-Ibarra, Alejandra Garcia-Gasca, Daniel Fregoso-Rueda |
| EPI_ISL_2264784, EPI_ISL_2264785 | Ospedale Vito Fazzi | Istituto Zooprofilattico Sperimentale della Puglia e della Basilicata | Parisi A., Bianco A., Capozzi L., Del Sambro L., Simone D., Difato L., Lobreglio G. |
| EPI_ISL_2264786, EPI_ISL_2264789, EPI_ISL_2264791, EPI_ISL_2264793, EPI_ISL_2264795, EPI_ISL_2264797 | Ospedale di Venere | Istituto Zooprofilattico Sperimentale della Puglia e della Basilicata | Parisi A., Bianco A., Capozzi L., Del Sambro L., Simone D., Iacobellis M., Ridolfi D., Giannico A. |
| EPI_ISL_2264799, EPI_ISL_2264802, EPI_ISL_2264804, EPI_ISL_2264810, EPI_ISL_2264812 | Ospedale Santa Caterina Novella | Istituto Zooprofilattico Sperimentale della Puglia e della Basilicata | Parisi A., Bianco A., Capozzi L., Del Sambro L., Simone D., Difato L., Bruno A. R. |
| EPI_ISL_2264814, EPI_ISL_2264815, EPI_ISL_2264817, EPI_ISL_2264819, EPI_ISL_2264821, EPI_ISL_2264823, EPI_ISL_2264824, EPI_ISL_2264825, EPI_ISL_2264826, EPI_ISL_2264828, EPI_ISL_2264831 | | | |
| see above | Presidio di Brindisi Di Summa - Perrino | Istituto Zooprofilattico Sperimentale della Puglia e della Basilicata | Parisi A., Bianco A., Capozzi L., Del Sambro L., Simone D., Difato L., Santoro A., Ridolfi D., Giannico A. |
| EPI_ISL_2264833, EPI_ISL_2264834, EPI_ISL_2264836, EPI_ISL_2264838, | Ospedale Vito Fazzi | Istituto Zooprofilattico Sperimentale della Puglia e della Basilicata | Parisi A., Bianco A., Capozzi L., Del Sambro L., Simone D., Difato L., Lobreglio G. |

| | | | |
|---|---|---|---|
| EPI_ISL_2264840, EPI_ISL_2264842, EPI_ISL_2264844, EPI_ISL_2264846, EPI_ISL_2264848, EPI_ISL_2264850 | | | |
| EPI_ISL_2264852, EPI_ISL_2264854, EPI_ISL_2264856, EPI_ISL_2264858 | I.R.C.C.S. "S. De Bellis" - Ente Ospedaliero | Istituto Zooprofilattico Sperimentale della Puglia e della Basilicata | Parisi A., Bianco A., Capozzi L., Del Sambro L., Simone D., Lippolis A., Notarnicola M. |
| EPI_ISL_2264859, EPI_ISL_2264861, EPI_ISL_2264863, EPI_ISL_2264865, EPI_ISL_2264871, EPI_ISL_2264873, EPI_ISL_2264877, EPI_ISL_2264879, EPI_ISL_2264880 | Ospedale Santissima Annunziata | Istituto Zooprofilattico Sperimentale della Puglia e della Basilicata | Parisi A., Bianco A., Capozzi L., Del Sambro L., Simone D., Difato L., Ridolfi D., Giannico A, Tampoia M. |
| EPI_ISL_2264884, EPI_ISL_2264886 | Ospedale Generale Regionale F. Miulli | Istituto Zooprofilattico Sperimentale della Puglia e della Basilicata | Parisi A., Bianco A., Capozzi L., Del Sambro L., Simone D., Ceci E. |
| EPI_ISL_2264888, EPI_ISL_2264893 | Ospedale Casa Sollievo della Sofferenza | Istituto Zooprofilattico Sperimentale della Puglia e della Basilicata | Parisi A., Bianco A., Capozzi L., Del Sambro L., Simone D., Difato L., Ridolfi D., Giannico A, Labonia M. |
| EPI_ISL_2270836 | DIP. PREV. AVEZZANO SERVIZIO DI IGIENE EPIDEMIOLOGIA E SANITA' PUBBLICA | Istituto Zooprofilattico Sperimentale dell'Abruzzo e Molise "G. Caporale" | Lorusso A, Marcacci M, Di Domenico M, Ancora M, Curini V, Di Lollo Valeria, Mangone I, Rinaldi A, Delli Compagni E, Scialabba S, Caporale M, Di Pasquale A, Cammà C, Puglia I, Calistri P, Savini G |
| EPI_ISL_2270837, EPI_ISL_2270838, EPI_ISL_2270839, EPI_ISL_2270840, EPI_ISL_2270841, EPI_ISL_2270842 | SIESP DIPARTIMENTO DI PREVENZIONE TERAMO | Istituto Zooprofilattico Sperimentale dell'Abruzzo e Molise "G. Caporale" | Lorusso A, Marcacci M, Di Domenico M, Ancora M, Curini V, Di Lollo Valeria, Mangone I, Rinaldi A, Delli Compagni E, Scialabba S, Caporale M, Di Pasquale A, Cammà C, Puglia I, Calistri P, Savini G |
| EPI_ISL_2270843, EPI_ISL_2270844 | DIP. PREV. AVEZZANO SERVIZIO DI IGIENE EPIDEMIOLOGIA E SANITA' PUBBLICA | Istituto Zooprofilattico Sperimentale dell'Abruzzo e Molise "G. Caporale" | Lorusso A, Marcacci M, Di Domenico M, Ancora M, Curini V, Di Lollo Valeria, Mangone I, Rinaldi A, Delli Compagni E, Scialabba S, Caporale M, Di Pasquale A, Cammà C, Puglia I, Calistri P, Savini G |
| EPI_ISL_2270845 | SIESP CHIETI - DRIVE IN LANCIANO | Istituto Zooprofilattico Sperimentale dell'Abruzzo e Molise "G. Caporale" | Lorusso A, Marcacci M, Di Domenico M, Ancora M, Curini V, Di Lollo Valeria, Mangone I, Rinaldi A, Delli Compagni E, Scialabba S, Caporale M, Di Pasquale A, Cammà C, Puglia I, Calistri P, Savini G |
| EPI_ISL_2270846 | PRESIDIO OSPEDALIERO TAGLIACOZZO | Istituto Zooprofilattico Sperimentale dell'Abruzzo e Molise "G. Caporale" | Lorusso A, Marcacci M, Di Domenico M, Ancora M, Curini V, Di Lollo Valeria, Mangone I, Rinaldi A, Delli Compagni E, Scialabba S, Caporale M, Di Pasquale A, Cammà C, Puglia I, Calistri P, Savini G |
| EPI_ISL_2270847, EPI_ISL_2270848 | SIESP CHIETI - DRIVE IN ORTONA | Istituto Zooprofilattico Sperimentale dell'Abruzzo e Molise "G. Caporale" | Lorusso A, Marcacci M, Di Domenico M, Ancora M, Curini V, Di Lollo Valeria, Mangone I, Rinaldi A, Delli Compagni E, Scialabba S, Caporale M, Di Pasquale A, Cammà C, Puglia I, Calistri P, Savini G |
| EPI_ISL_2270849 | SIESP CHIETI - DRIVE IN VASTO | Istituto Zooprofilattico Sperimentale dell'Abruzzo e Molise "G. Caporale" | Lorusso A, Marcacci M, Di Domenico M, Ancora M, Curini V, Di Lollo Valeria, Mangone I, Rinaldi A, Delli Compagni E, Scialabba S, Caporale M, Di Pasquale A, Cammà C, Puglia I, Calistri P, Savini G |
| EPI_ISL_2270850 | SIESP DIPARTIMENTO DI PREVENZIONE TERAMO | Istituto Zooprofilattico Sperimentale dell'Abruzzo e Molise "G. Caporale" | Lorusso A, Marcacci M, Di Domenico M, Ancora M, Curini V, Di Lollo Valeria, Mangone I, Rinaldi A, Delli Compagni E, Scialabba S, Caporale M, Di Pasquale A, Cammà C, Puglia I, Calistri P, Savini G |
| EPI_ISL_2270851 | USCA Castel di Sangro | Istituto Zooprofilattico Sperimentale dell'Abruzzo e Molise "G. Caporale" | Lorusso A, Marcacci M, Di Domenico M, Ancora M, Curini V, Di Lollo Valeria, Mangone I, Rinaldi A, Delli Compagni E, Scialabba S, Caporale M, Di Pasquale A, Cammà C, Puglia I, Calistri P, Savini G |
| EPI_ISL_2270852, EPI_ISL_2270853 | USCA Pescina | Istituto Zooprofilattico Sperimentale dell'Abruzzo e Molise "G. Caporale" | Lorusso A, Marcacci M, Di Domenico M, Ancora M, Curini V, Di Lollo Valeria, Mangone I, Rinaldi A, Delli Compagni E, Scialabba S, Caporale M, Di Pasquale A, Cammà C, Puglia I, Calistri P, Savini G |
| EPI_ISL_2270854 | SIESP CHIETI - DRIVE IN VASTO | Istituto Zooprofilattico Sperimentale dell'Abruzzo e Molise "G. Caporale" | Lorusso A, Marcacci M, Di Domenico M, Ancora M, Curini V, Di Lollo Valeria, Mangone I, Rinaldi A, Delli Compagni E, Scialabba S, Caporale M, Di Pasquale A, Cammà C, Puglia I, Calistri P, Savini G |
| EPI_ISL_2270855, EPI_ISL_2270856 | SIESP CHIETI - DRIVE IN CHIETI | Istituto Zooprofilattico Sperimentale dell'Abruzzo e Molise "G. Caporale" | Lorusso A, Marcacci M, Di Domenico M, Ancora M, Curini V, Di Lollo Valeria, Mangone I, Rinaldi A, Delli Compagni E, Scialabba S, Caporale M, Di Pasquale A, Cammà C, Puglia I, Calistri P, Savini G |
| EPI_ISL_2270857, EPI_ISL_2270858 | SIESP DIPARTIMENTO DI PREVENZIONE CHIETI | Istituto Zooprofilattico Sperimentale dell'Abruzzo e Molise "G. Caporale" | Lorusso A, Marcacci M, Di Domenico M, Ancora M, Curini V, Di Lollo Valeria, Mangone I, Rinaldi A, Delli Compagni E, Scialabba S, Caporale M, Di Pasquale A, Cammà C, Puglia I, Calistri P, Savini G |
| EPI_ISL_2270859, EPI_ISL_2270860, EPI_ISL_2270861 | SIESP CHIETI - DRIVE IN VASTO | Istituto Zooprofilattico Sperimentale dell'Abruzzo e Molise "G. Caporale" | Lorusso A, Marcacci M, Di Domenico M, Ancora M, Curini V, Di Lollo Valeria, Mangone I, Rinaldi A, Delli Compagni E, Scialabba S, Caporale M, Di Pasquale A, Cammà C, Puglia I, Calistri P, Savini G |
| EPI_ISL_2270862, EPI_ISL_2270863, EPI_ISL_2270864 | SIESP DIPARTIMENTO DI PREVENZIONE TERAMO | Istituto Zooprofilattico Sperimentale dell'Abruzzo e Molise "G. Caporale" | Lorusso A, Marcacci M, Di Domenico M, Ancora M, Curini V, Di Lollo Valeria, Mangone I, Rinaldi A, Delli Compagni E, Scialabba S, Caporale M, Di Pasquale A, Cammà C, Puglia I, Calistri P, Savini G |
| EPI_ISL_2270865, EPI_ISL_2270866, EPI_ISL_2270867 | SIESP DIPARTIMENTO DI PREVENZIONE CHIETI | Istituto Zooprofilattico Sperimentale dell'Abruzzo e Molise "G. Caporale" | Lorusso A, Marcacci M, Di Domenico M, Ancora M, Curini V, Di Lollo Valeria, Mangone I, Rinaldi A, Delli Compagni E, Scialabba S, Caporale M, Di Pasquale A, Cammà C, Puglia I, Calistri P, Savini G |
| EPI_ISL_2270868 | SIESP SULMONA | Istituto Zooprofilattico Sperimentale dell'Abruzzo e Molise "G. Caporale" | Lorusso A, Marcacci M, Di Domenico M, Ancora M, Curini V, Di Lollo Valeria, Mangone I, Rinaldi A, Delli Compagni E, Scialabba S, Caporale M, Di Pasquale A, Cammà C, Puglia I, Calistri P, Savini G |
| EPI_ISL_2270869, EPI_ISL_2270870, EPI_ISL_2270871, EPI_ISL_2270872, EPI_ISL_2270873 | DIP. PREV. AVEZZANO SERVIZIO DI IGIENE EPIDEMIOLOGIA E SANITA' PUBBLICA | Istituto Zooprofilattico Sperimentale dell'Abruzzo e Molise "G. Caporale" | Lorusso A, Marcacci M, Di Domenico M, Ancora M, Curini V, Di Lollo Valeria, Mangone I, Rinaldi A, Delli Compagni E, Scialabba S, Caporale M, Di Pasquale A, Cammà C, Puglia I, Calistri P, Savini G |
| EPI_ISL_2270874, EPI_ISL_2270875 | SIESP CHIETI - DRIVE IN LANCIANO | Istituto Zooprofilattico Sperimentale dell'Abruzzo e Molise "G. Caporale" | Lorusso A, Marcacci M, Di Domenico M, Ancora M, Curini V, Di Lollo Valeria, Mangone I, Rinaldi A, Delli Compagni E, Scialabba S, Caporale M, Di Pasquale A, Cammà C, Puglia I, Calistri P, Savini G |
| EPI_ISL_2270876, EPI_ISL_2270877, EPI_ISL_2270878, EPI_ISL_2270879, EPI_ISL_2270880 | SIESP DIPARTIMENTO DI PREVENZIONE TERAMO | Istituto Zooprofilattico Sperimentale dell'Abruzzo e Molise "G. Caporale" | Lorusso A, Marcacci M, Di Domenico M, Ancora M, Curini V, Di Lollo Valeria, Mangone I, Rinaldi A, Delli Compagni E, Scialabba S, Caporale M, Di Pasquale A, Cammà C, Puglia I, Calistri P, Savini G |
| EPI_ISL_2270881 | DIP. PREV. AVEZZANO SERVIZIO DI IGIENE EPIDEMIOLOGIA E SANITA' PUBBLICA | Istituto Zooprofilattico Sperimentale dell'Abruzzo e Molise "G. Caporale" | Lorusso A, Marcacci M, Di Domenico M, Ancora M, Curini V, Di Lollo Valeria, Mangone I, Rinaldi A, Delli Compagni E, Scialabba S, Caporale M, Di Pasquale A, Cammà C, Puglia I, Calistri P, Savini G |
| EPI_ISL_2270882, EPI_ISL_2270883 | SIESP CHIETI - DRIVE IN CHIETI | Istituto Zooprofilattico Sperimentale dell'Abruzzo e Molise "G. Caporale" | Lorusso A, Marcacci M, Di Domenico M, Ancora M, Curini V, Di Lollo Valeria, Mangone I, Rinaldi A, Delli Compagni E, Scialabba S, Caporale M, Di Pasquale A, Cammà C, Puglia I, Calistri P, Savini G |
| EPI_ISL_2270884 | USCA Tagliacozzo | Istituto Zooprofilattico Sperimentale dell'Abruzzo e Molise "G. Caporale" | Lorusso A, Marcacci M, Di Domenico M, Ancora M, Curini V, Di Lollo Valeria, Mangone I, Rinaldi A, Delli Compagni E, Scialabba S, Caporale M, Di Pasquale A, Cammà C, Puglia I, Calistri P, Savini G |
| EPI_ISL_2270885, EPI_ISL_2270886 | SIESP CHIETI - DRIVE IN ORTONA | Istituto Zooprofilattico Sperimentale dell'Abruzzo e Molise "G. Caporale" | Lorusso A, Marcacci M, Di Domenico M, Ancora M, Curini V, Di Lollo Valeria, Mangone I, Rinaldi A, Delli Compagni E, Scialabba S, Caporale M, Di Pasquale A, Cammà C, Puglia I, Calistri P, Savini G |
| EPI_ISL_2270887 | SIESP DIPARTIMENTO DI PREVENZIONE CHIETI | Istituto Zooprofilattico Sperimentale dell'Abruzzo e Molise "G. Caporale" | Lorusso A, Marcacci M, Di Domenico M, Ancora M, Curini V, Di Lollo Valeria, Mangone I, Rinaldi A, Delli Compagni E, Scialabba S, Caporale M, Di Pasquale A, Cammà C, Puglia I, Calistri P, Savini G |
| EPI_ISL_2270888, EPI_ISL_2270889 | SIESP DIPARTIMENTO DI PREVENZIONE TERAMO | Istituto Zooprofilattico Sperimentale dell'Abruzzo e Molise "G. Caporale" | Lorusso A, Marcacci M, Di Domenico M, Ancora M, Curini V, Di Lollo Valeria, Mangone I, Rinaldi A, Delli Compagni E, Scialabba S, Caporale M, Di Pasquale A, Cammà C, Puglia I, Calistri P, Savini G |
| EPI_ISL_2270890 | SIESP CHIETI - DRIVE IN LANCIANO | Istituto Zooprofilattico Sperimentale dell'Abruzzo e Molise "G. Caporale" | Lorusso A, Marcacci M, Di Domenico M, Ancora M, Curini V, Di Lollo Valeria, Mangone I, Rinaldi A, Delli Compagni E, Scialabba S, Caporale M, Di Pasquale A, Cammà C, Puglia I, Calistri P, Savini G |
| EPI_ISL_2270891 | SIESP DIPARTIMENTO DI PREVENZIONE CHIETI | Istituto Zooprofilattico Sperimentale dell'Abruzzo e Molise "G. | Lorusso A, Marcacci M, Di Domenico M, Ancora M, Curini V, Di Lollo Valeria, Mangone I, Rinaldi A, Delli Compagni E, Scialabba S, Caporale M, Di |

| | | | |
|---|---|---|---|
| | | Caporale" | Pasquale A, Cammà C, Puglia I, Calistri P, Savini G |
| EPI_ISL_2270892, EPI_ISL_2270893, EPI_ISL_2270894 | DIP. PREV. AVEZZANO SERVIZIO DI IGIENE EPIDEMIOLOGIA E SANITA' PUBBLICA | Istituto Zooprofilattico Sperimentale dell'Abruzzo e Molise "G. Caporale" | Lorusso A, Marcacci M, Di Domenico M, Ancora M, Curini V, Di Lollo Valeria, Mangone I, Rinaldi A, Delli Compagni E, Scialabba S, Caporale M, Di Pasquale A, Cammà C, Puglia I, Calistri P, Savini G |
| EPI_ISL_2270895, EPI_ISL_2270896, EPI_ISL_2270897, EPI_ISL_2270898, EPI_ISL_2270899 | SIESP DIPARTIMENTO DI PREVENZIONE TERAMO | Istituto Zooprofilattico Sperimentale dell'Abruzzo e Molise "G. Caporale" | Lorusso A, Marcacci M, Di Domenico M, Ancora M, Curini V, Di Lollo Valeria, Mangone I, Rinaldi A, Delli Compagni E, Scialabba S, Caporale M, Di Pasquale A, Cammà C, Puglia I, Calistri P, Savini G |
| EPI_ISL_2270900, EPI_ISL_2270901 | SIESP CHIETI - DRIVE IN LANCIANO | Istituto Zooprofilattico Sperimentale dell'Abruzzo e Molise "G. Caporale" | Lorusso A, Marcacci M, Di Domenico M, Ancora M, Curini V, Di Lollo Valeria, Mangone I, Rinaldi A, Delli Compagni E, Scialabba S, Caporale M, Di Pasquale A, Cammà C, Puglia I, Calistri P, Savini G |
| EPI_ISL_2270902 | DIP. PREV. AVEZZANO SERVIZIO DI IGIENE EPIDEMIOLOGIA E SANITA' PUBBLICA | Istituto Zooprofilattico Sperimentale dell'Abruzzo e Molise "G. Caporale" | Lorusso A, Marcacci M, Di Domenico M, Ancora M, Curini V, Di Lollo Valeria, Mangone I, Rinaldi A, Delli Compagni E, Scialabba S, Caporale M, Di Pasquale A, Cammà C, Puglia I, Calistri P, Savini G |
| EPI_ISL_2270903 | SIESP SULMONA | Istituto Zooprofilattico Sperimentale dell'Abruzzo e Molise "G. Caporale" | Lorusso A, Marcacci M, Di Domenico M, Ancora M, Curini V, Di Lollo Valeria, Mangone I, Rinaldi A, Delli Compagni E, Scialabba S, Caporale M, Di Pasquale A, Cammà C, Puglia I, Calistri P, Savini G |
| EPI_ISL_2270904 | DIP. PREV. AVEZZANO SERVIZIO DI IGIENE EPIDEMIOLOGIA E SANITA' PUBBLICA | Istituto Zooprofilattico Sperimentale dell'Abruzzo e Molise "G. Caporale" | Lorusso A, Marcacci M, Di Domenico M, Ancora M, Curini V, Di Lollo Valeria, Mangone I, Rinaldi A, Delli Compagni E, Scialabba S, Caporale M, Di Pasquale A, Cammà C, Puglia I, Calistri P, Savini G |
| EPI_ISL_2270905, EPI_ISL_2270906 | SIESP CHIETI - DRIVE IN LANCIANO | Istituto Zooprofilattico Sperimentale dell'Abruzzo e Molise "G. Caporale" | Lorusso A, Marcacci M, Di Domenico M, Ancora M, Curini V, Di Lollo Valeria, Mangone I, Rinaldi A, Delli Compagni E, Scialabba S, Caporale M, Di Pasquale A, Cammà C, Puglia I, Calistri P, Savini G |
| EPI_ISL_2270907, EPI_ISL_2270908, EPI_ISL_2270909, EPI_ISL_2270910 | SIESP DIPARTIMENTO DI PREVENZIONE CHIETI | Istituto Zooprofilattico Sperimentale dell'Abruzzo e Molise "G. Caporale" | Lorusso A, Marcacci M, Di Domenico M, Ancora M, Curini V, Di Lollo Valeria, Mangone I, Rinaldi A, Delli Compagni E, Scialabba S, Caporale M, Di Pasquale A, Cammà C, Puglia I, Calistri P, Savini G |
| EPI_ISL_2270911 | SIESP CHIETI - DRIVE IN ORTONA | Istituto Zooprofilattico Sperimentale dell'Abruzzo e Molise "G. Caporale" | Lorusso A, Marcacci M, Di Domenico M, Ancora M, Curini V, Di Lollo Valeria, Mangone I, Rinaldi A, Delli Compagni E, Scialabba S, Caporale M, Di Pasquale A, Cammà C, Puglia I, Calistri P, Savini G |
| EPI_ISL_2270912 | SIESP DIPARTIMENTO DI PREVENZIONE CHIETI | Istituto Zooprofilattico Sperimentale dell'Abruzzo e Molise "G. Caporale" | Lorusso A, Marcacci M, Di Domenico M, Ancora M, Curini V, Di Lollo Valeria, Mangone I, Rinaldi A, Delli Compagni E, Scialabba S, Caporale M, Di Pasquale A, Cammà C, Puglia I, Calistri P, Savini G |
| EPI_ISL_2270913 | SIESP CHIETI - DRIVE IN GISSI | Istituto Zooprofilattico Sperimentale dell'Abruzzo e Molise "G. Caporale" | Lorusso A, Marcacci M, Di Domenico M, Ancora M, Curini V, Di Lollo Valeria, Mangone I, Rinaldi A, Delli Compagni E, Scialabba S, Caporale M, Di Pasquale A, Cammà C, Puglia I, Calistri P, Savini G |
| EPI_ISL_2270914 | SIESP CHIETI - DRIVE IN VASTO | Istituto Zooprofilattico Sperimentale dell'Abruzzo e Molise "G. Caporale" | Lorusso A, Marcacci M, Di Domenico M, Ancora M, Curini V, Di Lollo Valeria, Mangone I, Rinaldi A, Delli Compagni E, Scialabba S, Caporale M, Di Pasquale A, Cammà C, Puglia I, Calistri P, Savini G |
| EPI_ISL_2270915 | DIP. PREV. AVEZZANO SERVIZIO DI IGIENE EPIDEMIOLOGIA E SANITA' PUBBLICA | Istituto Zooprofilattico Sperimentale dell'Abruzzo e Molise "G. Caporale" | Lorusso A, Marcacci M, Di Domenico M, Ancora M, Curini V, Di Lollo Valeria, Mangone I, Rinaldi A, Delli Compagni E, Scialabba S, Caporale M, Di Pasquale A, Cammà C, Puglia I, Calistri P, Savini G |
| EPI_ISL_2270917, EPI_ISL_2270918 | SIESP DIPARTIMENTO DI PREVENZIONE TERAMO | Istituto Zooprofilattico Sperimentale dell'Abruzzo e Molise "G. Caporale" | Lorusso A, Marcacci M, Di Domenico M, Ancora M, Curini V, Di Lollo Valeria, Mangone I, Rinaldi A, Delli Compagni E, Scialabba S, Caporale M, Di Pasquale A, Cammà C, Puglia I, Calistri P, Savini G |
| EPI_ISL_2270919 | SIESP CHIETI - DRIVE IN ORTONA | Istituto Zooprofilattico Sperimentale dell'Abruzzo e Molise "G. Caporale" | Lorusso A, Marcacci M, Di Domenico M, Ancora M, Curini V, Di Lollo Valeria, Mangone I, Rinaldi A, Delli Compagni E, Scialabba S, Caporale M, Di Pasquale A, Cammà C, Puglia I, Calistri P, Savini G |
| EPI_ISL_2270920, EPI_ISL_2270921 | SIESP CHIETI - DRIVE IN LANCIANO | Istituto Zooprofilattico Sperimentale dell'Abruzzo e Molise "G. Caporale" | Lorusso A, Marcacci M, Di Domenico M, Ancora M, Curini V, Di Lollo Valeria, Mangone I, Rinaldi A, Delli Compagni E, Scialabba S, Caporale M, Di Pasquale A, Cammà C, Puglia I, Calistri P, Savini G |
| EPI_ISL_2270922, EPI_ISL_2270923, EPI_ISL_2270924 | SIESP SULMONA | Istituto Zooprofilattico Sperimentale dell'Abruzzo e Molise "G. Caporale" | Lorusso A, Marcacci M, Di Domenico M, Ancora M, Curini V, Di Lollo Valeria, Mangone I, Rinaldi A, Delli Compagni E, Scialabba S, Caporale M, Di Pasquale A, Cammà C, Puglia I, Calistri P, Savini G |
| EPI_ISL_2270925, EPI_ISL_2270926 | SIESP DIPARTIMENTO DI PREVENZIONE TERAMO | Istituto Zooprofilattico Sperimentale dell'Abruzzo e Molise "G. Caporale" | Lorusso A, Marcacci M, Di Domenico M, Ancora M, Curini V, Di Lollo Valeria, Mangone I, Rinaldi A, Delli Compagni E, Scialabba S, Caporale M, Di Pasquale A, Cammà C, Puglia I, Calistri P, Savini G |
| EPI_ISL_2270927 | DIP. PREV. AVEZZANO SERVIZIO DI IGIENE EPIDEMIOLOGIA E SANITA' PUBBLICA | Istituto Zooprofilattico Sperimentale dell'Abruzzo e Molise "G. Caporale" | Lorusso A, Marcacci M, Di Domenico M, Ancora M, Curini V, Di Lollo Valeria, Mangone I, Rinaldi A, Delli Compagni E, Scialabba S, Caporale M, Di Pasquale A, Cammà C, Puglia I, Calistri P, Savini G |
| EPI_ISL_2270928 | SIESP DIPARTIMENTO DI PREVENZIONE TERAMO | Istituto Zooprofilattico Sperimentale dell'Abruzzo e Molise "G. Caporale" | Lorusso A, Marcacci M, Di Domenico M, Ancora M, Curini V, Di Lollo Valeria, Mangone I, Rinaldi A, Delli Compagni E, Scialabba S, Caporale M, Di Pasquale A, Cammà C, Puglia I, Calistri P, Savini G |
| EPI_ISL_2270929 | DIP. PREV. AVEZZANO SERVIZIO DI IGIENE EPIDEMIOLOGIA E SANITA' PUBBLICA | Istituto Zooprofilattico Sperimentale dell'Abruzzo e Molise "G. Caporale" | Lorusso A, Marcacci M, Di Domenico M, Ancora M, Curini V, Di Lollo Valeria, Mangone I, Rinaldi A, Delli Compagni E, Scialabba S, Caporale M, Di Pasquale A, Cammà C, Puglia I, Calistri P, Savini G |
| EPI_ISL_2274977 | Virology Unit, AOUP | Virology Unit, AOUP | Marialinda Vatteroni, Susi Frateschi, Mauro Pistello |
| EPI_ISL_2274979 | Aeroporto Internacional de Guarulhos | Instituto Adolfo Lutz, Interdiciplinary Procedures Center, Strategic Laboratory | Claudio Tavares Sacchi, Claudia Regina Gonçalves, Erica Valessa Ramos Gomes, Karoline Rodrigues Campos, Caio Vinicius Dias Lopes, Leonardo Jose Tadeu de Araujo |
| EPI_ISL_2283694 | LESP Oaxaca | Instituto de Diagnostico y Referencia Epidemiologicos (INDRE) | Claudia Wong-Arambula, Abril Rodriguez-Maldonado, Vanessa Rivero-Arredondo, Ariadna Medina-Benitez, Joaquin Quiroz-Mercado, Sergio Rangel-Guerrero, Natividad Cruz-Ortiz, Tatiana Nunez-Garcia, Gisela Barrera-Badillo, Lucia Hernandez-Rivas, Irma Lopez-Martinez, Ernesto Ramirez-Gonzalez. |
| EPI_ISL_2283697 | LESP Colima | Instituto de Diagnostico y Referencia Epidemiologicos (INDRE) | Claudia Wong-Arambula, Abril Rodriguez-Maldonado, Vanessa Rivero-Arredondo, Ariadna Medina-Benitez, Joaquin Quiroz-Mercado, Sergio Rangel-Guerrero, Natividad Cruz-Ortiz, Tatiana Nunez-Garcia, Gisela Barrera-Badillo, Lucia Hernandez-Rivas, Irma Lopez-Martinez, Ernesto Ramirez-Gonzalez. |
| EPI_ISL_2283706, EPI_ISL_2283707, EPI_ISL_2283708 | LESP Ciudad de Mexico | Instituto de Diagnostico y Referencia Epidemiologicos (INDRE) | Claudia Wong-Arambula, Abril Rodriguez-Maldonado, Vanessa Rivero-Arredondo, Ariadna Medina-Benitez, Joaquin Quiroz-Mercado, Sergio Rangel-Guerrero, Natividad Cruz-Ortiz, Tatiana Nunez-Garcia, Gisela Barrera-Badillo, Lucia Hernandez-Rivas, Irma Lopez-Martinez, Ernesto Ramirez-Gonzalez. |
| EPI_ISL_2283722 | LESP Guanajuato | Instituto de Diagnostico y Referencia Epidemiologicos (INDRE) | Claudia Wong-Arambula, Abril Rodriguez-Maldonado, Vanessa Rivero-Arredondo, Ariadna Medina-Benitez, Joaquin Quiroz-Mercado, Sergio Rangel-Guerrero, Natividad Cruz-Ortiz, Tatiana Nunez-Garcia, Gisela Barrera-Badillo, Lucia Hernandez-Rivas, Irma Lopez-Martinez, Ernesto Ramirez-Gonzalez. |
| EPI_ISL_2285112, EPI_ISL_2285113, EPI_ISL_2285114, EPI_ISL_2285115, EPI_ISL_2285116, EPI_ISL_2285117, EPI_ISL_2285118, EPI_ISL_2285119, EPI_ISL_2285120, EPI_ISL_2285121, EPI_ISL_2285122 | | | |
| see above | Microbiology Department. Complexo Hospitalario Universitario de Vigo | Microbiology Department. Complexo Hospitalario Universitario de Vigo | Alfaya N, Alonso I, Alvarez M, Cabrera JJ, Carballo R, Cores O, Cortizo S, del-Campo V, Martinez L, Mediero G, Perez S, Potel C, Regueiro B, Rey S, Vasallo FJ |
| EPI_ISL_2285125 | Complejo Hospitalario Universitario de Pontevedra | Microbiology Department. Complexo Hospitalario Universitario de Vigo | Campello M, Perez S, Regueiro B, Trigo M |
| EPI_ISL_2285135, EPI_ISL_2285136, EPI_ISL_2285137, EPI_ISL_2285138, EPI_ISL_2285139, EPI_ISL_2285142, EPI_ISL_2285144, EPI_ISL_2285145, EPI_ISL_2285146, EPI_ISL_2285147, EPI_ISL_2285151, EPI_ISL_2285153 | | | |
| see above | Microbiology Department. Complexo Hospitalario Universitario de Vigo | Microbiology Department. Complexo Hospitalario Universitario de Vigo | Alfaya N, Alonso I, Alvarez M, Cabrera JJ, Carballo R, Cores O, Cortizo S, del-Campo V, Martinez L, Mediero G, Perez S, Potel C, Regueiro B, Rey S, Vasallo FJ |
| EPI_ISL_2285157, EPI_ISL_2285158, EPI_ISL_2285159, EPI_ISL_2285160, EPI_ISL_2285161 | VITHAS Lab | Microbiology Department. Complexo Hospitalario Universitario de Vigo | Camacho AT, Perez S, Regueiro B |
| EPI_ISL_2285364, EPI_ISL_2285365, | University of Rome Tor Vergata: Departm Experim Medicine | University of Rome Tor Vergata: Departm Experim Medicine | Maria Concetta Bellocchi, Luca Carioti, Maria Botticelli, Lorenzo Piermatteo, Rossana Scutari, Marco Iannetta, Loredana Sarmati, Massimo Andreoni, |

| | | | |
|---|---|---|---|
| EPI_ISL_2285370, EPI_ISL_2285371, EPI_ISL_2285372, EPI_ISL_2285373, EPI_ISL_2285374, EPI_ISL_2285375 | Chair of Virology | Chair of Virology | Francesca Ceccherini-Silberstein |
| EPI_ISL_2285870, EPI_ISL_2285871 | Laboratorio Biologia Molecolare Sars Cov2 - UOC Laboratorio Analisi - Servizio Medicina di Laboratorio, Ospedale "San Francesco" - ATS-ASSL Nuoro | Laboratorio specialistico UOC Ematologia - Ospedale "San Francesco" - ATS-ASSL Nuoro | Asproni Rosanna, Pira Giovanna Pira, Fancello Patrizia, Rosu Valentina, Fiamma Maura,,Garau Maria Cristina,Mameli Giuseppe |
| EPI_ISL_2285872, EPI_ISL_2285873 | Laboratorio Biologia Molecolare Sars Cov2 - UOC Laboratorio Analisi - Servizio Medicina di Laboratorio, Ospedale "San Francesco" - ATS-ASSL Nuoro | Laboratorio specialistico UOC Ematologia - Ospedale "San Francesco" - ATS-ASSL Nuoro | Asproni Rosanna, Monne Maria Itria, Piras Giovanna, Palmas Angelo Domenico |
| EPI_ISL_2293254 | Imelda Ziekenhuis | Imelda Ziekenhuis | Johan Frans, Hanne Valgaeren, Dagmar Obbels |
| EPI_ISL_2293274 | Hospital | National Reference Center for Viruses of Respiratory Infections, Institut Pasteur, Paris | Marion Barbet, Sylvie Behillil, Méline Bizard, Angela Brisebarre, Camille Capel, Vincent Enouf, Louise Lefrançois, Frédéric Lemoine, Christophe Malabat, Corinne Maufrais, Pierre Lechat, Etienne Simon-Lorière, Maud Vanpeene, Sylvie Van der Werf ,Jacques Fourgeaud |
| EPI_ISL_2293284 | Hospital | National Reference Center for Viruses of Respiratory Infections, Institut Pasteur, Paris | Marion Barbet, Sylvie Behillil, Méline Bizard, Angela Brisebarre, Camille Capel, Vincent Enouf, Louise Lefrançois, Frédéric Lemoine, Christophe Malabat, Corinne Maufrais, Pierre Lechat, Etienne Simon-Lorière, Maud Vanpeene, Sylvie Van der Werf ,Laura Djamdjian |
| EPI_ISL_2293285 | Hospital | National Reference Center for Viruses of Respiratory Infections, Institut Pasteur, Paris | Marion Barbet, Sylvie Behillil, Méline Bizard, Angela Brisebarre, Camille Capel, Vincent Enouf, Louise Lefrançois, Frédéric Lemoine, Christophe Malabat, Corinne Maufrais, Etienne Simon-Lorière, Maud Vanpeene, Sylvie Van der Werf ,Laura Djamdjian |
| EPI_ISL_2293303 | Hospital | National Reference Center for Viruses of Respiratory Infections, Institut Pasteur, Paris | Marion Barbet, Sylvie Behillil, Méline Bizard, Angela Brisebarre, Camille Capel, Vincent Enouf, Louise Lefrançois, Frédéric Lemoine, Christophe Malabat, Corinne Maufrais, Pierre Lechat, Etienne Simon-Lorière, Maud Vanpeene, Sylvie Van der Werf ,Elodie Gleize |
| EPI_ISL_2293304, EPI_ISL_2293305, EPI_ISL_2293306 | Hospital | National Reference Center for Viruses of Respiratory Infections, Institut Pasteur, Paris | Marion Barbet, Sylvie Behillil, Méline Bizard, Angela Brisebarre, Camille Capel, Vincent Enouf, Louise Lefrançois, Frédéric Lemoine, Christophe Malabat, Corinne Maufrais, Etienne Simon-Lorière, Maud Vanpeene, Sylvie Van der Werf ,Elodie Gleize |
| EPI_ISL_2293307 | Hospital | National Reference Center for Viruses of Respiratory Infections, Institut Pasteur, Paris | Marion Barbet, Sylvie Behillil, Méline Bizard, Angela Brisebarre, Camille Capel, Vincent Enouf, Louise Lefrançois, Frédéric Lemoine, Christophe Malabat, Corinne Maufrais, Pierre Lechat, Etienne Simon-Lorière, Maud Vanpeene, Sylvie Van der Werf ,Elodie Gleize |
| EPI_ISL_2293308, EPI_ISL_2293309 | Hospital | National Reference Center for Viruses of Respiratory Infections, Institut Pasteur, Paris | Marion Barbet, Sylvie Behillil, Méline Bizard, Angela Brisebarre, Camille Capel, Vincent Enouf, Louise Lefrançois, Frédéric Lemoine, Christophe Malabat, Corinne Maufrais, Etienne Simon-Lorière, Maud Vanpeene, Sylvie Van der Werf ,Elodie Gleize |
| EPI_ISL_2293313, EPI_ISL_2293314 | Hospital | National Reference Center for Viruses of Respiratory Infections, Institut Pasteur, Paris | Marion Barbet, Sylvie Behillil, Méline Bizard, Angela Brisebarre, Camille Capel, Vincent Enouf, Louise Lefrançois, Frédéric Lemoine, Christophe Malabat, Corinne Maufrais, Etienne Simon-Lorière, Maud Vanpeene, Sylvie Van der Werf ,JéRome Guinard |
| EPI_ISL_2293315 | Hospital | National Reference Center for Viruses of Respiratory Infections, Institut Pasteur, Paris | Marion Barbet, Sylvie Behillil, Méline Bizard, Angela Brisebarre, Camille Capel, Vincent Enouf, Louise Lefrançois, Frédéric Lemoine, Christophe Malabat, Corinne Maufrais, Pierre Lechat, Etienne Simon-Lorière, Maud Vanpeene, Sylvie Van der Werf ,JéRome Guinard |
| EPI_ISL_2293316 | Hospital | National Reference Center for Viruses of Respiratory Infections, Institut Pasteur, Paris | Marion Barbet, Sylvie Behillil, Méline Bizard, Angela Brisebarre, Camille Capel, Vincent Enouf, Louise Lefrançois, Frédéric Lemoine, Christophe Malabat, Corinne Maufrais, Etienne Simon-Lorière, Maud Vanpeene, Sylvie Van der Werf ,JéRome Guinard |
| EPI_ISL_2293317 | Hospital | National Reference Center for Viruses of Respiratory Infections, Institut Pasteur, Paris | Marion Barbet, Sylvie Behillil, Méline Bizard, Angela Brisebarre, Camille Capel, Vincent Enouf, Louise Lefrançois, Frédéric Lemoine, Christophe Malabat, Corinne Maufrais, Pierre Lechat, Etienne Simon-Lorière, Maud Vanpeene, Sylvie Van der Werf ,JéRome Guinard |
| EPI_ISL_2293318, EPI_ISL_2293319 | Hospital | National Reference Center for Viruses of Respiratory Infections, Institut Pasteur, Paris | Marion Barbet, Sylvie Behillil, Méline Bizard, Angela Brisebarre, Camille Capel, Vincent Enouf, Louise Lefrançois, Frédéric Lemoine, Christophe Malabat, Corinne Maufrais, Etienne Simon-Lorière, Maud Vanpeene, Sylvie Van der Werf ,JéRome Guinard |
| EPI_ISL_2293324, EPI_ISL_2293325 | Armies | National Reference Center for Viruses of Respiratory Infections, Institut Pasteur, Paris | Marion Barbet, Sylvie Behillil, Méline Bizard, Angela Brisebarre, Camille Capel, Vincent Enouf, Louise Lefrançois, Frédéric Lemoine, Christophe Malabat, Corinne Maufrais, Etienne Simon-Lorière, Maud Vanpeene, Sylvie Van der Werf ,Marine Desroches |
| EPI_ISL_2293326, EPI_ISL_2293327 | Hospital | National Reference Center for Viruses of Respiratory Infections, Institut Pasteur, Paris | Marion Barbet, Sylvie Behillil, Méline Bizard, Angela Brisebarre, Camille Capel, Vincent Enouf, Louise Lefrançois, Frédéric Lemoine, Christophe Malabat, Corinne Maufrais, Etienne Simon-Lorière, Maud Vanpeene, Sylvie Van der Werf ,CéLine Ramanantsoa |
| EPI_ISL_2293383 | Hospital | National Reference Center for Viruses of Respiratory Infections, Institut Pasteur, Paris | Marion Barbet, Sylvie Behillil, Méline Bizard, Angela Brisebarre, Camille Capel, Vincent Enouf, Louise Lefrançois, Frédéric Lemoine, Christophe Malabat, Corinne Maufrais, Etienne Simon-Lorière, Maud Vanpeene, Sylvie Van der Werf ,ValéRie Serazin |
| EPI_ISL_2293384, EPI_ISL_2293385 | Hospital | National Reference Center for Viruses of Respiratory Infections, Institut Pasteur, Paris | Marion Barbet, Sylvie Behillil, Méline Bizard, Angela Brisebarre, Camille Capel, Vincent Enouf, Louise Lefrançois, Frédéric Lemoine, Christophe Malabat, Corinne Maufrais, Pierre Lechat, Etienne Simon-Lorière, Maud Vanpeene, Sylvie Van der Werf ,ValéRie Serazin |
| EPI_ISL_2293386, EPI_ISL_2293434, EPI_ISL_2293435, EPI_ISL_2293436, EPI_ISL_2293437, EPI_ISL_2293438, EPI_ISL_2293439, EPI_ISL_2293440, EPI_ISL_2293441 | Labo Analyses Med | National Reference Center for Viruses of Respiratory Infections, Institut Pasteur, Paris | Marion Barbet, Sylvie Behillil, Méline Bizard, Angela Brisebarre, Camille Capel, Vincent Enouf, Louise Lefrançois, Frédéric Lemoine, Christophe Malabat, Corinne Maufrais, Etienne Simon-Lorière, Maud Vanpeene, Sylvie Van der Werf ,Nouri Taleb |
| EPI_ISL_2293442, EPI_ISL_2293443 | Labo Analyses Med | National Reference Center for Viruses of Respiratory Infections, Institut Pasteur, Paris | Marion Barbet, Sylvie Behillil, Méline Bizard, Angela Brisebarre, Camille Capel, Vincent Enouf, Louise Lefrançois, Frédéric Lemoine, Christophe Malabat, Corinne Maufrais, Etienne Simon-Lorière, Maud Vanpeene, Sylvie Van der Werf ,Ronan Pichard |
| EPI_ISL_2293444, EPI_ISL_2293445, EPI_ISL_2293447, EPI_ISL_2293448, EPI_ISL_2293449, EPI_ISL_2293450, EPI_ISL_2293451, EPI_ISL_2293452, EPI_ISL_2293453, EPI_ISL_2293454, EPI_ISL_2293455, EPI_ISL_2293456, EPI_ISL_2293457, EPI_ISL_2293458, EPI_ISL_2293459, EPI_ISL_2293460, EPI_ISL_2293461, EPI_ISL_2293462, EPI_ISL_2293463 | | | |
| see above | Labo Analyses Med | National Reference Center for Viruses of Respiratory Infections, Institut Pasteur, Paris | Marion Barbet, Sylvie Behillil, Méline Bizard, Angela Brisebarre, Camille Capel, Vincent Enouf, Louise Lefrançois, Frédéric Lemoine, Christophe Malabat, Corinne Maufrais, Etienne Simon-Lorière, Maud Vanpeene, Sylvie Van der Werf ,Philippe Miara |
| EPI_ISL_2293464, EPI_ISL_2293465 | Hospital | National Reference Center for Viruses of Respiratory Infections, Institut Pasteur, Paris | Marion Barbet, Sylvie Behillil, Méline Bizard, Angela Brisebarre, Camille Capel, Vincent Enouf, Louise Lefrançois, Frédéric Lemoine, Christophe Malabat, Corinne Maufrais, Etienne Simon-Lorière, Maud Vanpeene, Sylvie Van der Werf ,Evelyne Schvoerer |
| EPI_ISL_2293466 | Hospital | National Reference Center for Viruses of Respiratory Infections, Institut Pasteur, Paris | Marion Barbet, Sylvie Behillil, Méline Bizard, Angela Brisebarre, Camille Capel, Vincent Enouf, Louise Lefrançois, Frédéric Lemoine, Christophe Malabat, Corinne Maufrais, Etienne Simon-Lorière, Maud Vanpeene, Sylvie Van der Werf ,Laurence Louvet |
| EPI_ISL_2293467, EPI_ISL_2293468, EPI_ISL_2293469, EPI_ISL_2293470, EPI_ISL_2293471, EPI_ISL_2293472 | Hospital | National Reference Center for Viruses of Respiratory Infections, Institut Pasteur, Paris | Marion Barbet, Sylvie Behillil, Méline Bizard, Angela Brisebarre, Camille Capel, Vincent Enouf, Louise Lefrançois, Frédéric Lemoine, Christophe Malabat, Corinne Maufrais, Etienne Simon-Lorière, Maud Vanpeene, Sylvie Van der Werf ,Ilka Engelmann |
| EPI_ISL_2293473 | Hospital | National Reference Center for Viruses of Respiratory Infections, Institut Pasteur, Paris | Marion Barbet, Sylvie Behillil, Méline Bizard, Angela Brisebarre, Camille Capel, Vincent Enouf, Louise Lefrançois, Frédéric Lemoine, Christophe Malabat, Corinne Maufrais, Etienne Simon-Lorière, Maud Vanpeene, Sylvie Van der Werf ,Sylvie Ledru |
| EPI_ISL_2293474, EPI_ISL_2293475, EPI_ISL_2293476, EPI_ISL_2293477, EPI_ISL_2293478, EPI_ISL_2293479, EPI_ISL_2293480 | Labo Analyses Med | National Reference Center for Viruses of Respiratory Infections, Institut Pasteur, Paris | Marion Barbet, Sylvie Behillil, Méline Bizard, Angela Brisebarre, Camille Capel, Vincent Enouf, Louise Lefrançois, Frédéric Lemoine, Christophe Malabat, Corinne Maufrais, Etienne Simon-Lorière, Maud Vanpeene, Sylvie Van der Werf ,Domitille Leman |
| EPI_ISL_2293481, EPI_ISL_2293482, EPI_ISL_2293484, EPI_ISL_2293485, EPI_ISL_2293486 | Hospital | National Reference Center for Viruses of Respiratory Infections, Institut Pasteur, Paris | Marion Barbet, Sylvie Behillil, Méline Bizard, Angela Brisebarre, Camille Capel, Vincent Enouf, Louise Lefrançois, Frédéric Lemoine, Christophe Malabat, Corinne Maufrais, Etienne Simon-Lorière, Maud Vanpeene, Sylvie Van der Werf ,Anne Brocard |
| EPI_ISL_2293487, EPI_ISL_2293488, EPI_ISL_2293489, EPI_ISL_2293490, | Hospital | National Reference Center for Viruses of Respiratory Infections, Institut Pasteur, Paris | Marion Barbet, Sylvie Behillil, Méline Bizard, Angela Brisebarre, Camille Capel, Vincent Enouf, Louise Lefrançois, Frédéric Lemoine, Christophe Malabat, Corinne Maufrais, Etienne Simon-Lorière, Maud Vanpeene, Sylvie Van der Werf ,FréDéRique Canis |

| | | | |
|---|---|---|---|
| EPI_ISL_2293491, EPI_ISL_2293492, EPI_ISL_2293493, EPI_ISL_2293494, EPI_ISL_2293495, EPI_ISL_2293496 | | | |
| EPI_ISL_2293497, EPI_ISL_2293501 | Hospital | National Reference Center for Viruses of Respiratory Infections, Institut Pasteur, Paris | Marion Barbet, Sylvie Behillil, Méline Bizard, Angela Brisebarre, Camille Capel, Vincent Enouf, Louise Lefrançois, Frédéric Lemoine, Christophe Malabat, Corinne Maufrais, Etienne Simon-Lorière, Maud Vanpeene, Sylvie Van der Werf ,Thibault Guinoiseau |
| EPI_ISL_2293502, EPI_ISL_2293503 | Hospital | National Reference Center for Viruses of Respiratory Infections, Institut Pasteur, Paris | Marion Barbet, Sylvie Behillil, Méline Bizard, Angela Brisebarre, Camille Capel, Vincent Enouf, Louise Lefrançois, Frédéric Lemoine, Christophe Malabat, Corinne Maufrais, Etienne Simon-Lorière, Maud Vanpeene, Sylvie Van der Werf ,Dominique Descamps |
| EPI_ISL_2293504, EPI_ISL_2293505 | Hospital | National Reference Center for Viruses of Respiratory Infections, Institut Pasteur, Paris | Marion Barbet, Sylvie Behillil, Méline Bizard, Angela Brisebarre, Camille Capel, Vincent Enouf, Louise Lefrançois, Frédéric Lemoine, Christophe Malabat, Corinne Maufrais, Etienne Simon-Lorière, Maud Vanpeene, Sylvie Van der Werf ,Salma Ben Hadj Yahia |
| EPI_ISL_2293506 | Hospital | National Reference Center for Viruses of Respiratory Infections, Institut Pasteur, Paris | Marion Barbet, Sylvie Behillil, Méline Bizard, Angela Brisebarre, Camille Capel, Vincent Enouf, Louise Lefrançois, Frédéric Lemoine, Christophe Malabat, Corinne Maufrais, Etienne Simon-Lorière, Maud Vanpeene, Sylvie Van der Werf ,Martine Escure |
| EPI_ISL_2293507, EPI_ISL_2293508, EPI_ISL_2293509, EPI_ISL_2293510, EPI_ISL_2293511, EPI_ISL_2293512, EPI_ISL_2293513, EPI_ISL_2293514, EPI_ISL_2293515, EPI_ISL_2293516, EPI_ISL_2293517, EPI_ISL_2293518, EPI_ISL_2293519, EPI_ISL_2293520, EPI_ISL_2293521 | | | |
| see above | Hospital | National Reference Center for Viruses of Respiratory Infections, Institut Pasteur, Paris | Marion Barbet, Sylvie Behillil, Méline Bizard, Angela Brisebarre, Camille Capel, Vincent Enouf, Louise Lefrançois, Frédéric Lemoine, Christophe Malabat, Corinne Maufrais, Etienne Simon-Lorière, Maud Vanpeene, Sylvie Van der Werf ,L. Courdavault |
| EPI_ISL_2293522, EPI_ISL_2293523, EPI_ISL_2293524, EPI_ISL_2293525, EPI_ISL_2293526, EPI_ISL_2293528, EPI_ISL_2293529, EPI_ISL_2293530, EPI_ISL_2293531, EPI_ISL_2293532, EPI_ISL_2293533, EPI_ISL_2293534, EPI_ISL_2293536, EPI_ISL_2293537, EPI_ISL_2293538, EPI_ISL_2293539, EPI_ISL_2293540, EPI_ISL_2293541, EPI_ISL_2293542, EPI_ISL_2293543, EPI_ISL_2293544, EPI_ISL_2293545, EPI_ISL_2293546, EPI_ISL_2293547, EPI_ISL_2293548, EPI_ISL_2293549, EPI_ISL_2293550, EPI_ISL_2293551, EPI_ISL_2293552, EPI_ISL_2293553, EPI_ISL_2293554, EPI_ISL_2293555, EPI_ISL_2293556, EPI_ISL_2293557, EPI_ISL_2293558, EPI_ISL_2293559, EPI_ISL_2293560, EPI_ISL_2293561, EPI_ISL_2293562, EPI_ISL_2293563, EPI_ISL_2293564, EPI_ISL_2293565 | | | |
| see above | Labo Analyses Med | National Reference Center for Viruses of Respiratory Infections, Institut Pasteur, Paris | Marion Barbet, Sylvie Behillil, Méline Bizard, Angela Brisebarre, Camille Capel, Vincent Enouf, Louise Lefrançois, Frédéric Lemoine, Christophe Malabat, Corinne Maufrais, Etienne Simon-Lorière, Maud Vanpeene, Sylvie Van der Werf ,Anne Holstein |
| EPI_ISL_2293567, EPI_ISL_2293568, EPI_ISL_2293569 | Labo Analyses Med | National Reference Center for Viruses of Respiratory Infections, Institut Pasteur, Paris | Marion Barbet, Sylvie Behillil, Méline Bizard, Angela Brisebarre, Camille Capel, Vincent Enouf, Louise Lefrançois, Frédéric Lemoine, Christophe Malabat, Corinne Maufrais, Etienne Simon-Lorière, Maud Vanpeene, Sylvie Van der Werf ,SéVerine Haouisee |
| EPI_ISL_2293570, EPI_ISL_2293571, EPI_ISL_2293572 | Hospital | National Reference Center for Viruses of Respiratory Infections, Institut Pasteur, Paris | Marion Barbet, Sylvie Behillil, Méline Bizard, Angela Brisebarre, Camille Capel, Vincent Enouf, Louise Lefrançois, Frédéric Lemoine, Christophe Malabat, Corinne Maufrais, Etienne Simon-Lorière, Maud Vanpeene, Sylvie Van der Werf ,Marianne Burgard |
| EPI_ISL_2293573, EPI_ISL_2293574, EPI_ISL_2293575, EPI_ISL_2293576, EPI_ISL_2293577, EPI_ISL_2293578, EPI_ISL_2293579, EPI_ISL_2293580, EPI_ISL_2293581, EPI_ISL_2293582, EPI_ISL_2293583, EPI_ISL_2293584, EPI_ISL_2293585, EPI_ISL_2293586, EPI_ISL_2293587, EPI_ISL_2293588, EPI_ISL_2293589, EPI_ISL_2293590, EPI_ISL_2293591, EPI_ISL_2293592, EPI_ISL_2293593, EPI_ISL_2293594, EPI_ISL_2293595, EPI_ISL_2293596, EPI_ISL_2293597, EPI_ISL_2293598, EPI_ISL_2293599, EPI_ISL_2293600, EPI_ISL_2293603, EPI_ISL_2293604, EPI_ISL_2293605, EPI_ISL_2293606, EPI_ISL_2293607 | | | |
| see above | Labo Analyses Med | National Reference Center for Viruses of Respiratory Infections, Institut Pasteur, Paris | Marion Barbet, Sylvie Behillil, Méline Bizard, Angela Brisebarre, Camille Capel, Vincent Enouf, Louise Lefrançois, Frédéric Lemoine, Christophe Malabat, Corinne Maufrais, Etienne Simon-Lorière, Maud Vanpeene, Sylvie Van der Werf , Pierre-Yves Leonard |
| EPI_ISL_2293608, EPI_ISL_2293609, EPI_ISL_2293610, EPI_ISL_2293611, EPI_ISL_2293612, EPI_ISL_2293613, EPI_ISL_2293614, EPI_ISL_2293615, EPI_ISL_2293616, EPI_ISL_2293617, EPI_ISL_2293618, EPI_ISL_2293619, EPI_ISL_2293620, EPI_ISL_2293621, EPI_ISL_2293622, EPI_ISL_2293623, EPI_ISL_2293624, EPI_ISL_2293625, EPI_ISL_2293629, EPI_ISL_2293630, EPI_ISL_2293631 | | | |
| see above | Labo Analyses Med | National Reference Center for Viruses of Respiratory Infections, Institut Pasteur, Paris | Marion Barbet, Sylvie Behillil, Méline Bizard, Angela Brisebarre, Camille Capel, Vincent Enouf, Louise Lefrançois, Frédéric Lemoine, Christophe Malabat, Corinne Maufrais, Etienne Simon-Lorière, Maud Vanpeene, Sylvie Van der Werf ,Jonas Amzalag |
| EPI_ISL_2294377, EPI_ISL_2294574, EPI_ISL_2294575 | S.C. Microbiologia e Virologia Laboratorio Virologia -Speciale Centro Influenza - AOU di Sassari - Viale san Pietro 43/B Palazzo Infettivologia | Laboratorio specialistico UOC Ematologia - Ospedale "San Francesco" - ATS-ASSL Nuoro | Rosanna Asproni, Maura Fiamma, Maria Cristina Garau,Giuseppe Mameli , Caterina Serra, Elena Rimini, Salvatore Rubino |
| EPI_ISL_2296064 | LESP Baja California | Instituto de Diagnostico y Referencia Epidemiologicos (INDRE) | Claudia Wong-Arambula, Abril Rodriguez-Maldonado, Vanessa Rivero-Arredondo, Ariadna Medina-Benitez, Joaquin Quiroz-Mercado, Sergio Rangel-Guerrero, Natividad Cruz-Ortiz, Tatiana Nunez-Garcia, Gisela Barrera-Badillo, Lucia Hernandez-Rivas, Irma Lopez-Martinez, Ernesto Ramirez-Gonzalez. |
| EPI_ISL_2296067 | LESP Colima | Instituto de Diagnostico y Referencia Epidemiologicos (INDRE) | Claudia Wong-Arambula, Abril Rodriguez-Maldonado, Vanessa Rivero-Arredondo, Ariadna Medina-Benitez, Joaquin Quiroz-Mercado, Sergio Rangel-Guerrero, Natividad Cruz-Ortiz, Tatiana Nunez-Garcia, Gisela Barrera-Badillo, Lucia Hernandez-Rivas, Irma Lopez-Martinez, Ernesto Ramirez-Gonzalez. |
| EPI_ISL_2296071 | LESP Sinaloa | Instituto de Diagnostico y Referencia Epidemiologicos (INDRE) | Claudia Wong-Arambula, Abril Rodriguez-Maldonado, Vanessa Rivero-Arredondo, Ariadna Medina-Benitez, Joaquin Quiroz-Mercado, Sergio Rangel-Guerrero, Natividad Cruz-Ortiz, Tatiana Nunez-Garcia, Gisela Barrera-Badillo, Lucia Hernandez-Rivas, Irma Lopez-Martinez, Ernesto Ramirez-Gonzalez. |
| EPI_ISL_2296073 | LESP Estado de Mexico | Instituto de Diagnostico y Referencia Epidemiologicos (INDRE) | Claudia Wong-Arambula, Abril Rodriguez-Maldonado, Vanessa Rivero-Arredondo, Ariadna Medina-Benitez, Joaquin Quiroz-Mercado, Sergio Rangel-Guerrero, Natividad Cruz-Ortiz, Tatiana Nunez-Garcia, Gisela Barrera-Badillo, Lucia Hernandez-Rivas, Irma Lopez-Martinez, Ernesto Ramirez-Gonzalez. |
| EPI_ISL_2296074, EPI_ISL_2296076, EPI_ISL_2296077, EPI_ISL_2296078 | LESP Ciudad de Mexico | Instituto de Diagnostico y Referencia Epidemiologicos (INDRE) | Claudia Wong-Arambula, Abril Rodriguez-Maldonado, Vanessa Rivero-Arredondo, Ariadna Medina-Benitez, Joaquin Quiroz-Mercado, Sergio Rangel-Guerrero, Natividad Cruz-Ortiz, Tatiana Nunez-Garcia, Gisela Barrera-Badillo, Lucia Hernandez-Rivas, Irma Lopez-Martinez, Ernesto Ramirez-Gonzalez. |
| EPI_ISL_2296096 | LESP Guanajuato | Instituto de Diagnostico y Referencia Epidemiologicos (INDRE) | Claudia Wong-Arambula, Abril Rodriguez-Maldonado, Vanessa Rivero-Arredondo, Ariadna Medina-Benitez, Joaquin Quiroz-Mercado, Sergio Rangel-Guerrero, Natividad Cruz-Ortiz, Tatiana Nunez-Garcia, Gisela Barrera-Badillo, Lucia Hernandez-Rivas, Irma Lopez-Martinez, Ernesto Ramirez-Gonzalez. |
| EPI_ISL_2296103 | LESP Sinaloa | Instituto de Diagnostico y Referencia Epidemiologicos (INDRE) | Claudia Wong-Arambula, Abril Rodriguez-Maldonado, Vanessa Rivero-Arredondo, Ariadna Medina-Benitez, Joaquin Quiroz-Mercado, Sergio Rangel-Guerrero, Natividad Cruz-Ortiz, Tatiana Nunez-Garcia, Gisela Barrera-Badillo, Lucia Hernandez-Rivas, Irma Lopez-Martinez, Ernesto Ramirez-Gonzalez. |
| EPI_ISL_2296104 | LESP Baja California | Instituto de Diagnostico y Referencia Epidemiologicos (INDRE) | Claudia Wong-Arambula, Abril Rodriguez-Maldonado, Vanessa Rivero-Arredondo, Ariadna Medina-Benitez, Joaquin Quiroz-Mercado, Sergio Rangel-Guerrero, Natividad Cruz-Ortiz, Tatiana Nunez-Garcia, Gisela Barrera-Badillo, Lucia Hernandez-Rivas, Irma Lopez-Martinez, Ernesto Ramirez-Gonzalez. |
| EPI_ISL_2304002, EPI_ISL_2304008, EPI_ISL_2304009, EPI_ISL_2304010, EPI_ISL_2304013, EPI_ISL_2304014, EPI_ISL_2304015, EPI_ISL_2304016, EPI_ISL_2304017, EPI_ISL_2304022, EPI_ISL_2304024, EPI_ISL_2304026, EPI_ISL_2304027, EPI_ISL_2304028 | | | |
| see above | University of Oregon COVID-19 MAP Laboratory | University of Oregon Genomics and Cell Characterczation Core Facility (GC3F) | Douglas Turnbull, Ariana White, Jeff Bishop, Jason Carriere, Jason Sydes, Peter Batzel |
| EPI_ISL_2308239, EPI_ISL_2308244, EPI_ISL_2308249 | National Institute of Infectious Diseases-Prof. Dr. Matei Bals Molecular Diagnostics Laboratory | National Institute of Infectious Diseases-Prof. Dr. Matei Bals Molecular Diagnostics Laboratory | Corina Casangiu, Leontina Banica, Marius Surleac, Ovidiu Vlaicu, Andreea Tudor, Simona Paraschiv, Dan Otelea |
| EPI_ISL_2308557, EPI_ISL_2308558, EPI_ISL_2308559, EPI_ISL_2308560, EPI_ISL_2308561, EPI_ISL_2308562, EPI_ISL_2308563, EPI_ISL_2308564, EPI_ISL_2308565 | SIESP DIPARTIMENTO DI PREVENZIONE TERAMO | Istituto Zooprofilattico Sperimentale dell'Abruzzo e Molise "G. Caporale" | Lorusso A, Marcacci M, Di Domenico M, Ancora M, Curini V, Di Lollo Valeria, Mangone I, Rinaldi A, Delli Compagni E, Scialabba S, Caporale M, Di Pasquale A, Cammà C, Puglia I, Calistri P, Savini G |
| EPI_ISL_2308566 | SIESP DIPARTIMENTO DI PREVENZIONE CHIETI | Istituto Zooprofilattico Sperimentale dell'Abruzzo e Molise "G. Caporale" | Lorusso A, Marcacci M, Di Domenico M, Ancora M, Curini V, Di Lollo Valeria, Mangone I, Rinaldi A, Delli Compagni E, Scialabba S, Caporale M, Di Pasquale A, Cammà C, Puglia I, Calistri P, Savini G |
| EPI_ISL_2308567 | SIESP DIPARTIMENTO DI PREVENZIONE TERAMO | Istituto Zooprofilattico Sperimentale dell'Abruzzo e Molise "G. Caporale" | Lorusso A, Marcacci M, Di Domenico M, Ancora M, Curini V, Di Lollo Valeria, Mangone I, Rinaldi A, Delli Compagni E, Scialabba S, Caporale M, Di Pasquale A, Cammà C, Puglia I, Calistri P, Savini G |
| EPI_ISL_2308568, EPI_ISL_2308569, | SIESP DIPARTIMENTO DI PREVENZIONE CHIETI | Istituto Zooprofilattico Sperimentale dell'Abruzzo e Molise "G. | Lorusso A, Marcacci M, Di Domenico M, Ancora M, Curini V, Di Lollo Valeria, Mangone I, Rinaldi A, Delli Compagni E, Scialabba S, Caporale M, Di |

| | | | |
|---|---|---|---|
| EPI_ISL_2308570 | | Caporale" | Pasquale A, Cammà C, Puglia I, Calistri P, Savini G |
| EPI_ISL_2308571, EPI_ISL_2308572, EPI_ISL_2308573 | SIESP DIPARTIMENTO DI PREVENZIONE TERAMO | Istituto Zooprofilattico Sperimentale dell'Abruzzo e Molise "G. Caporale" | Lorusso A, Marcacci M, Di Domenico M, Ancora M, Curini V, Di Lollo Valeria, Mangone I, Rinaldi A, Delli Compagni E, Scialabba S, Caporale M, Di Pasquale A, Cammà C, Puglia I, Calistri P, Savini G |
| EPI_ISL_2308574 | SIESP CHIETI - DRIVE IN LANCIANO | Istituto Zooprofilattico Sperimentale dell'Abruzzo e Molise "G. Caporale" | Lorusso A, Marcacci M, Di Domenico M, Ancora M, Curini V, Di Lollo Valeria, Mangone I, Rinaldi A, Delli Compagni E, Scialabba S, Caporale M, Di Pasquale A, Cammà C, Puglia I, Calistri P, Savini G |
| EPI_ISL_2308575 | SIESP CHIETI - DRIVE IN ORTONA | Istituto Zooprofilattico Sperimentale dell'Abruzzo e Molise "G. Caporale" | Lorusso A, Marcacci M, Di Domenico M, Ancora M, Curini V, Di Lollo Valeria, Mangone I, Rinaldi A, Delli Compagni E, Scialabba S, Caporale M, Di Pasquale A, Cammà C, Puglia I, Calistri P, Savini G |
| EPI_ISL_2308576, EPI_ISL_2308577 | SIESP CHIETI - DRIVE IN LANCIANO | Istituto Zooprofilattico Sperimentale dell'Abruzzo e Molise "G. Caporale" | Lorusso A, Marcacci M, Di Domenico M, Ancora M, Curini V, Di Lollo Valeria, Mangone I, Rinaldi A, Delli Compagni E, Scialabba S, Caporale M, Di Pasquale A, Cammà C, Puglia I, Calistri P, Savini G |
| EPI_ISL_2308578 | SIESP DIPARTIMENTO DI PREVENZIONE TERAMO | Istituto Zooprofilattico Sperimentale dell'Abruzzo e Molise "G. Caporale" | Lorusso A, Marcacci M, Di Domenico M, Ancora M, Curini V, Di Lollo Valeria, Mangone I, Rinaldi A, Delli Compagni E, Scialabba S, Caporale M, Di Pasquale A, Cammà C, Puglia I, Calistri P, Savini G |
| EPI_ISL_2308579 | PRESIDIO TERRITORIALE CASOLI | Istituto Zooprofilattico Sperimentale dell'Abruzzo e Molise "G. Caporale" | Lorusso A, Marcacci M, Di Domenico M, Ancora M, Curini V, Di Lollo Valeria, Mangone I, Rinaldi A, Delli Compagni E, Scialabba S, Caporale M, Di Pasquale A, Cammà C, Puglia I, Calistri P, Savini G |
| EPI_ISL_2308580, EPI_ISL_2308581 | SIESP DIPARTIMENTO DI PREVENZIONE CHIETI | Istituto Zooprofilattico Sperimentale dell'Abruzzo e Molise "G. Caporale" | Lorusso A, Marcacci M, Di Domenico M, Ancora M, Curini V, Di Lollo Valeria, Mangone I, Rinaldi A, Delli Compagni E, Scialabba S, Caporale M, Di Pasquale A, Cammà C, Puglia I, Calistri P, Savini G |
| EPI_ISL_2308582 | PRESIDIO TERRITORIALE CASOLI | Istituto Zooprofilattico Sperimentale dell'Abruzzo e Molise "G. Caporale" | Lorusso A, Marcacci M, Di Domenico M, Ancora M, Curini V, Di Lollo Valeria, Mangone I, Rinaldi A, Delli Compagni E, Scialabba S, Caporale M, Di Pasquale A, Cammà C, Puglia I, Calistri P, Savini G |
| EPI_ISL_2308583 | SIESP DIPARTIMENTO DI PREVENZIONE CHIETI | Istituto Zooprofilattico Sperimentale dell'Abruzzo e Molise "G. Caporale" | Lorusso A, Marcacci M, Di Domenico M, Ancora M, Curini V, Di Lollo Valeria, Mangone I, Rinaldi A, Delli Compagni E, Scialabba S, Caporale M, Di Pasquale A, Cammà C, Puglia I, Calistri P, Savini G |
| EPI_ISL_2308584 | SIESP CHIETI - DRIVE IN VASTO | Istituto Zooprofilattico Sperimentale dell'Abruzzo e Molise "G. Caporale" | Lorusso A, Marcacci M, Di Domenico M, Ancora M, Curini V, Di Lollo Valeria, Mangone I, Rinaldi A, Delli Compagni E, Scialabba S, Caporale M, Di Pasquale A, Cammà C, Puglia I, Calistri P, Savini G |
| EPI_ISL_2308585, EPI_ISL_2308586 | SIESP DIPARTIMENTO DI PREVENZIONE CHIETI | Istituto Zooprofilattico Sperimentale dell'Abruzzo e Molise "G. Caporale" | Lorusso A, Marcacci M, Di Domenico M, Ancora M, Curini V, Di Lollo Valeria, Mangone I, Rinaldi A, Delli Compagni E, Scialabba S, Caporale M, Di Pasquale A, Cammà C, Puglia I, Calistri P, Savini G |
| EPI_ISL_2308587, EPI_ISL_2308588 | SIESP DIPARTIMENTO DI PREVENZIONE TERAMO | Istituto Zooprofilattico Sperimentale dell'Abruzzo e Molise "G. Caporale" | Lorusso A, Marcacci M, Di Domenico M, Ancora M, Curini V, Di Lollo Valeria, Mangone I, Rinaldi A, Delli Compagni E, Scialabba S, Caporale M, Di Pasquale A, Cammà C, Puglia I, Calistri P, Savini G |
| EPI_ISL_2308589, EPI_ISL_2308590 | SIESP CHIETI - DRIVE IN VASTO | Istituto Zooprofilattico Sperimentale dell'Abruzzo e Molise "G. Caporale" | Lorusso A, Marcacci M, Di Domenico M, Ancora M, Curini V, Di Lollo Valeria, Mangone I, Rinaldi A, Delli Compagni E, Scialabba S, Caporale M, Di Pasquale A, Cammà C, Puglia I, Calistri P, Savini G |
| EPI_ISL_2308591 | SIESP DIPARTIMENTO DI PREVENZIONE CHIETI | Istituto Zooprofilattico Sperimentale dell'Abruzzo e Molise "G. Caporale" | Lorusso A, Marcacci M, Di Domenico M, Ancora M, Curini V, Di Lollo Valeria, Mangone I, Rinaldi A, Delli Compagni E, Scialabba S, Caporale M, Di Pasquale A, Cammà C, Puglia I, Calistri P, Savini G |
| EPI_ISL_2308592 | OSPEDALE CIVILE SANT'OMERO - PRONTO SOCCORSO | Istituto Zooprofilattico Sperimentale dell'Abruzzo e Molise "G. Caporale" | Lorusso A, Marcacci M, Di Domenico M, Ancora M, Curini V, Di Lollo Valeria, Mangone I, Rinaldi A, Delli Compagni E, Scialabba S, Caporale M, Di Pasquale A, Cammà C, Puglia I, Calistri P, Savini G |
| EPI_ISL_2308593 | SIESP DIPARTIMENTO DI PREVENZIONE TERAMO | Istituto Zooprofilattico Sperimentale dell'Abruzzo e Molise "G. Caporale" | Lorusso A, Marcacci M, Di Domenico M, Ancora M, Curini V, Di Lollo Valeria, Mangone I, Rinaldi A, Delli Compagni E, Scialabba S, Caporale M, Di Pasquale A, Cammà C, Puglia I, Calistri P, Savini G |
| EPI_ISL_2308594 | SIESP DIPARTIMENTO DI PREVENZIONE CHIETI | Istituto Zooprofilattico Sperimentale dell'Abruzzo e Molise "G. Caporale" | Lorusso A, Marcacci M, Di Domenico M, Ancora M, Curini V, Di Lollo Valeria, Mangone I, Rinaldi A, Delli Compagni E, Scialabba S, Caporale M, Di Pasquale A, Cammà C, Puglia I, Calistri P, Savini G |
| EPI_ISL_2308595, EPI_ISL_2308596, EPI_ISL_2308597 | SIESP DIPARTIMENTO DI PREVENZIONE TERAMO | Istituto Zooprofilattico Sperimentale dell'Abruzzo e Molise "G. Caporale" | Lorusso A, Marcacci M, Di Domenico M, Ancora M, Curini V, Di Lollo Valeria, Mangone I, Rinaldi A, Delli Compagni E, Scialabba S, Caporale M, Di Pasquale A, Cammà C, Puglia I, Calistri P, Savini G |
| EPI_ISL_2308598 | SIESP DIPARTIMENTO DI PREVENZIONE CHIETI | Istituto Zooprofilattico Sperimentale dell'Abruzzo e Molise "G. Caporale" | Lorusso A, Marcacci M, Di Domenico M, Ancora M, Curini V, Di Lollo Valeria, Mangone I, Rinaldi A, Delli Compagni E, Scialabba S, Caporale M, Di Pasquale A, Cammà C, Puglia I, Calistri P, Savini G |
| EPI_ISL_2308599, EPI_ISL_2308600, EPI_ISL_2308601 | SIESP DIPARTIMENTO DI PREVENZIONE TERAMO | Istituto Zooprofilattico Sperimentale dell'Abruzzo e Molise "G. Caporale" | Lorusso A, Marcacci M, Di Domenico M, Ancora M, Curini V, Di Lollo Valeria, Mangone I, Rinaldi A, Delli Compagni E, Scialabba S, Caporale M, Di Pasquale A, Cammà C, Puglia I, Calistri P, Savini G |
| EPI_ISL_2308602, EPI_ISL_2308603 | SIESP CHIETI - DRIVE IN LANCIANO | Istituto Zooprofilattico Sperimentale dell'Abruzzo e Molise "G. Caporale" | Lorusso A, Marcacci M, Di Domenico M, Ancora M, Curini V, Di Lollo Valeria, Mangone I, Rinaldi A, Delli Compagni E, Scialabba S, Caporale M, Di Pasquale A, Cammà C, Puglia I, Calistri P, Savini G |
| EPI_ISL_2308604 | SIESP CHIETI - DRIVE IN CHIETI | Istituto Zooprofilattico Sperimentale dell'Abruzzo e Molise "G. Caporale" | Lorusso A, Marcacci M, Di Domenico M, Ancora M, Curini V, Di Lollo Valeria, Mangone I, Rinaldi A, Delli Compagni E, Scialabba S, Caporale M, Di Pasquale A, Cammà C, Puglia I, Calistri P, Savini G |
| EPI_ISL_2308605 | SIESP DIPARTIMENTO DI PREVENZIONE CHIETI | Istituto Zooprofilattico Sperimentale dell'Abruzzo e Molise "G. Caporale" | Lorusso A, Marcacci M, Di Domenico M, Ancora M, Curini V, Di Lollo Valeria, Mangone I, Rinaldi A, Delli Compagni E, Scialabba S, Caporale M, Di Pasquale A, Cammà C, Puglia I, Calistri P, Savini G |
| EPI_ISL_2308606 | SIESP CHIETI - DRIVE IN ORTONA | Istituto Zooprofilattico Sperimentale dell'Abruzzo e Molise "G. Caporale" | Lorusso A, Marcacci M, Di Domenico M, Ancora M, Curini V, Di Lollo Valeria, Mangone I, Rinaldi A, Delli Compagni E, Scialabba S, Caporale M, Di Pasquale A, Cammà C, Puglia I, Calistri P, Savini G |
| EPI_ISL_2308607 | SIESP CHIETI - DRIVE IN GISSI | Istituto Zooprofilattico Sperimentale dell'Abruzzo e Molise "G. Caporale" | Lorusso A, Marcacci M, Di Domenico M, Ancora M, Curini V, Di Lollo Valeria, Mangone I, Rinaldi A, Delli Compagni E, Scialabba S, Caporale M, Di Pasquale A, Cammà C, Puglia I, Calistri P, Savini G |
| EPI_ISL_2308608 | OSPEDALE CIVILE SANT'OMERO - PRONTO SOCCORSO | Istituto Zooprofilattico Sperimentale dell'Abruzzo e Molise "G. Caporale" | Lorusso A, Marcacci M, Di Domenico M, Ancora M, Curini V, Di Lollo Valeria, Mangone I, Rinaldi A, Delli Compagni E, Scialabba S, Caporale M, Di Pasquale A, Cammà C, Puglia I, Calistri P, Savini G |
| EPI_ISL_2308611, EPI_ISL_2308612, EPI_ISL_2308613, EPI_ISL_2308614, EPI_ISL_2308615, EPI_ISL_2308616, EPI_ISL_2308617 | Azienda USL Umbria 2 | Istituto Zooprofilattico Sperimentale dell'Abruzzo e Molise "G. Caporale" | Proietti A, Pistoni E, Lorusso A, Marcacci M, Di Domenico M, Ancora M, Curini V, Mangone I, Rinaldi A, Delli Compagni E, Scialabba S, Di Pasquale A, Cammà C, Puglia I, Calistri P, Savini G |
| EPI_ISL_2308618, EPI_ISL_2308619, EPI_ISL_2308620, EPI_ISL_2308621, EPI_ISL_2308623, EPI_ISL_2308624, EPI_ISL_2308625, EPI_ISL_2308626, EPI_ISL_2308627, EPI_ISL_2308628, EPI_ISL_2308629, EPI_ISL_2308630, EPI_ISL_2308632, EPI_ISL_2308633, EPI_ISL_2308634, EPI_ISL_2308635 | | | |
| see above | Università degli Studi di Perugia | Istituto Zooprofilattico Sperimentale dell'Abruzzo e Molise "G. Caporale" | Mencacci A, Camilloni B, Biagetti M, Giammarioli M, Lorusso A, Marcacci M, Di Domenico M, Ancora M, Curini V, Mangone I, Rinaldi A, Delli Compagni E, Scialabba S, Di Pasquale A, Cammà C, Puglia I, Calistri P, Savini G |
| EPI_ISL_2308636 | Laboratorio Analisi Osp. Città di Castello - Azienda USL Umbria1 | Istituto Zooprofilattico Sperimentale dell'Abruzzo e Molise "G. Caporale" | Malagigi V, Taccani P,Lorusso A, Marcacci M, Di Domenico M, Ancora M, Curini V, Mangone I, Rinaldi A, Delli Compagni E, Scialabba S, Di Pasquale A, Cammà C, Puglia I, Calistri P, Savini G |
| EPI_ISL_2308637 | Laboratorio Analisi Osp. Città di Castello - Azienda USL Umbria2 | Istituto Zooprofilattico Sperimentale dell'Abruzzo e Molise "G. Caporale" | Malagigi V, Taccani P,Lorusso A, Marcacci M, Di Domenico M, Ancora M, Curini V, Mangone I, Rinaldi A, Delli Compagni E, Scialabba S, Di Pasquale A, Cammà C, Puglia I, Calistri P, Savini G |
| EPI_ISL_2308638 | Laboratorio Analisi Osp. Città di Castello - Azienda USL Umbria3 | Istituto Zooprofilattico Sperimentale dell'Abruzzo e Molise "G. Caporale" | Malagigi V, Taccani P,Lorusso A, Marcacci M, Di Domenico M, Ancora M, Curini V, Mangone I, Rinaldi A, Delli Compagni E, Scialabba S, Di Pasquale A, Cammà C, Puglia I, Calistri P, Savini G |
| EPI_ISL_2308639 | Laboratorio Analisi Osp. Città di Castello - Azienda USL Umbria4 | Istituto Zooprofilattico Sperimentale dell'Abruzzo e Molise "G. Caporale" | Malagigi V, Taccani P,Lorusso A, Marcacci M, Di Domenico M, Ancora M, Curini V, Mangone I, Rinaldi A, Delli Compagni E, Scialabba S, Di Pasquale A, Cammà C, Puglia I, Calistri P, Savini G |
| EPI_ISL_2308640 | Laboratorio Analisi Osp. Città di Castello - Azienda USL Umbria5 | Istituto Zooprofilattico Sperimentale dell'Abruzzo e Molise "G. Caporale" | Malagigi V, Taccani P,Lorusso A, Marcacci M, Di Domenico M, Ancora M, Curini V, Mangone I, Rinaldi A, Delli Compagni E, Scialabba S, Di Pasquale A, Cammà C, Puglia I, Calistri P, Savini G |

| | | | |
|---|---|---|---|
| EPI_ISL_2308641 | Laboratorio Analisi Osp. Città di Castello - Azienda USL Umbria6 | Istituto Zooprofilattico Sperimentale dell'Abruzzo e Molise "G. Caporale" | Malagigi V, Taccani P,Lorusso A, Marcacci M, Di Domenico M, Ancora M, Curini V, Mangone I, Rinaldi A, Delli Compagni E, Scialabba S, Di Pasquale A, Cammà C, Puglia I, Calistri P, Savini G |
| EPI_ISL_2308642 | Laboratorio Analisi Osp. Città di Castello - Azienda USL Umbria7 | Istituto Zooprofilattico Sperimentale dell'Abruzzo e Molise "G. Caporale" | Malagigi V, Taccani P,Lorusso A, Marcacci M, Di Domenico M, Ancora M, Curini V, Mangone I, Rinaldi A, Delli Compagni E, Scialabba S, Di Pasquale A, Cammà C, Puglia I, Calistri P, Savini G |
| EPI_ISL_2308643 | Laboratorio Analisi Osp. Città di Castello - Azienda USL Umbria8 | Istituto Zooprofilattico Sperimentale dell'Abruzzo e Molise "G. Caporale" | Malagigi V, Taccani P,Lorusso A, Marcacci M, Di Domenico M, Ancora M, Curini V, Mangone I, Rinaldi A, Delli Compagni E, Scialabba S, Di Pasquale A, Cammà C, Puglia I, Calistri P, Savini G |
| EPI_ISL_2308644 | Laboratorio Analisi Osp. Città di Castello - Azienda USL Umbria9 | Istituto Zooprofilattico Sperimentale dell'Abruzzo e Molise "G. Caporale" | Malagigi V, Taccani P,Lorusso A, Marcacci M, Di Domenico M, Ancora M, Curini V, Mangone I, Rinaldi A, Delli Compagni E, Scialabba S, Di Pasquale A, Cammà C, Puglia I, Calistri P, Savini G |
| EPI_ISL_2308684, EPI_ISL_2308685, EPI_ISL_2308686, EPI_ISL_2308687, EPI_ISL_2308688, EPI_ISL_2308689, EPI_ISL_2308690, EPI_ISL_2308691, EPI_ISL_2308692, EPI_ISL_2308693, EPI_ISL_2308694, EPI_ISL_2308695, EPI_ISL_2308696, EPI_ISL_2308697, EPI_ISL_2308698, EPI_ISL_2308699 | | | |
| see above | Azienda Ospedaliera Terni | Istituto Zooprofilattico Sperimentale dell'Abruzzo e Molise "G. Caporale" | Palumbo M, Scaccetti A, Lorusso A, Marcacci M, Di Domenico M, Ancora M, Curini V, Mangone I, Rinaldi A, Delli Compagni E, Scialabba S, Di Pasquale A, Cammà C, Puglia I, Calistri P, Savini G |
| EPI_ISL_2308700, EPI_ISL_2308702, EPI_ISL_2308703, EPI_ISL_2308704 | Azienda USL Umbria 2 | Istituto Zooprofilattico Sperimentale dell'Abruzzo e Molise "G. Caporale" | Proietti A, Pistoni E, Lorusso A, Marcacci M, Di Domenico M, Ancora M, Curini V, Mangone I, Rinaldi A, Delli Compagni E, Scialabba S, Di Pasquale A, Cammà C, Puglia I, Calistri P, Savini G |
| EPI_ISL_2308706, EPI_ISL_2308707, EPI_ISL_2308708, EPI_ISL_2308709 | Università degli Studi di Perugia | Istituto Zooprofilattico Sperimentale dell'Abruzzo e Molise "G. Caporale" | Mencacci A, Camilloni B, Biagetti M, GiammarioliM, Lorusso A, Marcacci M, Di Domenico M, Ancora M, Curini V, Mangone I, Rinaldi A, Delli Compagni E, Scialabba S, Di Pasquale A, Cammà C, Puglia I, Calistri P, Savini G |
| EPI_ISL_2308710 | Laboratorio Analisi Osp. Città di Castello - Azienda USL Umbria1 | Istituto Zooprofilattico Sperimentale dell'Abruzzo e Molise "G. Caporale" | Malagigi V, Tacconi P, Lorusso A, Marcacci M, Di Domenico M, Ancora M, Curini V, Mangone I, Rinaldi A, Delli Compagni E, Scialabba S, Di Pasquale A, Cammà C, Puglia I, Calistri P, Savini G |
| EPI_ISL_2308711 | Laboratorio Analisi Osp. Città di Castello - Azienda USL Umbria2 | Istituto Zooprofilattico Sperimentale dell'Abruzzo e Molise "G. Caporale" | Malagigi V, Tacconi P, Lorusso A, Marcacci M, Di Domenico M, Ancora M, Curini V, Mangone I, Rinaldi A, Delli Compagni E, Scialabba S, Di Pasquale A, Cammà C, Puglia I, Calistri P, Savini G |
| EPI_ISL_2308712 | Laboratorio Analisi Osp. Città di Castello - Azienda USL Umbria3 | Istituto Zooprofilattico Sperimentale dell'Abruzzo e Molise "G. Caporale" | Malagigi V, Tacconi P, Lorusso A, Marcacci M, Di Domenico M, Ancora M, Curini V, Mangone I, Rinaldi A, Delli Compagni E, Scialabba S, Di Pasquale A, Cammà C, Puglia I, Calistri P, Savini G |
| EPI_ISL_2308713 | Laboratorio Analisi Osp. Città di Castello - Azienda USL Umbria4 | Istituto Zooprofilattico Sperimentale dell'Abruzzo e Molise "G. Caporale" | Malagigi V, Tacconi P, Lorusso A, Marcacci M, Di Domenico M, Ancora M, Curini V, Mangone I, Rinaldi A, Delli Compagni E, Scialabba S, Di Pasquale A, Cammà C, Puglia I, Calistri P, Savini G |
| EPI_ISL_2308714 | Laboratorio Analisi Osp. Città di Castello - Azienda USL Umbria5 | Istituto Zooprofilattico Sperimentale dell'Abruzzo e Molise "G. Caporale" | Malagigi V, Tacconi P, Lorusso A, Marcacci M, Di Domenico M, Ancora M, Curini V, Mangone I, Rinaldi A, Delli Compagni E, Scialabba S, Di Pasquale A, Cammà C, Puglia I, Calistri P, Savini G |
| EPI_ISL_2308716 | Laboratorio Analisi Osp. Città di Castello - Azienda USL Umbria7 | Istituto Zooprofilattico Sperimentale dell'Abruzzo e Molise "G. Caporale" | Malagigi V, Tacconi P, Lorusso A, Marcacci M, Di Domenico M, Ancora M, Curini V, Mangone I, Rinaldi A, Delli Compagni E, Scialabba S, Di Pasquale A, Cammà C, Puglia I, Calistri P, Savini G |
| EPI_ISL_2308717 | Laboratorio Analisi Osp. Città di Castello - Azienda USL Umbria8 | Istituto Zooprofilattico Sperimentale dell'Abruzzo e Molise "G. Caporale" | Malagigi V, Tacconi P, Lorusso A, Marcacci M, Di Domenico M, Ancora M, Curini V, Mangone I, Rinaldi A, Delli Compagni E, Scialabba S, Di Pasquale A, Cammà C, Puglia I, Calistri P, Savini G |
| EPI_ISL_2308718 | Laboratorio Analisi Osp. Città di Castello - Azienda USL Umbria9 | Istituto Zooprofilattico Sperimentale dell'Abruzzo e Molise "G. Caporale" | Malagigi V, Tacconi P, Lorusso A, Marcacci M, Di Domenico M, Ancora M, Curini V, Mangone I, Rinaldi A, Delli Compagni E, Scialabba S, Di Pasquale A, Cammà C, Puglia I, Calistri P, Savini G |
| EPI_ISL_2308719 | Laboratorio Analisi Osp. Città di Castello - Azienda USL Umbria10 | Istituto Zooprofilattico Sperimentale dell'Abruzzo e Molise "G. Caporale" | Malagigi V, Tacconi P, Lorusso A, Marcacci M, Di Domenico M, Ancora M, Curini V, Mangone I, Rinaldi A, Delli Compagni E, Scialabba S, Di Pasquale A, Cammà C, Puglia I, Calistri P, Savini G |
| EPI_ISL_2308720 | Laboratorio Analisi Osp. Città di Castello - Azienda USL Umbria11 | Istituto Zooprofilattico Sperimentale dell'Abruzzo e Molise "G. Caporale" | Malagigi V, Tacconi P, Lorusso A, Marcacci M, Di Domenico M, Ancora M, Curini V, Mangone I, Rinaldi A, Delli Compagni E, Scialabba S, Di Pasquale A, Cammà C, Puglia I, Calistri P, Savini G |
| EPI_ISL_2311861 | Fundação Ezequiel Dias | Coordenação Geral de Laboratórios de Saúde Pública (CGLAB/DAEVS/SVS/MS) | Vagner Fonseca, et al. |
| EPI_ISL_2312665 | Centre Hospitalier Billom | CHU Clermont-Ferrand, service de virologie | Bisseux Maxime, Mirand Audrey, Combes Patricia, Henquell Cécile |
| EPI_ISL_2312666, EPI_ISL_2312667 | Centre Hospitalier Universitaire Clermont-Ferrand | CHU Clermont-Ferrand, service de virologie | Bisseux Maxime, Mirand Audrey, Combes Patricia, Henquell Cécile |
| EPI_ISL_2312668, EPI_ISL_2312669 | HOPITAL DE VICHY | CHU Clermont-Ferrand, service de virologie | Bisseux Maxime, Mirand Audrey, Combes Patricia, Henquell Cécile |
| EPI_ISL_2312670 | Centre Hospitalier Universitaire Clermont-Ferrand | CHU Clermont-Ferrand, service de virologie | Bisseux Maxime, Mirand Audrey, Combes Patricia, Henquell Cécile |
| EPI_ISL_2312671 | Centre Hospitalier Paul Ardier | CHU Clermont-Ferrand, service de virologie | Bisseux Maxime, Mirand Audrey, Combes Patricia, Henquell Cécile |
| EPI_ISL_2312673, EPI_ISL_2312674 | Centre Hospitalier Universitaire Clermont-Ferrand | CHU Clermont-Ferrand, service de virologie | Bisseux Maxime, Mirand Audrey, Combes Patricia, Henquell Cécile |
| EPI_ISL_2312678 | EHPAD | CHU Clermont-Ferrand, service de virologie | Bisseux Maxime, Mirand Audrey, Combes Patricia, Henquell Cécile |
| EPI_ISL_2312679 | HOPITAL DE MONTLUCON | CHU Clermont-Ferrand, service de virologie | Bisseux Maxime, Mirand Audrey, Combes Patricia, Henquell Cécile |
| EPI_ISL_2312680 | Centre Hospitalier Universitaire Clermont-Ferrand | CHU Clermont-Ferrand, service de virologie | Bisseux Maxime, Mirand Audrey, Combes Patricia, Henquell Cécile |
| EPI_ISL_2312681, EPI_ISL_2312682 | Centre Hospitalier Billom | CHU Clermont-Ferrand, service de virologie | Bisseux Maxime, Mirand Audrey, Combes Patricia, Henquell Cécile |
| EPI_ISL_2312683 | HOPITAL DE MOULINS | CHU Clermont-Ferrand, service de virologie | Bisseux Maxime, Mirand Audrey, Combes Patricia, Henquell Cécile |
| EPI_ISL_2312685 | Centre hospitalier D'AURILLAC | CHU Clermont-Ferrand, service de virologie | Bisseux Maxime, Mirand Audrey, Combes Patricia, Henquell Cécile |
| EPI_ISL_2312686 | HOPITAL DE VICHY | CHU Clermont-Ferrand, service de virologie | Bisseux Maxime, Mirand Audrey, Combes Patricia, Henquell Cécile |
| EPI_ISL_2312688 | Centre hospitalier D'AURILLAC | CHU Clermont-Ferrand, service de virologie | Bisseux Maxime, Mirand Audrey, Combes Patricia, Henquell Cécile |
| EPI_ISL_2312689, EPI_ISL_2312690, EPI_ISL_2312692, EPI_ISL_2312693 | Centre Hospitalier Universitaire Clermont-Ferrand | CHU Clermont-Ferrand, service de virologie | Bisseux Maxime, Mirand Audrey, Combes Patricia, Henquell Cécile |
| EPI_ISL_2312694 | EHPAD | CHU Clermont-Ferrand, service de virologie | Bisseux Maxime, Mirand Audrey, Combes Patricia, Henquell Cécile |
| EPI_ISL_2312695, EPI_ISL_2312696, EPI_ISL_2312697 | Centre Hospitalier Universitaire Clermont-Ferrand | CHU Clermont-Ferrand, service de virologie | Bisseux Maxime, Mirand Audrey, Combes Patricia, Henquell Cécile |
| EPI_ISL_2312699 | EHPAD | CHU Clermont-Ferrand, service de virologie | Bisseux Maxime, Mirand Audrey, Combes Patricia, Henquell Cécile |
| EPI_ISL_2312700 | laboratoire maymat | CHU Clermont-Ferrand, service de virologie | Bisseux Maxime, Mirand Audrey, Combes Patricia, Henquell Cécile |
| EPI_ISL_2312702, EPI_ISL_2312705, EPI_ISL_2312706, EPI_ISL_2312707, EPI_ISL_2312708, EPI_ISL_2312709, EPI_ISL_2312710 | Centre Hospitalier Universitaire Clermont-Ferrand | CHU Clermont-Ferrand, service de virologie | Bisseux Maxime, Mirand Audrey, Combes Patricia, Henquell Cécile |
| EPI_ISL_2312711 | Centre De Prélévement COVID VICHY | CHU Clermont-Ferrand, service de virologie | Bisseux Maxime, Mirand Audrey, Combes Patricia, Henquell Cécile |
| EPI_ISL_2312713 | HOPITAL DU MONT DORE | CHU Clermont-Ferrand, service de virologie | Bisseux Maxime, Mirand Audrey, Combes Patricia, Henquell Cécile |
| EPI_ISL_2312714, EPI_ISL_2312715, EPI_ISL_2312716 | Centre De Prelevement COVID RIOM | CHU Clermont-Ferrand, service de virologie | Bisseux Maxime, Mirand Audrey, Combes Patricia, Henquell Cécile |
| EPI_ISL_2312718 | Centre Hospitalier Universitaire Clermont-Ferrand | CHU Clermont-Ferrand, service de virologie | Bisseux Maxime, Mirand Audrey, Combes Patricia, Henquell Cécile |

| | | | |
|---|---|---|---|
| EPI_ISL_2312719 | Centre De Prélévement COVID VICHY | CHU Clermont-Ferrand, service de virologie | Bisseux Maxime, Mirand Audrey, Combes Patricia, Henquell Cécile |
| EPI_ISL_2312720, EPI_ISL_2312721 | CDP COVID MONTLUCON | CHU Clermont-Ferrand, service de virologie | Bisseux Maxime, Mirand Audrey, Combes Patricia, Henquell Cécile |
| EPI_ISL_2312722 | HOPITAL DU MONT DORE | CHU Clermont-Ferrand, service de virologie | Bisseux Maxime, Mirand Audrey, Combes Patricia, Henquell Cécile |
| EPI_ISL_2312723, EPI_ISL_2312725, EPI_ISL_2312726 | Centre Hospitalier Universitaire Clermont-Ferrand | CHU Clermont-Ferrand, service de virologie | Bisseux Maxime, Mirand Audrey, Combes Patricia, Henquell Cécile |
| EPI_ISL_2319604, EPI_ISL_2319605, EPI_ISL_2319606, EPI_ISL_2319607, EPI_ISL_2319608, EPI_ISL_2319609, EPI_ISL_2319610, EPI_ISL_2319611, EPI_ISL_2319612, EPI_ISL_2319613, EPI_ISL_2319614, EPI_ISL_2319615, EPI_ISL_2319616, EPI_ISL_2319617, EPI_ISL_2319618, EPI_ISL_2319619, EPI_ISL_2319620, EPI_ISL_2319621, EPI_ISL_2319622, EPI_ISL_2319623, EPI_ISL_2319624, EPI_ISL_2319625, EPI_ISL_2319626, EPI_ISL_2319627 | | | |
| see above | Cliniques universitaires Saint-Luc | UCLouvain/IREC/MBLG | Jean Ruelle, Ophélie Simon, Nicolas Pinte, Benoit Kabamba Mukadi |
| EPI_ISL_2322448 | National Institute of Infectious Diseases-Prof. Dr. Matei Bals Molecular Diagnostics Laboratory | National Institute of Infectious Diseases-Prof. Dr. Matei Bals Molecular Diagnostics Laboratory | Corina Casangiu, Leontina Banica, Marius Surleac, Ovidiu Vlaicu, Andreea Tudor, Simona Paraschiv, Dan Otelea |
| EPI_ISL_2322998 | Imeda Hospital | Imeda Hospital | Johan Frans, Dagmar Obbels, Hanne Valgaeren |
| EPI_ISL_2323001, EPI_ISL_2323004 | AZ Sint Maarten | Imeda Hospital | Johan Frans, Dagmar Obbels, Hanne Valgaeren |
| EPI_ISL_2323142, EPI_ISL_2323143, EPI_ISL_2323144, EPI_ISL_2323145, EPI_ISL_2323146, EPI_ISL_2323147, EPI_ISL_2323148, EPI_ISL_2323149 | Laboratorio Microbiologia PO Cardarelli | Laboratorio Microbiologia PO Cardarelli | Felice V., Niro G. Scutella' M. |
| EPI_ISL_2324301, EPI_ISL_2324302, EPI_ISL_2324303, EPI_ISL_2324304, EPI_ISL_2324305, EPI_ISL_2324306, EPI_ISL_2324307, EPI_ISL_2324308, EPI_ISL_2324309, EPI_ISL_2324310, EPI_ISL_2324312, EPI_ISL_2324317, EPI_ISL_2324318, EPI_ISL_2324319, EPI_ISL_2324320, EPI_ISL_2324321, EPI_ISL_2324322, EPI_ISL_2324323, EPI_ISL_2324324, EPI_ISL_2324325, EPI_ISL_2324326, EPI_ISL_2324327, EPI_ISL_2324330, EPI_ISL_2324335, EPI_ISL_2324336, EPI_ISL_2324337, EPI_ISL_2324354, EPI_ISL_2324355, EPI_ISL_2324356, EPI_ISL_2324357, EPI_ISL_2324358, EPI_ISL_2324359, EPI_ISL_2324361, EPI_ISL_2324364, EPI_ISL_2324365, EPI_ISL_2324366, EPI_ISL_2324367, EPI_ISL_2324368, EPI_ISL_2324369, EPI_ISL_2324370, EPI_ISL_2324371, EPI_ISL_2324372, EPI_ISL_2324383, EPI_ISL_2324385, EPI_ISL_2324387, EPI_ISL_2324388, EPI_ISL_2324389 | | | |
| see above | Austrian Agency for Health and Food Safety (AGES) | Bergthaler laboratory, CeMM Research Center for Molecular Medicine of the Austrian Academy of Sciences | Lukas Endler, Anna Schedl, Fabian Amman, Petr Triska, Thomas Penz, Benedikt Agerer, Maelle Le Moing, Michael Schuster, Bekir Erguner, Jan Laine, Martin Senekowitsch, Christoph Bock, Andreas Bergthaler |
| EPI_ISL_2324405, EPI_ISL_2324406, EPI_ISL_2324408, EPI_ISL_2324409, EPI_ISL_2324410 | Laboratorio di Genetica Medica, Matera, Italy | Laboratorio di Genetica Medica, Matera, Italy | Domenico Dell'Edera, Arianna Allegretti |
| EPI_ISL_2324411, EPI_ISL_2324412, EPI_ISL_2324413, EPI_ISL_2324414, EPI_ISL_2324415, EPI_ISL_2324416, EPI_ISL_2324417, EPI_ISL_2324419, EPI_ISL_2324420, EPI_ISL_2324422, EPI_ISL_2324423, EPI_ISL_2324424 | | | |
| see above | SC (UCO) Igiene e Sanità Pubblica, ASUGI, Trieste | ARGO Laboratorio Genomica ed Epigenomica | Licastro D, Dal Monego S, Degasperi M, Marcello A, Segat L, Piscianz E, D'Agaro P |
| EPI_ISL_2324425, EPI_ISL_2324426, EPI_ISL_2324427 | SC (UCO) Igiene e Sanità Pubblica, ASUGI, Trieste and SOC Igiene ed Epidemiologia Clinica, laboratorio di Virologia ,ASUFC, Udine | ARGO Laboratorio Genomica ed Epigenomica | Licastro D, Dal Monego S, Degasperi M, Marcello A, Segat L, Pipan C, D'Agaro P |
| EPI_ISL_2324428, EPI_ISL_2324429, EPI_ISL_2324430, EPI_ISL_2324431 | SC (UCO) Igiene e Sanità Pubblica, ASUGI, Trieste and SC Microbiologia e Virologia, ASFO, Pordenone | ARGO Laboratorio Genomica ed Epigenomica | Licastro D, Dal Monego S, Degasperi M, Marcello A, Segat L, De Rosa R, D'Agaro P |
| EPI_ISL_2324432 | SC (UCO) Igiene e Sanità Pubblica, ASUGI, Trieste and SOC Igiene ed Epidemiologia Clinica, laboratorio di Virologia ,ASUFC, Udine | ARGO Laboratorio Genomica ed Epigenomica | Licastro D, Dal Monego S, Degasperi M, Marcello A, Segat L, Pipan C, D'Agaro P |
| EPI_ISL_2324433, EPI_ISL_2324434, EPI_ISL_2324435, EPI_ISL_2324436, EPI_ISL_2324437, EPI_ISL_2324438, EPI_ISL_2324439 | SC (UCO) Igiene e Sanità Pubblica, ASUGI, Trieste | ARGO Laboratorio Genomica ed Epigenomica | Licastro D, Dal Monego S, Degasperi M, Marcello A, Segat L, Piscianz E, D'Agaro P |
| EPI_ISL_2332791, EPI_ISL_2332792, EPI_ISL_2332793, EPI_ISL_2332794, EPI_ISL_2332795, EPI_ISL_2332796, EPI_ISL_2332797, EPI_ISL_2332798, EPI_ISL_2332799, EPI_ISL_2332800, EPI_ISL_2332801, EPI_ISL_2332802, EPI_ISL_2332803, EPI_ISL_2332804, EPI_ISL_2332805, EPI_ISL_2332806, EPI_ISL_2332807, EPI_ISL_2332808, EPI_ISL_2332809, EPI_ISL_2332810, EPI_ISL_2332811 | | | |
| see above | OLVZ Aalst | OLVZ Aalst | Anne Vankeerberghen |
| EPI_ISL_2333654 | RSUD Cilacap | Genetics Working Group (Pokja Genetik) Faculty of Medicine, Public Health and Nursing Universitas Gadjah Mada (FK-KMK UGM); Disease Investigation Center Wates Ministry of Agriculture Indonesia; Department of Microbiology FK-KMK UGM; Laboratorium Diagnostik Yayasan Tahija World Mosquito Program (WMP) Yogyakarta Center for Tropical Medicine FK-KMK UGM; Integrated Research Center FK-KMK UGM; Department of Computer Science and Electronics FMIPA UGM; RSUP Dr. Sardjito | Gunadi, Hendra Wibawa, Marcellus, Mohamad S. Hakim, Edwin W. Daniwijaya, Ludhang P. Rizki, Endah Supriyati, Eggi Arguni, Titik Nuryastuti, Tri Wibawa, Dwi AA Nugrahaningsih, Afiahayati, Siswanto,Kristy Iskandar,Nungki Anggorowati, Pramesti G Dewi, Dwi Indaryati,Indarto Sulistiyono, Susan Simanjaya, Dyah A Puspitarani |
| EPI_ISL_2333790 | GH A.CHENEVIER-H.MONDOR | Department of Virology, Henri Mondor University Hospital, Assistance Publique Hôpitaux de Paris, Université Paris-Est Créteil, INSERM U955 | Christophe Rodriguez, Slim Fourati, Vanessa Demontant, Guillaume Gricourt, Melissa N'Debi, Alexandre Soulier, Elisabeth Trawinski, Jean-Michel Pawlotsky |
| EPI_ISL_2333810 | RSUD Cilacap | Genetics Working Group (Pokja Genetik) Faculty of Medicine, Public Health and Nursing Universitas Gadjah Mada (FK-KMK UGM); Disease Investigation Center Wates Ministry of Agriculture Indonesia; Department of Microbiology FK-KMK UGM; Laboratorium Diagnostik Yayasan Tahija World Mosquito Program (WMP) Yogyakarta Center for Tropical Medicine FK-KMK UGM; Integrated Research Center FK-KMK UGM; Department of Computer Science and Electronics FMIPA UGM; RSUP Dr. Sardjito | Gunadi, Hendra Wibawa, Marcellus, Mohamad S. Hakim, Edwin W. Daniwijaya, Ludhang P. Rizki, Endah Supriyati, Eggi Arguni, Titik Nuryastuti, Tri Wibawa, Dwi AA Nugrahaningsih, Afiahayati, Siswanto,Kristy Iskandar,Nungki Anggorowati, Pramesti G Dewi, Dwi Indaryati,Indarto Sulistiyono, Untung Riawan, Maria P Inggriani |
| EPI_ISL_2333811, EPI_ISL_2333827 | GH A.CHENEVIER-H.MONDOR | Department of Virology, Henri Mondor University Hospital, Assistance Publique Hôpitaux de Paris, Université Paris-Est Créteil, INSERM U955 | Christophe Rodriguez, Slim Fourati, Vanessa Demontant, Guillaume Gricourt, Melissa N'Debi, Alexandre Soulier, Elisabeth Trawinski, Jean-Michel Pawlotsky |
| EPI_ISL_2334094 | RSUD Cilacap | Genetics Working Group (Pokja Genetik) Faculty of Medicine, Public Health and Nursing Universitas Gadjah Mada (FK-KMK UGM); Disease Investigation Center Wates Ministry of Agriculture Indonesia; Department of Microbiology FK-KMK UGM; Laboratorium Diagnostik Yayasan Tahija World Mosquito Program (WMP) Yogyakarta Center for Tropical Medicine FK-KMK UGM; Integrated Research Center FK-KMK UGM; Department of Computer Science and Electronics FMIPA UGM; RSUP Dr. Sardjito | Gunadi, Hendra Wibawa, Marcellus, Mohamad S. Hakim, Edwin W. Daniwijaya, Ludhang P. Rizki, Endah Supriyati, Eggi Arguni, Titik Nuryastuti, Tri Wibawa, Dwi AA Nugrahaningsih, Afiahayati, Siswanto,Kristy Iskandar,Nungki Anggorowati, Pramesti G Dewi, Dwi Indaryati,Indarto Sulistiyono, Cita S Amalia, Alvina A Setiawan |

| | | | |
|---|---|---|---|
| EPI_ISL_2334474 | RSUD Cilacap | Genetics Working Group (Pokja Genetik) Faculty of Medicine, Public Health and Nursing Universitas Gadjah Mada (FK-KMK UGM); Disease Investigation Center Wates Ministry of Agriculture Indonesia; Department of Microbiology FK-KMK UGM; Laboratorium Diagnostik Yayasan Tahija World Mosquito Program (WMP) Yogyakarta Center for Tropical Medicine FK-KMK UGM; Integrated Research Center FK-KMK UGM; Department of Computer Science and Electronics FMIPA UGM; RSUP Dr. Sardjito | Gunadi, Hendra Wibawa, Marcellus, Mohamad S. Hakim, Edwin W. Daniwijaya, Ludhang P. Rizki, Endah Supriyati, Eggi Arguni, Titik Nuryastuti, Tri Wibawa, Dwi AA Nugrahaningsih, Afiahayati, Siswanto,Kristy Iskandar,Nungki Anggorowati, Pramesti G Dewi, Dwi Indaryati,Indarto Sulistiyono, Susan Simanjaya, I Putu Aditio A |
| EPI_ISL_2334587 | RSUD Cilacap | Genetics Working Group (Pokja Genetik) Faculty of Medicine, Public Health and Nursing Universitas Gadjah Mada (FK-KMK UGM); Disease Investigation Center Wates Ministry of Agriculture Indonesia; Department of Microbiology FK-KMK UGM; Laboratorium Diagnostik Yayasan Tahija World Mosquito Program (WMP) Yogyakarta Center for Tropical Medicine FK-KMK UGM; Integrated Research Center FK-KMK UGM; Department of Computer Science and Electronics FMIPA UGM; RSUP Dr. Sardjito | Gunadi, Hendra Wibawa, Marcellus, Mohamad S. Hakim, Edwin W. Daniwijaya, Ludhang P. Rizki, Endah Supriyati, Eggi Arguni, Titik Nuryastuti, Tri Wibawa, Dwi AA Nugrahaningsih, Afiahayati, Siswanto,Kristy Iskandar,Nungki Anggorowati, Pramesti G Dewi, Dwi Indaryati,Indarto Sulistiyono, Susan Simanjaya, Dyah A Puspitarani |
| EPI_ISL_2335021 | RSUD Cilacap | Genetics Working Group (Pokja Genetik) Faculty of Medicine, Public Health and Nursing Universitas Gadjah Mada (FK-KMK UGM); Disease Investigation Center Wates Ministry of Agriculture Indonesia; Department of Microbiology FK-KMK UGM; Laboratorium Diagnostik Yayasan Tahija World Mosquito Program (WMP) Yogyakarta Center for Tropical Medicine FK-KMK UGM; Integrated Research Center FK-KMK UGM; Department of Computer Science and Electronics FMIPA UGM; RSUP Dr. Sardjito | Gunadi, Hendra Wibawa, Marcellus, Mohamad S. Hakim, Edwin W. Daniwijaya, Ludhang P. Rizki, Endah Supriyati, Eggi Arguni, Titik Nuryastuti, Tri Wibawa, Dwi AA Nugrahaningsih, Afiahayati, Siswanto,Kristy Iskandar,Nungki Anggorowati, Pramesti G Dewi, Dwi Indaryati,Indarto Sulistiyono, Untung Riawan, Maria P Inggriani |
| EPI_ISL_2335090 | RSUD CILACAP | Genetics Working Group (Pokja Genetik) Faculty of Medicine, Public Health and Nursing Universitas Gadjah Mada (FK-KMK UGM); Disease Investigation Center Wates Ministry of Agriculture Indonesia; Department of Microbiology FK-KMK UGM; Laboratorium Diagnostik Yayasan Tahija World Mosquito Program (WMP) Yogyakarta Center for Tropical Medicine FK-KMK UGM; Integrated Research Center FK-KMK UGM; Department of Computer Science and Electronics FMIPA UGM; RSUP Dr. Sardjito | Gunadi, Hendra Wibawa, Marcellus, Mohamad S. Hakim, Edwin W. Daniwijaya, Ludhang P. Rizki, Endah Supriyati, Eggi Arguni, Titik Nuryastuti, Tri Wibawa, Dwi AA Nugrahaningsih, Afiahayati, Siswanto,Kristy Iskandar,Nungki Anggorowati, Pramesti G Dewi, Dwi Indaryati,Indarto Sulistiyono, Irene Tania, Hana F Hanifin |
| EPI_ISL_2335146 | RSUD CILACAP | Genetics Working Group (Pokja Genetik) Faculty of Medicine, Public Health and Nursing Universitas Gadjah Mada (FK-KMK UGM); Disease Investigation Center Wates Ministry of Agriculture Indonesia; Department of Microbiology FK-KMK UGM; Laboratorium Diagnostik Yayasan Tahija World Mosquito Program (WMP) Yogyakarta Center for Tropical Medicine FK-KMK UGM; Integrated Research Center FK-KMK UGM; Department of Computer Science and Electronics FMIPA UGM; RSUP Dr. Sardjito | Gunadi, Hendra Wibawa, Marcellus, Mohamad S. Hakim, Edwin W. Daniwijaya, Ludhang P. Rizki, Endah Supriyati, Eggi Arguni, Titik Nuryastuti, Tri Wibawa, Dwi AA Nugrahaningsih, Afiahayati, Siswanto,Kristy Iskandar,Nungki Anggorowati, Pramesti G Dewi, Dwi Indaryati,Indarto Sulistiyono, Cita S Amalia, Alvina A Setiawan |
| EPI_ISL_2335456 | RSUD CILACAP | Genetics Working Group (Pokja Genetik) Faculty of Medicine, Public Health and Nursing Universitas Gadjah Mada (FK-KMK UGM); Disease Investigation Center Wates Ministry of Agriculture Indonesia; Department of Microbiology FK-KMK UGM; Laboratorium Diagnostik Yayasan Tahija World Mosquito Program (WMP) Yogyakarta Center for Tropical Medicine FK-KMK UGM; Integrated Research Center FK-KMK UGM; Department of Computer Science and Electronics FMIPA UGM; RSUP Dr. Sardjito | Gunadi, Hendra Wibawa, Marcellus, Mohamad S. Hakim, Edwin W. Daniwijaya, Ludhang P. Rizki, Endah Supriyati, Eggi Arguni, Titik Nuryastuti, Tri Wibawa, Dwi AA Nugrahaningsih, Afiahayati, Siswanto,Kristy Iskandar,Nungki Anggorowati, Pramesti G Dewi, Dwi Indaryati,Indarto Sulistiyono, Susan Simanjaya, I Putu Aditio A |
| EPI_ISL_2335872 | RSUD CILACAP | Genetics Working Group (Pokja Genetik) Faculty of Medicine, Public Health and Nursing Universitas Gadjah Mada (FK-KMK UGM); Disease Investigation Center Wates Ministry of Agriculture Indonesia; Department of Microbiology FK-KMK UGM; Laboratorium Diagnostik Yayasan Tahija World Mosquito Program (WMP) Yogyakarta Center for Tropical Medicine FK-KMK UGM; Integrated Research Center FK-KMK UGM; Department of Computer Science and Electronics FMIPA UGM; RSUP Dr. Sardjito | Gunadi, Hendra Wibawa, Marcellus, Mohamad S. Hakim, Edwin W. Daniwijaya, Ludhang P. Rizki, Endah Supriyati, Eggi Arguni, Titik Nuryastuti, Tri Wibawa, Dwi AA Nugrahaningsih, Afiahayati, Siswanto,Kristy Iskandar,Nungki Anggorowati, Pramesti G Dewi, Dwi Indaryati,Indarto Sulistiyono, Susan Simanjaya, Dyah A Puspitarani |
| EPI_ISL_2336149 | RSUD CILACAP | Genetics Working Group (Pokja Genetik) Faculty of Medicine, Public Health and Nursing Universitas Gadjah Mada (FK-KMK UGM); Disease Investigation Center Wates Ministry of Agriculture Indonesia; Department of Microbiology FK-KMK UGM; Laboratorium Diagnostik Yayasan Tahija World Mosquito Program (WMP) Yogyakarta Center for Tropical Medicine FK-KMK UGM; Integrated Research Center FK-KMK UGM; Department of Computer Science and Electronics FMIPA UGM; RSUP Dr. Sardjito | Gunadi, Hendra Wibawa, Marcellus, Mohamad S. Hakim, Edwin W. Daniwijaya, Ludhang P. Rizki, Endah Supriyati, Eggi Arguni, Titik Nuryastuti, Tri Wibawa, Dwi AA Nugrahaningsih, Afiahayati, Siswanto,Kristy Iskandar,Nungki Anggorowati, Pramesti G Dewi, Dwi Indaryati,Indarto Sulistiyono, Untung Riawan, Maria P Inggriani |
| EPI_ISL_2337254 | RS Dr Oen Solo Baru | Genetics Working Group (Pokja Genetik) Faculty of Medicine, Public Health and Nursing Universitas Gadjah Mada (FK-KMK UGM); Disease Investigation Center Wates Ministry of Agriculture Indonesia; Department of Microbiology FK-KMK UGM; Laboratorium Diagnostik Yayasan Tahija World Mosquito Program (WMP) Yogyakarta Center for Tropical Medicine FK-KMK UGM; Integrated Research Center FK-KMK UGM; Department of Computer Science and Electronics FMIPA UGM; RSUP Dr. Sardjito | Gunadi, Hendra Wibawa, Marcellus, Mohamad S. Hakim, Edwin W. Daniwijaya, Ludhang P. Rizki, Endah Supriyati, Eggi Arguni, Titik Nuryastuti, Tri Wibawa, Dwi AA Nugrahaningsih, Afiahayati, Siswanto,Kristy Iskandar,Nungki Anggorowati,Irene Tania, Hana F Hanifin |
| EPI_ISL_2338534 | RSUP DR SARDJITO | Genetics Working Group (Pokja Genetik) Faculty of Medicine, Public Health and Nursing Universitas Gadjah Mada (FK-KMK | Gunadi, Hendra Wibawa, . Marcellus, Mohamad S. Hakim, Edwin W. Daniwijaya, Ludhang P. Rizki, Endah Supriyati, Eggi Arguni, Titik Nuryastuti, Tri Wibawa, Dwi AA Nugrahaningsih, Afiahayati, Siswanto, Kristy Iskandar, Nungki Anggorowati, Ika Trisnawati, Sumardi, Eko Budiono, Bambang Sigit |

| | | | |
|---|---|---|---|
| | | UGM); Disease Investigation Center Wates Ministry of Agriculture Indonesia; Department of Microbiology FK-KMK UGM; Laboratorium Diagnostik Yayasan Tahija World Mosquito Program (WMP) Yogyakarta Center for Tropical Medicine FK-KMK UGM; Integrated Research Center FK-KMK UGM; Department of Computer Science and Electronics FMIPA UGM; RSUP Dr. Sardjito | Riyanto, Heni Retnowulan, Munawar Gani, Satria Maulana, Nur Rahmi Ananda, Riat El Khair, Yunika Puspadewi, Osman Sianipar, Umi Solekhah Intansari, Elizabeth Henny Herningtiyas, Ira Puspitawati, Nur Imma Fatimah Harahap, Dyah A Puspitarani, Untung Riawan |
| EPI_ISL_2340887 | LESP Campeche | Instituto de Diagnostico y Referencia Epidemiologicos (INDRE) | Claudia Wong-Arambula, Abril Rodriguez-Maldonado, Vanessa Rivero-Arredondo, Ariadna Medina-Benitez, Joaquin Quiroz-Mercado, Sergio Rangel-Guerrero, Natividad Cruz-Ortiz, Tatiana Nunez-Garcia, Gisela Barrera-Badillo, Lucia Hernandez-Rivas, Irma Lopez-Martinez, Ernesto Ramirez-Gonzalez. |
| EPI_ISL_2341014, EPI_ISL_2341015 | LESP Morelos | Instituto de Diagnostico y Referencia Epidemiologicos (INDRE) | Claudia Wong-Arambula, Abril Rodriguez-Maldonado, Vanessa Rivero-Arredondo, Ariadna Medina-Benitez, Joaquin Quiroz-Mercado, Sergio Rangel-Guerrero, Natividad Cruz-Ortiz, Tatiana Nunez-Garcia, Gisela Barrera-Badillo, Lucia Hernandez-Rivas, Irma Lopez-Martinez, Ernesto Ramirez-Gonzalez. |
| EPI_ISL_2341020 | LESP Guanajuato | Instituto de Diagnostico y Referencia Epidemiologicos (INDRE) | Claudia Wong-Arambula, Abril Rodriguez-Maldonado, Vanessa Rivero-Arredondo, Ariadna Medina-Benitez, Joaquin Quiroz-Mercado, Sergio Rangel-Guerrero, Natividad Cruz-Ortiz, Tatiana Nunez-Garcia, Gisela Barrera-Badillo, Lucia Hernandez-Rivas, Irma Lopez-Martinez, Ernesto Ramirez-Gonzalez. |
| EPI_ISL_2341022 | LESP Nuevo Leon | Instituto de Diagnostico y Referencia Epidemiologicos (INDRE) | Claudia Wong-Arambula, Abril Rodriguez-Maldonado, Vanessa Rivero-Arredondo, Ariadna Medina-Benitez, Joaquin Quiroz-Mercado, Sergio Rangel-Guerrero, Natividad Cruz-Ortiz, Tatiana Nunez-Garcia, Gisela Barrera-Badillo, Lucia Hernandez-Rivas, Irma Lopez-Martinez, Ernesto Ramirez-Gonzalez. |
| EPI_ISL_2343017 | LESP Colima | Instituto de Diagnostico y Referencia Epidemiologicos (INDRE) | Claudia Wong-Arambula, Abril Rodriguez-Maldonado, Vanessa Rivero-Arredondo, Ariadna Medina-Benitez, Joaquin Quiroz-Mercado, Sergio Rangel-Guerrero, Natividad Cruz-Ortiz, Tatiana Nunez-Garcia, Gisela Barrera-Badillo, Lucia Hernandez-Rivas, Irma Lopez-Martinez, Ernesto Ramirez-Gonzalez. |
| EPI_ISL_2343208, EPI_ISL_2343209, EPI_ISL_2343210, EPI_ISL_2343211, EPI_ISL_2343212, EPI_ISL_2343213, EPI_ISL_2343214, EPI_ISL_2343215, EPI_ISL_2343216, EPI_ISL_2343217, EPI_ISL_2343218, EPI_ISL_2343219, EPI_ISL_2343220, EPI_ISL_2343221, EPI_ISL_2343222, EPI_ISL_2343223, EPI_ISL_2343224 | | | |
| see above | Banteay Meanchey Rapid Response Team | Virology Unit, Institut Pasteur du Cambodge | Jurre Y Siegers, Cecile Troupin, Leakhena Pum, Ly Sovann, Kraing Sidonn, Yi Sengdoeurn, Chin Savuth, Chau Darapheak, Veasna Duong, Erik A Karlsson |
| EPI_ISL_2343225, EPI_ISL_2343226 | Battambang Provincial Laboratory | Virology Unit, Institut Pasteur du Cambodge | Jurre Y Siegers, Cecile Troupin, Leakhena Pum, Ly Sovann, Kraing Sidonn, Yi Sengdoeurn, Chin Savuth, Chau Darapheak, Veasna Duong, Erik A Karlsson |
| EPI_ISL_2343227 | Kampong Cham Rapid Response Team | Virology Unit, Institut Pasteur du Cambodge | Jurre Y Siegers, Cecile Troupin, Leakhena Pum, Ly Sovann, Kraing Sidonn, Yi Sengdoeurn, Chin Savuth, Chau Darapheak, Veasna Duong, Erik A Karlsson |
| EPI_ISL_2343228 | Kampong Thom Rapid Response Team | Virology Unit, Institut Pasteur du Cambodge | Jurre Y Siegers, Cecile Troupin, Leakhena Pum, Ly Sovann, Kraing Sidonn, Yi Sengdoeurn, Chin Savuth, Chau Darapheak, Veasna Duong, Erik A Karlsson |
| EPI_ISL_2343229, EPI_ISL_2343230 | Oddar Meanchey Rapid Response Team | Virology Unit, Institut Pasteur du Cambodge | Jurre Y Siegers, Cecile Troupin, Leakhena Pum, Ly Sovann, Kraing Sidonn, Yi Sengdoeurn, Chin Savuth, Chau Darapheak, Veasna Duong, Erik A Karlsson |
| EPI_ISL_2343231, EPI_ISL_2343232, EPI_ISL_2343233, EPI_ISL_2343234, EPI_ISL_2343235, EPI_ISL_2343236, EPI_ISL_2343237, EPI_ISL_2343238, EPI_ISL_2343239, EPI_ISL_2343240, EPI_ISL_2343241, EPI_ISL_2343242, EPI_ISL_2343243, EPI_ISL_2343244, EPI_ISL_2343245, EPI_ISL_2343246, EPI_ISL_2343247, EPI_ISL_2343248 | | | |
| see above | Virology Unit, Institut Pasteur du Cambodge | Virology Unit, Institut Pasteur du Cambodge | Jurre Y Siegers, Cecile Troupin, Leakhena Pum, Ly Sovann, Kraing Sidonn, Yi Sengdoeurn, Chin Savuth, Chau Darapheak, Veasna Duong, Erik A Karlsson |
| EPI_ISL_2343249 | Kandal Rapid Response Team | Virology Unit, Institut Pasteur du Cambodge | Jurre Y Siegers, Cecile Troupin, Leakhena Pum, Ly Sovann, Kraing Sidonn, Yi Sengdoeurn, Chin Savuth, Chau Darapheak, Veasna Duong, Erik A Karlsson |
| EPI_ISL_2343250 | Kampot Rapid Response Team | Virology Unit, Institut Pasteur du Cambodge | Jurre Y Siegers, Cecile Troupin, Leakhena Pum, Ly Sovann, Kraing Sidonn, Yi Sengdoeurn, Chin Savuth, Chau Darapheak, Veasna Duong, Erik A Karlsson |
| EPI_ISL_2343251, EPI_ISL_2343252 | Banteay Meanchey Rapid Response Team | Virology Unit, Institut Pasteur du Cambodge | Jurre Y Siegers, Cecile Troupin, Leakhena Pum, Ly Sovann, Kraing Sidonn, Yi Sengdoeurn, Chin Savuth, Chau Darapheak, Veasna Duong, Erik A Karlsson |
| EPI_ISL_2346341, EPI_ISL_2346342, EPI_ISL_2346343, EPI_ISL_2346344, EPI_ISL_2346345, EPI_ISL_2346346, EPI_ISL_2346347, EPI_ISL_2346348, EPI_ISL_2346349, EPI_ISL_2346352, EPI_ISL_2346353, EPI_ISL_2346354, EPI_ISL_2346355, EPI_ISL_2346356, EPI_ISL_2346357, EPI_ISL_2346358, EPI_ISL_2346359, EPI_ISL_2346360, EPI_ISL_2346361, EPI_ISL_2346362, EPI_ISL_2346363, EPI_ISL_2346364, EPI_ISL_2346366, EPI_ISL_2346367, EPI_ISL_2346368, EPI_ISL_2346369, EPI_ISL_2346370, EPI_ISL_2346371, EPI_ISL_2346374, EPI_ISL_2346375, EPI_ISL_2346376, EPI_ISL_2346377, EPI_ISL_2346378, EPI_ISL_2346379, EPI_ISL_2346380 | | | |
| see above | Institute of Molecular and Translational Medicine / Laboratory of Experimental Medicine, Faculty of Medicine and Dentistry, Palacky University and University Hospital Olomouc | Institute of Molecular and Translational Medicine / Laboratory of Experimental Medicine, Faculty of Medicine and Dentistry, Palacky University | Rastislav Slavkovský, Hana Jaworek, Vladimíra Koudeláková, Barbora Blumová, Marián Hajdúch |
| EPI_ISL_2348840, EPI_ISL_2348841, EPI_ISL_2348842, EPI_ISL_2348843, EPI_ISL_2348844, EPI_ISL_2348845, EPI_ISL_2348846, EPI_ISL_2348847, EPI_ISL_2348848, EPI_ISL_2348849, EPI_ISL_2348850, EPI_ISL_2348851, EPI_ISL_2348852, EPI_ISL_2348853, EPI_ISL_2348854, EPI_ISL_2348855, EPI_ISL_2348856, EPI_ISL_2348857, EPI_ISL_2348858, EPI_ISL_2348859, EPI_ISL_2348860, EPI_ISL_2348861, EPI_ISL_2348862, EPI_ISL_2348863, EPI_ISL_2348864, EPI_ISL_2348865, EPI_ISL_2348866, EPI_ISL_2348867, EPI_ISL_2348868, EPI_ISL_2348869, EPI_ISL_2348870, EPI_ISL_2348871, EPI_ISL_2348872, EPI_ISL_2348873, EPI_ISL_2348874, EPI_ISL_2348875, EPI_ISL_2348876, EPI_ISL_2348877, EPI_ISL_2348878, EPI_ISL_2348879, EPI_ISL_2348880, EPI_ISL_2348881, EPI_ISL_2348882, EPI_ISL_2348883, EPI_ISL_2348884, EPI_ISL_2348885, EPI_ISL_2348886, EPI_ISL_2348887, EPI_ISL_2348889, EPI_ISL_2348890, EPI_ISL_2348891, EPI_ISL_2348893, EPI_ISL_2348894, EPI_ISL_2348895, EPI_ISL_2348896, EPI_ISL_2348897, EPI_ISL_2348898, EPI_ISL_2348899, EPI_ISL_2348900, EPI_ISL_2348901, EPI_ISL_2348902, EPI_ISL_2348903, EPI_ISL_2348904, EPI_ISL_2348905, EPI_ISL_2348906, EPI_ISL_2348907, EPI_ISL_2348908, EPI_ISL_2348910, EPI_ISL_2348911, EPI_ISL_2348912, EPI_ISL_2348915, EPI_ISL_2348916, EPI_ISL_2348917, EPI_ISL_2348918, EPI_ISL_2348919 | | | |
| see above | Cliniques universitaires Saint-Luc | UCLouvain/IREC/MBLG | Jean Ruelle, Ophélie Simon, Nicolas Pinte, Benoit Kabamba Mukadi |
| EPI_ISL_2348923, EPI_ISL_2348924, EPI_ISL_2348925 | Hôpitaux Iris Sud Etterbeek-Ixelles | UCLouvain/IREC/MBLG | Jean Ruelle, Ophélie Simon, Nicolas Pinte, Benoit Kabamba Mukadi |
| EPI_ISL_2349697, EPI_ISL_2349705 | Genome Analysis Center, Yamanashi Central Hospital | Genome Analysis Center, Yamanashi Central Hospital | Yosuke Hirotsu |
| EPI_ISL_2354481, EPI_ISL_2354499, EPI_ISL_2354806, EPI_ISL_2355213 | Dr. Gernot Walder GmbH | Dr. Gernot Walder GmbH | Sissy T. Sonnleitner |
| EPI_ISL_2357733, EPI_ISL_2357784, EPI_ISL_2357785, EPI_ISL_2357813, EPI_ISL_2357816, EPI_ISL_2357817, EPI_ISL_2357897, EPI_ISL_2358032 | Institut de Virologie du CHU de Strasbourg | Institut de Virologie du CHU de Strasbourg | Fafi-Kremer Samira, Gallais Floriane, Gantner Pierre, Laugel Elodie, Solis Morgane, Velay Aurélie, Wendling Marie-Josée |
| EPI_ISL_2358037 | Laboratoire Hôpital Sainte Musse | Laboratoire Hôpital Sainte Musse | Lionel Chollet |
| EPI_ISL_2360250 | Afzalipoor Hospital | National Influenza Center | V Salimi,NZ Shafiei Jandaghi, J Yavarian, A Nejati, K Sadeghi, N Ghavvami,F Ajaminejad and T Mokhtari Azad |
| EPI_ISL_2360251 | Afzalipoor Hospital | National Influenza Center | NZ Shafiei Jandaghi, V Salimi, A Nejati, K Sadeghi, J Yavarian, N Ghavvami,F Ajaminejad and T Mokhtari Azad |
| EPI_ISL_2361089, EPI_ISL_2361090, EPI_ISL_2361091 | Institute of Molecular and Translational Medicine / Laboratory of Experimental Medicine, Faculty of Medicine and Dentistry, Palacky University and University Hospital Olomouc | Institute of Molecular and Translational Medicine / Laboratory of Experimental Medicine, Faculty of Medicine and Dentistry, Palacky University | Rastislav Slavkovský, Hana Jaworek, Vladimíra Koudeláková, Barbora Blumová, Marián Hajdúch |
| EPI_ISL_2361523, EPI_ISL_2361524 | CH ROUBAIX | CHU Lille - Laboratoire de Virologie | AIT YAHYA Emilie, ALIDJINOU Enagnon Kazali, BOCKET Laurence, CREPIN Michel, DEMAY Christophe, ENGELMANN Ilka, GEFFROY Sandrine, GUIGON Aurélie, LAMBERT Valérie, LAZREK Mouna, NOBILLIAUX Florian, PREVOST Brigitte, THUILLIER Caroline, TINEZ Claire ,TCHANTCHOU NJOSSE YANICK |

| | | | |
|---|---|---|---|
| EPI_ISL_2361525 | CHU LILLE | CHU Lille - Laboratoire de Virologie | AIT YAHYA Emilie, ALIDJINOU Enagnon Kazali, BOCKET Laurence, CREPIN Michel, DEMAY Christophe, ENGELMANN Ilka, GEFFROY Sandrine, GUIGON Aurélie, LAMBERT Valérie, LAZREK Mouna, NOBILLIAUX Florian, PREVOST Brigitte, THUILLIER Caroline, TINEZ Claire ,TCHANTCHOU NJOSSE YANICK |
| EPI_ISL_2361526, EPI_ISL_2361527 | CH ROUBAIX | CHU Lille - Laboratoire de Virologie | AIT YAHYA Emilie, ALIDJINOU Enagnon Kazali, BOCKET Laurence, CREPIN Michel, DEMAY Christophe, ENGELMANN Ilka, GEFFROY Sandrine, GUIGON Aurélie, LAMBERT Valérie, LAZREK Mouna, NOBILLIAUX Florian, PREVOST Brigitte, THUILLIER Caroline, TINEZ Claire ,TCHANTCHOU NJOSSE YANICK |
| EPI_ISL_2361528 | CH CALAIS | CHU Lille - Laboratoire de Virologie | AIT YAHYA Emilie, ALIDJINOU Enagnon Kazali, BOCKET Laurence, CREPIN Michel, DEMAY Christophe, ENGELMANN Ilka, GEFFROY Sandrine, GUIGON Aurélie, LAMBERT Valérie, LAZREK Mouna, NOBILLIAUX Florian, PREVOST Brigitte, THUILLIER Caroline, TINEZ Claire ,TCHANTCHOU NJOSSE YANICK |
| EPI_ISL_2361529, EPI_ISL_2361530, EPI_ISL_2361532, EPI_ISL_2361533, EPI_ISL_2361534, EPI_ISL_2361535, EPI_ISL_2361536, EPI_ISL_2361537, EPI_ISL_2361538, EPI_ISL_2361540, EPI_ISL_2361541, EPI_ISL_2361542, EPI_ISL_2361543, EPI_ISL_2361544, EPI_ISL_2361545, EPI_ISL_2361546, EPI_ISL_2361547, EPI_ISL_2361548, EPI_ISL_2361549, EPI_ISL_2361550, EPI_ISL_2361551, EPI_ISL_2361552, EPI_ISL_2361553, EPI_ISL_2361554, EPI_ISL_2361555 | | | |
| see above | CHU LILLE | CHU Lille - Laboratoire de Virologie | AIT YAHYA Emilie, ALIDJINOU Enagnon Kazali, BOCKET Laurence, CREPIN Michel, DEMAY Christophe, ENGELMANN Ilka, GEFFROY Sandrine, GUIGON Aurélie, LAMBERT Valérie, LAZREK Mouna, NOBILLIAUX Florian, PREVOST Brigitte, THUILLIER Caroline, TINEZ Claire ,TCHANTCHOU NJOSSE YANICK |
| EPI_ISL_2361556 | CH SAINT-PHILIBERT | CHU Lille - Laboratoire de Virologie | AIT YAHYA Emilie, ALIDJINOU Enagnon Kazali, BOCKET Laurence, CREPIN Michel, DEMAY Christophe, ENGELMANN Ilka, GEFFROY Sandrine, GUIGON Aurélie, LAMBERT Valérie, LAZREK Mouna, NOBILLIAUX Florian, PREVOST Brigitte, THUILLIER Caroline, TINEZ Claire ,TCHANTCHOU NJOSSE YANICK |
| EPI_ISL_2361557 | CH ARMENTIERES | CHU Lille - Laboratoire de Virologie | AIT YAHYA Emilie, ALIDJINOU Enagnon Kazali, BOCKET Laurence, CREPIN Michel, DEMAY Christophe, ENGELMANN Ilka, GEFFROY Sandrine, GUIGON Aurélie, LAMBERT Valérie, LAZREK Mouna, NOBILLIAUX Florian, PREVOST Brigitte, THUILLIER Caroline, TINEZ Claire ,TCHANTCHOU NJOSSE YANICK |
| EPI_ISL_2361558, EPI_ISL_2361559 | CH BETHUNE | CHU Lille - Laboratoire de Virologie | AIT YAHYA Emilie, ALIDJINOU Enagnon Kazali, BOCKET Laurence, CREPIN Michel, DEMAY Christophe, ENGELMANN Ilka, GEFFROY Sandrine, GUIGON Aurélie, LAMBERT Valérie, LAZREK Mouna, NOBILLIAUX Florian, PREVOST Brigitte, THUILLIER Caroline, TINEZ Claire ,TCHANTCHOU NJOSSE YANICK |
| EPI_ISL_2361560, EPI_ISL_2361561 | CH BOULOGNE-SUR-MER | CHU Lille - Laboratoire de Virologie | AIT YAHYA Emilie, ALIDJINOU Enagnon Kazali, BOCKET Laurence, CREPIN Michel, DEMAY Christophe, ENGELMANN Ilka, GEFFROY Sandrine, GUIGON Aurélie, LAMBERT Valérie, LAZREK Mouna, NOBILLIAUX Florian, PREVOST Brigitte, THUILLIER Caroline, TINEZ Claire ,TCHANTCHOU NJOSSE YANICK |
| EPI_ISL_2361562, EPI_ISL_2361563 | CH ARRAS | CHU Lille - Laboratoire de Virologie | AIT YAHYA Emilie, ALIDJINOU Enagnon Kazali, BOCKET Laurence, CREPIN Michel, DEMAY Christophe, ENGELMANN Ilka, GEFFROY Sandrine, GUIGON Aurélie, LAMBERT Valérie, LAZREK Mouna, NOBILLIAUX Florian, PREVOST Brigitte, THUILLIER Caroline, TINEZ Claire ,TCHANTCHOU NJOSSE YANICK |
| EPI_ISL_2361564, EPI_ISL_2361565 | CH CALAIS | CHU Lille - Laboratoire de Virologie | AIT YAHYA Emilie, ALIDJINOU Enagnon Kazali, BOCKET Laurence, CREPIN Michel, DEMAY Christophe, ENGELMANN Ilka, GEFFROY Sandrine, GUIGON Aurélie, LAMBERT Valérie, LAZREK Mouna, NOBILLIAUX Florian, PREVOST Brigitte, THUILLIER Caroline, TINEZ Claire ,TCHANTCHOU NJOSSE YANICK |
| EPI_ISL_2361566 | CH DOUAI | CHU Lille - Laboratoire de Virologie | AIT YAHYA Emilie, ALIDJINOU Enagnon Kazali, BOCKET Laurence, CREPIN Michel, DEMAY Christophe, ENGELMANN Ilka, GEFFROY Sandrine, GUIGON Aurélie, LAMBERT Valérie, LAZREK Mouna, NOBILLIAUX Florian, PREVOST Brigitte, THUILLIER Caroline, TINEZ Claire ,TCHANTCHOU NJOSSE YANICK |
| EPI_ISL_2361567 | CHU LILLE | CHU Lille - Laboratoire de Virologie | AIT YAHYA Emilie, ALIDJINOU Enagnon Kazali, BOCKET Laurence, CREPIN Michel, DEMAY Christophe, ENGELMANN Ilka, GEFFROY Sandrine, GUIGON Aurélie, LAMBERT Valérie, LAZREK Mouna, NOBILLIAUX Florian, PREVOST Brigitte, THUILLIER Caroline, TINEZ Claire ,TCHANTCHOU NJOSSE YANICK |
| EPI_ISL_2361568 | CH LENS | CHU Lille - Laboratoire de Virologie | AIT YAHYA Emilie, ALIDJINOU Enagnon Kazali, BOCKET Laurence, CREPIN Michel, DEMAY Christophe, ENGELMANN Ilka, GEFFROY Sandrine, GUIGON Aurélie, LAMBERT Valérie, LAZREK Mouna, NOBILLIAUX Florian, PREVOST Brigitte, THUILLIER Caroline, TINEZ Claire ,TCHANTCHOU NJOSSE YANICK |
| EPI_ISL_2361569, EPI_ISL_2361570 | CH ROUBAIX | CHU Lille - Laboratoire de Virologie | AIT YAHYA Emilie, ALIDJINOU Enagnon Kazali, BOCKET Laurence, CREPIN Michel, DEMAY Christophe, ENGELMANN Ilka, GEFFROY Sandrine, GUIGON Aurélie, LAMBERT Valérie, LAZREK Mouna, NOBILLIAUX Florian, PREVOST Brigitte, THUILLIER Caroline, TINEZ Claire ,TCHANTCHOU NJOSSE YANICK |
| EPI_ISL_2361571, EPI_ISL_2361572, EPI_ISL_2361573, EPI_ISL_2361574, EPI_ISL_2361575, EPI_ISL_2361576, EPI_ISL_2361577, EPI_ISL_2361578, EPI_ISL_2361579, EPI_ISL_2361580 | CHU LILLE | CHU Lille - Laboratoire de Virologie | AIT YAHYA Emilie, ALIDJINOU Enagnon Kazali, BOCKET Laurence, CREPIN Michel, DEMAY Christophe, ENGELMANN Ilka, GEFFROY Sandrine, GUIGON Aurélie, LAMBERT Valérie, LAZREK Mouna, NOBILLIAUX Florian, PREVOST Brigitte, THUILLIER Caroline, TINEZ Claire ,TCHANTCHOU NJOSSE YANICK |
| EPI_ISL_2361581, EPI_ISL_2361582 | CH SAINT-PHILIBERT | CHU Lille - Laboratoire de Virologie | AIT YAHYA Emilie, ALIDJINOU Enagnon Kazali, BOCKET Laurence, CREPIN Michel, DEMAY Christophe, ENGELMANN Ilka, GEFFROY Sandrine, GUIGON Aurélie, LAMBERT Valérie, LAZREK Mouna, NOBILLIAUX Florian, PREVOST Brigitte, THUILLIER Caroline, TINEZ Claire ,TCHANTCHOU NJOSSE YANICK |
| EPI_ISL_2361583, EPI_ISL_2361584 | CH TOURCOING | CHU Lille - Laboratoire de Virologie | AIT YAHYA Emilie, ALIDJINOU Enagnon Kazali, BOCKET Laurence, CREPIN Michel, DEMAY Christophe, ENGELMANN Ilka, GEFFROY Sandrine, GUIGON Aurélie, LAMBERT Valérie, LAZREK Mouna, NOBILLIAUX Florian, PREVOST Brigitte, THUILLIER Caroline, TINEZ Claire ,TCHANTCHOU NJOSSE YANICK |
| EPI_ISL_2361585, EPI_ISL_2361586 | CH BOULOGNE-SUR-MER | CHU Lille - Laboratoire de Virologie | AIT YAHYA Emilie, ALIDJINOU Enagnon Kazali, BOCKET Laurence, CREPIN Michel, DEMAY Christophe, ENGELMANN Ilka, GEFFROY Sandrine, GUIGON Aurélie, LAMBERT Valérie, LAZREK Mouna, NOBILLIAUX Florian, PREVOST Brigitte, THUILLIER Caroline, TINEZ Claire ,TCHANTCHOU NJOSSE YANICK |
| EPI_ISL_2361587, EPI_ISL_2361588 | CH ARRAS | CHU Lille - Laboratoire de Virologie | AIT YAHYA Emilie, ALIDJINOU Enagnon Kazali, BOCKET Laurence, CREPIN Michel, DEMAY Christophe, ENGELMANN Ilka, GEFFROY Sandrine, GUIGON Aurélie, LAMBERT Valérie, LAZREK Mouna, NOBILLIAUX Florian, PREVOST Brigitte, THUILLIER Caroline, TINEZ Claire ,TCHANTCHOU NJOSSE YANICK |
| EPI_ISL_2361590 | CH CALAIS | CHU Lille - Laboratoire de Virologie | AIT YAHYA Emilie, ALIDJINOU Enagnon Kazali, BOCKET Laurence, CREPIN Michel, DEMAY Christophe, ENGELMANN Ilka, GEFFROY Sandrine, GUIGON Aurélie, LAMBERT Valérie, LAZREK Mouna, NOBILLIAUX Florian, PREVOST Brigitte, THUILLIER Caroline, TINEZ Claire ,TCHANTCHOU NJOSSE YANICK |
| EPI_ISL_2361591, EPI_ISL_2361592, EPI_ISL_2361593, EPI_ISL_2361596, EPI_ISL_2361597, EPI_ISL_2361598, EPI_ISL_2361599, EPI_ISL_2361600 | CHU LILLE | CHU Lille - Laboratoire de Virologie | AIT YAHYA Emilie, ALIDJINOU Enagnon Kazali, BOCKET Laurence, CREPIN Michel, DEMAY Christophe, ENGELMANN Ilka, GEFFROY Sandrine, GUIGON Aurélie, LAMBERT Valérie, LAZREK Mouna, NOBILLIAUX Florian, PREVOST Brigitte, THUILLIER Caroline, TINEZ Claire ,TCHANTCHOU NJOSSE YANICK |
| EPI_ISL_2362528, EPI_ISL_2362529, EPI_ISL_2362530 | St James's Hospital, Dublin 8, Ireland. | ELDA biotech | Deirdre M. Broderick, Peter R. Flanagan, Brendan Crowley |
| EPI_ISL_2362532, EPI_ISL_2362537 | Laboratoire Carage | Institut Pasteur de la Guyane | Anne Lavergne, Dominique Rousset, Antoine Enfissi, Arielle Salmier |
| EPI_ISL_2362548 | CNR Institut Pasteur de la Guyane | Institut Pasteur de la Guyane | Anne Lavergne, Dominique Rousset, Antoine Enfissi, Arielle Salmier |
| EPI_ISL_2362553 | Laboratoire Carage | Institut Pasteur de la Guyane | Anne Lavergne, Dominique Rousset, Antoine Enfissi, Arielle Salmier |
| EPI_ISL_2362685 | Microbiology and Virology Unit, Florence Careggi University | Microbiology and Virology Unit, Florence Careggi University | Vincenzo Di Pilato, Marco Coppi, Fabio Morecchiato, Noemi Aiezza, Ilaria Baccani, Nicla Giovacchini, Alberto Antonelli, Emanuele Gori, Gian Maria |

| | | | |
|---|---|---|---|
| | Hospital | Hospital | Rossolini |
| EPI_ISL_2365888, EPI_ISL_2365889, EPI_ISL_2365890, EPI_ISL_2365891, EPI_ISL_2365892, EPI_ISL_2365893, EPI_ISL_2365894, EPI_ISL_2365895 | Microbiology Department. Complexo Hospitalario Universitario de Vigo | Microbiology Department. Complexo Hospitalario Universitario de Vigo | Alfaya N, Alonso I, Alvarez M, Cabrera JJ, Carballo R, Cores O, Cortizo S, del-Campo V, Martinez L, Mediero G, Perez S, Potel C, Regueiro B, Rey S, Vasallo FJ |
| EPI_ISL_2367762 | CH BETHUNE | CHU Lille - Laboratoire de Virologie | AIT YAHYA Emilie, ALIDJINOU Enagnon Kazali, BOCKET Laurence, CREPIN Michel, DEMAY Christophe, ENGELMANN Ilka, GEFFROY Sandrine, GUIGON Aurélie, LAMBERT Valérie, LAZREK Mouna, NOBILLIAUX Florian, PREVOST Brigitte, THUILLIER Caroline, TINEZ Claire ,TCHANTCHOU NJOSSE YANICK |
| EPI_ISL_2367763, EPI_ISL_2367765, EPI_ISL_2367767, EPI_ISL_2367770, EPI_ISL_2367771, EPI_ISL_2367774, EPI_ISL_2367775, EPI_ISL_2367777, EPI_ISL_2367779, EPI_ISL_2367781, EPI_ISL_2367783 | | | |
| see above | CHU LILLE | CHU Lille - Laboratoire de Virologie | AIT YAHYA Emilie, ALIDJINOU Enagnon Kazali, BOCKET Laurence, CREPIN Michel, DEMAY Christophe, ENGELMANN Ilka, GEFFROY Sandrine, GUIGON Aurélie, LAMBERT Valérie, LAZREK Mouna, NOBILLIAUX Florian, PREVOST Brigitte, THUILLIER Caroline, TINEZ Claire ,TCHANTCHOU NJOSSE YANICK |
| EPI_ISL_2367785 | CH BETHUNE | CHU Lille - Laboratoire de Virologie | AIT YAHYA Emilie, ALIDJINOU Enagnon Kazali, BOCKET Laurence, CREPIN Michel, DEMAY Christophe, ENGELMANN Ilka, GEFFROY Sandrine, GUIGON Aurélie, LAMBERT Valérie, LAZREK Mouna, NOBILLIAUX Florian, PREVOST Brigitte, THUILLIER Caroline, TINEZ Claire ,TCHANTCHOU NJOSSE YANICK |
| EPI_ISL_2367787 | CH ARRAS | CHU Lille - Laboratoire de Virologie | AIT YAHYA Emilie, ALIDJINOU Enagnon Kazali, BOCKET Laurence, CREPIN Michel, DEMAY Christophe, ENGELMANN Ilka, GEFFROY Sandrine, GUIGON Aurélie, LAMBERT Valérie, LAZREK Mouna, NOBILLIAUX Florian, PREVOST Brigitte, THUILLIER Caroline, TINEZ Claire ,TCHANTCHOU NJOSSE YANICK |
| EPI_ISL_2367790 | CH CAMBRAI | CHU Lille - Laboratoire de Virologie | AIT YAHYA Emilie, ALIDJINOU Enagnon Kazali, BOCKET Laurence, CREPIN Michel, DEMAY Christophe, ENGELMANN Ilka, GEFFROY Sandrine, GUIGON Aurélie, LAMBERT Valérie, LAZREK Mouna, NOBILLIAUX Florian, PREVOST Brigitte, THUILLIER Caroline, TINEZ Claire ,TCHANTCHOU NJOSSE YANICK |
| EPI_ISL_2367791 | CH LENS | CHU Lille - Laboratoire de Virologie | AIT YAHYA Emilie, ALIDJINOU Enagnon Kazali, BOCKET Laurence, CREPIN Michel, DEMAY Christophe, ENGELMANN Ilka, GEFFROY Sandrine, GUIGON Aurélie, LAMBERT Valérie, LAZREK Mouna, NOBILLIAUX Florian, PREVOST Brigitte, THUILLIER Caroline, TINEZ Claire ,TCHANTCHOU NJOSSE YANICK |
| EPI_ISL_2372266, EPI_ISL_2372267 | OUCRU | OUCRU | Nguyen Van Vinh Chau, Nguyen To Anh, Nguyen Thi thu Hong, Nghiem My Ngoc, Huynh Trung Trieu, Le Nguyen Truc Nhu, Nguyen Thi Han Ny, Lam Minh Yen, Ngo Ngoc Quang Minh, Nguyen Thanh Phong, Nguyen Thanh Truong, Le Thi Thu Huong, Tran Nguyen Hoang Tu, Le Manh Hung, Tran Tan Thanh, Nguyen Thanh Dung, Nguyen Tri Dung, Guy Thwaites, Le Van Tan |
| EPI_ISL_2374170, EPI_ISL_2374171 | Hospital | National Reference Center for Viruses of Respiratory Infections, Institut Pasteur, Paris | Marion Barbet, Sylvie Behillil, Méline Bizard, Angela Brisebarre, Camille Capel, Vincent Enouf, Louise Lefrançois, Frédéric Lemoine, Christophe Malabat, Corinne Maufrais, Etienne Simon-Lorière, Maud Vanpeene, Sylvie Van der Werf ,Pierre Alain Billy |
| EPI_ISL_2374172 | Hospital | National Reference Center for Viruses of Respiratory Infections, Institut Pasteur, Paris | Marion Barbet, Sylvie Behillil, Méline Bizard, Angela Brisebarre, Camille Capel, Vincent Enouf, Louise Lefrançois, Frédéric Lemoine, Christophe Malabat, Corinne Maufrais, Emmanuelle Permal, Etienne Simon-Lorière, Maud Vanpeene, Sylvie Van der Werf ,Pierre Alain Billy |
| EPI_ISL_2374173, EPI_ISL_2374174, EPI_ISL_2374175, EPI_ISL_2374176, EPI_ISL_2374179 | Hospital | National Reference Center for Viruses of Respiratory Infections, Institut Pasteur, Paris | Marion Barbet, Sylvie Behillil, Méline Bizard, Angela Brisebarre, Camille Capel, Vincent Enouf, Louise Lefrançois, Frédéric Lemoine, Christophe Malabat, Corinne Maufrais, Etienne Simon-Lorière, Maud Vanpeene, Sylvie Van der Werf ,Sandrine Castelain |
| EPI_ISL_2374185, EPI_ISL_2374186, EPI_ISL_2374187 | Hospital | National Reference Center for Viruses of Respiratory Infections, Institut Pasteur, Paris | Marion Barbet, Sylvie Behillil, Méline Bizard, Angela Brisebarre, Camille Capel, Vincent Enouf, Louise Lefrançois, Frédéric Lemoine, Christophe Malabat, Corinne Maufrais, Etienne Simon-Lorière, Maud Vanpeene, Sylvie Van der Werf ,StéPhanie Marque Juillet |
| EPI_ISL_2374189, EPI_ISL_2374190, EPI_ISL_2374191 | Hospital | National Reference Center for Viruses of Respiratory Infections, Institut Pasteur, Paris | Marion Barbet, Sylvie Behillil, Méline Bizard, Angela Brisebarre, Camille Capel, Vincent Enouf, Louise Lefrançois, Frédéric Lemoine, Christophe Malabat, Corinne Maufrais, Etienne Simon-Lorière, Maud Vanpeene, Sylvie Van der Werf ,Pascale Martres |
| EPI_ISL_2374192, EPI_ISL_2374193 | Labo Analyses Med | National Reference Center for Viruses of Respiratory Infections, Institut Pasteur, Paris | Marion Barbet, Sylvie Behillil, Méline Bizard, Angela Brisebarre, Camille Capel, Vincent Enouf, Louise Lefrançois, Frédéric Lemoine, Christophe Malabat, Corinne Maufrais, Etienne Simon-Lorière, Maud Vanpeene, Sylvie Van der Werf ,Philippe Miara |
| EPI_ISL_2374194 | Labo Analyses Med | National Reference Center for Viruses of Respiratory Infections, Institut Pasteur, Paris | Marion Barbet, Sylvie Behillil, Méline Bizard, Angela Brisebarre, Camille Capel, Vincent Enouf, Louise Lefrançois, Frédéric Lemoine, Christophe Malabat, Corinne Maufrais, Emmanuelle Permal, Etienne Simon-Lorière, Maud Vanpeene, Sylvie Van der Werf ,Jonas Amzalag |
| EPI_ISL_2374195, EPI_ISL_2374196, EPI_ISL_2374197, EPI_ISL_2374198, EPI_ISL_2374199, EPI_ISL_2374200, EPI_ISL_2374201, EPI_ISL_2374202, EPI_ISL_2374203, EPI_ISL_2374204, EPI_ISL_2374205, EPI_ISL_2374206, EPI_ISL_2374207, EPI_ISL_2374208, EPI_ISL_2374209, EPI_ISL_2374210, EPI_ISL_2374211, EPI_ISL_2374212, EPI_ISL_2374213, EPI_ISL_2374214, EPI_ISL_2374215, EPI_ISL_2374216, EPI_ISL_2374217 | | | |
| see above | Labo Analyses Med | National Reference Center for Viruses of Respiratory Infections, Institut Pasteur, Paris | Marion Barbet, Sylvie Behillil, Méline Bizard, Angela Brisebarre, Camille Capel, Vincent Enouf, Louise Lefrançois, Frédéric Lemoine, Christophe Malabat, Corinne Maufrais, Etienne Simon-Lorière, Maud Vanpeene, Sylvie Van der Werf ,Jonas Amzalag |
| EPI_ISL_2374218 | Labo Analyses Med | National Reference Center for Viruses of Respiratory Infections, Institut Pasteur, Paris | Marion Barbet, Sylvie Behillil, Méline Bizard, Angela Brisebarre, Camille Capel, Vincent Enouf, Louise Lefrançois, Frédéric Lemoine, Christophe Malabat, Corinne Maufrais, Emmanuelle Permal, Etienne Simon-Lorière, Maud Vanpeene, Sylvie Van der Werf ,Jonas Amzalag |
| EPI_ISL_2374219, EPI_ISL_2374220, EPI_ISL_2374221 | Labo Analyses Med | National Reference Center for Viruses of Respiratory Infections, Institut Pasteur, Paris | Marion Barbet, Sylvie Behillil, Méline Bizard, Angela Brisebarre, Camille Capel, Vincent Enouf, Louise Lefrançois, Frédéric Lemoine, Christophe Malabat, Corinne Maufrais, Etienne Simon-Lorière, Maud Vanpeene, Sylvie Van der Werf ,Jonas Amzalag |
| EPI_ISL_2374223 | Labo Analyses Med | National Reference Center for Viruses of Respiratory Infections, Institut Pasteur, Paris | Marion Barbet, Sylvie Behillil, Méline Bizard, Angela Brisebarre, Camille Capel, Vincent Enouf, Louise Lefrançois, Frédéric Lemoine, Christophe Malabat, Corinne Maufrais, Emmanuelle Permal, Etienne Simon-Lorière, Maud Vanpeene, Sylvie Van der Werf ,Jonas Amzalag |
| EPI_ISL_2374224, EPI_ISL_2374225, EPI_ISL_2374226, EPI_ISL_2374227, EPI_ISL_2374228, EPI_ISL_2374229, EPI_ISL_2374230, EPI_ISL_2374231, EPI_ISL_2374232, EPI_ISL_2374233, EPI_ISL_2374234, EPI_ISL_2374235, EPI_ISL_2374236, EPI_ISL_2374237, EPI_ISL_2374238, EPI_ISL_2374239 | | | |
| see above | Labo Analyses Med | National Reference Center for Viruses of Respiratory Infections, Institut Pasteur, Paris | Marion Barbet, Sylvie Behillil, Méline Bizard, Angela Brisebarre, Camille Capel, Vincent Enouf, Louise Lefrançois, Frédéric Lemoine, Christophe Malabat, Corinne Maufrais, Etienne Simon-Lorière, Maud Vanpeene, Sylvie Van der Werf ,Jonas Amzalag |
| EPI_ISL_2374240 | Labo Analyses Med | National Reference Center for Viruses of Respiratory Infections, Institut Pasteur, Paris | Marion Barbet, Sylvie Behillil, Méline Bizard, Angela Brisebarre, Camille Capel, Vincent Enouf, Louise Lefrançois, Frédéric Lemoine, Christophe Malabat, Corinne Maufrais, Emmanuelle Permal, Etienne Simon-Lorière, Maud Vanpeene, Sylvie Van der Werf ,Jonas Amzalag |
| EPI_ISL_2374242, EPI_ISL_2374243, EPI_ISL_2374244, EPI_ISL_2374246, EPI_ISL_2374247, EPI_ISL_2374248, EPI_ISL_2374249, EPI_ISL_2374250 | Labo Analyses Med | National Reference Center for Viruses of Respiratory Infections, Institut Pasteur, Paris | Marion Barbet, Sylvie Behillil, Méline Bizard, Angela Brisebarre, Camille Capel, Vincent Enouf, Louise Lefrançois, Frédéric Lemoine, Christophe Malabat, Corinne Maufrais, Etienne Simon-Lorière, Maud Vanpeene, Sylvie Van der Werf ,Pierre-Yves Leonard |
| EPI_ISL_2374251 | Labo Analyses Med | National Reference Center for Viruses of Respiratory Infections, Institut Pasteur, Paris | Marion Barbet, Sylvie Behillil, Méline Bizard, Angela Brisebarre, Camille Capel, Vincent Enouf, Louise Lefrançois, Frédéric Lemoine, Christophe Malabat, Corinne Maufrais, Emmanuelle Permal, Etienne Simon-Lorière, Maud Vanpeene, Sylvie Van der Werf ,Pierre-Yves Leonard |
| EPI_ISL_2374252, EPI_ISL_2374253, EPI_ISL_2374254, EPI_ISL_2374255, EPI_ISL_2374256 | Labo Analyses Med | National Reference Center for Viruses of Respiratory Infections, Institut Pasteur, Paris | Marion Barbet, Sylvie Behillil, Méline Bizard, Angela Brisebarre, Camille Capel, Vincent Enouf, Louise Lefrançois, Frédéric Lemoine, Christophe Malabat, Corinne Maufrais, Etienne Simon-Lorière, Maud Vanpeene, Sylvie Van der Werf ,Pierre-Yves Leonard |
| EPI_ISL_2374257, EPI_ISL_2374258, EPI_ISL_2374259, EPI_ISL_2374260, EPI_ISL_2374261, EPI_ISL_2374262, EPI_ISL_2374263, EPI_ISL_2374264, EPI_ISL_2374265, EPI_ISL_2374266, EPI_ISL_2374267, EPI_ISL_2374268 | | | |
| see above | Labo Analyses Med | National Reference Center for Viruses of Respiratory Infections, Institut Pasteur, Paris | Marion Barbet, Sylvie Behillil, Méline Bizard, Angela Brisebarre, Camille Capel, Vincent Enouf, Louise Lefrançois, Frédéric Lemoine, Christophe Malabat, Corinne Maufrais, Etienne Simon-Lorière, Maud Vanpeene, Sylvie Van der Werf ,GréGoire Potiron |
| EPI_ISL_2374269, EPI_ISL_2374270, EPI_ISL_2374271, EPI_ISL_2374272, EPI_ISL_2374273 | Hospital | National Reference Center for Viruses of Respiratory Infections, Institut Pasteur, Paris | Marion Barbet, Sylvie Behillil, Méline Bizard, Angela Brisebarre, Camille Capel, Vincent Enouf, Louise Lefrançois, Frédéric Lemoine, Christophe Malabat, Corinne Maufrais, Etienne Simon-Lorière, Maud Vanpeene, Sylvie Van der Werf ,Anne Cady |

| | | | |
|---|---|---|---|
| EPI_ISL_2374274, EPI_ISL_2374275, EPI_ISL_2374276, EPI_ISL_2374277, EPI_ISL_2374278, EPI_ISL_2374279, EPI_ISL_2374280, EPI_ISL_2374281, EPI_ISL_2374282, EPI_ISL_2374283, EPI_ISL_2374284, EPI_ISL_2374285, EPI_ISL_2374286, EPI_ISL_2374287 | | | |
| see above | Labo Analyses Med | National Reference Center for Viruses of Respiratory Infections, Institut Pasteur, Paris | Marion Barbet, Sylvie Behillil, Méline Bizard, Angela Brisebarre, Camille Capel, Vincent Enouf, Louise Lefrançois, Frédéric Lemoine, Christophe Malabat, Corinne Maufrais, Etienne Simon-Lorière, Maud Vanpeene, Sylvie Van der Werf ,Edouard Lamandin |
| EPI_ISL_2374288 | Labo Analyses Med | National Reference Center for Viruses of Respiratory Infections, Institut Pasteur, Paris | Marion Barbet, Sylvie Behillil, Méline Bizard, Angela Brisebarre, Camille Capel, Vincent Enouf, Louise Lefrançois, Frédéric Lemoine, Christophe Malabat, Corinne Maufrais, Amaury Vaysse, Etienne Simon-Lorière, Maud Vanpeene, Sylvie Van der Werf ,Edouard Lamandin |
| EPI_ISL_2374289 | Labo Analyses Med | National Reference Center for Viruses of Respiratory Infections, Institut Pasteur, Paris | Marion Barbet, Sylvie Behillil, Méline Bizard, Angela Brisebarre, Camille Capel, Vincent Enouf, Louise Lefrançois, Frédéric Lemoine, Christophe Malabat, Corinne Maufrais, Etienne Simon-Lorière, Maud Vanpeene, Sylvie Van der Werf ,Arditi Cocco |
| EPI_ISL_2374290 | Labo Analyses Med | National Reference Center for Viruses of Respiratory Infections, Institut Pasteur, Paris | Marion Barbet, Sylvie Behillil, Méline Bizard, Angela Brisebarre, Camille Capel, Vincent Enouf, Louise Lefrançois, Frédéric Lemoine, Christophe Malabat, Corinne Maufrais, Amaury Vaysse, Etienne Simon-Lorière, Maud Vanpeene, Sylvie Van der Werf ,Arditi Cocco |
| EPI_ISL_2374291, EPI_ISL_2374292, EPI_ISL_2374293, EPI_ISL_2374294, EPI_ISL_2374295, EPI_ISL_2374296 | Labo Analyses Med | National Reference Center for Viruses of Respiratory Infections, Institut Pasteur, Paris | Marion Barbet, Sylvie Behillil, Méline Bizard, Angela Brisebarre, Camille Capel, Vincent Enouf, Louise Lefrançois, Frédéric Lemoine, Christophe Malabat, Corinne Maufrais, Etienne Simon-Lorière, Maud Vanpeene, Sylvie Van der Werf ,Arditi Cocco |
| EPI_ISL_2374297, EPI_ISL_2374298, EPI_ISL_2374299, EPI_ISL_2374300, EPI_ISL_2374302, EPI_ISL_2374305, EPI_ISL_2374306, EPI_ISL_2374307 | Labo Analyses Med | National Reference Center for Viruses of Respiratory Infections, Institut Pasteur, Paris | Marion Barbet, Sylvie Behillil, Méline Bizard, Angela Brisebarre, Camille Capel, Vincent Enouf, Louise Lefrançois, Frédéric Lemoine, Christophe Malabat, Corinne Maufrais, Emmanuelle Permal, Etienne Simon-Lorière, Maud Vanpeene, Sylvie Van der Werf ,Arditi Cocco |
| EPI_ISL_2374309, EPI_ISL_2374310, EPI_ISL_2374311, EPI_ISL_2374312, EPI_ISL_2374313, EPI_ISL_2374314, EPI_ISL_2374315, EPI_ISL_2374316, EPI_ISL_2374317, EPI_ISL_2374318, EPI_ISL_2374319, EPI_ISL_2374321, EPI_ISL_2374322, EPI_ISL_2374323, EPI_ISL_2374324, EPI_ISL_2374325, EPI_ISL_2374326, EPI_ISL_2374327 | | | |
| see above | Labo Analyses Med | National Reference Center for Viruses of Respiratory Infections, Institut Pasteur, Paris | Marion Barbet, Sylvie Behillil, Méline Bizard, Angela Brisebarre, Camille Capel, Vincent Enouf, Louise Lefrançois, Frédéric Lemoine, Christophe Malabat, Corinne Maufrais, Emmanuelle Permal, Etienne Simon-Lorière, Maud Vanpeene, Sylvie Van der Werf ,Claire Felloni |
| EPI_ISL_2374328, EPI_ISL_2374329, EPI_ISL_2374330, EPI_ISL_2374331, EPI_ISL_2374332 | Labo Analyses Med | National Reference Center for Viruses of Respiratory Infections, Institut Pasteur, Paris | Marion Barbet, Sylvie Behillil, Méline Bizard, Angela Brisebarre, Camille Capel, Vincent Enouf, Louise Lefrançois, Frédéric Lemoine, Christophe Malabat, Corinne Maufrais, Emmanuelle Permal, Etienne Simon-Lorière, Maud Vanpeene, Sylvie Van der Werf ,Christine Heurtin |
| EPI_ISL_2374333, EPI_ISL_2374334, EPI_ISL_2374335, EPI_ISL_2374336, EPI_ISL_2374337, EPI_ISL_2374338 | Hospital | National Reference Center for Viruses of Respiratory Infections, Institut Pasteur, Paris | Marion Barbet, Sylvie Behillil, Méline Bizard, Angela Brisebarre, Camille Capel, Vincent Enouf, Louise Lefrançois, Frédéric Lemoine, Christophe Malabat, Corinne Maufrais, Emmanuelle Permal, Etienne Simon-Lorière, Maud Vanpeene, Sylvie Van der Werf ,Vincent Thibault |
| EPI_ISL_2374339 | Hospital | National Reference Center for Viruses of Respiratory Infections, Institut Pasteur, Paris | Marion Barbet, Sylvie Behillil, Méline Bizard, Angela Brisebarre, Camille Capel, Vincent Enouf, Louise Lefrançois, Frédéric Lemoine, Christophe Malabat, Corinne Maufrais, Emmanuelle Permal, Etienne Simon-Lorière, Maud Vanpeene, Sylvie Van der Werf ,Pascale Martres |
| EPI_ISL_2374340, EPI_ISL_2374341, EPI_ISL_2374342, EPI_ISL_2374343, EPI_ISL_2374344, EPI_ISL_2374345, EPI_ISL_2374346, EPI_ISL_2374347 | Labo Analyses Med | National Reference Center for Viruses of Respiratory Infections, Institut Pasteur, Paris | Marion Barbet, Sylvie Behillil, Méline Bizard, Angela Brisebarre, Camille Capel, Vincent Enouf, Louise Lefrançois, Frédéric Lemoine, Christophe Malabat, Corinne Maufrais, Emmanuelle Permal, Etienne Simon-Lorière, Maud Vanpeene, Sylvie Van der Werf ,Armand Saaba |
| EPI_ISL_2374348 | Labo Analyses Med | National Reference Center for Viruses of Respiratory Infections, Institut Pasteur, Paris | Marion Barbet, Sylvie Behillil, Méline Bizard, Angela Brisebarre, Camille Capel, Vincent Enouf, Louise Lefrançois, Frédéric Lemoine, Christophe Malabat, Corinne Maufrais, Emmanuelle Permal, Etienne Simon-Lorière, Maud Vanpeene, Sylvie Van der Werf ,M. El Hamri |
| EPI_ISL_2374349, EPI_ISL_2374350, EPI_ISL_2374351, EPI_ISL_2374352, EPI_ISL_2374353, EPI_ISL_2374354, EPI_ISL_2374355, EPI_ISL_2374357, EPI_ISL_2374358, EPI_ISL_2374359, EPI_ISL_2374360, EPI_ISL_2374361, EPI_ISL_2374362, EPI_ISL_2374363, EPI_ISL_2374364, EPI_ISL_2374365, EPI_ISL_2374366, EPI_ISL_2374367, EPI_ISL_2374368, EPI_ISL_2374369, EPI_ISL_2374370 | | | |
| see above | Labo Analyses med | National Reference Center for Viruses of Respiratory Infections, Institut Pasteur, Paris | Marion Barbet, Sylvie Behillil, Méline Bizard, Angela Brisebarre, Camille Capel, Vincent Enouf, Louise Lefrançois, Frédéric Lemoine, Christophe Malabat, Corinne Maufrais, Emmanuelle Permal, Etienne Simon-Lorière, Maud Vanpeene, Sylvie Van der Werf ,Thierry Gussond |
| EPI_ISL_2374371 | Hospital | National Reference Center for Viruses of Respiratory Infections, Institut Pasteur, Paris | Marion Barbet, Sylvie Behillil, Méline Bizard, Angela Brisebarre, Camille Capel, Vincent Enouf, Louise Lefrançois, Frédéric Lemoine, Christophe Malabat, Corinne Maufrais, Emmanuelle Permal, Etienne Simon-Lorière, Maud Vanpeene, Sylvie Van der Werf ,Eric Farfour |
| EPI_ISL_2374372, EPI_ISL_2374373, EPI_ISL_2374374, EPI_ISL_2374375, EPI_ISL_2374376, EPI_ISL_2374377, EPI_ISL_2374378 | Labo Analyses Med | National Reference Center for Viruses of Respiratory Infections, Institut Pasteur, Paris | Marion Barbet, Sylvie Behillil, Méline Bizard, Angela Brisebarre, Camille Capel, Vincent Enouf, Louise Lefrançois, Frédéric Lemoine, Christophe Malabat, Corinne Maufrais, Emmanuelle Permal, Etienne Simon-Lorière, Maud Vanpeene, Sylvie Van der Werf ,Pierre-Yves Leonard |
| EPI_ISL_2374380, EPI_ISL_2374381, EPI_ISL_2374383 | Hospital | National Reference Center for Viruses of Respiratory Infections, Institut Pasteur, Paris | Marion Barbet, Sylvie Behillil, Méline Bizard, Angela Brisebarre, Camille Capel, Vincent Enouf, Louise Lefrançois, Frédéric Lemoine, Christophe Malabat, Corinne Maufrais, Emmanuelle Permal, Etienne Simon-Lorière, Maud Vanpeene, Sylvie Van der Werf ,Eric Farfour |
| EPI_ISL_2374384, EPI_ISL_2374385, EPI_ISL_2374386, EPI_ISL_2374387 | Labo Analyses Med | National Reference Center for Viruses of Respiratory Infections, Institut Pasteur, Paris | Marion Barbet, Sylvie Behillil, Méline Bizard, Angela Brisebarre, Camille Capel, Vincent Enouf, Louise Lefrançois, Frédéric Lemoine, Christophe Malabat, Corinne Maufrais, Emmanuelle Permal, Etienne Simon-Lorière, Maud Vanpeene, Sylvie Van der Werf ,Labm Mine Site De Thiant |
| EPI_ISL_2374388 | Hospital | National Reference Center for Viruses of Respiratory Infections, Institut Pasteur, Paris | Marion Barbet, Sylvie Behillil, Méline Bizard, Angela Brisebarre, Camille Capel, Vincent Enouf, Louise Lefrançois, Frédéric Lemoine, Christophe Malabat, Corinne Maufrais, Emmanuelle Permal, Etienne Simon-Lorière, Maud Vanpeene, Sylvie Van der Werf ,M. El Hamri |
| EPI_ISL_2374389, EPI_ISL_2374392 | Hospital | National Reference Center for Viruses of Respiratory Infections, Institut Pasteur, Paris | Marion Barbet, Sylvie Behillil, Méline Bizard, Angela Brisebarre, Camille Capel, Vincent Enouf, Louise Lefrançois, Frédéric Lemoine, Christophe Malabat, Corinne Maufrais, Emmanuelle Permal, Etienne Simon-Lorière, Maud Vanpeene, Sylvie Van der Werf ,Alexandra Ducancelle |
| EPI_ISL_2374395 | Labo Analyses Med | National Reference Center for Viruses of Respiratory Infections, Institut Pasteur, Paris | Marion Barbet, Sylvie Behillil, Méline Bizard, Angela Brisebarre, Camille Capel, Vincent Enouf, Louise Lefrançois, Frédéric Lemoine, Christophe Malabat, Corinne Maufrais, Emmanuelle Permal, Etienne Simon-Lorière, Maud Vanpeene, Sylvie Van der Werf |
| EPI_ISL_2374397, EPI_ISL_2374398, EPI_ISL_2374399, EPI_ISL_2374400 | Labo Analyses med | National Reference Center for Viruses of Respiratory Infections, Institut Pasteur, Paris | Marion Barbet, Sylvie Behillil, Méline Bizard, Angela Brisebarre, Camille Capel, Vincent Enouf, Louise Lefrançois, Frédéric Lemoine, Christophe Malabat, Corinne Maufrais, Emmanuelle Permal, Etienne Simon-Lorière, Maud Vanpeene, Sylvie Van der Werf ,CéCile Farrugia |
| EPI_ISL_2374401, EPI_ISL_2374402, EPI_ISL_2374403, EPI_ISL_2374404, EPI_ISL_2374405, EPI_ISL_2374407 | Labo Analyses Med | National Reference Center for Viruses of Respiratory Infections, Institut Pasteur, Paris | Marion Barbet, Sylvie Behillil, Méline Bizard, Angela Brisebarre, Camille Capel, Vincent Enouf, Louise Lefrançois, Frédéric Lemoine, Christophe Malabat, Corinne Maufrais, Emmanuelle Permal, Etienne Simon-Lorière, Maud Vanpeene, Sylvie Van der Werf ,Arnaud Jean |
| EPI_ISL_2374408, EPI_ISL_2374409 | Hospital | National Reference Center for Viruses of Respiratory Infections, Institut Pasteur, Paris | Marion Barbet, Sylvie Behillil, Méline Bizard, Angela Brisebarre, Camille Capel, Vincent Enouf, Louise Lefrançois, Frédéric Lemoine, Christophe Malabat, Corinne Maufrais, Emmanuelle Permal, Etienne Simon-Lorière, Maud Vanpeene, Sylvie Van der Werf ,CléMence Guillaume |
| EPI_ISL_2374413, EPI_ISL_2374414, EPI_ISL_2374415 | Hospital | National Reference Center for Viruses of Respiratory Infections, Institut Pasteur, Paris | Marion Barbet, Sylvie Behillil, Méline Bizard, Angela Brisebarre, Camille Capel, Vincent Enouf, Louise Lefrançois, Frédéric Lemoine, Christophe Malabat, Corinne Maufrais, Emmanuelle Permal, Etienne Simon-Lorière, Maud Vanpeene, Sylvie Van der Werf ,Esther Gyde |
| EPI_ISL_2374416, EPI_ISL_2374417, EPI_ISL_2374418, EPI_ISL_2374419, EPI_ISL_2374420, EPI_ISL_2374421, EPI_ISL_2374422, EPI_ISL_2374423, EPI_ISL_2374424, EPI_ISL_2374425, EPI_ISL_2374426, EPI_ISL_2374428, EPI_ISL_2374429, EPI_ISL_2374430, EPI_ISL_2374431, EPI_ISL_2374432, EPI_ISL_2374433, EPI_ISL_2374434, EPI_ISL_2374435, EPI_ISL_2374436, EPI_ISL_2374437, EPI_ISL_2374438, EPI_ISL_2374439 | | | |
| see above | Labo Analyses Med | National Reference Center for Viruses of Respiratory Infections, Institut Pasteur, Paris | Marion Barbet, Sylvie Behillil, Méline Bizard, Angela Brisebarre, Camille Capel, Vincent Enouf, Louise Lefrançois, Frédéric Lemoine, Christophe Malabat, Corinne Maufrais, Emmanuelle Permal, Etienne Simon-Lorière, Maud Vanpeene, Sylvie Van der Werf ,Benedicte Baccouch |
| EPI_ISL_2374440, EPI_ISL_2374441, EPI_ISL_2374442, EPI_ISL_2374443, EPI_ISL_2374444, EPI_ISL_2374445, EPI_ISL_2374446 | Hospital | National Reference Center for Viruses of Respiratory Infections, Institut Pasteur, Paris | Marion Barbet, Sylvie Behillil, Méline Bizard, Angela Brisebarre, Camille Capel, Vincent Enouf, Louise Lefrançois, Frédéric Lemoine, Christophe Malabat, Corinne Maufrais, Emmanuelle Permal, Etienne Simon-Lorière, Maud Vanpeene, Sylvie Van der Werf ,Axelle Paquin |
| EPI_ISL_2374447, EPI_ISL_2374448, EPI_ISL_2374449, EPI_ISL_2374450, | Hospital | National Reference Center for Viruses of Respiratory Infections, Institut Pasteur, Paris | Marion Barbet, Sylvie Behillil, Méline Bizard, Angela Brisebarre, Camille Capel, Vincent Enouf, Louise Lefrançois, Frédéric Lemoine, Christophe Malabat, Corinne Maufrais, Emmanuelle Permal, Etienne Simon-Lorière, Maud Vanpeene, Sylvie Van der Werf ,CéLine Ramanantsoa |

| | | | |
|---|---|---|---|
| EPI_ISL_2374451 | | | |
| EPI_ISL_2374452, EPI_ISL_2374454, EPI_ISL_2374455, EPI_ISL_2374456 | Hospital | National Reference Center for Viruses of Respiratory Infections, Institut Pasteur, Paris | Marion Barbet, Sylvie Behillil, Méline Bizard, Angela Brisebarre, Camille Capel, Vincent Enouf, Louise Lefrançois, Frédéric Lemoine, Christophe Malabat, Corinne Maufrais, Emmanuelle Permal, Etienne Simon-Lorière, Maud Vanpeene, Sylvie Van der Werf ,Axelle Paquin |
| EPI_ISL_2374457, EPI_ISL_2374458, EPI_ISL_2374459, EPI_ISL_2374460, EPI_ISL_2374461, EPI_ISL_2374462 | Labo Analyses Med | National Reference Center for Viruses of Respiratory Infections, Institut Pasteur, Paris | Marion Barbet, Sylvie Behillil, Méline Bizard, Angela Brisebarre, Camille Capel, Vincent Enouf, Louise Lefrançois, Frédéric Lemoine, Christophe Malabat, Corinne Maufrais, Emmanuelle Permal, Etienne Simon-Lorière, Maud Vanpeene, Sylvie Van der Werf ,Delphine MammÈS |
| EPI_ISL_2374463, EPI_ISL_2374464, EPI_ISL_2374465, EPI_ISL_2374466, EPI_ISL_2374467, EPI_ISL_2374468, EPI_ISL_2374469, EPI_ISL_2374470, EPI_ISL_2374471 | Labo Analyses med | National Reference Center for Viruses of Respiratory Infections, Institut Pasteur, Paris | Marion Barbet, Sylvie Behillil, Méline Bizard, Angela Brisebarre, Camille Capel, Vincent Enouf, Louise Lefrançois, Frédéric Lemoine, Christophe Malabat, Corinne Maufrais, Emmanuelle Permal, Etienne Simon-Lorière, Maud Vanpeene, Sylvie Van der Werf ,CéLine Bressollette |
| EPI_ISL_2374474 | Hospital | National Reference Center for Viruses of Respiratory Infections, Institut Pasteur, Paris | Marion Barbet, Sylvie Behillil, Méline Bizard, Angela Brisebarre, Camille Capel, Vincent Enouf, Louise Lefrançois, Frédéric Lemoine, Christophe Malabat, Corinne Maufrais, Emmanuelle Permal, Etienne Simon-Lorière, Maud Vanpeene, Sylvie Van der Werf ,Marie-Christine Legrand-Quillien |
| EPI_ISL_2374475, EPI_ISL_2374476, EPI_ISL_2374477 | Hospital | National Reference Center for Viruses of Respiratory Infections, Institut Pasteur, Paris | Marion Barbet, Sylvie Behillil, Méline Bizard, Angela Brisebarre, Camille Capel, Vincent Enouf, Louise Lefrançois, Frédéric Lemoine, Christophe Malabat, Corinne Maufrais, Emmanuelle Permal, Etienne Simon-Lorière, Maud Vanpeene, Sylvie Van der Werf ,E. Bichier |
| EPI_ISL_2374480 | Hospital | National Reference Center for Viruses of Respiratory Infections, Institut Pasteur, Paris | Marion Barbet, Sylvie Behillil, Méline Bizard, Angela Brisebarre, Camille Capel, Vincent Enouf, Louise Lefrançois, Frédéric Lemoine, Christophe Malabat, Corinne Maufrais, Emmanuelle Permal, Etienne Simon-Lorière, Maud Vanpeene, Sylvie Van der Werf ,FréDéRick Garel |
| EPI_ISL_2374481, EPI_ISL_2374483, EPI_ISL_2374484, EPI_ISL_2374485, EPI_ISL_2374486, EPI_ISL_2374487, EPI_ISL_2374489, EPI_ISL_2374490, EPI_ISL_2374491, EPI_ISL_2374492 | Labo Analyses med | National Reference Center for Viruses of Respiratory Infections, Institut Pasteur, Paris | Marion Barbet, Sylvie Behillil, Méline Bizard, Angela Brisebarre, Camille Capel, Vincent Enouf, Louise Lefrançois, Frédéric Lemoine, Christophe Malabat, Corinne Maufrais, Emmanuelle Permal, Etienne Simon-Lorière, Maud Vanpeene, Sylvie Van der Werf ,Jean-FrançOis Comes |
| EPI_ISL_2379461 | Hospital Melaka | Institute for Medical Research, Infectious Disease Research Centre, National Inst | Suppiah J, Kamel K, Mohd Zawawi Z, Azizan MA, Ramly N, Robert F, Thayan R |
| EPI_ISL_2379748 | National Heart Institute | Institute for Medical Research, Infectious Disease Research Centre, National Institutes of Health, Ministry of Health Malaysia | Suppiah J, Kamel K, Mohd Zawawi Z, Azizan MA, Ramly N, Robert F, Thayan R |
| EPI_ISL_2379749 | Tawau Hospital | Institute for Medical Research, Infectious Disease Research Centre, National Institutes of Health, Ministry of Health Malaysia | Suppiah J, Kamel K, Mohd Zawawi Z, Azizan MA, Ramly N, Robert F, Thayan R |
| EPI_ISL_2381459, EPI_ISL_2381503, EPI_ISL_2381504 | Institut de Virologie du CHU de Strasbourg | Institut de Virologie du CHU de Strasbourg | Fafi-Kremer Samira, Gallais Floriane, Gantner Pierre, Laugel Elodie, Solis Morgane, Velay Aurélie, Wendling Marie-Josée |
| EPI_ISL_2382000 | AZ Zeno Knokke-Blankenberge | AZ Sint Jan | Marijke Reynders, Patrick Descheemaeker, Laurien Hoornaert, Thomas Van Landschoot, Marie Madeleine Chabert-Consen, Jorn Hellemans |
| EPI_ISL_2382001 | AZ St.Jan campus Oostende | AZ Sint Jan | Marijke Reynders, Patrick Descheemaeker, Laurien Hoornaert, Thomas Van Landschoot, Marie Madeleine Chabert-Consen, Jorn Hellemans |
| EPI_ISL_2382003, EPI_ISL_2382004 | AZ Zeno Knokke-Blankenberge | AZ Sint Jan | Marijke Reynders, Patrick Descheemaeker, Laurien Hoornaert, Thomas Van Landschoot, Marie Madeleine Chabert-Consen, Jorn Hellemans |
| EPI_ISL_2382007, EPI_ISL_2382008 | AZ St.Jan campus Brugge | AZ Sint Jan | Marijke Reynders, Patrick Descheemaeker, Laurien Hoornaert, Thomas Van Landschoot, Marie Madeleine Chabert-Consen, Jorn Hellemans |
| EPI_ISL_2382036, EPI_ISL_2382040 | Institute for Laboratory Diagnostics and Microbiology, Klinikum Klagenfurt am Wörthersee | Bergthaler laboratory, CeMM Research Center for Molecular Medicine of the Austrian Academy of Sciences | Lukas Endler, Anna Schedl, Fabian Amman, Petr Triska, Thomas Penz, Benedikt Agerer, Maelle Le Moing, Michael Schuster, Bekir Erguner, Jan Laine, Martin Senekowitsch, Christoph Bock, Andreas Bergthaler |
| EPI_ISL_2382054 | Klinikum Wels-Grieskirchen | Bergthaler laboratory, CeMM Research Center for Molecular Medicine of the Austrian Academy of Sciences | Lukas Endler, Anna Schedl, Fabian Amman, Petr Triska, Thomas Penz, Benedikt Agerer, Maelle Le Moing, Michael Schuster, Bekir Erguner, Jan Laine, Martin Senekowitsch, Christoph Bock, Andreas Bergthaler |
| EPI_ISL_2382065, EPI_ISL_2382066, EPI_ISL_2382067, EPI_ISL_2382068, EPI_ISL_2382069, EPI_ISL_2382070, EPI_ISL_2382071, EPI_ISL_2382073, EPI_ISL_2382074 | Center for Virology, Medical University of Vienna | Bergthaler laboratory, CeMM Research Center for Molecular Medicine of the Austrian Academy of Sciences | Lukas Endler, Anna Schedl, Fabian Amman, Petr Triska, Thomas Penz, Benedikt Agerer, Maelle Le Moing, Michael Schuster, Bekir Erguner, Jan Laine, Martin Senekowitsch, Christoph Bock, Andreas Bergthaler |
| EPI_ISL_2391271, EPI_ISL_2391288, EPI_ISL_2391362 | Institut de Virologie du CHU de Strasbourg | Institut de Virologie du CHU de Strasbourg | Fafi-Kremer Samira, Gallais Floriane, Gantner Pierre, Laugel Elodie, Solis Morgane, Velay Aurélie, Wendling Marie-Josée |
| EPI_ISL_2391557, EPI_ISL_2391580, EPI_ISL_2391586, EPI_ISL_2391594, EPI_ISL_2391617, EPI_ISL_2391623, EPI_ISL_2391643, EPI_ISL_2391647 | Laboratorio Central de Epidemiología (LCE) | Instituto de Biotecnología de la UNAM | Consorcio Mexicano de Vigilancia Genómica (CoViGen-Mex). Authors (in alphabetical order): Julio Elias Alvarado-Yaah, Carlos F. Arias, Santiago Ávila-Ríos, Eduardo Becerril-Vargas, Víctor Hugo Borja-Aburto, Celia Boukadida, Cristóbal Cháidez-Quiróz, Juan Bautista Chale-Dzul, Ricardo Ciria Merce, Andreu Comas-García, Célida Duque Molina, Julissa Enciso-Ibarra, José Antonio Enciso-Moreno, Gloria Elena Espinosa-Ayala, Fernando Fontove-Herrera, Daniel Fregoso-Rueda, Victor Eduardo Garcia-Arias, Alejandra Garcia-Gasca, Bruno Gomez-Gil, Jean Pierre González, Irvin González-López, Concepción Grajales-Muñiz, Ricardo Grande, Rosa María Gutierrez Rios, Alejandra Hernández-Terán, Alfredo Herrera-Estrella, Carla Ivón Herrera-Najera, Pavel Isa, Daniel Lira Morales, Susana Lopez, Antonio Loza Román, Brenda Irasema Maldonado-Meza, Bernardo Martínez-Miguel, José Arturo Martínez-Orozco, Célida Martínez- Rodríguez, Margarita Matías-Florentino, Fidencio Mejía-Nepomuceno, María Guadalupe de Jesús Mireles-Rivera, Gloria María Molina-Salinas, Hector Montoya-Fuentes, , Mario Mújica-Sánchez, José Esteban Muñoz-Medina, José de Jesús Nuñez-Contreras, Alicia Ocaña-Mondragón, Luis Alberto Ochoa-Carrera, Hector Esteban Paz-Juárez, Francisco Pulido, Helen Haydee Fernanda Ramírez-Plascencia, Jorge Salas-Hernández, Angel Gustavo Salas-Lais, Alejandro Sanchez-Flores, Clara Esperanza Santacruz-Tinoco, María Guadalupe Santiago-Mauricio, Selene Zárate, Nelly Sélem-Mojica, Blanca Taboada, Gloria Vazquez, Joel Armando Vázquez-Pérez. |
| EPI_ISL_2392160, EPI_ISL_2392195 | Institut de Virologie du CHU de Strasbourg | Institut de Virologie du CHU de Strasbourg | Fafi-Kremer Samira, Gallais Floriane, Gantner Pierre, Laugel Elodie, Solis Morgane, Velay Aurélie, Wendling Marie-Josée |
| EPI_ISL_2396877, EPI_ISL_2396878, EPI_ISL_2396879, EPI_ISL_2396880, EPI_ISL_2396881, EPI_ISL_2396882, EPI_ISL_2396883, EPI_ISL_2396884, EPI_ISL_2396885 | Azienda Ospedaliera Pugliese Ciaccio di Catanzaro SOC Microbiologia e Virologia | Azienda Ospedaliera Pugliese Ciaccio di Catanzaro SOC Microbiologia e Virologia | Rossana Tallerico Federica Pasceri Marco De Fazio Ilenia Talotta Giuseppina Panduri Pasquale Minchella |
| EPI_ISL_2396888 | Institut de Virologie du CHU de Strasbourg | Institut de Virologie du CHU de Strasbourg | Fafi-Kremer Samira, Gallais Floriane, Gantner Pierre, Laugel Elodie, Solis Morgane, Velay Aurélie, Wendling Marie-Josée |
| EPI_ISL_2397166, EPI_ISL_2397167, EPI_ISL_2397168, EPI_ISL_2397169, EPI_ISL_2397170, EPI_ISL_2397171, EPI_ISL_2397172, EPI_ISL_2397173, EPI_ISL_2397174, EPI_ISL_2397175, EPI_ISL_2397176, EPI_ISL_2397177, EPI_ISL_2397178, EPI_ISL_2397179, EPI_ISL_2397180, EPI_ISL_2397181, EPI_ISL_2397182, EPI_ISL_2397183, EPI_ISL_2397184, EPI_ISL_2397185, EPI_ISL_2397186, EPI_ISL_2397187, EPI_ISL_2397188, EPI_ISL_2397189, EPI_ISL_2397190, EPI_ISL_2397191, EPI_ISL_2397192, EPI_ISL_2397193, EPI_ISL_2397194, EPI_ISL_2397195, EPI_ISL_2397196, EPI_ISL_2397197, EPI_ISL_2397198, EPI_ISL_2397199, EPI_ISL_2397200, EPI_ISL_2397201, EPI_ISL_2397202 | | | |
| see above | Klinisch Laboratorium ZNA | Klinisch Laboratorium ZNA | Verstrepen et al. |
| EPI_ISL_2397314, EPI_ISL_2397315, EPI_ISL_2397316, EPI_ISL_2397317, EPI_ISL_2397318, EPI_ISL_2397319, EPI_ISL_2397320, EPI_ISL_2397321, EPI_ISL_2397322, EPI_ISL_2397323, EPI_ISL_2399426 | | | |
| see above | Laboratori de Referencia de Catalunya | Laboratori de Referencia de Catalunya | Ramirez A., Bellosillo B., Padilla E., Vilas A., Hernandez JJ., Canal M. |
| EPI_ISL_2400376, EPI_ISL_2400393, EPI_ISL_2400394, EPI_ISL_2400395, EPI_ISL_2400396, EPI_ISL_2400398, EPI_ISL_2400399, EPI_ISL_2400400, EPI_ISL_2400401, EPI_ISL_2400402, EPI_ISL_2400403, EPI_ISL_2400404, EPI_ISL_2400405, EPI_ISL_2400406, EPI_ISL_2400407, EPI_ISL_2400408, EPI_ISL_2400409, EPI_ISL_2400410, EPI_ISL_2400411, EPI_ISL_2400412, EPI_ISL_2400413, EPI_ISL_2400414, EPI_ISL_2400415 | | | |

| | | | |
|---|---|---|---|
| see above | Klinisch Laboratorium ZNA | Klinisch Laboratorium ZNA | Verstrepen et al. |
| EPI_ISL_2400445, EPI_ISL_2400446, EPI_ISL_2400448 | Molecular Biology Laboratory, Westchester Medical Center | Genomics Core Laboratory, New York Medical College | Weihua Huang, Utsav Pandey, Jian Zhuge, Changhong Yin |
| EPI_ISL_2402535, EPI_ISL_2402536, EPI_ISL_2402537, EPI_ISL_2402540, EPI_ISL_2402541, EPI_ISL_2402543, EPI_ISL_2402544, EPI_ISL_2402545, EPI_ISL_2402546, EPI_ISL_2402547, EPI_ISL_2402548, EPI_ISL_2402549 | | | |
| see above | OLVZ Aalst | OLVZ Aalst | Astrid Holderbeke |
| EPI_ISL_2403193, EPI_ISL_2403196, EPI_ISL_2403197, EPI_ISL_2403201, EPI_ISL_2403202, EPI_ISL_2403205, EPI_ISL_2403207, EPI_ISL_2403209, EPI_ISL_2403211, EPI_ISL_2403217, EPI_ISL_2403221, EPI_ISL_2403224, EPI_ISL_2403228 | | | |
| see above | Institute of Molecular and Translational Medicine / Laboratory of Experimental Medicine, Faculty of Medicine and Dentistry, Palacky University and University Hospital Olomouc | Institute of Molecular and Translational Medicine / Laboratory of Experimental Medicine, Faculty of Medicine and Dentistry, Palacky University | Rastislav Slavkovský, Hana Jaworek, Vladimíra Koudeláková, Barbora Blumová, Marián Hajdúch |
| EPI_ISL_2403233, EPI_ISL_2403237 | CHU Pontchaillou | CHU Pontchaillou | GROLHIER Claire, DENOUAL Florent, ETCHEVERRY Amandine, SASSI Mohamed, JAGLINE Steven, FEBREAU Christine, PRONIER Charlotte, GALIBERT Marie Dominique, DE TAYRAC Marie, THIBAULT Vincent |
| EPI_ISL_2404840, EPI_ISL_2404842, EPI_ISL_2404843, EPI_ISL_2404845, EPI_ISL_2404846 | University of Oregon COVID-19 MAP Laboratory | University of Oregon Genomics and Cell Charactericzation Core Facility (GC3F) | Douglas Turnbull, Ariana White, Jeff Bishop, Jason Carriere, Jason Sydes, Peter Batzel |
| EPI_ISL_2405147, EPI_ISL_2405151, EPI_ISL_2405154, EPI_ISL_2405156, EPI_ISL_2405158, EPI_ISL_2405160, EPI_ISL_2405164, EPI_ISL_2405166 | Hospital Luis Vernaza | Omics Sciences Laboratory | Derly Andrade Molina, Gabriel Morey León, Darlyn Amaya, Rubén Armas González, Juan Carlos Fernández Cadena |
| EPI_ISL_2406458, EPI_ISL_2406459, EPI_ISL_2406460 | Battambang Provincial Laboratory | Virology Unit, Institut Pasteur du Cambodge | Cecile Troupin, Leakhena Pum, Jurre Y Siegers, Ly Sovann, Kraing Sidonn, Yi Sengdoeurn, Chin Savuth, Chau Darapheak, Veasna Duong, Erik A Karlsson |
| EPI_ISL_2406461 | Oddar Meanchey Rapid Response Team | Virology Unit, Institut Pasteur du Cambodge | Cecile Troupin, Leakhena Pum, Jurre Y Siegers, Ly Sovann, Kraing Sidonn, Yi Sengdoeurn, Chin Savuth, Chau Darapheak, Veasna Duong, Erik A Karlsson |
| EPI_ISL_2406462, EPI_ISL_2406463, EPI_ISL_2406464 | Virology Unit, Institut Pasteur du Cambodge | Virology Unit, Institut Pasteur du Cambodge | Cecile Troupin, Leakhena Pum, Jurre Y Siegers, Ly Sovann, Kraing Sidonn, Yi Sengdoeurn, Chin Savuth, Chau Darapheak, Veasna Duong, Erik A Karlsson |
| EPI_ISL_2406465, EPI_ISL_2406466, EPI_ISL_2406467 | Svay Rieng Rapid Response Team | Virology Unit, Institut Pasteur du Cambodge | Cecile Troupin, Leakhena Pum, Jurre Y Siegers, Ly Sovann, Kraing Sidonn, Yi Sengdoeurn, Chin Savuth, Chau Darapheak, Veasna Duong, Erik A Karlsson |
| EPI_ISL_2406468, EPI_ISL_2406469, EPI_ISL_2406470, EPI_ISL_2406471, EPI_ISL_2406472, EPI_ISL_2406473, EPI_ISL_2406474, EPI_ISL_2406475, EPI_ISL_2406476, EPI_ISL_2406477, EPI_ISL_2406478, EPI_ISL_2406479, EPI_ISL_2406480, EPI_ISL_2406481 | | | |
| see above | Virology Unit, Institut Pasteur du Cambodge | Virology Unit, Institut Pasteur du Cambodge | Cecile Troupin, Leakhena Pum, Jurre Y Siegers, Ly Sovann, Kraing Sidonn, Yi Sengdoeurn, Chin Savuth, Chau Darapheak, Veasna Duong, Erik A Karlsson |
| EPI_ISL_2408063 | Usansolo-Galdakao University Hospital | Cruces University Hospital | Ana Gual-de-Torrella, Izaskun Alejo-Cancho, Mikel Gallego, Ana Belén de la Hoz |
| EPI_ISL_2408116 | AZ Klina | AZ Klina | Carl Vael - Lynsey Berckmans |
| EPI_ISL_2420637, EPI_ISL_2420638, EPI_ISL_2420639, EPI_ISL_2420640, EPI_ISL_2420641, EPI_ISL_2420642, EPI_ISL_2420643, EPI_ISL_2420644, EPI_ISL_2420645, EPI_ISL_2420646, EPI_ISL_2420647, EPI_ISL_2420648, EPI_ISL_2420649 | | | |
| see above | Department of Virology, Istituto Zooprofilattico Sperimentale del Lazio e della Toscana (IZSLT) | Department of General Diagnostics; Department of Virology; Istituto Zooprofilattico Sperimentale del Lazio e della Toscana (IZSLT) | Patricia Alba, Giuseppe Manna, Elena L. Diaconu, Fabiola Feltrin, Raffaela Conti, Teresa Scicluna, Virginia Carfora, Antonella Cersini, Alessia Franco, Antonio Battisti. |
| EPI_ISL_2421279, EPI_ISL_2421280, EPI_ISL_2421281, EPI_ISL_2421282, EPI_ISL_2421284 | Laboratori de Referencia de Catalunya | Laboratori de Referencia de Catalunya | Ramirez A., Bellosillo B., Padilla E., Vilas A., Hernandez JJ., Canal M. |
| EPI_ISL_2422815, EPI_ISL_2422816, EPI_ISL_2422817, EPI_ISL_2422818, EPI_ISL_2422819, EPI_ISL_2422820, EPI_ISL_2422821, EPI_ISL_2422822, EPI_ISL_2422823, EPI_ISL_2422824, EPI_ISL_2422825, EPI_ISL_2422826, EPI_ISL_2422831, EPI_ISL_2422832 | | | |
| see above | Sharp HealthCare Laboratory | Andersen lab at Scripps Research | SEARCH Alliance San Diego with Aaron Harding, Jacquelyn Berumen, Cathy Woerle, Liam McGinnis, Art Mendoza, Omid Bakhtar |
| EPI_ISL_2426976, EPI_ISL_2426977, EPI_ISL_2426978, EPI_ISL_2426979, EPI_ISL_2426980, EPI_ISL_2426981 | Laboratoire Carage | Institut Pasteur de la Guyane | Anne Lavergne, Dominique Rousset, Antoine Enfissi, Arielle Salmier |
| EPI_ISL_2426982 | CHC Andrée Rosemon | Institut Pasteur de la Guyane | Anne Lavergne, Dominique Rousset, Antoine Enfissi, Arielle Salmier |
| EPI_ISL_2426983, EPI_ISL_2426984 | CNR Institut Pasteur de la Guyane | Institut Pasteur de la Guyane | Anne Lavergne, Dominique Rousset, Antoine Enfissi, Arielle Salmier |
| EPI_ISL_2426995, EPI_ISL_2426996, EPI_ISL_2426997, EPI_ISL_2426998, EPI_ISL_2426999, EPI_ISL_2427000, EPI_ISL_2427001, EPI_ISL_2427003, EPI_ISL_2427004, EPI_ISL_2427005, EPI_ISL_2427006, EPI_ISL_2427007, EPI_ISL_2427008, EPI_ISL_2427009, EPI_ISL_2427011, EPI_ISL_2427012, EPI_ISL_2427013, EPI_ISL_2427014, EPI_ISL_2427015, EPI_ISL_2427016, EPI_ISL_2427017, EPI_ISL_2427018, EPI_ISL_2427019, EPI_ISL_2427020, EPI_ISL_2427021, EPI_ISL_2427022, EPI_ISL_2427023, EPI_ISL_2427024, EPI_ISL_2427025, EPI_ISL_2427026, EPI_ISL_2427027, EPI_ISL_2427028, EPI_ISL_2427029, EPI_ISL_2427031, EPI_ISL_2427032, EPI_ISL_2427033, EPI_ISL_2427034, EPI_ISL_2427035, EPI_ISL_2427036, EPI_ISL_2427037, EPI_ISL_2427038, EPI_ISL_2427039, EPI_ISL_2427040, EPI_ISL_2427041, EPI_ISL_2427042, EPI_ISL_2427043, EPI_ISL_2427044, EPI_ISL_2427045, EPI_ISL_2427046, EPI_ISL_2427047, EPI_ISL_2427048, EPI_ISL_2427049, EPI_ISL_2427051, EPI_ISL_2427052, EPI_ISL_2427053, EPI_ISL_2427054, EPI_ISL_2427055, EPI_ISL_2427056, EPI_ISL_2427057, EPI_ISL_2427058, EPI_ISL_2427062, EPI_ISL_2427063, EPI_ISL_2427064, EPI_ISL_2427065, EPI_ISL_2427066, EPI_ISL_2427067, EPI_ISL_2427069, EPI_ISL_2427070, EPI_ISL_2427074, EPI_ISL_2427075, EPI_ISL_2427077, EPI_ISL_2427078, EPI_ISL_2427079, EPI_ISL_2427080, EPI_ISL_2427081, EPI_ISL_2427082, EPI_ISL_2427083, EPI_ISL_2427085, EPI_ISL_2427086, EPI_ISL_2427087, EPI_ISL_2427088, EPI_ISL_2427089, EPI_ISL_2427090, EPI_ISL_2427091, EPI_ISL_2427092, EPI_ISL_2427094, EPI_ISL_2427097, EPI_ISL_2427099, EPI_ISL_2427100, EPI_ISL_2427101, EPI_ISL_2427102, EPI_ISL_2427103, EPI_ISL_2427104, EPI_ISL_2427105, EPI_ISL_2427106, EPI_ISL_2427107, EPI_ISL_2427108, EPI_ISL_2427109, EPI_ISL_2427110, EPI_ISL_2427111, EPI_ISL_2427112, EPI_ISL_2427113, EPI_ISL_2427114, EPI_ISL_2427115, EPI_ISL_2427116, EPI_ISL_2427117, EPI_ISL_2427118, EPI_ISL_2427119, EPI_ISL_2427120, EPI_ISL_2427121, EPI_ISL_2427122, EPI_ISL_2427123, EPI_ISL_2427125, EPI_ISL_2427126, EPI_ISL_2427127, EPI_ISL_2427128, EPI_ISL_2427129, EPI_ISL_2427130, EPI_ISL_2427131, EPI_ISL_2427132, EPI_ISL_2427133, EPI_ISL_2427134, EPI_ISL_2427135, EPI_ISL_2427136, EPI_ISL_2427137, EPI_ISL_2427138, EPI_ISL_2427139, EPI_ISL_2427140, EPI_ISL_2427141, EPI_ISL_2427142, EPI_ISL_2427143, EPI_ISL_2427144, EPI_ISL_2427146, EPI_ISL_2427147, EPI_ISL_2427148, EPI_ISL_2427149, EPI_ISL_2427150, EPI_ISL_2427151, EPI_ISL_2427152, EPI_ISL_2427153, EPI_ISL_2427154, EPI_ISL_2427155, EPI_ISL_2427156, EPI_ISL_2427157, EPI_ISL_2427158, EPI_ISL_2427159, EPI_ISL_2427160, EPI_ISL_2427161, EPI_ISL_2427162, EPI_ISL_2427163, EPI_ISL_2427164, EPI_ISL_2427165, EPI_ISL_2427166, EPI_ISL_2427167, EPI_ISL_2427169, EPI_ISL_2427170, EPI_ISL_2427171, EPI_ISL_2427173, EPI_ISL_2427174, EPI_ISL_2427176, EPI_ISL_2427177, EPI_ISL_2427178, EPI_ISL_2427179, EPI_ISL_2427180, EPI_ISL_2427181, EPI_ISL_2427182, EPI_ISL_2427183, EPI_ISL_2427184, EPI_ISL_2427185, EPI_ISL_2427186, EPI_ISL_2427187, EPI_ISL_2427188, EPI_ISL_2427189, EPI_ISL_2427190, EPI_ISL_2427191, EPI_ISL_2427192, EPI_ISL_2427193, EPI_ISL_2427194, EPI_ISL_2427195, EPI_ISL_2427196, EPI_ISL_2427197, EPI_ISL_2427198, EPI_ISL_2427199, EPI_ISL_2427200, EPI_ISL_2427201, EPI_ISL_2427202, EPI_ISL_2427203, EPI_ISL_2427204, EPI_ISL_2427205, EPI_ISL_2427207, EPI_ISL_2427208, EPI_ISL_2427212, EPI_ISL_2427214, EPI_ISL_2427215, EPI_ISL_2427216, EPI_ISL_2427217, EPI_ISL_2427219, EPI_ISL_2427220, EPI_ISL_2427221, EPI_ISL_2427222, EPI_ISL_2427223, EPI_ISL_2427225, EPI_ISL_2427227, EPI_ISL_2427228 | | | |
| see above | Austrian Agency for Health and Food Safety (AGES) | Bergthaler laboratory, CeMM Research Center for Molecular Medicine of the Austrian Academy of Sciences | Lukas Endler, Anna Schedl, Fabian Amman, Petr Triska, Thomas Penz, Benedikt Agerer, Maelle Le Moing, Michael Schuster, Bekir Erguner, Jan Laine, Martin Senekowitsch, Christoph Bock, Andreas Bergthaler |
| EPI_ISL_2427322, EPI_ISL_2427323, EPI_ISL_2427324, EPI_ISL_2427325, EPI_ISL_2427326, EPI_ISL_2427327, EPI_ISL_2427328, EPI_ISL_2427329, EPI_ISL_2427330, EPI_ISL_2427331, EPI_ISL_2427332, EPI_ISL_2427333, EPI_ISL_2427334, EPI_ISL_2427335, EPI_ISL_2427336, EPI_ISL_2427338, EPI_ISL_2427339, EPI_ISL_2427340, EPI_ISL_2427341, EPI_ISL_2427342, EPI_ISL_2427343, EPI_ISL_2427344, EPI_ISL_2427345, EPI_ISL_2427346, EPI_ISL_2427347, EPI_ISL_2427348, EPI_ISL_2427349, EPI_ISL_2427350, EPI_ISL_2427351, EPI_ISL_2427352, EPI_ISL_2427353, EPI_ISL_2427354, EPI_ISL_2427355, EPI_ISL_2427356, EPI_ISL_2427357, EPI_ISL_2427358, EPI_ISL_2427359, EPI_ISL_2427360, EPI_ISL_2427361, EPI_ISL_2427362, EPI_ISL_2427363, EPI_ISL_2427366, EPI_ISL_2427367, EPI_ISL_2427368, EPI_ISL_2427369, EPI_ISL_2427370, EPI_ISL_2427371, EPI_ISL_2427372, EPI_ISL_2427373, EPI_ISL_2427375, EPI_ISL_2427377 | | | |
| see above | Cliniques universitaires Saint-Luc | UCLouvain/IREC/MBLG | Jean Ruelle, Ophélie Simon, Bertrand Bearzatto, Benoit Kabamba Mukadi |
| EPI_ISL_2427378, EPI_ISL_2427379, EPI_ISL_2427380, EPI_ISL_2427381, EPI_ISL_2427382, EPI_ISL_2427383, EPI_ISL_2427384, EPI_ISL_2427385, EPI_ISL_2427386, EPI_ISL_2427387, EPI_ISL_2427388, EPI_ISL_2427389, EPI_ISL_2427390, EPI_ISL_2427391, EPI_ISL_2427392, EPI_ISL_2427393, EPI_ISL_2427394, EPI_ISL_2427395, EPI_ISL_2427396, EPI_ISL_2427397, EPI_ISL_2427398, EPI_ISL_2427399, EPI_ISL_2427400, EPI_ISL_2427401, EPI_ISL_2427402 | | | |
| see above | Clinique Saint-Pierre Ottignies | UCLouvain/IREC/MBLG | Jean Ruelle, Ophélie Simon, Bertrand Bearzatto, Benoit Kabamba Mukadi |
| EPI_ISL_2429134, EPI_ISL_2429136, EPI_ISL_2429137, EPI_ISL_2429138, EPI_ISL_2429141, EPI_ISL_2429143, EPI_ISL_2429145 | Institute for Medical Research, Infectious Disease Research Centre, National Institutes of Health, Ministry of Health Malaysia | Institute for Medical Research, Infectious Disease Research Centre, National Institutes of Health, Ministry of Health Malaysia | Suppiah J, Kamel K, Mohd Zawawi Z, Azizan MA, Ramly N, Robert F, Thayan R |
| EPI_ISL_2434131, EPI_ISL_2434132, EPI_ISL_2434133, EPI_ISL_2434134, | Cliniques universitaires Saint-Luc | UCLouvain/IREC/MBLG | Jean Ruelle, Ophélie Simon, Bertrand Bearzatto, Benoit Kabamba Mukadi |

| | | | |
|---|---|---|---|
| EPI_ISL_2434135, EPI_ISL_2434136, EPI_ISL_2434137 | | | |
| EPI_ISL_2434138 | Clinique Saint-Pierre Ottignies | UCLouvain/IREC/MBLG | Jean Ruelle, Ophélie Simon, Bertrand Bearzatto, Benoit Kabamba Mukadi |
| EPI_ISL_2434144, EPI_ISL_2434145, EPI_ISL_2434146, EPI_ISL_2434147, EPI_ISL_2434148, EPI_ISL_2434149, EPI_ISL_2434150, EPI_ISL_2434151, EPI_ISL_2434154, EPI_ISL_2434155 | AZ Klina | AZ Klina | Carl Vael - Lynsey Berckmans |
| EPI_ISL_2434156 | AZ Klina | AZ Klina | Carl Vael- Lynsey Berckmans |
| EPI_ISL_2434159, EPI_ISL_2434160, EPI_ISL_2434162, EPI_ISL_2434163, EPI_ISL_2434164, EPI_ISL_2434165, EPI_ISL_2434166, EPI_ISL_2434167, EPI_ISL_2434170, EPI_ISL_2434171, EPI_ISL_2434172 | | | |
| see above | AZ Klina | AZ Klina | Carl Vael - Lynsey Berckmans |
| EPI_ISL_2434981, EPI_ISL_2434983, EPI_ISL_2434984 | Institut de Virologie du CHU de Strasbourg | Institut de Virologie du CHU de Strasbourg | Fafi-Kremer Samira, Gallais Floriane, Gantner Pierre, Laugel Elodie, Solis Morgane, Velay Aurélie, Wendling Marie-Josée |
| EPI_ISL_2439237 | Imeda Hospital | Imeda Hospital | Johan Frans, Dagmar Obbels, Hanne Valgaeren |
| EPI_ISL_2441781 | CHU de la Réunion - Nord | Laboratoire de virologie, CNR arbovirus Associé, Chu de la Réunion | Etienne Frumence, Marie-Christine Jaffar Bandjee, Nicolas Traversier, Sabrina Petit Genet |
| EPI_ISL_2441804, EPI_ISL_2441805, EPI_ISL_2441807 | CHU de la Réunion - Sud | Laboratoire de virologie, CNR arbovirus Associé, Chu de la Réunion | Etienne Frumence, Marie-Christine Jaffar Bandjee, Nicolas Traversier, Sabrina Petit Genet |
| EPI_ISL_2441808 | CHU de la Réunion - Nord | Laboratoire de virologie, CNR arbovirus Associé, Chu de la Réunion | Etienne Frumence, Marie-Christine Jaffar Bandjee, Nicolas Traversier, Sabrina Petit Genet |
| EPI_ISL_2441809, EPI_ISL_2441810, EPI_ISL_2441811 | CHU de la Réunion - Sud | Laboratoire de virologie, CNR arbovirus Associé, Chu de la Réunion | Etienne Frumence, Marie-Christine Jaffar Bandjee, Nicolas Traversier, Sabrina Petit Genet |
| EPI_ISL_2442278 | Laboratoire Hôpital Sainte Musse | Laboratoire Hôpital Sainte Musse | Lionel Chollet |
| EPI_ISL_2443048 | LESP Ciudad de Mexico | Instituto de Diagnostico y Referencia Epidemiologicos (INDRE) | Claudia Wong-Arambula, Abril Rodriguez-Maldonado, Vanessa Rivero-Arredondo, Ariadna Medina-Benitez, Joaquin Quiroz-Mercado, Sergio Rangel-Guerrero, Natividad Cruz-Ortiz, Tatiana Nunez-Garcia, Gisela Barrera-Badillo, Lucia Hernandez-Rivas, Irma Lopez-Martinez, Ernesto Ramirez-Gonzalez. |
| EPI_ISL_2443049 | LESP Quintana Roo | Instituto de Diagnostico y Referencia Epidemiologicos (INDRE) | Claudia Wong-Arambula, Abril Rodriguez-Maldonado, Vanessa Rivero-Arredondo, Ariadna Medina-Benitez, Joaquin Quiroz-Mercado, Sergio Rangel-Guerrero, Natividad Cruz-Ortiz, Tatiana Nunez-Garcia, Gisela Barrera-Badillo, Lucia Hernandez-Rivas, Irma Lopez-Martinez, Ernesto Ramirez-Gonzalez. |
| EPI_ISL_2443050 | LESP Michoacan | Instituto de Diagnostico y Referencia Epidemiologicos (INDRE) | Claudia Wong-Arambula, Abril Rodriguez-Maldonado, Vanessa Rivero-Arredondo, Ariadna Medina-Benitez, Joaquin Quiroz-Mercado, Sergio Rangel-Guerrero, Natividad Cruz-Ortiz, Tatiana Nunez-Garcia, Gisela Barrera-Badillo, Lucia Hernandez-Rivas, Irma Lopez-Martinez, Ernesto Ramirez-Gonzalez. |
| EPI_ISL_2443057 | Department of Health Technology and Informatics, The Hong Kong Polytechnic University | Department of Health Technology and Informatics, The Hong Kong Polytechnic University | Gilman Kit-Hang Siu, Lam-Kwong Lee, Kenneth Siu-Sing Leung, Jake Siu-Lun Leung, Wing-Hei Lo, Timothy Ting-Leung Ng, Chloe Toi-Mei Chan, Hoi-Ching Jim, Kingsley King-Gee Tam, Hiu-Yin Lao, Denise Sze-Hang Wong, Alan Ka-Lun Wu, Miranda Chong-Yee Yau, Yvette Wai-Man Lai, Kitty Sau-Chun Fung, Sandy Ka-Yee Chau, Barry Kin-Chung Wong, Wing-Kin To, Kristine Luk, Alex Yat-Man Ho, Tak-Lun Que, Kam-Tong Yip, Wing Cheong Yam, David Ho-Keung Shum, Shea Ping Yip |
| EPI_ISL_2443209, EPI_ISL_2443210, EPI_ISL_2443211, EPI_ISL_2443212, EPI_ISL_2443214, EPI_ISL_2443217, EPI_ISL_2443218, EPI_ISL_2443219, EPI_ISL_2443221, EPI_ISL_2443226, EPI_ISL_2443362 | | | |
| see above | Laboratoire Hôpital Sainte Musse | Laboratoire Hôpital Sainte Musse | Lionel Chollet |
| EPI_ISL_2443466, EPI_ISL_2443467, EPI_ISL_2443468, EPI_ISL_2443469, EPI_ISL_2443472, EPI_ISL_2443475, EPI_ISL_2443476, EPI_ISL_2443479, EPI_ISL_2443481, EPI_ISL_2443485, EPI_ISL_2443486, EPI_ISL_2443487, EPI_ISL_2443488, EPI_ISL_2443489, EPI_ISL_2443490, EPI_ISL_2443493, EPI_ISL_2443495, EPI_ISL_2443496, EPI_ISL_2443497, EPI_ISL_2443498, EPI_ISL_2443500, EPI_ISL_2443501, EPI_ISL_2443504, EPI_ISL_2443505, EPI_ISL_2443506, EPI_ISL_2443507, EPI_ISL_2443508, EPI_ISL_2443509, EPI_ISL_2443510, EPI_ISL_2443512, EPI_ISL_2443514, EPI_ISL_2443516, EPI_ISL_2443517, EPI_ISL_2443519 | | | |
| see above | Baylor Scott & White-Temple | Baylor Scott & White-Temple | Ari Rao, Linden Morales, Kimberly Walker, Marcus Volz, Shelby Johnson |
| EPI_ISL_2444318 | CHU LILLE | CHU Lille - Laboratoire de Virologie | AIT YAHYA Emilie, ALIDJINOU Enagnon Kazali, BOCKET Laurence, CREPIN Michel, DEMAY Christophe, ENGELMANN Ilka, GEFFROY Sandrine, GUIGON Aurélie, LAMBERT Valérie, LAZREK Mouna, NOBILLIAUX Florian, PREVOST Brigitte, THUILLIER Caroline, TINEZ Claire |
| EPI_ISL_2444319 | CH ARRAS | CHU Lille - Laboratoire de Virologie | AIT YAHYA Emilie, ALIDJINOU Enagnon Kazali, BOCKET Laurence, CREPIN Michel, DEMAY Christophe, ENGELMANN Ilka, GEFFROY Sandrine, GUIGON Aurélie, LAMBERT Valérie, LAZREK Mouna, NOBILLIAUX Florian, PREVOST Brigitte, THUILLIER Caroline, TINEZ Claire |
| EPI_ISL_2444320 | CH ROUBAIX | CHU Lille - Laboratoire de Virologie | AIT YAHYA Emilie, ALIDJINOU Enagnon Kazali, BOCKET Laurence, CREPIN Michel, DEMAY Christophe, ENGELMANN Ilka, GEFFROY Sandrine, GUIGON Aurélie, LAMBERT Valérie, LAZREK Mouna, NOBILLIAUX Florian, PREVOST Brigitte, THUILLIER Caroline, TINEZ Claire |
| EPI_ISL_2444321, EPI_ISL_2444322 | CH ARRAS | CHU Lille - Laboratoire de Virologie | AIT YAHYA Emilie, ALIDJINOU Enagnon Kazali, BOCKET Laurence, CREPIN Michel, DEMAY Christophe, ENGELMANN Ilka, GEFFROY Sandrine, GUIGON Aurélie, LAMBERT Valérie, LAZREK Mouna, NOBILLIAUX Florian, PREVOST Brigitte, THUILLIER Caroline, TINEZ Claire |
| EPI_ISL_2444323, EPI_ISL_2444324 | CH BETHUNE | CHU Lille - Laboratoire de Virologie | AIT YAHYA Emilie, ALIDJINOU Enagnon Kazali, BOCKET Laurence, CREPIN Michel, DEMAY Christophe, ENGELMANN Ilka, GEFFROY Sandrine, GUIGON Aurélie, LAMBERT Valérie, LAZREK Mouna, NOBILLIAUX Florian, PREVOST Brigitte, THUILLIER Caroline, TINEZ Claire |
| EPI_ISL_2444325, EPI_ISL_2444326 | CH LENS | CHU Lille - Laboratoire de Virologie | AIT YAHYA Emilie, ALIDJINOU Enagnon Kazali, BOCKET Laurence, CREPIN Michel, DEMAY Christophe, ENGELMANN Ilka, GEFFROY Sandrine, GUIGON Aurélie, LAMBERT Valérie, LAZREK Mouna, NOBILLIAUX Florian, PREVOST Brigitte, THUILLIER Caroline, TINEZ Claire |
| EPI_ISL_2444327, EPI_ISL_2444328, EPI_ISL_2444329 | CH CALAIS | CHU Lille - Laboratoire de Virologie | AIT YAHYA Emilie, ALIDJINOU Enagnon Kazali, BOCKET Laurence, CREPIN Michel, DEMAY Christophe, ENGELMANN Ilka, GEFFROY Sandrine, GUIGON Aurélie, LAMBERT Valérie, LAZREK Mouna, NOBILLIAUX Florian, PREVOST Brigitte, THUILLIER Caroline, TINEZ Claire |
| EPI_ISL_2444331 | CHU LILLE | CHU Lille - Laboratoire de Virologie | AIT YAHYA Emilie, ALIDJINOU Enagnon Kazali, BOCKET Laurence, CREPIN Michel, DEMAY Christophe, ENGELMANN Ilka, GEFFROY Sandrine, GUIGON Aurélie, LAMBERT Valérie, LAZREK Mouna, NOBILLIAUX Florian, PREVOST Brigitte, THUILLIER Caroline, TINEZ Claire |
| EPI_ISL_2444333 | CH CAMBRAI | CHU Lille - Laboratoire de Virologie | AIT YAHYA Emilie, ALIDJINOU Enagnon Kazali, BOCKET Laurence, CREPIN Michel, DEMAY Christophe, ENGELMANN Ilka, GEFFROY Sandrine, GUIGON Aurélie, LAMBERT Valérie, LAZREK Mouna, NOBILLIAUX Florian, PREVOST Brigitte, THUILLIER Caroline, TINEZ Claire |
| EPI_ISL_2444334, EPI_ISL_2444336, EPI_ISL_2444337, EPI_ISL_2444338 | CHU LILLE | CHU Lille - Laboratoire de Virologie | AIT YAHYA Emilie, ALIDJINOU Enagnon Kazali, BOCKET Laurence, CREPIN Michel, DEMAY Christophe, ENGELMANN Ilka, GEFFROY Sandrine, GUIGON Aurélie, LAMBERT Valérie, LAZREK Mouna, NOBILLIAUX Florian, PREVOST Brigitte, THUILLIER Caroline, TINEZ Claire |
| EPI_ISL_2444340 | CH BETHUNE | CHU Lille - Laboratoire de Virologie | AIT YAHYA Emilie, ALIDJINOU Enagnon Kazali, BOCKET Laurence, CREPIN Michel, DEMAY Christophe, ENGELMANN Ilka, GEFFROY Sandrine, GUIGON Aurélie, LAMBERT Valérie, LAZREK Mouna, NOBILLIAUX Florian, PREVOST Brigitte, THUILLIER Caroline, TINEZ Claire |
| EPI_ISL_2444341, EPI_ISL_2444342 | CH SAINT-PHILIBERT | CHU Lille - Laboratoire de Virologie | AIT YAHYA Emilie, ALIDJINOU Enagnon Kazali, BOCKET Laurence, CREPIN Michel, DEMAY Christophe, ENGELMANN Ilka, GEFFROY Sandrine, GUIGON Aurélie, LAMBERT Valérie, LAZREK Mouna, NOBILLIAUX Florian, PREVOST Brigitte, THUILLIER Caroline, TINEZ Claire |
| EPI_ISL_2444343 | CH TOURCOING | CHU Lille - Laboratoire de Virologie | AIT YAHYA Emilie, ALIDJINOU Enagnon Kazali, BOCKET Laurence, CREPIN Michel, DEMAY Christophe, ENGELMANN Ilka, GEFFROY Sandrine, GUIGON Aurélie, LAMBERT Valérie, LAZREK Mouna, NOBILLIAUX Florian, PREVOST Brigitte, THUILLIER Caroline, TINEZ Claire |
| EPI_ISL_2444344 | CH BETHUNE | CHU Lille - Laboratoire de Virologie | AIT YAHYA Emilie, ALIDJINOU Enagnon Kazali, BOCKET Laurence, CREPIN Michel, DEMAY Christophe, ENGELMANN Ilka, GEFFROY Sandrine, GUIGON Aurélie, LAMBERT Valérie, LAZREK Mouna, NOBILLIAUX Florian, PREVOST Brigitte, THUILLIER Caroline, TINEZ Claire |
| EPI_ISL_2444345 | CH DUNKERQUE | CHU Lille - Laboratoire de Virologie | AIT YAHYA Emilie, ALIDJINOU Enagnon Kazali, BOCKET Laurence, CREPIN Michel, DEMAY Christophe, ENGELMANN Ilka, GEFFROY Sandrine, |

| | | | |
|---|---|---|---|
| | | | GUIGON Aurélie, LAMBERT Valérie, LAZREK Mouna, NOBILLIAUX Florian, PREVOST Brigitte, THUILLIER Caroline, TINEZ Claire |
| EPI_ISL_2444346 | CH SECLIN | CHU Lille - Laboratoire de Virologie | AIT YAHYA Emilie, ALIDJINOU Enagnon Kazali, BOCKET Laurence, CREPIN Michel, DEMAY Christophe, ENGELMANN Ilka, GEFFROY Sandrine, GUIGON Aurélie, LAMBERT Valérie, LAZREK Mouna, NOBILLIAUX Florian, PREVOST Brigitte, THUILLIER Caroline, TINEZ Claire |
| EPI_ISL_2444347, EPI_ISL_2444348 | CH BOULOGNE-SUR-MER | CHU Lille - Laboratoire de Virologie | AIT YAHYA Emilie, ALIDJINOU Enagnon Kazali, BOCKET Laurence, CREPIN Michel, DEMAY Christophe, ENGELMANN Ilka, GEFFROY Sandrine, GUIGON Aurélie, LAMBERT Valérie, LAZREK Mouna, NOBILLIAUX Florian, PREVOST Brigitte, THUILLIER Caroline, TINEZ Claire |
| EPI_ISL_2444349 | CH SECLIN | CHU Lille - Laboratoire de Virologie | AIT YAHYA Emilie, ALIDJINOU Enagnon Kazali, BOCKET Laurence, CREPIN Michel, DEMAY Christophe, ENGELMANN Ilka, GEFFROY Sandrine, GUIGON Aurélie, LAMBERT Valérie, LAZREK Mouna, NOBILLIAUX Florian, PREVOST Brigitte, THUILLIER Caroline, TINEZ Claire |
| EPI_ISL_2444350, EPI_ISL_2444351, EPI_ISL_2444352 | CH DOUAI | CHU Lille - Laboratoire de Virologie | AIT YAHYA Emilie, ALIDJINOU Enagnon Kazali, BOCKET Laurence, CREPIN Michel, DEMAY Christophe, ENGELMANN Ilka, GEFFROY Sandrine, GUIGON Aurélie, LAMBERT Valérie, LAZREK Mouna, NOBILLIAUX Florian, PREVOST Brigitte, THUILLIER Caroline, TINEZ Claire |
| EPI_ISL_2444353, EPI_ISL_2444354 | CHU LILLE | CHU Lille - Laboratoire de Virologie | AIT YAHYA Emilie, ALIDJINOU Enagnon Kazali, BOCKET Laurence, CREPIN Michel, DEMAY Christophe, ENGELMANN Ilka, GEFFROY Sandrine, GUIGON Aurélie, LAMBERT Valérie, LAZREK Mouna, NOBILLIAUX Florian, PREVOST Brigitte, THUILLIER Caroline, TINEZ Claire |
| EPI_ISL_2444355, EPI_ISL_2444356 | CH ROUBAIX | CHU Lille - Laboratoire de Virologie | AIT YAHYA Emilie, ALIDJINOU Enagnon Kazali, BOCKET Laurence, CREPIN Michel, DEMAY Christophe, ENGELMANN Ilka, GEFFROY Sandrine, GUIGON Aurélie, LAMBERT Valérie, LAZREK Mouna, NOBILLIAUX Florian, PREVOST Brigitte, THUILLIER Caroline, TINEZ Claire |
| EPI_ISL_2444357 | CH BETHUNE | CHU Lille - Laboratoire de Virologie | AIT YAHYA Emilie, ALIDJINOU Enagnon Kazali, BOCKET Laurence, CREPIN Michel, DEMAY Christophe, ENGELMANN Ilka, GEFFROY Sandrine, GUIGON Aurélie, LAMBERT Valérie, LAZREK Mouna, NOBILLIAUX Florian, PREVOST Brigitte, THUILLIER Caroline, TINEZ Claire |
| EPI_ISL_2444358 | CH CAMBRAI | CHU Lille - Laboratoire de Virologie | AIT YAHYA Emilie, ALIDJINOU Enagnon Kazali, BOCKET Laurence, CREPIN Michel, DEMAY Christophe, ENGELMANN Ilka, GEFFROY Sandrine, GUIGON Aurélie, LAMBERT Valérie, LAZREK Mouna, NOBILLIAUX Florian, PREVOST Brigitte, THUILLIER Caroline, TINEZ Claire |
| EPI_ISL_2444359, EPI_ISL_2444360, EPI_ISL_2444361 | CHU LILLE | CHU Lille - Laboratoire de Virologie | AIT YAHYA Emilie, ALIDJINOU Enagnon Kazali, BOCKET Laurence, CREPIN Michel, DEMAY Christophe, ENGELMANN Ilka, GEFFROY Sandrine, GUIGON Aurélie, LAMBERT Valérie, LAZREK Mouna, NOBILLIAUX Florian, PREVOST Brigitte, THUILLIER Caroline, TINEZ Claire |
| EPI_ISL_2444362 | CH DUNKERQUE | CHU Lille - Laboratoire de Virologie | AIT YAHYA Emilie, ALIDJINOU Enagnon Kazali, BOCKET Laurence, CREPIN Michel, DEMAY Christophe, ENGELMANN Ilka, GEFFROY Sandrine, GUIGON Aurélie, LAMBERT Valérie, LAZREK Mouna, NOBILLIAUX Florian, PREVOST Brigitte, THUILLIER Caroline, TINEZ Claire |
| EPI_ISL_2444363 | CHU LILLE | CHU Lille - Laboratoire de Virologie | AIT YAHYA Emilie, ALIDJINOU Enagnon Kazali, BOCKET Laurence, CREPIN Michel, DEMAY Christophe, ENGELMANN Ilka, GEFFROY Sandrine, GUIGON Aurélie, LAMBERT Valérie, LAZREK Mouna, NOBILLIAUX Florian, PREVOST Brigitte, THUILLIER Caroline, TINEZ Claire |
| EPI_ISL_2444364, EPI_ISL_2444365 | CH ROUBAIX | CHU Lille - Laboratoire de Virologie | AIT YAHYA Emilie, ALIDJINOU Enagnon Kazali, BOCKET Laurence, CREPIN Michel, DEMAY Christophe, ENGELMANN Ilka, GEFFROY Sandrine, GUIGON Aurélie, LAMBERT Valérie, LAZREK Mouna, NOBILLIAUX Florian, PREVOST Brigitte, THUILLIER Caroline, TINEZ Claire |
| EPI_ISL_2444366 | CH SAINT-PHILIBERT | CHU Lille - Laboratoire de Virologie | AIT YAHYA Emilie, ALIDJINOU Enagnon Kazali, BOCKET Laurence, CREPIN Michel, DEMAY Christophe, ENGELMANN Ilka, GEFFROY Sandrine, GUIGON Aurélie, LAMBERT Valérie, LAZREK Mouna, NOBILLIAUX Florian, PREVOST Brigitte, THUILLIER Caroline, TINEZ Claire |
| EPI_ISL_2444367 | CH BOULOGNE-SUR-MER | CHU Lille - Laboratoire de Virologie | AIT YAHYA Emilie, ALIDJINOU Enagnon Kazali, BOCKET Laurence, CREPIN Michel, DEMAY Christophe, ENGELMANN Ilka, GEFFROY Sandrine, GUIGON Aurélie, LAMBERT Valérie, LAZREK Mouna, NOBILLIAUX Florian, PREVOST Brigitte, THUILLIER Caroline, TINEZ Claire |
| EPI_ISL_2444368 | CH ARMENTIERES | CHU Lille - Laboratoire de Virologie | AIT YAHYA Emilie, ALIDJINOU Enagnon Kazali, BOCKET Laurence, CREPIN Michel, DEMAY Christophe, ENGELMANN Ilka, GEFFROY Sandrine, GUIGON Aurélie, LAMBERT Valérie, LAZREK Mouna, NOBILLIAUX Florian, PREVOST Brigitte, THUILLIER Caroline, TINEZ Claire |
| EPI_ISL_2444369, EPI_ISL_2444370, EPI_ISL_2444371, EPI_ISL_2444372 | CHU LILLE | CHU Lille - Laboratoire de Virologie | AIT YAHYA Emilie, ALIDJINOU Enagnon Kazali, BOCKET Laurence, CREPIN Michel, DEMAY Christophe, ENGELMANN Ilka, GEFFROY Sandrine, GUIGON Aurélie, LAMBERT Valérie, LAZREK Mouna, NOBILLIAUX Florian, PREVOST Brigitte, THUILLIER Caroline, TINEZ Claire |
| EPI_ISL_2444373 | CH ARRAS | CHU Lille - Laboratoire de Virologie | AIT YAHYA Emilie, ALIDJINOU Enagnon Kazali, BOCKET Laurence, CREPIN Michel, DEMAY Christophe, ENGELMANN Ilka, GEFFROY Sandrine, GUIGON Aurélie, LAMBERT Valérie, LAZREK Mouna, NOBILLIAUX Florian, PREVOST Brigitte, THUILLIER Caroline, TINEZ Claire |
| EPI_ISL_2444374 | CH TOURCOING | CHU Lille - Laboratoire de Virologie | AIT YAHYA Emilie, ALIDJINOU Enagnon Kazali, BOCKET Laurence, CREPIN Michel, DEMAY Christophe, ENGELMANN Ilka, GEFFROY Sandrine, GUIGON Aurélie, LAMBERT Valérie, LAZREK Mouna, NOBILLIAUX Florian, PREVOST Brigitte, THUILLIER Caroline, TINEZ Claire |
| EPI_ISL_2444376, EPI_ISL_2444377 | CH LENS | CHU Lille - Laboratoire de Virologie | AIT YAHYA Emilie, ALIDJINOU Enagnon Kazali, BOCKET Laurence, CREPIN Michel, DEMAY Christophe, ENGELMANN Ilka, GEFFROY Sandrine, GUIGON Aurélie, LAMBERT Valérie, LAZREK Mouna, NOBILLIAUX Florian, PREVOST Brigitte, THUILLIER Caroline, TINEZ Claire |
| EPI_ISL_2444378 | CH BETHUNE | CHU Lille - Laboratoire de Virologie | AIT YAHYA Emilie, ALIDJINOU Enagnon Kazali, BOCKET Laurence, CREPIN Michel, DEMAY Christophe, ENGELMANN Ilka, GEFFROY Sandrine, GUIGON Aurélie, LAMBERT Valérie, LAZREK Mouna, NOBILLIAUX Florian, PREVOST Brigitte, THUILLIER Caroline, TINEZ Claire |
| EPI_ISL_2444379, EPI_ISL_2444380 | CH SAINT-PHILIBERT | CHU Lille - Laboratoire de Virologie | AIT YAHYA Emilie, ALIDJINOU Enagnon Kazali, BOCKET Laurence, CREPIN Michel, DEMAY Christophe, ENGELMANN Ilka, GEFFROY Sandrine, GUIGON Aurélie, LAMBERT Valérie, LAZREK Mouna, NOBILLIAUX Florian, PREVOST Brigitte, THUILLIER Caroline, TINEZ Claire |
| EPI_ISL_2444381 | CHU LILLE | CHU Lille - Laboratoire de Virologie | AIT YAHYA Emilie, ALIDJINOU Enagnon Kazali, BOCKET Laurence, CREPIN Michel, DEMAY Christophe, ENGELMANN Ilka, GEFFROY Sandrine, GUIGON Aurélie, LAMBERT Valérie, LAZREK Mouna, NOBILLIAUX Florian, PREVOST Brigitte, THUILLIER Caroline, TINEZ Claire |
| EPI_ISL_2444382 | CH ARRAS | CHU Lille - Laboratoire de Virologie | AIT YAHYA Emilie, ALIDJINOU Enagnon Kazali, BOCKET Laurence, CREPIN Michel, DEMAY Christophe, ENGELMANN Ilka, GEFFROY Sandrine, GUIGON Aurélie, LAMBERT Valérie, LAZREK Mouna, NOBILLIAUX Florian, PREVOST Brigitte, THUILLIER Caroline, TINEZ Claire |
| EPI_ISL_2444383, EPI_ISL_2444384 | CH ARMENTIERES | CHU Lille - Laboratoire de Virologie | AIT YAHYA Emilie, ALIDJINOU Enagnon Kazali, BOCKET Laurence, CREPIN Michel, DEMAY Christophe, ENGELMANN Ilka, GEFFROY Sandrine, GUIGON Aurélie, LAMBERT Valérie, LAZREK Mouna, NOBILLIAUX Florian, PREVOST Brigitte, THUILLIER Caroline, TINEZ Claire |
| EPI_ISL_2444386, EPI_ISL_2444387 | CHU LILLE | CHU Lille - Laboratoire de Virologie | AIT YAHYA Emilie, ALIDJINOU Enagnon Kazali, BOCKET Laurence, CREPIN Michel, DEMAY Christophe, ENGELMANN Ilka, GEFFROY Sandrine, GUIGON Aurélie, LAMBERT Valérie, LAZREK Mouna, NOBILLIAUX Florian, PREVOST Brigitte, THUILLIER Caroline, TINEZ Claire |
| EPI_ISL_2444388 | CH TOURCOING | CHU Lille - Laboratoire de Virologie | AIT YAHYA Emilie, ALIDJINOU Enagnon Kazali, BOCKET Laurence, CREPIN Michel, DEMAY Christophe, ENGELMANN Ilka, GEFFROY Sandrine, GUIGON Aurélie, LAMBERT Valérie, LAZREK Mouna, NOBILLIAUX Florian, PREVOST Brigitte, THUILLIER Caroline, TINEZ Claire |
| EPI_ISL_2444389 | CH ROUBAIX | CHU Lille - Laboratoire de Virologie | AIT YAHYA Emilie, ALIDJINOU Enagnon Kazali, BOCKET Laurence, CREPIN Michel, DEMAY Christophe, ENGELMANN Ilka, GEFFROY Sandrine, GUIGON Aurélie, LAMBERT Valérie, LAZREK Mouna, NOBILLIAUX Florian, PREVOST Brigitte, THUILLIER Caroline, TINEZ Claire |
| EPI_ISL_2444390 | CH SAINT-OMER | CHU Lille - Laboratoire de Virologie | AIT YAHYA Emilie, ALIDJINOU Enagnon Kazali, BOCKET Laurence, CREPIN Michel, DEMAY Christophe, ENGELMANN Ilka, GEFFROY Sandrine, GUIGON Aurélie, LAMBERT Valérie, LAZREK Mouna, NOBILLIAUX Florian, PREVOST Brigitte, THUILLIER Caroline, TINEZ Claire |
| EPI_ISL_2444391 | CH ARMENTIERES | CHU Lille - Laboratoire de Virologie | AIT YAHYA Emilie, ALIDJINOU Enagnon Kazali, BOCKET Laurence, CREPIN Michel, DEMAY Christophe, ENGELMANN Ilka, GEFFROY Sandrine, GUIGON Aurélie, LAMBERT Valérie, LAZREK Mouna, NOBILLIAUX Florian, PREVOST Brigitte, THUILLIER Caroline, TINEZ Claire |
| EPI_ISL_2444392 | CH TOURCOING | CHU Lille - Laboratoire de Virologie | AIT YAHYA Emilie, ALIDJINOU Enagnon Kazali, BOCKET Laurence, CREPIN Michel, DEMAY Christophe, ENGELMANN Ilka, GEFFROY Sandrine, GUIGON Aurélie, LAMBERT Valérie, LAZREK Mouna, NOBILLIAUX Florian, PREVOST Brigitte, THUILLIER Caroline, TINEZ Claire |
| EPI_ISL_2444393 | CH LENS | CHU Lille - Laboratoire de Virologie | AIT YAHYA Emilie, ALIDJINOU Enagnon Kazali, BOCKET Laurence, CREPIN Michel, DEMAY Christophe, ENGELMANN Ilka, GEFFROY Sandrine, GUIGON Aurélie, LAMBERT Valérie, LAZREK Mouna, NOBILLIAUX Florian, PREVOST Brigitte, THUILLIER Caroline, TINEZ Claire |
| EPI_ISL_2447989, EPI_ISL_2447990, EPI_ISL_2448109, EPI_ISL_2448110, EPI_ISL_2448160, EPI_ISL_2448297 | Institut de Virologie du CHU de Strasbourg | Institut de Virologie du CHU de Strasbourg | Fafi-Kremer Samira, Gallais Floriane, Gantner Pierre, Laugel Elodie, Solis Morgane, Velay Aurélie, Wendling Marie-Josée |
| EPI_ISL_2448394, EPI_ISL_2448395, EPI_ISL_2448396, EPI_ISL_2448397, EPI_ISL_2448398 | Klinisch Laboratorium ZNA | Klinisch Laboratorium ZNA | Verstrepen et al. |
| EPI_ISL_2448623 | CH Angoulême | CHU Poitiers | Caroline MICHAUD, Maxime PICHON, Manon PRAT, Valentin BON-BARET, Birama N'DIAYE, Magali GARCIA, Agnes BEBY-DEFAUX, Nicolas LEVEQUE |

| | | | |
|---|---|---|---|
| EPI_ISL_2448625 | Laboffice | CHU Poitiers | Caroline MICHAUD, Maxime PICHON, Manon PRAT, Valentin BON-BARET, Birama N'DIAYE, Magali GARCIA, Agnes BEBY-DEFAUX, Nicolas LEVEQUE |
| EPI_ISL_2448627 | CMSI | CHU Poitiers | Caroline MICHAUD, Maxime PICHON, Manon PRAT, Valentin BON-BARET, Birama N'DIAYE, Magali GARCIA, Agnes BEBY-DEFAUX, Nicolas LEVEQUE |
| EPI_ISL_2448628 | Centre REB | CHU Poitiers | Caroline MICHAUD, Maxime PICHON, Manon PRAT, Valentin BON-BARET, Birama N'DIAYE, Magali GARCIA, Agnes BEBY-DEFAUX, Nicolas LEVEQUE |
| EPI_ISL_2448629 | Dépistage Drive Montmorillon | CHU Poitiers | Caroline MICHAUD, Maxime PICHON, Manon PRAT, Valentin BON-BARET, Birama N'DIAYE, Magali GARCIA, Agnes BEBY-DEFAUX, Nicolas LEVEQUE |
| EPI_ISL_2448630 | Depistage Drive Poitiers | CHU Poitiers | Caroline MICHAUD, Maxime PICHON, Manon PRAT, Valentin BON-BARET, Birama N'DIAYE, Magali GARCIA, Agnes BEBY-DEFAUX, Nicolas LEVEQUE |
| EPI_ISL_2448631 | CHU Poitiers | CHU Poitiers | Caroline MICHAUD, Maxime PICHON, Manon PRAT, Valentin BON-BARET, Birama N'DIAYE, Magali GARCIA, Agnes BEBY-DEFAUX, Nicolas LEVEQUE |
| EPI_ISL_2448633 | Depistage Drive Poitiers | CHU Poitiers | Caroline MICHAUD, Maxime PICHON, Manon PRAT, Valentin BON-BARET, Birama N'DIAYE, Magali GARCIA, Agnes BEBY-DEFAUX, Nicolas LEVEQUE |
| EPI_ISL_2448634 | CH Niort | CHU Poitiers | Caroline MICHAUD, Maxime PICHON, Manon PRAT, Valentin BON-BARET, Birama N'DIAYE, Magali GARCIA, Agnes BEBY-DEFAUX, Nicolas LEVEQUE |
| EPI_ISL_2448635 | BIO86 | CHU Poitiers | Caroline MICHAUD, Maxime PICHON, Manon PRAT, Valentin BON-BARET, Birama N'DIAYE, Magali GARCIA, Agnes BEBY-DEFAUX, Nicolas LEVEQUE |
| EPI_ISL_2448636 | Laboffice | CHU Poitiers | Caroline MICHAUD, Maxime PICHON, Manon PRAT, Valentin BON-BARET, Birama N'DIAYE, Magali GARCIA, Agnes BEBY-DEFAUX, Nicolas LEVEQUE |
| EPI_ISL_2448637 | CH Niort | CHU Poitiers | Caroline MICHAUD, Maxime PICHON, Manon PRAT, Valentin BON-BARET, Birama N'DIAYE, Magali GARCIA, Agnes BEBY-DEFAUX, Nicolas LEVEQUE |
| EPI_ISL_2448638 | CHNDS | CHU Poitiers | Caroline MICHAUD, Maxime PICHON, Manon PRAT, Valentin BON-BARET, Birama N'DIAYE, Magali GARCIA, Agnes BEBY-DEFAUX, Nicolas LEVEQUE |
| EPI_ISL_2448639 | BIO86 | CHU Poitiers | Caroline MICHAUD, Maxime PICHON, Manon PRAT, Valentin BON-BARET, Birama N'DIAYE, Magali GARCIA, Agnes BEBY-DEFAUX, Nicolas LEVEQUE |
| EPI_ISL_2448640 | Depistage Drive Montmorillon | CHU Poitiers | Caroline MICHAUD, Maxime PICHON, Manon PRAT, Valentin BON-BARET, Birama N'DIAYE, Magali GARCIA, Agnes BEBY-DEFAUX, Nicolas LEVEQUE |
| EPI_ISL_2448642 | BIO86 | CHU Poitiers | Caroline MICHAUD, Maxime PICHON, Manon PRAT, Valentin BON-BARET, Birama N'DIAYE, Magali GARCIA, Agnes BEBY-DEFAUX, Nicolas LEVEQUE |
| EPI_ISL_2448643 | Medilab | CHU Poitiers | Caroline MICHAUD, Maxime PICHON, Manon PRAT, Valentin BON-BARET, Birama N'DIAYE, Magali GARCIA, Agnes BEBY-DEFAUX, Nicolas LEVEQUE |
| EPI_ISL_2448741 | Public Health Authority of the Slovak Republic | Laboratory of Genomics and Bioinformatics, Comenius University Science Park | Tatiana Sedláková, Diana Rusáková, Miroslav Böhmer, Anna Giová, Jaroslav Budiš, Tomáš Szemes, Jakub Styk |
| EPI_ISL_2455329, EPI_ISL_2455330 | Centers for Disease Control, R.O.C. (Taiwan) | Centers for Disease Control, R.O.C. (Taiwan) | Ji-Rong Yang, Yu-Chi-Lin, Jung-Jung Mu, Ming-Tsan-Liu |
| EPI_ISL_2455909, EPI_ISL_2455910, EPI_ISL_2455911, EPI_ISL_2455912 | LESP Queretaro | Instituto de Diagnostico y Referencia Epidemiologicos (INDRE) | Claudia Wong-Arambula, Abril Rodriguez-Maldonado, Vanessa Rivero-Arredondo, Ariadna Medina-Benitez, Joaquin Quiroz-Mercado, Sergio Rangel-Guerrero, Natividad Cruz-Ortiz, Tatiana Nunez-Garcia, Gisela Barrera-Badillo, Lucia Hernandez-Rivas, Irma Lopez-Martinez, Ernesto Ramirez-Gonzalez. |
| EPI_ISL_2455917, EPI_ISL_2455919, EPI_ISL_2455920 | LESP Chihuahua | Instituto de Diagnostico y Referencia Epidemiologicos (INDRE) | Claudia Wong-Arambula, Abril Rodriguez-Maldonado, Vanessa Rivero-Arredondo, Ariadna Medina-Benitez, Joaquin Quiroz-Mercado, Sergio Rangel-Guerrero, Natividad Cruz-Ortiz, Tatiana Nunez-Garcia, Gisela Barrera-Badillo, Lucia Hernandez-Rivas, Irma Lopez-Martinez, Ernesto Ramirez-Gonzalez. |
| EPI_ISL_2455922 | LESP Oaxaca | Instituto de Diagnostico y Referencia Epidemiologicos (INDRE) | Claudia Wong-Arambula, Abril Rodriguez-Maldonado, Vanessa Rivero-Arredondo, Ariadna Medina-Benitez, Joaquin Quiroz-Mercado, Sergio Rangel-Guerrero, Natividad Cruz-Ortiz, Tatiana Nunez-Garcia, Gisela Barrera-Badillo, Lucia Hernandez-Rivas, Irma Lopez-Martinez, Ernesto Ramirez-Gonzalez. |
| EPI_ISL_2455923, EPI_ISL_2455924, EPI_ISL_2455925, EPI_ISL_2455930, EPI_ISL_2455931, EPI_ISL_2455932, EPI_ISL_2455933, EPI_ISL_2455935, EPI_ISL_2455937, EPI_ISL_2455939, EPI_ISL_2455940, EPI_ISL_2455941, EPI_ISL_2455943, EPI_ISL_2455944, EPI_ISL_2455946, EPI_ISL_2455947, EPI_ISL_2455949, EPI_ISL_2455950, EPI_ISL_2455951, EPI_ISL_2455952 | | | |
| see above | LESP Tamaulipas | Instituto de Diagnostico y Referencia Epidemiologicos (INDRE) | Claudia Wong-Arambula, Abril Rodriguez-Maldonado, Vanessa Rivero-Arredondo, Ariadna Medina-Benitez, Joaquin Quiroz-Mercado, Sergio Rangel-Guerrero, Natividad Cruz-Ortiz, Tatiana Nunez-Garcia, Gisela Barrera-Badillo, Lucia Hernandez-Rivas, Irma Lopez-Martinez, Ernesto Ramirez-Gonzalez. |
| EPI_ISL_2455953 | LESP San Luis Potosi | Instituto de Diagnostico y Referencia Epidemiologicos (INDRE) | Claudia Wong-Arambula, Abril Rodriguez-Maldonado, Vanessa Rivero-Arredondo, Ariadna Medina-Benitez, Joaquin Quiroz-Mercado, Sergio Rangel-Guerrero, Natividad Cruz-Ortiz, Tatiana Nunez-Garcia, Gisela Barrera-Badillo, Lucia Hernandez-Rivas, Irma Lopez-Martinez, Ernesto Ramirez-Gonzalez. |
| EPI_ISL_2455954, EPI_ISL_2455956 | LESP Tamaulipas | Instituto de Diagnostico y Referencia Epidemiologicos (INDRE) | Claudia Wong-Arambula, Abril Rodriguez-Maldonado, Vanessa Rivero-Arredondo, Ariadna Medina-Benitez, Joaquin Quiroz-Mercado, Sergio Rangel-Guerrero, Natividad Cruz-Ortiz, Tatiana Nunez-Garcia, Gisela Barrera-Badillo, Lucia Hernandez-Rivas, Irma Lopez-Martinez, Ernesto Ramirez-Gonzalez. |
| EPI_ISL_2455959 | LESP Ciudad de Mexico | Instituto de Diagnostico y Referencia Epidemiologicos (INDRE) | Claudia Wong-Arambula, Abril Rodriguez-Maldonado, Vanessa Rivero-Arredondo, Ariadna Medina-Benitez, Joaquin Quiroz-Mercado, Sergio Rangel-Guerrero, Natividad Cruz-Ortiz, Tatiana Nunez-Garcia, Gisela Barrera-Badillo, Lucia Hernandez-Rivas, Irma Lopez-Martinez, Ernesto Ramirez-Gonzalez. |
| EPI_ISL_2455960, EPI_ISL_2455961, EPI_ISL_2455962 | LESP Queretaro | Instituto de Diagnostico y Referencia Epidemiologicos (INDRE) | Claudia Wong-Arambula, Abril Rodriguez-Maldonado, Vanessa Rivero-Arredondo, Ariadna Medina-Benitez, Joaquin Quiroz-Mercado, Sergio Rangel-Guerrero, Natividad Cruz-Ortiz, Tatiana Nunez-Garcia, Gisela Barrera-Badillo, Lucia Hernandez-Rivas, Irma Lopez-Martinez, Ernesto Ramirez-Gonzalez. |
| EPI_ISL_2455980 | LESP Colima | Instituto de Diagnostico y Referencia Epidemiologicos (INDRE) | Claudia Wong-Arambula, Abril Rodriguez-Maldonado, Vanessa Rivero-Arredondo, Ariadna Medina-Benitez, Joaquin Quiroz-Mercado, Sergio Rangel-Guerrero, Natividad Cruz-Ortiz, Tatiana Nunez-Garcia, Gisela Barrera-Badillo, Lucia Hernandez-Rivas, Irma Lopez-Martinez, Ernesto Ramirez-Gonzalez. |
| EPI_ISL_2455981 | LESP Queretaro | Instituto de Diagnostico y Referencia Epidemiologicos (INDRE) | Claudia Wong-Arambula, Abril Rodriguez-Maldonado, Vanessa Rivero-Arredondo, Ariadna Medina-Benitez, Joaquin Quiroz-Mercado, Sergio Rangel-Guerrero, Natividad Cruz-Ortiz, Tatiana Nunez-Garcia, Gisela Barrera-Badillo, Lucia Hernandez-Rivas, Irma Lopez-Martinez, Ernesto Ramirez-Gonzalez. |
| EPI_ISL_2455982, EPI_ISL_2455983, EPI_ISL_2455984, EPI_ISL_2455985 | LESP Nuevo Leon | Instituto de Diagnostico y Referencia Epidemiologicos (INDRE) | Claudia Wong-Arambula, Abril Rodriguez-Maldonado, Vanessa Rivero-Arredondo, Ariadna Medina-Benitez, Joaquin Quiroz-Mercado, Sergio Rangel-Guerrero, Natividad Cruz-Ortiz, Tatiana Nunez-Garcia, Gisela Barrera-Badillo, Lucia Hernandez-Rivas, Irma Lopez-Martinez, Ernesto Ramirez-Gonzalez. |
| EPI_ISL_2455986, EPI_ISL_2455987, EPI_ISL_2455988 | LESP Guanajuato | Instituto de Diagnostico y Referencia Epidemiologicos (INDRE) | Claudia Wong-Arambula, Abril Rodriguez-Maldonado, Vanessa Rivero-Arredondo, Ariadna Medina-Benitez, Joaquin Quiroz-Mercado, Sergio Rangel-Guerrero, Natividad Cruz-Ortiz, Tatiana Nunez-Garcia, Gisela Barrera-Badillo, Lucia Hernandez-Rivas, Irma Lopez-Martinez, Ernesto |

| | | | |
|---|---|---|---|
| | | | Ramirez-Gonzalez. |
| EPI_ISL_2455992 | LESP Estado de Mexico | Instituto de Diagnostico y Referencia Epidemiologicos (INDRE) | Claudia Wong-Arambula, Abril Rodriguez-Maldonado, Vanessa Rivero-Arredondo, Ariadna Medina-Benitez, Joaquin Quiroz-Mercado, Sergio Rangel-Guerrero, Natividad Cruz-Ortiz, Tatiana Nunez-Garcia, Gisela Barrera-Badillo, Lucia Hernandez-Rivas, Irma Lopez-Martinez, Ernesto Ramirez-Gonzalez. |
| EPI_ISL_2462052, EPI_ISL_2462054, EPI_ISL_2462057, EPI_ISL_2462058 | Genome Analysis Center, Yamanashi Central Hospital | Genome Analysis Center, Yamanashi Central Hospital | Yosuke Hirotsu |
| EPI_ISL_2462868, EPI_ISL_2462869, EPI_ISL_2462870, EPI_ISL_2462871 | AZ Sint-Lucas Gent | AZ Sint-Lucas Gent | Jos Van Acker, Charlotte Verfaillie, Elke Vanlaere, Klara De Rauw |
| EPI_ISL_2462886 | Hospital | National Reference Center for Viruses of Respiratory Infections, Institut Pasteur, Paris | Marion Barbet, Sylvie Behillil, Méline Bizard, Angela Brisebarre, Camille Capel, Vincent Enouf, Louise Lefrançois, Frédéric Lemoine, Christophe Malabat, Corinne Maufrais, Etienne Simon-Lorière, Maud Vanpeene, Sylvie Van der Werf ,Olivia Raulin |
| EPI_ISL_2462887 | Hospital | National Reference Center for Viruses of Respiratory Infections, Institut Pasteur, Paris | Marion Barbet, Sylvie Behillil, Méline Bizard, Angela Brisebarre, Camille Capel, Vincent Enouf, Louise Lefrançois, Frédéric Lemoine, Christophe Malabat, Corinne Maufrais, Etienne Simon-Lorière, Maud Vanpeene, Sylvie Van der Werf ,Thomas |
| EPI_ISL_2462888 | Labo Analyses med | National Reference Center for Viruses of Respiratory Infections, Institut Pasteur, Paris | Marion Barbet, Sylvie Behillil, Méline Bizard, Angela Brisebarre, Camille Capel, Vincent Enouf, Louise Lefrançois, Frédéric Lemoine, Christophe Malabat, Corinne Maufrais, Etienne Simon-Lorière, Maud Vanpeene, Sylvie Van der Werf ,Jean-FrançOis Comes |
| EPI_ISL_2462889 | Labo Analyses med | National Reference Center for Viruses of Respiratory Infections, Institut Pasteur, Paris | Marion Barbet, Sylvie Behillil, Méline Bizard, Angela Brisebarre, Camille Capel, Vincent Enouf, Louise Lefrançois, Frédéric Lemoine, Christophe Malabat, Corinne Maufrais, Amaury Vaysse, Etienne Simon-Lorière, Maud Vanpeene, Sylvie Van der Werf ,Jean-FrançOis Comes |
| EPI_ISL_2462890, EPI_ISL_2462891, EPI_ISL_2462892, EPI_ISL_2462893 | Labo Analyses med | National Reference Center for Viruses of Respiratory Infections, Institut Pasteur, Paris | Marion Barbet, Sylvie Behillil, Méline Bizard, Angela Brisebarre, Camille Capel, Vincent Enouf, Louise Lefrançois, Frédéric Lemoine, Christophe Malabat, Corinne Maufrais, Etienne Simon-Lorière, Maud Vanpeene, Sylvie Van der Werf ,Jean-FrançOis Comes |
| EPI_ISL_2462894 | Hospital | National Reference Center for Viruses of Respiratory Infections, Institut Pasteur, Paris | Marion Barbet, Sylvie Behillil, Méline Bizard, Angela Brisebarre, Camille Capel, Vincent Enouf, Louise Lefrançois, Frédéric Lemoine, Christophe Malabat, Corinne Maufrais, Etienne Simon-Lorière, Maud Vanpeene, Sylvie Van der Werf ,H. Seraphin |
| EPI_ISL_2462896, EPI_ISL_2462897, EPI_ISL_2462898, EPI_ISL_2462899, EPI_ISL_2462900, EPI_ISL_2462901 | Hospital | National Reference Center for Viruses of Respiratory Infections, Institut Pasteur, Paris | Marion Barbet, Sylvie Behillil, Méline Bizard, Angela Brisebarre, Camille Capel, Vincent Enouf, Louise Lefrançois, Frédéric Lemoine, Christophe Malabat, Corinne Maufrais, Etienne Simon-Lorière, Maud Vanpeene, Sylvie Van der Werf ,Olivia Raulin |
| EPI_ISL_2462902 | Hospital | National Reference Center for Viruses of Respiratory Infections, Institut Pasteur, Paris | Marion Barbet, Sylvie Behillil, Méline Bizard, Angela Brisebarre, Camille Capel, Vincent Enouf, Louise Lefrançois, Frédéric Lemoine, Christophe Malabat, Corinne Maufrais, Etienne Simon-Lorière, Maud Vanpeene, Sylvie Van der Werf ,Esther Gyde |
| EPI_ISL_2462904, EPI_ISL_2462905, EPI_ISL_2462906 | Hospital | National Reference Center for Viruses of Respiratory Infections, Institut Pasteur, Paris | Marion Barbet, Sylvie Behillil, Méline Bizard, Angela Brisebarre, Camille Capel, Vincent Enouf, Louise Lefrançois, Frédéric Lemoine, Christophe Malabat, Corinne Maufrais, Etienne Simon-Lorière, Maud Vanpeene, Sylvie Van der Werf ,Antoine Talarmin |
| EPI_ISL_2462908 | Labo Analyses Med | National Reference Center for Viruses of Respiratory Infections, Institut Pasteur, Paris | Marion Barbet, Sylvie Behillil, Méline Bizard, Angela Brisebarre, Camille Capel, Vincent Enouf, Louise Lefrançois, Frédéric Lemoine, Christophe Malabat, Corinne Maufrais, Etienne Simon-Lorière, Maud Vanpeene, Sylvie Van der Werf ,AuréLie Dupuis |
| EPI_ISL_2462909, EPI_ISL_2462910, EPI_ISL_2462911, EPI_ISL_2462912, EPI_ISL_2462913, EPI_ISL_2462914 | Labo Analyses med | National Reference Center for Viruses of Respiratory Infections, Institut Pasteur, Paris | Marion Barbet, Sylvie Behillil, Méline Bizard, Angela Brisebarre, Camille Capel, Vincent Enouf, Louise Lefrançois, Frédéric Lemoine, Christophe Malabat, Corinne Maufrais, Etienne Simon-Lorière, Maud Vanpeene, Sylvie Van der Werf ,Jean-FrançOis Comes |
| EPI_ISL_2462915, EPI_ISL_2462916 | Hospital | National Reference Center for Viruses of Respiratory Infections, Institut Pasteur, Paris | Marion Barbet, Sylvie Behillil, Méline Bizard, Angela Brisebarre, Camille Capel, Vincent Enouf, Louise Lefrançois, Frédéric Lemoine, Christophe Malabat, Corinne Maufrais, Etienne Simon-Lorière, Maud Vanpeene, Sylvie Van der Werf ,H. Seraphin |
| EPI_ISL_2462917, EPI_ISL_2462918 | Labo Analyses Med | National Reference Center for Viruses of Respiratory Infections, Institut Pasteur, Paris | Marion Barbet, Sylvie Behillil, Méline Bizard, Angela Brisebarre, Camille Capel, Vincent Enouf, Louise Lefrançois, Frédéric Lemoine, Christophe Malabat, Corinne Maufrais, Etienne Simon-Lorière, Maud Vanpeene, Sylvie Van der Werf ,Nouredine Sadeg |
| EPI_ISL_2462919, EPI_ISL_2462920 | Hospital | National Reference Center for Viruses of Respiratory Infections, Institut Pasteur, Paris | Marion Barbet, Sylvie Behillil, Méline Bizard, Angela Brisebarre, Camille Capel, Vincent Enouf, Louise Lefrançois, Frédéric Lemoine, Christophe Malabat, Corinne Maufrais, Etienne Simon-Lorière, Maud Vanpeene, Sylvie Van der Werf ,Olivia Raulin |
| EPI_ISL_2462921 | Labo Analyses Med | National Reference Center for Viruses of Respiratory Infections, Institut Pasteur, Paris | Marion Barbet, Sylvie Behillil, Méline Bizard, Angela Brisebarre, Camille Capel, Vincent Enouf, Louise Lefrançois, Frédéric Lemoine, Christophe Malabat, Corinne Maufrais, Etienne Simon-Lorière, Maud Vanpeene, Sylvie Van der Werf ,Vicky Le |
| EPI_ISL_2462922, EPI_ISL_2462923 | Hospital | National Reference Center for Viruses of Respiratory Infections, Institut Pasteur, Paris | Marion Barbet, Sylvie Behillil, Méline Bizard, Angela Brisebarre, Camille Capel, Vincent Enouf, Louise Lefrançois, Frédéric Lemoine, Christophe Malabat, Corinne Maufrais, Etienne Simon-Lorière, Maud Vanpeene, Sylvie Van der Werf ,Nabil Gastli |
| EPI_ISL_2462925 | Hospital | National Reference Center for Viruses of Respiratory Infections, Institut Pasteur, Paris | Marion Barbet, Sylvie Behillil, Méline Bizard, Angela Brisebarre, Camille Capel, Vincent Enouf, Louise Lefrançois, Frédéric Lemoine, Christophe Malabat, Corinne Maufrais, Etienne Simon-Lorière, Maud Vanpeene, Sylvie Van der Werf ,Marianne Burgard |
| EPI_ISL_2462926 | Labo Analyses Med | National Reference Center for Viruses of Respiratory Infections, Institut Pasteur, Paris | Marion Barbet, Sylvie Behillil, Méline Bizard, Angela Brisebarre, Camille Capel, Vincent Enouf, Louise Lefrançois, Frédéric Lemoine, Christophe Malabat, Corinne Maufrais, Etienne Simon-Lorière, Maud Vanpeene, Sylvie Van der Werf ,GréGoire Potiron |
| EPI_ISL_2462927 | Hospital | National Reference Center for Viruses of Respiratory Infections, Institut Pasteur, Paris | Marion Barbet, Sylvie Behillil, Méline Bizard, Angela Brisebarre, Camille Capel, Vincent Enouf, Louise Lefrançois, Frédéric Lemoine, Christophe Malabat, Corinne Maufrais, Etienne Simon-Lorière, Maud Vanpeene, Sylvie Van der Werf ,Didier Mattera |
| EPI_ISL_2462928, EPI_ISL_2462929, EPI_ISL_2462930, EPI_ISL_2462931 | Hospital | National Reference Center for Viruses of Respiratory Infections, Institut Pasteur, Paris | Marion Barbet, Sylvie Behillil, Méline Bizard, Angela Brisebarre, Camille Capel, Vincent Enouf, Louise Lefrançois, Frédéric Lemoine, Christophe Malabat, Corinne Maufrais, Etienne Simon-Lorière, Maud Vanpeene, Sylvie Van der Werf ,Antoine Talarmin |
| EPI_ISL_2462932, EPI_ISL_2462933, EPI_ISL_2462934, EPI_ISL_2462935 | Hospital | National Reference Center for Viruses of Respiratory Infections, Institut Pasteur, Paris | Marion Barbet, Sylvie Behillil, Méline Bizard, Angela Brisebarre, Camille Capel, Vincent Enouf, Louise Lefrançois, Frédéric Lemoine, Christophe Malabat, Corinne Maufrais, Etienne Simon-Lorière, Maud Vanpeene, Sylvie Van der Werf ,Sylvaine Bastian |
| EPI_ISL_2462936, EPI_ISL_2462937, EPI_ISL_2462938 | Hospital | National Reference Center for Viruses of Respiratory Infections, Institut Pasteur, Paris | Marion Barbet, Sylvie Behillil, Méline Bizard, Angela Brisebarre, Camille Capel, Vincent Enouf, Louise Lefrançois, Frédéric Lemoine, Christophe Malabat, Corinne Maufrais, Etienne Simon-Lorière, Maud Vanpeene, Sylvie Van der Werf ,Laurence Fagour |
| EPI_ISL_2462939 | Labo Analyses Med | National Reference Center for Viruses of Respiratory Infections, Institut Pasteur, Paris | Marion Barbet, Sylvie Behillil, Méline Bizard, Angela Brisebarre, Camille Capel, Vincent Enouf, Louise Lefrançois, Frédéric Lemoine, Christophe Malabat, Corinne Maufrais, Etienne Simon-Lorière, Maud Vanpeene, Sylvie Van der Werf |
| EPI_ISL_2462940 | Hospital | National Reference Center for Viruses of Respiratory Infections, Institut Pasteur, Paris | Marion Barbet, Sylvie Behillil, Méline Bizard, Angela Brisebarre, Camille Capel, Vincent Enouf, Louise Lefrançois, Frédéric Lemoine, Christophe Malabat, Corinne Maufrais, Etienne Simon-Lorière, Maud Vanpeene, Sylvie Van der Werf ,H. Seraphin |
| EPI_ISL_2462941 | Hospital | National Reference Center for Viruses of Respiratory Infections, Institut Pasteur, Paris | Marion Barbet, Sylvie Behillil, Méline Bizard, Angela Brisebarre, Camille Capel, Vincent Enouf, Louise Lefrançois, Frédéric Lemoine, Christophe Malabat, Corinne Maufrais, Amaury Vaysse, Etienne Simon-Lorière, Maud Vanpeene, Sylvie Van der Werf ,Axelle Paquin |
| EPI_ISL_2462949 | Labo Analyses Med | National Reference Center for Viruses of Respiratory Infections, Institut Pasteur, Paris | Marion Barbet, Sylvie Behillil, Méline Bizard, Angela Brisebarre, Camille Capel, Vincent Enouf, Louise Lefrançois, Frédéric Lemoine, Christophe Malabat, Corinne Maufrais, Etienne Simon-Lorière, Maud Vanpeene, Sylvie Van der Werf ,Jean-FrançOis Comes |
| EPI_ISL_2462950, EPI_ISL_2462951 | Labo Analyses med | National Reference Center for Viruses of Respiratory Infections, Institut Pasteur, Paris | Marion Barbet, Sylvie Behillil, Méline Bizard, Angela Brisebarre, Camille Capel, Vincent Enouf, Louise Lefrançois, Frédéric Lemoine, Christophe Malabat, Corinne Maufrais, Etienne Simon-Lorière, Maud Vanpeene, Sylvie Van der Werf ,Jean-FrançOis Comes |
| EPI_ISL_2462952 | Labo Analyses Med | National Reference Center for Viruses of Respiratory Infections, Institut Pasteur, Paris | Marion Barbet, Sylvie Behillil, Méline Bizard, Angela Brisebarre, Camille Capel, Vincent Enouf, Louise Lefrançois, Frédéric Lemoine, Christophe Malabat, Corinne Maufrais, Etienne Simon-Lorière, Maud Vanpeene, Sylvie Van der Werf ,Patricia Stoessel-Thouvenin |
| EPI_ISL_2462953, EPI_ISL_2462954 | Hospital | National Reference Center for Viruses of Respiratory Infections, Institut Pasteur, Paris | Marion Barbet, Sylvie Behillil, Méline Bizard, Angela Brisebarre, Camille Capel, Vincent Enouf, Louise Lefrançois, Frédéric Lemoine, Christophe Malabat, Corinne Maufrais, Etienne Simon-Lorière, Maud Vanpeene, Sylvie Van der Werf ,Pascale Martres |
| EPI_ISL_2462955 | Labo Analyses Med | National Reference Center for Viruses of Respiratory Infections, Institut Pasteur, Paris | Marion Barbet, Sylvie Behillil, Méline Bizard, Angela Brisebarre, Camille Capel, Vincent Enouf, Louise Lefrançois, Frédéric Lemoine, Christophe Malabat, Corinne Maufrais, Etienne Simon-Lorière, Maud Vanpeene, Sylvie Van der Werf ,Vincent Vieillefond |
| EPI_ISL_2462956, EPI_ISL_2462957 | Hospital | National Reference Center for Viruses of Respiratory | Marion Barbet, Sylvie Behillil, Méline Bizard, Angela Brisebarre, Camille Capel, Vincent Enouf, Louise Lefrançois, Frédéric Lemoine, Christophe Malabat, |

| | | | |
|---|---|---|---|
| | | Infections, Institut Pasteur, Paris | Corinne Maufrais, Etienne Simon-Lorière, Maud Vanpeene, Sylvie Van der Werf ,Nabil Gastli |
| EPI_ISL_2462958 | Hospital | National Reference Center for Viruses of Respiratory Infections, Institut Pasteur, Paris | Marion Barbet, Sylvie Behillil, Méline Bizard, Angela Brisebarre, Camille Capel, Vincent Enouf, Louise Lefrançois, Frédéric Lemoine, Christophe Malabat, Corinne Maufrais, Etienne Simon-Lorière, Maud Vanpeene, Sylvie Van der Werf ,AgnèS Scanvic |
| EPI_ISL_2462959 | Hospital | National Reference Center for Viruses of Respiratory Infections, Institut Pasteur, Paris | Marion Barbet, Sylvie Behillil, Méline Bizard, Angela Brisebarre, Camille Capel, Vincent Enouf, Louise Lefrançois, Frédéric Lemoine, Christophe Malabat, Corinne Maufrais, Etienne Simon-Lorière, Maud Vanpeene, Sylvie Van der Werf ,Franck Griscelli |
| EPI_ISL_2462960, EPI_ISL_2462962 | Hospital | National Reference Center for Viruses of Respiratory Infections, Institut Pasteur, Paris | Marion Barbet, Sylvie Behillil, Méline Bizard, Angela Brisebarre, Camille Capel, Vincent Enouf, Louise Lefrançois, Frédéric Lemoine, Christophe Malabat, Corinne Maufrais, Etienne Simon-Lorière, Maud Vanpeene, Sylvie Van der Werf ,Axelle Paquin |
| EPI_ISL_2462963 | Hospital | National Reference Center for Viruses of Respiratory Infections, Institut Pasteur, Paris | Marion Barbet, Sylvie Behillil, Méline Bizard, Angela Brisebarre, Camille Capel, Vincent Enouf, Louise Lefrançois, Frédéric Lemoine, Christophe Malabat, Corinne Maufrais, Etienne Simon-Lorière, Maud Vanpeene, Sylvie Van der Werf ,CéLine Ramanantsoa |
| EPI_ISL_2462964 | Labo Analyses Med | National Reference Center for Viruses of Respiratory Infections, Institut Pasteur, Paris | Marion Barbet, Sylvie Behillil, Méline Bizard, Angela Brisebarre, Camille Capel, Vincent Enouf, Louise Lefrançois, Frédéric Lemoine, Christophe Malabat, Corinne Maufrais, Etienne Simon-Lorière, Maud Vanpeene, Sylvie Van der Werf ,Nouredine Sadeg |
| EPI_ISL_2462965 | Hospital | National Reference Center for Viruses of Respiratory Infections, Institut Pasteur, Paris | Marion Barbet, Sylvie Behillil, Méline Bizard, Angela Brisebarre, Camille Capel, Vincent Enouf, Louise Lefrançois, Frédéric Lemoine, Christophe Malabat, Corinne Maufrais, Amaury Vaysse, Etienne Simon-Lorière, Maud Vanpeene, Sylvie Van der Werf ,Nabil Gastli |
| EPI_ISL_2462966, EPI_ISL_2462967 | Labo Analyses Med | National Reference Center for Viruses of Respiratory Infections, Institut Pasteur, Paris | Marion Barbet, Sylvie Behillil, Méline Bizard, Angela Brisebarre, Camille Capel, Vincent Enouf, Louise Lefrançois, Frédéric Lemoine, Christophe Malabat, Corinne Maufrais, Etienne Simon-Lorière, Maud Vanpeene, Sylvie Van der Werf ,Franck Ennouchi |
| EPI_ISL_2462968 | Hospital | National Reference Center for Viruses of Respiratory Infections, Institut Pasteur, Paris | Marion Barbet, Sylvie Behillil, Méline Bizard, Angela Brisebarre, Camille Capel, Vincent Enouf, Louise Lefrançois, Frédéric Lemoine, Christophe Malabat, Corinne Maufrais, Etienne Simon-Lorière, Maud Vanpeene, Sylvie Van der Werf ,Patricia Stoessel-Thouvenin |
| EPI_ISL_2462969, EPI_ISL_2462970 | Labo Analyses Med | National Reference Center for Viruses of Respiratory Infections, Institut Pasteur, Paris | Marion Barbet, Sylvie Behillil, Méline Bizard, Angela Brisebarre, Camille Capel, Vincent Enouf, Louise Lefrançois, Frédéric Lemoine, Christophe Malabat, Corinne Maufrais, Etienne Simon-Lorière, Maud Vanpeene, Sylvie Van der Werf ,Franck Ennouchi |
| EPI_ISL_2463291 | AZ Sint-Jan Brugge | AZ Sint Jan | Marijke Reynders, Patrick Descheemaeker, Laurien Hoornaert, Thomas Van Landschoot, Marie Madeleine Chabert-Consen, Jorn Hellemans |
| EPI_ISL_2463292, EPI_ISL_2463294, EPI_ISL_2463295 | AZ Zeno Knokke-Blankenberge | AZ Sint Jan | Marijke Reynders, Patrick Descheemaeker, Laurien Hoornaert, Thomas Van Landschoot, Marie Madeleine Chabert-Consen, Jorn Hellemans |
| EPI_ISL_2464025, EPI_ISL_2464026, EPI_ISL_2464027, EPI_ISL_2464028, EPI_ISL_2464029, EPI_ISL_2464030, EPI_ISL_2464031, EPI_ISL_2464032, EPI_ISL_2464033, EPI_ISL_2464034, EPI_ISL_2464035 | | | |
| see above | Hospital | National Reference Center for Viruses of Respiratory Infections, Institut Pasteur, Paris | Marion Barbet, Sylvie Behillil, Méline Bizard, Angela Brisebarre, Camille Capel, Vincent Enouf, Louise Lefrançois, Frédéric Lemoine, Christophe Malabat, Corinne Maufrais, Etienne Simon-Lorière, Maud Vanpeene, Sylvie Van der Werf ,Nabil Gastli |
| EPI_ISL_2464036 | Hospital | National Reference Center for Viruses of Respiratory Infections, Institut Pasteur, Paris | Marion Barbet, Sylvie Behillil, Méline Bizard, Angela Brisebarre, Camille Capel, Vincent Enouf, Louise Lefrançois, Frédéric Lemoine, Christophe Malabat, Corinne Maufrais, Amaury Vaysse, Etienne Simon-Lorière, Maud Vanpeene, Sylvie Van der Werf ,Nabil Gastli |
| EPI_ISL_2464038 | Hospital | National Reference Center for Viruses of Respiratory Infections, Institut Pasteur, Paris | Marion Barbet, Sylvie Behillil, Méline Bizard, Angela Brisebarre, Camille Capel, Vincent Enouf, Louise Lefrançois, Frédéric Lemoine, Christophe Malabat, Corinne Maufrais, Etienne Simon-Lorière, Maud Vanpeene, Sylvie Van der Werf ,Nabil Gastli |
| EPI_ISL_2464039 | Hospital | National Reference Center for Viruses of Respiratory Infections, Institut Pasteur, Paris | Marion Barbet, Sylvie Behillil, Méline Bizard, Angela Brisebarre, Camille Capel, Vincent Enouf, Louise Lefrançois, Frédéric Lemoine, Christophe Malabat, Corinne Maufrais, Amaury Vaysse, Etienne Simon-Lorière, Maud Vanpeene, Sylvie Van der Werf ,Nabil Gastli |
| EPI_ISL_2464040, EPI_ISL_2464041, EPI_ISL_2464042, EPI_ISL_2464044, EPI_ISL_2464045 | Hospital | National Reference Center for Viruses of Respiratory Infections, Institut Pasteur, Paris | Marion Barbet, Sylvie Behillil, Méline Bizard, Angela Brisebarre, Camille Capel, Vincent Enouf, Louise Lefrançois, Frédéric Lemoine, Christophe Malabat, Corinne Maufrais, Etienne Simon-Lorière, Maud Vanpeene, Sylvie Van der Werf ,Nabil Gastli |
| EPI_ISL_2464046 | Hospital | National Reference Center for Viruses of Respiratory Infections, Institut Pasteur, Paris | Marion Barbet, Sylvie Behillil, Méline Bizard, Angela Brisebarre, Camille Capel, Vincent Enouf, Louise Lefrançois, Frédéric Lemoine, Christophe Malabat, Corinne Maufrais, Amaury Vaysse, Etienne Simon-Lorière, Maud Vanpeene, Sylvie Van der Werf ,Nabil Gastli |
| EPI_ISL_2464047, EPI_ISL_2464048, EPI_ISL_2464050, EPI_ISL_2464052, EPI_ISL_2464053, EPI_ISL_2464055 | Hospital | National Reference Center for Viruses of Respiratory Infections, Institut Pasteur, Paris | Marion Barbet, Sylvie Behillil, Méline Bizard, Angela Brisebarre, Camille Capel, Vincent Enouf, Louise Lefrançois, Frédéric Lemoine, Christophe Malabat, Corinne Maufrais, Etienne Simon-Lorière, Maud Vanpeene, Sylvie Van der Werf ,Nabil Gastli |
| EPI_ISL_2464056 | Hospital | National Reference Center for Viruses of Respiratory Infections, Institut Pasteur, Paris | Marion Barbet, Sylvie Behillil, Méline Bizard, Angela Brisebarre, Camille Capel, Vincent Enouf, Louise Lefrançois, Frédéric Lemoine, Christophe Malabat, Corinne Maufrais, Amaury Vaysse, Etienne Simon-Lorière, Maud Vanpeene, Sylvie Van der Werf ,Nabil Gastli |
| EPI_ISL_2464061, EPI_ISL_2464062, EPI_ISL_2464063, EPI_ISL_2464064, EPI_ISL_2464065 | Hospital | National Reference Center for Viruses of Respiratory Infections, Institut Pasteur, Paris | Marion Barbet, Sylvie Behillil, Méline Bizard, Angela Brisebarre, Camille Capel, Vincent Enouf, Louise Lefrançois, Frédéric Lemoine, Christophe Malabat, Corinne Maufrais, Etienne Simon-Lorière, Maud Vanpeene, Sylvie Van der Werf ,Nabil Gastli |
| EPI_ISL_2464066 | Hospital | National Reference Center for Viruses of Respiratory Infections, Institut Pasteur, Paris | Marion Barbet, Sylvie Behillil, Méline Bizard, Angela Brisebarre, Camille Capel, Vincent Enouf, Louise Lefrançois, Frédéric Lemoine, Christophe Malabat, Corinne Maufrais, Amaury Vaysse, Etienne Simon-Lorière, Maud Vanpeene, Sylvie Van der Werf ,Nabil Gastli |
| EPI_ISL_2464067, EPI_ISL_2464068, EPI_ISL_2464069, EPI_ISL_2464071, EPI_ISL_2464072, EPI_ISL_2464073, EPI_ISL_2464074, EPI_ISL_2464075 | Hospital | National Reference Center for Viruses of Respiratory Infections, Institut Pasteur, Paris | Marion Barbet, Sylvie Behillil, Méline Bizard, Angela Brisebarre, Camille Capel, Vincent Enouf, Louise Lefrançois, Frédéric Lemoine, Christophe Malabat, Corinne Maufrais, Etienne Simon-Lorière, Maud Vanpeene, Sylvie Van der Werf ,Nabil Gastli |
| EPI_ISL_2464077, EPI_ISL_2464078 | Labo Analyses Med | National Reference Center for Viruses of Respiratory Infections, Institut Pasteur, Paris | Marion Barbet, Sylvie Behillil, Méline Bizard, Angela Brisebarre, Camille Capel, Vincent Enouf, Louise Lefrançois, Frédéric Lemoine, Christophe Malabat, Corinne Maufrais, Emmanuelle Permal, Etienne Simon-Lorière, Maud Vanpeene, Sylvie Van der Werf ,Anne Lise Bravetti |
| EPI_ISL_2464080 | Hospital | National Reference Center for Viruses of Respiratory Infections, Institut Pasteur, Paris | Marion Barbet, Sylvie Behillil, Méline Bizard, Angela Brisebarre, Camille Capel, Vincent Enouf, Louise Lefrançois, Frédéric Lemoine, Christophe Malabat, Corinne Maufrais, Emmanuelle Permal, Etienne Simon-Lorière, Maud Vanpeene, Sylvie Van der Werf ,Patricia Stoessel-Thouvenin |
| EPI_ISL_2464081, EPI_ISL_2464082 | Hospital | National Reference Center for Viruses of Respiratory Infections, Institut Pasteur, Paris | Marion Barbet, Sylvie Behillil, Méline Bizard, Angela Brisebarre, Camille Capel, Vincent Enouf, Louise Lefrançois, Frédéric Lemoine, Christophe Malabat, Corinne Maufrais, Etienne Simon-Lorière, Maud Vanpeene, Sylvie Van der Werf ,Marianne Burgard |
| EPI_ISL_2464085 | Hospital | National Reference Center for Viruses of Respiratory Infections, Institut Pasteur, Paris | Marion Barbet, Sylvie Behillil, Méline Bizard, Angela Brisebarre, Camille Capel, Vincent Enouf, Louise Lefrançois, Frédéric Lemoine, Christophe Malabat, Corinne Maufrais, Etienne Simon-Lorière, Maud Vanpeene, Sylvie Van der Werf ,JéRome Guinard |
| EPI_ISL_2464086 | Hospital | National Reference Center for Viruses of Respiratory Infections, Institut Pasteur, Paris | Marion Barbet, Sylvie Behillil, Méline Bizard, Angela Brisebarre, Camille Capel, Vincent Enouf, Louise Lefrançois, Frédéric Lemoine, Christophe Malabat, Corinne Maufrais, Amaury Vaysse, Etienne Simon-Lorière, Maud Vanpeene, Sylvie Van der Werf ,JéRome Guinard |
| EPI_ISL_2464087 | Hospital | National Reference Center for Viruses of Respiratory Infections, Institut Pasteur, Paris | Marion Barbet, Sylvie Behillil, Méline Bizard, Angela Brisebarre, Camille Capel, Vincent Enouf, Louise Lefrançois, Frédéric Lemoine, Christophe Malabat, Corinne Maufrais, Etienne Simon-Lorière, Maud Vanpeene, Sylvie Van der Werf ,JéRome Guinard |
| EPI_ISL_2464088, EPI_ISL_2464089 | Hospital | National Reference Center for Viruses of Respiratory Infections, Institut Pasteur, Paris | Marion Barbet, Sylvie Behillil, Méline Bizard, Angela Brisebarre, Camille Capel, Vincent Enouf, Louise Lefrançois, Frédéric Lemoine, Christophe Malabat, Corinne Maufrais, Etienne Simon-Lorière, Maud Vanpeene, Sylvie Van der Werf ,AgnèS Scanvic |
| EPI_ISL_2464090 | Hospital | National Reference Center for Viruses of Respiratory Infections, Institut Pasteur, Paris | Marion Barbet, Sylvie Behillil, Méline Bizard, Angela Brisebarre, Camille Capel, Vincent Enouf, Louise Lefrançois, Frédéric Lemoine, Christophe Malabat, Corinne Maufrais, Etienne Simon-Lorière, Maud Vanpeene, Sylvie Van der Werf ,Franck Griscelli |
| EPI_ISL_2464091 | Hospital | National Reference Center for Viruses of Respiratory Infections, Institut Pasteur, Paris | Marion Barbet, Sylvie Behillil, Méline Bizard, Angela Brisebarre, Camille Capel, Vincent Enouf, Louise Lefrançois, Frédéric Lemoine, Christophe Malabat, Corinne Maufrais, Etienne Simon-Lorière, Maud Vanpeene, Sylvie Van der Werf ,Isabelle Gros |
| EPI_ISL_2464092, EPI_ISL_2464093, EPI_ISL_2464094, EPI_ISL_2464095, EPI_ISL_2464096, EPI_ISL_2464097 | Hospital | National Reference Center for Viruses of Respiratory Infections, Institut Pasteur, Paris | Marion Barbet, Sylvie Behillil, Méline Bizard, Angela Brisebarre, Camille Capel, Vincent Enouf, Louise Lefrançois, Frédéric Lemoine, Christophe Malabat, Corinne Maufrais, Etienne Simon-Lorière, Maud Vanpeene, Sylvie Van der Werf ,Laura Djamdjian |

| | | | |
|---|---|---|---|
| EPI_ISL_2464098 | Armies | National Reference Center for Viruses of Respiratory Infections, Institut Pasteur, Paris | Marion Barbet, Sylvie Behillil, Méline Bizard, Angela Brisebarre, Camille Capel, Vincent Enouf, Louise Lefrançois, Frédéric Lemoine, Christophe Malabat, Corinne Maufrais, Etienne Simon-Lorière, Maud Vanpeene, Sylvie Van der Werf ,Marine Desroches |
| EPI_ISL_2464099, EPI_ISL_2464100, EPI_ISL_2464101, EPI_ISL_2464102, EPI_ISL_2464103, EPI_ISL_2464104, EPI_ISL_2464105, EPI_ISL_2464106, EPI_ISL_2464107 | Hospital | National Reference Center for Viruses of Respiratory Infections, Institut Pasteur, Paris | Marion Barbet, Sylvie Behillil, Méline Bizard, Angela Brisebarre, Camille Capel, Vincent Enouf, Louise Lefrançois, Frédéric Lemoine, Christophe Malabat, Corinne Maufrais, Etienne Simon-Lorière, Maud Vanpeene, Sylvie Van der Werf ,Ilka Engelmann |
| EPI_ISL_2464108 | Hospital | National Reference Center for Viruses of Respiratory Infections, Institut Pasteur, Paris | Marion Barbet, Sylvie Behillil, Méline Bizard, Angela Brisebarre, Camille Capel, Vincent Enouf, Louise Lefrançois, Frédéric Lemoine, Christophe Malabat, Corinne Maufrais, Etienne Simon-Lorière, Maud Vanpeene, Sylvie Van der Werf |
| EPI_ISL_2464109, EPI_ISL_2464110 | Hospital | National Reference Center for Viruses of Respiratory Infections, Institut Pasteur, Paris | Marion Barbet, Sylvie Behillil, Méline Bizard, Angela Brisebarre, Camille Capel, Vincent Enouf, Louise Lefrançois, Frédéric Lemoine, Christophe Malabat, Corinne Maufrais, Etienne Simon-Lorière, Maud Vanpeene, Sylvie Van der Werf ,Anne Vachee |
| EPI_ISL_2464111, EPI_ISL_2464112 | Hospital | National Reference Center for Viruses of Respiratory Infections, Institut Pasteur, Paris | Marion Barbet, Sylvie Behillil, Méline Bizard, Angela Brisebarre, Camille Capel, Vincent Enouf, Louise Lefrançois, Frédéric Lemoine, Christophe Malabat, Corinne Maufrais, Etienne Simon-Lorière, Maud Vanpeene, Sylvie Van der Werf ,Sylvie Ledru |
| EPI_ISL_2464113, EPI_ISL_2464114, EPI_ISL_2464115, EPI_ISL_2464117, EPI_ISL_2464118, EPI_ISL_2464119, EPI_ISL_2464122, EPI_ISL_2464123, EPI_ISL_2464124, EPI_ISL_2464125, EPI_ISL_2464127, EPI_ISL_2464130, EPI_ISL_2464131, EPI_ISL_2464132, EPI_ISL_2464134, EPI_ISL_2464135, EPI_ISL_2464136, EPI_ISL_2464137 | | | |
| see above | Labo Analyses Med | National Reference Center for Viruses of Respiratory Infections, Institut Pasteur, Paris | Marion Barbet, Sylvie Behillil, Méline Bizard, Angela Brisebarre, Camille Capel, Vincent Enouf, Louise Lefrançois, Frédéric Lemoine, Christophe Malabat, Corinne Maufrais, Etienne Simon-Lorière, Maud Vanpeene, Sylvie Van der Werf ,Philippe Miara |
| EPI_ISL_2464138, EPI_ISL_2464139 | Labo Analyses Med | National Reference Center for Viruses of Respiratory Infections, Institut Pasteur, Paris | Marion Barbet, Sylvie Behillil, Méline Bizard, Angela Brisebarre, Camille Capel, Vincent Enouf, Louise Lefrançois, Frédéric Lemoine, Christophe Malabat, Corinne Maufrais, Etienne Simon-Lorière, Maud Vanpeene, Sylvie Van der Werf ,Nicolas Capron |
| EPI_ISL_2464140 | Labo Analyses Med | National Reference Center for Viruses of Respiratory Infections, Institut Pasteur, Paris | Marion Barbet, Sylvie Behillil, Méline Bizard, Angela Brisebarre, Camille Capel, Vincent Enouf, Louise Lefrançois, Frédéric Lemoine, Christophe Malabat, Corinne Maufrais, Emmanuelle Permal, Etienne Simon-Lorière, Maud Vanpeene, Sylvie Van der Werf ,Melanie Caron |
| EPI_ISL_2464141, EPI_ISL_2464142, EPI_ISL_2464143, EPI_ISL_2464144, EPI_ISL_2464145, EPI_ISL_2464147, EPI_ISL_2464148, EPI_ISL_2464149, EPI_ISL_2464150, EPI_ISL_2464151, EPI_ISL_2464152, EPI_ISL_2464153 | | | |
| see above | Labo Analyses Med | National Reference Center for Viruses of Respiratory Infections, Institut Pasteur, Paris | Marion Barbet, Sylvie Behillil, Méline Bizard, Angela Brisebarre, Camille Capel, Vincent Enouf, Louise Lefrançois, Frédéric Lemoine, Christophe Malabat, Corinne Maufrais, Etienne Simon-Lorière, Maud Vanpeene, Sylvie Van der Werf ,Melanie Caron |
| EPI_ISL_2464154 | Labo Analyses Med | National Reference Center for Viruses of Respiratory Infections, Institut Pasteur, Paris | Marion Barbet, Sylvie Behillil, Méline Bizard, Angela Brisebarre, Camille Capel, Vincent Enouf, Louise Lefrançois, Frédéric Lemoine, Christophe Malabat, Corinne Maufrais, Emmanuelle Permal, Etienne Simon-Lorière, Maud Vanpeene, Sylvie Van der Werf ,Melanie Caron |
| EPI_ISL_2464155 | Labo Analyses Med | National Reference Center for Viruses of Respiratory Infections, Institut Pasteur, Paris | Marion Barbet, Sylvie Behillil, Méline Bizard, Angela Brisebarre, Camille Capel, Vincent Enouf, Louise Lefrançois, Frédéric Lemoine, Christophe Malabat, Corinne Maufrais, Etienne Simon-Lorière, Maud Vanpeene, Sylvie Van der Werf ,Arditi Cocco |
| EPI_ISL_2464156 | Labo Analyses Med | National Reference Center for Viruses of Respiratory Infections, Institut Pasteur, Paris | Marion Barbet, Sylvie Behillil, Méline Bizard, Angela Brisebarre, Camille Capel, Vincent Enouf, Louise Lefrançois, Frédéric Lemoine, Christophe Malabat, Corinne Maufrais, Emmanuelle Permal, Etienne Simon-Lorière, Maud Vanpeene, Sylvie Van der Werf ,Arditi Cocco |
| EPI_ISL_2464157, EPI_ISL_2464159, EPI_ISL_2464160, EPI_ISL_2464161, EPI_ISL_2464162, EPI_ISL_2464163, EPI_ISL_2464164, EPI_ISL_2464165, EPI_ISL_2464168, EPI_ISL_2464169, EPI_ISL_2464170, EPI_ISL_2464171, EPI_ISL_2464172, EPI_ISL_2464174, EPI_ISL_2464175, EPI_ISL_2464176, EPI_ISL_2464177, EPI_ISL_2464178, EPI_ISL_2464179, EPI_ISL_2464180, EPI_ISL_2464181, EPI_ISL_2464182, EPI_ISL_2464183, EPI_ISL_2464184, EPI_ISL_2464185 | | | |
| see above | Labo Analyses Med | National Reference Center for Viruses of Respiratory Infections, Institut Pasteur, Paris | Marion Barbet, Sylvie Behillil, Méline Bizard, Angela Brisebarre, Camille Capel, Vincent Enouf, Louise Lefrançois, Frédéric Lemoine, Christophe Malabat, Corinne Maufrais, Etienne Simon-Lorière, Maud Vanpeene, Sylvie Van der Werf ,Arditi Cocco |
| EPI_ISL_2464186 | Labo Analyses Med | National Reference Center for Viruses of Respiratory Infections, Institut Pasteur, Paris | Marion Barbet, Sylvie Behillil, Méline Bizard, Angela Brisebarre, Camille Capel, Vincent Enouf, Louise Lefrançois, Frédéric Lemoine, Christophe Malabat, Corinne Maufrais, Amaury Vaysse, Etienne Simon-Lorière, Maud Vanpeene, Sylvie Van der Werf ,Arditi Cocco |
| EPI_ISL_2464187, EPI_ISL_2464188 | Labo Analyses Med | National Reference Center for Viruses of Respiratory Infections, Institut Pasteur, Paris | Marion Barbet, Sylvie Behillil, Méline Bizard, Angela Brisebarre, Camille Capel, Vincent Enouf, Louise Lefrançois, Frédéric Lemoine, Christophe Malabat, Corinne Maufrais, Etienne Simon-Lorière, Maud Vanpeene, Sylvie Van der Werf ,Arditi Cocco |
| EPI_ISL_2464189, EPI_ISL_2464190, EPI_ISL_2464191 | Hospital | National Reference Center for Viruses of Respiratory Infections, Institut Pasteur, Paris | Marion Barbet, Sylvie Behillil, Méline Bizard, Angela Brisebarre, Camille Capel, Vincent Enouf, Louise Lefrançois, Frédéric Lemoine, Christophe Malabat, Corinne Maufrais, Etienne Simon-Lorière, Maud Vanpeene, Sylvie Van der Werf ,Anne Cady |
| EPI_ISL_2464192 | Labo Analyses Med | National Reference Center for Viruses of Respiratory Infections, Institut Pasteur, Paris | Marion Barbet, Sylvie Behillil, Méline Bizard, Angela Brisebarre, Camille Capel, Vincent Enouf, Louise Lefrançois, Frédéric Lemoine, Christophe Malabat, Corinne Maufrais, Amaury Vaysse, Etienne Simon-Lorière, Maud Vanpeene, Sylvie Van der Werf ,GréGoire Potiron |
| EPI_ISL_2464193, EPI_ISL_2464194, EPI_ISL_2464195, EPI_ISL_2464196, EPI_ISL_2464198, EPI_ISL_2464199, EPI_ISL_2464200 | Labo Analyses Med | National Reference Center for Viruses of Respiratory Infections, Institut Pasteur, Paris | Marion Barbet, Sylvie Behillil, Méline Bizard, Angela Brisebarre, Camille Capel, Vincent Enouf, Louise Lefrançois, Frédéric Lemoine, Christophe Malabat, Corinne Maufrais, Etienne Simon-Lorière, Maud Vanpeene, Sylvie Van der Werf ,GréGoire Potiron |
| EPI_ISL_2464201, EPI_ISL_2464202, EPI_ISL_2464203, EPI_ISL_2464204 | Labo Analyses Med | National Reference Center for Viruses of Respiratory Infections, Institut Pasteur, Paris | Marion Barbet, Sylvie Behillil, Méline Bizard, Angela Brisebarre, Camille Capel, Vincent Enouf, Louise Lefrançois, Frédéric Lemoine, Christophe Malabat, Corinne Maufrais, Etienne Simon-Lorière, Maud Vanpeene, Sylvie Van der Werf |
| EPI_ISL_2464205 | Labo Analyses Med | National Reference Center for Viruses of Respiratory Infections, Institut Pasteur, Paris | Marion Barbet, Sylvie Behillil, Méline Bizard, Angela Brisebarre, Camille Capel, Vincent Enouf, Louise Lefrançois, Frédéric Lemoine, Christophe Malabat, Corinne Maufrais, Etienne Simon-Lorière, Maud Vanpeene, Sylvie Van der Werf ,Edouard Lamandin |
| EPI_ISL_2464206 | Labo Analyses Med | National Reference Center for Viruses of Respiratory Infections, Institut Pasteur, Paris | Marion Barbet, Sylvie Behillil, Méline Bizard, Angela Brisebarre, Camille Capel, Vincent Enouf, Louise Lefrançois, Frédéric Lemoine, Christophe Malabat, Corinne Maufrais, Amaury Vaysse, Etienne Simon-Lorière, Maud Vanpeene, Sylvie Van der Werf ,Edouard Lamandin |
| EPI_ISL_2464207, EPI_ISL_2464208, EPI_ISL_2464210, EPI_ISL_2464211 | Labo Analyses Med | National Reference Center for Viruses of Respiratory Infections, Institut Pasteur, Paris | Marion Barbet, Sylvie Behillil, Méline Bizard, Angela Brisebarre, Camille Capel, Vincent Enouf, Louise Lefrançois, Frédéric Lemoine, Christophe Malabat, Corinne Maufrais, Etienne Simon-Lorière, Maud Vanpeene, Sylvie Van der Werf ,Edouard Lamandin |
| EPI_ISL_2464212, EPI_ISL_2464213, EPI_ISL_2464214, EPI_ISL_2464215, EPI_ISL_2464216, EPI_ISL_2464217, EPI_ISL_2464218, EPI_ISL_2464219, EPI_ISL_2464220, EPI_ISL_2464221, EPI_ISL_2464222, EPI_ISL_2464223, EPI_ISL_2464224, EPI_ISL_2464225, EPI_ISL_2464226, EPI_ISL_2464227 | | | |
| see above | Labo Analyses Med | National Reference Center for Viruses of Respiratory Infections, Institut Pasteur, Paris | Marion Barbet, Sylvie Behillil, Méline Bizard, Angela Brisebarre, Camille Capel, Vincent Enouf, Louise Lefrançois, Frédéric Lemoine, Christophe Malabat, Corinne Maufrais, Etienne Simon-Lorière, Maud Vanpeene, Sylvie Van der Werf ,Anne Holstein |
| EPI_ISL_2464228, EPI_ISL_2464230, EPI_ISL_2464231 | Labo Analyses Med | National Reference Center for Viruses of Respiratory Infections, Institut Pasteur, Paris | Marion Barbet, Sylvie Behillil, Méline Bizard, Angela Brisebarre, Camille Capel, Vincent Enouf, Louise Lefrançois, Frédéric Lemoine, Christophe Malabat, Corinne Maufrais, Amaury Vaysse, Etienne Simon-Lorière, Maud Vanpeene, Sylvie Van der Werf ,Anne Holstein |
| EPI_ISL_2464232, EPI_ISL_2464233 | Labo Analyses Med | National Reference Center for Viruses of Respiratory Infections, Institut Pasteur, Paris | Marion Barbet, Sylvie Behillil, Méline Bizard, Angela Brisebarre, Camille Capel, Vincent Enouf, Louise Lefrançois, Frédéric Lemoine, Christophe Malabat, Corinne Maufrais, Etienne Simon-Lorière, Maud Vanpeene, Sylvie Van der Werf ,Anne Holstein |
| EPI_ISL_2464235 | Labo Analyses Med | National Reference Center for Viruses of Respiratory Infections, Institut Pasteur, Paris | Marion Barbet, Sylvie Behillil, Méline Bizard, Angela Brisebarre, Camille Capel, Vincent Enouf, Louise Lefrançois, Frédéric Lemoine, Christophe Malabat, Corinne Maufrais, Amaury Vaysse, Etienne Simon-Lorière, Maud Vanpeene, Sylvie Van der Werf ,Anne Holstein |
| EPI_ISL_2464236, EPI_ISL_2464237, EPI_ISL_2464238 | Labo Analyses Med | National Reference Center for Viruses of Respiratory Infections, Institut Pasteur, Paris | Marion Barbet, Sylvie Behillil, Méline Bizard, Angela Brisebarre, Camille Capel, Vincent Enouf, Louise Lefrançois, Frédéric Lemoine, Christophe Malabat, Corinne Maufrais, Etienne Simon-Lorière, Maud Vanpeene, Sylvie Van der Werf ,Anne Holstein |
| EPI_ISL_2464239 | Hospital | National Reference Center for Viruses of Respiratory Infections, Institut Pasteur, Paris | Marion Barbet, Sylvie Behillil, Méline Bizard, Angela Brisebarre, Camille Capel, Vincent Enouf, Louise Lefrançois, Frédéric Lemoine, Christophe Malabat, Corinne Maufrais, Etienne Simon-Lorière, Maud Vanpeene, Sylvie Van der Werf ,Vincent Thibault |
| EPI_ISL_2464240 | Hospital | National Reference Center for Viruses of Respiratory Infections, Institut Pasteur, Paris | Marion Barbet, Sylvie Behillil, Méline Bizard, Angela Brisebarre, Camille Capel, Vincent Enouf, Louise Lefrançois, Frédéric Lemoine, Christophe Malabat, Corinne Maufrais, Etienne Simon-Lorière, Maud Vanpeene, Sylvie Van der Werf ,Pauline Morvan |
| EPI_ISL_2464241 | Hospital | National Reference Center for Viruses of Respiratory Infections, Institut Pasteur, Paris | Marion Barbet, Sylvie Behillil, Méline Bizard, Angela Brisebarre, Camille Capel, Vincent Enouf, Louise Lefrançois, Frédéric Lemoine, Christophe Malabat, Corinne Maufrais, Amaury Vaysse, Etienne Simon-Lorière, Maud Vanpeene, Sylvie Van der Werf ,A Hochart |

| | | | |
|---|---|---|---|
| EPI_ISL_2464242 | Hospital | National Reference Center for Viruses of Respiratory Infections, Institut Pasteur, Paris | Marion Barbet, Sylvie Behillil, Méline Bizard, Angela Brisebarre, Camille Capel, Vincent Enouf, Louise Lefrançois, Frédéric Lemoine, Christophe Malabat, Corinne Maufrais, Etienne Simon-Lorière, Maud Vanpeene, Sylvie Van der Werf ,Marianne Burgard |
| EPI_ISL_2464243, EPI_ISL_2464244 | Hospital | National Reference Center for Viruses of Respiratory Infections, Institut Pasteur, Paris | Marion Barbet, Sylvie Behillil, Méline Bizard, Angela Brisebarre, Camille Capel, Vincent Enouf, Louise Lefrançois, Frédéric Lemoine, Christophe Malabat, Corinne Maufrais, Etienne Simon-Lorière, Maud Vanpeene, Sylvie Van der Werf ,Jean Baptiste Bour |
| EPI_ISL_2464248 | Hospital | National Reference Center for Viruses of Respiratory Infections, Institut Pasteur, Paris | Marion Barbet, Sylvie Behillil, Méline Bizard, Angela Brisebarre, Camille Capel, Vincent Enouf, Louise Lefrançois, Frédéric Lemoine, Christophe Malabat, Corinne Maufrais, Etienne Simon-Lorière, Maud Vanpeene, Sylvie Van der Werf ,AgnèS Scanvic |
| EPI_ISL_2464249 | Hospital | National Reference Center for Viruses of Respiratory Infections, Institut Pasteur, Paris | Marion Barbet, Sylvie Behillil, Méline Bizard, Angela Brisebarre, Camille Capel, Vincent Enouf, Louise Lefrançois, Frédéric Lemoine, Christophe Malabat, Corinne Maufrais, Amaury Vaysse, Etienne Simon-Lorière, Maud Vanpeene, Sylvie Van der Werf ,Sandrine Castelain |
| EPI_ISL_2464250, EPI_ISL_2464252 | Hospital | National Reference Center for Viruses of Respiratory Infections, Institut Pasteur, Paris | Marion Barbet, Sylvie Behillil, Méline Bizard, Angela Brisebarre, Camille Capel, Vincent Enouf, Louise Lefrançois, Frédéric Lemoine, Christophe Malabat, Corinne Maufrais, Etienne Simon-Lorière, Maud Vanpeene, Sylvie Van der Werf ,Sandrine Castelain |
| EPI_ISL_2464253 | Hospital | National Reference Center for Viruses of Respiratory Infections, Institut Pasteur, Paris | Marion Barbet, Sylvie Behillil, Méline Bizard, Angela Brisebarre, Camille Capel, Vincent Enouf, Louise Lefrançois, Frédéric Lemoine, Christophe Malabat, Corinne Maufrais, Amaury Vaysse, Etienne Simon-Lorière, Maud Vanpeene, Sylvie Van der Werf ,Sandrine Castelain |
| EPI_ISL_2464254 | Hospital | National Reference Center for Viruses of Respiratory Infections, Institut Pasteur, Paris | Marion Barbet, Sylvie Behillil, Méline Bizard, Angela Brisebarre, Camille Capel, Vincent Enouf, Louise Lefrançois, Frédéric Lemoine, Christophe Malabat, Corinne Maufrais, Etienne Simon-Lorière, Maud Vanpeene, Sylvie Van der Werf ,Sandrine Castelain |
| EPI_ISL_2464256 | Hospital | National Reference Center for Viruses of Respiratory Infections, Institut Pasteur, Paris | Marion Barbet, Sylvie Behillil, Méline Bizard, Angela Brisebarre, Camille Capel, Vincent Enouf, Louise Lefrançois, Frédéric Lemoine, Christophe Malabat, Corinne Maufrais, Etienne Simon-Lorière, Maud Vanpeene, Sylvie Van der Werf ,Dominique Descamps |
| EPI_ISL_2464257 | Labo Analyses Med | National Reference Center for Viruses of Respiratory Infections, Institut Pasteur, Paris | Marion Barbet, Sylvie Behillil, Méline Bizard, Angela Brisebarre, Camille Capel, Vincent Enouf, Louise Lefrançois, Frédéric Lemoine, Christophe Malabat, Corinne Maufrais, Amaury Vaysse, Etienne Simon-Lorière, Maud Vanpeene, Sylvie Van der Werf ,Domitille Leman |
| EPI_ISL_2464258, EPI_ISL_2464259 | Labo Analyses Med | National Reference Center for Viruses of Respiratory Infections, Institut Pasteur, Paris | Marion Barbet, Sylvie Behillil, Méline Bizard, Angela Brisebarre, Camille Capel, Vincent Enouf, Louise Lefrançois, Frédéric Lemoine, Christophe Malabat, Corinne Maufrais, Etienne Simon-Lorière, Maud Vanpeene, Sylvie Van der Werf ,Domitille Leman |
| EPI_ISL_2464260, EPI_ISL_2464261 | Labo Analyses Med | National Reference Center for Viruses of Respiratory Infections, Institut Pasteur, Paris | Marion Barbet, Sylvie Behillil, Méline Bizard, Angela Brisebarre, Camille Capel, Vincent Enouf, Louise Lefrançois, Frédéric Lemoine, Christophe Malabat, Corinne Maufrais, Amaury Vaysse, Etienne Simon-Lorière, Maud Vanpeene, Sylvie Van der Werf ,Domitille Leman |
| EPI_ISL_2464262, EPI_ISL_2464263, EPI_ISL_2464264, EPI_ISL_2464265, EPI_ISL_2464266 | Labo Analyses Med | National Reference Center for Viruses of Respiratory Infections, Institut Pasteur, Paris | Marion Barbet, Sylvie Behillil, Méline Bizard, Angela Brisebarre, Camille Capel, Vincent Enouf, Louise Lefrançois, Frédéric Lemoine, Christophe Malabat, Corinne Maufrais, Etienne Simon-Lorière, Maud Vanpeene, Sylvie Van der Werf ,Domitille Leman |
| EPI_ISL_2464267, EPI_ISL_2464268 | Hospital | National Reference Center for Viruses of Respiratory Infections, Institut Pasteur, Paris | Marion Barbet, Sylvie Behillil, Méline Bizard, Angela Brisebarre, Camille Capel, Vincent Enouf, Louise Lefrançois, Frédéric Lemoine, Christophe Malabat, Corinne Maufrais, Etienne Simon-Lorière, Maud Vanpeene, Sylvie Van der Werf ,Laurence Courdavault |
| EPI_ISL_2464269, EPI_ISL_2464271, EPI_ISL_2464272, EPI_ISL_2464273 | Hospital | National Reference Center for Viruses of Respiratory Infections, Institut Pasteur, Paris | Marion Barbet, Sylvie Behillil, Méline Bizard, Angela Brisebarre, Camille Capel, Vincent Enouf, Louise Lefrançois, Frédéric Lemoine, Christophe Malabat, Corinne Maufrais, Etienne Simon-Lorière, Maud Vanpeene, Sylvie Van der Werf ,Nabil Gastli |
| EPI_ISL_2464274 | Hospital | National Reference Center for Viruses of Respiratory Infections, Institut Pasteur, Paris | Marion Barbet, Sylvie Behillil, Méline Bizard, Angela Brisebarre, Camille Capel, Vincent Enouf, Louise Lefrançois, Frédéric Lemoine, Christophe Malabat, Corinne Maufrais, Amaury Vaysse, Etienne Simon-Lorière, Maud Vanpeene, Sylvie Van der Werf ,Nabil Gastli |
| EPI_ISL_2464275, EPI_ISL_2464276 | Hospital | National Reference Center for Viruses of Respiratory Infections, Institut Pasteur, Paris | Marion Barbet, Sylvie Behillil, Méline Bizard, Angela Brisebarre, Camille Capel, Vincent Enouf, Louise Lefrançois, Frédéric Lemoine, Christophe Malabat, Corinne Maufrais, Etienne Simon-Lorière, Maud Vanpeene, Sylvie Van der Werf ,Nabil Gastli |
| EPI_ISL_2464277, EPI_ISL_2464278 | Hospital | National Reference Center for Viruses of Respiratory Infections, Institut Pasteur, Paris | Marion Barbet, Sylvie Behillil, Méline Bizard, Angela Brisebarre, Camille Capel, Vincent Enouf, Louise Lefrançois, Frédéric Lemoine, Christophe Malabat, Corinne Maufrais, Emmanuelle Permal, Etienne Simon-Lorière, Maud Vanpeene, Sylvie Van der Werf ,Nabil Gastli |
| EPI_ISL_2464499, EPI_ISL_2464503 | GH A.CHENEVIER-H.MONDOR | Department of Virology, Henri Mondor University Hospital, Assistance Publique Hôpitaux de Paris, Université Paris-Est Créteil, INSERM U955 | Christophe Rodriguez, Slim Fourati, Vanessa Demontant, Guillaume Gricourt, Melissa N'Debi, Alexandre Soulier, Elisabeth Trawinski, Jean-Michel Pawlotsky |
| EPI_ISL_2464504 | HOPITAL G. CLEMENCEAU | Department of Virology, Henri Mondor University Hospital, Assistance Publique Hôpitaux de Paris, Université Paris-Est Créteil, INSERM U955 | Christophe Rodriguez, Slim Fourati, Vanessa Demontant, Guillaume Gricourt, Melissa N'Debi, Alexandre Soulier, Elisabeth Trawinski, Jean-Michel Pawlotsky |
| EPI_ISL_2464507, EPI_ISL_2464512, EPI_ISL_2464513, EPI_ISL_2464516 | GH A.CHENEVIER-H.MONDOR | Department of Virology, Henri Mondor University Hospital, Assistance Publique Hôpitaux de Paris, Université Paris-Est Créteil, INSERM U955 | Christophe Rodriguez, Slim Fourati, Vanessa Demontant, Guillaume Gricourt, Melissa N'Debi, Alexandre Soulier, Elisabeth Trawinski, Jean-Michel Pawlotsky |
| EPI_ISL_2464521 | HOPITAL EMILE ROUX | Department of Virology, Henri Mondor University Hospital, Assistance Publique Hôpitaux de Paris, Université Paris-Est Créteil, INSERM U955 | Christophe Rodriguez, Slim Fourati, Vanessa Demontant, Guillaume Gricourt, Melissa N'Debi, Alexandre Soulier, Elisabeth Trawinski, Jean-Michel Pawlotsky |
| EPI_ISL_2464555, EPI_ISL_2464556, EPI_ISL_2464557, EPI_ISL_2464558, EPI_ISL_2464559 | Singapore General Hospital | Department of Microbiology | Nurdyana Abdul Rahman, Kenneth Xin Long Chan, Chayaporn Suphavilai, Kun Lee Lim, Sui Sin Goh, Kian Sing Chan, Lynette Oon, Kern Rei Chng, James Sim Heng Chiak, Niranjan Nagarajan, Karrie Ko |
| EPI_ISL_2464560, EPI_ISL_2464561, EPI_ISL_2464562, EPI_ISL_2464563, EPI_ISL_2464564 | Singapore General Hospital | Singapore General Hospital | Nurdyana Abdul Rahman, Kenneth Xin Long Chan, Chayaporn Suphavilai, Kun Lee Lim, Sui Sin Goh, Kian Sing Chan, Lynette Oon, Kern Rei Chng, James Sim Heng Chiak, Niranjan Nagarajan, Karrie Ko |
| EPI_ISL_2466276, EPI_ISL_2466277, EPI_ISL_2466280, EPI_ISL_2466281, EPI_ISL_2466282, EPI_ISL_2466283, EPI_ISL_2466284, EPI_ISL_2466285 | Presidio di Brindisi Di Summa - Perrino | Istituto Zooprofilattico Sperimentale della Puglia e della Basilicata | Parisi A., Bianco A., Capozzi L., Del Sambro L., Simone D., Difato L., Pennuzzi G., Santoro A. |
| EPI_ISL_2466286, EPI_ISL_2466287, EPI_ISL_2466288, EPI_ISL_2466289, EPI_ISL_2466290, EPI_ISL_2466291 | Ospedale Vito Fazzi | Istituto Zooprofilattico Sperimentale della Puglia e della Basilicata | Parisi A., Bianco A., Capozzi L., Del Sambro L., Simone D., Difato L., Bruno A. R. |
| EPI_ISL_2466292, EPI_ISL_2466293, EPI_ISL_2466294, EPI_ISL_2466295, EPI_ISL_2466296, EPI_ISL_2466297, EPI_ISL_2466298, EPI_ISL_2466299, EPI_ISL_2466300, EPI_ISL_2466301, EPI_ISL_2466302, EPI_ISL_2466303, EPI_ISL_2466304, EPI_ISL_2466305, EPI_ISL_2466306, EPI_ISL_2466307, EPI_ISL_2466308, EPI_ISL_2466309, EPI_ISL_2466310, EPI_ISL_2466311 | | | |
| see above | Ospedale Santa Caterina Novella | Istituto Zooprofilattico Sperimentale della Puglia e della Basilicata | Parisi A., Bianco A., Capozzi L., Del Sambro L., Simone D., Difato L., Lobreglio G. |
| EPI_ISL_2466312 | I.R.C.C.S. "S. De Bellis" - Ente Ospedaliero | Istituto Zooprofilattico Sperimentale della Puglia e della Basilicata | Parisi A., Bianco A., Capozzi L., Del Sambro L., Simone D., Difato L., Notarnicola M., Lippolis A. |
| EPI_ISL_2466340, EPI_ISL_2466346, EPI_ISL_2466347, EPI_ISL_2466348, EPI_ISL_2466349 | AZ Zeno Knokke-Blankenberge | AZ Sint Jan | Marijke Reynders, Patrick Descheemaeker, Laurien Hoornaert, Thomas Van Landschoot, Marie Madeleine Chabert-Consen, Jorn Hellemans |
| EPI_ISL_2466350 | AZ St.Jan campus Oostende | AZ Sint Jan | Marijke Reynders, Patrick Descheemaeker, Laurien Hoornaert, Thomas Van Landschoot, Marie Madeleine Chabert-Consen, Jorn Hellemans |
| EPI_ISL_2466352 | AZ Zeno Knokke-Blankenberge | AZ Sint Jan | Marijke Reynders, Patrick Descheemaeker, Laurien Hoornaert, Thomas Van Landschoot, Marie Madeleine Chabert-Consen, Jorn Hellemans |
| EPI_ISL_2466353 | AZ St.Jan campus Oostende | AZ Sint Jan | Marijke Reynders, Patrick Descheemaeker, Laurien Hoornaert, Thomas Van Landschoot, Marie Madeleine Chabert-Consen, Jorn Hellemans |

| | | | |
|---|---|---|---|
| EPI_ISL_2466354, EPI_ISL_2466355 | AZ Zeno Knokke-Blankenberge | AZ Sint Jan | Marijke Reynders, Patrick Descheemaeker, Laurien Hoornaert, Thomas Van Landschoot, Marie Madeleine Chabert-Consen, Jorn Hellemans |
| EPI_ISL_2467834, EPI_ISL_2467835, EPI_ISL_2467865 | Usansolo-Galdakao University Hospital | Cruces University Hospital | Ana Gual-de-Torrella, Izaskun Alejo-Cancho, Mikel Gallego, Ana Belén de la Hoz |
| EPI_ISL_2476409, EPI_ISL_2476410, EPI_ISL_2476411, EPI_ISL_2476412 | LESP Ciudad de Mexico | Instituto de Diagnostico y Referencia Epidemiologicos (INDRE) | Claudia Wong-Arambula, Abril Rodriguez-Maldonado, Vanessa Rivero-Arredondo, Ariadna Medina-Benitez, Joaquin Quiroz-Mercado, Sergio Rangel-Guerrero, Natividad Cruz-Ortiz, Tatiana Nunez-Garcia, Gisela Barrera-Badillo, Lucia Hernandez-Rivas, Irma Lopez-Martinez, Ernesto Ramirez-Gonzalez. |
| EPI_ISL_2476414 | LESP Yucatan | Instituto de Diagnostico y Referencia Epidemiologicos (INDRE) | Claudia Wong-Arambula, Abril Rodriguez-Maldonado, Vanessa Rivero-Arredondo, Ariadna Medina-Benitez, Joaquin Quiroz-Mercado, Sergio Rangel-Guerrero, Natividad Cruz-Ortiz, Tatiana Nunez-Garcia, Gisela Barrera-Badillo, Lucia Hernandez-Rivas, Irma Lopez-Martinez, Ernesto Ramirez-Gonzalez. |
| EPI_ISL_2476417 | LESP Zacatecas | Instituto de Diagnostico y Referencia Epidemiologicos (INDRE) | Claudia Wong-Arambula, Abril Rodriguez-Maldonado, Vanessa Rivero-Arredondo, Ariadna Medina-Benitez, Joaquin Quiroz-Mercado, Sergio Rangel-Guerrero, Natividad Cruz-Ortiz, Tatiana Nunez-Garcia, Gisela Barrera-Badillo, Lucia Hernandez-Rivas, Irma Lopez-Martinez, Ernesto Ramirez-Gonzalez. |
| EPI_ISL_2476418, EPI_ISL_2476419, EPI_ISL_2476420, EPI_ISL_2476421, EPI_ISL_2476422, EPI_ISL_2476426 | LESP Guanajuato | Instituto de Diagnostico y Referencia Epidemiologicos (INDRE) | Claudia Wong-Arambula, Abril Rodriguez-Maldonado, Vanessa Rivero-Arredondo, Ariadna Medina-Benitez, Joaquin Quiroz-Mercado, Sergio Rangel-Guerrero, Natividad Cruz-Ortiz, Tatiana Nunez-Garcia, Gisela Barrera-Badillo, Lucia Hernandez-Rivas, Irma Lopez-Martinez, Ernesto Ramirez-Gonzalez. |
| EPI_ISL_2476427, EPI_ISL_2476428, EPI_ISL_2476429 | LESP Ciudad de Mexico | Instituto de Diagnostico y Referencia Epidemiologicos (INDRE) | Claudia Wong-Arambula, Abril Rodriguez-Maldonado, Vanessa Rivero-Arredondo, Ariadna Medina-Benitez, Joaquin Quiroz-Mercado, Sergio Rangel-Guerrero, Natividad Cruz-Ortiz, Tatiana Nunez-Garcia, Gisela Barrera-Badillo, Lucia Hernandez-Rivas, Irma Lopez-Martinez, Ernesto Ramirez-Gonzalez. |
| EPI_ISL_2476430 | LESP Oaxaca | Instituto de Diagnostico y Referencia Epidemiologicos (INDRE) | Claudia Wong-Arambula, Abril Rodriguez-Maldonado, Vanessa Rivero-Arredondo, Ariadna Medina-Benitez, Joaquin Quiroz-Mercado, Sergio Rangel-Guerrero, Natividad Cruz-Ortiz, Tatiana Nunez-Garcia, Gisela Barrera-Badillo, Lucia Hernandez-Rivas, Irma Lopez-Martinez, Ernesto Ramirez-Gonzalez. |
| EPI_ISL_2476431, EPI_ISL_2476432, EPI_ISL_2476434, EPI_ISL_2476435, EPI_ISL_2476436 | LESP Tamaulipas | Instituto de Diagnostico y Referencia Epidemiologicos (INDRE) | Claudia Wong-Arambula, Abril Rodriguez-Maldonado, Vanessa Rivero-Arredondo, Ariadna Medina-Benitez, Joaquin Quiroz-Mercado, Sergio Rangel-Guerrero, Natividad Cruz-Ortiz, Tatiana Nunez-Garcia, Gisela Barrera-Badillo, Lucia Hernandez-Rivas, Irma Lopez-Martinez, Ernesto Ramirez-Gonzalez. |
| EPI_ISL_2476437 | LESP Sinaloa | Instituto de Diagnostico y Referencia Epidemiologicos (INDRE) | Claudia Wong-Arambula, Abril Rodriguez-Maldonado, Vanessa Rivero-Arredondo, Ariadna Medina-Benitez, Joaquin Quiroz-Mercado, Sergio Rangel-Guerrero, Natividad Cruz-Ortiz, Tatiana Nunez-Garcia, Gisela Barrera-Badillo, Lucia Hernandez-Rivas, Irma Lopez-Martinez, Ernesto Ramirez-Gonzalez. |
| EPI_ISL_2476438 | LESP Colima | Instituto de Diagnostico y Referencia Epidemiologicos (INDRE) | Claudia Wong-Arambula, Abril Rodriguez-Maldonado, Vanessa Rivero-Arredondo, Ariadna Medina-Benitez, Joaquin Quiroz-Mercado, Sergio Rangel-Guerrero, Natividad Cruz-Ortiz, Tatiana Nunez-Garcia, Gisela Barrera-Badillo, Lucia Hernandez-Rivas, Irma Lopez-Martinez, Ernesto Ramirez-Gonzalez. |
| EPI_ISL_2476439 | LESP Baja California | Instituto de Diagnostico y Referencia Epidemiologicos (INDRE) | Claudia Wong-Arambula, Abril Rodriguez-Maldonado, Vanessa Rivero-Arredondo, Ariadna Medina-Benitez, Joaquin Quiroz-Mercado, Sergio Rangel-Guerrero, Natividad Cruz-Ortiz, Tatiana Nunez-Garcia, Gisela Barrera-Badillo, Lucia Hernandez-Rivas, Irma Lopez-Martinez, Ernesto Ramirez-Gonzalez. |
| EPI_ISL_2476440 | LESP Morelos | Instituto de Diagnostico y Referencia Epidemiologicos (INDRE) | Claudia Wong-Arambula, Abril Rodriguez-Maldonado, Vanessa Rivero-Arredondo, Ariadna Medina-Benitez, Joaquin Quiroz-Mercado, Sergio Rangel-Guerrero, Natividad Cruz-Ortiz, Tatiana Nunez-Garcia, Gisela Barrera-Badillo, Lucia Hernandez-Rivas, Irma Lopez-Martinez, Ernesto Ramirez-Gonzalez. |
| EPI_ISL_2476441 | LESP Chihuahua | Instituto de Diagnostico y Referencia Epidemiologicos (INDRE) | Claudia Wong-Arambula, Abril Rodriguez-Maldonado, Vanessa Rivero-Arredondo, Ariadna Medina-Benitez, Joaquin Quiroz-Mercado, Sergio Rangel-Guerrero, Natividad Cruz-Ortiz, Tatiana Nunez-Garcia, Gisela Barrera-Badillo, Lucia Hernandez-Rivas, Irma Lopez-Martinez, Ernesto Ramirez-Gonzalez. |
| EPI_ISL_2476442 | LESP Campeche | Instituto de Diagnostico y Referencia Epidemiologicos (INDRE) | Claudia Wong-Arambula, Abril Rodriguez-Maldonado, Vanessa Rivero-Arredondo, Ariadna Medina-Benitez, Joaquin Quiroz-Mercado, Sergio Rangel-Guerrero, Natividad Cruz-Ortiz, Tatiana Nunez-Garcia, Gisela Barrera-Badillo, Lucia Hernandez-Rivas, Irma Lopez-Martinez, Ernesto Ramirez-Gonzalez. |
| EPI_ISL_2476443 | LESP Baja California | Instituto de Diagnostico y Referencia Epidemiologicos (INDRE) | Claudia Wong-Arambula, Abril Rodriguez-Maldonado, Vanessa Rivero-Arredondo, Ariadna Medina-Benitez, Joaquin Quiroz-Mercado, Sergio Rangel-Guerrero, Natividad Cruz-Ortiz, Tatiana Nunez-Garcia, Gisela Barrera-Badillo, Lucia Hernandez-Rivas, Irma Lopez-Martinez, Ernesto Ramirez-Gonzalez. |
| EPI_ISL_2476444 | LESP Sinaloa | Instituto de Diagnostico y Referencia Epidemiologicos (INDRE) | Claudia Wong-Arambula, Abril Rodriguez-Maldonado, Vanessa Rivero-Arredondo, Ariadna Medina-Benitez, Joaquin Quiroz-Mercado, Sergio Rangel-Guerrero, Natividad Cruz-Ortiz, Tatiana Nunez-Garcia, Gisela Barrera-Badillo, Lucia Hernandez-Rivas, Irma Lopez-Martinez, Ernesto Ramirez-Gonzalez. |
| EPI_ISL_2478137, EPI_ISL_2478140, EPI_ISL_2478142, EPI_ISL_2478144, EPI_ISL_2478145, EPI_ISL_2478149, EPI_ISL_2478152 | Laboratori de Referencia de Catalunya | Laboratori de Referencia de Catalunya | Ramirez A., Bellosillo B., Padilla E., Vilas A., Hernandez JJ., Canal M. |
| EPI_ISL_2479880 | LESP Queretaro | Instituto de Diagnostico y Referencia Epidemiologicos (INDRE) | Claudia Wong-Arambula, Abril Rodriguez-Maldonado, Vanessa Rivero-Arredondo, Ariadna Medina-Benitez, Joaquin Quiroz-Mercado, Sergio Rangel-Guerrero, Natividad Cruz-Ortiz, Tatiana Nunez-Garcia, Gisela Barrera-Badillo, Lucia Hernandez-Rivas, Irma Lopez-Martinez, Ernesto Ramirez-Gonzalez. |
| EPI_ISL_2479882 | LESP Ciudad de Mexico | Instituto de Diagnostico y Referencia Epidemiologicos (INDRE) | Claudia Wong-Arambula, Abril Rodriguez-Maldonado, Vanessa Rivero-Arredondo, Ariadna Medina-Benitez, Joaquin Quiroz-Mercado, Sergio Rangel-Guerrero, Natividad Cruz-Ortiz, Tatiana Nunez-Garcia, Gisela Barrera-Badillo, Lucia Hernandez-Rivas, Irma Lopez-Martinez, Ernesto Ramirez-Gonzalez. |
| EPI_ISL_2479887 | LESP Yucatan | Instituto de Diagnostico y Referencia Epidemiologicos (INDRE) | Claudia Wong-Arambula, Abril Rodriguez-Maldonado, Vanessa Rivero-Arredondo, Ariadna Medina-Benitez, Joaquin Quiroz-Mercado, Sergio Rangel-Guerrero, Natividad Cruz-Ortiz, Tatiana Nunez-Garcia, Gisela Barrera-Badillo, Lucia Hernandez-Rivas, Irma Lopez-Martinez, Ernesto Ramirez-Gonzalez. |
| EPI_ISL_2479888 | LESP Baja California | Instituto de Diagnostico y Referencia Epidemiologicos (INDRE) | Claudia Wong-Arambula, Abril Rodriguez-Maldonado, Vanessa Rivero-Arredondo, Ariadna Medina-Benitez, Joaquin Quiroz-Mercado, Sergio Rangel-Guerrero, Natividad Cruz-Ortiz, Tatiana Nunez-Garcia, Gisela Barrera-Badillo, Lucia Hernandez-Rivas, Irma Lopez-Martinez, Ernesto Ramirez-Gonzalez. |
| EPI_ISL_2479889, EPI_ISL_2479890 | LESP Quintana Roo | Instituto de Diagnostico y Referencia Epidemiologicos (INDRE) | Claudia Wong-Arambula, Abril Rodriguez-Maldonado, Vanessa Rivero-Arredondo, Ariadna Medina-Benitez, Joaquin Quiroz-Mercado, Sergio Rangel-Guerrero, Natividad Cruz-Ortiz, Tatiana Nunez-Garcia, Gisela Barrera-Badillo, Lucia Hernandez-Rivas, Irma Lopez-Martinez, Ernesto Ramirez-Gonzalez. |
| EPI_ISL_2479893 | LESP Queretaro | Instituto de Diagnostico y Referencia Epidemiologicos (INDRE) | Claudia Wong-Arambula, Abril Rodriguez-Maldonado, Vanessa Rivero-Arredondo, Ariadna Medina-Benitez, Joaquin Quiroz-Mercado, Sergio Rangel-Guerrero, Natividad Cruz-Ortiz, Tatiana Nunez-Garcia, Gisela Barrera-Badillo, Lucia Hernandez-Rivas, Irma Lopez-Martinez, Ernesto Ramirez-Gonzalez. |
| EPI_ISL_2479894, EPI_ISL_2479895, EPI_ISL_2479896, EPI_ISL_2479898 | LESP Nuevo Leon | Instituto de Diagnostico y Referencia Epidemiologicos (INDRE) | Claudia Wong-Arambula, Abril Rodriguez-Maldonado, Vanessa Rivero-Arredondo, Ariadna Medina-Benitez, Joaquin Quiroz-Mercado, Sergio Rangel-Guerrero, Natividad Cruz-Ortiz, Tatiana Nunez-Garcia, Gisela Barrera-Badillo, Lucia Hernandez-Rivas, Irma Lopez-Martinez, Ernesto Ramirez-Gonzalez. |
| EPI_ISL_2479899 | LESP Guanajuato | Instituto de Diagnostico y Referencia Epidemiologicos (INDRE) | Claudia Wong-Arambula, Abril Rodriguez-Maldonado, Vanessa Rivero-Arredondo, Ariadna Medina-Benitez, Joaquin Quiroz-Mercado, Sergio Rangel-Guerrero, Natividad Cruz-Ortiz, Tatiana Nunez-Garcia, Gisela Barrera-Badillo, Lucia Hernandez-Rivas, Irma Lopez-Martinez, Ernesto |

| | | | |
|---|---|---|---|
| | | | Ramirez-Gonzalez. |
| EPI_ISL_2479903, EPI_ISL_2479905 | LESP Puebla | Instituto de Diagnostico y Referencia Epidemiologicos (INDRE) | Claudia Wong-Arambula, Abril Rodriguez-Maldonado, Vanessa Rivero-Arredondo, Ariadna Medina-Benitez, Joaquin Quiroz-Mercado, Sergio Rangel-Guerrero, Natividad Cruz-Ortiz, Tatiana Nunez-Garcia, Gisela Barrera-Badillo, Lucia Hernandez-Rivas, Irma Lopez-Martinez, Ernesto Ramirez-Gonzalez. |
| EPI_ISL_2479906 | LESP Tamaulipas | Instituto de Diagnostico y Referencia Epidemiologicos (INDRE) | Claudia Wong-Arambula, Abril Rodriguez-Maldonado, Vanessa Rivero-Arredondo, Ariadna Medina-Benitez, Joaquin Quiroz-Mercado, Sergio Rangel-Guerrero, Natividad Cruz-Ortiz, Tatiana Nunez-Garcia, Gisela Barrera-Badillo, Lucia Hernandez-Rivas, Irma Lopez-Martinez, Ernesto Ramirez-Gonzalez. |
| EPI_ISL_2479907 | LESP Sonora | Instituto de Diagnostico y Referencia Epidemiologicos (INDRE) | Claudia Wong-Arambula, Abril Rodriguez-Maldonado, Vanessa Rivero-Arredondo, Ariadna Medina-Benitez, Joaquin Quiroz-Mercado, Sergio Rangel-Guerrero, Natividad Cruz-Ortiz, Tatiana Nunez-Garcia, Gisela Barrera-Badillo, Lucia Hernandez-Rivas, Irma Lopez-Martinez, Ernesto Ramirez-Gonzalez. |
| EPI_ISL_2479910 | LESP Colima | Instituto de Diagnostico y Referencia Epidemiologicos (INDRE) | Claudia Wong-Arambula, Abril Rodriguez-Maldonado, Vanessa Rivero-Arredondo, Ariadna Medina-Benitez, Joaquin Quiroz-Mercado, Sergio Rangel-Guerrero, Natividad Cruz-Ortiz, Tatiana Nunez-Garcia, Gisela Barrera-Badillo, Lucia Hernandez-Rivas, Irma Lopez-Martinez, Ernesto Ramirez-Gonzalez. |
| EPI_ISL_2479912 | LESP Hidalgo | Instituto de Diagnostico y Referencia Epidemiologicos (INDRE) | Claudia Wong-Arambula, Abril Rodriguez-Maldonado, Vanessa Rivero-Arredondo, Ariadna Medina-Benitez, Joaquin Quiroz-Mercado, Sergio Rangel-Guerrero, Natividad Cruz-Ortiz, Tatiana Nunez-Garcia, Gisela Barrera-Badillo, Lucia Hernandez-Rivas, Irma Lopez-Martinez, Ernesto Ramirez-Gonzalez. |
| EPI_ISL_2479913 | LESP Queretaro | Instituto de Diagnostico y Referencia Epidemiologicos (INDRE) | Claudia Wong-Arambula, Abril Rodriguez-Maldonado, Vanessa Rivero-Arredondo, Ariadna Medina-Benitez, Joaquin Quiroz-Mercado, Sergio Rangel-Guerrero, Natividad Cruz-Ortiz, Tatiana Nunez-Garcia, Gisela Barrera-Badillo, Lucia Hernandez-Rivas, Irma Lopez-Martinez, Ernesto Ramirez-Gonzalez. |
| EPI_ISL_2479914, EPI_ISL_2479915 | LESP Quintana Roo | Instituto de Diagnostico y Referencia Epidemiologicos (INDRE) | Claudia Wong-Arambula, Abril Rodriguez-Maldonado, Vanessa Rivero-Arredondo, Ariadna Medina-Benitez, Joaquin Quiroz-Mercado, Sergio Rangel-Guerrero, Natividad Cruz-Ortiz, Tatiana Nunez-Garcia, Gisela Barrera-Badillo, Lucia Hernandez-Rivas, Irma Lopez-Martinez, Ernesto Ramirez-Gonzalez. |
| EPI_ISL_2479916 | LESP Jalisco | Instituto de Diagnostico y Referencia Epidemiologicos (INDRE) | Claudia Wong-Arambula, Abril Rodriguez-Maldonado, Vanessa Rivero-Arredondo, Ariadna Medina-Benitez, Joaquin Quiroz-Mercado, Sergio Rangel-Guerrero, Natividad Cruz-Ortiz, Tatiana Nunez-Garcia, Gisela Barrera-Badillo, Lucia Hernandez-Rivas, Irma Lopez-Martinez, Ernesto Ramirez-Gonzalez. |
| EPI_ISL_2483058 | CHU Nice - Hopital Pasteur | Eurofins Biomnis | Xavier Naudot, Véronique Jacomo |
| EPI_ISL_2483060 | CH Philippe Le Bon | Eurofins Biomnis | Xavier Naudot, Véronique Jacomo |
| EPI_ISL_2483328, EPI_ISL_2483459 | Usansolo-Galdakao University Hospital | Cruces University Hospital | Ana Gual-de-Torrella, Izaskun Alejo-Cancho, Mikel Gallego, Ana Belén de la Hoz |
| EPI_ISL_2483563 | SIESP CHIETI - DRIVE IN LANCIANO | Istituto Zooprofilattico Sperimentale dell'Abruzzo e Molise "G. Caporale" | Lorusso A, Marcacci M, Di Domenico M, Ancora M, Curini V, Di Lollo Valeria, Mangone I, Rinaldi A, Delli Compagni E, Scialabba S, Caporale M, Di Pasquale A, Cammà C, Puglia I, Calistri P, Savini G |
| EPI_ISL_2483564 | SIESP CHIETI - DRIVE IN ORTONA | Istituto Zooprofilattico Sperimentale dell'Abruzzo e Molise "G. Caporale" | Lorusso A, Marcacci M, Di Domenico M, Ancora M, Curini V, Di Lollo Valeria, Mangone I, Rinaldi A, Delli Compagni E, Scialabba S, Caporale M, Di Pasquale A, Cammà C, Puglia I, Calistri P, Savini G |
| EPI_ISL_2483565 | SIESP CHIETI - DRIVE IN GISSI | Istituto Zooprofilattico Sperimentale dell'Abruzzo e Molise "G. Caporale" | Lorusso A, Marcacci M, Di Domenico M, Ancora M, Curini V, Di Lollo Valeria, Mangone I, Rinaldi A, Delli Compagni E, Scialabba S, Caporale M, Di Pasquale A, Cammà C, Puglia I, Calistri P, Savini G |
| EPI_ISL_2483566, EPI_ISL_2483567, EPI_ISL_2483568, EPI_ISL_2483569, EPI_ISL_2483570 | SIESP CHIETI - DRIVE IN LANCIANO | Istituto Zooprofilattico Sperimentale dell'Abruzzo e Molise "G. Caporale" | Lorusso A, Marcacci M, Di Domenico M, Ancora M, Curini V, Di Lollo Valeria, Mangone I, Rinaldi A, Delli Compagni E, Scialabba S, Caporale M, Di Pasquale A, Cammà C, Puglia I, Calistri P, Savini G |
| EPI_ISL_2483571, EPI_ISL_2483573 | SIESP DIPARTIMENTO DI PREVENZIONE CHIETI | Istituto Zooprofilattico Sperimentale dell'Abruzzo e Molise "G. Caporale" | Lorusso A, Marcacci M, Di Domenico M, Ancora M, Curini V, Di Lollo Valeria, Mangone I, Rinaldi A, Delli Compagni E, Scialabba S, Caporale M, Di Pasquale A, Cammà C, Puglia I, Calistri P, Savini G |
| EPI_ISL_2483574 | SIESP CHIETI - DRIVE IN CHIETI | Istituto Zooprofilattico Sperimentale dell'Abruzzo e Molise "G. Caporale" | Lorusso A, Marcacci M, Di Domenico M, Ancora M, Curini V, Di Lollo Valeria, Mangone I, Rinaldi A, Delli Compagni E, Scialabba S, Caporale M, Di Pasquale A, Cammà C, Puglia I, Calistri P, Savini G |
| EPI_ISL_2483575, EPI_ISL_2483576, EPI_ISL_2483577, EPI_ISL_2483578, EPI_ISL_2483579, EPI_ISL_2483580, EPI_ISL_2483581, EPI_ISL_2483582 | SIESP DIPARTIMENTO DI PREVENZIONE TERAMO | Istituto Zooprofilattico Sperimentale dell'Abruzzo e Molise "G. Caporale" | Lorusso A, Marcacci M, Di Domenico M, Ancora M, Curini V, Di Lollo Valeria, Mangone I, Rinaldi A, Delli Compagni E, Scialabba S, Caporale M, Di Pasquale A, Cammà C, Puglia I, Calistri P, Savini G |
| EPI_ISL_2483583 | SIESP CHIETI - DRIVE IN ORTONA | Istituto Zooprofilattico Sperimentale dell'Abruzzo e Molise "G. Caporale" | Lorusso A, Marcacci M, Di Domenico M, Ancora M, Curini V, Di Lollo Valeria, Mangone I, Rinaldi A, Delli Compagni E, Scialabba S, Caporale M, Di Pasquale A, Cammà C, Puglia I, Calistri P, Savini G |
| EPI_ISL_2483584, EPI_ISL_2483585, EPI_ISL_2483586, EPI_ISL_2483587 | SIESP DIPARTIMENTO DI PREVENZIONE TERAMO | Istituto Zooprofilattico Sperimentale dell'Abruzzo e Molise "G. Caporale" | Lorusso A, Marcacci M, Di Domenico M, Ancora M, Curini V, Di Lollo Valeria, Mangone I, Rinaldi A, Delli Compagni E, Scialabba S, Caporale M, Di Pasquale A, Cammà C, Puglia I, Calistri P, Savini G |
| EPI_ISL_2483588 | SIESP CHIETI - DRIVE IN ORTONA | Istituto Zooprofilattico Sperimentale dell'Abruzzo e Molise "G. Caporale" | Lorusso A, Marcacci M, Di Domenico M, Ancora M, Curini V, Di Lollo Valeria, Mangone I, Rinaldi A, Delli Compagni E, Scialabba S, Caporale M, Di Pasquale A, Cammà C, Puglia I, Calistri P, Savini G |
| EPI_ISL_2483589 | SIESP DIPARTIMENTO DI PREVENZIONE TERAMO | Istituto Zooprofilattico Sperimentale dell'Abruzzo e Molise "G. Caporale" | Lorusso A, Marcacci M, Di Domenico M, Ancora M, Curini V, Di Lollo Valeria, Mangone I, Rinaldi A, Delli Compagni E, Scialabba S, Caporale M, Di Pasquale A, Cammà C, Puglia I, Calistri P, Savini G |
| EPI_ISL_2483590 | SIESP DIPARTIMENTO DI PREVENZIONE CHIETI | Istituto Zooprofilattico Sperimentale dell'Abruzzo e Molise "G. Caporale" | Lorusso A, Marcacci M, Di Domenico M, Ancora M, Curini V, Di Lollo Valeria, Mangone I, Rinaldi A, Delli Compagni E, Scialabba S, Caporale M, Di Pasquale A, Cammà C, Puglia I, Calistri P, Savini G |
| EPI_ISL_2483591 | SIESP DIPARTIMENTO DI PREVENZIONE TERAMO | Istituto Zooprofilattico Sperimentale dell'Abruzzo e Molise "G. Caporale" | Lorusso A, Marcacci M, Di Domenico M, Ancora M, Curini V, Di Lollo Valeria, Mangone I, Rinaldi A, Delli Compagni E, Scialabba S, Caporale M, Di Pasquale A, Cammà C, Puglia I, Calistri P, Savini G |
| EPI_ISL_2483592 | SIESP DIPARTIMENTO DI PREVENZIONE CHIETI | Istituto Zooprofilattico Sperimentale dell'Abruzzo e Molise "G. Caporale" | Lorusso A, Marcacci M, Di Domenico M, Ancora M, Curini V, Di Lollo Valeria, Mangone I, Rinaldi A, Delli Compagni E, Scialabba S, Caporale M, Di Pasquale A, Cammà C, Puglia I, Calistri P, Savini G |
| EPI_ISL_2483593 | SIESP DIPARTIMENTO DI PREVENZIONE TERAMO | Istituto Zooprofilattico Sperimentale dell'Abruzzo e Molise "G. Caporale" | Lorusso A, Marcacci M, Di Domenico M, Ancora M, Curini V, Di Lollo Valeria, Mangone I, Rinaldi A, Delli Compagni E, Scialabba S, Caporale M, Di Pasquale A, Cammà C, Puglia I, Calistri P, Savini G |
| EPI_ISL_2483594 | SIESP CHIETI - DRIVE IN LANCIANO | Istituto Zooprofilattico Sperimentale dell'Abruzzo e Molise "G. Caporale" | Lorusso A, Marcacci M, Di Domenico M, Ancora M, Curini V, Di Lollo Valeria, Mangone I, Rinaldi A, Delli Compagni E, Scialabba S, Caporale M, Di Pasquale A, Cammà C, Puglia I, Calistri P, Savini G |
| EPI_ISL_2483595 | SIESP DIPARTIMENTO DI PREVENZIONE CHIETI | Istituto Zooprofilattico Sperimentale dell'Abruzzo e Molise "G. Caporale" | Lorusso A, Marcacci M, Di Domenico M, Ancora M, Curini V, Di Lollo Valeria, Mangone I, Rinaldi A, Delli Compagni E, Scialabba S, Caporale M, Di Pasquale A, Cammà C, Puglia I, Calistri P, Savini G |
| EPI_ISL_2483596 | DIP. PREV. AVEZZANO SERVIZIO DI IGIENE EPIDEMIOLOGIA E SANITA' PUBBLICA | Istituto Zooprofilattico Sperimentale dell'Abruzzo e Molise "G. Caporale" | Lorusso A, Marcacci M, Di Domenico M, Ancora M, Curini V, Di Lollo Valeria, Mangone I, Rinaldi A, Delli Compagni E, Scialabba S, Caporale M, Di Pasquale A, Cammà C, Puglia I, Calistri P, Savini G |
| EPI_ISL_2483597 | USCA Tagliacozzo | Istituto Zooprofilattico Sperimentale dell'Abruzzo e Molise "G. Caporale" | Lorusso A, Marcacci M, Di Domenico M, Ancora M, Curini V, Di Lollo Valeria, Mangone I, Rinaldi A, Delli Compagni E, Scialabba S, Caporale M, Di Pasquale A, Cammà C, Puglia I, Calistri P, Savini G |
| EPI_ISL_2483598 | SIESP CHIETI - DRIVE IN CHIETI | Istituto Zooprofilattico Sperimentale dell'Abruzzo e Molise "G. Caporale" | Lorusso A, Marcacci M, Di Domenico M, Ancora M, Curini V, Di Lollo Valeria, Mangone I, Rinaldi A, Delli Compagni E, Scialabba S, Caporale M, Di Pasquale A, Cammà C, Puglia I, Calistri P, Savini G |
| EPI_ISL_2483599 | SIESP DIPARTIMENTO DI PREVENZIONE TERAMO | Istituto Zooprofilattico Sperimentale dell'Abruzzo e Molise "G. | Lorusso A, Marcacci M, Di Domenico M, Ancora M, Curini V, Di Lollo Valeria, Mangone I, Rinaldi A, Delli Compagni E, Scialabba S, Caporale M, Di |

| | | | |
|---|---|---|---|
| | | Caporale" | Pasquale A, Cammà C, Puglia I, Calistri P, Savini G |
| EPI_ISL_2483600 | SIESP SULMONA | Istituto Zooprofilattico Sperimentale dell'Abruzzo e Molise "G. Caporale" | Lorusso A, Marcacci M, Di Domenico M, Ancora M, Curini V, Di Lollo Valeria, Mangone I, Rinaldi A, Delli Compagni E, Scialabba S, Caporale M, Di Pasquale A, Cammà C, Puglia I, Calistri P, Savini G |
| EPI_ISL_2483601 | SIESP DIPARTIMENTO DI PREVENZIONE CHIETI | Istituto Zooprofilattico Sperimentale dell'Abruzzo e Molise "G. Caporale" | Lorusso A, Marcacci M, Di Domenico M, Ancora M, Curini V, Di Lollo Valeria, Mangone I, Rinaldi A, Delli Compagni E, Scialabba S, Caporale M, Di Pasquale A, Cammà C, Puglia I, Calistri P, Savini G |
| EPI_ISL_2483602 | SIESP DIPARTIMENTO DI PREVENZIONE TERAMO | Istituto Zooprofilattico Sperimentale dell'Abruzzo e Molise "G. Caporale" | Lorusso A, Marcacci M, Di Domenico M, Ancora M, Curini V, Di Lollo Valeria, Mangone I, Rinaldi A, Delli Compagni E, Scialabba S, Caporale M, Di Pasquale A, Cammà C, Puglia I, Calistri P, Savini G |
| EPI_ISL_2483603 | SIESP CHIETI - DRIVE IN CHIETI | Istituto Zooprofilattico Sperimentale dell'Abruzzo e Molise "G. Caporale" | Lorusso A, Marcacci M, Di Domenico M, Ancora M, Curini V, Di Lollo Valeria, Mangone I, Rinaldi A, Delli Compagni E, Scialabba S, Caporale M, Di Pasquale A, Cammà C, Puglia I, Calistri P, Savini G |
| EPI_ISL_2483604 | DIP. PREV. AVEZZANO SERVIZIO DI IGIENE EPIDEMIOLOGIA E SANITA' PUBBLICA | Istituto Zooprofilattico Sperimentale dell'Abruzzo e Molise "G. Caporale" | Lorusso A, Marcacci M, Di Domenico M, Ancora M, Curini V, Di Lollo Valeria, Mangone I, Rinaldi A, Delli Compagni E, Scialabba S, Caporale M, Di Pasquale A, Cammà C, Puglia I, Calistri P, Savini G |
| EPI_ISL_2483605 | SIESP DIPARTIMENTO DI PREVENZIONE TERAMO | Istituto Zooprofilattico Sperimentale dell'Abruzzo e Molise "G. Caporale" | Lorusso A, Marcacci M, Di Domenico M, Ancora M, Curini V, Di Lollo Valeria, Mangone I, Rinaldi A, Delli Compagni E, Scialabba S, Caporale M, Di Pasquale A, Cammà C, Puglia I, Calistri P, Savini G |
| EPI_ISL_2483606, EPI_ISL_2483607 | SIESP CHIETI - DRIVE IN VASTO | Istituto Zooprofilattico Sperimentale dell'Abruzzo e Molise "G. Caporale" | Lorusso A, Marcacci M, Di Domenico M, Ancora M, Curini V, Di Lollo Valeria, Mangone I, Rinaldi A, Delli Compagni E, Scialabba S, Caporale M, Di Pasquale A, Cammà C, Puglia I, Calistri P, Savini G |
| EPI_ISL_2483608, EPI_ISL_2483609 | SIESP CHIETI - DRIVE IN ORTONA | Istituto Zooprofilattico Sperimentale dell'Abruzzo e Molise "G. Caporale" | Lorusso A, Marcacci M, Di Domenico M, Ancora M, Curini V, Di Lollo Valeria, Mangone I, Rinaldi A, Delli Compagni E, Scialabba S, Caporale M, Di Pasquale A, Cammà C, Puglia I, Calistri P, Savini G |
| EPI_ISL_2483610 | SIESP DIPARTIMENTO DI PREVENZIONE TERAMO | Istituto Zooprofilattico Sperimentale dell'Abruzzo e Molise "G. Caporale" | Lorusso A, Marcacci M, Di Domenico M, Ancora M, Curini V, Di Lollo Valeria, Mangone I, Rinaldi A, Delli Compagni E, Scialabba S, Caporale M, Di Pasquale A, Cammà C, Puglia I, Calistri P, Savini G |
| EPI_ISL_2483611, EPI_ISL_2483612 | SIESP CHIETI - DRIVE IN LANCIANO | Istituto Zooprofilattico Sperimentale dell'Abruzzo e Molise "G. Caporale" | Lorusso A, Marcacci M, Di Domenico M, Ancora M, Curini V, Di Lollo Valeria, Mangone I, Rinaldi A, Delli Compagni E, Scialabba S, Caporale M, Di Pasquale A, Cammà C, Puglia I, Calistri P, Savini G |
| EPI_ISL_2483613 | SIESP DIPARTIMENTO DI PREVENZIONE CHIETI | Istituto Zooprofilattico Sperimentale dell'Abruzzo e Molise "G. Caporale" | Lorusso A, Marcacci M, Di Domenico M, Ancora M, Curini V, Di Lollo Valeria, Mangone I, Rinaldi A, Delli Compagni E, Scialabba S, Caporale M, Di Pasquale A, Cammà C, Puglia I, Calistri P, Savini G |
| EPI_ISL_2483614 | SIESP CHIETI - DRIVE IN ORTONA | Istituto Zooprofilattico Sperimentale dell'Abruzzo e Molise "G. Caporale" | Lorusso A, Marcacci M, Di Domenico M, Ancora M, Curini V, Di Lollo Valeria, Mangone I, Rinaldi A, Delli Compagni E, Scialabba S, Caporale M, Di Pasquale A, Cammà C, Puglia I, Calistri P, Savini G |
| EPI_ISL_2483615, EPI_ISL_2483616 | SIESP DIPARTIMENTO DI PREVENZIONE TERAMO | Istituto Zooprofilattico Sperimentale dell'Abruzzo e Molise "G. Caporale" | Lorusso A, Marcacci M, Di Domenico M, Ancora M, Curini V, Di Lollo Valeria, Mangone I, Rinaldi A, Delli Compagni E, Scialabba S, Caporale M, Di Pasquale A, Cammà C, Puglia I, Calistri P, Savini G |
| EPI_ISL_2483617 | SIESP DIPARTIMENTO DI PREVENZIONE CHIETI | Istituto Zooprofilattico Sperimentale dell'Abruzzo e Molise "G. Caporale" | Lorusso A, Marcacci M, Di Domenico M, Ancora M, Curini V, Di Lollo Valeria, Mangone I, Rinaldi A, Delli Compagni E, Scialabba S, Caporale M, Di Pasquale A, Cammà C, Puglia I, Calistri P, Savini G |
| EPI_ISL_2483618 | SIESP CHIETI - DRIVE IN ORTONA | Istituto Zooprofilattico Sperimentale dell'Abruzzo e Molise "G. Caporale" | Lorusso A, Marcacci M, Di Domenico M, Ancora M, Curini V, Di Lollo Valeria, Mangone I, Rinaldi A, Delli Compagni E, Scialabba S, Caporale M, Di Pasquale A, Cammà C, Puglia I, Calistri P, Savini G |
| EPI_ISL_2483619 | SIESP DIPARTIMENTO DI PREVENZIONE TERAMO | Istituto Zooprofilattico Sperimentale dell'Abruzzo e Molise "G. Caporale" | Lorusso A, Marcacci M, Di Domenico M, Ancora M, Curini V, Di Lollo Valeria, Mangone I, Rinaldi A, Delli Compagni E, Scialabba S, Caporale M, Di Pasquale A, Cammà C, Puglia I, Calistri P, Savini G |
| EPI_ISL_2483620, EPI_ISL_2483621 | SIESP CHIETI - DRIVE IN CHIETI | Istituto Zooprofilattico Sperimentale dell'Abruzzo e Molise "G. Caporale" | Lorusso A, Marcacci M, Di Domenico M, Ancora M, Curini V, Di Lollo Valeria, Mangone I, Rinaldi A, Delli Compagni E, Scialabba S, Caporale M, Di Pasquale A, Cammà C, Puglia I, Calistri P, Savini G |
| EPI_ISL_2483622 | SIESP CHIETI - DRIVE IN ORTONA | Istituto Zooprofilattico Sperimentale dell'Abruzzo e Molise "G. Caporale" | Lorusso A, Marcacci M, Di Domenico M, Ancora M, Curini V, Di Lollo Valeria, Mangone I, Rinaldi A, Delli Compagni E, Scialabba S, Caporale M, Di Pasquale A, Cammà C, Puglia I, Calistri P, Savini G |
| EPI_ISL_2483623 | USCA Castel di Sangro | Istituto Zooprofilattico Sperimentale dell'Abruzzo e Molise "G. Caporale" | Lorusso A, Marcacci M, Di Domenico M, Ancora M, Curini V, Di Lollo Valeria, Mangone I, Rinaldi A, Delli Compagni E, Scialabba S, Caporale M, Di Pasquale A, Cammà C, Puglia I, Calistri P, Savini G |
| EPI_ISL_2483624 | SIESP CHIETI - DRIVE IN CHIETI | Istituto Zooprofilattico Sperimentale dell'Abruzzo e Molise "G. Caporale" | Lorusso A, Marcacci M, Di Domenico M, Ancora M, Curini V, Di Lollo Valeria, Mangone I, Rinaldi A, Delli Compagni E, Scialabba S, Caporale M, Di Pasquale A, Cammà C, Puglia I, Calistri P, Savini G |
| EPI_ISL_2483625 | SIESP CHIETI - DRIVE IN LANCIANO | Istituto Zooprofilattico Sperimentale dell'Abruzzo e Molise "G. Caporale" | Lorusso A, Marcacci M, Di Domenico M, Ancora M, Curini V, Di Lollo Valeria, Mangone I, Rinaldi A, Delli Compagni E, Scialabba S, Caporale M, Di Pasquale A, Cammà C, Puglia I, Calistri P, Savini G |
| EPI_ISL_2483626 | SIESP CHIETI - DRIVE IN VASTO | Istituto Zooprofilattico Sperimentale dell'Abruzzo e Molise "G. Caporale" | Lorusso A, Marcacci M, Di Domenico M, Ancora M, Curini V, Di Lollo Valeria, Mangone I, Rinaldi A, Delli Compagni E, Scialabba S, Caporale M, Di Pasquale A, Cammà C, Puglia I, Calistri P, Savini G |
| EPI_ISL_2483627, EPI_ISL_2483628 | SIESP CHIETI - DRIVE IN LANCIANO | Istituto Zooprofilattico Sperimentale dell'Abruzzo e Molise "G. Caporale" | Lorusso A, Marcacci M, Di Domenico M, Ancora M, Curini V, Di Lollo Valeria, Mangone I, Rinaldi A, Delli Compagni E, Scialabba S, Caporale M, Di Pasquale A, Cammà C, Puglia I, Calistri P, Savini G |
| EPI_ISL_2483629 | SIESP SULMONA | Istituto Zooprofilattico Sperimentale dell'Abruzzo e Molise "G. Caporale" | Lorusso A, Marcacci M, Di Domenico M, Ancora M, Curini V, Di Lollo Valeria, Mangone I, Rinaldi A, Delli Compagni E, Scialabba S, Caporale M, Di Pasquale A, Cammà C, Puglia I, Calistri P, Savini G |
| EPI_ISL_2483630 | SIESP DIPARTIMENTO DI PREVENZIONE TERAMO | Istituto Zooprofilattico Sperimentale dell'Abruzzo e Molise "G. Caporale" | Lorusso A, Marcacci M, Di Domenico M, Ancora M, Curini V, Di Lollo Valeria, Mangone I, Rinaldi A, Delli Compagni E, Scialabba S, Caporale M, Di Pasquale A, Cammà C, Puglia I, Calistri P, Savini G |
| EPI_ISL_2483631 | USCA Castel di Sangro | Istituto Zooprofilattico Sperimentale dell'Abruzzo e Molise "G. Caporale" | Lorusso A, Marcacci M, Di Domenico M, Ancora M, Curini V, Di Lollo Valeria, Mangone I, Rinaldi A, Delli Compagni E, Scialabba S, Caporale M, Di Pasquale A, Cammà C, Puglia I, Calistri P, Savini G |
| EPI_ISL_2483632 | SIESP CHIETI - DRIVE IN GISSI | Istituto Zooprofilattico Sperimentale dell'Abruzzo e Molise "G. Caporale" | Lorusso A, Marcacci M, Di Domenico M, Ancora M, Curini V, Di Lollo Valeria, Mangone I, Rinaldi A, Delli Compagni E, Scialabba S, Caporale M, Di Pasquale A, Cammà C, Puglia I, Calistri P, Savini G |
| EPI_ISL_2483633 | SIESP DIPARTIMENTO DI PREVENZIONE CHIETI | Istituto Zooprofilattico Sperimentale dell'Abruzzo e Molise "G. Caporale" | Lorusso A, Marcacci M, Di Domenico M, Ancora M, Curini V, Di Lollo Valeria, Mangone I, Rinaldi A, Delli Compagni E, Scialabba S, Caporale M, Di Pasquale A, Cammà C, Puglia I, Calistri P, Savini G |
| EPI_ISL_2483634 | RSA GIULIANOVA | Istituto Zooprofilattico Sperimentale dell'Abruzzo e Molise "G. Caporale" | Lorusso A, Marcacci M, Di Domenico M, Ancora M, Curini V, Di Lollo Valeria, Mangone I, Rinaldi A, Delli Compagni E, Scialabba S, Caporale M, Di Pasquale A, Cammà C, Puglia I, Calistri P, Savini G |
| EPI_ISL_2483635 | OSPEDALE CIVILE SANT'OMERO - PRONTO SOCCORSO | Istituto Zooprofilattico Sperimentale dell'Abruzzo e Molise "G. Caporale" | Lorusso A, Marcacci M, Di Domenico M, Ancora M, Curini V, Di Lollo Valeria, Mangone I, Rinaldi A, Delli Compagni E, Scialabba S, Caporale M, Di Pasquale A, Cammà C, Puglia I, Calistri P, Savini G |
| EPI_ISL_2483636 | RSA GIULIANOVA | Istituto Zooprofilattico Sperimentale dell'Abruzzo e Molise "G. Caporale" | Lorusso A, Marcacci M, Di Domenico M, Ancora M, Curini V, Di Lollo Valeria, Mangone I, Rinaldi A, Delli Compagni E, Scialabba S, Caporale M, Di Pasquale A, Cammà C, Puglia I, Calistri P, Savini G |
| EPI_ISL_2483637 | SIESP SULMONA | Istituto Zooprofilattico Sperimentale dell'Abruzzo e Molise "G. Caporale" | Lorusso A, Marcacci M, Di Domenico M, Ancora M, Curini V, Di Lollo Valeria, Mangone I, Rinaldi A, Delli Compagni E, Scialabba S, Caporale M, Di Pasquale A, Cammà C, Puglia I, Calistri P, Savini G |
| EPI_ISL_2483638 | SIESP DIPARTIMENTO DI PREVENZIONE TERAMO | Istituto Zooprofilattico Sperimentale dell'Abruzzo e Molise "G. Caporale" | Lorusso A, Marcacci M, Di Domenico M, Ancora M, Curini V, Di Lollo Valeria, Mangone I, Rinaldi A, Delli Compagni E, Scialabba S, Caporale M, Di Pasquale A, Cammà C, Puglia I, Calistri P, Savini G |
| EPI_ISL_2483639 | SIESP CHIETI - DRIVE IN ORTONA | Istituto Zooprofilattico Sperimentale dell'Abruzzo e Molise "G. Caporale" | Lorusso A, Marcacci M, Di Domenico M, Ancora M, Curini V, Di Lollo Valeria, Mangone I, Rinaldi A, Delli Compagni E, Scialabba S, Caporale M, Di Pasquale A, Cammà C, Puglia I, Calistri P, Savini G |
| EPI_ISL_2483640, EPI_ISL_2483641, | SIESP DIPARTIMENTO DI PREVENZIONE TERAMO | Istituto Zooprofilattico Sperimentale dell'Abruzzo e Molise "G. | Lorusso A, Marcacci M, Di Domenico M, Ancora M, Curini V, Di Lollo Valeria, Mangone I, Rinaldi A, Delli Compagni E, Scialabba S, Caporale M, Di |

| | | | |
|---|---|---|---|
| EPI_ISL_2483642 | | Caporale" | Pasquale A, Cammà C, Puglia I, Calistri P, Savini G |
| EPI_ISL_2483643 | SIESP DIPARTIMENTO DI PREVENZIONE CHIETI | Istituto Zooprofilattico Sperimentale dell'Abruzzo e Molise "G. Caporale" | Lorusso A, Marcacci M, Di Domenico M, Ancora M, Curini V, Di Lollo Valeria, Mangone I, Rinaldi A, Delli Compagni E, Scialabba S, Caporale M, Di Pasquale A, Cammà C, Puglia I, Calistri P, Savini G |
| EPI_ISL_2483644 | USCA Castel di Sangro | Istituto Zooprofilattico Sperimentale dell'Abruzzo e Molise "G. Caporale" | Lorusso A, Marcacci M, Di Domenico M, Ancora M, Curini V, Di Lollo Valeria, Mangone I, Rinaldi A, Delli Compagni E, Scialabba S, Caporale M, Di Pasquale A, Cammà C, Puglia I, Calistri P, Savini G |
| EPI_ISL_2483645 | SIESP DIPARTIMENTO DI PREVENZIONE TERAMO | Istituto Zooprofilattico Sperimentale dell'Abruzzo e Molise "G. Caporale" | Lorusso A, Marcacci M, Di Domenico M, Ancora M, Curini V, Di Lollo Valeria, Mangone I, Rinaldi A, Delli Compagni E, Scialabba S, Caporale M, Di Pasquale A, Cammà C, Puglia I, Calistri P, Savini G |
| EPI_ISL_2483646, EPI_ISL_2483647 | SIESP CHIETI - DRIVE IN ORTONA | Istituto Zooprofilattico Sperimentale dell'Abruzzo e Molise "G. Caporale" | Lorusso A, Marcacci M, Di Domenico M, Ancora M, Curini V, Di Lollo Valeria, Mangone I, Rinaldi A, Delli Compagni E, Scialabba S, Caporale M, Di Pasquale A, Cammà C, Puglia I, Calistri P, Savini G |
| EPI_ISL_2483648 | SIESP CHIETI - DRIVE IN GISSI | Istituto Zooprofilattico Sperimentale dell'Abruzzo e Molise "G. Caporale" | Lorusso A, Marcacci M, Di Domenico M, Ancora M, Curini V, Di Lollo Valeria, Mangone I, Rinaldi A, Delli Compagni E, Scialabba S, Caporale M, Di Pasquale A, Cammà C, Puglia I, Calistri P, Savini G |
| EPI_ISL_2483649, EPI_ISL_2483650 | SIESP CHIETI - DRIVE IN LANCIANO | Istituto Zooprofilattico Sperimentale dell'Abruzzo e Molise "G. Caporale" | Lorusso A, Marcacci M, Di Domenico M, Ancora M, Curini V, Di Lollo Valeria, Mangone I, Rinaldi A, Delli Compagni E, Scialabba S, Caporale M, Di Pasquale A, Cammà C, Puglia I, Calistri P, Savini G |
| EPI_ISL_2483651 | SIESP CHIETI - DRIVE IN CHIETI | Istituto Zooprofilattico Sperimentale dell'Abruzzo e Molise "G. Caporale" | Lorusso A, Marcacci M, Di Domenico M, Ancora M, Curini V, Di Lollo Valeria, Mangone I, Rinaldi A, Delli Compagni E, Scialabba S, Caporale M, Di Pasquale A, Cammà C, Puglia I, Calistri P, Savini G |
| EPI_ISL_2483652 | SIESP CHIETI - DRIVE IN LANCIANO | Istituto Zooprofilattico Sperimentale dell'Abruzzo e Molise "G. Caporale" | Lorusso A, Marcacci M, Di Domenico M, Ancora M, Curini V, Di Lollo Valeria, Mangone I, Rinaldi A, Delli Compagni E, Scialabba S, Caporale M, Di Pasquale A, Cammà C, Puglia I, Calistri P, Savini G |
| EPI_ISL_2483653, EPI_ISL_2483654, EPI_ISL_2483655, EPI_ISL_2483656 | SIESP DIPARTIMENTO DI PREVENZIONE TERAMO | Istituto Zooprofilattico Sperimentale dell'Abruzzo e Molise "G. Caporale" | Lorusso A, Marcacci M, Di Domenico M, Ancora M, Curini V, Di Lollo Valeria, Mangone I, Rinaldi A, Delli Compagni E, Scialabba S, Caporale M, Di Pasquale A, Cammà C, Puglia I, Calistri P, Savini G |
| EPI_ISL_2483657 | SIESP DIPARTIMENTO DI PREVENZIONE CHIETI | Istituto Zooprofilattico Sperimentale dell'Abruzzo e Molise "G. Caporale" | Lorusso A, Marcacci M, Di Domenico M, Ancora M, Curini V, Di Lollo Valeria, Mangone I, Rinaldi A, Delli Compagni E, Scialabba S, Caporale M, Di Pasquale A, Cammà C, Puglia I, Calistri P, Savini G |
| EPI_ISL_2483658 | SIESP CHIETI - DRIVE IN LANCIANO | Istituto Zooprofilattico Sperimentale dell'Abruzzo e Molise "G. Caporale" | Lorusso A, Marcacci M, Di Domenico M, Ancora M, Curini V, Di Lollo Valeria, Mangone I, Rinaldi A, Delli Compagni E, Scialabba S, Caporale M, Di Pasquale A, Cammà C, Puglia I, Calistri P, Savini G |
| EPI_ISL_2483659 | SIESP DIPARTIMENTO DI PREVENZIONE CHIETI | Istituto Zooprofilattico Sperimentale dell'Abruzzo e Molise "G. Caporale" | Lorusso A, Marcacci M, Di Domenico M, Ancora M, Curini V, Di Lollo Valeria, Mangone I, Rinaldi A, Delli Compagni E, Scialabba S, Caporale M, Di Pasquale A, Cammà C, Puglia I, Calistri P, Savini G |
| EPI_ISL_2483660 | SIESP DIPARTIMENTO DI PREVENZIONE TERAMO | Istituto Zooprofilattico Sperimentale dell'Abruzzo e Molise "G. Caporale" | Lorusso A, Marcacci M, Di Domenico M, Ancora M, Curini V, Di Lollo Valeria, Mangone I, Rinaldi A, Delli Compagni E, Scialabba S, Caporale M, Di Pasquale A, Cammà C, Puglia I, Calistri P, Savini G |
| EPI_ISL_2483661 | SIESP CHIETI - DRIVE IN LANCIANO | Istituto Zooprofilattico Sperimentale dell'Abruzzo e Molise "G. Caporale" | Lorusso A, Marcacci M, Di Domenico M, Ancora M, Curini V, Di Lollo Valeria, Mangone I, Rinaldi A, Delli Compagni E, Scialabba S, Caporale M, Di Pasquale A, Cammà C, Puglia I, Calistri P, Savini G |
| EPI_ISL_2483662, EPI_ISL_2483663 | SIESP DIPARTIMENTO DI PREVENZIONE CHIETI | Istituto Zooprofilattico Sperimentale dell'Abruzzo e Molise "G. Caporale" | Lorusso A, Marcacci M, Di Domenico M, Ancora M, Curini V, Di Lollo Valeria, Mangone I, Rinaldi A, Delli Compagni E, Scialabba S, Caporale M, Di Pasquale A, Cammà C, Puglia I, Calistri P, Savini G |
| EPI_ISL_2483664 | SIESP CHIETI - DRIVE IN LANCIANO | Istituto Zooprofilattico Sperimentale dell'Abruzzo e Molise "G. Caporale" | Lorusso A, Marcacci M, Di Domenico M, Ancora M, Curini V, Di Lollo Valeria, Mangone I, Rinaldi A, Delli Compagni E, Scialabba S, Caporale M, Di Pasquale A, Cammà C, Puglia I, Calistri P, Savini G |
| EPI_ISL_2483665 | SIESP DIPARTIMENTO DI PREVENZIONE TERAMO | Istituto Zooprofilattico Sperimentale dell'Abruzzo e Molise "G. Caporale" | Lorusso A, Marcacci M, Di Domenico M, Ancora M, Curini V, Di Lollo Valeria, Mangone I, Rinaldi A, Delli Compagni E, Scialabba S, Caporale M, Di Pasquale A, Cammà C, Puglia I, Calistri P, Savini G |
| EPI_ISL_2483666, EPI_ISL_2483667 | SIESP CHIETI - DRIVE IN CHIETI | Istituto Zooprofilattico Sperimentale dell'Abruzzo e Molise "G. Caporale" | Lorusso A, Marcacci M, Di Domenico M, Ancora M, Curini V, Di Lollo Valeria, Mangone I, Rinaldi A, Delli Compagni E, Scialabba S, Caporale M, Di Pasquale A, Cammà C, Puglia I, Calistri P, Savini G |
| EPI_ISL_2483668 | SIESP CHIETI - DRIVE IN LANCIANO | Istituto Zooprofilattico Sperimentale dell'Abruzzo e Molise "G. Caporale" | Lorusso A, Marcacci M, Di Domenico M, Ancora M, Curini V, Di Lollo Valeria, Mangone I, Rinaldi A, Delli Compagni E, Scialabba S, Caporale M, Di Pasquale A, Cammà C, Puglia I, Calistri P, Savini G |
| EPI_ISL_2483669, EPI_ISL_2483670 | SIESP CHIETI - DRIVE IN VASTO | Istituto Zooprofilattico Sperimentale dell'Abruzzo e Molise "G. Caporale" | Lorusso A, Marcacci M, Di Domenico M, Ancora M, Curini V, Di Lollo Valeria, Mangone I, Rinaldi A, Delli Compagni E, Scialabba S, Caporale M, Di Pasquale A, Cammà C, Puglia I, Calistri P, Savini G |
| EPI_ISL_2483671 | RSA GIULIANOVA | Istituto Zooprofilattico Sperimentale dell'Abruzzo e Molise "G. Caporale" | Lorusso A, Marcacci M, Di Domenico M, Ancora M, Curini V, Di Lollo Valeria, Mangone I, Rinaldi A, Delli Compagni E, Scialabba S, Caporale M, Di Pasquale A, Cammà C, Puglia I, Calistri P, Savini G |
| EPI_ISL_2483672 | SIESP CHIETI - DRIVE IN ORTONA | Istituto Zooprofilattico Sperimentale dell'Abruzzo e Molise "G. Caporale" | Lorusso A, Marcacci M, Di Domenico M, Ancora M, Curini V, Di Lollo Valeria, Mangone I, Rinaldi A, Delli Compagni E, Scialabba S, Caporale M, Di Pasquale A, Cammà C, Puglia I, Calistri P, Savini G |
| EPI_ISL_2483673, EPI_ISL_2483674 | SIESP CHIETI - DRIVE IN VASTO | Istituto Zooprofilattico Sperimentale dell'Abruzzo e Molise "G. Caporale" | Lorusso A, Marcacci M, Di Domenico M, Ancora M, Curini V, Di Lollo Valeria, Mangone I, Rinaldi A, Delli Compagni E, Scialabba S, Caporale M, Di Pasquale A, Cammà C, Puglia I, Calistri P, Savini G |
| EPI_ISL_2483675 | SIESP DIPARTIMENTO DI PREVENZIONE CHIETI | Istituto Zooprofilattico Sperimentale dell'Abruzzo e Molise "G. Caporale" | Lorusso A, Marcacci M, Di Domenico M, Ancora M, Curini V, Di Lollo Valeria, Mangone I, Rinaldi A, Delli Compagni E, Scialabba S, Caporale M, Di Pasquale A, Cammà C, Puglia I, Calistri P, Savini G |
| EPI_ISL_2483676 | SIESP CHIETI - DRIVE IN VASTO | Istituto Zooprofilattico Sperimentale dell'Abruzzo e Molise "G. Caporale" | Lorusso A, Marcacci M, Di Domenico M, Ancora M, Curini V, Di Lollo Valeria, Mangone I, Rinaldi A, Delli Compagni E, Scialabba S, Caporale M, Di Pasquale A, Cammà C, Puglia I, Calistri P, Savini G |
| EPI_ISL_2483677 | SIESP DIPARTIMENTO DI PREVENZIONE CHIETI | Istituto Zooprofilattico Sperimentale dell'Abruzzo e Molise "G. Caporale" | Lorusso A, Marcacci M, Di Domenico M, Ancora M, Curini V, Di Lollo Valeria, Mangone I, Rinaldi A, Delli Compagni E, Scialabba S, Caporale M, Di Pasquale A, Cammà C, Puglia I, Calistri P, Savini G |
| EPI_ISL_2483678 | SIESP CHIETI - DRIVE IN CHIETI | Istituto Zooprofilattico Sperimentale dell'Abruzzo e Molise "G. Caporale" | Lorusso A, Marcacci M, Di Domenico M, Ancora M, Curini V, Di Lollo Valeria, Mangone I, Rinaldi A, Delli Compagni E, Scialabba S, Caporale M, Di Pasquale A, Cammà C, Puglia I, Calistri P, Savini G |
| EPI_ISL_2483679 | SIESP DIPARTIMENTO DI PREVENZIONE TERAMO | Istituto Zooprofilattico Sperimentale dell'Abruzzo e Molise "G. Caporale" | Lorusso A, Marcacci M, Di Domenico M, Ancora M, Curini V, Di Lollo Valeria, Mangone I, Rinaldi A, Delli Compagni E, Scialabba S, Caporale M, Di Pasquale A, Cammà C, Puglia I, Calistri P, Savini G |
| EPI_ISL_2483680 | PRESIDIO OSPEDALIERO SULMONA | Istituto Zooprofilattico Sperimentale dell'Abruzzo e Molise "G. Caporale" | Lorusso A, Marcacci M, Di Domenico M, Ancora M, Curini V, Di Lollo Valeria, Mangone I, Rinaldi A, Delli Compagni E, Scialabba S, Caporale M, Di Pasquale A, Cammà C, Puglia I, Calistri P, Savini G |
| EPI_ISL_2483681 | USCA Sulmona | Istituto Zooprofilattico Sperimentale dell'Abruzzo e Molise "G. Caporale" | Lorusso A, Marcacci M, Di Domenico M, Ancora M, Curini V, Di Lollo Valeria, Mangone I, Rinaldi A, Delli Compagni E, Scialabba S, Caporale M, Di Pasquale A, Cammà C, Puglia I, Calistri P, Savini G |
| EPI_ISL_2483682, EPI_ISL_2483683, EPI_ISL_2483684, EPI_ISL_2483685 | SIESP DIPARTIMENTO DI PREVENZIONE CHIETI | Istituto Zooprofilattico Sperimentale dell'Abruzzo e Molise "G. Caporale" | Lorusso A, Marcacci M, Di Domenico M, Ancora M, Curini V, Di Lollo Valeria, Mangone I, Rinaldi A, Delli Compagni E, Scialabba S, Caporale M, Di Pasquale A, Cammà C, Puglia I, Calistri P, Savini G |
| EPI_ISL_2483686 | SIESP DIPARTIMENTO DI PREVENZIONE TERAMO | Istituto Zooprofilattico Sperimentale dell'Abruzzo e Molise "G. Caporale" | Lorusso A, Marcacci M, Di Domenico M, Ancora M, Curini V, Di Lollo Valeria, Mangone I, Rinaldi A, Delli Compagni E, Scialabba S, Caporale M, Di Pasquale A, Cammà C, Puglia I, Calistri P, Savini G |
| EPI_ISL_2483687, EPI_ISL_2483688, EPI_ISL_2483689 | SIESP CHIETI - DRIVE IN VASTO | Istituto Zooprofilattico Sperimentale dell'Abruzzo e Molise "G. Caporale" | Lorusso A, Marcacci M, Di Domenico M, Ancora M, Curini V, Di Lollo Valeria, Mangone I, Rinaldi A, Delli Compagni E, Scialabba S, Caporale M, Di Pasquale A, Cammà C, Puglia I, Calistri P, Savini G |
| EPI_ISL_2483690 | RSA GIULIANOVA | Istituto Zooprofilattico Sperimentale dell'Abruzzo e Molise "G. Caporale" | Lorusso A, Marcacci M, Di Domenico M, Ancora M, Curini V, Di Lollo Valeria, Mangone I, Rinaldi A, Delli Compagni E, Scialabba S, Caporale M, Di Pasquale A, Cammà C, Puglia I, Calistri P, Savini G |
| EPI_ISL_2483691 | SIESP CHIETI - DRIVE IN VASTO | Istituto Zooprofilattico Sperimentale dell'Abruzzo e Molise "G. | Lorusso A, Marcacci M, Di Domenico M, Ancora M, Curini V, Di Lollo Valeria, Mangone I, Rinaldi A, Delli Compagni E, Scialabba S, Caporale M, Di |

| | | | |
|---|---|---|---|
| | | Caporale" | Pasquale A, Cammà C, Puglia I, Calistri P, Savini G |
| EPI_ISL_2483692, EPI_ISL_2483693 | SIESP CHIETI - DRIVE IN CHIETI | Istituto Zooprofilattico Sperimentale dell'Abruzzo e Molise "G. Caporale" | Lorusso A, Marcacci M, Di Domenico M, Ancora M, Curini V, Di Lollo Valeria, Mangone I, Rinaldi A, Delli Compagni E, Scialabba S, Caporale M, Di Pasquale A, Cammà C, Puglia I, Calistri P, Savini G |
| EPI_ISL_2483694 | SIESP CHIETI - DRIVE IN VASTO | Istituto Zooprofilattico Sperimentale dell'Abruzzo e Molise "G. Caporale" | Lorusso A, Marcacci M, Di Domenico M, Ancora M, Curini V, Di Lollo Valeria, Mangone I, Rinaldi A, Delli Compagni E, Scialabba S, Caporale M, Di Pasquale A, Cammà C, Puglia I, Calistri P, Savini G |
| EPI_ISL_2483695 | SIESP CHIETI - DRIVE IN ORTONA | Istituto Zooprofilattico Sperimentale dell'Abruzzo e Molise "G. Caporale" | Lorusso A, Marcacci M, Di Domenico M, Ancora M, Curini V, Di Lollo Valeria, Mangone I, Rinaldi A, Delli Compagni E, Scialabba S, Caporale M, Di Pasquale A, Cammà C, Puglia I, Calistri P, Savini G |
| EPI_ISL_2483696, EPI_ISL_2483697 | SIESP DIPARTIMENTO DI PREVENZIONE CHIETI | Istituto Zooprofilattico Sperimentale dell'Abruzzo e Molise "G. Caporale" | Lorusso A, Marcacci M, Di Domenico M, Ancora M, Curini V, Di Lollo Valeria, Mangone I, Rinaldi A, Delli Compagni E, Scialabba S, Caporale M, Di Pasquale A, Cammà C, Puglia I, Calistri P, Savini G |
| EPI_ISL_2483698, EPI_ISL_2483699 | SIESP CHIETI - DRIVE IN VASTO | Istituto Zooprofilattico Sperimentale dell'Abruzzo e Molise "G. Caporale" | Lorusso A, Marcacci M, Di Domenico M, Ancora M, Curini V, Di Lollo Valeria, Mangone I, Rinaldi A, Delli Compagni E, Scialabba S, Caporale M, Di Pasquale A, Cammà C, Puglia I, Calistri P, Savini G |
| EPI_ISL_2483701, EPI_ISL_2483702 | SIESP DIPARTIMENTO DI PREVENZIONE TERAMO | Istituto Zooprofilattico Sperimentale dell'Abruzzo e Molise "G. Caporale" | Lorusso A, Marcacci M, Di Domenico M, Ancora M, Curini V, Di Lollo Valeria, Mangone I, Rinaldi A, Delli Compagni E, Scialabba S, Caporale M, Di Pasquale A, Cammà C, Puglia I, Calistri P, Savini G |
| EPI_ISL_2483813, EPI_ISL_2483814, EPI_ISL_2483815, EPI_ISL_2483816, EPI_ISL_2483817, EPI_ISL_2483818, EPI_ISL_2483819, EPI_ISL_2483820, EPI_ISL_2483821, EPI_ISL_2483822, EPI_ISL_2483823, EPI_ISL_2483824, EPI_ISL_2483825, EPI_ISL_2483826, EPI_ISL_2483827, EPI_ISL_2483828, EPI_ISL_2483829, EPI_ISL_2483830, EPI_ISL_2483831, EPI_ISL_2483832, EPI_ISL_2483833 | | | |
| see above | Klinisch Laboratorium ZNA | Klinisch Laboratorium ZNA | Verstrepen et al. |
| EPI_ISL_2484166 | LESP Zacatecas | Instituto de Diagnostico y Referencia Epidemiologicos (INDRE) | Claudia Wong-Arambula, Abril Rodriguez-Maldonado, Vanessa Rivero-Arredondo, Ariadna Medina-Benitez, Joaquin Quiroz-Mercado, Sergio Rangel-Guerrero, Natividad Cruz-Ortiz, Tatiana Nunez-Garcia, Gisela Barrera-Badillo, Lucia Hernandez-Rivas, Irma Lopez-Martinez, Ernesto Ramirez-Gonzalez. |
| EPI_ISL_2484276 | AZ Zeno Knokke-Blankenberge | AZ Sint Jan | Marijke Reynders, Patrick Descheemaeker, Laurien Hoornaert, Thomas Van Landschoot, Marie Madeleine Chabert-Consen, Jorn Hellemans |
| EPI_ISL_2484602, EPI_ISL_2484616, EPI_ISL_2484617, EPI_ISL_2484632, EPI_ISL_2484633, EPI_ISL_2484635, EPI_ISL_2484636, EPI_ISL_2484637, EPI_ISL_2484641, EPI_ISL_2484642, EPI_ISL_2484643 | | | |
| see above | NL-Dr. Leonard A. Miller Centre for Health Services | National Microbiology Laboratory (NML) | Anna Majer, Shari Tyson, Grace Seo, Philip Mabon, Elsie Grudeski, Rhiannon Huzarewich, Russell Mandes, Anneliese Landgraff, Jennifer Tanner, Natalie Knox, Morag Graham, Gary Van Domselaar, Robert Needle, Yang Yu, Adel Malek, Laura Gilbert, George Zahariadis, Nathalie Bastien, Yan Li, Timothy Booth, Darian Hole, Madison Chapel, Kirsten Biggar, Kerri Smith, CanCOGeN's metadata curation team, Public Health Agency of Canada CanCOGeN team |
| EPI_ISL_2484967, EPI_ISL_2484968, EPI_ISL_2484969, EPI_ISL_2484970, EPI_ISL_2484972, EPI_ISL_2484973, EPI_ISL_2484974, EPI_ISL_2484975, EPI_ISL_2484976, EPI_ISL_2484977, EPI_ISL_2484978, EPI_ISL_2484979, EPI_ISL_2484980, EPI_ISL_2484983, EPI_ISL_2484984, EPI_ISL_2484985, EPI_ISL_2484986, EPI_ISL_2484988, EPI_ISL_2484989, EPI_ISL_2484990, EPI_ISL_2484991, EPI_ISL_2484992, EPI_ISL_2484993, EPI_ISL_2484994, EPI_ISL_2484996, EPI_ISL_2484997, EPI_ISL_2484998, EPI_ISL_2485000, EPI_ISL_2485001, EPI_ISL_2485002, EPI_ISL_2485003, EPI_ISL_2485004, EPI_ISL_2485005, EPI_ISL_2485007, EPI_ISL_2485008, EPI_ISL_2485009, EPI_ISL_2485010, EPI_ISL_2485011, EPI_ISL_2485012, EPI_ISL_2485013, EPI_ISL_2485014, EPI_ISL_2485015, EPI_ISL_2485016, EPI_ISL_2485017, EPI_ISL_2485018, EPI_ISL_2485019, EPI_ISL_2485020, EPI_ISL_2485022, EPI_ISL_2485024, EPI_ISL_2485025, EPI_ISL_2485026, EPI_ISL_2485027, EPI_ISL_2485028, EPI_ISL_2485029, EPI_ISL_2485030, EPI_ISL_2485031, EPI_ISL_2485032, EPI_ISL_2485033, EPI_ISL_2485034, EPI_ISL_2485038, EPI_ISL_2485039, EPI_ISL_2485040, EPI_ISL_2485041, EPI_ISL_2485042, EPI_ISL_2485043, EPI_ISL_2485044, EPI_ISL_2485045, EPI_ISL_2485046, EPI_ISL_2485047, EPI_ISL_2485048, EPI_ISL_2485049, EPI_ISL_2485050, EPI_ISL_2485051, EPI_ISL_2485052, EPI_ISL_2485055, EPI_ISL_2485056, EPI_ISL_2485057, EPI_ISL_2485059, EPI_ISL_2485062, EPI_ISL_2485063, EPI_ISL_2485064, EPI_ISL_2485065, EPI_ISL_2485066, EPI_ISL_2485067, EPI_ISL_2485068, EPI_ISL_2485069, EPI_ISL_2485070, EPI_ISL_2485071, EPI_ISL_2485072, EPI_ISL_2485073, EPI_ISL_2485074, EPI_ISL_2485075, EPI_ISL_2485077, EPI_ISL_2485079, EPI_ISL_2485080, EPI_ISL_2485081, EPI_ISL_2485083, EPI_ISL_2485084, EPI_ISL_2485085, EPI_ISL_2485087, EPI_ISL_2485088, EPI_ISL_2485089, EPI_ISL_2485090, EPI_ISL_2485091, EPI_ISL_2485092, EPI_ISL_2485093, EPI_ISL_2485094, EPI_ISL_2485095, EPI_ISL_2485096, EPI_ISL_2485097, EPI_ISL_2485098, EPI_ISL_2485099, EPI_ISL_2485100, EPI_ISL_2485101, EPI_ISL_2485102, EPI_ISL_2485103, EPI_ISL_2485104, EPI_ISL_2485105, EPI_ISL_2485106, EPI_ISL_2485107, EPI_ISL_2485108, EPI_ISL_2485109, EPI_ISL_2485110, EPI_ISL_2485111, EPI_ISL_2485112, EPI_ISL_2485113, EPI_ISL_2485114, EPI_ISL_2485115, EPI_ISL_2485116, EPI_ISL_2485117, EPI_ISL_2485118, EPI_ISL_2485119, EPI_ISL_2485120, EPI_ISL_2485121, EPI_ISL_2485122, EPI_ISL_2485123, EPI_ISL_2485124, EPI_ISL_2485125, EPI_ISL_2485126, EPI_ISL_2485128, EPI_ISL_2485129, EPI_ISL_2485132, EPI_ISL_2485134, EPI_ISL_2485135, EPI_ISL_2485136, EPI_ISL_2485137, EPI_ISL_2485138, EPI_ISL_2485140, EPI_ISL_2485141, EPI_ISL_2485142, EPI_ISL_2485143, EPI_ISL_2485144, EPI_ISL_2485145, EPI_ISL_2485146, EPI_ISL_2485147, EPI_ISL_2485148, EPI_ISL_2485149, EPI_ISL_2485150, EPI_ISL_2485151, EPI_ISL_2485152, EPI_ISL_2485153, EPI_ISL_2485154, EPI_ISL_2485155, EPI_ISL_2485157, EPI_ISL_2485158, EPI_ISL_2485160, EPI_ISL_2485163, EPI_ISL_2485165, EPI_ISL_2485166, EPI_ISL_2485167, EPI_ISL_2485168 | | | |
| see above | Austrian Agency for Health and Food Safety (AGES) | Bergthaler laboratory, CeMM Research Center for Molecular Medicine of the Austrian Academy of Sciences | Lukas Endler, Anna Schedl, Fabian Amman, Petr Triska, Thomas Penz, Benedikt Agerer, Maelle Le Moing, Michael Schuster, Bekir Erguner, Jan Laine, Martin Senekowitsch, Christoph Bock, Andreas Bergthaler |
| EPI_ISL_2485169 | 3. Medizinische Abteilung, Hanusch Krankenhaus | Bergthaler laboratory, CeMM Research Center for Molecular Medicine of the Austrian Academy of Sciences | Lukas Endler, Anna Schedl, Fabian Amman, Petr Triska, Thomas Penz, Benedikt Agerer, Maelle Le Moing, Michael Schuster, Bekir Erguner, Jan Laine, Martin Senekowitsch, Christoph Bock, Andreas Bergthaler |
| EPI_ISL_2488813 | Evandro Chagas Institute | Evandro Chagas Institute | Santos, M.C.; Silva, A.M.; Junior, W.D.C.; Barbagelata, L.S.; Ferreira, J.A.; Sousa, E.M.A.; da Silva, P.S.; Pinheiro, K.C.; L.C.; Sousa Junior, E.C. |
| EPI_ISL_2490307, EPI_ISL_2490308, EPI_ISL_2490309 | Basurto University Hospital: Clinical Microbiology Laboratory | Biocruces Bizkaia | Mikel Urrutikoetxea-Gutiérrez, Mª Carmen Nieto Toboso, Estibaliz Ugalde Zarraga, Mikel Gallego Rodrigo, Ana de la Hoz, José Luis Díaz de Tuesta del Arco |
| EPI_ISL_2490313, EPI_ISL_2490315, EPI_ISL_2490316, EPI_ISL_2490320, EPI_ISL_2490321, EPI_ISL_2490322, EPI_ISL_2490325, EPI_ISL_2490326 | Centro de Investigación Biomédica del Noreste (CIBIN) | Centro de Investigación en Enfermedades Infecciosas (CIENI), Instituto Nacional de Enfermedades Respiratorias (INER) | Consorcio Mexicano de Vigilancia Genómica (CoViGen-Mex). Authors (in alphabetical order): Julio Elias Alvarado-Yaah, Carlos F. Arias, Santiago Ávila-Ríos, Eduardo Becerril-Vargas, Víctor Hugo Borja-Aburto, Celia Boukadida, Cristóbal Cháidez-Quiróz, Juan Bautista Chale-Dzul, Ricardo Ciria Merce, Andreu Comas-García, Célida Duque Molina, Julissa Enciso-Ibarra, José Antonio Enciso-Moreno, Gloria Elena Espinosa-Ayala, Fernando Fontove-Herrera, Daniel Fregoso-Rueda, Victor Eduardo Garcia-Arias, Alejandra Garcia-Gasca, Bruno Gomez-Gil, Jean Pierre González, Irvin González-López, Concepción Grajales-Muñiz, Ricardo Grande, Rosa María Gutierrez Rios, Alejandra Hernández-Terán, Alfredo Herrera-Estrella, Carla Ivón Herrera-Najera, Pavel Isa, Daniel Lira Morales, Susana Lopez, Antonio Loza Román, Brenda Irasema Maldonado-Meza, Bernardo Martínez-Miguel, José Arturo Martínez-Orozco, Célida Martínez- Rodríguez, Margarita Matías-Florentino, Fidencio Mejía-Nepomuceno, María Guadalupe de Jesús Mireles-Rivera, Gloria María Molina-Salinas, Hector Montoya-Fuentes, , Mario Mújica-Sánchez, José Esteban Muñoz-Medina, José de Jesús Nuñez-Contreras, Alicia Ocaña-Mondragón, Luis Alberto Ochoa-Carrera, Hector Esteban Paz-Juárez, Marissa Perez-Garcia, Francisco Pulido, Helen Haydee Fernanda Ramírez-Plascencia, Jorge Salas-Hernández, Angel Gustavo Salas-Lais, Alejandro Sanchez-Flores, Clara Esperanza Santacruz-Tinoco, María Guadalupe Santiago-Mauricio, Selene Zárate, Nelly Sélem-Mojica, Blanca Taboada, Gloria Vazquez, Joel Armando Vázquez-Pérez. |
| EPI_ISL_2490328, EPI_ISL_2490329, EPI_ISL_2490330 | Unidad de Investigación Médica de Yucatán (UIMY) | Centro de Investigación en Enfermedades Infecciosas (CIENI), Instituto Nacional de Enfermedades Respiratorias (INER) | Consorcio Mexicano de Vigilancia Genómica (CoViGen-Mex). Authors (in alphabetical order): Julio Elias Alvarado-Yaah, Carlos F. Arias, Santiago Ávila-Ríos, Eduardo Becerril-Vargas, Víctor Hugo Borja-Aburto, Celia Boukadida, Cristóbal Cháidez-Quiróz, Juan Bautista Chale-Dzul, Ricardo Ciria Merce, Andreu Comas-García, Célida Duque Molina, Julissa Enciso-Ibarra, José Antonio Enciso-Moreno, Gloria Elena Espinosa-Ayala, Fernando Fontove-Herrera, Daniel Fregoso-Rueda, Victor Eduardo Garcia-Arias, Alejandra Garcia-Gasca, Bruno Gomez-Gil, Jean Pierre González, Irvin González-López, Concepción Grajales-Muñiz, Ricardo Grande, Rosa María Gutierrez Rios, Alejandra Hernández-Terán, Alfredo Herrera-Estrella, Carla Ivón Herrera-Najera, Pavel Isa, Daniel Lira Morales, Susana Lopez, Antonio Loza Román, Brenda Irasema Maldonado-Meza, Bernardo Martínez-Miguel, José Arturo Martínez-Orozco, Célida Martínez- Rodríguez, Margarita Matías-Florentino, Fidencio Mejía-Nepomuceno, María Guadalupe de Jesús Mireles-Rivera, Gloria María Molina-Salinas, Hector Montoya-Fuentes, , Mario Mújica-Sánchez, José Esteban Muñoz-Medina, José de Jesús Nuñez-Contreras, Alicia Ocaña-Mondragón, Luis Alberto Ochoa-Carrera, Hector Esteban Paz-Juárez, Marissa Perez-Garcia, Francisco Pulido, Helen Haydee Fernanda Ramírez-Plascencia, Jorge Salas-Hernández, Angel Gustavo Salas-Lais, Alejandro Sanchez-Flores, Clara Esperanza Santacruz-Tinoco, María Guadalupe Santiago-Mauricio, Selene Zárate, Nelly Sélem-Mojica, Blanca Taboada, Gloria Vazquez, Joel Armando Vázquez-Pérez. |
| EPI_ISL_2490331, EPI_ISL_2490332 | Centro de Investigación Biomédica de Occidente (CIBO) | Centro de Investigación en Enfermedades Infecciosas (CIENI), Instituto Nacional de Enfermedades Respiratorias (INER) | Consorcio Mexicano de Vigilancia Genómica (CoViGen-Mex). Authors (in alphabetical order): Julio Elias Alvarado-Yaah, Carlos F. Arias, Santiago Ávila-Ríos, Eduardo Becerril-Vargas, Víctor Hugo Borja-Aburto, Celia Boukadida, Cristóbal Cháidez-Quiróz, Juan Bautista Chale-Dzul, Ricardo Ciria Merce, Andreu Comas-García, Célida Duque Molina, Julissa Enciso-Ibarra, José Antonio Enciso-Moreno, Gloria Elena Espinosa-Ayala, Fernando Fontove-Herrera, Daniel Fregoso-Rueda, Victor Eduardo Garcia-Arias, Alejandra Garcia-Gasca, Bruno Gomez-Gil, Jean Pierre González, Irvin González-López, Concepción Grajales-Muñiz, Ricardo Grande, Rosa María Gutierrez Rios, Alejandra Hernández-Terán, Alfredo Herrera-Estrella, Carla Ivón Herrera-Najera, Pavel Isa, Daniel Lira Morales, Susana Lopez, Antonio Loza Román, Brenda Irasema Maldonado-Meza, Bernardo Martínez-Miguel, José Arturo Martínez-Orozco, Célida Martínez- Rodríguez, Margarita Matías-Florentino, Fidencio Mejía-Nepomuceno, María Guadalupe de Jesús Mireles-Rivera, Gloria María Molina-Salinas, Hector Montoya-Fuentes, , Mario Mújica-Sánchez, José Esteban Muñoz-Medina, José de Jesús Nuñez-Contreras, Alicia Ocaña-Mondragón, Luis Alberto Ochoa-Carrera, Hector Esteban Paz-Juárez, Marissa Perez-Garcia, Francisco Pulido, Helen Haydee Fernanda Ramírez-Plascencia, Jorge Salas-Hernández, Angel Gustavo Salas-Lais, Alejandro Sanchez-Flores, Clara Esperanza |

| | | | |
|---|---|---|---|
| | | | Santacruz-Tinoco, María Guadalupe Santiago-Mauricio, Selene Zárate, Nelly Sélem-Mojica, Blanca Taboada, Gloria Vazquez, Joel Armando Vázquez-Pérez. |
| EPI_ISL_2490333, EPI_ISL_2490335 | Unidad de Investigación Biomédica de Zacatecas (UIBZ) | Centro de Investigación en Enfermedades Infecciosas (CIENI), Instituto Nacional de Enfermedades Respiratorias (INER) | Consorcio Mexicano de Vigilancia Genómica (CoViGen-Mex). Authors (in alphabetical order): Julio Elias Alvarado-Yaah, Carlos F. Arias, Santiago Ávila-Ríos, Eduardo Becerril-Vargas, Víctor Hugo Borja-Aburto, Celia Boukadida, Cristóbal Cháidez-Quiróz, Juan Bautista Chale-Dzul, Ricardo Ciria Merce, Andreu Comas-García, Célida Duque Molina, Julissa Enciso-Ibarra, José Antonio Enciso-Moreno, Gloria Elena Espinosa-Ayala, Fernando Fontove-Herrera, Daniel Fregoso-Rueda, Victor Eduardo Garcia-Arias, Alejandra Garcia-Gasca, Bruno Gomez-Gil, Jean Pierre González, Irvin González-López, Concepción Grajales-Muñiz, Ricardo Grande, Rosa María Gutierrez Rios, Alejandra Hernández-Terán, Alfredo Herrera-Estrella, Carla Ivón Herrera-Najera, Pavel Isa, Daniel Lira Morales, Susana Lopez, Antonio Loza Román, Brenda Irasema Maldonado-Meza, Bernardo Martínez-Miguel, José Arturo Martínez-Orozco, Célida Martínez- Rodríguez, Margarita Matías-Florentino, Fidencio Mejía-Nepomuceno, María Guadalupe de Jesús Mireles-Rivera, Gloria María Molina-Salinas, Hector Montoya-Fuentes, , Mario Mújica-Sánchez, José Esteban Muñoz-Medina, José de Jesús Nuñez-Contreras, Alicia Ocaña-Mondragón, Luis Alberto Ochoa-Carrera, Hector Esteban Paz-Juárez, Marissa Perez-Garcia, Francisco Pulido, Helen Haydee Fernanda Ramírez-Plascencia, Jorge Salas-Hernández, Angel Gustavo Salas-Lais, Alejandro Sanchez-Flores, Clara Esperanza Santacruz-Tinoco, María Guadalupe Santiago-Mauricio, Selene Zárate, Nelly Sélem-Mojica, Blanca Taboada, Gloria Vazquez, Joel Armando Vázquez-Pérez. |
| EPI_ISL_2490336 | Laboratorio Central de Epidemiología (LCE) | Centro de Investigación en Enfermedades Infecciosas (CIENI), Instituto Nacional de Enfermedades Respiratorias (INER) | Consorcio Mexicano de Vigilancia Genómica (CoViGen-Mex). Authors (in alphabetical order): Julio Elias Alvarado-Yaah, Carlos F. Arias, Santiago Ávila-Ríos, Eduardo Becerril-Vargas, Víctor Hugo Borja-Aburto, Celia Boukadida, Cristóbal Cháidez-Quiróz, Juan Bautista Chale-Dzul, Ricardo Ciria Merce, Andreu Comas-García, Célida Duque Molina, Julissa Enciso-Ibarra, José Antonio Enciso-Moreno, Gloria Elena Espinosa-Ayala, Fernando Fontove-Herrera, Daniel Fregoso-Rueda, Victor Eduardo Garcia-Arias, Alejandra Garcia-Gasca, Bruno Gomez-Gil, Jean Pierre González, Irvin González-López, Concepción Grajales-Muñiz, Ricardo Grande, Rosa María Gutierrez Rios, Alejandra Hernández-Terán, Alfredo Herrera-Estrella, Carla Ivón Herrera-Najera, Pavel Isa, Daniel Lira Morales, Susana Lopez, Antonio Loza Román, Brenda Irasema Maldonado-Meza, Bernardo Martínez-Miguel, José Arturo Martínez-Orozco, Célida Martínez- Rodríguez, Margarita Matías-Florentino, Fidencio Mejía-Nepomuceno, María Guadalupe de Jesús Mireles-Rivera, Gloria María Molina-Salinas, Hector Montoya-Fuentes, , Mario Mújica-Sánchez, José Esteban Muñoz-Medina, José de Jesús Nuñez-Contreras, Alicia Ocaña-Mondragón, Luis Alberto Ochoa-Carrera, Hector Esteban Paz-Juárez, Marissa Perez-Garcia, Francisco Pulido, Helen Haydee Fernanda Ramírez-Plascencia, Jorge Salas-Hernández, Angel Gustavo Salas-Lais, Alejandro Sanchez-Flores, Clara Esperanza Santacruz-Tinoco, María Guadalupe Santiago-Mauricio, Selene Zárate, Nelly Sélem-Mojica, Blanca Taboada, Gloria Vazquez, Joel Armando Vázquez-Pérez. |
| EPI_ISL_2490339, EPI_ISL_2490340, EPI_ISL_2490341, EPI_ISL_2490342 | Centro de Investigación Biomédica del Noreste (CIBIN) | Centro de Investigación en Enfermedades Infecciosas (CIENI), Instituto Nacional de Enfermedades Respiratorias (INER) | Consorcio Mexicano de Vigilancia Genómica (CoViGen-Mex). Authors (in alphabetical order): Julio Elias Alvarado-Yaah, Carlos F. Arias, Santiago Ávila-Ríos, Eduardo Becerril-Vargas, Víctor Hugo Borja-Aburto, Celia Boukadida, Cristóbal Cháidez-Quiróz, Juan Bautista Chale-Dzul, Ricardo Ciria Merce, Andreu Comas-García, Célida Duque Molina, Julissa Enciso-Ibarra, José Antonio Enciso-Moreno, Gloria Elena Espinosa-Ayala, Fernando Fontove-Herrera, Daniel Fregoso-Rueda, Victor Eduardo Garcia-Arias, Alejandra Garcia-Gasca, Bruno Gomez-Gil, Jean Pierre González, Irvin González-López, Concepción Grajales-Muñiz, Ricardo Grande, Rosa María Gutierrez Rios, Alejandra Hernández-Terán, Alfredo Herrera-Estrella, Carla Ivón Herrera-Najera, Pavel Isa, Daniel Lira Morales, Susana Lopez, Antonio Loza Román, Brenda Irasema Maldonado-Meza, Bernardo Martínez-Miguel, José Arturo Martínez-Orozco, Célida Martínez- Rodríguez, Margarita Matías-Florentino, Fidencio Mejía-Nepomuceno, María Guadalupe de Jesús Mireles-Rivera, Gloria María Molina-Salinas, Hector Montoya-Fuentes, , Mario Mújica-Sánchez, José Esteban Muñoz-Medina, José de Jesús Nuñez-Contreras, Alicia Ocaña-Mondragón, Luis Alberto Ochoa-Carrera, Hector Esteban Paz-Juárez, Marissa Perez-Garcia, Francisco Pulido, Helen Haydee Fernanda Ramírez-Plascencia, Jorge Salas-Hernández, Angel Gustavo Salas-Lais, Alejandro Sanchez-Flores, Clara Esperanza Santacruz-Tinoco, María Guadalupe Santiago-Mauricio, Selene Zárate, Nelly Sélem-Mojica, Blanca Taboada, Gloria Vazquez, Joel Armando Vázquez-Pérez. |
| EPI_ISL_2490347 | Unidad de Investigación Médica de Yucatán (UIMY) | Centro de Investigación en Enfermedades Infecciosas (CIENI), Instituto Nacional de Enfermedades Respiratorias (INER) | Consorcio Mexicano de Vigilancia Genómica (CoViGen-Mex). Authors (in alphabetical order): Julio Elias Alvarado-Yaah, Carlos F. Arias, Santiago Ávila-Ríos, Eduardo Becerril-Vargas, Víctor Hugo Borja-Aburto, Celia Boukadida, Cristóbal Cháidez-Quiróz, Juan Bautista Chale-Dzul, Ricardo Ciria Merce, Andreu Comas-García, Célida Duque Molina, Julissa Enciso-Ibarra, José Antonio Enciso-Moreno, Gloria Elena Espinosa-Ayala, Fernando Fontove-Herrera, Daniel Fregoso-Rueda, Victor Eduardo Garcia-Arias, Alejandra Garcia-Gasca, Bruno Gomez-Gil, Jean Pierre González, Irvin González-López, Concepción Grajales-Muñiz, Ricardo Grande, Rosa María Gutierrez Rios, Alejandra Hernández-Terán, Alfredo Herrera-Estrella, Carla Ivón Herrera-Najera, Pavel Isa, Daniel Lira Morales, Susana Lopez, Antonio Loza Román, Brenda Irasema Maldonado-Meza, Bernardo Martínez-Miguel, José Arturo Martínez-Orozco, Célida Martínez- Rodríguez, Margarita Matías-Florentino, Fidencio Mejía-Nepomuceno, María Guadalupe de Jesús Mireles-Rivera, Gloria María Molina-Salinas, Hector Montoya-Fuentes, , Mario Mújica-Sánchez, José Esteban Muñoz-Medina, José de Jesús Nuñez-Contreras, Alicia Ocaña-Mondragón, Luis Alberto Ochoa-Carrera, Hector Esteban Paz-Juárez, Marissa Perez-Garcia, Francisco Pulido, Helen Haydee Fernanda Ramírez-Plascencia, Jorge Salas-Hernández, Angel Gustavo Salas-Lais, Alejandro Sanchez-Flores, Clara Esperanza Santacruz-Tinoco, María Guadalupe Santiago-Mauricio, Selene Zárate, Nelly Sélem-Mojica, Blanca Taboada, Gloria Vazquez, Joel Armando Vázquez-Pérez. |
| EPI_ISL_2490348 | Unidad de Investigación Biomédica de Zacatecas (UIBZ) | Centro de Investigación en Enfermedades Infecciosas (CIENI), Instituto Nacional de Enfermedades Respiratorias (INER) | Consorcio Mexicano de Vigilancia Genómica (CoViGen-Mex). Authors (in alphabetical order): Julio Elias Alvarado-Yaah, Carlos F. Arias, Santiago Ávila-Ríos, Eduardo Becerril-Vargas, Víctor Hugo Borja-Aburto, Celia Boukadida, Cristóbal Cháidez-Quiróz, Juan Bautista Chale-Dzul, Ricardo Ciria Merce, Andreu Comas-García, Célida Duque Molina, Julissa Enciso-Ibarra, José Antonio Enciso-Moreno, Gloria Elena Espinosa-Ayala, Fernando Fontove-Herrera, Daniel Fregoso-Rueda, Victor Eduardo Garcia-Arias, Alejandra Garcia-Gasca, Bruno Gomez-Gil, Jean Pierre González, Irvin González-López, Concepción Grajales-Muñiz, Ricardo Grande, Rosa María Gutierrez Rios, Alejandra Hernández-Terán, Alfredo Herrera-Estrella, Carla Ivón Herrera-Najera, Pavel Isa, Daniel Lira Morales, Susana Lopez, Antonio Loza Román, Brenda Irasema Maldonado-Meza, Bernardo Martínez-Miguel, José Arturo Martínez-Orozco, Célida Martínez- Rodríguez, Margarita Matías-Florentino, Fidencio Mejía-Nepomuceno, María Guadalupe de Jesús Mireles-Rivera, Gloria María Molina-Salinas, Hector Montoya-Fuentes, , Mario Mújica-Sánchez, José Esteban Muñoz-Medina, José de Jesús Nuñez-Contreras, Alicia Ocaña-Mondragón, Luis Alberto Ochoa-Carrera, Hector Esteban Paz-Juárez, Marissa Perez-Garcia, Francisco Pulido, Helen Haydee Fernanda Ramírez-Plascencia, Jorge Salas-Hernández, Angel Gustavo Salas-Lais, Alejandro Sanchez-Flores, Clara Esperanza Santacruz-Tinoco, María Guadalupe Santiago-Mauricio, Selene Zárate, Nelly Sélem-Mojica, Blanca Taboada, Gloria Vazquez, Joel Armando Vázquez-Pérez. |
| EPI_ISL_2490349, EPI_ISL_2490350, EPI_ISL_2490351, EPI_ISL_2490352, EPI_ISL_2490353, EPI_ISL_2490354, EPI_ISL_2490355 | Laboratorio Central de Epidemiología (LCE) | Centro de Investigación en Enfermedades Infecciosas (CIENI), Instituto Nacional de Enfermedades Respiratorias (INER) | Consorcio Mexicano de Vigilancia Genómica (CoViGen-Mex). Authors (in alphabetical order): Julio Elias Alvarado-Yaah, Carlos F. Arias, Santiago Ávila-Ríos, Eduardo Becerril-Vargas, Víctor Hugo Borja-Aburto, Celia Boukadida, Cristóbal Cháidez-Quiróz, Juan Bautista Chale-Dzul, Ricardo Ciria Merce, Andreu Comas-García, Célida Duque Molina, Julissa Enciso-Ibarra, José Antonio Enciso-Moreno, Gloria Elena Espinosa-Ayala, Fernando Fontove-Herrera, Daniel Fregoso-Rueda, Victor Eduardo Garcia-Arias, Alejandra Garcia-Gasca, Bruno Gomez-Gil, Jean Pierre González, Irvin González-López, Concepción Grajales-Muñiz, Ricardo Grande, Rosa María Gutierrez Rios, Alejandra Hernández-Terán, Alfredo Herrera-Estrella, Carla Ivón Herrera-Najera, Pavel Isa, Daniel Lira Morales, Susana Lopez, Antonio Loza Román, Brenda Irasema Maldonado-Meza, Bernardo Martínez-Miguel, José Arturo Martínez-Orozco, Célida Martínez- Rodríguez, Margarita Matías-Florentino, Fidencio Mejía-Nepomuceno, María Guadalupe de Jesús Mireles-Rivera, Gloria María Molina-Salinas, Hector Montoya-Fuentes, , Mario Mújica-Sánchez, José Esteban Muñoz-Medina, José de Jesús Nuñez-Contreras, Alicia Ocaña-Mondragón, Luis Alberto Ochoa-Carrera, Hector Esteban Paz-Juárez, Marissa Perez-Garcia, Francisco Pulido, Helen Haydee Fernanda Ramírez-Plascencia, Jorge Salas-Hernández, Angel Gustavo Salas-Lais, Alejandro Sanchez-Flores, Clara Esperanza Santacruz-Tinoco, María Guadalupe Santiago-Mauricio, Selene Zárate, Nelly Sélem-Mojica, Blanca Taboada, Gloria Vazquez, Joel Armando Vázquez-Pérez. |
| EPI_ISL_2490356, EPI_ISL_2490362, EPI_ISL_2490363, EPI_ISL_2490366 | Centro de Investigación Biomédica del Noreste (CIBIN) | Centro de Investigación en Enfermedades Infecciosas (CIENI), Instituto Nacional de Enfermedades Respiratorias (INER) | Consorcio Mexicano de Vigilancia Genómica (CoViGen-Mex). Authors (in alphabetical order): Julio Elias Alvarado-Yaah, Carlos F. Arias, Santiago Ávila-Ríos, Eduardo Becerril-Vargas, Víctor Hugo Borja-Aburto, Celia Boukadida, Cristóbal Cháidez-Quiróz, Juan Bautista Chale-Dzul, Ricardo Ciria Merce, Andreu Comas-García, Célida Duque Molina, Julissa Enciso-Ibarra, José Antonio Enciso-Moreno, Gloria Elena Espinosa-Ayala, Fernando Fontove-Herrera, Daniel Fregoso-Rueda, Victor Eduardo Garcia-Arias, Alejandra Garcia-Gasca, Bruno Gomez-Gil, Jean Pierre González, Irvin González-López, Concepción Grajales-Muñiz, Ricardo Grande, Rosa María Gutierrez Rios, Alejandra Hernández-Terán, Alfredo Herrera-Estrella, Carla Ivón Herrera-Najera, Pavel Isa, Daniel Lira Morales, Susana Lopez, Antonio Loza Román, Brenda Irasema Maldonado-Meza, Bernardo Martínez-Miguel, José Arturo Martínez-Orozco, Célida Martínez- Rodríguez, Margarita Matías-Florentino, Fidencio Mejía-Nepomuceno, María Guadalupe de Jesús Mireles-Rivera, Gloria María Molina-Salinas, Hector Montoya-Fuentes, , Mario Mújica-Sánchez, José Esteban Muñoz-Medina, José de Jesús Nuñez-Contreras, Alicia Ocaña-Mondragón, Luis Alberto Ochoa-Carrera, Hector Esteban Paz-Juárez, Marissa Perez-Garcia, Francisco Pulido, Helen Haydee Fernanda Ramírez-Plascencia, Jorge Salas-Hernández, Angel Gustavo Salas-Lais, Alejandro Sanchez-Flores, Clara Esperanza |

| | | | |
|---|---|---|---|
| | | | Santacruz-Tinoco, María Guadalupe Santiago-Mauricio, Selene Zárate, Nelly Sélem-Mojica, Blanca Taboada, Gloria Vazquez, Joel Armando Vázquez-Pérez. |
| EPI_ISL_2490373, EPI_ISL_2490374, EPI_ISL_2490375, EPI_ISL_2490376, EPI_ISL_2490377, EPI_ISL_2490378, EPI_ISL_2490379, EPI_ISL_2490380, EPI_ISL_2490381 | Unidad de Investigación Médica de Yucatán (UIMY) | Centro de Investigación en Enfermedades Infecciosas (CIENI), Instituto Nacional de Enfermedades Respiratorias (INER) | Consorcio Mexicano de Vigilancia Genómica (CoViGen-Mex). Authors (in alphabetical order): Julio Elias Alvarado-Yaah, Carlos F. Arias, Santiago Ávila-Ríos, Eduardo Becerril-Vargas, Víctor Hugo Borja-Aburto, Celia Boukadida, Cristóbal Cháidez-Quiróz, Juan Bautista Chale-Dzul, Ricardo Ciria Merce, Andreu Comas-García, Célida Duque Molina, Julissa Enciso-Ibarra, José Antonio Enciso-Moreno, Gloria Elena Espinosa-Ayala, Fernando Fontove-Herrera, Daniel Fregoso-Rueda, Victor Eduardo Garcia-Arias, Alejandra Garcia-Gasca, Bruno Gomez-Gil, Jean Pierre González, Irvin González-López, Concepción Grajales-Muñiz, Ricardo Grande, Rosa María Gutierrez Rios, Alejandra Hernández-Terán, Alfredo Herrera-Estrella, Carla Ivón Herrera-Najera, Pavel Isa, Daniel Lira Morales, Susana Lopez, Antonio Loza Román, Brenda Irasema Maldonado-Meza, Bernardo Martínez-Miguel, José Arturo Martínez-Orozco, Célida Martínez- Rodríguez, Margarita Matías-Florentino, Fidencio Mejía-Nepomuceno, María Guadalupe de Jesús Mireles-Rivera, Gloria María Molina-Salinas, Hector Montoya-Fuentes, , Mario Mújica-Sánchez, José Esteban Muñoz-Medina, José de Jesús Nuñez-Contreras, Alicia Ocaña-Mondragón, Luis Alberto Ochoa-Carrera, Hector Esteban Paz-Juárez, Marissa Perez-Garcia, Francisco Pulido, Helen Haydee Fernanda Ramírez-Plascencia, Jorge Salas-Hernández, Angel Gustavo Salas-Lais, Alejandro Sanchez-Flores, Clara Esperanza Santacruz-Tinoco, María Guadalupe Santiago-Mauricio, Selene Zárate, Nelly Sélem-Mojica, Blanca Taboada, Gloria Vazquez, Joel Armando Vázquez-Pérez. |
| EPI_ISL_2490382, EPI_ISL_2490383, EPI_ISL_2490384 | Centro de Investigación Biomédica de Occidente (CIBO) | Centro de Investigación en Enfermedades Infecciosas (CIENI), Instituto Nacional de Enfermedades Respiratorias (INER) | Consorcio Mexicano de Vigilancia Genómica (CoViGen-Mex). Authors (in alphabetical order): Julio Elias Alvarado-Yaah, Carlos F. Arias, Santiago Ávila-Ríos, Eduardo Becerril-Vargas, Víctor Hugo Borja-Aburto, Celia Boukadida, Cristóbal Cháidez-Quiróz, Juan Bautista Chale-Dzul, Ricardo Ciria Merce, Andreu Comas-García, Célida Duque Molina, Julissa Enciso-Ibarra, José Antonio Enciso-Moreno, Gloria Elena Espinosa-Ayala, Fernando Fontove-Herrera, Daniel Fregoso-Rueda, Victor Eduardo Garcia-Arias, Alejandra Garcia-Gasca, Bruno Gomez-Gil, Jean Pierre González, Irvin González-López, Concepción Grajales-Muñiz, Ricardo Grande, Rosa María Gutierrez Rios, Alejandra Hernández-Terán, Alfredo Herrera-Estrella, Carla Ivón Herrera-Najera, Pavel Isa, Daniel Lira Morales, Susana Lopez, Antonio Loza Román, Brenda Irasema Maldonado-Meza, Bernardo Martínez-Miguel, José Arturo Martínez-Orozco, Célida Martínez- Rodríguez, Margarita Matías-Florentino, Fidencio Mejía-Nepomuceno, María Guadalupe de Jesús Mireles-Rivera, Gloria María Molina-Salinas, Hector Montoya-Fuentes, , Mario Mújica-Sánchez, José Esteban Muñoz-Medina, José de Jesús Nuñez-Contreras, Alicia Ocaña-Mondragón, Luis Alberto Ochoa-Carrera, Hector Esteban Paz-Juárez, Marissa Perez-Garcia, Francisco Pulido, Helen Haydee Fernanda Ramírez-Plascencia, Jorge Salas-Hernández, Angel Gustavo Salas-Lais, Alejandro Sanchez-Flores, Clara Esperanza Santacruz-Tinoco, María Guadalupe Santiago-Mauricio, Selene Zárate, Nelly Sélem-Mojica, Blanca Taboada, Gloria Vazquez, Joel Armando Vázquez-Pérez. |
| EPI_ISL_2490386 | Unidad de Investigación Biomédica de Zacatecas (UIBZ) | Centro de Investigación en Enfermedades Infecciosas (CIENI), Instituto Nacional de Enfermedades Respiratorias (INER) | Consorcio Mexicano de Vigilancia Genómica (CoViGen-Mex). Authors (in alphabetical order): Julio Elias Alvarado-Yaah, Carlos F. Arias, Santiago Ávila-Ríos, Eduardo Becerril-Vargas, Víctor Hugo Borja-Aburto, Celia Boukadida, Cristóbal Cháidez-Quiróz, Juan Bautista Chale-Dzul, Ricardo Ciria Merce, Andreu Comas-García, Célida Duque Molina, Julissa Enciso-Ibarra, José Antonio Enciso-Moreno, Gloria Elena Espinosa-Ayala, Fernando Fontove-Herrera, Daniel Fregoso-Rueda, Victor Eduardo Garcia-Arias, Alejandra Garcia-Gasca, Bruno Gomez-Gil, Jean Pierre González, Irvin González-López, Concepción Grajales-Muñiz, Ricardo Grande, Rosa María Gutierrez Rios, Alejandra Hernández-Terán, Alfredo Herrera-Estrella, Carla Ivón Herrera-Najera, Pavel Isa, Daniel Lira Morales, Susana Lopez, Antonio Loza Román, Brenda Irasema Maldonado-Meza, Bernardo Martínez-Miguel, José Arturo Martínez-Orozco, Célida Martínez- Rodríguez, Margarita Matías-Florentino, Fidencio Mejía-Nepomuceno, María Guadalupe de Jesús Mireles-Rivera, Gloria María Molina-Salinas, Hector Montoya-Fuentes, , Mario Mújica-Sánchez, José Esteban Muñoz-Medina, José de Jesús Nuñez-Contreras, Alicia Ocaña-Mondragón, Luis Alberto Ochoa-Carrera, Hector Esteban Paz-Juárez, Marissa Perez-Garcia, Francisco Pulido, Helen Haydee Fernanda Ramírez-Plascencia, Jorge Salas-Hernández, Angel Gustavo Salas-Lais, Alejandro Sanchez-Flores, Clara Esperanza Santacruz-Tinoco, María Guadalupe Santiago-Mauricio, Selene Zárate, Nelly Sélem-Mojica, Blanca Taboada, Gloria Vazquez, Joel Armando Vázquez-Pérez. |
| EPI_ISL_2490387, EPI_ISL_2490388, EPI_ISL_2490389, EPI_ISL_2490390, EPI_ISL_2490391, EPI_ISL_2490392, EPI_ISL_2490393, EPI_ISL_2490396 | Laboratorio Central de Epidemiología (LCE) | Centro de Investigación en Enfermedades Infecciosas (CIENI), Instituto Nacional de Enfermedades Respiratorias (INER) | Consorcio Mexicano de Vigilancia Genómica (CoViGen-Mex). Authors (in alphabetical order): Julio Elias Alvarado-Yaah, Carlos F. Arias, Santiago Ávila-Ríos, Eduardo Becerril-Vargas, Víctor Hugo Borja-Aburto, Celia Boukadida, Cristóbal Cháidez-Quiróz, Juan Bautista Chale-Dzul, Ricardo Ciria Merce, Andreu Comas-García, Célida Duque Molina, Julissa Enciso-Ibarra, José Antonio Enciso-Moreno, Gloria Elena Espinosa-Ayala, Fernando Fontove-Herrera, Daniel Fregoso-Rueda, Victor Eduardo Garcia-Arias, Alejandra Garcia-Gasca, Bruno Gomez-Gil, Jean Pierre González, Irvin González-López, Concepción Grajales-Muñiz, Ricardo Grande, Rosa María Gutierrez Rios, Alejandra Hernández-Terán, Alfredo Herrera-Estrella, Carla Ivón Herrera-Najera, Pavel Isa, Daniel Lira Morales, Susana Lopez, Antonio Loza Román, Brenda Irasema Maldonado-Meza, Bernardo Martínez-Miguel, José Arturo Martínez-Orozco, Célida Martínez- Rodríguez, Margarita Matías-Florentino, Fidencio Mejía-Nepomuceno, María Guadalupe de Jesús Mireles-Rivera, Gloria María Molina-Salinas, Hector Montoya-Fuentes, , Mario Mújica-Sánchez, José Esteban Muñoz-Medina, José de Jesús Nuñez-Contreras, Alicia Ocaña-Mondragón, Luis Alberto Ochoa-Carrera, Hector Esteban Paz-Juárez, Marissa Perez-Garcia, Francisco Pulido, Helen Haydee Fernanda Ramírez-Plascencia, Jorge Salas-Hernández, Angel Gustavo Salas-Lais, Alejandro Sanchez-Flores, Clara Esperanza Santacruz-Tinoco, María Guadalupe Santiago-Mauricio, Selene Zárate, Nelly Sélem-Mojica, Blanca Taboada, Gloria Vazquez, Joel Armando Vázquez-Pérez. |
| EPI_ISL_2490397, EPI_ISL_2490398, EPI_ISL_2490399, EPI_ISL_2490400 | Centro de Investigación Biomédica de Occidente (CIBO) | Centro de Investigación en Enfermedades Infecciosas (CIENI), Instituto Nacional de Enfermedades Respiratorias (INER) | Consorcio Mexicano de Vigilancia Genómica (CoViGen-Mex). Authors (in alphabetical order): Julio Elias Alvarado-Yaah, Carlos F. Arias, Santiago Ávila-Ríos, Eduardo Becerril-Vargas, Víctor Hugo Borja-Aburto, Celia Boukadida, Cristóbal Cháidez-Quiróz, Juan Bautista Chale-Dzul, Ricardo Ciria Merce, Andreu Comas-García, Célida Duque Molina, Julissa Enciso-Ibarra, José Antonio Enciso-Moreno, Gloria Elena Espinosa-Ayala, Fernando Fontove-Herrera, Daniel Fregoso-Rueda, Victor Eduardo Garcia-Arias, Alejandra Garcia-Gasca, Bruno Gomez-Gil, Jean Pierre González, Irvin González-López, Concepción Grajales-Muñiz, Ricardo Grande, Rosa María Gutierrez Rios, Alejandra Hernández-Terán, Alfredo Herrera-Estrella, Carla Ivón Herrera-Najera, Pavel Isa, Daniel Lira Morales, Susana Lopez, Antonio Loza Román, Brenda Irasema Maldonado-Meza, Bernardo Martínez-Miguel, José Arturo Martínez-Orozco, Célida Martínez- Rodríguez, Margarita Matías-Florentino, Fidencio Mejía-Nepomuceno, María Guadalupe de Jesús Mireles-Rivera, Gloria María Molina-Salinas, Hector Montoya-Fuentes, , Mario Mújica-Sánchez, José Esteban Muñoz-Medina, José de Jesús Nuñez-Contreras, Alicia Ocaña-Mondragón, Luis Alberto Ochoa-Carrera, Hector Esteban Paz-Juárez, Marissa Perez-Garcia, Francisco Pulido, Helen Haydee Fernanda Ramírez-Plascencia, Jorge Salas-Hernández, Angel Gustavo Salas-Lais, Alejandro Sanchez-Flores, Clara Esperanza Santacruz-Tinoco, María Guadalupe Santiago-Mauricio, Selene Zárate, Nelly Sélem-Mojica, Blanca Taboada, Gloria Vazquez, Joel Armando Vázquez-Pérez. |
| EPI_ISL_2490410, EPI_ISL_2490411 | Centro de Investigación Biomédica del Noreste (CIBIN) | Centro de Investigación en Enfermedades Infecciosas (CIENI), Instituto Nacional de Enfermedades Respiratorias (INER) | Consorcio Mexicano de Vigilancia Genómica (CoViGen-Mex). Authors (in alphabetical order): Julio Elias Alvarado-Yaah, Carlos F. Arias, Santiago Ávila-Ríos, Eduardo Becerril-Vargas, Víctor Hugo Borja-Aburto, Celia Boukadida, Cristóbal Cháidez-Quiróz, Juan Bautista Chale-Dzul, Ricardo Ciria Merce, Andreu Comas-García, Célida Duque Molina, Julissa Enciso-Ibarra, José Antonio Enciso-Moreno, Gloria Elena Espinosa-Ayala, Fernando Fontove-Herrera, Daniel Fregoso-Rueda, Victor Eduardo Garcia-Arias, Alejandra Garcia-Gasca, Bruno Gomez-Gil, Jean Pierre González, Irvin González-López, Concepción Grajales-Muñiz, Ricardo Grande, Rosa María Gutierrez Rios, Alejandra Hernández-Terán, Alfredo Herrera-Estrella, Carla Ivón Herrera-Najera, Pavel Isa, Daniel Lira Morales, Susana Lopez, Antonio Loza Román, Brenda Irasema Maldonado-Meza, Bernardo Martínez-Miguel, José Arturo Martínez-Orozco, Célida Martínez- Rodríguez, Margarita Matías-Florentino, Fidencio Mejía-Nepomuceno, María Guadalupe de Jesús Mireles-Rivera, Gloria María Molina-Salinas, Hector Montoya-Fuentes, , Mario Mújica-Sánchez, José Esteban Muñoz-Medina, José de Jesús Nuñez-Contreras, Alicia Ocaña-Mondragón, Luis Alberto Ochoa-Carrera, Hector Esteban Paz-Juárez, Marissa Perez-Garcia, Francisco Pulido, Helen Haydee Fernanda Ramírez-Plascencia, Jorge Salas-Hernández, Angel Gustavo Salas-Lais, Alejandro Sanchez-Flores, Clara Esperanza Santacruz-Tinoco, María Guadalupe Santiago-Mauricio, Selene Zárate, Nelly Sélem-Mojica, Blanca Taboada, Gloria Vazquez, Joel Armando Vázquez-Pérez. |
| EPI_ISL_2490412, EPI_ISL_2490413, EPI_ISL_2490414, EPI_ISL_2490415, EPI_ISL_2490416 | Laboratorio Central de Epidemiología (LCE) | Centro de Investigación en Enfermedades Infecciosas (CIENI), Instituto Nacional de Enfermedades Respiratorias (INER) | Consorcio Mexicano de Vigilancia Genómica (CoViGen-Mex). Authors (in alphabetical order): Julio Elias Alvarado-Yaah, Carlos F. Arias, Santiago Ávila-Ríos, Eduardo Becerril-Vargas, Víctor Hugo Borja-Aburto, Celia Boukadida, Cristóbal Cháidez-Quiróz, Juan Bautista Chale-Dzul, Ricardo Ciria Merce, Andreu Comas-García, Célida Duque Molina, Julissa Enciso-Ibarra, José Antonio Enciso-Moreno, Gloria Elena Espinosa-Ayala, Fernando Fontove-Herrera, Daniel Fregoso-Rueda, Victor Eduardo Garcia-Arias, Alejandra Garcia-Gasca, Bruno Gomez-Gil, Jean Pierre González, Irvin González-López, Concepción Grajales-Muñiz, Ricardo Grande, Rosa María Gutierrez Rios, Alejandra Hernández-Terán, Alfredo Herrera-Estrella, Carla Ivón Herrera-Najera, Pavel Isa, Daniel Lira Morales, Susana Lopez, Antonio Loza Román, Brenda Irasema Maldonado-Meza, Bernardo Martínez-Miguel, José Arturo Martínez-Orozco, Célida Martínez- Rodríguez, Margarita Matías-Florentino, Fidencio Mejía-Nepomuceno, María Guadalupe de Jesús Mireles-Rivera, Gloria María Molina-Salinas, Hector Montoya-Fuentes, , Mario Mújica-Sánchez, José Esteban Muñoz-Medina, José de Jesús Nuñez-Contreras, Alicia Ocaña-Mondragón, Luis Alberto Ochoa-Carrera, Hector Esteban Paz-Juárez, Marissa Perez-Garcia, Francisco Pulido, Helen Haydee Fernanda Ramírez-Plascencia, Jorge Salas-Hernández, Angel Gustavo Salas-Lais, Alejandro Sanchez-Flores, Clara Esperanza |

| | | | |
|---|---|---|---|
| | | | Santacruz-Tinoco, María Guadalupe Santiago-Mauricio, Selene Zárate, Nelly Sélem-Mojica, Blanca Taboada, Gloria Vazquez, Joel Armando Vázquez-Pérez. |
| EPI_ISL_2490421, EPI_ISL_2490422 | Unidad de Investigación Médica de Yucatán (UIMY) | Centro de Investigación en Enfermedades Infecciosas (CIENI), Instituto Nacional de Enfermedades Respiratorias (INER) | Consorcio Mexicano de Vigilancia Genómica (CoViGen-Mex). Authors (in alphabetical order): Julio Elias Alvarado-Yaah, Carlos F. Arias, Santiago Ávila-Ríos, Eduardo Becerril-Vargas, Víctor Hugo Borja-Aburto, Celia Boukadida, Cristóbal Cháidez-Quiróz, Juan Bautista Chale-Dzul, Ricardo Ciria Merce, Andreu Comas-García, Célida Duque Molina, Julissa Enciso-Ibarra, José Antonio Enciso-Moreno, Gloria Elena Espinosa-Ayala, Fernando Fontove-Herrera, Daniel Fregoso-Rueda, Victor Eduardo Garcia-Arias, Alejandra Garcia-Gasca, Bruno Gomez-Gil, Jean Pierre González, Irvin González-López, Concepción Grajales-Muñiz, Ricardo Grande, Rosa María Gutierrez Rios, Alejandra Hernández-Terán, Alfredo Herrera-Estrella, Carla Ivón Herrera-Najera, Pavel Isa, Daniel Lira Morales, Susana Lopez, Antonio Loza Román, Brenda Irasema Maldonado-Meza, Bernardo Martínez-Miguel, José Arturo Martínez-Orozco, Célida Martínez- Rodríguez, Margarita Matías-Florentino, Fidencio Mejía-Nepomuceno, María Guadalupe de Jesús Mireles-Rivera, Gloria María Molina-Salinas, Hector Montoya-Fuentes, , Mario Mújica-Sánchez, José Esteban Muñoz-Medina, José de Jesús Nuñez-Contreras, Alicia Ocaña-Mondragón, Luis Alberto Ochoa-Carrera, Hector Esteban Paz-Juárez, Marissa Perez-Garcia, Francisco Pulido, Helen Haydee Fernanda Ramírez-Plascencia, Jorge Salas-Hernández, Angel Gustavo Salas-Lais, Alejandro Sanchez-Flores, Clara Esperanza Santacruz-Tinoco, María Guadalupe Santiago-Mauricio, Selene Zárate, Nelly Sélem-Mojica, Blanca Taboada, Gloria Vazquez, Joel Armando Vázquez-Pérez. |
| EPI_ISL_2490423 | Laboratorio Central de Epidemiología (LCE) | Centro de Investigación en Enfermedades Infecciosas (CIENI), Instituto Nacional de Enfermedades Respiratorias (INER) | Consorcio Mexicano de Vigilancia Genómica (CoViGen-Mex). Authors (in alphabetical order): Julio Elias Alvarado-Yaah, Carlos F. Arias, Santiago Ávila-Ríos, Eduardo Becerril-Vargas, Víctor Hugo Borja-Aburto, Celia Boukadida, Cristóbal Cháidez-Quiróz, Juan Bautista Chale-Dzul, Ricardo Ciria Merce, Andreu Comas-García, Célida Duque Molina, Julissa Enciso-Ibarra, José Antonio Enciso-Moreno, Gloria Elena Espinosa-Ayala, Fernando Fontove-Herrera, Daniel Fregoso-Rueda, Victor Eduardo Garcia-Arias, Alejandra Garcia-Gasca, Bruno Gomez-Gil, Jean Pierre González, Irvin González-López, Concepción Grajales-Muñiz, Ricardo Grande, Rosa María Gutierrez Rios, Alejandra Hernández-Terán, Alfredo Herrera-Estrella, Carla Ivón Herrera-Najera, Pavel Isa, Daniel Lira Morales, Susana Lopez, Antonio Loza Román, Brenda Irasema Maldonado-Meza, Bernardo Martínez-Miguel, José Arturo Martínez-Orozco, Célida Martínez- Rodríguez, Margarita Matías-Florentino, Fidencio Mejía-Nepomuceno, María Guadalupe de Jesús Mireles-Rivera, Gloria María Molina-Salinas, Hector Montoya-Fuentes, , Mario Mújica-Sánchez, José Esteban Muñoz-Medina, José de Jesús Nuñez-Contreras, Alicia Ocaña-Mondragón, Luis Alberto Ochoa-Carrera, Hector Esteban Paz-Juárez, Marissa Perez-Garcia, Francisco Pulido, Helen Haydee Fernanda Ramírez-Plascencia, Jorge Salas-Hernández, Angel Gustavo Salas-Lais, Alejandro Sanchez-Flores, Clara Esperanza Santacruz-Tinoco, María Guadalupe Santiago-Mauricio, Selene Zárate, Nelly Sélem-Mojica, Blanca Taboada, Gloria Vazquez, Joel Armando Vázquez-Pérez. |
| EPI_ISL_2490424 | Unidad de Investigación Biomédica de Zacatecas (UIBZ) | Centro de Investigación en Enfermedades Infecciosas (CIENI), Instituto Nacional de Enfermedades Respiratorias (INER) | Consorcio Mexicano de Vigilancia Genómica (CoViGen-Mex). Authors (in alphabetical order): Julio Elias Alvarado-Yaah, Carlos F. Arias, Santiago Ávila-Ríos, Eduardo Becerril-Vargas, Víctor Hugo Borja-Aburto, Celia Boukadida, Cristóbal Cháidez-Quiróz, Juan Bautista Chale-Dzul, Ricardo Ciria Merce, Andreu Comas-García, Célida Duque Molina, Julissa Enciso-Ibarra, José Antonio Enciso-Moreno, Gloria Elena Espinosa-Ayala, Fernando Fontove-Herrera, Daniel Fregoso-Rueda, Victor Eduardo Garcia-Arias, Alejandra Garcia-Gasca, Bruno Gomez-Gil, Jean Pierre González, Irvin González-López, Concepción Grajales-Muñiz, Ricardo Grande, Rosa María Gutierrez Rios, Alejandra Hernández-Terán, Alfredo Herrera-Estrella, Carla Ivón Herrera-Najera, Pavel Isa, Daniel Lira Morales, Susana Lopez, Antonio Loza Román, Brenda Irasema Maldonado-Meza, Bernardo Martínez-Miguel, José Arturo Martínez-Orozco, Célida Martínez- Rodríguez, Margarita Matías-Florentino, Fidencio Mejía-Nepomuceno, María Guadalupe de Jesús Mireles-Rivera, Gloria María Molina-Salinas, Hector Montoya-Fuentes, , Mario Mújica-Sánchez, José Esteban Muñoz-Medina, José de Jesús Nuñez-Contreras, Alicia Ocaña-Mondragón, Luis Alberto Ochoa-Carrera, Hector Esteban Paz-Juárez, Marissa Perez-Garcia, Francisco Pulido, Helen Haydee Fernanda Ramírez-Plascencia, Jorge Salas-Hernández, Angel Gustavo Salas-Lais, Alejandro Sanchez-Flores, Clara Esperanza Santacruz-Tinoco, María Guadalupe Santiago-Mauricio, Selene Zárate, Nelly Sélem-Mojica, Blanca Taboada, Gloria Vazquez, Joel Armando Vázquez-Pérez. |
| EPI_ISL_2490425 | Laboratorio Central de Epidemiología (LCE) | Centro de Investigación en Enfermedades Infecciosas (CIENI), Instituto Nacional de Enfermedades Respiratorias (INER) | Consorcio Mexicano de Vigilancia Genómica (CoViGen-Mex). Authors (in alphabetical order): Julio Elias Alvarado-Yaah, Carlos F. Arias, Santiago Ávila-Ríos, Eduardo Becerril-Vargas, Víctor Hugo Borja-Aburto, Celia Boukadida, Cristóbal Cháidez-Quiróz, Juan Bautista Chale-Dzul, Ricardo Ciria Merce, Andreu Comas-García, Célida Duque Molina, Julissa Enciso-Ibarra, José Antonio Enciso-Moreno, Gloria Elena Espinosa-Ayala, Fernando Fontove-Herrera, Daniel Fregoso-Rueda, Victor Eduardo Garcia-Arias, Alejandra Garcia-Gasca, Bruno Gomez-Gil, Jean Pierre González, Irvin González-López, Concepción Grajales-Muñiz, Ricardo Grande, Rosa María Gutierrez Rios, Alejandra Hernández-Terán, Alfredo Herrera-Estrella, Carla Ivón Herrera-Najera, Pavel Isa, Daniel Lira Morales, Susana Lopez, Antonio Loza Román, Brenda Irasema Maldonado-Meza, Bernardo Martínez-Miguel, José Arturo Martínez-Orozco, Célida Martínez- Rodríguez, Margarita Matías-Florentino, Fidencio Mejía-Nepomuceno, María Guadalupe de Jesús Mireles-Rivera, Gloria María Molina-Salinas, Hector Montoya-Fuentes, , Mario Mújica-Sánchez, José Esteban Muñoz-Medina, José de Jesús Nuñez-Contreras, Alicia Ocaña-Mondragón, Luis Alberto Ochoa-Carrera, Hector Esteban Paz-Juárez, Marissa Perez-Garcia, Francisco Pulido, Helen Haydee Fernanda Ramírez-Plascencia, Jorge Salas-Hernández, Angel Gustavo Salas-Lais, Alejandro Sanchez-Flores, Clara Esperanza Santacruz-Tinoco, María Guadalupe Santiago-Mauricio, Selene Zárate, Nelly Sélem-Mojica, Blanca Taboada, Gloria Vazquez, Joel Armando Vázquez-Pérez. |
| EPI_ISL_2490426 | Centro de Investigación Biomédica del Noreste (CIBIN) | Centro de Investigación en Enfermedades Infecciosas (CIENI), Instituto Nacional de Enfermedades Respiratorias (INER) | Consorcio Mexicano de Vigilancia Genómica (CoViGen-Mex). Authors (in alphabetical order): Julio Elias Alvarado-Yaah, Carlos F. Arias, Santiago Ávila-Ríos, Eduardo Becerril-Vargas, Víctor Hugo Borja-Aburto, Celia Boukadida, Cristóbal Cháidez-Quiróz, Juan Bautista Chale-Dzul, Ricardo Ciria Merce, Andreu Comas-García, Célida Duque Molina, Julissa Enciso-Ibarra, José Antonio Enciso-Moreno, Gloria Elena Espinosa-Ayala, Fernando Fontove-Herrera, Daniel Fregoso-Rueda, Victor Eduardo Garcia-Arias, Alejandra Garcia-Gasca, Bruno Gomez-Gil, Jean Pierre González, Irvin González-López, Concepción Grajales-Muñiz, Ricardo Grande, Rosa María Gutierrez Rios, Alejandra Hernández-Terán, Alfredo Herrera-Estrella, Carla Ivón Herrera-Najera, Pavel Isa, Daniel Lira Morales, Susana Lopez, Antonio Loza Román, Brenda Irasema Maldonado-Meza, Bernardo Martínez-Miguel, José Arturo Martínez-Orozco, Célida Martínez- Rodríguez, Margarita Matías-Florentino, Fidencio Mejía-Nepomuceno, María Guadalupe de Jesús Mireles-Rivera, Gloria María Molina-Salinas, Hector Montoya-Fuentes, , Mario Mújica-Sánchez, José Esteban Muñoz-Medina, José de Jesús Nuñez-Contreras, Alicia Ocaña-Mondragón, Luis Alberto Ochoa-Carrera, Hector Esteban Paz-Juárez, Marissa Perez-Garcia, Francisco Pulido, Helen Haydee Fernanda Ramírez-Plascencia, Jorge Salas-Hernández, Angel Gustavo Salas-Lais, Alejandro Sanchez-Flores, Clara Esperanza Santacruz-Tinoco, María Guadalupe Santiago-Mauricio, Selene Zárate, Nelly Sélem-Mojica, Blanca Taboada, Gloria Vazquez, Joel Armando Vázquez-Pérez. |
| EPI_ISL_2490433, EPI_ISL_2490434, EPI_ISL_2490435, EPI_ISL_2490437, EPI_ISL_2490438 | Laboratorio Central de Epidemiología (LCE) | Centro de Investigación en Enfermedades Infecciosas (CIENI), Instituto Nacional de Enfermedades Respiratorias (INER) | Consorcio Mexicano de Vigilancia Genómica (CoViGen-Mex). Authors (in alphabetical order): Julio Elias Alvarado-Yaah, Carlos F. Arias, Santiago Ávila-Ríos, Eduardo Becerril-Vargas, Víctor Hugo Borja-Aburto, Celia Boukadida, Cristóbal Cháidez-Quiróz, Juan Bautista Chale-Dzul, Ricardo Ciria Merce, Andreu Comas-García, Célida Duque Molina, Julissa Enciso-Ibarra, José Antonio Enciso-Moreno, Gloria Elena Espinosa-Ayala, Fernando Fontove-Herrera, Daniel Fregoso-Rueda, Victor Eduardo Garcia-Arias, Alejandra Garcia-Gasca, Bruno Gomez-Gil, Jean Pierre González, Irvin González-López, Concepción Grajales-Muñiz, Ricardo Grande, Rosa María Gutierrez Rios, Alejandra Hernández-Terán, Alfredo Herrera-Estrella, Carla Ivón Herrera-Najera, Pavel Isa, Daniel Lira Morales, Susana Lopez, Antonio Loza Román, Brenda Irasema Maldonado-Meza, Bernardo Martínez-Miguel, José Arturo Martínez-Orozco, Célida Martínez- Rodríguez, Margarita Matías-Florentino, Fidencio Mejía-Nepomuceno, María Guadalupe de Jesús Mireles-Rivera, Gloria María Molina-Salinas, Hector Montoya-Fuentes, , Mario Mújica-Sánchez, José Esteban Muñoz-Medina, José de Jesús Nuñez-Contreras, Alicia Ocaña-Mondragón, Luis Alberto Ochoa-Carrera, Hector Esteban Paz-Juárez, Marissa Perez-Garcia, Francisco Pulido, Helen Haydee Fernanda Ramírez-Plascencia, Jorge Salas-Hernández, Angel Gustavo Salas-Lais, Alejandro Sanchez-Flores, Clara Esperanza Santacruz-Tinoco, María Guadalupe Santiago-Mauricio, Selene Zárate, Nelly Sélem-Mojica, Blanca Taboada, Gloria Vazquez, Joel Armando Vázquez-Pérez. |
| EPI_ISL_2490452, EPI_ISL_2490454, EPI_ISL_2490455, EPI_ISL_2490456, EPI_ISL_2490457, EPI_ISL_2490458, EPI_ISL_2490459, EPI_ISL_2490460, EPI_ISL_2490461, EPI_ISL_2490462, EPI_ISL_2490463, EPI_ISL_2490464, EPI_ISL_2490465 | | | |
| see above | Centro de Investigación Biomédica del Noreste (CIBIN) | Centro de Investigación en Enfermedades Infecciosas (CIENI), Instituto Nacional de Enfermedades Respiratorias (INER) | Consorcio Mexicano de Vigilancia Genómica (CoViGen-Mex). Authors (in alphabetical order): Julio Elias Alvarado-Yaah, Carlos F. Arias, Santiago Ávila-Ríos, Eduardo Becerril-Vargas, Víctor Hugo Borja-Aburto, Celia Boukadida, Cristóbal Cháidez-Quiróz, Juan Bautista Chale-Dzul, Ricardo Ciria Merce, Andreu Comas-García, Célida Duque Molina, Julissa Enciso-Ibarra, José Antonio Enciso-Moreno, Gloria Elena Espinosa-Ayala, Fernando Fontove-Herrera, Daniel Fregoso-Rueda, Victor Eduardo Garcia-Arias, Alejandra Garcia-Gasca, Bruno Gomez-Gil, Jean Pierre González, Irvin González-López, Concepción Grajales-Muñiz, Ricardo Grande, Rosa María Gutierrez Rios, Alejandra Hernández-Terán, Alfredo Herrera-Estrella, Carla Ivón Herrera-Najera, Pavel Isa, Daniel Lira Morales, Susana Lopez, Antonio Loza Román, Brenda Irasema Maldonado-Meza, Bernardo Martínez-Miguel, José Arturo Martínez-Orozco, Célida Martínez- Rodríguez, Margarita Matías-Florentino, Fidencio Mejía-Nepomuceno, María Guadalupe de Jesús Mireles-Rivera, Gloria María Molina-Salinas, Hector Montoya-Fuentes, , Mario Mújica-Sánchez, José Esteban Muñoz-Medina, José de Jesús |

| | | | |
|---|---|---|---|
| | | | Nuñez-Contreras, Alicia Ocaña-Mondragón, Luis Alberto Ochoa-Carrera, Hector Esteban Paz-Juárez, Marissa Perez-Garcia, Francisco Pulido, Helen Haydee Fernanda Ramírez-Plascencia, Jorge Salas-Hernández, Angel Gustavo Salas-Lais, Alejandro Sanchez-Flores, Clara Esperanza Santacruz-Tinoco, María Guadalupe Santiago-Mauricio, Selene Zárate, Nelly Sélem-Mojica, Blanca Taboada, Gloria Vazquez, Joel Armando Vázquez-Pérez. |
| EPI_ISL_2490479, EPI_ISL_2490480, EPI_ISL_2490481, EPI_ISL_2490482, EPI_ISL_2490483, EPI_ISL_2490484, EPI_ISL_2490485, EPI_ISL_2490486, EPI_ISL_2490487, EPI_ISL_2490488, EPI_ISL_2490489, EPI_ISL_2490490, EPI_ISL_2490491, EPI_ISL_2490492 | | | |
| see above | Unidad de Investigación Médica de Yucatán (UIMY) | Centro de Investigación en Enfermedades Infecciosas (CIENI), Instituto Nacional de Enfermedades Respiratorias (INER) | Consorcio Mexicano de Vigilancia Genómica (CoViGen-Mex). Authors (in alphabetical order): Julio Elias Alvarado-Yaah, Carlos F. Arias, Santiago Ávila-Ríos, Eduardo Becerril-Vargas, Víctor Hugo Borja-Aburto, Celia Boukadida, Cristóbal Cháidez-Quiróz, Juan Bautista Chale-Dzul, Ricardo Ciria Merce, Andreu Comas-García, Célida Duque Molina, Julissa Enciso-Ibarra, José Antonio Enciso-Moreno, Gloria Elena Espinosa-Ayala, Fernando Fontove-Herrera, Daniel Fregoso-Rueda, Victor Eduardo Garcia-Arias, Alejandra Garcia-Gasca, Bruno Gomez-Gil, Jean Pierre González, Irvin González-López, Concepción Grajales-Muñiz, Ricardo Grande, Rosa María Gutierrez Rios, Alejandra Hernández-Terán, Alfredo Herrera-Estrella, Carla Ivón Herrera-Najera, Pavel Isa, Daniel Lira Morales, Susana Lopez, Antonio Loza Román, Brenda Irasema Maldonado-Meza, Bernardo Martínez-Miguel, José Arturo Martínez-Orozco, Célida Martínez- Rodríguez, Margarita Matías-Florentino, Fidencio Mejía-Nepomuceno, María Guadalupe de Jesús Mireles-Rivera, Gloria María Molina-Salinas, Hector Montoya-Fuentes, , Mario Mújica-Sánchez, José Esteban Muñoz-Medina, José de Jesús Nuñez-Contreras, Alicia Ocaña-Mondragón, Luis Alberto Ochoa-Carrera, Hector Esteban Paz-Juárez, Marissa Perez-Garcia, Francisco Pulido, Helen Haydee Fernanda Ramírez-Plascencia, Jorge Salas-Hernández, Angel Gustavo Salas-Lais, Alejandro Sanchez-Flores, Clara Esperanza Santacruz-Tinoco, María Guadalupe Santiago-Mauricio, Selene Zárate, Nelly Sélem-Mojica, Blanca Taboada, Gloria Vazquez, Joel Armando Vázquez-Pérez. |
| EPI_ISL_2490496, EPI_ISL_2490498 | Centro de Investigación Biomédica de Occidente (CIBO) | Centro de Investigación en Enfermedades Infecciosas (CIENI), Instituto Nacional de Enfermedades Respiratorias (INER) | Consorcio Mexicano de Vigilancia Genómica (CoViGen-Mex). Authors (in alphabetical order): Julio Elias Alvarado-Yaah, Carlos F. Arias, Santiago Ávila-Ríos, Eduardo Becerril-Vargas, Víctor Hugo Borja-Aburto, Celia Boukadida, Cristóbal Cháidez-Quiróz, Juan Bautista Chale-Dzul, Ricardo Ciria Merce, Andreu Comas-García, Célida Duque Molina, Julissa Enciso-Ibarra, José Antonio Enciso-Moreno, Gloria Elena Espinosa-Ayala, Fernando Fontove-Herrera, Daniel Fregoso-Rueda, Victor Eduardo Garcia-Arias, Alejandra Garcia-Gasca, Bruno Gomez-Gil, Jean Pierre González, Irvin González-López, Concepción Grajales-Muñiz, Ricardo Grande, Rosa María Gutierrez Rios, Alejandra Hernández-Terán, Alfredo Herrera-Estrella, Carla Ivón Herrera-Najera, Pavel Isa, Daniel Lira Morales, Susana Lopez, Antonio Loza Román, Brenda Irasema Maldonado-Meza, Bernardo Martínez-Miguel, José Arturo Martínez-Orozco, Célida Martínez- Rodríguez, Margarita Matías-Florentino, Fidencio Mejía-Nepomuceno, María Guadalupe de Jesús Mireles-Rivera, Gloria María Molina-Salinas, Hector Montoya-Fuentes, , Mario Mújica-Sánchez, José Esteban Muñoz-Medina, José de Jesús Nuñez-Contreras, Alicia Ocaña-Mondragón, Luis Alberto Ochoa-Carrera, Hector Esteban Paz-Juárez, Marissa Perez-Garcia, Francisco Pulido, Helen Haydee Fernanda Ramírez-Plascencia, Jorge Salas-Hernández, Angel Gustavo Salas-Lais, Alejandro Sanchez-Flores, Clara Esperanza Santacruz-Tinoco, María Guadalupe Santiago-Mauricio, Selene Zárate, Nelly Sélem-Mojica, Blanca Taboada, Gloria Vazquez, Joel Armando Vázquez-Pérez. |
| EPI_ISL_2490511, EPI_ISL_2490512, EPI_ISL_2490513, EPI_ISL_2490514, EPI_ISL_2490515, EPI_ISL_2490516, EPI_ISL_2490517, EPI_ISL_2490518, EPI_ISL_2490519, EPI_ISL_2490520, EPI_ISL_2490521, EPI_ISL_2490522, EPI_ISL_2490523, EPI_ISL_2490524, EPI_ISL_2490525, EPI_ISL_2490526, EPI_ISL_2490527, EPI_ISL_2490528, EPI_ISL_2490529, EPI_ISL_2490530, EPI_ISL_2490531, EPI_ISL_2490532, EPI_ISL_2490533 | | | |
| see above | Unidad de Investigación Biomédica de Zacatecas (UIBZ) | Centro de Investigación en Enfermedades Infecciosas (CIENI), Instituto Nacional de Enfermedades Respiratorias (INER) | Consorcio Mexicano de Vigilancia Genómica (CoViGen-Mex). Authors (in alphabetical order): Julio Elias Alvarado-Yaah, Carlos F. Arias, Santiago Ávila-Ríos, Eduardo Becerril-Vargas, Víctor Hugo Borja-Aburto, Celia Boukadida, Cristóbal Cháidez-Quiróz, Juan Bautista Chale-Dzul, Ricardo Ciria Merce, Andreu Comas-García, Célida Duque Molina, Julissa Enciso-Ibarra, José Antonio Enciso-Moreno, Gloria Elena Espinosa-Ayala, Fernando Fontove-Herrera, Daniel Fregoso-Rueda, Victor Eduardo Garcia-Arias, Alejandra Garcia-Gasca, Bruno Gomez-Gil, Jean Pierre González, Irvin González-López, Concepción Grajales-Muñiz, Ricardo Grande, Rosa María Gutierrez Rios, Alejandra Hernández-Terán, Alfredo Herrera-Estrella, Carla Ivón Herrera-Najera, Pavel Isa, Daniel Lira Morales, Susana Lopez, Antonio Loza Román, Brenda Irasema Maldonado-Meza, Bernardo Martínez-Miguel, José Arturo Martínez-Orozco, Célida Martínez- Rodríguez, Margarita Matías-Florentino, Fidencio Mejía-Nepomuceno, María Guadalupe de Jesús Mireles-Rivera, Gloria María Molina-Salinas, Hector Montoya-Fuentes, , Mario Mújica-Sánchez, José Esteban Muñoz-Medina, José de Jesús Nuñez-Contreras, Alicia Ocaña-Mondragón, Luis Alberto Ochoa-Carrera, Hector Esteban Paz-Juárez, Marissa Perez-Garcia, Francisco Pulido, Helen Haydee Fernanda Ramírez-Plascencia, Jorge Salas-Hernández, Angel Gustavo Salas-Lais, Alejandro Sanchez-Flores, Clara Esperanza Santacruz-Tinoco, María Guadalupe Santiago-Mauricio, Selene Zárate, Nelly Sélem-Mojica, Blanca Taboada, Gloria Vazquez, Joel Armando Vázquez-Pérez. |
| EPI_ISL_2490534 | Basurto University Hospital: Clinical Microbiology Laboratory | Biocruces Bizkaia | Mikel Urrutikoetxea-Gutiérrez, Mª Carmen Nieto Toboso, Estibaliz Ugalde Zarraga, Mikel Gallego Rodrigo, Ana de la Hoz, José Luis Díaz de Tuesta del Arco |
| EPI_ISL_2490535, EPI_ISL_2490536, EPI_ISL_2490537, EPI_ISL_2490538, EPI_ISL_2490539, EPI_ISL_2490540, EPI_ISL_2490541 | Unidad de Investigación Biomédica de Zacatecas (UIBZ) | Centro de Investigación en Enfermedades Infecciosas (CIENI), Instituto Nacional de Enfermedades Respiratorias (INER) | Consorcio Mexicano de Vigilancia Genómica (CoViGen-Mex). Authors (in alphabetical order): Julio Elias Alvarado-Yaah, Carlos F. Arias, Santiago Ávila-Ríos, Eduardo Becerril-Vargas, Víctor Hugo Borja-Aburto, Celia Boukadida, Cristóbal Cháidez-Quiróz, Juan Bautista Chale-Dzul, Ricardo Ciria Merce, Andreu Comas-García, Célida Duque Molina, Julissa Enciso-Ibarra, José Antonio Enciso-Moreno, Gloria Elena Espinosa-Ayala, Fernando Fontove-Herrera, Daniel Fregoso-Rueda, Victor Eduardo Garcia-Arias, Alejandra Garcia-Gasca, Bruno Gomez-Gil, Jean Pierre González, Irvin González-López, Concepción Grajales-Muñiz, Ricardo Grande, Rosa María Gutierrez Rios, Alejandra Hernández-Terán, Alfredo Herrera-Estrella, Carla Ivón Herrera-Najera, Pavel Isa, Daniel Lira Morales, Susana Lopez, Antonio Loza Román, Brenda Irasema Maldonado-Meza, Bernardo Martínez-Miguel, José Arturo Martínez-Orozco, Célida Martínez- Rodríguez, Margarita Matías-Florentino, Fidencio Mejía-Nepomuceno, María Guadalupe de Jesús Mireles-Rivera, Gloria María Molina-Salinas, Hector Montoya-Fuentes, , Mario Mújica-Sánchez, José Esteban Muñoz-Medina, José de Jesús Nuñez-Contreras, Alicia Ocaña-Mondragón, Luis Alberto Ochoa-Carrera, Hector Esteban Paz-Juárez, Marissa Perez-Garcia, Francisco Pulido, Helen Haydee Fernanda Ramírez-Plascencia, Jorge Salas-Hernández, Angel Gustavo Salas-Lais, Alejandro Sanchez-Flores, Clara Esperanza Santacruz-Tinoco, María Guadalupe Santiago-Mauricio, Selene Zárate, Nelly Sélem-Mojica, Blanca Taboada, Gloria Vazquez, Joel Armando Vázquez-Pérez. |
| EPI_ISL_2490542, EPI_ISL_2490543, EPI_ISL_2490544, EPI_ISL_2490545, EPI_ISL_2490546, EPI_ISL_2490547, EPI_ISL_2490548, EPI_ISL_2490549, EPI_ISL_2490550, EPI_ISL_2490551, EPI_ISL_2490552, EPI_ISL_2490553, EPI_ISL_2490554, EPI_ISL_2490555, EPI_ISL_2490556, EPI_ISL_2490557, EPI_ISL_2490562, EPI_ISL_2490564, EPI_ISL_2490565, EPI_ISL_2490566, EPI_ISL_2490567, EPI_ISL_2490568, EPI_ISL_2490569, EPI_ISL_2490570, EPI_ISL_2490571, EPI_ISL_2490572, EPI_ISL_2490573, EPI_ISL_2490574, EPI_ISL_2490575, EPI_ISL_2490576, EPI_ISL_2490577, EPI_ISL_2490578, EPI_ISL_2490579, EPI_ISL_2490580, EPI_ISL_2490581, EPI_ISL_2490582, EPI_ISL_2490584 | | | |
| see above | Laboratorio Central de Epidemiología (LCE) | Centro de Investigación en Enfermedades Infecciosas (CIENI), Instituto Nacional de Enfermedades Respiratorias (INER) | Consorcio Mexicano de Vigilancia Genómica (CoViGen-Mex). Authors (in alphabetical order): Julio Elias Alvarado-Yaah, Carlos F. Arias, Santiago Ávila-Ríos, Eduardo Becerril-Vargas, Víctor Hugo Borja-Aburto, Celia Boukadida, Cristóbal Cháidez-Quiróz, Juan Bautista Chale-Dzul, Ricardo Ciria Merce, Andreu Comas-García, Célida Duque Molina, Julissa Enciso-Ibarra, José Antonio Enciso-Moreno, Gloria Elena Espinosa-Ayala, Fernando Fontove-Herrera, Daniel Fregoso-Rueda, Victor Eduardo Garcia-Arias, Alejandra Garcia-Gasca, Bruno Gomez-Gil, Jean Pierre González, Irvin González-López, Concepción Grajales-Muñiz, Ricardo Grande, Rosa María Gutierrez Rios, Alejandra Hernández-Terán, Alfredo Herrera-Estrella, Carla Ivón Herrera-Najera, Pavel Isa, Daniel Lira Morales, Susana Lopez, Antonio Loza Román, Brenda Irasema Maldonado-Meza, Bernardo Martínez-Miguel, José Arturo Martínez-Orozco, Célida Martínez- Rodríguez, Margarita Matías-Florentino, Fidencio Mejía-Nepomuceno, María Guadalupe de Jesús Mireles-Rivera, Gloria María Molina-Salinas, Hector Montoya-Fuentes, , Mario Mújica-Sánchez, José Esteban Muñoz-Medina, José de Jesús Nuñez-Contreras, Alicia Ocaña-Mondragón, Luis Alberto Ochoa-Carrera, Hector Esteban Paz-Juárez, Marissa Perez-Garcia, Francisco Pulido, Helen Haydee Fernanda Ramírez-Plascencia, Jorge Salas-Hernández, Angel Gustavo Salas-Lais, Alejandro Sanchez-Flores, Clara Esperanza Santacruz-Tinoco, María Guadalupe Santiago-Mauricio, Selene Zárate, Nelly Sélem-Mojica, Blanca Taboada, Gloria Vazquez, Joel Armando Vázquez-Pérez. |
| EPI_ISL_2490585, EPI_ISL_2490586, EPI_ISL_2490587, EPI_ISL_2490588, EPI_ISL_2490589, EPI_ISL_2490590, EPI_ISL_2490595, EPI_ISL_2490596, EPI_ISL_2490597, EPI_ISL_2490598, EPI_ISL_2490599, EPI_ISL_2490600, EPI_ISL_2490601, EPI_ISL_2490602, EPI_ISL_2490603, EPI_ISL_2490604, EPI_ISL_2490605, EPI_ISL_2490606, EPI_ISL_2490607, EPI_ISL_2490608 | | | |
| see above | Instituto Nacional de Enfermedades Respiratorias (INER) | Centro de Investigación en Enfermedades Infecciosas (CIENI), Instituto Nacional de Enfermedades Respiratorias (INER) | Consorcio Mexicano de Vigilancia Genómica (CoViGen-Mex). Authors (in alphabetical order): Julio Elias Alvarado-Yaah, Carlos F. Arias, Santiago Ávila-Ríos, Eduardo Becerril-Vargas, Víctor Hugo Borja-Aburto, Celia Boukadida, Cristóbal Cháidez-Quiróz, Juan Bautista Chale-Dzul, Ricardo Ciria Merce, Andreu Comas-García, Célida Duque Molina, Julissa Enciso-Ibarra, José Antonio Enciso-Moreno, Gloria Elena Espinosa-Ayala, Fernando Fontove-Herrera, Daniel Fregoso-Rueda, Victor Eduardo Garcia-Arias, Alejandra Garcia-Gasca, Bruno Gomez-Gil, Jean Pierre González, Irvin González-López, Concepción Grajales-Muñiz, Ricardo Grande, Rosa María Gutierrez Rios, Alejandra Hernández-Terán, Alfredo Herrera-Estrella, Carla Ivón Herrera-Najera, Pavel Isa, Daniel Lira Morales, Susana Lopez, Antonio Loza Román, Brenda Irasema Maldonado-Meza, Bernardo Martínez-Miguel, José Arturo Martínez-Orozco, Célida Martínez- Rodríguez, Margarita Matías-Florentino, Fidencio Mejía-Nepomuceno, María Guadalupe de Jesús Mireles-Rivera, Gloria María Molina-Salinas, Hector Montoya-Fuentes, , Mario Mújica-Sánchez, José Esteban Muñoz-Medina, José de Jesús |

| | | | |
|---|---|---|---|
| | | | Nuñez-Contreras, Alicia Ocaña-Mondragón, Luis Alberto Ochoa-Carrera, Hector Esteban Paz-Juárez, Marissa Perez-Garcia, Francisco Pulido, Helen Haydee Fernanda Ramírez-Plascencia, Jorge Salas-Hernández, Angel Gustavo Salas-Lais, Alejandro Sanchez-Flores, Clara Esperanza Santacruz-Tinoco, María Guadalupe Santiago-Mauricio, Selene Zárate, Nelly Sélem-Mojica, Blanca Taboada, Gloria Vazquez, Joel Armando Vázquez-Pérez. |
| EPI_ISL_2491608 | Centre Hospitalier Universitaire Clermont-Ferrand | CHU Clermont-Ferrand, service de virologie | Bisseux Maxime, Mirand Audrey, Combes Patricia, Henquell Cécile |
| EPI_ISL_2491609 | HOPITAL DE MOULINS | CHU Clermont-Ferrand, service de virologie | Bisseux Maxime, Mirand Audrey, Combes Patricia, Henquell Cécile |
| EPI_ISL_2491611, EPI_ISL_2491612 | Centre Hospitalier Universitaire Clermont-Ferrand | CHU Clermont-Ferrand, service de virologie | Bisseux Maxime, Mirand Audrey, Combes Patricia, Henquell Cécile |
| EPI_ISL_2491613 | Centre Hospitalier RIOM | CHU Clermont-Ferrand, service de virologie | Bisseux Maxime, Mirand Audrey, Combes Patricia, Henquell Cécile |
| EPI_ISL_2491614, EPI_ISL_2491615, EPI_ISL_2491616, EPI_ISL_2491617, EPI_ISL_2491618 | Centre Hospitalier Universitaire Clermont-Ferrand | CHU Clermont-Ferrand, service de virologie | Bisseux Maxime, Mirand Audrey, Combes Patricia, Henquell Cécile |
| EPI_ISL_2491619 | Centre De Prélévement COVID VICHY | CHU Clermont-Ferrand, service de virologie | Bisseux Maxime, Mirand Audrey, Combes Patricia, Henquell Cécile |
| EPI_ISL_2491620, EPI_ISL_2491621 | HOPITAL DU MONT DORE | CHU Clermont-Ferrand, service de virologie | Bisseux Maxime, Mirand Audrey, Combes Patricia, Henquell Cécile |
| EPI_ISL_2491622, EPI_ISL_2491623, EPI_ISL_2491624, EPI_ISL_2491625, EPI_ISL_2491626, EPI_ISL_2491627 | Centre Hospitalier Universitaire Clermont-Ferrand | CHU Clermont-Ferrand, service de virologie | Bisseux Maxime, Mirand Audrey, Combes Patricia, Henquell Cécile |
| EPI_ISL_2491628 | HOPITAL DU MONT DORE | CHU Clermont-Ferrand, service de virologie | Bisseux Maxime, Mirand Audrey, Combes Patricia, Henquell Cécile |
| EPI_ISL_2491629 | Centre Hospitalier Universitaire Clermont-Ferrand | CHU Clermont-Ferrand, service de virologie | Bisseux Maxime, Mirand Audrey, Combes Patricia, Henquell Cécile |
| EPI_ISL_2491633 | laboratoire Maymat | CHU Clermont-Ferrand, service de virologie | Bisseux Maxime, Mirand Audrey, Combes Patricia, Henquell Cécile |
| EPI_ISL_2491634, EPI_ISL_2491635, EPI_ISL_2491636 | Centre Hospitalier Universitaire Clermont-Ferrand | CHU Clermont-Ferrand, service de virologie | Bisseux Maxime, Mirand Audrey, Combes Patricia, Henquell Cécile |
| EPI_ISL_2491637, EPI_ISL_2491638 | EHPAD | CHU Clermont-Ferrand, service de virologie | Bisseux Maxime, Mirand Audrey, Combes Patricia, Henquell Cécile |
| EPI_ISL_2491639 | HOPITAL DE MONTLUCON | CHU Clermont-Ferrand, service de virologie | Bisseux Maxime, Mirand Audrey, Combes Patricia, Henquell Cécile |
| EPI_ISL_2491640 | Centre Hospitalier Universitaire Clermont-Ferrand | CHU Clermont-Ferrand, service de virologie | Bisseux Maxime, Mirand Audrey, Combes Patricia, Henquell Cécile |
| EPI_ISL_2491641 | HOPITAL DE MONTLUCON | CHU Clermont-Ferrand, service de virologie | Bisseux Maxime, Mirand Audrey, Combes Patricia, Henquell Cécile |
| EPI_ISL_2491642 | HOPITAL DE VICHY | CHU Clermont-Ferrand, service de virologie | Bisseux Maxime, Mirand Audrey, Combes Patricia, Henquell Cécile |
| EPI_ISL_2491643, EPI_ISL_2491644, EPI_ISL_2491645 | Centre Hospitalier Universitaire Clermont-Ferrand | CHU Clermont-Ferrand, service de virologie | Bisseux Maxime, Mirand Audrey, Combes Patricia, Henquell Cécile |
| EPI_ISL_2491646 | CDP COVID MONTLUCON | CHU Clermont-Ferrand, service de virologie | Bisseux Maxime, Mirand Audrey, Combes Patricia, Henquell Cécile |
| EPI_ISL_2491647 | Centre Hospitalier Universitaire Clermont-Ferrand | CHU Clermont-Ferrand, service de virologie | Bisseux Maxime, Mirand Audrey, Combes Patricia, Henquell Cécile |
| EPI_ISL_2491648 | center de prélévement COVID ISSOIRE | CHU Clermont-Ferrand, service de virologie | Bisseux Maxime, Mirand Audrey, Combes Patricia, Henquell Cécile |
| EPI_ISL_2491649, EPI_ISL_2491650, EPI_ISL_2491651, EPI_ISL_2491652, EPI_ISL_2491653 | Centre Hospitalier Universitaire Clermont-Ferrand | CHU Clermont-Ferrand, service de virologie | Bisseux Maxime, Mirand Audrey, Combes Patricia, Henquell Cécile |
| EPI_ISL_2491654, EPI_ISL_2491655, EPI_ISL_2491656, EPI_ISL_2491657 | CDP COVID MONTLUCON | CHU Clermont-Ferrand, service de virologie | Bisseux Maxime, Mirand Audrey, Combes Patricia, Henquell Cécile |
| EPI_ISL_2491658, EPI_ISL_2491659, EPI_ISL_2491660, EPI_ISL_2491661 | Centre Hospitalier Universitaire Clermont-Ferrand | CHU Clermont-Ferrand, service de virologie | Bisseux Maxime, Mirand Audrey, Combes Patricia, Henquell Cécile |
| EPI_ISL_2491662 | Centre De Prelevement COVID RIOM | CHU Clermont-Ferrand, service de virologie | Bisseux Maxime, Mirand Audrey, Combes Patricia, Henquell Cécile |
| EPI_ISL_2491663 | Centre Hospitalier Universitaire Clermont-Ferrand | CHU Clermont-Ferrand, service de virologie | Bisseux Maxime, Mirand Audrey, Combes Patricia, Henquell Cécile |
| EPI_ISL_2491664 | Centre hospitalier D'AURILLAC | CHU Clermont-Ferrand, service de virologie | Bisseux Maxime, Mirand Audrey, Combes Patricia, Henquell Cécile |
| EPI_ISL_2492366, EPI_ISL_2492367, EPI_ISL_2492368, EPI_ISL_2492369, EPI_ISL_2492370, EPI_ISL_2492371, EPI_ISL_2492372, EPI_ISL_2492373, EPI_ISL_2492374, EPI_ISL_2492375, EPI_ISL_2492376, EPI_ISL_2492377, EPI_ISL_2492378, EPI_ISL_2492379, EPI_ISL_2492380, EPI_ISL_2492381, EPI_ISL_2492382, EPI_ISL_2492383, EPI_ISL_2492384, EPI_ISL_2492385, EPI_ISL_2492386, EPI_ISL_2492387, EPI_ISL_2492388, EPI_ISL_2492389, EPI_ISL_2492390, EPI_ISL_2492391, EPI_ISL_2492392 | | | |
| see above | Cliniques universitaires Saint-Luc | UCLouvain/IREC/MBLG | Jean Ruelle, Benoit Kabamba Mukadi |
| EPI_ISL_2492393, EPI_ISL_2492394, EPI_ISL_2492395, EPI_ISL_2492396, EPI_ISL_2492397, EPI_ISL_2492398, EPI_ISL_2492399, EPI_ISL_2492400, EPI_ISL_2492401, EPI_ISL_2492402, EPI_ISL_2492403 | | | |
| see above | Clinique Saint-Pierre Ottignies | UCLouvain/IREC/MBLG | Jean Ruelle, Benoit Kabamba Mukadi |
| EPI_ISL_2492404, EPI_ISL_2492405, EPI_ISL_2492406, EPI_ISL_2492407, EPI_ISL_2492408, EPI_ISL_2492409, EPI_ISL_2492410, EPI_ISL_2492411, EPI_ISL_2492412, EPI_ISL_2492413 | Cliniques universitaires Saint-Luc | UCLouvain/IREC/MBLG | Jean Ruelle, Benoit Kabamba Mukadi |
| EPI_ISL_2492414, EPI_ISL_2492415, EPI_ISL_2492416 | Clinique Saint-Pierre Ottignies | UCLouvain/IREC/MBLG | Jean Ruelle, Benoit Kabamba Mukadi |
| EPI_ISL_2492417, EPI_ISL_2492418, EPI_ISL_2492419, EPI_ISL_2492420, EPI_ISL_2492421, EPI_ISL_2492422, EPI_ISL_2492423, EPI_ISL_2492424, EPI_ISL_2492425, EPI_ISL_2492426, EPI_ISL_2492427, EPI_ISL_2492428, EPI_ISL_2492429, EPI_ISL_2492430, EPI_ISL_2492431, EPI_ISL_2492432, EPI_ISL_2492433, EPI_ISL_2492434, EPI_ISL_2492435, EPI_ISL_2492436 | | | |
| see above | Cliniques universitaires Saint-Luc | UCLouvain/IREC/MBLG | Jean Ruelle, Benoit Kabamba Mukadi |
| EPI_ISL_2492437, EPI_ISL_2492438, EPI_ISL_2492439 | Clinique Saint-Pierre Ottignies | UCLouvain/IREC/MBLG | Jean Ruelle, Benoit Kabamba Mukadi |
| EPI_ISL_2492440 | Cliniques universitaires Saint-Luc | UCLouvain/IREC/MBLG | Jean Ruelle, Benoit Kabamba Mukadi |
| EPI_ISL_2492462, EPI_ISL_2492463 | LESP Queretaro | Instituto de Diagnostico y Referencia Epidemiologicos (INDRE) | Claudia Wong-Arambula, Abril Rodriguez-Maldonado, Vanessa Rivero-Arredondo, Ariadna Medina-Benitez, Joaquin Quiroz-Mercado, Sergio Rangel-Guerrero, Natividad Cruz-Ortiz, Tatiana Nunez-Garcia, Gisela Barrera-Badillo, Lucia Hernandez-Rivas, Irma Lopez-Martinez, Ernesto Ramirez-Gonzalez. |
| EPI_ISL_2492466 | LESP Ciudad de Mexico | Instituto de Diagnostico y Referencia Epidemiologicos (INDRE) | Claudia Wong-Arambula, Abril Rodriguez-Maldonado, Vanessa Rivero-Arredondo, Ariadna Medina-Benitez, Joaquin Quiroz-Mercado, Sergio Rangel-Guerrero, Natividad Cruz-Ortiz, Tatiana Nunez-Garcia, Gisela Barrera-Badillo, Lucia Hernandez-Rivas, Irma Lopez-Martinez, Ernesto Ramirez-Gonzalez. |
| EPI_ISL_2492468, EPI_ISL_2492469, EPI_ISL_2492470 | LESP Queretaro | Instituto de Diagnostico y Referencia Epidemiologicos (INDRE) | Claudia Wong-Arambula, Abril Rodriguez-Maldonado, Vanessa Rivero-Arredondo, Ariadna Medina-Benitez, Joaquin Quiroz-Mercado, Sergio Rangel-Guerrero, Natividad Cruz-Ortiz, Tatiana Nunez-Garcia, Gisela Barrera-Badillo, Lucia Hernandez-Rivas, Irma Lopez-Martinez, Ernesto Ramirez-Gonzalez. |

| | | | |
|---|---|---|---|
| EPI_ISL_2492471 | LESP Guanajuato | Instituto de Diagnostico y Referencia Epidemiologicos (INDRE) | Claudia Wong-Arambula, Abril Rodriguez-Maldonado, Vanessa Rivero-Arredondo, Ariadna Medina-Benitez, Joaquin Quiroz-Mercado, Sergio Rangel-Guerrero, Natividad Cruz-Ortiz, Tatiana Nunez-Garcia, Gisela Barrera-Badillo, Lucia Hernandez-Rivas, Irma Lopez-Martinez, Ernesto Ramirez-Gonzalez. |
| EPI_ISL_2492474 | LESP Queretaro | Instituto de Diagnostico y Referencia Epidemiologicos (INDRE) | Claudia Wong-Arambula, Abril Rodriguez-Maldonado, Vanessa Rivero-Arredondo, Ariadna Medina-Benitez, Joaquin Quiroz-Mercado, Sergio Rangel-Guerrero, Natividad Cruz-Ortiz, Tatiana Nunez-Garcia, Gisela Barrera-Badillo, Lucia Hernandez-Rivas, Irma Lopez-Martinez, Ernesto Ramirez-Gonzalez. |
| EPI_ISL_2492475, EPI_ISL_2492476, EPI_ISL_2492477 | LESP Nuevo Leon | Instituto de Diagnostico y Referencia Epidemiologicos (INDRE) | Claudia Wong-Arambula, Abril Rodriguez-Maldonado, Vanessa Rivero-Arredondo, Ariadna Medina-Benitez, Joaquin Quiroz-Mercado, Sergio Rangel-Guerrero, Natividad Cruz-Ortiz, Tatiana Nunez-Garcia, Gisela Barrera-Badillo, Lucia Hernandez-Rivas, Irma Lopez-Martinez, Ernesto Ramirez-Gonzalez. |
| EPI_ISL_2492478, EPI_ISL_2492479, EPI_ISL_2492480, EPI_ISL_2492481, EPI_ISL_2492482, EPI_ISL_2492483 | LESP Guanajuato | Instituto de Diagnostico y Referencia Epidemiologicos (INDRE) | Claudia Wong-Arambula, Abril Rodriguez-Maldonado, Vanessa Rivero-Arredondo, Ariadna Medina-Benitez, Joaquin Quiroz-Mercado, Sergio Rangel-Guerrero, Natividad Cruz-Ortiz, Tatiana Nunez-Garcia, Gisela Barrera-Badillo, Lucia Hernandez-Rivas, Irma Lopez-Martinez, Ernesto Ramirez-Gonzalez. |
| EPI_ISL_2492484 | LESP Ciudad de Mexico | Instituto de Diagnostico y Referencia Epidemiologicos (INDRE) | Claudia Wong-Arambula, Abril Rodriguez-Maldonado, Vanessa Rivero-Arredondo, Ariadna Medina-Benitez, Joaquin Quiroz-Mercado, Sergio Rangel-Guerrero, Natividad Cruz-Ortiz, Tatiana Nunez-Garcia, Gisela Barrera-Badillo, Lucia Hernandez-Rivas, Irma Lopez-Martinez, Ernesto Ramirez-Gonzalez. |
| EPI_ISL_2492489 | LESP Chiapas | Instituto de Diagnostico y Referencia Epidemiologicos (INDRE) | Claudia Wong-Arambula, Abril Rodriguez-Maldonado, Vanessa Rivero-Arredondo, Ariadna Medina-Benitez, Joaquin Quiroz-Mercado, Sergio Rangel-Guerrero, Natividad Cruz-Ortiz, Tatiana Nunez-Garcia, Gisela Barrera-Badillo, Lucia Hernandez-Rivas, Irma Lopez-Martinez, Ernesto Ramirez-Gonzalez. |
| EPI_ISL_2492490, EPI_ISL_2492491 | LESP Ciudad de Mexico | Instituto de Diagnostico y Referencia Epidemiologicos (INDRE) | Claudia Wong-Arambula, Abril Rodriguez-Maldonado, Vanessa Rivero-Arredondo, Ariadna Medina-Benitez, Joaquin Quiroz-Mercado, Sergio Rangel-Guerrero, Natividad Cruz-Ortiz, Tatiana Nunez-Garcia, Gisela Barrera-Badillo, Lucia Hernandez-Rivas, Irma Lopez-Martinez, Ernesto Ramirez-Gonzalez. |
| EPI_ISL_2492492, EPI_ISL_2492493 | LESP Tamaulipas | Instituto de Diagnostico y Referencia Epidemiologicos (INDRE) | Claudia Wong-Arambula, Abril Rodriguez-Maldonado, Vanessa Rivero-Arredondo, Ariadna Medina-Benitez, Joaquin Quiroz-Mercado, Sergio Rangel-Guerrero, Natividad Cruz-Ortiz, Tatiana Nunez-Garcia, Gisela Barrera-Badillo, Lucia Hernandez-Rivas, Irma Lopez-Martinez, Ernesto Ramirez-Gonzalez. |
| EPI_ISL_2492494 | LESP Chiapas | Instituto de Diagnostico y Referencia Epidemiologicos (INDRE) | Claudia Wong-Arambula, Abril Rodriguez-Maldonado, Vanessa Rivero-Arredondo, Ariadna Medina-Benitez, Joaquin Quiroz-Mercado, Sergio Rangel-Guerrero, Natividad Cruz-Ortiz, Tatiana Nunez-Garcia, Gisela Barrera-Badillo, Lucia Hernandez-Rivas, Irma Lopez-Martinez, Ernesto Ramirez-Gonzalez. |
| EPI_ISL_2492495, EPI_ISL_2492496 | LESP Quintana Roo | Instituto de Diagnostico y Referencia Epidemiologicos (INDRE) | Claudia Wong-Arambula, Abril Rodriguez-Maldonado, Vanessa Rivero-Arredondo, Ariadna Medina-Benitez, Joaquin Quiroz-Mercado, Sergio Rangel-Guerrero, Natividad Cruz-Ortiz, Tatiana Nunez-Garcia, Gisela Barrera-Badillo, Lucia Hernandez-Rivas, Irma Lopez-Martinez, Ernesto Ramirez-Gonzalez. |
| EPI_ISL_2492497, EPI_ISL_2492498 | LESP Queretaro | Instituto de Diagnostico y Referencia Epidemiologicos (INDRE) | Claudia Wong-Arambula, Abril Rodriguez-Maldonado, Vanessa Rivero-Arredondo, Ariadna Medina-Benitez, Joaquin Quiroz-Mercado, Sergio Rangel-Guerrero, Natividad Cruz-Ortiz, Tatiana Nunez-Garcia, Gisela Barrera-Badillo, Lucia Hernandez-Rivas, Irma Lopez-Martinez, Ernesto Ramirez-Gonzalez. |
| EPI_ISL_2495586, EPI_ISL_2495587, EPI_ISL_2495588, EPI_ISL_2495589, EPI_ISL_2495590, EPI_ISL_2495591, EPI_ISL_2495592, EPI_ISL_2495597, EPI_ISL_2495598, EPI_ISL_2495599, EPI_ISL_2495600, EPI_ISL_2495603, EPI_ISL_2495617, EPI_ISL_2495618, EPI_ISL_2495622, EPI_ISL_2495623, EPI_ISL_2495624, EPI_ISL_2495625, EPI_ISL_2495626, EPI_ISL_2495628, EPI_ISL_2495629, EPI_ISL_2495630, EPI_ISL_2495631, EPI_ISL_2495632, EPI_ISL_2495633, EPI_ISL_2495634, EPI_ISL_2495635, EPI_ISL_2495636, EPI_ISL_2495637, EPI_ISL_2495638, EPI_ISL_2495639, EPI_ISL_2495640, EPI_ISL_2495641, EPI_ISL_2495642, EPI_ISL_2495643, EPI_ISL_2495644, EPI_ISL_2495645, EPI_ISL_2495646, EPI_ISL_2495647, EPI_ISL_2495648, EPI_ISL_2495649, EPI_ISL_2495650, EPI_ISL_2495651, EPI_ISL_2495652, EPI_ISL_2495653, EPI_ISL_2495654, EPI_ISL_2495655, EPI_ISL_2495656, EPI_ISL_2495657, EPI_ISL_2495658, EPI_ISL_2495659, EPI_ISL_2495660, EPI_ISL_2495661, EPI_ISL_2495662, EPI_ISL_2495663, EPI_ISL_2495664, EPI_ISL_2495665, EPI_ISL_2495666, EPI_ISL_2495683, EPI_ISL_2495684, EPI_ISL_2495685, EPI_ISL_2495686, EPI_ISL_2495687, EPI_ISL_2495688, EPI_ISL_2495689, EPI_ISL_2495690, EPI_ISL_2495691, EPI_ISL_2495692, EPI_ISL_2495693, EPI_ISL_2495694, EPI_ISL_2495695, EPI_ISL_2495696, EPI_ISL_2495697, EPI_ISL_2495698, EPI_ISL_2495699, EPI_ISL_2495700, EPI_ISL_2495701, EPI_ISL_2495702, EPI_ISL_2495703, EPI_ISL_2495704, EPI_ISL_2495705, EPI_ISL_2495706, EPI_ISL_2495707, EPI_ISL_2495708, EPI_ISL_2495709, EPI_ISL_2495710, EPI_ISL_2495711, EPI_ISL_2495712, EPI_ISL_2495743, EPI_ISL_2495744, EPI_ISL_2495745, EPI_ISL_2495746, EPI_ISL_2495747, EPI_ISL_2495748, EPI_ISL_2495749, EPI_ISL_2495750, EPI_ISL_2495751, EPI_ISL_2495752, EPI_ISL_2495753, EPI_ISL_2495754, EPI_ISL_2495755, EPI_ISL_2495756 | | | |
| see above | MB-Cadham Provincial laboratory | National Microbiology Laboratory (NML) | Anna Majer, Shari Tyson, Grace Seo, Philip Mabon, Elsie Grudeski, Rhiannon Huzarewich, Russell Mandes, Anneliese Landgraff, Jennifer Tanner, Natalie Knox, Morag Graham, Gary Van Domselaar, Paul Van Caeseele, Jared Bullard, David Alexander, Kerry Dust, Nathalie Bastien, Yan Li, Timothy Booth, Darian Hole, Madison Chapel, Kirsten Biggar, CanCOGeN's metadata curation team, Public Health Agency of Canada CanCOGeN team |
| EPI_ISL_2495930 | LESP Jalisco | Instituto de Diagnostico y Referencia Epidemiologicos (INDRE) | Claudia Wong-Arambula, Abril Rodriguez-Maldonado, Vanessa Rivero-Arredondo, Ariadna Medina-Benitez, Joaquin Quiroz-Mercado, Sergio Rangel-Guerrero, Natividad Cruz-Ortiz, Tatiana Nunez-Garcia, Gisela Barrera-Badillo, Lucia Hernandez-Rivas, Irma Lopez-Martinez, Ernesto Ramirez-Gonzalez. |
| EPI_ISL_2495934 | LESP Nuevo Leon | Instituto de Diagnostico y Referencia Epidemiologicos (INDRE) | Claudia Wong-Arambula, Abril Rodriguez-Maldonado, Vanessa Rivero-Arredondo, Ariadna Medina-Benitez, Joaquin Quiroz-Mercado, Sergio Rangel-Guerrero, Natividad Cruz-Ortiz, Tatiana Nunez-Garcia, Gisela Barrera-Badillo, Lucia Hernandez-Rivas, Irma Lopez-Martinez, Ernesto Ramirez-Gonzalez. |
| EPI_ISL_2495938, EPI_ISL_2495939, EPI_ISL_2495948, EPI_ISL_2495949, EPI_ISL_2495950, EPI_ISL_2495951, EPI_ISL_2495952, EPI_ISL_2495953, EPI_ISL_2495954, EPI_ISL_2495955, EPI_ISL_2495956, EPI_ISL_2495957, EPI_ISL_2495958, EPI_ISL_2495959, EPI_ISL_2495960, EPI_ISL_2495961, EPI_ISL_2495962 | | | |
| see above | LESP Queretaro | Instituto de Diagnostico y Referencia Epidemiologicos (INDRE) | Claudia Wong-Arambula, Abril Rodriguez-Maldonado, Vanessa Rivero-Arredondo, Ariadna Medina-Benitez, Joaquin Quiroz-Mercado, Sergio Rangel-Guerrero, Natividad Cruz-Ortiz, Tatiana Nunez-Garcia, Gisela Barrera-Badillo, Lucia Hernandez-Rivas, Irma Lopez-Martinez, Ernesto Ramirez-Gonzalez. |
| EPI_ISL_2495965 | LESP Yucatan | Instituto de Diagnostico y Referencia Epidemiologicos (INDRE) | Claudia Wong-Arambula, Abril Rodriguez-Maldonado, Vanessa Rivero-Arredondo, Ariadna Medina-Benitez, Joaquin Quiroz-Mercado, Sergio Rangel-Guerrero, Natividad Cruz-Ortiz, Tatiana Nunez-Garcia, Gisela Barrera-Badillo, Lucia Hernandez-Rivas, Irma Lopez-Martinez, Ernesto Ramirez-Gonzalez. |
| EPI_ISL_2495966 | LESP Quintana Roo | Instituto de Diagnostico y Referencia Epidemiologicos (INDRE) | Claudia Wong-Arambula, Abril Rodriguez-Maldonado, Vanessa Rivero-Arredondo, Ariadna Medina-Benitez, Joaquin Quiroz-Mercado, Sergio Rangel-Guerrero, Natividad Cruz-Ortiz, Tatiana Nunez-Garcia, Gisela Barrera-Badillo, Lucia Hernandez-Rivas, Irma Lopez-Martinez, Ernesto Ramirez-Gonzalez. |
| EPI_ISL_2495967, EPI_ISL_2495968 | LESP Queretaro | Instituto de Diagnostico y Referencia Epidemiologicos (INDRE) | Claudia Wong-Arambula, Abril Rodriguez-Maldonado, Vanessa Rivero-Arredondo, Ariadna Medina-Benitez, Joaquin Quiroz-Mercado, Sergio Rangel-Guerrero, Natividad Cruz-Ortiz, Tatiana Nunez-Garcia, Gisela Barrera-Badillo, Lucia Hernandez-Rivas, Irma Lopez-Martinez, Ernesto Ramirez-Gonzalez. |
| EPI_ISL_2495971, EPI_ISL_2495972, EPI_ISL_2495973, EPI_ISL_2495974, EPI_ISL_2495975 | LESP Zacatecas | Instituto de Diagnostico y Referencia Epidemiologicos (INDRE) | Claudia Wong-Arambula, Abril Rodriguez-Maldonado, Vanessa Rivero-Arredondo, Ariadna Medina-Benitez, Joaquin Quiroz-Mercado, Sergio Rangel-Guerrero, Natividad Cruz-Ortiz, Tatiana Nunez-Garcia, Gisela Barrera-Badillo, Lucia Hernandez-Rivas, Irma Lopez-Martinez, Ernesto Ramirez-Gonzalez. |
| EPI_ISL_2495976 | LESP Hidalgo | Instituto de Diagnostico y Referencia Epidemiologicos (INDRE) | Claudia Wong-Arambula, Abril Rodriguez-Maldonado, Vanessa Rivero-Arredondo, Ariadna Medina-Benitez, Joaquin Quiroz-Mercado, Sergio Rangel-Guerrero, Natividad Cruz-Ortiz, Tatiana Nunez-Garcia, Gisela Barrera-Badillo, Lucia Hernandez-Rivas, Irma Lopez-Martinez, Ernesto Ramirez-Gonzalez. |
| EPI_ISL_2495977, EPI_ISL_2495978 | LESP Queretaro | Instituto de Diagnostico y Referencia Epidemiologicos (INDRE) | Claudia Wong-Arambula, Abril Rodriguez-Maldonado, Vanessa Rivero-Arredondo, Ariadna Medina-Benitez, Joaquin Quiroz-Mercado, Sergio Rangel-Guerrero, Natividad Cruz-Ortiz, Tatiana Nunez-Garcia, Gisela Barrera-Badillo, Lucia Hernandez-Rivas, Irma Lopez-Martinez, Ernesto Ramirez-Gonzalez. |
| EPI_ISL_2495979 | LESP Tabasco | Instituto de Diagnostico y Referencia Epidemiologicos (INDRE) | Claudia Wong-Arambula, Abril Rodriguez-Maldonado, Vanessa Rivero-Arredondo, Ariadna Medina-Benitez, Joaquin Quiroz-Mercado, Sergio Rangel-Guerrero, Natividad Cruz-Ortiz, Tatiana Nunez-Garcia, Gisela Barrera-Badillo, Lucia Hernandez-Rivas, Irma Lopez-Martinez, Ernesto Ramirez-Gonzalez. |
| EPI_ISL_2495980, EPI_ISL_2495981 | LESP Nuevo Leon | Instituto de Diagnostico y Referencia Epidemiologicos (INDRE) | Claudia Wong-Arambula, Abril Rodriguez-Maldonado, Vanessa Rivero-Arredondo, Ariadna Medina-Benitez, Joaquin Quiroz-Mercado, Sergio Rangel-Guerrero, Natividad Cruz-Ortiz, Tatiana Nunez-Garcia, Gisela Barrera-Badillo, Lucia Hernandez-Rivas, Irma Lopez-Martinez, Ernesto |

| | | | |
|---|---|---|---|
| | | | Ramirez-Gonzalez. |
| EPI_ISL_2495982, EPI_ISL_2495983, EPI_ISL_2495984, EPI_ISL_2495985, EPI_ISL_2495986, EPI_ISL_2495987 | LESP Guanajuato | Instituto de Diagnostico y Referencia Epidemiologicos (INDRE) | Claudia Wong-Arambula, Abril Rodriguez-Maldonado, Vanessa Rivero-Arredondo, Ariadna Medina-Benitez, Joaquin Quiroz-Mercado, Sergio Rangel-Guerrero, Natividad Cruz-Ortiz, Tatiana Nunez-Garcia, Gisela Barrera-Badillo, Lucia Hernandez-Rivas, Irma Lopez-Martinez, Ernesto Ramirez-Gonzalez. |
| EPI_ISL_2495988 | LESP Guerrero | Instituto de Diagnostico y Referencia Epidemiologicos (INDRE) | Claudia Wong-Arambula, Abril Rodriguez-Maldonado, Vanessa Rivero-Arredondo, Ariadna Medina-Benitez, Joaquin Quiroz-Mercado, Sergio Rangel-Guerrero, Natividad Cruz-Ortiz, Tatiana Nunez-Garcia, Gisela Barrera-Badillo, Lucia Hernandez-Rivas, Irma Lopez-Martinez, Ernesto Ramirez-Gonzalez. |
| EPI_ISL_2495989, EPI_ISL_2495990, EPI_ISL_2495991 | LESP Guanajuato | Instituto de Diagnostico y Referencia Epidemiologicos (INDRE) | Claudia Wong-Arambula, Abril Rodriguez-Maldonado, Vanessa Rivero-Arredondo, Ariadna Medina-Benitez, Joaquin Quiroz-Mercado, Sergio Rangel-Guerrero, Natividad Cruz-Ortiz, Tatiana Nunez-Garcia, Gisela Barrera-Badillo, Lucia Hernandez-Rivas, Irma Lopez-Martinez, Ernesto Ramirez-Gonzalez. |
| EPI_ISL_2495992, EPI_ISL_2495994 | LESP Morelos | Instituto de Diagnostico y Referencia Epidemiologicos (INDRE) | Claudia Wong-Arambula, Abril Rodriguez-Maldonado, Vanessa Rivero-Arredondo, Ariadna Medina-Benitez, Joaquin Quiroz-Mercado, Sergio Rangel-Guerrero, Natividad Cruz-Ortiz, Tatiana Nunez-Garcia, Gisela Barrera-Badillo, Lucia Hernandez-Rivas, Irma Lopez-Martinez, Ernesto Ramirez-Gonzalez. |
| EPI_ISL_2495999 | LESP Guanajuato | Instituto de Diagnostico y Referencia Epidemiologicos (INDRE) | Claudia Wong-Arambula, Abril Rodriguez-Maldonado, Vanessa Rivero-Arredondo, Ariadna Medina-Benitez, Joaquin Quiroz-Mercado, Sergio Rangel-Guerrero, Natividad Cruz-Ortiz, Tatiana Nunez-Garcia, Gisela Barrera-Badillo, Lucia Hernandez-Rivas, Irma Lopez-Martinez, Ernesto Ramirez-Gonzalez. |
| EPI_ISL_2496003 | LESP Chihuahua | Instituto de Diagnostico y Referencia Epidemiologicos (INDRE) | Claudia Wong-Arambula, Abril Rodriguez-Maldonado, Vanessa Rivero-Arredondo, Ariadna Medina-Benitez, Joaquin Quiroz-Mercado, Sergio Rangel-Guerrero, Natividad Cruz-Ortiz, Tatiana Nunez-Garcia, Gisela Barrera-Badillo, Lucia Hernandez-Rivas, Irma Lopez-Martinez, Ernesto Ramirez-Gonzalez. |
| EPI_ISL_2496004 | LESP Tamaulipas | Instituto de Diagnostico y Referencia Epidemiologicos (INDRE) | Claudia Wong-Arambula, Abril Rodriguez-Maldonado, Vanessa Rivero-Arredondo, Ariadna Medina-Benitez, Joaquin Quiroz-Mercado, Sergio Rangel-Guerrero, Natividad Cruz-Ortiz, Tatiana Nunez-Garcia, Gisela Barrera-Badillo, Lucia Hernandez-Rivas, Irma Lopez-Martinez, Ernesto Ramirez-Gonzalez. |
| EPI_ISL_2496005 | LESP Campeche | Instituto de Diagnostico y Referencia Epidemiologicos (INDRE) | Claudia Wong-Arambula, Abril Rodriguez-Maldonado, Vanessa Rivero-Arredondo, Ariadna Medina-Benitez, Joaquin Quiroz-Mercado, Sergio Rangel-Guerrero, Natividad Cruz-Ortiz, Tatiana Nunez-Garcia, Gisela Barrera-Badillo, Lucia Hernandez-Rivas, Irma Lopez-Martinez, Ernesto Ramirez-Gonzalez. |
| EPI_ISL_2496008, EPI_ISL_2496009, EPI_ISL_2496012 | LESP Michoacan | Instituto de Diagnostico y Referencia Epidemiologicos (INDRE) | Claudia Wong-Arambula, Abril Rodriguez-Maldonado, Vanessa Rivero-Arredondo, Ariadna Medina-Benitez, Joaquin Quiroz-Mercado, Sergio Rangel-Guerrero, Natividad Cruz-Ortiz, Tatiana Nunez-Garcia, Gisela Barrera-Badillo, Lucia Hernandez-Rivas, Irma Lopez-Martinez, Ernesto Ramirez-Gonzalez. |
| EPI_ISL_2496013, EPI_ISL_2496014 | LESP Sinaloa | Instituto de Diagnostico y Referencia Epidemiologicos (INDRE) | Claudia Wong-Arambula, Abril Rodriguez-Maldonado, Vanessa Rivero-Arredondo, Ariadna Medina-Benitez, Joaquin Quiroz-Mercado, Sergio Rangel-Guerrero, Natividad Cruz-Ortiz, Tatiana Nunez-Garcia, Gisela Barrera-Badillo, Lucia Hernandez-Rivas, Irma Lopez-Martinez, Ernesto Ramirez-Gonzalez. |
| EPI_ISL_2496015, EPI_ISL_2496016, EPI_ISL_2496017, EPI_ISL_2496018, EPI_ISL_2496019 | LESP Ciudad de Mexico | Instituto de Diagnostico y Referencia Epidemiologicos (INDRE) | Claudia Wong-Arambula, Abril Rodriguez-Maldonado, Vanessa Rivero-Arredondo, Ariadna Medina-Benitez, Joaquin Quiroz-Mercado, Sergio Rangel-Guerrero, Natividad Cruz-Ortiz, Tatiana Nunez-Garcia, Gisela Barrera-Badillo, Lucia Hernandez-Rivas, Irma Lopez-Martinez, Ernesto Ramirez-Gonzalez. |
| EPI_ISL_2496020, EPI_ISL_2496021, EPI_ISL_2496022 | LESP Queretaro | Instituto de Diagnostico y Referencia Epidemiologicos (INDRE) | Claudia Wong-Arambula, Abril Rodriguez-Maldonado, Vanessa Rivero-Arredondo, Ariadna Medina-Benitez, Joaquin Quiroz-Mercado, Sergio Rangel-Guerrero, Natividad Cruz-Ortiz, Tatiana Nunez-Garcia, Gisela Barrera-Badillo, Lucia Hernandez-Rivas, Irma Lopez-Martinez, Ernesto Ramirez-Gonzalez. |
| EPI_ISL_2496023, EPI_ISL_2496024 | LESP Jalisco | Instituto de Diagnostico y Referencia Epidemiologicos (INDRE) | Claudia Wong-Arambula, Abril Rodriguez-Maldonado, Vanessa Rivero-Arredondo, Ariadna Medina-Benitez, Joaquin Quiroz-Mercado, Sergio Rangel-Guerrero, Natividad Cruz-Ortiz, Tatiana Nunez-Garcia, Gisela Barrera-Badillo, Lucia Hernandez-Rivas, Irma Lopez-Martinez, Ernesto Ramirez-Gonzalez. |
| EPI_ISL_2496025 | LESP Quintana Roo | Instituto de Diagnostico y Referencia Epidemiologicos (INDRE) | Claudia Wong-Arambula, Abril Rodriguez-Maldonado, Vanessa Rivero-Arredondo, Ariadna Medina-Benitez, Joaquin Quiroz-Mercado, Sergio Rangel-Guerrero, Natividad Cruz-Ortiz, Tatiana Nunez-Garcia, Gisela Barrera-Badillo, Lucia Hernandez-Rivas, Irma Lopez-Martinez, Ernesto Ramirez-Gonzalez. |
| EPI_ISL_2497483, EPI_ISL_2497486 | Genome Analysis Center, Yamanashi Central Hospital | Genome Analysis Center, Yamanashi Central Hospital | Yosuke Hirotsu |
| EPI_ISL_2497870, EPI_ISL_2497872, EPI_ISL_2497873, EPI_ISL_2497875, EPI_ISL_2497876, EPI_ISL_2497877, EPI_ISL_2497878, EPI_ISL_2497879 | Laboratori de Referencia de Catalunya | Laboratori de Referencia de Catalunya | Ramirez A., Bellosillo B., Padilla E., Vilas A., Hernandez JJ., Canal M. |
| EPI_ISL_2499966 | LESP Sinaloa | Instituto de Diagnostico y Referencia Epidemiologicos (INDRE) | Claudia Wong-Arambula, Abril Rodriguez-Maldonado, Vanessa Rivero-Arredondo, Ariadna Medina-Benitez, Joaquin Quiroz-Mercado, Sergio Rangel-Guerrero, Natividad Cruz-Ortiz, Tatiana Nunez-Garcia, Gisela Barrera-Badillo, Lucia Hernandez-Rivas, Irma Lopez-Martinez, Ernesto Ramirez-Gonzalez. |
| EPI_ISL_2499967 | LESP Michoacan | Instituto de Diagnostico y Referencia Epidemiologicos (INDRE) | Claudia Wong-Arambula, Abril Rodriguez-Maldonado, Vanessa Rivero-Arredondo, Ariadna Medina-Benitez, Joaquin Quiroz-Mercado, Sergio Rangel-Guerrero, Natividad Cruz-Ortiz, Tatiana Nunez-Garcia, Gisela Barrera-Badillo, Lucia Hernandez-Rivas, Irma Lopez-Martinez, Ernesto Ramirez-Gonzalez. |
| EPI_ISL_2500966 | Instituto Nacional de Salud | Instituto Nacional de Salud- Dirección de Investigación en Salud Pública | Katherine Laiton-Donato, Diego A. Álvarez-Díaz, Carlos Franco-Muñoz, Hector Alejandro Ruiz-Moreno, Paola Rojas, Maria T. Herrera-Sepúlveda, Diego Andrés Prada, Jhonnatan Reales-González, Sheryll Corchuelo, Julian Naizaque, Jorge Rivera, Gerardo Santamaría, Sergio Gomez, Lisseth Pardo, Juan Camilo Martinez, Marta Lopez Blanco, Ángela Alarcon Cruz, Diana Malo, Carmen Osorio, Magdalena Wiesner, Martha Lucia Ospina Martinez, Marcela Mercado-Reyes |
| EPI_ISL_2501020, EPI_ISL_2501021, EPI_ISL_2501022, EPI_ISL_2501023, EPI_ISL_2501024, EPI_ISL_2501025, EPI_ISL_2501026 | Banteay Meanchey Rapid Response Team | Virology Unit, Institut Pasteur du Cambodge | Cecile Troupin, Leakhena Pum, Jurre Y Siegers, Ly Sovann, Kraing Sidonn, Yi Sengdoeurn, Chin Savuth, Chau Darapheak, Veasna Duong, Erik A Karlsson |
| EPI_ISL_2501027, EPI_ISL_2501028 | Battambang Provincial Laboratory | Virology Unit, Institut Pasteur du Cambodge | Cecile Troupin, Leakhena Pum, Jurre Y Siegers, Ly Sovann, Kraing Sidonn, Yi Sengdoeurn, Chin Savuth, Chau Darapheak, Veasna Duong, Erik A Karlsson |
| EPI_ISL_2501029, EPI_ISL_2501030 | Siem Reap Provinical Laboratory | Virology Unit, Institut Pasteur du Cambodge | Cecile Troupin, Leakhena Pum, Jurre Y Siegers, Ly Sovann, Kraing Sidonn, Yi Sengdoeurn, Chin Savuth, Chau Darapheak, Veasna Duong, Erik A Karlsson |
| EPI_ISL_2501031, EPI_ISL_2501032 | Oddar Meanchey Rapid Response Team | Virology Unit, Institut Pasteur du Cambodge | Cecile Troupin, Leakhena Pum, Jurre Y Siegers, Ly Sovann, Kraing Sidonn, Yi Sengdoeurn, Chin Savuth, Chau Darapheak, Veasna Duong, Erik A Karlsson |
| EPI_ISL_2501033, EPI_ISL_2501034, EPI_ISL_2501035 | Tboung Khmum Rapid Response Team | Virology Unit, Institut Pasteur du Cambodge | Cecile Troupin, Leakhena Pum, Jurre Y Siegers, Ly Sovann, Kraing Sidonn, Yi Sengdoeurn, Chin Savuth, Chau Darapheak, Veasna Duong, Erik A Karlsson |
| EPI_ISL_2501036, EPI_ISL_2501037, EPI_ISL_2501038, EPI_ISL_2501039, EPI_ISL_2501040, EPI_ISL_2501041, EPI_ISL_2501042, EPI_ISL_2501043, EPI_ISL_2501044, EPI_ISL_2501045, EPI_ISL_2501046, EPI_ISL_2501047, EPI_ISL_2501048, EPI_ISL_2501049, EPI_ISL_2501050, EPI_ISL_2501051, EPI_ISL_2501052, EPI_ISL_2501053, EPI_ISL_2501054, EPI_ISL_2501055, EPI_ISL_2501056, EPI_ISL_2501057, EPI_ISL_2501058, EPI_ISL_2501059, EPI_ISL_2501060, EPI_ISL_2501061, EPI_ISL_2501062, EPI_ISL_2501063 | | | |
| see above | Virology Unit, Institut Pasteur du Cambodge | Virology Unit, Institut Pasteur du Cambodge | Cecile Troupin, Leakhena Pum, Jurre Y Siegers, Ly Sovann, Kraing Sidonn, Yi Sengdoeurn, Chin Savuth, Chau Darapheak, Veasna Duong, Erik A Karlsson |

| | | | |
|---|---|---|---|
| EPI_ISL_2501225, EPI_ISL_2501226 | CHU LILLE | CHU Lille - Laboratoire de Virologie | AIT YAHYA Emilie, ALIDJINOU Enagnon Kazali, BOCKET Laurence, CREPIN Michel, DEMAY Christophe, ENGELMANN Ilka, GEFFROY Sandrine, GUIGON Aurélie, LAMBERT Valérie, LAZREK Mouna, NOBILLIAUX Florian, PREVOST Brigitte, THUILLIER Caroline, TINEZ Claire ,TCHANTCHOU NJOSSE YANICK |
| EPI_ISL_2501227 | CH ROUBAIX | CHU Lille - Laboratoire de Virologie | AIT YAHYA Emilie, ALIDJINOU Enagnon Kazali, BOCKET Laurence, CREPIN Michel, DEMAY Christophe, ENGELMANN Ilka, GEFFROY Sandrine, GUIGON Aurélie, LAMBERT Valérie, LAZREK Mouna, NOBILLIAUX Florian, PREVOST Brigitte, THUILLIER Caroline, TINEZ Claire ,TCHANTCHOU NJOSSE YANICK |
| EPI_ISL_2501228 | CHU LILLE | CHU Lille - Laboratoire de Virologie | AIT YAHYA Emilie, ALIDJINOU Enagnon Kazali, BOCKET Laurence, CREPIN Michel, DEMAY Christophe, ENGELMANN Ilka, GEFFROY Sandrine, GUIGON Aurélie, LAMBERT Valérie, LAZREK Mouna, NOBILLIAUX Florian, PREVOST Brigitte, THUILLIER Caroline, TINEZ Claire ,TCHANTCHOU NJOSSE YANICK |
| EPI_ISL_2501229, EPI_ISL_2501230 | CH ARRAS | CHU Lille - Laboratoire de Virologie | AIT YAHYA Emilie, ALIDJINOU Enagnon Kazali, BOCKET Laurence, CREPIN Michel, DEMAY Christophe, ENGELMANN Ilka, GEFFROY Sandrine, GUIGON Aurélie, LAMBERT Valérie, LAZREK Mouna, NOBILLIAUX Florian, PREVOST Brigitte, THUILLIER Caroline, TINEZ Claire ,TCHANTCHOU NJOSSE YANICK |
| EPI_ISL_2501231 | CH SAINT-PHILIBERT | CHU Lille - Laboratoire de Virologie | AIT YAHYA Emilie, ALIDJINOU Enagnon Kazali, BOCKET Laurence, CREPIN Michel, DEMAY Christophe, ENGELMANN Ilka, GEFFROY Sandrine, GUIGON Aurélie, LAMBERT Valérie, LAZREK Mouna, NOBILLIAUX Florian, PREVOST Brigitte, THUILLIER Caroline, TINEZ Claire ,TCHANTCHOU NJOSSE YANICK |
| EPI_ISL_2501232 | CH CAMBRAI | CHU Lille - Laboratoire de Virologie | AIT YAHYA Emilie, ALIDJINOU Enagnon Kazali, BOCKET Laurence, CREPIN Michel, DEMAY Christophe, ENGELMANN Ilka, GEFFROY Sandrine, GUIGON Aurélie, LAMBERT Valérie, LAZREK Mouna, NOBILLIAUX Florian, PREVOST Brigitte, THUILLIER Caroline, TINEZ Claire ,TCHANTCHOU NJOSSE YANICK |
| EPI_ISL_2501233 | CHU LILLE | CHU Lille - Laboratoire de Virologie | AIT YAHYA Emilie, ALIDJINOU Enagnon Kazali, BOCKET Laurence, CREPIN Michel, DEMAY Christophe, ENGELMANN Ilka, GEFFROY Sandrine, GUIGON Aurélie, LAMBERT Valérie, LAZREK Mouna, NOBILLIAUX Florian, PREVOST Brigitte, THUILLIER Caroline, TINEZ Claire ,TCHANTCHOU NJOSSE YANICK |
| EPI_ISL_2501234, EPI_ISL_2501235, EPI_ISL_2501236 | CH ROUBAIX | CHU Lille - Laboratoire de Virologie | AIT YAHYA Emilie, ALIDJINOU Enagnon Kazali, BOCKET Laurence, CREPIN Michel, DEMAY Christophe, ENGELMANN Ilka, GEFFROY Sandrine, GUIGON Aurélie, LAMBERT Valérie, LAZREK Mouna, NOBILLIAUX Florian, PREVOST Brigitte, THUILLIER Caroline, TINEZ Claire ,TCHANTCHOU NJOSSE YANICK |
| EPI_ISL_2501237, EPI_ISL_2501238, EPI_ISL_2501239 | CH TOURCOING | CHU Lille - Laboratoire de Virologie | AIT YAHYA Emilie, ALIDJINOU Enagnon Kazali, BOCKET Laurence, CREPIN Michel, DEMAY Christophe, ENGELMANN Ilka, GEFFROY Sandrine, GUIGON Aurélie, LAMBERT Valérie, LAZREK Mouna, NOBILLIAUX Florian, PREVOST Brigitte, THUILLIER Caroline, TINEZ Claire ,TCHANTCHOU NJOSSE YANICK |
| EPI_ISL_2501240 | CHU LILLE | CHU Lille - Laboratoire de Virologie | AIT YAHYA Emilie, ALIDJINOU Enagnon Kazali, BOCKET Laurence, CREPIN Michel, DEMAY Christophe, ENGELMANN Ilka, GEFFROY Sandrine, GUIGON Aurélie, LAMBERT Valérie, LAZREK Mouna, NOBILLIAUX Florian, PREVOST Brigitte, THUILLIER Caroline, TINEZ Claire ,TCHANTCHOU NJOSSE YANICK |
| EPI_ISL_2501241 | CH BOULOGNE-SUR-MER | CHU Lille - Laboratoire de Virologie | AIT YAHYA Emilie, ALIDJINOU Enagnon Kazali, BOCKET Laurence, CREPIN Michel, DEMAY Christophe, ENGELMANN Ilka, GEFFROY Sandrine, GUIGON Aurélie, LAMBERT Valérie, LAZREK Mouna, NOBILLIAUX Florian, PREVOST Brigitte, THUILLIER Caroline, TINEZ Claire ,TCHANTCHOU NJOSSE YANICK |
| EPI_ISL_2501242, EPI_ISL_2501243 | CHU LILLE | CHU Lille - Laboratoire de Virologie | AIT YAHYA Emilie, ALIDJINOU Enagnon Kazali, BOCKET Laurence, CREPIN Michel, DEMAY Christophe, ENGELMANN Ilka, GEFFROY Sandrine, GUIGON Aurélie, LAMBERT Valérie, LAZREK Mouna, NOBILLIAUX Florian, PREVOST Brigitte, THUILLIER Caroline, TINEZ Claire ,TCHANTCHOU NJOSSE YANICK |
| EPI_ISL_2501244, EPI_ISL_2501245, EPI_ISL_2501246 | CH ROUBAIX | CHU Lille - Laboratoire de Virologie | AIT YAHYA Emilie, ALIDJINOU Enagnon Kazali, BOCKET Laurence, CREPIN Michel, DEMAY Christophe, ENGELMANN Ilka, GEFFROY Sandrine, GUIGON Aurélie, LAMBERT Valérie, LAZREK Mouna, NOBILLIAUX Florian, PREVOST Brigitte, THUILLIER Caroline, TINEZ Claire ,TCHANTCHOU NJOSSE YANICK |
| EPI_ISL_2501247 | CH BOULOGNE-SUR-MER | CHU Lille - Laboratoire de Virologie | AIT YAHYA Emilie, ALIDJINOU Enagnon Kazali, BOCKET Laurence, CREPIN Michel, DEMAY Christophe, ENGELMANN Ilka, GEFFROY Sandrine, GUIGON Aurélie, LAMBERT Valérie, LAZREK Mouna, NOBILLIAUX Florian, PREVOST Brigitte, THUILLIER Caroline, TINEZ Claire ,TCHANTCHOU NJOSSE YANICK |
| EPI_ISL_2501248, EPI_ISL_2501249 | CHU LILLE | CHU Lille - Laboratoire de Virologie | AIT YAHYA Emilie, ALIDJINOU Enagnon Kazali, BOCKET Laurence, CREPIN Michel, DEMAY Christophe, ENGELMANN Ilka, GEFFROY Sandrine, GUIGON Aurélie, LAMBERT Valérie, LAZREK Mouna, NOBILLIAUX Florian, PREVOST Brigitte, THUILLIER Caroline, TINEZ Claire ,TCHANTCHOU NJOSSE YANICK |
| EPI_ISL_2501250 | CH SECLIN | CHU Lille - Laboratoire de Virologie | AIT YAHYA Emilie, ALIDJINOU Enagnon Kazali, BOCKET Laurence, CREPIN Michel, DEMAY Christophe, ENGELMANN Ilka, GEFFROY Sandrine, GUIGON Aurélie, LAMBERT Valérie, LAZREK Mouna, NOBILLIAUX Florian, PREVOST Brigitte, THUILLIER Caroline, TINEZ Claire ,TCHANTCHOU NJOSSE YANICK |
| EPI_ISL_2501251, EPI_ISL_2501252, EPI_ISL_2501253, EPI_ISL_2501254, EPI_ISL_2501255, EPI_ISL_2501256 | CHU LILLE | CHU Lille - Laboratoire de Virologie | AIT YAHYA Emilie, ALIDJINOU Enagnon Kazali, BOCKET Laurence, CREPIN Michel, DEMAY Christophe, ENGELMANN Ilka, GEFFROY Sandrine, GUIGON Aurélie, LAMBERT Valérie, LAZREK Mouna, NOBILLIAUX Florian, PREVOST Brigitte, THUILLIER Caroline, TINEZ Claire ,TCHANTCHOU NJOSSE YANICK |
| EPI_ISL_2501257 | CH SAINT-PHILIBERT | CHU Lille - Laboratoire de Virologie | AIT YAHYA Emilie, ALIDJINOU Enagnon Kazali, BOCKET Laurence, CREPIN Michel, DEMAY Christophe, ENGELMANN Ilka, GEFFROY Sandrine, GUIGON Aurélie, LAMBERT Valérie, LAZREK Mouna, NOBILLIAUX Florian, PREVOST Brigitte, THUILLIER Caroline, TINEZ Claire ,TCHANTCHOU NJOSSE YANICK |
| EPI_ISL_2501258, EPI_ISL_2501259, EPI_ISL_2501260, EPI_ISL_2501261, EPI_ISL_2501262 | CH ROUBAIX | CHU Lille - Laboratoire de Virologie | AIT YAHYA Emilie, ALIDJINOU Enagnon Kazali, BOCKET Laurence, CREPIN Michel, DEMAY Christophe, ENGELMANN Ilka, GEFFROY Sandrine, GUIGON Aurélie, LAMBERT Valérie, LAZREK Mouna, NOBILLIAUX Florian, PREVOST Brigitte, THUILLIER Caroline, TINEZ Claire ,TCHANTCHOU NJOSSE YANICK |
| EPI_ISL_2501263, EPI_ISL_2501264 | CHU LILLE | CHU Lille - Laboratoire de Virologie | AIT YAHYA Emilie, ALIDJINOU Enagnon Kazali, BOCKET Laurence, CREPIN Michel, DEMAY Christophe, ENGELMANN Ilka, GEFFROY Sandrine, GUIGON Aurélie, LAMBERT Valérie, LAZREK Mouna, NOBILLIAUX Florian, PREVOST Brigitte, THUILLIER Caroline, TINEZ Claire ,TCHANTCHOU NJOSSE YANICK |
| EPI_ISL_2501265 | CH ARRAS | CHU Lille - Laboratoire de Virologie | AIT YAHYA Emilie, ALIDJINOU Enagnon Kazali, BOCKET Laurence, CREPIN Michel, DEMAY Christophe, ENGELMANN Ilka, GEFFROY Sandrine, GUIGON Aurélie, LAMBERT Valérie, LAZREK Mouna, NOBILLIAUX Florian, PREVOST Brigitte, THUILLIER Caroline, TINEZ Claire ,TCHANTCHOU NJOSSE YANICK |
| EPI_ISL_2501266 | CH TOURCOING | CHU Lille - Laboratoire de Virologie | AIT YAHYA Emilie, ALIDJINOU Enagnon Kazali, BOCKET Laurence, CREPIN Michel, DEMAY Christophe, ENGELMANN Ilka, GEFFROY Sandrine, GUIGON Aurélie, LAMBERT Valérie, LAZREK Mouna, NOBILLIAUX Florian, PREVOST Brigitte, THUILLIER Caroline, TINEZ Claire ,TCHANTCHOU NJOSSE YANICK |
| EPI_ISL_2501267 | CH SAINT-PHILIBERT | CHU Lille - Laboratoire de Virologie | AIT YAHYA Emilie, ALIDJINOU Enagnon Kazali, BOCKET Laurence, CREPIN Michel, DEMAY Christophe, ENGELMANN Ilka, GEFFROY Sandrine, GUIGON Aurélie, LAMBERT Valérie, LAZREK Mouna, NOBILLIAUX Florian, PREVOST Brigitte, THUILLIER Caroline, TINEZ Claire ,TCHANTCHOU NJOSSE YANICK |
| EPI_ISL_2501268, EPI_ISL_2501269, EPI_ISL_2501270 | CHU LILLE | CHU Lille - Laboratoire de Virologie | AIT YAHYA Emilie, ALIDJINOU Enagnon Kazali, BOCKET Laurence, CREPIN Michel, DEMAY Christophe, ENGELMANN Ilka, GEFFROY Sandrine, GUIGON Aurélie, LAMBERT Valérie, LAZREK Mouna, NOBILLIAUX Florian, PREVOST Brigitte, THUILLIER Caroline, TINEZ Claire ,TCHANTCHOU NJOSSE YANICK |
| EPI_ISL_2501271 | CH ARRAS | CHU Lille - Laboratoire de Virologie | AIT YAHYA Emilie, ALIDJINOU Enagnon Kazali, BOCKET Laurence, CREPIN Michel, DEMAY Christophe, ENGELMANN Ilka, GEFFROY Sandrine, GUIGON Aurélie, LAMBERT Valérie, LAZREK Mouna, NOBILLIAUX Florian, PREVOST Brigitte, THUILLIER Caroline, TINEZ Claire ,TCHANTCHOU NJOSSE YANICK |

| | | | |
|---|---|---|---|
| EPI_ISL_2501272 | CHU LILLE | CHU Lille - Laboratoire de Virologie | AIT YAHYA Emilie, ALIDJINOU Enagnon Kazali, BOCKET Laurence, CREPIN Michel, DEMAY Christophe, ENGELMANN Ilka, GEFFROY Sandrine, GUIGON Aurélie, LAMBERT Valérie, LAZREK Mouna, NOBILLIAUX Florian, PREVOST Brigitte, THUILLIER Caroline, TINEZ Claire ,TCHANTCHOU NJOSSE YANICK |
| EPI_ISL_2501273 | CH ARRAS | CHU Lille - Laboratoire de Virologie | AIT YAHYA Emilie, ALIDJINOU Enagnon Kazali, BOCKET Laurence, CREPIN Michel, DEMAY Christophe, ENGELMANN Ilka, GEFFROY Sandrine, GUIGON Aurélie, LAMBERT Valérie, LAZREK Mouna, NOBILLIAUX Florian, PREVOST Brigitte, THUILLIER Caroline, TINEZ Claire ,TCHANTCHOU NJOSSE YANICK |
| EPI_ISL_2501274, EPI_ISL_2501275, EPI_ISL_2501276, EPI_ISL_2501277 | CH TOURCOING | CHU Lille - Laboratoire de Virologie | AIT YAHYA Emilie, ALIDJINOU Enagnon Kazali, BOCKET Laurence, CREPIN Michel, DEMAY Christophe, ENGELMANN Ilka, GEFFROY Sandrine, GUIGON Aurélie, LAMBERT Valérie, LAZREK Mouna, NOBILLIAUX Florian, PREVOST Brigitte, THUILLIER Caroline, TINEZ Claire ,TCHANTCHOU NJOSSE YANICK |
| EPI_ISL_2501278, EPI_ISL_2501279 | CHU LILLE | CHU Lille - Laboratoire de Virologie | AIT YAHYA Emilie, ALIDJINOU Enagnon Kazali, BOCKET Laurence, CREPIN Michel, DEMAY Christophe, ENGELMANN Ilka, GEFFROY Sandrine, GUIGON Aurélie, LAMBERT Valérie, LAZREK Mouna, NOBILLIAUX Florian, PREVOST Brigitte, THUILLIER Caroline, TINEZ Claire ,TCHANTCHOU NJOSSE YANICK |
| EPI_ISL_2501280, EPI_ISL_2501281, EPI_ISL_2501282 | CH ROUBAIX | CHU Lille - Laboratoire de Virologie | AIT YAHYA Emilie, ALIDJINOU Enagnon Kazali, BOCKET Laurence, CREPIN Michel, DEMAY Christophe, ENGELMANN Ilka, GEFFROY Sandrine, GUIGON Aurélie, LAMBERT Valérie, LAZREK Mouna, NOBILLIAUX Florian, PREVOST Brigitte, THUILLIER Caroline, TINEZ Claire ,TCHANTCHOU NJOSSE YANICK |
| EPI_ISL_2501283, EPI_ISL_2501284 | CH LENS | CHU Lille - Laboratoire de Virologie | AIT YAHYA Emilie, ALIDJINOU Enagnon Kazali, BOCKET Laurence, CREPIN Michel, DEMAY Christophe, ENGELMANN Ilka, GEFFROY Sandrine, GUIGON Aurélie, LAMBERT Valérie, LAZREK Mouna, NOBILLIAUX Florian, PREVOST Brigitte, THUILLIER Caroline, TINEZ Claire ,TCHANTCHOU NJOSSE YANICK |
| EPI_ISL_2501285 | CH TOURCOING | CHU Lille - Laboratoire de Virologie | AIT YAHYA Emilie, ALIDJINOU Enagnon Kazali, BOCKET Laurence, CREPIN Michel, DEMAY Christophe, ENGELMANN Ilka, GEFFROY Sandrine, GUIGON Aurélie, LAMBERT Valérie, LAZREK Mouna, NOBILLIAUX Florian, PREVOST Brigitte, THUILLIER Caroline, TINEZ Claire ,TCHANTCHOU NJOSSE YANICK |
| EPI_ISL_2501286 | CHU LILLE | CHU Lille - Laboratoire de Virologie | AIT YAHYA Emilie, ALIDJINOU Enagnon Kazali, BOCKET Laurence, CREPIN Michel, DEMAY Christophe, ENGELMANN Ilka, GEFFROY Sandrine, GUIGON Aurélie, LAMBERT Valérie, LAZREK Mouna, NOBILLIAUX Florian, PREVOST Brigitte, THUILLIER Caroline, TINEZ Claire ,TCHANTCHOU NJOSSE YANICK |
| EPI_ISL_2501287 | CH ARMENTIERES | CHU Lille - Laboratoire de Virologie | AIT YAHYA Emilie, ALIDJINOU Enagnon Kazali, BOCKET Laurence, CREPIN Michel, DEMAY Christophe, ENGELMANN Ilka, GEFFROY Sandrine, GUIGON Aurélie, LAMBERT Valérie, LAZREK Mouna, NOBILLIAUX Florian, PREVOST Brigitte, THUILLIER Caroline, TINEZ Claire ,TCHANTCHOU NJOSSE YANICK |
| EPI_ISL_2501288 | CH BETHUNE | CHU Lille - Laboratoire de Virologie | AIT YAHYA Emilie, ALIDJINOU Enagnon Kazali, BOCKET Laurence, CREPIN Michel, DEMAY Christophe, ENGELMANN Ilka, GEFFROY Sandrine, GUIGON Aurélie, LAMBERT Valérie, LAZREK Mouna, NOBILLIAUX Florian, PREVOST Brigitte, THUILLIER Caroline, TINEZ Claire ,TCHANTCHOU NJOSSE YANICK |
| EPI_ISL_2501289, EPI_ISL_2501290 | CH SAINT-PHILIBERT | CHU Lille - Laboratoire de Virologie | AIT YAHYA Emilie, ALIDJINOU Enagnon Kazali, BOCKET Laurence, CREPIN Michel, DEMAY Christophe, ENGELMANN Ilka, GEFFROY Sandrine, GUIGON Aurélie, LAMBERT Valérie, LAZREK Mouna, NOBILLIAUX Florian, PREVOST Brigitte, THUILLIER Caroline, TINEZ Claire ,TCHANTCHOU NJOSSE YANICK |
| EPI_ISL_2501291, EPI_ISL_2501292, EPI_ISL_2501293, EPI_ISL_2501294 | CHU LILLE | CHU Lille - Laboratoire de Virologie | AIT YAHYA Emilie, ALIDJINOU Enagnon Kazali, BOCKET Laurence, CREPIN Michel, DEMAY Christophe, ENGELMANN Ilka, GEFFROY Sandrine, GUIGON Aurélie, LAMBERT Valérie, LAZREK Mouna, NOBILLIAUX Florian, PREVOST Brigitte, THUILLIER Caroline, TINEZ Claire ,TCHANTCHOU NJOSSE YANICK |
| EPI_ISL_2501295 | CH TOURCOING | CHU Lille - Laboratoire de Virologie | AIT YAHYA Emilie, ALIDJINOU Enagnon Kazali, BOCKET Laurence, CREPIN Michel, DEMAY Christophe, ENGELMANN Ilka, GEFFROY Sandrine, GUIGON Aurélie, LAMBERT Valérie, LAZREK Mouna, NOBILLIAUX Florian, PREVOST Brigitte, THUILLIER Caroline, TINEZ Claire ,TCHANTCHOU NJOSSE YANICK |
| EPI_ISL_2501296 | CHU LILLE | CHU Lille - Laboratoire de Virologie | AIT YAHYA Emilie, ALIDJINOU Enagnon Kazali, BOCKET Laurence, CREPIN Michel, DEMAY Christophe, ENGELMANN Ilka, GEFFROY Sandrine, GUIGON Aurélie, LAMBERT Valérie, LAZREK Mouna, NOBILLIAUX Florian, PREVOST Brigitte, THUILLIER Caroline, TINEZ Claire ,TCHANTCHOU NJOSSE YANICK |
| EPI_ISL_2501297 | CH CAMBRAI | CHU Lille - Laboratoire de Virologie | AIT YAHYA Emilie, ALIDJINOU Enagnon Kazali, BOCKET Laurence, CREPIN Michel, DEMAY Christophe, ENGELMANN Ilka, GEFFROY Sandrine, GUIGON Aurélie, LAMBERT Valérie, LAZREK Mouna, NOBILLIAUX Florian, PREVOST Brigitte, THUILLIER Caroline, TINEZ Claire ,TCHANTCHOU NJOSSE YANICK |
| EPI_ISL_2501299, EPI_ISL_2501300 | CHU LILLE | CHU Lille - Laboratoire de Virologie | AIT YAHYA Emilie, ALIDJINOU Enagnon Kazali, BOCKET Laurence, CREPIN Michel, DEMAY Christophe, ENGELMANN Ilka, GEFFROY Sandrine, GUIGON Aurélie, LAMBERT Valérie, LAZREK Mouna, NOBILLIAUX Florian, PREVOST Brigitte, THUILLIER Caroline, TINEZ Claire ,TCHANTCHOU NJOSSE YANICK |
| EPI_ISL_2501301 | CH LENS | CHU Lille - Laboratoire de Virologie | AIT YAHYA Emilie, ALIDJINOU Enagnon Kazali, BOCKET Laurence, CREPIN Michel, DEMAY Christophe, ENGELMANN Ilka, GEFFROY Sandrine, GUIGON Aurélie, LAMBERT Valérie, LAZREK Mouna, NOBILLIAUX Florian, PREVOST Brigitte, THUILLIER Caroline, TINEZ Claire ,TCHANTCHOU NJOSSE YANICK |
| EPI_ISL_2501304, EPI_ISL_2501305, EPI_ISL_2501307 | Heilig hart Lier | Imelda Ziekenhuis | Johan Frans, Hanne Valgaeren, Dagmar Obbels |

EPI_ISL_2501313, EPI_ISL_2501314, EPI_ISL_2501315, EPI_ISL_2501316, EPI_ISL_2501317, EPI_ISL_2501318, EPI_ISL_2501319, EPI_ISL_2501320, EPI_ISL_2501321, EPI_ISL_2501322, EPI_ISL_2501323, EPI_ISL_2501324, EPI_ISL_2501325, EPI_ISL_2501326, EPI_ISL_2501327, EPI_ISL_2501328, EPI_ISL_2501329, EPI_ISL_2501330, EPI_ISL_2501331, EPI_ISL_2501332, EPI_ISL_2501335, EPI_ISL_2501336, EPI_ISL_2501337, EPI_ISL_2501338, EPI_ISL_2501339, EPI_ISL_2501340, EPI_ISL_2501341, EPI_ISL_2501342, EPI_ISL_2501343, EPI_ISL_2501344, EPI_ISL_2501345, EPI_ISL_2501346, EPI_ISL_2501347, EPI_ISL_2501363, EPI_ISL_2501364, EPI_ISL_2501365, EPI_ISL_2501366, EPI_ISL_2501367, EPI_ISL_2501368, EPI_ISL_2501369, EPI_ISL_2501370, EPI_ISL_2501371, EPI_ISL_2501372, EPI_ISL_2501373, EPI_ISL_2501374, EPI_ISL_2501375, EPI_ISL_2501376, EPI_ISL_2501377, EPI_ISL_2501378

| | | | |
|---|---|---|---|
| see above | Microbiology Department. Complexo Hospitalario Universitario de Vigo | Microbiology Department. Complexo Hospitalario Universitario de Vigo | Alfaya N, Alonso I, Alvarez M, Cabrera JJ, Carballo R, Cores O, Cortizo S, del-Campo V, Martinez L, Mediero G, Perez S, Potel C, Regueiro B, Rey S, Vasallo FJ |
| EPI_ISL_2502615 | RSUD dr Loekmono Hadi | Genetics Working Group (Pokja Genetik) Faculty of Medicine, Public Health and Nursing Universitas Gadjah Mada (FK-KMK UGM); Disease Investigation Center Wates Ministry of Agriculture Indonesia; Department of Microbiology FK-KMK UGM; Laboratorium Diagnostik Yayasan Tahija World Mosquito Program (WMP) Yogyakarta Center for Tropical Medicine FK-KMK UGM; Integrated Research Center FK-KMK UGM; Department of Computer Science and Electronics FMIPA UGM; RSUP Dr. Sardjito | Gunadi, Hendra Wibawa, Marcellus, Mohamad S. Hakim, Edwin W. Daniwijaya, Ludhang P. Rizki, Endah Supriyati, Eggi Arguni, Titik Nuryastuti, Tri Wibawa, Dwi AA Nugrahaningsih, Afiahayati, Siswanto,Kristy Iskandar,Nungki Anggorowati, Pramesti G Dewi, Dwi Indaryati,Indarto Sulistiyono, Susan Simanjaya, Dyah A Puspitarani |
| EPI_ISL_2502687 | RSUD dr. LOEKMONO HADI | Genetics Working Group (Pokja Genetik) Faculty of Medicine, Public Health and Nursing Universitas Gadjah Mada (FK-KMK UGM); Disease Investigation Center Wates Ministry of Agriculture Indonesia; Department of Microbiology FK-KMK UGM; Laboratorium Diagnostik Yayasan Tahija World Mosquito Program (WMP) Yogyakarta Center for Tropical Medicine FK-KMK UGM; Integrated Research Center FK-KMK UGM; Department of Computer Science and Electronics FMIPA UGM; RSUP Dr. Sardjito | Gunadi, Hendra Wibawa, Marcellus, Mohamad S. Hakim, Edwin W. Daniwijaya, Ludhang P. Rizki, Endah Supriyati, Eggi Arguni, Titik Nuryastuti, Tri Wibawa, Dwi AA Nugrahaningsih, Afiahayati, Siswanto,Kristy Iskandar,Nungki Anggorowati, Pramesti G Dewi, Dwi Indaryati,Indarto Sulistiyono, Cita S Amalia, Alvina A Setiawan |

| | | | |
|---|---|---|---|
| EPI_ISL_2502735 | AREA DE SALUD CATEDRAL NORESTE | Inciensa, Instituto Costarricense de InvestigaciOn y Enseñanza en NutriciOn y Salud | Francisco Duarte, Hebleen Porras, Claudio Soto-Garita, Estela Cordero, Adriana Godínez, Melany CalderOn, Jose Luis Vargas, Mariela Gutierrez, Joselyn Prado & Mariel LOpez |
| EPI_ISL_2502738 | HOSPITAL DR. MAX TERAN VALLS | Inciensa, Instituto Costarricense de InvestigaciOn y Enseñanza en NutriciOn y Salud | Francisco Duarte, Hebleen Porras, Claudio Soto-Garita, Estela Cordero, Adriana Godínez, Melany CalderOn, Jose Luis Vargas, Mariela Gutierrez, Joselyn Prado & María Jose GOmez-Umaña |
| EPI_ISL_2502739 | AREA DE SALUD MATA REDONDA -HOSPITAL -CLINICA DR. MORENO CAÑAS | Inciensa, Instituto Costarricense de InvestigaciOn y Enseñanza en NutriciOn y Salud | Francisco Duarte, Hebleen Porras, Claudio Soto-Garita, Estela Cordero, Adriana Godínez, Melany CalderOn, Jose Luis Vargas, Mariela Gutierrez, Joselyn Prado & Carolina Duarte |
| EPI_ISL_2502740 | LABORATORIO CLINICO LABIN | Inciensa, Instituto Costarricense de InvestigaciOn y Enseñanza en NutriciOn y Salud | Francisco Duarte, Hebleen Porras, Claudio Soto-Garita, Estela Cordero, Adriana Godínez, Melany CalderOn, Jose Luis Vargas, Mariela Gutierrez, Joselyn Prado & Pei Chan |
| EPI_ISL_2502741 | HOSPITAL SAN FRANCISCO DE ASIS | Inciensa, Instituto Costarricense de InvestigaciOn y Enseñanza en NutriciOn y Salud | Francisco Duarte, Hebleen Porras, Claudio Soto-Garita, Estela Cordero, Adriana Godínez, Melany CalderOn, Jose Luis Vargas, Mariela Gutierrez, Joselyn Prado & Adrián Fallas-Mora |
| EPI_ISL_2502742 | AREA DE SALUD OSA | Inciensa, Instituto Costarricense de InvestigaciOn y Enseñanza en NutriciOn y Salud | Francisco Duarte, Hebleen Porras, Claudio Soto-Garita, Estela Cordero, Adriana Godínez, Melany CalderOn, Jose Luis Vargas, Mariela Gutierrez, Joselyn Prado & María Ramírez |
| EPI_ISL_2502743 | HOSPITAL DR. FERNANDO ESCALANTE PRADILLA | Inciensa, Instituto Costarricense de InvestigaciOn y Enseñanza en NutriciOn y Salud | Francisco Duarte, Hebleen Porras, Claudio Soto-Garita, Estela Cordero, Adriana Godínez, Melany CalderOn, Jose Luis Vargas, Mariela Gutierrez, Joselyn Prado & María Matamoros-Madrigal |
| EPI_ISL_2502744 | AREA DE SALUD SIQUIRRES | Inciensa, Instituto Costarricense de InvestigaciOn y Enseñanza en NutriciOn y Salud | Francisco Duarte, Hebleen Porras, Claudio Soto-Garita, Estela Cordero, Adriana Godínez, Melany CalderOn, Jose Luis Vargas, Mariela Gutierrez, Joselyn Prado & Karla LOpez |
| EPI_ISL_2502745 | AREA DE SALUD CARTAGO | Inciensa, Instituto Costarricense de InvestigaciOn y Enseñanza en NutriciOn y Salud | Francisco Duarte, Hebleen Porras, Claudio Soto-Garita, Estela Cordero, Adriana Godínez, Melany CalderOn, Jose Luis Vargas, Mariela Gutierrez, Joselyn Prado & Tania Garro-Vargas |
| EPI_ISL_2502746 | HOSPITAL MEXICO | Inciensa, Instituto Costarricense de InvestigaciOn y Enseñanza en NutriciOn y Salud | Francisco Duarte, Hebleen Porras, Claudio Soto-Garita, Estela Cordero, Adriana Godínez, Melany CalderOn, Jose Luis Vargas, Mariela Gutierrez, Joselyn Prado & Juan Vilallobos |
| EPI_ISL_2502747 | AREA DE SALUD PARAISO-CERVANTES | Inciensa, Instituto Costarricense de InvestigaciOn y Enseñanza en NutriciOn y Salud | Francisco Duarte, Hebleen Porras, Claudio Soto-Garita, Estela Cordero, Adriana Godínez, Melany CalderOn, Jose Luis Vargas, Mariela Gutierrez & Joselyn Prado |
| EPI_ISL_2502748 | AREA DE SALUD DESAMPARADOS 2 (COOPESALUD) | Inciensa, Instituto Costarricense de InvestigaciOn y Enseñanza en NutriciOn y Salud | Francisco Duarte, Hebleen Porras, Claudio Soto-Garita, Estela Cordero, Adriana Godínez, Melany CalderOn, Jose Luis Vargas, Mariela Gutierrez, Joselyn Prado & Paola Sánchez-Traña |
| EPI_ISL_2502749 | HOSPITAL SAN JUAN DE DIOS | Inciensa, Instituto Costarricense de InvestigaciOn y Enseñanza en NutriciOn y Salud | Francisco Duarte, Hebleen Porras, Claudio Soto-Garita, Estela Cordero, Adriana Godínez, Melany CalderOn, Jose Luis Vargas, Mariela Gutierrez, Joselyn Prado & Paola Sánchez-Traña |
| EPI_ISL_2502750 | AREA DE SALUD PEREZ ZELEDON | Inciensa, Instituto Costarricense de InvestigaciOn y Enseñanza en NutriciOn y Salud | Francisco Duarte, Hebleen Porras, Claudio Soto-Garita, Estela Cordero, Adriana Godínez, Melany CalderOn, Jose Luis Vargas, Mariela Gutierrez, Joselyn Prado & Fabricio Aguilar |
| EPI_ISL_2502751 | AREA DE SALUD PAQUERA | Inciensa, Instituto Costarricense de InvestigaciOn y Enseñanza en NutriciOn y Salud | Francisco Duarte, Hebleen Porras, Claudio Soto-Garita, Estela Cordero, Adriana Godínez, Melany CalderOn, Jose Luis Vargas, Mariela Gutierrez, Joselyn Prado & Ronald Núñez-Barrientos |
| EPI_ISL_2502752 | AREA DE SALUD ALAJUELITA | Inciensa, Instituto Costarricense de InvestigaciOn y Enseñanza en NutriciOn y Salud | Francisco Duarte, Hebleen Porras, Claudio Soto-Garita, Estela Cordero, Adriana Godínez, Melany CalderOn, Jose Luis Vargas, Mariela Gutierrez & Joselyn Prado |
| EPI_ISL_2502753, EPI_ISL_2502754 | HOSPITAL SAN JUAN DE DIOS | Inciensa, Instituto Costarricense de InvestigaciOn y Enseñanza en NutriciOn y Salud | Francisco Duarte, Hebleen Porras, Claudio Soto-Garita, Estela Cordero, Adriana Godínez, Melany CalderOn, Jose Luis Vargas, Mariela Gutierrez, Joselyn Prado & Marco Cháves-Otárola |
| EPI_ISL_2502755 | AREA DE SALUD GOICOECHEA 2 - CLINICA DR. JIMENEZ NUÑEZ | Inciensa, Instituto Costarricense de InvestigaciOn y Enseñanza en NutriciOn y Salud | Francisco Duarte, Hebleen Porras, Claudio Soto-Garita, Estela Cordero, Adriana Godínez, Melany CalderOn, Jose Luis Vargas, Mariela Gutierrez, Joselyn Prado & Alberto Madrigal-Vega |
| EPI_ISL_2502756 | AREA DE SALUD LA UNION | Inciensa, Instituto Costarricense de InvestigaciOn y Enseñanza en NutriciOn y Salud | Francisco Duarte, Hebleen Porras, Claudio Soto-Garita, Estela Cordero, Adriana Godínez, Melany CalderOn, Jose Luis Vargas, Mariela Gutierrez, Joselyn Prado & MOnica Charpentier |
| EPI_ISL_2502757 | AREA DE SALUD PARAISO-CERVANTES | Inciensa, Instituto Costarricense de InvestigaciOn y Enseñanza en NutriciOn y Salud | Francisco Duarte, Hebleen Porras, Claudio Soto-Garita, Estela Cordero, Adriana Godínez, Melany CalderOn, Jose Luis Vargas, Mariela Gutierrez, Joselyn Prado & MOnica Charpentier |
| EPI_ISL_2502758, EPI_ISL_2502759 | AREA DE SALUD ZAPOTE-CATEDRAL - CLINICA DR. CARLOS DURAN | Inciensa, Instituto Costarricense de InvestigaciOn y Enseñanza en NutriciOn y Salud | Francisco Duarte, Hebleen Porras, Claudio Soto-Garita, Estela Cordero, Adriana Godínez, Melany CalderOn, Jose Luis Vargas, Mariela Gutierrez, Joselyn Prado & Marco Cháves-Otárola |
| EPI_ISL_2502760, EPI_ISL_2502761, EPI_ISL_2502762 | HOSPITAL SAN JUAN DE DIOS | Inciensa, Instituto Costarricense de InvestigaciOn y Enseñanza en NutriciOn y Salud | Francisco Duarte, Hebleen Porras, Claudio Soto-Garita, Estela Cordero, Adriana Godínez, Melany CalderOn, Jose Luis Vargas, Mariela Gutierrez, Joselyn Prado & Marco Cháves-Otárola |
| EPI_ISL_2502764 | RSUD dr. LOEKMONO HADI | Genetics Working Group (Pokja Genetik) Faculty of Medicine, Public Health and Nursing Universitas Gadjah Mada (FK-KMK UGM); Disease Investigation Center Wates Ministry of Agriculture Indonesia; Department of Microbiology FK-KMK UGM; Laboratorium Diagnostik Yayasan Tahija World Mosquito Program (WMP) Yogyakarta Center for Tropical Medicine FK-KMK UGM; Integrated Research Center FK-KMK UGM; Department of Computer Science and Electronics FMIPA UGM; RSUP Dr. Sardjito | Gunadi, Hendra Wibawa, Marcellus, Mohamad S. Hakim, Edwin W. Daniwijaya, Ludhang P. Rizki, Endah Supriyati, Eggi Arguni, Titik Nuryastuti, Tri Wibawa, Dwi AA Nugrahaningsih, Afiahayati, Siswanto,Kristy Iskandar,Nungki Anggorowati, Pramesti G Dewi, Dwi Indaryati,Indarto Sulistiyono, Irene Tania, Hana F Hanifin |
| EPI_ISL_2503135, EPI_ISL_2503137, EPI_ISL_2503138, EPI_ISL_2503139 | Laboratorio Biologia Molecolare Sars Cov2 - UOC Laboratorio Analisi - Servizio Medicina di Laboratorio, Ospedale "San Francesco" - ATS-ASSL Nuoro | Laboratorio specialistico UOC Ematologia - Ospedale "San Francesco" - ATS-ASSL Nuoro | Asproni Rosanna,Piras Giovanna, Pira Giovanna, Fancello Patrizia, Rosu Valentina, Carta Franco,Fiamma Maura, Mameli Giuseppe, Maria Cristina Garau |
| EPI_ISL_2504068, EPI_ISL_2504071, EPI_ISL_2504072, EPI_ISL_2504073, EPI_ISL_2504088 | Botswana Harvard HIV Reference Laboratory | Botswana Harvard HIV Reference Laboratory | Sikhulile Moyo, Dorcas Maruapula, Wonderful T. Choga, Thongbotho Mphoyakgosi, Boitumelo Zuze, Botshelo Radibe, Legodile Kooepile, Ontlametse T. Bareng, Keoratile Ntshambiwa, Modisa Motswaledi, Madisa Mine, Joseph Makhema, Roger Shapiro, Shahin Lockman, Mosepele Mosepele, Simani Gaseitsiwe |
| EPI_ISL_2504114 | Palapye Primary Hospital Laboratory | Botswana Harvard HIV Reference Laboratory | Sikhulile Moyo, Dorcas Maruapula, Wonderful T. Choga, Thongbotho Mphoyakgosi, Boitumelo Zuze, Botshelo Radibe, Legodile Kooepile, Ontlametse T. Bareng, Letsibogo Gaoraelwe, Thela Tefelo, Keoratile Ntshambiwa, Modisa Motswaledi, Madisa Mine, Joseph Makhema, Roger Shapiro, Shahin Lockman, Mosepele Mosepele, Simani Gaseitsiwe |
| EPI_ISL_2506041 | RSUD dr. LOEKMONO HADI | Genetics Working Group (Pokja Genetik) Faculty of Medicine, Public Health and Nursing Universitas Gadjah Mada (FK-KMK UGM); Disease Investigation Center Wates Ministry of Agriculture Indonesia; Department of Microbiology FK-KMK UGM; Laboratorium Diagnostik Yayasan Tahija World Mosquito Program (WMP) Yogyakarta Center for Tropical Medicine FK-KMK UGM; Integrated Research Center FK-KMK UGM; Department of Computer Science and Electronics FMIPA UGM; RSUP Dr. Sardjito | Gunadi, Hendra Wibawa, Marcellus, Mohamad S. Hakim, Edwin W. Daniwijaya, Ludhang P. Rizki, Endah Supriyati, Eggi Arguni, Titik Nuryastuti, Tri Wibawa, Dwi AA Nugrahaningsih, Afiahayati, Siswanto,Kristy Iskandar,Nungki Anggorowati, Pramesti G Dewi, Dwi Indaryati,Indarto Sulistiyono, Irene Tania, Hana F Hanifin |
| EPI_ISL_2509275, EPI_ISL_2509326, EPI_ISL_2509327, EPI_ISL_2509328, EPI_ISL_2509330 | Institut de Virologie du CHU de Strasbourg | Institut de Virologie du CHU de Strasbourg | Fafi-Kremer Samira, Gallais Floriane, Gantner Pierre, Laugel Elodie, Solis Morgane, Velay Aurélie, Wendling Marie-Josée |

| | | | |
|---|---|---|---|
| EPI_ISL_2509334 | Palapye Primary Hospital Laboratory | Botswana Harvard HIV Reference Laboratory | Sikhulile Moyo, Dorcas Maruapula, Wonderful T. Choga, Thongbotho Mphoyakgosi, Boitumelo Zuze, Botshelo Radibe, Legodile Kooepile, Ontlametse T. Bareng, Letsibogo Gaoraelwe, Thela Tefelo, Keoratile Ntshambiwa, Modisa Motswaledi, Madisa Mine, Joseph Makhema, Roger Shapiro, Shahin Lockman, Mosepele Mosepele, Simani Gaseitsiwe |
| EPI_ISL_2509336 | Institut de Virologie du CHU de Strasbourg | Institut de Virologie du CHU de Strasbourg | Fafi-Kremer Samira, Gallais Floriane, Gantner Pierre, Laugel Elodie, Solis Morgane, Velay Aurélie, Wendling Marie-Josée |
| EPI_ISL_2510687 | Surabaya Health Office | Institute of Tropical Disease, Universitas Airlangga | Jezzy R Dewantari, Rima R Prasetya, Krisnoadi Rahardjo, Aldise M Nastri, Febria Rachmanita, Gatot Soegiarto, Laksmi Wulandari, Resti Yudhawati, Yasuko Mori, Soetjipto, Kazufumi Shimizu, Maria I Lusida |
| EPI_ISL_2517046, EPI_ISL_2517047, EPI_ISL_2517048, EPI_ISL_2517049, EPI_ISL_2517050, EPI_ISL_2517051, EPI_ISL_2517052, EPI_ISL_2517053, EPI_ISL_2517054, EPI_ISL_2517055, EPI_ISL_2517056, EPI_ISL_2517057, EPI_ISL_2517058, EPI_ISL_2517059, EPI_ISL_2517060, EPI_ISL_2517061, EPI_ISL_2517062 | | | |
| see above | Althaia. Xarxa Assistencial Universitària de Manresa | IrsiCaixa | Marc Noguera-Julian, Mariona Parera, Maria Casadellà, Pilar Armengol, Francesc Catala-Moll, Roger Paredes, Bonaventura Clotet Gloria Trujillo, Rafel Perez Vidal, Jaume Trapé, Carolina Gonzalez-Fernandez, Miquel Micó, Antonia Flor, Roger Paredes, Eulalia Grau, Bonaventura Clotet |
| EPI_ISL_2530256, EPI_ISL_2530258, EPI_ISL_2530259, EPI_ISL_2530261, EPI_ISL_2530262, EPI_ISL_2530290, EPI_ISL_2530384, EPI_ISL_2530388, EPI_ISL_2530390, EPI_ISL_2530391, EPI_ISL_2530392, EPI_ISL_2530394 | | | |
| see above | Hematopathology Laboratory, ACTREC, TMC | Hematopathology Laboratory, ACTREC, TMC | Hematopathology Laboratory, ACTREC |
| EPI_ISL_2531827 | Hospital | National Reference Center for Viruses of Respiratory Infections, Institut Pasteur, Paris | Marion Barbet, Sylvie Behillil, Méline Bizard, Angela Brisebarre, Camille Capel, Vincent Enouf, Louise Lefrançois, Frédéric Lemoine, Christophe Malabat, Corinne Maufrais, Etienne Simon-Lorière, Maud Vanpeene, Sylvie Van der Werf ,Pascale Martres |
| EPI_ISL_2531834, EPI_ISL_2531839, EPI_ISL_2531843, EPI_ISL_2531851 | Hospital | National Reference Center for Viruses of Respiratory Infections, Institut Pasteur, Paris | Marion Barbet, Sylvie Behillil, Méline Bizard, Angela Brisebarre, Camille Capel, Vincent Enouf, Louise Lefrançois, Frédéric Lemoine, Christophe Malabat, Corinne Maufrais, Etienne Simon-Lorière, Maud Vanpeene, Sylvie Van der Werf ,Raphael Guiheneuf |
| EPI_ISL_2531877 | Hospital | National Reference Center for Viruses of Respiratory Infections, Institut Pasteur, Paris | Marion Barbet, Sylvie Behillil, Méline Bizard, Angela Brisebarre, Camille Capel, Vincent Enouf, Louise Lefrançois, Frédéric Lemoine, Christophe Malabat, Corinne Maufrais, Etienne Simon-Lorière, Maud Vanpeene, Sylvie Van der Werf ,Marianne Burgard |
| EPI_ISL_2531881, EPI_ISL_2531885, EPI_ISL_2531889, EPI_ISL_2531894, EPI_ISL_2531897, EPI_ISL_2531901, EPI_ISL_2531905, EPI_ISL_2531908, EPI_ISL_2531912, EPI_ISL_2531917, EPI_ISL_2531922, EPI_ISL_2531926, EPI_ISL_2531930, EPI_ISL_2531934, EPI_ISL_2531939, EPI_ISL_2531940, EPI_ISL_2531942, EPI_ISL_2531943 | | | |
| see above | RIIP | National Reference Center for Viruses of Respiratory Infections, Institut Pasteur, Paris | Marion Barbet, Sylvie Behillil, Méline Bizard, Angela Brisebarre, Camille Capel, Vincent Enouf, Louise Lefrançois, Frédéric Lemoine, Christophe Malabat, Corinne Maufrais, Etienne Simon-Lorière, Maud Vanpeene, Sylvie Van der Werf ,StéPhanie Guyomard-Rabenirina |
| EPI_ISL_2531945, EPI_ISL_2531947, EPI_ISL_2531948, EPI_ISL_2531953 | Labo Analyses Med | National Reference Center for Viruses of Respiratory Infections, Institut Pasteur, Paris | Marion Barbet, Sylvie Behillil, Méline Bizard, Angela Brisebarre, Camille Capel, Vincent Enouf, Louise Lefrançois, Frédéric Lemoine, Christophe Malabat, Corinne Maufrais, Etienne Simon-Lorière, Maud Vanpeene, Sylvie Van der Werf ,Hugues Leroy |
| EPI_ISL_2531958, EPI_ISL_2531962 | Labo Analyses Med | National Reference Center for Viruses of Respiratory Infections, Institut Pasteur, Paris | Marion Barbet, Sylvie Behillil, Méline Bizard, Angela Brisebarre, Camille Capel, Vincent Enouf, Louise Lefrançois, Frédéric Lemoine, Christophe Malabat, Corinne Maufrais, Etienne Simon-Lorière, Maud Vanpeene, Sylvie Van der Werf ,Vicky Le |
| EPI_ISL_2532193 | Hospital | National Reference Center for Viruses of Respiratory Infections, Institut Pasteur, Paris | Marion Barbet, Sylvie Behillil, Méline Bizard, Angela Brisebarre, Camille Capel, Vincent Enouf, Louise Lefrançois, Frédéric Lemoine, Christophe Malabat, Corinne Maufrais, Emmanuelle Permal, Etienne Simon-Lorière, Maud Vanpeene, Sylvie Van der Werf ,Eric Farfour |
| EPI_ISL_2532279, EPI_ISL_2532280, EPI_ISL_2532281, EPI_ISL_2532282, EPI_ISL_2532283 | Labo Analyses Med | National Reference Center for Viruses of Respiratory Infections, Institut Pasteur, Paris | Marion Barbet, Sylvie Behillil, Méline Bizard, Angela Brisebarre, Camille Capel, Vincent Enouf, Louise Lefrançois, Frédéric Lemoine, Christophe Malabat, Corinne Maufrais, Etienne Simon-Lorière, Maud Vanpeene, Sylvie Van der Werf ,OphéLie Said-Delattre |
| EPI_ISL_2532284, EPI_ISL_2532285, EPI_ISL_2532286 | Labo Analyses Med | National Reference Center for Viruses of Respiratory Infections, Institut Pasteur, Paris | Marion Barbet, Sylvie Behillil, Méline Bizard, Angela Brisebarre, Camille Capel, Vincent Enouf, Louise Lefrançois, Frédéric Lemoine, Christophe Malabat, Corinne Maufrais, Etienne Simon-Lorière, Maud Vanpeene, Sylvie Van der Werf ,Eve Haguenoer |
| EPI_ISL_2532287, EPI_ISL_2532288 | Labo Analyses Med | National Reference Center for Viruses of Respiratory Infections, Institut Pasteur, Paris | Marion Barbet, Sylvie Behillil, Méline Bizard, Angela Brisebarre, Camille Capel, Vincent Enouf, Louise Lefrançois, Frédéric Lemoine, Christophe Malabat, Corinne Maufrais, Etienne Simon-Lorière, Maud Vanpeene, Sylvie Van der Werf ,GréGoire Potiron |
| EPI_ISL_2532291 | Labo Analyses Med | National Reference Center for Viruses of Respiratory Infections, Institut Pasteur, Paris | Marion Barbet, Sylvie Behillil, Méline Bizard, Angela Brisebarre, Camille Capel, Vincent Enouf, Louise Lefrançois, Frédéric Lemoine, Christophe Malabat, Corinne Maufrais, Emmanuelle Permal, Etienne Simon-Lorière, Maud Vanpeene, Sylvie Van der Werf ,BéNéDicte Roquebert |
| EPI_ISL_2532292 | Hospital | National Reference Center for Viruses of Respiratory Infections, Institut Pasteur, Paris | Marion Barbet, Sylvie Behillil, Méline Bizard, Angela Brisebarre, Camille Capel, Vincent Enouf, Louise Lefrançois, Frédéric Lemoine, Christophe Malabat, Corinne Maufrais, Emmanuelle Permal, Etienne Simon-Lorière, Maud Vanpeene, Sylvie Van der Werf ,CéLine Ramanantsoa |
| EPI_ISL_2532293 | Hospital | National Reference Center for Viruses of Respiratory Infections, Institut Pasteur, Paris | Marion Barbet, Sylvie Behillil, Méline Bizard, Angela Brisebarre, Camille Capel, Vincent Enouf, Louise Lefrançois, Frédéric Lemoine, Christophe Malabat, Corinne Maufrais, Amaury Vaysse, Etienne Simon-Lorière, Maud Vanpeene, Sylvie Van der Werf ,JéRome Guinard |
| EPI_ISL_2532295, EPI_ISL_2532296 | Hospital | National Reference Center for Viruses of Respiratory Infections, Institut Pasteur, Paris | Marion Barbet, Sylvie Behillil, Méline Bizard, Angela Brisebarre, Camille Capel, Vincent Enouf, Louise Lefrançois, Frédéric Lemoine, Christophe Malabat, Corinne Maufrais, Etienne Simon-Lorière, Maud Vanpeene, Sylvie Van der Werf ,Thibault Guinoiseau |
| EPI_ISL_2532300 | Hospital | National Reference Center for Viruses of Respiratory Infections, Institut Pasteur, Paris | Marion Barbet, Sylvie Behillil, Méline Bizard, Angela Brisebarre, Camille Capel, Vincent Enouf, Louise Lefrançois, Frédéric Lemoine, Christophe Malabat, Corinne Maufrais, Amaury Vaysse, Etienne Simon-Lorière, Maud Vanpeene, Sylvie Van der Werf ,Sylvaine Bastian |
| EPI_ISL_2532301 | Hospital | National Reference Center for Viruses of Respiratory Infections, Institut Pasteur, Paris | Marion Barbet, Sylvie Behillil, Méline Bizard, Angela Brisebarre, Camille Capel, Vincent Enouf, Louise Lefrançois, Frédéric Lemoine, Christophe Malabat, Corinne Maufrais, Etienne Simon-Lorière, Maud Vanpeene, Sylvie Van der Werf ,Sylvaine Bastian |
| EPI_ISL_2532303, EPI_ISL_2532304, EPI_ISL_2532305, EPI_ISL_2532307, EPI_ISL_2532308, EPI_ISL_2532309 | Hospital | National Reference Center for Viruses of Respiratory Infections, Institut Pasteur, Paris | Marion Barbet, Sylvie Behillil, Méline Bizard, Angela Brisebarre, Camille Capel, Vincent Enouf, Louise Lefrançois, Frédéric Lemoine, Christophe Malabat, Corinne Maufrais, Etienne Simon-Lorière, Maud Vanpeene, Sylvie Van der Werf ,FréDéRique Canis |
| EPI_ISL_2532311, EPI_ISL_2532312, EPI_ISL_2532313 | Hospital | National Reference Center for Viruses of Respiratory Infections, Institut Pasteur, Paris | Marion Barbet, Sylvie Behillil, Méline Bizard, Angela Brisebarre, Camille Capel, Vincent Enouf, Louise Lefrançois, Frédéric Lemoine, Christophe Malabat, Corinne Maufrais, Etienne Simon-Lorière, Maud Vanpeene, Sylvie Van der Werf ,Pascale Martres |
| EPI_ISL_2532315 | Hospital | National Reference Center for Viruses of Respiratory Infections, Institut Pasteur, Paris | Marion Barbet, Sylvie Behillil, Méline Bizard, Angela Brisebarre, Camille Capel, Vincent Enouf, Louise Lefrançois, Frédéric Lemoine, Christophe Malabat, Corinne Maufrais, Amaury Vaysse, Etienne Simon-Lorière, Maud Vanpeene, Sylvie Van der Werf ,Pascale Martres |
| EPI_ISL_2532339, EPI_ISL_2532340, EPI_ISL_2532341, EPI_ISL_2532342, EPI_ISL_2532343, EPI_ISL_2532344, EPI_ISL_2532345, EPI_ISL_2532346, EPI_ISL_2532347, EPI_ISL_2532348, EPI_ISL_2532349, EPI_ISL_2532350, EPI_ISL_2532351, EPI_ISL_2532352 | | | |
| see above | Labo Analyses Med | National Reference Center for Viruses of Respiratory Infections, Institut Pasteur, Paris | Marion Barbet, Sylvie Behillil, Méline Bizard, Angela Brisebarre, Camille Capel, Vincent Enouf, Louise Lefrançois, Frédéric Lemoine, Christophe Malabat, Corinne Maufrais, Etienne Simon-Lorière, Maud Vanpeene, Sylvie Van der Werf ,Ronan Pichard |
| EPI_ISL_2532353, EPI_ISL_2532354, EPI_ISL_2532355, EPI_ISL_2532356, EPI_ISL_2532357, EPI_ISL_2532358, EPI_ISL_2532359, EPI_ISL_2532360 | Hospital | National Reference Center for Viruses of Respiratory Infections, Institut Pasteur, Paris | Marion Barbet, Sylvie Behillil, Méline Bizard, Angela Brisebarre, Camille Capel, Vincent Enouf, Louise Lefrançois, Frédéric Lemoine, Christophe Malabat, Corinne Maufrais, Etienne Simon-Lorière, Maud Vanpeene, Sylvie Van der Werf ,CéLine Bressollette |
| EPI_ISL_2532361, EPI_ISL_2532362, EPI_ISL_2532363, EPI_ISL_2532364, EPI_ISL_2532365, EPI_ISL_2532366 | Labo Analyses Med | National Reference Center for Viruses of Respiratory Infections, Institut Pasteur, Paris | Marion Barbet, Sylvie Behillil, Méline Bizard, Angela Brisebarre, Camille Capel, Vincent Enouf, Louise Lefrançois, Frédéric Lemoine, Christophe Malabat, Corinne Maufrais, Etienne Simon-Lorière, Maud Vanpeene, Sylvie Van der Werf ,Hervé Dupont |
| EPI_ISL_2532367 | Labo Analyses Med | National Reference Center for Viruses of Respiratory Infections, Institut Pasteur, Paris | Marion Barbet, Sylvie Behillil, Méline Bizard, Angela Brisebarre, Camille Capel, Vincent Enouf, Louise Lefrançois, Frédéric Lemoine, Christophe Malabat, Corinne Maufrais, Victoire Baillet, Etienne Simon-Lorière, Maud Vanpeene, Sylvie Van der Werf ,Hervé Dupont |
| EPI_ISL_2532368, EPI_ISL_2532370, EPI_ISL_2532371, EPI_ISL_2532372, EPI_ISL_2532373, EPI_ISL_2532374, EPI_ISL_2532375, EPI_ISL_2532376, EPI_ISL_2532377, EPI_ISL_2532378 | Labo Analyses Med | National Reference Center for Viruses of Respiratory Infections, Institut Pasteur, Paris | Marion Barbet, Sylvie Behillil, Méline Bizard, Angela Brisebarre, Camille Capel, Vincent Enouf, Louise Lefrançois, Frédéric Lemoine, Christophe Malabat, Corinne Maufrais, Etienne Simon-Lorière, Maud Vanpeene, Sylvie Van der Werf ,Hervé Dupont |
| EPI_ISL_2532388, EPI_ISL_2532389 | Hospital | National Reference Center for Viruses of Respiratory | Marion Barbet, Sylvie Behillil, Méline Bizard, Angela Brisebarre, Camille Capel, Vincent Enouf, Louise Lefrançois, Frédéric Lemoine, Christophe Malabat, |

| | | | |
|---|---|---|---|
| | | Infections, Institut Pasteur, Paris | Corinne Maufrais, Etienne Simon-Lorière, Maud Vanpeene, Sylvie Van der Werf ,AgnèS Scanvic |
| EPI_ISL_2532390 | Labo Analyses Med | National Reference Center for Viruses of Respiratory Infections, Institut Pasteur, Paris | Marion Barbet, Sylvie Behillil, Méline Bizard, Angela Brisebarre, Camille Capel, Vincent Enouf, Louise Lefrançois, Frédéric Lemoine, Christophe Malabat, Corinne Maufrais, Etienne Simon-Lorière, Maud Vanpeene, Sylvie Van der Werf ,Christine Heurtin |
| EPI_ISL_2532392 | Hospital | National Reference Center for Viruses of Respiratory Infections, Institut Pasteur, Paris | Marion Barbet, Sylvie Behillil, Méline Bizard, Angela Brisebarre, Camille Capel, Vincent Enouf, Louise Lefrançois, Frédéric Lemoine, Christophe Malabat, Corinne Maufrais, Etienne Simon-Lorière, Maud Vanpeene, Sylvie Van der Werf ,Patrice Combe |
| EPI_ISL_2532397, EPI_ISL_2532398, EPI_ISL_2532399, EPI_ISL_2532400, EPI_ISL_2532401, EPI_ISL_2532402, EPI_ISL_2532403, EPI_ISL_2532404, EPI_ISL_2532405, EPI_ISL_2532406, EPI_ISL_2532408, EPI_ISL_2532409, EPI_ISL_2532410, EPI_ISL_2532411, EPI_ISL_2532412, EPI_ISL_2532413, EPI_ISL_2532414, EPI_ISL_2532415, EPI_ISL_2532416, EPI_ISL_2532417, EPI_ISL_2532418, EPI_ISL_2532419, EPI_ISL_2532420, EPI_ISL_2532421, EPI_ISL_2532422, EPI_ISL_2532423, EPI_ISL_2532424, EPI_ISL_2532425, EPI_ISL_2532427, EPI_ISL_2532428, EPI_ISL_2532429, EPI_ISL_2532430, EPI_ISL_2532433, EPI_ISL_2532434, EPI_ISL_2532435, EPI_ISL_2532436, EPI_ISL_2532437, EPI_ISL_2532438, EPI_ISL_2532440, EPI_ISL_2532441, EPI_ISL_2532442, EPI_ISL_2532443, EPI_ISL_2532444, EPI_ISL_2532445, EPI_ISL_2532446, EPI_ISL_2532447, EPI_ISL_2532448, EPI_ISL_2532449, EPI_ISL_2532450, EPI_ISL_2532451, EPI_ISL_2532452 | | | |
| see above | Labo Analyses Med | National Reference Center for Viruses of Respiratory Infections, Institut Pasteur, Paris | Marion Barbet, Sylvie Behillil, Méline Bizard, Angela Brisebarre, Camille Capel, Vincent Enouf, Louise Lefrançois, Frédéric Lemoine, Christophe Malabat, Corinne Maufrais, Etienne Simon-Lorière, Maud Vanpeene, Sylvie Van der Werf ,Pierre-Yves Leonard |
| EPI_ISL_2532453, EPI_ISL_2532454, EPI_ISL_2532456, EPI_ISL_2532457, EPI_ISL_2532458, EPI_ISL_2532459, EPI_ISL_2532460, EPI_ISL_2532461, EPI_ISL_2532462, EPI_ISL_2532463 | Labo Analyses Med | National Reference Center for Viruses of Respiratory Infections, Institut Pasteur, Paris | Marion Barbet, Sylvie Behillil, Méline Bizard, Angela Brisebarre, Camille Capel, Vincent Enouf, Louise Lefrançois, Frédéric Lemoine, Christophe Malabat, Corinne Maufrais, Etienne Simon-Lorière, Maud Vanpeene, Sylvie Van der Werf |
| EPI_ISL_2532464 | Labo Analyses Med | National Reference Center for Viruses of Respiratory Infections, Institut Pasteur, Paris | Marion Barbet, Sylvie Behillil, Méline Bizard, Angela Brisebarre, Camille Capel, Vincent Enouf, Louise Lefrançois, Frédéric Lemoine, Christophe Malabat, Corinne Maufrais, Adrien Pain, Etienne Simon-Lorière, Maud Vanpeene, Sylvie Van der Werf |
| EPI_ISL_2532465, EPI_ISL_2532466, EPI_ISL_2532468, EPI_ISL_2532469, EPI_ISL_2532471 | Labo Analyses Med | National Reference Center for Viruses of Respiratory Infections, Institut Pasteur, Paris | Marion Barbet, Sylvie Behillil, Méline Bizard, Angela Brisebarre, Camille Capel, Vincent Enouf, Louise Lefrançois, Frédéric Lemoine, Christophe Malabat, Corinne Maufrais, Etienne Simon-Lorière, Maud Vanpeene, Sylvie Van der Werf |
| EPI_ISL_2532472 | Presidio di Brindisi Di Summa - Perrino | Istituto Zooprofilattico Sperimentale della Puglia e della Basilicata | Parisi A., Bianco A., Capozzi L., Del Sambro L., Simone D., Difato L., Pennuzzi G., Santoro A. |
| EPI_ISL_2532729 | SIESP DIPARTIMENTO DI PREVENZIONE TERAMO | Istituto Zooprofilattico Sperimentale dell'Abruzzo e Molise "G. Caporale" | Lorusso A, Marcacci M, Di Domenico M, Ancora M, Curini V, Di Lollo Valeria, Mangone I, Rinaldi A, Delli Compagni E, Scialabba S, Caporale M, Di Pasquale A, Cammà C, Puglia I, Calistri P, Savini G |
| EPI_ISL_2533759, EPI_ISL_2533760, EPI_ISL_2533761, EPI_ISL_2533762 | LESP Chihuahua | Instituto de Diagnostico y Referencia Epidemiologicos (INDRE) | Claudia Wong-Arambula, Abril Rodriguez-Maldonado, Vanessa Rivero-Arredondo, Ariadna Medina-Benitez, Joaquin Quiroz-Mercado, Sergio Rangel-Guerrero, Natividad Cruz-Ortiz, Tatiana Nunez-Garcia, Gisela Barrera-Badillo, Lucia Hernandez-Rivas, Irma Lopez-Martinez, Ernesto Ramirez-Gonzalez. |
| EPI_ISL_2533763 | LESP Coahuila | Instituto de Diagnostico y Referencia Epidemiologicos (INDRE) | Claudia Wong-Arambula, Abril Rodriguez-Maldonado, Vanessa Rivero-Arredondo, Ariadna Medina-Benitez, Joaquin Quiroz-Mercado, Sergio Rangel-Guerrero, Natividad Cruz-Ortiz, Tatiana Nunez-Garcia, Gisela Barrera-Badillo, Lucia Hernandez-Rivas, Irma Lopez-Martinez, Ernesto Ramirez-Gonzalez. |
| EPI_ISL_2533764, EPI_ISL_2533765, EPI_ISL_2533766 | LESP Queretaro | Instituto de Diagnostico y Referencia Epidemiologicos (INDRE) | Claudia Wong-Arambula, Abril Rodriguez-Maldonado, Vanessa Rivero-Arredondo, Ariadna Medina-Benitez, Joaquin Quiroz-Mercado, Sergio Rangel-Guerrero, Natividad Cruz-Ortiz, Tatiana Nunez-Garcia, Gisela Barrera-Badillo, Lucia Hernandez-Rivas, Irma Lopez-Martinez, Ernesto Ramirez-Gonzalez. |
| EPI_ISL_2533767 | LESP Jalisco | Instituto de Diagnostico y Referencia Epidemiologicos (INDRE) | Claudia Wong-Arambula, Abril Rodriguez-Maldonado, Vanessa Rivero-Arredondo, Ariadna Medina-Benitez, Joaquin Quiroz-Mercado, Sergio Rangel-Guerrero, Natividad Cruz-Ortiz, Tatiana Nunez-Garcia, Gisela Barrera-Badillo, Lucia Hernandez-Rivas, Irma Lopez-Martinez, Ernesto Ramirez-Gonzalez. |
| EPI_ISL_2533768, EPI_ISL_2533769 | LESP Tabasco | Instituto de Diagnostico y Referencia Epidemiologicos (INDRE) | Claudia Wong-Arambula, Abril Rodriguez-Maldonado, Vanessa Rivero-Arredondo, Ariadna Medina-Benitez, Joaquin Quiroz-Mercado, Sergio Rangel-Guerrero, Natividad Cruz-Ortiz, Tatiana Nunez-Garcia, Gisela Barrera-Badillo, Lucia Hernandez-Rivas, Irma Lopez-Martinez, Ernesto Ramirez-Gonzalez. |
| EPI_ISL_2533770, EPI_ISL_2533771, EPI_ISL_2533772, EPI_ISL_2533773, EPI_ISL_2533774, EPI_ISL_2533775, EPI_ISL_2533776, EPI_ISL_2533777, EPI_ISL_2533778, EPI_ISL_2533779, EPI_ISL_2533780, EPI_ISL_2533781, EPI_ISL_2533782, EPI_ISL_2533783, EPI_ISL_2533784, EPI_ISL_2533785, EPI_ISL_2533786, EPI_ISL_2533787, EPI_ISL_2533788, EPI_ISL_2533789, EPI_ISL_2533790, EPI_ISL_2533791, EPI_ISL_2533792, EPI_ISL_2533793, EPI_ISL_2533794, EPI_ISL_2533795, EPI_ISL_2533796 | | | |
| see above | LESP Yucatan | Instituto de Diagnostico y Referencia Epidemiologicos (INDRE) | Claudia Wong-Arambula, Abril Rodriguez-Maldonado, Vanessa Rivero-Arredondo, Ariadna Medina-Benitez, Joaquin Quiroz-Mercado, Sergio Rangel-Guerrero, Natividad Cruz-Ortiz, Tatiana Nunez-Garcia, Gisela Barrera-Badillo, Lucia Hernandez-Rivas, Irma Lopez-Martinez, Ernesto Ramirez-Gonzalez. |
| EPI_ISL_2533797, EPI_ISL_2533798, EPI_ISL_2533799, EPI_ISL_2533800, EPI_ISL_2533801 | LESP Tamaulipas | Instituto de Diagnostico y Referencia Epidemiologicos (INDRE) | Claudia Wong-Arambula, Abril Rodriguez-Maldonado, Vanessa Rivero-Arredondo, Ariadna Medina-Benitez, Joaquin Quiroz-Mercado, Sergio Rangel-Guerrero, Natividad Cruz-Ortiz, Tatiana Nunez-Garcia, Gisela Barrera-Badillo, Lucia Hernandez-Rivas, Irma Lopez-Martinez, Ernesto Ramirez-Gonzalez. |
| EPI_ISL_2533803 | LESP Coahuila | Instituto de Diagnostico y Referencia Epidemiologicos (INDRE) | Claudia Wong-Arambula, Abril Rodriguez-Maldonado, Vanessa Rivero-Arredondo, Ariadna Medina-Benitez, Joaquin Quiroz-Mercado, Sergio Rangel-Guerrero, Natividad Cruz-Ortiz, Tatiana Nunez-Garcia, Gisela Barrera-Badillo, Lucia Hernandez-Rivas, Irma Lopez-Martinez, Ernesto Ramirez-Gonzalez. |
| EPI_ISL_2533804, EPI_ISL_2533805 | LESP Quintana Roo | Instituto de Diagnostico y Referencia Epidemiologicos (INDRE) | Claudia Wong-Arambula, Abril Rodriguez-Maldonado, Vanessa Rivero-Arredondo, Ariadna Medina-Benitez, Joaquin Quiroz-Mercado, Sergio Rangel-Guerrero, Natividad Cruz-Ortiz, Tatiana Nunez-Garcia, Gisela Barrera-Badillo, Lucia Hernandez-Rivas, Irma Lopez-Martinez, Ernesto Ramirez-Gonzalez. |
| EPI_ISL_2533806 | LESP San Luis Potosi | Instituto de Diagnostico y Referencia Epidemiologicos (INDRE) | Claudia Wong-Arambula, Abril Rodriguez-Maldonado, Vanessa Rivero-Arredondo, Ariadna Medina-Benitez, Joaquin Quiroz-Mercado, Sergio Rangel-Guerrero, Natividad Cruz-Ortiz, Tatiana Nunez-Garcia, Gisela Barrera-Badillo, Lucia Hernandez-Rivas, Irma Lopez-Martinez, Ernesto Ramirez-Gonzalez. |
| EPI_ISL_2533807 | LESP Ciudad de Mexico | Instituto de Diagnostico y Referencia Epidemiologicos (INDRE) | Claudia Wong-Arambula, Abril Rodriguez-Maldonado, Vanessa Rivero-Arredondo, Ariadna Medina-Benitez, Joaquin Quiroz-Mercado, Sergio Rangel-Guerrero, Natividad Cruz-Ortiz, Tatiana Nunez-Garcia, Gisela Barrera-Badillo, Lucia Hernandez-Rivas, Irma Lopez-Martinez, Ernesto Ramirez-Gonzalez. |
| EPI_ISL_2533809 | LESP Estado de Mexico | Instituto de Diagnostico y Referencia Epidemiologicos (INDRE) | Claudia Wong-Arambula, Abril Rodriguez-Maldonado, Vanessa Rivero-Arredondo, Ariadna Medina-Benitez, Joaquin Quiroz-Mercado, Sergio Rangel-Guerrero, Natividad Cruz-Ortiz, Tatiana Nunez-Garcia, Gisela Barrera-Badillo, Lucia Hernandez-Rivas, Irma Lopez-Martinez, Ernesto Ramirez-Gonzalez. |
| EPI_ISL_2533810, EPI_ISL_2533811, EPI_ISL_2533812, EPI_ISL_2533813, EPI_ISL_2533814, EPI_ISL_2533815, EPI_ISL_2533817 | LESP Ciudad de Mexico | Instituto de Diagnostico y Referencia Epidemiologicos (INDRE) | Claudia Wong-Arambula, Abril Rodriguez-Maldonado, Vanessa Rivero-Arredondo, Ariadna Medina-Benitez, Joaquin Quiroz-Mercado, Sergio Rangel-Guerrero, Natividad Cruz-Ortiz, Tatiana Nunez-Garcia, Gisela Barrera-Badillo, Lucia Hernandez-Rivas, Irma Lopez-Martinez, Ernesto Ramirez-Gonzalez. |
| EPI_ISL_2533818 | LESP Baja California Sur | Instituto de Diagnostico y Referencia Epidemiologicos (INDRE) | Claudia Wong-Arambula, Abril Rodriguez-Maldonado, Vanessa Rivero-Arredondo, Ariadna Medina-Benitez, Joaquin Quiroz-Mercado, Sergio Rangel-Guerrero, Natividad Cruz-Ortiz, Tatiana Nunez-Garcia, Gisela Barrera-Badillo, Lucia Hernandez-Rivas, Irma Lopez-Martinez, Ernesto Ramirez-Gonzalez. |
| EPI_ISL_2533819 | LESP Yucatan | Instituto de Diagnostico y Referencia Epidemiologicos (INDRE) | Claudia Wong-Arambula, Abril Rodriguez-Maldonado, Vanessa Rivero-Arredondo, Ariadna Medina-Benitez, Joaquin Quiroz-Mercado, Sergio Rangel-Guerrero, Natividad Cruz-Ortiz, Tatiana Nunez-Garcia, Gisela Barrera-Badillo, Lucia Hernandez-Rivas, Irma Lopez-Martinez, Ernesto Ramirez-Gonzalez. |
| EPI_ISL_2533820 | LESP Quintana Roo | Instituto de Diagnostico y Referencia Epidemiologicos (INDRE) | Claudia Wong-Arambula, Abril Rodriguez-Maldonado, Vanessa Rivero-Arredondo, Ariadna Medina-Benitez, Joaquin Quiroz-Mercado, Sergio Rangel-Guerrero, Natividad Cruz-Ortiz, Tatiana Nunez-Garcia, Gisela Barrera-Badillo, Lucia Hernandez-Rivas, Irma Lopez-Martinez, Ernesto |

| | | | |
|---|---|---|---|
| | | | Ramirez-Gonzalez. |
| EPI_ISL_2533932 | CIAD-Hermosillo | Immunology Laboratory | Consorcio Mexicano de Vigilancia Genómica (CoViGen-Mex). Authors (in alphabetical order): Julio Elias Alvarado-Yaah, Carlos F. Arias, Santiago Ávila-Ríos, Eduardo Becerril-Vargas, Víctor Hugo Borja-Aburto, Celia Boukadida, Cristóbal Chaidez-Quiróz, Juan Bautista Chale-Dzul, Ricardo Ciria Merce, Andreu Comas-García, Célida Duque Molina, Julissa Enciso-Ibarra, José Antonio Enciso-Moreno, Gloria Elena Espinosa-Ayala, Fernando Fontove-Herrera, Daniel Fregoso-Rueda, Victor Eduardo García-Arias, Alejandra García-Gasca, Bruno Gómez-Gil, Jean Pierre González, Irvin González-López, Concepción Grajales-Muñiz, Ricardo Grande, Rosa María Gutiérrez Rios, Jesús Hernández, Alejandra Hernández-Terán, Alfredo Herrera-Estrella, Carla Ivón Herrera-Najera, Pavel Isa, Verónica Mata-Haro, Daniel Lira Morales, Susana Lopez, Antonio Loza Román, Brenda Irasema Maldonado-Meza, Bernardo Martínez-Miguel, José Arturo Martínez-Orozco, Célida Martínez- Rodríguez, Margarita Matías-Florentino, Fidencio Mejía-Nepomuceno, María Guadalupe de Jesús Mireles-Rivera, Gloria María Molina-Salinas, Hector Montoya-Fuentes, , Mario Mújica-Sánchez, José Esteban Muñoz-Medina, José de Jesús Nuñez-Contreras, Alicia Ocaña-Mondragón, Luis Alberto Ochoa-Carrera, Hector Esteban Paz-Juárez, Marissa Perez-Garcia, Francisco Pulido, Helen Haydee Fernanda Ramírez-Plascencia, Jorge Salas-Hernández, Angel Gustavo Salas-Lais, Alejandro Sánchez-Flores, Clara Esperanza Santacruz-Tinoco, María Guadalupe Santiago-Mauricio, Selene Zárate, Nelly Sélem-Mojica, Blanca Taboada, Gloria Vazquez, Joel Armando Vázquez-Pérez. |
| EPI_ISL_2534079 | RSUD dr. LOEKMONO HADI | Genetics Working Group (Pokja Genetik) Faculty of Medicine, Public Health and Nursing Universitas Gadjah Mada (FK-KMK UGM); Disease Investigation Center Wates Ministry of Agriculture Indonesia; Department of Microbiology FK-KMK UGM; Laboratorium Diagnostik Yayasan Tahija World Mosquito Program (WMP) Yogyakarta Center for Tropical Medicine FK-KMK UGM; Integrated Research Center FK-KMK UGM; Department of Computer Science and Electronics FMIPA UGM; RSUP Dr. Sardjito | Gunadi, Hendra Wibawa, Marcellus, Mohamad S. Hakim, Edwin W. Daniwijaya, Ludhang P. Rizki, Endah Supriyati, Eggi Arguni, Titik Nuryastuti, Tri Wibawa, Dwi AA Nugrahaningsih, Afiahayati, Siswanto,Kristy Iskandar,Nungki Anggorowati, Pramesti G Dewi, Dwi Indaryati,Indarto Sulistiyono, Susan Simanjaya, Dyah A Puspitarani |
| EPI_ISL_2534310, EPI_ISL_2534311, EPI_ISL_2534313, EPI_ISL_2534314, EPI_ISL_2534315 | The Ohio State University Applied Microbiology Services Laboratory | The Ohio State University Applied Microbiology Services Laboratory | Seth A. Faith PhD |
| EPI_ISL_2534379, EPI_ISL_2534457, EPI_ISL_2534489 | RSUD dr. LOEKMONO HADI | Genetics Working Group (Pokja Genetik) Faculty of Medicine, Public Health and Nursing Universitas Gadjah Mada (FK-KMK UGM); Disease Investigation Center Wates Ministry of Agriculture Indonesia; Department of Microbiology FK-KMK UGM; Laboratorium Diagnostik Yayasan Tahija World Mosquito Program (WMP) Yogyakarta Center for Tropical Medicine FK-KMK UGM; Integrated Research Center FK-KMK UGM; Department of Computer Science and Electronics FMIPA UGM; RSUP Dr. Sardjito | Gunadi, Hendra Wibawa, Marcellus, Mohamad S. Hakim, Edwin W. Daniwijaya, Ludhang P. Rizki, Endah Supriyati, Eggi Arguni, Titik Nuryastuti, Tri Wibawa, Dwi AA Nugrahaningsih, Afiahayati, Siswanto,Kristy Iskandar,Nungki Anggorowati, Pramesti G Dewi, Dwi Indaryati,Indarto Sulistiyono, Cita S Amalia, Alvina A Setiawan |
| EPI_ISL_2534490 | RSUD dr. LOEKMONO HADI | Genetics Working Group (Pokja Genetik) Faculty of Medicine, Public Health and Nursing Universitas Gadjah Mada (FK-KMK UGM); Disease Investigation Center Wates Ministry of Agriculture Indonesia; Department of Microbiology FK-KMK UGM; Laboratorium Diagnostik Yayasan Tahija World Mosquito Program (WMP) Yogyakarta Center for Tropical Medicine FK-KMK UGM; Integrated Research Center FK-KMK UGM; Department of Computer Science and Electronics FMIPA UGM; RSUP Dr. Sardjito | Gunadi, Hendra Wibawa, Marcellus, Mohamad S. Hakim, Edwin W. Daniwijaya, Ludhang P. Rizki, Endah Supriyati, Eggi Arguni, Titik Nuryastuti, Tri Wibawa, Dwi AA Nugrahaningsih, Afiahayati, Siswanto,Kristy Iskandar,Nungki Anggorowati, Pramesti G Dewi, Dwi Indaryati,Indarto Sulistiyono, Khanza a Vujira, Alvina A Setiawan |
| EPI_ISL_2534518 | RSUD dr. LOEKMONO HADI | Genetics Working Group (Pokja Genetik) Faculty of Medicine, Public Health and Nursing Universitas Gadjah Mada (FK-KMK UGM); Disease Investigation Center Wates Ministry of Agriculture Indonesia; Department of Microbiology FK-KMK UGM; Laboratorium Diagnostik Yayasan Tahija World Mosquito Program (WMP) Yogyakarta Center for Tropical Medicine FK-KMK UGM; Integrated Research Center FK-KMK UGM; Department of Computer Science and Electronics FMIPA UGM; RSUP Dr. Sardjito | Gunadi, Hendra Wibawa, Marcellus, Mohamad S. Hakim, Edwin W. Daniwijaya, Ludhang P. Rizki, Endah Supriyati, Eggi Arguni, Titik Nuryastuti, Tri Wibawa, Dwi AA Nugrahaningsih, Afiahayati, Siswanto,Kristy Iskandar,Nungki Anggorowati, Pramesti G Dewi, Dwi Indaryati,Indarto Sulistiyono, Susan Simanjaya, Dyah A Puspitarani |
| EPI_ISL_2535237, EPI_ISL_2535238, EPI_ISL_2535239, EPI_ISL_2535240, EPI_ISL_2535241 | Labo Analyses Med | National Reference Center for Viruses of Respiratory Infections, Institut Pasteur, Paris | Marion Barbet, Sylvie Behillil, Méline Bizard, Angela Brisebarre, Camille Capel, Vincent Enouf, Louise Lefrançois, Frédéric Lemoine, Christophe Malabat, Corinne Maufrais, Etienne Simon-Lorière, Maud Vanpeene, Sylvie Van der Werf |
| EPI_ISL_2535242 | Hospital | National Reference Center for Viruses of Respiratory Infections, Institut Pasteur, Paris | Marion Barbet, Sylvie Behillil, Méline Bizard, Angela Brisebarre, Camille Capel, Vincent Enouf, Louise Lefrançois, Frédéric Lemoine, Christophe Malabat, Corinne Maufrais, Etienne Simon-Lorière, Maud Vanpeene, Sylvie Van der Werf ,JéRome Guinard |
| EPI_ISL_2535243 | Labo Analyses Med | National Reference Center for Viruses of Respiratory Infections, Institut Pasteur, Paris | Marion Barbet, Sylvie Behillil, Méline Bizard, Angela Brisebarre, Camille Capel, Vincent Enouf, Louise Lefrançois, Frédéric Lemoine, Christophe Malabat, Corinne Maufrais, Etienne Simon-Lorière, Maud Vanpeene, Sylvie Van der Werf ,Anne Holstein |
| EPI_ISL_2535244, EPI_ISL_2535245 | Labo Analyses Med | National Reference Center for Viruses of Respiratory Infections, Institut Pasteur, Paris | Marion Barbet, Sylvie Behillil, Méline Bizard, Angela Brisebarre, Camille Capel, Vincent Enouf, Louise Lefrançois, Frédéric Lemoine, Christophe Malabat, Corinne Maufrais, Etienne Simon-Lorière, Maud Vanpeene, Sylvie Van der Werf ,Nouredine Sadeg |
| EPI_ISL_2535247, EPI_ISL_2535248, EPI_ISL_2535249, EPI_ISL_2535250, EPI_ISL_2535251 | Labo Analyses Med | National Reference Center for Viruses of Respiratory Infections, Institut Pasteur, Paris | Marion Barbet, Sylvie Behillil, Méline Bizard, Angela Brisebarre, Camille Capel, Vincent Enouf, Louise Lefrançois, Frédéric Lemoine, Christophe Malabat, Corinne Maufrais, Etienne Simon-Lorière, Maud Vanpeene, Sylvie Van der Werf ,V. Bourdrel |
| EPI_ISL_2535252, EPI_ISL_2535253, EPI_ISL_2535254, EPI_ISL_2535255, EPI_ISL_2535256, EPI_ISL_2535257, EPI_ISL_2535258, EPI_ISL_2535259 | Labo Analyses Med | National Reference Center for Viruses of Respiratory Infections, Institut Pasteur, Paris | Marion Barbet, Sylvie Behillil, Méline Bizard, Angela Brisebarre, Camille Capel, Vincent Enouf, Louise Lefrançois, Frédéric Lemoine, Christophe Malabat, Corinne Maufrais, Etienne Simon-Lorière, Maud Vanpeene, Sylvie Van der Werf |
| EPI_ISL_2535260, EPI_ISL_2535262, EPI_ISL_2535264 | Labo Analyses Med | National Reference Center for Viruses of Respiratory Infections, Institut Pasteur, Paris | Marion Barbet, Sylvie Behillil, Méline Bizard, Angela Brisebarre, Camille Capel, Vincent Enouf, Louise Lefrançois, Frédéric Lemoine, Christophe Malabat, Corinne Maufrais, Etienne Simon-Lorière, Maud Vanpeene, Sylvie Van der Werf ,GréGoire Potiron |
| EPI_ISL_2535265 | Labo Analyses Med | National Reference Center for Viruses of Respiratory Infections, Institut Pasteur, Paris | Marion Barbet, Sylvie Behillil, Méline Bizard, Angela Brisebarre, Camille Capel, Vincent Enouf, Louise Lefrançois, Frédéric Lemoine, Christophe Malabat, Corinne Maufrais, Etienne Simon-Lorière, Maud Vanpeene, Sylvie Van der Werf ,Hugues Leroy |
| EPI_ISL_2535266, EPI_ISL_2535267, EPI_ISL_2535269, EPI_ISL_2535270 | Hospital | National Reference Center for Viruses of Respiratory Infections, Institut Pasteur, Paris | Marion Barbet, Sylvie Behillil, Méline Bizard, Angela Brisebarre, Camille Capel, Vincent Enouf, Louise Lefrançois, Frédéric Lemoine, Christophe Malabat, Corinne Maufrais, Etienne Simon-Lorière, Maud Vanpeene, Sylvie Van der Werf ,Olivia Raulin |
| EPI_ISL_2535271 | Hospital | National Reference Center for Viruses of Respiratory Infections, Institut Pasteur, Paris | Marion Barbet, Sylvie Behillil, Méline Bizard, Angela Brisebarre, Camille Capel, Vincent Enouf, Louise Lefrançois, Frédéric Lemoine, Christophe Malabat, Corinne Maufrais, Damien Mornico, Etienne Simon-Lorière, Maud Vanpeene, Sylvie Van der Werf ,Pascale Martres |
| EPI_ISL_2535272, EPI_ISL_2535274 | Hospital | National Reference Center for Viruses of Respiratory Infections, Institut Pasteur, Paris | Marion Barbet, Sylvie Behillil, Méline Bizard, Angela Brisebarre, Camille Capel, Vincent Enouf, Louise Lefrançois, Frédéric Lemoine, Christophe Malabat, Corinne Maufrais, Etienne Simon-Lorière, Maud Vanpeene, Sylvie Van der Werf ,Pascale Martres |

| | | | |
|---|---|---|---|
| EPI_ISL_2535275 | Labo Analyses Med | National Reference Center for Viruses of Respiratory Infections, Institut Pasteur, Paris | Marion Barbet, Sylvie Behillil, Méline Bizard, Angela Brisebarre, Camille Capel, Vincent Enouf, Louise Lefrançois, Frédéric Lemoine, Christophe Malabat, Corinne Maufrais, Etienne Simon-Lorière, Maud Vanpeene, Sylvie Van der Werf ,OphéLie Said-Delattre |
| EPI_ISL_2535276 | Labo Analyses Med | National Reference Center for Viruses of Respiratory Infections, Institut Pasteur, Paris | Marion Barbet, Sylvie Behillil, Méline Bizard, Angela Brisebarre, Camille Capel, Vincent Enouf, Louise Lefrançois, Frédéric Lemoine, Christophe Malabat, Corinne Maufrais, Victoire Baillet, Etienne Simon-Lorière, Maud Vanpeene, Sylvie Van der Werf ,OphéLie Said-Delattre |
| EPI_ISL_2535277, EPI_ISL_2535278, EPI_ISL_2535279, EPI_ISL_2535280, EPI_ISL_2535281, EPI_ISL_2535282, EPI_ISL_2535283, EPI_ISL_2535284, EPI_ISL_2535285, EPI_ISL_2535286, EPI_ISL_2535287, EPI_ISL_2535288, EPI_ISL_2535289, EPI_ISL_2535290, EPI_ISL_2535291, EPI_ISL_2535295 | | | |
| see above | Labo Analyses Med | National Reference Center for Viruses of Respiratory Infections, Institut Pasteur, Paris | Marion Barbet, Sylvie Behillil, Méline Bizard, Angela Brisebarre, Camille Capel, Vincent Enouf, Louise Lefrançois, Frédéric Lemoine, Christophe Malabat, Corinne Maufrais, Etienne Simon-Lorière, Maud Vanpeene, Sylvie Van der Werf ,OphéLie Said-Delattre |
| EPI_ISL_2535296 | Labo Analyses Med | National Reference Center for Viruses of Respiratory Infections, Institut Pasteur, Paris | Marion Barbet, Sylvie Behillil, Méline Bizard, Angela Brisebarre, Camille Capel, Vincent Enouf, Louise Lefrançois, Frédéric Lemoine, Christophe Malabat, Corinne Maufrais, Victoire Baillet, Etienne Simon-Lorière, Maud Vanpeene, Sylvie Van der Werf ,OphéLie Said-Delattre |
| EPI_ISL_2535297, EPI_ISL_2535299, EPI_ISL_2535300, EPI_ISL_2535301 | Labo Analyses Med | National Reference Center for Viruses of Respiratory Infections, Institut Pasteur, Paris | Marion Barbet, Sylvie Behillil, Méline Bizard, Angela Brisebarre, Camille Capel, Vincent Enouf, Louise Lefrançois, Frédéric Lemoine, Christophe Malabat, Corinne Maufrais, Etienne Simon-Lorière, Maud Vanpeene, Sylvie Van der Werf ,OphéLie Said-Delattre |
| EPI_ISL_2535436, EPI_ISL_2535437, EPI_ISL_2535438, EPI_ISL_2535439, EPI_ISL_2535440, EPI_ISL_2535441, EPI_ISL_2535442, EPI_ISL_2535443, EPI_ISL_2535444 | Hospital | National Reference Center for Viruses of Respiratory Infections, Institut Pasteur, Paris | Marion Barbet, Sylvie Behillil, Méline Bizard, Angela Brisebarre, Camille Capel, Vincent Enouf, Louise Lefrançois, Frédéric Lemoine, Christophe Malabat, Corinne Maufrais, Etienne Simon-Lorière, Maud Vanpeene, Sylvie Van der Werf ,Nabil Gastli |
| EPI_ISL_2535445, EPI_ISL_2535446, EPI_ISL_2535447, EPI_ISL_2535448, EPI_ISL_2535449, EPI_ISL_2535450, EPI_ISL_2535451, EPI_ISL_2535452, EPI_ISL_2535453, EPI_ISL_2535454, EPI_ISL_2535455, EPI_ISL_2535456, EPI_ISL_2535457, EPI_ISL_2535458, EPI_ISL_2535459, EPI_ISL_2535460, EPI_ISL_2535461, EPI_ISL_2535462 | | | |
| see above | Labo Analyses Med | National Reference Center for Viruses of Respiratory Infections, Institut Pasteur, Paris | Marion Barbet, Sylvie Behillil, Méline Bizard, Angela Brisebarre, Camille Capel, Vincent Enouf, Louise Lefrançois, Frédéric Lemoine, Christophe Malabat, Corinne Maufrais, Etienne Simon-Lorière, Maud Vanpeene, Sylvie Van der Werf ,Claire Felloni |
| EPI_ISL_2535463 | Labo Analyses Med | National Reference Center for Viruses of Respiratory Infections, Institut Pasteur, Paris | Marion Barbet, Sylvie Behillil, Méline Bizard, Angela Brisebarre, Camille Capel, Vincent Enouf, Louise Lefrançois, Frédéric Lemoine, Christophe Malabat, Corinne Maufrais, Etienne Simon-Lorière, Maud Vanpeene, Sylvie Van der Werf ,Jean-Baptiste Foulquier |
| EPI_ISL_2535464, EPI_ISL_2535466, EPI_ISL_2535467, EPI_ISL_2535468, EPI_ISL_2535469, EPI_ISL_2535470 | Labo Analyses Med | National Reference Center for Viruses of Respiratory Infections, Institut Pasteur, Paris | Marion Barbet, Sylvie Behillil, Méline Bizard, Angela Brisebarre, Camille Capel, Vincent Enouf, Louise Lefrançois, Frédéric Lemoine, Christophe Malabat, Corinne Maufrais, Etienne Simon-Lorière, Maud Vanpeene, Sylvie Van der Werf ,Benedicte Baccouch |
| EPI_ISL_2535476, EPI_ISL_2535477, EPI_ISL_2535478 | Hospital | National Reference Center for Viruses of Respiratory Infections, Institut Pasteur, Paris | Marion Barbet, Sylvie Behillil, Méline Bizard, Angela Brisebarre, Camille Capel, Vincent Enouf, Louise Lefrançois, Frédéric Lemoine, Christophe Malabat, Corinne Maufrais, Etienne Simon-Lorière, Maud Vanpeene, Sylvie Van der Werf ,LéA Pilorge |
| EPI_ISL_2535480, EPI_ISL_2535481 | Hospital | National Reference Center for Viruses of Respiratory Infections, Institut Pasteur, Paris | Marion Barbet, Sylvie Behillil, Méline Bizard, Angela Brisebarre, Camille Capel, Vincent Enouf, Louise Lefrançois, Frédéric Lemoine, Christophe Malabat, Corinne Maufrais, Etienne Simon-Lorière, Maud Vanpeene, Sylvie Van der Werf ,Alexandra Ducancelle |
| EPI_ISL_2535483, EPI_ISL_2535484, EPI_ISL_2535485, EPI_ISL_2535486, EPI_ISL_2535487, EPI_ISL_2535488, EPI_ISL_2535489, EPI_ISL_2535490, EPI_ISL_2535491, EPI_ISL_2535492, EPI_ISL_2535493, EPI_ISL_2535495, EPI_ISL_2535496 | | | |
| see above | Labo Analyses Med | National Reference Center for Viruses of Respiratory Infections, Institut Pasteur, Paris | Marion Barbet, Sylvie Behillil, Méline Bizard, Angela Brisebarre, Camille Capel, Vincent Enouf, Louise Lefrançois, Frédéric Lemoine, Christophe Malabat, Corinne Maufrais, Etienne Simon-Lorière, Maud Vanpeene, Sylvie Van der Werf , Hugues Leroy |
| EPI_ISL_2535497 | Labo Analyses Med | National Reference Center for Viruses of Respiratory Infections, Institut Pasteur, Paris | Marion Barbet, Sylvie Behillil, Méline Bizard, Angela Brisebarre, Camille Capel, Vincent Enouf, Louise Lefrançois, Frédéric Lemoine, Christophe Malabat, Corinne Maufrais, Emmanulelle Permal, Etienne Simon-Lorière, Maud Vanpeene, Sylvie Van der Werf , Hugues Leroy |
| EPI_ISL_2535498, EPI_ISL_2535499, EPI_ISL_2535500, EPI_ISL_2535501, EPI_ISL_2535502, EPI_ISL_2535503, EPI_ISL_2535504, EPI_ISL_2535505, EPI_ISL_2535506, EPI_ISL_2535507, EPI_ISL_2535508, EPI_ISL_2535509, EPI_ISL_2535510, EPI_ISL_2535511, EPI_ISL_2535512, EPI_ISL_2535513, EPI_ISL_2535514, EPI_ISL_2535515, EPI_ISL_2535516, EPI_ISL_2535517, EPI_ISL_2535518, EPI_ISL_2535520, EPI_ISL_2535521, EPI_ISL_2535522, EPI_ISL_2535523, EPI_ISL_2535524, EPI_ISL_2535525, EPI_ISL_2535526, EPI_ISL_2535527, EPI_ISL_2535528, EPI_ISL_2535529, EPI_ISL_2535530, EPI_ISL_2535532, EPI_ISL_2535533, EPI_ISL_2535534, EPI_ISL_2535535, EPI_ISL_2535536, EPI_ISL_2535537, EPI_ISL_2535538, EPI_ISL_2535539, EPI_ISL_2535540, EPI_ISL_2535541, EPI_ISL_2535542, EPI_ISL_2535543, EPI_ISL_2535544, EPI_ISL_2535545, EPI_ISL_2535546, EPI_ISL_2535547, EPI_ISL_2535548, EPI_ISL_2535549, EPI_ISL_2535550, EPI_ISL_2535551, EPI_ISL_2535552, EPI_ISL_2535553, EPI_ISL_2535554 | | | |
| see above | Labo Analyses Med | National Reference Center for Viruses of Respiratory Infections, Institut Pasteur, Paris | Marion Barbet, Sylvie Behillil, Méline Bizard, Angela Brisebarre, Camille Capel, Vincent Enouf, Louise Lefrançois, Frédéric Lemoine, Christophe Malabat, Corinne Maufrais, Etienne Simon-Lorière, Maud Vanpeene, Sylvie Van der Werf , Hugues Leroy |
| EPI_ISL_2535555 | Hospital | National Reference Center for Viruses of Respiratory Infections, Institut Pasteur, Paris | Marion Barbet, Sylvie Behillil, Méline Bizard, Angela Brisebarre, Camille Capel, Vincent Enouf, Louise Lefrançois, Frédéric Lemoine, Christophe Malabat, Corinne Maufrais, Etienne Simon-Lorière, Maud Vanpeene, Sylvie Van der Werf ,Isabelle Woercel |
| EPI_ISL_2535556 | Labo Analyses Med | National Reference Center for Viruses of Respiratory Infections, Institut Pasteur, Paris | Marion Barbet, Sylvie Behillil, Méline Bizard, Angela Brisebarre, Camille Capel, Vincent Enouf, Louise Lefrançois, Frédéric Lemoine, Christophe Malabat, Corinne Maufrais, Etienne Simon-Lorière, Maud Vanpeene, Sylvie Van der Werf ,Isabelle Woercel |
| EPI_ISL_2535557, EPI_ISL_2535558, EPI_ISL_2535559 | Hospital | National Reference Center for Viruses of Respiratory Infections, Institut Pasteur, Paris | Marion Barbet, Sylvie Behillil, Méline Bizard, Angela Brisebarre, Camille Capel, Vincent Enouf, Louise Lefrançois, Frédéric Lemoine, Christophe Malabat, Corinne Maufrais, Etienne Simon-Lorière, Maud Vanpeene, Sylvie Van der Werf ,CéCile Farrugia |
| EPI_ISL_2535560, EPI_ISL_2535561, EPI_ISL_2535562, EPI_ISL_2535563, EPI_ISL_2535564 | Labo Analyses Med | National Reference Center for Viruses of Respiratory Infections, Institut Pasteur, Paris | Marion Barbet, Sylvie Behillil, Méline Bizard, Angela Brisebarre, Camille Capel, Vincent Enouf, Louise Lefrançois, Frédéric Lemoine, Christophe Malabat, Corinne Maufrais, Etienne Simon-Lorière, Maud Vanpeene, Sylvie Van der Werf ,Benedicte Baccouch |
| EPI_ISL_2535566, EPI_ISL_2535567, EPI_ISL_2535568, EPI_ISL_2535569, EPI_ISL_2535570, EPI_ISL_2535571, EPI_ISL_2535572, EPI_ISL_2535573, EPI_ISL_2535574, EPI_ISL_2535575, EPI_ISL_2535576, EPI_ISL_2535577, EPI_ISL_2535578, EPI_ISL_2535579, EPI_ISL_2535580, EPI_ISL_2535581, EPI_ISL_2535582, EPI_ISL_2535583, EPI_ISL_2535584, EPI_ISL_2535585, EPI_ISL_2535586, EPI_ISL_2535587, EPI_ISL_2535588 | | | |
| see above | Labo Analyses Med | National Reference Center for Viruses of Respiratory Infections, Institut Pasteur, Paris | Marion Barbet, Sylvie Behillil, Méline Bizard, Angela Brisebarre, Camille Capel, Vincent Enouf, Louise Lefrançois, Frédéric Lemoine, Christophe Malabat, Corinne Maufrais, Etienne Simon-Lorière, Maud Vanpeene, Sylvie Van der Werf ,Arnaud Serpentini |
| EPI_ISL_2535589, EPI_ISL_2535590, EPI_ISL_2535591, EPI_ISL_2535592, EPI_ISL_2535593, EPI_ISL_2535594, EPI_ISL_2535595, EPI_ISL_2535596, EPI_ISL_2535597, EPI_ISL_2535598, EPI_ISL_2535599, EPI_ISL_2535600, EPI_ISL_2535601 | | | |
| see above | Labo Analyses Med | National Reference Center for Viruses of Respiratory Infections, Institut Pasteur, Paris | Marion Barbet, Sylvie Behillil, Méline Bizard, Angela Brisebarre, Camille Capel, Vincent Enouf, Louise Lefrançois, Frédéric Lemoine, Christophe Malabat, Corinne Maufrais, Etienne Simon-Lorière, Maud Vanpeene, Sylvie Van der Werf ,Ronan Pichard |
| EPI_ISL_2537480 | Laboratori de Referencia de Catalunya | Laboratori de Referencia de Catalunya | Ramirez A., Bellosillo B., Padilla E., Vilas A., Hernandez JJ., Canal M. |
| EPI_ISL_2543433 | Provincial Public Health Reference Laboratory (PPHRL) | Provincial Public Health Reference Laboratory | ANDLEEB HANIF |
| EPI_ISL_2543474 | Provincial Public Health Reference Laboratory (PPHRL) | Provincial Public Health Reference Laboratory (PPHRL) | ANDLEEB HANIF, MARIAM JAVED |
| EPI_ISL_2543612 | Provincial Public Health Reference Laboratory (PPHRL) P& SHD | Provincial Public Health Reference Laboratory (PPHRL) P& SHD | ANDLEEB HANIF, MARIAM JAVED |
| EPI_ISL_2543613 | Provincial Public Health Reference Laboratory (PPHRL) P& SHD | Provincial Public Health Reference Laboratory (PPHRL) P& SHD | ANDLEEB HANIF |
| EPI_ISL_2543899 | Sterling Hospital, Memnagar, Ahmedabad | Gujarat Biotechnology Research Centre | Atulbhai Patel, Sonal Sharma ,Zuber Saiyed,Ramesh Pandit,Janvi Raval,Zarna Patel,Nitin Savaliya,Dinesh Kumar,Twinkle Soni,Umang Mishra,Nitesh Shah,Chaitanya Joshi,Madhvi Joshi |
| EPI_ISL_2543900 | N.H.L Municipal Medical College, Ahmedabad | Gujarat Biotechnology Research Centre | Zuber Saiyed,Ramesh Pandit,Janvi Raval,Zarna Patel,Nitin Savaliya,Dinesh Kumar,Twinkle Soni,Sonal Sharma ,Jayshri Pethani,Umang Mishra,Nitesh Shah,Chaitanya Joshi,Madhvi Joshi |
| EPI_ISL_2543902 | N.H.L Municipal Medical College, Ahmedabad | Gujarat Biotechnology Research Centre | Janvi Raval,Zarna Patel,Nitin Savaliya,Dinesh Kumar,Twinkle Soni,Sonal Sharma ,Zuber Saiyed,Ramesh Pandit,Jayshri Pethani,Umang Mishra,Nitesh Shah,Chaitanya Joshi,Madhvi Joshi |
| EPI_ISL_2543903 | N.H.L Municipal Medical College, Ahmedabad | Gujarat Biotechnology Research Centre | Zarna Patel,Nitin Savaliya,Dinesh Kumar,Twinkle Soni,Sonal Sharma ,Zuber Saiyed,Ramesh Pandit,Janvi Raval,Jayshri Pethani,Umang Mishra,Nitesh Shah,Chaitanya Joshi,Madhvi Joshi |

| | | | |
|---|---|---|---|
| EPI_ISL_2543904 | N.H.L Municipal Medical College, Ahmedabad | Gujarat Biotechnology Research Centre | Nitin Savaliya,Dinesh Kumar,Twinkle Soni,Sonal Sharma ,Zuber Saiyed,Ramesh Pandit,Janvi Raval,Zarna Patel,Jayshri Pethani,Umang Mishra,Nitesh Shah,Chaitanya Joshi,Madhvi Joshi |
| EPI_ISL_2543905 | N.H.L Municipal Medical College, Ahmedabad | Gujarat Biotechnology Research Centre | Dinesh Kumar,Twinkle Soni,Sonal Sharma ,Zuber Saiyed,Ramesh Pandit,Janvi Raval,Zarna Patel,Nitin Savaliya,Jayshri Pethani,Umang Mishra,Nitesh Shah,Chaitanya Joshi,Madhvi Joshi |
| EPI_ISL_2543906 | N.H.L Municipal Medical College, Ahmedabad | Gujarat Biotechnology Research Centre | Twinkle Soni,Sonal Sharma ,Zuber Saiyed,Ramesh Pandit,Janvi Raval,Zarna Patel,Nitin Savaliya,Dinesh Kumar,Jayshri Pethani,Umang Mishra,Nitesh Shah,Chaitanya Joshi,Madhvi Joshi |
| EPI_ISL_2543907 | N.H.L Municipal Medical College, Ahmedabad | Gujarat Biotechnology Research Centre | Sonal Sharma ,Zuber Saiyed,Ramesh Pandit,Janvi Raval,Zarna Patel,Nitin Savaliya,Dinesh Kumar,Twinkle Soni,Jayshri Pethani,Umang Mishra,Nitesh Shah,Chaitanya Joshi,Madhvi Joshi |
| EPI_ISL_2543908 | N.H.L Municipal Medical College, Ahmedabad | Gujarat Biotechnology Research Centre | Zuber Saiyed,Ramesh Pandit,Janvi Raval,Zarna Patel,Nitin Savaliya,Dinesh Kumar,Twinkle Soni,Sonal Sharma ,Jayshri Pethani,Umang Mishra,Nitesh Shah,Chaitanya Joshi,Madhvi Joshi |
| EPI_ISL_2543909 | N.H.L Municipal Medical College, Ahmedabad | Gujarat Biotechnology Research Centre | Ramesh Pandit,Janvi Raval,Zarna Patel,Nitin Savaliya,Dinesh Kumar,Twinkle Soni,Sonal Sharma ,Zuber Saiyed,Jayshri Pethani,Umang Mishra,Nitesh Shah,Chaitanya Joshi,Madhvi Joshi |
| EPI_ISL_2543910 | N.H.L Municipal Medical College, Ahmedabad | Gujarat Biotechnology Research Centre | Janvi Raval,Zarna Patel,Nitin Savaliya,Dinesh Kumar,Twinkle Soni,Sonal Sharma ,Zuber Saiyed,Ramesh Pandit,Jayshri Pethani,Umang Mishra,Nitesh Shah,Chaitanya Joshi,Madhvi Joshi |
| EPI_ISL_2543911 | N.H.L Municipal Medical College, Ahmedabad | Gujarat Biotechnology Research Centre | Zarna Patel,Nitin Savaliya,Dinesh Kumar,Twinkle Soni,Sonal Sharma ,Zuber Saiyed,Ramesh Pandit,Janvi Raval,Jayshri Pethani,Umang Mishra,Nitesh Shah,Chaitanya Joshi,Madhvi Joshi |
| EPI_ISL_2543912 | N.H.L Municipal Medical College, Ahmedabad | Gujarat Biotechnology Research Centre | Nitin Savaliya,Dinesh Kumar,Twinkle Soni,Sonal Sharma ,Zuber Saiyed,Ramesh Pandit,Janvi Raval,Zarna Patel,Jayshri Pethani,Umang Mishra,Nitesh Shah,Chaitanya Joshi,Madhvi Joshi |
| EPI_ISL_2543913 | N.H.L Municipal Medical College, Ahmedabad | Gujarat Biotechnology Research Centre | Dinesh Kumar,Twinkle Soni,Sonal Sharma ,Zuber Saiyed,Ramesh Pandit,Janvi Raval,Zarna Patel,Nitin Savaliya,Jayshri Pethani,Umang Mishra,Nitesh Shah,Chaitanya Joshi,Madhvi Joshi |
| EPI_ISL_2543914 | N.H.L Municipal Medical College, Ahmedabad | Gujarat Biotechnology Research Centre | Twinkle Soni,Sonal Sharma ,Zuber Saiyed,Ramesh Pandit,Janvi Raval,Zarna Patel,Nitin Savaliya,Dinesh Kumar,Jayshri Pethani,Umang Mishra,Nitesh Shah,Chaitanya Joshi,Madhvi Joshi |
| EPI_ISL_2543915 | N.H.L Municipal Medical College, Ahmedabad | Gujarat Biotechnology Research Centre | Sonal Sharma ,Zuber Saiyed,Ramesh Pandit,Janvi Raval,Zarna Patel,Nitin Savaliya,Dinesh Kumar,Twinkle Soni,Jayshri Pethani,Umang Mishra,Nitesh Shah,Chaitanya Joshi,Madhvi Joshi |
| EPI_ISL_2544514, EPI_ISL_2544515, EPI_ISL_2544516, EPI_ISL_2544517, EPI_ISL_2544519, EPI_ISL_2544520, EPI_ISL_2544522, EPI_ISL_2544523 | Department of Virology, Istituto Zooprofilattico Sperimentale del Lazio e della Toscana (IZSLT) | Department of General Diagnostics; Department of Virology; Istituto Zooprofilattico Sperimentale del Lazio e della Toscana (IZSLT) | Patricia Alba, Giuseppe Manna, Elena L. Diaconu, Fabiola Feltrin, Raffaela Conti, Teresa Scicluna, Virginia Carfora, Antonella Cersini, Alessia Franco, Antonio Battisti. |
| EPI_ISL_2544528, EPI_ISL_2544529 | Basurto University Hospital: Clinical Microbiology Laboratory | Biocruces Bizkaia | Mikel Urrutikoetxea-Gutiérrez, Mª Carmen Nieto Toboso, Estibaliz Ugalde Zarraga, Mikel Gallego Rodrigo, Ana de la Hoz, José Luis Díaz de Tuesta del Arco |
| EPI_ISL_2544744, EPI_ISL_2544745, EPI_ISL_2544747 | Laboratoire Hôpital Sainte Musse | Laboratoire Hôpital Sainte Musse | Lionel Chollet |
| EPI_ISL_2544820 | GMERS Government Medical College (GMC), Gandhinagar | Gujarat Biotechnology Research Centre | Zuber Saiyed,Ramesh Pandit,Janvi Raval,Zarna Patel,Nitin Savaliya,Dinesh Kumar,Twinkle Soni,Sonal Sharma ,Gaurishankar Shrimali,Umang Mishra,Nitesh Shah,Chaitanya Joshi,Madhvi Joshi |
| EPI_ISL_2544821 | GMERS Government Medical College (GMC), Gandhinagar | Gujarat Biotechnology Research Centre | Ramesh Pandit,Janvi Raval,Zarna Patel,Nitin Savaliya,Dinesh Kumar,Twinkle Soni,Sonal Sharma ,Zuber Saiyed,Gaurishankar Shrimali,Umang Mishra,Nitesh Shah,Chaitanya Joshi,Madhvi Joshi |
| EPI_ISL_2544823 | GMERS Government Medical College (GMC), Gotri | Gujarat Biotechnology Research Centre | Zarna Patel,Nitin Savaliya,Dinesh Kumar,Twinkle Soni,Sonal Sharma ,Zuber Saiyed,Ramesh Pandit,Janvi Raval,Bithika Duttaroy,Umang Mishra,Nitesh Shah,Chaitanya Joshi,Madhvi Joshi |
| EPI_ISL_2544825 | GMERS Government Medical College (GMC), Gotri | Gujarat Biotechnology Research Centre | Dinesh Kumar,Twinkle Soni,Sonal Sharma ,Zuber Saiyed,Ramesh Pandit,Janvi Raval,Zarna Patel,Nitin Savaliya,Bithika Duttaroy,Umang Mishra,Nitesh Shah,Chaitanya Joshi,Madhvi Joshi |
| EPI_ISL_2544827 | GMERS Government Medical College (GMC), Gotri | Gujarat Biotechnology Research Centre | Sonal Sharma ,Zuber Saiyed,Ramesh Pandit,Janvi Raval,Zarna Patel,Nitin Savaliya,Dinesh Kumar,Twinkle Soni,Bithika Duttaroy,Umang Mishra,Nitesh Shah,Chaitanya Joshi,Madhvi Joshi |
| EPI_ISL_2544829 | GMERS Government Medical College (GMC), Gotri | Gujarat Biotechnology Research Centre | Ramesh Pandit,Janvi Raval,Zarna Patel,Nitin Savaliya,Dinesh Kumar,Twinkle Soni,Sonal Sharma ,Zuber Saiyed,Bithika Duttaroy,Umang Mishra,Nitesh Shah,Chaitanya Joshi,Madhvi Joshi |
| EPI_ISL_2544830 | GMERS Government Medical College (GMC), Gotri | Gujarat Biotechnology Research Centre | Janvi Raval,Zarna Patel,Nitin Savaliya,Dinesh Kumar,Twinkle Soni,Sonal Sharma ,Zuber Saiyed,Ramesh Pandit,Bithika Duttaroy,Umang Mishra,Nitesh Shah,Chaitanya Joshi,Madhvi Joshi |
| EPI_ISL_2544831 | GMERS Government Medical College (GMC), Gotri | Gujarat Biotechnology Research Centre | Zarna Patel,Nitin Savaliya,Dinesh Kumar,Twinkle Soni,Sonal Sharma ,Zuber Saiyed,Ramesh Pandit,Janvi Raval,Bithika Duttaroy,Umang Mishra,Nitesh Shah,Chaitanya Joshi,Madhvi Joshi |
| EPI_ISL_2544834 | GMERS Government Medical College (GMC), Gotri | Gujarat Biotechnology Research Centre | Twinkle Soni,Sonal Sharma ,Zuber Saiyed,Ramesh Pandit,Janvi Raval,Zarna Patel,Nitin Savaliya,Dinesh Kumar,Bithika Duttaroy,Umang Mishra,Nitesh Shah,Chaitanya Joshi,Madhvi Joshi |
| EPI_ISL_2544835 | GMERS Government Medical College (GMC), Gotri | Gujarat Biotechnology Research Centre | Sonal Sharma ,Zuber Saiyed,Ramesh Pandit,Janvi Raval,Zarna Patel,Nitin Savaliya,Dinesh Kumar,Twinkle Soni,Bithika Duttaroy,Umang Mishra,Nitesh Shah,Chaitanya Joshi,Madhvi Joshi |
| EPI_ISL_2544899, EPI_ISL_2544900, EPI_ISL_2544901, EPI_ISL_2544902, EPI_ISL_2544903, EPI_ISL_2544904, EPI_ISL_2544905, EPI_ISL_2544906, EPI_ISL_2544907, EPI_ISL_2544908, EPI_ISL_2544909, EPI_ISL_2544910, EPI_ISL_2544912 | | | |
| see above | OLVZ Aalst | OLVZ Aalst | Anne Vankeerberghen |
| EPI_ISL_2545667 | Tripler Army Medical Center | Tripler Army Medical Center | Rachael L. Downham, Jonathan A. D'Ambrozio, Kendra N. Carter, Lindsey L. O'Neal, Zakariae Bram, and Catherine F. Uyehara |
| EPI_ISL_2545683 | LESP Jalisco | Instituto de Diagnostico y Referencia Epidemiologicos (INDRE) | Claudia Wong-Arambula, Abril Rodriguez-Maldonado, Vanessa Rivero-Arredondo, Ariadna Medina-Benitez, Joaquin Quiroz-Mercado, Sergio Rangel-Guerrero, Natividad Cruz-Ortiz, Tatiana Nunez-Garcia, Gisela Barrera-Badillo, Lucia Hernandez-Rivas, Irma Lopez-Martinez, Ernesto Ramirez-Gonzalez. |
| EPI_ISL_2545684, EPI_ISL_2545685, EPI_ISL_2545686, EPI_ISL_2545687, EPI_ISL_2545688 | LESP Yucatan | Instituto de Diagnostico y Referencia Epidemiologicos (INDRE) | Claudia Wong-Arambula, Abril Rodriguez-Maldonado, Vanessa Rivero-Arredondo, Ariadna Medina-Benitez, Joaquin Quiroz-Mercado, Sergio Rangel-Guerrero, Natividad Cruz-Ortiz, Tatiana Nunez-Garcia, Gisela Barrera-Badillo, Lucia Hernandez-Rivas, Irma Lopez-Martinez, Ernesto Ramirez-Gonzalez. |
| EPI_ISL_2545690, EPI_ISL_2545691 | LESP Tamaulipas | Instituto de Diagnostico y Referencia Epidemiologicos (INDRE) | Claudia Wong-Arambula, Abril Rodriguez-Maldonado, Vanessa Rivero-Arredondo, Ariadna Medina-Benitez, Joaquin Quiroz-Mercado, Sergio Rangel-Guerrero, Natividad Cruz-Ortiz, Tatiana Nunez-Garcia, Gisela Barrera-Badillo, Lucia Hernandez-Rivas, Irma Lopez-Martinez, Ernesto Ramirez-Gonzalez. |
| EPI_ISL_2545692 | LESP Sinaloa | Instituto de Diagnostico y Referencia Epidemiologicos (INDRE) | Claudia Wong-Arambula, Abril Rodriguez-Maldonado, Vanessa Rivero-Arredondo, Ariadna Medina-Benitez, Joaquin Quiroz-Mercado, Sergio Rangel-Guerrero, Natividad Cruz-Ortiz, Tatiana Nunez-Garcia, Gisela Barrera-Badillo, Lucia Hernandez-Rivas, Irma Lopez-Martinez, Ernesto Ramirez-Gonzalez. |
| EPI_ISL_2545693, EPI_ISL_2545694 | LESP Quintana Roo | Instituto de Diagnostico y Referencia Epidemiologicos (INDRE) | Claudia Wong-Arambula, Abril Rodriguez-Maldonado, Vanessa Rivero-Arredondo, Ariadna Medina-Benitez, Joaquin Quiroz-Mercado, Sergio Rangel-Guerrero, Natividad Cruz-Ortiz, Tatiana Nunez-Garcia, Gisela Barrera-Badillo, Lucia Hernandez-Rivas, Irma Lopez-Martinez, Ernesto Ramirez-Gonzalez. |
| EPI_ISL_2545695, EPI_ISL_2545696, | LESP Ciudad de Mexico | Instituto de Diagnostico y Referencia Epidemiologicos | Claudia Wong-Arambula, Abril Rodriguez-Maldonado, Vanessa Rivero-Arredondo, Ariadna Medina-Benitez, Joaquin Quiroz-Mercado, Sergio |

| | | | |
|---|---|---|---|
| EPI_ISL_2545697 | | (INDRE) | Rangel-Guerrero, Natividad Cruz-Ortiz, Tatiana Nunez-Garcia, Gisela Barrera-Badillo, Lucia Hernandez-Rivas, Irma Lopez-Martinez, Ernesto Ramirez-Gonzalez. |
| EPI_ISL_2545698 | LESP Estado de Mexico | Instituto de Diagnostico y Referencia Epidemiologicos (INDRE) | Claudia Wong-Arambula, Abril Rodriguez-Maldonado, Vanessa Rivero-Arredondo, Ariadna Medina-Benitez, Joaquin Quiroz-Mercado, Sergio Rangel-Guerrero, Natividad Cruz-Ortiz, Tatiana Nunez-Garcia, Gisela Barrera-Badillo, Lucia Hernandez-Rivas, Irma Lopez-Martinez, Ernesto Ramirez-Gonzalez. |
| EPI_ISL_2545703, EPI_ISL_2545704 | LESP Sinaloa | Instituto de Diagnostico y Referencia Epidemiologicos (INDRE) | Claudia Wong-Arambula, Abril Rodriguez-Maldonado, Vanessa Rivero-Arredondo, Ariadna Medina-Benitez, Joaquin Quiroz-Mercado, Sergio Rangel-Guerrero, Natividad Cruz-Ortiz, Tatiana Nunez-Garcia, Gisela Barrera-Badillo, Lucia Hernandez-Rivas, Irma Lopez-Martinez, Ernesto Ramirez-Gonzalez. |
| EPI_ISL_2545708, EPI_ISL_2545709 | LESP Chihuahua | Instituto de Diagnostico y Referencia Epidemiologicos (INDRE) | Claudia Wong-Arambula, Abril Rodriguez-Maldonado, Vanessa Rivero-Arredondo, Ariadna Medina-Benitez, Joaquin Quiroz-Mercado, Sergio Rangel-Guerrero, Natividad Cruz-Ortiz, Tatiana Nunez-Garcia, Gisela Barrera-Badillo, Lucia Hernandez-Rivas, Irma Lopez-Martinez, Ernesto Ramirez-Gonzalez. |
| EPI_ISL_2545710, EPI_ISL_2545711 | LESP Veracruz | Instituto de Diagnostico y Referencia Epidemiologicos (INDRE) | Claudia Wong-Arambula, Abril Rodriguez-Maldonado, Vanessa Rivero-Arredondo, Ariadna Medina-Benitez, Joaquin Quiroz-Mercado, Sergio Rangel-Guerrero, Natividad Cruz-Ortiz, Tatiana Nunez-Garcia, Gisela Barrera-Badillo, Lucia Hernandez-Rivas, Irma Lopez-Martinez, Ernesto Ramirez-Gonzalez. |
| EPI_ISL_2545722 | LESP Ciudad de Mexico | Instituto de Diagnostico y Referencia Epidemiologicos (INDRE) | Claudia Wong-Arambula, Abril Rodriguez-Maldonado, Vanessa Rivero-Arredondo, Ariadna Medina-Benitez, Joaquin Quiroz-Mercado, Sergio Rangel-Guerrero, Natividad Cruz-Ortiz, Tatiana Nunez-Garcia, Gisela Barrera-Badillo, Lucia Hernandez-Rivas, Irma Lopez-Martinez, Ernesto Ramirez-Gonzalez. |
| EPI_ISL_2545723 | LESP Estado de Mexico | Instituto de Diagnostico y Referencia Epidemiologicos (INDRE) | Claudia Wong-Arambula, Abril Rodriguez-Maldonado, Vanessa Rivero-Arredondo, Ariadna Medina-Benitez, Joaquin Quiroz-Mercado, Sergio Rangel-Guerrero, Natividad Cruz-Ortiz, Tatiana Nunez-Garcia, Gisela Barrera-Badillo, Lucia Hernandez-Rivas, Irma Lopez-Martinez, Ernesto Ramirez-Gonzalez. |
| EPI_ISL_2545724, EPI_ISL_2545725, EPI_ISL_2545726, EPI_ISL_2545727 | LESP Campeche | Instituto de Diagnostico y Referencia Epidemiologicos (INDRE) | Claudia Wong-Arambula, Abril Rodriguez-Maldonado, Vanessa Rivero-Arredondo, Ariadna Medina-Benitez, Joaquin Quiroz-Mercado, Sergio Rangel-Guerrero, Natividad Cruz-Ortiz, Tatiana Nunez-Garcia, Gisela Barrera-Badillo, Lucia Hernandez-Rivas, Irma Lopez-Martinez, Ernesto Ramirez-Gonzalez. |
| EPI_ISL_2545728 | LESP Coahuila | Instituto de Diagnostico y Referencia Epidemiologicos (INDRE) | Claudia Wong-Arambula, Abril Rodriguez-Maldonado, Vanessa Rivero-Arredondo, Ariadna Medina-Benitez, Joaquin Quiroz-Mercado, Sergio Rangel-Guerrero, Natividad Cruz-Ortiz, Tatiana Nunez-Garcia, Gisela Barrera-Badillo, Lucia Hernandez-Rivas, Irma Lopez-Martinez, Ernesto Ramirez-Gonzalez. |
| EPI_ISL_2545729, EPI_ISL_2545730 | LESP Baja California Sur | Instituto de Diagnostico y Referencia Epidemiologicos (INDRE) | Claudia Wong-Arambula, Abril Rodriguez-Maldonado, Vanessa Rivero-Arredondo, Ariadna Medina-Benitez, Joaquin Quiroz-Mercado, Sergio Rangel-Guerrero, Natividad Cruz-Ortiz, Tatiana Nunez-Garcia, Gisela Barrera-Badillo, Lucia Hernandez-Rivas, Irma Lopez-Martinez, Ernesto Ramirez-Gonzalez. |
| EPI_ISL_2545732 | LESP Queretaro | Instituto de Diagnostico y Referencia Epidemiologicos (INDRE) | Claudia Wong-Arambula, Abril Rodriguez-Maldonado, Vanessa Rivero-Arredondo, Ariadna Medina-Benitez, Joaquin Quiroz-Mercado, Sergio Rangel-Guerrero, Natividad Cruz-Ortiz, Tatiana Nunez-Garcia, Gisela Barrera-Badillo, Lucia Hernandez-Rivas, Irma Lopez-Martinez, Ernesto Ramirez-Gonzalez. |
| EPI_ISL_2545735, EPI_ISL_2545736, EPI_ISL_2545737, EPI_ISL_2545738, EPI_ISL_2545739, EPI_ISL_2545740, EPI_ISL_2545741, EPI_ISL_2545742, EPI_ISL_2545743, EPI_ISL_2545744, EPI_ISL_2545745, EPI_ISL_2545746, EPI_ISL_2545747, EPI_ISL_2545748, EPI_ISL_2545749, EPI_ISL_2545750, EPI_ISL_2545751, EPI_ISL_2545752, EPI_ISL_2545753, EPI_ISL_2545754, EPI_ISL_2545755, EPI_ISL_2545756, EPI_ISL_2545757, EPI_ISL_2545758, EPI_ISL_2545759, EPI_ISL_2545760, EPI_ISL_2545761, EPI_ISL_2545762, EPI_ISL_2545763, EPI_ISL_2545764, EPI_ISL_2545765, EPI_ISL_2545766, EPI_ISL_2545767, EPI_ISL_2545768, EPI_ISL_2545769, EPI_ISL_2545770, EPI_ISL_2545771, EPI_ISL_2545772, EPI_ISL_2545773, EPI_ISL_2545774, EPI_ISL_2545775, EPI_ISL_2545776, EPI_ISL_2545777, EPI_ISL_2545778, EPI_ISL_2545779, EPI_ISL_2545780, EPI_ISL_2545781, EPI_ISL_2545782, EPI_ISL_2545783, EPI_ISL_2545784, EPI_ISL_2545785, EPI_ISL_2545786, EPI_ISL_2545787, EPI_ISL_2545788, EPI_ISL_2545789, EPI_ISL_2545790, EPI_ISL_2545791, EPI_ISL_2545792, EPI_ISL_2545793, EPI_ISL_2545794, EPI_ISL_2545795, EPI_ISL_2545796, EPI_ISL_2545797, EPI_ISL_2545798, EPI_ISL_2545799, EPI_ISL_2545800, EPI_ISL_2545801, EPI_ISL_2545802, EPI_ISL_2545803 | | | |
| see above | LESP Yucatan | Instituto de Diagnostico y Referencia Epidemiologicos (INDRE) | Claudia Wong-Arambula, Abril Rodriguez-Maldonado, Vanessa Rivero-Arredondo, Ariadna Medina-Benitez, Joaquin Quiroz-Mercado, Sergio Rangel-Guerrero, Natividad Cruz-Ortiz, Tatiana Nunez-Garcia, Gisela Barrera-Badillo, Lucia Hernandez-Rivas, Irma Lopez-Martinez, Ernesto Ramirez-Gonzalez. |
| EPI_ISL_2545804 | LESP Michoacan | Instituto de Diagnostico y Referencia Epidemiologicos (INDRE) | Claudia Wong-Arambula, Abril Rodriguez-Maldonado, Vanessa Rivero-Arredondo, Ariadna Medina-Benitez, Joaquin Quiroz-Mercado, Sergio Rangel-Guerrero, Natividad Cruz-Ortiz, Tatiana Nunez-Garcia, Gisela Barrera-Badillo, Lucia Hernandez-Rivas, Irma Lopez-Martinez, Ernesto Ramirez-Gonzalez. |
| EPI_ISL_2545805 | LESP Durango | Instituto de Diagnostico y Referencia Epidemiologicos (INDRE) | Claudia Wong-Arambula, Abril Rodriguez-Maldonado, Vanessa Rivero-Arredondo, Ariadna Medina-Benitez, Joaquin Quiroz-Mercado, Sergio Rangel-Guerrero, Natividad Cruz-Ortiz, Tatiana Nunez-Garcia, Gisela Barrera-Badillo, Lucia Hernandez-Rivas, Irma Lopez-Martinez, Ernesto Ramirez-Gonzalez. |
| EPI_ISL_2545806 | LESP Campeche | Instituto de Diagnostico y Referencia Epidemiologicos (INDRE) | Claudia Wong-Arambula, Abril Rodriguez-Maldonado, Vanessa Rivero-Arredondo, Ariadna Medina-Benitez, Joaquin Quiroz-Mercado, Sergio Rangel-Guerrero, Natividad Cruz-Ortiz, Tatiana Nunez-Garcia, Gisela Barrera-Badillo, Lucia Hernandez-Rivas, Irma Lopez-Martinez, Ernesto Ramirez-Gonzalez. |
| EPI_ISL_2545808, EPI_ISL_2545809, EPI_ISL_2545810, EPI_ISL_2545811 | LESP Quintana Roo | Instituto de Diagnostico y Referencia Epidemiologicos (INDRE) | Claudia Wong-Arambula, Abril Rodriguez-Maldonado, Vanessa Rivero-Arredondo, Ariadna Medina-Benitez, Joaquin Quiroz-Mercado, Sergio Rangel-Guerrero, Natividad Cruz-Ortiz, Tatiana Nunez-Garcia, Gisela Barrera-Badillo, Lucia Hernandez-Rivas, Irma Lopez-Martinez, Ernesto Ramirez-Gonzalez. |
| EPI_ISL_2545812 | LESP Sonora | Instituto de Diagnostico y Referencia Epidemiologicos (INDRE) | Claudia Wong-Arambula, Abril Rodriguez-Maldonado, Vanessa Rivero-Arredondo, Ariadna Medina-Benitez, Joaquin Quiroz-Mercado, Sergio Rangel-Guerrero, Natividad Cruz-Ortiz, Tatiana Nunez-Garcia, Gisela Barrera-Badillo, Lucia Hernandez-Rivas, Irma Lopez-Martinez, Ernesto Ramirez-Gonzalez. |
| EPI_ISL_2545813, EPI_ISL_2545814 | LESP San Luis Potosi | Instituto de Diagnostico y Referencia Epidemiologicos (INDRE) | Claudia Wong-Arambula, Abril Rodriguez-Maldonado, Vanessa Rivero-Arredondo, Ariadna Medina-Benitez, Joaquin Quiroz-Mercado, Sergio Rangel-Guerrero, Natividad Cruz-Ortiz, Tatiana Nunez-Garcia, Gisela Barrera-Badillo, Lucia Hernandez-Rivas, Irma Lopez-Martinez, Ernesto Ramirez-Gonzalez. |
| EPI_ISL_2545815, EPI_ISL_2545818 | LESP Ciudad de Mexico | Instituto de Diagnostico y Referencia Epidemiologicos (INDRE) | Claudia Wong-Arambula, Abril Rodriguez-Maldonado, Vanessa Rivero-Arredondo, Ariadna Medina-Benitez, Joaquin Quiroz-Mercado, Sergio Rangel-Guerrero, Natividad Cruz-Ortiz, Tatiana Nunez-Garcia, Gisela Barrera-Badillo, Lucia Hernandez-Rivas, Irma Lopez-Martinez, Ernesto Ramirez-Gonzalez. |
| EPI_ISL_2545819 | LESP San Luis Potosi | Instituto de Diagnostico y Referencia Epidemiologicos (INDRE) | Claudia Wong-Arambula, Abril Rodriguez-Maldonado, Vanessa Rivero-Arredondo, Ariadna Medina-Benitez, Joaquin Quiroz-Mercado, Sergio Rangel-Guerrero, Natividad Cruz-Ortiz, Tatiana Nunez-Garcia, Gisela Barrera-Badillo, Lucia Hernandez-Rivas, Irma Lopez-Martinez, Ernesto Ramirez-Gonzalez. |
| EPI_ISL_2546901 | Laboratory of Communicable Diseases | 1. Laboratory of Communicable Diseases (Estonia); 2. Eurofins Genomics Europe Sequencing GmbH | Liidia Dotsenko et al. |
| EPI_ISL_2547525 | INT Fondazione Pascale | INT Fondazione Pascale | INT Fondazione Pascale |
| EPI_ISL_2549198, EPI_ISL_2549199 | Virology Department, Central Health Laboratory | The Francis Crick Institute | Ramuth M, Manraj SS, Sonoo J, Bahadoor BS, Crawford M, Patel H, Harvey R, Goldstone R, Daniels RS, Nicod J |
| EPI_ISL_2549253 | Institut de Virologie du CHU de Strasbourg | Institut de Virologie du CHU de Strasbourg | Fafi-Kremer Samira, Gallais Floriane, Gantner Pierre, Laugel Elodie, Solis Morgane, Velay Aurélie, Wendling Marie-Josée |
| EPI_ISL_2549576 | Hospital Tuanku Fauziah | Institute for Medical Research, Infectious Disease Research Centre, National Institutes of Health, Ministry of Health Malaysia | Suppiah J, Kamel K, Mohd Zawawi Z, Azizan MA, Ramly N, Robert F, Thayan R |
| EPI_ISL_2550714, EPI_ISL_2550731 | Johor Bahru Public Health Laboratory | Institute for Medical Research, Infectious Disease Research Centre, National Institutes of Health, Ministry of Health | Suppiah J, Kamel K, Mohd Zawawi Z, Azizan MA, Ramly N, Robert F, Thayan R |

| | | | |
|---|---|---|---|
| | | Malaysia | |
| EPI_ISL_2550732 | UOC Microbiologia e Virologia, Azienda Ospedaliera Universitaria Senese, Siena, Italy | Dipartimento di Biotecnologie Mediche | Maria Grazia Cusi, Gianni Gori Savellini, Gabriele Anichini, Claudia Gandolfo |
| EPI_ISL_2550733 | Hospital Labuan | Institute for Medical Research, Infectious Disease Research Centre, National Institutes of Health, Ministry of Health Malaysia | Suppiah J, Kamel K, Mohd Zawawi Z, Azizan MA, Ramly N, Robert F, Thayan R |
| EPI_ISL_2550758, EPI_ISL_2550759 | Port Dickson Hospital | Institute for Medical Research, Infectious Disease Research Centre, National Institutes of Health, Ministry of Health Malaysia | Suppiah J, Kamel K, Mohd Zawawi Z, Azizan MA, Ramly N, Robert F, Thayan R |
| EPI_ISL_2556117, EPI_ISL_2556119, EPI_ISL_2557251 | MOH - Jaber Al-Ahmad Hospital (Innovation Research Laboratory) | MOH - Jaber Al-Ahmad Hospital (Innovation Research Laboratory) | Salman Al-Sabah , Mohammad Alghounaim |
| EPI_ISL_2557364, EPI_ISL_2557366, EPI_ISL_2557375 | Lboratorio Central de Saude Publica do Estado do Parana (LACEN/PR) | Laboratory of Respiratory Viruses and Measles, Oswaldo Cruz Institute, FIOCRUZ | Paola Resende, Luciana Appolinario, Fernando Motta, Anna Carolina Paixao, Ana Carolina Mendonca, Alice Sampaio Rocha, Taina Venas, Elisa Cavalcante Pereira, Renata Serrano Lopes, Irina Riediger, Marilda Siqueira on behalf of the Fiocruz COVID-19 Genomic Surveillance Network |
| EPI_ISL_2557459 | MOH - Jaber Al-Ahmad Hospital (Innovation Research Laboratory) | MOH - Jaber Al-Ahmad Hospital (Innovation Research Laboratory) | Salman Al-Sabah , Mohammad Alghounaim |
| EPI_ISL_2557840 | University Hospital Limerick | University Hospital Limerick | Carolyn Meaney, Colm McDonnell, Patrick Stapleton |
| EPI_ISL_2559271, EPI_ISL_2559273, EPI_ISL_2559275 | LESP Chihuahua | Instituto de Diagnostico y Referencia Epidemiologicos (INDRE) | Claudia Wong-Arambula, Abril Rodriguez-Maldonado, Vanessa Rivero-Arredondo, Ariadna Medina-Benitez, Joaquin Quiroz-Mercado, Sergio Rangel-Guerrero, Natividad Cruz-Ortiz, Tatiana Nunez-Garcia, Gisela Barrera-Badillo, Lucia Hernandez-Rivas, Irma Lopez-Martinez, Ernesto Ramirez-Gonzalez. |
| EPI_ISL_2559277 | LESP Veracruz | Instituto de Diagnostico y Referencia Epidemiologicos (INDRE) | Claudia Wong-Arambula, Abril Rodriguez-Maldonado, Vanessa Rivero-Arredondo, Ariadna Medina-Benitez, Joaquin Quiroz-Mercado, Sergio Rangel-Guerrero, Natividad Cruz-Ortiz, Tatiana Nunez-Garcia, Gisela Barrera-Badillo, Lucia Hernandez-Rivas, Irma Lopez-Martinez, Ernesto Ramirez-Gonzalez. |
| EPI_ISL_2559354 | LESP Morelos | Instituto de Diagnostico y Referencia Epidemiologicos (INDRE) | Claudia Wong-Arambula, Abril Rodriguez-Maldonado, Vanessa Rivero-Arredondo, Ariadna Medina-Benitez, Joaquin Quiroz-Mercado, Sergio Rangel-Guerrero, Natividad Cruz-Ortiz, Tatiana Nunez-Garcia, Gisela Barrera-Badillo, Lucia Hernandez-Rivas, Irma Lopez-Martinez, Ernesto Ramirez-Gonzalez. |
| EPI_ISL_2559356, EPI_ISL_2559358 | LESP Chihuahua | Instituto de Diagnostico y Referencia Epidemiologicos (INDRE) | Claudia Wong-Arambula, Abril Rodriguez-Maldonado, Vanessa Rivero-Arredondo, Ariadna Medina-Benitez, Joaquin Quiroz-Mercado, Sergio Rangel-Guerrero, Natividad Cruz-Ortiz, Tatiana Nunez-Garcia, Gisela Barrera-Badillo, Lucia Hernandez-Rivas, Irma Lopez-Martinez, Ernesto Ramirez-Gonzalez. |
| EPI_ISL_2559361 | LESP Campeche | Instituto de Diagnostico y Referencia Epidemiologicos (INDRE) | Claudia Wong-Arambula, Abril Rodriguez-Maldonado, Vanessa Rivero-Arredondo, Ariadna Medina-Benitez, Joaquin Quiroz-Mercado, Sergio Rangel-Guerrero, Natividad Cruz-Ortiz, Tatiana Nunez-Garcia, Gisela Barrera-Badillo, Lucia Hernandez-Rivas, Irma Lopez-Martinez, Ernesto Ramirez-Gonzalez. |
| EPI_ISL_2559363, EPI_ISL_2559365 | LESP Puebla | Instituto de Diagnostico y Referencia Epidemiologicos (INDRE) | Claudia Wong-Arambula, Abril Rodriguez-Maldonado, Vanessa Rivero-Arredondo, Ariadna Medina-Benitez, Joaquin Quiroz-Mercado, Sergio Rangel-Guerrero, Natividad Cruz-Ortiz, Tatiana Nunez-Garcia, Gisela Barrera-Badillo, Lucia Hernandez-Rivas, Irma Lopez-Martinez, Ernesto Ramirez-Gonzalez. |
| EPI_ISL_2559367, EPI_ISL_2559369, EPI_ISL_2559371, EPI_ISL_2559373 | LESP Queretaro | Instituto de Diagnostico y Referencia Epidemiologicos (INDRE) | Claudia Wong-Arambula, Abril Rodriguez-Maldonado, Vanessa Rivero-Arredondo, Ariadna Medina-Benitez, Joaquin Quiroz-Mercado, Sergio Rangel-Guerrero, Natividad Cruz-Ortiz, Tatiana Nunez-Garcia, Gisela Barrera-Badillo, Lucia Hernandez-Rivas, Irma Lopez-Martinez, Ernesto Ramirez-Gonzalez. |
| EPI_ISL_2559375, EPI_ISL_2559377 | LESP Tabasco | Instituto de Diagnostico y Referencia Epidemiologicos (INDRE) | Claudia Wong-Arambula, Abril Rodriguez-Maldonado, Vanessa Rivero-Arredondo, Ariadna Medina-Benitez, Joaquin Quiroz-Mercado, Sergio Rangel-Guerrero, Natividad Cruz-Ortiz, Tatiana Nunez-Garcia, Gisela Barrera-Badillo, Lucia Hernandez-Rivas, Irma Lopez-Martinez, Ernesto Ramirez-Gonzalez. |
| EPI_ISL_2559379 | LESP Veracruz | Instituto de Diagnostico y Referencia Epidemiologicos (INDRE) | Claudia Wong-Arambula, Abril Rodriguez-Maldonado, Vanessa Rivero-Arredondo, Ariadna Medina-Benitez, Joaquin Quiroz-Mercado, Sergio Rangel-Guerrero, Natividad Cruz-Ortiz, Tatiana Nunez-Garcia, Gisela Barrera-Badillo, Lucia Hernandez-Rivas, Irma Lopez-Martinez, Ernesto Ramirez-Gonzalez. |
| EPI_ISL_2559381, EPI_ISL_2559384, EPI_ISL_2559386, EPI_ISL_2559388, EPI_ISL_2559389, EPI_ISL_2559391, EPI_ISL_2559393, EPI_ISL_2559395, EPI_ISL_2559397, EPI_ISL_2559399, EPI_ISL_2559402, EPI_ISL_2559404, EPI_ISL_2559406, EPI_ISL_2559408, EPI_ISL_2559410, EPI_ISL_2559412, EPI_ISL_2559414, EPI_ISL_2559416, EPI_ISL_2559419, EPI_ISL_2559421, EPI_ISL_2559423 | | | |
| see above | LESP Yucatan | Instituto de Diagnostico y Referencia Epidemiologicos (INDRE) | Claudia Wong-Arambula, Abril Rodriguez-Maldonado, Vanessa Rivero-Arredondo, Ariadna Medina-Benitez, Joaquin Quiroz-Mercado, Sergio Rangel-Guerrero, Natividad Cruz-Ortiz, Tatiana Nunez-Garcia, Gisela Barrera-Badillo, Lucia Hernandez-Rivas, Irma Lopez-Martinez, Ernesto Ramirez-Gonzalez. |
| EPI_ISL_2559425, EPI_ISL_2559427, EPI_ISL_2559429, EPI_ISL_2559435, EPI_ISL_2559437, EPI_ISL_2559439, EPI_ISL_2559442, EPI_ISL_2559444 | LESP Michoacan | Instituto de Diagnostico y Referencia Epidemiologicos (INDRE) | Claudia Wong-Arambula, Abril Rodriguez-Maldonado, Vanessa Rivero-Arredondo, Ariadna Medina-Benitez, Joaquin Quiroz-Mercado, Sergio Rangel-Guerrero, Natividad Cruz-Ortiz, Tatiana Nunez-Garcia, Gisela Barrera-Badillo, Lucia Hernandez-Rivas, Irma Lopez-Martinez, Ernesto Ramirez-Gonzalez. |
| EPI_ISL_2559448 | LESP Durango | Instituto de Diagnostico y Referencia Epidemiologicos (INDRE) | Claudia Wong-Arambula, Abril Rodriguez-Maldonado, Vanessa Rivero-Arredondo, Ariadna Medina-Benitez, Joaquin Quiroz-Mercado, Sergio Rangel-Guerrero, Natividad Cruz-Ortiz, Tatiana Nunez-Garcia, Gisela Barrera-Badillo, Lucia Hernandez-Rivas, Irma Lopez-Martinez, Ernesto Ramirez-Gonzalez. |
| EPI_ISL_2559450 | LESP Sinaloa | Instituto de Diagnostico y Referencia Epidemiologicos (INDRE) | Claudia Wong-Arambula, Abril Rodriguez-Maldonado, Vanessa Rivero-Arredondo, Ariadna Medina-Benitez, Joaquin Quiroz-Mercado, Sergio Rangel-Guerrero, Natividad Cruz-Ortiz, Tatiana Nunez-Garcia, Gisela Barrera-Badillo, Lucia Hernandez-Rivas, Irma Lopez-Martinez, Ernesto Ramirez-Gonzalez. |
| EPI_ISL_2559452 | LESP Colima | Instituto de Diagnostico y Referencia Epidemiologicos (INDRE) | Claudia Wong-Arambula, Abril Rodriguez-Maldonado, Vanessa Rivero-Arredondo, Ariadna Medina-Benitez, Joaquin Quiroz-Mercado, Sergio Rangel-Guerrero, Natividad Cruz-Ortiz, Tatiana Nunez-Garcia, Gisela Barrera-Badillo, Lucia Hernandez-Rivas, Irma Lopez-Martinez, Ernesto Ramirez-Gonzalez. |
| EPI_ISL_2559456, EPI_ISL_2559458 | LESP Ciudad de Mexico | Instituto de Diagnostico y Referencia Epidemiologicos (INDRE) | Claudia Wong-Arambula, Abril Rodriguez-Maldonado, Vanessa Rivero-Arredondo, Ariadna Medina-Benitez, Joaquin Quiroz-Mercado, Sergio Rangel-Guerrero, Natividad Cruz-Ortiz, Tatiana Nunez-Garcia, Gisela Barrera-Badillo, Lucia Hernandez-Rivas, Irma Lopez-Martinez, Ernesto Ramirez-Gonzalez. |
| EPI_ISL_2559460 | LESP Estado de Mexico | Instituto de Diagnostico y Referencia Epidemiologicos (INDRE) | Claudia Wong-Arambula, Abril Rodriguez-Maldonado, Vanessa Rivero-Arredondo, Ariadna Medina-Benitez, Joaquin Quiroz-Mercado, Sergio Rangel-Guerrero, Natividad Cruz-Ortiz, Tatiana Nunez-Garcia, Gisela Barrera-Badillo, Lucia Hernandez-Rivas, Irma Lopez-Martinez, Ernesto Ramirez-Gonzalez. |
| EPI_ISL_2559463, EPI_ISL_2559465, EPI_ISL_2559467 | LESP Ciudad de Mexico | Instituto de Diagnostico y Referencia Epidemiologicos (INDRE) | Claudia Wong-Arambula, Abril Rodriguez-Maldonado, Vanessa Rivero-Arredondo, Ariadna Medina-Benitez, Joaquin Quiroz-Mercado, Sergio Rangel-Guerrero, Natividad Cruz-Ortiz, Tatiana Nunez-Garcia, Gisela Barrera-Badillo, Lucia Hernandez-Rivas, Irma Lopez-Martinez, Ernesto Ramirez-Gonzalez. |
| EPI_ISL_2559469 | LESP Estado de Mexico | Instituto de Diagnostico y Referencia Epidemiologicos (INDRE) | Claudia Wong-Arambula, Abril Rodriguez-Maldonado, Vanessa Rivero-Arredondo, Ariadna Medina-Benitez, Joaquin Quiroz-Mercado, Sergio Rangel-Guerrero, Natividad Cruz-Ortiz, Tatiana Nunez-Garcia, Gisela Barrera-Badillo, Lucia Hernandez-Rivas, Irma Lopez-Martinez, Ernesto Ramirez-Gonzalez. |
| EPI_ISL_2559471, EPI_ISL_2559472, | LESP Ciudad de Mexico | Instituto de Diagnostico y Referencia Epidemiologicos | Claudia Wong-Arambula, Abril Rodriguez-Maldonado, Vanessa Rivero-Arredondo, Ariadna Medina-Benitez, Joaquin Quiroz-Mercado, Sergio |

| | | | |
|---|---|---|---|
| EPI_ISL_2559473, EPI_ISL_2559475, EPI_ISL_2559477, EPI_ISL_2559479 | | (INDRE) | Rangel-Guerrero, Natividad Cruz-Ortiz, Tatiana Nunez-Garcia, Gisela Barrera-Badillo, Lucia Hernandez-Rivas, Irma Lopez-Martinez, Ernesto Ramirez-Gonzalez. |
| EPI_ISL_2562085 | GMERS Government Medical College (GMC), Gotri | Gujarat Biotechnology Research Centre | Ramesh Pandit,Janvi Raval,Zarna Patel,Nitin Savaliya,Dinesh Kumar,Twinkle Soni,Sonal Sharma ,Zuber Saiyed,Bithika Duttaroy,Umang Mishra,Nitesh Shah,Chaitanya Joshi,Madhvi Joshi |
| EPI_ISL_2562086 | GMERS Government Medical College (GMC), Gotri | Gujarat Biotechnology Research Centre | Janvi Raval,Zarna Patel,Nitin Savaliya,Dinesh Kumar,Twinkle Soni,Sonal Sharma ,Zuber Saiyed,Ramesh Pandit,Bithika Duttaroy,Umang Mishra,Nitesh Shah,Chaitanya Joshi,Madhvi Joshi |
| EPI_ISL_2562101 | GMERS Government Medical College (GMC), Gotri | Gujarat Biotechnology Research Centre | Ramesh Pandit,Janvi Raval,Zarna Patel,Nitin Savaliya,Dinesh Kumar,Twinkle Soni,Sonal Sharma ,Zuber Saiyed,Bithika Duttaroy,Umang Mishra,Nitesh Shah,Chaitanya Joshi,Madhvi Joshi |
| EPI_ISL_2562102 | GMERS Government Medical College (GMC), Gotri | Gujarat Biotechnology Research Centre | Janvi Raval,Zarna Patel,Nitin Savaliya,Dinesh Kumar,Twinkle Soni,Sonal Sharma ,Zuber Saiyed,Ramesh Pandit,Bithika Duttaroy,Umang Mishra,Nitesh Shah,Chaitanya Joshi,Madhvi Joshi |
| EPI_ISL_2562106 | GMERS Government Medical College (GMC), Gotri | Gujarat Biotechnology Research Centre | Twinkle Soni,Sonal Sharma ,Zuber Saiyed,Ramesh Pandit,Janvi Raval,Zarna Patel,Nitin Savaliya,Dinesh Kumar,Bithika Duttaroy,Umang Mishra,Nitesh Shah,Chaitanya Joshi,Madhvi Joshi |
| EPI_ISL_2562107 | GMERS Government Medical College (GMC), Gotri | Gujarat Biotechnology Research Centre | Sonal Sharma ,Zuber Saiyed,Ramesh Pandit,Janvi Raval,Zarna Patel,Nitin Savaliya,Dinesh Kumar,Twinkle Soni,Bithika Duttaroy,Umang Mishra,Nitesh Shah,Chaitanya Joshi,Madhvi Joshi |
| EPI_ISL_2562108 | GMERS Government Medical College (GMC), Gotri | Gujarat Biotechnology Research Centre | Zuber Saiyed,Ramesh Pandit,Janvi Raval,Zarna Patel,Nitin Savaliya,Dinesh Kumar,Twinkle Soni,Sonal Sharma ,Bithika Duttaroy,Umang Mishra,Nitesh Shah,Chaitanya Joshi,Madhvi Joshi |
| EPI_ISL_2562110 | GMERS Government Medical College (GMC), Gotri | Gujarat Biotechnology Research Centre | Janvi Raval,Zarna Patel,Nitin Savaliya,Dinesh Kumar,Twinkle Soni,Sonal Sharma ,Zuber Saiyed,Ramesh Pandit,Bithika Duttaroy,Umang Mishra,Nitesh Shah,Chaitanya Joshi,Madhvi Joshi |
| EPI_ISL_2562111 | GMERS Government Medical College (GMC), Gotri | Gujarat Biotechnology Research Centre | Zarna Patel,Nitin Savaliya,Dinesh Kumar,Twinkle Soni,Sonal Sharma ,Zuber Saiyed,Ramesh Pandit,Janvi Raval,Bithika Duttaroy,Umang Mishra,Nitesh Shah,Chaitanya Joshi,Madhvi Joshi |
| EPI_ISL_2565970, EPI_ISL_2565977, EPI_ISL_2565981 | Institute of Molecular and Translational Medicine / Laboratory of Experimental Medicine, Faculty of Medicine and Dentistry, Palacky University and University Hospital Olomouc | Institute of Molecular and Translational Medicine / Laboratory of Experimental Medicine, Faculty of Medicine and Dentistry, Palacky University | Rastislav Slavkovský, Hana Dímalová, Vladimíra Koudeláková, Barbora Blumová, Marián Hajdúch |
| EPI_ISL_2565984, EPI_ISL_2565986, EPI_ISL_2565992, EPI_ISL_2565993, EPI_ISL_2565994, EPI_ISL_2565996 | Genome Analysis Center, Yamanashi Central Hospital | Genome Analysis Center, Yamanashi Central Hospital | Yosuke Hirotsu |
| EPI_ISL_2566164, EPI_ISL_2566165, EPI_ISL_2566166, EPI_ISL_2566169, EPI_ISL_2566170, EPI_ISL_2566171, EPI_ISL_2566172, EPI_ISL_2566173, EPI_ISL_2566174, EPI_ISL_2566175, EPI_ISL_2566176, EPI_ISL_2566177, EPI_ISL_2566178, EPI_ISL_2566179, EPI_ISL_2566180, EPI_ISL_2566181, EPI_ISL_2566183, EPI_ISL_2566186, EPI_ISL_2566188, EPI_ISL_2566189, EPI_ISL_2566191, EPI_ISL_2566193 | | | |
| see above | Microbiology Department. Complexo Hospitalario Universitario de Vigo | Microbiology Department. Complexo Hospitalario Universitario de Vigo | Alfaya N, Alonso I, Alvarez M, Cabrera JJ, Carballo R, Cores O, Cortizo S, del-Campo V, Martinez L, Mediero G, Perez S, Potel C, Regueiro B, Rey S, Vasallo FJ |
| EPI_ISL_2566444 | Singapore General Hospital | Department of Microbiology | Nurdyana Abdul Rahman, Kenneth Xin Long Chan, Chayaporn Suphavilai, Kun Lee Lim, Sui Sin Goh, Kian Sing Chan, Lynette Oon, Kern Rei Chng, James Sim Heng Chiak, Niranjan Nagarajan, Karrie Ko |
| EPI_ISL_2566447, EPI_ISL_2566448, EPI_ISL_2566459, EPI_ISL_2566460, EPI_ISL_2566466 | National Institute of Infectious Diseases-Prof. Dr. Matei Bals Molecular Diagnostics Laboratory | National Institute of Infectious Diseases-Prof. Dr. Matei Bals Molecular Diagnostics Laboratory | Corina Casangiu, Leontina Banica, Marius Surleac, Petre Milu, Robert Hohan, Simona Paraschiv, Dan Otelea |
| EPI_ISL_2566676, EPI_ISL_2566677, EPI_ISL_2566678, EPI_ISL_2566679, EPI_ISL_2566873 | Institut de Virologie du CHU de Strasbourg | Institut de Virologie du CHU de Strasbourg | Fafi-Kremer Samira, Gallais Floriane, Gantner Pierre, Laugel Elodie, Solis Morgane, Velay Aurélie, Wendling Marie-Josée |
| EPI_ISL_2566919, EPI_ISL_2566920, EPI_ISL_2566921, EPI_ISL_2566922, EPI_ISL_2566923, EPI_ISL_2566924 | AZ Klina | AZ Klina | Carl Vael - Lynsey Berckmans |
| EPI_ISL_2566925 | AZ Klina | AZ Klina | Carl Vael -Lynsey Berckmans |
| EPI_ISL_2566927, EPI_ISL_2566928, EPI_ISL_2566929, EPI_ISL_2566930, EPI_ISL_2566931, EPI_ISL_2566932, EPI_ISL_2567001, EPI_ISL_2567002, EPI_ISL_2567003, EPI_ISL_2567075 | AZ Klina | AZ Klina | Carl Vael - Lynsey Berckmans |
| EPI_ISL_2567620, EPI_ISL_2567622 | Institut de Virologie du CHU de Strasbourg | Institut de Virologie du CHU de Strasbourg | Fafi-Kremer Samira, Gallais Floriane, Gantner Pierre, Laugel Elodie, Solis Morgane, Velay Aurélie, Wendling Marie-Josée |
| EPI_ISL_2570809 | Surabaya Health Office | Institute of Tropical Disease, Universitas Airlangga | Aldise M Nastri, Jezzy R Dewantari, Rima R Prasetya, Krisnoadi Rahardjo, Febria Rachmanita, Herlin Ferliana, Gatot Soegiarto, Laksmi Wulandari, Resti Yudhawati, Yasuko Mori, Soetjipto, Kazufumi Shimizu, Maria I Lusida |
| EPI_ISL_2570811 | Surabaya Health Office | Institute of Tropical Disease, Universitas Airlangga | Jezzy R Dewantari, Rima R Prasetya, Krisnoadi Rahardjo, Aldise M Nastri, Febria Rachmanita, Herlin Ferliana, Gatot Soegiarto, Laksmi Wulandari, Resti Yudhawati, Yasuko Mori, Soetjipto, Kazufumi Shimizu, Maria I Lusida |
| EPI_ISL_2592630, EPI_ISL_2593764, EPI_ISL_2595725, EPI_ISL_2597228, EPI_ISL_2597229, EPI_ISL_2597312 | DSMRC | DSMRC | Khine Zaw Oo, Nay Myo Aung, Ko Ko Win, Phyo Kyaw Aung, Pai Khant Kyaw, Thet Wai Oo, Zaw Win Htun, Kyee Myint, Ko Ko Lwin |
| EPI_ISL_2600536, EPI_ISL_2600537, EPI_ISL_2600538, EPI_ISL_2600539, EPI_ISL_2600541, EPI_ISL_2600542, EPI_ISL_2600543 | Università degli Studi di Perugia | Istituto Zooprofilattico Sperimentale dell'Abruzzo e Molise "G. Caporale" | Mencacci A, Camilloni B, Biagetti M, Giammarioli M, Lorusso A, Marcacci M, Di Domenico M, Ancora M, Curini V, Mangone I, Rinaldi A, Delli Compagni E, Scialabba S, Di Pasquale A, Cammà C, Puglia I, Calistri P, Savini G |
| EPI_ISL_2600544, EPI_ISL_2600545, EPI_ISL_2600546, EPI_ISL_2600547, EPI_ISL_2600548, EPI_ISL_2600549 | Laboratorio Analisi Osp. Città di Castello - Azienda USL Umbria1 | Istituto Zooprofilattico Sperimentale dell'Abruzzo e Molise "G. Caporale" | Malagigi V, Taccani P,Lorusso A, Marcacci M, Di Domenico M, Ancora M, Curini V, Mangone I, Rinaldi A, Delli Compagni E, Scialabba S, Di Pasquale A, Cammà C, Puglia I, Calistri P, Savini G |
| EPI_ISL_2600550, EPI_ISL_2600551, EPI_ISL_2600552, EPI_ISL_2600553, EPI_ISL_2600554, EPI_ISL_2600555, EPI_ISL_2600556, EPI_ISL_2600557, EPI_ISL_2600558, EPI_ISL_2600559, EPI_ISL_2600560, EPI_ISL_2600561 | | | |
| see above | Azienda Ospedaliera Terni | Istituto Zooprofilattico Sperimentale dell'Abruzzo e Molise "G. Caporale" | Palumbo M, Scaccetti A, Lorusso A, Marcacci M, Di Domenico M, Ancora M, Curini V, Mangone I, Rinaldi A, Delli Compagni E, Scialabba S, Di Pasquale A, Cammà C, Puglia I, Calistri P, Savini G |
| EPI_ISL_2600733, EPI_ISL_2600734, EPI_ISL_2600735, EPI_ISL_2600736, EPI_ISL_2600738, EPI_ISL_2600739, EPI_ISL_2600740, EPI_ISL_2600741, EPI_ISL_2600742, EPI_ISL_2600743, EPI_ISL_2600744, EPI_ISL_2600745, EPI_ISL_2600746, EPI_ISL_2600747, EPI_ISL_2600748, EPI_ISL_2600749, EPI_ISL_2600750, EPI_ISL_2600751, EPI_ISL_2600752, EPI_ISL_2600753, EPI_ISL_2600754, EPI_ISL_2600755, EPI_ISL_2600756, EPI_ISL_2600758, EPI_ISL_2600759, EPI_ISL_2600760, EPI_ISL_2600761, EPI_ISL_2600762, EPI_ISL_2600766, EPI_ISL_2600767, EPI_ISL_2600768, EPI_ISL_2600769, EPI_ISL_2600771, EPI_ISL_2600772, EPI_ISL_2600773, EPI_ISL_2600774, EPI_ISL_2600775, EPI_ISL_2600778, EPI_ISL_2600779 | | | |
| see above | Università degli Studi di Perugia | Istituto Zooprofilattico Sperimentale dell'Abruzzo e Molise "G. Caporale" | Mencacci A, Camilloni B, Biagetti M, Giammarioli M, Lorusso A, Marcacci M, Di Domenico M, Ancora M, Curini V, Mangone I, Rinaldi A, Delli Compagni E, Scialabba S, Di Pasquale A, Cammà C, Puglia I, Calistri P, Savini G |
| EPI_ISL_2600784, EPI_ISL_2600785, EPI_ISL_2600786, EPI_ISL_2600787, EPI_ISL_2600788, EPI_ISL_2600789, | Azienda Ospedaliera Terni | Istituto Zooprofilattico Sperimentale dell'Abruzzo e Molise "G. Caporale" | Palumbo M, Scaccetti A, Lorusso A, Marcacci M, Di Domenico M, Ancora M, Curini V, Mangone I, Rinaldi A, Delli Compagni E, Scialabba S, Di Pasquale A, Cammà C, Puglia I, Calistri P, Savini G |

| | | | |
|---|---|---|---|
| EPI_ISL_2600790, EPI_ISL_2600791, EPI_ISL_2600792 | | | |
| EPI_ISL_2600793, EPI_ISL_2600795, EPI_ISL_2600796, EPI_ISL_2600797, EPI_ISL_2600798, EPI_ISL_2600799 | Azienda USL Umbria 2 | Istituto Zooprofilattico Sperimentale dell'Abruzzo e Molise "G. Caporale" | Proietti A, Pistoni E, Lorusso A, Marcacci M, Di Domenico M, Ancora M, Curini V, Mangone I, Rinaldi A, Delli Compagni E, Scialabba S, Di Pasquale A, Cammà C, Puglia I, Calistri P, Savini G |
| EPI_ISL_2600800 | PRESIDIO OSPEDALIERO SULMONA | Istituto Zooprofilattico Sperimentale dell'Abruzzo e Molise "G. Caporale" | Lorusso A, Marcacci M, Di Domenico M, Ancora M, Curini V, Di Lollo Valeria, Mangone I, Rinaldi A, Delli Compagni E, Scialabba S, Caporale M, Di Pasquale A, Cammà C, Puglia I, Calistri P, Savini G |
| EPI_ISL_2600801 | SIESP CHIETI - DRIVE IN VASTO | Istituto Zooprofilattico Sperimentale dell'Abruzzo e Molise "G. Caporale" | Lorusso A, Marcacci M, Di Domenico M, Ancora M, Curini V, Di Lollo Valeria, Mangone I, Rinaldi A, Delli Compagni E, Scialabba S, Caporale M, Di Pasquale A, Cammà C, Puglia I, Calistri P, Savini G |
| EPI_ISL_2600802 | SIESP DIPARTIMENTO DI PREVENZIONE CHIETI | Istituto Zooprofilattico Sperimentale dell'Abruzzo e Molise "G. Caporale" | Lorusso A, Marcacci M, Di Domenico M, Ancora M, Curini V, Di Lollo Valeria, Mangone I, Rinaldi A, Delli Compagni E, Scialabba S, Caporale M, Di Pasquale A, Cammà C, Puglia I, Calistri P, Savini G |
| EPI_ISL_2600803, EPI_ISL_2600804, EPI_ISL_2600805 | SIESP CHIETI - DRIVE IN LANCIANO | Istituto Zooprofilattico Sperimentale dell'Abruzzo e Molise "G. Caporale" | Lorusso A, Marcacci M, Di Domenico M, Ancora M, Curini V, Di Lollo Valeria, Mangone I, Rinaldi A, Delli Compagni E, Scialabba S, Caporale M, Di Pasquale A, Cammà C, Puglia I, Calistri P, Savini G |
| EPI_ISL_2600806, EPI_ISL_2600807 | PO AVEZZANO - U.O.C. Laboratorio Analisi | Istituto Zooprofilattico Sperimentale dell'Abruzzo e Molise "G. Caporale" | Lorusso A, Marcacci M, Di Domenico M, Ancora M, Curini V, Di Lollo Valeria, Mangone I, Rinaldi A, Delli Compagni E, Scialabba S, Caporale M, Di Pasquale A, Cammà C, Puglia I, Calistri P, Savini G |
| EPI_ISL_2600808, EPI_ISL_2600809 | SIESP DIPARTIMENTO DI PREVENZIONE TERAMO | Istituto Zooprofilattico Sperimentale dell'Abruzzo e Molise "G. Caporale" | Lorusso A, Marcacci M, Di Domenico M, Ancora M, Curini V, Di Lollo Valeria, Mangone I, Rinaldi A, Delli Compagni E, Scialabba S, Caporale M, Di Pasquale A, Cammà C, Puglia I, Calistri P, Savini G |
| EPI_ISL_2600810 | SIESP CHIETI - DRIVE IN LANCIANO | Istituto Zooprofilattico Sperimentale dell'Abruzzo e Molise "G. Caporale" | Lorusso A, Marcacci M, Di Domenico M, Ancora M, Curini V, Di Lollo Valeria, Mangone I, Rinaldi A, Delli Compagni E, Scialabba S, Caporale M, Di Pasquale A, Cammà C, Puglia I, Calistri P, Savini G |
| EPI_ISL_2600811, EPI_ISL_2600812 | SIESP DIPARTIMENTO DI PREVENZIONE CHIETI | Istituto Zooprofilattico Sperimentale dell'Abruzzo e Molise "G. Caporale" | Lorusso A, Marcacci M, Di Domenico M, Ancora M, Curini V, Di Lollo Valeria, Mangone I, Rinaldi A, Delli Compagni E, Scialabba S, Caporale M, Di Pasquale A, Cammà C, Puglia I, Calistri P, Savini G |
| EPI_ISL_2600813, EPI_ISL_2600814 | SIESP CHIETI - DRIVE IN LANCIANO | Istituto Zooprofilattico Sperimentale dell'Abruzzo e Molise "G. Caporale" | Lorusso A, Marcacci M, Di Domenico M, Ancora M, Curini V, Di Lollo Valeria, Mangone I, Rinaldi A, Delli Compagni E, Scialabba S, Caporale M, Di Pasquale A, Cammà C, Puglia I, Calistri P, Savini G |
| EPI_ISL_2600815 | RSA GIULIANOVA | Istituto Zooprofilattico Sperimentale dell'Abruzzo e Molise "G. Caporale" | Lorusso A, Marcacci M, Di Domenico M, Ancora M, Curini V, Di Lollo Valeria, Mangone I, Rinaldi A, Delli Compagni E, Scialabba S, Caporale M, Di Pasquale A, Cammà C, Puglia I, Calistri P, Savini G |
| EPI_ISL_2600816 | SIESP CHIETI - DRIVE IN CHIETI | Istituto Zooprofilattico Sperimentale dell'Abruzzo e Molise "G. Caporale" | Lorusso A, Marcacci M, Di Domenico M, Ancora M, Curini V, Di Lollo Valeria, Mangone I, Rinaldi A, Delli Compagni E, Scialabba S, Caporale M, Di Pasquale A, Cammà C, Puglia I, Calistri P, Savini G |
| EPI_ISL_2600817, EPI_ISL_2600818, EPI_ISL_2600819, EPI_ISL_2600821 | SIESP CHIETI - DRIVE IN VASTO | Istituto Zooprofilattico Sperimentale dell'Abruzzo e Molise "G. Caporale" | Lorusso A, Marcacci M, Di Domenico M, Ancora M, Curini V, Di Lollo Valeria, Mangone I, Rinaldi A, Delli Compagni E, Scialabba S, Caporale M, Di Pasquale A, Cammà C, Puglia I, Calistri P, Savini G |
| EPI_ISL_2600822 | SIESP DIPARTIMENTO DI PREVENZIONE CHIETI | Istituto Zooprofilattico Sperimentale dell'Abruzzo e Molise "G. Caporale" | Lorusso A, Marcacci M, Di Domenico M, Ancora M, Curini V, Di Lollo Valeria, Mangone I, Rinaldi A, Delli Compagni E, Scialabba S, Caporale M, Di Pasquale A, Cammà C, Puglia I, Calistri P, Savini G |
| EPI_ISL_2600823 | SIESP CHIETI - DRIVE IN LANCIANO | Istituto Zooprofilattico Sperimentale dell'Abruzzo e Molise "G. Caporale" | Lorusso A, Marcacci M, Di Domenico M, Ancora M, Curini V, Di Lollo Valeria, Mangone I, Rinaldi A, Delli Compagni E, Scialabba S, Caporale M, Di Pasquale A, Cammà C, Puglia I, Calistri P, Savini G |
| EPI_ISL_2600824, EPI_ISL_2600825 | USCA Sulmona | Istituto Zooprofilattico Sperimentale dell'Abruzzo e Molise "G. Caporale" | Lorusso A, Marcacci M, Di Domenico M, Ancora M, Curini V, Di Lollo Valeria, Mangone I, Rinaldi A, Delli Compagni E, Scialabba S, Caporale M, Di Pasquale A, Cammà C, Puglia I, Calistri P, Savini G |
| EPI_ISL_2600826 | SIESP DIPARTIMENTO DI PREVENZIONE CHIETI | Istituto Zooprofilattico Sperimentale dell'Abruzzo e Molise "G. Caporale" | Lorusso A, Marcacci M, Di Domenico M, Ancora M, Curini V, Di Lollo Valeria, Mangone I, Rinaldi A, Delli Compagni E, Scialabba S, Caporale M, Di Pasquale A, Cammà C, Puglia I, Calistri P, Savini G |
| EPI_ISL_2600827, EPI_ISL_2600828 | SIESP CHIETI - DRIVE IN VASTO | Istituto Zooprofilattico Sperimentale dell'Abruzzo e Molise "G. Caporale" | Lorusso A, Marcacci M, Di Domenico M, Ancora M, Curini V, Di Lollo Valeria, Mangone I, Rinaldi A, Delli Compagni E, Scialabba S, Caporale M, Di Pasquale A, Cammà C, Puglia I, Calistri P, Savini G |
| EPI_ISL_2600830 | SIESP DIPARTIMENTO DI PREVENZIONE TERAMO | Istituto Zooprofilattico Sperimentale dell'Abruzzo e Molise "G. Caporale" | Lorusso A, Marcacci M, Di Domenico M, Ancora M, Curini V, Di Lollo Valeria, Mangone I, Rinaldi A, Delli Compagni E, Scialabba S, Caporale M, Di Pasquale A, Cammà C, Puglia I, Calistri P, Savini G |
| EPI_ISL_2600831, EPI_ISL_2600832 | SIESP CHIETI - DRIVE IN VASTO | Istituto Zooprofilattico Sperimentale dell'Abruzzo e Molise "G. Caporale" | Lorusso A, Marcacci M, Di Domenico M, Ancora M, Curini V, Di Lollo Valeria, Mangone I, Rinaldi A, Delli Compagni E, Scialabba S, Caporale M, Di Pasquale A, Cammà C, Puglia I, Calistri P, Savini G |
| EPI_ISL_2600833 | SIESP CHIETI - DRIVE IN CHIETI | Istituto Zooprofilattico Sperimentale dell'Abruzzo e Molise "G. Caporale" | Lorusso A, Marcacci M, Di Domenico M, Ancora M, Curini V, Di Lollo Valeria, Mangone I, Rinaldi A, Delli Compagni E, Scialabba S, Caporale M, Di Pasquale A, Cammà C, Puglia I, Calistri P, Savini G |
| EPI_ISL_2600834 | OSPEDALE CIVILE TERAMO | Istituto Zooprofilattico Sperimentale dell'Abruzzo e Molise "G. Caporale" | Lorusso A, Marcacci M, Di Domenico M, Ancora M, Curini V, Di Lollo Valeria, Mangone I, Rinaldi A, Delli Compagni E, Scialabba S, Caporale M, Di Pasquale A, Cammà C, Puglia I, Calistri P, Savini G |
| EPI_ISL_2600964 | Molecular Hepatology Laboratory AOU Messina | Molecular Hepatology Laboratory AOU Messina | Giuseppina Raffa, Cristina Musolino, Valeria Chines, Daniele Lombardo, Teresa Pollicino. |
| EPI_ISL_2601037 | Instituto de Medicina Tropical & Salud Global, Universidad Iberoamericana (UNIBE) | Instituto de Medicina Tropical & Salud Global, Universidad Iberoamericana (UNIBE) | Victor Virgilio Calderon, Alejandro Vallejo Degaudenzi & Robert Paulino-Ramirez |
| EPI_ISL_2603432, EPI_ISL_2603433, EPI_ISL_2603441, EPI_ISL_2603442, EPI_ISL_2603443, EPI_ISL_2603444, EPI_ISL_2603446, EPI_ISL_2603447, EPI_ISL_2603448 | Laboratorio Central de Saude Publica do Estado do Rio Grande do Sul (LACEN-RS) | Laboratory of Respiratory Viruses and Measles, Oswaldo Cruz Institute, FIOCRUZ | Paola Resende, Luciana Appolinario, Fernando Motta, Anna Carolina Paixao, Ana Carolina Mendonca, Alice Sampaio Rocha, Taina Venas, Elisa Cavalcante Pereira, Renata Serrano Lopes, Anderson Brandao Leite, Marilda Siqueira on behalf of the Fiocruz COVID-19 Genomic Surveillance Network |
| EPI_ISL_2603840, EPI_ISL_2603842, EPI_ISL_2603856, EPI_ISL_2603898 | Institut de Virologie du CHU de Strasbourg | Institut de Virologie du CHU de Strasbourg | Fafi-Kremer Samira, Gallais Floriane, Gantner Pierre, Laugel Elodie, Solis Morgane, Velay Aurélie, Wendling Marie-Josée |
| EPI_ISL_2603899 | PRESIDIO OSPEDALIERO TAGLIACOZZO | Istituto Zooprofilattico Sperimentale dell'Abruzzo e Molise "G. Caporale" | Lorusso A, Marcacci M, Di Domenico M, Ancora M, Curini V, Di Lollo Valeria, Mangone I, Rinaldi A, Delli Compagni E, Scialabba S, Caporale M, Di Pasquale A, Cammà C, Puglia I, Calistri P, Savini G |
| EPI_ISL_2604033, EPI_ISL_2604102 | Institut de Virologie du CHU de Strasbourg | Institut de Virologie du CHU de Strasbourg | Fafi-Kremer Samira, Gallais Floriane, Gantner Pierre, Laugel Elodie, Solis Morgane, Velay Aurélie, Wendling Marie-Josée |
| EPI_ISL_2604147 | Azienda Ospedaliera Terni | Istituto Zooprofilattico Sperimentale dell'Abruzzo e Molise "G. Caporale" | Palumbo M, Scaccetti A, Lorusso A, Marcacci M, Di Domenico M, Ancora M, Curini V, Mangone I, Rinaldi A, Delli Compagni E, Scialabba S, Di Pasquale A, Cammà C, Puglia I, Calistri P, Savini G |
| EPI_ISL_2604216, EPI_ISL_2604251 | Ministry of Health, Jaber Al-Ahmad Hospital | Virology Unit, Department of Microbiology, Faculty of Medicine, Kuwait University | Nada Madi, Hussain Safar, Ebaa Al-Awadhi, Anfal Al-Adwani |
| EPI_ISL_2604822 | Laboratorio di Riferimento Regionale della Sicilia Occidentale per l'Emergenza COVID-19 | Laboratorio di Riferimento Regionale della Sicilia Occidentale per l'Emergenza COVID-19 | Fabio Tramuto, Carmelo Massimo Maida, Daniela Di Naro, Giulia Randazzo, Walter Mazzucco, Giorgio Graziano, Vincenzo Restivo, Claudio Costantino, Francesco Vitale |
| EPI_ISL_2604869, EPI_ISL_2604882, EPI_ISL_2604888 | Omics Sciences Laboratory | Omics Sciences Laboratory | Derly Andrade Molina, Gabriel Morey León, Darlyn Amaya, Rubén Armas González, Juan Carlos Fernández Cadena |
| EPI_ISL_2608421 | Kampong Cham Rapid Response Team | Virology Unit, Institut Pasteur du Cambodge | Cecile Troupin, Leakhena Pum, Jurre Y Siegers, Ly Sovann, Kraing Sidonn, Yi Sengdoeurn, Chin Savuth, Chau Darapheak, Veasna Duong, Erik A Karlsson |
| EPI_ISL_2608422, EPI_ISL_2608423 | Siem Reap Provinical Laboratory | Virology Unit, Institut Pasteur du Cambodge | Cecile Troupin, Leakhena Pum, Jurre Y Siegers, Ly Sovann, Kraing Sidonn, Yi Sengdoeurn, Chin Savuth, Chau Darapheak, Veasna Duong, Erik A |

| | | | |
|---|---|---|---|
| | | | Karlsson |
| EPI_ISL_2608424, EPI_ISL_2608425, EPI_ISL_2608426, EPI_ISL_2608427, EPI_ISL_2608428, EPI_ISL_2608429, EPI_ISL_2608430, EPI_ISL_2608431, EPI_ISL_2608432, EPI_ISL_2608433, EPI_ISL_2608434, EPI_ISL_2608435, EPI_ISL_2608436, EPI_ISL_2608437, EPI_ISL_2608438, EPI_ISL_2608439, EPI_ISL_2608440, EPI_ISL_2608441, EPI_ISL_2608442, EPI_ISL_2608443 | | | |
| see above | Virology Unit, Institut Pasteur du Cambodge | Virology Unit, Institut Pasteur du Cambodge | Cecile Troupin, Leakhena Pum, Jurre Y Siegers, Ly Sovann, Kraing Sidonn, Yi Sengdoeurn, Chin Savuth, Chau Darapheak, Veasna Duong, Erik A Karlsson |
| EPI_ISL_2608444 | Kampot Rapid Response Team | Virology Unit, Institut Pasteur du Cambodge | Cecile Troupin, Leakhena Pum, Jurre Y Siegers, Ly Sovann, Kraing Sidonn, Yi Sengdoeurn, Chin Savuth, Chau Darapheak, Veasna Duong, Erik A Karlsson |
| EPI_ISL_2611661 | Temporary Specimen Collection Centre at the AsiaWorld-Expo | Hong Kong Department of Health | Alan K.L. Tsang, Peter C.W. Yip, Ken H.L. Ng, Edman T.K. Lam, Rickjason C.W. Chan, Dominic N.C. Tsang |
| EPI_ISL_2611662 | PRINCESS MARGARET HOSPITAL | Hong Kong Department of Health | Alan K.L. Tsang, Peter C.W. Yip, Ken H.L. Ng, Edman T.K. Lam, Rickjason C.W. Chan, Dominic N.C. Tsang |
| EPI_ISL_2611984, EPI_ISL_2611985 | I.R.C.C.S. "S. De Bellis" - Ente Ospedaliero | Istituto Zooprofilattico Sperimentale della Puglia e della Basilicata | Parisi A., Bianco A., Capozzi L., Del Sambro L., Simone D., Difato L., Notarnicola M., Lippolis A. |
| EPI_ISL_2611988, EPI_ISL_2611989, EPI_ISL_2611990, EPI_ISL_2611991, EPI_ISL_2611992, EPI_ISL_2611994, EPI_ISL_2611995, EPI_ISL_2611996 | Presidio di Brindisi Di Summa - Perrino | Istituto Zooprofilattico Sperimentale della Puglia e della Basilicata | Parisi A., Bianco A., Capozzi L., Del Sambro L., Simone D., Difato L., Pennuzzi G., Santoro A. |
| EPI_ISL_2611997, EPI_ISL_2611998 | Istituto Zooprofilattico Sperimentale della Puglia e della Basilicata | Istituto Zooprofilattico Sperimentale della Puglia e della Basilicata | Parisi A., Bianco A., Capozzi L., Del Sambro L., Simone D., Difato L., Pennuzzi G., Pace L., Galante D. |
| EPI_ISL_2612000, EPI_ISL_2612001, EPI_ISL_2612002, EPI_ISL_2612003 | Ospedale Vito Fazzi | Istituto Zooprofilattico Sperimentale della Puglia e della Basilicata | Parisi A., Bianco A., Capozzi L., Del Sambro L., Simone D., Difato L., Lobreglio G. |
| EPI_ISL_2612004 | Ospedale Santa Caterina Novella | Istituto Zooprofilattico Sperimentale della Puglia e della Basilicata | Parisi A., Bianco A., Capozzi L., Del Sambro L., Simone D., Difato L., Bruno A. R. |
| EPI_ISL_2612005, EPI_ISL_2612006, EPI_ISL_2612007 | Ospedale Vito Fazzi | Istituto Zooprofilattico Sperimentale della Puglia e della Basilicata | Parisi A., Bianco A., Capozzi L., Del Sambro L., Simone D., Difato L., Lobreglio G. |
| EPI_ISL_2612008, EPI_ISL_2612009, EPI_ISL_2612010 | Ospedale Santa Caterina Novella | Istituto Zooprofilattico Sperimentale della Puglia e della Basilicata | Parisi A., Bianco A., Capozzi L., Del Sambro L., Simone D., Difato L., Bruno A. R. |
| EPI_ISL_2612011, EPI_ISL_2612012 | Ospedale Cardinale G. Panico | Istituto Zooprofilattico Sperimentale della Puglia e della Basilicata | Parisi A., Bianco A., Capozzi L., Del Sambro L., Simone D., Difato L., Bruno A. R. |
| EPI_ISL_2612013, EPI_ISL_2612014, EPI_ISL_2612015, EPI_ISL_2612016, EPI_ISL_2612017, EPI_ISL_2612019, EPI_ISL_2612020, EPI_ISL_2612021, EPI_ISL_2612022 | Ospedale Santissima Annunziata | Istituto Zooprofilattico Sperimentale della Puglia e della Basilicata | Parisi A., Bianco A., Capozzi L., Del Sambro L., Simone D., Difato L., Ridolfi D., Giannico A, Tampoia M. |
| EPI_ISL_2612023, EPI_ISL_2612024, EPI_ISL_2612025, EPI_ISL_2612026, EPI_ISL_2612027, EPI_ISL_2612028, EPI_ISL_2612029, EPI_ISL_2612030, EPI_ISL_2612031, EPI_ISL_2612033, EPI_ISL_2612034, EPI_ISL_2612035, EPI_ISL_2612036, EPI_ISL_2612037, EPI_ISL_2612038, EPI_ISL_2612039, EPI_ISL_2612040, EPI_ISL_2612041, EPI_ISL_2612042, EPI_ISL_2612043, EPI_ISL_2612044 | | | |
| see above | Klinisch Laboratorium ZNA | Klinisch Laboratorium ZNA | Verstrepen et al. |
| EPI_ISL_2612301, EPI_ISL_2612302, EPI_ISL_2612303, EPI_ISL_2612304, EPI_ISL_2612305, EPI_ISL_2612306, EPI_ISL_2612307, EPI_ISL_2612308, EPI_ISL_2612309, EPI_ISL_2612310, EPI_ISL_2612311, EPI_ISL_2612312, EPI_ISL_2612313, EPI_ISL_2612314 | | | |
| see above | Università degli Studi di Perugia | Istituto Zooprofilattico Sperimentale dell'Abruzzo e Molise "G. Caporale" | Mencacci A, Camilloni B, Biagetti M, Giammarioli M, Lorusso A, Marcacci M, Di Domenico M, Ancora M, Curini V, Mangone I, Rinaldi A, Delli Compagni E, Scialabba S, Di Pasquale A, Cammà C, Puglia I, Calistri P, Savini G |
| EPI_ISL_2612563 | OCME Office Of Chief Medical Examiner | New York City Public Health Laboratory | Jade Wang, et al. |
| EPI_ISL_2612564 | Labo Analyses Med | National Reference Center for Viruses of Respiratory Infections, Institut Pasteur, Paris | Marion Barbet, Sylvie Behillil, Méline Bizard, Angela Brisebarre, Camille Capel, Vincent Enouf, Louise Lefrançois, Frédéric Lemoine, Christophe Malabat, Corinne Maufrais, Etienne Simon-Lorière, Maud Vanpeene, Sylvie Van der Werf ,Sophie Chalmin |
| EPI_ISL_2612565 | Labo Analyses Med | National Reference Center for Viruses of Respiratory Infections, Institut Pasteur, Paris | Marion Barbet, Sylvie Behillil, Méline Bizard, Angela Brisebarre, Camille Capel, Vincent Enouf, Louise Lefrançois, Frédéric Lemoine, Christophe Malabat, Corinne Maufrais, Adrien Pain, Etienne Simon-Lorière, Maud Vanpeene, Sylvie Van der Werf ,Sophie Chalmin |
| EPI_ISL_2612566 | Labo Analyses Med | National Reference Center for Viruses of Respiratory Infections, Institut Pasteur, Paris | Marion Barbet, Sylvie Behillil, Méline Bizard, Angela Brisebarre, Camille Capel, Vincent Enouf, Louise Lefrançois, Frédéric Lemoine, Christophe Malabat, Corinne Maufrais, Etienne Simon-Lorière, Maud Vanpeene, Sylvie Van der Werf ,Sophie Chalmin |
| EPI_ISL_2612567 | Labo Analyses Med | National Reference Center for Viruses of Respiratory Infections, Institut Pasteur, Paris | Marion Barbet, Sylvie Behillil, Méline Bizard, Angela Brisebarre, Camille Capel, Vincent Enouf, Louise Lefrançois, Frédéric Lemoine, Christophe Malabat, Corinne Maufrais, Adrien Pain, Etienne Simon-Lorière, Maud Vanpeene, Sylvie Van der Werf ,Sophie Chalmin |
| EPI_ISL_2612568, EPI_ISL_2612569, EPI_ISL_2612570, EPI_ISL_2612571, EPI_ISL_2612572, EPI_ISL_2612573, EPI_ISL_2612574, EPI_ISL_2612575, EPI_ISL_2612576, EPI_ISL_2612577, EPI_ISL_2612578 | | | |
| see above | Labo Analyses Med | National Reference Center for Viruses of Respiratory Infections, Institut Pasteur, Paris | Marion Barbet, Sylvie Behillil, Méline Bizard, Angela Brisebarre, Camille Capel, Vincent Enouf, Louise Lefrançois, Frédéric Lemoine, Christophe Malabat, Corinne Maufrais, Etienne Simon-Lorière, Maud Vanpeene, Sylvie Van der Werf ,Sophie Chalmin |
| EPI_ISL_2612579, EPI_ISL_2612580, EPI_ISL_2612581, EPI_ISL_2612582, EPI_ISL_2612583, EPI_ISL_2612584, EPI_ISL_2612585, EPI_ISL_2612586 | Labo Analyses Med | National Reference Center for Viruses of Respiratory Infections, Institut Pasteur, Paris | Marion Barbet, Sylvie Behillil, Méline Bizard, Angela Brisebarre, Camille Capel, Vincent Enouf, Louise Lefrançois, Frédéric Lemoine, Christophe Malabat, Corinne Maufrais, Etienne Simon-Lorière, Maud Vanpeene, Sylvie Van der Werf ,Melanie Caron |
| EPI_ISL_2612588, EPI_ISL_2612589 | Labo Analyses Med | National Reference Center for Viruses of Respiratory Infections, Institut Pasteur, Paris | Marion Barbet, Sylvie Behillil, Méline Bizard, Angela Brisebarre, Camille Capel, Vincent Enouf, Louise Lefrançois, Frédéric Lemoine, Christophe Malabat, Corinne Maufrais, Etienne Simon-Lorière, Maud Vanpeene, Sylvie Van der Werf ,Nicolas Capron |
| EPI_ISL_2612590, EPI_ISL_2612591, EPI_ISL_2612592, EPI_ISL_2612593, EPI_ISL_2612594 | Labo Analyses Med | National Reference Center for Viruses of Respiratory Infections, Institut Pasteur, Paris | Marion Barbet, Sylvie Behillil, Méline Bizard, Angela Brisebarre, Camille Capel, Vincent Enouf, Louise Lefrançois, Frédéric Lemoine, Christophe Malabat, Corinne Maufrais, Etienne Simon-Lorière, Maud Vanpeene, Sylvie Van der Werf ,GréGoire Potiron |
| EPI_ISL_2612595, EPI_ISL_2612596, EPI_ISL_2612597 | Labo Analyses Med | National Reference Center for Viruses of Respiratory Infections, Institut Pasteur, Paris | Marion Barbet, Sylvie Behillil, Méline Bizard, Angela Brisebarre, Camille Capel, Vincent Enouf, Louise Lefrançois, Frédéric Lemoine, Christophe Malabat, Corinne Maufrais, Etienne Simon-Lorière, Maud Vanpeene, Sylvie Van der Werf ,Delphine MammÈS |
| EPI_ISL_2612598, EPI_ISL_2612599, EPI_ISL_2612600 | Hospital | National Reference Center for Viruses of Respiratory Infections, Institut Pasteur, Paris | Marion Barbet, Sylvie Behillil, Méline Bizard, Angela Brisebarre, Camille Capel, Vincent Enouf, Louise Lefrançois, Frédéric Lemoine, Christophe Malabat, Corinne Maufrais, Etienne Simon-Lorière, Maud Vanpeene, Sylvie Van der Werf ,Christine Febreau |
| EPI_ISL_2612602, EPI_ISL_2612603, EPI_ISL_2612604, EPI_ISL_2612605 | Hospital | National Reference Center for Viruses of Respiratory Infections, Institut Pasteur, Paris | Marion Barbet, Sylvie Behillil, Méline Bizard, Angela Brisebarre, Camille Capel, Vincent Enouf, Louise Lefrançois, Frédéric Lemoine, Christophe Malabat, Corinne Maufrais, Etienne Simon-Lorière, Maud Vanpeene, Sylvie Van der Werf ,Pierre Alain Billy |
| EPI_ISL_2612606, EPI_ISL_2612608, EPI_ISL_2612609, EPI_ISL_2612610, EPI_ISL_2612611, EPI_ISL_2612613, EPI_ISL_2612614, EPI_ISL_2612615 | Labo Analyses Med | National Reference Center for Viruses of Respiratory Infections, Institut Pasteur, Paris | Marion Barbet, Sylvie Behillil, Méline Bizard, Angela Brisebarre, Camille Capel, Vincent Enouf, Louise Lefrançois, Frédéric Lemoine, Christophe Malabat, Corinne Maufrais, Etienne Simon-Lorière, Maud Vanpeene, Sylvie Van der Werf ,Pierre-Yves Leonard |
| EPI_ISL_2612616 | Labo Analyses Med | National Reference Center for Viruses of Respiratory Infections, Institut Pasteur, Paris | Marion Barbet, Sylvie Behillil, Méline Bizard, Angela Brisebarre, Camille Capel, Vincent Enouf, Louise Lefrançois, Frédéric Lemoine, Christophe Malabat, Corinne Maufrais, Emmanuelle Permal, Etienne Simon-Lorière, Maud Vanpeene, Sylvie Van der Werf ,Pierre-Yves Leonard |

| | | | |
|---|---|---|---|
| EPI_ISL_2612617 | Labo Analyses Med | National Reference Center for Viruses of Respiratory Infections, Institut Pasteur, Paris | Marion Barbet, Sylvie Behillil, Méline Bizard, Angela Brisebarre, Camille Capel, Vincent Enouf, Louise Lefrançois, Frédéric Lemoine, Christophe Malabat, Corinne Maufrais, Etienne Simon-Lorière, Maud Vanpeene, Sylvie Van der Werf ,Pierre-Yves Leonard |
| EPI_ISL_2612618 | Labo Analyses Med | National Reference Center for Viruses of Respiratory Infections, Institut Pasteur, Paris | Marion Barbet, Sylvie Behillil, Méline Bizard, Angela Brisebarre, Camille Capel, Vincent Enouf, Louise Lefrançois, Frédéric Lemoine, Christophe Malabat, Corinne Maufrais, Amaury Vaysse, Etienne Simon-Lorière, Maud Vanpeene, Sylvie Van der Werf ,Pierre-Yves Leonard |
| EPI_ISL_2612619, EPI_ISL_2612620, EPI_ISL_2612621 | Labo Analyses Med | National Reference Center for Viruses of Respiratory Infections, Institut Pasteur, Paris | Marion Barbet, Sylvie Behillil, Méline Bizard, Angela Brisebarre, Camille Capel, Vincent Enouf, Louise Lefrançois, Frédéric Lemoine, Christophe Malabat, Corinne Maufrais, Etienne Simon-Lorière, Maud Vanpeene, Sylvie Van der Werf ,Pierre-Yves Leonard |
| EPI_ISL_2612622, EPI_ISL_2612623, EPI_ISL_2612624 | Hospital | National Reference Center for Viruses of Respiratory Infections, Institut Pasteur, Paris | Marion Barbet, Sylvie Behillil, Méline Bizard, Angela Brisebarre, Camille Capel, Vincent Enouf, Louise Lefrançois, Frédéric Lemoine, Christophe Malabat, Corinne Maufrais, Etienne Simon-Lorière, Maud Vanpeene, Sylvie Van der Werf ,Axelle Paquin |
| EPI_ISL_2612625 | Hospital | National Reference Center for Viruses of Respiratory Infections, Institut Pasteur, Paris | Marion Barbet, Sylvie Behillil, Méline Bizard, Angela Brisebarre, Camille Capel, Vincent Enouf, Louise Lefrançois, Frédéric Lemoine, Christophe Malabat, Corinne Maufrais, Etienne Simon-Lorière, Maud Vanpeene, Sylvie Van der Werf ,Alexandra Ducancelle |
| EPI_ISL_2612629 | Hospital | National Reference Center for Viruses of Respiratory Infections, Institut Pasteur, Paris | Marion Barbet, Sylvie Behillil, Méline Bizard, Angela Brisebarre, Camille Capel, Vincent Enouf, Louise Lefrançois, Frédéric Lemoine, Christophe Malabat, Corinne Maufrais, Etienne Simon-Lorière, Maud Vanpeene, Sylvie Van der Werf ,Caroline Lefeuvre |
| EPI_ISL_2612630 | Hospital | National Reference Center for Viruses of Respiratory Infections, Institut Pasteur, Paris | Marion Barbet, Sylvie Behillil, Méline Bizard, Angela Brisebarre, Camille Capel, Vincent Enouf, Louise Lefrançois, Frédéric Lemoine, Christophe Malabat, Corinne Maufrais, Etienne Simon-Lorière, Maud Vanpeene, Sylvie Van der Werf ,Franck Labbe |
| EPI_ISL_2612631 | Labo Analyses Med | National Reference Center for Viruses of Respiratory Infections, Institut Pasteur, Paris | Marion Barbet, Sylvie Behillil, Méline Bizard, Angela Brisebarre, Camille Capel, Vincent Enouf, Louise Lefrançois, Frédéric Lemoine, Christophe Malabat, Corinne Maufrais, Amaury Vaysse, Etienne Simon-Lorière, Maud Vanpeene, Sylvie Van der Werf ,Anne Holstein |
| EPI_ISL_2612632, EPI_ISL_2612633 | Labo Analyses Med | National Reference Center for Viruses of Respiratory Infections, Institut Pasteur, Paris | Marion Barbet, Sylvie Behillil, Méline Bizard, Angela Brisebarre, Camille Capel, Vincent Enouf, Louise Lefrançois, Frédéric Lemoine, Christophe Malabat, Corinne Maufrais, Etienne Simon-Lorière, Maud Vanpeene, Sylvie Van der Werf ,Anne Holstein |
| EPI_ISL_2612634 | Labo Analyses Med | National Reference Center for Viruses of Respiratory Infections, Institut Pasteur, Paris | Marion Barbet, Sylvie Behillil, Méline Bizard, Angela Brisebarre, Camille Capel, Vincent Enouf, Louise Lefrançois, Frédéric Lemoine, Christophe Malabat, Corinne Maufrais, Amaury Vaysse, Etienne Simon-Lorière, Maud Vanpeene, Sylvie Van der Werf ,Anne Holstein |
| EPI_ISL_2612635, EPI_ISL_2612636, EPI_ISL_2612637, EPI_ISL_2612638, EPI_ISL_2612639, EPI_ISL_2612640, EPI_ISL_2612641, EPI_ISL_2612642, EPI_ISL_2612643, EPI_ISL_2612644, EPI_ISL_2612645, EPI_ISL_2612646, EPI_ISL_2612647, EPI_ISL_2612648, EPI_ISL_2612649, EPI_ISL_2612650, EPI_ISL_2612651, EPI_ISL_2612652, EPI_ISL_2612653, EPI_ISL_2612654, EPI_ISL_2612655, EPI_ISL_2612656, EPI_ISL_2612657 | | | |
| see above | Labo Analyses Med | National Reference Center for Viruses of Respiratory Infections, Institut Pasteur, Paris | Marion Barbet, Sylvie Behillil, Méline Bizard, Angela Brisebarre, Camille Capel, Vincent Enouf, Louise Lefrançois, Frédéric Lemoine, Christophe Malabat, Corinne Maufrais, Etienne Simon-Lorière, Maud Vanpeene, Sylvie Van der Werf ,Anne Holstein |
| EPI_ISL_2612658, EPI_ISL_2612659, EPI_ISL_2612660, EPI_ISL_2612661, EPI_ISL_2612662, EPI_ISL_2612663, EPI_ISL_2612664, EPI_ISL_2612665, EPI_ISL_2612666, EPI_ISL_2612667, EPI_ISL_2612668 | | | |
| see above | Labo Analyses Med | National Reference Center for Viruses of Respiratory Infections, Institut Pasteur, Paris | Marion Barbet, Sylvie Behillil, Méline Bizard, Angela Brisebarre, Camille Capel, Vincent Enouf, Louise Lefrançois, Frédéric Lemoine, Christophe Malabat, Corinne Maufrais, Etienne Simon-Lorière, Maud Vanpeene, Sylvie Van der Werf ,Jean-FrançOis Comes |
| EPI_ISL_2612669, EPI_ISL_2612670 | Hospital | National Reference Center for Viruses of Respiratory Infections, Institut Pasteur, Paris | Marion Barbet, Sylvie Behillil, Méline Bizard, Angela Brisebarre, Camille Capel, Vincent Enouf, Louise Lefrançois, Frédéric Lemoine, Christophe Malabat, Corinne Maufrais, Etienne Simon-Lorière, Maud Vanpeene, Sylvie Van der Werf ,Anne Cady |
| EPI_ISL_2612671, EPI_ISL_2612672, EPI_ISL_2612673, EPI_ISL_2612674, EPI_ISL_2612675, EPI_ISL_2612676, EPI_ISL_2612677, EPI_ISL_2612680, EPI_ISL_2612681 | Labo Analyses Med | National Reference Center for Viruses of Respiratory Infections, Institut Pasteur, Paris | Marion Barbet, Sylvie Behillil, Méline Bizard, Angela Brisebarre, Camille Capel, Vincent Enouf, Louise Lefrançois, Frédéric Lemoine, Christophe Malabat, Corinne Maufrais, Etienne Simon-Lorière, Maud Vanpeene, Sylvie Van der Werf ,Arditi Cocco |
| EPI_ISL_2613609, EPI_ISL_2613610, EPI_ISL_2613611 | Dipartimento di Medicina di Laboratorio, Azienda sanitaria universitaria Friuli Centrale (ASU FC) | Dipartimento di Medicina di Laboratorio, Azienda sanitaria universitaria Friuli Centrale (ASU FC) | Corrado Pipan, Catia Mio, Chiara Dal Secco, Stefania Marzinotto, Francesco Curcio |
| EPI_ISL_2613612 | CH SAINT-PHILIBERT | CHU Lille - Laboratoire de Virologie | AIT YAHYA Emilie, ALIDJINOU Enagnon Kazali, BOCKET Laurence, CREPIN Michel, DEMAY Christophe, ENGELMANN Ilka, GEFFROY Sandrine, GUIGON Aurélie, LAMBERT Valérie, LAZREK Mouna, NOBILLIAUX Florian, PREVOST Brigitte, THUILLIER Caroline, TINEZ Claire |
| EPI_ISL_2613613 | CH SECLIN | CHU Lille - Laboratoire de Virologie | AIT YAHYA Emilie, ALIDJINOU Enagnon Kazali, BOCKET Laurence, CREPIN Michel, DEMAY Christophe, ENGELMANN Ilka, GEFFROY Sandrine, GUIGON Aurélie, LAMBERT Valérie, LAZREK Mouna, NOBILLIAUX Florian, PREVOST Brigitte, THUILLIER Caroline, TINEZ Claire |
| EPI_ISL_2613614 | CHU LILLE | CHU Lille - Laboratoire de Virologie | AIT YAHYA Emilie, ALIDJINOU Enagnon Kazali, BOCKET Laurence, CREPIN Michel, DEMAY Christophe, ENGELMANN Ilka, GEFFROY Sandrine, GUIGON Aurélie, LAMBERT Valérie, LAZREK Mouna, NOBILLIAUX Florian, PREVOST Brigitte, THUILLIER Caroline, TINEZ Claire |
| EPI_ISL_2613615 | CH BETHUNE | CHU Lille - Laboratoire de Virologie | AIT YAHYA Emilie, ALIDJINOU Enagnon Kazali, BOCKET Laurence, CREPIN Michel, DEMAY Christophe, ENGELMANN Ilka, GEFFROY Sandrine, GUIGON Aurélie, LAMBERT Valérie, LAZREK Mouna, NOBILLIAUX Florian, PREVOST Brigitte, THUILLIER Caroline, TINEZ Claire |
| EPI_ISL_2613616 | CH LENS | CHU Lille - Laboratoire de Virologie | AIT YAHYA Emilie, ALIDJINOU Enagnon Kazali, BOCKET Laurence, CREPIN Michel, DEMAY Christophe, ENGELMANN Ilka, GEFFROY Sandrine, GUIGON Aurélie, LAMBERT Valérie, LAZREK Mouna, NOBILLIAUX Florian, PREVOST Brigitte, THUILLIER Caroline, TINEZ Claire |
| EPI_ISL_2613617 | CHU LILLE | CHU Lille - Laboratoire de Virologie | AIT YAHYA Emilie, ALIDJINOU Enagnon Kazali, BOCKET Laurence, CREPIN Michel, DEMAY Christophe, ENGELMANN Ilka, GEFFROY Sandrine, GUIGON Aurélie, LAMBERT Valérie, LAZREK Mouna, NOBILLIAUX Florian, PREVOST Brigitte, THUILLIER Caroline, TINEZ Claire |
| EPI_ISL_2613618 | CH TOURCOING | CHU Lille - Laboratoire de Virologie | AIT YAHYA Emilie, ALIDJINOU Enagnon Kazali, BOCKET Laurence, CREPIN Michel, DEMAY Christophe, ENGELMANN Ilka, GEFFROY Sandrine, GUIGON Aurélie, LAMBERT Valérie, LAZREK Mouna, NOBILLIAUX Florian, PREVOST Brigitte, THUILLIER Caroline, TINEZ Claire |
| EPI_ISL_2613619, EPI_ISL_2613620 | CHU LILLE | CHU Lille - Laboratoire de Virologie | AIT YAHYA Emilie, ALIDJINOU Enagnon Kazali, BOCKET Laurence, CREPIN Michel, DEMAY Christophe, ENGELMANN Ilka, GEFFROY Sandrine, GUIGON Aurélie, LAMBERT Valérie, LAZREK Mouna, NOBILLIAUX Florian, PREVOST Brigitte, THUILLIER Caroline, TINEZ Claire |
| EPI_ISL_2613621 | CH SECLIN | CHU Lille - Laboratoire de Virologie | AIT YAHYA Emilie, ALIDJINOU Enagnon Kazali, BOCKET Laurence, CREPIN Michel, DEMAY Christophe, ENGELMANN Ilka, GEFFROY Sandrine, GUIGON Aurélie, LAMBERT Valérie, LAZREK Mouna, NOBILLIAUX Florian, PREVOST Brigitte, THUILLIER Caroline, TINEZ Claire |
| EPI_ISL_2613622, EPI_ISL_2613623 | CHU LILLE | CHU Lille - Laboratoire de Virologie | AIT YAHYA Emilie, ALIDJINOU Enagnon Kazali, BOCKET Laurence, CREPIN Michel, DEMAY Christophe, ENGELMANN Ilka, GEFFROY Sandrine, GUIGON Aurélie, LAMBERT Valérie, LAZREK Mouna, NOBILLIAUX Florian, PREVOST Brigitte, THUILLIER Caroline, TINEZ Claire |
| EPI_ISL_2613624 | CH TOURCOING | CHU Lille - Laboratoire de Virologie | AIT YAHYA Emilie, ALIDJINOU Enagnon Kazali, BOCKET Laurence, CREPIN Michel, DEMAY Christophe, ENGELMANN Ilka, GEFFROY Sandrine, GUIGON Aurélie, LAMBERT Valérie, LAZREK Mouna, NOBILLIAUX Florian, PREVOST Brigitte, THUILLIER Caroline, TINEZ Claire |
| EPI_ISL_2613810, EPI_ISL_2613811, EPI_ISL_2613815 | Lboratorio Central de Saude Publica do Estado do Parana (LACEN/PR) | Laboratory of Respiratory Viruses and Measles, Oswaldo Cruz Institute, FIOCRUZ | Paola Resende, Luciana Appolinario, Fernando Motta, Anna Carolina Paixao, Ana Carolina Mendonca, Alice Sampaio Rocha, Taina Venas, Elisa Cavalcante Pereira, Renata Serrano Lopes, Irina Riediger, Marilda Siqueira on behalf of the Fiocruz COVID-19 Genomic Surveillance Network |
| EPI_ISL_2613825 | CH SAINT-PHILIBERT | CHU Lille - Laboratoire de Virologie | AIT YAHYA Emilie, ALIDJINOU Enagnon Kazali, BOCKET Laurence, CREPIN Michel, DEMAY Christophe, ENGELMANN Ilka, GEFFROY Sandrine, GUIGON Aurélie, LAMBERT Valérie, LAZREK Mouna, NOBILLIAUX Florian, PREVOST Brigitte, THUILLIER Caroline, TINEZ Claire ,TCHANTCHOU NJOSSE YANICK |
| EPI_ISL_2613826 | CH ROUBAIX | CHU Lille - Laboratoire de Virologie | AIT YAHYA Emilie, ALIDJINOU Enagnon Kazali, BOCKET Laurence, CREPIN Michel, DEMAY Christophe, ENGELMANN Ilka, GEFFROY Sandrine, GUIGON Aurélie, LAMBERT Valérie, LAZREK Mouna, NOBILLIAUX Florian, PREVOST Brigitte, THUILLIER Caroline, TINEZ Claire ,TCHANTCHOU NJOSSE YANICK |
| EPI_ISL_2613827, EPI_ISL_2613828 | CHU LILLE | CHU Lille - Laboratoire de Virologie | AIT YAHYA Emilie, ALIDJINOU Enagnon Kazali, BOCKET Laurence, CREPIN Michel, DEMAY Christophe, ENGELMANN Ilka, GEFFROY Sandrine, GUIGON Aurélie, LAMBERT Valérie, LAZREK Mouna, NOBILLIAUX Florian, PREVOST Brigitte, THUILLIER Caroline, TINEZ Claire ,TCHANTCHOU NJOSSE YANICK |
| EPI_ISL_2613829 | CH LENS | CHU Lille - Laboratoire de Virologie | AIT YAHYA Emilie, ALIDJINOU Enagnon Kazali, BOCKET Laurence, CREPIN Michel, DEMAY Christophe, ENGELMANN Ilka, GEFFROY Sandrine, GUIGON Aurélie, LAMBERT Valérie, LAZREK Mouna, NOBILLIAUX Florian, PREVOST Brigitte, THUILLIER Caroline, TINEZ Claire ,TCHANTCHOU NJOSSE YANICK |

| | | | |
|---|---|---|---|
| EPI_ISL_2613830 | CH BETHUNE | CHU Lille - Laboratoire de Virologie | AIT YAHYA Emilie, ALIDJINOU Enagnon Kazali, BOCKET Laurence, CREPIN Michel, DEMAY Christophe, ENGELMANN Ilka, GEFFROY Sandrine, GUIGON Aurélie, LAMBERT Valérie, LAZREK Mouna, NOBILLIAUX Florian, PREVOST Brigitte, THUILLIER Caroline, TINEZ Claire ,TCHANTCHOU NJOSSE YANICK |
| EPI_ISL_2613831, EPI_ISL_2613832 | CH VALENCIENNES | CHU Lille - Laboratoire de Virologie | AIT YAHYA Emilie, ALIDJINOU Enagnon Kazali, BOCKET Laurence, CREPIN Michel, DEMAY Christophe, ENGELMANN Ilka, GEFFROY Sandrine, GUIGON Aurélie, LAMBERT Valérie, LAZREK Mouna, NOBILLIAUX Florian, PREVOST Brigitte, THUILLIER Caroline, TINEZ Claire ,TCHANTCHOU NJOSSE YANICK |
| EPI_ISL_2613833 | CH ROUBAIX | CHU Lille - Laboratoire de Virologie | AIT YAHYA Emilie, ALIDJINOU Enagnon Kazali, BOCKET Laurence, CREPIN Michel, DEMAY Christophe, ENGELMANN Ilka, GEFFROY Sandrine, GUIGON Aurélie, LAMBERT Valérie, LAZREK Mouna, NOBILLIAUX Florian, PREVOST Brigitte, THUILLIER Caroline, TINEZ Claire ,TCHANTCHOU NJOSSE YANICK |
| EPI_ISL_2613834 | CHU LILLE | CHU Lille - Laboratoire de Virologie | AIT YAHYA Emilie, ALIDJINOU Enagnon Kazali, BOCKET Laurence, CREPIN Michel, DEMAY Christophe, ENGELMANN Ilka, GEFFROY Sandrine, GUIGON Aurélie, LAMBERT Valérie, LAZREK Mouna, NOBILLIAUX Florian, PREVOST Brigitte, THUILLIER Caroline, TINEZ Claire ,TCHANTCHOU NJOSSE YANICK |
| EPI_ISL_2613835 | CH ROUBAIX | CHU Lille - Laboratoire de Virologie | AIT YAHYA Emilie, ALIDJINOU Enagnon Kazali, BOCKET Laurence, CREPIN Michel, DEMAY Christophe, ENGELMANN Ilka, GEFFROY Sandrine, GUIGON Aurélie, LAMBERT Valérie, LAZREK Mouna, NOBILLIAUX Florian, PREVOST Brigitte, THUILLIER Caroline, TINEZ Claire ,TCHANTCHOU NJOSSE YANICK |
| EPI_ISL_2614419, EPI_ISL_2614420, EPI_ISL_2614421, EPI_ISL_2614422, EPI_ISL_2614423, EPI_ISL_2614424, EPI_ISL_2614425 | Klinisch Laboratorium ZNA | Klinisch Laboratorium ZNA | Verstrepen et al. |
| EPI_ISL_2614617, EPI_ISL_2614619, EPI_ISL_2614621, EPI_ISL_2614623, EPI_ISL_2614624, EPI_ISL_2614625 | Instituto Adolfo Lutz - Regional de Presidente Prudente | Instituto Adolfo Lutz, Interdiciplinary Procedures Center, Strategic Laboratory | Claudio Tavares Sacchi, Claudia Regina Gonçalves, Erica Valessa Ramos Gomes, Karoline Rodrigues Campos, Caio Vinicius Dias Lopes, Leonardo Jose Tadeu de Araujo |
| EPI_ISL_2614634, EPI_ISL_2614635, EPI_ISL_2614636, EPI_ISL_2614637, EPI_ISL_2614638, EPI_ISL_2614641, EPI_ISL_2614642 | Dr. Yuichiro Oba Department of General Medicine, Osaka General Medical Center | Department of Developmental Medicine, Research Institute, Osaka Women's and Children's Hospital | Itaru Yanagihara, Yukiko Nakura |
| EPI_ISL_2614721 | RS Mitra Keluarga Bintaro | Eijkman Institute for Molecular Biology, National Research and Innovation Agency | Lidwina Priliani, Edison Johar, Frilasita A Yudhaputri, Muhammad Rezki Rasyak, Willy Agustine, Hidayat Trimarsanto, Lydia V. Panggalo, Iskandar Adnan, Sukma Oktavianthi, Safarina G Malik, Khin Saw Myint, Amin Soebandrio |
| EPI_ISL_2614723 | US Embassy Health Unit | Eijkman Institute for Molecular Biology, National Research and Innovation Agency | Frilasita A Yudhaputri, Muhammad Rezki Rasyak, Willy Agustine, Hidayat Trimarsanto, Lydia V. Panggalo, Iskandar Adnan, Sukma Oktavianthi, Lidwina Priliani, Edison Johar, Safarina G Malik, Khin Saw Myint, Amin Soebandrio |
| EPI_ISL_2614724 | Excellent Beneficial Diagnostic Center | Eijkman Institute for Molecular Biology, National Research and Innovation Agency | Muhammad Rezki Rasyak, Willy Agustine, Hidayat Trimarsanto, Lydia V. Panggalo, Iskandar Adnan, Sukma Oktavianthi, Lidwina Priliani, Edison Johar, Frilasita A Yudhaputri, Safarina G Malik, Khin Saw Myint, Amin Soebandrio |
| EPI_ISL_2614725 | RS Pelni | Eijkman Institute for Molecular Biology, National Research and Innovation Agency | Willy Agustine, Hidayat Trimarsanto, Lydia V. Panggalo, Iskandar Adnan, Sukma Oktavianthi, Lidwina Priliani, Edison Johar, Frilasita A Yudhaputri, Muhammad Rezki Rasyak, Safarina G Malik, Khin Saw Myint, Amin Soebandrio |
| EPI_ISL_2614726 | RS Pelni | Eijkman Institute for Molecular Biology, National Research and Innovation Agency | Hidayat Trimarsanto, Lydia V. Panggalo, Iskandar Adnan, Sukma Oktavianthi, Lidwina Priliani, Edison Johar, Frilasita A Yudhaputri, Muhammad Rezki Rasyak, Willy Agustine, Safarina G Malik, Khin Saw Myint, Amin Soebandrio |
| EPI_ISL_2614727 | RS Pelni | Eijkman Institute for Molecular Biology, National Research and Innovation Agency | Lydia V. Panggalo, Iskandar Adnan, Sukma Oktavianthi, Lidwina Priliani, Edison Johar, Frilasita A Yudhaputri, Muhammad Rezki Rasyak, Willy Agustine, Hidayat Trimarsanto, Safarina G Malik, Khin Saw Myint, Amin Soebandrio |
| EPI_ISL_2614728 | RS Grha Kedoya | Eijkman Institute for Molecular Biology, National Research and Innovation Agency | Iskandar Adnan, Sukma Oktavianthi, Lidwina Priliani, Edison Johar, Frilasita A Yudhaputri, Muhammad Rezki Rasyak, Willy Agustine, Hidayat Trimarsanto, Lydia V. Panggalo, Safarina G Malik, Khin Saw Myint, Amin Soebandrio |
| EPI_ISL_2614730 | EKA Hospital BSD | Eijkman Institute for Molecular Biology, National Research and Innovation Agency | Lidwina Priliani, Edison Johar, Frilasita A Yudhaputri, Muhammad Rezki Rasyak, Willy Agustine, Hidayat Trimarsanto, Lydia V. Panggalo, Iskandar Adnan, Sukma Oktavianthi, Safarina G Malik, Khin Saw Myint, Amin Soebandrio |
| EPI_ISL_2614731 | RS Pelni | Eijkman Institute for Molecular Biology, National Research and Innovation Agency | Iskandar Adnan, Sukma Oktavianthi, Lidwina Priliani, Edison Johar, Frilasita A Yudhaputri, Muhammad Rezki Rasyak, Willy Agustine, Hidayat Trimarsanto, Lydia V. Panggalo, Safarina G Malik, Khin Saw Myint, Amin Soebandrio |
| EPI_ISL_2614732 | RS Sumber Waras | Eijkman Institute for Molecular Biology, National Research and Innovation Agency | Sukma Oktavianthi, Lidwina Priliani, Edison Johar, Frilasita A Yudhaputri, Muhammad Rezki Rasyak, Willy Agustine, Hidayat Trimarsanto, Lydia V. Panggalo, Iskandar Adnan, Safarina G Malik, Khin Saw Myint, Amin Soebandrio |
| EPI_ISL_2614733 | RS Pusat Pertamina | Eijkman Institute for Molecular Biology, National Research and Innovation Agency | Lidwina Priliani, Edison Johar, Frilasita A Yudhaputri, Muhammad Rezki Rasyak, Willy Agustine, Hidayat Trimarsanto, Lydia V. Panggalo, Iskandar Adnan, Sukma Oktavianthi, Safarina G Malik, Khin Saw Myint, Amin Soebandrio |
| EPI_ISL_2614734 | RS Pusat Pertamina | Eijkman Institute for Molecular Biology, National Research and Innovation Agency | Edison Johar, Frilasita A Yudhaputri, Muhammad Rezki Rasyak, Willy Agustine, Hidayat Trimarsanto, Lydia V. Panggalo, Iskandar Adnan, Sukma Oktavianthi, Lidwina Priliani, Safarina G Malik, Khin Saw Myint, Amin Soebandrio |
| EPI_ISL_2614735 | RS Pondok Indah Pondok Indah | Eijkman Institute for Molecular Biology, National Research and Innovation Agency | Frilasita A Yudhaputri, Muhammad Rezki Rasyak, Willy Agustine, Hidayat Trimarsanto, Lydia V. Panggalo, Iskandar Adnan, Sukma Oktavianthi, Lidwina Priliani, Edison Johar, Safarina G Malik, Khin Saw Myint, Amin Soebandrio |
| EPI_ISL_2614736 | RS Hermina Depok | Eijkman Institute for Molecular Biology, National Research and Innovation Agency | Muhammad Rezki Rasyak, Willy Agustine, Hidayat Trimarsanto, Lydia V. Panggalo, Iskandar Adnan, Sukma Oktavianthi, Lidwina Priliani, Edison Johar, Frilasita A Yudhaputri, Safarina G Malik, Khin Saw Myint, Amin Soebandrio |
| EPI_ISL_2614737 | RS Pondok Indah Pondok Indah | Eijkman Institute for Molecular Biology, National Research and Innovation Agency | Willy Agustine, Hidayat Trimarsanto, Lydia V. Panggalo, Iskandar Adnan, Sukma Oktavianthi, Lidwina Priliani, Edison Johar, Frilasita A Yudhaputri, Muhammad Rezki Rasyak, Safarina G Malik, Khin Saw Myint, Amin Soebandrio |
| EPI_ISL_2614738 | RS Pondok Indah Pondok Indah | Eijkman Institute for Molecular Biology, National Research and Innovation Agency | Hidayat Trimarsanto, Lydia V. Panggalo, Iskandar Adnan, Sukma Oktavianthi, Lidwina Priliani, Edison Johar, Frilasita A Yudhaputri, Muhammad Rezki Rasyak, Willy Agustine, Safarina G Malik, Khin Saw Myint, Amin Soebandrio |
| EPI_ISL_2614743 | Sungai Buloh Hospital | Institute for Medical Research, Infectious Disease Research Centre, National Institutes of Health, Ministry of Health Malaysia | Suppiah J, Kamel K, Mohd Zawawi Z, Azizan MA, Ramly N, Robert F, Thayan R |
| EPI_ISL_2614744 | Tuanku Ampuan Najihah Hospital, Kuala Pilah | Institute for Medical Research, Infectious Disease Research Centre, National Institutes of Health, Ministry of Health Malaysia | Suppiah J, Kamel K, Mohd Zawawi Z, Azizan MA, Ramly N, Robert F, Thayan R |
| EPI_ISL_2614745 | Sarawak General Hospital | Institute for Medical Research, Infectious Disease Research Centre, National Institutes of Health, Ministry of Health Malaysia | Suppiah J, Kamel K, Mohd Zawawi Z, Azizan MA, Ramly N, Robert F, Thayan R |
| EPI_ISL_2615399 | MOH - Jaber Al-Ahmad Hospital (Innovation Research Laboratory) | MOH - Jaber Al-Ahmad Hospital (Innovation Research Laboratory) | Salman Al-Sabah , Mohammad Alghounaim |

EPI_ISL_2616217, EPI_ISL_2616218, EPI_ISL_2616219, EPI_ISL_2616220, EPI_ISL_2616224, EPI_ISL_2616225, EPI_ISL_2616226, EPI_ISL_2616227, EPI_ISL_2616228, EPI_ISL_2616229, EPI_ISL_2616230, EPI_ISL_2616231, EPI_ISL_2616232, EPI_ISL_2616233, EPI_ISL_2616234, EPI_ISL_2616235, EPI_ISL_2616236, EPI_ISL_2616237, EPI_ISL_2616238, EPI_ISL_2616239, EPI_ISL_2616240, EPI_ISL_2616241, EPI_ISL_2616242, EPI_ISL_2616243, EPI_ISL_2616244, EPI_ISL_2616245, EPI_ISL_2616246, EPI_ISL_2616247, EPI_ISL_2616248, EPI_ISL_2616249, EPI_ISL_2616250, EPI_ISL_2616251, EPI_ISL_2616252, EPI_ISL_2616253, EPI_ISL_2616254, EPI_ISL_2616255, EPI_ISL_2616256, EPI_ISL_2616257, EPI_ISL_2616258, EPI_ISL_2616259, EPI_ISL_2616260, EPI_ISL_2616261, EPI_ISL_2616262, EPI_ISL_2616265, EPI_ISL_2616266, EPI_ISL_2616267, EPI_ISL_2616268, EPI_ISL_2616269, EPI_ISL_2616270, EPI_ISL_2616271, EPI_ISL_2616272, EPI_ISL_2616273, EPI_ISL_2616274, EPI_ISL_2616275, EPI_ISL_2616276, EPI_ISL_2616277, EPI_ISL_2616278, EPI_ISL_2616279, EPI_ISL_2616280, EPI_ISL_2616281, EPI_ISL_2616282, EPI_ISL_2616283, EPI_ISL_2616284, EPI_ISL_2616285, EPI_ISL_2616286, EPI_ISL_2616287, EPI_ISL_2616288, EPI_ISL_2616289,

| | | | |
|---|---|---|---|
| EPI_ISL_2616290, EPI_ISL_2616291, EPI_ISL_2616292, EPI_ISL_2616293, EPI_ISL_2616294, EPI_ISL_2616295, EPI_ISL_2616296, EPI_ISL_2616297, EPI_ISL_2616298, EPI_ISL_2616299, EPI_ISL_2616300, EPI_ISL_2616301, EPI_ISL_2616302, EPI_ISL_2616303, EPI_ISL_2616304, EPI_ISL_2616305, EPI_ISL_2616306, EPI_ISL_2616307, EPI_ISL_2616308, EPI_ISL_2616309, EPI_ISL_2616310, EPI_ISL_2616311, EPI_ISL_2616312, EPI_ISL_2616313, EPI_ISL_2616314, EPI_ISL_2616315, EPI_ISL_2616316, EPI_ISL_2616317, EPI_ISL_2616318 | | | |
| see above | Austrian Agency for Health and Food Safety (AGES) | Bergthaler laboratory, CeMM Research Center for Molecular Medicine of the Austrian Academy of Sciences | Lukas Endler, Anna Schedl, Fabian Amman, Petr Triska, Matthew Thornton, Thomas Penz, Benedikt Agerer, Maelle Le Moing, Michael Schuster, Bekir Erguner, Jan Laine, Martin Senekowitsch, Christoph Bock, Andreas Bergthaler |
| EPI_ISL_2616319 | Center for Virology, Medical University of Vienna | Bergthaler laboratory, CeMM Research Center for Molecular Medicine of the Austrian Academy of Sciences | Lukas Endler, Anna Schedl, Fabian Amman, Petr Triska, Matthew Thornton, Thomas Penz, Benedikt Agerer, Maelle Le Moing, Michael Schuster, Bekir Erguner, Jan Laine, Martin Senekowitsch, Christoph Bock, Andreas Bergthaler |
| EPI_ISL_2616320, EPI_ISL_2616321, EPI_ISL_2616322, EPI_ISL_2616323, EPI_ISL_2616324, EPI_ISL_2616325, EPI_ISL_2616326, EPI_ISL_2616327, EPI_ISL_2616328, EPI_ISL_2616329, EPI_ISL_2616330, EPI_ISL_2616331, EPI_ISL_2616332, EPI_ISL_2616333 | | | |
| see above | Austrian Agency for Health and Food Safety (AGES) | Bergthaler laboratory, CeMM Research Center for Molecular Medicine of the Austrian Academy of Sciences | Lukas Endler, Anna Schedl, Fabian Amman, Petr Triska, Matthew Thornton, Thomas Penz, Benedikt Agerer, Maelle Le Moing, Michael Schuster, Bekir Erguner, Jan Laine, Martin Senekowitsch, Christoph Bock, Andreas Bergthaler |
| EPI_ISL_2616334, EPI_ISL_2616335, EPI_ISL_2616336, EPI_ISL_2616342 | Elling group, Institute of Molecular Biotechnology (IMBA) | Bergthaler laboratory, CeMM Research Center for Molecular Medicine of the Austrian Academy of Sciences | Lukas Endler, Anna Schedl, Fabian Amman, Petr Triska, Matthew Thornton, Thomas Penz, Benedikt Agerer, Maelle Le Moing, Michael Schuster, Bekir Erguner, Jan Laine, Martin Senekowitsch, Christoph Bock, Andreas Bergthaler |
| EPI_ISL_2616343, EPI_ISL_2616344, EPI_ISL_2616345, EPI_ISL_2616346, EPI_ISL_2616362, EPI_ISL_2616364, EPI_ISL_2616365 | Austrian Agency for Health and Food Safety (AGES) | Bergthaler laboratory, CeMM Research Center for Molecular Medicine of the Austrian Academy of Sciences | Lukas Endler, Anna Schedl, Fabian Amman, Petr Triska, Matthew Thornton, Thomas Penz, Benedikt Agerer, Maelle Le Moing, Michael Schuster, Bekir Erguner, Jan Laine, Martin Senekowitsch, Christoph Bock, Andreas Bergthaler |
| EPI_ISL_2617440, EPI_ISL_2617441, EPI_ISL_2617442, EPI_ISL_2617443, EPI_ISL_2617445 | laboratorio Microbiologia PO Cardarelli | laboratorio Microbiologia PO Cardarelli | Felice V., Niro G. Scutella' M. |
| EPI_ISL_2617632, EPI_ISL_2617633, EPI_ISL_2617634, EPI_ISL_2617635, EPI_ISL_2617636, EPI_ISL_2617637, EPI_ISL_2617638, EPI_ISL_2617639, EPI_ISL_2617640, EPI_ISL_2617641, EPI_ISL_2617642, EPI_ISL_2617643, EPI_ISL_2617644, EPI_ISL_2617645, EPI_ISL_2617646, EPI_ISL_2617647, EPI_ISL_2617648, EPI_ISL_2617649, EPI_ISL_2617650, EPI_ISL_2617651, EPI_ISL_2617652, EPI_ISL_2617653, EPI_ISL_2617654, EPI_ISL_2617655, EPI_ISL_2617656, EPI_ISL_2617657, EPI_ISL_2617658, EPI_ISL_2617659 | | | |
| see above | Austrian Agency for Health and Food Safety (AGES) | Bergthaler laboratory, CeMM Research Center for Molecular Medicine of the Austrian Academy of Sciences | Lukas Endler, Anna Schedl, Fabian Amman, Petr Triska, Matthew Thornton, Thomas Penz, Benedikt Agerer, Maelle Le Moing, Michael Schuster, Bekir Erguner, Jan Laine, Martin Senekowitsch, Christoph Bock, Andreas Bergthaler |
| EPI_ISL_2617660, EPI_ISL_2617661, EPI_ISL_2617662, EPI_ISL_2617663, EPI_ISL_2617664, EPI_ISL_2617665, EPI_ISL_2617666, EPI_ISL_2617667, EPI_ISL_2617668, EPI_ISL_2617669, EPI_ISL_2617670, EPI_ISL_2617671, EPI_ISL_2617672, EPI_ISL_2617673, EPI_ISL_2617674, EPI_ISL_2617675, EPI_ISL_2617676, EPI_ISL_2617677, EPI_ISL_2617678, EPI_ISL_2617679 | | | |
| see above | Elling group, Institute of Molecular Biotechnology (IMBA) | Bergthaler laboratory, CeMM Research Center for Molecular Medicine of the Austrian Academy of Sciences | Lukas Endler, Anna Schedl, Fabian Amman, Petr Triska, Matthew Thornton, Thomas Penz, Benedikt Agerer, Maelle Le Moing, Michael Schuster, Bekir Erguner, Jan Laine, Martin Senekowitsch, Christoph Bock, Andreas Bergthaler |
| EPI_ISL_2620714 | BBTKLPP Jakarta | National Institute of Health Research and Development | Kartika Dewi Puspa, Arie Ardiansyah Nugraha, Hana Apsari Pawestri, Hartanti Dian Ikawati, Subangkit,Triyani Soekarso, Krisna Pangesti, Nelly Puspandari, Vivi Setiawaty |
| EPI_ISL_2620715 | Hospital Tuanku Fauziah, Kangar | Institute for Medical Research, Infectious Disease Research Centre, National Institutes of Health, Ministry of Health Malaysia | Suppiah J, Kamel K, Mohd Zawawi Z, Azizan MA, Ramly N, Robert F, Thayan R |
| EPI_ISL_2621148, EPI_ISL_2621149 | Synlab | Universidad Nacional de Colombia - Laboratorio Genómico One Health | Andres F. Cardona-Rios, Maria Stella López, Daniel O. Maldonado-Perez, Simón Villegas Velásquez, Lina M. Hurtado, Laura Silvana Perez, Karl A Ciuoderis, Maria Angélica Maya, Carolina Muñoz-Arango, Celeny Ortiz, Rita Almanza Payares, Idabely Betancur Ortiz, Sandra Ines Cano, Diego A. Álvarez-Díaz, Carlos Franco-Muñoz, Katherine Laiton-Donato, Hector Alejandro Ruiz-Moreno, Marcela Mercado-Reyes, Jorge E. Osorio, Juan P. Hernandez-Ortiz |
| EPI_ISL_2621161 | ICMT-APARTADO | Universidad Nacional de Colombia - Laboratorio Genómico One Health | Simón Villegas Velásquez, Andres F. Cardona-Rios, Maria Stella López, Daniel O. Maldonado-Perez, Lina M. Hurtado, Laura Silvana Perez, Karl A Ciuoderis, Maria Angélica Maya, Carolina Muñoz-Arango, Celeny Ortiz, Rita Almanza Payares, Idabely Betancur Ortiz, Sandra Ines Cano, Diego A. Álvarez-Díaz, Carlos Franco-Muñoz, Katherine Laiton-Donato, Hector Alejandro Ruiz-Moreno, Marcela Mercado-Reyes, Jorge E. Osorio, Juan P. Hernandez-Ortiz |
| EPI_ISL_2621167, EPI_ISL_2621168 | ICMT-APARTADO | Universidad Nacional de Colombia - Laboratorio Genómico One Health | Andres F. Cardona-Rios, Maria Stella López, Daniel O. Maldonado-Perez, Simón Villegas Velásquez, Lina M. Hurtado, Laura Silvana Perez, Karl A Ciuoderis, Maria Angélica Maya, Carolina Muñoz-Arango, Celeny Ortiz, Rita Almanza Payares, Idabely Betancur Ortiz, Sandra Ines Cano, Diego A. Álvarez-Díaz, Carlos Franco-Muñoz, Katherine Laiton-Donato, Hector Alejandro Ruiz-Moreno, Marcela Mercado-Reyes, Jorge E. Osorio, Juan P. Hernandez-Ortiz |
| EPI_ISL_2621177, EPI_ISL_2621178, EPI_ISL_2621179 | PRIME DIAGNOSTICS | Universidad Nacional de Colombia - Laboratorio Genómico One Health | Andres F. Cardona-Rios, Maria Stella López, Daniel O. Maldonado-Perez, Simón Villegas Velásquez, Lina M. Hurtado, Laura Silvana Perez, Karl A Ciuoderis, Maria Angélica Maya, Carolina Muñoz-Arango, Celeny Ortiz, Rita Almanza Payares, Idabely Betancur Ortiz, Sandra Ines Cano, Diego A. Álvarez-Díaz, Carlos Franco-Muñoz, Katherine Laiton-Donato, Hector Alejandro Ruiz-Moreno, Marcela Mercado-Reyes, Jorge E. Osorio, Juan P. Hernandez-Ortiz |
| EPI_ISL_2621189 | Somer | Universidad Nacional de Colombia - Laboratorio Genómico One Health | Maria Stella López, Andres F. Cardona-Rios, Daniel O. Maldonado-Perez, Simón Villegas Velásquez, Lina M. Hurtado, Laura Silvana Perez, Karl A Ciuoderis, Maria Angélica Maya, Carolina Muñoz-Arango, Celeny Ortiz, Rita Almanza Payares, Idabely Betancur Ortiz, Sandra Ines Cano, Diego A. Álvarez-Díaz, Carlos Franco-Muñoz, Katherine Laiton-Donato, Hector Alejandro Ruiz-Moreno, Marcela Mercado-Reyes, Jorge E. Osorio, Juan P. Hernandez-Ortiz |
| EPI_ISL_2621202 | Synlab | Universidad Nacional de Colombia - Laboratorio Genómico One Health | Simón Villegas Velásquez, Andres F. Cardona-Rios, Maria Stella López, Daniel O. Maldonado-Perez, Lina M. Hurtado, Laura Silvana Perez, Karl A Ciuoderis, Maria Angélica Maya, Carolina Muñoz-Arango, Celeny Ortiz, Rita Almanza Payares, Idabely Betancur Ortiz, Sandra Ines Cano, Diego A. Álvarez-Díaz, Carlos Franco-Muñoz, Katherine Laiton-Donato, Hector Alejandro Ruiz-Moreno, Marcela Mercado-Reyes, Jorge E. Osorio, Juan P. Hernandez-Ortiz |
| EPI_ISL_2621211 | SAN VICENTE FUNDACION | Universidad Nacional de Colombia - Laboratorio Genómico One Health | Andres F. Cardona-Rios, Maria Stella López, Daniel O. Maldonado-Perez, Simón Villegas Velásquez, Lina M. Hurtado, Laura Silvana Perez, Karl A Ciuoderis, Maria Angélica Maya, Carolina Muñoz-Arango, Celeny Ortiz, Rita Almanza Payares, Idabely Betancur Ortiz, Sandra Ines Cano, Diego A. Álvarez-Díaz, Carlos Franco-Muñoz, Katherine Laiton-Donato, Hector Alejandro Ruiz-Moreno, Marcela Mercado-Reyes, Jorge E. Osorio, Juan P. Hernandez-Ortiz |
| EPI_ISL_2621218 | ADILAB | Universidad Nacional de Colombia - Laboratorio Genómico One Health | Simón Villegas Velásquez, Andres F. Cardona-Rios, Maria Stella López, Daniel O. Maldonado-Perez, Lina M. Hurtado, Laura Silvana Perez, Karl A Ciuoderis, Maria Angélica Maya, Carolina Muñoz-Arango, Celeny Ortiz, Rita Almanza Payares, Idabely Betancur Ortiz, Sandra Ines Cano, Diego A. Álvarez-Díaz, Carlos Franco-Muñoz, Katherine Laiton-Donato, Hector Alejandro Ruiz-Moreno, Marcela Mercado-Reyes, Jorge E. Osorio, Juan P. Hernandez-Ortiz |
| EPI_ISL_2621454, EPI_ISL_2621460, EPI_ISL_2621461, EPI_ISL_2621464, EPI_ISL_2621465, EPI_ISL_2621466, EPI_ISL_2621467, EPI_ISL_2621469, EPI_ISL_2621475, EPI_ISL_2621476, EPI_ISL_2621477, EPI_ISL_2621478, EPI_ISL_2621479, EPI_ISL_2621480, EPI_ISL_2621481, EPI_ISL_2621482, EPI_ISL_2621483, EPI_ISL_2621485, EPI_ISL_2621486, EPI_ISL_2621487, EPI_ISL_2621488, EPI_ISL_2621489, EPI_ISL_2621490, EPI_ISL_2621491, EPI_ISL_2621492, EPI_ISL_2621494, EPI_ISL_2621495, EPI_ISL_2621496, EPI_ISL_2621497 | | | |
| see above | National Center of Infectious and Parasitic Diseases | National Center of Infectious and Parasitic Diseases | Alexiev et al |
| EPI_ISL_2621731, EPI_ISL_2621736 | Genome Analysis Center, Yamanashi Central Hospital | Genome Analysis Center, Yamanashi Central Hospital | Yosuke Hirotsu |
| EPI_ISL_2622160, EPI_ISL_2622161, EPI_ISL_2622162, EPI_ISL_2622164, EPI_ISL_2622165, EPI_ISL_2622166, EPI_ISL_2622167, EPI_ISL_2622169, EPI_ISL_2622171, EPI_ISL_2622172, EPI_ISL_2622174, EPI_ISL_2622175 | | | |
| see above | Azienda Ospedaliera Pugliese Ciaccio di Catanzaro SOC Microbiologia e Virologia | Azienda Ospedaliera Pugliese Ciaccio di Catanzaro SOC Microbiologia e Virologia | Rossana Tallerico Federica Pasceri Marco De Fazio Ilenia Talotta Giuseppina Panduri Pasquale Minchella |
| EPI_ISL_2625622 | Usansolo-Galdakao University Hospital | Cruces University Hospital | Ana Gual-de-Torrella, Izaskun Alejo-Cancho, Mikel Gallego, Ana Belén de la Hoz |
| EPI_ISL_2625972, EPI_ISL_2625973, EPI_ISL_2625974, EPI_ISL_2625975, EPI_ISL_2625977, EPI_ISL_2625978, EPI_ISL_2625981, EPI_ISL_2625982, EPI_ISL_2625983, EPI_ISL_2625984, EPI_ISL_2625987, EPI_ISL_2625988, EPI_ISL_2625989, EPI_ISL_2625990, EPI_ISL_2625991, EPI_ISL_2625992, EPI_ISL_2625993, EPI_ISL_2625994, EPI_ISL_2625995, EPI_ISL_2625996, EPI_ISL_2625997, EPI_ISL_2625998, EPI_ISL_2625999, EPI_ISL_2626000, EPI_ISL_2626001, EPI_ISL_2626002, EPI_ISL_2626003, EPI_ISL_2626004, EPI_ISL_2626005, EPI_ISL_2626006, EPI_ISL_2626007, EPI_ISL_2626008 | | | |

| | | | |
|---|---|---|---|
| see above | Microbiology Department, University Hospital Donostia | Microbiology Department, University Hospital Donostia | Marimon JM, Montes M, Piñeiro L, Martin-Peñaranda T, Gomez M, Sorarrain A, Cilla G. |
| EPI_ISL_2626010, EPI_ISL_2626011, EPI_ISL_2626012, EPI_ISL_2626013, EPI_ISL_2626014, EPI_ISL_2626015, EPI_ISL_2626016, EPI_ISL_2626017, EPI_ISL_2626018, EPI_ISL_2626019, EPI_ISL_2626020, EPI_ISL_2626021, EPI_ISL_2626022, EPI_ISL_2626023, EPI_ISL_2626024, EPI_ISL_2626025, EPI_ISL_2626026, EPI_ISL_2626027, EPI_ISL_2626028, EPI_ISL_2626029, EPI_ISL_2626030, EPI_ISL_2626031, EPI_ISL_2626032, EPI_ISL_2626033, EPI_ISL_2626034, EPI_ISL_2626035, EPI_ISL_2626036, EPI_ISL_2626037, EPI_ISL_2626038, EPI_ISL_2626039, EPI_ISL_2626040, EPI_ISL_2626041, EPI_ISL_2626042, EPI_ISL_2626043, EPI_ISL_2626044, EPI_ISL_2626045, EPI_ISL_2626046, EPI_ISL_2626047, EPI_ISL_2626048, EPI_ISL_2626049, EPI_ISL_2626050, EPI_ISL_2626051, EPI_ISL_2626053, EPI_ISL_2626054, EPI_ISL_2626055, EPI_ISL_2626056, EPI_ISL_2626057, EPI_ISL_2626058, EPI_ISL_2626059, EPI_ISL_2626060, EPI_ISL_2626061 | | | |
| see above | Cliniques universitaires Saint-Luc | UCLouvain/IREC/MBLG | Jean Ruelle, Bertrand Bearzatto, Benoit Kabamba Mukadi |
| EPI_ISL_2626062, EPI_ISL_2626063, EPI_ISL_2626064, EPI_ISL_2626065, EPI_ISL_2626066, EPI_ISL_2626067, EPI_ISL_2626068, EPI_ISL_2626069, EPI_ISL_2626070, EPI_ISL_2626071, EPI_ISL_2626072, EPI_ISL_2626073, EPI_ISL_2626074, EPI_ISL_2626075, EPI_ISL_2626077, EPI_ISL_2626078, EPI_ISL_2626079, EPI_ISL_2626080, EPI_ISL_2626081 | | | |
| see above | Clinique Saint-Pierre Ottignies | UCLouvain/IREC/MBLG | Jean Ruelle, Bertrand Bearzatto, Benoit Kabamba Mukadi |
| EPI_ISL_2626082, EPI_ISL_2626083, EPI_ISL_2626084, EPI_ISL_2626085, EPI_ISL_2626086, EPI_ISL_2626087, EPI_ISL_2626088, EPI_ISL_2626089, EPI_ISL_2626090, EPI_ISL_2626091, EPI_ISL_2626092, EPI_ISL_2626093, EPI_ISL_2626094, EPI_ISL_2626095, EPI_ISL_2626096 | | | |
| see above | Cliniques universitaires Saint-Luc | UCLouvain/IREC/MBLG | Jean Ruelle, Bertrand Bearzatto, Benoit Kabamba Mukadi |
| EPI_ISL_2626097 | Hôpitaux Iris Sud Etterbeek-Ixelles | UCLouvain/IREC/MBLG | Jean Ruelle, Bertrand Bearzatto, Benoit Kabamba Mukadi |
| EPI_ISL_2626180, EPI_ISL_2626181 | Usansolo-Galdakao University Hospital | Cruces University Hospital | Ana Gual-de-Torrella, Izaskun Alejo-Cancho, Mikel Gallego, Ana Belén de la Hoz |
| EPI_ISL_2626182, EPI_ISL_2626183 | Virology Unit, AOUP | Virology Unit, AOUP | Marialinda Vatteroni, Susi Frateschi, Mauro Pistello |
| EPI_ISL_2626433 | Laboratori de Referencia de Catalunya | Laboratori de Referencia de Catalunya | Ramirez A., Bellosillo B., Padilla E., Vilas A., Hernandez JJ., Canal M. |
| EPI_ISL_2626441 | Microbiology Department, University Hospital Donostia | Microbiology Department, University Hospital Donostia | Marimon JM, Montes M, Piñeiro L, Martin-Peñaranda T, Gomez M, Sorarrain A, Cilla G. |
| EPI_ISL_2626770, EPI_ISL_2626771, EPI_ISL_2626772, EPI_ISL_2626774, EPI_ISL_2626777, EPI_ISL_2626778, EPI_ISL_2626779, EPI_ISL_2626782, EPI_ISL_2626783, EPI_ISL_2626784, EPI_ISL_2626785, EPI_ISL_2626786, EPI_ISL_2626787, EPI_ISL_2626788, EPI_ISL_2626789, EPI_ISL_2626790, EPI_ISL_2626794, EPI_ISL_2626795, EPI_ISL_2626797, EPI_ISL_2626799, EPI_ISL_2626800, EPI_ISL_2626803, EPI_ISL_2626807, EPI_ISL_2626808, EPI_ISL_2626809, EPI_ISL_2626810, EPI_ISL_2626811, EPI_ISL_2626812, EPI_ISL_2626813, EPI_ISL_2626814, EPI_ISL_2626816, EPI_ISL_2626818, EPI_ISL_2626819, EPI_ISL_2626820, EPI_ISL_2626821, EPI_ISL_2626822, EPI_ISL_2626824, EPI_ISL_2626825, EPI_ISL_2626826, EPI_ISL_2626827, EPI_ISL_2626831, EPI_ISL_2626835, EPI_ISL_2626836, EPI_ISL_2626837, EPI_ISL_2626838 | | | |
| see above | Baylor Scott & White-Temple | Baylor Scott & White-Temple | Ari Rao, Linden Morales, Kimberly Walker, Marcus Volz, Shelby Johnson |
| EPI_ISL_2627491, EPI_ISL_2627492, EPI_ISL_2627493, EPI_ISL_2627494, EPI_ISL_2627495, EPI_ISL_2627496, EPI_ISL_2627497, EPI_ISL_2627498 | Klinisch Laboratorium ZNA | Klinisch Laboratorium ZNA | Verstrepen et al. |
| EPI_ISL_2627846 | CHC Andrée Rosemon | Institut Pasteur de la Guyane | Anne Lavergne, Dominique Rousset, Antoine Enfissi, Arielle Salmier |
| EPI_ISL_2627940, EPI_ISL_2627941 | Hospital | National Reference Center for Viruses of Respiratory Infections, Institut Pasteur, Paris | Marion Barbet, Sylvie Behillil, Méline Bizard, Angela Brisebarre, Camille Capel, Vincent Enouf, Louise Lefrançois, Frédéric Lemoine, Christophe Malabat, Corinne Maufrais, Etienne Simon-Lorière, Maud Vanpeene, Sylvie Van der Werf ,Patricia Stoessel-Thouvenin |
| EPI_ISL_2627942 | Hospital | National Reference Center for Viruses of Respiratory Infections, Institut Pasteur, Paris | Marion Barbet, Sylvie Behillil, Méline Bizard, Angela Brisebarre, Camille Capel, Vincent Enouf, Louise Lefrançois, Frédéric Lemoine, Christophe Malabat, Corinne Maufrais, Pierre Lechat, Etienne Simon-Lorière, Maud Vanpeene, Sylvie Van der Werf ,Patricia Stoessel-Thouvenin |
| EPI_ISL_2627943, EPI_ISL_2627944, EPI_ISL_2627945 | Hospital | National Reference Center for Viruses of Respiratory Infections, Institut Pasteur, Paris | Marion Barbet, Sylvie Behillil, Méline Bizard, Angela Brisebarre, Camille Capel, Vincent Enouf, Louise Lefrançois, Frédéric Lemoine, Christophe Malabat, Corinne Maufrais, Etienne Simon-Lorière, Maud Vanpeene, Sylvie Van der Werf ,Patricia Stoessel-Thouvenin |
| EPI_ISL_2627947, EPI_ISL_2627948 | Hospital | National Reference Center for Viruses of Respiratory Infections, Institut Pasteur, Paris | Marion Barbet, Sylvie Behillil, Méline Bizard, Angela Brisebarre, Camille Capel, Vincent Enouf, Louise Lefrançois, Frédéric Lemoine, Christophe Malabat, Corinne Maufrais, Etienne Simon-Lorière, Maud Vanpeene, Sylvie Van der Werf ,ValéRie Serazin |
| EPI_ISL_2627949, EPI_ISL_2627950 | Labo Analyses Med | National Reference Center for Viruses of Respiratory Infections, Institut Pasteur, Paris | Marion Barbet, Sylvie Behillil, Méline Bizard, Angela Brisebarre, Camille Capel, Vincent Enouf, Louise Lefrançois, Frédéric Lemoine, Christophe Malabat, Corinne Maufrais, Etienne Simon-Lorière, Maud Vanpeene, Sylvie Van der Werf ,GréGoire Potiron |
| EPI_ISL_2627952 | Hospital | National Reference Center for Viruses of Respiratory Infections, Institut Pasteur, Paris | Marion Barbet, Sylvie Behillil, Méline Bizard, Angela Brisebarre, Camille Capel, Vincent Enouf, Louise Lefrançois, Frédéric Lemoine, Christophe Malabat, Corinne Maufrais, Etienne Simon-Lorière, Maud Vanpeene, Sylvie Van der Werf ,Anne-CéCile Hochart |
| EPI_ISL_2627953, EPI_ISL_2627954, EPI_ISL_2627955 | Hospital | National Reference Center for Viruses of Respiratory Infections, Institut Pasteur, Paris | Marion Barbet, Sylvie Behillil, Méline Bizard, Angela Brisebarre, Camille Capel, Vincent Enouf, Louise Lefrançois, Frédéric Lemoine, Christophe Malabat, Corinne Maufrais, Etienne Simon-Lorière, Maud Vanpeene, Sylvie Van der Werf ,Pascale Martres |
| EPI_ISL_2627956 | Hospital | National Reference Center for Viruses of Respiratory Infections, Institut Pasteur, Paris | Marion Barbet, Sylvie Behillil, Méline Bizard, Angela Brisebarre, Camille Capel, Vincent Enouf, Louise Lefrançois, Frédéric Lemoine, Christophe Malabat, Corinne Maufrais, Etienne Simon-Lorière, Maud Vanpeene, Sylvie Van der Werf ,StéPhanie Marque Juillet |
| EPI_ISL_2627958, EPI_ISL_2627959 | Hospital | National Reference Center for Viruses of Respiratory Infections, Institut Pasteur, Paris | Marion Barbet, Sylvie Behillil, Méline Bizard, Angela Brisebarre, Camille Capel, Vincent Enouf, Louise Lefrançois, Frédéric Lemoine, Christophe Malabat, Corinne Maufrais, Etienne Simon-Lorière, Maud Vanpeene, Sylvie Van der Werf ,JéRome Guinard |
| EPI_ISL_2627960 | Hospital | National Reference Center for Viruses of Respiratory Infections, Institut Pasteur, Paris | Marion Barbet, Sylvie Behillil, Méline Bizard, Angela Brisebarre, Camille Capel, Vincent Enouf, Louise Lefrançois, Frédéric Lemoine, Christophe Malabat, Corinne Maufrais, Pierre Lechat, Etienne Simon-Lorière, Maud Vanpeene, Sylvie Van der Werf ,JéRome Guinard |
| EPI_ISL_2627961, EPI_ISL_2627963 | Hospital | National Reference Center for Viruses of Respiratory Infections, Institut Pasteur, Paris | Marion Barbet, Sylvie Behillil, Méline Bizard, Angela Brisebarre, Camille Capel, Vincent Enouf, Louise Lefrançois, Frédéric Lemoine, Christophe Malabat, Corinne Maufrais, Etienne Simon-Lorière, Maud Vanpeene, Sylvie Van der Werf ,JéRome Guinard |
| EPI_ISL_2627965 | Hospital | National Reference Center for Viruses of Respiratory Infections, Institut Pasteur, Paris | Marion Barbet, Sylvie Behillil, Méline Bizard, Angela Brisebarre, Camille Capel, Vincent Enouf, Louise Lefrançois, Frédéric Lemoine, Christophe Malabat, Corinne Maufrais, Etienne Simon-Lorière, Maud Vanpeene, Sylvie Van der Werf ,Laura Djamdjian |
| EPI_ISL_2627966, EPI_ISL_2627967, EPI_ISL_2627968, EPI_ISL_2627969 | Hospital | National Reference Center for Viruses of Respiratory Infections, Institut Pasteur, Paris | Marion Barbet, Sylvie Behillil, Méline Bizard, Angela Brisebarre, Camille Capel, Vincent Enouf, Louise Lefrançois, Frédéric Lemoine, Christophe Malabat, Corinne Maufrais, Etienne Simon-Lorière, Maud Vanpeene, Sylvie Van der Werf ,Laurence Bocket |
| EPI_ISL_2627970 | Hospital | National Reference Center for Viruses of Respiratory Infections, Institut Pasteur, Paris | Marion Barbet, Sylvie Behillil, Méline Bizard, Angela Brisebarre, Camille Capel, Vincent Enouf, Louise Lefrançois, Frédéric Lemoine, Christophe Malabat, Corinne Maufrais, Etienne Simon-Lorière, Maud Vanpeene, Sylvie Van der Werf ,Sylvie Ledru |
| EPI_ISL_2627971, EPI_ISL_2627972, EPI_ISL_2627973 | Labo Analyses Med | National Reference Center for Viruses of Respiratory Infections, Institut Pasteur, Paris | Marion Barbet, Sylvie Behillil, Méline Bizard, Angela Brisebarre, Camille Capel, Vincent Enouf, Louise Lefrançois, Frédéric Lemoine, Christophe Malabat, Corinne Maufrais, Etienne Simon-Lorière, Maud Vanpeene, Sylvie Van der Werf ,Sophie Chalmin |
| EPI_ISL_2627975, EPI_ISL_2627976, EPI_ISL_2627978 | Hospital | National Reference Center for Viruses of Respiratory Infections, Institut Pasteur, Paris | Marion Barbet, Sylvie Behillil, Méline Bizard, Angela Brisebarre, Camille Capel, Vincent Enouf, Louise Lefrançois, Frédéric Lemoine, Christophe Malabat, Corinne Maufrais, Etienne Simon-Lorière, Maud Vanpeene, Sylvie Van der Werf ,Pascale Martres |
| EPI_ISL_2627979, EPI_ISL_2627980, EPI_ISL_2627981, EPI_ISL_2627982 | Hospital | National Reference Center for Viruses of Respiratory Infections, Institut Pasteur, Paris | Marion Barbet, Sylvie Behillil, Méline Bizard, Angela Brisebarre, Camille Capel, Vincent Enouf, Louise Lefrançois, Frédéric Lemoine, Christophe Malabat, Corinne Maufrais, Etienne Simon-Lorière, Maud Vanpeene, Sylvie Van der Werf ,Sandrine Castelain |
| EPI_ISL_2627983, EPI_ISL_2627984 | Hospital | National Reference Center for Viruses of Respiratory Infections, Institut Pasteur, Paris | Marion Barbet, Sylvie Behillil, Méline Bizard, Angela Brisebarre, Camille Capel, Vincent Enouf, Louise Lefrançois, Frédéric Lemoine, Christophe Malabat, Corinne Maufrais, Etienne Simon-Lorière, Maud Vanpeene, Sylvie Van der Werf ,L. Courdavault |
| EPI_ISL_2627988 | Hospital | National Reference Center for Viruses of Respiratory Infections, Institut Pasteur, Paris | Marion Barbet, Sylvie Behillil, Méline Bizard, Angela Brisebarre, Camille Capel, Vincent Enouf, Louise Lefrançois, Frédéric Lemoine, Christophe Malabat, Corinne Maufrais, Etienne Simon-Lorière, Maud Vanpeene, Sylvie Van der Werf ,Eric Farfour |
| EPI_ISL_2627989, EPI_ISL_2627990 | Armies | National Reference Center for Viruses of Respiratory Infections, Institut Pasteur, Paris | Marion Barbet, Sylvie Behillil, Méline Bizard, Angela Brisebarre, Camille Capel, Vincent Enouf, Louise Lefrançois, Frédéric Lemoine, Christophe Malabat, Corinne Maufrais, Etienne Simon-Lorière, Maud Vanpeene, Sylvie Van der Werf ,Marine Desroches |
| EPI_ISL_2627991, EPI_ISL_2627992, EPI_ISL_2627993, EPI_ISL_2627994 | Hospital | National Reference Center for Viruses of Respiratory Infections, Institut Pasteur, Paris | Marion Barbet, Sylvie Behillil, Méline Bizard, Angela Brisebarre, Camille Capel, Vincent Enouf, Louise Lefrançois, Frédéric Lemoine, Christophe Malabat, Corinne Maufrais, Etienne Simon-Lorière, Maud Vanpeene, Sylvie Van der Werf ,Nabil Gastli |
| EPI_ISL_2627996 | Hospital | National Reference Center for Viruses of Respiratory Infections, Institut Pasteur, Paris | Marion Barbet, Sylvie Behillil, Méline Bizard, Angela Brisebarre, Camille Capel, Vincent Enouf, Louise Lefrançois, Frédéric Lemoine, Christophe Malabat, Corinne Maufrais, Etienne Simon-Lorière, Maud Vanpeene, Sylvie Van der Werf ,Catherine Gaudy Graffin |
| EPI_ISL_2627997 | Labo Analyses Med | National Reference Center for Viruses of Respiratory | Marion Barbet, Sylvie Behillil, Méline Bizard, Angela Brisebarre, Camille Capel, Vincent Enouf, Louise Lefrançois, Frédéric Lemoine, Christophe Malabat, |

| | | | |
|---|---|---|---|
| | | Infections, Institut Pasteur, Paris | Corinne Maufrais, Pierre Lechat, Etienne Simon-Lorière, Maud Vanpeene, Sylvie Van der Werf ,Caroline Devaux |
| EPI_ISL_2627998, EPI_ISL_2627999, EPI_ISL_2628000, EPI_ISL_2628001, EPI_ISL_2628002 | Labo Analyses Med | National Reference Center for Viruses of Respiratory Infections, Institut Pasteur, Paris | Marion Barbet, Sylvie Behillil, Méline Bizard, Angela Brisebarre, Camille Capel, Vincent Enouf, Louise Lefrançois, Frédéric Lemoine, Christophe Malabat, Corinne Maufrais, Etienne Simon-Lorière, Maud Vanpeene, Sylvie Van der Werf ,Caroline Devaux |
| EPI_ISL_2628004 | Labo Analyses Med | National Reference Center for Viruses of Respiratory Infections, Institut Pasteur, Paris | Marion Barbet, Sylvie Behillil, Méline Bizard, Angela Brisebarre, Camille Capel, Vincent Enouf, Louise Lefrançois, Frédéric Lemoine, Christophe Malabat, Corinne Maufrais, Etienne Simon-Lorière, Maud Vanpeene, Sylvie Van der Werf ,Arditi Cocco |
| EPI_ISL_2628272, EPI_ISL_2628273 | Labo Analyses Med | National Reference Center for Viruses of Respiratory Infections, Institut Pasteur, Paris | Marion Barbet, Sylvie Behillil, Méline Bizard, Angela Brisebarre, Camille Capel, Vincent Enouf, Louise Lefrançois, Frédéric Lemoine, Christophe Malabat, Corinne Maufrais, Etienne Simon-Lorière, Maud Vanpeene, Sylvie Van der Werf ,Caroline Devaux |
| EPI_ISL_2628274 | Hospital | National Reference Center for Viruses of Respiratory Infections, Institut Pasteur, Paris | Marion Barbet, Sylvie Behillil, Méline Bizard, Angela Brisebarre, Camille Capel, Vincent Enouf, Louise Lefrançois, Frédéric Lemoine, Christophe Malabat, Corinne Maufrais, Etienne Simon-Lorière, Maud Vanpeene, Sylvie Van der Werf ,E. Bichier |
| EPI_ISL_2628275, EPI_ISL_2628276 | Hospital | National Reference Center for Viruses of Respiratory Infections, Institut Pasteur, Paris | Marion Barbet, Sylvie Behillil, Méline Bizard, Angela Brisebarre, Camille Capel, Vincent Enouf, Louise Lefrançois, Frédéric Lemoine, Christophe Malabat, Corinne Maufrais, Etienne Simon-Lorière, Maud Vanpeene, Sylvie Van der Werf ,CéCile Hermann |
| EPI_ISL_2628277 | Hospital | National Reference Center for Viruses of Respiratory Infections, Institut Pasteur, Paris | Marion Barbet, Sylvie Behillil, Méline Bizard, Angela Brisebarre, Camille Capel, Vincent Enouf, Louise Lefrançois, Frédéric Lemoine, Christophe Malabat, Corinne Maufrais, Etienne Simon-Lorière, Maud Vanpeene, Sylvie Van der Werf ,Marianne Burgard |
| EPI_ISL_2628283, EPI_ISL_2628284, EPI_ISL_2628285, EPI_ISL_2628286 | Hospital | National Reference Center for Viruses of Respiratory Infections, Institut Pasteur, Paris | Marion Barbet, Sylvie Behillil, Méline Bizard, Angela Brisebarre, Camille Capel, Vincent Enouf, Louise Lefrançois, Frédéric Lemoine, Christophe Malabat, Corinne Maufrais, Etienne Simon-Lorière, Maud Vanpeene, Sylvie Van der Werf ,Marianne Coste-Burel |
| EPI_ISL_2628289, EPI_ISL_2628292, EPI_ISL_2628294 | Hospital | National Reference Center for Viruses of Respiratory Infections, Institut Pasteur, Paris | Marion Barbet, Sylvie Behillil, Méline Bizard, Angela Brisebarre, Camille Capel, Vincent Enouf, Louise Lefrançois, Frédéric Lemoine, Christophe Malabat, Corinne Maufrais, Etienne Simon-Lorière, Maud Vanpeene, Sylvie Van der Werf ,LéA Pilorge |
| EPI_ISL_2628295, EPI_ISL_2628296 | Labo Analyses Med | National Reference Center for Viruses of Respiratory Infections, Institut Pasteur, Paris | Marion Barbet, Sylvie Behillil, Méline Bizard, Angela Brisebarre, Camille Capel, Vincent Enouf, Louise Lefrançois, Frédéric Lemoine, Christophe Malabat, Corinne Maufrais, Etienne Simon-Lorière, Maud Vanpeene, Sylvie Van der Werf ,Claire Felloni |
| EPI_ISL_2637227 | Genome Analysis Center, Yamanashi Central Hospital | Genome Analysis Center, Yamanashi Central Hospital | Yosuke Hirotsu |
| EPI_ISL_2637418, EPI_ISL_2637495, EPI_ISL_2637510, EPI_ISL_2637569, EPI_ISL_2637659, EPI_ISL_2637700, EPI_ISL_2637735 | GH A.CHENEVIER-H.MONDOR | Department of Virology, Henri Mondor University Hospital, Assistance Publique Hôpitaux de Paris, Université Paris-Est Créteil, INSERM U955 | Christophe Rodriguez, Slim Fourati, Vanessa Demontant, Guillaume Gricourt, Melissa N'Debi, Alexandre Soulier, Elisabeth Trawinski, Jean-Michel Pawlotsky |
| EPI_ISL_2638046 | Clinique Saint-Pierre Ottignies | UCLouvain/IREC/MBLG | Jean Ruelle, Bertrand Bearzatto, Benoit Kabamba Mukadi |
| EPI_ISL_2642158 | AOU Policlinico " G. Rodolico- S. Marco" PO Rodolico | AOU Policlinico " G. Rodolico- S. Marco" PO Rodolico and BIOMETEC department UNICT | Stefani and Scalia |
| EPI_ISL_2642542 | AOU Policlinico "G. Rodolico-S. Marco", P.O. "G. Rodolico" | AOU Policlinico "G. Rodolico-S. Marco", P.O. "G. Rodolico" | Stefani and Scalia |
| EPI_ISL_2642543 | Laboratory of Microbiology, Varese | Laboratory of Microbiology, Varese | Federica Novazzi, Francesca Dragoferrante, Angelo Genoni, Daniele Focosi, Andreina Baj, Fabrizio Maggi |
| EPI_ISL_2642545, EPI_ISL_2642549 | AOP Policlinico " G. Rodolico- S. Marco" PO Rodolico Catania | AOP Policlinico " G. Rodolico- S. Marco" PO Rodolico Catania and BIOMETEC UNICT | Stefani and Scalia |
| EPI_ISL_2642552 | Laboratorio "L.C. Campisi" Avola | AOU Policlinico "G. Rodolico-S. Marco", P.O. "G. Rodolico" | Stefani and Scalia |
| EPI_ISL_2642654, EPI_ISL_2642898 | AOP Policlinico " G. Rodolico- S. Marco" PO Rodolico Catania | AOP Policlinico " G. Rodolico- S. Marco" PO Rodolico Catania and BIOMETEC UNICT | Stefani and Scalia |
| EPI_ISL_2643119 | Laboratorio "L.C. Campisi" Avola (Siracusa) | AOU Policlinico "G. Rodolico-S. Marco", P.O. "G. Rodolico" | Stefani and Scalia |
| EPI_ISL_2643671 | AOU Policlinico "G. Rodolico-S. Marco", P.O. "G. Rodolico" | AOU Policlinico "G. Rodolico-S. Marco", P.O. "G. Rodolico" | Stefani and Scalia |
| EPI_ISL_2644656, EPI_ISL_2644664 | Genome Analysis Center, Yamanashi Central Hospital | Genome Analysis Center, Yamanashi Central Hospital | Yosuke Hirotsu |
| EPI_ISL_2644834 | LESP HIDALGO | LESP HIDALGO GENE2LIFE | HERRERA-GOMEZ CONNIE ALEJANDRA; CASTILLO-ORTEGA LAURA SOFIA; GARCIA-REYES MELITO; VIGUERAS-MORALES YAJAIRA SULIM; GARCIA-GONZALEZ OCTAVIO; VILLAFUENTES-TELLEZ HECTOR ALBERTO |
| EPI_ISL_2645445, EPI_ISL_2645446, EPI_ISL_2645447, EPI_ISL_2645448, EPI_ISL_2645449, EPI_ISL_2645450, EPI_ISL_2645452, EPI_ISL_2645453, EPI_ISL_2645455, EPI_ISL_2645457, EPI_ISL_2645458, EPI_ISL_2645459, EPI_ISL_2645460, EPI_ISL_2645461, EPI_ISL_2645462, EPI_ISL_2645463, EPI_ISL_2645464, EPI_ISL_2645465, EPI_ISL_2645466, EPI_ISL_2645467, EPI_ISL_2645468, EPI_ISL_2645469, EPI_ISL_2645470, EPI_ISL_2645471, EPI_ISL_2645472, EPI_ISL_2645473, EPI_ISL_2645474, EPI_ISL_2645475, EPI_ISL_2645476, EPI_ISL_2645477, EPI_ISL_2645478, EPI_ISL_2645479, EPI_ISL_2645481, EPI_ISL_2645482, EPI_ISL_2645483, EPI_ISL_2645484, EPI_ISL_2645486, EPI_ISL_2645491, EPI_ISL_2645492, EPI_ISL_2645493, EPI_ISL_2645494, EPI_ISL_2645495, EPI_ISL_2645496, EPI_ISL_2645499, EPI_ISL_2645500, EPI_ISL_2645501, EPI_ISL_2645503, EPI_ISL_2645504 | | | |
| see above | Laboratorio Central de Saude Publica do Estado do Parana (LACEN/PR) | Laboratory of Respiratory Viruses and Measles, Oswaldo Cruz Institute, FIOCRUZ | Paola Resende, Luciana Appolinario, Fernando Motta, Anna Carolina Paixao, Ana Carolina Mendonca, Alice Sampaio Rocha, Taina Venas, Elisa Cavalcante Pereira, Renata Serrano Lopes, Irina Riediger, Marilda Siqueira on behalf of the Fiocruz COVID-19 Genomic Surveillance Network |
| EPI_ISL_2646140 | CH ARMENTIERES | CHU Lille - Laboratoire de Virologie | AIT YAHYA Emilie, ALIDJINOU Enagnon Kazali, BOCKET Laurence, CREPIN Michel, DEMAY Christophe, ENGELMANN Ilka, GEFFROY Sandrine, GUIGON Aurélie, LAMBERT Valérie, LAZREK Mouna, NOBILLIAUX Florian, PREVOST Brigitte, THUILLIER Caroline, TINEZ Claire ,TCHANTCHOU NJOSSE YANICK |
| EPI_ISL_2646141 | CH BETHUNE | CHU Lille - Laboratoire de Virologie | AIT YAHYA Emilie, ALIDJINOU Enagnon Kazali, BOCKET Laurence, CREPIN Michel, DEMAY Christophe, ENGELMANN Ilka, GEFFROY Sandrine, GUIGON Aurélie, LAMBERT Valérie, LAZREK Mouna, NOBILLIAUX Florian, PREVOST Brigitte, THUILLIER Caroline, TINEZ Claire ,TCHANTCHOU NJOSSE YANICK |
| EPI_ISL_2646142, EPI_ISL_2646143 | CH ROUBAIX | CHU Lille - Laboratoire de Virologie | AIT YAHYA Emilie, ALIDJINOU Enagnon Kazali, BOCKET Laurence, CREPIN Michel, DEMAY Christophe, ENGELMANN Ilka, GEFFROY Sandrine, GUIGON Aurélie, LAMBERT Valérie, LAZREK Mouna, NOBILLIAUX Florian, PREVOST Brigitte, THUILLIER Caroline, TINEZ Claire ,TCHANTCHOU NJOSSE YANICK |
| EPI_ISL_2646144 | CH ARMENTIERES | CHU Lille - Laboratoire de Virologie | AIT YAHYA Emilie, ALIDJINOU Enagnon Kazali, BOCKET Laurence, CREPIN Michel, DEMAY Christophe, ENGELMANN Ilka, GEFFROY Sandrine, GUIGON Aurélie, LAMBERT Valérie, LAZREK Mouna, NOBILLIAUX Florian, PREVOST Brigitte, THUILLIER Caroline, TINEZ Claire ,TCHANTCHOU NJOSSE YANICK |
| EPI_ISL_2646145, EPI_ISL_2646146 | CH CALAIS | CHU Lille - Laboratoire de Virologie | AIT YAHYA Emilie, ALIDJINOU Enagnon Kazali, BOCKET Laurence, CREPIN Michel, DEMAY Christophe, ENGELMANN Ilka, GEFFROY Sandrine, GUIGON Aurélie, LAMBERT Valérie, LAZREK Mouna, NOBILLIAUX Florian, PREVOST Brigitte, THUILLIER Caroline, TINEZ Claire ,TCHANTCHOU NJOSSE YANICK |
| EPI_ISL_2646147 | CH ROUBAIX | CHU Lille - Laboratoire de Virologie | AIT YAHYA Emilie, ALIDJINOU Enagnon Kazali, BOCKET Laurence, CREPIN Michel, DEMAY Christophe, ENGELMANN Ilka, GEFFROY Sandrine, GUIGON Aurélie, LAMBERT Valérie, LAZREK Mouna, NOBILLIAUX Florian, PREVOST Brigitte, THUILLIER Caroline, TINEZ Claire ,TCHANTCHOU NJOSSE YANICK |
| EPI_ISL_2646148 | CH ARMENTIERES | CHU Lille - Laboratoire de Virologie | AIT YAHYA Emilie, ALIDJINOU Enagnon Kazali, BOCKET Laurence, CREPIN Michel, DEMAY Christophe, ENGELMANN Ilka, GEFFROY Sandrine, GUIGON Aurélie, LAMBERT Valérie, LAZREK Mouna, NOBILLIAUX Florian, PREVOST Brigitte, THUILLIER Caroline, TINEZ Claire ,TCHANTCHOU NJOSSE YANICK |
| EPI_ISL_2646149 | CH LENS | CHU Lille - Laboratoire de Virologie | AIT YAHYA Emilie, ALIDJINOU Enagnon Kazali, BOCKET Laurence, CREPIN Michel, DEMAY Christophe, ENGELMANN Ilka, GEFFROY Sandrine, GUIGON Aurélie, LAMBERT Valérie, LAZREK Mouna, NOBILLIAUX Florian, PREVOST Brigitte, THUILLIER Caroline, TINEZ Claire ,TCHANTCHOU NJOSSE YANICK |
| EPI_ISL_2646150, EPI_ISL_2646151 | CH ARMENTIERES | CHU Lille - Laboratoire de Virologie | AIT YAHYA Emilie, ALIDJINOU Enagnon Kazali, BOCKET Laurence, CREPIN Michel, DEMAY Christophe, ENGELMANN Ilka, GEFFROY Sandrine, GUIGON Aurélie, LAMBERT Valérie, LAZREK Mouna, NOBILLIAUX Florian, PREVOST Brigitte, THUILLIER Caroline, TINEZ Claire ,TCHANTCHOU |

| | | | |
|---|---|---|---|
| | | | NJOSSE YANICK |
| EPI_ISL_2646152, EPI_ISL_2646153, EPI_ISL_2646154 | CHU LILLE | CHU Lille - Laboratoire de Virologie | AIT YAHYA Emilie, ALIDJINOU Enagnon Kazali, BOCKET Laurence, CREPIN Michel, DEMAY Christophe, ENGELMANN Ilka, GEFFROY Sandrine, GUIGON Aurélie, LAMBERT Valérie, LAZREK Mouna, NOBILLIAUX Florian, PREVOST Brigitte, THUILLIER Caroline, TINEZ Claire ,TCHANTCHOU NJOSSE YANICK |
| EPI_ISL_2646155 | CH CALAIS | CHU Lille - Laboratoire de Virologie | AIT YAHYA Emilie, ALIDJINOU Enagnon Kazali, BOCKET Laurence, CREPIN Michel, DEMAY Christophe, ENGELMANN Ilka, GEFFROY Sandrine, GUIGON Aurélie, LAMBERT Valérie, LAZREK Mouna, NOBILLIAUX Florian, PREVOST Brigitte, THUILLIER Caroline, TINEZ Claire ,TCHANTCHOU NJOSSE YANICK |
| EPI_ISL_2646156 | CH ARRAS | CHU Lille - Laboratoire de Virologie | AIT YAHYA Emilie, ALIDJINOU Enagnon Kazali, BOCKET Laurence, CREPIN Michel, DEMAY Christophe, ENGELMANN Ilka, GEFFROY Sandrine, GUIGON Aurélie, LAMBERT Valérie, LAZREK Mouna, NOBILLIAUX Florian, PREVOST Brigitte, THUILLIER Caroline, TINEZ Claire ,TCHANTCHOU NJOSSE YANICK |
| EPI_ISL_2646157 | CH ROUBAIX | CHU Lille - Laboratoire de Virologie | AIT YAHYA Emilie, ALIDJINOU Enagnon Kazali, BOCKET Laurence, CREPIN Michel, DEMAY Christophe, ENGELMANN Ilka, GEFFROY Sandrine, GUIGON Aurélie, LAMBERT Valérie, LAZREK Mouna, NOBILLIAUX Florian, PREVOST Brigitte, THUILLIER Caroline, TINEZ Claire ,TCHANTCHOU NJOSSE YANICK |
| EPI_ISL_2646158, EPI_ISL_2646160 | CHU LILLE | CHU Lille - Laboratoire de Virologie | AIT YAHYA Emilie, ALIDJINOU Enagnon Kazali, BOCKET Laurence, CREPIN Michel, DEMAY Christophe, ENGELMANN Ilka, GEFFROY Sandrine, GUIGON Aurélie, LAMBERT Valérie, LAZREK Mouna, NOBILLIAUX Florian, PREVOST Brigitte, THUILLIER Caroline, TINEZ Claire ,TCHANTCHOU NJOSSE YANICK |
| EPI_ISL_2646161 | CH ROUBAIX | CHU Lille - Laboratoire de Virologie | AIT YAHYA Emilie, ALIDJINOU Enagnon Kazali, BOCKET Laurence, CREPIN Michel, DEMAY Christophe, ENGELMANN Ilka, GEFFROY Sandrine, GUIGON Aurélie, LAMBERT Valérie, LAZREK Mouna, NOBILLIAUX Florian, PREVOST Brigitte, THUILLIER Caroline, TINEZ Claire ,TCHANTCHOU NJOSSE YANICK |
| EPI_ISL_2646162, EPI_ISL_2646163 | CH BETHUNE | CHU Lille - Laboratoire de Virologie | AIT YAHYA Emilie, ALIDJINOU Enagnon Kazali, BOCKET Laurence, CREPIN Michel, DEMAY Christophe, ENGELMANN Ilka, GEFFROY Sandrine, GUIGON Aurélie, LAMBERT Valérie, LAZREK Mouna, NOBILLIAUX Florian, PREVOST Brigitte, THUILLIER Caroline, TINEZ Claire ,TCHANTCHOU NJOSSE YANICK |
| EPI_ISL_2646164, EPI_ISL_2646165, EPI_ISL_2646166, EPI_ISL_2646167, EPI_ISL_2646168 | CHU LILLE | CHU Lille - Laboratoire de Virologie | AIT YAHYA Emilie, ALIDJINOU Enagnon Kazali, BOCKET Laurence, CREPIN Michel, DEMAY Christophe, ENGELMANN Ilka, GEFFROY Sandrine, GUIGON Aurélie, LAMBERT Valérie, LAZREK Mouna, NOBILLIAUX Florian, PREVOST Brigitte, THUILLIER Caroline, TINEZ Claire ,TCHANTCHOU NJOSSE YANICK |
| EPI_ISL_2646169 | CH LENS | CHU Lille - Laboratoire de Virologie | AIT YAHYA Emilie, ALIDJINOU Enagnon Kazali, BOCKET Laurence, CREPIN Michel, DEMAY Christophe, ENGELMANN Ilka, GEFFROY Sandrine, GUIGON Aurélie, LAMBERT Valérie, LAZREK Mouna, NOBILLIAUX Florian, PREVOST Brigitte, THUILLIER Caroline, TINEZ Claire ,TCHANTCHOU NJOSSE YANICK |
| EPI_ISL_2646170, EPI_ISL_2646171 | CHU LILLE | CHU Lille - Laboratoire de Virologie | AIT YAHYA Emilie, ALIDJINOU Enagnon Kazali, BOCKET Laurence, CREPIN Michel, DEMAY Christophe, ENGELMANN Ilka, GEFFROY Sandrine, GUIGON Aurélie, LAMBERT Valérie, LAZREK Mouna, NOBILLIAUX Florian, PREVOST Brigitte, THUILLIER Caroline, TINEZ Claire ,TCHANTCHOU NJOSSE YANICK |
| EPI_ISL_2646172, EPI_ISL_2646173 | CH ROUBAIX | CHU Lille - Laboratoire de Virologie | AIT YAHYA Emilie, ALIDJINOU Enagnon Kazali, BOCKET Laurence, CREPIN Michel, DEMAY Christophe, ENGELMANN Ilka, GEFFROY Sandrine, GUIGON Aurélie, LAMBERT Valérie, LAZREK Mouna, NOBILLIAUX Florian, PREVOST Brigitte, THUILLIER Caroline, TINEZ Claire ,TCHANTCHOU NJOSSE YANICK |
| EPI_ISL_2646174 | CH TOURCOING | CHU Lille - Laboratoire de Virologie | AIT YAHYA Emilie, ALIDJINOU Enagnon Kazali, BOCKET Laurence, CREPIN Michel, DEMAY Christophe, ENGELMANN Ilka, GEFFROY Sandrine, GUIGON Aurélie, LAMBERT Valérie, LAZREK Mouna, NOBILLIAUX Florian, PREVOST Brigitte, THUILLIER Caroline, TINEZ Claire ,TCHANTCHOU NJOSSE YANICK |
| EPI_ISL_2646175 | CH BETHUNE | CHU Lille - Laboratoire de Virologie | AIT YAHYA Emilie, ALIDJINOU Enagnon Kazali, BOCKET Laurence, CREPIN Michel, DEMAY Christophe, ENGELMANN Ilka, GEFFROY Sandrine, GUIGON Aurélie, LAMBERT Valérie, LAZREK Mouna, NOBILLIAUX Florian, PREVOST Brigitte, THUILLIER Caroline, TINEZ Claire ,TCHANTCHOU NJOSSE YANICK |
| EPI_ISL_2646177, EPI_ISL_2646178 | CHU LILLE | CHU Lille - Laboratoire de Virologie | AIT YAHYA Emilie, ALIDJINOU Enagnon Kazali, BOCKET Laurence, CREPIN Michel, DEMAY Christophe, ENGELMANN Ilka, GEFFROY Sandrine, GUIGON Aurélie, LAMBERT Valérie, LAZREK Mouna, NOBILLIAUX Florian, PREVOST Brigitte, THUILLIER Caroline, TINEZ Claire ,TCHANTCHOU NJOSSE YANICK |
| EPI_ISL_2646179 | CH ARMENTIERES | CHU Lille - Laboratoire de Virologie | AIT YAHYA Emilie, ALIDJINOU Enagnon Kazali, BOCKET Laurence, CREPIN Michel, DEMAY Christophe, ENGELMANN Ilka, GEFFROY Sandrine, GUIGON Aurélie, LAMBERT Valérie, LAZREK Mouna, NOBILLIAUX Florian, PREVOST Brigitte, THUILLIER Caroline, TINEZ Claire ,TCHANTCHOU NJOSSE YANICK |
| EPI_ISL_2646180, EPI_ISL_2646181, EPI_ISL_2646182, EPI_ISL_2646183, EPI_ISL_2646184, EPI_ISL_2646185, EPI_ISL_2646186 | CHU LILLE | CHU Lille - Laboratoire de Virologie | AIT YAHYA Emilie, ALIDJINOU Enagnon Kazali, BOCKET Laurence, CREPIN Michel, DEMAY Christophe, ENGELMANN Ilka, GEFFROY Sandrine, GUIGON Aurélie, LAMBERT Valérie, LAZREK Mouna, NOBILLIAUX Florian, PREVOST Brigitte, THUILLIER Caroline, TINEZ Claire ,TCHANTCHOU NJOSSE YANICK |
| EPI_ISL_2646187, EPI_ISL_2646188 | CH ROUBAIX | CHU Lille - Laboratoire de Virologie | AIT YAHYA Emilie, ALIDJINOU Enagnon Kazali, BOCKET Laurence, CREPIN Michel, DEMAY Christophe, ENGELMANN Ilka, GEFFROY Sandrine, GUIGON Aurélie, LAMBERT Valérie, LAZREK Mouna, NOBILLIAUX Florian, PREVOST Brigitte, THUILLIER Caroline, TINEZ Claire ,TCHANTCHOU NJOSSE YANICK |
| EPI_ISL_2646189, EPI_ISL_2646190 | CHU LILLE | CHU Lille - Laboratoire de Virologie | AIT YAHYA Emilie, ALIDJINOU Enagnon Kazali, BOCKET Laurence, CREPIN Michel, DEMAY Christophe, ENGELMANN Ilka, GEFFROY Sandrine, GUIGON Aurélie, LAMBERT Valérie, LAZREK Mouna, NOBILLIAUX Florian, PREVOST Brigitte, THUILLIER Caroline, TINEZ Claire ,TCHANTCHOU NJOSSE YANICK |
| EPI_ISL_2646192, EPI_ISL_2646193 | CH ARRAS | CHU Lille - Laboratoire de Virologie | AIT YAHYA Emilie, ALIDJINOU Enagnon Kazali, BOCKET Laurence, CREPIN Michel, DEMAY Christophe, ENGELMANN Ilka, GEFFROY Sandrine, GUIGON Aurélie, LAMBERT Valérie, LAZREK Mouna, NOBILLIAUX Florian, PREVOST Brigitte, THUILLIER Caroline, TINEZ Claire ,TCHANTCHOU NJOSSE YANICK |
| EPI_ISL_2646195, EPI_ISL_2646196 | CH ROUBAIX | CHU Lille - Laboratoire de Virologie | AIT YAHYA Emilie, ALIDJINOU Enagnon Kazali, BOCKET Laurence, CREPIN Michel, DEMAY Christophe, ENGELMANN Ilka, GEFFROY Sandrine, GUIGON Aurélie, LAMBERT Valérie, LAZREK Mouna, NOBILLIAUX Florian, PREVOST Brigitte, THUILLIER Caroline, TINEZ Claire ,TCHANTCHOU NJOSSE YANICK |
| EPI_ISL_2646197 | CH BETHUNE | CHU Lille - Laboratoire de Virologie | AIT YAHYA Emilie, ALIDJINOU Enagnon Kazali, BOCKET Laurence, CREPIN Michel, DEMAY Christophe, ENGELMANN Ilka, GEFFROY Sandrine, GUIGON Aurélie, LAMBERT Valérie, LAZREK Mouna, NOBILLIAUX Florian, PREVOST Brigitte, THUILLIER Caroline, TINEZ Claire ,TCHANTCHOU NJOSSE YANICK |
| EPI_ISL_2646198, EPI_ISL_2646199 | CH ARMENTIERES | CHU Lille - Laboratoire de Virologie | AIT YAHYA Emilie, ALIDJINOU Enagnon Kazali, BOCKET Laurence, CREPIN Michel, DEMAY Christophe, ENGELMANN Ilka, GEFFROY Sandrine, GUIGON Aurélie, LAMBERT Valérie, LAZREK Mouna, NOBILLIAUX Florian, PREVOST Brigitte, THUILLIER Caroline, TINEZ Claire ,TCHANTCHOU NJOSSE YANICK |
| EPI_ISL_2646200 | CH TOURCOING | CHU Lille - Laboratoire de Virologie | AIT YAHYA Emilie, ALIDJINOU Enagnon Kazali, BOCKET Laurence, CREPIN Michel, DEMAY Christophe, ENGELMANN Ilka, GEFFROY Sandrine, GUIGON Aurélie, LAMBERT Valérie, LAZREK Mouna, NOBILLIAUX Florian, PREVOST Brigitte, THUILLIER Caroline, TINEZ Claire ,TCHANTCHOU NJOSSE YANICK |
| EPI_ISL_2646201 | CH ARMENTIERES | CHU Lille - Laboratoire de Virologie | AIT YAHYA Emilie, ALIDJINOU Enagnon Kazali, BOCKET Laurence, CREPIN Michel, DEMAY Christophe, ENGELMANN Ilka, GEFFROY Sandrine, GUIGON Aurélie, LAMBERT Valérie, LAZREK Mouna, NOBILLIAUX Florian, PREVOST Brigitte, THUILLIER Caroline, TINEZ Claire ,TCHANTCHOU NJOSSE YANICK |
| EPI_ISL_2646202 | CH CALAIS | CHU Lille - Laboratoire de Virologie | AIT YAHYA Emilie, ALIDJINOU Enagnon Kazali, BOCKET Laurence, CREPIN Michel, DEMAY Christophe, ENGELMANN Ilka, GEFFROY Sandrine, |

| | | | |
|---|---|---|---|
| | | | GUIGON Aurélie, LAMBERT Valérie, LAZREK Mouna, NOBILLIAUX Florian, PREVOST Brigitte, THUILLIER Caroline, TINEZ Claire ,TCHANTCHOU NJOSSE YANICK |
| EPI_ISL_2646203 | CHU LILLE | CHU Lille - Laboratoire de Virologie | AIT YAHYA Emilie, ALIDJINOU Enagnon Kazali, BOCKET Laurence, CREPIN Michel, DEMAY Christophe, ENGELMANN Ilka, GEFFROY Sandrine, GUIGON Aurélie, LAMBERT Valérie, LAZREK Mouna, NOBILLIAUX Florian, PREVOST Brigitte, THUILLIER Caroline, TINEZ Claire ,TCHANTCHOU NJOSSE YANICK |
| EPI_ISL_2646204 | CH CAMBRAI | CHU Lille - Laboratoire de Virologie | AIT YAHYA Emilie, ALIDJINOU Enagnon Kazali, BOCKET Laurence, CREPIN Michel, DEMAY Christophe, ENGELMANN Ilka, GEFFROY Sandrine, GUIGON Aurélie, LAMBERT Valérie, LAZREK Mouna, NOBILLIAUX Florian, PREVOST Brigitte, THUILLIER Caroline, TINEZ Claire ,TCHANTCHOU NJOSSE YANICK |
| EPI_ISL_2646205, EPI_ISL_2646206, EPI_ISL_2646207 | CHU LILLE | CHU Lille - Laboratoire de Virologie | AIT YAHYA Emilie, ALIDJINOU Enagnon Kazali, BOCKET Laurence, CREPIN Michel, DEMAY Christophe, ENGELMANN Ilka, GEFFROY Sandrine, GUIGON Aurélie, LAMBERT Valérie, LAZREK Mouna, NOBILLIAUX Florian, PREVOST Brigitte, THUILLIER Caroline, TINEZ Claire ,TCHANTCHOU NJOSSE YANICK |
| EPI_ISL_2646208 | CH TOURCOING | CHU Lille - Laboratoire de Virologie | AIT YAHYA Emilie, ALIDJINOU Enagnon Kazali, BOCKET Laurence, CREPIN Michel, DEMAY Christophe, ENGELMANN Ilka, GEFFROY Sandrine, GUIGON Aurélie, LAMBERT Valérie, LAZREK Mouna, NOBILLIAUX Florian, PREVOST Brigitte, THUILLIER Caroline, TINEZ Claire ,TCHANTCHOU NJOSSE YANICK |
| EPI_ISL_2646210 | CH Valenciennes | CHU Lille - Laboratoire de Virologie | AIT YAHYA Emilie, ALIDJINOU Enagnon Kazali, BOCKET Laurence, CREPIN Michel, DEMAY Christophe, ENGELMANN Ilka, GEFFROY Sandrine, GUIGON Aurélie, LAMBERT Valérie, LAZREK Mouna, NOBILLIAUX Florian, PREVOST Brigitte, THUILLIER Caroline, TINEZ Claire ,TCHANTCHOU NJOSSE YANICK |
| EPI_ISL_2646211 | CHU LILLE | CHU Lille - Laboratoire de Virologie | AIT YAHYA Emilie, ALIDJINOU Enagnon Kazali, BOCKET Laurence, CREPIN Michel, DEMAY Christophe, ENGELMANN Ilka, GEFFROY Sandrine, GUIGON Aurélie, LAMBERT Valérie, LAZREK Mouna, NOBILLIAUX Florian, PREVOST Brigitte, THUILLIER Caroline, TINEZ Claire ,TCHANTCHOU NJOSSE YANICK |
| EPI_ISL_2646212 | CH Valenciennes | CHU Lille - Laboratoire de Virologie | AIT YAHYA Emilie, ALIDJINOU Enagnon Kazali, BOCKET Laurence, CREPIN Michel, DEMAY Christophe, ENGELMANN Ilka, GEFFROY Sandrine, GUIGON Aurélie, LAMBERT Valérie, LAZREK Mouna, NOBILLIAUX Florian, PREVOST Brigitte, THUILLIER Caroline, TINEZ Claire ,TCHANTCHOU NJOSSE YANICK |
| EPI_ISL_2646213 | CHU LILLE | CHU Lille - Laboratoire de Virologie | AIT YAHYA Emilie, ALIDJINOU Enagnon Kazali, BOCKET Laurence, CREPIN Michel, DEMAY Christophe, ENGELMANN Ilka, GEFFROY Sandrine, GUIGON Aurélie, LAMBERT Valérie, LAZREK Mouna, NOBILLIAUX Florian, PREVOST Brigitte, THUILLIER Caroline, TINEZ Claire ,TCHANTCHOU NJOSSE YANICK |
| EPI_ISL_2646214 | CH ARRAS | CHU Lille - Laboratoire de Virologie | AIT YAHYA Emilie, ALIDJINOU Enagnon Kazali, BOCKET Laurence, CREPIN Michel, DEMAY Christophe, ENGELMANN Ilka, GEFFROY Sandrine, GUIGON Aurélie, LAMBERT Valérie, LAZREK Mouna, NOBILLIAUX Florian, PREVOST Brigitte, THUILLIER Caroline, TINEZ Claire ,TCHANTCHOU NJOSSE YANICK |
| EPI_ISL_2646215 | CHU LILLE | CHU Lille - Laboratoire de Virologie | AIT YAHYA Emilie, ALIDJINOU Enagnon Kazali, BOCKET Laurence, CREPIN Michel, DEMAY Christophe, ENGELMANN Ilka, GEFFROY Sandrine, GUIGON Aurélie, LAMBERT Valérie, LAZREK Mouna, NOBILLIAUX Florian, PREVOST Brigitte, THUILLIER Caroline, TINEZ Claire ,TCHANTCHOU NJOSSE YANICK |
| EPI_ISL_2646216 | CH CAMBRAI | CHU Lille - Laboratoire de Virologie | AIT YAHYA Emilie, ALIDJINOU Enagnon Kazali, BOCKET Laurence, CREPIN Michel, DEMAY Christophe, ENGELMANN Ilka, GEFFROY Sandrine, GUIGON Aurélie, LAMBERT Valérie, LAZREK Mouna, NOBILLIAUX Florian, PREVOST Brigitte, THUILLIER Caroline, TINEZ Claire ,TCHANTCHOU NJOSSE YANICK |
| EPI_ISL_2646217, EPI_ISL_2646218 | CHU LILLE | CHU Lille - Laboratoire de Virologie | AIT YAHYA Emilie, ALIDJINOU Enagnon Kazali, BOCKET Laurence, CREPIN Michel, DEMAY Christophe, ENGELMANN Ilka, GEFFROY Sandrine, GUIGON Aurélie, LAMBERT Valérie, LAZREK Mouna, NOBILLIAUX Florian, PREVOST Brigitte, THUILLIER Caroline, TINEZ Claire ,TCHANTCHOU NJOSSE YANICK |
| EPI_ISL_2646219 | cerballiance-IDF | Cerba lab | Emmanuel Lecorche, Sabine Trombert-Paolantoni, Aude Lessenne, Kader Merah, Thierry Collin, Laura Verdurme, Patrice Herisson, Stéphanie Haïm-Boukobza, Bénédicte Roquebert |
| EPI_ISL_2646220, EPI_ISL_2646221, EPI_ISL_2646222, EPI_ISL_2646223 | Exalab | Cerba lab | Emmanuel Lecorche, Sabine Trombert-Paolantoni, Aude Lessenne, Kader Merah, Thierry Collin, Laura Verdurme, Patrice Herisson, Stéphanie Haïm-Boukobza, Bénédicte Roquebert |
| EPI_ISL_2646224 | cerballiance-IDF | Cerba lab | Emmanuel Lecorche, Sabine Trombert-Paolantoni, Aude Lessenne, Kader Merah, Thierry Collin, Laura Verdurme, Patrice Herisson, Stéphanie Haïm-Boukobza, Bénédicte Roquebert |
| EPI_ISL_2646225, EPI_ISL_2646226, EPI_ISL_2646227, EPI_ISL_2646228, EPI_ISL_2646229, EPI_ISL_2646230, EPI_ISL_2646231 | Exalab | Cerba lab | Emmanuel Lecorche, Sabine Trombert-Paolantoni, Aude Lessenne, Kader Merah, Thierry Collin, Laura Verdurme, Patrice Herisson, Stéphanie Haïm-Boukobza, Bénédicte Roquebert |
| EPI_ISL_2646232 | Clinique de Meudon | Cerba lab | Emmanuel Lecorche, Sabine Trombert-Paolantoni, Aude Lessenne, Kader Merah, Thierry Collin, Laura Verdurme, Patrice Herisson, Stéphanie Haïm-Boukobza, Bénédicte Roquebert |
| EPI_ISL_2646233, EPI_ISL_2646234, EPI_ISL_2646235, EPI_ISL_2646236, EPI_ISL_2646237, EPI_ISL_2646238 | Cerballiance-IDF | Cerba lab | Emmanuel Lecorche, Sabine Trombert-Paolantoni, Aude Lessenne, Kader Merah, Thierry Collin, Laura Verdurme, Patrice Herisson, Stéphanie Haïm-Boukobza, Bénédicte Roquebert |
| EPI_ISL_2646239, EPI_ISL_2646240 | Cerballiance-Provence | Cerba lab | Emmanuel Lecorche, Sabine Trombert-Paolantoni, Aude Lessenne, Kader Merah, Thierry Collin, Laura Verdurme, Patrice Herisson, Stéphanie Haïm-Boukobza, Bénédicte Roquebert |
| EPI_ISL_2646241 | Cerballiance-Normandie | Cerba lab | Emmanuel Lecorche, Sabine Trombert-Paolantoni, Aude Lessenne, Kader Merah, Thierry Collin, Laura Verdurme, Patrice Herisson, Stéphanie Haïm-Boukobza, Bénédicte Roquebert |
| EPI_ISL_2646242 | Cerballiance-IDF | Cerba lab | Emmanuel Lecorche, Sabine Trombert-Paolantoni, Aude Lessenne, Kader Merah, Thierry Collin, Laura Verdurme, Patrice Herisson, Stéphanie Haïm-Boukobza, Bénédicte Roquebert |
| EPI_ISL_2646243 | Cerballiance-Provence | Cerba lab | Emmanuel Lecorche, Sabine Trombert-Paolantoni, Aude Lessenne, Kader Merah, Thierry Collin, Laura Verdurme, Patrice Herisson, Stéphanie Haïm-Boukobza, Bénédicte Roquebert |
| EPI_ISL_2646245, EPI_ISL_2646246 | Cerballiance-Paris | Cerba lab | Emmanuel Lecorche, Sabine Trombert-Paolantoni, Aude Lessenne, Kader Merah, Thierry Collin, Laura Verdurme, Patrice Herisson, Stéphanie Haïm-Boukobza, Bénédicte Roquebert |
| EPI_ISL_2646247 | Cerballiance-Normandie | Cerba lab | Emmanuel Lecorche, Sabine Trombert-Paolantoni, Aude Lessenne, Kader Merah, Thierry Collin, Laura Verdurme, Patrice Herisson, Stéphanie Haïm-Boukobza, Bénédicte Roquebert |
| EPI_ISL_2646248, EPI_ISL_2646249, EPI_ISL_2646250 | Cerballiance-IDF | Cerba lab | Emmanuel Lecorche, Sabine Trombert-Paolantoni, Aude Lessenne, Kader Merah, Thierry Collin, Laura Verdurme, Patrice Herisson, Stéphanie Haïm-Boukobza, Bénédicte Roquebert |
| EPI_ISL_2646251 | Synlab | Cerba lab | Emmanuel Lecorche, Sabine Trombert-Paolantoni, Aude Lessenne, Kader Merah, Thierry Collin, Laura Verdurme, Patrice Herisson, Stéphanie Haïm-Boukobza, Bénédicte Roquebert |
| EPI_ISL_2646252 | Cerballiance-Paris | Cerba lab | Emmanuel Lecorche, Sabine Trombert-Paolantoni, Aude Lessenne, Kader Merah, Thierry Collin, Laura Verdurme, Patrice Herisson, Stéphanie Haïm-Boukobza, Bénédicte Roquebert |
| EPI_ISL_2646253, EPI_ISL_2646254 | Cerballiance-Pyrénées | Cerba lab | Emmanuel Lecorche, Sabine Trombert-Paolantoni, Aude Lessenne, Kader Merah, Thierry Collin, Laura Verdurme, Patrice Herisson, Stéphanie Haïm-Boukobza, Bénédicte Roquebert |

| | | | |
|---|---|---|---|
| EPI_ISL_2646255 | Cerballiance-Paris | Cerba lab | Emmanuel Lecorche, Sabine Trombert-Paolantoni, Aude Lessenne, Kader Merah, Thierry Collin, Laura Verdurme, Patrice Herisson, Stéphanie Haïm-Boukobza, Bénédicte Roquebert |
| EPI_ISL_2646256 | Cerballiance-Centre val de Loire | Cerba lab | Emmanuel Lecorche, Sabine Trombert-Paolantoni, Aude Lessenne, Kader Merah, Thierry Collin, Laura Verdurme, Patrice Herisson, Stéphanie Haïm-Boukobza, Bénédicte Roquebert |
| EPI_ISL_2646257 | Cerballiance-Pyrénées | Cerba lab | Emmanuel Lecorche, Sabine Trombert-Paolantoni, Aude Lessenne, Kader Merah, Thierry Collin, Laura Verdurme, Patrice Herisson, Stéphanie Haïm-Boukobza, Bénédicte Roquebert |
| EPI_ISL_2646258 | Cerballiance-Paris | Cerba lab | Emmanuel Lecorche, Sabine Trombert-Paolantoni, Aude Lessenne, Kader Merah, Thierry Collin, Laura Verdurme, Patrice Herisson, Stéphanie Haïm-Boukobza, Bénédicte Roquebert |
| EPI_ISL_2646259 | COSEM | Cerba lab | Emmanuel Lecorche, Sabine Trombert-Paolantoni, Aude Lessenne, Kader Merah, Thierry Collin, Laura Verdurme, Patrice Herisson, Stéphanie Haïm-Boukobza, Bénédicte Roquebert |
| EPI_ISL_2646260 | Cerballiance-IDF | Cerba lab | Emmanuel Lecorche, Sabine Trombert-Paolantoni, Aude Lessenne, Kader Merah, Thierry Collin, Laura Verdurme, Patrice Herisson, Stéphanie Haïm-Boukobza, Bénédicte Roquebert |
| EPI_ISL_2646261 | Cerballiance-Paris | Cerba lab | Emmanuel Lecorche, Sabine Trombert-Paolantoni, Aude Lessenne, Kader Merah, Thierry Collin, Laura Verdurme, Patrice Herisson, Stéphanie Haïm-Boukobza, Bénédicte Roquebert |
| EPI_ISL_2646262, EPI_ISL_2646263, EPI_ISL_2646264 | Cerballiance-IDF | Cerba lab | Emmanuel Lecorche, Sabine Trombert-Paolantoni, Aude Lessenne, Kader Merah, Thierry Collin, Laura Verdurme, Patrice Herisson, Stéphanie Haïm-Boukobza, Bénédicte Roquebert |
| EPI_ISL_2646265 | Cerballiance-Paris | Cerba lab | Emmanuel Lecorche, Sabine Trombert-Paolantoni, Aude Lessenne, Kader Merah, Thierry Collin, Laura Verdurme, Patrice Herisson, Stéphanie Haïm-Boukobza, Bénédicte Roquebert |
| EPI_ISL_2646266 | Cerballiance-IDF | Cerba lab | Emmanuel Lecorche, Sabine Trombert-Paolantoni, Aude Lessenne, Kader Merah, Thierry Collin, Laura Verdurme, Patrice Herisson, Stéphanie Haïm-Boukobza, Bénédicte Roquebert |
| EPI_ISL_2646267 | Cerballiance-Paris | Cerba lab | Emmanuel Lecorche, Sabine Trombert-Paolantoni, Aude Lessenne, Kader Merah, Thierry Collin, Laura Verdurme, Patrice Herisson, Stéphanie Haïm-Boukobza, Bénédicte Roquebert |
| EPI_ISL_2646268 | Unilabs | Cerba lab | Emmanuel Lecorche, Sabine Trombert-Paolantoni, Aude Lessenne, Kader Merah, Thierry Collin, Laura Verdurme, Patrice Herisson, Stéphanie Haïm-Boukobza, Bénédicte Roquebert |
| EPI_ISL_2646269 | Cerballiance-IDF | Cerba lab | Emmanuel Lecorche, Sabine Trombert-Paolantoni, Aude Lessenne, Kader Merah, Thierry Collin, Laura Verdurme, Patrice Herisson, Stéphanie Haïm-Boukobza, Bénédicte Roquebert |
| EPI_ISL_2646270 | Cerballiance-HDF | Cerba lab | Emmanuel Lecorche, Sabine Trombert-Paolantoni, Aude Lessenne, Kader Merah, Thierry Collin, Laura Verdurme, Patrice Herisson, Stéphanie Haïm-Boukobza, Bénédicte Roquebert |
| EPI_ISL_2646271 | Kuate | Cerba lab | Emmanuel Lecorche, Sabine Trombert-Paolantoni, Aude Lessenne, Kader Merah, Thierry Collin, Laura Verdurme, Patrice Herisson, Stéphanie Haïm-Boukobza, Bénédicte Roquebert |
| EPI_ISL_2646272 | Hopital Max Fourestier | Cerba lab | Emmanuel Lecorche, Sabine Trombert-Paolantoni, Aude Lessenne, Kader Merah, Thierry Collin, Laura Verdurme, Patrice Herisson, Stéphanie Haïm-Boukobza, Bénédicte Roquebert |
| EPI_ISL_2646273 | Laboratoire Frechin | Cerba lab | Emmanuel Lecorche, Sabine Trombert-Paolantoni, Aude Lessenne, Kader Merah, Thierry Collin, Laura Verdurme, Patrice Herisson, Stéphanie Haïm-Boukobza, Bénédicte Roquebert |
| EPI_ISL_2646274 | Cerballiance-Provence | Cerba lab | Emmanuel Lecorche, Sabine Trombert-Paolantoni, Aude Lessenne, Kader Merah, Thierry Collin, Laura Verdurme, Patrice Herisson, Stéphanie Haïm-Boukobza, Bénédicte Roquebert |
| EPI_ISL_2646275, EPI_ISL_2646276 | Cerballiance-Paris | Cerba lab | Emmanuel Lecorche, Sabine Trombert-Paolantoni, Aude Lessenne, Kader Merah, Thierry Collin, Laura Verdurme, Patrice Herisson, Stéphanie Haïm-Boukobza, Bénédicte Roquebert |
| EPI_ISL_2646277 | Cerballiance-Provence | Cerba lab | Emmanuel Lecorche, Sabine Trombert-Paolantoni, Aude Lessenne, Kader Merah, Thierry Collin, Laura Verdurme, Patrice Herisson, Stéphanie Haïm-Boukobza, Bénédicte Roquebert |
| EPI_ISL_2646278 | Cerballiance-Paris | Cerba lab | Emmanuel Lecorche, Sabine Trombert-Paolantoni, Aude Lessenne, Kader Merah, Thierry Collin, Laura Verdurme, Patrice Herisson, Stéphanie Haïm-Boukobza, Bénédicte Roquebert |
| EPI_ISL_2646279 | Cerballiance-IDF | Cerba lab | Emmanuel Lecorche, Sabine Trombert-Paolantoni, Aude Lessenne, Kader Merah, Thierry Collin, Laura Verdurme, Patrice Herisson, Stéphanie Haïm-Boukobza, Bénédicte Roquebert |
| EPI_ISL_2646280 | Ch Dourdan | Cerba lab | Emmanuel Lecorche, Sabine Trombert-Paolantoni, Aude Lessenne, Kader Merah, Thierry Collin, Laura Verdurme, Patrice Herisson, Stéphanie Haïm-Boukobza, Bénédicte Roquebert |
| EPI_ISL_2646281 | Hopital Max Fourestier | Cerba lab | Emmanuel Lecorche, Sabine Trombert-Paolantoni, Aude Lessenne, Kader Merah, Thierry Collin, Laura Verdurme, Patrice Herisson, Stéphanie Haïm-Boukobza, Bénédicte Roquebert |
| EPI_ISL_2646285 | CHU de la Réunion - Sud | Laboratoire de virologie, CNR arbovirus Associé, Chu de la Réunion | Etienne Frumence, Marie-Christine Jaffar Bandjee, Nicolas Traversier, Sabrina Petit Genet |
| EPI_ISL_2646288 | CHU de la Réunion - Nord | Laboratoire de virologie, CNR arbovirus Associé, Chu de la Réunion | Etienne Frumence, Marie-Christine Jaffar Bandjee, Nicolas Traversier, Sabrina Petit Genet |
| EPI_ISL_2646289, EPI_ISL_2646292, EPI_ISL_2646293, EPI_ISL_2646297 | CHU de la Réunion - Sud | Laboratoire de virologie, CNR arbovirus Associé, Chu de la Réunion | Etienne Frumence, Marie-Christine Jaffar Bandjee, Nicolas Traversier, Sabrina Petit Genet |
| EPI_ISL_2646298 | CHU de la Réunion - Nord | Laboratoire de virologie, CNR arbovirus Associé, Chu de la Réunion | Etienne Frumence, Marie-Christine Jaffar Bandjee, Nicolas Traversier, Sabrina Petit Genet |
| EPI_ISL_2648066, EPI_ISL_2648067, EPI_ISL_2648084, EPI_ISL_2648085, EPI_ISL_2648086, EPI_ISL_2648087, EPI_ISL_2648088, EPI_ISL_2648089, EPI_ISL_2648090, EPI_ISL_2648091, EPI_ISL_2648092, EPI_ISL_2648093, EPI_ISL_2648094, EPI_ISL_2648095, EPI_ISL_2648096, EPI_ISL_2648097, EPI_ISL_2648098, EPI_ISL_2648099, EPI_ISL_2648100, EPI_ISL_2648101, EPI_ISL_2648102, EPI_ISL_2648103, EPI_ISL_2648104, EPI_ISL_2648105, EPI_ISL_2648106, EPI_ISL_2648107, EPI_ISL_2648108, EPI_ISL_2648109 | | | |
| see above | National Center of Infectious and Parasitic Diseases | National Center of Infectious and Parasitic Diseases | Alexiev et al |
| EPI_ISL_2648115, EPI_ISL_2648116, EPI_ISL_2648117, EPI_ISL_2648118 | Instituto Nacional de Enfermedades Respiratorias (INER) | Centro de Investigación en Enfermedades Infecciosas (CIENI), Instituto Nacional de Enfermedades Respiratorias (INER) | Consorcio Mexicano de Vigilancia Genómica (CoViGen-Mex). Authors (in alphabetical order): Julio Elias Alvarado-Yaah, Carlos F. Arias, Santiago Ávila-Ríos, Eduardo Becerril-Vargas, Víctor Hugo Borja-Aburto, Celia Boukadida, Cristóbal Cháidez-Quiróz, Juan Bautista Chale-Dzul, Ricardo Ciria Merce, Andreu Comas-García, Célida Duque Molina, Julissa Enciso-Ibarra, José Antonio Enciso-Moreno, Gloria Elena Espinosa-Ayala, Fernando Fontove-Herrera, Daniel Fregoso-Rueda, Victor Eduardo Garcia-Arias, Alejandra Garcia-Gasca, Bruno Gomez-Gil, Jean Pierre González, Irvin González-López, Concepción Grajales-Muñiz, Ricardo Grande, Rosa María Gutierrez Rios, Alejandra Hernández-Terán, Alfredo Herrera-Estrella, Carla Ivón Herrera-Najera, Pavel Isa, Daniel Lira Morales, Susana Lopez, Antonio Loza Román, Brenda Irasema Maldonado-Meza, Bernardo Martínez-Miguel, José Arturo Martínez-Orozco, Célida Martínez- Rodríguez, Margarita Matías-Florentino, Fidencio Mejía-Nepomuceno, María Guadalupe de Jesús Mireles-Rivera, Gloria María Molina-Salinas, Hector Montoya-Fuentes, , Mario Mújica-Sánchez, José Esteban Muñoz-Medina, José de Jesús Nuñez-Contreras, Alicia Ocaña-Mondragón, Luis Alberto Ochoa-Carrera, Hector Esteban Paz-Juárez, Marissa Pérez-García, Francisco Pulido, Helen Haydee Fernanda Ramírez-Plascencia, Jorge Salas-Hernández, Angel Gustavo Salas-Lais, Alejandro Sanchez-Flores, Clara Esperanza Santacruz-Tinoco, María Guadalupe Santiago-Mauricio, Selene Zárate, Nelly Sélem-Mojica, Blanca Taboada, Gloria Vazquez, Joel Armando Vázquez-Pérez. |
| EPI_ISL_2648206 | Debswana Orapa Mine Hospital Laboratory | Botswana Harvard HIV Reference Laboratory | Sikhulile Moyo, Wonderful T. Choga, Dorcas Maruapula, Keoratile Ntshambiwa, Lesedi Magama, Mpo Molapisi, Koketso Maotwe, Thongbotho |

| | | | |
|---|---|---|---|
| | | | Mphoyakgosi, Boitumelo Zuze, Botshelo Radibe, Legodile Kooepile, Ontlametse T. Bareng, Roger Shapiro, Shahin Lockman, Joseph Makhema, Madisa Mine, Mosepele Mosepele, Simani Gaseitsiwe |
| EPI_ISL_2648221, EPI_ISL_2648223, EPI_ISL_2648224, EPI_ISL_2648226, EPI_ISL_2648235, EPI_ISL_2648236, EPI_ISL_2648237, EPI_ISL_2648239 | Kasane Primary Hospital Laboratory | Botswana Harvard HIV Reference Laboratory | Sikhulile Moyo, Wonderful T. Choga, Dorcas Maruapula, Agnes Karutwaeng, Rose Munyere, Keoratile Ntshambiwa, Thongbotho Mphoyakgosi, Boitumelo Zuze, Botshelo Radibe, Legodile Kooepile, Ontlametse T. Bareng, Roger Shapiro, Shahin Lockman, Joseph Makhema, Madisa Mine, Mosepele Mosepele, Simani Gaseitsiwe |
| EPI_ISL_2649991 | Kudat Hospital | Institute for Medical Research, Infectious Disease Research Centre, National Institutes of Health, Ministry of Health Malaysia | Suppiah J, Kamel K, Mohd Zawawi Z, Azizan MA, Ramly N, Robert F, Thayan R |
| EPI_ISL_2649992 | Tuanku Ampuan Najihah Hospital, Kuala Pilah | Institute for Medical Research, Infectious Disease Research Centre, National Institutes of Health, Ministry of Health Malaysia | Suppiah J, Kamel K, Mohd Zawawi Z, Azizan MA, Ramly N, Robert F, Thayan R |
| EPI_ISL_2649994 | Tuanku Fauziah Hopsital | Institute for Medical Research, Infectious Disease Research Centre, National Institutes of Health, Ministry of Health Malaysia | Suppiah J, Kamel K, Mohd Zawawi Z, Azizan MA, Ramly N, Robert F, Thayan R |
| EPI_ISL_2649995 | Segamat Hospital | Institute for Medical Research, Infectious Disease Research Centre, National Institutes of Health, Ministry of Health Malaysia | Suppiah J, Kamel K, Mohd Zawawi Z, Azizan MA, Ramly N, Robert F, Thayan R |
| EPI_ISL_2649996 | Johor Bahru Public Health Laboratory | Institute for Medical Research, Infectious Disease Research Centre, National Institutes of Health, Ministry of Health Malaysia | Suppiah J, Kamel K, Mohd Zawawi Z, Azizan MA, Ramly N, Robert F, Thayan R |
| EPI_ISL_2649997 | Sultan Ismail Hospital | Institute for Medical Research, Infectious Disease Research Centre, National Institutes of Health, Ministry of Health Malaysia | Suppiah J, Kamel K, Mohd Zawawi Z, Azizan MA, Ramly N, Robert F, Thayan R |
| EPI_ISL_2649998, EPI_ISL_2649999 | Penang General Hospital | Institute for Medical Research, Infectious Disease Research Centre, National Institutes of Health, Ministry of Health Malaysia | Suppiah J, Kamel K, Mohd Zawawi Z, Azizan MA, Ramly N, Robert F, Thayan R |
| EPI_ISL_2650001 | Hospital Sultanah Nur Zahirah Kuala Terengganu | Institute for Medical Research, Infectious Disease Research Centre, National Institutes of Health, Ministry of Health Malaysia | Suppiah J, Kamel K, Mohd Zawawi Z, Azizan MA, Ramly N, Robert F, Thayan R |
| EPI_ISL_2650003 | Hospital Wanita Dan Kanak-Kanak Sabah | Institute for Medical Research, Infectious Disease Research Centre, National Institutes of Health, Ministry of Health Malaysia | Jalan Setia Murni U13/52, Bandar Setia Alam, 40170 Selangor, Malaysia |
| EPI_ISL_2650004, EPI_ISL_2650005 | Sungai Buloh Hospital | Institute for Medical Research, Infectious Disease Research Centre, National Institutes of Health, Ministry of Health Malaysia | Suppiah J, Kamel K, Mohd Zawawi Z, Azizan MA, Ramly N, Robert F, Thayan R |
| EPI_ISL_2650006 | Tumpat Hospital | Institute for Medical Research, Infectious Disease Research Centre, National Institutes of Health, Ministry of Health Malaysia | Suppiah J, Kamel K, Mohd Zawawi Z, Azizan MA, Ramly N, Robert F, Thayan R |
| EPI_ISL_2650007 | Ipoh Public Health Laboratory (MKAI), Ministry of Health Malaysia | Institute for Medical Research, Infectious Disease Research Centre, National Institutes of Health, Ministry of Health Malaysia | Suppiah J, Kamel K, Mohd Zawawi Z, Azizan MA, Ramly N, Robert F, Thayan R |
| EPI_ISL_2650010 | Sungai Buloh Hospital | Institute for Medical Research, Infectious Disease Research Centre, National Institutes of Health, Ministry of Health Malaysia | Suppiah J, Kamel K, Mohd Zawawi Z, Azizan MA, Ramly N, Robert F, Thayan R |
| EPI_ISL_2650015 | Sultanah Bahiyah Hospital | Institute for Medical Research, Infectious Disease Research Centre, National Institutes of Health, Ministry of Health Malaysia | Suppiah J, Kamel K, Mohd Zawawi Z, Azizan MA, Ramly N, Robert F, Thayan R |
| EPI_ISL_2650027 | Queen Elizabeth Hospital | Institute for Medical Research, Infectious Disease Research Centre, National Institutes of Health, Ministry of Health Malaysia | Suppiah J, Kamel K, Mohd Zawawi Z, Azizan MA, Ramly N, Robert F, Thayan R |
| EPI_ISL_2650256 | Johor Bahru Public Health Laboratory | Institute for Medical Research, Infectious Disease Research Centre, National Institutes of Health, Ministry of Health Malaysia | Suppiah J, Kamel K, Mohd Zawawi Z, Azizan MA, Ramly N, Robert F, Thayan R |
| EPI_ISL_2650430, EPI_ISL_2650432, EPI_ISL_2650499, EPI_ISL_2650505 | Queen Elizabeth Hospital | Institute for Medical Research, Infectious Disease Research Centre, National Institutes of Health, Ministry of Health Malaysia | Suppiah J, Kamel K, Mohd Zawawi Z, Azizan MA, Ramly N, Robert F, Thayan R |
| EPI_ISL_2650506 | Tawau Hospital | Institute for Medical Research, Infectious Disease Research Centre, National Institutes of Health, Ministry of Health Malaysia | Jalan Setia Murni U13/52, Bandar Setia Alam, 40170 Selangor, Malaysia |
| EPI_ISL_2650507, EPI_ISL_2650508, EPI_ISL_2650510, EPI_ISL_2650511, EPI_ISL_2650512, EPI_ISL_2650513, EPI_ISL_2650514, EPI_ISL_2650515, EPI_ISL_2650516, EPI_ISL_2650517, EPI_ISL_2650518, EPI_ISL_2650519, EPI_ISL_2650521 | | | |
| see above | University of Oregon COVID-19 MAP Laboratory | University of Oregon Genomics and Cell Characterization Core Facility (GC3F) | Douglas Turnbull, Ariana White, Jeff Bishop, Jason Carriere, Jason Sydes, Peter Batzel |
| EPI_ISL_2651115, EPI_ISL_2651116 | Temporary Specimen Collection Centre at the AsiaWorld-Expo | Hong Kong Department of Health | Alan K.L. Tsang, Peter C.W. Yip, Ken H.L. Ng, Edman T.K. Lam, Rickjason C.W. Chan, Dominic N.C. Tsang |
| EPI_ISL_2651117 | North Lantau Hospital | Hong Kong Department of Health | Alan K.L. Tsang, Peter C.W. Yip, Ken H.L. Ng, Edman T.K. Lam, Rickjason C.W. Chan, Dominic N.C. Tsang |
| EPI_ISL_2651118 | HOME QUARANTINE TASKFORCE | Hong Kong Department of Health | Alan K.L. Tsang, Peter C.W. Yip, Ken H.L. Ng, Edman T.K. Lam, Rickjason C.W. Chan, Dominic N.C. Tsang |
| EPI_ISL_2651119, EPI_ISL_2651120, EPI_ISL_2651121 | Temporary Specimen Collection Centre at the AsiaWorld-Expo | Hong Kong Department of Health | Alan K.L. Tsang, Peter C.W. Yip, Ken H.L. Ng, Edman T.K. Lam, Rickjason C.W. Chan, Dominic N.C. Tsang |
| EPI_ISL_2651122 | QUARANTINE CAMP | Hong Kong Department of Health | Alan K.L. Tsang, Peter C.W. Yip, Ken H.L. Ng, Edman T.K. Lam, Rickjason C.W. Chan, Dominic N.C. Tsang |
| EPI_ISL_2651123 | Temporary Specimen Collection Centre at the AsiaWorld-Expo | Hong Kong Department of Health | Alan K.L. Tsang, Peter C.W. Yip, Ken H.L. Ng, Edman T.K. Lam, Rickjason C.W. Chan, Dominic N.C. Tsang |
| EPI_ISL_2651124 | North Lantau Hospital | Hong Kong Department of Health | Alan K.L. Tsang, Peter C.W. Yip, Ken H.L. Ng, Edman T.K. Lam, Rickjason C.W. Chan, Dominic N.C. Tsang |
| EPI_ISL_2651125 | YAN CHAI HOSPITAL | Hong Kong Department of Health | Alan K.L. Tsang, Peter C.W. Yip, Ken H.L. Ng, Edman T.K. Lam, Rickjason C.W. Chan, Dominic N.C. Tsang |
| EPI_ISL_2651127 | TUEN MUN HOSPITAL | Hong Kong Department of Health | Alan K.L. Tsang, Peter C.W. Yip, Ken H.L. Ng, Edman T.K. Lam, Rickjason C.W. Chan, Dominic N.C. Tsang |

| | | | |
|---|---|---|---|
| EPI_ISL_2651128, EPI_ISL_2651129, EPI_ISL_2651130 | North Lantau Hospital | Hong Kong Department of Health | Alan K.L. Tsang, Peter C.W. Yip, Ken H.L. Ng, Edman T.K. Lam, Rickjason C.W. Chan, Dominic N.C. Tsang |
| EPI_ISL_2651131 | HOME QUARANTINE TASKFORCE | Hong Kong Department of Health | Alan K.L. Tsang, Peter C.W. Yip, Ken H.L. Ng, Edman T.K. Lam, Rickjason C.W. Chan, Dominic N.C. Tsang |
| EPI_ISL_2651229 | LABORATORIO HEMATOLOGICO | Universidad Nacional de Colombia - Laboratorio Genómico One Health | Simón Villegas Velásquez, Andres F. Cardona-Rios, Maria Stella López, Daniel O. Maldonado-Perez, Lina M. Hurtado, Laura Silvana Perez, Karl A Ciuoderis, Maria Angélica Maya, Carolina Muñoz-Arango, Celeny Ortiz, Rita Almanza Payares, Idabely Betancur Ortiz, Sandra Ines Cano, Diego A. Álvarez-Díaz, Carlos Franco-Muñoz, Katherine Laiton-Donato, Hector Alejandro Ruiz-Moreno, Marcela Mercado-Reyes, Jorge E. Osorio, Juan P. Hernandez-Ortiz |
| EPI_ISL_2651236 | AOU Policlinico "G. Rodolico- S. Marco" PO G. Rodolico | AOU Policlinico "G. Rodolico- S. Marco" PO G. Rodolico and Biometec UNICT | Stefani and Scalia |
| EPI_ISL_2651266, EPI_ISL_2651267 | CHU de la Réunion - Nord | Laboratoire de virologie, CNR arbovirus Associé, Chu de la Réunion | Etienne Frumence, Marie-Christine Jaffar Bandjee, Nicolas Traversier, Sabrina Petit Genet |
| EPI_ISL_2651270 | CHU de la Réunion - Sud | Laboratoire de virologie, CNR arbovirus Associé, Chu de la Réunion | Etienne Frumence, Marie-Christine Jaffar Bandjee, Nicolas Traversier, Sabrina Petit Genet |
| EPI_ISL_2651272 | CHU de la Réunion - Nord | Laboratoire de virologie, CNR arbovirus Associé, Chu de la Réunion | Etienne Frumence, Marie-Christine Jaffar Bandjee, Nicolas Traversier, Sabrina Petit Genet |
| EPI_ISL_2651273, EPI_ISL_2651274, EPI_ISL_2651275, EPI_ISL_2651276, EPI_ISL_2651277 | CHU de la Réunion - Sud | Laboratoire de virologie, CNR arbovirus Associé, Chu de la Réunion | Etienne Frumence, Marie-Christine Jaffar Bandjee, Nicolas Traversier, Sabrina Petit Genet |
| EPI_ISL_2651279 | CHU de la Réunion - Nord | Laboratoire de virologie, CNR arbovirus Associé, Chu de la Réunion | Etienne Frumence, Marie-Christine Jaffar Bandjee, Nicolas Traversier, Sabrina Petit Genet |
| EPI_ISL_2651365, EPI_ISL_2651366 | CHU de la Réunion - Sud | Laboratoire de virologie, CNR arbovirus Associé, Chu de la Réunion | Etienne Frumence, Marie-Christine Jaffar Bandjee, Nicolas Traversier, Sabrina Petit Genet |
| EPI_ISL_2651367, EPI_ISL_2651368 | CHU de la Réunion - Nord | Laboratoire de virologie, CNR arbovirus Associé, Chu de la Réunion | Etienne Frumence, Marie-Christine Jaffar Bandjee, Nicolas Traversier, Sabrina Petit Genet |
| EPI_ISL_2651370, EPI_ISL_2651371, EPI_ISL_2651372, EPI_ISL_2651373, EPI_ISL_2651374 | CHU de la Réunion - Sud | Laboratoire de virologie, CNR arbovirus Associé, Chu de la Réunion | Etienne Frumence, Marie-Christine Jaffar Bandjee, Nicolas Traversier, Sabrina Petit Genet |
| EPI_ISL_2651376 | CHU de la Réunion - Nord | Laboratoire de virologie, CNR arbovirus Associé, Chu de la Réunion | Etienne Frumence, Marie-Christine Jaffar Bandjee, Nicolas Traversier, Sabrina Petit Genet |
| EPI_ISL_2651378, EPI_ISL_2651379, EPI_ISL_2651380, EPI_ISL_2651381, EPI_ISL_2651382 | CHU de la Réunion - Sud | Laboratoire de virologie, CNR arbovirus Associé, Chu de la Réunion | Etienne Frumence, Marie-Christine Jaffar Bandjee, Nicolas Traversier, Sabrina Petit Genet |
| EPI_ISL_2651386, EPI_ISL_2651387, EPI_ISL_2651388, EPI_ISL_2651389, EPI_ISL_2651390 | CHU de la Réunion - Nord | Laboratoire de virologie, CNR arbovirus Associé, Chu de la Réunion | Etienne Frumence, Marie-Christine Jaffar Bandjee, Nicolas Traversier, Sabrina Petit Genet |
| EPI_ISL_2651391, EPI_ISL_2651392 | CHU de la Réunion - Sud | Laboratoire de virologie, CNR arbovirus Associé, Chu de la Réunion | Etienne Frumence, Marie-Christine Jaffar Bandjee, Nicolas Traversier, Sabrina Petit Genet |
| EPI_ISL_2651395, EPI_ISL_2651396, EPI_ISL_2651397, EPI_ISL_2651399 | CHU de la Réunion - Nord | Laboratoire de virologie, CNR arbovirus Associé, Chu de la Réunion | Etienne Frumence, Marie-Christine Jaffar Bandjee, Nicolas Traversier, Sabrina Petit Genet |
| EPI_ISL_2651402, EPI_ISL_2651403, EPI_ISL_2651404, EPI_ISL_2651405, EPI_ISL_2651406 | CHU de la Réunion - Sud | Laboratoire de virologie, CNR arbovirus Associé, Chu de la Réunion | Etienne Frumence, Marie-Christine Jaffar Bandjee, Nicolas Traversier, Sabrina Petit Genet |
| EPI_ISL_2652114, EPI_ISL_2652115, EPI_ISL_2652116 | Cerballiance-IDF | Cerba lab | Emmanuel Lecorche, Sabine Trombert-Paolantoni, Aude Lessenne, Kader Merah, Thierry Collin, Laura Verdurme, Patrice Herisson, Stéphanie Haïm-Boukobza, Bénédicte Roquebert |
| EPI_ISL_2652120, EPI_ISL_2652122, EPI_ISL_2652151, EPI_ISL_2652152, EPI_ISL_2652154, EPI_ISL_2652155, EPI_ISL_2652157, EPI_ISL_2652158, EPI_ISL_2652160, EPI_ISL_2652161, EPI_ISL_2652162, EPI_ISL_2652163 | | | |
| see above | AZ Klina | AZ Klina | Carl Vael - Lynsey Berckmans |
| EPI_ISL_2652165, EPI_ISL_2652166, EPI_ISL_2652167, EPI_ISL_2652168, EPI_ISL_2652169, EPI_ISL_2652170, EPI_ISL_2652171 | Cantacuzino National Military-Medical Institute, Viral Respiratory Infections Laboratory | Cantacuzino Institute Virology | Carmen Cherciu, Luiza Ustea, Nicoleta Paraschiv, Sorin Dinu, Mihaela Oprea, Mihaela Lazar |
| EPI_ISL_2652172, EPI_ISL_2652173, EPI_ISL_2652174, EPI_ISL_2652175, EPI_ISL_2652176, EPI_ISL_2652177, EPI_ISL_2652178, EPI_ISL_2652179, EPI_ISL_2652180, EPI_ISL_2652181, EPI_ISL_2652182, EPI_ISL_2652183, EPI_ISL_2652184, EPI_ISL_2652185, EPI_ISL_2652186, EPI_ISL_2652187, EPI_ISL_2652188, EPI_ISL_2652189, EPI_ISL_2652190, EPI_ISL_2652191, EPI_ISL_2652192 | | | |
| see above | Infectious diseases Clinical Hospital, Cluj Napoca | Cantacuzino Institute Virology | Carmen Cherciu, Luiza Ustea, Nicoleta Paraschiv, Sorin Dinu, Mihaela Oprea, Mihaela Lazar |
| EPI_ISL_2652194, EPI_ISL_2652195, EPI_ISL_2652196, EPI_ISL_2652197, EPI_ISL_2652198, EPI_ISL_2652199, EPI_ISL_2652200, EPI_ISL_2652201, EPI_ISL_2652202, EPI_ISL_2652203 | Cantacuzino National Military-Medical Institute, Viral Respiratory Infections Laboratory | Cantacuzino Institute Virology | Carmen Cherciu, Luiza Ustea, Nicoleta Paraschiv, Sorin Dinu, Mihaela Oprea, Mihaela Lazar |
| EPI_ISL_2652204, EPI_ISL_2652205, EPI_ISL_2652206, EPI_ISL_2652207, EPI_ISL_2652208, EPI_ISL_2652209, EPI_ISL_2652210, EPI_ISL_2652211, EPI_ISL_2652212, EPI_ISL_2652213 | Clinical Emergency County Hospital, Craiova | Cantacuzino Institute Virology | Carmen Cherciu, Luiza Ustea, Nicoleta Paraschiv, Sorin Dinu, Mihaela Oprea, Mihaela Lazar |
| EPI_ISL_2652214 | Cantacuzino National Military-Medical Institute, Viral Respiratory Infections Laboratory | Cantacuzino Institute Virology | Carmen Cherciu, Luiza Ustea, Nicoleta Paraschiv, Sorin Dinu, Mihaela Oprea, Mihaela Lazar |
| EPI_ISL_2652215 | Clinical Emergency County Hospital, Craiova | Cantacuzino Institute Virology | Carmen Cherciu, Luiza Ustea, Nicoleta Paraschiv, Sorin Dinu, Mihaela Oprea, Mihaela Lazar |
| EPI_ISL_2652216, EPI_ISL_2652217 | Cantacuzino National Military-Medical Institute, Viral Respiratory Infections Laboratory | Cantacuzino Institute Virology | Carmen Cherciu, Luiza Ustea, Nicoleta Paraschiv, Sorin Dinu, Mihaela Oprea, Mihaela Lazar |
| EPI_ISL_2652218, EPI_ISL_2652219, EPI_ISL_2652242, EPI_ISL_2652243, EPI_ISL_2652244, EPI_ISL_2652245, EPI_ISL_2652246, EPI_ISL_2652247, EPI_ISL_2652248 | AZ Klina | AZ Klina | Carl Vael - Lynsey Berckmans |

| | | | |
|---|---|---|---|
| EPI_ISL_2657376, EPI_ISL_2657377, EPI_ISL_2657379, EPI_ISL_2657381, EPI_ISL_2657382, EPI_ISL_2657385, EPI_ISL_2657386, EPI_ISL_2657387, EPI_ISL_2657390, EPI_ISL_2657391, EPI_ISL_2657395, EPI_ISL_2657396, EPI_ISL_2657399, EPI_ISL_2657400, EPI_ISL_2657401, EPI_ISL_2657403, EPI_ISL_2657404, EPI_ISL_2657406, EPI_ISL_2657407, EPI_ISL_2657408, EPI_ISL_2657409, EPI_ISL_2657410, EPI_ISL_2657411, EPI_ISL_2657412, EPI_ISL_2657413, EPI_ISL_2657414, EPI_ISL_2657415, EPI_ISL_2657416, EPI_ISL_2657417, EPI_ISL_2657419, EPI_ISL_2657420, EPI_ISL_2657421, EPI_ISL_2657422, EPI_ISL_2657423, EPI_ISL_2657424, EPI_ISL_2657425, EPI_ISL_2657426, EPI_ISL_2657429, EPI_ISL_2657430, EPI_ISL_2657431, EPI_ISL_2657432, EPI_ISL_2657433, EPI_ISL_2657434, EPI_ISL_2657436, EPI_ISL_2657437, EPI_ISL_2657438, EPI_ISL_2657439, EPI_ISL_2657440, EPI_ISL_2657441, EPI_ISL_2657442, EPI_ISL_2657443, EPI_ISL_2657444, EPI_ISL_2657445, EPI_ISL_2657447, EPI_ISL_2657448, EPI_ISL_2657449, EPI_ISL_2657450, EPI_ISL_2657451, EPI_ISL_2657452, EPI_ISL_2657454, EPI_ISL_2657461, EPI_ISL_2657462, EPI_ISL_2657463, EPI_ISL_2657464, EPI_ISL_2657465, EPI_ISL_2657466, EPI_ISL_2657467, EPI_ISL_2657468, EPI_ISL_2657469, EPI_ISL_2657470, EPI_ISL_2657471, EPI_ISL_2657474, EPI_ISL_2657475, EPI_ISL_2657482, EPI_ISL_2657485, EPI_ISL_2657486, EPI_ISL_2657487, EPI_ISL_2657488, EPI_ISL_2657489, EPI_ISL_2657490, EPI_ISL_2657492, EPI_ISL_2657495, EPI_ISL_2657498, EPI_ISL_2657501, EPI_ISL_2657502, EPI_ISL_2657503, EPI_ISL_2657504, EPI_ISL_2657506, EPI_ISL_2657507, EPI_ISL_2657508, EPI_ISL_2657509, EPI_ISL_2657510, EPI_ISL_2657511 | | | |
| see above | Austrian Agency for Health and Food Safety (AGES) | Bergthaler laboratory, CeMM Research Center for Molecular Medicine of the Austrian Academy of Sciences | Lukas Endler, Anna Schedl, Fabian Amman, Petr Triska, Matthew Thornton, Thomas Penz, Benedikt Agerer, Maelle Le Moing, Michael Schuster, Bekir Erguner, Jan Laine, Martin Senekowitsch, Christoph Bock, Andreas Bergthaler |
| EPI_ISL_2657517, EPI_ISL_2657518, EPI_ISL_2657519, EPI_ISL_2657520, EPI_ISL_2657521, EPI_ISL_2657522, EPI_ISL_2657523, EPI_ISL_2657524, EPI_ISL_2657525 | Dr. Gernot Walder GmbH | Bergthaler laboratory, CeMM Research Center for Molecular Medicine of the Austrian Academy of Sciences | Lukas Endler, Anna Schedl, Fabian Amman, Petr Triska, Matthew Thornton, Thomas Penz, Benedikt Agerer, Maelle Le Moing, Michael Schuster, Bekir Erguner, Jan Laine, Martin Senekowitsch, Christoph Bock, Andreas Bergthaler |
| EPI_ISL_2657526 | ILV Kärnten | Bergthaler laboratory, CeMM Research Center for Molecular Medicine of the Austrian Academy of Sciences | Lukas Endler, Anna Schedl, Fabian Amman, Petr Triska, Matthew Thornton, Thomas Penz, Benedikt Agerer, Maelle Le Moing, Michael Schuster, Bekir Erguner, Jan Laine, Martin Senekowitsch, Christoph Bock, Andreas Bergthaler |
| EPI_ISL_2657527, EPI_ISL_2657531, EPI_ISL_2657532, EPI_ISL_2657533, EPI_ISL_2657534, EPI_ISL_2657535, EPI_ISL_2657537, EPI_ISL_2657538, EPI_ISL_2657539, EPI_ISL_2657540, EPI_ISL_2657542, EPI_ISL_2657543, EPI_ISL_2657544, EPI_ISL_2657546, EPI_ISL_2657547, EPI_ISL_2657548, EPI_ISL_2657550, EPI_ISL_2657551, EPI_ISL_2657552, EPI_ISL_2657554, EPI_ISL_2657555 | | | |
| see above | Elling group, Institute of Molecular Biotechnology (IMBA) | Bergthaler laboratory, CeMM Research Center for Molecular Medicine of the Austrian Academy of Sciences | Lukas Endler, Anna Schedl, Fabian Amman, Petr Triska, Matthew Thornton, Thomas Penz, Benedikt Agerer, Maelle Le Moing, Michael Schuster, Bekir Erguner, Jan Laine, Martin Senekowitsch, Christoph Bock, Andreas Bergthaler |
| EPI_ISL_2657570 | Medizinische Universität Graz | Bergthaler laboratory, CeMM Research Center for Molecular Medicine of the Austrian Academy of Sciences | Lukas Endler, Anna Schedl, Fabian Amman, Petr Triska, Matthew Thornton, Thomas Penz, Benedikt Agerer, Maelle Le Moing, Michael Schuster, Bekir Erguner, Jan Laine, Martin Senekowitsch, Christoph Bock, Andreas Bergthaler |
| EPI_ISL_2657573, EPI_ISL_2657574 | Vienna COVID-19 Detection Initiative (VCDI) | Bergthaler laboratory, CeMM Research Center for Molecular Medicine of the Austrian Academy of Sciences | Lukas Endler, Anna Schedl, Fabian Amman, Petr Triska, Matthew Thornton, Thomas Penz, Benedikt Agerer, Maelle Le Moing, Michael Schuster, Bekir Erguner, Jan Laine, Martin Senekowitsch, Christoph Bock, Andreas Bergthaler |
| EPI_ISL_2657687 | HOSPITAL SAN JUAN DE DIOS | Inciensa, Instituto Costarricense de Investigación y Enseñanza en Nutrición y Salud | Francisco Duarte, Hebleen Porras, Claudio Soto-Garita, Estela Cordero, Adriana Godínez, Melany Calderón, José Luis Vargas, Mariela Gutiérrez, Joselyn Prado & Marco Cháves-Otárola |
| EPI_ISL_2657688 | HOSPITAL DR. CARLOS LUIS VALVERDE VEGA | Inciensa, Instituto Costarricense de Investigación y Enseñanza en Nutrición y Salud | Francisco Duarte, Hebleen Porras, Claudio Soto-Garita, Estela Cordero, Adriana Godínez, Melany Calderón, José Luis Vargas, Mariela Gutiérrez, Joselyn Prado & Yahaira Ramos |
| EPI_ISL_2657858 | Laboratorios del Valle SAS | Instituto Nacional de Salud- Dirección de Investigación en Salud Pública | Katherine Laiton-Donato, Diego A. Álvarez-Díaz, Carlos Franco-Muñoz, Hector Alejandro Ruiz-Moreno, Paola Rojas, Maria T. Herrera-Sepúlveda, Diego Andrés Prada, Jhonnatan Reales-González, Sheryll Corchuelo, Julian Naizaque, Jorge Rivera, Gerardo Santamaría, Sergio Gomez, Lisseth Pardo, Juan Camilo Martinez, Marta Lopez Blanco, Ángela Alarcon Cruz, Diana Malo, Carmen Osorio, Magdalena Wiesner, Martha Lucia Ospina Martinez, Marcela Mercado-Reyes |
| EPI_ISL_2657859, EPI_ISL_2657862, EPI_ISL_2657863, EPI_ISL_2657864 | Laboratorio de Salud Pública Fronterizo de Arauca | Instituto Nacional de Salud- Dirección de Investigación en Salud Pública | Katherine Laiton-Donato, Diego A. Álvarez-Díaz, Carlos Franco-Muñoz, Hector Alejandro Ruiz-Moreno, Paola Rojas, Maria T. Herrera-Sepúlveda, Diego Andrés Prada, Jhonnatan Reales-González, Sheryll Corchuelo, Julian Naizaque, Jorge Rivera, Gerardo Santamaría, Sergio Gomez, Lisseth Pardo, Juan Camilo Martinez, Marta Lopez Blanco, Ángela Alarcon Cruz, Diana Malo, Carmen Osorio, Magdalena Wiesner, Martha Lucia Ospina Martinez, Marcela Mercado-Reyes |
| EPI_ISL_2657867, EPI_ISL_2657868 | COVID 19- EPS FAMISANAR | Instituto Nacional de Salud- Dirección de Investigación en Salud Pública | Katherine Laiton-Donato, Diego A. Álvarez-Díaz, Carlos Franco-Muñoz, Hector Alejandro Ruiz-Moreno, Paola Rojas, Maria T. Herrera-Sepúlveda, Diego Andrés Prada, Jhonnatan Reales-González, Sheryll Corchuelo, Julian Naizaque, Jorge Rivera, Gerardo Santamaría, Sergio Gomez, Lisseth Pardo, Juan Camilo Martinez, Marta Lopez Blanco, Ángela Alarcon Cruz, Diana Malo, Carmen Osorio, Magdalena Wiesner, Martha Lucia Ospina Martinez, Marcela Mercado-Reyes |
| EPI_ISL_2657873, EPI_ISL_2657874 | LABORATORIO BILOGIA MOLECULAR IMAT SAS | Instituto Nacional de Salud- Dirección de Investigación en Salud Pública | Katherine Laiton-Donato, Diego A. Álvarez-Díaz, Carlos Franco-Muñoz, Hector Alejandro Ruiz-Moreno, Paola Rojas, Maria T. Herrera-Sepúlveda, Diego Andrés Prada, Jhonnatan Reales-González, Sheryll Corchuelo, Julian Naizaque, Jorge Rivera, Gerardo Santamaría, Sergio Gomez, Lisseth Pardo, Juan Camilo Martinez, Marta Lopez Blanco, Ángela Alarcon Cruz, Diana Malo, Carmen Osorio, Magdalena Wiesner, Martha Lucia Ospina Martinez, Marcela Mercado-Reyes |
| EPI_ISL_2657875 | laboratorio de salud publica de cundinamarca | Instituto Nacional de Salud- Dirección de Investigación en Salud Pública | Katherine Laiton-Donato, Diego A. Álvarez-Díaz, Carlos Franco-Muñoz, Hector Alejandro Ruiz-Moreno, Paola Rojas, Maria T. Herrera-Sepúlveda, Diego Andrés Prada, Jhonnatan Reales-González, Sheryll Corchuelo, Julian Naizaque, Jorge Rivera, Gerardo Santamaría, Sergio Gomez, Lisseth Pardo, Juan Camilo Martinez, Marta Lopez Blanco, Ángela Alarcon Cruz, Diana Malo, Carmen Osorio, Magdalena Wiesner, Martha Lucia Ospina Martinez, Marcela Mercado-Reyes |
| EPI_ISL_2657876 | Lab microbiologia FVL | Instituto Nacional de Salud- Dirección de Investigación en Salud Pública | Katherine Laiton-Donato, Diego A. Álvarez-Díaz, Carlos Franco-Muñoz, Hector Alejandro Ruiz-Moreno, Paola Rojas, Maria T. Herrera-Sepúlveda, Diego Andrés Prada, Jhonnatan Reales-González, Sheryll Corchuelo, Julian Naizaque, Jorge Rivera, Gerardo Santamaría, Sergio Gomez, Lisseth Pardo, Juan Camilo Martinez, Marta Lopez Blanco, Ángela Alarcon Cruz, Diana Malo, Carmen Osorio, Magdalena Wiesner, Martha Lucia Ospina Martinez, Marcela Mercado-Reyes |
| EPI_ISL_2657918, EPI_ISL_2657936 | Public Health Authority of the Slovak Republic | Laboratory of Genomics and Bioinformatics, Comenius University Science Park | Tatiana Sedláková, Diana Rusáková, Miroslav Böhmer, Anna Giová, Jaroslav Budiš, Tomáš Szemes, Jakub Styk |
| EPI_ISL_2658282 | AREA DE SALUD EL GUARCO | Inciensa, Instituto Costarricense de Investigación y Enseñanza en Nutrición y Salud | Francisco Duarte, Hebleen Porras, Claudio Soto-Garita, Estela Cordero, Adriana Godínez, Melany Calderón, José Luis Vargas, Mariela Gutiérrez, Joselyn Prado & Pamela Serrano Valerín |
| EPI_ISL_2658283 | AREA DE SALUD ALAJUELITA | Inciensa, Instituto Costarricense de Investigación y Enseñanza en Nutrición y Salud | Francisco Duarte, Hebleen Porras, Claudio Soto-Garita, Estela Cordero, Adriana Godínez, Melany Calderón, José Luis Vargas, Mariela Gutiérrez & Joselyn Prado |
| EPI_ISL_2658284 | AREA DE SALUD MATA REDONDA-HOSPITAL - CLINICA DR. MORENO CAÑAS | Inciensa, Instituto Costarricense de Investigación y Enseñanza en Nutrición y Salud | Francisco Duarte, Hebleen Porras, Claudio Soto-Garita, Estela Cordero, Adriana Godínez, Melany Calderón, José Luis Vargas, Mariela Gutiérrez & Joselyn Prado |
| EPI_ISL_2658285 | AREA DE SALUD MORAVIA | Inciensa, Instituto Costarricense de Investigación y Enseñanza en Nutrición y Salud | Francisco Duarte, Hebleen Porras, Claudio Soto-Garita, Estela Cordero, Adriana Godínez, Melany Calderón, José Luis Vargas, Mariela Gutiérrez, Joselyn Prado & Mariel López |
| EPI_ISL_2658454, EPI_ISL_2658455, EPI_ISL_2658457 | Laboratori de Referencia de Catalunya | Laboratori de Referencia de Catalunya | Ramirez A., Bellosillo B., Padilla E., Vilas A., Hernandez JJ., Canal M. |
| EPI_ISL_2658950 | Basurto University Hospital: Clinical Microbiology Laboratory | Biocruces Bizkaia | Mikel Urrutikoetxea-Gutiérrez, Mª Carmen Nieto Toboso, Estibaliz Ugalde Zarraga, Mikel Gallego Rodrigo, Ana de la Hoz, José Luis Díaz de Tuesta del Arco |
| EPI_ISL_2658951, EPI_ISL_2658952 | LESP San Luis Potosi | Instituto de Diagnostico y Referencia Epidemiologicos (INDRE) | Claudia Wong-Arambula, Abril Rodriguez-Maldonado, Vanessa Rivero-Arredondo, Ariadna Medina-Benitez, Joaquin Quiroz-Mercado, Sergio Rangel-Guerrero, Natividad Cruz-Ortiz, Tatiana Nunez-Garcia, Gisela Barrera-Badillo, Lucia Hernandez-Rivas, Irma Lopez-Martinez, Ernesto Ramirez-Gonzalez. |
| EPI_ISL_2658953 | LESP Nuevo Leon | Instituto de Diagnostico y Referencia Epidemiologicos (INDRE) | Claudia Wong-Arambula, Abril Rodriguez-Maldonado, Vanessa Rivero-Arredondo, Ariadna Medina-Benitez, Joaquin Quiroz-Mercado, Sergio Rangel-Guerrero, Natividad Cruz-Ortiz, Tatiana Nunez-Garcia, Gisela Barrera-Badillo, Lucia Hernandez-Rivas, Irma Lopez-Martinez, Ernesto Ramirez-Gonzalez. |
| EPI_ISL_2662825 | Laboratorio de Virología HUCA | Laboratorio de Virología HUCA | Sandoval M, Castelló C, Gómez de Oña J, Boga JA, Rojo S, Alvarez-Arguelles ME, Abreu F, Costales I, Perez-Martínez Z, Martín-Rodríguez G, Coto E, Melón S |
| EPI_ISL_2663128, EPI_ISL_2663131, | Ochsner Health | Bioinfoexperts, LLC | Judy Crabtree, Amy Feehan, Rebecca Rose, David J. Nolan, Tessa LaFleur, Julia-Garcia-Diaz, Lucio Miele, Susanna L. Lamers |

| | | | |
|---|---|---|---|
| EPI_ISL_2663132, EPI_ISL_2663134, EPI_ISL_2663135, EPI_ISL_2663139, EPI_ISL_2663140, EPI_ISL_2663142, EPI_ISL_2663143, EPI_ISL_2663146 | | | |
| EPI_ISL_2663327 | LESP Nuevo Leon | Instituto de Diagnostico y Referencia Epidemiologicos (INDRE) | Claudia Wong-Arambula, Abril Rodriguez-Maldonado, Vanessa Rivero-Arredondo, Ariadna Medina-Benitez, Joaquin Quiroz-Mercado, Sergio Rangel-Guerrero, Natividad Cruz-Ortiz, Tatiana Nunez-Garcia, Gisela Barrera-Badillo, Lucia Hernandez-Rivas, Irma Lopez-Martinez, Ernesto Ramirez-Gonzalez. |
| EPI_ISL_2663328, EPI_ISL_2663329, EPI_ISL_2663330 | LESP Ciudad de Mexico | Instituto de Diagnostico y Referencia Epidemiologicos (INDRE) | Claudia Wong-Arambula, Abril Rodriguez-Maldonado, Vanessa Rivero-Arredondo, Ariadna Medina-Benitez, Joaquin Quiroz-Mercado, Sergio Rangel-Guerrero, Natividad Cruz-Ortiz, Tatiana Nunez-Garcia, Gisela Barrera-Badillo, Lucia Hernandez-Rivas, Irma Lopez-Martinez, Ernesto Ramirez-Gonzalez. |
| EPI_ISL_2663331 | LESP Queretaro | Instituto de Diagnostico y Referencia Epidemiologicos (INDRE) | Claudia Wong-Arambula, Abril Rodriguez-Maldonado, Vanessa Rivero-Arredondo, Ariadna Medina-Benitez, Joaquin Quiroz-Mercado, Sergio Rangel-Guerrero, Natividad Cruz-Ortiz, Tatiana Nunez-Garcia, Gisela Barrera-Badillo, Lucia Hernandez-Rivas, Irma Lopez-Martinez, Ernesto Ramirez-Gonzalez. |
| EPI_ISL_2663333 | LESP Sinaloa | Instituto de Diagnostico y Referencia Epidemiologicos (INDRE) | Claudia Wong-Arambula, Abril Rodriguez-Maldonado, Vanessa Rivero-Arredondo, Ariadna Medina-Benitez, Joaquin Quiroz-Mercado, Sergio Rangel-Guerrero, Natividad Cruz-Ortiz, Tatiana Nunez-Garcia, Gisela Barrera-Badillo, Lucia Hernandez-Rivas, Irma Lopez-Martinez, Ernesto Ramirez-Gonzalez. |
| EPI_ISL_2663336, EPI_ISL_2663337, EPI_ISL_2663338, EPI_ISL_2663339, EPI_ISL_2663340, EPI_ISL_2663341 | LESP Tabasco | Instituto de Diagnostico y Referencia Epidemiologicos (INDRE) | Claudia Wong-Arambula, Abril Rodriguez-Maldonado, Vanessa Rivero-Arredondo, Ariadna Medina-Benitez, Joaquin Quiroz-Mercado, Sergio Rangel-Guerrero, Natividad Cruz-Ortiz, Tatiana Nunez-Garcia, Gisela Barrera-Badillo, Lucia Hernandez-Rivas, Irma Lopez-Martinez, Ernesto Ramirez-Gonzalez. |
| EPI_ISL_2663342 | LESP Ciudad de Mexico | Instituto de Diagnostico y Referencia Epidemiologicos (INDRE) | Claudia Wong-Arambula, Abril Rodriguez-Maldonado, Vanessa Rivero-Arredondo, Ariadna Medina-Benitez, Joaquin Quiroz-Mercado, Sergio Rangel-Guerrero, Natividad Cruz-Ortiz, Tatiana Nunez-Garcia, Gisela Barrera-Badillo, Lucia Hernandez-Rivas, Irma Lopez-Martinez, Ernesto Ramirez-Gonzalez. |
| EPI_ISL_2663345 | LESP Colima | Instituto de Diagnostico y Referencia Epidemiologicos (INDRE) | Claudia Wong-Arambula, Abril Rodriguez-Maldonado, Vanessa Rivero-Arredondo, Ariadna Medina-Benitez, Joaquin Quiroz-Mercado, Sergio Rangel-Guerrero, Natividad Cruz-Ortiz, Tatiana Nunez-Garcia, Gisela Barrera-Badillo, Lucia Hernandez-Rivas, Irma Lopez-Martinez, Ernesto Ramirez-Gonzalez. |
| EPI_ISL_2663346, EPI_ISL_2663347, EPI_ISL_2663348, EPI_ISL_2663349 | LESP Tabasco | Instituto de Diagnostico y Referencia Epidemiologicos (INDRE) | Claudia Wong-Arambula, Abril Rodriguez-Maldonado, Vanessa Rivero-Arredondo, Ariadna Medina-Benitez, Joaquin Quiroz-Mercado, Sergio Rangel-Guerrero, Natividad Cruz-Ortiz, Tatiana Nunez-Garcia, Gisela Barrera-Badillo, Lucia Hernandez-Rivas, Irma Lopez-Martinez, Ernesto Ramirez-Gonzalez. |
| EPI_ISL_2663350 | LESP Tamaulipas | Instituto de Diagnostico y Referencia Epidemiologicos (INDRE) | Claudia Wong-Arambula, Abril Rodriguez-Maldonado, Vanessa Rivero-Arredondo, Ariadna Medina-Benitez, Joaquin Quiroz-Mercado, Sergio Rangel-Guerrero, Natividad Cruz-Ortiz, Tatiana Nunez-Garcia, Gisela Barrera-Badillo, Lucia Hernandez-Rivas, Irma Lopez-Martinez, Ernesto Ramirez-Gonzalez. |
| EPI_ISL_2663352 | LESP Oaxaca | Instituto de Diagnostico y Referencia Epidemiologicos (INDRE) | Claudia Wong-Arambula, Abril Rodriguez-Maldonado, Vanessa Rivero-Arredondo, Ariadna Medina-Benitez, Joaquin Quiroz-Mercado, Sergio Rangel-Guerrero, Natividad Cruz-Ortiz, Tatiana Nunez-Garcia, Gisela Barrera-Badillo, Lucia Hernandez-Rivas, Irma Lopez-Martinez, Ernesto Ramirez-Gonzalez. |
| EPI_ISL_2663353 | LESP Baja California Sur | Instituto de Diagnostico y Referencia Epidemiologicos (INDRE) | Claudia Wong-Arambula, Abril Rodriguez-Maldonado, Vanessa Rivero-Arredondo, Ariadna Medina-Benitez, Joaquin Quiroz-Mercado, Sergio Rangel-Guerrero, Natividad Cruz-Ortiz, Tatiana Nunez-Garcia, Gisela Barrera-Badillo, Lucia Hernandez-Rivas, Irma Lopez-Martinez, Ernesto Ramirez-Gonzalez. |
| EPI_ISL_2663354, EPI_ISL_2663355 | LESP Sonora | Instituto de Diagnostico y Referencia Epidemiologicos (INDRE) | Claudia Wong-Arambula, Abril Rodriguez-Maldonado, Vanessa Rivero-Arredondo, Ariadna Medina-Benitez, Joaquin Quiroz-Mercado, Sergio Rangel-Guerrero, Natividad Cruz-Ortiz, Tatiana Nunez-Garcia, Gisela Barrera-Badillo, Lucia Hernandez-Rivas, Irma Lopez-Martinez, Ernesto Ramirez-Gonzalez. |
| EPI_ISL_2663356, EPI_ISL_2663357 | LESP Quintana Roo | Instituto de Diagnostico y Referencia Epidemiologicos (INDRE) | Claudia Wong-Arambula, Abril Rodriguez-Maldonado, Vanessa Rivero-Arredondo, Ariadna Medina-Benitez, Joaquin Quiroz-Mercado, Sergio Rangel-Guerrero, Natividad Cruz-Ortiz, Tatiana Nunez-Garcia, Gisela Barrera-Badillo, Lucia Hernandez-Rivas, Irma Lopez-Martinez, Ernesto Ramirez-Gonzalez. |
| EPI_ISL_2663358 | LESP Ciudad de Mexico | Instituto de Diagnostico y Referencia Epidemiologicos (INDRE) | Claudia Wong-Arambula, Abril Rodriguez-Maldonado, Vanessa Rivero-Arredondo, Ariadna Medina-Benitez, Joaquin Quiroz-Mercado, Sergio Rangel-Guerrero, Natividad Cruz-Ortiz, Tatiana Nunez-Garcia, Gisela Barrera-Badillo, Lucia Hernandez-Rivas, Irma Lopez-Martinez, Ernesto Ramirez-Gonzalez. |
| EPI_ISL_2663359 | LESP Sonora | Instituto de Diagnostico y Referencia Epidemiologicos (INDRE) | Claudia Wong-Arambula, Abril Rodriguez-Maldonado, Vanessa Rivero-Arredondo, Ariadna Medina-Benitez, Joaquin Quiroz-Mercado, Sergio Rangel-Guerrero, Natividad Cruz-Ortiz, Tatiana Nunez-Garcia, Gisela Barrera-Badillo, Lucia Hernandez-Rivas, Irma Lopez-Martinez, Ernesto Ramirez-Gonzalez. |
| EPI_ISL_2663360, EPI_ISL_2663361, EPI_ISL_2663362, EPI_ISL_2663363, EPI_ISL_2663364, EPI_ISL_2663365, EPI_ISL_2663366, EPI_ISL_2663367, EPI_ISL_2663368, EPI_ISL_2663369, EPI_ISL_2663370, EPI_ISL_2663371, EPI_ISL_2663372, EPI_ISL_2663373, EPI_ISL_2663374, EPI_ISL_2663375 | | | |
| see above | LESP Hidalgo | Instituto de Diagnostico y Referencia Epidemiologicos (INDRE) | Claudia Wong-Arambula, Abril Rodriguez-Maldonado, Vanessa Rivero-Arredondo, Ariadna Medina-Benitez, Joaquin Quiroz-Mercado, Sergio Rangel-Guerrero, Natividad Cruz-Ortiz, Tatiana Nunez-Garcia, Gisela Barrera-Badillo, Lucia Hernandez-Rivas, Irma Lopez-Martinez, Ernesto Ramirez-Gonzalez. |
| EPI_ISL_2663377 | LESP Queretaro | Instituto de Diagnostico y Referencia Epidemiologicos (INDRE) | Claudia Wong-Arambula, Abril Rodriguez-Maldonado, Vanessa Rivero-Arredondo, Ariadna Medina-Benitez, Joaquin Quiroz-Mercado, Sergio Rangel-Guerrero, Natividad Cruz-Ortiz, Tatiana Nunez-Garcia, Gisela Barrera-Badillo, Lucia Hernandez-Rivas, Irma Lopez-Martinez, Ernesto Ramirez-Gonzalez. |
| EPI_ISL_2663378, EPI_ISL_2663379 | LESP Yucatan | Instituto de Diagnostico y Referencia Epidemiologicos (INDRE) | Claudia Wong-Arambula, Abril Rodriguez-Maldonado, Vanessa Rivero-Arredondo, Ariadna Medina-Benitez, Joaquin Quiroz-Mercado, Sergio Rangel-Guerrero, Natividad Cruz-Ortiz, Tatiana Nunez-Garcia, Gisela Barrera-Badillo, Lucia Hernandez-Rivas, Irma Lopez-Martinez, Ernesto Ramirez-Gonzalez. |
| EPI_ISL_2663380, EPI_ISL_2663381 | LESP Queretaro | Instituto de Diagnostico y Referencia Epidemiologicos (INDRE) | Claudia Wong-Arambula, Abril Rodriguez-Maldonado, Vanessa Rivero-Arredondo, Ariadna Medina-Benitez, Joaquin Quiroz-Mercado, Sergio Rangel-Guerrero, Natividad Cruz-Ortiz, Tatiana Nunez-Garcia, Gisela Barrera-Badillo, Lucia Hernandez-Rivas, Irma Lopez-Martinez, Ernesto Ramirez-Gonzalez. |
| EPI_ISL_2663382 | LESP Morelos | Instituto de Diagnostico y Referencia Epidemiologicos (INDRE) | Claudia Wong-Arambula, Abril Rodriguez-Maldonado, Vanessa Rivero-Arredondo, Ariadna Medina-Benitez, Joaquin Quiroz-Mercado, Sergio Rangel-Guerrero, Natividad Cruz-Ortiz, Tatiana Nunez-Garcia, Gisela Barrera-Badillo, Lucia Hernandez-Rivas, Irma Lopez-Martinez, Ernesto Ramirez-Gonzalez. |
| EPI_ISL_2663383, EPI_ISL_2663384, EPI_ISL_2663385, EPI_ISL_2663386, EPI_ISL_2663387, EPI_ISL_2663388 | LESP Guanajuato | Instituto de Diagnostico y Referencia Epidemiologicos (INDRE) | Claudia Wong-Arambula, Abril Rodriguez-Maldonado, Vanessa Rivero-Arredondo, Ariadna Medina-Benitez, Joaquin Quiroz-Mercado, Sergio Rangel-Guerrero, Natividad Cruz-Ortiz, Tatiana Nunez-Garcia, Gisela Barrera-Badillo, Lucia Hernandez-Rivas, Irma Lopez-Martinez, Ernesto Ramirez-Gonzalez. |
| EPI_ISL_2663389, EPI_ISL_2663390, EPI_ISL_2663391, EPI_ISL_2663392 | LESP Nuevo Leon | Instituto de Diagnostico y Referencia Epidemiologicos (INDRE) | Claudia Wong-Arambula, Abril Rodriguez-Maldonado, Vanessa Rivero-Arredondo, Ariadna Medina-Benitez, Joaquin Quiroz-Mercado, Sergio Rangel-Guerrero, Natividad Cruz-Ortiz, Tatiana Nunez-Garcia, Gisela Barrera-Badillo, Lucia Hernandez-Rivas, Irma Lopez-Martinez, Ernesto Ramirez-Gonzalez. |
| EPI_ISL_2663393 | LESP Ciudad de Mexico | Instituto de Diagnostico y Referencia Epidemiologicos (INDRE) | Claudia Wong-Arambula, Abril Rodriguez-Maldonado, Vanessa Rivero-Arredondo, Ariadna Medina-Benitez, Joaquin Quiroz-Mercado, Sergio Rangel-Guerrero, Natividad Cruz-Ortiz, Tatiana Nunez-Garcia, Gisela Barrera-Badillo, Lucia Hernandez-Rivas, Irma Lopez-Martinez, Ernesto Ramirez-Gonzalez. |
| EPI_ISL_2663394, EPI_ISL_2663395, | LESP Chihuahua | Instituto de Diagnostico y Referencia Epidemiologicos | Claudia Wong-Arambula, Abril Rodriguez-Maldonado, Vanessa Rivero-Arredondo, Ariadna Medina-Benitez, Joaquin Quiroz-Mercado, Sergio |

| | | | |
|---|---|---|---|
| EPI_ISL_2663396, EPI_ISL_2663397, EPI_ISL_2663398 | | (INDRE) | Rangel-Guerrero, Natividad Cruz-Ortiz, Tatiana Nunez-Garcia, Gisela Barrera-Badillo, Lucia Hernandez-Rivas, Irma Lopez-Martinez, Ernesto Ramirez-Gonzalez. |
| EPI_ISL_2663399 | LESP Puebla | Instituto de Diagnostico y Referencia Epidemiologicos (INDRE) | Claudia Wong-Arambula, Abril Rodriguez-Maldonado, Vanessa Rivero-Arredondo, Ariadna Medina-Benitez, Joaquin Quiroz-Mercado, Sergio Rangel-Guerrero, Natividad Cruz-Ortiz, Tatiana Nunez-Garcia, Gisela Barrera-Badillo, Lucia Hernandez-Rivas, Irma Lopez-Martinez, Ernesto Ramirez-Gonzalez. |
| EPI_ISL_2663400, EPI_ISL_2663401 | LESP Tamaulipas | Instituto de Diagnostico y Referencia Epidemiologicos (INDRE) | Claudia Wong-Arambula, Abril Rodriguez-Maldonado, Vanessa Rivero-Arredondo, Ariadna Medina-Benitez, Joaquin Quiroz-Mercado, Sergio Rangel-Guerrero, Natividad Cruz-Ortiz, Tatiana Nunez-Garcia, Gisela Barrera-Badillo, Lucia Hernandez-Rivas, Irma Lopez-Martinez, Ernesto Ramirez-Gonzalez. |
| EPI_ISL_2663402 | LESP Yucatan | Instituto de Diagnostico y Referencia Epidemiologicos (INDRE) | Claudia Wong-Arambula, Abril Rodriguez-Maldonado, Vanessa Rivero-Arredondo, Ariadna Medina-Benitez, Joaquin Quiroz-Mercado, Sergio Rangel-Guerrero, Natividad Cruz-Ortiz, Tatiana Nunez-Garcia, Gisela Barrera-Badillo, Lucia Hernandez-Rivas, Irma Lopez-Martinez, Ernesto Ramirez-Gonzalez. |
| EPI_ISL_2663403 | LESP Sonora | Instituto de Diagnostico y Referencia Epidemiologicos (INDRE) | Claudia Wong-Arambula, Abril Rodriguez-Maldonado, Vanessa Rivero-Arredondo, Ariadna Medina-Benitez, Joaquin Quiroz-Mercado, Sergio Rangel-Guerrero, Natividad Cruz-Ortiz, Tatiana Nunez-Garcia, Gisela Barrera-Badillo, Lucia Hernandez-Rivas, Irma Lopez-Martinez, Ernesto Ramirez-Gonzalez. |
| EPI_ISL_2671484 | Centro de Investigacion Biomedica de Occidente (CIBO) | Unidad de Genomica Avanzada | Consorcio Mexicano de Vigilancia Genomica (CoViGen-Mex). Authors (in alphabetical order): Julio Elias Alvarado-Yaah, Carlos F. Arias, Santiago avila-Rios, Eduardo Becerril-Vargas, Victor Hugo Borja-Aburto, Celia Boukadida, Cristobal Chaidez-Quiroz, Juan Bautista Chale-Dzul, Ricardo Ciria Merce, Andreu Comas-Garcia, Celida Duque Molina, Julissa Enciso-Ibarra, Jose Antonio Enciso-Moreno, Gloria Elena Espinosa-Ayala, Fernando Fontove-Herrera, Daniel Fregoso-Rueda, Victor Eduardo Garcia-Arias, Alejandra Garcia-Gasca, Bruno Gomez-Gil, Jean Pierre Gonzalez, Irvin Gonzalez-Lopez, Concepcion Grajales-Muñiz, Ricardo Grande, Rosa Maria Gutierrez Rios, Jesus Hernandez, Alejandra Hernandez-Teran, Alfredo Herrera-Estrella, Carla Ivon Herrera-Najera, Pavel Isa, Veronica Mata-Haro, Daniel Lira Morales, Susana Lopez, Antonio Loza Roman, Brenda Irasema Maldonado-Meza, Bernardo Martinez-Miguel, Jose Arturo Martinez-Orozco, Celida Martinez- Rodriguez, Margarita Matias-Florentino, Fidencio Mejia-Nepomuceno, Maria Guadalupe de Jesus Mireles-Rivera, Gloria Maria Molina-Salinas, Hector Montoya-Fuentes, , Mario Mujica-Sanchez, Jose Esteban Muñoz-Medina, Jose de Jesus Nuñez-Contreras, Alicia Ocaña-Mondragon, Luis Alberto Ochoa-Carrera, Hector Esteban Paz-Juarez, Marissa Perez-Garcia, Francisco Pulido, Helen Haydee Fernanda Ramirez-Plascencia, Jorge Salas-Hernandez, Angel Gustavo Salas-Lais, Alejandro Sanchez-Flores, Clara Esperanza Santacruz-Tinoco, Maria Guadalupe Santiago-Mauricio, Selene Zarate, Nelly Selem-Mojica, Blanca Taboada, Gloria Vazquez, Joel Armando Vazquez-Perez. |
| EPI_ISL_2671485, EPI_ISL_2671486, EPI_ISL_2671488, EPI_ISL_2671489, EPI_ISL_2671490, EPI_ISL_2671492 | Laboratorio Central de Epidemiologi-a (LCE) | Unidad de Genomica Avanzada | Consorcio Mexicano de Vigilancia Genomica (CoViGen-Mex). Authors (in alphabetical order): Julio Elias Alvarado-Yaah, Carlos F. Arias, Santiago avila-Rios, Eduardo Becerril-Vargas, Victor Hugo Borja-Aburto, Celia Boukadida, Cristobal Chaidez-Quiroz, Juan Bautista Chale-Dzul, Ricardo Ciria Merce, Andreu Comas-Garcia, Celida Duque Molina, Julissa Enciso-Ibarra, Jose Antonio Enciso-Moreno, Gloria Elena Espinosa-Ayala, Fernando Fontove-Herrera, Daniel Fregoso-Rueda, Victor Eduardo Garcia-Arias, Alejandra Garcia-Gasca, Bruno Gomez-Gil, Jean Pierre Gonzalez, Irvin Gonzalez-Lopez, Concepcion Grajales-Muñiz, Ricardo Grande, Rosa Maria Gutierrez Rios, Jesus Hernandez, Alejandra Hernandez-Teran, Alfredo Herrera-Estrella, Carla Ivon Herrera-Najera, Pavel Isa, Veronica Mata-Haro, Daniel Lira Morales, Susana Lopez, Antonio Loza Roman, Brenda Irasema Maldonado-Meza, Bernardo Martinez-Miguel, Jose Arturo Martinez-Orozco, Celida Martinez- Rodriguez, Margarita Matias-Florentino, Fidencio Mejia-Nepomuceno, Maria Guadalupe de Jesus Mireles-Rivera, Gloria Maria Molina-Salinas, Hector Montoya-Fuentes, , Mario Mujica-Sanchez, Jose Esteban Muñoz-Medina, Jose de Jesus Nuñez-Contreras, Alicia Ocaña-Mondragon, Luis Alberto Ochoa-Carrera, Hector Esteban Paz-Juarez, Marissa Perez-Garcia, Francisco Pulido, Helen Haydee Fernanda Ramirez-Plascencia, Jorge Salas-Hernandez, Angel Gustavo Salas-Lais, Alejandro Sanchez-Flores, Clara Esperanza Santacruz-Tinoco, Maria Guadalupe Santiago-Mauricio, Selene Zarate, Nelly Selem-Mojica, Blanca Taboada, Gloria Vazquez, Joel Armando Vazquez-Perez. |
| EPI_ISL_2671494, EPI_ISL_2671495, EPI_ISL_2671496, EPI_ISL_2671497, EPI_ISL_2671498, EPI_ISL_2671499, EPI_ISL_2671500, EPI_ISL_2671501, EPI_ISL_2671502, EPI_ISL_2671503, EPI_ISL_2671504 | | | |
| see above | Unidad de Investigacion Medica de Yucatan (UIMY) | Unidad de Genomica Avanzada | Consorcio Mexicano de Vigilancia Genomica (CoViGen-Mex). Authors (in alphabetical order): Julio Elias Alvarado-Yaah, Carlos F. Arias, Santiago avila-Rios, Eduardo Becerril-Vargas, Victor Hugo Borja-Aburto, Celia Boukadida, Cristobal Chaidez-Quiroz, Juan Bautista Chale-Dzul, Ricardo Ciria Merce, Andreu Comas-Garcia, Celida Duque Molina, Julissa Enciso-Ibarra, Jose Antonio Enciso-Moreno, Gloria Elena Espinosa-Ayala, Fernando Fontove-Herrera, Daniel Fregoso-Rueda, Victor Eduardo Garcia-Arias, Alejandra Garcia-Gasca, Bruno Gomez-Gil, Jean Pierre Gonzalez, Irvin Gonzalez-Lopez, Concepcion Grajales-Muñiz, Ricardo Grande, Rosa Maria Gutierrez Rios, Jesus Hernandez, Alejandra Hernandez-Teran, Alfredo Herrera-Estrella, Carla Ivon Herrera-Najera, Pavel Isa, Veronica Mata-Haro, Daniel Lira Morales, Susana Lopez, Antonio Loza Roman, Brenda Irasema Maldonado-Meza, Bernardo Martinez-Miguel, Jose Arturo Martinez-Orozco, Celida Martinez- Rodriguez, Margarita Matias-Florentino, Fidencio Mejia-Nepomuceno, Maria Guadalupe de Jesus Mireles-Rivera, Gloria Maria Molina-Salinas, Hector Montoya-Fuentes, , Mario Mujica-Sanchez, Jose Esteban Muñoz-Medina, Jose de Jesus Nuñez-Contreras, Alicia Ocaña-Mondragon, Luis Alberto Ochoa-Carrera, Hector Esteban Paz-Juarez, Marissa Perez-Garcia, Francisco Pulido, Helen Haydee Fernanda Ramirez-Plascencia, Jorge Salas-Hernandez, Angel Gustavo Salas-Lais, Alejandro Sanchez-Flores, Clara Esperanza Santacruz-Tinoco, Maria Guadalupe Santiago-Mauricio, Selene Zarate, Nelly Selem-Mojica, Blanca Taboada, Gloria Vazquez, Joel Armando Vazquez-Perez. |
| EPI_ISL_2671505, EPI_ISL_2671506, EPI_ISL_2671509, EPI_ISL_2671510, EPI_ISL_2671513, EPI_ISL_2671515, EPI_ISL_2671516, EPI_ISL_2671517, EPI_ISL_2671518, EPI_ISL_2671519, EPI_ISL_2671520, EPI_ISL_2671521, EPI_ISL_2671522, EPI_ISL_2671523, EPI_ISL_2671524, EPI_ISL_2671525 | | | |
| see above | Centro de Investigacion Biomedica del Noreste (CIBIN) | Unidad de Genomica Avanzada | Consorcio Mexicano de Vigilancia Genomica (CoViGen-Mex). Authors (in alphabetical order): Julio Elias Alvarado-Yaah, Carlos F. Arias, Santiago avila-Rios, Eduardo Becerril-Vargas, Victor Hugo Borja-Aburto, Celia Boukadida, Cristobal Chaidez-Quiroz, Juan Bautista Chale-Dzul, Ricardo Ciria Merce, Andreu Comas-Garcia, Celida Duque Molina, Julissa Enciso-Ibarra, Jose Antonio Enciso-Moreno, Gloria Elena Espinosa-Ayala, Fernando Fontove-Herrera, Daniel Fregoso-Rueda, Victor Eduardo Garcia-Arias, Alejandra Garcia-Gasca, Bruno Gomez-Gil, Jean Pierre Gonzalez, Irvin Gonzalez-Lopez, Concepcion Grajales-Muñiz, Ricardo Grande, Rosa Maria Gutierrez Rios, Jesus Hernandez, Alejandra Hernandez-Teran, Alfredo Herrera-Estrella, Carla Ivon Herrera-Najera, Pavel Isa, Veronica Mata-Haro, Daniel Lira Morales, Susana Lopez, Antonio Loza Roman, Brenda Irasema Maldonado-Meza, Bernardo Martinez-Miguel, Jose Arturo Martinez-Orozco, Celida Martinez- Rodriguez, Margarita Matias-Florentino, Fidencio Mejia-Nepomuceno, Maria Guadalupe de Jesus Mireles-Rivera, Gloria Maria Molina-Salinas, Hector Montoya-Fuentes, , Mario Mujica-Sanchez, Jose Esteban Muñoz-Medina, Jose de Jesus Nuñez-Contreras, Alicia Ocaña-Mondragon, Luis Alberto Ochoa-Carrera, Hector Esteban Paz-Juarez, Marissa Perez-Garcia, Francisco Pulido, Helen Haydee Fernanda Ramirez-Plascencia, Jorge Salas-Hernandez, Angel Gustavo Salas-Lais, Alejandro Sanchez-Flores, Clara Esperanza Santacruz-Tinoco, Maria Guadalupe Santiago-Mauricio, Selene Zarate, Nelly Selem-Mojica, Blanca Taboada, Gloria Vazquez, Joel Armando Vazquez-Perez. |
| EPI_ISL_2671526, EPI_ISL_2671528, EPI_ISL_2671529, EPI_ISL_2671530, EPI_ISL_2671531, EPI_ISL_2671532, EPI_ISL_2671533, EPI_ISL_2671535, EPI_ISL_2671537, EPI_ISL_2671538, EPI_ISL_2671539, EPI_ISL_2671541 | | | |
| see above | Laboratorio Central de Epidemiologi-a (LCE) | Unidad de Genomica Avanzada | Consorcio Mexicano de Vigilancia Genomica (CoViGen-Mex). Authors (in alphabetical order): Julio Elias Alvarado-Yaah, Carlos F. Arias, Santiago avila-Rios, Eduardo Becerril-Vargas, Victor Hugo Borja-Aburto, Celia Boukadida, Cristobal Chaidez-Quiroz, Juan Bautista Chale-Dzul, Ricardo Ciria Merce, Andreu Comas-Garcia, Celida Duque Molina, Julissa Enciso-Ibarra, Jose Antonio Enciso-Moreno, Gloria Elena Espinosa-Ayala, Fernando Fontove-Herrera, Daniel Fregoso-Rueda, Victor Eduardo Garcia-Arias, Alejandra Garcia-Gasca, Bruno Gomez-Gil, Jean Pierre Gonzalez, Irvin Gonzalez-Lopez, Concepcion Grajales-Muñiz, Ricardo Grande, Rosa Maria Gutierrez Rios, Jesus Hernandez, Alejandra Hernandez-Teran, Alfredo Herrera-Estrella, Carla Ivon Herrera-Najera, Pavel Isa, Veronica Mata-Haro, Daniel Lira Morales, Susana Lopez, Antonio Loza Roman, Brenda Irasema Maldonado-Meza, Bernardo Martinez-Miguel, Jose Arturo Martinez-Orozco, Celida Martinez- Rodriguez, Margarita Matias-Florentino, Fidencio Mejia-Nepomuceno, Maria Guadalupe de Jesus Mireles-Rivera, Gloria Maria Molina-Salinas, Hector Montoya-Fuentes, , Mario Mujica-Sanchez, Jose Esteban Muñoz-Medina, Jose de Jesus Nuñez-Contreras, Alicia Ocaña-Mondragon, Luis Alberto Ochoa-Carrera, Hector Esteban Paz-Juarez, Marissa Perez-Garcia, Francisco Pulido, Helen Haydee Fernanda Ramirez-Plascencia, Jorge Salas-Hernandez, Angel Gustavo Salas-Lais, Alejandro Sanchez-Flores, Clara Esperanza Santacruz-Tinoco, Maria Guadalupe Santiago-Mauricio, Selene Zarate, Nelly Selem-Mojica, Blanca Taboada, Gloria Vazquez, Joel Armando Vazquez-Perez. |
| EPI_ISL_2671543, EPI_ISL_2671544, EPI_ISL_2671545, EPI_ISL_2671547 | Centro de Investigacion Biomedica del Noreste (CIBIN) | Unidad de Genomica Avanzada | Consorcio Mexicano de Vigilancia Genomica (CoViGen-Mex). Authors (in alphabetical order): Julio Elias Alvarado-Yaah, Carlos F. Arias, Santiago avila-Rios, Eduardo Becerril-Vargas, Victor Hugo Borja-Aburto, Celia Boukadida, Cristobal Chaidez-Quiroz, Juan Bautista Chale-Dzul, Ricardo Ciria Merce, Andreu Comas-Garcia, Celida Duque Molina, Julissa Enciso-Ibarra, Jose Antonio Enciso-Moreno, Gloria Elena Espinosa-Ayala, Fernando Fontove-Herrera, Daniel Fregoso-Rueda, Victor Eduardo Garcia-Arias, Alejandra Garcia-Gasca, Bruno Gomez-Gil, Jean Pierre Gonzalez, Irvin |

| | | | |
|---|---|---|---|
| | | | Gonzalez-Lopez, Concepcion Grajales-Muñiz, Ricardo Grande, Rosa Maria Gutierrez Rios, Jesus Hernandez, Alejandra Hernandez-Teran, Alfredo Herrera-Estrella, Carla Ivon Herrera-Najera, Pavel Isa, Veronica Mata-Haro, Daniel Lira Morales, Susana Lopez, Antonio Loza Roman, Brenda Irasema Maldonado-Meza, Bernardo Martinez-Miguel, Jose Arturo Martinez-Orozco, Celida Martinez- Rodriguez, Margarita Matias-Florentino, Fidencio Mejia-Nepomuceno, Maria Guadalupe de Jesus Mireles-Rivera, Gloria Maria Molina-Salinas, Hector Montoya-Fuentes, , Mario Mujica-Sanchez, Jose Esteban Muñoz-Medina, Jose de Jesus Nuñez-Contreras, Alicia Ocaña-Mondragon, Luis Alberto Ochoa-Carrera, Hector Esteban Paz-Juarez, Marissa Perez-Garcia, Francisco Pulido, Helen Haydee Fernanda Ramirez-Plascencia, Jorge Salas-Hernandez, Angel Gustavo Salas-Lais, Alejandro Sanchez-Flores, Clara Esperanza Santacruz-Tinoco, Maria Guadalupe Santiago-Mauricio, Selene Zarate, Nelly Selem-Mojica, Blanca Taboada, Gloria Vazquez, Joel Armando Vazquez-Perez. |
| EPI_ISL_2671548, EPI_ISL_2671549, EPI_ISL_2671550, EPI_ISL_2671551, EPI_ISL_2671552, EPI_ISL_2671553, EPI_ISL_2671554, EPI_ISL_2671555, EPI_ISL_2671556, EPI_ISL_2671557 | Centro de Investigacion Biomedica de Occidente (CIBO) | Unidad de Genomica Avanzada | Consorcio Mexicano de Vigilancia Genomica (CoViGen-Mex). Authors (in alphabetical order): Julio Elias Alvarado-Yaah, Carlos F. Arias, Santiago avila-Rios, Eduardo Becerril-Vargas, Victor Hugo Borja-Aburto, Celia Boukadida, Cristobal Chaidez-Quiroz, Juan Bautista Chale-Dzul, Ricardo Ciria Merce, Andreu Comas-Garcia, Celida Duque Molina, Julissa Enciso-Ibarra, Jose Antonio Enciso-Moreno, Gloria Elena Espinosa-Ayala, Fernando Fontove-Herrera, Daniel Fregoso-Rueda, Victor Eduardo Garcia-Arias, Alejandra Garcia-Gasca, Bruno Gomez-Gil, Jean Pierre Gonzalez, Irvin Gonzalez-Lopez, Concepcion Grajales-Muñiz, Ricardo Grande, Rosa Maria Gutierrez Rios, Jesus Hernandez, Alejandra Hernandez-Teran, Alfredo Herrera-Estrella, Carla Ivon Herrera-Najera, Pavel Isa, Veronica Mata-Haro, Daniel Lira Morales, Susana Lopez, Antonio Loza Roman, Brenda Irasema Maldonado-Meza, Bernardo Martinez-Miguel, Jose Arturo Martinez-Orozco, Celida Martinez- Rodriguez, Margarita Matias-Florentino, Fidencio Mejia-Nepomuceno, Maria Guadalupe de Jesus Mireles-Rivera, Gloria Maria Molina-Salinas, Hector Montoya-Fuentes, , Mario Mujica-Sanchez, Jose Esteban Muñoz-Medina, Jose de Jesus Nuñez-Contreras, Alicia Ocaña-Mondragon, Luis Alberto Ochoa-Carrera, Hector Esteban Paz-Juarez, Marissa Perez-Garcia, Francisco Pulido, Helen Haydee Fernanda Ramirez-Plascencia, Jorge Salas-Hernandez, Angel Gustavo Salas-Lais, Alejandro Sanchez-Flores, Clara Esperanza Santacruz-Tinoco, Maria Guadalupe Santiago-Mauricio, Selene Zarate, Nelly Selem-Mojica, Blanca Taboada, Gloria Vazquez, Joel Armando Vazquez-Perez. |
| EPI_ISL_2671558, EPI_ISL_2671559, EPI_ISL_2671560 | Centro de Investigacion Biomedica del Noreste (CIBIN) | Unidad de Genomica Avanzada | Consorcio Mexicano de Vigilancia Genomica (CoViGen-Mex). Authors (in alphabetical order): Julio Elias Alvarado-Yaah, Carlos F. Arias, Santiago avila-Rios, Eduardo Becerril-Vargas, Victor Hugo Borja-Aburto, Celia Boukadida, Cristobal Chaidez-Quiroz, Juan Bautista Chale-Dzul, Ricardo Ciria Merce, Andreu Comas-Garcia, Celida Duque Molina, Julissa Enciso-Ibarra, Jose Antonio Enciso-Moreno, Gloria Elena Espinosa-Ayala, Fernando Fontove-Herrera, Daniel Fregoso-Rueda, Victor Eduardo Garcia-Arias, Alejandra Garcia-Gasca, Bruno Gomez-Gil, Jean Pierre Gonzalez, Irvin Gonzalez-Lopez, Concepcion Grajales-Muñiz, Ricardo Grande, Rosa Maria Gutierrez Rios, Jesus Hernandez, Alejandra Hernandez-Teran, Alfredo Herrera-Estrella, Carla Ivon Herrera-Najera, Pavel Isa, Veronica Mata-Haro, Daniel Lira Morales, Susana Lopez, Antonio Loza Roman, Brenda Irasema Maldonado-Meza, Bernardo Martinez-Miguel, Jose Arturo Martinez-Orozco, Celida Martinez- Rodriguez, Margarita Matias-Florentino, Fidencio Mejia-Nepomuceno, Maria Guadalupe de Jesus Mireles-Rivera, Gloria Maria Molina-Salinas, Hector Montoya-Fuentes, , Mario Mujica-Sanchez, Jose Esteban Muñoz-Medina, Jose de Jesus Nuñez-Contreras, Alicia Ocaña-Mondragon, Luis Alberto Ochoa-Carrera, Hector Esteban Paz-Juarez, Marissa Perez-Garcia, Francisco Pulido, Helen Haydee Fernanda Ramirez-Plascencia, Jorge Salas-Hernandez, Angel Gustavo Salas-Lais, Alejandro Sanchez-Flores, Clara Esperanza Santacruz-Tinoco, Maria Guadalupe Santiago-Mauricio, Selene Zarate, Nelly Selem-Mojica, Blanca Taboada, Gloria Vazquez, Joel Armando Vazquez-Perez. |
| EPI_ISL_2671561, EPI_ISL_2671562, EPI_ISL_2671563 | Laboratorio Central de Epidemiologi-a (LCE) | Unidad de Genomica Avanzada | Consorcio Mexicano de Vigilancia Genomica (CoViGen-Mex). Authors (in alphabetical order): Julio Elias Alvarado-Yaah, Carlos F. Arias, Santiago avila-Rios, Eduardo Becerril-Vargas, Victor Hugo Borja-Aburto, Celia Boukadida, Cristobal Chaidez-Quiroz, Juan Bautista Chale-Dzul, Ricardo Ciria Merce, Andreu Comas-Garcia, Celida Duque Molina, Julissa Enciso-Ibarra, Jose Antonio Enciso-Moreno, Gloria Elena Espinosa-Ayala, Fernando Fontove-Herrera, Daniel Fregoso-Rueda, Victor Eduardo Garcia-Arias, Alejandra Garcia-Gasca, Bruno Gomez-Gil, Jean Pierre Gonzalez, Irvin Gonzalez-Lopez, Concepcion Grajales-Muñiz, Ricardo Grande, Rosa Maria Gutierrez Rios, Jesus Hernandez, Alejandra Hernandez-Teran, Alfredo Herrera-Estrella, Carla Ivon Herrera-Najera, Pavel Isa, Veronica Mata-Haro, Daniel Lira Morales, Susana Lopez, Antonio Loza Roman, Brenda Irasema Maldonado-Meza, Bernardo Martinez-Miguel, Jose Arturo Martinez-Orozco, Celida Martinez- Rodriguez, Margarita Matias-Florentino, Fidencio Mejia-Nepomuceno, Maria Guadalupe de Jesus Mireles-Rivera, Gloria Maria Molina-Salinas, Hector Montoya-Fuentes, , Mario Mujica-Sanchez, Jose Esteban Muñoz-Medina, Jose de Jesus Nuñez-Contreras, Alicia Ocaña-Mondragon, Luis Alberto Ochoa-Carrera, Hector Esteban Paz-Juarez, Marissa Perez-Garcia, Francisco Pulido, Helen Haydee Fernanda Ramirez-Plascencia, Jorge Salas-Hernandez, Angel Gustavo Salas-Lais, Alejandro Sanchez-Flores, Clara Esperanza Santacruz-Tinoco, Maria Guadalupe Santiago-Mauricio, Selene Zarate, Nelly Selem-Mojica, Blanca Taboada, Gloria Vazquez, Joel Armando Vazquez-Perez. |
| EPI_ISL_2671564, EPI_ISL_2671565, EPI_ISL_2671566, EPI_ISL_2671567, EPI_ISL_2671568, EPI_ISL_2671570, EPI_ISL_2671571, EPI_ISL_2671572, EPI_ISL_2671573, EPI_ISL_2671574, EPI_ISL_2671575, EPI_ISL_2671576, EPI_ISL_2671577, EPI_ISL_2671578, EPI_ISL_2671579, EPI_ISL_2671580, EPI_ISL_2671581, EPI_ISL_2671582, EPI_ISL_2671583, EPI_ISL_2671584, EPI_ISL_2671585, EPI_ISL_2671586 | | | |
| see above | Centro de Investigacion Biomedica de Occidente (CIBO) | Unidad de Genomica Avanzada | Consorcio Mexicano de Vigilancia Genomica (CoViGen-Mex). Authors (in alphabetical order): Julio Elias Alvarado-Yaah, Carlos F. Arias, Santiago avila-Rios, Eduardo Becerril-Vargas, Victor Hugo Borja-Aburto, Celia Boukadida, Cristobal Chaidez-Quiroz, Juan Bautista Chale-Dzul, Ricardo Ciria Merce, Andreu Comas-Garcia, Celida Duque Molina, Julissa Enciso-Ibarra, Jose Antonio Enciso-Moreno, Gloria Elena Espinosa-Ayala, Fernando Fontove-Herrera, Daniel Fregoso-Rueda, Victor Eduardo Garcia-Arias, Alejandra Garcia-Gasca, Bruno Gomez-Gil, Jean Pierre Gonzalez, Irvin Gonzalez-Lopez, Concepcion Grajales-Muñiz, Ricardo Grande, Rosa Maria Gutierrez Rios, Jesus Hernandez, Alejandra Hernandez-Teran, Alfredo Herrera-Estrella, Carla Ivon Herrera-Najera, Pavel Isa, Veronica Mata-Haro, Daniel Lira Morales, Susana Lopez, Antonio Loza Roman, Brenda Irasema Maldonado-Meza, Bernardo Martinez-Miguel, Jose Arturo Martinez-Orozco, Celida Martinez- Rodriguez, Margarita Matias-Florentino, Fidencio Mejia-Nepomuceno, Maria Guadalupe de Jesus Mireles-Rivera, Gloria Maria Molina-Salinas, Hector Montoya-Fuentes, , Mario Mujica-Sanchez, Jose Esteban Muñoz-Medina, Jose de Jesus Nuñez-Contreras, Alicia Ocaña-Mondragon, Luis Alberto Ochoa-Carrera, Hector Esteban Paz-Juarez, Marissa Perez-Garcia, Francisco Pulido, Helen Haydee Fernanda Ramirez-Plascencia, Jorge Salas-Hernandez, Angel Gustavo Salas-Lais, Alejandro Sanchez-Flores, Clara Esperanza Santacruz-Tinoco, Maria Guadalupe Santiago-Mauricio, Selene Zarate, Nelly Selem-Mojica, Blanca Taboada, Gloria Vazquez, Joel Armando Vazquez-Perez. |
| EPI_ISL_2671589, EPI_ISL_2671590, EPI_ISL_2671591, EPI_ISL_2671593, EPI_ISL_2671594, EPI_ISL_2671595, EPI_ISL_2671596, EPI_ISL_2671597, EPI_ISL_2671598, EPI_ISL_2671599, EPI_ISL_2671600, EPI_ISL_2671601, EPI_ISL_2671602, EPI_ISL_2671603, EPI_ISL_2671604, EPI_ISL_2671605, EPI_ISL_2671606, EPI_ISL_2671607 | | | |
| see above | Laboratorio Central de Epidemiologi-a (LCE) | Unidad de Genomica Avanzada | Consorcio Mexicano de Vigilancia Genomica (CoViGen-Mex). Authors (in alphabetical order): Julio Elias Alvarado-Yaah, Carlos F. Arias, Santiago avila-Rios, Eduardo Becerril-Vargas, Victor Hugo Borja-Aburto, Celia Boukadida, Cristobal Chaidez-Quiroz, Juan Bautista Chale-Dzul, Ricardo Ciria Merce, Andreu Comas-Garcia, Celida Duque Molina, Julissa Enciso-Ibarra, Jose Antonio Enciso-Moreno, Gloria Elena Espinosa-Ayala, Fernando Fontove-Herrera, Daniel Fregoso-Rueda, Victor Eduardo Garcia-Arias, Alejandra Garcia-Gasca, Bruno Gomez-Gil, Jean Pierre Gonzalez, Irvin Gonzalez-Lopez, Concepcion Grajales-Muñiz, Ricardo Grande, Rosa Maria Gutierrez Rios, Jesus Hernandez, Alejandra Hernandez-Teran, Alfredo Herrera-Estrella, Carla Ivon Herrera-Najera, Pavel Isa, Veronica Mata-Haro, Daniel Lira Morales, Susana Lopez, Antonio Loza Roman, Brenda Irasema Maldonado-Meza, Bernardo Martinez-Miguel, Jose Arturo Martinez-Orozco, Celida Martinez- Rodriguez, Margarita Matias-Florentino, Fidencio Mejia-Nepomuceno, Maria Guadalupe de Jesus Mireles-Rivera, Gloria Maria Molina-Salinas, Hector Montoya-Fuentes, , Mario Mujica-Sanchez, Jose Esteban Muñoz-Medina, Jose de Jesus Nuñez-Contreras, Alicia Ocaña-Mondragon, Luis Alberto Ochoa-Carrera, Hector Esteban Paz-Juarez, Marissa Perez-Garcia, Francisco Pulido, Helen Haydee Fernanda Ramirez-Plascencia, Jorge Salas-Hernandez, Angel Gustavo Salas-Lais, Alejandro Sanchez-Flores, Clara Esperanza Santacruz-Tinoco, Maria Guadalupe Santiago-Mauricio, Selene Zarate, Nelly Selem-Mojica, Blanca Taboada, Gloria Vazquez, Joel Armando Vazquez-Perez. |
| EPI_ISL_2671608, EPI_ISL_2671609, EPI_ISL_2671610, EPI_ISL_2671611, EPI_ISL_2671612 | Centro de Investigacion Biomedica de Occidente (CIBO) | Unidad de Genomica Avanzada | Consorcio Mexicano de Vigilancia Genomica (CoViGen-Mex). Authors (in alphabetical order): Julio Elias Alvarado-Yaah, Carlos F. Arias, Santiago avila-Rios, Eduardo Becerril-Vargas, Victor Hugo Borja-Aburto, Celia Boukadida, Cristobal Chaidez-Quiroz, Juan Bautista Chale-Dzul, Ricardo Ciria Merce, Andreu Comas-Garcia, Celida Duque Molina, Julissa Enciso-Ibarra, Jose Antonio Enciso-Moreno, Gloria Elena Espinosa-Ayala, Fernando Fontove-Herrera, Daniel Fregoso-Rueda, Victor Eduardo Garcia-Arias, Alejandra Garcia-Gasca, Bruno Gomez-Gil, Jean Pierre Gonzalez, Irvin Gonzalez-Lopez, Concepcion Grajales-Muñiz, Ricardo Grande, Rosa Maria Gutierrez Rios, Jesus Hernandez, Alejandra Hernandez-Teran, Alfredo Herrera-Estrella, Carla Ivon Herrera-Najera, Pavel Isa, Veronica Mata-Haro, Daniel Lira Morales, Susana Lopez, Antonio Loza Roman, Brenda Irasema Maldonado-Meza, Bernardo Martinez-Miguel, Jose Arturo Martinez-Orozco, Celida Martinez- Rodriguez, Margarita Matias-Florentino, Fidencio Mejia-Nepomuceno, Maria Guadalupe de Jesus Mireles-Rivera, Gloria Maria Molina-Salinas, Hector Montoya-Fuentes, , Mario Mujica-Sanchez, Jose Esteban Muñoz-Medina, Jose de Jesus Nuñez-Contreras, Alicia Ocaña-Mondragon, Luis Alberto Ochoa-Carrera, Hector Esteban Paz-Juarez, Marissa |

| | | | |
|---|---|---|---|
| | | | Perez-Garcia, Francisco Pulido, Helen Haydee Fernanda Ramirez-Plascencia, Jorge Salas-Hernandez, Angel Gustavo Salas-Lais, Alejandro Sanchez-Flores, Clara Esperanza Santacruz-Tinoco, Maria Guadalupe Santiago-Mauricio, Selene Zarate, Nelly Selem-Mojica, Blanca Taboada, Gloria Vazquez, Joel Armando Vazquez-Perez. |
| EPI_ISL_2671613, EPI_ISL_2671614, EPI_ISL_2671615 | Laboratorio Central de Epidemiologi-a (LCE) | Unidad de Genomica Avanzada | Consorcio Mexicano de Vigilancia Genomica (CoViGen-Mex). Authors (in alphabetical order): Julio Elias Alvarado-Yaah, Carlos F. Arias, Santiago avila-Rios, Eduardo Becerril-Vargas, Victor Hugo Borja-Aburto, Celia Boukadida, Cristobal Chaidez-Quiroz, Juan Bautista Chale-Dzul, Ricardo Ciria Merce, Andreu Comas-Garcia, Celida Duque Molina, Julissa Enciso-Ibarra, Jose Antonio Enciso-Moreno, Gloria Elena Espinosa-Ayala, Fernando Fontove-Herrera, Daniel Fregoso-Rueda, Victor Eduardo Garcia-Arias, Alejandra Garcia-Gasca, Bruno Gomez-Gil, Jean Pierre Gonzalez, Irvin Gonzalez-Lopez, Concepcion Grajales-Muñiz, Ricardo Grande, Rosa Maria Gutierrez Rios, Jesus Hernandez, Alejandra Hernandez-Teran, Alfredo Herrera-Estrella, Carla Ivon Herrera-Najera, Pavel Isa, Veronica Mata-Haro, Daniel Lira Morales, Susana Lopez, Antonio Loza Roman, Brenda Irasema Maldonado-Meza, Bernardo Martinez-Miguel, Jose Arturo Martinez-Orozco, Celida Martinez- Rodriguez, Margarita Matias-Florentino, Fidencio Mejia-Nepomuceno, Maria Guadalupe de Jesus Mireles-Rivera, Gloria Maria Molina-Salinas, Hector Montoya-Fuentes, , Mario Mujica-Sanchez, Jose Esteban Muñoz-Medina, Jose de Jesus Nuñez-Contreras, Alicia Ocaña-Mondragon, Luis Alberto Ochoa-Carrera, Hector Esteban Paz-Juarez, Marissa Perez-Garcia, Francisco Pulido, Helen Haydee Fernanda Ramirez-Plascencia, Jorge Salas-Hernandez, Angel Gustavo Salas-Lais, Alejandro Sanchez-Flores, Clara Esperanza Santacruz-Tinoco, Maria Guadalupe Santiago-Mauricio, Selene Zarate, Nelly Selem-Mojica, Blanca Taboada, Gloria Vazquez, Joel Armando Vazquez-Perez. |
| EPI_ISL_2671616, EPI_ISL_2671617, EPI_ISL_2671618 | Centro de Investigacion Biomedica de Occidente (CIBO) | Unidad de Genomica Avanzada | Consorcio Mexicano de Vigilancia Genomica (CoViGen-Mex). Authors (in alphabetical order): Julio Elias Alvarado-Yaah, Carlos F. Arias, Santiago avila-Rios, Eduardo Becerril-Vargas, Victor Hugo Borja-Aburto, Celia Boukadida, Cristobal Chaidez-Quiroz, Juan Bautista Chale-Dzul, Ricardo Ciria Merce, Andreu Comas-Garcia, Celida Duque Molina, Julissa Enciso-Ibarra, Jose Antonio Enciso-Moreno, Gloria Elena Espinosa-Ayala, Fernando Fontove-Herrera, Daniel Fregoso-Rueda, Victor Eduardo Garcia-Arias, Alejandra Garcia-Gasca, Bruno Gomez-Gil, Jean Pierre Gonzalez, Irvin Gonzalez-Lopez, Concepcion Grajales-Muñiz, Ricardo Grande, Rosa Maria Gutierrez Rios, Jesus Hernandez, Alejandra Hernandez-Teran, Alfredo Herrera-Estrella, Carla Ivon Herrera-Najera, Pavel Isa, Veronica Mata-Haro, Daniel Lira Morales, Susana Lopez, Antonio Loza Roman, Brenda Irasema Maldonado-Meza, Bernardo Martinez-Miguel, Jose Arturo Martinez-Orozco, Celida Martinez- Rodriguez, Margarita Matias-Florentino, Fidencio Mejia-Nepomuceno, Maria Guadalupe de Jesus Mireles-Rivera, Gloria Maria Molina-Salinas, Hector Montoya-Fuentes, , Mario Mujica-Sanchez, Jose Esteban Muñoz-Medina, Jose de Jesus Nuñez-Contreras, Alicia Ocaña-Mondragon, Luis Alberto Ochoa-Carrera, Hector Esteban Paz-Juarez, Marissa Perez-Garcia, Francisco Pulido, Helen Haydee Fernanda Ramirez-Plascencia, Jorge Salas-Hernandez, Angel Gustavo Salas-Lais, Alejandro Sanchez-Flores, Clara Esperanza Santacruz-Tinoco, Maria Guadalupe Santiago-Mauricio, Selene Zarate, Nelly Selem-Mojica, Blanca Taboada, Gloria Vazquez, Joel Armando Vazquez-Perez. |
| EPI_ISL_2671619, EPI_ISL_2671620, EPI_ISL_2671621, EPI_ISL_2671622, EPI_ISL_2671623, EPI_ISL_2671624, EPI_ISL_2671625, EPI_ISL_2671627, EPI_ISL_2671628, EPI_ISL_2671629, EPI_ISL_2671630, EPI_ISL_2671631, EPI_ISL_2671633, EPI_ISL_2671634, EPI_ISL_2671635, EPI_ISL_2671636 | | | |
| see above | Centro de Investigacion Biomedica del Noreste (CIBIN) | Unidad de Genomica Avanzada | Consorcio Mexicano de Vigilancia Genomica (CoViGen-Mex). Authors (in alphabetical order): Julio Elias Alvarado-Yaah, Carlos F. Arias, Santiago avila-Rios, Eduardo Becerril-Vargas, Victor Hugo Borja-Aburto, Celia Boukadida, Cristobal Chaidez-Quiroz, Juan Bautista Chale-Dzul, Ricardo Ciria Merce, Andreu Comas-Garcia, Celida Duque Molina, Julissa Enciso-Ibarra, Jose Antonio Enciso-Moreno, Gloria Elena Espinosa-Ayala, Fernando Fontove-Herrera, Daniel Fregoso-Rueda, Victor Eduardo Garcia-Arias, Alejandra Garcia-Gasca, Bruno Gomez-Gil, Jean Pierre Gonzalez, Irvin Gonzalez-Lopez, Concepcion Grajales-Muñiz, Ricardo Grande, Rosa Maria Gutierrez Rios, Jesus Hernandez, Alejandra Hernandez-Teran, Alfredo Herrera-Estrella, Carla Ivon Herrera-Najera, Pavel Isa, Veronica Mata-Haro, Daniel Lira Morales, Susana Lopez, Antonio Loza Roman, Brenda Irasema Maldonado-Meza, Bernardo Martinez-Miguel, Jose Arturo Martinez-Orozco, Celida Martinez- Rodriguez, Margarita Matias-Florentino, Fidencio Mejia-Nepomuceno, Maria Guadalupe de Jesus Mireles-Rivera, Gloria Maria Molina-Salinas, Hector Montoya-Fuentes, , Mario Mujica-Sanchez, Jose Esteban Muñoz-Medina, Jose de Jesus Nuñez-Contreras, Alicia Ocaña-Mondragon, Luis Alberto Ochoa-Carrera, Hector Esteban Paz-Juarez, Marissa Perez-Garcia, Francisco Pulido, Helen Haydee Fernanda Ramirez-Plascencia, Jorge Salas-Hernandez, Angel Gustavo Salas-Lais, Alejandro Sanchez-Flores, Clara Esperanza Santacruz-Tinoco, Maria Guadalupe Santiago-Mauricio, Selene Zarate, Nelly Selem-Mojica, Blanca Taboada, Gloria Vazquez, Joel Armando Vazquez-Perez. |
| EPI_ISL_2671637, EPI_ISL_2671639, EPI_ISL_2671640, EPI_ISL_2671641, EPI_ISL_2671642, EPI_ISL_2671643, EPI_ISL_2671644, EPI_ISL_2671645, EPI_ISL_2671646, EPI_ISL_2671647, EPI_ISL_2671648, EPI_ISL_2671649, EPI_ISL_2671650, EPI_ISL_2671651, EPI_ISL_2671652 | | | |
| see above | Laboratorio Central de Epidemiologi-a (LCE) | Unidad de Genomica Avanzada | Consorcio Mexicano de Vigilancia Genomica (CoViGen-Mex). Authors (in alphabetical order): Julio Elias Alvarado-Yaah, Carlos F. Arias, Santiago avila-Rios, Eduardo Becerril-Vargas, Victor Hugo Borja-Aburto, Celia Boukadida, Cristobal Chaidez-Quiroz, Juan Bautista Chale-Dzul, Ricardo Ciria Merce, Andreu Comas-Garcia, Celida Duque Molina, Julissa Enciso-Ibarra, Jose Antonio Enciso-Moreno, Gloria Elena Espinosa-Ayala, Fernando Fontove-Herrera, Daniel Fregoso-Rueda, Victor Eduardo Garcia-Arias, Alejandra Garcia-Gasca, Bruno Gomez-Gil, Jean Pierre Gonzalez, Irvin Gonzalez-Lopez, Concepcion Grajales-Muñiz, Ricardo Grande, Rosa Maria Gutierrez Rios, Jesus Hernandez, Alejandra Hernandez-Teran, Alfredo Herrera-Estrella, Carla Ivon Herrera-Najera, Pavel Isa, Veronica Mata-Haro, Daniel Lira Morales, Susana Lopez, Antonio Loza Roman, Brenda Irasema Maldonado-Meza, Bernardo Martinez-Miguel, Jose Arturo Martinez-Orozco, Celida Martinez- Rodriguez, Margarita Matias-Florentino, Fidencio Mejia-Nepomuceno, Maria Guadalupe de Jesus Mireles-Rivera, Gloria Maria Molina-Salinas, Hector Montoya-Fuentes, , Mario Mujica-Sanchez, Jose Esteban Muñoz-Medina, Jose de Jesus Nuñez-Contreras, Alicia Ocaña-Mondragon, Luis Alberto Ochoa-Carrera, Hector Esteban Paz-Juarez, Marissa Perez-Garcia, Francisco Pulido, Helen Haydee Fernanda Ramirez-Plascencia, Jorge Salas-Hernandez, Angel Gustavo Salas-Lais, Alejandro Sanchez-Flores, Clara Esperanza Santacruz-Tinoco, Maria Guadalupe Santiago-Mauricio, Selene Zarate, Nelly Selem-Mojica, Blanca Taboada, Gloria Vazquez, Joel Armando Vazquez-Perez. |
| EPI_ISL_2671653, EPI_ISL_2671654, EPI_ISL_2671655, EPI_ISL_2671656, EPI_ISL_2671657, EPI_ISL_2671658, EPI_ISL_2671659, EPI_ISL_2671660, EPI_ISL_2671663, EPI_ISL_2671664, EPI_ISL_2671666, EPI_ISL_2671667, EPI_ISL_2671669, EPI_ISL_2671670, EPI_ISL_2671671, EPI_ISL_2671672, EPI_ISL_2671673, EPI_ISL_2671674, EPI_ISL_2671676, EPI_ISL_2671677, EPI_ISL_2671678, EPI_ISL_2671679, EPI_ISL_2671680, EPI_ISL_2671681, EPI_ISL_2671683, EPI_ISL_2671684, EPI_ISL_2671685, EPI_ISL_2671686, EPI_ISL_2671687, EPI_ISL_2671688, EPI_ISL_2671689, EPI_ISL_2671690, EPI_ISL_2671692, EPI_ISL_2671694, EPI_ISL_2671695, EPI_ISL_2671696, EPI_ISL_2671697 | | | |
| see above | Unidad de Investigacion Medica de Yucatan (UIMY) | Unidad de Genomica Avanzada | Consorcio Mexicano de Vigilancia Genomica (CoViGen-Mex). Authors (in alphabetical order): Julio Elias Alvarado-Yaah, Carlos F. Arias, Santiago avila-Rios, Eduardo Becerril-Vargas, Victor Hugo Borja-Aburto, Celia Boukadida, Cristobal Chaidez-Quiroz, Juan Bautista Chale-Dzul, Ricardo Ciria Merce, Andreu Comas-Garcia, Celida Duque Molina, Julissa Enciso-Ibarra, Jose Antonio Enciso-Moreno, Gloria Elena Espinosa-Ayala, Fernando Fontove-Herrera, Daniel Fregoso-Rueda, Victor Eduardo Garcia-Arias, Alejandra Garcia-Gasca, Bruno Gomez-Gil, Jean Pierre Gonzalez, Irvin Gonzalez-Lopez, Concepcion Grajales-Muñiz, Ricardo Grande, Rosa Maria Gutierrez Rios, Jesus Hernandez, Alejandra Hernandez-Teran, Alfredo Herrera-Estrella, Carla Ivon Herrera-Najera, Pavel Isa, Veronica Mata-Haro, Daniel Lira Morales, Susana Lopez, Antonio Loza Roman, Brenda Irasema Maldonado-Meza, Bernardo Martinez-Miguel, Jose Arturo Martinez-Orozco, Celida Martinez- Rodriguez, Margarita Matias-Florentino, Fidencio Mejia-Nepomuceno, Maria Guadalupe de Jesus Mireles-Rivera, Gloria Maria Molina-Salinas, Hector Montoya-Fuentes, , Mario Mujica-Sanchez, Jose Esteban Muñoz-Medina, Jose de Jesus Nuñez-Contreras, Alicia Ocaña-Mondragon, Luis Alberto Ochoa-Carrera, Hector Esteban Paz-Juarez, Marissa Perez-Garcia, Francisco Pulido, Helen Haydee Fernanda Ramirez-Plascencia, Jorge Salas-Hernandez, Angel Gustavo Salas-Lais, Alejandro Sanchez-Flores, Clara Esperanza Santacruz-Tinoco, Maria Guadalupe Santiago-Mauricio, Selene Zarate, Nelly Selem-Mojica, Blanca Taboada, Gloria Vazquez, Joel Armando Vazquez-Perez. |
| EPI_ISL_2671698, EPI_ISL_2671699 | Centro de Investigacion Biomedica de Occidente (CIBO) | Unidad de Genomica Avanzada | Consorcio Mexicano de Vigilancia Genomica (CoViGen-Mex). Authors (in alphabetical order): Julio Elias Alvarado-Yaah, Carlos F. Arias, Santiago avila-Rios, Eduardo Becerril-Vargas, Victor Hugo Borja-Aburto, Celia Boukadida, Cristobal Chaidez-Quiroz, Juan Bautista Chale-Dzul, Ricardo Ciria Merce, Andreu Comas-Garcia, Celida Duque Molina, Julissa Enciso-Ibarra, Jose Antonio Enciso-Moreno, Gloria Elena Espinosa-Ayala, Fernando Fontove-Herrera, Daniel Fregoso-Rueda, Victor Eduardo Garcia-Arias, Alejandra Garcia-Gasca, Bruno Gomez-Gil, Jean Pierre Gonzalez, Irvin Gonzalez-Lopez, Concepcion Grajales-Muñiz, Ricardo Grande, Rosa Maria Gutierrez Rios, Jesus Hernandez, Alejandra Hernandez-Teran, Alfredo Herrera-Estrella, Carla Ivon Herrera-Najera, Pavel Isa, Veronica Mata-Haro, Daniel Lira Morales, Susana Lopez, Antonio Loza Roman, Brenda Irasema Maldonado-Meza, Bernardo Martinez-Miguel, Jose Arturo Martinez-Orozco, Celida Martinez- Rodriguez, Margarita Matias-Florentino, Fidencio Mejia-Nepomuceno, Maria Guadalupe de Jesus Mireles-Rivera, Gloria Maria Molina-Salinas, Hector Montoya-Fuentes, , Mario Mujica-Sanchez, Jose Esteban Muñoz-Medina, Jose de Jesus Nuñez-Contreras, Alicia Ocaña-Mondragon, Luis Alberto Ochoa-Carrera, Hector Esteban Paz-Juarez, Marissa Perez-Garcia, Francisco Pulido, Helen Haydee Fernanda Ramirez-Plascencia, Jorge Salas-Hernandez, Angel Gustavo Salas-Lais, Alejandro Sanchez-Flores, Clara Esperanza Santacruz-Tinoco, Maria Guadalupe Santiago-Mauricio, Selene Zarate, Nelly Selem-Mojica, Blanca Taboada, Gloria Vazquez, Joel Armando Vazquez-Perez. |
| EPI_ISL_2671700, EPI_ISL_2671701, EPI_ISL_2671702 | Unidad de Investigacion Biomedica de Zacatecas (UIBZ) | Unidad de Genomica Avanzada | Consorcio Mexicano de Vigilancia Genomica (CoViGen-Mex). Authors (in alphabetical order): Julio Elias Alvarado-Yaah, Carlos F. Arias, Santiago avila-Rios, Eduardo Becerril-Vargas, Victor Hugo Borja-Aburto, Celia Boukadida, Cristobal Chaidez-Quiroz, Juan Bautista Chale-Dzul, Ricardo Ciria |

| | | | |
|---|---|---|---|
| | | | Merce, Andreu Comas-Garcia, Celida Duque Molina, Julissa Enciso-Ibarra, Jose Antonio Enciso-Moreno, Gloria Elena Espinosa-Ayala, Fernando Fontove-Herrera, Daniel Fregoso-Rueda, Victor Eduardo Garcia-Arias, Alejandra Garcia-Gasca, Bruno Gomez-Gil, Jean Pierre Gonzalez, Irvin Gonzalez-Lopez, Concepcion Grajales-Muñiz, Ricardo Grande, Rosa Maria Gutierrez Rios, Jesus Hernandez, Alejandra Hernandez-Teran, Alfredo Herrera-Estrella, Carla Ivon Herrera-Najera, Pavel Isa, Veronica Mata-Haro, Daniel Lira Morales, Susana Lopez, Antonio Loza Roman, Brenda Irasema Maldonado-Meza, Bernardo Martinez-Miguel, Jose Arturo Martinez-Orozco, Celida Martinez- Rodriguez, Margarita Matias-Florentino, Fidencio Mejia-Nepomuceno, Maria Guadalupe de Jesus Mireles-Rivera, Gloria Maria Molina-Salinas, Hector Montoya-Fuentes, , Mario Mujica-Sanchez, Jose Esteban Muñoz-Medina, Jose de Jesus Nuñez-Contreras, Alicia Ocaña-Mondragon, Luis Alberto Ochoa-Carrera, Hector Esteban Paz-Juarez, Marissa Perez-Garcia, Francisco Pulido, Helen Haydee Fernanda Ramirez-Plascencia, Jorge Salas-Hernandez, Angel Gustavo Salas-Lais, Alejandro Sanchez-Flores, Clara Esperanza Santacruz-Tinoco, Maria Guadalupe Santiago-Mauricio, Selene Zarate, Nelly Selem-Mojica, Blanca Taboada, Gloria Vazquez, Joel Armando Vazquez-Perez. |
| EPI_ISL_2671703 | Centro de Investigacion Biomedica de Occidente (CIBO) | Unidad de Genomica Avanzada | Consorcio Mexicano de Vigilancia Genomica (CoViGen-Mex). Authors (in alphabetical order): Julio Elias Alvarado-Yaah, Carlos F. Arias, Santiago avila-Rios, Eduardo Becerril-Vargas, Victor Hugo Borja-Aburto, Celia Boukadida, Cristobal Chaidez-Quiroz, Juan Bautista Chale-Dzul, Ricardo Ciria Merce, Andreu Comas-Garcia, Celida Duque Molina, Julissa Enciso-Ibarra, Jose Antonio Enciso-Moreno, Gloria Elena Espinosa-Ayala, Fernando Fontove-Herrera, Daniel Fregoso-Rueda, Victor Eduardo Garcia-Arias, Alejandra Garcia-Gasca, Bruno Gomez-Gil, Jean Pierre Gonzalez, Irvin Gonzalez-Lopez, Concepcion Grajales-Muñiz, Ricardo Grande, Rosa Maria Gutierrez Rios, Jesus Hernandez, Alejandra Hernandez-Teran, Alfredo Herrera-Estrella, Carla Ivon Herrera-Najera, Pavel Isa, Veronica Mata-Haro, Daniel Lira Morales, Susana Lopez, Antonio Loza Roman, Brenda Irasema Maldonado-Meza, Bernardo Martinez-Miguel, Jose Arturo Martinez-Orozco, Celida Martinez- Rodriguez, Margarita Matias-Florentino, Fidencio Mejia-Nepomuceno, Maria Guadalupe de Jesus Mireles-Rivera, Gloria Maria Molina-Salinas, Hector Montoya-Fuentes, , Mario Mujica-Sanchez, Jose Esteban Muñoz-Medina, Jose de Jesus Nuñez-Contreras, Alicia Ocaña-Mondragon, Luis Alberto Ochoa-Carrera, Hector Esteban Paz-Juarez, Marissa Perez-Garcia, Francisco Pulido, Helen Haydee Fernanda Ramirez-Plascencia, Jorge Salas-Hernandez, Angel Gustavo Salas-Lais, Alejandro Sanchez-Flores, Clara Esperanza Santacruz-Tinoco, Maria Guadalupe Santiago-Mauricio, Selene Zarate, Nelly Selem-Mojica, Blanca Taboada, Gloria Vazquez, Joel Armando Vazquez-Perez. |
| EPI_ISL_2671704, EPI_ISL_2671706, EPI_ISL_2671707, EPI_ISL_2671708, EPI_ISL_2671709, EPI_ISL_2671710, EPI_ISL_2671712, EPI_ISL_2671714, EPI_ISL_2671715 | Unidad de Investigacion Medica de Yucatan (UIMY) | Unidad de Genomica Avanzada | Consorcio Mexicano de Vigilancia Genomica (CoViGen-Mex). Authors (in alphabetical order): Julio Elias Alvarado-Yaah, Carlos F. Arias, Santiago avila-Rios, Eduardo Becerril-Vargas, Victor Hugo Borja-Aburto, Celia Boukadida, Cristobal Chaidez-Quiroz, Juan Bautista Chale-Dzul, Ricardo Ciria Merce, Andreu Comas-Garcia, Celida Duque Molina, Julissa Enciso-Ibarra, Jose Antonio Enciso-Moreno, Gloria Elena Espinosa-Ayala, Fernando Fontove-Herrera, Daniel Fregoso-Rueda, Victor Eduardo Garcia-Arias, Alejandra Garcia-Gasca, Bruno Gomez-Gil, Jean Pierre Gonzalez, Irvin Gonzalez-Lopez, Concepcion Grajales-Muñiz, Ricardo Grande, Rosa Maria Gutierrez Rios, Jesus Hernandez, Alejandra Hernandez-Teran, Alfredo Herrera-Estrella, Carla Ivon Herrera-Najera, Pavel Isa, Veronica Mata-Haro, Daniel Lira Morales, Susana Lopez, Antonio Loza Roman, Brenda Irasema Maldonado-Meza, Bernardo Martinez-Miguel, Jose Arturo Martinez-Orozco, Celida Martinez- Rodriguez, Margarita Matias-Florentino, Fidencio Mejia-Nepomuceno, Maria Guadalupe de Jesus Mireles-Rivera, Gloria Maria Molina-Salinas, Hector Montoya-Fuentes, , Mario Mujica-Sanchez, Jose Esteban Muñoz-Medina, Jose de Jesus Nuñez-Contreras, Alicia Ocaña-Mondragon, Luis Alberto Ochoa-Carrera, Hector Esteban Paz-Juarez, Marissa Perez-Garcia, Francisco Pulido, Helen Haydee Fernanda Ramirez-Plascencia, Jorge Salas-Hernandez, Angel Gustavo Salas-Lais, Alejandro Sanchez-Flores, Clara Esperanza Santacruz-Tinoco, Maria Guadalupe Santiago-Mauricio, Selene Zarate, Nelly Selem-Mojica, Blanca Taboada, Gloria Vazquez, Joel Armando Vazquez-Perez. |
| EPI_ISL_2671716, EPI_ISL_2671717, EPI_ISL_2671718, EPI_ISL_2671719, EPI_ISL_2671721, EPI_ISL_2671722 | Centro de Investigacion Biomedica del Noreste (CIBIN) | Unidad de Genomica Avanzada | Consorcio Mexicano de Vigilancia Genomica (CoViGen-Mex). Authors (in alphabetical order): Julio Elias Alvarado-Yaah, Carlos F. Arias, Santiago avila-Rios, Eduardo Becerril-Vargas, Victor Hugo Borja-Aburto, Celia Boukadida, Cristobal Chaidez-Quiroz, Juan Bautista Chale-Dzul, Ricardo Ciria Merce, Andreu Comas-Garcia, Celida Duque Molina, Julissa Enciso-Ibarra, Jose Antonio Enciso-Moreno, Gloria Elena Espinosa-Ayala, Fernando Fontove-Herrera, Daniel Fregoso-Rueda, Victor Eduardo Garcia-Arias, Alejandra Garcia-Gasca, Bruno Gomez-Gil, Jean Pierre Gonzalez, Irvin Gonzalez-Lopez, Concepcion Grajales-Muñiz, Ricardo Grande, Rosa Maria Gutierrez Rios, Jesus Hernandez, Alejandra Hernandez-Teran, Alfredo Herrera-Estrella, Carla Ivon Herrera-Najera, Pavel Isa, Veronica Mata-Haro, Daniel Lira Morales, Susana Lopez, Antonio Loza Roman, Brenda Irasema Maldonado-Meza, Bernardo Martinez-Miguel, Jose Arturo Martinez-Orozco, Celida Martinez- Rodriguez, Margarita Matias-Florentino, Fidencio Mejia-Nepomuceno, Maria Guadalupe de Jesus Mireles-Rivera, Gloria Maria Molina-Salinas, Hector Montoya-Fuentes, , Mario Mujica-Sanchez, Jose Esteban Muñoz-Medina, Jose de Jesus Nuñez-Contreras, Alicia Ocaña-Mondragon, Luis Alberto Ochoa-Carrera, Hector Esteban Paz-Juarez, Marissa Perez-Garcia, Francisco Pulido, Helen Haydee Fernanda Ramirez-Plascencia, Jorge Salas-Hernandez, Angel Gustavo Salas-Lais, Alejandro Sanchez-Flores, Clara Esperanza Santacruz-Tinoco, Maria Guadalupe Santiago-Mauricio, Selene Zarate, Nelly Selem-Mojica, Blanca Taboada, Gloria Vazquez, Joel Armando Vazquez-Perez. |
| EPI_ISL_2671723, EPI_ISL_2671726, EPI_ISL_2671728, EPI_ISL_2671729, EPI_ISL_2671730, EPI_ISL_2671731, EPI_ISL_2671732, EPI_ISL_2671733, EPI_ISL_2671734, EPI_ISL_2671735, EPI_ISL_2671736, EPI_ISL_2671737, EPI_ISL_2671739, EPI_ISL_2671740, EPI_ISL_2671741, EPI_ISL_2671742, EPI_ISL_2671743, EPI_ISL_2671744, EPI_ISL_2671745 | | | |
| see above | Laboratorio Central de Epidemiologi-a (LCE) | Unidad de Genomica Avanzada | Consorcio Mexicano de Vigilancia Genomica (CoViGen-Mex). Authors (in alphabetical order): Julio Elias Alvarado-Yaah, Carlos F. Arias, Santiago avila-Rios, Eduardo Becerril-Vargas, Victor Hugo Borja-Aburto, Celia Boukadida, Cristobal Chaidez-Quiroz, Juan Bautista Chale-Dzul, Ricardo Ciria Merce, Andreu Comas-Garcia, Celida Duque Molina, Julissa Enciso-Ibarra, Jose Antonio Enciso-Moreno, Gloria Elena Espinosa-Ayala, Fernando Fontove-Herrera, Daniel Fregoso-Rueda, Victor Eduardo Garcia-Arias, Alejandra Garcia-Gasca, Bruno Gomez-Gil, Jean Pierre Gonzalez, Irvin Gonzalez-Lopez, Concepcion Grajales-Muñiz, Ricardo Grande, Rosa Maria Gutierrez Rios, Jesus Hernandez, Alejandra Hernandez-Teran, Alfredo Herrera-Estrella, Carla Ivon Herrera-Najera, Pavel Isa, Veronica Mata-Haro, Daniel Lira Morales, Susana Lopez, Antonio Loza Roman, Brenda Irasema Maldonado-Meza, Bernardo Martinez-Miguel, Jose Arturo Martinez-Orozco, Celida Martinez- Rodriguez, Margarita Matias-Florentino, Fidencio Mejia-Nepomuceno, Maria Guadalupe de Jesus Mireles-Rivera, Gloria Maria Molina-Salinas, Hector Montoya-Fuentes, , Mario Mujica-Sanchez, Jose Esteban Muñoz-Medina, Jose de Jesus Nuñez-Contreras, Alicia Ocaña-Mondragon, Luis Alberto Ochoa-Carrera, Hector Esteban Paz-Juarez, Marissa Perez-Garcia, Francisco Pulido, Helen Haydee Fernanda Ramirez-Plascencia, Jorge Salas-Hernandez, Angel Gustavo Salas-Lais, Alejandro Sanchez-Flores, Clara Esperanza Santacruz-Tinoco, Maria Guadalupe Santiago-Mauricio, Selene Zarate, Nelly Selem-Mojica, Blanca Taboada, Gloria Vazquez, Joel Armando Vazquez-Perez. |
| EPI_ISL_2671746, EPI_ISL_2671747, EPI_ISL_2671748, EPI_ISL_2671749, EPI_ISL_2671750, EPI_ISL_2671752, EPI_ISL_2671753, EPI_ISL_2671754, EPI_ISL_2671755, EPI_ISL_2671756, EPI_ISL_2671757 | | | |
| see above | Unidad de Investigacion Medica de Yucatan (UIMY) | Unidad de Genomica Avanzada | Consorcio Mexicano de Vigilancia Genomica (CoViGen-Mex). Authors (in alphabetical order): Julio Elias Alvarado-Yaah, Carlos F. Arias, Santiago avila-Rios, Eduardo Becerril-Vargas, Victor Hugo Borja-Aburto, Celia Boukadida, Cristobal Chaidez-Quiroz, Juan Bautista Chale-Dzul, Ricardo Ciria Merce, Andreu Comas-Garcia, Celida Duque Molina, Julissa Enciso-Ibarra, Jose Antonio Enciso-Moreno, Gloria Elena Espinosa-Ayala, Fernando Fontove-Herrera, Daniel Fregoso-Rueda, Victor Eduardo Garcia-Arias, Alejandra Garcia-Gasca, Bruno Gomez-Gil, Jean Pierre Gonzalez, Irvin Gonzalez-Lopez, Concepcion Grajales-Muñiz, Ricardo Grande, Rosa Maria Gutierrez Rios, Jesus Hernandez, Alejandra Hernandez-Teran, Alfredo Herrera-Estrella, Carla Ivon Herrera-Najera, Pavel Isa, Veronica Mata-Haro, Daniel Lira Morales, Susana Lopez, Antonio Loza Roman, Brenda Irasema Maldonado-Meza, Bernardo Martinez-Miguel, Jose Arturo Martinez-Orozco, Celida Martinez- Rodriguez, Margarita Matias-Florentino, Fidencio Mejia-Nepomuceno, Maria Guadalupe de Jesus Mireles-Rivera, Gloria Maria Molina-Salinas, Hector Montoya-Fuentes, , Mario Mujica-Sanchez, Jose Esteban Muñoz-Medina, Jose de Jesus Nuñez-Contreras, Alicia Ocaña-Mondragon, Luis Alberto Ochoa-Carrera, Hector Esteban Paz-Juarez, Marissa Perez-Garcia, Francisco Pulido, Helen Haydee Fernanda Ramirez-Plascencia, Jorge Salas-Hernandez, Angel Gustavo Salas-Lais, Alejandro Sanchez-Flores, Clara Esperanza Santacruz-Tinoco, Maria Guadalupe Santiago-Mauricio, Selene Zarate, Nelly Selem-Mojica, Blanca Taboada, Gloria Vazquez, Joel Armando Vazquez-Perez. |
| EPI_ISL_2671758, EPI_ISL_2671760, EPI_ISL_2671761, EPI_ISL_2671762, EPI_ISL_2671764, EPI_ISL_2671766, EPI_ISL_2671767, EPI_ISL_2671768, EPI_ISL_2671769, EPI_ISL_2671770, EPI_ISL_2671771, EPI_ISL_2671772, EPI_ISL_2671774, EPI_ISL_2671775, EPI_ISL_2671776, EPI_ISL_2671777, EPI_ISL_2671779, EPI_ISL_2671780 | | | |
| see above | Unidad de Investigacion Biomedica de Zacatecas (UIBZ) | Unidad de Genomica Avanzada | Consorcio Mexicano de Vigilancia Genomica (CoViGen-Mex). Authors (in alphabetical order): Julio Elias Alvarado-Yaah, Carlos F. Arias, Santiago avila-Rios, Eduardo Becerril-Vargas, Victor Hugo Borja-Aburto, Celia Boukadida, Cristobal Chaidez-Quiroz, Juan Bautista Chale-Dzul, Ricardo Ciria Merce, Andreu Comas-Garcia, Celida Duque Molina, Julissa Enciso-Ibarra, Jose Antonio Enciso-Moreno, Gloria Elena Espinosa-Ayala, Fernando Fontove-Herrera, Daniel Fregoso-Rueda, Victor Eduardo Garcia-Arias, Alejandra Garcia-Gasca, Bruno Gomez-Gil, Jean Pierre Gonzalez, Irvin Gonzalez-Lopez, Concepcion Grajales-Muñiz, Ricardo Grande, Rosa Maria Gutierrez Rios, Jesus Hernandez, Alejandra Hernandez-Teran, Alfredo Herrera-Estrella, Carla Ivon Herrera-Najera, Pavel Isa, Veronica Mata-Haro, Daniel Lira Morales, Susana Lopez, Antonio Loza Roman, Brenda Irasema Maldonado-Meza, Bernardo Martinez-Miguel, Jose Arturo Martinez-Orozco, Celida Martinez- Rodriguez, Margarita Matias-Florentino, Fidencio Mejia-Nepomuceno, Maria Guadalupe de Jesus Mireles-Rivera, Gloria Maria Molina-Salinas, Hector Montoya-Fuentes, , Mario Mujica-Sanchez, Jose |

| | | | |
|---|---|---|---|
| | | | Esteban Muñoz-Medina, Jose de Jesus Nuñez-Contreras, Alicia Ocaña-Mondragon, Luis Alberto Ochoa-Carrera, Hector Esteban Paz-Juarez, Marissa Perez-Garcia, Francisco Pulido, Helen Haydee Fernanda Ramirez-Plascencia, Jorge Salas-Hernandez, Angel Gustavo Salas-Lais, Alejandro Sanchez-Flores, Clara Esperanza Santacruz-Tinoco, Maria Guadalupe Santiago-Mauricio, Selene Zarate, Nelly Selem-Mojica, Blanca Taboada, Gloria Vazquez, Joel Armando Vazquez-Perez. |
| EPI_ISL_2671782, EPI_ISL_2671783, EPI_ISL_2671785 | Unidad de Investigacion Medica de Yucatan (UIMY) | Unidad de Genomica Avanzada | Consorcio Mexicano de Vigilancia Genomica (CoViGen-Mex). Authors (in alphabetical order): Julio Elias Alvarado-Yaah, Carlos F. Arias, Santiago avila-Rios, Eduardo Becerril-Vargas, Victor Hugo Borja-Aburto, Celia Boukadida, Cristobal Chaidez-Quiroz, Juan Bautista Chale-Dzul, Ricardo Ciria Merce, Andreu Comas-Garcia, Celida Duque Molina, Julissa Enciso-Ibarra, Jose Antonio Enciso-Moreno, Gloria Elena Espinosa-Ayala, Fernando Fontove-Herrera, Daniel Fregoso-Rueda, Victor Eduardo Garcia-Arias, Alejandra Garcia-Gasca, Bruno Gomez-Gil, Jean Pierre Gonzalez, Irvin Gonzalez-Lopez, Concepcion Grajales-Muñiz, Ricardo Grande, Rosa Maria Gutierrez Rios, Jesus Hernandez, Alejandra Hernandez-Teran, Alfredo Herrera-Estrella, Carla Ivon Herrera-Najera, Pavel Isa, Veronica Mata-Haro, Daniel Lira Morales, Susana Lopez, Antonio Loza Roman, Brenda Irasema Maldonado-Meza, Bernardo Martinez-Miguel, Jose Arturo Martinez-Orozco, Celida Martinez- Rodriguez, Margarita Matias-Florentino, Fidencio Mejia-Nepomuceno, Maria Guadalupe de Jesus Mireles-Rivera, Gloria Maria Molina-Salinas, Hector Montoya-Fuentes, , Mario Mujica-Sanchez, Jose Esteban Muñoz-Medina, Jose de Jesus Nuñez-Contreras, Alicia Ocaña-Mondragon, Luis Alberto Ochoa-Carrera, Hector Esteban Paz-Juarez, Marissa Perez-Garcia, Francisco Pulido, Helen Haydee Fernanda Ramirez-Plascencia, Jorge Salas-Hernandez, Angel Gustavo Salas-Lais, Alejandro Sanchez-Flores, Clara Esperanza Santacruz-Tinoco, Maria Guadalupe Santiago-Mauricio, Selene Zarate, Nelly Selem-Mojica, Blanca Taboada, Gloria Vazquez, Joel Armando Vazquez-Perez. |
| EPI_ISL_2671834 | COVID LAB | Cayman Islands Forensic Science Lab | Jonathan Smellie, Karla Montes, Brittany Balcewich, Allen Grolla |
| EPI_ISL_2673506, EPI_ISL_2673507, EPI_ISL_2673508, EPI_ISL_2673509, EPI_ISL_2673510, EPI_ISL_2673511, EPI_ISL_2673512, EPI_ISL_2673513, EPI_ISL_2673514, EPI_ISL_2673515, EPI_ISL_2673516, EPI_ISL_2673517, EPI_ISL_2673518 | | | |
| see above | Department of Virology, Istituto Zooprofilattico Sperimentale del Lazio e della Toscana (IZSLT) | Department of General Diagnostics; Department of Virology; Istituto Zooprofilattico Sperimentale del Lazio e della Toscana (IZSLT) | Patricia Alba, Giuseppe Manna, Elena L. Diaconu, Fabiola Feltrin, Raffaela Conti, Teresa Scicluna, Virginia Carfora, Antonella Cersini, Alessia Franco, Antonio Battisti. |
| EPI_ISL_2673648, EPI_ISL_2673649, EPI_ISL_2673650, EPI_ISL_2673651 | SIESP CHIETI - DRIVE IN VASTO | Istituto Zooprofilattico Sperimentale dell'Abruzzo e Molise "G. Caporale" | Lorusso A, Marcacci M, Di Domenico M, Ancora M, Curini V, Di Lollo Valeria, Mangone I, Rinaldi A, Delli Compagni E, Scialabba S, Caporale M, Di Pasquale A, Cammà C, Puglia I, Calistri P, Savini G |
| EPI_ISL_2673652 | SIESP DIPARTIMENTO DI PREVENZIONE TERAMO | Istituto Zooprofilattico Sperimentale dell'Abruzzo e Molise "G. Caporale" | Lorusso A, Marcacci M, Di Domenico M, Ancora M, Curini V, Di Lollo Valeria, Mangone I, Rinaldi A, Delli Compagni E, Scialabba S, Caporale M, Di Pasquale A, Cammà C, Puglia I, Calistri P, Savini G |
| EPI_ISL_2673653 | SIESP CHIETI - DRIVE IN CHIETI | Istituto Zooprofilattico Sperimentale dell'Abruzzo e Molise "G. Caporale" | Lorusso A, Marcacci M, Di Domenico M, Ancora M, Curini V, Di Lollo Valeria, Mangone I, Rinaldi A, Delli Compagni E, Scialabba S, Caporale M, Di Pasquale A, Cammà C, Puglia I, Calistri P, Savini G |
| EPI_ISL_2673654 | SIESP CHIETI - DRIVE IN VASTO | Istituto Zooprofilattico Sperimentale dell'Abruzzo e Molise "G. Caporale" | Lorusso A, Marcacci M, Di Domenico M, Ancora M, Curini V, Di Lollo Valeria, Mangone I, Rinaldi A, Delli Compagni E, Scialabba S, Caporale M, Di Pasquale A, Cammà C, Puglia I, Calistri P, Savini G |
| EPI_ISL_2673655 | SIESP CHIETI - DRIVE IN CHIETI | Istituto Zooprofilattico Sperimentale dell'Abruzzo e Molise "G. Caporale" | Lorusso A, Marcacci M, Di Domenico M, Ancora M, Curini V, Di Lollo Valeria, Mangone I, Rinaldi A, Delli Compagni E, Scialabba S, Caporale M, Di Pasquale A, Cammà C, Puglia I, Calistri P, Savini G |
| EPI_ISL_2673656 | SIESP DIPARTIMENTO DI PREVENZIONE CHIETI | Istituto Zooprofilattico Sperimentale dell'Abruzzo e Molise "G. Caporale" | Lorusso A, Marcacci M, Di Domenico M, Ancora M, Curini V, Di Lollo Valeria, Mangone I, Rinaldi A, Delli Compagni E, Scialabba S, Caporale M, Di Pasquale A, Cammà C, Puglia I, Calistri P, Savini G |
| EPI_ISL_2673657, EPI_ISL_2673658 | SIESP CHIETI - DRIVE IN LANCIANO | Istituto Zooprofilattico Sperimentale dell'Abruzzo e Molise "G. Caporale" | Lorusso A, Marcacci M, Di Domenico M, Ancora M, Curini V, Di Lollo Valeria, Mangone I, Rinaldi A, Delli Compagni E, Scialabba S, Caporale M, Di Pasquale A, Cammà C, Puglia I, Calistri P, Savini G |
| EPI_ISL_2673659 | SIESP DIPARTIMENTO DI PREVENZIONE CHIETI | Istituto Zooprofilattico Sperimentale dell'Abruzzo e Molise "G. Caporale" | Lorusso A, Marcacci M, Di Domenico M, Ancora M, Curini V, Di Lollo Valeria, Mangone I, Rinaldi A, Delli Compagni E, Scialabba S, Caporale M, Di Pasquale A, Cammà C, Puglia I, Calistri P, Savini G |
| EPI_ISL_2673660, EPI_ISL_2673661, EPI_ISL_2673662, EPI_ISL_2673663, EPI_ISL_2673664 | SIESP CHIETI - DRIVE IN VASTO | Istituto Zooprofilattico Sperimentale dell'Abruzzo e Molise "G. Caporale" | Lorusso A, Marcacci M, Di Domenico M, Ancora M, Curini V, Di Lollo Valeria, Mangone I, Rinaldi A, Delli Compagni E, Scialabba S, Caporale M, Di Pasquale A, Cammà C, Puglia I, Calistri P, Savini G |
| EPI_ISL_2673665 | SIESP CHIETI - DRIVE IN ORTONA | Istituto Zooprofilattico Sperimentale dell'Abruzzo e Molise "G. Caporale" | Lorusso A, Marcacci M, Di Domenico M, Ancora M, Curini V, Di Lollo Valeria, Mangone I, Rinaldi A, Delli Compagni E, Scialabba S, Caporale M, Di Pasquale A, Cammà C, Puglia I, Calistri P, Savini G |
| EPI_ISL_2673666, EPI_ISL_2673667 | SIESP CHIETI - DRIVE IN LANCIANO | Istituto Zooprofilattico Sperimentale dell'Abruzzo e Molise "G. Caporale" | Lorusso A, Marcacci M, Di Domenico M, Ancora M, Curini V, Di Lollo Valeria, Mangone I, Rinaldi A, Delli Compagni E, Scialabba S, Caporale M, Di Pasquale A, Cammà C, Puglia I, Calistri P, Savini G |
| EPI_ISL_2673668, EPI_ISL_2673669, EPI_ISL_2673670 | SIESP CHIETI - DRIVE IN VASTO | Istituto Zooprofilattico Sperimentale dell'Abruzzo e Molise "G. Caporale" | Lorusso A, Marcacci M, Di Domenico M, Ancora M, Curini V, Di Lollo Valeria, Mangone I, Rinaldi A, Delli Compagni E, Scialabba S, Caporale M, Di Pasquale A, Cammà C, Puglia I, Calistri P, Savini G |
| EPI_ISL_2673671 | SIESP CHIETI - DRIVE IN LANCIANO | Istituto Zooprofilattico Sperimentale dell'Abruzzo e Molise "G. Caporale" | Lorusso A, Marcacci M, Di Domenico M, Ancora M, Curini V, Di Lollo Valeria, Mangone I, Rinaldi A, Delli Compagni E, Scialabba S, Caporale M, Di Pasquale A, Cammà C, Puglia I, Calistri P, Savini G |
| EPI_ISL_2673672 | SIESP CHIETI - DRIVE IN ORTONA | Istituto Zooprofilattico Sperimentale dell'Abruzzo e Molise "G. Caporale" | Lorusso A, Marcacci M, Di Domenico M, Ancora M, Curini V, Di Lollo Valeria, Mangone I, Rinaldi A, Delli Compagni E, Scialabba S, Caporale M, Di Pasquale A, Cammà C, Puglia I, Calistri P, Savini G |
| EPI_ISL_2673673 | SIESP CHIETI - DRIVE IN VASTO | Istituto Zooprofilattico Sperimentale dell'Abruzzo e Molise "G. Caporale" | Lorusso A, Marcacci M, Di Domenico M, Ancora M, Curini V, Di Lollo Valeria, Mangone I, Rinaldi A, Delli Compagni E, Scialabba S, Caporale M, Di Pasquale A, Cammà C, Puglia I, Calistri P, Savini G |
| EPI_ISL_2673674 | SIESP CHIETI - DRIVE IN LANCIANO | Istituto Zooprofilattico Sperimentale dell'Abruzzo e Molise "G. Caporale" | Lorusso A, Marcacci M, Di Domenico M, Ancora M, Curini V, Di Lollo Valeria, Mangone I, Rinaldi A, Delli Compagni E, Scialabba S, Caporale M, Di Pasquale A, Cammà C, Puglia I, Calistri P, Savini G |
| EPI_ISL_2673675 | SIESP CHIETI - DRIVE IN CHIETI | Istituto Zooprofilattico Sperimentale dell'Abruzzo e Molise "G. Caporale" | Lorusso A, Marcacci M, Di Domenico M, Ancora M, Curini V, Di Lollo Valeria, Mangone I, Rinaldi A, Delli Compagni E, Scialabba S, Caporale M, Di Pasquale A, Cammà C, Puglia I, Calistri P, Savini G |
| EPI_ISL_2673676, EPI_ISL_2673677, EPI_ISL_2673678 | OSPEDALE CIVILE TERAMO - CENTRO TRASFUSIONALE | Istituto Zooprofilattico Sperimentale dell'Abruzzo e Molise "G. Caporale" | Lorusso A, Marcacci M, Di Domenico M, Ancora M, Curini V, Di Lollo Valeria, Mangone I, Rinaldi A, Delli Compagni E, Scialabba S, Caporale M, Di Pasquale A, Cammà C, Puglia I, Calistri P, Savini G |
| EPI_ISL_2673679, EPI_ISL_2673680 | PRESIDIO OSPEDALIERO TAGLIACOZZO | Istituto Zooprofilattico Sperimentale dell'Abruzzo e Molise "G. Caporale" | Lorusso A, Marcacci M, Di Domenico M, Ancora M, Curini V, Di Lollo Valeria, Mangone I, Rinaldi A, Delli Compagni E, Scialabba S, Caporale M, Di Pasquale A, Cammà C, Puglia I, Calistri P, Savini G |
| EPI_ISL_2673681, EPI_ISL_2673682 | SIESP CHIETI - DRIVE IN LANCIANO | Istituto Zooprofilattico Sperimentale dell'Abruzzo e Molise "G. Caporale" | Lorusso A, Marcacci M, Di Domenico M, Ancora M, Curini V, Di Lollo Valeria, Mangone I, Rinaldi A, Delli Compagni E, Scialabba S, Caporale M, Di Pasquale A, Cammà C, Puglia I, Calistri P, Savini G |
| EPI_ISL_2673696 | Instituto Nacional de Enfermedades Respiratorias (INER) | Centro de Investigación en Enfermedades Infecciosas (CIENI), Instituto Nacional de Enfermedades Respiratorias (INER) | Consorcio Mexicano de Vigilancia Genómica (CoViGen-Mex). Authors (in alphabetical order): Julio Elias Alvarado-Yaah, Carlos F. Arias, Santiago Ávila-Ríos, Eduardo Becerril-Vargas, Víctor Hugo Borja-Aburto, Celia Boukadida, Cristóbal Cháidez-Quiróz, Juan Bautista Chale-Dzul, Ricardo Ciria Merce, Andreu Comas-García, Célida Duque Molina, Julissa Enciso-Ibarra, José Antonio Enciso-Moreno, Gloria Elena Espinosa-Ayala, Fernando Fontove-Herrera, Daniel Fregoso-Rueda, Victor Eduardo Garcia-Arias, Alejandra Garcia-Gasca, Bruno Gomez-Gil, Jean Pierre González, Irvin González-López, Concepción Grajales-Muñiz, Ricardo Grande, Rosa María Gutierrez Rios, Alejandra Hernández-Terán, Alfredo Herrera-Estrella, Carla Ivón Herrera-Najera, Pavel Isa, Daniel Lira Morales, Susana Lopez, Antonio Loza Román, Brenda Irasema Maldonado-Meza, Bernardo Martínez-Miguel, José Arturo Martínez-Orozco, Célida Martínez- Rodríguez, Margarita Matías-Florentino, Fidencio Mejía-Nepomuceno, María Guadalupe de Jesús Mireles-Rivera, Gloria María Molina-Salinas, Hector Montoya-Fuentes, , Mario Mújica-Sánchez, José Esteban Muñoz-Medina, José de Jesús Nuñez-Contreras, Alicia Ocaña-Mondragón, Luis Alberto Ochoa-Carrera, Hector Esteban Paz-Juárez, Marissa Pérez-García, Francisco Pulido, Helen Haydee Fernanda Ramírez-Plascencia, Jorge Salas-Hernández, Angel Gustavo Salas-Lais, Alejandro Sanchez-Flores, Clara Esperanza Santacruz-Tinoco, María Guadalupe Santiago-Mauricio, Selene Zárate, Nelly Sélem-Mojica, Blanca Taboada, Gloria Vazquez, Joel Armando Vázquez-Pérez. |

| | | | |
|---|---|---|---|
| EPI_ISL_2674297, EPI_ISL_2674300 | LABORATORIO CARVAJAL | Instituto Nacional de Salud- Dirección de Investigación en Salud Pública | Katherine Laiton-Donato, Diego A. Álvarez-Díaz, Carlos Franco-Muñoz, Hector Alejandro Ruiz-Moreno, Paola Rojas, Maria T. Herrera-Sepúlveda, Diego Andrés Prada, Jhonnatan Reales-González, Sheryll Corchuelo, Julian Naizaque, Jorge Rivera, Gerardo Santamaría, Sergio Gomez, Lisseth Pardo, Juan Camilo Martinez, Marta Lopez Blanco, Ángela Alarcon Cruz, Diana Malo, Carmen Osorio, Magdalena Wiesner, Martha Lucia Ospina Martinez, Marcela Mercado-Reyes |
| EPI_ISL_2674306, EPI_ISL_2674307, EPI_ISL_2674308 | SURA | Instituto Nacional de Salud- Dirección de Investigación en Salud Pública | Katherine Laiton-Donato, Diego A. Álvarez-Díaz, Carlos Franco-Muñoz, Hector Alejandro Ruiz-Moreno, Paola Rojas, Maria T. Herrera-Sepúlveda, Diego Andrés Prada, Jhonnatan Reales-González, Sheryll Corchuelo, Julian Naizaque, Jorge Rivera, Gerardo Santamaría, Sergio Gomez, Lisseth Pardo, Juan Camilo Martinez, Marta Lopez Blanco, Ángela Alarcon Cruz, Diana Malo, Carmen Osorio, Magdalena Wiesner, Martha Lucia Ospina Martinez, Marcela Mercado-Reyes |
| EPI_ISL_2674318 | LDSP TOLIMA | Instituto Nacional de Salud- Dirección de Investigación en Salud Pública | Katherine Laiton-Donato, Diego A. Álvarez-Díaz, Carlos Franco-Muñoz, Hector Alejandro Ruiz-Moreno, Paola Rojas, Maria T. Herrera-Sepúlveda, Diego Andrés Prada, Jhonnatan Reales-González, Sheryll Corchuelo, Julian Naizaque, Jorge Rivera, Gerardo Santamaría, Sergio Gomez, Lisseth Pardo, Juan Camilo Martinez, Marta Lopez Blanco, Ángela Alarcon Cruz, Diana Malo, Carmen Osorio, Magdalena Wiesner, Martha Lucia Ospina Martinez, Marcela Mercado-Reyes |
| EPI_ISL_2674320, EPI_ISL_2674321, EPI_ISL_2674322 | CARVAJAL LABORATORIOS | Instituto Nacional de Salud- Dirección de Investigación en Salud Pública | Katherine Laiton-Donato, Diego A. Álvarez-Díaz, Carlos Franco-Muñoz, Hector Alejandro Ruiz-Moreno, Paola Rojas, Maria T. Herrera-Sepúlveda, Diego Andrés Prada, Jhonnatan Reales-González, Sheryll Corchuelo, Julian Naizaque, Jorge Rivera, Gerardo Santamaría, Sergio Gomez, Lisseth Pardo, Juan Camilo Martinez, Marta Lopez Blanco, Ángela Alarcon Cruz, Diana Malo, Carmen Osorio, Magdalena Wiesner, Martha Lucia Ospina Martinez, Marcela Mercado-Reyes |
| EPI_ISL_2674323, EPI_ISL_2674324 | LDSP HUILA | Instituto Nacional de Salud- Dirección de Investigación en Salud Pública | Katherine Laiton-Donato, Diego A. Álvarez-Díaz, Carlos Franco-Muñoz, Hector Alejandro Ruiz-Moreno, Paola Rojas, Maria T. Herrera-Sepúlveda, Diego Andrés Prada, Jhonnatan Reales-González, Sheryll Corchuelo, Julian Naizaque, Jorge Rivera, Gerardo Santamaría, Sergio Gomez, Lisseth Pardo, Juan Camilo Martinez, Marta Lopez Blanco, Ángela Alarcon Cruz, Diana Malo, Carmen Osorio, Magdalena Wiesner, Martha Lucia Ospina Martinez, Marcela Mercado-Reyes |
| EPI_ISL_2674330 | Laboratorio de Inmunología y Biología Molecular | Instituto Nacional de Salud- Dirección de Investigación en Salud Pública | Katherine Laiton-Donato, Diego A. Álvarez-Díaz, Carlos Franco-Muñoz, Hector Alejandro Ruiz-Moreno, Paola Rojas, Maria T. Herrera-Sepúlveda, Diego Andrés Prada, Jhonnatan Reales-González, Sheryll Corchuelo, Julian Naizaque, Jorge Rivera, Gerardo Santamaría, Sergio Gomez, Lisseth Pardo, Juan Camilo Martinez, Marta Lopez Blanco, Ángela Alarcon Cruz, Diana Malo, Carmen Osorio, Magdalena Wiesner, Martha Lucia Ospina Martinez, Marcela Mercado-Reyes |
| EPI_ISL_2674344 | HOSPITAL PEDRO LEON ALVAREZ DIAZ | Instituto Nacional de Salud- Dirección de Investigación en Salud Pública | Katherine Laiton-Donato, Diego A. Álvarez-Díaz, Carlos Franco-Muñoz, Hector Alejandro Ruiz-Moreno, Paola Rojas, Maria T. Herrera-Sepúlveda, Diego Andrés Prada, Jhonnatan Reales-González, Sheryll Corchuelo, Julian Naizaque, Jorge Rivera, Gerardo Santamaría, Sergio Gomez, Lisseth Pardo, Juan Camilo Martinez, Marta Lopez Blanco, Ángela Alarcon Cruz, Diana Malo, Carmen Osorio, Magdalena Wiesner, Martha Lucia Ospina Martinez, Marcela Mercado-Reyes |
| EPI_ISL_2674346 | HOSPITAL REGIONAL DE LA ORINOQUIA | Instituto Nacional de Salud- Dirección de Investigación en Salud Pública | Katherine Laiton-Donato, Diego A. Álvarez-Díaz, Carlos Franco-Muñoz, Hector Alejandro Ruiz-Moreno, Paola Rojas, Maria T. Herrera-Sepúlveda, Diego Andrés Prada, Jhonnatan Reales-González, Sheryll Corchuelo, Julian Naizaque, Jorge Rivera, Gerardo Santamaría, Sergio Gomez, Lisseth Pardo, Juan Camilo Martinez, Marta Lopez Blanco, Ángela Alarcon Cruz, Diana Malo, Carmen Osorio, Magdalena Wiesner, Martha Lucia Ospina Martinez, Marcela Mercado-Reyes |
| EPI_ISL_2674347 | HOSPITAL REGIONAL ORINOQUIA | Instituto Nacional de Salud- Dirección de Investigación en Salud Pública | Katherine Laiton-Donato, Diego A. Álvarez-Díaz, Carlos Franco-Muñoz, Hector Alejandro Ruiz-Moreno, Paola Rojas, Maria T. Herrera-Sepúlveda, Diego Andrés Prada, Jhonnatan Reales-González, Sheryll Corchuelo, Julian Naizaque, Jorge Rivera, Gerardo Santamaría, Sergio Gomez, Lisseth Pardo, Juan Camilo Martinez, Marta Lopez Blanco, Ángela Alarcon Cruz, Diana Malo, Carmen Osorio, Magdalena Wiesner, Martha Lucia Ospina Martinez, Marcela Mercado-Reyes |
| EPI_ISL_2674348 | HOSPITAL REGIONAL DE LA ORINOQUIA | Instituto Nacional de Salud- Dirección de Investigación en Salud Pública | Katherine Laiton-Donato, Diego A. Álvarez-Díaz, Carlos Franco-Muñoz, Hector Alejandro Ruiz-Moreno, Paola Rojas, Maria T. Herrera-Sepúlveda, Diego Andrés Prada, Jhonnatan Reales-González, Sheryll Corchuelo, Julian Naizaque, Jorge Rivera, Gerardo Santamaría, Sergio Gomez, Lisseth Pardo, Juan Camilo Martinez, Marta Lopez Blanco, Ángela Alarcon Cruz, Diana Malo, Carmen Osorio, Magdalena Wiesner, Martha Lucia Ospina Martinez, Marcela Mercado-Reyes |
| EPI_ISL_2674349 | EMPRESA SOCIAL DEL ESTADO HOSPITAL UNIVERSITARIO DE SANTANDER | Instituto Nacional de Salud- Dirección de Investigación en Salud Pública | Katherine Laiton-Donato, Diego A. Álvarez-Díaz, Carlos Franco-Muñoz, Hector Alejandro Ruiz-Moreno, Paola Rojas, Maria T. Herrera-Sepúlveda, Diego Andrés Prada, Jhonnatan Reales-González, Sheryll Corchuelo, Julian Naizaque, Jorge Rivera, Gerardo Santamaría, Sergio Gomez, Lisseth Pardo, Juan Camilo Martinez, Marta Lopez Blanco, Ángela Alarcon Cruz, Diana Malo, Carmen Osorio, Magdalena Wiesner, Martha Lucia Ospina Martinez, Marcela Mercado-Reyes |
| EPI_ISL_2674353, EPI_ISL_2674354 | Laboratorio de Salud Publica San Andrés Islas | Instituto Nacional de Salud- Dirección de Investigación en Salud Pública | Katherine Laiton-Donato, Diego A. Álvarez-Díaz, Carlos Franco-Muñoz, Hector Alejandro Ruiz-Moreno, Paola Rojas, Maria T. Herrera-Sepúlveda, Diego Andrés Prada, Jhonnatan Reales-González, Sheryll Corchuelo, Julian Naizaque, Jorge Rivera, Gerardo Santamaría, Sergio Gomez, Lisseth Pardo, Juan Camilo Martinez, Marta Lopez Blanco, Ángela Alarcon Cruz, Diana Malo, Carmen Osorio, Magdalena Wiesner, Martha Lucia Ospina Martinez, Marcela Mercado-Reyes |
| EPI_ISL_2674355, EPI_ISL_2674358, EPI_ISL_2674359, EPI_ISL_2674360 | Laboratorio Salud Publica San Andres Islas | Instituto Nacional de Salud- Dirección de Investigación en Salud Pública | Katherine Laiton-Donato, Diego A. Álvarez-Díaz, Carlos Franco-Muñoz, Hector Alejandro Ruiz-Moreno, Paola Rojas, Maria T. Herrera-Sepúlveda, Diego Andrés Prada, Jhonnatan Reales-González, Sheryll Corchuelo, Julian Naizaque, Jorge Rivera, Gerardo Santamaría, Sergio Gomez, Lisseth Pardo, Juan Camilo Martinez, Marta Lopez Blanco, Ángela Alarcon Cruz, Diana Malo, Carmen Osorio, Magdalena Wiesner, Martha Lucia Ospina Martinez, Marcela Mercado-Reyes |
| EPI_ISL_2674361 | Laboratorio de Salud Publica San Andrés Islas | Instituto Nacional de Salud- Dirección de Investigación en Salud Pública | Katherine Laiton-Donato, Diego A. Álvarez-Díaz, Carlos Franco-Muñoz, Hector Alejandro Ruiz-Moreno, Paola Rojas, Maria T. Herrera-Sepúlveda, Diego Andrés Prada, Jhonnatan Reales-González, Sheryll Corchuelo, Julian Naizaque, Jorge Rivera, Gerardo Santamaría, Sergio Gomez, Lisseth Pardo, Juan Camilo Martinez, Marta Lopez Blanco, Ángela Alarcon Cruz, Diana Malo, Carmen Osorio, Magdalena Wiesner, Martha Lucia Ospina Martinez, Marcela Mercado-Reyes |
| EPI_ISL_2674364, EPI_ISL_2674365 | Laboratorio de Salud Publica | Instituto Nacional de Salud- Dirección de Investigación en Salud Pública | Katherine Laiton-Donato, Diego A. Álvarez-Díaz, Carlos Franco-Muñoz, Hector Alejandro Ruiz-Moreno, Paola Rojas, Maria T. Herrera-Sepúlveda, Diego Andrés Prada, Jhonnatan Reales-González, Sheryll Corchuelo, Julian Naizaque, Jorge Rivera, Gerardo Santamaría, Sergio Gomez, Lisseth Pardo, Juan Camilo Martinez, Marta Lopez Blanco, Ángela Alarcon Cruz, Diana Malo, Carmen Osorio, Magdalena Wiesner, Martha Lucia Ospina Martinez, Marcela Mercado-Reyes |
| EPI_ISL_2674366, EPI_ISL_2674367 | Laboratorio Salud Publica San Andres Islas | Instituto Nacional de Salud- Dirección de Investigación en Salud Pública | Katherine Laiton-Donato, Diego A. Álvarez-Díaz, Carlos Franco-Muñoz, Hector Alejandro Ruiz-Moreno, Paola Rojas, Maria T. Herrera-Sepúlveda, Diego Andrés Prada, Jhonnatan Reales-González, Sheryll Corchuelo, Julian Naizaque, Jorge Rivera, Gerardo Santamaría, Sergio Gomez, Lisseth Pardo, Juan Camilo Martinez, Marta Lopez Blanco, Ángela Alarcon Cruz, Diana Malo, Carmen Osorio, Magdalena Wiesner, Martha Lucia Ospina Martinez, Marcela Mercado-Reyes |
| EPI_ISL_2674629, EPI_ISL_2674631 | LESP Nuevo Leon | Instituto de Diagnostico y Referencia Epidemiologicos (INDRE) | Claudia Wong-Arambula, Abril Rodriguez-Maldonado, Vanessa Rivero-Arredondo, Ariadna Medina-Benitez, Joaquin Quiroz-Mercado, Sergio Rangel-Guerrero, Natividad Cruz-Ortiz, Tatiana Nunez-Garcia, Gisela Barrera-Badillo, Lucia Hernandez-Rivas, Irma Lopez-Martinez, Ernesto Ramirez-Gonzalez. |
| EPI_ISL_2674632 | LESP Ciudad de Mexico | Instituto de Diagnostico y Referencia Epidemiologicos (INDRE) | Claudia Wong-Arambula, Abril Rodriguez-Maldonado, Vanessa Rivero-Arredondo, Ariadna Medina-Benitez, Joaquin Quiroz-Mercado, Sergio Rangel-Guerrero, Natividad Cruz-Ortiz, Tatiana Nunez-Garcia, Gisela Barrera-Badillo, Lucia Hernandez-Rivas, Irma Lopez-Martinez, Ernesto Ramirez-Gonzalez. |
| EPI_ISL_2674633, EPI_ISL_2674634 | LESP Sinaloa | Instituto de Diagnostico y Referencia Epidemiologicos (INDRE) | Claudia Wong-Arambula, Abril Rodriguez-Maldonado, Vanessa Rivero-Arredondo, Ariadna Medina-Benitez, Joaquin Quiroz-Mercado, Sergio Rangel-Guerrero, Natividad Cruz-Ortiz, Tatiana Nunez-Garcia, Gisela Barrera-Badillo, Lucia Hernandez-Rivas, Irma Lopez-Martinez, Ernesto Ramirez-Gonzalez. |
| EPI_ISL_2674635, EPI_ISL_2674636, EPI_ISL_2674637, EPI_ISL_2674638, EPI_ISL_2674639, EPI_ISL_2674640 | LESP Tabasco | Instituto de Diagnostico y Referencia Epidemiologicos (INDRE) | Claudia Wong-Arambula, Abril Rodriguez-Maldonado, Vanessa Rivero-Arredondo, Ariadna Medina-Benitez, Joaquin Quiroz-Mercado, Sergio Rangel-Guerrero, Natividad Cruz-Ortiz, Tatiana Nunez-Garcia, Gisela Barrera-Badillo, Lucia Hernandez-Rivas, Irma Lopez-Martinez, Ernesto Ramirez-Gonzalez. |
| EPI_ISL_2674641 | LESP Michoacan | Instituto de Diagnostico y Referencia Epidemiologicos | Claudia Wong-Arambula, Abril Rodriguez-Maldonado, Vanessa Rivero-Arredondo, Ariadna Medina-Benitez, Joaquin Quiroz-Mercado, Sergio |

| | | | |
|---|---|---|---|
| | | (INDRE) | Rangel-Guerrero, Natividad Cruz-Ortiz, Tatiana Nunez-Garcia, Gisela Barrera-Badillo, Lucia Hernandez-Rivas, Irma Lopez-Martinez, Ernesto Ramirez-Gonzalez. |
| EPI_ISL_2674642 | LESP Durango | Instituto de Diagnostico y Referencia Epidemiologicos (INDRE) | Claudia Wong-Arambula, Abril Rodriguez-Maldonado, Vanessa Rivero-Arredondo, Ariadna Medina-Benitez, Joaquin Quiroz-Mercado, Sergio Rangel-Guerrero, Natividad Cruz-Ortiz, Tatiana Nunez-Garcia, Gisela Barrera-Badillo, Lucia Hernandez-Rivas, Irma Lopez-Martinez, Ernesto Ramirez-Gonzalez. |
| EPI_ISL_2674643, EPI_ISL_2674644, EPI_ISL_2674645, EPI_ISL_2674646, EPI_ISL_2674647, EPI_ISL_2674648, EPI_ISL_2674649 | LESP Sonora | Instituto de Diagnostico y Referencia Epidemiologicos (INDRE) | Claudia Wong-Arambula, Abril Rodriguez-Maldonado, Vanessa Rivero-Arredondo, Ariadna Medina-Benitez, Joaquin Quiroz-Mercado, Sergio Rangel-Guerrero, Natividad Cruz-Ortiz, Tatiana Nunez-Garcia, Gisela Barrera-Badillo, Lucia Hernandez-Rivas, Irma Lopez-Martinez, Ernesto Ramirez-Gonzalez. |
| EPI_ISL_2674650 | LESP Jalisco | Instituto de Diagnostico y Referencia Epidemiologicos (INDRE) | Claudia Wong-Arambula, Abril Rodriguez-Maldonado, Vanessa Rivero-Arredondo, Ariadna Medina-Benitez, Joaquin Quiroz-Mercado, Sergio Rangel-Guerrero, Natividad Cruz-Ortiz, Tatiana Nunez-Garcia, Gisela Barrera-Badillo, Lucia Hernandez-Rivas, Irma Lopez-Martinez, Ernesto Ramirez-Gonzalez. |
| EPI_ISL_2674651, EPI_ISL_2674652, EPI_ISL_2674653, EPI_ISL_2674654, EPI_ISL_2674655 | LESP Quintana Roo | Instituto de Diagnostico y Referencia Epidemiologicos (INDRE) | Claudia Wong-Arambula, Abril Rodriguez-Maldonado, Vanessa Rivero-Arredondo, Ariadna Medina-Benitez, Joaquin Quiroz-Mercado, Sergio Rangel-Guerrero, Natividad Cruz-Ortiz, Tatiana Nunez-Garcia, Gisela Barrera-Badillo, Lucia Hernandez-Rivas, Irma Lopez-Martinez, Ernesto Ramirez-Gonzalez. |
| EPI_ISL_2674658, EPI_ISL_2674659, EPI_ISL_2674660, EPI_ISL_2674661, EPI_ISL_2674662, EPI_ISL_2674663 | LESP Hidalgo | Instituto de Diagnostico y Referencia Epidemiologicos (INDRE) | Claudia Wong-Arambula, Abril Rodriguez-Maldonado, Vanessa Rivero-Arredondo, Ariadna Medina-Benitez, Joaquin Quiroz-Mercado, Sergio Rangel-Guerrero, Natividad Cruz-Ortiz, Tatiana Nunez-Garcia, Gisela Barrera-Badillo, Lucia Hernandez-Rivas, Irma Lopez-Martinez, Ernesto Ramirez-Gonzalez. |
| EPI_ISL_2674664 | LESP Estado de Mexico | Instituto de Diagnostico y Referencia Epidemiologicos (INDRE) | Claudia Wong-Arambula, Abril Rodriguez-Maldonado, Vanessa Rivero-Arredondo, Ariadna Medina-Benitez, Joaquin Quiroz-Mercado, Sergio Rangel-Guerrero, Natividad Cruz-Ortiz, Tatiana Nunez-Garcia, Gisela Barrera-Badillo, Lucia Hernandez-Rivas, Irma Lopez-Martinez, Ernesto Ramirez-Gonzalez. |
| EPI_ISL_2674665, EPI_ISL_2674666, EPI_ISL_2674667 | LESP Hidalgo | Instituto de Diagnostico y Referencia Epidemiologicos (INDRE) | Claudia Wong-Arambula, Abril Rodriguez-Maldonado, Vanessa Rivero-Arredondo, Ariadna Medina-Benitez, Joaquin Quiroz-Mercado, Sergio Rangel-Guerrero, Natividad Cruz-Ortiz, Tatiana Nunez-Garcia, Gisela Barrera-Badillo, Lucia Hernandez-Rivas, Irma Lopez-Martinez, Ernesto Ramirez-Gonzalez. |
| EPI_ISL_2674668, EPI_ISL_2674669, EPI_ISL_2674670 | LESP Estado de Mexico | Instituto de Diagnostico y Referencia Epidemiologicos (INDRE) | Claudia Wong-Arambula, Abril Rodriguez-Maldonado, Vanessa Rivero-Arredondo, Ariadna Medina-Benitez, Joaquin Quiroz-Mercado, Sergio Rangel-Guerrero, Natividad Cruz-Ortiz, Tatiana Nunez-Garcia, Gisela Barrera-Badillo, Lucia Hernandez-Rivas, Irma Lopez-Martinez, Ernesto Ramirez-Gonzalez. |
| EPI_ISL_2674672 | LESP Hidalgo | Instituto de Diagnostico y Referencia Epidemiologicos (INDRE) | Claudia Wong-Arambula, Abril Rodriguez-Maldonado, Vanessa Rivero-Arredondo, Ariadna Medina-Benitez, Joaquin Quiroz-Mercado, Sergio Rangel-Guerrero, Natividad Cruz-Ortiz, Tatiana Nunez-Garcia, Gisela Barrera-Badillo, Lucia Hernandez-Rivas, Irma Lopez-Martinez, Ernesto Ramirez-Gonzalez. |
| EPI_ISL_2674673 | LESP Puebla | Instituto de Diagnostico y Referencia Epidemiologicos (INDRE) | Claudia Wong-Arambula, Abril Rodriguez-Maldonado, Vanessa Rivero-Arredondo, Ariadna Medina-Benitez, Joaquin Quiroz-Mercado, Sergio Rangel-Guerrero, Natividad Cruz-Ortiz, Tatiana Nunez-Garcia, Gisela Barrera-Badillo, Lucia Hernandez-Rivas, Irma Lopez-Martinez, Ernesto Ramirez-Gonzalez. |
| EPI_ISL_2674674, EPI_ISL_2674675, EPI_ISL_2674676, EPI_ISL_2674677, EPI_ISL_2674678, EPI_ISL_2674679, EPI_ISL_2674680, EPI_ISL_2674681, EPI_ISL_2674682 | LESP Hidalgo | Instituto de Diagnostico y Referencia Epidemiologicos (INDRE) | Claudia Wong-Arambula, Abril Rodriguez-Maldonado, Vanessa Rivero-Arredondo, Ariadna Medina-Benitez, Joaquin Quiroz-Mercado, Sergio Rangel-Guerrero, Natividad Cruz-Ortiz, Tatiana Nunez-Garcia, Gisela Barrera-Badillo, Lucia Hernandez-Rivas, Irma Lopez-Martinez, Ernesto Ramirez-Gonzalez. |
| EPI_ISL_2674683 | LESP Queretaro | Instituto de Diagnostico y Referencia Epidemiologicos (INDRE) | Claudia Wong-Arambula, Abril Rodriguez-Maldonado, Vanessa Rivero-Arredondo, Ariadna Medina-Benitez, Joaquin Quiroz-Mercado, Sergio Rangel-Guerrero, Natividad Cruz-Ortiz, Tatiana Nunez-Garcia, Gisela Barrera-Badillo, Lucia Hernandez-Rivas, Irma Lopez-Martinez, Ernesto Ramirez-Gonzalez. |
| EPI_ISL_2674684, EPI_ISL_2674685, EPI_ISL_2674686, EPI_ISL_2674687, EPI_ISL_2674688 | LESP Yucatan | Instituto de Diagnostico y Referencia Epidemiologicos (INDRE) | Claudia Wong-Arambula, Abril Rodriguez-Maldonado, Vanessa Rivero-Arredondo, Ariadna Medina-Benitez, Joaquin Quiroz-Mercado, Sergio Rangel-Guerrero, Natividad Cruz-Ortiz, Tatiana Nunez-Garcia, Gisela Barrera-Badillo, Lucia Hernandez-Rivas, Irma Lopez-Martinez, Ernesto Ramirez-Gonzalez. |
| EPI_ISL_2674689 | LESP Coahuila | Instituto de Diagnostico y Referencia Epidemiologicos (INDRE) | Claudia Wong-Arambula, Abril Rodriguez-Maldonado, Vanessa Rivero-Arredondo, Ariadna Medina-Benitez, Joaquin Quiroz-Mercado, Sergio Rangel-Guerrero, Natividad Cruz-Ortiz, Tatiana Nunez-Garcia, Gisela Barrera-Badillo, Lucia Hernandez-Rivas, Irma Lopez-Martinez, Ernesto Ramirez-Gonzalez. |
| EPI_ISL_2674690 | LESP Hidalgo | Instituto de Diagnostico y Referencia Epidemiologicos (INDRE) | Claudia Wong-Arambula, Abril Rodriguez-Maldonado, Vanessa Rivero-Arredondo, Ariadna Medina-Benitez, Joaquin Quiroz-Mercado, Sergio Rangel-Guerrero, Natividad Cruz-Ortiz, Tatiana Nunez-Garcia, Gisela Barrera-Badillo, Lucia Hernandez-Rivas, Irma Lopez-Martinez, Ernesto Ramirez-Gonzalez. |
| EPI_ISL_2674691, EPI_ISL_2674692, EPI_ISL_2674693, EPI_ISL_2674694, EPI_ISL_2674695, EPI_ISL_2674696 | LESP Guanajuato | Instituto de Diagnostico y Referencia Epidemiologicos (INDRE) | Claudia Wong-Arambula, Abril Rodriguez-Maldonado, Vanessa Rivero-Arredondo, Ariadna Medina-Benitez, Joaquin Quiroz-Mercado, Sergio Rangel-Guerrero, Natividad Cruz-Ortiz, Tatiana Nunez-Garcia, Gisela Barrera-Badillo, Lucia Hernandez-Rivas, Irma Lopez-Martinez, Ernesto Ramirez-Gonzalez. |
| EPI_ISL_2674697 | LESP Nuevo Leon | Instituto de Diagnostico y Referencia Epidemiologicos (INDRE) | Claudia Wong-Arambula, Abril Rodriguez-Maldonado, Vanessa Rivero-Arredondo, Ariadna Medina-Benitez, Joaquin Quiroz-Mercado, Sergio Rangel-Guerrero, Natividad Cruz-Ortiz, Tatiana Nunez-Garcia, Gisela Barrera-Badillo, Lucia Hernandez-Rivas, Irma Lopez-Martinez, Ernesto Ramirez-Gonzalez. |
| EPI_ISL_2674698 | LESP Ciudad de Mexico | Instituto de Diagnostico y Referencia Epidemiologicos (INDRE) | Claudia Wong-Arambula, Abril Rodriguez-Maldonado, Vanessa Rivero-Arredondo, Ariadna Medina-Benitez, Joaquin Quiroz-Mercado, Sergio Rangel-Guerrero, Natividad Cruz-Ortiz, Tatiana Nunez-Garcia, Gisela Barrera-Badillo, Lucia Hernandez-Rivas, Irma Lopez-Martinez, Ernesto Ramirez-Gonzalez. |
| EPI_ISL_2674699 | LESP Chihuahua | Instituto de Diagnostico y Referencia Epidemiologicos (INDRE) | Claudia Wong-Arambula, Abril Rodriguez-Maldonado, Vanessa Rivero-Arredondo, Ariadna Medina-Benitez, Joaquin Quiroz-Mercado, Sergio Rangel-Guerrero, Natividad Cruz-Ortiz, Tatiana Nunez-Garcia, Gisela Barrera-Badillo, Lucia Hernandez-Rivas, Irma Lopez-Martinez, Ernesto Ramirez-Gonzalez. |
| EPI_ISL_2674700 | LESP Puebla | Instituto de Diagnostico y Referencia Epidemiologicos (INDRE) | Claudia Wong-Arambula, Abril Rodriguez-Maldonado, Vanessa Rivero-Arredondo, Ariadna Medina-Benitez, Joaquin Quiroz-Mercado, Sergio Rangel-Guerrero, Natividad Cruz-Ortiz, Tatiana Nunez-Garcia, Gisela Barrera-Badillo, Lucia Hernandez-Rivas, Irma Lopez-Martinez, Ernesto Ramirez-Gonzalez. |
| EPI_ISL_2674701, EPI_ISL_2674703, EPI_ISL_2674704, EPI_ISL_2674705, EPI_ISL_2674706, EPI_ISL_2674707, EPI_ISL_2674708, EPI_ISL_2674709, EPI_ISL_2674710, EPI_ISL_2674711, EPI_ISL_2674712, EPI_ISL_2674713 | | | |
| see above | LESP Yucatan | Instituto de Diagnostico y Referencia Epidemiologicos (INDRE) | Claudia Wong-Arambula, Abril Rodriguez-Maldonado, Vanessa Rivero-Arredondo, Ariadna Medina-Benitez, Joaquin Quiroz-Mercado, Sergio Rangel-Guerrero, Natividad Cruz-Ortiz, Tatiana Nunez-Garcia, Gisela Barrera-Badillo, Lucia Hernandez-Rivas, Irma Lopez-Martinez, Ernesto Ramirez-Gonzalez. |
| EPI_ISL_2674714 | LESP Quintana Roo | Instituto de Diagnostico y Referencia Epidemiologicos (INDRE) | Claudia Wong-Arambula, Abril Rodriguez-Maldonado, Vanessa Rivero-Arredondo, Ariadna Medina-Benitez, Joaquin Quiroz-Mercado, Sergio Rangel-Guerrero, Natividad Cruz-Ortiz, Tatiana Nunez-Garcia, Gisela Barrera-Badillo, Lucia Hernandez-Rivas, Irma Lopez-Martinez, Ernesto Ramirez-Gonzalez. |
| EPI_ISL_2674715 | LESP Yucatan | Instituto de Diagnostico y Referencia Epidemiologicos (INDRE) | Claudia Wong-Arambula, Abril Rodriguez-Maldonado, Vanessa Rivero-Arredondo, Ariadna Medina-Benitez, Joaquin Quiroz-Mercado, Sergio Rangel-Guerrero, Natividad Cruz-Ortiz, Tatiana Nunez-Garcia, Gisela Barrera-Badillo, Lucia Hernandez-Rivas, Irma Lopez-Martinez, Ernesto Ramirez-Gonzalez. |

| | | | |
|---|---|---|---|
| EPI_ISL_2674716 | LESP San Luis Potosi | Instituto de Diagnostico y Referencia Epidemiologicos (INDRE) | Claudia Wong-Arambula, Abril Rodriguez-Maldonado, Vanessa Rivero-Arredondo, Ariadna Medina-Benitez, Joaquin Quiroz-Mercado, Sergio Rangel-Guerrero, Natividad Cruz-Ortiz, Tatiana Nunez-Garcia, Gisela Barrera-Badillo, Lucia Hernandez-Rivas, Irma Lopez-Martinez, Ernesto Ramirez-Gonzalez. |
| EPI_ISL_2674718, EPI_ISL_2674719 | LESP Sonora | Instituto de Diagnostico y Referencia Epidemiologicos (INDRE) | Claudia Wong-Arambula, Abril Rodriguez-Maldonado, Vanessa Rivero-Arredondo, Ariadna Medina-Benitez, Joaquin Quiroz-Mercado, Sergio Rangel-Guerrero, Natividad Cruz-Ortiz, Tatiana Nunez-Garcia, Gisela Barrera-Badillo, Lucia Hernandez-Rivas, Irma Lopez-Martinez, Ernesto Ramirez-Gonzalez. |
| EPI_ISL_2674900, EPI_ISL_2674901, EPI_ISL_2674902 | LESP Ciudad de Mexico | Instituto de Diagnostico y Referencia Epidemiologicos (INDRE) | Claudia Wong-Arambula, Abril Rodriguez-Maldonado, Vanessa Rivero-Arredondo, Ariadna Medina-Benitez, Joaquin Quiroz-Mercado, Sergio Rangel-Guerrero, Natividad Cruz-Ortiz, Tatiana Nunez-Garcia, Gisela Barrera-Badillo, Lucia Hernandez-Rivas, Irma Lopez-Martinez, Ernesto Ramirez-Gonzalez. |
| EPI_ISL_2674903 | LESP Jalisco | Instituto de Diagnostico y Referencia Epidemiologicos (INDRE) | Claudia Wong-Arambula, Abril Rodriguez-Maldonado, Vanessa Rivero-Arredondo, Ariadna Medina-Benitez, Joaquin Quiroz-Mercado, Sergio Rangel-Guerrero, Natividad Cruz-Ortiz, Tatiana Nunez-Garcia, Gisela Barrera-Badillo, Lucia Hernandez-Rivas, Irma Lopez-Martinez, Ernesto Ramirez-Gonzalez. |
| EPI_ISL_2674904 | LESP Ciudad de Mexico | Instituto de Diagnostico y Referencia Epidemiologicos (INDRE) | Claudia Wong-Arambula, Abril Rodriguez-Maldonado, Vanessa Rivero-Arredondo, Ariadna Medina-Benitez, Joaquin Quiroz-Mercado, Sergio Rangel-Guerrero, Natividad Cruz-Ortiz, Tatiana Nunez-Garcia, Gisela Barrera-Badillo, Lucia Hernandez-Rivas, Irma Lopez-Martinez, Ernesto Ramirez-Gonzalez. |
| EPI_ISL_2674905 | LESP Yucatan | Instituto de Diagnostico y Referencia Epidemiologicos (INDRE) | Claudia Wong-Arambula, Abril Rodriguez-Maldonado, Vanessa Rivero-Arredondo, Ariadna Medina-Benitez, Joaquin Quiroz-Mercado, Sergio Rangel-Guerrero, Natividad Cruz-Ortiz, Tatiana Nunez-Garcia, Gisela Barrera-Badillo, Lucia Hernandez-Rivas, Irma Lopez-Martinez, Ernesto Ramirez-Gonzalez. |
| EPI_ISL_2680931 | Hospital Margarita Maza de Juárez | Microbial Genomics Laboratory | Consorcio Mexicano de Vigilancia Genómica (CoViGen-Mex). Authors (in alphabetical order): Julio Elias Alvarado-Yaah, Carlos F. Arias, Santiago Ávila-Ríos, Eduardo Becerril-Vargas, Víctor Hugo Borja-Aburto, Celia Boukadida, Cristóbal Cháidez-Quiróz, Juan Bautista Chale-Dzul, Ricardo Ciria Merce, Andreu Comas-García, Célida Duque Molina, Julissa Enciso-Ibarra, José Antonio Enciso-Moreno, Gloria Elena Espinosa-Ayala, Fernando Fontove-Herrera, Daniel Fregoso-Rueda, Victor Eduardo García-Arias, Alejandra García-Gasca, Bruno Gómez-Gil, Jean Pierre González, Irvin González-López, Concepción Grajales-Muñiz, Ricardo Grande, Rosa María Gutiérrez Rios, Jesús Hernández, Alejandra Hernández-Terán, Alfredo Herrera-Estrella, Carla Ivón Herrera-Najera, Pavel Isa, Verónica Mata-Haro, Daniel Lira Morales, Susana Lopez, Antonio Loza Román, Brenda Irasema Maldonado-Meza, Bernardo Martínez-Miguel, José Arturo Martínez-Orozco, Célida Martínez- Rodríguez, Margarita Matías-Florentino, Fidencio Mejía-Nepomuceno, María Guadalupe de Jesús Mireles-Rivera, Gloria María Molina-Salinas, Hector Montoya-Fuentes, , Mario Mújica-Sánchez, José Esteban Muñoz-Medina, José de Jesús Nuñez-Contreras, Alicia Ocaña-Mondragón, Luis Alberto Ochoa-Carrera, Hector Esteban Paz-Juárez, Marissa Perez-Garcia, Francisco Pulido, Helen Haydee Fernanda Ramírez-Plascencia, Jorge Salas-Hernández, Angel Gustavo Salas-Lais, Alejandro Sánchez-Flores, Clara Esperanza Santacruz-Tinoco, María Guadalupe Santiago-Mauricio, Selene Zárate, Nelly Sélem-Mojica, Blanca Taboada, Gloria Vazquez, Joel Armando Vázquez-Pérez. |
| EPI_ISL_2680934 | Imágenes Diagnósticas de Mazatlán | Microbial Genomics Laboratory | Consorcio Mexicano de Vigilancia Genómica (CoViGen-Mex). Authors (in alphabetical order): Julio Elias Alvarado-Yaah, Carlos F. Arias, Santiago Ávila-Ríos, Eduardo Becerril-Vargas, Víctor Hugo Borja-Aburto, Celia Boukadida, Cristóbal Cháidez-Quiróz, Juan Bautista Chale-Dzul, Ricardo Ciria Merce, Andreu Comas-García, Célida Duque Molina, Julissa Enciso-Ibarra, José Antonio Enciso-Moreno, Gloria Elena Espinosa-Ayala, Fernando Fontove-Herrera, Daniel Fregoso-Rueda, Victor Eduardo García-Arias, Alejandra García-Gasca, Bruno Gómez-Gil, Jean Pierre González, Irvin González-López, Concepción Grajales-Muñiz, Ricardo Grande, Rosa María Gutiérrez Rios, Jesús Hernández, Alejandra Hernández-Terán, Alfredo Herrera-Estrella, Carla Ivón Herrera-Najera, Pavel Isa, Verónica Mata-Haro, Daniel Lira Morales, Susana Lopez, Antonio Loza Román, Brenda Irasema Maldonado-Meza, Bernardo Martínez-Miguel, José Arturo Martínez-Orozco, Célida Martínez- Rodríguez, Margarita Matías-Florentino, Fidencio Mejía-Nepomuceno, María Guadalupe de Jesús Mireles-Rivera, Gloria María Molina-Salinas, Hector Montoya-Fuentes, , Mario Mújica-Sánchez, José Esteban Muñoz-Medina, José de Jesús Nuñez-Contreras, Alicia Ocaña-Mondragón, Luis Alberto Ochoa-Carrera, Hector Esteban Paz-Juárez, Marissa Perez-Garcia, Francisco Pulido, Helen Haydee Fernanda Ramírez-Plascencia, Jorge Salas-Hernández, Angel Gustavo Salas-Lais, Alejandro Sánchez-Flores, Clara Esperanza Santacruz-Tinoco, María Guadalupe Santiago-Mauricio, Selene Zárate, Nelly Sélem-Mojica, Blanca Taboada, Gloria Vazquez, Joel Armando Vázquez-Pérez. |
| EPI_ISL_2680935 | Hospital Sharp | Microbial Genomics Laboratory | Consorcio Mexicano de Vigilancia Genómica (CoViGen-Mex). Authors (in alphabetical order): Julio Elias Alvarado-Yaah, Carlos F. Arias, Santiago Ávila-Ríos, Eduardo Becerril-Vargas, Víctor Hugo Borja-Aburto, Celia Boukadida, Cristóbal Cháidez-Quiróz, Juan Bautista Chale-Dzul, Ricardo Ciria Merce, Andreu Comas-García, Célida Duque Molina, Julissa Enciso-Ibarra, José Antonio Enciso-Moreno, Gloria Elena Espinosa-Ayala, Fernando Fontove-Herrera, Daniel Fregoso-Rueda, Victor Eduardo García-Arias, Alejandra García-Gasca, Bruno Gómez-Gil, Jean Pierre González, Irvin González-López, Concepción Grajales-Muñiz, Ricardo Grande, Rosa María Gutiérrez Rios, Jesús Hernández, Alejandra Hernández-Terán, Alfredo Herrera-Estrella, Carla Ivón Herrera-Najera, Pavel Isa, Verónica Mata-Haro, Daniel Lira Morales, Susana Lopez, Antonio Loza Román, Brenda Irasema Maldonado-Meza, Bernardo Martínez-Miguel, José Arturo Martínez-Orozco, Célida Martínez- Rodríguez, Margarita Matías-Florentino, Fidencio Mejía-Nepomuceno, María Guadalupe de Jesús Mireles-Rivera, Gloria María Molina-Salinas, Hector Montoya-Fuentes, , Mario Mújica-Sánchez, José Esteban Muñoz-Medina, José de Jesús Nuñez-Contreras, Alicia Ocaña-Mondragón, Luis Alberto Ochoa-Carrera, Hector Esteban Paz-Juárez, Marissa Perez-Garcia, Francisco Pulido, Helen Haydee Fernanda Ramírez-Plascencia, Jorge Salas-Hernández, Angel Gustavo Salas-Lais, Alejandro Sánchez-Flores, Clara Esperanza Santacruz-Tinoco, María Guadalupe Santiago-Mauricio, Selene Zárate, Nelly Sélem-Mojica, Blanca Taboada, Gloria Vazquez, Joel Armando Vázquez-Pérez. |
| EPI_ISL_2680936, EPI_ISL_2680937 | Hospital Margarita Maza de Juárez | Microbial Genomics Laboratory | Consorcio Mexicano de Vigilancia Genómica (CoViGen-Mex). Authors (in alphabetical order): Julio Elias Alvarado-Yaah, Carlos F. Arias, Santiago Ávila-Ríos, Eduardo Becerril-Vargas, Víctor Hugo Borja-Aburto, Celia Boukadida, Cristóbal Cháidez-Quiróz, Juan Bautista Chale-Dzul, Ricardo Ciria Merce, Andreu Comas-García, Célida Duque Molina, Julissa Enciso-Ibarra, José Antonio Enciso-Moreno, Gloria Elena Espinosa-Ayala, Fernando Fontove-Herrera, Daniel Fregoso-Rueda, Victor Eduardo García-Arias, Alejandra García-Gasca, Bruno Gómez-Gil, Jean Pierre González, Irvin González-López, Concepción Grajales-Muñiz, Ricardo Grande, Rosa María Gutiérrez Rios, Jesús Hernández, Alejandra Hernández-Terán, Alfredo Herrera-Estrella, Carla Ivón Herrera-Najera, Pavel Isa, Verónica Mata-Haro, Daniel Lira Morales, Susana Lopez, Antonio Loza Román, Brenda Irasema Maldonado-Meza, Bernardo Martínez-Miguel, José Arturo Martínez-Orozco, Célida Martínez- Rodríguez, Margarita Matías-Florentino, Fidencio Mejía-Nepomuceno, María Guadalupe de Jesús Mireles-Rivera, Gloria María Molina-Salinas, Hector Montoya-Fuentes, , Mario Mújica-Sánchez, José Esteban Muñoz-Medina, José de Jesús Nuñez-Contreras, Alicia Ocaña-Mondragón, Luis Alberto Ochoa-Carrera, Hector Esteban Paz-Juárez, Marissa Perez-Garcia, Francisco Pulido, Helen Haydee Fernanda Ramírez-Plascencia, Jorge Salas-Hernández, Angel Gustavo Salas-Lais, Alejandro Sánchez-Flores, Clara Esperanza Santacruz-Tinoco, María Guadalupe Santiago-Mauricio, Selene Zárate, Nelly Sélem-Mojica, Blanca Taboada, Gloria Vazquez, Joel Armando Vázquez-Pérez. |
| EPI_ISL_2680938, EPI_ISL_2680939, EPI_ISL_2680940, EPI_ISL_2680942 | Hospital Sharp | Microbial Genomics Laboratory | Consorcio Mexicano de Vigilancia Genómica (CoViGen-Mex). Authors (in alphabetical order): Julio Elias Alvarado-Yaah, Carlos F. Arias, Santiago Ávila-Ríos, Eduardo Becerril-Vargas, Víctor Hugo Borja-Aburto, Celia Boukadida, Cristóbal Cháidez-Quiróz, Juan Bautista Chale-Dzul, Ricardo Ciria Merce, Andreu Comas-García, Célida Duque Molina, Julissa Enciso-Ibarra, José Antonio Enciso-Moreno, Gloria Elena Espinosa-Ayala, Fernando Fontove-Herrera, Daniel Fregoso-Rueda, Victor Eduardo García-Arias, Alejandra García-Gasca, Bruno Gómez-Gil, Jean Pierre González, Irvin González-López, Concepción Grajales-Muñiz, Ricardo Grande, Rosa María Gutiérrez Rios, Jesús Hernández, Alejandra Hernández-Terán, Alfredo Herrera-Estrella, Carla Ivón Herrera-Najera, Pavel Isa, Verónica Mata-Haro, Daniel Lira Morales, Susana Lopez, Antonio Loza Román, Brenda Irasema Maldonado-Meza, Bernardo Martínez-Miguel, José Arturo Martínez-Orozco, Célida Martínez- Rodríguez, Margarita Matías-Florentino, Fidencio Mejía-Nepomuceno, María Guadalupe de Jesús Mireles-Rivera, Gloria María Molina-Salinas, Hector Montoya-Fuentes, , Mario Mújica-Sánchez, José Esteban Muñoz-Medina, José de Jesús Nuñez-Contreras, Alicia Ocaña-Mondragón, Luis Alberto Ochoa-Carrera, Hector Esteban Paz-Juárez, Marissa Perez-Garcia, Francisco Pulido, Helen Haydee Fernanda Ramírez-Plascencia, Jorge Salas-Hernández, Angel Gustavo Salas-Lais, Alejandro Sánchez-Flores, Clara Esperanza Santacruz-Tinoco, María Guadalupe Santiago-Mauricio, Selene Zárate, Nelly Sélem-Mojica, Blanca Taboada, Gloria Vazquez, Joel Armando Vázquez-Pérez. |
| EPI_ISL_2681046 | Laboratorio Central de Epidemiología (LCE) | Instituto de Biotecnología de la UNAM | Consorcio Mexicano de Vigilancia Genómica (CoViGen-Mex). Authors (in alphabetical order): Julio Elias Alvarado-Yaah, Carlos F. Arias, Santiago Ávila-Ríos, Eduardo Becerril-Vargas, Víctor Hugo Borja-Aburto, Celia Boukadida, Cristóbal Cháidez-Quiróz, Juan Bautista Chale-Dzul, Ricardo Ciria Merce, Andreu Comas-García, Célida Duque Molina, Julissa Enciso-Ibarra, José Antonio Enciso-Moreno, Gloria Elena Espinosa-Ayala, Fernando Fontove-Herrera, Daniel Fregoso-Rueda, Victor Eduardo García-Arias, Alejandra García-Gasca, Bruno Gómez-Gil, Jean Pierre González, Irvin |

| | | | |
|---|---|---|---|
| | | | González-López, Concepción Grajales-Muñiz, Ricardo Grande, Rosa María Gutiérrez Rios, Jesús Hernández, Alejandra Hernández-Terán, Alfredo Herrera-Estrella, Carla Ivón Herrera-Najera, Pavel Isa, Verónica Mata-Haro, Daniel Lira Morales, Susana Lopez, Antonio Loza Román, Brenda Irasema Maldonado-Meza, Bernardo Martínez-Miguel, José Arturo Martínez-Orozco, Célida Martínez- Rodríguez, Margarita Matías-Florentino, Fidencio Mejía-Nepomuceno, María Guadalupe de Jesús Mireles-Rivera, Gloria María Molina-Salinas, Hector Montoya-Fuentes, , Mario Mújica-Sánchez, José Esteban Muñoz-Medina, José de Jesús Nuñez-Contreras, Alicia Ocaña-Mondragón, Luis Alberto Ochoa-Carrera, Hector Esteban Paz-Juárez, Marissa Perez-Garcia, Francisco Pulido, Helen Haydee Fernanda Ramírez-Plascencia, Jorge Salas-Hernández, Angel Gustavo Salas-Lais, Alejandro Sánchez-Flores, Clara Esperanza Santacruz-Tinoco, María Guadalupe Santiago-Mauricio, Selene Zárate, Nelly Sélem-Mojica, Blanca Taboada, Gloria Vazquez, Joel Armando Vázquez-Pérez. |
| EPI_ISL_2681047, EPI_ISL_2681048, EPI_ISL_2681049 | Unidad de Investigación Biomédica de Zacatecas (UIBZ) | Instituto de Biotecnología de la UNAM | Consorcio Mexicano de Vigilancia Genómica (CoViGen-Mex). Authors (in alphabetical order): Julio Elias Alvarado-Yaah, Carlos F. Arias, Santiago Ávila-Ríos, Eduardo Becerril-Vargas, Víctor Hugo Borja-Aburto, Celia Boukadida, Cristóbal Cháidez-Quiróz, Juan Bautista Chale-Dzul, Ricardo Ciria Merce, Andreu Comas-García, Célida Duque Molina, Julissa Enciso-Ibarra, José Antonio Enciso-Moreno, Gloria Elena Espinosa-Ayala, Fernando Fontove-Herrera, Daniel Fregoso-Rueda, Victor Eduardo García-Arias, Alejandra García-Gasca, Bruno Gómez-Gil, Jean Pierre González, Irvin González-López, Concepción Grajales-Muñiz, Ricardo Grande, Rosa María Gutiérrez Rios, Jesús Hernández, Alejandra Hernández-Terán, Alfredo Herrera-Estrella, Carla Ivón Herrera-Najera, Pavel Isa, Verónica Mata-Haro, Daniel Lira Morales, Susana Lopez, Antonio Loza Román, Brenda Irasema Maldonado-Meza, Bernardo Martínez-Miguel, José Arturo Martínez-Orozco, Célida Martínez- Rodríguez, Margarita Matías-Florentino, Fidencio Mejía-Nepomuceno, María Guadalupe de Jesús Mireles-Rivera, Gloria María Molina-Salinas, Hector Montoya-Fuentes, , Mario Mújica-Sánchez, José Esteban Muñoz-Medina, José de Jesús Nuñez-Contreras, Alicia Ocaña-Mondragón, Luis Alberto Ochoa-Carrera, Hector Esteban Paz-Juárez, Marissa Perez-Garcia, Francisco Pulido, Helen Haydee Fernanda Ramírez-Plascencia, Jorge Salas-Hernández, Angel Gustavo Salas-Lais, Alejandro Sánchez-Flores, Clara Esperanza Santacruz-Tinoco, María Guadalupe Santiago-Mauricio, Selene Zárate, Nelly Sélem-Mojica, Blanca Taboada, Gloria Vazquez, Joel Armando Vázquez-Pérez. |
| EPI_ISL_2681050 | Unidad de Investigación Médica de Yucatán (UIMY) | Instituto de Biotecnología de la UNAM | Consorcio Mexicano de Vigilancia Genómica (CoViGen-Mex). Authors (in alphabetical order): Julio Elias Alvarado-Yaah, Carlos F. Arias, Santiago Ávila-Ríos, Eduardo Becerril-Vargas, Víctor Hugo Borja-Aburto, Celia Boukadida, Cristóbal Cháidez-Quiróz, Juan Bautista Chale-Dzul, Ricardo Ciria Merce, Andreu Comas-García, Célida Duque Molina, Julissa Enciso-Ibarra, José Antonio Enciso-Moreno, Gloria Elena Espinosa-Ayala, Fernando Fontove-Herrera, Daniel Fregoso-Rueda, Victor Eduardo García-Arias, Alejandra García-Gasca, Bruno Gómez-Gil, Jean Pierre González, Irvin González-López, Concepción Grajales-Muñiz, Ricardo Grande, Rosa María Gutiérrez Rios, Jesús Hernández, Alejandra Hernández-Terán, Alfredo Herrera-Estrella, Carla Ivón Herrera-Najera, Pavel Isa, Verónica Mata-Haro, Daniel Lira Morales, Susana Lopez, Antonio Loza Román, Brenda Irasema Maldonado-Meza, Bernardo Martínez-Miguel, José Arturo Martínez-Orozco, Célida Martínez- Rodríguez, Margarita Matías-Florentino, Fidencio Mejía-Nepomuceno, María Guadalupe de Jesús Mireles-Rivera, Gloria María Molina-Salinas, Hector Montoya-Fuentes, , Mario Mújica-Sánchez, José Esteban Muñoz-Medina, José de Jesús Nuñez-Contreras, Alicia Ocaña-Mondragón, Luis Alberto Ochoa-Carrera, Hector Esteban Paz-Juárez, Marissa Perez-Garcia, Francisco Pulido, Helen Haydee Fernanda Ramírez-Plascencia, Jorge Salas-Hernández, Angel Gustavo Salas-Lais, Alejandro Sánchez-Flores, Clara Esperanza Santacruz-Tinoco, María Guadalupe Santiago-Mauricio, Selene Zárate, Nelly Sélem-Mojica, Blanca Taboada, Gloria Vazquez, Joel Armando Vázquez-Pérez. |
| EPI_ISL_2681051 | Laboratorio Central de Epidemiología (LCE) | Instituto de Biotecnología de la UNAM | Consorcio Mexicano de Vigilancia Genómica (CoViGen-Mex). Authors (in alphabetical order): Julio Elias Alvarado-Yaah, Carlos F. Arias, Santiago Ávila-Ríos, Eduardo Becerril-Vargas, Víctor Hugo Borja-Aburto, Celia Boukadida, Cristóbal Cháidez-Quiróz, Juan Bautista Chale-Dzul, Ricardo Ciria Merce, Andreu Comas-García, Célida Duque Molina, Julissa Enciso-Ibarra, José Antonio Enciso-Moreno, Gloria Elena Espinosa-Ayala, Fernando Fontove-Herrera, Daniel Fregoso-Rueda, Victor Eduardo García-Arias, Alejandra García-Gasca, Bruno Gómez-Gil, Jean Pierre González, Irvin González-López, Concepción Grajales-Muñiz, Ricardo Grande, Rosa María Gutiérrez Rios, Jesús Hernández, Alejandra Hernández-Terán, Alfredo Herrera-Estrella, Carla Ivón Herrera-Najera, Pavel Isa, Verónica Mata-Haro, Daniel Lira Morales, Susana Lopez, Antonio Loza Román, Brenda Irasema Maldonado-Meza, Bernardo Martínez-Miguel, José Arturo Martínez-Orozco, Célida Martínez- Rodríguez, Margarita Matías-Florentino, Fidencio Mejía-Nepomuceno, María Guadalupe de Jesús Mireles-Rivera, Gloria María Molina-Salinas, Hector Montoya-Fuentes, , Mario Mújica-Sánchez, José Esteban Muñoz-Medina, José de Jesús Nuñez-Contreras, Alicia Ocaña-Mondragón, Luis Alberto Ochoa-Carrera, Hector Esteban Paz-Juárez, Marissa Perez-Garcia, Francisco Pulido, Helen Haydee Fernanda Ramírez-Plascencia, Jorge Salas-Hernández, Angel Gustavo Salas-Lais, Alejandro Sánchez-Flores, Clara Esperanza Santacruz-Tinoco, María Guadalupe Santiago-Mauricio, Selene Zárate, Nelly Sélem-Mojica, Blanca Taboada, Gloria Vazquez, Joel Armando Vázquez-Pérez. |
| EPI_ISL_2681052 | Unidad de Investigación Biomédica de Zacatecas (UIBZ) | Instituto de Biotecnología de la UNAM | Consorcio Mexicano de Vigilancia Genómica (CoViGen-Mex). Authors (in alphabetical order): Julio Elias Alvarado-Yaah, Carlos F. Arias, Santiago Ávila-Ríos, Eduardo Becerril-Vargas, Víctor Hugo Borja-Aburto, Celia Boukadida, Cristóbal Cháidez-Quiróz, Juan Bautista Chale-Dzul, Ricardo Ciria Merce, Andreu Comas-García, Célida Duque Molina, Julissa Enciso-Ibarra, José Antonio Enciso-Moreno, Gloria Elena Espinosa-Ayala, Fernando Fontove-Herrera, Daniel Fregoso-Rueda, Victor Eduardo García-Arias, Alejandra García-Gasca, Bruno Gómez-Gil, Jean Pierre González, Irvin González-López, Concepción Grajales-Muñiz, Ricardo Grande, Rosa María Gutiérrez Rios, Jesús Hernández, Alejandra Hernández-Terán, Alfredo Herrera-Estrella, Carla Ivón Herrera-Najera, Pavel Isa, Verónica Mata-Haro, Daniel Lira Morales, Susana Lopez, Antonio Loza Román, Brenda Irasema Maldonado-Meza, Bernardo Martínez-Miguel, José Arturo Martínez-Orozco, Célida Martínez- Rodríguez, Margarita Matías-Florentino, Fidencio Mejía-Nepomuceno, María Guadalupe de Jesús Mireles-Rivera, Gloria María Molina-Salinas, Hector Montoya-Fuentes, , Mario Mújica-Sánchez, José Esteban Muñoz-Medina, José de Jesús Nuñez-Contreras, Alicia Ocaña-Mondragón, Luis Alberto Ochoa-Carrera, Hector Esteban Paz-Juárez, Marissa Perez-Garcia, Francisco Pulido, Helen Haydee Fernanda Ramírez-Plascencia, Jorge Salas-Hernández, Angel Gustavo Salas-Lais, Alejandro Sánchez-Flores, Clara Esperanza Santacruz-Tinoco, María Guadalupe Santiago-Mauricio, Selene Zárate, Nelly Sélem-Mojica, Blanca Taboada, Gloria Vazquez, Joel Armando Vázquez-Pérez. |
| EPI_ISL_2681054 | Laboratorio Central de Epidemiología (LCE) | Instituto de Biotecnología de la UNAM | Consorcio Mexicano de Vigilancia Genómica (CoViGen-Mex). Authors (in alphabetical order): Julio Elias Alvarado-Yaah, Carlos F. Arias, Santiago Ávila-Ríos, Eduardo Becerril-Vargas, Víctor Hugo Borja-Aburto, Celia Boukadida, Cristóbal Cháidez-Quiróz, Juan Bautista Chale-Dzul, Ricardo Ciria Merce, Andreu Comas-García, Célida Duque Molina, Julissa Enciso-Ibarra, José Antonio Enciso-Moreno, Gloria Elena Espinosa-Ayala, Fernando Fontove-Herrera, Daniel Fregoso-Rueda, Victor Eduardo García-Arias, Alejandra García-Gasca, Bruno Gómez-Gil, Jean Pierre González, Irvin González-López, Concepción Grajales-Muñiz, Ricardo Grande, Rosa María Gutiérrez Rios, Jesús Hernández, Alejandra Hernández-Terán, Alfredo Herrera-Estrella, Carla Ivón Herrera-Najera, Pavel Isa, Verónica Mata-Haro, Daniel Lira Morales, Susana Lopez, Antonio Loza Román, Brenda Irasema Maldonado-Meza, Bernardo Martínez-Miguel, José Arturo Martínez-Orozco, Célida Martínez- Rodríguez, Margarita Matías-Florentino, Fidencio Mejía-Nepomuceno, María Guadalupe de Jesús Mireles-Rivera, Gloria María Molina-Salinas, Hector Montoya-Fuentes, , Mario Mújica-Sánchez, José Esteban Muñoz-Medina, José de Jesús Nuñez-Contreras, Alicia Ocaña-Mondragón, Luis Alberto Ochoa-Carrera, Hector Esteban Paz-Juárez, Marissa Perez-Garcia, Francisco Pulido, Helen Haydee Fernanda Ramírez-Plascencia, Jorge Salas-Hernández, Angel Gustavo Salas-Lais, Alejandro Sánchez-Flores, Clara Esperanza Santacruz-Tinoco, María Guadalupe Santiago-Mauricio, Selene Zárate, Nelly Sélem-Mojica, Blanca Taboada, Gloria Vazquez, Joel Armando Vázquez-Pérez. |
| EPI_ISL_2681055 | Unidad de Investigación Médica de Yucatán (UIMY) | Instituto de Biotecnología de la UNAM | Consorcio Mexicano de Vigilancia Genómica (CoViGen-Mex). Authors (in alphabetical order): Julio Elias Alvarado-Yaah, Carlos F. Arias, Santiago Ávila-Ríos, Eduardo Becerril-Vargas, Víctor Hugo Borja-Aburto, Celia Boukadida, Cristóbal Cháidez-Quiróz, Juan Bautista Chale-Dzul, Ricardo Ciria Merce, Andreu Comas-García, Célida Duque Molina, Julissa Enciso-Ibarra, José Antonio Enciso-Moreno, Gloria Elena Espinosa-Ayala, Fernando Fontove-Herrera, Daniel Fregoso-Rueda, Victor Eduardo García-Arias, Alejandra García-Gasca, Bruno Gómez-Gil, Jean Pierre González, Irvin González-López, Concepción Grajales-Muñiz, Ricardo Grande, Rosa María Gutiérrez Rios, Jesús Hernández, Alejandra Hernández-Terán, Alfredo Herrera-Estrella, Carla Ivón Herrera-Najera, Pavel Isa, Verónica Mata-Haro, Daniel Lira Morales, Susana Lopez, Antonio Loza Román, Brenda Irasema Maldonado-Meza, Bernardo Martínez-Miguel, José Arturo Martínez-Orozco, Célida Martínez- Rodríguez, Margarita Matías-Florentino, Fidencio Mejía-Nepomuceno, María Guadalupe de Jesús Mireles-Rivera, Gloria María Molina-Salinas, Hector Montoya-Fuentes, , Mario Mújica-Sánchez, José Esteban Muñoz-Medina, José de Jesús Nuñez-Contreras, Alicia Ocaña-Mondragón, Luis Alberto Ochoa-Carrera, Hector Esteban Paz-Juárez, Marissa Perez-Garcia, Francisco Pulido, Helen Haydee Fernanda Ramírez-Plascencia, Jorge Salas-Hernández, Angel Gustavo Salas-Lais, Alejandro Sánchez-Flores, Clara Esperanza Santacruz-Tinoco, María Guadalupe Santiago-Mauricio, Selene Zárate, Nelly Sélem-Mojica, Blanca Taboada, Gloria Vazquez, Joel Armando Vázquez-Pérez. |
| EPI_ISL_2681056 | Laboratorio Central de Epidemiología (LCE) | Instituto de Biotecnología de la UNAM | Consorcio Mexicano de Vigilancia Genómica (CoViGen-Mex). Authors (in alphabetical order): Julio Elias Alvarado-Yaah, Carlos F. Arias, Santiago Ávila-Ríos, Eduardo Becerril-Vargas, Víctor Hugo Borja-Aburto, Celia Boukadida, Cristóbal Cháidez-Quiróz, Juan Bautista Chale-Dzul, Ricardo Ciria Merce, Andreu Comas-García, Célida Duque Molina, Julissa Enciso-Ibarra, José Antonio Enciso-Moreno, Gloria Elena Espinosa-Ayala, Fernando Fontove-Herrera, Daniel Fregoso-Rueda, Victor Eduardo García-Arias, Alejandra García-Gasca, Bruno Gómez-Gil, Jean Pierre González, Irvin |

| | | | |
|---|---|---|---|
| | | | González-López, Concepción Grajales-Muñiz, Ricardo Grande, Rosa María Gutiérrez Rios, Jesús Hernández, Alejandra Hernández-Terán, Alfredo Herrera-Estrella, Carla Ivón Herrera-Najera, Pavel Isa, Verónica Mata-Haro, Daniel Lira Morales, Susana Lopez, Antonio Loza Román, Brenda Irasema Maldonado-Meza, Bernardo Martínez-Miguel, José Arturo Martínez-Orozco, Célida Martínez- Rodríguez, Margarita Matías-Florentino, Fidencio Mejía-Nepomuceno, María Guadalupe de Jesús Mireles-Rivera, Gloria María Molina-Salinas, Hector Montoya-Fuentes, , Mario Mújica-Sánchez, José Esteban Muñoz-Medina, José de Jesús Nuñez-Contreras, Alicia Ocaña-Mondragón, Luis Alberto Ochoa-Carrera, Hector Esteban Paz-Juárez, Marissa Perez-Garcia, Francisco Pulido, Helen Haydee Fernanda Ramírez-Plascencia, Jorge Salas-Hernández, Angel Gustavo Salas-Lais, Alejandro Sánchez-Flores, Clara Esperanza Santacruz-Tinoco, María Guadalupe Santiago-Mauricio, Selene Zárate, Nelly Sélem-Mojica, Blanca Taboada, Gloria Vazquez, Joel Armando Vázquez-Pérez. |
| EPI_ISL_2681057 | Unidad de Investigación Biomédica de Zacatecas (UIBZ) | Instituto de Biotecnología de la UNAM | Consorcio Mexicano de Vigilancia Genómica (CoViGen-Mex). Authors (in alphabetical order): Julio Elias Alvarado-Yaah, Carlos F. Arias, Santiago Ávila-Ríos, Eduardo Becerril-Vargas, Víctor Hugo Borja-Aburto, Celia Boukadida, Cristóbal Cháidez-Quiróz, Juan Bautista Chale-Dzul, Ricardo Ciria Merce, Andreu Comas-García, Célida Duque Molina, Julissa Enciso-Ibarra, José Antonio Enciso-Moreno, Gloria Elena Espinosa-Ayala, Fernando Fontove-Herrera, Daniel Fregoso-Rueda, Victor Eduardo García-Arias, Alejandra García-Gasca, Bruno Gómez-Gil, Jean Pierre González, Irvin González-López, Concepción Grajales-Muñiz, Ricardo Grande, Rosa María Gutiérrez Rios, Jesús Hernández, Alejandra Hernández-Terán, Alfredo Herrera-Estrella, Carla Ivón Herrera-Najera, Pavel Isa, Verónica Mata-Haro, Daniel Lira Morales, Susana Lopez, Antonio Loza Román, Brenda Irasema Maldonado-Meza, Bernardo Martínez-Miguel, José Arturo Martínez-Orozco, Célida Martínez- Rodríguez, Margarita Matías-Florentino, Fidencio Mejía-Nepomuceno, María Guadalupe de Jesús Mireles-Rivera, Gloria María Molina-Salinas, Hector Montoya-Fuentes, , Mario Mújica-Sánchez, José Esteban Muñoz-Medina, José de Jesús Nuñez-Contreras, Alicia Ocaña-Mondragón, Luis Alberto Ochoa-Carrera, Hector Esteban Paz-Juárez, Marissa Perez-Garcia, Francisco Pulido, Helen Haydee Fernanda Ramírez-Plascencia, Jorge Salas-Hernández, Angel Gustavo Salas-Lais, Alejandro Sánchez-Flores, Clara Esperanza Santacruz-Tinoco, María Guadalupe Santiago-Mauricio, Selene Zárate, Nelly Sélem-Mojica, Blanca Taboada, Gloria Vazquez, Joel Armando Vázquez-Pérez. |
| EPI_ISL_2681058, EPI_ISL_2681059, EPI_ISL_2681060, EPI_ISL_2681061, EPI_ISL_2681062, EPI_ISL_2681063, EPI_ISL_2681064, EPI_ISL_2681065, EPI_ISL_2681066 | Laboratorio Central de Epidemiología (LCE) | Instituto de Biotecnología de la UNAM | Consorcio Mexicano de Vigilancia Genómica (CoViGen-Mex). Authors (in alphabetical order): Julio Elias Alvarado-Yaah, Carlos F. Arias, Santiago Ávila-Ríos, Eduardo Becerril-Vargas, Víctor Hugo Borja-Aburto, Celia Boukadida, Cristóbal Cháidez-Quiróz, Juan Bautista Chale-Dzul, Ricardo Ciria Merce, Andreu Comas-García, Célida Duque Molina, Julissa Enciso-Ibarra, José Antonio Enciso-Moreno, Gloria Elena Espinosa-Ayala, Fernando Fontove-Herrera, Daniel Fregoso-Rueda, Victor Eduardo García-Arias, Alejandra García-Gasca, Bruno Gómez-Gil, Jean Pierre González, Irvin González-López, Concepción Grajales-Muñiz, Ricardo Grande, Rosa María Gutiérrez Rios, Jesús Hernández, Alejandra Hernández-Terán, Alfredo Herrera-Estrella, Carla Ivón Herrera-Najera, Pavel Isa, Verónica Mata-Haro, Daniel Lira Morales, Susana Lopez, Antonio Loza Román, Brenda Irasema Maldonado-Meza, Bernardo Martínez-Miguel, José Arturo Martínez-Orozco, Célida Martínez- Rodríguez, Margarita Matías-Florentino, Fidencio Mejía-Nepomuceno, María Guadalupe de Jesús Mireles-Rivera, Gloria María Molina-Salinas, Hector Montoya-Fuentes, , Mario Mújica-Sánchez, José Esteban Muñoz-Medina, José de Jesús Nuñez-Contreras, Alicia Ocaña-Mondragón, Luis Alberto Ochoa-Carrera, Hector Esteban Paz-Juárez, Marissa Perez-Garcia, Francisco Pulido, Helen Haydee Fernanda Ramírez-Plascencia, Jorge Salas-Hernández, Angel Gustavo Salas-Lais, Alejandro Sánchez-Flores, Clara Esperanza Santacruz-Tinoco, María Guadalupe Santiago-Mauricio, Selene Zárate, Nelly Sélem-Mojica, Blanca Taboada, Gloria Vazquez, Joel Armando Vázquez-Pérez. |
| EPI_ISL_2681067 | Unidad de Investigación Médica de Yucatán (UIMY) | Instituto de Biotecnología de la UNAM | Consorcio Mexicano de Vigilancia Genómica (CoViGen-Mex). Authors (in alphabetical order): Julio Elias Alvarado-Yaah, Carlos F. Arias, Santiago Ávila-Ríos, Eduardo Becerril-Vargas, Víctor Hugo Borja-Aburto, Celia Boukadida, Cristóbal Cháidez-Quiróz, Juan Bautista Chale-Dzul, Ricardo Ciria Merce, Andreu Comas-García, Célida Duque Molina, Julissa Enciso-Ibarra, José Antonio Enciso-Moreno, Gloria Elena Espinosa-Ayala, Fernando Fontove-Herrera, Daniel Fregoso-Rueda, Victor Eduardo García-Arias, Alejandra García-Gasca, Bruno Gómez-Gil, Jean Pierre González, Irvin González-López, Concepción Grajales-Muñiz, Ricardo Grande, Rosa María Gutiérrez Rios, Jesús Hernández, Alejandra Hernández-Terán, Alfredo Herrera-Estrella, Carla Ivón Herrera-Najera, Pavel Isa, Verónica Mata-Haro, Daniel Lira Morales, Susana Lopez, Antonio Loza Román, Brenda Irasema Maldonado-Meza, Bernardo Martínez-Miguel, José Arturo Martínez-Orozco, Célida Martínez- Rodríguez, Margarita Matías-Florentino, Fidencio Mejía-Nepomuceno, María Guadalupe de Jesús Mireles-Rivera, Gloria María Molina-Salinas, Hector Montoya-Fuentes, , Mario Mújica-Sánchez, José Esteban Muñoz-Medina, José de Jesús Nuñez-Contreras, Alicia Ocaña-Mondragón, Luis Alberto Ochoa-Carrera, Hector Esteban Paz-Juárez, Marissa Perez-Garcia, Francisco Pulido, Helen Haydee Fernanda Ramírez-Plascencia, Jorge Salas-Hernández, Angel Gustavo Salas-Lais, Alejandro Sánchez-Flores, Clara Esperanza Santacruz-Tinoco, María Guadalupe Santiago-Mauricio, Selene Zárate, Nelly Sélem-Mojica, Blanca Taboada, Gloria Vazquez, Joel Armando Vázquez-Pérez. |
| EPI_ISL_2681068 | Unidad de Investigación Biomédica de Zacatecas (UIBZ) | Instituto de Biotecnología de la UNAM | Consorcio Mexicano de Vigilancia Genómica (CoViGen-Mex). Authors (in alphabetical order): Julio Elias Alvarado-Yaah, Carlos F. Arias, Santiago Ávila-Ríos, Eduardo Becerril-Vargas, Víctor Hugo Borja-Aburto, Celia Boukadida, Cristóbal Cháidez-Quiróz, Juan Bautista Chale-Dzul, Ricardo Ciria Merce, Andreu Comas-García, Célida Duque Molina, Julissa Enciso-Ibarra, José Antonio Enciso-Moreno, Gloria Elena Espinosa-Ayala, Fernando Fontove-Herrera, Daniel Fregoso-Rueda, Victor Eduardo García-Arias, Alejandra García-Gasca, Bruno Gómez-Gil, Jean Pierre González, Irvin González-López, Concepción Grajales-Muñiz, Ricardo Grande, Rosa María Gutiérrez Rios, Jesús Hernández, Alejandra Hernández-Terán, Alfredo Herrera-Estrella, Carla Ivón Herrera-Najera, Pavel Isa, Verónica Mata-Haro, Daniel Lira Morales, Susana Lopez, Antonio Loza Román, Brenda Irasema Maldonado-Meza, Bernardo Martínez-Miguel, José Arturo Martínez-Orozco, Célida Martínez- Rodríguez, Margarita Matías-Florentino, Fidencio Mejía-Nepomuceno, María Guadalupe de Jesús Mireles-Rivera, Gloria María Molina-Salinas, Hector Montoya-Fuentes, , Mario Mújica-Sánchez, José Esteban Muñoz-Medina, José de Jesús Nuñez-Contreras, Alicia Ocaña-Mondragón, Luis Alberto Ochoa-Carrera, Hector Esteban Paz-Juárez, Marissa Perez-Garcia, Francisco Pulido, Helen Haydee Fernanda Ramírez-Plascencia, Jorge Salas-Hernández, Angel Gustavo Salas-Lais, Alejandro Sánchez-Flores, Clara Esperanza Santacruz-Tinoco, María Guadalupe Santiago-Mauricio, Selene Zárate, Nelly Sélem-Mojica, Blanca Taboada, Gloria Vazquez, Joel Armando Vázquez-Pérez. |
| EPI_ISL_2681069, EPI_ISL_2681070, EPI_ISL_2681071, EPI_ISL_2681072, EPI_ISL_2681073 | Laboratorio Central de Epidemiología (LCE) | Instituto de Biotecnología de la UNAM | Consorcio Mexicano de Vigilancia Genómica (CoViGen-Mex). Authors (in alphabetical order): Julio Elias Alvarado-Yaah, Carlos F. Arias, Santiago Ávila-Ríos, Eduardo Becerril-Vargas, Víctor Hugo Borja-Aburto, Celia Boukadida, Cristóbal Cháidez-Quiróz, Juan Bautista Chale-Dzul, Ricardo Ciria Merce, Andreu Comas-García, Célida Duque Molina, Julissa Enciso-Ibarra, José Antonio Enciso-Moreno, Gloria Elena Espinosa-Ayala, Fernando Fontove-Herrera, Daniel Fregoso-Rueda, Victor Eduardo García-Arias, Alejandra García-Gasca, Bruno Gómez-Gil, Jean Pierre González, Irvin González-López, Concepción Grajales-Muñiz, Ricardo Grande, Rosa María Gutiérrez Rios, Jesús Hernández, Alejandra Hernández-Terán, Alfredo Herrera-Estrella, Carla Ivón Herrera-Najera, Pavel Isa, Verónica Mata-Haro, Daniel Lira Morales, Susana Lopez, Antonio Loza Román, Brenda Irasema Maldonado-Meza, Bernardo Martínez-Miguel, José Arturo Martínez-Orozco, Célida Martínez- Rodríguez, Margarita Matías-Florentino, Fidencio Mejía-Nepomuceno, María Guadalupe de Jesús Mireles-Rivera, Gloria María Molina-Salinas, Hector Montoya-Fuentes, , Mario Mújica-Sánchez, José Esteban Muñoz-Medina, José de Jesús Nuñez-Contreras, Alicia Ocaña-Mondragón, Luis Alberto Ochoa-Carrera, Hector Esteban Paz-Juárez, Marissa Perez-Garcia, Francisco Pulido, Helen Haydee Fernanda Ramírez-Plascencia, Jorge Salas-Hernández, Angel Gustavo Salas-Lais, Alejandro Sánchez-Flores, Clara Esperanza Santacruz-Tinoco, María Guadalupe Santiago-Mauricio, Selene Zárate, Nelly Sélem-Mojica, Blanca Taboada, Gloria Vazquez, Joel Armando Vázquez-Pérez. |
| EPI_ISL_2681074, EPI_ISL_2681075 | Unidad de Investigación Médica de Yucatán (UIMY) | Instituto de Biotecnología de la UNAM | Consorcio Mexicano de Vigilancia Genómica (CoViGen-Mex). Authors (in alphabetical order): Julio Elias Alvarado-Yaah, Carlos F. Arias, Santiago Ávila-Ríos, Eduardo Becerril-Vargas, Víctor Hugo Borja-Aburto, Celia Boukadida, Cristóbal Cháidez-Quiróz, Juan Bautista Chale-Dzul, Ricardo Ciria Merce, Andreu Comas-García, Célida Duque Molina, Julissa Enciso-Ibarra, José Antonio Enciso-Moreno, Gloria Elena Espinosa-Ayala, Fernando Fontove-Herrera, Daniel Fregoso-Rueda, Victor Eduardo García-Arias, Alejandra García-Gasca, Bruno Gómez-Gil, Jean Pierre González, Irvin González-López, Concepción Grajales-Muñiz, Ricardo Grande, Rosa María Gutiérrez Rios, Jesús Hernández, Alejandra Hernández-Terán, Alfredo Herrera-Estrella, Carla Ivón Herrera-Najera, Pavel Isa, Verónica Mata-Haro, Daniel Lira Morales, Susana Lopez, Antonio Loza Román, Brenda Irasema Maldonado-Meza, Bernardo Martínez-Miguel, José Arturo Martínez-Orozco, Célida Martínez- Rodríguez, Margarita Matías-Florentino, Fidencio Mejía-Nepomuceno, María Guadalupe de Jesús Mireles-Rivera, Gloria María Molina-Salinas, Hector Montoya-Fuentes, , Mario Mújica-Sánchez, José Esteban Muñoz-Medina, José de Jesús Nuñez-Contreras, Alicia Ocaña-Mondragón, Luis Alberto Ochoa-Carrera, Hector Esteban Paz-Juárez, Marissa Perez-Garcia, Francisco Pulido, Helen Haydee Fernanda Ramírez-Plascencia, Jorge Salas-Hernández, Angel Gustavo Salas-Lais, Alejandro Sánchez-Flores, Clara Esperanza Santacruz-Tinoco, María Guadalupe Santiago-Mauricio, Selene Zárate, Nelly Sélem-Mojica, Blanca Taboada, Gloria Vazquez, Joel Armando Vázquez-Pérez. |
| EPI_ISL_2681076 | Laboratorio Central de Epidemiología (LCE) | Instituto de Biotecnología de la UNAM | Consorcio Mexicano de Vigilancia Genómica (CoViGen-Mex). Authors (in alphabetical order): Julio Elias Alvarado-Yaah, Carlos F. Arias, Santiago Ávila-Ríos, Eduardo Becerril-Vargas, Víctor Hugo Borja-Aburto, Celia Boukadida, Cristóbal Cháidez-Quiróz, Juan Bautista Chale-Dzul, Ricardo Ciria Merce, Andreu Comas-García, Célida Duque Molina, Julissa Enciso-Ibarra, José Antonio Enciso-Moreno, Gloria Elena Espinosa-Ayala, Fernando Fontove-Herrera, Daniel Fregoso-Rueda, Victor Eduardo García-Arias, Alejandra García-Gasca, Bruno Gómez-Gil, Jean Pierre González, Irvin |

| | | | |
|---|---|---|---|
| | | | González-López, Concepción Grajales-Muñiz, Ricardo Grande, Rosa María Gutiérrez Rios, Jesús Hernández, Alejandra Hernández-Terán, Alfredo Herrera-Estrella, Carla Ivón Herrera-Najera, Pavel Isa, Verónica Mata-Haro, Daniel Lira Morales, Susana Lopez, Antonio Loza Román, Brenda Irasema Maldonado-Meza, Bernardo Martínez-Miguel, José Arturo Martínez-Orozco, Célida Martínez- Rodríguez, Margarita Matías-Florentino, Fidencio Mejía-Nepomuceno, María Guadalupe de Jesús Mireles-Rivera, Gloria María Molina-Salinas, Hector Montoya-Fuentes, , Mario Mújica-Sánchez, José Esteban Muñoz-Medina, José de Jesús Nuñez-Contreras, Alicia Ocaña-Mondragón, Luis Alberto Ochoa-Carrera, Hector Esteban Paz-Juárez, Marissa Perez-Garcia, Francisco Pulido, Helen Haydee Fernanda Ramírez-Plascencia, Jorge Salas-Hernández, Angel Gustavo Salas-Lais, Alejandro Sánchez-Flores, Clara Esperanza Santacruz-Tinoco, María Guadalupe Santiago-Mauricio, Selene Zárate, Nelly Sélem-Mojica, Blanca Taboada, Gloria Vazquez, Joel Armando Vázquez-Pérez. |
| EPI_ISL_2681077, EPI_ISL_2681078, EPI_ISL_2681079, EPI_ISL_2681080 | Unidad de Investigación Médica de Yucatán (UIMY) | Instituto de Biotecnología de la UNAM | Consorcio Mexicano de Vigilancia Genómica (CoViGen-Mex). Authors (in alphabetical order): Julio Elias Alvarado-Yaah, Carlos F. Arias, Santiago Ávila-Ríos, Eduardo Becerril-Vargas, Víctor Hugo Borja-Aburto, Celia Boukadida, Cristóbal Cháidez-Quiróz, Juan Bautista Chale-Dzul, Ricardo Ciria Merce, Andreu Comas-García, Célida Duque Molina, Julissa Enciso-Ibarra, José Antonio Enciso-Moreno, Gloria Elena Espinosa-Ayala, Fernando Fontove-Herrera, Daniel Fregoso-Rueda, Victor Eduardo García-Arias, Alejandra García-Gasca, Bruno Gómez-Gil, Jean Pierre González, Irvin González-López, Concepción Grajales-Muñiz, Ricardo Grande, Rosa María Gutiérrez Rios, Jesús Hernández, Alejandra Hernández-Terán, Alfredo Herrera-Estrella, Carla Ivón Herrera-Najera, Pavel Isa, Verónica Mata-Haro, Daniel Lira Morales, Susana Lopez, Antonio Loza Román, Brenda Irasema Maldonado-Meza, Bernardo Martínez-Miguel, José Arturo Martínez-Orozco, Célida Martínez- Rodríguez, Margarita Matías-Florentino, Fidencio Mejía-Nepomuceno, María Guadalupe de Jesús Mireles-Rivera, Gloria María Molina-Salinas, Hector Montoya-Fuentes, , Mario Mújica-Sánchez, José Esteban Muñoz-Medina, José de Jesús Nuñez-Contreras, Alicia Ocaña-Mondragón, Luis Alberto Ochoa-Carrera, Hector Esteban Paz-Juárez, Marissa Perez-Garcia, Francisco Pulido, Helen Haydee Fernanda Ramírez-Plascencia, Jorge Salas-Hernández, Angel Gustavo Salas-Lais, Alejandro Sánchez-Flores, Clara Esperanza Santacruz-Tinoco, María Guadalupe Santiago-Mauricio, Selene Zárate, Nelly Sélem-Mojica, Blanca Taboada, Gloria Vazquez, Joel Armando Vázquez-Pérez. |
| EPI_ISL_2681081 | Laboratorio Central de Epidemiología (LCE) | Instituto de Biotecnología de la UNAM | Consorcio Mexicano de Vigilancia Genómica (CoViGen-Mex). Authors (in alphabetical order): Julio Elias Alvarado-Yaah, Carlos F. Arias, Santiago Ávila-Ríos, Eduardo Becerril-Vargas, Víctor Hugo Borja-Aburto, Celia Boukadida, Cristóbal Cháidez-Quiróz, Juan Bautista Chale-Dzul, Ricardo Ciria Merce, Andreu Comas-García, Célida Duque Molina, Julissa Enciso-Ibarra, José Antonio Enciso-Moreno, Gloria Elena Espinosa-Ayala, Fernando Fontove-Herrera, Daniel Fregoso-Rueda, Victor Eduardo García-Arias, Alejandra García-Gasca, Bruno Gómez-Gil, Jean Pierre González, Irvin González-López, Concepción Grajales-Muñiz, Ricardo Grande, Rosa María Gutiérrez Rios, Jesús Hernández, Alejandra Hernández-Terán, Alfredo Herrera-Estrella, Carla Ivón Herrera-Najera, Pavel Isa, Verónica Mata-Haro, Daniel Lira Morales, Susana Lopez, Antonio Loza Román, Brenda Irasema Maldonado-Meza, Bernardo Martínez-Miguel, José Arturo Martínez-Orozco, Célida Martínez- Rodríguez, Margarita Matías-Florentino, Fidencio Mejía-Nepomuceno, María Guadalupe de Jesús Mireles-Rivera, Gloria María Molina-Salinas, Hector Montoya-Fuentes, , Mario Mújica-Sánchez, José Esteban Muñoz-Medina, José de Jesús Nuñez-Contreras, Alicia Ocaña-Mondragón, Luis Alberto Ochoa-Carrera, Hector Esteban Paz-Juárez, Marissa Perez-Garcia, Francisco Pulido, Helen Haydee Fernanda Ramírez-Plascencia, Jorge Salas-Hernández, Angel Gustavo Salas-Lais, Alejandro Sánchez-Flores, Clara Esperanza Santacruz-Tinoco, María Guadalupe Santiago-Mauricio, Selene Zárate, Nelly Sélem-Mojica, Blanca Taboada, Gloria Vazquez, Joel Armando Vázquez-Pérez. |
| EPI_ISL_2681082, EPI_ISL_2681083, EPI_ISL_2681084, EPI_ISL_2681085 | Unidad de Investigación Médica de Yucatán (UIMY) | Instituto de Biotecnología de la UNAM | Consorcio Mexicano de Vigilancia Genómica (CoViGen-Mex). Authors (in alphabetical order): Julio Elias Alvarado-Yaah, Carlos F. Arias, Santiago Ávila-Ríos, Eduardo Becerril-Vargas, Víctor Hugo Borja-Aburto, Celia Boukadida, Cristóbal Cháidez-Quiróz, Juan Bautista Chale-Dzul, Ricardo Ciria Merce, Andreu Comas-García, Célida Duque Molina, Julissa Enciso-Ibarra, José Antonio Enciso-Moreno, Gloria Elena Espinosa-Ayala, Fernando Fontove-Herrera, Daniel Fregoso-Rueda, Victor Eduardo García-Arias, Alejandra García-Gasca, Bruno Gómez-Gil, Jean Pierre González, Irvin González-López, Concepción Grajales-Muñiz, Ricardo Grande, Rosa María Gutiérrez Rios, Jesús Hernández, Alejandra Hernández-Terán, Alfredo Herrera-Estrella, Carla Ivón Herrera-Najera, Pavel Isa, Verónica Mata-Haro, Daniel Lira Morales, Susana Lopez, Antonio Loza Román, Brenda Irasema Maldonado-Meza, Bernardo Martínez-Miguel, José Arturo Martínez-Orozco, Célida Martínez- Rodríguez, Margarita Matías-Florentino, Fidencio Mejía-Nepomuceno, María Guadalupe de Jesús Mireles-Rivera, Gloria María Molina-Salinas, Hector Montoya-Fuentes, , Mario Mújica-Sánchez, José Esteban Muñoz-Medina, José de Jesús Nuñez-Contreras, Alicia Ocaña-Mondragón, Luis Alberto Ochoa-Carrera, Hector Esteban Paz-Juárez, Marissa Perez-Garcia, Francisco Pulido, Helen Haydee Fernanda Ramírez-Plascencia, Jorge Salas-Hernández, Angel Gustavo Salas-Lais, Alejandro Sánchez-Flores, Clara Esperanza Santacruz-Tinoco, María Guadalupe Santiago-Mauricio, Selene Zárate, Nelly Sélem-Mojica, Blanca Taboada, Gloria Vazquez, Joel Armando Vázquez-Pérez. |
| EPI_ISL_2681086 | Centro de Investigación Biomédica del Noreste (CIBIN) | Instituto de Biotecnología de la UNAM | Consorcio Mexicano de Vigilancia Genómica (CoViGen-Mex). Authors (in alphabetical order): Julio Elias Alvarado-Yaah, Carlos F. Arias, Santiago Ávila-Ríos, Eduardo Becerril-Vargas, Víctor Hugo Borja-Aburto, Celia Boukadida, Cristóbal Cháidez-Quiróz, Juan Bautista Chale-Dzul, Ricardo Ciria Merce, Andreu Comas-García, Célida Duque Molina, Julissa Enciso-Ibarra, José Antonio Enciso-Moreno, Gloria Elena Espinosa-Ayala, Fernando Fontove-Herrera, Daniel Fregoso-Rueda, Victor Eduardo García-Arias, Alejandra García-Gasca, Bruno Gómez-Gil, Jean Pierre González, Irvin González-López, Concepción Grajales-Muñiz, Ricardo Grande, Rosa María Gutiérrez Rios, Jesús Hernández, Alejandra Hernández-Terán, Alfredo Herrera-Estrella, Carla Ivón Herrera-Najera, Pavel Isa, Verónica Mata-Haro, Daniel Lira Morales, Susana Lopez, Antonio Loza Román, Brenda Irasema Maldonado-Meza, Bernardo Martínez-Miguel, José Arturo Martínez-Orozco, Célida Martínez- Rodríguez, Margarita Matías-Florentino, Fidencio Mejía-Nepomuceno, María Guadalupe de Jesús Mireles-Rivera, Gloria María Molina-Salinas, Hector Montoya-Fuentes, , Mario Mújica-Sánchez, José Esteban Muñoz-Medina, José de Jesús Nuñez-Contreras, Alicia Ocaña-Mondragón, Luis Alberto Ochoa-Carrera, Hector Esteban Paz-Juárez, Marissa Perez-Garcia, Francisco Pulido, Helen Haydee Fernanda Ramírez-Plascencia, Jorge Salas-Hernández, Angel Gustavo Salas-Lais, Alejandro Sánchez-Flores, Clara Esperanza Santacruz-Tinoco, María Guadalupe Santiago-Mauricio, Selene Zárate, Nelly Sélem-Mojica, Blanca Taboada, Gloria Vazquez, Joel Armando Vázquez-Pérez. |
| EPI_ISL_2681087 | Laboratorio Central de Epidemiología (LCE) | Instituto de Biotecnología de la UNAM | Consorcio Mexicano de Vigilancia Genómica (CoViGen-Mex). Authors (in alphabetical order): Julio Elias Alvarado-Yaah, Carlos F. Arias, Santiago Ávila-Ríos, Eduardo Becerril-Vargas, Víctor Hugo Borja-Aburto, Celia Boukadida, Cristóbal Cháidez-Quiróz, Juan Bautista Chale-Dzul, Ricardo Ciria Merce, Andreu Comas-García, Célida Duque Molina, Julissa Enciso-Ibarra, José Antonio Enciso-Moreno, Gloria Elena Espinosa-Ayala, Fernando Fontove-Herrera, Daniel Fregoso-Rueda, Victor Eduardo García-Arias, Alejandra García-Gasca, Bruno Gómez-Gil, Jean Pierre González, Irvin González-López, Concepción Grajales-Muñiz, Ricardo Grande, Rosa María Gutiérrez Rios, Jesús Hernández, Alejandra Hernández-Terán, Alfredo Herrera-Estrella, Carla Ivón Herrera-Najera, Pavel Isa, Verónica Mata-Haro, Daniel Lira Morales, Susana Lopez, Antonio Loza Román, Brenda Irasema Maldonado-Meza, Bernardo Martínez-Miguel, José Arturo Martínez-Orozco, Célida Martínez- Rodríguez, Margarita Matías-Florentino, Fidencio Mejía-Nepomuceno, María Guadalupe de Jesús Mireles-Rivera, Gloria María Molina-Salinas, Hector Montoya-Fuentes, , Mario Mújica-Sánchez, José Esteban Muñoz-Medina, José de Jesús Nuñez-Contreras, Alicia Ocaña-Mondragón, Luis Alberto Ochoa-Carrera, Hector Esteban Paz-Juárez, Marissa Perez-Garcia, Francisco Pulido, Helen Haydee Fernanda Ramírez-Plascencia, Jorge Salas-Hernández, Angel Gustavo Salas-Lais, Alejandro Sánchez-Flores, Clara Esperanza Santacruz-Tinoco, María Guadalupe Santiago-Mauricio, Selene Zárate, Nelly Sélem-Mojica, Blanca Taboada, Gloria Vazquez, Joel Armando Vázquez-Pérez. |
| EPI_ISL_2681088, EPI_ISL_2681089, EPI_ISL_2681090, EPI_ISL_2681091 | Unidad de Investigación Médica de Yucatán (UIMY) | Instituto de Biotecnología de la UNAM | Consorcio Mexicano de Vigilancia Genómica (CoViGen-Mex). Authors (in alphabetical order): Julio Elias Alvarado-Yaah, Carlos F. Arias, Santiago Ávila-Ríos, Eduardo Becerril-Vargas, Víctor Hugo Borja-Aburto, Celia Boukadida, Cristóbal Cháidez-Quiróz, Juan Bautista Chale-Dzul, Ricardo Ciria Merce, Andreu Comas-García, Célida Duque Molina, Julissa Enciso-Ibarra, José Antonio Enciso-Moreno, Gloria Elena Espinosa-Ayala, Fernando Fontove-Herrera, Daniel Fregoso-Rueda, Victor Eduardo García-Arias, Alejandra García-Gasca, Bruno Gómez-Gil, Jean Pierre González, Irvin González-López, Concepción Grajales-Muñiz, Ricardo Grande, Rosa María Gutiérrez Rios, Jesús Hernández, Alejandra Hernández-Terán, Alfredo Herrera-Estrella, Carla Ivón Herrera-Najera, Pavel Isa, Verónica Mata-Haro, Daniel Lira Morales, Susana Lopez, Antonio Loza Román, Brenda Irasema Maldonado-Meza, Bernardo Martínez-Miguel, José Arturo Martínez-Orozco, Célida Martínez- Rodríguez, Margarita Matías-Florentino, Fidencio Mejía-Nepomuceno, María Guadalupe de Jesús Mireles-Rivera, Gloria María Molina-Salinas, Hector Montoya-Fuentes, , Mario Mújica-Sánchez, José Esteban Muñoz-Medina, José de Jesús Nuñez-Contreras, Alicia Ocaña-Mondragón, Luis Alberto Ochoa-Carrera, Hector Esteban Paz-Juárez, Marissa Perez-Garcia, Francisco Pulido, Helen Haydee Fernanda Ramírez-Plascencia, Jorge Salas-Hernández, Angel Gustavo Salas-Lais, Alejandro Sánchez-Flores, Clara Esperanza Santacruz-Tinoco, María Guadalupe Santiago-Mauricio, Selene Zárate, Nelly Sélem-Mojica, Blanca Taboada, Gloria Vazquez, Joel Armando Vázquez-Pérez. |
| EPI_ISL_2681092 | Unidad de Investigación Biomédica de Zacatecas (UIBZ) | Instituto de Biotecnología de la UNAM | Consorcio Mexicano de Vigilancia Genómica (CoViGen-Mex). Authors (in alphabetical order): Julio Elias Alvarado-Yaah, Carlos F. Arias, Santiago Ávila-Ríos, Eduardo Becerril-Vargas, Víctor Hugo Borja-Aburto, Celia Boukadida, Cristóbal Cháidez-Quiróz, Juan Bautista Chale-Dzul, Ricardo Ciria Merce, Andreu Comas-García, Célida Duque Molina, Julissa Enciso-Ibarra, José Antonio Enciso-Moreno, Gloria Elena Espinosa-Ayala, Fernando Fontove-Herrera, Daniel Fregoso-Rueda, Victor Eduardo García-Arias, Alejandra García-Gasca, Bruno Gómez-Gil, Jean Pierre González, Irvin |

| | | | |
|---|---|---|---|
| | | | González-López, Concepción Grajales-Muñiz, Ricardo Grande, Rosa María Gutiérrez Rios, Jesús Hernández, Alejandra Hernández-Terán, Alfredo Herrera-Estrella, Carla Ivón Herrera-Najera, Pavel Isa, Verónica Mata-Haro, Daniel Lira Morales, Susana Lopez, Antonio Loza Román, Brenda Irasema Maldonado-Meza, Bernardo Martínez-Miguel, José Arturo Martínez-Orozco, Célida Martínez- Rodríguez, Margarita Matías-Florentino, Fidencio Mejía-Nepomuceno, María Guadalupe de Jesús Mireles-Rivera, Gloria María Molina-Salinas, Hector Montoya-Fuentes, , Mario Mújica-Sánchez, José Esteban Muñoz-Medina, José de Jesús Nuñez-Contreras, Alicia Ocaña-Mondragón, Luis Alberto Ochoa-Carrera, Hector Esteban Paz-Juárez, Marissa Perez-Garcia, Francisco Pulido, Helen Haydee Fernanda Ramírez-Plascencia, Jorge Salas-Hernández, Angel Gustavo Salas-Lais, Alejandro Sánchez-Flores, Clara Esperanza Santacruz-Tinoco, María Guadalupe Santiago-Mauricio, Selene Zárate, Nelly Sélem-Mojica, Blanca Taboada, Gloria Vazquez, Joel Armando Vázquez-Pérez. |
| EPI_ISL_2681093 | Laboratorio Central de Epidemiología (LCE) | Instituto de Biotecnología de la UNAM | Consorcio Mexicano de Vigilancia Genómica (CoViGen-Mex). Authors (in alphabetical order): Julio Elias Alvarado-Yaah, Carlos F. Arias, Santiago Ávila-Ríos, Eduardo Becerril-Vargas, Víctor Hugo Borja-Aburto, Celia Boukadida, Cristóbal Cháidez-Quiróz, Juan Bautista Chale-Dzul, Ricardo Ciria Merce, Andreu Comas-García, Célida Duque Molina, Julissa Enciso-Ibarra, José Antonio Enciso-Moreno, Gloria Elena Espinosa-Ayala, Fernando Fontove-Herrera, Daniel Fregoso-Rueda, Victor Eduardo García-Arias, Alejandra García-Gasca, Bruno Gómez-Gil, Jean Pierre González, Irvin González-López, Concepción Grajales-Muñiz, Ricardo Grande, Rosa María Gutiérrez Rios, Jesús Hernández, Alejandra Hernández-Terán, Alfredo Herrera-Estrella, Carla Ivón Herrera-Najera, Pavel Isa, Verónica Mata-Haro, Daniel Lira Morales, Susana Lopez, Antonio Loza Román, Brenda Irasema Maldonado-Meza, Bernardo Martínez-Miguel, José Arturo Martínez-Orozco, Célida Martínez- Rodríguez, Margarita Matías-Florentino, Fidencio Mejía-Nepomuceno, María Guadalupe de Jesús Mireles-Rivera, Gloria María Molina-Salinas, Hector Montoya-Fuentes, , Mario Mújica-Sánchez, José Esteban Muñoz-Medina, José de Jesús Nuñez-Contreras, Alicia Ocaña-Mondragón, Luis Alberto Ochoa-Carrera, Hector Esteban Paz-Juárez, Marissa Perez-Garcia, Francisco Pulido, Helen Haydee Fernanda Ramírez-Plascencia, Jorge Salas-Hernández, Angel Gustavo Salas-Lais, Alejandro Sánchez-Flores, Clara Esperanza Santacruz-Tinoco, María Guadalupe Santiago-Mauricio, Selene Zárate, Nelly Sélem-Mojica, Blanca Taboada, Gloria Vazquez, Joel Armando Vázquez-Pérez. |
| EPI_ISL_2681094 | Unidad de Investigación Médica de Yucatán (UIMY) | Instituto de Biotecnología de la UNAM | Consorcio Mexicano de Vigilancia Genómica (CoViGen-Mex). Authors (in alphabetical order): Julio Elias Alvarado-Yaah, Carlos F. Arias, Santiago Ávila-Ríos, Eduardo Becerril-Vargas, Víctor Hugo Borja-Aburto, Celia Boukadida, Cristóbal Cháidez-Quiróz, Juan Bautista Chale-Dzul, Ricardo Ciria Merce, Andreu Comas-García, Célida Duque Molina, Julissa Enciso-Ibarra, José Antonio Enciso-Moreno, Gloria Elena Espinosa-Ayala, Fernando Fontove-Herrera, Daniel Fregoso-Rueda, Victor Eduardo García-Arias, Alejandra García-Gasca, Bruno Gómez-Gil, Jean Pierre González, Irvin González-López, Concepción Grajales-Muñiz, Ricardo Grande, Rosa María Gutiérrez Rios, Jesús Hernández, Alejandra Hernández-Terán, Alfredo Herrera-Estrella, Carla Ivón Herrera-Najera, Pavel Isa, Verónica Mata-Haro, Daniel Lira Morales, Susana Lopez, Antonio Loza Román, Brenda Irasema Maldonado-Meza, Bernardo Martínez-Miguel, José Arturo Martínez-Orozco, Célida Martínez- Rodríguez, Margarita Matías-Florentino, Fidencio Mejía-Nepomuceno, María Guadalupe de Jesús Mireles-Rivera, Gloria María Molina-Salinas, Hector Montoya-Fuentes, , Mario Mújica-Sánchez, José Esteban Muñoz-Medina, José de Jesús Nuñez-Contreras, Alicia Ocaña-Mondragón, Luis Alberto Ochoa-Carrera, Hector Esteban Paz-Juárez, Marissa Perez-Garcia, Francisco Pulido, Helen Haydee Fernanda Ramírez-Plascencia, Jorge Salas-Hernández, Angel Gustavo Salas-Lais, Alejandro Sánchez-Flores, Clara Esperanza Santacruz-Tinoco, María Guadalupe Santiago-Mauricio, Selene Zárate, Nelly Sélem-Mojica, Blanca Taboada, Gloria Vazquez, Joel Armando Vázquez-Pérez. |
| EPI_ISL_2681095, EPI_ISL_2681096 | Centro de Investigación Biomédica del Noreste (CIBIN) | Instituto de Biotecnología de la UNAM | Consorcio Mexicano de Vigilancia Genómica (CoViGen-Mex). Authors (in alphabetical order): Julio Elias Alvarado-Yaah, Carlos F. Arias, Santiago Ávila-Ríos, Eduardo Becerril-Vargas, Víctor Hugo Borja-Aburto, Celia Boukadida, Cristóbal Cháidez-Quiróz, Juan Bautista Chale-Dzul, Ricardo Ciria Merce, Andreu Comas-García, Célida Duque Molina, Julissa Enciso-Ibarra, José Antonio Enciso-Moreno, Gloria Elena Espinosa-Ayala, Fernando Fontove-Herrera, Daniel Fregoso-Rueda, Victor Eduardo García-Arias, Alejandra García-Gasca, Bruno Gómez-Gil, Jean Pierre González, Irvin González-López, Concepción Grajales-Muñiz, Ricardo Grande, Rosa María Gutiérrez Rios, Jesús Hernández, Alejandra Hernández-Terán, Alfredo Herrera-Estrella, Carla Ivón Herrera-Najera, Pavel Isa, Verónica Mata-Haro, Daniel Lira Morales, Susana Lopez, Antonio Loza Román, Brenda Irasema Maldonado-Meza, Bernardo Martínez-Miguel, José Arturo Martínez-Orozco, Célida Martínez- Rodríguez, Margarita Matías-Florentino, Fidencio Mejía-Nepomuceno, María Guadalupe de Jesús Mireles-Rivera, Gloria María Molina-Salinas, Hector Montoya-Fuentes, , Mario Mújica-Sánchez, José Esteban Muñoz-Medina, José de Jesús Nuñez-Contreras, Alicia Ocaña-Mondragón, Luis Alberto Ochoa-Carrera, Hector Esteban Paz-Juárez, Marissa Perez-Garcia, Francisco Pulido, Helen Haydee Fernanda Ramírez-Plascencia, Jorge Salas-Hernández, Angel Gustavo Salas-Lais, Alejandro Sánchez-Flores, Clara Esperanza Santacruz-Tinoco, María Guadalupe Santiago-Mauricio, Selene Zárate, Nelly Sélem-Mojica, Blanca Taboada, Gloria Vazquez, Joel Armando Vázquez-Pérez. |
| EPI_ISL_2681097 | Unidad de Investigación Biomédica de Zacatecas (UIBZ) | Instituto de Biotecnología de la UNAM | Consorcio Mexicano de Vigilancia Genómica (CoViGen-Mex). Authors (in alphabetical order): Julio Elias Alvarado-Yaah, Carlos F. Arias, Santiago Ávila-Ríos, Eduardo Becerril-Vargas, Víctor Hugo Borja-Aburto, Celia Boukadida, Cristóbal Cháidez-Quiróz, Juan Bautista Chale-Dzul, Ricardo Ciria Merce, Andreu Comas-García, Célida Duque Molina, Julissa Enciso-Ibarra, José Antonio Enciso-Moreno, Gloria Elena Espinosa-Ayala, Fernando Fontove-Herrera, Daniel Fregoso-Rueda, Victor Eduardo García-Arias, Alejandra García-Gasca, Bruno Gómez-Gil, Jean Pierre González, Irvin González-López, Concepción Grajales-Muñiz, Ricardo Grande, Rosa María Gutiérrez Rios, Jesús Hernández, Alejandra Hernández-Terán, Alfredo Herrera-Estrella, Carla Ivón Herrera-Najera, Pavel Isa, Verónica Mata-Haro, Daniel Lira Morales, Susana Lopez, Antonio Loza Román, Brenda Irasema Maldonado-Meza, Bernardo Martínez-Miguel, José Arturo Martínez-Orozco, Célida Martínez- Rodríguez, Margarita Matías-Florentino, Fidencio Mejía-Nepomuceno, María Guadalupe de Jesús Mireles-Rivera, Gloria María Molina-Salinas, Hector Montoya-Fuentes, , Mario Mújica-Sánchez, José Esteban Muñoz-Medina, José de Jesús Nuñez-Contreras, Alicia Ocaña-Mondragón, Luis Alberto Ochoa-Carrera, Hector Esteban Paz-Juárez, Marissa Perez-Garcia, Francisco Pulido, Helen Haydee Fernanda Ramírez-Plascencia, Jorge Salas-Hernández, Angel Gustavo Salas-Lais, Alejandro Sánchez-Flores, Clara Esperanza Santacruz-Tinoco, María Guadalupe Santiago-Mauricio, Selene Zárate, Nelly Sélem-Mojica, Blanca Taboada, Gloria Vazquez, Joel Armando Vázquez-Pérez. |
| EPI_ISL_2681098, EPI_ISL_2681099, EPI_ISL_2681100, EPI_ISL_2681101 | Unidad de Investigación Médica de Yucatán (UIMY) | Instituto de Biotecnología de la UNAM | Consorcio Mexicano de Vigilancia Genómica (CoViGen-Mex). Authors (in alphabetical order): Julio Elias Alvarado-Yaah, Carlos F. Arias, Santiago Ávila-Ríos, Eduardo Becerril-Vargas, Víctor Hugo Borja-Aburto, Celia Boukadida, Cristóbal Cháidez-Quiróz, Juan Bautista Chale-Dzul, Ricardo Ciria Merce, Andreu Comas-García, Célida Duque Molina, Julissa Enciso-Ibarra, José Antonio Enciso-Moreno, Gloria Elena Espinosa-Ayala, Fernando Fontove-Herrera, Daniel Fregoso-Rueda, Victor Eduardo García-Arias, Alejandra García-Gasca, Bruno Gómez-Gil, Jean Pierre González, Irvin González-López, Concepción Grajales-Muñiz, Ricardo Grande, Rosa María Gutiérrez Rios, Jesús Hernández, Alejandra Hernández-Terán, Alfredo Herrera-Estrella, Carla Ivón Herrera-Najera, Pavel Isa, Verónica Mata-Haro, Daniel Lira Morales, Susana Lopez, Antonio Loza Román, Brenda Irasema Maldonado-Meza, Bernardo Martínez-Miguel, José Arturo Martínez-Orozco, Célida Martínez- Rodríguez, Margarita Matías-Florentino, Fidencio Mejía-Nepomuceno, María Guadalupe de Jesús Mireles-Rivera, Gloria María Molina-Salinas, Hector Montoya-Fuentes, , Mario Mújica-Sánchez, José Esteban Muñoz-Medina, José de Jesús Nuñez-Contreras, Alicia Ocaña-Mondragón, Luis Alberto Ochoa-Carrera, Hector Esteban Paz-Juárez, Marissa Perez-Garcia, Francisco Pulido, Helen Haydee Fernanda Ramírez-Plascencia, Jorge Salas-Hernández, Angel Gustavo Salas-Lais, Alejandro Sánchez-Flores, Clara Esperanza Santacruz-Tinoco, María Guadalupe Santiago-Mauricio, Selene Zárate, Nelly Sélem-Mojica, Blanca Taboada, Gloria Vazquez, Joel Armando Vázquez-Pérez. |
| EPI_ISL_2681102, EPI_ISL_2681103, EPI_ISL_2681104 | Centro de Investigación Biomédica del Noreste (CIBIN) | Instituto de Biotecnología de la UNAM | Consorcio Mexicano de Vigilancia Genómica (CoViGen-Mex). Authors (in alphabetical order): Julio Elias Alvarado-Yaah, Carlos F. Arias, Santiago Ávila-Ríos, Eduardo Becerril-Vargas, Víctor Hugo Borja-Aburto, Celia Boukadida, Cristóbal Cháidez-Quiróz, Juan Bautista Chale-Dzul, Ricardo Ciria Merce, Andreu Comas-García, Célida Duque Molina, Julissa Enciso-Ibarra, José Antonio Enciso-Moreno, Gloria Elena Espinosa-Ayala, Fernando Fontove-Herrera, Daniel Fregoso-Rueda, Victor Eduardo García-Arias, Alejandra García-Gasca, Bruno Gómez-Gil, Jean Pierre González, Irvin González-López, Concepción Grajales-Muñiz, Ricardo Grande, Rosa María Gutiérrez Rios, Jesús Hernández, Alejandra Hernández-Terán, Alfredo Herrera-Estrella, Carla Ivón Herrera-Najera, Pavel Isa, Verónica Mata-Haro, Daniel Lira Morales, Susana Lopez, Antonio Loza Román, Brenda Irasema Maldonado-Meza, Bernardo Martínez-Miguel, José Arturo Martínez-Orozco, Célida Martínez- Rodríguez, Margarita Matías-Florentino, Fidencio Mejía-Nepomuceno, María Guadalupe de Jesús Mireles-Rivera, Gloria María Molina-Salinas, Hector Montoya-Fuentes, , Mario Mújica-Sánchez, José Esteban Muñoz-Medina, José de Jesús Nuñez-Contreras, Alicia Ocaña-Mondragón, Luis Alberto Ochoa-Carrera, Hector Esteban Paz-Juárez, Marissa Perez-Garcia, Francisco Pulido, Helen Haydee Fernanda Ramírez-Plascencia, Jorge Salas-Hernández, Angel Gustavo Salas-Lais, Alejandro Sánchez-Flores, Clara Esperanza Santacruz-Tinoco, María Guadalupe Santiago-Mauricio, Selene Zárate, Nelly Sélem-Mojica, Blanca Taboada, Gloria Vazquez, Joel Armando Vázquez-Pérez. |
| EPI_ISL_2681105 | Laboratorio Central de Epidemiología (LCE) | Instituto de Biotecnología de la UNAM | Consorcio Mexicano de Vigilancia Genómica (CoViGen-Mex). Authors (in alphabetical order): Julio Elias Alvarado-Yaah, Carlos F. Arias, Santiago Ávila-Ríos, Eduardo Becerril-Vargas, Víctor Hugo Borja-Aburto, Celia Boukadida, Cristóbal Cháidez-Quiróz, Juan Bautista Chale-Dzul, Ricardo Ciria Merce, Andreu Comas-García, Célida Duque Molina, Julissa Enciso-Ibarra, José Antonio Enciso-Moreno, Gloria Elena Espinosa-Ayala, Fernando Fontove-Herrera, Daniel Fregoso-Rueda, Victor Eduardo García-Arias, Alejandra García-Gasca, Bruno Gómez-Gil, Jean Pierre González, Irvin |

| | | | |
|---|---|---|---|
| | | | González-López, Concepción Grajales-Muñiz, Ricardo Grande, Rosa María Gutiérrez Rios, Jesús Hernández, Alejandra Hernández-Terán, Alfredo Herrera-Estrella, Carla Ivón Herrera-Najera, Pavel Isa, Verónica Mata-Haro, Daniel Lira Morales, Susana Lopez, Antonio Loza Román, Brenda Irasema Maldonado-Meza, Bernardo Martínez-Miguel, José Arturo Martínez-Orozco, Célida Martínez- Rodríguez, Margarita Matías-Florentino, Fidencio Mejía-Nepomuceno, María Guadalupe de Jesús Mireles-Rivera, Gloria María Molina-Salinas, Hector Montoya-Fuentes, , Mario Mújica-Sánchez, José Esteban Muñoz-Medina, José de Jesús Nuñez-Contreras, Alicia Ocaña-Mondragón, Luis Alberto Ochoa-Carrera, Hector Esteban Paz-Juárez, Marissa Perez-Garcia, Francisco Pulido, Helen Haydee Fernanda Ramírez-Plascencia, Jorge Salas-Hernández, Angel Gustavo Salas-Lais, Alejandro Sánchez-Flores, Clara Esperanza Santacruz-Tinoco, María Guadalupe Santiago-Mauricio, Selene Zárate, Nelly Sélem-Mojica, Blanca Taboada, Gloria Vazquez, Joel Armando Vázquez-Pérez. |
| EPI_ISL_2681106, EPI_ISL_2681107, EPI_ISL_2681108, EPI_ISL_2681109 | Unidad de Investigación Médica de Yucatán (UIMY) | Instituto de Biotecnología de la UNAM | Consorcio Mexicano de Vigilancia Genómica (CoViGen-Mex). Authors (in alphabetical order): Julio Elias Alvarado-Yaah, Carlos F. Arias, Santiago Ávila-Ríos, Eduardo Becerril-Vargas, Víctor Hugo Borja-Aburto, Celia Boukadida, Cristóbal Cháidez-Quiróz, Juan Bautista Chale-Dzul, Ricardo Ciria Merce, Andreu Comas-García, Célida Duque Molina, Julissa Enciso-Ibarra, José Antonio Enciso-Moreno, Gloria Elena Espinosa-Ayala, Fernando Fontove-Herrera, Daniel Fregoso-Rueda, Victor Eduardo García-Arias, Alejandra García-Gasca, Bruno Gómez-Gil, Jean Pierre González, Irvin González-López, Concepción Grajales-Muñiz, Ricardo Grande, Rosa María Gutiérrez Rios, Jesús Hernández, Alejandra Hernández-Terán, Alfredo Herrera-Estrella, Carla Ivón Herrera-Najera, Pavel Isa, Verónica Mata-Haro, Daniel Lira Morales, Susana Lopez, Antonio Loza Román, Brenda Irasema Maldonado-Meza, Bernardo Martínez-Miguel, José Arturo Martínez-Orozco, Célida Martínez- Rodríguez, Margarita Matías-Florentino, Fidencio Mejía-Nepomuceno, María Guadalupe de Jesús Mireles-Rivera, Gloria María Molina-Salinas, Hector Montoya-Fuentes, , Mario Mújica-Sánchez, José Esteban Muñoz-Medina, José de Jesús Nuñez-Contreras, Alicia Ocaña-Mondragón, Luis Alberto Ochoa-Carrera, Hector Esteban Paz-Juárez, Marissa Perez-Garcia, Francisco Pulido, Helen Haydee Fernanda Ramírez-Plascencia, Jorge Salas-Hernández, Angel Gustavo Salas-Lais, Alejandro Sánchez-Flores, Clara Esperanza Santacruz-Tinoco, María Guadalupe Santiago-Mauricio, Selene Zárate, Nelly Sélem-Mojica, Blanca Taboada, Gloria Vazquez, Joel Armando Vázquez-Pérez. |
| EPI_ISL_2681110 | Centro de Investigación Biomédica del Noreste (CIBIN) | Instituto de Biotecnología de la UNAM | Consorcio Mexicano de Vigilancia Genómica (CoViGen-Mex). Authors (in alphabetical order): Julio Elias Alvarado-Yaah, Carlos F. Arias, Santiago Ávila-Ríos, Eduardo Becerril-Vargas, Víctor Hugo Borja-Aburto, Celia Boukadida, Cristóbal Cháidez-Quiróz, Juan Bautista Chale-Dzul, Ricardo Ciria Merce, Andreu Comas-García, Célida Duque Molina, Julissa Enciso-Ibarra, José Antonio Enciso-Moreno, Gloria Elena Espinosa-Ayala, Fernando Fontove-Herrera, Daniel Fregoso-Rueda, Victor Eduardo García-Arias, Alejandra García-Gasca, Bruno Gómez-Gil, Jean Pierre González, Irvin González-López, Concepción Grajales-Muñiz, Ricardo Grande, Rosa María Gutiérrez Rios, Jesús Hernández, Alejandra Hernández-Terán, Alfredo Herrera-Estrella, Carla Ivón Herrera-Najera, Pavel Isa, Verónica Mata-Haro, Daniel Lira Morales, Susana Lopez, Antonio Loza Román, Brenda Irasema Maldonado-Meza, Bernardo Martínez-Miguel, José Arturo Martínez-Orozco, Célida Martínez- Rodríguez, Margarita Matías-Florentino, Fidencio Mejía-Nepomuceno, María Guadalupe de Jesús Mireles-Rivera, Gloria María Molina-Salinas, Hector Montoya-Fuentes, , Mario Mújica-Sánchez, José Esteban Muñoz-Medina, José de Jesús Nuñez-Contreras, Alicia Ocaña-Mondragón, Luis Alberto Ochoa-Carrera, Hector Esteban Paz-Juárez, Marissa Perez-Garcia, Francisco Pulido, Helen Haydee Fernanda Ramírez-Plascencia, Jorge Salas-Hernández, Angel Gustavo Salas-Lais, Alejandro Sánchez-Flores, Clara Esperanza Santacruz-Tinoco, María Guadalupe Santiago-Mauricio, Selene Zárate, Nelly Sélem-Mojica, Blanca Taboada, Gloria Vazquez, Joel Armando Vázquez-Pérez. |
| EPI_ISL_2681111, EPI_ISL_2681112, EPI_ISL_2681113 | Laboratorio Central de Epidemiología (LCE) | Instituto de Biotecnología de la UNAM | Consorcio Mexicano de Vigilancia Genómica (CoViGen-Mex). Authors (in alphabetical order): Julio Elias Alvarado-Yaah, Carlos F. Arias, Santiago Ávila-Ríos, Eduardo Becerril-Vargas, Víctor Hugo Borja-Aburto, Celia Boukadida, Cristóbal Cháidez-Quiróz, Juan Bautista Chale-Dzul, Ricardo Ciria Merce, Andreu Comas-García, Célida Duque Molina, Julissa Enciso-Ibarra, José Antonio Enciso-Moreno, Gloria Elena Espinosa-Ayala, Fernando Fontove-Herrera, Daniel Fregoso-Rueda, Victor Eduardo García-Arias, Alejandra García-Gasca, Bruno Gómez-Gil, Jean Pierre González, Irvin González-López, Concepción Grajales-Muñiz, Ricardo Grande, Rosa María Gutiérrez Rios, Jesús Hernández, Alejandra Hernández-Terán, Alfredo Herrera-Estrella, Carla Ivón Herrera-Najera, Pavel Isa, Verónica Mata-Haro, Daniel Lira Morales, Susana Lopez, Antonio Loza Román, Brenda Irasema Maldonado-Meza, Bernardo Martínez-Miguel, José Arturo Martínez-Orozco, Célida Martínez- Rodríguez, Margarita Matías-Florentino, Fidencio Mejía-Nepomuceno, María Guadalupe de Jesús Mireles-Rivera, Gloria María Molina-Salinas, Hector Montoya-Fuentes, , Mario Mújica-Sánchez, José Esteban Muñoz-Medina, José de Jesús Nuñez-Contreras, Alicia Ocaña-Mondragón, Luis Alberto Ochoa-Carrera, Hector Esteban Paz-Juárez, Marissa Perez-Garcia, Francisco Pulido, Helen Haydee Fernanda Ramírez-Plascencia, Jorge Salas-Hernández, Angel Gustavo Salas-Lais, Alejandro Sánchez-Flores, Clara Esperanza Santacruz-Tinoco, María Guadalupe Santiago-Mauricio, Selene Zárate, Nelly Sélem-Mojica, Blanca Taboada, Gloria Vazquez, Joel Armando Vázquez-Pérez. |
| EPI_ISL_2681114, EPI_ISL_2681115, EPI_ISL_2681116, EPI_ISL_2681117, EPI_ISL_2681118 | Unidad de Investigación Médica de Yucatán (UIMY) | Instituto de Biotecnología de la UNAM | Consorcio Mexicano de Vigilancia Genómica (CoViGen-Mex). Authors (in alphabetical order): Julio Elias Alvarado-Yaah, Carlos F. Arias, Santiago Ávila-Ríos, Eduardo Becerril-Vargas, Víctor Hugo Borja-Aburto, Celia Boukadida, Cristóbal Cháidez-Quiróz, Juan Bautista Chale-Dzul, Ricardo Ciria Merce, Andreu Comas-García, Célida Duque Molina, Julissa Enciso-Ibarra, José Antonio Enciso-Moreno, Gloria Elena Espinosa-Ayala, Fernando Fontove-Herrera, Daniel Fregoso-Rueda, Victor Eduardo García-Arias, Alejandra García-Gasca, Bruno Gómez-Gil, Jean Pierre González, Irvin González-López, Concepción Grajales-Muñiz, Ricardo Grande, Rosa María Gutiérrez Rios, Jesús Hernández, Alejandra Hernández-Terán, Alfredo Herrera-Estrella, Carla Ivón Herrera-Najera, Pavel Isa, Verónica Mata-Haro, Daniel Lira Morales, Susana Lopez, Antonio Loza Román, Brenda Irasema Maldonado-Meza, Bernardo Martínez-Miguel, José Arturo Martínez-Orozco, Célida Martínez- Rodríguez, Margarita Matías-Florentino, Fidencio Mejía-Nepomuceno, María Guadalupe de Jesús Mireles-Rivera, Gloria María Molina-Salinas, Hector Montoya-Fuentes, , Mario Mújica-Sánchez, José Esteban Muñoz-Medina, José de Jesús Nuñez-Contreras, Alicia Ocaña-Mondragón, Luis Alberto Ochoa-Carrera, Hector Esteban Paz-Juárez, Marissa Perez-Garcia, Francisco Pulido, Helen Haydee Fernanda Ramírez-Plascencia, Jorge Salas-Hernández, Angel Gustavo Salas-Lais, Alejandro Sánchez-Flores, Clara Esperanza Santacruz-Tinoco, María Guadalupe Santiago-Mauricio, Selene Zárate, Nelly Sélem-Mojica, Blanca Taboada, Gloria Vazquez, Joel Armando Vázquez-Pérez. |
| EPI_ISL_2681119, EPI_ISL_2681120, EPI_ISL_2681121, EPI_ISL_2681122, EPI_ISL_2681123 | Unidad de Investigación Biomédica de Zacatecas (UIBZ) | Instituto de Biotecnología de la UNAM | Consorcio Mexicano de Vigilancia Genómica (CoViGen-Mex). Authors (in alphabetical order): Julio Elias Alvarado-Yaah, Carlos F. Arias, Santiago Ávila-Ríos, Eduardo Becerril-Vargas, Víctor Hugo Borja-Aburto, Celia Boukadida, Cristóbal Cháidez-Quiróz, Juan Bautista Chale-Dzul, Ricardo Ciria Merce, Andreu Comas-García, Célida Duque Molina, Julissa Enciso-Ibarra, José Antonio Enciso-Moreno, Gloria Elena Espinosa-Ayala, Fernando Fontove-Herrera, Daniel Fregoso-Rueda, Victor Eduardo García-Arias, Alejandra García-Gasca, Bruno Gómez-Gil, Jean Pierre González, Irvin González-López, Concepción Grajales-Muñiz, Ricardo Grande, Rosa María Gutiérrez Rios, Jesús Hernández, Alejandra Hernández-Terán, Alfredo Herrera-Estrella, Carla Ivón Herrera-Najera, Pavel Isa, Verónica Mata-Haro, Daniel Lira Morales, Susana Lopez, Antonio Loza Román, Brenda Irasema Maldonado-Meza, Bernardo Martínez-Miguel, José Arturo Martínez-Orozco, Célida Martínez- Rodríguez, Margarita Matías-Florentino, Fidencio Mejía-Nepomuceno, María Guadalupe de Jesús Mireles-Rivera, Gloria María Molina-Salinas, Hector Montoya-Fuentes, , Mario Mújica-Sánchez, José Esteban Muñoz-Medina, José de Jesús Nuñez-Contreras, Alicia Ocaña-Mondragón, Luis Alberto Ochoa-Carrera, Hector Esteban Paz-Juárez, Marissa Perez-Garcia, Francisco Pulido, Helen Haydee Fernanda Ramírez-Plascencia, Jorge Salas-Hernández, Angel Gustavo Salas-Lais, Alejandro Sánchez-Flores, Clara Esperanza Santacruz-Tinoco, María Guadalupe Santiago-Mauricio, Selene Zárate, Nelly Sélem-Mojica, Blanca Taboada, Gloria Vazquez, Joel Armando Vázquez-Pérez. |
| EPI_ISL_2681124, EPI_ISL_2681125 | Laboratorio Central de Epidemiología (LCE) | Instituto de Biotecnología de la UNAM | Consorcio Mexicano de Vigilancia Genómica (CoViGen-Mex). Authors (in alphabetical order): Julio Elias Alvarado-Yaah, Carlos F. Arias, Santiago Ávila-Ríos, Eduardo Becerril-Vargas, Víctor Hugo Borja-Aburto, Celia Boukadida, Cristóbal Cháidez-Quiróz, Juan Bautista Chale-Dzul, Ricardo Ciria Merce, Andreu Comas-García, Célida Duque Molina, Julissa Enciso-Ibarra, José Antonio Enciso-Moreno, Gloria Elena Espinosa-Ayala, Fernando Fontove-Herrera, Daniel Fregoso-Rueda, Victor Eduardo García-Arias, Alejandra García-Gasca, Bruno Gómez-Gil, Jean Pierre González, Irvin González-López, Concepción Grajales-Muñiz, Ricardo Grande, Rosa María Gutiérrez Rios, Jesús Hernández, Alejandra Hernández-Terán, Alfredo Herrera-Estrella, Carla Ivón Herrera-Najera, Pavel Isa, Verónica Mata-Haro, Daniel Lira Morales, Susana Lopez, Antonio Loza Román, Brenda Irasema Maldonado-Meza, Bernardo Martínez-Miguel, José Arturo Martínez-Orozco, Célida Martínez- Rodríguez, Margarita Matías-Florentino, Fidencio Mejía-Nepomuceno, María Guadalupe de Jesús Mireles-Rivera, Gloria María Molina-Salinas, Hector Montoya-Fuentes, , Mario Mújica-Sánchez, José Esteban Muñoz-Medina, José de Jesús Nuñez-Contreras, Alicia Ocaña-Mondragón, Luis Alberto Ochoa-Carrera, Hector Esteban Paz-Juárez, Marissa Perez-Garcia, Francisco Pulido, Helen Haydee Fernanda Ramírez-Plascencia, Jorge Salas-Hernández, Angel Gustavo Salas-Lais, Alejandro Sánchez-Flores, Clara Esperanza Santacruz-Tinoco, María Guadalupe Santiago-Mauricio, Selene Zárate, Nelly Sélem-Mojica, Blanca Taboada, Gloria Vazquez, Joel Armando Vázquez-Pérez. |
| EPI_ISL_2681126 | Unidad de Investigación Médica de Yucatán (UIMY) | Instituto de Biotecnología de la UNAM | Consorcio Mexicano de Vigilancia Genómica (CoViGen-Mex). Authors (in alphabetical order): Julio Elias Alvarado-Yaah, Carlos F. Arias, Santiago Ávila-Ríos, Eduardo Becerril-Vargas, Víctor Hugo Borja-Aburto, Celia Boukadida, Cristóbal Cháidez-Quiróz, Juan Bautista Chale-Dzul, Ricardo Ciria Merce, Andreu Comas-García, Célida Duque Molina, Julissa Enciso-Ibarra, José Antonio Enciso-Moreno, Gloria Elena Espinosa-Ayala, Fernando Fontove-Herrera, Daniel Fregoso-Rueda, Victor Eduardo García-Arias, Alejandra García-Gasca, Bruno Gómez-Gil, Jean Pierre González, Irvin |

| | | | |
|---|---|---|---|
| | | | González-López, Concepción Grajales-Muñiz, Ricardo Grande, Rosa María Gutiérrez Rios, Jesús Hernández, Alejandra Hernández-Terán, Alfredo Herrera-Estrella, Carla Ivón Herrera-Najera, Pavel Isa, Verónica Mata-Haro, Daniel Lira Morales, Susana Lopez, Antonio Loza Román, Brenda Irasema Maldonado-Meza, Bernardo Martínez-Miguel, José Arturo Martínez-Orozco, Célida Martínez- Rodríguez, Margarita Matías-Florentino, Fidencio Mejía-Nepomuceno, María Guadalupe de Jesús Mireles-Rivera, Gloria María Molina-Salinas, Hector Montoya-Fuentes, , Mario Mújica-Sánchez, José Esteban Muñoz-Medina, José de Jesús Nuñez-Contreras, Alicia Ocaña-Mondragón, Luis Alberto Ochoa-Carrera, Hector Esteban Paz-Juárez, Marissa Perez-Garcia, Francisco Pulido, Helen Haydee Fernanda Ramírez-Plascencia, Jorge Salas-Hernández, Angel Gustavo Salas-Lais, Alejandro Sánchez-Flores, Clara Esperanza Santacruz-Tinoco, María Guadalupe Santiago-Mauricio, Selene Zárate, Nelly Sélem-Mojica, Blanca Taboada, Gloria Vazquez, Joel Armando Vázquez-Pérez. |
| EPI_ISL_2681127 | Centro de Investigación Biomédica del Noreste (CIBIN) | Instituto de Biotecnología de la UNAM | Consorcio Mexicano de Vigilancia Genómica (CoViGen-Mex). Authors (in alphabetical order): Julio Elias Alvarado-Yaah, Carlos F. Arias, Santiago Ávila-Ríos, Eduardo Becerril-Vargas, Víctor Hugo Borja-Aburto, Celia Boukadida, Cristóbal Cháidez-Quiróz, Juan Bautista Chale-Dzul, Ricardo Ciria Merce, Andreu Comas-García, Célida Duque Molina, Julissa Enciso-Ibarra, José Antonio Enciso-Moreno, Gloria Elena Espinosa-Ayala, Fernando Fontove-Herrera, Daniel Fregoso-Rueda, Victor Eduardo García-Arias, Alejandra García-Gasca, Bruno Gómez-Gil, Jean Pierre González, Irvin González-López, Concepción Grajales-Muñiz, Ricardo Grande, Rosa María Gutiérrez Rios, Jesús Hernández, Alejandra Hernández-Terán, Alfredo Herrera-Estrella, Carla Ivón Herrera-Najera, Pavel Isa, Verónica Mata-Haro, Daniel Lira Morales, Susana Lopez, Antonio Loza Román, Brenda Irasema Maldonado-Meza, Bernardo Martínez-Miguel, José Arturo Martínez-Orozco, Célida Martínez- Rodríguez, Margarita Matías-Florentino, Fidencio Mejía-Nepomuceno, María Guadalupe de Jesús Mireles-Rivera, Gloria María Molina-Salinas, Hector Montoya-Fuentes, , Mario Mújica-Sánchez, José Esteban Muñoz-Medina, José de Jesús Nuñez-Contreras, Alicia Ocaña-Mondragón, Luis Alberto Ochoa-Carrera, Hector Esteban Paz-Juárez, Marissa Perez-Garcia, Francisco Pulido, Helen Haydee Fernanda Ramírez-Plascencia, Jorge Salas-Hernández, Angel Gustavo Salas-Lais, Alejandro Sánchez-Flores, Clara Esperanza Santacruz-Tinoco, María Guadalupe Santiago-Mauricio, Selene Zárate, Nelly Sélem-Mojica, Blanca Taboada, Gloria Vazquez, Joel Armando Vázquez-Pérez. |
| EPI_ISL_2681128, EPI_ISL_2681129 | Laboratorio Central de Epidemiología (LCE) | Instituto de Biotecnología de la UNAM | Consorcio Mexicano de Vigilancia Genómica (CoViGen-Mex). Authors (in alphabetical order): Julio Elias Alvarado-Yaah, Carlos F. Arias, Santiago Ávila-Ríos, Eduardo Becerril-Vargas, Víctor Hugo Borja-Aburto, Celia Boukadida, Cristóbal Cháidez-Quiróz, Juan Bautista Chale-Dzul, Ricardo Ciria Merce, Andreu Comas-García, Célida Duque Molina, Julissa Enciso-Ibarra, José Antonio Enciso-Moreno, Gloria Elena Espinosa-Ayala, Fernando Fontove-Herrera, Daniel Fregoso-Rueda, Victor Eduardo García-Arias, Alejandra García-Gasca, Bruno Gómez-Gil, Jean Pierre González, Irvin González-López, Concepción Grajales-Muñiz, Ricardo Grande, Rosa María Gutiérrez Rios, Jesús Hernández, Alejandra Hernández-Terán, Alfredo Herrera-Estrella, Carla Ivón Herrera-Najera, Pavel Isa, Verónica Mata-Haro, Daniel Lira Morales, Susana Lopez, Antonio Loza Román, Brenda Irasema Maldonado-Meza, Bernardo Martínez-Miguel, José Arturo Martínez-Orozco, Célida Martínez- Rodríguez, Margarita Matías-Florentino, Fidencio Mejía-Nepomuceno, María Guadalupe de Jesús Mireles-Rivera, Gloria María Molina-Salinas, Hector Montoya-Fuentes, , Mario Mújica-Sánchez, José Esteban Muñoz-Medina, José de Jesús Nuñez-Contreras, Alicia Ocaña-Mondragón, Luis Alberto Ochoa-Carrera, Hector Esteban Paz-Juárez, Marissa Perez-Garcia, Francisco Pulido, Helen Haydee Fernanda Ramírez-Plascencia, Jorge Salas-Hernández, Angel Gustavo Salas-Lais, Alejandro Sánchez-Flores, Clara Esperanza Santacruz-Tinoco, María Guadalupe Santiago-Mauricio, Selene Zárate, Nelly Sélem-Mojica, Blanca Taboada, Gloria Vazquez, Joel Armando Vázquez-Pérez. |
| EPI_ISL_2681130 | Unidad de Investigación Médica de Yucatán (UIMY) | Instituto de Biotecnología de la UNAM | Consorcio Mexicano de Vigilancia Genómica (CoViGen-Mex). Authors (in alphabetical order): Julio Elias Alvarado-Yaah, Carlos F. Arias, Santiago Ávila-Ríos, Eduardo Becerril-Vargas, Víctor Hugo Borja-Aburto, Celia Boukadida, Cristóbal Cháidez-Quiróz, Juan Bautista Chale-Dzul, Ricardo Ciria Merce, Andreu Comas-García, Célida Duque Molina, Julissa Enciso-Ibarra, José Antonio Enciso-Moreno, Gloria Elena Espinosa-Ayala, Fernando Fontove-Herrera, Daniel Fregoso-Rueda, Victor Eduardo García-Arias, Alejandra García-Gasca, Bruno Gómez-Gil, Jean Pierre González, Irvin González-López, Concepción Grajales-Muñiz, Ricardo Grande, Rosa María Gutiérrez Rios, Jesús Hernández, Alejandra Hernández-Terán, Alfredo Herrera-Estrella, Carla Ivón Herrera-Najera, Pavel Isa, Verónica Mata-Haro, Daniel Lira Morales, Susana Lopez, Antonio Loza Román, Brenda Irasema Maldonado-Meza, Bernardo Martínez-Miguel, José Arturo Martínez-Orozco, Célida Martínez- Rodríguez, Margarita Matías-Florentino, Fidencio Mejía-Nepomuceno, María Guadalupe de Jesús Mireles-Rivera, Gloria María Molina-Salinas, Hector Montoya-Fuentes, , Mario Mújica-Sánchez, José Esteban Muñoz-Medina, José de Jesús Nuñez-Contreras, Alicia Ocaña-Mondragón, Luis Alberto Ochoa-Carrera, Hector Esteban Paz-Juárez, Marissa Perez-Garcia, Francisco Pulido, Helen Haydee Fernanda Ramírez-Plascencia, Jorge Salas-Hernández, Angel Gustavo Salas-Lais, Alejandro Sánchez-Flores, Clara Esperanza Santacruz-Tinoco, María Guadalupe Santiago-Mauricio, Selene Zárate, Nelly Sélem-Mojica, Blanca Taboada, Gloria Vazquez, Joel Armando Vázquez-Pérez. |
| EPI_ISL_2681131, EPI_ISL_2681132 | Laboratorio Central de Epidemiología (LCE) | Instituto de Biotecnología de la UNAM | Consorcio Mexicano de Vigilancia Genómica (CoViGen-Mex). Authors (in alphabetical order): Julio Elias Alvarado-Yaah, Carlos F. Arias, Santiago Ávila-Ríos, Eduardo Becerril-Vargas, Víctor Hugo Borja-Aburto, Celia Boukadida, Cristóbal Cháidez-Quiróz, Juan Bautista Chale-Dzul, Ricardo Ciria Merce, Andreu Comas-García, Célida Duque Molina, Julissa Enciso-Ibarra, José Antonio Enciso-Moreno, Gloria Elena Espinosa-Ayala, Fernando Fontove-Herrera, Daniel Fregoso-Rueda, Victor Eduardo García-Arias, Alejandra García-Gasca, Bruno Gómez-Gil, Jean Pierre González, Irvin González-López, Concepción Grajales-Muñiz, Ricardo Grande, Rosa María Gutiérrez Rios, Jesús Hernández, Alejandra Hernández-Terán, Alfredo Herrera-Estrella, Carla Ivón Herrera-Najera, Pavel Isa, Verónica Mata-Haro, Daniel Lira Morales, Susana Lopez, Antonio Loza Román, Brenda Irasema Maldonado-Meza, Bernardo Martínez-Miguel, José Arturo Martínez-Orozco, Célida Martínez- Rodríguez, Margarita Matías-Florentino, Fidencio Mejía-Nepomuceno, María Guadalupe de Jesús Mireles-Rivera, Gloria María Molina-Salinas, Hector Montoya-Fuentes, , Mario Mújica-Sánchez, José Esteban Muñoz-Medina, José de Jesús Nuñez-Contreras, Alicia Ocaña-Mondragón, Luis Alberto Ochoa-Carrera, Hector Esteban Paz-Juárez, Marissa Perez-Garcia, Francisco Pulido, Helen Haydee Fernanda Ramírez-Plascencia, Jorge Salas-Hernández, Angel Gustavo Salas-Lais, Alejandro Sánchez-Flores, Clara Esperanza Santacruz-Tinoco, María Guadalupe Santiago-Mauricio, Selene Zárate, Nelly Sélem-Mojica, Blanca Taboada, Gloria Vazquez, Joel Armando Vázquez-Pérez. |
| EPI_ISL_2681133 | Unidad de Investigación Médica de Yucatán (UIMY) | Instituto de Biotecnología de la UNAM | Consorcio Mexicano de Vigilancia Genómica (CoViGen-Mex). Authors (in alphabetical order): Julio Elias Alvarado-Yaah, Carlos F. Arias, Santiago Ávila-Ríos, Eduardo Becerril-Vargas, Víctor Hugo Borja-Aburto, Celia Boukadida, Cristóbal Cháidez-Quiróz, Juan Bautista Chale-Dzul, Ricardo Ciria Merce, Andreu Comas-García, Célida Duque Molina, Julissa Enciso-Ibarra, José Antonio Enciso-Moreno, Gloria Elena Espinosa-Ayala, Fernando Fontove-Herrera, Daniel Fregoso-Rueda, Victor Eduardo García-Arias, Alejandra García-Gasca, Bruno Gómez-Gil, Jean Pierre González, Irvin González-López, Concepción Grajales-Muñiz, Ricardo Grande, Rosa María Gutiérrez Rios, Jesús Hernández, Alejandra Hernández-Terán, Alfredo Herrera-Estrella, Carla Ivón Herrera-Najera, Pavel Isa, Verónica Mata-Haro, Daniel Lira Morales, Susana Lopez, Antonio Loza Román, Brenda Irasema Maldonado-Meza, Bernardo Martínez-Miguel, José Arturo Martínez-Orozco, Célida Martínez- Rodríguez, Margarita Matías-Florentino, Fidencio Mejía-Nepomuceno, María Guadalupe de Jesús Mireles-Rivera, Gloria María Molina-Salinas, Hector Montoya-Fuentes, , Mario Mújica-Sánchez, José Esteban Muñoz-Medina, José de Jesús Nuñez-Contreras, Alicia Ocaña-Mondragón, Luis Alberto Ochoa-Carrera, Hector Esteban Paz-Juárez, Marissa Perez-Garcia, Francisco Pulido, Helen Haydee Fernanda Ramírez-Plascencia, Jorge Salas-Hernández, Angel Gustavo Salas-Lais, Alejandro Sánchez-Flores, Clara Esperanza Santacruz-Tinoco, María Guadalupe Santiago-Mauricio, Selene Zárate, Nelly Sélem-Mojica, Blanca Taboada, Gloria Vazquez, Joel Armando Vázquez-Pérez. |
| EPI_ISL_2681134, EPI_ISL_2681135 | Laboratorio Central de Epidemiología (LCE) | Instituto de Biotecnología de la UNAM | Consorcio Mexicano de Vigilancia Genómica (CoViGen-Mex). Authors (in alphabetical order): Julio Elias Alvarado-Yaah, Carlos F. Arias, Santiago Ávila-Ríos, Eduardo Becerril-Vargas, Víctor Hugo Borja-Aburto, Celia Boukadida, Cristóbal Cháidez-Quiróz, Juan Bautista Chale-Dzul, Ricardo Ciria Merce, Andreu Comas-García, Célida Duque Molina, Julissa Enciso-Ibarra, José Antonio Enciso-Moreno, Gloria Elena Espinosa-Ayala, Fernando Fontove-Herrera, Daniel Fregoso-Rueda, Victor Eduardo García-Arias, Alejandra García-Gasca, Bruno Gómez-Gil, Jean Pierre González, Irvin González-López, Concepción Grajales-Muñiz, Ricardo Grande, Rosa María Gutiérrez Rios, Jesús Hernández, Alejandra Hernández-Terán, Alfredo Herrera-Estrella, Carla Ivón Herrera-Najera, Pavel Isa, Verónica Mata-Haro, Daniel Lira Morales, Susana Lopez, Antonio Loza Román, Brenda Irasema Maldonado-Meza, Bernardo Martínez-Miguel, José Arturo Martínez-Orozco, Célida Martínez- Rodríguez, Margarita Matías-Florentino, Fidencio Mejía-Nepomuceno, María Guadalupe de Jesús Mireles-Rivera, Gloria María Molina-Salinas, Hector Montoya-Fuentes, , Mario Mújica-Sánchez, José Esteban Muñoz-Medina, José de Jesús Nuñez-Contreras, Alicia Ocaña-Mondragón, Luis Alberto Ochoa-Carrera, Hector Esteban Paz-Juárez, Marissa Perez-Garcia, Francisco Pulido, Helen Haydee Fernanda Ramírez-Plascencia, Jorge Salas-Hernández, Angel Gustavo Salas-Lais, Alejandro Sánchez-Flores, Clara Esperanza Santacruz-Tinoco, María Guadalupe Santiago-Mauricio, Selene Zárate, Nelly Sélem-Mojica, Blanca Taboada, Gloria Vazquez, Joel Armando Vázquez-Pérez. |
| EPI_ISL_2681136 | Unidad de Investigación Médica de Yucatán (UIMY) | Instituto de Biotecnología de la UNAM | Consorcio Mexicano de Vigilancia Genómica (CoViGen-Mex). Authors (in alphabetical order): Julio Elias Alvarado-Yaah, Carlos F. Arias, Santiago Ávila-Ríos, Eduardo Becerril-Vargas, Víctor Hugo Borja-Aburto, Celia Boukadida, Cristóbal Cháidez-Quiróz, Juan Bautista Chale-Dzul, Ricardo Ciria Merce, Andreu Comas-García, Célida Duque Molina, Julissa Enciso-Ibarra, José Antonio Enciso-Moreno, Gloria Elena Espinosa-Ayala, Fernando Fontove-Herrera, Daniel Fregoso-Rueda, Victor Eduardo García-Arias, Alejandra García-Gasca, Bruno Gómez-Gil, Jean Pierre González, Irvin |

| | | | |
|---|---|---|---|
| | | | González-López, Concepción Grajales-Muñiz, Ricardo Grande, Rosa María Gutiérrez Rios, Jesús Hernández, Alejandra Hernández-Terán, Alfredo Herrera-Estrella, Carla Ivón Herrera-Najera, Pavel Isa, Verónica Mata-Haro, Daniel Lira Morales, Susana Lopez, Antonio Loza Román, Brenda Irasema Maldonado-Meza, Bernardo Martínez-Miguel, José Arturo Martínez-Orozco, Célida Martínez- Rodríguez, Margarita Matías-Florentino, Fidencio Mejía-Nepomuceno, María Guadalupe de Jesús Mireles-Rivera, Gloria María Molina-Salinas, Hector Montoya-Fuentes, , Mario Mújica-Sánchez, José Esteban Muñoz-Medina, José de Jesús Nuñez-Contreras, Alicia Ocaña-Mondragón, Luis Alberto Ochoa-Carrera, Hector Esteban Paz-Juárez, Marissa Perez-Garcia, Francisco Pulido, Helen Haydee Fernanda Ramírez-Plascencia, Jorge Salas-Hernández, Angel Gustavo Salas-Lais, Alejandro Sánchez-Flores, Clara Esperanza Santacruz-Tinoco, María Guadalupe Santiago-Mauricio, Selene Zárate, Nelly Sélem-Mojica, Blanca Taboada, Gloria Vazquez, Joel Armando Vázquez-Pérez. |
| EPI_ISL_2681137 | Centro de Investigación Biomédica del Noreste (CIBIN) | Instituto de Biotecnología de la UNAM | Consorcio Mexicano de Vigilancia Genómica (CoViGen-Mex). Authors (in alphabetical order): Julio Elias Alvarado-Yaah, Carlos F. Arias, Santiago Ávila-Ríos, Eduardo Becerril-Vargas, Víctor Hugo Borja-Aburto, Celia Boukadida, Cristóbal Cháidez-Quiróz, Juan Bautista Chale-Dzul, Ricardo Ciria Merce, Andreu Comas-García, Célida Duque Molina, Julissa Enciso-Ibarra, José Antonio Enciso-Moreno, Gloria Elena Espinosa-Ayala, Fernando Fontove-Herrera, Daniel Fregoso-Rueda, Victor Eduardo García-Arias, Alejandra García-Gasca, Bruno Gómez-Gil, Jean Pierre González, Irvin González-López, Concepción Grajales-Muñiz, Ricardo Grande, Rosa María Gutiérrez Rios, Jesús Hernández, Alejandra Hernández-Terán, Alfredo Herrera-Estrella, Carla Ivón Herrera-Najera, Pavel Isa, Verónica Mata-Haro, Daniel Lira Morales, Susana Lopez, Antonio Loza Román, Brenda Irasema Maldonado-Meza, Bernardo Martínez-Miguel, José Arturo Martínez-Orozco, Célida Martínez- Rodríguez, Margarita Matías-Florentino, Fidencio Mejía-Nepomuceno, María Guadalupe de Jesús Mireles-Rivera, Gloria María Molina-Salinas, Hector Montoya-Fuentes, , Mario Mújica-Sánchez, José Esteban Muñoz-Medina, José de Jesús Nuñez-Contreras, Alicia Ocaña-Mondragón, Luis Alberto Ochoa-Carrera, Hector Esteban Paz-Juárez, Marissa Perez-Garcia, Francisco Pulido, Helen Haydee Fernanda Ramírez-Plascencia, Jorge Salas-Hernández, Angel Gustavo Salas-Lais, Alejandro Sánchez-Flores, Clara Esperanza Santacruz-Tinoco, María Guadalupe Santiago-Mauricio, Selene Zárate, Nelly Sélem-Mojica, Blanca Taboada, Gloria Vazquez, Joel Armando Vázquez-Pérez. |
| EPI_ISL_2681139, EPI_ISL_2681140 | Laboratorio Central de Epidemiología (LCE) | Instituto de Biotecnología de la UNAM | Consorcio Mexicano de Vigilancia Genómica (CoViGen-Mex). Authors (in alphabetical order): Julio Elias Alvarado-Yaah, Carlos F. Arias, Santiago Ávila-Ríos, Eduardo Becerril-Vargas, Víctor Hugo Borja-Aburto, Celia Boukadida, Cristóbal Cháidez-Quiróz, Juan Bautista Chale-Dzul, Ricardo Ciria Merce, Andreu Comas-García, Célida Duque Molina, Julissa Enciso-Ibarra, José Antonio Enciso-Moreno, Gloria Elena Espinosa-Ayala, Fernando Fontove-Herrera, Daniel Fregoso-Rueda, Victor Eduardo García-Arias, Alejandra García-Gasca, Bruno Gómez-Gil, Jean Pierre González, Irvin González-López, Concepción Grajales-Muñiz, Ricardo Grande, Rosa María Gutiérrez Rios, Jesús Hernández, Alejandra Hernández-Terán, Alfredo Herrera-Estrella, Carla Ivón Herrera-Najera, Pavel Isa, Verónica Mata-Haro, Daniel Lira Morales, Susana Lopez, Antonio Loza Román, Brenda Irasema Maldonado-Meza, Bernardo Martínez-Miguel, José Arturo Martínez-Orozco, Célida Martínez- Rodríguez, Margarita Matías-Florentino, Fidencio Mejía-Nepomuceno, María Guadalupe de Jesús Mireles-Rivera, Gloria María Molina-Salinas, Hector Montoya-Fuentes, , Mario Mújica-Sánchez, José Esteban Muñoz-Medina, José de Jesús Nuñez-Contreras, Alicia Ocaña-Mondragón, Luis Alberto Ochoa-Carrera, Hector Esteban Paz-Juárez, Marissa Perez-Garcia, Francisco Pulido, Helen Haydee Fernanda Ramírez-Plascencia, Jorge Salas-Hernández, Angel Gustavo Salas-Lais, Alejandro Sánchez-Flores, Clara Esperanza Santacruz-Tinoco, María Guadalupe Santiago-Mauricio, Selene Zárate, Nelly Sélem-Mojica, Blanca Taboada, Gloria Vazquez, Joel Armando Vázquez-Pérez. |
| EPI_ISL_2681141 | Unidad de Investigación Médica de Yucatán (UIMY) | Instituto de Biotecnología de la UNAM | Consorcio Mexicano de Vigilancia Genómica (CoViGen-Mex). Authors (in alphabetical order): Julio Elias Alvarado-Yaah, Carlos F. Arias, Santiago Ávila-Ríos, Eduardo Becerril-Vargas, Víctor Hugo Borja-Aburto, Celia Boukadida, Cristóbal Cháidez-Quiróz, Juan Bautista Chale-Dzul, Ricardo Ciria Merce, Andreu Comas-García, Célida Duque Molina, Julissa Enciso-Ibarra, José Antonio Enciso-Moreno, Gloria Elena Espinosa-Ayala, Fernando Fontove-Herrera, Daniel Fregoso-Rueda, Victor Eduardo García-Arias, Alejandra García-Gasca, Bruno Gómez-Gil, Jean Pierre González, Irvin González-López, Concepción Grajales-Muñiz, Ricardo Grande, Rosa María Gutiérrez Rios, Jesús Hernández, Alejandra Hernández-Terán, Alfredo Herrera-Estrella, Carla Ivón Herrera-Najera, Pavel Isa, Verónica Mata-Haro, Daniel Lira Morales, Susana Lopez, Antonio Loza Román, Brenda Irasema Maldonado-Meza, Bernardo Martínez-Miguel, José Arturo Martínez-Orozco, Célida Martínez- Rodríguez, Margarita Matías-Florentino, Fidencio Mejía-Nepomuceno, María Guadalupe de Jesús Mireles-Rivera, Gloria María Molina-Salinas, Hector Montoya-Fuentes, , Mario Mújica-Sánchez, José Esteban Muñoz-Medina, José de Jesús Nuñez-Contreras, Alicia Ocaña-Mondragón, Luis Alberto Ochoa-Carrera, Hector Esteban Paz-Juárez, Marissa Perez-Garcia, Francisco Pulido, Helen Haydee Fernanda Ramírez-Plascencia, Jorge Salas-Hernández, Angel Gustavo Salas-Lais, Alejandro Sánchez-Flores, Clara Esperanza Santacruz-Tinoco, María Guadalupe Santiago-Mauricio, Selene Zárate, Nelly Sélem-Mojica, Blanca Taboada, Gloria Vazquez, Joel Armando Vázquez-Pérez. |
| EPI_ISL_2681142 | Unidad de Investigación Biomédica de Zacatecas (UIBZ) | Instituto de Biotecnología de la UNAM | Consorcio Mexicano de Vigilancia Genómica (CoViGen-Mex). Authors (in alphabetical order): Julio Elias Alvarado-Yaah, Carlos F. Arias, Santiago Ávila-Ríos, Eduardo Becerril-Vargas, Víctor Hugo Borja-Aburto, Celia Boukadida, Cristóbal Cháidez-Quiróz, Juan Bautista Chale-Dzul, Ricardo Ciria Merce, Andreu Comas-García, Célida Duque Molina, Julissa Enciso-Ibarra, José Antonio Enciso-Moreno, Gloria Elena Espinosa-Ayala, Fernando Fontove-Herrera, Daniel Fregoso-Rueda, Victor Eduardo García-Arias, Alejandra García-Gasca, Bruno Gómez-Gil, Jean Pierre González, Irvin González-López, Concepción Grajales-Muñiz, Ricardo Grande, Rosa María Gutiérrez Rios, Jesús Hernández, Alejandra Hernández-Terán, Alfredo Herrera-Estrella, Carla Ivón Herrera-Najera, Pavel Isa, Verónica Mata-Haro, Daniel Lira Morales, Susana Lopez, Antonio Loza Román, Brenda Irasema Maldonado-Meza, Bernardo Martínez-Miguel, José Arturo Martínez-Orozco, Célida Martínez- Rodríguez, Margarita Matías-Florentino, Fidencio Mejía-Nepomuceno, María Guadalupe de Jesús Mireles-Rivera, Gloria María Molina-Salinas, Hector Montoya-Fuentes, , Mario Mújica-Sánchez, José Esteban Muñoz-Medina, José de Jesús Nuñez-Contreras, Alicia Ocaña-Mondragón, Luis Alberto Ochoa-Carrera, Hector Esteban Paz-Juárez, Marissa Perez-Garcia, Francisco Pulido, Helen Haydee Fernanda Ramírez-Plascencia, Jorge Salas-Hernández, Angel Gustavo Salas-Lais, Alejandro Sánchez-Flores, Clara Esperanza Santacruz-Tinoco, María Guadalupe Santiago-Mauricio, Selene Zárate, Nelly Sélem-Mojica, Blanca Taboada, Gloria Vazquez, Joel Armando Vázquez-Pérez. |
| EPI_ISL_2681143, EPI_ISL_2681144, EPI_ISL_2681145, EPI_ISL_2681146, EPI_ISL_2681147, EPI_ISL_2681148, EPI_ISL_2681149, EPI_ISL_2681150, EPI_ISL_2681151, EPI_ISL_2681152, EPI_ISL_2681153, EPI_ISL_2681154, EPI_ISL_2681155, EPI_ISL_2681156, EPI_ISL_2681157, EPI_ISL_2681158, EPI_ISL_2681159 | | | |
| see above | Laboratorio Central de Epidemiología (LCE) | Instituto de Biotecnología de la UNAM | Consorcio Mexicano de Vigilancia Genómica (CoViGen-Mex). Authors (in alphabetical order): Julio Elias Alvarado-Yaah, Carlos F. Arias, Santiago Ávila-Ríos, Eduardo Becerril-Vargas, Víctor Hugo Borja-Aburto, Celia Boukadida, Cristóbal Cháidez-Quiróz, Juan Bautista Chale-Dzul, Ricardo Ciria Merce, Andreu Comas-García, Célida Duque Molina, Julissa Enciso-Ibarra, José Antonio Enciso-Moreno, Gloria Elena Espinosa-Ayala, Fernando Fontove-Herrera, Daniel Fregoso-Rueda, Victor Eduardo García-Arias, Alejandra García-Gasca, Bruno Gómez-Gil, Jean Pierre González, Irvin González-López, Concepción Grajales-Muñiz, Ricardo Grande, Rosa María Gutiérrez Rios, Jesús Hernández, Alejandra Hernández-Terán, Alfredo Herrera-Estrella, Carla Ivón Herrera-Najera, Pavel Isa, Verónica Mata-Haro, Daniel Lira Morales, Susana Lopez, Antonio Loza Román, Brenda Irasema Maldonado-Meza, Bernardo Martínez-Miguel, José Arturo Martínez-Orozco, Célida Martínez- Rodríguez, Margarita Matías-Florentino, Fidencio Mejía-Nepomuceno, María Guadalupe de Jesús Mireles-Rivera, Gloria María Molina-Salinas, Hector Montoya-Fuentes, , Mario Mújica-Sánchez, José Esteban Muñoz-Medina, José de Jesús Nuñez-Contreras, Alicia Ocaña-Mondragón, Luis Alberto Ochoa-Carrera, Hector Esteban Paz-Juárez, Marissa Perez-Garcia, Francisco Pulido, Helen Haydee Fernanda Ramírez-Plascencia, Jorge Salas-Hernández, Angel Gustavo Salas-Lais, Alejandro Sánchez-Flores, Clara Esperanza Santacruz-Tinoco, María Guadalupe Santiago-Mauricio, Selene Zárate, Nelly Sélem-Mojica, Blanca Taboada, Gloria Vazquez, Joel Armando Vázquez-Pérez. |
| EPI_ISL_2681160 | Unidad de Investigación Biomédica de Zacatecas (UIBZ) | Instituto de Biotecnología de la UNAM | Consorcio Mexicano de Vigilancia Genómica (CoViGen-Mex). Authors (in alphabetical order): Julio Elias Alvarado-Yaah, Carlos F. Arias, Santiago Ávila-Ríos, Eduardo Becerril-Vargas, Víctor Hugo Borja-Aburto, Celia Boukadida, Cristóbal Cháidez-Quiróz, Juan Bautista Chale-Dzul, Ricardo Ciria Merce, Andreu Comas-García, Célida Duque Molina, Julissa Enciso-Ibarra, José Antonio Enciso-Moreno, Gloria Elena Espinosa-Ayala, Fernando Fontove-Herrera, Daniel Fregoso-Rueda, Victor Eduardo García-Arias, Alejandra García-Gasca, Bruno Gómez-Gil, Jean Pierre González, Irvin González-López, Concepción Grajales-Muñiz, Ricardo Grande, Rosa María Gutiérrez Rios, Jesús Hernández, Alejandra Hernández-Terán, Alfredo Herrera-Estrella, Carla Ivón Herrera-Najera, Pavel Isa, Verónica Mata-Haro, Daniel Lira Morales, Susana Lopez, Antonio Loza Román, Brenda Irasema Maldonado-Meza, Bernardo Martínez-Miguel, José Arturo Martínez-Orozco, Célida Martínez- Rodríguez, Margarita Matías-Florentino, Fidencio Mejía-Nepomuceno, María Guadalupe de Jesús Mireles-Rivera, Gloria María Molina-Salinas, Hector Montoya-Fuentes, , Mario Mújica-Sánchez, José Esteban Muñoz-Medina, José de Jesús Nuñez-Contreras, Alicia Ocaña-Mondragón, Luis Alberto Ochoa-Carrera, Hector Esteban Paz-Juárez, Marissa Perez-Garcia, Francisco Pulido, Helen Haydee Fernanda Ramírez-Plascencia, Jorge Salas-Hernández, Angel Gustavo Salas-Lais, Alejandro Sánchez-Flores, Clara Esperanza Santacruz-Tinoco, María Guadalupe Santiago-Mauricio, Selene Zárate, Nelly Sélem-Mojica, Blanca Taboada, Gloria Vazquez, Joel Armando Vázquez-Pérez. |
| EPI_ISL_2681161, EPI_ISL_2681162, EPI_ISL_2681163, EPI_ISL_2681164, EPI_ISL_2681165, EPI_ISL_2681166, EPI_ISL_2681167, EPI_ISL_2681168, EPI_ISL_2681169, EPI_ISL_2681170, EPI_ISL_2681171, EPI_ISL_2681172, EPI_ISL_2681173, EPI_ISL_2681174, EPI_ISL_2681175 | | | |
| see above | Laboratorio Central de Epidemiología (LCE) | Instituto de Biotecnología de la UNAM | Consorcio Mexicano de Vigilancia Genómica (CoViGen-Mex). Authors (in alphabetical order): Julio Elias Alvarado-Yaah, Carlos F. Arias, Santiago |

| | | | |
|---|---|---|---|
| | | | Ávila-Ríos, Eduardo Becerril-Vargas, Víctor Hugo Borja-Aburto, Celia Boukadida, Cristóbal Cháidez-Quiróz, Juan Bautista Chale-Dzul, Ricardo Ciria Merce, Andreu Comas-García, Célida Duque Molina, Julissa Enciso-Ibarra, José Antonio Enciso-Moreno, Gloria Elena Espinosa-Ayala, Fernando Fontove-Herrera, Daniel Fregoso-Rueda, Victor Eduardo García-Arias, Alejandra García-Gasca, Bruno Gómez-Gil, Jean Pierre González, Irvin González-López, Concepción Grajales-Muñiz, Ricardo Grande, Rosa María Gutiérrez Rios, Jesús Hernández, Alejandra Hernández-Terán, Alfredo Herrera-Estrella, Carla Ivón Herrera-Najera, Pavel Isa, Verónica Mata-Haro, Daniel Lira Morales, Susana Lopez, Antonio Loza Román, Brenda Irasema Maldonado-Meza, Bernardo Martínez-Miguel, José Arturo Martínez-Orozco, Célida Martínez- Rodríguez, Margarita Matías-Florentino, Fidencio Mejía-Nepomuceno, María Guadalupe de Jesús Mireles-Rivera, Gloria María Molina-Salinas, Hector Montoya-Fuentes, , Mario Mújica-Sánchez, José Esteban Muñoz-Medina, José de Jesús Nuñez-Contreras, Alicia Ocaña-Mondragón, Luis Alberto Ochoa-Carrera, Hector Esteban Paz-Juárez, Marissa Perez-Garcia, Francisco Pulido, Helen Haydee Fernanda Ramírez-Plascencia, Jorge Salas-Hernández, Angel Gustavo Salas-Lais, Alejandro Sánchez-Flores, Clara Esperanza Santacruz-Tinoco, María Guadalupe Santiago-Mauricio, Selene Zárate, Nelly Sélem-Mojica, Blanca Taboada, Gloria Vazquez, Joel Armando Vázquez-Pérez. |
| EPI_ISL_2681176 | Unidad de Investigación Médica de Yucatán (UIMY) | Instituto de Biotecnología de la UNAM | Consorcio Mexicano de Vigilancia Genómica (CoViGen-Mex). Authors (in alphabetical order): Julio Elias Alvarado-Yaah, Carlos F. Arias, Santiago Ávila-Ríos, Eduardo Becerril-Vargas, Víctor Hugo Borja-Aburto, Celia Boukadida, Cristóbal Cháidez-Quiróz, Juan Bautista Chale-Dzul, Ricardo Ciria Merce, Andreu Comas-García, Célida Duque Molina, Julissa Enciso-Ibarra, José Antonio Enciso-Moreno, Gloria Elena Espinosa-Ayala, Fernando Fontove-Herrera, Daniel Fregoso-Rueda, Victor Eduardo García-Arias, Alejandra García-Gasca, Bruno Gómez-Gil, Jean Pierre González, Irvin González-López, Concepción Grajales-Muñiz, Ricardo Grande, Rosa María Gutiérrez Rios, Jesús Hernández, Alejandra Hernández-Terán, Alfredo Herrera-Estrella, Carla Ivón Herrera-Najera, Pavel Isa, Verónica Mata-Haro, Daniel Lira Morales, Susana Lopez, Antonio Loza Román, Brenda Irasema Maldonado-Meza, Bernardo Martínez-Miguel, José Arturo Martínez-Orozco, Célida Martínez- Rodríguez, Margarita Matías-Florentino, Fidencio Mejía-Nepomuceno, María Guadalupe de Jesús Mireles-Rivera, Gloria María Molina-Salinas, Hector Montoya-Fuentes, , Mario Mújica-Sánchez, José Esteban Muñoz-Medina, José de Jesús Nuñez-Contreras, Alicia Ocaña-Mondragón, Luis Alberto Ochoa-Carrera, Hector Esteban Paz-Juárez, Marissa Perez-Garcia, Francisco Pulido, Helen Haydee Fernanda Ramírez-Plascencia, Jorge Salas-Hernández, Angel Gustavo Salas-Lais, Alejandro Sánchez-Flores, Clara Esperanza Santacruz-Tinoco, María Guadalupe Santiago-Mauricio, Selene Zárate, Nelly Sélem-Mojica, Blanca Taboada, Gloria Vazquez, Joel Armando Vázquez-Pérez. |
| EPI_ISL_2681177, EPI_ISL_2681178, EPI_ISL_2681179, EPI_ISL_2681180, EPI_ISL_2681181, EPI_ISL_2681182, EPI_ISL_2681183, EPI_ISL_2681184, EPI_ISL_2681185, EPI_ISL_2681186, EPI_ISL_2681187, EPI_ISL_2681188, EPI_ISL_2681189 | | | |
| see above | Laboratorio Central de Epidemiología (LCE) | Instituto de Biotecnología de la UNAM | Consorcio Mexicano de Vigilancia Genómica (CoViGen-Mex). Authors (in alphabetical order): Julio Elias Alvarado-Yaah, Carlos F. Arias, Santiago Ávila-Ríos, Eduardo Becerril-Vargas, Víctor Hugo Borja-Aburto, Celia Boukadida, Cristóbal Cháidez-Quiróz, Juan Bautista Chale-Dzul, Ricardo Ciria Merce, Andreu Comas-García, Célida Duque Molina, Julissa Enciso-Ibarra, José Antonio Enciso-Moreno, Gloria Elena Espinosa-Ayala, Fernando Fontove-Herrera, Daniel Fregoso-Rueda, Victor Eduardo García-Arias, Alejandra García-Gasca, Bruno Gómez-Gil, Jean Pierre González, Irvin González-López, Concepción Grajales-Muñiz, Ricardo Grande, Rosa María Gutiérrez Rios, Jesús Hernández, Alejandra Hernández-Terán, Alfredo Herrera-Estrella, Carla Ivón Herrera-Najera, Pavel Isa, Verónica Mata-Haro, Daniel Lira Morales, Susana Lopez, Antonio Loza Román, Brenda Irasema Maldonado-Meza, Bernardo Martínez-Miguel, José Arturo Martínez-Orozco, Célida Martínez- Rodríguez, Margarita Matías-Florentino, Fidencio Mejía-Nepomuceno, María Guadalupe de Jesús Mireles-Rivera, Gloria María Molina-Salinas, Hector Montoya-Fuentes, , Mario Mújica-Sánchez, José Esteban Muñoz-Medina, José de Jesús Nuñez-Contreras, Alicia Ocaña-Mondragón, Luis Alberto Ochoa-Carrera, Hector Esteban Paz-Juárez, Marissa Perez-Garcia, Francisco Pulido, Helen Haydee Fernanda Ramírez-Plascencia, Jorge Salas-Hernández, Angel Gustavo Salas-Lais, Alejandro Sánchez-Flores, Clara Esperanza Santacruz-Tinoco, María Guadalupe Santiago-Mauricio, Selene Zárate, Nelly Sélem-Mojica, Blanca Taboada, Gloria Vazquez, Joel Armando Vázquez-Pérez. |
| EPI_ISL_2681190 | Unidad de Investigación Biomédica de Zacatecas (UIBZ) | Instituto de Biotecnología de la UNAM | Consorcio Mexicano de Vigilancia Genómica (CoViGen-Mex). Authors (in alphabetical order): Julio Elias Alvarado-Yaah, Carlos F. Arias, Santiago Ávila-Ríos, Eduardo Becerril-Vargas, Víctor Hugo Borja-Aburto, Celia Boukadida, Cristóbal Cháidez-Quiróz, Juan Bautista Chale-Dzul, Ricardo Ciria Merce, Andreu Comas-García, Célida Duque Molina, Julissa Enciso-Ibarra, José Antonio Enciso-Moreno, Gloria Elena Espinosa-Ayala, Fernando Fontove-Herrera, Daniel Fregoso-Rueda, Victor Eduardo García-Arias, Alejandra García-Gasca, Bruno Gómez-Gil, Jean Pierre González, Irvin González-López, Concepción Grajales-Muñiz, Ricardo Grande, Rosa María Gutiérrez Rios, Jesús Hernández, Alejandra Hernández-Terán, Alfredo Herrera-Estrella, Carla Ivón Herrera-Najera, Pavel Isa, Verónica Mata-Haro, Daniel Lira Morales, Susana Lopez, Antonio Loza Román, Brenda Irasema Maldonado-Meza, Bernardo Martínez-Miguel, José Arturo Martínez-Orozco, Célida Martínez- Rodríguez, Margarita Matías-Florentino, Fidencio Mejía-Nepomuceno, María Guadalupe de Jesús Mireles-Rivera, Gloria María Molina-Salinas, Hector Montoya-Fuentes, , Mario Mújica-Sánchez, José Esteban Muñoz-Medina, José de Jesús Nuñez-Contreras, Alicia Ocaña-Mondragón, Luis Alberto Ochoa-Carrera, Hector Esteban Paz-Juárez, Marissa Perez-Garcia, Francisco Pulido, Helen Haydee Fernanda Ramírez-Plascencia, Jorge Salas-Hernández, Angel Gustavo Salas-Lais, Alejandro Sánchez-Flores, Clara Esperanza Santacruz-Tinoco, María Guadalupe Santiago-Mauricio, Selene Zárate, Nelly Sélem-Mojica, Blanca Taboada, Gloria Vazquez, Joel Armando Vázquez-Pérez. |
| EPI_ISL_2681191, EPI_ISL_2681192, EPI_ISL_2681193, EPI_ISL_2681194, EPI_ISL_2681195, EPI_ISL_2681196 | Laboratorio Central de Epidemiología (LCE) | Instituto de Biotecnología de la UNAM | Consorcio Mexicano de Vigilancia Genómica (CoViGen-Mex). Authors (in alphabetical order): Julio Elias Alvarado-Yaah, Carlos F. Arias, Santiago Ávila-Ríos, Eduardo Becerril-Vargas, Víctor Hugo Borja-Aburto, Celia Boukadida, Cristóbal Cháidez-Quiróz, Juan Bautista Chale-Dzul, Ricardo Ciria Merce, Andreu Comas-García, Célida Duque Molina, Julissa Enciso-Ibarra, José Antonio Enciso-Moreno, Gloria Elena Espinosa-Ayala, Fernando Fontove-Herrera, Daniel Fregoso-Rueda, Victor Eduardo García-Arias, Alejandra García-Gasca, Bruno Gómez-Gil, Jean Pierre González, Irvin González-López, Concepción Grajales-Muñiz, Ricardo Grande, Rosa María Gutiérrez Rios, Jesús Hernández, Alejandra Hernández-Terán, Alfredo Herrera-Estrella, Carla Ivón Herrera-Najera, Pavel Isa, Verónica Mata-Haro, Daniel Lira Morales, Susana Lopez, Antonio Loza Román, Brenda Irasema Maldonado-Meza, Bernardo Martínez-Miguel, José Arturo Martínez-Orozco, Célida Martínez- Rodríguez, Margarita Matías-Florentino, Fidencio Mejía-Nepomuceno, María Guadalupe de Jesús Mireles-Rivera, Gloria María Molina-Salinas, Hector Montoya-Fuentes, , Mario Mújica-Sánchez, José Esteban Muñoz-Medina, José de Jesús Nuñez-Contreras, Alicia Ocaña-Mondragón, Luis Alberto Ochoa-Carrera, Hector Esteban Paz-Juárez, Marissa Perez-Garcia, Francisco Pulido, Helen Haydee Fernanda Ramírez-Plascencia, Jorge Salas-Hernández, Angel Gustavo Salas-Lais, Alejandro Sánchez-Flores, Clara Esperanza Santacruz-Tinoco, María Guadalupe Santiago-Mauricio, Selene Zárate, Nelly Sélem-Mojica, Blanca Taboada, Gloria Vazquez, Joel Armando Vázquez-Pérez. |
| EPI_ISL_2681197 | Centro de Investigación Biomédica del Noreste (CIBIN) | Instituto de Biotecnología de la UNAM | Consorcio Mexicano de Vigilancia Genómica (CoViGen-Mex). Authors (in alphabetical order): Julio Elias Alvarado-Yaah, Carlos F. Arias, Santiago Ávila-Ríos, Eduardo Becerril-Vargas, Víctor Hugo Borja-Aburto, Celia Boukadida, Cristóbal Cháidez-Quiróz, Juan Bautista Chale-Dzul, Ricardo Ciria Merce, Andreu Comas-García, Célida Duque Molina, Julissa Enciso-Ibarra, José Antonio Enciso-Moreno, Gloria Elena Espinosa-Ayala, Fernando Fontove-Herrera, Daniel Fregoso-Rueda, Victor Eduardo García-Arias, Alejandra García-Gasca, Bruno Gómez-Gil, Jean Pierre González, Irvin González-López, Concepción Grajales-Muñiz, Ricardo Grande, Rosa María Gutiérrez Rios, Jesús Hernández, Alejandra Hernández-Terán, Alfredo Herrera-Estrella, Carla Ivón Herrera-Najera, Pavel Isa, Verónica Mata-Haro, Daniel Lira Morales, Susana Lopez, Antonio Loza Román, Brenda Irasema Maldonado-Meza, Bernardo Martínez-Miguel, José Arturo Martínez-Orozco, Célida Martínez- Rodríguez, Margarita Matías-Florentino, Fidencio Mejía-Nepomuceno, María Guadalupe de Jesús Mireles-Rivera, Gloria María Molina-Salinas, Hector Montoya-Fuentes, , Mario Mújica-Sánchez, José Esteban Muñoz-Medina, José de Jesús Nuñez-Contreras, Alicia Ocaña-Mondragón, Luis Alberto Ochoa-Carrera, Hector Esteban Paz-Juárez, Marissa Perez-Garcia, Francisco Pulido, Helen Haydee Fernanda Ramírez-Plascencia, Jorge Salas-Hernández, Angel Gustavo Salas-Lais, Alejandro Sánchez-Flores, Clara Esperanza Santacruz-Tinoco, María Guadalupe Santiago-Mauricio, Selene Zárate, Nelly Sélem-Mojica, Blanca Taboada, Gloria Vazquez, Joel Armando Vázquez-Pérez. |
| EPI_ISL_2681198, EPI_ISL_2681199, EPI_ISL_2681200 | Laboratorio Central de Epidemiología (LCE) | Instituto de Biotecnología de la UNAM | Consorcio Mexicano de Vigilancia Genómica (CoViGen-Mex). Authors (in alphabetical order): Julio Elias Alvarado-Yaah, Carlos F. Arias, Santiago Ávila-Ríos, Eduardo Becerril-Vargas, Víctor Hugo Borja-Aburto, Celia Boukadida, Cristóbal Cháidez-Quiróz, Juan Bautista Chale-Dzul, Ricardo Ciria Merce, Andreu Comas-García, Célida Duque Molina, Julissa Enciso-Ibarra, José Antonio Enciso-Moreno, Gloria Elena Espinosa-Ayala, Fernando Fontove-Herrera, Daniel Fregoso-Rueda, Victor Eduardo García-Arias, Alejandra García-Gasca, Bruno Gómez-Gil, Jean Pierre González, Irvin González-López, Concepción Grajales-Muñiz, Ricardo Grande, Rosa María Gutiérrez Rios, Jesús Hernández, Alejandra Hernández-Terán, Alfredo Herrera-Estrella, Carla Ivón Herrera-Najera, Pavel Isa, Verónica Mata-Haro, Daniel Lira Morales, Susana Lopez, Antonio Loza Román, Brenda Irasema Maldonado-Meza, Bernardo Martínez-Miguel, José Arturo Martínez-Orozco, Célida Martínez- Rodríguez, Margarita Matías-Florentino, Fidencio Mejía-Nepomuceno, María Guadalupe de Jesús Mireles-Rivera, Gloria María Molina-Salinas, Hector Montoya-Fuentes, , Mario Mújica-Sánchez, José Esteban Muñoz-Medina, José de Jesús Nuñez-Contreras, Alicia Ocaña-Mondragón, Luis Alberto Ochoa-Carrera, Hector Esteban Paz-Juárez, Marissa Perez-Garcia, Francisco Pulido, Helen Haydee Fernanda Ramírez-Plascencia, Jorge Salas-Hernández, Angel Gustavo Salas-Lais, Alejandro Sánchez-Flores, Clara Esperanza Santacruz-Tinoco, María Guadalupe Santiago-Mauricio, Selene Zárate, Nelly Sélem-Mojica, Blanca Taboada, Gloria Vazquez, Joel Armando Vázquez-Pérez. |

| | | | |
|---|---|---|---|
| EPI_ISL_2681201 | Centro de Investigación Biomédica del Noreste (CIBIN) | Instituto de Biotecnología de la UNAM | Consorcio Mexicano de Vigilancia Genómica (CoViGen-Mex). Authors (in alphabetical order): Julio Elias Alvarado-Yaah, Carlos F. Arias, Santiago Ávila-Ríos, Eduardo Becerril-Vargas, Víctor Hugo Borja-Aburto, Celia Boukadida, Cristóbal Cháidez-Quiróz, Juan Bautista Chale-Dzul, Ricardo Ciria Merce, Andreu Comas-García, Célida Duque Molina, Julissa Enciso-Ibarra, José Antonio Enciso-Moreno, Gloria Elena Espinosa-Ayala, Fernando Fontove-Herrera, Daniel Fregoso-Rueda, Victor Eduardo García-Arias, Alejandra García-Gasca, Bruno Gómez-Gil, Jean Pierre González, Irvin González-López, Concepción Grajales-Muñiz, Ricardo Grande, Rosa María Gutiérrez Rios, Jesús Hernández, Alejandra Hernández-Terán, Alfredo Herrera-Estrella, Carla Ivón Herrera-Najera, Pavel Isa, Verónica Mata-Haro, Daniel Lira Morales, Susana Lopez, Antonio Loza Román, Brenda Irasema Maldonado-Meza, Bernardo Martínez-Miguel, José Arturo Martínez-Orozco, Célida Martínez- Rodríguez, Margarita Matías-Florentino, Fidencio Mejía-Nepomuceno, María Guadalupe de Jesús Mireles-Rivera, Gloria María Molina-Salinas, Hector Montoya-Fuentes, , Mario Mújica-Sánchez, José Esteban Muñoz-Medina, José de Jesús Nuñez-Contreras, Alicia Ocaña-Mondragón, Luis Alberto Ochoa-Carrera, Hector Esteban Paz-Juárez, Marissa Perez-Garcia, Francisco Pulido, Helen Haydee Fernanda Ramírez-Plascencia, Jorge Salas-Hernández, Angel Gustavo Salas-Lais, Alejandro Sánchez-Flores, Clara Esperanza Santacruz-Tinoco, María Guadalupe Santiago-Mauricio, Selene Zárate, Nelly Sélem-Mojica, Blanca Taboada, Gloria Vazquez, Joel Armando Vázquez-Pérez. |
| EPI_ISL_2681202, EPI_ISL_2681203, EPI_ISL_2681204, EPI_ISL_2681205, EPI_ISL_2681206 | Laboratorio Central de Epidemiología (LCE) | Instituto de Biotecnología de la UNAM | Consorcio Mexicano de Vigilancia Genómica (CoViGen-Mex). Authors (in alphabetical order): Julio Elias Alvarado-Yaah, Carlos F. Arias, Santiago Ávila-Ríos, Eduardo Becerril-Vargas, Víctor Hugo Borja-Aburto, Celia Boukadida, Cristóbal Cháidez-Quiróz, Juan Bautista Chale-Dzul, Ricardo Ciria Merce, Andreu Comas-García, Célida Duque Molina, Julissa Enciso-Ibarra, José Antonio Enciso-Moreno, Gloria Elena Espinosa-Ayala, Fernando Fontove-Herrera, Daniel Fregoso-Rueda, Victor Eduardo García-Arias, Alejandra García-Gasca, Bruno Gómez-Gil, Jean Pierre González, Irvin González-López, Concepción Grajales-Muñiz, Ricardo Grande, Rosa María Gutiérrez Rios, Jesús Hernández, Alejandra Hernández-Terán, Alfredo Herrera-Estrella, Carla Ivón Herrera-Najera, Pavel Isa, Verónica Mata-Haro, Daniel Lira Morales, Susana Lopez, Antonio Loza Román, Brenda Irasema Maldonado-Meza, Bernardo Martínez-Miguel, José Arturo Martínez-Orozco, Célida Martínez- Rodríguez, Margarita Matías-Florentino, Fidencio Mejía-Nepomuceno, María Guadalupe de Jesús Mireles-Rivera, Gloria María Molina-Salinas, Hector Montoya-Fuentes, , Mario Mújica-Sánchez, José Esteban Muñoz-Medina, José de Jesús Nuñez-Contreras, Alicia Ocaña-Mondragón, Luis Alberto Ochoa-Carrera, Hector Esteban Paz-Juárez, Marissa Perez-Garcia, Francisco Pulido, Helen Haydee Fernanda Ramírez-Plascencia, Jorge Salas-Hernández, Angel Gustavo Salas-Lais, Alejandro Sánchez-Flores, Clara Esperanza Santacruz-Tinoco, María Guadalupe Santiago-Mauricio, Selene Zárate, Nelly Sélem-Mojica, Blanca Taboada, Gloria Vazquez, Joel Armando Vázquez-Pérez. |
| EPI_ISL_2681207 | Unidad de Investigación Biomédica de Zacatecas (UIBZ) | Instituto de Biotecnología de la UNAM | Consorcio Mexicano de Vigilancia Genómica (CoViGen-Mex). Authors (in alphabetical order): Julio Elias Alvarado-Yaah, Carlos F. Arias, Santiago Ávila-Ríos, Eduardo Becerril-Vargas, Víctor Hugo Borja-Aburto, Celia Boukadida, Cristóbal Cháidez-Quiróz, Juan Bautista Chale-Dzul, Ricardo Ciria Merce, Andreu Comas-García, Célida Duque Molina, Julissa Enciso-Ibarra, José Antonio Enciso-Moreno, Gloria Elena Espinosa-Ayala, Fernando Fontove-Herrera, Daniel Fregoso-Rueda, Victor Eduardo García-Arias, Alejandra García-Gasca, Bruno Gómez-Gil, Jean Pierre González, Irvin González-López, Concepción Grajales-Muñiz, Ricardo Grande, Rosa María Gutiérrez Rios, Jesús Hernández, Alejandra Hernández-Terán, Alfredo Herrera-Estrella, Carla Ivón Herrera-Najera, Pavel Isa, Verónica Mata-Haro, Daniel Lira Morales, Susana Lopez, Antonio Loza Román, Brenda Irasema Maldonado-Meza, Bernardo Martínez-Miguel, José Arturo Martínez-Orozco, Célida Martínez- Rodríguez, Margarita Matías-Florentino, Fidencio Mejía-Nepomuceno, María Guadalupe de Jesús Mireles-Rivera, Gloria María Molina-Salinas, Hector Montoya-Fuentes, , Mario Mújica-Sánchez, José Esteban Muñoz-Medina, José de Jesús Nuñez-Contreras, Alicia Ocaña-Mondragón, Luis Alberto Ochoa-Carrera, Hector Esteban Paz-Juárez, Marissa Perez-Garcia, Francisco Pulido, Helen Haydee Fernanda Ramírez-Plascencia, Jorge Salas-Hernández, Angel Gustavo Salas-Lais, Alejandro Sánchez-Flores, Clara Esperanza Santacruz-Tinoco, María Guadalupe Santiago-Mauricio, Selene Zárate, Nelly Sélem-Mojica, Blanca Taboada, Gloria Vazquez, Joel Armando Vázquez-Pérez. |
| EPI_ISL_2681208 | Laboratorio Central de Epidemiología (LCE) | Instituto de Biotecnología de la UNAM | Consorcio Mexicano de Vigilancia Genómica (CoViGen-Mex). Authors (in alphabetical order): Julio Elias Alvarado-Yaah, Carlos F. Arias, Santiago Ávila-Ríos, Eduardo Becerril-Vargas, Víctor Hugo Borja-Aburto, Celia Boukadida, Cristóbal Cháidez-Quiróz, Juan Bautista Chale-Dzul, Ricardo Ciria Merce, Andreu Comas-García, Célida Duque Molina, Julissa Enciso-Ibarra, José Antonio Enciso-Moreno, Gloria Elena Espinosa-Ayala, Fernando Fontove-Herrera, Daniel Fregoso-Rueda, Victor Eduardo García-Arias, Alejandra García-Gasca, Bruno Gómez-Gil, Jean Pierre González, Irvin González-López, Concepción Grajales-Muñiz, Ricardo Grande, Rosa María Gutiérrez Rios, Jesús Hernández, Alejandra Hernández-Terán, Alfredo Herrera-Estrella, Carla Ivón Herrera-Najera, Pavel Isa, Verónica Mata-Haro, Daniel Lira Morales, Susana Lopez, Antonio Loza Román, Brenda Irasema Maldonado-Meza, Bernardo Martínez-Miguel, José Arturo Martínez-Orozco, Célida Martínez- Rodríguez, Margarita Matías-Florentino, Fidencio Mejía-Nepomuceno, María Guadalupe de Jesús Mireles-Rivera, Gloria María Molina-Salinas, Hector Montoya-Fuentes, , Mario Mújica-Sánchez, José Esteban Muñoz-Medina, José de Jesús Nuñez-Contreras, Alicia Ocaña-Mondragón, Luis Alberto Ochoa-Carrera, Hector Esteban Paz-Juárez, Marissa Perez-Garcia, Francisco Pulido, Helen Haydee Fernanda Ramírez-Plascencia, Jorge Salas-Hernández, Angel Gustavo Salas-Lais, Alejandro Sánchez-Flores, Clara Esperanza Santacruz-Tinoco, María Guadalupe Santiago-Mauricio, Selene Zárate, Nelly Sélem-Mojica, Blanca Taboada, Gloria Vazquez, Joel Armando Vázquez-Pérez. |
| EPI_ISL_2681209 | Unidad de Investigación Biomédica de Zacatecas (UIBZ) | Instituto de Biotecnología de la UNAM | Consorcio Mexicano de Vigilancia Genómica (CoViGen-Mex). Authors (in alphabetical order): Julio Elias Alvarado-Yaah, Carlos F. Arias, Santiago Ávila-Ríos, Eduardo Becerril-Vargas, Víctor Hugo Borja-Aburto, Celia Boukadida, Cristóbal Cháidez-Quiróz, Juan Bautista Chale-Dzul, Ricardo Ciria Merce, Andreu Comas-García, Célida Duque Molina, Julissa Enciso-Ibarra, José Antonio Enciso-Moreno, Gloria Elena Espinosa-Ayala, Fernando Fontove-Herrera, Daniel Fregoso-Rueda, Victor Eduardo García-Arias, Alejandra García-Gasca, Bruno Gómez-Gil, Jean Pierre González, Irvin González-López, Concepción Grajales-Muñiz, Ricardo Grande, Rosa María Gutiérrez Rios, Jesús Hernández, Alejandra Hernández-Terán, Alfredo Herrera-Estrella, Carla Ivón Herrera-Najera, Pavel Isa, Verónica Mata-Haro, Daniel Lira Morales, Susana Lopez, Antonio Loza Román, Brenda Irasema Maldonado-Meza, Bernardo Martínez-Miguel, José Arturo Martínez-Orozco, Célida Martínez- Rodríguez, Margarita Matías-Florentino, Fidencio Mejía-Nepomuceno, María Guadalupe de Jesús Mireles-Rivera, Gloria María Molina-Salinas, Hector Montoya-Fuentes, , Mario Mújica-Sánchez, José Esteban Muñoz-Medina, José de Jesús Nuñez-Contreras, Alicia Ocaña-Mondragón, Luis Alberto Ochoa-Carrera, Hector Esteban Paz-Juárez, Marissa Perez-Garcia, Francisco Pulido, Helen Haydee Fernanda Ramírez-Plascencia, Jorge Salas-Hernández, Angel Gustavo Salas-Lais, Alejandro Sánchez-Flores, Clara Esperanza Santacruz-Tinoco, María Guadalupe Santiago-Mauricio, Selene Zárate, Nelly Sélem-Mojica, Blanca Taboada, Gloria Vazquez, Joel Armando Vázquez-Pérez. |
| EPI_ISL_2681210 | Laboratorio Central de Epidemiología (LCE) | Instituto de Biotecnología de la UNAM | Consorcio Mexicano de Vigilancia Genómica (CoViGen-Mex). Authors (in alphabetical order): Julio Elias Alvarado-Yaah, Carlos F. Arias, Santiago Ávila-Ríos, Eduardo Becerril-Vargas, Víctor Hugo Borja-Aburto, Celia Boukadida, Cristóbal Cháidez-Quiróz, Juan Bautista Chale-Dzul, Ricardo Ciria Merce, Andreu Comas-García, Célida Duque Molina, Julissa Enciso-Ibarra, José Antonio Enciso-Moreno, Gloria Elena Espinosa-Ayala, Fernando Fontove-Herrera, Daniel Fregoso-Rueda, Victor Eduardo García-Arias, Alejandra García-Gasca, Bruno Gómez-Gil, Jean Pierre González, Irvin González-López, Concepción Grajales-Muñiz, Ricardo Grande, Rosa María Gutiérrez Rios, Jesús Hernández, Alejandra Hernández-Terán, Alfredo Herrera-Estrella, Carla Ivón Herrera-Najera, Pavel Isa, Verónica Mata-Haro, Daniel Lira Morales, Susana Lopez, Antonio Loza Román, Brenda Irasema Maldonado-Meza, Bernardo Martínez-Miguel, José Arturo Martínez-Orozco, Célida Martínez- Rodríguez, Margarita Matías-Florentino, Fidencio Mejía-Nepomuceno, María Guadalupe de Jesús Mireles-Rivera, Gloria María Molina-Salinas, Hector Montoya-Fuentes, , Mario Mújica-Sánchez, José Esteban Muñoz-Medina, José de Jesús Nuñez-Contreras, Alicia Ocaña-Mondragón, Luis Alberto Ochoa-Carrera, Hector Esteban Paz-Juárez, Marissa Perez-Garcia, Francisco Pulido, Helen Haydee Fernanda Ramírez-Plascencia, Jorge Salas-Hernández, Angel Gustavo Salas-Lais, Alejandro Sánchez-Flores, Clara Esperanza Santacruz-Tinoco, María Guadalupe Santiago-Mauricio, Selene Zárate, Nelly Sélem-Mojica, Blanca Taboada, Gloria Vazquez, Joel Armando Vázquez-Pérez. |
| EPI_ISL_2681211 | Unidad de Investigación Médica de Yucatán (UIMY) | Instituto de Biotecnología de la UNAM | Consorcio Mexicano de Vigilancia Genómica (CoViGen-Mex). Authors (in alphabetical order): Julio Elias Alvarado-Yaah, Carlos F. Arias, Santiago Ávila-Ríos, Eduardo Becerril-Vargas, Víctor Hugo Borja-Aburto, Celia Boukadida, Cristóbal Cháidez-Quiróz, Juan Bautista Chale-Dzul, Ricardo Ciria Merce, Andreu Comas-García, Célida Duque Molina, Julissa Enciso-Ibarra, José Antonio Enciso-Moreno, Gloria Elena Espinosa-Ayala, Fernando Fontove-Herrera, Daniel Fregoso-Rueda, Victor Eduardo García-Arias, Alejandra García-Gasca, Bruno Gómez-Gil, Jean Pierre González, Irvin González-López, Concepción Grajales-Muñiz, Ricardo Grande, Rosa María Gutiérrez Rios, Jesús Hernández, Alejandra Hernández-Terán, Alfredo Herrera-Estrella, Carla Ivón Herrera-Najera, Pavel Isa, Verónica Mata-Haro, Daniel Lira Morales, Susana Lopez, Antonio Loza Román, Brenda Irasema Maldonado-Meza, Bernardo Martínez-Miguel, José Arturo Martínez-Orozco, Célida Martínez- Rodríguez, Margarita Matías-Florentino, Fidencio Mejía-Nepomuceno, María Guadalupe de Jesús Mireles-Rivera, Gloria María Molina-Salinas, Hector Montoya-Fuentes, , Mario Mújica-Sánchez, José Esteban Muñoz-Medina, José de Jesús Nuñez-Contreras, Alicia Ocaña-Mondragón, Luis Alberto Ochoa-Carrera, Hector Esteban Paz-Juárez, Marissa Perez-Garcia, Francisco Pulido, Helen Haydee Fernanda Ramírez-Plascencia, Jorge Salas-Hernández, Angel Gustavo Salas-Lais, Alejandro Sánchez-Flores, Clara Esperanza Santacruz-Tinoco, María Guadalupe Santiago-Mauricio, Selene Zárate, Nelly Sélem-Mojica, Blanca Taboada, Gloria Vazquez, Joel Armando Vázquez-Pérez. |

| | | | |
|---|---|---|---|
| EPI_ISL_2681212 | Unidad de Investigación Biomédica de Zacatecas (UIBZ) | Instituto de Biotecnología de la UNAM | Consorcio Mexicano de Vigilancia Genómica (CoViGen-Mex). Authors (in alphabetical order): Julio Elias Alvarado-Yaah, Carlos F. Arias, Santiago Ávila-Ríos, Eduardo Becerril-Vargas, Víctor Hugo Borja-Aburto, Celia Boukadida, Cristóbal Cháidez-Quiróz, Juan Bautista Chale-Dzul, Ricardo Ciria Merce, Andreu Comas-García, Célida Duque Molina, Julissa Enciso-Ibarra, José Antonio Enciso-Moreno, Gloria Elena Espinosa-Ayala, Fernando Fontove-Herrera, Daniel Fregoso-Rueda, Victor Eduardo García-Arias, Alejandra García-Gasca, Bruno Gómez-Gil, Jean Pierre González, Irvin González-López, Concepción Grajales-Muñiz, Ricardo Grande, Rosa María Gutiérrez Rios, Jesús Hernández, Alejandra Hernández-Terán, Alfredo Herrera-Estrella, Carla Ivón Herrera-Najera, Pavel Isa, Verónica Mata-Haro, Daniel Lira Morales, Susana Lopez, Antonio Loza Román, Brenda Irasema Maldonado-Meza, Bernardo Martínez-Miguel, José Arturo Martínez-Orozco, Célida Martínez- Rodríguez, Margarita Matías-Florentino, Fidencio Mejía-Nepomuceno, María Guadalupe de Jesús Mireles-Rivera, Gloria María Molina-Salinas, Hector Montoya-Fuentes, , Mario Mújica-Sánchez, José Esteban Muñoz-Medina, José de Jesús Nuñez-Contreras, Alicia Ocaña-Mondragón, Luis Alberto Ochoa-Carrera, Hector Esteban Paz-Juárez, Marissa Perez-Garcia, Francisco Pulido, Helen Haydee Fernanda Ramírez-Plascencia, Jorge Salas-Hernández, Angel Gustavo Salas-Lais, Alejandro Sánchez-Flores, Clara Esperanza Santacruz-Tinoco, María Guadalupe Santiago-Mauricio, Selene Zárate, Nelly Sélem-Mojica, Blanca Taboada, Gloria Vazquez, Joel Armando Vázquez-Pérez. |
| EPI_ISL_2681213, EPI_ISL_2681214, EPI_ISL_2681215, EPI_ISL_2681216, EPI_ISL_2681217 | Laboratorio Central de Epidemiología (LCE) | Instituto de Biotecnología de la UNAM | Consorcio Mexicano de Vigilancia Genómica (CoViGen-Mex). Authors (in alphabetical order): Julio Elias Alvarado-Yaah, Carlos F. Arias, Santiago Ávila-Ríos, Eduardo Becerril-Vargas, Víctor Hugo Borja-Aburto, Celia Boukadida, Cristóbal Cháidez-Quiróz, Juan Bautista Chale-Dzul, Ricardo Ciria Merce, Andreu Comas-García, Célida Duque Molina, Julissa Enciso-Ibarra, José Antonio Enciso-Moreno, Gloria Elena Espinosa-Ayala, Fernando Fontove-Herrera, Daniel Fregoso-Rueda, Victor Eduardo García-Arias, Alejandra García-Gasca, Bruno Gómez-Gil, Jean Pierre González, Irvin González-López, Concepción Grajales-Muñiz, Ricardo Grande, Rosa María Gutiérrez Rios, Jesús Hernández, Alejandra Hernández-Terán, Alfredo Herrera-Estrella, Carla Ivón Herrera-Najera, Pavel Isa, Verónica Mata-Haro, Daniel Lira Morales, Susana Lopez, Antonio Loza Román, Brenda Irasema Maldonado-Meza, Bernardo Martínez-Miguel, José Arturo Martínez-Orozco, Célida Martínez- Rodríguez, Margarita Matías-Florentino, Fidencio Mejía-Nepomuceno, María Guadalupe de Jesús Mireles-Rivera, Gloria María Molina-Salinas, Hector Montoya-Fuentes, , Mario Mújica-Sánchez, José Esteban Muñoz-Medina, José de Jesús Nuñez-Contreras, Alicia Ocaña-Mondragón, Luis Alberto Ochoa-Carrera, Hector Esteban Paz-Juárez, Marissa Perez-Garcia, Francisco Pulido, Helen Haydee Fernanda Ramírez-Plascencia, Jorge Salas-Hernández, Angel Gustavo Salas-Lais, Alejandro Sánchez-Flores, Clara Esperanza Santacruz-Tinoco, María Guadalupe Santiago-Mauricio, Selene Zárate, Nelly Sélem-Mojica, Blanca Taboada, Gloria Vazquez, Joel Armando Vázquez-Pérez. |
| EPI_ISL_2681218 | Unidad de Investigación Médica de Yucatán (UIMY) | Instituto de Biotecnología de la UNAM | Consorcio Mexicano de Vigilancia Genómica (CoViGen-Mex). Authors (in alphabetical order): Julio Elias Alvarado-Yaah, Carlos F. Arias, Santiago Ávila-Ríos, Eduardo Becerril-Vargas, Víctor Hugo Borja-Aburto, Celia Boukadida, Cristóbal Cháidez-Quiróz, Juan Bautista Chale-Dzul, Ricardo Ciria Merce, Andreu Comas-García, Célida Duque Molina, Julissa Enciso-Ibarra, José Antonio Enciso-Moreno, Gloria Elena Espinosa-Ayala, Fernando Fontove-Herrera, Daniel Fregoso-Rueda, Victor Eduardo García-Arias, Alejandra García-Gasca, Bruno Gómez-Gil, Jean Pierre González, Irvin González-López, Concepción Grajales-Muñiz, Ricardo Grande, Rosa María Gutiérrez Rios, Jesús Hernández, Alejandra Hernández-Terán, Alfredo Herrera-Estrella, Carla Ivón Herrera-Najera, Pavel Isa, Verónica Mata-Haro, Daniel Lira Morales, Susana Lopez, Antonio Loza Román, Brenda Irasema Maldonado-Meza, Bernardo Martínez-Miguel, José Arturo Martínez-Orozco, Célida Martínez- Rodríguez, Margarita Matías-Florentino, Fidencio Mejía-Nepomuceno, María Guadalupe de Jesús Mireles-Rivera, Gloria María Molina-Salinas, Hector Montoya-Fuentes, , Mario Mújica-Sánchez, José Esteban Muñoz-Medina, José de Jesús Nuñez-Contreras, Alicia Ocaña-Mondragón, Luis Alberto Ochoa-Carrera, Hector Esteban Paz-Juárez, Marissa Perez-Garcia, Francisco Pulido, Helen Haydee Fernanda Ramírez-Plascencia, Jorge Salas-Hernández, Angel Gustavo Salas-Lais, Alejandro Sánchez-Flores, Clara Esperanza Santacruz-Tinoco, María Guadalupe Santiago-Mauricio, Selene Zárate, Nelly Sélem-Mojica, Blanca Taboada, Gloria Vazquez, Joel Armando Vázquez-Pérez. |
| EPI_ISL_2681219 | Centro de Investigación Biomédica del Noreste (CIBIN) | Instituto de Biotecnología de la UNAM | Consorcio Mexicano de Vigilancia Genómica (CoViGen-Mex). Authors (in alphabetical order): Julio Elias Alvarado-Yaah, Carlos F. Arias, Santiago Ávila-Ríos, Eduardo Becerril-Vargas, Víctor Hugo Borja-Aburto, Celia Boukadida, Cristóbal Cháidez-Quiróz, Juan Bautista Chale-Dzul, Ricardo Ciria Merce, Andreu Comas-García, Célida Duque Molina, Julissa Enciso-Ibarra, José Antonio Enciso-Moreno, Gloria Elena Espinosa-Ayala, Fernando Fontove-Herrera, Daniel Fregoso-Rueda, Victor Eduardo García-Arias, Alejandra García-Gasca, Bruno Gómez-Gil, Jean Pierre González, Irvin González-López, Concepción Grajales-Muñiz, Ricardo Grande, Rosa María Gutiérrez Rios, Jesús Hernández, Alejandra Hernández-Terán, Alfredo Herrera-Estrella, Carla Ivón Herrera-Najera, Pavel Isa, Verónica Mata-Haro, Daniel Lira Morales, Susana Lopez, Antonio Loza Román, Brenda Irasema Maldonado-Meza, Bernardo Martínez-Miguel, José Arturo Martínez-Orozco, Célida Martínez- Rodríguez, Margarita Matías-Florentino, Fidencio Mejía-Nepomuceno, María Guadalupe de Jesús Mireles-Rivera, Gloria María Molina-Salinas, Hector Montoya-Fuentes, , Mario Mújica-Sánchez, José Esteban Muñoz-Medina, José de Jesús Nuñez-Contreras, Alicia Ocaña-Mondragón, Luis Alberto Ochoa-Carrera, Hector Esteban Paz-Juárez, Marissa Perez-Garcia, Francisco Pulido, Helen Haydee Fernanda Ramírez-Plascencia, Jorge Salas-Hernández, Angel Gustavo Salas-Lais, Alejandro Sánchez-Flores, Clara Esperanza Santacruz-Tinoco, María Guadalupe Santiago-Mauricio, Selene Zárate, Nelly Sélem-Mojica, Blanca Taboada, Gloria Vazquez, Joel Armando Vázquez-Pérez. |
| EPI_ISL_2681220 | Laboratorio Central de Epidemiología (LCE) | Instituto de Biotecnología de la UNAM | Consorcio Mexicano de Vigilancia Genómica (CoViGen-Mex). Authors (in alphabetical order): Julio Elias Alvarado-Yaah, Carlos F. Arias, Santiago Ávila-Ríos, Eduardo Becerril-Vargas, Víctor Hugo Borja-Aburto, Celia Boukadida, Cristóbal Cháidez-Quiróz, Juan Bautista Chale-Dzul, Ricardo Ciria Merce, Andreu Comas-García, Célida Duque Molina, Julissa Enciso-Ibarra, José Antonio Enciso-Moreno, Gloria Elena Espinosa-Ayala, Fernando Fontove-Herrera, Daniel Fregoso-Rueda, Victor Eduardo García-Arias, Alejandra García-Gasca, Bruno Gómez-Gil, Jean Pierre González, Irvin González-López, Concepción Grajales-Muñiz, Ricardo Grande, Rosa María Gutiérrez Rios, Jesús Hernández, Alejandra Hernández-Terán, Alfredo Herrera-Estrella, Carla Ivón Herrera-Najera, Pavel Isa, Verónica Mata-Haro, Daniel Lira Morales, Susana Lopez, Antonio Loza Román, Brenda Irasema Maldonado-Meza, Bernardo Martínez-Miguel, José Arturo Martínez-Orozco, Célida Martínez- Rodríguez, Margarita Matías-Florentino, Fidencio Mejía-Nepomuceno, María Guadalupe de Jesús Mireles-Rivera, Gloria María Molina-Salinas, Hector Montoya-Fuentes, , Mario Mújica-Sánchez, José Esteban Muñoz-Medina, José de Jesús Nuñez-Contreras, Alicia Ocaña-Mondragón, Luis Alberto Ochoa-Carrera, Hector Esteban Paz-Juárez, Marissa Perez-Garcia, Francisco Pulido, Helen Haydee Fernanda Ramírez-Plascencia, Jorge Salas-Hernández, Angel Gustavo Salas-Lais, Alejandro Sánchez-Flores, Clara Esperanza Santacruz-Tinoco, María Guadalupe Santiago-Mauricio, Selene Zárate, Nelly Sélem-Mojica, Blanca Taboada, Gloria Vazquez, Joel Armando Vázquez-Pérez. |
| EPI_ISL_2681221 | Unidad de Investigación Biomédica de Zacatecas (UIBZ) | Instituto de Biotecnología de la UNAM | Consorcio Mexicano de Vigilancia Genómica (CoViGen-Mex). Authors (in alphabetical order): Julio Elias Alvarado-Yaah, Carlos F. Arias, Santiago Ávila-Ríos, Eduardo Becerril-Vargas, Víctor Hugo Borja-Aburto, Celia Boukadida, Cristóbal Cháidez-Quiróz, Juan Bautista Chale-Dzul, Ricardo Ciria Merce, Andreu Comas-García, Célida Duque Molina, Julissa Enciso-Ibarra, José Antonio Enciso-Moreno, Gloria Elena Espinosa-Ayala, Fernando Fontove-Herrera, Daniel Fregoso-Rueda, Victor Eduardo García-Arias, Alejandra García-Gasca, Bruno Gómez-Gil, Jean Pierre González, Irvin González-López, Concepción Grajales-Muñiz, Ricardo Grande, Rosa María Gutiérrez Rios, Jesús Hernández, Alejandra Hernández-Terán, Alfredo Herrera-Estrella, Carla Ivón Herrera-Najera, Pavel Isa, Verónica Mata-Haro, Daniel Lira Morales, Susana Lopez, Antonio Loza Román, Brenda Irasema Maldonado-Meza, Bernardo Martínez-Miguel, José Arturo Martínez-Orozco, Célida Martínez- Rodríguez, Margarita Matías-Florentino, Fidencio Mejía-Nepomuceno, María Guadalupe de Jesús Mireles-Rivera, Gloria María Molina-Salinas, Hector Montoya-Fuentes, , Mario Mújica-Sánchez, José Esteban Muñoz-Medina, José de Jesús Nuñez-Contreras, Alicia Ocaña-Mondragón, Luis Alberto Ochoa-Carrera, Hector Esteban Paz-Juárez, Marissa Perez-Garcia, Francisco Pulido, Helen Haydee Fernanda Ramírez-Plascencia, Jorge Salas-Hernández, Angel Gustavo Salas-Lais, Alejandro Sánchez-Flores, Clara Esperanza Santacruz-Tinoco, María Guadalupe Santiago-Mauricio, Selene Zárate, Nelly Sélem-Mojica, Blanca Taboada, Gloria Vazquez, Joel Armando Vázquez-Pérez. |
| EPI_ISL_2681222 | Laboratorio Central de Epidemiología (LCE) | Instituto de Biotecnología de la UNAM | Consorcio Mexicano de Vigilancia Genómica (CoViGen-Mex). Authors (in alphabetical order): Julio Elias Alvarado-Yaah, Carlos F. Arias, Santiago Ávila-Ríos, Eduardo Becerril-Vargas, Víctor Hugo Borja-Aburto, Celia Boukadida, Cristóbal Cháidez-Quiróz, Juan Bautista Chale-Dzul, Ricardo Ciria Merce, Andreu Comas-García, Célida Duque Molina, Julissa Enciso-Ibarra, José Antonio Enciso-Moreno, Gloria Elena Espinosa-Ayala, Fernando Fontove-Herrera, Daniel Fregoso-Rueda, Victor Eduardo García-Arias, Alejandra García-Gasca, Bruno Gómez-Gil, Jean Pierre González, Irvin González-López, Concepción Grajales-Muñiz, Ricardo Grande, Rosa María Gutiérrez Rios, Jesús Hernández, Alejandra Hernández-Terán, Alfredo Herrera-Estrella, Carla Ivón Herrera-Najera, Pavel Isa, Verónica Mata-Haro, Daniel Lira Morales, Susana Lopez, Antonio Loza Román, Brenda Irasema Maldonado-Meza, Bernardo Martínez-Miguel, José Arturo Martínez-Orozco, Célida Martínez- Rodríguez, Margarita Matías-Florentino, Fidencio Mejía-Nepomuceno, María Guadalupe de Jesús Mireles-Rivera, Gloria María Molina-Salinas, Hector Montoya-Fuentes, , Mario Mújica-Sánchez, José Esteban Muñoz-Medina, José de Jesús Nuñez-Contreras, Alicia Ocaña-Mondragón, Luis Alberto Ochoa-Carrera, Hector Esteban Paz-Juárez, Marissa Perez-Garcia, Francisco Pulido, Helen Haydee Fernanda Ramírez-Plascencia, Jorge Salas-Hernández, Angel Gustavo Salas-Lais, Alejandro Sánchez-Flores, Clara Esperanza Santacruz-Tinoco, María Guadalupe Santiago-Mauricio, Selene Zárate, Nelly Sélem-Mojica, Blanca Taboada, Gloria Vazquez, Joel Armando Vázquez-Pérez. |

| | | | |
|---|---|---|---|
| EPI_ISL_2681223 | Unidad de Investigación Biomédica de Zacatecas (UIBZ) | Instituto de Biotecnología de la UNAM | Consorcio Mexicano de Vigilancia Genómica (CoViGen-Mex). Authors (in alphabetical order): Julio Elias Alvarado-Yaah, Carlos F. Arias, Santiago Ávila-Ríos, Eduardo Becerril-Vargas, Víctor Hugo Borja-Aburto, Celia Boukadida, Cristóbal Cháidez-Quiróz, Juan Bautista Chale-Dzul, Ricardo Ciria Merce, Andreu Comas-García, Célida Duque Molina, Julissa Enciso-Ibarra, José Antonio Enciso-Moreno, Gloria Elena Espinosa-Ayala, Fernando Fontove-Herrera, Daniel Fregoso-Rueda, Victor Eduardo García-Arias, Alejandra García-Gasca, Bruno Gómez-Gil, Jean Pierre González, Irvin González-López, Concepción Grajales-Muñiz, Ricardo Grande, Rosa María Gutiérrez Rios, Jesús Hernández, Alejandra Hernández-Terán, Alfredo Herrera-Estrella, Carla Ivón Herrera-Najera, Pavel Isa, Verónica Mata-Haro, Daniel Lira Morales, Susana Lopez, Antonio Loza Román, Brenda Irasema Maldonado-Meza, Bernardo Martínez-Miguel, José Arturo Martínez-Orozco, Célida Martínez- Rodríguez, Margarita Matías-Florentino, Fidencio Mejía-Nepomuceno, María Guadalupe de Jesús Mireles-Rivera, Gloria María Molina-Salinas, Hector Montoya-Fuentes, , Mario Mújica-Sánchez, José Esteban Muñoz-Medina, José de Jesús Nuñez-Contreras, Alicia Ocaña-Mondragón, Luis Alberto Ochoa-Carrera, Hector Esteban Paz-Juárez, Marissa Perez-Garcia, Francisco Pulido, Helen Haydee Fernanda Ramírez-Plascencia, Jorge Salas-Hernández, Angel Gustavo Salas-Lais, Alejandro Sánchez-Flores, Clara Esperanza Santacruz-Tinoco, María Guadalupe Santiago-Mauricio, Selene Zárate, Nelly Sélem-Mojica, Blanca Taboada, Gloria Vazquez, Joel Armando Vázquez-Pérez. |
| EPI_ISL_2681224, EPI_ISL_2681225, EPI_ISL_2681226 | Laboratorio Central de Epidemiología (LCE) | Instituto de Biotecnología de la UNAM | Consorcio Mexicano de Vigilancia Genómica (CoViGen-Mex). Authors (in alphabetical order): Julio Elias Alvarado-Yaah, Carlos F. Arias, Santiago Ávila-Ríos, Eduardo Becerril-Vargas, Víctor Hugo Borja-Aburto, Celia Boukadida, Cristóbal Cháidez-Quiróz, Juan Bautista Chale-Dzul, Ricardo Ciria Merce, Andreu Comas-García, Célida Duque Molina, Julissa Enciso-Ibarra, José Antonio Enciso-Moreno, Gloria Elena Espinosa-Ayala, Fernando Fontove-Herrera, Daniel Fregoso-Rueda, Victor Eduardo García-Arias, Alejandra García-Gasca, Bruno Gómez-Gil, Jean Pierre González, Irvin González-López, Concepción Grajales-Muñiz, Ricardo Grande, Rosa María Gutiérrez Rios, Jesús Hernández, Alejandra Hernández-Terán, Alfredo Herrera-Estrella, Carla Ivón Herrera-Najera, Pavel Isa, Verónica Mata-Haro, Daniel Lira Morales, Susana Lopez, Antonio Loza Román, Brenda Irasema Maldonado-Meza, Bernardo Martínez-Miguel, José Arturo Martínez-Orozco, Célida Martínez- Rodríguez, Margarita Matías-Florentino, Fidencio Mejía-Nepomuceno, María Guadalupe de Jesús Mireles-Rivera, Gloria María Molina-Salinas, Hector Montoya-Fuentes, , Mario Mújica-Sánchez, José Esteban Muñoz-Medina, José de Jesús Nuñez-Contreras, Alicia Ocaña-Mondragón, Luis Alberto Ochoa-Carrera, Hector Esteban Paz-Juárez, Marissa Perez-Garcia, Francisco Pulido, Helen Haydee Fernanda Ramírez-Plascencia, Jorge Salas-Hernández, Angel Gustavo Salas-Lais, Alejandro Sánchez-Flores, Clara Esperanza Santacruz-Tinoco, María Guadalupe Santiago-Mauricio, Selene Zárate, Nelly Sélem-Mojica, Blanca Taboada, Gloria Vazquez, Joel Armando Vázquez-Pérez. |
| EPI_ISL_2681227, EPI_ISL_2681228 | Centro de Investigación Biomédica del Noreste (CIBIN) | Instituto de Biotecnología de la UNAM | Consorcio Mexicano de Vigilancia Genómica (CoViGen-Mex). Authors (in alphabetical order): Julio Elias Alvarado-Yaah, Carlos F. Arias, Santiago Ávila-Ríos, Eduardo Becerril-Vargas, Víctor Hugo Borja-Aburto, Celia Boukadida, Cristóbal Cháidez-Quiróz, Juan Bautista Chale-Dzul, Ricardo Ciria Merce, Andreu Comas-García, Célida Duque Molina, Julissa Enciso-Ibarra, José Antonio Enciso-Moreno, Gloria Elena Espinosa-Ayala, Fernando Fontove-Herrera, Daniel Fregoso-Rueda, Victor Eduardo García-Arias, Alejandra García-Gasca, Bruno Gómez-Gil, Jean Pierre González, Irvin González-López, Concepción Grajales-Muñiz, Ricardo Grande, Rosa María Gutiérrez Rios, Jesús Hernández, Alejandra Hernández-Terán, Alfredo Herrera-Estrella, Carla Ivón Herrera-Najera, Pavel Isa, Verónica Mata-Haro, Daniel Lira Morales, Susana Lopez, Antonio Loza Román, Brenda Irasema Maldonado-Meza, Bernardo Martínez-Miguel, José Arturo Martínez-Orozco, Célida Martínez- Rodríguez, Margarita Matías-Florentino, Fidencio Mejía-Nepomuceno, María Guadalupe de Jesús Mireles-Rivera, Gloria María Molina-Salinas, Hector Montoya-Fuentes, , Mario Mújica-Sánchez, José Esteban Muñoz-Medina, José de Jesús Nuñez-Contreras, Alicia Ocaña-Mondragón, Luis Alberto Ochoa-Carrera, Hector Esteban Paz-Juárez, Marissa Perez-Garcia, Francisco Pulido, Helen Haydee Fernanda Ramírez-Plascencia, Jorge Salas-Hernández, Angel Gustavo Salas-Lais, Alejandro Sánchez-Flores, Clara Esperanza Santacruz-Tinoco, María Guadalupe Santiago-Mauricio, Selene Zárate, Nelly Sélem-Mojica, Blanca Taboada, Gloria Vazquez, Joel Armando Vázquez-Pérez. |
| EPI_ISL_2681229, EPI_ISL_2681230, EPI_ISL_2681231 | Laboratorio Central de Epidemiología (LCE) | Instituto de Biotecnología de la UNAM | Consorcio Mexicano de Vigilancia Genómica (CoViGen-Mex). Authors (in alphabetical order): Julio Elias Alvarado-Yaah, Carlos F. Arias, Santiago Ávila-Ríos, Eduardo Becerril-Vargas, Víctor Hugo Borja-Aburto, Celia Boukadida, Cristóbal Cháidez-Quiróz, Juan Bautista Chale-Dzul, Ricardo Ciria Merce, Andreu Comas-García, Célida Duque Molina, Julissa Enciso-Ibarra, José Antonio Enciso-Moreno, Gloria Elena Espinosa-Ayala, Fernando Fontove-Herrera, Daniel Fregoso-Rueda, Victor Eduardo García-Arias, Alejandra García-Gasca, Bruno Gómez-Gil, Jean Pierre González, Irvin González-López, Concepción Grajales-Muñiz, Ricardo Grande, Rosa María Gutiérrez Rios, Jesús Hernández, Alejandra Hernández-Terán, Alfredo Herrera-Estrella, Carla Ivón Herrera-Najera, Pavel Isa, Verónica Mata-Haro, Daniel Lira Morales, Susana Lopez, Antonio Loza Román, Brenda Irasema Maldonado-Meza, Bernardo Martínez-Miguel, José Arturo Martínez-Orozco, Célida Martínez- Rodríguez, Margarita Matías-Florentino, Fidencio Mejía-Nepomuceno, María Guadalupe de Jesús Mireles-Rivera, Gloria María Molina-Salinas, Hector Montoya-Fuentes, , Mario Mújica-Sánchez, José Esteban Muñoz-Medina, José de Jesús Nuñez-Contreras, Alicia Ocaña-Mondragón, Luis Alberto Ochoa-Carrera, Hector Esteban Paz-Juárez, Marissa Perez-Garcia, Francisco Pulido, Helen Haydee Fernanda Ramírez-Plascencia, Jorge Salas-Hernández, Angel Gustavo Salas-Lais, Alejandro Sánchez-Flores, Clara Esperanza Santacruz-Tinoco, María Guadalupe Santiago-Mauricio, Selene Zárate, Nelly Sélem-Mojica, Blanca Taboada, Gloria Vazquez, Joel Armando Vázquez-Pérez. |
| EPI_ISL_2681232, EPI_ISL_2681233 | Unidad de Investigación Médica de Yucatán (UIMY) | Instituto de Biotecnología de la UNAM | Consorcio Mexicano de Vigilancia Genómica (CoViGen-Mex). Authors (in alphabetical order): Julio Elias Alvarado-Yaah, Carlos F. Arias, Santiago Ávila-Ríos, Eduardo Becerril-Vargas, Víctor Hugo Borja-Aburto, Celia Boukadida, Cristóbal Cháidez-Quiróz, Juan Bautista Chale-Dzul, Ricardo Ciria Merce, Andreu Comas-García, Célida Duque Molina, Julissa Enciso-Ibarra, José Antonio Enciso-Moreno, Gloria Elena Espinosa-Ayala, Fernando Fontove-Herrera, Daniel Fregoso-Rueda, Victor Eduardo García-Arias, Alejandra García-Gasca, Bruno Gómez-Gil, Jean Pierre González, Irvin González-López, Concepción Grajales-Muñiz, Ricardo Grande, Rosa María Gutiérrez Rios, Jesús Hernández, Alejandra Hernández-Terán, Alfredo Herrera-Estrella, Carla Ivón Herrera-Najera, Pavel Isa, Verónica Mata-Haro, Daniel Lira Morales, Susana Lopez, Antonio Loza Román, Brenda Irasema Maldonado-Meza, Bernardo Martínez-Miguel, José Arturo Martínez-Orozco, Célida Martínez- Rodríguez, Margarita Matías-Florentino, Fidencio Mejía-Nepomuceno, María Guadalupe de Jesús Mireles-Rivera, Gloria María Molina-Salinas, Hector Montoya-Fuentes, , Mario Mújica-Sánchez, José Esteban Muñoz-Medina, José de Jesús Nuñez-Contreras, Alicia Ocaña-Mondragón, Luis Alberto Ochoa-Carrera, Hector Esteban Paz-Juárez, Marissa Perez-Garcia, Francisco Pulido, Helen Haydee Fernanda Ramírez-Plascencia, Jorge Salas-Hernández, Angel Gustavo Salas-Lais, Alejandro Sánchez-Flores, Clara Esperanza Santacruz-Tinoco, María Guadalupe Santiago-Mauricio, Selene Zárate, Nelly Sélem-Mojica, Blanca Taboada, Gloria Vazquez, Joel Armando Vázquez-Pérez. |
| EPI_ISL_2681234, EPI_ISL_2681235 | Unidad de Investigación Biomédica de Zacatecas (UIBZ) | Instituto de Biotecnología de la UNAM | Consorcio Mexicano de Vigilancia Genómica (CoViGen-Mex). Authors (in alphabetical order): Julio Elias Alvarado-Yaah, Carlos F. Arias, Santiago Ávila-Ríos, Eduardo Becerril-Vargas, Víctor Hugo Borja-Aburto, Celia Boukadida, Cristóbal Cháidez-Quiróz, Juan Bautista Chale-Dzul, Ricardo Ciria Merce, Andreu Comas-García, Célida Duque Molina, Julissa Enciso-Ibarra, José Antonio Enciso-Moreno, Gloria Elena Espinosa-Ayala, Fernando Fontove-Herrera, Daniel Fregoso-Rueda, Victor Eduardo García-Arias, Alejandra García-Gasca, Bruno Gómez-Gil, Jean Pierre González, Irvin González-López, Concepción Grajales-Muñiz, Ricardo Grande, Rosa María Gutiérrez Rios, Jesús Hernández, Alejandra Hernández-Terán, Alfredo Herrera-Estrella, Carla Ivón Herrera-Najera, Pavel Isa, Verónica Mata-Haro, Daniel Lira Morales, Susana Lopez, Antonio Loza Román, Brenda Irasema Maldonado-Meza, Bernardo Martínez-Miguel, José Arturo Martínez-Orozco, Célida Martínez- Rodríguez, Margarita Matías-Florentino, Fidencio Mejía-Nepomuceno, María Guadalupe de Jesús Mireles-Rivera, Gloria María Molina-Salinas, Hector Montoya-Fuentes, , Mario Mújica-Sánchez, José Esteban Muñoz-Medina, José de Jesús Nuñez-Contreras, Alicia Ocaña-Mondragón, Luis Alberto Ochoa-Carrera, Hector Esteban Paz-Juárez, Marissa Perez-Garcia, Francisco Pulido, Helen Haydee Fernanda Ramírez-Plascencia, Jorge Salas-Hernández, Angel Gustavo Salas-Lais, Alejandro Sánchez-Flores, Clara Esperanza Santacruz-Tinoco, María Guadalupe Santiago-Mauricio, Selene Zárate, Nelly Sélem-Mojica, Blanca Taboada, Gloria Vazquez, Joel Armando Vázquez-Pérez. |
| EPI_ISL_2681236 | Unidad de Investigación Médica de Yucatán (UIMY) | Instituto de Biotecnología de la UNAM | Consorcio Mexicano de Vigilancia Genómica (CoViGen-Mex). Authors (in alphabetical order): Julio Elias Alvarado-Yaah, Carlos F. Arias, Santiago Ávila-Ríos, Eduardo Becerril-Vargas, Víctor Hugo Borja-Aburto, Celia Boukadida, Cristóbal Cháidez-Quiróz, Juan Bautista Chale-Dzul, Ricardo Ciria Merce, Andreu Comas-García, Célida Duque Molina, Julissa Enciso-Ibarra, José Antonio Enciso-Moreno, Gloria Elena Espinosa-Ayala, Fernando Fontove-Herrera, Daniel Fregoso-Rueda, Victor Eduardo García-Arias, Alejandra García-Gasca, Bruno Gómez-Gil, Jean Pierre González, Irvin González-López, Concepción Grajales-Muñiz, Ricardo Grande, Rosa María Gutiérrez Rios, Jesús Hernández, Alejandra Hernández-Terán, Alfredo Herrera-Estrella, Carla Ivón Herrera-Najera, Pavel Isa, Verónica Mata-Haro, Daniel Lira Morales, Susana Lopez, Antonio Loza Román, Brenda Irasema Maldonado-Meza, Bernardo Martínez-Miguel, José Arturo Martínez-Orozco, Célida Martínez- Rodríguez, Margarita Matías-Florentino, Fidencio Mejía-Nepomuceno, María Guadalupe de Jesús Mireles-Rivera, Gloria María Molina-Salinas, Hector Montoya-Fuentes, , Mario Mújica-Sánchez, José Esteban Muñoz-Medina, José de Jesús Nuñez-Contreras, Alicia Ocaña-Mondragón, Luis Alberto Ochoa-Carrera, Hector Esteban Paz-Juárez, Marissa Perez-Garcia, Francisco Pulido, Helen Haydee Fernanda Ramírez-Plascencia, Jorge Salas-Hernández, Angel Gustavo Salas-Lais, Alejandro Sánchez-Flores, Clara Esperanza Santacruz-Tinoco, María Guadalupe Santiago-Mauricio, Selene Zárate, Nelly Sélem-Mojica, Blanca Taboada, Gloria Vazquez, Joel Armando Vázquez-Pérez. |

| | | | |
|---|---|---|---|
| EPI_ISL_2681238 | Laboratorio Central de Epidemiología (LCE) | Instituto de Biotecnología de la UNAM | Consorcio Mexicano de Vigilancia Genómica (CoViGen-Mex). Authors (in alphabetical order): Julio Elias Alvarado-Yaah, Carlos F. Arias, Santiago Ávila-Ríos, Eduardo Becerril-Vargas, Víctor Hugo Borja-Aburto, Celia Boukadida, Cristóbal Cháidez-Quiróz, Juan Bautista Chale-Dzul, Ricardo Ciria Merce, Andreu Comas-García, Célida Duque Molina, Julissa Enciso-Ibarra, José Antonio Enciso-Moreno, Gloria Elena Espinosa-Ayala, Fernando Fontove-Herrera, Daniel Fregoso-Rueda, Victor Eduardo García-Arias, Alejandra García-Gasca, Bruno Gómez-Gil, Jean Pierre González, Irvin González-López, Concepción Grajales-Muñiz, Ricardo Grande, Rosa María Gutiérrez Rios, Jesús Hernández, Alejandra Hernández-Terán, Alfredo Herrera-Estrella, Carla Ivón Herrera-Najera, Pavel Isa, Verónica Mata-Haro, Daniel Lira Morales, Susana Lopez, Antonio Loza Román, Brenda Irasema Maldonado-Meza, Bernardo Martínez-Miguel, José Arturo Martínez-Orozco, Célida Martínez- Rodríguez, Margarita Matías-Florentino, Fidencio Mejía-Nepomuceno, María Guadalupe de Jesús Mireles-Rivera, Gloria María Molina-Salinas, Hector Montoya-Fuentes, , Mario Mújica-Sánchez, José Esteban Muñoz-Medina, José de Jesús Nuñez-Contreras, Alicia Ocaña-Mondragón, Luis Alberto Ochoa-Carrera, Hector Esteban Paz-Juárez, Marissa Perez-Garcia, Francisco Pulido, Helen Haydee Fernanda Ramírez-Plascencia, Jorge Salas-Hernández, Angel Gustavo Salas-Lais, Alejandro Sánchez-Flores, Clara Esperanza Santacruz-Tinoco, María Guadalupe Santiago-Mauricio, Selene Zárate, Nelly Sélem-Mojica, Blanca Taboada, Gloria Vazquez, Joel Armando Vázquez-Pérez. |
| EPI_ISL_2681239 | Unidad de Investigación Biomédica de Zacatecas (UIBZ) | Instituto de Biotecnología de la UNAM | Consorcio Mexicano de Vigilancia Genómica (CoViGen-Mex). Authors (in alphabetical order): Julio Elias Alvarado-Yaah, Carlos F. Arias, Santiago Ávila-Ríos, Eduardo Becerril-Vargas, Víctor Hugo Borja-Aburto, Celia Boukadida, Cristóbal Cháidez-Quiróz, Juan Bautista Chale-Dzul, Ricardo Ciria Merce, Andreu Comas-García, Célida Duque Molina, Julissa Enciso-Ibarra, José Antonio Enciso-Moreno, Gloria Elena Espinosa-Ayala, Fernando Fontove-Herrera, Daniel Fregoso-Rueda, Victor Eduardo García-Arias, Alejandra García-Gasca, Bruno Gómez-Gil, Jean Pierre González, Irvin González-López, Concepción Grajales-Muñiz, Ricardo Grande, Rosa María Gutiérrez Rios, Jesús Hernández, Alejandra Hernández-Terán, Alfredo Herrera-Estrella, Carla Ivón Herrera-Najera, Pavel Isa, Verónica Mata-Haro, Daniel Lira Morales, Susana Lopez, Antonio Loza Román, Brenda Irasema Maldonado-Meza, Bernardo Martínez-Miguel, José Arturo Martínez-Orozco, Célida Martínez- Rodríguez, Margarita Matías-Florentino, Fidencio Mejía-Nepomuceno, María Guadalupe de Jesús Mireles-Rivera, Gloria María Molina-Salinas, Hector Montoya-Fuentes, , Mario Mújica-Sánchez, José Esteban Muñoz-Medina, José de Jesús Nuñez-Contreras, Alicia Ocaña-Mondragón, Luis Alberto Ochoa-Carrera, Hector Esteban Paz-Juárez, Marissa Perez-Garcia, Francisco Pulido, Helen Haydee Fernanda Ramírez-Plascencia, Jorge Salas-Hernández, Angel Gustavo Salas-Lais, Alejandro Sánchez-Flores, Clara Esperanza Santacruz-Tinoco, María Guadalupe Santiago-Mauricio, Selene Zárate, Nelly Sélem-Mojica, Blanca Taboada, Gloria Vazquez, Joel Armando Vázquez-Pérez. |
| EPI_ISL_2681240 | Laboratorio Central de Epidemiología (LCE) | Instituto de Biotecnología de la UNAM | Consorcio Mexicano de Vigilancia Genómica (CoViGen-Mex). Authors (in alphabetical order): Julio Elias Alvarado-Yaah, Carlos F. Arias, Santiago Ávila-Ríos, Eduardo Becerril-Vargas, Víctor Hugo Borja-Aburto, Celia Boukadida, Cristóbal Cháidez-Quiróz, Juan Bautista Chale-Dzul, Ricardo Ciria Merce, Andreu Comas-García, Célida Duque Molina, Julissa Enciso-Ibarra, José Antonio Enciso-Moreno, Gloria Elena Espinosa-Ayala, Fernando Fontove-Herrera, Daniel Fregoso-Rueda, Victor Eduardo García-Arias, Alejandra García-Gasca, Bruno Gómez-Gil, Jean Pierre González, Irvin González-López, Concepción Grajales-Muñiz, Ricardo Grande, Rosa María Gutiérrez Rios, Jesús Hernández, Alejandra Hernández-Terán, Alfredo Herrera-Estrella, Carla Ivón Herrera-Najera, Pavel Isa, Verónica Mata-Haro, Daniel Lira Morales, Susana Lopez, Antonio Loza Román, Brenda Irasema Maldonado-Meza, Bernardo Martínez-Miguel, José Arturo Martínez-Orozco, Célida Martínez- Rodríguez, Margarita Matías-Florentino, Fidencio Mejía-Nepomuceno, María Guadalupe de Jesús Mireles-Rivera, Gloria María Molina-Salinas, Hector Montoya-Fuentes, , Mario Mújica-Sánchez, José Esteban Muñoz-Medina, José de Jesús Nuñez-Contreras, Alicia Ocaña-Mondragón, Luis Alberto Ochoa-Carrera, Hector Esteban Paz-Juárez, Marissa Perez-Garcia, Francisco Pulido, Helen Haydee Fernanda Ramírez-Plascencia, Jorge Salas-Hernández, Angel Gustavo Salas-Lais, Alejandro Sánchez-Flores, Clara Esperanza Santacruz-Tinoco, María Guadalupe Santiago-Mauricio, Selene Zárate, Nelly Sélem-Mojica, Blanca Taboada, Gloria Vazquez, Joel Armando Vázquez-Pérez. |
| EPI_ISL_2681241 | Unidad de Investigación Médica de Yucatán (UIMY) | Instituto de Biotecnología de la UNAM | Consorcio Mexicano de Vigilancia Genómica (CoViGen-Mex). Authors (in alphabetical order): Julio Elias Alvarado-Yaah, Carlos F. Arias, Santiago Ávila-Ríos, Eduardo Becerril-Vargas, Víctor Hugo Borja-Aburto, Celia Boukadida, Cristóbal Cháidez-Quiróz, Juan Bautista Chale-Dzul, Ricardo Ciria Merce, Andreu Comas-García, Célida Duque Molina, Julissa Enciso-Ibarra, José Antonio Enciso-Moreno, Gloria Elena Espinosa-Ayala, Fernando Fontove-Herrera, Daniel Fregoso-Rueda, Victor Eduardo García-Arias, Alejandra García-Gasca, Bruno Gómez-Gil, Jean Pierre González, Irvin González-López, Concepción Grajales-Muñiz, Ricardo Grande, Rosa María Gutiérrez Rios, Jesús Hernández, Alejandra Hernández-Terán, Alfredo Herrera-Estrella, Carla Ivón Herrera-Najera, Pavel Isa, Verónica Mata-Haro, Daniel Lira Morales, Susana Lopez, Antonio Loza Román, Brenda Irasema Maldonado-Meza, Bernardo Martínez-Miguel, José Arturo Martínez-Orozco, Célida Martínez- Rodríguez, Margarita Matías-Florentino, Fidencio Mejía-Nepomuceno, María Guadalupe de Jesús Mireles-Rivera, Gloria María Molina-Salinas, Hector Montoya-Fuentes, , Mario Mújica-Sánchez, José Esteban Muñoz-Medina, José de Jesús Nuñez-Contreras, Alicia Ocaña-Mondragón, Luis Alberto Ochoa-Carrera, Hector Esteban Paz-Juárez, Marissa Perez-Garcia, Francisco Pulido, Helen Haydee Fernanda Ramírez-Plascencia, Jorge Salas-Hernández, Angel Gustavo Salas-Lais, Alejandro Sánchez-Flores, Clara Esperanza Santacruz-Tinoco, María Guadalupe Santiago-Mauricio, Selene Zárate, Nelly Sélem-Mojica, Blanca Taboada, Gloria Vazquez, Joel Armando Vázquez-Pérez. |
| EPI_ISL_2681242 | Unidad de Investigación Biomédica de Zacatecas (UIBZ) | Instituto de Biotecnología de la UNAM | Consorcio Mexicano de Vigilancia Genómica (CoViGen-Mex). Authors (in alphabetical order): Julio Elias Alvarado-Yaah, Carlos F. Arias, Santiago Ávila-Ríos, Eduardo Becerril-Vargas, Víctor Hugo Borja-Aburto, Celia Boukadida, Cristóbal Cháidez-Quiróz, Juan Bautista Chale-Dzul, Ricardo Ciria Merce, Andreu Comas-García, Célida Duque Molina, Julissa Enciso-Ibarra, José Antonio Enciso-Moreno, Gloria Elena Espinosa-Ayala, Fernando Fontove-Herrera, Daniel Fregoso-Rueda, Victor Eduardo García-Arias, Alejandra García-Gasca, Bruno Gómez-Gil, Jean Pierre González, Irvin González-López, Concepción Grajales-Muñiz, Ricardo Grande, Rosa María Gutiérrez Rios, Jesús Hernández, Alejandra Hernández-Terán, Alfredo Herrera-Estrella, Carla Ivón Herrera-Najera, Pavel Isa, Verónica Mata-Haro, Daniel Lira Morales, Susana Lopez, Antonio Loza Román, Brenda Irasema Maldonado-Meza, Bernardo Martínez-Miguel, José Arturo Martínez-Orozco, Célida Martínez- Rodríguez, Margarita Matías-Florentino, Fidencio Mejía-Nepomuceno, María Guadalupe de Jesús Mireles-Rivera, Gloria María Molina-Salinas, Hector Montoya-Fuentes, , Mario Mújica-Sánchez, José Esteban Muñoz-Medina, José de Jesús Nuñez-Contreras, Alicia Ocaña-Mondragón, Luis Alberto Ochoa-Carrera, Hector Esteban Paz-Juárez, Marissa Perez-Garcia, Francisco Pulido, Helen Haydee Fernanda Ramírez-Plascencia, Jorge Salas-Hernández, Angel Gustavo Salas-Lais, Alejandro Sánchez-Flores, Clara Esperanza Santacruz-Tinoco, María Guadalupe Santiago-Mauricio, Selene Zárate, Nelly Sélem-Mojica, Blanca Taboada, Gloria Vazquez, Joel Armando Vázquez-Pérez. |
| EPI_ISL_2681243 | Laboratorio Central de Epidemiología (LCE) | Instituto de Biotecnología de la UNAM | Consorcio Mexicano de Vigilancia Genómica (CoViGen-Mex). Authors (in alphabetical order): Julio Elias Alvarado-Yaah, Carlos F. Arias, Santiago Ávila-Ríos, Eduardo Becerril-Vargas, Víctor Hugo Borja-Aburto, Celia Boukadida, Cristóbal Cháidez-Quiróz, Juan Bautista Chale-Dzul, Ricardo Ciria Merce, Andreu Comas-García, Célida Duque Molina, Julissa Enciso-Ibarra, José Antonio Enciso-Moreno, Gloria Elena Espinosa-Ayala, Fernando Fontove-Herrera, Daniel Fregoso-Rueda, Victor Eduardo García-Arias, Alejandra García-Gasca, Bruno Gómez-Gil, Jean Pierre González, Irvin González-López, Concepción Grajales-Muñiz, Ricardo Grande, Rosa María Gutiérrez Rios, Jesús Hernández, Alejandra Hernández-Terán, Alfredo Herrera-Estrella, Carla Ivón Herrera-Najera, Pavel Isa, Verónica Mata-Haro, Daniel Lira Morales, Susana Lopez, Antonio Loza Román, Brenda Irasema Maldonado-Meza, Bernardo Martínez-Miguel, José Arturo Martínez-Orozco, Célida Martínez- Rodríguez, Margarita Matías-Florentino, Fidencio Mejía-Nepomuceno, María Guadalupe de Jesús Mireles-Rivera, Gloria María Molina-Salinas, Hector Montoya-Fuentes, , Mario Mújica-Sánchez, José Esteban Muñoz-Medina, José de Jesús Nuñez-Contreras, Alicia Ocaña-Mondragón, Luis Alberto Ochoa-Carrera, Hector Esteban Paz-Juárez, Marissa Perez-Garcia, Francisco Pulido, Helen Haydee Fernanda Ramírez-Plascencia, Jorge Salas-Hernández, Angel Gustavo Salas-Lais, Alejandro Sánchez-Flores, Clara Esperanza Santacruz-Tinoco, María Guadalupe Santiago-Mauricio, Selene Zárate, Nelly Sélem-Mojica, Blanca Taboada, Gloria Vazquez, Joel Armando Vázquez-Pérez. |
| EPI_ISL_2681244 | Unidad de Investigación Médica de Yucatán (UIMY) | Instituto de Biotecnología de la UNAM | Consorcio Mexicano de Vigilancia Genómica (CoViGen-Mex). Authors (in alphabetical order): Julio Elias Alvarado-Yaah, Carlos F. Arias, Santiago Ávila-Ríos, Eduardo Becerril-Vargas, Víctor Hugo Borja-Aburto, Celia Boukadida, Cristóbal Cháidez-Quiróz, Juan Bautista Chale-Dzul, Ricardo Ciria Merce, Andreu Comas-García, Célida Duque Molina, Julissa Enciso-Ibarra, José Antonio Enciso-Moreno, Gloria Elena Espinosa-Ayala, Fernando Fontove-Herrera, Daniel Fregoso-Rueda, Victor Eduardo García-Arias, Alejandra García-Gasca, Bruno Gómez-Gil, Jean Pierre González, Irvin González-López, Concepción Grajales-Muñiz, Ricardo Grande, Rosa María Gutiérrez Rios, Jesús Hernández, Alejandra Hernández-Terán, Alfredo Herrera-Estrella, Carla Ivón Herrera-Najera, Pavel Isa, Verónica Mata-Haro, Daniel Lira Morales, Susana Lopez, Antonio Loza Román, Brenda Irasema Maldonado-Meza, Bernardo Martínez-Miguel, José Arturo Martínez-Orozco, Célida Martínez- Rodríguez, Margarita Matías-Florentino, Fidencio Mejía-Nepomuceno, María Guadalupe de Jesús Mireles-Rivera, Gloria María Molina-Salinas, Hector Montoya-Fuentes, , Mario Mújica-Sánchez, José Esteban Muñoz-Medina, José de Jesús Nuñez-Contreras, Alicia Ocaña-Mondragón, Luis Alberto Ochoa-Carrera, Hector Esteban Paz-Juárez, Marissa Perez-Garcia, Francisco Pulido, Helen Haydee Fernanda Ramírez-Plascencia, Jorge Salas-Hernández, Angel Gustavo Salas-Lais, Alejandro Sánchez-Flores, Clara Esperanza Santacruz-Tinoco, María Guadalupe Santiago-Mauricio, Selene Zárate, Nelly Sélem-Mojica, Blanca Taboada, Gloria Vazquez, Joel Armando Vázquez-Pérez. |

| | | | |
|---|---|---|---|
| EPI_ISL_2681245, EPI_ISL_2681247 | Laboratorio Central de Epidemiología (LCE) | Instituto de Biotecnología de la UNAM | Consorcio Mexicano de Vigilancia Genómica (CoViGen-Mex). Authors (in alphabetical order): Julio Elias Alvarado-Yaah, Carlos F. Arias, Santiago Ávila-Ríos, Eduardo Becerril-Vargas, Víctor Hugo Borja-Aburto, Celia Boukadida, Cristóbal Cháidez-Quiróz, Juan Bautista Chale-Dzul, Ricardo Ciria Merce, Andreu Comas-García, Célida Duque Molina, Julissa Enciso-Ibarra, José Antonio Enciso-Moreno, Gloria Elena Espinosa-Ayala, Fernando Fontove-Herrera, Daniel Fregoso-Rueda, Victor Eduardo García-Arias, Alejandra García-Gasca, Bruno Gómez-Gil, Jean Pierre González, Irvin González-López, Concepción Grajales-Muñiz, Ricardo Grande, Rosa María Gutiérrez Rios, Jesús Hernández, Alejandra Hernández-Terán, Alfredo Herrera-Estrella, Carla Ivón Herrera-Najera, Pavel Isa, Verónica Mata-Haro, Daniel Lira Morales, Susana Lopez, Antonio Loza Román, Brenda Irasema Maldonado-Meza, Bernardo Martínez-Miguel, José Arturo Martínez-Orozco, Célida Martínez- Rodríguez, Margarita Matías-Florentino, Fidencio Mejía-Nepomuceno, María Guadalupe de Jesús Mireles-Rivera, Gloria María Molina-Salinas, Hector Montoya-Fuentes, , Mario Mújica-Sánchez, José Esteban Muñoz-Medina, José de Jesús Nuñez-Contreras, Alicia Ocaña-Mondragón, Luis Alberto Ochoa-Carrera, Hector Esteban Paz-Juárez, Marissa Perez-Garcia, Francisco Pulido, Helen Haydee Fernanda Ramírez-Plascencia, Jorge Salas-Hernández, Angel Gustavo Salas-Lais, Alejandro Sánchez-Flores, Clara Esperanza Santacruz-Tinoco, María Guadalupe Santiago-Mauricio, Selene Zárate, Nelly Sélem-Mojica, Blanca Taboada, Gloria Vazquez, Joel Armando Vázquez-Pérez. |
| EPI_ISL_2681248 | Centro de Investigación Biomédica del Noreste (CIBIN) | Instituto de Biotecnología de la UNAM | Consorcio Mexicano de Vigilancia Genómica (CoViGen-Mex). Authors (in alphabetical order): Julio Elias Alvarado-Yaah, Carlos F. Arias, Santiago Ávila-Ríos, Eduardo Becerril-Vargas, Víctor Hugo Borja-Aburto, Celia Boukadida, Cristóbal Cháidez-Quiróz, Juan Bautista Chale-Dzul, Ricardo Ciria Merce, Andreu Comas-García, Célida Duque Molina, Julissa Enciso-Ibarra, José Antonio Enciso-Moreno, Gloria Elena Espinosa-Ayala, Fernando Fontove-Herrera, Daniel Fregoso-Rueda, Victor Eduardo García-Arias, Alejandra García-Gasca, Bruno Gómez-Gil, Jean Pierre González, Irvin González-López, Concepción Grajales-Muñiz, Ricardo Grande, Rosa María Gutiérrez Rios, Jesús Hernández, Alejandra Hernández-Terán, Alfredo Herrera-Estrella, Carla Ivón Herrera-Najera, Pavel Isa, Verónica Mata-Haro, Daniel Lira Morales, Susana Lopez, Antonio Loza Román, Brenda Irasema Maldonado-Meza, Bernardo Martínez-Miguel, José Arturo Martínez-Orozco, Célida Martínez- Rodríguez, Margarita Matías-Florentino, Fidencio Mejía-Nepomuceno, María Guadalupe de Jesús Mireles-Rivera, Gloria María Molina-Salinas, Hector Montoya-Fuentes, , Mario Mújica-Sánchez, José Esteban Muñoz-Medina, José de Jesús Nuñez-Contreras, Alicia Ocaña-Mondragón, Luis Alberto Ochoa-Carrera, Hector Esteban Paz-Juárez, Marissa Perez-Garcia, Francisco Pulido, Helen Haydee Fernanda Ramírez-Plascencia, Jorge Salas-Hernández, Angel Gustavo Salas-Lais, Alejandro Sánchez-Flores, Clara Esperanza Santacruz-Tinoco, María Guadalupe Santiago-Mauricio, Selene Zárate, Nelly Sélem-Mojica, Blanca Taboada, Gloria Vazquez, Joel Armando Vázquez-Pérez. |
| EPI_ISL_2681249, EPI_ISL_2681250, EPI_ISL_2681251, EPI_ISL_2681252, EPI_ISL_2681253, EPI_ISL_2681254, EPI_ISL_2681255, EPI_ISL_2681256, EPI_ISL_2681257, EPI_ISL_2681258, EPI_ISL_2681259 | | | |
| see above | Laboratorio Central de Epidemiología (LCE) | Instituto de Biotecnología de la UNAM | Consorcio Mexicano de Vigilancia Genómica (CoViGen-Mex). Authors (in alphabetical order): Julio Elias Alvarado-Yaah, Carlos F. Arias, Santiago Ávila-Ríos, Eduardo Becerril-Vargas, Víctor Hugo Borja-Aburto, Celia Boukadida, Cristóbal Cháidez-Quiróz, Juan Bautista Chale-Dzul, Ricardo Ciria Merce, Andreu Comas-García, Célida Duque Molina, Julissa Enciso-Ibarra, José Antonio Enciso-Moreno, Gloria Elena Espinosa-Ayala, Fernando Fontove-Herrera, Daniel Fregoso-Rueda, Victor Eduardo García-Arias, Alejandra García-Gasca, Bruno Gómez-Gil, Jean Pierre González, Irvin González-López, Concepción Grajales-Muñiz, Ricardo Grande, Rosa María Gutiérrez Rios, Jesús Hernández, Alejandra Hernández-Terán, Alfredo Herrera-Estrella, Carla Ivón Herrera-Najera, Pavel Isa, Verónica Mata-Haro, Daniel Lira Morales, Susana Lopez, Antonio Loza Román, Brenda Irasema Maldonado-Meza, Bernardo Martínez-Miguel, José Arturo Martínez-Orozco, Célida Martínez- Rodríguez, Margarita Matías-Florentino, Fidencio Mejía-Nepomuceno, María Guadalupe de Jesús Mireles-Rivera, Gloria María Molina-Salinas, Hector Montoya-Fuentes, , Mario Mújica-Sánchez, José Esteban Muñoz-Medina, José de Jesús Nuñez-Contreras, Alicia Ocaña-Mondragón, Luis Alberto Ochoa-Carrera, Hector Esteban Paz-Juárez, Marissa Perez-Garcia, Francisco Pulido, Helen Haydee Fernanda Ramírez-Plascencia, Jorge Salas-Hernández, Angel Gustavo Salas-Lais, Alejandro Sánchez-Flores, Clara Esperanza Santacruz-Tinoco, María Guadalupe Santiago-Mauricio, Selene Zárate, Nelly Sélem-Mojica, Blanca Taboada, Gloria Vazquez, Joel Armando Vázquez-Pérez. |
| EPI_ISL_2681261 | Unidad de Investigación Médica de Yucatán (UIMY) | Instituto de Biotecnología de la UNAM | Consorcio Mexicano de Vigilancia Genómica (CoViGen-Mex). Authors (in alphabetical order): Julio Elias Alvarado-Yaah, Carlos F. Arias, Santiago Ávila-Ríos, Eduardo Becerril-Vargas, Víctor Hugo Borja-Aburto, Celia Boukadida, Cristóbal Cháidez-Quiróz, Juan Bautista Chale-Dzul, Ricardo Ciria Merce, Andreu Comas-García, Célida Duque Molina, Julissa Enciso-Ibarra, José Antonio Enciso-Moreno, Gloria Elena Espinosa-Ayala, Fernando Fontove-Herrera, Daniel Fregoso-Rueda, Victor Eduardo García-Arias, Alejandra García-Gasca, Bruno Gómez-Gil, Jean Pierre González, Irvin González-López, Concepción Grajales-Muñiz, Ricardo Grande, Rosa María Gutiérrez Rios, Jesús Hernández, Alejandra Hernández-Terán, Alfredo Herrera-Estrella, Carla Ivón Herrera-Najera, Pavel Isa, Verónica Mata-Haro, Daniel Lira Morales, Susana Lopez, Antonio Loza Román, Brenda Irasema Maldonado-Meza, Bernardo Martínez-Miguel, José Arturo Martínez-Orozco, Célida Martínez- Rodríguez, Margarita Matías-Florentino, Fidencio Mejía-Nepomuceno, María Guadalupe de Jesús Mireles-Rivera, Gloria María Molina-Salinas, Hector Montoya-Fuentes, , Mario Mújica-Sánchez, José Esteban Muñoz-Medina, José de Jesús Nuñez-Contreras, Alicia Ocaña-Mondragón, Luis Alberto Ochoa-Carrera, Hector Esteban Paz-Juárez, Marissa Perez-Garcia, Francisco Pulido, Helen Haydee Fernanda Ramírez-Plascencia, Jorge Salas-Hernández, Angel Gustavo Salas-Lais, Alejandro Sánchez-Flores, Clara Esperanza Santacruz-Tinoco, María Guadalupe Santiago-Mauricio, Selene Zárate, Nelly Sélem-Mojica, Blanca Taboada, Gloria Vazquez, Joel Armando Vázquez-Pérez. |
| EPI_ISL_2681262, EPI_ISL_2681264 | Laboratorio Central de Epidemiología (LCE) | Instituto de Biotecnología de la UNAM | Consorcio Mexicano de Vigilancia Genómica (CoViGen-Mex). Authors (in alphabetical order): Julio Elias Alvarado-Yaah, Carlos F. Arias, Santiago Ávila-Ríos, Eduardo Becerril-Vargas, Víctor Hugo Borja-Aburto, Celia Boukadida, Cristóbal Cháidez-Quiróz, Juan Bautista Chale-Dzul, Ricardo Ciria Merce, Andreu Comas-García, Célida Duque Molina, Julissa Enciso-Ibarra, José Antonio Enciso-Moreno, Gloria Elena Espinosa-Ayala, Fernando Fontove-Herrera, Daniel Fregoso-Rueda, Victor Eduardo García-Arias, Alejandra García-Gasca, Bruno Gómez-Gil, Jean Pierre González, Irvin González-López, Concepción Grajales-Muñiz, Ricardo Grande, Rosa María Gutiérrez Rios, Jesús Hernández, Alejandra Hernández-Terán, Alfredo Herrera-Estrella, Carla Ivón Herrera-Najera, Pavel Isa, Verónica Mata-Haro, Daniel Lira Morales, Susana Lopez, Antonio Loza Román, Brenda Irasema Maldonado-Meza, Bernardo Martínez-Miguel, José Arturo Martínez-Orozco, Célida Martínez- Rodríguez, Margarita Matías-Florentino, Fidencio Mejía-Nepomuceno, María Guadalupe de Jesús Mireles-Rivera, Gloria María Molina-Salinas, Hector Montoya-Fuentes, , Mario Mújica-Sánchez, José Esteban Muñoz-Medina, José de Jesús Nuñez-Contreras, Alicia Ocaña-Mondragón, Luis Alberto Ochoa-Carrera, Hector Esteban Paz-Juárez, Marissa Perez-Garcia, Francisco Pulido, Helen Haydee Fernanda Ramírez-Plascencia, Jorge Salas-Hernández, Angel Gustavo Salas-Lais, Alejandro Sánchez-Flores, Clara Esperanza Santacruz-Tinoco, María Guadalupe Santiago-Mauricio, Selene Zárate, Nelly Sélem-Mojica, Blanca Taboada, Gloria Vazquez, Joel Armando Vázquez-Pérez. |
| EPI_ISL_2681265 | Centro de Investigación Biomédica del Noreste (CIBIN) | Instituto de Biotecnología de la UNAM | Consorcio Mexicano de Vigilancia Genómica (CoViGen-Mex). Authors (in alphabetical order): Julio Elias Alvarado-Yaah, Carlos F. Arias, Santiago Ávila-Ríos, Eduardo Becerril-Vargas, Víctor Hugo Borja-Aburto, Celia Boukadida, Cristóbal Cháidez-Quiróz, Juan Bautista Chale-Dzul, Ricardo Ciria Merce, Andreu Comas-García, Célida Duque Molina, Julissa Enciso-Ibarra, José Antonio Enciso-Moreno, Gloria Elena Espinosa-Ayala, Fernando Fontove-Herrera, Daniel Fregoso-Rueda, Victor Eduardo García-Arias, Alejandra García-Gasca, Bruno Gómez-Gil, Jean Pierre González, Irvin González-López, Concepción Grajales-Muñiz, Ricardo Grande, Rosa María Gutiérrez Rios, Jesús Hernández, Alejandra Hernández-Terán, Alfredo Herrera-Estrella, Carla Ivón Herrera-Najera, Pavel Isa, Verónica Mata-Haro, Daniel Lira Morales, Susana Lopez, Antonio Loza Román, Brenda Irasema Maldonado-Meza, Bernardo Martínez-Miguel, José Arturo Martínez-Orozco, Célida Martínez- Rodríguez, Margarita Matías-Florentino, Fidencio Mejía-Nepomuceno, María Guadalupe de Jesús Mireles-Rivera, Gloria María Molina-Salinas, Hector Montoya-Fuentes, , Mario Mújica-Sánchez, José Esteban Muñoz-Medina, José de Jesús Nuñez-Contreras, Alicia Ocaña-Mondragón, Luis Alberto Ochoa-Carrera, Hector Esteban Paz-Juárez, Marissa Perez-Garcia, Francisco Pulido, Helen Haydee Fernanda Ramírez-Plascencia, Jorge Salas-Hernández, Angel Gustavo Salas-Lais, Alejandro Sánchez-Flores, Clara Esperanza Santacruz-Tinoco, María Guadalupe Santiago-Mauricio, Selene Zárate, Nelly Sélem-Mojica, Blanca Taboada, Gloria Vazquez, Joel Armando Vázquez-Pérez. |
| EPI_ISL_2681267, EPI_ISL_2681268 | Laboratorio Central de Epidemiología (LCE) | Instituto de Biotecnología de la UNAM | Consorcio Mexicano de Vigilancia Genómica (CoViGen-Mex). Authors (in alphabetical order): Julio Elias Alvarado-Yaah, Carlos F. Arias, Santiago Ávila-Ríos, Eduardo Becerril-Vargas, Víctor Hugo Borja-Aburto, Celia Boukadida, Cristóbal Cháidez-Quiróz, Juan Bautista Chale-Dzul, Ricardo Ciria Merce, Andreu Comas-García, Célida Duque Molina, Julissa Enciso-Ibarra, José Antonio Enciso-Moreno, Gloria Elena Espinosa-Ayala, Fernando Fontove-Herrera, Daniel Fregoso-Rueda, Victor Eduardo García-Arias, Alejandra García-Gasca, Bruno Gómez-Gil, Jean Pierre González, Irvin González-López, Concepción Grajales-Muñiz, Ricardo Grande, Rosa María Gutiérrez Rios, Jesús Hernández, Alejandra Hernández-Terán, Alfredo Herrera-Estrella, Carla Ivón Herrera-Najera, Pavel Isa, Verónica Mata-Haro, Daniel Lira Morales, Susana Lopez, Antonio Loza Román, Brenda Irasema Maldonado-Meza, Bernardo Martínez-Miguel, José Arturo Martínez-Orozco, Célida Martínez- Rodríguez, Margarita Matías-Florentino, Fidencio Mejía-Nepomuceno, María Guadalupe de Jesús Mireles-Rivera, Gloria María Molina-Salinas, Hector Montoya-Fuentes, , Mario Mújica-Sánchez, José Esteban Muñoz-Medina, José de Jesús Nuñez-Contreras, Alicia Ocaña-Mondragón, Luis Alberto Ochoa-Carrera, Hector Esteban Paz-Juárez, Marissa |

| | | | |
|---|---|---|---|
| | | | Perez-Garcia, Francisco Pulido, Helen Haydee Fernanda Ramírez-Plascencia, Jorge Salas-Hernández, Angel Gustavo Salas-Lais, Alejandro Sánchez-Flores, Clara Esperanza Santacruz-Tinoco, María Guadalupe Santiago-Mauricio, Selene Zárate, Nelly Sélem-Mojica, Blanca Taboada, Gloria Vazquez, Joel Armando Vázquez-Pérez. |
| EPI_ISL_2681271 | Centro de Investigación Biomédica del Noreste (CIBIN) | Instituto de Biotecnología de la UNAM | Consorcio Mexicano de Vigilancia Genómica (CoViGen-Mex). Authors (in alphabetical order): Julio Elias Alvarado-Yaah, Carlos F. Arias, Santiago Ávila-Ríos, Eduardo Becerril-Vargas, Víctor Hugo Borja-Aburto, Celia Boukadida, Cristóbal Cháidez-Quiróz, Juan Bautista Chale-Dzul, Ricardo Ciria Merce, Andreu Comas-García, Célida Duque Molina, Julissa Enciso-Ibarra, José Antonio Enciso-Moreno, Gloria Elena Espinosa-Ayala, Fernando Fontove-Herrera, Daniel Fregoso-Rueda, Victor Eduardo García-Arias, Alejandra García-Gasca, Bruno Gómez-Gil, Jean Pierre González, Irvin González-López, Concepción Grajales-Muñiz, Ricardo Grande, Rosa María Gutiérrez Rios, Jesús Hernández, Alejandra Hernández-Terán, Alfredo Herrera-Estrella, Carla Ivón Herrera-Najera, Pavel Isa, Verónica Mata-Haro, Daniel Lira Morales, Susana Lopez, Antonio Loza Román, Brenda Irasema Maldonado-Meza, Bernardo Martínez-Miguel, José Arturo Martínez-Orozco, Célida Martínez- Rodríguez, Margarita Matías-Florentino, Fidencio Mejía-Nepomuceno, María Guadalupe de Jesús Mireles-Rivera, Gloria María Molina-Salinas, Hector Montoya-Fuentes, , Mario Mújica-Sánchez, José Esteban Muñoz-Medina, José de Jesús Nuñez-Contreras, Alicia Ocaña-Mondragón, Luis Alberto Ochoa-Carrera, Hector Esteban Paz-Juárez, Marissa Perez-Garcia, Francisco Pulido, Helen Haydee Fernanda Ramírez-Plascencia, Jorge Salas-Hernández, Angel Gustavo Salas-Lais, Alejandro Sánchez-Flores, Clara Esperanza Santacruz-Tinoco, María Guadalupe Santiago-Mauricio, Selene Zárate, Nelly Sélem-Mojica, Blanca Taboada, Gloria Vazquez, Joel Armando Vázquez-Pérez. |
| EPI_ISL_2681275 | Unidad de Investigación Médica de Yucatán (UIMY) | Instituto de Biotecnología de la UNAM | Consorcio Mexicano de Vigilancia Genómica (CoViGen-Mex). Authors (in alphabetical order): Julio Elias Alvarado-Yaah, Carlos F. Arias, Santiago Ávila-Ríos, Eduardo Becerril-Vargas, Víctor Hugo Borja-Aburto, Celia Boukadida, Cristóbal Cháidez-Quiróz, Juan Bautista Chale-Dzul, Ricardo Ciria Merce, Andreu Comas-García, Célida Duque Molina, Julissa Enciso-Ibarra, José Antonio Enciso-Moreno, Gloria Elena Espinosa-Ayala, Fernando Fontove-Herrera, Daniel Fregoso-Rueda, Victor Eduardo García-Arias, Alejandra García-Gasca, Bruno Gómez-Gil, Jean Pierre González, Irvin González-López, Concepción Grajales-Muñiz, Ricardo Grande, Rosa María Gutiérrez Rios, Jesús Hernández, Alejandra Hernández-Terán, Alfredo Herrera-Estrella, Carla Ivón Herrera-Najera, Pavel Isa, Verónica Mata-Haro, Daniel Lira Morales, Susana Lopez, Antonio Loza Román, Brenda Irasema Maldonado-Meza, Bernardo Martínez-Miguel, José Arturo Martínez-Orozco, Célida Martínez- Rodríguez, Margarita Matías-Florentino, Fidencio Mejía-Nepomuceno, María Guadalupe de Jesús Mireles-Rivera, Gloria María Molina-Salinas, Hector Montoya-Fuentes, , Mario Mújica-Sánchez, José Esteban Muñoz-Medina, José de Jesús Nuñez-Contreras, Alicia Ocaña-Mondragón, Luis Alberto Ochoa-Carrera, Hector Esteban Paz-Juárez, Marissa Perez-Garcia, Francisco Pulido, Helen Haydee Fernanda Ramírez-Plascencia, Jorge Salas-Hernández, Angel Gustavo Salas-Lais, Alejandro Sánchez-Flores, Clara Esperanza Santacruz-Tinoco, María Guadalupe Santiago-Mauricio, Selene Zárate, Nelly Sélem-Mojica, Blanca Taboada, Gloria Vazquez, Joel Armando Vázquez-Pérez. |
| EPI_ISL_2681276, EPI_ISL_2681278, EPI_ISL_2681279, EPI_ISL_2681281, EPI_ISL_2681282, EPI_ISL_2681283, EPI_ISL_2681284, EPI_ISL_2681288 | Laboratorio Central de Epidemiología (LCE) | Instituto de Biotecnología de la UNAM | Consorcio Mexicano de Vigilancia Genómica (CoViGen-Mex). Authors (in alphabetical order): Julio Elias Alvarado-Yaah, Carlos F. Arias, Santiago Ávila-Ríos, Eduardo Becerril-Vargas, Víctor Hugo Borja-Aburto, Celia Boukadida, Cristóbal Cháidez-Quiróz, Juan Bautista Chale-Dzul, Ricardo Ciria Merce, Andreu Comas-García, Célida Duque Molina, Julissa Enciso-Ibarra, José Antonio Enciso-Moreno, Gloria Elena Espinosa-Ayala, Fernando Fontove-Herrera, Daniel Fregoso-Rueda, Victor Eduardo García-Arias, Alejandra García-Gasca, Bruno Gómez-Gil, Jean Pierre González, Irvin González-López, Concepción Grajales-Muñiz, Ricardo Grande, Rosa María Gutiérrez Rios, Jesús Hernández, Alejandra Hernández-Terán, Alfredo Herrera-Estrella, Carla Ivón Herrera-Najera, Pavel Isa, Verónica Mata-Haro, Daniel Lira Morales, Susana Lopez, Antonio Loza Román, Brenda Irasema Maldonado-Meza, Bernardo Martínez-Miguel, José Arturo Martínez-Orozco, Célida Martínez- Rodríguez, Margarita Matías-Florentino, Fidencio Mejía-Nepomuceno, María Guadalupe de Jesús Mireles-Rivera, Gloria María Molina-Salinas, Hector Montoya-Fuentes, , Mario Mújica-Sánchez, José Esteban Muñoz-Medina, José de Jesús Nuñez-Contreras, Alicia Ocaña-Mondragón, Luis Alberto Ochoa-Carrera, Hector Esteban Paz-Juárez, Marissa Perez-Garcia, Francisco Pulido, Helen Haydee Fernanda Ramírez-Plascencia, Jorge Salas-Hernández, Angel Gustavo Salas-Lais, Alejandro Sánchez-Flores, Clara Esperanza Santacruz-Tinoco, María Guadalupe Santiago-Mauricio, Selene Zárate, Nelly Sélem-Mojica, Blanca Taboada, Gloria Vazquez, Joel Armando Vázquez-Pérez. |
| EPI_ISL_2681289 | Centro de Investigación Biomédica del Noreste (CIBIN) | Instituto de Biotecnología de la UNAM | Consorcio Mexicano de Vigilancia Genómica (CoViGen-Mex). Authors (in alphabetical order): Julio Elias Alvarado-Yaah, Carlos F. Arias, Santiago Ávila-Ríos, Eduardo Becerril-Vargas, Víctor Hugo Borja-Aburto, Celia Boukadida, Cristóbal Cháidez-Quiróz, Juan Bautista Chale-Dzul, Ricardo Ciria Merce, Andreu Comas-García, Célida Duque Molina, Julissa Enciso-Ibarra, José Antonio Enciso-Moreno, Gloria Elena Espinosa-Ayala, Fernando Fontove-Herrera, Daniel Fregoso-Rueda, Victor Eduardo García-Arias, Alejandra García-Gasca, Bruno Gómez-Gil, Jean Pierre González, Irvin González-López, Concepción Grajales-Muñiz, Ricardo Grande, Rosa María Gutiérrez Rios, Jesús Hernández, Alejandra Hernández-Terán, Alfredo Herrera-Estrella, Carla Ivón Herrera-Najera, Pavel Isa, Verónica Mata-Haro, Daniel Lira Morales, Susana Lopez, Antonio Loza Román, Brenda Irasema Maldonado-Meza, Bernardo Martínez-Miguel, José Arturo Martínez-Orozco, Célida Martínez- Rodríguez, Margarita Matías-Florentino, Fidencio Mejía-Nepomuceno, María Guadalupe de Jesús Mireles-Rivera, Gloria María Molina-Salinas, Hector Montoya-Fuentes, , Mario Mújica-Sánchez, José Esteban Muñoz-Medina, José de Jesús Nuñez-Contreras, Alicia Ocaña-Mondragón, Luis Alberto Ochoa-Carrera, Hector Esteban Paz-Juárez, Marissa Perez-Garcia, Francisco Pulido, Helen Haydee Fernanda Ramírez-Plascencia, Jorge Salas-Hernández, Angel Gustavo Salas-Lais, Alejandro Sánchez-Flores, Clara Esperanza Santacruz-Tinoco, María Guadalupe Santiago-Mauricio, Selene Zárate, Nelly Sélem-Mojica, Blanca Taboada, Gloria Vazquez, Joel Armando Vázquez-Pérez. |
| EPI_ISL_2681386, EPI_ISL_2681388 | Laboratorio Central de Epidemiología (LCE) | Instituto de Biotecnología de la UNAM | Consorcio Mexicano de Vigilancia Genómica (CoViGen-Mex). Authors (in alphabetical order): Julio Elias Alvarado-Yaah, Carlos F. Arias, Santiago Ávila-Ríos, Eduardo Becerril-Vargas, Víctor Hugo Borja-Aburto, Celia Boukadida, Cristóbal Cháidez-Quiróz, Juan Bautista Chale-Dzul, Ricardo Ciria Merce, Andreu Comas-García, Célida Duque Molina, Julissa Enciso-Ibarra, José Antonio Enciso-Moreno, Gloria Elena Espinosa-Ayala, Fernando Fontove-Herrera, Daniel Fregoso-Rueda, Victor Eduardo García-Arias, Alejandra García-Gasca, Bruno Gómez-Gil, Jean Pierre González, Irvin González-López, Concepción Grajales-Muñiz, Ricardo Grande, Rosa María Gutiérrez Rios, Jesús Hernández, Alejandra Hernández-Terán, Alfredo Herrera-Estrella, Carla Ivón Herrera-Najera, Pavel Isa, Verónica Mata-Haro, Daniel Lira Morales, Susana Lopez, Antonio Loza Román, Brenda Irasema Maldonado-Meza, Bernardo Martínez-Miguel, José Arturo Martínez-Orozco, Célida Martínez- Rodríguez, Margarita Matías-Florentino, Fidencio Mejía-Nepomuceno, María Guadalupe de Jesús Mireles-Rivera, Gloria María Molina-Salinas, Hector Montoya-Fuentes, , Mario Mújica-Sánchez, José Esteban Muñoz-Medina, José de Jesús Nuñez-Contreras, Alicia Ocaña-Mondragón, Luis Alberto Ochoa-Carrera, Hector Esteban Paz-Juárez, Marissa Perez-Garcia, Francisco Pulido, Helen Haydee Fernanda Ramírez-Plascencia, Jorge Salas-Hernández, Angel Gustavo Salas-Lais, Alejandro Sánchez-Flores, Clara Esperanza Santacruz-Tinoco, María Guadalupe Santiago-Mauricio, Selene Zárate, Nelly Sélem-Mojica, Blanca Taboada, Gloria Vazquez, Joel Armando Vázquez-Pérez. |
| EPI_ISL_2681441, EPI_ISL_2681442, EPI_ISL_2681443, EPI_ISL_2681444, EPI_ISL_2681445, EPI_ISL_2681446, EPI_ISL_2681447, EPI_ISL_2681448, EPI_ISL_2681449 | Althaia. Xarxa Assistencial Universitària de Manresa | IrsiCaixa | Marc Noguera-Julian, Mariona Parera, Maria Casadellà, Pilar Armengol, Francesc Catala-Moll, Roger Paredes, Bonaventura Clotet Gloria Trujillo, Rafel Perez Vidal, Jaume Trapé, Carolina Gonzalez-Fernandez, Miquel Micó, Antonia Flor, Roger Paredes, Eulalia Grau, Bonaventura Clotet |
| EPI_ISL_2683247, EPI_ISL_2683496 | Sultanah Bahiyah Hospital, Alor Setar | Institute for Medical Research, Infectious Disease Research Centre, National Institutes of Health, Ministry of Health Malaysia | Suppiah J, Kamel K, Mohd Zawawi Z, Azizan MA, Ramly N, Robert F, Thayan R |
| EPI_ISL_2683498 | Ipoh Public Health Laboratory (MKAI), Ministry of Health Malaysia | Institute for Medical Research, Infectious Disease Research Centre, National Institutes of Health, Ministry of Health Malaysia | Suppiah J, Kamel K, Mohd Zawawi Z, Azizan MA, Ramly N, Robert F, Thayan R |
| EPI_ISL_2684231, EPI_ISL_2684232, EPI_ISL_2684233 | Institute for Medical Research, Infectious Disease Research Centre, National Institutes of Health, Ministry of Health Malaysia | Institute for Medical Research, Infectious Disease Research Centre, National Institutes of Health, Ministry of Health Malaysia | Suppiah J, Kamel K, Mohd Zawawi Z, Azizan MA, Ramly N, Robert F, Thayan R |
| EPI_ISL_2684234, EPI_ISL_2684235 | Melaka Hospital | Institute for Medical Research, Infectious Disease Research Centre, National Institutes of Health, Ministry of Health Malaysia | Suppiah J, Kamel K, Mohd Zawawi Z, Azizan MA, Ramly N, Robert F, Thayan R |
| EPI_ISL_2684236 | Sultanah Bahiyah Hospital, Alor Setar | Institute for Medical Research, Infectious Disease Research Centre, National Institutes of Health, Ministry of Health | Suppiah J, Kamel K, Mohd Zawawi Z, Azizan MA, Ramly N, Robert F, Thayan R |

| | | | |
|---|---|---|---|
| | | Malaysia | |
| EPI_ISL_2684545, EPI_ISL_2685359, EPI_ISL_2685766, EPI_ISL_2685835, EPI_ISL_2685836, EPI_ISL_2685837, EPI_ISL_2685838, EPI_ISL_2685839, EPI_ISL_2685840, EPI_ISL_2685841, EPI_ISL_2685843, EPI_ISL_2685844, EPI_ISL_2685845 | | | |
| see above | Institute for Medical Research, Infectious Disease Research Centre, National Institutes of Health, Ministry of Health Malaysia | Institute for Medical Research, Infectious Disease Research Centre, National Institutes of Health, Ministry of Health Malaysia | Suppiah J, Kamel K, Mohd Zawawi Z, Azizan MA, Ramly N, Robert F, Thayan R |
| EPI_ISL_2686028, EPI_ISL_2686029, EPI_ISL_2686030, EPI_ISL_2686031, EPI_ISL_2686032, EPI_ISL_2686033, EPI_ISL_2686034, EPI_ISL_2686035, EPI_ISL_2686036, EPI_ISL_2686037, EPI_ISL_2686038, EPI_ISL_2686039, EPI_ISL_2686040, EPI_ISL_2686041, EPI_ISL_2686042, EPI_ISL_2686043, EPI_ISL_2686044, EPI_ISL_2686045, EPI_ISL_2686046, EPI_ISL_2686047, EPI_ISL_2686048, EPI_ISL_2686049, EPI_ISL_2686050, EPI_ISL_2686051, EPI_ISL_2686052, EPI_ISL_2686053 | | | |
| see above | Università degli Studi di Perugia | Istituto Zooprofilattico Sperimentale dell'Abruzzo e Molise "G. Caporale" | Mencacci A, Camilloni B, Biagetti M, Giammarioli M, Lorusso A, Marcacci M, Di Domenico M, Ancora M, Curini V, Mangone I, Rinaldi A, Delli Compagni E, Scialabba S, Di Pasquale A, Cammà C, Puglia I, Calistri P, Savini G |
| EPI_ISL_2686054, EPI_ISL_2686055, EPI_ISL_2686057, EPI_ISL_2686059 | Laboratorio Analisi Osp. Città di Castello - Azienda USL Umbria1 | Istituto Zooprofilattico Sperimentale dell'Abruzzo e Molise "G. Caporale" | Malagigi V, Taccani P,Lorusso A, Marcacci M, Di Domenico M, Ancora M, Curini V, Mangone I, Rinaldi A, Delli Compagni E, Scialabba S, Di Pasquale A, Cammà C, Puglia I, Calistri P, Savini G |
| EPI_ISL_2686060, EPI_ISL_2686061 | Università degli Studi di Perugia | Istituto Zooprofilattico Sperimentale dell'Abruzzo e Molise "G. Caporale" | Mencacci A, Camilloni B, Biagetti M, Giammarioli M, Lorusso A, Marcacci M, Di Domenico M, Ancora M, Curini V, Mangone I, Rinaldi A, Delli Compagni E, Scialabba S, Di Pasquale A, Cammà C, Puglia I, Calistri P, Savini G |
| EPI_ISL_2686062 | Azienda Ospedaliera Terni | Istituto Zooprofilattico Sperimentale dell'Abruzzo e Molise "G. Caporale" | Palumbo M, Scaccetti A, Lorusso A, Marcacci M, Di Domenico M, Ancora M, Curini V, Mangone I, Rinaldi A, Delli Compagni E, Scialabba S, Di Pasquale A, Cammà C, Puglia I, Calistri P, Savini G |
| EPI_ISL_2686063, EPI_ISL_2686064, EPI_ISL_2686065 | Azienda USL Umbria 2 | Istituto Zooprofilattico Sperimentale dell'Abruzzo e Molise "G. Caporale" | Proietti A, Pistoni E, Lorusso A, Marcacci M, Di Domenico M, Ancora M, Curini V, Mangone I, Rinaldi A, Delli Compagni E, Scialabba S, Di Pasquale A, Cammà C, Puglia I, Calistri P, Savini G |
| EPI_ISL_2686066, EPI_ISL_2686067, EPI_ISL_2686068, EPI_ISL_2686069 | Università degli Studi di Perugia | Istituto Zooprofilattico Sperimentale dell'Abruzzo e Molise "G. Caporale" | Mencacci A, Camilloni B, Biagetti M, Giammarioli M, Lorusso A, Marcacci M, Di Domenico M, Ancora M, Curini V, Mangone I, Rinaldi A, Delli Compagni E, Scialabba S, Di Pasquale A, Cammà C, Puglia I, Calistri P, Savini G |
| EPI_ISL_2686070, EPI_ISL_2686071, EPI_ISL_2686072, EPI_ISL_2686073 | Laboratorio Analisi Osp. Città di Castello - Azienda USL Umbria1 | Istituto Zooprofilattico Sperimentale dell'Abruzzo e Molise "G. Caporale" | Malagigi V, Taccani P,Lorusso A, Marcacci M, Di Domenico M, Ancora M, Curini V, Mangone I, Rinaldi A, Delli Compagni E, Scialabba S, Di Pasquale A, Cammà C, Puglia I, Calistri P, Savini G |
| EPI_ISL_2686074, EPI_ISL_2686075, EPI_ISL_2686076, EPI_ISL_2686077, EPI_ISL_2686078, EPI_ISL_2686079, EPI_ISL_2686080 | Azienda USL Umbria 2 | Istituto Zooprofilattico Sperimentale dell'Abruzzo e Molise "G. Caporale" | Proietti A, Pistoni E, Lorusso A, Marcacci M, Di Domenico M, Ancora M, Curini V, Mangone I, Rinaldi A, Delli Compagni E, Scialabba S, Di Pasquale A, Cammà C, Puglia I, Calistri P, Savini G |
| EPI_ISL_2686081, EPI_ISL_2686082, EPI_ISL_2686083, EPI_ISL_2686084 | Azienda Ospedaliera Terni | Istituto Zooprofilattico Sperimentale dell'Abruzzo e Molise "G. Caporale" | Palumbo M, Scaccetti A, Lorusso A, Marcacci M, Di Domenico M, Ancora M, Curini V, Mangone I, Rinaldi A, Delli Compagni E, Scialabba S, Di Pasquale A, Cammà C, Puglia I, Calistri P, Savini G |
| EPI_ISL_2686390 | AZ Zeno | AZ Sint Jan | Marijke Reynders, Patrick Descheemaeker, Laurien Hoornaert, Thomas Van Landschoot, Marie Madeleine Chabert-Consen, Jorn Hellemans |
| EPI_ISL_2687705, EPI_ISL_2687765, EPI_ISL_2687787 | Public Health Authority of the Slovak Republic | Public Health Authority of the Slovak Republic | Barbora Kotvasová, Lucia Ševíková, Terézia Vrabová, Anna Giová, Elena Tichá, Miroslav Böhmer, Tomáš Szemes, Pavol Mišenko |
| EPI_ISL_2687993 | National Institute of Public Health | National Institute of Infectious Diseases-Prof. Dr. Matei Bals Molecular Diagnostics Laboratory | Corina Casangiu, Leontina Banica, Marius Surleac, Petre Milu, Robert Hohan, Simona Paraschiv, Dan Otelea |
| EPI_ISL_2687995, EPI_ISL_2687996, EPI_ISL_2688000 | National Institute of Infectious Diseases-Prof. Dr. Matei Bals Molecular Diagnostics Laboratory | National Institute of Infectious Diseases-Prof. Dr. Matei Bals Molecular Diagnostics Laboratory | Corina Casangiu, Leontina Banica, Marius Surleac, Petre Milu, Robert Hohan, Simona Paraschiv, Dan Otelea |
| EPI_ISL_2688259, EPI_ISL_2688260, EPI_ISL_2688271, EPI_ISL_2688274 | Centers for Disease Control and Prevention, Dengue Branch | Centers for Disease Control and Prevention, Dengue Branch | Gilberto A. Santiago, Glenda Gonzalez, Betzabel Flores, Keyla Charriez, Gabriela Paz-Bailey, Jorge L. Munoz-Jordan |
| EPI_ISL_2688304, EPI_ISL_2688305, EPI_ISL_2688306, EPI_ISL_2688307, EPI_ISL_2688308, EPI_ISL_2688309, EPI_ISL_2688310, EPI_ISL_2688311, EPI_ISL_2688312, EPI_ISL_2688313, EPI_ISL_2688314, EPI_ISL_2688315, EPI_ISL_2688316, EPI_ISL_2688317, EPI_ISL_2688318, EPI_ISL_2688319 | | | |
| see above | Puerto Rico Department of Health | Centers for Disease Control and Prevention, Dengue Branch | Gilberto A. Santiago, Glenda Gonzalez, Betzabel Flores, Keyla Charriez, Gabriela Paz-Bailey, Jorge L. Munoz-Jordan |
| EPI_ISL_2688911 | LESP Ciudad de Mexico | Instituto de Diagnostico y Referencia Epidemiologicos (INDRE) | Claudia Wong-Arambula, Abril Rodriguez-Maldonado, Vanessa Rivero-Arredondo, Ariadna Medina-Benitez, Joaquin Quiroz-Mercado, Sergio Rangel-Guerrero, Natividad Cruz-Ortiz, Tatiana Nunez-Garcia, Gisela Barrera-Badillo, Lucia Hernandez-Rivas, Irma Lopez-Martinez, Ernesto Ramirez-Gonzalez. |
| EPI_ISL_2688912 | LESP Nuevo Leon | Instituto de Diagnostico y Referencia Epidemiologicos (INDRE) | Claudia Wong-Arambula, Abril Rodriguez-Maldonado, Vanessa Rivero-Arredondo, Ariadna Medina-Benitez, Joaquin Quiroz-Mercado, Sergio Rangel-Guerrero, Natividad Cruz-Ortiz, Tatiana Nunez-Garcia, Gisela Barrera-Badillo, Lucia Hernandez-Rivas, Irma Lopez-Martinez, Ernesto Ramirez-Gonzalez. |
| EPI_ISL_2688913, EPI_ISL_2688914 | LESP Jalisco | Instituto de Diagnostico y Referencia Epidemiologicos (INDRE) | Claudia Wong-Arambula, Abril Rodriguez-Maldonado, Vanessa Rivero-Arredondo, Ariadna Medina-Benitez, Joaquin Quiroz-Mercado, Sergio Rangel-Guerrero, Natividad Cruz-Ortiz, Tatiana Nunez-Garcia, Gisela Barrera-Badillo, Lucia Hernandez-Rivas, Irma Lopez-Martinez, Ernesto Ramirez-Gonzalez. |
| EPI_ISL_2688915, EPI_ISL_2688916, EPI_ISL_2688917, EPI_ISL_2688918 | LESP Morelos | Instituto de Diagnostico y Referencia Epidemiologicos (INDRE) | Claudia Wong-Arambula, Abril Rodriguez-Maldonado, Vanessa Rivero-Arredondo, Ariadna Medina-Benitez, Joaquin Quiroz-Mercado, Sergio Rangel-Guerrero, Natividad Cruz-Ortiz, Tatiana Nunez-Garcia, Gisela Barrera-Badillo, Lucia Hernandez-Rivas, Irma Lopez-Martinez, Ernesto Ramirez-Gonzalez. |
| EPI_ISL_2688919, EPI_ISL_2688921, EPI_ISL_2688922, EPI_ISL_2688923 | LESP Ciudad de Mexico | Instituto de Diagnostico y Referencia Epidemiologicos (INDRE) | Claudia Wong-Arambula, Abril Rodriguez-Maldonado, Vanessa Rivero-Arredondo, Ariadna Medina-Benitez, Joaquin Quiroz-Mercado, Sergio Rangel-Guerrero, Natividad Cruz-Ortiz, Tatiana Nunez-Garcia, Gisela Barrera-Badillo, Lucia Hernandez-Rivas, Irma Lopez-Martinez, Ernesto Ramirez-Gonzalez. |
| EPI_ISL_2688924, EPI_ISL_2688925 | LESP Queretaro | Instituto de Diagnostico y Referencia Epidemiologicos (INDRE) | Claudia Wong-Arambula, Abril Rodriguez-Maldonado, Vanessa Rivero-Arredondo, Ariadna Medina-Benitez, Joaquin Quiroz-Mercado, Sergio Rangel-Guerrero, Natividad Cruz-Ortiz, Tatiana Nunez-Garcia, Gisela Barrera-Badillo, Lucia Hernandez-Rivas, Irma Lopez-Martinez, Ernesto Ramirez-Gonzalez. |
| EPI_ISL_2688926 | LESP Ciudad de Mexico | Instituto de Diagnostico y Referencia Epidemiologicos (INDRE) | Claudia Wong-Arambula, Abril Rodriguez-Maldonado, Vanessa Rivero-Arredondo, Ariadna Medina-Benitez, Joaquin Quiroz-Mercado, Sergio Rangel-Guerrero, Natividad Cruz-Ortiz, Tatiana Nunez-Garcia, Gisela Barrera-Badillo, Lucia Hernandez-Rivas, Irma Lopez-Martinez, Ernesto Ramirez-Gonzalez. |
| EPI_ISL_2688927, EPI_ISL_2688928, EPI_ISL_2688929, EPI_ISL_2688930, EPI_ISL_2688931, EPI_ISL_2688932, EPI_ISL_2688933, EPI_ISL_2688934, EPI_ISL_2688935, EPI_ISL_2688936, EPI_ISL_2688937, EPI_ISL_2688938, EPI_ISL_2688939, EPI_ISL_2688940, EPI_ISL_2688941, EPI_ISL_2688942 | | | |
| see above | LESP Tabasco | Instituto de Diagnostico y Referencia Epidemiologicos (INDRE) | Claudia Wong-Arambula, Abril Rodriguez-Maldonado, Vanessa Rivero-Arredondo, Ariadna Medina-Benitez, Joaquin Quiroz-Mercado, Sergio Rangel-Guerrero, Natividad Cruz-Ortiz, Tatiana Nunez-Garcia, Gisela Barrera-Badillo, Lucia Hernandez-Rivas, Irma Lopez-Martinez, Ernesto Ramirez-Gonzalez. |
| EPI_ISL_2688943, EPI_ISL_2688944, EPI_ISL_2688945, EPI_ISL_2688946, EPI_ISL_2688947 | LESP Michoacan | Instituto de Diagnostico y Referencia Epidemiologicos (INDRE) | Claudia Wong-Arambula, Abril Rodriguez-Maldonado, Vanessa Rivero-Arredondo, Ariadna Medina-Benitez, Joaquin Quiroz-Mercado, Sergio Rangel-Guerrero, Natividad Cruz-Ortiz, Tatiana Nunez-Garcia, Gisela Barrera-Badillo, Lucia Hernandez-Rivas, Irma Lopez-Martinez, Ernesto Ramirez-Gonzalez. |
| EPI_ISL_2688948, EPI_ISL_2688949, EPI_ISL_2688950, EPI_ISL_2688951, EPI_ISL_2688952, EPI_ISL_2688953, EPI_ISL_2688954, EPI_ISL_2688955, EPI_ISL_2688956, EPI_ISL_2688958 | LESP Tabasco | Instituto de Diagnostico y Referencia Epidemiologicos (INDRE) | Claudia Wong-Arambula, Abril Rodriguez-Maldonado, Vanessa Rivero-Arredondo, Ariadna Medina-Benitez, Joaquin Quiroz-Mercado, Sergio Rangel-Guerrero, Natividad Cruz-Ortiz, Tatiana Nunez-Garcia, Gisela Barrera-Badillo, Lucia Hernandez-Rivas, Irma Lopez-Martinez, Ernesto Ramirez-Gonzalez. |
| EPI_ISL_2688959 | LESP Estado de Mexico | Instituto de Diagnostico y Referencia Epidemiologicos | Claudia Wong-Arambula, Abril Rodriguez-Maldonado, Vanessa Rivero-Arredondo, Ariadna Medina-Benitez, Joaquin Quiroz-Mercado, Sergio |

| | | | |
|---|---|---|---|
| | | (INDRE) | Rangel-Guerrero, Natividad Cruz-Ortiz, Tatiana Nunez-Garcia, Gisela Barrera-Badillo, Lucia Hernandez-Rivas, Irma Lopez-Martinez, Ernesto Ramirez-Gonzalez. |
| EPI_ISL_2688960, EPI_ISL_2688962, EPI_ISL_2688963, EPI_ISL_2688964 | LESP Ciudad de Mexico | Instituto de Diagnostico y Referencia Epidemiologicos (INDRE) | Claudia Wong-Arambula, Abril Rodriguez-Maldonado, Vanessa Rivero-Arredondo, Ariadna Medina-Benitez, Joaquin Quiroz-Mercado, Sergio Rangel-Guerrero, Natividad Cruz-Ortiz, Tatiana Nunez-Garcia, Gisela Barrera-Badillo, Lucia Hernandez-Rivas, Irma Lopez-Martinez, Ernesto Ramirez-Gonzalez. |
| EPI_ISL_2688965 | LESP Estado de Mexico | Instituto de Diagnostico y Referencia Epidemiologicos (INDRE) | Claudia Wong-Arambula, Abril Rodriguez-Maldonado, Vanessa Rivero-Arredondo, Ariadna Medina-Benitez, Joaquin Quiroz-Mercado, Sergio Rangel-Guerrero, Natividad Cruz-Ortiz, Tatiana Nunez-Garcia, Gisela Barrera-Badillo, Lucia Hernandez-Rivas, Irma Lopez-Martinez, Ernesto Ramirez-Gonzalez. |
| EPI_ISL_2688966, EPI_ISL_2688967, EPI_ISL_2688968 | LESP Ciudad de Mexico | Instituto de Diagnostico y Referencia Epidemiologicos (INDRE) | Claudia Wong-Arambula, Abril Rodriguez-Maldonado, Vanessa Rivero-Arredondo, Ariadna Medina-Benitez, Joaquin Quiroz-Mercado, Sergio Rangel-Guerrero, Natividad Cruz-Ortiz, Tatiana Nunez-Garcia, Gisela Barrera-Badillo, Lucia Hernandez-Rivas, Irma Lopez-Martinez, Ernesto Ramirez-Gonzalez. |
| EPI_ISL_2688969 | LESP Queretaro | Instituto de Diagnostico y Referencia Epidemiologicos (INDRE) | Claudia Wong-Arambula, Abril Rodriguez-Maldonado, Vanessa Rivero-Arredondo, Ariadna Medina-Benitez, Joaquin Quiroz-Mercado, Sergio Rangel-Guerrero, Natividad Cruz-Ortiz, Tatiana Nunez-Garcia, Gisela Barrera-Badillo, Lucia Hernandez-Rivas, Irma Lopez-Martinez, Ernesto Ramirez-Gonzalez. |
| EPI_ISL_2688970 | LESP Michoacan | Instituto de Diagnostico y Referencia Epidemiologicos (INDRE) | Claudia Wong-Arambula, Abril Rodriguez-Maldonado, Vanessa Rivero-Arredondo, Ariadna Medina-Benitez, Joaquin Quiroz-Mercado, Sergio Rangel-Guerrero, Natividad Cruz-Ortiz, Tatiana Nunez-Garcia, Gisela Barrera-Badillo, Lucia Hernandez-Rivas, Irma Lopez-Martinez, Ernesto Ramirez-Gonzalez. |
| EPI_ISL_2688971 | LESP Zacatecas | Instituto de Diagnostico y Referencia Epidemiologicos (INDRE) | Claudia Wong-Arambula, Abril Rodriguez-Maldonado, Vanessa Rivero-Arredondo, Ariadna Medina-Benitez, Joaquin Quiroz-Mercado, Sergio Rangel-Guerrero, Natividad Cruz-Ortiz, Tatiana Nunez-Garcia, Gisela Barrera-Badillo, Lucia Hernandez-Rivas, Irma Lopez-Martinez, Ernesto Ramirez-Gonzalez. |
| EPI_ISL_2688972 | LESP Hidalgo | Instituto de Diagnostico y Referencia Epidemiologicos (INDRE) | Claudia Wong-Arambula, Abril Rodriguez-Maldonado, Vanessa Rivero-Arredondo, Ariadna Medina-Benitez, Joaquin Quiroz-Mercado, Sergio Rangel-Guerrero, Natividad Cruz-Ortiz, Tatiana Nunez-Garcia, Gisela Barrera-Badillo, Lucia Hernandez-Rivas, Irma Lopez-Martinez, Ernesto Ramirez-Gonzalez. |
| EPI_ISL_2688973 | LESP Morelos | Instituto de Diagnostico y Referencia Epidemiologicos (INDRE) | Claudia Wong-Arambula, Abril Rodriguez-Maldonado, Vanessa Rivero-Arredondo, Ariadna Medina-Benitez, Joaquin Quiroz-Mercado, Sergio Rangel-Guerrero, Natividad Cruz-Ortiz, Tatiana Nunez-Garcia, Gisela Barrera-Badillo, Lucia Hernandez-Rivas, Irma Lopez-Martinez, Ernesto Ramirez-Gonzalez. |
| EPI_ISL_2688974 | LESP Guanajuato | Instituto de Diagnostico y Referencia Epidemiologicos (INDRE) | Claudia Wong-Arambula, Abril Rodriguez-Maldonado, Vanessa Rivero-Arredondo, Ariadna Medina-Benitez, Joaquin Quiroz-Mercado, Sergio Rangel-Guerrero, Natividad Cruz-Ortiz, Tatiana Nunez-Garcia, Gisela Barrera-Badillo, Lucia Hernandez-Rivas, Irma Lopez-Martinez, Ernesto Ramirez-Gonzalez. |
| EPI_ISL_2688975 | LESP Jalisco | Instituto de Diagnostico y Referencia Epidemiologicos (INDRE) | Claudia Wong-Arambula, Abril Rodriguez-Maldonado, Vanessa Rivero-Arredondo, Ariadna Medina-Benitez, Joaquin Quiroz-Mercado, Sergio Rangel-Guerrero, Natividad Cruz-Ortiz, Tatiana Nunez-Garcia, Gisela Barrera-Badillo, Lucia Hernandez-Rivas, Irma Lopez-Martinez, Ernesto Ramirez-Gonzalez. |
| EPI_ISL_2688976, EPI_ISL_2688977, EPI_ISL_2688978, EPI_ISL_2688979 | LESP Guanajuato | Instituto de Diagnostico y Referencia Epidemiologicos (INDRE) | Claudia Wong-Arambula, Abril Rodriguez-Maldonado, Vanessa Rivero-Arredondo, Ariadna Medina-Benitez, Joaquin Quiroz-Mercado, Sergio Rangel-Guerrero, Natividad Cruz-Ortiz, Tatiana Nunez-Garcia, Gisela Barrera-Badillo, Lucia Hernandez-Rivas, Irma Lopez-Martinez, Ernesto Ramirez-Gonzalez. |
| EPI_ISL_2688980, EPI_ISL_2688981, EPI_ISL_2688982, EPI_ISL_2688983, EPI_ISL_2688984 | LESP Chihuahua | Instituto de Diagnostico y Referencia Epidemiologicos (INDRE) | Claudia Wong-Arambula, Abril Rodriguez-Maldonado, Vanessa Rivero-Arredondo, Ariadna Medina-Benitez, Joaquin Quiroz-Mercado, Sergio Rangel-Guerrero, Natividad Cruz-Ortiz, Tatiana Nunez-Garcia, Gisela Barrera-Badillo, Lucia Hernandez-Rivas, Irma Lopez-Martinez, Ernesto Ramirez-Gonzalez. |
| EPI_ISL_2688985 | LESP Coahuila | Instituto de Diagnostico y Referencia Epidemiologicos (INDRE) | Claudia Wong-Arambula, Abril Rodriguez-Maldonado, Vanessa Rivero-Arredondo, Ariadna Medina-Benitez, Joaquin Quiroz-Mercado, Sergio Rangel-Guerrero, Natividad Cruz-Ortiz, Tatiana Nunez-Garcia, Gisela Barrera-Badillo, Lucia Hernandez-Rivas, Irma Lopez-Martinez, Ernesto Ramirez-Gonzalez. |
| EPI_ISL_2688986, EPI_ISL_2688987 | LESP Puebla | Instituto de Diagnostico y Referencia Epidemiologicos (INDRE) | Claudia Wong-Arambula, Abril Rodriguez-Maldonado, Vanessa Rivero-Arredondo, Ariadna Medina-Benitez, Joaquin Quiroz-Mercado, Sergio Rangel-Guerrero, Natividad Cruz-Ortiz, Tatiana Nunez-Garcia, Gisela Barrera-Badillo, Lucia Hernandez-Rivas, Irma Lopez-Martinez, Ernesto Ramirez-Gonzalez. |
| EPI_ISL_2688988, EPI_ISL_2688989 | LESP Veracruz | Instituto de Diagnostico y Referencia Epidemiologicos (INDRE) | Claudia Wong-Arambula, Abril Rodriguez-Maldonado, Vanessa Rivero-Arredondo, Ariadna Medina-Benitez, Joaquin Quiroz-Mercado, Sergio Rangel-Guerrero, Natividad Cruz-Ortiz, Tatiana Nunez-Garcia, Gisela Barrera-Badillo, Lucia Hernandez-Rivas, Irma Lopez-Martinez, Ernesto Ramirez-Gonzalez. |
| EPI_ISL_2688990 | LESP Yucatan | Instituto de Diagnostico y Referencia Epidemiologicos (INDRE) | Claudia Wong-Arambula, Abril Rodriguez-Maldonado, Vanessa Rivero-Arredondo, Ariadna Medina-Benitez, Joaquin Quiroz-Mercado, Sergio Rangel-Guerrero, Natividad Cruz-Ortiz, Tatiana Nunez-Garcia, Gisela Barrera-Badillo, Lucia Hernandez-Rivas, Irma Lopez-Martinez, Ernesto Ramirez-Gonzalez. |
| EPI_ISL_2689794 | LESP Sonora | Instituto de Diagnostico y Referencia Epidemiologicos (INDRE) | Claudia Wong-Arambula, Abril Rodriguez-Maldonado, Vanessa Rivero-Arredondo, Ariadna Medina-Benitez, Joaquin Quiroz-Mercado, Sergio Rangel-Guerrero, Natividad Cruz-Ortiz, Tatiana Nunez-Garcia, Gisela Barrera-Badillo, Lucia Hernandez-Rivas, Irma Lopez-Martinez, Ernesto Ramirez-Gonzalez. |
| EPI_ISL_2689796, EPI_ISL_2689798, EPI_ISL_2689801, EPI_ISL_2689802, EPI_ISL_2689803, EPI_ISL_2689804 | LESP Ciudad de Mexico | Instituto de Diagnostico y Referencia Epidemiologicos (INDRE) | Claudia Wong-Arambula, Abril Rodriguez-Maldonado, Vanessa Rivero-Arredondo, Ariadna Medina-Benitez, Joaquin Quiroz-Mercado, Sergio Rangel-Guerrero, Natividad Cruz-Ortiz, Tatiana Nunez-Garcia, Gisela Barrera-Badillo, Lucia Hernandez-Rivas, Irma Lopez-Martinez, Ernesto Ramirez-Gonzalez. |
| EPI_ISL_2689806 | LESP Sinaloa | Instituto de Diagnostico y Referencia Epidemiologicos (INDRE) | Claudia Wong-Arambula, Abril Rodriguez-Maldonado, Vanessa Rivero-Arredondo, Ariadna Medina-Benitez, Joaquin Quiroz-Mercado, Sergio Rangel-Guerrero, Natividad Cruz-Ortiz, Tatiana Nunez-Garcia, Gisela Barrera-Badillo, Lucia Hernandez-Rivas, Irma Lopez-Martinez, Ernesto Ramirez-Gonzalez. |
| EPI_ISL_2689808 | LESP Tabasco | Instituto de Diagnostico y Referencia Epidemiologicos (INDRE) | Claudia Wong-Arambula, Abril Rodriguez-Maldonado, Vanessa Rivero-Arredondo, Ariadna Medina-Benitez, Joaquin Quiroz-Mercado, Sergio Rangel-Guerrero, Natividad Cruz-Ortiz, Tatiana Nunez-Garcia, Gisela Barrera-Badillo, Lucia Hernandez-Rivas, Irma Lopez-Martinez, Ernesto Ramirez-Gonzalez. |
| EPI_ISL_2689812 | LESP Michoacan | Instituto de Diagnostico y Referencia Epidemiologicos (INDRE) | Claudia Wong-Arambula, Abril Rodriguez-Maldonado, Vanessa Rivero-Arredondo, Ariadna Medina-Benitez, Joaquin Quiroz-Mercado, Sergio Rangel-Guerrero, Natividad Cruz-Ortiz, Tatiana Nunez-Garcia, Gisela Barrera-Badillo, Lucia Hernandez-Rivas, Irma Lopez-Martinez, Ernesto Ramirez-Gonzalez. |
| EPI_ISL_2689818, EPI_ISL_2689819, EPI_ISL_2689822, EPI_ISL_2689823 | LESP Ciudad de Mexico | Instituto de Diagnostico y Referencia Epidemiologicos (INDRE) | Claudia Wong-Arambula, Abril Rodriguez-Maldonado, Vanessa Rivero-Arredondo, Ariadna Medina-Benitez, Joaquin Quiroz-Mercado, Sergio Rangel-Guerrero, Natividad Cruz-Ortiz, Tatiana Nunez-Garcia, Gisela Barrera-Badillo, Lucia Hernandez-Rivas, Irma Lopez-Martinez, Ernesto Ramirez-Gonzalez. |
| EPI_ISL_2689824 | LESP Queretaro | Instituto de Diagnostico y Referencia Epidemiologicos (INDRE) | Claudia Wong-Arambula, Abril Rodriguez-Maldonado, Vanessa Rivero-Arredondo, Ariadna Medina-Benitez, Joaquin Quiroz-Mercado, Sergio Rangel-Guerrero, Natividad Cruz-Ortiz, Tatiana Nunez-Garcia, Gisela Barrera-Badillo, Lucia Hernandez-Rivas, Irma Lopez-Martinez, Ernesto Ramirez-Gonzalez. |
| EPI_ISL_2689825 | LESP Michoacan | Instituto de Diagnostico y Referencia Epidemiologicos (INDRE) | Claudia Wong-Arambula, Abril Rodriguez-Maldonado, Vanessa Rivero-Arredondo, Ariadna Medina-Benitez, Joaquin Quiroz-Mercado, Sergio Rangel-Guerrero, Natividad Cruz-Ortiz, Tatiana Nunez-Garcia, Gisela Barrera-Badillo, Lucia Hernandez-Rivas, Irma Lopez-Martinez, Ernesto Ramirez-Gonzalez. |
| EPI_ISL_2689826 | LESP Veracruz | Instituto de Diagnostico y Referencia Epidemiologicos | Claudia Wong-Arambula, Abril Rodriguez-Maldonado, Vanessa Rivero-Arredondo, Ariadna Medina-Benitez, Joaquin Quiroz-Mercado, Sergio |

| | | | |
|---|---|---|---|
| | | (INDRE) | Rangel-Guerrero, Natividad Cruz-Ortiz, Tatiana Nunez-Garcia, Gisela Barrera-Badillo, Lucia Hernandez-Rivas, Irma Lopez-Martinez, Ernesto Ramirez-Gonzalez. |
| EPI_ISL_2689827, EPI_ISL_2689828 | LESP Chihuahua | Instituto de Diagnostico y Referencia Epidemiologicos (INDRE) | Claudia Wong-Arambula, Abril Rodriguez-Maldonado, Vanessa Rivero-Arredondo, Ariadna Medina-Benitez, Joaquin Quiroz-Mercado, Sergio Rangel-Guerrero, Natividad Cruz-Ortiz, Tatiana Nunez-Garcia, Gisela Barrera-Badillo, Lucia Hernandez-Rivas, Irma Lopez-Martinez, Ernesto Ramirez-Gonzalez. |
| EPI_ISL_2689829 | LESP Sonora | Instituto de Diagnostico y Referencia Epidemiologicos (INDRE) | Claudia Wong-Arambula, Abril Rodriguez-Maldonado, Vanessa Rivero-Arredondo, Ariadna Medina-Benitez, Joaquin Quiroz-Mercado, Sergio Rangel-Guerrero, Natividad Cruz-Ortiz, Tatiana Nunez-Garcia, Gisela Barrera-Badillo, Lucia Hernandez-Rivas, Irma Lopez-Martinez, Ernesto Ramirez-Gonzalez. |
| EPI_ISL_2690133, EPI_ISL_2690135, EPI_ISL_2690136 | Ochsner Health | Bioinfoexperts, LLC | Judy Crabtree, Amy Feehan, Rebecca Rose, David J. Nolan, Tessa LaFleur, Julia-Garcia-Diaz, Lucio Miele, Susanna L. Lamers |
| EPI_ISL_2690698 | MSK Microbiology Lab | MSK Microbiology Lab | Esther Babady, Tracy McMillen, Krupa Jani |
| EPI_ISL_2691110, EPI_ISL_2691111, EPI_ISL_2691112 | Instituto Adolfo Lutz - Regional de Bauru | Instituto Adolfo Lutz, Interdiciplinary Procedures Center, Strategic Laboratory | Claudio Tavares Sacchi, Claudia Regina Gonçalves, Erica Valessa Ramos Gomes, Karoline Rodrigues Campos, Caio Vinicius Dias Lopes, Leonardo Jose Tadeu de Araujo |
| EPI_ISL_2691145 | Instituto Adolfo Lutz - Regional de Sao Jose do Rio Preto | Instituto Adolfo Lutz, Interdiciplinary Procedures Center, Strategic Laboratory | Claudio Tavares Sacchi, Claudia Regina Gonçalves, Erica Valessa Ramos Gomes, Karoline Rodrigues Campos, Caio Vinicius Dias Lopes, Leonardo Jose Tadeu de Araujo |
| EPI_ISL_2691151, EPI_ISL_2691152, EPI_ISL_2691153, EPI_ISL_2691154, EPI_ISL_2691155, EPI_ISL_2691156, EPI_ISL_2691159, EPI_ISL_2691160, EPI_ISL_2691161, EPI_ISL_2691162, EPI_ISL_2691163, EPI_ISL_2691164, EPI_ISL_2691165, EPI_ISL_2691166, EPI_ISL_2691167, EPI_ISL_2691168, EPI_ISL_2691169, EPI_ISL_2691170 | | | |
| see above | Instituo Adolfo Lutz Central | Instituto Adolfo Lutz, Interdiciplinary Procedures Center, Strategic Laboratory | Claudio Tavares Sacchi, Claudia Regina Gonçalves, Erica Valessa Ramos Gomes, Karoline Rodrigues Campos, Caio Vinicius Dias Lopes, Leonardo Jose Tadeu de Araujo |
| EPI_ISL_2691171, EPI_ISL_2691172, EPI_ISL_2691173, EPI_ISL_2691174, EPI_ISL_2691175, EPI_ISL_2691176, EPI_ISL_2691177, EPI_ISL_2691178, EPI_ISL_2691179, EPI_ISL_2691180, EPI_ISL_2691181, EPI_ISL_2691182, EPI_ISL_2691183, EPI_ISL_2691184, EPI_ISL_2691185, EPI_ISL_2691186, EPI_ISL_2691187, EPI_ISL_2691188, EPI_ISL_2691189, EPI_ISL_2691190, EPI_ISL_2691191, EPI_ISL_2691192, EPI_ISL_2691193, EPI_ISL_2691194, EPI_ISL_2691195, EPI_ISL_2691196, EPI_ISL_2691197, EPI_ISL_2691198, EPI_ISL_2691199, EPI_ISL_2691200, EPI_ISL_2691201, EPI_ISL_2691202, EPI_ISL_2691203 | | | |
| see above | Instituto Adolfo Lutz - Reginal de Sao José do Rio Preto | Instituto Adolfo Lutz, Interdiciplinary Procedures Center, Strategic Laboratory | Claudio Tavares Sacchi, Claudia Regina Gonçalves, Erica Valessa Ramos Gomes, Karoline Rodrigues Campos, Caio Vinicius Dias Lopes, Leonardo Jose Tadeu de Araujo |
| EPI_ISL_2691209, EPI_ISL_2691210, EPI_ISL_2691211 | LACEN do Estado de Goias | Instituto Adolfo Lutz, Interdiciplinary Procedures Center, Strategic Laboratory | Claudio Tavares Sacchi, Claudia Regina Gonçalves, Erica Valessa Ramos Gomes, Karoline Rodrigues Campos, Caio Vinicius Dias Lopes, Leonardo Jose Tadeu de Araujo |
| EPI_ISL_2692533, EPI_ISL_2692905 | Clinical Microbiology Lab RSPTN Universitas Hasanuddin | Clinical Microbiology Lab RSPTN Universitas Hasanuddin | Muhammad Nasrum Massi, Handayani Halik, Rizalinda Sjahril, Gita Vita Soraya, Triwibawa, Muhammad Yusuf Usman, Rufika Shari |
| EPI_ISL_2693006 | TSGH-CP molecular lab | TSGH-CP molecular lab | Cherng-Lih Perng, Ming-Jr JIAN, Chih-Kai Chang, Jung-Chung Lin, Kuo-Ming Yeh, Chien-Wen Chen, Sheng-Kang Chiu, Hsing-Yi Chung, Shih-Hung Tsai, Kuo-Sheng Hung, Tien-Yao Chang, Feng-Yee Chang, Hung-Sheng Shang |
| EPI_ISL_2693109, EPI_ISL_2693110 | St James's Hospital, Dublin 8, Ireland. | ELDA biotech | Deirdre M. Broderick, Peter R. Flanagan, Brendan Crowley |
| EPI_ISL_2693909, EPI_ISL_2693910, EPI_ISL_2693911, EPI_ISL_2693912, EPI_ISL_2693913, EPI_ISL_2693914, EPI_ISL_2693915, EPI_ISL_2693916, EPI_ISL_2693917, EPI_ISL_2693918, EPI_ISL_2693919, EPI_ISL_2693920, EPI_ISL_2693921, EPI_ISL_2693922, EPI_ISL_2693923, EPI_ISL_2693924, EPI_ISL_2693925, EPI_ISL_2693926, EPI_ISL_2693927, EPI_ISL_2693928, EPI_ISL_2693929, EPI_ISL_2693930, EPI_ISL_2693931, EPI_ISL_2693932, EPI_ISL_2693933, EPI_ISL_2693934, EPI_ISL_2693935, EPI_ISL_2693936, EPI_ISL_2693937, EPI_ISL_2693938, EPI_ISL_2693939, EPI_ISL_2693940, EPI_ISL_2693941, EPI_ISL_2693942, EPI_ISL_2693943, EPI_ISL_2693944, EPI_ISL_2693945, EPI_ISL_2693946, EPI_ISL_2693947, EPI_ISL_2693948, EPI_ISL_2693949, EPI_ISL_2693950, EPI_ISL_2693951 | | | |
| see above | Virology Unit, Institut Pasteur du Cambodge | Virology Unit, Institut Pasteur du Cambodge | Cecile Troupin, Leakhena Pum, Jurre Y Siegers, Ly Sovann, Kraing Sidonn, Yi Sengdoeurn, Chin Savuth, Chau Darapheak, Veasna Duong, Erik A Karlsson |
| EPI_ISL_2693960, EPI_ISL_2693963, EPI_ISL_2693964, EPI_ISL_2693965, EPI_ISL_2693966, EPI_ISL_2693967, EPI_ISL_2693968, EPI_ISL_2693969, EPI_ISL_2693970 | laboratorio Microbiologia PO Cardarelli | laboratorio Microbiologia PO Cardarelli | Felice V., Niro G. Scutella' M. |
| EPI_ISL_2695878, EPI_ISL_2695879, EPI_ISL_2695880, EPI_ISL_2695883 | Osaka General Medical Center | Department of Developmental Medicine, Research Institute, Osaka Women's and Children's Hospital | Itaru Yanagihara, Yukiko Nakura |
| EPI_ISL_2695962 | Department of Developmental Medicine, Research Institute, Osaka Women's and Children's Hospital | Department of Developmental Medicine, Research Institute, Osaka Women's and Children's Hospital | Itaru Yanagihara, Yukiko Nakura |
| EPI_ISL_2705575 | Centers for Disease Control, R.O.C. (Taiwan) | Centers for Disease Control, R.O.C. (Taiwan) | Ji-Rong Yang, Yu-Chi-Lin, Jung-Jung Mu, Ming-Tsan-Liu |
| EPI_ISL_2708444, EPI_ISL_2708957, EPI_ISL_2708958, EPI_ISL_2708959, EPI_ISL_2708978, EPI_ISL_2708979, EPI_ISL_2708980, EPI_ISL_2708981 | Institut de Virologie du CHU de Strasbourg | Institut de Virologie du CHU de Strasbourg | Fafi-Kremer Samira, Gallais Floriane, Gantner Pierre, Laugel Elodie, Solis Morgane, Velay Aurélie, Wendling Marie-Josée |
| EPI_ISL_2709251, EPI_ISL_2709256 | CHU Pontchaillou | CHU Pontchaillou | GROLHIER Claire, DENOUAL Florent, ETCHEVERRY Amandine, SASSI Mohamed, JAGLINE Steven, FEBREAU Christine, PRONIER Charlotte, GALIBERT Marie Dominique, DE TAYRAC Marie, THIBAULT Vincent |
| EPI_ISL_2709257 | Labo Analyses Med | National Reference Center for Viruses of Respiratory Infections, Institut Pasteur, Paris | Marion Barbet, Sylvie Behillil, Méline Bizard, Angela Brisebarre, Camille Capel, Vincent Enouf, Louise Lefrançois, Frédéric Lemoine, Christophe Malabat, Corinne Maufrais, Etienne Simon-Lorière, Maud Vanpeene, Sylvie Van der Werf ,Franck Ennouchi |
| EPI_ISL_2709258 | Hospital | National Reference Center for Viruses of Respiratory Infections, Institut Pasteur, Paris | Marion Barbet, Sylvie Behillil, Méline Bizard, Angela Brisebarre, Camille Capel, Vincent Enouf, Louise Lefrançois, Frédéric Lemoine, Christophe Malabat, Corinne Maufrais, Etienne Simon-Lorière, Maud Vanpeene, Sylvie Van der Werf ,Salma Ben Hadj Yahia |
| EPI_ISL_2709259 | Hospital | National Reference Center for Viruses of Respiratory Infections, Institut Pasteur, Paris | Marion Barbet, Sylvie Behillil, Méline Bizard, Angela Brisebarre, Camille Capel, Vincent Enouf, Louise Lefrançois, Frédéric Lemoine, Christophe Malabat, Corinne Maufrais, Damien Mornico, Etienne Simon-Lorière, Maud Vanpeene, Sylvie Van der Werf ,Laurent Bret |
| EPI_ISL_2709261 | Labo Analyses Med | National Reference Center for Viruses of Respiratory Infections, Institut Pasteur, Paris | Marion Barbet, Sylvie Behillil, Méline Bizard, Angela Brisebarre, Camille Capel, Vincent Enouf, Louise Lefrançois, Frédéric Lemoine, Christophe Malabat, Corinne Maufrais, Etienne Simon-Lorière, Maud Vanpeene, Sylvie Van der Werf ,Patricia Tamby |
| EPI_ISL_2709262 | Labo Analyses Med | National Reference Center for Viruses of Respiratory Infections, Institut Pasteur, Paris | Marion Barbet, Sylvie Behillil, Méline Bizard, Angela Brisebarre, Camille Capel, Vincent Enouf, Louise Lefrançois, Frédéric Lemoine, Christophe Malabat, Corinne Maufrais, Etienne Simon-Lorière, Maud Vanpeene, Sylvie Van der Werf ,Vicky Le |
| EPI_ISL_2709263 | Labo Analyses Med | National Reference Center for Viruses of Respiratory Infections, Institut Pasteur, Paris | Marion Barbet, Sylvie Behillil, Méline Bizard, Angela Brisebarre, Camille Capel, Vincent Enouf, Louise Lefrançois, Frédéric Lemoine, Christophe Malabat, Corinne Maufrais, Damien Mornico, Etienne Simon-Lorière, Maud Vanpeene, Sylvie Van der Werf ,Vicky Le |
| EPI_ISL_2709264 | Labo Analyses Med | National Reference Center for Viruses of Respiratory Infections, Institut Pasteur, Paris | Marion Barbet, Sylvie Behillil, Méline Bizard, Angela Brisebarre, Camille Capel, Vincent Enouf, Louise Lefrançois, Frédéric Lemoine, Christophe Malabat, Corinne Maufrais, Etienne Simon-Lorière, Maud Vanpeene, Sylvie Van der Werf ,GréGoire Potiron |
| EPI_ISL_2709265 | Labo Analyses Med | National Reference Center for Viruses of Respiratory Infections, Institut Pasteur, Paris | Marion Barbet, Sylvie Behillil, Méline Bizard, Angela Brisebarre, Camille Capel, Vincent Enouf, Louise Lefrançois, Frédéric Lemoine, Christophe Malabat, Corinne Maufrais, Etienne Simon-Lorière, Maud Vanpeene, Sylvie Van der Werf ,Hugues Leroy |
| EPI_ISL_2709266 | Hospital | National Reference Center for Viruses of Respiratory Infections, Institut Pasteur, Paris | Marion Barbet, Sylvie Behillil, Méline Bizard, Angela Brisebarre, Camille Capel, Vincent Enouf, Louise Lefrançois, Frédéric Lemoine, Christophe Malabat, Corinne Maufrais, Etienne Simon-Lorière, Maud Vanpeene, Sylvie Van der Werf ,Farida Benouadi |
| EPI_ISL_2709267 | Hospital | National Reference Center for Viruses of Respiratory Infections, Institut Pasteur, Paris | Marion Barbet, Sylvie Behillil, Méline Bizard, Angela Brisebarre, Camille Capel, Vincent Enouf, Louise Lefrançois, Frédéric Lemoine, Christophe Malabat, Corinne Maufrais, Etienne Simon-Lorière, Maud Vanpeene, Sylvie Van der Werf ,Laurence Courdavault |
| EPI_ISL_2709268, EPI_ISL_2709269, | Labo Analyses Med | National Reference Center for Viruses of Respiratory | Marion Barbet, Sylvie Behillil, Méline Bizard, Angela Brisebarre, Camille Capel, Vincent Enouf, Louise Lefrançois, Frédéric Lemoine, Christophe Malabat, |

| | | | |
|---|---|---|---|
| EPI_ISL_2709270, EPI_ISL_2709271, EPI_ISL_2709273 | | Infections, Institut Pasteur, Paris | Corinne Maufrais, Etienne Simon-Lorière, Maud Vanpeene, Sylvie Van der Werf ,Anne-Laure Garand |
| EPI_ISL_2709274 | Labo Analyses Med | National Reference Center for Viruses of Respiratory Infections, Institut Pasteur, Paris | Marion Barbet, Sylvie Behillil, Méline Bizard, Angela Brisebarre, Camille Capel, Vincent Enouf, Louise Lefrançois, Frédéric Lemoine, Christophe Malabat, Corinne Maufrais, Damien Mornico, Etienne Simon-Lorière, Maud Vanpeene, Sylvie Van der Werf ,Anne-Laure Garand |
| EPI_ISL_2709275, EPI_ISL_2709276 | Labo Analyses Med | National Reference Center for Viruses of Respiratory Infections, Institut Pasteur, Paris | Marion Barbet, Sylvie Behillil, Méline Bizard, Angela Brisebarre, Camille Capel, Vincent Enouf, Louise Lefrançois, Frédéric Lemoine, Christophe Malabat, Corinne Maufrais, Etienne Simon-Lorière, Maud Vanpeene, Sylvie Van der Werf ,Anne-Laure Garand |
| EPI_ISL_2709277 | Labo Analyses Med | National Reference Center for Viruses of Respiratory Infections, Institut Pasteur, Paris | Marion Barbet, Sylvie Behillil, Méline Bizard, Angela Brisebarre, Camille Capel, Vincent Enouf, Louise Lefrançois, Frédéric Lemoine, Christophe Malabat, Corinne Maufrais, Damien Mornico, Etienne Simon-Lorière, Maud Vanpeene, Sylvie Van der Werf ,Hugues Leroy |
| EPI_ISL_2709279, EPI_ISL_2709280, EPI_ISL_2709281, EPI_ISL_2709282, EPI_ISL_2709283, EPI_ISL_2709284, EPI_ISL_2709285, EPI_ISL_2709286, EPI_ISL_2709287, EPI_ISL_2709288, EPI_ISL_2709289, EPI_ISL_2709290, EPI_ISL_2709291, EPI_ISL_2709292, EPI_ISL_2709293 | | | |
| see above | Labo Analyses med | National Reference Center for Viruses of Respiratory Infections, Institut Pasteur, Paris | Marion Barbet, Sylvie Behillil, Méline Bizard, Angela Brisebarre, Camille Capel, Vincent Enouf, Louise Lefrançois, Frédéric Lemoine, Christophe Malabat, Corinne Maufrais, Etienne Simon-Lorière, Maud Vanpeene, Sylvie Van der Werf ,OphéLie Said-Delattre |
| EPI_ISL_2709294 | Labo Analyses med | National Reference Center for Viruses of Respiratory Infections, Institut Pasteur, Paris | Marion Barbet, Sylvie Behillil, Méline Bizard, Angela Brisebarre, Camille Capel, Vincent Enouf, Louise Lefrançois, Frédéric Lemoine, Christophe Malabat, Corinne Maufrais, Damien Mornico, Etienne Simon-Lorière, Maud Vanpeene, Sylvie Van der Werf ,OphéLie Said-Delattre |
| EPI_ISL_2709295 | Labo Analyses med | National Reference Center for Viruses of Respiratory Infections, Institut Pasteur, Paris | Marion Barbet, Sylvie Behillil, Méline Bizard, Angela Brisebarre, Camille Capel, Vincent Enouf, Louise Lefrançois, Frédéric Lemoine, Christophe Malabat, Corinne Maufrais, Etienne Simon-Lorière, Maud Vanpeene, Sylvie Van der Werf ,OphéLie Said-Delattre |
| EPI_ISL_2709296 | Labo Analyses med | National Reference Center for Viruses of Respiratory Infections, Institut Pasteur, Paris | Marion Barbet, Sylvie Behillil, Méline Bizard, Angela Brisebarre, Camille Capel, Vincent Enouf, Louise Lefrançois, Frédéric Lemoine, Christophe Malabat, Corinne Maufrais, Damien Mornico, Etienne Simon-Lorière, Maud Vanpeene, Sylvie Van der Werf ,OphéLie Said-Delattre |
| EPI_ISL_2709297, EPI_ISL_2709298, EPI_ISL_2709299, EPI_ISL_2709301, EPI_ISL_2709302, EPI_ISL_2709303, EPI_ISL_2709304, EPI_ISL_2709305, EPI_ISL_2709306, EPI_ISL_2709307, EPI_ISL_2709308, EPI_ISL_2709309, EPI_ISL_2709310, EPI_ISL_2709311, EPI_ISL_2709312 | | | |
| see above | Labo Analyses med | National Reference Center for Viruses of Respiratory Infections, Institut Pasteur, Paris | Marion Barbet, Sylvie Behillil, Méline Bizard, Angela Brisebarre, Camille Capel, Vincent Enouf, Louise Lefrançois, Frédéric Lemoine, Christophe Malabat, Corinne Maufrais, Etienne Simon-Lorière, Maud Vanpeene, Sylvie Van der Werf ,OphéLie Said-Delattre |
| EPI_ISL_2709603 | AZ Sint-Lucas Gent | AZ Sint-Lucas Gent | Jos Van Acker, Charlotte Verfaillie, Elke Vanlaere, Klara De Rauw |
| EPI_ISL_2709686 | LESP Queretaro | Instituto de Diagnostico y Referencia Epidemiologicos (INDRE) | Claudia Wong-Arambula, Abril Rodriguez-Maldonado, Vanessa Rivero-Arredondo, Ariadna Medina-Benitez, Joaquin Quiroz-Mercado, Sergio Rangel-Guerrero, Natividad Cruz-Ortiz, Tatiana Nunez-Garcia, Gisela Barrera-Badillo, Lucia Hernandez-Rivas, Irma Lopez-Martinez, Ernesto Ramirez-Gonzalez. |
| EPI_ISL_2709687 | LESP Baja California | Instituto de Diagnostico y Referencia Epidemiologicos (INDRE) | Claudia Wong-Arambula, Abril Rodriguez-Maldonado, Vanessa Rivero-Arredondo, Ariadna Medina-Benitez, Joaquin Quiroz-Mercado, Sergio Rangel-Guerrero, Natividad Cruz-Ortiz, Tatiana Nunez-Garcia, Gisela Barrera-Badillo, Lucia Hernandez-Rivas, Irma Lopez-Martinez, Ernesto Ramirez-Gonzalez. |
| EPI_ISL_2709688 | LESP Michoacan | Instituto de Diagnostico y Referencia Epidemiologicos (INDRE) | Claudia Wong-Arambula, Abril Rodriguez-Maldonado, Vanessa Rivero-Arredondo, Ariadna Medina-Benitez, Joaquin Quiroz-Mercado, Sergio Rangel-Guerrero, Natividad Cruz-Ortiz, Tatiana Nunez-Garcia, Gisela Barrera-Badillo, Lucia Hernandez-Rivas, Irma Lopez-Martinez, Ernesto Ramirez-Gonzalez. |
| EPI_ISL_2709690, EPI_ISL_2709691, EPI_ISL_2709692, EPI_ISL_2709693, EPI_ISL_2709694, EPI_ISL_2709695, EPI_ISL_2709696, EPI_ISL_2709697, EPI_ISL_2709698 | LESP Nuevo Leon | Instituto de Diagnostico y Referencia Epidemiologicos (INDRE) | Claudia Wong-Arambula, Abril Rodriguez-Maldonado, Vanessa Rivero-Arredondo, Ariadna Medina-Benitez, Joaquin Quiroz-Mercado, Sergio Rangel-Guerrero, Natividad Cruz-Ortiz, Tatiana Nunez-Garcia, Gisela Barrera-Badillo, Lucia Hernandez-Rivas, Irma Lopez-Martinez, Ernesto Ramirez-Gonzalez. |
| EPI_ISL_2709699 | LESP Jalisco | Instituto de Diagnostico y Referencia Epidemiologicos (INDRE) | Claudia Wong-Arambula, Abril Rodriguez-Maldonado, Vanessa Rivero-Arredondo, Ariadna Medina-Benitez, Joaquin Quiroz-Mercado, Sergio Rangel-Guerrero, Natividad Cruz-Ortiz, Tatiana Nunez-Garcia, Gisela Barrera-Badillo, Lucia Hernandez-Rivas, Irma Lopez-Martinez, Ernesto Ramirez-Gonzalez. |
| EPI_ISL_2709700, EPI_ISL_2709701, EPI_ISL_2709702 | LESP Morelos | Instituto de Diagnostico y Referencia Epidemiologicos (INDRE) | Claudia Wong-Arambula, Abril Rodriguez-Maldonado, Vanessa Rivero-Arredondo, Ariadna Medina-Benitez, Joaquin Quiroz-Mercado, Sergio Rangel-Guerrero, Natividad Cruz-Ortiz, Tatiana Nunez-Garcia, Gisela Barrera-Badillo, Lucia Hernandez-Rivas, Irma Lopez-Martinez, Ernesto Ramirez-Gonzalez. |
| EPI_ISL_2709703 | LESP Ciudad de Mexico | Instituto de Diagnostico y Referencia Epidemiologicos (INDRE) | Claudia Wong-Arambula, Abril Rodriguez-Maldonado, Vanessa Rivero-Arredondo, Ariadna Medina-Benitez, Joaquin Quiroz-Mercado, Sergio Rangel-Guerrero, Natividad Cruz-Ortiz, Tatiana Nunez-Garcia, Gisela Barrera-Badillo, Lucia Hernandez-Rivas, Irma Lopez-Martinez, Ernesto Ramirez-Gonzalez. |
| EPI_ISL_2709704, EPI_ISL_2709705 | LESP Estado de Mexico | Instituto de Diagnostico y Referencia Epidemiologicos (INDRE) | Claudia Wong-Arambula, Abril Rodriguez-Maldonado, Vanessa Rivero-Arredondo, Ariadna Medina-Benitez, Joaquin Quiroz-Mercado, Sergio Rangel-Guerrero, Natividad Cruz-Ortiz, Tatiana Nunez-Garcia, Gisela Barrera-Badillo, Lucia Hernandez-Rivas, Irma Lopez-Martinez, Ernesto Ramirez-Gonzalez. |
| EPI_ISL_2709706, EPI_ISL_2709707, EPI_ISL_2709708 | LESP Queretaro | Instituto de Diagnostico y Referencia Epidemiologicos (INDRE) | Claudia Wong-Arambula, Abril Rodriguez-Maldonado, Vanessa Rivero-Arredondo, Ariadna Medina-Benitez, Joaquin Quiroz-Mercado, Sergio Rangel-Guerrero, Natividad Cruz-Ortiz, Tatiana Nunez-Garcia, Gisela Barrera-Badillo, Lucia Hernandez-Rivas, Irma Lopez-Martinez, Ernesto Ramirez-Gonzalez. |
| EPI_ISL_2709709, EPI_ISL_2709710, EPI_ISL_2709711 | LESP Tabasco | Instituto de Diagnostico y Referencia Epidemiologicos (INDRE) | Claudia Wong-Arambula, Abril Rodriguez-Maldonado, Vanessa Rivero-Arredondo, Ariadna Medina-Benitez, Joaquin Quiroz-Mercado, Sergio Rangel-Guerrero, Natividad Cruz-Ortiz, Tatiana Nunez-Garcia, Gisela Barrera-Badillo, Lucia Hernandez-Rivas, Irma Lopez-Martinez, Ernesto Ramirez-Gonzalez. |
| EPI_ISL_2709712, EPI_ISL_2709713, EPI_ISL_2709714, EPI_ISL_2709715, EPI_ISL_2709716 | LESP Ciudad de Mexico | Instituto de Diagnostico y Referencia Epidemiologicos (INDRE) | Claudia Wong-Arambula, Abril Rodriguez-Maldonado, Vanessa Rivero-Arredondo, Ariadna Medina-Benitez, Joaquin Quiroz-Mercado, Sergio Rangel-Guerrero, Natividad Cruz-Ortiz, Tatiana Nunez-Garcia, Gisela Barrera-Badillo, Lucia Hernandez-Rivas, Irma Lopez-Martinez, Ernesto Ramirez-Gonzalez. |
| EPI_ISL_2709717 | LESP Jalisco | Instituto de Diagnostico y Referencia Epidemiologicos (INDRE) | Claudia Wong-Arambula, Abril Rodriguez-Maldonado, Vanessa Rivero-Arredondo, Ariadna Medina-Benitez, Joaquin Quiroz-Mercado, Sergio Rangel-Guerrero, Natividad Cruz-Ortiz, Tatiana Nunez-Garcia, Gisela Barrera-Badillo, Lucia Hernandez-Rivas, Irma Lopez-Martinez, Ernesto Ramirez-Gonzalez. |
| EPI_ISL_2709718 | LESP Sonora | Instituto de Diagnostico y Referencia Epidemiologicos (INDRE) | Claudia Wong-Arambula, Abril Rodriguez-Maldonado, Vanessa Rivero-Arredondo, Ariadna Medina-Benitez, Joaquin Quiroz-Mercado, Sergio Rangel-Guerrero, Natividad Cruz-Ortiz, Tatiana Nunez-Garcia, Gisela Barrera-Badillo, Lucia Hernandez-Rivas, Irma Lopez-Martinez, Ernesto Ramirez-Gonzalez. |
| EPI_ISL_2709719 | LESP Estado de Mexico | Instituto de Diagnostico y Referencia Epidemiologicos (INDRE) | Claudia Wong-Arambula, Abril Rodriguez-Maldonado, Vanessa Rivero-Arredondo, Ariadna Medina-Benitez, Joaquin Quiroz-Mercado, Sergio Rangel-Guerrero, Natividad Cruz-Ortiz, Tatiana Nunez-Garcia, Gisela Barrera-Badillo, Lucia Hernandez-Rivas, Irma Lopez-Martinez, Ernesto Ramirez-Gonzalez. |
| EPI_ISL_2709720 | LESP Ciudad de Mexico | Instituto de Diagnostico y Referencia Epidemiologicos (INDRE) | Claudia Wong-Arambula, Abril Rodriguez-Maldonado, Vanessa Rivero-Arredondo, Ariadna Medina-Benitez, Joaquin Quiroz-Mercado, Sergio Rangel-Guerrero, Natividad Cruz-Ortiz, Tatiana Nunez-Garcia, Gisela Barrera-Badillo, Lucia Hernandez-Rivas, Irma Lopez-Martinez, Ernesto Ramirez-Gonzalez. |
| EPI_ISL_2709721 | LESP Estado de Mexico | Instituto de Diagnostico y Referencia Epidemiologicos (INDRE) | Claudia Wong-Arambula, Abril Rodriguez-Maldonado, Vanessa Rivero-Arredondo, Ariadna Medina-Benitez, Joaquin Quiroz-Mercado, Sergio Rangel-Guerrero, Natividad Cruz-Ortiz, Tatiana Nunez-Garcia, Gisela Barrera-Badillo, Lucia Hernandez-Rivas, Irma Lopez-Martinez, Ernesto Ramirez-Gonzalez. |
| EPI_ISL_2709722 | LESP Hidalgo | Instituto de Diagnostico y Referencia Epidemiologicos (INDRE) | Claudia Wong-Arambula, Abril Rodriguez-Maldonado, Vanessa Rivero-Arredondo, Ariadna Medina-Benitez, Joaquin Quiroz-Mercado, Sergio Rangel-Guerrero, Natividad Cruz-Ortiz, Tatiana Nunez-Garcia, Gisela Barrera-Badillo, Lucia Hernandez-Rivas, Irma Lopez-Martinez, Ernesto |

| | | | |
|---|---|---|---|
| | | | Ramirez-Gonzalez. |
| EPI_ISL_2709723 | LESP Veracruz | Instituto de Diagnostico y Referencia Epidemiologicos (INDRE) | Claudia Wong-Arambula, Abril Rodriguez-Maldonado, Vanessa Rivero-Arredondo, Ariadna Medina-Benitez, Joaquin Quiroz-Mercado, Sergio Rangel-Guerrero, Natividad Cruz-Ortiz, Tatiana Nunez-Garcia, Gisela Barrera-Badillo, Lucia Hernandez-Rivas, Irma Lopez-Martinez, Ernesto Ramirez-Gonzalez. |
| EPI_ISL_2709724, EPI_ISL_2709725 | LESP Guanajuato | Instituto de Diagnostico y Referencia Epidemiologicos (INDRE) | Claudia Wong-Arambula, Abril Rodriguez-Maldonado, Vanessa Rivero-Arredondo, Ariadna Medina-Benitez, Joaquin Quiroz-Mercado, Sergio Rangel-Guerrero, Natividad Cruz-Ortiz, Tatiana Nunez-Garcia, Gisela Barrera-Badillo, Lucia Hernandez-Rivas, Irma Lopez-Martinez, Ernesto Ramirez-Gonzalez. |
| EPI_ISL_2709726, EPI_ISL_2709727, EPI_ISL_2709728, EPI_ISL_2709729 | LESP Nuevo Leon | Instituto de Diagnostico y Referencia Epidemiologicos (INDRE) | Claudia Wong-Arambula, Abril Rodriguez-Maldonado, Vanessa Rivero-Arredondo, Ariadna Medina-Benitez, Joaquin Quiroz-Mercado, Sergio Rangel-Guerrero, Natividad Cruz-Ortiz, Tatiana Nunez-Garcia, Gisela Barrera-Badillo, Lucia Hernandez-Rivas, Irma Lopez-Martinez, Ernesto Ramirez-Gonzalez. |
| EPI_ISL_2709730 | LESP Ciudad de Mexico | Instituto de Diagnostico y Referencia Epidemiologicos (INDRE) | Claudia Wong-Arambula, Abril Rodriguez-Maldonado, Vanessa Rivero-Arredondo, Ariadna Medina-Benitez, Joaquin Quiroz-Mercado, Sergio Rangel-Guerrero, Natividad Cruz-Ortiz, Tatiana Nunez-Garcia, Gisela Barrera-Badillo, Lucia Hernandez-Rivas, Irma Lopez-Martinez, Ernesto Ramirez-Gonzalez. |
| EPI_ISL_2709731 | LESP Coahuila | Instituto de Diagnostico y Referencia Epidemiologicos (INDRE) | Claudia Wong-Arambula, Abril Rodriguez-Maldonado, Vanessa Rivero-Arredondo, Ariadna Medina-Benitez, Joaquin Quiroz-Mercado, Sergio Rangel-Guerrero, Natividad Cruz-Ortiz, Tatiana Nunez-Garcia, Gisela Barrera-Badillo, Lucia Hernandez-Rivas, Irma Lopez-Martinez, Ernesto Ramirez-Gonzalez. |
| EPI_ISL_2709732 | LESP Chiapas | Instituto de Diagnostico y Referencia Epidemiologicos (INDRE) | Claudia Wong-Arambula, Abril Rodriguez-Maldonado, Vanessa Rivero-Arredondo, Ariadna Medina-Benitez, Joaquin Quiroz-Mercado, Sergio Rangel-Guerrero, Natividad Cruz-Ortiz, Tatiana Nunez-Garcia, Gisela Barrera-Badillo, Lucia Hernandez-Rivas, Irma Lopez-Martinez, Ernesto Ramirez-Gonzalez. |
| EPI_ISL_2709733 | LESP Veracruz | Instituto de Diagnostico y Referencia Epidemiologicos (INDRE) | Claudia Wong-Arambula, Abril Rodriguez-Maldonado, Vanessa Rivero-Arredondo, Ariadna Medina-Benitez, Joaquin Quiroz-Mercado, Sergio Rangel-Guerrero, Natividad Cruz-Ortiz, Tatiana Nunez-Garcia, Gisela Barrera-Badillo, Lucia Hernandez-Rivas, Irma Lopez-Martinez, Ernesto Ramirez-Gonzalez. |
| EPI_ISL_2709750, EPI_ISL_2709751 | Department of Virology, Istituto Zooprofilattico Sperimentale del Lazio e della Toscana (IZSLT) | Department of General Diagnostics; Department of Virology; Istituto Zooprofilattico Sperimentale del Lazio e della Toscana (IZSLT) | Patricia Alba, Giuseppe Manna, Elena L. Diaconu, Fabiola Feltrin, Raffaela Conti, Teresa Scicluna, Virginia Carfora, Antonella Cersini, Alessia Franco, Antonio Battisti. |
| EPI_ISL_2709771, EPI_ISL_2709772, EPI_ISL_2709773, EPI_ISL_2709774, EPI_ISL_2709775, EPI_ISL_2709776, EPI_ISL_2709777, EPI_ISL_2709778 | Institute of Molecular and Translational Medicine / Laboratory of Experimental Medicine, Faculty of Medicine and Dentistry, Palacky University and University Hospital Olomouc | Institute of Molecular and Translational Medicine / Laboratory of Experimental Medicine, Faculty of Medicine and Dentistry, Palacky University | Rastislav Slavkovský, Hana Dímalová, Vladimíra Koudeláková, Barbora Blumová, Marián Hajdúch |
| EPI_ISL_2709779 | Department of Microbiology, Faculty of Medicine and Dentistry, Palacky University and University Hospital Olomouc | Institute of Molecular and Translational Medicine / Laboratory of Experimental Medicine, Faculty of Medicine and Dentistry, Palacky University | Rastislav Slavkovský, Hana Dímalová, Pavel Sauer, Vladimíra Koudeláková, Barbora Blumová, Marián Hajdúch |
| EPI_ISL_2709780, EPI_ISL_2709781, EPI_ISL_2709782, EPI_ISL_2709783, EPI_ISL_2709784, EPI_ISL_2709785, EPI_ISL_2709786, EPI_ISL_2709787, EPI_ISL_2709788, EPI_ISL_2709789, EPI_ISL_2709790, EPI_ISL_2709791, EPI_ISL_2709792, EPI_ISL_2709794, EPI_ISL_2709795, EPI_ISL_2709796, EPI_ISL_2709797, EPI_ISL_2709798, EPI_ISL_2709799 | | | |
| see above | Institute of Molecular and Translational Medicine / Laboratory of Experimental Medicine, Faculty of Medicine and Dentistry, Palacky University and University Hospital Olomouc | Institute of Molecular and Translational Medicine / Laboratory of Experimental Medicine, Faculty of Medicine and Dentistry, Palacky University | Rastislav Slavkovský, Hana Dímalová, Vladimíra Koudeláková, Barbora Blumová, Marián Hajdúch |
| EPI_ISL_2710017, EPI_ISL_2710018 | Ochsner Health | Bioinfoexperts, LLC | Judy Crabtree, Amy Feehan, Rebecca Rose, David J. Nolan, Tessa LaFleur, Julia-Garcia-Diaz, Lucio Miele, Susanna L. Lamers |
| EPI_ISL_2713259, EPI_ISL_2713262, EPI_ISL_2713263, EPI_ISL_2713267, EPI_ISL_2713303, EPI_ISL_2713304, EPI_ISL_2713305, EPI_ISL_2713306, EPI_ISL_2713307, EPI_ISL_2713308, EPI_ISL_2713309, EPI_ISL_2713310, EPI_ISL_2713311, EPI_ISL_2713312, EPI_ISL_2713313 | | | |
| see above | Department of Health Technology and Informatics, The Hong Kong Polytechnic University | Department of Health Technology and Informatics, The Hong Kong Polytechnic University | Gilman Kit-Hang Siu, Lam-Kwong Lee, Kenneth Siu-Sing Leung, Jake Siu-Lun Leung, Wing-Hei Lo, Timothy Ting-Leung Ng, Chloe Toi-Mei Chan, Hoi-Ching Jim, Ivan Tak-Fai Wong, Kingsley King-Gee Tam, Hiu-Yin Lao, Denise Sze-Hang Wong, Alan Ka-Lun Wu, Miranda Chong-Yee Yau, Yvette Wai-Man Lai, Kitty Sau-Chun Fung, Sandy Ka-Yee Chau, Barry Kin-Chung Wong, Wing-Kin To, Kristine Luk, Alex Yat-Man Ho, Tak-Lun Que, Kam-Tong Yip, Wing Cheong Yam, David Ho-Keung Shum, Shea Ping Yip |
| EPI_ISL_2723570, EPI_ISL_2723571, EPI_ISL_2723572, EPI_ISL_2723573, EPI_ISL_2723574, EPI_ISL_2723575, EPI_ISL_2723576, EPI_ISL_2723577, EPI_ISL_2723578, EPI_ISL_2723579, EPI_ISL_2723580, EPI_ISL_2723581, EPI_ISL_2723582 | | | |
| see above | SC (UCO) Igiene e Sanità Pubblica, ASUGI, Trieste | ARGO Laboratorio Genomica ed Epigenomica | Licastro D, Dal Monego S, Degasperi M, Marcello A, Segat L, Piscianz E, Busetti M, Fontana F, Braida C, Barbone F, D'Agaro P |
| EPI_ISL_2723583, EPI_ISL_2723584 | SC (UCO) Igiene e Sanità Pubblica, ASUGI, Trieste | ARGO Laboratorio Genomica ed Epigenomica | Licastro D, Dal Monego S, Degasperi M, Marcello A, Segat L, Piscianz E, Pipan C, Braida C, Barbone F, D'Agaro P |
| EPI_ISL_2723585, EPI_ISL_2723586, EPI_ISL_2723587, EPI_ISL_2723588, EPI_ISL_2723589, EPI_ISL_2723590, EPI_ISL_2723591 | SC (UCO) Igiene e Sanità Pubblica, ASUGI, Trieste | ARGO Laboratorio Genomica ed Epigenomica | Licastro D, Dal Monego S, Degasperi M, Marcello A, Segat L, Piscianz E, Busetti M, Fontana F, Braida C, Barbone F, D'Agaro P |
| EPI_ISL_2723592 | SC (UCO) Igiene e Sanità Pubblica, ASUGI, Trieste | ARGO Laboratorio Genomica ed Epigenomica | Licastro D, Dal Monego S, Degasperi M, Marcello A, Segat L, De Rosa R, Braida C, Barbone F, D'Agaro P |
| EPI_ISL_2723593 | SC (UCO) Igiene e Sanità Pubblica, ASUGI, Trieste | ARGO Laboratorio Genomica ed Epigenomica | Licastro D, Dal Monego S, Degasperi M, Marcello A, Segat L, Piscianz E, Busetti M, Fontana F, Braida C, Barbone F, D'Agaro P |
| EPI_ISL_2723594 | SC (UCO) Igiene e Sanità Pubblica, ASUGI, Trieste | ARGO Laboratorio Genomica ed Epigenomica | Licastro D, Dal Monego S, Degasperi M, Marcello A, Segat L, De Rosa R, Braida C, Barbone F, D'Agaro P |
| EPI_ISL_2723595, EPI_ISL_2723596 | SC (UCO) Igiene e Sanità Pubblica, ASUGI, Trieste | ARGO Laboratorio Genomica ed Epigenomica | Licastro D, Dal Monego S, Degasperi M, Marcello A, Segat L, Piscianz E, Busetti M, Fontana F, Braida C, Barbone F, D'Agaro P |
| EPI_ISL_2723597, EPI_ISL_2723598 | SC (UCO) Igiene e Sanità Pubblica, ASUGI, Trieste | ARGO Laboratorio Genomica ed Epigenomica | Licastro D, Dal Monego S, Degasperi M, Marcello A, Segat L, De Rosa R, Braida C, Barbone F, D'Agaro P |
| EPI_ISL_2723599 | SC (UCO) Igiene e Sanità Pubblica, ASUGI, Trieste | ARGO Laboratorio Genomica ed Epigenomica | Licastro D, Dal Monego S, Degasperi M, Marcello A, Segat L, Piscianz E, Busetti M, Fontana F, Braida C, Barbone F, D'Agaro P |
| EPI_ISL_2724009 | Molecular Biology Laboratory, Infectious Diseases Department, Instituto Nacional de Ciencias Medicas y Nutricion Salvador Zubiran | Instituto Nacional de Ciencias Medicas y Nutricion Infectious Diseases | Pilar Ramos Cervantes, Fernando Ledesma Barrientos, Violeta Ibarra Gonzalez, Patricia Leal, Eric Ochoa, Arturo Galindo, Guillermo Ruiz-Palacios y Santos |
| EPI_ISL_2725106 | Laboratori de Referencia de Catalunya | Laboratori de Referencia de Catalunya | Ramirez A., Bellosillo B., Padilla E., Vilas A., Hernandez JJ., Canal M. |
| EPI_ISL_2725107, EPI_ISL_2725108, EPI_ISL_2725109 | Laboratorio di Genetica Medica, Matera, Italy | Laboratorio di Genetica Medica, Matera, Italy | Domenico Dell'Edera |
| EPI_ISL_2727499, EPI_ISL_2727500, EPI_ISL_2727501 | Cantacuzino National Military-Medical Institute, Viral Respiratory Infections Laboratory | Cantacuzino Institute Virology | Carmen Cherciu, Luiza Ustea, Nicoleta Paraschiv, Sorin Dinu, Mihaela Oprea, Mihaela Lazar |
| EPI_ISL_2727502, EPI_ISL_2727503, EPI_ISL_2727504, EPI_ISL_2727505, EPI_ISL_2727506, EPI_ISL_2727507, EPI_ISL_2727508, EPI_ISL_2727511, EPI_ISL_2727512, EPI_ISL_2727515, EPI_ISL_2727516, EPI_ISL_2727517, EPI_ISL_2727518, EPI_ISL_2727519, EPI_ISL_2727521 | | | |
| see above | Institute of Molecular and Translational Medicine / Laboratory of Experimental Medicine, Faculty of Medicine and Dentistry, Palacky University and University Hospital Olomouc | Institute of Molecular and Translational Medicine / Laboratory of Experimental Medicine, Faculty of Medicine and Dentistry, Palacky University | Rastislav Slavkovský, Hana Dímalová, Vladimíra Koudeláková, Barbora Blumová, Marián Hajdúch |
| EPI_ISL_2727565, EPI_ISL_2727566, EPI_ISL_2727567, EPI_ISL_2727568 | Cantacuzino National Military-Medical Institute, Viral Respiratory Infections Laboratory | Cantacuzino Institute Virology | Carmen Cherciu, Luiza Ustea, Nicoleta Paraschiv, Sorin Dinu, Mihaela Oprea, Mihaela Lazar |
| EPI_ISL_2727836 | Hospital Angkatan Tentera Kota Kinabalu | Institute for Medical Research, Infectious Disease Research | Suppiah J, Kamel K, Mohd Zawawi Z, Azizan MA, Ramly N, Robert F, Thayan R |

| | | | |
|---|---|---|---|
| | | Centre, National Institutes of Health, Ministry of Health Malaysia | |
| EPI_ISL_2727837 | Hospital Angkatan Tentera Wilayah Kota Kinabalu | Institute for Medical Research, Infectious Disease Research Centre, National Institutes of Health, Ministry of Health Malaysia | Suppiah J, Kamel K, Mohd Zawawi Z, Azizan MA, Ramly N, Robert F, Thayan R |
| EPI_ISL_2728560, EPI_ISL_2728561, EPI_ISL_2728562, EPI_ISL_2728572, EPI_ISL_2728577, EPI_ISL_2728578 | Laboratorio de Infectologia y Virologia Molecular | Laboratory of Molecular Virology, School of Medicine, Pontificia Universidad Catolica de Chile | Catalina Pardo-Roa, Constanza Martinez-Valdevenito, Jennifer Angulo, Tamara Garcia-Salum, Estefany Poblete, Maria Jose Avendano, Leonardo I. Almonacid, Ana Maria Contreras, Carlos Palma, Jorge Levican, Eileen Serrano, Constanza Maldonado, M. Belen Leyton, Erick Salinas, Andres E. Munoz-Marcos, Francisco Melo, Marcela Ferres, Alejandro Bhrun, Magdalena Vera, Rafael A. Medina |
| EPI_ISL_2731271 | S.C Microbiologia e virologia - Speciale Centro influenza -AOU di Sassari | Laboratorio Biologia Molecolare Sars Cov2 - UOC Laboratorio Analisi - Servizio Medicina di Laboratorio, Ospedale "San Francesco" - ATS-ASSL Nuoro and Laboratorio specialistico UOC Ematologia - Ospedale "San Francesco" - ATS-ASSL Nuoro | Asproni Rosanna, Piras Giovanna, Malune Paolo, Pira Giovanna, Fancello Patrizia, Rosu Valentina, Carta Franco, Fiamma Maura, Mameli Giuseppe, Garau Maria Cristina , Palmas Angelo Domenico,Rimini Elena ,Serra Caterina, Salvatore Rubino |
| EPI_ISL_2732231, EPI_ISL_2732235, EPI_ISL_2732238 | Department of Virology, Istituto Zooprofilattico Sperimentale del Lazio e della Toscana (IZSLT) | Department of General Diagnostics; Department of Virology; Istituto Zooprofilattico Sperimentale del Lazio e della Toscana (IZSLT) | Patricia Alba, Giuseppe Manna, Elena L. Diaconu, Fabiola Feltrin, Raffaela Conti, Teresa Scicluna, Virginia Carfora, Antonella Cersini, Alessia Franco, Antonio Battisti. |
| EPI_ISL_2736780, EPI_ISL_2736781, EPI_ISL_2736783 | LESP Sinaloa | Instituto de Diagnostico y Referencia Epidemiologicos (INDRE) | Claudia Wong-Arambula, Abril Rodriguez-Maldonado, Vanessa Rivero-Arredondo, Ariadna Medina-Benitez, Joaquin Quiroz-Mercado, Sergio Rangel-Guerrero, Natividad Cruz-Ortiz, Tatiana Nunez-Garcia, Gisela Barrera-Badillo, Lucia Hernandez-Rivas, Irma Lopez-Martinez, Ernesto Ramirez-Gonzalez. |
| EPI_ISL_2736785 | LESP Puebla | Instituto de Diagnostico y Referencia Epidemiologicos (INDRE) | Claudia Wong-Arambula, Abril Rodriguez-Maldonado, Vanessa Rivero-Arredondo, Ariadna Medina-Benitez, Joaquin Quiroz-Mercado, Sergio Rangel-Guerrero, Natividad Cruz-Ortiz, Tatiana Nunez-Garcia, Gisela Barrera-Badillo, Lucia Hernandez-Rivas, Irma Lopez-Martinez, Ernesto Ramirez-Gonzalez. |
| EPI_ISL_2736787 | LESP Guanajuato | Instituto de Diagnostico y Referencia Epidemiologicos (INDRE) | Claudia Wong-Arambula, Abril Rodriguez-Maldonado, Vanessa Rivero-Arredondo, Ariadna Medina-Benitez, Joaquin Quiroz-Mercado, Sergio Rangel-Guerrero, Natividad Cruz-Ortiz, Tatiana Nunez-Garcia, Gisela Barrera-Badillo, Lucia Hernandez-Rivas, Irma Lopez-Martinez, Ernesto Ramirez-Gonzalez. |
| EPI_ISL_2736825 | LESP Michoacan | Instituto de Diagnostico y Referencia Epidemiologicos (INDRE) | Claudia Wong-Arambula, Abril Rodriguez-Maldonado, Vanessa Rivero-Arredondo, Ariadna Medina-Benitez, Joaquin Quiroz-Mercado, Sergio Rangel-Guerrero, Natividad Cruz-Ortiz, Tatiana Nunez-Garcia, Gisela Barrera-Badillo, Lucia Hernandez-Rivas, Irma Lopez-Martinez, Ernesto Ramirez-Gonzalez. |
| EPI_ISL_2736827 | LESP Guanajuato | Instituto de Diagnostico y Referencia Epidemiologicos (INDRE) | Claudia Wong-Arambula, Abril Rodriguez-Maldonado, Vanessa Rivero-Arredondo, Ariadna Medina-Benitez, Joaquin Quiroz-Mercado, Sergio Rangel-Guerrero, Natividad Cruz-Ortiz, Tatiana Nunez-Garcia, Gisela Barrera-Badillo, Lucia Hernandez-Rivas, Irma Lopez-Martinez, Ernesto Ramirez-Gonzalez. |
| EPI_ISL_2736829 | LESP Coahuila | Instituto de Diagnostico y Referencia Epidemiologicos (INDRE) | Claudia Wong-Arambula, Abril Rodriguez-Maldonado, Vanessa Rivero-Arredondo, Ariadna Medina-Benitez, Joaquin Quiroz-Mercado, Sergio Rangel-Guerrero, Natividad Cruz-Ortiz, Tatiana Nunez-Garcia, Gisela Barrera-Badillo, Lucia Hernandez-Rivas, Irma Lopez-Martinez, Ernesto Ramirez-Gonzalez. |
| EPI_ISL_2736832 | LESP Baja California | Instituto de Diagnostico y Referencia Epidemiologicos (INDRE) | Claudia Wong-Arambula, Abril Rodriguez-Maldonado, Vanessa Rivero-Arredondo, Ariadna Medina-Benitez, Joaquin Quiroz-Mercado, Sergio Rangel-Guerrero, Natividad Cruz-Ortiz, Tatiana Nunez-Garcia, Gisela Barrera-Badillo, Lucia Hernandez-Rivas, Irma Lopez-Martinez, Ernesto Ramirez-Gonzalez. |
| EPI_ISL_2736837, EPI_ISL_2736839 | LESP Tabasco | Instituto de Diagnostico y Referencia Epidemiologicos (INDRE) | Claudia Wong-Arambula, Abril Rodriguez-Maldonado, Vanessa Rivero-Arredondo, Ariadna Medina-Benitez, Joaquin Quiroz-Mercado, Sergio Rangel-Guerrero, Natividad Cruz-Ortiz, Tatiana Nunez-Garcia, Gisela Barrera-Badillo, Lucia Hernandez-Rivas, Irma Lopez-Martinez, Ernesto Ramirez-Gonzalez. |
| EPI_ISL_2736841 | LESP Guerrero | Instituto de Diagnostico y Referencia Epidemiologicos (INDRE) | Claudia Wong-Arambula, Abril Rodriguez-Maldonado, Vanessa Rivero-Arredondo, Ariadna Medina-Benitez, Joaquin Quiroz-Mercado, Sergio Rangel-Guerrero, Natividad Cruz-Ortiz, Tatiana Nunez-Garcia, Gisela Barrera-Badillo, Lucia Hernandez-Rivas, Irma Lopez-Martinez, Ernesto Ramirez-Gonzalez. |
| EPI_ISL_2736845 | LESP Yucatan | Instituto de Diagnostico y Referencia Epidemiologicos (INDRE) | Claudia Wong-Arambula, Abril Rodriguez-Maldonado, Vanessa Rivero-Arredondo, Ariadna Medina-Benitez, Joaquin Quiroz-Mercado, Sergio Rangel-Guerrero, Natividad Cruz-Ortiz, Tatiana Nunez-Garcia, Gisela Barrera-Badillo, Lucia Hernandez-Rivas, Irma Lopez-Martinez, Ernesto Ramirez-Gonzalez. |
| EPI_ISL_2736848, EPI_ISL_2736849, EPI_ISL_2736851 | LESP Michoacan | Instituto de Diagnostico y Referencia Epidemiologicos (INDRE) | Claudia Wong-Arambula, Abril Rodriguez-Maldonado, Vanessa Rivero-Arredondo, Ariadna Medina-Benitez, Joaquin Quiroz-Mercado, Sergio Rangel-Guerrero, Natividad Cruz-Ortiz, Tatiana Nunez-Garcia, Gisela Barrera-Badillo, Lucia Hernandez-Rivas, Irma Lopez-Martinez, Ernesto Ramirez-Gonzalez. |
| EPI_ISL_2736856 | LESP Sonora | Instituto de Diagnostico y Referencia Epidemiologicos (INDRE) | Claudia Wong-Arambula, Abril Rodriguez-Maldonado, Vanessa Rivero-Arredondo, Ariadna Medina-Benitez, Joaquin Quiroz-Mercado, Sergio Rangel-Guerrero, Natividad Cruz-Ortiz, Tatiana Nunez-Garcia, Gisela Barrera-Badillo, Lucia Hernandez-Rivas, Irma Lopez-Martinez, Ernesto Ramirez-Gonzalez. |
| EPI_ISL_2736857 | LESP Campeche | Instituto de Diagnostico y Referencia Epidemiologicos (INDRE) | Claudia Wong-Arambula, Abril Rodriguez-Maldonado, Vanessa Rivero-Arredondo, Ariadna Medina-Benitez, Joaquin Quiroz-Mercado, Sergio Rangel-Guerrero, Natividad Cruz-Ortiz, Tatiana Nunez-Garcia, Gisela Barrera-Badillo, Lucia Hernandez-Rivas, Irma Lopez-Martinez, Ernesto Ramirez-Gonzalez. |
| EPI_ISL_2736861, EPI_ISL_2736864 | LESP Guanajuato | Instituto de Diagnostico y Referencia Epidemiologicos (INDRE) | Claudia Wong-Arambula, Abril Rodriguez-Maldonado, Vanessa Rivero-Arredondo, Ariadna Medina-Benitez, Joaquin Quiroz-Mercado, Sergio Rangel-Guerrero, Natividad Cruz-Ortiz, Tatiana Nunez-Garcia, Gisela Barrera-Badillo, Lucia Hernandez-Rivas, Irma Lopez-Martinez, Ernesto Ramirez-Gonzalez. |
| EPI_ISL_2754138 | Genome Analysis Center, Yamanashi Central Hospital | Genome Analysis Center, Yamanashi Central Hospital | Yosuke Hirotsu |
| EPI_ISL_2754196 | S.C Microbiologia e virologia - Speciale Centro influenza -AOU di Sassari | Laboratorio Biologia Molecolare Sars Cov2 - UOC Laboratorio Analisi - Servizio Medicina di Laboratorio, Ospedale "San Francesco" - ATS-ASSL Nuoro and Laboratorio specialistico UOC Ematologia - Ospedale "San Francesco" - ATS-ASSL Nuoro | Asproni Rosanna, Piras Giovanna, Malune Paolo, Pira Giovanna, Fancello Patrizia, Rosu Valentina, Carta Franco, Fiamma Maura, Mameli Giuseppe, Garau Maria Cristina , Palmas Angelo Domenico,Rimini Elena ,Serra Caterina, Salvatore Rubino |
| EPI_ISL_2754197 | S.C Microbiologia e virologia - Speciale Centro influenza -AOU di Sassari | Laboratorio Biologia Molecolare Sars Cov2 - UOC Laboratorio Analisi - Servizio Medicina di Laboratorio, Ospedale "San Francesco" - ATS-ASSL Nuoro and Laboratorio specialistico UOC Ematologia - Ospedale "San Francesco" - ATS-ASSL Nuoro | Asproni Rosanna, Piras Giovanna, Malune Paolo, Pira Giovanna, Fancello Patrizia, Rosu Valentina, Carta Franco, Fiamma Maura, Mameli Giuseppe, Garau Maria Cristina , Palmas Angelo Domenico,Rimini Elena ,Serra Caterina, Rubino Salvatore |
| EPI_ISL_2754199, EPI_ISL_2754201, EPI_ISL_2754202 | S.C Microbiologia e virologia - Speciale Centro influenza -AOU di Sassari | Laboratorio Biologia Molecolare Sars Cov2 - UOC Laboratorio Analisi - Servizio Medicina di Laboratorio, Ospedale "San Francesco" - ATS-ASSL Nuoro and Laboratorio specialistico UOC Ematologia - Ospedale "San Francesco" - ATS-ASSL Nuoro | Asproni Rosanna, Piras Giovanna, Malune Paolo, Pira Giovanna, Fancello Patrizia, Rosu Valentina, Carta Franco, Fiamma Maura, Mameli Giuseppe, Garau Maria Cristina , Palmas Angelo Domenico,Rimini Elena ,Serra Caterina, Salvatore Rubino |

EPI_ISL_2756364, EPI_ISL_2756365, EPI_ISL_2756366, EPI_ISL_2756367, EPI_ISL_2756368, EPI_ISL_2756369, EPI_ISL_2756370, EPI_ISL_2756371, EPI_ISL_2756372, EPI_ISL_2756373, EPI_ISL_2756374, EPI_ISL_2756375, EPI_ISL_2756376, EPI_ISL_2756377, EPI_ISL_2756379, EPI_ISL_2756381, EPI_ISL_2756382, EPI_ISL_2756384, EPI_ISL_2756385, EPI_ISL_2756386, EPI_ISL_2756387, EPI_ISL_2756389, EPI_ISL_2756390, EPI_ISL_2756391, EPI_ISL_2756392, EPI_ISL_2756393, EPI_ISL_2756394, EPI_ISL_2756395, EPI_ISL_2756396, EPI_ISL_2756397, EPI_ISL_2756398, EPI_ISL_2756400, EPI_ISL_2756401, EPI_ISL_2756402,

| | | | |
|---|---|---|---|
| EPI_ISL_2756403, EPI_ISL_2756404, EPI_ISL_2756405, EPI_ISL_2756406, EPI_ISL_2756407, EPI_ISL_2756408, EPI_ISL_2756409, EPI_ISL_2756411, EPI_ISL_2756414, EPI_ISL_2756415, EPI_ISL_2756416, EPI_ISL_2756418, EPI_ISL_2756419, EPI_ISL_2756420, EPI_ISL_2756421 | | | |
| see above | Microbiology Department, University Hospital Donostia | Microbiology Department, University Hospital Donostia | Marimon JM, Montes M, Piñeiro L, Martin-Peñaranda T, Gomez M, Sorarrain A, Cilla G. |
| EPI_ISL_2756431 | Instituto Adolfo Lutz Central | Instituto Adolfo Lutz, Interdiciplinary Procedures Center, Strategic Laboratory | Claudio Tavares Sacchi, Claudia Regina Gonçalves, Erica Valessa Ramos Gomes, Karoline Rodrigues Campos, Caio Vinicius Dias Lopes, Leonardo Jose Tadeu de Araujo |
| EPI_ISL_2756557 | Centers for Disease Control and Prevention, Dengue Branch | Centers for Disease Control and Prevention, Dengue Branch | Gilberto A. Santiago, Glenda Gonzalez, Betzabel Flores, Keyla Charriez, Gabriela Paz-Bailey, Jorge L. Munoz-Jordan |
| EPI_ISL_2756605, EPI_ISL_2756606, EPI_ISL_2756607, EPI_ISL_2756608, EPI_ISL_2756609 | Dipartimento di Medicina di Laboratorio, Azienda sanitaria universitaria Friuli Centrale (ASU FC) | Dipartimento di Medicina di Laboratorio, Azienda sanitaria universitaria Friuli Centrale (ASU FC) | Corrado Pipan, Catia Mio, Chiara Dal Secco, Stefania Marzinotto, Francesco Curcio |
| EPI_ISL_2757658 | Hospital | National Reference Center for Viruses of Respiratory Infections, Institut Pasteur, Paris | Marion Barbet, Sylvie Behillil, Méline Bizard, Angela Brisebarre, Camille Capel, Vincent Enouf, Louise Lefrançois, Frédéric Lemoine, Christophe Malabat, Corinne Maufrais, Etienne Simon-Lorière, Maud Vanpeene, Sylvie Van der Werf ,JéRome Guinard |
| EPI_ISL_2757659 | Labo Analyses Med | National Reference Center for Viruses of Respiratory Infections, Institut Pasteur, Paris | Marion Barbet, Sylvie Behillil, Méline Bizard, Angela Brisebarre, Camille Capel, Vincent Enouf, Louise Lefrançois, Frédéric Lemoine, Christophe Malabat, Corinne Maufrais, Emmanuelle Permal, Etienne Simon-Lorière, Maud Vanpeene, Sylvie Van der Werf ,GréGoire Potiron |
| EPI_ISL_2757660 | Hospital | National Reference Center for Viruses of Respiratory Infections, Institut Pasteur, Paris | Marion Barbet, Sylvie Behillil, Méline Bizard, Angela Brisebarre, Camille Capel, Vincent Enouf, Louise Lefrançois, Frédéric Lemoine, Christophe Malabat, Corinne Maufrais, Etienne Simon-Lorière, Maud Vanpeene, Sylvie Van der Werf ,Eric Farfour |
| EPI_ISL_2757661 | Hospital | National Reference Center for Viruses of Respiratory Infections, Institut Pasteur, Paris | Marion Barbet, Sylvie Behillil, Méline Bizard, Angela Brisebarre, Camille Capel, Vincent Enouf, Louise Lefrançois, Frédéric Lemoine, Christophe Malabat, Corinne Maufrais, Etienne Simon-Lorière, Maud Vanpeene, Sylvie Van der Werf ,Laurence Courdavault |
| EPI_ISL_2757662 | Hospital | National Reference Center for Viruses of Respiratory Infections, Institut Pasteur, Paris | Marion Barbet, Sylvie Behillil, Méline Bizard, Angela Brisebarre, Camille Capel, Vincent Enouf, Louise Lefrançois, Frédéric Lemoine, Christophe Malabat, Corinne Maufrais, Adrien Pain, Etienne Simon-Lorière, Maud Vanpeene, Sylvie Van der Werf ,Laurence Courdavault |
| EPI_ISL_2757663, EPI_ISL_2757664, EPI_ISL_2757665 | Hospital | National Reference Center for Viruses of Respiratory Infections, Institut Pasteur, Paris | Marion Barbet, Sylvie Behillil, Méline Bizard, Angela Brisebarre, Camille Capel, Vincent Enouf, Louise Lefrançois, Frédéric Lemoine, Christophe Malabat, Corinne Maufrais, Etienne Simon-Lorière, Maud Vanpeene, Sylvie Van der Werf ,JéRôMe Durivault |
| EPI_ISL_2757666 | Hospital | National Reference Center for Viruses of Respiratory Infections, Institut Pasteur, Paris | Marion Barbet, Sylvie Behillil, Méline Bizard, Angela Brisebarre, Camille Capel, Vincent Enouf, Louise Lefrançois, Frédéric Lemoine, Christophe Malabat, Corinne Maufrais, Adrien Pain, Etienne Simon-Lorière, Maud Vanpeene, Sylvie Van der Werf ,JéRôMe Durivault |
| EPI_ISL_2757667, EPI_ISL_2757668, EPI_ISL_2757669, EPI_ISL_2757670, EPI_ISL_2757671, EPI_ISL_2757672, EPI_ISL_2757673 | Hospital | National Reference Center for Viruses of Respiratory Infections, Institut Pasteur, Paris | Marion Barbet, Sylvie Behillil, Méline Bizard, Angela Brisebarre, Camille Capel, Vincent Enouf, Louise Lefrançois, Frédéric Lemoine, Christophe Malabat, Corinne Maufrais, Etienne Simon-Lorière, Maud Vanpeene, Sylvie Van der Werf ,JéRôMe Durivault |
| EPI_ISL_2757674, EPI_ISL_2757675 | Hospital | National Reference Center for Viruses of Respiratory Infections, Institut Pasteur, Paris | Marion Barbet, Sylvie Behillil, Méline Bizard, Angela Brisebarre, Camille Capel, Vincent Enouf, Louise Lefrançois, Frédéric Lemoine, Christophe Malabat, Corinne Maufrais, Amaury Vaysse, Etienne Simon-Lorière, Maud Vanpeene, Sylvie Van der Werf ,JéRôMe Durivault |
| EPI_ISL_2757676, EPI_ISL_2757677, EPI_ISL_2757678 | Hospital | National Reference Center for Viruses of Respiratory Infections, Institut Pasteur, Paris | Marion Barbet, Sylvie Behillil, Méline Bizard, Angela Brisebarre, Camille Capel, Vincent Enouf, Louise Lefrançois, Frédéric Lemoine, Christophe Malabat, Corinne Maufrais, Etienne Simon-Lorière, Maud Vanpeene, Sylvie Van der Werf ,JéRôMe Durivault |
| EPI_ISL_2757679 | Armies | National Reference Center for Viruses of Respiratory Infections, Institut Pasteur, Paris | Marion Barbet, Sylvie Behillil, Méline Bizard, Angela Brisebarre, Camille Capel, Vincent Enouf, Louise Lefrançois, Frédéric Lemoine, Christophe Malabat, Corinne Maufrais, Etienne Simon-Lorière, Maud Vanpeene, Sylvie Van der Werf ,Marine Desroches |
| EPI_ISL_2757680, EPI_ISL_2757681 | Labo Analyses Med | National Reference Center for Viruses of Respiratory Infections, Institut Pasteur, Paris | Marion Barbet, Sylvie Behillil, Méline Bizard, Angela Brisebarre, Camille Capel, Vincent Enouf, Louise Lefrançois, Frédéric Lemoine, Christophe Malabat, Corinne Maufrais, Etienne Simon-Lorière, Maud Vanpeene, Sylvie Van der Werf ,Nader Kherbek |
| EPI_ISL_2757682 | Labo Analyses Med | National Reference Center for Viruses of Respiratory Infections, Institut Pasteur, Paris | Marion Barbet, Sylvie Behillil, Méline Bizard, Angela Brisebarre, Camille Capel, Vincent Enouf, Louise Lefrançois, Frédéric Lemoine, Christophe Malabat, Corinne Maufrais, Amaury Vaysse, Etienne Simon-Lorière, Maud Vanpeene, Sylvie Van der Werf ,GréGoire Potiron |
| EPI_ISL_2757683, EPI_ISL_2757684, EPI_ISL_2757685, EPI_ISL_2757686, EPI_ISL_2757687, EPI_ISL_2757689 | Presidio di Brindisi Di Summa - Perrino | Istituto Zooprofilattico Sperimentale della Puglia e della Basilicata | Parisi A., Bianco A., Capozzi L., Del Sambro L., Simone D., Difato L., Santoro A., Ridolfi D., Giannico A. |
| EPI_ISL_2757692, EPI_ISL_2757693, EPI_ISL_2757694 | Ospedale Santa Caterina Novella | Istituto Zooprofilattico Sperimentale della Puglia e della Basilicata | Parisi A., Bianco A., Capozzi L., Del Sambro L., Simone D., Difato L., Bruno A. R. |
| EPI_ISL_2757695, EPI_ISL_2757696 | Ospedale Cardinale G. Panico | Istituto Zooprofilattico Sperimentale della Puglia e della Basilicata | Parisi A., Bianco A., Capozzi L., Del Sambro L., Simone D., Difato L., Bruno A. R. |
| EPI_ISL_2757697, EPI_ISL_2757698, EPI_ISL_2757699, EPI_ISL_2757700, EPI_ISL_2757701 | Ospedale Vito Fazzi | Istituto Zooprofilattico Sperimentale della Puglia e della Basilicata | Parisi A., Bianco A., Capozzi L., Del Sambro L., Simone D., Difato L., Lobreglio G. |
| EPI_ISL_2757702, EPI_ISL_2757703, EPI_ISL_2757704, EPI_ISL_2757705 | Ospedale Santa Caterina Novella | Istituto Zooprofilattico Sperimentale della Puglia e della Basilicata | Parisi A., Bianco A., Capozzi L., Del Sambro L., Simone D., Difato L., Bruno A. R. |
| EPI_ISL_2757706, EPI_ISL_2757707, EPI_ISL_2757708, EPI_ISL_2757709, EPI_ISL_2757710, EPI_ISL_2757711, EPI_ISL_2757712 | Presidio di Brindisi Di Summa - Perrino | Istituto Zooprofilattico Sperimentale della Puglia e della Basilicata | Parisi A., Bianco A., Capozzi L., Del Sambro L., Simone D., Difato L., Santoro A., Ridolfi D., Giannico A. |
| EPI_ISL_2757713 | Ospedale Santissima Annunziata | Istituto Zooprofilattico Sperimentale della Puglia e della Basilicata | Parisi A., Bianco A., Capozzi L., Del Sambro L., Simone D., Difato L., Ridolfi D., Giannico A, Tampoia M. |
| EPI_ISL_2757714, EPI_ISL_2757716 | Istituto Zooprofilattico Sperimentale della Puglia e della Basilicata | Istituto Zooprofilattico Sperimentale della Puglia e della Basilicata | Parisi A., Bianco A., Capozzi L., Del Sambro L., Simone D., Difato L., Pennuzzi G., Pace L., Galante D. |
| EPI_ISL_2757718, EPI_ISL_2757719, EPI_ISL_2757720, EPI_ISL_2757721 | Ospedale Santa Caterina Novella | Istituto Zooprofilattico Sperimentale della Puglia e della Basilicata | Parisi A., Bianco A., Capozzi L., Del Sambro L., Simone D., Difato L., Bruno A. R. |
| EPI_ISL_2757722 | Dipartimento di prevenzione Bari | Istituto Zooprofilattico Sperimentale della Puglia e della Basilicata | Parisi A., Bianco A., Capozzi L., Del Sambro L., Simone D., Difato L., Ridolfi D., Giannico A., Lagravinese D. |
| EPI_ISL_2757723, EPI_ISL_2757724, EPI_ISL_2757725, EPI_ISL_2757726, EPI_ISL_2757728, EPI_ISL_2757729, EPI_ISL_2757731 | Ospedale Santissima Annunziata | Istituto Zooprofilattico Sperimentale della Puglia e della Basilicata | Parisi A., Bianco A., Capozzi L., Del Sambro L., Simone D., Difato L., Ridolfi D., Giannico A, Tampoia M. |
| EPI_ISL_2757766, EPI_ISL_2757768 | Laboratoire Hôpital Sainte Musse | Laboratoire Hôpital Sainte Musse | Lionel Chollet |
| EPI_ISL_2757772, EPI_ISL_2757773, EPI_ISL_2757774, EPI_ISL_2757775, EPI_ISL_2757776, EPI_ISL_2757777, EPI_ISL_2757778, EPI_ISL_2757779, EPI_ISL_2757780, EPI_ISL_2757781, EPI_ISL_2757782, EPI_ISL_2757783, EPI_ISL_2757785, EPI_ISL_2757786, EPI_ISL_2757787, EPI_ISL_2757788, EPI_ISL_2757789, EPI_ISL_2757790, EPI_ISL_2757791, EPI_ISL_2757793, EPI_ISL_2757794, EPI_ISL_2757796, EPI_ISL_2757797 | | | |
| see above | Microbiology Department, University Hospital Donostia | Microbiology Department, University Hospital Donostia | Marimon JM, Montes M, Piñeiro L, Martin-Peñaranda T, Gomez M, Sorarrain A, Cilla G. |
| EPI_ISL_2757798 | SIESP CHIETI - DRIVE IN LANCIANO | Istituto Zooprofilattico Sperimentale dell'Abruzzo e Molise "G. Caporale" | Lorusso A, Marcacci M, Di Domenico M, Ancora M, Curini V, Di Lollo Valeria, Mangone I, Rinaldi A, Delli Compagni E, Scialabba S, Caporale M, Di Pasquale A, Cammà C, Puglia I, Calistri P, Savini G |
| EPI_ISL_2757799 | SIESP CHIETI - DRIVE IN VASTO | Istituto Zooprofilattico Sperimentale dell'Abruzzo e Molise "G. | Lorusso A, Marcacci M, Di Domenico M, Ancora M, Curini V, Di Lollo Valeria, Mangone I, Rinaldi A, Delli Compagni E, Scialabba S, Caporale M, Di |

| | | | |
|---|---|---|---|
| | | Caporale" | Pasquale A, Cammà C, Puglia I, Calistri P, Savini G |
| EPI_ISL_2757800, EPI_ISL_2757801, EPI_ISL_2757802 | SIESP CHIETI - DRIVE IN LANCIANO | Istituto Zooprofilattico Sperimentale dell'Abruzzo e Molise "G. Caporale" | Lorusso A, Marcacci M, Di Domenico M, Ancora M, Curini V, Di Lollo Valeria, Mangone I, Rinaldi A, Delli Compagni E, Scialabba S, Caporale M, Di Pasquale A, Cammà C, Puglia I, Calistri P, Savini G |
| EPI_ISL_2757803 | PRESIDIO OSPEDALIERO TAGLIACOZZO | Istituto Zooprofilattico Sperimentale dell'Abruzzo e Molise "G. Caporale" | Lorusso A, Marcacci M, Di Domenico M, Ancora M, Curini V, Di Lollo Valeria, Mangone I, Rinaldi A, Delli Compagni E, Scialabba S, Caporale M, Di Pasquale A, Cammà C, Puglia I, Calistri P, Savini G |
| EPI_ISL_2757804, EPI_ISL_2757805, EPI_ISL_2757806 | SIESP DIPARTIMENTO DI PREVENZIONE CHIETI | Istituto Zooprofilattico Sperimentale dell'Abruzzo e Molise "G. Caporale" | Lorusso A, Marcacci M, Di Domenico M, Ancora M, Curini V, Di Lollo Valeria, Mangone I, Rinaldi A, Delli Compagni E, Scialabba S, Caporale M, Di Pasquale A, Cammà C, Puglia I, Calistri P, Savini G |
| EPI_ISL_2757807, EPI_ISL_2757808 | SIESP CHIETI - DRIVE IN LANCIANO | Istituto Zooprofilattico Sperimentale dell'Abruzzo e Molise "G. Caporale" | Lorusso A, Marcacci M, Di Domenico M, Ancora M, Curini V, Di Lollo Valeria, Mangone I, Rinaldi A, Delli Compagni E, Scialabba S, Caporale M, Di Pasquale A, Cammà C, Puglia I, Calistri P, Savini G |
| EPI_ISL_2757809 | SIESP DIPARTIMENTO DI PREVENZIONE CHIETI | Istituto Zooprofilattico Sperimentale dell'Abruzzo e Molise "G. Caporale" | Lorusso A, Marcacci M, Di Domenico M, Ancora M, Curini V, Di Lollo Valeria, Mangone I, Rinaldi A, Delli Compagni E, Scialabba S, Caporale M, Di Pasquale A, Cammà C, Puglia I, Calistri P, Savini G |
| EPI_ISL_2757810 | SIESP CHIETI - DRIVE IN LANCIANO | Istituto Zooprofilattico Sperimentale dell'Abruzzo e Molise "G. Caporale" | Lorusso A, Marcacci M, Di Domenico M, Ancora M, Curini V, Di Lollo Valeria, Mangone I, Rinaldi A, Delli Compagni E, Scialabba S, Caporale M, Di Pasquale A, Cammà C, Puglia I, Calistri P, Savini G |
| EPI_ISL_2757811, EPI_ISL_2757812, EPI_ISL_2757813, EPI_ISL_2757814, EPI_ISL_2757815, EPI_ISL_2757816, EPI_ISL_2757817 | OSPEDALE CIVILE TERAMO - CENTRO TRASFUSIONALE | Istituto Zooprofilattico Sperimentale dell'Abruzzo e Molise "G. Caporale" | Lorusso A, Marcacci M, Di Domenico M, Ancora M, Curini V, Di Lollo Valeria, Mangone I, Rinaldi A, Delli Compagni E, Scialabba S, Caporale M, Di Pasquale A, Cammà C, Puglia I, Calistri P, Savini G |
| EPI_ISL_2757818, EPI_ISL_2757819, EPI_ISL_2757820, EPI_ISL_2757821 | SIESP CHIETI - DRIVE IN LANCIANO | Istituto Zooprofilattico Sperimentale dell'Abruzzo e Molise "G. Caporale" | Lorusso A, Marcacci M, Di Domenico M, Ancora M, Curini V, Di Lollo Valeria, Mangone I, Rinaldi A, Delli Compagni E, Scialabba S, Caporale M, Di Pasquale A, Cammà C, Puglia I, Calistri P, Savini G |
| EPI_ISL_2757822, EPI_ISL_2757823 | OSPEDALE CIVILE TERAMO - CENTRO TRASFUSIONALE | Istituto Zooprofilattico Sperimentale dell'Abruzzo e Molise "G. Caporale" | Lorusso A, Marcacci M, Di Domenico M, Ancora M, Curini V, Di Lollo Valeria, Mangone I, Rinaldi A, Delli Compagni E, Scialabba S, Caporale M, Di Pasquale A, Cammà C, Puglia I, Calistri P, Savini G |
| EPI_ISL_2757824, EPI_ISL_2757825, EPI_ISL_2757826 | SIESP CHIETI - DRIVE IN VASTO | Istituto Zooprofilattico Sperimentale dell'Abruzzo e Molise "G. Caporale" | Lorusso A, Marcacci M, Di Domenico M, Ancora M, Curini V, Di Lollo Valeria, Mangone I, Rinaldi A, Delli Compagni E, Scialabba S, Caporale M, Di Pasquale A, Cammà C, Puglia I, Calistri P, Savini G |
| EPI_ISL_2757827, EPI_ISL_2757828, EPI_ISL_2757829, EPI_ISL_2757830, EPI_ISL_2757831 | SIESP CHIETI - DRIVE IN LANCIANO | Istituto Zooprofilattico Sperimentale dell'Abruzzo e Molise "G. Caporale" | Lorusso A, Marcacci M, Di Domenico M, Ancora M, Curini V, Di Lollo Valeria, Mangone I, Rinaldi A, Delli Compagni E, Scialabba S, Caporale M, Di Pasquale A, Cammà C, Puglia I, Calistri P, Savini G |
| EPI_ISL_2757832 | OSPEDALE CIVILE TERAMO - CENTRO TRASFUSIONALE | Istituto Zooprofilattico Sperimentale dell'Abruzzo e Molise "G. Caporale" | Lorusso A, Marcacci M, Di Domenico M, Ancora M, Curini V, Di Lollo Valeria, Mangone I, Rinaldi A, Delli Compagni E, Scialabba S, Caporale M, Di Pasquale A, Cammà C, Puglia I, Calistri P, Savini G |
| EPI_ISL_2757930 | Johor State Health Department | Institute for Medical Research, Infectious Disease Research Centre, National Institutes of Health, Ministry of Health Malaysia | Suppiah J, Kamel K, Mohd Zawawi Z, Azizan MA, Ramly N, Robert F, Thayan R |
| EPI_ISL_2757941, EPI_ISL_2757942 | Queen Elizabeth Hospital, Kota Kinabalu | Institute for Medical Research, Infectious Disease Research Centre, National Institutes of Health, Ministry of Health Malaysia | Suppiah J, Kamel K, Mohd Zawawi Z, Azizan MA, Ramly N, Robert F, Thayan R |
| EPI_ISL_2757951 | Johor State Health Department | Institute for Medical Research, Infectious Disease Research Centre, National Institutes of Health, Ministry of Health Malaysia | Suppiah J, Kamel K, Mohd Zawawi Z, Azizan MA, Ramly N, Robert F, Thayan R |
| EPI_ISL_2758216, EPI_ISL_2758217, EPI_ISL_2758219, EPI_ISL_2758220, EPI_ISL_2758221, EPI_ISL_2758222, EPI_ISL_2758223, EPI_ISL_2758224 | Laboratoire Hôpital Sainte Musse | Laboratoire Hôpital Sainte Musse | Lionel Chollet |
| EPI_ISL_2758645 | OCME Office Of Chief Medical Examiner | New York City Public Health Laboratory | Jade Wang, et al. |
| EPI_ISL_2759090 | Virology Unit, AOUP | Virology Unit, AOUP | Marialinda Vatteroni, Susi Frateschi, Mauro Pistello |
| EPI_ISL_2759091 | Virology Unit, AOUP | Virology Unit | Marialinda Vatteroni, Susi Frateschi, Mauro Pistello |
| EPI_ISL_2759092 | Virology Unit, AOUP | Virology Unit, AOUP | Marialinda Vatteroni, Susi Frateschi |
| EPI_ISL_2759093 | MOH - Jaber Al-Ahmad Hospital (Innovation Research Laboratory) | MOH - Jaber Al-Ahmad Hospital (Innovation Research Laboratory) | Salman Al-Sabah , Mohammad Alghounaim |
| EPI_ISL_2759095 | Virology Unit, AOUP | Virology Unit | Marialinda Vatteroni, Susi Frateschi |
| EPI_ISL_2759096 | Virology Unit, AOUP | Virology Unit, AOUP | Marialinda Vatteroni, Susi Frateschi |
| EPI_ISL_2763624 | MOH - Jaber Al-Ahmad Hospital (Innovation Research Laboratory) | MOH - Jaber Al-Ahmad Hospital (Innovation Research Laboratory) | Salman Al-Sabah , Mohammad Alghounaim |
| EPI_ISL_2773986, EPI_ISL_2773987, EPI_ISL_2773988, EPI_ISL_2773989, EPI_ISL_2773990, EPI_ISL_2773991, EPI_ISL_2773993, EPI_ISL_2773994, EPI_ISL_2773995, EPI_ISL_2773996, EPI_ISL_2773997, EPI_ISL_2773998, EPI_ISL_2773999 | | | |
| see above | OLVZ Aalst | OLVZ Aalst | Astrid Holderbeke |
| EPI_ISL_2774004 | Cantacuzino National Military-Medical Institute, Viral Respiratory Infections Laboratory | Cantacuzino Institute Virology | Luiza Ustea, Nicoleta Paraschiv, Sorin Dinu, Mihaela Oprea, Mihaela Lazar |
| EPI_ISL_2774005 | Clinical Emergency County Hospital, Craiova | Cantacuzino Institute Virology | Luiza Ustea, Nicoleta Paraschiv, Sorin Dinu, Mihaela Oprea, Mihaela Lazar |
| EPI_ISL_2774006, EPI_ISL_2774007, EPI_ISL_2774008, EPI_ISL_2774010, EPI_ISL_2774011, EPI_ISL_2774012, EPI_ISL_2774013, EPI_ISL_2774014, EPI_ISL_2774015 | Cantacuzino National Military-Medical Institute, Viral Respiratory Infections Laboratory | Cantacuzino Institute Virology | Luiza Ustea, Nicoleta Paraschiv, Sorin Dinu, Mihaela Oprea, Mihaela Lazar |
| EPI_ISL_2774017, EPI_ISL_2774018, EPI_ISL_2774019, EPI_ISL_2774020, EPI_ISL_2774021, EPI_ISL_2774022, EPI_ISL_2774023, EPI_ISL_2774024, EPI_ISL_2774025, EPI_ISL_2774026, EPI_ISL_2774027, EPI_ISL_2774028, EPI_ISL_2774029, EPI_ISL_2774030, EPI_ISL_2774031, EPI_ISL_2774032, EPI_ISL_2774033, EPI_ISL_2774034 | | | |
| see above | Clinical Emergency County Hospital, Craiova | Cantacuzino Institute Virology | Luiza Ustea, Nicoleta Paraschiv, Sorin Dinu, Mihaela Oprea, Mihaela Lazar |
| EPI_ISL_2774035 | Cantacuzino National Military-Medical Institute, Viral Respiratory Infections Laboratory | Cantacuzino Institute Virology | Luiza Ustea, Nicoleta Paraschiv, Sorin Dinu, Mihaela Oprea, Mihaela Lazar |
| EPI_ISL_2774037 | Clinical Emergency County Hospital, Craiova | Cantacuzino Institute Virology | Luiza Ustea, Nicoleta Paraschiv, Sorin Dinu, Mihaela Oprea, Mihaela Lazar |
| EPI_ISL_2774094, EPI_ISL_2774096, EPI_ISL_2774097, EPI_ISL_2774098, EPI_ISL_2774099, EPI_ISL_2774100, EPI_ISL_2774101, EPI_ISL_2774102, EPI_ISL_2774104, EPI_ISL_2774105, EPI_ISL_2774106, EPI_ISL_2774107, EPI_ISL_2774108, EPI_ISL_2774109, EPI_ISL_2774110, EPI_ISL_2774111, EPI_ISL_2774112, EPI_ISL_2774113, EPI_ISL_2774114, EPI_ISL_2774115, EPI_ISL_2774116, EPI_ISL_2774117, EPI_ISL_2774118, EPI_ISL_2774119, EPI_ISL_2774120, EPI_ISL_2774121, EPI_ISL_2774122, EPI_ISL_2774123, EPI_ISL_2774124, EPI_ISL_2774125, EPI_ISL_2774126, EPI_ISL_2774127, EPI_ISL_2774128, EPI_ISL_2774129, EPI_ISL_2774130, EPI_ISL_2774131, EPI_ISL_2774132 | | | |

| | | | |
|---|---|---|---|
| see above | Microbiology Department, University Hospital Donostia | Microbiology Department, University Hospital Donostia | Marimon JM, Montes M, Piñeiro L, Martin-Peñaranda T, Gomez M, Sorarrain A, Cilla G. |
| EPI_ISL_2774241, EPI_ISL_2774242, EPI_ISL_2774244, EPI_ISL_2774245, EPI_ISL_2774246, EPI_ISL_2774247, EPI_ISL_2774248, EPI_ISL_2774249, EPI_ISL_2774252, EPI_ISL_2774257, EPI_ISL_2774259, EPI_ISL_2774260, EPI_ISL_2774261, EPI_ISL_2774265, EPI_ISL_2774266, EPI_ISL_2774270, EPI_ISL_2774271, EPI_ISL_2774272, EPI_ISL_2774274, EPI_ISL_2774276, EPI_ISL_2774277, EPI_ISL_2774278, EPI_ISL_2774279, EPI_ISL_2774281, EPI_ISL_2774285, EPI_ISL_2774286, EPI_ISL_2774287, EPI_ISL_2774288, EPI_ISL_2774289, EPI_ISL_2774290, EPI_ISL_2774291, EPI_ISL_2774294, EPI_ISL_2774295, EPI_ISL_2774296, EPI_ISL_2774297, EPI_ISL_2774298, EPI_ISL_2774303, EPI_ISL_2774304 | | | |
| see above | Baylor Scott & White-Temple | Baylor Scott & White-Temple | Ari Rao, Linden Morales, Kimberly Walker, Marcus Volz, Shelby Johnson |
| EPI_ISL_2774306 | Baylor Scott & White-Irving | Baylor Scott & White-Temple | Ari Rao, Linden Morales, Kimberly Walker, Marcus Volz, Shelby Johnson |
| EPI_ISL_2776369 | Ministry of Health, Jaber Al-Ahmad Hospital | Virology Unit, Department of Microbiology, Faulty of Medicine, Kuwait University | Nada Madi, Hussain Safar, Ebaa Al-Awadhi, Anfal Al-Adwani |
| EPI_ISL_2776430 | Ministry of Health, Jaber Al-Ahmad Hospital | Virology Unit, Department of Microbiology, Faculty of Medicine, Kuwait University | Nada Madi, Hussain Safar, Ebaa Al-Awadhi, Anfal Al-Adwani |
| EPI_ISL_2777211, EPI_ISL_2777212, EPI_ISL_2777213, EPI_ISL_2777214 | Instituto Nacional de Enfermedades Respiratorias (INER) | Centro de Investigación en Enfermedades Infecciosas (CIENI), Instituto Nacional de Enfermedades Respiratorias (INER) | Consorcio Mexicano de Vigilancia Genómica (CoViGen-Mex). Authors (in alphabetical order): Julio Elias Alvarado-Yaah, Carlos F. Arias, Santiago Ávila-Ríos, Eduardo Becerril-Vargas, Víctor Hugo Borja-Aburto, Celia Boukadida, Cristóbal Cháidez-Quiróz, Juan Bautista Chale-Dzul, Ricardo Ciria Merce, Andreu Comas-García, Célida Duque Molina, Julissa Enciso-Ibarra, José Antonio Enciso-Moreno, Gloria Elena Espinosa-Ayala, Fernando Fontove-Herrera, Daniel Fregoso-Rueda, Victor Eduardo Garcia-Arias, Alejandra Garcia-Gasca, Bruno Gomez-Gil, Jean Pierre González, Irvin González-López, Concepción Grajales-Muñiz, Ricardo Grande, Rosa María Gutierrez Rios, Alejandra Hernández-Terán, Alfredo Herrera-Estrella, Carla Ivón Herrera-Najera, Pavel Isa, Daniel Lira Morales, Susana Lopez, Antonio Loza Román, Brenda Irasema Maldonado-Meza, Bernardo Martínez-Miguel, José Arturo Martínez-Orozco, Célida Martínez- Rodríguez, Margarita Matías-Florentino, Fidencio Mejía-Nepomuceno, María Guadalupe de Jesús Mireles-Rivera, Gloria María Molina-Salinas, Hector Montoya-Fuentes, Mario Mújica-Sánchez, José Esteban Muñoz-Medina, José de Jesús Nuñez-Contreras, Alicia Ocaña-Mondragón, Luis Alberto Ochoa-Carrera, Hector Esteban Paz-Juárez, Marissa Pérez-García, Francisco Pulido, Helen Haydee Fernanda Ramírez-Plascencia, Jorge Salas-Hernández, Angel Gustavo Salas-Lais, Alejandro Sanchez-Flores, Clara Esperanza Santacruz-Tinoco, María Guadalupe Santiago-Mauricio, Selene Zárate, Nelly Sélem-Mojica, Blanca Taboada, Kathia Elizabeth Tapia-Diaz, Gloria Vazquez, Joel Armando Vázquez-Pérez. |
| EPI_ISL_2777215 | Instituto Nacional de Cancerología | Centro de Investigación en Enfermedades Infecciosas (CIENI), Instituto Nacional de Enfermedades Respiratorias (INER) | Consorcio Mexicano de Vigilancia Genómica (CoViGen-Mex). Authors (in alphabetical order): Julio Elias Alvarado-Yaah, Carlos F. Arias, Santiago Ávila-Ríos, Eduardo Becerril-Vargas, Víctor Hugo Borja-Aburto, Celia Boukadida, Cristóbal Cháidez-Quiróz, Juan Bautista Chale-Dzul, Ricardo Ciria Merce, Andreu Comas-García, Célida Duque Molina, Julissa Enciso-Ibarra, José Antonio Enciso-Moreno, Gloria Elena Espinosa-Ayala, Fernando Fontove-Herrera, Daniel Fregoso-Rueda, Victor Eduardo Garcia-Arias, Alejandra Garcia-Gasca, Bruno Gomez-Gil, Jean Pierre González, Irvin González-López, Concepción Grajales-Muñiz, Ricardo Grande, Rosa María Gutierrez Rios, Alejandra Hernández-Terán, Alfredo Herrera-Estrella, Carla Ivón Herrera-Najera, Pavel Isa, Daniel Lira Morales, Susana Lopez, Antonio Loza Román, Brenda Irasema Maldonado-Meza, Bernardo Martínez-Miguel, José Arturo Martínez-Orozco, Célida Martínez- Rodríguez, Margarita Matías-Florentino, Fidencio Mejía-Nepomuceno, María Guadalupe de Jesús Mireles-Rivera, Gloria María Molina-Salinas, Hector Montoya-Fuentes, Mario Mújica-Sánchez, José Esteban Muñoz-Medina, José de Jesús Nuñez-Contreras, Alicia Ocaña-Mondragón, Luis Alberto Ochoa-Carrera, Hector Esteban Paz-Juárez, Marissa Pérez-García, Francisco Pulido, Helen Haydee Fernanda Ramírez-Plascencia, Jorge Salas-Hernández, Angel Gustavo Salas-Lais, Alejandro Sanchez-Flores, Clara Esperanza Santacruz-Tinoco, María Guadalupe Santiago-Mauricio, Selene Zárate, Nelly Sélem-Mojica, Blanca Taboada, Kathia Elizabeth Tapia-Diaz, Gloria Vazquez, Joel Armando Vázquez-Pérez. |
| EPI_ISL_2777216, EPI_ISL_2777217, EPI_ISL_2777218, EPI_ISL_2777219, EPI_ISL_2777220 | Instituto Nacional de Enfermedades Respiratorias (INER) | Centro de Investigación en Enfermedades Infecciosas (CIENI), Instituto Nacional de Enfermedades Respiratorias (INER) | Consorcio Mexicano de Vigilancia Genómica (CoViGen-Mex). Authors (in alphabetical order): Julio Elias Alvarado-Yaah, Carlos F. Arias, Santiago Ávila-Ríos, Eduardo Becerril-Vargas, Víctor Hugo Borja-Aburto, Celia Boukadida, Cristóbal Cháidez-Quiróz, Juan Bautista Chale-Dzul, Ricardo Ciria Merce, Andreu Comas-García, Célida Duque Molina, Julissa Enciso-Ibarra, José Antonio Enciso-Moreno, Gloria Elena Espinosa-Ayala, Fernando Fontove-Herrera, Daniel Fregoso-Rueda, Victor Eduardo Garcia-Arias, Alejandra Garcia-Gasca, Bruno Gomez-Gil, Jean Pierre González, Irvin González-López, Concepción Grajales-Muñiz, Ricardo Grande, Rosa María Gutierrez Rios, Alejandra Hernández-Terán, Alfredo Herrera-Estrella, Carla Ivón Herrera-Najera, Pavel Isa, Daniel Lira Morales, Susana Lopez, Antonio Loza Román, Brenda Irasema Maldonado-Meza, Bernardo Martínez-Miguel, José Arturo Martínez-Orozco, Célida Martínez- Rodríguez, Margarita Matías-Florentino, Fidencio Mejía-Nepomuceno, María Guadalupe de Jesús Mireles-Rivera, Gloria María Molina-Salinas, Hector Montoya-Fuentes, Mario Mújica-Sánchez, José Esteban Muñoz-Medina, José de Jesús Nuñez-Contreras, Alicia Ocaña-Mondragón, Luis Alberto Ochoa-Carrera, Hector Esteban Paz-Juárez, Marissa Pérez-García, Francisco Pulido, Helen Haydee Fernanda Ramírez-Plascencia, Jorge Salas-Hernández, Angel Gustavo Salas-Lais, Alejandro Sanchez-Flores, Clara Esperanza Santacruz-Tinoco, María Guadalupe Santiago-Mauricio, Selene Zárate, Nelly Sélem-Mojica, Blanca Taboada, Kathia Elizabeth Tapia-Diaz, Gloria Vazquez, Joel Armando Vázquez-Pérez. |
| EPI_ISL_2777222, EPI_ISL_2777223, EPI_ISL_2777224, EPI_ISL_2777225, EPI_ISL_2777226, EPI_ISL_2777227, EPI_ISL_2777229, EPI_ISL_2777230, EPI_ISL_2777231, EPI_ISL_2777232, EPI_ISL_2777233, EPI_ISL_2777234 | | | |
| see above | Instituto Nacional de Cancerología | Centro de Investigación en Enfermedades Infecciosas (CIENI), Instituto Nacional de Enfermedades Respiratorias (INER) | Consorcio Mexicano de Vigilancia Genómica (CoViGen-Mex). Authors (in alphabetical order): Julio Elias Alvarado-Yaah, Carlos F. Arias, Santiago Ávila-Ríos, Eduardo Becerril-Vargas, Víctor Hugo Borja-Aburto, Celia Boukadida, Cristóbal Cháidez-Quiróz, Juan Bautista Chale-Dzul, Ricardo Ciria Merce, Andreu Comas-García, Célida Duque Molina, Julissa Enciso-Ibarra, José Antonio Enciso-Moreno, Gloria Elena Espinosa-Ayala, Fernando Fontove-Herrera, Daniel Fregoso-Rueda, Victor Eduardo Garcia-Arias, Alejandra Garcia-Gasca, Bruno Gomez-Gil, Jean Pierre González, Irvin González-López, Concepción Grajales-Muñiz, Ricardo Grande, Rosa María Gutierrez Rios, Alejandra Hernández-Terán, Alfredo Herrera-Estrella, Carla Ivón Herrera-Najera, Pavel Isa, Daniel Lira Morales, Susana Lopez, Antonio Loza Román, Brenda Irasema Maldonado-Meza, Bernardo Martínez-Miguel, José Arturo Martínez-Orozco, Célida Martínez- Rodríguez, Margarita Matías-Florentino, Fidencio Mejía-Nepomuceno, María Guadalupe de Jesús Mireles-Rivera, Gloria María Molina-Salinas, Hector Montoya-Fuentes, Mario Mújica-Sánchez, José Esteban Muñoz-Medina, José de Jesús Nuñez-Contreras, Alicia Ocaña-Mondragón, Luis Alberto Ochoa-Carrera, Hector Esteban Paz-Juárez, Marissa Pérez-García, Francisco Pulido, Helen Haydee Fernanda Ramírez-Plascencia, Jorge Salas-Hernández, Angel Gustavo Salas-Lais, Alejandro Sanchez-Flores, Clara Esperanza Santacruz-Tinoco, María Guadalupe Santiago-Mauricio, Selene Zárate, Nelly Sélem-Mojica, Blanca Taboada, Kathia Elizabeth Tapia-Diaz, Gloria Vazquez, Joel Armando Vázquez-Pérez. |
| EPI_ISL_2778018, EPI_ISL_2778019, EPI_ISL_2778020, EPI_ISL_2778021, EPI_ISL_2778022, EPI_ISL_2778023, EPI_ISL_2778024, EPI_ISL_2778025, EPI_ISL_2778026, EPI_ISL_2778027, EPI_ISL_2778028, EPI_ISL_2778029, EPI_ISL_2778030, EPI_ISL_2778031, EPI_ISL_2778032, EPI_ISL_2778033, EPI_ISL_2778034, EPI_ISL_2778035, EPI_ISL_2778036, EPI_ISL_2778037, EPI_ISL_2778038, EPI_ISL_2778039, EPI_ISL_2778040, EPI_ISL_2778041, EPI_ISL_2778042, EPI_ISL_2778043, EPI_ISL_2778045, EPI_ISL_2778046, EPI_ISL_2778047, EPI_ISL_2778048, EPI_ISL_2778049, EPI_ISL_2778050, EPI_ISL_2778051, EPI_ISL_2778052, EPI_ISL_2778053, EPI_ISL_2778054, EPI_ISL_2778055, EPI_ISL_2778056, EPI_ISL_2778057, EPI_ISL_2778058, EPI_ISL_2778059, EPI_ISL_2778060, EPI_ISL_2778061, EPI_ISL_2778062, EPI_ISL_2778063, EPI_ISL_2778064, EPI_ISL_2778065, EPI_ISL_2778066, EPI_ISL_2778067, EPI_ISL_2778068, EPI_ISL_2778069, EPI_ISL_2778070, EPI_ISL_2778071, EPI_ISL_2778072, EPI_ISL_2778073, EPI_ISL_2778074, EPI_ISL_2778075, EPI_ISL_2778076, EPI_ISL_2778077, EPI_ISL_2778078, EPI_ISL_2778079, EPI_ISL_2778080, EPI_ISL_2778081, EPI_ISL_2778082, EPI_ISL_2778083, EPI_ISL_2778084, EPI_ISL_2778085, EPI_ISL_2778086, EPI_ISL_2778087, EPI_ISL_2778088, EPI_ISL_2778089, EPI_ISL_2778090, EPI_ISL_2778091, EPI_ISL_2778092, EPI_ISL_2778093, EPI_ISL_2778094, EPI_ISL_2778095, EPI_ISL_2778096 | | | |
| see above | Cliniques universitaires Saint-Luc | UCLouvain/IREC/MBLG | Jean Ruelle, Bertrand Bearzatto, Benoit Kabamba Mukadi |
| EPI_ISL_2778097, EPI_ISL_2778098, EPI_ISL_2778099 | Clinique Saint-Pierre Ottignies | UCLouvain/IREC/MBLG | Jean Ruelle, Bertrand Bearzatto, Benoit Kabamba Mukadi |
| EPI_ISL_2778101, EPI_ISL_2778103 | Cliniques universitaires Saint-Luc | UCLouvain/IREC/MBLG | Jean Ruelle, Bertrand Bearzatto, Benoit Kabamba Mukadi |
| EPI_ISL_2778984, EPI_ISL_2778985, EPI_ISL_2778986, EPI_ISL_2778987, EPI_ISL_2778988 | LESP Tabasco | Instituto de Diagnostico y Referencia Epidemiologicos (INDRE) | Claudia Wong-Arambula, Abril Rodriguez-Maldonado, Vanessa Rivero-Arredondo, Ariadna Medina-Benitez, Joaquin Quiroz-Mercado, Sergio Rangel-Guerrero, Maribel Gonzalez-Villa, Natividad Cruz-Ortiz, Tatiana Nunez-Garcia, Gisela Barrera-Badillo, Lucia Hernandez-Rivas, Irma Lopez-Martinez, Ernesto Ramirez-Gonzalez. |
| EPI_ISL_2778989 | LESP Sonora | Instituto de Diagnostico y Referencia Epidemiologicos (INDRE) | Claudia Wong-Arambula, Abril Rodriguez-Maldonado, Vanessa Rivero-Arredondo, Ariadna Medina-Benitez, Joaquin Quiroz-Mercado, Sergio Rangel-Guerrero, Maribel Gonzalez-Villa, Natividad Cruz-Ortiz, Tatiana Nunez-Garcia, Gisela Barrera-Badillo, Lucia Hernandez-Rivas, Irma Lopez-Martinez, Ernesto Ramirez-Gonzalez. |
| EPI_ISL_2778990, EPI_ISL_2778991 | LESP Guanajuato | Instituto de Diagnostico y Referencia Epidemiologicos (INDRE) | Claudia Wong-Arambula, Abril Rodriguez-Maldonado, Vanessa Rivero-Arredondo, Ariadna Medina-Benitez, Joaquin Quiroz-Mercado, Sergio Rangel-Guerrero, Maribel Gonzalez-Villa, Natividad Cruz-Ortiz, Tatiana Nunez-Garcia, Gisela Barrera-Badillo, Lucia Hernandez-Rivas, Irma Lopez-Martinez, Ernesto Ramirez-Gonzalez. |
| EPI_ISL_2778993, EPI_ISL_2778994 | LESP Durango | Instituto de Diagnostico y Referencia Epidemiologicos | Claudia Wong-Arambula, Abril Rodriguez-Maldonado, Vanessa Rivero-Arredondo, Ariadna Medina-Benitez, Joaquin Quiroz-Mercado, Sergio |

| | | | |
|---|---|---|---|
| | | (INDRE) | Rangel-Guerrero, Maribel Gonzalez-Villa, Natividad Cruz-Ortiz, Tatiana Nunez-Garcia, Gisela Barrera-Badillo, Lucia Hernandez-Rivas, Irma Lopez-Martinez, Ernesto Ramirez-Gonzalez. |
| EPI_ISL_2778995 | LESP Nayarit | Instituto de Diagnostico y Referencia Epidemiologicos (INDRE) | Claudia Wong-Arambula, Abril Rodriguez-Maldonado, Vanessa Rivero-Arredondo, Ariadna Medina-Benitez, Joaquin Quiroz-Mercado, Sergio Rangel-Guerrero, Maribel Gonzalez-Villa, Natividad Cruz-Ortiz, Tatiana Nunez-Garcia, Gisela Barrera-Badillo, Lucia Hernandez-Rivas, Irma Lopez-Martinez, Ernesto Ramirez-Gonzalez. |
| EPI_ISL_2778996 | LESP Durango | Instituto de Diagnostico y Referencia Epidemiologicos (INDRE) | Claudia Wong-Arambula, Abril Rodriguez-Maldonado, Vanessa Rivero-Arredondo, Ariadna Medina-Benitez, Joaquin Quiroz-Mercado, Sergio Rangel-Guerrero, Maribel Gonzalez-Villa, Natividad Cruz-Ortiz, Tatiana Nunez-Garcia, Gisela Barrera-Badillo, Lucia Hernandez-Rivas, Irma Lopez-Martinez, Ernesto Ramirez-Gonzalez. |
| EPI_ISL_2778997 | LESP Chiapas | Instituto de Diagnostico y Referencia Epidemiologicos (INDRE) | Claudia Wong-Arambula, Abril Rodriguez-Maldonado, Vanessa Rivero-Arredondo, Ariadna Medina-Benitez, Joaquin Quiroz-Mercado, Sergio Rangel-Guerrero, Maribel Gonzalez-Villa, Natividad Cruz-Ortiz, Tatiana Nunez-Garcia, Gisela Barrera-Badillo, Lucia Hernandez-Rivas, Irma Lopez-Martinez, Ernesto Ramirez-Gonzalez. |
| EPI_ISL_2778998 | LESP Sinaloa | Instituto de Diagnostico y Referencia Epidemiologicos (INDRE) | Claudia Wong-Arambula, Abril Rodriguez-Maldonado, Vanessa Rivero-Arredondo, Ariadna Medina-Benitez, Joaquin Quiroz-Mercado, Sergio Rangel-Guerrero, Maribel Gonzalez-Villa, Natividad Cruz-Ortiz, Tatiana Nunez-Garcia, Gisela Barrera-Badillo, Lucia Hernandez-Rivas, Irma Lopez-Martinez, Ernesto Ramirez-Gonzalez. |
| EPI_ISL_2778999 | LESP Estado de Mexico | Instituto de Diagnostico y Referencia Epidemiologicos (INDRE) | Claudia Wong-Arambula, Abril Rodriguez-Maldonado, Vanessa Rivero-Arredondo, Ariadna Medina-Benitez, Joaquin Quiroz-Mercado, Sergio Rangel-Guerrero, Maribel Gonzalez-Villa, Natividad Cruz-Ortiz, Tatiana Nunez-Garcia, Gisela Barrera-Badillo, Lucia Hernandez-Rivas, Irma Lopez-Martinez, Ernesto Ramirez-Gonzalez. |
| EPI_ISL_2779000 | LESP Durango | Instituto de Diagnostico y Referencia Epidemiologicos (INDRE) | Claudia Wong-Arambula, Abril Rodriguez-Maldonado, Vanessa Rivero-Arredondo, Ariadna Medina-Benitez, Joaquin Quiroz-Mercado, Sergio Rangel-Guerrero, Maribel Gonzalez-Villa, Natividad Cruz-Ortiz, Tatiana Nunez-Garcia, Gisela Barrera-Badillo, Lucia Hernandez-Rivas, Irma Lopez-Martinez, Ernesto Ramirez-Gonzalez. |
| EPI_ISL_2779001 | LESP Campeche | Instituto de Diagnostico y Referencia Epidemiologicos (INDRE) | Claudia Wong-Arambula, Abril Rodriguez-Maldonado, Vanessa Rivero-Arredondo, Ariadna Medina-Benitez, Joaquin Quiroz-Mercado, Sergio Rangel-Guerrero, Maribel Gonzalez-Villa, Natividad Cruz-Ortiz, Tatiana Nunez-Garcia, Gisela Barrera-Badillo, Lucia Hernandez-Rivas, Irma Lopez-Martinez, Ernesto Ramirez-Gonzalez. |
| EPI_ISL_2779002, EPI_ISL_2779003, EPI_ISL_2779004 | LESP Tamaulipas | Instituto de Diagnostico y Referencia Epidemiologicos (INDRE) | Claudia Wong-Arambula, Abril Rodriguez-Maldonado, Vanessa Rivero-Arredondo, Ariadna Medina-Benitez, Joaquin Quiroz-Mercado, Sergio Rangel-Guerrero, Maribel Gonzalez-Villa, Natividad Cruz-Ortiz, Tatiana Nunez-Garcia, Gisela Barrera-Badillo, Lucia Hernandez-Rivas, Irma Lopez-Martinez, Ernesto Ramirez-Gonzalez. |
| EPI_ISL_2779005, EPI_ISL_2779006, EPI_ISL_2779007 | LESP Sinaloa | Instituto de Diagnostico y Referencia Epidemiologicos (INDRE) | Claudia Wong-Arambula, Abril Rodriguez-Maldonado, Vanessa Rivero-Arredondo, Ariadna Medina-Benitez, Joaquin Quiroz-Mercado, Sergio Rangel-Guerrero, Maribel Gonzalez-Villa, Natividad Cruz-Ortiz, Tatiana Nunez-Garcia, Gisela Barrera-Badillo, Lucia Hernandez-Rivas, Irma Lopez-Martinez, Ernesto Ramirez-Gonzalez. |
| EPI_ISL_2779008, EPI_ISL_2779009, EPI_ISL_2779010, EPI_ISL_2779011, EPI_ISL_2779012, EPI_ISL_2779013, EPI_ISL_2779014, EPI_ISL_2779015, EPI_ISL_2779016, EPI_ISL_2779017, EPI_ISL_2779018 | | | |
| see above | LESP Yucatan | Instituto de Diagnostico y Referencia Epidemiologicos (INDRE) | Claudia Wong-Arambula, Abril Rodriguez-Maldonado, Vanessa Rivero-Arredondo, Ariadna Medina-Benitez, Joaquin Quiroz-Mercado, Sergio Rangel-Guerrero, Maribel Gonzalez-Villa, Natividad Cruz-Ortiz, Tatiana Nunez-Garcia, Gisela Barrera-Badillo, Lucia Hernandez-Rivas, Irma Lopez-Martinez, Ernesto Ramirez-Gonzalez. |
| EPI_ISL_2779019 | LESP Guanajuato | Instituto de Diagnostico y Referencia Epidemiologicos (INDRE) | Claudia Wong-Arambula, Abril Rodriguez-Maldonado, Vanessa Rivero-Arredondo, Ariadna Medina-Benitez, Joaquin Quiroz-Mercado, Sergio Rangel-Guerrero, Maribel Gonzalez-Villa, Natividad Cruz-Ortiz, Tatiana Nunez-Garcia, Gisela Barrera-Badillo, Lucia Hernandez-Rivas, Irma Lopez-Martinez, Ernesto Ramirez-Gonzalez. |
| EPI_ISL_2779020, EPI_ISL_2779021, EPI_ISL_2779022, EPI_ISL_2779023 | LESP Yucatan | Instituto de Diagnostico y Referencia Epidemiologicos (INDRE) | Claudia Wong-Arambula, Abril Rodriguez-Maldonado, Vanessa Rivero-Arredondo, Ariadna Medina-Benitez, Joaquin Quiroz-Mercado, Sergio Rangel-Guerrero, Maribel Gonzalez-Villa, Natividad Cruz-Ortiz, Tatiana Nunez-Garcia, Gisela Barrera-Badillo, Lucia Hernandez-Rivas, Irma Lopez-Martinez, Ernesto Ramirez-Gonzalez. |
| EPI_ISL_2779024, EPI_ISL_2779025, EPI_ISL_2779026, EPI_ISL_2779027, EPI_ISL_2779028, EPI_ISL_2779029, EPI_ISL_2779030, EPI_ISL_2779031, EPI_ISL_2779032, EPI_ISL_2779033 | LESP Tabasco | Instituto de Diagnostico y Referencia Epidemiologicos (INDRE) | Claudia Wong-Arambula, Abril Rodriguez-Maldonado, Vanessa Rivero-Arredondo, Ariadna Medina-Benitez, Joaquin Quiroz-Mercado, Sergio Rangel-Guerrero, Maribel Gonzalez-Villa, Natividad Cruz-Ortiz, Tatiana Nunez-Garcia, Gisela Barrera-Badillo, Lucia Hernandez-Rivas, Irma Lopez-Martinez, Ernesto Ramirez-Gonzalez. |
| EPI_ISL_2779034 | LESP Coahuila | Instituto de Diagnostico y Referencia Epidemiologicos (INDRE) | Claudia Wong-Arambula, Abril Rodriguez-Maldonado, Vanessa Rivero-Arredondo, Ariadna Medina-Benitez, Joaquin Quiroz-Mercado, Sergio Rangel-Guerrero, Maribel Gonzalez-Villa, Natividad Cruz-Ortiz, Tatiana Nunez-Garcia, Gisela Barrera-Badillo, Lucia Hernandez-Rivas, Irma Lopez-Martinez, Ernesto Ramirez-Gonzalez. |
| EPI_ISL_2779035, EPI_ISL_2779036, EPI_ISL_2779037, EPI_ISL_2779038, EPI_ISL_2779039, EPI_ISL_2779040, EPI_ISL_2779041, EPI_ISL_2779042 | LESP Puebla | Instituto de Diagnostico y Referencia Epidemiologicos (INDRE) | Claudia Wong-Arambula, Abril Rodriguez-Maldonado, Vanessa Rivero-Arredondo, Ariadna Medina-Benitez, Joaquin Quiroz-Mercado, Sergio Rangel-Guerrero, Maribel Gonzalez-Villa, Natividad Cruz-Ortiz, Tatiana Nunez-Garcia, Gisela Barrera-Badillo, Lucia Hernandez-Rivas, Irma Lopez-Martinez, Ernesto Ramirez-Gonzalez. |
| EPI_ISL_2779043, EPI_ISL_2779044, EPI_ISL_2779045, EPI_ISL_2779046 | LESP Tabasco | Instituto de Diagnostico y Referencia Epidemiologicos (INDRE) | Claudia Wong-Arambula, Abril Rodriguez-Maldonado, Vanessa Rivero-Arredondo, Ariadna Medina-Benitez, Joaquin Quiroz-Mercado, Sergio Rangel-Guerrero, Maribel Gonzalez-Villa, Natividad Cruz-Ortiz, Tatiana Nunez-Garcia, Gisela Barrera-Badillo, Lucia Hernandez-Rivas, Irma Lopez-Martinez, Ernesto Ramirez-Gonzalez. |
| EPI_ISL_2779047 | LESP Guanajuato | Instituto de Diagnostico y Referencia Epidemiologicos (INDRE) | Claudia Wong-Arambula, Abril Rodriguez-Maldonado, Vanessa Rivero-Arredondo, Ariadna Medina-Benitez, Joaquin Quiroz-Mercado, Sergio Rangel-Guerrero, Maribel Gonzalez-Villa, Natividad Cruz-Ortiz, Tatiana Nunez-Garcia, Gisela Barrera-Badillo, Lucia Hernandez-Rivas, Irma Lopez-Martinez, Ernesto Ramirez-Gonzalez. |
| EPI_ISL_2779050 | LESP Guerrero | Instituto de Diagnostico y Referencia Epidemiologicos (INDRE) | Claudia Wong-Arambula, Abril Rodriguez-Maldonado, Vanessa Rivero-Arredondo, Ariadna Medina-Benitez, Joaquin Quiroz-Mercado, Sergio Rangel-Guerrero, Maribel Gonzalez-Villa, Natividad Cruz-Ortiz, Tatiana Nunez-Garcia, Gisela Barrera-Badillo, Lucia Hernandez-Rivas, Irma Lopez-Martinez, Ernesto Ramirez-Gonzalez. |
| EPI_ISL_2779051, EPI_ISL_2779052 | LESP Quintana Roo | Instituto de Diagnostico y Referencia Epidemiologicos (INDRE) | Claudia Wong-Arambula, Abril Rodriguez-Maldonado, Vanessa Rivero-Arredondo, Ariadna Medina-Benitez, Joaquin Quiroz-Mercado, Sergio Rangel-Guerrero, Maribel Gonzalez-Villa, Natividad Cruz-Ortiz, Tatiana Nunez-Garcia, Gisela Barrera-Badillo, Lucia Hernandez-Rivas, Irma Lopez-Martinez, Ernesto Ramirez-Gonzalez. |
| EPI_ISL_2779053, EPI_ISL_2779054 | LESP Tabasco | Instituto de Diagnostico y Referencia Epidemiologicos (INDRE) | Claudia Wong-Arambula, Abril Rodriguez-Maldonado, Vanessa Rivero-Arredondo, Ariadna Medina-Benitez, Joaquin Quiroz-Mercado, Sergio Rangel-Guerrero, Maribel Gonzalez-Villa, Natividad Cruz-Ortiz, Tatiana Nunez-Garcia, Gisela Barrera-Badillo, Lucia Hernandez-Rivas, Irma Lopez-Martinez, Ernesto Ramirez-Gonzalez. |
| EPI_ISL_2779055 | LESP Sonora | Instituto de Diagnostico y Referencia Epidemiologicos (INDRE) | Claudia Wong-Arambula, Abril Rodriguez-Maldonado, Vanessa Rivero-Arredondo, Ariadna Medina-Benitez, Joaquin Quiroz-Mercado, Sergio Rangel-Guerrero, Maribel Gonzalez-Villa, Natividad Cruz-Ortiz, Tatiana Nunez-Garcia, Gisela Barrera-Badillo, Lucia Hernandez-Rivas, Irma Lopez-Martinez, Ernesto Ramirez-Gonzalez. |
| EPI_ISL_2779056, EPI_ISL_2779057, EPI_ISL_2779058 | LESP Guanajuato | Instituto de Diagnostico y Referencia Epidemiologicos (INDRE) | Claudia Wong-Arambula, Abril Rodriguez-Maldonado, Vanessa Rivero-Arredondo, Ariadna Medina-Benitez, Joaquin Quiroz-Mercado, Sergio Rangel-Guerrero, Maribel Gonzalez-Villa, Natividad Cruz-Ortiz, Tatiana Nunez-Garcia, Gisela Barrera-Badillo, Lucia Hernandez-Rivas, Irma Lopez-Martinez, Ernesto Ramirez-Gonzalez. |
| EPI_ISL_2779059 | LESP Campeche | Instituto de Diagnostico y Referencia Epidemiologicos (INDRE) | Claudia Wong-Arambula, Abril Rodriguez-Maldonado, Vanessa Rivero-Arredondo, Ariadna Medina-Benitez, Joaquin Quiroz-Mercado, Sergio Rangel-Guerrero, Maribel Gonzalez-Villa, Natividad Cruz-Ortiz, Tatiana Nunez-Garcia, Gisela Barrera-Badillo, Lucia Hernandez-Rivas, Irma Lopez-Martinez, Ernesto Ramirez-Gonzalez. |

| | | | |
|---|---|---|---|
| EPI_ISL_2779060, EPI_ISL_2779061, EPI_ISL_2779062, EPI_ISL_2779063, EPI_ISL_2779064, EPI_ISL_2779065 | LESP Tamaulipas | Instituto de Diagnostico y Referencia Epidemiologicos (INDRE) | Claudia Wong-Arambula, Abril Rodriguez-Maldonado, Vanessa Rivero-Arredondo, Ariadna Medina-Benitez, Joaquin Quiroz-Mercado, Sergio Rangel-Guerrero, Maribel Gonzalez-Villa, Natividad Cruz-Ortiz, Tatiana Nunez-Garcia, Gisela Barrera-Badillo, Lucia Hernandez-Rivas, Irma Lopez-Martinez, Ernesto Ramirez-Gonzalez. |
| EPI_ISL_2779066, EPI_ISL_2779067, EPI_ISL_2779068, EPI_ISL_2779069, EPI_ISL_2779070, EPI_ISL_2779071, EPI_ISL_2779072, EPI_ISL_2779073, EPI_ISL_2779074, EPI_ISL_2779075, EPI_ISL_2779076, EPI_ISL_2779077, EPI_ISL_2779078, EPI_ISL_2779079, EPI_ISL_2779080, EPI_ISL_2779081, EPI_ISL_2779082, EPI_ISL_2779083, EPI_ISL_2779084, EPI_ISL_2779085, EPI_ISL_2779086, EPI_ISL_2779087, EPI_ISL_2779088, EPI_ISL_2779089, EPI_ISL_2779090, EPI_ISL_2779091, EPI_ISL_2779092, EPI_ISL_2779093, EPI_ISL_2779094, EPI_ISL_2779095, EPI_ISL_2779096, EPI_ISL_2779097, EPI_ISL_2779098, EPI_ISL_2779099, EPI_ISL_2779100, EPI_ISL_2779101, EPI_ISL_2779102, EPI_ISL_2779103, EPI_ISL_2779104, EPI_ISL_2779105, EPI_ISL_2779106, EPI_ISL_2779107, EPI_ISL_2779108, EPI_ISL_2779109, EPI_ISL_2779110, EPI_ISL_2779111, EPI_ISL_2779112, EPI_ISL_2779113, EPI_ISL_2779114, EPI_ISL_2779115, EPI_ISL_2779116, EPI_ISL_2779117, EPI_ISL_2779118, EPI_ISL_2779119 | | | |
| see above | LESP Yucatan | Instituto de Diagnostico y Referencia Epidemiologicos (INDRE) | Claudia Wong-Arambula, Abril Rodriguez-Maldonado, Vanessa Rivero-Arredondo, Ariadna Medina-Benitez, Joaquin Quiroz-Mercado, Sergio Rangel-Guerrero, Maribel Gonzalez-Villa, Natividad Cruz-Ortiz, Tatiana Nunez-Garcia, Gisela Barrera-Badillo, Lucia Hernandez-Rivas, Irma Lopez-Martinez, Ernesto Ramirez-Gonzalez. |
| EPI_ISL_2779120 | LESP Guanajuato | Instituto de Diagnostico y Referencia Epidemiologicos (INDRE) | Claudia Wong-Arambula, Abril Rodriguez-Maldonado, Vanessa Rivero-Arredondo, Ariadna Medina-Benitez, Joaquin Quiroz-Mercado, Sergio Rangel-Guerrero, Maribel Gonzalez-Villa, Natividad Cruz-Ortiz, Tatiana Nunez-Garcia, Gisela Barrera-Badillo, Lucia Hernandez-Rivas, Irma Lopez-Martinez, Ernesto Ramirez-Gonzalez. |
| EPI_ISL_2779121, EPI_ISL_2779122, EPI_ISL_2779123, EPI_ISL_2779124, EPI_ISL_2779125, EPI_ISL_2779126, EPI_ISL_2779127, EPI_ISL_2779128, EPI_ISL_2779130, EPI_ISL_2779131, EPI_ISL_2779132, EPI_ISL_2779133, EPI_ISL_2779134, EPI_ISL_2779135, EPI_ISL_2779136, EPI_ISL_2779137, EPI_ISL_2779138, EPI_ISL_2779139, EPI_ISL_2779140, EPI_ISL_2779141, EPI_ISL_2779142, EPI_ISL_2779143, EPI_ISL_2779144, EPI_ISL_2779145, EPI_ISL_2779146, EPI_ISL_2779147, EPI_ISL_2779148, EPI_ISL_2779149, EPI_ISL_2779150, EPI_ISL_2779151, EPI_ISL_2779152, EPI_ISL_2779153, EPI_ISL_2779154, EPI_ISL_2779155, EPI_ISL_2779156, EPI_ISL_2779157, EPI_ISL_2779158 | | | |
| see above | LESP Yucatan | Instituto de Diagnostico y Referencia Epidemiologicos (INDRE) | Claudia Wong-Arambula, Abril Rodriguez-Maldonado, Vanessa Rivero-Arredondo, Ariadna Medina-Benitez, Joaquin Quiroz-Mercado, Sergio Rangel-Guerrero, Maribel Gonzalez-Villa, Natividad Cruz-Ortiz, Tatiana Nunez-Garcia, Gisela Barrera-Badillo, Lucia Hernandez-Rivas, Irma Lopez-Martinez, Ernesto Ramirez-Gonzalez. |
| EPI_ISL_2779159 | LESP San Luis Potosi | Instituto de Diagnostico y Referencia Epidemiologicos (INDRE) | Claudia Wong-Arambula, Abril Rodriguez-Maldonado, Vanessa Rivero-Arredondo, Ariadna Medina-Benitez, Joaquin Quiroz-Mercado, Sergio Rangel-Guerrero, Maribel Gonzalez-Villa, Natividad Cruz-Ortiz, Tatiana Nunez-Garcia, Gisela Barrera-Badillo, Lucia Hernandez-Rivas, Irma Lopez-Martinez, Ernesto Ramirez-Gonzalez. |
| EPI_ISL_2779160, EPI_ISL_2779161, EPI_ISL_2779162, EPI_ISL_2779163, EPI_ISL_2779164, EPI_ISL_2779165, EPI_ISL_2779166, EPI_ISL_2779167, EPI_ISL_2779168, EPI_ISL_2779169, EPI_ISL_2779170, EPI_ISL_2779171, EPI_ISL_2779172 | | | |
| see above | LESP Tabasco | Instituto de Diagnostico y Referencia Epidemiologicos (INDRE) | Claudia Wong-Arambula, Abril Rodriguez-Maldonado, Vanessa Rivero-Arredondo, Ariadna Medina-Benitez, Joaquin Quiroz-Mercado, Sergio Rangel-Guerrero, Maribel Gonzalez-Villa, Natividad Cruz-Ortiz, Tatiana Nunez-Garcia, Gisela Barrera-Badillo, Lucia Hernandez-Rivas, Irma Lopez-Martinez, Ernesto Ramirez-Gonzalez. |
| EPI_ISL_2779173 | LESP Ciudad de Mexico | Instituto de Diagnostico y Referencia Epidemiologicos (INDRE) | Claudia Wong-Arambula, Abril Rodriguez-Maldonado, Vanessa Rivero-Arredondo, Ariadna Medina-Benitez, Joaquin Quiroz-Mercado, Sergio Rangel-Guerrero, Maribel Gonzalez-Villa, Natividad Cruz-Ortiz, Tatiana Nunez-Garcia, Gisela Barrera-Badillo, Lucia Hernandez-Rivas, Irma Lopez-Martinez, Ernesto Ramirez-Gonzalez. |
| EPI_ISL_2779174 | LESP Puebla | Instituto de Diagnostico y Referencia Epidemiologicos (INDRE) | Claudia Wong-Arambula, Abril Rodriguez-Maldonado, Vanessa Rivero-Arredondo, Ariadna Medina-Benitez, Joaquin Quiroz-Mercado, Sergio Rangel-Guerrero, Maribel Gonzalez-Villa, Natividad Cruz-Ortiz, Tatiana Nunez-Garcia, Gisela Barrera-Badillo, Lucia Hernandez-Rivas, Irma Lopez-Martinez, Ernesto Ramirez-Gonzalez. |
| EPI_ISL_2779175 | LESP Jalisco | Instituto de Diagnostico y Referencia Epidemiologicos (INDRE) | Claudia Wong-Arambula, Abril Rodriguez-Maldonado, Vanessa Rivero-Arredondo, Ariadna Medina-Benitez, Joaquin Quiroz-Mercado, Sergio Rangel-Guerrero, Maribel Gonzalez-Villa, Natividad Cruz-Ortiz, Tatiana Nunez-Garcia, Gisela Barrera-Badillo, Lucia Hernandez-Rivas, Irma Lopez-Martinez, Ernesto Ramirez-Gonzalez. |
| EPI_ISL_2779176, EPI_ISL_2779177, EPI_ISL_2779178, EPI_ISL_2779179, EPI_ISL_2779180, EPI_ISL_2779181 | LESP Quintana Roo | Instituto de Diagnostico y Referencia Epidemiologicos (INDRE) | Claudia Wong-Arambula, Abril Rodriguez-Maldonado, Vanessa Rivero-Arredondo, Ariadna Medina-Benitez, Joaquin Quiroz-Mercado, Sergio Rangel-Guerrero, Maribel Gonzalez-Villa, Natividad Cruz-Ortiz, Tatiana Nunez-Garcia, Gisela Barrera-Badillo, Lucia Hernandez-Rivas, Irma Lopez-Martinez, Ernesto Ramirez-Gonzalez. |
| EPI_ISL_2779182, EPI_ISL_2779183 | LESP Tabasco | Instituto de Diagnostico y Referencia Epidemiologicos (INDRE) | Claudia Wong-Arambula, Abril Rodriguez-Maldonado, Vanessa Rivero-Arredondo, Ariadna Medina-Benitez, Joaquin Quiroz-Mercado, Sergio Rangel-Guerrero, Maribel Gonzalez-Villa, Natividad Cruz-Ortiz, Tatiana Nunez-Garcia, Gisela Barrera-Badillo, Lucia Hernandez-Rivas, Irma Lopez-Martinez, Ernesto Ramirez-Gonzalez. |
| EPI_ISL_2779184, EPI_ISL_2779185 | LESP Sonora | Instituto de Diagnostico y Referencia Epidemiologicos (INDRE) | Claudia Wong-Arambula, Abril Rodriguez-Maldonado, Vanessa Rivero-Arredondo, Ariadna Medina-Benitez, Joaquin Quiroz-Mercado, Sergio Rangel-Guerrero, Maribel Gonzalez-Villa, Natividad Cruz-Ortiz, Tatiana Nunez-Garcia, Gisela Barrera-Badillo, Lucia Hernandez-Rivas, Irma Lopez-Martinez, Ernesto Ramirez-Gonzalez. |
| EPI_ISL_2779186, EPI_ISL_2779187, EPI_ISL_2779188, EPI_ISL_2779190, EPI_ISL_2779191, EPI_ISL_2779192, EPI_ISL_2779193 | LESP Guanajuato | Instituto de Diagnostico y Referencia Epidemiologicos (INDRE) | Claudia Wong-Arambula, Abril Rodriguez-Maldonado, Vanessa Rivero-Arredondo, Ariadna Medina-Benitez, Joaquin Quiroz-Mercado, Sergio Rangel-Guerrero, Maribel Gonzalez-Villa, Natividad Cruz-Ortiz, Tatiana Nunez-Garcia, Gisela Barrera-Badillo, Lucia Hernandez-Rivas, Irma Lopez-Martinez, Ernesto Ramirez-Gonzalez. |
| EPI_ISL_2779194 | LESP Tabasco | Instituto de Diagnostico y Referencia Epidemiologicos (INDRE) | Claudia Wong-Arambula, Abril Rodriguez-Maldonado, Vanessa Rivero-Arredondo, Ariadna Medina-Benitez, Joaquin Quiroz-Mercado, Sergio Rangel-Guerrero, Maribel Gonzalez-Villa, Natividad Cruz-Ortiz, Tatiana Nunez-Garcia, Gisela Barrera-Badillo, Lucia Hernandez-Rivas, Irma Lopez-Martinez, Ernesto Ramirez-Gonzalez. |
| EPI_ISL_2779195 | LESP Campeche | Instituto de Diagnostico y Referencia Epidemiologicos (INDRE) | Claudia Wong-Arambula, Abril Rodriguez-Maldonado, Vanessa Rivero-Arredondo, Ariadna Medina-Benitez, Joaquin Quiroz-Mercado, Sergio Rangel-Guerrero, Maribel Gonzalez-Villa, Natividad Cruz-Ortiz, Tatiana Nunez-Garcia, Gisela Barrera-Badillo, Lucia Hernandez-Rivas, Irma Lopez-Martinez, Ernesto Ramirez-Gonzalez. |
| EPI_ISL_2779196 | LESP Guerrero | Instituto de Diagnostico y Referencia Epidemiologicos (INDRE) | Claudia Wong-Arambula, Abril Rodriguez-Maldonado, Vanessa Rivero-Arredondo, Ariadna Medina-Benitez, Joaquin Quiroz-Mercado, Sergio Rangel-Guerrero, Maribel Gonzalez-Villa, Natividad Cruz-Ortiz, Tatiana Nunez-Garcia, Gisela Barrera-Badillo, Lucia Hernandez-Rivas, Irma Lopez-Martinez, Ernesto Ramirez-Gonzalez. |
| EPI_ISL_2779197 | LESP Baja California Sur | Instituto de Diagnostico y Referencia Epidemiologicos (INDRE) | Claudia Wong-Arambula, Abril Rodriguez-Maldonado, Vanessa Rivero-Arredondo, Ariadna Medina-Benitez, Joaquin Quiroz-Mercado, Sergio Rangel-Guerrero, Maribel Gonzalez-Villa, Natividad Cruz-Ortiz, Tatiana Nunez-Garcia, Gisela Barrera-Badillo, Lucia Hernandez-Rivas, Irma Lopez-Martinez, Ernesto Ramirez-Gonzalez. |
| EPI_ISL_2779198, EPI_ISL_2779199, EPI_ISL_2779200, EPI_ISL_2779201, EPI_ISL_2779202, EPI_ISL_2779203, EPI_ISL_2779204, EPI_ISL_2779205, EPI_ISL_2779206, EPI_ISL_2779207, EPI_ISL_2779208, EPI_ISL_2779209, EPI_ISL_2779210 | | | |
| see above | LESP Tabasco | Instituto de Diagnostico y Referencia Epidemiologicos (INDRE) | Claudia Wong-Arambula, Abril Rodriguez-Maldonado, Vanessa Rivero-Arredondo, Ariadna Medina-Benitez, Joaquin Quiroz-Mercado, Sergio Rangel-Guerrero, Maribel Gonzalez-Villa, Natividad Cruz-Ortiz, Tatiana Nunez-Garcia, Gisela Barrera-Badillo, Lucia Hernandez-Rivas, Irma Lopez-Martinez, Ernesto Ramirez-Gonzalez. |
| EPI_ISL_2779211, EPI_ISL_2779212 | LESP Puebla | Instituto de Diagnostico y Referencia Epidemiologicos (INDRE) | Claudia Wong-Arambula, Abril Rodriguez-Maldonado, Vanessa Rivero-Arredondo, Ariadna Medina-Benitez, Joaquin Quiroz-Mercado, Sergio Rangel-Guerrero, Maribel Gonzalez-Villa, Natividad Cruz-Ortiz, Tatiana Nunez-Garcia, Gisela Barrera-Badillo, Lucia Hernandez-Rivas, Irma Lopez-Martinez, Ernesto Ramirez-Gonzalez. |
| EPI_ISL_2779213 | LESP Sonora | Instituto de Diagnostico y Referencia Epidemiologicos (INDRE) | Claudia Wong-Arambula, Abril Rodriguez-Maldonado, Vanessa Rivero-Arredondo, Ariadna Medina-Benitez, Joaquin Quiroz-Mercado, Sergio Rangel-Guerrero, Maribel Gonzalez-Villa, Natividad Cruz-Ortiz, Tatiana Nunez-Garcia, Gisela Barrera-Badillo, Lucia Hernandez-Rivas, Irma Lopez-Martinez, Ernesto Ramirez-Gonzalez. |
| EPI_ISL_2779214 | LESP Guerrero | Instituto de Diagnostico y Referencia Epidemiologicos (INDRE) | Claudia Wong-Arambula, Abril Rodriguez-Maldonado, Vanessa Rivero-Arredondo, Ariadna Medina-Benitez, Joaquin Quiroz-Mercado, Sergio Rangel-Guerrero, Maribel Gonzalez-Villa, Natividad Cruz-Ortiz, Tatiana Nunez-Garcia, Gisela Barrera-Badillo, Lucia Hernandez-Rivas, Irma Lopez-Martinez, Ernesto Ramirez-Gonzalez. |
| EPI_ISL_2779215, EPI_ISL_2779217, EPI_ISL_2779218 | LESP Guanajuato | Instituto de Diagnostico y Referencia Epidemiologicos (INDRE) | Claudia Wong-Arambula, Abril Rodriguez-Maldonado, Vanessa Rivero-Arredondo, Ariadna Medina-Benitez, Joaquin Quiroz-Mercado, Sergio Rangel-Guerrero, Maribel Gonzalez-Villa, Natividad Cruz-Ortiz, Tatiana Nunez-Garcia, Gisela Barrera-Badillo, Lucia Hernandez-Rivas, Irma |

| | | | |
|---|---|---|---|
| | | | Lopez-Martinez, Ernesto Ramirez-Gonzalez. |
| EPI_ISL_2779219 | LESP Tamaulipas | Instituto de Diagnostico y Referencia Epidemiologicos (INDRE) | Claudia Wong-Arambula, Abril Rodriguez-Maldonado, Vanessa Rivero-Arredondo, Ariadna Medina-Benitez, Joaquin Quiroz-Mercado, Sergio Rangel-Guerrero, Maribel Gonzalez-Villa, Natividad Cruz-Ortiz, Tatiana Nunez-Garcia, Gisela Barrera-Badillo, Lucia Hernandez-Rivas, Irma Lopez-Martinez, Ernesto Ramirez-Gonzalez. |
| EPI_ISL_2779220 | LESP Sinaloa | Instituto de Diagnostico y Referencia Epidemiologicos (INDRE) | Claudia Wong-Arambula, Abril Rodriguez-Maldonado, Vanessa Rivero-Arredondo, Ariadna Medina-Benitez, Joaquin Quiroz-Mercado, Sergio Rangel-Guerrero, Maribel Gonzalez-Villa, Natividad Cruz-Ortiz, Tatiana Nunez-Garcia, Gisela Barrera-Badillo, Lucia Hernandez-Rivas, Irma Lopez-Martinez, Ernesto Ramirez-Gonzalez. |
| EPI_ISL_2779221, EPI_ISL_2779222, EPI_ISL_2779223, EPI_ISL_2779224, EPI_ISL_2779225 | LESP Yucatan | Instituto de Diagnostico y Referencia Epidemiologicos (INDRE) | Claudia Wong-Arambula, Abril Rodriguez-Maldonado, Vanessa Rivero-Arredondo, Ariadna Medina-Benitez, Joaquin Quiroz-Mercado, Sergio Rangel-Guerrero, Maribel Gonzalez-Villa, Natividad Cruz-Ortiz, Tatiana Nunez-Garcia, Gisela Barrera-Badillo, Lucia Hernandez-Rivas, Irma Lopez-Martinez, Ernesto Ramirez-Gonzalez. |
| EPI_ISL_2779226 | LESP Michoacan | Instituto de Diagnostico y Referencia Epidemiologicos (INDRE) | Claudia Wong-Arambula, Abril Rodriguez-Maldonado, Vanessa Rivero-Arredondo, Ariadna Medina-Benitez, Joaquin Quiroz-Mercado, Sergio Rangel-Guerrero, Maribel Gonzalez-Villa, Natividad Cruz-Ortiz, Tatiana Nunez-Garcia, Gisela Barrera-Badillo, Lucia Hernandez-Rivas, Irma Lopez-Martinez, Ernesto Ramirez-Gonzalez. |
| EPI_ISL_2779227 | LESP Guerrero | Instituto de Diagnostico y Referencia Epidemiologicos (INDRE) | Claudia Wong-Arambula, Abril Rodriguez-Maldonado, Vanessa Rivero-Arredondo, Ariadna Medina-Benitez, Joaquin Quiroz-Mercado, Sergio Rangel-Guerrero, Maribel Gonzalez-Villa, Natividad Cruz-Ortiz, Tatiana Nunez-Garcia, Gisela Barrera-Badillo, Lucia Hernandez-Rivas, Irma Lopez-Martinez, Ernesto Ramirez-Gonzalez. |
| EPI_ISL_2779228, EPI_ISL_2779229, EPI_ISL_2779230, EPI_ISL_2779231 | LESP Yucatan | Instituto de Diagnostico y Referencia Epidemiologicos (INDRE) | Claudia Wong-Arambula, Abril Rodriguez-Maldonado, Vanessa Rivero-Arredondo, Ariadna Medina-Benitez, Joaquin Quiroz-Mercado, Sergio Rangel-Guerrero, Maribel Gonzalez-Villa, Natividad Cruz-Ortiz, Tatiana Nunez-Garcia, Gisela Barrera-Badillo, Lucia Hernandez-Rivas, Irma Lopez-Martinez, Ernesto Ramirez-Gonzalez. |
| EPI_ISL_2779232, EPI_ISL_2779233 | LESP Michoacan | Instituto de Diagnostico y Referencia Epidemiologicos (INDRE) | Claudia Wong-Arambula, Abril Rodriguez-Maldonado, Vanessa Rivero-Arredondo, Ariadna Medina-Benitez, Joaquin Quiroz-Mercado, Sergio Rangel-Guerrero, Maribel Gonzalez-Villa, Natividad Cruz-Ortiz, Tatiana Nunez-Garcia, Gisela Barrera-Badillo, Lucia Hernandez-Rivas, Irma Lopez-Martinez, Ernesto Ramirez-Gonzalez. |
| EPI_ISL_2779234, EPI_ISL_2779235, EPI_ISL_2779236, EPI_ISL_2779237, EPI_ISL_2779238, EPI_ISL_2779239, EPI_ISL_2779240, EPI_ISL_2779241, EPI_ISL_2779242, EPI_ISL_2779243, EPI_ISL_2779244, EPI_ISL_2779245, EPI_ISL_2779246, EPI_ISL_2779247, EPI_ISL_2779248, EPI_ISL_2779249, EPI_ISL_2779250, EPI_ISL_2779251, EPI_ISL_2779252, EPI_ISL_2779253, EPI_ISL_2779254, EPI_ISL_2779255, EPI_ISL_2779256 | | | |
| see above | LESP Tabasco | Instituto de Diagnostico y Referencia Epidemiologicos (INDRE) | Claudia Wong-Arambula, Abril Rodriguez-Maldonado, Vanessa Rivero-Arredondo, Ariadna Medina-Benitez, Joaquin Quiroz-Mercado, Sergio Rangel-Guerrero, Maribel Gonzalez-Villa, Natividad Cruz-Ortiz, Tatiana Nunez-Garcia, Gisela Barrera-Badillo, Lucia Hernandez-Rivas, Irma Lopez-Martinez, Ernesto Ramirez-Gonzalez. |
| EPI_ISL_2779257, EPI_ISL_2779258, EPI_ISL_2779259, EPI_ISL_2779260, EPI_ISL_2779261, EPI_ISL_2779262, EPI_ISL_2779263, EPI_ISL_2779264, EPI_ISL_2779265, EPI_ISL_2779266, EPI_ISL_2779267, EPI_ISL_2779268, EPI_ISL_2779269 | | | |
| see above | LESP Puebla | Instituto de Diagnostico y Referencia Epidemiologicos (INDRE) | Claudia Wong-Arambula, Abril Rodriguez-Maldonado, Vanessa Rivero-Arredondo, Ariadna Medina-Benitez, Joaquin Quiroz-Mercado, Sergio Rangel-Guerrero, Maribel Gonzalez-Villa, Natividad Cruz-Ortiz, Tatiana Nunez-Garcia, Gisela Barrera-Badillo, Lucia Hernandez-Rivas, Irma Lopez-Martinez, Ernesto Ramirez-Gonzalez. |
| EPI_ISL_2779270, EPI_ISL_2779271, EPI_ISL_2779272, EPI_ISL_2779273, EPI_ISL_2779274, EPI_ISL_2779275 | LESP Tabasco | Instituto de Diagnostico y Referencia Epidemiologicos (INDRE) | Claudia Wong-Arambula, Abril Rodriguez-Maldonado, Vanessa Rivero-Arredondo, Ariadna Medina-Benitez, Joaquin Quiroz-Mercado, Sergio Rangel-Guerrero, Maribel Gonzalez-Villa, Natividad Cruz-Ortiz, Tatiana Nunez-Garcia, Gisela Barrera-Badillo, Lucia Hernandez-Rivas, Irma Lopez-Martinez, Ernesto Ramirez-Gonzalez. |
| EPI_ISL_2779276 | LESP Guanajuato | Instituto de Diagnostico y Referencia Epidemiologicos (INDRE) | Claudia Wong-Arambula, Abril Rodriguez-Maldonado, Vanessa Rivero-Arredondo, Ariadna Medina-Benitez, Joaquin Quiroz-Mercado, Sergio Rangel-Guerrero, Maribel Gonzalez-Villa, Natividad Cruz-Ortiz, Tatiana Nunez-Garcia, Gisela Barrera-Badillo, Lucia Hernandez-Rivas, Irma Lopez-Martinez, Ernesto Ramirez-Gonzalez. |
| EPI_ISL_2779277, EPI_ISL_2779278, EPI_ISL_2779279 | LESP Tabasco | Instituto de Diagnostico y Referencia Epidemiologicos (INDRE) | Claudia Wong-Arambula, Abril Rodriguez-Maldonado, Vanessa Rivero-Arredondo, Ariadna Medina-Benitez, Joaquin Quiroz-Mercado, Sergio Rangel-Guerrero, Maribel Gonzalez-Villa, Natividad Cruz-Ortiz, Tatiana Nunez-Garcia, Gisela Barrera-Badillo, Lucia Hernandez-Rivas, Irma Lopez-Martinez, Ernesto Ramirez-Gonzalez. |
| EPI_ISL_2779280 | LESP Puebla | Instituto de Diagnostico y Referencia Epidemiologicos (INDRE) | Claudia Wong-Arambula, Abril Rodriguez-Maldonado, Vanessa Rivero-Arredondo, Ariadna Medina-Benitez, Joaquin Quiroz-Mercado, Sergio Rangel-Guerrero, Maribel Gonzalez-Villa, Natividad Cruz-Ortiz, Tatiana Nunez-Garcia, Gisela Barrera-Badillo, Lucia Hernandez-Rivas, Irma Lopez-Martinez, Ernesto Ramirez-Gonzalez. |
| EPI_ISL_2779281 | LESP Guanajuato | Instituto de Diagnostico y Referencia Epidemiologicos (INDRE) | Claudia Wong-Arambula, Abril Rodriguez-Maldonado, Vanessa Rivero-Arredondo, Ariadna Medina-Benitez, Joaquin Quiroz-Mercado, Sergio Rangel-Guerrero, Maribel Gonzalez-Villa, Natividad Cruz-Ortiz, Tatiana Nunez-Garcia, Gisela Barrera-Badillo, Lucia Hernandez-Rivas, Irma Lopez-Martinez, Ernesto Ramirez-Gonzalez. |
| EPI_ISL_2785968 | Hospital Clínico Universitario Virgen de la Arrixaca | Hospital Clínico Universitario Virgen de la Arrixaca | Marina Simón Páez y Laura Moreno Parrado |
| EPI_ISL_2786928 | Inovie AS LABOSUD | Inovie AS GenBio | Florian VERCRUYSSEN / Pauline LAURENT |
| EPI_ISL_2786968, EPI_ISL_2786969 | Microbiology Department, University Hospital Donostia | Microbiology Department, University Hospital Donostia | Marimon JM, Montes M, Piñeiro L, Martin-Peñaranda T, Gomez M, Sorarrain A, Cilla G. |
| EPI_ISL_2786994 | private medical practitioners | Hong Kong Department of Health | Alan K.L. Tsang, Peter C.W. Yip, Ken H.L. Ng, Edman T.K. Lam, Rickjason C.W. Chan |
| EPI_ISL_2786995 | VICTORIA PUBLIC MORTUARY | Hong Kong Department of Health | Alan K.L. Tsang, Peter C.W. Yip, Ken H.L. Ng, Edman T.K. Lam, Rickjason C.W. Chan |
| EPI_ISL_2786996 | Queen Elizabeth Hospital | Hong Kong Department of Health | Alan K.L. Tsang, Peter C.W. Yip, Ken H.L. Ng, Edman T.K. Lam, Rickjason C.W. Chan |
| EPI_ISL_2786997, EPI_ISL_2786998, EPI_ISL_2786999, EPI_ISL_2787000, EPI_ISL_2787001 | North Lantau Hospital | Hong Kong Department of Health | Alan K.L. Tsang, Peter C.W. Yip, Ken H.L. Ng, Edman T.K. Lam, Rickjason C.W. Chan |
| EPI_ISL_2787002 | Queen Elizabeth Hospital | Hong Kong Department of Health | Alan K.L. Tsang, Peter C.W. Yip, Ken H.L. Ng, Edman T.K. Lam, Rickjason C.W. Chan |
| EPI_ISL_2787004, EPI_ISL_2787005, EPI_ISL_2787006, EPI_ISL_2787007, EPI_ISL_2787008, EPI_ISL_2787009 | Temporary Specimen Collection Centre at the AsiaWorld-Expo | Hong Kong Department of Health | Alan K.L. Tsang, Peter C.W. Yip, Ken H.L. Ng, Edman T.K. Lam, Rickjason C.W. Chan |
| EPI_ISL_2787010 | HOME QUARANTINE TASKFORCE | Hong Kong Department of Health | Alan K.L. Tsang, Peter C.W. Yip, Ken H.L. Ng, Edman T.K. Lam, Rickjason C.W. Chan |
| EPI_ISL_2787011, EPI_ISL_2787012 | QUARANTINE CAMP | Hong Kong Department of Health | Alan K.L. Tsang, Peter C.W. Yip, Ken H.L. Ng, Edman T.K. Lam, Rickjason C.W. Chan |
| EPI_ISL_2787013, EPI_ISL_2787014 | HOME QUARANTINE TASKFORCE | Hong Kong Department of Health | Alan K.L. Tsang, Peter C.W. Yip, Ken H.L. Ng, Edman T.K. Lam, Rickjason C.W. Chan |
| EPI_ISL_2787015 | Queen Elizabeth Hospital | Hong Kong Department of Health | Alan K.L. Tsang, Peter C.W. Yip, Ken H.L. Ng, Edman T.K. Lam, Rickjason C.W. Chan |
| EPI_ISL_2788723, EPI_ISL_2788724, EPI_ISL_2788725, EPI_ISL_2788726, EPI_ISL_2788731, EPI_ISL_2788733, EPI_ISL_2788736, EPI_ISL_2788737, EPI_ISL_2788738, EPI_ISL_2788740, EPI_ISL_2788741, EPI_ISL_2788742, EPI_ISL_2788743, EPI_ISL_2788744, EPI_ISL_2788745, EPI_ISL_2788746, EPI_ISL_2788752, EPI_ISL_2788753, EPI_ISL_2788754, EPI_ISL_2788755, EPI_ISL_2788756, EPI_ISL_2788757, EPI_ISL_2788761, EPI_ISL_2788762, EPI_ISL_2788763, EPI_ISL_2788778, EPI_ISL_2788779, EPI_ISL_2788780, EPI_ISL_2788781, EPI_ISL_2788782, EPI_ISL_2788784, EPI_ISL_2788785, EPI_ISL_2788786, EPI_ISL_2788787, EPI_ISL_2788788, EPI_ISL_2788789, EPI_ISL_2788790, EPI_ISL_2788791, EPI_ISL_2788792, EPI_ISL_2788793, EPI_ISL_2788794, EPI_ISL_2788796, EPI_ISL_2788797, EPI_ISL_2788799, EPI_ISL_2788800, EPI_ISL_2788801, EPI_ISL_2788803, EPI_ISL_2788804, EPI_ISL_2788805, EPI_ISL_2788806, EPI_ISL_2788807, EPI_ISL_2788808, EPI_ISL_2788809, EPI_ISL_2788811, EPI_ISL_2788815, EPI_ISL_2788816, EPI_ISL_2788817, EPI_ISL_2788818, EPI_ISL_2788821, EPI_ISL_2788822, EPI_ISL_2788826, EPI_ISL_2788827, EPI_ISL_2788828, EPI_ISL_2788829, EPI_ISL_2788830, EPI_ISL_2788831, EPI_ISL_2788833, EPI_ISL_2788834, EPI_ISL_2788835, EPI_ISL_2788836, EPI_ISL_2788837, EPI_ISL_2788842, EPI_ISL_2788843, EPI_ISL_2788844, EPI_ISL_2788845, EPI_ISL_2788847, EPI_ISL_2788848, EPI_ISL_2788849, EPI_ISL_2788850, EPI_ISL_2788851, EPI_ISL_2788855, EPI_ISL_2788857, EPI_ISL_2788858, EPI_ISL_2788859, EPI_ISL_2788860, EPI_ISL_2788865, EPI_ISL_2788866, EPI_ISL_2788867, EPI_ISL_2788868, EPI_ISL_2788870, EPI_ISL_2788871, EPI_ISL_2788874, EPI_ISL_2788876 | | | |
| see above | Synlab Eesti OÜ | 1. Laboratory of Communicable Diseases (Estonia); 2. Eurofins Genomics Europe Sequencing GmbH | Liidia Dotsenko et al. |
| EPI_ISL_2789815, EPI_ISL_2789816, | Centre Hospitalier Universitaire Clermont-Ferrand | CHU Clermont-Ferrand, service de virologie | Bisseux Maxime, Mirand Audrey, Combes Patricia, Henquell Cécile |

| | | | |
|---|---|---|---|
| EPI_ISL_2789817, EPI_ISL_2789818, EPI_ISL_2789835, EPI_ISL_2789836 | | | |
| EPI_ISL_2790132, EPI_ISL_2790133, EPI_ISL_2790134, EPI_ISL_2790135, EPI_ISL_2790136 | Instituto Nacional de Cancerología | Centro de Investigación en Enfermedades Infecciosas (CIENI), Instituto Nacional de Enfermedades Respiratorias (INER) | Consorcio Mexicano de Vigilancia Genómica (CoViGen-Mex). Authors (in alphabetical order): Julio Elias Alvarado-Yaah, Carlos F. Arias, Santiago Ávila-Ríos, Eduardo Becerril-Vargas, Víctor Hugo Borja-Aburto, Celia Boukadida, Cristóbal Cháidez-Quiróz, Juan Bautista Chale-Dzul, Ricardo Ciria Merce, Andreu Comas-García, Célida Duque Molina, Julissa Enciso-Ibarra, José Antonio Enciso-Moreno, Gloria Elena Espinosa-Ayala, Fernando Fontove-Herrera, Daniel Fregoso-Rueda, Victor Eduardo Garcia-Arias, Alejandra Garcia-Gasca, Bruno Gomez-Gil, Jean Pierre González, Irvin González-López, Concepción Grajales-Muñiz, Ricardo Grande, Rosa María Gutierrez Rios, Alejandra Hernández-Terán, Alfredo Herrera-Estrella, Carla Ivón Herrera-Najera, Pavel Isa, Daniel Lira Morales, Susana Lopez, Antonio Loza Román, Brenda Irasema Maldonado-Meza, Bernardo Martínez-Miguel, José Arturo Martínez-Orozco, Célida Martínez- Rodríguez, Margarita Matías-Florentino, Fidencio Mejía-Nepomuceno, María Guadalupe de Jesús Mireles-Rivera, Gloria María Molina-Salinas, Hector Montoya-Fuentes, Mario Mújica-Sánchez, José Esteban Muñoz-Medina, José de Jesús Nuñez-Contreras, Alicia Ocaña-Mondragón, Luis Alberto Ochoa-Carrera, Hector Esteban Paz-Juárez, Marissa Pérez-García, Francisco Pulido, Helen Haydee Fernanda Ramírez-Plascencia, Jorge Salas-Hernández, Angel Gustavo Salas-Lais, Alejandro Sanchez-Flores, Clara Esperanza Santacruz-Tinoco, María Guadalupe Santiago-Mauricio, Selene Zárate, Nelly Sélem-Mojica, Blanca Taboada, Kathia Elizabeth Tapia-Diaz, Gloria Vazquez, Joel Armando Vázquez-Pérez. |
| EPI_ISL_2790175, EPI_ISL_2790176 | Instituto Nacional de Enfermedades Respiratorias (INER) | Centro de Investigación en Enfermedades Infecciosas (CIENI), Instituto Nacional de Enfermedades Respiratorias (INER) | Consorcio Mexicano de Vigilancia Genómica (CoViGen-Mex). Authors (in alphabetical order): Julio Elias Alvarado-Yaah, Carlos F. Arias, Santiago Ávila-Ríos, Eduardo Becerril-Vargas, Víctor Hugo Borja-Aburto, Celia Boukadida, Cristóbal Cháidez-Quiróz, Juan Bautista Chale-Dzul, Ricardo Ciria Merce, Andreu Comas-García, Célida Duque Molina, Julissa Enciso-Ibarra, José Antonio Enciso-Moreno, Gloria Elena Espinosa-Ayala, Fernando Fontove-Herrera, Daniel Fregoso-Rueda, Victor Eduardo Garcia-Arias, Alejandra Garcia-Gasca, Bruno Gomez-Gil, Jean Pierre González, Irvin González-López, Concepción Grajales-Muñiz, Ricardo Grande, Rosa María Gutierrez Rios, Alejandra Hernández-Terán, Alfredo Herrera-Estrella, Carla Ivón Herrera-Najera, Pavel Isa, Daniel Lira Morales, Susana Lopez, Antonio Loza Román, Brenda Irasema Maldonado-Meza, Bernardo Martínez-Miguel, José Arturo Martínez-Orozco, Célida Martínez- Rodríguez, Margarita Matías-Florentino, Fidencio Mejía-Nepomuceno, María Guadalupe de Jesús Mireles-Rivera, Gloria María Molina-Salinas, Hector Montoya-Fuentes, , Mario Mújica-Sánchez, José Esteban Muñoz-Medina, José de Jesús Nuñez-Contreras, Alicia Ocaña-Mondragón, Luis Alberto Ochoa-Carrera, Hector Esteban Paz-Juárez, Marissa Perez-Garcia, Francisco Pulido, Helen Haydee Fernanda Ramírez-Plascencia, Jorge Salas-Hernández, Angel Gustavo Salas-Lais, Alejandro Sanchez-Flores, Clara Esperanza Santacruz-Tinoco, María Guadalupe Santiago-Mauricio, Selene Zárate, Nelly Sélem-Mojica, Blanca Taboada, Gloria Vazquez, Joel Armando Vázquez-Pérez. |
| EPI_ISL_2790236, EPI_ISL_2790239, EPI_ISL_2790242, EPI_ISL_2790243, EPI_ISL_2790244, EPI_ISL_2790245 | CHWAPI - SITE NOTRE DAME | Institut de Pathologie et Genetique (IPG) | Pascale Hilbert, Jérémie Gras |
| EPI_ISL_2790249, EPI_ISL_2790250, EPI_ISL_2790251, EPI_ISL_2790252, EPI_ISL_2790253, EPI_ISL_2790254, EPI_ISL_2790255 | Università degli Studi di Perugia | Istituto Zooprofilattico Sperimentale dell'Abruzzo e Molise "G. Caporale" | Mencacci A, Camilloni B, Biagetti M, Giammarioli M, Lorusso A, Marcacci M, Di Domenico M, Ancora M, Curini V, Mangone I, Rinaldi A, Delli Compagni E, Scialabba S, Di Pasquale A, Cammà C, Puglia I, Calistri P, Savini G |
| EPI_ISL_2790256, EPI_ISL_2790257, EPI_ISL_2790258, EPI_ISL_2790259, EPI_ISL_2790260, EPI_ISL_2790261 | Laboratorio Analisi Osp. Città di Castello - Azienda USL Umbria1 | Istituto Zooprofilattico Sperimentale dell'Abruzzo e Molise "G. Caporale" | Malagigi V, Taccani P,Lorusso A, Marcacci M, Di Domenico M, Ancora M, Curini V, Mangone I, Rinaldi A, Delli Compagni E, Scialabba S, Di Pasquale A, Cammà C, Puglia I, Calistri P, Savini G |
| EPI_ISL_2790262 | Università degli Studi di Perugia | Istituto Zooprofilattico Sperimentale dell'Abruzzo e Molise "G. Caporale" | Mencacci A, Camilloni B, Biagetti M, Giammarioli M, Lorusso A, Marcacci M, Di Domenico M, Ancora M, Curini V, Mangone I, Rinaldi A, Delli Compagni E, Scialabba S, Di Pasquale A, Cammà C, Puglia I, Calistri P, Savini G |
| EPI_ISL_2790263 | Laboratorio Analisi Osp. Città di Castello - Azienda USL Umbria1 | Istituto Zooprofilattico Sperimentale dell'Abruzzo e Molise "G. Caporale" | Malagigi V, Taccani P,Lorusso A, Marcacci M, Di Domenico M, Ancora M, Curini V, Mangone I, Rinaldi A, Delli Compagni E, Scialabba S, Di Pasquale A, Cammà C, Puglia I, Calistri P, Savini G |
| EPI_ISL_2790264, EPI_ISL_2790265, EPI_ISL_2790266, EPI_ISL_2790267, EPI_ISL_2790268, EPI_ISL_2790269, EPI_ISL_2790270, EPI_ISL_2790271, EPI_ISL_2790272, EPI_ISL_2790273, EPI_ISL_2790274, EPI_ISL_2790275, EPI_ISL_2790277 | | | |
| see above | Azienda USL Umbria 2 | Istituto Zooprofilattico Sperimentale dell'Abruzzo e Molise "G. Caporale" | Proietti A, Pistoni E, Lorusso A, Marcacci M, Di Domenico M, Ancora M, Curini V, Mangone I, Rinaldi A, Delli Compagni E, Scialabba S, Di Pasquale A, Cammà C, Puglia I, Calistri P, Savini G |
| EPI_ISL_2790280, EPI_ISL_2790281, EPI_ISL_2790282, EPI_ISL_2790283, EPI_ISL_2790284, EPI_ISL_2790285, EPI_ISL_2790286, EPI_ISL_2790287, EPI_ISL_2790288 | Azienda Ospedaliera Terni | Istituto Zooprofilattico Sperimentale dell'Abruzzo e Molise "G. Caporale" | Palumbo M, Scaccetti A, Lorusso A, Marcacci M, Di Domenico M, Ancora M, Curini V, Mangone I, Rinaldi A, Delli Compagni E, Scialabba S, Di Pasquale A, Cammà C, Puglia I, Calistri P, Savini G |
| EPI_ISL_2790310 | Genome Analysis Center, Yamanashi Central Hospital | Genome Analysis Center, Yamanashi Central Hospital | Yosuke Hirotsu |
| EPI_ISL_2790622, EPI_ISL_2790623, EPI_ISL_2790624, EPI_ISL_2790625, EPI_ISL_2790627, EPI_ISL_2790630, EPI_ISL_2790631, EPI_ISL_2790632, EPI_ISL_2790633, EPI_ISL_2790634, EPI_ISL_2790635, EPI_ISL_2790639, EPI_ISL_2790640, EPI_ISL_2790642, EPI_ISL_2790643, EPI_ISL_2790644, EPI_ISL_2790645, EPI_ISL_2790647, EPI_ISL_2790648, EPI_ISL_2790652, EPI_ISL_2790654, EPI_ISL_2790655, EPI_ISL_2790656, EPI_ISL_2790657, EPI_ISL_2790660, EPI_ISL_2790661, EPI_ISL_2790662, EPI_ISL_2790663, EPI_ISL_2790666, EPI_ISL_2790667, EPI_ISL_2790668, EPI_ISL_2790671, EPI_ISL_2790673, EPI_ISL_2790675, EPI_ISL_2790676, EPI_ISL_2790677, EPI_ISL_2790679, EPI_ISL_2790680, EPI_ISL_2790681, EPI_ISL_2790682, EPI_ISL_2790683, EPI_ISL_2790684, EPI_ISL_2790685, EPI_ISL_2790688, EPI_ISL_2790689, EPI_ISL_2790691, EPI_ISL_2790692, EPI_ISL_2790693 | | | |
| see above | Colorado Department of Public Health and Environment | Colorado Department of Public Health and Environment | Laura Bankers, Molly C. Hetherington-Rauth, Diana Ir, Alexandria Rossheim, Michael Martin, Mandy Waters, Shannon R. Matzinger, Sarah Elizabeth Totten, Emily A. Travanty |
| EPI_ISL_2790862, EPI_ISL_2790863 | Labo Analyses Med | National Reference Center for Viruses of Respiratory Infections, Institut Pasteur, Paris | Marion Barbet, Sylvie Behillil, Méline Bizard, Angela Brisebarre, Camille Capel, Vincent Enouf, Louise Lefrançois, Frédéric Lemoine, Christophe Malabat, Corinne Maufrais, Emmanuelle Permal, Etienne Simon-Lorière, Maud Vanpeene, Sylvie Van der Werf ,GréGoire Potiron |
| EPI_ISL_2790864 | Labo Analyses Med | National Reference Center for Viruses of Respiratory Infections, Institut Pasteur, Paris | Marion Barbet, Sylvie Behillil, Méline Bizard, Angela Brisebarre, Camille Capel, Vincent Enouf, Louise Lefrançois, Frédéric Lemoine, Christophe Malabat, Corinne Maufrais, Emmanuelle Permal, Etienne Simon-Lorière, Maud Vanpeene, Sylvie Van der Werf ,Sabine Trombert |
| EPI_ISL_2790865, EPI_ISL_2790866, EPI_ISL_2790867, EPI_ISL_2790868, EPI_ISL_2790869 | Labo Analyses Med | National Reference Center for Viruses of Respiratory Infections, Institut Pasteur, Paris | Marion Barbet, Sylvie Behillil, Méline Bizard, Angela Brisebarre, Camille Capel, Vincent Enouf, Louise Lefrançois, Frédéric Lemoine, Christophe Malabat, Corinne Maufrais, Emmanuelle Permal, Etienne Simon-Lorière, Maud Vanpeene, Sylvie Van der Werf ,BéNéDicte Roquebert |
| EPI_ISL_2790870, EPI_ISL_2790871, EPI_ISL_2790872 | Labo Analyses Med | National Reference Center for Viruses of Respiratory Infections, Institut Pasteur, Paris | Marion Barbet, Sylvie Behillil, Méline Bizard, Angela Brisebarre, Camille Capel, Vincent Enouf, Louise Lefrançois, Frédéric Lemoine, Christophe Malabat, Corinne Maufrais, Emmanuelle Permal, Etienne Simon-Lorière, Maud Vanpeene, Sylvie Van der Werf ,Sabine Trombert |
| EPI_ISL_2790910, EPI_ISL_2790911 | Hôpital | National Reference Center for Viruses of Respiratory Infections, Institut Pasteur, Paris | Marion Barbet, Sylvie Behillil, Méline Bizard, Angela Brisebarre, Camille Capel, Vincent Enouf, Louise Lefrançois, Frédéric Lemoine, Christophe Malabat, Corinne Maufrais, Emmanuelle Permal, Etienne Simon-Lorière, Maud Vanpeene, Sylvie Van der Werf ,Nabil Gastli |
| EPI_ISL_2790912, EPI_ISL_2790913, EPI_ISL_2790914, EPI_ISL_2790915 | Labo Analyses Med | National Reference Center for Viruses of Respiratory Infections, Institut Pasteur, Paris | Marion Barbet, Sylvie Behillil, Méline Bizard, Angela Brisebarre, Camille Capel, Vincent Enouf, Louise Lefrançois, Frédéric Lemoine, Christophe Malabat, Corinne Maufrais, Emmanuelle Permal, Etienne Simon-Lorière, Maud Vanpeene, Sylvie Van der Werf ,Nicolas Capron |
| EPI_ISL_2790924 | Labo Analyses Med | National Reference Center for Viruses of Respiratory Infections, Institut Pasteur, Paris | Marion Barbet, Sylvie Behillil, Méline Bizard, Angela Brisebarre, Camille Capel, Vincent Enouf, Louise Lefrançois, Frédéric Lemoine, Christophe Malabat, Corinne Maufrais, Emmanuelle Permal, Etienne Simon-Lorière, Maud Vanpeene, Sylvie Van der Werf ,Brieuc Gestin |
| EPI_ISL_2790925 | Hospital | National Reference Center for Viruses of Respiratory Infections, Institut Pasteur, Paris | Marion Barbet, Sylvie Behillil, Méline Bizard, Angela Brisebarre, Camille Capel, Vincent Enouf, Louise Lefrançois, Frédéric Lemoine, Christophe Malabat, Corinne Maufrais, Emmanuelle Permal, Etienne Simon-Lorière, Maud Vanpeene, Sylvie Van der Werf ,Nabil Gastli |
| EPI_ISL_2790927 | Hospital | National Reference Center for Viruses of Respiratory Infections, Institut Pasteur, Paris | Marion Barbet, Sylvie Behillil, Méline Bizard, Angela Brisebarre, Camille Capel, Vincent Enouf, Louise Lefrançois, Frédéric Lemoine, Christophe Malabat, Corinne Maufrais, Emmanuelle Permal, Etienne Simon-Lorière, Maud Vanpeene, Sylvie Van der Werf ,Laurence Louvet |
| EPI_ISL_2790929 | Labo Analyses Med | National Reference Center for Viruses of Respiratory Infections, Institut Pasteur, Paris | Marion Barbet, Sylvie Behillil, Méline Bizard, Angela Brisebarre, Camille Capel, Vincent Enouf, Louise Lefrançois, Frédéric Lemoine, Christophe Malabat, Corinne Maufrais, Emmanuelle Permal, Etienne Simon-Lorière, Maud Vanpeene, Sylvie Van der Werf ,BéNéDicte Roquebert |

| | | | |
|---|---|---|---|
| EPI_ISL_2790930, EPI_ISL_2790931, EPI_ISL_2790932, EPI_ISL_2790933, EPI_ISL_2790935, EPI_ISL_2790936, EPI_ISL_2790937, EPI_ISL_2790938, EPI_ISL_2790939, EPI_ISL_2790940, EPI_ISL_2790941, EPI_ISL_2790942, EPI_ISL_2790943, EPI_ISL_2790944, EPI_ISL_2790945, EPI_ISL_2790946, EPI_ISL_2790947, EPI_ISL_2790948, EPI_ISL_2790949, EPI_ISL_2790950, EPI_ISL_2790952, EPI_ISL_2790953, EPI_ISL_2790954, EPI_ISL_2790955, EPI_ISL_2790956, EPI_ISL_2790957, EPI_ISL_2790959, EPI_ISL_2790960, EPI_ISL_2790961, EPI_ISL_2790963, EPI_ISL_2790964, EPI_ISL_2790965, EPI_ISL_2790966 | | | |
| see above | Hôpital | National Reference Center for Viruses of Respiratory Infections, Institut Pasteur, Paris | Marion Barbet, Sylvie Behillil, Méline Bizard, Angela Brisebarre, Camille Capel, Vincent Enouf, Louise Lefrançois, Frédéric Lemoine, Christophe Malabat, Corinne Maufrais, Emmanuelle Permal, Etienne Simon-Lorière, Maud Vanpeene, Sylvie Van der Werf ,BéNéDicte Roquebert |
| EPI_ISL_2790967, EPI_ISL_2790968, EPI_ISL_2790969, EPI_ISL_2790970, EPI_ISL_2790971, EPI_ISL_2790972, EPI_ISL_2790973 | Labo Analyses Med | National Reference Center for Viruses of Respiratory Infections, Institut Pasteur, Paris | Marion Barbet, Sylvie Behillil, Méline Bizard, Angela Brisebarre, Camille Capel, Vincent Enouf, Louise Lefrançois, Frédéric Lemoine, Christophe Malabat, Corinne Maufrais, Emmanuelle Permal, Etienne Simon-Lorière, Maud Vanpeene, Sylvie Van der Werf ,Sabine Trombert |
| EPI_ISL_2790974 | Labo Analyses Med | National Reference Center for Viruses of Respiratory Infections, Institut Pasteur, Paris | Marion Barbet, Sylvie Behillil, Méline Bizard, Angela Brisebarre, Camille Capel, Vincent Enouf, Louise Lefrançois, Frédéric Lemoine, Christophe Malabat, Corinne Maufrais, Emmanuelle Permal, Etienne Simon-Lorière, Maud Vanpeene, Sylvie Van der Werf ,GréGoire Potiron |
| EPI_ISL_2790975, EPI_ISL_2790976, EPI_ISL_2790977, EPI_ISL_2790978 | Labo Analyses Med | National Reference Center for Viruses of Respiratory Infections, Institut Pasteur, Paris | Marion Barbet, Sylvie Behillil, Méline Bizard, Angela Brisebarre, Camille Capel, Vincent Enouf, Louise Lefrançois, Frédéric Lemoine, Christophe Malabat, Corinne Maufrais, Emmanuelle Permal, Etienne Simon-Lorière, Maud Vanpeene, Sylvie Van der Werf ,Eve Haguenoer |
| EPI_ISL_2790979, EPI_ISL_2790980 | Labo Analyses Med | National Reference Center for Viruses of Respiratory Infections, Institut Pasteur, Paris | Marion Barbet, Sylvie Behillil, Méline Bizard, Angela Brisebarre, Camille Capel, Vincent Enouf, Louise Lefrançois, Frédéric Lemoine, Christophe Malabat, Corinne Maufrais, Emmanuelle Permal, Etienne Simon-Lorière, Maud Vanpeene, Sylvie Van der Werf ,Nicolas Capron |
| EPI_ISL_2790981, EPI_ISL_2790982, EPI_ISL_2790983, EPI_ISL_2790984 | Hospital | National Reference Center for Viruses of Respiratory Infections, Institut Pasteur, Paris | Marion Barbet, Sylvie Behillil, Méline Bizard, Angela Brisebarre, Camille Capel, Vincent Enouf, Louise Lefrançois, Frédéric Lemoine, Christophe Malabat, Corinne Maufrais, Emmanuelle Permal, Etienne Simon-Lorière, Maud Vanpeene, Sylvie Van der Werf ,JéRôMe Durivault |
| EPI_ISL_2790985, EPI_ISL_2790986 | Hospital | National Reference Center for Viruses of Respiratory Infections, Institut Pasteur, Paris | Marion Barbet, Sylvie Behillil, Méline Bizard, Angela Brisebarre, Camille Capel, Vincent Enouf, Louise Lefrançois, Frédéric Lemoine, Christophe Malabat, Corinne Maufrais, Emmanuelle Permal, Etienne Simon-Lorière, Maud Vanpeene, Sylvie Van der Werf ,Patricia Stoessel-Thouvenin |
| EPI_ISL_2790988, EPI_ISL_2790989, EPI_ISL_2790990, EPI_ISL_2790991 | Labo Analyses Med | National Reference Center for Viruses of Respiratory Infections, Institut Pasteur, Paris | Marion Barbet, Sylvie Behillil, Méline Bizard, Angela Brisebarre, Camille Capel, Vincent Enouf, Louise Lefrançois, Frédéric Lemoine, Christophe Malabat, Corinne Maufrais, Emmanuelle Permal, Etienne Simon-Lorière, Maud Vanpeene, Sylvie Van der Werf ,BéNéDicte Roquebert |
| EPI_ISL_2791278, EPI_ISL_2791298, EPI_ISL_2791300, EPI_ISL_2791321 | GH A.CHENEVIER-H.MONDOR | Department of Virology, Henri Mondor University Hospital, Assistance Publique Hôpitaux de Paris, Université Paris-Est Créteil, INSERM U955 | Christophe Rodriguez, Slim Fourati, Vanessa Demontant, Guillaume Gricourt, Melissa N'Debi, Alexandre Soulier, Elisabeth Trawinski, Jean-Michel Pawlotsky |
| EPI_ISL_2800916 | LESP Yucatan | Instituto de Diagnostico y Referencia Epidemiologicos (INDRE) | Claudia Wong-Arambula, Abril Rodriguez-Maldonado, Vanessa Rivero-Arredondo, Ariadna Medina-Benitez, Joaquin Quiroz-Mercado, Sergio Rangel-Guerrero, Maribel Gonzalez-Villa, Natividad Cruz-Ortiz, Tatiana Nunez-Garcia, Gisela Barrera-Badillo, Lucia Hernandez-Rivas, Irma Lopez-Martinez, Ernesto Ramirez-Gonzalez. |
| EPI_ISL_2800917 | LESP Guanajuato | Instituto de Diagnostico y Referencia Epidemiologicos (INDRE) | Claudia Wong-Arambula, Abril Rodriguez-Maldonado, Vanessa Rivero-Arredondo, Ariadna Medina-Benitez, Joaquin Quiroz-Mercado, Sergio Rangel-Guerrero, Maribel Gonzalez-Villa, Natividad Cruz-Ortiz, Tatiana Nunez-Garcia, Gisela Barrera-Badillo, Lucia Hernandez-Rivas, Irma Lopez-Martinez, Ernesto Ramirez-Gonzalez. |
| EPI_ISL_2800918, EPI_ISL_2800919, EPI_ISL_2800920, EPI_ISL_2800921, EPI_ISL_2800922, EPI_ISL_2800923 | LESP Yucatan | Instituto de Diagnostico y Referencia Epidemiologicos (INDRE) | Claudia Wong-Arambula, Abril Rodriguez-Maldonado, Vanessa Rivero-Arredondo, Ariadna Medina-Benitez, Joaquin Quiroz-Mercado, Sergio Rangel-Guerrero, Maribel Gonzalez-Villa, Natividad Cruz-Ortiz, Tatiana Nunez-Garcia, Gisela Barrera-Badillo, Lucia Hernandez-Rivas, Irma Lopez-Martinez, Ernesto Ramirez-Gonzalez. |
| EPI_ISL_2800924 | LESP Guanajuato | Instituto de Diagnostico y Referencia Epidemiologicos (INDRE) | Claudia Wong-Arambula, Abril Rodriguez-Maldonado, Vanessa Rivero-Arredondo, Ariadna Medina-Benitez, Joaquin Quiroz-Mercado, Sergio Rangel-Guerrero, Maribel Gonzalez-Villa, Natividad Cruz-Ortiz, Tatiana Nunez-Garcia, Gisela Barrera-Badillo, Lucia Hernandez-Rivas, Irma Lopez-Martinez, Ernesto Ramirez-Gonzalez. |
| EPI_ISL_2800925 | LESP Tabasco | Instituto de Diagnostico y Referencia Epidemiologicos (INDRE) | Claudia Wong-Arambula, Abril Rodriguez-Maldonado, Vanessa Rivero-Arredondo, Ariadna Medina-Benitez, Joaquin Quiroz-Mercado, Sergio Rangel-Guerrero, Natividad Cruz-Ortiz, Tatiana Nunez-Garcia, Gisela Barrera-Badillo, Lucia Hernandez-Rivas, Irma Lopez-Martinez, Ernesto Ramirez-Gonzalez. |
| EPI_ISL_2800926 | LESP Guanajuato | Instituto de Diagnostico y Referencia Epidemiologicos (INDRE) | Claudia Wong-Arambula, Abril Rodriguez-Maldonado, Vanessa Rivero-Arredondo, Ariadna Medina-Benitez, Joaquin Quiroz-Mercado, Sergio Rangel-Guerrero, Natividad Cruz-Ortiz, Tatiana Nunez-Garcia, Gisela Barrera-Badillo, Lucia Hernandez-Rivas, Irma Lopez-Martinez, Ernesto Ramirez-Gonzalez. |
| EPI_ISL_2800927, EPI_ISL_2800928, EPI_ISL_2800929 | LESP Tabasco | Instituto de Diagnostico y Referencia Epidemiologicos (INDRE) | Claudia Wong-Arambula, Abril Rodriguez-Maldonado, Vanessa Rivero-Arredondo, Ariadna Medina-Benitez, Joaquin Quiroz-Mercado, Sergio Rangel-Guerrero, Natividad Cruz-Ortiz, Tatiana Nunez-Garcia, Gisela Barrera-Badillo, Lucia Hernandez-Rivas, Irma Lopez-Martinez, Ernesto Ramirez-Gonzalez. |
| EPI_ISL_2800930, EPI_ISL_2800931 | LESP Sonora | Instituto de Diagnostico y Referencia Epidemiologicos (INDRE) | Claudia Wong-Arambula, Abril Rodriguez-Maldonado, Vanessa Rivero-Arredondo, Ariadna Medina-Benitez, Joaquin Quiroz-Mercado, Sergio Rangel-Guerrero, Natividad Cruz-Ortiz, Tatiana Nunez-Garcia, Gisela Barrera-Badillo, Lucia Hernandez-Rivas, Irma Lopez-Martinez, Ernesto Ramirez-Gonzalez. |
| EPI_ISL_2800932 | LESP Guanajuato | Instituto de Diagnostico y Referencia Epidemiologicos (INDRE) | Claudia Wong-Arambula, Abril Rodriguez-Maldonado, Vanessa Rivero-Arredondo, Ariadna Medina-Benitez, Joaquin Quiroz-Mercado, Sergio Rangel-Guerrero, Natividad Cruz-Ortiz, Tatiana Nunez-Garcia, Gisela Barrera-Badillo, Lucia Hernandez-Rivas, Irma Lopez-Martinez, Ernesto Ramirez-Gonzalez. |
| EPI_ISL_2800933, EPI_ISL_2800934 | LESP Durango | Instituto de Diagnostico y Referencia Epidemiologicos (INDRE) | Claudia Wong-Arambula, Abril Rodriguez-Maldonado, Vanessa Rivero-Arredondo, Ariadna Medina-Benitez, Joaquin Quiroz-Mercado, Sergio Rangel-Guerrero, Natividad Cruz-Ortiz, Tatiana Nunez-Garcia, Gisela Barrera-Badillo, Lucia Hernandez-Rivas, Irma Lopez-Martinez, Ernesto Ramirez-Gonzalez. |
| EPI_ISL_2800935 | LESP Tamaulipas | Instituto de Diagnostico y Referencia Epidemiologicos (INDRE) | Claudia Wong-Arambula, Abril Rodriguez-Maldonado, Vanessa Rivero-Arredondo, Ariadna Medina-Benitez, Joaquin Quiroz-Mercado, Sergio Rangel-Guerrero, Natividad Cruz-Ortiz, Tatiana Nunez-Garcia, Gisela Barrera-Badillo, Lucia Hernandez-Rivas, Irma Lopez-Martinez, Ernesto Ramirez-Gonzalez. |
| EPI_ISL_2800936 | LESP Puebla | Instituto de Diagnostico y Referencia Epidemiologicos (INDRE) | Claudia Wong-Arambula, Abril Rodriguez-Maldonado, Vanessa Rivero-Arredondo, Ariadna Medina-Benitez, Joaquin Quiroz-Mercado, Sergio Rangel-Guerrero, Natividad Cruz-Ortiz, Tatiana Nunez-Garcia, Gisela Barrera-Badillo, Lucia Hernandez-Rivas, Irma Lopez-Martinez, Ernesto Ramirez-Gonzalez. |
| EPI_ISL_2800937 | LESP Sonora | Instituto de Diagnostico y Referencia Epidemiologicos (INDRE) | Claudia Wong-Arambula, Abril Rodriguez-Maldonado, Vanessa Rivero-Arredondo, Ariadna Medina-Benitez, Joaquin Quiroz-Mercado, Sergio Rangel-Guerrero, Natividad Cruz-Ortiz, Tatiana Nunez-Garcia, Gisela Barrera-Badillo, Lucia Hernandez-Rivas, Irma Lopez-Martinez, Ernesto Ramirez-Gonzalez. |
| EPI_ISL_2800938 | LESP Quintana Roo | Instituto de Diagnostico y Referencia Epidemiologicos (INDRE) | Claudia Wong-Arambula, Abril Rodriguez-Maldonado, Vanessa Rivero-Arredondo, Ariadna Medina-Benitez, Joaquin Quiroz-Mercado, Sergio Rangel-Guerrero, Natividad Cruz-Ortiz, Tatiana Nunez-Garcia, Gisela Barrera-Badillo, Lucia Hernandez-Rivas, Irma Lopez-Martinez, Ernesto Ramirez-Gonzalez. |
| EPI_ISL_2800939, EPI_ISL_2800940 | LESP Tabasco | Instituto de Diagnostico y Referencia Epidemiologicos (INDRE) | Claudia Wong-Arambula, Abril Rodriguez-Maldonado, Vanessa Rivero-Arredondo, Ariadna Medina-Benitez, Joaquin Quiroz-Mercado, Sergio Rangel-Guerrero, Natividad Cruz-Ortiz, Tatiana Nunez-Garcia, Gisela Barrera-Badillo, Lucia Hernandez-Rivas, Irma Lopez-Martinez, Ernesto Ramirez-Gonzalez. |
| EPI_ISL_2800941 | LESP Guanajuato | Instituto de Diagnostico y Referencia Epidemiologicos (INDRE) | Claudia Wong-Arambula, Abril Rodriguez-Maldonado, Vanessa Rivero-Arredondo, Ariadna Medina-Benitez, Joaquin Quiroz-Mercado, Sergio Rangel-Guerrero, Natividad Cruz-Ortiz, Tatiana Nunez-Garcia, Gisela Barrera-Badillo, Lucia Hernandez-Rivas, Irma Lopez-Martinez, Ernesto Ramirez-Gonzalez. |
| EPI_ISL_2800942 | LESP Tabasco | Instituto de Diagnostico y Referencia Epidemiologicos (INDRE) | Claudia Wong-Arambula, Abril Rodriguez-Maldonado, Vanessa Rivero-Arredondo, Ariadna Medina-Benitez, Joaquin Quiroz-Mercado, Sergio Rangel-Guerrero, Maribel Gonzalez-Villa, Natividad Cruz-Ortiz, Tatiana Nunez-Garcia, Gisela Barrera-Badillo, Lucia Hernandez-Rivas, Irma Lopez-Martinez, Ernesto Ramirez-Gonzalez. |

| | | | |
|---|---|---|---|
| EPI_ISL_2800943 | LESP Guanajuato | Instituto de Diagnostico y Referencia Epidemiologicos (INDRE) | Claudia Wong-Arambula, Abril Rodriguez-Maldonado, Vanessa Rivero-Arredondo, Ariadna Medina-Benitez, Joaquin Quiroz-Mercado, Sergio Rangel-Guerrero, Natividad Cruz-Ortiz, Tatiana Nunez-Garcia, Gisela Barrera-Badillo, Lucia Hernandez-Rivas, Irma Lopez-Martinez, Ernesto Ramirez-Gonzalez. |
| EPI_ISL_2800944 | LESP Yucatan | Instituto de Diagnostico y Referencia Epidemiologicos (INDRE) | Claudia Wong-Arambula, Abril Rodriguez-Maldonado, Vanessa Rivero-Arredondo, Ariadna Medina-Benitez, Joaquin Quiroz-Mercado, Sergio Rangel-Guerrero, Natividad Cruz-Ortiz, Tatiana Nunez-Garcia, Gisela Barrera-Badillo, Lucia Hernandez-Rivas, Irma Lopez-Martinez, Ernesto Ramirez-Gonzalez. |
| EPI_ISL_2800945 | LESP Michoacan | Instituto de Diagnostico y Referencia Epidemiologicos (INDRE) | Claudia Wong-Arambula, Abril Rodriguez-Maldonado, Vanessa Rivero-Arredondo, Ariadna Medina-Benitez, Joaquin Quiroz-Mercado, Sergio Rangel-Guerrero, Natividad Cruz-Ortiz, Tatiana Nunez-Garcia, Gisela Barrera-Badillo, Lucia Hernandez-Rivas, Irma Lopez-Martinez, Ernesto Ramirez-Gonzalez. |
| EPI_ISL_2801565, EPI_ISL_2801566, EPI_ISL_2801571, EPI_ISL_2801576, EPI_ISL_2801578, EPI_ISL_2801579, EPI_ISL_2801580, EPI_ISL_2801581, EPI_ISL_2801583, EPI_ISL_2801584, EPI_ISL_2801585, EPI_ISL_2801586, EPI_ISL_2801587, EPI_ISL_2801588, EPI_ISL_2801591, EPI_ISL_2801592, EPI_ISL_2801593, EPI_ISL_2801595, EPI_ISL_2801596, EPI_ISL_2801597, EPI_ISL_2801598, EPI_ISL_2801599, EPI_ISL_2801600, EPI_ISL_2801603, EPI_ISL_2801604, EPI_ISL_2801606, EPI_ISL_2801609, EPI_ISL_2801610, EPI_ISL_2801611, EPI_ISL_2801612, EPI_ISL_2801617, EPI_ISL_2801618, EPI_ISL_2801619, EPI_ISL_2801622, EPI_ISL_2801623, EPI_ISL_2801624, EPI_ISL_2801625, EPI_ISL_2801628, EPI_ISL_2801629, EPI_ISL_2801631, EPI_ISL_2801637, EPI_ISL_2801641, EPI_ISL_2801642, EPI_ISL_2801643, EPI_ISL_2801644, EPI_ISL_2801645, EPI_ISL_2801646, EPI_ISL_2801648, EPI_ISL_2801649, EPI_ISL_2801652, EPI_ISL_2801653, EPI_ISL_2801654, EPI_ISL_2801655, EPI_ISL_2801656, EPI_ISL_2801657 | | | |
| see above | Unidad de Investigación Médica de Yucatán (UIMY) | Instituto de Biotecnología de la UNAM | Consorcio Mexicano de Vigilancia Genómica (CoViGen-Mex). Authors (in alphabetical order): Julio Elias Alvarado-Yaah, Carlos F. Arias, Santiago Ávila-Ríos, Eduardo Becerril-Vargas, Víctor Hugo Borja-Aburto, Celia Boukadida, Cristóbal Cháidez-Quiróz, Juan Bautista Chale-Dzul, Ricardo Ciria Merce, Andreu Comas-García, Célida Duque Molina, Julissa Enciso-Ibarra, José Antonio Enciso-Moreno, Gloria Elena Espinosa-Ayala, Fernando Fontove-Herrera, Daniel Fregoso-Rueda, Victor Eduardo García-Arias, Alejandra García-Gasca, Bruno Gómez-Gil, Jean Pierre González, Irvin González-López, Concepción Grajales-Muñiz, Ricardo Grande, Rosa María Gutiérrez Rios, Jesús Hernández, Alejandra Hernández-Terán, Alfredo Herrera-Estrella, Carla Ivón Herrera-Najera, Pavel Isa, Verónica Mata-Haro, Daniel Lira Morales, Susana Lopez, Antonio Loza Román, Brenda Irasema Maldonado-Meza, Bernardo Martínez-Miguel, José Arturo Martínez-Orozco, Célida Martínez- Rodríguez, Margarita Matías-Florentino, Fidencio Mejía-Nepomuceno, María Guadalupe de Jesús Mireles-Rivera, Gloria María Molina-Salinas, Hector Montoya-Fuentes, , Mario Mújica-Sánchez, José Esteban Muñoz-Medina, José de Jesús Nuñez-Contreras, Alicia Ocaña-Mondragón, Luis Alberto Ochoa-Carrera, Hector Esteban Paz-Juárez, Marissa Perez-Garcia, Francisco Pulido, Helen Haydee Fernanda Ramírez-Plascencia, Jorge Salas-Hernández, Angel Gustavo Salas-Lais, Alejandro Sánchez-Flores, Clara Esperanza Santacruz-Tinoco, María Guadalupe Santiago-Mauricio, Selene Zárate, Nelly Sélem-Mojica, Blanca Taboada, Gloria Vazquez, Joel Armando Vázquez-Pérez. |
| EPI_ISL_2801658, EPI_ISL_2801664, EPI_ISL_2801667, EPI_ISL_2801668, EPI_ISL_2801669, EPI_ISL_2801670, EPI_ISL_2801673, EPI_ISL_2801674, EPI_ISL_2801675, EPI_ISL_2801680, EPI_ISL_2801681, EPI_ISL_2801683, EPI_ISL_2801685, EPI_ISL_2801692, EPI_ISL_2801695 | | | |
| see above | Centro de Investigación Biomédica del Noreste (CIBIN) | Instituto de Biotecnología de la UNAM | Consorcio Mexicano de Vigilancia Genómica (CoViGen-Mex). Authors (in alphabetical order): Julio Elias Alvarado-Yaah, Carlos F. Arias, Santiago Ávila-Ríos, Eduardo Becerril-Vargas, Víctor Hugo Borja-Aburto, Celia Boukadida, Cristóbal Cháidez-Quiróz, Juan Bautista Chale-Dzul, Ricardo Ciria Merce, Andreu Comas-García, Célida Duque Molina, Julissa Enciso-Ibarra, José Antonio Enciso-Moreno, Gloria Elena Espinosa-Ayala, Fernando Fontove-Herrera, Daniel Fregoso-Rueda, Victor Eduardo García-Arias, Alejandra García-Gasca, Bruno Gómez-Gil, Jean Pierre González, Irvin González-López, Concepción Grajales-Muñiz, Ricardo Grande, Rosa María Gutiérrez Rios, Jesús Hernández, Alejandra Hernández-Terán, Alfredo Herrera-Estrella, Carla Ivón Herrera-Najera, Pavel Isa, Verónica Mata-Haro, Daniel Lira Morales, Susana Lopez, Antonio Loza Román, Brenda Irasema Maldonado-Meza, Bernardo Martínez-Miguel, José Arturo Martínez-Orozco, Célida Martínez- Rodríguez, Margarita Matías-Florentino, Fidencio Mejía-Nepomuceno, María Guadalupe de Jesús Mireles-Rivera, Gloria María Molina-Salinas, Hector Montoya-Fuentes, , Mario Mújica-Sánchez, José Esteban Muñoz-Medina, José de Jesús Nuñez-Contreras, Alicia Ocaña-Mondragón, Luis Alberto Ochoa-Carrera, Hector Esteban Paz-Juárez, Marissa Perez-Garcia, Francisco Pulido, Helen Haydee Fernanda Ramírez-Plascencia, Jorge Salas-Hernández, Angel Gustavo Salas-Lais, Alejandro Sánchez-Flores, Clara Esperanza Santacruz-Tinoco, María Guadalupe Santiago-Mauricio, Selene Zárate, Nelly Sélem-Mojica, Blanca Taboada, Gloria Vazquez, Joel Armando Vázquez-Pérez. |
| EPI_ISL_2801698, EPI_ISL_2801700, EPI_ISL_2801701, EPI_ISL_2801702, EPI_ISL_2801704, EPI_ISL_2801705 | Unidad de Investigación Biomédica de Zacatecas (UIBZ) | Instituto de Biotecnología de la UNAM | Consorcio Mexicano de Vigilancia Genómica (CoViGen-Mex). Authors (in alphabetical order): Julio Elias Alvarado-Yaah, Carlos F. Arias, Santiago Ávila-Ríos, Eduardo Becerril-Vargas, Víctor Hugo Borja-Aburto, Celia Boukadida, Cristóbal Cháidez-Quiróz, Juan Bautista Chale-Dzul, Ricardo Ciria Merce, Andreu Comas-García, Célida Duque Molina, Julissa Enciso-Ibarra, José Antonio Enciso-Moreno, Gloria Elena Espinosa-Ayala, Fernando Fontove-Herrera, Daniel Fregoso-Rueda, Victor Eduardo García-Arias, Alejandra García-Gasca, Bruno Gómez-Gil, Jean Pierre González, Irvin González-López, Concepción Grajales-Muñiz, Ricardo Grande, Rosa María Gutiérrez Rios, Jesús Hernández, Alejandra Hernández-Terán, Alfredo Herrera-Estrella, Carla Ivón Herrera-Najera, Pavel Isa, Verónica Mata-Haro, Daniel Lira Morales, Susana Lopez, Antonio Loza Román, Brenda Irasema Maldonado-Meza, Bernardo Martínez-Miguel, José Arturo Martínez-Orozco, Célida Martínez- Rodríguez, Margarita Matías-Florentino, Fidencio Mejía-Nepomuceno, María Guadalupe de Jesús Mireles-Rivera, Gloria María Molina-Salinas, Hector Montoya-Fuentes, , Mario Mújica-Sánchez, José Esteban Muñoz-Medina, José de Jesús Nuñez-Contreras, Alicia Ocaña-Mondragón, Luis Alberto Ochoa-Carrera, Hector Esteban Paz-Juárez, Marissa Perez-Garcia, Francisco Pulido, Helen Haydee Fernanda Ramírez-Plascencia, Jorge Salas-Hernández, Angel Gustavo Salas-Lais, Alejandro Sánchez-Flores, Clara Esperanza Santacruz-Tinoco, María Guadalupe Santiago-Mauricio, Selene Zárate, Nelly Sélem-Mojica, Blanca Taboada, Gloria Vazquez, Joel Armando Vázquez-Pérez. |
| EPI_ISL_2801707, EPI_ISL_2801708, EPI_ISL_2801709, EPI_ISL_2801710, EPI_ISL_2801713, EPI_ISL_2801718, EPI_ISL_2801719, EPI_ISL_2801720, EPI_ISL_2801721, EPI_ISL_2801722, EPI_ISL_2801723, EPI_ISL_2801724, EPI_ISL_2801725, EPI_ISL_2801726, EPI_ISL_2801727, EPI_ISL_2801729, EPI_ISL_2801730, EPI_ISL_2801731, EPI_ISL_2801732, EPI_ISL_2801733, EPI_ISL_2801734, EPI_ISL_2801735, EPI_ISL_2801736, EPI_ISL_2801737, EPI_ISL_2801738, EPI_ISL_2801739, EPI_ISL_2801740, EPI_ISL_2801742, EPI_ISL_2801745, EPI_ISL_2801746, EPI_ISL_2801748, EPI_ISL_2801750 | | | |
| see above | Centro de Investigación Biomédica de Occidente (CIBO) | Instituto de Biotecnología de la UNAM | Consorcio Mexicano de Vigilancia Genómica (CoViGen-Mex). Authors (in alphabetical order): Julio Elias Alvarado-Yaah, Carlos F. Arias, Santiago Ávila-Ríos, Eduardo Becerril-Vargas, Víctor Hugo Borja-Aburto, Celia Boukadida, Cristóbal Cháidez-Quiróz, Juan Bautista Chale-Dzul, Ricardo Ciria Merce, Andreu Comas-García, Célida Duque Molina, Julissa Enciso-Ibarra, José Antonio Enciso-Moreno, Gloria Elena Espinosa-Ayala, Fernando Fontove-Herrera, Daniel Fregoso-Rueda, Victor Eduardo García-Arias, Alejandra García-Gasca, Bruno Gómez-Gil, Jean Pierre González, Irvin González-López, Concepción Grajales-Muñiz, Ricardo Grande, Rosa María Gutiérrez Rios, Jesús Hernández, Alejandra Hernández-Terán, Alfredo Herrera-Estrella, Carla Ivón Herrera-Najera, Pavel Isa, Verónica Mata-Haro, Daniel Lira Morales, Susana Lopez, Antonio Loza Román, Brenda Irasema Maldonado-Meza, Bernardo Martínez-Miguel, José Arturo Martínez-Orozco, Célida Martínez- Rodríguez, Margarita Matías-Florentino, Fidencio Mejía-Nepomuceno, María Guadalupe de Jesús Mireles-Rivera, Gloria María Molina-Salinas, Hector Montoya-Fuentes, , Mario Mújica-Sánchez, José Esteban Muñoz-Medina, José de Jesús Nuñez-Contreras, Alicia Ocaña-Mondragón, Luis Alberto Ochoa-Carrera, Hector Esteban Paz-Juárez, Marissa Perez-Garcia, Francisco Pulido, Helen Haydee Fernanda Ramírez-Plascencia, Jorge Salas-Hernández, Angel Gustavo Salas-Lais, Alejandro Sánchez-Flores, Clara Esperanza Santacruz-Tinoco, María Guadalupe Santiago-Mauricio, Selene Zárate, Nelly Sélem-Mojica, Blanca Taboada, Gloria Vazquez, Joel Armando Vázquez-Pérez. |
| EPI_ISL_2801758, EPI_ISL_2801759, EPI_ISL_2801762, EPI_ISL_2801763, EPI_ISL_2801764, EPI_ISL_2801765, EPI_ISL_2801766, EPI_ISL_2801767, EPI_ISL_2801768, EPI_ISL_2801769, EPI_ISL_2801772, EPI_ISL_2801774, EPI_ISL_2801775, EPI_ISL_2801776, EPI_ISL_2801779, EPI_ISL_2801780, EPI_ISL_2801783, EPI_ISL_2801786, EPI_ISL_2801787, EPI_ISL_2801789, EPI_ISL_2801795, EPI_ISL_2801796, EPI_ISL_2801797, EPI_ISL_2801802, EPI_ISL_2801804, EPI_ISL_2801806, EPI_ISL_2801807, EPI_ISL_2801809, EPI_ISL_2801810, EPI_ISL_2801811, EPI_ISL_2801812, EPI_ISL_2801814, EPI_ISL_2801815, EPI_ISL_2801816, EPI_ISL_2801817, EPI_ISL_2801819, EPI_ISL_2801822, EPI_ISL_2801823, EPI_ISL_2801825, EPI_ISL_2801826, EPI_ISL_2801827, EPI_ISL_2801829, EPI_ISL_2801832, EPI_ISL_2801834, EPI_ISL_2801837, EPI_ISL_2801838, EPI_ISL_2801839, EPI_ISL_2801841, EPI_ISL_2801842, EPI_ISL_2801845, EPI_ISL_2801846, EPI_ISL_2801847, EPI_ISL_2801848, EPI_ISL_2801849, EPI_ISL_2801850, EPI_ISL_2801851, EPI_ISL_2801854, EPI_ISL_2801855, EPI_ISL_2801856, EPI_ISL_2801858, EPI_ISL_2801860, EPI_ISL_2801861, EPI_ISL_2801862, EPI_ISL_2801864, EPI_ISL_2801865, EPI_ISL_2801866, EPI_ISL_2801867, EPI_ISL_2801869, EPI_ISL_2801870, EPI_ISL_2801871, EPI_ISL_2801872, EPI_ISL_2801873, EPI_ISL_2801874, EPI_ISL_2801875, EPI_ISL_2801876, EPI_ISL_2801877, EPI_ISL_2801878, EPI_ISL_2801880, EPI_ISL_2801881, EPI_ISL_2801883, EPI_ISL_2801884, EPI_ISL_2801887, EPI_ISL_2801888, EPI_ISL_2801889, EPI_ISL_2801890 | | | |
| see above | Laboratorio Central de Epidemiología (LCE) | Instituto de Biotecnología de la UNAM | Consorcio Mexicano de Vigilancia Genómica (CoViGen-Mex). Authors (in alphabetical order): Julio Elias Alvarado-Yaah, Carlos F. Arias, Santiago Ávila-Ríos, Eduardo Becerril-Vargas, Víctor Hugo Borja-Aburto, Celia Boukadida, Cristóbal Cháidez-Quiróz, Juan Bautista Chale-Dzul, Ricardo Ciria Merce, Andreu Comas-García, Célida Duque Molina, Julissa Enciso-Ibarra, José Antonio Enciso-Moreno, Gloria Elena Espinosa-Ayala, Fernando Fontove-Herrera, Daniel Fregoso-Rueda, Victor Eduardo García-Arias, Alejandra García-Gasca, Bruno Gómez-Gil, Jean Pierre González, Irvin González-López, Concepción Grajales-Muñiz, Ricardo Grande, Rosa María Gutiérrez Rios, Jesús Hernández, Alejandra Hernández-Terán, Alfredo Herrera-Estrella, Carla Ivón Herrera-Najera, Pavel Isa, Verónica Mata-Haro, Daniel Lira Morales, Susana Lopez, Antonio Loza Román, Brenda Irasema Maldonado-Meza, Bernardo Martínez-Miguel, José Arturo Martínez-Orozco, Célida Martínez- Rodríguez, Margarita Matías-Florentino, Fidencio Mejía-Nepomuceno, María Guadalupe de Jesús Mireles-Rivera, Gloria María Molina-Salinas, Hector Montoya-Fuentes, , Mario Mújica-Sánchez, José Esteban Muñoz-Medina, José de Jesús Nuñez-Contreras, Alicia Ocaña-Mondragón, Luis Alberto Ochoa-Carrera, Hector Esteban Paz-Juárez, Marissa Perez-Garcia, Francisco Pulido, Helen Haydee Fernanda Ramírez-Plascencia, Jorge Salas-Hernández, Angel Gustavo Salas-Lais, Alejandro Sánchez-Flores, Clara Esperanza Santacruz-Tinoco, María Guadalupe Santiago-Mauricio, Selene Zárate, Nelly Sélem-Mojica, Blanca Taboada, Gloria Vazquez, Joel Armando Vázquez-Pérez. |

| | | | |
|---|---|---|---|
| EPI_ISL_2801892, EPI_ISL_2801893, EPI_ISL_2801894 | Banteay Meanchey Regional Laboratory | Virology Unit, Institut Pasteur du Cambodge | Cecile Troupin, Leakhena Pum, Jurre Y Siegers, Ly Sovann, Kraing Sidonn, Yi Sengdoeurn, Chin Savuth, Chau Darapheak, Veasna Duong, Erik A Karlsson |
| EPI_ISL_2801895 | Battambang Rapid Response Team | Virology Unit, Institut Pasteur du Cambodge | Cecile Troupin, Leakhena Pum, Jurre Y Siegers, Ly Sovann, Kraing Sidonn, Yi Sengdoeurn, Chin Savuth, Chau Darapheak, Veasna Duong, Erik A Karlsson |
| EPI_ISL_2801896, EPI_ISL_2801897 | Kampong Cham Rapid Response Team | Virology Unit, Institut Pasteur du Cambodge | Cecile Troupin, Leakhena Pum, Jurre Y Siegers, Ly Sovann, Kraing Sidonn, Yi Sengdoeurn, Chin Savuth, Chau Darapheak, Veasna Duong, Erik A Karlsson |
| EPI_ISL_2801898 | Pursat Rapid Response Team | Virology Unit, Institut Pasteur du Cambodge | Cecile Troupin, Leakhena Pum, Jurre Y Siegers, Ly Sovann, Kraing Sidonn, Yi Sengdoeurn, Chin Savuth, Chau Darapheak, Veasna Duong, Erik A Karlsson |
| EPI_ISL_2801899, EPI_ISL_2801900, EPI_ISL_2801901, EPI_ISL_2801902, EPI_ISL_2801903 | Oddar Meanchey Rapid Response Team | Virology Unit, Institut Pasteur du Cambodge | Cecile Troupin, Leakhena Pum, Jurre Y Siegers, Ly Sovann, Kraing Sidonn, Yi Sengdoeurn, Chin Savuth, Chau Darapheak, Veasna Duong, Erik A Karlsson |
| EPI_ISL_2801904, EPI_ISL_2801905, EPI_ISL_2801906 | Tboung Khmum Rapid Response Team | Virology Unit, Institut Pasteur du Cambodge | Cecile Troupin, Leakhena Pum, Jurre Y Siegers, Ly Sovann, Kraing Sidonn, Yi Sengdoeurn, Chin Savuth, Chau Darapheak, Veasna Duong, Erik A Karlsson |
| EPI_ISL_2801907, EPI_ISL_2801908, EPI_ISL_2801909, EPI_ISL_2801910, EPI_ISL_2801911, EPI_ISL_2801912, EPI_ISL_2801913, EPI_ISL_2801914, EPI_ISL_2801915, EPI_ISL_2801916, EPI_ISL_2801917, EPI_ISL_2801918, EPI_ISL_2801919, EPI_ISL_2801920, EPI_ISL_2801921, EPI_ISL_2801922, EPI_ISL_2801923, EPI_ISL_2801924, EPI_ISL_2801925, EPI_ISL_2801926, EPI_ISL_2801927 | | | |
| see above | Virology Unit, Institut Pasteur du Cambodge | Virology Unit, Institut Pasteur du Cambodge | Cecile Troupin, Leakhena Pum, Jurre Y Siegers, Ly Sovann, Kraing Sidonn, Yi Sengdoeurn, Chin Savuth, Chau Darapheak, Veasna Duong, Erik A Karlsson |
| EPI_ISL_2811883 | Sultanah Nora Ismail Hospital, Batu Pahat | Institute for Medical Research, Infectious Disease Research Centre, National Institutes of Health, Ministry of Health Malaysia | Suppiah J, Kamel K, Mohd Zawawi Z, Azizan MA, Ramly N, Robert F, Thayan R |
| EPI_ISL_2811886 | Raja Perempuan Zainab II Hospital, Kota Bharu | Institute for Medical Research, Infectious Disease Research Centre, National Institutes of Health, Ministry of Health Malaysia | Suppiah J, Kamel K, Mohd Zawawi Z, Azizan MA, Ramly N, Robert F, Thayan R |
| EPI_ISL_2811887 | Institute for Medical Research, Infectious Disease Research Centre, National Institutes of Health, Ministry of Health Malaysia | Institute for Medical Research, Infectious Disease Research Centre, National Institutes of Health, Ministry of Health Malaysia | Suppiah J, Kamel K, Mohd Zawawi Z, Azizan MA, Ramly N, Robert F, Thayan R |
| EPI_ISL_2811959 | Tengku Ampuan Afzan Hospital (HTAA) | Institute for Medical Research, Infectious Disease Research Centre, National Institutes of Health, Ministry of Health Malaysia | Suppiah J, Kamel K, Mohd Zawawi Z, Azizan MA, Ramly N, Robert F, Thayan R |
| EPI_ISL_2811984 | Pekan Tajau Health Clinic | Institute for Medical Research, Infectious Disease Research Centre, National Institutes of Health, Ministry of Health Malaysia | Suppiah J, Kamel K, Mohd Zawawi Z, Azizan MA, Ramly N, Robert F, Thayan R |
| EPI_ISL_2811998 | Bandar Kuantan Health Clinic | Institute for Medical Research, Infectious Disease Research Centre, National Institutes of Health, Ministry of Health Malaysia | Suppiah J, Kamel K, Mohd Zawawi Z, Azizan MA, Ramly N, Robert F, Thayan R |
| EPI_ISL_2812026, EPI_ISL_2812027 | Rembau Health Clinic | Institute for Medical Research, Infectious Disease Research Centre, National Institutes of Health, Ministry of Health Malaysia | Suppiah J, Kamel K, Mohd Zawawi Z, Azizan MA, Ramly N, Robert F, Thayan R |
| EPI_ISL_2812146, EPI_ISL_2812147, EPI_ISL_2812148, EPI_ISL_2812149, EPI_ISL_2812150, EPI_ISL_2812151, EPI_ISL_2812152, EPI_ISL_2812153, EPI_ISL_2812154, EPI_ISL_2812156 | Singapore General Hospital | Department of Microbiology | Nurdyana Abdul Rahman, Kenneth Xin Long Chan, Chayaporn Suphavilai, Kun Lee Lim, Sui Sin Goh, Kian Sing Chan, Lynette Oon, Kern Rei Chng, James Sim Heng Chiak, Niranjan Nagarajan, Karrie Ko |
| EPI_ISL_2812441, EPI_ISL_2812537 | Laboratorio de Virología HUCA | Laboratorio de Virología HUCA | Sandoval M, Castelló C, Gómez de Oña J, Boga JA, Rojo S, Alvarez-Arguelles ME, Abreu F, Costales I, Perez-Martínez Z, Martín-Rodríguez G, Coto E, Melón S |
| EPI_ISL_2812576 | Kemaman Hospital | Institute for Medical Research, Infectious Disease Research Centre, National Institutes of Health, Ministry of Health Malaysia | Suppiah J, Kamel K, Mohd Zawawi Z, Azizan MA, Ramly N, Robert F, Thayan R |
| EPI_ISL_2812654, EPI_ISL_2812704 | Sultanah Bahiyah Hospital, Alor Setar | Institute for Medical Research, Infectious Disease Research Centre, National Institutes of Health, Ministry of Health Malaysia | Suppiah J, Kamel K, Mohd Zawawi Z, Azizan MA, Ramly N, Robert F, Thayan R |
| EPI_ISL_2812706 | Sultan Haji Ahmad Shah Hospital | Institute for Medical Research, Infectious Disease Research Centre, National Institutes of Health, Ministry of Health Malaysia | Suppiah J, Kamel K, Mohd Zawawi Z, Azizan MA, Ramly N, Robert F, Thayan R |
| EPI_ISL_2812958, EPI_ISL_2812959, EPI_ISL_2812960, EPI_ISL_2812961, EPI_ISL_2812962, EPI_ISL_2812963, EPI_ISL_2812964, EPI_ISL_2812965, EPI_ISL_2812966, EPI_ISL_2812967, EPI_ISL_2812968 | | | |
| see above | Department of Health Technology and Informatics, The Hong Kong Polytechnic University | Department of Health Technology and Informatics, The Hong Kong Polytechnic University | Gilman Kit-Hang Siu, Lam-Kwong Lee, Kenneth Siu-Sing Leung, Jake Siu-Lun Leung, Wing-Hei Lo, Timothy Ting-Leung Ng, Chloe Toi-Mei Chan, Hoi-Ching Jim, Kingsley King-Gee Tam, Hiu-Yin Lao, Denise Sze-Hang Wong, Alan Ka-Lun Wu, Miranda Chong-Yee Yau, Yvette Wai-Man Lai, Kitty Sau-Chun Fung, Sandy Ka-Yee Chau, Barry Kin-Chung Wong, Wing-Kin To, Kristine Luk, Alex Yat-Man Ho, Tak-Lun Que, Kam-Tong Yip, Wing Cheong Yam, David Ho-Keung Shum, Shea Ping Yip |
| EPI_ISL_2812971 | Labuan Hospital | Institute for Medical Research, Infectious Disease Research Centre, National Institutes of Health, Ministry of Health Malaysia | Suppiah J, Kamel K, Mohd Zawawi Z, Azizan MA, Ramly N, Robert F, Thayan R |
| EPI_ISL_2812981 | Johor State Health Department | Institute for Medical Research, Infectious Disease Research Centre, National Institutes of Health, Ministry of Health Malaysia | Suppiah J, Kamel K, Mohd Zawawi Z, Azizan MA, Ramly N, Robert F, Thayan R |
| EPI_ISL_2812982, EPI_ISL_2812983, EPI_ISL_2814168 | L. C. Laboratori Campisi Group | AOU Polclinico "G Rodolico-S.Marco" PO Rodolico and BIOMETEC UNICT CATANIA | STEFANI AND GUIDO SCALIA |
| EPI_ISL_2814916, EPI_ISL_2815327, EPI_ISL_2815328 | Melaka Hospital | Institute for Medical Research, Infectious Disease Research Centre, National Institutes of Health, Ministry of Health Malaysia | Suppiah J, Kamel K, Mohd Zawawi Z, Azizan MA, Ramly N, Robert F, Thayan R |
| EPI_ISL_2815329, EPI_ISL_2815330 | Ipoh Public Health Laboratory (MKAI), Ministry of Health Malaysia | Institute for Medical Research, Infectious Disease Research Centre, National Institutes of Health, Ministry of Health Malaysia | Suppiah J, Kamel K, Mohd Zawawi Z, Azizan MA, Ramly N, Robert F, Thayan R |
| EPI_ISL_2815331 | Kuala Lumpur General Hospital | Institute for Medical Research, Infectious Disease Research | Suppiah J, Kamel K, Mohd Zawawi Z, Azizan MA, Ramly N, Robert F, Thayan R |

| | | | |
|---|---|---|---|
| | | Centre, National Institutes of Health, Ministry of Health Malaysia | |
| EPI_ISL_2815375 | Institute for Medical Research, Infectious Disease Research Centre, National Institutes of Health, Ministry of Health Malaysia | Institute for Medical Research, Infectious Disease Research Centre, National Institutes of Health, Ministry of Health Malaysia | Suppiah J, Kamel K, Mohd Zawawi Z, Azizan MA, Ramly N, Robert F, Thayan R |
| EPI_ISL_2815971 | Sultanah Bahiyah Hospital, Alor Setar | Institute for Medical Research, Infectious Disease Research Centre, National Institutes of Health, Ministry of Health Malaysia | Suppiah J, Kamel K, Mohd Zawawi Z, Azizan MA, Ramly N, Robert F, Thayan R |
| EPI_ISL_2819024, EPI_ISL_2819571, EPI_ISL_2819919 | Institute for Medical Research, Infectious Disease Research Centre, National Institutes of Health, Ministry of Health Malaysia | Institute for Medical Research, Infectious Disease Research Centre, National Institutes of Health, Ministry of Health Malaysia | Suppiah J, Kamel K, Mohd Zawawi Z, Azizan MA, Ramly N, Robert F, Thayan R |
| EPI_ISL_2820290, EPI_ISL_2820296, EPI_ISL_2820300, EPI_ISL_2820303, EPI_ISL_2820304, EPI_ISL_2820307, EPI_ISL_2820308, EPI_ISL_2820310, EPI_ISL_2820312, EPI_ISL_2820321, EPI_ISL_2820322, EPI_ISL_2820325, EPI_ISL_2820326, EPI_ISL_2820330, EPI_ISL_2820331, EPI_ISL_2820333 | | | |
| see above | Kasane Primary Hospital Laboratory | Botswana Harvard HIV Reference Laboratory | Sikhulile Moyo, Wonderful T. Choga, Dorcas Maruapula, Agnes Karutwaeng, Rose Munyere, Keoratile Ntshambiwa, Thongbotho Mphoyakgosi, Boitumelo Zuze, Botshelo Radibe, Legodile Kooepile, Ontlametse T. Bareng, Roger Shapiro, Shahin Lockman, Joseph Makhema, Madisa Mine, Mosepele Mosepele, Simani Gaseitsiwe |
| EPI_ISL_2820339, EPI_ISL_2820340, EPI_ISL_2820341, EPI_ISL_2820342, EPI_ISL_2820343, EPI_ISL_2820345, EPI_ISL_2820350, EPI_ISL_2820353, EPI_ISL_2820359, EPI_ISL_2820366, EPI_ISL_2820374 | | | |
| see above | Botswana Harvard HIV Reference Laboratory | Botswana Harvard HIV Reference Laboratory | Sikhulile Moyo, Wonderful T. Choga, Dorcas Maruapula, Thongbotho Mphoyakgosi, Boitumelo Zuze, Botshelo Radibe, Legodile Kooepile, Ontlametse T. Bareng, Motlalepule L. Pone, Godfrey Simoonga, Kereng Mphoyakgosi, Keoratile Ntshambiwa, Modisa Motswaledi, Madisa Mine, Joseph Makhema, Roger Shapiro, Shahin Lockman, Mosepele Mosepele, Simani Gaseitsiwe |
| EPI_ISL_2820386, EPI_ISL_2820388, EPI_ISL_2820392, EPI_ISL_2820400 | Kasane Primary Hospital Laboratory | Botswana Harvard HIV Reference Laboratory | Sikhulile Moyo, Wonderful T. Choga, Dorcas Maruapula, Agnes Karutwaeng, Rose Munyere, Keoratile Ntshambiwa, Thongbotho Mphoyakgosi, Boitumelo Zuze, Botshelo Radibe, Legodile Kooepile, Ontlametse T. Bareng, Roger Shapiro, Shahin Lockman, Joseph Makhema, Madisa Mine, Mosepele Mosepele, Simani Gaseitsiwe |
| EPI_ISL_2820448, EPI_ISL_2820451, EPI_ISL_2820459, EPI_ISL_2820462 | Botswana Harvard HIV Reference Laboratory | Botswana Harvard HIV Reference Laboratory | Sikhulile Moyo, Wonderful T. Choga, Dorcas Maruapula, Thongbotho Mphoyakgosi, Boitumelo Zuze, Botshelo Radibe, Legodile Kooepile, Ontlametse T. Bareng, Keoratile Ntshambiwa, Modisa Motswaledi, Madisa Mine, Joseph Makhema, Roger Shapiro, Shahin Lockman, Mosepele Mosepele, Simani Gaseitsiwe |
| EPI_ISL_2820677, EPI_ISL_2820678, EPI_ISL_2820679, EPI_ISL_2820683, EPI_ISL_2820684 | Università degli Studi di Perugia | Istituto Zooprofilattico Sperimentale dell'Abruzzo e Molise "G. Caporale" | Mencacci A, Camilloni B, Biagetti M, Giammarioli M, Lorusso A, Marcacci M, Di Domenico M, Ancora M, Curini V, Mangone I, Rinaldi A, Delli Compagni E, Scialabba S, Di Pasquale A, Cammà C, Puglia I, Calistri P, Savini G |
| EPI_ISL_2820707 | Presidio di Brindisi Di Summa - Perrino | Istituto Zooprofilattico Sperimentale della Puglia e della Basilicata | Parisi A., Bianco A., Capozzi L., Del Sambro L., Simone D., Difato L., Santoro A., Ridolfi D., Giannico A., Manzari C., Pesole G. |
| EPI_ISL_2820715, EPI_ISL_2820716, EPI_ISL_2820717 | I.R.C.C.S. "S. De Bellis" - Ente Ospedaliero | Istituto Zooprofilattico Sperimentale della Puglia e della Basilicata | Parisi A., Bianco A., Capozzi L., Del Sambro L., Simone D., Difato L., Notarnicola M., Lippolis A., Manzari C., Pesole G. |
| EPI_ISL_2820718, EPI_ISL_2820719, EPI_ISL_2820720, EPI_ISL_2820721, EPI_ISL_2820722, EPI_ISL_2820723 | Ospedale Santa Caterina Novella | Istituto Zooprofilattico Sperimentale della Puglia e della Basilicata | Parisi A., Bianco A., Capozzi L., Del Sambro L., Simone D., Difato L., Bruno A. R., Manzari C., Pesole G. |
| EPI_ISL_2820724, EPI_ISL_2820726 | Ospedale Vito Fazzi | Istituto Zooprofilattico Sperimentale della Puglia e della Basilicata | Parisi A., Bianco A., Capozzi L., Del Sambro L., Simone D., Difato L., Lobreglio G., Manzari C., Pesole G. |
| EPI_ISL_2820729, EPI_ISL_2820730, EPI_ISL_2820741, EPI_ISL_2820742, EPI_ISL_2820743, EPI_ISL_2820744 | Presidio di Brindisi Di Summa - Perrino | Istituto Zooprofilattico Sperimentale della Puglia e della Basilicata | Parisi A., Bianco A., Capozzi L., Del Sambro L., Simone D., Difato L., Santoro A., Ridolfi D., Giannico A., Manzari C., Pesole G. |
| EPI_ISL_2820746, EPI_ISL_2820747, EPI_ISL_2820748, EPI_ISL_2820749, EPI_ISL_2820750, EPI_ISL_2820751, EPI_ISL_2820752, EPI_ISL_2820753, EPI_ISL_2820754 | Ospedale Santissima Annunziata | Istituto Zooprofilattico Sperimentale della Puglia e della Basilicata | Parisi A., Bianco A., Capozzi L., Del Sambro L., Simone D., Difato L., Ridolfi D., Giannico A, Tampoia M., Manzari C., Pesole G. |
| EPI_ISL_2827779 | LDSP - CAQUETA HOSPITAL MARIA INMACULADA | Instituto Nacional de Salud- Dirección de Investigación en Salud Pública | Katherine Laiton-Donato, Diego A. Álvarez-Díaz, Carlos Franco-Muñoz, Hector Alejandro Ruiz-Moreno, Paola Rojas, Maria T. Herrera-Sepúlveda, Diego Andrés Prada, Jhonnatan Reales-González, Sheryll Corchuelo, Julian Naizaque, Jorge Rivera, Gerardo Santamaría, Sergio Gomez, Lisseth Pardo, Juan Camilo Martinez, Marta Lopez Blanco, Ángela Alarcon Cruz, Diana Malo, Carmen Osorio, Magdalena Wiesner, Martha Lucia Ospina Martinez, Marcela Mercado-Reyes |
| EPI_ISL_2827783 | LDSP - CAQUETA HOSPITAL MALVINAS | Instituto Nacional de Salud- Dirección de Investigación en Salud Pública | Katherine Laiton-Donato, Diego A. Álvarez-Díaz, Carlos Franco-Muñoz, Hector Alejandro Ruiz-Moreno, Paola Rojas, Maria T. Herrera-Sepúlveda, Diego Andrés Prada, Jhonnatan Reales-González, Sheryll Corchuelo, Julian Naizaque, Jorge Rivera, Gerardo Santamaría, Sergio Gomez, Lisseth Pardo, Juan Camilo Martinez, Marta Lopez Blanco, Ángela Alarcon Cruz, Diana Malo, Carmen Osorio, Magdalena Wiesner, Martha Lucia Ospina Martinez, Marcela Mercado-Reyes |
| EPI_ISL_2827961, EPI_ISL_2827962 | AREA DE SALUD HATILLO - CLINICA DR. SOLON NUÑEZ | Inciensa, Instituto Costarricense de Investigación y Enseñanza en Nutrición y Salud | Francisco Duarte, Hebleen Porras, Claudio Soto-Garita, Estela Cordero, Adriana Godínez, Melany Calderón, José Luis Vargas, Mariela Gutiérrez, Joselyn Prado & Roberto Brilla |
| EPI_ISL_2827963, EPI_ISL_2827964, EPI_ISL_2827965 | HOSPITAL DR. WILLIAM ALLEN | Inciensa, Instituto Costarricense de Investigación y Enseñanza en Nutrición y Salud | Francisco Duarte, Hebleen Porras, Claudio Soto-Garita, Estela Cordero, Adriana Godínez, Melany Calderón, José Luis Vargas, Mariela Gutiérrez & Joselyn Prado |
| EPI_ISL_2827966 | AREA DE SALUD CARIARI | Inciensa, Instituto Costarricense de Investigación y Enseñanza en Nutrición y Salud | Francisco Duarte, Hebleen Porras, Claudio Soto-Garita, Estela Cordero, Adriana Godínez, Melany Calderón, José Luis Vargas, Mariela Gutiérrez, Joselyn Prado & Vanessa Jiménez |
| EPI_ISL_2827967, EPI_ISL_2827968 | HOSPITAL DR. RAFAEL A. CALDERON GUARDIA | Inciensa, Instituto Costarricense de Investigación y Enseñanza en Nutrición y Salud | Francisco Duarte, Hebleen Porras, Claudio Soto-Garita, Estela Cordero, Adriana Godínez, Melany Calderón, José Luis Vargas, Mariela Gutiérrez, Joselyn Prado & Fabian Salas |
| EPI_ISL_2827970 | AREA DE SALUD GUATUSO | Inciensa, Instituto Costarricense de Investigación y Enseñanza en Nutrición y Salud | Francisco Duarte, Hebleen Porras, Claudio Soto-Garita, Estela Cordero, Adriana Godínez, Melany Calderón, José Luis Vargas, Mariela Gutiérrez, Joselyn Prado & Francisco Chacón |
| EPI_ISL_2827971 | HOSPITAL CIUDAD NEILY | Inciensa, Instituto Costarricense de Investigación y Enseñanza en Nutrición y Salud | Francisco Duarte, Hebleen Porras, Claudio Soto-Garita, Estela Cordero, Adriana Godínez, Melany Calderón, José Luis Vargas, Mariela Gutiérrez, Joselyn Prado & María Isabel Victoria |
| EPI_ISL_2827972 | AREA DE SALUD PAVAS (COOPESALUD) | Inciensa, Instituto Costarricense de Investigación y Enseñanza en Nutrición y Salud | Francisco Duarte, Hebleen Porras, Claudio Soto-Garita, Estela Cordero, Adriana Godínez, Melany Calderón, José Luis Vargas, Mariela Gutiérrez, Joselyn Prado & Paola Sánchez |
| EPI_ISL_2827973, EPI_ISL_2827974 | HOSPITAL SAN JUAN DE DIOS | Inciensa, Instituto Costarricense de Investigación y Enseñanza en Nutrición y Salud | Francisco Duarte, Hebleen Porras, Claudio Soto-Garita, Estela Cordero, Adriana Godínez, Melany Calderón, José Luis Vargas, Mariela Gutiérrez, Joselyn Prado & Paola Sánchez |
| EPI_ISL_2827975 | HOSPITAL LA ANEXION | Inciensa, Instituto Costarricense de Investigación y Enseñanza en Nutrición y Salud | Francisco Duarte, Hebleen Porras, Claudio Soto-Garita, Estela Cordero, Adriana Godínez, Melany Calderón, José Luis Vargas, Mariela Gutiérrez, Joselyn Prado & Adriana Bermúdez |
| EPI_ISL_2827976, EPI_ISL_2827977 | HOSPITAL DR. ENRIQUE BALTODANO BRICEÑO | Inciensa, Instituto Costarricense de Investigación y Enseñanza en Nutrición y Salud | Francisco Duarte, Hebleen Porras, Claudio Soto-Garita, Estela Cordero, Adriana Godínez, Melany Calderón, José Luis Vargas, Mariela Gutiérrez, Joselyn Prado & Adriana Bermúdez |

| | | | |
|---|---|---|---|
| EPI_ISL_2827978 | AREA DE SALUD SANTA CRUZ | Inciensa, Instituto Costarricense de Investigación y Enseñanza en Nutrición y Salud | Francisco Duarte, Hebleen Porras, Claudio Soto-Garita, Estela Cordero, Adriana Godínez, Melany Calderón, José Luis Vargas, Mariela Gutiérrez, Joselyn Prado & Adriana Bermúdez |
| EPI_ISL_2827979, EPI_ISL_2827980 | AREA DE SALUD CARTAGO | Inciensa, Instituto Costarricense de Investigación y Enseñanza en Nutrición y Salud | Francisco Duarte, Hebleen Porras, Claudio Soto-Garita, Estela Cordero, Adriana Godínez, Melany Calderón, José Luis Vargas, Mariela Gutiérrez, Joselyn Prado & Tania Garro |
| EPI_ISL_2827981 | HOSPITAL MONSEÑOR SANABRIA | Inciensa, Instituto Costarricense de Investigación y Enseñanza en Nutrición y Salud | Francisco Duarte, Hebleen Porras, Claudio Soto-Garita, Estela Cordero, Adriana Godínez, Melany Calderón, José Luis Vargas, Mariela Gutiérrez, Joselyn Prado & María José Gómez |
| EPI_ISL_2827982 | AREA DE SALUD COBANO | Inciensa, Instituto Costarricense de Investigación y Enseñanza en Nutrición y Salud | Francisco Duarte, Hebleen Porras, Claudio Soto-Garita, Estela Cordero, Adriana Godínez, Melany Calderón, José Luis Vargas, Mariela Gutiérrez, Joselyn Prado & María José Gómez |
| EPI_ISL_2827983 | AREA DE SALUD BARRANCA - CLINICA DR. ROBERTO SOTO | Inciensa, Instituto Costarricense de Investigación y Enseñanza en Nutrición y Salud | Francisco Duarte, Hebleen Porras, Claudio Soto-Garita, Estela Cordero, Adriana Godínez, Melany Calderón, José Luis Vargas, Mariela Gutiérrez, Joselyn Prado & María José Gómez |
| EPI_ISL_2827984 | AREA DE SALUD COBANO | Inciensa, Instituto Costarricense de Investigación y Enseñanza en Nutrición y Salud | Francisco Duarte, Hebleen Porras, Claudio Soto-Garita, Estela Cordero, Adriana Godínez, Melany Calderón, José Luis Vargas, Mariela Gutiérrez, Joselyn Prado & María José Gómez |
| EPI_ISL_2827985 | AREA DE SALUD ESPARZA | Inciensa, Instituto Costarricense de Investigación y Enseñanza en Nutrición y Salud | Francisco Duarte, Hebleen Porras, Claudio Soto-Garita, Estela Cordero, Adriana Godínez, Melany Calderón, José Luis Vargas, Mariela Gutiérrez, Joselyn Prado & María José Gómez |
| EPI_ISL_2827986 | HOSPITAL SAN VICENTE DE PAUL | Inciensa, Instituto Costarricense de Investigación y Enseñanza en Nutrición y Salud | Francisco Duarte, Hebleen Porras, Claudio Soto-Garita, Estela Cordero, Adriana Godínez, Melany Calderón, José Luis Vargas, Mariela Gutiérrez, Joselyn Prado & Josiany Arce |
| EPI_ISL_2827987 | HOSPITAL DR. CARLOS LUIS VALVERDE VEGA | Inciensa, Instituto Costarricense de Investigación y Enseñanza en Nutrición y Salud | Francisco Duarte, Hebleen Porras, Claudio Soto-Garita, Estela Cordero, Adriana Godínez, Melany Calderón, José Luis Vargas, Mariela Gutiérrez, Joselyn Prado & Yahaira Ramos |
| EPI_ISL_2827988, EPI_ISL_2827989, EPI_ISL_2827990 | AREA DE SALUD CIUDAD QUESADA | Inciensa, Instituto Costarricense de Investigación y Enseñanza en Nutrición y Salud | Francisco Duarte, Hebleen Porras, Claudio Soto-Garita, Estela Cordero, Adriana Godínez, Melany Calderón, José Luis Vargas, Mariela Gutiérrez, Joselyn Prado & Juan Carlos Villalobos |
| EPI_ISL_2827991, EPI_ISL_2827992, EPI_ISL_2827993 | HOSPITAL MEXICO | Inciensa, Instituto Costarricense de Investigación y Enseñanza en Nutrición y Salud | Francisco Duarte, Hebleen Porras, Claudio Soto-Garita, Estela Cordero, Adriana Godínez, Melany Calderón, José Luis Vargas, Mariela Gutiérrez, Joselyn Prado & Juan Carlos Villalobos |
| EPI_ISL_2827994 | AREA DE SALUD CARIARI | Inciensa, Instituto Costarricense de Investigación y Enseñanza en Nutrición y Salud | Francisco Duarte, Hebleen Porras, Claudio Soto-Garita, Estela Cordero, Adriana Godínez, Melany Calderón, José Luis Vargas, Mariela Gutiérrez, Joselyn Prado & Blanca Gallegos |
| EPI_ISL_2827995, EPI_ISL_2827996 | AREA DE SALUD CARTAGO | Inciensa, Instituto Costarricense de Investigación y Enseñanza en Nutrición y Salud | Francisco Duarte, Hebleen Porras, Claudio Soto-Garita, Estela Cordero, Adriana Godínez, Melany Calderón, José Luis Vargas, Mariela Gutiérrez, Joselyn Prado & Monserrat Segura |
| EPI_ISL_2827997, EPI_ISL_2827998 | AREA DE SALUD BAGACES | Inciensa, Instituto Costarricense de Investigación y Enseñanza en Nutrición y Salud | Francisco Duarte, Hebleen Porras, Claudio Soto-Garita, Estela Cordero, Adriana Godínez, Melany Calderón, José Luis Vargas, Mariela Gutiérrez, Joselyn Prado & Karol Matamoros |
| EPI_ISL_2827999 | AREA DE SALUD DESAMPARADOS 2 (COOPESALUD) | Inciensa, Instituto Costarricense de Investigación y Enseñanza en Nutrición y Salud | Francisco Duarte, Hebleen Porras, Claudio Soto-Garita, Estela Cordero, Adriana Godínez, Melany Calderón, José Luis Vargas, Mariela Gutiérrez, Joselyn Prado & Alberto Quirós |
| EPI_ISL_2828000 | AREA DE SALUD GARABITO | Inciensa, Instituto Costarricense de Investigación y Enseñanza en Nutrición y Salud | Francisco Duarte, Hebleen Porras, Claudio Soto-Garita, Estela Cordero, Adriana Godínez, Melany Calderón, José Luis Vargas, Mariela Gutiérrez, Joselyn Prado & María José Gómez |
| EPI_ISL_2828001 | HOSPITAL MONSEÑOR SANABRIA | Inciensa, Instituto Costarricense de Investigación y Enseñanza en Nutrición y Salud | Francisco Duarte, Hebleen Porras, Claudio Soto-Garita, Estela Cordero, Adriana Godínez, Melany Calderón, José Luis Vargas, Mariela Gutiérrez, Joselyn Prado & María José Gómez |
| EPI_ISL_2828002 | AREA DE SALUD GOICOECHEA 2 - CLINICA DR. JIMENEZ NUÑEZ | Inciensa, Instituto Costarricense de Investigación y Enseñanza en Nutrición y Salud | Francisco Duarte, Hebleen Porras, Claudio Soto-Garita, Estela Cordero, Adriana Godínez, Melany Calderón, José Luis Vargas, Mariela Gutiérrez, Joselyn Prado & Juan Carlos Cartes |
| EPI_ISL_2828003 | AREA DE SALUD ACOSTA | Inciensa, Instituto Costarricense de Investigación y Enseñanza en Nutrición y Salud | Francisco Duarte, Hebleen Porras, Claudio Soto-Garita, Estela Cordero, Adriana Godínez, Melany Calderón, José Luis Vargas, Mariela Gutiérrez & Joselyn Prado |
| EPI_ISL_2828004 | AREA DE SALUD MATA REDONDA-HOSPITAL - CLINICA DR. MORENO CAÑAS | Inciensa, Instituto Costarricense de Investigación y Enseñanza en Nutrición y Salud | Francisco Duarte, Hebleen Porras, Claudio Soto-Garita, Estela Cordero, Adriana Godínez, Melany Calderón, José Luis Vargas, Mariela Gutiérrez, Joselyn Prado & Ricardo González |
| EPI_ISL_2828006 | AREA DE SALUD ESCAZU (COOPESANA) | Inciensa, Instituto Costarricense de Investigación y Enseñanza en Nutrición y Salud | Francisco Duarte, Hebleen Porras, Claudio Soto-Garita, Estela Cordero, Adriana Godínez, Melany Calderón, José Luis Vargas, Mariela Gutiérrez, Joselyn Prado & Juliana Mora |
| EPI_ISL_2828007, EPI_ISL_2828008 | AREA DE SALUD SAN RAMON | Inciensa, Instituto Costarricense de Investigación y Enseñanza en Nutrición y Salud | Francisco Duarte, Hebleen Porras, Claudio Soto-Garita, Estela Cordero, Adriana Godínez, Melany Calderón, José Luis Vargas, Mariela Gutiérrez & Joselyn Prado |
| EPI_ISL_2828009 | AREA DE SALUD SIQUIRRES | Inciensa, Instituto Costarricense de Investigación y Enseñanza en Nutrición y Salud | Francisco Duarte, Hebleen Porras, Claudio Soto-Garita, Estela Cordero, Adriana Godínez, Melany Calderón, José Luis Vargas, Mariela Gutiérrez, Joselyn Prado & Ileana Chavez |
| EPI_ISL_2828010 | HOSPITAL DR. RAFAEL A. CALDERON GUARDIA | Inciensa, Instituto Costarricense de Investigación y Enseñanza en Nutrición y Salud | Francisco Duarte, Hebleen Porras, Claudio Soto-Garita, Estela Cordero, Adriana Godínez, Melany Calderón, José Luis Vargas, Mariela Gutiérrez, Joselyn Prado & Fabian Salas |
| EPI_ISL_2828011 | AREA DE SALUD SAN RAFAEL | Inciensa, Instituto Costarricense de Investigación y Enseñanza en Nutrición y Salud | Francisco Duarte, Hebleen Porras, Claudio Soto-Garita, Estela Cordero, Adriana Godínez, Melany Calderón, José Luis Vargas, Mariela Gutiérrez, Joselyn Prado & María José Gómez |
| EPI_ISL_2828012 | AREA DE SALUD GARABITO | Inciensa, Instituto Costarricense de Investigación y Enseñanza en Nutrición y Salud | Francisco Duarte, Hebleen Porras, Claudio Soto-Garita, Estela Cordero, Adriana Godínez, Melany Calderón, José Luis Vargas, Mariela Gutiérrez, Joselyn Prado & María José Gómez |
| EPI_ISL_2828013 | HOSPITAL DR. MAX TERAN VALLS | Inciensa, Instituto Costarricense de Investigación y Enseñanza en Nutrición y Salud | Francisco Duarte, Hebleen Porras, Claudio Soto-Garita, Estela Cordero, Adriana Godínez, Melany Calderón, José Luis Vargas, Mariela Gutiérrez, Joselyn Prado & María José Gómez |
| EPI_ISL_2828014 | HOSPITAL MONSEÑOR SANABRIA | Inciensa, Instituto Costarricense de Investigación y Enseñanza en Nutrición y Salud | Francisco Duarte, Hebleen Porras, Claudio Soto-Garita, Estela Cordero, Adriana Godínez, Melany Calderón, José Luis Vargas, Mariela Gutiérrez, Joselyn Prado & María José Gómez |
| EPI_ISL_2828015 | HOSPITAL MEXICO | Inciensa, Instituto Costarricense de Investigación y Enseñanza en Nutrición y Salud | Francisco Duarte, Hebleen Porras, Claudio Soto-Garita, Estela Cordero, Adriana Godínez, Melany Calderón, José Luis Vargas, Mariela Gutiérrez, Joselyn Prado & Juan Carlos Villalobos |
| EPI_ISL_2828327 | Hospital Clínico Universitario Virgen de la Arrixaca | Hospital Clínico Universitario Virgen de la Arrixaca | Marina Simón Páez y Laura Moreno Parrado |
| EPI_ISL_2828383 | SC (UCO) Igiene e Sanità Pubblica, ASUGI, Trieste | ARGO Laboratorio Genomica ed Epigenomica | Licastro D, Dal Monego S, Degasperi M, Marcello A, Segat L, Piscianz E, Busetti M, Fontana F, Breda C, Barbone F, D'Agaro P |
| EPI_ISL_2828384 | SC (UCO) Igiene e Sanità Pubblica, ASUGI, Trieste and SC Microbiologia e Virologia, ASFO, Pordenone | ARGO Laboratorio Genomica ed Epigenomica | Licastro D, Dal Monego S, Degasperi M, Marcello A, Segat L, De Rosa R, Breda C, Barbone F, D'Agaro P |
| EPI_ISL_2828385, EPI_ISL_2828386, EPI_ISL_2828387, EPI_ISL_2828388, EPI_ISL_2828389 | SC (UCO) Igiene e Sanità Pubblica, ASUGI, Trieste | ARGO Laboratorio Genomica ed Epigenomica | Licastro D, Dal Monego S, Degasperi M, Marcello A, Segat L, Piscianz E, Busetti M, Fontana F, Breda C, Barbone F, D'Agaro P |
| EPI_ISL_2828390 | SC (UCO) Igiene e Sanità Pubblica, ASUGI, Trieste and SC Microbiologia e Virologia, ASFO, Pordenone | ARGO Laboratorio Genomica ed Epigenomica | Licastro D, Dal Monego S, Degasperi M, Marcello A, Segat L, De Rosa R, Breda C, Barbone F, D'Agaro P |
| EPI_ISL_2828391, EPI_ISL_2828392, EPI_ISL_2828393, EPI_ISL_2828394, EPI_ISL_2828395, EPI_ISL_2828396, EPI_ISL_2828397, EPI_ISL_2828398, EPI_ISL_2828399, EPI_ISL_2828400, EPI_ISL_2828401, EPI_ISL_2828402, EPI_ISL_2828403 | | | |
| see above | SC (UCO) Igiene e Sanità Pubblica, ASUGI, Trieste | ARGO Laboratorio Genomica ed Epigenomica | Licastro D, Dal Monego S, Degasperi M, Marcello A, Segat L, Piscianz E, Busetti M, Fontana F, Breda C, Barbone F, D'Agaro P |
| EPI_ISL_2828404 | SC (UCO) Igiene e Sanità Pubblica, ASUGI, Trieste and SC | ARGO Laboratorio Genomica ed Epigenomica | Licastro D, Dal Monego S, Degasperi M, Marcello A, Segat L, De Rosa R, Breda C, Barbone F, D'Agaro P |

| | | | |
|---|---|---|---|
| | Microbiologia e Virologia, ASFO, Pordenone | | |
| EPI_ISL_2828405, EPI_ISL_2828406, EPI_ISL_2828407, EPI_ISL_2828408, EPI_ISL_2828409, EPI_ISL_2828410, EPI_ISL_2828411, EPI_ISL_2828412, EPI_ISL_2828413, EPI_ISL_2828414 | SC (UCO) Igiene e Sanità Pubblica, ASUGI, Trieste | ARGO Laboratorio Genomica ed Epigenomica | Licastro D, Dal Monego S, Degasperi M, Marcello A, Segat L, Piscianz E, Busetti M, Fontana F, Breda C, Barbone F, D'Agaro P |
| EPI_ISL_2828415, EPI_ISL_2828416, EPI_ISL_2828417, EPI_ISL_2828418 | SC (UCO) Igiene e Sanità Pubblica, ASUGI, Trieste and SC Microbiologia e Virologia, ASFO, Pordenone | ARGO Laboratorio Genomica ed Epigenomica | Licastro D, Dal Monego S, Degasperi M, Marcello A, Segat L, De Rosa R, Breda C, Barbone F, D'Agaro P |
| EPI_ISL_2828419, EPI_ISL_2828420, EPI_ISL_2828421, EPI_ISL_2828422, EPI_ISL_2828423, EPI_ISL_2828424, EPI_ISL_2828425, EPI_ISL_2828426, EPI_ISL_2828427, EPI_ISL_2828428, EPI_ISL_2828429, EPI_ISL_2828430, EPI_ISL_2828431, EPI_ISL_2828432, EPI_ISL_2828433, EPI_ISL_2828434, EPI_ISL_2828435, EPI_ISL_2828436, EPI_ISL_2828437, EPI_ISL_2828438, EPI_ISL_2828439, EPI_ISL_2828440, EPI_ISL_2828441, EPI_ISL_2828442, EPI_ISL_2828443, EPI_ISL_2828444 | | | |
| see above | SC (UCO) Igiene e Sanità Pubblica, ASUGI, Trieste | ARGO Laboratorio Genomica ed Epigenomica | Licastro D, Dal Monego S, Degasperi M, Marcello A, Segat L, Piscianz E, Busetti M, Fontana F, Breda C, Barbone F, D'Agaro P |
| EPI_ISL_2828445, EPI_ISL_2828446, EPI_ISL_2828447, EPI_ISL_2828448, EPI_ISL_2828449, EPI_ISL_2828450, EPI_ISL_2828451, EPI_ISL_2828452, EPI_ISL_2828453, EPI_ISL_2828454, EPI_ISL_2828455, EPI_ISL_2828456, EPI_ISL_2828457, EPI_ISL_2828458 | | | |
| see above | SC (UCO) Igiene e Sanità Pubblica, ASUGI, Trieste and SC Microbiologia e Virologia, ASFO, Pordenone | ARGO Laboratorio Genomica ed Epigenomica | Licastro D, Dal Monego S, Degasperi M, Marcello A, Segat L, De Rosa R, Breda C, Barbone F, D'Agaro P |
| EPI_ISL_2828459, EPI_ISL_2828460, EPI_ISL_2828461, EPI_ISL_2828462, EPI_ISL_2828463, EPI_ISL_2828464, EPI_ISL_2828465, EPI_ISL_2828466, EPI_ISL_2828467, EPI_ISL_2828468, EPI_ISL_2828469, EPI_ISL_2828470, EPI_ISL_2828471 | | | |
| see above | SC (UCO) Igiene e Sanità Pubblica, ASUGI, Trieste | ARGO Laboratorio Genomica ed Epigenomica | Licastro D, Dal Monego S, Degasperi M, Marcello A, Segat L, Piscianz E, Busetti M, Fontana F, Breda C, Barbone F, D'Agaro P |
| EPI_ISL_2828647, EPI_ISL_2828648, EPI_ISL_2828649, EPI_ISL_2828650, EPI_ISL_2828651, EPI_ISL_2828652, EPI_ISL_2828653 | Instituto Adolfo Lutz - Reginal de Sao José do Rio Preto | Instituto Adolfo Lutz, Interdiciplinary Procedures Center, Strategic Laboratory | Claudio Tavares Sacchi, Claudia Regina Gonçalves, Erica Valessa Ramos Gomes, Karoline Rodrigues Campos, Caio Vinicius Dias Lopes |
| EPI_ISL_2828654, EPI_ISL_2828655, EPI_ISL_2828656, EPI_ISL_2828657, EPI_ISL_2828658, EPI_ISL_2828659, EPI_ISL_2828660 | Instituo Adolfo Lutz Central | Instituto Adolfo Lutz, Interdiciplinary Procedures Center, Strategic Laboratory | Claudio Tavares Sacchi, Claudia Regina Gonçalves, Erica Valessa Ramos Gomes, Karoline Rodrigues Campos, Caio Vinicius Dias Lopes |
| EPI_ISL_2828663, EPI_ISL_2828664, EPI_ISL_2828665 | Instituo Adolfo Lutz - Reginal de Ribeirao Preto | Instituto Adolfo Lutz, Interdiciplinary Procedures Center, Strategic Laboratory | Claudio Tavares Sacchi, Claudia Regina Gonçalves, Erica Valessa Ramos Gomes, Karoline Rodrigues Campos, Caio Vinicius Dias Lopes |
| EPI_ISL_2828668 | FoxEs Enterprise Solutions | Instituto Adolfo Lutz, Interdiciplinary Procedures Center, Strategic Laboratory | Claudio Tavares Sacchi, Claudia Regina Gonçalves, Erica Valessa Ramos Gomes, Karoline Rodrigues Campos, Caio Vinicius Dias Lopes |
| EPI_ISL_2828669, EPI_ISL_2828670, EPI_ISL_2828671, EPI_ISL_2828672, EPI_ISL_2828673, EPI_ISL_2828674, EPI_ISL_2828675, EPI_ISL_2828676, EPI_ISL_2828677, EPI_ISL_2828678, EPI_ISL_2828679, EPI_ISL_2828680, EPI_ISL_2828681, EPI_ISL_2828682, EPI_ISL_2828683 | | | |
| see above | Instituto Adolfo Lutz - Regional de Santo Andre | Instituto Adolfo Lutz, Interdiciplinary Procedures Center, Strategic Laboratory | Claudio Tavares Sacchi, Claudia Regina Gonçalves, Erica Valessa Ramos Gomes, Karoline Rodrigues Campos, Caio Vinicius Dias Lopes |
| EPI_ISL_2828684, EPI_ISL_2828685 | Instituto Adolfo Lutz - Regional de Campinas | Instituto Adolfo Lutz, Interdiciplinary Procedures Center, Strategic Laboratory | Claudio Tavares Sacchi, Claudia Regina Gonçalves, Erica Valessa Ramos Gomes, Karoline Rodrigues Campos, Caio Vinicius Dias Lopes |
| EPI_ISL_2828698 | Instituto Adolfo Lutz - Regional de Santos | Instituto Adolfo Lutz, Interdiciplinary Procedures Center, Strategic Laboratory | Claudio Tavares Sacchi, Claudia Regina Gonçalves, Erica Valessa Ramos Gomes, Karoline Rodrigues Campos, Caio Vinicius Dias Lopes |
| EPI_ISL_2832354, EPI_ISL_2832356, EPI_ISL_2832358, EPI_ISL_2832360 | Department of Virology, Istituto Zooprofilattico Sperimentale del Lazio e della Toscana (IZSLT) | Department of General Diagnostics; Department of Virology; Istituto Zooprofilattico Sperimentale del Lazio e della Toscana (IZSLT) | Patricia Alba, Giuseppe Manna, Elena L. Diaconu, Fabiola Feltrin, Raffaela Conti, Teresa Scicluna, Virginia Carfora, Antonella Cersini, Alessia Franco, Antonio Battisti. |
| EPI_ISL_2833731, EPI_ISL_2833734, EPI_ISL_2833735, EPI_ISL_2833738, EPI_ISL_2833739 | MOH - Jaber Al-Ahmad Hospital (Innovation Research Laboratory) | MOH - Jaber Al-Ahmad Hospital (Innovation Research Laboratory) | Salman Al-Sabah , Mohammad Alghounaim |
| EPI_ISL_2833785, EPI_ISL_2833788, EPI_ISL_2833789, EPI_ISL_2833790 | University of Oregon COVID-19 MAP Laboratory | University of Oregon Genomics and Cell Charactericzation Core Facility (GC3F) | Douglas Turnbull, Ariana White, Jeff Bishop, Jason Carriere, Jason Sydes, Peter Batzel |
| EPI_ISL_2834159, EPI_ISL_2834161, EPI_ISL_2834162, EPI_ISL_2834163, EPI_ISL_2834165, EPI_ISL_2834169, EPI_ISL_2834170, EPI_ISL_2834171, EPI_ISL_2834173, EPI_ISL_2834174, EPI_ISL_2834178, EPI_ISL_2834184, EPI_ISL_2834185, EPI_ISL_2834186, EPI_ISL_2834188, EPI_ISL_2834192, EPI_ISL_2834193 | | | |
| see above | Baylor Scott & White-Temple | Baylor Scott & White-Temple | Ari Rao, Linden Morales, Kimberly Walker, Marcus Volz, Shelby Johnson |
| EPI_ISL_2834733, EPI_ISL_2834734, EPI_ISL_2834735, EPI_ISL_2834736, EPI_ISL_2834737, EPI_ISL_2834738, EPI_ISL_2834739, EPI_ISL_2834740, EPI_ISL_2834741, EPI_ISL_2834742, EPI_ISL_2834743, EPI_ISL_2834744 | | | |
| see above | Ochsner Health | Bioinfoexperts, LLC | Judy Crabtree, Amy Feehan, Rebecca Rose, David J. Nolan, Tessa LaFleur, Julia-Garcia-Diaz, Lucio Miele, Susanna L. Lamers |
| EPI_ISL_2834925, EPI_ISL_2834926, EPI_ISL_2834927 | Central Health Laboratory/Airport Health Laboratory | Central Health Laboratory ,Victoria Hospital, Candos,Ministry of Health and Wellness, Mauritius | Manraj SS, Sonoo J, Patoo M, Bahadoor BS, Mathur H, Sujeewon C, Jannoo N, Ramuth M |
| EPI_ISL_2835005, EPI_ISL_2835006 | LESP Ciudad de Mexico | Instituto de Diagnostico y Referencia Epidemiologicos (INDRE) | Claudia Wong-Arambula, Abril Rodriguez-Maldonado, Vanessa Rivero-Arredondo, Ariadna Medina-Benitez, Joaquin Quiroz-Mercado, Sergio Rangel-Guerrero, Natividad Cruz-Ortiz, Tatiana Nunez-Garcia, Gisela Barrera-Badillo, Lucia Hernandez-Rivas, Irma Lopez-Martinez, Ernesto Ramirez-Gonzalez. |
| EPI_ISL_2835007 | LESP Veracruz | Instituto de Diagnostico y Referencia Epidemiologicos (INDRE) | Claudia Wong-Arambula, Abril Rodriguez-Maldonado, Vanessa Rivero-Arredondo, Ariadna Medina-Benitez, Joaquin Quiroz-Mercado, Sergio Rangel-Guerrero, Natividad Cruz-Ortiz, Tatiana Nunez-Garcia, Gisela Barrera-Badillo, Lucia Hernandez-Rivas, Irma Lopez-Martinez, Ernesto Ramirez-Gonzalez. |
| EPI_ISL_2835008 | LESP Morelos | Instituto de Diagnostico y Referencia Epidemiologicos (INDRE) | Claudia Wong-Arambula, Abril Rodriguez-Maldonado, Vanessa Rivero-Arredondo, Ariadna Medina-Benitez, Joaquin Quiroz-Mercado, Sergio Rangel-Guerrero, Natividad Cruz-Ortiz, Tatiana Nunez-Garcia, Gisela Barrera-Badillo, Lucia Hernandez-Rivas, Irma Lopez-Martinez, Ernesto Ramirez-Gonzalez. |
| EPI_ISL_2835009, EPI_ISL_2835010, EPI_ISL_2835011, EPI_ISL_2835012, EPI_ISL_2835013 | LESP Ciudad de Mexico | Instituto de Diagnostico y Referencia Epidemiologicos (INDRE) | Claudia Wong-Arambula, Abril Rodriguez-Maldonado, Vanessa Rivero-Arredondo, Ariadna Medina-Benitez, Joaquin Quiroz-Mercado, Sergio Rangel-Guerrero, Natividad Cruz-Ortiz, Tatiana Nunez-Garcia, Gisela Barrera-Badillo, Lucia Hernandez-Rivas, Irma Lopez-Martinez, Ernesto Ramirez-Gonzalez. |
| EPI_ISL_2835014 | LESP Queretaro | Instituto de Diagnostico y Referencia Epidemiologicos (INDRE) | Claudia Wong-Arambula, Abril Rodriguez-Maldonado, Vanessa Rivero-Arredondo, Ariadna Medina-Benitez, Joaquin Quiroz-Mercado, Sergio Rangel-Guerrero, Natividad Cruz-Ortiz, Tatiana Nunez-Garcia, Gisela Barrera-Badillo, Lucia Hernandez-Rivas, Irma Lopez-Martinez, Ernesto Ramirez-Gonzalez. |
| EPI_ISL_2835016 | LESP Jalisco | Instituto de Diagnostico y Referencia Epidemiologicos (INDRE) | Claudia Wong-Arambula, Abril Rodriguez-Maldonado, Vanessa Rivero-Arredondo, Ariadna Medina-Benitez, Joaquin Quiroz-Mercado, Sergio Rangel-Guerrero, Natividad Cruz-Ortiz, Tatiana Nunez-Garcia, Gisela Barrera-Badillo, Lucia Hernandez-Rivas, Irma Lopez-Martinez, Ernesto Ramirez-Gonzalez. |
| EPI_ISL_2835017, EPI_ISL_2835018, EPI_ISL_2835019, EPI_ISL_2835020, EPI_ISL_2835021 | LESP Sinaloa | Instituto de Diagnostico y Referencia Epidemiologicos (INDRE) | Claudia Wong-Arambula, Abril Rodriguez-Maldonado, Vanessa Rivero-Arredondo, Ariadna Medina-Benitez, Joaquin Quiroz-Mercado, Sergio Rangel-Guerrero, Natividad Cruz-Ortiz, Tatiana Nunez-Garcia, Gisela Barrera-Badillo, Lucia Hernandez-Rivas, Irma Lopez-Martinez, Ernesto Ramirez-Gonzalez. |
| EPI_ISL_2835022 | LESP Aguascalientes | Instituto de Diagnostico y Referencia Epidemiologicos (INDRE) | Claudia Wong-Arambula, Abril Rodriguez-Maldonado, Vanessa Rivero-Arredondo, Ariadna Medina-Benitez, Joaquin Quiroz-Mercado, Sergio Rangel-Guerrero, Natividad Cruz-Ortiz, Tatiana Nunez-Garcia, Gisela Barrera-Badillo, Lucia Hernandez-Rivas, Irma Lopez-Martinez, Ernesto |

| | | | |
|---|---|---|---|
| | | | Ramirez-Gonzalez. |
| EPI_ISL_2835023 | LESP Baja California | Instituto de Diagnostico y Referencia Epidemiologicos (INDRE) | Claudia Wong-Arambula, Abril Rodriguez-Maldonado, Vanessa Rivero-Arredondo, Ariadna Medina-Benitez, Joaquin Quiroz-Mercado, Sergio Rangel-Guerrero, Natividad Cruz-Ortiz, Tatiana Nunez-Garcia, Gisela Barrera-Badillo, Lucia Hernandez-Rivas, Irma Lopez-Martinez, Ernesto Ramirez-Gonzalez. |
| EPI_ISL_2835024 | LESP Campeche | Instituto de Diagnostico y Referencia Epidemiologicos (INDRE) | Claudia Wong-Arambula, Abril Rodriguez-Maldonado, Vanessa Rivero-Arredondo, Ariadna Medina-Benitez, Joaquin Quiroz-Mercado, Sergio Rangel-Guerrero, Natividad Cruz-Ortiz, Tatiana Nunez-Garcia, Gisela Barrera-Badillo, Lucia Hernandez-Rivas, Irma Lopez-Martinez, Ernesto Ramirez-Gonzalez. |
| EPI_ISL_2835025 | LESP Sonora | Instituto de Diagnostico y Referencia Epidemiologicos (INDRE) | Claudia Wong-Arambula, Abril Rodriguez-Maldonado, Vanessa Rivero-Arredondo, Ariadna Medina-Benitez, Joaquin Quiroz-Mercado, Sergio Rangel-Guerrero, Maribel Gonzalez-Villa, Natividad Cruz-Ortiz, Tatiana Nunez-Garcia, Gisela Barrera-Badillo, Lucia Hernandez-Rivas, Irma Lopez-Martinez, Ernesto Ramirez-Gonzalez. |
| EPI_ISL_2835026, EPI_ISL_2835027, EPI_ISL_2835028, EPI_ISL_2835029, EPI_ISL_2835030, EPI_ISL_2835031, EPI_ISL_2835032, EPI_ISL_2835033, EPI_ISL_2835034, EPI_ISL_2835035, EPI_ISL_2835036, EPI_ISL_2835037, EPI_ISL_2835038, EPI_ISL_2835039 | | | |
| see above | LESP Ciudad de Mexico | Instituto de Diagnostico y Referencia Epidemiologicos (INDRE) | Claudia Wong-Arambula, Abril Rodriguez-Maldonado, Vanessa Rivero-Arredondo, Ariadna Medina-Benitez, Joaquin Quiroz-Mercado, Sergio Rangel-Guerrero, Maribel Gonzalez-Villa, Natividad Cruz-Ortiz, Tatiana Nunez-Garcia, Gisela Barrera-Badillo, Lucia Hernandez-Rivas, Irma Lopez-Martinez, Ernesto Ramirez-Gonzalez. |
| EPI_ISL_2835040 | LESP Estado de Mexico | Instituto de Diagnostico y Referencia Epidemiologicos (INDRE) | Claudia Wong-Arambula, Abril Rodriguez-Maldonado, Vanessa Rivero-Arredondo, Ariadna Medina-Benitez, Joaquin Quiroz-Mercado, Sergio Rangel-Guerrero, Maribel Gonzalez-Villa, Natividad Cruz-Ortiz, Tatiana Nunez-Garcia, Gisela Barrera-Badillo, Lucia Hernandez-Rivas, Irma Lopez-Martinez, Ernesto Ramirez-Gonzalez. |
| EPI_ISL_2835041, EPI_ISL_2835042, EPI_ISL_2835043 | LESP Ciudad de Mexico | Instituto de Diagnostico y Referencia Epidemiologicos (INDRE) | Claudia Wong-Arambula, Abril Rodriguez-Maldonado, Vanessa Rivero-Arredondo, Ariadna Medina-Benitez, Joaquin Quiroz-Mercado, Sergio Rangel-Guerrero, Maribel Gonzalez-Villa, Natividad Cruz-Ortiz, Tatiana Nunez-Garcia, Gisela Barrera-Badillo, Lucia Hernandez-Rivas, Irma Lopez-Martinez, Ernesto Ramirez-Gonzalez. |
| EPI_ISL_2835044 | LESP Nuevo Leon | Instituto de Diagnostico y Referencia Epidemiologicos (INDRE) | Claudia Wong-Arambula, Abril Rodriguez-Maldonado, Vanessa Rivero-Arredondo, Ariadna Medina-Benitez, Joaquin Quiroz-Mercado, Sergio Rangel-Guerrero, Maribel Gonzalez-Villa, Natividad Cruz-Ortiz, Tatiana Nunez-Garcia, Gisela Barrera-Badillo, Lucia Hernandez-Rivas, Irma Lopez-Martinez, Ernesto Ramirez-Gonzalez. |
| EPI_ISL_2835045 | LESP Chihuahua | Instituto de Diagnostico y Referencia Epidemiologicos (INDRE) | Claudia Wong-Arambula, Abril Rodriguez-Maldonado, Vanessa Rivero-Arredondo, Ariadna Medina-Benitez, Joaquin Quiroz-Mercado, Sergio Rangel-Guerrero, Maribel Gonzalez-Villa, Natividad Cruz-Ortiz, Tatiana Nunez-Garcia, Gisela Barrera-Badillo, Lucia Hernandez-Rivas, Irma Lopez-Martinez, Ernesto Ramirez-Gonzalez. |
| EPI_ISL_2835047 | LESP Veracruz | Instituto de Diagnostico y Referencia Epidemiologicos (INDRE) | Claudia Wong-Arambula, Abril Rodriguez-Maldonado, Vanessa Rivero-Arredondo, Ariadna Medina-Benitez, Joaquin Quiroz-Mercado, Sergio Rangel-Guerrero, Maribel Gonzalez-Villa, Natividad Cruz-Ortiz, Tatiana Nunez-Garcia, Gisela Barrera-Badillo, Lucia Hernandez-Rivas, Irma Lopez-Martinez, Ernesto Ramirez-Gonzalez. |
| EPI_ISL_2835048 | LESP Tamaulipas | Instituto de Diagnostico y Referencia Epidemiologicos (INDRE) | Claudia Wong-Arambula, Abril Rodriguez-Maldonado, Vanessa Rivero-Arredondo, Ariadna Medina-Benitez, Joaquin Quiroz-Mercado, Sergio Rangel-Guerrero, Maribel Gonzalez-Villa, Natividad Cruz-Ortiz, Tatiana Nunez-Garcia, Gisela Barrera-Badillo, Lucia Hernandez-Rivas, Irma Lopez-Martinez, Ernesto Ramirez-Gonzalez. |
| EPI_ISL_2835053 | LESP Ciudad de Mexico | Instituto de Diagnostico y Referencia Epidemiologicos (INDRE) | Claudia Wong-Arambula, Abril Rodriguez-Maldonado, Vanessa Rivero-Arredondo, Ariadna Medina-Benitez, Joaquin Quiroz-Mercado, Sergio Rangel-Guerrero, Maribel Gonzalez-Villa, Natividad Cruz-Ortiz, Tatiana Nunez-Garcia, Gisela Barrera-Badillo, Lucia Hernandez-Rivas, Irma Lopez-Martinez, Ernesto Ramirez-Gonzalez. |
| EPI_ISL_2835054 | LESP Sinaloa | Instituto de Diagnostico y Referencia Epidemiologicos (INDRE) | Claudia Wong-Arambula, Abril Rodriguez-Maldonado, Vanessa Rivero-Arredondo, Ariadna Medina-Benitez, Joaquin Quiroz-Mercado, Sergio Rangel-Guerrero, Maribel Gonzalez-Villa, Natividad Cruz-Ortiz, Tatiana Nunez-Garcia, Gisela Barrera-Badillo, Lucia Hernandez-Rivas, Irma Lopez-Martinez, Ernesto Ramirez-Gonzalez. |
| EPI_ISL_2835056 | LESP Chihuahua | Instituto de Diagnostico y Referencia Epidemiologicos (INDRE) | Claudia Wong-Arambula, Abril Rodriguez-Maldonado, Vanessa Rivero-Arredondo, Ariadna Medina-Benitez, Joaquin Quiroz-Mercado, Sergio Rangel-Guerrero, Maribel Gonzalez-Villa, Natividad Cruz-Ortiz, Tatiana Nunez-Garcia, Gisela Barrera-Badillo, Lucia Hernandez-Rivas, Irma Lopez-Martinez, Ernesto Ramirez-Gonzalez. |
| EPI_ISL_2835057, EPI_ISL_2835058, EPI_ISL_2835059, EPI_ISL_2835060, EPI_ISL_2835061, EPI_ISL_2835062, EPI_ISL_2835063, EPI_ISL_2835064, EPI_ISL_2835065, EPI_ISL_2835066, EPI_ISL_2835067, EPI_ISL_2835068, EPI_ISL_2835069, EPI_ISL_2835070 | | | |
| see above | LESP Tabasco | Instituto de Diagnostico y Referencia Epidemiologicos (INDRE) | Claudia Wong-Arambula, Abril Rodriguez-Maldonado, Vanessa Rivero-Arredondo, Ariadna Medina-Benitez, Joaquin Quiroz-Mercado, Sergio Rangel-Guerrero, Maribel Gonzalez-Villa, Natividad Cruz-Ortiz, Tatiana Nunez-Garcia, Gisela Barrera-Badillo, Lucia Hernandez-Rivas, Irma Lopez-Martinez, Ernesto Ramirez-Gonzalez. |
| EPI_ISL_2835071 | LESP Aguascalientes | Instituto de Diagnostico y Referencia Epidemiologicos (INDRE) | Claudia Wong-Arambula, Abril Rodriguez-Maldonado, Vanessa Rivero-Arredondo, Ariadna Medina-Benitez, Joaquin Quiroz-Mercado, Sergio Rangel-Guerrero, Maribel Gonzalez-Villa, Natividad Cruz-Ortiz, Tatiana Nunez-Garcia, Gisela Barrera-Badillo, Lucia Hernandez-Rivas, Irma Lopez-Martinez, Ernesto Ramirez-Gonzalez. |
| EPI_ISL_2835072, EPI_ISL_2835073 | LESP Sonora | Instituto de Diagnostico y Referencia Epidemiologicos (INDRE) | Claudia Wong-Arambula, Abril Rodriguez-Maldonado, Vanessa Rivero-Arredondo, Ariadna Medina-Benitez, Joaquin Quiroz-Mercado, Sergio Rangel-Guerrero, Maribel Gonzalez-Villa, Natividad Cruz-Ortiz, Tatiana Nunez-Garcia, Gisela Barrera-Badillo, Lucia Hernandez-Rivas, Irma Lopez-Martinez, Ernesto Ramirez-Gonzalez. |
| EPI_ISL_2835074, EPI_ISL_2835075, EPI_ISL_2835076 | LESP Nuevo Leon | Instituto de Diagnostico y Referencia Epidemiologicos (INDRE) | Claudia Wong-Arambula, Abril Rodriguez-Maldonado, Vanessa Rivero-Arredondo, Ariadna Medina-Benitez, Joaquin Quiroz-Mercado, Sergio Rangel-Guerrero, Maribel Gonzalez-Villa, Natividad Cruz-Ortiz, Tatiana Nunez-Garcia, Gisela Barrera-Badillo, Lucia Hernandez-Rivas, Irma Lopez-Martinez, Ernesto Ramirez-Gonzalez. |
| EPI_ISL_2835077 | LESP Sinaloa | Instituto de Diagnostico y Referencia Epidemiologicos (INDRE) | Claudia Wong-Arambula, Abril Rodriguez-Maldonado, Vanessa Rivero-Arredondo, Ariadna Medina-Benitez, Joaquin Quiroz-Mercado, Sergio Rangel-Guerrero, Maribel Gonzalez-Villa, Natividad Cruz-Ortiz, Tatiana Nunez-Garcia, Gisela Barrera-Badillo, Lucia Hernandez-Rivas, Irma Lopez-Martinez, Ernesto Ramirez-Gonzalez. |
| EPI_ISL_2835078, EPI_ISL_2835079, EPI_ISL_2835080, EPI_ISL_2835081, EPI_ISL_2835082, EPI_ISL_2835083, EPI_ISL_2835084 | LESP Tabasco | Instituto de Diagnostico y Referencia Epidemiologicos (INDRE) | Claudia Wong-Arambula, Abril Rodriguez-Maldonado, Vanessa Rivero-Arredondo, Ariadna Medina-Benitez, Joaquin Quiroz-Mercado, Sergio Rangel-Guerrero, Maribel Gonzalez-Villa, Natividad Cruz-Ortiz, Tatiana Nunez-Garcia, Gisela Barrera-Badillo, Lucia Hernandez-Rivas, Irma Lopez-Martinez, Ernesto Ramirez-Gonzalez. |
| EPI_ISL_2835085 | LESP Tamaulipas | Instituto de Diagnostico y Referencia Epidemiologicos (INDRE) | Claudia Wong-Arambula, Abril Rodriguez-Maldonado, Vanessa Rivero-Arredondo, Ariadna Medina-Benitez, Joaquin Quiroz-Mercado, Sergio Rangel-Guerrero, Maribel Gonzalez-Villa, Natividad Cruz-Ortiz, Tatiana Nunez-Garcia, Gisela Barrera-Badillo, Lucia Hernandez-Rivas, Irma Lopez-Martinez, Ernesto Ramirez-Gonzalez. |
| EPI_ISL_2835086, EPI_ISL_2835087 | LESP Campeche | Instituto de Diagnostico y Referencia Epidemiologicos (INDRE) | Claudia Wong-Arambula, Abril Rodriguez-Maldonado, Vanessa Rivero-Arredondo, Ariadna Medina-Benitez, Joaquin Quiroz-Mercado, Sergio Rangel-Guerrero, Maribel Gonzalez-Villa, Natividad Cruz-Ortiz, Tatiana Nunez-Garcia, Gisela Barrera-Badillo, Lucia Hernandez-Rivas, Irma Lopez-Martinez, Ernesto Ramirez-Gonzalez. |
| EPI_ISL_2835088 | LESP Tabasco | Instituto de Diagnostico y Referencia Epidemiologicos (INDRE) | Claudia Wong-Arambula, Abril Rodriguez-Maldonado, Vanessa Rivero-Arredondo, Ariadna Medina-Benitez, Joaquin Quiroz-Mercado, Sergio Rangel-Guerrero, Maribel Gonzalez-Villa, Natividad Cruz-Ortiz, Tatiana Nunez-Garcia, Gisela Barrera-Badillo, Lucia Hernandez-Rivas, Irma Lopez-Martinez, Ernesto Ramirez-Gonzalez. |
| EPI_ISL_2835089 | LESP Queretaro | Instituto de Diagnostico y Referencia Epidemiologicos (INDRE) | Claudia Wong-Arambula, Abril Rodriguez-Maldonado, Vanessa Rivero-Arredondo, Ariadna Medina-Benitez, Joaquin Quiroz-Mercado, Sergio Rangel-Guerrero, Maribel Gonzalez-Villa, Natividad Cruz-Ortiz, Tatiana Nunez-Garcia, Gisela Barrera-Badillo, Lucia Hernandez-Rivas, Irma Lopez-Martinez, Ernesto Ramirez-Gonzalez. |
| EPI_ISL_2835552, EPI_ISL_2835553, EPI_ISL_2835554, EPI_ISL_2835555, EPI_ISL_2835556, EPI_ISL_2835557, EPI_ISL_2835558, EPI_ISL_2835559, EPI_ISL_2835560, EPI_ISL_2835561, EPI_ISL_2835562, EPI_ISL_2835563, EPI_ISL_2835564, EPI_ISL_2835565 | | | |

| | | | |
|---|---|---|---|
| see above | Sharp HealthCare Laboratory | Andersen lab at Scripps Research | SEARCH Alliance San Diego with Aaron Harding, Jacquelyn Berumen, Cathy Woerle, Liam McGinnis, Art Mendoza, Omid Bakhtar |
| EPI_ISL_2835685 | Clinical Microbiology Laboratory, Faculty of Medicine, Universitas Indonesia | Faculty of Medicine, Universitas Indonesia | Fera Ibrahim, Fadilah, Andi Yasmon, Yulia Rosa, Rafika Indah Paramita, Khaerunissa Anbar Istiadi, Linda Erlina, Hana Apsari Pawestri, Vivi Setiawaty, Fithriyah, Beti Ernawati, Pratiwi Sudarmono, Budi Wiweko, Badriul Hegar, Anis Karuniawati, Ari Fahrial Syam |
| EPI_ISL_2835704, EPI_ISL_2835705, EPI_ISL_2835706, EPI_ISL_2835707, EPI_ISL_2835708, EPI_ISL_2835709, EPI_ISL_2835710, EPI_ISL_2835711, EPI_ISL_2835712, EPI_ISL_2835714, EPI_ISL_2835715, EPI_ISL_2835716, EPI_ISL_2835717, EPI_ISL_2835718, EPI_ISL_2835719, EPI_ISL_2835720, EPI_ISL_2835721, EPI_ISL_2835722, EPI_ISL_2835723, EPI_ISL_2835724, EPI_ISL_2835726, EPI_ISL_2835727, EPI_ISL_2835728, EPI_ISL_2835729, EPI_ISL_2835730, EPI_ISL_2835731, EPI_ISL_2835732, EPI_ISL_2835734, EPI_ISL_2835735, EPI_ISL_2835736 | | | |
| see above | Labo Analyses Med | National Reference Center for Viruses of Respiratory Infections, Institut Pasteur, Paris | Marion Barbet, Sylvie Behillil, Méline Bizard, Angela Brisebarre, Camille Capel, Vincent Enouf, Louise Lefrançois, Frédéric Lemoine, Christophe Malabat, Corinne Maufrais, Etienne Simon-Lorière, Maud Vanpeene, Sylvie Van der Werf ,BéNéDicte Roquebert |
| EPI_ISL_2835737 | Hospital | National Reference Center for Viruses of Respiratory Infections, Institut Pasteur, Paris | Marion Barbet, Sylvie Behillil, Méline Bizard, Angela Brisebarre, Camille Capel, Vincent Enouf, Louise Lefrançois, Frédéric Lemoine, Christophe Malabat, Corinne Maufrais, Etienne Simon-Lorière, Maud Vanpeene, Sylvie Van der Werf ,JéRome Guinard |
| EPI_ISL_2835739 | Hospital | National Reference Center for Viruses of Respiratory Infections, Institut Pasteur, Paris | Marion Barbet, Sylvie Behillil, Méline Bizard, Angela Brisebarre, Camille Capel, Vincent Enouf, Louise Lefrançois, Frédéric Lemoine, Christophe Malabat, Corinne Maufrais, Etienne Simon-Lorière, Maud Vanpeene, Sylvie Van der Werf ,Laurent Bret |
| EPI_ISL_2835740 | Hospital | National Reference Center for Viruses of Respiratory Infections, Institut Pasteur, Paris | Marion Barbet, Sylvie Behillil, Méline Bizard, Angela Brisebarre, Camille Capel, Vincent Enouf, Louise Lefrançois, Frédéric Lemoine, Christophe Malabat, Corinne Maufrais, Etienne Simon-Lorière, Maud Vanpeene, Sylvie Van der Werf ,Benjamin Maneglie |
| EPI_ISL_2835741 | Hospital | National Reference Center for Viruses of Respiratory Infections, Institut Pasteur, Paris | Marion Barbet, Sylvie Behillil, Méline Bizard, Angela Brisebarre, Camille Capel, Vincent Enouf, Louise Lefrançois, Frédéric Lemoine, Christophe Malabat, Corinne Maufrais, Etienne Simon-Lorière, Maud Vanpeene, Sylvie Van der Werf ,ValéRie Serazin |
| EPI_ISL_2835742 | Hospital | National Reference Center for Viruses of Respiratory Infections, Institut Pasteur, Paris | Marion Barbet, Sylvie Behillil, Méline Bizard, Angela Brisebarre, Camille Capel, Vincent Enouf, Louise Lefrançois, Frédéric Lemoine, Christophe Malabat, Corinne Maufrais, Etienne Simon-Lorière, Maud Vanpeene, Sylvie Van der Werf ,Laura Djamdjian |
| EPI_ISL_2835743 | Labo Analyses Med | National Reference Center for Viruses of Respiratory Infections, Institut Pasteur, Paris | Marion Barbet, Sylvie Behillil, Méline Bizard, Angela Brisebarre, Camille Capel, Vincent Enouf, Louise Lefrançois, Frédéric Lemoine, Christophe Malabat, Corinne Maufrais, Etienne Simon-Lorière, Maud Vanpeene, Sylvie Van der Werf ,Eve Haguenoer |
| EPI_ISL_2835744, EPI_ISL_2835745, EPI_ISL_2835748 | Labo Analyses Med | National Reference Center for Viruses of Respiratory Infections, Institut Pasteur, Paris | Marion Barbet, Sylvie Behillil, Méline Bizard, Angela Brisebarre, Camille Capel, Vincent Enouf, Louise Lefrançois, Frédéric Lemoine, Christophe Malabat, Corinne Maufrais, Etienne Simon-Lorière, Maud Vanpeene, Sylvie Van der Werf ,Sabine Trombert |
| EPI_ISL_2835749, EPI_ISL_2835752, EPI_ISL_2835754, EPI_ISL_2835755, EPI_ISL_2835756, EPI_ISL_2835757, EPI_ISL_2835758, EPI_ISL_2835760 | Labo Analyses Med | National Reference Center for Viruses of Respiratory Infections, Institut Pasteur, Paris | Marion Barbet, Sylvie Behillil, Méline Bizard, Angela Brisebarre, Camille Capel, Vincent Enouf, Louise Lefrançois, Frédéric Lemoine, Christophe Malabat, Corinne Maufrais, Etienne Simon-Lorière, Maud Vanpeene, Sylvie Van der Werf ,BéNéDicte Roquebert |
| EPI_ISL_2835761, EPI_ISL_2835762, EPI_ISL_2835763, EPI_ISL_2835764 | Labo Analyses Med | National Reference Center for Viruses of Respiratory Infections, Institut Pasteur, Paris | Marion Barbet, Sylvie Behillil, Méline Bizard, Angela Brisebarre, Camille Capel, Vincent Enouf, Louise Lefrançois, Frédéric Lemoine, Christophe Malabat, Corinne Maufrais, Etienne Simon-Lorière, Maud Vanpeene, Sylvie Van der Werf ,Caroline Devaux |
| EPI_ISL_2835765, EPI_ISL_2835766, EPI_ISL_2835767, EPI_ISL_2835768 | Labo Analyses Med | National Reference Center for Viruses of Respiratory Infections, Institut Pasteur, Paris | Marion Barbet, Sylvie Behillil, Méline Bizard, Angela Brisebarre, Camille Capel, Vincent Enouf, Louise Lefrançois, Frédéric Lemoine, Christophe Malabat, Corinne Maufrais, Etienne Simon-Lorière, Maud Vanpeene, Sylvie Van der Werf ,Stephanie Hainos |
| EPI_ISL_2835769 | Labo Analyses Med | National Reference Center for Viruses of Respiratory Infections, Institut Pasteur, Paris | Marion Barbet, Sylvie Behillil, Méline Bizard, Angela Brisebarre, Camille Capel, Vincent Enouf, Louise Lefrançois, Frédéric Lemoine, Christophe Malabat, Corinne Maufrais, Etienne Simon-Lorière, Maud Vanpeene, Sylvie Van der Werf ,Brieuc Gestin |
| EPI_ISL_2835770 | Labo Analyses Med | National Reference Center for Viruses of Respiratory Infections, Institut Pasteur, Paris | Marion Barbet, Sylvie Behillil, Méline Bizard, Angela Brisebarre, Camille Capel, Vincent Enouf, Louise Lefrançois, Frédéric Lemoine, Christophe Malabat, Corinne Maufrais, Etienne Simon-Lorière, Maud Vanpeene, Sylvie Van der Werf ,GréGoire Potiron |
| EPI_ISL_2835771, EPI_ISL_2835772, EPI_ISL_2835773, EPI_ISL_2835774, EPI_ISL_2835775 | Labo Analyses Med | National Reference Center for Viruses of Respiratory Infections, Institut Pasteur, Paris | Marion Barbet, Sylvie Behillil, Méline Bizard, Angela Brisebarre, Camille Capel, Vincent Enouf, Louise Lefrançois, Frédéric Lemoine, Christophe Malabat, Corinne Maufrais, Etienne Simon-Lorière, Maud Vanpeene, Sylvie Van der Werf ,BéAtrice Ricard |
| EPI_ISL_2835776, EPI_ISL_2835777, EPI_ISL_2835779, EPI_ISL_2835780, EPI_ISL_2835781, EPI_ISL_2835782, EPI_ISL_2835783 | Labo Analyses Med | National Reference Center for Viruses of Respiratory Infections, Institut Pasteur, Paris | Marion Barbet, Sylvie Behillil, Méline Bizard, Angela Brisebarre, Camille Capel, Vincent Enouf, Louise Lefrançois, Frédéric Lemoine, Christophe Malabat, Corinne Maufrais, Etienne Simon-Lorière, Maud Vanpeene, Sylvie Van der Werf ,Jean-FrançOis Comes |
| EPI_ISL_2835784, EPI_ISL_2835785, EPI_ISL_2835787, EPI_ISL_2835788, EPI_ISL_2835789, EPI_ISL_2835790, EPI_ISL_2835791, EPI_ISL_2835792, EPI_ISL_2835793, EPI_ISL_2835794, EPI_ISL_2835795, EPI_ISL_2835796, EPI_ISL_2835797, EPI_ISL_2835798, EPI_ISL_2835799, EPI_ISL_2835800, EPI_ISL_2835801, EPI_ISL_2835802, EPI_ISL_2835804, EPI_ISL_2835805, EPI_ISL_2835806, EPI_ISL_2835807 | | | |
| see above | Labo Analyses Med | National Reference Center for Viruses of Respiratory Infections, Institut Pasteur, Paris | Marion Barbet, Sylvie Behillil, Méline Bizard, Angela Brisebarre, Camille Capel, Vincent Enouf, Louise Lefrançois, Frédéric Lemoine, Christophe Malabat, Corinne Maufrais, Etienne Simon-Lorière, Maud Vanpeene, Sylvie Van der Werf ,Anne Holstein |
| EPI_ISL_2835809, EPI_ISL_2835810, EPI_ISL_2835811, EPI_ISL_2835812, EPI_ISL_2835813, EPI_ISL_2835815, EPI_ISL_2835816, EPI_ISL_2835817, EPI_ISL_2835818, EPI_ISL_2835819, EPI_ISL_2835820, EPI_ISL_2835821, EPI_ISL_2835822, EPI_ISL_2835823, EPI_ISL_2835824, EPI_ISL_2835825, EPI_ISL_2835826, EPI_ISL_2835827, EPI_ISL_2835828, EPI_ISL_2835829, EPI_ISL_2835830, EPI_ISL_2835831, EPI_ISL_2835832, EPI_ISL_2835833 | | | |
| see above | Labo Analyses Med | National Reference Center for Viruses of Respiratory Infections, Institut Pasteur, Paris | Marion Barbet, Sylvie Behillil, Méline Bizard, Angela Brisebarre, Camille Capel, Vincent Enouf, Louise Lefrançois, Frédéric Lemoine, Christophe Malabat, Corinne Maufrais, Etienne Simon-Lorière, Maud Vanpeene, Sylvie Van der Werf ,Vincent Vieillefond |
| EPI_ISL_2835835 | Hospital | National Reference Center for Viruses of Respiratory Infections, Institut Pasteur, Paris | Marion Barbet, Sylvie Behillil, Méline Bizard, Angela Brisebarre, Camille Capel, Vincent Enouf, Louise Lefrançois, Frédéric Lemoine, Christophe Malabat, Corinne Maufrais, Etienne Simon-Lorière, Maud Vanpeene, Sylvie Van der Werf ,Axelle Paquin |
| EPI_ISL_2835836, EPI_ISL_2835837 | Labo Analyses Med | National Reference Center for Viruses of Respiratory Infections, Institut Pasteur, Paris | Marion Barbet, Sylvie Behillil, Méline Bizard, Angela Brisebarre, Camille Capel, Vincent Enouf, Louise Lefrançois, Frédéric Lemoine, Christophe Malabat, Corinne Maufrais, Etienne Simon-Lorière, Maud Vanpeene, Sylvie Van der Werf ,GwéNolé Prigent |
| EPI_ISL_2835838, EPI_ISL_2835839 | Labo Analyses Med | National Reference Center for Viruses of Respiratory Infections, Institut Pasteur, Paris | Marion Barbet, Sylvie Behillil, Méline Bizard, Angela Brisebarre, Camille Capel, Vincent Enouf, Louise Lefrançois, Frédéric Lemoine, Christophe Malabat, Corinne Maufrais, Etienne Simon-Lorière, Maud Vanpeene, Sylvie Van der Werf ,Eric Grandsire |
| EPI_ISL_2835840, EPI_ISL_2835841, EPI_ISL_2835842 | Labo Analyses Med | National Reference Center for Viruses of Respiratory Infections, Institut Pasteur, Paris | Marion Barbet, Sylvie Behillil, Méline Bizard, Angela Brisebarre, Camille Capel, Vincent Enouf, Louise Lefrançois, Frédéric Lemoine, Christophe Malabat, Corinne Maufrais, Etienne Simon-Lorière, Maud Vanpeene, Sylvie Van der Werf |
| EPI_ISL_2835843, EPI_ISL_2835844, EPI_ISL_2835845 | Hospital | National Reference Center for Viruses of Respiratory Infections, Institut Pasteur, Paris | Marion Barbet, Sylvie Behillil, Méline Bizard, Angela Brisebarre, Camille Capel, Vincent Enouf, Louise Lefrançois, Frédéric Lemoine, Christophe Malabat, Corinne Maufrais, Etienne Simon-Lorière, Maud Vanpeene, Sylvie Van der Werf ,Pascale Martres |
| EPI_ISL_2835849, EPI_ISL_2835850, EPI_ISL_2835851, EPI_ISL_2835852, EPI_ISL_2835853, EPI_ISL_2835854, EPI_ISL_2835855, EPI_ISL_2835856, EPI_ISL_2835857, EPI_ISL_2835858, EPI_ISL_2835860, EPI_ISL_2835861, EPI_ISL_2835863, EPI_ISL_2835864, EPI_ISL_2835866, EPI_ISL_2835868, EPI_ISL_2835870, EPI_ISL_2835871, EPI_ISL_2835873, EPI_ISL_2835874, EPI_ISL_2835875, EPI_ISL_2835876 | | | |
| see above | Labo Analyses Med | National Reference Center for Viruses of Respiratory Infections, Institut Pasteur, Paris | Marion Barbet, Sylvie Behillil, Méline Bizard, Angela Brisebarre, Camille Capel, Vincent Enouf, Louise Lefrançois, Frédéric Lemoine, Christophe Malabat, Corinne Maufrais, Etienne Simon-Lorière, Maud Vanpeene, Sylvie Van der Werf ,Sabine Trombert |
| EPI_ISL_2835877, EPI_ISL_2835878, EPI_ISL_2835879, EPI_ISL_2835880 | Hospital | National Reference Center for Viruses of Respiratory Infections, Institut Pasteur, Paris | Marion Barbet, Sylvie Behillil, Méline Bizard, Angela Brisebarre, Camille Capel, Vincent Enouf, Louise Lefrançois, Frédéric Lemoine, Christophe Malabat, Corinne Maufrais, Etienne Simon-Lorière, Maud Vanpeene, Sylvie Van der Werf ,Sylvaine Bastian |
| EPI_ISL_2835881 | Hospital | National Reference Center for Viruses of Respiratory Infections, Institut Pasteur, Paris | Marion Barbet, Sylvie Behillil, Méline Bizard, Angela Brisebarre, Camille Capel, Vincent Enouf, Louise Lefrançois, Frédéric Lemoine, Christophe Malabat, Corinne Maufrais, Etienne Simon-Lorière, Maud Vanpeene, Sylvie Van der Werf ,Alexandra Ducancelle |
| EPI_ISL_2835882, EPI_ISL_2835883 | Labo Analyses Med | National Reference Center for Viruses of Respiratory Infections, Institut Pasteur, Paris | Marion Barbet, Sylvie Behillil, Méline Bizard, Angela Brisebarre, Camille Capel, Vincent Enouf, Louise Lefrançois, Frédéric Lemoine, Christophe Malabat, Corinne Maufrais, Etienne Simon-Lorière, Maud Vanpeene, Sylvie Van der Werf ,Claire Felloni |
| EPI_ISL_2835884 | Hospital | National Reference Center for Viruses of Respiratory Infections, Institut Pasteur, Paris | Marion Barbet, Sylvie Behillil, Méline Bizard, Angela Brisebarre, Camille Capel, Vincent Enouf, Louise Lefrançois, Frédéric Lemoine, Christophe Malabat, Corinne Maufrais, Etienne Simon-Lorière, Maud Vanpeene, Sylvie Van der Werf ,CéLine Bressollette |
| EPI_ISL_2835886, EPI_ISL_2835887, | Labo Analyses Med | National Reference Center for Viruses of Respiratory | Marion Barbet, Sylvie Behillil, Méline Bizard, Angela Brisebarre, Camille Capel, Vincent Enouf, Louise Lefrançois, Frédéric Lemoine, Christophe Malabat, |

| | | | |
|---|---|---|---|
| EPI_ISL_2835888, EPI_ISL_2835889, EPI_ISL_2835890, EPI_ISL_2835891, EPI_ISL_2835892, EPI_ISL_2835893, EPI_ISL_2835894, EPI_ISL_2835895 | | Infections, Institut Pasteur, Paris | Corinne Maufrais, Etienne Simon-Lorière, Maud Vanpeene, Sylvie Van der Werf ,Pierre-Yves Leonard |
| EPI_ISL_2835897 | Hospital | National Reference Center for Viruses of Respiratory Infections, Institut Pasteur, Paris | Marion Barbet, Sylvie Behillil, Méline Bizard, Angela Brisebarre, Camille Capel, Vincent Enouf, Louise Lefrançois, Frédéric Lemoine, Christophe Malabat, Corinne Maufrais, Etienne Simon-Lorière, Maud Vanpeene, Sylvie Van der Werf ,LéA Pilorge |
| EPI_ISL_2835898 | Hospital | National Reference Center for Viruses of Respiratory Infections, Institut Pasteur, Paris | Marion Barbet, Sylvie Behillil, Méline Bizard, Angela Brisebarre, Camille Capel, Vincent Enouf, Louise Lefrançois, Frédéric Lemoine, Christophe Malabat, Corinne Maufrais, Etienne Simon-Lorière, Maud Vanpeene, Sylvie Van der Werf ,Laurence Louvet |
| EPI_ISL_2835899, EPI_ISL_2835900, EPI_ISL_2835901, EPI_ISL_2835902, EPI_ISL_2835903, EPI_ISL_2835904 | Hospital | National Reference Center for Viruses of Respiratory Infections, Institut Pasteur, Paris | Marion Barbet, Sylvie Behillil, Méline Bizard, Angela Brisebarre, Camille Capel, Vincent Enouf, Louise Lefrançois, Frédéric Lemoine, Christophe Malabat, Corinne Maufrais, Etienne Simon-Lorière, Maud Vanpeene, Sylvie Van der Werf ,Sandrine Castelain |
| EPI_ISL_2835905 | Hospital | National Reference Center for Viruses of Respiratory Infections, Institut Pasteur, Paris | Marion Barbet, Sylvie Behillil, Méline Bizard, Angela Brisebarre, Camille Capel, Vincent Enouf, Louise Lefrançois, Frédéric Lemoine, Christophe Malabat, Corinne Maufrais, Etienne Simon-Lorière, Maud Vanpeene, Sylvie Van der Werf ,Hammas |
| EPI_ISL_2835906, EPI_ISL_2835907 | Hospital | National Reference Center for Viruses of Respiratory Infections, Institut Pasteur, Paris | Marion Barbet, Sylvie Behillil, Méline Bizard, Angela Brisebarre, Camille Capel, Vincent Enouf, Louise Lefrançois, Frédéric Lemoine, Christophe Malabat, Corinne Maufrais, Etienne Simon-Lorière, Maud Vanpeene, Sylvie Van der Werf ,Didier Mattera |
| EPI_ISL_2835908, EPI_ISL_2835909, EPI_ISL_2835910 | Hospital | National Reference Center for Viruses of Respiratory Infections, Institut Pasteur, Paris | Marion Barbet, Sylvie Behillil, Méline Bizard, Angela Brisebarre, Camille Capel, Vincent Enouf, Louise Lefrançois, Frédéric Lemoine, Christophe Malabat, Corinne Maufrais, Etienne Simon-Lorière, Maud Vanpeene, Sylvie Van der Werf ,Odile Petsaris |
| EPI_ISL_2835911, EPI_ISL_2835912 | Labo Analyses Med | National Reference Center for Viruses of Respiratory Infections, Institut Pasteur, Paris | Marion Barbet, Sylvie Behillil, Méline Bizard, Angela Brisebarre, Camille Capel, Vincent Enouf, Louise Lefrançois, Frédéric Lemoine, Christophe Malabat, Corinne Maufrais, Etienne Simon-Lorière, Maud Vanpeene, Sylvie Van der Werf ,Sabine Trombert |
| EPI_ISL_2835977 | CHC Andrée Rosemon | Institut Pasteur de la Guyane | Anne Lavergne, Dominique Rousset, Antoine Enfissi, Arielle Salmier |
| EPI_ISL_2835979, EPI_ISL_2835980, EPI_ISL_2835981, EPI_ISL_2835982, EPI_ISL_2835984 | CNR Institut Pasteur de la Guyane | Institut Pasteur de la Guyane | Anne Lavergne, Dominique Rousset, Antoine Enfissi, Arielle Salmier |
| EPI_ISL_2835986, EPI_ISL_2835987, EPI_ISL_2835988, EPI_ISL_2835989 | Laboratoire Carage | Institut Pasteur de la Guyane | Anne Lavergne, Dominique Rousset, Antoine Enfissi, Arielle Salmier |
| EPI_ISL_2838050 | UMR190-Unité des virus émergents | UMR190-Unité des virus émergents | C.Baronti |
| EPI_ISL_2838051 | Hospital Clínico Universitario Virgen de la Arrixaca | Hospital Clínico Universitario Virgen de la Arrixaca | Marina Simón y Laura Moreno |
| EPI_ISL_2838052 | Hospital Clínico Universitario Virgen de la Arrixaca | Hospital Clínico Universitario Virgen de la Arrixaca | Laura Moreno y Marina Simón |
| EPI_ISL_2840609, EPI_ISL_2840611, EPI_ISL_2840612, EPI_ISL_2840613, EPI_ISL_2840617, EPI_ISL_2840618, EPI_ISL_2840619, EPI_ISL_2840620, EPI_ISL_2840622, EPI_ISL_2840623, EPI_ISL_2840624 | | | |
| see above | S.C. Laboratorio Analisi, ASL 3 Liguria | U.O. Igiene, Ospedale Policlinico San Martino | Bruzzone Bianca, Caligiuri Patrizia, De Pace Vanessa, Domnich Alexander, Orsi Andrea, Ricucci Valentina, Spitaleri Antonino, Icardi Giancarlo on behalf of SARS-CoV-2 ITALIAN RESEARCH ENTERPRISE-(SCIRE) Collaborative Group |
| EPI_ISL_2840625 | U.O. Igiene, Ospedale Policlinico San Martino IRCCS Genova | U.O. Igiene, Ospedale Policlinico San Martino | Bruzzone Bianca, Caligiuri Patrizia, De Pace Vanessa, Domnich Alexander, Orsi Andrea, Ricucci Valentina, Icardi Giancarlo on behalf of SARS-CoV-2 ITALIAN RESEARCH ENTERPRISE-(SCIRE) Collaborative Group |
| EPI_ISL_2840626, EPI_ISL_2840627, EPI_ISL_2840628, EPI_ISL_2840629, EPI_ISL_2840630 | S.C. Laboratorio Analisi, ASL 3 Liguria | U.O. Igiene, Ospedale Policlinico San Martino | Bruzzone Bianca, Caligiuri Patrizia, De Pace Vanessa, Domnich Alexander, Orsi Andrea, Ricucci Valentina, Spitaleri Antonino, Icardi Giancarlo on behalf of SARS-CoV-2 ITALIAN RESEARCH ENTERPRISE-(SCIRE) Collaborative Group |
| EPI_ISL_2840631, EPI_ISL_2840632, EPI_ISL_2840633 | S.S.D. Microbiologia, Ospedale Santa Corona di Pietra Ligure, ASL 2 Liguria | U.O. Igiene, Ospedale Policlinico San Martino | Bruzzone Bianca, Caligiuri Patrizia, De Pace Vanessa, Domnich Alexander, Orsi Andrea, Ricucci Valentina, Valle Caterina, Icardi Giancarlo on behalf of SARS-CoV-2 ITALIAN RESEARCH ENTERPRISE-(SCIRE) Collaborative Group |
| EPI_ISL_2840634, EPI_ISL_2840635 | S.C. Patologia Clinica, Ospedale Sant'Andrea, ASL 5 | U.O. Igiene, Ospedale Policlinico San Martino | Bruzzone Bianca, Battolla Enrico, Caligiuri Patrizia, De Pace Vanessa, Domnich Alexander, Orsi Andrea, Ricucci Valentina, Icardi Giancarlo on behalf of SARS-CoV-2 ITALIAN RESEARCH ENTERPRISE-(SCIRE) Collaborative Group |
| EPI_ISL_2840636 | S.C. Laboratorio Analisi, Ospedale di Lavagna, ASL 4 Liguria | U.O. Igiene, Ospedale Policlinico San Martino | Bruzzone Bianca, Bandettini Roberto, Caligiuri Patrizia, De Pace Vanessa, Domnich Alexander, Orsi Andrea, Ricucci Valentina, Icardi Giancarlo on behalf of SARS-CoV-2 ITALIAN RESEARCH ENTERPRISE-(SCIRE) Collaborative Group |
| EPI_ISL_2840637 | U.O. Igiene, Ospedale Policlinico San Martino IRCCS Genova | U.O. Igiene, Ospedale Policlinico San Martino | Bruzzone Bianca, Caligiuri Patrizia, De Pace Vanessa, Domnich Alexander, Orsi Andrea, Ricucci Valentina, Icardi Giancarlo on behalf of SARS-CoV-2 ITALIAN RESEARCH ENTERPRISE-(SCIRE) Collaborative Group |
| EPI_ISL_2840643, EPI_ISL_2840644, EPI_ISL_2840645, EPI_ISL_2840647, EPI_ISL_2840648, EPI_ISL_2840649 | Università degli Studi di Perugia | Istituto Zooprofilattico Sperimentale dell'Abruzzo e Molise "G. Caporale" | Mencacci A, Camilloni B, Biagetti M, Giammarioli M, Lorusso A, Marcacci M, Di Domenico M, Ancora M, Curini V, Mangone I, Rinaldi A, Delli Compagni E, Scialabba S, Di Pasquale A, Cammà C, Puglia I, Calistri P, Savini G |
| EPI_ISL_2840655 | Laboratorio Analisi Osp. Città di Castello - Azienda USL Umbria1 | Istituto Zooprofilattico Sperimentale dell'Abruzzo e Molise "G. Caporale" | Malagigi V, Taccani P,Lorusso A, Marcacci M, Di Domenico M, Ancora M, Curini V, Mangone I, Rinaldi A, Delli Compagni E, Scialabba S, Di Pasquale A, Cammà C, Puglia I, Calistri P, Savini G |
| EPI_ISL_2840656 | Azienda Ospedaliera Terni | Istituto Zooprofilattico Sperimentale dell'Abruzzo e Molise "G. Caporale" | Palumbo M, Scaccetti A, Lorusso A, Marcacci M, Di Domenico M, Ancora M, Curini V, Mangone I, Rinaldi A, Delli Compagni E, Scialabba S, Di Pasquale A, Cammà C, Puglia I, Calistri P, Savini G |
| EPI_ISL_2840657, EPI_ISL_2840658, EPI_ISL_2840659, EPI_ISL_2840660, EPI_ISL_2840661, EPI_ISL_2840662, EPI_ISL_2840663, EPI_ISL_2840664, EPI_ISL_2840665, EPI_ISL_2840666, EPI_ISL_2840667, EPI_ISL_2840668, EPI_ISL_2840669 | | | |
| see above | Azienda USL Umbria 2 | Istituto Zooprofilattico Sperimentale dell'Abruzzo e Molise "G. Caporale" | Proietti A, Pistoni E, Lorusso A, Marcacci M, Di Domenico M, Ancora M, Curini V, Mangone I, Rinaldi A, Delli Compagni E, Scialabba S, Di Pasquale A, Cammà C, Puglia I, Calistri P, Savini G |
| EPI_ISL_2840672 | SIESP CHIETI - DRIVE IN VASTO | Istituto Zooprofilattico Sperimentale dell'Abruzzo e Molise "G. Caporale" | Lorusso A, Marcacci M, Di Domenico M, Ancora M, Curini V, Di Lollo Valeria, Mangone I, Rinaldi A, Delli Compagni E, Scialabba S, Caporale M, Di Pasquale A, Cammà C, Puglia I, Calistri P, Savini G |
| EPI_ISL_2840673, EPI_ISL_2840674, EPI_ISL_2840675, EPI_ISL_2840676 | SIESP CHIETI - DRIVE IN CHIETI | Istituto Zooprofilattico Sperimentale dell'Abruzzo e Molise "G. Caporale" | Lorusso A, Marcacci M, Di Domenico M, Ancora M, Curini V, Di Lollo Valeria, Mangone I, Rinaldi A, Delli Compagni E, Scialabba S, Caporale M, Di Pasquale A, Cammà C, Puglia I, Calistri P, Savini G |
| EPI_ISL_2840678 | OSPEDALE CIVILE TERAMO - CENTRO TRASFUSIONALE | Istituto Zooprofilattico Sperimentale dell'Abruzzo e Molise "G. Caporale" | Lorusso A, Marcacci M, Di Domenico M, Ancora M, Curini V, Di Lollo Valeria, Mangone I, Rinaldi A, Delli Compagni E, Scialabba S, Caporale M, Di Pasquale A, Cammà C, Puglia I, Calistri P, Savini G |
| EPI_ISL_2840679 | SIESP DIPARTIMENTO DI PREVENZIONE CHIETI | Istituto Zooprofilattico Sperimentale dell'Abruzzo e Molise "G. Caporale" | Lorusso A, Marcacci M, Di Domenico M, Ancora M, Curini V, Di Lollo Valeria, Mangone I, Rinaldi A, Delli Compagni E, Scialabba S, Caporale M, Di Pasquale A, Cammà C, Puglia I, Calistri P, Savini G |
| EPI_ISL_2840680, EPI_ISL_2840681, EPI_ISL_2840682 | SIESP CHIETI - DRIVE IN LANCIANO | Istituto Zooprofilattico Sperimentale dell'Abruzzo e Molise "G. Caporale" | Lorusso A, Marcacci M, Di Domenico M, Ancora M, Curini V, Di Lollo Valeria, Mangone I, Rinaldi A, Delli Compagni E, Scialabba S, Caporale M, Di Pasquale A, Cammà C, Puglia I, Calistri P, Savini G |
| EPI_ISL_2840683, EPI_ISL_2840684 | DIP. PREV. AVEZZANO SERVIZIO DI IGIENE EPIDEMIOLOGIA E SANITA' PUBBLICA | Istituto Zooprofilattico Sperimentale dell'Abruzzo e Molise "G. Caporale" | Lorusso A, Marcacci M, Di Domenico M, Ancora M, Curini V, Di Lollo Valeria, Mangone I, Rinaldi A, Delli Compagni E, Scialabba S, Caporale M, Di Pasquale A, Cammà C, Puglia I, Calistri P, Savini G |
| EPI_ISL_2840685 | SIESP CHIETI - DRIVE IN CHIETI | Istituto Zooprofilattico Sperimentale dell'Abruzzo e Molise "G. Caporale" | Lorusso A, Marcacci M, Di Domenico M, Ancora M, Curini V, Di Lollo Valeria, Mangone I, Rinaldi A, Delli Compagni E, Scialabba S, Caporale M, Di Pasquale A, Cammà C, Puglia I, Calistri P, Savini G |
| EPI_ISL_2840686, EPI_ISL_2840687, EPI_ISL_2840688, EPI_ISL_2840689, EPI_ISL_2840690, EPI_ISL_2840691, EPI_ISL_2840692, EPI_ISL_2840693, EPI_ISL_2840694, EPI_ISL_2840695, EPI_ISL_2840696, EPI_ISL_2840697, EPI_ISL_2840698, EPI_ISL_2840699, EPI_ISL_2840700, EPI_ISL_2840701, EPI_ISL_2840702, EPI_ISL_2840703 | | | |

| | | | |
|---|---|---|---|
| see above | OSPEDALE CIVILE TERAMO - CENTRO TRASFUSIONALE | Istituto Zooprofilattico Sperimentale dell'Abruzzo e Molise "G. Caporale" | Lorusso A, Marcacci M, Di Domenico M, Ancora M, Curini V, Di Lollo Valeria, Mangone I, Rinaldi A, Delli Compagni E, Scialabba S, Caporale M, Di Pasquale A, Cammà C, Puglia I, Calistri P, Savini G |
| EPI_ISL_2841556 | Department of Virology, Istituto Zooprofilattico Sperimentale del Lazio e della Toscana (IZSLT) | Department of General Diagnostics; Department of Virology; Istituto Zooprofilattico Sperimentale del Lazio e della Toscana (IZSLT) | Patricia Alba, Giuseppe Manna, Elena L. Diaconu, Fabiola Feltrin, Raffaela Conti, Teresa Scicluna, Virginia Carfora, Antonella Cersini, Alessia Franco, Antonio Battisti. |
| EPI_ISL_2841869, EPI_ISL_2841870, EPI_ISL_2841871, EPI_ISL_2841872, EPI_ISL_2841875, EPI_ISL_2841876, EPI_ISL_2841877, EPI_ISL_2841878, EPI_ISL_2841881, EPI_ISL_2841882, EPI_ISL_2841883, EPI_ISL_2841884, EPI_ISL_2841886, EPI_ISL_2841887, EPI_ISL_2841888, EPI_ISL_2841889, EPI_ISL_2841890, EPI_ISL_2841891, EPI_ISL_2841892, EPI_ISL_2841893, EPI_ISL_2841895, EPI_ISL_2841896, EPI_ISL_2841897, EPI_ISL_2841898, EPI_ISL_2841899, EPI_ISL_2841900, EPI_ISL_2841901, EPI_ISL_2841903, EPI_ISL_2841904, EPI_ISL_2841905, EPI_ISL_2841906, EPI_ISL_2841907, EPI_ISL_2841908, EPI_ISL_2841909, EPI_ISL_2841910, EPI_ISL_2841911, EPI_ISL_2841912, EPI_ISL_2841913, EPI_ISL_2841914, EPI_ISL_2841915, EPI_ISL_2841916, EPI_ISL_2841917, EPI_ISL_2841918, EPI_ISL_2841919, EPI_ISL_2841920, EPI_ISL_2841921, EPI_ISL_2841942, EPI_ISL_2841943, EPI_ISL_2841944, EPI_ISL_2841945, EPI_ISL_2841946, EPI_ISL_2841947, EPI_ISL_2841948, EPI_ISL_2841949, EPI_ISL_2841953, EPI_ISL_2841954, EPI_ISL_2841955, EPI_ISL_2841956, EPI_ISL_2841957, EPI_ISL_2841958, EPI_ISL_2841959, EPI_ISL_2841960, EPI_ISL_2841961, EPI_ISL_2841962, EPI_ISL_2841964, EPI_ISL_2841965, EPI_ISL_2841966, EPI_ISL_2841967, EPI_ISL_2841968, EPI_ISL_2841969, EPI_ISL_2841970, EPI_ISL_2841973, EPI_ISL_2841975, EPI_ISL_2841976, EPI_ISL_2841977, EPI_ISL_2841978, EPI_ISL_2841979, EPI_ISL_2841982, EPI_ISL_2841983, EPI_ISL_2841984, EPI_ISL_2841985, EPI_ISL_2841986, EPI_ISL_2841987, EPI_ISL_2841988, EPI_ISL_2841989, EPI_ISL_2841991, EPI_ISL_2841992, EPI_ISL_2841993, EPI_ISL_2841994, EPI_ISL_2841995, EPI_ISL_2841996, EPI_ISL_2841997, EPI_ISL_2841998, EPI_ISL_2841999 | | | |
| see above | National Center of Infectious and Parasitic Diseases | National Center of Infectious and Parasitic Diseases | Alexiev et al |
| EPI_ISL_2842625 | Università di Parma, Laboratorio di Igiene e Sanità Pubblica | Istituto Zooprofilattico Sperimentale della Lombardia e dell'Emilia Romagna (IZSLER), Risk Analysis and Genomic Epidemiology Unit | Maria Eugenia Colucci, Licia Veronesi, Paola Affanni, Marina Morganti, Ilaria Menozzi, Erika Scaltriti, Stefano Pongolini |
| EPI_ISL_2846010, EPI_ISL_2846011, EPI_ISL_2846012, EPI_ISL_2846013, EPI_ISL_2846014, EPI_ISL_2846015 | AZ Klina | AZ Klina | Carl Vael - Lynsey Berckmans |
| EPI_ISL_2854048 | Labuan Hospital | Institute for Medical Research, Infectious Disease Research Centre, National Institutes of Health, Ministry of Health Malaysia | Suppiah J, Kamel K, Mohd Zawawi Z, Azizan MA, Ramly N, Robert F, Anasir MI, Thayan R |
| EPI_ISL_2854052 | Ampang Hospital | Institute for Medical Research, Infectious Disease Research Centre, National Institutes of Health, Ministry of Health Malaysia | Suppiah J, Kamel K, Mohd Zawawi Z, Azizan MA, Ramly N, Robert F, Anasir MI, Thayan R |
| EPI_ISL_2854053 | Kemaman Hospital | Institute for Medical Research, Infectious Disease Research Centre, National Institutes of Health, Ministry of Health Malaysia | Suppiah J, Kamel K, Mohd Zawawi Z, Azizan MA, Ramly N, Robert F, Anasir MI, Thayan R |
| EPI_ISL_2854069 | Bayan Lepas Health Clinic | Institute for Medical Research, Infectious Disease Research Centre, National Institutes of Health, Ministry of Health Malaysia | Suppiah J, Kamel K, Mohd Zawawi Z, Azizan MA, Ramly N, Robert F, Anasir MI, Thayan R |
| EPI_ISL_2854070, EPI_ISL_2854103, EPI_ISL_2854104, EPI_ISL_2854105 | Tawau Hospital | Institute for Medical Research, Infectious Disease Research Centre, National Institutes of Health, Ministry of Health Malaysia | Suppiah J, Kamel K, Mohd Zawawi Z, Azizan MA, Ramly N, Robert F, Anasir MI, Thayan R |
| EPI_ISL_2854106 | Queen Elizabeth Hospital | Institute for Medical Research, Infectious Disease Research Centre, National Institutes of Health, Ministry of Health Malaysia | Suppiah J, Kamel K, Mohd Zawawi Z, Azizan MA, Ramly N, Robert F, Anasir MI, Thayan R |
| EPI_ISL_2854107 | Sultan Ismail Hospital | Institute for Medical Research, Infectious Disease Research Centre, National Institutes of Health, Ministry of Health Malaysia | Suppiah J, Kamel K, Mohd Zawawi Z, Azizan MA, Ramly N, Robert F, Anasir MI, Thayan R |
| EPI_ISL_2854108, EPI_ISL_2854109, EPI_ISL_2854174, EPI_ISL_2854186 | Tawau Hospital | Institute for Medical Research, Infectious Disease Research Centre, National Institutes of Health, Ministry of Health Malaysia | Suppiah J, Kamel K, Mohd Zawawi Z, Azizan MA, Ramly N, Robert F, Anasir MI, Thayan R |
| EPI_ISL_2854187 | Kuala Terengganu District Health Office | Institute for Medical Research, Infectious Disease Research Centre, National Institutes of Health, Ministry of Health Malaysia | Suppiah J, Kamel K, Mohd Zawawi Z, Azizan MA, Ramly N, Robert F, Anasir MI, Thayan R |
| EPI_ISL_2854188 | Marang District Health Office | Institute for Medical Research, Infectious Disease Research Centre, National Institutes of Health, Ministry of Health Malaysia | Suppiah J, Kamel K, Mohd Zawawi Z, Azizan MA, Ramly N, Robert F, Anasir MI, Thayan R |
| EPI_ISL_2854747 | RSUD Ciawi | National Institute of Health Research and Development | Subangkit, Hana Apsari Pawestri, , Kartika Dewi Puspa, Arie Ardiansyah Nugraha, Hartanti Dian Ikawati, Krisna Pangesti, Triyani Soekarso, Nelly Puspandari, Vivi Setiawaty |
| EPI_ISL_2854779 | KKP Kelas II Cilacap | National Institute of Health Research and Development | Hana Apsari Pawestri, Subangkit, Kartika Dewi Puspa, Arie Ardiansyah Nugraha, Hartanti Dian Ikawati, Krisna Pangesti, Triyani Soekarso, Nelly Puspandari, Vivi Setiawaty |
| EPI_ISL_2854794 | Kantor Kesehatan Pelabuhan Kelas I Medan | National Institute of Health Research and Development | Hana Apsari Pawestri, Subangkit, Kartika Dewi Puspa, Arie Ardiansyah Nugraha, Hartanti Dian Ikawati, Krisna Pangesti, Triyani Soekarso, Nelly Puspandari, Vivi Setiawaty |
| EPI_ISL_2858935, EPI_ISL_2858936, EPI_ISL_2858937 | LESP Nuevo Leon | Instituto de Diagnostico y Referencia Epidemiologicos (INDRE) | Claudia Wong-Arambula, Abril Rodriguez-Maldonado, Vanessa Rivero-Arredondo, Ariadna Medina-Benitez, Joaquin Quiroz-Mercado, Sergio Rangel-Guerrero, Natividad Cruz-Ortiz, Tatiana Nunez-Garcia, Gisela Barrera-Badillo, Lucia Hernandez-Rivas, Irma Lopez-Martinez, Ernesto Ramirez-Gonzalez. |
| EPI_ISL_2858939, EPI_ISL_2858940 | LESP Chihuahua | Instituto de Diagnostico y Referencia Epidemiologicos (INDRE) | Claudia Wong-Arambula, Abril Rodriguez-Maldonado, Vanessa Rivero-Arredondo, Ariadna Medina-Benitez, Joaquin Quiroz-Mercado, Sergio Rangel-Guerrero, Natividad Cruz-Ortiz, Tatiana Nunez-Garcia, Gisela Barrera-Badillo, Lucia Hernandez-Rivas, Irma Lopez-Martinez, Ernesto Ramirez-Gonzalez. |
| EPI_ISL_2858941, EPI_ISL_2858942, EPI_ISL_2858943 | LESP Tamaulipas | Instituto de Diagnostico y Referencia Epidemiologicos (INDRE) | Claudia Wong-Arambula, Abril Rodriguez-Maldonado, Vanessa Rivero-Arredondo, Ariadna Medina-Benitez, Joaquin Quiroz-Mercado, Sergio Rangel-Guerrero, Natividad Cruz-Ortiz, Tatiana Nunez-Garcia, Gisela Barrera-Badillo, Lucia Hernandez-Rivas, Irma Lopez-Martinez, Ernesto Ramirez-Gonzalez. |
| EPI_ISL_2858944, EPI_ISL_2858945 | LESP Veracruz | Instituto de Diagnostico y Referencia Epidemiologicos (INDRE) | Claudia Wong-Arambula, Abril Rodriguez-Maldonado, Vanessa Rivero-Arredondo, Ariadna Medina-Benitez, Joaquin Quiroz-Mercado, Sergio Rangel-Guerrero, Natividad Cruz-Ortiz, Tatiana Nunez-Garcia, Gisela Barrera-Badillo, Lucia Hernandez-Rivas, Irma Lopez-Martinez, Ernesto Ramirez-Gonzalez. |
| EPI_ISL_2858946 | LESP Yucatan | Instituto de Diagnostico y Referencia Epidemiologicos (INDRE) | Claudia Wong-Arambula, Abril Rodriguez-Maldonado, Vanessa Rivero-Arredondo, Ariadna Medina-Benitez, Joaquin Quiroz-Mercado, Sergio Rangel-Guerrero, Natividad Cruz-Ortiz, Tatiana Nunez-Garcia, Gisela Barrera-Badillo, Lucia Hernandez-Rivas, Irma Lopez-Martinez, Ernesto Ramirez-Gonzalez. |
| EPI_ISL_2858948, EPI_ISL_2858949, EPI_ISL_2858950, EPI_ISL_2858951 | LESP Ciudad de Mexico | Instituto de Diagnostico y Referencia Epidemiologicos (INDRE) | Claudia Wong-Arambula, Abril Rodriguez-Maldonado, Vanessa Rivero-Arredondo, Ariadna Medina-Benitez, Joaquin Quiroz-Mercado, Sergio Rangel-Guerrero, Natividad Cruz-Ortiz, Tatiana Nunez-Garcia, Gisela Barrera-Badillo, Lucia Hernandez-Rivas, Irma Lopez-Martinez, Ernesto Ramirez-Gonzalez. |
| EPI_ISL_2858952, EPI_ISL_2858953 | LESP Tamaulipas | Instituto de Diagnostico y Referencia Epidemiologicos (INDRE) | Claudia Wong-Arambula, Abril Rodriguez-Maldonado, Vanessa Rivero-Arredondo, Ariadna Medina-Benitez, Joaquin Quiroz-Mercado, Sergio Rangel-Guerrero, Natividad Cruz-Ortiz, Tatiana Nunez-Garcia, Gisela Barrera-Badillo, Lucia Hernandez-Rivas, Irma Lopez-Martinez, Ernesto Ramirez-Gonzalez. |

| | | | |
|---|---|---|---|
| EPI_ISL_2858956 | LESP Chihuahua | Instituto de Diagnostico y Referencia Epidemiologicos (INDRE) | Claudia Wong-Arambula, Abril Rodriguez-Maldonado, Vanessa Rivero-Arredondo, Ariadna Medina-Benitez, Joaquin Quiroz-Mercado, Sergio Rangel-Guerrero, Maribel Gonzalez-Villa, Natividad Cruz-Ortiz, Tatiana Nunez-Garcia, Gisela Barrera-Badillo, Lucia Hernandez-Rivas, Irma Lopez-Martinez, Ernesto Ramirez-Gonzalez. |
| EPI_ISL_2858957, EPI_ISL_2858958, EPI_ISL_2858959 | LESP Tabasco | Instituto de Diagnostico y Referencia Epidemiologicos (INDRE) | Claudia Wong-Arambula, Abril Rodriguez-Maldonado, Vanessa Rivero-Arredondo, Ariadna Medina-Benitez, Joaquin Quiroz-Mercado, Sergio Rangel-Guerrero, Maribel Gonzalez-Villa, Natividad Cruz-Ortiz, Tatiana Nunez-Garcia, Gisela Barrera-Badillo, Lucia Hernandez-Rivas, Irma Lopez-Martinez, Ernesto Ramirez-Gonzalez. |
| EPI_ISL_2858960 | LESP Michoacan | Instituto de Diagnostico y Referencia Epidemiologicos (INDRE) | Claudia Wong-Arambula, Abril Rodriguez-Maldonado, Vanessa Rivero-Arredondo, Ariadna Medina-Benitez, Joaquin Quiroz-Mercado, Sergio Rangel-Guerrero, Maribel Gonzalez-Villa, Natividad Cruz-Ortiz, Tatiana Nunez-Garcia, Gisela Barrera-Badillo, Lucia Hernandez-Rivas, Irma Lopez-Martinez, Ernesto Ramirez-Gonzalez. |
| EPI_ISL_2858961 | LESP Sonora | Instituto de Diagnostico y Referencia Epidemiologicos (INDRE) | Claudia Wong-Arambula, Abril Rodriguez-Maldonado, Vanessa Rivero-Arredondo, Ariadna Medina-Benitez, Joaquin Quiroz-Mercado, Sergio Rangel-Guerrero, Maribel Gonzalez-Villa, Natividad Cruz-Ortiz, Tatiana Nunez-Garcia, Gisela Barrera-Badillo, Lucia Hernandez-Rivas, Irma Lopez-Martinez, Ernesto Ramirez-Gonzalez. |
| EPI_ISL_2858962, EPI_ISL_2858963, EPI_ISL_2858964 | LESP Tabasco | Instituto de Diagnostico y Referencia Epidemiologicos (INDRE) | Claudia Wong-Arambula, Abril Rodriguez-Maldonado, Vanessa Rivero-Arredondo, Ariadna Medina-Benitez, Joaquin Quiroz-Mercado, Sergio Rangel-Guerrero, Maribel Gonzalez-Villa, Natividad Cruz-Ortiz, Tatiana Nunez-Garcia, Gisela Barrera-Badillo, Lucia Hernandez-Rivas, Irma Lopez-Martinez, Ernesto Ramirez-Gonzalez. |
| EPI_ISL_2858965, EPI_ISL_2858966, EPI_ISL_2858967, EPI_ISL_2858968, EPI_ISL_2858969, EPI_ISL_2858970, EPI_ISL_2858971 | LESP Tamaulipas | Instituto de Diagnostico y Referencia Epidemiologicos (INDRE) | Claudia Wong-Arambula, Abril Rodriguez-Maldonado, Vanessa Rivero-Arredondo, Ariadna Medina-Benitez, Joaquin Quiroz-Mercado, Sergio Rangel-Guerrero, Maribel Gonzalez-Villa, Natividad Cruz-Ortiz, Tatiana Nunez-Garcia, Gisela Barrera-Badillo, Lucia Hernandez-Rivas, Irma Lopez-Martinez, Ernesto Ramirez-Gonzalez. |
| EPI_ISL_2858973 | LESP Sinaloa | Instituto de Diagnostico y Referencia Epidemiologicos (INDRE) | Claudia Wong-Arambula, Abril Rodriguez-Maldonado, Vanessa Rivero-Arredondo, Ariadna Medina-Benitez, Joaquin Quiroz-Mercado, Sergio Rangel-Guerrero, Maribel Gonzalez-Villa, Natividad Cruz-Ortiz, Tatiana Nunez-Garcia, Gisela Barrera-Badillo, Lucia Hernandez-Rivas, Irma Lopez-Martinez, Ernesto Ramirez-Gonzalez. |
| EPI_ISL_2858974, EPI_ISL_2858975, EPI_ISL_2858976, EPI_ISL_2858977, EPI_ISL_2858978 | LESP Tabasco | Instituto de Diagnostico y Referencia Epidemiologicos (INDRE) | Claudia Wong-Arambula, Abril Rodriguez-Maldonado, Vanessa Rivero-Arredondo, Ariadna Medina-Benitez, Joaquin Quiroz-Mercado, Sergio Rangel-Guerrero, Maribel Gonzalez-Villa, Natividad Cruz-Ortiz, Tatiana Nunez-Garcia, Gisela Barrera-Badillo, Lucia Hernandez-Rivas, Irma Lopez-Martinez, Ernesto Ramirez-Gonzalez. |
| EPI_ISL_2858979 | LESP Chihuahua | Instituto de Diagnostico y Referencia Epidemiologicos (INDRE) | Claudia Wong-Arambula, Abril Rodriguez-Maldonado, Vanessa Rivero-Arredondo, Ariadna Medina-Benitez, Joaquin Quiroz-Mercado, Sergio Rangel-Guerrero, Maribel Gonzalez-Villa, Natividad Cruz-Ortiz, Tatiana Nunez-Garcia, Gisela Barrera-Badillo, Lucia Hernandez-Rivas, Irma Lopez-Martinez, Ernesto Ramirez-Gonzalez. |
| EPI_ISL_2858980 | LESP Baja California | Instituto de Diagnostico y Referencia Epidemiologicos (INDRE) | Claudia Wong-Arambula, Abril Rodriguez-Maldonado, Vanessa Rivero-Arredondo, Ariadna Medina-Benitez, Joaquin Quiroz-Mercado, Sergio Rangel-Guerrero, Maribel Gonzalez-Villa, Natividad Cruz-Ortiz, Tatiana Nunez-Garcia, Gisela Barrera-Badillo, Lucia Hernandez-Rivas, Irma Lopez-Martinez, Ernesto Ramirez-Gonzalez. |
| EPI_ISL_2858981, EPI_ISL_2858982 | LESP Sonora | Instituto de Diagnostico y Referencia Epidemiologicos (INDRE) | Claudia Wong-Arambula, Abril Rodriguez-Maldonado, Vanessa Rivero-Arredondo, Ariadna Medina-Benitez, Joaquin Quiroz-Mercado, Sergio Rangel-Guerrero, Maribel Gonzalez-Villa, Natividad Cruz-Ortiz, Tatiana Nunez-Garcia, Gisela Barrera-Badillo, Lucia Hernandez-Rivas, Irma Lopez-Martinez, Ernesto Ramirez-Gonzalez. |
| EPI_ISL_2858983 | LESP Nuevo Leon | Instituto de Diagnostico y Referencia Epidemiologicos (INDRE) | Claudia Wong-Arambula, Abril Rodriguez-Maldonado, Vanessa Rivero-Arredondo, Ariadna Medina-Benitez, Joaquin Quiroz-Mercado, Sergio Rangel-Guerrero, Maribel Gonzalez-Villa, Natividad Cruz-Ortiz, Tatiana Nunez-Garcia, Gisela Barrera-Badillo, Lucia Hernandez-Rivas, Irma Lopez-Martinez, Ernesto Ramirez-Gonzalez. |
| EPI_ISL_2858984, EPI_ISL_2858985, EPI_ISL_2858986 | LESP Tabasco | Instituto de Diagnostico y Referencia Epidemiologicos (INDRE) | Claudia Wong-Arambula, Abril Rodriguez-Maldonado, Vanessa Rivero-Arredondo, Ariadna Medina-Benitez, Joaquin Quiroz-Mercado, Sergio Rangel-Guerrero, Maribel Gonzalez-Villa, Natividad Cruz-Ortiz, Tatiana Nunez-Garcia, Gisela Barrera-Badillo, Lucia Hernandez-Rivas, Irma Lopez-Martinez, Ernesto Ramirez-Gonzalez. |
| EPI_ISL_2858987, EPI_ISL_2858988, EPI_ISL_2858989, EPI_ISL_2858990 | LESP San Luis Potosi | Instituto de Diagnostico y Referencia Epidemiologicos (INDRE) | Claudia Wong-Arambula, Abril Rodriguez-Maldonado, Vanessa Rivero-Arredondo, Ariadna Medina-Benitez, Joaquin Quiroz-Mercado, Sergio Rangel-Guerrero, Maribel Gonzalez-Villa, Natividad Cruz-Ortiz, Tatiana Nunez-Garcia, Gisela Barrera-Badillo, Lucia Hernandez-Rivas, Irma Lopez-Martinez, Ernesto Ramirez-Gonzalez. |
| EPI_ISL_2858991, EPI_ISL_2858992, EPI_ISL_2858993 | LESP Guanajuato | Instituto de Diagnostico y Referencia Epidemiologicos (INDRE) | Claudia Wong-Arambula, Abril Rodriguez-Maldonado, Vanessa Rivero-Arredondo, Ariadna Medina-Benitez, Joaquin Quiroz-Mercado, Sergio Rangel-Guerrero, Maribel Gonzalez-Villa, Natividad Cruz-Ortiz, Tatiana Nunez-Garcia, Gisela Barrera-Badillo, Lucia Hernandez-Rivas, Irma Lopez-Martinez, Ernesto Ramirez-Gonzalez. |
| EPI_ISL_2858994, EPI_ISL_2858995 | LESP Jalisco | Instituto de Diagnostico y Referencia Epidemiologicos (INDRE) | Claudia Wong-Arambula, Abril Rodriguez-Maldonado, Vanessa Rivero-Arredondo, Ariadna Medina-Benitez, Joaquin Quiroz-Mercado, Sergio Rangel-Guerrero, Maribel Gonzalez-Villa, Natividad Cruz-Ortiz, Tatiana Nunez-Garcia, Gisela Barrera-Badillo, Lucia Hernandez-Rivas, Irma Lopez-Martinez, Ernesto Ramirez-Gonzalez. |
| EPI_ISL_2858996, EPI_ISL_2858997, EPI_ISL_2858998, EPI_ISL_2858999 | LESP Tamaulipas | Instituto de Diagnostico y Referencia Epidemiologicos (INDRE) | Claudia Wong-Arambula, Abril Rodriguez-Maldonado, Vanessa Rivero-Arredondo, Ariadna Medina-Benitez, Joaquin Quiroz-Mercado, Sergio Rangel-Guerrero, Maribel Gonzalez-Villa, Natividad Cruz-Ortiz, Tatiana Nunez-Garcia, Gisela Barrera-Badillo, Lucia Hernandez-Rivas, Irma Lopez-Martinez, Ernesto Ramirez-Gonzalez. |
| EPI_ISL_2859000 | LESP Veracruz | Instituto de Diagnostico y Referencia Epidemiologicos (INDRE) | Claudia Wong-Arambula, Abril Rodriguez-Maldonado, Vanessa Rivero-Arredondo, Ariadna Medina-Benitez, Joaquin Quiroz-Mercado, Sergio Rangel-Guerrero, Maribel Gonzalez-Villa, Natividad Cruz-Ortiz, Tatiana Nunez-Garcia, Gisela Barrera-Badillo, Lucia Hernandez-Rivas, Irma Lopez-Martinez, Ernesto Ramirez-Gonzalez. |
| EPI_ISL_2859001, EPI_ISL_2859002, EPI_ISL_2859003, EPI_ISL_2859005, EPI_ISL_2859006, EPI_ISL_2859007 | LESP Yucatan | Instituto de Diagnostico y Referencia Epidemiologicos (INDRE) | Claudia Wong-Arambula, Abril Rodriguez-Maldonado, Vanessa Rivero-Arredondo, Ariadna Medina-Benitez, Joaquin Quiroz-Mercado, Sergio Rangel-Guerrero, Maribel Gonzalez-Villa, Natividad Cruz-Ortiz, Tatiana Nunez-Garcia, Gisela Barrera-Badillo, Lucia Hernandez-Rivas, Irma Lopez-Martinez, Ernesto Ramirez-Gonzalez. |
| EPI_ISL_2859008, EPI_ISL_2859009, EPI_ISL_2859010, EPI_ISL_2859011 | LESP Chiapas | Instituto de Diagnostico y Referencia Epidemiologicos (INDRE) | Claudia Wong-Arambula, Abril Rodriguez-Maldonado, Vanessa Rivero-Arredondo, Ariadna Medina-Benitez, Joaquin Quiroz-Mercado, Sergio Rangel-Guerrero, Maribel Gonzalez-Villa, Natividad Cruz-Ortiz, Tatiana Nunez-Garcia, Gisela Barrera-Badillo, Lucia Hernandez-Rivas, Irma Lopez-Martinez, Ernesto Ramirez-Gonzalez. |
| EPI_ISL_2859012 | LESP Hidalgo | Instituto de Diagnostico y Referencia Epidemiologicos (INDRE) | Claudia Wong-Arambula, Abril Rodriguez-Maldonado, Vanessa Rivero-Arredondo, Ariadna Medina-Benitez, Joaquin Quiroz-Mercado, Sergio Rangel-Guerrero, Maribel Gonzalez-Villa, Natividad Cruz-Ortiz, Tatiana Nunez-Garcia, Gisela Barrera-Badillo, Lucia Hernandez-Rivas, Irma Lopez-Martinez, Ernesto Ramirez-Gonzalez. |
| EPI_ISL_2859013, EPI_ISL_2859015 | LESP Sinaloa | Instituto de Diagnostico y Referencia Epidemiologicos (INDRE) | Claudia Wong-Arambula, Abril Rodriguez-Maldonado, Vanessa Rivero-Arredondo, Ariadna Medina-Benitez, Joaquin Quiroz-Mercado, Sergio Rangel-Guerrero, Maribel Gonzalez-Villa, Natividad Cruz-Ortiz, Tatiana Nunez-Garcia, Gisela Barrera-Badillo, Lucia Hernandez-Rivas, Irma Lopez-Martinez, Ernesto Ramirez-Gonzalez. |
| EPI_ISL_2859016, EPI_ISL_2859017, EPI_ISL_2859018, EPI_ISL_2859020, EPI_ISL_2859021, EPI_ISL_2859022, EPI_ISL_2859023, EPI_ISL_2859024, EPI_ISL_2859025, EPI_ISL_2859026, EPI_ISL_2859027 | | | |
| see above | LESP Tabasco | Instituto de Diagnostico y Referencia Epidemiologicos (INDRE) | Claudia Wong-Arambula, Abril Rodriguez-Maldonado, Vanessa Rivero-Arredondo, Ariadna Medina-Benitez, Joaquin Quiroz-Mercado, Sergio Rangel-Guerrero, Maribel Gonzalez-Villa, Natividad Cruz-Ortiz, Tatiana Nunez-Garcia, Gisela Barrera-Badillo, Lucia Hernandez-Rivas, Irma Lopez-Martinez, Ernesto Ramirez-Gonzalez. |
| EPI_ISL_2859028 | LESP Jalisco | Instituto de Diagnostico y Referencia Epidemiologicos (INDRE) | Claudia Wong-Arambula, Abril Rodriguez-Maldonado, Vanessa Rivero-Arredondo, Ariadna Medina-Benitez, Joaquin Quiroz-Mercado, Sergio Rangel-Guerrero, Maribel Gonzalez-Villa, Natividad Cruz-Ortiz, Tatiana Nunez-Garcia, Gisela Barrera-Badillo, Lucia Hernandez-Rivas, Irma Lopez-Martinez, Ernesto Ramirez-Gonzalez. |
| EPI_ISL_2859029, EPI_ISL_2859030, | LESP Tamaulipas | Instituto de Diagnostico y Referencia Epidemiologicos | Claudia Wong-Arambula, Abril Rodriguez-Maldonado, Vanessa Rivero-Arredondo, Ariadna Medina-Benitez, Joaquin Quiroz-Mercado, Sergio |

| | | | |
|---|---|---|---|
| EPI_ISL_2859031, EPI_ISL_2859032, EPI_ISL_2859033 | | (INDRE) | Rangel-Guerrero, Maribel Gonzalez-Villa, Natividad Cruz-Ortiz, Tatiana Nunez-Garcia, Gisela Barrera-Badillo, Lucia Hernandez-Rivas, Irma Lopez-Martinez, Ernesto Ramirez-Gonzalez. |
| EPI_ISL_2859034 | LESP Sinaloa | Instituto de Diagnostico y Referencia Epidemiologicos (INDRE) | Claudia Wong-Arambula, Abril Rodriguez-Maldonado, Vanessa Rivero-Arredondo, Ariadna Medina-Benitez, Joaquin Quiroz-Mercado, Sergio Rangel-Guerrero, Maribel Gonzalez-Villa, Natividad Cruz-Ortiz, Tatiana Nunez-Garcia, Gisela Barrera-Badillo, Lucia Hernandez-Rivas, Irma Lopez-Martinez, Ernesto Ramirez-Gonzalez. |
| EPI_ISL_2859035, EPI_ISL_2859036, EPI_ISL_2859037, EPI_ISL_2859038 | LESP Campeche | Instituto de Diagnostico y Referencia Epidemiologicos (INDRE) | Claudia Wong-Arambula, Abril Rodriguez-Maldonado, Vanessa Rivero-Arredondo, Ariadna Medina-Benitez, Joaquin Quiroz-Mercado, Sergio Rangel-Guerrero, Maribel Gonzalez-Villa, Natividad Cruz-Ortiz, Tatiana Nunez-Garcia, Gisela Barrera-Badillo, Lucia Hernandez-Rivas, Irma Lopez-Martinez, Ernesto Ramirez-Gonzalez. |
| EPI_ISL_2859039, EPI_ISL_2859040, EPI_ISL_2859041, EPI_ISL_2859042, EPI_ISL_2859043, EPI_ISL_2859044, EPI_ISL_2859045, EPI_ISL_2859046, EPI_ISL_2859047, EPI_ISL_2859048 | LESP Tabasco | Instituto de Diagnostico y Referencia Epidemiologicos (INDRE) | Claudia Wong-Arambula, Abril Rodriguez-Maldonado, Vanessa Rivero-Arredondo, Ariadna Medina-Benitez, Joaquin Quiroz-Mercado, Sergio Rangel-Guerrero, Maribel Gonzalez-Villa, Natividad Cruz-Ortiz, Tatiana Nunez-Garcia, Gisela Barrera-Badillo, Lucia Hernandez-Rivas, Irma Lopez-Martinez, Ernesto Ramirez-Gonzalez. |
| EPI_ISL_2859049, EPI_ISL_2859050 | LESP Veracruz | Instituto de Diagnostico y Referencia Epidemiologicos (INDRE) | Claudia Wong-Arambula, Abril Rodriguez-Maldonado, Vanessa Rivero-Arredondo, Ariadna Medina-Benitez, Joaquin Quiroz-Mercado, Sergio Rangel-Guerrero, Maribel Gonzalez-Villa, Natividad Cruz-Ortiz, Tatiana Nunez-Garcia, Gisela Barrera-Badillo, Lucia Hernandez-Rivas, Irma Lopez-Martinez, Ernesto Ramirez-Gonzalez. |
| EPI_ISL_2859051 | LESP Sonora | Instituto de Diagnostico y Referencia Epidemiologicos (INDRE) | Claudia Wong-Arambula, Abril Rodriguez-Maldonado, Vanessa Rivero-Arredondo, Ariadna Medina-Benitez, Joaquin Quiroz-Mercado, Sergio Rangel-Guerrero, Maribel Gonzalez-Villa, Natividad Cruz-Ortiz, Tatiana Nunez-Garcia, Gisela Barrera-Badillo, Lucia Hernandez-Rivas, Irma Lopez-Martinez, Ernesto Ramirez-Gonzalez. |
| EPI_ISL_2859052, EPI_ISL_2859053 | LESP Quintana Roo | Instituto de Diagnostico y Referencia Epidemiologicos (INDRE) | Claudia Wong-Arambula, Abril Rodriguez-Maldonado, Vanessa Rivero-Arredondo, Ariadna Medina-Benitez, Joaquin Quiroz-Mercado, Sergio Rangel-Guerrero, Maribel Gonzalez-Villa, Natividad Cruz-Ortiz, Tatiana Nunez-Garcia, Gisela Barrera-Badillo, Lucia Hernandez-Rivas, Irma Lopez-Martinez, Ernesto Ramirez-Gonzalez. |
| EPI_ISL_2859054 | LESP Nuevo Leon | Instituto de Diagnostico y Referencia Epidemiologicos (INDRE) | Claudia Wong-Arambula, Abril Rodriguez-Maldonado, Vanessa Rivero-Arredondo, Ariadna Medina-Benitez, Joaquin Quiroz-Mercado, Sergio Rangel-Guerrero, Maribel Gonzalez-Villa, Natividad Cruz-Ortiz, Tatiana Nunez-Garcia, Gisela Barrera-Badillo, Lucia Hernandez-Rivas, Irma Lopez-Martinez, Ernesto Ramirez-Gonzalez. |
| EPI_ISL_2859055, EPI_ISL_2859056, EPI_ISL_2859057 | LESP Tabasco | Instituto de Diagnostico y Referencia Epidemiologicos (INDRE) | Claudia Wong-Arambula, Abril Rodriguez-Maldonado, Vanessa Rivero-Arredondo, Ariadna Medina-Benitez, Joaquin Quiroz-Mercado, Sergio Rangel-Guerrero, Maribel Gonzalez-Villa, Natividad Cruz-Ortiz, Tatiana Nunez-Garcia, Gisela Barrera-Badillo, Lucia Hernandez-Rivas, Irma Lopez-Martinez, Ernesto Ramirez-Gonzalez. |
| EPI_ISL_2859059 | LESP San Luis Potosi | Instituto de Diagnostico y Referencia Epidemiologicos (INDRE) | Claudia Wong-Arambula, Abril Rodriguez-Maldonado, Vanessa Rivero-Arredondo, Ariadna Medina-Benitez, Joaquin Quiroz-Mercado, Sergio Rangel-Guerrero, Maribel Gonzalez-Villa, Natividad Cruz-Ortiz, Tatiana Nunez-Garcia, Gisela Barrera-Badillo, Lucia Hernandez-Rivas, Irma Lopez-Martinez, Ernesto Ramirez-Gonzalez. |
| EPI_ISL_2859061, EPI_ISL_2859062 | LESP Ciudad de Mexico | Instituto de Diagnostico y Referencia Epidemiologicos (INDRE) | Claudia Wong-Arambula, Abril Rodriguez-Maldonado, Vanessa Rivero-Arredondo, Ariadna Medina-Benitez, Joaquin Quiroz-Mercado, Sergio Rangel-Guerrero, Maribel Gonzalez-Villa, Natividad Cruz-Ortiz, Tatiana Nunez-Garcia, Gisela Barrera-Badillo, Lucia Hernandez-Rivas, Irma Lopez-Martinez, Ernesto Ramirez-Gonzalez. |
| EPI_ISL_2859063 | LESP Yucatan | Instituto de Diagnostico y Referencia Epidemiologicos (INDRE) | Claudia Wong-Arambula, Abril Rodriguez-Maldonado, Vanessa Rivero-Arredondo, Ariadna Medina-Benitez, Joaquin Quiroz-Mercado, Sergio Rangel-Guerrero, Maribel Gonzalez-Villa, Natividad Cruz-Ortiz, Tatiana Nunez-Garcia, Gisela Barrera-Badillo, Lucia Hernandez-Rivas, Irma Lopez-Martinez, Ernesto Ramirez-Gonzalez. |
| EPI_ISL_2859065 | LESP Michoacan | Instituto de Diagnostico y Referencia Epidemiologicos (INDRE) | Claudia Wong-Arambula, Abril Rodriguez-Maldonado, Vanessa Rivero-Arredondo, Ariadna Medina-Benitez, Joaquin Quiroz-Mercado, Sergio Rangel-Guerrero, Maribel Gonzalez-Villa, Natividad Cruz-Ortiz, Tatiana Nunez-Garcia, Gisela Barrera-Badillo, Lucia Hernandez-Rivas, Irma Lopez-Martinez, Ernesto Ramirez-Gonzalez. |
| EPI_ISL_2859072, EPI_ISL_2859073 | LESP Ciudad de Mexico | Instituto de Diagnostico y Referencia Epidemiologicos (INDRE) | Claudia Wong-Arambula, Abril Rodriguez-Maldonado, Vanessa Rivero-Arredondo, Ariadna Medina-Benitez, Joaquin Quiroz-Mercado, Sergio Rangel-Guerrero, Maribel Gonzalez-Villa, Natividad Cruz-Ortiz, Tatiana Nunez-Garcia, Gisela Barrera-Badillo, Lucia Hernandez-Rivas, Irma Lopez-Martinez, Ernesto Ramirez-Gonzalez. |
| EPI_ISL_2859077, EPI_ISL_2859078, EPI_ISL_2859079, EPI_ISL_2859081, EPI_ISL_2859082, EPI_ISL_2859083, EPI_ISL_2859084, EPI_ISL_2859085, EPI_ISL_2859086, EPI_ISL_2859087, EPI_ISL_2859088, EPI_ISL_2859089, EPI_ISL_2859090, EPI_ISL_2859091, EPI_ISL_2859092, EPI_ISL_2859093, EPI_ISL_2859094, EPI_ISL_2859095, EPI_ISL_2859096, EPI_ISL_2859097, EPI_ISL_2859098, EPI_ISL_2859099, EPI_ISL_2859100, EPI_ISL_2859101, EPI_ISL_2859102, EPI_ISL_2859103, EPI_ISL_2859104, EPI_ISL_2859105, EPI_ISL_2859106 | | | |
| see above | LESP Tabasco | Instituto de Diagnostico y Referencia Epidemiologicos (INDRE) | Claudia Wong-Arambula, Abril Rodriguez-Maldonado, Vanessa Rivero-Arredondo, Ariadna Medina-Benitez, Joaquin Quiroz-Mercado, Sergio Rangel-Guerrero, Maribel Gonzalez-Villa, Natividad Cruz-Ortiz, Tatiana Nunez-Garcia, Gisela Barrera-Badillo, Lucia Hernandez-Rivas, Irma Lopez-Martinez, Ernesto Ramirez-Gonzalez. |
| EPI_ISL_2859107 | LESP Jalisco | Instituto de Diagnostico y Referencia Epidemiologicos (INDRE) | Claudia Wong-Arambula, Abril Rodriguez-Maldonado, Vanessa Rivero-Arredondo, Ariadna Medina-Benitez, Joaquin Quiroz-Mercado, Sergio Rangel-Guerrero, Maribel Gonzalez-Villa, Natividad Cruz-Ortiz, Tatiana Nunez-Garcia, Gisela Barrera-Badillo, Lucia Hernandez-Rivas, Irma Lopez-Martinez, Ernesto Ramirez-Gonzalez. |
| EPI_ISL_2859108 | LESP San Luis Potosi | Instituto de Diagnostico y Referencia Epidemiologicos (INDRE) | Claudia Wong-Arambula, Abril Rodriguez-Maldonado, Vanessa Rivero-Arredondo, Ariadna Medina-Benitez, Joaquin Quiroz-Mercado, Sergio Rangel-Guerrero, Maribel Gonzalez-Villa, Natividad Cruz-Ortiz, Tatiana Nunez-Garcia, Gisela Barrera-Badillo, Lucia Hernandez-Rivas, Irma Lopez-Martinez, Ernesto Ramirez-Gonzalez. |
| EPI_ISL_2859109, EPI_ISL_2859110, EPI_ISL_2859111, EPI_ISL_2859112, EPI_ISL_2859113, EPI_ISL_2859114, EPI_ISL_2859115, EPI_ISL_2859116, EPI_ISL_2859117, EPI_ISL_2859118, EPI_ISL_2859119, EPI_ISL_2859120 | | | |
| see above | LESP Tabasco | Instituto de Diagnostico y Referencia Epidemiologicos (INDRE) | Claudia Wong-Arambula, Abril Rodriguez-Maldonado, Vanessa Rivero-Arredondo, Ariadna Medina-Benitez, Joaquin Quiroz-Mercado, Sergio Rangel-Guerrero, Maribel Gonzalez-Villa, Natividad Cruz-Ortiz, Tatiana Nunez-Garcia, Gisela Barrera-Badillo, Lucia Hernandez-Rivas, Irma Lopez-Martinez, Ernesto Ramirez-Gonzalez. |
| EPI_ISL_2859121 | LESP Nuevo Leon | Instituto de Diagnostico y Referencia Epidemiologicos (INDRE) | Claudia Wong-Arambula, Abril Rodriguez-Maldonado, Vanessa Rivero-Arredondo, Ariadna Medina-Benitez, Joaquin Quiroz-Mercado, Sergio Rangel-Guerrero, Maribel Gonzalez-Villa, Natividad Cruz-Ortiz, Tatiana Nunez-Garcia, Gisela Barrera-Badillo, Lucia Hernandez-Rivas, Irma Lopez-Martinez, Ernesto Ramirez-Gonzalez. |
| EPI_ISL_2859122, EPI_ISL_2859123, EPI_ISL_2859124, EPI_ISL_2859125 | LESP Tabasco | Instituto de Diagnostico y Referencia Epidemiologicos (INDRE) | Claudia Wong-Arambula, Abril Rodriguez-Maldonado, Vanessa Rivero-Arredondo, Ariadna Medina-Benitez, Joaquin Quiroz-Mercado, Sergio Rangel-Guerrero, Maribel Gonzalez-Villa, Natividad Cruz-Ortiz, Tatiana Nunez-Garcia, Gisela Barrera-Badillo, Lucia Hernandez-Rivas, Irma Lopez-Martinez, Ernesto Ramirez-Gonzalez. |
| EPI_ISL_2860824, EPI_ISL_2860825, EPI_ISL_2860826 | AZ Sint-Lucas Gent | AZ Sint-Lucas Gent | Jos Van Acker, Charlotte Verfaillie, Elke Vanlaere, Klara De Rauw |
| EPI_ISL_2861158 | LESP Ciudad de Mexico | Instituto de Diagnostico y Referencia Epidemiologicos (INDRE) | Claudia Wong-Arambula, Abril Rodriguez-Maldonado, Vanessa Rivero-Arredondo, Ariadna Medina-Benitez, Joaquin Quiroz-Mercado, Sergio Rangel-Guerrero, Maribel Gonzalez-Villa, Natividad Cruz-Ortiz, Tatiana Nunez-Garcia, Gisela Barrera-Badillo, Lucia Hernandez-Rivas, Irma Lopez-Martinez, Ernesto Ramirez-Gonzalez. |
| EPI_ISL_2861159, EPI_ISL_2861160, EPI_ISL_2861161 | LESP Nuevo Leon | Instituto de Diagnostico y Referencia Epidemiologicos (INDRE) | Claudia Wong-Arambula, Abril Rodriguez-Maldonado, Vanessa Rivero-Arredondo, Ariadna Medina-Benitez, Joaquin Quiroz-Mercado, Sergio Rangel-Guerrero, Maribel Gonzalez-Villa, Natividad Cruz-Ortiz, Tatiana Nunez-Garcia, Gisela Barrera-Badillo, Lucia Hernandez-Rivas, Irma Lopez-Martinez, Ernesto Ramirez-Gonzalez. |
| EPI_ISL_2861162 | LESP Tamaulipas | Instituto de Diagnostico y Referencia Epidemiologicos (INDRE) | Claudia Wong-Arambula, Abril Rodriguez-Maldonado, Vanessa Rivero-Arredondo, Ariadna Medina-Benitez, Joaquin Quiroz-Mercado, Sergio Rangel-Guerrero, Maribel Gonzalez-Villa, Natividad Cruz-Ortiz, Tatiana Nunez-Garcia, Gisela Barrera-Badillo, Lucia Hernandez-Rivas, Irma Lopez-Martinez, Ernesto Ramirez-Gonzalez. |

| | | | |
|---|---|---|---|
| EPI_ISL_2861163 | LESP Nuevo Leon | Instituto de Diagnostico y Referencia Epidemiologicos (INDRE) | Claudia Wong-Arambula, Abril Rodriguez-Maldonado, Vanessa Rivero-Arredondo, Ariadna Medina-Benitez, Joaquin Quiroz-Mercado, Sergio Rangel-Guerrero, Maribel Gonzalez-Villa, Natividad Cruz-Ortiz, Tatiana Nunez-Garcia, Gisela Barrera-Badillo, Lucia Hernandez-Rivas, Irma Lopez-Martinez, Ernesto Ramirez-Gonzalez. |
| EPI_ISL_2861164, EPI_ISL_2861165 | LESP Tamaulipas | Instituto de Diagnostico y Referencia Epidemiologicos (INDRE) | Claudia Wong-Arambula, Abril Rodriguez-Maldonado, Vanessa Rivero-Arredondo, Ariadna Medina-Benitez, Joaquin Quiroz-Mercado, Sergio Rangel-Guerrero, Maribel Gonzalez-Villa, Natividad Cruz-Ortiz, Tatiana Nunez-Garcia, Gisela Barrera-Badillo, Lucia Hernandez-Rivas, Irma Lopez-Martinez, Ernesto Ramirez-Gonzalez. |
| EPI_ISL_2861166 | LESP Veracruz | Instituto de Diagnostico y Referencia Epidemiologicos (INDRE) | Claudia Wong-Arambula, Abril Rodriguez-Maldonado, Vanessa Rivero-Arredondo, Ariadna Medina-Benitez, Joaquin Quiroz-Mercado, Sergio Rangel-Guerrero, Maribel Gonzalez-Villa, Natividad Cruz-Ortiz, Tatiana Nunez-Garcia, Gisela Barrera-Badillo, Lucia Hernandez-Rivas, Irma Lopez-Martinez, Ernesto Ramirez-Gonzalez. |
| EPI_ISL_2861167 | LESP Guanajuato | Instituto de Diagnostico y Referencia Epidemiologicos (INDRE) | Claudia Wong-Arambula, Abril Rodriguez-Maldonado, Vanessa Rivero-Arredondo, Ariadna Medina-Benitez, Joaquin Quiroz-Mercado, Sergio Rangel-Guerrero, Maribel Gonzalez-Villa, Natividad Cruz-Ortiz, Tatiana Nunez-Garcia, Gisela Barrera-Badillo, Lucia Hernandez-Rivas, Irma Lopez-Martinez, Ernesto Ramirez-Gonzalez. |
| EPI_ISL_2861168, EPI_ISL_2861169, EPI_ISL_2861170 | LESP Durango | Instituto de Diagnostico y Referencia Epidemiologicos (INDRE) | Claudia Wong-Arambula, Abril Rodriguez-Maldonado, Vanessa Rivero-Arredondo, Ariadna Medina-Benitez, Joaquin Quiroz-Mercado, Sergio Rangel-Guerrero, Maribel Gonzalez-Villa, Natividad Cruz-Ortiz, Tatiana Nunez-Garcia, Gisela Barrera-Badillo, Lucia Hernandez-Rivas, Irma Lopez-Martinez, Ernesto Ramirez-Gonzalez. |
| EPI_ISL_2861171 | LESP Chiapas | Instituto de Diagnostico y Referencia Epidemiologicos (INDRE) | Claudia Wong-Arambula, Abril Rodriguez-Maldonado, Vanessa Rivero-Arredondo, Ariadna Medina-Benitez, Joaquin Quiroz-Mercado, Sergio Rangel-Guerrero, Maribel Gonzalez-Villa, Natividad Cruz-Ortiz, Tatiana Nunez-Garcia, Gisela Barrera-Badillo, Lucia Hernandez-Rivas, Irma Lopez-Martinez, Ernesto Ramirez-Gonzalez. |
| EPI_ISL_2861172 | LESP Michoacan | Instituto de Diagnostico y Referencia Epidemiologicos (INDRE) | Claudia Wong-Arambula, Abril Rodriguez-Maldonado, Vanessa Rivero-Arredondo, Ariadna Medina-Benitez, Joaquin Quiroz-Mercado, Sergio Rangel-Guerrero, Maribel Gonzalez-Villa, Natividad Cruz-Ortiz, Tatiana Nunez-Garcia, Gisela Barrera-Badillo, Lucia Hernandez-Rivas, Irma Lopez-Martinez, Ernesto Ramirez-Gonzalez. |
| EPI_ISL_2861173 | LESP Hidalgo | Instituto de Diagnostico y Referencia Epidemiologicos (INDRE) | Claudia Wong-Arambula, Abril Rodriguez-Maldonado, Vanessa Rivero-Arredondo, Ariadna Medina-Benitez, Joaquin Quiroz-Mercado, Sergio Rangel-Guerrero, Maribel Gonzalez-Villa, Natividad Cruz-Ortiz, Tatiana Nunez-Garcia, Gisela Barrera-Badillo, Lucia Hernandez-Rivas, Irma Lopez-Martinez, Ernesto Ramirez-Gonzalez. |
| EPI_ISL_2861174, EPI_ISL_2861175 | LESP Queretaro | Instituto de Diagnostico y Referencia Epidemiologicos (INDRE) | Claudia Wong-Arambula, Abril Rodriguez-Maldonado, Vanessa Rivero-Arredondo, Ariadna Medina-Benitez, Joaquin Quiroz-Mercado, Sergio Rangel-Guerrero, Maribel Gonzalez-Villa, Natividad Cruz-Ortiz, Tatiana Nunez-Garcia, Gisela Barrera-Badillo, Lucia Hernandez-Rivas, Irma Lopez-Martinez, Ernesto Ramirez-Gonzalez. |
| EPI_ISL_2861176, EPI_ISL_2861177, EPI_ISL_2861178, EPI_ISL_2861179 | LESP Tabasco | Instituto de Diagnostico y Referencia Epidemiologicos (INDRE) | Claudia Wong-Arambula, Abril Rodriguez-Maldonado, Vanessa Rivero-Arredondo, Ariadna Medina-Benitez, Joaquin Quiroz-Mercado, Sergio Rangel-Guerrero, Maribel Gonzalez-Villa, Natividad Cruz-Ortiz, Tatiana Nunez-Garcia, Gisela Barrera-Badillo, Lucia Hernandez-Rivas, Irma Lopez-Martinez, Ernesto Ramirez-Gonzalez. |
| EPI_ISL_2861180 | LESP Coahuila | Instituto de Diagnostico y Referencia Epidemiologicos (INDRE) | Claudia Wong-Arambula, Abril Rodriguez-Maldonado, Vanessa Rivero-Arredondo, Ariadna Medina-Benitez, Joaquin Quiroz-Mercado, Sergio Rangel-Guerrero, Maribel Gonzalez-Villa, Natividad Cruz-Ortiz, Tatiana Nunez-Garcia, Gisela Barrera-Badillo, Lucia Hernandez-Rivas, Irma Lopez-Martinez, Ernesto Ramirez-Gonzalez. |
| EPI_ISL_2861181, EPI_ISL_2861182 | LESP Tamaulipas | Instituto de Diagnostico y Referencia Epidemiologicos (INDRE) | Claudia Wong-Arambula, Abril Rodriguez-Maldonado, Vanessa Rivero-Arredondo, Ariadna Medina-Benitez, Joaquin Quiroz-Mercado, Sergio Rangel-Guerrero, Maribel Gonzalez-Villa, Natividad Cruz-Ortiz, Tatiana Nunez-Garcia, Gisela Barrera-Badillo, Lucia Hernandez-Rivas, Irma Lopez-Martinez, Ernesto Ramirez-Gonzalez. |
| EPI_ISL_2861183, EPI_ISL_2861184, EPI_ISL_2861185 | LESP Tabasco | Instituto de Diagnostico y Referencia Epidemiologicos (INDRE) | Claudia Wong-Arambula, Abril Rodriguez-Maldonado, Vanessa Rivero-Arredondo, Ariadna Medina-Benitez, Joaquin Quiroz-Mercado, Sergio Rangel-Guerrero, Maribel Gonzalez-Villa, Natividad Cruz-Ortiz, Tatiana Nunez-Garcia, Gisela Barrera-Badillo, Lucia Hernandez-Rivas, Irma Lopez-Martinez, Ernesto Ramirez-Gonzalez. |
| EPI_ISL_2861186, EPI_ISL_2861187 | LESP Aguascalientes | Instituto de Diagnostico y Referencia Epidemiologicos (INDRE) | Claudia Wong-Arambula, Abril Rodriguez-Maldonado, Vanessa Rivero-Arredondo, Ariadna Medina-Benitez, Joaquin Quiroz-Mercado, Sergio Rangel-Guerrero, Maribel Gonzalez-Villa, Natividad Cruz-Ortiz, Tatiana Nunez-Garcia, Gisela Barrera-Badillo, Lucia Hernandez-Rivas, Irma Lopez-Martinez, Ernesto Ramirez-Gonzalez. |
| EPI_ISL_2861188, EPI_ISL_2861189, EPI_ISL_2861190, EPI_ISL_2861191 | LESP Tabasco | Instituto de Diagnostico y Referencia Epidemiologicos (INDRE) | Claudia Wong-Arambula, Abril Rodriguez-Maldonado, Vanessa Rivero-Arredondo, Ariadna Medina-Benitez, Joaquin Quiroz-Mercado, Sergio Rangel-Guerrero, Maribel Gonzalez-Villa, Natividad Cruz-Ortiz, Tatiana Nunez-Garcia, Gisela Barrera-Badillo, Lucia Hernandez-Rivas, Irma Lopez-Martinez, Ernesto Ramirez-Gonzalez. |
| EPI_ISL_2861192, EPI_ISL_2861193, EPI_ISL_2861194, EPI_ISL_2861195, EPI_ISL_2861196 | LESP Ciudad de Mexico | Instituto de Diagnostico y Referencia Epidemiologicos (INDRE) | Claudia Wong-Arambula, Abril Rodriguez-Maldonado, Vanessa Rivero-Arredondo, Ariadna Medina-Benitez, Joaquin Quiroz-Mercado, Sergio Rangel-Guerrero, Maribel Gonzalez-Villa, Natividad Cruz-Ortiz, Tatiana Nunez-Garcia, Gisela Barrera-Badillo, Lucia Hernandez-Rivas, Irma Lopez-Martinez, Ernesto Ramirez-Gonzalez. |
| EPI_ISL_2861262, EPI_ISL_2861263, EPI_ISL_2861264, EPI_ISL_2861265 | Dr. Yoshitaka Tamura Department of Clinical Laboratory, Osaka Habikino Medical Center | Department of Developmental Medicine, Research Institute, Osaka Women's and Children's Hospital | Itaru Yanagihara, Yukiko Nakura |
| EPI_ISL_2861267, EPI_ISL_2861268 | Osaka General Medical Center | Department of Developmental Medicine, Research Institute, Osaka Women's and Children's Hospital | Itaru Yanagihara, Yukiko Nakura |
| EPI_ISL_2861280 | Azienda Ospedaliera Terni | Istituto Zooprofilattico Sperimentale dell'Abruzzo e Molise "G. Caporale" | Palumbo M, Scaccetti A, Lorusso A, Marcacci M, Di Domenico M, Ancora M, Curini V, Mangone I, Rinaldi A, Delli Compagni E, Scialabba S, Di Pasquale A, Cammà C, Puglia I, Calistri P, Savini G |
| EPI_ISL_2861288, EPI_ISL_2861309 | Microbiology Department. Complexo Hospitalario Universitario de Vigo | Microbiology Department. Complexo Hospitalario Universitario de Vigo | Alfaya N, Alonso I, Alvarez M, Cabrera JJ, Carballo R, Cores O, Cortizo S, del-Campo V, Martinez L, Mediero G, Perez S, Davina C, Potel C, Regueiro B, Rey S, Vasallo FJ |
| EPI_ISL_2861318, EPI_ISL_2861319, EPI_ISL_2861320, EPI_ISL_2861321, EPI_ISL_2861322, EPI_ISL_2861323 | Microbiology Department, University Hospital Donostia | Microbiology Department, University Hospital Donostia | Marimon JM, Montes M, Piñeiro L, Martin-Peñaranda T, Gomez M, Sorarrain A, Cilla G. |
| EPI_ISL_2861339, EPI_ISL_2861340 | Istituto Zooprofilattico Sperimentale della Puglia e della Basilicata | Istituto Zooprofilattico Sperimentale della Puglia e della Basilicata | Parisi A., Bianco A., Capozzi L., Del Sambro L., Simone D., Difato L., Santoro A., Ridolfi D., Giannico A., Manzari C., Pesole G., Pace L., Galante D. |
| EPI_ISL_2861341 | Ospedale Santissima Annunziata | Istituto Zooprofilattico Sperimentale della Puglia e della Basilicata | Parisi A., Bianco A., Capozzi L., Del Sambro L., Simone D., Difato L., Ridolfi D., Giannico A, Tampoia M., Manzari C., Pesole G. |
| EPI_ISL_2863571, EPI_ISL_2863576 | Lboratorio Central de Saude Publica do Estado do Parana (LACEN/PR) | Laboratory of Respiratory Viruses and Measles, Oswaldo Cruz Institute, FIOCRUZ | Paola Resende, Luciana Appolinario, Fernando Motta, Anna Carolina Paixao, Ana Carolina Mendonca, Ighor Leonardo Arantes Gomes, Agatha Cristinne Prudencio, Alice Sampaio Rocha, Taina Venas, Elisa Cavalcante Pereira, Renata Serrano Lopes, Irina Riediger, Marilda Siqueira on behalf of the Fiocruz COVID-19 Genomic Surveillance Network |
| EPI_ISL_2863666 | Laboratorio Central de Saude Publica do Estado de Santa Catarina (LACEN-SC) | Laboratory of Respiratory Viruses and Measles, Oswaldo Cruz Institute, FIOCRUZ | Paola Resende, Luciana Appolinario, Fernando Motta, Anna Carolina Paixao, Ana Carolina Mendonca, Alice Sampaio Rocha, Taina Venas, Elisa Cavalcante Pereira, Renata Serrano Lopes, Darcita Buerger Rovaris, Sandra Bianchini Fernandes, Marilda Siqueira on behalf of the Fiocruz COVID-19 Genomic Surveillance Network |
| EPI_ISL_2863682, EPI_ISL_2863684, EPI_ISL_2863685, EPI_ISL_2863725 | Lboratorio Central de Saude Publica do Estado do Parana (LACEN/PR) | Laboratory of Respiratory Viruses and Measles, Oswaldo Cruz Institute, FIOCRUZ | Paola Resende, Luciana Appolinario, Fernando Motta, Anna Carolina Paixao, Ana Carolina Mendonca, Ighor Leonardo Arantes Gomes, Agatha Cristinne Prudencio, Alice Sampaio Rocha, Taina Venas, Elisa Cavalcante Pereira, Renata Serrano Lopes, Irina Riediger, Marilda Siqueira on behalf of the Fiocruz COVID-19 Genomic Surveillance Network |
| EPI_ISL_2863889 | Lboratorio Central de Saude Publica do Estado do Parana (LACEN/PR) | Laboratory of Respiratory Viruses and Measles, Oswaldo Cruz Institute, FIOCRUZ | Paola Resende, Luciana Appolinario, Fernando Motta, Anna Carolina Paixao, Ana Carolina Mendonca, Alice Sampaio Rocha, Taina Venas, Elisa Cavalcante Pereira, Renata Serrano Lopes, Agatha Cristinne Prudencio Soares, Ighor Leonardo Arantes Gomes, Irina Riediger, Marilda Siqueira on behalf |

| | | | |
|---|---|---|---|
| | | | of the Fiocruz COVID-19 Genomic Surveillance Network |
| EPI_ISL_2864141, EPI_ISL_2864142, EPI_ISL_2864144, EPI_ISL_2864145, EPI_ISL_2864146, EPI_ISL_2864147 | Microbiology Department. Complexo Hospitalario Universitario de Vigo | Microbiology Department. Complexo Hospitalario Universitario de Vigo | Alfaya N, Alonso I, Alvarez M, Cabrera JJ, Carballo R, Cores O, Cortizo S, del-Campo V, Martinez L, Mediero G, Perez S, Davina C, Potel C, Regueiro B, Rey S, Vasallo FJ |
| EPI_ISL_2868892 | BTKLPP Kelas I Makassar | National Institute of Health Research and Development | Vivi Setiawaty, Hana Apsari Pawestri, Subangkit, Kartika Dewi Puspa, Arie Ardiansyah Nugraha, Hartanti Dian Ikawati, Krisna Pangesti, Triyani Soekarso, Nelly Puspandari |
| EPI_ISL_2868913 | RS Royal Progress | National Institute of Health Research and Development | Vivi Setiawaty, Hana Apsari Pawestri, Subangkit, Kartika Dewi Puspa, Arie Ardiansyah Nugraha, Hartanti Dian Ikawati, Krisna Pangesti, Triyani Soekarso, Nelly Puspandari |
| EPI_ISL_2876292, EPI_ISL_2876296 | Genome Analysis Center, Yamanashi Central Hospital | Genome Analysis Center, Yamanashi Central Hospital | Yosuke Hirotsu |
| EPI_ISL_2885350, EPI_ISL_2885351 | CHU de la Réunion - Nord | Laboratoire de virologie, CNR arbovirus Associé, Chu de la Réunion | Etienne Frumence, Marie-Christine Jaffar Bandjee, Nicolas Traversier, Sabrina Petit Genet |
| EPI_ISL_2885355, EPI_ISL_2885356, EPI_ISL_2885357 | CHU de la Réunion - Sud | Laboratoire de virologie, CNR arbovirus Associé, Chu de la Réunion | Etienne Frumence, Marie-Christine Jaffar Bandjee, Nicolas Traversier, Sabrina Petit Genet |
| EPI_ISL_2885358, EPI_ISL_2885359, EPI_ISL_2885361 | CHU de la Réunion - Nord | Laboratoire de virologie, CNR arbovirus Associé, Chu de la Réunion | Etienne Frumence, Marie-Christine Jaffar Bandjee, Nicolas Traversier, Sabrina Petit Genet |
| EPI_ISL_2885362, EPI_ISL_2885363 | CHU de la Réunion - Sud | Laboratoire de virologie, CNR arbovirus Associé, Chu de la Réunion | Etienne Frumence, Marie-Christine Jaffar Bandjee, Nicolas Traversier, Sabrina Petit Genet |
| EPI_ISL_2885364 | CHU de la Réunion - Nord | Laboratoire de virologie, CNR arbovirus Associé, Chu de la Réunion | Etienne Frumence, Marie-Christine Jaffar Bandjee, Nicolas Traversier, Sabrina Petit Genet |
| EPI_ISL_2885365, EPI_ISL_2885366 | CHU de la Réunion - Sud | Laboratoire de virologie, CNR arbovirus Associé, Chu de la Réunion | Etienne Frumence, Marie-Christine Jaffar Bandjee, Nicolas Traversier, Sabrina Petit Genet |
| EPI_ISL_2885367, EPI_ISL_2885368 | CHU de la Réunion - Nord | Laboratoire de virologie, CNR arbovirus Associé, Chu de la Réunion | Etienne Frumence, Marie-Christine Jaffar Bandjee, Nicolas Traversier, Sabrina Petit Genet |
| EPI_ISL_2885369, EPI_ISL_2885370, EPI_ISL_2885371, EPI_ISL_2885372 | CHU de la Réunion - Sud | Laboratoire de virologie, CNR arbovirus Associé, Chu de la Réunion | Etienne Frumence, Marie-Christine Jaffar Bandjee, Nicolas Traversier, Sabrina Petit Genet |
| EPI_ISL_2885375, EPI_ISL_2885376, EPI_ISL_2885378, EPI_ISL_2885379, EPI_ISL_2885380, EPI_ISL_2885381, EPI_ISL_2885382, EPI_ISL_2885383 | CHU de la Réunion - Nord | Laboratoire de virologie, CNR arbovirus Associé, Chu de la Réunion | Etienne Frumence, Marie-Christine Jaffar Bandjee, Nicolas Traversier, Sabrina Petit Genet |
| EPI_ISL_2885622, EPI_ISL_2885634, EPI_ISL_2885637, EPI_ISL_2885697, EPI_ISL_2885698 | Cliniques universitaires Saint-Luc | UCLouvain/IREC/MBLG | Jean Ruelle, Bertrand Bearzatto, Benoit Kabamba Mukadi |
| EPI_ISL_2885901 | LBM Bonhoure | Cerba lab | e.g. Jane Doe, John Doe |
| EPI_ISL_2886157, EPI_ISL_2886159, EPI_ISL_2886165 | Institute for Vector and Reservoir Control, Research and Development (IVRCRD)- National Institute of Health Research and Development (NIHRD), Indonesia | Institute for Vector and Reservoir Control, Research and Development (IVRCRD)- National Institute of Health Research and Development (NIHRD), Indonesia | Arief Mulyono, Triwibowo Ambar Garjito, Restu Choirul Saban, Ayu Pradipta Pratiwi, Dwi Susilo, Yusnita Mirna Anggraeni, Mujiyanto, Mega Tyas Prihatin, Nur Hidayati, Arif Suryo Prasetyo, Tika Fiona Sari, Sri Wahyuni Handayani, Aryo Ardanto, Wimbi Kartika Ratnasari, Kusumaningtyas Sekar Negari, Aprilia Safitri, Dimas Bagus Wicaksono Putro, Esti Rahardianingtyas, Sidiq Setyo Nugroho, Marjianto, Muhidin, Subangkit, Hana Apsari Pawestri, Siti Alfiah, Ristiyanto, Evi sulistyorini, Lulus Susanti, Vivi Setiawaty, Joko Waluyo |

We gratefully acknowledge the following Authors from the Originating laboratories responsible for obtaining the specimens, as well as the Submitting laboratories where the genome data were generated and shared via GISAID, on which this research is based.

All Submitters of data may be contacted directly via www.gisaid.org

Authors are sorted alphabetically.

| Accession ID | Originating Laboratory | Submitting Laboratory | Authors |
|---|---|---|---|
| EPI_ISL_2774000, EPI_ISL_2774001, EPI_ISL_2774002 | OLVZ Aalst | OLVZ Aalst | Astrid Holderbeke |
| EPI_ISL_2803757 | The Ohio State University Applied Microbiology Services Laboratory | The Ohio State University Applied Microbiology Services Laboratory | Seth A. Faith PhD |
| EPI_ISL_2812155 | Singapore General Hospital | Department of Microbiology | Nurdyana Abdul Rahman, Kenneth Xin Long Chan, Chayaporn Suphavilai, Kun Lee Lim, Sui Sin Goh, Kian Sing Chan, Lynette Oon, Kern Rei Chng, James Sim Heng Chiak, Niranjan Nagarajan, Karrie Ko |
| EPI_ISL_2812969 | Department of Health Technology and Informatics, The Hong Kong Polytechnic University | Department of Health Technology and Informatics, The Hong Kong Polytechnic University | Gilman Kit-Hang Siu, Lam-Kwong Lee, Kenneth Siu-Sing Leung, Jake Siu-Lun Leung, Wing-Hei Lo, Timothy Ting-Leung Ng, Chloe Toi-Mei Chan, Hoi-Ching Jim, Kingsley King-Gee Tam, Hiu-Yin Lao, Denise Sze-Hang Wong, Alan Ka-Lun Wu, Miranda Chong-Yee Yau, Yvette Wai-Man Lai, Kitty Sau-Chun Fung, Sandy Ka-Yee Chau, Barry Kin-Chung Wong, Wing-Kin To, Kristine Luk, Alex Yat-Man Ho, Tak-Lun Que, Kam-Tong Yip, Wing Cheong Yam, David Ho-Keung Shum, Shea Ping Yip |
| EPI_ISL_2820430 | Botswana Harvard HIV Reference Laboratory | Botswana Harvard HIV Reference Laboratory | Sikhulile Moyo, Wonderful T. Choga, Dorcas Maruapula, Thongbotho Mphoyakgosi, Boitumelo Zuze, Botshelo Radibe, Legodile Kooepile, Ontlametse T. Bareng, Keoratile Ntshambiwa, Modisa Motswaledi, Madisa Mine, Joseph Makhema, Roger Shapiro, Shahin Lockman, Mosepele Mosepele, Simani Gaseitsiwe |
| EPI_ISL_2820731, EPI_ISL_2820733, EPI_ISL_2820735, EPI_ISL_2820738, EPI_ISL_2820739 | Presidio di Brindisi Di Summa - Perrino | Istituto Zooprofilattico Sperimentale della Puglia e della Basilicata | Parisi A., Bianco A., Capozzi L., Del Sambro L., Simone D., Difato L., Santoro A., Ridolfi D., Giannico A., Manzari C., Pesole G. |
| EPI_ISL_2824964 | Laboratoire Hôpital Sainte Musse | Laboratoire Hôpital Sainte Musse | Lionel Chollet |
| EPI_ISL_2834141 | St James's Hospital, Dublin 8, Ireland. | ELDA biotech | Deirdre M. Broderick, Peter R. Flanagan, Brendan Crowley |
| EPI_ISL_2834181, EPI_ISL_2834194, EPI_ISL_2834195, EPI_ISL_2834196, EPI_ISL_2834197, EPI_ISL_2834198, EPI_ISL_2834199, EPI_ISL_2834202, EPI_ISL_2834204, EPI_ISL_2834205, EPI_ISL_2834206 | | | |
| see above | Baylor Scott & White-Temple | Baylor Scott & White-Temple | Ari Rao, Linden Morales, Kimberly Walker, Marcus Volz, Shelby Johnson |
| EPI_ISL_2835862 | Labo Analyses Med | National Reference Center for Viruses of Respiratory Infections, Institut Pasteur, Paris | Marion Barbet, Sylvie Behillil, Méline Bizard, Angela Brisebarre, Camille Capel, Vincent Enouf, Louise Lefrançois, Frédéric Lemoine, Christophe Malabat, Corinne Maufrais, Etienne Simon-Lorière, Maud Vanpeene, Sylvie Van der Werf ,Sabine Trombert |
| EPI_ISL_2840652, EPI_ISL_2840653 | Università degli Studi di Perugia | Istituto Zooprofilattico Sperimentale dell'Abruzzo e Molise "G. Caporale" | Mencacci A, Camilloni B, Biagetti M, Giammarioli M, Lorusso A, Marcacci M, Di Domenico M, Ancora M, Curini V, Mangone I, Rinaldi A, Delli Compagni E, Scialabba S, Di Pasquale A, Cammà C, Puglia I, Calistri P, Savini G |
| EPI_ISL_2840704, EPI_ISL_2840705, EPI_ISL_2840706, EPI_ISL_2840707, EPI_ISL_2840708 | PO AVEZZANO - U.O.C. Laboratorio Analisi | Istituto Zooprofilattico Sperimentale dell'Abruzzo e Molise "G. Caporale" | Lorusso A, Marcacci M, Di Domenico M, Ancora M, Curini V, Di Lollo Valeria, Mangone I, Rinaldi A, Delli Compagni E, Scialabba S, Caporale M, Di Pasquale A, Cammà C, Puglia I, Calistri P, Savini G |
| EPI_ISL_2840709, EPI_ISL_2840710, EPI_ISL_2840711, EPI_ISL_2840712, EPI_ISL_2840713, EPI_ISL_2840714, EPI_ISL_2840715, EPI_ISL_2840716, EPI_ISL_2840717, EPI_ISL_2840718, EPI_ISL_2840719, EPI_ISL_2840720, EPI_ISL_2840721, EPI_ISL_2840722, EPI_ISL_2840723, EPI_ISL_2840724, EPI_ISL_2840725, EPI_ISL_2840726, EPI_ISL_2840727, EPI_ISL_2840728, EPI_ISL_2840729, EPI_ISL_2840730, EPI_ISL_2840731, EPI_ISL_2840732, EPI_ISL_2840733, EPI_ISL_2840734, EPI_ISL_2840735, EPI_ISL_2840736, EPI_ISL_2840737, EPI_ISL_2840738, EPI_ISL_2840739, EPI_ISL_2840740, EPI_ISL_2840741, EPI_ISL_2840742, EPI_ISL_2840743 | | | |
| see above | OSPEDALE CIVILE TERAMO - CENTRO TRASFUSIONALE | Istituto Zooprofilattico Sperimentale dell'Abruzzo e Molise "G. Caporale" | Lorusso A, Marcacci M, Di Domenico M, Ancora M, Curini V, Di Lollo Valeria, Mangone I, Rinaldi A, Delli Compagni E, Scialabba S, Caporale M, Di Pasquale A, Cammà C, Puglia I, Calistri P, Savini G |
| EPI_ISL_2846016, EPI_ISL_2846017, EPI_ISL_2846018, EPI_ISL_2846019, EPI_ISL_2846020, EPI_ISL_2846021, EPI_ISL_2846342, EPI_ISL_2847115, EPI_ISL_2847710, EPI_ISL_2848203, EPI_ISL_2848709, EPI_ISL_2849133, EPI_ISL_2849677, EPI_ISL_2850193, EPI_ISL_2850671, EPI_ISL_2851133, EPI_ISL_2851616, EPI_ISL_2852110 | | | |
| see above | AZ Klina | AZ Klina | Carl Vael - Lynsey Berckmans |
| EPI_ISL_2858872 | Genomic Research Laboratory, Bangabandhu Sheikh Mujib Medical University | Genomic Research Laboratory, BSMMU | Laila Anjuman Banu, Md.Saydur Rahman, Sharadindu Kanti Sinha, Arifa Akram, Md.Sharfuddin Ahmed |
| EPI_ISL_2859060 | LESP San Luis Potosi | Instituto de Diagnostico y Referencia Epidemiologicos (INDRE) | Claudia Wong-Arambula, Abril Rodriguez-Maldonado, Vanessa Rivero-Arredondo, Ariadna Medina-Benitez, Joaquin Quiroz-Mercado, Sergio Rangel-Guerrero, Maribel Gonzalez-Villa, Natividad Cruz-Ortiz, Tatiana Nunez-Garcia, Gisela Barrera-Badillo, Lucia Hernandez-Rivas, Irma Lopez-Martinez, Ernesto Ramirez-Gonzalez. |
| EPI_ISL_2861290, EPI_ISL_2861292, EPI_ISL_2861293, EPI_ISL_2861294, EPI_ISL_2861295, EPI_ISL_2861296, EPI_ISL_2861297, EPI_ISL_2861298, EPI_ISL_2861299, EPI_ISL_2861303, EPI_ISL_2861304, EPI_ISL_2861305, EPI_ISL_2861306, EPI_ISL_2861307, EPI_ISL_2861308, EPI_ISL_2861310, EPI_ISL_2861311, EPI_ISL_2861314 | | | |
| see above | Microbiology Department. Complexo Hospitalario Universitario de Vigo | Microbiology Department. Complexo Hospitalario Universitario de Vigo | Alfaya N, Alonso I, Alvarez M, Cabrera JJ, Carballo R, Cores O, Cortizo S, del-Campo V, Martinez L, Mediero G, Perez S, Davina C, Potel C, Regueiro B, Rey S, Vasallo FJ |
| EPI_ISL_2861376 | St James's Hospital, Dublin 8, Ireland. | ELDA biotech | Deirdre M. Broderick, Peter R. Flanagan, Brendan Crowley |
| EPI_ISL_2862010 | Genomic Research Laboratory, Bangabandhu Sheikh Mujib Medical University | Genomic Research Laboratory, BSMMU | Laila Anjuman Banu, Md.Saydur Rahman, Sharadindu Kanti Sinha, Arifa Akram, Md.Sharfuddin Ahmed |
| EPI_ISL_2862015 | Genomic Research Laboratory, Bangabandhu Sheikh Mujib Medical University (BSMMU) Shahbag, Dhaka_1000 | Genomic Research Laboratory, BSMMU | Laila Anjuman Banu, Md.Saydur Rahman, Sharadindu Kanti Sinha, Arifa Akram, Md.Sharfuddin Ahmed |
| EPI_ISL_2862016, EPI_ISL_2862048 | Genomic Research Laboratory, Bangabandhu Sheikh Mujib Medical University | Genomic Research Laboratory, BSMMU | Laila Anjuman Banu, Md.Saydur Rahman, Sharadindu Kanti Sinha, Arifa Akram, Md.Sharfuddin Ahmed |
| EPI_ISL_2868394, EPI_ISL_2868395, EPI_ISL_2868396, EPI_ISL_2868397, EPI_ISL_2868398, EPI_ISL_2868399, EPI_ISL_2868400, EPI_ISL_2868401, EPI_ISL_2868402, EPI_ISL_2868403, EPI_ISL_2868404, EPI_ISL_2868405, EPI_ISL_2868406, EPI_ISL_2868407, EPI_ISL_2868408, EPI_ISL_2868409, EPI_ISL_2868410, EPI_ISL_2868411, EPI_ISL_2868412, EPI_ISL_2868413 | | | |
| see above | Botswana Harvard HIV Reference Laboratory | Botswana Harvard HIV Reference Laboratory | Sikhulile Moyo, Wonderful T. Choga, Dorcas Maruapula, Thongbotho Mphoyakgosi, Boitumelo Zuze, Botshelo Radibe, Legodile Kooepile, Ontlametse T. Bareng, Keoratile Ntshambiwa, Modisa Motswaledi, Madisa Mine, Joseph Makhema, Roger Shapiro, Shahin Lockman, Mosepele Mosepele, Simani Gaseitsiwe |
| EPI_ISL_2876298 | Genome Analysis Center, Yamanashi Central Hospital | Genome Analysis Center, Yamanashi Central Hospital | Yosuke Hirotsu |
| EPI_ISL_2885610, EPI_ISL_2885611, EPI_ISL_2885612, EPI_ISL_2885613, EPI_ISL_2885614, EPI_ISL_2885615, EPI_ISL_2885616, EPI_ISL_2885617, EPI_ISL_2885618, EPI_ISL_2885619, EPI_ISL_2885620, EPI_ISL_2885621, EPI_ISL_2885623, EPI_ISL_2885624, EPI_ISL_2885625, EPI_ISL_2885626, EPI_ISL_2885627, EPI_ISL_2885628, EPI_ISL_2885629, EPI_ISL_2885630, EPI_ISL_2885631, EPI_ISL_2885632, EPI_ISL_2885633, EPI_ISL_2885635, EPI_ISL_2885636, EPI_ISL_2885638, EPI_ISL_2885639, EPI_ISL_2885640, EPI_ISL_2885641, EPI_ISL_2885642, EPI_ISL_2885643, EPI_ISL_2885644, EPI_ISL_2885645, EPI_ISL_2885646, EPI_ISL_2885647, EPI_ISL_2885648, EPI_ISL_2885649, EPI_ISL_2885650, EPI_ISL_2885651, EPI_ISL_2885652, EPI_ISL_2885653, EPI_ISL_2885654, EPI_ISL_2885655, EPI_ISL_2885656, EPI_ISL_2885657, EPI_ISL_2885658, EPI_ISL_2885659, EPI_ISL_2885660, EPI_ISL_2885661, EPI_ISL_2885662, EPI_ISL_2885663, EPI_ISL_2885664, EPI_ISL_2885665, EPI_ISL_2885666, EPI_ISL_2885667, EPI_ISL_2885668, EPI_ISL_2885669, EPI_ISL_2885670, EPI_ISL_2885671, EPI_ISL_2885672, EPI_ISL_2885673, EPI_ISL_2885674, EPI_ISL_2885675, EPI_ISL_2885676, EPI_ISL_2885677, EPI_ISL_2885678, EPI_ISL_2885679 | | | |

| | | | |
|---|---|---|---|
| see above | Cliniques universitaires Saint-Luc | UCLouvain/IREC/MBLG | Jean Ruelle, Bertrand Bearzatto, Benoit Kabamba Mukadi |

EPI_ISL_2885680, EPI_ISL_2885681, EPI_ISL_2885682, EPI_ISL_2885684, EPI_ISL_2885685, EPI_ISL_2885686, EPI_ISL_2885687, EPI_ISL_2885688, EPI_ISL_2885689, EPI_ISL_2885690, EPI_ISL_2885691, EPI_ISL_2885692, EPI_ISL_2885693, EPI_ISL_2885694, EPI_ISL_2885695, EPI_ISL_2885696

| | | | |
|---|---|---|---|
| see above | Clinique Saint-Pierre Ottignies | UCLouvain/IREC/MBLG | Jean Ruelle, Bertrand Bearzatto, Benoit Kabamba Mukadi |

We gratefully acknowledge the following Authors from the Originating laboratories responsible for obtaining the specimens, as well as the Submitting laboratories where the genome data were generated and shared via GISAID, on which this research is based.

All Submitters of data may be contacted directly via www.gisaid.org

Authors are sorted alphabetically.

| Accession ID | Originating Laboratory | Submitting Laboratory | Authors |
|---|---|---|---|
| EPI_ISL_2332805 | OLVZ Aalst | OLVZ Aalst | Anne Vankeerberghen |
| EPI_ISL_2402535, EPI_ISL_2402536, EPI_ISL_2402537, EPI_ISL_2402540, EPI_ISL_2402541, EPI_ISL_2402543, EPI_ISL_2402544, EPI_ISL_2402545, EPI_ISL_2402546, EPI_ISL_2402547, EPI_ISL_2402548, EPI_ISL_2402549 | | | |
| see above | OLVZ Aalst | OLVZ Aalst | Astrid Holderbeke |
| EPI_ISL_2405147, EPI_ISL_2405151, EPI_ISL_2405154, EPI_ISL_2405156, EPI_ISL_2405158, EPI_ISL_2405160, EPI_ISL_2405164, EPI_ISL_2405166 | Hospital Luis Vernaza | Omics Sciences Laboratory | Derly Andrade Molina, Gabriel Morey León, Darlyn Amaya, Rubén Armas González, Juan Carlos Fernández Cadena |
| EPI_ISL_2406458, EPI_ISL_2406459, EPI_ISL_2406460 | Battambang Provincial Laboratory | Virology Unit, Institut Pasteur du Cambodge | Cecile Troupin, Leakhena Pum, Jurre Y Siegers, Ly Sovann, Kraing Sidonn, Yi Sengdoeurn, Chin Savuth, Chau Darapheak, Veasna Duong, Erik A Karlsson |
| EPI_ISL_2406461 | Oddar Meanchey Rapid Response Team | Virology Unit, Institut Pasteur du Cambodge | Cecile Troupin, Leakhena Pum, Jurre Y Siegers, Ly Sovann, Kraing Sidonn, Yi Sengdoeurn, Chin Savuth, Chau Darapheak, Veasna Duong, Erik A Karlsson |
| EPI_ISL_2406462, EPI_ISL_2406463, EPI_ISL_2406464 | Virology Unit, Institut Pasteur du Cambodge | Virology Unit, Institut Pasteur du Cambodge | Cecile Troupin, Leakhena Pum, Jurre Y Siegers, Ly Sovann, Kraing Sidonn, Yi Sengdoeurn, Chin Savuth, Chau Darapheak, Veasna Duong, Erik A Karlsson |
| EPI_ISL_2406465, EPI_ISL_2406466, EPI_ISL_2406467 | Svay Rieng Rapid Response Team | Virology Unit, Institut Pasteur du Cambodge | Cecile Troupin, Leakhena Pum, Jurre Y Siegers, Ly Sovann, Kraing Sidonn, Yi Sengdoeurn, Chin Savuth, Chau Darapheak, Veasna Duong, Erik A Karlsson |
| EPI_ISL_2406468, EPI_ISL_2406469, EPI_ISL_2406470, EPI_ISL_2406471, EPI_ISL_2406472, EPI_ISL_2406473, EPI_ISL_2406474, EPI_ISL_2406475, EPI_ISL_2406476, EPI_ISL_2406477, EPI_ISL_2406478, EPI_ISL_2406479, EPI_ISL_2406480, EPI_ISL_2406481 | | | |
| see above | Virology Unit, Institut Pasteur du Cambodge | Virology Unit, Institut Pasteur du Cambodge | Cecile Troupin, Leakhena Pum, Jurre Y Siegers, Ly Sovann, Kraing Sidonn, Yi Sengdoeurn, Chin Savuth, Chau Darapheak, Veasna Duong, Erik A Karlsson |
| EPI_ISL_2426981 | Laboratoire Carage | Institut Pasteur de la Guyane | Anne Lavergne, Dominique Rousset, Antoine Enfissi, Arielle Salmier |
| EPI_ISL_2426982 | CHC Andrée Rosemon | Institut Pasteur de la Guyane | Anne Lavergne, Dominique Rousset, Antoine Enfissi, Arielle Salmier |
| EPI_ISL_2426983, EPI_ISL_2426984 | CNR Institut Pasteur de la Guyane | Institut Pasteur de la Guyane | Anne Lavergne, Dominique Rousset, Antoine Enfissi, Arielle Salmier |
| EPI_ISL_2427322, EPI_ISL_2427324, EPI_ISL_2427325, EPI_ISL_2427327, EPI_ISL_2427328, EPI_ISL_2427330, EPI_ISL_2427331, EPI_ISL_2427334, EPI_ISL_2427338, EPI_ISL_2427340, EPI_ISL_2427342, EPI_ISL_2427346, EPI_ISL_2427348, EPI_ISL_2427350, EPI_ISL_2427351, EPI_ISL_2427352, EPI_ISL_2427353, EPI_ISL_2427359, EPI_ISL_2427362, EPI_ISL_2427363, EPI_ISL_2427367, EPI_ISL_2427368, EPI_ISL_2427369, EPI_ISL_2427370, EPI_ISL_2427371, EPI_ISL_2427373, EPI_ISL_2427377, EPI_ISL_2434131, EPI_ISL_2434132, EPI_ISL_2434133, EPI_ISL_2434134, EPI_ISL_2434136 | | | |
| see above | Cliniques universitaires Saint-Luc | UCLouvain/IREC/MBLG | Jean Ruelle, Ophélie Simon, Bertrand Bearzatto, Benoit Kabamba Mukadi |
| EPI_ISL_2439237 | Imeda Hospital | Imeda Hospital | Johan Frans, Dagmar Obbels, Hanne Valgaeren |
| EPI_ISL_2443057 | Department of Health Technology and Informatics, The Hong Kong Polytechnic University | Department of Health Technology and Informatics, The Hong Kong Polytechnic University | Gilman Kit-Hang Siu, Lam-Kwong Lee, Kenneth Siu-Sing Leung, Jake Siu-Lun Leung, Wing-Hei Lo, Timothy Ting-Leung Ng, Chloe Toi-Mei Chan, Hoi-Ching Jim, Kingsley King-Gee Tam, Hiu-Yin Lao, Denise Sze-Hang Wong, Alan Ka-Lun Wu, Miranda Chong-Yee Yau, Yvette Wai-Man Lai, Kitty Sau-Chun Fung, Sandy Ka-Yee Chau, Barry Kin-Chung Wong, Wing-Kin To, Kristine Luk, Alex Yat-Man Ho, Tak-Lun Que, Kam-Tong Yip, Wing Cheong Yam, David Ho-Keung Shum, Shea Ping Yip |
| EPI_ISL_2443210, EPI_ISL_2443211, EPI_ISL_2443212, EPI_ISL_2443214, EPI_ISL_2443217, EPI_ISL_2443218, EPI_ISL_2443219, EPI_ISL_2443221, EPI_ISL_2443226 | Laboratoire Hôpital Sainte Musse | Laboratoire Hôpital Sainte Musse | Lionel Chollet |
| EPI_ISL_2444386, EPI_ISL_2444387 | CHU LILLE | CHU Lille - Laboratoire de Virologie | AIT YAHYA Emilie, ALIDJINOU Enagnon Kazali, BOCKET Laurence, CREPIN Michel, DEMAY Christophe, ENGELMANN Ilka, GEFFROY Sandrine, GUIGON Aurélie, LAMBERT Valérie, LAZREK Mouna, NOBILLIAUX Florian, PREVOST Brigitte, THUILLIER Caroline, TINEZ Claire |
| EPI_ISL_2444389 | CH ROUBAIX | CHU Lille - Laboratoire de Virologie | AIT YAHYA Emilie, ALIDJINOU Enagnon Kazali, BOCKET Laurence, CREPIN Michel, DEMAY Christophe, ENGELMANN Ilka, GEFFROY Sandrine, GUIGON Aurélie, LAMBERT Valérie, LAZREK Mouna, NOBILLIAUX Florian, PREVOST Brigitte, THUILLIER Caroline, TINEZ Claire |
| EPI_ISL_2444390 | CH SAINT-OMER | CHU Lille - Laboratoire de Virologie | AIT YAHYA Emilie, ALIDJINOU Enagnon Kazali, BOCKET Laurence, CREPIN Michel, DEMAY Christophe, ENGELMANN Ilka, GEFFROY Sandrine, GUIGON Aurélie, LAMBERT Valérie, LAZREK Mouna, NOBILLIAUX Florian, PREVOST Brigitte, THUILLIER Caroline, TINEZ Claire |
| EPI_ISL_2444392 | CH TOURCOING | CHU Lille - Laboratoire de Virologie | AIT YAHYA Emilie, ALIDJINOU Enagnon Kazali, BOCKET Laurence, CREPIN Michel, DEMAY Christophe, ENGELMANN Ilka, GEFFROY Sandrine, GUIGON Aurélie, LAMBERT Valérie, LAZREK Mouna, NOBILLIAUX Florian, PREVOST Brigitte, THUILLIER Caroline, TINEZ Claire |
| EPI_ISL_2444393 | CH LENS | CHU Lille - Laboratoire de Virologie | AIT YAHYA Emilie, ALIDJINOU Enagnon Kazali, BOCKET Laurence, CREPIN Michel, DEMAY Christophe, ENGELMANN Ilka, GEFFROY Sandrine, GUIGON Aurélie, LAMBERT Valérie, LAZREK Mouna, NOBILLIAUX Florian, PREVOST Brigitte, THUILLIER Caroline, TINEZ Claire |
| EPI_ISL_2448638 | CHNDS | CHU Poitiers | Caroline MICHAUD, Maxime PICHON, Manon PRAT, Valentin BON-BARET, Birama N'DIAYE, Magali GARCIA, Agnes BEBY-DEFAUX, Nicolas LEVEQUE |
| EPI_ISL_2448640 | Depistage Drive Montmorillon | CHU Poitiers | Caroline MICHAUD, Maxime PICHON, Manon PRAT, Valentin BON-BARET, Birama N'DIAYE, Magali GARCIA, Agnes BEBY-DEFAUX, Nicolas LEVEQUE |
| EPI_ISL_2455329, EPI_ISL_2455330 | Centers for Disease Control, R.O.C. (Taiwan) | Centers for Disease Control, R.O.C. (Taiwan) | Ji-Rong Yang, Yu-Chi-Lin, Jung-Jung Mu, Ming-Tsan-Liu |
| EPI_ISL_2462057, EPI_ISL_2462058 | Genome Analysis Center, Yamanashi Central Hospital | Genome Analysis Center, Yamanashi Central Hospital | Yosuke Hirotsu |
| EPI_ISL_2462871 | AZ Sint-Lucas Gent | AZ Sint-Lucas Gent | Jos Van Acker, Charlotte Verfaillie, Elke Vanlaere, Klara De Rauw |
| EPI_ISL_2463295 | AZ Zeno Knokke-Blankenberge | AZ Sint Jan | Marijke Reynders, Patrick Descheemaeker, Laurien Hoornaert, Thomas Van Landschoot, Marie Madeleine Chabert-Consen, Jorn Hellemans |
| EPI_ISL_2464085 | Hospital | National Reference Center for Viruses of Respiratory Infections, Institut Pasteur, Paris | Marion Barbet, Sylvie Behillil, Méline Bizard, Angela Brisebarre, Camille Capel, Vincent Enouf, Louise Lefrançois, Frédéric Lemoine, Christophe Malabat, Corinne Maufrais, Etienne Simon-Lorière, Maud Vanpeene, Sylvie Van der Werf ,JéRome Guinard |
| EPI_ISL_2464086 | Hospital | National Reference Center for Viruses of Respiratory Infections, Institut Pasteur, Paris | Marion Barbet, Sylvie Behillil, Méline Bizard, Angela Brisebarre, Camille Capel, Vincent Enouf, Louise Lefrançois, Frédéric Lemoine, Christophe Malabat, Corinne Maufrais, Amaury Vaysse, Etienne Simon-Lorière, Maud Vanpeene, Sylvie Van der Werf ,JéRome Guinard |
| EPI_ISL_2464087 | Hospital | National Reference Center for Viruses of Respiratory Infections, Institut Pasteur, Paris | Marion Barbet, Sylvie Behillil, Méline Bizard, Angela Brisebarre, Camille Capel, Vincent Enouf, Louise Lefrançois, Frédéric Lemoine, Christophe Malabat, Corinne Maufrais, Etienne Simon-Lorière, Maud Vanpeene, Sylvie Van der Werf ,JéRome Guinard |
| EPI_ISL_2464090 | Hospital | National Reference Center for Viruses of Respiratory | Marion Barbet, Sylvie Behillil, Méline Bizard, Angela Brisebarre, Camille Capel, Vincent Enouf, Louise Lefrançois, Frédéric Lemoine, Christophe Malabat, |

| | | | |
|---|---|---|---|
| | | Infections, Institut Pasteur, Paris | Corinne Maufrais, Etienne Simon-Lorière, Maud Vanpeene, Sylvie Van der Werf ,Franck Griscelli |
| EPI_ISL_2464091 | Hospital | National Reference Center for Viruses of Respiratory Infections, Institut Pasteur, Paris | Marion Barbet, Sylvie Behillil, Méline Bizard, Angela Brisebarre, Camille Capel, Vincent Enouf, Louise Lefrançois, Frédéric Lemoine, Christophe Malabat, Corinne Maufrais, Etienne Simon-Lorière, Maud Vanpeene, Sylvie Van der Werf ,Isabelle Gros |
| EPI_ISL_2464092, EPI_ISL_2464093, EPI_ISL_2464094, EPI_ISL_2464095, EPI_ISL_2464096, EPI_ISL_2464097 | Hospital | National Reference Center for Viruses of Respiratory Infections, Institut Pasteur, Paris | Marion Barbet, Sylvie Behillil, Méline Bizard, Angela Brisebarre, Camille Capel, Vincent Enouf, Louise Lefrançois, Frédéric Lemoine, Christophe Malabat, Corinne Maufrais, Etienne Simon-Lorière, Maud Vanpeene, Sylvie Van der Werf ,Laura Djamdjian |
| EPI_ISL_2464098 | Armies | National Reference Center for Viruses of Respiratory Infections, Institut Pasteur, Paris | Marion Barbet, Sylvie Behillil, Méline Bizard, Angela Brisebarre, Camille Capel, Vincent Enouf, Louise Lefrançois, Frédéric Lemoine, Christophe Malabat, Corinne Maufrais, Etienne Simon-Lorière, Maud Vanpeene, Sylvie Van der Werf ,Marine Desroches |
| EPI_ISL_2464099, EPI_ISL_2464100, EPI_ISL_2464101, EPI_ISL_2464102, EPI_ISL_2464103, EPI_ISL_2464104, EPI_ISL_2464105, EPI_ISL_2464106, EPI_ISL_2464107 | Hospital | National Reference Center for Viruses of Respiratory Infections, Institut Pasteur, Paris | Marion Barbet, Sylvie Behillil, Méline Bizard, Angela Brisebarre, Camille Capel, Vincent Enouf, Louise Lefrançois, Frédéric Lemoine, Christophe Malabat, Corinne Maufrais, Etienne Simon-Lorière, Maud Vanpeene, Sylvie Van der Werf ,Ilka Engelmann |
| EPI_ISL_2464108 | Hospital | National Reference Center for Viruses of Respiratory Infections, Institut Pasteur, Paris | Marion Barbet, Sylvie Behillil, Méline Bizard, Angela Brisebarre, Camille Capel, Vincent Enouf, Louise Lefrançois, Frédéric Lemoine, Christophe Malabat, Corinne Maufrais, Etienne Simon-Lorière, Maud Vanpeene, Sylvie Van der Werf |
| EPI_ISL_2464109, EPI_ISL_2464110 | Hospital | National Reference Center for Viruses of Respiratory Infections, Institut Pasteur, Paris | Marion Barbet, Sylvie Behillil, Méline Bizard, Angela Brisebarre, Camille Capel, Vincent Enouf, Louise Lefrançois, Frédéric Lemoine, Christophe Malabat, Corinne Maufrais, Etienne Simon-Lorière, Maud Vanpeene, Sylvie Van der Werf ,Anne Vachee |
| EPI_ISL_2464111, EPI_ISL_2464112 | Hospital | National Reference Center for Viruses of Respiratory Infections, Institut Pasteur, Paris | Marion Barbet, Sylvie Behillil, Méline Bizard, Angela Brisebarre, Camille Capel, Vincent Enouf, Louise Lefrançois, Frédéric Lemoine, Christophe Malabat, Corinne Maufrais, Etienne Simon-Lorière, Maud Vanpeene, Sylvie Van der Werf ,Sylvie Ledru |
| EPI_ISL_2464113, EPI_ISL_2464114, EPI_ISL_2464115, EPI_ISL_2464117, EPI_ISL_2464118, EPI_ISL_2464119, EPI_ISL_2464122, EPI_ISL_2464123, EPI_ISL_2464124, EPI_ISL_2464125, EPI_ISL_2464127, EPI_ISL_2464130, EPI_ISL_2464131, EPI_ISL_2464132, EPI_ISL_2464134, EPI_ISL_2464135, EPI_ISL_2464136, EPI_ISL_2464137 | | | |
| see above | Labo Analyses Med | National Reference Center for Viruses of Respiratory Infections, Institut Pasteur, Paris | Marion Barbet, Sylvie Behillil, Méline Bizard, Angela Brisebarre, Camille Capel, Vincent Enouf, Louise Lefrançois, Frédéric Lemoine, Christophe Malabat, Corinne Maufrais, Etienne Simon-Lorière, Maud Vanpeene, Sylvie Van der Werf ,Philippe Miara |
| EPI_ISL_2464138, EPI_ISL_2464139 | Labo Analyses Med | National Reference Center for Viruses of Respiratory Infections, Institut Pasteur, Paris | Marion Barbet, Sylvie Behillil, Méline Bizard, Angela Brisebarre, Camille Capel, Vincent Enouf, Louise Lefrançois, Frédéric Lemoine, Christophe Malabat, Corinne Maufrais, Etienne Simon-Lorière, Maud Vanpeene, Sylvie Van der Werf ,Nicolas Capron |
| EPI_ISL_2464140 | Labo Analyses Med | National Reference Center for Viruses of Respiratory Infections, Institut Pasteur, Paris | Marion Barbet, Sylvie Behillil, Méline Bizard, Angela Brisebarre, Camille Capel, Vincent Enouf, Louise Lefrançois, Frédéric Lemoine, Christophe Malabat, Corinne Maufrais, Emmanuelle Permal, Etienne Simon-Lorière, Maud Vanpeene, Sylvie Van der Werf ,Melanie Caron |
| EPI_ISL_2464141, EPI_ISL_2464142, EPI_ISL_2464143, EPI_ISL_2464144, EPI_ISL_2464145, EPI_ISL_2464147, EPI_ISL_2464148, EPI_ISL_2464149, EPI_ISL_2464150, EPI_ISL_2464151, EPI_ISL_2464152, EPI_ISL_2464153 | | | |
| see above | Labo Analyses Med | National Reference Center for Viruses of Respiratory Infections, Institut Pasteur, Paris | Marion Barbet, Sylvie Behillil, Méline Bizard, Angela Brisebarre, Camille Capel, Vincent Enouf, Louise Lefrançois, Frédéric Lemoine, Christophe Malabat, Corinne Maufrais, Etienne Simon-Lorière, Maud Vanpeene, Sylvie Van der Werf ,Melanie Caron |
| EPI_ISL_2464154 | Labo Analyses Med | National Reference Center for Viruses of Respiratory Infections, Institut Pasteur, Paris | Marion Barbet, Sylvie Behillil, Méline Bizard, Angela Brisebarre, Camille Capel, Vincent Enouf, Louise Lefrançois, Frédéric Lemoine, Christophe Malabat, Corinne Maufrais, Emmanuelle Permal, Etienne Simon-Lorière, Maud Vanpeene, Sylvie Van der Werf ,Melanie Caron |
| EPI_ISL_2464155 | Labo Analyses Med | National Reference Center for Viruses of Respiratory Infections, Institut Pasteur, Paris | Marion Barbet, Sylvie Behillil, Méline Bizard, Angela Brisebarre, Camille Capel, Vincent Enouf, Louise Lefrançois, Frédéric Lemoine, Christophe Malabat, Corinne Maufrais, Etienne Simon-Lorière, Maud Vanpeene, Sylvie Van der Werf ,Arditi Cocco |
| EPI_ISL_2464156 | Labo Analyses Med | National Reference Center for Viruses of Respiratory Infections, Institut Pasteur, Paris | Marion Barbet, Sylvie Behillil, Méline Bizard, Angela Brisebarre, Camille Capel, Vincent Enouf, Louise Lefrançois, Frédéric Lemoine, Christophe Malabat, Corinne Maufrais, Emmanuelle Permal, Etienne Simon-Lorière, Maud Vanpeene, Sylvie Van der Werf ,Arditi Cocco |
| EPI_ISL_2464157, EPI_ISL_2464159, EPI_ISL_2464160, EPI_ISL_2464161, EPI_ISL_2464162, EPI_ISL_2464163, EPI_ISL_2464164, EPI_ISL_2464165, EPI_ISL_2464168, EPI_ISL_2464169, EPI_ISL_2464170, EPI_ISL_2464171, EPI_ISL_2464172, EPI_ISL_2464174, EPI_ISL_2464175, EPI_ISL_2464176, EPI_ISL_2464177, EPI_ISL_2464178, EPI_ISL_2464179, EPI_ISL_2464180, EPI_ISL_2464181, EPI_ISL_2464182, EPI_ISL_2464183, EPI_ISL_2464184, EPI_ISL_2464185 | | | |
| see above | Labo Analyses Med | National Reference Center for Viruses of Respiratory Infections, Institut Pasteur, Paris | Marion Barbet, Sylvie Behillil, Méline Bizard, Angela Brisebarre, Camille Capel, Vincent Enouf, Louise Lefrançois, Frédéric Lemoine, Christophe Malabat, Corinne Maufrais, Etienne Simon-Lorière, Maud Vanpeene, Sylvie Van der Werf ,Arditi Cocco |
| EPI_ISL_2464186 | Labo Analyses Med | National Reference Center for Viruses of Respiratory Infections, Institut Pasteur, Paris | Marion Barbet, Sylvie Behillil, Méline Bizard, Angela Brisebarre, Camille Capel, Vincent Enouf, Louise Lefrançois, Frédéric Lemoine, Christophe Malabat, Corinne Maufrais, Amaury Vaysse, Etienne Simon-Lorière, Maud Vanpeene, Sylvie Van der Werf ,Arditi Cocco |
| EPI_ISL_2464187, EPI_ISL_2464188 | Labo Analyses Med | National Reference Center for Viruses of Respiratory Infections, Institut Pasteur, Paris | Marion Barbet, Sylvie Behillil, Méline Bizard, Angela Brisebarre, Camille Capel, Vincent Enouf, Louise Lefrançois, Frédéric Lemoine, Christophe Malabat, Corinne Maufrais, Etienne Simon-Lorière, Maud Vanpeene, Sylvie Van der Werf ,Arditi Cocco |
| EPI_ISL_2464189, EPI_ISL_2464190, EPI_ISL_2464191 | Hospital | National Reference Center for Viruses of Respiratory Infections, Institut Pasteur, Paris | Marion Barbet, Sylvie Behillil, Méline Bizard, Angela Brisebarre, Camille Capel, Vincent Enouf, Louise Lefrançois, Frédéric Lemoine, Christophe Malabat, Corinne Maufrais, Etienne Simon-Lorière, Maud Vanpeene, Sylvie Van der Werf ,Anne Cady |
| EPI_ISL_2464192 | Labo Analyses Med | National Reference Center for Viruses of Respiratory Infections, Institut Pasteur, Paris | Marion Barbet, Sylvie Behillil, Méline Bizard, Angela Brisebarre, Camille Capel, Vincent Enouf, Louise Lefrançois, Frédéric Lemoine, Christophe Malabat, Corinne Maufrais, Amaury Vaysse, Etienne Simon-Lorière, Maud Vanpeene, Sylvie Van der Werf ,GréGoire Potiron |
| EPI_ISL_2464193, EPI_ISL_2464194, EPI_ISL_2464195, EPI_ISL_2464196, EPI_ISL_2464198, EPI_ISL_2464199, EPI_ISL_2464200 | Labo Analyses Med | National Reference Center for Viruses of Respiratory Infections, Institut Pasteur, Paris | Marion Barbet, Sylvie Behillil, Méline Bizard, Angela Brisebarre, Camille Capel, Vincent Enouf, Louise Lefrançois, Frédéric Lemoine, Christophe Malabat, Corinne Maufrais, Etienne Simon-Lorière, Maud Vanpeene, Sylvie Van der Werf ,GréGoire Potiron |
| EPI_ISL_2464201, EPI_ISL_2464202, EPI_ISL_2464203, EPI_ISL_2464204 | Labo Analyses Med | National Reference Center for Viruses of Respiratory Infections, Institut Pasteur, Paris | Marion Barbet, Sylvie Behillil, Méline Bizard, Angela Brisebarre, Camille Capel, Vincent Enouf, Louise Lefrançois, Frédéric Lemoine, Christophe Malabat, Corinne Maufrais, Etienne Simon-Lorière, Maud Vanpeene, Sylvie Van der Werf |
| EPI_ISL_2464205 | Labo Analyses Med | National Reference Center for Viruses of Respiratory Infections, Institut Pasteur, Paris | Marion Barbet, Sylvie Behillil, Méline Bizard, Angela Brisebarre, Camille Capel, Vincent Enouf, Louise Lefrançois, Frédéric Lemoine, Christophe Malabat, Corinne Maufrais, Etienne Simon-Lorière, Maud Vanpeene, Sylvie Van der Werf ,Edouard Lamandin |
| EPI_ISL_2464206 | Labo Analyses Med | National Reference Center for Viruses of Respiratory Infections, Institut Pasteur, Paris | Marion Barbet, Sylvie Behillil, Méline Bizard, Angela Brisebarre, Camille Capel, Vincent Enouf, Louise Lefrançois, Frédéric Lemoine, Christophe Malabat, Corinne Maufrais, Amaury Vaysse, Etienne Simon-Lorière, Maud Vanpeene, Sylvie Van der Werf ,Edouard Lamandin |
| EPI_ISL_2464207, EPI_ISL_2464208, EPI_ISL_2464210, EPI_ISL_2464211 | Labo Analyses Med | National Reference Center for Viruses of Respiratory Infections, Institut Pasteur, Paris | Marion Barbet, Sylvie Behillil, Méline Bizard, Angela Brisebarre, Camille Capel, Vincent Enouf, Louise Lefrançois, Frédéric Lemoine, Christophe Malabat, Corinne Maufrais, Etienne Simon-Lorière, Maud Vanpeene, Sylvie Van der Werf ,Edouard Lamandin |
| EPI_ISL_2464212, EPI_ISL_2464213, EPI_ISL_2464214, EPI_ISL_2464215, EPI_ISL_2464216, EPI_ISL_2464217, EPI_ISL_2464218, EPI_ISL_2464219, EPI_ISL_2464220, EPI_ISL_2464221, EPI_ISL_2464222, EPI_ISL_2464223, EPI_ISL_2464224, EPI_ISL_2464225, EPI_ISL_2464226, EPI_ISL_2464227 | | | |
| see above | Labo Analyses Med | National Reference Center for Viruses of Respiratory Infections, Institut Pasteur, Paris | Marion Barbet, Sylvie Behillil, Méline Bizard, Angela Brisebarre, Camille Capel, Vincent Enouf, Louise Lefrançois, Frédéric Lemoine, Christophe Malabat, Corinne Maufrais, Etienne Simon-Lorière, Maud Vanpeene, Sylvie Van der Werf ,Anne Holstein |
| EPI_ISL_2464228, EPI_ISL_2464230, EPI_ISL_2464231 | Labo Analyses Med | National Reference Center for Viruses of Respiratory Infections, Institut Pasteur, Paris | Marion Barbet, Sylvie Behillil, Méline Bizard, Angela Brisebarre, Camille Capel, Vincent Enouf, Louise Lefrançois, Frédéric Lemoine, Christophe Malabat, Corinne Maufrais, Amaury Vaysse, Etienne Simon-Lorière, Maud Vanpeene, Sylvie Van der Werf ,Anne Holstein |
| EPI_ISL_2464232, EPI_ISL_2464233 | Labo Analyses Med | National Reference Center for Viruses of Respiratory Infections, Institut Pasteur, Paris | Marion Barbet, Sylvie Behillil, Méline Bizard, Angela Brisebarre, Camille Capel, Vincent Enouf, Louise Lefrançois, Frédéric Lemoine, Christophe Malabat, Corinne Maufrais, Etienne Simon-Lorière, Maud Vanpeene, Sylvie Van der Werf ,Anne Holstein |
| EPI_ISL_2464235 | Labo Analyses Med | National Reference Center for Viruses of Respiratory Infections, Institut Pasteur, Paris | Marion Barbet, Sylvie Behillil, Méline Bizard, Angela Brisebarre, Camille Capel, Vincent Enouf, Louise Lefrançois, Frédéric Lemoine, Christophe Malabat, Corinne Maufrais, Amaury Vaysse, Etienne Simon-Lorière, Maud Vanpeene, Sylvie Van der Werf ,Anne Holstein |
| EPI_ISL_2464236, EPI_ISL_2464237, EPI_ISL_2464238 | Labo Analyses Med | National Reference Center for Viruses of Respiratory Infections, Institut Pasteur, Paris | Marion Barbet, Sylvie Behillil, Méline Bizard, Angela Brisebarre, Camille Capel, Vincent Enouf, Louise Lefrançois, Frédéric Lemoine, Christophe Malabat, Corinne Maufrais, Etienne Simon-Lorière, Maud Vanpeene, Sylvie Van der Werf ,Anne Holstein |

| | | | |
|---|---|---|---|
| EPI_ISL_2464239 | Hospital | National Reference Center for Viruses of Respiratory Infections, Institut Pasteur, Paris | Marion Barbet, Sylvie Behillil, Méline Bizard, Angela Brisebarre, Camille Capel, Vincent Enouf, Louise Lefrançois, Frédéric Lemoine, Christophe Malabat, Corinne Maufrais, Etienne Simon-Lorière, Maud Vanpeene, Sylvie Van der Werf ,Vincent Thibault |
| EPI_ISL_2464240 | Hospital | National Reference Center for Viruses of Respiratory Infections, Institut Pasteur, Paris | Marion Barbet, Sylvie Behillil, Méline Bizard, Angela Brisebarre, Camille Capel, Vincent Enouf, Louise Lefrançois, Frédéric Lemoine, Christophe Malabat, Corinne Maufrais, Etienne Simon-Lorière, Maud Vanpeene, Sylvie Van der Werf ,Pauline Morvan |
| EPI_ISL_2464241 | Hospital | National Reference Center for Viruses of Respiratory Infections, Institut Pasteur, Paris | Marion Barbet, Sylvie Behillil, Méline Bizard, Angela Brisebarre, Camille Capel, Vincent Enouf, Louise Lefrançois, Frédéric Lemoine, Christophe Malabat, Corinne Maufrais, Amaury Vaysse, Etienne Simon-Lorière, Maud Vanpeene, Sylvie Van der Werf ,A Hochart |
| EPI_ISL_2464242 | Hospital | National Reference Center for Viruses of Respiratory Infections, Institut Pasteur, Paris | Marion Barbet, Sylvie Behillil, Méline Bizard, Angela Brisebarre, Camille Capel, Vincent Enouf, Louise Lefrançois, Frédéric Lemoine, Christophe Malabat, Corinne Maufrais, Etienne Simon-Lorière, Maud Vanpeene, Sylvie Van der Werf ,Marianne Burgard |
| EPI_ISL_2464243, EPI_ISL_2464244 | Hospital | National Reference Center for Viruses of Respiratory Infections, Institut Pasteur, Paris | Marion Barbet, Sylvie Behillil, Méline Bizard, Angela Brisebarre, Camille Capel, Vincent Enouf, Louise Lefrançois, Frédéric Lemoine, Christophe Malabat, Corinne Maufrais, Etienne Simon-Lorière, Maud Vanpeene, Sylvie Van der Werf ,Jean Baptiste Bour |
| EPI_ISL_2464248 | Hospital | National Reference Center for Viruses of Respiratory Infections, Institut Pasteur, Paris | Marion Barbet, Sylvie Behillil, Méline Bizard, Angela Brisebarre, Camille Capel, Vincent Enouf, Louise Lefrançois, Frédéric Lemoine, Christophe Malabat, Corinne Maufrais, Etienne Simon-Lorière, Maud Vanpeene, Sylvie Van der Werf ,AgnèS Scanvic |
| EPI_ISL_2464249 | Hospital | National Reference Center for Viruses of Respiratory Infections, Institut Pasteur, Paris | Marion Barbet, Sylvie Behillil, Méline Bizard, Angela Brisebarre, Camille Capel, Vincent Enouf, Louise Lefrançois, Frédéric Lemoine, Christophe Malabat, Corinne Maufrais, Amaury Vaysse, Etienne Simon-Lorière, Maud Vanpeene, Sylvie Van der Werf ,Sandrine Castelain |
| EPI_ISL_2464250, EPI_ISL_2464252 | Hospital | National Reference Center for Viruses of Respiratory Infections, Institut Pasteur, Paris | Marion Barbet, Sylvie Behillil, Méline Bizard, Angela Brisebarre, Camille Capel, Vincent Enouf, Louise Lefrançois, Frédéric Lemoine, Christophe Malabat, Corinne Maufrais, Etienne Simon-Lorière, Maud Vanpeene, Sylvie Van der Werf ,Sandrine Castelain |
| EPI_ISL_2464253 | Hospital | National Reference Center for Viruses of Respiratory Infections, Institut Pasteur, Paris | Marion Barbet, Sylvie Behillil, Méline Bizard, Angela Brisebarre, Camille Capel, Vincent Enouf, Louise Lefrançois, Frédéric Lemoine, Christophe Malabat, Corinne Maufrais, Amaury Vaysse, Etienne Simon-Lorière, Maud Vanpeene, Sylvie Van der Werf ,Sandrine Castelain |
| EPI_ISL_2464254 | Hospital | National Reference Center for Viruses of Respiratory Infections, Institut Pasteur, Paris | Marion Barbet, Sylvie Behillil, Méline Bizard, Angela Brisebarre, Camille Capel, Vincent Enouf, Louise Lefrançois, Frédéric Lemoine, Christophe Malabat, Corinne Maufrais, Etienne Simon-Lorière, Maud Vanpeene, Sylvie Van der Werf ,Sandrine Castelain |
| EPI_ISL_2464256 | Hospital | National Reference Center for Viruses of Respiratory Infections, Institut Pasteur, Paris | Marion Barbet, Sylvie Behillil, Méline Bizard, Angela Brisebarre, Camille Capel, Vincent Enouf, Louise Lefrançois, Frédéric Lemoine, Christophe Malabat, Corinne Maufrais, Etienne Simon-Lorière, Maud Vanpeene, Sylvie Van der Werf ,Dominique Descamps |
| EPI_ISL_2464257 | Labo Analyses Med | National Reference Center for Viruses of Respiratory Infections, Institut Pasteur, Paris | Marion Barbet, Sylvie Behillil, Méline Bizard, Angela Brisebarre, Camille Capel, Vincent Enouf, Louise Lefrançois, Frédéric Lemoine, Christophe Malabat, Corinne Maufrais, Amaury Vaysse, Etienne Simon-Lorière, Maud Vanpeene, Sylvie Van der Werf ,Domitille Leman |
| EPI_ISL_2464258, EPI_ISL_2464259 | Labo Analyses Med | National Reference Center for Viruses of Respiratory Infections, Institut Pasteur, Paris | Marion Barbet, Sylvie Behillil, Méline Bizard, Angela Brisebarre, Camille Capel, Vincent Enouf, Louise Lefrançois, Frédéric Lemoine, Christophe Malabat, Corinne Maufrais, Etienne Simon-Lorière, Maud Vanpeene, Sylvie Van der Werf ,Domitille Leman |
| EPI_ISL_2464260, EPI_ISL_2464261 | Labo Analyses Med | National Reference Center for Viruses of Respiratory Infections, Institut Pasteur, Paris | Marion Barbet, Sylvie Behillil, Méline Bizard, Angela Brisebarre, Camille Capel, Vincent Enouf, Louise Lefrançois, Frédéric Lemoine, Christophe Malabat, Corinne Maufrais, Amaury Vaysse, Etienne Simon-Lorière, Maud Vanpeene, Sylvie Van der Werf ,Domitille Leman |
| EPI_ISL_2464262, EPI_ISL_2464263, EPI_ISL_2464264, EPI_ISL_2464265, EPI_ISL_2464266 | Labo Analyses Med | National Reference Center for Viruses of Respiratory Infections, Institut Pasteur, Paris | Marion Barbet, Sylvie Behillil, Méline Bizard, Angela Brisebarre, Camille Capel, Vincent Enouf, Louise Lefrançois, Frédéric Lemoine, Christophe Malabat, Corinne Maufrais, Etienne Simon-Lorière, Maud Vanpeene, Sylvie Van der Werf ,Domitille Leman |
| EPI_ISL_2464267, EPI_ISL_2464268 | Hospital | National Reference Center for Viruses of Respiratory Infections, Institut Pasteur, Paris | Marion Barbet, Sylvie Behillil, Méline Bizard, Angela Brisebarre, Camille Capel, Vincent Enouf, Louise Lefrançois, Frédéric Lemoine, Christophe Malabat, Corinne Maufrais, Etienne Simon-Lorière, Maud Vanpeene, Sylvie Van der Werf ,Laurence Courdavault |
| EPI_ISL_2464269, EPI_ISL_2464271, EPI_ISL_2464272, EPI_ISL_2464273 | Hospital | National Reference Center for Viruses of Respiratory Infections, Institut Pasteur, Paris | Marion Barbet, Sylvie Behillil, Méline Bizard, Angela Brisebarre, Camille Capel, Vincent Enouf, Louise Lefrançois, Frédéric Lemoine, Christophe Malabat, Corinne Maufrais, Etienne Simon-Lorière, Maud Vanpeene, Sylvie Van der Werf ,Nabil Gastli |
| EPI_ISL_2464274 | Hospital | National Reference Center for Viruses of Respiratory Infections, Institut Pasteur, Paris | Marion Barbet, Sylvie Behillil, Méline Bizard, Angela Brisebarre, Camille Capel, Vincent Enouf, Louise Lefrançois, Frédéric Lemoine, Christophe Malabat, Corinne Maufrais, Amaury Vaysse, Etienne Simon-Lorière, Maud Vanpeene, Sylvie Van der Werf ,Nabil Gastli |
| EPI_ISL_2464275, EPI_ISL_2464276 | Hospital | National Reference Center for Viruses of Respiratory Infections, Institut Pasteur, Paris | Marion Barbet, Sylvie Behillil, Méline Bizard, Angela Brisebarre, Camille Capel, Vincent Enouf, Louise Lefrançois, Frédéric Lemoine, Christophe Malabat, Corinne Maufrais, Etienne Simon-Lorière, Maud Vanpeene, Sylvie Van der Werf ,Nabil Gastli |
| EPI_ISL_2464277, EPI_ISL_2464278 | Hospital | National Reference Center for Viruses of Respiratory Infections, Institut Pasteur, Paris | Marion Barbet, Sylvie Behillil, Méline Bizard, Angela Brisebarre, Camille Capel, Vincent Enouf, Louise Lefrançois, Frédéric Lemoine, Christophe Malabat, Corinne Maufrais, Emmanuelle Permal, Etienne Simon-Lorière, Maud Vanpeene, Sylvie Van der Werf ,Nabil Gastli |
| EPI_ISL_2464563, EPI_ISL_2464564 | Singapore General Hospital | Singapore General Hospital | Nurdyana Abdul Rahman, Kenneth Xin Long Chan, Chayaporn Suphavilai, Kun Lee Lim, Sui Sin Goh, Kian Sing Chan, Lynette Oon, Kern Rei Chng, James Sim Heng Chiak, Niranjan Nagarajan, Karrie Ko |
| EPI_ISL_2466276, EPI_ISL_2466277, EPI_ISL_2466285 | Presidio di Brindisi Di Summa - Perrino | Istituto Zooprofilattico Sperimentale della Puglia e della Basilicata | Parisi A., Bianco A., Capozzi L., Del Sambro L., Simone D., Difato L., Pennuzzi G., Santoro A. |
| EPI_ISL_2466286, EPI_ISL_2466287, EPI_ISL_2466288, EPI_ISL_2466289, EPI_ISL_2466290, EPI_ISL_2466291 | Ospedale Vito Fazzi | Istituto Zooprofilattico Sperimentale della Puglia e della Basilicata | Parisi A., Bianco A., Capozzi L., Del Sambro L., Simone D., Difato L., Bruno A. R. |
| EPI_ISL_2466292, EPI_ISL_2466293, EPI_ISL_2466294, EPI_ISL_2466295, EPI_ISL_2466296, EPI_ISL_2466297, EPI_ISL_2466298, EPI_ISL_2466299, EPI_ISL_2466300, EPI_ISL_2466301, EPI_ISL_2466302, EPI_ISL_2466303, EPI_ISL_2466304, EPI_ISL_2466305, EPI_ISL_2466306, EPI_ISL_2466307, EPI_ISL_2466308, EPI_ISL_2466309, EPI_ISL_2466310, EPI_ISL_2466311 | | | |
| see above | Ospedale Santa Caterina Novella | Istituto Zooprofilattico Sperimentale della Puglia e della Basilicata | Parisi A., Bianco A., Capozzi L., Del Sambro L., Simone D., Difato L., Lobreglio G. |
| EPI_ISL_2466312 | I.R.C.C.S. "S. De Bellis" - Ente Ospedaliero | Istituto Zooprofilattico Sperimentale della Puglia e della Basilicata | Parisi A., Bianco A., Capozzi L., Del Sambro L., Simone D., Difato L., Notarnicola M., Lippolis A. |
| EPI_ISL_2467835, EPI_ISL_2467865 | Usansolo-Galdakao University Hospital | Cruces University Hospital | Ana Gual-de-Torrella, Izaskun Alejo-Cancho, Mikel Gallego, Ana Belén de la Hoz |
| EPI_ISL_2479916 | LESP Jalisco | Instituto de Diagnostico y Referencia Epidemiologicos (INDRE) | Claudia Wong-Arambula, Abril Rodriguez-Maldonado, Vanessa Rivero-Arredondo, Ariadna Medina-Benitez, Joaquin Quiroz-Mercado, Sergio Rangel-Guerrero, Natividad Cruz-Ortiz, Tatiana Nunez-Garcia, Gisela Barrera-Badillo, Lucia Hernandez-Rivas, Irma Lopez-Martinez, Ernesto Ramirez-Gonzalez. |
| EPI_ISL_2483568, EPI_ISL_2483569, EPI_ISL_2483570 | SIESP CHIETI - DRIVE IN LANCIANO | Istituto Zooprofilattico Sperimentale dell'Abruzzo e Molise "G. Caporale" | Lorusso A, Marcacci M, Di Domenico M, Ancora M, Curini V, Di Lollo Valeria, Mangone I, Rinaldi A, Delli Compagni E, Scialabba S, Caporale M, Di Pasquale A, Cammà C, Puglia I, Calistri P, Savini G |
| EPI_ISL_2483577, EPI_ISL_2483591 | SIESP DIPARTIMENTO DI PREVENZIONE TERAMO | Istituto Zooprofilattico Sperimentale dell'Abruzzo e Molise "G. Caporale" | Lorusso A, Marcacci M, Di Domenico M, Ancora M, Curini V, Di Lollo Valeria, Mangone I, Rinaldi A, Delli Compagni E, Scialabba S, Caporale M, Di Pasquale A, Cammà C, Puglia I, Calistri P, Savini G |
| EPI_ISL_2483592 | SIESP DIPARTIMENTO DI PREVENZIONE CHIETI | Istituto Zooprofilattico Sperimentale dell'Abruzzo e Molise "G. Caporale" | Lorusso A, Marcacci M, Di Domenico M, Ancora M, Curini V, Di Lollo Valeria, Mangone I, Rinaldi A, Delli Compagni E, Scialabba S, Caporale M, Di Pasquale A, Cammà C, Puglia I, Calistri P, Savini G |
| EPI_ISL_2483593, EPI_ISL_2483653, EPI_ISL_2483654, EPI_ISL_2483655, EPI_ISL_2483656 | SIESP DIPARTIMENTO DI PREVENZIONE TERAMO | Istituto Zooprofilattico Sperimentale dell'Abruzzo e Molise "G. Caporale" | Lorusso A, Marcacci M, Di Domenico M, Ancora M, Curini V, Di Lollo Valeria, Mangone I, Rinaldi A, Delli Compagni E, Scialabba S, Caporale M, Di Pasquale A, Cammà C, Puglia I, Calistri P, Savini G |
| EPI_ISL_2483657 | SIESP DIPARTIMENTO DI PREVENZIONE CHIETI | Istituto Zooprofilattico Sperimentale dell'Abruzzo e Molise "G. Caporale" | Lorusso A, Marcacci M, Di Domenico M, Ancora M, Curini V, Di Lollo Valeria, Mangone I, Rinaldi A, Delli Compagni E, Scialabba S, Caporale M, Di Pasquale A, Cammà C, Puglia I, Calistri P, Savini G |
| EPI_ISL_2483661 | SIESP CHIETI - DRIVE IN LANCIANO | Istituto Zooprofilattico Sperimentale dell'Abruzzo e Molise "G. Caporale" | Lorusso A, Marcacci M, Di Domenico M, Ancora M, Curini V, Di Lollo Valeria, Mangone I, Rinaldi A, Delli Compagni E, Scialabba S, Caporale M, Di Pasquale A, Cammà C, Puglia I, Calistri P, Savini G |

| | | | |
|---|---|---|---|
| EPI_ISL_2483662, EPI_ISL_2483663 | SIESP DIPARTIMENTO DI PREVENZIONE CHIETI | Istituto Zooprofilattico Sperimentale dell'Abruzzo e Molise "G. Caporale" | Lorusso A, Marcacci M, Di Domenico M, Ancora M, Curini V, Di Lollo Valeria, Mangone I, Rinaldi A, Delli Compagni E, Scialabba S, Caporale M, Di Pasquale A, Cammà C, Puglia I, Calistri P, Savini G |
| EPI_ISL_2483664 | SIESP CHIETI - DRIVE IN LANCIANO | Istituto Zooprofilattico Sperimentale dell'Abruzzo e Molise "G. Caporale" | Lorusso A, Marcacci M, Di Domenico M, Ancora M, Curini V, Di Lollo Valeria, Mangone I, Rinaldi A, Delli Compagni E, Scialabba S, Caporale M, Di Pasquale A, Cammà C, Puglia I, Calistri P, Savini G |
| EPI_ISL_2483665 | SIESP DIPARTIMENTO DI PREVENZIONE TERAMO | Istituto Zooprofilattico Sperimentale dell'Abruzzo e Molise "G. Caporale" | Lorusso A, Marcacci M, Di Domenico M, Ancora M, Curini V, Di Lollo Valeria, Mangone I, Rinaldi A, Delli Compagni E, Scialabba S, Caporale M, Di Pasquale A, Cammà C, Puglia I, Calistri P, Savini G |
| EPI_ISL_2483666, EPI_ISL_2483667 | SIESP CHIETI - DRIVE IN CHIETI | Istituto Zooprofilattico Sperimentale dell'Abruzzo e Molise "G. Caporale" | Lorusso A, Marcacci M, Di Domenico M, Ancora M, Curini V, Di Lollo Valeria, Mangone I, Rinaldi A, Delli Compagni E, Scialabba S, Caporale M, Di Pasquale A, Cammà C, Puglia I, Calistri P, Savini G |
| EPI_ISL_2483668 | SIESP CHIETI - DRIVE IN LANCIANO | Istituto Zooprofilattico Sperimentale dell'Abruzzo e Molise "G. Caporale" | Lorusso A, Marcacci M, Di Domenico M, Ancora M, Curini V, Di Lollo Valeria, Mangone I, Rinaldi A, Delli Compagni E, Scialabba S, Caporale M, Di Pasquale A, Cammà C, Puglia I, Calistri P, Savini G |
| EPI_ISL_2483669, EPI_ISL_2483670 | SIESP CHIETI - DRIVE IN VASTO | Istituto Zooprofilattico Sperimentale dell'Abruzzo e Molise "G. Caporale" | Lorusso A, Marcacci M, Di Domenico M, Ancora M, Curini V, Di Lollo Valeria, Mangone I, Rinaldi A, Delli Compagni E, Scialabba S, Caporale M, Di Pasquale A, Cammà C, Puglia I, Calistri P, Savini G |
| EPI_ISL_2483671 | RSA GIULIANOVA | Istituto Zooprofilattico Sperimentale dell'Abruzzo e Molise "G. Caporale" | Lorusso A, Marcacci M, Di Domenico M, Ancora M, Curini V, Di Lollo Valeria, Mangone I, Rinaldi A, Delli Compagni E, Scialabba S, Caporale M, Di Pasquale A, Cammà C, Puglia I, Calistri P, Savini G |
| EPI_ISL_2483672 | SIESP CHIETI - DRIVE IN ORTONA | Istituto Zooprofilattico Sperimentale dell'Abruzzo e Molise "G. Caporale" | Lorusso A, Marcacci M, Di Domenico M, Ancora M, Curini V, Di Lollo Valeria, Mangone I, Rinaldi A, Delli Compagni E, Scialabba S, Caporale M, Di Pasquale A, Cammà C, Puglia I, Calistri P, Savini G |
| EPI_ISL_2483673, EPI_ISL_2483674 | SIESP CHIETI - DRIVE IN VASTO | Istituto Zooprofilattico Sperimentale dell'Abruzzo e Molise "G. Caporale" | Lorusso A, Marcacci M, Di Domenico M, Ancora M, Curini V, Di Lollo Valeria, Mangone I, Rinaldi A, Delli Compagni E, Scialabba S, Caporale M, Di Pasquale A, Cammà C, Puglia I, Calistri P, Savini G |
| EPI_ISL_2483675 | SIESP DIPARTIMENTO DI PREVENZIONE CHIETI | Istituto Zooprofilattico Sperimentale dell'Abruzzo e Molise "G. Caporale" | Lorusso A, Marcacci M, Di Domenico M, Ancora M, Curini V, Di Lollo Valeria, Mangone I, Rinaldi A, Delli Compagni E, Scialabba S, Caporale M, Di Pasquale A, Cammà C, Puglia I, Calistri P, Savini G |
| EPI_ISL_2483676 | SIESP CHIETI - DRIVE IN VASTO | Istituto Zooprofilattico Sperimentale dell'Abruzzo e Molise "G. Caporale" | Lorusso A, Marcacci M, Di Domenico M, Ancora M, Curini V, Di Lollo Valeria, Mangone I, Rinaldi A, Delli Compagni E, Scialabba S, Caporale M, Di Pasquale A, Cammà C, Puglia I, Calistri P, Savini G |
| EPI_ISL_2483677 | SIESP DIPARTIMENTO DI PREVENZIONE CHIETI | Istituto Zooprofilattico Sperimentale dell'Abruzzo e Molise "G. Caporale" | Lorusso A, Marcacci M, Di Domenico M, Ancora M, Curini V, Di Lollo Valeria, Mangone I, Rinaldi A, Delli Compagni E, Scialabba S, Caporale M, Di Pasquale A, Cammà C, Puglia I, Calistri P, Savini G |
| EPI_ISL_2483678 | SIESP CHIETI - DRIVE IN CHIETI | Istituto Zooprofilattico Sperimentale dell'Abruzzo e Molise "G. Caporale" | Lorusso A, Marcacci M, Di Domenico M, Ancora M, Curini V, Di Lollo Valeria, Mangone I, Rinaldi A, Delli Compagni E, Scialabba S, Caporale M, Di Pasquale A, Cammà C, Puglia I, Calistri P, Savini G |
| EPI_ISL_2483679 | SIESP DIPARTIMENTO DI PREVENZIONE TERAMO | Istituto Zooprofilattico Sperimentale dell'Abruzzo e Molise "G. Caporale" | Lorusso A, Marcacci M, Di Domenico M, Ancora M, Curini V, Di Lollo Valeria, Mangone I, Rinaldi A, Delli Compagni E, Scialabba S, Caporale M, Di Pasquale A, Cammà C, Puglia I, Calistri P, Savini G |
| EPI_ISL_2483680 | PRESIDIO OSPEDALIERO SULMONA | Istituto Zooprofilattico Sperimentale dell'Abruzzo e Molise "G. Caporale" | Lorusso A, Marcacci M, Di Domenico M, Ancora M, Curini V, Di Lollo Valeria, Mangone I, Rinaldi A, Delli Compagni E, Scialabba S, Caporale M, Di Pasquale A, Cammà C, Puglia I, Calistri P, Savini G |
| EPI_ISL_2483681 | USCA Sulmona | Istituto Zooprofilattico Sperimentale dell'Abruzzo e Molise "G. Caporale" | Lorusso A, Marcacci M, Di Domenico M, Ancora M, Curini V, Di Lollo Valeria, Mangone I, Rinaldi A, Delli Compagni E, Scialabba S, Caporale M, Di Pasquale A, Cammà C, Puglia I, Calistri P, Savini G |
| EPI_ISL_2483682, EPI_ISL_2483683, EPI_ISL_2483684, EPI_ISL_2483685 | SIESP DIPARTIMENTO DI PREVENZIONE CHIETI | Istituto Zooprofilattico Sperimentale dell'Abruzzo e Molise "G. Caporale" | Lorusso A, Marcacci M, Di Domenico M, Ancora M, Curini V, Di Lollo Valeria, Mangone I, Rinaldi A, Delli Compagni E, Scialabba S, Caporale M, Di Pasquale A, Cammà C, Puglia I, Calistri P, Savini G |
| EPI_ISL_2483686 | SIESP DIPARTIMENTO DI PREVENZIONE TERAMO | Istituto Zooprofilattico Sperimentale dell'Abruzzo e Molise "G. Caporale" | Lorusso A, Marcacci M, Di Domenico M, Ancora M, Curini V, Di Lollo Valeria, Mangone I, Rinaldi A, Delli Compagni E, Scialabba S, Caporale M, Di Pasquale A, Cammà C, Puglia I, Calistri P, Savini G |
| EPI_ISL_2483687, EPI_ISL_2483688, EPI_ISL_2483689 | SIESP CHIETI - DRIVE IN VASTO | Istituto Zooprofilattico Sperimentale dell'Abruzzo e Molise "G. Caporale" | Lorusso A, Marcacci M, Di Domenico M, Ancora M, Curini V, Di Lollo Valeria, Mangone I, Rinaldi A, Delli Compagni E, Scialabba S, Caporale M, Di Pasquale A, Cammà C, Puglia I, Calistri P, Savini G |
| EPI_ISL_2483690 | RSA GIULIANOVA | Istituto Zooprofilattico Sperimentale dell'Abruzzo e Molise "G. Caporale" | Lorusso A, Marcacci M, Di Domenico M, Ancora M, Curini V, Di Lollo Valeria, Mangone I, Rinaldi A, Delli Compagni E, Scialabba S, Caporale M, Di Pasquale A, Cammà C, Puglia I, Calistri P, Savini G |
| EPI_ISL_2483691 | SIESP CHIETI - DRIVE IN VASTO | Istituto Zooprofilattico Sperimentale dell'Abruzzo e Molise "G. Caporale" | Lorusso A, Marcacci M, Di Domenico M, Ancora M, Curini V, Di Lollo Valeria, Mangone I, Rinaldi A, Delli Compagni E, Scialabba S, Caporale M, Di Pasquale A, Cammà C, Puglia I, Calistri P, Savini G |
| EPI_ISL_2483692, EPI_ISL_2483693 | SIESP CHIETI - DRIVE IN CHIETI | Istituto Zooprofilattico Sperimentale dell'Abruzzo e Molise "G. Caporale" | Lorusso A, Marcacci M, Di Domenico M, Ancora M, Curini V, Di Lollo Valeria, Mangone I, Rinaldi A, Delli Compagni E, Scialabba S, Caporale M, Di Pasquale A, Cammà C, Puglia I, Calistri P, Savini G |
| EPI_ISL_2483694 | SIESP CHIETI - DRIVE IN VASTO | Istituto Zooprofilattico Sperimentale dell'Abruzzo e Molise "G. Caporale" | Lorusso A, Marcacci M, Di Domenico M, Ancora M, Curini V, Di Lollo Valeria, Mangone I, Rinaldi A, Delli Compagni E, Scialabba S, Caporale M, Di Pasquale A, Cammà C, Puglia I, Calistri P, Savini G |
| EPI_ISL_2483695 | SIESP CHIETI - DRIVE IN ORTONA | Istituto Zooprofilattico Sperimentale dell'Abruzzo e Molise "G. Caporale" | Lorusso A, Marcacci M, Di Domenico M, Ancora M, Curini V, Di Lollo Valeria, Mangone I, Rinaldi A, Delli Compagni E, Scialabba S, Caporale M, Di Pasquale A, Cammà C, Puglia I, Calistri P, Savini G |
| EPI_ISL_2483696, EPI_ISL_2483697 | SIESP DIPARTIMENTO DI PREVENZIONE CHIETI | Istituto Zooprofilattico Sperimentale dell'Abruzzo e Molise "G. Caporale" | Lorusso A, Marcacci M, Di Domenico M, Ancora M, Curini V, Di Lollo Valeria, Mangone I, Rinaldi A, Delli Compagni E, Scialabba S, Caporale M, Di Pasquale A, Cammà C, Puglia I, Calistri P, Savini G |
| EPI_ISL_2483698, EPI_ISL_2483699 | SIESP CHIETI - DRIVE IN VASTO | Istituto Zooprofilattico Sperimentale dell'Abruzzo e Molise "G. Caporale" | Lorusso A, Marcacci M, Di Domenico M, Ancora M, Curini V, Di Lollo Valeria, Mangone I, Rinaldi A, Delli Compagni E, Scialabba S, Caporale M, Di Pasquale A, Cammà C, Puglia I, Calistri P, Savini G |
| EPI_ISL_2483701, EPI_ISL_2483702 | SIESP DIPARTIMENTO DI PREVENZIONE TERAMO | Istituto Zooprofilattico Sperimentale dell'Abruzzo e Molise "G. Caporale" | Lorusso A, Marcacci M, Di Domenico M, Ancora M, Curini V, Di Lollo Valeria, Mangone I, Rinaldi A, Delli Compagni E, Scialabba S, Caporale M, Di Pasquale A, Cammà C, Puglia I, Calistri P, Savini G |
| EPI_ISL_2483822, EPI_ISL_2483823, EPI_ISL_2483824, EPI_ISL_2483825, EPI_ISL_2483826, EPI_ISL_2483827, EPI_ISL_2483828, EPI_ISL_2483829, EPI_ISL_2483830, EPI_ISL_2483831, EPI_ISL_2483832, EPI_ISL_2483833 | | | |
| see above | Klinisch Laboratorium ZNA | Klinisch Laboratorium ZNA | Verstrepen et al. |
| EPI_ISL_2484967, EPI_ISL_2484968, EPI_ISL_2484969, EPI_ISL_2484970, EPI_ISL_2484972, EPI_ISL_2484973, EPI_ISL_2484974, EPI_ISL_2484975, EPI_ISL_2484977, EPI_ISL_2484983, EPI_ISL_2484984, EPI_ISL_2484985, EPI_ISL_2485091, EPI_ISL_2485103, EPI_ISL_2485117, EPI_ISL_2485118, EPI_ISL_2485119, EPI_ISL_2485120, EPI_ISL_2485121, EPI_ISL_2485122, EPI_ISL_2485123, EPI_ISL_2485124, EPI_ISL_2485125, EPI_ISL_2485126, EPI_ISL_2485134 | | | |
| see above | Austrian Agency for Health and Food Safety (AGES) | Bergthaler laboratory, CeMM Research Center for Molecular Medicine of the Austrian Academy of Sciences | Lukas Endler, Anna Schedl, Fabian Amman, Petr Triska, Thomas Penz, Benedikt Agerer, Maelle Le Moing, Michael Schuster, Bekir Erguner, Jan Laine, Martin Senekowitsch, Christoph Bock, Andreas Bergthaler |
| EPI_ISL_2485169 | 3. Medizinische Abteilung, Hanusch Krankenhaus | Bergthaler laboratory, CeMM Research Center for Molecular Medicine of the Austrian Academy of Sciences | Lukas Endler, Anna Schedl, Fabian Amman, Petr Triska, Thomas Penz, Benedikt Agerer, Maelle Le Moing, Michael Schuster, Bekir Erguner, Jan Laine, Martin Senekowitsch, Christoph Bock, Andreas Bergthaler |
| EPI_ISL_2490307, EPI_ISL_2490308, EPI_ISL_2490309, EPI_ISL_2490534 | Basurto University Hospital: Clinical Microbiology Laboratory | Biocruces Bizkaia | Mikel Urrutikoetxea-Gutiérrez, Mª Carmen Nieto Toboso, Estibaliz Ugalde Zarraga, Mikel Gallego Rodrigo, Ana de la Hoz, José Luis Díaz de Tuesta del Arco |
| EPI_ISL_2491608 | Centre Hospitalier Universitaire Clermont-Ferrand | CHU Clermont-Ferrand, service de virologie | Bisseux Maxime, Mirand Audrey, Combes Patricia, Henquell Cécile |
| EPI_ISL_2491609 | HOPITAL DE MOULINS | CHU Clermont-Ferrand, service de virologie | Bisseux Maxime, Mirand Audrey, Combes Patricia, Henquell Cécile |
| EPI_ISL_2491611, EPI_ISL_2491612 | Centre Hospitalier Universitaire Clermont-Ferrand | CHU Clermont-Ferrand, service de virologie | Bisseux Maxime, Mirand Audrey, Combes Patricia, Henquell Cécile |
| EPI_ISL_2491613 | Centre Hospitalier RIOM | CHU Clermont-Ferrand, service de virologie | Bisseux Maxime, Mirand Audrey, Combes Patricia, Henquell Cécile |
| EPI_ISL_2491614, EPI_ISL_2491615, | Centre Hospitalier Universitaire Clermont-Ferrand | CHU Clermont-Ferrand, service de virologie | Bisseux Maxime, Mirand Audrey, Combes Patricia, Henquell Cécile |

| | | | |
|---|---|---|---|
| EPI_ISL_2491616, EPI_ISL_2491617, EPI_ISL_2491618 | | | |
| EPI_ISL_2491619 | Centre De Prélévement COVID VICHY | CHU Clermont-Ferrand, service de virologie | Bisseux Maxime, Mirand Audrey, Combes Patricia, Henquell Cécile |
| EPI_ISL_2491620, EPI_ISL_2491621 | HOPITAL DU MONT DORE | CHU Clermont-Ferrand, service de virologie | Bisseux Maxime, Mirand Audrey, Combes Patricia, Henquell Cécile |
| EPI_ISL_2491622, EPI_ISL_2491623, EPI_ISL_2491624, EPI_ISL_2491625, EPI_ISL_2491626, EPI_ISL_2491627 | Centre Hospitalier Universitaire Clermont-Ferrand | CHU Clermont-Ferrand, service de virologie | Bisseux Maxime, Mirand Audrey, Combes Patricia, Henquell Cécile |
| EPI_ISL_2491628 | HOPITAL DU MONT DORE | CHU Clermont-Ferrand, service de virologie | Bisseux Maxime, Mirand Audrey, Combes Patricia, Henquell Cécile |
| EPI_ISL_2491629 | Centre Hospitalier Universitaire Clermont-Ferrand | CHU Clermont-Ferrand, service de virologie | Bisseux Maxime, Mirand Audrey, Combes Patricia, Henquell Cécile |
| EPI_ISL_2491633 | laboratoire Maymat | CHU Clermont-Ferrand, service de virologie | Bisseux Maxime, Mirand Audrey, Combes Patricia, Henquell Cécile |
| EPI_ISL_2491634, EPI_ISL_2491635, EPI_ISL_2491636 | Centre Hospitalier Universitaire Clermont-Ferrand | CHU Clermont-Ferrand, service de virologie | Bisseux Maxime, Mirand Audrey, Combes Patricia, Henquell Cécile |
| EPI_ISL_2491637, EPI_ISL_2491638 | EHPAD | CHU Clermont-Ferrand, service de virologie | Bisseux Maxime, Mirand Audrey, Combes Patricia, Henquell Cécile |
| EPI_ISL_2491639 | HOPITAL DE MONTLUCON | CHU Clermont-Ferrand, service de virologie | Bisseux Maxime, Mirand Audrey, Combes Patricia, Henquell Cécile |
| EPI_ISL_2491640 | Centre Hospitalier Universitaire Clermont-Ferrand | CHU Clermont-Ferrand, service de virologie | Bisseux Maxime, Mirand Audrey, Combes Patricia, Henquell Cécile |
| EPI_ISL_2491641 | HOPITAL DE MONTLUCON | CHU Clermont-Ferrand, service de virologie | Bisseux Maxime, Mirand Audrey, Combes Patricia, Henquell Cécile |
| EPI_ISL_2491642 | HOPITAL DE VICHY | CHU Clermont-Ferrand, service de virologie | Bisseux Maxime, Mirand Audrey, Combes Patricia, Henquell Cécile |
| EPI_ISL_2491643, EPI_ISL_2491644, EPI_ISL_2491645 | Centre Hospitalier Universitaire Clermont-Ferrand | CHU Clermont-Ferrand, service de virologie | Bisseux Maxime, Mirand Audrey, Combes Patricia, Henquell Cécile |
| EPI_ISL_2491646 | CDP COVID MONTLUCON | CHU Clermont-Ferrand, service de virologie | Bisseux Maxime, Mirand Audrey, Combes Patricia, Henquell Cécile |
| EPI_ISL_2491647 | Centre Hospitalier Universitaire Clermont-Ferrand | CHU Clermont-Ferrand, service de virologie | Bisseux Maxime, Mirand Audrey, Combes Patricia, Henquell Cécile |
| EPI_ISL_2491648 | center de prélévement COVID ISSOIRE | CHU Clermont-Ferrand, service de virologie | Bisseux Maxime, Mirand Audrey, Combes Patricia, Henquell Cécile |
| EPI_ISL_2491649, EPI_ISL_2491650, EPI_ISL_2491651, EPI_ISL_2491652, EPI_ISL_2491653 | Centre Hospitalier Universitaire Clermont-Ferrand | CHU Clermont-Ferrand, service de virologie | Bisseux Maxime, Mirand Audrey, Combes Patricia, Henquell Cécile |
| EPI_ISL_2491654, EPI_ISL_2491655, EPI_ISL_2491656, EPI_ISL_2491657 | CDP COVID MONTLUCON | CHU Clermont-Ferrand, service de virologie | Bisseux Maxime, Mirand Audrey, Combes Patricia, Henquell Cécile |
| EPI_ISL_2491658, EPI_ISL_2491659, EPI_ISL_2491660, EPI_ISL_2491661 | Centre Hospitalier Universitaire Clermont-Ferrand | CHU Clermont-Ferrand, service de virologie | Bisseux Maxime, Mirand Audrey, Combes Patricia, Henquell Cécile |
| EPI_ISL_2491662 | Centre De Prelevement COVID RIOM | CHU Clermont-Ferrand, service de virologie | Bisseux Maxime, Mirand Audrey, Combes Patricia, Henquell Cécile |
| EPI_ISL_2491663 | Centre Hospitalier Universitaire Clermont-Ferrand | CHU Clermont-Ferrand, service de virologie | Bisseux Maxime, Mirand Audrey, Combes Patricia, Henquell Cécile |
| EPI_ISL_2491664 | Centre hospitalier D'AURILLAC | CHU Clermont-Ferrand, service de virologie | Bisseux Maxime, Mirand Audrey, Combes Patricia, Henquell Cécile |
| EPI_ISL_2492366, EPI_ISL_2492367, EPI_ISL_2492368, EPI_ISL_2492369, EPI_ISL_2492370, EPI_ISL_2492371, EPI_ISL_2492372, EPI_ISL_2492373, EPI_ISL_2492374, EPI_ISL_2492375, EPI_ISL_2492376, EPI_ISL_2492377, EPI_ISL_2492378, EPI_ISL_2492379, EPI_ISL_2492380, EPI_ISL_2492381, EPI_ISL_2492382, EPI_ISL_2492383, EPI_ISL_2492384, EPI_ISL_2492385, EPI_ISL_2492386, EPI_ISL_2492387, EPI_ISL_2492388, EPI_ISL_2492389, EPI_ISL_2492390, EPI_ISL_2492391, EPI_ISL_2492392 | | | |
| see above | Cliniques universitaires Saint-Luc | UCLouvain/IREC/MBLG | Jean Ruelle, Benoit Kabamba Mukadi |
| EPI_ISL_2492393, EPI_ISL_2492394, EPI_ISL_2492395, EPI_ISL_2492396, EPI_ISL_2492397, EPI_ISL_2492398, EPI_ISL_2492399, EPI_ISL_2492400, EPI_ISL_2492401, EPI_ISL_2492402, EPI_ISL_2492403 | | | |
| see above | Clinique Saint-Pierre Ottignies | UCLouvain/IREC/MBLG | Jean Ruelle, Benoit Kabamba Mukadi |
| EPI_ISL_2492404, EPI_ISL_2492405, EPI_ISL_2492406, EPI_ISL_2492407, EPI_ISL_2492408, EPI_ISL_2492409, EPI_ISL_2492410, EPI_ISL_2492411, EPI_ISL_2492412, EPI_ISL_2492413 | Cliniques universitaires Saint-Luc | UCLouvain/IREC/MBLG | Jean Ruelle, Benoit Kabamba Mukadi |
| EPI_ISL_2492414, EPI_ISL_2492415, EPI_ISL_2492416 | Clinique Saint-Pierre Ottignies | UCLouvain/IREC/MBLG | Jean Ruelle, Benoit Kabamba Mukadi |
| EPI_ISL_2492417, EPI_ISL_2492418, EPI_ISL_2492419, EPI_ISL_2492420, EPI_ISL_2492421, EPI_ISL_2492422, EPI_ISL_2492423, EPI_ISL_2492424, EPI_ISL_2492425, EPI_ISL_2492426, EPI_ISL_2492427, EPI_ISL_2492428, EPI_ISL_2492429, EPI_ISL_2492430, EPI_ISL_2492431, EPI_ISL_2492432, EPI_ISL_2492433, EPI_ISL_2492434, EPI_ISL_2492435, EPI_ISL_2492436 | | | |
| see above | Cliniques universitaires Saint-Luc | UCLouvain/IREC/MBLG | Jean Ruelle, Benoit Kabamba Mukadi |
| EPI_ISL_2492437, EPI_ISL_2492438, EPI_ISL_2492439 | Clinique Saint-Pierre Ottignies | UCLouvain/IREC/MBLG | Jean Ruelle, Benoit Kabamba Mukadi |
| EPI_ISL_2492440 | Cliniques universitaires Saint-Luc | UCLouvain/IREC/MBLG | Jean Ruelle, Benoit Kabamba Mukadi |
| EPI_ISL_2492498 | LESP Queretaro | Instituto de Diagnostico y Referencia Epidemiologicos (INDRE) | Claudia Wong-Arambula, Abril Rodriguez-Maldonado, Vanessa Rivero-Arredondo, Ariadna Medina-Benitez, Joaquin Quiroz-Mercado, Sergio Rangel-Guerrero, Natividad Cruz-Ortiz, Tatiana Nunez-Garcia, Gisela Barrera-Badillo, Lucia Hernandez-Rivas, Irma Lopez-Martinez, Ernesto Ramirez-Gonzalez. |
| EPI_ISL_2501020, EPI_ISL_2501021, EPI_ISL_2501022, EPI_ISL_2501023, EPI_ISL_2501024, EPI_ISL_2501025, EPI_ISL_2501026 | Banteay Meanchey Rapid Response Team | Virology Unit, Institut Pasteur du Cambodge | Cecile Troupin, Leakhena Pum, Jurre Y Siegers, Ly Sovann, Kraing Sidonn, Yi Sengdoeurn, Chin Savuth, Chau Darapheak, Veasna Duong, Erik A Karlsson |
| EPI_ISL_2501027, EPI_ISL_2501028 | Battambang Provincial Laboratory | Virology Unit, Institut Pasteur du Cambodge | Cecile Troupin, Leakhena Pum, Jurre Y Siegers, Ly Sovann, Kraing Sidonn, Yi Sengdoeurn, Chin Savuth, Chau Darapheak, Veasna Duong, Erik A Karlsson |
| EPI_ISL_2501029, EPI_ISL_2501030 | Siem Reap Provinical Laboratory | Virology Unit, Institut Pasteur du Cambodge | Cecile Troupin, Leakhena Pum, Jurre Y Siegers, Ly Sovann, Kraing Sidonn, Yi Sengdoeurn, Chin Savuth, Chau Darapheak, Veasna Duong, Erik A Karlsson |
| EPI_ISL_2501031, EPI_ISL_2501032 | Oddar Meanchey Rapid Response Team | Virology Unit, Institut Pasteur du Cambodge | Cecile Troupin, Leakhena Pum, Jurre Y Siegers, Ly Sovann, Kraing Sidonn, Yi Sengdoeurn, Chin Savuth, Chau Darapheak, Veasna Duong, Erik A Karlsson |
| EPI_ISL_2501033, EPI_ISL_2501034, EPI_ISL_2501035 | Tboung Khmum Rapid Response Team | Virology Unit, Institut Pasteur du Cambodge | Cecile Troupin, Leakhena Pum, Jurre Y Siegers, Ly Sovann, Kraing Sidonn, Yi Sengdoeurn, Chin Savuth, Chau Darapheak, Veasna Duong, Erik A Karlsson |
| EPI_ISL_2501036, EPI_ISL_2501037, EPI_ISL_2501038, EPI_ISL_2501039, EPI_ISL_2501040, EPI_ISL_2501041, EPI_ISL_2501042, EPI_ISL_2501043, EPI_ISL_2501044, EPI_ISL_2501045, EPI_ISL_2501046, EPI_ISL_2501047, EPI_ISL_2501048, EPI_ISL_2501049, EPI_ISL_2501050, EPI_ISL_2501051, EPI_ISL_2501052, EPI_ISL_2501053, EPI_ISL_2501054, EPI_ISL_2501055, EPI_ISL_2501056, EPI_ISL_2501057, EPI_ISL_2501058, EPI_ISL_2501059, EPI_ISL_2501060, EPI_ISL_2501061, EPI_ISL_2501062, EPI_ISL_2501063 | | | |
| see above | Virology Unit, Institut Pasteur du Cambodge | Virology Unit, Institut Pasteur du Cambodge | Cecile Troupin, Leakhena Pum, Jurre Y Siegers, Ly Sovann, Kraing Sidonn, Yi Sengdoeurn, Chin Savuth, Chau Darapheak, Veasna Duong, Erik A |

| | | | |
|---|---|---|---|
| | | | Karlsson |
| EPI_ISL_2501225, EPI_ISL_2501226, EPI_ISL_2501228 | CHU LILLE | CHU Lille - Laboratoire de Virologie | AIT YAHYA Emilie, ALIDJINOU Enagnon Kazali, BOCKET Laurence, CREPIN Michel, DEMAY Christophe, ENGELMANN Ilka, GEFFROY Sandrine, GUIGON Aurélie, LAMBERT Valérie, LAZREK Mouna, NOBILLIAUX Florian, PREVOST Brigitte, THUILLIER Caroline, TINEZ Claire ,TCHANTCHOU NJOSSE YANICK |
| EPI_ISL_2501229, EPI_ISL_2501230 | CH ARRAS | CHU Lille - Laboratoire de Virologie | AIT YAHYA Emilie, ALIDJINOU Enagnon Kazali, BOCKET Laurence, CREPIN Michel, DEMAY Christophe, ENGELMANN Ilka, GEFFROY Sandrine, GUIGON Aurélie, LAMBERT Valérie, LAZREK Mouna, NOBILLIAUX Florian, PREVOST Brigitte, THUILLIER Caroline, TINEZ Claire ,TCHANTCHOU NJOSSE YANICK |
| EPI_ISL_2501232 | CH CAMBRAI | CHU Lille - Laboratoire de Virologie | AIT YAHYA Emilie, ALIDJINOU Enagnon Kazali, BOCKET Laurence, CREPIN Michel, DEMAY Christophe, ENGELMANN Ilka, GEFFROY Sandrine, GUIGON Aurélie, LAMBERT Valérie, LAZREK Mouna, NOBILLIAUX Florian, PREVOST Brigitte, THUILLIER Caroline, TINEZ Claire ,TCHANTCHOU NJOSSE YANICK |
| EPI_ISL_2501233 | CHU LILLE | CHU Lille - Laboratoire de Virologie | AIT YAHYA Emilie, ALIDJINOU Enagnon Kazali, BOCKET Laurence, CREPIN Michel, DEMAY Christophe, ENGELMANN Ilka, GEFFROY Sandrine, GUIGON Aurélie, LAMBERT Valérie, LAZREK Mouna, NOBILLIAUX Florian, PREVOST Brigitte, THUILLIER Caroline, TINEZ Claire ,TCHANTCHOU NJOSSE YANICK |
| EPI_ISL_2501234, EPI_ISL_2501235, EPI_ISL_2501236 | CH ROUBAIX | CHU Lille - Laboratoire de Virologie | AIT YAHYA Emilie, ALIDJINOU Enagnon Kazali, BOCKET Laurence, CREPIN Michel, DEMAY Christophe, ENGELMANN Ilka, GEFFROY Sandrine, GUIGON Aurélie, LAMBERT Valérie, LAZREK Mouna, NOBILLIAUX Florian, PREVOST Brigitte, THUILLIER Caroline, TINEZ Claire ,TCHANTCHOU NJOSSE YANICK |
| EPI_ISL_2501237, EPI_ISL_2501238, EPI_ISL_2501239 | CH TOURCOING | CHU Lille - Laboratoire de Virologie | AIT YAHYA Emilie, ALIDJINOU Enagnon Kazali, BOCKET Laurence, CREPIN Michel, DEMAY Christophe, ENGELMANN Ilka, GEFFROY Sandrine, GUIGON Aurélie, LAMBERT Valérie, LAZREK Mouna, NOBILLIAUX Florian, PREVOST Brigitte, THUILLIER Caroline, TINEZ Claire ,TCHANTCHOU NJOSSE YANICK |
| EPI_ISL_2501240 | CHU LILLE | CHU Lille - Laboratoire de Virologie | AIT YAHYA Emilie, ALIDJINOU Enagnon Kazali, BOCKET Laurence, CREPIN Michel, DEMAY Christophe, ENGELMANN Ilka, GEFFROY Sandrine, GUIGON Aurélie, LAMBERT Valérie, LAZREK Mouna, NOBILLIAUX Florian, PREVOST Brigitte, THUILLIER Caroline, TINEZ Claire ,TCHANTCHOU NJOSSE YANICK |
| EPI_ISL_2501241 | CH BOULOGNE-SUR-MER | CHU Lille - Laboratoire de Virologie | AIT YAHYA Emilie, ALIDJINOU Enagnon Kazali, BOCKET Laurence, CREPIN Michel, DEMAY Christophe, ENGELMANN Ilka, GEFFROY Sandrine, GUIGON Aurélie, LAMBERT Valérie, LAZREK Mouna, NOBILLIAUX Florian, PREVOST Brigitte, THUILLIER Caroline, TINEZ Claire ,TCHANTCHOU NJOSSE YANICK |
| EPI_ISL_2501242, EPI_ISL_2501243 | CHU LILLE | CHU Lille - Laboratoire de Virologie | AIT YAHYA Emilie, ALIDJINOU Enagnon Kazali, BOCKET Laurence, CREPIN Michel, DEMAY Christophe, ENGELMANN Ilka, GEFFROY Sandrine, GUIGON Aurélie, LAMBERT Valérie, LAZREK Mouna, NOBILLIAUX Florian, PREVOST Brigitte, THUILLIER Caroline, TINEZ Claire ,TCHANTCHOU NJOSSE YANICK |
| EPI_ISL_2501244, EPI_ISL_2501245, EPI_ISL_2501246 | CH ROUBAIX | CHU Lille - Laboratoire de Virologie | AIT YAHYA Emilie, ALIDJINOU Enagnon Kazali, BOCKET Laurence, CREPIN Michel, DEMAY Christophe, ENGELMANN Ilka, GEFFROY Sandrine, GUIGON Aurélie, LAMBERT Valérie, LAZREK Mouna, NOBILLIAUX Florian, PREVOST Brigitte, THUILLIER Caroline, TINEZ Claire ,TCHANTCHOU NJOSSE YANICK |
| EPI_ISL_2501247 | CH BOULOGNE-SUR-MER | CHU Lille - Laboratoire de Virologie | AIT YAHYA Emilie, ALIDJINOU Enagnon Kazali, BOCKET Laurence, CREPIN Michel, DEMAY Christophe, ENGELMANN Ilka, GEFFROY Sandrine, GUIGON Aurélie, LAMBERT Valérie, LAZREK Mouna, NOBILLIAUX Florian, PREVOST Brigitte, THUILLIER Caroline, TINEZ Claire ,TCHANTCHOU NJOSSE YANICK |
| EPI_ISL_2501248, EPI_ISL_2501249 | CHU LILLE | CHU Lille - Laboratoire de Virologie | AIT YAHYA Emilie, ALIDJINOU Enagnon Kazali, BOCKET Laurence, CREPIN Michel, DEMAY Christophe, ENGELMANN Ilka, GEFFROY Sandrine, GUIGON Aurélie, LAMBERT Valérie, LAZREK Mouna, NOBILLIAUX Florian, PREVOST Brigitte, THUILLIER Caroline, TINEZ Claire ,TCHANTCHOU NJOSSE YANICK |
| EPI_ISL_2501250 | CH SECLIN | CHU Lille - Laboratoire de Virologie | AIT YAHYA Emilie, ALIDJINOU Enagnon Kazali, BOCKET Laurence, CREPIN Michel, DEMAY Christophe, ENGELMANN Ilka, GEFFROY Sandrine, GUIGON Aurélie, LAMBERT Valérie, LAZREK Mouna, NOBILLIAUX Florian, PREVOST Brigitte, THUILLIER Caroline, TINEZ Claire ,TCHANTCHOU NJOSSE YANICK |
| EPI_ISL_2501251, EPI_ISL_2501252, EPI_ISL_2501253, EPI_ISL_2501254, EPI_ISL_2501255, EPI_ISL_2501256 | CHU LILLE | CHU Lille - Laboratoire de Virologie | AIT YAHYA Emilie, ALIDJINOU Enagnon Kazali, BOCKET Laurence, CREPIN Michel, DEMAY Christophe, ENGELMANN Ilka, GEFFROY Sandrine, GUIGON Aurélie, LAMBERT Valérie, LAZREK Mouna, NOBILLIAUX Florian, PREVOST Brigitte, THUILLIER Caroline, TINEZ Claire ,TCHANTCHOU NJOSSE YANICK |
| EPI_ISL_2501257 | CH SAINT-PHILIBERT | CHU Lille - Laboratoire de Virologie | AIT YAHYA Emilie, ALIDJINOU Enagnon Kazali, BOCKET Laurence, CREPIN Michel, DEMAY Christophe, ENGELMANN Ilka, GEFFROY Sandrine, GUIGON Aurélie, LAMBERT Valérie, LAZREK Mouna, NOBILLIAUX Florian, PREVOST Brigitte, THUILLIER Caroline, TINEZ Claire ,TCHANTCHOU NJOSSE YANICK |
| EPI_ISL_2501258, EPI_ISL_2501259, EPI_ISL_2501260, EPI_ISL_2501261, EPI_ISL_2501262 | CH ROUBAIX | CHU Lille - Laboratoire de Virologie | AIT YAHYA Emilie, ALIDJINOU Enagnon Kazali, BOCKET Laurence, CREPIN Michel, DEMAY Christophe, ENGELMANN Ilka, GEFFROY Sandrine, GUIGON Aurélie, LAMBERT Valérie, LAZREK Mouna, NOBILLIAUX Florian, PREVOST Brigitte, THUILLIER Caroline, TINEZ Claire ,TCHANTCHOU NJOSSE YANICK |
| EPI_ISL_2501263, EPI_ISL_2501264 | CHU LILLE | CHU Lille - Laboratoire de Virologie | AIT YAHYA Emilie, ALIDJINOU Enagnon Kazali, BOCKET Laurence, CREPIN Michel, DEMAY Christophe, ENGELMANN Ilka, GEFFROY Sandrine, GUIGON Aurélie, LAMBERT Valérie, LAZREK Mouna, NOBILLIAUX Florian, PREVOST Brigitte, THUILLIER Caroline, TINEZ Claire ,TCHANTCHOU NJOSSE YANICK |
| EPI_ISL_2501267 | CH SAINT-PHILIBERT | CHU Lille - Laboratoire de Virologie | AIT YAHYA Emilie, ALIDJINOU Enagnon Kazali, BOCKET Laurence, CREPIN Michel, DEMAY Christophe, ENGELMANN Ilka, GEFFROY Sandrine, GUIGON Aurélie, LAMBERT Valérie, LAZREK Mouna, NOBILLIAUX Florian, PREVOST Brigitte, THUILLIER Caroline, TINEZ Claire ,TCHANTCHOU NJOSSE YANICK |
| EPI_ISL_2501268, EPI_ISL_2501269, EPI_ISL_2501270 | CHU LILLE | CHU Lille - Laboratoire de Virologie | AIT YAHYA Emilie, ALIDJINOU Enagnon Kazali, BOCKET Laurence, CREPIN Michel, DEMAY Christophe, ENGELMANN Ilka, GEFFROY Sandrine, GUIGON Aurélie, LAMBERT Valérie, LAZREK Mouna, NOBILLIAUX Florian, PREVOST Brigitte, THUILLIER Caroline, TINEZ Claire ,TCHANTCHOU NJOSSE YANICK |
| EPI_ISL_2501271 | CH ARRAS | CHU Lille - Laboratoire de Virologie | AIT YAHYA Emilie, ALIDJINOU Enagnon Kazali, BOCKET Laurence, CREPIN Michel, DEMAY Christophe, ENGELMANN Ilka, GEFFROY Sandrine, GUIGON Aurélie, LAMBERT Valérie, LAZREK Mouna, NOBILLIAUX Florian, PREVOST Brigitte, THUILLIER Caroline, TINEZ Claire ,TCHANTCHOU NJOSSE YANICK |
| EPI_ISL_2501272 | CHU LILLE | CHU Lille - Laboratoire de Virologie | AIT YAHYA Emilie, ALIDJINOU Enagnon Kazali, BOCKET Laurence, CREPIN Michel, DEMAY Christophe, ENGELMANN Ilka, GEFFROY Sandrine, GUIGON Aurélie, LAMBERT Valérie, LAZREK Mouna, NOBILLIAUX Florian, PREVOST Brigitte, THUILLIER Caroline, TINEZ Claire ,TCHANTCHOU NJOSSE YANICK |
| EPI_ISL_2501275, EPI_ISL_2501276 | CH TOURCOING | CHU Lille - Laboratoire de Virologie | AIT YAHYA Emilie, ALIDJINOU Enagnon Kazali, BOCKET Laurence, CREPIN Michel, DEMAY Christophe, ENGELMANN Ilka, GEFFROY Sandrine, GUIGON Aurélie, LAMBERT Valérie, LAZREK Mouna, NOBILLIAUX Florian, PREVOST Brigitte, THUILLIER Caroline, TINEZ Claire ,TCHANTCHOU NJOSSE YANICK |
| EPI_ISL_2501278, EPI_ISL_2501279 | CHU LILLE | CHU Lille - Laboratoire de Virologie | AIT YAHYA Emilie, ALIDJINOU Enagnon Kazali, BOCKET Laurence, CREPIN Michel, DEMAY Christophe, ENGELMANN Ilka, GEFFROY Sandrine, GUIGON Aurélie, LAMBERT Valérie, LAZREK Mouna, NOBILLIAUX Florian, PREVOST Brigitte, THUILLIER Caroline, TINEZ Claire ,TCHANTCHOU NJOSSE YANICK |
| EPI_ISL_2501280, EPI_ISL_2501281 | CH ROUBAIX | CHU Lille - Laboratoire de Virologie | AIT YAHYA Emilie, ALIDJINOU Enagnon Kazali, BOCKET Laurence, CREPIN Michel, DEMAY Christophe, ENGELMANN Ilka, GEFFROY Sandrine, GUIGON Aurélie, LAMBERT Valérie, LAZREK Mouna, NOBILLIAUX Florian, PREVOST Brigitte, THUILLIER Caroline, TINEZ Claire ,TCHANTCHOU NJOSSE YANICK |
| EPI_ISL_2501283, EPI_ISL_2501284 | CH LENS | CHU Lille - Laboratoire de Virologie | AIT YAHYA Emilie, ALIDJINOU Enagnon Kazali, BOCKET Laurence, CREPIN Michel, DEMAY Christophe, ENGELMANN Ilka, GEFFROY Sandrine, GUIGON Aurélie, LAMBERT Valérie, LAZREK Mouna, NOBILLIAUX Florian, PREVOST Brigitte, THUILLIER Caroline, TINEZ Claire ,TCHANTCHOU |

| | | | |
|---|---|---|---|
| | | | NJOSSE YANICK |
| EPI_ISL_2501285 | CH TOURCOING | CHU Lille - Laboratoire de Virologie | AIT YAHYA Emilie, ALIDJINOU Enagnon Kazali, BOCKET Laurence, CREPIN Michel, DEMAY Christophe, ENGELMANN Ilka, GEFFROY Sandrine, GUIGON Aurélie, LAMBERT Valérie, LAZREK Mouna, NOBILLIAUX Florian, PREVOST Brigitte, THUILLIER Caroline, TINEZ Claire ,TCHANTCHOU NJOSSE YANICK |
| EPI_ISL_2501286 | CHU LILLE | CHU Lille - Laboratoire de Virologie | AIT YAHYA Emilie, ALIDJINOU Enagnon Kazali, BOCKET Laurence, CREPIN Michel, DEMAY Christophe, ENGELMANN Ilka, GEFFROY Sandrine, GUIGON Aurélie, LAMBERT Valérie, LAZREK Mouna, NOBILLIAUX Florian, PREVOST Brigitte, THUILLIER Caroline, TINEZ Claire ,TCHANTCHOU NJOSSE YANICK |
| EPI_ISL_2501287 | CH ARMENTIERES | CHU Lille - Laboratoire de Virologie | AIT YAHYA Emilie, ALIDJINOU Enagnon Kazali, BOCKET Laurence, CREPIN Michel, DEMAY Christophe, ENGELMANN Ilka, GEFFROY Sandrine, GUIGON Aurélie, LAMBERT Valérie, LAZREK Mouna, NOBILLIAUX Florian, PREVOST Brigitte, THUILLIER Caroline, TINEZ Claire ,TCHANTCHOU NJOSSE YANICK |
| EPI_ISL_2501288 | CH BETHUNE | CHU Lille - Laboratoire de Virologie | AIT YAHYA Emilie, ALIDJINOU Enagnon Kazali, BOCKET Laurence, CREPIN Michel, DEMAY Christophe, ENGELMANN Ilka, GEFFROY Sandrine, GUIGON Aurélie, LAMBERT Valérie, LAZREK Mouna, NOBILLIAUX Florian, PREVOST Brigitte, THUILLIER Caroline, TINEZ Claire ,TCHANTCHOU NJOSSE YANICK |
| EPI_ISL_2501289, EPI_ISL_2501290 | CH SAINT-PHILIBERT | CHU Lille - Laboratoire de Virologie | AIT YAHYA Emilie, ALIDJINOU Enagnon Kazali, BOCKET Laurence, CREPIN Michel, DEMAY Christophe, ENGELMANN Ilka, GEFFROY Sandrine, GUIGON Aurélie, LAMBERT Valérie, LAZREK Mouna, NOBILLIAUX Florian, PREVOST Brigitte, THUILLIER Caroline, TINEZ Claire ,TCHANTCHOU NJOSSE YANICK |
| EPI_ISL_2501291, EPI_ISL_2501292, EPI_ISL_2501293, EPI_ISL_2501294 | CHU LILLE | CHU Lille - Laboratoire de Virologie | AIT YAHYA Emilie, ALIDJINOU Enagnon Kazali, BOCKET Laurence, CREPIN Michel, DEMAY Christophe, ENGELMANN Ilka, GEFFROY Sandrine, GUIGON Aurélie, LAMBERT Valérie, LAZREK Mouna, NOBILLIAUX Florian, PREVOST Brigitte, THUILLIER Caroline, TINEZ Claire ,TCHANTCHOU NJOSSE YANICK |
| EPI_ISL_2501295 | CH TOURCOING | CHU Lille - Laboratoire de Virologie | AIT YAHYA Emilie, ALIDJINOU Enagnon Kazali, BOCKET Laurence, CREPIN Michel, DEMAY Christophe, ENGELMANN Ilka, GEFFROY Sandrine, GUIGON Aurélie, LAMBERT Valérie, LAZREK Mouna, NOBILLIAUX Florian, PREVOST Brigitte, THUILLIER Caroline, TINEZ Claire ,TCHANTCHOU NJOSSE YANICK |
| EPI_ISL_2501296 | CHU LILLE | CHU Lille - Laboratoire de Virologie | AIT YAHYA Emilie, ALIDJINOU Enagnon Kazali, BOCKET Laurence, CREPIN Michel, DEMAY Christophe, ENGELMANN Ilka, GEFFROY Sandrine, GUIGON Aurélie, LAMBERT Valérie, LAZREK Mouna, NOBILLIAUX Florian, PREVOST Brigitte, THUILLIER Caroline, TINEZ Claire ,TCHANTCHOU NJOSSE YANICK |
| EPI_ISL_2501297 | CH CAMBRAI | CHU Lille - Laboratoire de Virologie | AIT YAHYA Emilie, ALIDJINOU Enagnon Kazali, BOCKET Laurence, CREPIN Michel, DEMAY Christophe, ENGELMANN Ilka, GEFFROY Sandrine, GUIGON Aurélie, LAMBERT Valérie, LAZREK Mouna, NOBILLIAUX Florian, PREVOST Brigitte, THUILLIER Caroline, TINEZ Claire ,TCHANTCHOU NJOSSE YANICK |
| EPI_ISL_2501299, EPI_ISL_2501300 | CHU LILLE | CHU Lille - Laboratoire de Virologie | AIT YAHYA Emilie, ALIDJINOU Enagnon Kazali, BOCKET Laurence, CREPIN Michel, DEMAY Christophe, ENGELMANN Ilka, GEFFROY Sandrine, GUIGON Aurélie, LAMBERT Valérie, LAZREK Mouna, NOBILLIAUX Florian, PREVOST Brigitte, THUILLIER Caroline, TINEZ Claire ,TCHANTCHOU NJOSSE YANICK |
| EPI_ISL_2501301 | CH LENS | CHU Lille - Laboratoire de Virologie | AIT YAHYA Emilie, ALIDJINOU Enagnon Kazali, BOCKET Laurence, CREPIN Michel, DEMAY Christophe, ENGELMANN Ilka, GEFFROY Sandrine, GUIGON Aurélie, LAMBERT Valérie, LAZREK Mouna, NOBILLIAUX Florian, PREVOST Brigitte, THUILLIER Caroline, TINEZ Claire ,TCHANTCHOU NJOSSE YANICK |
| EPI_ISL_2501304, EPI_ISL_2501305, EPI_ISL_2501307 | Heilig hart Lier | Imelda Ziekenhuis | Johan Frans, Hanne Valgaeren, Dagmar Obbels |
| EPI_ISL_2501316, EPI_ISL_2501317, EPI_ISL_2501322, EPI_ISL_2501323, EPI_ISL_2501324, EPI_ISL_2501325, EPI_ISL_2501326, EPI_ISL_2501327, EPI_ISL_2501328, EPI_ISL_2501329, EPI_ISL_2501330, EPI_ISL_2501331, EPI_ISL_2501332, EPI_ISL_2501337, EPI_ISL_2501338, EPI_ISL_2501339, EPI_ISL_2501342, EPI_ISL_2501343, EPI_ISL_2501344, EPI_ISL_2501345, EPI_ISL_2501346, EPI_ISL_2501347, EPI_ISL_2501364, EPI_ISL_2501366, EPI_ISL_2501367, EPI_ISL_2501368, EPI_ISL_2501370, EPI_ISL_2501377, EPI_ISL_2501378 | | | |
| see above | Microbiology Department. Complexo Hospitalario Universitario de Vigo | Microbiology Department. Complexo Hospitalario Universitario de Vigo | Alfaya N, Alonso I, Alvarez M, Cabrera JJ, Carballo R, Cores O, Cortizo S, del-Campo V, Martinez L, Mediero G, Perez S, Potel C, Regueiro B, Rey S, Vasallo FJ |
| EPI_ISL_2502615 | RSUD dr Loekmono Hadi | Genetics Working Group (Pokja Genetik) Faculty of Medicine, Public Health and Nursing Universitas Gadjah Mada (FK-KMK UGM); Disease Investigation Center Wates Ministry of Agriculture Indonesia; Department of Microbiology FK-KMK UGM; Laboratorium Diagnostik Yayasan Tahija World Mosquito Program (WMP) Yogyakarta Center for Tropical Medicine FK-KMK UGM; Integrated Research Center FK-KMK UGM; Department of Computer Science and Electronics FMIPA UGM; RSUP Dr. Sardjito | Gunadi, Hendra Wibawa, Marcellus, Mohamad S. Hakim, Edwin W. Daniwijaya, Ludhang P. Rizki, Endah Supriyati, Eggi Arguni, Titik Nuryastuti, Tri Wibawa, Dwi AA Nugrahaningsih, Afiahayati, Siswanto,Kristy Iskandar,Nungki Anggorowati, Pramesti G Dewi, Dwi Indaryati,Indarto Sulistiyono, Susan Simanjaya, Dyah A Puspitarani |
| EPI_ISL_2502687 | RSUD dr. LOEKMONO HADI | Genetics Working Group (Pokja Genetik) Faculty of Medicine, Public Health and Nursing Universitas Gadjah Mada (FK-KMK UGM); Disease Investigation Center Wates Ministry of Agriculture Indonesia; Department of Microbiology FK-KMK UGM; Laboratorium Diagnostik Yayasan Tahija World Mosquito Program (WMP) Yogyakarta Center for Tropical Medicine FK-KMK UGM; Integrated Research Center FK-KMK UGM; Department of Computer Science and Electronics FMIPA UGM; RSUP Dr. Sardjito | Gunadi, Hendra Wibawa, Marcellus, Mohamad S. Hakim, Edwin W. Daniwijaya, Ludhang P. Rizki, Endah Supriyati, Eggi Arguni, Titik Nuryastuti, Tri Wibawa, Dwi AA Nugrahaningsih, Afiahayati, Siswanto,Kristy Iskandar,Nungki Anggorowati, Pramesti G Dewi, Dwi Indaryati,Indarto Sulistiyono, Cita S Amalia, Alvina A Setiawan |
| EPI_ISL_2502756 | AREA DE SALUD LA UNION | Inciensa, Instituto Costarricense de InvestigaciOn y Enseñanza en NutriciOn y Salud | Francisco Duarte, Hebleen Porras, Claudio Soto-Garita, Estela Cordero, Adriana Godínez, Melany CalderOn, Jose Luis Vargas, Mariela Gutierrez, Joselyn Prado & MOnica Charpentier |
| EPI_ISL_2502757 | AREA DE SALUD PARAISO-CERVANTES | Inciensa, Instituto Costarricense de InvestigaciOn y Enseñanza en NutriciOn y Salud | Francisco Duarte, Hebleen Porras, Claudio Soto-Garita, Estela Cordero, Adriana Godínez, Melany CalderOn, Jose Luis Vargas, Mariela Gutierrez, Joselyn Prado & MOnica Charpentier |
| EPI_ISL_2502764 | RSUD dr. LOEKMONO HADI | Genetics Working Group (Pokja Genetik) Faculty of Medicine, Public Health and Nursing Universitas Gadjah Mada (FK-KMK UGM); Disease Investigation Center Wates Ministry of Agriculture Indonesia; Department of Microbiology FK-KMK UGM; Laboratorium Diagnostik Yayasan Tahija World Mosquito Program (WMP) Yogyakarta Center for Tropical Medicine FK-KMK UGM; Integrated Research Center FK-KMK UGM; Department of Computer Science and Electronics FMIPA UGM; RSUP Dr. Sardjito | Gunadi, Hendra Wibawa, Marcellus, Mohamad S. Hakim, Edwin W. Daniwijaya, Ludhang P. Rizki, Endah Supriyati, Eggi Arguni, Titik Nuryastuti, Tri Wibawa, Dwi AA Nugrahaningsih, Afiahayati, Siswanto,Kristy Iskandar,Nungki Anggorowati, Pramesti G Dewi, Dwi Indaryati,Indarto Sulistiyono, Irene Tania, Hana F Hanifin |
| EPI_ISL_2503137, EPI_ISL_2503138, EPI_ISL_2503139 | Laboratorio Biologia Molecolare Sars Cov2 - UOC Laboratorio Analisi - Servizio Medicina di Laboratorio, Ospedale "San Francesco" - ATS-ASSL Nuoro | Laboratorio specialistico UOC Ematologia - Ospedale "San Francesco" - ATS-ASSL Nuoro | Aspenoni Rosanna,Piras Giovanna, Pira Giovanna, Fancello Patrizia, Rosu Valentina, Carta Franco,Fiamma Maura, Mameli Giuseppe, Maria Cristina Garau |
| EPI_ISL_2504068, EPI_ISL_2504071, EPI_ISL_2504072, EPI_ISL_2504073, EPI_ISL_2504088 | Botswana Harvard HIV Reference Laboratory | Botswana Harvard HIV Reference Laboratory | Sikhulile Moyo, Dorcas Maruapula, Wonderful T. Choga, Thongbotho Mphoyakgosi, Boitumelo Zuze, Botshelo Radibe, Legodile Kooepile, Ontlametse T. Bareng, Keoratile Ntshambiwa, Modisa Motswaledi, Madisa Mine, Joseph Makhema, Roger Shapiro, Shahin Lockman, Mosepele Mosepele, Simani Gaseitsiwe |

| | | | |
|---|---|---|---|
| EPI_ISL_2504114 | Palapye Primary Hospital Laboratory | Botswana Harvard HIV Reference Laboratory | Sikhulile Moyo, Dorcas Maruapula, Wonderful T. Choga, Thongbotho Mphoyakgosi, Boitumelo Zuze, Botshelo Radibe, Legodile Kooepile, Ontlametse T. Bareng, Letsibogo Gaoraelwe, Thela Tefelo, Keoratile Ntshambiwa, Modisa Motswaledi, Madisa Mine, Joseph Makhema, Roger Shapiro, Shahin Lockman, Mosepele Mosepele, Simani Gaseitsiwe |
| EPI_ISL_2506041 | RSUD dr. LOEKMONO HADI | Genetics Working Group (Pokja Genetik) Faculty of Medicine, Public Health and Nursing Universitas Gadjah Mada (FK-KMK UGM); Disease Investigation Center Wates Ministry of Agriculture Indonesia; Department of Microbiology FK-KMK UGM; Laboratorium Diagnostik Yayasan Tahija World Mosquito Program (WMP) Yogyakarta Center for Tropical Medicine FK-KMK UGM; Integrated Research Center FK-KMK UGM; Department of Computer Science and Electronics FMIPA UGM; RSUP Dr. Sardjito | Gunadi, Hendra Wibawa, Marcellus, Mohamad S. Hakim, Edwin W. Daniwijaya, Ludhang P. Rizki, Endah Supriyati, Eggi Arguni, Titik Nuryastuti, Tri Wibawa, Dwi AA Nugrahaningsih, Afiahayati, Siswanto,Kristy Iskandar,Nungki Anggorowati, Pramesti G Dewi, Dwi Indaryati,Indarto Sulistiyono, Irene Tania, Hana F Hanifin |
| EPI_ISL_2509334 | Palapye Primary Hospital Laboratory | Botswana Harvard HIV Reference Laboratory | Sikhulile Moyo, Dorcas Maruapula, Wonderful T. Choga, Thongbotho Mphoyakgosi, Boitumelo Zuze, Botshelo Radibe, Legodile Kooepile, Ontlametse T. Bareng, Letsibogo Gaoraelwe, Thela Tefelo, Keoratile Ntshambiwa, Modisa Motswaledi, Madisa Mine, Joseph Makhema, Roger Shapiro, Shahin Lockman, Mosepele Mosepele, Simani Gaseitsiwe |
| EPI_ISL_2510687 | Surabaya Health Office | Institute of Tropical Disease, Universitas Airlangga | Jezzy R Dewantari, Rima R Prasetya, Krisnoadi Rahardjo, Aldise M Nastri, Febria Rachmanita, Gatot Soegiarto, Laksmi Wulandari, Resti Yudhawati, Yasuko Mori, Soetjipto, Kazufumi Shimizu, Maria I Lusida |
| EPI_ISL_2517047, EPI_ISL_2517048, EPI_ISL_2517051, EPI_ISL_2517052, EPI_ISL_2517053, EPI_ISL_2517054, EPI_ISL_2517055, EPI_ISL_2517056, EPI_ISL_2517058, EPI_ISL_2517059, EPI_ISL_2517060, EPI_ISL_2517061, EPI_ISL_2517062 | | | |
| see above | Althaia. Xarxa Assistencial Universitària de Manresa | IrsiCaixa | Marc Noguera-Julian, Mariona Parera, Maria Casadellà, Pilar Armengol, Francesc Catala-Moll, Roger Paredes, Bonaventura Clotet Gloria Trujillo, Rafel Perez Vidal, Jaume Trapé, Carolina Gonzalez-Fernandez, Miquel Micó, Antonia Flor, Roger Paredes, Eulalia Grau, Bonaventura Clotet |
| EPI_ISL_2530391, EPI_ISL_2530392, EPI_ISL_2530394 | Hematopathology Laboratory, ACTREC, TMC | Hematopathology Laboratory, ACTREC, TMC | Hematopathology Laboratory, ACTREC |
| EPI_ISL_2531827 | Hospital | National Reference Center for Viruses of Respiratory Infections, Institut Pasteur, Paris | Marion Barbet, Sylvie Behillil, Méline Bizard, Angela Brisebarre, Camille Capel, Vincent Enouf, Louise Lefrançois, Frédéric Lemoine, Christophe Malabat, Corinne Maufrais, Etienne Simon-Lorière, Maud Vanpeene, Sylvie Van der Werf ,Pascale Martres |
| EPI_ISL_2531834, EPI_ISL_2531839, EPI_ISL_2531843, EPI_ISL_2531851 | Hospital | National Reference Center for Viruses of Respiratory Infections, Institut Pasteur, Paris | Marion Barbet, Sylvie Behillil, Méline Bizard, Angela Brisebarre, Camille Capel, Vincent Enouf, Louise Lefrançois, Frédéric Lemoine, Christophe Malabat, Corinne Maufrais, Etienne Simon-Lorière, Maud Vanpeene, Sylvie Van der Werf ,Raphael Guiheneuf |
| EPI_ISL_2531877 | Hospital | National Reference Center for Viruses of Respiratory Infections, Institut Pasteur, Paris | Marion Barbet, Sylvie Behillil, Méline Bizard, Angela Brisebarre, Camille Capel, Vincent Enouf, Louise Lefrançois, Frédéric Lemoine, Christophe Malabat, Corinne Maufrais, Etienne Simon-Lorière, Maud Vanpeene, Sylvie Van der Werf ,Marianne Burgard |
| EPI_ISL_2531881, EPI_ISL_2531885, EPI_ISL_2531889, EPI_ISL_2531894, EPI_ISL_2531897, EPI_ISL_2531901, EPI_ISL_2531905, EPI_ISL_2531908, EPI_ISL_2531912, EPI_ISL_2531917, EPI_ISL_2531922, EPI_ISL_2531926, EPI_ISL_2531930, EPI_ISL_2531934, EPI_ISL_2531939, EPI_ISL_2531940, EPI_ISL_2531942, EPI_ISL_2531943 | | | |
| see above | RIIP | National Reference Center for Viruses of Respiratory Infections, Institut Pasteur, Paris | Marion Barbet, Sylvie Behillil, Méline Bizard, Angela Brisebarre, Camille Capel, Vincent Enouf, Louise Lefrançois, Frédéric Lemoine, Christophe Malabat, Corinne Maufrais, Etienne Simon-Lorière, Maud Vanpeene, Sylvie Van der Werf ,StéPhanie Guyomard-Rabenirina |
| EPI_ISL_2531945, EPI_ISL_2531947, EPI_ISL_2531948, EPI_ISL_2531953 | Labo Analyses Med | National Reference Center for Viruses of Respiratory Infections, Institut Pasteur, Paris | Marion Barbet, Sylvie Behillil, Méline Bizard, Angela Brisebarre, Camille Capel, Vincent Enouf, Louise Lefrançois, Frédéric Lemoine, Christophe Malabat, Corinne Maufrais, Etienne Simon-Lorière, Maud Vanpeene, Sylvie Van der Werf ,Hugues Leroy |
| EPI_ISL_2531958, EPI_ISL_2531962 | Labo Analyses Med | National Reference Center for Viruses of Respiratory Infections, Institut Pasteur, Paris | Marion Barbet, Sylvie Behillil, Méline Bizard, Angela Brisebarre, Camille Capel, Vincent Enouf, Louise Lefrançois, Frédéric Lemoine, Christophe Malabat, Corinne Maufrais, Etienne Simon-Lorière, Maud Vanpeene, Sylvie Van der Werf ,Vicky Le |
| EPI_ISL_2532193 | Hospital | National Reference Center for Viruses of Respiratory Infections, Institut Pasteur, Paris | Marion Barbet, Sylvie Behillil, Méline Bizard, Angela Brisebarre, Camille Capel, Vincent Enouf, Louise Lefrançois, Frédéric Lemoine, Christophe Malabat, Corinne Maufrais, Emmanuelle Permal, Etienne Simon-Lorière, Maud Vanpeene, Sylvie Van der Werf ,Eric Farfour |
| EPI_ISL_2532279, EPI_ISL_2532280, EPI_ISL_2532281, EPI_ISL_2532282, EPI_ISL_2532283 | Labo Analyses Med | National Reference Center for Viruses of Respiratory Infections, Institut Pasteur, Paris | Marion Barbet, Sylvie Behillil, Méline Bizard, Angela Brisebarre, Camille Capel, Vincent Enouf, Louise Lefrançois, Frédéric Lemoine, Christophe Malabat, Corinne Maufrais, Etienne Simon-Lorière, Maud Vanpeene, Sylvie Van der Werf ,OphéLie Said-Delattre |
| EPI_ISL_2532284, EPI_ISL_2532285, EPI_ISL_2532286 | Labo Analyses Med | National Reference Center for Viruses of Respiratory Infections, Institut Pasteur, Paris | Marion Barbet, Sylvie Behillil, Méline Bizard, Angela Brisebarre, Camille Capel, Vincent Enouf, Louise Lefrançois, Frédéric Lemoine, Christophe Malabat, Corinne Maufrais, Etienne Simon-Lorière, Maud Vanpeene, Sylvie Van der Werf ,Eve Haguenoer |
| EPI_ISL_2532287, EPI_ISL_2532288 | Labo Analyses Med | National Reference Center for Viruses of Respiratory Infections, Institut Pasteur, Paris | Marion Barbet, Sylvie Behillil, Méline Bizard, Angela Brisebarre, Camille Capel, Vincent Enouf, Louise Lefrançois, Frédéric Lemoine, Christophe Malabat, Corinne Maufrais, Etienne Simon-Lorière, Maud Vanpeene, Sylvie Van der Werf ,GréGoire Potiron |
| EPI_ISL_2532291 | Labo Analyses Med | National Reference Center for Viruses of Respiratory Infections, Institut Pasteur, Paris | Marion Barbet, Sylvie Behillil, Méline Bizard, Angela Brisebarre, Camille Capel, Vincent Enouf, Louise Lefrançois, Frédéric Lemoine, Christophe Malabat, Corinne Maufrais, Emmanuelle Permal, Etienne Simon-Lorière, Maud Vanpeene, Sylvie Van der Werf ,BéNéDicte Roquebert |
| EPI_ISL_2532292 | Hospital | National Reference Center for Viruses of Respiratory Infections, Institut Pasteur, Paris | Marion Barbet, Sylvie Behillil, Méline Bizard, Angela Brisebarre, Camille Capel, Vincent Enouf, Louise Lefrançois, Frédéric Lemoine, Christophe Malabat, Corinne Maufrais, Emmanuelle Permal, Etienne Simon-Lorière, Maud Vanpeene, Sylvie Van der Werf ,CéLine Ramanantsoa |
| EPI_ISL_2532293 | Hospital | National Reference Center for Viruses of Respiratory Infections, Institut Pasteur, Paris | Marion Barbet, Sylvie Behillil, Méline Bizard, Angela Brisebarre, Camille Capel, Vincent Enouf, Louise Lefrançois, Frédéric Lemoine, Christophe Malabat, Corinne Maufrais, Amaury Vaysse, Etienne Simon-Lorière, Maud Vanpeene, Sylvie Van der Werf ,JéRome Guinard |
| EPI_ISL_2532295, EPI_ISL_2532296 | Hospital | National Reference Center for Viruses of Respiratory Infections, Institut Pasteur, Paris | Marion Barbet, Sylvie Behillil, Méline Bizard, Angela Brisebarre, Camille Capel, Vincent Enouf, Louise Lefrançois, Frédéric Lemoine, Christophe Malabat, Corinne Maufrais, Etienne Simon-Lorière, Maud Vanpeene, Sylvie Van der Werf ,Thibault Guinoiseau |
| EPI_ISL_2532300 | Hospital | National Reference Center for Viruses of Respiratory Infections, Institut Pasteur, Paris | Marion Barbet, Sylvie Behillil, Méline Bizard, Angela Brisebarre, Camille Capel, Vincent Enouf, Louise Lefrançois, Frédéric Lemoine, Christophe Malabat, Corinne Maufrais, Amaury Vaysse, Etienne Simon-Lorière, Maud Vanpeene, Sylvie Van der Werf ,Sylvaine Bastian |
| EPI_ISL_2532301 | Hospital | National Reference Center for Viruses of Respiratory Infections, Institut Pasteur, Paris | Marion Barbet, Sylvie Behillil, Méline Bizard, Angela Brisebarre, Camille Capel, Vincent Enouf, Louise Lefrançois, Frédéric Lemoine, Christophe Malabat, Corinne Maufrais, Etienne Simon-Lorière, Maud Vanpeene, Sylvie Van der Werf ,Sylvaine Bastian |
| EPI_ISL_2532303, EPI_ISL_2532304, EPI_ISL_2532305, EPI_ISL_2532307, EPI_ISL_2532308, EPI_ISL_2532309 | Hospital | National Reference Center for Viruses of Respiratory Infections, Institut Pasteur, Paris | Marion Barbet, Sylvie Behillil, Méline Bizard, Angela Brisebarre, Camille Capel, Vincent Enouf, Louise Lefrançois, Frédéric Lemoine, Christophe Malabat, Corinne Maufrais, Etienne Simon-Lorière, Maud Vanpeene, Sylvie Van der Werf ,FréDéRique Canis |
| EPI_ISL_2532311, EPI_ISL_2532312, EPI_ISL_2532313 | Hospital | National Reference Center for Viruses of Respiratory Infections, Institut Pasteur, Paris | Marion Barbet, Sylvie Behillil, Méline Bizard, Angela Brisebarre, Camille Capel, Vincent Enouf, Louise Lefrançois, Frédéric Lemoine, Christophe Malabat, Corinne Maufrais, Etienne Simon-Lorière, Maud Vanpeene, Sylvie Van der Werf ,Pascale Martres |
| EPI_ISL_2532315 | Hospital | National Reference Center for Viruses of Respiratory Infections, Institut Pasteur, Paris | Marion Barbet, Sylvie Behillil, Méline Bizard, Angela Brisebarre, Camille Capel, Vincent Enouf, Louise Lefrançois, Frédéric Lemoine, Christophe Malabat, Corinne Maufrais, Amaury Vaysse, Etienne Simon-Lorière, Maud Vanpeene, Sylvie Van der Werf ,Pascale Martres |
| EPI_ISL_2532339, EPI_ISL_2532347, EPI_ISL_2532350, EPI_ISL_2532351, EPI_ISL_2532352 | Labo Analyses Med | National Reference Center for Viruses of Respiratory Infections, Institut Pasteur, Paris | Marion Barbet, Sylvie Behillil, Méline Bizard, Angela Brisebarre, Camille Capel, Vincent Enouf, Louise Lefrançois, Frédéric Lemoine, Christophe Malabat, Corinne Maufrais, Etienne Simon-Lorière, Maud Vanpeene, Sylvie Van der Werf ,Ronan Pichard |
| EPI_ISL_2532353, EPI_ISL_2532354, EPI_ISL_2532355, EPI_ISL_2532356, EPI_ISL_2532357, EPI_ISL_2532358, EPI_ISL_2532359, EPI_ISL_2532360 | Hospital | National Reference Center for Viruses of Respiratory Infections, Institut Pasteur, Paris | Marion Barbet, Sylvie Behillil, Méline Bizard, Angela Brisebarre, Camille Capel, Vincent Enouf, Louise Lefrançois, Frédéric Lemoine, Christophe Malabat, Corinne Maufrais, Etienne Simon-Lorière, Maud Vanpeene, Sylvie Van der Werf ,CéLine Bressollette |
| EPI_ISL_2532361, EPI_ISL_2532362, EPI_ISL_2532363, EPI_ISL_2532364, | Labo Analyses Med | National Reference Center for Viruses of Respiratory Infections, Institut Pasteur, Paris | Marion Barbet, Sylvie Behillil, Méline Bizard, Angela Brisebarre, Camille Capel, Vincent Enouf, Louise Lefrançois, Frédéric Lemoine, Christophe Malabat, Corinne Maufrais, Etienne Simon-Lorière, Maud Vanpeene, Sylvie Van der Werf ,Hervé Dupont |

| | | | |
|---|---|---|---|
| EPI_ISL_2532365, EPI_ISL_2532366 | | | |
| EPI_ISL_2532367 | Labo Analyses Med | National Reference Center for Viruses of Respiratory Infections, Institut Pasteur, Paris | Marion Barbet, Sylvie Behillil, Méline Bizard, Angela Brisebarre, Camille Capel, Vincent Enouf, Louise Lefrançois, Frédéric Lemoine, Christophe Malabat, Corinne Maufrais, Victoire Baillet, Etienne Simon-Lorière, Maud Vanpeene, Sylvie Van der Werf ,Hervé Dupont |
| EPI_ISL_2532368, EPI_ISL_2532370, EPI_ISL_2532371, EPI_ISL_2532372, EPI_ISL_2532373, EPI_ISL_2532374, EPI_ISL_2532375, EPI_ISL_2532376, EPI_ISL_2532377, EPI_ISL_2532378 | Labo Analyses Med | National Reference Center for Viruses of Respiratory Infections, Institut Pasteur, Paris | Marion Barbet, Sylvie Behillil, Méline Bizard, Angela Brisebarre, Camille Capel, Vincent Enouf, Louise Lefrançois, Frédéric Lemoine, Christophe Malabat, Corinne Maufrais, Etienne Simon-Lorière, Maud Vanpeene, Sylvie Van der Werf ,Hervé Dupont |
| EPI_ISL_2532388, EPI_ISL_2532389 | Hospital | National Reference Center for Viruses of Respiratory Infections, Institut Pasteur, Paris | Marion Barbet, Sylvie Behillil, Méline Bizard, Angela Brisebarre, Camille Capel, Vincent Enouf, Louise Lefrançois, Frédéric Lemoine, Christophe Malabat, Corinne Maufrais, Etienne Simon-Lorière, Maud Vanpeene, Sylvie Van der Werf ,AgnèS Scanvic |
| EPI_ISL_2532397, EPI_ISL_2532398, EPI_ISL_2532399, EPI_ISL_2532400, EPI_ISL_2532401, EPI_ISL_2532402, EPI_ISL_2532403, EPI_ISL_2532404, EPI_ISL_2532405, EPI_ISL_2532406, EPI_ISL_2532408, EPI_ISL_2532409, EPI_ISL_2532410, EPI_ISL_2532411, EPI_ISL_2532412, EPI_ISL_2532413, EPI_ISL_2532414, EPI_ISL_2532415, EPI_ISL_2532416, EPI_ISL_2532417, EPI_ISL_2532418, EPI_ISL_2532419, EPI_ISL_2532420, EPI_ISL_2532421, EPI_ISL_2532422, EPI_ISL_2532423, EPI_ISL_2532424, EPI_ISL_2532425, EPI_ISL_2532427, EPI_ISL_2532428, EPI_ISL_2532429, EPI_ISL_2532430, EPI_ISL_2532433, EPI_ISL_2532434, EPI_ISL_2532435, EPI_ISL_2532436, EPI_ISL_2532437, EPI_ISL_2532438, EPI_ISL_2532440, EPI_ISL_2532441, EPI_ISL_2532442, EPI_ISL_2532443, EPI_ISL_2532444, EPI_ISL_2532445, EPI_ISL_2532446, EPI_ISL_2532447, EPI_ISL_2532448, EPI_ISL_2532449, EPI_ISL_2532450, EPI_ISL_2532451, EPI_ISL_2532452 | | | |
| see above | Labo Analyses Med | National Reference Center for Viruses of Respiratory Infections, Institut Pasteur, Paris | Marion Barbet, Sylvie Behillil, Méline Bizard, Angela Brisebarre, Camille Capel, Vincent Enouf, Louise Lefrançois, Frédéric Lemoine, Christophe Malabat, Corinne Maufrais, Etienne Simon-Lorière, Maud Vanpeene, Sylvie Van der Werf ,Pierre-Yves Leonard |
| EPI_ISL_2532453, EPI_ISL_2532454, EPI_ISL_2532456, EPI_ISL_2532457, EPI_ISL_2532458, EPI_ISL_2532459, EPI_ISL_2532460, EPI_ISL_2532461, EPI_ISL_2532462, EPI_ISL_2532463 | Labo Analyses Med | National Reference Center for Viruses of Respiratory Infections, Institut Pasteur, Paris | Marion Barbet, Sylvie Behillil, Méline Bizard, Angela Brisebarre, Camille Capel, Vincent Enouf, Louise Lefrançois, Frédéric Lemoine, Christophe Malabat, Corinne Maufrais, Etienne Simon-Lorière, Maud Vanpeene, Sylvie Van der Werf |
| EPI_ISL_2532464 | Labo Analyses Med | National Reference Center for Viruses of Respiratory Infections, Institut Pasteur, Paris | Marion Barbet, Sylvie Behillil, Méline Bizard, Angela Brisebarre, Camille Capel, Vincent Enouf, Louise Lefrançois, Frédéric Lemoine, Christophe Malabat, Corinne Maufrais, Adrien Pain, Etienne Simon-Lorière, Maud Vanpeene, Sylvie Van der Werf |
| EPI_ISL_2532465, EPI_ISL_2532466, EPI_ISL_2532468, EPI_ISL_2532469, EPI_ISL_2532471 | Labo Analyses Med | National Reference Center for Viruses of Respiratory Infections, Institut Pasteur, Paris | Marion Barbet, Sylvie Behillil, Méline Bizard, Angela Brisebarre, Camille Capel, Vincent Enouf, Louise Lefrançois, Frédéric Lemoine, Christophe Malabat, Corinne Maufrais, Etienne Simon-Lorière, Maud Vanpeene, Sylvie Van der Werf |
| EPI_ISL_2532729 | SIESP DIPARTIMENTO DI PREVENZIONE TERAMO | Istituto Zooprofilattico Sperimentale dell'Abruzzo e Molise "G. Caporale" | Lorusso A, Marcacci M, Di Domenico M, Ancora M, Curini V, Di Lollo Valeria, Mangone I, Rinaldi A, Delli Compagni E, Scialabba S, Caporale M, Di Pasquale A, Cammà C, Puglia I, Calistri P, Savini G |
| EPI_ISL_2533764, EPI_ISL_2533766 | LESP Queretaro | Instituto de Diagnostico y Referencia Epidemiologicos (INDRE) | Claudia Wong-Arambula, Abril Rodriguez-Maldonado, Vanessa Rivero-Arredondo, Ariadna Medina-Benitez, Joaquin Quiroz-Mercado, Sergio Rangel-Guerrero, Natividad Cruz-Ortiz, Tatiana Nunez-Garcia, Gisela Barrera-Badillo, Lucia Hernandez-Rivas, Irma Lopez-Martinez, Ernesto Ramirez-Gonzalez. |
| EPI_ISL_2533767 | LESP Jalisco | Instituto de Diagnostico y Referencia Epidemiologicos (INDRE) | Claudia Wong-Arambula, Abril Rodriguez-Maldonado, Vanessa Rivero-Arredondo, Ariadna Medina-Benitez, Joaquin Quiroz-Mercado, Sergio Rangel-Guerrero, Natividad Cruz-Ortiz, Tatiana Nunez-Garcia, Gisela Barrera-Badillo, Lucia Hernandez-Rivas, Irma Lopez-Martinez, Ernesto Ramirez-Gonzalez. |
| EPI_ISL_2533768, EPI_ISL_2533769 | LESP Tabasco | Instituto de Diagnostico y Referencia Epidemiologicos (INDRE) | Claudia Wong-Arambula, Abril Rodriguez-Maldonado, Vanessa Rivero-Arredondo, Ariadna Medina-Benitez, Joaquin Quiroz-Mercado, Sergio Rangel-Guerrero, Natividad Cruz-Ortiz, Tatiana Nunez-Garcia, Gisela Barrera-Badillo, Lucia Hernandez-Rivas, Irma Lopez-Martinez, Ernesto Ramirez-Gonzalez. |
| EPI_ISL_2533784, EPI_ISL_2533785, EPI_ISL_2533786, EPI_ISL_2533787, EPI_ISL_2533788, EPI_ISL_2533789, EPI_ISL_2533790, EPI_ISL_2533791, EPI_ISL_2533792, EPI_ISL_2533793, EPI_ISL_2533794, EPI_ISL_2533795, EPI_ISL_2533796 | | | |
| see above | LESP Yucatan | Instituto de Diagnostico y Referencia Epidemiologicos (INDRE) | Claudia Wong-Arambula, Abril Rodriguez-Maldonado, Vanessa Rivero-Arredondo, Ariadna Medina-Benitez, Joaquin Quiroz-Mercado, Sergio Rangel-Guerrero, Natividad Cruz-Ortiz, Tatiana Nunez-Garcia, Gisela Barrera-Badillo, Lucia Hernandez-Rivas, Irma Lopez-Martinez, Ernesto Ramirez-Gonzalez. |
| EPI_ISL_2533801 | LESP Tamaulipas | Instituto de Diagnostico y Referencia Epidemiologicos (INDRE) | Claudia Wong-Arambula, Abril Rodriguez-Maldonado, Vanessa Rivero-Arredondo, Ariadna Medina-Benitez, Joaquin Quiroz-Mercado, Sergio Rangel-Guerrero, Natividad Cruz-Ortiz, Tatiana Nunez-Garcia, Gisela Barrera-Badillo, Lucia Hernandez-Rivas, Irma Lopez-Martinez, Ernesto Ramirez-Gonzalez. |
| EPI_ISL_2533803 | LESP Coahuila | Instituto de Diagnostico y Referencia Epidemiologicos (INDRE) | Claudia Wong-Arambula, Abril Rodriguez-Maldonado, Vanessa Rivero-Arredondo, Ariadna Medina-Benitez, Joaquin Quiroz-Mercado, Sergio Rangel-Guerrero, Natividad Cruz-Ortiz, Tatiana Nunez-Garcia, Gisela Barrera-Badillo, Lucia Hernandez-Rivas, Irma Lopez-Martinez, Ernesto Ramirez-Gonzalez. |
| EPI_ISL_2533804, EPI_ISL_2533805 | LESP Quintana Roo | Instituto de Diagnostico y Referencia Epidemiologicos (INDRE) | Claudia Wong-Arambula, Abril Rodriguez-Maldonado, Vanessa Rivero-Arredondo, Ariadna Medina-Benitez, Joaquin Quiroz-Mercado, Sergio Rangel-Guerrero, Natividad Cruz-Ortiz, Tatiana Nunez-Garcia, Gisela Barrera-Badillo, Lucia Hernandez-Rivas, Irma Lopez-Martinez, Ernesto Ramirez-Gonzalez. |
| EPI_ISL_2533817 | LESP Ciudad de Mexico | Instituto de Diagnostico y Referencia Epidemiologicos (INDRE) | Claudia Wong-Arambula, Abril Rodriguez-Maldonado, Vanessa Rivero-Arredondo, Ariadna Medina-Benitez, Joaquin Quiroz-Mercado, Sergio Rangel-Guerrero, Natividad Cruz-Ortiz, Tatiana Nunez-Garcia, Gisela Barrera-Badillo, Lucia Hernandez-Rivas, Irma Lopez-Martinez, Ernesto Ramirez-Gonzalez. |
| EPI_ISL_2533932 | CIAD-Hermosillo | Immunology Laboratory | Consorcio Mexicano de Vigilancia Genómica (CoViGen-Mex). Authors (in alphabetical order): Julio Elias Alvarado-Yaah, Carlos F. Arias, Santiago Ávila-Ríos, Eduardo Becerril-Vargas, Víctor Hugo Borja-Aburto, Celia Boukadida, Cristóbal Cháidez-Quiróz, Juan Bautista Chale-Dzul, Ricardo Ciria Merce, Andreu Comas-García, Célida Duque Molina, Julissa Enciso-Ibarra, José Antonio Enciso-Moreno, Gloria Elena Espinosa-Ayala, Fernando Fontove-Herrera, Daniel Fregoso-Rueda, Victor Eduardo García-Arias, Alejandra García-Gasca, Bruno Gómez-Gil, Jean Pierre González, Irvin González-López, Concepción Grajales-Muñiz, Ricardo Grande, Rosa María Gutiérrez Rios, Jesús Hernández, Alejandra Hernández-Terán, Alfredo Herrera-Estrella, Carla Ivón Herrera-Najera, Pavel Isa, Verónica Mata-Haro, Daniel Lira Morales, Susana Lopez, Antonio Loza Román, Brenda Irasema Maldonado-Meza, Bernardo Martínez-Miguel, José Arturo Martínez-Orozco, Célida Martínez- Rodríguez, Margarita Matías-Florentino, Fidencio Mejía-Nepomuceno, María Guadalupe de Jesús Mireles-Rivera, Gloria María Molina-Salinas, Hector Montoya-Fuentes, , Mario Mújica-Sánchez, José Esteban Muñoz-Medina, José de Jesús Nuñez-Contreras, Alicia Ocaña-Mondragón, Luis Alberto Ochoa-Carrera, Hector Esteban Paz-Juárez, Marissa Perez-Garcia, Francisco Pulido, Helen Haydee Fernanda Ramírez-Plascencia, Jorge Salas-Hernández, Angel Gustavo Salas-Lais, Alejandro Sánchez-Flores, Clara Esperanza Santacruz-Tinoco, María Guadalupe Santiago-Mauricio, Selene Zárate, Nelly Sélem-Mojica, Blanca Taboada, Gloria Vazquez, Joel Armando Vázquez-Pérez. |
| EPI_ISL_2534079 | RSUD dr. LOEKMONO HADI | Genetics Working Group (Pokja Genetik) Faculty of Medicine, Public Health and Nursing Universitas Gadjah Mada (FK-KMK UGM); Disease Investigation Center Wates Ministry of Agriculture Indonesia; Department of Microbiology FK-KMK UGM; Laboratorium Diagnostik Yayasan Tahija World Mosquito Program (WMP) Yogyakarta Center for Tropical Medicine FK-KMK UGM; Integrated Research Center FK-KMK UGM; Department of Computer Science and Electronics FMIPA UGM; RSUP Dr. Sardjito | Gunadi, Hendra Wibawa, Marcellus, Mohamad S. Hakim, Edwin W. Daniwijaya, Ludhang P. Rizki, Endah Supriyati, Eggi Arguni, Titik Nuryastuti, Tri Wibawa, Dwi AA Nugrahaningsih, Afiahayati, Siswanto,Kristy Iskandar,Nungki Anggorowati, Pramesti G Dewi, Dwi Indaryati,Indarto Sulistiyono, Susan Simanjaya, Dyah A Puspitarani |
| EPI_ISL_2534313, EPI_ISL_2534314, EPI_ISL_2534315 | The Ohio State University Applied Microbiology Services Laboratory | The Ohio State University Applied Microbiology Services Laboratory | Seth A. Faith PhD |
| EPI_ISL_2534379, EPI_ISL_2534457, | RSUD dr. LOEKMONO HADI | Genetics Working Group (Pokja Genetik) Faculty of Medicine, | Gunadi, Hendra Wibawa, Marcellus, Mohamad S. Hakim, Edwin W. Daniwijaya, Ludhang P. Rizki, Endah Supriyati, Eggi Arguni, Titik Nuryastuti, Tri |

| | | | |
|---|---|---|---|
| EPI_ISL_2534489 | | Public Health and Nursing Universitas Gadjah Mada (FK-KMK UGM); Disease Investigation Center Wates Ministry of Agriculture Indonesia; Department of Microbiology FK-KMK UGM; Laboratorium Diagnostik Yayasan Tahija World Mosquito Program (WMP) Yogyakarta Center for Tropical Medicine FK-KMK UGM; Integrated Research Center FK-KMK UGM; Department of Computer Science and Electronics FMIPA UGM; RSUP Dr. Sardjito | Wibawa, Dwi AA Nugrahaningsih, Afiahayati, Siswanto,Kristy Iskandar,Nungki Anggorowati, Pramesti G Dewi, Dwi Indaryati,Indarto Sulistiyono, Cita S Amalia, Alvina A Setiawan |
| EPI_ISL_2534490 | RSUD dr. LOEKMONO HADI | Genetics Working Group (Pokja Genetik) Faculty of Medicine, Public Health and Nursing Universitas Gadjah Mada (FK-KMK UGM); Disease Investigation Center Wates Ministry of Agriculture Indonesia; Department of Microbiology FK-KMK UGM; Laboratorium Diagnostik Yayasan Tahija World Mosquito Program (WMP) Yogyakarta Center for Tropical Medicine FK-KMK UGM; Integrated Research Center FK-KMK UGM; Department of Computer Science and Electronics FMIPA UGM; RSUP Dr. Sardjito | Gunadi, Hendra Wibawa, Marcellus, Mohamad S. Hakim, Edwin W. Daniwijaya, Ludhang P. Rizki, Endah Supriyati, Eggi Arguni, Titik Nuryastuti, Tri Wibawa, Dwi AA Nugrahaningsih, Afiahayati, Siswanto,Kristy Iskandar,Nungki Anggorowati, Pramesti G Dewi, Dwi Indaryati,Indarto Sulistiyono, Khanza a Vujira, Alvina A Setiawan |
| EPI_ISL_2534518 | RSUD dr. LOEKMONO HADI | Genetics Working Group (Pokja Genetik) Faculty of Medicine, Public Health and Nursing Universitas Gadjah Mada (FK-KMK UGM); Disease Investigation Center Wates Ministry of Agriculture Indonesia; Department of Microbiology FK-KMK UGM; Laboratorium Diagnostik Yayasan Tahija World Mosquito Program (WMP) Yogyakarta Center for Tropical Medicine FK-KMK UGM; Integrated Research Center FK-KMK UGM; Department of Computer Science and Electronics FMIPA UGM; RSUP Dr. Sardjito | Gunadi, Hendra Wibawa, Marcellus, Mohamad S. Hakim, Edwin W. Daniwijaya, Ludhang P. Rizki, Endah Supriyati, Eggi Arguni, Titik Nuryastuti, Tri Wibawa, Dwi AA Nugrahaningsih, Afiahayati, Siswanto,Kristy Iskandar,Nungki Anggorowati, Pramesti G Dewi, Dwi Indaryati,Indarto Sulistiyono, Susan Simanjaya, Dyah A Puspitarani |
| EPI_ISL_2535237, EPI_ISL_2535238, EPI_ISL_2535239, EPI_ISL_2535240, EPI_ISL_2535241 | Labo Analyses Med | National Reference Center for Viruses of Respiratory Infections, Institut Pasteur, Paris | Marion Barbet, Sylvie Behillil, Méline Bizard, Angela Brisebarre, Camille Capel, Vincent Enouf, Louise Lefrançois, Frédéric Lemoine, Christophe Malabat, Corinne Maufrais, Etienne Simon-Lorière, Maud Vanpeene, Sylvie Van der Werf |
| EPI_ISL_2535242 | Hospital | National Reference Center for Viruses of Respiratory Infections, Institut Pasteur, Paris | Marion Barbet, Sylvie Behillil, Méline Bizard, Angela Brisebarre, Camille Capel, Vincent Enouf, Louise Lefrançois, Frédéric Lemoine, Christophe Malabat, Corinne Maufrais, Etienne Simon-Lorière, Maud Vanpeene, Sylvie Van der Werf ,JéRome Guinard |
| EPI_ISL_2535244, EPI_ISL_2535245 | Labo Analyses Med | National Reference Center for Viruses of Respiratory Infections, Institut Pasteur, Paris | Marion Barbet, Sylvie Behillil, Méline Bizard, Angela Brisebarre, Camille Capel, Vincent Enouf, Louise Lefrançois, Frédéric Lemoine, Christophe Malabat, Corinne Maufrais, Etienne Simon-Lorière, Maud Vanpeene, Sylvie Van der Werf ,Nouredine Sadeg |
| EPI_ISL_2535247, EPI_ISL_2535248, EPI_ISL_2535249, EPI_ISL_2535250, EPI_ISL_2535251 | Labo Analyses Med | National Reference Center for Viruses of Respiratory Infections, Institut Pasteur, Paris | Marion Barbet, Sylvie Behillil, Méline Bizard, Angela Brisebarre, Camille Capel, Vincent Enouf, Louise Lefrançois, Frédéric Lemoine, Christophe Malabat, Corinne Maufrais, Etienne Simon-Lorière, Maud Vanpeene, Sylvie Van der Werf ,V. Bourdrel |
| EPI_ISL_2535252, EPI_ISL_2535253, EPI_ISL_2535254, EPI_ISL_2535255, EPI_ISL_2535256, EPI_ISL_2535257, EPI_ISL_2535258, EPI_ISL_2535259 | Labo Analyses Med | National Reference Center for Viruses of Respiratory Infections, Institut Pasteur, Paris | Marion Barbet, Sylvie Behillil, Méline Bizard, Angela Brisebarre, Camille Capel, Vincent Enouf, Louise Lefrançois, Frédéric Lemoine, Christophe Malabat, Corinne Maufrais, Etienne Simon-Lorière, Maud Vanpeene, Sylvie Van der Werf |
| EPI_ISL_2535260, EPI_ISL_2535262, EPI_ISL_2535264 | Labo Analyses Med | National Reference Center for Viruses of Respiratory Infections, Institut Pasteur, Paris | Marion Barbet, Sylvie Behillil, Méline Bizard, Angela Brisebarre, Camille Capel, Vincent Enouf, Louise Lefrançois, Frédéric Lemoine, Christophe Malabat, Corinne Maufrais, Etienne Simon-Lorière, Maud Vanpeene, Sylvie Van der Werf ,GréGoire Potiron |
| EPI_ISL_2535265 | Labo Analyses Med | National Reference Center for Viruses of Respiratory Infections, Institut Pasteur, Paris | Marion Barbet, Sylvie Behillil, Méline Bizard, Angela Brisebarre, Camille Capel, Vincent Enouf, Louise Lefrançois, Frédéric Lemoine, Christophe Malabat, Corinne Maufrais, Etienne Simon-Lorière, Maud Vanpeene, Sylvie Van der Werf ,Hugues Leroy |
| EPI_ISL_2535266, EPI_ISL_2535267, EPI_ISL_2535269, EPI_ISL_2535270 | Hospital | National Reference Center for Viruses of Respiratory Infections, Institut Pasteur, Paris | Marion Barbet, Sylvie Behillil, Méline Bizard, Angela Brisebarre, Camille Capel, Vincent Enouf, Louise Lefrançois, Frédéric Lemoine, Christophe Malabat, Corinne Maufrais, Etienne Simon-Lorière, Maud Vanpeene, Sylvie Van der Werf ,Olivia Raulin |
| EPI_ISL_2535271 | Hospital | National Reference Center for Viruses of Respiratory Infections, Institut Pasteur, Paris | Marion Barbet, Sylvie Behillil, Méline Bizard, Angela Brisebarre, Camille Capel, Vincent Enouf, Louise Lefrançois, Frédéric Lemoine, Christophe Malabat, Corinne Maufrais, Damien Mornico, Etienne Simon-Lorière, Maud Vanpeene, Sylvie Van der Werf ,Pascale Martres |
| EPI_ISL_2535272, EPI_ISL_2535274 | Hospital | National Reference Center for Viruses of Respiratory Infections, Institut Pasteur, Paris | Marion Barbet, Sylvie Behillil, Méline Bizard, Angela Brisebarre, Camille Capel, Vincent Enouf, Louise Lefrançois, Frédéric Lemoine, Christophe Malabat, Corinne Maufrais, Etienne Simon-Lorière, Maud Vanpeene, Sylvie Van der Werf ,Pascale Martres |
| EPI_ISL_2535275 | Labo Analyses Med | National Reference Center for Viruses of Respiratory Infections, Institut Pasteur, Paris | Marion Barbet, Sylvie Behillil, Méline Bizard, Angela Brisebarre, Camille Capel, Vincent Enouf, Louise Lefrançois, Frédéric Lemoine, Christophe Malabat, Corinne Maufrais, Etienne Simon-Lorière, Maud Vanpeene, Sylvie Van der Werf ,OphéLie Said-Delattre |
| EPI_ISL_2535276 | Labo Analyses Med | National Reference Center for Viruses of Respiratory Infections, Institut Pasteur, Paris | Marion Barbet, Sylvie Behillil, Méline Bizard, Angela Brisebarre, Camille Capel, Vincent Enouf, Louise Lefrançois, Frédéric Lemoine, Christophe Malabat, Corinne Maufrais, Victoire Baillet, Etienne Simon-Lorière, Maud Vanpeene, Sylvie Van der Werf ,OphéLie Said-Delattre |
| EPI_ISL_2535277, EPI_ISL_2535278, EPI_ISL_2535279, EPI_ISL_2535280, EPI_ISL_2535281, EPI_ISL_2535282, EPI_ISL_2535283, EPI_ISL_2535284, EPI_ISL_2535285, EPI_ISL_2535286, EPI_ISL_2535287, EPI_ISL_2535288, EPI_ISL_2535289, EPI_ISL_2535290, EPI_ISL_2535291, EPI_ISL_2535295 | | | |
| see above | Labo Analyses Med | National Reference Center for Viruses of Respiratory Infections, Institut Pasteur, Paris | Marion Barbet, Sylvie Behillil, Méline Bizard, Angela Brisebarre, Camille Capel, Vincent Enouf, Louise Lefrançois, Frédéric Lemoine, Christophe Malabat, Corinne Maufrais, Etienne Simon-Lorière, Maud Vanpeene, Sylvie Van der Werf ,OphéLie Said-Delattre |
| EPI_ISL_2535296 | Labo Analyses Med | National Reference Center for Viruses of Respiratory Infections, Institut Pasteur, Paris | Marion Barbet, Sylvie Behillil, Méline Bizard, Angela Brisebarre, Camille Capel, Vincent Enouf, Louise Lefrançois, Frédéric Lemoine, Christophe Malabat, Corinne Maufrais, Victoire Baillet, Etienne Simon-Lorière, Maud Vanpeene, Sylvie Van der Werf ,OphéLie Said-Delattre |
| EPI_ISL_2535297, EPI_ISL_2535299, EPI_ISL_2535300, EPI_ISL_2535301 | Labo Analyses Med | National Reference Center for Viruses of Respiratory Infections, Institut Pasteur, Paris | Marion Barbet, Sylvie Behillil, Méline Bizard, Angela Brisebarre, Camille Capel, Vincent Enouf, Louise Lefrançois, Frédéric Lemoine, Christophe Malabat, Corinne Maufrais, Etienne Simon-Lorière, Maud Vanpeene, Sylvie Van der Werf ,OphéLie Said-Delattre |
| EPI_ISL_2535436, EPI_ISL_2535437, EPI_ISL_2535438, EPI_ISL_2535439, EPI_ISL_2535440, EPI_ISL_2535441, EPI_ISL_2535442, EPI_ISL_2535443, EPI_ISL_2535444 | Hospital | National Reference Center for Viruses of Respiratory Infections, Institut Pasteur, Paris | Marion Barbet, Sylvie Behillil, Méline Bizard, Angela Brisebarre, Camille Capel, Vincent Enouf, Louise Lefrançois, Frédéric Lemoine, Christophe Malabat, Corinne Maufrais, Etienne Simon-Lorière, Maud Vanpeene, Sylvie Van der Werf ,Nabil Gastli |
| EPI_ISL_2535445, EPI_ISL_2535446, EPI_ISL_2535447, EPI_ISL_2535448, EPI_ISL_2535449, EPI_ISL_2535450, EPI_ISL_2535451, EPI_ISL_2535452, EPI_ISL_2535453, EPI_ISL_2535454, EPI_ISL_2535455, EPI_ISL_2535456, EPI_ISL_2535457, EPI_ISL_2535458, EPI_ISL_2535459, EPI_ISL_2535460, EPI_ISL_2535461, EPI_ISL_2535462 | | | |
| see above | Labo Analyses Med | National Reference Center for Viruses of Respiratory Infections, Institut Pasteur, Paris | Marion Barbet, Sylvie Behillil, Méline Bizard, Angela Brisebarre, Camille Capel, Vincent Enouf, Louise Lefrançois, Frédéric Lemoine, Christophe Malabat, Corinne Maufrais, Etienne Simon-Lorière, Maud Vanpeene, Sylvie Van der Werf ,Claire Felloni |
| EPI_ISL_2535463 | Labo Analyses Med | National Reference Center for Viruses of Respiratory Infections, Institut Pasteur, Paris | Marion Barbet, Sylvie Behillil, Méline Bizard, Angela Brisebarre, Camille Capel, Vincent Enouf, Louise Lefrançois, Frédéric Lemoine, Christophe Malabat, Corinne Maufrais, Etienne Simon-Lorière, Maud Vanpeene, Sylvie Van der Werf ,Jean-Baptiste Foulquier |
| EPI_ISL_2535464, EPI_ISL_2535466, EPI_ISL_2535467, EPI_ISL_2535468, EPI_ISL_2535469, EPI_ISL_2535470 | Labo Analyses Med | National Reference Center for Viruses of Respiratory Infections, Institut Pasteur, Paris | Marion Barbet, Sylvie Behillil, Méline Bizard, Angela Brisebarre, Camille Capel, Vincent Enouf, Louise Lefrançois, Frédéric Lemoine, Christophe Malabat, Corinne Maufrais, Etienne Simon-Lorière, Maud Vanpeene, Sylvie Van der Werf ,Benedicte Baccouch |

| | | | |
|---|---|---|---|
| EPI_ISL_2535476, EPI_ISL_2535477, EPI_ISL_2535478 | Hospital | National Reference Center for Viruses of Respiratory Infections, Institut Pasteur, Paris | Marion Barbet, Sylvie Behillil, Méline Bizard, Angela Brisebarre, Camille Capel, Vincent Enouf, Louise Lefrançois, Frédéric Lemoine, Christophe Malabat, Corinne Maufrais, Etienne Simon-Lorière, Maud Vanpeene, Sylvie Van der Werf ,LéA Pilorge |
| EPI_ISL_2535480, EPI_ISL_2535481 | Hospital | National Reference Center for Viruses of Respiratory Infections, Institut Pasteur, Paris | Marion Barbet, Sylvie Behillil, Méline Bizard, Angela Brisebarre, Camille Capel, Vincent Enouf, Louise Lefrançois, Frédéric Lemoine, Christophe Malabat, Corinne Maufrais, Etienne Simon-Lorière, Maud Vanpeene, Sylvie Van der Werf ,Alexandra Ducancelle |
| EPI_ISL_2535483, EPI_ISL_2535484, EPI_ISL_2535485, EPI_ISL_2535486, EPI_ISL_2535487, EPI_ISL_2535488, EPI_ISL_2535489, EPI_ISL_2535490, EPI_ISL_2535491, EPI_ISL_2535492, EPI_ISL_2535493, EPI_ISL_2535495, EPI_ISL_2535496 | | | |
| see above | Labo Analyses Med | National Reference Center for Viruses of Respiratory Infections, Institut Pasteur, Paris | Marion Barbet, Sylvie Behillil, Méline Bizard, Angela Brisebarre, Camille Capel, Vincent Enouf, Louise Lefrançois, Frédéric Lemoine, Christophe Malabat, Corinne Maufrais, Etienne Simon-Lorière, Maud Vanpeene, Sylvie Van der Werf , Hugues Leroy |
| EPI_ISL_2535497 | Labo Analyses Med | National Reference Center for Viruses of Respiratory Infections, Institut Pasteur, Paris | Marion Barbet, Sylvie Behillil, Méline Bizard, Angela Brisebarre, Camille Capel, Vincent Enouf, Louise Lefrançois, Frédéric Lemoine, Christophe Malabat, Corinne Maufrais, Emmanulelle Permal, Etienne Simon-Lorière, Maud Vanpeene, Sylvie Van der Werf , Hugues Leroy |
| EPI_ISL_2535498, EPI_ISL_2535499, EPI_ISL_2535500, EPI_ISL_2535501, EPI_ISL_2535502, EPI_ISL_2535503, EPI_ISL_2535504, EPI_ISL_2535505, EPI_ISL_2535506, EPI_ISL_2535507, EPI_ISL_2535508, EPI_ISL_2535509, EPI_ISL_2535510, EPI_ISL_2535511, EPI_ISL_2535512, EPI_ISL_2535513, EPI_ISL_2535514, EPI_ISL_2535515, EPI_ISL_2535516, EPI_ISL_2535517, EPI_ISL_2535518, EPI_ISL_2535520, EPI_ISL_2535521, EPI_ISL_2535522, EPI_ISL_2535523, EPI_ISL_2535524, EPI_ISL_2535525, EPI_ISL_2535526, EPI_ISL_2535527, EPI_ISL_2535528, EPI_ISL_2535529, EPI_ISL_2535530, EPI_ISL_2535532, EPI_ISL_2535533, EPI_ISL_2535534, EPI_ISL_2535535, EPI_ISL_2535536, EPI_ISL_2535537, EPI_ISL_2535538, EPI_ISL_2535539, EPI_ISL_2535540, EPI_ISL_2535541, EPI_ISL_2535542, EPI_ISL_2535543, EPI_ISL_2535544, EPI_ISL_2535545, EPI_ISL_2535546, EPI_ISL_2535547, EPI_ISL_2535548, EPI_ISL_2535549, EPI_ISL_2535550, EPI_ISL_2535551, EPI_ISL_2535552, EPI_ISL_2535553, EPI_ISL_2535554 | | | |
| see above | Labo Analyses Med | National Reference Center for Viruses of Respiratory Infections, Institut Pasteur, Paris | Marion Barbet, Sylvie Behillil, Méline Bizard, Angela Brisebarre, Camille Capel, Vincent Enouf, Louise Lefrançois, Frédéric Lemoine, Christophe Malabat, Corinne Maufrais, Etienne Simon-Lorière, Maud Vanpeene, Sylvie Van der Werf , Hugues Leroy |
| EPI_ISL_2535555 | Hospital | National Reference Center for Viruses of Respiratory Infections, Institut Pasteur, Paris | Marion Barbet, Sylvie Behillil, Méline Bizard, Angela Brisebarre, Camille Capel, Vincent Enouf, Louise Lefrançois, Frédéric Lemoine, Christophe Malabat, Corinne Maufrais, Etienne Simon-Lorière, Maud Vanpeene, Sylvie Van der Werf ,Isabelle Woercel |
| EPI_ISL_2535556 | Labo Analyses Med | National Reference Center for Viruses of Respiratory Infections, Institut Pasteur, Paris | Marion Barbet, Sylvie Behillil, Méline Bizard, Angela Brisebarre, Camille Capel, Vincent Enouf, Louise Lefrançois, Frédéric Lemoine, Christophe Malabat, Corinne Maufrais, Etienne Simon-Lorière, Maud Vanpeene, Sylvie Van der Werf ,Isabelle Woercel |
| EPI_ISL_2535557, EPI_ISL_2535558, EPI_ISL_2535559 | Hospital | National Reference Center for Viruses of Respiratory Infections, Institut Pasteur, Paris | Marion Barbet, Sylvie Behillil, Méline Bizard, Angela Brisebarre, Camille Capel, Vincent Enouf, Louise Lefrançois, Frédéric Lemoine, Christophe Malabat, Corinne Maufrais, Etienne Simon-Lorière, Maud Vanpeene, Sylvie Van der Werf ,CéCile Farrugia |
| EPI_ISL_2535560, EPI_ISL_2535561, EPI_ISL_2535562, EPI_ISL_2535563, EPI_ISL_2535564 | Labo Analyses Med | National Reference Center for Viruses of Respiratory Infections, Institut Pasteur, Paris | Marion Barbet, Sylvie Behillil, Méline Bizard, Angela Brisebarre, Camille Capel, Vincent Enouf, Louise Lefrançois, Frédéric Lemoine, Christophe Malabat, Corinne Maufrais, Etienne Simon-Lorière, Maud Vanpeene, Sylvie Van der Werf ,Benedicte Baccouch |
| EPI_ISL_2535566, EPI_ISL_2535567, EPI_ISL_2535568, EPI_ISL_2535569, EPI_ISL_2535570, EPI_ISL_2535571, EPI_ISL_2535572, EPI_ISL_2535573, EPI_ISL_2535574, EPI_ISL_2535575, EPI_ISL_2535576, EPI_ISL_2535577, EPI_ISL_2535578, EPI_ISL_2535579, EPI_ISL_2535580, EPI_ISL_2535581, EPI_ISL_2535582, EPI_ISL_2535583, EPI_ISL_2535584, EPI_ISL_2535585, EPI_ISL_2535586, EPI_ISL_2535587, EPI_ISL_2535588 | | | |
| see above | Labo Analyses Med | National Reference Center for Viruses of Respiratory Infections, Institut Pasteur, Paris | Marion Barbet, Sylvie Behillil, Méline Bizard, Angela Brisebarre, Camille Capel, Vincent Enouf, Louise Lefrançois, Frédéric Lemoine, Christophe Malabat, Corinne Maufrais, Etienne Simon-Lorière, Maud Vanpeene, Sylvie Van der Werf ,Arnaud Serpentini |
| EPI_ISL_2535589, EPI_ISL_2535590, EPI_ISL_2535591, EPI_ISL_2535592, EPI_ISL_2535593, EPI_ISL_2535594, EPI_ISL_2535595, EPI_ISL_2535596, EPI_ISL_2535597, EPI_ISL_2535598, EPI_ISL_2535599, EPI_ISL_2535600, EPI_ISL_2535601 | | | |
| see above | Labo Analyses Med | National Reference Center for Viruses of Respiratory Infections, Institut Pasteur, Paris | Marion Barbet, Sylvie Behillil, Méline Bizard, Angela Brisebarre, Camille Capel, Vincent Enouf, Louise Lefrançois, Frédéric Lemoine, Christophe Malabat, Corinne Maufrais, Etienne Simon-Lorière, Maud Vanpeene, Sylvie Van der Werf ,Ronan Pichard |
| EPI_ISL_2544528, EPI_ISL_2544529 | Basurto University Hospital: Clinical Microbiology Laboratory | Biocruces Bizkaia | Mikel Urrutikoetxea-Gutiérrez, Mª Carmen Nieto Toboso, Estibaliz Ugalde Zarraga, Mikel Gallego Rodrigo, Ana de la Hoz, José Luis Díaz de Tuesta del Arco |
| EPI_ISL_2544744, EPI_ISL_2544745, EPI_ISL_2544747 | Laboratoire Hôpital Sainte Musse | Laboratoire Hôpital Sainte Musse | Lionel Chollet |
| EPI_ISL_2544899, EPI_ISL_2544900, EPI_ISL_2544901, EPI_ISL_2544902, EPI_ISL_2544903, EPI_ISL_2544904, EPI_ISL_2544905, EPI_ISL_2544906, EPI_ISL_2544907, EPI_ISL_2544908, EPI_ISL_2544909, EPI_ISL_2544910, EPI_ISL_2544912 | | | |
| see above | OLVZ Aalst | OLVZ Aalst | Anne Vankeerberghen |
| EPI_ISL_2545667 | Tripler Army Medical Center | Tripler Army Medical Center | Rachael L. Downham, Jonathan A. D'Ambrozio, Kendra N. Carter, Lindsey L. O'Neal, Zakariae Bram, and Catherine F. Uyehara |
| EPI_ISL_2545683 | LESP Jalisco | Instituto de Diagnostico y Referencia Epidemiologicos (INDRE) | Claudia Wong-Arambula, Abril Rodriguez-Maldonado, Vanessa Rivero-Arredondo, Ariadna Medina-Benitez, Joaquin Quiroz-Mercado, Sergio Rangel-Guerrero, Natividad Cruz-Ortiz, Tatiana Nunez-Garcia, Gisela Barrera-Badillo, Lucia Hernandez-Rivas, Irma Lopez-Martinez, Ernesto Ramirez-Gonzalez. |
| EPI_ISL_2545692 | LESP Sinaloa | Instituto de Diagnostico y Referencia Epidemiologicos (INDRE) | Claudia Wong-Arambula, Abril Rodriguez-Maldonado, Vanessa Rivero-Arredondo, Ariadna Medina-Benitez, Joaquin Quiroz-Mercado, Sergio Rangel-Guerrero, Natividad Cruz-Ortiz, Tatiana Nunez-Garcia, Gisela Barrera-Badillo, Lucia Hernandez-Rivas, Irma Lopez-Martinez, Ernesto Ramirez-Gonzalez. |
| EPI_ISL_2545693, EPI_ISL_2545694 | LESP Quintana Roo | Instituto de Diagnostico y Referencia Epidemiologicos (INDRE) | Claudia Wong-Arambula, Abril Rodriguez-Maldonado, Vanessa Rivero-Arredondo, Ariadna Medina-Benitez, Joaquin Quiroz-Mercado, Sergio Rangel-Guerrero, Natividad Cruz-Ortiz, Tatiana Nunez-Garcia, Gisela Barrera-Badillo, Lucia Hernandez-Rivas, Irma Lopez-Martinez, Ernesto Ramirez-Gonzalez. |
| EPI_ISL_2545703, EPI_ISL_2545704 | LESP Sinaloa | Instituto de Diagnostico y Referencia Epidemiologicos (INDRE) | Claudia Wong-Arambula, Abril Rodriguez-Maldonado, Vanessa Rivero-Arredondo, Ariadna Medina-Benitez, Joaquin Quiroz-Mercado, Sergio Rangel-Guerrero, Natividad Cruz-Ortiz, Tatiana Nunez-Garcia, Gisela Barrera-Badillo, Lucia Hernandez-Rivas, Irma Lopez-Martinez, Ernesto Ramirez-Gonzalez. |
| EPI_ISL_2545723 | LESP Estado de Mexico | Instituto de Diagnostico y Referencia Epidemiologicos (INDRE) | Claudia Wong-Arambula, Abril Rodriguez-Maldonado, Vanessa Rivero-Arredondo, Ariadna Medina-Benitez, Joaquin Quiroz-Mercado, Sergio Rangel-Guerrero, Natividad Cruz-Ortiz, Tatiana Nunez-Garcia, Gisela Barrera-Badillo, Lucia Hernandez-Rivas, Irma Lopez-Martinez, Ernesto Ramirez-Gonzalez. |
| EPI_ISL_2545732 | LESP Queretaro | Instituto de Diagnostico y Referencia Epidemiologicos (INDRE) | Claudia Wong-Arambula, Abril Rodriguez-Maldonado, Vanessa Rivero-Arredondo, Ariadna Medina-Benitez, Joaquin Quiroz-Mercado, Sergio Rangel-Guerrero, Natividad Cruz-Ortiz, Tatiana Nunez-Garcia, Gisela Barrera-Badillo, Lucia Hernandez-Rivas, Irma Lopez-Martinez, Ernesto Ramirez-Gonzalez. |
| EPI_ISL_2545782, EPI_ISL_2545783, EPI_ISL_2545784, EPI_ISL_2545785, EPI_ISL_2545786, EPI_ISL_2545787, EPI_ISL_2545788, EPI_ISL_2545790, EPI_ISL_2545791, EPI_ISL_2545792, EPI_ISL_2545793, EPI_ISL_2545794, EPI_ISL_2545795, EPI_ISL_2545796, EPI_ISL_2545797, EPI_ISL_2545798, EPI_ISL_2545799, EPI_ISL_2545800, EPI_ISL_2545801, EPI_ISL_2545802, EPI_ISL_2545803 | | | |
| see above | LESP Yucatan | Instituto de Diagnostico y Referencia Epidemiologicos (INDRE) | Claudia Wong-Arambula, Abril Rodriguez-Maldonado, Vanessa Rivero-Arredondo, Ariadna Medina-Benitez, Joaquin Quiroz-Mercado, Sergio Rangel-Guerrero, Natividad Cruz-Ortiz, Tatiana Nunez-Garcia, Gisela Barrera-Badillo, Lucia Hernandez-Rivas, Irma Lopez-Martinez, Ernesto Ramirez-Gonzalez. |
| EPI_ISL_2545805 | LESP Durango | Instituto de Diagnostico y Referencia Epidemiologicos (INDRE) | Claudia Wong-Arambula, Abril Rodriguez-Maldonado, Vanessa Rivero-Arredondo, Ariadna Medina-Benitez, Joaquin Quiroz-Mercado, Sergio Rangel-Guerrero, Natividad Cruz-Ortiz, Tatiana Nunez-Garcia, Gisela Barrera-Badillo, Lucia Hernandez-Rivas, Irma Lopez-Martinez, Ernesto Ramirez-Gonzalez. |
| EPI_ISL_2545806 | LESP Campeche | Instituto de Diagnostico y Referencia Epidemiologicos (INDRE) | Claudia Wong-Arambula, Abril Rodriguez-Maldonado, Vanessa Rivero-Arredondo, Ariadna Medina-Benitez, Joaquin Quiroz-Mercado, Sergio Rangel-Guerrero, Natividad Cruz-Ortiz, Tatiana Nunez-Garcia, Gisela Barrera-Badillo, Lucia Hernandez-Rivas, Irma Lopez-Martinez, Ernesto Ramirez-Gonzalez. |
| EPI_ISL_2545808, EPI_ISL_2545809, EPI_ISL_2545810, EPI_ISL_2545811 | LESP Quintana Roo | Instituto de Diagnostico y Referencia Epidemiologicos (INDRE) | Claudia Wong-Arambula, Abril Rodriguez-Maldonado, Vanessa Rivero-Arredondo, Ariadna Medina-Benitez, Joaquin Quiroz-Mercado, Sergio Rangel-Guerrero, Natividad Cruz-Ortiz, Tatiana Nunez-Garcia, Gisela Barrera-Badillo, Lucia Hernandez-Rivas, Irma Lopez-Martinez, Ernesto Ramirez-Gonzalez. |
| EPI_ISL_2556117, EPI_ISL_2556119, EPI_ISL_2557251, EPI_ISL_2557459 | MOH - Jaber Al-Ahmad Hospital (Innovation Research Laboratory) | MOH - Jaber Al-Ahmad Hospital (Innovation Research Laboratory) | Salman Al-Sabah , Mohammad Alghounaim |

| | | | |
|---|---|---|---|
| EPI_ISL_2559367, EPI_ISL_2559369, EPI_ISL_2559371, EPI_ISL_2559373 | LESP Queretaro | Instituto de Diagnostico y Referencia Epidemiologicos (INDRE) | Claudia Wong-Arambula, Abril Rodriguez-Maldonado, Vanessa Rivero-Arredondo, Ariadna Medina-Benitez, Joaquin Quiroz-Mercado, Sergio Rangel-Guerrero, Natividad Cruz-Ortiz, Tatiana Nunez-Garcia, Gisela Barrera-Badillo, Lucia Hernandez-Rivas, Irma Lopez-Martinez, Ernesto Ramirez-Gonzalez. |
| EPI_ISL_2559375, EPI_ISL_2559377 | LESP Tabasco | Instituto de Diagnostico y Referencia Epidemiologicos (INDRE) | Claudia Wong-Arambula, Abril Rodriguez-Maldonado, Vanessa Rivero-Arredondo, Ariadna Medina-Benitez, Joaquin Quiroz-Mercado, Sergio Rangel-Guerrero, Natividad Cruz-Ortiz, Tatiana Nunez-Garcia, Gisela Barrera-Badillo, Lucia Hernandez-Rivas, Irma Lopez-Martinez, Ernesto Ramirez-Gonzalez. |
| EPI_ISL_2559379 | LESP Veracruz | Instituto de Diagnostico y Referencia Epidemiologicos (INDRE) | Claudia Wong-Arambula, Abril Rodriguez-Maldonado, Vanessa Rivero-Arredondo, Ariadna Medina-Benitez, Joaquin Quiroz-Mercado, Sergio Rangel-Guerrero, Natividad Cruz-Ortiz, Tatiana Nunez-Garcia, Gisela Barrera-Badillo, Lucia Hernandez-Rivas, Irma Lopez-Martinez, Ernesto Ramirez-Gonzalez. |
| EPI_ISL_2559416, EPI_ISL_2559419, EPI_ISL_2559421, EPI_ISL_2559423 | LESP Yucatan | Instituto de Diagnostico y Referencia Epidemiologicos (INDRE) | Claudia Wong-Arambula, Abril Rodriguez-Maldonado, Vanessa Rivero-Arredondo, Ariadna Medina-Benitez, Joaquin Quiroz-Mercado, Sergio Rangel-Guerrero, Natividad Cruz-Ortiz, Tatiana Nunez-Garcia, Gisela Barrera-Badillo, Lucia Hernandez-Rivas, Irma Lopez-Martinez, Ernesto Ramirez-Gonzalez. |
| EPI_ISL_2559450 | LESP Sinaloa | Instituto de Diagnostico y Referencia Epidemiologicos (INDRE) | Claudia Wong-Arambula, Abril Rodriguez-Maldonado, Vanessa Rivero-Arredondo, Ariadna Medina-Benitez, Joaquin Quiroz-Mercado, Sergio Rangel-Guerrero, Natividad Cruz-Ortiz, Tatiana Nunez-Garcia, Gisela Barrera-Badillo, Lucia Hernandez-Rivas, Irma Lopez-Martinez, Ernesto Ramirez-Gonzalez. |
| EPI_ISL_2559452 | LESP Colima | Instituto de Diagnostico y Referencia Epidemiologicos (INDRE) | Claudia Wong-Arambula, Abril Rodriguez-Maldonado, Vanessa Rivero-Arredondo, Ariadna Medina-Benitez, Joaquin Quiroz-Mercado, Sergio Rangel-Guerrero, Natividad Cruz-Ortiz, Tatiana Nunez-Garcia, Gisela Barrera-Badillo, Lucia Hernandez-Rivas, Irma Lopez-Martinez, Ernesto Ramirez-Gonzalez. |
| EPI_ISL_2565970, EPI_ISL_2565977, EPI_ISL_2565981 | Institute of Molecular and Translational Medicine / Laboratory of Experimental Medicine, Faculty of Medicine and Dentistry, Palacky University and University Hospital Olomouc | Institute of Molecular and Translational Medicine / Laboratory of Experimental Medicine, Faculty of Medicine and Dentistry, Palacky University | Rastislav Slavkovský, Hana Dímalová, Vladimíra Koudeláková, Barbora Blumová, Marián Hajdúch |
| EPI_ISL_2565984, EPI_ISL_2565986, EPI_ISL_2565992, EPI_ISL_2565993, EPI_ISL_2565994, EPI_ISL_2565996 | Genome Analysis Center, Yamanashi Central Hospital | Genome Analysis Center, Yamanashi Central Hospital | Yosuke Hirotsu |
| EPI_ISL_2566164, EPI_ISL_2566165, EPI_ISL_2566166, EPI_ISL_2566169, EPI_ISL_2566170, EPI_ISL_2566171, EPI_ISL_2566172, EPI_ISL_2566173, EPI_ISL_2566174, EPI_ISL_2566175, EPI_ISL_2566176, EPI_ISL_2566177, EPI_ISL_2566178, EPI_ISL_2566179, EPI_ISL_2566180, EPI_ISL_2566181, EPI_ISL_2566183, EPI_ISL_2566186, EPI_ISL_2566188, EPI_ISL_2566189, EPI_ISL_2566191, EPI_ISL_2566193 | | | |
| see above | Microbiology Department. Complexo Hospitalario Universitario de Vigo | Microbiology Department. Complexo Hospitalario Universitario de Vigo | Alfaya N, Alonso I, Alvarez M, Cabrera JJ, Carballo R, Cores O, Cortizo S, del-Campo V, Martinez L, Mediero G, Perez S, Potel C, Regueiro B, Rey S, Vasallo FJ |
| EPI_ISL_2566444 | Singapore General Hospital | Department of Microbiology | Nurdyana Abdul Rahman, Kenneth Xin Long Chan, Chayaporn Suphavilai, Kun Lee Lim, Sui Sin Goh, Kian Sing Chan, Lynette Oon, Kern Rei Chng, James Sim Heng Chiak, Niranjan Nagarajan, Karrie Ko |
| EPI_ISL_2566448, EPI_ISL_2566459, EPI_ISL_2566460 | National Institute of Infectious Diseases-Prof. Dr. Matei Bals Molecular Diagnostics Laboratory | National Institute of Infectious Diseases-Prof. Dr. Matei Bals Molecular Diagnostics Laboratory | Corina Casangiu, Leontina Banica, Marius Surleac, Petre Milu, Robert Hohan, Simona Paraschiv, Dan Otelea |
| EPI_ISL_2566919, EPI_ISL_2566920, EPI_ISL_2566921, EPI_ISL_2566922, EPI_ISL_2566923, EPI_ISL_2566924 | AZ Klina | AZ Klina | Carl Vael - Lynsey Berckmans |
| EPI_ISL_2566925 | AZ Klina | AZ Klina | Carl Vael -Lynsey Berckmans |
| EPI_ISL_2566927, EPI_ISL_2566928, EPI_ISL_2566929, EPI_ISL_2566930, EPI_ISL_2566931, EPI_ISL_2566932, EPI_ISL_2567001, EPI_ISL_2567002, EPI_ISL_2567003, EPI_ISL_2567075 | AZ Klina | AZ Klina | Carl Vael - Lynsey Berckmans |
| EPI_ISL_2570809 | Surabaya Health Office | Institute of Tropical Disease, Universitas Airlangga | Aldise M Nastri, Jezzy R Dewantari, Rima R Prasetya, Krisnoadi Rahardjo, Febria Rachmanita, Herlin Ferliana, Gatot Soegiarto, Laksmi Wulandari, Resti Yudhawati, Yasuko Mori, Soetjipto, Kazufumi Shimizu, Maria I Lusida |
| EPI_ISL_2570811 | Surabaya Health Office | Institute of Tropical Disease, Universitas Airlangga | Jezzy R Dewantari, Rima R Prasetya, Krisnoadi Rahardjo, Aldise M Nastri, Febria Rachmanita, Herlin Ferliana, Gatot Soegiarto, Laksmi Wulandari, Resti Yudhawati, Yasuko Mori, Soetjipto, Kazufumi Shimizu, Maria I Lusida |
| EPI_ISL_2592630, EPI_ISL_2593764, EPI_ISL_2595725, EPI_ISL_2597228, EPI_ISL_2597229, EPI_ISL_2597312 | DSMRC | DSMRC | Khine Zaw Oo, Nay Myo Aung, Ko Ko Win, Phyo Kyaw Aung, Pai Khant Kyaw, Thet Wai Oo, Zaw Win Htun, Kyee Myint, Ko Ko Lwin |
| EPI_ISL_2600538, EPI_ISL_2600539, EPI_ISL_2600541, EPI_ISL_2600542, EPI_ISL_2600543, EPI_ISL_2600733, EPI_ISL_2600734, EPI_ISL_2600735, EPI_ISL_2600736, EPI_ISL_2600738, EPI_ISL_2600739, EPI_ISL_2600740, EPI_ISL_2600741, EPI_ISL_2600742, EPI_ISL_2600743, EPI_ISL_2600744, EPI_ISL_2600745, EPI_ISL_2600746, EPI_ISL_2600747, EPI_ISL_2600748, EPI_ISL_2600749, EPI_ISL_2600750, EPI_ISL_2600751, EPI_ISL_2600752, EPI_ISL_2600753, EPI_ISL_2600754, EPI_ISL_2600755, EPI_ISL_2600756, EPI_ISL_2600758, EPI_ISL_2600759, EPI_ISL_2600760, EPI_ISL_2600761, EPI_ISL_2600762, EPI_ISL_2600766, EPI_ISL_2600767, EPI_ISL_2600768, EPI_ISL_2600769, EPI_ISL_2600771, EPI_ISL_2600772, EPI_ISL_2600773, EPI_ISL_2600774, EPI_ISL_2600775, EPI_ISL_2600778, EPI_ISL_2600779 | | | |
| see above | Università degli Studi di Perugia | Istituto Zooprofilattico Sperimentale dell'Abruzzo e Molise "G. Caporale" | Mencacci A, Camilloni B, Biagetti M, Giammarioli M, Lorusso A, Marcacci M, Di Domenico M, Ancora M, Curini V, Mangone I, Rinaldi A, Delli Compagni E, Scialabba S, Di Pasquale A, Cammà C, Puglia I, Calistri P, Savini G |
| EPI_ISL_2600784, EPI_ISL_2600785, EPI_ISL_2600786, EPI_ISL_2600787, EPI_ISL_2600788, EPI_ISL_2600789, EPI_ISL_2600790, EPI_ISL_2600791, EPI_ISL_2600792 | Azienda Ospedaliera Terni | Istituto Zooprofilattico Sperimentale dell'Abruzzo e Molise "G. Caporale" | Palumbo M, Scaccetti A, Lorusso A, Marcacci M, Di Domenico M, Ancora M, Curini V, Mangone I, Rinaldi A, Delli Compagni E, Scialabba S, Di Pasquale A, Cammà C, Puglia I, Calistri P, Savini G |
| EPI_ISL_2600793, EPI_ISL_2600795, EPI_ISL_2600796, EPI_ISL_2600797, EPI_ISL_2600798, EPI_ISL_2600799 | Azienda USL Umbria 2 | Istituto Zooprofilattico Sperimentale dell'Abruzzo e Molise "G. Caporale" | Proietti A, Pistoni E, Lorusso A, Marcacci M, Di Domenico M, Ancora M, Curini V, Mangone I, Rinaldi A, Delli Compagni E, Scialabba S, Di Pasquale A, Cammà C, Puglia I, Calistri P, Savini G |
| EPI_ISL_2600800 | PRESIDIO OSPEDALIERO SULMONA | Istituto Zooprofilattico Sperimentale dell'Abruzzo e Molise "G. Caporale" | Lorusso A, Marcacci M, Di Domenico M, Ancora M, Curini V, Di Lollo Valeria, Mangone I, Rinaldi A, Delli Compagni E, Scialabba S, Caporale M, Di Pasquale A, Cammà C, Puglia I, Calistri P, Savini G |
| EPI_ISL_2600801 | SIESP CHIETI - DRIVE IN VASTO | Istituto Zooprofilattico Sperimentale dell'Abruzzo e Molise "G. Caporale" | Lorusso A, Marcacci M, Di Domenico M, Ancora M, Curini V, Di Lollo Valeria, Mangone I, Rinaldi A, Delli Compagni E, Scialabba S, Caporale M, Di Pasquale A, Cammà C, Puglia I, Calistri P, Savini G |
| EPI_ISL_2600802 | SIESP DIPARTIMENTO DI PREVENZIONE CHIETI | Istituto Zooprofilattico Sperimentale dell'Abruzzo e Molise "G. Caporale" | Lorusso A, Marcacci M, Di Domenico M, Ancora M, Curini V, Di Lollo Valeria, Mangone I, Rinaldi A, Delli Compagni E, Scialabba S, Caporale M, Di Pasquale A, Cammà C, Puglia I, Calistri P, Savini G |
| EPI_ISL_2600803, EPI_ISL_2600804, EPI_ISL_2600805 | SIESP CHIETI - DRIVE IN LANCIANO | Istituto Zooprofilattico Sperimentale dell'Abruzzo e Molise "G. Caporale" | Lorusso A, Marcacci M, Di Domenico M, Ancora M, Curini V, Di Lollo Valeria, Mangone I, Rinaldi A, Delli Compagni E, Scialabba S, Caporale M, Di Pasquale A, Cammà C, Puglia I, Calistri P, Savini G |
| EPI_ISL_2600806, EPI_ISL_2600807 | PO AVEZZANO - U.O.C. Laboratorio Analisi | Istituto Zooprofilattico Sperimentale dell'Abruzzo e Molise "G. Caporale" | Lorusso A, Marcacci M, Di Domenico M, Ancora M, Curini V, Di Lollo Valeria, Mangone I, Rinaldi A, Delli Compagni E, Scialabba S, Caporale M, Di Pasquale A, Cammà C, Puglia I, Calistri P, Savini G |
| EPI_ISL_2600808, EPI_ISL_2600809 | SIESP DIPARTIMENTO DI PREVENZIONE TERAMO | Istituto Zooprofilattico Sperimentale dell'Abruzzo e Molise "G. Caporale" | Lorusso A, Marcacci M, Di Domenico M, Ancora M, Curini V, Di Lollo Valeria, Mangone I, Rinaldi A, Delli Compagni E, Scialabba S, Caporale M, Di Pasquale A, Cammà C, Puglia I, Calistri P, Savini G |

| | | | |
|---|---|---|---|
| EPI_ISL_2600810 | SIESP CHIETI - DRIVE IN LANCIANO | Istituto Zooprofilattico Sperimentale dell'Abruzzo e Molise "G. Caporale" | Lorusso A, Marcacci M, Di Domenico M, Ancora M, Curini V, Di Lollo Valeria, Mangone I, Rinaldi A, Delli Compagni E, Scialabba S, Caporale M, Di Pasquale A, Cammà C, Puglia I, Calistri P, Savini G |
| EPI_ISL_2600811, EPI_ISL_2600812 | SIESP DIPARTIMENTO DI PREVENZIONE CHIETI | Istituto Zooprofilattico Sperimentale dell'Abruzzo e Molise "G. Caporale" | Lorusso A, Marcacci M, Di Domenico M, Ancora M, Curini V, Di Lollo Valeria, Mangone I, Rinaldi A, Delli Compagni E, Scialabba S, Caporale M, Di Pasquale A, Cammà C, Puglia I, Calistri P, Savini G |
| EPI_ISL_2600813, EPI_ISL_2600814 | SIESP CHIETI - DRIVE IN LANCIANO | Istituto Zooprofilattico Sperimentale dell'Abruzzo e Molise "G. Caporale" | Lorusso A, Marcacci M, Di Domenico M, Ancora M, Curini V, Di Lollo Valeria, Mangone I, Rinaldi A, Delli Compagni E, Scialabba S, Caporale M, Di Pasquale A, Cammà C, Puglia I, Calistri P, Savini G |
| EPI_ISL_2600815 | RSA GIULIANOVA | Istituto Zooprofilattico Sperimentale dell'Abruzzo e Molise "G. Caporale" | Lorusso A, Marcacci M, Di Domenico M, Ancora M, Curini V, Di Lollo Valeria, Mangone I, Rinaldi A, Delli Compagni E, Scialabba S, Caporale M, Di Pasquale A, Cammà C, Puglia I, Calistri P, Savini G |
| EPI_ISL_2600816 | SIESP CHIETI - DRIVE IN CHIETI | Istituto Zooprofilattico Sperimentale dell'Abruzzo e Molise "G. Caporale" | Lorusso A, Marcacci M, Di Domenico M, Ancora M, Curini V, Di Lollo Valeria, Mangone I, Rinaldi A, Delli Compagni E, Scialabba S, Caporale M, Di Pasquale A, Cammà C, Puglia I, Calistri P, Savini G |
| EPI_ISL_2600817, EPI_ISL_2600818, EPI_ISL_2600819, EPI_ISL_2600821 | SIESP CHIETI - DRIVE IN VASTO | Istituto Zooprofilattico Sperimentale dell'Abruzzo e Molise "G. Caporale" | Lorusso A, Marcacci M, Di Domenico M, Ancora M, Curini V, Di Lollo Valeria, Mangone I, Rinaldi A, Delli Compagni E, Scialabba S, Caporale M, Di Pasquale A, Cammà C, Puglia I, Calistri P, Savini G |
| EPI_ISL_2600822 | SIESP DIPARTIMENTO DI PREVENZIONE CHIETI | Istituto Zooprofilattico Sperimentale dell'Abruzzo e Molise "G. Caporale" | Lorusso A, Marcacci M, Di Domenico M, Ancora M, Curini V, Di Lollo Valeria, Mangone I, Rinaldi A, Delli Compagni E, Scialabba S, Caporale M, Di Pasquale A, Cammà C, Puglia I, Calistri P, Savini G |
| EPI_ISL_2600823 | SIESP CHIETI - DRIVE IN LANCIANO | Istituto Zooprofilattico Sperimentale dell'Abruzzo e Molise "G. Caporale" | Lorusso A, Marcacci M, Di Domenico M, Ancora M, Curini V, Di Lollo Valeria, Mangone I, Rinaldi A, Delli Compagni E, Scialabba S, Caporale M, Di Pasquale A, Cammà C, Puglia I, Calistri P, Savini G |
| EPI_ISL_2600824, EPI_ISL_2600825 | USCA Sulmona | Istituto Zooprofilattico Sperimentale dell'Abruzzo e Molise "G. Caporale" | Lorusso A, Marcacci M, Di Domenico M, Ancora M, Curini V, Di Lollo Valeria, Mangone I, Rinaldi A, Delli Compagni E, Scialabba S, Caporale M, Di Pasquale A, Cammà C, Puglia I, Calistri P, Savini G |
| EPI_ISL_2600826 | SIESP DIPARTIMENTO DI PREVENZIONE CHIETI | Istituto Zooprofilattico Sperimentale dell'Abruzzo e Molise "G. Caporale" | Lorusso A, Marcacci M, Di Domenico M, Ancora M, Curini V, Di Lollo Valeria, Mangone I, Rinaldi A, Delli Compagni E, Scialabba S, Caporale M, Di Pasquale A, Cammà C, Puglia I, Calistri P, Savini G |
| EPI_ISL_2600827, EPI_ISL_2600828 | SIESP CHIETI - DRIVE IN VASTO | Istituto Zooprofilattico Sperimentale dell'Abruzzo e Molise "G. Caporale" | Lorusso A, Marcacci M, Di Domenico M, Ancora M, Curini V, Di Lollo Valeria, Mangone I, Rinaldi A, Delli Compagni E, Scialabba S, Caporale M, Di Pasquale A, Cammà C, Puglia I, Calistri P, Savini G |
| EPI_ISL_2600830 | SIESP DIPARTIMENTO DI PREVENZIONE TERAMO | Istituto Zooprofilattico Sperimentale dell'Abruzzo e Molise "G. Caporale" | Lorusso A, Marcacci M, Di Domenico M, Ancora M, Curini V, Di Lollo Valeria, Mangone I, Rinaldi A, Delli Compagni E, Scialabba S, Caporale M, Di Pasquale A, Cammà C, Puglia I, Calistri P, Savini G |
| EPI_ISL_2600831, EPI_ISL_2600832 | SIESP CHIETI - DRIVE IN VASTO | Istituto Zooprofilattico Sperimentale dell'Abruzzo e Molise "G. Caporale" | Lorusso A, Marcacci M, Di Domenico M, Ancora M, Curini V, Di Lollo Valeria, Mangone I, Rinaldi A, Delli Compagni E, Scialabba S, Caporale M, Di Pasquale A, Cammà C, Puglia I, Calistri P, Savini G |
| EPI_ISL_2600833 | SIESP CHIETI - DRIVE IN CHIETI | Istituto Zooprofilattico Sperimentale dell'Abruzzo e Molise "G. Caporale" | Lorusso A, Marcacci M, Di Domenico M, Ancora M, Curini V, Di Lollo Valeria, Mangone I, Rinaldi A, Delli Compagni E, Scialabba S, Caporale M, Di Pasquale A, Cammà C, Puglia I, Calistri P, Savini G |
| EPI_ISL_2600834 | OSPEDALE CIVILE TERAMO | Istituto Zooprofilattico Sperimentale dell'Abruzzo e Molise "G. Caporale" | Lorusso A, Marcacci M, Di Domenico M, Ancora M, Curini V, Di Lollo Valeria, Mangone I, Rinaldi A, Delli Compagni E, Scialabba S, Caporale M, Di Pasquale A, Cammà C, Puglia I, Calistri P, Savini G |
| EPI_ISL_2600964 | Molecular Hepatology Laboratory AOU Messina | Molecular Hepatology Laboratory AOU Messina | Giuseppina Raffa, Cristina Musolino, Valeria Chines, Daniele Lombardo, Teresa Pollicino. |
| EPI_ISL_2601037 | Instituto de Medicina Tropical & Salud Global, Universidad Iberoamericana (UNIBE) | Instituto de Medicina Tropical & Salud Global, Universidad Iberoamericana (UNIBE) | Victor Virgilio Calderon, Alejandro Vallejo Degaudenzi & Robert Paulino-Ramirez |
| EPI_ISL_2603441 | Laboratorio Central de Saude Publica do Estado do Rio Grande do Sul (LACEN-RS) | Laboratory of Respiratory Viruses and Measles, Oswaldo Cruz Institute, FIOCRUZ | Paola Resende, Luciana Appolinario, Fernando Motta, Anna Carolina Paixao, Ana Carolina Mendonca, Alice Sampaio Rocha, Taina Venas, Elisa Cavalcante Pereira, Renata Serrano Lopes, Anderson Brandao Leite, Marilda Siqueira on behalf of the Fiocruz COVID-19 Genomic Surveillance Network |
| EPI_ISL_2603899 | PRESIDIO OSPEDALIERO TAGLIACOZZO | Istituto Zooprofilattico Sperimentale dell'Abruzzo e Molise "G. Caporale" | Lorusso A, Marcacci M, Di Domenico M, Ancora M, Curini V, Di Lollo Valeria, Mangone I, Rinaldi A, Delli Compagni E, Scialabba S, Caporale M, Di Pasquale A, Cammà C, Puglia I, Calistri P, Savini G |
| EPI_ISL_2604216, EPI_ISL_2604251 | Ministry of Health, Jaber Al-Ahmad Hospital | Virology Unit, Department of Microbiology, Faculty of Medicine, Kuwait University | Nada Madi, Hussain Safar, Ebaa Al-Awadhi, Anfal Al-Adwani |
| EPI_ISL_2604822 | Laboratorio di Riferimento Regionale della Sicilia Occidentale per l'Emergenza COVID-19 | Laboratorio di Riferimento Regionale della Sicilia Occidentale per l'Emergenza COVID-19 | Fabio Tramuto, Carmelo Massimo Maida, Daniela Di Naro, Giulia Randazzo, Walter Mazzucco, Giorgio Graziano, Vincenzo Restivo, Claudio Costantino, Francesco Vitale |
| EPI_ISL_2604869, EPI_ISL_2604882, EPI_ISL_2604888 | Omics Sciences Laboratory | Omics Sciences Laboratory | Derly Andrade Molina, Gabriel Morey León, Darlyn Amaya, Rubén Armas González, Juan Carlos Fernández Cadena |
| EPI_ISL_2608421 | Kampong Cham Rapid Response Team | Virology Unit, Institut Pasteur du Cambodge | Cecile Troupin, Leakhena Pum, Jurre Y Siegers, Ly Sovann, Kraing Sidonn, Yi Sengdoeurn, Chin Savuth, Chau Darapheak, Veasna Duong, Erik A Karlsson |
| EPI_ISL_2608422, EPI_ISL_2608423 | Siem Reap Provinical Laboratory | Virology Unit, Institut Pasteur du Cambodge | Cecile Troupin, Leakhena Pum, Jurre Y Siegers, Ly Sovann, Kraing Sidonn, Yi Sengdoeurn, Chin Savuth, Chau Darapheak, Veasna Duong, Erik A Karlsson |
| EPI_ISL_2608424, EPI_ISL_2608425, EPI_ISL_2608426, EPI_ISL_2608427, EPI_ISL_2608428, EPI_ISL_2608429, EPI_ISL_2608430, EPI_ISL_2608431, EPI_ISL_2608432, EPI_ISL_2608433, EPI_ISL_2608434, EPI_ISL_2608435, EPI_ISL_2608436, EPI_ISL_2608437, EPI_ISL_2608438, EPI_ISL_2608439, EPI_ISL_2608440, EPI_ISL_2608441, EPI_ISL_2608442, EPI_ISL_2608443 | | | |
| see above | Virology Unit, Institut Pasteur du Cambodge | Virology Unit, Institut Pasteur du Cambodge | Cecile Troupin, Leakhena Pum, Jurre Y Siegers, Ly Sovann, Kraing Sidonn, Yi Sengdoeurn, Chin Savuth, Chau Darapheak, Veasna Duong, Erik A Karlsson |
| EPI_ISL_2608444 | Kampot Rapid Response Team | Virology Unit, Institut Pasteur du Cambodge | Cecile Troupin, Leakhena Pum, Jurre Y Siegers, Ly Sovann, Kraing Sidonn, Yi Sengdoeurn, Chin Savuth, Chau Darapheak, Veasna Duong, Erik A Karlsson |
| EPI_ISL_2611661 | Temporary Specimen Collection Centre at the AsiaWorld-Expo | Hong Kong Department of Health | Alan K.L. Tsang, Peter C.W. Yip, Ken H.L. Ng, Edman T.K. Lam, Rickjason C.W. Chan, Dominic N.C. Tsang |
| EPI_ISL_2611984, EPI_ISL_2611985 | I.R.C.C.S. "S. De Bellis" - Ente Ospedaliero | Istituto Zooprofilattico Sperimentale della Puglia e della Basilicata | Parisi A., Bianco A., Capozzi L., Del Sambro L., Simone D., Difato L., Notarnicola M., Lippolis A. |
| EPI_ISL_2611990, EPI_ISL_2611991, EPI_ISL_2611994, EPI_ISL_2611995 | Presidio di Brindisi Di Summa - Perrino | Istituto Zooprofilattico Sperimentale della Puglia e della Basilicata | Parisi A., Bianco A., Capozzi L., Del Sambro L., Simone D., Difato L., Pennuzzi G., Santoro A. |
| EPI_ISL_2611997, EPI_ISL_2611998 | Istituto Zooprofilattico Sperimentale della Puglia e della Basilicata | Istituto Zooprofilattico Sperimentale della Puglia e della Basilicata | Parisi A., Bianco A., Capozzi L., Del Sambro L., Simone D., Difato L., Pennuzzi G., Pace L., Galante D. |
| EPI_ISL_2612000, EPI_ISL_2612001, EPI_ISL_2612002, EPI_ISL_2612003 | Ospedale Vito Fazzi | Istituto Zooprofilattico Sperimentale della Puglia e della Basilicata | Parisi A., Bianco A., Capozzi L., Del Sambro L., Simone D., Difato L., Lobreglio G. |
| EPI_ISL_2612004 | Ospedale Santa Caterina Novella | Istituto Zooprofilattico Sperimentale della Puglia e della Basilicata | Parisi A., Bianco A., Capozzi L., Del Sambro L., Simone D., Difato L., Bruno A. R. |
| EPI_ISL_2612005, EPI_ISL_2612006, EPI_ISL_2612007 | Ospedale Vito Fazzi | Istituto Zooprofilattico Sperimentale della Puglia e della Basilicata | Parisi A., Bianco A., Capozzi L., Del Sambro L., Simone D., Difato L., Lobreglio G. |
| EPI_ISL_2612008, EPI_ISL_2612009, EPI_ISL_2612010 | Ospedale Santa Caterina Novella | Istituto Zooprofilattico Sperimentale della Puglia e della Basilicata | Parisi A., Bianco A., Capozzi L., Del Sambro L., Simone D., Difato L., Bruno A. R. |
| EPI_ISL_2612011, EPI_ISL_2612012 | Ospedale Cardinale G. Panico | Istituto Zooprofilattico Sperimentale della Puglia e della Basilicata | Parisi A., Bianco A., Capozzi L., Del Sambro L., Simone D., Difato L., Bruno A. R. |

| | | | |
|---|---|---|---|
| EPI_ISL_2612013, EPI_ISL_2612014, EPI_ISL_2612015, EPI_ISL_2612016, EPI_ISL_2612017, EPI_ISL_2612019, EPI_ISL_2612020, EPI_ISL_2612021, EPI_ISL_2612022 | Ospedale Santissima Annunziata | Istituto Zooprofilattico Sperimentale della Puglia e della Basilicata | Parisi A., Bianco A., Capozzi L., Del Sambro L., Simone D., Difato L., Ridolfi D., Giannico A, Tampoia M. |
| EPI_ISL_2612025, EPI_ISL_2612026, EPI_ISL_2612027, EPI_ISL_2612028, EPI_ISL_2612029, EPI_ISL_2612030, EPI_ISL_2612031, EPI_ISL_2612033, EPI_ISL_2612034, EPI_ISL_2612035, EPI_ISL_2612036, EPI_ISL_2612037, EPI_ISL_2612038, EPI_ISL_2612039, EPI_ISL_2612040, EPI_ISL_2612041, EPI_ISL_2612042, EPI_ISL_2612043, EPI_ISL_2612044 | | | |
| see above | Klinisch Laboratorium ZNA | Klinisch Laboratorium ZNA | Verstrepen et al. |
| EPI_ISL_2612301, EPI_ISL_2612302, EPI_ISL_2612303, EPI_ISL_2612304, EPI_ISL_2612305, EPI_ISL_2612306, EPI_ISL_2612307, EPI_ISL_2612308, EPI_ISL_2612309, EPI_ISL_2612310, EPI_ISL_2612311, EPI_ISL_2612312, EPI_ISL_2612313, EPI_ISL_2612314 | | | |
| see above | Università degli Studi di Perugia | Istituto Zooprofilattico Sperimentale dell'Abruzzo e Molise "G. Caporale" | Mencacci A, Camilloni B, Biagetti M, Giammarioli M, Lorusso A, Marcacci M, Di Domenico M, Ancora M, Curini V, Mangone I, Rinaldi A, Delli Compagni E, Scialabba S, Di Pasquale A, Cammà C, Puglia I, Calistri P, Savini G |
| EPI_ISL_2612563 | OCME Office Of Chief Medical Examiner | New York City Public Health Laboratory | Jade Wang, et al. |
| EPI_ISL_2612564 | Labo Analyses Med | National Reference Center for Viruses of Respiratory Infections, Institut Pasteur, Paris | Marion Barbet, Sylvie Behillil, Méline Bizard, Angela Brisebarre, Camille Capel, Vincent Enouf, Louise Lefrançois, Frédéric Lemoine, Christophe Malabat, Corinne Maufrais, Etienne Simon-Lorière, Maud Vanpeene, Sylvie Van der Werf ,Sophie Chalmin |
| EPI_ISL_2612565 | Labo Analyses Med | National Reference Center for Viruses of Respiratory Infections, Institut Pasteur, Paris | Marion Barbet, Sylvie Behillil, Méline Bizard, Angela Brisebarre, Camille Capel, Vincent Enouf, Louise Lefrançois, Frédéric Lemoine, Christophe Malabat, Corinne Maufrais, Adrien Pain, Etienne Simon-Lorière, Maud Vanpeene, Sylvie Van der Werf ,Sophie Chalmin |
| EPI_ISL_2612566 | Labo Analyses Med | National Reference Center for Viruses of Respiratory Infections, Institut Pasteur, Paris | Marion Barbet, Sylvie Behillil, Méline Bizard, Angela Brisebarre, Camille Capel, Vincent Enouf, Louise Lefrançois, Frédéric Lemoine, Christophe Malabat, Corinne Maufrais, Etienne Simon-Lorière, Maud Vanpeene, Sylvie Van der Werf ,Sophie Chalmin |
| EPI_ISL_2612567 | Labo Analyses Med | National Reference Center for Viruses of Respiratory Infections, Institut Pasteur, Paris | Marion Barbet, Sylvie Behillil, Méline Bizard, Angela Brisebarre, Camille Capel, Vincent Enouf, Louise Lefrançois, Frédéric Lemoine, Christophe Malabat, Corinne Maufrais, Adrien Pain, Etienne Simon-Lorière, Maud Vanpeene, Sylvie Van der Werf ,Sophie Chalmin |
| EPI_ISL_2612568, EPI_ISL_2612569, EPI_ISL_2612570, EPI_ISL_2612571, EPI_ISL_2612572, EPI_ISL_2612573, EPI_ISL_2612574, EPI_ISL_2612575, EPI_ISL_2612576, EPI_ISL_2612577, EPI_ISL_2612578 | | | |
| see above | Labo Analyses Med | National Reference Center for Viruses of Respiratory Infections, Institut Pasteur, Paris | Marion Barbet, Sylvie Behillil, Méline Bizard, Angela Brisebarre, Camille Capel, Vincent Enouf, Louise Lefrançois, Frédéric Lemoine, Christophe Malabat, Corinne Maufrais, Etienne Simon-Lorière, Maud Vanpeene, Sylvie Van der Werf ,Sophie Chalmin |
| EPI_ISL_2612579, EPI_ISL_2612580, EPI_ISL_2612581, EPI_ISL_2612582, EPI_ISL_2612583, EPI_ISL_2612584, EPI_ISL_2612585, EPI_ISL_2612586 | Labo Analyses Med | National Reference Center for Viruses of Respiratory Infections, Institut Pasteur, Paris | Marion Barbet, Sylvie Behillil, Méline Bizard, Angela Brisebarre, Camille Capel, Vincent Enouf, Louise Lefrançois, Frédéric Lemoine, Christophe Malabat, Corinne Maufrais, Etienne Simon-Lorière, Maud Vanpeene, Sylvie Van der Werf ,Melanie Caron |
| EPI_ISL_2612588, EPI_ISL_2612589 | Labo Analyses Med | National Reference Center for Viruses of Respiratory Infections, Institut Pasteur, Paris | Marion Barbet, Sylvie Behillil, Méline Bizard, Angela Brisebarre, Camille Capel, Vincent Enouf, Louise Lefrançois, Frédéric Lemoine, Christophe Malabat, Corinne Maufrais, Etienne Simon-Lorière, Maud Vanpeene, Sylvie Van der Werf ,Nicolas Capron |
| EPI_ISL_2612590, EPI_ISL_2612591, EPI_ISL_2612592, EPI_ISL_2612593, EPI_ISL_2612594 | Labo Analyses Med | National Reference Center for Viruses of Respiratory Infections, Institut Pasteur, Paris | Marion Barbet, Sylvie Behillil, Méline Bizard, Angela Brisebarre, Camille Capel, Vincent Enouf, Louise Lefrançois, Frédéric Lemoine, Christophe Malabat, Corinne Maufrais, Etienne Simon-Lorière, Maud Vanpeene, Sylvie Van der Werf ,GréGoire Potiron |
| EPI_ISL_2612595, EPI_ISL_2612596, EPI_ISL_2612597 | Labo Analyses Med | National Reference Center for Viruses of Respiratory Infections, Institut Pasteur, Paris | Marion Barbet, Sylvie Behillil, Méline Bizard, Angela Brisebarre, Camille Capel, Vincent Enouf, Louise Lefrançois, Frédéric Lemoine, Christophe Malabat, Corinne Maufrais, Etienne Simon-Lorière, Maud Vanpeene, Sylvie Van der Werf ,Delphine MammÈS |
| EPI_ISL_2612598, EPI_ISL_2612599, EPI_ISL_2612600 | Hospital | National Reference Center for Viruses of Respiratory Infections, Institut Pasteur, Paris | Marion Barbet, Sylvie Behillil, Méline Bizard, Angela Brisebarre, Camille Capel, Vincent Enouf, Louise Lefrançois, Frédéric Lemoine, Christophe Malabat, Corinne Maufrais, Etienne Simon-Lorière, Maud Vanpeene, Sylvie Van der Werf ,Christine Febreau |
| EPI_ISL_2612602, EPI_ISL_2612603, EPI_ISL_2612604, EPI_ISL_2612605 | Hospital | National Reference Center for Viruses of Respiratory Infections, Institut Pasteur, Paris | Marion Barbet, Sylvie Behillil, Méline Bizard, Angela Brisebarre, Camille Capel, Vincent Enouf, Louise Lefrançois, Frédéric Lemoine, Christophe Malabat, Corinne Maufrais, Etienne Simon-Lorière, Maud Vanpeene, Sylvie Van der Werf ,Pierre Alain Billy |
| EPI_ISL_2612606, EPI_ISL_2612608, EPI_ISL_2612609, EPI_ISL_2612610, EPI_ISL_2612611, EPI_ISL_2612613, EPI_ISL_2612614, EPI_ISL_2612615 | Labo Analyses Med | National Reference Center for Viruses of Respiratory Infections, Institut Pasteur, Paris | Marion Barbet, Sylvie Behillil, Méline Bizard, Angela Brisebarre, Camille Capel, Vincent Enouf, Louise Lefrançois, Frédéric Lemoine, Christophe Malabat, Corinne Maufrais, Etienne Simon-Lorière, Maud Vanpeene, Sylvie Van der Werf ,Pierre-Yves Leonard |
| EPI_ISL_2612616 | Labo Analyses Med | National Reference Center for Viruses of Respiratory Infections, Institut Pasteur, Paris | Marion Barbet, Sylvie Behillil, Méline Bizard, Angela Brisebarre, Camille Capel, Vincent Enouf, Louise Lefrançois, Frédéric Lemoine, Christophe Malabat, Corinne Maufrais, Emmanuelle Permal, Etienne Simon-Lorière, Maud Vanpeene, Sylvie Van der Werf ,Pierre-Yves Leonard |
| EPI_ISL_2612617 | Labo Analyses Med | National Reference Center for Viruses of Respiratory Infections, Institut Pasteur, Paris | Marion Barbet, Sylvie Behillil, Méline Bizard, Angela Brisebarre, Camille Capel, Vincent Enouf, Louise Lefrançois, Frédéric Lemoine, Christophe Malabat, Corinne Maufrais, Etienne Simon-Lorière, Maud Vanpeene, Sylvie Van der Werf ,Pierre-Yves Leonard |
| EPI_ISL_2612618 | Labo Analyses Med | National Reference Center for Viruses of Respiratory Infections, Institut Pasteur, Paris | Marion Barbet, Sylvie Behillil, Méline Bizard, Angela Brisebarre, Camille Capel, Vincent Enouf, Louise Lefrançois, Frédéric Lemoine, Christophe Malabat, Corinne Maufrais, Amaury Vaysse, Etienne Simon-Lorière, Maud Vanpeene, Sylvie Van der Werf ,Pierre-Yves Leonard |
| EPI_ISL_2612619, EPI_ISL_2612620, EPI_ISL_2612621 | Labo Analyses Med | National Reference Center for Viruses of Respiratory Infections, Institut Pasteur, Paris | Marion Barbet, Sylvie Behillil, Méline Bizard, Angela Brisebarre, Camille Capel, Vincent Enouf, Louise Lefrançois, Frédéric Lemoine, Christophe Malabat, Corinne Maufrais, Etienne Simon-Lorière, Maud Vanpeene, Sylvie Van der Werf ,Pierre-Yves Leonard |
| EPI_ISL_2612622, EPI_ISL_2612623, EPI_ISL_2612624 | Hospital | National Reference Center for Viruses of Respiratory Infections, Institut Pasteur, Paris | Marion Barbet, Sylvie Behillil, Méline Bizard, Angela Brisebarre, Camille Capel, Vincent Enouf, Louise Lefrançois, Frédéric Lemoine, Christophe Malabat, Corinne Maufrais, Etienne Simon-Lorière, Maud Vanpeene, Sylvie Van der Werf ,Axelle Paquin |
| EPI_ISL_2612625 | Hospital | National Reference Center for Viruses of Respiratory Infections, Institut Pasteur, Paris | Marion Barbet, Sylvie Behillil, Méline Bizard, Angela Brisebarre, Camille Capel, Vincent Enouf, Louise Lefrançois, Frédéric Lemoine, Christophe Malabat, Corinne Maufrais, Etienne Simon-Lorière, Maud Vanpeene, Sylvie Van der Werf ,Alexandra Ducancelle |
| EPI_ISL_2612629 | Hospital | National Reference Center for Viruses of Respiratory Infections, Institut Pasteur, Paris | Marion Barbet, Sylvie Behillil, Méline Bizard, Angela Brisebarre, Camille Capel, Vincent Enouf, Louise Lefrançois, Frédéric Lemoine, Christophe Malabat, Corinne Maufrais, Etienne Simon-Lorière, Maud Vanpeene, Sylvie Van der Werf ,Caroline Lefeuvre |
| EPI_ISL_2612630 | Hospital | National Reference Center for Viruses of Respiratory Infections, Institut Pasteur, Paris | Marion Barbet, Sylvie Behillil, Méline Bizard, Angela Brisebarre, Camille Capel, Vincent Enouf, Louise Lefrançois, Frédéric Lemoine, Christophe Malabat, Corinne Maufrais, Etienne Simon-Lorière, Maud Vanpeene, Sylvie Van der Werf ,Franck Labbe |
| EPI_ISL_2612631 | Labo Analyses Med | National Reference Center for Viruses of Respiratory Infections, Institut Pasteur, Paris | Marion Barbet, Sylvie Behillil, Méline Bizard, Angela Brisebarre, Camille Capel, Vincent Enouf, Louise Lefrançois, Frédéric Lemoine, Christophe Malabat, Corinne Maufrais, Amaury Vaysse, Etienne Simon-Lorière, Maud Vanpeene, Sylvie Van der Werf ,Anne Holstein |
| EPI_ISL_2612632, EPI_ISL_2612633 | Labo Analyses Med | National Reference Center for Viruses of Respiratory Infections, Institut Pasteur, Paris | Marion Barbet, Sylvie Behillil, Méline Bizard, Angela Brisebarre, Camille Capel, Vincent Enouf, Louise Lefrançois, Frédéric Lemoine, Christophe Malabat, Corinne Maufrais, Etienne Simon-Lorière, Maud Vanpeene, Sylvie Van der Werf ,Anne Holstein |
| EPI_ISL_2612634 | Labo Analyses Med | National Reference Center for Viruses of Respiratory Infections, Institut Pasteur, Paris | Marion Barbet, Sylvie Behillil, Méline Bizard, Angela Brisebarre, Camille Capel, Vincent Enouf, Louise Lefrançois, Frédéric Lemoine, Christophe Malabat, Corinne Maufrais, Amaury Vaysse, Etienne Simon-Lorière, Maud Vanpeene, Sylvie Van der Werf ,Anne Holstein |
| EPI_ISL_2612635, EPI_ISL_2612636, EPI_ISL_2612637, EPI_ISL_2612638, EPI_ISL_2612639, EPI_ISL_2612640, EPI_ISL_2612641, EPI_ISL_2612642, EPI_ISL_2612643, EPI_ISL_2612644, EPI_ISL_2612645, EPI_ISL_2612646, EPI_ISL_2612647, EPI_ISL_2612648, EPI_ISL_2612649, EPI_ISL_2612650, EPI_ISL_2612651, EPI_ISL_2612652, EPI_ISL_2612653, EPI_ISL_2612654, EPI_ISL_2612655, EPI_ISL_2612656, EPI_ISL_2612657 | | | |
| see above | Labo Analyses Med | National Reference Center for Viruses of Respiratory Infections, Institut Pasteur, Paris | Marion Barbet, Sylvie Behillil, Méline Bizard, Angela Brisebarre, Camille Capel, Vincent Enouf, Louise Lefrançois, Frédéric Lemoine, Christophe Malabat, Corinne Maufrais, Etienne Simon-Lorière, Maud Vanpeene, Sylvie Van der Werf ,Anne Holstein |
| EPI_ISL_2612658, EPI_ISL_2612659, EPI_ISL_2612660, EPI_ISL_2612661, EPI_ISL_2612662, EPI_ISL_2612663, EPI_ISL_2612664, EPI_ISL_2612665, EPI_ISL_2612666, EPI_ISL_2612667, EPI_ISL_2612668 | | | |
| see above | Labo Analyses Med | National Reference Center for Viruses of Respiratory Infections, Institut Pasteur, Paris | Marion Barbet, Sylvie Behillil, Méline Bizard, Angela Brisebarre, Camille Capel, Vincent Enouf, Louise Lefrançois, Frédéric Lemoine, Christophe Malabat, Corinne Maufrais, Etienne Simon-Lorière, Maud Vanpeene, Sylvie Van der Werf ,Jean-FrançOis Comes |

| | | | |
|---|---|---|---|
| EPI_ISL_2612669, EPI_ISL_2612670 | Hospital | National Reference Center for Viruses of Respiratory Infections, Institut Pasteur, Paris | Marion Barbet, Sylvie Behillil, Méline Bizard, Angela Brisebarre, Camille Capel, Vincent Enouf, Louise Lefrançois, Frédéric Lemoine, Christophe Malabat, Corinne Maufrais, Etienne Simon-Lorière, Maud Vanpeene, Sylvie Van der Werf ,Anne Cady |
| EPI_ISL_2612671, EPI_ISL_2612672, EPI_ISL_2612673, EPI_ISL_2612674, EPI_ISL_2612675, EPI_ISL_2612676, EPI_ISL_2612677, EPI_ISL_2612680, EPI_ISL_2612681 | Labo Analyses Med | National Reference Center for Viruses of Respiratory Infections, Institut Pasteur, Paris | Marion Barbet, Sylvie Behillil, Méline Bizard, Angela Brisebarre, Camille Capel, Vincent Enouf, Louise Lefrançois, Frédéric Lemoine, Christophe Malabat, Corinne Maufrais, Etienne Simon-Lorière, Maud Vanpeene, Sylvie Van der Werf ,Arditi Cocco |
| EPI_ISL_2613623 | CHU LILLE | CHU Lille - Laboratoire de Virologie | AIT YAHYA Emilie, ALIDJINOU Enagnon Kazali, BOCKET Laurence, CREPIN Michel, DEMAY Christophe, ENGELMANN Ilka, GEFFROY Sandrine, GUIGON Aurélie, LAMBERT Valérie, LAZREK Mouna, NOBILLIAUX Florian, PREVOST Brigitte, THUILLIER Caroline, TINEZ Claire |
| EPI_ISL_2613624 | CH TOURCOING | CHU Lille - Laboratoire de Virologie | AIT YAHYA Emilie, ALIDJINOU Enagnon Kazali, BOCKET Laurence, CREPIN Michel, DEMAY Christophe, ENGELMANN Ilka, GEFFROY Sandrine, GUIGON Aurélie, LAMBERT Valérie, LAZREK Mouna, NOBILLIAUX Florian, PREVOST Brigitte, THUILLIER Caroline, TINEZ Claire |
| EPI_ISL_2613810, EPI_ISL_2613811, EPI_ISL_2613815 | Lboratorio Central de Saude Publica do Estado do Parana (LACEN/PR) | Laboratory of Respiratory Viruses and Measles, Oswaldo Cruz Institute, FIOCRUZ | Paola Resende, Luciana Appolinario, Fernando Motta, Anna Carolina Paixao, Ana Carolina Mendonca, Alice Sampaio Rocha, Taina Venas, Elisa Cavalcante Pereira, Renata Serrano Lopes, Irina Riediger, Marilda Siqueira on behalf of the Fiocruz COVID-19 Genomic Surveillance Network |
| EPI_ISL_2613826 | CH ROUBAIX | CHU Lille - Laboratoire de Virologie | AIT YAHYA Emilie, ALIDJINOU Enagnon Kazali, BOCKET Laurence, CREPIN Michel, DEMAY Christophe, ENGELMANN Ilka, GEFFROY Sandrine, GUIGON Aurélie, LAMBERT Valérie, LAZREK Mouna, NOBILLIAUX Florian, PREVOST Brigitte, THUILLIER Caroline, TINEZ Claire ,TCHANTCHOU NJOSSE YANICK |
| EPI_ISL_2613827, EPI_ISL_2613828 | CHU LILLE | CHU Lille - Laboratoire de Virologie | AIT YAHYA Emilie, ALIDJINOU Enagnon Kazali, BOCKET Laurence, CREPIN Michel, DEMAY Christophe, ENGELMANN Ilka, GEFFROY Sandrine, GUIGON Aurélie, LAMBERT Valérie, LAZREK Mouna, NOBILLIAUX Florian, PREVOST Brigitte, THUILLIER Caroline, TINEZ Claire ,TCHANTCHOU NJOSSE YANICK |
| EPI_ISL_2613829 | CH LENS | CHU Lille - Laboratoire de Virologie | AIT YAHYA Emilie, ALIDJINOU Enagnon Kazali, BOCKET Laurence, CREPIN Michel, DEMAY Christophe, ENGELMANN Ilka, GEFFROY Sandrine, GUIGON Aurélie, LAMBERT Valérie, LAZREK Mouna, NOBILLIAUX Florian, PREVOST Brigitte, THUILLIER Caroline, TINEZ Claire ,TCHANTCHOU NJOSSE YANICK |
| EPI_ISL_2613830 | CH BETHUNE | CHU Lille - Laboratoire de Virologie | AIT YAHYA Emilie, ALIDJINOU Enagnon Kazali, BOCKET Laurence, CREPIN Michel, DEMAY Christophe, ENGELMANN Ilka, GEFFROY Sandrine, GUIGON Aurélie, LAMBERT Valérie, LAZREK Mouna, NOBILLIAUX Florian, PREVOST Brigitte, THUILLIER Caroline, TINEZ Claire ,TCHANTCHOU NJOSSE YANICK |
| EPI_ISL_2613831 | CH VALENCIENNES | CHU Lille - Laboratoire de Virologie | AIT YAHYA Emilie, ALIDJINOU Enagnon Kazali, BOCKET Laurence, CREPIN Michel, DEMAY Christophe, ENGELMANN Ilka, GEFFROY Sandrine, GUIGON Aurélie, LAMBERT Valérie, LAZREK Mouna, NOBILLIAUX Florian, PREVOST Brigitte, THUILLIER Caroline, TINEZ Claire ,TCHANTCHOU NJOSSE YANICK |
| EPI_ISL_2613834 | CHU LILLE | CHU Lille - Laboratoire de Virologie | AIT YAHYA Emilie, ALIDJINOU Enagnon Kazali, BOCKET Laurence, CREPIN Michel, DEMAY Christophe, ENGELMANN Ilka, GEFFROY Sandrine, GUIGON Aurélie, LAMBERT Valérie, LAZREK Mouna, NOBILLIAUX Florian, PREVOST Brigitte, THUILLIER Caroline, TINEZ Claire ,TCHANTCHOU NJOSSE YANICK |
| EPI_ISL_2613835 | CH ROUBAIX | CHU Lille - Laboratoire de Virologie | AIT YAHYA Emilie, ALIDJINOU Enagnon Kazali, BOCKET Laurence, CREPIN Michel, DEMAY Christophe, ENGELMANN Ilka, GEFFROY Sandrine, GUIGON Aurélie, LAMBERT Valérie, LAZREK Mouna, NOBILLIAUX Florian, PREVOST Brigitte, THUILLIER Caroline, TINEZ Claire ,TCHANTCHOU NJOSSE YANICK |
| EPI_ISL_2614425 | Klinisch Laboratorium ZNA | Klinisch Laboratorium ZNA | Verstrepen et al. |
| EPI_ISL_2614634, EPI_ISL_2614635, EPI_ISL_2614636, EPI_ISL_2614637, EPI_ISL_2614638, EPI_ISL_2614641, EPI_ISL_2614642 | Dr. Yuichiro Oba Department of General Medicine, Osaka General Medical Center | Department of Developmental Medicine, Research Institute, Osaka Women's and Children's Hospital | Itaru Yanagihara, Yukiko Nakura |
| EPI_ISL_2614725 | RS Pelni | Eijkman Institute for Molecular Biology, National Research and Innovation Agency | Willy Agustine, Hidayat Trimarsanto, Lydia V. Panggalo, Iskandar Adnan, Sukma Oktavianthi, Lidwina Priliani, Edison Johar, Frilasita A Yudhaputri, Muhammad Rezki Rasyak, Safarina G Malik, Khin Saw Myint, Amin Soebandrio |
| EPI_ISL_2614726 | RS Pelni | Eijkman Institute for Molecular Biology, National Research and Innovation Agency | Hidayat Trimarsanto, Lydia V. Panggalo, Iskandar Adnan, Sukma Oktavianthi, Lidwina Priliani, Edison Johar, Frilasita A Yudhaputri, Muhammad Rezki Rasyak, Willy Agustine, Safarina G Malik, Khin Saw Myint, Amin Soebandrio |
| EPI_ISL_2614727 | RS Pelni | Eijkman Institute for Molecular Biology, National Research and Innovation Agency | Lydia V. Panggalo, Iskandar Adnan, Sukma Oktavianthi, Lidwina Priliani, Edison Johar, Frilasita A Yudhaputri, Muhammad Rezki Rasyak, Willy Agustine, Hidayat Trimarsanto, Safarina G Malik, Khin Saw Myint, Amin Soebandrio |
| EPI_ISL_2614728 | RS Grha Kedoya | Eijkman Institute for Molecular Biology, National Research and Innovation Agency | Iskandar Adnan, Sukma Oktavianthi, Lidwina Priliani, Edison Johar, Frilasita A Yudhaputri, Muhammad Rezki Rasyak, Willy Agustine, Hidayat Trimarsanto, Lydia V. Panggalo, Safarina G Malik, Khin Saw Myint, Amin Soebandrio |
| EPI_ISL_2614730 | EKA Hospital BSD | Eijkman Institute for Molecular Biology, National Research and Innovation Agency | Lidwina Priliani, Edison Johar, Frilasita A Yudhaputri, Muhammad Rezki Rasyak, Willy Agustine, Hidayat Trimarsanto, Lydia V. Panggalo, Iskandar Adnan, Sukma Oktavianthi, Safarina G Malik, Khin Saw Myint, Amin Soebandrio |
| EPI_ISL_2614731 | RS Pelni | Eijkman Institute for Molecular Biology, National Research and Innovation Agency | Iskandar Adnan, Sukma Oktavianthi, Lidwina Priliani, Edison Johar, Frilasita A Yudhaputri, Muhammad Rezki Rasyak, Willy Agustine, Hidayat Trimarsanto, Lydia V. Panggalo, Safarina G Malik, Khin Saw Myint, Amin Soebandrio |
| EPI_ISL_2614732 | RS Sumber Waras | Eijkman Institute for Molecular Biology, National Research and Innovation Agency | Sukma Oktavianthi, Lidwina Priliani, Edison Johar, Frilasita A Yudhaputri, Muhammad Rezki Rasyak, Willy Agustine, Hidayat Trimarsanto, Lydia V. Panggalo, Iskandar Adnan, Safarina G Malik, Khin Saw Myint, Amin Soebandrio |
| EPI_ISL_2614733 | RS Pusat Pertamina | Eijkman Institute for Molecular Biology, National Research and Innovation Agency | Lidwina Priliani, Edison Johar, Frilasita A Yudhaputri, Muhammad Rezki Rasyak, Willy Agustine, Hidayat Trimarsanto, Lydia V. Panggalo, Iskandar Adnan, Sukma Oktavianthi, Safarina G Malik, Khin Saw Myint, Amin Soebandrio |
| EPI_ISL_2614734 | RS Pusat Pertamina | Eijkman Institute for Molecular Biology, National Research and Innovation Agency | Edison Johar, Frilasita A Yudhaputri, Muhammad Rezki Rasyak, Willy Agustine, Hidayat Trimarsanto, Lydia V. Panggalo, Iskandar Adnan, Sukma Oktavianthi, Lidwina Priliani, Safarina G Malik, Khin Saw Myint, Amin Soebandrio |
| EPI_ISL_2614735 | RS Pondok Indah Pondok Indah | Eijkman Institute for Molecular Biology, National Research and Innovation Agency | Frilasita A Yudhaputri, Muhammad Rezki Rasyak, Willy Agustine, Hidayat Trimarsanto, Lydia V. Panggalo, Iskandar Adnan, Sukma Oktavianthi, Lidwina Priliani, Edison Johar, Safarina G Malik, Khin Saw Myint, Amin Soebandrio |
| EPI_ISL_2614736 | RS Hermina Depok | Eijkman Institute for Molecular Biology, National Research and Innovation Agency | Muhammad Rezki Rasyak, Willy Agustine, Hidayat Trimarsanto, Lydia V. Panggalo, Iskandar Adnan, Sukma Oktavianthi, Lidwina Priliani, Edison Johar, Frilasita A Yudhaputri, Safarina G Malik, Khin Saw Myint, Amin Soebandrio |
| EPI_ISL_2614737 | RS Pondok Indah Pondok Indah | Eijkman Institute for Molecular Biology, National Research and Innovation Agency | Willy Agustine, Hidayat Trimarsanto, Lydia V. Panggalo, Iskandar Adnan, Sukma Oktavianthi, Lidwina Priliani, Edison Johar, Frilasita A Yudhaputri, Muhammad Rezki Rasyak, Safarina G Malik, Khin Saw Myint, Amin Soebandrio |
| EPI_ISL_2614738 | RS Pondok Indah Pondok Indah | Eijkman Institute for Molecular Biology, National Research and Innovation Agency | Hidayat Trimarsanto, Lydia V. Panggalo, Iskandar Adnan, Sukma Oktavianthi, Lidwina Priliani, Edison Johar, Frilasita A Yudhaputri, Muhammad Rezki Rasyak, Willy Agustine, Safarina G Malik, Khin Saw Myint, Amin Soebandrio |
| EPI_ISL_2614743 | Sungai Buloh Hospital | Institute for Medical Research, Infectious Disease Research Centre, National Institutes of Health, Ministry of Health Malaysia | Suppiah J, Kamel K, Mohd Zawawi Z, Azizan MA, Ramly N, Robert F, Thayan R |
| EPI_ISL_2614744 | Tuanku Ampuan Najihah Hospital, Kuala Pilah | Institute for Medical Research, Infectious Disease Research Centre, National Institutes of Health, Ministry of Health Malaysia | Suppiah J, Kamel K, Mohd Zawawi Z, Azizan MA, Ramly N, Robert F, Thayan R |
| EPI_ISL_2614745 | Sarawak General Hospital | Institute for Medical Research, Infectious Disease Research Centre, National Institutes of Health, Ministry of Health | Suppiah J, Kamel K, Mohd Zawawi Z, Azizan MA, Ramly N, Robert F, Thayan R |

| | | | |
|---|---|---|---|
| | | Malaysia | |
| EPI_ISL_2615399 | MOH - Jaber Al-Ahmad Hospital (Innovation Research Laboratory) | MOH - Jaber Al-Ahmad Hospital (Innovation Research Laboratory) | Salman Al-Sabah , Mohammad Alghounaim |
| EPI_ISL_2616219, EPI_ISL_2616220, EPI_ISL_2616232, EPI_ISL_2616234, EPI_ISL_2616235, EPI_ISL_2616236, EPI_ISL_2616237, EPI_ISL_2616238, EPI_ISL_2616239, EPI_ISL_2616240, EPI_ISL_2616241, EPI_ISL_2616242, EPI_ISL_2616243, EPI_ISL_2616244, EPI_ISL_2616245, EPI_ISL_2616261, EPI_ISL_2616262, EPI_ISL_2616293, EPI_ISL_2616299, EPI_ISL_2616313, EPI_ISL_2616314, EPI_ISL_2616315, EPI_ISL_2616316, EPI_ISL_2616317, EPI_ISL_2616318 | | | |
| see above | Austrian Agency for Health and Food Safety (AGES) | Bergthaler laboratory, CeMM Research Center for Molecular Medicine of the Austrian Academy of Sciences | Lukas Endler, Anna Schedl, Fabian Amman, Petr Triska, Matthew Thornton, Thomas Penz, Benedikt Agerer, Maelle Le Moing, Michael Schuster, Bekir Erguner, Jan Laine, Martin Senekowitsch, Christoph Bock, Andreas Bergthaler |
| EPI_ISL_2616319 | Center for Virology, Medical University of Vienna | Bergthaler laboratory, CeMM Research Center for Molecular Medicine of the Austrian Academy of Sciences | Lukas Endler, Anna Schedl, Fabian Amman, Petr Triska, Matthew Thornton, Thomas Penz, Benedikt Agerer, Maelle Le Moing, Michael Schuster, Bekir Erguner, Jan Laine, Martin Senekowitsch, Christoph Bock, Andreas Bergthaler |
| EPI_ISL_2616320, EPI_ISL_2616321, EPI_ISL_2616322, EPI_ISL_2616323, EPI_ISL_2616325, EPI_ISL_2616326, EPI_ISL_2616327, EPI_ISL_2616328, EPI_ISL_2616329, EPI_ISL_2616330, EPI_ISL_2616331, EPI_ISL_2616332, EPI_ISL_2616333, EPI_ISL_2616343, EPI_ISL_2616344, EPI_ISL_2616345, EPI_ISL_2616346, EPI_ISL_2616365 | | | |
| see above | Austrian Agency for Health and Food Safety (AGES) | Bergthaler laboratory, CeMM Research Center for Molecular Medicine of the Austrian Academy of Sciences | Lukas Endler, Anna Schedl, Fabian Amman, Petr Triska, Matthew Thornton, Thomas Penz, Benedikt Agerer, Maelle Le Moing, Michael Schuster, Bekir Erguner, Jan Laine, Martin Senekowitsch, Christoph Bock, Andreas Bergthaler |
| EPI_ISL_2617440, EPI_ISL_2617441, EPI_ISL_2617442, EPI_ISL_2617443, EPI_ISL_2617445 | laboratorio Microbiologia PO Cardarelli | laboratorio Microbiologia PO Cardarelli | Felice V., Niro G. Scutella' M. |
| EPI_ISL_2621736 | Genome Analysis Center, Yamanashi Central Hospital | Genome Analysis Center, Yamanashi Central Hospital | Yosuke Hirotsu |
| EPI_ISL_2622160, EPI_ISL_2622161, EPI_ISL_2622162, EPI_ISL_2622164, EPI_ISL_2622165, EPI_ISL_2622166, EPI_ISL_2622167, EPI_ISL_2622169, EPI_ISL_2622171, EPI_ISL_2622172, EPI_ISL_2622174, EPI_ISL_2622175 | | | |
| see above | Azienda Ospedaliera Pugliese Ciaccio di Catanzaro SOC Microbiologia e Virologia | Azienda Ospedaliera Pugliese Ciaccio di Catanzaro SOC Microbiologia e Virologia | Rossana Tallerico Federica Pasceri Marco De Fazio Ilenia Talotta Giuseppina Panduri Pasquale Minchella |
| EPI_ISL_2625622 | Usansolo-Galdakao University Hospital | Cruces University Hospital | Ana Gual-de-Torrella, Izaskun Alejo-Cancho, Mikel Gallego, Ana Belén de la Hoz |
| EPI_ISL_2626010, EPI_ISL_2626011, EPI_ISL_2626012, EPI_ISL_2626013, EPI_ISL_2626014, EPI_ISL_2626015, EPI_ISL_2626016, EPI_ISL_2626017, EPI_ISL_2626018, EPI_ISL_2626019, EPI_ISL_2626020, EPI_ISL_2626021, EPI_ISL_2626022, EPI_ISL_2626023, EPI_ISL_2626024, EPI_ISL_2626025, EPI_ISL_2626026, EPI_ISL_2626027, EPI_ISL_2626028, EPI_ISL_2626029, EPI_ISL_2626030, EPI_ISL_2626031, EPI_ISL_2626032, EPI_ISL_2626033, EPI_ISL_2626034, EPI_ISL_2626035, EPI_ISL_2626036, EPI_ISL_2626037, EPI_ISL_2626038, EPI_ISL_2626039, EPI_ISL_2626040, EPI_ISL_2626041, EPI_ISL_2626042, EPI_ISL_2626043, EPI_ISL_2626044, EPI_ISL_2626045, EPI_ISL_2626046, EPI_ISL_2626047, EPI_ISL_2626048, EPI_ISL_2626049, EPI_ISL_2626050, EPI_ISL_2626051, EPI_ISL_2626053, EPI_ISL_2626054, EPI_ISL_2626055, EPI_ISL_2626056, EPI_ISL_2626057, EPI_ISL_2626058, EPI_ISL_2626059, EPI_ISL_2626060, EPI_ISL_2626061 | | | |
| see above | Cliniques universitaires Saint-Luc | UCLouvain/IREC/MBLG | Jean Ruelle, Bertrand Bearzatto, Benoit Kabamba Mukadi |
| EPI_ISL_2626062, EPI_ISL_2626063, EPI_ISL_2626064, EPI_ISL_2626065, EPI_ISL_2626066, EPI_ISL_2626067, EPI_ISL_2626068, EPI_ISL_2626069, EPI_ISL_2626070, EPI_ISL_2626071, EPI_ISL_2626072, EPI_ISL_2626073, EPI_ISL_2626074, EPI_ISL_2626075, EPI_ISL_2626077, EPI_ISL_2626078, EPI_ISL_2626079, EPI_ISL_2626080, EPI_ISL_2626081 | | | |
| see above | Clinique Saint-Pierre Ottignies | UCLouvain/IREC/MBLG | Jean Ruelle, Bertrand Bearzatto, Benoit Kabamba Mukadi |
| EPI_ISL_2626082, EPI_ISL_2626083, EPI_ISL_2626084, EPI_ISL_2626085, EPI_ISL_2626086, EPI_ISL_2626087, EPI_ISL_2626088, EPI_ISL_2626089, EPI_ISL_2626090, EPI_ISL_2626091, EPI_ISL_2626092, EPI_ISL_2626093, EPI_ISL_2626096 | | | |
| see above | Cliniques universitaires Saint-Luc | UCLouvain/IREC/MBLG | Jean Ruelle, Bertrand Bearzatto, Benoit Kabamba Mukadi |
| EPI_ISL_2626180, EPI_ISL_2626181 | Usansolo-Galdakao University Hospital | Cruces University Hospital | Ana Gual-de-Torrella, Izaskun Alejo-Cancho, Mikel Gallego, Ana Belén de la Hoz |
| EPI_ISL_2626770, EPI_ISL_2626771, EPI_ISL_2626772, EPI_ISL_2626774, EPI_ISL_2626777, EPI_ISL_2626778, EPI_ISL_2626779, EPI_ISL_2626782, EPI_ISL_2626783, EPI_ISL_2626784, EPI_ISL_2626785, EPI_ISL_2626786, EPI_ISL_2626787, EPI_ISL_2626788, EPI_ISL_2626789, EPI_ISL_2626790, EPI_ISL_2626794, EPI_ISL_2626795, EPI_ISL_2626797, EPI_ISL_2626799, EPI_ISL_2626800, EPI_ISL_2626803, EPI_ISL_2626807, EPI_ISL_2626808, EPI_ISL_2626809, EPI_ISL_2626810, EPI_ISL_2626811, EPI_ISL_2626812, EPI_ISL_2626813, EPI_ISL_2626814, EPI_ISL_2626816, EPI_ISL_2626818, EPI_ISL_2626819, EPI_ISL_2626820, EPI_ISL_2626821, EPI_ISL_2626822, EPI_ISL_2626824, EPI_ISL_2626825, EPI_ISL_2626826, EPI_ISL_2626827, EPI_ISL_2626831, EPI_ISL_2626835, EPI_ISL_2626836, EPI_ISL_2626837, EPI_ISL_2626838 | | | |
| see above | Baylor Scott & White-Temple | Baylor Scott & White-Temple | Ari Rao, Linden Morales, Kimberly Walker, Marcus Volz, Shelby Johnson |
| EPI_ISL_2627491, EPI_ISL_2627492, EPI_ISL_2627493, EPI_ISL_2627494, EPI_ISL_2627495, EPI_ISL_2627496, EPI_ISL_2627497, EPI_ISL_2627498 | Klinisch Laboratorium ZNA | Klinisch Laboratorium ZNA | Verstrepen et al. |
| EPI_ISL_2627940, EPI_ISL_2627941 | Hospital | National Reference Center for Viruses of Respiratory Infections, Institut Pasteur, Paris | Marion Barbet, Sylvie Behillil, Méline Bizard, Angela Brisebarre, Camille Capel, Vincent Enouf, Louise Lefrançois, Frédéric Lemoine, Christophe Malabat, Corinne Maufrais, Etienne Simon-Lorière, Maud Vanpeene, Sylvie Van der Werf ,Patricia Stoessel-Thouvenin |
| EPI_ISL_2627942 | Hospital | National Reference Center for Viruses of Respiratory Infections, Institut Pasteur, Paris | Marion Barbet, Sylvie Behillil, Méline Bizard, Angela Brisebarre, Camille Capel, Vincent Enouf, Louise Lefrançois, Frédéric Lemoine, Christophe Malabat, Corinne Maufrais, Pierre Lechat, Etienne Simon-Lorière, Maud Vanpeene, Sylvie Van der Werf ,Patricia Stoessel-Thouvenin |
| EPI_ISL_2627943, EPI_ISL_2627944, EPI_ISL_2627945 | Hospital | National Reference Center for Viruses of Respiratory Infections, Institut Pasteur, Paris | Marion Barbet, Sylvie Behillil, Méline Bizard, Angela Brisebarre, Camille Capel, Vincent Enouf, Louise Lefrançois, Frédéric Lemoine, Christophe Malabat, Corinne Maufrais, Etienne Simon-Lorière, Maud Vanpeene, Sylvie Van der Werf ,Patricia Stoessel-Thouvenin |
| EPI_ISL_2627947, EPI_ISL_2627948 | Hospital | National Reference Center for Viruses of Respiratory Infections, Institut Pasteur, Paris | Marion Barbet, Sylvie Behillil, Méline Bizard, Angela Brisebarre, Camille Capel, Vincent Enouf, Louise Lefrançois, Frédéric Lemoine, Christophe Malabat, Corinne Maufrais, Etienne Simon-Lorière, Maud Vanpeene, Sylvie Van der Werf ,ValéRie Serazin |
| EPI_ISL_2627949, EPI_ISL_2627950 | Labo Analyses Med | National Reference Center for Viruses of Respiratory Infections, Institut Pasteur, Paris | Marion Barbet, Sylvie Behillil, Méline Bizard, Angela Brisebarre, Camille Capel, Vincent Enouf, Louise Lefrançois, Frédéric Lemoine, Christophe Malabat, Corinne Maufrais, Etienne Simon-Lorière, Maud Vanpeene, Sylvie Van der Werf ,GréGoire Potiron |
| EPI_ISL_2627952 | Hospital | National Reference Center for Viruses of Respiratory Infections, Institut Pasteur, Paris | Marion Barbet, Sylvie Behillil, Méline Bizard, Angela Brisebarre, Camille Capel, Vincent Enouf, Louise Lefrançois, Frédéric Lemoine, Christophe Malabat, Corinne Maufrais, Etienne Simon-Lorière, Maud Vanpeene, Sylvie Van der Werf ,Anne-CéCile Hochart |
| EPI_ISL_2627953, EPI_ISL_2627954, EPI_ISL_2627955 | Hospital | National Reference Center for Viruses of Respiratory Infections, Institut Pasteur, Paris | Marion Barbet, Sylvie Behillil, Méline Bizard, Angela Brisebarre, Camille Capel, Vincent Enouf, Louise Lefrançois, Frédéric Lemoine, Christophe Malabat, Corinne Maufrais, Etienne Simon-Lorière, Maud Vanpeene, Sylvie Van der Werf ,Pascale Martres |
| EPI_ISL_2627956 | Hospital | National Reference Center for Viruses of Respiratory Infections, Institut Pasteur, Paris | Marion Barbet, Sylvie Behillil, Méline Bizard, Angela Brisebarre, Camille Capel, Vincent Enouf, Louise Lefrançois, Frédéric Lemoine, Christophe Malabat, Corinne Maufrais, Etienne Simon-Lorière, Maud Vanpeene, Sylvie Van der Werf ,StéPhanie Marque Juillet |
| EPI_ISL_2627958, EPI_ISL_2627959 | Hospital | National Reference Center for Viruses of Respiratory Infections, Institut Pasteur, Paris | Marion Barbet, Sylvie Behillil, Méline Bizard, Angela Brisebarre, Camille Capel, Vincent Enouf, Louise Lefrançois, Frédéric Lemoine, Christophe Malabat, Corinne Maufrais, Etienne Simon-Lorière, Maud Vanpeene, Sylvie Van der Werf ,JéRome Guinard |
| EPI_ISL_2627960 | Hospital | National Reference Center for Viruses of Respiratory Infections, Institut Pasteur, Paris | Marion Barbet, Sylvie Behillil, Méline Bizard, Angela Brisebarre, Camille Capel, Vincent Enouf, Louise Lefrançois, Frédéric Lemoine, Christophe Malabat, Corinne Maufrais, Pierre Lechat, Etienne Simon-Lorière, Maud Vanpeene, Sylvie Van der Werf ,JéRome Guinard |
| EPI_ISL_2627961, EPI_ISL_2627963 | Hospital | National Reference Center for Viruses of Respiratory Infections, Institut Pasteur, Paris | Marion Barbet, Sylvie Behillil, Méline Bizard, Angela Brisebarre, Camille Capel, Vincent Enouf, Louise Lefrançois, Frédéric Lemoine, Christophe Malabat, Corinne Maufrais, Etienne Simon-Lorière, Maud Vanpeene, Sylvie Van der Werf ,JéRome Guinard |
| EPI_ISL_2627965 | Hospital | National Reference Center for Viruses of Respiratory Infections, Institut Pasteur, Paris | Marion Barbet, Sylvie Behillil, Méline Bizard, Angela Brisebarre, Camille Capel, Vincent Enouf, Louise Lefrançois, Frédéric Lemoine, Christophe Malabat, Corinne Maufrais, Etienne Simon-Lorière, Maud Vanpeene, Sylvie Van der Werf ,Laura Djamdjian |
| EPI_ISL_2627966, EPI_ISL_2627967, EPI_ISL_2627968, EPI_ISL_2627969 | Hospital | National Reference Center for Viruses of Respiratory Infections, Institut Pasteur, Paris | Marion Barbet, Sylvie Behillil, Méline Bizard, Angela Brisebarre, Camille Capel, Vincent Enouf, Louise Lefrançois, Frédéric Lemoine, Christophe Malabat, Corinne Maufrais, Etienne Simon-Lorière, Maud Vanpeene, Sylvie Van der Werf ,Laurence Bocket |
| EPI_ISL_2627970 | Hospital | National Reference Center for Viruses of Respiratory Infections, Institut Pasteur, Paris | Marion Barbet, Sylvie Behillil, Méline Bizard, Angela Brisebarre, Camille Capel, Vincent Enouf, Louise Lefrançois, Frédéric Lemoine, Christophe Malabat, Corinne Maufrais, Etienne Simon-Lorière, Maud Vanpeene, Sylvie Van der Werf ,Sylvie Ledru |
| EPI_ISL_2627971, EPI_ISL_2627972, EPI_ISL_2627973 | Labo Analyses Med | National Reference Center for Viruses of Respiratory Infections, Institut Pasteur, Paris | Marion Barbet, Sylvie Behillil, Méline Bizard, Angela Brisebarre, Camille Capel, Vincent Enouf, Louise Lefrançois, Frédéric Lemoine, Christophe Malabat, Corinne Maufrais, Etienne Simon-Lorière, Maud Vanpeene, Sylvie Van der Werf ,Sophie Chalmin |

| | | | |
|---|---|---|---|
| EPI_ISL_2627975, EPI_ISL_2627976, EPI_ISL_2627978 | Hospital | National Reference Center for Viruses of Respiratory Infections, Institut Pasteur, Paris | Marion Barbet, Sylvie Behillil, Méline Bizard, Angela Brisebarre, Camille Capel, Vincent Enouf, Louise Lefrançois, Frédéric Lemoine, Christophe Malabat, Corinne Maufrais, Etienne Simon-Lorière, Maud Vanpeene, Sylvie Van der Werf ,Pascale Martres |
| EPI_ISL_2627979, EPI_ISL_2627980, EPI_ISL_2627981 | Hospital | National Reference Center for Viruses of Respiratory Infections, Institut Pasteur, Paris | Marion Barbet, Sylvie Behillil, Méline Bizard, Angela Brisebarre, Camille Capel, Vincent Enouf, Louise Lefrançois, Frédéric Lemoine, Christophe Malabat, Corinne Maufrais, Etienne Simon-Lorière, Maud Vanpeene, Sylvie Van der Werf ,Sandrine Castelain |
| EPI_ISL_2627983, EPI_ISL_2627984 | Hospital | National Reference Center for Viruses of Respiratory Infections, Institut Pasteur, Paris | Marion Barbet, Sylvie Behillil, Méline Bizard, Angela Brisebarre, Camille Capel, Vincent Enouf, Louise Lefrançois, Frédéric Lemoine, Christophe Malabat, Corinne Maufrais, Etienne Simon-Lorière, Maud Vanpeene, Sylvie Van der Werf ,L. Courdavault |
| EPI_ISL_2627988 | Hospital | National Reference Center for Viruses of Respiratory Infections, Institut Pasteur, Paris | Marion Barbet, Sylvie Behillil, Méline Bizard, Angela Brisebarre, Camille Capel, Vincent Enouf, Louise Lefrançois, Frédéric Lemoine, Christophe Malabat, Corinne Maufrais, Etienne Simon-Lorière, Maud Vanpeene, Sylvie Van der Werf ,Eric Farfour |
| EPI_ISL_2627989, EPI_ISL_2627990 | Armies | National Reference Center for Viruses of Respiratory Infections, Institut Pasteur, Paris | Marion Barbet, Sylvie Behillil, Méline Bizard, Angela Brisebarre, Camille Capel, Vincent Enouf, Louise Lefrançois, Frédéric Lemoine, Christophe Malabat, Corinne Maufrais, Etienne Simon-Lorière, Maud Vanpeene, Sylvie Van der Werf ,Marine Desroches |
| EPI_ISL_2627991, EPI_ISL_2627992, EPI_ISL_2627993, EPI_ISL_2627994 | Hospital | National Reference Center for Viruses of Respiratory Infections, Institut Pasteur, Paris | Marion Barbet, Sylvie Behillil, Méline Bizard, Angela Brisebarre, Camille Capel, Vincent Enouf, Louise Lefrançois, Frédéric Lemoine, Christophe Malabat, Corinne Maufrais, Etienne Simon-Lorière, Maud Vanpeene, Sylvie Van der Werf ,Nabil Gastli |
| EPI_ISL_2627996 | Hospital | National Reference Center for Viruses of Respiratory Infections, Institut Pasteur, Paris | Marion Barbet, Sylvie Behillil, Méline Bizard, Angela Brisebarre, Camille Capel, Vincent Enouf, Louise Lefrançois, Frédéric Lemoine, Christophe Malabat, Corinne Maufrais, Etienne Simon-Lorière, Maud Vanpeene, Sylvie Van der Werf ,Catherine Gaudy Graffin |
| EPI_ISL_2627997 | Labo Analyses Med | National Reference Center for Viruses of Respiratory Infections, Institut Pasteur, Paris | Marion Barbet, Sylvie Behillil, Méline Bizard, Angela Brisebarre, Camille Capel, Vincent Enouf, Louise Lefrançois, Frédéric Lemoine, Christophe Malabat, Corinne Maufrais, Pierre Lechat, Etienne Simon-Lorière, Maud Vanpeene, Sylvie Van der Werf ,Caroline Devaux |
| EPI_ISL_2627998, EPI_ISL_2627999, EPI_ISL_2628000, EPI_ISL_2628001, EPI_ISL_2628002 | Labo Analyses Med | National Reference Center for Viruses of Respiratory Infections, Institut Pasteur, Paris | Marion Barbet, Sylvie Behillil, Méline Bizard, Angela Brisebarre, Camille Capel, Vincent Enouf, Louise Lefrançois, Frédéric Lemoine, Christophe Malabat, Corinne Maufrais, Etienne Simon-Lorière, Maud Vanpeene, Sylvie Van der Werf ,Caroline Devaux |
| EPI_ISL_2628004 | Labo Analyses Med | National Reference Center for Viruses of Respiratory Infections, Institut Pasteur, Paris | Marion Barbet, Sylvie Behillil, Méline Bizard, Angela Brisebarre, Camille Capel, Vincent Enouf, Louise Lefrançois, Frédéric Lemoine, Christophe Malabat, Corinne Maufrais, Etienne Simon-Lorière, Maud Vanpeene, Sylvie Van der Werf ,Arditi Cocco |
| EPI_ISL_2628272, EPI_ISL_2628273 | Labo Analyses Med | National Reference Center for Viruses of Respiratory Infections, Institut Pasteur, Paris | Marion Barbet, Sylvie Behillil, Méline Bizard, Angela Brisebarre, Camille Capel, Vincent Enouf, Louise Lefrançois, Frédéric Lemoine, Christophe Malabat, Corinne Maufrais, Etienne Simon-Lorière, Maud Vanpeene, Sylvie Van der Werf ,Caroline Devaux |
| EPI_ISL_2628274 | Hospital | National Reference Center for Viruses of Respiratory Infections, Institut Pasteur, Paris | Marion Barbet, Sylvie Behillil, Méline Bizard, Angela Brisebarre, Camille Capel, Vincent Enouf, Louise Lefrançois, Frédéric Lemoine, Christophe Malabat, Corinne Maufrais, Etienne Simon-Lorière, Maud Vanpeene, Sylvie Van der Werf ,E. Bichier |
| EPI_ISL_2628275, EPI_ISL_2628276 | Hospital | National Reference Center for Viruses of Respiratory Infections, Institut Pasteur, Paris | Marion Barbet, Sylvie Behillil, Méline Bizard, Angela Brisebarre, Camille Capel, Vincent Enouf, Louise Lefrançois, Frédéric Lemoine, Christophe Malabat, Corinne Maufrais, Etienne Simon-Lorière, Maud Vanpeene, Sylvie Van der Werf ,CéCile Hermann |
| EPI_ISL_2628277 | Hospital | National Reference Center for Viruses of Respiratory Infections, Institut Pasteur, Paris | Marion Barbet, Sylvie Behillil, Méline Bizard, Angela Brisebarre, Camille Capel, Vincent Enouf, Louise Lefrançois, Frédéric Lemoine, Christophe Malabat, Corinne Maufrais, Etienne Simon-Lorière, Maud Vanpeene, Sylvie Van der Werf ,Marianne Burgard |
| EPI_ISL_2628283, EPI_ISL_2628284, EPI_ISL_2628285, EPI_ISL_2628286 | Hospital | National Reference Center for Viruses of Respiratory Infections, Institut Pasteur, Paris | Marion Barbet, Sylvie Behillil, Méline Bizard, Angela Brisebarre, Camille Capel, Vincent Enouf, Louise Lefrançois, Frédéric Lemoine, Christophe Malabat, Corinne Maufrais, Etienne Simon-Lorière, Maud Vanpeene, Sylvie Van der Werf ,Marianne Coste-Burel |
| EPI_ISL_2628289, EPI_ISL_2628292, EPI_ISL_2628294 | Hospital | National Reference Center for Viruses of Respiratory Infections, Institut Pasteur, Paris | Marion Barbet, Sylvie Behillil, Méline Bizard, Angela Brisebarre, Camille Capel, Vincent Enouf, Louise Lefrançois, Frédéric Lemoine, Christophe Malabat, Corinne Maufrais, Etienne Simon-Lorière, Maud Vanpeene, Sylvie Van der Werf ,LéA Pilorge |
| EPI_ISL_2628295, EPI_ISL_2628296 | Labo Analyses Med | National Reference Center for Viruses of Respiratory Infections, Institut Pasteur, Paris | Marion Barbet, Sylvie Behillil, Méline Bizard, Angela Brisebarre, Camille Capel, Vincent Enouf, Louise Lefrançois, Frédéric Lemoine, Christophe Malabat, Corinne Maufrais, Etienne Simon-Lorière, Maud Vanpeene, Sylvie Van der Werf ,Claire Felloni |
| EPI_ISL_2637227 | Genome Analysis Center, Yamanashi Central Hospital | Genome Analysis Center, Yamanashi Central Hospital | Yosuke Hirotsu |
| EPI_ISL_2637418, EPI_ISL_2637495, EPI_ISL_2637510, EPI_ISL_2637569, EPI_ISL_2637659, EPI_ISL_2637700, EPI_ISL_2637735 | GH A.CHENEVIER-H.MONDOR | Department of Virology, Henri Mondor University Hospital, Assistance Publique Hôpitaux de Paris, Université Paris-Est Créteil, INSERM U955 | Christophe Rodriguez, Slim Fourati, Vanessa Demontant, Guillaume Gricourt, Melissa N'Debi, Alexandre Soulier, Elisabeth Trawinski, Jean-Michel Pawlotsky |
| EPI_ISL_2638046 | Clinique Saint-Pierre Ottignies | UCLouvain/IREC/MBLG | Jean Ruelle, Bertrand Bearzatto, Benoit Kabamba Mukadi |
| EPI_ISL_2644664 | Genome Analysis Center, Yamanashi Central Hospital | Genome Analysis Center, Yamanashi Central Hospital | Yosuke Hirotsu |
| EPI_ISL_2645445, EPI_ISL_2645446, EPI_ISL_2645447, EPI_ISL_2645448, EPI_ISL_2645449, EPI_ISL_2645450, EPI_ISL_2645452, EPI_ISL_2645453, EPI_ISL_2645455, EPI_ISL_2645457, EPI_ISL_2645459, EPI_ISL_2645460, EPI_ISL_2645462, EPI_ISL_2645464, EPI_ISL_2645467, EPI_ISL_2645468, EPI_ISL_2645469, EPI_ISL_2645473, EPI_ISL_2645475, EPI_ISL_2645476, EPI_ISL_2645477, EPI_ISL_2645478, EPI_ISL_2645479, EPI_ISL_2645481, EPI_ISL_2645483, EPI_ISL_2645486, EPI_ISL_2645493 | | | |
| see above | Laboratorio Central de Saude Publica do Estado do Parana (LACEN/PR) | Laboratory of Respiratory Viruses and Measles, Oswaldo Cruz Institute, FIOCRUZ | Paola Resende, Luciana Appolinario, Fernando Motta, Anna Carolina Paixao, Ana Carolina Mendonca, Alice Sampaio Rocha, Taina Venas, Elisa Cavalcante Pereira, Renata Serrano Lopes, Irina Riediger, Marilda Siqueira on behalf of the Fiocruz COVID-19 Genomic Surveillance Network |
| EPI_ISL_2646140 | CH ARMENTIERES | CHU Lille - Laboratoire de Virologie | AIT YAHYA Emilie, ALIDJINOU Enagnon Kazali, BOCKET Laurence, CREPIN Michel, DEMAY Christophe, ENGELMANN Ilka, GEFFROY Sandrine, GUIGON Aurélie, LAMBERT Valérie, LAZREK Mouna, NOBILLIAUX Florian, PREVOST Brigitte, THUILLIER Caroline, TINEZ Claire ,TCHANTCHOU NJOSSE YANICK |
| EPI_ISL_2646141 | CH BETHUNE | CHU Lille - Laboratoire de Virologie | AIT YAHYA Emilie, ALIDJINOU Enagnon Kazali, BOCKET Laurence, CREPIN Michel, DEMAY Christophe, ENGELMANN Ilka, GEFFROY Sandrine, GUIGON Aurélie, LAMBERT Valérie, LAZREK Mouna, NOBILLIAUX Florian, PREVOST Brigitte, THUILLIER Caroline, TINEZ Claire ,TCHANTCHOU NJOSSE YANICK |
| EPI_ISL_2646142, EPI_ISL_2646143 | CH ROUBAIX | CHU Lille - Laboratoire de Virologie | AIT YAHYA Emilie, ALIDJINOU Enagnon Kazali, BOCKET Laurence, CREPIN Michel, DEMAY Christophe, ENGELMANN Ilka, GEFFROY Sandrine, GUIGON Aurélie, LAMBERT Valérie, LAZREK Mouna, NOBILLIAUX Florian, PREVOST Brigitte, THUILLIER Caroline, TINEZ Claire ,TCHANTCHOU NJOSSE YANICK |
| EPI_ISL_2646144 | CH ARMENTIERES | CHU Lille - Laboratoire de Virologie | AIT YAHYA Emilie, ALIDJINOU Enagnon Kazali, BOCKET Laurence, CREPIN Michel, DEMAY Christophe, ENGELMANN Ilka, GEFFROY Sandrine, GUIGON Aurélie, LAMBERT Valérie, LAZREK Mouna, NOBILLIAUX Florian, PREVOST Brigitte, THUILLIER Caroline, TINEZ Claire ,TCHANTCHOU NJOSSE YANICK |
| EPI_ISL_2646145, EPI_ISL_2646146 | CH CALAIS | CHU Lille - Laboratoire de Virologie | AIT YAHYA Emilie, ALIDJINOU Enagnon Kazali, BOCKET Laurence, CREPIN Michel, DEMAY Christophe, ENGELMANN Ilka, GEFFROY Sandrine, GUIGON Aurélie, LAMBERT Valérie, LAZREK Mouna, NOBILLIAUX Florian, PREVOST Brigitte, THUILLIER Caroline, TINEZ Claire ,TCHANTCHOU NJOSSE YANICK |
| EPI_ISL_2646147 | CH ROUBAIX | CHU Lille - Laboratoire de Virologie | AIT YAHYA Emilie, ALIDJINOU Enagnon Kazali, BOCKET Laurence, CREPIN Michel, DEMAY Christophe, ENGELMANN Ilka, GEFFROY Sandrine, GUIGON Aurélie, LAMBERT Valérie, LAZREK Mouna, NOBILLIAUX Florian, PREVOST Brigitte, THUILLIER Caroline, TINEZ Claire ,TCHANTCHOU NJOSSE YANICK |
| EPI_ISL_2646148 | CH ARMENTIERES | CHU Lille - Laboratoire de Virologie | AIT YAHYA Emilie, ALIDJINOU Enagnon Kazali, BOCKET Laurence, CREPIN Michel, DEMAY Christophe, ENGELMANN Ilka, GEFFROY Sandrine, GUIGON Aurélie, LAMBERT Valérie, LAZREK Mouna, NOBILLIAUX Florian, PREVOST Brigitte, THUILLIER Caroline, TINEZ Claire ,TCHANTCHOU NJOSSE YANICK |
| EPI_ISL_2646149 | CH LENS | CHU Lille - Laboratoire de Virologie | AIT YAHYA Emilie, ALIDJINOU Enagnon Kazali, BOCKET Laurence, CREPIN Michel, DEMAY Christophe, ENGELMANN Ilka, GEFFROY Sandrine, GUIGON Aurélie, LAMBERT Valérie, LAZREK Mouna, NOBILLIAUX Florian, PREVOST Brigitte, THUILLIER Caroline, TINEZ Claire ,TCHANTCHOU NJOSSE YANICK |
| EPI_ISL_2646150, EPI_ISL_2646151 | CH ARMENTIERES | CHU Lille - Laboratoire de Virologie | AIT YAHYA Emilie, ALIDJINOU Enagnon Kazali, BOCKET Laurence, CREPIN Michel, DEMAY Christophe, ENGELMANN Ilka, GEFFROY Sandrine, GUIGON Aurélie, LAMBERT Valérie, LAZREK Mouna, NOBILLIAUX Florian, PREVOST Brigitte, THUILLIER Caroline, TINEZ Claire ,TCHANTCHOU |

| | | | |
|---|---|---|---|
| | | | NJOSSE YANICK |
| EPI_ISL_2646152, EPI_ISL_2646153, EPI_ISL_2646154 | CHU LILLE | CHU Lille - Laboratoire de Virologie | AIT YAHYA Emilie, ALIDJINOU Enagnon Kazali, BOCKET Laurence, CREPIN Michel, DEMAY Christophe, ENGELMANN Ilka, GEFFROY Sandrine, GUIGON Aurélie, LAMBERT Valérie, LAZREK Mouna, NOBILLIAUX Florian, PREVOST Brigitte, THUILLIER Caroline, TINEZ Claire ,TCHANTCHOU NJOSSE YANICK |
| EPI_ISL_2646155 | CH CALAIS | CHU Lille - Laboratoire de Virologie | AIT YAHYA Emilie, ALIDJINOU Enagnon Kazali, BOCKET Laurence, CREPIN Michel, DEMAY Christophe, ENGELMANN Ilka, GEFFROY Sandrine, GUIGON Aurélie, LAMBERT Valérie, LAZREK Mouna, NOBILLIAUX Florian, PREVOST Brigitte, THUILLIER Caroline, TINEZ Claire ,TCHANTCHOU NJOSSE YANICK |
| EPI_ISL_2646156 | CH ARRAS | CHU Lille - Laboratoire de Virologie | AIT YAHYA Emilie, ALIDJINOU Enagnon Kazali, BOCKET Laurence, CREPIN Michel, DEMAY Christophe, ENGELMANN Ilka, GEFFROY Sandrine, GUIGON Aurélie, LAMBERT Valérie, LAZREK Mouna, NOBILLIAUX Florian, PREVOST Brigitte, THUILLIER Caroline, TINEZ Claire ,TCHANTCHOU NJOSSE YANICK |
| EPI_ISL_2646157 | CH ROUBAIX | CHU Lille - Laboratoire de Virologie | AIT YAHYA Emilie, ALIDJINOU Enagnon Kazali, BOCKET Laurence, CREPIN Michel, DEMAY Christophe, ENGELMANN Ilka, GEFFROY Sandrine, GUIGON Aurélie, LAMBERT Valérie, LAZREK Mouna, NOBILLIAUX Florian, PREVOST Brigitte, THUILLIER Caroline, TINEZ Claire ,TCHANTCHOU NJOSSE YANICK |
| EPI_ISL_2646158, EPI_ISL_2646160 | CHU LILLE | CHU Lille - Laboratoire de Virologie | AIT YAHYA Emilie, ALIDJINOU Enagnon Kazali, BOCKET Laurence, CREPIN Michel, DEMAY Christophe, ENGELMANN Ilka, GEFFROY Sandrine, GUIGON Aurélie, LAMBERT Valérie, LAZREK Mouna, NOBILLIAUX Florian, PREVOST Brigitte, THUILLIER Caroline, TINEZ Claire ,TCHANTCHOU NJOSSE YANICK |
| EPI_ISL_2646161 | CH ROUBAIX | CHU Lille - Laboratoire de Virologie | AIT YAHYA Emilie, ALIDJINOU Enagnon Kazali, BOCKET Laurence, CREPIN Michel, DEMAY Christophe, ENGELMANN Ilka, GEFFROY Sandrine, GUIGON Aurélie, LAMBERT Valérie, LAZREK Mouna, NOBILLIAUX Florian, PREVOST Brigitte, THUILLIER Caroline, TINEZ Claire ,TCHANTCHOU NJOSSE YANICK |
| EPI_ISL_2646162, EPI_ISL_2646163 | CH BETHUNE | CHU Lille - Laboratoire de Virologie | AIT YAHYA Emilie, ALIDJINOU Enagnon Kazali, BOCKET Laurence, CREPIN Michel, DEMAY Christophe, ENGELMANN Ilka, GEFFROY Sandrine, GUIGON Aurélie, LAMBERT Valérie, LAZREK Mouna, NOBILLIAUX Florian, PREVOST Brigitte, THUILLIER Caroline, TINEZ Claire ,TCHANTCHOU NJOSSE YANICK |
| EPI_ISL_2646164, EPI_ISL_2646165, EPI_ISL_2646166, EPI_ISL_2646167, EPI_ISL_2646168 | CHU LILLE | CHU Lille - Laboratoire de Virologie | AIT YAHYA Emilie, ALIDJINOU Enagnon Kazali, BOCKET Laurence, CREPIN Michel, DEMAY Christophe, ENGELMANN Ilka, GEFFROY Sandrine, GUIGON Aurélie, LAMBERT Valérie, LAZREK Mouna, NOBILLIAUX Florian, PREVOST Brigitte, THUILLIER Caroline, TINEZ Claire ,TCHANTCHOU NJOSSE YANICK |
| EPI_ISL_2646169 | CH LENS | CHU Lille - Laboratoire de Virologie | AIT YAHYA Emilie, ALIDJINOU Enagnon Kazali, BOCKET Laurence, CREPIN Michel, DEMAY Christophe, ENGELMANN Ilka, GEFFROY Sandrine, GUIGON Aurélie, LAMBERT Valérie, LAZREK Mouna, NOBILLIAUX Florian, PREVOST Brigitte, THUILLIER Caroline, TINEZ Claire ,TCHANTCHOU NJOSSE YANICK |
| EPI_ISL_2646170, EPI_ISL_2646171 | CHU LILLE | CHU Lille - Laboratoire de Virologie | AIT YAHYA Emilie, ALIDJINOU Enagnon Kazali, BOCKET Laurence, CREPIN Michel, DEMAY Christophe, ENGELMANN Ilka, GEFFROY Sandrine, GUIGON Aurélie, LAMBERT Valérie, LAZREK Mouna, NOBILLIAUX Florian, PREVOST Brigitte, THUILLIER Caroline, TINEZ Claire ,TCHANTCHOU NJOSSE YANICK |
| EPI_ISL_2646172, EPI_ISL_2646173 | CH ROUBAIX | CHU Lille - Laboratoire de Virologie | AIT YAHYA Emilie, ALIDJINOU Enagnon Kazali, BOCKET Laurence, CREPIN Michel, DEMAY Christophe, ENGELMANN Ilka, GEFFROY Sandrine, GUIGON Aurélie, LAMBERT Valérie, LAZREK Mouna, NOBILLIAUX Florian, PREVOST Brigitte, THUILLIER Caroline, TINEZ Claire ,TCHANTCHOU NJOSSE YANICK |
| EPI_ISL_2646174 | CH TOURCOING | CHU Lille - Laboratoire de Virologie | AIT YAHYA Emilie, ALIDJINOU Enagnon Kazali, BOCKET Laurence, CREPIN Michel, DEMAY Christophe, ENGELMANN Ilka, GEFFROY Sandrine, GUIGON Aurélie, LAMBERT Valérie, LAZREK Mouna, NOBILLIAUX Florian, PREVOST Brigitte, THUILLIER Caroline, TINEZ Claire ,TCHANTCHOU NJOSSE YANICK |
| EPI_ISL_2646175 | CH BETHUNE | CHU Lille - Laboratoire de Virologie | AIT YAHYA Emilie, ALIDJINOU Enagnon Kazali, BOCKET Laurence, CREPIN Michel, DEMAY Christophe, ENGELMANN Ilka, GEFFROY Sandrine, GUIGON Aurélie, LAMBERT Valérie, LAZREK Mouna, NOBILLIAUX Florian, PREVOST Brigitte, THUILLIER Caroline, TINEZ Claire ,TCHANTCHOU NJOSSE YANICK |
| EPI_ISL_2646177, EPI_ISL_2646178 | CHU LILLE | CHU Lille - Laboratoire de Virologie | AIT YAHYA Emilie, ALIDJINOU Enagnon Kazali, BOCKET Laurence, CREPIN Michel, DEMAY Christophe, ENGELMANN Ilka, GEFFROY Sandrine, GUIGON Aurélie, LAMBERT Valérie, LAZREK Mouna, NOBILLIAUX Florian, PREVOST Brigitte, THUILLIER Caroline, TINEZ Claire ,TCHANTCHOU NJOSSE YANICK |
| EPI_ISL_2646179 | CH ARMENTIERES | CHU Lille - Laboratoire de Virologie | AIT YAHYA Emilie, ALIDJINOU Enagnon Kazali, BOCKET Laurence, CREPIN Michel, DEMAY Christophe, ENGELMANN Ilka, GEFFROY Sandrine, GUIGON Aurélie, LAMBERT Valérie, LAZREK Mouna, NOBILLIAUX Florian, PREVOST Brigitte, THUILLIER Caroline, TINEZ Claire ,TCHANTCHOU NJOSSE YANICK |
| EPI_ISL_2646180, EPI_ISL_2646181, EPI_ISL_2646182, EPI_ISL_2646183, EPI_ISL_2646184, EPI_ISL_2646185, EPI_ISL_2646186 | CHU LILLE | CHU Lille - Laboratoire de Virologie | AIT YAHYA Emilie, ALIDJINOU Enagnon Kazali, BOCKET Laurence, CREPIN Michel, DEMAY Christophe, ENGELMANN Ilka, GEFFROY Sandrine, GUIGON Aurélie, LAMBERT Valérie, LAZREK Mouna, NOBILLIAUX Florian, PREVOST Brigitte, THUILLIER Caroline, TINEZ Claire ,TCHANTCHOU NJOSSE YANICK |
| EPI_ISL_2646187, EPI_ISL_2646188 | CH ROUBAIX | CHU Lille - Laboratoire de Virologie | AIT YAHYA Emilie, ALIDJINOU Enagnon Kazali, BOCKET Laurence, CREPIN Michel, DEMAY Christophe, ENGELMANN Ilka, GEFFROY Sandrine, GUIGON Aurélie, LAMBERT Valérie, LAZREK Mouna, NOBILLIAUX Florian, PREVOST Brigitte, THUILLIER Caroline, TINEZ Claire ,TCHANTCHOU NJOSSE YANICK |
| EPI_ISL_2646189, EPI_ISL_2646190 | CHU LILLE | CHU Lille - Laboratoire de Virologie | AIT YAHYA Emilie, ALIDJINOU Enagnon Kazali, BOCKET Laurence, CREPIN Michel, DEMAY Christophe, ENGELMANN Ilka, GEFFROY Sandrine, GUIGON Aurélie, LAMBERT Valérie, LAZREK Mouna, NOBILLIAUX Florian, PREVOST Brigitte, THUILLIER Caroline, TINEZ Claire ,TCHANTCHOU NJOSSE YANICK |
| EPI_ISL_2646192, EPI_ISL_2646193 | CH ARRAS | CHU Lille - Laboratoire de Virologie | AIT YAHYA Emilie, ALIDJINOU Enagnon Kazali, BOCKET Laurence, CREPIN Michel, DEMAY Christophe, ENGELMANN Ilka, GEFFROY Sandrine, GUIGON Aurélie, LAMBERT Valérie, LAZREK Mouna, NOBILLIAUX Florian, PREVOST Brigitte, THUILLIER Caroline, TINEZ Claire ,TCHANTCHOU NJOSSE YANICK |
| EPI_ISL_2646195, EPI_ISL_2646196 | CH ROUBAIX | CHU Lille - Laboratoire de Virologie | AIT YAHYA Emilie, ALIDJINOU Enagnon Kazali, BOCKET Laurence, CREPIN Michel, DEMAY Christophe, ENGELMANN Ilka, GEFFROY Sandrine, GUIGON Aurélie, LAMBERT Valérie, LAZREK Mouna, NOBILLIAUX Florian, PREVOST Brigitte, THUILLIER Caroline, TINEZ Claire ,TCHANTCHOU NJOSSE YANICK |
| EPI_ISL_2646197 | CH BETHUNE | CHU Lille - Laboratoire de Virologie | AIT YAHYA Emilie, ALIDJINOU Enagnon Kazali, BOCKET Laurence, CREPIN Michel, DEMAY Christophe, ENGELMANN Ilka, GEFFROY Sandrine, GUIGON Aurélie, LAMBERT Valérie, LAZREK Mouna, NOBILLIAUX Florian, PREVOST Brigitte, THUILLIER Caroline, TINEZ Claire ,TCHANTCHOU NJOSSE YANICK |
| EPI_ISL_2646198, EPI_ISL_2646199 | CH ARMENTIERES | CHU Lille - Laboratoire de Virologie | AIT YAHYA Emilie, ALIDJINOU Enagnon Kazali, BOCKET Laurence, CREPIN Michel, DEMAY Christophe, ENGELMANN Ilka, GEFFROY Sandrine, GUIGON Aurélie, LAMBERT Valérie, LAZREK Mouna, NOBILLIAUX Florian, PREVOST Brigitte, THUILLIER Caroline, TINEZ Claire ,TCHANTCHOU NJOSSE YANICK |
| EPI_ISL_2646200 | CH TOURCOING | CHU Lille - Laboratoire de Virologie | AIT YAHYA Emilie, ALIDJINOU Enagnon Kazali, BOCKET Laurence, CREPIN Michel, DEMAY Christophe, ENGELMANN Ilka, GEFFROY Sandrine, GUIGON Aurélie, LAMBERT Valérie, LAZREK Mouna, NOBILLIAUX Florian, PREVOST Brigitte, THUILLIER Caroline, TINEZ Claire ,TCHANTCHOU NJOSSE YANICK |
| EPI_ISL_2646201 | CH ARMENTIERES | CHU Lille - Laboratoire de Virologie | AIT YAHYA Emilie, ALIDJINOU Enagnon Kazali, BOCKET Laurence, CREPIN Michel, DEMAY Christophe, ENGELMANN Ilka, GEFFROY Sandrine, GUIGON Aurélie, LAMBERT Valérie, LAZREK Mouna, NOBILLIAUX Florian, PREVOST Brigitte, THUILLIER Caroline, TINEZ Claire ,TCHANTCHOU NJOSSE YANICK |
| EPI_ISL_2646202 | CH CALAIS | CHU Lille - Laboratoire de Virologie | AIT YAHYA Emilie, ALIDJINOU Enagnon Kazali, BOCKET Laurence, CREPIN Michel, DEMAY Christophe, ENGELMANN Ilka, GEFFROY Sandrine, |

| | | | |
|---|---|---|---|
| | | | GUIGON Aurélie, LAMBERT Valérie, LAZREK Mouna, NOBILLIAUX Florian, PREVOST Brigitte, THUILLIER Caroline, TINEZ Claire ,TCHANTCHOU NJOSSE YANICK |
| EPI_ISL_2646203 | CHU LILLE | CHU Lille - Laboratoire de Virologie | AIT YAHYA Emilie, ALIDJINOU Enagnon Kazali, BOCKET Laurence, CREPIN Michel, DEMAY Christophe, ENGELMANN Ilka, GEFFROY Sandrine, GUIGON Aurélie, LAMBERT Valérie, LAZREK Mouna, NOBILLIAUX Florian, PREVOST Brigitte, THUILLIER Caroline, TINEZ Claire ,TCHANTCHOU NJOSSE YANICK |
| EPI_ISL_2646204 | CH CAMBRAI | CHU Lille - Laboratoire de Virologie | AIT YAHYA Emilie, ALIDJINOU Enagnon Kazali, BOCKET Laurence, CREPIN Michel, DEMAY Christophe, ENGELMANN Ilka, GEFFROY Sandrine, GUIGON Aurélie, LAMBERT Valérie, LAZREK Mouna, NOBILLIAUX Florian, PREVOST Brigitte, THUILLIER Caroline, TINEZ Claire ,TCHANTCHOU NJOSSE YANICK |
| EPI_ISL_2646205, EPI_ISL_2646206, EPI_ISL_2646207 | CHU LILLE | CHU Lille - Laboratoire de Virologie | AIT YAHYA Emilie, ALIDJINOU Enagnon Kazali, BOCKET Laurence, CREPIN Michel, DEMAY Christophe, ENGELMANN Ilka, GEFFROY Sandrine, GUIGON Aurélie, LAMBERT Valérie, LAZREK Mouna, NOBILLIAUX Florian, PREVOST Brigitte, THUILLIER Caroline, TINEZ Claire ,TCHANTCHOU NJOSSE YANICK |
| EPI_ISL_2646208 | CH TOURCOING | CHU Lille - Laboratoire de Virologie | AIT YAHYA Emilie, ALIDJINOU Enagnon Kazali, BOCKET Laurence, CREPIN Michel, DEMAY Christophe, ENGELMANN Ilka, GEFFROY Sandrine, GUIGON Aurélie, LAMBERT Valérie, LAZREK Mouna, NOBILLIAUX Florian, PREVOST Brigitte, THUILLIER Caroline, TINEZ Claire ,TCHANTCHOU NJOSSE YANICK |
| EPI_ISL_2646210 | CH Valenciennes | CHU Lille - Laboratoire de Virologie | AIT YAHYA Emilie, ALIDJINOU Enagnon Kazali, BOCKET Laurence, CREPIN Michel, DEMAY Christophe, ENGELMANN Ilka, GEFFROY Sandrine, GUIGON Aurélie, LAMBERT Valérie, LAZREK Mouna, NOBILLIAUX Florian, PREVOST Brigitte, THUILLIER Caroline, TINEZ Claire ,TCHANTCHOU NJOSSE YANICK |
| EPI_ISL_2646211 | CHU LILLE | CHU Lille - Laboratoire de Virologie | AIT YAHYA Emilie, ALIDJINOU Enagnon Kazali, BOCKET Laurence, CREPIN Michel, DEMAY Christophe, ENGELMANN Ilka, GEFFROY Sandrine, GUIGON Aurélie, LAMBERT Valérie, LAZREK Mouna, NOBILLIAUX Florian, PREVOST Brigitte, THUILLIER Caroline, TINEZ Claire ,TCHANTCHOU NJOSSE YANICK |
| EPI_ISL_2646212 | CH Valenciennes | CHU Lille - Laboratoire de Virologie | AIT YAHYA Emilie, ALIDJINOU Enagnon Kazali, BOCKET Laurence, CREPIN Michel, DEMAY Christophe, ENGELMANN Ilka, GEFFROY Sandrine, GUIGON Aurélie, LAMBERT Valérie, LAZREK Mouna, NOBILLIAUX Florian, PREVOST Brigitte, THUILLIER Caroline, TINEZ Claire ,TCHANTCHOU NJOSSE YANICK |
| EPI_ISL_2646213 | CHU LILLE | CHU Lille - Laboratoire de Virologie | AIT YAHYA Emilie, ALIDJINOU Enagnon Kazali, BOCKET Laurence, CREPIN Michel, DEMAY Christophe, ENGELMANN Ilka, GEFFROY Sandrine, GUIGON Aurélie, LAMBERT Valérie, LAZREK Mouna, NOBILLIAUX Florian, PREVOST Brigitte, THUILLIER Caroline, TINEZ Claire ,TCHANTCHOU NJOSSE YANICK |
| EPI_ISL_2646214 | CH ARRAS | CHU Lille - Laboratoire de Virologie | AIT YAHYA Emilie, ALIDJINOU Enagnon Kazali, BOCKET Laurence, CREPIN Michel, DEMAY Christophe, ENGELMANN Ilka, GEFFROY Sandrine, GUIGON Aurélie, LAMBERT Valérie, LAZREK Mouna, NOBILLIAUX Florian, PREVOST Brigitte, THUILLIER Caroline, TINEZ Claire ,TCHANTCHOU NJOSSE YANICK |
| EPI_ISL_2646215 | CHU LILLE | CHU Lille - Laboratoire de Virologie | AIT YAHYA Emilie, ALIDJINOU Enagnon Kazali, BOCKET Laurence, CREPIN Michel, DEMAY Christophe, ENGELMANN Ilka, GEFFROY Sandrine, GUIGON Aurélie, LAMBERT Valérie, LAZREK Mouna, NOBILLIAUX Florian, PREVOST Brigitte, THUILLIER Caroline, TINEZ Claire ,TCHANTCHOU NJOSSE YANICK |
| EPI_ISL_2646216 | CH CAMBRAI | CHU Lille - Laboratoire de Virologie | AIT YAHYA Emilie, ALIDJINOU Enagnon Kazali, BOCKET Laurence, CREPIN Michel, DEMAY Christophe, ENGELMANN Ilka, GEFFROY Sandrine, GUIGON Aurélie, LAMBERT Valérie, LAZREK Mouna, NOBILLIAUX Florian, PREVOST Brigitte, THUILLIER Caroline, TINEZ Claire ,TCHANTCHOU NJOSSE YANICK |
| EPI_ISL_2646217, EPI_ISL_2646218 | CHU LILLE | CHU Lille - Laboratoire de Virologie | AIT YAHYA Emilie, ALIDJINOU Enagnon Kazali, BOCKET Laurence, CREPIN Michel, DEMAY Christophe, ENGELMANN Ilka, GEFFROY Sandrine, GUIGON Aurélie, LAMBERT Valérie, LAZREK Mouna, NOBILLIAUX Florian, PREVOST Brigitte, THUILLIER Caroline, TINEZ Claire ,TCHANTCHOU NJOSSE YANICK |
| EPI_ISL_2646219 | cerballiance-IDF | Cerba lab | Emmanuel Lecorche, Sabine Trombert-Paolantoni, Aude Lessenne, Kader Merah, Thierry Collin, Laura Verdurme, Patrice Herisson, Stéphanie Haïm-Boukobza, Bénédicte Roquebert |
| EPI_ISL_2646220, EPI_ISL_2646221, EPI_ISL_2646222, EPI_ISL_2646223 | Exalab | Cerba lab | Emmanuel Lecorche, Sabine Trombert-Paolantoni, Aude Lessenne, Kader Merah, Thierry Collin, Laura Verdurme, Patrice Herisson, Stéphanie Haïm-Boukobza, Bénédicte Roquebert |
| EPI_ISL_2646224 | cerballiance-IDF | Cerba lab | Emmanuel Lecorche, Sabine Trombert-Paolantoni, Aude Lessenne, Kader Merah, Thierry Collin, Laura Verdurme, Patrice Herisson, Stéphanie Haïm-Boukobza, Bénédicte Roquebert |
| EPI_ISL_2646225, EPI_ISL_2646226, EPI_ISL_2646227, EPI_ISL_2646228, EPI_ISL_2646229, EPI_ISL_2646230, EPI_ISL_2646231 | Exalab | Cerba lab | Emmanuel Lecorche, Sabine Trombert-Paolantoni, Aude Lessenne, Kader Merah, Thierry Collin, Laura Verdurme, Patrice Herisson, Stéphanie Haïm-Boukobza, Bénédicte Roquebert |
| EPI_ISL_2646232 | Clinique de Meudon | Cerba lab | Emmanuel Lecorche, Sabine Trombert-Paolantoni, Aude Lessenne, Kader Merah, Thierry Collin, Laura Verdurme, Patrice Herisson, Stéphanie Haïm-Boukobza, Bénédicte Roquebert |
| EPI_ISL_2646233, EPI_ISL_2646234, EPI_ISL_2646235, EPI_ISL_2646236, EPI_ISL_2646237, EPI_ISL_2646238 | Cerballiance-IDF | Cerba lab | Emmanuel Lecorche, Sabine Trombert-Paolantoni, Aude Lessenne, Kader Merah, Thierry Collin, Laura Verdurme, Patrice Herisson, Stéphanie Haïm-Boukobza, Bénédicte Roquebert |
| EPI_ISL_2646239, EPI_ISL_2646240 | Cerballiance-Provence | Cerba lab | Emmanuel Lecorche, Sabine Trombert-Paolantoni, Aude Lessenne, Kader Merah, Thierry Collin, Laura Verdurme, Patrice Herisson, Stéphanie Haïm-Boukobza, Bénédicte Roquebert |
| EPI_ISL_2646241 | Cerballiance-Normandie | Cerba lab | Emmanuel Lecorche, Sabine Trombert-Paolantoni, Aude Lessenne, Kader Merah, Thierry Collin, Laura Verdurme, Patrice Herisson, Stéphanie Haïm-Boukobza, Bénédicte Roquebert |
| EPI_ISL_2646242 | Cerballiance-IDF | Cerba lab | Emmanuel Lecorche, Sabine Trombert-Paolantoni, Aude Lessenne, Kader Merah, Thierry Collin, Laura Verdurme, Patrice Herisson, Stéphanie Haïm-Boukobza, Bénédicte Roquebert |
| EPI_ISL_2646243 | Cerballiance-Provence | Cerba lab | Emmanuel Lecorche, Sabine Trombert-Paolantoni, Aude Lessenne, Kader Merah, Thierry Collin, Laura Verdurme, Patrice Herisson, Stéphanie Haïm-Boukobza, Bénédicte Roquebert |
| EPI_ISL_2646245, EPI_ISL_2646246 | Cerballiance-Paris | Cerba lab | Emmanuel Lecorche, Sabine Trombert-Paolantoni, Aude Lessenne, Kader Merah, Thierry Collin, Laura Verdurme, Patrice Herisson, Stéphanie Haïm-Boukobza, Bénédicte Roquebert |
| EPI_ISL_2646247 | Cerballiance-Normandie | Cerba lab | Emmanuel Lecorche, Sabine Trombert-Paolantoni, Aude Lessenne, Kader Merah, Thierry Collin, Laura Verdurme, Patrice Herisson, Stéphanie Haïm-Boukobza, Bénédicte Roquebert |
| EPI_ISL_2646248, EPI_ISL_2646249, EPI_ISL_2646250 | Cerballiance-IDF | Cerba lab | Emmanuel Lecorche, Sabine Trombert-Paolantoni, Aude Lessenne, Kader Merah, Thierry Collin, Laura Verdurme, Patrice Herisson, Stéphanie Haïm-Boukobza, Bénédicte Roquebert |
| EPI_ISL_2646251 | Synlab | Cerba lab | Emmanuel Lecorche, Sabine Trombert-Paolantoni, Aude Lessenne, Kader Merah, Thierry Collin, Laura Verdurme, Patrice Herisson, Stéphanie Haïm-Boukobza, Bénédicte Roquebert |
| EPI_ISL_2646252 | Cerballiance-Paris | Cerba lab | Emmanuel Lecorche, Sabine Trombert-Paolantoni, Aude Lessenne, Kader Merah, Thierry Collin, Laura Verdurme, Patrice Herisson, Stéphanie Haïm-Boukobza, Bénédicte Roquebert |
| EPI_ISL_2646253, EPI_ISL_2646254 | Cerballiance-Pyrénées | Cerba lab | Emmanuel Lecorche, Sabine Trombert-Paolantoni, Aude Lessenne, Kader Merah, Thierry Collin, Laura Verdurme, Patrice Herisson, Stéphanie Haïm-Boukobza, Bénédicte Roquebert |

| | | | |
|---|---|---|---|
| EPI_ISL_2646255 | Cerballiance-Paris | Cerba lab | Emmanuel Lecorche, Sabine Trombert-Paolantoni, Aude Lessenne, Kader Merah, Thierry Collin, Laura Verdurme, Patrice Herisson, Stéphanie Haïm-Boukobza, Bénédicte Roquebert |
| EPI_ISL_2646256 | Cerballiance-Centre val de Loire | Cerba lab | Emmanuel Lecorche, Sabine Trombert-Paolantoni, Aude Lessenne, Kader Merah, Thierry Collin, Laura Verdurme, Patrice Herisson, Stéphanie Haïm-Boukobza, Bénédicte Roquebert |
| EPI_ISL_2646257 | Cerballiance-Pyrénées | Cerba lab | Emmanuel Lecorche, Sabine Trombert-Paolantoni, Aude Lessenne, Kader Merah, Thierry Collin, Laura Verdurme, Patrice Herisson, Stéphanie Haïm-Boukobza, Bénédicte Roquebert |
| EPI_ISL_2646258 | Cerballiance-Paris | Cerba lab | Emmanuel Lecorche, Sabine Trombert-Paolantoni, Aude Lessenne, Kader Merah, Thierry Collin, Laura Verdurme, Patrice Herisson, Stéphanie Haïm-Boukobza, Bénédicte Roquebert |
| EPI_ISL_2646259 | COSEM | Cerba lab | Emmanuel Lecorche, Sabine Trombert-Paolantoni, Aude Lessenne, Kader Merah, Thierry Collin, Laura Verdurme, Patrice Herisson, Stéphanie Haïm-Boukobza, Bénédicte Roquebert |
| EPI_ISL_2646260 | Cerballiance-IDF | Cerba lab | Emmanuel Lecorche, Sabine Trombert-Paolantoni, Aude Lessenne, Kader Merah, Thierry Collin, Laura Verdurme, Patrice Herisson, Stéphanie Haïm-Boukobza, Bénédicte Roquebert |
| EPI_ISL_2646261 | Cerballiance-Paris | Cerba lab | Emmanuel Lecorche, Sabine Trombert-Paolantoni, Aude Lessenne, Kader Merah, Thierry Collin, Laura Verdurme, Patrice Herisson, Stéphanie Haïm-Boukobza, Bénédicte Roquebert |
| EPI_ISL_2646262, EPI_ISL_2646263, EPI_ISL_2646264 | Cerballiance-IDF | Cerba lab | Emmanuel Lecorche, Sabine Trombert-Paolantoni, Aude Lessenne, Kader Merah, Thierry Collin, Laura Verdurme, Patrice Herisson, Stéphanie Haïm-Boukobza, Bénédicte Roquebert |
| EPI_ISL_2646265 | Cerballiance-Paris | Cerba lab | Emmanuel Lecorche, Sabine Trombert-Paolantoni, Aude Lessenne, Kader Merah, Thierry Collin, Laura Verdurme, Patrice Herisson, Stéphanie Haïm-Boukobza, Bénédicte Roquebert |
| EPI_ISL_2646266 | Cerballiance-IDF | Cerba lab | Emmanuel Lecorche, Sabine Trombert-Paolantoni, Aude Lessenne, Kader Merah, Thierry Collin, Laura Verdurme, Patrice Herisson, Stéphanie Haïm-Boukobza, Bénédicte Roquebert |
| EPI_ISL_2646267 | Cerballiance-Paris | Cerba lab | Emmanuel Lecorche, Sabine Trombert-Paolantoni, Aude Lessenne, Kader Merah, Thierry Collin, Laura Verdurme, Patrice Herisson, Stéphanie Haïm-Boukobza, Bénédicte Roquebert |
| EPI_ISL_2646268 | Unilabs | Cerba lab | Emmanuel Lecorche, Sabine Trombert-Paolantoni, Aude Lessenne, Kader Merah, Thierry Collin, Laura Verdurme, Patrice Herisson, Stéphanie Haïm-Boukobza, Bénédicte Roquebert |
| EPI_ISL_2646269 | Cerballiance-IDF | Cerba lab | Emmanuel Lecorche, Sabine Trombert-Paolantoni, Aude Lessenne, Kader Merah, Thierry Collin, Laura Verdurme, Patrice Herisson, Stéphanie Haïm-Boukobza, Bénédicte Roquebert |
| EPI_ISL_2646270 | Cerballiance-HDF | Cerba lab | Emmanuel Lecorche, Sabine Trombert-Paolantoni, Aude Lessenne, Kader Merah, Thierry Collin, Laura Verdurme, Patrice Herisson, Stéphanie Haïm-Boukobza, Bénédicte Roquebert |
| EPI_ISL_2646271 | Kuate | Cerba lab | Emmanuel Lecorche, Sabine Trombert-Paolantoni, Aude Lessenne, Kader Merah, Thierry Collin, Laura Verdurme, Patrice Herisson, Stéphanie Haïm-Boukobza, Bénédicte Roquebert |
| EPI_ISL_2646272 | Hopital Max Fourestier | Cerba lab | Emmanuel Lecorche, Sabine Trombert-Paolantoni, Aude Lessenne, Kader Merah, Thierry Collin, Laura Verdurme, Patrice Herisson, Stéphanie Haïm-Boukobza, Bénédicte Roquebert |
| EPI_ISL_2646273 | Laboratoire Frechin | Cerba lab | Emmanuel Lecorche, Sabine Trombert-Paolantoni, Aude Lessenne, Kader Merah, Thierry Collin, Laura Verdurme, Patrice Herisson, Stéphanie Haïm-Boukobza, Bénédicte Roquebert |
| EPI_ISL_2646274 | Cerballiance-Provence | Cerba lab | Emmanuel Lecorche, Sabine Trombert-Paolantoni, Aude Lessenne, Kader Merah, Thierry Collin, Laura Verdurme, Patrice Herisson, Stéphanie Haïm-Boukobza, Bénédicte Roquebert |
| EPI_ISL_2646275, EPI_ISL_2646276 | Cerballiance-Paris | Cerba lab | Emmanuel Lecorche, Sabine Trombert-Paolantoni, Aude Lessenne, Kader Merah, Thierry Collin, Laura Verdurme, Patrice Herisson, Stéphanie Haïm-Boukobza, Bénédicte Roquebert |
| EPI_ISL_2646277 | Cerballiance-Provence | Cerba lab | Emmanuel Lecorche, Sabine Trombert-Paolantoni, Aude Lessenne, Kader Merah, Thierry Collin, Laura Verdurme, Patrice Herisson, Stéphanie Haïm-Boukobza, Bénédicte Roquebert |
| EPI_ISL_2646278 | Cerballiance-Paris | Cerba lab | Emmanuel Lecorche, Sabine Trombert-Paolantoni, Aude Lessenne, Kader Merah, Thierry Collin, Laura Verdurme, Patrice Herisson, Stéphanie Haïm-Boukobza, Bénédicte Roquebert |
| EPI_ISL_2646279 | Cerballiance-IDF | Cerba lab | Emmanuel Lecorche, Sabine Trombert-Paolantoni, Aude Lessenne, Kader Merah, Thierry Collin, Laura Verdurme, Patrice Herisson, Stéphanie Haïm-Boukobza, Bénédicte Roquebert |
| EPI_ISL_2646280 | Ch Dourdan | Cerba lab | Emmanuel Lecorche, Sabine Trombert-Paolantoni, Aude Lessenne, Kader Merah, Thierry Collin, Laura Verdurme, Patrice Herisson, Stéphanie Haïm-Boukobza, Bénédicte Roquebert |
| EPI_ISL_2646281 | Hopital Max Fourestier | Cerba lab | Emmanuel Lecorche, Sabine Trombert-Paolantoni, Aude Lessenne, Kader Merah, Thierry Collin, Laura Verdurme, Patrice Herisson, Stéphanie Haïm-Boukobza, Bénédicte Roquebert |
| EPI_ISL_2646285 | CHU de la Réunion - Sud | Laboratoire de virologie, CNR arbovirus Associé, Chu de la Réunion | Etienne Frumence, Marie-Christine Jaffar Bandjee, Nicolas Traversier, Sabrina Petit Genet |
| EPI_ISL_2646288 | CHU de la Réunion - Nord | Laboratoire de virologie, CNR arbovirus Associé, Chu de la Réunion | Etienne Frumence, Marie-Christine Jaffar Bandjee, Nicolas Traversier, Sabrina Petit Genet |
| EPI_ISL_2646289, EPI_ISL_2646292, EPI_ISL_2646293, EPI_ISL_2646297 | CHU de la Réunion - Sud | Laboratoire de virologie, CNR arbovirus Associé, Chu de la Réunion | Etienne Frumence, Marie-Christine Jaffar Bandjee, Nicolas Traversier, Sabrina Petit Genet |
| EPI_ISL_2646298 | CHU de la Réunion - Nord | Laboratoire de virologie, CNR arbovirus Associé, Chu de la Réunion | Etienne Frumence, Marie-Christine Jaffar Bandjee, Nicolas Traversier, Sabrina Petit Genet |
| EPI_ISL_2648115, EPI_ISL_2648116, EPI_ISL_2648117, EPI_ISL_2648118 | Instituto Nacional de Enfermedades Respiratorias (INER) | Centro de Investigación en Enfermedades Infecciosas (CIENI), Instituto Nacional de Enfermedades Respiratorias (INER) | Consorcio Mexicano de Vigilancia Genómica (CoViGen-Mex). Authors (in alphabetical order): Julio Elias Alvarado-Yaah, Carlos F. Arias, Santiago Ávila-Ríos, Eduardo Becerril-Vargas, Víctor Hugo Borja-Aburto, Celia Boukadida, Cristóbal Cháidez-Quiróz, Juan Bautista Chale-Dzul, Ricardo Ciria Merce, Andreu Comas-García, Célida Duque Molina, Julissa Enciso-Ibarra, José Antonio Enciso-Moreno, Gloria Elena Espinosa-Ayala, Fernando Fontove-Herrera, Daniel Fregoso-Rueda, Victor Eduardo Garcia-Arias, Alejandra Garcia-Gasca, Bruno Gomez-Gil, Jean Pierre González, Irvin González-López, Concepción Grajales-Muñiz, Ricardo Grande, Rosa María Gutierrez Rios, Alejandra Hernández-Terán, Alfredo Herrera-Estrella, Carla Ivón Herrera-Najera, Pavel Isa, Daniel Lira Morales, Susana Lopez, Antonio Loza Román, Brenda Irasema Maldonado-Meza, Bernardo Martínez-Miguel, José Arturo Martínez-Orozco, Célida Martínez- Rodríguez, Margarita Matías-Florentino, Fidencio Mejía-Nepomuceno, María Guadalupe de Jesús Mireles-Rivera, Gloria María Molina-Salinas, Hector Montoya-Fuentes, , Mario Mújica-Sánchez, José Esteban Muñoz-Medina, José de Jesús Nuñez-Contreras, Alicia Ocaña-Mondragón, Luis Alberto Ochoa-Carrera, Hector Esteban Paz-Juárez, Marissa Pérez-García, Francisco Pulido, Helen Haydee Fernanda Ramírez-Plascencia, Jorge Salas-Hernández, Angel Gustavo Salas-Lais, Alejandro Sanchez-Flores, Clara Esperanza Santacruz-Tinoco, María Guadalupe Santiago-Mauricio, Selene Zárate, Nelly Sélem-Mojica, Blanca Taboada, Gloria Vazquez, Joel Armando Vázquez-Pérez. |
| EPI_ISL_2648221, EPI_ISL_2648223, EPI_ISL_2648224, EPI_ISL_2648226, EPI_ISL_2648235, EPI_ISL_2648236, EPI_ISL_2648237, EPI_ISL_2648239 | Kasane Primary Hospital Laboratory | Botswana Harvard HIV Reference Laboratory | Sikhulile Moyo, Wonderful T. Choga, Dorcas Maruapula, Agnes Karutwaeng, Rose Munyere, Keoratile Ntshambiwa, Thongbotho Mphoyakgosi, Boitumelo Zuze, Botshelo Radibe, Legodile Kooepile, Ontlametse T. Bareng, Roger Shapiro, Shahin Lockman, Joseph Makhema, Madisa Mine, Mosepele Mosepele, Simani Gaseitsiwe |
| EPI_ISL_2649992 | Tuanku Ampuan Najihah Hospital, Kuala Pilah | Institute for Medical Research, Infectious Disease Research | Suppiah J, Kamel K, Mohd Zawawi Z, Azizan MA, Ramly N, Robert F, Thayan R |

| | | | |
|---|---|---|---|
| | | Centre, National Institutes of Health, Ministry of Health Malaysia | |
| EPI_ISL_2650001 | Hospital Sultanah Nur Zahirah Kuala Terengganu | Institute for Medical Research, Infectious Disease Research Centre, National Institutes of Health, Ministry of Health Malaysia | Suppiah J, Kamel K, Mohd Zawawi Z, Azizan MA, Ramly N, Robert F, Thayan R |
| EPI_ISL_2650003 | Hospital Wanita Dan Kanak-Kanak Sabah | Institute for Medical Research, Infectious Disease Research Centre, National Institutes of Health, Ministry of Health Malaysia | Jalan Setia Murni U13/52, Bandar Setia Alam, 40170 Selangor, Malaysia |
| EPI_ISL_2650004 | Sungai Buloh Hospital | Institute for Medical Research, Infectious Disease Research Centre, National Institutes of Health, Ministry of Health Malaysia | Suppiah J, Kamel K, Mohd Zawawi Z, Azizan MA, Ramly N, Robert F, Thayan R |
| EPI_ISL_2650256 | Johor Bahru Public Health Laboratory | Institute for Medical Research, Infectious Disease Research Centre, National Institutes of Health, Ministry of Health Malaysia | Suppiah J, Kamel K, Mohd Zawawi Z, Azizan MA, Ramly N, Robert F, Thayan R |
| EPI_ISL_2650499, EPI_ISL_2650505 | Queen Elizabeth Hospital | Institute for Medical Research, Infectious Disease Research Centre, National Institutes of Health, Ministry of Health Malaysia | Suppiah J, Kamel K, Mohd Zawawi Z, Azizan MA, Ramly N, Robert F, Thayan R |
| EPI_ISL_2650507, EPI_ISL_2650508, EPI_ISL_2650510, EPI_ISL_2650511, EPI_ISL_2650512, EPI_ISL_2650513, EPI_ISL_2650514, EPI_ISL_2650515, EPI_ISL_2650516, EPI_ISL_2650517, EPI_ISL_2650519, EPI_ISL_2650521 | | | |
| see above | University of Oregon COVID-19 MAP Laboratory | University of Oregon Genomics and Cell Characterczation Core Facility (GC3F) | Douglas Turnbull, Ariana White, Jeff Bishop, Jason Carriere, Jason Sydes, Peter Batzel |
| EPI_ISL_2651115, EPI_ISL_2651116 | Temporary Specimen Collection Centre at the AsiaWorld-Expo | Hong Kong Department of Health | Alan K.L. Tsang, Peter C.W. Yip, Ken H.L. Ng, Edman T.K. Lam, Rickjason C.W. Chan, Dominic N.C. Tsang |
| EPI_ISL_2651117 | North Lantau Hospital | Hong Kong Department of Health | Alan K.L. Tsang, Peter C.W. Yip, Ken H.L. Ng, Edman T.K. Lam, Rickjason C.W. Chan, Dominic N.C. Tsang |
| EPI_ISL_2651118 | HOME QUARANTINE TASKFORCE | Hong Kong Department of Health | Alan K.L. Tsang, Peter C.W. Yip, Ken H.L. Ng, Edman T.K. Lam, Rickjason C.W. Chan, Dominic N.C. Tsang |
| EPI_ISL_2651119, EPI_ISL_2651120, EPI_ISL_2651121 | Temporary Specimen Collection Centre at the AsiaWorld-Expo | Hong Kong Department of Health | Alan K.L. Tsang, Peter C.W. Yip, Ken H.L. Ng, Edman T.K. Lam, Rickjason C.W. Chan, Dominic N.C. Tsang |
| EPI_ISL_2651122 | QUARANTINE CAMP | Hong Kong Department of Health | Alan K.L. Tsang, Peter C.W. Yip, Ken H.L. Ng, Edman T.K. Lam, Rickjason C.W. Chan, Dominic N.C. Tsang |
| EPI_ISL_2651123 | Temporary Specimen Collection Centre at the AsiaWorld-Expo | Hong Kong Department of Health | Alan K.L. Tsang, Peter C.W. Yip, Ken H.L. Ng, Edman T.K. Lam, Rickjason C.W. Chan, Dominic N.C. Tsang |
| EPI_ISL_2651124 | North Lantau Hospital | Hong Kong Department of Health | Alan K.L. Tsang, Peter C.W. Yip, Ken H.L. Ng, Edman T.K. Lam, Rickjason C.W. Chan, Dominic N.C. Tsang |
| EPI_ISL_2651125 | YAN CHAI HOSPITAL | Hong Kong Department of Health | Alan K.L. Tsang, Peter C.W. Yip, Ken H.L. Ng, Edman T.K. Lam, Rickjason C.W. Chan, Dominic N.C. Tsang |
| EPI_ISL_2651127 | TUEN MUN HOSPITAL | Hong Kong Department of Health | Alan K.L. Tsang, Peter C.W. Yip, Ken H.L. Ng, Edman T.K. Lam, Rickjason C.W. Chan, Dominic N.C. Tsang |
| EPI_ISL_2651128, EPI_ISL_2651129, EPI_ISL_2651130 | North Lantau Hospital | Hong Kong Department of Health | Alan K.L. Tsang, Peter C.W. Yip, Ken H.L. Ng, Edman T.K. Lam, Rickjason C.W. Chan, Dominic N.C. Tsang |
| EPI_ISL_2651131 | HOME QUARANTINE TASKFORCE | Hong Kong Department of Health | Alan K.L. Tsang, Peter C.W. Yip, Ken H.L. Ng, Edman T.K. Lam, Rickjason C.W. Chan, Dominic N.C. Tsang |
| EPI_ISL_2651236 | AOU Policlinico "G. Rodolico- S. Marco" PO G. Rodolico | AOU Policlinico "G. Rodolico- S. Marco" PO G. Rodolico and Biometec UNICT | Stefani and Scalia |
| EPI_ISL_2651266, EPI_ISL_2651267 | CHU de la Réunion - Nord | Laboratoire de virologie, CNR arbovirus Associé, Chu de la Réunion | Etienne Frumence, Marie-Christine Jaffar Bandjee, Nicolas Traversier, Sabrina Petit Genet |
| EPI_ISL_2651270 | CHU de la Réunion - Sud | Laboratoire de virologie, CNR arbovirus Associé, Chu de la Réunion | Etienne Frumence, Marie-Christine Jaffar Bandjee, Nicolas Traversier, Sabrina Petit Genet |
| EPI_ISL_2651272 | CHU de la Réunion - Nord | Laboratoire de virologie, CNR arbovirus Associé, Chu de la Réunion | Etienne Frumence, Marie-Christine Jaffar Bandjee, Nicolas Traversier, Sabrina Petit Genet |
| EPI_ISL_2651273, EPI_ISL_2651274, EPI_ISL_2651275, EPI_ISL_2651276, EPI_ISL_2651277 | CHU de la Réunion - Sud | Laboratoire de virologie, CNR arbovirus Associé, Chu de la Réunion | Etienne Frumence, Marie-Christine Jaffar Bandjee, Nicolas Traversier, Sabrina Petit Genet |
| EPI_ISL_2651279 | CHU de la Réunion - Nord | Laboratoire de virologie, CNR arbovirus Associé, Chu de la Réunion | Etienne Frumence, Marie-Christine Jaffar Bandjee, Nicolas Traversier, Sabrina Petit Genet |
| EPI_ISL_2651365, EPI_ISL_2651366 | CHU de la Réunion - Sud | Laboratoire de virologie, CNR arbovirus Associé, Chu de la Réunion | Etienne Frumence, Marie-Christine Jaffar Bandjee, Nicolas Traversier, Sabrina Petit Genet |
| EPI_ISL_2651367, EPI_ISL_2651368 | CHU de la Réunion - Nord | Laboratoire de virologie, CNR arbovirus Associé, Chu de la Réunion | Etienne Frumence, Marie-Christine Jaffar Bandjee, Nicolas Traversier, Sabrina Petit Genet |
| EPI_ISL_2651370, EPI_ISL_2651371, EPI_ISL_2651372, EPI_ISL_2651373, EPI_ISL_2651374 | CHU de la Réunion - Sud | Laboratoire de virologie, CNR arbovirus Associé, Chu de la Réunion | Etienne Frumence, Marie-Christine Jaffar Bandjee, Nicolas Traversier, Sabrina Petit Genet |
| EPI_ISL_2651376 | CHU de la Réunion - Nord | Laboratoire de virologie, CNR arbovirus Associé, Chu de la Réunion | Etienne Frumence, Marie-Christine Jaffar Bandjee, Nicolas Traversier, Sabrina Petit Genet |
| EPI_ISL_2651378, EPI_ISL_2651379, EPI_ISL_2651380, EPI_ISL_2651381, EPI_ISL_2651382 | CHU de la Réunion - Sud | Laboratoire de virologie, CNR arbovirus Associé, Chu de la Réunion | Etienne Frumence, Marie-Christine Jaffar Bandjee, Nicolas Traversier, Sabrina Petit Genet |
| EPI_ISL_2651386, EPI_ISL_2651387, EPI_ISL_2651388, EPI_ISL_2651389, EPI_ISL_2651390 | CHU de la Réunion - Nord | Laboratoire de virologie, CNR arbovirus Associé, Chu de la Réunion | Etienne Frumence, Marie-Christine Jaffar Bandjee, Nicolas Traversier, Sabrina Petit Genet |
| EPI_ISL_2651391, EPI_ISL_2651392 | CHU de la Réunion - Sud | Laboratoire de virologie, CNR arbovirus Associé, Chu de la Réunion | Etienne Frumence, Marie-Christine Jaffar Bandjee, Nicolas Traversier, Sabrina Petit Genet |
| EPI_ISL_2651395, EPI_ISL_2651396, EPI_ISL_2651397, EPI_ISL_2651399 | CHU de la Réunion - Nord | Laboratoire de virologie, CNR arbovirus Associé, Chu de la Réunion | Etienne Frumence, Marie-Christine Jaffar Bandjee, Nicolas Traversier, Sabrina Petit Genet |
| EPI_ISL_2651402, EPI_ISL_2651403, EPI_ISL_2651404, EPI_ISL_2651405, EPI_ISL_2651406 | CHU de la Réunion - Sud | Laboratoire de virologie, CNR arbovirus Associé, Chu de la Réunion | Etienne Frumence, Marie-Christine Jaffar Bandjee, Nicolas Traversier, Sabrina Petit Genet |
| EPI_ISL_2652114, EPI_ISL_2652115, EPI_ISL_2652116 | Cerballiance-IDF | Cerba lab | Emmanuel Lecorche, Sabine Trombert-Paolantoni, Aude Lessenne, Kader Merah, Thierry Collin, Laura Verdurme, Patrice Herisson, Stéphanie Haïm-Boukobza, Bénédicte Roquebert |

| | | | |
|---|---|---|---|
| EPI_ISL_2652120, EPI_ISL_2652122, EPI_ISL_2652151, EPI_ISL_2652152, EPI_ISL_2652154, EPI_ISL_2652155, EPI_ISL_2652157, EPI_ISL_2652158, EPI_ISL_2652160, EPI_ISL_2652161, EPI_ISL_2652162, EPI_ISL_2652163 | | | |
| see above | AZ Klina | AZ Klina | Carl Vael - Lynsey Berckmans |
| EPI_ISL_2652165, EPI_ISL_2652166, EPI_ISL_2652167, EPI_ISL_2652168, EPI_ISL_2652170, EPI_ISL_2652171 | Cantacuzino National Military-Medical Institute, Viral Respiratory Infections Laboratory | Cantacuzino Institute Virology | Carmen Cherciu, Luiza Ustea, Nicoleta Paraschiv, Sorin Dinu, Mihaela Oprea, Mihaela Lazar |
| EPI_ISL_2652172, EPI_ISL_2652173, EPI_ISL_2652174, EPI_ISL_2652175, EPI_ISL_2652176, EPI_ISL_2652177, EPI_ISL_2652178, EPI_ISL_2652179, EPI_ISL_2652180, EPI_ISL_2652181, EPI_ISL_2652182, EPI_ISL_2652183, EPI_ISL_2652184, EPI_ISL_2652185, EPI_ISL_2652186, EPI_ISL_2652187, EPI_ISL_2652188, EPI_ISL_2652189, EPI_ISL_2652190, EPI_ISL_2652191, EPI_ISL_2652192 | | | |
| see above | Infectious diseases Clinical Hospital, Cluj Napoca | Cantacuzino Institute Virology | Carmen Cherciu, Luiza Ustea, Nicoleta Paraschiv, Sorin Dinu, Mihaela Oprea, Mihaela Lazar |
| EPI_ISL_2652194, EPI_ISL_2652195, EPI_ISL_2652196, EPI_ISL_2652197, EPI_ISL_2652198, EPI_ISL_2652199, EPI_ISL_2652200, EPI_ISL_2652201, EPI_ISL_2652202, EPI_ISL_2652203 | Cantacuzino National Military-Medical Institute, Viral Respiratory Infections Laboratory | Cantacuzino Institute Virology | Carmen Cherciu, Luiza Ustea, Nicoleta Paraschiv, Sorin Dinu, Mihaela Oprea, Mihaela Lazar |
| EPI_ISL_2652204, EPI_ISL_2652205, EPI_ISL_2652206, EPI_ISL_2652207, EPI_ISL_2652208, EPI_ISL_2652209, EPI_ISL_2652210, EPI_ISL_2652211, EPI_ISL_2652212, EPI_ISL_2652213 | Clinical Emergency County Hospital, Craiova | Cantacuzino Institute Virology | Carmen Cherciu, Luiza Ustea, Nicoleta Paraschiv, Sorin Dinu, Mihaela Oprea, Mihaela Lazar |
| EPI_ISL_2652214 | Cantacuzino National Military-Medical Institute, Viral Respiratory Infections Laboratory | Cantacuzino Institute Virology | Carmen Cherciu, Luiza Ustea, Nicoleta Paraschiv, Sorin Dinu, Mihaela Oprea, Mihaela Lazar |
| EPI_ISL_2652215 | Clinical Emergency County Hospital, Craiova | Cantacuzino Institute Virology | Carmen Cherciu, Luiza Ustea, Nicoleta Paraschiv, Sorin Dinu, Mihaela Oprea, Mihaela Lazar |
| EPI_ISL_2652216, EPI_ISL_2652217 | Cantacuzino National Military-Medical Institute, Viral Respiratory Infections Laboratory | Cantacuzino Institute Virology | Carmen Cherciu, Luiza Ustea, Nicoleta Paraschiv, Sorin Dinu, Mihaela Oprea, Mihaela Lazar |
| EPI_ISL_2652218, EPI_ISL_2652219, EPI_ISL_2652242, EPI_ISL_2652243, EPI_ISL_2652244, EPI_ISL_2652245, EPI_ISL_2652246, EPI_ISL_2652247, EPI_ISL_2652248 | AZ Klina | AZ Klina | Carl Vael - Lynsey Berckmans |
| EPI_ISL_2657376, EPI_ISL_2657377, EPI_ISL_2657379, EPI_ISL_2657381, EPI_ISL_2657382, EPI_ISL_2657385, EPI_ISL_2657386, EPI_ISL_2657387, EPI_ISL_2657390, EPI_ISL_2657391, EPI_ISL_2657395, EPI_ISL_2657396, EPI_ISL_2657399, EPI_ISL_2657400, EPI_ISL_2657401, EPI_ISL_2657403, EPI_ISL_2657404, EPI_ISL_2657406, EPI_ISL_2657407, EPI_ISL_2657408, EPI_ISL_2657409, EPI_ISL_2657410, EPI_ISL_2657411, EPI_ISL_2657412, EPI_ISL_2657413, EPI_ISL_2657414, EPI_ISL_2657420, EPI_ISL_2657421, EPI_ISL_2657422, EPI_ISL_2657423, EPI_ISL_2657424, EPI_ISL_2657425, EPI_ISL_2657426, EPI_ISL_2657429, EPI_ISL_2657430, EPI_ISL_2657431, EPI_ISL_2657432, EPI_ISL_2657433, EPI_ISL_2657434, EPI_ISL_2657436, EPI_ISL_2657437, EPI_ISL_2657438, EPI_ISL_2657439, EPI_ISL_2657441, EPI_ISL_2657442, EPI_ISL_2657443, EPI_ISL_2657444, EPI_ISL_2657445, EPI_ISL_2657447, EPI_ISL_2657448, EPI_ISL_2657449, EPI_ISL_2657450, EPI_ISL_2657451, EPI_ISL_2657452, EPI_ISL_2657454, EPI_ISL_2657461, EPI_ISL_2657462, EPI_ISL_2657463, EPI_ISL_2657464, EPI_ISL_2657465, EPI_ISL_2657466, EPI_ISL_2657467, EPI_ISL_2657468, EPI_ISL_2657469, EPI_ISL_2657470, EPI_ISL_2657471, EPI_ISL_2657474, EPI_ISL_2657475, EPI_ISL_2657482, EPI_ISL_2657485, EPI_ISL_2657486, EPI_ISL_2657487, EPI_ISL_2657488, EPI_ISL_2657489, EPI_ISL_2657490, EPI_ISL_2657492, EPI_ISL_2657495, EPI_ISL_2657498, EPI_ISL_2657501, EPI_ISL_2657502, EPI_ISL_2657504, EPI_ISL_2657506, EPI_ISL_2657507, EPI_ISL_2657508, EPI_ISL_2657509, EPI_ISL_2657510, EPI_ISL_2657511 | | | |
| see above | Austrian Agency for Health and Food Safety (AGES) | Bergthaler laboratory, CeMM Research Center for Molecular Medicine of the Austrian Academy of Sciences | Lukas Endler, Anna Schedl, Fabian Amman, Petr Triska, Matthew Thornton, Thomas Penz, Benedikt Agerer, Maelle Le Moing, Michael Schuster, Bekir Erguner, Jan Laine, Martin Senekowitsch, Christoph Bock, Andreas Bergthaler |
| EPI_ISL_2657526 | ILV Kärnten | Bergthaler laboratory, CeMM Research Center for Molecular Medicine of the Austrian Academy of Sciences | Lukas Endler, Anna Schedl, Fabian Amman, Petr Triska, Matthew Thornton, Thomas Penz, Benedikt Agerer, Maelle Le Moing, Michael Schuster, Bekir Erguner, Jan Laine, Martin Senekowitsch, Christoph Bock, Andreas Bergthaler |
| EPI_ISL_2657574 | Vienna COVID-19 Detection Initiative (VCDI) | Bergthaler laboratory, CeMM Research Center for Molecular Medicine of the Austrian Academy of Sciences | Lukas Endler, Anna Schedl, Fabian Amman, Petr Triska, Matthew Thornton, Thomas Penz, Benedikt Agerer, Maelle Le Moing, Michael Schuster, Bekir Erguner, Jan Laine, Martin Senekowitsch, Christoph Bock, Andreas Bergthaler |
| EPI_ISL_2657867, EPI_ISL_2657868 | COVID 19- EPS FAMISANAR | Instituto Nacional de Salud- Dirección de Investigación en Salud Pública | Katherine Laiton-Donato, Diego A. Álvarez-Díaz, Carlos Franco-Muñoz, Hector Alejandro Ruiz-Moreno, Paola Rojas, Maria T. Herrera-Sepúlveda, Diego Andrés Prada, Jhonnatan Reales-González, Sheryll Corchuelo, Julian Naizaque, Jorge Rivera, Gerardo Santamaría, Sergio Gomez, Lisseth Pardo, Juan Camilo Martinez, Marta Lopez Blanco, Ángela Alarcon Cruz, Diana Malo, Carmen Osorio, Magdalena Wiesner, Martha Lucia Ospina Martinez, Marcela Mercado-Reyes |
| EPI_ISL_2657873, EPI_ISL_2657874 | LABORATORIO BILOGIA MOLECULAR IMAT SAS | Instituto Nacional de Salud- Dirección de Investigación en Salud Pública | Katherine Laiton-Donato, Diego A. Álvarez-Díaz, Carlos Franco-Muñoz, Hector Alejandro Ruiz-Moreno, Paola Rojas, Maria T. Herrera-Sepúlveda, Diego Andrés Prada, Jhonnatan Reales-González, Sheryll Corchuelo, Julian Naizaque, Jorge Rivera, Gerardo Santamaría, Sergio Gomez, Lisseth Pardo, Juan Camilo Martinez, Marta Lopez Blanco, Ángela Alarcon Cruz, Diana Malo, Carmen Osorio, Magdalena Wiesner, Martha Lucia Ospina Martinez, Marcela Mercado-Reyes |
| EPI_ISL_2657918, EPI_ISL_2657936 | Public Health Authority of the Slovak Republic | Laboratory of Genomics and Bioinformatics, Comenius University Science Park | Tatiana Sedláková, Diana Rusáková, Miroslav Böhmer, Anna Giová, Jaroslav Budiš, Tomáš Szemes, Jakub Styk |
| EPI_ISL_2658950 | Basurto University Hospital: Clinical Microbiology Laboratory | Biocruces Bizkaia | Mikel Urrutikoetxea-Gutiérrez, Mª Carmen Nieto Toboso, Estibaliz Ugalde Zarraga, Mikel Gallego Rodrigo, Ana de la Hoz, José Luis Díaz de Tuesta del Arco |
| EPI_ISL_2658951, EPI_ISL_2658952 | LESP San Luis Potosi | Instituto de Diagnostico y Referencia Epidemiologicos (INDRE) | Claudia Wong-Arambula, Abril Rodriguez-Maldonado, Vanessa Rivero-Arredondo, Ariadna Medina-Benitez, Joaquin Quiroz-Mercado, Sergio Rangel-Guerrero, Natividad Cruz-Ortiz, Tatiana Nunez-Garcia, Gisela Barrera-Badillo, Lucia Hernandez-Rivas, Irma Lopez-Martinez, Ernesto Ramirez-Gonzalez. |
| EPI_ISL_2658953 | LESP Nuevo Leon | Instituto de Diagnostico y Referencia Epidemiologicos (INDRE) | Claudia Wong-Arambula, Abril Rodriguez-Maldonado, Vanessa Rivero-Arredondo, Ariadna Medina-Benitez, Joaquin Quiroz-Mercado, Sergio Rangel-Guerrero, Natividad Cruz-Ortiz, Tatiana Nunez-Garcia, Gisela Barrera-Badillo, Lucia Hernandez-Rivas, Irma Lopez-Martinez, Ernesto Ramirez-Gonzalez. |
| EPI_ISL_2663131, EPI_ISL_2663132, EPI_ISL_2663139, EPI_ISL_2663140, EPI_ISL_2663142, EPI_ISL_2663143 | Ochsner Health | Bioinfoexperts, LLC | Judy Crabtree, Amy Feehan, Rebecca Rose, David J. Nolan, Tessa LaFleur, Julia-Garcia-Diaz, Lucio Miele, Susanna L. Lamers |
| EPI_ISL_2663327 | LESP Nuevo Leon | Instituto de Diagnostico y Referencia Epidemiologicos (INDRE) | Claudia Wong-Arambula, Abril Rodriguez-Maldonado, Vanessa Rivero-Arredondo, Ariadna Medina-Benitez, Joaquin Quiroz-Mercado, Sergio Rangel-Guerrero, Natividad Cruz-Ortiz, Tatiana Nunez-Garcia, Gisela Barrera-Badillo, Lucia Hernandez-Rivas, Irma Lopez-Martinez, Ernesto Ramirez-Gonzalez. |
| EPI_ISL_2663328, EPI_ISL_2663329, EPI_ISL_2663330 | LESP Ciudad de Mexico | Instituto de Diagnostico y Referencia Epidemiologicos (INDRE) | Claudia Wong-Arambula, Abril Rodriguez-Maldonado, Vanessa Rivero-Arredondo, Ariadna Medina-Benitez, Joaquin Quiroz-Mercado, Sergio Rangel-Guerrero, Natividad Cruz-Ortiz, Tatiana Nunez-Garcia, Gisela Barrera-Badillo, Lucia Hernandez-Rivas, Irma Lopez-Martinez, Ernesto Ramirez-Gonzalez. |
| EPI_ISL_2663331 | LESP Queretaro | Instituto de Diagnostico y Referencia Epidemiologicos (INDRE) | Claudia Wong-Arambula, Abril Rodriguez-Maldonado, Vanessa Rivero-Arredondo, Ariadna Medina-Benitez, Joaquin Quiroz-Mercado, Sergio Rangel-Guerrero, Natividad Cruz-Ortiz, Tatiana Nunez-Garcia, Gisela Barrera-Badillo, Lucia Hernandez-Rivas, Irma Lopez-Martinez, Ernesto Ramirez-Gonzalez. |
| EPI_ISL_2663333 | LESP Sinaloa | Instituto de Diagnostico y Referencia Epidemiologicos (INDRE) | Claudia Wong-Arambula, Abril Rodriguez-Maldonado, Vanessa Rivero-Arredondo, Ariadna Medina-Benitez, Joaquin Quiroz-Mercado, Sergio Rangel-Guerrero, Natividad Cruz-Ortiz, Tatiana Nunez-Garcia, Gisela Barrera-Badillo, Lucia Hernandez-Rivas, Irma Lopez-Martinez, Ernesto Ramirez-Gonzalez. |
| EPI_ISL_2663336, EPI_ISL_2663337, | LESP Tabasco | Instituto de Diagnostico y Referencia Epidemiologicos | Claudia Wong-Arambula, Abril Rodriguez-Maldonado, Vanessa Rivero-Arredondo, Ariadna Medina-Benitez, Joaquin Quiroz-Mercado, Sergio |

| | | | |
|---|---|---|---|
| EPI_ISL_2663338, EPI_ISL_2663339, EPI_ISL_2663340, EPI_ISL_2663341, EPI_ISL_2663346, EPI_ISL_2663347, EPI_ISL_2663348, EPI_ISL_2663349 | | (INDRE) | Rangel-Guerrero, Natividad Cruz-Ortiz, Tatiana Nunez-Garcia, Gisela Barrera-Badillo, Lucia Hernandez-Rivas, Irma Lopez-Martinez, Ernesto Ramirez-Gonzalez. |
| EPI_ISL_2663350 | LESP Tamaulipas | Instituto de Diagnostico y Referencia Epidemiologicos (INDRE) | Claudia Wong-Arambula, Abril Rodriguez-Maldonado, Vanessa Rivero-Arredondo, Ariadna Medina-Benitez, Joaquin Quiroz-Mercado, Sergio Rangel-Guerrero, Natividad Cruz-Ortiz, Tatiana Nunez-Garcia, Gisela Barrera-Badillo, Lucia Hernandez-Rivas, Irma Lopez-Martinez, Ernesto Ramirez-Gonzalez. |
| EPI_ISL_2663353 | LESP Baja California Sur | Instituto de Diagnostico y Referencia Epidemiologicos (INDRE) | Claudia Wong-Arambula, Abril Rodriguez-Maldonado, Vanessa Rivero-Arredondo, Ariadna Medina-Benitez, Joaquin Quiroz-Mercado, Sergio Rangel-Guerrero, Natividad Cruz-Ortiz, Tatiana Nunez-Garcia, Gisela Barrera-Badillo, Lucia Hernandez-Rivas, Irma Lopez-Martinez, Ernesto Ramirez-Gonzalez. |
| EPI_ISL_2663354, EPI_ISL_2663355 | LESP Sonora | Instituto de Diagnostico y Referencia Epidemiologicos (INDRE) | Claudia Wong-Arambula, Abril Rodriguez-Maldonado, Vanessa Rivero-Arredondo, Ariadna Medina-Benitez, Joaquin Quiroz-Mercado, Sergio Rangel-Guerrero, Natividad Cruz-Ortiz, Tatiana Nunez-Garcia, Gisela Barrera-Badillo, Lucia Hernandez-Rivas, Irma Lopez-Martinez, Ernesto Ramirez-Gonzalez. |
| EPI_ISL_2663356, EPI_ISL_2663357 | LESP Quintana Roo | Instituto de Diagnostico y Referencia Epidemiologicos (INDRE) | Claudia Wong-Arambula, Abril Rodriguez-Maldonado, Vanessa Rivero-Arredondo, Ariadna Medina-Benitez, Joaquin Quiroz-Mercado, Sergio Rangel-Guerrero, Natividad Cruz-Ortiz, Tatiana Nunez-Garcia, Gisela Barrera-Badillo, Lucia Hernandez-Rivas, Irma Lopez-Martinez, Ernesto Ramirez-Gonzalez. |
| EPI_ISL_2663359 | LESP Sonora | Instituto de Diagnostico y Referencia Epidemiologicos (INDRE) | Claudia Wong-Arambula, Abril Rodriguez-Maldonado, Vanessa Rivero-Arredondo, Ariadna Medina-Benitez, Joaquin Quiroz-Mercado, Sergio Rangel-Guerrero, Natividad Cruz-Ortiz, Tatiana Nunez-Garcia, Gisela Barrera-Badillo, Lucia Hernandez-Rivas, Irma Lopez-Martinez, Ernesto Ramirez-Gonzalez. |
| EPI_ISL_2663361, EPI_ISL_2663362, EPI_ISL_2663363, EPI_ISL_2663364, EPI_ISL_2663365, EPI_ISL_2663366, EPI_ISL_2663367, EPI_ISL_2663368, EPI_ISL_2663369, EPI_ISL_2663370, EPI_ISL_2663371, EPI_ISL_2663372, EPI_ISL_2663373, EPI_ISL_2663374, EPI_ISL_2663375 | | | |
| see above | LESP Hidalgo | Instituto de Diagnostico y Referencia Epidemiologicos (INDRE) | Claudia Wong-Arambula, Abril Rodriguez-Maldonado, Vanessa Rivero-Arredondo, Ariadna Medina-Benitez, Joaquin Quiroz-Mercado, Sergio Rangel-Guerrero, Natividad Cruz-Ortiz, Tatiana Nunez-Garcia, Gisela Barrera-Badillo, Lucia Hernandez-Rivas, Irma Lopez-Martinez, Ernesto Ramirez-Gonzalez. |
| EPI_ISL_2663377 | LESP Queretaro | Instituto de Diagnostico y Referencia Epidemiologicos (INDRE) | Claudia Wong-Arambula, Abril Rodriguez-Maldonado, Vanessa Rivero-Arredondo, Ariadna Medina-Benitez, Joaquin Quiroz-Mercado, Sergio Rangel-Guerrero, Natividad Cruz-Ortiz, Tatiana Nunez-Garcia, Gisela Barrera-Badillo, Lucia Hernandez-Rivas, Irma Lopez-Martinez, Ernesto Ramirez-Gonzalez. |
| EPI_ISL_2663378, EPI_ISL_2663379 | LESP Yucatan | Instituto de Diagnostico y Referencia Epidemiologicos (INDRE) | Claudia Wong-Arambula, Abril Rodriguez-Maldonado, Vanessa Rivero-Arredondo, Ariadna Medina-Benitez, Joaquin Quiroz-Mercado, Sergio Rangel-Guerrero, Natividad Cruz-Ortiz, Tatiana Nunez-Garcia, Gisela Barrera-Badillo, Lucia Hernandez-Rivas, Irma Lopez-Martinez, Ernesto Ramirez-Gonzalez. |
| EPI_ISL_2663380, EPI_ISL_2663381 | LESP Queretaro | Instituto de Diagnostico y Referencia Epidemiologicos (INDRE) | Claudia Wong-Arambula, Abril Rodriguez-Maldonado, Vanessa Rivero-Arredondo, Ariadna Medina-Benitez, Joaquin Quiroz-Mercado, Sergio Rangel-Guerrero, Natividad Cruz-Ortiz, Tatiana Nunez-Garcia, Gisela Barrera-Badillo, Lucia Hernandez-Rivas, Irma Lopez-Martinez, Ernesto Ramirez-Gonzalez. |
| EPI_ISL_2663382 | LESP Morelos | Instituto de Diagnostico y Referencia Epidemiologicos (INDRE) | Claudia Wong-Arambula, Abril Rodriguez-Maldonado, Vanessa Rivero-Arredondo, Ariadna Medina-Benitez, Joaquin Quiroz-Mercado, Sergio Rangel-Guerrero, Natividad Cruz-Ortiz, Tatiana Nunez-Garcia, Gisela Barrera-Badillo, Lucia Hernandez-Rivas, Irma Lopez-Martinez, Ernesto Ramirez-Gonzalez. |
| EPI_ISL_2663383, EPI_ISL_2663384, EPI_ISL_2663385, EPI_ISL_2663386, EPI_ISL_2663387, EPI_ISL_2663388 | LESP Guanajuato | Instituto de Diagnostico y Referencia Epidemiologicos (INDRE) | Claudia Wong-Arambula, Abril Rodriguez-Maldonado, Vanessa Rivero-Arredondo, Ariadna Medina-Benitez, Joaquin Quiroz-Mercado, Sergio Rangel-Guerrero, Natividad Cruz-Ortiz, Tatiana Nunez-Garcia, Gisela Barrera-Badillo, Lucia Hernandez-Rivas, Irma Lopez-Martinez, Ernesto Ramirez-Gonzalez. |
| EPI_ISL_2663389, EPI_ISL_2663390, EPI_ISL_2663391, EPI_ISL_2663392 | LESP Nuevo Leon | Instituto de Diagnostico y Referencia Epidemiologicos (INDRE) | Claudia Wong-Arambula, Abril Rodriguez-Maldonado, Vanessa Rivero-Arredondo, Ariadna Medina-Benitez, Joaquin Quiroz-Mercado, Sergio Rangel-Guerrero, Natividad Cruz-Ortiz, Tatiana Nunez-Garcia, Gisela Barrera-Badillo, Lucia Hernandez-Rivas, Irma Lopez-Martinez, Ernesto Ramirez-Gonzalez. |
| EPI_ISL_2663393 | LESP Ciudad de Mexico | Instituto de Diagnostico y Referencia Epidemiologicos (INDRE) | Claudia Wong-Arambula, Abril Rodriguez-Maldonado, Vanessa Rivero-Arredondo, Ariadna Medina-Benitez, Joaquin Quiroz-Mercado, Sergio Rangel-Guerrero, Natividad Cruz-Ortiz, Tatiana Nunez-Garcia, Gisela Barrera-Badillo, Lucia Hernandez-Rivas, Irma Lopez-Martinez, Ernesto Ramirez-Gonzalez. |
| EPI_ISL_2663394, EPI_ISL_2663395, EPI_ISL_2663396, EPI_ISL_2663397, EPI_ISL_2663398 | LESP Chihuahua | Instituto de Diagnostico y Referencia Epidemiologicos (INDRE) | Claudia Wong-Arambula, Abril Rodriguez-Maldonado, Vanessa Rivero-Arredondo, Ariadna Medina-Benitez, Joaquin Quiroz-Mercado, Sergio Rangel-Guerrero, Natividad Cruz-Ortiz, Tatiana Nunez-Garcia, Gisela Barrera-Badillo, Lucia Hernandez-Rivas, Irma Lopez-Martinez, Ernesto Ramirez-Gonzalez. |
| EPI_ISL_2663399 | LESP Puebla | Instituto de Diagnostico y Referencia Epidemiologicos (INDRE) | Claudia Wong-Arambula, Abril Rodriguez-Maldonado, Vanessa Rivero-Arredondo, Ariadna Medina-Benitez, Joaquin Quiroz-Mercado, Sergio Rangel-Guerrero, Natividad Cruz-Ortiz, Tatiana Nunez-Garcia, Gisela Barrera-Badillo, Lucia Hernandez-Rivas, Irma Lopez-Martinez, Ernesto Ramirez-Gonzalez. |
| EPI_ISL_2663400, EPI_ISL_2663401 | LESP Tamaulipas | Instituto de Diagnostico y Referencia Epidemiologicos (INDRE) | Claudia Wong-Arambula, Abril Rodriguez-Maldonado, Vanessa Rivero-Arredondo, Ariadna Medina-Benitez, Joaquin Quiroz-Mercado, Sergio Rangel-Guerrero, Natividad Cruz-Ortiz, Tatiana Nunez-Garcia, Gisela Barrera-Badillo, Lucia Hernandez-Rivas, Irma Lopez-Martinez, Ernesto Ramirez-Gonzalez. |
| EPI_ISL_2663402 | LESP Yucatan | Instituto de Diagnostico y Referencia Epidemiologicos (INDRE) | Claudia Wong-Arambula, Abril Rodriguez-Maldonado, Vanessa Rivero-Arredondo, Ariadna Medina-Benitez, Joaquin Quiroz-Mercado, Sergio Rangel-Guerrero, Natividad Cruz-Ortiz, Tatiana Nunez-Garcia, Gisela Barrera-Badillo, Lucia Hernandez-Rivas, Irma Lopez-Martinez, Ernesto Ramirez-Gonzalez. |
| EPI_ISL_2663403 | LESP Sonora | Instituto de Diagnostico y Referencia Epidemiologicos (INDRE) | Claudia Wong-Arambula, Abril Rodriguez-Maldonado, Vanessa Rivero-Arredondo, Ariadna Medina-Benitez, Joaquin Quiroz-Mercado, Sergio Rangel-Guerrero, Natividad Cruz-Ortiz, Tatiana Nunez-Garcia, Gisela Barrera-Badillo, Lucia Hernandez-Rivas, Irma Lopez-Martinez, Ernesto Ramirez-Gonzalez. |
| EPI_ISL_2671484 | Centro de Investigacion Biomedica de Occidente (CIBO) | Unidad de Genomica Avanzada | Consorcio Mexicano de Vigilancia Genomica (CoViGen-Mex). Authors (in alphabetical order): Julio Elias Alvarado-Yaah, Carlos F. Arias, Santiago avila-Rios, Eduardo Becerril-Vargas, Victor Hugo Borja-Aburto, Celia Boukadida, Cristobal Chaidez-Quiroz, Juan Bautista Chale-Dzul, Ricardo Ciria Merce, Andreu Comas-Garcia, Celida Duque Molina, Julissa Enciso-Ibarra, Jose Antonio Enciso-Moreno, Gloria Elena Espinosa-Ayala, Fernando Fontove-Herrera, Daniel Fregoso-Rueda, Victor Eduardo Garcia-Arias, Alejandra Garcia-Gasca, Bruno Gomez-Gil, Jean Pierre Gonzalez, Irvin Gonzalez-Lopez, Concepcion Grajales-Muñiz, Ricardo Grande, Rosa Maria Gutierrez Rios, Jesus Hernandez, Alejandra Hernandez-Teran, Alfredo Herrera-Estrella, Carla Ivon Herrera-Najera, Pavel Isa, Veronica Mata-Haro, Daniel Lira Morales, Susana Lopez, Antonio Loza Roman, Brenda Irasema Maldonado-Meza, Bernardo Martinez-Miguel, Jose Arturo Martinez-Orozco, Celida Martinez- Rodriguez, Margarita Matias-Florentino, Fidencio Mejia-Nepomuceno, Maria Guadalupe de Jesus Mireles-Rivera, Gloria Maria Molina-Salinas, Hector Montoya-Fuentes, , Mario Mujica-Sanchez, Jose Esteban Muñoz-Medina, Jose de Jesus Nuñez-Contreras, Alicia Ocaña-Mondragon, Luis Alberto Ochoa-Carrera, Hector Esteban Paz-Juarez, Marissa Perez-Garcia, Francisco Pulido, Helen Haydee Fernanda Ramirez-Plascencia, Jorge Salas-Hernandez, Angel Gustavo Salas-Lais, Alejandro Sanchez-Flores, Clara Esperanza Santacruz-Tinoco, Maria Guadalupe Santiago-Mauricio, Selene Zarate, Nelly Selem-Mojica, Blanca Taboada, Gloria Vazquez, Joel Armando Vazquez-Perez. |
| EPI_ISL_2671502, EPI_ISL_2671503, EPI_ISL_2671504 | Unidad de Investigacion Medica de Yucatan (UIMY) | Unidad de Genomica Avanzada | Consorcio Mexicano de Vigilancia Genomica (CoViGen-Mex). Authors (in alphabetical order): Julio Elias Alvarado-Yaah, Carlos F. Arias, Santiago avila-Rios, Eduardo Becerril-Vargas, Victor Hugo Borja-Aburto, Celia Boukadida, Cristobal Chaidez-Quiroz, Juan Bautista Chale-Dzul, Ricardo Ciria Merce, Andreu Comas-Garcia, Celida Duque Molina, Julissa Enciso-Ibarra, Jose Antonio Enciso-Moreno, Gloria Elena Espinosa-Ayala, Fernando Fontove-Herrera, Daniel Fregoso-Rueda, Victor Eduardo Garcia-Arias, Alejandra Garcia-Gasca, Bruno Gomez-Gil, Jean Pierre Gonzalez, Irvin Gonzalez-Lopez, Concepcion Grajales-Muñiz, Ricardo Grande, Rosa Maria Gutierrez Rios, Jesus Hernandez, Alejandra Hernandez-Teran, Alfredo Herrera-Estrella, Carla Ivon Herrera-Najera, Pavel Isa, Veronica Mata-Haro, Daniel Lira Morales, Susana Lopez, Antonio Loza Roman, Brenda Irasema |

| | | | |
|---|---|---|---|
| | | | Maldonado-Meza, Bernardo Martinez-Miguel, Jose Arturo Martinez-Orozco, Celida Martinez- Rodriguez, Margarita Matias-Florentino, Fidencio Mejia-Nepomuceno, Maria Guadalupe de Jesus Mireles-Rivera, Gloria Maria Molina-Salinas, Hector Montoya-Fuentes, , Mario Mujica-Sanchez, Jose Esteban Muñoz-Medina, Jose de Jesus Nuñez-Contreras, Alicia Ocaña-Mondragon, Luis Alberto Ochoa-Carrera, Hector Esteban Paz-Juarez, Marissa Perez-Garcia, Francisco Pulido, Helen Haydee Fernanda Ramirez-Plascencia, Jorge Salas-Hernandez, Angel Gustavo Salas-Lais, Alejandro Sanchez-Flores, Clara Esperanza Santacruz-Tinoco, Maria Guadalupe Santiago-Mauricio, Selene Zarate, Nelly Selem-Mojica, Blanca Taboada, Gloria Vazquez, Joel Armando Vazquez-Perez. |
| EPI_ISL_2671525, EPI_ISL_2671547 | Centro de Investigacion Biomedica del Noreste (CIBIN) | Unidad de Genomica Avanzada | Consorcio Mexicano de Vigilancia Genomica (CoViGen-Mex). Authors (in alphabetical order): Julio Elias Alvarado-Yaah, Carlos F. Arias, Santiago avila-Rios, Eduardo Becerril-Vargas, Victor Hugo Borja-Aburto, Celia Boukadida, Cristobal Chaidez-Quiroz, Juan Bautista Chale-Dzul, Ricardo Ciria Merce, Andreu Comas-Garcia, Celida Duque Molina, Julissa Enciso-Ibarra, Jose Antonio Enciso-Moreno, Gloria Elena Espinosa-Ayala, Fernando Fontove-Herrera, Daniel Fregoso-Rueda, Victor Eduardo Garcia-Arias, Alejandra Garcia-Gasca, Bruno Gomez-Gil, Jean Pierre Gonzalez, Irvin Gonzalez-Lopez, Concepcion Grajales-Muñiz, Ricardo Grande, Rosa Maria Gutierrez Rios, Jesus Hernandez, Alejandra Hernandez-Teran, Alfredo Herrera-Estrella, Carla Ivon Herrera-Najera, Pavel Isa, Veronica Mata-Haro, Daniel Lira Morales, Susana Lopez, Antonio Loza Roman, Brenda Irasema Maldonado-Meza, Bernardo Martinez-Miguel, Jose Arturo Martinez-Orozco, Celida Martinez- Rodriguez, Margarita Matias-Florentino, Fidencio Mejia-Nepomuceno, Maria Guadalupe de Jesus Mireles-Rivera, Gloria Maria Molina-Salinas, Hector Montoya-Fuentes, , Mario Mujica-Sanchez, Jose Esteban Muñoz-Medina, Jose de Jesus Nuñez-Contreras, Alicia Ocaña-Mondragon, Luis Alberto Ochoa-Carrera, Hector Esteban Paz-Juarez, Marissa Perez-Garcia, Francisco Pulido, Helen Haydee Fernanda Ramirez-Plascencia, Jorge Salas-Hernandez, Angel Gustavo Salas-Lais, Alejandro Sanchez-Flores, Clara Esperanza Santacruz-Tinoco, Maria Guadalupe Santiago-Mauricio, Selene Zarate, Nelly Selem-Mojica, Blanca Taboada, Gloria Vazquez, Joel Armando Vazquez-Perez. |
| EPI_ISL_2671554, EPI_ISL_2671555, EPI_ISL_2671556, EPI_ISL_2671557 | Centro de Investigacion Biomedica de Occidente (CIBO) | Unidad de Genomica Avanzada | Consorcio Mexicano de Vigilancia Genomica (CoViGen-Mex). Authors (in alphabetical order): Julio Elias Alvarado-Yaah, Carlos F. Arias, Santiago avila-Rios, Eduardo Becerril-Vargas, Victor Hugo Borja-Aburto, Celia Boukadida, Cristobal Chaidez-Quiroz, Juan Bautista Chale-Dzul, Ricardo Ciria Merce, Andreu Comas-Garcia, Celida Duque Molina, Julissa Enciso-Ibarra, Jose Antonio Enciso-Moreno, Gloria Elena Espinosa-Ayala, Fernando Fontove-Herrera, Daniel Fregoso-Rueda, Victor Eduardo Garcia-Arias, Alejandra Garcia-Gasca, Bruno Gomez-Gil, Jean Pierre Gonzalez, Irvin Gonzalez-Lopez, Concepcion Grajales-Muñiz, Ricardo Grande, Rosa Maria Gutierrez Rios, Jesus Hernandez, Alejandra Hernandez-Teran, Alfredo Herrera-Estrella, Carla Ivon Herrera-Najera, Pavel Isa, Veronica Mata-Haro, Daniel Lira Morales, Susana Lopez, Antonio Loza Roman, Brenda Irasema Maldonado-Meza, Bernardo Martinez-Miguel, Jose Arturo Martinez-Orozco, Celida Martinez- Rodriguez, Margarita Matias-Florentino, Fidencio Mejia-Nepomuceno, Maria Guadalupe de Jesus Mireles-Rivera, Gloria Maria Molina-Salinas, Hector Montoya-Fuentes, , Mario Mujica-Sanchez, Jose Esteban Muñoz-Medina, Jose de Jesus Nuñez-Contreras, Alicia Ocaña-Mondragon, Luis Alberto Ochoa-Carrera, Hector Esteban Paz-Juarez, Marissa Perez-Garcia, Francisco Pulido, Helen Haydee Fernanda Ramirez-Plascencia, Jorge Salas-Hernandez, Angel Gustavo Salas-Lais, Alejandro Sanchez-Flores, Clara Esperanza Santacruz-Tinoco, Maria Guadalupe Santiago-Mauricio, Selene Zarate, Nelly Selem-Mojica, Blanca Taboada, Gloria Vazquez, Joel Armando Vazquez-Perez. |
| EPI_ISL_2671560 | Centro de Investigacion Biomedica del Noreste (CIBIN) | Unidad de Genomica Avanzada | Consorcio Mexicano de Vigilancia Genomica (CoViGen-Mex). Authors (in alphabetical order): Julio Elias Alvarado-Yaah, Carlos F. Arias, Santiago avila-Rios, Eduardo Becerril-Vargas, Victor Hugo Borja-Aburto, Celia Boukadida, Cristobal Chaidez-Quiroz, Juan Bautista Chale-Dzul, Ricardo Ciria Merce, Andreu Comas-Garcia, Celida Duque Molina, Julissa Enciso-Ibarra, Jose Antonio Enciso-Moreno, Gloria Elena Espinosa-Ayala, Fernando Fontove-Herrera, Daniel Fregoso-Rueda, Victor Eduardo Garcia-Arias, Alejandra Garcia-Gasca, Bruno Gomez-Gil, Jean Pierre Gonzalez, Irvin Gonzalez-Lopez, Concepcion Grajales-Muñiz, Ricardo Grande, Rosa Maria Gutierrez Rios, Jesus Hernandez, Alejandra Hernandez-Teran, Alfredo Herrera-Estrella, Carla Ivon Herrera-Najera, Pavel Isa, Veronica Mata-Haro, Daniel Lira Morales, Susana Lopez, Antonio Loza Roman, Brenda Irasema Maldonado-Meza, Bernardo Martinez-Miguel, Jose Arturo Martinez-Orozco, Celida Martinez- Rodriguez, Margarita Matias-Florentino, Fidencio Mejia-Nepomuceno, Maria Guadalupe de Jesus Mireles-Rivera, Gloria Maria Molina-Salinas, Hector Montoya-Fuentes, , Mario Mujica-Sanchez, Jose Esteban Muñoz-Medina, Jose de Jesus Nuñez-Contreras, Alicia Ocaña-Mondragon, Luis Alberto Ochoa-Carrera, Hector Esteban Paz-Juarez, Marissa Perez-Garcia, Francisco Pulido, Helen Haydee Fernanda Ramirez-Plascencia, Jorge Salas-Hernandez, Angel Gustavo Salas-Lais, Alejandro Sanchez-Flores, Clara Esperanza Santacruz-Tinoco, Maria Guadalupe Santiago-Mauricio, Selene Zarate, Nelly Selem-Mojica, Blanca Taboada, Gloria Vazquez, Joel Armando Vazquez-Perez. |
| EPI_ISL_2671568, EPI_ISL_2671575, EPI_ISL_2671576, EPI_ISL_2671577, EPI_ISL_2671578, EPI_ISL_2671579, EPI_ISL_2671580, EPI_ISL_2671581, EPI_ISL_2671582, EPI_ISL_2671583, EPI_ISL_2671584, EPI_ISL_2671585, EPI_ISL_2671586, EPI_ISL_2671611, EPI_ISL_2671612 | | | |
| see above | Centro de Investigacion Biomedica de Occidente (CIBO) | Unidad de Genomica Avanzada | Consorcio Mexicano de Vigilancia Genomica (CoViGen-Mex). Authors (in alphabetical order): Julio Elias Alvarado-Yaah, Carlos F. Arias, Santiago avila-Rios, Eduardo Becerril-Vargas, Victor Hugo Borja-Aburto, Celia Boukadida, Cristobal Chaidez-Quiroz, Juan Bautista Chale-Dzul, Ricardo Ciria Merce, Andreu Comas-Garcia, Celida Duque Molina, Julissa Enciso-Ibarra, Jose Antonio Enciso-Moreno, Gloria Elena Espinosa-Ayala, Fernando Fontove-Herrera, Daniel Fregoso-Rueda, Victor Eduardo Garcia-Arias, Alejandra Garcia-Gasca, Bruno Gomez-Gil, Jean Pierre Gonzalez, Irvin Gonzalez-Lopez, Concepcion Grajales-Muñiz, Ricardo Grande, Rosa Maria Gutierrez Rios, Jesus Hernandez, Alejandra Hernandez-Teran, Alfredo Herrera-Estrella, Carla Ivon Herrera-Najera, Pavel Isa, Veronica Mata-Haro, Daniel Lira Morales, Susana Lopez, Antonio Loza Roman, Brenda Irasema Maldonado-Meza, Bernardo Martinez-Miguel, Jose Arturo Martinez-Orozco, Celida Martinez- Rodriguez, Margarita Matias-Florentino, Fidencio Mejia-Nepomuceno, Maria Guadalupe de Jesus Mireles-Rivera, Gloria Maria Molina-Salinas, Hector Montoya-Fuentes, , Mario Mujica-Sanchez, Jose Esteban Muñoz-Medina, Jose de Jesus Nuñez-Contreras, Alicia Ocaña-Mondragon, Luis Alberto Ochoa-Carrera, Hector Esteban Paz-Juarez, Marissa Perez-Garcia, Francisco Pulido, Helen Haydee Fernanda Ramirez-Plascencia, Jorge Salas-Hernandez, Angel Gustavo Salas-Lais, Alejandro Sanchez-Flores, Clara Esperanza Santacruz-Tinoco, Maria Guadalupe Santiago-Mauricio, Selene Zarate, Nelly Selem-Mojica, Blanca Taboada, Gloria Vazquez, Joel Armando Vazquez-Perez. |
| EPI_ISL_2671615 | Laboratorio Central de Epidemiologi-a (LCE) | Unidad de Genomica Avanzada | Consorcio Mexicano de Vigilancia Genomica (CoViGen-Mex). Authors (in alphabetical order): Julio Elias Alvarado-Yaah, Carlos F. Arias, Santiago avila-Rios, Eduardo Becerril-Vargas, Victor Hugo Borja-Aburto, Celia Boukadida, Cristobal Chaidez-Quiroz, Juan Bautista Chale-Dzul, Ricardo Ciria Merce, Andreu Comas-Garcia, Celida Duque Molina, Julissa Enciso-Ibarra, Jose Antonio Enciso-Moreno, Gloria Elena Espinosa-Ayala, Fernando Fontove-Herrera, Daniel Fregoso-Rueda, Victor Eduardo Garcia-Arias, Alejandra Garcia-Gasca, Bruno Gomez-Gil, Jean Pierre Gonzalez, Irvin Gonzalez-Lopez, Concepcion Grajales-Muñiz, Ricardo Grande, Rosa Maria Gutierrez Rios, Jesus Hernandez, Alejandra Hernandez-Teran, Alfredo Herrera-Estrella, Carla Ivon Herrera-Najera, Pavel Isa, Veronica Mata-Haro, Daniel Lira Morales, Susana Lopez, Antonio Loza Roman, Brenda Irasema Maldonado-Meza, Bernardo Martinez-Miguel, Jose Arturo Martinez-Orozco, Celida Martinez- Rodriguez, Margarita Matias-Florentino, Fidencio Mejia-Nepomuceno, Maria Guadalupe de Jesus Mireles-Rivera, Gloria Maria Molina-Salinas, Hector Montoya-Fuentes, , Mario Mujica-Sanchez, Jose Esteban Muñoz-Medina, Jose de Jesus Nuñez-Contreras, Alicia Ocaña-Mondragon, Luis Alberto Ochoa-Carrera, Hector Esteban Paz-Juarez, Marissa Perez-Garcia, Francisco Pulido, Helen Haydee Fernanda Ramirez-Plascencia, Jorge Salas-Hernandez, Angel Gustavo Salas-Lais, Alejandro Sanchez-Flores, Clara Esperanza Santacruz-Tinoco, Maria Guadalupe Santiago-Mauricio, Selene Zarate, Nelly Selem-Mojica, Blanca Taboada, Gloria Vazquez, Joel Armando Vazquez-Perez. |
| EPI_ISL_2671617, EPI_ISL_2671618 | Centro de Investigacion Biomedica de Occidente (CIBO) | Unidad de Genomica Avanzada | Consorcio Mexicano de Vigilancia Genomica (CoViGen-Mex). Authors (in alphabetical order): Julio Elias Alvarado-Yaah, Carlos F. Arias, Santiago avila-Rios, Eduardo Becerril-Vargas, Victor Hugo Borja-Aburto, Celia Boukadida, Cristobal Chaidez-Quiroz, Juan Bautista Chale-Dzul, Ricardo Ciria Merce, Andreu Comas-Garcia, Celida Duque Molina, Julissa Enciso-Ibarra, Jose Antonio Enciso-Moreno, Gloria Elena Espinosa-Ayala, Fernando Fontove-Herrera, Daniel Fregoso-Rueda, Victor Eduardo Garcia-Arias, Alejandra Garcia-Gasca, Bruno Gomez-Gil, Jean Pierre Gonzalez, Irvin Gonzalez-Lopez, Concepcion Grajales-Muñiz, Ricardo Grande, Rosa Maria Gutierrez Rios, Jesus Hernandez, Alejandra Hernandez-Teran, Alfredo Herrera-Estrella, Carla Ivon Herrera-Najera, Pavel Isa, Veronica Mata-Haro, Daniel Lira Morales, Susana Lopez, Antonio Loza Roman, Brenda Irasema Maldonado-Meza, Bernardo Martinez-Miguel, Jose Arturo Martinez-Orozco, Celida Martinez- Rodriguez, Margarita Matias-Florentino, Fidencio Mejia-Nepomuceno, Maria Guadalupe de Jesus Mireles-Rivera, Gloria Maria Molina-Salinas, Hector Montoya-Fuentes, , Mario Mujica-Sanchez, Jose Esteban Muñoz-Medina, Jose de Jesus Nuñez-Contreras, Alicia Ocaña-Mondragon, Luis Alberto Ochoa-Carrera, Hector Esteban Paz-Juarez, Marissa Perez-Garcia, Francisco Pulido, Helen Haydee Fernanda Ramirez-Plascencia, Jorge Salas-Hernandez, Angel Gustavo Salas-Lais, Alejandro Sanchez-Flores, Clara Esperanza Santacruz-Tinoco, Maria Guadalupe Santiago-Mauricio, Selene Zarate, Nelly Selem-Mojica, Blanca Taboada, Gloria Vazquez, Joel Armando Vazquez-Perez. |
| EPI_ISL_2671651 | Laboratorio Central de Epidemiologi-a (LCE) | Unidad de Genomica Avanzada | Consorcio Mexicano de Vigilancia Genomica (CoViGen-Mex). Authors (in alphabetical order): Julio Elias Alvarado-Yaah, Carlos F. Arias, Santiago avila-Rios, Eduardo Becerril-Vargas, Victor Hugo Borja-Aburto, Celia Boukadida, Cristobal Chaidez-Quiroz, Juan Bautista Chale-Dzul, Ricardo Ciria Merce, Andreu Comas-Garcia, Celida Duque Molina, Julissa Enciso-Ibarra, Jose Antonio Enciso-Moreno, Gloria Elena Espinosa-Ayala, Fernando Fontove-Herrera, Daniel Fregoso-Rueda, Victor Eduardo Garcia-Arias, Alejandra Garcia-Gasca, Bruno Gomez-Gil, Jean Pierre Gonzalez, Irvin |

| | | | |
|---|---|---|---|
| | | | Gonzalez-Lopez, Concepcion Grajales-Muñiz, Ricardo Grande, Rosa Maria Gutierrez Rios, Jesus Hernandez, Alejandra Hernandez-Teran, Alfredo Herrera-Estrella, Carla Ivon Herrera-Najera, Pavel Isa, Veronica Mata-Haro, Daniel Lira Morales, Susana Lopez, Antonio Loza Roman, Brenda Irasema Maldonado-Meza, Bernardo Martinez-Miguel, Jose Arturo Martinez-Orozco, Celida Martinez- Rodriguez, Margarita Matias-Florentino, Fidencio Mejia-Nepomuceno, Maria Guadalupe de Jesus Mireles-Rivera, Gloria Maria Molina-Salinas, Hector Montoya-Fuentes, , Mario Mujica-Sanchez, Jose Esteban Muñoz-Medina, Jose de Jesus Nuñez-Contreras, Alicia Ocaña-Mondragon, Luis Alberto Ochoa-Carrera, Hector Esteban Paz-Juarez, Marissa Perez-Garcia, Francisco Pulido, Helen Haydee Fernanda Ramirez-Plascencia, Jorge Salas-Hernandez, Angel Gustavo Salas-Lais, Alejandro Sanchez-Flores, Clara Esperanza Santacruz-Tinoco, Maria Guadalupe Santiago-Mauricio, Selene Zarate, Nelly Selem-Mojica, Blanca Taboada, Gloria Vazquez, Joel Armando Vazquez-Perez. |
| EPI_ISL_2671685, EPI_ISL_2671687, EPI_ISL_2671688, EPI_ISL_2671689, EPI_ISL_2671690, EPI_ISL_2671692, EPI_ISL_2671694, EPI_ISL_2671695, EPI_ISL_2671696, EPI_ISL_2671697 | Unidad de Investigacion Medica de Yucatan (UIMY) | Unidad de Genomica Avanzada | Consorcio Mexicano de Vigilancia Genomica (CoViGen-Mex). Authors (in alphabetical order): Julio Elias Alvarado-Yaah, Carlos F. Arias, Santiago avila-Rios, Eduardo Becerril-Vargas, Victor Hugo Borja-Aburto, Celia Boukadida, Cristobal Chaidez-Quiroz, Juan Bautista Chale-Dzul, Ricardo Ciria Merce, Andreu Comas-Garcia, Celida Duque Molina, Julissa Enciso-Ibarra, Jose Antonio Enciso-Moreno, Gloria Elena Espinosa-Ayala, Fernando Fontove-Herrera, Daniel Fregoso-Rueda, Victor Eduardo Garcia-Arias, Alejandra Garcia-Gasca, Bruno Gomez-Gil, Jean Pierre Gonzalez, Irvin Gonzalez-Lopez, Concepcion Grajales-Muñiz, Ricardo Grande, Rosa Maria Gutierrez Rios, Jesus Hernandez, Alejandra Hernandez-Teran, Alfredo Herrera-Estrella, Carla Ivon Herrera-Najera, Pavel Isa, Veronica Mata-Haro, Daniel Lira Morales, Susana Lopez, Antonio Loza Roman, Brenda Irasema Maldonado-Meza, Bernardo Martinez-Miguel, Jose Arturo Martinez-Orozco, Celida Martinez- Rodriguez, Margarita Matias-Florentino, Fidencio Mejia-Nepomuceno, Maria Guadalupe de Jesus Mireles-Rivera, Gloria Maria Molina-Salinas, Hector Montoya-Fuentes, , Mario Mujica-Sanchez, Jose Esteban Muñoz-Medina, Jose de Jesus Nuñez-Contreras, Alicia Ocaña-Mondragon, Luis Alberto Ochoa-Carrera, Hector Esteban Paz-Juarez, Marissa Perez-Garcia, Francisco Pulido, Helen Haydee Fernanda Ramirez-Plascencia, Jorge Salas-Hernandez, Angel Gustavo Salas-Lais, Alejandro Sanchez-Flores, Clara Esperanza Santacruz-Tinoco, Maria Guadalupe Santiago-Mauricio, Selene Zarate, Nelly Selem-Mojica, Blanca Taboada, Gloria Vazquez, Joel Armando Vazquez-Perez. |
| EPI_ISL_2671703 | Centro de Investigacion Biomedica de Occidente (CIBO) | Unidad de Genomica Avanzada | Consorcio Mexicano de Vigilancia Genomica (CoViGen-Mex). Authors (in alphabetical order): Julio Elias Alvarado-Yaah, Carlos F. Arias, Santiago avila-Rios, Eduardo Becerril-Vargas, Victor Hugo Borja-Aburto, Celia Boukadida, Cristobal Chaidez-Quiroz, Juan Bautista Chale-Dzul, Ricardo Ciria Merce, Andreu Comas-Garcia, Celida Duque Molina, Julissa Enciso-Ibarra, Jose Antonio Enciso-Moreno, Gloria Elena Espinosa-Ayala, Fernando Fontove-Herrera, Daniel Fregoso-Rueda, Victor Eduardo Garcia-Arias, Alejandra Garcia-Gasca, Bruno Gomez-Gil, Jean Pierre Gonzalez, Irvin Gonzalez-Lopez, Concepcion Grajales-Muñiz, Ricardo Grande, Rosa Maria Gutierrez Rios, Jesus Hernandez, Alejandra Hernandez-Teran, Alfredo Herrera-Estrella, Carla Ivon Herrera-Najera, Pavel Isa, Veronica Mata-Haro, Daniel Lira Morales, Susana Lopez, Antonio Loza Roman, Brenda Irasema Maldonado-Meza, Bernardo Martinez-Miguel, Jose Arturo Martinez-Orozco, Celida Martinez- Rodriguez, Margarita Matias-Florentino, Fidencio Mejia-Nepomuceno, Maria Guadalupe de Jesus Mireles-Rivera, Gloria Maria Molina-Salinas, Hector Montoya-Fuentes, , Mario Mujica-Sanchez, Jose Esteban Muñoz-Medina, Jose de Jesus Nuñez-Contreras, Alicia Ocaña-Mondragon, Luis Alberto Ochoa-Carrera, Hector Esteban Paz-Juarez, Marissa Perez-Garcia, Francisco Pulido, Helen Haydee Fernanda Ramirez-Plascencia, Jorge Salas-Hernandez, Angel Gustavo Salas-Lais, Alejandro Sanchez-Flores, Clara Esperanza Santacruz-Tinoco, Maria Guadalupe Santiago-Mauricio, Selene Zarate, Nelly Selem-Mojica, Blanca Taboada, Gloria Vazquez, Joel Armando Vazquez-Perez. |
| EPI_ISL_2671721, EPI_ISL_2671722 | Centro de Investigacion Biomedica del Noreste (CIBIN) | Unidad de Genomica Avanzada | Consorcio Mexicano de Vigilancia Genomica (CoViGen-Mex). Authors (in alphabetical order): Julio Elias Alvarado-Yaah, Carlos F. Arias, Santiago avila-Rios, Eduardo Becerril-Vargas, Victor Hugo Borja-Aburto, Celia Boukadida, Cristobal Chaidez-Quiroz, Juan Bautista Chale-Dzul, Ricardo Ciria Merce, Andreu Comas-Garcia, Celida Duque Molina, Julissa Enciso-Ibarra, Jose Antonio Enciso-Moreno, Gloria Elena Espinosa-Ayala, Fernando Fontove-Herrera, Daniel Fregoso-Rueda, Victor Eduardo Garcia-Arias, Alejandra Garcia-Gasca, Bruno Gomez-Gil, Jean Pierre Gonzalez, Irvin Gonzalez-Lopez, Concepcion Grajales-Muñiz, Ricardo Grande, Rosa Maria Gutierrez Rios, Jesus Hernandez, Alejandra Hernandez-Teran, Alfredo Herrera-Estrella, Carla Ivon Herrera-Najera, Pavel Isa, Veronica Mata-Haro, Daniel Lira Morales, Susana Lopez, Antonio Loza Roman, Brenda Irasema Maldonado-Meza, Bernardo Martinez-Miguel, Jose Arturo Martinez-Orozco, Celida Martinez- Rodriguez, Margarita Matias-Florentino, Fidencio Mejia-Nepomuceno, Maria Guadalupe de Jesus Mireles-Rivera, Gloria Maria Molina-Salinas, Hector Montoya-Fuentes, , Mario Mujica-Sanchez, Jose Esteban Muñoz-Medina, Jose de Jesus Nuñez-Contreras, Alicia Ocaña-Mondragon, Luis Alberto Ochoa-Carrera, Hector Esteban Paz-Juarez, Marissa Perez-Garcia, Francisco Pulido, Helen Haydee Fernanda Ramirez-Plascencia, Jorge Salas-Hernandez, Angel Gustavo Salas-Lais, Alejandro Sanchez-Flores, Clara Esperanza Santacruz-Tinoco, Maria Guadalupe Santiago-Mauricio, Selene Zarate, Nelly Selem-Mojica, Blanca Taboada, Gloria Vazquez, Joel Armando Vazquez-Perez. |
| EPI_ISL_2671750, EPI_ISL_2671752, EPI_ISL_2671753, EPI_ISL_2671754, EPI_ISL_2671755, EPI_ISL_2671756, EPI_ISL_2671757, EPI_ISL_2671785 | Unidad de Investigacion Medica de Yucatan (UIMY) | Unidad de Genomica Avanzada | Consorcio Mexicano de Vigilancia Genomica (CoViGen-Mex). Authors (in alphabetical order): Julio Elias Alvarado-Yaah, Carlos F. Arias, Santiago avila-Rios, Eduardo Becerril-Vargas, Victor Hugo Borja-Aburto, Celia Boukadida, Cristobal Chaidez-Quiroz, Juan Bautista Chale-Dzul, Ricardo Ciria Merce, Andreu Comas-Garcia, Celida Duque Molina, Julissa Enciso-Ibarra, Jose Antonio Enciso-Moreno, Gloria Elena Espinosa-Ayala, Fernando Fontove-Herrera, Daniel Fregoso-Rueda, Victor Eduardo Garcia-Arias, Alejandra Garcia-Gasca, Bruno Gomez-Gil, Jean Pierre Gonzalez, Irvin Gonzalez-Lopez, Concepcion Grajales-Muñiz, Ricardo Grande, Rosa Maria Gutierrez Rios, Jesus Hernandez, Alejandra Hernandez-Teran, Alfredo Herrera-Estrella, Carla Ivon Herrera-Najera, Pavel Isa, Veronica Mata-Haro, Daniel Lira Morales, Susana Lopez, Antonio Loza Roman, Brenda Irasema Maldonado-Meza, Bernardo Martinez-Miguel, Jose Arturo Martinez-Orozco, Celida Martinez- Rodriguez, Margarita Matias-Florentino, Fidencio Mejia-Nepomuceno, Maria Guadalupe de Jesus Mireles-Rivera, Gloria Maria Molina-Salinas, Hector Montoya-Fuentes, , Mario Mujica-Sanchez, Jose Esteban Muñoz-Medina, Jose de Jesus Nuñez-Contreras, Alicia Ocaña-Mondragon, Luis Alberto Ochoa-Carrera, Hector Esteban Paz-Juarez, Marissa Perez-Garcia, Francisco Pulido, Helen Haydee Fernanda Ramirez-Plascencia, Jorge Salas-Hernandez, Angel Gustavo Salas-Lais, Alejandro Sanchez-Flores, Clara Esperanza Santacruz-Tinoco, Maria Guadalupe Santiago-Mauricio, Selene Zarate, Nelly Selem-Mojica, Blanca Taboada, Gloria Vazquez, Joel Armando Vazquez-Perez. |
| EPI_ISL_2671834 | COVID LAB | Cayman Islands Forensic Science Lab | Jonathan Smellie, Karla Montes, Brittany Balcewich, Allen Grolla |
| EPI_ISL_2673506, EPI_ISL_2673507, EPI_ISL_2673508, EPI_ISL_2673509, EPI_ISL_2673510, EPI_ISL_2673511, EPI_ISL_2673512, EPI_ISL_2673513, EPI_ISL_2673514, EPI_ISL_2673515, EPI_ISL_2673516, EPI_ISL_2673517, EPI_ISL_2673518 | | | |
| see above | Department of Virology, Istituto Zooprofilattico Sperimentale del Lazio e della Toscana (IZSLT) | Department of General Diagnostics; Department of Virology; Istituto Zooprofilattico Sperimentale del Lazio e della Toscana (IZSLT) | Patricia Alba, Giuseppe Manna, Elena L. Diaconu, Fabiola Feltrin, Raffaela Conti, Teresa Scicluna, Virginia Carfora, Antonella Cersini, Alessia Franco, Antonio Battisti. |
| EPI_ISL_2673648, EPI_ISL_2673649, EPI_ISL_2673650, EPI_ISL_2673651 | SIESP CHIETI - DRIVE IN VASTO | Istituto Zooprofilattico Sperimentale dell'Abruzzo e Molise "G. Caporale" | Lorusso A, Marcacci M, Di Domenico M, Ancora M, Curini V, Di Lollo Valeria, Mangone I, Rinaldi A, Delli Compagni E, Scialabba S, Caporale M, Di Pasquale A, Cammà C, Puglia I, Calistri P, Savini G |
| EPI_ISL_2673652 | SIESP DIPARTIMENTO DI PREVENZIONE TERAMO | Istituto Zooprofilattico Sperimentale dell'Abruzzo e Molise "G. Caporale" | Lorusso A, Marcacci M, Di Domenico M, Ancora M, Curini V, Di Lollo Valeria, Mangone I, Rinaldi A, Delli Compagni E, Scialabba S, Caporale M, Di Pasquale A, Cammà C, Puglia I, Calistri P, Savini G |
| EPI_ISL_2673653 | SIESP CHIETI - DRIVE IN CHIETI | Istituto Zooprofilattico Sperimentale dell'Abruzzo e Molise "G. Caporale" | Lorusso A, Marcacci M, Di Domenico M, Ancora M, Curini V, Di Lollo Valeria, Mangone I, Rinaldi A, Delli Compagni E, Scialabba S, Caporale M, Di Pasquale A, Cammà C, Puglia I, Calistri P, Savini G |
| EPI_ISL_2673654 | SIESP CHIETI - DRIVE IN VASTO | Istituto Zooprofilattico Sperimentale dell'Abruzzo e Molise "G. Caporale" | Lorusso A, Marcacci M, Di Domenico M, Ancora M, Curini V, Di Lollo Valeria, Mangone I, Rinaldi A, Delli Compagni E, Scialabba S, Caporale M, Di Pasquale A, Cammà C, Puglia I, Calistri P, Savini G |
| EPI_ISL_2673655 | SIESP CHIETI - DRIVE IN CHIETI | Istituto Zooprofilattico Sperimentale dell'Abruzzo e Molise "G. Caporale" | Lorusso A, Marcacci M, Di Domenico M, Ancora M, Curini V, Di Lollo Valeria, Mangone I, Rinaldi A, Delli Compagni E, Scialabba S, Caporale M, Di Pasquale A, Cammà C, Puglia I, Calistri P, Savini G |
| EPI_ISL_2673656 | SIESP DIPARTIMENTO DI PREVENZIONE CHIETI | Istituto Zooprofilattico Sperimentale dell'Abruzzo e Molise "G. Caporale" | Lorusso A, Marcacci M, Di Domenico M, Ancora M, Curini V, Di Lollo Valeria, Mangone I, Rinaldi A, Delli Compagni E, Scialabba S, Caporale M, Di Pasquale A, Cammà C, Puglia I, Calistri P, Savini G |
| EPI_ISL_2673657, EPI_ISL_2673658 | SIESP CHIETI - DRIVE IN LANCIANO | Istituto Zooprofilattico Sperimentale dell'Abruzzo e Molise "G. Caporale" | Lorusso A, Marcacci M, Di Domenico M, Ancora M, Curini V, Di Lollo Valeria, Mangone I, Rinaldi A, Delli Compagni E, Scialabba S, Caporale M, Di Pasquale A, Cammà C, Puglia I, Calistri P, Savini G |
| EPI_ISL_2673659 | SIESP DIPARTIMENTO DI PREVENZIONE CHIETI | Istituto Zooprofilattico Sperimentale dell'Abruzzo e Molise "G. Caporale" | Lorusso A, Marcacci M, Di Domenico M, Ancora M, Curini V, Di Lollo Valeria, Mangone I, Rinaldi A, Delli Compagni E, Scialabba S, Caporale M, Di Pasquale A, Cammà C, Puglia I, Calistri P, Savini G |
| EPI_ISL_2673660, EPI_ISL_2673661, EPI_ISL_2673662, EPI_ISL_2673663, | SIESP CHIETI - DRIVE IN VASTO | Istituto Zooprofilattico Sperimentale dell'Abruzzo e Molise "G. Caporale" | Lorusso A, Marcacci M, Di Domenico M, Ancora M, Curini V, Di Lollo Valeria, Mangone I, Rinaldi A, Delli Compagni E, Scialabba S, Caporale M, Di Pasquale A, Cammà C, Puglia I, Calistri P, Savini G |

| | | | |
|---|---|---|---|
| EPI_ISL_2673664 | | | |
| EPI_ISL_2673665 | SIESP CHIETI - DRIVE IN ORTONA | Istituto Zooprofilattico Sperimentale dell'Abruzzo e Molise "G. Caporale" | Lorusso A, Marcacci M, Di Domenico M, Ancora M, Curini V, Di Lollo Valeria, Mangone I, Rinaldi A, Delli Compagni E, Scialabba S, Caporale M, Di Pasquale A, Cammà C, Puglia I, Calistri P, Savini G |
| EPI_ISL_2673666, EPI_ISL_2673667 | SIESP CHIETI - DRIVE IN LANCIANO | Istituto Zooprofilattico Sperimentale dell'Abruzzo e Molise "G. Caporale" | Lorusso A, Marcacci M, Di Domenico M, Ancora M, Curini V, Di Lollo Valeria, Mangone I, Rinaldi A, Delli Compagni E, Scialabba S, Caporale M, Di Pasquale A, Cammà C, Puglia I, Calistri P, Savini G |
| EPI_ISL_2673668, EPI_ISL_2673669, EPI_ISL_2673670 | SIESP CHIETI - DRIVE IN VASTO | Istituto Zooprofilattico Sperimentale dell'Abruzzo e Molise "G. Caporale" | Lorusso A, Marcacci M, Di Domenico M, Ancora M, Curini V, Di Lollo Valeria, Mangone I, Rinaldi A, Delli Compagni E, Scialabba S, Caporale M, Di Pasquale A, Cammà C, Puglia I, Calistri P, Savini G |
| EPI_ISL_2673671 | SIESP CHIETI - DRIVE IN LANCIANO | Istituto Zooprofilattico Sperimentale dell'Abruzzo e Molise "G. Caporale" | Lorusso A, Marcacci M, Di Domenico M, Ancora M, Curini V, Di Lollo Valeria, Mangone I, Rinaldi A, Delli Compagni E, Scialabba S, Caporale M, Di Pasquale A, Cammà C, Puglia I, Calistri P, Savini G |
| EPI_ISL_2673672 | SIESP CHIETI - DRIVE IN ORTONA | Istituto Zooprofilattico Sperimentale dell'Abruzzo e Molise "G. Caporale" | Lorusso A, Marcacci M, Di Domenico M, Ancora M, Curini V, Di Lollo Valeria, Mangone I, Rinaldi A, Delli Compagni E, Scialabba S, Caporale M, Di Pasquale A, Cammà C, Puglia I, Calistri P, Savini G |
| EPI_ISL_2673673 | SIESP CHIETI - DRIVE IN VASTO | Istituto Zooprofilattico Sperimentale dell'Abruzzo e Molise "G. Caporale" | Lorusso A, Marcacci M, Di Domenico M, Ancora M, Curini V, Di Lollo Valeria, Mangone I, Rinaldi A, Delli Compagni E, Scialabba S, Caporale M, Di Pasquale A, Cammà C, Puglia I, Calistri P, Savini G |
| EPI_ISL_2673674 | SIESP CHIETI - DRIVE IN LANCIANO | Istituto Zooprofilattico Sperimentale dell'Abruzzo e Molise "G. Caporale" | Lorusso A, Marcacci M, Di Domenico M, Ancora M, Curini V, Di Lollo Valeria, Mangone I, Rinaldi A, Delli Compagni E, Scialabba S, Caporale M, Di Pasquale A, Cammà C, Puglia I, Calistri P, Savini G |
| EPI_ISL_2673675 | SIESP CHIETI - DRIVE IN CHIETI | Istituto Zooprofilattico Sperimentale dell'Abruzzo e Molise "G. Caporale" | Lorusso A, Marcacci M, Di Domenico M, Ancora M, Curini V, Di Lollo Valeria, Mangone I, Rinaldi A, Delli Compagni E, Scialabba S, Caporale M, Di Pasquale A, Cammà C, Puglia I, Calistri P, Savini G |
| EPI_ISL_2673676, EPI_ISL_2673677, EPI_ISL_2673678 | OSPEDALE CIVILE TERAMO - CENTRO TRASFUSIONALE | Istituto Zooprofilattico Sperimentale dell'Abruzzo e Molise "G. Caporale" | Lorusso A, Marcacci M, Di Domenico M, Ancora M, Curini V, Di Lollo Valeria, Mangone I, Rinaldi A, Delli Compagni E, Scialabba S, Caporale M, Di Pasquale A, Cammà C, Puglia I, Calistri P, Savini G |
| EPI_ISL_2673679, EPI_ISL_2673680 | PRESIDIO OSPEDALIERO TAGLIACOZZO | Istituto Zooprofilattico Sperimentale dell'Abruzzo e Molise "G. Caporale" | Lorusso A, Marcacci M, Di Domenico M, Ancora M, Curini V, Di Lollo Valeria, Mangone I, Rinaldi A, Delli Compagni E, Scialabba S, Caporale M, Di Pasquale A, Cammà C, Puglia I, Calistri P, Savini G |
| EPI_ISL_2673681, EPI_ISL_2673682 | SIESP CHIETI - DRIVE IN LANCIANO | Istituto Zooprofilattico Sperimentale dell'Abruzzo e Molise "G. Caporale" | Lorusso A, Marcacci M, Di Domenico M, Ancora M, Curini V, Di Lollo Valeria, Mangone I, Rinaldi A, Delli Compagni E, Scialabba S, Caporale M, Di Pasquale A, Cammà C, Puglia I, Calistri P, Savini G |
| EPI_ISL_2674629, EPI_ISL_2674631 | LESP Nuevo Leon | Instituto de Diagnostico y Referencia Epidemiologicos (INDRE) | Claudia Wong-Arambula, Abril Rodriguez-Maldonado, Vanessa Rivero-Arredondo, Ariadna Medina-Benitez, Joaquin Quiroz-Mercado, Sergio Rangel-Guerrero, Natividad Cruz-Ortiz, Tatiana Nunez-Garcia, Gisela Barrera-Badillo, Lucia Hernandez-Rivas, Irma Lopez-Martinez, Ernesto Ramirez-Gonzalez. |
| EPI_ISL_2674633, EPI_ISL_2674634 | LESP Sinaloa | Instituto de Diagnostico y Referencia Epidemiologicos (INDRE) | Claudia Wong-Arambula, Abril Rodriguez-Maldonado, Vanessa Rivero-Arredondo, Ariadna Medina-Benitez, Joaquin Quiroz-Mercado, Sergio Rangel-Guerrero, Natividad Cruz-Ortiz, Tatiana Nunez-Garcia, Gisela Barrera-Badillo, Lucia Hernandez-Rivas, Irma Lopez-Martinez, Ernesto Ramirez-Gonzalez. |
| EPI_ISL_2674635, EPI_ISL_2674636, EPI_ISL_2674637, EPI_ISL_2674638, EPI_ISL_2674639, EPI_ISL_2674640 | LESP Tabasco | Instituto de Diagnostico y Referencia Epidemiologicos (INDRE) | Claudia Wong-Arambula, Abril Rodriguez-Maldonado, Vanessa Rivero-Arredondo, Ariadna Medina-Benitez, Joaquin Quiroz-Mercado, Sergio Rangel-Guerrero, Natividad Cruz-Ortiz, Tatiana Nunez-Garcia, Gisela Barrera-Badillo, Lucia Hernandez-Rivas, Irma Lopez-Martinez, Ernesto Ramirez-Gonzalez. |
| EPI_ISL_2674642 | LESP Durango | Instituto de Diagnostico y Referencia Epidemiologicos (INDRE) | Claudia Wong-Arambula, Abril Rodriguez-Maldonado, Vanessa Rivero-Arredondo, Ariadna Medina-Benitez, Joaquin Quiroz-Mercado, Sergio Rangel-Guerrero, Natividad Cruz-Ortiz, Tatiana Nunez-Garcia, Gisela Barrera-Badillo, Lucia Hernandez-Rivas, Irma Lopez-Martinez, Ernesto Ramirez-Gonzalez. |
| EPI_ISL_2674643, EPI_ISL_2674644, EPI_ISL_2674645, EPI_ISL_2674646, EPI_ISL_2674647, EPI_ISL_2674648, EPI_ISL_2674649 | LESP Sonora | Instituto de Diagnostico y Referencia Epidemiologicos (INDRE) | Claudia Wong-Arambula, Abril Rodriguez-Maldonado, Vanessa Rivero-Arredondo, Ariadna Medina-Benitez, Joaquin Quiroz-Mercado, Sergio Rangel-Guerrero, Natividad Cruz-Ortiz, Tatiana Nunez-Garcia, Gisela Barrera-Badillo, Lucia Hernandez-Rivas, Irma Lopez-Martinez, Ernesto Ramirez-Gonzalez. |
| EPI_ISL_2674650 | LESP Jalisco | Instituto de Diagnostico y Referencia Epidemiologicos (INDRE) | Claudia Wong-Arambula, Abril Rodriguez-Maldonado, Vanessa Rivero-Arredondo, Ariadna Medina-Benitez, Joaquin Quiroz-Mercado, Sergio Rangel-Guerrero, Natividad Cruz-Ortiz, Tatiana Nunez-Garcia, Gisela Barrera-Badillo, Lucia Hernandez-Rivas, Irma Lopez-Martinez, Ernesto Ramirez-Gonzalez. |
| EPI_ISL_2674651, EPI_ISL_2674652, EPI_ISL_2674653, EPI_ISL_2674654, EPI_ISL_2674655 | LESP Quintana Roo | Instituto de Diagnostico y Referencia Epidemiologicos (INDRE) | Claudia Wong-Arambula, Abril Rodriguez-Maldonado, Vanessa Rivero-Arredondo, Ariadna Medina-Benitez, Joaquin Quiroz-Mercado, Sergio Rangel-Guerrero, Natividad Cruz-Ortiz, Tatiana Nunez-Garcia, Gisela Barrera-Badillo, Lucia Hernandez-Rivas, Irma Lopez-Martinez, Ernesto Ramirez-Gonzalez. |
| EPI_ISL_2674661, EPI_ISL_2674662, EPI_ISL_2674663 | LESP Hidalgo | Instituto de Diagnostico y Referencia Epidemiologicos (INDRE) | Claudia Wong-Arambula, Abril Rodriguez-Maldonado, Vanessa Rivero-Arredondo, Ariadna Medina-Benitez, Joaquin Quiroz-Mercado, Sergio Rangel-Guerrero, Natividad Cruz-Ortiz, Tatiana Nunez-Garcia, Gisela Barrera-Badillo, Lucia Hernandez-Rivas, Irma Lopez-Martinez, Ernesto Ramirez-Gonzalez. |
| EPI_ISL_2674664 | LESP Estado de Mexico | Instituto de Diagnostico y Referencia Epidemiologicos (INDRE) | Claudia Wong-Arambula, Abril Rodriguez-Maldonado, Vanessa Rivero-Arredondo, Ariadna Medina-Benitez, Joaquin Quiroz-Mercado, Sergio Rangel-Guerrero, Natividad Cruz-Ortiz, Tatiana Nunez-Garcia, Gisela Barrera-Badillo, Lucia Hernandez-Rivas, Irma Lopez-Martinez, Ernesto Ramirez-Gonzalez. |
| EPI_ISL_2674665, EPI_ISL_2674666, EPI_ISL_2674667 | LESP Hidalgo | Instituto de Diagnostico y Referencia Epidemiologicos (INDRE) | Claudia Wong-Arambula, Abril Rodriguez-Maldonado, Vanessa Rivero-Arredondo, Ariadna Medina-Benitez, Joaquin Quiroz-Mercado, Sergio Rangel-Guerrero, Natividad Cruz-Ortiz, Tatiana Nunez-Garcia, Gisela Barrera-Badillo, Lucia Hernandez-Rivas, Irma Lopez-Martinez, Ernesto Ramirez-Gonzalez. |
| EPI_ISL_2674668, EPI_ISL_2674669, EPI_ISL_2674670 | LESP Estado de Mexico | Instituto de Diagnostico y Referencia Epidemiologicos (INDRE) | Claudia Wong-Arambula, Abril Rodriguez-Maldonado, Vanessa Rivero-Arredondo, Ariadna Medina-Benitez, Joaquin Quiroz-Mercado, Sergio Rangel-Guerrero, Natividad Cruz-Ortiz, Tatiana Nunez-Garcia, Gisela Barrera-Badillo, Lucia Hernandez-Rivas, Irma Lopez-Martinez, Ernesto Ramirez-Gonzalez. |
| EPI_ISL_2674672 | LESP Hidalgo | Instituto de Diagnostico y Referencia Epidemiologicos (INDRE) | Claudia Wong-Arambula, Abril Rodriguez-Maldonado, Vanessa Rivero-Arredondo, Ariadna Medina-Benitez, Joaquin Quiroz-Mercado, Sergio Rangel-Guerrero, Natividad Cruz-Ortiz, Tatiana Nunez-Garcia, Gisela Barrera-Badillo, Lucia Hernandez-Rivas, Irma Lopez-Martinez, Ernesto Ramirez-Gonzalez. |
| EPI_ISL_2674673 | LESP Puebla | Instituto de Diagnostico y Referencia Epidemiologicos (INDRE) | Claudia Wong-Arambula, Abril Rodriguez-Maldonado, Vanessa Rivero-Arredondo, Ariadna Medina-Benitez, Joaquin Quiroz-Mercado, Sergio Rangel-Guerrero, Natividad Cruz-Ortiz, Tatiana Nunez-Garcia, Gisela Barrera-Badillo, Lucia Hernandez-Rivas, Irma Lopez-Martinez, Ernesto Ramirez-Gonzalez. |
| EPI_ISL_2674674, EPI_ISL_2674675, EPI_ISL_2674676, EPI_ISL_2674677, EPI_ISL_2674678, EPI_ISL_2674679, EPI_ISL_2674680, EPI_ISL_2674681, EPI_ISL_2674682 | LESP Hidalgo | Instituto de Diagnostico y Referencia Epidemiologicos (INDRE) | Claudia Wong-Arambula, Abril Rodriguez-Maldonado, Vanessa Rivero-Arredondo, Ariadna Medina-Benitez, Joaquin Quiroz-Mercado, Sergio Rangel-Guerrero, Natividad Cruz-Ortiz, Tatiana Nunez-Garcia, Gisela Barrera-Badillo, Lucia Hernandez-Rivas, Irma Lopez-Martinez, Ernesto Ramirez-Gonzalez. |
| EPI_ISL_2674683 | LESP Queretaro | Instituto de Diagnostico y Referencia Epidemiologicos (INDRE) | Claudia Wong-Arambula, Abril Rodriguez-Maldonado, Vanessa Rivero-Arredondo, Ariadna Medina-Benitez, Joaquin Quiroz-Mercado, Sergio Rangel-Guerrero, Natividad Cruz-Ortiz, Tatiana Nunez-Garcia, Gisela Barrera-Badillo, Lucia Hernandez-Rivas, Irma Lopez-Martinez, Ernesto Ramirez-Gonzalez. |
| EPI_ISL_2674684, EPI_ISL_2674685, EPI_ISL_2674686, EPI_ISL_2674687, EPI_ISL_2674688 | LESP Yucatan | Instituto de Diagnostico y Referencia Epidemiologicos (INDRE) | Claudia Wong-Arambula, Abril Rodriguez-Maldonado, Vanessa Rivero-Arredondo, Ariadna Medina-Benitez, Joaquin Quiroz-Mercado, Sergio Rangel-Guerrero, Natividad Cruz-Ortiz, Tatiana Nunez-Garcia, Gisela Barrera-Badillo, Lucia Hernandez-Rivas, Irma Lopez-Martinez, Ernesto Ramirez-Gonzalez. |

| | | | |
|---|---|---|---|
| EPI_ISL_2674689 | LESP Coahuila | Instituto de Diagnostico y Referencia Epidemiologicos (INDRE) | Claudia Wong-Arambula, Abril Rodriguez-Maldonado, Vanessa Rivero-Arredondo, Ariadna Medina-Benitez, Joaquin Quiroz-Mercado, Sergio Rangel-Guerrero, Natividad Cruz-Ortiz, Tatiana Nunez-Garcia, Gisela Barrera-Badillo, Lucia Hernandez-Rivas, Irma Lopez-Martinez, Ernesto Ramirez-Gonzalez. |
| EPI_ISL_2674690 | LESP Hidalgo | Instituto de Diagnostico y Referencia Epidemiologicos (INDRE) | Claudia Wong-Arambula, Abril Rodriguez-Maldonado, Vanessa Rivero-Arredondo, Ariadna Medina-Benitez, Joaquin Quiroz-Mercado, Sergio Rangel-Guerrero, Natividad Cruz-Ortiz, Tatiana Nunez-Garcia, Gisela Barrera-Badillo, Lucia Hernandez-Rivas, Irma Lopez-Martinez, Ernesto Ramirez-Gonzalez. |
| EPI_ISL_2674691, EPI_ISL_2674692, EPI_ISL_2674693, EPI_ISL_2674694, EPI_ISL_2674695, EPI_ISL_2674696 | LESP Guanajuato | Instituto de Diagnostico y Referencia Epidemiologicos (INDRE) | Claudia Wong-Arambula, Abril Rodriguez-Maldonado, Vanessa Rivero-Arredondo, Ariadna Medina-Benitez, Joaquin Quiroz-Mercado, Sergio Rangel-Guerrero, Natividad Cruz-Ortiz, Tatiana Nunez-Garcia, Gisela Barrera-Badillo, Lucia Hernandez-Rivas, Irma Lopez-Martinez, Ernesto Ramirez-Gonzalez. |
| EPI_ISL_2674697 | LESP Nuevo Leon | Instituto de Diagnostico y Referencia Epidemiologicos (INDRE) | Claudia Wong-Arambula, Abril Rodriguez-Maldonado, Vanessa Rivero-Arredondo, Ariadna Medina-Benitez, Joaquin Quiroz-Mercado, Sergio Rangel-Guerrero, Natividad Cruz-Ortiz, Tatiana Nunez-Garcia, Gisela Barrera-Badillo, Lucia Hernandez-Rivas, Irma Lopez-Martinez, Ernesto Ramirez-Gonzalez. |
| EPI_ISL_2674698 | LESP Ciudad de Mexico | Instituto de Diagnostico y Referencia Epidemiologicos (INDRE) | Claudia Wong-Arambula, Abril Rodriguez-Maldonado, Vanessa Rivero-Arredondo, Ariadna Medina-Benitez, Joaquin Quiroz-Mercado, Sergio Rangel-Guerrero, Natividad Cruz-Ortiz, Tatiana Nunez-Garcia, Gisela Barrera-Badillo, Lucia Hernandez-Rivas, Irma Lopez-Martinez, Ernesto Ramirez-Gonzalez. |
| EPI_ISL_2674699 | LESP Chihuahua | Instituto de Diagnostico y Referencia Epidemiologicos (INDRE) | Claudia Wong-Arambula, Abril Rodriguez-Maldonado, Vanessa Rivero-Arredondo, Ariadna Medina-Benitez, Joaquin Quiroz-Mercado, Sergio Rangel-Guerrero, Natividad Cruz-Ortiz, Tatiana Nunez-Garcia, Gisela Barrera-Badillo, Lucia Hernandez-Rivas, Irma Lopez-Martinez, Ernesto Ramirez-Gonzalez. |
| EPI_ISL_2674700 | LESP Puebla | Instituto de Diagnostico y Referencia Epidemiologicos (INDRE) | Claudia Wong-Arambula, Abril Rodriguez-Maldonado, Vanessa Rivero-Arredondo, Ariadna Medina-Benitez, Joaquin Quiroz-Mercado, Sergio Rangel-Guerrero, Natividad Cruz-Ortiz, Tatiana Nunez-Garcia, Gisela Barrera-Badillo, Lucia Hernandez-Rivas, Irma Lopez-Martinez, Ernesto Ramirez-Gonzalez. |
| EPI_ISL_2674701, EPI_ISL_2674703, EPI_ISL_2674704, EPI_ISL_2674705, EPI_ISL_2674706, EPI_ISL_2674707, EPI_ISL_2674708, EPI_ISL_2674709, EPI_ISL_2674710, EPI_ISL_2674711, EPI_ISL_2674712, EPI_ISL_2674713 | | | |
| see above | LESP Yucatan | Instituto de Diagnostico y Referencia Epidemiologicos (INDRE) | Claudia Wong-Arambula, Abril Rodriguez-Maldonado, Vanessa Rivero-Arredondo, Ariadna Medina-Benitez, Joaquin Quiroz-Mercado, Sergio Rangel-Guerrero, Natividad Cruz-Ortiz, Tatiana Nunez-Garcia, Gisela Barrera-Badillo, Lucia Hernandez-Rivas, Irma Lopez-Martinez, Ernesto Ramirez-Gonzalez. |
| EPI_ISL_2674714 | LESP Quintana Roo | Instituto de Diagnostico y Referencia Epidemiologicos (INDRE) | Claudia Wong-Arambula, Abril Rodriguez-Maldonado, Vanessa Rivero-Arredondo, Ariadna Medina-Benitez, Joaquin Quiroz-Mercado, Sergio Rangel-Guerrero, Natividad Cruz-Ortiz, Tatiana Nunez-Garcia, Gisela Barrera-Badillo, Lucia Hernandez-Rivas, Irma Lopez-Martinez, Ernesto Ramirez-Gonzalez. |
| EPI_ISL_2674715 | LESP Yucatan | Instituto de Diagnostico y Referencia Epidemiologicos (INDRE) | Claudia Wong-Arambula, Abril Rodriguez-Maldonado, Vanessa Rivero-Arredondo, Ariadna Medina-Benitez, Joaquin Quiroz-Mercado, Sergio Rangel-Guerrero, Natividad Cruz-Ortiz, Tatiana Nunez-Garcia, Gisela Barrera-Badillo, Lucia Hernandez-Rivas, Irma Lopez-Martinez, Ernesto Ramirez-Gonzalez. |
| EPI_ISL_2674716 | LESP San Luis Potosi | Instituto de Diagnostico y Referencia Epidemiologicos (INDRE) | Claudia Wong-Arambula, Abril Rodriguez-Maldonado, Vanessa Rivero-Arredondo, Ariadna Medina-Benitez, Joaquin Quiroz-Mercado, Sergio Rangel-Guerrero, Natividad Cruz-Ortiz, Tatiana Nunez-Garcia, Gisela Barrera-Badillo, Lucia Hernandez-Rivas, Irma Lopez-Martinez, Ernesto Ramirez-Gonzalez. |
| EPI_ISL_2674718, EPI_ISL_2674719 | LESP Sonora | Instituto de Diagnostico y Referencia Epidemiologicos (INDRE) | Claudia Wong-Arambula, Abril Rodriguez-Maldonado, Vanessa Rivero-Arredondo, Ariadna Medina-Benitez, Joaquin Quiroz-Mercado, Sergio Rangel-Guerrero, Natividad Cruz-Ortiz, Tatiana Nunez-Garcia, Gisela Barrera-Badillo, Lucia Hernandez-Rivas, Irma Lopez-Martinez, Ernesto Ramirez-Gonzalez. |
| EPI_ISL_2674900, EPI_ISL_2674901, EPI_ISL_2674902 | LESP Ciudad de Mexico | Instituto de Diagnostico y Referencia Epidemiologicos (INDRE) | Claudia Wong-Arambula, Abril Rodriguez-Maldonado, Vanessa Rivero-Arredondo, Ariadna Medina-Benitez, Joaquin Quiroz-Mercado, Sergio Rangel-Guerrero, Natividad Cruz-Ortiz, Tatiana Nunez-Garcia, Gisela Barrera-Badillo, Lucia Hernandez-Rivas, Irma Lopez-Martinez, Ernesto Ramirez-Gonzalez. |
| EPI_ISL_2674903 | LESP Jalisco | Instituto de Diagnostico y Referencia Epidemiologicos (INDRE) | Claudia Wong-Arambula, Abril Rodriguez-Maldonado, Vanessa Rivero-Arredondo, Ariadna Medina-Benitez, Joaquin Quiroz-Mercado, Sergio Rangel-Guerrero, Natividad Cruz-Ortiz, Tatiana Nunez-Garcia, Gisela Barrera-Badillo, Lucia Hernandez-Rivas, Irma Lopez-Martinez, Ernesto Ramirez-Gonzalez. |
| EPI_ISL_2674904 | LESP Ciudad de Mexico | Instituto de Diagnostico y Referencia Epidemiologicos (INDRE) | Claudia Wong-Arambula, Abril Rodriguez-Maldonado, Vanessa Rivero-Arredondo, Ariadna Medina-Benitez, Joaquin Quiroz-Mercado, Sergio Rangel-Guerrero, Natividad Cruz-Ortiz, Tatiana Nunez-Garcia, Gisela Barrera-Badillo, Lucia Hernandez-Rivas, Irma Lopez-Martinez, Ernesto Ramirez-Gonzalez. |
| EPI_ISL_2674905 | LESP Yucatan | Instituto de Diagnostico y Referencia Epidemiologicos (INDRE) | Claudia Wong-Arambula, Abril Rodriguez-Maldonado, Vanessa Rivero-Arredondo, Ariadna Medina-Benitez, Joaquin Quiroz-Mercado, Sergio Rangel-Guerrero, Natividad Cruz-Ortiz, Tatiana Nunez-Garcia, Gisela Barrera-Badillo, Lucia Hernandez-Rivas, Irma Lopez-Martinez, Ernesto Ramirez-Gonzalez. |
| EPI_ISL_2680931 | Hospital Margarita Maza de Juárez | Microbial Genomics Laboratory | Consorcio Mexicano de Vigilancia Genómica (CoViGen-Mex). Authors (in alphabetical order): Julio Elias Alvarado-Yaah, Carlos F. Arias, Santiago Ávila-Ríos, Eduardo Becerril-Vargas, Víctor Hugo Borja-Aburto, Celia Boukadida, Cristóbal Cháidez-Quiróz, Juan Bautista Chale-Dzul, Ricardo Ciria Merce, Andreu Comas-García, Célida Duque Molina, Julissa Enciso-Ibarra, José Antonio Enciso-Moreno, Gloria Elena Espinosa-Ayala, Fernando Fontove-Herrera, Daniel Fregoso-Rueda, Victor Eduardo García-Arias, Alejandra García-Gasca, Bruno Gómez-Gil, Jean Pierre González, Irvin González-López, Concepción Grajales-Muñiz, Ricardo Grande, Rosa María Gutiérrez Rios, Jesús Hernández, Alejandra Hernández-Terán, Alfredo Herrera-Estrella, Carla Ivón Herrera-Najera, Pavel Isa, Verónica Mata-Haro, Daniel Lira Morales, Susana Lopez, Antonio Loza Román, Brenda Irasema Maldonado-Meza, Bernardo Martínez-Miguel, José Arturo Martínez-Orozco, Célida Martínez- Rodríguez, Margarita Matías-Florentino, Fidencio Mejía-Nepomuceno, María Guadalupe de Jesús Mireles-Rivera, Gloria María Molina-Salinas, Hector Montoya-Fuentes, , Mario Mújica-Sánchez, José Esteban Muñoz-Medina, José de Jesús Nuñez-Contreras, Alicia Ocaña-Mondragón, Luis Alberto Ochoa-Carrera, Hector Esteban Paz-Juárez, Marissa Perez-Garcia, Francisco Pulido, Helen Haydee Fernanda Ramírez-Plascencia, Jorge Salas-Hernández, Angel Gustavo Salas-Lais, Alejandro Sánchez-Flores, Clara Esperanza Santacruz-Tinoco, María Guadalupe Santiago-Mauricio, Selene Zárate, Nelly Sélem-Mojica, Blanca Taboada, Gloria Vazquez, Joel Armando Vázquez-Pérez. |
| EPI_ISL_2680934 | Imágenes Diagnósticas de Mazatlán | Microbial Genomics Laboratory | Consorcio Mexicano de Vigilancia Genómica (CoViGen-Mex). Authors (in alphabetical order): Julio Elias Alvarado-Yaah, Carlos F. Arias, Santiago Ávila-Ríos, Eduardo Becerril-Vargas, Víctor Hugo Borja-Aburto, Celia Boukadida, Cristóbal Cháidez-Quiróz, Juan Bautista Chale-Dzul, Ricardo Ciria Merce, Andreu Comas-García, Célida Duque Molina, Julissa Enciso-Ibarra, José Antonio Enciso-Moreno, Gloria Elena Espinosa-Ayala, Fernando Fontove-Herrera, Daniel Fregoso-Rueda, Victor Eduardo García-Arias, Alejandra García-Gasca, Bruno Gómez-Gil, Jean Pierre González, Irvin González-López, Concepción Grajales-Muñiz, Ricardo Grande, Rosa María Gutiérrez Rios, Jesús Hernández, Alejandra Hernández-Terán, Alfredo Herrera-Estrella, Carla Ivón Herrera-Najera, Pavel Isa, Verónica Mata-Haro, Daniel Lira Morales, Susana Lopez, Antonio Loza Román, Brenda Irasema Maldonado-Meza, Bernardo Martínez-Miguel, José Arturo Martínez-Orozco, Célida Martínez- Rodríguez, Margarita Matías-Florentino, Fidencio Mejía-Nepomuceno, María Guadalupe de Jesús Mireles-Rivera, Gloria María Molina-Salinas, Hector Montoya-Fuentes, , Mario Mújica-Sánchez, José Esteban Muñoz-Medina, José de Jesús Nuñez-Contreras, Alicia Ocaña-Mondragón, Luis Alberto Ochoa-Carrera, Hector Esteban Paz-Juárez, Marissa Perez-Garcia, Francisco Pulido, Helen Haydee Fernanda Ramírez-Plascencia, Jorge Salas-Hernández, Angel Gustavo Salas-Lais, Alejandro Sánchez-Flores, Clara Esperanza Santacruz-Tinoco, María Guadalupe Santiago-Mauricio, Selene Zárate, Nelly Sélem-Mojica, Blanca Taboada, Gloria Vazquez, Joel Armando Vázquez-Pérez. |
| EPI_ISL_2680935 | Hospital Sharp | Microbial Genomics Laboratory | Consorcio Mexicano de Vigilancia Genómica (CoViGen-Mex). Authors (in alphabetical order): Julio Elias Alvarado-Yaah, Carlos F. Arias, Santiago Ávila-Ríos, Eduardo Becerril-Vargas, Víctor Hugo Borja-Aburto, Celia Boukadida, Cristóbal Cháidez-Quiróz, Juan Bautista Chale-Dzul, Ricardo Ciria Merce, Andreu Comas-García, Célida Duque Molina, Julissa Enciso-Ibarra, José Antonio Enciso-Moreno, Gloria Elena Espinosa-Ayala, Fernando Fontove-Herrera, Daniel Fregoso-Rueda, Victor Eduardo García-Arias, Alejandra García-Gasca, Bruno Gómez-Gil, Jean Pierre González, Irvin González-López, Concepción Grajales-Muñiz, Ricardo Grande, Rosa María Gutiérrez Rios, Jesús Hernández, Alejandra Hernández-Terán, Alfredo |

| | | | |
|---|---|---|---|
| | | | Herrera-Estrella, Carla Ivón Herrera-Najera, Pavel Isa, Verónica Mata-Haro, Daniel Lira Morales, Susana Lopez, Antonio Loza Román, Brenda Irasema Maldonado-Meza, Bernardo Martínez-Miguel, José Arturo Martínez-Orozco, Célida Martínez- Rodríguez, Margarita Matías-Florentino, Fidencio Mejía-Nepomuceno, María Guadalupe de Jesús Mireles-Rivera, Gloria María Molina-Salinas, Hector Montoya-Fuentes, , Mario Mújica-Sánchez, José Esteban Muñoz-Medina, José de Jesús Nuñez-Contreras, Alicia Ocaña-Mondragón, Luis Alberto Ochoa-Carrera, Hector Esteban Paz-Juárez, Marissa Perez-Garcia, Francisco Pulido, Helen Haydee Fernanda Ramírez-Plascencia, Jorge Salas-Hernández, Angel Gustavo Salas-Lais, Alejandro Sánchez-Flores, Clara Esperanza Santacruz-Tinoco, María Guadalupe Santiago-Mauricio, Selene Zárate, Nelly Sélem-Mojica, Blanca Taboada, Gloria Vazquez, Joel Armando Vázquez-Pérez. |
| EPI_ISL_2680936, EPI_ISL_2680937 | Hospital Margarita Maza de Juárez | Microbial Genomics Laboratory | Consorcio Mexicano de Vigilancia Genómica (CoViGen-Mex). Authors (in alphabetical order): Julio Elias Alvarado-Yaah, Carlos F. Arias, Santiago Ávila-Ríos, Eduardo Becerril-Vargas, Víctor Hugo Borja-Aburto, Cristóbal Cháidez-Quiróz, Juan Bautista Chale-Dzul, Ricardo Ciria Merce, Andreu Comas-García, Célida Duque Molina, Julissa Enciso-Ibarra, José Antonio Enciso-Moreno, Gloria Elena Espinosa-Ayala, Fernando Fontove-Herrera, Daniel Fregoso-Rueda, Victor Eduardo García-Arias, Alejandra García-Gasca, Bruno Gómez-Gil, Jean Pierre González, Irvin González-López, Concepción Grajales-Muñiz, Ricardo Grande, Rosa María Gutiérrez Rios, Jesús Hernández, Alejandra Hernández-Terán, Alfredo Herrera-Estrella, Carla Ivón Herrera-Najera, Pavel Isa, Verónica Mata-Haro, Daniel Lira Morales, Susana Lopez, Antonio Loza Román, Brenda Irasema Maldonado-Meza, Bernardo Martínez-Miguel, José Arturo Martínez-Orozco, Célida Martínez- Rodríguez, Margarita Matías-Florentino, Fidencio Mejía-Nepomuceno, María Guadalupe de Jesús Mireles-Rivera, Gloria María Molina-Salinas, Hector Montoya-Fuentes, , Mario Mújica-Sánchez, José Esteban Muñoz-Medina, José de Jesús Nuñez-Contreras, Alicia Ocaña-Mondragón, Luis Alberto Ochoa-Carrera, Hector Esteban Paz-Juárez, Marissa Perez-Garcia, Francisco Pulido, Helen Haydee Fernanda Ramírez-Plascencia, Jorge Salas-Hernández, Angel Gustavo Salas-Lais, Alejandro Sánchez-Flores, Clara Esperanza Santacruz-Tinoco, María Guadalupe Santiago-Mauricio, Selene Zárate, Nelly Sélem-Mojica, Blanca Taboada, Gloria Vazquez, Joel Armando Vázquez-Pérez. |
| EPI_ISL_2680938, EPI_ISL_2680939, EPI_ISL_2680940, EPI_ISL_2680942 | Hospital Sharp | Microbial Genomics Laboratory | Consorcio Mexicano de Vigilancia Genómica (CoViGen-Mex). Authors (in alphabetical order): Julio Elias Alvarado-Yaah, Carlos F. Arias, Santiago Ávila-Ríos, Eduardo Becerril-Vargas, Víctor Hugo Borja-Aburto, Celia Boukadida, Cristóbal Cháidez-Quiróz, Juan Bautista Chale-Dzul, Ricardo Ciria Merce, Andreu Comas-García, Célida Duque Molina, Julissa Enciso-Ibarra, José Antonio Enciso-Moreno, Gloria Elena Espinosa-Ayala, Fernando Fontove-Herrera, Daniel Fregoso-Rueda, Victor Eduardo García-Arias, Alejandra García-Gasca, Bruno Gómez-Gil, Jean Pierre González, Irvin González-López, Concepción Grajales-Muñiz, Ricardo Grande, Rosa María Gutiérrez Rios, Jesús Hernández, Alejandra Hernández-Terán, Alfredo Herrera-Estrella, Carla Ivón Herrera-Najera, Pavel Isa, Verónica Mata-Haro, Daniel Lira Morales, Susana Lopez, Antonio Loza Román, Brenda Irasema Maldonado-Meza, Bernardo Martínez-Miguel, José Arturo Martínez-Orozco, Célida Martínez- Rodríguez, Margarita Matías-Florentino, Fidencio Mejía-Nepomuceno, María Guadalupe de Jesús Mireles-Rivera, Gloria María Molina-Salinas, Hector Montoya-Fuentes, , Mario Mújica-Sánchez, José Esteban Muñoz-Medina, José de Jesús Nuñez-Contreras, Alicia Ocaña-Mondragón, Luis Alberto Ochoa-Carrera, Hector Esteban Paz-Juárez, Marissa Perez-Garcia, Francisco Pulido, Helen Haydee Fernanda Ramírez-Plascencia, Jorge Salas-Hernández, Angel Gustavo Salas-Lais, Alejandro Sánchez-Flores, Clara Esperanza Santacruz-Tinoco, María Guadalupe Santiago-Mauricio, Selene Zárate, Nelly Sélem-Mojica, Blanca Taboada, Gloria Vazquez, Joel Armando Vázquez-Pérez. |
| EPI_ISL_2681047, EPI_ISL_2681048, EPI_ISL_2681049 | Unidad de Investigación Biomédica de Zacatecas (UIBZ) | Instituto de Biotecnología de la UNAM | Consorcio Mexicano de Vigilancia Genómica (CoViGen-Mex). Authors (in alphabetical order): Julio Elias Alvarado-Yaah, Carlos F. Arias, Santiago Ávila-Ríos, Eduardo Becerril-Vargas, Víctor Hugo Borja-Aburto, Celia Boukadida, Cristóbal Cháidez-Quiróz, Juan Bautista Chale-Dzul, Ricardo Ciria Merce, Andreu Comas-García, Célida Duque Molina, Julissa Enciso-Ibarra, José Antonio Enciso-Moreno, Gloria Elena Espinosa-Ayala, Fernando Fontove-Herrera, Daniel Fregoso-Rueda, Victor Eduardo García-Arias, Alejandra García-Gasca, Bruno Gómez-Gil, Jean Pierre González, Irvin González-López, Concepción Grajales-Muñiz, Ricardo Grande, Rosa María Gutiérrez Rios, Jesús Hernández, Alejandra Hernández-Terán, Alfredo Herrera-Estrella, Carla Ivón Herrera-Najera, Pavel Isa, Verónica Mata-Haro, Daniel Lira Morales, Susana Lopez, Antonio Loza Román, Brenda Irasema Maldonado-Meza, Bernardo Martínez-Miguel, José Arturo Martínez-Orozco, Célida Martínez- Rodríguez, Margarita Matías-Florentino, Fidencio Mejía-Nepomuceno, María Guadalupe de Jesús Mireles-Rivera, Gloria María Molina-Salinas, Hector Montoya-Fuentes, , Mario Mújica-Sánchez, José Esteban Muñoz-Medina, José de Jesús Nuñez-Contreras, Alicia Ocaña-Mondragón, Luis Alberto Ochoa-Carrera, Hector Esteban Paz-Juárez, Marissa Perez-Garcia, Francisco Pulido, Helen Haydee Fernanda Ramírez-Plascencia, Jorge Salas-Hernández, Angel Gustavo Salas-Lais, Alejandro Sánchez-Flores, Clara Esperanza Santacruz-Tinoco, María Guadalupe Santiago-Mauricio, Selene Zárate, Nelly Sélem-Mojica, Blanca Taboada, Gloria Vazquez, Joel Armando Vázquez-Pérez. |
| EPI_ISL_2681050 | Unidad de Investigación Médica de Yucatán (UIMY) | Instituto de Biotecnología de la UNAM | Consorcio Mexicano de Vigilancia Genómica (CoViGen-Mex). Authors (in alphabetical order): Julio Elias Alvarado-Yaah, Carlos F. Arias, Santiago Ávila-Ríos, Eduardo Becerril-Vargas, Víctor Hugo Borja-Aburto, Celia Boukadida, Cristóbal Cháidez-Quiróz, Juan Bautista Chale-Dzul, Ricardo Ciria Merce, Andreu Comas-García, Célida Duque Molina, Julissa Enciso-Ibarra, José Antonio Enciso-Moreno, Gloria Elena Espinosa-Ayala, Fernando Fontove-Herrera, Daniel Fregoso-Rueda, Victor Eduardo García-Arias, Alejandra García-Gasca, Bruno Gómez-Gil, Jean Pierre González, Irvin González-López, Concepción Grajales-Muñiz, Ricardo Grande, Rosa María Gutiérrez Rios, Jesús Hernández, Alejandra Hernández-Terán, Alfredo Herrera-Estrella, Carla Ivón Herrera-Najera, Pavel Isa, Verónica Mata-Haro, Daniel Lira Morales, Susana Lopez, Antonio Loza Román, Brenda Irasema Maldonado-Meza, Bernardo Martínez-Miguel, José Arturo Martínez-Orozco, Célida Martínez- Rodríguez, Margarita Matías-Florentino, Fidencio Mejía-Nepomuceno, María Guadalupe de Jesús Mireles-Rivera, Gloria María Molina-Salinas, Hector Montoya-Fuentes, , Mario Mújica-Sánchez, José Esteban Muñoz-Medina, José de Jesús Nuñez-Contreras, Alicia Ocaña-Mondragón, Luis Alberto Ochoa-Carrera, Hector Esteban Paz-Juárez, Marissa Perez-Garcia, Francisco Pulido, Helen Haydee Fernanda Ramírez-Plascencia, Jorge Salas-Hernández, Angel Gustavo Salas-Lais, Alejandro Sánchez-Flores, Clara Esperanza Santacruz-Tinoco, María Guadalupe Santiago-Mauricio, Selene Zárate, Nelly Sélem-Mojica, Blanca Taboada, Gloria Vazquez, Joel Armando Vázquez-Pérez. |
| EPI_ISL_2681051 | Laboratorio Central de Epidemiología (LCE) | Instituto de Biotecnología de la UNAM | Consorcio Mexicano de Vigilancia Genómica (CoViGen-Mex). Authors (in alphabetical order): Julio Elias Alvarado-Yaah, Carlos F. Arias, Santiago Ávila-Ríos, Eduardo Becerril-Vargas, Víctor Hugo Borja-Aburto, Celia Boukadida, Cristóbal Cháidez-Quiróz, Juan Bautista Chale-Dzul, Ricardo Ciria Merce, Andreu Comas-García, Célida Duque Molina, Julissa Enciso-Ibarra, José Antonio Enciso-Moreno, Gloria Elena Espinosa-Ayala, Fernando Fontove-Herrera, Daniel Fregoso-Rueda, Victor Eduardo García-Arias, Alejandra García-Gasca, Bruno Gómez-Gil, Jean Pierre González, Irvin González-López, Concepción Grajales-Muñiz, Ricardo Grande, Rosa María Gutiérrez Rios, Jesús Hernández, Alejandra Hernández-Terán, Alfredo Herrera-Estrella, Carla Ivón Herrera-Najera, Pavel Isa, Verónica Mata-Haro, Daniel Lira Morales, Susana Lopez, Antonio Loza Román, Brenda Irasema Maldonado-Meza, Bernardo Martínez-Miguel, José Arturo Martínez-Orozco, Célida Martínez- Rodríguez, Margarita Matías-Florentino, Fidencio Mejía-Nepomuceno, María Guadalupe de Jesús Mireles-Rivera, Gloria María Molina-Salinas, Hector Montoya-Fuentes, , Mario Mújica-Sánchez, José Esteban Muñoz-Medina, José de Jesús Nuñez-Contreras, Alicia Ocaña-Mondragón, Luis Alberto Ochoa-Carrera, Hector Esteban Paz-Juárez, Marissa Perez-Garcia, Francisco Pulido, Helen Haydee Fernanda Ramírez-Plascencia, Jorge Salas-Hernández, Angel Gustavo Salas-Lais, Alejandro Sánchez-Flores, Clara Esperanza Santacruz-Tinoco, María Guadalupe Santiago-Mauricio, Selene Zárate, Nelly Sélem-Mojica, Blanca Taboada, Gloria Vazquez, Joel Armando Vázquez-Pérez. |
| EPI_ISL_2681052 | Unidad de Investigación Biomédica de Zacatecas (UIBZ) | Instituto de Biotecnología de la UNAM | Consorcio Mexicano de Vigilancia Genómica (CoViGen-Mex). Authors (in alphabetical order): Julio Elias Alvarado-Yaah, Carlos F. Arias, Santiago Ávila-Ríos, Eduardo Becerril-Vargas, Víctor Hugo Borja-Aburto, Celia Boukadida, Cristóbal Cháidez-Quiróz, Juan Bautista Chale-Dzul, Ricardo Ciria Merce, Andreu Comas-García, Célida Duque Molina, Julissa Enciso-Ibarra, José Antonio Enciso-Moreno, Gloria Elena Espinosa-Ayala, Fernando Fontove-Herrera, Daniel Fregoso-Rueda, Victor Eduardo García-Arias, Alejandra García-Gasca, Bruno Gómez-Gil, Jean Pierre González, Irvin González-López, Concepción Grajales-Muñiz, Ricardo Grande, Rosa María Gutiérrez Rios, Jesús Hernández, Alejandra Hernández-Terán, Alfredo Herrera-Estrella, Carla Ivón Herrera-Najera, Pavel Isa, Verónica Mata-Haro, Daniel Lira Morales, Susana Lopez, Antonio Loza Román, Brenda Irasema Maldonado-Meza, Bernardo Martínez-Miguel, José Arturo Martínez-Orozco, Célida Martínez- Rodríguez, Margarita Matías-Florentino, Fidencio Mejía-Nepomuceno, María Guadalupe de Jesús Mireles-Rivera, Gloria María Molina-Salinas, Hector Montoya-Fuentes, , Mario Mújica-Sánchez, José Esteban Muñoz-Medina, José de Jesús Nuñez-Contreras, Alicia Ocaña-Mondragón, Luis Alberto Ochoa-Carrera, Hector Esteban Paz-Juárez, Marissa Perez-Garcia, Francisco Pulido, Helen Haydee Fernanda Ramírez-Plascencia, Jorge Salas-Hernández, Angel Gustavo Salas-Lais, Alejandro Sánchez-Flores, Clara Esperanza Santacruz-Tinoco, María Guadalupe Santiago-Mauricio, Selene Zárate, Nelly Sélem-Mojica, Blanca Taboada, Gloria Vazquez, Joel Armando Vázquez-Pérez. |
| EPI_ISL_2681054 | Laboratorio Central de Epidemiología (LCE) | Instituto de Biotecnología de la UNAM | Consorcio Mexicano de Vigilancia Genómica (CoViGen-Mex). Authors (in alphabetical order): Julio Elias Alvarado-Yaah, Carlos F. Arias, Santiago Ávila-Ríos, Eduardo Becerril-Vargas, Víctor Hugo Borja-Aburto, Celia Boukadida, Cristóbal Cháidez-Quiróz, Juan Bautista Chale-Dzul, Ricardo Ciria Merce, Andreu Comas-García, Célida Duque Molina, Julissa Enciso-Ibarra, José Antonio Enciso-Moreno, Gloria Elena Espinosa-Ayala, Fernando Fontove-Herrera, Daniel Fregoso-Rueda, Victor Eduardo García-Arias, Alejandra García-Gasca, Bruno Gómez-Gil, Jean Pierre González, Irvin González-López, Concepción Grajales-Muñiz, Ricardo Grande, Rosa María Gutiérrez Rios, Jesús Hernández, Alejandra Hernández-Terán, Alfredo |

| | | | |
|---|---|---|---|
| | | | Herrera-Estrella, Carla Ivón Herrera-Najera, Pavel Isa, Verónica Mata-Haro, Daniel Lira Morales, Susana Lopez, Antonio Loza Román, Brenda Irasema Maldonado-Meza, Bernardo Martínez-Miguel, José Arturo Martínez-Orozco, Célida Martínez- Rodríguez, Margarita Matías-Florentino, Fidencio Mejía-Nepomuceno, María Guadalupe de Jesús Mireles-Rivera, Gloria María Molina-Salinas, Hector Montoya-Fuentes, , Mario Mújica-Sánchez, José Esteban Muñoz-Medina, José de Jesús Nuñez-Contreras, Alicia Ocaña-Mondragón, Luis Alberto Ochoa-Carrera, Hector Esteban Paz-Juárez, Marissa Perez-Garcia, Francisco Pulido, Helen Haydee Fernanda Ramírez-Plascencia, Jorge Salas-Hernández, Angel Gustavo Salas-Lais, Alejandro Sánchez-Flores, Clara Esperanza Santacruz-Tinoco, María Guadalupe Santiago-Mauricio, Selene Zárate, Nelly Sélem-Mojica, Blanca Taboada, Gloria Vazquez, Joel Armando Vázquez-Pérez. |
| EPI_ISL_2681055 | Unidad de Investigación Médica de Yucatán (UIMY) | Instituto de Biotecnología de la UNAM | Consorcio Mexicano de Vigilancia Genómica (CoViGen-Mex). Authors (in alphabetical order): Julio Elias Alvarado-Yaah, Carlos F. Arias, Santiago Ávila-Ríos, Eduardo Becerril-Vargas, Víctor Hugo Borja-Aburto, Celia Boukadida, Cristóbal Cháidez-Quiróz, Juan Bautista Chale-Dzul, Ricardo Ciria Merce, Andreu Comas-García, Célida Duque Molina, Julissa Enciso-Ibarra, José Antonio Enciso-Moreno, Gloria Elena Espinosa-Ayala, Fernando Fontove-Herrera, Daniel Fregoso-Rueda, Victor Eduardo García-Arias, Alejandra García-Gasca, Bruno Gómez-Gil, Jean Pierre González, Irvin González-López, Concepción Grajales-Muñiz, Ricardo Grande, Rosa María Gutiérrez Rios, Jesús Hernández, Alejandra Hernández-Terán, Alfredo Herrera-Estrella, Carla Ivón Herrera-Najera, Pavel Isa, Verónica Mata-Haro, Daniel Lira Morales, Susana Lopez, Antonio Loza Román, Brenda Irasema Maldonado-Meza, Bernardo Martínez-Miguel, José Arturo Martínez-Orozco, Célida Martínez- Rodríguez, Margarita Matías-Florentino, Fidencio Mejía-Nepomuceno, María Guadalupe de Jesús Mireles-Rivera, Gloria María Molina-Salinas, Hector Montoya-Fuentes, , Mario Mújica-Sánchez, José Esteban Muñoz-Medina, José de Jesús Nuñez-Contreras, Alicia Ocaña-Mondragón, Luis Alberto Ochoa-Carrera, Hector Esteban Paz-Juárez, Marissa Perez-Garcia, Francisco Pulido, Helen Haydee Fernanda Ramírez-Plascencia, Jorge Salas-Hernández, Angel Gustavo Salas-Lais, Alejandro Sánchez-Flores, Clara Esperanza Santacruz-Tinoco, María Guadalupe Santiago-Mauricio, Selene Zárate, Nelly Sélem-Mojica, Blanca Taboada, Gloria Vazquez, Joel Armando Vázquez-Pérez. |
| EPI_ISL_2681056 | Laboratorio Central de Epidemiología (LCE) | Instituto de Biotecnología de la UNAM | Consorcio Mexicano de Vigilancia Genómica (CoViGen-Mex). Authors (in alphabetical order): Julio Elias Alvarado-Yaah, Carlos F. Arias, Santiago Ávila-Ríos, Eduardo Becerril-Vargas, Víctor Hugo Borja-Aburto, Celia Boukadida, Cristóbal Cháidez-Quiróz, Juan Bautista Chale-Dzul, Ricardo Ciria Merce, Andreu Comas-García, Célida Duque Molina, Julissa Enciso-Ibarra, José Antonio Enciso-Moreno, Gloria Elena Espinosa-Ayala, Fernando Fontove-Herrera, Daniel Fregoso-Rueda, Victor Eduardo García-Arias, Alejandra García-Gasca, Bruno Gómez-Gil, Jean Pierre González, Irvin González-López, Concepción Grajales-Muñiz, Ricardo Grande, Rosa María Gutiérrez Rios, Jesús Hernández, Alejandra Hernández-Terán, Alfredo Herrera-Estrella, Carla Ivón Herrera-Najera, Pavel Isa, Verónica Mata-Haro, Daniel Lira Morales, Susana Lopez, Antonio Loza Román, Brenda Irasema Maldonado-Meza, Bernardo Martínez-Miguel, José Arturo Martínez-Orozco, Célida Martínez- Rodríguez, Margarita Matías-Florentino, Fidencio Mejía-Nepomuceno, María Guadalupe de Jesús Mireles-Rivera, Gloria María Molina-Salinas, Hector Montoya-Fuentes, , Mario Mújica-Sánchez, José Esteban Muñoz-Medina, José de Jesús Nuñez-Contreras, Alicia Ocaña-Mondragón, Luis Alberto Ochoa-Carrera, Hector Esteban Paz-Juárez, Marissa Perez-Garcia, Francisco Pulido, Helen Haydee Fernanda Ramírez-Plascencia, Jorge Salas-Hernández, Angel Gustavo Salas-Lais, Alejandro Sánchez-Flores, Clara Esperanza Santacruz-Tinoco, María Guadalupe Santiago-Mauricio, Selene Zárate, Nelly Sélem-Mojica, Blanca Taboada, Gloria Vazquez, Joel Armando Vázquez-Pérez. |
| EPI_ISL_2681057 | Unidad de Investigación Biomédica de Zacatecas (UIBZ) | Instituto de Biotecnología de la UNAM | Consorcio Mexicano de Vigilancia Genómica (CoViGen-Mex). Authors (in alphabetical order): Julio Elias Alvarado-Yaah, Carlos F. Arias, Santiago Ávila-Ríos, Eduardo Becerril-Vargas, Víctor Hugo Borja-Aburto, Celia Boukadida, Cristóbal Cháidez-Quiróz, Juan Bautista Chale-Dzul, Ricardo Ciria Merce, Andreu Comas-García, Célida Duque Molina, Julissa Enciso-Ibarra, José Antonio Enciso-Moreno, Gloria Elena Espinosa-Ayala, Fernando Fontove-Herrera, Daniel Fregoso-Rueda, Victor Eduardo García-Arias, Alejandra García-Gasca, Bruno Gómez-Gil, Jean Pierre González, Irvin González-López, Concepción Grajales-Muñiz, Ricardo Grande, Rosa María Gutiérrez Rios, Jesús Hernández, Alejandra Hernández-Terán, Alfredo Herrera-Estrella, Carla Ivón Herrera-Najera, Pavel Isa, Verónica Mata-Haro, Daniel Lira Morales, Susana Lopez, Antonio Loza Román, Brenda Irasema Maldonado-Meza, Bernardo Martínez-Miguel, José Arturo Martínez-Orozco, Célida Martínez- Rodríguez, Margarita Matías-Florentino, Fidencio Mejía-Nepomuceno, María Guadalupe de Jesús Mireles-Rivera, Gloria María Molina-Salinas, Hector Montoya-Fuentes, , Mario Mújica-Sánchez, José Esteban Muñoz-Medina, José de Jesús Nuñez-Contreras, Alicia Ocaña-Mondragón, Luis Alberto Ochoa-Carrera, Hector Esteban Paz-Juárez, Marissa Perez-Garcia, Francisco Pulido, Helen Haydee Fernanda Ramírez-Plascencia, Jorge Salas-Hernández, Angel Gustavo Salas-Lais, Alejandro Sánchez-Flores, Clara Esperanza Santacruz-Tinoco, María Guadalupe Santiago-Mauricio, Selene Zárate, Nelly Sélem-Mojica, Blanca Taboada, Gloria Vazquez, Joel Armando Vázquez-Pérez. |
| EPI_ISL_2681058, EPI_ISL_2681059, EPI_ISL_2681060, EPI_ISL_2681062, EPI_ISL_2681064, EPI_ISL_2681065, EPI_ISL_2681066 | Laboratorio Central de Epidemiología (LCE) | Instituto de Biotecnología de la UNAM | Consorcio Mexicano de Vigilancia Genómica (CoViGen-Mex). Authors (in alphabetical order): Julio Elias Alvarado-Yaah, Carlos F. Arias, Santiago Ávila-Ríos, Eduardo Becerril-Vargas, Víctor Hugo Borja-Aburto, Celia Boukadida, Cristóbal Cháidez-Quiróz, Juan Bautista Chale-Dzul, Ricardo Ciria Merce, Andreu Comas-García, Célida Duque Molina, Julissa Enciso-Ibarra, José Antonio Enciso-Moreno, Gloria Elena Espinosa-Ayala, Fernando Fontove-Herrera, Daniel Fregoso-Rueda, Victor Eduardo García-Arias, Alejandra García-Gasca, Bruno Gómez-Gil, Jean Pierre González, Irvin González-López, Concepción Grajales-Muñiz, Ricardo Grande, Rosa María Gutiérrez Rios, Jesús Hernández, Alejandra Hernández-Terán, Alfredo Herrera-Estrella, Carla Ivón Herrera-Najera, Pavel Isa, Verónica Mata-Haro, Daniel Lira Morales, Susana Lopez, Antonio Loza Román, Brenda Irasema Maldonado-Meza, Bernardo Martínez-Miguel, José Arturo Martínez-Orozco, Célida Martínez- Rodríguez, Margarita Matías-Florentino, Fidencio Mejía-Nepomuceno, María Guadalupe de Jesús Mireles-Rivera, Gloria María Molina-Salinas, Hector Montoya-Fuentes, , Mario Mújica-Sánchez, José Esteban Muñoz-Medina, José de Jesús Nuñez-Contreras, Alicia Ocaña-Mondragón, Luis Alberto Ochoa-Carrera, Hector Esteban Paz-Juárez, Marissa Perez-Garcia, Francisco Pulido, Helen Haydee Fernanda Ramírez-Plascencia, Jorge Salas-Hernández, Angel Gustavo Salas-Lais, Alejandro Sánchez-Flores, Clara Esperanza Santacruz-Tinoco, María Guadalupe Santiago-Mauricio, Selene Zárate, Nelly Sélem-Mojica, Blanca Taboada, Gloria Vazquez, Joel Armando Vázquez-Pérez. |
| EPI_ISL_2681067 | Unidad de Investigación Médica de Yucatán (UIMY) | Instituto de Biotecnología de la UNAM | Consorcio Mexicano de Vigilancia Genómica (CoViGen-Mex). Authors (in alphabetical order): Julio Elias Alvarado-Yaah, Carlos F. Arias, Santiago Ávila-Ríos, Eduardo Becerril-Vargas, Víctor Hugo Borja-Aburto, Celia Boukadida, Cristóbal Cháidez-Quiróz, Juan Bautista Chale-Dzul, Ricardo Ciria Merce, Andreu Comas-García, Célida Duque Molina, Julissa Enciso-Ibarra, José Antonio Enciso-Moreno, Gloria Elena Espinosa-Ayala, Fernando Fontove-Herrera, Daniel Fregoso-Rueda, Victor Eduardo García-Arias, Alejandra García-Gasca, Bruno Gómez-Gil, Jean Pierre González, Irvin González-López, Concepción Grajales-Muñiz, Ricardo Grande, Rosa María Gutiérrez Rios, Jesús Hernández, Alejandra Hernández-Terán, Alfredo Herrera-Estrella, Carla Ivón Herrera-Najera, Pavel Isa, Verónica Mata-Haro, Daniel Lira Morales, Susana Lopez, Antonio Loza Román, Brenda Irasema Maldonado-Meza, Bernardo Martínez-Miguel, José Arturo Martínez-Orozco, Célida Martínez- Rodríguez, Margarita Matías-Florentino, Fidencio Mejía-Nepomuceno, María Guadalupe de Jesús Mireles-Rivera, Gloria María Molina-Salinas, Hector Montoya-Fuentes, , Mario Mújica-Sánchez, José Esteban Muñoz-Medina, José de Jesús Nuñez-Contreras, Alicia Ocaña-Mondragón, Luis Alberto Ochoa-Carrera, Hector Esteban Paz-Juárez, Marissa Perez-Garcia, Francisco Pulido, Helen Haydee Fernanda Ramírez-Plascencia, Jorge Salas-Hernández, Angel Gustavo Salas-Lais, Alejandro Sánchez-Flores, Clara Esperanza Santacruz-Tinoco, María Guadalupe Santiago-Mauricio, Selene Zárate, Nelly Sélem-Mojica, Blanca Taboada, Gloria Vazquez, Joel Armando Vázquez-Pérez. |
| EPI_ISL_2681068 | Unidad de Investigación Biomédica de Zacatecas (UIBZ) | Instituto de Biotecnología de la UNAM | Consorcio Mexicano de Vigilancia Genómica (CoViGen-Mex). Authors (in alphabetical order): Julio Elias Alvarado-Yaah, Carlos F. Arias, Santiago Ávila-Ríos, Eduardo Becerril-Vargas, Víctor Hugo Borja-Aburto, Celia Boukadida, Cristóbal Cháidez-Quiróz, Juan Bautista Chale-Dzul, Ricardo Ciria Merce, Andreu Comas-García, Célida Duque Molina, Julissa Enciso-Ibarra, José Antonio Enciso-Moreno, Gloria Elena Espinosa-Ayala, Fernando Fontove-Herrera, Daniel Fregoso-Rueda, Victor Eduardo García-Arias, Alejandra García-Gasca, Bruno Gómez-Gil, Jean Pierre González, Irvin González-López, Concepción Grajales-Muñiz, Ricardo Grande, Rosa María Gutiérrez Rios, Jesús Hernández, Alejandra Hernández-Terán, Alfredo Herrera-Estrella, Carla Ivón Herrera-Najera, Pavel Isa, Verónica Mata-Haro, Daniel Lira Morales, Susana Lopez, Antonio Loza Román, Brenda Irasema Maldonado-Meza, Bernardo Martínez-Miguel, José Arturo Martínez-Orozco, Célida Martínez- Rodríguez, Margarita Matías-Florentino, Fidencio Mejía-Nepomuceno, María Guadalupe de Jesús Mireles-Rivera, Gloria María Molina-Salinas, Hector Montoya-Fuentes, , Mario Mújica-Sánchez, José Esteban Muñoz-Medina, José de Jesús Nuñez-Contreras, Alicia Ocaña-Mondragón, Luis Alberto Ochoa-Carrera, Hector Esteban Paz-Juárez, Marissa Perez-Garcia, Francisco Pulido, Helen Haydee Fernanda Ramírez-Plascencia, Jorge Salas-Hernández, Angel Gustavo Salas-Lais, Alejandro Sánchez-Flores, Clara Esperanza Santacruz-Tinoco, María Guadalupe Santiago-Mauricio, Selene Zárate, Nelly Sélem-Mojica, Blanca Taboada, Gloria Vazquez, Joel Armando Vázquez-Pérez. |
| EPI_ISL_2681069, EPI_ISL_2681070, EPI_ISL_2681071, EPI_ISL_2681072, EPI_ISL_2681073 | Laboratorio Central de Epidemiología (LCE) | Instituto de Biotecnología de la UNAM | Consorcio Mexicano de Vigilancia Genómica (CoViGen-Mex). Authors (in alphabetical order): Julio Elias Alvarado-Yaah, Carlos F. Arias, Santiago Ávila-Ríos, Eduardo Becerril-Vargas, Víctor Hugo Borja-Aburto, Celia Boukadida, Cristóbal Cháidez-Quiróz, Juan Bautista Chale-Dzul, Ricardo Ciria Merce, Andreu Comas-García, Célida Duque Molina, Julissa Enciso-Ibarra, José Antonio Enciso-Moreno, Gloria Elena Espinosa-Ayala, Fernando Fontove-Herrera, Daniel Fregoso-Rueda, Victor Eduardo García-Arias, Alejandra García-Gasca, Bruno Gómez-Gil, Jean Pierre González, Irvin González-López, Concepción Grajales-Muñiz, Ricardo Grande, Rosa María Gutiérrez Rios, Jesús Hernández, Alejandra Hernández-Terán, Alfredo |

| | | | |
|---|---|---|---|
| | | | Herrera-Estrella, Carla Ivón Herrera-Najera, Pavel Isa, Verónica Mata-Haro, Daniel Lira Morales, Susana Lopez, Antonio Loza Román, Brenda Irasema Maldonado-Meza, Bernardo Martínez-Miguel, José Arturo Martínez-Orozco, Célida Martínez- Rodríguez, Margarita Matías-Florentino, Fidencio Mejía-Nepomuceno, María Guadalupe de Jesús Mireles-Rivera, Gloria María Molina-Salinas, Hector Montoya-Fuentes, , Mario Mújica-Sánchez, José Esteban Muñoz-Medina, José de Jesús Nuñez-Contreras, Alicia Ocaña-Mondragón, Luis Alberto Ochoa-Carrera, Hector Esteban Paz-Juárez, Marissa Perez-Garcia, Francisco Pulido, Helen Haydee Fernanda Ramírez-Plascencia, Jorge Salas-Hernández, Angel Gustavo Salas-Lais, Alejandro Sánchez-Flores, Clara Esperanza Santacruz-Tinoco, María Guadalupe Santiago-Mauricio, Selene Zárate, Nelly Sélem-Mojica, Blanca Taboada, Gloria Vazquez, Joel Armando Vázquez-Pérez. |
| EPI_ISL_2681074, EPI_ISL_2681075 | Unidad de Investigación Médica de Yucatán (UIMY) | Instituto de Biotecnología de la UNAM | Consorcio Mexicano de Vigilancia Genómica (CoViGen-Mex). Authors (in alphabetical order): Julio Elias Alvarado-Yaah, Carlos F. Arias, Santiago Ávila-Ríos, Eduardo Becerril-Vargas, Víctor Hugo Borja-Aburto, Celia Boukadida, Cristóbal Cháidez-Quiróz, Juan Bautista Chale-Dzul, Ricardo Ciria Merce, Andreu Comas-García, Célida Duque Molina, Julissa Enciso-Ibarra, José Antonio Enciso-Moreno, Gloria Elena Espinosa-Ayala, Fernando Fontove-Herrera, Daniel Fregoso-Rueda, Victor Eduardo García-Arias, Alejandra García-Gasca, Bruno Gómez-Gil, Jean Pierre González, Irvin González-López, Concepción Grajales-Muñiz, Ricardo Grande, Rosa María Gutiérrez Rios, Jesús Hernández, Alejandra Hernández-Terán, Alfredo Herrera-Estrella, Carla Ivón Herrera-Najera, Pavel Isa, Verónica Mata-Haro, Daniel Lira Morales, Susana Lopez, Antonio Loza Román, Brenda Irasema Maldonado-Meza, Bernardo Martínez-Miguel, José Arturo Martínez-Orozco, Célida Martínez- Rodríguez, Margarita Matías-Florentino, Fidencio Mejía-Nepomuceno, María Guadalupe de Jesús Mireles-Rivera, Gloria María Molina-Salinas, Hector Montoya-Fuentes, , Mario Mújica-Sánchez, José Esteban Muñoz-Medina, José de Jesús Nuñez-Contreras, Alicia Ocaña-Mondragón, Luis Alberto Ochoa-Carrera, Hector Esteban Paz-Juárez, Marissa Perez-Garcia, Francisco Pulido, Helen Haydee Fernanda Ramírez-Plascencia, Jorge Salas-Hernández, Angel Gustavo Salas-Lais, Alejandro Sánchez-Flores, Clara Esperanza Santacruz-Tinoco, María Guadalupe Santiago-Mauricio, Selene Zárate, Nelly Sélem-Mojica, Blanca Taboada, Gloria Vazquez, Joel Armando Vázquez-Pérez. |
| EPI_ISL_2681076 | Laboratorio Central de Epidemiología (LCE) | Instituto de Biotecnología de la UNAM | Consorcio Mexicano de Vigilancia Genómica (CoViGen-Mex). Authors (in alphabetical order): Julio Elias Alvarado-Yaah, Carlos F. Arias, Santiago Ávila-Ríos, Eduardo Becerril-Vargas, Víctor Hugo Borja-Aburto, Celia Boukadida, Cristóbal Cháidez-Quiróz, Juan Bautista Chale-Dzul, Ricardo Ciria Merce, Andreu Comas-García, Célida Duque Molina, Julissa Enciso-Ibarra, José Antonio Enciso-Moreno, Gloria Elena Espinosa-Ayala, Fernando Fontove-Herrera, Daniel Fregoso-Rueda, Victor Eduardo García-Arias, Alejandra García-Gasca, Bruno Gómez-Gil, Jean Pierre González, Irvin González-López, Concepción Grajales-Muñiz, Ricardo Grande, Rosa María Gutiérrez Rios, Jesús Hernández, Alejandra Hernández-Terán, Alfredo Herrera-Estrella, Carla Ivón Herrera-Najera, Pavel Isa, Verónica Mata-Haro, Daniel Lira Morales, Susana Lopez, Antonio Loza Román, Brenda Irasema Maldonado-Meza, Bernardo Martínez-Miguel, José Arturo Martínez-Orozco, Célida Martínez- Rodríguez, Margarita Matías-Florentino, Fidencio Mejía-Nepomuceno, María Guadalupe de Jesús Mireles-Rivera, Gloria María Molina-Salinas, Hector Montoya-Fuentes, , Mario Mújica-Sánchez, José Esteban Muñoz-Medina, José de Jesús Nuñez-Contreras, Alicia Ocaña-Mondragón, Luis Alberto Ochoa-Carrera, Hector Esteban Paz-Juárez, Marissa Perez-Garcia, Francisco Pulido, Helen Haydee Fernanda Ramírez-Plascencia, Jorge Salas-Hernández, Angel Gustavo Salas-Lais, Alejandro Sánchez-Flores, Clara Esperanza Santacruz-Tinoco, María Guadalupe Santiago-Mauricio, Selene Zárate, Nelly Sélem-Mojica, Blanca Taboada, Gloria Vazquez, Joel Armando Vázquez-Pérez. |
| EPI_ISL_2681077, EPI_ISL_2681078, EPI_ISL_2681079, EPI_ISL_2681080, EPI_ISL_2681082, EPI_ISL_2681083, EPI_ISL_2681084, EPI_ISL_2681085 | Unidad de Investigación Médica de Yucatán (UIMY) | Instituto de Biotecnología de la UNAM | Consorcio Mexicano de Vigilancia Genómica (CoViGen-Mex). Authors (in alphabetical order): Julio Elias Alvarado-Yaah, Carlos F. Arias, Santiago Ávila-Ríos, Eduardo Becerril-Vargas, Víctor Hugo Borja-Aburto, Celia Boukadida, Cristóbal Cháidez-Quiróz, Juan Bautista Chale-Dzul, Ricardo Ciria Merce, Andreu Comas-García, Célida Duque Molina, Julissa Enciso-Ibarra, José Antonio Enciso-Moreno, Gloria Elena Espinosa-Ayala, Fernando Fontove-Herrera, Daniel Fregoso-Rueda, Victor Eduardo García-Arias, Alejandra García-Gasca, Bruno Gómez-Gil, Jean Pierre González, Irvin González-López, Concepción Grajales-Muñiz, Ricardo Grande, Rosa María Gutiérrez Rios, Jesús Hernández, Alejandra Hernández-Terán, Alfredo Herrera-Estrella, Carla Ivón Herrera-Najera, Pavel Isa, Verónica Mata-Haro, Daniel Lira Morales, Susana Lopez, Antonio Loza Román, Brenda Irasema Maldonado-Meza, Bernardo Martínez-Miguel, José Arturo Martínez-Orozco, Célida Martínez- Rodríguez, Margarita Matías-Florentino, Fidencio Mejía-Nepomuceno, María Guadalupe de Jesús Mireles-Rivera, Gloria María Molina-Salinas, Hector Montoya-Fuentes, , Mario Mújica-Sánchez, José Esteban Muñoz-Medina, José de Jesús Nuñez-Contreras, Alicia Ocaña-Mondragón, Luis Alberto Ochoa-Carrera, Hector Esteban Paz-Juárez, Marissa Perez-Garcia, Francisco Pulido, Helen Haydee Fernanda Ramírez-Plascencia, Jorge Salas-Hernández, Angel Gustavo Salas-Lais, Alejandro Sánchez-Flores, Clara Esperanza Santacruz-Tinoco, María Guadalupe Santiago-Mauricio, Selene Zárate, Nelly Sélem-Mojica, Blanca Taboada, Gloria Vazquez, Joel Armando Vázquez-Pérez. |
| EPI_ISL_2681086 | Centro de Investigación Biomédica del Noreste (CIBIN) | Instituto de Biotecnología de la UNAM | Consorcio Mexicano de Vigilancia Genómica (CoViGen-Mex). Authors (in alphabetical order): Julio Elias Alvarado-Yaah, Carlos F. Arias, Santiago Ávila-Ríos, Eduardo Becerril-Vargas, Víctor Hugo Borja-Aburto, Celia Boukadida, Cristóbal Cháidez-Quiróz, Juan Bautista Chale-Dzul, Ricardo Ciria Merce, Andreu Comas-García, Célida Duque Molina, Julissa Enciso-Ibarra, José Antonio Enciso-Moreno, Gloria Elena Espinosa-Ayala, Fernando Fontove-Herrera, Daniel Fregoso-Rueda, Victor Eduardo García-Arias, Alejandra García-Gasca, Bruno Gómez-Gil, Jean Pierre González, Irvin González-López, Concepción Grajales-Muñiz, Ricardo Grande, Rosa María Gutiérrez Rios, Jesús Hernández, Alejandra Hernández-Terán, Alfredo Herrera-Estrella, Carla Ivón Herrera-Najera, Pavel Isa, Verónica Mata-Haro, Daniel Lira Morales, Susana Lopez, Antonio Loza Román, Brenda Irasema Maldonado-Meza, Bernardo Martínez-Miguel, José Arturo Martínez-Orozco, Célida Martínez- Rodríguez, Margarita Matías-Florentino, Fidencio Mejía-Nepomuceno, María Guadalupe de Jesús Mireles-Rivera, Gloria María Molina-Salinas, Hector Montoya-Fuentes, , Mario Mújica-Sánchez, José Esteban Muñoz-Medina, José de Jesús Nuñez-Contreras, Alicia Ocaña-Mondragón, Luis Alberto Ochoa-Carrera, Hector Esteban Paz-Juárez, Marissa Perez-Garcia, Francisco Pulido, Helen Haydee Fernanda Ramírez-Plascencia, Jorge Salas-Hernández, Angel Gustavo Salas-Lais, Alejandro Sánchez-Flores, Clara Esperanza Santacruz-Tinoco, María Guadalupe Santiago-Mauricio, Selene Zárate, Nelly Sélem-Mojica, Blanca Taboada, Gloria Vazquez, Joel Armando Vázquez-Pérez. |
| EPI_ISL_2681087 | Laboratorio Central de Epidemiología (LCE) | Instituto de Biotecnología de la UNAM | Consorcio Mexicano de Vigilancia Genómica (CoViGen-Mex). Authors (in alphabetical order): Julio Elias Alvarado-Yaah, Carlos F. Arias, Santiago Ávila-Ríos, Eduardo Becerril-Vargas, Víctor Hugo Borja-Aburto, Celia Boukadida, Cristóbal Cháidez-Quiróz, Juan Bautista Chale-Dzul, Ricardo Ciria Merce, Andreu Comas-García, Célida Duque Molina, Julissa Enciso-Ibarra, José Antonio Enciso-Moreno, Gloria Elena Espinosa-Ayala, Fernando Fontove-Herrera, Daniel Fregoso-Rueda, Victor Eduardo García-Arias, Alejandra García-Gasca, Bruno Gómez-Gil, Jean Pierre González, Irvin González-López, Concepción Grajales-Muñiz, Ricardo Grande, Rosa María Gutiérrez Rios, Jesús Hernández, Alejandra Hernández-Terán, Alfredo Herrera-Estrella, Carla Ivón Herrera-Najera, Pavel Isa, Verónica Mata-Haro, Daniel Lira Morales, Susana Lopez, Antonio Loza Román, Brenda Irasema Maldonado-Meza, Bernardo Martínez-Miguel, José Arturo Martínez-Orozco, Célida Martínez- Rodríguez, Margarita Matías-Florentino, Fidencio Mejía-Nepomuceno, María Guadalupe de Jesús Mireles-Rivera, Gloria María Molina-Salinas, Hector Montoya-Fuentes, , Mario Mújica-Sánchez, José Esteban Muñoz-Medina, José de Jesús Nuñez-Contreras, Alicia Ocaña-Mondragón, Luis Alberto Ochoa-Carrera, Hector Esteban Paz-Juárez, Marissa Perez-Garcia, Francisco Pulido, Helen Haydee Fernanda Ramírez-Plascencia, Jorge Salas-Hernández, Angel Gustavo Salas-Lais, Alejandro Sánchez-Flores, Clara Esperanza Santacruz-Tinoco, María Guadalupe Santiago-Mauricio, Selene Zárate, Nelly Sélem-Mojica, Blanca Taboada, Gloria Vazquez, Joel Armando Vázquez-Pérez. |
| EPI_ISL_2681088, EPI_ISL_2681089, EPI_ISL_2681090, EPI_ISL_2681091 | Unidad de Investigación Médica de Yucatán (UIMY) | Instituto de Biotecnología de la UNAM | Consorcio Mexicano de Vigilancia Genómica (CoViGen-Mex). Authors (in alphabetical order): Julio Elias Alvarado-Yaah, Carlos F. Arias, Santiago Ávila-Ríos, Eduardo Becerril-Vargas, Víctor Hugo Borja-Aburto, Celia Boukadida, Cristóbal Cháidez-Quiróz, Juan Bautista Chale-Dzul, Ricardo Ciria Merce, Andreu Comas-García, Célida Duque Molina, Julissa Enciso-Ibarra, José Antonio Enciso-Moreno, Gloria Elena Espinosa-Ayala, Fernando Fontove-Herrera, Daniel Fregoso-Rueda, Victor Eduardo García-Arias, Alejandra García-Gasca, Bruno Gómez-Gil, Jean Pierre González, Irvin González-López, Concepción Grajales-Muñiz, Ricardo Grande, Rosa María Gutiérrez Rios, Jesús Hernández, Alejandra Hernández-Terán, Alfredo Herrera-Estrella, Carla Ivón Herrera-Najera, Pavel Isa, Verónica Mata-Haro, Daniel Lira Morales, Susana Lopez, Antonio Loza Román, Brenda Irasema Maldonado-Meza, Bernardo Martínez-Miguel, José Arturo Martínez-Orozco, Célida Martínez- Rodríguez, Margarita Matías-Florentino, Fidencio Mejía-Nepomuceno, María Guadalupe de Jesús Mireles-Rivera, Gloria María Molina-Salinas, Hector Montoya-Fuentes, , Mario Mújica-Sánchez, José Esteban Muñoz-Medina, José de Jesús Nuñez-Contreras, Alicia Ocaña-Mondragón, Luis Alberto Ochoa-Carrera, Hector Esteban Paz-Juárez, Marissa Perez-Garcia, Francisco Pulido, Helen Haydee Fernanda Ramírez-Plascencia, Jorge Salas-Hernández, Angel Gustavo Salas-Lais, Alejandro Sánchez-Flores, Clara Esperanza Santacruz-Tinoco, María Guadalupe Santiago-Mauricio, Selene Zárate, Nelly Sélem-Mojica, Blanca Taboada, Gloria Vazquez, Joel Armando Vázquez-Pérez. |
| EPI_ISL_2681092 | Unidad de Investigación Biomédica de Zacatecas (UIBZ) | Instituto de Biotecnología de la UNAM | Consorcio Mexicano de Vigilancia Genómica (CoViGen-Mex). Authors (in alphabetical order): Julio Elias Alvarado-Yaah, Carlos F. Arias, Santiago Ávila-Ríos, Eduardo Becerril-Vargas, Víctor Hugo Borja-Aburto, Celia Boukadida, Cristóbal Cháidez-Quiróz, Juan Bautista Chale-Dzul, Ricardo Ciria Merce, Andreu Comas-García, Célida Duque Molina, Julissa Enciso-Ibarra, José Antonio Enciso-Moreno, Gloria Elena Espinosa-Ayala, Fernando Fontove-Herrera, Daniel Fregoso-Rueda, Victor Eduardo García-Arias, Alejandra García-Gasca, Bruno Gómez-Gil, Jean Pierre González, Irvin González-López, Concepción Grajales-Muñiz, Ricardo Grande, Rosa María Gutiérrez Rios, Jesús Hernández, Alejandra Hernández-Terán, Alfredo |

| | | | |
|---|---|---|---|
| | | | Herrera-Estrella, Carla Ivón Herrera-Najera, Pavel Isa, Verónica Mata-Haro, Daniel Lira Morales, Susana Lopez, Antonio Loza Román, Brenda Irasema Maldonado-Meza, Bernardo Martínez-Miguel, José Arturo Martínez-Orozco, Célida Martínez- Rodríguez, Margarita Matías-Florentino, Fidencio Mejía-Nepomuceno, María Guadalupe de Jesús Mireles-Rivera, Gloria María Molina-Salinas, Hector Montoya-Fuentes, , Mario Mújica-Sánchez, José Esteban Muñoz-Medina, José de Jesús Nuñez-Contreras, Alicia Ocaña-Mondragón, Luis Alberto Ochoa-Carrera, Hector Esteban Paz-Juárez, Marissa Perez-Garcia, Francisco Pulido, Helen Haydee Fernanda Ramírez-Plascencia, Jorge Salas-Hernández, Angel Gustavo Salas-Lais, Alejandro Sánchez-Flores, Clara Esperanza Santacruz-Tinoco, María Guadalupe Santiago-Mauricio, Selene Zárate, Nelly Sélem-Mojica, Blanca Taboada, Gloria Vazquez, Joel Armando Vázquez-Pérez. |
| EPI_ISL_2681093 | Laboratorio Central de Epidemiología (LCE) | Instituto de Biotecnología de la UNAM | Consorcio Mexicano de Vigilancia Genómica (CoViGen-Mex). Authors (in alphabetical order): Julio Elias Alvarado-Yaah, Carlos F. Arias, Santiago Ávila-Ríos, Eduardo Becerril-Vargas, Víctor Hugo Borja-Aburto, Celia Boukadida, Cristóbal Cháidez-Quiróz, Juan Bautista Chale-Dzul, Ricardo Ciria Merce, Andreu Comas-García, Célida Duque Molina, Julissa Enciso-Ibarra, José Antonio Enciso-Moreno, Gloria Elena Espinosa-Ayala, Fernando Fontove-Herrera, Daniel Fregoso-Rueda, Victor Eduardo García-Arias, Alejandra García-Gasca, Bruno Gómez-Gil, Jean Pierre González, Irvin González-López, Concepción Grajales-Muñiz, Ricardo Grande, Rosa María Gutiérrez Rios, Jesús Hernández, Alejandra Hernández-Terán, Alfredo Herrera-Estrella, Carla Ivón Herrera-Najera, Pavel Isa, Verónica Mata-Haro, Daniel Lira Morales, Susana Lopez, Antonio Loza Román, Brenda Irasema Maldonado-Meza, Bernardo Martínez-Miguel, José Arturo Martínez-Orozco, Célida Martínez- Rodríguez, Margarita Matías-Florentino, Fidencio Mejía-Nepomuceno, María Guadalupe de Jesús Mireles-Rivera, Gloria María Molina-Salinas, Hector Montoya-Fuentes, , Mario Mújica-Sánchez, José Esteban Muñoz-Medina, José de Jesús Nuñez-Contreras, Alicia Ocaña-Mondragón, Luis Alberto Ochoa-Carrera, Hector Esteban Paz-Juárez, Marissa Perez-Garcia, Francisco Pulido, Helen Haydee Fernanda Ramírez-Plascencia, Jorge Salas-Hernández, Angel Gustavo Salas-Lais, Alejandro Sánchez-Flores, Clara Esperanza Santacruz-Tinoco, María Guadalupe Santiago-Mauricio, Selene Zárate, Nelly Sélem-Mojica, Blanca Taboada, Gloria Vazquez, Joel Armando Vázquez-Pérez. |
| EPI_ISL_2681094 | Unidad de Investigación Médica de Yucatán (UIMY) | Instituto de Biotecnología de la UNAM | Consorcio Mexicano de Vigilancia Genómica (CoViGen-Mex). Authors (in alphabetical order): Julio Elias Alvarado-Yaah, Carlos F. Arias, Santiago Ávila-Ríos, Eduardo Becerril-Vargas, Víctor Hugo Borja-Aburto, Celia Boukadida, Cristóbal Cháidez-Quiróz, Juan Bautista Chale-Dzul, Ricardo Ciria Merce, Andreu Comas-García, Célida Duque Molina, Julissa Enciso-Ibarra, José Antonio Enciso-Moreno, Gloria Elena Espinosa-Ayala, Fernando Fontove-Herrera, Daniel Fregoso-Rueda, Victor Eduardo García-Arias, Alejandra García-Gasca, Bruno Gómez-Gil, Jean Pierre González, Irvin González-López, Concepción Grajales-Muñiz, Ricardo Grande, Rosa María Gutiérrez Rios, Jesús Hernández, Alejandra Hernández-Terán, Alfredo Herrera-Estrella, Carla Ivón Herrera-Najera, Pavel Isa, Verónica Mata-Haro, Daniel Lira Morales, Susana Lopez, Antonio Loza Román, Brenda Irasema Maldonado-Meza, Bernardo Martínez-Miguel, José Arturo Martínez-Orozco, Célida Martínez- Rodríguez, Margarita Matías-Florentino, Fidencio Mejía-Nepomuceno, María Guadalupe de Jesús Mireles-Rivera, Gloria María Molina-Salinas, Hector Montoya-Fuentes, , Mario Mújica-Sánchez, José Esteban Muñoz-Medina, José de Jesús Nuñez-Contreras, Alicia Ocaña-Mondragón, Luis Alberto Ochoa-Carrera, Hector Esteban Paz-Juárez, Marissa Perez-Garcia, Francisco Pulido, Helen Haydee Fernanda Ramírez-Plascencia, Jorge Salas-Hernández, Angel Gustavo Salas-Lais, Alejandro Sánchez-Flores, Clara Esperanza Santacruz-Tinoco, María Guadalupe Santiago-Mauricio, Selene Zárate, Nelly Sélem-Mojica, Blanca Taboada, Gloria Vazquez, Joel Armando Vázquez-Pérez. |
| EPI_ISL_2681095, EPI_ISL_2681096 | Centro de Investigación Biomédica del Noreste (CIBIN) | Instituto de Biotecnología de la UNAM | Consorcio Mexicano de Vigilancia Genómica (CoViGen-Mex). Authors (in alphabetical order): Julio Elias Alvarado-Yaah, Carlos F. Arias, Santiago Ávila-Ríos, Eduardo Becerril-Vargas, Víctor Hugo Borja-Aburto, Celia Boukadida, Cristóbal Cháidez-Quiróz, Juan Bautista Chale-Dzul, Ricardo Ciria Merce, Andreu Comas-García, Célida Duque Molina, Julissa Enciso-Ibarra, José Antonio Enciso-Moreno, Gloria Elena Espinosa-Ayala, Fernando Fontove-Herrera, Daniel Fregoso-Rueda, Victor Eduardo García-Arias, Alejandra García-Gasca, Bruno Gómez-Gil, Jean Pierre González, Irvin González-López, Concepción Grajales-Muñiz, Ricardo Grande, Rosa María Gutiérrez Rios, Jesús Hernández, Alejandra Hernández-Terán, Alfredo Herrera-Estrella, Carla Ivón Herrera-Najera, Pavel Isa, Verónica Mata-Haro, Daniel Lira Morales, Susana Lopez, Antonio Loza Román, Brenda Irasema Maldonado-Meza, Bernardo Martínez-Miguel, José Arturo Martínez-Orozco, Célida Martínez- Rodríguez, Margarita Matías-Florentino, Fidencio Mejía-Nepomuceno, María Guadalupe de Jesús Mireles-Rivera, Gloria María Molina-Salinas, Hector Montoya-Fuentes, , Mario Mújica-Sánchez, José Esteban Muñoz-Medina, José de Jesús Nuñez-Contreras, Alicia Ocaña-Mondragón, Luis Alberto Ochoa-Carrera, Hector Esteban Paz-Juárez, Marissa Perez-Garcia, Francisco Pulido, Helen Haydee Fernanda Ramírez-Plascencia, Jorge Salas-Hernández, Angel Gustavo Salas-Lais, Alejandro Sánchez-Flores, Clara Esperanza Santacruz-Tinoco, María Guadalupe Santiago-Mauricio, Selene Zárate, Nelly Sélem-Mojica, Blanca Taboada, Gloria Vazquez, Joel Armando Vázquez-Pérez. |
| EPI_ISL_2681098, EPI_ISL_2681099, EPI_ISL_2681100, EPI_ISL_2681101 | Unidad de Investigación Médica de Yucatán (UIMY) | Instituto de Biotecnología de la UNAM | Consorcio Mexicano de Vigilancia Genómica (CoViGen-Mex). Authors (in alphabetical order): Julio Elias Alvarado-Yaah, Carlos F. Arias, Santiago Ávila-Ríos, Eduardo Becerril-Vargas, Víctor Hugo Borja-Aburto, Celia Boukadida, Cristóbal Cháidez-Quiróz, Juan Bautista Chale-Dzul, Ricardo Ciria Merce, Andreu Comas-García, Célida Duque Molina, Julissa Enciso-Ibarra, José Antonio Enciso-Moreno, Gloria Elena Espinosa-Ayala, Fernando Fontove-Herrera, Daniel Fregoso-Rueda, Victor Eduardo García-Arias, Alejandra García-Gasca, Bruno Gómez-Gil, Jean Pierre González, Irvin González-López, Concepción Grajales-Muñiz, Ricardo Grande, Rosa María Gutiérrez Rios, Jesús Hernández, Alejandra Hernández-Terán, Alfredo Herrera-Estrella, Carla Ivón Herrera-Najera, Pavel Isa, Verónica Mata-Haro, Daniel Lira Morales, Susana Lopez, Antonio Loza Román, Brenda Irasema Maldonado-Meza, Bernardo Martínez-Miguel, José Arturo Martínez-Orozco, Célida Martínez- Rodríguez, Margarita Matías-Florentino, Fidencio Mejía-Nepomuceno, María Guadalupe de Jesús Mireles-Rivera, Gloria María Molina-Salinas, Hector Montoya-Fuentes, , Mario Mújica-Sánchez, José Esteban Muñoz-Medina, José de Jesús Nuñez-Contreras, Alicia Ocaña-Mondragón, Luis Alberto Ochoa-Carrera, Hector Esteban Paz-Juárez, Marissa Perez-Garcia, Francisco Pulido, Helen Haydee Fernanda Ramírez-Plascencia, Jorge Salas-Hernández, Angel Gustavo Salas-Lais, Alejandro Sánchez-Flores, Clara Esperanza Santacruz-Tinoco, María Guadalupe Santiago-Mauricio, Selene Zárate, Nelly Sélem-Mojica, Blanca Taboada, Gloria Vazquez, Joel Armando Vázquez-Pérez. |
| EPI_ISL_2681102, EPI_ISL_2681103, EPI_ISL_2681104 | Centro de Investigación Biomédica del Noreste (CIBIN) | Instituto de Biotecnología de la UNAM | Consorcio Mexicano de Vigilancia Genómica (CoViGen-Mex). Authors (in alphabetical order): Julio Elias Alvarado-Yaah, Carlos F. Arias, Santiago Ávila-Ríos, Eduardo Becerril-Vargas, Víctor Hugo Borja-Aburto, Celia Boukadida, Cristóbal Cháidez-Quiróz, Juan Bautista Chale-Dzul, Ricardo Ciria Merce, Andreu Comas-García, Célida Duque Molina, Julissa Enciso-Ibarra, José Antonio Enciso-Moreno, Gloria Elena Espinosa-Ayala, Fernando Fontove-Herrera, Daniel Fregoso-Rueda, Victor Eduardo García-Arias, Alejandra García-Gasca, Bruno Gómez-Gil, Jean Pierre González, Irvin González-López, Concepción Grajales-Muñiz, Ricardo Grande, Rosa María Gutiérrez Rios, Jesús Hernández, Alejandra Hernández-Terán, Alfredo Herrera-Estrella, Carla Ivón Herrera-Najera, Pavel Isa, Verónica Mata-Haro, Daniel Lira Morales, Susana Lopez, Antonio Loza Román, Brenda Irasema Maldonado-Meza, Bernardo Martínez-Miguel, José Arturo Martínez-Orozco, Célida Martínez- Rodríguez, Margarita Matías-Florentino, Fidencio Mejía-Nepomuceno, María Guadalupe de Jesús Mireles-Rivera, Gloria María Molina-Salinas, Hector Montoya-Fuentes, , Mario Mújica-Sánchez, José Esteban Muñoz-Medina, José de Jesús Nuñez-Contreras, Alicia Ocaña-Mondragón, Luis Alberto Ochoa-Carrera, Hector Esteban Paz-Juárez, Marissa Perez-Garcia, Francisco Pulido, Helen Haydee Fernanda Ramírez-Plascencia, Jorge Salas-Hernández, Angel Gustavo Salas-Lais, Alejandro Sánchez-Flores, Clara Esperanza Santacruz-Tinoco, María Guadalupe Santiago-Mauricio, Selene Zárate, Nelly Sélem-Mojica, Blanca Taboada, Gloria Vazquez, Joel Armando Vázquez-Pérez. |
| EPI_ISL_2681105 | Laboratorio Central de Epidemiología (LCE) | Instituto de Biotecnología de la UNAM | Consorcio Mexicano de Vigilancia Genómica (CoViGen-Mex). Authors (in alphabetical order): Julio Elias Alvarado-Yaah, Carlos F. Arias, Santiago Ávila-Ríos, Eduardo Becerril-Vargas, Víctor Hugo Borja-Aburto, Celia Boukadida, Cristóbal Cháidez-Quiróz, Juan Bautista Chale-Dzul, Ricardo Ciria Merce, Andreu Comas-García, Célida Duque Molina, Julissa Enciso-Ibarra, José Antonio Enciso-Moreno, Gloria Elena Espinosa-Ayala, Fernando Fontove-Herrera, Daniel Fregoso-Rueda, Victor Eduardo García-Arias, Alejandra García-Gasca, Bruno Gómez-Gil, Jean Pierre González, Irvin González-López, Concepción Grajales-Muñiz, Ricardo Grande, Rosa María Gutiérrez Rios, Jesús Hernández, Alejandra Hernández-Terán, Alfredo Herrera-Estrella, Carla Ivón Herrera-Najera, Pavel Isa, Verónica Mata-Haro, Daniel Lira Morales, Susana Lopez, Antonio Loza Román, Brenda Irasema Maldonado-Meza, Bernardo Martínez-Miguel, José Arturo Martínez-Orozco, Célida Martínez- Rodríguez, Margarita Matías-Florentino, Fidencio Mejía-Nepomuceno, María Guadalupe de Jesús Mireles-Rivera, Gloria María Molina-Salinas, Hector Montoya-Fuentes, , Mario Mújica-Sánchez, José Esteban Muñoz-Medina, José de Jesús Nuñez-Contreras, Alicia Ocaña-Mondragón, Luis Alberto Ochoa-Carrera, Hector Esteban Paz-Juárez, Marissa Perez-Garcia, Francisco Pulido, Helen Haydee Fernanda Ramírez-Plascencia, Jorge Salas-Hernández, Angel Gustavo Salas-Lais, Alejandro Sánchez-Flores, Clara Esperanza Santacruz-Tinoco, María Guadalupe Santiago-Mauricio, Selene Zárate, Nelly Sélem-Mojica, Blanca Taboada, Gloria Vazquez, Joel Armando Vázquez-Pérez. |
| EPI_ISL_2681106, EPI_ISL_2681107, EPI_ISL_2681108, EPI_ISL_2681109 | Unidad de Investigación Médica de Yucatán (UIMY) | Instituto de Biotecnología de la UNAM | Consorcio Mexicano de Vigilancia Genómica (CoViGen-Mex). Authors (in alphabetical order): Julio Elias Alvarado-Yaah, Carlos F. Arias, Santiago Ávila-Ríos, Eduardo Becerril-Vargas, Víctor Hugo Borja-Aburto, Celia Boukadida, Cristóbal Cháidez-Quiróz, Juan Bautista Chale-Dzul, Ricardo Ciria Merce, Andreu Comas-García, Célida Duque Molina, Julissa Enciso-Ibarra, José Antonio Enciso-Moreno, Gloria Elena Espinosa-Ayala, Fernando Fontove-Herrera, Daniel Fregoso-Rueda, Victor Eduardo García-Arias, Alejandra García-Gasca, Bruno Gómez-Gil, Jean Pierre González, Irvin González-López, Concepción Grajales-Muñiz, Ricardo Grande, Rosa María Gutiérrez Rios, Jesús Hernández, Alejandra Hernández-Terán, Alfredo |

| | | | |
|---|---|---|---|
| | | | Herrera-Estrella, Carla Ivón Herrera-Najera, Pavel Isa, Verónica Mata-Haro, Daniel Lira Morales, Susana Lopez, Antonio Loza Román, Brenda Irasema Maldonado-Meza, Bernardo Martínez-Miguel, José Arturo Martínez-Orozco, Célida Martínez- Rodríguez, Margarita Matías-Florentino, Fidencio Mejía-Nepomuceno, María Guadalupe de Jesús Mireles-Rivera, Gloria María Molina-Salinas, Hector Montoya-Fuentes, , Mario Mújica-Sánchez, José Esteban Muñoz-Medina, José de Jesús Nuñez-Contreras, Alicia Ocaña-Mondragón, Luis Alberto Ochoa-Carrera, Hector Esteban Paz-Juárez, Marissa Perez-Garcia, Francisco Pulido, Helen Haydee Fernanda Ramírez-Plascencia, Jorge Salas-Hernández, Angel Gustavo Salas-Lais, Alejandro Sánchez-Flores, Clara Esperanza Santacruz-Tinoco, María Guadalupe Santiago-Mauricio, Selene Zárate, Nelly Sélem-Mojica, Blanca Taboada, Gloria Vazquez, Joel Armando Vázquez-Pérez. |
| EPI_ISL_2681110 | Centro de Investigación Biomédica del Noreste (CIBIN) | Instituto de Biotecnología de la UNAM | Consorcio Mexicano de Vigilancia Genómica (CoViGen-Mex). Authors (in alphabetical order): Julio Elias Alvarado-Yaah, Carlos F. Arias, Santiago Ávila-Ríos, Eduardo Becerril-Vargas, Víctor Hugo Borja-Aburto, Celia Boukadida, Cristóbal Cháidez-Quiróz, Juan Bautista Chale-Dzul, Ricardo Ciria Merce, Andreu Comas-García, Célida Duque Molina, Julissa Enciso-Ibarra, José Antonio Enciso-Moreno, Gloria Elena Espinosa-Ayala, Fernando Fontove-Herrera, Daniel Fregoso-Rueda, Victor Eduardo García-Arias, Alejandra García-Gasca, Bruno Gómez-Gil, Jean Pierre González, Irvin González-López, Concepción Grajales-Muñiz, Ricardo Grande, Rosa María Gutiérrez Rios, Jesús Hernández, Alejandra Hernández-Terán, Alfredo Herrera-Estrella, Carla Ivón Herrera-Najera, Pavel Isa, Verónica Mata-Haro, Daniel Lira Morales, Susana Lopez, Antonio Loza Román, Brenda Irasema Maldonado-Meza, Bernardo Martínez-Miguel, José Arturo Martínez-Orozco, Célida Martínez- Rodríguez, Margarita Matías-Florentino, Fidencio Mejía-Nepomuceno, María Guadalupe de Jesús Mireles-Rivera, Gloria María Molina-Salinas, Hector Montoya-Fuentes, , Mario Mújica-Sánchez, José Esteban Muñoz-Medina, José de Jesús Nuñez-Contreras, Alicia Ocaña-Mondragón, Luis Alberto Ochoa-Carrera, Hector Esteban Paz-Juárez, Marissa Perez-Garcia, Francisco Pulido, Helen Haydee Fernanda Ramírez-Plascencia, Jorge Salas-Hernández, Angel Gustavo Salas-Lais, Alejandro Sánchez-Flores, Clara Esperanza Santacruz-Tinoco, María Guadalupe Santiago-Mauricio, Selene Zárate, Nelly Sélem-Mojica, Blanca Taboada, Gloria Vazquez, Joel Armando Vázquez-Pérez. |
| EPI_ISL_2681111, EPI_ISL_2681112, EPI_ISL_2681113 | Laboratorio Central de Epidemiología (LCE) | Instituto de Biotecnología de la UNAM | Consorcio Mexicano de Vigilancia Genómica (CoViGen-Mex). Authors (in alphabetical order): Julio Elias Alvarado-Yaah, Carlos F. Arias, Santiago Ávila-Ríos, Eduardo Becerril-Vargas, Víctor Hugo Borja-Aburto, Celia Boukadida, Cristóbal Cháidez-Quiróz, Juan Bautista Chale-Dzul, Ricardo Ciria Merce, Andreu Comas-García, Célida Duque Molina, Julissa Enciso-Ibarra, José Antonio Enciso-Moreno, Gloria Elena Espinosa-Ayala, Fernando Fontove-Herrera, Daniel Fregoso-Rueda, Victor Eduardo García-Arias, Alejandra García-Gasca, Bruno Gómez-Gil, Jean Pierre González, Irvin González-López, Concepción Grajales-Muñiz, Ricardo Grande, Rosa María Gutiérrez Rios, Jesús Hernández, Alejandra Hernández-Terán, Alfredo Herrera-Estrella, Carla Ivón Herrera-Najera, Pavel Isa, Verónica Mata-Haro, Daniel Lira Morales, Susana Lopez, Antonio Loza Román, Brenda Irasema Maldonado-Meza, Bernardo Martínez-Miguel, José Arturo Martínez-Orozco, Célida Martínez- Rodríguez, Margarita Matías-Florentino, Fidencio Mejía-Nepomuceno, María Guadalupe de Jesús Mireles-Rivera, Gloria María Molina-Salinas, Hector Montoya-Fuentes, , Mario Mújica-Sánchez, José Esteban Muñoz-Medina, José de Jesús Nuñez-Contreras, Alicia Ocaña-Mondragón, Luis Alberto Ochoa-Carrera, Hector Esteban Paz-Juárez, Marissa Perez-Garcia, Francisco Pulido, Helen Haydee Fernanda Ramírez-Plascencia, Jorge Salas-Hernández, Angel Gustavo Salas-Lais, Alejandro Sánchez-Flores, Clara Esperanza Santacruz-Tinoco, María Guadalupe Santiago-Mauricio, Selene Zárate, Nelly Sélem-Mojica, Blanca Taboada, Gloria Vazquez, Joel Armando Vázquez-Pérez. |
| EPI_ISL_2681114, EPI_ISL_2681115, EPI_ISL_2681116, EPI_ISL_2681117, EPI_ISL_2681118 | Unidad de Investigación Médica de Yucatán (UIMY) | Instituto de Biotecnología de la UNAM | Consorcio Mexicano de Vigilancia Genómica (CoViGen-Mex). Authors (in alphabetical order): Julio Elias Alvarado-Yaah, Carlos F. Arias, Santiago Ávila-Ríos, Eduardo Becerril-Vargas, Víctor Hugo Borja-Aburto, Celia Boukadida, Cristóbal Cháidez-Quiróz, Juan Bautista Chale-Dzul, Ricardo Ciria Merce, Andreu Comas-García, Célida Duque Molina, Julissa Enciso-Ibarra, José Antonio Enciso-Moreno, Gloria Elena Espinosa-Ayala, Fernando Fontove-Herrera, Daniel Fregoso-Rueda, Victor Eduardo García-Arias, Alejandra García-Gasca, Bruno Gómez-Gil, Jean Pierre González, Irvin González-López, Concepción Grajales-Muñiz, Ricardo Grande, Rosa María Gutiérrez Rios, Jesús Hernández, Alejandra Hernández-Terán, Alfredo Herrera-Estrella, Carla Ivón Herrera-Najera, Pavel Isa, Verónica Mata-Haro, Daniel Lira Morales, Susana Lopez, Antonio Loza Román, Brenda Irasema Maldonado-Meza, Bernardo Martínez-Miguel, José Arturo Martínez-Orozco, Célida Martínez- Rodríguez, Margarita Matías-Florentino, Fidencio Mejía-Nepomuceno, María Guadalupe de Jesús Mireles-Rivera, Gloria María Molina-Salinas, Hector Montoya-Fuentes, , Mario Mújica-Sánchez, José Esteban Muñoz-Medina, José de Jesús Nuñez-Contreras, Alicia Ocaña-Mondragón, Luis Alberto Ochoa-Carrera, Hector Esteban Paz-Juárez, Marissa Perez-Garcia, Francisco Pulido, Helen Haydee Fernanda Ramírez-Plascencia, Jorge Salas-Hernández, Angel Gustavo Salas-Lais, Alejandro Sánchez-Flores, Clara Esperanza Santacruz-Tinoco, María Guadalupe Santiago-Mauricio, Selene Zárate, Nelly Sélem-Mojica, Blanca Taboada, Gloria Vazquez, Joel Armando Vázquez-Pérez. |
| EPI_ISL_2681120, EPI_ISL_2681121, EPI_ISL_2681122, EPI_ISL_2681123 | Unidad de Investigación Biomédica de Zacatecas (UIBZ) | Instituto de Biotecnología de la UNAM | Consorcio Mexicano de Vigilancia Genómica (CoViGen-Mex). Authors (in alphabetical order): Julio Elias Alvarado-Yaah, Carlos F. Arias, Santiago Ávila-Ríos, Eduardo Becerril-Vargas, Víctor Hugo Borja-Aburto, Celia Boukadida, Cristóbal Cháidez-Quiróz, Juan Bautista Chale-Dzul, Ricardo Ciria Merce, Andreu Comas-García, Célida Duque Molina, Julissa Enciso-Ibarra, José Antonio Enciso-Moreno, Gloria Elena Espinosa-Ayala, Fernando Fontove-Herrera, Daniel Fregoso-Rueda, Victor Eduardo García-Arias, Alejandra García-Gasca, Bruno Gómez-Gil, Jean Pierre González, Irvin González-López, Concepción Grajales-Muñiz, Ricardo Grande, Rosa María Gutiérrez Rios, Jesús Hernández, Alejandra Hernández-Terán, Alfredo Herrera-Estrella, Carla Ivón Herrera-Najera, Pavel Isa, Verónica Mata-Haro, Daniel Lira Morales, Susana Lopez, Antonio Loza Román, Brenda Irasema Maldonado-Meza, Bernardo Martínez-Miguel, José Arturo Martínez-Orozco, Célida Martínez- Rodríguez, Margarita Matías-Florentino, Fidencio Mejía-Nepomuceno, María Guadalupe de Jesús Mireles-Rivera, Gloria María Molina-Salinas, Hector Montoya-Fuentes, , Mario Mújica-Sánchez, José Esteban Muñoz-Medina, José de Jesús Nuñez-Contreras, Alicia Ocaña-Mondragón, Luis Alberto Ochoa-Carrera, Hector Esteban Paz-Juárez, Marissa Perez-Garcia, Francisco Pulido, Helen Haydee Fernanda Ramírez-Plascencia, Jorge Salas-Hernández, Angel Gustavo Salas-Lais, Alejandro Sánchez-Flores, Clara Esperanza Santacruz-Tinoco, María Guadalupe Santiago-Mauricio, Selene Zárate, Nelly Sélem-Mojica, Blanca Taboada, Gloria Vazquez, Joel Armando Vázquez-Pérez. |
| EPI_ISL_2681124, EPI_ISL_2681125 | Laboratorio Central de Epidemiología (LCE) | Instituto de Biotecnología de la UNAM | Consorcio Mexicano de Vigilancia Genómica (CoViGen-Mex). Authors (in alphabetical order): Julio Elias Alvarado-Yaah, Carlos F. Arias, Santiago Ávila-Ríos, Eduardo Becerril-Vargas, Víctor Hugo Borja-Aburto, Celia Boukadida, Cristóbal Cháidez-Quiróz, Juan Bautista Chale-Dzul, Ricardo Ciria Merce, Andreu Comas-García, Célida Duque Molina, Julissa Enciso-Ibarra, José Antonio Enciso-Moreno, Gloria Elena Espinosa-Ayala, Fernando Fontove-Herrera, Daniel Fregoso-Rueda, Victor Eduardo García-Arias, Alejandra García-Gasca, Bruno Gómez-Gil, Jean Pierre González, Irvin González-López, Concepción Grajales-Muñiz, Ricardo Grande, Rosa María Gutiérrez Rios, Jesús Hernández, Alejandra Hernández-Terán, Alfredo Herrera-Estrella, Carla Ivón Herrera-Najera, Pavel Isa, Verónica Mata-Haro, Daniel Lira Morales, Susana Lopez, Antonio Loza Román, Brenda Irasema Maldonado-Meza, Bernardo Martínez-Miguel, José Arturo Martínez-Orozco, Célida Martínez- Rodríguez, Margarita Matías-Florentino, Fidencio Mejía-Nepomuceno, María Guadalupe de Jesús Mireles-Rivera, Gloria María Molina-Salinas, Hector Montoya-Fuentes, , Mario Mújica-Sánchez, José Esteban Muñoz-Medina, José de Jesús Nuñez-Contreras, Alicia Ocaña-Mondragón, Luis Alberto Ochoa-Carrera, Hector Esteban Paz-Juárez, Marissa Perez-Garcia, Francisco Pulido, Helen Haydee Fernanda Ramírez-Plascencia, Jorge Salas-Hernández, Angel Gustavo Salas-Lais, Alejandro Sánchez-Flores, Clara Esperanza Santacruz-Tinoco, María Guadalupe Santiago-Mauricio, Selene Zárate, Nelly Sélem-Mojica, Blanca Taboada, Gloria Vazquez, Joel Armando Vázquez-Pérez. |
| EPI_ISL_2681126 | Unidad de Investigación Médica de Yucatán (UIMY) | Instituto de Biotecnología de la UNAM | Consorcio Mexicano de Vigilancia Genómica (CoViGen-Mex). Authors (in alphabetical order): Julio Elias Alvarado-Yaah, Carlos F. Arias, Santiago Ávila-Ríos, Eduardo Becerril-Vargas, Víctor Hugo Borja-Aburto, Celia Boukadida, Cristóbal Cháidez-Quiróz, Juan Bautista Chale-Dzul, Ricardo Ciria Merce, Andreu Comas-García, Célida Duque Molina, Julissa Enciso-Ibarra, José Antonio Enciso-Moreno, Gloria Elena Espinosa-Ayala, Fernando Fontove-Herrera, Daniel Fregoso-Rueda, Victor Eduardo García-Arias, Alejandra García-Gasca, Bruno Gómez-Gil, Jean Pierre González, Irvin González-López, Concepción Grajales-Muñiz, Ricardo Grande, Rosa María Gutiérrez Rios, Jesús Hernández, Alejandra Hernández-Terán, Alfredo Herrera-Estrella, Carla Ivón Herrera-Najera, Pavel Isa, Verónica Mata-Haro, Daniel Lira Morales, Susana Lopez, Antonio Loza Román, Brenda Irasema Maldonado-Meza, Bernardo Martínez-Miguel, José Arturo Martínez-Orozco, Célida Martínez- Rodríguez, Margarita Matías-Florentino, Fidencio Mejía-Nepomuceno, María Guadalupe de Jesús Mireles-Rivera, Gloria María Molina-Salinas, Hector Montoya-Fuentes, , Mario Mújica-Sánchez, José Esteban Muñoz-Medina, José de Jesús Nuñez-Contreras, Alicia Ocaña-Mondragón, Luis Alberto Ochoa-Carrera, Hector Esteban Paz-Juárez, Marissa Perez-Garcia, Francisco Pulido, Helen Haydee Fernanda Ramírez-Plascencia, Jorge Salas-Hernández, Angel Gustavo Salas-Lais, Alejandro Sánchez-Flores, Clara Esperanza Santacruz-Tinoco, María Guadalupe Santiago-Mauricio, Selene Zárate, Nelly Sélem-Mojica, Blanca Taboada, Gloria Vazquez, Joel Armando Vázquez-Pérez. |
| EPI_ISL_2681127 | Centro de Investigación Biomédica del Noreste (CIBIN) | Instituto de Biotecnología de la UNAM | Consorcio Mexicano de Vigilancia Genómica (CoViGen-Mex). Authors (in alphabetical order): Julio Elias Alvarado-Yaah, Carlos F. Arias, Santiago Ávila-Ríos, Eduardo Becerril-Vargas, Víctor Hugo Borja-Aburto, Celia Boukadida, Cristóbal Cháidez-Quiróz, Juan Bautista Chale-Dzul, Ricardo Ciria Merce, Andreu Comas-García, Célida Duque Molina, Julissa Enciso-Ibarra, José Antonio Enciso-Moreno, Gloria Elena Espinosa-Ayala, Fernando Fontove-Herrera, Daniel Fregoso-Rueda, Victor Eduardo García-Arias, Alejandra García-Gasca, Bruno Gómez-Gil, Jean Pierre González, Irvin González-López, Concepción Grajales-Muñiz, Ricardo Grande, Rosa María Gutiérrez Rios, Jesús Hernández, Alejandra Hernández-Terán, Alfredo |

| | | | |
|---|---|---|---|
| | | | Herrera-Estrella, Carla Ivón Herrera-Najera, Pavel Isa, Verónica Mata-Haro, Daniel Lira Morales, Susana Lopez, Antonio Loza Román, Brenda Irasema Maldonado-Meza, Bernardo Martínez-Miguel, José Arturo Martínez-Orozco, Célida Martínez- Rodríguez, Margarita Matías-Florentino, Fidencio Mejía-Nepomuceno, María Guadalupe de Jesús Mireles-Rivera, Gloria María Molina-Salinas, Hector Montoya-Fuentes, , Mario Mújica-Sánchez, José Esteban Muñoz-Medina, José de Jesús Nuñez-Contreras, Alicia Ocaña-Mondragón, Luis Alberto Ochoa-Carrera, Hector Esteban Paz-Juárez, Marissa Perez-Garcia, Francisco Pulido, Helen Haydee Fernanda Ramírez-Plascencia, Jorge Salas-Hernández, Angel Gustavo Salas-Lais, Alejandro Sánchez-Flores, Clara Esperanza Santacruz-Tinoco, María Guadalupe Santiago-Mauricio, Selene Zárate, Nelly Sélem-Mojica, Blanca Taboada, Gloria Vazquez, Joel Armando Vázquez-Pérez. |
| EPI_ISL_2681128, EPI_ISL_2681129 | Laboratorio Central de Epidemiología (LCE) | Instituto de Biotecnología de la UNAM | Consorcio Mexicano de Vigilancia Genómica (CoViGen-Mex). Authors (in alphabetical order): Julio Elias Alvarado-Yaah, Carlos F. Arias, Santiago Ávila-Ríos, Eduardo Becerril-Vargas, Víctor Hugo Borja-Aburto, Celia Boukadida, Cristóbal Cháidez-Quiróz, Juan Bautista Chale-Dzul, Ricardo Ciria Merce, Andreu Comas-García, Célida Duque Molina, Julissa Enciso-Ibarra, José Antonio Enciso-Moreno, Gloria Elena Espinosa-Ayala, Fernando Fontove-Herrera, Daniel Fregoso-Rueda, Victor Eduardo García-Arias, Alejandra García-Gasca, Bruno Gómez-Gil, Jean Pierre González, Irvin González-López, Concepción Grajales-Muñiz, Ricardo Grande, Rosa María Gutiérrez Rios, Jesús Hernández, Alejandra Hernández-Terán, Alfredo Herrera-Estrella, Carla Ivón Herrera-Najera, Pavel Isa, Verónica Mata-Haro, Daniel Lira Morales, Susana Lopez, Antonio Loza Román, Brenda Irasema Maldonado-Meza, Bernardo Martínez-Miguel, José Arturo Martínez-Orozco, Célida Martínez- Rodríguez, Margarita Matías-Florentino, Fidencio Mejía-Nepomuceno, María Guadalupe de Jesús Mireles-Rivera, Gloria María Molina-Salinas, Hector Montoya-Fuentes, , Mario Mújica-Sánchez, José Esteban Muñoz-Medina, José de Jesús Nuñez-Contreras, Alicia Ocaña-Mondragón, Luis Alberto Ochoa-Carrera, Hector Esteban Paz-Juárez, Marissa Perez-Garcia, Francisco Pulido, Helen Haydee Fernanda Ramírez-Plascencia, Jorge Salas-Hernández, Angel Gustavo Salas-Lais, Alejandro Sánchez-Flores, Clara Esperanza Santacruz-Tinoco, María Guadalupe Santiago-Mauricio, Selene Zárate, Nelly Sélem-Mojica, Blanca Taboada, Gloria Vazquez, Joel Armando Vázquez-Pérez. |
| EPI_ISL_2681130 | Unidad de Investigación Médica de Yucatán (UIMY) | Instituto de Biotecnología de la UNAM | Consorcio Mexicano de Vigilancia Genómica (CoViGen-Mex). Authors (in alphabetical order): Julio Elias Alvarado-Yaah, Carlos F. Arias, Santiago Ávila-Ríos, Eduardo Becerril-Vargas, Víctor Hugo Borja-Aburto, Celia Boukadida, Cristóbal Cháidez-Quiróz, Juan Bautista Chale-Dzul, Ricardo Ciria Merce, Andreu Comas-García, Célida Duque Molina, Julissa Enciso-Ibarra, José Antonio Enciso-Moreno, Gloria Elena Espinosa-Ayala, Fernando Fontove-Herrera, Daniel Fregoso-Rueda, Victor Eduardo García-Arias, Alejandra García-Gasca, Bruno Gómez-Gil, Jean Pierre González, Irvin González-López, Concepción Grajales-Muñiz, Ricardo Grande, Rosa María Gutiérrez Rios, Jesús Hernández, Alejandra Hernández-Terán, Alfredo Herrera-Estrella, Carla Ivón Herrera-Najera, Pavel Isa, Verónica Mata-Haro, Daniel Lira Morales, Susana Lopez, Antonio Loza Román, Brenda Irasema Maldonado-Meza, Bernardo Martínez-Miguel, José Arturo Martínez-Orozco, Célida Martínez- Rodríguez, Margarita Matías-Florentino, Fidencio Mejía-Nepomuceno, María Guadalupe de Jesús Mireles-Rivera, Gloria María Molina-Salinas, Hector Montoya-Fuentes, , Mario Mújica-Sánchez, José Esteban Muñoz-Medina, José de Jesús Nuñez-Contreras, Alicia Ocaña-Mondragón, Luis Alberto Ochoa-Carrera, Hector Esteban Paz-Juárez, Marissa Perez-Garcia, Francisco Pulido, Helen Haydee Fernanda Ramírez-Plascencia, Jorge Salas-Hernández, Angel Gustavo Salas-Lais, Alejandro Sánchez-Flores, Clara Esperanza Santacruz-Tinoco, María Guadalupe Santiago-Mauricio, Selene Zárate, Nelly Sélem-Mojica, Blanca Taboada, Gloria Vazquez, Joel Armando Vázquez-Pérez. |
| EPI_ISL_2681132 | Laboratorio Central de Epidemiología (LCE) | Instituto de Biotecnología de la UNAM | Consorcio Mexicano de Vigilancia Genómica (CoViGen-Mex). Authors (in alphabetical order): Julio Elias Alvarado-Yaah, Carlos F. Arias, Santiago Ávila-Ríos, Eduardo Becerril-Vargas, Víctor Hugo Borja-Aburto, Celia Boukadida, Cristóbal Cháidez-Quiróz, Juan Bautista Chale-Dzul, Ricardo Ciria Merce, Andreu Comas-García, Célida Duque Molina, Julissa Enciso-Ibarra, José Antonio Enciso-Moreno, Gloria Elena Espinosa-Ayala, Fernando Fontove-Herrera, Daniel Fregoso-Rueda, Victor Eduardo García-Arias, Alejandra García-Gasca, Bruno Gómez-Gil, Jean Pierre González, Irvin González-López, Concepción Grajales-Muñiz, Ricardo Grande, Rosa María Gutiérrez Rios, Jesús Hernández, Alejandra Hernández-Terán, Alfredo Herrera-Estrella, Carla Ivón Herrera-Najera, Pavel Isa, Verónica Mata-Haro, Daniel Lira Morales, Susana Lopez, Antonio Loza Román, Brenda Irasema Maldonado-Meza, Bernardo Martínez-Miguel, José Arturo Martínez-Orozco, Célida Martínez- Rodríguez, Margarita Matías-Florentino, Fidencio Mejía-Nepomuceno, María Guadalupe de Jesús Mireles-Rivera, Gloria María Molina-Salinas, Hector Montoya-Fuentes, , Mario Mújica-Sánchez, José Esteban Muñoz-Medina, José de Jesús Nuñez-Contreras, Alicia Ocaña-Mondragón, Luis Alberto Ochoa-Carrera, Hector Esteban Paz-Juárez, Marissa Perez-Garcia, Francisco Pulido, Helen Haydee Fernanda Ramírez-Plascencia, Jorge Salas-Hernández, Angel Gustavo Salas-Lais, Alejandro Sánchez-Flores, Clara Esperanza Santacruz-Tinoco, María Guadalupe Santiago-Mauricio, Selene Zárate, Nelly Sélem-Mojica, Blanca Taboada, Gloria Vazquez, Joel Armando Vázquez-Pérez. |
| EPI_ISL_2681133 | Unidad de Investigación Médica de Yucatán (UIMY) | Instituto de Biotecnología de la UNAM | Consorcio Mexicano de Vigilancia Genómica (CoViGen-Mex). Authors (in alphabetical order): Julio Elias Alvarado-Yaah, Carlos F. Arias, Santiago Ávila-Ríos, Eduardo Becerril-Vargas, Víctor Hugo Borja-Aburto, Celia Boukadida, Cristóbal Cháidez-Quiróz, Juan Bautista Chale-Dzul, Ricardo Ciria Merce, Andreu Comas-García, Célida Duque Molina, Julissa Enciso-Ibarra, José Antonio Enciso-Moreno, Gloria Elena Espinosa-Ayala, Fernando Fontove-Herrera, Daniel Fregoso-Rueda, Victor Eduardo García-Arias, Alejandra García-Gasca, Bruno Gómez-Gil, Jean Pierre González, Irvin González-López, Concepción Grajales-Muñiz, Ricardo Grande, Rosa María Gutiérrez Rios, Jesús Hernández, Alejandra Hernández-Terán, Alfredo Herrera-Estrella, Carla Ivón Herrera-Najera, Pavel Isa, Verónica Mata-Haro, Daniel Lira Morales, Susana Lopez, Antonio Loza Román, Brenda Irasema Maldonado-Meza, Bernardo Martínez-Miguel, José Arturo Martínez-Orozco, Célida Martínez- Rodríguez, Margarita Matías-Florentino, Fidencio Mejía-Nepomuceno, María Guadalupe de Jesús Mireles-Rivera, Gloria María Molina-Salinas, Hector Montoya-Fuentes, , Mario Mújica-Sánchez, José Esteban Muñoz-Medina, José de Jesús Nuñez-Contreras, Alicia Ocaña-Mondragón, Luis Alberto Ochoa-Carrera, Hector Esteban Paz-Juárez, Marissa Perez-Garcia, Francisco Pulido, Helen Haydee Fernanda Ramírez-Plascencia, Jorge Salas-Hernández, Angel Gustavo Salas-Lais, Alejandro Sánchez-Flores, Clara Esperanza Santacruz-Tinoco, María Guadalupe Santiago-Mauricio, Selene Zárate, Nelly Sélem-Mojica, Blanca Taboada, Gloria Vazquez, Joel Armando Vázquez-Pérez. |
| EPI_ISL_2681134, EPI_ISL_2681135 | Laboratorio Central de Epidemiología (LCE) | Instituto de Biotecnología de la UNAM | Consorcio Mexicano de Vigilancia Genómica (CoViGen-Mex). Authors (in alphabetical order): Julio Elias Alvarado-Yaah, Carlos F. Arias, Santiago Ávila-Ríos, Eduardo Becerril-Vargas, Víctor Hugo Borja-Aburto, Celia Boukadida, Cristóbal Cháidez-Quiróz, Juan Bautista Chale-Dzul, Ricardo Ciria Merce, Andreu Comas-García, Célida Duque Molina, Julissa Enciso-Ibarra, José Antonio Enciso-Moreno, Gloria Elena Espinosa-Ayala, Fernando Fontove-Herrera, Daniel Fregoso-Rueda, Victor Eduardo García-Arias, Alejandra García-Gasca, Bruno Gómez-Gil, Jean Pierre González, Irvin González-López, Concepción Grajales-Muñiz, Ricardo Grande, Rosa María Gutiérrez Rios, Jesús Hernández, Alejandra Hernández-Terán, Alfredo Herrera-Estrella, Carla Ivón Herrera-Najera, Pavel Isa, Verónica Mata-Haro, Daniel Lira Morales, Susana Lopez, Antonio Loza Román, Brenda Irasema Maldonado-Meza, Bernardo Martínez-Miguel, José Arturo Martínez-Orozco, Célida Martínez- Rodríguez, Margarita Matías-Florentino, Fidencio Mejía-Nepomuceno, María Guadalupe de Jesús Mireles-Rivera, Gloria María Molina-Salinas, Hector Montoya-Fuentes, , Mario Mújica-Sánchez, José Esteban Muñoz-Medina, José de Jesús Nuñez-Contreras, Alicia Ocaña-Mondragón, Luis Alberto Ochoa-Carrera, Hector Esteban Paz-Juárez, Marissa Perez-Garcia, Francisco Pulido, Helen Haydee Fernanda Ramírez-Plascencia, Jorge Salas-Hernández, Angel Gustavo Salas-Lais, Alejandro Sánchez-Flores, Clara Esperanza Santacruz-Tinoco, María Guadalupe Santiago-Mauricio, Selene Zárate, Nelly Sélem-Mojica, Blanca Taboada, Gloria Vazquez, Joel Armando Vázquez-Pérez. |
| EPI_ISL_2681136 | Unidad de Investigación Médica de Yucatán (UIMY) | Instituto de Biotecnología de la UNAM | Consorcio Mexicano de Vigilancia Genómica (CoViGen-Mex). Authors (in alphabetical order): Julio Elias Alvarado-Yaah, Carlos F. Arias, Santiago Ávila-Ríos, Eduardo Becerril-Vargas, Víctor Hugo Borja-Aburto, Celia Boukadida, Cristóbal Cháidez-Quiróz, Juan Bautista Chale-Dzul, Ricardo Ciria Merce, Andreu Comas-García, Célida Duque Molina, Julissa Enciso-Ibarra, José Antonio Enciso-Moreno, Gloria Elena Espinosa-Ayala, Fernando Fontove-Herrera, Daniel Fregoso-Rueda, Victor Eduardo García-Arias, Alejandra García-Gasca, Bruno Gómez-Gil, Jean Pierre González, Irvin González-López, Concepción Grajales-Muñiz, Ricardo Grande, Rosa María Gutiérrez Rios, Jesús Hernández, Alejandra Hernández-Terán, Alfredo Herrera-Estrella, Carla Ivón Herrera-Najera, Pavel Isa, Verónica Mata-Haro, Daniel Lira Morales, Susana Lopez, Antonio Loza Román, Brenda Irasema Maldonado-Meza, Bernardo Martínez-Miguel, José Arturo Martínez-Orozco, Célida Martínez- Rodríguez, Margarita Matías-Florentino, Fidencio Mejía-Nepomuceno, María Guadalupe de Jesús Mireles-Rivera, Gloria María Molina-Salinas, Hector Montoya-Fuentes, , Mario Mújica-Sánchez, José Esteban Muñoz-Medina, José de Jesús Nuñez-Contreras, Alicia Ocaña-Mondragón, Luis Alberto Ochoa-Carrera, Hector Esteban Paz-Juárez, Marissa Perez-Garcia, Francisco Pulido, Helen Haydee Fernanda Ramírez-Plascencia, Jorge Salas-Hernández, Angel Gustavo Salas-Lais, Alejandro Sánchez-Flores, Clara Esperanza Santacruz-Tinoco, María Guadalupe Santiago-Mauricio, Selene Zárate, Nelly Sélem-Mojica, Blanca Taboada, Gloria Vazquez, Joel Armando Vázquez-Pérez. |
| EPI_ISL_2681137 | Centro de Investigación Biomédica del Noreste (CIBIN) | Instituto de Biotecnología de la UNAM | Consorcio Mexicano de Vigilancia Genómica (CoViGen-Mex). Authors (in alphabetical order): Julio Elias Alvarado-Yaah, Carlos F. Arias, Santiago Ávila-Ríos, Eduardo Becerril-Vargas, Víctor Hugo Borja-Aburto, Celia Boukadida, Cristóbal Cháidez-Quiróz, Juan Bautista Chale-Dzul, Ricardo Ciria Merce, Andreu Comas-García, Célida Duque Molina, Julissa Enciso-Ibarra, José Antonio Enciso-Moreno, Gloria Elena Espinosa-Ayala, Fernando Fontove-Herrera, Daniel Fregoso-Rueda, Victor Eduardo García-Arias, Alejandra García-Gasca, Bruno Gómez-Gil, Jean Pierre González, Irvin González-López, Concepción Grajales-Muñiz, Ricardo Grande, Rosa María Gutiérrez Rios, Jesús Hernández, Alejandra Hernández-Terán, Alfredo |

| | | | |
|---|---|---|---|
| | | | Herrera-Estrella, Carla Ivón Herrera-Najera, Pavel Isa, Verónica Mata-Haro, Daniel Lira Morales, Susana Lopez, Antonio Loza Román, Brenda Irasema Maldonado-Meza, Bernardo Martínez-Miguel, José Arturo Martínez-Orozco, Célida Martínez- Rodríguez, Margarita Matías-Florentino, Fidencio Mejía-Nepomuceno, María Guadalupe de Jesús Mireles-Rivera, Gloria María Molina-Salinas, Hector Montoya-Fuentes, , Mario Mújica-Sánchez, José Esteban Muñoz-Medina, José de Jesús Nuñez-Contreras, Alicia Ocaña-Mondragón, Luis Alberto Ochoa-Carrera, Hector Esteban Paz-Juárez, Marissa Perez-Garcia, Francisco Pulido, Helen Haydee Fernanda Ramírez-Plascencia, Jorge Salas-Hernández, Angel Gustavo Salas-Lais, Alejandro Sánchez-Flores, Clara Esperanza Santacruz-Tinoco, María Guadalupe Santiago-Mauricio, Selene Zárate, Nelly Sélem-Mojica, Blanca Taboada, Gloria Vazquez, Joel Armando Vázquez-Pérez. |
| EPI_ISL_2681139, EPI_ISL_2681140 | Laboratorio Central de Epidemiología (LCE) | Instituto de Biotecnología de la UNAM | Consorcio Mexicano de Vigilancia Genómica (CoViGen-Mex). Authors (in alphabetical order): Julio Elias Alvarado-Yaah, Carlos F. Arias, Santiago Ávila-Ríos, Eduardo Becerril-Vargas, Víctor Hugo Borja-Aburto, Celia Boukadida, Cristóbal Cháidez-Quiróz, Juan Bautista Chale-Dzul, Ricardo Ciria Merce, Andreu Comas-García, Célida Duque Molina, Julissa Enciso-Ibarra, José Antonio Enciso-Moreno, Gloria Elena Espinosa-Ayala, Fernando Fontove-Herrera, Daniel Fregoso-Rueda, Victor Eduardo García-Arias, Alejandra García-Gasca, Bruno Gómez-Gil, Jean Pierre González, Irvin González-López, Concepción Grajales-Muñiz, Ricardo Grande, Rosa María Gutiérrez Rios, Jesús Hernández, Alejandra Hernández-Terán, Alfredo Herrera-Estrella, Carla Ivón Herrera-Najera, Pavel Isa, Verónica Mata-Haro, Daniel Lira Morales, Susana Lopez, Antonio Loza Román, Brenda Irasema Maldonado-Meza, Bernardo Martínez-Miguel, José Arturo Martínez-Orozco, Célida Martínez- Rodríguez, Margarita Matías-Florentino, Fidencio Mejía-Nepomuceno, María Guadalupe de Jesús Mireles-Rivera, Gloria María Molina-Salinas, Hector Montoya-Fuentes, , Mario Mújica-Sánchez, José Esteban Muñoz-Medina, José de Jesús Nuñez-Contreras, Alicia Ocaña-Mondragón, Luis Alberto Ochoa-Carrera, Hector Esteban Paz-Juárez, Marissa Perez-Garcia, Francisco Pulido, Helen Haydee Fernanda Ramírez-Plascencia, Jorge Salas-Hernández, Angel Gustavo Salas-Lais, Alejandro Sánchez-Flores, Clara Esperanza Santacruz-Tinoco, María Guadalupe Santiago-Mauricio, Selene Zárate, Nelly Sélem-Mojica, Blanca Taboada, Gloria Vazquez, Joel Armando Vázquez-Pérez. |
| EPI_ISL_2681141 | Unidad de Investigación Médica de Yucatán (UIMY) | Instituto de Biotecnología de la UNAM | Consorcio Mexicano de Vigilancia Genómica (CoViGen-Mex). Authors (in alphabetical order): Julio Elias Alvarado-Yaah, Carlos F. Arias, Santiago Ávila-Ríos, Eduardo Becerril-Vargas, Víctor Hugo Borja-Aburto, Celia Boukadida, Cristóbal Cháidez-Quiróz, Juan Bautista Chale-Dzul, Ricardo Ciria Merce, Andreu Comas-García, Célida Duque Molina, Julissa Enciso-Ibarra, José Antonio Enciso-Moreno, Gloria Elena Espinosa-Ayala, Fernando Fontove-Herrera, Daniel Fregoso-Rueda, Victor Eduardo García-Arias, Alejandra García-Gasca, Bruno Gómez-Gil, Jean Pierre González, Irvin González-López, Concepción Grajales-Muñiz, Ricardo Grande, Rosa María Gutiérrez Rios, Jesús Hernández, Alejandra Hernández-Terán, Alfredo Herrera-Estrella, Carla Ivón Herrera-Najera, Pavel Isa, Verónica Mata-Haro, Daniel Lira Morales, Susana Lopez, Antonio Loza Román, Brenda Irasema Maldonado-Meza, Bernardo Martínez-Miguel, José Arturo Martínez-Orozco, Célida Martínez- Rodríguez, Margarita Matías-Florentino, Fidencio Mejía-Nepomuceno, María Guadalupe de Jesús Mireles-Rivera, Gloria María Molina-Salinas, Hector Montoya-Fuentes, , Mario Mújica-Sánchez, José Esteban Muñoz-Medina, José de Jesús Nuñez-Contreras, Alicia Ocaña-Mondragón, Luis Alberto Ochoa-Carrera, Hector Esteban Paz-Juárez, Marissa Perez-Garcia, Francisco Pulido, Helen Haydee Fernanda Ramírez-Plascencia, Jorge Salas-Hernández, Angel Gustavo Salas-Lais, Alejandro Sánchez-Flores, Clara Esperanza Santacruz-Tinoco, María Guadalupe Santiago-Mauricio, Selene Zárate, Nelly Sélem-Mojica, Blanca Taboada, Gloria Vazquez, Joel Armando Vázquez-Pérez. |
| EPI_ISL_2681142 | Unidad de Investigación Biomédica de Zacatecas (UIBZ) | Instituto de Biotecnología de la UNAM | Consorcio Mexicano de Vigilancia Genómica (CoViGen-Mex). Authors (in alphabetical order): Julio Elias Alvarado-Yaah, Carlos F. Arias, Santiago Ávila-Ríos, Eduardo Becerril-Vargas, Víctor Hugo Borja-Aburto, Celia Boukadida, Cristóbal Cháidez-Quiróz, Juan Bautista Chale-Dzul, Ricardo Ciria Merce, Andreu Comas-García, Célida Duque Molina, Julissa Enciso-Ibarra, José Antonio Enciso-Moreno, Gloria Elena Espinosa-Ayala, Fernando Fontove-Herrera, Daniel Fregoso-Rueda, Victor Eduardo García-Arias, Alejandra García-Gasca, Bruno Gómez-Gil, Jean Pierre González, Irvin González-López, Concepción Grajales-Muñiz, Ricardo Grande, Rosa María Gutiérrez Rios, Jesús Hernández, Alejandra Hernández-Terán, Alfredo Herrera-Estrella, Carla Ivón Herrera-Najera, Pavel Isa, Verónica Mata-Haro, Daniel Lira Morales, Susana Lopez, Antonio Loza Román, Brenda Irasema Maldonado-Meza, Bernardo Martínez-Miguel, José Arturo Martínez-Orozco, Célida Martínez- Rodríguez, Margarita Matías-Florentino, Fidencio Mejía-Nepomuceno, María Guadalupe de Jesús Mireles-Rivera, Gloria María Molina-Salinas, Hector Montoya-Fuentes, , Mario Mújica-Sánchez, José Esteban Muñoz-Medina, José de Jesús Nuñez-Contreras, Alicia Ocaña-Mondragón, Luis Alberto Ochoa-Carrera, Hector Esteban Paz-Juárez, Marissa Perez-Garcia, Francisco Pulido, Helen Haydee Fernanda Ramírez-Plascencia, Jorge Salas-Hernández, Angel Gustavo Salas-Lais, Alejandro Sánchez-Flores, Clara Esperanza Santacruz-Tinoco, María Guadalupe Santiago-Mauricio, Selene Zárate, Nelly Sélem-Mojica, Blanca Taboada, Gloria Vazquez, Joel Armando Vázquez-Pérez. |
| EPI_ISL_2681144, EPI_ISL_2681145, EPI_ISL_2681146, EPI_ISL_2681147, EPI_ISL_2681148, EPI_ISL_2681149, EPI_ISL_2681150, EPI_ISL_2681151, EPI_ISL_2681154, EPI_ISL_2681155, EPI_ISL_2681156, EPI_ISL_2681157, EPI_ISL_2681158, EPI_ISL_2681159 | | | |
| see above | Laboratorio Central de Epidemiología (LCE) | Instituto de Biotecnología de la UNAM | Consorcio Mexicano de Vigilancia Genómica (CoViGen-Mex). Authors (in alphabetical order): Julio Elias Alvarado-Yaah, Carlos F. Arias, Santiago Ávila-Ríos, Eduardo Becerril-Vargas, Víctor Hugo Borja-Aburto, Celia Boukadida, Cristóbal Cháidez-Quiróz, Juan Bautista Chale-Dzul, Ricardo Ciria Merce, Andreu Comas-García, Célida Duque Molina, Julissa Enciso-Ibarra, José Antonio Enciso-Moreno, Gloria Elena Espinosa-Ayala, Fernando Fontove-Herrera, Daniel Fregoso-Rueda, Victor Eduardo García-Arias, Alejandra García-Gasca, Bruno Gómez-Gil, Jean Pierre González, Irvin González-López, Concepción Grajales-Muñiz, Ricardo Grande, Rosa María Gutiérrez Rios, Jesús Hernández, Alejandra Hernández-Terán, Alfredo Herrera-Estrella, Carla Ivón Herrera-Najera, Pavel Isa, Verónica Mata-Haro, Daniel Lira Morales, Susana Lopez, Antonio Loza Román, Brenda Irasema Maldonado-Meza, Bernardo Martínez-Miguel, José Arturo Martínez-Orozco, Célida Martínez- Rodríguez, Margarita Matías-Florentino, Fidencio Mejía-Nepomuceno, María Guadalupe de Jesús Mireles-Rivera, Gloria María Molina-Salinas, Hector Montoya-Fuentes, , Mario Mújica-Sánchez, José Esteban Muñoz-Medina, José de Jesús Nuñez-Contreras, Alicia Ocaña-Mondragón, Luis Alberto Ochoa-Carrera, Hector Esteban Paz-Juárez, Marissa Perez-Garcia, Francisco Pulido, Helen Haydee Fernanda Ramírez-Plascencia, Jorge Salas-Hernández, Angel Gustavo Salas-Lais, Alejandro Sánchez-Flores, Clara Esperanza Santacruz-Tinoco, María Guadalupe Santiago-Mauricio, Selene Zárate, Nelly Sélem-Mojica, Blanca Taboada, Gloria Vazquez, Joel Armando Vázquez-Pérez. |
| EPI_ISL_2681160 | Unidad de Investigación Biomédica de Zacatecas (UIBZ) | Instituto de Biotecnología de la UNAM | Consorcio Mexicano de Vigilancia Genómica (CoViGen-Mex). Authors (in alphabetical order): Julio Elias Alvarado-Yaah, Carlos F. Arias, Santiago Ávila-Ríos, Eduardo Becerril-Vargas, Víctor Hugo Borja-Aburto, Celia Boukadida, Cristóbal Cháidez-Quiróz, Juan Bautista Chale-Dzul, Ricardo Ciria Merce, Andreu Comas-García, Célida Duque Molina, Julissa Enciso-Ibarra, José Antonio Enciso-Moreno, Gloria Elena Espinosa-Ayala, Fernando Fontove-Herrera, Daniel Fregoso-Rueda, Victor Eduardo García-Arias, Alejandra García-Gasca, Bruno Gómez-Gil, Jean Pierre González, Irvin González-López, Concepción Grajales-Muñiz, Ricardo Grande, Rosa María Gutiérrez Rios, Jesús Hernández, Alejandra Hernández-Terán, Alfredo Herrera-Estrella, Carla Ivón Herrera-Najera, Pavel Isa, Verónica Mata-Haro, Daniel Lira Morales, Susana Lopez, Antonio Loza Román, Brenda Irasema Maldonado-Meza, Bernardo Martínez-Miguel, José Arturo Martínez-Orozco, Célida Martínez- Rodríguez, Margarita Matías-Florentino, Fidencio Mejía-Nepomuceno, María Guadalupe de Jesús Mireles-Rivera, Gloria María Molina-Salinas, Hector Montoya-Fuentes, , Mario Mújica-Sánchez, José Esteban Muñoz-Medina, José de Jesús Nuñez-Contreras, Alicia Ocaña-Mondragón, Luis Alberto Ochoa-Carrera, Hector Esteban Paz-Juárez, Marissa Perez-Garcia, Francisco Pulido, Helen Haydee Fernanda Ramírez-Plascencia, Jorge Salas-Hernández, Angel Gustavo Salas-Lais, Alejandro Sánchez-Flores, Clara Esperanza Santacruz-Tinoco, María Guadalupe Santiago-Mauricio, Selene Zárate, Nelly Sélem-Mojica, Blanca Taboada, Gloria Vazquez, Joel Armando Vázquez-Pérez. |
| EPI_ISL_2681163, EPI_ISL_2681164, EPI_ISL_2681165, EPI_ISL_2681166, EPI_ISL_2681167, EPI_ISL_2681168, EPI_ISL_2681169, EPI_ISL_2681170, EPI_ISL_2681171, EPI_ISL_2681172, EPI_ISL_2681173, EPI_ISL_2681174, EPI_ISL_2681175 | | | |
| see above | Laboratorio Central de Epidemiología (LCE) | Instituto de Biotecnología de la UNAM | Consorcio Mexicano de Vigilancia Genómica (CoViGen-Mex). Authors (in alphabetical order): Julio Elias Alvarado-Yaah, Carlos F. Arias, Santiago Ávila-Ríos, Eduardo Becerril-Vargas, Víctor Hugo Borja-Aburto, Celia Boukadida, Cristóbal Cháidez-Quiróz, Juan Bautista Chale-Dzul, Ricardo Ciria Merce, Andreu Comas-García, Célida Duque Molina, Julissa Enciso-Ibarra, José Antonio Enciso-Moreno, Gloria Elena Espinosa-Ayala, Fernando Fontove-Herrera, Daniel Fregoso-Rueda, Victor Eduardo García-Arias, Alejandra García-Gasca, Bruno Gómez-Gil, Jean Pierre González, Irvin González-López, Concepción Grajales-Muñiz, Ricardo Grande, Rosa María Gutiérrez Rios, Jesús Hernández, Alejandra Hernández-Terán, Alfredo Herrera-Estrella, Carla Ivón Herrera-Najera, Pavel Isa, Verónica Mata-Haro, Daniel Lira Morales, Susana Lopez, Antonio Loza Román, Brenda Irasema Maldonado-Meza, Bernardo Martínez-Miguel, José Arturo Martínez-Orozco, Célida Martínez- Rodríguez, Margarita Matías-Florentino, Fidencio Mejía-Nepomuceno, María Guadalupe de Jesús Mireles-Rivera, Gloria María Molina-Salinas, Hector Montoya-Fuentes, , Mario Mújica-Sánchez, José Esteban Muñoz-Medina, José de Jesús Nuñez-Contreras, Alicia Ocaña-Mondragón, Luis Alberto Ochoa-Carrera, Hector Esteban Paz-Juárez, Marissa Perez-Garcia, Francisco Pulido, Helen Haydee Fernanda Ramírez-Plascencia, Jorge Salas-Hernández, Angel Gustavo Salas-Lais, Alejandro Sánchez-Flores, Clara Esperanza Santacruz-Tinoco, María Guadalupe Santiago-Mauricio, Selene Zárate, Nelly Sélem-Mojica, Blanca Taboada, Gloria Vazquez, Joel Armando Vázquez-Pérez. |
| EPI_ISL_2681176 | Unidad de Investigación Médica de Yucatán (UIMY) | Instituto de Biotecnología de la UNAM | Consorcio Mexicano de Vigilancia Genómica (CoViGen-Mex). Authors (in alphabetical order): Julio Elias Alvarado-Yaah, Carlos F. Arias, Santiago Ávila-Ríos, Eduardo Becerril-Vargas, Víctor Hugo Borja-Aburto, Celia Boukadida, Cristóbal Cháidez-Quiróz, Juan Bautista Chale-Dzul, Ricardo Ciria |

| | | | |
|---|---|---|---|
| | | | Merce, Andreu Comas-García, Célida Duque Molina, Julissa Enciso-Ibarra, José Antonio Enciso-Moreno, Gloria Elena Espinosa-Ayala, Fernando Fontove-Herrera, Daniel Fregoso-Rueda, Victor Eduardo García-Arias, Alejandra García-Gasca, Bruno Gómez-Gil, Jean Pierre González, Irvin González-López, Concepción Grajales-Muñiz, Ricardo Grande, Rosa María Gutiérrez Rios, Jesús Hernández, Alejandra Hernández-Terán, Alfredo Herrera-Estrella, Carla Ivón Herrera-Najera, Pavel Isa, Verónica Mata-Haro, Daniel Lira Morales, Susana Lopez, Antonio Loza Román, Brenda Irasema Maldonado-Meza, Bernardo Martínez-Miguel, José Arturo Martínez-Orozco, Célida Martínez- Rodríguez, Margarita Matías-Florentino, Fidencio Mejía-Nepomuceno, María Guadalupe de Jesús Mireles-Rivera, Gloria María Molina-Salinas, Hector Montoya-Fuentes, , Mario Mújica-Sánchez, José Esteban Muñoz-Medina, José de Jesús Nuñez-Contreras, Alicia Ocaña-Mondragón, Luis Alberto Ochoa-Carrera, Hector Esteban Paz-Juárez, Marissa Perez-Garcia, Francisco Pulido, Helen Haydee Fernanda Ramírez-Plascencia, Jorge Salas-Hernández, Angel Gustavo Salas-Lais, Alejandro Sánchez-Flores, Clara Esperanza Santacruz-Tinoco, María Guadalupe Santiago-Mauricio, Selene Zárate, Nelly Sélem-Mojica, Blanca Taboada, Gloria Vazquez, Joel Armando Vázquez-Pérez. |
| EPI_ISL_2681178, EPI_ISL_2681179, EPI_ISL_2681180, EPI_ISL_2681181, EPI_ISL_2681182, EPI_ISL_2681183, EPI_ISL_2681184, EPI_ISL_2681185, EPI_ISL_2681186, EPI_ISL_2681187, EPI_ISL_2681188, EPI_ISL_2681189 | | | |
| see above | Laboratorio Central de Epidemiología (LCE) | Instituto de Biotecnología de la UNAM | Consorcio Mexicano de Vigilancia Genómica (CoViGen-Mex). Authors (in alphabetical order): Julio Elias Alvarado-Yaah, Carlos F. Arias, Santiago Ávila-Ríos, Eduardo Becerril-Vargas, Víctor Hugo Borja-Aburto, Celia Boukadida, Cristóbal Cháidez-Quiróz, Juan Bautista Chale-Dzul, Ricardo Ciria Merce, Andreu Comas-García, Célida Duque Molina, Julissa Enciso-Ibarra, José Antonio Enciso-Moreno, Gloria Elena Espinosa-Ayala, Fernando Fontove-Herrera, Daniel Fregoso-Rueda, Victor Eduardo García-Arias, Alejandra García-Gasca, Bruno Gómez-Gil, Jean Pierre González, Irvin González-López, Concepción Grajales-Muñiz, Ricardo Grande, Rosa María Gutiérrez Rios, Jesús Hernández, Alejandra Hernández-Terán, Alfredo Herrera-Estrella, Carla Ivón Herrera-Najera, Pavel Isa, Verónica Mata-Haro, Daniel Lira Morales, Susana Lopez, Antonio Loza Román, Brenda Irasema Maldonado-Meza, Bernardo Martínez-Miguel, José Arturo Martínez-Orozco, Célida Martínez- Rodríguez, Margarita Matías-Florentino, Fidencio Mejía-Nepomuceno, María Guadalupe de Jesús Mireles-Rivera, Gloria María Molina-Salinas, Hector Montoya-Fuentes, , Mario Mújica-Sánchez, José Esteban Muñoz-Medina, José de Jesús Nuñez-Contreras, Alicia Ocaña-Mondragón, Luis Alberto Ochoa-Carrera, Hector Esteban Paz-Juárez, Marissa Perez-Garcia, Francisco Pulido, Helen Haydee Fernanda Ramírez-Plascencia, Jorge Salas-Hernández, Angel Gustavo Salas-Lais, Alejandro Sánchez-Flores, Clara Esperanza Santacruz-Tinoco, María Guadalupe Santiago-Mauricio, Selene Zárate, Nelly Sélem-Mojica, Blanca Taboada, Gloria Vazquez, Joel Armando Vázquez-Pérez. |
| EPI_ISL_2681190 | Unidad de Investigación Biomédica de Zacatecas (UIBZ) | Instituto de Biotecnología de la UNAM | Consorcio Mexicano de Vigilancia Genómica (CoViGen-Mex). Authors (in alphabetical order): Julio Elias Alvarado-Yaah, Carlos F. Arias, Santiago Ávila-Ríos, Eduardo Becerril-Vargas, Víctor Hugo Borja-Aburto, Celia Boukadida, Cristóbal Cháidez-Quiróz, Juan Bautista Chale-Dzul, Ricardo Ciria Merce, Andreu Comas-García, Célida Duque Molina, Julissa Enciso-Ibarra, José Antonio Enciso-Moreno, Gloria Elena Espinosa-Ayala, Fernando Fontove-Herrera, Daniel Fregoso-Rueda, Victor Eduardo García-Arias, Alejandra García-Gasca, Bruno Gómez-Gil, Jean Pierre González, Irvin González-López, Concepción Grajales-Muñiz, Ricardo Grande, Rosa María Gutiérrez Rios, Jesús Hernández, Alejandra Hernández-Terán, Alfredo Herrera-Estrella, Carla Ivón Herrera-Najera, Pavel Isa, Verónica Mata-Haro, Daniel Lira Morales, Susana Lopez, Antonio Loza Román, Brenda Irasema Maldonado-Meza, Bernardo Martínez-Miguel, José Arturo Martínez-Orozco, Célida Martínez- Rodríguez, Margarita Matías-Florentino, Fidencio Mejía-Nepomuceno, María Guadalupe de Jesús Mireles-Rivera, Gloria María Molina-Salinas, Hector Montoya-Fuentes, , Mario Mújica-Sánchez, José Esteban Muñoz-Medina, José de Jesús Nuñez-Contreras, Alicia Ocaña-Mondragón, Luis Alberto Ochoa-Carrera, Hector Esteban Paz-Juárez, Marissa Perez-Garcia, Francisco Pulido, Helen Haydee Fernanda Ramírez-Plascencia, Jorge Salas-Hernández, Angel Gustavo Salas-Lais, Alejandro Sánchez-Flores, Clara Esperanza Santacruz-Tinoco, María Guadalupe Santiago-Mauricio, Selene Zárate, Nelly Sélem-Mojica, Blanca Taboada, Gloria Vazquez, Joel Armando Vázquez-Pérez. |
| EPI_ISL_2681192, EPI_ISL_2681193, EPI_ISL_2681194, EPI_ISL_2681195, EPI_ISL_2681196 | Laboratorio Central de Epidemiología (LCE) | Instituto de Biotecnología de la UNAM | Consorcio Mexicano de Vigilancia Genómica (CoViGen-Mex). Authors (in alphabetical order): Julio Elias Alvarado-Yaah, Carlos F. Arias, Santiago Ávila-Ríos, Eduardo Becerril-Vargas, Víctor Hugo Borja-Aburto, Celia Boukadida, Cristóbal Cháidez-Quiróz, Juan Bautista Chale-Dzul, Ricardo Ciria Merce, Andreu Comas-García, Célida Duque Molina, Julissa Enciso-Ibarra, José Antonio Enciso-Moreno, Gloria Elena Espinosa-Ayala, Fernando Fontove-Herrera, Daniel Fregoso-Rueda, Victor Eduardo García-Arias, Alejandra García-Gasca, Bruno Gómez-Gil, Jean Pierre González, Irvin González-López, Concepción Grajales-Muñiz, Ricardo Grande, Rosa María Gutiérrez Rios, Jesús Hernández, Alejandra Hernández-Terán, Alfredo Herrera-Estrella, Carla Ivón Herrera-Najera, Pavel Isa, Verónica Mata-Haro, Daniel Lira Morales, Susana Lopez, Antonio Loza Román, Brenda Irasema Maldonado-Meza, Bernardo Martínez-Miguel, José Arturo Martínez-Orozco, Célida Martínez- Rodríguez, Margarita Matías-Florentino, Fidencio Mejía-Nepomuceno, María Guadalupe de Jesús Mireles-Rivera, Gloria María Molina-Salinas, Hector Montoya-Fuentes, , Mario Mújica-Sánchez, José Esteban Muñoz-Medina, José de Jesús Nuñez-Contreras, Alicia Ocaña-Mondragón, Luis Alberto Ochoa-Carrera, Hector Esteban Paz-Juárez, Marissa Perez-Garcia, Francisco Pulido, Helen Haydee Fernanda Ramírez-Plascencia, Jorge Salas-Hernández, Angel Gustavo Salas-Lais, Alejandro Sánchez-Flores, Clara Esperanza Santacruz-Tinoco, María Guadalupe Santiago-Mauricio, Selene Zárate, Nelly Sélem-Mojica, Blanca Taboada, Gloria Vazquez, Joel Armando Vázquez-Pérez. |
| EPI_ISL_2681197 | Centro de Investigación Biomédica del Noreste (CIBIN) | Instituto de Biotecnología de la UNAM | Consorcio Mexicano de Vigilancia Genómica (CoViGen-Mex). Authors (in alphabetical order): Julio Elias Alvarado-Yaah, Carlos F. Arias, Santiago Ávila-Ríos, Eduardo Becerril-Vargas, Víctor Hugo Borja-Aburto, Celia Boukadida, Cristóbal Cháidez-Quiróz, Juan Bautista Chale-Dzul, Ricardo Ciria Merce, Andreu Comas-García, Célida Duque Molina, Julissa Enciso-Ibarra, José Antonio Enciso-Moreno, Gloria Elena Espinosa-Ayala, Fernando Fontove-Herrera, Daniel Fregoso-Rueda, Victor Eduardo García-Arias, Alejandra García-Gasca, Bruno Gómez-Gil, Jean Pierre González, Irvin González-López, Concepción Grajales-Muñiz, Ricardo Grande, Rosa María Gutiérrez Rios, Jesús Hernández, Alejandra Hernández-Terán, Alfredo Herrera-Estrella, Carla Ivón Herrera-Najera, Pavel Isa, Verónica Mata-Haro, Daniel Lira Morales, Susana Lopez, Antonio Loza Román, Brenda Irasema Maldonado-Meza, Bernardo Martínez-Miguel, José Arturo Martínez-Orozco, Célida Martínez- Rodríguez, Margarita Matías-Florentino, Fidencio Mejía-Nepomuceno, María Guadalupe de Jesús Mireles-Rivera, Gloria María Molina-Salinas, Hector Montoya-Fuentes, , Mario Mújica-Sánchez, José Esteban Muñoz-Medina, José de Jesús Nuñez-Contreras, Alicia Ocaña-Mondragón, Luis Alberto Ochoa-Carrera, Hector Esteban Paz-Juárez, Marissa Perez-Garcia, Francisco Pulido, Helen Haydee Fernanda Ramírez-Plascencia, Jorge Salas-Hernández, Angel Gustavo Salas-Lais, Alejandro Sánchez-Flores, Clara Esperanza Santacruz-Tinoco, María Guadalupe Santiago-Mauricio, Selene Zárate, Nelly Sélem-Mojica, Blanca Taboada, Gloria Vazquez, Joel Armando Vázquez-Pérez. |
| EPI_ISL_2681198, EPI_ISL_2681199, EPI_ISL_2681200 | Laboratorio Central de Epidemiología (LCE) | Instituto de Biotecnología de la UNAM | Consorcio Mexicano de Vigilancia Genómica (CoViGen-Mex). Authors (in alphabetical order): Julio Elias Alvarado-Yaah, Carlos F. Arias, Santiago Ávila-Ríos, Eduardo Becerril-Vargas, Víctor Hugo Borja-Aburto, Celia Boukadida, Cristóbal Cháidez-Quiróz, Juan Bautista Chale-Dzul, Ricardo Ciria Merce, Andreu Comas-García, Célida Duque Molina, Julissa Enciso-Ibarra, José Antonio Enciso-Moreno, Gloria Elena Espinosa-Ayala, Fernando Fontove-Herrera, Daniel Fregoso-Rueda, Victor Eduardo García-Arias, Alejandra García-Gasca, Bruno Gómez-Gil, Jean Pierre González, Irvin González-López, Concepción Grajales-Muñiz, Ricardo Grande, Rosa María Gutiérrez Rios, Jesús Hernández, Alejandra Hernández-Terán, Alfredo Herrera-Estrella, Carla Ivón Herrera-Najera, Pavel Isa, Verónica Mata-Haro, Daniel Lira Morales, Susana Lopez, Antonio Loza Román, Brenda Irasema Maldonado-Meza, Bernardo Martínez-Miguel, José Arturo Martínez-Orozco, Célida Martínez- Rodríguez, Margarita Matías-Florentino, Fidencio Mejía-Nepomuceno, María Guadalupe de Jesús Mireles-Rivera, Gloria María Molina-Salinas, Hector Montoya-Fuentes, , Mario Mújica-Sánchez, José Esteban Muñoz-Medina, José de Jesús Nuñez-Contreras, Alicia Ocaña-Mondragón, Luis Alberto Ochoa-Carrera, Hector Esteban Paz-Juárez, Marissa Perez-Garcia, Francisco Pulido, Helen Haydee Fernanda Ramírez-Plascencia, Jorge Salas-Hernández, Angel Gustavo Salas-Lais, Alejandro Sánchez-Flores, Clara Esperanza Santacruz-Tinoco, María Guadalupe Santiago-Mauricio, Selene Zárate, Nelly Sélem-Mojica, Blanca Taboada, Gloria Vazquez, Joel Armando Vázquez-Pérez. |
| EPI_ISL_2681201 | Centro de Investigación Biomédica del Noreste (CIBIN) | Instituto de Biotecnología de la UNAM | Consorcio Mexicano de Vigilancia Genómica (CoViGen-Mex). Authors (in alphabetical order): Julio Elias Alvarado-Yaah, Carlos F. Arias, Santiago Ávila-Ríos, Eduardo Becerril-Vargas, Víctor Hugo Borja-Aburto, Celia Boukadida, Cristóbal Cháidez-Quiróz, Juan Bautista Chale-Dzul, Ricardo Ciria Merce, Andreu Comas-García, Célida Duque Molina, Julissa Enciso-Ibarra, José Antonio Enciso-Moreno, Gloria Elena Espinosa-Ayala, Fernando Fontove-Herrera, Daniel Fregoso-Rueda, Victor Eduardo García-Arias, Alejandra García-Gasca, Bruno Gómez-Gil, Jean Pierre González, Irvin González-López, Concepción Grajales-Muñiz, Ricardo Grande, Rosa María Gutiérrez Rios, Jesús Hernández, Alejandra Hernández-Terán, Alfredo Herrera-Estrella, Carla Ivón Herrera-Najera, Pavel Isa, Verónica Mata-Haro, Daniel Lira Morales, Susana Lopez, Antonio Loza Román, Brenda Irasema Maldonado-Meza, Bernardo Martínez-Miguel, José Arturo Martínez-Orozco, Célida Martínez- Rodríguez, Margarita Matías-Florentino, Fidencio Mejía-Nepomuceno, María Guadalupe de Jesús Mireles-Rivera, Gloria María Molina-Salinas, Hector Montoya-Fuentes, , Mario Mújica-Sánchez, José Esteban Muñoz-Medina, José de Jesús Nuñez-Contreras, Alicia Ocaña-Mondragón, Luis Alberto Ochoa-Carrera, Hector Esteban Paz-Juárez, Marissa Perez-Garcia, Francisco Pulido, Helen Haydee Fernanda Ramírez-Plascencia, Jorge Salas-Hernández, Angel Gustavo Salas-Lais, Alejandro Sánchez-Flores, Clara Esperanza Santacruz-Tinoco, María Guadalupe Santiago-Mauricio, Selene Zárate, Nelly Sélem-Mojica, Blanca Taboada, Gloria Vazquez, Joel Armando Vázquez-Pérez. |

| | | | |
|---|---|---|---|
| EPI_ISL_2681202, EPI_ISL_2681204, EPI_ISL_2681205, EPI_ISL_2681206 | Laboratorio Central de Epidemiología (LCE) | Instituto de Biotecnología de la UNAM | Consorcio Mexicano de Vigilancia Genómica (CoViGen-Mex). Authors (in alphabetical order): Julio Elias Alvarado-Yaah, Carlos F. Arias, Santiago Ávila-Ríos, Eduardo Becerril-Vargas, Víctor Hugo Borja-Aburto, Celia Boukadida, Cristóbal Cháidez-Quiróz, Juan Bautista Chale-Dzul, Ricardo Ciria Merce, Andreu Comas-García, Célida Duque Molina, Julissa Enciso-Ibarra, José Antonio Enciso-Moreno, Gloria Elena Espinosa-Ayala, Fernando Fontove-Herrera, Daniel Fregoso-Rueda, Victor Eduardo García-Arias, Alejandra García-Gasca, Bruno Gómez-Gil, Jean Pierre González, Irvin González-López, Concepción Grajales-Muñiz, Ricardo Grande, Rosa María Gutiérrez Rios, Jesús Hernández, Alejandra Hernández-Terán, Alfredo Herrera-Estrella, Carla Ivón Herrera-Najera, Pavel Isa, Verónica Mata-Haro, Daniel Lira Morales, Susana Lopez, Antonio Loza Román, Brenda Irasema Maldonado-Meza, Bernardo Martínez-Miguel, José Arturo Martínez-Orozco, Célida Martínez- Rodríguez, Margarita Matías-Florentino, Fidencio Mejía-Nepomuceno, María Guadalupe de Jesús Mireles-Rivera, Gloria María Molina-Salinas, Hector Montoya-Fuentes, , Mario Mújica-Sánchez, José Esteban Muñoz-Medina, José de Jesús Nuñez-Contreras, Alicia Ocaña-Mondragón, Luis Alberto Ochoa-Carrera, Hector Esteban Paz-Juárez, Marissa Perez-Garcia, Francisco Pulido, Helen Haydee Fernanda Ramírez-Plascencia, Jorge Salas-Hernández, Angel Gustavo Salas-Lais, Alejandro Sánchez-Flores, Clara Esperanza Santacruz-Tinoco, María Guadalupe Santiago-Mauricio, Selene Zárate, Nelly Sélem-Mojica, Blanca Taboada, Gloria Vazquez, Joel Armando Vázquez-Pérez. |
| EPI_ISL_2681207 | Unidad de Investigación Biomédica de Zacatecas (UIBZ) | Instituto de Biotecnología de la UNAM | Consorcio Mexicano de Vigilancia Genómica (CoViGen-Mex). Authors (in alphabetical order): Julio Elias Alvarado-Yaah, Carlos F. Arias, Santiago Ávila-Ríos, Eduardo Becerril-Vargas, Víctor Hugo Borja-Aburto, Celia Boukadida, Cristóbal Cháidez-Quiróz, Juan Bautista Chale-Dzul, Ricardo Ciria Merce, Andreu Comas-García, Célida Duque Molina, Julissa Enciso-Ibarra, José Antonio Enciso-Moreno, Gloria Elena Espinosa-Ayala, Fernando Fontove-Herrera, Daniel Fregoso-Rueda, Victor Eduardo García-Arias, Alejandra García-Gasca, Bruno Gómez-Gil, Jean Pierre González, Irvin González-López, Concepción Grajales-Muñiz, Ricardo Grande, Rosa María Gutiérrez Rios, Jesús Hernández, Alejandra Hernández-Terán, Alfredo Herrera-Estrella, Carla Ivón Herrera-Najera, Pavel Isa, Verónica Mata-Haro, Daniel Lira Morales, Susana Lopez, Antonio Loza Román, Brenda Irasema Maldonado-Meza, Bernardo Martínez-Miguel, José Arturo Martínez-Orozco, Célida Martínez- Rodríguez, Margarita Matías-Florentino, Fidencio Mejía-Nepomuceno, María Guadalupe de Jesús Mireles-Rivera, Gloria María Molina-Salinas, Hector Montoya-Fuentes, , Mario Mújica-Sánchez, José Esteban Muñoz-Medina, José de Jesús Nuñez-Contreras, Alicia Ocaña-Mondragón, Luis Alberto Ochoa-Carrera, Hector Esteban Paz-Juárez, Marissa Perez-Garcia, Francisco Pulido, Helen Haydee Fernanda Ramírez-Plascencia, Jorge Salas-Hernández, Angel Gustavo Salas-Lais, Alejandro Sánchez-Flores, Clara Esperanza Santacruz-Tinoco, María Guadalupe Santiago-Mauricio, Selene Zárate, Nelly Sélem-Mojica, Blanca Taboada, Gloria Vazquez, Joel Armando Vázquez-Pérez. |
| EPI_ISL_2681208 | Laboratorio Central de Epidemiología (LCE) | Instituto de Biotecnología de la UNAM | Consorcio Mexicano de Vigilancia Genómica (CoViGen-Mex). Authors (in alphabetical order): Julio Elias Alvarado-Yaah, Carlos F. Arias, Santiago Ávila-Ríos, Eduardo Becerril-Vargas, Víctor Hugo Borja-Aburto, Celia Boukadida, Cristóbal Cháidez-Quiróz, Juan Bautista Chale-Dzul, Ricardo Ciria Merce, Andreu Comas-García, Célida Duque Molina, Julissa Enciso-Ibarra, José Antonio Enciso-Moreno, Gloria Elena Espinosa-Ayala, Fernando Fontove-Herrera, Daniel Fregoso-Rueda, Victor Eduardo García-Arias, Alejandra García-Gasca, Bruno Gómez-Gil, Jean Pierre González, Irvin González-López, Concepción Grajales-Muñiz, Ricardo Grande, Rosa María Gutiérrez Rios, Jesús Hernández, Alejandra Hernández-Terán, Alfredo Herrera-Estrella, Carla Ivón Herrera-Najera, Pavel Isa, Verónica Mata-Haro, Daniel Lira Morales, Susana Lopez, Antonio Loza Román, Brenda Irasema Maldonado-Meza, Bernardo Martínez-Miguel, José Arturo Martínez-Orozco, Célida Martínez- Rodríguez, Margarita Matías-Florentino, Fidencio Mejía-Nepomuceno, María Guadalupe de Jesús Mireles-Rivera, Gloria María Molina-Salinas, Hector Montoya-Fuentes, , Mario Mújica-Sánchez, José Esteban Muñoz-Medina, José de Jesús Nuñez-Contreras, Alicia Ocaña-Mondragón, Luis Alberto Ochoa-Carrera, Hector Esteban Paz-Juárez, Marissa Perez-Garcia, Francisco Pulido, Helen Haydee Fernanda Ramírez-Plascencia, Jorge Salas-Hernández, Angel Gustavo Salas-Lais, Alejandro Sánchez-Flores, Clara Esperanza Santacruz-Tinoco, María Guadalupe Santiago-Mauricio, Selene Zárate, Nelly Sélem-Mojica, Blanca Taboada, Gloria Vazquez, Joel Armando Vázquez-Pérez. |
| EPI_ISL_2681209 | Unidad de Investigación Biomédica de Zacatecas (UIBZ) | Instituto de Biotecnología de la UNAM | Consorcio Mexicano de Vigilancia Genómica (CoViGen-Mex). Authors (in alphabetical order): Julio Elias Alvarado-Yaah, Carlos F. Arias, Santiago Ávila-Ríos, Eduardo Becerril-Vargas, Víctor Hugo Borja-Aburto, Celia Boukadida, Cristóbal Cháidez-Quiróz, Juan Bautista Chale-Dzul, Ricardo Ciria Merce, Andreu Comas-García, Célida Duque Molina, Julissa Enciso-Ibarra, José Antonio Enciso-Moreno, Gloria Elena Espinosa-Ayala, Fernando Fontove-Herrera, Daniel Fregoso-Rueda, Victor Eduardo García-Arias, Alejandra García-Gasca, Bruno Gómez-Gil, Jean Pierre González, Irvin González-López, Concepción Grajales-Muñiz, Ricardo Grande, Rosa María Gutiérrez Rios, Jesús Hernández, Alejandra Hernández-Terán, Alfredo Herrera-Estrella, Carla Ivón Herrera-Najera, Pavel Isa, Verónica Mata-Haro, Daniel Lira Morales, Susana Lopez, Antonio Loza Román, Brenda Irasema Maldonado-Meza, Bernardo Martínez-Miguel, José Arturo Martínez-Orozco, Célida Martínez- Rodríguez, Margarita Matías-Florentino, Fidencio Mejía-Nepomuceno, María Guadalupe de Jesús Mireles-Rivera, Gloria María Molina-Salinas, Hector Montoya-Fuentes, , Mario Mújica-Sánchez, José Esteban Muñoz-Medina, José de Jesús Nuñez-Contreras, Alicia Ocaña-Mondragón, Luis Alberto Ochoa-Carrera, Hector Esteban Paz-Juárez, Marissa Perez-Garcia, Francisco Pulido, Helen Haydee Fernanda Ramírez-Plascencia, Jorge Salas-Hernández, Angel Gustavo Salas-Lais, Alejandro Sánchez-Flores, Clara Esperanza Santacruz-Tinoco, María Guadalupe Santiago-Mauricio, Selene Zárate, Nelly Sélem-Mojica, Blanca Taboada, Gloria Vazquez, Joel Armando Vázquez-Pérez. |
| EPI_ISL_2681210 | Laboratorio Central de Epidemiología (LCE) | Instituto de Biotecnología de la UNAM | Consorcio Mexicano de Vigilancia Genómica (CoViGen-Mex). Authors (in alphabetical order): Julio Elias Alvarado-Yaah, Carlos F. Arias, Santiago Ávila-Ríos, Eduardo Becerril-Vargas, Víctor Hugo Borja-Aburto, Celia Boukadida, Cristóbal Cháidez-Quiróz, Juan Bautista Chale-Dzul, Ricardo Ciria Merce, Andreu Comas-García, Célida Duque Molina, Julissa Enciso-Ibarra, José Antonio Enciso-Moreno, Gloria Elena Espinosa-Ayala, Fernando Fontove-Herrera, Daniel Fregoso-Rueda, Victor Eduardo García-Arias, Alejandra García-Gasca, Bruno Gómez-Gil, Jean Pierre González, Irvin González-López, Concepción Grajales-Muñiz, Ricardo Grande, Rosa María Gutiérrez Rios, Jesús Hernández, Alejandra Hernández-Terán, Alfredo Herrera-Estrella, Carla Ivón Herrera-Najera, Pavel Isa, Verónica Mata-Haro, Daniel Lira Morales, Susana Lopez, Antonio Loza Román, Brenda Irasema Maldonado-Meza, Bernardo Martínez-Miguel, José Arturo Martínez-Orozco, Célida Martínez- Rodríguez, Margarita Matías-Florentino, Fidencio Mejía-Nepomuceno, María Guadalupe de Jesús Mireles-Rivera, Gloria María Molina-Salinas, Hector Montoya-Fuentes, , Mario Mújica-Sánchez, José Esteban Muñoz-Medina, José de Jesús Nuñez-Contreras, Alicia Ocaña-Mondragón, Luis Alberto Ochoa-Carrera, Hector Esteban Paz-Juárez, Marissa Perez-Garcia, Francisco Pulido, Helen Haydee Fernanda Ramírez-Plascencia, Jorge Salas-Hernández, Angel Gustavo Salas-Lais, Alejandro Sánchez-Flores, Clara Esperanza Santacruz-Tinoco, María Guadalupe Santiago-Mauricio, Selene Zárate, Nelly Sélem-Mojica, Blanca Taboada, Gloria Vazquez, Joel Armando Vázquez-Pérez. |
| EPI_ISL_2681211 | Unidad de Investigación Médica de Yucatán (UIMY) | Instituto de Biotecnología de la UNAM | Consorcio Mexicano de Vigilancia Genómica (CoViGen-Mex). Authors (in alphabetical order): Julio Elias Alvarado-Yaah, Carlos F. Arias, Santiago Ávila-Ríos, Eduardo Becerril-Vargas, Víctor Hugo Borja-Aburto, Celia Boukadida, Cristóbal Cháidez-Quiróz, Juan Bautista Chale-Dzul, Ricardo Ciria Merce, Andreu Comas-García, Célida Duque Molina, Julissa Enciso-Ibarra, José Antonio Enciso-Moreno, Gloria Elena Espinosa-Ayala, Fernando Fontove-Herrera, Daniel Fregoso-Rueda, Victor Eduardo García-Arias, Alejandra García-Gasca, Bruno Gómez-Gil, Jean Pierre González, Irvin González-López, Concepción Grajales-Muñiz, Ricardo Grande, Rosa María Gutiérrez Rios, Jesús Hernández, Alejandra Hernández-Terán, Alfredo Herrera-Estrella, Carla Ivón Herrera-Najera, Pavel Isa, Verónica Mata-Haro, Daniel Lira Morales, Susana Lopez, Antonio Loza Román, Brenda Irasema Maldonado-Meza, Bernardo Martínez-Miguel, José Arturo Martínez-Orozco, Célida Martínez- Rodríguez, Margarita Matías-Florentino, Fidencio Mejía-Nepomuceno, María Guadalupe de Jesús Mireles-Rivera, Gloria María Molina-Salinas, Hector Montoya-Fuentes, , Mario Mújica-Sánchez, José Esteban Muñoz-Medina, José de Jesús Nuñez-Contreras, Alicia Ocaña-Mondragón, Luis Alberto Ochoa-Carrera, Hector Esteban Paz-Juárez, Marissa Perez-Garcia, Francisco Pulido, Helen Haydee Fernanda Ramírez-Plascencia, Jorge Salas-Hernández, Angel Gustavo Salas-Lais, Alejandro Sánchez-Flores, Clara Esperanza Santacruz-Tinoco, María Guadalupe Santiago-Mauricio, Selene Zárate, Nelly Sélem-Mojica, Blanca Taboada, Gloria Vazquez, Joel Armando Vázquez-Pérez. |
| EPI_ISL_2681212 | Unidad de Investigación Biomédica de Zacatecas (UIBZ) | Instituto de Biotecnología de la UNAM | Consorcio Mexicano de Vigilancia Genómica (CoViGen-Mex). Authors (in alphabetical order): Julio Elias Alvarado-Yaah, Carlos F. Arias, Santiago Ávila-Ríos, Eduardo Becerril-Vargas, Víctor Hugo Borja-Aburto, Celia Boukadida, Cristóbal Cháidez-Quiróz, Juan Bautista Chale-Dzul, Ricardo Ciria Merce, Andreu Comas-García, Célida Duque Molina, Julissa Enciso-Ibarra, José Antonio Enciso-Moreno, Gloria Elena Espinosa-Ayala, Fernando Fontove-Herrera, Daniel Fregoso-Rueda, Victor Eduardo García-Arias, Alejandra García-Gasca, Bruno Gómez-Gil, Jean Pierre González, Irvin González-López, Concepción Grajales-Muñiz, Ricardo Grande, Rosa María Gutiérrez Rios, Jesús Hernández, Alejandra Hernández-Terán, Alfredo Herrera-Estrella, Carla Ivón Herrera-Najera, Pavel Isa, Verónica Mata-Haro, Daniel Lira Morales, Susana Lopez, Antonio Loza Román, Brenda Irasema Maldonado-Meza, Bernardo Martínez-Miguel, José Arturo Martínez-Orozco, Célida Martínez- Rodríguez, Margarita Matías-Florentino, Fidencio Mejía-Nepomuceno, María Guadalupe de Jesús Mireles-Rivera, Gloria María Molina-Salinas, Hector Montoya-Fuentes, , Mario Mújica-Sánchez, José Esteban Muñoz-Medina, José de Jesús Nuñez-Contreras, Alicia Ocaña-Mondragón, Luis Alberto Ochoa-Carrera, Hector Esteban Paz-Juárez, Marissa Perez-Garcia, Francisco Pulido, Helen Haydee Fernanda Ramírez-Plascencia, Jorge Salas-Hernández, Angel Gustavo Salas-Lais, Alejandro Sánchez-Flores, Clara Esperanza Santacruz-Tinoco, María Guadalupe Santiago-Mauricio, Selene Zárate, Nelly Sélem-Mojica, Blanca Taboada, Gloria Vazquez, Joel Armando Vázquez-Pérez. |

| | | | |
|---|---|---|---|
| EPI_ISL_2681213, EPI_ISL_2681214, EPI_ISL_2681216, EPI_ISL_2681217 | Laboratorio Central de Epidemiología (LCE) | Instituto de Biotecnología de la UNAM | Consorcio Mexicano de Vigilancia Genómica (CoViGen-Mex). Authors (in alphabetical order): Julio Elias Alvarado-Yaah, Carlos F. Arias, Santiago Ávila-Ríos, Eduardo Becerril-Vargas, Víctor Hugo Borja-Aburto, Celia Boukadida, Cristóbal Cháidez-Quiróz, Juan Bautista Chale-Dzul, Ricardo Ciria Merce, Andreu Comas-García, Célida Duque Molina, Julissa Enciso-Ibarra, José Antonio Enciso-Moreno, Gloria Elena Espinosa-Ayala, Fernando Fontove-Herrera, Daniel Fregoso-Rueda, Victor Eduardo García-Arias, Alejandra García-Gasca, Bruno Gómez-Gil, Jean Pierre González, Irvin González-López, Concepción Grajales-Muñiz, Ricardo Grande, Rosa María Gutiérrez Rios, Jesús Hernández, Alejandra Hernández-Terán, Alfredo Herrera-Estrella, Carla Ivón Herrera-Najera, Pavel Isa, Verónica Mata-Haro, Daniel Lira Morales, Susana Lopez, Antonio Loza Román, Brenda Irasema Maldonado-Meza, Bernardo Martínez-Miguel, José Arturo Martínez-Orozco, Célida Martínez- Rodríguez, Margarita Matías-Florentino, Fidencio Mejía-Nepomuceno, María Guadalupe de Jesús Mireles-Rivera, Gloria María Molina-Salinas, Hector Montoya-Fuentes, , Mario Mújica-Sánchez, José Esteban Muñoz-Medina, José de Jesús Nuñez-Contreras, Alicia Ocaña-Mondragón, Luis Alberto Ochoa-Carrera, Hector Esteban Paz-Juárez, Marissa Perez-Garcia, Francisco Pulido, Helen Haydee Fernanda Ramírez-Plascencia, Jorge Salas-Hernández, Angel Gustavo Salas-Lais, Alejandro Sánchez-Flores, Clara Esperanza Santacruz-Tinoco, María Guadalupe Santiago-Mauricio, Selene Zárate, Nelly Sélem-Mojica, Blanca Taboada, Gloria Vazquez, Joel Armando Vázquez-Pérez. |
| EPI_ISL_2681218 | Unidad de Investigación Médica de Yucatán (UIMY) | Instituto de Biotecnología de la UNAM | Consorcio Mexicano de Vigilancia Genómica (CoViGen-Mex). Authors (in alphabetical order): Julio Elias Alvarado-Yaah, Carlos F. Arias, Santiago Ávila-Ríos, Eduardo Becerril-Vargas, Víctor Hugo Borja-Aburto, Celia Boukadida, Cristóbal Cháidez-Quiróz, Juan Bautista Chale-Dzul, Ricardo Ciria Merce, Andreu Comas-García, Célida Duque Molina, Julissa Enciso-Ibarra, José Antonio Enciso-Moreno, Gloria Elena Espinosa-Ayala, Fernando Fontove-Herrera, Daniel Fregoso-Rueda, Victor Eduardo García-Arias, Alejandra García-Gasca, Bruno Gómez-Gil, Jean Pierre González, Irvin González-López, Concepción Grajales-Muñiz, Ricardo Grande, Rosa María Gutiérrez Rios, Jesús Hernández, Alejandra Hernández-Terán, Alfredo Herrera-Estrella, Carla Ivón Herrera-Najera, Pavel Isa, Verónica Mata-Haro, Daniel Lira Morales, Susana Lopez, Antonio Loza Román, Brenda Irasema Maldonado-Meza, Bernardo Martínez-Miguel, José Arturo Martínez-Orozco, Célida Martínez- Rodríguez, Margarita Matías-Florentino, Fidencio Mejía-Nepomuceno, María Guadalupe de Jesús Mireles-Rivera, Gloria María Molina-Salinas, Hector Montoya-Fuentes, , Mario Mújica-Sánchez, José Esteban Muñoz-Medina, José de Jesús Nuñez-Contreras, Alicia Ocaña-Mondragón, Luis Alberto Ochoa-Carrera, Hector Esteban Paz-Juárez, Marissa Perez-Garcia, Francisco Pulido, Helen Haydee Fernanda Ramírez-Plascencia, Jorge Salas-Hernández, Angel Gustavo Salas-Lais, Alejandro Sánchez-Flores, Clara Esperanza Santacruz-Tinoco, María Guadalupe Santiago-Mauricio, Selene Zárate, Nelly Sélem-Mojica, Blanca Taboada, Gloria Vazquez, Joel Armando Vázquez-Pérez. |
| EPI_ISL_2681219 | Centro de Investigación Biomédica del Noreste (CIBIN) | Instituto de Biotecnología de la UNAM | Consorcio Mexicano de Vigilancia Genómica (CoViGen-Mex). Authors (in alphabetical order): Julio Elias Alvarado-Yaah, Carlos F. Arias, Santiago Ávila-Ríos, Eduardo Becerril-Vargas, Víctor Hugo Borja-Aburto, Celia Boukadida, Cristóbal Cháidez-Quiróz, Juan Bautista Chale-Dzul, Ricardo Ciria Merce, Andreu Comas-García, Célida Duque Molina, Julissa Enciso-Ibarra, José Antonio Enciso-Moreno, Gloria Elena Espinosa-Ayala, Fernando Fontove-Herrera, Daniel Fregoso-Rueda, Victor Eduardo García-Arias, Alejandra García-Gasca, Bruno Gómez-Gil, Jean Pierre González, Irvin González-López, Concepción Grajales-Muñiz, Ricardo Grande, Rosa María Gutiérrez Rios, Jesús Hernández, Alejandra Hernández-Terán, Alfredo Herrera-Estrella, Carla Ivón Herrera-Najera, Pavel Isa, Verónica Mata-Haro, Daniel Lira Morales, Susana Lopez, Antonio Loza Román, Brenda Irasema Maldonado-Meza, Bernardo Martínez-Miguel, José Arturo Martínez-Orozco, Célida Martínez- Rodríguez, Margarita Matías-Florentino, Fidencio Mejía-Nepomuceno, María Guadalupe de Jesús Mireles-Rivera, Gloria María Molina-Salinas, Hector Montoya-Fuentes, , Mario Mújica-Sánchez, José Esteban Muñoz-Medina, José de Jesús Nuñez-Contreras, Alicia Ocaña-Mondragón, Luis Alberto Ochoa-Carrera, Hector Esteban Paz-Juárez, Marissa Perez-Garcia, Francisco Pulido, Helen Haydee Fernanda Ramírez-Plascencia, Jorge Salas-Hernández, Angel Gustavo Salas-Lais, Alejandro Sánchez-Flores, Clara Esperanza Santacruz-Tinoco, María Guadalupe Santiago-Mauricio, Selene Zárate, Nelly Sélem-Mojica, Blanca Taboada, Gloria Vazquez, Joel Armando Vázquez-Pérez. |
| EPI_ISL_2681220 | Laboratorio Central de Epidemiología (LCE) | Instituto de Biotecnología de la UNAM | Consorcio Mexicano de Vigilancia Genómica (CoViGen-Mex). Authors (in alphabetical order): Julio Elias Alvarado-Yaah, Carlos F. Arias, Santiago Ávila-Ríos, Eduardo Becerril-Vargas, Víctor Hugo Borja-Aburto, Celia Boukadida, Cristóbal Cháidez-Quiróz, Juan Bautista Chale-Dzul, Ricardo Ciria Merce, Andreu Comas-García, Célida Duque Molina, Julissa Enciso-Ibarra, José Antonio Enciso-Moreno, Gloria Elena Espinosa-Ayala, Fernando Fontove-Herrera, Daniel Fregoso-Rueda, Victor Eduardo García-Arias, Alejandra García-Gasca, Bruno Gómez-Gil, Jean Pierre González, Irvin González-López, Concepción Grajales-Muñiz, Ricardo Grande, Rosa María Gutiérrez Rios, Jesús Hernández, Alejandra Hernández-Terán, Alfredo Herrera-Estrella, Carla Ivón Herrera-Najera, Pavel Isa, Verónica Mata-Haro, Daniel Lira Morales, Susana Lopez, Antonio Loza Román, Brenda Irasema Maldonado-Meza, Bernardo Martínez-Miguel, José Arturo Martínez-Orozco, Célida Martínez- Rodríguez, Margarita Matías-Florentino, Fidencio Mejía-Nepomuceno, María Guadalupe de Jesús Mireles-Rivera, Gloria María Molina-Salinas, Hector Montoya-Fuentes, , Mario Mújica-Sánchez, José Esteban Muñoz-Medina, José de Jesús Nuñez-Contreras, Alicia Ocaña-Mondragón, Luis Alberto Ochoa-Carrera, Hector Esteban Paz-Juárez, Marissa Perez-Garcia, Francisco Pulido, Helen Haydee Fernanda Ramírez-Plascencia, Jorge Salas-Hernández, Angel Gustavo Salas-Lais, Alejandro Sánchez-Flores, Clara Esperanza Santacruz-Tinoco, María Guadalupe Santiago-Mauricio, Selene Zárate, Nelly Sélem-Mojica, Blanca Taboada, Gloria Vazquez, Joel Armando Vázquez-Pérez. |
| EPI_ISL_2681221 | Unidad de Investigación Biomédica de Zacatecas (UIBZ) | Instituto de Biotecnología de la UNAM | Consorcio Mexicano de Vigilancia Genómica (CoViGen-Mex). Authors (in alphabetical order): Julio Elias Alvarado-Yaah, Carlos F. Arias, Santiago Ávila-Ríos, Eduardo Becerril-Vargas, Víctor Hugo Borja-Aburto, Celia Boukadida, Cristóbal Cháidez-Quiróz, Juan Bautista Chale-Dzul, Ricardo Ciria Merce, Andreu Comas-García, Célida Duque Molina, Julissa Enciso-Ibarra, José Antonio Enciso-Moreno, Gloria Elena Espinosa-Ayala, Fernando Fontove-Herrera, Daniel Fregoso-Rueda, Victor Eduardo García-Arias, Alejandra García-Gasca, Bruno Gómez-Gil, Jean Pierre González, Irvin González-López, Concepción Grajales-Muñiz, Ricardo Grande, Rosa María Gutiérrez Rios, Jesús Hernández, Alejandra Hernández-Terán, Alfredo Herrera-Estrella, Carla Ivón Herrera-Najera, Pavel Isa, Verónica Mata-Haro, Daniel Lira Morales, Susana Lopez, Antonio Loza Román, Brenda Irasema Maldonado-Meza, Bernardo Martínez-Miguel, José Arturo Martínez-Orozco, Célida Martínez- Rodríguez, Margarita Matías-Florentino, Fidencio Mejía-Nepomuceno, María Guadalupe de Jesús Mireles-Rivera, Gloria María Molina-Salinas, Hector Montoya-Fuentes, , Mario Mújica-Sánchez, José Esteban Muñoz-Medina, José de Jesús Nuñez-Contreras, Alicia Ocaña-Mondragón, Luis Alberto Ochoa-Carrera, Hector Esteban Paz-Juárez, Marissa Perez-Garcia, Francisco Pulido, Helen Haydee Fernanda Ramírez-Plascencia, Jorge Salas-Hernández, Angel Gustavo Salas-Lais, Alejandro Sánchez-Flores, Clara Esperanza Santacruz-Tinoco, María Guadalupe Santiago-Mauricio, Selene Zárate, Nelly Sélem-Mojica, Blanca Taboada, Gloria Vazquez, Joel Armando Vázquez-Pérez. |
| EPI_ISL_2681222 | Laboratorio Central de Epidemiología (LCE) | Instituto de Biotecnología de la UNAM | Consorcio Mexicano de Vigilancia Genómica (CoViGen-Mex). Authors (in alphabetical order): Julio Elias Alvarado-Yaah, Carlos F. Arias, Santiago Ávila-Ríos, Eduardo Becerril-Vargas, Víctor Hugo Borja-Aburto, Celia Boukadida, Cristóbal Cháidez-Quiróz, Juan Bautista Chale-Dzul, Ricardo Ciria Merce, Andreu Comas-García, Célida Duque Molina, Julissa Enciso-Ibarra, José Antonio Enciso-Moreno, Gloria Elena Espinosa-Ayala, Fernando Fontove-Herrera, Daniel Fregoso-Rueda, Victor Eduardo García-Arias, Alejandra García-Gasca, Bruno Gómez-Gil, Jean Pierre González, Irvin González-López, Concepción Grajales-Muñiz, Ricardo Grande, Rosa María Gutiérrez Rios, Jesús Hernández, Alejandra Hernández-Terán, Alfredo Herrera-Estrella, Carla Ivón Herrera-Najera, Pavel Isa, Verónica Mata-Haro, Daniel Lira Morales, Susana Lopez, Antonio Loza Román, Brenda Irasema Maldonado-Meza, Bernardo Martínez-Miguel, José Arturo Martínez-Orozco, Célida Martínez- Rodríguez, Margarita Matías-Florentino, Fidencio Mejía-Nepomuceno, María Guadalupe de Jesús Mireles-Rivera, Gloria María Molina-Salinas, Hector Montoya-Fuentes, , Mario Mújica-Sánchez, José Esteban Muñoz-Medina, José de Jesús Nuñez-Contreras, Alicia Ocaña-Mondragón, Luis Alberto Ochoa-Carrera, Hector Esteban Paz-Juárez, Marissa Perez-Garcia, Francisco Pulido, Helen Haydee Fernanda Ramírez-Plascencia, Jorge Salas-Hernández, Angel Gustavo Salas-Lais, Alejandro Sánchez-Flores, Clara Esperanza Santacruz-Tinoco, María Guadalupe Santiago-Mauricio, Selene Zárate, Nelly Sélem-Mojica, Blanca Taboada, Gloria Vazquez, Joel Armando Vázquez-Pérez. |
| EPI_ISL_2681223 | Unidad de Investigación Biomédica de Zacatecas (UIBZ) | Instituto de Biotecnología de la UNAM | Consorcio Mexicano de Vigilancia Genómica (CoViGen-Mex). Authors (in alphabetical order): Julio Elias Alvarado-Yaah, Carlos F. Arias, Santiago Ávila-Ríos, Eduardo Becerril-Vargas, Víctor Hugo Borja-Aburto, Celia Boukadida, Cristóbal Cháidez-Quiróz, Juan Bautista Chale-Dzul, Ricardo Ciria Merce, Andreu Comas-García, Célida Duque Molina, Julissa Enciso-Ibarra, José Antonio Enciso-Moreno, Gloria Elena Espinosa-Ayala, Fernando Fontove-Herrera, Daniel Fregoso-Rueda, Victor Eduardo García-Arias, Alejandra García-Gasca, Bruno Gómez-Gil, Jean Pierre González, Irvin González-López, Concepción Grajales-Muñiz, Ricardo Grande, Rosa María Gutiérrez Rios, Jesús Hernández, Alejandra Hernández-Terán, Alfredo Herrera-Estrella, Carla Ivón Herrera-Najera, Pavel Isa, Verónica Mata-Haro, Daniel Lira Morales, Susana Lopez, Antonio Loza Román, Brenda Irasema Maldonado-Meza, Bernardo Martínez-Miguel, José Arturo Martínez-Orozco, Célida Martínez- Rodríguez, Margarita Matías-Florentino, Fidencio Mejía-Nepomuceno, María Guadalupe de Jesús Mireles-Rivera, Gloria María Molina-Salinas, Hector Montoya-Fuentes, , Mario Mújica-Sánchez, José Esteban Muñoz-Medina, José de Jesús Nuñez-Contreras, Alicia Ocaña-Mondragón, Luis Alberto Ochoa-Carrera, Hector Esteban Paz-Juárez, Marissa Perez-Garcia, Francisco Pulido, Helen Haydee Fernanda Ramírez-Plascencia, Jorge Salas-Hernández, Angel Gustavo Salas-Lais, Alejandro Sánchez-Flores, Clara Esperanza Santacruz-Tinoco, María Guadalupe Santiago-Mauricio, Selene Zárate, Nelly Sélem-Mojica, Blanca Taboada, Gloria Vazquez, Joel Armando Vázquez-Pérez. |

| | | | |
|---|---|---|---|
| EPI_ISL_2681225, EPI_ISL_2681226 | Laboratorio Central de Epidemiología (LCE) | Instituto de Biotecnología de la UNAM | Consorcio Mexicano de Vigilancia Genómica (CoViGen-Mex). Authors (in alphabetical order): Julio Elias Alvarado-Yaah, Carlos F. Arias, Santiago Ávila-Ríos, Eduardo Becerril-Vargas, Víctor Hugo Borja-Aburto, Celia Boukadida, Cristóbal Cháidez-Quiróz, Juan Bautista Chale-Dzul, Ricardo Ciria Merce, Andreu Comas-García, Célida Duque Molina, Julissa Enciso-Ibarra, José Antonio Enciso-Moreno, Gloria Elena Espinosa-Ayala, Fernando Fontove-Herrera, Daniel Fregoso-Rueda, Victor Eduardo García-Arias, Alejandra García-Gasca, Bruno Gómez-Gil, Jean Pierre González, Irvin González-López, Concepción Grajales-Muñiz, Ricardo Grande, Rosa María Gutiérrez Rios, Jesús Hernández, Alejandra Hernández-Terán, Alfredo Herrera-Estrella, Carla Ivón Herrera-Najera, Pavel Isa, Verónica Mata-Haro, Daniel Lira Morales, Susana Lopez, Antonio Loza Román, Brenda Irasema Maldonado-Meza, Bernardo Martínez-Miguel, José Arturo Martínez-Orozco, Célida Martínez- Rodríguez, Margarita Matías-Florentino, Fidencio Mejía-Nepomuceno, María Guadalupe de Jesús Mireles-Rivera, Gloria María Molina-Salinas, Hector Montoya-Fuentes, , Mario Mújica-Sánchez, José Esteban Muñoz-Medina, José de Jesús Nuñez-Contreras, Alicia Ocaña-Mondragón, Luis Alberto Ochoa-Carrera, Hector Esteban Paz-Juárez, Marissa Perez-Garcia, Francisco Pulido, Helen Haydee Fernanda Ramírez-Plascencia, Jorge Salas-Hernández, Angel Gustavo Salas-Lais, Alejandro Sánchez-Flores, Clara Esperanza Santacruz-Tinoco, María Guadalupe Santiago-Mauricio, Selene Zárate, Nelly Sélem-Mojica, Blanca Taboada, Gloria Vazquez, Joel Armando Vázquez-Pérez. |
| EPI_ISL_2681227, EPI_ISL_2681228 | Centro de Investigación Biomédica del Noreste (CIBIN) | Instituto de Biotecnología de la UNAM | Consorcio Mexicano de Vigilancia Genómica (CoViGen-Mex). Authors (in alphabetical order): Julio Elias Alvarado-Yaah, Carlos F. Arias, Santiago Ávila-Ríos, Eduardo Becerril-Vargas, Víctor Hugo Borja-Aburto, Celia Boukadida, Cristóbal Cháidez-Quiróz, Juan Bautista Chale-Dzul, Ricardo Ciria Merce, Andreu Comas-García, Célida Duque Molina, Julissa Enciso-Ibarra, José Antonio Enciso-Moreno, Gloria Elena Espinosa-Ayala, Fernando Fontove-Herrera, Daniel Fregoso-Rueda, Victor Eduardo García-Arias, Alejandra García-Gasca, Bruno Gómez-Gil, Jean Pierre González, Irvin González-López, Concepción Grajales-Muñiz, Ricardo Grande, Rosa María Gutiérrez Rios, Jesús Hernández, Alejandra Hernández-Terán, Alfredo Herrera-Estrella, Carla Ivón Herrera-Najera, Pavel Isa, Verónica Mata-Haro, Daniel Lira Morales, Susana Lopez, Antonio Loza Román, Brenda Irasema Maldonado-Meza, Bernardo Martínez-Miguel, José Arturo Martínez-Orozco, Célida Martínez- Rodríguez, Margarita Matías-Florentino, Fidencio Mejía-Nepomuceno, María Guadalupe de Jesús Mireles-Rivera, Gloria María Molina-Salinas, Hector Montoya-Fuentes, , Mario Mújica-Sánchez, José Esteban Muñoz-Medina, José de Jesús Nuñez-Contreras, Alicia Ocaña-Mondragón, Luis Alberto Ochoa-Carrera, Hector Esteban Paz-Juárez, Marissa Perez-Garcia, Francisco Pulido, Helen Haydee Fernanda Ramírez-Plascencia, Jorge Salas-Hernández, Angel Gustavo Salas-Lais, Alejandro Sánchez-Flores, Clara Esperanza Santacruz-Tinoco, María Guadalupe Santiago-Mauricio, Selene Zárate, Nelly Sélem-Mojica, Blanca Taboada, Gloria Vazquez, Joel Armando Vázquez-Pérez. |
| EPI_ISL_2681229, EPI_ISL_2681230, EPI_ISL_2681231 | Laboratorio Central de Epidemiología (LCE) | Instituto de Biotecnología de la UNAM | Consorcio Mexicano de Vigilancia Genómica (CoViGen-Mex). Authors (in alphabetical order): Julio Elias Alvarado-Yaah, Carlos F. Arias, Santiago Ávila-Ríos, Eduardo Becerril-Vargas, Víctor Hugo Borja-Aburto, Celia Boukadida, Cristóbal Cháidez-Quiróz, Juan Bautista Chale-Dzul, Ricardo Ciria Merce, Andreu Comas-García, Célida Duque Molina, Julissa Enciso-Ibarra, José Antonio Enciso-Moreno, Gloria Elena Espinosa-Ayala, Fernando Fontove-Herrera, Daniel Fregoso-Rueda, Victor Eduardo García-Arias, Alejandra García-Gasca, Bruno Gómez-Gil, Jean Pierre González, Irvin González-López, Concepción Grajales-Muñiz, Ricardo Grande, Rosa María Gutiérrez Rios, Jesús Hernández, Alejandra Hernández-Terán, Alfredo Herrera-Estrella, Carla Ivón Herrera-Najera, Pavel Isa, Verónica Mata-Haro, Daniel Lira Morales, Susana Lopez, Antonio Loza Román, Brenda Irasema Maldonado-Meza, Bernardo Martínez-Miguel, José Arturo Martínez-Orozco, Célida Martínez- Rodríguez, Margarita Matías-Florentino, Fidencio Mejía-Nepomuceno, María Guadalupe de Jesús Mireles-Rivera, Gloria María Molina-Salinas, Hector Montoya-Fuentes, , Mario Mújica-Sánchez, José Esteban Muñoz-Medina, José de Jesús Nuñez-Contreras, Alicia Ocaña-Mondragón, Luis Alberto Ochoa-Carrera, Hector Esteban Paz-Juárez, Marissa Perez-Garcia, Francisco Pulido, Helen Haydee Fernanda Ramírez-Plascencia, Jorge Salas-Hernández, Angel Gustavo Salas-Lais, Alejandro Sánchez-Flores, Clara Esperanza Santacruz-Tinoco, María Guadalupe Santiago-Mauricio, Selene Zárate, Nelly Sélem-Mojica, Blanca Taboada, Gloria Vazquez, Joel Armando Vázquez-Pérez. |
| EPI_ISL_2681232, EPI_ISL_2681233 | Unidad de Investigación Médica de Yucatán (UIMY) | Instituto de Biotecnología de la UNAM | Consorcio Mexicano de Vigilancia Genómica (CoViGen-Mex). Authors (in alphabetical order): Julio Elias Alvarado-Yaah, Carlos F. Arias, Santiago Ávila-Ríos, Eduardo Becerril-Vargas, Víctor Hugo Borja-Aburto, Celia Boukadida, Cristóbal Cháidez-Quiróz, Juan Bautista Chale-Dzul, Ricardo Ciria Merce, Andreu Comas-García, Célida Duque Molina, Julissa Enciso-Ibarra, José Antonio Enciso-Moreno, Gloria Elena Espinosa-Ayala, Fernando Fontove-Herrera, Daniel Fregoso-Rueda, Victor Eduardo García-Arias, Alejandra García-Gasca, Bruno Gómez-Gil, Jean Pierre González, Irvin González-López, Concepción Grajales-Muñiz, Ricardo Grande, Rosa María Gutiérrez Rios, Jesús Hernández, Alejandra Hernández-Terán, Alfredo Herrera-Estrella, Carla Ivón Herrera-Najera, Pavel Isa, Verónica Mata-Haro, Daniel Lira Morales, Susana Lopez, Antonio Loza Román, Brenda Irasema Maldonado-Meza, Bernardo Martínez-Miguel, José Arturo Martínez-Orozco, Célida Martínez- Rodríguez, Margarita Matías-Florentino, Fidencio Mejía-Nepomuceno, María Guadalupe de Jesús Mireles-Rivera, Gloria María Molina-Salinas, Hector Montoya-Fuentes, , Mario Mújica-Sánchez, José Esteban Muñoz-Medina, José de Jesús Nuñez-Contreras, Alicia Ocaña-Mondragón, Luis Alberto Ochoa-Carrera, Hector Esteban Paz-Juárez, Marissa Perez-Garcia, Francisco Pulido, Helen Haydee Fernanda Ramírez-Plascencia, Jorge Salas-Hernández, Angel Gustavo Salas-Lais, Alejandro Sánchez-Flores, Clara Esperanza Santacruz-Tinoco, María Guadalupe Santiago-Mauricio, Selene Zárate, Nelly Sélem-Mojica, Blanca Taboada, Gloria Vazquez, Joel Armando Vázquez-Pérez. |
| EPI_ISL_2681234, EPI_ISL_2681235 | Unidad de Investigación Biomédica de Zacatecas (UIBZ) | Instituto de Biotecnología de la UNAM | Consorcio Mexicano de Vigilancia Genómica (CoViGen-Mex). Authors (in alphabetical order): Julio Elias Alvarado-Yaah, Carlos F. Arias, Santiago Ávila-Ríos, Eduardo Becerril-Vargas, Víctor Hugo Borja-Aburto, Celia Boukadida, Cristóbal Cháidez-Quiróz, Juan Bautista Chale-Dzul, Ricardo Ciria Merce, Andreu Comas-García, Célida Duque Molina, Julissa Enciso-Ibarra, José Antonio Enciso-Moreno, Gloria Elena Espinosa-Ayala, Fernando Fontove-Herrera, Daniel Fregoso-Rueda, Victor Eduardo García-Arias, Alejandra García-Gasca, Bruno Gómez-Gil, Jean Pierre González, Irvin González-López, Concepción Grajales-Muñiz, Ricardo Grande, Rosa María Gutiérrez Rios, Jesús Hernández, Alejandra Hernández-Terán, Alfredo Herrera-Estrella, Carla Ivón Herrera-Najera, Pavel Isa, Verónica Mata-Haro, Daniel Lira Morales, Susana Lopez, Antonio Loza Román, Brenda Irasema Maldonado-Meza, Bernardo Martínez-Miguel, José Arturo Martínez-Orozco, Célida Martínez- Rodríguez, Margarita Matías-Florentino, Fidencio Mejía-Nepomuceno, María Guadalupe de Jesús Mireles-Rivera, Gloria María Molina-Salinas, Hector Montoya-Fuentes, , Mario Mújica-Sánchez, José Esteban Muñoz-Medina, José de Jesús Nuñez-Contreras, Alicia Ocaña-Mondragón, Luis Alberto Ochoa-Carrera, Hector Esteban Paz-Juárez, Marissa Perez-Garcia, Francisco Pulido, Helen Haydee Fernanda Ramírez-Plascencia, Jorge Salas-Hernández, Angel Gustavo Salas-Lais, Alejandro Sánchez-Flores, Clara Esperanza Santacruz-Tinoco, María Guadalupe Santiago-Mauricio, Selene Zárate, Nelly Sélem-Mojica, Blanca Taboada, Gloria Vazquez, Joel Armando Vázquez-Pérez. |
| EPI_ISL_2681236 | Unidad de Investigación Médica de Yucatán (UIMY) | Instituto de Biotecnología de la UNAM | Consorcio Mexicano de Vigilancia Genómica (CoViGen-Mex). Authors (in alphabetical order): Julio Elias Alvarado-Yaah, Carlos F. Arias, Santiago Ávila-Ríos, Eduardo Becerril-Vargas, Víctor Hugo Borja-Aburto, Celia Boukadida, Cristóbal Cháidez-Quiróz, Juan Bautista Chale-Dzul, Ricardo Ciria Merce, Andreu Comas-García, Célida Duque Molina, Julissa Enciso-Ibarra, José Antonio Enciso-Moreno, Gloria Elena Espinosa-Ayala, Fernando Fontove-Herrera, Daniel Fregoso-Rueda, Victor Eduardo García-Arias, Alejandra García-Gasca, Bruno Gómez-Gil, Jean Pierre González, Irvin González-López, Concepción Grajales-Muñiz, Ricardo Grande, Rosa María Gutiérrez Rios, Jesús Hernández, Alejandra Hernández-Terán, Alfredo Herrera-Estrella, Carla Ivón Herrera-Najera, Pavel Isa, Verónica Mata-Haro, Daniel Lira Morales, Susana Lopez, Antonio Loza Román, Brenda Irasema Maldonado-Meza, Bernardo Martínez-Miguel, José Arturo Martínez-Orozco, Célida Martínez- Rodríguez, Margarita Matías-Florentino, Fidencio Mejía-Nepomuceno, María Guadalupe de Jesús Mireles-Rivera, Gloria María Molina-Salinas, Hector Montoya-Fuentes, , Mario Mújica-Sánchez, José Esteban Muñoz-Medina, José de Jesús Nuñez-Contreras, Alicia Ocaña-Mondragón, Luis Alberto Ochoa-Carrera, Hector Esteban Paz-Juárez, Marissa Perez-Garcia, Francisco Pulido, Helen Haydee Fernanda Ramírez-Plascencia, Jorge Salas-Hernández, Angel Gustavo Salas-Lais, Alejandro Sánchez-Flores, Clara Esperanza Santacruz-Tinoco, María Guadalupe Santiago-Mauricio, Selene Zárate, Nelly Sélem-Mojica, Blanca Taboada, Gloria Vazquez, Joel Armando Vázquez-Pérez. |
| EPI_ISL_2681238 | Laboratorio Central de Epidemiología (LCE) | Instituto de Biotecnología de la UNAM | Consorcio Mexicano de Vigilancia Genómica (CoViGen-Mex). Authors (in alphabetical order): Julio Elias Alvarado-Yaah, Carlos F. Arias, Santiago Ávila-Ríos, Eduardo Becerril-Vargas, Víctor Hugo Borja-Aburto, Celia Boukadida, Cristóbal Cháidez-Quiróz, Juan Bautista Chale-Dzul, Ricardo Ciria Merce, Andreu Comas-García, Célida Duque Molina, Julissa Enciso-Ibarra, José Antonio Enciso-Moreno, Gloria Elena Espinosa-Ayala, Fernando Fontove-Herrera, Daniel Fregoso-Rueda, Victor Eduardo García-Arias, Alejandra García-Gasca, Bruno Gómez-Gil, Jean Pierre González, Irvin González-López, Concepción Grajales-Muñiz, Ricardo Grande, Rosa María Gutiérrez Rios, Jesús Hernández, Alejandra Hernández-Terán, Alfredo Herrera-Estrella, Carla Ivón Herrera-Najera, Pavel Isa, Verónica Mata-Haro, Daniel Lira Morales, Susana Lopez, Antonio Loza Román, Brenda Irasema Maldonado-Meza, Bernardo Martínez-Miguel, José Arturo Martínez-Orozco, Célida Martínez- Rodríguez, Margarita Matías-Florentino, Fidencio Mejía-Nepomuceno, María Guadalupe de Jesús Mireles-Rivera, Gloria María Molina-Salinas, Hector Montoya-Fuentes, , Mario Mújica-Sánchez, José Esteban Muñoz-Medina, José de Jesús Nuñez-Contreras, Alicia Ocaña-Mondragón, Luis Alberto Ochoa-Carrera, Hector Esteban Paz-Juárez, Marissa Perez-Garcia, Francisco Pulido, Helen Haydee Fernanda Ramírez-Plascencia, Jorge Salas-Hernández, Angel Gustavo Salas-Lais, Alejandro Sánchez-Flores, Clara Esperanza Santacruz-Tinoco, María Guadalupe Santiago-Mauricio, Selene Zárate, Nelly Sélem-Mojica, Blanca Taboada, Gloria Vazquez, Joel Armando Vázquez-Pérez. |

| | | | |
|---|---|---|---|
| EPI_ISL_2681239 | Unidad de Investigación Biomédica de Zacatecas (UIBZ) | Instituto de Biotecnología de la UNAM | Consorcio Mexicano de Vigilancia Genómica (CoViGen-Mex). Authors (in alphabetical order): Julio Elias Alvarado-Yaah, Carlos F. Arias, Santiago Ávila-Ríos, Eduardo Becerril-Vargas, Víctor Hugo Borja-Aburto, Celia Boukadida, Cristóbal Cháidez-Quiróz, Juan Bautista Chale-Dzul, Ricardo Ciria Merce, Andreu Comas-García, Célida Duque Molina, Julissa Enciso-Ibarra, José Antonio Enciso-Moreno, Gloria Elena Espinosa-Ayala, Fernando Fontove-Herrera, Daniel Fregoso-Rueda, Victor Eduardo García-Arias, Alejandra García-Gasca, Bruno Gómez-Gil, Jean Pierre González, Irvin González-López, Concepción Grajales-Muñiz, Ricardo Grande, Rosa María Gutiérrez Rios, Jesús Hernández, Alejandra Hernández-Terán, Alfredo Herrera-Estrella, Carla Ivón Herrera-Najera, Pavel Isa, Verónica Mata-Haro, Daniel Lira Morales, Susana Lopez, Antonio Loza Román, Brenda Irasema Maldonado-Meza, Bernardo Martínez-Miguel, José Arturo Martínez-Orozco, Célida Martínez- Rodríguez, Margarita Matías-Florentino, Fidencio Mejía-Nepomuceno, María Guadalupe de Jesús Mireles-Rivera, Gloria María Molina-Salinas, Hector Montoya-Fuentes, , Mario Mújica-Sánchez, José Esteban Muñoz-Medina, José de Jesús Nuñez-Contreras, Alicia Ocaña-Mondragón, Luis Alberto Ochoa-Carrera, Hector Esteban Paz-Juárez, Marissa Perez-Garcia, Francisco Pulido, Helen Haydee Fernanda Ramírez-Plascencia, Jorge Salas-Hernández, Angel Gustavo Salas-Lais, Alejandro Sánchez-Flores, Clara Esperanza Santacruz-Tinoco, María Guadalupe Santiago-Mauricio, Selene Zárate, Nelly Sélem-Mojica, Blanca Taboada, Gloria Vazquez, Joel Armando Vázquez-Pérez. |
| EPI_ISL_2681240 | Laboratorio Central de Epidemiología (LCE) | Instituto de Biotecnología de la UNAM | Consorcio Mexicano de Vigilancia Genómica (CoViGen-Mex). Authors (in alphabetical order): Julio Elias Alvarado-Yaah, Carlos F. Arias, Santiago Ávila-Ríos, Eduardo Becerril-Vargas, Víctor Hugo Borja-Aburto, Celia Boukadida, Cristóbal Cháidez-Quiróz, Juan Bautista Chale-Dzul, Ricardo Ciria Merce, Andreu Comas-García, Célida Duque Molina, Julissa Enciso-Ibarra, José Antonio Enciso-Moreno, Gloria Elena Espinosa-Ayala, Fernando Fontove-Herrera, Daniel Fregoso-Rueda, Victor Eduardo García-Arias, Alejandra García-Gasca, Bruno Gómez-Gil, Jean Pierre González, Irvin González-López, Concepción Grajales-Muñiz, Ricardo Grande, Rosa María Gutiérrez Rios, Jesús Hernández, Alejandra Hernández-Terán, Alfredo Herrera-Estrella, Carla Ivón Herrera-Najera, Pavel Isa, Verónica Mata-Haro, Daniel Lira Morales, Susana Lopez, Antonio Loza Román, Brenda Irasema Maldonado-Meza, Bernardo Martínez-Miguel, José Arturo Martínez-Orozco, Célida Martínez- Rodríguez, Margarita Matías-Florentino, Fidencio Mejía-Nepomuceno, María Guadalupe de Jesús Mireles-Rivera, Gloria María Molina-Salinas, Hector Montoya-Fuentes, , Mario Mújica-Sánchez, José Esteban Muñoz-Medina, José de Jesús Nuñez-Contreras, Alicia Ocaña-Mondragón, Luis Alberto Ochoa-Carrera, Hector Esteban Paz-Juárez, Marissa Perez-Garcia, Francisco Pulido, Helen Haydee Fernanda Ramírez-Plascencia, Jorge Salas-Hernández, Angel Gustavo Salas-Lais, Alejandro Sánchez-Flores, Clara Esperanza Santacruz-Tinoco, María Guadalupe Santiago-Mauricio, Selene Zárate, Nelly Sélem-Mojica, Blanca Taboada, Gloria Vazquez, Joel Armando Vázquez-Pérez. |
| EPI_ISL_2681241 | Unidad de Investigación Médica de Yucatán (UIMY) | Instituto de Biotecnología de la UNAM | Consorcio Mexicano de Vigilancia Genómica (CoViGen-Mex). Authors (in alphabetical order): Julio Elias Alvarado-Yaah, Carlos F. Arias, Santiago Ávila-Ríos, Eduardo Becerril-Vargas, Víctor Hugo Borja-Aburto, Celia Boukadida, Cristóbal Cháidez-Quiróz, Juan Bautista Chale-Dzul, Ricardo Ciria Merce, Andreu Comas-García, Célida Duque Molina, Julissa Enciso-Ibarra, José Antonio Enciso-Moreno, Gloria Elena Espinosa-Ayala, Fernando Fontove-Herrera, Daniel Fregoso-Rueda, Victor Eduardo García-Arias, Alejandra García-Gasca, Bruno Gómez-Gil, Jean Pierre González, Irvin González-López, Concepción Grajales-Muñiz, Ricardo Grande, Rosa María Gutiérrez Rios, Jesús Hernández, Alejandra Hernández-Terán, Alfredo Herrera-Estrella, Carla Ivón Herrera-Najera, Pavel Isa, Verónica Mata-Haro, Daniel Lira Morales, Susana Lopez, Antonio Loza Román, Brenda Irasema Maldonado-Meza, Bernardo Martínez-Miguel, José Arturo Martínez-Orozco, Célida Martínez- Rodríguez, Margarita Matías-Florentino, Fidencio Mejía-Nepomuceno, María Guadalupe de Jesús Mireles-Rivera, Gloria María Molina-Salinas, Hector Montoya-Fuentes, , Mario Mújica-Sánchez, José Esteban Muñoz-Medina, José de Jesús Nuñez-Contreras, Alicia Ocaña-Mondragón, Luis Alberto Ochoa-Carrera, Hector Esteban Paz-Juárez, Marissa Perez-Garcia, Francisco Pulido, Helen Haydee Fernanda Ramírez-Plascencia, Jorge Salas-Hernández, Angel Gustavo Salas-Lais, Alejandro Sánchez-Flores, Clara Esperanza Santacruz-Tinoco, María Guadalupe Santiago-Mauricio, Selene Zárate, Nelly Sélem-Mojica, Blanca Taboada, Gloria Vazquez, Joel Armando Vázquez-Pérez. |
| EPI_ISL_2681243 | Laboratorio Central de Epidemiología (LCE) | Instituto de Biotecnología de la UNAM | Consorcio Mexicano de Vigilancia Genómica (CoViGen-Mex). Authors (in alphabetical order): Julio Elias Alvarado-Yaah, Carlos F. Arias, Santiago Ávila-Ríos, Eduardo Becerril-Vargas, Víctor Hugo Borja-Aburto, Celia Boukadida, Cristóbal Cháidez-Quiróz, Juan Bautista Chale-Dzul, Ricardo Ciria Merce, Andreu Comas-García, Célida Duque Molina, Julissa Enciso-Ibarra, José Antonio Enciso-Moreno, Gloria Elena Espinosa-Ayala, Fernando Fontove-Herrera, Daniel Fregoso-Rueda, Victor Eduardo García-Arias, Alejandra García-Gasca, Bruno Gómez-Gil, Jean Pierre González, Irvin González-López, Concepción Grajales-Muñiz, Ricardo Grande, Rosa María Gutiérrez Rios, Jesús Hernández, Alejandra Hernández-Terán, Alfredo Herrera-Estrella, Carla Ivón Herrera-Najera, Pavel Isa, Verónica Mata-Haro, Daniel Lira Morales, Susana Lopez, Antonio Loza Román, Brenda Irasema Maldonado-Meza, Bernardo Martínez-Miguel, José Arturo Martínez-Orozco, Célida Martínez- Rodríguez, Margarita Matías-Florentino, Fidencio Mejía-Nepomuceno, María Guadalupe de Jesús Mireles-Rivera, Gloria María Molina-Salinas, Hector Montoya-Fuentes, , Mario Mújica-Sánchez, José Esteban Muñoz-Medina, José de Jesús Nuñez-Contreras, Alicia Ocaña-Mondragón, Luis Alberto Ochoa-Carrera, Hector Esteban Paz-Juárez, Marissa Perez-Garcia, Francisco Pulido, Helen Haydee Fernanda Ramírez-Plascencia, Jorge Salas-Hernández, Angel Gustavo Salas-Lais, Alejandro Sánchez-Flores, Clara Esperanza Santacruz-Tinoco, María Guadalupe Santiago-Mauricio, Selene Zárate, Nelly Sélem-Mojica, Blanca Taboada, Gloria Vazquez, Joel Armando Vázquez-Pérez. |
| EPI_ISL_2681244 | Unidad de Investigación Médica de Yucatán (UIMY) | Instituto de Biotecnología de la UNAM | Consorcio Mexicano de Vigilancia Genómica (CoViGen-Mex). Authors (in alphabetical order): Julio Elias Alvarado-Yaah, Carlos F. Arias, Santiago Ávila-Ríos, Eduardo Becerril-Vargas, Víctor Hugo Borja-Aburto, Celia Boukadida, Cristóbal Cháidez-Quiróz, Juan Bautista Chale-Dzul, Ricardo Ciria Merce, Andreu Comas-García, Célida Duque Molina, Julissa Enciso-Ibarra, José Antonio Enciso-Moreno, Gloria Elena Espinosa-Ayala, Fernando Fontove-Herrera, Daniel Fregoso-Rueda, Victor Eduardo García-Arias, Alejandra García-Gasca, Bruno Gómez-Gil, Jean Pierre González, Irvin González-López, Concepción Grajales-Muñiz, Ricardo Grande, Rosa María Gutiérrez Rios, Jesús Hernández, Alejandra Hernández-Terán, Alfredo Herrera-Estrella, Carla Ivón Herrera-Najera, Pavel Isa, Verónica Mata-Haro, Daniel Lira Morales, Susana Lopez, Antonio Loza Román, Brenda Irasema Maldonado-Meza, Bernardo Martínez-Miguel, José Arturo Martínez-Orozco, Célida Martínez- Rodríguez, Margarita Matías-Florentino, Fidencio Mejía-Nepomuceno, María Guadalupe de Jesús Mireles-Rivera, Gloria María Molina-Salinas, Hector Montoya-Fuentes, , Mario Mújica-Sánchez, José Esteban Muñoz-Medina, José de Jesús Nuñez-Contreras, Alicia Ocaña-Mondragón, Luis Alberto Ochoa-Carrera, Hector Esteban Paz-Juárez, Marissa Perez-Garcia, Francisco Pulido, Helen Haydee Fernanda Ramírez-Plascencia, Jorge Salas-Hernández, Angel Gustavo Salas-Lais, Alejandro Sánchez-Flores, Clara Esperanza Santacruz-Tinoco, María Guadalupe Santiago-Mauricio, Selene Zárate, Nelly Sélem-Mojica, Blanca Taboada, Gloria Vazquez, Joel Armando Vázquez-Pérez. |
| EPI_ISL_2681245 | Laboratorio Central de Epidemiología (LCE) | Instituto de Biotecnología de la UNAM | Consorcio Mexicano de Vigilancia Genómica (CoViGen-Mex). Authors (in alphabetical order): Julio Elias Alvarado-Yaah, Carlos F. Arias, Santiago Ávila-Ríos, Eduardo Becerril-Vargas, Víctor Hugo Borja-Aburto, Celia Boukadida, Cristóbal Cháidez-Quiróz, Juan Bautista Chale-Dzul, Ricardo Ciria Merce, Andreu Comas-García, Célida Duque Molina, Julissa Enciso-Ibarra, José Antonio Enciso-Moreno, Gloria Elena Espinosa-Ayala, Fernando Fontove-Herrera, Daniel Fregoso-Rueda, Victor Eduardo García-Arias, Alejandra García-Gasca, Bruno Gómez-Gil, Jean Pierre González, Irvin González-López, Concepción Grajales-Muñiz, Ricardo Grande, Rosa María Gutiérrez Rios, Jesús Hernández, Alejandra Hernández-Terán, Alfredo Herrera-Estrella, Carla Ivón Herrera-Najera, Pavel Isa, Verónica Mata-Haro, Daniel Lira Morales, Susana Lopez, Antonio Loza Román, Brenda Irasema Maldonado-Meza, Bernardo Martínez-Miguel, José Arturo Martínez-Orozco, Célida Martínez- Rodríguez, Margarita Matías-Florentino, Fidencio Mejía-Nepomuceno, María Guadalupe de Jesús Mireles-Rivera, Gloria María Molina-Salinas, Hector Montoya-Fuentes, , Mario Mújica-Sánchez, José Esteban Muñoz-Medina, José de Jesús Nuñez-Contreras, Alicia Ocaña-Mondragón, Luis Alberto Ochoa-Carrera, Hector Esteban Paz-Juárez, Marissa Perez-Garcia, Francisco Pulido, Helen Haydee Fernanda Ramírez-Plascencia, Jorge Salas-Hernández, Angel Gustavo Salas-Lais, Alejandro Sánchez-Flores, Clara Esperanza Santacruz-Tinoco, María Guadalupe Santiago-Mauricio, Selene Zárate, Nelly Sélem-Mojica, Blanca Taboada, Gloria Vazquez, Joel Armando Vázquez-Pérez. |
| EPI_ISL_2681248 | Centro de Investigación Biomédica del Noreste (CIBIN) | Instituto de Biotecnología de la UNAM | Consorcio Mexicano de Vigilancia Genómica (CoViGen-Mex). Authors (in alphabetical order): Julio Elias Alvarado-Yaah, Carlos F. Arias, Santiago Ávila-Ríos, Eduardo Becerril-Vargas, Víctor Hugo Borja-Aburto, Celia Boukadida, Cristóbal Cháidez-Quiróz, Juan Bautista Chale-Dzul, Ricardo Ciria Merce, Andreu Comas-García, Célida Duque Molina, Julissa Enciso-Ibarra, José Antonio Enciso-Moreno, Gloria Elena Espinosa-Ayala, Fernando Fontove-Herrera, Daniel Fregoso-Rueda, Victor Eduardo García-Arias, Alejandra García-Gasca, Bruno Gómez-Gil, Jean Pierre González, Irvin González-López, Concepción Grajales-Muñiz, Ricardo Grande, Rosa María Gutiérrez Rios, Jesús Hernández, Alejandra Hernández-Terán, Alfredo Herrera-Estrella, Carla Ivón Herrera-Najera, Pavel Isa, Verónica Mata-Haro, Daniel Lira Morales, Susana Lopez, Antonio Loza Román, Brenda Irasema Maldonado-Meza, Bernardo Martínez-Miguel, José Arturo Martínez-Orozco, Célida Martínez- Rodríguez, Margarita Matías-Florentino, Fidencio Mejía-Nepomuceno, María Guadalupe de Jesús Mireles-Rivera, Gloria María Molina-Salinas, Hector Montoya-Fuentes, , Mario Mújica-Sánchez, José Esteban Muñoz-Medina, José de Jesús Nuñez-Contreras, Alicia Ocaña-Mondragón, Luis Alberto Ochoa-Carrera, Hector Esteban Paz-Juárez, Marissa Perez-Garcia, Francisco Pulido, Helen Haydee Fernanda Ramírez-Plascencia, Jorge Salas-Hernández, Angel Gustavo Salas-Lais, Alejandro Sánchez-Flores, Clara Esperanza Santacruz-Tinoco, María Guadalupe Santiago-Mauricio, Selene Zárate, Nelly Sélem-Mojica, Blanca Taboada, Gloria Vazquez, Joel Armando Vázquez-Pérez. |

EPI_ISL_2681249, EPI_ISL_2681250, EPI_ISL_2681251, EPI_ISL_2681252, EPI_ISL_2681253, EPI_ISL_2681254, EPI_ISL_2681255, EPI_ISL_2681256, EPI_ISL_2681257, EPI_ISL_2681258, EPI_ISL_2681259

| | | | |
|---|---|---|---|
| see above | Laboratorio Central de Epidemiología (LCE) | Instituto de Biotecnología de la UNAM | Consorcio Mexicano de Vigilancia Genómica (CoViGen-Mex). Authors (in alphabetical order): Julio Elias Alvarado-Yaah, Carlos F. Arias, Santiago Ávila-Ríos, Eduardo Becerril-Vargas, Víctor Hugo Borja-Aburto, Celia Boukadida, Cristóbal Cháidez-Quiróz, Juan Bautista Chale-Dzul, Ricardo Ciria Merce, Andreu Comas-García, Célida Duque Molina, Julissa Enciso-Ibarra, José Antonio Enciso-Moreno, Gloria Elena Espinosa-Ayala, Fernando Fontove-Herrera, Daniel Fregoso-Rueda, Victor Eduardo García-Arias, Alejandra García-Gasca, Bruno Gómez-Gil, Jean Pierre González, Irvin González-López, Concepción Grajales-Muñiz, Ricardo Grande, Rosa María Gutiérrez Rios, Jesús Hernández, Alejandra Hernández-Terán, Alfredo Herrera-Estrella, Carla Ivón Herrera-Najera, Pavel Isa, Verónica Mata-Haro, Daniel Lira Morales, Susana Lopez, Antonio Loza Román, Brenda Irasema Maldonado-Meza, Bernardo Martínez-Miguel, José Arturo Martínez-Orozco, Célida Martínez- Rodríguez, Margarita Matías-Florentino, Fidencio Mejía-Nepomuceno, María Guadalupe de Jesús Mireles-Rivera, Gloria María Molina-Salinas, Hector Montoya-Fuentes, , Mario Mújica-Sánchez, José Esteban Muñoz-Medina, José de Jesús Nuñez-Contreras, Alicia Ocaña-Mondragón, Luis Alberto Ochoa-Carrera, Hector Esteban Paz-Juárez, Marissa Perez-Garcia, Francisco Pulido, Helen Haydee Fernanda Ramírez-Plascencia, Jorge Salas-Hernández, Angel Gustavo Salas-Lais, Alejandro Sánchez-Flores, Clara Esperanza Santacruz-Tinoco, María Guadalupe Santiago-Mauricio, Selene Zárate, Nelly Sélem-Mojica, Blanca Taboada, Gloria Vazquez, Joel Armando Vázquez-Pérez. |
| EPI_ISL_2681261 | Unidad de Investigación Médica de Yucatán (UIMY) | Instituto de Biotecnología de la UNAM | Consorcio Mexicano de Vigilancia Genómica (CoViGen-Mex). Authors (in alphabetical order): Julio Elias Alvarado-Yaah, Carlos F. Arias, Santiago Ávila-Ríos, Eduardo Becerril-Vargas, Víctor Hugo Borja-Aburto, Celia Boukadida, Cristóbal Cháidez-Quiróz, Juan Bautista Chale-Dzul, Ricardo Ciria Merce, Andreu Comas-García, Célida Duque Molina, Julissa Enciso-Ibarra, José Antonio Enciso-Moreno, Gloria Elena Espinosa-Ayala, Fernando Fontove-Herrera, Daniel Fregoso-Rueda, Victor Eduardo García-Arias, Alejandra García-Gasca, Bruno Gómez-Gil, Jean Pierre González, Irvin González-López, Concepción Grajales-Muñiz, Ricardo Grande, Rosa María Gutiérrez Rios, Jesús Hernández, Alejandra Hernández-Terán, Alfredo Herrera-Estrella, Carla Ivón Herrera-Najera, Pavel Isa, Verónica Mata-Haro, Daniel Lira Morales, Susana Lopez, Antonio Loza Román, Brenda Irasema Maldonado-Meza, Bernardo Martínez-Miguel, José Arturo Martínez-Orozco, Célida Martínez- Rodríguez, Margarita Matías-Florentino, Fidencio Mejía-Nepomuceno, María Guadalupe de Jesús Mireles-Rivera, Gloria María Molina-Salinas, Hector Montoya-Fuentes, , Mario Mújica-Sánchez, José Esteban Muñoz-Medina, José de Jesús Nuñez-Contreras, Alicia Ocaña-Mondragón, Luis Alberto Ochoa-Carrera, Hector Esteban Paz-Juárez, Marissa Perez-Garcia, Francisco Pulido, Helen Haydee Fernanda Ramírez-Plascencia, Jorge Salas-Hernández, Angel Gustavo Salas-Lais, Alejandro Sánchez-Flores, Clara Esperanza Santacruz-Tinoco, María Guadalupe Santiago-Mauricio, Selene Zárate, Nelly Sélem-Mojica, Blanca Taboada, Gloria Vazquez, Joel Armando Vázquez-Pérez. |
| EPI_ISL_2681262, EPI_ISL_2681264 | Laboratorio Central de Epidemiología (LCE) | Instituto de Biotecnología de la UNAM | Consorcio Mexicano de Vigilancia Genómica (CoViGen-Mex). Authors (in alphabetical order): Julio Elias Alvarado-Yaah, Carlos F. Arias, Santiago Ávila-Ríos, Eduardo Becerril-Vargas, Víctor Hugo Borja-Aburto, Celia Boukadida, Cristóbal Cháidez-Quiróz, Juan Bautista Chale-Dzul, Ricardo Ciria Merce, Andreu Comas-García, Célida Duque Molina, Julissa Enciso-Ibarra, José Antonio Enciso-Moreno, Gloria Elena Espinosa-Ayala, Fernando Fontove-Herrera, Daniel Fregoso-Rueda, Victor Eduardo García-Arias, Alejandra García-Gasca, Bruno Gómez-Gil, Jean Pierre González, Irvin González-López, Concepción Grajales-Muñiz, Ricardo Grande, Rosa María Gutiérrez Rios, Jesús Hernández, Alejandra Hernández-Terán, Alfredo Herrera-Estrella, Carla Ivón Herrera-Najera, Pavel Isa, Verónica Mata-Haro, Daniel Lira Morales, Susana Lopez, Antonio Loza Román, Brenda Irasema Maldonado-Meza, Bernardo Martínez-Miguel, José Arturo Martínez-Orozco, Célida Martínez- Rodríguez, Margarita Matías-Florentino, Fidencio Mejía-Nepomuceno, María Guadalupe de Jesús Mireles-Rivera, Gloria María Molina-Salinas, Hector Montoya-Fuentes, , Mario Mújica-Sánchez, José Esteban Muñoz-Medina, José de Jesús Nuñez-Contreras, Alicia Ocaña-Mondragón, Luis Alberto Ochoa-Carrera, Hector Esteban Paz-Juárez, Marissa Perez-Garcia, Francisco Pulido, Helen Haydee Fernanda Ramírez-Plascencia, Jorge Salas-Hernández, Angel Gustavo Salas-Lais, Alejandro Sánchez-Flores, Clara Esperanza Santacruz-Tinoco, María Guadalupe Santiago-Mauricio, Selene Zárate, Nelly Sélem-Mojica, Blanca Taboada, Gloria Vazquez, Joel Armando Vázquez-Pérez. |
| EPI_ISL_2681265 | Centro de Investigación Biomédica del Noreste (CIBIN) | Instituto de Biotecnología de la UNAM | Consorcio Mexicano de Vigilancia Genómica (CoViGen-Mex). Authors (in alphabetical order): Julio Elias Alvarado-Yaah, Carlos F. Arias, Santiago Ávila-Ríos, Eduardo Becerril-Vargas, Víctor Hugo Borja-Aburto, Celia Boukadida, Cristóbal Cháidez-Quiróz, Juan Bautista Chale-Dzul, Ricardo Ciria Merce, Andreu Comas-García, Célida Duque Molina, Julissa Enciso-Ibarra, José Antonio Enciso-Moreno, Gloria Elena Espinosa-Ayala, Fernando Fontove-Herrera, Daniel Fregoso-Rueda, Victor Eduardo García-Arias, Alejandra García-Gasca, Bruno Gómez-Gil, Jean Pierre González, Irvin González-López, Concepción Grajales-Muñiz, Ricardo Grande, Rosa María Gutiérrez Rios, Jesús Hernández, Alejandra Hernández-Terán, Alfredo Herrera-Estrella, Carla Ivón Herrera-Najera, Pavel Isa, Verónica Mata-Haro, Daniel Lira Morales, Susana Lopez, Antonio Loza Román, Brenda Irasema Maldonado-Meza, Bernardo Martínez-Miguel, José Arturo Martínez-Orozco, Célida Martínez- Rodríguez, Margarita Matías-Florentino, Fidencio Mejía-Nepomuceno, María Guadalupe de Jesús Mireles-Rivera, Gloria María Molina-Salinas, Hector Montoya-Fuentes, , Mario Mújica-Sánchez, José Esteban Muñoz-Medina, José de Jesús Nuñez-Contreras, Alicia Ocaña-Mondragón, Luis Alberto Ochoa-Carrera, Hector Esteban Paz-Juárez, Marissa Perez-Garcia, Francisco Pulido, Helen Haydee Fernanda Ramírez-Plascencia, Jorge Salas-Hernández, Angel Gustavo Salas-Lais, Alejandro Sánchez-Flores, Clara Esperanza Santacruz-Tinoco, María Guadalupe Santiago-Mauricio, Selene Zárate, Nelly Sélem-Mojica, Blanca Taboada, Gloria Vazquez, Joel Armando Vázquez-Pérez. |
| EPI_ISL_2681267, EPI_ISL_2681268 | Laboratorio Central de Epidemiología (LCE) | Instituto de Biotecnología de la UNAM | Consorcio Mexicano de Vigilancia Genómica (CoViGen-Mex). Authors (in alphabetical order): Julio Elias Alvarado-Yaah, Carlos F. Arias, Santiago Ávila-Ríos, Eduardo Becerril-Vargas, Víctor Hugo Borja-Aburto, Celia Boukadida, Cristóbal Cháidez-Quiróz, Juan Bautista Chale-Dzul, Ricardo Ciria Merce, Andreu Comas-García, Célida Duque Molina, Julissa Enciso-Ibarra, José Antonio Enciso-Moreno, Gloria Elena Espinosa-Ayala, Fernando Fontove-Herrera, Daniel Fregoso-Rueda, Victor Eduardo García-Arias, Alejandra García-Gasca, Bruno Gómez-Gil, Jean Pierre González, Irvin González-López, Concepción Grajales-Muñiz, Ricardo Grande, Rosa María Gutiérrez Rios, Jesús Hernández, Alejandra Hernández-Terán, Alfredo Herrera-Estrella, Carla Ivón Herrera-Najera, Pavel Isa, Verónica Mata-Haro, Daniel Lira Morales, Susana Lopez, Antonio Loza Román, Brenda Irasema Maldonado-Meza, Bernardo Martínez-Miguel, José Arturo Martínez-Orozco, Célida Martínez- Rodríguez, Margarita Matías-Florentino, Fidencio Mejía-Nepomuceno, María Guadalupe de Jesús Mireles-Rivera, Gloria María Molina-Salinas, Hector Montoya-Fuentes, , Mario Mújica-Sánchez, José Esteban Muñoz-Medina, José de Jesús Nuñez-Contreras, Alicia Ocaña-Mondragón, Luis Alberto Ochoa-Carrera, Hector Esteban Paz-Juárez, Marissa Perez-Garcia, Francisco Pulido, Helen Haydee Fernanda Ramírez-Plascencia, Jorge Salas-Hernández, Angel Gustavo Salas-Lais, Alejandro Sánchez-Flores, Clara Esperanza Santacruz-Tinoco, María Guadalupe Santiago-Mauricio, Selene Zárate, Nelly Sélem-Mojica, Blanca Taboada, Gloria Vazquez, Joel Armando Vázquez-Pérez. |
| EPI_ISL_2681271 | Centro de Investigación Biomédica del Noreste (CIBIN) | Instituto de Biotecnología de la UNAM | Consorcio Mexicano de Vigilancia Genómica (CoViGen-Mex). Authors (in alphabetical order): Julio Elias Alvarado-Yaah, Carlos F. Arias, Santiago Ávila-Ríos, Eduardo Becerril-Vargas, Víctor Hugo Borja-Aburto, Celia Boukadida, Cristóbal Cháidez-Quiróz, Juan Bautista Chale-Dzul, Ricardo Ciria Merce, Andreu Comas-García, Célida Duque Molina, Julissa Enciso-Ibarra, José Antonio Enciso-Moreno, Gloria Elena Espinosa-Ayala, Fernando Fontove-Herrera, Daniel Fregoso-Rueda, Victor Eduardo García-Arias, Alejandra García-Gasca, Bruno Gómez-Gil, Jean Pierre González, Irvin González-López, Concepción Grajales-Muñiz, Ricardo Grande, Rosa María Gutiérrez Rios, Jesús Hernández, Alejandra Hernández-Terán, Alfredo Herrera-Estrella, Carla Ivón Herrera-Najera, Pavel Isa, Verónica Mata-Haro, Daniel Lira Morales, Susana Lopez, Antonio Loza Román, Brenda Irasema Maldonado-Meza, Bernardo Martínez-Miguel, José Arturo Martínez-Orozco, Célida Martínez- Rodríguez, Margarita Matías-Florentino, Fidencio Mejía-Nepomuceno, María Guadalupe de Jesús Mireles-Rivera, Gloria María Molina-Salinas, Hector Montoya-Fuentes, , Mario Mújica-Sánchez, José Esteban Muñoz-Medina, José de Jesús Nuñez-Contreras, Alicia Ocaña-Mondragón, Luis Alberto Ochoa-Carrera, Hector Esteban Paz-Juárez, Marissa Perez-Garcia, Francisco Pulido, Helen Haydee Fernanda Ramírez-Plascencia, Jorge Salas-Hernández, Angel Gustavo Salas-Lais, Alejandro Sánchez-Flores, Clara Esperanza Santacruz-Tinoco, María Guadalupe Santiago-Mauricio, Selene Zárate, Nelly Sélem-Mojica, Blanca Taboada, Gloria Vazquez, Joel Armando Vázquez-Pérez. |
| EPI_ISL_2681275 | Unidad de Investigación Médica de Yucatán (UIMY) | Instituto de Biotecnología de la UNAM | Consorcio Mexicano de Vigilancia Genómica (CoViGen-Mex). Authors (in alphabetical order): Julio Elias Alvarado-Yaah, Carlos F. Arias, Santiago Ávila-Ríos, Eduardo Becerril-Vargas, Víctor Hugo Borja-Aburto, Celia Boukadida, Cristóbal Cháidez-Quiróz, Juan Bautista Chale-Dzul, Ricardo Ciria Merce, Andreu Comas-García, Célida Duque Molina, Julissa Enciso-Ibarra, José Antonio Enciso-Moreno, Gloria Elena Espinosa-Ayala, Fernando Fontove-Herrera, Daniel Fregoso-Rueda, Victor Eduardo García-Arias, Alejandra García-Gasca, Bruno Gómez-Gil, Jean Pierre González, Irvin González-López, Concepción Grajales-Muñiz, Ricardo Grande, Rosa María Gutiérrez Rios, Jesús Hernández, Alejandra Hernández-Terán, Alfredo Herrera-Estrella, Carla Ivón Herrera-Najera, Pavel Isa, Verónica Mata-Haro, Daniel Lira Morales, Susana Lopez, Antonio Loza Román, Brenda Irasema Maldonado-Meza, Bernardo Martínez-Miguel, José Arturo Martínez-Orozco, Célida Martínez- Rodríguez, Margarita Matías-Florentino, Fidencio Mejía-Nepomuceno, María Guadalupe de Jesús Mireles-Rivera, Gloria María Molina-Salinas, Hector Montoya-Fuentes, , Mario Mújica-Sánchez, José Esteban Muñoz-Medina, José de Jesús Nuñez-Contreras, Alicia Ocaña-Mondragón, Luis Alberto Ochoa-Carrera, Hector Esteban Paz-Juárez, Marissa |

| | | | |
|---|---|---|---|
| | | | Perez-Garcia, Francisco Pulido, Helen Haydee Fernanda Ramírez-Plascencia, Jorge Salas-Hernández, Angel Gustavo Salas-Lais, Alejandro Sánchez-Flores, Clara Esperanza Santacruz-Tinoco, María Guadalupe Santiago-Mauricio, Selene Zárate, Nelly Sélem-Mojica, Blanca Taboada, Gloria Vazquez, Joel Armando Vázquez-Pérez. |
| EPI_ISL_2681276, EPI_ISL_2681278, EPI_ISL_2681279, EPI_ISL_2681281, EPI_ISL_2681282, EPI_ISL_2681283, EPI_ISL_2681284, EPI_ISL_2681288 | Laboratorio Central de Epidemiología (LCE) | Instituto de Biotecnología de la UNAM | Consorcio Mexicano de Vigilancia Genómica (CoViGen-Mex). Authors (in alphabetical order): Julio Elias Alvarado-Yaah, Carlos F. Arias, Santiago Ávila-Ríos, Eduardo Becerril-Vargas, Víctor Hugo Borja-Aburto, Celia Boukadida, Cristóbal Cháidez-Quiróz, Juan Bautista Chale-Dzul, Ricardo Ciria Merce, Andreu Comas-García, Célida Duque Molina, Julissa Enciso-Ibarra, José Antonio Enciso-Moreno, Gloria Elena Espinosa-Ayala, Fernando Fontove-Herrera, Daniel Fregoso-Rueda, Victor Eduardo García-Arias, Alejandra García-Gasca, Bruno Gómez-Gil, Jean Pierre González, Irvin González-López, Concepción Grajales-Muñiz, Ricardo Grande, Rosa María Gutiérrez Rios, Jesús Hernández, Alejandra Hernández-Terán, Alfredo Herrera-Estrella, Carla Ivón Herrera-Najera, Pavel Isa, Verónica Mata-Haro, Daniel Lira Morales, Susana Lopez, Antonio Loza Román, Brenda Irasema Maldonado-Meza, Bernardo Martínez-Miguel, José Arturo Martínez-Orozco, Célida Martínez- Rodríguez, Margarita Matías-Florentino, Fidencio Mejía-Nepomuceno, María Guadalupe de Jesús Mireles-Rivera, Gloria María Molina-Salinas, Hector Montoya-Fuentes, , Mario Mújica-Sánchez, José Esteban Muñoz-Medina, José de Jesús Nuñez-Contreras, Alicia Ocaña-Mondragón, Luis Alberto Ochoa-Carrera, Hector Esteban Paz-Juárez, Marissa Perez-Garcia, Francisco Pulido, Helen Haydee Fernanda Ramírez-Plascencia, Jorge Salas-Hernández, Angel Gustavo Salas-Lais, Alejandro Sánchez-Flores, Clara Esperanza Santacruz-Tinoco, María Guadalupe Santiago-Mauricio, Selene Zárate, Nelly Sélem-Mojica, Blanca Taboada, Gloria Vazquez, Joel Armando Vázquez-Pérez. |
| EPI_ISL_2681289 | Centro de Investigación Biomédica del Noreste (CIBIN) | Instituto de Biotecnología de la UNAM | Consorcio Mexicano de Vigilancia Genómica (CoViGen-Mex). Authors (in alphabetical order): Julio Elias Alvarado-Yaah, Carlos F. Arias, Santiago Ávila-Ríos, Eduardo Becerril-Vargas, Víctor Hugo Borja-Aburto, Celia Boukadida, Cristóbal Cháidez-Quiróz, Juan Bautista Chale-Dzul, Ricardo Ciria Merce, Andreu Comas-García, Célida Duque Molina, Julissa Enciso-Ibarra, José Antonio Enciso-Moreno, Gloria Elena Espinosa-Ayala, Fernando Fontove-Herrera, Daniel Fregoso-Rueda, Victor Eduardo García-Arias, Alejandra García-Gasca, Bruno Gómez-Gil, Jean Pierre González, Irvin González-López, Concepción Grajales-Muñiz, Ricardo Grande, Rosa María Gutiérrez Rios, Jesús Hernández, Alejandra Hernández-Terán, Alfredo Herrera-Estrella, Carla Ivón Herrera-Najera, Pavel Isa, Verónica Mata-Haro, Daniel Lira Morales, Susana Lopez, Antonio Loza Román, Brenda Irasema Maldonado-Meza, Bernardo Martínez-Miguel, José Arturo Martínez-Orozco, Célida Martínez- Rodríguez, Margarita Matías-Florentino, Fidencio Mejía-Nepomuceno, María Guadalupe de Jesús Mireles-Rivera, Gloria María Molina-Salinas, Hector Montoya-Fuentes, , Mario Mújica-Sánchez, José Esteban Muñoz-Medina, José de Jesús Nuñez-Contreras, Alicia Ocaña-Mondragón, Luis Alberto Ochoa-Carrera, Hector Esteban Paz-Juárez, Marissa Perez-Garcia, Francisco Pulido, Helen Haydee Fernanda Ramírez-Plascencia, Jorge Salas-Hernández, Angel Gustavo Salas-Lais, Alejandro Sánchez-Flores, Clara Esperanza Santacruz-Tinoco, María Guadalupe Santiago-Mauricio, Selene Zárate, Nelly Sélem-Mojica, Blanca Taboada, Gloria Vazquez, Joel Armando Vázquez-Pérez. |
| EPI_ISL_2681388 | Laboratorio Central de Epidemiología (LCE) | Instituto de Biotecnología de la UNAM | Consorcio Mexicano de Vigilancia Genómica (CoViGen-Mex). Authors (in alphabetical order): Julio Elias Alvarado-Yaah, Carlos F. Arias, Santiago Ávila-Ríos, Eduardo Becerril-Vargas, Víctor Hugo Borja-Aburto, Celia Boukadida, Cristóbal Cháidez-Quiróz, Juan Bautista Chale-Dzul, Ricardo Ciria Merce, Andreu Comas-García, Célida Duque Molina, Julissa Enciso-Ibarra, José Antonio Enciso-Moreno, Gloria Elena Espinosa-Ayala, Fernando Fontove-Herrera, Daniel Fregoso-Rueda, Victor Eduardo García-Arias, Alejandra García-Gasca, Bruno Gómez-Gil, Jean Pierre González, Irvin González-López, Concepción Grajales-Muñiz, Ricardo Grande, Rosa María Gutiérrez Rios, Jesús Hernández, Alejandra Hernández-Terán, Alfredo Herrera-Estrella, Carla Ivón Herrera-Najera, Pavel Isa, Verónica Mata-Haro, Daniel Lira Morales, Susana Lopez, Antonio Loza Román, Brenda Irasema Maldonado-Meza, Bernardo Martínez-Miguel, José Arturo Martínez-Orozco, Célida Martínez- Rodríguez, Margarita Matías-Florentino, Fidencio Mejía-Nepomuceno, María Guadalupe de Jesús Mireles-Rivera, Gloria María Molina-Salinas, Hector Montoya-Fuentes, , Mario Mújica-Sánchez, José Esteban Muñoz-Medina, José de Jesús Nuñez-Contreras, Alicia Ocaña-Mondragón, Luis Alberto Ochoa-Carrera, Hector Esteban Paz-Juárez, Marissa Perez-Garcia, Francisco Pulido, Helen Haydee Fernanda Ramírez-Plascencia, Jorge Salas-Hernández, Angel Gustavo Salas-Lais, Alejandro Sánchez-Flores, Clara Esperanza Santacruz-Tinoco, María Guadalupe Santiago-Mauricio, Selene Zárate, Nelly Sélem-Mojica, Blanca Taboada, Gloria Vazquez, Joel Armando Vázquez-Pérez. |
| EPI_ISL_2681442, EPI_ISL_2681443, EPI_ISL_2681444, EPI_ISL_2681445, EPI_ISL_2681446, EPI_ISL_2681447, EPI_ISL_2681448, EPI_ISL_2681449 | Althaia. Xarxa Assistencial Universitària de Manresa | IrsiCaixa | Marc Noguera-Julian, Mariona Parera, Maria Casadellà, Pilar Armengol, Francesc Catala-Moll, Roger Paredes, Bonaventura Clotet Gloria Trujillo, Rafel Perez Vidal, Jaume Trapé, Carolina Gonzalez-Fernandez, Miquel Micó, Antonia Flor, Roger Paredes, Eulalia Grau, Bonaventura Clotet |
| EPI_ISL_2686028, EPI_ISL_2686029, EPI_ISL_2686030, EPI_ISL_2686031, EPI_ISL_2686032, EPI_ISL_2686033, EPI_ISL_2686034, EPI_ISL_2686035, EPI_ISL_2686036, EPI_ISL_2686037, EPI_ISL_2686038, EPI_ISL_2686039, EPI_ISL_2686040, EPI_ISL_2686041, EPI_ISL_2686042, EPI_ISL_2686043, EPI_ISL_2686044, EPI_ISL_2686045, EPI_ISL_2686046, EPI_ISL_2686047, EPI_ISL_2686048, EPI_ISL_2686049, EPI_ISL_2686050, EPI_ISL_2686051, EPI_ISL_2686052, EPI_ISL_2686053 | | | |
| see above | Università degli Studi di Perugia | Istituto Zooprofilattico Sperimentale dell'Abruzzo e Molise "G. Caporale" | Mencacci A, Camilloni B, Biagetti M, Giammarioli M, Lorusso A, Marcacci M, Di Domenico M, Ancora M, Curini V, Mangone I, Rinaldi A, Delli Compagni E, Scialabba S, Di Pasquale A, Cammà C, Puglia I, Calistri P, Savini G |
| EPI_ISL_2686054, EPI_ISL_2686057, EPI_ISL_2686059 | Laboratorio Analisi Osp. Città di Castello - Azienda USL Umbria1 | Istituto Zooprofilattico Sperimentale dell'Abruzzo e Molise "G. Caporale" | Malagigi V, Taccani P,Lorusso A, Marcacci M, Di Domenico M, Ancora M, Curini V, Mangone I, Rinaldi A, Delli Compagni E, Scialabba S, Di Pasquale A, Cammà C, Puglia I, Calistri P, Savini G |
| EPI_ISL_2686060, EPI_ISL_2686061 | Università degli Studi di Perugia | Istituto Zooprofilattico Sperimentale dell'Abruzzo e Molise "G. Caporale" | Mencacci A, Camilloni B, Biagetti M, Giammarioli M, Lorusso A, Marcacci M, Di Domenico M, Ancora M, Curini V, Mangone I, Rinaldi A, Delli Compagni E, Scialabba S, Di Pasquale A, Cammà C, Puglia I, Calistri P, Savini G |
| EPI_ISL_2686062 | Azienda Ospedaliera Terni | Istituto Zooprofilattico Sperimentale dell'Abruzzo e Molise "G. Caporale" | Palumbo M, Scaccetti A, Lorusso A, Marcacci M, Di Domenico M, Ancora M, Curini V, Mangone I, Rinaldi A, Delli Compagni E, Scialabba S, Di Pasquale A, Cammà C, Puglia I, Calistri P, Savini G |
| EPI_ISL_2686063, EPI_ISL_2686064, EPI_ISL_2686065 | Azienda USL Umbria 2 | Istituto Zooprofilattico Sperimentale dell'Abruzzo e Molise "G. Caporale" | Proietti A, Pistoni E, Lorusso A, Marcacci M, Di Domenico M, Ancora M, Curini V, Mangone I, Rinaldi A, Delli Compagni E, Scialabba S, Di Pasquale A, Cammà C, Puglia I, Calistri P, Savini G |
| EPI_ISL_2686066, EPI_ISL_2686067, EPI_ISL_2686068, EPI_ISL_2686069 | Università degli Studi di Perugia | Istituto Zooprofilattico Sperimentale dell'Abruzzo e Molise "G. Caporale" | Mencacci A, Camilloni B, Biagetti M, Giammarioli M, Lorusso A, Marcacci M, Di Domenico M, Ancora M, Curini V, Mangone I, Rinaldi A, Delli Compagni E, Scialabba S, Di Pasquale A, Cammà C, Puglia I, Calistri P, Savini G |
| EPI_ISL_2686070, EPI_ISL_2686071, EPI_ISL_2686072, EPI_ISL_2686073 | Laboratorio Analisi Osp. Città di Castello - Azienda USL Umbria1 | Istituto Zooprofilattico Sperimentale dell'Abruzzo e Molise "G. Caporale" | Malagigi V, Taccani P,Lorusso A, Marcacci M, Di Domenico M, Ancora M, Curini V, Mangone I, Rinaldi A, Delli Compagni E, Scialabba S, Di Pasquale A, Cammà C, Puglia I, Calistri P, Savini G |
| EPI_ISL_2686074, EPI_ISL_2686075, EPI_ISL_2686076, EPI_ISL_2686077, EPI_ISL_2686078, EPI_ISL_2686079, EPI_ISL_2686080 | Azienda USL Umbria 2 | Istituto Zooprofilattico Sperimentale dell'Abruzzo e Molise "G. Caporale" | Proietti A, Pistoni E, Lorusso A, Marcacci M, Di Domenico M, Ancora M, Curini V, Mangone I, Rinaldi A, Delli Compagni E, Scialabba S, Di Pasquale A, Cammà C, Puglia I, Calistri P, Savini G |
| EPI_ISL_2686081, EPI_ISL_2686082, EPI_ISL_2686083, EPI_ISL_2686084 | Azienda Ospedaliera Terni | Istituto Zooprofilattico Sperimentale dell'Abruzzo e Molise "G. Caporale" | Palumbo M, Scaccetti A, Lorusso A, Marcacci M, Di Domenico M, Ancora M, Curini V, Mangone I, Rinaldi A, Delli Compagni E, Scialabba S, Di Pasquale A, Cammà C, Puglia I, Calistri P, Savini G |
| EPI_ISL_2686390 | AZ Zeno | AZ Sint Jan | Marijke Reynders, Patrick Descheemaeker, Laurien Hoornaert, Thomas Van Landschoot, Marie Madeleine Chabert-Consen, Jorn Hellemans |
| EPI_ISL_2687993 | National Institute of Public Health | National Institute of Infectious Diseases-Prof. Dr. Matei Bals Molecular Diagnostics Laboratory | Corina Casangiu, Leontina Banica, Marius Surleac, Petre Milu, Robert Hohan, Simona Paraschiv, Dan Otelea |
| EPI_ISL_2687995, EPI_ISL_2687996, EPI_ISL_2688000 | National Institute of Infectious Diseases-Prof. Dr. Matei Bals Molecular Diagnostics Laboratory | National Institute of Infectious Diseases-Prof. Dr. Matei Bals Molecular Diagnostics Laboratory | Corina Casangiu, Leontina Banica, Marius Surleac, Petre Milu, Robert Hohan, Simona Paraschiv, Dan Otelea |
| EPI_ISL_2688319 | Puerto Rico Department of Health | Centers for Disease Control and Prevention, Dengue Branch | Gilberto A. Santiago, Glenda Gonzalez, Betzabel Flores, Keyla Charriez, Gabriela Paz-Bailey, Jorge L. Munoz-Jordan |
| EPI_ISL_2688913, EPI_ISL_2688914 | LESP Jalisco | Instituto de Diagnostico y Referencia Epidemiologicos (INDRE) | Claudia Wong-Arambula, Abril Rodriguez-Maldonado, Vanessa Rivero-Arredondo, Ariadna Medina-Benitez, Joaquin Quiroz-Mercado, Sergio Rangel-Guerrero, Natividad Cruz-Ortiz, Tatiana Nunez-Garcia, Gisela Barrera-Badillo, Lucia Hernandez-Rivas, Irma Lopez-Martinez, Ernesto Ramirez-Gonzalez. |
| EPI_ISL_2688915, EPI_ISL_2688916, EPI_ISL_2688917, EPI_ISL_2688918 | LESP Morelos | Instituto de Diagnostico y Referencia Epidemiologicos (INDRE) | Claudia Wong-Arambula, Abril Rodriguez-Maldonado, Vanessa Rivero-Arredondo, Ariadna Medina-Benitez, Joaquin Quiroz-Mercado, Sergio Rangel-Guerrero, Natividad Cruz-Ortiz, Tatiana Nunez-Garcia, Gisela Barrera-Badillo, Lucia Hernandez-Rivas, Irma Lopez-Martinez, Ernesto Ramirez-Gonzalez. |

| | | | |
|---|---|---|---|
| EPI_ISL_2688924, EPI_ISL_2688925 | LESP Queretaro | Instituto de Diagnostico y Referencia Epidemiologicos (INDRE) | Claudia Wong-Arambula, Abril Rodriguez-Maldonado, Vanessa Rivero-Arredondo, Ariadna Medina-Benitez, Joaquin Quiroz-Mercado, Sergio Rangel-Guerrero, Natividad Cruz-Ortiz, Tatiana Nunez-Garcia, Gisela Barrera-Badillo, Lucia Hernandez-Rivas, Irma Lopez-Martinez, Ernesto Ramirez-Gonzalez. |
| EPI_ISL_2688927, EPI_ISL_2688928, EPI_ISL_2688929, EPI_ISL_2688930, EPI_ISL_2688931, EPI_ISL_2688932, EPI_ISL_2688933, EPI_ISL_2688934, EPI_ISL_2688935, EPI_ISL_2688936, EPI_ISL_2688937, EPI_ISL_2688938, EPI_ISL_2688939, EPI_ISL_2688940, EPI_ISL_2688941, EPI_ISL_2688942 | | | |
| see above | LESP Tabasco | Instituto de Diagnostico y Referencia Epidemiologicos (INDRE) | Claudia Wong-Arambula, Abril Rodriguez-Maldonado, Vanessa Rivero-Arredondo, Ariadna Medina-Benitez, Joaquin Quiroz-Mercado, Sergio Rangel-Guerrero, Natividad Cruz-Ortiz, Tatiana Nunez-Garcia, Gisela Barrera-Badillo, Lucia Hernandez-Rivas, Irma Lopez-Martinez, Ernesto Ramirez-Gonzalez. |
| EPI_ISL_2688945, EPI_ISL_2688946, EPI_ISL_2688947 | LESP Michoacan | Instituto de Diagnostico y Referencia Epidemiologicos (INDRE) | Claudia Wong-Arambula, Abril Rodriguez-Maldonado, Vanessa Rivero-Arredondo, Ariadna Medina-Benitez, Joaquin Quiroz-Mercado, Sergio Rangel-Guerrero, Natividad Cruz-Ortiz, Tatiana Nunez-Garcia, Gisela Barrera-Badillo, Lucia Hernandez-Rivas, Irma Lopez-Martinez, Ernesto Ramirez-Gonzalez. |
| EPI_ISL_2688948, EPI_ISL_2688949, EPI_ISL_2688950, EPI_ISL_2688951, EPI_ISL_2688952, EPI_ISL_2688953, EPI_ISL_2688954, EPI_ISL_2688955, EPI_ISL_2688956, EPI_ISL_2688958 | LESP Tabasco | Instituto de Diagnostico y Referencia Epidemiologicos (INDRE) | Claudia Wong-Arambula, Abril Rodriguez-Maldonado, Vanessa Rivero-Arredondo, Ariadna Medina-Benitez, Joaquin Quiroz-Mercado, Sergio Rangel-Guerrero, Natividad Cruz-Ortiz, Tatiana Nunez-Garcia, Gisela Barrera-Badillo, Lucia Hernandez-Rivas, Irma Lopez-Martinez, Ernesto Ramirez-Gonzalez. |
| EPI_ISL_2688969 | LESP Queretaro | Instituto de Diagnostico y Referencia Epidemiologicos (INDRE) | Claudia Wong-Arambula, Abril Rodriguez-Maldonado, Vanessa Rivero-Arredondo, Ariadna Medina-Benitez, Joaquin Quiroz-Mercado, Sergio Rangel-Guerrero, Natividad Cruz-Ortiz, Tatiana Nunez-Garcia, Gisela Barrera-Badillo, Lucia Hernandez-Rivas, Irma Lopez-Martinez, Ernesto Ramirez-Gonzalez. |
| EPI_ISL_2688970 | LESP Michoacan | Instituto de Diagnostico y Referencia Epidemiologicos (INDRE) | Claudia Wong-Arambula, Abril Rodriguez-Maldonado, Vanessa Rivero-Arredondo, Ariadna Medina-Benitez, Joaquin Quiroz-Mercado, Sergio Rangel-Guerrero, Natividad Cruz-Ortiz, Tatiana Nunez-Garcia, Gisela Barrera-Badillo, Lucia Hernandez-Rivas, Irma Lopez-Martinez, Ernesto Ramirez-Gonzalez. |
| EPI_ISL_2688971 | LESP Zacatecas | Instituto de Diagnostico y Referencia Epidemiologicos (INDRE) | Claudia Wong-Arambula, Abril Rodriguez-Maldonado, Vanessa Rivero-Arredondo, Ariadna Medina-Benitez, Joaquin Quiroz-Mercado, Sergio Rangel-Guerrero, Natividad Cruz-Ortiz, Tatiana Nunez-Garcia, Gisela Barrera-Badillo, Lucia Hernandez-Rivas, Irma Lopez-Martinez, Ernesto Ramirez-Gonzalez. |
| EPI_ISL_2688972 | LESP Hidalgo | Instituto de Diagnostico y Referencia Epidemiologicos (INDRE) | Claudia Wong-Arambula, Abril Rodriguez-Maldonado, Vanessa Rivero-Arredondo, Ariadna Medina-Benitez, Joaquin Quiroz-Mercado, Sergio Rangel-Guerrero, Natividad Cruz-Ortiz, Tatiana Nunez-Garcia, Gisela Barrera-Badillo, Lucia Hernandez-Rivas, Irma Lopez-Martinez, Ernesto Ramirez-Gonzalez. |
| EPI_ISL_2688973 | LESP Morelos | Instituto de Diagnostico y Referencia Epidemiologicos (INDRE) | Claudia Wong-Arambula, Abril Rodriguez-Maldonado, Vanessa Rivero-Arredondo, Ariadna Medina-Benitez, Joaquin Quiroz-Mercado, Sergio Rangel-Guerrero, Natividad Cruz-Ortiz, Tatiana Nunez-Garcia, Gisela Barrera-Badillo, Lucia Hernandez-Rivas, Irma Lopez-Martinez, Ernesto Ramirez-Gonzalez. |
| EPI_ISL_2688974 | LESP Guanajuato | Instituto de Diagnostico y Referencia Epidemiologicos (INDRE) | Claudia Wong-Arambula, Abril Rodriguez-Maldonado, Vanessa Rivero-Arredondo, Ariadna Medina-Benitez, Joaquin Quiroz-Mercado, Sergio Rangel-Guerrero, Natividad Cruz-Ortiz, Tatiana Nunez-Garcia, Gisela Barrera-Badillo, Lucia Hernandez-Rivas, Irma Lopez-Martinez, Ernesto Ramirez-Gonzalez. |
| EPI_ISL_2688975 | LESP Jalisco | Instituto de Diagnostico y Referencia Epidemiologicos (INDRE) | Claudia Wong-Arambula, Abril Rodriguez-Maldonado, Vanessa Rivero-Arredondo, Ariadna Medina-Benitez, Joaquin Quiroz-Mercado, Sergio Rangel-Guerrero, Natividad Cruz-Ortiz, Tatiana Nunez-Garcia, Gisela Barrera-Badillo, Lucia Hernandez-Rivas, Irma Lopez-Martinez, Ernesto Ramirez-Gonzalez. |
| EPI_ISL_2688976, EPI_ISL_2688977, EPI_ISL_2688978, EPI_ISL_2688979 | LESP Guanajuato | Instituto de Diagnostico y Referencia Epidemiologicos (INDRE) | Claudia Wong-Arambula, Abril Rodriguez-Maldonado, Vanessa Rivero-Arredondo, Ariadna Medina-Benitez, Joaquin Quiroz-Mercado, Sergio Rangel-Guerrero, Natividad Cruz-Ortiz, Tatiana Nunez-Garcia, Gisela Barrera-Badillo, Lucia Hernandez-Rivas, Irma Lopez-Martinez, Ernesto Ramirez-Gonzalez. |
| EPI_ISL_2688980, EPI_ISL_2688981, EPI_ISL_2688982, EPI_ISL_2688983, EPI_ISL_2688984 | LESP Chihuahua | Instituto de Diagnostico y Referencia Epidemiologicos (INDRE) | Claudia Wong-Arambula, Abril Rodriguez-Maldonado, Vanessa Rivero-Arredondo, Ariadna Medina-Benitez, Joaquin Quiroz-Mercado, Sergio Rangel-Guerrero, Natividad Cruz-Ortiz, Tatiana Nunez-Garcia, Gisela Barrera-Badillo, Lucia Hernandez-Rivas, Irma Lopez-Martinez, Ernesto Ramirez-Gonzalez. |
| EPI_ISL_2688985 | LESP Coahuila | Instituto de Diagnostico y Referencia Epidemiologicos (INDRE) | Claudia Wong-Arambula, Abril Rodriguez-Maldonado, Vanessa Rivero-Arredondo, Ariadna Medina-Benitez, Joaquin Quiroz-Mercado, Sergio Rangel-Guerrero, Natividad Cruz-Ortiz, Tatiana Nunez-Garcia, Gisela Barrera-Badillo, Lucia Hernandez-Rivas, Irma Lopez-Martinez, Ernesto Ramirez-Gonzalez. |
| EPI_ISL_2688986, EPI_ISL_2688987 | LESP Puebla | Instituto de Diagnostico y Referencia Epidemiologicos (INDRE) | Claudia Wong-Arambula, Abril Rodriguez-Maldonado, Vanessa Rivero-Arredondo, Ariadna Medina-Benitez, Joaquin Quiroz-Mercado, Sergio Rangel-Guerrero, Natividad Cruz-Ortiz, Tatiana Nunez-Garcia, Gisela Barrera-Badillo, Lucia Hernandez-Rivas, Irma Lopez-Martinez, Ernesto Ramirez-Gonzalez. |
| EPI_ISL_2688988, EPI_ISL_2688989 | LESP Veracruz | Instituto de Diagnostico y Referencia Epidemiologicos (INDRE) | Claudia Wong-Arambula, Abril Rodriguez-Maldonado, Vanessa Rivero-Arredondo, Ariadna Medina-Benitez, Joaquin Quiroz-Mercado, Sergio Rangel-Guerrero, Natividad Cruz-Ortiz, Tatiana Nunez-Garcia, Gisela Barrera-Badillo, Lucia Hernandez-Rivas, Irma Lopez-Martinez, Ernesto Ramirez-Gonzalez. |
| EPI_ISL_2688990 | LESP Yucatan | Instituto de Diagnostico y Referencia Epidemiologicos (INDRE) | Claudia Wong-Arambula, Abril Rodriguez-Maldonado, Vanessa Rivero-Arredondo, Ariadna Medina-Benitez, Joaquin Quiroz-Mercado, Sergio Rangel-Guerrero, Natividad Cruz-Ortiz, Tatiana Nunez-Garcia, Gisela Barrera-Badillo, Lucia Hernandez-Rivas, Irma Lopez-Martinez, Ernesto Ramirez-Gonzalez. |
| EPI_ISL_2689796 | LESP Ciudad de Mexico | Instituto de Diagnostico y Referencia Epidemiologicos (INDRE) | Claudia Wong-Arambula, Abril Rodriguez-Maldonado, Vanessa Rivero-Arredondo, Ariadna Medina-Benitez, Joaquin Quiroz-Mercado, Sergio Rangel-Guerrero, Natividad Cruz-Ortiz, Tatiana Nunez-Garcia, Gisela Barrera-Badillo, Lucia Hernandez-Rivas, Irma Lopez-Martinez, Ernesto Ramirez-Gonzalez. |
| EPI_ISL_2689806 | LESP Sinaloa | Instituto de Diagnostico y Referencia Epidemiologicos (INDRE) | Claudia Wong-Arambula, Abril Rodriguez-Maldonado, Vanessa Rivero-Arredondo, Ariadna Medina-Benitez, Joaquin Quiroz-Mercado, Sergio Rangel-Guerrero, Natividad Cruz-Ortiz, Tatiana Nunez-Garcia, Gisela Barrera-Badillo, Lucia Hernandez-Rivas, Irma Lopez-Martinez, Ernesto Ramirez-Gonzalez. |
| EPI_ISL_2689808 | LESP Tabasco | Instituto de Diagnostico y Referencia Epidemiologicos (INDRE) | Claudia Wong-Arambula, Abril Rodriguez-Maldonado, Vanessa Rivero-Arredondo, Ariadna Medina-Benitez, Joaquin Quiroz-Mercado, Sergio Rangel-Guerrero, Natividad Cruz-Ortiz, Tatiana Nunez-Garcia, Gisela Barrera-Badillo, Lucia Hernandez-Rivas, Irma Lopez-Martinez, Ernesto Ramirez-Gonzalez. |
| EPI_ISL_2689812 | LESP Michoacan | Instituto de Diagnostico y Referencia Epidemiologicos (INDRE) | Claudia Wong-Arambula, Abril Rodriguez-Maldonado, Vanessa Rivero-Arredondo, Ariadna Medina-Benitez, Joaquin Quiroz-Mercado, Sergio Rangel-Guerrero, Natividad Cruz-Ortiz, Tatiana Nunez-Garcia, Gisela Barrera-Badillo, Lucia Hernandez-Rivas, Irma Lopez-Martinez, Ernesto Ramirez-Gonzalez. |
| EPI_ISL_2689824 | LESP Queretaro | Instituto de Diagnostico y Referencia Epidemiologicos (INDRE) | Claudia Wong-Arambula, Abril Rodriguez-Maldonado, Vanessa Rivero-Arredondo, Ariadna Medina-Benitez, Joaquin Quiroz-Mercado, Sergio Rangel-Guerrero, Natividad Cruz-Ortiz, Tatiana Nunez-Garcia, Gisela Barrera-Badillo, Lucia Hernandez-Rivas, Irma Lopez-Martinez, Ernesto Ramirez-Gonzalez. |
| EPI_ISL_2689827, EPI_ISL_2689828 | LESP Chihuahua | Instituto de Diagnostico y Referencia Epidemiologicos (INDRE) | Claudia Wong-Arambula, Abril Rodriguez-Maldonado, Vanessa Rivero-Arredondo, Ariadna Medina-Benitez, Joaquin Quiroz-Mercado, Sergio Rangel-Guerrero, Natividad Cruz-Ortiz, Tatiana Nunez-Garcia, Gisela Barrera-Badillo, Lucia Hernandez-Rivas, Irma Lopez-Martinez, Ernesto Ramirez-Gonzalez. |
| EPI_ISL_2689829 | LESP Sonora | Instituto de Diagnostico y Referencia Epidemiologicos (INDRE) | Claudia Wong-Arambula, Abril Rodriguez-Maldonado, Vanessa Rivero-Arredondo, Ariadna Medina-Benitez, Joaquin Quiroz-Mercado, Sergio Rangel-Guerrero, Natividad Cruz-Ortiz, Tatiana Nunez-Garcia, Gisela Barrera-Badillo, Lucia Hernandez-Rivas, Irma Lopez-Martinez, Ernesto Ramirez-Gonzalez. |

| | | | |
|---|---|---|---|
| EPI_ISL_2690135 | Ochsner Health | Bioinfoexperts, LLC | Judy Crabtree, Amy Feehan, Rebecca Rose, David J. Nolan, Tessa LaFleur, Julia-Garcia-Diaz, Lucio Miele, Susanna L. Lamers |
| EPI_ISL_2692533, EPI_ISL_2692905 | Clinical Microbiology Lab RSPTN Universitas Hasanuddin | Clinical Microbiology Lab RSPTN Universitas Hasanuddin | Muhammad Nasrum Massi, Handayani Halik, Rizalinda Sjahril, Gita Vita Soraya, Triwibawa, Muhammad Yusuf Usman, Rufika Shari |
| EPI_ISL_2693109, EPI_ISL_2693110 | St James's Hospital, Dublin 8, Ireland. | ELDA biotech | Deirdre M. Broderick, Peter R. Flanagan, Brendan Crowley |
| EPI_ISL_2693909, EPI_ISL_2693910, EPI_ISL_2693911, EPI_ISL_2693912, EPI_ISL_2693913, EPI_ISL_2693914, EPI_ISL_2693915, EPI_ISL_2693916, EPI_ISL_2693917, EPI_ISL_2693918, EPI_ISL_2693919, EPI_ISL_2693920, EPI_ISL_2693921, EPI_ISL_2693922, EPI_ISL_2693923, EPI_ISL_2693924, EPI_ISL_2693925, EPI_ISL_2693926, EPI_ISL_2693927, EPI_ISL_2693928, EPI_ISL_2693929, EPI_ISL_2693930, EPI_ISL_2693931, EPI_ISL_2693932, EPI_ISL_2693933, EPI_ISL_2693934, EPI_ISL_2693935, EPI_ISL_2693936, EPI_ISL_2693937, EPI_ISL_2693938, EPI_ISL_2693939, EPI_ISL_2693940, EPI_ISL_2693941, EPI_ISL_2693942, EPI_ISL_2693943, EPI_ISL_2693944, EPI_ISL_2693945, EPI_ISL_2693946, EPI_ISL_2693947, EPI_ISL_2693948, EPI_ISL_2693949, EPI_ISL_2693950, EPI_ISL_2693951 | | | |
| see above | Virology Unit, Institut Pasteur du Cambodge | Virology Unit, Institut Pasteur du Cambodge | Cecile Troupin, Leakhena Pum, Jurre Y Siegers, Ly Sovann, Kraing Sidonn, Yi Sengdoeurn, Chin Savuth, Chau Darapheak, Veasna Duong, Erik A Karlsson |
| EPI_ISL_2693960, EPI_ISL_2693963, EPI_ISL_2693964, EPI_ISL_2693965, EPI_ISL_2693966, EPI_ISL_2693967, EPI_ISL_2693968, EPI_ISL_2693969, EPI_ISL_2693970 | laboratorio Microbiologia PO Cardarelli | laboratorio Microbiologia PO Cardarelli | Felice V., Niro G. Scutella' M. |
| EPI_ISL_2695878, EPI_ISL_2695879, EPI_ISL_2695880, EPI_ISL_2695883 | Osaka General Medical Center | Department of Developmental Medicine, Research Institute, Osaka Women's and Children's Hospital | Itaru Yanagihara, Yukiko Nakura |
| EPI_ISL_2695962 | Department of Developmental Medicine, Research Institute, Osaka Women's and Children's Hospital | Department of Developmental Medicine, Research Institute, Osaka Women's and Children's Hospital | Itaru Yanagihara, Yukiko Nakura |
| EPI_ISL_2705575 | Centers for Disease Control, R.O.C. (Taiwan) | Centers for Disease Control, R.O.C. (Taiwan) | Ji-Rong Yang, Yu-Chi-Lin, Jung-Jung Mu, Ming-Tsan-Liu |
| EPI_ISL_2709256 | CHU Pontchaillou | CHU Pontchaillou | GROLHIER Claire, DENOUAL Florent, ETCHEVERRY Amandine, SASSI Mohamed, JAGLINE Steven, FEBREAU Christine, PRONIER Charlotte, GALIBERT Marie Dominique, DE TAYRAC Marie, THIBAULT Vincent |
| EPI_ISL_2709257 | Labo Analyses Med | National Reference Center for Viruses of Respiratory Infections, Institut Pasteur, Paris | Marion Barbet, Sylvie Behillil, Méline Bizard, Angela Brisebarre, Camille Capel, Vincent Enouf, Louise Lefrançois, Frédéric Lemoine, Christophe Malabat, Corinne Maufrais, Etienne Simon-Lorière, Maud Vanpeene, Sylvie Van der Werf ,Franck Ennouchi |
| EPI_ISL_2709258 | Hospital | National Reference Center for Viruses of Respiratory Infections, Institut Pasteur, Paris | Marion Barbet, Sylvie Behillil, Méline Bizard, Angela Brisebarre, Camille Capel, Vincent Enouf, Louise Lefrançois, Frédéric Lemoine, Christophe Malabat, Corinne Maufrais, Etienne Simon-Lorière, Maud Vanpeene, Sylvie Van der Werf ,Salma Ben Hadj Yahia |
| EPI_ISL_2709259 | Hospital | National Reference Center for Viruses of Respiratory Infections, Institut Pasteur, Paris | Marion Barbet, Sylvie Behillil, Méline Bizard, Angela Brisebarre, Camille Capel, Vincent Enouf, Louise Lefrançois, Frédéric Lemoine, Christophe Malabat, Corinne Maufrais, Damien Mornico, Etienne Simon-Lorière, Maud Vanpeene, Sylvie Van der Werf ,Laurent Bret |
| EPI_ISL_2709261 | Labo Analyses Med | National Reference Center for Viruses of Respiratory Infections, Institut Pasteur, Paris | Marion Barbet, Sylvie Behillil, Méline Bizard, Angela Brisebarre, Camille Capel, Vincent Enouf, Louise Lefrançois, Frédéric Lemoine, Christophe Malabat, Corinne Maufrais, Etienne Simon-Lorière, Maud Vanpeene, Sylvie Van der Werf ,Patricia Tamby |
| EPI_ISL_2709262 | Labo Analyses Med | National Reference Center for Viruses of Respiratory Infections, Institut Pasteur, Paris | Marion Barbet, Sylvie Behillil, Méline Bizard, Angela Brisebarre, Camille Capel, Vincent Enouf, Louise Lefrançois, Frédéric Lemoine, Christophe Malabat, Corinne Maufrais, Etienne Simon-Lorière, Maud Vanpeene, Sylvie Van der Werf ,Vicky Le |
| EPI_ISL_2709263 | Labo Analyses Med | National Reference Center for Viruses of Respiratory Infections, Institut Pasteur, Paris | Marion Barbet, Sylvie Behillil, Méline Bizard, Angela Brisebarre, Camille Capel, Vincent Enouf, Louise Lefrançois, Frédéric Lemoine, Christophe Malabat, Corinne Maufrais, Damien Mornico, Etienne Simon-Lorière, Maud Vanpeene, Sylvie Van der Werf ,Vicky Le |
| EPI_ISL_2709264 | Labo Analyses Med | National Reference Center for Viruses of Respiratory Infections, Institut Pasteur, Paris | Marion Barbet, Sylvie Behillil, Méline Bizard, Angela Brisebarre, Camille Capel, Vincent Enouf, Louise Lefrançois, Frédéric Lemoine, Christophe Malabat, Corinne Maufrais, Etienne Simon-Lorière, Maud Vanpeene, Sylvie Van der Werf ,GréGoire Potiron |
| EPI_ISL_2709265 | Labo Analyses Med | National Reference Center for Viruses of Respiratory Infections, Institut Pasteur, Paris | Marion Barbet, Sylvie Behillil, Méline Bizard, Angela Brisebarre, Camille Capel, Vincent Enouf, Louise Lefrançois, Frédéric Lemoine, Christophe Malabat, Corinne Maufrais, Etienne Simon-Lorière, Maud Vanpeene, Sylvie Van der Werf ,Hugues Leroy |
| EPI_ISL_2709266 | Hospital | National Reference Center for Viruses of Respiratory Infections, Institut Pasteur, Paris | Marion Barbet, Sylvie Behillil, Méline Bizard, Angela Brisebarre, Camille Capel, Vincent Enouf, Louise Lefrançois, Frédéric Lemoine, Christophe Malabat, Corinne Maufrais, Etienne Simon-Lorière, Maud Vanpeene, Sylvie Van der Werf ,Farida Benouadi |
| EPI_ISL_2709267 | Hospital | National Reference Center for Viruses of Respiratory Infections, Institut Pasteur, Paris | Marion Barbet, Sylvie Behillil, Méline Bizard, Angela Brisebarre, Camille Capel, Vincent Enouf, Louise Lefrançois, Frédéric Lemoine, Christophe Malabat, Corinne Maufrais, Etienne Simon-Lorière, Maud Vanpeene, Sylvie Van der Werf ,Laurence Courdavault |
| EPI_ISL_2709268, EPI_ISL_2709270, EPI_ISL_2709271 | Labo Analyses Med | National Reference Center for Viruses of Respiratory Infections, Institut Pasteur, Paris | Marion Barbet, Sylvie Behillil, Méline Bizard, Angela Brisebarre, Camille Capel, Vincent Enouf, Louise Lefrançois, Frédéric Lemoine, Christophe Malabat, Corinne Maufrais, Etienne Simon-Lorière, Maud Vanpeene, Sylvie Van der Werf ,Anne-Laure Garand |
| EPI_ISL_2709274 | Labo Analyses Med | National Reference Center for Viruses of Respiratory Infections, Institut Pasteur, Paris | Marion Barbet, Sylvie Behillil, Méline Bizard, Angela Brisebarre, Camille Capel, Vincent Enouf, Louise Lefrançois, Frédéric Lemoine, Christophe Malabat, Corinne Maufrais, Damien Mornico, Etienne Simon-Lorière, Maud Vanpeene, Sylvie Van der Werf ,Anne-Laure Garand |
| EPI_ISL_2709275 | Labo Analyses Med | National Reference Center for Viruses of Respiratory Infections, Institut Pasteur, Paris | Marion Barbet, Sylvie Behillil, Méline Bizard, Angela Brisebarre, Camille Capel, Vincent Enouf, Louise Lefrançois, Frédéric Lemoine, Christophe Malabat, Corinne Maufrais, Etienne Simon-Lorière, Maud Vanpeene, Sylvie Van der Werf ,Anne-Laure Garand |
| EPI_ISL_2709277 | Labo Analyses Med | National Reference Center for Viruses of Respiratory Infections, Institut Pasteur, Paris | Marion Barbet, Sylvie Behillil, Méline Bizard, Angela Brisebarre, Camille Capel, Vincent Enouf, Louise Lefrançois, Frédéric Lemoine, Christophe Malabat, Corinne Maufrais, Damien Mornico, Etienne Simon-Lorière, Maud Vanpeene, Sylvie Van der Werf ,Hugues Leroy |
| EPI_ISL_2709279, EPI_ISL_2709280, EPI_ISL_2709281, EPI_ISL_2709282, EPI_ISL_2709283, EPI_ISL_2709284, EPI_ISL_2709285, EPI_ISL_2709286, EPI_ISL_2709287, EPI_ISL_2709288, EPI_ISL_2709289, EPI_ISL_2709290, EPI_ISL_2709291, EPI_ISL_2709292, EPI_ISL_2709293 | | | |
| see above | Labo Analyses med | National Reference Center for Viruses of Respiratory Infections, Institut Pasteur, Paris | Marion Barbet, Sylvie Behillil, Méline Bizard, Angela Brisebarre, Camille Capel, Vincent Enouf, Louise Lefrançois, Frédéric Lemoine, Christophe Malabat, Corinne Maufrais, Etienne Simon-Lorière, Maud Vanpeene, Sylvie Van der Werf ,OphéLie Said-Delattre |
| EPI_ISL_2709294 | Labo Analyses med | National Reference Center for Viruses of Respiratory Infections, Institut Pasteur, Paris | Marion Barbet, Sylvie Behillil, Méline Bizard, Angela Brisebarre, Camille Capel, Vincent Enouf, Louise Lefrançois, Frédéric Lemoine, Christophe Malabat, Corinne Maufrais, Damien Mornico, Etienne Simon-Lorière, Maud Vanpeene, Sylvie Van der Werf ,OphéLie Said-Delattre |
| EPI_ISL_2709295 | Labo Analyses med | National Reference Center for Viruses of Respiratory Infections, Institut Pasteur, Paris | Marion Barbet, Sylvie Behillil, Méline Bizard, Angela Brisebarre, Camille Capel, Vincent Enouf, Louise Lefrançois, Frédéric Lemoine, Christophe Malabat, Corinne Maufrais, Etienne Simon-Lorière, Maud Vanpeene, Sylvie Van der Werf ,OphéLie Said-Delattre |
| EPI_ISL_2709296 | Labo Analyses med | National Reference Center for Viruses of Respiratory Infections, Institut Pasteur, Paris | Marion Barbet, Sylvie Behillil, Méline Bizard, Angela Brisebarre, Camille Capel, Vincent Enouf, Louise Lefrançois, Frédéric Lemoine, Christophe Malabat, Corinne Maufrais, Damien Mornico, Etienne Simon-Lorière, Maud Vanpeene, Sylvie Van der Werf ,OphéLie Said-Delattre |
| EPI_ISL_2709297, EPI_ISL_2709298, EPI_ISL_2709299, EPI_ISL_2709301, EPI_ISL_2709302, EPI_ISL_2709303, EPI_ISL_2709304, EPI_ISL_2709305, EPI_ISL_2709306, EPI_ISL_2709307, EPI_ISL_2709308, EPI_ISL_2709309, EPI_ISL_2709310, EPI_ISL_2709311, EPI_ISL_2709312 | | | |
| see above | Labo Analyses med | National Reference Center for Viruses of Respiratory Infections, Institut Pasteur, Paris | Marion Barbet, Sylvie Behillil, Méline Bizard, Angela Brisebarre, Camille Capel, Vincent Enouf, Louise Lefrançois, Frédéric Lemoine, Christophe Malabat, Corinne Maufrais, Etienne Simon-Lorière, Maud Vanpeene, Sylvie Van der Werf ,OphéLie Said-Delattre |
| EPI_ISL_2709603 | AZ Sint-Lucas Gent | AZ Sint-Lucas Gent | Jos Van Acker, Charlotte Verfaillie, Elke Vanlaere, Klara De Rauw |
| EPI_ISL_2709686 | LESP Queretaro | Instituto de Diagnostico y Referencia Epidemiologicos (INDRE) | Claudia Wong-Arambula, Abril Rodriguez-Maldonado, Vanessa Rivero-Arredondo, Ariadna Medina-Benitez, Joaquin Quiroz-Mercado, Sergio Rangel-Guerrero, Natividad Cruz-Ortiz, Tatiana Nunez-Garcia, Gisela Barrera-Badillo, Lucia Hernandez-Rivas, Irma Lopez-Martinez, Ernesto Ramirez-Gonzalez. |
| EPI_ISL_2709687 | LESP Baja California | Instituto de Diagnostico y Referencia Epidemiologicos (INDRE) | Claudia Wong-Arambula, Abril Rodriguez-Maldonado, Vanessa Rivero-Arredondo, Ariadna Medina-Benitez, Joaquin Quiroz-Mercado, Sergio Rangel-Guerrero, Natividad Cruz-Ortiz, Tatiana Nunez-Garcia, Gisela Barrera-Badillo, Lucia Hernandez-Rivas, Irma Lopez-Martinez, Ernesto Ramirez-Gonzalez. |
| EPI_ISL_2709688 | LESP Michoacan | Instituto de Diagnostico y Referencia Epidemiologicos (INDRE) | Claudia Wong-Arambula, Abril Rodriguez-Maldonado, Vanessa Rivero-Arredondo, Ariadna Medina-Benitez, Joaquin Quiroz-Mercado, Sergio Rangel-Guerrero, Natividad Cruz-Ortiz, Tatiana Nunez-Garcia, Gisela Barrera-Badillo, Lucia Hernandez-Rivas, Irma Lopez-Martinez, Ernesto Ramirez-Gonzalez. |

| | | | |
|---|---|---|---|
| EPI_ISL_2709690, EPI_ISL_2709691, EPI_ISL_2709692, EPI_ISL_2709693, EPI_ISL_2709694, EPI_ISL_2709695, EPI_ISL_2709696, EPI_ISL_2709697, EPI_ISL_2709698 | LESP Nuevo Leon | Instituto de Diagnostico y Referencia Epidemiologicos (INDRE) | Claudia Wong-Arambula, Abril Rodriguez-Maldonado, Vanessa Rivero-Arredondo, Ariadna Medina-Benitez, Joaquin Quiroz-Mercado, Sergio Rangel-Guerrero, Natividad Cruz-Ortiz, Tatiana Nunez-Garcia, Gisela Barrera-Badillo, Lucia Hernandez-Rivas, Irma Lopez-Martinez, Ernesto Ramirez-Gonzalez. |
| EPI_ISL_2709699 | LESP Jalisco | Instituto de Diagnostico y Referencia Epidemiologicos (INDRE) | Claudia Wong-Arambula, Abril Rodriguez-Maldonado, Vanessa Rivero-Arredondo, Ariadna Medina-Benitez, Joaquin Quiroz-Mercado, Sergio Rangel-Guerrero, Natividad Cruz-Ortiz, Tatiana Nunez-Garcia, Gisela Barrera-Badillo, Lucia Hernandez-Rivas, Irma Lopez-Martinez, Ernesto Ramirez-Gonzalez. |
| EPI_ISL_2709700, EPI_ISL_2709701, EPI_ISL_2709702 | LESP Morelos | Instituto de Diagnostico y Referencia Epidemiologicos (INDRE) | Claudia Wong-Arambula, Abril Rodriguez-Maldonado, Vanessa Rivero-Arredondo, Ariadna Medina-Benitez, Joaquin Quiroz-Mercado, Sergio Rangel-Guerrero, Natividad Cruz-Ortiz, Tatiana Nunez-Garcia, Gisela Barrera-Badillo, Lucia Hernandez-Rivas, Irma Lopez-Martinez, Ernesto Ramirez-Gonzalez. |
| EPI_ISL_2709703 | LESP Ciudad de Mexico | Instituto de Diagnostico y Referencia Epidemiologicos (INDRE) | Claudia Wong-Arambula, Abril Rodriguez-Maldonado, Vanessa Rivero-Arredondo, Ariadna Medina-Benitez, Joaquin Quiroz-Mercado, Sergio Rangel-Guerrero, Natividad Cruz-Ortiz, Tatiana Nunez-Garcia, Gisela Barrera-Badillo, Lucia Hernandez-Rivas, Irma Lopez-Martinez, Ernesto Ramirez-Gonzalez. |
| EPI_ISL_2709706, EPI_ISL_2709707, EPI_ISL_2709708 | LESP Queretaro | Instituto de Diagnostico y Referencia Epidemiologicos (INDRE) | Claudia Wong-Arambula, Abril Rodriguez-Maldonado, Vanessa Rivero-Arredondo, Ariadna Medina-Benitez, Joaquin Quiroz-Mercado, Sergio Rangel-Guerrero, Natividad Cruz-Ortiz, Tatiana Nunez-Garcia, Gisela Barrera-Badillo, Lucia Hernandez-Rivas, Irma Lopez-Martinez, Ernesto Ramirez-Gonzalez. |
| EPI_ISL_2709709, EPI_ISL_2709710, EPI_ISL_2709711 | LESP Tabasco | Instituto de Diagnostico y Referencia Epidemiologicos (INDRE) | Claudia Wong-Arambula, Abril Rodriguez-Maldonado, Vanessa Rivero-Arredondo, Ariadna Medina-Benitez, Joaquin Quiroz-Mercado, Sergio Rangel-Guerrero, Natividad Cruz-Ortiz, Tatiana Nunez-Garcia, Gisela Barrera-Badillo, Lucia Hernandez-Rivas, Irma Lopez-Martinez, Ernesto Ramirez-Gonzalez. |
| EPI_ISL_2709717 | LESP Jalisco | Instituto de Diagnostico y Referencia Epidemiologicos (INDRE) | Claudia Wong-Arambula, Abril Rodriguez-Maldonado, Vanessa Rivero-Arredondo, Ariadna Medina-Benitez, Joaquin Quiroz-Mercado, Sergio Rangel-Guerrero, Natividad Cruz-Ortiz, Tatiana Nunez-Garcia, Gisela Barrera-Badillo, Lucia Hernandez-Rivas, Irma Lopez-Martinez, Ernesto Ramirez-Gonzalez. |
| EPI_ISL_2709718 | LESP Sonora | Instituto de Diagnostico y Referencia Epidemiologicos (INDRE) | Claudia Wong-Arambula, Abril Rodriguez-Maldonado, Vanessa Rivero-Arredondo, Ariadna Medina-Benitez, Joaquin Quiroz-Mercado, Sergio Rangel-Guerrero, Natividad Cruz-Ortiz, Tatiana Nunez-Garcia, Gisela Barrera-Badillo, Lucia Hernandez-Rivas, Irma Lopez-Martinez, Ernesto Ramirez-Gonzalez. |
| EPI_ISL_2709723 | LESP Veracruz | Instituto de Diagnostico y Referencia Epidemiologicos (INDRE) | Claudia Wong-Arambula, Abril Rodriguez-Maldonado, Vanessa Rivero-Arredondo, Ariadna Medina-Benitez, Joaquin Quiroz-Mercado, Sergio Rangel-Guerrero, Natividad Cruz-Ortiz, Tatiana Nunez-Garcia, Gisela Barrera-Badillo, Lucia Hernandez-Rivas, Irma Lopez-Martinez, Ernesto Ramirez-Gonzalez. |
| EPI_ISL_2709724, EPI_ISL_2709725 | LESP Guanajuato | Instituto de Diagnostico y Referencia Epidemiologicos (INDRE) | Claudia Wong-Arambula, Abril Rodriguez-Maldonado, Vanessa Rivero-Arredondo, Ariadna Medina-Benitez, Joaquin Quiroz-Mercado, Sergio Rangel-Guerrero, Natividad Cruz-Ortiz, Tatiana Nunez-Garcia, Gisela Barrera-Badillo, Lucia Hernandez-Rivas, Irma Lopez-Martinez, Ernesto Ramirez-Gonzalez. |
| EPI_ISL_2709726, EPI_ISL_2709727, EPI_ISL_2709728, EPI_ISL_2709729 | LESP Nuevo Leon | Instituto de Diagnostico y Referencia Epidemiologicos (INDRE) | Claudia Wong-Arambula, Abril Rodriguez-Maldonado, Vanessa Rivero-Arredondo, Ariadna Medina-Benitez, Joaquin Quiroz-Mercado, Sergio Rangel-Guerrero, Natividad Cruz-Ortiz, Tatiana Nunez-Garcia, Gisela Barrera-Badillo, Lucia Hernandez-Rivas, Irma Lopez-Martinez, Ernesto Ramirez-Gonzalez. |
| EPI_ISL_2709730 | LESP Ciudad de Mexico | Instituto de Diagnostico y Referencia Epidemiologicos (INDRE) | Claudia Wong-Arambula, Abril Rodriguez-Maldonado, Vanessa Rivero-Arredondo, Ariadna Medina-Benitez, Joaquin Quiroz-Mercado, Sergio Rangel-Guerrero, Natividad Cruz-Ortiz, Tatiana Nunez-Garcia, Gisela Barrera-Badillo, Lucia Hernandez-Rivas, Irma Lopez-Martinez, Ernesto Ramirez-Gonzalez. |
| EPI_ISL_2709731 | LESP Coahuila | Instituto de Diagnostico y Referencia Epidemiologicos (INDRE) | Claudia Wong-Arambula, Abril Rodriguez-Maldonado, Vanessa Rivero-Arredondo, Ariadna Medina-Benitez, Joaquin Quiroz-Mercado, Sergio Rangel-Guerrero, Natividad Cruz-Ortiz, Tatiana Nunez-Garcia, Gisela Barrera-Badillo, Lucia Hernandez-Rivas, Irma Lopez-Martinez, Ernesto Ramirez-Gonzalez. |
| EPI_ISL_2709732 | LESP Chiapas | Instituto de Diagnostico y Referencia Epidemiologicos (INDRE) | Claudia Wong-Arambula, Abril Rodriguez-Maldonado, Vanessa Rivero-Arredondo, Ariadna Medina-Benitez, Joaquin Quiroz-Mercado, Sergio Rangel-Guerrero, Natividad Cruz-Ortiz, Tatiana Nunez-Garcia, Gisela Barrera-Badillo, Lucia Hernandez-Rivas, Irma Lopez-Martinez, Ernesto Ramirez-Gonzalez. |
| EPI_ISL_2709733 | LESP Veracruz | Instituto de Diagnostico y Referencia Epidemiologicos (INDRE) | Claudia Wong-Arambula, Abril Rodriguez-Maldonado, Vanessa Rivero-Arredondo, Ariadna Medina-Benitez, Joaquin Quiroz-Mercado, Sergio Rangel-Guerrero, Natividad Cruz-Ortiz, Tatiana Nunez-Garcia, Gisela Barrera-Badillo, Lucia Hernandez-Rivas, Irma Lopez-Martinez, Ernesto Ramirez-Gonzalez. |
| EPI_ISL_2709750, EPI_ISL_2709751 | Department of Virology, Istituto Zooprofilattico Sperimentale del Lazio e della Toscana (IZSLT) | Department of General Diagnostics; Department of Virology; Istituto Zooprofilattico Sperimentale del Lazio e della Toscana (IZSLT) | Patricia Alba, Giuseppe Manna, Elena L. Diaconu, Fabiola Feltrin, Raffaela Conti, Teresa Scicluna, Virginia Carfora, Antonella Cersini, Alessia Franco, Antonio Battisti. |
| EPI_ISL_2709771, EPI_ISL_2709772, EPI_ISL_2709773, EPI_ISL_2709774, EPI_ISL_2709775, EPI_ISL_2709776, EPI_ISL_2709777, EPI_ISL_2709778 | Institute of Molecular and Translational Medicine / Laboratory of Experimental Medicine, Faculty of Medicine and Dentistry, Palacky University and University Hospital Olomouc | Institute of Molecular and Translational Medicine / Laboratory of Experimental Medicine, Faculty of Medicine and Dentistry, Palacky University | Rastislav Slavkovský, Hana Dímalová, Vladimíra Koudeláková, Barbora Blumová, Marián Hajdúch |
| EPI_ISL_2709779 | Department of Microbiology, Faculty of Medicine and Dentistry, Palacky University and University Hospital Olomouc | Institute of Molecular and Translational Medicine / Laboratory of Experimental Medicine, Faculty of Medicine and Dentistry, Palacky University | Rastislav Slavkovský, Hana Dímalová, Pavel Sauer, Vladimíra Koudeláková, Barbora Blumová, Marián Hajdúch |
| EPI_ISL_2709780, EPI_ISL_2709781, EPI_ISL_2709782, EPI_ISL_2709783, EPI_ISL_2709784, EPI_ISL_2709785, EPI_ISL_2709786, EPI_ISL_2709787, EPI_ISL_2709788, EPI_ISL_2709789, EPI_ISL_2709790, EPI_ISL_2709791, EPI_ISL_2709792, EPI_ISL_2709794, EPI_ISL_2709795, EPI_ISL_2709796, EPI_ISL_2709797, EPI_ISL_2709798, EPI_ISL_2709799 | | | |
| see above | Institute of Molecular and Translational Medicine / Laboratory of Experimental Medicine, Faculty of Medicine and Dentistry, Palacky University and University Hospital Olomouc | Institute of Molecular and Translational Medicine / Laboratory of Experimental Medicine, Faculty of Medicine and Dentistry, Palacky University | Rastislav Slavkovský, Hana Dímalová, Vladimíra Koudeláková, Barbora Blumová, Marián Hajdúch |
| EPI_ISL_2713263, EPI_ISL_2713267, EPI_ISL_2713303, EPI_ISL_2713304, EPI_ISL_2713305, EPI_ISL_2713306, EPI_ISL_2713307, EPI_ISL_2713308, EPI_ISL_2713309, EPI_ISL_2713310, EPI_ISL_2713311, EPI_ISL_2713312, EPI_ISL_2713313 | | | |
| see above | Department of Health Technology and Informatics, The Hong Kong Polytechnic University | Department of Health Technology and Informatics, The Hong Kong Polytechnic University | Gilman Kit-Hang Siu, Lam-Kwong Lee, Kenneth Siu-Sing Leung, Jake Siu-Lun Leung, Wing-Hei Lo, Timothy Ting-Leung Ng, Chloe Toi-Mei Chan, Hoi-Ching Jim, Ivan Tak-Fai Wong, Kingsley King-Gee Tam, Hiu-Yin Lao, Denise Sze-Hang Wong, Alan Ka-Lun Wu, Miranda Chong-Yee Yau, Yvette Wai-Man Lai, Kitty Sau-Chun Fung, Sandy Ka-Yee Chau, Barry Kin-Chung Wong, Wing-Kin To, Kristine Luk, Alex Yat-Man Ho, Tak-Lun Que, Kam-Tong Yip, Wing Cheong Yam, David Ho-Keung Shum, Shea Ping Yip |
| EPI_ISL_2723570, EPI_ISL_2723571, EPI_ISL_2723572, EPI_ISL_2723573, EPI_ISL_2723574, EPI_ISL_2723575, EPI_ISL_2723576, EPI_ISL_2723577, EPI_ISL_2723578, EPI_ISL_2723579, EPI_ISL_2723580, EPI_ISL_2723581, EPI_ISL_2723582 | | | |
| see above | SC (UCO) Igiene e Sanità Pubblica, ASUGI, Trieste | ARGO Laboratorio Genomica ed Epigenomica | Licastro D, Dal Monego S, Degasperi M, Marcello A, Segat L, Piscianz E, Busetti M, Fontana F, Braida C, Barbone F, D'Agaro P |
| EPI_ISL_2723583, EPI_ISL_2723584 | SC (UCO) Igiene e Sanità Pubblica, ASUGI, Trieste | ARGO Laboratorio Genomica ed Epigenomica | Licastro D, Dal Monego S, Degasperi M, Marcello A, Segat L, Piscianz E, Pipan C, Braida C, Barbone F, D'Agaro P |
| EPI_ISL_2723585, EPI_ISL_2723586, EPI_ISL_2723587, EPI_ISL_2723588, EPI_ISL_2723589, EPI_ISL_2723590, EPI_ISL_2723591 | SC (UCO) Igiene e Sanità Pubblica, ASUGI, Trieste | ARGO Laboratorio Genomica ed Epigenomica | Licastro D, Dal Monego S, Degasperi M, Marcello A, Segat L, Piscianz E, Busetti M, Fontana F, Braida C, Barbone F, D'Agaro P |

| | | | |
|---|---|---|---|
| EPI_ISL_2723592 | SC (UCO) Igiene e Sanità Pubblica, ASUGI, Trieste | ARGO Laboratorio Genomica ed Epigenomica | Licastro D, Dal Monego S, Degasperi M, Marcello A, Segat L, De Rosa R, Braida C, Barbone F, D'Agaro P |
| EPI_ISL_2723593 | SC (UCO) Igiene e Sanità Pubblica, ASUGI, Trieste | ARGO Laboratorio Genomica ed Epigenomica | Licastro D, Dal Monego S, Degasperi M, Marcello A, Segat L, Piscianz E, Busetti M, Fontana F, Braida C, Barbone F, D'Agaro P |
| EPI_ISL_2723594 | SC (UCO) Igiene e Sanità Pubblica, ASUGI, Trieste | ARGO Laboratorio Genomica ed Epigenomica | Licastro D, Dal Monego S, Degasperi M, Marcello A, Segat L, De Rosa R, Braida C, Barbone F, D'Agaro P |
| EPI_ISL_2723595, EPI_ISL_2723596 | SC (UCO) Igiene e Sanità Pubblica, ASUGI, Trieste | ARGO Laboratorio Genomica ed Epigenomica | Licastro D, Dal Monego S, Degasperi M, Marcello A, Segat L, Piscianz E, Busetti M, Fontana F, Braida C, Barbone F, D'Agaro P |
| EPI_ISL_2723597, EPI_ISL_2723598 | SC (UCO) Igiene e Sanità Pubblica, ASUGI, Trieste | ARGO Laboratorio Genomica ed Epigenomica | Licastro D, Dal Monego S, Degasperi M, Marcello A, Segat L, De Rosa R, Braida C, Barbone F, D'Agaro P |
| EPI_ISL_2723599 | SC (UCO) Igiene e Sanità Pubblica, ASUGI, Trieste | ARGO Laboratorio Genomica ed Epigenomica | Licastro D, Dal Monego S, Degasperi M, Marcello A, Segat L, Piscianz E, Busetti M, Fontana F, Braida C, Barbone F, D'Agaro P |
| EPI_ISL_2724009 | Molecular Biology Laboratory, Infectious Diseases Department, Instituto Nacional de Ciencias Medicas y Nutricion Salvador Zubiran | Instituto Nacional de Ciencias Medicas y Nutricion Infectious Diseases | Pilar Ramos Cervantes, Fernando Ledesma Barrientos, Violeta Ibarra Gonzalez, Patricia Leal, Eric Ochoa, Arturo Galindo, Guillermo Ruiz-Palacios y Santos |
| EPI_ISL_2725107, EPI_ISL_2725108, EPI_ISL_2725109 | Laboratorio di Genetica Medica, Matera, Italy | Laboratorio di Genetica Medica, Matera, Italy | Domenico Dell'Edera |
| EPI_ISL_2727501 | Cantacuzino National Military-Medical Institute, Viral Respiratory Infections Laboratory | Cantacuzino Institute Virology | Carmen Cherciu, Luiza Ustea, Nicoleta Paraschiv, Sorin Dinu, Mihaela Oprea, Mihaela Lazar |
| EPI_ISL_2727502, EPI_ISL_2727503, EPI_ISL_2727504, EPI_ISL_2727505, EPI_ISL_2727506, EPI_ISL_2727507, EPI_ISL_2727508, EPI_ISL_2727511, EPI_ISL_2727512, EPI_ISL_2727515, EPI_ISL_2727516, EPI_ISL_2727517, EPI_ISL_2727518, EPI_ISL_2727519, EPI_ISL_2727521 | | | |
| see above | Institute of Molecular and Translational Medicine / Laboratory of Experimental Medicine, Faculty of Medicine and Dentistry, Palacky University and University Hospital Olomouc | Institute of Molecular and Translational Medicine / Laboratory of Experimental Medicine, Faculty of Medicine and Dentistry, Palacky University | Rastislav Slavkovský, Hana Dímalová, Vladimíra Koudeláková, Barbora Blumová, Marián Hajdúch |
| EPI_ISL_2727567, EPI_ISL_2727568 | Cantacuzino National Military-Medical Institute, Viral Respiratory Infections Laboratory | Cantacuzino Institute Virology | Carmen Cherciu, Luiza Ustea, Nicoleta Paraschiv, Sorin Dinu, Mihaela Oprea, Mihaela Lazar |
| EPI_ISL_2727836 | Hospital Angkatan Tentera Kota Kinabalu | Institute for Medical Research, Infectious Disease Research Centre, National Institutes of Health, Ministry of Health Malaysia | Suppiah J, Kamel K, Mohd Zawawi Z, Azizan MA, Ramly N, Robert F, Thayan R |
| EPI_ISL_2727837 | Hospital Angkatan Tentera Wilayah Kota Kinabalu | Institute for Medical Research, Infectious Disease Research Centre, National Institutes of Health, Ministry of Health Malaysia | Suppiah J, Kamel K, Mohd Zawawi Z, Azizan MA, Ramly N, Robert F, Thayan R |
| EPI_ISL_2728577, EPI_ISL_2728578 | Laboratorio de Infectologia y Virologia Molecular | Laboratory of Molecular Virology, School of Medicine, Pontificia Universidad Catolica de Chile | Catalina Pardo-Roa, Constanza Martinez-Valdevenito, Jennifer Angulo, Tamara Garcia-Salum, Estefany Poblete, Maria Jose Avendano, Leonardo I. Almonacid, Ana Maria Contreras, Carlos Palma, Jorge Levican, Eileen Serrano, Constanza Maldonado, M. Belen Leyton, Erick Salinas, Andres E. Munoz-Marcos, Francisco Melo, Marcela Ferres, Alejandro Bhrun, Magdalena Vera, Rafael A. Medina |
| EPI_ISL_2731271 | S.C Microbiologia e virologia - Speciale Centro influenza -AOU di Sassari | Laboratorio Biologia Molecolare Sars Cov2 - UOC Laboratorio Analisi - Servizio Medicina di Laboratorio, Ospedale "San Francesco" - ATS-ASSL Nuoro and Laboratorio specialistico UOC Ematologia - Ospedale "San Francesco" - ATS-ASSL Nuoro | Asproni Rosanna, Piras Giovanna, Malune Paolo, Pira Giovanna, Fancello Patrizia, Rosu Valentina, Carta Franco, Fiamma Maura, Mameli Giuseppe, Garau Maria Cristina , Palmas Angelo Domenico,Rimini Elena ,Serra Caterina, Salvatore Rubino |
| EPI_ISL_2732231, EPI_ISL_2732235, EPI_ISL_2732238 | Department of Virology, Istituto Zooprofilattico Sperimentale del Lazio e della Toscana (IZSLT) | Department of General Diagnostics; Department of Virology; Istituto Zooprofilattico Sperimentale del Lazio e della Toscana (IZSLT) | Patricia Alba, Giuseppe Manna, Elena L. Diaconu, Fabiola Feltrin, Raffaela Conti, Teresa Scicluna, Virginia Carfora, Antonella Cersini, Alessia Franco, Antonio Battisti. |
| EPI_ISL_2736780, EPI_ISL_2736781, EPI_ISL_2736783 | LESP Sinaloa | Instituto de Diagnostico y Referencia Epidemiologicos (INDRE) | Claudia Wong-Arambula, Abril Rodriguez-Maldonado, Vanessa Rivero-Arredondo, Ariadna Medina-Benitez, Joaquin Quiroz-Mercado, Sergio Rangel-Guerrero, Natividad Cruz-Ortiz, Tatiana Nunez-Garcia, Gisela Barrera-Badillo, Lucia Hernandez-Rivas, Irma Lopez-Martinez, Ernesto Ramirez-Gonzalez. |
| EPI_ISL_2736785 | LESP Puebla | Instituto de Diagnostico y Referencia Epidemiologicos (INDRE) | Claudia Wong-Arambula, Abril Rodriguez-Maldonado, Vanessa Rivero-Arredondo, Ariadna Medina-Benitez, Joaquin Quiroz-Mercado, Sergio Rangel-Guerrero, Natividad Cruz-Ortiz, Tatiana Nunez-Garcia, Gisela Barrera-Badillo, Lucia Hernandez-Rivas, Irma Lopez-Martinez, Ernesto Ramirez-Gonzalez. |
| EPI_ISL_2736787 | LESP Guanajuato | Instituto de Diagnostico y Referencia Epidemiologicos (INDRE) | Claudia Wong-Arambula, Abril Rodriguez-Maldonado, Vanessa Rivero-Arredondo, Ariadna Medina-Benitez, Joaquin Quiroz-Mercado, Sergio Rangel-Guerrero, Natividad Cruz-Ortiz, Tatiana Nunez-Garcia, Gisela Barrera-Badillo, Lucia Hernandez-Rivas, Irma Lopez-Martinez, Ernesto Ramirez-Gonzalez. |
| EPI_ISL_2736829 | LESP Coahuila | Instituto de Diagnostico y Referencia Epidemiologicos (INDRE) | Claudia Wong-Arambula, Abril Rodriguez-Maldonado, Vanessa Rivero-Arredondo, Ariadna Medina-Benitez, Joaquin Quiroz-Mercado, Sergio Rangel-Guerrero, Natividad Cruz-Ortiz, Tatiana Nunez-Garcia, Gisela Barrera-Badillo, Lucia Hernandez-Rivas, Irma Lopez-Martinez, Ernesto Ramirez-Gonzalez. |
| EPI_ISL_2736832 | LESP Baja California | Instituto de Diagnostico y Referencia Epidemiologicos (INDRE) | Claudia Wong-Arambula, Abril Rodriguez-Maldonado, Vanessa Rivero-Arredondo, Ariadna Medina-Benitez, Joaquin Quiroz-Mercado, Sergio Rangel-Guerrero, Natividad Cruz-Ortiz, Tatiana Nunez-Garcia, Gisela Barrera-Badillo, Lucia Hernandez-Rivas, Irma Lopez-Martinez, Ernesto Ramirez-Gonzalez. |
| EPI_ISL_2736837, EPI_ISL_2736839 | LESP Tabasco | Instituto de Diagnostico y Referencia Epidemiologicos (INDRE) | Claudia Wong-Arambula, Abril Rodriguez-Maldonado, Vanessa Rivero-Arredondo, Ariadna Medina-Benitez, Joaquin Quiroz-Mercado, Sergio Rangel-Guerrero, Natividad Cruz-Ortiz, Tatiana Nunez-Garcia, Gisela Barrera-Badillo, Lucia Hernandez-Rivas, Irma Lopez-Martinez, Ernesto Ramirez-Gonzalez. |
| EPI_ISL_2736841 | LESP Guerrero | Instituto de Diagnostico y Referencia Epidemiologicos (INDRE) | Claudia Wong-Arambula, Abril Rodriguez-Maldonado, Vanessa Rivero-Arredondo, Ariadna Medina-Benitez, Joaquin Quiroz-Mercado, Sergio Rangel-Guerrero, Natividad Cruz-Ortiz, Tatiana Nunez-Garcia, Gisela Barrera-Badillo, Lucia Hernandez-Rivas, Irma Lopez-Martinez, Ernesto Ramirez-Gonzalez. |
| EPI_ISL_2736845 | LESP Yucatan | Instituto de Diagnostico y Referencia Epidemiologicos (INDRE) | Claudia Wong-Arambula, Abril Rodriguez-Maldonado, Vanessa Rivero-Arredondo, Ariadna Medina-Benitez, Joaquin Quiroz-Mercado, Sergio Rangel-Guerrero, Natividad Cruz-Ortiz, Tatiana Nunez-Garcia, Gisela Barrera-Badillo, Lucia Hernandez-Rivas, Irma Lopez-Martinez, Ernesto Ramirez-Gonzalez. |
| EPI_ISL_2736848, EPI_ISL_2736849, EPI_ISL_2736851 | LESP Michoacan | Instituto de Diagnostico y Referencia Epidemiologicos (INDRE) | Claudia Wong-Arambula, Abril Rodriguez-Maldonado, Vanessa Rivero-Arredondo, Ariadna Medina-Benitez, Joaquin Quiroz-Mercado, Sergio Rangel-Guerrero, Natividad Cruz-Ortiz, Tatiana Nunez-Garcia, Gisela Barrera-Badillo, Lucia Hernandez-Rivas, Irma Lopez-Martinez, Ernesto Ramirez-Gonzalez. |
| EPI_ISL_2736856 | LESP Sonora | Instituto de Diagnostico y Referencia Epidemiologicos (INDRE) | Claudia Wong-Arambula, Abril Rodriguez-Maldonado, Vanessa Rivero-Arredondo, Ariadna Medina-Benitez, Joaquin Quiroz-Mercado, Sergio Rangel-Guerrero, Natividad Cruz-Ortiz, Tatiana Nunez-Garcia, Gisela Barrera-Badillo, Lucia Hernandez-Rivas, Irma Lopez-Martinez, Ernesto Ramirez-Gonzalez. |
| EPI_ISL_2736857 | LESP Campeche | Instituto de Diagnostico y Referencia Epidemiologicos (INDRE) | Claudia Wong-Arambula, Abril Rodriguez-Maldonado, Vanessa Rivero-Arredondo, Ariadna Medina-Benitez, Joaquin Quiroz-Mercado, Sergio Rangel-Guerrero, Natividad Cruz-Ortiz, Tatiana Nunez-Garcia, Gisela Barrera-Badillo, Lucia Hernandez-Rivas, Irma Lopez-Martinez, Ernesto Ramirez-Gonzalez. |
| EPI_ISL_2736861, EPI_ISL_2736864 | LESP Guanajuato | Instituto de Diagnostico y Referencia Epidemiologicos (INDRE) | Claudia Wong-Arambula, Abril Rodriguez-Maldonado, Vanessa Rivero-Arredondo, Ariadna Medina-Benitez, Joaquin Quiroz-Mercado, Sergio Rangel-Guerrero, Natividad Cruz-Ortiz, Tatiana Nunez-Garcia, Gisela Barrera-Badillo, Lucia Hernandez-Rivas, Irma Lopez-Martinez, Ernesto Ramirez-Gonzalez. |
| EPI_ISL_2754138 | Genome Analysis Center, Yamanashi Central Hospital | Genome Analysis Center, Yamanashi Central Hospital | Yosuke Hirotsu |

| | | | |
|---|---|---|---|
| EPI_ISL_2754196 | S.C Microbiologia e virologia - Speciale Centro influenza -AOU di Sassari | Laboratorio Biologia Molecolare Sars Cov2 - UOC Laboratorio Analisi - Servizio Medicina di Laboratorio, Ospedale "San Francesco" - ATS-ASSL Nuoro and Laboratorio specialistico UOC Ematologia - Ospedale "San Francesco" - ATS-ASSL Nuoro | Asproni Rosanna, Piras Giovanna, Malune Paolo, Pira Giovanna, Fancello Patrizia, Rosu Valentina, Carta Franco, Fiamma Maura, Mameli Giuseppe, Garau Maria Cristina , Palmas Angelo Domenico,Rimini Elena ,Serra Caterina, Salvatore Rubino |
| EPI_ISL_2754197 | S.C Microbiologia e virologia - Speciale Centro influenza -AOU di Sassari | Laboratorio Biologia Molecolare Sars Cov2 - UOC Laboratorio Analisi - Servizio Medicina di Laboratorio, Ospedale "San Francesco" - ATS-ASSL Nuoro and Laboratorio specialistico UOC Ematologia - Ospedale "San Francesco" - ATS-ASSL Nuoro | Asproni Rosanna, Piras Giovanna, Malune Paolo, Pira Giovanna, Fancello Patrizia, Rosu Valentina, Carta Franco, Fiamma Maura, Mameli Giuseppe, Garau Maria Cristina , Palmas Angelo Domenico,Rimini Elena ,Serra Caterina, Rubino Salvatore |
| EPI_ISL_2754199, EPI_ISL_2754201, EPI_ISL_2754202 | S.C Microbiologia e virologia - Speciale Centro influenza -AOU di Sassari | Laboratorio Biologia Molecolare Sars Cov2 - UOC Laboratorio Analisi - Servizio Medicina di Laboratorio, Ospedale "San Francesco" - ATS-ASSL Nuoro and Laboratorio specialistico UOC Ematologia - Ospedale "San Francesco" - ATS-ASSL Nuoro | Asproni Rosanna, Piras Giovanna, Malune Paolo, Pira Giovanna, Fancello Patrizia, Rosu Valentina, Carta Franco, Fiamma Maura, Mameli Giuseppe, Garau Maria Cristina , Palmas Angelo Domenico,Rimini Elena ,Serra Caterina, Salvatore Rubino |
| EPI_ISL_2756384, EPI_ISL_2756385, EPI_ISL_2756386, EPI_ISL_2756389, EPI_ISL_2756390, EPI_ISL_2756392, EPI_ISL_2756394, EPI_ISL_2756395, EPI_ISL_2756396, EPI_ISL_2756397, EPI_ISL_2756398, EPI_ISL_2756402, EPI_ISL_2756403, EPI_ISL_2756406, EPI_ISL_2756407, EPI_ISL_2756408, EPI_ISL_2756415, EPI_ISL_2756416, EPI_ISL_2756418, EPI_ISL_2756421 | | | |
| see above | Microbiology Department, University Hospital Donostia | Microbiology Department, University Hospital Donostia | Marimon JM, Montes M, Piñeiro L, Martin-Peñaranda T, Gomez M, Sorarrain A, Cilla G. |
| EPI_ISL_2756557 | Centers for Disease Control and Prevention, Dengue Branch | Centers for Disease Control and Prevention, Dengue Branch | Gilberto A. Santiago, Glenda Gonzalez, Betzabel Flores, Keyla Charriez, Gabriela Paz-Bailey, Jorge L. Munoz-Jordan |
| EPI_ISL_2756607, EPI_ISL_2756608, EPI_ISL_2756609 | Dipartimento di Medicina di Laboratorio, Azienda sanitaria universitaria Friuli Centrale (ASU FC) | Dipartimento di Medicina di Laboratorio, Azienda sanitaria universitaria Friuli Centrale (ASU FC) | Corrado Pipan, Catia Mio, Chiara Dal Secco, Stefania Marzinotto, Francesco Curcio |
| EPI_ISL_2757658 | Hospital | National Reference Center for Viruses of Respiratory Infections, Institut Pasteur, Paris | Marion Barbet, Sylvie Behillil, Méline Bizard, Angela Brisebarre, Camille Capel, Vincent Enouf, Louise Lefrançois, Frédéric Lemoine, Christophe Malabat, Corinne Maufrais, Etienne Simon-Lorière, Maud Vanpeene, Sylvie Van der Werf ,JéRome Guinard |
| EPI_ISL_2757659 | Labo Analyses Med | National Reference Center for Viruses of Respiratory Infections, Institut Pasteur, Paris | Marion Barbet, Sylvie Behillil, Méline Bizard, Angela Brisebarre, Camille Capel, Vincent Enouf, Louise Lefrançois, Frédéric Lemoine, Christophe Malabat, Corinne Maufrais, Emmanuelle Permal, Etienne Simon-Lorière, Maud Vanpeene, Sylvie Van der Werf ,GréGoire Potiron |
| EPI_ISL_2757660 | Hospital | National Reference Center for Viruses of Respiratory Infections, Institut Pasteur, Paris | Marion Barbet, Sylvie Behillil, Méline Bizard, Angela Brisebarre, Camille Capel, Vincent Enouf, Louise Lefrançois, Frédéric Lemoine, Christophe Malabat, Corinne Maufrais, Etienne Simon-Lorière, Maud Vanpeene, Sylvie Van der Werf ,Eric Farfour |
| EPI_ISL_2757661 | Hospital | National Reference Center for Viruses of Respiratory Infections, Institut Pasteur, Paris | Marion Barbet, Sylvie Behillil, Méline Bizard, Angela Brisebarre, Camille Capel, Vincent Enouf, Louise Lefrançois, Frédéric Lemoine, Christophe Malabat, Corinne Maufrais, Etienne Simon-Lorière, Maud Vanpeene, Sylvie Van der Werf ,Laurence Courdavault |
| EPI_ISL_2757662 | Hospital | National Reference Center for Viruses of Respiratory Infections, Institut Pasteur, Paris | Marion Barbet, Sylvie Behillil, Méline Bizard, Angela Brisebarre, Camille Capel, Vincent Enouf, Louise Lefrançois, Frédéric Lemoine, Christophe Malabat, Corinne Maufrais, Adrien Pain, Etienne Simon-Lorière, Maud Vanpeene, Sylvie Van der Werf ,Laurence Courdavault |
| EPI_ISL_2757663, EPI_ISL_2757664, EPI_ISL_2757665 | Hospital | National Reference Center for Viruses of Respiratory Infections, Institut Pasteur, Paris | Marion Barbet, Sylvie Behillil, Méline Bizard, Angela Brisebarre, Camille Capel, Vincent Enouf, Louise Lefrançois, Frédéric Lemoine, Christophe Malabat, Corinne Maufrais, Etienne Simon-Lorière, Maud Vanpeene, Sylvie Van der Werf ,JéRôMe Durivault |
| EPI_ISL_2757666 | Hospital | National Reference Center for Viruses of Respiratory Infections, Institut Pasteur, Paris | Marion Barbet, Sylvie Behillil, Méline Bizard, Angela Brisebarre, Camille Capel, Vincent Enouf, Louise Lefrançois, Frédéric Lemoine, Christophe Malabat, Corinne Maufrais, Adrien Pain, Etienne Simon-Lorière, Maud Vanpeene, Sylvie Van der Werf ,JéRôMe Durivault |
| EPI_ISL_2757667, EPI_ISL_2757668, EPI_ISL_2757669, EPI_ISL_2757670, EPI_ISL_2757671, EPI_ISL_2757672, EPI_ISL_2757673 | Hospital | National Reference Center for Viruses of Respiratory Infections, Institut Pasteur, Paris | Marion Barbet, Sylvie Behillil, Méline Bizard, Angela Brisebarre, Camille Capel, Vincent Enouf, Louise Lefrançois, Frédéric Lemoine, Christophe Malabat, Corinne Maufrais, Etienne Simon-Lorière, Maud Vanpeene, Sylvie Van der Werf ,JéRôMe Durivault |
| EPI_ISL_2757674, EPI_ISL_2757675 | Hospital | National Reference Center for Viruses of Respiratory Infections, Institut Pasteur, Paris | Marion Barbet, Sylvie Behillil, Méline Bizard, Angela Brisebarre, Camille Capel, Vincent Enouf, Louise Lefrançois, Frédéric Lemoine, Christophe Malabat, Corinne Maufrais, Amaury Vaysse, Etienne Simon-Lorière, Maud Vanpeene, Sylvie Van der Werf ,JéRôMe Durivault |
| EPI_ISL_2757676, EPI_ISL_2757677, EPI_ISL_2757678 | Hospital | National Reference Center for Viruses of Respiratory Infections, Institut Pasteur, Paris | Marion Barbet, Sylvie Behillil, Méline Bizard, Angela Brisebarre, Camille Capel, Vincent Enouf, Louise Lefrançois, Frédéric Lemoine, Christophe Malabat, Corinne Maufrais, Etienne Simon-Lorière, Maud Vanpeene, Sylvie Van der Werf ,JéRôMe Durivault |
| EPI_ISL_2757679 | Armies | National Reference Center for Viruses of Respiratory Infections, Institut Pasteur, Paris | Marion Barbet, Sylvie Behillil, Méline Bizard, Angela Brisebarre, Camille Capel, Vincent Enouf, Louise Lefrançois, Frédéric Lemoine, Christophe Malabat, Corinne Maufrais, Etienne Simon-Lorière, Maud Vanpeene, Sylvie Van der Werf ,Marine Desroches |
| EPI_ISL_2757680, EPI_ISL_2757681 | Labo Analyses Med | National Reference Center for Viruses of Respiratory Infections, Institut Pasteur, Paris | Marion Barbet, Sylvie Behillil, Méline Bizard, Angela Brisebarre, Camille Capel, Vincent Enouf, Louise Lefrançois, Frédéric Lemoine, Christophe Malabat, Corinne Maufrais, Etienne Simon-Lorière, Maud Vanpeene, Sylvie Van der Werf ,Nader Kherbek |
| EPI_ISL_2757682 | Labo Analyses Med | National Reference Center for Viruses of Respiratory Infections, Institut Pasteur, Paris | Marion Barbet, Sylvie Behillil, Méline Bizard, Angela Brisebarre, Camille Capel, Vincent Enouf, Louise Lefrançois, Frédéric Lemoine, Christophe Malabat, Corinne Maufrais, Amaury Vaysse, Etienne Simon-Lorière, Maud Vanpeene, Sylvie Van der Werf ,GréGoire Potiron |
| EPI_ISL_2757683, EPI_ISL_2757684, EPI_ISL_2757685, EPI_ISL_2757686, EPI_ISL_2757687, EPI_ISL_2757689 | Presidio di Brindisi Di Summa - Perrino | Istituto Zooprofilattico Sperimentale della Puglia e della Basilicata | Parisi A., Bianco A., Capozzi L., Del Sambro L., Simone D., Difato L., Santoro A., Ridolfi D., Giannico A. |
| EPI_ISL_2757692, EPI_ISL_2757693, EPI_ISL_2757694 | Ospedale Santa Caterina Novella | Istituto Zooprofilattico Sperimentale della Puglia e della Basilicata | Parisi A., Bianco A., Capozzi L., Del Sambro L., Simone D., Difato L., Bruno A. R. |
| EPI_ISL_2757695, EPI_ISL_2757696 | Ospedale Cardinale G. Panico | Istituto Zooprofilattico Sperimentale della Puglia e della Basilicata | Parisi A., Bianco A., Capozzi L., Del Sambro L., Simone D., Difato L., Bruno A. R. |
| EPI_ISL_2757697, EPI_ISL_2757698, EPI_ISL_2757699, EPI_ISL_2757700, EPI_ISL_2757701 | Ospedale Vito Fazzi | Istituto Zooprofilattico Sperimentale della Puglia e della Basilicata | Parisi A., Bianco A., Capozzi L., Del Sambro L., Simone D., Difato L., Lobreglio G. |
| EPI_ISL_2757702, EPI_ISL_2757703, EPI_ISL_2757704, EPI_ISL_2757705 | Ospedale Santa Caterina Novella | Istituto Zooprofilattico Sperimentale della Puglia e della Basilicata | Parisi A., Bianco A., Capozzi L., Del Sambro L., Simone D., Difato L., Bruno A. R. |
| EPI_ISL_2757706, EPI_ISL_2757707, EPI_ISL_2757708, EPI_ISL_2757709, EPI_ISL_2757710, EPI_ISL_2757711, EPI_ISL_2757712 | Presidio di Brindisi Di Summa - Perrino | Istituto Zooprofilattico Sperimentale della Puglia e della Basilicata | Parisi A., Bianco A., Capozzi L., Del Sambro L., Simone D., Difato L., Santoro A., Ridolfi D., Giannico A. |
| EPI_ISL_2757713 | Ospedale Santissima Annunziata | Istituto Zooprofilattico Sperimentale della Puglia e della Basilicata | Parisi A., Bianco A., Capozzi L., Del Sambro L., Simone D., Difato L., Ridolfi D., Giannico A, Tampoia M. |
| EPI_ISL_2757718, EPI_ISL_2757719, EPI_ISL_2757720, EPI_ISL_2757721 | Ospedale Santa Caterina Novella | Istituto Zooprofilattico Sperimentale della Puglia e della Basilicata | Parisi A., Bianco A., Capozzi L., Del Sambro L., Simone D., Difato L., Bruno A. R. |
| EPI_ISL_2757722 | Dipartimento di prevenzione Bari | Istituto Zooprofilattico Sperimentale della Puglia e della Basilicata | Parisi A., Bianco A., Capozzi L., Del Sambro L., Simone D., Difato L., Ridolfi D., Giannico A., Lagravinese D. |
| EPI_ISL_2757723, EPI_ISL_2757724, EPI_ISL_2757725, EPI_ISL_2757726, | Ospedale Santissima Annunziata | Istituto Zooprofilattico Sperimentale della Puglia e della Basilicata | Parisi A., Bianco A., Capozzi L., Del Sambro L., Simone D., Difato L., Ridolfi D., Giannico A, Tampoia M. |

| | | | |
|---|---|---|---|
| EPI_ISL_2757728, EPI_ISL_2757729, EPI_ISL_2757731 | | | |
| EPI_ISL_2757766, EPI_ISL_2757768 | Laboratoire Hôpital Sainte Musse | Laboratoire Hôpital Sainte Musse | Lionel Chollet |
| EPI_ISL_2757772, EPI_ISL_2757773, EPI_ISL_2757774, EPI_ISL_2757775, EPI_ISL_2757776, EPI_ISL_2757777, EPI_ISL_2757778, EPI_ISL_2757779, EPI_ISL_2757780, EPI_ISL_2757781, EPI_ISL_2757782, EPI_ISL_2757783, EPI_ISL_2757785, EPI_ISL_2757786, EPI_ISL_2757787, EPI_ISL_2757788, EPI_ISL_2757789, EPI_ISL_2757790, EPI_ISL_2757791, EPI_ISL_2757793, EPI_ISL_2757794, EPI_ISL_2757796, EPI_ISL_2757797 | | | |
| see above | Microbiology Department, University Hospital Donostia | Microbiology Department, University Hospital Donostia | Marimon JM, Montes M, Piñeiro L, Martin-Peñaranda T, Gomez M, Sorarrain A, Cilla G. |
| EPI_ISL_2757798 | SIESP CHIETI - DRIVE IN LANCIANO | Istituto Zooprofilattico Sperimentale dell'Abruzzo e Molise "G. Caporale" | Lorusso A, Marcacci M, Di Domenico M, Ancora M, Curini V, Di Lollo Valeria, Mangone I, Rinaldi A, Delli Compagni E, Scialabba S, Caporale M, Di Pasquale A, Cammà C, Puglia I, Calistri P, Savini G |
| EPI_ISL_2757799 | SIESP CHIETI - DRIVE IN VASTO | Istituto Zooprofilattico Sperimentale dell'Abruzzo e Molise "G. Caporale" | Lorusso A, Marcacci M, Di Domenico M, Ancora M, Curini V, Di Lollo Valeria, Mangone I, Rinaldi A, Delli Compagni E, Scialabba S, Caporale M, Di Pasquale A, Cammà C, Puglia I, Calistri P, Savini G |
| EPI_ISL_2757800, EPI_ISL_2757801, EPI_ISL_2757802 | SIESP CHIETI - DRIVE IN LANCIANO | Istituto Zooprofilattico Sperimentale dell'Abruzzo e Molise "G. Caporale" | Lorusso A, Marcacci M, Di Domenico M, Ancora M, Curini V, Di Lollo Valeria, Mangone I, Rinaldi A, Delli Compagni E, Scialabba S, Caporale M, Di Pasquale A, Cammà C, Puglia I, Calistri P, Savini G |
| EPI_ISL_2757803 | PRESIDIO OSPEDALIERO TAGLIACOZZO | Istituto Zooprofilattico Sperimentale dell'Abruzzo e Molise "G. Caporale" | Lorusso A, Marcacci M, Di Domenico M, Ancora M, Curini V, Di Lollo Valeria, Mangone I, Rinaldi A, Delli Compagni E, Scialabba S, Caporale M, Di Pasquale A, Cammà C, Puglia I, Calistri P, Savini G |
| EPI_ISL_2757804, EPI_ISL_2757805, EPI_ISL_2757806 | SIESP DIPARTIMENTO DI PREVENZIONE CHIETI | Istituto Zooprofilattico Sperimentale dell'Abruzzo e Molise "G. Caporale" | Lorusso A, Marcacci M, Di Domenico M, Ancora M, Curini V, Di Lollo Valeria, Mangone I, Rinaldi A, Delli Compagni E, Scialabba S, Caporale M, Di Pasquale A, Cammà C, Puglia I, Calistri P, Savini G |
| EPI_ISL_2757807, EPI_ISL_2757808 | SIESP CHIETI - DRIVE IN LANCIANO | Istituto Zooprofilattico Sperimentale dell'Abruzzo e Molise "G. Caporale" | Lorusso A, Marcacci M, Di Domenico M, Ancora M, Curini V, Di Lollo Valeria, Mangone I, Rinaldi A, Delli Compagni E, Scialabba S, Caporale M, Di Pasquale A, Cammà C, Puglia I, Calistri P, Savini G |
| EPI_ISL_2757809 | SIESP DIPARTIMENTO DI PREVENZIONE CHIETI | Istituto Zooprofilattico Sperimentale dell'Abruzzo e Molise "G. Caporale" | Lorusso A, Marcacci M, Di Domenico M, Ancora M, Curini V, Di Lollo Valeria, Mangone I, Rinaldi A, Delli Compagni E, Scialabba S, Caporale M, Di Pasquale A, Cammà C, Puglia I, Calistri P, Savini G |
| EPI_ISL_2757810 | SIESP CHIETI - DRIVE IN LANCIANO | Istituto Zooprofilattico Sperimentale dell'Abruzzo e Molise "G. Caporale" | Lorusso A, Marcacci M, Di Domenico M, Ancora M, Curini V, Di Lollo Valeria, Mangone I, Rinaldi A, Delli Compagni E, Scialabba S, Caporale M, Di Pasquale A, Cammà C, Puglia I, Calistri P, Savini G |
| EPI_ISL_2757811, EPI_ISL_2757812, EPI_ISL_2757813, EPI_ISL_2757814, EPI_ISL_2757815, EPI_ISL_2757816, EPI_ISL_2757817 | OSPEDALE CIVILE TERAMO - CENTRO TRASFUSIONALE | Istituto Zooprofilattico Sperimentale dell'Abruzzo e Molise "G. Caporale" | Lorusso A, Marcacci M, Di Domenico M, Ancora M, Curini V, Di Lollo Valeria, Mangone I, Rinaldi A, Delli Compagni E, Scialabba S, Caporale M, Di Pasquale A, Cammà C, Puglia I, Calistri P, Savini G |
| EPI_ISL_2757818, EPI_ISL_2757819, EPI_ISL_2757820, EPI_ISL_2757821 | SIESP CHIETI - DRIVE IN LANCIANO | Istituto Zooprofilattico Sperimentale dell'Abruzzo e Molise "G. Caporale" | Lorusso A, Marcacci M, Di Domenico M, Ancora M, Curini V, Di Lollo Valeria, Mangone I, Rinaldi A, Delli Compagni E, Scialabba S, Caporale M, Di Pasquale A, Cammà C, Puglia I, Calistri P, Savini G |
| EPI_ISL_2757822, EPI_ISL_2757823 | OSPEDALE CIVILE TERAMO - CENTRO TRASFUSIONALE | Istituto Zooprofilattico Sperimentale dell'Abruzzo e Molise "G. Caporale" | Lorusso A, Marcacci M, Di Domenico M, Ancora M, Curini V, Di Lollo Valeria, Mangone I, Rinaldi A, Delli Compagni E, Scialabba S, Caporale M, Di Pasquale A, Cammà C, Puglia I, Calistri P, Savini G |
| EPI_ISL_2757824, EPI_ISL_2757825, EPI_ISL_2757826 | SIESP CHIETI - DRIVE IN VASTO | Istituto Zooprofilattico Sperimentale dell'Abruzzo e Molise "G. Caporale" | Lorusso A, Marcacci M, Di Domenico M, Ancora M, Curini V, Di Lollo Valeria, Mangone I, Rinaldi A, Delli Compagni E, Scialabba S, Caporale M, Di Pasquale A, Cammà C, Puglia I, Calistri P, Savini G |
| EPI_ISL_2757827, EPI_ISL_2757828, EPI_ISL_2757829, EPI_ISL_2757830, EPI_ISL_2757831 | SIESP CHIETI - DRIVE IN LANCIANO | Istituto Zooprofilattico Sperimentale dell'Abruzzo e Molise "G. Caporale" | Lorusso A, Marcacci M, Di Domenico M, Ancora M, Curini V, Di Lollo Valeria, Mangone I, Rinaldi A, Delli Compagni E, Scialabba S, Caporale M, Di Pasquale A, Cammà C, Puglia I, Calistri P, Savini G |
| EPI_ISL_2757832 | OSPEDALE CIVILE TERAMO - CENTRO TRASFUSIONALE | Istituto Zooprofilattico Sperimentale dell'Abruzzo e Molise "G. Caporale" | Lorusso A, Marcacci M, Di Domenico M, Ancora M, Curini V, Di Lollo Valeria, Mangone I, Rinaldi A, Delli Compagni E, Scialabba S, Caporale M, Di Pasquale A, Cammà C, Puglia I, Calistri P, Savini G |
| EPI_ISL_2757930 | Johor State Health Department | Institute for Medical Research, Infectious Disease Research Centre, National Institutes of Health, Ministry of Health Malaysia | Suppiah J, Kamel K, Mohd Zawawi Z, Azizan MA, Ramly N, Robert F, Thayan R |
| EPI_ISL_2757941, EPI_ISL_2757942 | Queen Elizabeth Hospital, Kota Kinabalu | Institute for Medical Research, Infectious Disease Research Centre, National Institutes of Health, Ministry of Health Malaysia | Suppiah J, Kamel K, Mohd Zawawi Z, Azizan MA, Ramly N, Robert F, Thayan R |
| EPI_ISL_2757951 | Johor State Health Department | Institute for Medical Research, Infectious Disease Research Centre, National Institutes of Health, Ministry of Health Malaysia | Suppiah J, Kamel K, Mohd Zawawi Z, Azizan MA, Ramly N, Robert F, Thayan R |
| EPI_ISL_2758216, EPI_ISL_2758217, EPI_ISL_2758219, EPI_ISL_2758220, EPI_ISL_2758221, EPI_ISL_2758222, EPI_ISL_2758223, EPI_ISL_2758224 | Laboratoire Hôpital Sainte Musse | Laboratoire Hôpital Sainte Musse | Lionel Chollet |
| EPI_ISL_2758645 | OCME Office Of Chief Medical Examiner | New York City Public Health Laboratory | Jade Wang, et al. |
| EPI_ISL_2759093, EPI_ISL_2763624 | MOH - Jaber Al-Ahmad Hospital (Innovation Research Laboratory) | MOH - Jaber Al-Ahmad Hospital (Innovation Research Laboratory) | Salman Al-Sabah , Mohammad Alghounaim |
| EPI_ISL_2773986, EPI_ISL_2773987, EPI_ISL_2773988, EPI_ISL_2773989, EPI_ISL_2773990, EPI_ISL_2773991, EPI_ISL_2773993, EPI_ISL_2773994, EPI_ISL_2773995, EPI_ISL_2773996, EPI_ISL_2773997, EPI_ISL_2773998, EPI_ISL_2773999, EPI_ISL_2774000, EPI_ISL_2774001, EPI_ISL_2774002 | | | |
| see above | OLVZ Aalst | OLVZ Aalst | Astrid Holderbeke |
| EPI_ISL_2774010, EPI_ISL_2774011, EPI_ISL_2774012, EPI_ISL_2774013, EPI_ISL_2774014, EPI_ISL_2774015 | Cantacuzino National Military-Medical Institute, Viral Respiratory Infections Laboratory | Cantacuzino Institute Virology | Luiza Ustea, Nicoleta Paraschiv, Sorin Dinu, Mihaela Oprea, Mihaela Lazar |
| EPI_ISL_2774017, EPI_ISL_2774018, EPI_ISL_2774019, EPI_ISL_2774020, EPI_ISL_2774021, EPI_ISL_2774022, EPI_ISL_2774023, EPI_ISL_2774024, EPI_ISL_2774025, EPI_ISL_2774026, EPI_ISL_2774027, EPI_ISL_2774028, EPI_ISL_2774029, EPI_ISL_2774030, EPI_ISL_2774031, EPI_ISL_2774032, EPI_ISL_2774033, EPI_ISL_2774034 | | | |
| see above | Clinical Emergency County Hospital, Craiova | Cantacuzino Institute Virology | Luiza Ustea, Nicoleta Paraschiv, Sorin Dinu, Mihaela Oprea, Mihaela Lazar |
| EPI_ISL_2774035 | Cantacuzino National Military-Medical Institute, Viral Respiratory Infections Laboratory | Cantacuzino Institute Virology | Luiza Ustea, Nicoleta Paraschiv, Sorin Dinu, Mihaela Oprea, Mihaela Lazar |
| EPI_ISL_2774037 | Clinical Emergency County Hospital, Craiova | Cantacuzino Institute Virology | Luiza Ustea, Nicoleta Paraschiv, Sorin Dinu, Mihaela Oprea, Mihaela Lazar |
| EPI_ISL_2774094, EPI_ISL_2774096, EPI_ISL_2774097, EPI_ISL_2774098, EPI_ISL_2774099, EPI_ISL_2774100, EPI_ISL_2774101, EPI_ISL_2774102, EPI_ISL_2774104, EPI_ISL_2774105, EPI_ISL_2774106, EPI_ISL_2774107, EPI_ISL_2774108, EPI_ISL_2774109, EPI_ISL_2774110, EPI_ISL_2774111, EPI_ISL_2774112, EPI_ISL_2774113, EPI_ISL_2774114, EPI_ISL_2774115, EPI_ISL_2774116, EPI_ISL_2774117, EPI_ISL_2774118, EPI_ISL_2774119, EPI_ISL_2774120, EPI_ISL_2774121, EPI_ISL_2774122, EPI_ISL_2774123, EPI_ISL_2774124, EPI_ISL_2774125, EPI_ISL_2774126, EPI_ISL_2774127, EPI_ISL_2774128, EPI_ISL_2774129, EPI_ISL_2774130, EPI_ISL_2774131 | | | |
| see above | Microbiology Department, University Hospital Donostia | Microbiology Department, University Hospital Donostia | Marimon JM, Montes M, Piñeiro L, Martin-Peñaranda T, Gomez M, Sorarrain A, Cilla G. |
| EPI_ISL_2774241, EPI_ISL_2774242, EPI_ISL_2774244, EPI_ISL_2774245, EPI_ISL_2774246, EPI_ISL_2774247, EPI_ISL_2774248, EPI_ISL_2774249, EPI_ISL_2774252, EPI_ISL_2774257, EPI_ISL_2774259, EPI_ISL_2774260, EPI_ISL_2774261, EPI_ISL_2774265, EPI_ISL_2774266, EPI_ISL_2774270, EPI_ISL_2774271, EPI_ISL_2774272, EPI_ISL_2774274, EPI_ISL_2774276, EPI_ISL_2774277, EPI_ISL_2774278, EPI_ISL_2774279, EPI_ISL_2774281, EPI_ISL_2774285, EPI_ISL_2774286, EPI_ISL_2774287, EPI_ISL_2774288, EPI_ISL_2774289, EPI_ISL_2774290, EPI_ISL_2774291, EPI_ISL_2774294, EPI_ISL_2774295, EPI_ISL_2774296, EPI_ISL_2774297, EPI_ISL_2774298, EPI_ISL_2774303, EPI_ISL_2774304 | | | |

| | | | |
|---|---|---|---|
| see above | Baylor Scott & White-Temple | Baylor Scott & White-Temple | Ari Rao, Linden Morales, Kimberly Walker, Marcus Volz, Shelby Johnson |
| EPI_ISL_2774306 | Baylor Scott & White-Irving | Baylor Scott & White-Temple | Ari Rao, Linden Morales, Kimberly Walker, Marcus Volz, Shelby Johnson |
| EPI_ISL_2776369 | Ministry of Health, Jaber Al-Ahmad Hospital | Virology Unit, Department of Microbiology, Faulty of Medicine, Kuwait University | Nada Madi, Hussain Safar, Ebaa Al-Awadhi, Anfal Al-Adwani |
| EPI_ISL_2776430 | Ministry of Health, Jaber Al-Ahmad Hospital | Virology Unit, Department of Microbiology, Faculty of Medicine, Kuwait University | Nada Madi, Hussain Safar, Ebaa Al-Awadhi, Anfal Al-Adwani |
| EPI_ISL_2777211, EPI_ISL_2777212, EPI_ISL_2777213, EPI_ISL_2777214 | Instituto Nacional de Enfermedades Respiratorias (INER) | Centro de Investigación en Enfermedades Infecciosas (CIENI), Instituto Nacional de Enfermedades Respiratorias (INER) | Consorcio Mexicano de Vigilancia Genómica (CoViGen-Mex). Authors (in alphabetical order): Julio Elias Alvarado-Yaah, Carlos F. Arias, Santiago Ávila-Ríos, Eduardo Becerril-Vargas, Víctor Hugo Borja-Aburto, Celia Boukadida, Cristóbal Cháidez-Quiróz, Juan Bautista Chale-Dzul, Ricardo Ciria Merce, Andreu Comas-García, Célida Duque Molina, Julissa Enciso-Ibarra, José Antonio Enciso-Moreno, Gloria Elena Espinosa-Ayala, Fernando Fontove-Herrera, Daniel Fregoso-Rueda, Victor Eduardo Garcia-Arias, Alejandra Garcia-Gasca, Bruno Gomez-Gil, Jean Pierre González, Irvin González-López, Concepción Grajales-Muñiz, Ricardo Grande, Rosa María Gutierrez Rios, Alejandra Hernández-Terán, Alfredo Herrera-Estrella, Carla Ivón Herrera-Najera, Pavel Isa, Daniel Lira Morales, Susana Lopez, Antonio Loza Román, Brenda Irasema Maldonado-Meza, Bernardo Martínez-Miguel, José Arturo Martínez-Orozco, Célida Martínez- Rodríguez, Margarita Matías-Florentino, Fidencio Mejía-Nepomuceno, María Guadalupe de Jesús Mireles-Rivera, Gloria María Molina-Salinas, Hector Montoya-Fuentes, Mario Mújica-Sánchez, José Esteban Muñoz-Medina, José de Jesús Nuñez-Contreras, Alicia Ocaña-Mondragón, Luis Alberto Ochoa-Carrera, Hector Esteban Paz-Juárez, Marissa Pérez-García, Francisco Pulido, Helen Haydee Fernanda Ramírez-Plascencia, Jorge Salas-Hernández, Angel Gustavo Salas-Lais, Alejandro Sanchez-Flores, Clara Esperanza Santacruz-Tinoco, María Guadalupe Santiago-Mauricio, Selene Zárate, Nelly Sélem-Mojica, Blanca Taboada, Kathia Elizabeth Tapia-Diaz, Gloria Vazquez, Joel Armando Vázquez-Pérez. |
| EPI_ISL_2777215 | Instituto Nacional de Cancerología | Centro de Investigación en Enfermedades Infecciosas (CIENI), Instituto Nacional de Enfermedades Respiratorias (INER) | Consorcio Mexicano de Vigilancia Genómica (CoViGen-Mex). Authors (in alphabetical order): Julio Elias Alvarado-Yaah, Carlos F. Arias, Santiago Ávila-Ríos, Eduardo Becerril-Vargas, Víctor Hugo Borja-Aburto, Celia Boukadida, Cristóbal Cháidez-Quiróz, Juan Bautista Chale-Dzul, Ricardo Ciria Merce, Andreu Comas-García, Célida Duque Molina, Julissa Enciso-Ibarra, José Antonio Enciso-Moreno, Gloria Elena Espinosa-Ayala, Fernando Fontove-Herrera, Daniel Fregoso-Rueda, Victor Eduardo Garcia-Arias, Alejandra Garcia-Gasca, Bruno Gomez-Gil, Jean Pierre González, Irvin González-López, Concepción Grajales-Muñiz, Ricardo Grande, Rosa María Gutierrez Rios, Alejandra Hernández-Terán, Alfredo Herrera-Estrella, Carla Ivón Herrera-Najera, Pavel Isa, Daniel Lira Morales, Susana Lopez, Antonio Loza Román, Brenda Irasema Maldonado-Meza, Bernardo Martínez-Miguel, José Arturo Martínez-Orozco, Célida Martínez- Rodríguez, Margarita Matías-Florentino, Fidencio Mejía-Nepomuceno, María Guadalupe de Jesús Mireles-Rivera, Gloria María Molina-Salinas, Hector Montoya-Fuentes, Mario Mújica-Sánchez, José Esteban Muñoz-Medina, José de Jesús Nuñez-Contreras, Alicia Ocaña-Mondragón, Luis Alberto Ochoa-Carrera, Hector Esteban Paz-Juárez, Marissa Pérez-García, Francisco Pulido, Helen Haydee Fernanda Ramírez-Plascencia, Jorge Salas-Hernández, Angel Gustavo Salas-Lais, Alejandro Sanchez-Flores, Clara Esperanza Santacruz-Tinoco, María Guadalupe Santiago-Mauricio, Selene Zárate, Nelly Sélem-Mojica, Blanca Taboada, Kathia Elizabeth Tapia-Diaz, Gloria Vazquez, Joel Armando Vázquez-Pérez. |
| EPI_ISL_2777216, EPI_ISL_2777217, EPI_ISL_2777218, EPI_ISL_2777219, EPI_ISL_2777220 | Instituto Nacional de Enfermedades Respiratorias (INER) | Centro de Investigación en Enfermedades Infecciosas (CIENI), Instituto Nacional de Enfermedades Respiratorias (INER) | Consorcio Mexicano de Vigilancia Genómica (CoViGen-Mex). Authors (in alphabetical order): Julio Elias Alvarado-Yaah, Carlos F. Arias, Santiago Ávila-Ríos, Eduardo Becerril-Vargas, Víctor Hugo Borja-Aburto, Celia Boukadida, Cristóbal Cháidez-Quiróz, Juan Bautista Chale-Dzul, Ricardo Ciria Merce, Andreu Comas-García, Célida Duque Molina, Julissa Enciso-Ibarra, José Antonio Enciso-Moreno, Gloria Elena Espinosa-Ayala, Fernando Fontove-Herrera, Daniel Fregoso-Rueda, Victor Eduardo Garcia-Arias, Alejandra Garcia-Gasca, Bruno Gomez-Gil, Jean Pierre González, Irvin González-López, Concepción Grajales-Muñiz, Ricardo Grande, Rosa María Gutierrez Rios, Alejandra Hernández-Terán, Alfredo Herrera-Estrella, Carla Ivón Herrera-Najera, Pavel Isa, Daniel Lira Morales, Susana Lopez, Antonio Loza Román, Brenda Irasema Maldonado-Meza, Bernardo Martínez-Miguel, José Arturo Martínez-Orozco, Célida Martínez- Rodríguez, Margarita Matías-Florentino, Fidencio Mejía-Nepomuceno, María Guadalupe de Jesús Mireles-Rivera, Gloria María Molina-Salinas, Hector Montoya-Fuentes, Mario Mújica-Sánchez, José Esteban Muñoz-Medina, José de Jesús Nuñez-Contreras, Alicia Ocaña-Mondragón, Luis Alberto Ochoa-Carrera, Hector Esteban Paz-Juárez, Marissa Pérez-García, Francisco Pulido, Helen Haydee Fernanda Ramírez-Plascencia, Jorge Salas-Hernández, Angel Gustavo Salas-Lais, Alejandro Sanchez-Flores, Clara Esperanza Santacruz-Tinoco, María Guadalupe Santiago-Mauricio, Selene Zárate, Nelly Sélem-Mojica, Blanca Taboada, Kathia Elizabeth Tapia-Diaz, Gloria Vazquez, Joel Armando Vázquez-Pérez. |
| EPI_ISL_2777222, EPI_ISL_2777223, EPI_ISL_2777224, EPI_ISL_2777225, EPI_ISL_2777226, EPI_ISL_2777227, EPI_ISL_2777229, EPI_ISL_2777230, EPI_ISL_2777231, EPI_ISL_2777232, EPI_ISL_2777233, EPI_ISL_2777234 | | | |
| see above | Instituto Nacional de Cancerología | Centro de Investigación en Enfermedades Infecciosas (CIENI), Instituto Nacional de Enfermedades Respiratorias (INER) | Consorcio Mexicano de Vigilancia Genómica (CoViGen-Mex). Authors (in alphabetical order): Julio Elias Alvarado-Yaah, Carlos F. Arias, Santiago Ávila-Ríos, Eduardo Becerril-Vargas, Víctor Hugo Borja-Aburto, Celia Boukadida, Cristóbal Cháidez-Quiróz, Juan Bautista Chale-Dzul, Ricardo Ciria Merce, Andreu Comas-García, Célida Duque Molina, Julissa Enciso-Ibarra, José Antonio Enciso-Moreno, Gloria Elena Espinosa-Ayala, Fernando Fontove-Herrera, Daniel Fregoso-Rueda, Victor Eduardo Garcia-Arias, Alejandra Garcia-Gasca, Bruno Gomez-Gil, Jean Pierre González, Irvin González-López, Concepción Grajales-Muñiz, Ricardo Grande, Rosa María Gutierrez Rios, Alejandra Hernández-Terán, Alfredo Herrera-Estrella, Carla Ivón Herrera-Najera, Pavel Isa, Daniel Lira Morales, Susana Lopez, Antonio Loza Román, Brenda Irasema Maldonado-Meza, Bernardo Martínez-Miguel, José Arturo Martínez-Orozco, Célida Martínez- Rodríguez, Margarita Matías-Florentino, Fidencio Mejía-Nepomuceno, María Guadalupe de Jesús Mireles-Rivera, Gloria María Molina-Salinas, Hector Montoya-Fuentes, Mario Mújica-Sánchez, José Esteban Muñoz-Medina, José de Jesús Nuñez-Contreras, Alicia Ocaña-Mondragón, Luis Alberto Ochoa-Carrera, Hector Esteban Paz-Juárez, Marissa Pérez-García, Francisco Pulido, Helen Haydee Fernanda Ramírez-Plascencia, Jorge Salas-Hernández, Angel Gustavo Salas-Lais, Alejandro Sanchez-Flores, Clara Esperanza Santacruz-Tinoco, María Guadalupe Santiago-Mauricio, Selene Zárate, Nelly Sélem-Mojica, Blanca Taboada, Kathia Elizabeth Tapia-Diaz, Gloria Vazquez, Joel Armando Vázquez-Pérez. |
| EPI_ISL_2778018, EPI_ISL_2778019, EPI_ISL_2778020, EPI_ISL_2778021, EPI_ISL_2778022, EPI_ISL_2778023, EPI_ISL_2778024, EPI_ISL_2778025, EPI_ISL_2778026, EPI_ISL_2778027, EPI_ISL_2778028, EPI_ISL_2778029, EPI_ISL_2778030, EPI_ISL_2778031, EPI_ISL_2778032, EPI_ISL_2778033, EPI_ISL_2778035, EPI_ISL_2778036, EPI_ISL_2778037, EPI_ISL_2778040, EPI_ISL_2778041, EPI_ISL_2778043, EPI_ISL_2778045, EPI_ISL_2778046, EPI_ISL_2778047, EPI_ISL_2778048, EPI_ISL_2778049, EPI_ISL_2778050, EPI_ISL_2778051, EPI_ISL_2778052, EPI_ISL_2778053, EPI_ISL_2778054, EPI_ISL_2778055, EPI_ISL_2778057, EPI_ISL_2778060, EPI_ISL_2778062, EPI_ISL_2778063, EPI_ISL_2778064, EPI_ISL_2778065, EPI_ISL_2778066, EPI_ISL_2778067, EPI_ISL_2778069, EPI_ISL_2778071, EPI_ISL_2778072, EPI_ISL_2778073, EPI_ISL_2778074, EPI_ISL_2778076, EPI_ISL_2778077, EPI_ISL_2778078, EPI_ISL_2778079, EPI_ISL_2778080, EPI_ISL_2778081, EPI_ISL_2778085, EPI_ISL_2778086, EPI_ISL_2778087, EPI_ISL_2778088, EPI_ISL_2778090, EPI_ISL_2778093, EPI_ISL_2778095, EPI_ISL_2778096 | | | |
| see above | Cliniques universitaires Saint-Luc | UCLouvain/IREC/MBLG | Jean Ruelle, Bertrand Bearzatto, Benoit Kabamba Mukadi |
| EPI_ISL_2778097, EPI_ISL_2778098, EPI_ISL_2778099 | Clinique Saint-Pierre Ottignies | UCLouvain/IREC/MBLG | Jean Ruelle, Bertrand Bearzatto, Benoit Kabamba Mukadi |
| EPI_ISL_2778101, EPI_ISL_2778103 | Cliniques universitaires Saint-Luc | UCLouvain/IREC/MBLG | Jean Ruelle, Bertrand Bearzatto, Benoit Kabamba Mukadi |
| EPI_ISL_2778984, EPI_ISL_2778985, EPI_ISL_2778986, EPI_ISL_2778987, EPI_ISL_2778988 | LESP Tabasco | Instituto de Diagnostico y Referencia Epidemiologicos (INDRE) | Claudia Wong-Arambula, Abril Rodriguez-Maldonado, Vanessa Rivero-Arredondo, Ariadna Medina-Benitez, Joaquin Quiroz-Mercado, Sergio Rangel-Guerrero, Maribel Gonzalez-Villa, Natividad Cruz-Ortiz, Tatiana Nunez-Garcia, Gisela Barrera-Badillo, Lucia Hernandez-Rivas, Irma Lopez-Martinez, Ernesto Ramirez-Gonzalez. |
| EPI_ISL_2778990, EPI_ISL_2778991 | LESP Guanajuato | Instituto de Diagnostico y Referencia Epidemiologicos (INDRE) | Claudia Wong-Arambula, Abril Rodriguez-Maldonado, Vanessa Rivero-Arredondo, Ariadna Medina-Benitez, Joaquin Quiroz-Mercado, Sergio Rangel-Guerrero, Maribel Gonzalez-Villa, Natividad Cruz-Ortiz, Tatiana Nunez-Garcia, Gisela Barrera-Badillo, Lucia Hernandez-Rivas, Irma Lopez-Martinez, Ernesto Ramirez-Gonzalez. |
| EPI_ISL_2778993, EPI_ISL_2778994 | LESP Durango | Instituto de Diagnostico y Referencia Epidemiologicos (INDRE) | Claudia Wong-Arambula, Abril Rodriguez-Maldonado, Vanessa Rivero-Arredondo, Ariadna Medina-Benitez, Joaquin Quiroz-Mercado, Sergio Rangel-Guerrero, Maribel Gonzalez-Villa, Natividad Cruz-Ortiz, Tatiana Nunez-Garcia, Gisela Barrera-Badillo, Lucia Hernandez-Rivas, Irma Lopez-Martinez, Ernesto Ramirez-Gonzalez. |
| EPI_ISL_2778995 | LESP Nayarit | Instituto de Diagnostico y Referencia Epidemiologicos (INDRE) | Claudia Wong-Arambula, Abril Rodriguez-Maldonado, Vanessa Rivero-Arredondo, Ariadna Medina-Benitez, Joaquin Quiroz-Mercado, Sergio Rangel-Guerrero, Maribel Gonzalez-Villa, Natividad Cruz-Ortiz, Tatiana Nunez-Garcia, Gisela Barrera-Badillo, Lucia Hernandez-Rivas, Irma Lopez-Martinez, Ernesto Ramirez-Gonzalez. |
| EPI_ISL_2778996 | LESP Durango | Instituto de Diagnostico y Referencia Epidemiologicos (INDRE) | Claudia Wong-Arambula, Abril Rodriguez-Maldonado, Vanessa Rivero-Arredondo, Ariadna Medina-Benitez, Joaquin Quiroz-Mercado, Sergio Rangel-Guerrero, Maribel Gonzalez-Villa, Natividad Cruz-Ortiz, Tatiana Nunez-Garcia, Gisela Barrera-Badillo, Lucia Hernandez-Rivas, Irma Lopez-Martinez, Ernesto Ramirez-Gonzalez. |

| | | | |
|---|---|---|---|
| EPI_ISL_2778997 | LESP Chiapas | Instituto de Diagnostico y Referencia Epidemiologicos (INDRE) | Claudia Wong-Arambula, Abril Rodriguez-Maldonado, Vanessa Rivero-Arredondo, Ariadna Medina-Benitez, Joaquin Quiroz-Mercado, Sergio Rangel-Guerrero, Maribel Gonzalez-Villa, Natividad Cruz-Ortiz, Tatiana Nunez-Garcia, Gisela Barrera-Badillo, Lucia Hernandez-Rivas, Irma Lopez-Martinez, Ernesto Ramirez-Gonzalez. |
| EPI_ISL_2778998 | LESP Sinaloa | Instituto de Diagnostico y Referencia Epidemiologicos (INDRE) | Claudia Wong-Arambula, Abril Rodriguez-Maldonado, Vanessa Rivero-Arredondo, Ariadna Medina-Benitez, Joaquin Quiroz-Mercado, Sergio Rangel-Guerrero, Maribel Gonzalez-Villa, Natividad Cruz-Ortiz, Tatiana Nunez-Garcia, Gisela Barrera-Badillo, Lucia Hernandez-Rivas, Irma Lopez-Martinez, Ernesto Ramirez-Gonzalez. |
| EPI_ISL_2778999 | LESP Estado de Mexico | Instituto de Diagnostico y Referencia Epidemiologicos (INDRE) | Claudia Wong-Arambula, Abril Rodriguez-Maldonado, Vanessa Rivero-Arredondo, Ariadna Medina-Benitez, Joaquin Quiroz-Mercado, Sergio Rangel-Guerrero, Maribel Gonzalez-Villa, Natividad Cruz-Ortiz, Tatiana Nunez-Garcia, Gisela Barrera-Badillo, Lucia Hernandez-Rivas, Irma Lopez-Martinez, Ernesto Ramirez-Gonzalez. |
| EPI_ISL_2779000 | LESP Durango | Instituto de Diagnostico y Referencia Epidemiologicos (INDRE) | Claudia Wong-Arambula, Abril Rodriguez-Maldonado, Vanessa Rivero-Arredondo, Ariadna Medina-Benitez, Joaquin Quiroz-Mercado, Sergio Rangel-Guerrero, Maribel Gonzalez-Villa, Natividad Cruz-Ortiz, Tatiana Nunez-Garcia, Gisela Barrera-Badillo, Lucia Hernandez-Rivas, Irma Lopez-Martinez, Ernesto Ramirez-Gonzalez. |
| EPI_ISL_2779001 | LESP Campeche | Instituto de Diagnostico y Referencia Epidemiologicos (INDRE) | Claudia Wong-Arambula, Abril Rodriguez-Maldonado, Vanessa Rivero-Arredondo, Ariadna Medina-Benitez, Joaquin Quiroz-Mercado, Sergio Rangel-Guerrero, Maribel Gonzalez-Villa, Natividad Cruz-Ortiz, Tatiana Nunez-Garcia, Gisela Barrera-Badillo, Lucia Hernandez-Rivas, Irma Lopez-Martinez, Ernesto Ramirez-Gonzalez. |
| EPI_ISL_2779002, EPI_ISL_2779003, EPI_ISL_2779004 | LESP Tamaulipas | Instituto de Diagnostico y Referencia Epidemiologicos (INDRE) | Claudia Wong-Arambula, Abril Rodriguez-Maldonado, Vanessa Rivero-Arredondo, Ariadna Medina-Benitez, Joaquin Quiroz-Mercado, Sergio Rangel-Guerrero, Maribel Gonzalez-Villa, Natividad Cruz-Ortiz, Tatiana Nunez-Garcia, Gisela Barrera-Badillo, Lucia Hernandez-Rivas, Irma Lopez-Martinez, Ernesto Ramirez-Gonzalez. |
| EPI_ISL_2779005, EPI_ISL_2779006, EPI_ISL_2779007 | LESP Sinaloa | Instituto de Diagnostico y Referencia Epidemiologicos (INDRE) | Claudia Wong-Arambula, Abril Rodriguez-Maldonado, Vanessa Rivero-Arredondo, Ariadna Medina-Benitez, Joaquin Quiroz-Mercado, Sergio Rangel-Guerrero, Maribel Gonzalez-Villa, Natividad Cruz-Ortiz, Tatiana Nunez-Garcia, Gisela Barrera-Badillo, Lucia Hernandez-Rivas, Irma Lopez-Martinez, Ernesto Ramirez-Gonzalez. |
| EPI_ISL_2779008, EPI_ISL_2779009, EPI_ISL_2779010, EPI_ISL_2779011, EPI_ISL_2779012, EPI_ISL_2779013, EPI_ISL_2779014, EPI_ISL_2779015, EPI_ISL_2779016, EPI_ISL_2779017, EPI_ISL_2779018 | | | |
| see above | LESP Yucatan | Instituto de Diagnostico y Referencia Epidemiologicos (INDRE) | Claudia Wong-Arambula, Abril Rodriguez-Maldonado, Vanessa Rivero-Arredondo, Ariadna Medina-Benitez, Joaquin Quiroz-Mercado, Sergio Rangel-Guerrero, Maribel Gonzalez-Villa, Natividad Cruz-Ortiz, Tatiana Nunez-Garcia, Gisela Barrera-Badillo, Lucia Hernandez-Rivas, Irma Lopez-Martinez, Ernesto Ramirez-Gonzalez. |
| EPI_ISL_2779019 | LESP Guanajuato | Instituto de Diagnostico y Referencia Epidemiologicos (INDRE) | Claudia Wong-Arambula, Abril Rodriguez-Maldonado, Vanessa Rivero-Arredondo, Ariadna Medina-Benitez, Joaquin Quiroz-Mercado, Sergio Rangel-Guerrero, Maribel Gonzalez-Villa, Natividad Cruz-Ortiz, Tatiana Nunez-Garcia, Gisela Barrera-Badillo, Lucia Hernandez-Rivas, Irma Lopez-Martinez, Ernesto Ramirez-Gonzalez. |
| EPI_ISL_2779020, EPI_ISL_2779021, EPI_ISL_2779022, EPI_ISL_2779023 | LESP Yucatan | Instituto de Diagnostico y Referencia Epidemiologicos (INDRE) | Claudia Wong-Arambula, Abril Rodriguez-Maldonado, Vanessa Rivero-Arredondo, Ariadna Medina-Benitez, Joaquin Quiroz-Mercado, Sergio Rangel-Guerrero, Maribel Gonzalez-Villa, Natividad Cruz-Ortiz, Tatiana Nunez-Garcia, Gisela Barrera-Badillo, Lucia Hernandez-Rivas, Irma Lopez-Martinez, Ernesto Ramirez-Gonzalez. |
| EPI_ISL_2779024, EPI_ISL_2779025, EPI_ISL_2779026, EPI_ISL_2779027, EPI_ISL_2779028, EPI_ISL_2779029, EPI_ISL_2779030, EPI_ISL_2779031, EPI_ISL_2779032, EPI_ISL_2779033 | LESP Tabasco | Instituto de Diagnostico y Referencia Epidemiologicos (INDRE) | Claudia Wong-Arambula, Abril Rodriguez-Maldonado, Vanessa Rivero-Arredondo, Ariadna Medina-Benitez, Joaquin Quiroz-Mercado, Sergio Rangel-Guerrero, Maribel Gonzalez-Villa, Natividad Cruz-Ortiz, Tatiana Nunez-Garcia, Gisela Barrera-Badillo, Lucia Hernandez-Rivas, Irma Lopez-Martinez, Ernesto Ramirez-Gonzalez. |
| EPI_ISL_2779034 | LESP Coahuila | Instituto de Diagnostico y Referencia Epidemiologicos (INDRE) | Claudia Wong-Arambula, Abril Rodriguez-Maldonado, Vanessa Rivero-Arredondo, Ariadna Medina-Benitez, Joaquin Quiroz-Mercado, Sergio Rangel-Guerrero, Maribel Gonzalez-Villa, Natividad Cruz-Ortiz, Tatiana Nunez-Garcia, Gisela Barrera-Badillo, Lucia Hernandez-Rivas, Irma Lopez-Martinez, Ernesto Ramirez-Gonzalez. |
| EPI_ISL_2779035, EPI_ISL_2779036, EPI_ISL_2779037, EPI_ISL_2779038, EPI_ISL_2779039, EPI_ISL_2779040, EPI_ISL_2779041, EPI_ISL_2779042 | LESP Puebla | Instituto de Diagnostico y Referencia Epidemiologicos (INDRE) | Claudia Wong-Arambula, Abril Rodriguez-Maldonado, Vanessa Rivero-Arredondo, Ariadna Medina-Benitez, Joaquin Quiroz-Mercado, Sergio Rangel-Guerrero, Maribel Gonzalez-Villa, Natividad Cruz-Ortiz, Tatiana Nunez-Garcia, Gisela Barrera-Badillo, Lucia Hernandez-Rivas, Irma Lopez-Martinez, Ernesto Ramirez-Gonzalez. |
| EPI_ISL_2779043, EPI_ISL_2779044, EPI_ISL_2779045, EPI_ISL_2779046 | LESP Tabasco | Instituto de Diagnostico y Referencia Epidemiologicos (INDRE) | Claudia Wong-Arambula, Abril Rodriguez-Maldonado, Vanessa Rivero-Arredondo, Ariadna Medina-Benitez, Joaquin Quiroz-Mercado, Sergio Rangel-Guerrero, Maribel Gonzalez-Villa, Natividad Cruz-Ortiz, Tatiana Nunez-Garcia, Gisela Barrera-Badillo, Lucia Hernandez-Rivas, Irma Lopez-Martinez, Ernesto Ramirez-Gonzalez. |
| EPI_ISL_2779047 | LESP Guanajuato | Instituto de Diagnostico y Referencia Epidemiologicos (INDRE) | Claudia Wong-Arambula, Abril Rodriguez-Maldonado, Vanessa Rivero-Arredondo, Ariadna Medina-Benitez, Joaquin Quiroz-Mercado, Sergio Rangel-Guerrero, Maribel Gonzalez-Villa, Natividad Cruz-Ortiz, Tatiana Nunez-Garcia, Gisela Barrera-Badillo, Lucia Hernandez-Rivas, Irma Lopez-Martinez, Ernesto Ramirez-Gonzalez. |
| EPI_ISL_2779050 | LESP Guerrero | Instituto de Diagnostico y Referencia Epidemiologicos (INDRE) | Claudia Wong-Arambula, Abril Rodriguez-Maldonado, Vanessa Rivero-Arredondo, Ariadna Medina-Benitez, Joaquin Quiroz-Mercado, Sergio Rangel-Guerrero, Maribel Gonzalez-Villa, Natividad Cruz-Ortiz, Tatiana Nunez-Garcia, Gisela Barrera-Badillo, Lucia Hernandez-Rivas, Irma Lopez-Martinez, Ernesto Ramirez-Gonzalez. |
| EPI_ISL_2779051, EPI_ISL_2779052 | LESP Quintana Roo | Instituto de Diagnostico y Referencia Epidemiologicos (INDRE) | Claudia Wong-Arambula, Abril Rodriguez-Maldonado, Vanessa Rivero-Arredondo, Ariadna Medina-Benitez, Joaquin Quiroz-Mercado, Sergio Rangel-Guerrero, Maribel Gonzalez-Villa, Natividad Cruz-Ortiz, Tatiana Nunez-Garcia, Gisela Barrera-Badillo, Lucia Hernandez-Rivas, Irma Lopez-Martinez, Ernesto Ramirez-Gonzalez. |
| EPI_ISL_2779053, EPI_ISL_2779054 | LESP Tabasco | Instituto de Diagnostico y Referencia Epidemiologicos (INDRE) | Claudia Wong-Arambula, Abril Rodriguez-Maldonado, Vanessa Rivero-Arredondo, Ariadna Medina-Benitez, Joaquin Quiroz-Mercado, Sergio Rangel-Guerrero, Maribel Gonzalez-Villa, Natividad Cruz-Ortiz, Tatiana Nunez-Garcia, Gisela Barrera-Badillo, Lucia Hernandez-Rivas, Irma Lopez-Martinez, Ernesto Ramirez-Gonzalez. |
| EPI_ISL_2779055 | LESP Sonora | Instituto de Diagnostico y Referencia Epidemiologicos (INDRE) | Claudia Wong-Arambula, Abril Rodriguez-Maldonado, Vanessa Rivero-Arredondo, Ariadna Medina-Benitez, Joaquin Quiroz-Mercado, Sergio Rangel-Guerrero, Maribel Gonzalez-Villa, Natividad Cruz-Ortiz, Tatiana Nunez-Garcia, Gisela Barrera-Badillo, Lucia Hernandez-Rivas, Irma Lopez-Martinez, Ernesto Ramirez-Gonzalez. |
| EPI_ISL_2779057, EPI_ISL_2779058 | LESP Guanajuato | Instituto de Diagnostico y Referencia Epidemiologicos (INDRE) | Claudia Wong-Arambula, Abril Rodriguez-Maldonado, Vanessa Rivero-Arredondo, Ariadna Medina-Benitez, Joaquin Quiroz-Mercado, Sergio Rangel-Guerrero, Maribel Gonzalez-Villa, Natividad Cruz-Ortiz, Tatiana Nunez-Garcia, Gisela Barrera-Badillo, Lucia Hernandez-Rivas, Irma Lopez-Martinez, Ernesto Ramirez-Gonzalez. |
| EPI_ISL_2779059 | LESP Campeche | Instituto de Diagnostico y Referencia Epidemiologicos (INDRE) | Claudia Wong-Arambula, Abril Rodriguez-Maldonado, Vanessa Rivero-Arredondo, Ariadna Medina-Benitez, Joaquin Quiroz-Mercado, Sergio Rangel-Guerrero, Maribel Gonzalez-Villa, Natividad Cruz-Ortiz, Tatiana Nunez-Garcia, Gisela Barrera-Badillo, Lucia Hernandez-Rivas, Irma Lopez-Martinez, Ernesto Ramirez-Gonzalez. |
| EPI_ISL_2779060, EPI_ISL_2779061, EPI_ISL_2779062, EPI_ISL_2779063, EPI_ISL_2779064, EPI_ISL_2779065 | LESP Tamaulipas | Instituto de Diagnostico y Referencia Epidemiologicos (INDRE) | Claudia Wong-Arambula, Abril Rodriguez-Maldonado, Vanessa Rivero-Arredondo, Ariadna Medina-Benitez, Joaquin Quiroz-Mercado, Sergio Rangel-Guerrero, Maribel Gonzalez-Villa, Natividad Cruz-Ortiz, Tatiana Nunez-Garcia, Gisela Barrera-Badillo, Lucia Hernandez-Rivas, Irma Lopez-Martinez, Ernesto Ramirez-Gonzalez. |
| EPI_ISL_2779066, EPI_ISL_2779067, EPI_ISL_2779068, EPI_ISL_2779069, EPI_ISL_2779070, EPI_ISL_2779071, EPI_ISL_2779072, EPI_ISL_2779073, EPI_ISL_2779074, EPI_ISL_2779075, EPI_ISL_2779076, EPI_ISL_2779077, EPI_ISL_2779078, EPI_ISL_2779079, EPI_ISL_2779080, EPI_ISL_2779081, EPI_ISL_2779082, EPI_ISL_2779083, EPI_ISL_2779084, EPI_ISL_2779085, EPI_ISL_2779086, EPI_ISL_2779087, EPI_ISL_2779088, EPI_ISL_2779089, EPI_ISL_2779090, EPI_ISL_2779091, EPI_ISL_2779092, EPI_ISL_2779093, EPI_ISL_2779094, EPI_ISL_2779095, EPI_ISL_2779096, EPI_ISL_2779097, EPI_ISL_2779098, EPI_ISL_2779099, EPI_ISL_2779100, EPI_ISL_2779101, EPI_ISL_2779102, EPI_ISL_2779103, EPI_ISL_2779104, EPI_ISL_2779105, EPI_ISL_2779106, EPI_ISL_2779107, EPI_ISL_2779108, EPI_ISL_2779109, EPI_ISL_2779110, EPI_ISL_2779111, EPI_ISL_2779112, EPI_ISL_2779113, EPI_ISL_2779114, EPI_ISL_2779115, EPI_ISL_2779116, EPI_ISL_2779117, EPI_ISL_2779118, EPI_ISL_2779119 | | | |
| see above | LESP Yucatan | Instituto de Diagnostico y Referencia Epidemiologicos | Claudia Wong-Arambula, Abril Rodriguez-Maldonado, Vanessa Rivero-Arredondo, Ariadna Medina-Benitez, Joaquin Quiroz-Mercado, Sergio |

| | | | |
|---|---|---|---|
| | | (INDRE) | Rangel-Guerrero, Maribel Gonzalez-Villa, Natividad Cruz-Ortiz, Tatiana Nunez-Garcia, Gisela Barrera-Badillo, Lucia Hernandez-Rivas, Irma Lopez-Martinez, Ernesto Ramirez-Gonzalez. |
| EPI_ISL_2779120 | LESP Guanajuato | Instituto de Diagnostico y Referencia Epidemiologicos (INDRE) | Claudia Wong-Arambula, Abril Rodriguez-Maldonado, Vanessa Rivero-Arredondo, Ariadna Medina-Benitez, Joaquin Quiroz-Mercado, Sergio Rangel-Guerrero, Maribel Gonzalez-Villa, Natividad Cruz-Ortiz, Tatiana Nunez-Garcia, Gisela Barrera-Badillo, Lucia Hernandez-Rivas, Irma Lopez-Martinez, Ernesto Ramirez-Gonzalez. |
| EPI_ISL_2779121, EPI_ISL_2779122, EPI_ISL_2779123, EPI_ISL_2779124, EPI_ISL_2779125, EPI_ISL_2779126, EPI_ISL_2779127, EPI_ISL_2779128, EPI_ISL_2779130, EPI_ISL_2779131, EPI_ISL_2779132, EPI_ISL_2779133, EPI_ISL_2779134, EPI_ISL_2779135, EPI_ISL_2779136, EPI_ISL_2779137, EPI_ISL_2779138, EPI_ISL_2779139, EPI_ISL_2779140, EPI_ISL_2779141, EPI_ISL_2779142, EPI_ISL_2779143, EPI_ISL_2779144, EPI_ISL_2779145, EPI_ISL_2779146, EPI_ISL_2779147, EPI_ISL_2779148, EPI_ISL_2779149, EPI_ISL_2779150, EPI_ISL_2779151, EPI_ISL_2779152, EPI_ISL_2779153, EPI_ISL_2779154, EPI_ISL_2779155, EPI_ISL_2779156, EPI_ISL_2779157, EPI_ISL_2779158 | | | |
| see above | LESP Yucatan | Instituto de Diagnostico y Referencia Epidemiologicos (INDRE) | Claudia Wong-Arambula, Abril Rodriguez-Maldonado, Vanessa Rivero-Arredondo, Ariadna Medina-Benitez, Joaquin Quiroz-Mercado, Sergio Rangel-Guerrero, Maribel Gonzalez-Villa, Natividad Cruz-Ortiz, Tatiana Nunez-Garcia, Gisela Barrera-Badillo, Lucia Hernandez-Rivas, Irma Lopez-Martinez, Ernesto Ramirez-Gonzalez. |
| EPI_ISL_2779159 | LESP San Luis Potosi | Instituto de Diagnostico y Referencia Epidemiologicos (INDRE) | Claudia Wong-Arambula, Abril Rodriguez-Maldonado, Vanessa Rivero-Arredondo, Ariadna Medina-Benitez, Joaquin Quiroz-Mercado, Sergio Rangel-Guerrero, Maribel Gonzalez-Villa, Natividad Cruz-Ortiz, Tatiana Nunez-Garcia, Gisela Barrera-Badillo, Lucia Hernandez-Rivas, Irma Lopez-Martinez, Ernesto Ramirez-Gonzalez. |
| EPI_ISL_2779160, EPI_ISL_2779161, EPI_ISL_2779162, EPI_ISL_2779163, EPI_ISL_2779164, EPI_ISL_2779165, EPI_ISL_2779166, EPI_ISL_2779167, EPI_ISL_2779168, EPI_ISL_2779169, EPI_ISL_2779170, EPI_ISL_2779171, EPI_ISL_2779172 | | | |
| see above | LESP Tabasco | Instituto de Diagnostico y Referencia Epidemiologicos (INDRE) | Claudia Wong-Arambula, Abril Rodriguez-Maldonado, Vanessa Rivero-Arredondo, Ariadna Medina-Benitez, Joaquin Quiroz-Mercado, Sergio Rangel-Guerrero, Maribel Gonzalez-Villa, Natividad Cruz-Ortiz, Tatiana Nunez-Garcia, Gisela Barrera-Badillo, Lucia Hernandez-Rivas, Irma Lopez-Martinez, Ernesto Ramirez-Gonzalez. |
| EPI_ISL_2779173 | LESP Ciudad de Mexico | Instituto de Diagnostico y Referencia Epidemiologicos (INDRE) | Claudia Wong-Arambula, Abril Rodriguez-Maldonado, Vanessa Rivero-Arredondo, Ariadna Medina-Benitez, Joaquin Quiroz-Mercado, Sergio Rangel-Guerrero, Maribel Gonzalez-Villa, Natividad Cruz-Ortiz, Tatiana Nunez-Garcia, Gisela Barrera-Badillo, Lucia Hernandez-Rivas, Irma Lopez-Martinez, Ernesto Ramirez-Gonzalez. |
| EPI_ISL_2779174 | LESP Puebla | Instituto de Diagnostico y Referencia Epidemiologicos (INDRE) | Claudia Wong-Arambula, Abril Rodriguez-Maldonado, Vanessa Rivero-Arredondo, Ariadna Medina-Benitez, Joaquin Quiroz-Mercado, Sergio Rangel-Guerrero, Maribel Gonzalez-Villa, Natividad Cruz-Ortiz, Tatiana Nunez-Garcia, Gisela Barrera-Badillo, Lucia Hernandez-Rivas, Irma Lopez-Martinez, Ernesto Ramirez-Gonzalez. |
| EPI_ISL_2779175 | LESP Jalisco | Instituto de Diagnostico y Referencia Epidemiologicos (INDRE) | Claudia Wong-Arambula, Abril Rodriguez-Maldonado, Vanessa Rivero-Arredondo, Ariadna Medina-Benitez, Joaquin Quiroz-Mercado, Sergio Rangel-Guerrero, Maribel Gonzalez-Villa, Natividad Cruz-Ortiz, Tatiana Nunez-Garcia, Gisela Barrera-Badillo, Lucia Hernandez-Rivas, Irma Lopez-Martinez, Ernesto Ramirez-Gonzalez. |
| EPI_ISL_2779176, EPI_ISL_2779177, EPI_ISL_2779178, EPI_ISL_2779179, EPI_ISL_2779180, EPI_ISL_2779181 | LESP Quintana Roo | Instituto de Diagnostico y Referencia Epidemiologicos (INDRE) | Claudia Wong-Arambula, Abril Rodriguez-Maldonado, Vanessa Rivero-Arredondo, Ariadna Medina-Benitez, Joaquin Quiroz-Mercado, Sergio Rangel-Guerrero, Maribel Gonzalez-Villa, Natividad Cruz-Ortiz, Tatiana Nunez-Garcia, Gisela Barrera-Badillo, Lucia Hernandez-Rivas, Irma Lopez-Martinez, Ernesto Ramirez-Gonzalez. |
| EPI_ISL_2779182, EPI_ISL_2779183 | LESP Tabasco | Instituto de Diagnostico y Referencia Epidemiologicos (INDRE) | Claudia Wong-Arambula, Abril Rodriguez-Maldonado, Vanessa Rivero-Arredondo, Ariadna Medina-Benitez, Joaquin Quiroz-Mercado, Sergio Rangel-Guerrero, Maribel Gonzalez-Villa, Natividad Cruz-Ortiz, Tatiana Nunez-Garcia, Gisela Barrera-Badillo, Lucia Hernandez-Rivas, Irma Lopez-Martinez, Ernesto Ramirez-Gonzalez. |
| EPI_ISL_2779184, EPI_ISL_2779185 | LESP Sonora | Instituto de Diagnostico y Referencia Epidemiologicos (INDRE) | Claudia Wong-Arambula, Abril Rodriguez-Maldonado, Vanessa Rivero-Arredondo, Ariadna Medina-Benitez, Joaquin Quiroz-Mercado, Sergio Rangel-Guerrero, Maribel Gonzalez-Villa, Natividad Cruz-Ortiz, Tatiana Nunez-Garcia, Gisela Barrera-Badillo, Lucia Hernandez-Rivas, Irma Lopez-Martinez, Ernesto Ramirez-Gonzalez. |
| EPI_ISL_2779186, EPI_ISL_2779187, EPI_ISL_2779188, EPI_ISL_2779190, EPI_ISL_2779191, EPI_ISL_2779192, EPI_ISL_2779193 | LESP Guanajuato | Instituto de Diagnostico y Referencia Epidemiologicos (INDRE) | Claudia Wong-Arambula, Abril Rodriguez-Maldonado, Vanessa Rivero-Arredondo, Ariadna Medina-Benitez, Joaquin Quiroz-Mercado, Sergio Rangel-Guerrero, Maribel Gonzalez-Villa, Natividad Cruz-Ortiz, Tatiana Nunez-Garcia, Gisela Barrera-Badillo, Lucia Hernandez-Rivas, Irma Lopez-Martinez, Ernesto Ramirez-Gonzalez. |
| EPI_ISL_2779194 | LESP Tabasco | Instituto de Diagnostico y Referencia Epidemiologicos (INDRE) | Claudia Wong-Arambula, Abril Rodriguez-Maldonado, Vanessa Rivero-Arredondo, Ariadna Medina-Benitez, Joaquin Quiroz-Mercado, Sergio Rangel-Guerrero, Maribel Gonzalez-Villa, Natividad Cruz-Ortiz, Tatiana Nunez-Garcia, Gisela Barrera-Badillo, Lucia Hernandez-Rivas, Irma Lopez-Martinez, Ernesto Ramirez-Gonzalez. |
| EPI_ISL_2779196 | LESP Guerrero | Instituto de Diagnostico y Referencia Epidemiologicos (INDRE) | Claudia Wong-Arambula, Abril Rodriguez-Maldonado, Vanessa Rivero-Arredondo, Ariadna Medina-Benitez, Joaquin Quiroz-Mercado, Sergio Rangel-Guerrero, Maribel Gonzalez-Villa, Natividad Cruz-Ortiz, Tatiana Nunez-Garcia, Gisela Barrera-Badillo, Lucia Hernandez-Rivas, Irma Lopez-Martinez, Ernesto Ramirez-Gonzalez. |
| EPI_ISL_2779197 | LESP Baja California Sur | Instituto de Diagnostico y Referencia Epidemiologicos (INDRE) | Claudia Wong-Arambula, Abril Rodriguez-Maldonado, Vanessa Rivero-Arredondo, Ariadna Medina-Benitez, Joaquin Quiroz-Mercado, Sergio Rangel-Guerrero, Maribel Gonzalez-Villa, Natividad Cruz-Ortiz, Tatiana Nunez-Garcia, Gisela Barrera-Badillo, Lucia Hernandez-Rivas, Irma Lopez-Martinez, Ernesto Ramirez-Gonzalez. |
| EPI_ISL_2779198, EPI_ISL_2779199, EPI_ISL_2779200, EPI_ISL_2779201, EPI_ISL_2779202, EPI_ISL_2779203, EPI_ISL_2779204, EPI_ISL_2779205, EPI_ISL_2779206, EPI_ISL_2779207, EPI_ISL_2779208, EPI_ISL_2779209, EPI_ISL_2779210 | | | |
| see above | LESP Tabasco | Instituto de Diagnostico y Referencia Epidemiologicos (INDRE) | Claudia Wong-Arambula, Abril Rodriguez-Maldonado, Vanessa Rivero-Arredondo, Ariadna Medina-Benitez, Joaquin Quiroz-Mercado, Sergio Rangel-Guerrero, Maribel Gonzalez-Villa, Natividad Cruz-Ortiz, Tatiana Nunez-Garcia, Gisela Barrera-Badillo, Lucia Hernandez-Rivas, Irma Lopez-Martinez, Ernesto Ramirez-Gonzalez. |
| EPI_ISL_2779217, EPI_ISL_2779218 | LESP Guanajuato | Instituto de Diagnostico y Referencia Epidemiologicos (INDRE) | Claudia Wong-Arambula, Abril Rodriguez-Maldonado, Vanessa Rivero-Arredondo, Ariadna Medina-Benitez, Joaquin Quiroz-Mercado, Sergio Rangel-Guerrero, Maribel Gonzalez-Villa, Natividad Cruz-Ortiz, Tatiana Nunez-Garcia, Gisela Barrera-Badillo, Lucia Hernandez-Rivas, Irma Lopez-Martinez, Ernesto Ramirez-Gonzalez. |
| EPI_ISL_2779219 | LESP Tamaulipas | Instituto de Diagnostico y Referencia Epidemiologicos (INDRE) | Claudia Wong-Arambula, Abril Rodriguez-Maldonado, Vanessa Rivero-Arredondo, Ariadna Medina-Benitez, Joaquin Quiroz-Mercado, Sergio Rangel-Guerrero, Maribel Gonzalez-Villa, Natividad Cruz-Ortiz, Tatiana Nunez-Garcia, Gisela Barrera-Badillo, Lucia Hernandez-Rivas, Irma Lopez-Martinez, Ernesto Ramirez-Gonzalez. |
| EPI_ISL_2779220 | LESP Sinaloa | Instituto de Diagnostico y Referencia Epidemiologicos (INDRE) | Claudia Wong-Arambula, Abril Rodriguez-Maldonado, Vanessa Rivero-Arredondo, Ariadna Medina-Benitez, Joaquin Quiroz-Mercado, Sergio Rangel-Guerrero, Maribel Gonzalez-Villa, Natividad Cruz-Ortiz, Tatiana Nunez-Garcia, Gisela Barrera-Badillo, Lucia Hernandez-Rivas, Irma Lopez-Martinez, Ernesto Ramirez-Gonzalez. |
| EPI_ISL_2779221, EPI_ISL_2779222, EPI_ISL_2779223, EPI_ISL_2779224, EPI_ISL_2779225 | LESP Yucatan | Instituto de Diagnostico y Referencia Epidemiologicos (INDRE) | Claudia Wong-Arambula, Abril Rodriguez-Maldonado, Vanessa Rivero-Arredondo, Ariadna Medina-Benitez, Joaquin Quiroz-Mercado, Sergio Rangel-Guerrero, Maribel Gonzalez-Villa, Natividad Cruz-Ortiz, Tatiana Nunez-Garcia, Gisela Barrera-Badillo, Lucia Hernandez-Rivas, Irma Lopez-Martinez, Ernesto Ramirez-Gonzalez. |
| EPI_ISL_2779226 | LESP Michoacan | Instituto de Diagnostico y Referencia Epidemiologicos (INDRE) | Claudia Wong-Arambula, Abril Rodriguez-Maldonado, Vanessa Rivero-Arredondo, Ariadna Medina-Benitez, Joaquin Quiroz-Mercado, Sergio Rangel-Guerrero, Maribel Gonzalez-Villa, Natividad Cruz-Ortiz, Tatiana Nunez-Garcia, Gisela Barrera-Badillo, Lucia Hernandez-Rivas, Irma Lopez-Martinez, Ernesto Ramirez-Gonzalez. |
| EPI_ISL_2779227 | LESP Guerrero | Instituto de Diagnostico y Referencia Epidemiologicos (INDRE) | Claudia Wong-Arambula, Abril Rodriguez-Maldonado, Vanessa Rivero-Arredondo, Ariadna Medina-Benitez, Joaquin Quiroz-Mercado, Sergio Rangel-Guerrero, Maribel Gonzalez-Villa, Natividad Cruz-Ortiz, Tatiana Nunez-Garcia, Gisela Barrera-Badillo, Lucia Hernandez-Rivas, Irma Lopez-Martinez, Ernesto Ramirez-Gonzalez. |
| EPI_ISL_2779228, EPI_ISL_2779229, EPI_ISL_2779230, EPI_ISL_2779231 | LESP Yucatan | Instituto de Diagnostico y Referencia Epidemiologicos (INDRE) | Claudia Wong-Arambula, Abril Rodriguez-Maldonado, Vanessa Rivero-Arredondo, Ariadna Medina-Benitez, Joaquin Quiroz-Mercado, Sergio Rangel-Guerrero, Maribel Gonzalez-Villa, Natividad Cruz-Ortiz, Tatiana Nunez-Garcia, Gisela Barrera-Badillo, Lucia Hernandez-Rivas, Irma Lopez-Martinez, Ernesto Ramirez-Gonzalez. |

| | | | |
|---|---|---|---|
| EPI_ISL_2779232, EPI_ISL_2779233 | LESP Michoacan | Instituto de Diagnostico y Referencia Epidemiologicos (INDRE) | Claudia Wong-Arambula, Abril Rodriguez-Maldonado, Vanessa Rivero-Arredondo, Ariadna Medina-Benitez, Joaquin Quiroz-Mercado, Sergio Rangel-Guerrero, Maribel Gonzalez-Villa, Natividad Cruz-Ortiz, Tatiana Nunez-Garcia, Gisela Barrera-Badillo, Lucia Hernandez-Rivas, Irma Lopez-Martinez, Ernesto Ramirez-Gonzalez. |
| EPI_ISL_2779234, EPI_ISL_2779235, EPI_ISL_2779236, EPI_ISL_2779237, EPI_ISL_2779238, EPI_ISL_2779239, EPI_ISL_2779240, EPI_ISL_2779241, EPI_ISL_2779242, EPI_ISL_2779243, EPI_ISL_2779244, EPI_ISL_2779245, EPI_ISL_2779246, EPI_ISL_2779247, EPI_ISL_2779248, EPI_ISL_2779249, EPI_ISL_2779250, EPI_ISL_2779251, EPI_ISL_2779252, EPI_ISL_2779253, EPI_ISL_2779254, EPI_ISL_2779255, EPI_ISL_2779256 | | | |
| see above | LESP Tabasco | Instituto de Diagnostico y Referencia Epidemiologicos (INDRE) | Claudia Wong-Arambula, Abril Rodriguez-Maldonado, Vanessa Rivero-Arredondo, Ariadna Medina-Benitez, Joaquin Quiroz-Mercado, Sergio Rangel-Guerrero, Maribel Gonzalez-Villa, Natividad Cruz-Ortiz, Tatiana Nunez-Garcia, Gisela Barrera-Badillo, Lucia Hernandez-Rivas, Irma Lopez-Martinez, Ernesto Ramirez-Gonzalez. |
| EPI_ISL_2779257, EPI_ISL_2779258, EPI_ISL_2779259, EPI_ISL_2779260, EPI_ISL_2779261, EPI_ISL_2779262, EPI_ISL_2779263, EPI_ISL_2779264, EPI_ISL_2779265, EPI_ISL_2779266, EPI_ISL_2779267, EPI_ISL_2779268, EPI_ISL_2779269 | | | |
| see above | LESP Puebla | Instituto de Diagnostico y Referencia Epidemiologicos (INDRE) | Claudia Wong-Arambula, Abril Rodriguez-Maldonado, Vanessa Rivero-Arredondo, Ariadna Medina-Benitez, Joaquin Quiroz-Mercado, Sergio Rangel-Guerrero, Maribel Gonzalez-Villa, Natividad Cruz-Ortiz, Tatiana Nunez-Garcia, Gisela Barrera-Badillo, Lucia Hernandez-Rivas, Irma Lopez-Martinez, Ernesto Ramirez-Gonzalez. |
| EPI_ISL_2779271, EPI_ISL_2779272, EPI_ISL_2779273, EPI_ISL_2779274, EPI_ISL_2779275 | LESP Tabasco | Instituto de Diagnostico y Referencia Epidemiologicos (INDRE) | Claudia Wong-Arambula, Abril Rodriguez-Maldonado, Vanessa Rivero-Arredondo, Ariadna Medina-Benitez, Joaquin Quiroz-Mercado, Sergio Rangel-Guerrero, Maribel Gonzalez-Villa, Natividad Cruz-Ortiz, Tatiana Nunez-Garcia, Gisela Barrera-Badillo, Lucia Hernandez-Rivas, Irma Lopez-Martinez, Ernesto Ramirez-Gonzalez. |
| EPI_ISL_2779276 | LESP Guanajuato | Instituto de Diagnostico y Referencia Epidemiologicos (INDRE) | Claudia Wong-Arambula, Abril Rodriguez-Maldonado, Vanessa Rivero-Arredondo, Ariadna Medina-Benitez, Joaquin Quiroz-Mercado, Sergio Rangel-Guerrero, Maribel Gonzalez-Villa, Natividad Cruz-Ortiz, Tatiana Nunez-Garcia, Gisela Barrera-Badillo, Lucia Hernandez-Rivas, Irma Lopez-Martinez, Ernesto Ramirez-Gonzalez. |
| EPI_ISL_2779277, EPI_ISL_2779278, EPI_ISL_2779279 | LESP Tabasco | Instituto de Diagnostico y Referencia Epidemiologicos (INDRE) | Claudia Wong-Arambula, Abril Rodriguez-Maldonado, Vanessa Rivero-Arredondo, Ariadna Medina-Benitez, Joaquin Quiroz-Mercado, Sergio Rangel-Guerrero, Maribel Gonzalez-Villa, Natividad Cruz-Ortiz, Tatiana Nunez-Garcia, Gisela Barrera-Badillo, Lucia Hernandez-Rivas, Irma Lopez-Martinez, Ernesto Ramirez-Gonzalez. |
| EPI_ISL_2779280 | LESP Puebla | Instituto de Diagnostico y Referencia Epidemiologicos (INDRE) | Claudia Wong-Arambula, Abril Rodriguez-Maldonado, Vanessa Rivero-Arredondo, Ariadna Medina-Benitez, Joaquin Quiroz-Mercado, Sergio Rangel-Guerrero, Maribel Gonzalez-Villa, Natividad Cruz-Ortiz, Tatiana Nunez-Garcia, Gisela Barrera-Badillo, Lucia Hernandez-Rivas, Irma Lopez-Martinez, Ernesto Ramirez-Gonzalez. |
| EPI_ISL_2779281 | LESP Guanajuato | Instituto de Diagnostico y Referencia Epidemiologicos (INDRE) | Claudia Wong-Arambula, Abril Rodriguez-Maldonado, Vanessa Rivero-Arredondo, Ariadna Medina-Benitez, Joaquin Quiroz-Mercado, Sergio Rangel-Guerrero, Maribel Gonzalez-Villa, Natividad Cruz-Ortiz, Tatiana Nunez-Garcia, Gisela Barrera-Badillo, Lucia Hernandez-Rivas, Irma Lopez-Martinez, Ernesto Ramirez-Gonzalez. |
| EPI_ISL_2785968 | Hospital Clínico Universitario Virgen de la Arrixaca | Hospital Clínico Universitario Virgen de la Arrixaca | Marina Simón Páez y Laura Moreno Parrado |
| EPI_ISL_2786928 | Inovie AS LABOSUD | Inovie AS GenBio | Florian VERCRUYSSEN / Pauline LAURENT |
| EPI_ISL_2786968, EPI_ISL_2786969 | Microbiology Department, University Hospital Donostia | Microbiology Department, University Hospital Donostia | Marimon JM, Montes M, Piñeiro L, Martin-Peñaranda T, Gomez M, Sorarrain A, Cilla G. |
| EPI_ISL_2786994 | private medical practitioners | Hong Kong Department of Health | Alan K.L. Tsang, Peter C.W. Yip, Ken H.L. Ng, Edman T.K. Lam, Rickjason C.W. Chan |
| EPI_ISL_2786995 | VICTORIA PUBLIC MORTUARY | Hong Kong Department of Health | Alan K.L. Tsang, Peter C.W. Yip, Ken H.L. Ng, Edman T.K. Lam, Rickjason C.W. Chan |
| EPI_ISL_2786996 | Queen Elizabeth Hospital | Hong Kong Department of Health | Alan K.L. Tsang, Peter C.W. Yip, Ken H.L. Ng, Edman T.K. Lam, Rickjason C.W. Chan |
| EPI_ISL_2786997, EPI_ISL_2786998, EPI_ISL_2786999, EPI_ISL_2787000, EPI_ISL_2787001 | North Lantau Hospital | Hong Kong Department of Health | Alan K.L. Tsang, Peter C.W. Yip, Ken H.L. Ng, Edman T.K. Lam, Rickjason C.W. Chan |
| EPI_ISL_2787002 | Queen Elizabeth Hospital | Hong Kong Department of Health | Alan K.L. Tsang, Peter C.W. Yip, Ken H.L. Ng, Edman T.K. Lam, Rickjason C.W. Chan |
| EPI_ISL_2787004, EPI_ISL_2787005, EPI_ISL_2787006, EPI_ISL_2787007, EPI_ISL_2787008, EPI_ISL_2787009 | Temporary Specimen Collection Centre at the AsiaWorld-Expo | Hong Kong Department of Health | Alan K.L. Tsang, Peter C.W. Yip, Ken H.L. Ng, Edman T.K. Lam, Rickjason C.W. Chan |
| EPI_ISL_2787010 | HOME QUARANTINE TASKFORCE | Hong Kong Department of Health | Alan K.L. Tsang, Peter C.W. Yip, Ken H.L. Ng, Edman T.K. Lam, Rickjason C.W. Chan |
| EPI_ISL_2787011, EPI_ISL_2787012 | QUARANTINE CAMP | Hong Kong Department of Health | Alan K.L. Tsang, Peter C.W. Yip, Ken H.L. Ng, Edman T.K. Lam, Rickjason C.W. Chan |
| EPI_ISL_2787013, EPI_ISL_2787014 | HOME QUARANTINE TASKFORCE | Hong Kong Department of Health | Alan K.L. Tsang, Peter C.W. Yip, Ken H.L. Ng, Edman T.K. Lam, Rickjason C.W. Chan |
| EPI_ISL_2787015 | Queen Elizabeth Hospital | Hong Kong Department of Health | Alan K.L. Tsang, Peter C.W. Yip, Ken H.L. Ng, Edman T.K. Lam, Rickjason C.W. Chan |
| EPI_ISL_2789815, EPI_ISL_2789816, EPI_ISL_2789817, EPI_ISL_2789818, EPI_ISL_2789835, EPI_ISL_2789836 | Centre Hospitalier Universitaire Clermont-Ferrand | CHU Clermont-Ferrand, service de virologie | Bisseux Maxime, Mirand Audrey, Combes Patricia, Henquell Cécile |
| EPI_ISL_2790132, EPI_ISL_2790133, EPI_ISL_2790134, EPI_ISL_2790135, EPI_ISL_2790136 | Instituto Nacional de Cancerología | Centro de Investigación en Enfermedades Infecciosas (CIENI), Instituto Nacional de Enfermedades Respiratorias (INER) | Consorcio Mexicano de Vigilancia Genómica (CoViGen-Mex). Authors (in alphabetical order): Julio Elias Alvarado-Yaah, Carlos F. Arias, Santiago Ávila-Ríos, Eduardo Becerril-Vargas, Víctor Hugo Borja-Aburto, Celia Boukadida, Cristóbal Cháidez-Quiróz, Juan Bautista Chale-Dzul, Ricardo Ciria Merce, Andreu Comas-García, Célida Duque Molina, Julissa Enciso-Ibarra, José Antonio Enciso-Moreno, Gloria Elena Espinosa-Ayala, Fernando Fontove-Herrera, Daniel Fregoso-Rueda, Victor Eduardo Garcia-Arias, Alejandra Garcia-Gasca, Bruno Gomez-Gil, Jean Pierre González, Irvin González-López, Concepción Grajales-Muñiz, Ricardo Grande, Rosa María Gutierrez Rios, Alejandra Hernández-Terán, Alfredo Herrera-Estrella, Carla Ivón Herrera-Najera, Pavel Isa, Daniel Lira Morales, Susana Lopez, Antonio Loza Román, Brenda Irasema Maldonado-Meza, Bernardo Martínez-Miguel, José Arturo Martínez-Orozco, Célida Martínez- Rodríguez, Margarita Matías-Florentino, Fidencio Mejía-Nepomuceno, María Guadalupe de Jesús Mireles-Rivera, Gloria María Molina-Salinas, Hector Montoya-Fuentes, Mario Mújica-Sánchez, José Esteban Muñoz-Medina, José de Jesús Nuñez-Contreras, Alicia Ocaña-Mondragón, Luis Alberto Ochoa-Carrera, Hector Esteban Paz-Juárez, Marissa Pérez-García, Francisco Pulido, Helen Haydee Fernanda Ramírez-Plascencia, Jorge Salas-Hernández, Angel Gustavo Salas-Lais, Alejandro Sanchez-Flores, Clara Esperanza Santacruz-Tinoco, María Guadalupe Santiago-Mauricio, Selene Zárate, Nelly Sélem-Mojica, Blanca Taboada, Kathia Elizabeth Tapia-Diaz, Gloria Vazquez, Joel Armando Vázquez-Pérez. |
| EPI_ISL_2790236, EPI_ISL_2790239, EPI_ISL_2790242, EPI_ISL_2790243, EPI_ISL_2790244, EPI_ISL_2790245 | CHWAPI - SITE NOTRE DAME | Institut de Pathologie et Genetique (IPG) | Pascale Hilbert, Jérémie Gras |
| EPI_ISL_2790249, EPI_ISL_2790250, EPI_ISL_2790251, EPI_ISL_2790252, EPI_ISL_2790253, EPI_ISL_2790254, EPI_ISL_2790255 | Università degli Studi di Perugia | Istituto Zooprofilattico Sperimentale dell'Abruzzo e Molise "G. Caporale" | Mencacci A, Camilloni B, Biagetti M, Giammarioli M, Lorusso A, Marcacci M, Di Domenico M, Ancora M, Curini V, Mangone I, Rinaldi A, Delli Compagni E, Scialabba S, Di Pasquale A, Cammà C, Puglia I, Calistri P, Savini G |
| EPI_ISL_2790256, EPI_ISL_2790257, EPI_ISL_2790258, EPI_ISL_2790259, EPI_ISL_2790260, EPI_ISL_2790261 | Laboratorio Analisi Osp. Città di Castello - Azienda USL Umbria1 | Istituto Zooprofilattico Sperimentale dell'Abruzzo e Molise "G. Caporale" | Malagigi V, Taccani P,Lorusso A, Marcacci M, Di Domenico M, Ancora M, Curini V, Mangone I, Rinaldi A, Delli Compagni E, Scialabba S, Di Pasquale A, Cammà C, Puglia I, Calistri P, Savini G |
| EPI_ISL_2790262 | Università degli Studi di Perugia | Istituto Zooprofilattico Sperimentale dell'Abruzzo e Molise "G. Caporale" | Mencacci A, Camilloni B, Biagetti M, Giammarioli M, Lorusso A, Marcacci M, Di Domenico M, Ancora M, Curini V, Mangone I, Rinaldi A, Delli Compagni E, Scialabba S, Di Pasquale A, Cammà C, Puglia I, Calistri P, Savini G |
| EPI_ISL_2790263 | Laboratorio Analisi Osp. Città di Castello - Azienda USL Umbria1 | Istituto Zooprofilattico Sperimentale dell'Abruzzo e Molise "G. Caporale" | Malagigi V, Taccani P,Lorusso A, Marcacci M, Di Domenico M, Ancora M, Curini V, Mangone I, Rinaldi A, Delli Compagni E, Scialabba S, Di Pasquale A, Cammà C, Puglia I, Calistri P, Savini G |

| | | | |
|---|---|---|---|
| EPI_ISL_2790264, EPI_ISL_2790265, EPI_ISL_2790266, EPI_ISL_2790267, EPI_ISL_2790268, EPI_ISL_2790269, EPI_ISL_2790270, EPI_ISL_2790271, EPI_ISL_2790272, EPI_ISL_2790273, EPI_ISL_2790274, EPI_ISL_2790275, EPI_ISL_2790277 | | | |
| see above | Azienda USL Umbria 2 | Istituto Zooprofilattico Sperimentale dell'Abruzzo e Molise "G. Caporale" | Proietti A, Pistoni E, Lorusso A, Marcacci M, Di Domenico M, Ancora M, Curini V, Mangone I, Rinaldi A, Delli Compagni E, Scialabba S, Di Pasquale A, Cammà C, Puglia I, Calistri P, Savini G |
| EPI_ISL_2790280, EPI_ISL_2790281, EPI_ISL_2790282, EPI_ISL_2790283, EPI_ISL_2790284, EPI_ISL_2790285, EPI_ISL_2790286, EPI_ISL_2790287, EPI_ISL_2790288 | Azienda Ospedaliera Terni | Istituto Zooprofilattico Sperimentale dell'Abruzzo e Molise "G. Caporale" | Palumbo M, Scaccetti A, Lorusso A, Marcacci M, Di Domenico M, Ancora M, Curini V, Mangone I, Rinaldi A, Delli Compagni E, Scialabba S, Di Pasquale A, Cammà C, Puglia I, Calistri P, Savini G |
| EPI_ISL_2790310 | Genome Analysis Center, Yamanashi Central Hospital | Genome Analysis Center, Yamanashi Central Hospital | Yosuke Hirotsu |
| EPI_ISL_2790622, EPI_ISL_2790623, EPI_ISL_2790624, EPI_ISL_2790625, EPI_ISL_2790627, EPI_ISL_2790630, EPI_ISL_2790631, EPI_ISL_2790632, EPI_ISL_2790633, EPI_ISL_2790634, EPI_ISL_2790635, EPI_ISL_2790639, EPI_ISL_2790640, EPI_ISL_2790642, EPI_ISL_2790643, EPI_ISL_2790644, EPI_ISL_2790645, EPI_ISL_2790647, EPI_ISL_2790648, EPI_ISL_2790652, EPI_ISL_2790654, EPI_ISL_2790655, EPI_ISL_2790656, EPI_ISL_2790657, EPI_ISL_2790660, EPI_ISL_2790661, EPI_ISL_2790662, EPI_ISL_2790663, EPI_ISL_2790666, EPI_ISL_2790667, EPI_ISL_2790668, EPI_ISL_2790671, EPI_ISL_2790673, EPI_ISL_2790675, EPI_ISL_2790676, EPI_ISL_2790677, EPI_ISL_2790679, EPI_ISL_2790680, EPI_ISL_2790681, EPI_ISL_2790682, EPI_ISL_2790683, EPI_ISL_2790684, EPI_ISL_2790685, EPI_ISL_2790688, EPI_ISL_2790689, EPI_ISL_2790691, EPI_ISL_2790692, EPI_ISL_2790693 | | | |
| see above | Colorado Department of Public Health and Environment | Colorado Department of Public Health and Environment | Laura Bankers, Molly C. Hetherington-Rauth, Diana Ir, Alexandria Rossheim, Michael Martin, Mandy Waters, Shannon R. Matzinger, Sarah Elizabeth Totten, Emily A. Travanty |
| EPI_ISL_2790862, EPI_ISL_2790863 | Labo Analyses Med | National Reference Center for Viruses of Respiratory Infections, Institut Pasteur, Paris | Marion Barbet, Sylvie Behillil, Méline Bizard, Angela Brisebarre, Camille Capel, Vincent Enouf, Louise Lefrançois, Frédéric Lemoine, Christophe Malabat, Corinne Maufrais, Emmanuelle Permal, Etienne Simon-Lorière, Maud Vanpeene, Sylvie Van der Werf ,GréGoire Potiron |
| EPI_ISL_2790864 | Labo Analyses Med | National Reference Center for Viruses of Respiratory Infections, Institut Pasteur, Paris | Marion Barbet, Sylvie Behillil, Méline Bizard, Angela Brisebarre, Camille Capel, Vincent Enouf, Louise Lefrançois, Frédéric Lemoine, Christophe Malabat, Corinne Maufrais, Emmanuelle Permal, Etienne Simon-Lorière, Maud Vanpeene, Sylvie Van der Werf ,Sabine Trombert |
| EPI_ISL_2790865, EPI_ISL_2790866, EPI_ISL_2790867, EPI_ISL_2790868, EPI_ISL_2790869 | Labo Analyses Med | National Reference Center for Viruses of Respiratory Infections, Institut Pasteur, Paris | Marion Barbet, Sylvie Behillil, Méline Bizard, Angela Brisebarre, Camille Capel, Vincent Enouf, Louise Lefrançois, Frédéric Lemoine, Christophe Malabat, Corinne Maufrais, Emmanuelle Permal, Etienne Simon-Lorière, Maud Vanpeene, Sylvie Van der Werf ,BéNéDicte Roquebert |
| EPI_ISL_2790870, EPI_ISL_2790871, EPI_ISL_2790872 | Labo Analyses Med | National Reference Center for Viruses of Respiratory Infections, Institut Pasteur, Paris | Marion Barbet, Sylvie Behillil, Méline Bizard, Angela Brisebarre, Camille Capel, Vincent Enouf, Louise Lefrançois, Frédéric Lemoine, Christophe Malabat, Corinne Maufrais, Emmanuelle Permal, Etienne Simon-Lorière, Maud Vanpeene, Sylvie Van der Werf ,Sabine Trombert |
| EPI_ISL_2790910, EPI_ISL_2790911 | Hôpital | National Reference Center for Viruses of Respiratory Infections, Institut Pasteur, Paris | Marion Barbet, Sylvie Behillil, Méline Bizard, Angela Brisebarre, Camille Capel, Vincent Enouf, Louise Lefrançois, Frédéric Lemoine, Christophe Malabat, Corinne Maufrais, Emmanuelle Permal, Etienne Simon-Lorière, Maud Vanpeene, Sylvie Van der Werf ,Nabil Gastli |
| EPI_ISL_2790912, EPI_ISL_2790913, EPI_ISL_2790914, EPI_ISL_2790915 | Labo Analyses Med | National Reference Center for Viruses of Respiratory Infections, Institut Pasteur, Paris | Marion Barbet, Sylvie Behillil, Méline Bizard, Angela Brisebarre, Camille Capel, Vincent Enouf, Louise Lefrançois, Frédéric Lemoine, Christophe Malabat, Corinne Maufrais, Emmanuelle Permal, Etienne Simon-Lorière, Maud Vanpeene, Sylvie Van der Werf ,Nicolas Capron |
| EPI_ISL_2790924 | Labo Analyses Med | National Reference Center for Viruses of Respiratory Infections, Institut Pasteur, Paris | Marion Barbet, Sylvie Behillil, Méline Bizard, Angela Brisebarre, Camille Capel, Vincent Enouf, Louise Lefrançois, Frédéric Lemoine, Christophe Malabat, Corinne Maufrais, Emmanuelle Permal, Etienne Simon-Lorière, Maud Vanpeene, Sylvie Van der Werf ,Brieuc Gestin |
| EPI_ISL_2790925 | Hospital | National Reference Center for Viruses of Respiratory Infections, Institut Pasteur, Paris | Marion Barbet, Sylvie Behillil, Méline Bizard, Angela Brisebarre, Camille Capel, Vincent Enouf, Louise Lefrançois, Frédéric Lemoine, Christophe Malabat, Corinne Maufrais, Emmanuelle Permal, Etienne Simon-Lorière, Maud Vanpeene, Sylvie Van der Werf ,Nabil Gastli |
| EPI_ISL_2790927 | Hospital | National Reference Center for Viruses of Respiratory Infections, Institut Pasteur, Paris | Marion Barbet, Sylvie Behillil, Méline Bizard, Angela Brisebarre, Camille Capel, Vincent Enouf, Louise Lefrançois, Frédéric Lemoine, Christophe Malabat, Corinne Maufrais, Emmanuelle Permal, Etienne Simon-Lorière, Maud Vanpeene, Sylvie Van der Werf ,Laurence Louvet |
| EPI_ISL_2790929 | Labo Analyses Med | National Reference Center for Viruses of Respiratory Infections, Institut Pasteur, Paris | Marion Barbet, Sylvie Behillil, Méline Bizard, Angela Brisebarre, Camille Capel, Vincent Enouf, Louise Lefrançois, Frédéric Lemoine, Christophe Malabat, Corinne Maufrais, Emmanuelle Permal, Etienne Simon-Lorière, Maud Vanpeene, Sylvie Van der Werf ,BéNéDicte Roquebert |
| EPI_ISL_2790930, EPI_ISL_2790931, EPI_ISL_2790932, EPI_ISL_2790933, EPI_ISL_2790935, EPI_ISL_2790936, EPI_ISL_2790937, EPI_ISL_2790938, EPI_ISL_2790939, EPI_ISL_2790940, EPI_ISL_2790941, EPI_ISL_2790942, EPI_ISL_2790943, EPI_ISL_2790944, EPI_ISL_2790945, EPI_ISL_2790946, EPI_ISL_2790947, EPI_ISL_2790948, EPI_ISL_2790949, EPI_ISL_2790950, EPI_ISL_2790952, EPI_ISL_2790953, EPI_ISL_2790954, EPI_ISL_2790955, EPI_ISL_2790956, EPI_ISL_2790957, EPI_ISL_2790959, EPI_ISL_2790960, EPI_ISL_2790961, EPI_ISL_2790963, EPI_ISL_2790964, EPI_ISL_2790965, EPI_ISL_2790966 | | | |
| see above | Hôpital | National Reference Center for Viruses of Respiratory Infections, Institut Pasteur, Paris | Marion Barbet, Sylvie Behillil, Méline Bizard, Angela Brisebarre, Camille Capel, Vincent Enouf, Louise Lefrançois, Frédéric Lemoine, Christophe Malabat, Corinne Maufrais, Emmanuelle Permal, Etienne Simon-Lorière, Maud Vanpeene, Sylvie Van der Werf ,BéNéDicte Roquebert |
| EPI_ISL_2790969, EPI_ISL_2790970, EPI_ISL_2790971, EPI_ISL_2790972, EPI_ISL_2790973 | Labo Analyses Med | National Reference Center for Viruses of Respiratory Infections, Institut Pasteur, Paris | Marion Barbet, Sylvie Behillil, Méline Bizard, Angela Brisebarre, Camille Capel, Vincent Enouf, Louise Lefrançois, Frédéric Lemoine, Christophe Malabat, Corinne Maufrais, Emmanuelle Permal, Etienne Simon-Lorière, Maud Vanpeene, Sylvie Van der Werf ,Sabine Trombert |
| EPI_ISL_2790974 | Labo Analyses Med | National Reference Center for Viruses of Respiratory Infections, Institut Pasteur, Paris | Marion Barbet, Sylvie Behillil, Méline Bizard, Angela Brisebarre, Camille Capel, Vincent Enouf, Louise Lefrançois, Frédéric Lemoine, Christophe Malabat, Corinne Maufrais, Emmanuelle Permal, Etienne Simon-Lorière, Maud Vanpeene, Sylvie Van der Werf ,GréGoire Potiron |
| EPI_ISL_2790975, EPI_ISL_2790976, EPI_ISL_2790977, EPI_ISL_2790978 | Labo Analyses Med | National Reference Center for Viruses of Respiratory Infections, Institut Pasteur, Paris | Marion Barbet, Sylvie Behillil, Méline Bizard, Angela Brisebarre, Camille Capel, Vincent Enouf, Louise Lefrançois, Frédéric Lemoine, Christophe Malabat, Corinne Maufrais, Emmanuelle Permal, Etienne Simon-Lorière, Maud Vanpeene, Sylvie Van der Werf ,Eve Haguenoer |
| EPI_ISL_2790979, EPI_ISL_2790980 | Labo Analyses Med | National Reference Center for Viruses of Respiratory Infections, Institut Pasteur, Paris | Marion Barbet, Sylvie Behillil, Méline Bizard, Angela Brisebarre, Camille Capel, Vincent Enouf, Louise Lefrançois, Frédéric Lemoine, Christophe Malabat, Corinne Maufrais, Emmanuelle Permal, Etienne Simon-Lorière, Maud Vanpeene, Sylvie Van der Werf ,Nicolas Capron |
| EPI_ISL_2790981, EPI_ISL_2790982, EPI_ISL_2790983, EPI_ISL_2790984 | Hospital | National Reference Center for Viruses of Respiratory Infections, Institut Pasteur, Paris | Marion Barbet, Sylvie Behillil, Méline Bizard, Angela Brisebarre, Camille Capel, Vincent Enouf, Louise Lefrançois, Frédéric Lemoine, Christophe Malabat, Corinne Maufrais, Emmanuelle Permal, Etienne Simon-Lorière, Maud Vanpeene, Sylvie Van der Werf ,JéRôMe Durivault |
| EPI_ISL_2790985, EPI_ISL_2790986 | Hospital | National Reference Center for Viruses of Respiratory Infections, Institut Pasteur, Paris | Marion Barbet, Sylvie Behillil, Méline Bizard, Angela Brisebarre, Camille Capel, Vincent Enouf, Louise Lefrançois, Frédéric Lemoine, Christophe Malabat, Corinne Maufrais, Emmanuelle Permal, Etienne Simon-Lorière, Maud Vanpeene, Sylvie Van der Werf ,Patricia Stoessel-Thouvenin |
| EPI_ISL_2790988, EPI_ISL_2790989, EPI_ISL_2790990, EPI_ISL_2790991 | Labo Analyses Med | National Reference Center for Viruses of Respiratory Infections, Institut Pasteur, Paris | Marion Barbet, Sylvie Behillil, Méline Bizard, Angela Brisebarre, Camille Capel, Vincent Enouf, Louise Lefrançois, Frédéric Lemoine, Christophe Malabat, Corinne Maufrais, Emmanuelle Permal, Etienne Simon-Lorière, Maud Vanpeene, Sylvie Van der Werf ,BéNéDicte Roquebert |
| EPI_ISL_2791278, EPI_ISL_2791298, EPI_ISL_2791300, EPI_ISL_2791321 | GH A.CHENEVIER-H.MONDOR | Department of Virology, Henri Mondor University Hospital, Assistance Publique Hôpitaux de Paris, Université Paris-Est Créteil, INSERM U955 | Christophe Rodriguez, Slim Fourati, Vanessa Demontant, Guillaume Gricourt, Melissa N'Debi, Alexandre Soulier, Elisabeth Trawinski, Jean-Michel Pawlotsky |
| EPI_ISL_2800916 | LESP Yucatan | Instituto de Diagnostico y Referencia Epidemiologicos (INDRE) | Claudia Wong-Arambula, Abril Rodriguez-Maldonado, Vanessa Rivero-Arredondo, Ariadna Medina-Benitez, Joaquin Quiroz-Mercado, Sergio Rangel-Guerrero, Maribel Gonzalez-Villa, Natividad Cruz-Ortiz, Tatiana Nunez-Garcia, Gisela Barrera-Badillo, Lucia Hernandez-Rivas, Irma Lopez-Martinez, Ernesto Ramirez-Gonzalez. |
| EPI_ISL_2800917 | LESP Guanajuato | Instituto de Diagnostico y Referencia Epidemiologicos (INDRE) | Claudia Wong-Arambula, Abril Rodriguez-Maldonado, Vanessa Rivero-Arredondo, Ariadna Medina-Benitez, Joaquin Quiroz-Mercado, Sergio Rangel-Guerrero, Maribel Gonzalez-Villa, Natividad Cruz-Ortiz, Tatiana Nunez-Garcia, Gisela Barrera-Badillo, Lucia Hernandez-Rivas, Irma Lopez-Martinez, Ernesto Ramirez-Gonzalez. |
| EPI_ISL_2800918, EPI_ISL_2800919, EPI_ISL_2800920, EPI_ISL_2800921, EPI_ISL_2800922, EPI_ISL_2800923 | LESP Yucatan | Instituto de Diagnostico y Referencia Epidemiologicos (INDRE) | Claudia Wong-Arambula, Abril Rodriguez-Maldonado, Vanessa Rivero-Arredondo, Ariadna Medina-Benitez, Joaquin Quiroz-Mercado, Sergio Rangel-Guerrero, Maribel Gonzalez-Villa, Natividad Cruz-Ortiz, Tatiana Nunez-Garcia, Gisela Barrera-Badillo, Lucia Hernandez-Rivas, Irma Lopez-Martinez, Ernesto Ramirez-Gonzalez. |
| EPI_ISL_2800924 | LESP Guanajuato | Instituto de Diagnostico y Referencia Epidemiologicos (INDRE) | Claudia Wong-Arambula, Abril Rodriguez-Maldonado, Vanessa Rivero-Arredondo, Ariadna Medina-Benitez, Joaquin Quiroz-Mercado, Sergio Rangel-Guerrero, Maribel Gonzalez-Villa, Natividad Cruz-Ortiz, Tatiana Nunez-Garcia, Gisela Barrera-Badillo, Lucia Hernandez-Rivas, Irma Lopez-Martinez, Ernesto Ramirez-Gonzalez. |
| EPI_ISL_2800925 | LESP Tabasco | Instituto de Diagnostico y Referencia Epidemiologicos (INDRE) | Claudia Wong-Arambula, Abril Rodriguez-Maldonado, Vanessa Rivero-Arredondo, Ariadna Medina-Benitez, Joaquin Quiroz-Mercado, Sergio Rangel-Guerrero, Natividad Cruz-Ortiz, Tatiana Nunez-Garcia, Gisela Barrera-Badillo, Lucia Hernandez-Rivas, Irma Lopez-Martinez, Ernesto Ramirez-Gonzalez. |

| | | | |
|---|---|---|---|
| EPI_ISL_2800926 | LESP Guanajuato | Instituto de Diagnostico y Referencia Epidemiologicos (INDRE) | Claudia Wong-Arambula, Abril Rodriguez-Maldonado, Vanessa Rivero-Arredondo, Ariadna Medina-Benitez, Joaquin Quiroz-Mercado, Sergio Rangel-Guerrero, Natividad Cruz-Ortiz, Tatiana Nunez-Garcia, Gisela Barrera-Badillo, Lucia Hernandez-Rivas, Irma Lopez-Martinez, Ernesto Ramirez-Gonzalez. |
| EPI_ISL_2800927, EPI_ISL_2800928, EPI_ISL_2800929 | LESP Tabasco | Instituto de Diagnostico y Referencia Epidemiologicos (INDRE) | Claudia Wong-Arambula, Abril Rodriguez-Maldonado, Vanessa Rivero-Arredondo, Ariadna Medina-Benitez, Joaquin Quiroz-Mercado, Sergio Rangel-Guerrero, Natividad Cruz-Ortiz, Tatiana Nunez-Garcia, Gisela Barrera-Badillo, Lucia Hernandez-Rivas, Irma Lopez-Martinez, Ernesto Ramirez-Gonzalez. |
| EPI_ISL_2800930, EPI_ISL_2800931 | LESP Sonora | Instituto de Diagnostico y Referencia Epidemiologicos (INDRE) | Claudia Wong-Arambula, Abril Rodriguez-Maldonado, Vanessa Rivero-Arredondo, Ariadna Medina-Benitez, Joaquin Quiroz-Mercado, Sergio Rangel-Guerrero, Natividad Cruz-Ortiz, Tatiana Nunez-Garcia, Gisela Barrera-Badillo, Lucia Hernandez-Rivas, Irma Lopez-Martinez, Ernesto Ramirez-Gonzalez. |
| EPI_ISL_2800932 | LESP Guanajuato | Instituto de Diagnostico y Referencia Epidemiologicos (INDRE) | Claudia Wong-Arambula, Abril Rodriguez-Maldonado, Vanessa Rivero-Arredondo, Ariadna Medina-Benitez, Joaquin Quiroz-Mercado, Sergio Rangel-Guerrero, Natividad Cruz-Ortiz, Tatiana Nunez-Garcia, Gisela Barrera-Badillo, Lucia Hernandez-Rivas, Irma Lopez-Martinez, Ernesto Ramirez-Gonzalez. |
| EPI_ISL_2800933, EPI_ISL_2800934 | LESP Durango | Instituto de Diagnostico y Referencia Epidemiologicos (INDRE) | Claudia Wong-Arambula, Abril Rodriguez-Maldonado, Vanessa Rivero-Arredondo, Ariadna Medina-Benitez, Joaquin Quiroz-Mercado, Sergio Rangel-Guerrero, Natividad Cruz-Ortiz, Tatiana Nunez-Garcia, Gisela Barrera-Badillo, Lucia Hernandez-Rivas, Irma Lopez-Martinez, Ernesto Ramirez-Gonzalez. |
| EPI_ISL_2800935 | LESP Tamaulipas | Instituto de Diagnostico y Referencia Epidemiologicos (INDRE) | Claudia Wong-Arambula, Abril Rodriguez-Maldonado, Vanessa Rivero-Arredondo, Ariadna Medina-Benitez, Joaquin Quiroz-Mercado, Sergio Rangel-Guerrero, Natividad Cruz-Ortiz, Tatiana Nunez-Garcia, Gisela Barrera-Badillo, Lucia Hernandez-Rivas, Irma Lopez-Martinez, Ernesto Ramirez-Gonzalez. |
| EPI_ISL_2800936 | LESP Puebla | Instituto de Diagnostico y Referencia Epidemiologicos (INDRE) | Claudia Wong-Arambula, Abril Rodriguez-Maldonado, Vanessa Rivero-Arredondo, Ariadna Medina-Benitez, Joaquin Quiroz-Mercado, Sergio Rangel-Guerrero, Natividad Cruz-Ortiz, Tatiana Nunez-Garcia, Gisela Barrera-Badillo, Lucia Hernandez-Rivas, Irma Lopez-Martinez, Ernesto Ramirez-Gonzalez. |
| EPI_ISL_2800937 | LESP Sonora | Instituto de Diagnostico y Referencia Epidemiologicos (INDRE) | Claudia Wong-Arambula, Abril Rodriguez-Maldonado, Vanessa Rivero-Arredondo, Ariadna Medina-Benitez, Joaquin Quiroz-Mercado, Sergio Rangel-Guerrero, Natividad Cruz-Ortiz, Tatiana Nunez-Garcia, Gisela Barrera-Badillo, Lucia Hernandez-Rivas, Irma Lopez-Martinez, Ernesto Ramirez-Gonzalez. |
| EPI_ISL_2800938 | LESP Quintana Roo | Instituto de Diagnostico y Referencia Epidemiologicos (INDRE) | Claudia Wong-Arambula, Abril Rodriguez-Maldonado, Vanessa Rivero-Arredondo, Ariadna Medina-Benitez, Joaquin Quiroz-Mercado, Sergio Rangel-Guerrero, Natividad Cruz-Ortiz, Tatiana Nunez-Garcia, Gisela Barrera-Badillo, Lucia Hernandez-Rivas, Irma Lopez-Martinez, Ernesto Ramirez-Gonzalez. |
| EPI_ISL_2800939, EPI_ISL_2800940 | LESP Tabasco | Instituto de Diagnostico y Referencia Epidemiologicos (INDRE) | Claudia Wong-Arambula, Abril Rodriguez-Maldonado, Vanessa Rivero-Arredondo, Ariadna Medina-Benitez, Joaquin Quiroz-Mercado, Sergio Rangel-Guerrero, Natividad Cruz-Ortiz, Tatiana Nunez-Garcia, Gisela Barrera-Badillo, Lucia Hernandez-Rivas, Irma Lopez-Martinez, Ernesto Ramirez-Gonzalez. |
| EPI_ISL_2800941 | LESP Guanajuato | Instituto de Diagnostico y Referencia Epidemiologicos (INDRE) | Claudia Wong-Arambula, Abril Rodriguez-Maldonado, Vanessa Rivero-Arredondo, Ariadna Medina-Benitez, Joaquin Quiroz-Mercado, Sergio Rangel-Guerrero, Natividad Cruz-Ortiz, Tatiana Nunez-Garcia, Gisela Barrera-Badillo, Lucia Hernandez-Rivas, Irma Lopez-Martinez, Ernesto Ramirez-Gonzalez. |
| EPI_ISL_2800942 | LESP Tabasco | Instituto de Diagnostico y Referencia Epidemiologicos (INDRE) | Claudia Wong-Arambula, Abril Rodriguez-Maldonado, Vanessa Rivero-Arredondo, Ariadna Medina-Benitez, Joaquin Quiroz-Mercado, Sergio Rangel-Guerrero, Maribel Gonzalez-Villa, Natividad Cruz-Ortiz, Tatiana Nunez-Garcia, Gisela Barrera-Badillo, Lucia Hernandez-Rivas, Irma Lopez-Martinez, Ernesto Ramirez-Gonzalez. |
| EPI_ISL_2800943 | LESP Guanajuato | Instituto de Diagnostico y Referencia Epidemiologicos (INDRE) | Claudia Wong-Arambula, Abril Rodriguez-Maldonado, Vanessa Rivero-Arredondo, Ariadna Medina-Benitez, Joaquin Quiroz-Mercado, Sergio Rangel-Guerrero, Natividad Cruz-Ortiz, Tatiana Nunez-Garcia, Gisela Barrera-Badillo, Lucia Hernandez-Rivas, Irma Lopez-Martinez, Ernesto Ramirez-Gonzalez. |
| EPI_ISL_2800944 | LESP Yucatan | Instituto de Diagnostico y Referencia Epidemiologicos (INDRE) | Claudia Wong-Arambula, Abril Rodriguez-Maldonado, Vanessa Rivero-Arredondo, Ariadna Medina-Benitez, Joaquin Quiroz-Mercado, Sergio Rangel-Guerrero, Natividad Cruz-Ortiz, Tatiana Nunez-Garcia, Gisela Barrera-Badillo, Lucia Hernandez-Rivas, Irma Lopez-Martinez, Ernesto Ramirez-Gonzalez. |
| EPI_ISL_2800945 | LESP Michoacan | Instituto de Diagnostico y Referencia Epidemiologicos (INDRE) | Claudia Wong-Arambula, Abril Rodriguez-Maldonado, Vanessa Rivero-Arredondo, Ariadna Medina-Benitez, Joaquin Quiroz-Mercado, Sergio Rangel-Guerrero, Natividad Cruz-Ortiz, Tatiana Nunez-Garcia, Gisela Barrera-Badillo, Lucia Hernandez-Rivas, Irma Lopez-Martinez, Ernesto Ramirez-Gonzalez. |
| EPI_ISL_2801565, EPI_ISL_2801566, EPI_ISL_2801571, EPI_ISL_2801576, EPI_ISL_2801578, EPI_ISL_2801579, EPI_ISL_2801580, EPI_ISL_2801581, EPI_ISL_2801583, EPI_ISL_2801584, EPI_ISL_2801585, EPI_ISL_2801586, EPI_ISL_2801587, EPI_ISL_2801588, EPI_ISL_2801591, EPI_ISL_2801592, EPI_ISL_2801593, EPI_ISL_2801595, EPI_ISL_2801596, EPI_ISL_2801597, EPI_ISL_2801598, EPI_ISL_2801599, EPI_ISL_2801600, EPI_ISL_2801603, EPI_ISL_2801604, EPI_ISL_2801606, EPI_ISL_2801609, EPI_ISL_2801610, EPI_ISL_2801611, EPI_ISL_2801612, EPI_ISL_2801617, EPI_ISL_2801618, EPI_ISL_2801619, EPI_ISL_2801622, EPI_ISL_2801623, EPI_ISL_2801624, EPI_ISL_2801625, EPI_ISL_2801628, EPI_ISL_2801629, EPI_ISL_2801631, EPI_ISL_2801637, EPI_ISL_2801641, EPI_ISL_2801642, EPI_ISL_2801643, EPI_ISL_2801644, EPI_ISL_2801645, EPI_ISL_2801646, EPI_ISL_2801648, EPI_ISL_2801649, EPI_ISL_2801652, EPI_ISL_2801653, EPI_ISL_2801654, EPI_ISL_2801655, EPI_ISL_2801656, EPI_ISL_2801657 | | | |
| see above | Unidad de Investigación Médica de Yucatán (UIMY) | Instituto de Biotecnología de la UNAM | Consorcio Mexicano de Vigilancia Genómica (CoViGen-Mex). Authors (in alphabetical order): Julio Elias Alvarado-Yaah, Carlos F. Arias, Santiago Ávila-Ríos, Eduardo Becerril-Vargas, Víctor Hugo Borja-Aburto, Celia Boukadida, Cristóbal Cháidez-Quiróz, Juan Bautista Chale-Dzul, Ricardo Ciria Merce, Andreu Comas-García, Célida Duque Molina, Julissa Enciso-Ibarra, José Antonio Enciso-Moreno, Gloria Elena Espinosa-Ayala, Fernando Fontove-Herrera, Daniel Fregoso-Rueda, Victor Eduardo García-Arias, Alejandra García-Gasca, Bruno Gómez-Gil, Jean Pierre González, Irvin González-López, Concepción Grajales-Muñiz, Ricardo Grande, Rosa María Gutiérrez Rios, Jesús Hernández, Alejandra Hernández-Terán, Alfredo Herrera-Estrella, Carla Ivón Herrera-Najera, Pavel Isa, Verónica Mata-Haro, Daniel Lira Morales, Susana Lopez, Antonio Loza Román, Brenda Irasema Maldonado-Meza, Bernardo Martínez-Miguel, José Arturo Martínez-Orozco, Célida Martínez- Rodríguez, Margarita Matías-Florentino, Fidencio Mejía-Nepomuceno, María Guadalupe de Jesús Mireles-Rivera, Gloria María Molina-Salinas, Hector Montoya-Fuentes, , Mario Mújica-Sánchez, José Esteban Muñoz-Medina, José de Jesús Nuñez-Contreras, Alicia Ocaña-Mondragón, Luis Alberto Ochoa-Carrera, Hector Esteban Paz-Juárez, Marissa Perez-Garcia, Francisco Pulido, Helen Haydee Fernanda Ramírez-Plascencia, Jorge Salas-Hernández, Angel Gustavo Salas-Lais, Alejandro Sánchez-Flores, Clara Esperanza Santacruz-Tinoco, María Guadalupe Santiago-Mauricio, Selene Zárate, Nelly Sélem-Mojica, Blanca Taboada, Gloria Vazquez, Joel Armando Vázquez-Pérez. |
| EPI_ISL_2801658, EPI_ISL_2801664, EPI_ISL_2801667, EPI_ISL_2801668, EPI_ISL_2801669, EPI_ISL_2801670, EPI_ISL_2801673, EPI_ISL_2801674, EPI_ISL_2801675, EPI_ISL_2801680, EPI_ISL_2801681, EPI_ISL_2801683, EPI_ISL_2801685, EPI_ISL_2801692, EPI_ISL_2801695 | | | |
| see above | Centro de Investigación Biomédica del Noreste (CIBIN) | Instituto de Biotecnología de la UNAM | Consorcio Mexicano de Vigilancia Genómica (CoViGen-Mex). Authors (in alphabetical order): Julio Elias Alvarado-Yaah, Carlos F. Arias, Santiago Ávila-Ríos, Eduardo Becerril-Vargas, Víctor Hugo Borja-Aburto, Celia Boukadida, Cristóbal Cháidez-Quiróz, Juan Bautista Chale-Dzul, Ricardo Ciria Merce, Andreu Comas-García, Célida Duque Molina, Julissa Enciso-Ibarra, José Antonio Enciso-Moreno, Gloria Elena Espinosa-Ayala, Fernando Fontove-Herrera, Daniel Fregoso-Rueda, Victor Eduardo García-Arias, Alejandra García-Gasca, Bruno Gómez-Gil, Jean Pierre González, Irvin González-López, Concepción Grajales-Muñiz, Ricardo Grande, Rosa María Gutiérrez Rios, Jesús Hernández, Alejandra Hernández-Terán, Alfredo Herrera-Estrella, Carla Ivón Herrera-Najera, Pavel Isa, Verónica Mata-Haro, Daniel Lira Morales, Susana Lopez, Antonio Loza Román, Brenda Irasema Maldonado-Meza, Bernardo Martínez-Miguel, José Arturo Martínez-Orozco, Célida Martínez- Rodríguez, Margarita Matías-Florentino, Fidencio Mejía-Nepomuceno, María Guadalupe de Jesús Mireles-Rivera, Gloria María Molina-Salinas, Hector Montoya-Fuentes, , Mario Mújica-Sánchez, José Esteban Muñoz-Medina, José de Jesús Nuñez-Contreras, Alicia Ocaña-Mondragón, Luis Alberto Ochoa-Carrera, Hector Esteban Paz-Juárez, Marissa Perez-Garcia, Francisco Pulido, Helen Haydee Fernanda Ramírez-Plascencia, Jorge Salas-Hernández, Angel Gustavo Salas-Lais, Alejandro Sánchez-Flores, Clara Esperanza Santacruz-Tinoco, María Guadalupe Santiago-Mauricio, Selene Zárate, Nelly Sélem-Mojica, Blanca Taboada, Gloria Vazquez, Joel Armando Vázquez-Pérez. |
| EPI_ISL_2801698, EPI_ISL_2801700, EPI_ISL_2801701, EPI_ISL_2801702, EPI_ISL_2801704, EPI_ISL_2801705 | Unidad de Investigación Biomédica de Zacatecas (UIBZ) | Instituto de Biotecnología de la UNAM | Consorcio Mexicano de Vigilancia Genómica (CoViGen-Mex). Authors (in alphabetical order): Julio Elias Alvarado-Yaah, Carlos F. Arias, Santiago Ávila-Ríos, Eduardo Becerril-Vargas, Víctor Hugo Borja-Aburto, Celia Boukadida, Cristóbal Cháidez-Quiróz, Juan Bautista Chale-Dzul, Ricardo Ciria Merce, Andreu Comas-García, Célida Duque Molina, Julissa Enciso-Ibarra, José Antonio Enciso-Moreno, Gloria Elena Espinosa-Ayala, Fernando Fontove-Herrera, Daniel Fregoso-Rueda, Victor Eduardo García-Arias, Alejandra García-Gasca, Bruno Gómez-Gil, Jean Pierre González, Irvin |

| | | | |
|---|---|---|---|
| | | | González-López, Concepción Grajales-Muñiz, Ricardo Grande, Rosa María Gutiérrez Rios, Jesús Hernández, Alejandra Hernández-Terán, Alfredo Herrera-Estrella, Carla Ivón Herrera-Najera, Pavel Isa, Verónica Mata-Haro, Daniel Lira Morales, Susana Lopez, Antonio Loza Román, Brenda Irasema Maldonado-Meza, Bernardo Martínez-Miguel, José Arturo Martínez-Orozco, Célida Martínez- Rodríguez, Margarita Matías-Florentino, Fidencio Mejía-Nepomuceno, María Guadalupe de Jesús Mireles-Rivera, Gloria María Molina-Salinas, Hector Montoya-Fuentes, , Mario Mújica-Sánchez, José Esteban Muñoz-Medina, José de Jesús Nuñez-Contreras, Alicia Ocaña-Mondragón, Luis Alberto Ochoa-Carrera, Hector Esteban Paz-Juárez, Marissa Perez-Garcia, Francisco Pulido, Helen Haydee Fernanda Ramírez-Plascencia, Jorge Salas-Hernández, Angel Gustavo Salas-Lais, Alejandro Sánchez-Flores, Clara Esperanza Santacruz-Tinoco, María Guadalupe Santiago-Mauricio, Selene Zárate, Nelly Sélem-Mojica, Blanca Taboada, Gloria Vazquez, Joel Armando Vázquez-Pérez. |
| EPI_ISL_2801707, EPI_ISL_2801708, EPI_ISL_2801709, EPI_ISL_2801710, EPI_ISL_2801713, EPI_ISL_2801718, EPI_ISL_2801719, EPI_ISL_2801720, EPI_ISL_2801721, EPI_ISL_2801722, EPI_ISL_2801723, EPI_ISL_2801724, EPI_ISL_2801725, EPI_ISL_2801726, EPI_ISL_2801727, EPI_ISL_2801729, EPI_ISL_2801730, EPI_ISL_2801731, EPI_ISL_2801732, EPI_ISL_2801733, EPI_ISL_2801734, EPI_ISL_2801735, EPI_ISL_2801736, EPI_ISL_2801737, EPI_ISL_2801738, EPI_ISL_2801739, EPI_ISL_2801740, EPI_ISL_2801742, EPI_ISL_2801745, EPI_ISL_2801746, EPI_ISL_2801748, EPI_ISL_2801750 | | | |
| see above | Centro de Investigación Biomédica de Occidente (CIBO) | Instituto de Biotecnología de la UNAM | Consorcio Mexicano de Vigilancia Genómica (CoViGen-Mex). Authors (in alphabetical order): Julio Elias Alvarado-Yaah, Carlos F. Arias, Santiago Ávila-Ríos, Eduardo Becerril-Vargas, Víctor Hugo Borja-Aburto, Celia Boukadida, Cristóbal Cháidez-Quiróz, Juan Bautista Chale-Dzul, Ricardo Ciria Merce, Andreu Comas-García, Célida Duque Molina, Julissa Enciso-Ibarra, José Antonio Enciso-Moreno, Gloria Elena Espinosa-Ayala, Fernando Fontove-Herrera, Daniel Fregoso-Rueda, Victor Eduardo García-Arias, Alejandra García-Gasca, Bruno Gómez-Gil, Jean Pierre González, Irvin González-López, Concepción Grajales-Muñiz, Ricardo Grande, Rosa María Gutiérrez Rios, Jesús Hernández, Alejandra Hernández-Terán, Alfredo Herrera-Estrella, Carla Ivón Herrera-Najera, Pavel Isa, Verónica Mata-Haro, Daniel Lira Morales, Susana Lopez, Antonio Loza Román, Brenda Irasema Maldonado-Meza, Bernardo Martínez-Miguel, José Arturo Martínez-Orozco, Célida Martínez- Rodríguez, Margarita Matías-Florentino, Fidencio Mejía-Nepomuceno, María Guadalupe de Jesús Mireles-Rivera, Gloria María Molina-Salinas, Hector Montoya-Fuentes, , Mario Mújica-Sánchez, José Esteban Muñoz-Medina, José de Jesús Nuñez-Contreras, Alicia Ocaña-Mondragón, Luis Alberto Ochoa-Carrera, Hector Esteban Paz-Juárez, Marissa Perez-Garcia, Francisco Pulido, Helen Haydee Fernanda Ramírez-Plascencia, Jorge Salas-Hernández, Angel Gustavo Salas-Lais, Alejandro Sánchez-Flores, Clara Esperanza Santacruz-Tinoco, María Guadalupe Santiago-Mauricio, Selene Zárate, Nelly Sélem-Mojica, Blanca Taboada, Gloria Vazquez, Joel Armando Vázquez-Pérez. |
| EPI_ISL_2801758, EPI_ISL_2801759, EPI_ISL_2801762, EPI_ISL_2801763, EPI_ISL_2801764, EPI_ISL_2801765, EPI_ISL_2801766, EPI_ISL_2801767, EPI_ISL_2801768, EPI_ISL_2801769, EPI_ISL_2801772, EPI_ISL_2801774, EPI_ISL_2801775, EPI_ISL_2801776, EPI_ISL_2801779, EPI_ISL_2801780, EPI_ISL_2801783, EPI_ISL_2801786, EPI_ISL_2801787, EPI_ISL_2801789, EPI_ISL_2801795, EPI_ISL_2801796, EPI_ISL_2801797, EPI_ISL_2801802, EPI_ISL_2801804, EPI_ISL_2801806, EPI_ISL_2801807, EPI_ISL_2801809, EPI_ISL_2801810, EPI_ISL_2801811, EPI_ISL_2801812, EPI_ISL_2801814, EPI_ISL_2801815, EPI_ISL_2801816, EPI_ISL_2801817, EPI_ISL_2801819, EPI_ISL_2801822, EPI_ISL_2801823, EPI_ISL_2801825, EPI_ISL_2801826, EPI_ISL_2801827, EPI_ISL_2801829, EPI_ISL_2801832, EPI_ISL_2801834, EPI_ISL_2801837, EPI_ISL_2801838, EPI_ISL_2801839, EPI_ISL_2801841, EPI_ISL_2801842, EPI_ISL_2801845, EPI_ISL_2801846, EPI_ISL_2801847, EPI_ISL_2801848, EPI_ISL_2801849, EPI_ISL_2801850, EPI_ISL_2801851, EPI_ISL_2801854, EPI_ISL_2801855, EPI_ISL_2801856, EPI_ISL_2801858, EPI_ISL_2801860, EPI_ISL_2801861, EPI_ISL_2801862, EPI_ISL_2801864, EPI_ISL_2801865, EPI_ISL_2801866, EPI_ISL_2801867, EPI_ISL_2801869, EPI_ISL_2801870, EPI_ISL_2801871, EPI_ISL_2801872, EPI_ISL_2801873, EPI_ISL_2801874, EPI_ISL_2801875, EPI_ISL_2801876, EPI_ISL_2801877, EPI_ISL_2801878, EPI_ISL_2801880, EPI_ISL_2801881, EPI_ISL_2801883, EPI_ISL_2801884, EPI_ISL_2801887, EPI_ISL_2801888, EPI_ISL_2801889, EPI_ISL_2801890 | | | |
| see above | Laboratorio Central de Epidemiología (LCE) | Instituto de Biotecnología de la UNAM | Consorcio Mexicano de Vigilancia Genómica (CoViGen-Mex). Authors (in alphabetical order): Julio Elias Alvarado-Yaah, Carlos F. Arias, Santiago Ávila-Ríos, Eduardo Becerril-Vargas, Víctor Hugo Borja-Aburto, Celia Boukadida, Cristóbal Cháidez-Quiróz, Juan Bautista Chale-Dzul, Ricardo Ciria Merce, Andreu Comas-García, Célida Duque Molina, Julissa Enciso-Ibarra, José Antonio Enciso-Moreno, Gloria Elena Espinosa-Ayala, Fernando Fontove-Herrera, Daniel Fregoso-Rueda, Victor Eduardo García-Arias, Alejandra García-Gasca, Bruno Gómez-Gil, Jean Pierre González, Irvin González-López, Concepción Grajales-Muñiz, Ricardo Grande, Rosa María Gutiérrez Rios, Jesús Hernández, Alejandra Hernández-Terán, Alfredo Herrera-Estrella, Carla Ivón Herrera-Najera, Pavel Isa, Verónica Mata-Haro, Daniel Lira Morales, Susana Lopez, Antonio Loza Román, Brenda Irasema Maldonado-Meza, Bernardo Martínez-Miguel, José Arturo Martínez-Orozco, Célida Martínez- Rodríguez, Margarita Matías-Florentino, Fidencio Mejía-Nepomuceno, María Guadalupe de Jesús Mireles-Rivera, Gloria María Molina-Salinas, Hector Montoya-Fuentes, , Mario Mújica-Sánchez, José Esteban Muñoz-Medina, José de Jesús Nuñez-Contreras, Alicia Ocaña-Mondragón, Luis Alberto Ochoa-Carrera, Hector Esteban Paz-Juárez, Marissa Perez-Garcia, Francisco Pulido, Helen Haydee Fernanda Ramírez-Plascencia, Jorge Salas-Hernández, Angel Gustavo Salas-Lais, Alejandro Sánchez-Flores, Clara Esperanza Santacruz-Tinoco, María Guadalupe Santiago-Mauricio, Selene Zárate, Nelly Sélem-Mojica, Blanca Taboada, Gloria Vazquez, Joel Armando Vázquez-Pérez. |
| EPI_ISL_2801892, EPI_ISL_2801893, EPI_ISL_2801894 | Banteay Meanchey Regional Laboratory | Virology Unit, Institut Pasteur du Cambodge | Cecile Troupin, Leakhena Pum, Jurre Y Siegers, Ly Sovann, Kraing Sidonn, Yi Sengdoeurn, Chin Savuth, Chau Darapheak, Veasna Duong, Erik A Karlsson |
| EPI_ISL_2801895 | Battambang Rapid Response Team | Virology Unit, Institut Pasteur du Cambodge | Cecile Troupin, Leakhena Pum, Jurre Y Siegers, Ly Sovann, Kraing Sidonn, Yi Sengdoeurn, Chin Savuth, Chau Darapheak, Veasna Duong, Erik A Karlsson |
| EPI_ISL_2801896, EPI_ISL_2801897 | Kampong Cham Rapid Response Team | Virology Unit, Institut Pasteur du Cambodge | Cecile Troupin, Leakhena Pum, Jurre Y Siegers, Ly Sovann, Kraing Sidonn, Yi Sengdoeurn, Chin Savuth, Chau Darapheak, Veasna Duong, Erik A Karlsson |
| EPI_ISL_2801898 | Pursat Rapid Response Team | Virology Unit, Institut Pasteur du Cambodge | Cecile Troupin, Leakhena Pum, Jurre Y Siegers, Ly Sovann, Kraing Sidonn, Yi Sengdoeurn, Chin Savuth, Chau Darapheak, Veasna Duong, Erik A Karlsson |
| EPI_ISL_2801899, EPI_ISL_2801900, EPI_ISL_2801901, EPI_ISL_2801902, EPI_ISL_2801903 | Oddar Meanchey Rapid Response Team | Virology Unit, Institut Pasteur du Cambodge | Cecile Troupin, Leakhena Pum, Jurre Y Siegers, Ly Sovann, Kraing Sidonn, Yi Sengdoeurn, Chin Savuth, Chau Darapheak, Veasna Duong, Erik A Karlsson |
| EPI_ISL_2801904, EPI_ISL_2801905, EPI_ISL_2801906 | Tboung Khmum Rapid Response Team | Virology Unit, Institut Pasteur du Cambodge | Cecile Troupin, Leakhena Pum, Jurre Y Siegers, Ly Sovann, Kraing Sidonn, Yi Sengdoeurn, Chin Savuth, Chau Darapheak, Veasna Duong, Erik A Karlsson |
| EPI_ISL_2801907, EPI_ISL_2801908, EPI_ISL_2801909, EPI_ISL_2801910, EPI_ISL_2801911, EPI_ISL_2801912, EPI_ISL_2801913, EPI_ISL_2801914, EPI_ISL_2801915, EPI_ISL_2801916, EPI_ISL_2801917, EPI_ISL_2801918, EPI_ISL_2801919, EPI_ISL_2801920, EPI_ISL_2801921, EPI_ISL_2801922, EPI_ISL_2801923, EPI_ISL_2801924, EPI_ISL_2801925, EPI_ISL_2801926, EPI_ISL_2801927 | | | |
| see above | Virology Unit, Institut Pasteur du Cambodge | Virology Unit, Institut Pasteur du Cambodge | Cecile Troupin, Leakhena Pum, Jurre Y Siegers, Ly Sovann, Kraing Sidonn, Yi Sengdoeurn, Chin Savuth, Chau Darapheak, Veasna Duong, Erik A Karlsson |
| EPI_ISL_2803757 | The Ohio State University Applied Microbiology Services Laboratory | The Ohio State University Applied Microbiology Services Laboratory | Seth A. Faith PhD |
| EPI_ISL_2811959 | Tengku Ampuan Afzan Hospital (HTAA) | Institute for Medical Research, Infectious Disease Research Centre, National Institutes of Health, Ministry of Health Malaysia | Suppiah J, Kamel K, Mohd Zawawi Z, Azizan MA, Ramly N, Robert F, Thayan R |
| EPI_ISL_2811984 | Pekan Tajau Health Clinic | Institute for Medical Research, Infectious Disease Research Centre, National Institutes of Health, Ministry of Health Malaysia | Suppiah J, Kamel K, Mohd Zawawi Z, Azizan MA, Ramly N, Robert F, Thayan R |
| EPI_ISL_2811998 | Bandar Kuantan Health Clinic | Institute for Medical Research, Infectious Disease Research Centre, National Institutes of Health, Ministry of Health Malaysia | Suppiah J, Kamel K, Mohd Zawawi Z, Azizan MA, Ramly N, Robert F, Thayan R |
| EPI_ISL_2812026, EPI_ISL_2812027 | Rembau Health Clinic | Institute for Medical Research, Infectious Disease Research Centre, National Institutes of Health, Ministry of Health Malaysia | Suppiah J, Kamel K, Mohd Zawawi Z, Azizan MA, Ramly N, Robert F, Thayan R |
| EPI_ISL_2812146, EPI_ISL_2812147, EPI_ISL_2812148, EPI_ISL_2812149, EPI_ISL_2812150, EPI_ISL_2812151, EPI_ISL_2812152, EPI_ISL_2812153, EPI_ISL_2812154, EPI_ISL_2812155, EPI_ISL_2812156 | | | |
| see above | Singapore General Hospital | Department of Microbiology | Nurdyana Abdul Rahman, Kenneth Xin Long Chan, Chayaporn Suphavilai, Kun Lee Lim, Sui Sin Goh, Kian Sing Chan, Lynette Oon, Kern Rei Chng, James Sim Heng Chiak, Niranjan Nagarajan, Karrie Ko |
| EPI_ISL_2812537 | Laboratorio de Virología HUCA | Laboratorio de Virología HUCA | Sandoval M, Castelló C, Gómez de Oña J, Boga JA, Rojo S, Alvarez-Arguelles ME, Abreu F, Costales I, Perez-Martínez Z, Martín-Rodríguez G, Coto E, Melón S |
| EPI_ISL_2812576 | Kemaman Hospital | Institute for Medical Research, Infectious Disease Research Centre, National Institutes of Health, Ministry of Health | Suppiah J, Kamel K, Mohd Zawawi Z, Azizan MA, Ramly N, Robert F, Thayan R |

| | | | |
|---|---|---|---|
| | | Malaysia | |
| EPI_ISL_2812654 | Sultanah Bahiyah Hospital, Alor Setar | Institute for Medical Research, Infectious Disease Research Centre, National Institutes of Health, Ministry of Health Malaysia | Suppiah J, Kamel K, Mohd Zawawi Z, Azizan MA, Ramly N, Robert F, Thayan R |
| EPI_ISL_2812706 | Sultan Haji Ahmad Shah Hospital | Institute for Medical Research, Infectious Disease Research Centre, National Institutes of Health, Ministry of Health Malaysia | Suppiah J, Kamel K, Mohd Zawawi Z, Azizan MA, Ramly N, Robert F, Thayan R |
| EPI_ISL_2812958, EPI_ISL_2812959, EPI_ISL_2812960, EPI_ISL_2812961, EPI_ISL_2812962, EPI_ISL_2812963, EPI_ISL_2812964, EPI_ISL_2812965, EPI_ISL_2812966, EPI_ISL_2812967, EPI_ISL_2812968, EPI_ISL_2812969 | | | |
| see above | Department of Health Technology and Informatics, The Hong Kong Polytechnic University | Department of Health Technology and Informatics, The Hong Kong Polytechnic University | Gilman Kit-Hang Siu, Lam-Kwong Lee, Kenneth Siu-Sing Leung, Jake Siu-Lun Leung, Wing-Hei Lo, Timothy Ting-Leung Ng, Chloe Toi-Mei Chan, Hoi-Ching Jim, Kingsley King-Gee Tam, Hiu-Yin Lao, Denise Sze-Hang Wong, Alan Ka-Lun Wu, Miranda Chong-Yee Yau, Yvette Wai-Man Lai, Kitty Sau-Chun Fung, Sandy Ka-Yee Chau, Barry Kin-Chung Wong, Wing-Kin To, Kristine Luk, Alex Yat-Man Ho, Tak-Lun Que, Kam-Tong Yip, Wing Cheong Yam, David Ho-Keung Shum, Shea Ping Yip |
| EPI_ISL_2812971 | Labuan Hospital | Institute for Medical Research, Infectious Disease Research Centre, National Institutes of Health, Ministry of Health Malaysia | Suppiah J, Kamel K, Mohd Zawawi Z, Azizan MA, Ramly N, Robert F, Thayan R |
| EPI_ISL_2812981 | Johor State Health Department | Institute for Medical Research, Infectious Disease Research Centre, National Institutes of Health, Ministry of Health Malaysia | Suppiah J, Kamel K, Mohd Zawawi Z, Azizan MA, Ramly N, Robert F, Thayan R |
| EPI_ISL_2812982, EPI_ISL_2812983, EPI_ISL_2814168 | L. C. Laboratori Campisi Group | AOU Polclinico "G Rodolico-S.Marco" PO Rodolico and BIOMETEC UNICT CATANIA | STEFANI AND GUIDO SCALIA |
| EPI_ISL_2814916, EPI_ISL_2815327, EPI_ISL_2815328 | Melaka Hospital | Institute for Medical Research, Infectious Disease Research Centre, National Institutes of Health, Ministry of Health Malaysia | Suppiah J, Kamel K, Mohd Zawawi Z, Azizan MA, Ramly N, Robert F, Thayan R |
| EPI_ISL_2815329, EPI_ISL_2815330 | Ipoh Public Health Laboratory (MKAI), Ministry of Health Malaysia | Institute for Medical Research, Infectious Disease Research Centre, National Institutes of Health, Ministry of Health Malaysia | Suppiah J, Kamel K, Mohd Zawawi Z, Azizan MA, Ramly N, Robert F, Thayan R |
| EPI_ISL_2815331 | Kuala Lumpur General Hospital | Institute for Medical Research, Infectious Disease Research Centre, National Institutes of Health, Ministry of Health Malaysia | Suppiah J, Kamel K, Mohd Zawawi Z, Azizan MA, Ramly N, Robert F, Thayan R |
| EPI_ISL_2815375, EPI_ISL_2819024, EPI_ISL_2819571, EPI_ISL_2819919 | Institute for Medical Research, Infectious Disease Research Centre, National Institutes of Health, Ministry of Health Malaysia | Institute for Medical Research, Infectious Disease Research Centre, National Institutes of Health, Ministry of Health Malaysia | Suppiah J, Kamel K, Mohd Zawawi Z, Azizan MA, Ramly N, Robert F, Thayan R |
| EPI_ISL_2820290, EPI_ISL_2820296, EPI_ISL_2820300, EPI_ISL_2820303, EPI_ISL_2820304, EPI_ISL_2820307, EPI_ISL_2820308, EPI_ISL_2820310, EPI_ISL_2820312, EPI_ISL_2820321, EPI_ISL_2820322, EPI_ISL_2820325, EPI_ISL_2820326, EPI_ISL_2820330, EPI_ISL_2820331, EPI_ISL_2820333 | | | |
| see above | Kasane Primary Hospital Laboratory | Botswana Harvard HIV Reference Laboratory | Sikhulile Moyo, Wonderful T. Choga, Dorcas Maruapula, Agnes Karutwaeng, Rose Munyere, Keoratile Ntshambiwa, Thongbotho Mphoyakgosi, Boitumelo Zuze, Botshelo Radibe, Legodile Kooepile, Ontlametse T. Bareng, Roger Shapiro, Shahin Lockman, Joseph Makhema, Madisa Mine, Mosepele Mosepele, Simani Gaseitsiwe |
| EPI_ISL_2820339, EPI_ISL_2820340, EPI_ISL_2820341, EPI_ISL_2820342, EPI_ISL_2820343, EPI_ISL_2820345, EPI_ISL_2820350, EPI_ISL_2820353, EPI_ISL_2820359, EPI_ISL_2820366, EPI_ISL_2820374 | | | |
| see above | Botswana Harvard HIV Reference Laboratory | Botswana Harvard HIV Reference Laboratory | Sikhulile Moyo, Wonderful T. Choga, Dorcas Maruapula, Thongbotho Mphoyakgosi, Boitumelo Zuze, Botshelo Radibe, Legodile Kooepile, Ontlametse T. Bareng, Motlalepule L. Pone, Godfrey Simoonga, Kereng Mphoyakgosi, Keoratile Ntshambiwa, Modisa Motswaledi, Madisa Mine, Joseph Makhema, Roger Shapiro, Shahin Lockman, Mosepele Mosepele, Simani Gaseitsiwe |
| EPI_ISL_2820386, EPI_ISL_2820388, EPI_ISL_2820392, EPI_ISL_2820400 | Kasane Primary Hospital Laboratory | Botswana Harvard HIV Reference Laboratory | Sikhulile Moyo, Wonderful T. Choga, Dorcas Maruapula, Agnes Karutwaeng, Rose Munyere, Keoratile Ntshambiwa, Thongbotho Mphoyakgosi, Boitumelo Zuze, Botshelo Radibe, Legodile Kooepile, Ontlametse T. Bareng, Roger Shapiro, Shahin Lockman, Joseph Makhema, Madisa Mine, Mosepele Mosepele, Simani Gaseitsiwe |
| EPI_ISL_2820430, EPI_ISL_2820448, EPI_ISL_2820451, EPI_ISL_2820459, EPI_ISL_2820462 | Botswana Harvard HIV Reference Laboratory | Botswana Harvard HIV Reference Laboratory | Sikhulile Moyo, Wonderful T. Choga, Dorcas Maruapula, Thongbotho Mphoyakgosi, Boitumelo Zuze, Botshelo Radibe, Legodile Kooepile, Ontlametse T. Bareng, Keoratile Ntshambiwa, Modisa Motswaledi, Madisa Mine, Joseph Makhema, Roger Shapiro, Shahin Lockman, Mosepele Mosepele, Simani Gaseitsiwe |
| EPI_ISL_2820677, EPI_ISL_2820678, EPI_ISL_2820679, EPI_ISL_2820683, EPI_ISL_2820684 | Università degli Studi di Perugia | Istituto Zooprofilattico Sperimentale dell'Abruzzo e Molise "G. Caporale" | Mencacci A, Camilloni B, Biagetti M, Giammarioli M, Lorusso A, Marcacci M, Di Domenico M, Ancora M, Curini V, Mangone I, Rinaldi A, Delli Compagni E, Scialabba S, Di Pasquale A, Cammà C, Puglia I, Calistri P, Savini G |
| EPI_ISL_2820707 | Presidio di Brindisi Di Summa - Perrino | Istituto Zooprofilattico Sperimentale della Puglia e della Basilicata | Parisi A., Bianco A., Capozzi L., Del Sambro L., Simone D., Difato L., Santoro A., Ridolfi D., Giannico A., Manzari C., Pesole G. |
| EPI_ISL_2820715, EPI_ISL_2820716, EPI_ISL_2820717 | I.R.C.C.S. "S. De Bellis" - Ente Ospedaliero | Istituto Zooprofilattico Sperimentale della Puglia e della Basilicata | Parisi A., Bianco A., Capozzi L., Del Sambro L., Simone D., Difato L., Notarnicola M., Lippolis A., Manzari C., Pesole G. |
| EPI_ISL_2820718, EPI_ISL_2820719, EPI_ISL_2820720, EPI_ISL_2820721, EPI_ISL_2820722, EPI_ISL_2820723 | Ospedale Santa Caterina Novella | Istituto Zooprofilattico Sperimentale della Puglia e della Basilicata | Parisi A., Bianco A., Capozzi L., Del Sambro L., Simone D., Difato L., Bruno A. R., Manzari C., Pesole G. |
| EPI_ISL_2820724, EPI_ISL_2820726 | Ospedale Vito Fazzi | Istituto Zooprofilattico Sperimentale della Puglia e della Basilicata | Parisi A., Bianco A., Capozzi L., Del Sambro L., Simone D., Difato L., Lobreglio G., Manzari C., Pesole G. |
| EPI_ISL_2820729, EPI_ISL_2820730, EPI_ISL_2820731, EPI_ISL_2820733, EPI_ISL_2820735, EPI_ISL_2820738, EPI_ISL_2820739, EPI_ISL_2820741, EPI_ISL_2820742, EPI_ISL_2820743, EPI_ISL_2820744 | | | |
| see above | Presidio di Brindisi Di Summa - Perrino | Istituto Zooprofilattico Sperimentale della Puglia e della Basilicata | Parisi A., Bianco A., Capozzi L., Del Sambro L., Simone D., Difato L., Santoro A., Ridolfi D., Giannico A., Manzari C., Pesole G. |
| EPI_ISL_2820746, EPI_ISL_2820747, EPI_ISL_2820748, EPI_ISL_2820749, EPI_ISL_2820750, EPI_ISL_2820751, EPI_ISL_2820752, EPI_ISL_2820753, EPI_ISL_2820754 | Ospedale Santissima Annunziata | Istituto Zooprofilattico Sperimentale della Puglia e della Basilicata | Parisi A., Bianco A., Capozzi L., Del Sambro L., Simone D., Difato L., Ridolfi D., Giannico A, Tampoia M., Manzari C., Pesole G. |
| EPI_ISL_2827779 | LDSP - CAQUETA HOSPITAL MARIA INMACULADA | Instituto Nacional de Salud- Dirección de Investigación en Salud Pública | Katherine Laiton-Donato, Diego A. Álvarez-Díaz, Carlos Franco-Muñoz, Hector Alejandro Ruiz-Moreno, Paola Rojas, Maria T. Herrera-Sepúlveda, Diego Andrés Prada, Jhonnatan Reales-González, Sheryll Corchuelo, Julian Naizaque, Jorge Rivera, Gerardo Santamaría, Sergio Gomez, Lisseth Pardo, Juan Camilo Martinez, Marta Lopez Blanco, Ángela Alarcon Cruz, Diana Malo, Carmen Osorio, Magdalena Wiesner, Martha Lucia Ospina Martinez, Marcela Mercado-Reyes |
| EPI_ISL_2827783 | LDSP - CAQUETA HOSPITAL MALVINAS | Instituto Nacional de Salud- Dirección de Investigación en Salud Pública | Katherine Laiton-Donato, Diego A. Álvarez-Díaz, Carlos Franco-Muñoz, Hector Alejandro Ruiz-Moreno, Paola Rojas, Maria T. Herrera-Sepúlveda, Diego Andrés Prada, Jhonnatan Reales-González, Sheryll Corchuelo, Julian Naizaque, Jorge Rivera, Gerardo Santamaría, Sergio Gomez, Lisseth Pardo, Juan Camilo Martinez, Marta Lopez Blanco, Ángela Alarcon Cruz, Diana Malo, Carmen Osorio, Magdalena Wiesner, Martha Lucia Ospina Martinez, Marcela Mercado-Reyes |

| | | | |
|---|---|---|---|
| EPI_ISL_2827961, EPI_ISL_2827962 | AREA DE SALUD HATILLO - CLINICA DR. SOLON NUÑEZ | Inciensa, Instituto Costarricense de Investigación y Enseñanza en Nutrición y Salud | Francisco Duarte, Hebleen Porras, Claudio Soto-Garita, Estela Cordero, Adriana Godínez, Melany Calderón, José Luis Vargas, Mariela Gutiérrez, Joselyn Prado & Roberto Brilla |
| EPI_ISL_2827963, EPI_ISL_2827964, EPI_ISL_2827965 | HOSPITAL DR. WILLIAM ALLEN | Inciensa, Instituto Costarricense de Investigación y Enseñanza en Nutrición y Salud | Francisco Duarte, Hebleen Porras, Claudio Soto-Garita, Estela Cordero, Adriana Godínez, Melany Calderón, José Luis Vargas, Mariela Gutiérrez & Joselyn Prado |
| EPI_ISL_2827966 | AREA DE SALUD CARIARI | Inciensa, Instituto Costarricense de Investigación y Enseñanza en Nutrición y Salud | Francisco Duarte, Hebleen Porras, Claudio Soto-Garita, Estela Cordero, Adriana Godínez, Melany Calderón, José Luis Vargas, Mariela Gutiérrez, Joselyn Prado & Vanessa Jiménez |
| EPI_ISL_2827967, EPI_ISL_2827968 | HOSPITAL DR. RAFAEL A. CALDERON GUARDIA | Inciensa, Instituto Costarricense de Investigación y Enseñanza en Nutrición y Salud | Francisco Duarte, Hebleen Porras, Claudio Soto-Garita, Estela Cordero, Adriana Godínez, Melany Calderón, José Luis Vargas, Mariela Gutiérrez, Joselyn Prado & Fabian Salas |
| EPI_ISL_2827970 | AREA DE SALUD GUATUSO | Inciensa, Instituto Costarricense de Investigación y Enseñanza en Nutrición y Salud | Francisco Duarte, Hebleen Porras, Claudio Soto-Garita, Estela Cordero, Adriana Godínez, Melany Calderón, José Luis Vargas, Mariela Gutiérrez, Joselyn Prado & Francisco Chacón |
| EPI_ISL_2827971 | HOSPITAL CIUDAD NEILY | Inciensa, Instituto Costarricense de Investigación y Enseñanza en Nutrición y Salud | Francisco Duarte, Hebleen Porras, Claudio Soto-Garita, Estela Cordero, Adriana Godínez, Melany Calderón, José Luis Vargas, Mariela Gutiérrez, Joselyn Prado & María Isabel Victoria |
| EPI_ISL_2827972 | AREA DE SALUD PAVAS (COOPESALUD) | Inciensa, Instituto Costarricense de Investigación y Enseñanza en Nutrición y Salud | Francisco Duarte, Hebleen Porras, Claudio Soto-Garita, Estela Cordero, Adriana Godínez, Melany Calderón, José Luis Vargas, Mariela Gutiérrez, Joselyn Prado & Paola Sánchez |
| EPI_ISL_2827973, EPI_ISL_2827974 | HOSPITAL SAN JUAN DE DIOS | Inciensa, Instituto Costarricense de Investigación y Enseñanza en Nutrición y Salud | Francisco Duarte, Hebleen Porras, Claudio Soto-Garita, Estela Cordero, Adriana Godínez, Melany Calderón, José Luis Vargas, Mariela Gutiérrez, Joselyn Prado & Paola Sánchez |
| EPI_ISL_2827975 | HOSPITAL LA ANEXION | Inciensa, Instituto Costarricense de Investigación y Enseñanza en Nutrición y Salud | Francisco Duarte, Hebleen Porras, Claudio Soto-Garita, Estela Cordero, Adriana Godínez, Melany Calderón, José Luis Vargas, Mariela Gutiérrez, Joselyn Prado & Adriana Bermúdez |
| EPI_ISL_2827976, EPI_ISL_2827977 | HOSPITAL DR. ENRIQUE BALTODANO BRICEÑO | Inciensa, Instituto Costarricense de Investigación y Enseñanza en Nutrición y Salud | Francisco Duarte, Hebleen Porras, Claudio Soto-Garita, Estela Cordero, Adriana Godínez, Melany Calderón, José Luis Vargas, Mariela Gutiérrez, Joselyn Prado & Adriana Bermúdez |
| EPI_ISL_2827978 | AREA DE SALUD SANTA CRUZ | Inciensa, Instituto Costarricense de Investigación y Enseñanza en Nutrición y Salud | Francisco Duarte, Hebleen Porras, Claudio Soto-Garita, Estela Cordero, Adriana Godínez, Melany Calderón, José Luis Vargas, Mariela Gutiérrez, Joselyn Prado & Adriana Bermúdez |
| EPI_ISL_2827979, EPI_ISL_2827980 | AREA DE SALUD CARTAGO | Inciensa, Instituto Costarricense de Investigación y Enseñanza en Nutrición y Salud | Francisco Duarte, Hebleen Porras, Claudio Soto-Garita, Estela Cordero, Adriana Godínez, Melany Calderón, José Luis Vargas, Mariela Gutiérrez, Joselyn Prado & Tania Garro |
| EPI_ISL_2827981 | HOSPITAL MONSEÑOR SANABRIA | Inciensa, Instituto Costarricense de Investigación y Enseñanza en Nutrición y Salud | Francisco Duarte, Hebleen Porras, Claudio Soto-Garita, Estela Cordero, Adriana Godínez, Melany Calderón, José Luis Vargas, Mariela Gutiérrez, Joselyn Prado & María José Gómez |
| EPI_ISL_2827982 | AREA DE SALUD COBANO | Inciensa, Instituto Costarricense de Investigación y Enseñanza en Nutrición y Salud | Francisco Duarte, Hebleen Porras, Claudio Soto-Garita, Estela Cordero, Adriana Godínez, Melany Calderón, José Luis Vargas, Mariela Gutiérrez, Joselyn Prado & María José Gómez |
| EPI_ISL_2827983 | AREA DE SALUD BARRANCA - CLINICA DR. ROBERTO SOTO | Inciensa, Instituto Costarricense de Investigación y Enseñanza en Nutrición y Salud | Francisco Duarte, Hebleen Porras, Claudio Soto-Garita, Estela Cordero, Adriana Godínez, Melany Calderón, José Luis Vargas, Mariela Gutiérrez, Joselyn Prado & María José Gómez |
| EPI_ISL_2827984 | AREA DE SALUD COBANO | Inciensa, Instituto Costarricense de Investigación y Enseñanza en Nutrición y Salud | Francisco Duarte, Hebleen Porras, Claudio Soto-Garita, Estela Cordero, Adriana Godínez, Melany Calderón, José Luis Vargas, Mariela Gutiérrez, Joselyn Prado & María José Gómez |
| EPI_ISL_2827985 | AREA DE SALUD ESPARZA | Inciensa, Instituto Costarricense de Investigación y Enseñanza en Nutrición y Salud | Francisco Duarte, Hebleen Porras, Claudio Soto-Garita, Estela Cordero, Adriana Godínez, Melany Calderón, José Luis Vargas, Mariela Gutiérrez, Joselyn Prado & María José Gómez |
| EPI_ISL_2827986 | HOSPITAL SAN VICENTE DE PAUL | Inciensa, Instituto Costarricense de Investigación y Enseñanza en Nutrición y Salud | Francisco Duarte, Hebleen Porras, Claudio Soto-Garita, Estela Cordero, Adriana Godínez, Melany Calderón, José Luis Vargas, Mariela Gutiérrez, Joselyn Prado & Josiany Arce |
| EPI_ISL_2827987 | HOSPITAL DR. CARLOS LUIS VALVERDE VEGA | Inciensa, Instituto Costarricense de Investigación y Enseñanza en Nutrición y Salud | Francisco Duarte, Hebleen Porras, Claudio Soto-Garita, Estela Cordero, Adriana Godínez, Melany Calderón, José Luis Vargas, Mariela Gutiérrez, Joselyn Prado & Yahaira Ramos |
| EPI_ISL_2827988, EPI_ISL_2827989, EPI_ISL_2827990 | AREA DE SALUD CIUDAD QUESADA | Inciensa, Instituto Costarricense de Investigación y Enseñanza en Nutrición y Salud | Francisco Duarte, Hebleen Porras, Claudio Soto-Garita, Estela Cordero, Adriana Godínez, Melany Calderón, José Luis Vargas, Mariela Gutiérrez, Joselyn Prado & Juan Carlos Villalobos |
| EPI_ISL_2827991, EPI_ISL_2827992, EPI_ISL_2827993 | HOSPITAL MEXICO | Inciensa, Instituto Costarricense de Investigación y Enseñanza en Nutrición y Salud | Francisco Duarte, Hebleen Porras, Claudio Soto-Garita, Estela Cordero, Adriana Godínez, Melany Calderón, José Luis Vargas, Mariela Gutiérrez, Joselyn Prado & Juan Carlos Villalobos |
| EPI_ISL_2827994 | AREA DE SALUD CARIARI | Inciensa, Instituto Costarricense de Investigación y Enseñanza en Nutrición y Salud | Francisco Duarte, Hebleen Porras, Claudio Soto-Garita, Estela Cordero, Adriana Godínez, Melany Calderón, José Luis Vargas, Mariela Gutiérrez, Joselyn Prado & Blanca Gallegos |
| EPI_ISL_2827995, EPI_ISL_2827996 | AREA DE SALUD CARTAGO | Inciensa, Instituto Costarricense de Investigación y Enseñanza en Nutrición y Salud | Francisco Duarte, Hebleen Porras, Claudio Soto-Garita, Estela Cordero, Adriana Godínez, Melany Calderón, José Luis Vargas, Mariela Gutiérrez, Joselyn Prado & Monserrat Segura |
| EPI_ISL_2827997, EPI_ISL_2827998 | AREA DE SALUD BAGACES | Inciensa, Instituto Costarricense de Investigación y Enseñanza en Nutrición y Salud | Francisco Duarte, Hebleen Porras, Claudio Soto-Garita, Estela Cordero, Adriana Godínez, Melany Calderón, José Luis Vargas, Mariela Gutiérrez, Joselyn Prado & Karol Matamoros |
| EPI_ISL_2827999 | AREA DE SALUD DESAMPARADOS 2 (COOPESALUD) | Inciensa, Instituto Costarricense de Investigación y Enseñanza en Nutrición y Salud | Francisco Duarte, Hebleen Porras, Claudio Soto-Garita, Estela Cordero, Adriana Godínez, Melany Calderón, José Luis Vargas, Mariela Gutiérrez, Joselyn Prado & Alberto Quirós |
| EPI_ISL_2828000 | AREA DE SALUD GARABITO | Inciensa, Instituto Costarricense de Investigación y Enseñanza en Nutrición y Salud | Francisco Duarte, Hebleen Porras, Claudio Soto-Garita, Estela Cordero, Adriana Godínez, Melany Calderón, José Luis Vargas, Mariela Gutiérrez, Joselyn Prado & María José Gómez |
| EPI_ISL_2828001 | HOSPITAL MONSEÑOR SANABRIA | Inciensa, Instituto Costarricense de Investigación y Enseñanza en Nutrición y Salud | Francisco Duarte, Hebleen Porras, Claudio Soto-Garita, Estela Cordero, Adriana Godínez, Melany Calderón, José Luis Vargas, Mariela Gutiérrez, Joselyn Prado & María José Gómez |
| EPI_ISL_2828002 | AREA DE SALUD GOICOECHEA 2 - CLINICA DR. JIMENEZ NUÑEZ | Inciensa, Instituto Costarricense de Investigación y Enseñanza en Nutrición y Salud | Francisco Duarte, Hebleen Porras, Claudio Soto-Garita, Estela Cordero, Adriana Godínez, Melany Calderón, José Luis Vargas, Mariela Gutiérrez, Joselyn Prado & Juan Carlos Cartes |
| EPI_ISL_2828003 | AREA DE SALUD ACOSTA | Inciensa, Instituto Costarricense de Investigación y Enseñanza en Nutrición y Salud | Francisco Duarte, Hebleen Porras, Claudio Soto-Garita, Estela Cordero, Adriana Godínez, Melany Calderón, José Luis Vargas, Mariela Gutiérrez & Joselyn Prado |
| EPI_ISL_2828004 | AREA DE SALUD MATA REDONDA-HOSPITAL - CLINICA DR. MORENO CAÑAS | Inciensa, Instituto Costarricense de Investigación y Enseñanza en Nutrición y Salud | Francisco Duarte, Hebleen Porras, Claudio Soto-Garita, Estela Cordero, Adriana Godínez, Melany Calderón, José Luis Vargas, Mariela Gutiérrez, Joselyn Prado & Ricardo González |
| EPI_ISL_2828006 | AREA DE SALUD ESCAZU (COOPESANA) | Inciensa, Instituto Costarricense de Investigación y Enseñanza en Nutrición y Salud | Francisco Duarte, Hebleen Porras, Claudio Soto-Garita, Estela Cordero, Adriana Godínez, Melany Calderón, José Luis Vargas, Mariela Gutiérrez, Joselyn Prado & Juliana Mora |
| EPI_ISL_2828007, EPI_ISL_2828008 | AREA DE SALUD SAN RAMON | Inciensa, Instituto Costarricense de Investigación y Enseñanza en Nutrición y Salud | Francisco Duarte, Hebleen Porras, Claudio Soto-Garita, Estela Cordero, Adriana Godínez, Melany Calderón, José Luis Vargas, Mariela Gutiérrez & Joselyn Prado |
| EPI_ISL_2828009 | AREA DE SALUD SIQUIRRES | Inciensa, Instituto Costarricense de Investigación y Enseñanza en Nutrición y Salud | Francisco Duarte, Hebleen Porras, Claudio Soto-Garita, Estela Cordero, Adriana Godínez, Melany Calderón, José Luis Vargas, Mariela Gutiérrez, Joselyn Prado & Ileana Chavez |
| EPI_ISL_2828010 | HOSPITAL DR. RAFAEL A. CALDERON GUARDIA | Inciensa, Instituto Costarricense de Investigación y Enseñanza en Nutrición y Salud | Francisco Duarte, Hebleen Porras, Claudio Soto-Garita, Estela Cordero, Adriana Godínez, Melany Calderón, José Luis Vargas, Mariela Gutiérrez, Joselyn Prado & Fabian Salas |
| EPI_ISL_2828011 | AREA DE SALUD SAN RAFAEL | Inciensa, Instituto Costarricense de Investigación y Enseñanza en Nutrición y Salud | Francisco Duarte, Hebleen Porras, Claudio Soto-Garita, Estela Cordero, Adriana Godínez, Melany Calderón, José Luis Vargas, Mariela Gutiérrez, Joselyn Prado & María José Gómez |

| | | | |
|---|---|---|---|
| EPI_ISL_2828012 | AREA DE SALUD GARABITO | Inciensa, Instituto Costarricense de Investigación y Enseñanza en Nutrición y Salud | Francisco Duarte, Hebleen Porras, Claudio Soto-Garita, Estela Cordero, Adriana Godínez, Melany Calderón, José Luis Vargas, Mariela Gutiérrez, Joselyn Prado & María José Gómez |
| EPI_ISL_2828013 | HOSPITAL DR. MAX TERAN VALLS | Inciensa, Instituto Costarricense de Investigación y Enseñanza en Nutrición y Salud | Francisco Duarte, Hebleen Porras, Claudio Soto-Garita, Estela Cordero, Adriana Godínez, Melany Calderón, José Luis Vargas, Mariela Gutiérrez, Joselyn Prado & María José Gómez |
| EPI_ISL_2828014 | HOSPITAL MONSEÑOR SANABRIA | Inciensa, Instituto Costarricense de Investigación y Enseñanza en Nutrición y Salud | Francisco Duarte, Hebleen Porras, Claudio Soto-Garita, Estela Cordero, Adriana Godínez, Melany Calderón, José Luis Vargas, Mariela Gutiérrez, Joselyn Prado & María José Gómez |
| EPI_ISL_2828015 | HOSPITAL MEXICO | Inciensa, Instituto Costarricense de Investigación y Enseñanza en Nutrición y Salud | Francisco Duarte, Hebleen Porras, Claudio Soto-Garita, Estela Cordero, Adriana Godínez, Melany Calderón, José Luis Vargas, Mariela Gutiérrez, Joselyn Prado & Juan Carlos Villalobos |
| EPI_ISL_2828327 | Hospital Clínico Universitario Virgen de la Arrixaca | Hospital Clínico Universitario Virgen de la Arrixaca | Marina Simón Páez y Laura Moreno Parrado |
| EPI_ISL_2828383 | SC (UCO) Igiene e Sanità Pubblica, ASUGI, Trieste | ARGO Laboratorio Genomica ed Epigenomica | Licastro D, Dal Monego S, Degasperi M, Marcello A, Segat L, Piscianz E, Busetti M, Fontana F, Breda C, Barbone F, D'Agaro P |
| EPI_ISL_2828384 | SC (UCO) Igiene e Sanità Pubblica, ASUGI, Trieste and SC Microbiologia e Virologia, ASFO, Pordenone | ARGO Laboratorio Genomica ed Epigenomica | Licastro D, Dal Monego S, Degasperi M, Marcello A, Segat L, De Rosa R, Breda C, Barbone F, D'Agaro P |
| EPI_ISL_2828385, EPI_ISL_2828386, EPI_ISL_2828387, EPI_ISL_2828388, EPI_ISL_2828389 | SC (UCO) Igiene e Sanità Pubblica, ASUGI, Trieste | ARGO Laboratorio Genomica ed Epigenomica | Licastro D, Dal Monego S, Degasperi M, Marcello A, Segat L, Piscianz E, Busetti M, Fontana F, Breda C, Barbone F, D'Agaro P |
| EPI_ISL_2828390 | SC (UCO) Igiene e Sanità Pubblica, ASUGI, Trieste and SC Microbiologia e Virologia, ASFO, Pordenone | ARGO Laboratorio Genomica ed Epigenomica | Licastro D, Dal Monego S, Degasperi M, Marcello A, Segat L, De Rosa R, Breda C, Barbone F, D'Agaro P |
| EPI_ISL_2828391, EPI_ISL_2828392, EPI_ISL_2828393, EPI_ISL_2828394, EPI_ISL_2828395, EPI_ISL_2828396, EPI_ISL_2828397, EPI_ISL_2828398, EPI_ISL_2828399, EPI_ISL_2828400, EPI_ISL_2828401, EPI_ISL_2828402, EPI_ISL_2828403 | | | |
| see above | SC (UCO) Igiene e Sanità Pubblica, ASUGI, Trieste | ARGO Laboratorio Genomica ed Epigenomica | Licastro D, Dal Monego S, Degasperi M, Marcello A, Segat L, Piscianz E, Busetti M, Fontana F, Breda C, Barbone F, D'Agaro P |
| EPI_ISL_2828404 | SC (UCO) Igiene e Sanità Pubblica, ASUGI, Trieste and SC Microbiologia e Virologia, ASFO, Pordenone | ARGO Laboratorio Genomica ed Epigenomica | Licastro D, Dal Monego S, Degasperi M, Marcello A, Segat L, De Rosa R, Breda C, Barbone F, D'Agaro P |
| EPI_ISL_2828405, EPI_ISL_2828406, EPI_ISL_2828407, EPI_ISL_2828408, EPI_ISL_2828409, EPI_ISL_2828410, EPI_ISL_2828411, EPI_ISL_2828412, EPI_ISL_2828413, EPI_ISL_2828414 | SC (UCO) Igiene e Sanità Pubblica, ASUGI, Trieste | ARGO Laboratorio Genomica ed Epigenomica | Licastro D, Dal Monego S, Degasperi M, Marcello A, Segat L, Piscianz E, Busetti M, Fontana F, Breda C, Barbone F, D'Agaro P |
| EPI_ISL_2828415, EPI_ISL_2828416, EPI_ISL_2828417, EPI_ISL_2828418 | SC (UCO) Igiene e Sanità Pubblica, ASUGI, Trieste and SC Microbiologia e Virologia, ASFO, Pordenone | ARGO Laboratorio Genomica ed Epigenomica | Licastro D, Dal Monego S, Degasperi M, Marcello A, Segat L, De Rosa R, Breda C, Barbone F, D'Agaro P |
| EPI_ISL_2828419, EPI_ISL_2828420, EPI_ISL_2828421, EPI_ISL_2828422, EPI_ISL_2828423, EPI_ISL_2828424, EPI_ISL_2828425, EPI_ISL_2828426, EPI_ISL_2828427, EPI_ISL_2828428, EPI_ISL_2828429, EPI_ISL_2828430, EPI_ISL_2828431, EPI_ISL_2828432, EPI_ISL_2828433, EPI_ISL_2828434, EPI_ISL_2828435, EPI_ISL_2828436, EPI_ISL_2828437, EPI_ISL_2828438, EPI_ISL_2828439, EPI_ISL_2828440, EPI_ISL_2828441, EPI_ISL_2828442, EPI_ISL_2828443, EPI_ISL_2828444 | | | |
| see above | SC (UCO) Igiene e Sanità Pubblica, ASUGI, Trieste | ARGO Laboratorio Genomica ed Epigenomica | Licastro D, Dal Monego S, Degasperi M, Marcello A, Segat L, Piscianz E, Busetti M, Fontana F, Breda C, Barbone F, D'Agaro P |
| EPI_ISL_2828445, EPI_ISL_2828446, EPI_ISL_2828447, EPI_ISL_2828448, EPI_ISL_2828449, EPI_ISL_2828450, EPI_ISL_2828451, EPI_ISL_2828452, EPI_ISL_2828453, EPI_ISL_2828454, EPI_ISL_2828455, EPI_ISL_2828456, EPI_ISL_2828457, EPI_ISL_2828458 | | | |
| see above | SC (UCO) Igiene e Sanità Pubblica, ASUGI, Trieste and SC Microbiologia e Virologia, ASFO, Pordenone | ARGO Laboratorio Genomica ed Epigenomica | Licastro D, Dal Monego S, Degasperi M, Marcello A, Segat L, De Rosa R, Breda C, Barbone F, D'Agaro P |
| EPI_ISL_2828459, EPI_ISL_2828460, EPI_ISL_2828461, EPI_ISL_2828462, EPI_ISL_2828463, EPI_ISL_2828464, EPI_ISL_2828465, EPI_ISL_2828466, EPI_ISL_2828467, EPI_ISL_2828468, EPI_ISL_2828469, EPI_ISL_2828470, EPI_ISL_2828471 | | | |
| see above | SC (UCO) Igiene e Sanità Pubblica, ASUGI, Trieste | ARGO Laboratorio Genomica ed Epigenomica | Licastro D, Dal Monego S, Degasperi M, Marcello A, Segat L, Piscianz E, Busetti M, Fontana F, Breda C, Barbone F, D'Agaro P |
| EPI_ISL_2828668 | FoxEs Enterprise Solutions | Instituto Adolfo Lutz, Interdiciplinary Procedures Center, Strategic Laboratory | Claudio Tavares Sacchi, Claudia Regina Gonçalves, Erica Valessa Ramos Gomes, Karoline Rodrigues Campos, Caio Vinicius Dias Lopes |
| EPI_ISL_2828669, EPI_ISL_2828670, EPI_ISL_2828671, EPI_ISL_2828672, EPI_ISL_2828673, EPI_ISL_2828674, EPI_ISL_2828675, EPI_ISL_2828676, EPI_ISL_2828677, EPI_ISL_2828678, EPI_ISL_2828679, EPI_ISL_2828680, EPI_ISL_2828681, EPI_ISL_2828682, EPI_ISL_2828683 | | | |
| see above | Instituto Adolfo Lutz - Regional de Santo Andre | Instituto Adolfo Lutz, Interdiciplinary Procedures Center, Strategic Laboratory | Claudio Tavares Sacchi, Claudia Regina Gonçalves, Erica Valessa Ramos Gomes, Karoline Rodrigues Campos, Caio Vinicius Dias Lopes |
| EPI_ISL_2828684, EPI_ISL_2828685 | Instituto Adolfo Lutz - Regional de Campinas | Instituto Adolfo Lutz, Interdiciplinary Procedures Center, Strategic Laboratory | Claudio Tavares Sacchi, Claudia Regina Gonçalves, Erica Valessa Ramos Gomes, Karoline Rodrigues Campos, Caio Vinicius Dias Lopes |
| EPI_ISL_2832356, EPI_ISL_2832358, EPI_ISL_2832360 | Department of Virology, Istituto Zooprofilattico Sperimentale del Lazio e della Toscana (IZSLT) | Department of General Diagnostics; Department of Virology; Istituto Zooprofilattico Sperimentale del Lazio e della Toscana (IZSLT) | Patricia Alba, Giuseppe Manna, Elena L. Diaconu, Fabiola Feltrin, Raffaela Conti, Teresa Scicluna, Virginia Carfora, Antonella Cersini, Alessia Franco, Antonio Battisti. |
| EPI_ISL_2833731, EPI_ISL_2833734, EPI_ISL_2833735, EPI_ISL_2833738, EPI_ISL_2833739 | MOH - Jaber Al-Ahmad Hospital (Innovation Research Laboratory) | MOH - Jaber Al-Ahmad Hospital (Innovation Research Laboratory) | Salman Al-Sabah , Mohammad Alghounaim |
| EPI_ISL_2833785, EPI_ISL_2833788, EPI_ISL_2833789, EPI_ISL_2833790 | University of Oregon COVID-19 MAP Laboratory | University of Oregon Genomics and Cell Characterization Core Facility (GC3F) | Douglas Turnbull, Ariana White, Jeff Bishop, Jason Carriere, Jason Sydes, Peter Batzel |
| EPI_ISL_2834141 | St James's Hospital, Dublin 8, Ireland. | ELDA biotech | Deirdre M. Broderick, Peter R. Flanagan, Brendan Crowley |
| EPI_ISL_2834159, EPI_ISL_2834161, EPI_ISL_2834162, EPI_ISL_2834163, EPI_ISL_2834165, EPI_ISL_2834169, EPI_ISL_2834170, EPI_ISL_2834171, EPI_ISL_2834173, EPI_ISL_2834174, EPI_ISL_2834178, EPI_ISL_2834181, EPI_ISL_2834184, EPI_ISL_2834185, EPI_ISL_2834186, EPI_ISL_2834188, EPI_ISL_2834192, EPI_ISL_2834193, EPI_ISL_2834194, EPI_ISL_2834195, EPI_ISL_2834196, EPI_ISL_2834197, EPI_ISL_2834198, EPI_ISL_2834199, EPI_ISL_2834202, EPI_ISL_2834204, EPI_ISL_2834205, EPI_ISL_2834206 | | | |
| see above | Baylor Scott & White-Temple | Baylor Scott & White-Temple | Ari Rao, Linden Morales, Kimberly Walker, Marcus Volz, Shelby Johnson |
| EPI_ISL_2834733, EPI_ISL_2834734, EPI_ISL_2834735, EPI_ISL_2834736, EPI_ISL_2834737, EPI_ISL_2834738, EPI_ISL_2834739, EPI_ISL_2834740, EPI_ISL_2834741, EPI_ISL_2834742, EPI_ISL_2834743, EPI_ISL_2834744 | | | |
| see above | Ochsner Health | Bioinfoexperts, LLC | Judy Crabtree, Amy Feehan, Rebecca Rose, David J. Nolan, Tessa LaFleur, Julia-Garcia-Diaz, Lucio Miele, Susanna L. Lamers |
| EPI_ISL_2834925, EPI_ISL_2834926, EPI_ISL_2834927 | Central Health Laboratory/Airport Health Laboratory | Central Health Laboratory ,Victoria Hospital, Candos,Ministry of Health and Wellness, Mauritius | Manraj SS, Sonoo J, Patoo M, Bahadoor BS, Mathur H, Sujeewon C, Jannoo N, Ramuth M |
| EPI_ISL_2835005, EPI_ISL_2835006 | LESP Ciudad de Mexico | Instituto de Diagnostico y Referencia Epidemiologicos (INDRE) | Claudia Wong-Arambula, Abril Rodriguez-Maldonado, Vanessa Rivero-Arredondo, Ariadna Medina-Benitez, Joaquin Quiroz-Mercado, Sergio Rangel-Guerrero, Natividad Cruz-Ortiz, Tatiana Nunez-Garcia, Gisela Barrera-Badillo, Lucia Hernandez-Rivas, Irma Lopez-Martinez, Ernesto Ramirez-Gonzalez. |
| EPI_ISL_2835007 | LESP Veracruz | Instituto de Diagnostico y Referencia Epidemiologicos (INDRE) | Claudia Wong-Arambula, Abril Rodriguez-Maldonado, Vanessa Rivero-Arredondo, Ariadna Medina-Benitez, Joaquin Quiroz-Mercado, Sergio Rangel-Guerrero, Natividad Cruz-Ortiz, Tatiana Nunez-Garcia, Gisela Barrera-Badillo, Lucia Hernandez-Rivas, Irma Lopez-Martinez, Ernesto Ramirez-Gonzalez. |
| EPI_ISL_2835008 | LESP Morelos | Instituto de Diagnostico y Referencia Epidemiologicos (INDRE) | Claudia Wong-Arambula, Abril Rodriguez-Maldonado, Vanessa Rivero-Arredondo, Ariadna Medina-Benitez, Joaquin Quiroz-Mercado, Sergio Rangel-Guerrero, Natividad Cruz-Ortiz, Tatiana Nunez-Garcia, Gisela Barrera-Badillo, Lucia Hernandez-Rivas, Irma Lopez-Martinez, Ernesto Ramirez-Gonzalez. |
| EPI_ISL_2835009, EPI_ISL_2835010, EPI_ISL_2835011, EPI_ISL_2835012, | LESP Ciudad de Mexico | Instituto de Diagnostico y Referencia Epidemiologicos (INDRE) | Claudia Wong-Arambula, Abril Rodriguez-Maldonado, Vanessa Rivero-Arredondo, Ariadna Medina-Benitez, Joaquin Quiroz-Mercado, Sergio Rangel-Guerrero, Natividad Cruz-Ortiz, Tatiana Nunez-Garcia, Gisela Barrera-Badillo, Lucia Hernandez-Rivas, Irma Lopez-Martinez, Ernesto |

| | | | |
|---|---|---|---|
| EPI_ISL_2835013 | | | Ramirez-Gonzalez. |
| EPI_ISL_2835014 | LESP Queretaro | Instituto de Diagnostico y Referencia Epidemiologicos (INDRE) | Claudia Wong-Arambula, Abril Rodriguez-Maldonado, Vanessa Rivero-Arredondo, Ariadna Medina-Benitez, Joaquin Quiroz-Mercado, Sergio Rangel-Guerrero, Natividad Cruz-Ortiz, Tatiana Nunez-Garcia, Gisela Barrera-Badillo, Lucia Hernandez-Rivas, Irma Lopez-Martinez, Ernesto Ramirez-Gonzalez. |
| EPI_ISL_2835016 | LESP Jalisco | Instituto de Diagnostico y Referencia Epidemiologicos (INDRE) | Claudia Wong-Arambula, Abril Rodriguez-Maldonado, Vanessa Rivero-Arredondo, Ariadna Medina-Benitez, Joaquin Quiroz-Mercado, Sergio Rangel-Guerrero, Natividad Cruz-Ortiz, Tatiana Nunez-Garcia, Gisela Barrera-Badillo, Lucia Hernandez-Rivas, Irma Lopez-Martinez, Ernesto Ramirez-Gonzalez. |
| EPI_ISL_2835017, EPI_ISL_2835018, EPI_ISL_2835019, EPI_ISL_2835020, EPI_ISL_2835021 | LESP Sinaloa | Instituto de Diagnostico y Referencia Epidemiologicos (INDRE) | Claudia Wong-Arambula, Abril Rodriguez-Maldonado, Vanessa Rivero-Arredondo, Ariadna Medina-Benitez, Joaquin Quiroz-Mercado, Sergio Rangel-Guerrero, Natividad Cruz-Ortiz, Tatiana Nunez-Garcia, Gisela Barrera-Badillo, Lucia Hernandez-Rivas, Irma Lopez-Martinez, Ernesto Ramirez-Gonzalez. |
| EPI_ISL_2835022 | LESP Aguascalientes | Instituto de Diagnostico y Referencia Epidemiologicos (INDRE) | Claudia Wong-Arambula, Abril Rodriguez-Maldonado, Vanessa Rivero-Arredondo, Ariadna Medina-Benitez, Joaquin Quiroz-Mercado, Sergio Rangel-Guerrero, Natividad Cruz-Ortiz, Tatiana Nunez-Garcia, Gisela Barrera-Badillo, Lucia Hernandez-Rivas, Irma Lopez-Martinez, Ernesto Ramirez-Gonzalez. |
| EPI_ISL_2835023 | LESP Baja California | Instituto de Diagnostico y Referencia Epidemiologicos (INDRE) | Claudia Wong-Arambula, Abril Rodriguez-Maldonado, Vanessa Rivero-Arredondo, Ariadna Medina-Benitez, Joaquin Quiroz-Mercado, Sergio Rangel-Guerrero, Natividad Cruz-Ortiz, Tatiana Nunez-Garcia, Gisela Barrera-Badillo, Lucia Hernandez-Rivas, Irma Lopez-Martinez, Ernesto Ramirez-Gonzalez. |
| EPI_ISL_2835024 | LESP Campeche | Instituto de Diagnostico y Referencia Epidemiologicos (INDRE) | Claudia Wong-Arambula, Abril Rodriguez-Maldonado, Vanessa Rivero-Arredondo, Ariadna Medina-Benitez, Joaquin Quiroz-Mercado, Sergio Rangel-Guerrero, Natividad Cruz-Ortiz, Tatiana Nunez-Garcia, Gisela Barrera-Badillo, Lucia Hernandez-Rivas, Irma Lopez-Martinez, Ernesto Ramirez-Gonzalez. |
| EPI_ISL_2835025 | LESP Sonora | Instituto de Diagnostico y Referencia Epidemiologicos (INDRE) | Claudia Wong-Arambula, Abril Rodriguez-Maldonado, Vanessa Rivero-Arredondo, Ariadna Medina-Benitez, Joaquin Quiroz-Mercado, Sergio Rangel-Guerrero, Maribel Gonzalez-Villa, Natividad Cruz-Ortiz, Tatiana Nunez-Garcia, Gisela Barrera-Badillo, Lucia Hernandez-Rivas, Irma Lopez-Martinez, Ernesto Ramirez-Gonzalez. |
| EPI_ISL_2835026, EPI_ISL_2835027, EPI_ISL_2835028, EPI_ISL_2835029, EPI_ISL_2835030, EPI_ISL_2835031, EPI_ISL_2835032, EPI_ISL_2835033, EPI_ISL_2835034, EPI_ISL_2835035, EPI_ISL_2835036, EPI_ISL_2835037, EPI_ISL_2835038, EPI_ISL_2835039 | | | |
| see above | LESP Ciudad de Mexico | Instituto de Diagnostico y Referencia Epidemiologicos (INDRE) | Claudia Wong-Arambula, Abril Rodriguez-Maldonado, Vanessa Rivero-Arredondo, Ariadna Medina-Benitez, Joaquin Quiroz-Mercado, Sergio Rangel-Guerrero, Maribel Gonzalez-Villa, Natividad Cruz-Ortiz, Tatiana Nunez-Garcia, Gisela Barrera-Badillo, Lucia Hernandez-Rivas, Irma Lopez-Martinez, Ernesto Ramirez-Gonzalez. |
| EPI_ISL_2835040 | LESP Estado de Mexico | Instituto de Diagnostico y Referencia Epidemiologicos (INDRE) | Claudia Wong-Arambula, Abril Rodriguez-Maldonado, Vanessa Rivero-Arredondo, Ariadna Medina-Benitez, Joaquin Quiroz-Mercado, Sergio Rangel-Guerrero, Maribel Gonzalez-Villa, Natividad Cruz-Ortiz, Tatiana Nunez-Garcia, Gisela Barrera-Badillo, Lucia Hernandez-Rivas, Irma Lopez-Martinez, Ernesto Ramirez-Gonzalez. |
| EPI_ISL_2835041, EPI_ISL_2835042, EPI_ISL_2835043 | LESP Ciudad de Mexico | Instituto de Diagnostico y Referencia Epidemiologicos (INDRE) | Claudia Wong-Arambula, Abril Rodriguez-Maldonado, Vanessa Rivero-Arredondo, Ariadna Medina-Benitez, Joaquin Quiroz-Mercado, Sergio Rangel-Guerrero, Maribel Gonzalez-Villa, Natividad Cruz-Ortiz, Tatiana Nunez-Garcia, Gisela Barrera-Badillo, Lucia Hernandez-Rivas, Irma Lopez-Martinez, Ernesto Ramirez-Gonzalez. |
| EPI_ISL_2835044 | LESP Nuevo Leon | Instituto de Diagnostico y Referencia Epidemiologicos (INDRE) | Claudia Wong-Arambula, Abril Rodriguez-Maldonado, Vanessa Rivero-Arredondo, Ariadna Medina-Benitez, Joaquin Quiroz-Mercado, Sergio Rangel-Guerrero, Maribel Gonzalez-Villa, Natividad Cruz-Ortiz, Tatiana Nunez-Garcia, Gisela Barrera-Badillo, Lucia Hernandez-Rivas, Irma Lopez-Martinez, Ernesto Ramirez-Gonzalez. |
| EPI_ISL_2835045 | LESP Chihuahua | Instituto de Diagnostico y Referencia Epidemiologicos (INDRE) | Claudia Wong-Arambula, Abril Rodriguez-Maldonado, Vanessa Rivero-Arredondo, Ariadna Medina-Benitez, Joaquin Quiroz-Mercado, Sergio Rangel-Guerrero, Maribel Gonzalez-Villa, Natividad Cruz-Ortiz, Tatiana Nunez-Garcia, Gisela Barrera-Badillo, Lucia Hernandez-Rivas, Irma Lopez-Martinez, Ernesto Ramirez-Gonzalez. |
| EPI_ISL_2835048 | LESP Tamaulipas | Instituto de Diagnostico y Referencia Epidemiologicos (INDRE) | Claudia Wong-Arambula, Abril Rodriguez-Maldonado, Vanessa Rivero-Arredondo, Ariadna Medina-Benitez, Joaquin Quiroz-Mercado, Sergio Rangel-Guerrero, Maribel Gonzalez-Villa, Natividad Cruz-Ortiz, Tatiana Nunez-Garcia, Gisela Barrera-Badillo, Lucia Hernandez-Rivas, Irma Lopez-Martinez, Ernesto Ramirez-Gonzalez. |
| EPI_ISL_2835053 | LESP Ciudad de Mexico | Instituto de Diagnostico y Referencia Epidemiologicos (INDRE) | Claudia Wong-Arambula, Abril Rodriguez-Maldonado, Vanessa Rivero-Arredondo, Ariadna Medina-Benitez, Joaquin Quiroz-Mercado, Sergio Rangel-Guerrero, Maribel Gonzalez-Villa, Natividad Cruz-Ortiz, Tatiana Nunez-Garcia, Gisela Barrera-Badillo, Lucia Hernandez-Rivas, Irma Lopez-Martinez, Ernesto Ramirez-Gonzalez. |
| EPI_ISL_2835056 | LESP Chihuahua | Instituto de Diagnostico y Referencia Epidemiologicos (INDRE) | Claudia Wong-Arambula, Abril Rodriguez-Maldonado, Vanessa Rivero-Arredondo, Ariadna Medina-Benitez, Joaquin Quiroz-Mercado, Sergio Rangel-Guerrero, Maribel Gonzalez-Villa, Natividad Cruz-Ortiz, Tatiana Nunez-Garcia, Gisela Barrera-Badillo, Lucia Hernandez-Rivas, Irma Lopez-Martinez, Ernesto Ramirez-Gonzalez. |
| EPI_ISL_2835057, EPI_ISL_2835058, EPI_ISL_2835059, EPI_ISL_2835060, EPI_ISL_2835061, EPI_ISL_2835062, EPI_ISL_2835063, EPI_ISL_2835064, EPI_ISL_2835065, EPI_ISL_2835066, EPI_ISL_2835067, EPI_ISL_2835068, EPI_ISL_2835069, EPI_ISL_2835070 | | | |
| see above | LESP Tabasco | Instituto de Diagnostico y Referencia Epidemiologicos (INDRE) | Claudia Wong-Arambula, Abril Rodriguez-Maldonado, Vanessa Rivero-Arredondo, Ariadna Medina-Benitez, Joaquin Quiroz-Mercado, Sergio Rangel-Guerrero, Maribel Gonzalez-Villa, Natividad Cruz-Ortiz, Tatiana Nunez-Garcia, Gisela Barrera-Badillo, Lucia Hernandez-Rivas, Irma Lopez-Martinez, Ernesto Ramirez-Gonzalez. |
| EPI_ISL_2835071 | LESP Aguascalientes | Instituto de Diagnostico y Referencia Epidemiologicos (INDRE) | Claudia Wong-Arambula, Abril Rodriguez-Maldonado, Vanessa Rivero-Arredondo, Ariadna Medina-Benitez, Joaquin Quiroz-Mercado, Sergio Rangel-Guerrero, Maribel Gonzalez-Villa, Natividad Cruz-Ortiz, Tatiana Nunez-Garcia, Gisela Barrera-Badillo, Lucia Hernandez-Rivas, Irma Lopez-Martinez, Ernesto Ramirez-Gonzalez. |
| EPI_ISL_2835072, EPI_ISL_2835073 | LESP Sonora | Instituto de Diagnostico y Referencia Epidemiologicos (INDRE) | Claudia Wong-Arambula, Abril Rodriguez-Maldonado, Vanessa Rivero-Arredondo, Ariadna Medina-Benitez, Joaquin Quiroz-Mercado, Sergio Rangel-Guerrero, Maribel Gonzalez-Villa, Natividad Cruz-Ortiz, Tatiana Nunez-Garcia, Gisela Barrera-Badillo, Lucia Hernandez-Rivas, Irma Lopez-Martinez, Ernesto Ramirez-Gonzalez. |
| EPI_ISL_2835074, EPI_ISL_2835075, EPI_ISL_2835076 | LESP Nuevo Leon | Instituto de Diagnostico y Referencia Epidemiologicos (INDRE) | Claudia Wong-Arambula, Abril Rodriguez-Maldonado, Vanessa Rivero-Arredondo, Ariadna Medina-Benitez, Joaquin Quiroz-Mercado, Sergio Rangel-Guerrero, Maribel Gonzalez-Villa, Natividad Cruz-Ortiz, Tatiana Nunez-Garcia, Gisela Barrera-Badillo, Lucia Hernandez-Rivas, Irma Lopez-Martinez, Ernesto Ramirez-Gonzalez. |
| EPI_ISL_2835077 | LESP Sinaloa | Instituto de Diagnostico y Referencia Epidemiologicos (INDRE) | Claudia Wong-Arambula, Abril Rodriguez-Maldonado, Vanessa Rivero-Arredondo, Ariadna Medina-Benitez, Joaquin Quiroz-Mercado, Sergio Rangel-Guerrero, Maribel Gonzalez-Villa, Natividad Cruz-Ortiz, Tatiana Nunez-Garcia, Gisela Barrera-Badillo, Lucia Hernandez-Rivas, Irma Lopez-Martinez, Ernesto Ramirez-Gonzalez. |
| EPI_ISL_2835078, EPI_ISL_2835079, EPI_ISL_2835080, EPI_ISL_2835081, EPI_ISL_2835082 | LESP Tabasco | Instituto de Diagnostico y Referencia Epidemiologicos (INDRE) | Claudia Wong-Arambula, Abril Rodriguez-Maldonado, Vanessa Rivero-Arredondo, Ariadna Medina-Benitez, Joaquin Quiroz-Mercado, Sergio Rangel-Guerrero, Maribel Gonzalez-Villa, Natividad Cruz-Ortiz, Tatiana Nunez-Garcia, Gisela Barrera-Badillo, Lucia Hernandez-Rivas, Irma Lopez-Martinez, Ernesto Ramirez-Gonzalez. |
| EPI_ISL_2835085 | LESP Tamaulipas | Instituto de Diagnostico y Referencia Epidemiologicos (INDRE) | Claudia Wong-Arambula, Abril Rodriguez-Maldonado, Vanessa Rivero-Arredondo, Ariadna Medina-Benitez, Joaquin Quiroz-Mercado, Sergio Rangel-Guerrero, Maribel Gonzalez-Villa, Natividad Cruz-Ortiz, Tatiana Nunez-Garcia, Gisela Barrera-Badillo, Lucia Hernandez-Rivas, Irma Lopez-Martinez, Ernesto Ramirez-Gonzalez. |
| EPI_ISL_2835086, EPI_ISL_2835087 | LESP Campeche | Instituto de Diagnostico y Referencia Epidemiologicos (INDRE) | Claudia Wong-Arambula, Abril Rodriguez-Maldonado, Vanessa Rivero-Arredondo, Ariadna Medina-Benitez, Joaquin Quiroz-Mercado, Sergio Rangel-Guerrero, Maribel Gonzalez-Villa, Natividad Cruz-Ortiz, Tatiana Nunez-Garcia, Gisela Barrera-Badillo, Lucia Hernandez-Rivas, Irma Lopez-Martinez, Ernesto Ramirez-Gonzalez. |
| EPI_ISL_2835088 | LESP Tabasco | Instituto de Diagnostico y Referencia Epidemiologicos (INDRE) | Claudia Wong-Arambula, Abril Rodriguez-Maldonado, Vanessa Rivero-Arredondo, Ariadna Medina-Benitez, Joaquin Quiroz-Mercado, Sergio Rangel-Guerrero, Maribel Gonzalez-Villa, Natividad Cruz-Ortiz, Tatiana Nunez-Garcia, Gisela Barrera-Badillo, Lucia Hernandez-Rivas, Irma |

| | | | |
|---|---|---|---|
| | | | Lopez-Martinez, Ernesto Ramirez-Gonzalez. |
| EPI_ISL_2835089 | LESP Queretaro | Instituto de Diagnostico y Referencia Epidemiologicos (INDRE) | Claudia Wong-Arambula, Abril Rodriguez-Maldonado, Vanessa Rivero-Arredondo, Ariadna Medina-Benitez, Joaquin Quiroz-Mercado, Sergio Rangel-Guerrero, Maribel Gonzalez-Villa, Natividad Cruz-Ortiz, Tatiana Nunez-Garcia, Gisela Barrera-Badillo, Lucia Hernandez-Rivas, Irma Lopez-Martinez, Ernesto Ramirez-Gonzalez. |
| EPI_ISL_2835552, EPI_ISL_2835553, EPI_ISL_2835554, EPI_ISL_2835555, EPI_ISL_2835556, EPI_ISL_2835557, EPI_ISL_2835558, EPI_ISL_2835559, EPI_ISL_2835560, EPI_ISL_2835561, EPI_ISL_2835562, EPI_ISL_2835563, EPI_ISL_2835564, EPI_ISL_2835565 | | | |
| see above | Sharp HealthCare Laboratory | Andersen lab at Scripps Research | SEARCH Alliance San Diego with Aaron Harding, Jacquelyn Berumen, Cathy Woerle, Liam McGinnis, Art Mendoza, Omid Bakhtar |
| EPI_ISL_2835685 | Clinical Microbiology Laboratory, Faculty of Medicine, Universitas Indonesia | Faculty of Medicine, Universitas Indonesia | Fera Ibrahim, Fadilah, Andi Yasmon, Yulia Rosa, Rafika Indah Paramita, Khaerunissa Anbar Istiadi, Linda Erlina, Hana Apsari Pawestri, Vivi Setiawaty, Fithriyah, Beti Ernawati, Pratiwi Sudarmono, Budi Wiweko, Badriul Hegar, Anis Karuniawati, Ari Fahrial Syam |
| EPI_ISL_2835704, EPI_ISL_2835705, EPI_ISL_2835706, EPI_ISL_2835707, EPI_ISL_2835708, EPI_ISL_2835709, EPI_ISL_2835710, EPI_ISL_2835711, EPI_ISL_2835712, EPI_ISL_2835714, EPI_ISL_2835715, EPI_ISL_2835716, EPI_ISL_2835717, EPI_ISL_2835718, EPI_ISL_2835719, EPI_ISL_2835720, EPI_ISL_2835721, EPI_ISL_2835722, EPI_ISL_2835723, EPI_ISL_2835724, EPI_ISL_2835726, EPI_ISL_2835727, EPI_ISL_2835728, EPI_ISL_2835729, EPI_ISL_2835730, EPI_ISL_2835731, EPI_ISL_2835732, EPI_ISL_2835734, EPI_ISL_2835735, EPI_ISL_2835736 | | | |
| see above | Labo Analyses Med | National Reference Center for Viruses of Respiratory Infections, Institut Pasteur, Paris | Marion Barbet, Sylvie Behillil, Méline Bizard, Angela Brisebarre, Camille Capel, Vincent Enouf, Louise Lefrançois, Frédéric Lemoine, Christophe Malabat, Corinne Maufrais, Etienne Simon-Lorière, Maud Vanpeene, Sylvie Van der Werf ,BéNéDicte Roquebert |
| EPI_ISL_2835737 | Hospital | National Reference Center for Viruses of Respiratory Infections, Institut Pasteur, Paris | Marion Barbet, Sylvie Behillil, Méline Bizard, Angela Brisebarre, Camille Capel, Vincent Enouf, Louise Lefrançois, Frédéric Lemoine, Christophe Malabat, Corinne Maufrais, Etienne Simon-Lorière, Maud Vanpeene, Sylvie Van der Werf ,JéRome Guinard |
| EPI_ISL_2835739 | Hospital | National Reference Center for Viruses of Respiratory Infections, Institut Pasteur, Paris | Marion Barbet, Sylvie Behillil, Méline Bizard, Angela Brisebarre, Camille Capel, Vincent Enouf, Louise Lefrançois, Frédéric Lemoine, Christophe Malabat, Corinne Maufrais, Etienne Simon-Lorière, Maud Vanpeene, Sylvie Van der Werf ,Laurent Bret |
| EPI_ISL_2835740 | Hospital | National Reference Center for Viruses of Respiratory Infections, Institut Pasteur, Paris | Marion Barbet, Sylvie Behillil, Méline Bizard, Angela Brisebarre, Camille Capel, Vincent Enouf, Louise Lefrançois, Frédéric Lemoine, Christophe Malabat, Corinne Maufrais, Etienne Simon-Lorière, Maud Vanpeene, Sylvie Van der Werf ,Benjamin Maneglie |
| EPI_ISL_2835741 | Hospital | National Reference Center for Viruses of Respiratory Infections, Institut Pasteur, Paris | Marion Barbet, Sylvie Behillil, Méline Bizard, Angela Brisebarre, Camille Capel, Vincent Enouf, Louise Lefrançois, Frédéric Lemoine, Christophe Malabat, Corinne Maufrais, Etienne Simon-Lorière, Maud Vanpeene, Sylvie Van der Werf ,ValéRie Serazin |
| EPI_ISL_2835742 | Hospital | National Reference Center for Viruses of Respiratory Infections, Institut Pasteur, Paris | Marion Barbet, Sylvie Behillil, Méline Bizard, Angela Brisebarre, Camille Capel, Vincent Enouf, Louise Lefrançois, Frédéric Lemoine, Christophe Malabat, Corinne Maufrais, Etienne Simon-Lorière, Maud Vanpeene, Sylvie Van der Werf ,Laura Djamdjian |
| EPI_ISL_2835743 | Labo Analyses Med | National Reference Center for Viruses of Respiratory Infections, Institut Pasteur, Paris | Marion Barbet, Sylvie Behillil, Méline Bizard, Angela Brisebarre, Camille Capel, Vincent Enouf, Louise Lefrançois, Frédéric Lemoine, Christophe Malabat, Corinne Maufrais, Etienne Simon-Lorière, Maud Vanpeene, Sylvie Van der Werf ,Eve Haguenoer |
| EPI_ISL_2835744, EPI_ISL_2835745, EPI_ISL_2835748 | Labo Analyses Med | National Reference Center for Viruses of Respiratory Infections, Institut Pasteur, Paris | Marion Barbet, Sylvie Behillil, Méline Bizard, Angela Brisebarre, Camille Capel, Vincent Enouf, Louise Lefrançois, Frédéric Lemoine, Christophe Malabat, Corinne Maufrais, Etienne Simon-Lorière, Maud Vanpeene, Sylvie Van der Werf ,Sabine Trombert |
| EPI_ISL_2835749, EPI_ISL_2835752, EPI_ISL_2835754, EPI_ISL_2835755, EPI_ISL_2835756, EPI_ISL_2835757, EPI_ISL_2835758, EPI_ISL_2835760 | Labo Analyses Med | National Reference Center for Viruses of Respiratory Infections, Institut Pasteur, Paris | Marion Barbet, Sylvie Behillil, Méline Bizard, Angela Brisebarre, Camille Capel, Vincent Enouf, Louise Lefrançois, Frédéric Lemoine, Christophe Malabat, Corinne Maufrais, Etienne Simon-Lorière, Maud Vanpeene, Sylvie Van der Werf ,BéNéDicte Roquebert |
| EPI_ISL_2835761, EPI_ISL_2835762, EPI_ISL_2835763, EPI_ISL_2835764 | Labo Analyses Med | National Reference Center for Viruses of Respiratory Infections, Institut Pasteur, Paris | Marion Barbet, Sylvie Behillil, Méline Bizard, Angela Brisebarre, Camille Capel, Vincent Enouf, Louise Lefrançois, Frédéric Lemoine, Christophe Malabat, Corinne Maufrais, Etienne Simon-Lorière, Maud Vanpeene, Sylvie Van der Werf ,Caroline Devaux |
| EPI_ISL_2835765, EPI_ISL_2835766, EPI_ISL_2835767, EPI_ISL_2835768 | Labo Analyses Med | National Reference Center for Viruses of Respiratory Infections, Institut Pasteur, Paris | Marion Barbet, Sylvie Behillil, Méline Bizard, Angela Brisebarre, Camille Capel, Vincent Enouf, Louise Lefrançois, Frédéric Lemoine, Christophe Malabat, Corinne Maufrais, Etienne Simon-Lorière, Maud Vanpeene, Sylvie Van der Werf ,Stephanie Hainos |
| EPI_ISL_2835769 | Labo Analyses Med | National Reference Center for Viruses of Respiratory Infections, Institut Pasteur, Paris | Marion Barbet, Sylvie Behillil, Méline Bizard, Angela Brisebarre, Camille Capel, Vincent Enouf, Louise Lefrançois, Frédéric Lemoine, Christophe Malabat, Corinne Maufrais, Etienne Simon-Lorière, Maud Vanpeene, Sylvie Van der Werf ,Brieuc Gestin |
| EPI_ISL_2835770 | Labo Analyses Med | National Reference Center for Viruses of Respiratory Infections, Institut Pasteur, Paris | Marion Barbet, Sylvie Behillil, Méline Bizard, Angela Brisebarre, Camille Capel, Vincent Enouf, Louise Lefrançois, Frédéric Lemoine, Christophe Malabat, Corinne Maufrais, Etienne Simon-Lorière, Maud Vanpeene, Sylvie Van der Werf ,GréGoire Potiron |
| EPI_ISL_2835771, EPI_ISL_2835772, EPI_ISL_2835773, EPI_ISL_2835774, EPI_ISL_2835775 | Labo Analyses Med | National Reference Center for Viruses of Respiratory Infections, Institut Pasteur, Paris | Marion Barbet, Sylvie Behillil, Méline Bizard, Angela Brisebarre, Camille Capel, Vincent Enouf, Louise Lefrançois, Frédéric Lemoine, Christophe Malabat, Corinne Maufrais, Etienne Simon-Lorière, Maud Vanpeene, Sylvie Van der Werf ,BéAtrice Ricard |
| EPI_ISL_2835776, EPI_ISL_2835777, EPI_ISL_2835779, EPI_ISL_2835780, EPI_ISL_2835781, EPI_ISL_2835782, EPI_ISL_2835783 | Labo Analyses Med | National Reference Center for Viruses of Respiratory Infections, Institut Pasteur, Paris | Marion Barbet, Sylvie Behillil, Méline Bizard, Angela Brisebarre, Camille Capel, Vincent Enouf, Louise Lefrançois, Frédéric Lemoine, Christophe Malabat, Corinne Maufrais, Etienne Simon-Lorière, Maud Vanpeene, Sylvie Van der Werf ,Jean-FrançOis Comes |
| EPI_ISL_2835784, EPI_ISL_2835785, EPI_ISL_2835787, EPI_ISL_2835788, EPI_ISL_2835789, EPI_ISL_2835790, EPI_ISL_2835791, EPI_ISL_2835792, EPI_ISL_2835793, EPI_ISL_2835794, EPI_ISL_2835795, EPI_ISL_2835796, EPI_ISL_2835797, EPI_ISL_2835798, EPI_ISL_2835799, EPI_ISL_2835800, EPI_ISL_2835801, EPI_ISL_2835802, EPI_ISL_2835804, EPI_ISL_2835805, EPI_ISL_2835806, EPI_ISL_2835807 | | | |
| see above | Labo Analyses Med | National Reference Center for Viruses of Respiratory Infections, Institut Pasteur, Paris | Marion Barbet, Sylvie Behillil, Méline Bizard, Angela Brisebarre, Camille Capel, Vincent Enouf, Louise Lefrançois, Frédéric Lemoine, Christophe Malabat, Corinne Maufrais, Etienne Simon-Lorière, Maud Vanpeene, Sylvie Van der Werf ,Anne Holstein |
| EPI_ISL_2835809, EPI_ISL_2835810, EPI_ISL_2835811, EPI_ISL_2835812, EPI_ISL_2835813, EPI_ISL_2835815, EPI_ISL_2835816, EPI_ISL_2835817, EPI_ISL_2835818, EPI_ISL_2835819, EPI_ISL_2835820, EPI_ISL_2835821, EPI_ISL_2835822, EPI_ISL_2835823, EPI_ISL_2835824, EPI_ISL_2835825, EPI_ISL_2835826, EPI_ISL_2835827, EPI_ISL_2835828, EPI_ISL_2835829, EPI_ISL_2835830, EPI_ISL_2835831, EPI_ISL_2835832, EPI_ISL_2835833 | | | |
| see above | Labo Analyses Med | National Reference Center for Viruses of Respiratory Infections, Institut Pasteur, Paris | Marion Barbet, Sylvie Behillil, Méline Bizard, Angela Brisebarre, Camille Capel, Vincent Enouf, Louise Lefrançois, Frédéric Lemoine, Christophe Malabat, Corinne Maufrais, Etienne Simon-Lorière, Maud Vanpeene, Sylvie Van der Werf ,Vincent Vieillefond |
| EPI_ISL_2835835 | Hospital | National Reference Center for Viruses of Respiratory Infections, Institut Pasteur, Paris | Marion Barbet, Sylvie Behillil, Méline Bizard, Angela Brisebarre, Camille Capel, Vincent Enouf, Louise Lefrançois, Frédéric Lemoine, Christophe Malabat, Corinne Maufrais, Etienne Simon-Lorière, Maud Vanpeene, Sylvie Van der Werf ,Axelle Paquin |
| EPI_ISL_2835836, EPI_ISL_2835837 | Labo Analyses Med | National Reference Center for Viruses of Respiratory Infections, Institut Pasteur, Paris | Marion Barbet, Sylvie Behillil, Méline Bizard, Angela Brisebarre, Camille Capel, Vincent Enouf, Louise Lefrançois, Frédéric Lemoine, Christophe Malabat, Corinne Maufrais, Etienne Simon-Lorière, Maud Vanpeene, Sylvie Van der Werf ,GwéNolé Prigent |
| EPI_ISL_2835838, EPI_ISL_2835839 | Labo Analyses Med | National Reference Center for Viruses of Respiratory Infections, Institut Pasteur, Paris | Marion Barbet, Sylvie Behillil, Méline Bizard, Angela Brisebarre, Camille Capel, Vincent Enouf, Louise Lefrançois, Frédéric Lemoine, Christophe Malabat, Corinne Maufrais, Etienne Simon-Lorière, Maud Vanpeene, Sylvie Van der Werf ,Eric Grandsire |
| EPI_ISL_2835840, EPI_ISL_2835841, EPI_ISL_2835842 | Labo Analyses Med | National Reference Center for Viruses of Respiratory Infections, Institut Pasteur, Paris | Marion Barbet, Sylvie Behillil, Méline Bizard, Angela Brisebarre, Camille Capel, Vincent Enouf, Louise Lefrançois, Frédéric Lemoine, Christophe Malabat, Corinne Maufrais, Etienne Simon-Lorière, Maud Vanpeene, Sylvie Van der Werf |
| EPI_ISL_2835843, EPI_ISL_2835844, EPI_ISL_2835845 | Hospital | National Reference Center for Viruses of Respiratory Infections, Institut Pasteur, Paris | Marion Barbet, Sylvie Behillil, Méline Bizard, Angela Brisebarre, Camille Capel, Vincent Enouf, Louise Lefrançois, Frédéric Lemoine, Christophe Malabat, Corinne Maufrais, Etienne Simon-Lorière, Maud Vanpeene, Sylvie Van der Werf ,Pascale Martres |
| EPI_ISL_2835849, EPI_ISL_2835850, EPI_ISL_2835851, EPI_ISL_2835852, EPI_ISL_2835853, EPI_ISL_2835854, EPI_ISL_2835855, EPI_ISL_2835856, EPI_ISL_2835857, EPI_ISL_2835858, EPI_ISL_2835860, EPI_ISL_2835861, EPI_ISL_2835862, EPI_ISL_2835863, EPI_ISL_2835864, EPI_ISL_2835866, EPI_ISL_2835868, EPI_ISL_2835870, EPI_ISL_2835871, EPI_ISL_2835873, EPI_ISL_2835874, EPI_ISL_2835875, EPI_ISL_2835876 | | | |
| see above | Labo Analyses Med | National Reference Center for Viruses of Respiratory Infections, Institut Pasteur, Paris | Marion Barbet, Sylvie Behillil, Méline Bizard, Angela Brisebarre, Camille Capel, Vincent Enouf, Louise Lefrançois, Frédéric Lemoine, Christophe Malabat, Corinne Maufrais, Etienne Simon-Lorière, Maud Vanpeene, Sylvie Van der Werf ,Sabine Trombert |
| EPI_ISL_2835877, EPI_ISL_2835878, EPI_ISL_2835879, EPI_ISL_2835880 | Hospital | National Reference Center for Viruses of Respiratory Infections, Institut Pasteur, Paris | Marion Barbet, Sylvie Behillil, Méline Bizard, Angela Brisebarre, Camille Capel, Vincent Enouf, Louise Lefrançois, Frédéric Lemoine, Christophe Malabat, Corinne Maufrais, Etienne Simon-Lorière, Maud Vanpeene, Sylvie Van der Werf ,Sylvaine Bastian |
| EPI_ISL_2835881 | Hospital | National Reference Center for Viruses of Respiratory Infections, Institut Pasteur, Paris | Marion Barbet, Sylvie Behillil, Méline Bizard, Angela Brisebarre, Camille Capel, Vincent Enouf, Louise Lefrançois, Frédéric Lemoine, Christophe Malabat, Corinne Maufrais, Etienne Simon-Lorière, Maud Vanpeene, Sylvie Van der Werf ,Alexandra Ducancelle |

| | | | |
|---|---|---|---|
| EPI_ISL_2835882, EPI_ISL_2835883 | Labo Analyses Med | National Reference Center for Viruses of Respiratory Infections, Institut Pasteur, Paris | Marion Barbet, Sylvie Behillil, Méline Bizard, Angela Brisebarre, Camille Capel, Vincent Enouf, Louise Lefrançois, Frédéric Lemoine, Christophe Malabat, Corinne Maufrais, Etienne Simon-Lorière, Maud Vanpeene, Sylvie Van der Werf ,Claire Felloni |
| EPI_ISL_2835884 | Hospital | National Reference Center for Viruses of Respiratory Infections, Institut Pasteur, Paris | Marion Barbet, Sylvie Behillil, Méline Bizard, Angela Brisebarre, Camille Capel, Vincent Enouf, Louise Lefrançois, Frédéric Lemoine, Christophe Malabat, Corinne Maufrais, Etienne Simon-Lorière, Maud Vanpeene, Sylvie Van der Werf ,CéLine Bressollette |
| EPI_ISL_2835886, EPI_ISL_2835887, EPI_ISL_2835888, EPI_ISL_2835889, EPI_ISL_2835890, EPI_ISL_2835891, EPI_ISL_2835892, EPI_ISL_2835893, EPI_ISL_2835894, EPI_ISL_2835895 | Labo Analyses Med | National Reference Center for Viruses of Respiratory Infections, Institut Pasteur, Paris | Marion Barbet, Sylvie Behillil, Méline Bizard, Angela Brisebarre, Camille Capel, Vincent Enouf, Louise Lefrançois, Frédéric Lemoine, Christophe Malabat, Corinne Maufrais, Etienne Simon-Lorière, Maud Vanpeene, Sylvie Van der Werf ,Pierre-Yves Leonard |
| EPI_ISL_2835897 | Hospital | National Reference Center for Viruses of Respiratory Infections, Institut Pasteur, Paris | Marion Barbet, Sylvie Behillil, Méline Bizard, Angela Brisebarre, Camille Capel, Vincent Enouf, Louise Lefrançois, Frédéric Lemoine, Christophe Malabat, Corinne Maufrais, Etienne Simon-Lorière, Maud Vanpeene, Sylvie Van der Werf ,LéA Pilorge |
| EPI_ISL_2835898 | Hospital | National Reference Center for Viruses of Respiratory Infections, Institut Pasteur, Paris | Marion Barbet, Sylvie Behillil, Méline Bizard, Angela Brisebarre, Camille Capel, Vincent Enouf, Louise Lefrançois, Frédéric Lemoine, Christophe Malabat, Corinne Maufrais, Etienne Simon-Lorière, Maud Vanpeene, Sylvie Van der Werf ,Laurence Louvet |
| EPI_ISL_2835899, EPI_ISL_2835900, EPI_ISL_2835901, EPI_ISL_2835902, EPI_ISL_2835903, EPI_ISL_2835904 | Hospital | National Reference Center for Viruses of Respiratory Infections, Institut Pasteur, Paris | Marion Barbet, Sylvie Behillil, Méline Bizard, Angela Brisebarre, Camille Capel, Vincent Enouf, Louise Lefrançois, Frédéric Lemoine, Christophe Malabat, Corinne Maufrais, Etienne Simon-Lorière, Maud Vanpeene, Sylvie Van der Werf ,Sandrine Castelain |
| EPI_ISL_2835905 | Hospital | National Reference Center for Viruses of Respiratory Infections, Institut Pasteur, Paris | Marion Barbet, Sylvie Behillil, Méline Bizard, Angela Brisebarre, Camille Capel, Vincent Enouf, Louise Lefrançois, Frédéric Lemoine, Christophe Malabat, Corinne Maufrais, Etienne Simon-Lorière, Maud Vanpeene, Sylvie Van der Werf ,Hammas |
| EPI_ISL_2835906, EPI_ISL_2835907 | Hospital | National Reference Center for Viruses of Respiratory Infections, Institut Pasteur, Paris | Marion Barbet, Sylvie Behillil, Méline Bizard, Angela Brisebarre, Camille Capel, Vincent Enouf, Louise Lefrançois, Frédéric Lemoine, Christophe Malabat, Corinne Maufrais, Etienne Simon-Lorière, Maud Vanpeene, Sylvie Van der Werf ,Didier Mattera |
| EPI_ISL_2835908, EPI_ISL_2835909, EPI_ISL_2835910 | Hospital | National Reference Center for Viruses of Respiratory Infections, Institut Pasteur, Paris | Marion Barbet, Sylvie Behillil, Méline Bizard, Angela Brisebarre, Camille Capel, Vincent Enouf, Louise Lefrançois, Frédéric Lemoine, Christophe Malabat, Corinne Maufrais, Etienne Simon-Lorière, Maud Vanpeene, Sylvie Van der Werf ,Odile Petsaris |
| EPI_ISL_2835911, EPI_ISL_2835912 | Labo Analyses Med | National Reference Center for Viruses of Respiratory Infections, Institut Pasteur, Paris | Marion Barbet, Sylvie Behillil, Méline Bizard, Angela Brisebarre, Camille Capel, Vincent Enouf, Louise Lefrançois, Frédéric Lemoine, Christophe Malabat, Corinne Maufrais, Etienne Simon-Lorière, Maud Vanpeene, Sylvie Van der Werf ,Sabine Trombert |
| EPI_ISL_2835977 | CHC Andrée Rosemon | Institut Pasteur de la Guyane | Anne Lavergne, Dominique Rousset, Antoine Enfissi, Arielle Salmier |
| EPI_ISL_2835979, EPI_ISL_2835980, EPI_ISL_2835981, EPI_ISL_2835982, EPI_ISL_2835984 | CNR Institut Pasteur de la Guyane | Institut Pasteur de la Guyane | Anne Lavergne, Dominique Rousset, Antoine Enfissi, Arielle Salmier |
| EPI_ISL_2835986, EPI_ISL_2835987, EPI_ISL_2835988, EPI_ISL_2835989 | Laboratoire Carage | Institut Pasteur de la Guyane | Anne Lavergne, Dominique Rousset, Antoine Enfissi, Arielle Salmier |
| EPI_ISL_2838051 | Hospital Clínico Universitario Virgen de la Arrixaca | Hospital Clínico Universitario Virgen de la Arrixaca | Marina Simón y Laura Moreno |
| EPI_ISL_2838052 | Hospital Clínico Universitario Virgen de la Arrixaca | Hospital Clínico Universitario Virgen de la Arrixaca | Laura Moreno y Marina Simón |
| EPI_ISL_2840622, EPI_ISL_2840623, EPI_ISL_2840624 | S.C. Laboratorio Analisi, ASL 3 Liguria | U.O. Igiene, Ospedale Policlinico San Martino | Bruzzone Bianca, Caligiuri Patrizia, De Pace Vanessa, Domnich Alexander, Orsi Andrea, Ricucci Valentina, Spitaleri Antonino, Icardi Giancarlo on behalf of SARS-CoV-2 ITALIAN RESEARCH ENTERPRISE-(SCIRE) Collaborative Group |
| EPI_ISL_2840625 | U.O. Igiene, Ospedale Policlinico San Martino IRCCS Genova | U.O. Igiene, Ospedale Policlinico San Martino | Bruzzone Bianca, Caligiuri Patrizia, De Pace Vanessa, Domnich Alexander, Orsi Andrea, Ricucci Valentina, Icardi Giancarlo on behalf of SARS-CoV-2 ITALIAN RESEARCH ENTERPRISE-(SCIRE) Collaborative Group |
| EPI_ISL_2840626, EPI_ISL_2840627, EPI_ISL_2840628, EPI_ISL_2840629, EPI_ISL_2840630 | S.C. Laboratorio Analisi, ASL 3 Liguria | U.O. Igiene, Ospedale Policlinico San Martino | Bruzzone Bianca, Caligiuri Patrizia, De Pace Vanessa, Domnich Alexander, Orsi Andrea, Ricucci Valentina, Spitaleri Antonino, Icardi Giancarlo on behalf of SARS-CoV-2 ITALIAN RESEARCH ENTERPRISE-(SCIRE) Collaborative Group |
| EPI_ISL_2840631, EPI_ISL_2840632, EPI_ISL_2840633 | S.S.D. Microbiologia, Ospedale Santa Corona di Pietra Ligure, ASL 2 Liguria | U.O. Igiene, Ospedale Policlinico San Martino | Bruzzone Bianca, Caligiuri Patrizia, De Pace Vanessa, Domnich Alexander, Orsi Andrea, Ricucci Valentina, Valle Caterina, Icardi Giancarlo on behalf of SARS-CoV-2 ITALIAN RESEARCH ENTERPRISE-(SCIRE) Collaborative Group |
| EPI_ISL_2840634, EPI_ISL_2840635 | S.C. Patologia Clinica, Ospedale Sant'Andrea, ASL 5 | U.O. Igiene, Ospedale Policlinico San Martino | Bruzzone Bianca, Battolla Enrico, Caligiuri Patrizia, De Pace Vanessa, Domnich Alexander, Orsi Andrea, Ricucci Valentina, Icardi Giancarlo on behalf of SARS-CoV-2 ITALIAN RESEARCH ENTERPRISE-(SCIRE) Collaborative Group |
| EPI_ISL_2840636 | S.C. Laboratorio Analisi, Ospedale di Lavagna, ASL 4 Liguria | U.O. Igiene, Ospedale Policlinico San Martino | Bruzzone Bianca, Bandettini Roberto, Caligiuri Patrizia, De Pace Vanessa, Domnich Alexander, Orsi Andrea, Ricucci Valentina, Icardi Giancarlo on behalf of SARS-CoV-2 ITALIAN RESEARCH ENTERPRISE-(SCIRE) Collaborative Group |
| EPI_ISL_2840637 | U.O. Igiene, Ospedale Policlinico San Martino IRCCS Genova | U.O. Igiene, Ospedale Policlinico San Martino | Bruzzone Bianca, Caligiuri Patrizia, De Pace Vanessa, Domnich Alexander, Orsi Andrea, Ricucci Valentina, Icardi Giancarlo on behalf of SARS-CoV-2 ITALIAN RESEARCH ENTERPRISE-(SCIRE) Collaborative Group |
| EPI_ISL_2840643, EPI_ISL_2840644, EPI_ISL_2840645, EPI_ISL_2840647, EPI_ISL_2840648, EPI_ISL_2840649, EPI_ISL_2840652, EPI_ISL_2840653 | Università degli Studi di Perugia | Istituto Zooprofilattico Sperimentale dell'Abruzzo e Molise "G. Caporale" | Mencacci A, Camilloni B, Biagetti M, Giammarioli M, Lorusso A, Marcacci M, Di Domenico M, Ancora M, Curini V, Mangone I, Rinaldi A, Delli Compagni E, Scialabba S, Di Pasquale A, Cammà C, Puglia I, Calistri P, Savini G |
| EPI_ISL_2840655 | Laboratorio Analisi Osp. Città di Castello - Azienda USL Umbria1 | Istituto Zooprofilattico Sperimentale dell'Abruzzo e Molise "G. Caporale" | Malagigi V, Taccani P,Lorusso A, Marcacci M, Di Domenico M, Ancora M, Curini V, Mangone I, Rinaldi A, Delli Compagni E, Scialabba S, Di Pasquale A, Cammà C, Puglia I, Calistri P, Savini G |
| EPI_ISL_2840656 | Azienda Ospedaliera Terni | Istituto Zooprofilattico Sperimentale dell'Abruzzo e Molise "G. Caporale" | Palumbo M, Scaccetti A, Lorusso A, Marcacci M, Di Domenico M, Ancora M, Curini V, Mangone I, Rinaldi A, Delli Compagni E, Scialabba S, Di Pasquale A, Cammà C, Puglia I, Calistri P, Savini G |
| EPI_ISL_2840657, EPI_ISL_2840658, EPI_ISL_2840659, EPI_ISL_2840660, EPI_ISL_2840661, EPI_ISL_2840662, EPI_ISL_2840663, EPI_ISL_2840664, EPI_ISL_2840665, EPI_ISL_2840666, EPI_ISL_2840667, EPI_ISL_2840668, EPI_ISL_2840669 | | | |
| see above | Azienda USL Umbria 2 | Istituto Zooprofilattico Sperimentale dell'Abruzzo e Molise "G. Caporale" | Proietti A, Pistoni E, Lorusso A, Marcacci M, Di Domenico M, Ancora M, Curini V, Mangone I, Rinaldi A, Delli Compagni E, Scialabba S, Di Pasquale A, Cammà C, Puglia I, Calistri P, Savini G |
| EPI_ISL_2840672 | SIESP CHIETI - DRIVE IN VASTO | Istituto Zooprofilattico Sperimentale dell'Abruzzo e Molise "G. Caporale" | Lorusso A, Marcacci M, Di Domenico M, Ancora M, Curini V, Di Lollo Valeria, Mangone I, Rinaldi A, Delli Compagni E, Scialabba S, Caporale M, Di Pasquale A, Cammà C, Puglia I, Calistri P, Savini G |
| EPI_ISL_2840673, EPI_ISL_2840674, EPI_ISL_2840675, EPI_ISL_2840676 | SIESP CHIETI - DRIVE IN CHIETI | Istituto Zooprofilattico Sperimentale dell'Abruzzo e Molise "G. Caporale" | Lorusso A, Marcacci M, Di Domenico M, Ancora M, Curini V, Di Lollo Valeria, Mangone I, Rinaldi A, Delli Compagni E, Scialabba S, Caporale M, Di Pasquale A, Cammà C, Puglia I, Calistri P, Savini G |
| EPI_ISL_2840678 | OSPEDALE CIVILE TERAMO - CENTRO TRASFUSIONALE | Istituto Zooprofilattico Sperimentale dell'Abruzzo e Molise "G. Caporale" | Lorusso A, Marcacci M, Di Domenico M, Ancora M, Curini V, Di Lollo Valeria, Mangone I, Rinaldi A, Delli Compagni E, Scialabba S, Caporale M, Di Pasquale A, Cammà C, Puglia I, Calistri P, Savini G |
| EPI_ISL_2840679 | SIESP DIPARTIMENTO DI PREVENZIONE CHIETI | Istituto Zooprofilattico Sperimentale dell'Abruzzo e Molise "G. Caporale" | Lorusso A, Marcacci M, Di Domenico M, Ancora M, Curini V, Di Lollo Valeria, Mangone I, Rinaldi A, Delli Compagni E, Scialabba S, Caporale M, Di Pasquale A, Cammà C, Puglia I, Calistri P, Savini G |
| EPI_ISL_2840680, EPI_ISL_2840681, EPI_ISL_2840682 | SIESP CHIETI - DRIVE IN LANCIANO | Istituto Zooprofilattico Sperimentale dell'Abruzzo e Molise "G. Caporale" | Lorusso A, Marcacci M, Di Domenico M, Ancora M, Curini V, Di Lollo Valeria, Mangone I, Rinaldi A, Delli Compagni E, Scialabba S, Caporale M, Di Pasquale A, Cammà C, Puglia I, Calistri P, Savini G |
| EPI_ISL_2840683, EPI_ISL_2840684 | DIP. PREV. AVEZZANO SERVIZIO DI IGIENE EPIDEMIOLOGIA E SANITA' PUBBLICA | Istituto Zooprofilattico Sperimentale dell'Abruzzo e Molise "G. Caporale" | Lorusso A, Marcacci M, Di Domenico M, Ancora M, Curini V, Di Lollo Valeria, Mangone I, Rinaldi A, Delli Compagni E, Scialabba S, Caporale M, Di Pasquale A, Cammà C, Puglia I, Calistri P, Savini G |
| EPI_ISL_2840685 | SIESP CHIETI - DRIVE IN CHIETI | Istituto Zooprofilattico Sperimentale dell'Abruzzo e Molise "G. | Lorusso A, Marcacci M, Di Domenico M, Ancora M, Curini V, Di Lollo Valeria, Mangone I, Rinaldi A, Delli Compagni E, Scialabba S, Caporale M, Di |

| | | | |
|---|---|---|---|
| | | Caporale" | Pasquale A, Cammà C, Puglia I, Calistri P, Savini G |
| EPI_ISL_2840686, EPI_ISL_2840687, EPI_ISL_2840688, EPI_ISL_2840689, EPI_ISL_2840690, EPI_ISL_2840691, EPI_ISL_2840692, EPI_ISL_2840693, EPI_ISL_2840694, EPI_ISL_2840695, EPI_ISL_2840696, EPI_ISL_2840697, EPI_ISL_2840698, EPI_ISL_2840699, EPI_ISL_2840700, EPI_ISL_2840701, EPI_ISL_2840702, EPI_ISL_2840703 | | | |
| see above | OSPEDALE CIVILE TERAMO - CENTRO TRASFUSIONALE | Istituto Zooprofilattico Sperimentale dell'Abruzzo e Molise "G. Caporale" | Lorusso A, Marcacci M, Di Domenico M, Ancora M, Curini V, Di Lollo Valeria, Mangone I, Rinaldi A, Delli Compagni E, Scialabba S, Caporale M, Di Pasquale A, Cammà C, Puglia I, Calistri P, Savini G |
| EPI_ISL_2840704, EPI_ISL_2840705, EPI_ISL_2840706, EPI_ISL_2840707, EPI_ISL_2840708 | PO AVEZZANO - U.O.C. Laboratorio Analisi | Istituto Zooprofilattico Sperimentale dell'Abruzzo e Molise "G. Caporale" | Lorusso A, Marcacci M, Di Domenico M, Ancora M, Curini V, Di Lollo Valeria, Mangone I, Rinaldi A, Delli Compagni E, Scialabba S, Caporale M, Di Pasquale A, Cammà C, Puglia I, Calistri P, Savini G |
| EPI_ISL_2840709, EPI_ISL_2840710, EPI_ISL_2840711, EPI_ISL_2840712, EPI_ISL_2840713, EPI_ISL_2840714, EPI_ISL_2840715, EPI_ISL_2840716, EPI_ISL_2840717, EPI_ISL_2840718, EPI_ISL_2840719, EPI_ISL_2840720, EPI_ISL_2840721, EPI_ISL_2840722, EPI_ISL_2840723, EPI_ISL_2840724, EPI_ISL_2840725, EPI_ISL_2840726, EPI_ISL_2840727, EPI_ISL_2840728, EPI_ISL_2840729, EPI_ISL_2840730, EPI_ISL_2840731, EPI_ISL_2840732, EPI_ISL_2840733, EPI_ISL_2840734, EPI_ISL_2840735, EPI_ISL_2840736, EPI_ISL_2840737, EPI_ISL_2840738, EPI_ISL_2840739, EPI_ISL_2840740, EPI_ISL_2840741, EPI_ISL_2840742, EPI_ISL_2840743 | | | |
| see above | OSPEDALE CIVILE TERAMO - CENTRO TRASFUSIONALE | Istituto Zooprofilattico Sperimentale dell'Abruzzo e Molise "G. Caporale" | Lorusso A, Marcacci M, Di Domenico M, Ancora M, Curini V, Di Lollo Valeria, Mangone I, Rinaldi A, Delli Compagni E, Scialabba S, Caporale M, Di Pasquale A, Cammà C, Puglia I, Calistri P, Savini G |
| EPI_ISL_2841556 | Department of Virology, Istituto Zooprofilattico Sperimentale del Lazio e della Toscana (IZSLT) | Department of General Diagnostics; Department of Virology; Istituto Zooprofilattico Sperimentale del Lazio e della Toscana (IZSLT) | Patricia Alba, Giuseppe Manna, Elena L. Diaconu, Fabiola Feltrin, Raffaela Conti, Teresa Scicluna, Virginia Carfora, Antonella Cersini, Alessia Franco, Antonio Battisti. |
| EPI_ISL_2841872, EPI_ISL_2841875, EPI_ISL_2841876, EPI_ISL_2841877, EPI_ISL_2841878, EPI_ISL_2841881, EPI_ISL_2841882, EPI_ISL_2841883, EPI_ISL_2841884, EPI_ISL_2841886, EPI_ISL_2841887, EPI_ISL_2841888, EPI_ISL_2841889, EPI_ISL_2841892, EPI_ISL_2841893, EPI_ISL_2841895, EPI_ISL_2841896, EPI_ISL_2841897, EPI_ISL_2841898, EPI_ISL_2841899, EPI_ISL_2841900, EPI_ISL_2841901, EPI_ISL_2841903, EPI_ISL_2841904, EPI_ISL_2841905, EPI_ISL_2841906, EPI_ISL_2841907, EPI_ISL_2841908, EPI_ISL_2841909, EPI_ISL_2841910, EPI_ISL_2841911, EPI_ISL_2841912, EPI_ISL_2841920, EPI_ISL_2841921, EPI_ISL_2841942, EPI_ISL_2841943, EPI_ISL_2841944, EPI_ISL_2841945, EPI_ISL_2841946, EPI_ISL_2841947, EPI_ISL_2841948, EPI_ISL_2841949, EPI_ISL_2841966, EPI_ISL_2841967, EPI_ISL_2841968, EPI_ISL_2841969, EPI_ISL_2841970, EPI_ISL_2841973, EPI_ISL_2841975, EPI_ISL_2841976, EPI_ISL_2841977, EPI_ISL_2841978, EPI_ISL_2841979, EPI_ISL_2841982, EPI_ISL_2841983, EPI_ISL_2841984, EPI_ISL_2841985, EPI_ISL_2841986, EPI_ISL_2841987, EPI_ISL_2841988, EPI_ISL_2841989, EPI_ISL_2841991, EPI_ISL_2841992, EPI_ISL_2841993, EPI_ISL_2841994, EPI_ISL_2841995, EPI_ISL_2841996, EPI_ISL_2841997, EPI_ISL_2841998, EPI_ISL_2841999 | | | |
| see above | National Center of Infectious and Parasitic Diseases | National Center of Infectious and Parasitic Diseases | Alexiev et al |
| EPI_ISL_2842625 | Università di Parma, Laboratorio di Igiene e Sanità Pubblica | Istituto Zooprofilattico Sperimentale della Lombardia e dell'Emilia Romagna (IZSLER), Risk Analysis and Genomic Epidemiology Unit | Maria Eugenia Colucci, Licia Veronesi, Paola Affanni, Marina Morganti, Ilaria Menozzi, Erika Scaltriti, Stefano Pongolini |
| EPI_ISL_2846010, EPI_ISL_2846011, EPI_ISL_2846012, EPI_ISL_2846013, EPI_ISL_2846014, EPI_ISL_2846015, EPI_ISL_2846016, EPI_ISL_2846017, EPI_ISL_2846018, EPI_ISL_2846019, EPI_ISL_2846020, EPI_ISL_2846021, EPI_ISL_2846342, EPI_ISL_2847115, EPI_ISL_2847710, EPI_ISL_2848203, EPI_ISL_2848709, EPI_ISL_2849133, EPI_ISL_2849677, EPI_ISL_2850193, EPI_ISL_2850671, EPI_ISL_2851133, EPI_ISL_2851616, EPI_ISL_2852110 | | | |
| see above | AZ Klina | AZ Klina | Carl Vael - Lynsey Berckmans |
| EPI_ISL_2854048 | Labuan Hospital | Institute for Medical Research, Infectious Disease Research Centre, National Institutes of Health, Ministry of Health Malaysia | Suppiah J, Kamel K, Mohd Zawawi Z, Azizan MA, Ramly N, Robert F, Anasir MI, Thayan R |
| EPI_ISL_2854052 | Ampang Hospital | Institute for Medical Research, Infectious Disease Research Centre, National Institutes of Health, Ministry of Health Malaysia | Suppiah J, Kamel K, Mohd Zawawi Z, Azizan MA, Ramly N, Robert F, Anasir MI, Thayan R |
| EPI_ISL_2854053 | Kemaman Hospital | Institute for Medical Research, Infectious Disease Research Centre, National Institutes of Health, Ministry of Health Malaysia | Suppiah J, Kamel K, Mohd Zawawi Z, Azizan MA, Ramly N, Robert F, Anasir MI, Thayan R |
| EPI_ISL_2854069 | Bayan Lepas Health Clinic | Institute for Medical Research, Infectious Disease Research Centre, National Institutes of Health, Ministry of Health Malaysia | Suppiah J, Kamel K, Mohd Zawawi Z, Azizan MA, Ramly N, Robert F, Anasir MI, Thayan R |
| EPI_ISL_2854106 | Queen Elizabeth Hospital | Institute for Medical Research, Infectious Disease Research Centre, National Institutes of Health, Ministry of Health Malaysia | Suppiah J, Kamel K, Mohd Zawawi Z, Azizan MA, Ramly N, Robert F, Anasir MI, Thayan R |
| EPI_ISL_2854107 | Sultan Ismail Hospital | Institute for Medical Research, Infectious Disease Research Centre, National Institutes of Health, Ministry of Health Malaysia | Suppiah J, Kamel K, Mohd Zawawi Z, Azizan MA, Ramly N, Robert F, Anasir MI, Thayan R |
| EPI_ISL_2854109 | Tawau Hospital | Institute for Medical Research, Infectious Disease Research Centre, National Institutes of Health, Ministry of Health Malaysia | Suppiah J, Kamel K, Mohd Zawawi Z, Azizan MA, Ramly N, Robert F, Anasir MI, Thayan R |
| EPI_ISL_2854187 | Kuala Terengganu District Health Office | Institute for Medical Research, Infectious Disease Research Centre, National Institutes of Health, Ministry of Health Malaysia | Suppiah J, Kamel K, Mohd Zawawi Z, Azizan MA, Ramly N, Robert F, Anasir MI, Thayan R |
| EPI_ISL_2854188 | Marang District Health Office | Institute for Medical Research, Infectious Disease Research Centre, National Institutes of Health, Ministry of Health Malaysia | Suppiah J, Kamel K, Mohd Zawawi Z, Azizan MA, Ramly N, Robert F, Anasir MI, Thayan R |
| EPI_ISL_2854779 | KKP Kelas II Cilacap | National Institute of Health Research and Development | Hana Apsari Pawestri, Subangkit, Kartika Dewi Puspa, Arie Ardiansyah Nugraha, Hartanti Dian Ikawati, Krisna Pangesti, Triyani Soekarso, Nelly Puspandari, Vivi Setiawaty |
| EPI_ISL_2854794 | Kantor Kesehatan Pelabuhan Kelas I Medan | National Institute of Health Research and Development | Hana Apsari Pawestri, Subangkit, Kartika Dewi Puspa, Arie Ardiansyah Nugraha, Hartanti Dian Ikawati, Krisna Pangesti, Triyani Soekarso, Nelly Puspandari, Vivi Setiawaty |
| EPI_ISL_2858872 | Genomic Research Laboratory, Bangabandhu Sheikh Mujib Medical University | Genomic Research Laboratory, BSMMU | Laila Anjuman Banu, Md.Saydur Rahman, Sharadindu Kanti Sinha, Arifa Akram, Md.Sharfuddin Ahmed |
| EPI_ISL_2858935, EPI_ISL_2858936, EPI_ISL_2858937 | LESP Nuevo Leon | Instituto de Diagnostico y Referencia Epidemiologicos (INDRE) | Claudia Wong-Arambula, Abril Rodriguez-Maldonado, Vanessa Rivero-Arredondo, Ariadna Medina-Benitez, Joaquin Quiroz-Mercado, Sergio Rangel-Guerrero, Natividad Cruz-Ortiz, Tatiana Nunez-Garcia, Gisela Barrera-Badillo, Lucia Hernandez-Rivas, Irma Lopez-Martinez, Ernesto Ramirez-Gonzalez. |
| EPI_ISL_2858939, EPI_ISL_2858940 | LESP Chihuahua | Instituto de Diagnostico y Referencia Epidemiologicos (INDRE) | Claudia Wong-Arambula, Abril Rodriguez-Maldonado, Vanessa Rivero-Arredondo, Ariadna Medina-Benitez, Joaquin Quiroz-Mercado, Sergio Rangel-Guerrero, Natividad Cruz-Ortiz, Tatiana Nunez-Garcia, Gisela Barrera-Badillo, Lucia Hernandez-Rivas, Irma Lopez-Martinez, Ernesto Ramirez-Gonzalez. |
| EPI_ISL_2858941, EPI_ISL_2858942, EPI_ISL_2858943 | LESP Tamaulipas | Instituto de Diagnostico y Referencia Epidemiologicos (INDRE) | Claudia Wong-Arambula, Abril Rodriguez-Maldonado, Vanessa Rivero-Arredondo, Ariadna Medina-Benitez, Joaquin Quiroz-Mercado, Sergio Rangel-Guerrero, Natividad Cruz-Ortiz, Tatiana Nunez-Garcia, Gisela Barrera-Badillo, Lucia Hernandez-Rivas, Irma Lopez-Martinez, Ernesto Ramirez-Gonzalez. |
| EPI_ISL_2858944, EPI_ISL_2858945 | LESP Veracruz | Instituto de Diagnostico y Referencia Epidemiologicos (INDRE) | Claudia Wong-Arambula, Abril Rodriguez-Maldonado, Vanessa Rivero-Arredondo, Ariadna Medina-Benitez, Joaquin Quiroz-Mercado, Sergio Rangel-Guerrero, Natividad Cruz-Ortiz, Tatiana Nunez-Garcia, Gisela Barrera-Badillo, Lucia Hernandez-Rivas, Irma Lopez-Martinez, Ernesto Ramirez-Gonzalez. |

| | | | |
|---|---|---|---|
| EPI_ISL_2858946 | LESP Yucatan | Instituto de Diagnostico y Referencia Epidemiologicos (INDRE) | Claudia Wong-Arambula, Abril Rodriguez-Maldonado, Vanessa Rivero-Arredondo, Ariadna Medina-Benitez, Joaquin Quiroz-Mercado, Sergio Rangel-Guerrero, Natividad Cruz-Ortiz, Tatiana Nunez-Garcia, Gisela Barrera-Badillo, Lucia Hernandez-Rivas, Irma Lopez-Martinez, Ernesto Ramirez-Gonzalez. |
| EPI_ISL_2858948, EPI_ISL_2858949, EPI_ISL_2858950, EPI_ISL_2858951 | LESP Ciudad de Mexico | Instituto de Diagnostico y Referencia Epidemiologicos (INDRE) | Claudia Wong-Arambula, Abril Rodriguez-Maldonado, Vanessa Rivero-Arredondo, Ariadna Medina-Benitez, Joaquin Quiroz-Mercado, Sergio Rangel-Guerrero, Natividad Cruz-Ortiz, Tatiana Nunez-Garcia, Gisela Barrera-Badillo, Lucia Hernandez-Rivas, Irma Lopez-Martinez, Ernesto Ramirez-Gonzalez. |
| EPI_ISL_2858956 | LESP Chihuahua | Instituto de Diagnostico y Referencia Epidemiologicos (INDRE) | Claudia Wong-Arambula, Abril Rodriguez-Maldonado, Vanessa Rivero-Arredondo, Ariadna Medina-Benitez, Joaquin Quiroz-Mercado, Sergio Rangel-Guerrero, Maribel Gonzalez-Villa, Natividad Cruz-Ortiz, Tatiana Nunez-Garcia, Gisela Barrera-Badillo, Lucia Hernandez-Rivas, Irma Lopez-Martinez, Ernesto Ramirez-Gonzalez. |
| EPI_ISL_2858957, EPI_ISL_2858958, EPI_ISL_2858959 | LESP Tabasco | Instituto de Diagnostico y Referencia Epidemiologicos (INDRE) | Claudia Wong-Arambula, Abril Rodriguez-Maldonado, Vanessa Rivero-Arredondo, Ariadna Medina-Benitez, Joaquin Quiroz-Mercado, Sergio Rangel-Guerrero, Maribel Gonzalez-Villa, Natividad Cruz-Ortiz, Tatiana Nunez-Garcia, Gisela Barrera-Badillo, Lucia Hernandez-Rivas, Irma Lopez-Martinez, Ernesto Ramirez-Gonzalez. |
| EPI_ISL_2858960 | LESP Michoacan | Instituto de Diagnostico y Referencia Epidemiologicos (INDRE) | Claudia Wong-Arambula, Abril Rodriguez-Maldonado, Vanessa Rivero-Arredondo, Ariadna Medina-Benitez, Joaquin Quiroz-Mercado, Sergio Rangel-Guerrero, Maribel Gonzalez-Villa, Natividad Cruz-Ortiz, Tatiana Nunez-Garcia, Gisela Barrera-Badillo, Lucia Hernandez-Rivas, Irma Lopez-Martinez, Ernesto Ramirez-Gonzalez. |
| EPI_ISL_2858961 | LESP Sonora | Instituto de Diagnostico y Referencia Epidemiologicos (INDRE) | Claudia Wong-Arambula, Abril Rodriguez-Maldonado, Vanessa Rivero-Arredondo, Ariadna Medina-Benitez, Joaquin Quiroz-Mercado, Sergio Rangel-Guerrero, Maribel Gonzalez-Villa, Natividad Cruz-Ortiz, Tatiana Nunez-Garcia, Gisela Barrera-Badillo, Lucia Hernandez-Rivas, Irma Lopez-Martinez, Ernesto Ramirez-Gonzalez. |
| EPI_ISL_2858962, EPI_ISL_2858963, EPI_ISL_2858964 | LESP Tabasco | Instituto de Diagnostico y Referencia Epidemiologicos (INDRE) | Claudia Wong-Arambula, Abril Rodriguez-Maldonado, Vanessa Rivero-Arredondo, Ariadna Medina-Benitez, Joaquin Quiroz-Mercado, Sergio Rangel-Guerrero, Maribel Gonzalez-Villa, Natividad Cruz-Ortiz, Tatiana Nunez-Garcia, Gisela Barrera-Badillo, Lucia Hernandez-Rivas, Irma Lopez-Martinez, Ernesto Ramirez-Gonzalez. |
| EPI_ISL_2858965, EPI_ISL_2858966, EPI_ISL_2858967, EPI_ISL_2858968, EPI_ISL_2858969, EPI_ISL_2858970, EPI_ISL_2858971 | LESP Tamaulipas | Instituto de Diagnostico y Referencia Epidemiologicos (INDRE) | Claudia Wong-Arambula, Abril Rodriguez-Maldonado, Vanessa Rivero-Arredondo, Ariadna Medina-Benitez, Joaquin Quiroz-Mercado, Sergio Rangel-Guerrero, Maribel Gonzalez-Villa, Natividad Cruz-Ortiz, Tatiana Nunez-Garcia, Gisela Barrera-Badillo, Lucia Hernandez-Rivas, Irma Lopez-Martinez, Ernesto Ramirez-Gonzalez. |
| EPI_ISL_2858973 | LESP Sinaloa | Instituto de Diagnostico y Referencia Epidemiologicos (INDRE) | Claudia Wong-Arambula, Abril Rodriguez-Maldonado, Vanessa Rivero-Arredondo, Ariadna Medina-Benitez, Joaquin Quiroz-Mercado, Sergio Rangel-Guerrero, Maribel Gonzalez-Villa, Natividad Cruz-Ortiz, Tatiana Nunez-Garcia, Gisela Barrera-Badillo, Lucia Hernandez-Rivas, Irma Lopez-Martinez, Ernesto Ramirez-Gonzalez. |
| EPI_ISL_2858974, EPI_ISL_2858975, EPI_ISL_2858976, EPI_ISL_2858977, EPI_ISL_2858978 | LESP Tabasco | Instituto de Diagnostico y Referencia Epidemiologicos (INDRE) | Claudia Wong-Arambula, Abril Rodriguez-Maldonado, Vanessa Rivero-Arredondo, Ariadna Medina-Benitez, Joaquin Quiroz-Mercado, Sergio Rangel-Guerrero, Maribel Gonzalez-Villa, Natividad Cruz-Ortiz, Tatiana Nunez-Garcia, Gisela Barrera-Badillo, Lucia Hernandez-Rivas, Irma Lopez-Martinez, Ernesto Ramirez-Gonzalez. |
| EPI_ISL_2858979 | LESP Chihuahua | Instituto de Diagnostico y Referencia Epidemiologicos (INDRE) | Claudia Wong-Arambula, Abril Rodriguez-Maldonado, Vanessa Rivero-Arredondo, Ariadna Medina-Benitez, Joaquin Quiroz-Mercado, Sergio Rangel-Guerrero, Maribel Gonzalez-Villa, Natividad Cruz-Ortiz, Tatiana Nunez-Garcia, Gisela Barrera-Badillo, Lucia Hernandez-Rivas, Irma Lopez-Martinez, Ernesto Ramirez-Gonzalez. |
| EPI_ISL_2858980 | LESP Baja California | Instituto de Diagnostico y Referencia Epidemiologicos (INDRE) | Claudia Wong-Arambula, Abril Rodriguez-Maldonado, Vanessa Rivero-Arredondo, Ariadna Medina-Benitez, Joaquin Quiroz-Mercado, Sergio Rangel-Guerrero, Maribel Gonzalez-Villa, Natividad Cruz-Ortiz, Tatiana Nunez-Garcia, Gisela Barrera-Badillo, Lucia Hernandez-Rivas, Irma Lopez-Martinez, Ernesto Ramirez-Gonzalez. |
| EPI_ISL_2858981, EPI_ISL_2858982 | LESP Sonora | Instituto de Diagnostico y Referencia Epidemiologicos (INDRE) | Claudia Wong-Arambula, Abril Rodriguez-Maldonado, Vanessa Rivero-Arredondo, Ariadna Medina-Benitez, Joaquin Quiroz-Mercado, Sergio Rangel-Guerrero, Maribel Gonzalez-Villa, Natividad Cruz-Ortiz, Tatiana Nunez-Garcia, Gisela Barrera-Badillo, Lucia Hernandez-Rivas, Irma Lopez-Martinez, Ernesto Ramirez-Gonzalez. |
| EPI_ISL_2858983 | LESP Nuevo Leon | Instituto de Diagnostico y Referencia Epidemiologicos (INDRE) | Claudia Wong-Arambula, Abril Rodriguez-Maldonado, Vanessa Rivero-Arredondo, Ariadna Medina-Benitez, Joaquin Quiroz-Mercado, Sergio Rangel-Guerrero, Maribel Gonzalez-Villa, Natividad Cruz-Ortiz, Tatiana Nunez-Garcia, Gisela Barrera-Badillo, Lucia Hernandez-Rivas, Irma Lopez-Martinez, Ernesto Ramirez-Gonzalez. |
| EPI_ISL_2858987, EPI_ISL_2858988, EPI_ISL_2858989, EPI_ISL_2858990 | LESP San Luis Potosi | Instituto de Diagnostico y Referencia Epidemiologicos (INDRE) | Claudia Wong-Arambula, Abril Rodriguez-Maldonado, Vanessa Rivero-Arredondo, Ariadna Medina-Benitez, Joaquin Quiroz-Mercado, Sergio Rangel-Guerrero, Maribel Gonzalez-Villa, Natividad Cruz-Ortiz, Tatiana Nunez-Garcia, Gisela Barrera-Badillo, Lucia Hernandez-Rivas, Irma Lopez-Martinez, Ernesto Ramirez-Gonzalez. |
| EPI_ISL_2858991, EPI_ISL_2858992, EPI_ISL_2858993 | LESP Guanajuato | Instituto de Diagnostico y Referencia Epidemiologicos (INDRE) | Claudia Wong-Arambula, Abril Rodriguez-Maldonado, Vanessa Rivero-Arredondo, Ariadna Medina-Benitez, Joaquin Quiroz-Mercado, Sergio Rangel-Guerrero, Maribel Gonzalez-Villa, Natividad Cruz-Ortiz, Tatiana Nunez-Garcia, Gisela Barrera-Badillo, Lucia Hernandez-Rivas, Irma Lopez-Martinez, Ernesto Ramirez-Gonzalez. |
| EPI_ISL_2858994, EPI_ISL_2858995 | LESP Jalisco | Instituto de Diagnostico y Referencia Epidemiologicos (INDRE) | Claudia Wong-Arambula, Abril Rodriguez-Maldonado, Vanessa Rivero-Arredondo, Ariadna Medina-Benitez, Joaquin Quiroz-Mercado, Sergio Rangel-Guerrero, Maribel Gonzalez-Villa, Natividad Cruz-Ortiz, Tatiana Nunez-Garcia, Gisela Barrera-Badillo, Lucia Hernandez-Rivas, Irma Lopez-Martinez, Ernesto Ramirez-Gonzalez. |
| EPI_ISL_2858996, EPI_ISL_2858997, EPI_ISL_2858998, EPI_ISL_2858999 | LESP Tamaulipas | Instituto de Diagnostico y Referencia Epidemiologicos (INDRE) | Claudia Wong-Arambula, Abril Rodriguez-Maldonado, Vanessa Rivero-Arredondo, Ariadna Medina-Benitez, Joaquin Quiroz-Mercado, Sergio Rangel-Guerrero, Maribel Gonzalez-Villa, Natividad Cruz-Ortiz, Tatiana Nunez-Garcia, Gisela Barrera-Badillo, Lucia Hernandez-Rivas, Irma Lopez-Martinez, Ernesto Ramirez-Gonzalez. |
| EPI_ISL_2859000 | LESP Veracruz | Instituto de Diagnostico y Referencia Epidemiologicos (INDRE) | Claudia Wong-Arambula, Abril Rodriguez-Maldonado, Vanessa Rivero-Arredondo, Ariadna Medina-Benitez, Joaquin Quiroz-Mercado, Sergio Rangel-Guerrero, Maribel Gonzalez-Villa, Natividad Cruz-Ortiz, Tatiana Nunez-Garcia, Gisela Barrera-Badillo, Lucia Hernandez-Rivas, Irma Lopez-Martinez, Ernesto Ramirez-Gonzalez. |
| EPI_ISL_2859001, EPI_ISL_2859002, EPI_ISL_2859003, EPI_ISL_2859005, EPI_ISL_2859006, EPI_ISL_2859007 | LESP Yucatan | Instituto de Diagnostico y Referencia Epidemiologicos (INDRE) | Claudia Wong-Arambula, Abril Rodriguez-Maldonado, Vanessa Rivero-Arredondo, Ariadna Medina-Benitez, Joaquin Quiroz-Mercado, Sergio Rangel-Guerrero, Maribel Gonzalez-Villa, Natividad Cruz-Ortiz, Tatiana Nunez-Garcia, Gisela Barrera-Badillo, Lucia Hernandez-Rivas, Irma Lopez-Martinez, Ernesto Ramirez-Gonzalez. |
| EPI_ISL_2859008, EPI_ISL_2859009, EPI_ISL_2859010, EPI_ISL_2859011 | LESP Chiapas | Instituto de Diagnostico y Referencia Epidemiologicos (INDRE) | Claudia Wong-Arambula, Abril Rodriguez-Maldonado, Vanessa Rivero-Arredondo, Ariadna Medina-Benitez, Joaquin Quiroz-Mercado, Sergio Rangel-Guerrero, Maribel Gonzalez-Villa, Natividad Cruz-Ortiz, Tatiana Nunez-Garcia, Gisela Barrera-Badillo, Lucia Hernandez-Rivas, Irma Lopez-Martinez, Ernesto Ramirez-Gonzalez. |
| EPI_ISL_2859012 | LESP Hidalgo | Instituto de Diagnostico y Referencia Epidemiologicos (INDRE) | Claudia Wong-Arambula, Abril Rodriguez-Maldonado, Vanessa Rivero-Arredondo, Ariadna Medina-Benitez, Joaquin Quiroz-Mercado, Sergio Rangel-Guerrero, Maribel Gonzalez-Villa, Natividad Cruz-Ortiz, Tatiana Nunez-Garcia, Gisela Barrera-Badillo, Lucia Hernandez-Rivas, Irma Lopez-Martinez, Ernesto Ramirez-Gonzalez. |
| EPI_ISL_2859013, EPI_ISL_2859015 | LESP Sinaloa | Instituto de Diagnostico y Referencia Epidemiologicos (INDRE) | Claudia Wong-Arambula, Abril Rodriguez-Maldonado, Vanessa Rivero-Arredondo, Ariadna Medina-Benitez, Joaquin Quiroz-Mercado, Sergio Rangel-Guerrero, Maribel Gonzalez-Villa, Natividad Cruz-Ortiz, Tatiana Nunez-Garcia, Gisela Barrera-Badillo, Lucia Hernandez-Rivas, Irma Lopez-Martinez, Ernesto Ramirez-Gonzalez. |
| EPI_ISL_2859016, EPI_ISL_2859017, EPI_ISL_2859018, EPI_ISL_2859020, EPI_ISL_2859021, EPI_ISL_2859022, EPI_ISL_2859023, EPI_ISL_2859024, EPI_ISL_2859025, EPI_ISL_2859026, EPI_ISL_2859027 | | | |
| see above | LESP Tabasco | Instituto de Diagnostico y Referencia Epidemiologicos (INDRE) | Claudia Wong-Arambula, Abril Rodriguez-Maldonado, Vanessa Rivero-Arredondo, Ariadna Medina-Benitez, Joaquin Quiroz-Mercado, Sergio Rangel-Guerrero, Maribel Gonzalez-Villa, Natividad Cruz-Ortiz, Tatiana Nunez-Garcia, Gisela Barrera-Badillo, Lucia Hernandez-Rivas, Irma Lopez-Martinez, Ernesto Ramirez-Gonzalez. |
| EPI_ISL_2859028 | LESP Jalisco | Instituto de Diagnostico y Referencia Epidemiologicos | Claudia Wong-Arambula, Abril Rodriguez-Maldonado, Vanessa Rivero-Arredondo, Ariadna Medina-Benitez, Joaquin Quiroz-Mercado, Sergio |

| | | | |
|---|---|---|---|
| | | (INDRE) | Rangel-Guerrero, Maribel Gonzalez-Villa, Natividad Cruz-Ortiz, Tatiana Nunez-Garcia, Gisela Barrera-Badillo, Lucia Hernandez-Rivas, Irma Lopez-Martinez, Ernesto Ramirez-Gonzalez. |
| EPI_ISL_2859029, EPI_ISL_2859030, EPI_ISL_2859031, EPI_ISL_2859032, EPI_ISL_2859033 | LESP Tamaulipas | Instituto de Diagnostico y Referencia Epidemiologicos (INDRE) | Claudia Wong-Arambula, Abril Rodriguez-Maldonado, Vanessa Rivero-Arredondo, Ariadna Medina-Benitez, Joaquin Quiroz-Mercado, Sergio Rangel-Guerrero, Maribel Gonzalez-Villa, Natividad Cruz-Ortiz, Tatiana Nunez-Garcia, Gisela Barrera-Badillo, Lucia Hernandez-Rivas, Irma Lopez-Martinez, Ernesto Ramirez-Gonzalez. |
| EPI_ISL_2859034 | LESP Sinaloa | Instituto de Diagnostico y Referencia Epidemiologicos (INDRE) | Claudia Wong-Arambula, Abril Rodriguez-Maldonado, Vanessa Rivero-Arredondo, Ariadna Medina-Benitez, Joaquin Quiroz-Mercado, Sergio Rangel-Guerrero, Maribel Gonzalez-Villa, Natividad Cruz-Ortiz, Tatiana Nunez-Garcia, Gisela Barrera-Badillo, Lucia Hernandez-Rivas, Irma Lopez-Martinez, Ernesto Ramirez-Gonzalez. |
| EPI_ISL_2859035, EPI_ISL_2859036, EPI_ISL_2859037, EPI_ISL_2859038 | LESP Campeche | Instituto de Diagnostico y Referencia Epidemiologicos (INDRE) | Claudia Wong-Arambula, Abril Rodriguez-Maldonado, Vanessa Rivero-Arredondo, Ariadna Medina-Benitez, Joaquin Quiroz-Mercado, Sergio Rangel-Guerrero, Maribel Gonzalez-Villa, Natividad Cruz-Ortiz, Tatiana Nunez-Garcia, Gisela Barrera-Badillo, Lucia Hernandez-Rivas, Irma Lopez-Martinez, Ernesto Ramirez-Gonzalez. |
| EPI_ISL_2859039, EPI_ISL_2859040, EPI_ISL_2859041, EPI_ISL_2859042, EPI_ISL_2859043, EPI_ISL_2859044, EPI_ISL_2859045, EPI_ISL_2859046, EPI_ISL_2859047, EPI_ISL_2859048 | LESP Tabasco | Instituto de Diagnostico y Referencia Epidemiologicos (INDRE) | Claudia Wong-Arambula, Abril Rodriguez-Maldonado, Vanessa Rivero-Arredondo, Ariadna Medina-Benitez, Joaquin Quiroz-Mercado, Sergio Rangel-Guerrero, Maribel Gonzalez-Villa, Natividad Cruz-Ortiz, Tatiana Nunez-Garcia, Gisela Barrera-Badillo, Lucia Hernandez-Rivas, Irma Lopez-Martinez, Ernesto Ramirez-Gonzalez. |
| EPI_ISL_2859049, EPI_ISL_2859050 | LESP Veracruz | Instituto de Diagnostico y Referencia Epidemiologicos (INDRE) | Claudia Wong-Arambula, Abril Rodriguez-Maldonado, Vanessa Rivero-Arredondo, Ariadna Medina-Benitez, Joaquin Quiroz-Mercado, Sergio Rangel-Guerrero, Maribel Gonzalez-Villa, Natividad Cruz-Ortiz, Tatiana Nunez-Garcia, Gisela Barrera-Badillo, Lucia Hernandez-Rivas, Irma Lopez-Martinez, Ernesto Ramirez-Gonzalez. |
| EPI_ISL_2859051 | LESP Sonora | Instituto de Diagnostico y Referencia Epidemiologicos (INDRE) | Claudia Wong-Arambula, Abril Rodriguez-Maldonado, Vanessa Rivero-Arredondo, Ariadna Medina-Benitez, Joaquin Quiroz-Mercado, Sergio Rangel-Guerrero, Maribel Gonzalez-Villa, Natividad Cruz-Ortiz, Tatiana Nunez-Garcia, Gisela Barrera-Badillo, Lucia Hernandez-Rivas, Irma Lopez-Martinez, Ernesto Ramirez-Gonzalez. |
| EPI_ISL_2859054 | LESP Nuevo Leon | Instituto de Diagnostico y Referencia Epidemiologicos (INDRE) | Claudia Wong-Arambula, Abril Rodriguez-Maldonado, Vanessa Rivero-Arredondo, Ariadna Medina-Benitez, Joaquin Quiroz-Mercado, Sergio Rangel-Guerrero, Maribel Gonzalez-Villa, Natividad Cruz-Ortiz, Tatiana Nunez-Garcia, Gisela Barrera-Badillo, Lucia Hernandez-Rivas, Irma Lopez-Martinez, Ernesto Ramirez-Gonzalez. |
| EPI_ISL_2859055, EPI_ISL_2859056, EPI_ISL_2859057 | LESP Tabasco | Instituto de Diagnostico y Referencia Epidemiologicos (INDRE) | Claudia Wong-Arambula, Abril Rodriguez-Maldonado, Vanessa Rivero-Arredondo, Ariadna Medina-Benitez, Joaquin Quiroz-Mercado, Sergio Rangel-Guerrero, Maribel Gonzalez-Villa, Natividad Cruz-Ortiz, Tatiana Nunez-Garcia, Gisela Barrera-Badillo, Lucia Hernandez-Rivas, Irma Lopez-Martinez, Ernesto Ramirez-Gonzalez. |
| EPI_ISL_2859059, EPI_ISL_2859060 | LESP San Luis Potosi | Instituto de Diagnostico y Referencia Epidemiologicos (INDRE) | Claudia Wong-Arambula, Abril Rodriguez-Maldonado, Vanessa Rivero-Arredondo, Ariadna Medina-Benitez, Joaquin Quiroz-Mercado, Sergio Rangel-Guerrero, Maribel Gonzalez-Villa, Natividad Cruz-Ortiz, Tatiana Nunez-Garcia, Gisela Barrera-Badillo, Lucia Hernandez-Rivas, Irma Lopez-Martinez, Ernesto Ramirez-Gonzalez. |
| EPI_ISL_2859061, EPI_ISL_2859062 | LESP Ciudad de Mexico | Instituto de Diagnostico y Referencia Epidemiologicos (INDRE) | Claudia Wong-Arambula, Abril Rodriguez-Maldonado, Vanessa Rivero-Arredondo, Ariadna Medina-Benitez, Joaquin Quiroz-Mercado, Sergio Rangel-Guerrero, Maribel Gonzalez-Villa, Natividad Cruz-Ortiz, Tatiana Nunez-Garcia, Gisela Barrera-Badillo, Lucia Hernandez-Rivas, Irma Lopez-Martinez, Ernesto Ramirez-Gonzalez. |
| EPI_ISL_2859063 | LESP Yucatan | Instituto de Diagnostico y Referencia Epidemiologicos (INDRE) | Claudia Wong-Arambula, Abril Rodriguez-Maldonado, Vanessa Rivero-Arredondo, Ariadna Medina-Benitez, Joaquin Quiroz-Mercado, Sergio Rangel-Guerrero, Maribel Gonzalez-Villa, Natividad Cruz-Ortiz, Tatiana Nunez-Garcia, Gisela Barrera-Badillo, Lucia Hernandez-Rivas, Irma Lopez-Martinez, Ernesto Ramirez-Gonzalez. |
| EPI_ISL_2859065 | LESP Michoacan | Instituto de Diagnostico y Referencia Epidemiologicos (INDRE) | Claudia Wong-Arambula, Abril Rodriguez-Maldonado, Vanessa Rivero-Arredondo, Ariadna Medina-Benitez, Joaquin Quiroz-Mercado, Sergio Rangel-Guerrero, Maribel Gonzalez-Villa, Natividad Cruz-Ortiz, Tatiana Nunez-Garcia, Gisela Barrera-Badillo, Lucia Hernandez-Rivas, Irma Lopez-Martinez, Ernesto Ramirez-Gonzalez. |
| EPI_ISL_2859072, EPI_ISL_2859073 | LESP Ciudad de Mexico | Instituto de Diagnostico y Referencia Epidemiologicos (INDRE) | Claudia Wong-Arambula, Abril Rodriguez-Maldonado, Vanessa Rivero-Arredondo, Ariadna Medina-Benitez, Joaquin Quiroz-Mercado, Sergio Rangel-Guerrero, Maribel Gonzalez-Villa, Natividad Cruz-Ortiz, Tatiana Nunez-Garcia, Gisela Barrera-Badillo, Lucia Hernandez-Rivas, Irma Lopez-Martinez, Ernesto Ramirez-Gonzalez. |
| EPI_ISL_2859077, EPI_ISL_2859078, EPI_ISL_2859079, EPI_ISL_2859081, EPI_ISL_2859082, EPI_ISL_2859083, EPI_ISL_2859084, EPI_ISL_2859085, EPI_ISL_2859086, EPI_ISL_2859087, EPI_ISL_2859088, EPI_ISL_2859089, EPI_ISL_2859090, EPI_ISL_2859091, EPI_ISL_2859092, EPI_ISL_2859093, EPI_ISL_2859094, EPI_ISL_2859095, EPI_ISL_2859096, EPI_ISL_2859097, EPI_ISL_2859098, EPI_ISL_2859099, EPI_ISL_2859100, EPI_ISL_2859101, EPI_ISL_2859102, EPI_ISL_2859103, EPI_ISL_2859104, EPI_ISL_2859105, EPI_ISL_2859106 | | | |
| see above | LESP Tabasco | Instituto de Diagnostico y Referencia Epidemiologicos (INDRE) | Claudia Wong-Arambula, Abril Rodriguez-Maldonado, Vanessa Rivero-Arredondo, Ariadna Medina-Benitez, Joaquin Quiroz-Mercado, Sergio Rangel-Guerrero, Maribel Gonzalez-Villa, Natividad Cruz-Ortiz, Tatiana Nunez-Garcia, Gisela Barrera-Badillo, Lucia Hernandez-Rivas, Irma Lopez-Martinez, Ernesto Ramirez-Gonzalez. |
| EPI_ISL_2859107 | LESP Jalisco | Instituto de Diagnostico y Referencia Epidemiologicos (INDRE) | Claudia Wong-Arambula, Abril Rodriguez-Maldonado, Vanessa Rivero-Arredondo, Ariadna Medina-Benitez, Joaquin Quiroz-Mercado, Sergio Rangel-Guerrero, Maribel Gonzalez-Villa, Natividad Cruz-Ortiz, Tatiana Nunez-Garcia, Gisela Barrera-Badillo, Lucia Hernandez-Rivas, Irma Lopez-Martinez, Ernesto Ramirez-Gonzalez. |
| EPI_ISL_2859108 | LESP San Luis Potosi | Instituto de Diagnostico y Referencia Epidemiologicos (INDRE) | Claudia Wong-Arambula, Abril Rodriguez-Maldonado, Vanessa Rivero-Arredondo, Ariadna Medina-Benitez, Joaquin Quiroz-Mercado, Sergio Rangel-Guerrero, Maribel Gonzalez-Villa, Natividad Cruz-Ortiz, Tatiana Nunez-Garcia, Gisela Barrera-Badillo, Lucia Hernandez-Rivas, Irma Lopez-Martinez, Ernesto Ramirez-Gonzalez. |
| EPI_ISL_2859109, EPI_ISL_2859110, EPI_ISL_2859111, EPI_ISL_2859112, EPI_ISL_2859113, EPI_ISL_2859114, EPI_ISL_2859115, EPI_ISL_2859116, EPI_ISL_2859117, EPI_ISL_2859118, EPI_ISL_2859119, EPI_ISL_2859120 | | | |
| see above | LESP Tabasco | Instituto de Diagnostico y Referencia Epidemiologicos (INDRE) | Claudia Wong-Arambula, Abril Rodriguez-Maldonado, Vanessa Rivero-Arredondo, Ariadna Medina-Benitez, Joaquin Quiroz-Mercado, Sergio Rangel-Guerrero, Maribel Gonzalez-Villa, Natividad Cruz-Ortiz, Tatiana Nunez-Garcia, Gisela Barrera-Badillo, Lucia Hernandez-Rivas, Irma Lopez-Martinez, Ernesto Ramirez-Gonzalez. |
| EPI_ISL_2859121 | LESP Nuevo Leon | Instituto de Diagnostico y Referencia Epidemiologicos (INDRE) | Claudia Wong-Arambula, Abril Rodriguez-Maldonado, Vanessa Rivero-Arredondo, Ariadna Medina-Benitez, Joaquin Quiroz-Mercado, Sergio Rangel-Guerrero, Maribel Gonzalez-Villa, Natividad Cruz-Ortiz, Tatiana Nunez-Garcia, Gisela Barrera-Badillo, Lucia Hernandez-Rivas, Irma Lopez-Martinez, Ernesto Ramirez-Gonzalez. |
| EPI_ISL_2859122, EPI_ISL_2859123, EPI_ISL_2859124, EPI_ISL_2859125 | LESP Tabasco | Instituto de Diagnostico y Referencia Epidemiologicos (INDRE) | Claudia Wong-Arambula, Abril Rodriguez-Maldonado, Vanessa Rivero-Arredondo, Ariadna Medina-Benitez, Joaquin Quiroz-Mercado, Sergio Rangel-Guerrero, Maribel Gonzalez-Villa, Natividad Cruz-Ortiz, Tatiana Nunez-Garcia, Gisela Barrera-Badillo, Lucia Hernandez-Rivas, Irma Lopez-Martinez, Ernesto Ramirez-Gonzalez. |
| EPI_ISL_2860825, EPI_ISL_2860826 | AZ Sint-Lucas Gent | AZ Sint-Lucas Gent | Jos Van Acker, Charlotte Verfaillie, Elke Vanlaere, Klara De Rauw |
| EPI_ISL_2861158 | LESP Ciudad de Mexico | Instituto de Diagnostico y Referencia Epidemiologicos (INDRE) | Claudia Wong-Arambula, Abril Rodriguez-Maldonado, Vanessa Rivero-Arredondo, Ariadna Medina-Benitez, Joaquin Quiroz-Mercado, Sergio Rangel-Guerrero, Maribel Gonzalez-Villa, Natividad Cruz-Ortiz, Tatiana Nunez-Garcia, Gisela Barrera-Badillo, Lucia Hernandez-Rivas, Irma Lopez-Martinez, Ernesto Ramirez-Gonzalez. |
| EPI_ISL_2861159, EPI_ISL_2861160, EPI_ISL_2861161 | LESP Nuevo Leon | Instituto de Diagnostico y Referencia Epidemiologicos (INDRE) | Claudia Wong-Arambula, Abril Rodriguez-Maldonado, Vanessa Rivero-Arredondo, Ariadna Medina-Benitez, Joaquin Quiroz-Mercado, Sergio Rangel-Guerrero, Maribel Gonzalez-Villa, Natividad Cruz-Ortiz, Tatiana Nunez-Garcia, Gisela Barrera-Badillo, Lucia Hernandez-Rivas, Irma Lopez-Martinez, Ernesto Ramirez-Gonzalez. |
| EPI_ISL_2861162 | LESP Tamaulipas | Instituto de Diagnostico y Referencia Epidemiologicos (INDRE) | Claudia Wong-Arambula, Abril Rodriguez-Maldonado, Vanessa Rivero-Arredondo, Ariadna Medina-Benitez, Joaquin Quiroz-Mercado, Sergio Rangel-Guerrero, Maribel Gonzalez-Villa, Natividad Cruz-Ortiz, Tatiana Nunez-Garcia, Gisela Barrera-Badillo, Lucia Hernandez-Rivas, Irma Lopez-Martinez, Ernesto Ramirez-Gonzalez. |

| | | | |
|---|---|---|---|
| EPI_ISL_2861163 | LESP Nuevo Leon | Instituto de Diagnostico y Referencia Epidemiologicos (INDRE) | Claudia Wong-Arambula, Abril Rodriguez-Maldonado, Vanessa Rivero-Arredondo, Ariadna Medina-Benitez, Joaquin Quiroz-Mercado, Sergio Rangel-Guerrero, Maribel Gonzalez-Villa, Natividad Cruz-Ortiz, Tatiana Nunez-Garcia, Gisela Barrera-Badillo, Lucia Hernandez-Rivas, Irma Lopez-Martinez, Ernesto Ramirez-Gonzalez. |
| EPI_ISL_2861164, EPI_ISL_2861165 | LESP Tamaulipas | Instituto de Diagnostico y Referencia Epidemiologicos (INDRE) | Claudia Wong-Arambula, Abril Rodriguez-Maldonado, Vanessa Rivero-Arredondo, Ariadna Medina-Benitez, Joaquin Quiroz-Mercado, Sergio Rangel-Guerrero, Maribel Gonzalez-Villa, Natividad Cruz-Ortiz, Tatiana Nunez-Garcia, Gisela Barrera-Badillo, Lucia Hernandez-Rivas, Irma Lopez-Martinez, Ernesto Ramirez-Gonzalez. |
| EPI_ISL_2861166 | LESP Veracruz | Instituto de Diagnostico y Referencia Epidemiologicos (INDRE) | Claudia Wong-Arambula, Abril Rodriguez-Maldonado, Vanessa Rivero-Arredondo, Ariadna Medina-Benitez, Joaquin Quiroz-Mercado, Sergio Rangel-Guerrero, Maribel Gonzalez-Villa, Natividad Cruz-Ortiz, Tatiana Nunez-Garcia, Gisela Barrera-Badillo, Lucia Hernandez-Rivas, Irma Lopez-Martinez, Ernesto Ramirez-Gonzalez. |
| EPI_ISL_2861167 | LESP Guanajuato | Instituto de Diagnostico y Referencia Epidemiologicos (INDRE) | Claudia Wong-Arambula, Abril Rodriguez-Maldonado, Vanessa Rivero-Arredondo, Ariadna Medina-Benitez, Joaquin Quiroz-Mercado, Sergio Rangel-Guerrero, Maribel Gonzalez-Villa, Natividad Cruz-Ortiz, Tatiana Nunez-Garcia, Gisela Barrera-Badillo, Lucia Hernandez-Rivas, Irma Lopez-Martinez, Ernesto Ramirez-Gonzalez. |
| EPI_ISL_2861168, EPI_ISL_2861169, EPI_ISL_2861170 | LESP Durango | Instituto de Diagnostico y Referencia Epidemiologicos (INDRE) | Claudia Wong-Arambula, Abril Rodriguez-Maldonado, Vanessa Rivero-Arredondo, Ariadna Medina-Benitez, Joaquin Quiroz-Mercado, Sergio Rangel-Guerrero, Maribel Gonzalez-Villa, Natividad Cruz-Ortiz, Tatiana Nunez-Garcia, Gisela Barrera-Badillo, Lucia Hernandez-Rivas, Irma Lopez-Martinez, Ernesto Ramirez-Gonzalez. |
| EPI_ISL_2861171 | LESP Chiapas | Instituto de Diagnostico y Referencia Epidemiologicos (INDRE) | Claudia Wong-Arambula, Abril Rodriguez-Maldonado, Vanessa Rivero-Arredondo, Ariadna Medina-Benitez, Joaquin Quiroz-Mercado, Sergio Rangel-Guerrero, Maribel Gonzalez-Villa, Natividad Cruz-Ortiz, Tatiana Nunez-Garcia, Gisela Barrera-Badillo, Lucia Hernandez-Rivas, Irma Lopez-Martinez, Ernesto Ramirez-Gonzalez. |
| EPI_ISL_2861172 | LESP Michoacan | Instituto de Diagnostico y Referencia Epidemiologicos (INDRE) | Claudia Wong-Arambula, Abril Rodriguez-Maldonado, Vanessa Rivero-Arredondo, Ariadna Medina-Benitez, Joaquin Quiroz-Mercado, Sergio Rangel-Guerrero, Maribel Gonzalez-Villa, Natividad Cruz-Ortiz, Tatiana Nunez-Garcia, Gisela Barrera-Badillo, Lucia Hernandez-Rivas, Irma Lopez-Martinez, Ernesto Ramirez-Gonzalez. |
| EPI_ISL_2861173 | LESP Hidalgo | Instituto de Diagnostico y Referencia Epidemiologicos (INDRE) | Claudia Wong-Arambula, Abril Rodriguez-Maldonado, Vanessa Rivero-Arredondo, Ariadna Medina-Benitez, Joaquin Quiroz-Mercado, Sergio Rangel-Guerrero, Maribel Gonzalez-Villa, Natividad Cruz-Ortiz, Tatiana Nunez-Garcia, Gisela Barrera-Badillo, Lucia Hernandez-Rivas, Irma Lopez-Martinez, Ernesto Ramirez-Gonzalez. |
| EPI_ISL_2861174, EPI_ISL_2861175 | LESP Queretaro | Instituto de Diagnostico y Referencia Epidemiologicos (INDRE) | Claudia Wong-Arambula, Abril Rodriguez-Maldonado, Vanessa Rivero-Arredondo, Ariadna Medina-Benitez, Joaquin Quiroz-Mercado, Sergio Rangel-Guerrero, Maribel Gonzalez-Villa, Natividad Cruz-Ortiz, Tatiana Nunez-Garcia, Gisela Barrera-Badillo, Lucia Hernandez-Rivas, Irma Lopez-Martinez, Ernesto Ramirez-Gonzalez. |
| EPI_ISL_2861176, EPI_ISL_2861177, EPI_ISL_2861178, EPI_ISL_2861179 | LESP Tabasco | Instituto de Diagnostico y Referencia Epidemiologicos (INDRE) | Claudia Wong-Arambula, Abril Rodriguez-Maldonado, Vanessa Rivero-Arredondo, Ariadna Medina-Benitez, Joaquin Quiroz-Mercado, Sergio Rangel-Guerrero, Maribel Gonzalez-Villa, Natividad Cruz-Ortiz, Tatiana Nunez-Garcia, Gisela Barrera-Badillo, Lucia Hernandez-Rivas, Irma Lopez-Martinez, Ernesto Ramirez-Gonzalez. |
| EPI_ISL_2861180 | LESP Coahuila | Instituto de Diagnostico y Referencia Epidemiologicos (INDRE) | Claudia Wong-Arambula, Abril Rodriguez-Maldonado, Vanessa Rivero-Arredondo, Ariadna Medina-Benitez, Joaquin Quiroz-Mercado, Sergio Rangel-Guerrero, Maribel Gonzalez-Villa, Natividad Cruz-Ortiz, Tatiana Nunez-Garcia, Gisela Barrera-Badillo, Lucia Hernandez-Rivas, Irma Lopez-Martinez, Ernesto Ramirez-Gonzalez. |
| EPI_ISL_2861181, EPI_ISL_2861182 | LESP Tamaulipas | Instituto de Diagnostico y Referencia Epidemiologicos (INDRE) | Claudia Wong-Arambula, Abril Rodriguez-Maldonado, Vanessa Rivero-Arredondo, Ariadna Medina-Benitez, Joaquin Quiroz-Mercado, Sergio Rangel-Guerrero, Maribel Gonzalez-Villa, Natividad Cruz-Ortiz, Tatiana Nunez-Garcia, Gisela Barrera-Badillo, Lucia Hernandez-Rivas, Irma Lopez-Martinez, Ernesto Ramirez-Gonzalez. |
| EPI_ISL_2861183, EPI_ISL_2861184, EPI_ISL_2861185 | LESP Tabasco | Instituto de Diagnostico y Referencia Epidemiologicos (INDRE) | Claudia Wong-Arambula, Abril Rodriguez-Maldonado, Vanessa Rivero-Arredondo, Ariadna Medina-Benitez, Joaquin Quiroz-Mercado, Sergio Rangel-Guerrero, Maribel Gonzalez-Villa, Natividad Cruz-Ortiz, Tatiana Nunez-Garcia, Gisela Barrera-Badillo, Lucia Hernandez-Rivas, Irma Lopez-Martinez, Ernesto Ramirez-Gonzalez. |
| EPI_ISL_2861186, EPI_ISL_2861187 | LESP Aguascalientes | Instituto de Diagnostico y Referencia Epidemiologicos (INDRE) | Claudia Wong-Arambula, Abril Rodriguez-Maldonado, Vanessa Rivero-Arredondo, Ariadna Medina-Benitez, Joaquin Quiroz-Mercado, Sergio Rangel-Guerrero, Maribel Gonzalez-Villa, Natividad Cruz-Ortiz, Tatiana Nunez-Garcia, Gisela Barrera-Badillo, Lucia Hernandez-Rivas, Irma Lopez-Martinez, Ernesto Ramirez-Gonzalez. |
| EPI_ISL_2861188, EPI_ISL_2861189, EPI_ISL_2861190, EPI_ISL_2861191 | LESP Tabasco | Instituto de Diagnostico y Referencia Epidemiologicos (INDRE) | Claudia Wong-Arambula, Abril Rodriguez-Maldonado, Vanessa Rivero-Arredondo, Ariadna Medina-Benitez, Joaquin Quiroz-Mercado, Sergio Rangel-Guerrero, Maribel Gonzalez-Villa, Natividad Cruz-Ortiz, Tatiana Nunez-Garcia, Gisela Barrera-Badillo, Lucia Hernandez-Rivas, Irma Lopez-Martinez, Ernesto Ramirez-Gonzalez. |
| EPI_ISL_2861192, EPI_ISL_2861193, EPI_ISL_2861194, EPI_ISL_2861195, EPI_ISL_2861196 | LESP Ciudad de Mexico | Instituto de Diagnostico y Referencia Epidemiologicos (INDRE) | Claudia Wong-Arambula, Abril Rodriguez-Maldonado, Vanessa Rivero-Arredondo, Ariadna Medina-Benitez, Joaquin Quiroz-Mercado, Sergio Rangel-Guerrero, Maribel Gonzalez-Villa, Natividad Cruz-Ortiz, Tatiana Nunez-Garcia, Gisela Barrera-Badillo, Lucia Hernandez-Rivas, Irma Lopez-Martinez, Ernesto Ramirez-Gonzalez. |
| EPI_ISL_2861262, EPI_ISL_2861263, EPI_ISL_2861264, EPI_ISL_2861265 | Dr. Yoshitaka Tamura Department of Clinical Laboratory, Osaka Habikino Medical Center | Department of Developmental Medicine, Research Institute, Osaka Women's and Children's Hospital | Itaru Yanagihara, Yukiko Nakura |
| EPI_ISL_2861267, EPI_ISL_2861268 | Osaka General Medical Center | Department of Developmental Medicine, Research Institute, Osaka Women's and Children's Hospital | Itaru Yanagihara, Yukiko Nakura |
| EPI_ISL_2861280 | Azienda Ospedaliera Terni | Istituto Zooprofilattico Sperimentale dell'Abruzzo e Molise "G. Caporale" | Palumbo M, Scaccetti A, Lorusso A, Marcacci M, Di Domenico M, Ancora M, Curini V, Mangone I, Rinaldi A, Delli Compagni E, Scialabba S, Di Pasquale A, Cammà C, Puglia I, Calistri P, Savini G |
| EPI_ISL_2861288, EPI_ISL_2861290, EPI_ISL_2861292, EPI_ISL_2861293, EPI_ISL_2861294, EPI_ISL_2861295, EPI_ISL_2861296, EPI_ISL_2861297, EPI_ISL_2861298, EPI_ISL_2861299, EPI_ISL_2861303, EPI_ISL_2861304, EPI_ISL_2861305, EPI_ISL_2861306, EPI_ISL_2861307, EPI_ISL_2861308, EPI_ISL_2861309, EPI_ISL_2861310, EPI_ISL_2861311, EPI_ISL_2861314 | | | |
| see above | Microbiology Department. Complexo Hospitalario Universitario de Vigo | Microbiology Department. Complexo Hospitalario Universitario de Vigo | Alfaya N, Alonso I, Alvarez M, Cabrera JJ, Carballo R, Cores O, Cortizo S, del-Campo V, Martinez L, Mediero G, Perez S, Davina C, Potel C, Regueiro B, Rey S, Vasallo FJ |
| EPI_ISL_2861318, EPI_ISL_2861319, EPI_ISL_2861320, EPI_ISL_2861321, EPI_ISL_2861322, EPI_ISL_2861323 | Microbiology Department, University Hospital Donostia | Microbiology Department, University Hospital Donostia | Marimon JM, Montes M, Piñeiro L, Martin-Peñaranda T, Gomez M, Sorarrain A, Cilla G. |
| EPI_ISL_2861339, EPI_ISL_2861340 | Istituto Zooprofilattico Sperimentale della Puglia e della Basilicata | Istituto Zooprofilattico Sperimentale della Puglia e della Basilicata | Parisi A., Bianco A., Capozzi L., Del Sambro L., Simone D., Difato L., Santoro A., Ridolfi D., Giannico A., Manzari C., Pesole G., Pace L., Galante D. |
| EPI_ISL_2861341 | Ospedale Santissima Annunziata | Istituto Zooprofilattico Sperimentale della Puglia e della Basilicata | Parisi A., Bianco A., Capozzi L., Del Sambro L., Simone D., Difato L., Ridolfi D., Giannico A, Tampoia M., Manzari C., Pesole G. |
| EPI_ISL_2861376 | St James's Hospital, Dublin 8, Ireland. | ELDA biotech | Deirdre M. Broderick, Peter R. Flanagan, Brendan Crowley |
| EPI_ISL_2862010 | Genomic Research Laboratory, Bangabandhu Sheikh Mujib Medical University | Genomic Research Laboratory, BSMMU | Laila Anjuman Banu, Md.Saydur Rahman, Sharadindu Kanti Sinha, Arifa Akram, Md.Sharfuddin Ahmed |
| EPI_ISL_2862015 | Genomic Research Laboratory, Bangabandhu Sheikh Mujib Medical University (BSMMU) Shahbag, Dhaka_1000 | Genomic Research Laboratory, BSMMU | Laila Anjuman Banu, Md.Saydur Rahman, Sharadindu Kanti Sinha, Arifa Akram, Md.Sharfuddin Ahmed |
| EPI_ISL_2862016, EPI_ISL_2862048 | Genomic Research Laboratory, Bangabandhu Sheikh Mujib Medical University | Genomic Research Laboratory, BSMMU | Laila Anjuman Banu, Md.Saydur Rahman, Sharadindu Kanti Sinha, Arifa Akram, Md.Sharfuddin Ahmed |
| EPI_ISL_2863666 | Laboratorio Central de Saude Publica do Estado de Santa | Laboratory of Respiratory Viruses and Measles, Oswaldo Cruz | Paola Resende, Luciana Appolinario, Fernando Motta, Anna Carolina Paixao, Ana Carolina Mendonca, Alice Sampaio Rocha, Taina Venas, Elisa |

| | | | |
|---|---|---|---|
| | Catarina (LACEN-SC) | Institute, FIOCRUZ | Cavalcante Pereira, Renata Serrano Lopes, Darcita Buerger Rovaris, Sandra Bianchini Fernandes, Marilda Siqueira on behalf of the Fiocruz COVID-19 Genomic Surveillance Network |
| EPI_ISL_2864141, EPI_ISL_2864142, EPI_ISL_2864144, EPI_ISL_2864145, EPI_ISL_2864146, EPI_ISL_2864147 | Microbiology Department. Complexo Hospitalario Universitario de Vigo | Microbiology Department. Complexo Hospitalario Universitario de Vigo | Alfaya N, Alonso I, Alvarez M, Cabrera JJ, Carballo R, Cores O, Cortizo S, del-Campo V, Martinez L, Mediero G, Perez S, Davina C, Potel C, Regueiro B, Rey S, Vasallo FJ |
| EPI_ISL_2868394, EPI_ISL_2868395, EPI_ISL_2868396, EPI_ISL_2868397, EPI_ISL_2868398, EPI_ISL_2868399, EPI_ISL_2868400, EPI_ISL_2868401, EPI_ISL_2868402, EPI_ISL_2868403, EPI_ISL_2868404, EPI_ISL_2868405, EPI_ISL_2868406, EPI_ISL_2868407, EPI_ISL_2868408, EPI_ISL_2868409, EPI_ISL_2868410, EPI_ISL_2868411, EPI_ISL_2868412, EPI_ISL_2868413 | | | |
| see above | Botswana Harvard HIV Reference Laboratory | Botswana Harvard HIV Reference Laboratory | Sikhulile Moyo, Wonderful T. Choga, Dorcas Maruapula, Thongbotho Mphoyakgosi, Boitumelo Zuze, Botshelo Radibe, Legodile Kooepile, Ontlametse T. Bareng, Keoratile Ntshambiwa, Modisa Motswaledi, Madisa Mine, Joseph Makhema, Roger Shapiro, Shahin Lockman, Mosepele Mosepele, Simani Gaseitsiwe |
| EPI_ISL_2868892 | BTKLPP Kelas I Makassar | National Institute of Health Research and Development | Vivi Setiawaty, Hana Apsari Pawestri, Subangkit, Kartika Dewi Puspa, Arie Ardiansyah Nugraha, Hartanti Dian Ikawati, Krisna Pangesti, Triyani Soekarso, Nelly Puspandari |
| EPI_ISL_2868913 | RS Royal Progress | National Institute of Health Research and Development | Vivi Setiawaty, Hana Apsari Pawestri, Subangkit, Kartika Dewi Puspa, Arie Ardiansyah Nugraha, Hartanti Dian Ikawati, Krisna Pangesti, Triyani Soekarso, Nelly Puspandari |
| EPI_ISL_2876292, EPI_ISL_2876296, EPI_ISL_2876298 | Genome Analysis Center, Yamanashi Central Hospital | Genome Analysis Center, Yamanashi Central Hospital | Yosuke Hirotsu |
| EPI_ISL_2885350, EPI_ISL_2885351 | CHU de la Réunion - Nord | Laboratoire de virologie, CNR arbovirus Associé, Chu de la Réunion | Etienne Frumence, Marie-Christine Jaffar Bandjee, Nicolas Traversier, Sabrina Petit Genet |
| EPI_ISL_2885355, EPI_ISL_2885356, EPI_ISL_2885357 | CHU de la Réunion - Sud | Laboratoire de virologie, CNR arbovirus Associé, Chu de la Réunion | Etienne Frumence, Marie-Christine Jaffar Bandjee, Nicolas Traversier, Sabrina Petit Genet |
| EPI_ISL_2885358, EPI_ISL_2885359, EPI_ISL_2885361 | CHU de la Réunion - Nord | Laboratoire de virologie, CNR arbovirus Associé, Chu de la Réunion | Etienne Frumence, Marie-Christine Jaffar Bandjee, Nicolas Traversier, Sabrina Petit Genet |
| EPI_ISL_2885362, EPI_ISL_2885363 | CHU de la Réunion - Sud | Laboratoire de virologie, CNR arbovirus Associé, Chu de la Réunion | Etienne Frumence, Marie-Christine Jaffar Bandjee, Nicolas Traversier, Sabrina Petit Genet |
| EPI_ISL_2885364 | CHU de la Réunion - Nord | Laboratoire de virologie, CNR arbovirus Associé, Chu de la Réunion | Etienne Frumence, Marie-Christine Jaffar Bandjee, Nicolas Traversier, Sabrina Petit Genet |
| EPI_ISL_2885365, EPI_ISL_2885366 | CHU de la Réunion - Sud | Laboratoire de virologie, CNR arbovirus Associé, Chu de la Réunion | Etienne Frumence, Marie-Christine Jaffar Bandjee, Nicolas Traversier, Sabrina Petit Genet |
| EPI_ISL_2885367, EPI_ISL_2885368 | CHU de la Réunion - Nord | Laboratoire de virologie, CNR arbovirus Associé, Chu de la Réunion | Etienne Frumence, Marie-Christine Jaffar Bandjee, Nicolas Traversier, Sabrina Petit Genet |
| EPI_ISL_2885369, EPI_ISL_2885370, EPI_ISL_2885371, EPI_ISL_2885372 | CHU de la Réunion - Sud | Laboratoire de virologie, CNR arbovirus Associé, Chu de la Réunion | Etienne Frumence, Marie-Christine Jaffar Bandjee, Nicolas Traversier, Sabrina Petit Genet |
| EPI_ISL_2885375, EPI_ISL_2885376, EPI_ISL_2885378, EPI_ISL_2885379, EPI_ISL_2885380, EPI_ISL_2885381, EPI_ISL_2885382, EPI_ISL_2885383 | CHU de la Réunion - Nord | Laboratoire de virologie, CNR arbovirus Associé, Chu de la Réunion | Etienne Frumence, Marie-Christine Jaffar Bandjee, Nicolas Traversier, Sabrina Petit Genet |
| EPI_ISL_2885610, EPI_ISL_2885611, EPI_ISL_2885612, EPI_ISL_2885613, EPI_ISL_2885614, EPI_ISL_2885615, EPI_ISL_2885616, EPI_ISL_2885617, EPI_ISL_2885618, EPI_ISL_2885619, EPI_ISL_2885620, EPI_ISL_2885621, EPI_ISL_2885622, EPI_ISL_2885623, EPI_ISL_2885624, EPI_ISL_2885625, EPI_ISL_2885626, EPI_ISL_2885627, EPI_ISL_2885628, EPI_ISL_2885629, EPI_ISL_2885630, EPI_ISL_2885631, EPI_ISL_2885632, EPI_ISL_2885633, EPI_ISL_2885634, EPI_ISL_2885635, EPI_ISL_2885636, EPI_ISL_2885637, EPI_ISL_2885638, EPI_ISL_2885639, EPI_ISL_2885640, EPI_ISL_2885641, EPI_ISL_2885642, EPI_ISL_2885643, EPI_ISL_2885644, EPI_ISL_2885645, EPI_ISL_2885646, EPI_ISL_2885647, EPI_ISL_2885648, EPI_ISL_2885649, EPI_ISL_2885650, EPI_ISL_2885651, EPI_ISL_2885652, EPI_ISL_2885653, EPI_ISL_2885654, EPI_ISL_2885655, EPI_ISL_2885656, EPI_ISL_2885657, EPI_ISL_2885658, EPI_ISL_2885659, EPI_ISL_2885660, EPI_ISL_2885661, EPI_ISL_2885662, EPI_ISL_2885663, EPI_ISL_2885664, EPI_ISL_2885665, EPI_ISL_2885666, EPI_ISL_2885667, EPI_ISL_2885668, EPI_ISL_2885669, EPI_ISL_2885670, EPI_ISL_2885671, EPI_ISL_2885672, EPI_ISL_2885673, EPI_ISL_2885674, EPI_ISL_2885675, EPI_ISL_2885676, EPI_ISL_2885677, EPI_ISL_2885678, EPI_ISL_2885679 | | | |
| see above | Cliniques universitaires Saint-Luc | UCLouvain/IREC/MBLG | Jean Ruelle, Bertrand Bearzatto, Benoit Kabamba Mukadi |
| EPI_ISL_2885680, EPI_ISL_2885681, EPI_ISL_2885682, EPI_ISL_2885684, EPI_ISL_2885685, EPI_ISL_2885686, EPI_ISL_2885687, EPI_ISL_2885688, EPI_ISL_2885689, EPI_ISL_2885690, EPI_ISL_2885691, EPI_ISL_2885692, EPI_ISL_2885693, EPI_ISL_2885694, EPI_ISL_2885695, EPI_ISL_2885696 | | | |
| see above | Clinique Saint-Pierre Ottignies | UCLouvain/IREC/MBLG | Jean Ruelle, Bertrand Bearzatto, Benoit Kabamba Mukadi |
| EPI_ISL_2885697, EPI_ISL_2885698 | Cliniques universitaires Saint-Luc | UCLouvain/IREC/MBLG | Jean Ruelle, Bertrand Bearzatto, Benoit Kabamba Mukadi |
| EPI_ISL_2885901 | LBM Bonhoure | Cerba lab | e.g. Jane Doe, John Doe |
| EPI_ISL_2886157, EPI_ISL_2886159 | IVRCRD RSPAW | Institute for Vector and Reservoir Control, Research and Development (IVRCRD)- National Institute of Health Research and Development (NIHRD), Indonesia | Arief Mulyono, Triwibowo Ambar Garjito, Restu Choirul Saban, Ayu Pradipta Pratiwi, Dwi Susilo, Yusnita Mirna Anggraeni, Mujiyanto, Mega Tyas Prihatin, Nur Hidayati, Arif Suryo Prasetyo, Tika Fiona Sari, Sri Wahyuni Handayani, Aryo Ardanto, Wimbi Kartika Ratnasari, Kusumaningtyas Sekar Negari, Aprilia Safitri, Dimas Bagus Wicaksono Putro, Esti Rahardianingtyas, Sidiq Setyo Nugroho, Marjianto, Muhidin, Subangkit, Hana Apsari Pawestri, Siti Alfiah, Ristiyanto, Evi sulistyorini, Lulus Susanti, Vivi Setiawaty, Joko Waluyo |
| EPI_ISL_2886165 | IVRCRD RSUD Budi Rahayu | Institute for Vector and Reservoir Control, Research and Development (IVRCRD)- National Institute of Health Research and Development (NIHRD), Indonesia | Arief Mulyono, Triwibowo Ambar Garjito, Restu Choirul Saban, Ayu Pradipta Pratiwi, Dwi Susilo, Yusnita Mirna Anggraeni, Mujiyanto, Mega Tyas Prihatin, Nur Hidayati, Arif Suryo Prasetyo, Tika Fiona Sari, Sri Wahyuni Handayani, Aryo Ardanto, Wimbi Kartika Ratnasari, Kusumaningtyas Sekar Negari, Aprilia Safitri, Dimas Bagus Wicaksono Putro, Esti Rahardianingtyas, Sidiq Setyo Nugroho, Marjianto, Muhidin, Subangkit, Hana Apsari Pawestri, Siti Alfiah, Ristiyanto, Evi sulistyorini, Lulus Susanti, Vivi Setiawaty, Joko Waluyo |
| EPI_ISL_2886194 | Laboratorio di Riferimento Regionale della Sicilia Occidentale per l'Emergenza COVID-19 | Laboratorio di Riferimento Regionale della Sicilia Occidentale per l'Emergenza COVID-19 | Fabio Tramuto, Carmelo Massimo Maida, Daniela Di Naro, Giulia Randazzo, Walter Mazzucco, Giorgio Graziano, Vincenzo Restivo, Claudio Costantino, Francesco Vitale |
| EPI_ISL_2886437, EPI_ISL_2886438, EPI_ISL_2886439, EPI_ISL_2886441 | CHU de la Réunion - Nord | Laboratoire de virologie, CNR arbovirus Associé, Chu de la Réunion | Etienne Frumence, Marie-Christine Jaffar Bandjee, Nicolas Traversier, Sabrina Petit Genet |
| EPI_ISL_2886442 | CHU de la Réunion - Sud | Laboratoire de virologie, CNR arbovirus Associé, Chu de la Réunion | Etienne Frumence, Marie-Christine Jaffar Bandjee, Nicolas Traversier, Sabrina Petit Genet |
| EPI_ISL_2886444 | CHU de la Réunion - Nord | Laboratoire de virologie, CNR arbovirus Associé, Chu de la Réunion | Etienne Frumence, Marie-Christine Jaffar Bandjee, Nicolas Traversier, Sabrina Petit Genet |
| EPI_ISL_2886445, EPI_ISL_2886447, EPI_ISL_2886449 | CHU de la Réunion - Sud | Laboratoire de virologie, CNR arbovirus Associé, Chu de la Réunion | Etienne Frumence, Marie-Christine Jaffar Bandjee, Nicolas Traversier, Sabrina Petit Genet |
| EPI_ISL_2886450 | CHU de la Réunion - Nord | Laboratoire de virologie, CNR arbovirus Associé, Chu de la Réunion | Etienne Frumence, Marie-Christine Jaffar Bandjee, Nicolas Traversier, Sabrina Petit Genet |
| EPI_ISL_2886451, EPI_ISL_2886452 | CHU de la Réunion - Sud | Laboratoire de virologie, CNR arbovirus Associé, Chu de la Réunion | Etienne Frumence, Marie-Christine Jaffar Bandjee, Nicolas Traversier, Sabrina Petit Genet |
| EPI_ISL_2886453, EPI_ISL_2886454, EPI_ISL_2886455 | CHU de la Réunion - Nord | Laboratoire de virologie, CNR arbovirus Associé, Chu de la Réunion | Etienne Frumence, Marie-Christine Jaffar Bandjee, Nicolas Traversier, Sabrina Petit Genet |
| EPI_ISL_2887417, EPI_ISL_2887418, EPI_ISL_2887420, EPI_ISL_2887422, EPI_ISL_2887423, EPI_ISL_2887424, EPI_ISL_2887425, EPI_ISL_2887429, EPI_ISL_2887430, EPI_ISL_2887432, EPI_ISL_2887433, EPI_ISL_2887434, EPI_ISL_2887436, EPI_ISL_2887439, EPI_ISL_2887441, EPI_ISL_2887442, EPI_ISL_2887443, EPI_ISL_2887445, EPI_ISL_2887448, EPI_ISL_2887450, EPI_ISL_2887451, EPI_ISL_2887452, EPI_ISL_2887453, EPI_ISL_2887454, EPI_ISL_2887456, EPI_ISL_2887457, EPI_ISL_2887460, EPI_ISL_2887461, EPI_ISL_2887462, EPI_ISL_2887463, EPI_ISL_2887464, EPI_ISL_2887466, EPI_ISL_2887469, EPI_ISL_2887471, | | | |

| | | | |
|---|---|---|---|
| EPI_ISL_2887472, EPI_ISL_2887473, EPI_ISL_2887477, EPI_ISL_2887478, EPI_ISL_2887480, EPI_ISL_2887481, EPI_ISL_2887483, EPI_ISL_2887484, EPI_ISL_2887485, EPI_ISL_2887486, EPI_ISL_2887487, EPI_ISL_2887488, EPI_ISL_2887490, EPI_ISL_2887491, EPI_ISL_2887493, EPI_ISL_2887494, EPI_ISL_2887498, EPI_ISL_2887499, EPI_ISL_2887501, EPI_ISL_2887502, EPI_ISL_2887503, EPI_ISL_2887505, EPI_ISL_2887506, EPI_ISL_2887507, EPI_ISL_2887508, EPI_ISL_2887510, EPI_ISL_2887511, EPI_ISL_2887512, EPI_ISL_2887513, EPI_ISL_2887515, EPI_ISL_2887516, EPI_ISL_2887519, EPI_ISL_2887520, EPI_ISL_2887522, EPI_ISL_2887524, EPI_ISL_2887525, EPI_ISL_2887527, EPI_ISL_2887528, EPI_ISL_2887530, EPI_ISL_2887531, EPI_ISL_2887534, EPI_ISL_2887536, EPI_ISL_2887537, EPI_ISL_2887538, EPI_ISL_2887539, EPI_ISL_2887540, EPI_ISL_2887543, EPI_ISL_2887544, EPI_ISL_2887545, EPI_ISL_2887546, EPI_ISL_2887547, EPI_ISL_2887548, EPI_ISL_2887550, EPI_ISL_2887553, EPI_ISL_2887554, EPI_ISL_2887555, EPI_ISL_2887556, EPI_ISL_2887557, EPI_ISL_2887558, EPI_ISL_2887559, EPI_ISL_2887560, EPI_ISL_2887561, EPI_ISL_2887562, EPI_ISL_2887563, EPI_ISL_2887564, EPI_ISL_2887566, EPI_ISL_2887567, EPI_ISL_2887568, EPI_ISL_2887569, EPI_ISL_2887570, EPI_ISL_2887571, EPI_ISL_2887572, EPI_ISL_2887573, EPI_ISL_2887574, EPI_ISL_2887576, EPI_ISL_2887577, EPI_ISL_2887578, EPI_ISL_2887579, EPI_ISL_2887580, EPI_ISL_2887581, EPI_ISL_2887582, EPI_ISL_2887583, EPI_ISL_2887585, EPI_ISL_2887586, EPI_ISL_2887587, EPI_ISL_2887588, EPI_ISL_2887589, EPI_ISL_2887590, EPI_ISL_2887592, EPI_ISL_2887594, EPI_ISL_2887596, EPI_ISL_2887597, EPI_ISL_2887598, EPI_ISL_2887599, EPI_ISL_2887600, EPI_ISL_2887601, EPI_ISL_2887603, EPI_ISL_2887604, EPI_ISL_2887605, EPI_ISL_2887607, EPI_ISL_2887609, EPI_ISL_2887610, EPI_ISL_2887611, EPI_ISL_2887613, EPI_ISL_2887614, EPI_ISL_2887615, EPI_ISL_2887616, EPI_ISL_2887617, EPI_ISL_2887618, EPI_ISL_2887619, EPI_ISL_2887620, EPI_ISL_2887621, EPI_ISL_2887622, EPI_ISL_2887623, EPI_ISL_2887624, EPI_ISL_2887625, EPI_ISL_2887626, EPI_ISL_2887627, EPI_ISL_2887628, EPI_ISL_2887629, EPI_ISL_2887631, EPI_ISL_2887632, EPI_ISL_2887633, EPI_ISL_2887634, EPI_ISL_2887636, EPI_ISL_2887637, EPI_ISL_2887638, EPI_ISL_2887639, EPI_ISL_2887640, EPI_ISL_2887641, EPI_ISL_2887642, EPI_ISL_2887643, EPI_ISL_2887644, EPI_ISL_2887645, EPI_ISL_2887647, EPI_ISL_2887648, EPI_ISL_2887649, EPI_ISL_2887650, EPI_ISL_2887651, EPI_ISL_2887652, EPI_ISL_2887653, EPI_ISL_2887654, EPI_ISL_2887655, EPI_ISL_2887656, EPI_ISL_2887657, EPI_ISL_2887658, EPI_ISL_2887659, EPI_ISL_2887660, EPI_ISL_2887661, EPI_ISL_2887662, EPI_ISL_2887663, EPI_ISL_2887664, EPI_ISL_2887665, EPI_ISL_2887666, EPI_ISL_2887667, EPI_ISL_2887668, EPI_ISL_2887669, EPI_ISL_2887670, EPI_ISL_2887671, EPI_ISL_2887672, EPI_ISL_2887673, EPI_ISL_2887674, EPI_ISL_2887675, EPI_ISL_2887676, EPI_ISL_2887677, EPI_ISL_2887679, EPI_ISL_2887681, EPI_ISL_2887682, EPI_ISL_2887683 | | | |
| see above | Austrian Agency for Health and Food Safety (AGES) | Bergthaler laboratory, CeMM Research Center for Molecular Medicine of the Austrian Academy of Sciences | Lukas Endler, Anna Schedl, Fabian Amman, Petr Triska, Matthew Thornton, Thomas Penz, Benedikt Agerer, Maelle Le Moing, Michael Schuster, Bekir Erguner, Jan Laine, Martin Senekowitsch, Christoph Bock, Andreas Bergthaler |
| EPI_ISL_2887701 | Center for Virology, Medical University of Vienna | Bergthaler laboratory, CeMM Research Center for Molecular Medicine of the Austrian Academy of Sciences | Lukas Endler, Anna Schedl, Fabian Amman, Petr Triska, Matthew Thornton, Thomas Penz, Benedikt Agerer, Maelle Le Moing, Michael Schuster, Bekir Erguner, Jan Laine, Martin Senekowitsch, Christoph Bock, Andreas Bergthaler |
| EPI_ISL_2887703, EPI_ISL_2887704, EPI_ISL_2887705, EPI_ISL_2887706, EPI_ISL_2887707, EPI_ISL_2887708, EPI_ISL_2887709, EPI_ISL_2887710, EPI_ISL_2887711, EPI_ISL_2887712 | CHU Lille | CHU Lille - Laboratoire de Virologie | AIT YAHYA Emilie, ALIDJINOU Enagnon Kazali, BOCKET Laurence, CREPIN Michel, DEMAY Christophe, ENGELMANN Ilka, GEFFROY Sandrine, GUIGON Aurélie, LAMBERT Valérie, LAZREK Mouna, NOBILLIAUX Florian, PREVOST Brigitte, THUILLIER Caroline, TINEZ Claire ,TCHANTCHOU NJOSSE YANICK |
| EPI_ISL_2887713 | CH Bethune | CHU Lille - Laboratoire de Virologie | AIT YAHYA Emilie, ALIDJINOU Enagnon Kazali, BOCKET Laurence, CREPIN Michel, DEMAY Christophe, ENGELMANN Ilka, GEFFROY Sandrine, GUIGON Aurélie, LAMBERT Valérie, LAZREK Mouna, NOBILLIAUX Florian, PREVOST Brigitte, THUILLIER Caroline, TINEZ Claire ,TCHANTCHOU NJOSSE YANICK |
| EPI_ISL_2887714, EPI_ISL_2887715 | CH Boulogne sur Mer | CHU Lille - Laboratoire de Virologie | AIT YAHYA Emilie, ALIDJINOU Enagnon Kazali, BOCKET Laurence, CREPIN Michel, DEMAY Christophe, ENGELMANN Ilka, GEFFROY Sandrine, GUIGON Aurélie, LAMBERT Valérie, LAZREK Mouna, NOBILLIAUX Florian, PREVOST Brigitte, THUILLIER Caroline, TINEZ Claire ,TCHANTCHOU NJOSSE YANICK |
| EPI_ISL_2887716, EPI_ISL_2887717, EPI_ISL_2887718 | CH Roubaix | CHU Lille - Laboratoire de Virologie | AIT YAHYA Emilie, ALIDJINOU Enagnon Kazali, BOCKET Laurence, CREPIN Michel, DEMAY Christophe, ENGELMANN Ilka, GEFFROY Sandrine, GUIGON Aurélie, LAMBERT Valérie, LAZREK Mouna, NOBILLIAUX Florian, PREVOST Brigitte, THUILLIER Caroline, TINEZ Claire ,TCHANTCHOU NJOSSE YANICK |
| EPI_ISL_2887719, EPI_ISL_2887720, EPI_ISL_2887721 | CH Calais | CHU Lille - Laboratoire de Virologie | AIT YAHYA Emilie, ALIDJINOU Enagnon Kazali, BOCKET Laurence, CREPIN Michel, DEMAY Christophe, ENGELMANN Ilka, GEFFROY Sandrine, GUIGON Aurélie, LAMBERT Valérie, LAZREK Mouna, NOBILLIAUX Florian, PREVOST Brigitte, THUILLIER Caroline, TINEZ Claire ,TCHANTCHOU NJOSSE YANICK |
| EPI_ISL_2887722, EPI_ISL_2887723 | CH Cambrai | CHU Lille - Laboratoire de Virologie | AIT YAHYA Emilie, ALIDJINOU Enagnon Kazali, BOCKET Laurence, CREPIN Michel, DEMAY Christophe, ENGELMANN Ilka, GEFFROY Sandrine, GUIGON Aurélie, LAMBERT Valérie, LAZREK Mouna, NOBILLIAUX Florian, PREVOST Brigitte, THUILLIER Caroline, TINEZ Claire ,TCHANTCHOU NJOSSE YANICK |
| EPI_ISL_2887724, EPI_ISL_2887725 | CH Dunkerque | CHU Lille - Laboratoire de Virologie | AIT YAHYA Emilie, ALIDJINOU Enagnon Kazali, BOCKET Laurence, CREPIN Michel, DEMAY Christophe, ENGELMANN Ilka, GEFFROY Sandrine, GUIGON Aurélie, LAMBERT Valérie, LAZREK Mouna, NOBILLIAUX Florian, PREVOST Brigitte, THUILLIER Caroline, TINEZ Claire ,TCHANTCHOU NJOSSE YANICK |
| EPI_ISL_2887726, EPI_ISL_2887727, EPI_ISL_2887728, EPI_ISL_2887729 | CH Valenciennes | CHU Lille - Laboratoire de Virologie | AIT YAHYA Emilie, ALIDJINOU Enagnon Kazali, BOCKET Laurence, CREPIN Michel, DEMAY Christophe, ENGELMANN Ilka, GEFFROY Sandrine, GUIGON Aurélie, LAMBERT Valérie, LAZREK Mouna, NOBILLIAUX Florian, PREVOST Brigitte, THUILLIER Caroline, TINEZ Claire ,TCHANTCHOU NJOSSE YANICK |
| EPI_ISL_2887730 | CH Lens | CHU Lille - Laboratoire de Virologie | AIT YAHYA Emilie, ALIDJINOU Enagnon Kazali, BOCKET Laurence, CREPIN Michel, DEMAY Christophe, ENGELMANN Ilka, GEFFROY Sandrine, GUIGON Aurélie, LAMBERT Valérie, LAZREK Mouna, NOBILLIAUX Florian, PREVOST Brigitte, THUILLIER Caroline, TINEZ Claire ,TCHANTCHOU NJOSSE YANICK |
| EPI_ISL_2887732, EPI_ISL_2887733 | CH Roubaix | CHU Lille - Laboratoire de Virologie | AIT YAHYA Emilie, ALIDJINOU Enagnon Kazali, BOCKET Laurence, CREPIN Michel, DEMAY Christophe, ENGELMANN Ilka, GEFFROY Sandrine, GUIGON Aurélie, LAMBERT Valérie, LAZREK Mouna, NOBILLIAUX Florian, PREVOST Brigitte, THUILLIER Caroline, TINEZ Claire ,TCHANTCHOU NJOSSE YANICK |
| EPI_ISL_2887734, EPI_ISL_2887735, EPI_ISL_2887736, EPI_ISL_2887737, EPI_ISL_2887738, EPI_ISL_2887739, EPI_ISL_2887740, EPI_ISL_2887741, EPI_ISL_2887742, EPI_ISL_2887743, EPI_ISL_2887744 | | | |
| see above | CHU Lille | CHU Lille - Laboratoire de Virologie | AIT YAHYA Emilie, ALIDJINOU Enagnon Kazali, BOCKET Laurence, CREPIN Michel, DEMAY Christophe, ENGELMANN Ilka, GEFFROY Sandrine, GUIGON Aurélie, LAMBERT Valérie, LAZREK Mouna, NOBILLIAUX Florian, PREVOST Brigitte, THUILLIER Caroline, TINEZ Claire ,TCHANTCHOU NJOSSE YANICK |
| EPI_ISL_2887745, EPI_ISL_2887746, EPI_ISL_2887747, EPI_ISL_2887748 | CH Tourcoing | CHU Lille - Laboratoire de Virologie | AIT YAHYA Emilie, ALIDJINOU Enagnon Kazali, BOCKET Laurence, CREPIN Michel, DEMAY Christophe, ENGELMANN Ilka, GEFFROY Sandrine, GUIGON Aurélie, LAMBERT Valérie, LAZREK Mouna, NOBILLIAUX Florian, PREVOST Brigitte, THUILLIER Caroline, TINEZ Claire ,TCHANTCHOU NJOSSE YANICK |
| EPI_ISL_2888012, EPI_ISL_2888013, EPI_ISL_2888014, EPI_ISL_2888015, EPI_ISL_2888017, EPI_ISL_2888019, EPI_ISL_2888020 | Ospedale Sandro Pertini ASL Roma2 | Department of General Diagnostics; Department of Virology; Istituto Zooprofilattico Sperimentale del Lazio e della Toscana (IZSLT) | Patricia Alba, Giuseppe Manna, Elena L. Diaconu, Fabiola Feltrin, Raffaela Conti, Teresa Scicluna, Virginia Carfora, Antonella Cersini, Alessia Franco, Antonio Battisti. |
| EPI_ISL_2888022, EPI_ISL_2888023, EPI_ISL_2888024, EPI_ISL_2888025, EPI_ISL_2888026, EPI_ISL_2888028, EPI_ISL_2888029, EPI_ISL_2888032, EPI_ISL_2888035, EPI_ISL_2888037, EPI_ISL_2888038, EPI_ISL_2888040, EPI_ISL_2888041, EPI_ISL_2888042 | | | |
| see above | Ospedale San Filippo Neri | Department of General Diagnostics; Department of Virology; Istituto Zooprofilattico Sperimentale del Lazio e della Toscana (IZSLT) | Patricia Alba, Giuseppe Manna, Elena L. Diaconu, Fabiola Feltrin, Raffaela Conti, Teresa Scicluna, Virginia Carfora, Antonella Cersini, Alessia Franco, Antonio Battisti. |
| EPI_ISL_2893524, EPI_ISL_2893525, EPI_ISL_2893526, EPI_ISL_2893527, EPI_ISL_2893530, EPI_ISL_2893531, EPI_ISL_2893532 | Laboratoire Hôpital Sainte Musse | Laboratoire Hôpital Sainte Musse | Lionel Chollet |
| EPI_ISL_2894130 | MEDILAB | CHU Poitiers | Caroline MICHAUD, Maxime PICHON, Manon PRAT, Valentin BON-BARET, Birama N'DIAYE, Magali GARCIA, Agnes BEBY-DEFAUX, Nicolas LEVEQUE |
| EPI_ISL_2894131, EPI_ISL_2894132, EPI_ISL_2894133 | BIO86 GENCAY | CHU Poitiers | Caroline MICHAUD, Maxime PICHON, Manon PRAT, Valentin BON-BARET, Birama N'DIAYE, Magali GARCIA, Agnes BEBY-DEFAUX, Nicolas LEVEQUE |
| EPI_ISL_2894547, EPI_ISL_2894548 | LACEN do Estado de Mato Grosso | Instituto Adolfo Lutz, Interdiciplinary Procedures Center, Strategic Laboratory | Claudio Tavares Sacchi, Claudia Regina Gonçalves, Karoline Rodrigues Campos, Caio Vinicius Dias Lopes |
| EPI_ISL_2894553, EPI_ISL_2894554, EPI_ISL_2894555, EPI_ISL_2894556, EPI_ISL_2894557, EPI_ISL_2894558, EPI_ISL_2894559, EPI_ISL_2894560, EPI_ISL_2894561 | Hospital Margarita Maza de Juárez | Microbial Genomics Laboratory | Consorcio Mexicano de Vigilancia Genómica (CoViGen-Mex). Authors (in alphabetical order): Julio Elias Alvarado-Yaah, Carlos F. Arias, Santiago Ávila-Ríos, Eduardo Becerril-Vargas, Víctor Hugo Borja-Aburto, Celia Boukadida, Cristóbal Cháidez-Quiróz, Juan Bautista Chale-Dzul, Ricardo Ciria Merce, Andreu Comas-García, Célida Duque Molina, Julissa Enciso-Ibarra, José Antonio Enciso-Moreno, Gloria Elena Espinosa-Ayala, Fernando Fontove-Herrera, Daniel Fregoso-Rueda, Victor Eduardo García-Arias, Alejandra García-Gasca, Bruno Gómez-Gil, Jean Pierre González, Irvin González-López, Concepción Grajales-Muñiz, Ricardo Grande, Rosa María Gutiérrez Rios, Jesús Hernández, Alejandra Hernández-Terán, Alfredo |

| | | | |
|---|---|---|---|
| | | | Herrera-Estrella, Carla Ivón Herrera-Najera, Pavel Isa, Verónica Mata-Haro, Daniel Lira Morales, Susana Lopez, Antonio Loza Román, Brenda Irasema Maldonado-Meza, Bernardo Martínez-Miguel, José Arturo Martínez-Orozco, Célida Martínez- Rodríguez, Margarita Matías-Florentino, Fidencio Mejía-Nepomuceno, María Guadalupe de Jesús Mireles-Rivera, Gloria María Molina-Salinas, Hector Montoya-Fuentes, , Mario Mújica-Sánchez, José Esteban Muñoz-Medina, José de Jesús Nuñez-Contreras, Alicia Ocaña-Mondragón, Luis Alberto Ochoa-Carrera, Hector Esteban Paz-Juárez, Marissa Perez-Garcia, Francisco Pulido, Helen Haydee Fernanda Ramírez-Plascencia, Jorge Salas-Hernández, Angel Gustavo Salas-Lais, Alejandro Sánchez-Flores, Clara Esperanza Santacruz-Tinoco, María Guadalupe Santiago-Mauricio, Selene Zárate, Nelly Sélem-Mojica, Blanca Taboada, Gloria Vazquez, Joel Armando Vázquez-Pérez. |
| EPI_ISL_2894562, EPI_ISL_2894563, EPI_ISL_2894564 | Hospital Sharp | Microbial Genomics Laboratory | Consorcio Mexicano de Vigilancia Genómica (CoViGen-Mex). Authors (in alphabetical order): Julio Elias Alvarado-Yaah, Carlos F. Arias, Santiago Ávila-Ríos, Eduardo Becerril-Vargas, Víctor Hugo Borja-Aburto, Celia Boukadida, Cristóbal Cháidez-Quiróz, Juan Bautista Chale-Dzul, Ricardo Ciria Merce, Andreu Comas-García, Célida Duque Molina, Julissa Enciso-Ibarra, José Antonio Enciso-Moreno, Gloria Elena Espinosa-Ayala, Fernando Fontove-Herrera, Daniel Fregoso-Rueda, Victor Eduardo García-Arias, Alejandra García-Gasca, Bruno Gómez-Gil, Jean Pierre González, Irvin González-López, Concepción Grajales-Muñiz, Ricardo Grande, Rosa María Gutiérrez Rios, Jesús Hernández, Alejandra Hernández-Terán, Alfredo Herrera-Estrella, Carla Ivón Herrera-Najera, Pavel Isa, Verónica Mata-Haro, Daniel Lira Morales, Susana Lopez, Antonio Loza Román, Brenda Irasema Maldonado-Meza, Bernardo Martínez-Miguel, José Arturo Martínez-Orozco, Célida Martínez- Rodríguez, Margarita Matías-Florentino, Fidencio Mejía-Nepomuceno, María Guadalupe de Jesús Mireles-Rivera, Gloria María Molina-Salinas, Hector Montoya-Fuentes, , Mario Mújica-Sánchez, José Esteban Muñoz-Medina, José de Jesús Nuñez-Contreras, Alicia Ocaña-Mondragón, Luis Alberto Ochoa-Carrera, Hector Esteban Paz-Juárez, Marissa Perez-Garcia, Francisco Pulido, Helen Haydee Fernanda Ramírez-Plascencia, Jorge Salas-Hernández, Angel Gustavo Salas-Lais, Alejandro Sánchez-Flores, Clara Esperanza Santacruz-Tinoco, María Guadalupe Santiago-Mauricio, Selene Zárate, Nelly Sélem-Mojica, Blanca Taboada, Gloria Vazquez, Joel Armando Vázquez-Pérez. |
| EPI_ISL_2894566, EPI_ISL_2894568, EPI_ISL_2894569 | CIAD Hermosillo | CIAD Hermosillo | Consorcio Mexicano de Vigilancia Genómica (CoViGen-Mex). Authors (in alphabetical order): Julio Elias Alvarado-Yaah, Carlos F. Arias, Santiago Ávila-Ríos, Eduardo Becerril-Vargas, Víctor Hugo Borja-Aburto, Celia Boukadida, Cristóbal Cháidez-Quiróz, Juan Bautista Chale-Dzul, Ricardo Ciria Merce, Andreu Comas-García, Célida Duque Molina, Julissa Enciso-Ibarra, José Antonio Enciso-Moreno, Gloria Elena Espinosa-Ayala, Fernando Fontove-Herrera, Daniel Fregoso-Rueda, Victor Eduardo García-Arias, Alejandra García-Gasca, Bruno Gómez-Gil, Jean Pierre González, Irvin González-López, Concepción Grajales-Muñiz, Ricardo Grande, Rosa María Gutiérrez Rios, Jesús Hernández, Alejandra Hernández-Terán, Alfredo Herrera-Estrella, Carla Ivón Herrera-Najera, Pavel Isa, Verónica Mata-Haro, Daniel Lira Morales, Susana Lopez, Antonio Loza Román, Brenda Irasema Maldonado-Meza, Bernardo Martínez-Miguel, José Arturo Martínez-Orozco, Célida Martínez- Rodríguez, Margarita Matías-Florentino, Fidencio Mejía-Nepomuceno, María Guadalupe de Jesús Mireles-Rivera, Gloria María Molina-Salinas, Hector Montoya-Fuentes, , Mario Mújica-Sánchez, José Esteban Muñoz-Medina, José de Jesús Nuñez-Contreras, Alicia Ocaña-Mondragón, Luis Alberto Ochoa-Carrera, Hector Esteban Paz-Juárez, Marissa Perez-Garcia, Francisco Pulido, Helen Haydee Fernanda Ramírez-Plascencia, Jorge Salas-Hernández, Angel Gustavo Salas-Lais, Alejandro Sánchez-Flores, Clara Esperanza Santacruz-Tinoco, María Guadalupe Santiago-Mauricio, Selene Zárate, Nelly Sélem-Mojica, Blanca Taboada, Gloria Vazquez, Joel Armando Vázquez-Pérez. |
| EPI_ISL_2896232, EPI_ISL_2896233, EPI_ISL_2896234, EPI_ISL_2896235, EPI_ISL_2896236, EPI_ISL_2896237, EPI_ISL_2896238, EPI_ISL_2896239, EPI_ISL_2896240, EPI_ISL_2896241 | LACEN do Estado de Mato Grosso | Instituto Adolfo Lutz, Interdiciplinary Procedures Center, Strategic Laboratory | Claudio Tavares Sacchi, Claudia Regina Gonçalves, Karoline Rodrigues Campos, Caio Vinicius Dias Lopes |
| EPI_ISL_2896258 | Osaka General Medical Center | Department of Developmental Medicine, Research Institute, Osaka Women's and Children's Hospital | Itaru Yanagihara, Yukiko Nakura |
| EPI_ISL_2897094 | Althaia. Xarxa Assistencial Universitària de Manresa | IrsiCaixa | Marc Noguera-Julian, Mariona Parera, Maria Casadellà, Pilar Armengol, Francesc Catala-Moll, Roger Paredes, Bonaventura Clotet Gloria Trujillo, Rafel Perez Vidal, Jaume Trapé, Carolina Gonzalez-Fernandez, Miquel Micó, Antonia Flor, Roger Paredes, Eulalia Grau, Bonaventura Clotet |
| EPI_ISL_2918940 | Biomédicos de Mérida | Biomédicos de Mérida | CANCHE-PECH JOSE REYES, MENDOZA-OLIVEROS TAHALI, CHIN-HAU LETICIA, SAURI-MEDINA EMILY, CAUICH-VERA KRISTIAN, CEBALLOS-LOPEZ ADRIAN, LOPEZ-NOVELO MARIA E, GARCIA-GONZALEZ IGRID |
| EPI_ISL_2918997 | Laboratório de Microbiologia Molecular - Universidade FEEVALE | Laboratório de Microbiologia Molecular - Universidade FEEVALE | Alana Witt Hansen, Fágner Henrique Heldt, Fernando Rosado Spilki, Flávio Silveira, Juliana Schons Gulart, Juliane Deise Fleck, Mariana Soares da Silva, Meriane Demoliner, Matheus Nunes Weber, Paula Rodrigues de Almeida, Michele Filippi. |
| EPI_ISL_2918998 | Laboratório de Microbiologia Molecular - Universidade FEEVALE | Laboratório de Microbiologia Molecular - Universidade FEEVALE | Alana Witt Hansen, Fágner Henrique Heldt, Fernando Rosado Spilki, Flávio Silveira, Juliana Schons Gularte, Juliane Deise Fleck, Mariana Soares da Silva, Meriane Demoliner, Matheus Nunes Weber, Paula Rodrigues de Almeida, Micheli Filippi. |
| EPI_ISL_2918999, EPI_ISL_2919001 | Laboratório de Microbiologia Molecular - Universidade FEEVALE | Laboratório de Microbiologia Molecular - Universidade FEEVALE | Alana Witt Hansen, Fágner Henrique Heldt, Fernando Rosado Spilki, Flávio Silveira, Juliana Schons Gulart, Juliane Deise Fleck, Mariana Soares da Silva, Meriane Demoliner, Matheus Nunes Weber, Paula Rodrigues de Almeida, Michele Filippi. |
| EPI_ISL_2919002 | Laboratório de Microbiologia Molecular - Universidade FEEVALE | Laboratório de Microbiologia Molecular - Universidade FEEVALE | Alana Witt Hansen, Fágner Henrique Heldt, Fernando Rosado Spilki, Flávio Silveira, Juliana Schons Gularte, Juliane Deise Fleck, Mariana Soares da Silva, Meriane Demoliner, Matheus Nunes Weber, Paula Rodrigues de Almeida, Micheli Filippi. |
| EPI_ISL_2919003 | Laboratório de Microbiologia Molecular - Universidade FEEVALE | Laboratório de Microbiologia Molecular - Universidade FEEVALE | Alana Witt Hansen, Fágner Henrique Heldt, Fernando Rosado Spilki, Flávio Silveira, Juliana Schons Gulart, Juliane Deise Fleck, Mariana Soares da Silva, Meriane Demoliner, Matheus Nunes Weber, Paula Rodrigues de Almeida, Michele Filippi. |
| EPI_ISL_2919004 | Laboratório de Microbiologia Molecular - Universidade FEEVALE | Laboratório de Microbiologia Molecular - Universidade FEEVALE | Alana Witt Hansen, Fágner Henrique Heldt, Fernando Rosado Spilki, Flávio Silveira, Juliana Schons Gularte, Juliane Deise Fleck, Mariana Soares da Silva, Meriane Demoliner, Matheus Nunes Weber, Paula Rodrigues de Almeida, Micheli Filippi. |
| EPI_ISL_2919005 | Laboratório de Microbiologia Molecular - Universidade FEEVALE | Laboratório de Microbiologia Molecular - Universidade FEEVALE | Alana Witt Hansen, Fágner Henrique Heldt, Fernando Rosado Spilki, Flávio Silveira, Juliana Schons Gulart, Juliane Deise Fleck, Mariana Soares da Silva, Meriane Demoliner, Matheus Nunes Weber, Paula Rodrigues de Almeida, Michele Filippi. |
| EPI_ISL_2919006 | Laboratório de Microbiologia Molecular - Universidade FEEVALE | Laboratório de Microbiologia Molecular - Universidade FEEVALE | Alana Witt Hansen, Fágner Henrique Heldt, Fernando Rosado Spilki, Flávio Silveira, Juliana Schons Gularte, Juliane Deise Fleck, Mariana Soares da Silva, Meriane Demoliner, Matheus Nunes Weber, Paula Rodrigues de Almeida, Micheli Filippi. |
| EPI_ISL_2919013 | Secretaria Municipal da Saúde- Porto Alegre | Laboratório de Microbiologia Molecular - Universidade FEEVALE | Alana Witt Hansen, Fágner Henrique Heldt, Fernando Rosado Spilki, Flávio Silveira, Juliana Schons Gularte, Juliane Deise Fleck, Mariana Soares da Silva, Meriane Demoliner, Matheus Nunes Weber, Paula Rodrigues de Almeida, Micheli Filippi. |
| EPI_ISL_2919014 | Secretaria Municipal da Saúde- Porto Alegre | Laboratório de Microbiologia Molecular - Universidade FEEVALE | Alana Witt Hansen, Fágner Henrique Heldt, Fernando Rosado Spilki, Flávio Silveira, Juliana Schons Gulart, Juliane Deise Fleck, Mariana Soares da Silva, Meriane Demoliner, Matheus Nunes Weber, Paula Rodrigues de Almeida, Michele Filippi. |
| EPI_ISL_2919104, EPI_ISL_2919105, EPI_ISL_2919106, EPI_ISL_2919107, EPI_ISL_2919108, EPI_ISL_2919109, EPI_ISL_2919111, EPI_ISL_2919114, EPI_ISL_2919115, EPI_ISL_2919116 | Università degli Studi di Perugia | Istituto Zooprofilattico Sperimentale dell'Abruzzo e Molise "G. Caporale" | Mencacci A, Camilloni B, Biagetti M, Giammarioli M, Lorusso A, Marcacci M, Di Domenico M, Ancora M, Curini V, Mangone I, Rinaldi A, Delli Compagni E, Scialabba S, Di Pasquale A, Cammà C, Puglia I, Calistri P, Savini G |
| EPI_ISL_2919118, EPI_ISL_2919119 | Laboratorio Analisi Osp. Città di Castello - Azienda USL Umbria1 | Istituto Zooprofilattico Sperimentale dell'Abruzzo e Molise "G. Caporale" | Malagigi V, Tacconi P,Lorusso A, Marcacci M, Di Domenico M, Ancora M, Curini V, Mangone I, Rinaldi A, Delli Compagni E, Scialabba S, Di Pasquale A, Cammà C, Puglia I, Calistri P, Savini G |
| EPI_ISL_2919120 | Università degli Studi di Perugia | Istituto Zooprofilattico Sperimentale dell'Abruzzo e Molise "G. Caporale" | Mencacci A, Camilloni B, Biagetti M, Giammarioli M, Lorusso A, Marcacci M, Di Domenico M, Ancora M, Curini V, Mangone I, Rinaldi A, Delli Compagni E, Scialabba S, Di Pasquale A, Cammà C, Puglia I, Calistri P, Savini G |
| EPI_ISL_2919122, EPI_ISL_2919123, EPI_ISL_2919125, EPI_ISL_2919126, EPI_ISL_2919127, EPI_ISL_2919128, EPI_ISL_2919129 | Azienda USL Umbria 2 | Istituto Zooprofilattico Sperimentale dell'Abruzzo e Molise "G. Caporale" | Proietti A, Pistoni E, Lorusso A, Marcacci M, Di Domenico M, Ancora M, Curini V, Mangone I, Rinaldi A, Delli Compagni E, Scialabba S, Di Pasquale A, Cammà C, Puglia I, Calistri P, Savini G |
| EPI_ISL_2919141, EPI_ISL_2919142, EPI_ISL_2919148, EPI_ISL_2919154, | Fakultní nemocnice Hradec Králové | Fakultní nemocnice Hradec Králové | Marketa Gancarcikova, Helena Kovarikova |

| | | | |
|---|---|---|---|
| EPI_ISL_2919157, EPI_ISL_2919158, EPI_ISL_2919162 | | | |
| EPI_ISL_2919233, EPI_ISL_2919234 | Instituto Adolfo Lutz Central | Instituto Adolfo Lutz, Interdiciplinary Procedures Center, Strategic Laboratory | Claudio Tavares Sacchi, Claudia Regina Gonçalves, Erica Valessa Ramos Gomes, Karoline Rodrigues Campos, Caio Vinicius Dias Lopes |
| EPI_ISL_2919255, EPI_ISL_2919256, EPI_ISL_2919257 | Instituto Adolfo Lutz - Regional de Bauru | Instituto Adolfo Lutz, Interdiciplinary Procedures Center, Strategic Laboratory | Claudio Tavares Sacchi, Claudia Regina Gonçalves, Erica Valessa Ramos Gomes, Karoline Rodrigues Campos, Caio Vinicius Dias Lopes |
| EPI_ISL_2919263 | Instituto Adolfo Lutz - Regional de Campinas | Instituto Adolfo Lutz, Interdiciplinary Procedures Center, Strategic Laboratory | Claudio Tavares Sacchi, Claudia Regina Gonçalves, Erica Valessa Ramos Gomes, Karoline Rodrigues Campos, Caio Vinicius Dias Lopes |
| EPI_ISL_2919265, EPI_ISL_2919266, EPI_ISL_2919267 | Instituto Adolfo Lutz - Regional de Sao Jose do Rio Preto | Instituto Adolfo Lutz, Interdiciplinary Procedures Center, Strategic Laboratory | Claudio Tavares Sacchi, Claudia Regina Gonçalves, Erica Valessa Ramos Gomes, Karoline Rodrigues Campos, Caio Vinicius Dias Lopes |
| EPI_ISL_2919268, EPI_ISL_2919269, EPI_ISL_2919270, EPI_ISL_2919271, EPI_ISL_2919272, EPI_ISL_2919273, EPI_ISL_2919274, EPI_ISL_2919275, EPI_ISL_2919276, EPI_ISL_2919277, EPI_ISL_2919278, EPI_ISL_2919279, EPI_ISL_2919280, EPI_ISL_2919281, EPI_ISL_2919282, EPI_ISL_2919283, EPI_ISL_2919284, EPI_ISL_2919285, EPI_ISL_2919286, EPI_ISL_2919287, EPI_ISL_2919288, EPI_ISL_2919289, EPI_ISL_2919290, EPI_ISL_2919291, EPI_ISL_2919292, EPI_ISL_2919293, EPI_ISL_2919294 | | | |
| see above | Instituto Adolfo Lutz Central | Instituto Adolfo Lutz, Interdiciplinary Procedures Center, Strategic Laboratory | Claudio Tavares Sacchi, Claudia Regina Gonçalves, Erica Valessa Ramos Gomes, Karoline Rodrigues Campos, Caio Vinicius Dias Lopes |
| EPI_ISL_2920683 | LESP Coahuila | Instituto de Diagnostico y Referencia Epidemiologicos (INDRE) | Claudia Wong-Arambula, Abril Rodriguez-Maldonado, Vanessa Rivero-Arredondo, Ariadna Medina-Benitez, Joaquin Quiroz-Mercado, Sergio Rangel-Guerrero, Natividad Cruz-Ortiz, Tatiana Nunez-Garcia, Gisela Barrera-Badillo, Lucia Hernandez-Rivas, Irma Lopez-Martinez, Ernesto Ramirez-Gonzalez. |
| EPI_ISL_2920684 | LESP Zacatecas | Instituto de Diagnostico y Referencia Epidemiologicos (INDRE) | Claudia Wong-Arambula, Abril Rodriguez-Maldonado, Vanessa Rivero-Arredondo, Ariadna Medina-Benitez, Joaquin Quiroz-Mercado, Sergio Rangel-Guerrero, Natividad Cruz-Ortiz, Tatiana Nunez-Garcia, Gisela Barrera-Badillo, Lucia Hernandez-Rivas, Irma Lopez-Martinez, Ernesto Ramirez-Gonzalez. |
| EPI_ISL_2920685 | LESP Baja California | Instituto de Diagnostico y Referencia Epidemiologicos (INDRE) | Claudia Wong-Arambula, Abril Rodriguez-Maldonado, Vanessa Rivero-Arredondo, Ariadna Medina-Benitez, Joaquin Quiroz-Mercado, Sergio Rangel-Guerrero, Natividad Cruz-Ortiz, Tatiana Nunez-Garcia, Gisela Barrera-Badillo, Lucia Hernandez-Rivas, Irma Lopez-Martinez, Ernesto Ramirez-Gonzalez. |
| EPI_ISL_2920686 | LESP Colima | Instituto de Diagnostico y Referencia Epidemiologicos (INDRE) | Claudia Wong-Arambula, Abril Rodriguez-Maldonado, Vanessa Rivero-Arredondo, Ariadna Medina-Benitez, Joaquin Quiroz-Mercado, Sergio Rangel-Guerrero, Natividad Cruz-Ortiz, Tatiana Nunez-Garcia, Gisela Barrera-Badillo, Lucia Hernandez-Rivas, Irma Lopez-Martinez, Ernesto Ramirez-Gonzalez. |
| EPI_ISL_2920687, EPI_ISL_2920688, EPI_ISL_2920689, EPI_ISL_2920690, EPI_ISL_2920691, EPI_ISL_2920692 | LESP Nuevo Leon | Instituto de Diagnostico y Referencia Epidemiologicos (INDRE) | Claudia Wong-Arambula, Abril Rodriguez-Maldonado, Vanessa Rivero-Arredondo, Ariadna Medina-Benitez, Joaquin Quiroz-Mercado, Sergio Rangel-Guerrero, Natividad Cruz-Ortiz, Tatiana Nunez-Garcia, Gisela Barrera-Badillo, Lucia Hernandez-Rivas, Irma Lopez-Martinez, Ernesto Ramirez-Gonzalez. |
| EPI_ISL_2920693, EPI_ISL_2920694, EPI_ISL_2920695, EPI_ISL_2920696, EPI_ISL_2920697, EPI_ISL_2920698, EPI_ISL_2920699, EPI_ISL_2920700, EPI_ISL_2920701 | LESP Tamaulipas | Instituto de Diagnostico y Referencia Epidemiologicos (INDRE) | Claudia Wong-Arambula, Abril Rodriguez-Maldonado, Vanessa Rivero-Arredondo, Ariadna Medina-Benitez, Joaquin Quiroz-Mercado, Sergio Rangel-Guerrero, Natividad Cruz-Ortiz, Tatiana Nunez-Garcia, Gisela Barrera-Badillo, Lucia Hernandez-Rivas, Irma Lopez-Martinez, Ernesto Ramirez-Gonzalez. |
| EPI_ISL_2920702 | LESP Chihuahua | Instituto de Diagnostico y Referencia Epidemiologicos (INDRE) | Claudia Wong-Arambula, Abril Rodriguez-Maldonado, Vanessa Rivero-Arredondo, Ariadna Medina-Benitez, Joaquin Quiroz-Mercado, Sergio Rangel-Guerrero, Natividad Cruz-Ortiz, Tatiana Nunez-Garcia, Gisela Barrera-Badillo, Lucia Hernandez-Rivas, Irma Lopez-Martinez, Ernesto Ramirez-Gonzalez. |
| EPI_ISL_2920703, EPI_ISL_2920704 | LESP Chihuahua | Instituto de Diagnostico y Referencia Epidemiologicos (INDRE) | Claudia Wong-Arambula, Abril Rodriguez-Maldonado, Vanessa Rivero-Arredondo, Ariadna Medina-Benitez, Joaquin Quiroz-Mercado, Sergio Rangel-Guerrero, Maribel Gonzalez-Villa, Natividad Cruz-Ortiz, Tatiana Nunez-Garcia, Gisela Barrera-Badillo, Lucia Hernandez-Rivas, Irma Lopez-Martinez, Ernesto Ramirez-Gonzalez. |
| EPI_ISL_2920705 | LESP Coahuila | Instituto de Diagnostico y Referencia Epidemiologicos (INDRE) | Claudia Wong-Arambula, Abril Rodriguez-Maldonado, Vanessa Rivero-Arredondo, Ariadna Medina-Benitez, Joaquin Quiroz-Mercado, Sergio Rangel-Guerrero, Maribel Gonzalez-Villa, Natividad Cruz-Ortiz, Tatiana Nunez-Garcia, Gisela Barrera-Badillo, Lucia Hernandez-Rivas, Irma Lopez-Martinez, Ernesto Ramirez-Gonzalez. |
| EPI_ISL_2920706 | LESP Sonora | Instituto de Diagnostico y Referencia Epidemiologicos (INDRE) | Claudia Wong-Arambula, Abril Rodriguez-Maldonado, Vanessa Rivero-Arredondo, Ariadna Medina-Benitez, Joaquin Quiroz-Mercado, Sergio Rangel-Guerrero, Maribel Gonzalez-Villa, Natividad Cruz-Ortiz, Tatiana Nunez-Garcia, Gisela Barrera-Badillo, Lucia Hernandez-Rivas, Irma Lopez-Martinez, Ernesto Ramirez-Gonzalez. |
| EPI_ISL_2920707, EPI_ISL_2920708, EPI_ISL_2920709, EPI_ISL_2920710, EPI_ISL_2920711, EPI_ISL_2920712 | LESP Ciudad de Mexico | Instituto de Diagnostico y Referencia Epidemiologicos (INDRE) | Claudia Wong-Arambula, Abril Rodriguez-Maldonado, Vanessa Rivero-Arredondo, Ariadna Medina-Benitez, Joaquin Quiroz-Mercado, Sergio Rangel-Guerrero, Maribel Gonzalez-Villa, Natividad Cruz-Ortiz, Tatiana Nunez-Garcia, Gisela Barrera-Badillo, Lucia Hernandez-Rivas, Irma Lopez-Martinez, Ernesto Ramirez-Gonzalez. |
| EPI_ISL_2920713 | LESP Queretaro | Instituto de Diagnostico y Referencia Epidemiologicos (INDRE) | Claudia Wong-Arambula, Abril Rodriguez-Maldonado, Vanessa Rivero-Arredondo, Ariadna Medina-Benitez, Joaquin Quiroz-Mercado, Sergio Rangel-Guerrero, Maribel Gonzalez-Villa, Natividad Cruz-Ortiz, Tatiana Nunez-Garcia, Gisela Barrera-Badillo, Lucia Hernandez-Rivas, Irma Lopez-Martinez, Ernesto Ramirez-Gonzalez. |
| EPI_ISL_2920714, EPI_ISL_2920715, EPI_ISL_2920716, EPI_ISL_2920717, EPI_ISL_2920719, EPI_ISL_2920720, EPI_ISL_2920721, EPI_ISL_2920722, EPI_ISL_2920723 | LESP Tabasco | Instituto de Diagnostico y Referencia Epidemiologicos (INDRE) | Claudia Wong-Arambula, Abril Rodriguez-Maldonado, Vanessa Rivero-Arredondo, Ariadna Medina-Benitez, Joaquin Quiroz-Mercado, Sergio Rangel-Guerrero, Maribel Gonzalez-Villa, Natividad Cruz-Ortiz, Tatiana Nunez-Garcia, Gisela Barrera-Badillo, Lucia Hernandez-Rivas, Irma Lopez-Martinez, Ernesto Ramirez-Gonzalez. |
| EPI_ISL_2920724 | LESP Oaxaca | Instituto de Diagnostico y Referencia Epidemiologicos (INDRE) | Claudia Wong-Arambula, Abril Rodriguez-Maldonado, Vanessa Rivero-Arredondo, Ariadna Medina-Benitez, Joaquin Quiroz-Mercado, Sergio Rangel-Guerrero, Maribel Gonzalez-Villa, Natividad Cruz-Ortiz, Tatiana Nunez-Garcia, Gisela Barrera-Badillo, Lucia Hernandez-Rivas, Irma Lopez-Martinez, Ernesto Ramirez-Gonzalez. |
| EPI_ISL_2920725, EPI_ISL_2920726, EPI_ISL_2920727, EPI_ISL_2920728, EPI_ISL_2920729, EPI_ISL_2920730 | LESP Tamaulipas | Instituto de Diagnostico y Referencia Epidemiologicos (INDRE) | Claudia Wong-Arambula, Abril Rodriguez-Maldonado, Vanessa Rivero-Arredondo, Ariadna Medina-Benitez, Joaquin Quiroz-Mercado, Sergio Rangel-Guerrero, Maribel Gonzalez-Villa, Natividad Cruz-Ortiz, Tatiana Nunez-Garcia, Gisela Barrera-Badillo, Lucia Hernandez-Rivas, Irma Lopez-Martinez, Ernesto Ramirez-Gonzalez. |
| EPI_ISL_2920739 | LESP Nuevo Leon | Instituto de Diagnostico y Referencia Epidemiologicos (INDRE) | Claudia Wong-Arambula, Abril Rodriguez-Maldonado, Vanessa Rivero-Arredondo, Ariadna Medina-Benitez, Joaquin Quiroz-Mercado, Sergio Rangel-Guerrero, Maribel Gonzalez-Villa, Natividad Cruz-Ortiz, Tatiana Nunez-Garcia, Gisela Barrera-Badillo, Lucia Hernandez-Rivas, Irma Lopez-Martinez, Ernesto Ramirez-Gonzalez. |
| EPI_ISL_2920740 | LESP Veracruz | Instituto de Diagnostico y Referencia Epidemiologicos (INDRE) | Claudia Wong-Arambula, Abril Rodriguez-Maldonado, Vanessa Rivero-Arredondo, Ariadna Medina-Benitez, Joaquin Quiroz-Mercado, Sergio Rangel-Guerrero, Maribel Gonzalez-Villa, Natividad Cruz-Ortiz, Tatiana Nunez-Garcia, Gisela Barrera-Badillo, Lucia Hernandez-Rivas, Irma Lopez-Martinez, Ernesto Ramirez-Gonzalez. |
| EPI_ISL_2920741 | LESP Tabasco | Instituto de Diagnostico y Referencia Epidemiologicos (INDRE) | Claudia Wong-Arambula, Abril Rodriguez-Maldonado, Vanessa Rivero-Arredondo, Ariadna Medina-Benitez, Joaquin Quiroz-Mercado, Sergio Rangel-Guerrero, Maribel Gonzalez-Villa, Natividad Cruz-Ortiz, Tatiana Nunez-Garcia, Gisela Barrera-Badillo, Lucia Hernandez-Rivas, Irma Lopez-Martinez, Ernesto Ramirez-Gonzalez. |
| EPI_ISL_2920742 | LESP Estado de Mexico | Instituto de Diagnostico y Referencia Epidemiologicos (INDRE) | Claudia Wong-Arambula, Abril Rodriguez-Maldonado, Vanessa Rivero-Arredondo, Ariadna Medina-Benitez, Joaquin Quiroz-Mercado, Sergio Rangel-Guerrero, Maribel Gonzalez-Villa, Natividad Cruz-Ortiz, Tatiana Nunez-Garcia, Gisela Barrera-Badillo, Lucia Hernandez-Rivas, Irma Lopez-Martinez, Ernesto Ramirez-Gonzalez. |

| | | | |
|---|---|---|---|
| EPI_ISL_2920743, EPI_ISL_2920744, EPI_ISL_2920745 | LESP Ciudad de Mexico | Instituto de Diagnostico y Referencia Epidemiologicos (INDRE) | Claudia Wong-Arambula, Abril Rodriguez-Maldonado, Vanessa Rivero-Arredondo, Ariadna Medina-Benitez, Joaquin Quiroz-Mercado, Sergio Rangel-Guerrero, Maribel Gonzalez-Villa, Natividad Cruz-Ortiz, Tatiana Nunez-Garcia, Gisela Barrera-Badillo, Lucia Hernandez-Rivas, Irma Lopez-Martinez, Ernesto Ramirez-Gonzalez. |
| EPI_ISL_2920746 | LESP Tabasco | Instituto de Diagnostico y Referencia Epidemiologicos (INDRE) | Claudia Wong-Arambula, Abril Rodriguez-Maldonado, Vanessa Rivero-Arredondo, Ariadna Medina-Benitez, Joaquin Quiroz-Mercado, Sergio Rangel-Guerrero, Maribel Gonzalez-Villa, Natividad Cruz-Ortiz, Tatiana Nunez-Garcia, Gisela Barrera-Badillo, Lucia Hernandez-Rivas, Irma Lopez-Martinez, Ernesto Ramirez-Gonzalez. |
| EPI_ISL_2920749, EPI_ISL_2920750 | LESP Guerrero | Instituto de Diagnostico y Referencia Epidemiologicos (INDRE) | Claudia Wong-Arambula, Abril Rodriguez-Maldonado, Vanessa Rivero-Arredondo, Ariadna Medina-Benitez, Joaquin Quiroz-Mercado, Sergio Rangel-Guerrero, Maribel Gonzalez-Villa, Natividad Cruz-Ortiz, Tatiana Nunez-Garcia, Gisela Barrera-Badillo, Lucia Hernandez-Rivas, Irma Lopez-Martinez, Ernesto Ramirez-Gonzalez. |
| EPI_ISL_2920751 | LESP Estado de Mexico | Instituto de Diagnostico y Referencia Epidemiologicos (INDRE) | Claudia Wong-Arambula, Abril Rodriguez-Maldonado, Vanessa Rivero-Arredondo, Ariadna Medina-Benitez, Joaquin Quiroz-Mercado, Sergio Rangel-Guerrero, Maribel Gonzalez-Villa, Natividad Cruz-Ortiz, Tatiana Nunez-Garcia, Gisela Barrera-Badillo, Lucia Hernandez-Rivas, Irma Lopez-Martinez, Ernesto Ramirez-Gonzalez. |
| EPI_ISL_2920752 | LESP Ciudad de Mexico | Instituto de Diagnostico y Referencia Epidemiologicos (INDRE) | Claudia Wong-Arambula, Abril Rodriguez-Maldonado, Vanessa Rivero-Arredondo, Ariadna Medina-Benitez, Joaquin Quiroz-Mercado, Sergio Rangel-Guerrero, Maribel Gonzalez-Villa, Natividad Cruz-Ortiz, Tatiana Nunez-Garcia, Gisela Barrera-Badillo, Lucia Hernandez-Rivas, Irma Lopez-Martinez, Ernesto Ramirez-Gonzalez. |
| EPI_ISL_2920753 | LESP Michoacan | Instituto de Diagnostico y Referencia Epidemiologicos (INDRE) | Claudia Wong-Arambula, Abril Rodriguez-Maldonado, Vanessa Rivero-Arredondo, Ariadna Medina-Benitez, Joaquin Quiroz-Mercado, Sergio Rangel-Guerrero, Maribel Gonzalez-Villa, Natividad Cruz-Ortiz, Tatiana Nunez-Garcia, Gisela Barrera-Badillo, Lucia Hernandez-Rivas, Irma Lopez-Martinez, Ernesto Ramirez-Gonzalez. |
| EPI_ISL_2920754 | LESP Queretaro | Instituto de Diagnostico y Referencia Epidemiologicos (INDRE) | Claudia Wong-Arambula, Abril Rodriguez-Maldonado, Vanessa Rivero-Arredondo, Ariadna Medina-Benitez, Joaquin Quiroz-Mercado, Sergio Rangel-Guerrero, Maribel Gonzalez-Villa, Natividad Cruz-Ortiz, Tatiana Nunez-Garcia, Gisela Barrera-Badillo, Lucia Hernandez-Rivas, Irma Lopez-Martinez, Ernesto Ramirez-Gonzalez. |
| EPI_ISL_2920755, EPI_ISL_2920756, EPI_ISL_2920757, EPI_ISL_2920758, EPI_ISL_2920759, EPI_ISL_2920760, EPI_ISL_2920761, EPI_ISL_2920762, EPI_ISL_2920763, EPI_ISL_2920764, EPI_ISL_2920765, EPI_ISL_2920766, EPI_ISL_2920767, EPI_ISL_2920768, EPI_ISL_2920769, EPI_ISL_2920770, EPI_ISL_2920771, EPI_ISL_2920772, EPI_ISL_2920773, EPI_ISL_2920774, EPI_ISL_2920775, EPI_ISL_2920776, EPI_ISL_2920777, EPI_ISL_2920778, EPI_ISL_2920780, EPI_ISL_2920781, EPI_ISL_2920782, EPI_ISL_2920783, EPI_ISL_2920784, EPI_ISL_2920785, EPI_ISL_2920786, EPI_ISL_2920787, EPI_ISL_2920788, EPI_ISL_2920789, EPI_ISL_2920790 | | | |
| see above | LESP Tabasco | Instituto de Diagnostico y Referencia Epidemiologicos (INDRE) | Claudia Wong-Arambula, Abril Rodriguez-Maldonado, Vanessa Rivero-Arredondo, Ariadna Medina-Benitez, Joaquin Quiroz-Mercado, Sergio Rangel-Guerrero, Maribel Gonzalez-Villa, Natividad Cruz-Ortiz, Tatiana Nunez-Garcia, Gisela Barrera-Badillo, Lucia Hernandez-Rivas, Irma Lopez-Martinez, Ernesto Ramirez-Gonzalez. |
| EPI_ISL_2920791 | LESP Aguascalientes | Instituto de Diagnostico y Referencia Epidemiologicos (INDRE) | Claudia Wong-Arambula, Abril Rodriguez-Maldonado, Vanessa Rivero-Arredondo, Ariadna Medina-Benitez, Joaquin Quiroz-Mercado, Sergio Rangel-Guerrero, Maribel Gonzalez-Villa, Natividad Cruz-Ortiz, Tatiana Nunez-Garcia, Gisela Barrera-Badillo, Lucia Hernandez-Rivas, Irma Lopez-Martinez, Ernesto Ramirez-Gonzalez. |
| EPI_ISL_2921202, EPI_ISL_2921203 | SIESP CHIETI - DRIVE IN CHIETI | Istituto Zooprofilattico Sperimentale dell'Abruzzo e Molise "G. Caporale" | Lorusso A, Marcacci M, Di Domenico M, Ancora M, Curini V, Di Lollo Valeria, Mangone I, Rinaldi A, Delli Compagni E, Scialabba S, Caporale M, Di Pasquale A, Cammà C, Puglia I, Calistri P, Savini G |
| EPI_ISL_2921204 | SIESP CHIETI - DRIVE IN ORTONA | Istituto Zooprofilattico Sperimentale dell'Abruzzo e Molise "G. Caporale" | Lorusso A, Marcacci M, Di Domenico M, Ancora M, Curini V, Di Lollo Valeria, Mangone I, Rinaldi A, Delli Compagni E, Scialabba S, Caporale M, Di Pasquale A, Cammà C, Puglia I, Calistri P, Savini G |
| EPI_ISL_2921205, EPI_ISL_2921206 | SIESP CHIETI - DRIVE IN CHIETI | Istituto Zooprofilattico Sperimentale dell'Abruzzo e Molise "G. Caporale" | Lorusso A, Marcacci M, Di Domenico M, Ancora M, Curini V, Di Lollo Valeria, Mangone I, Rinaldi A, Delli Compagni E, Scialabba S, Caporale M, Di Pasquale A, Cammà C, Puglia I, Calistri P, Savini G |
| EPI_ISL_2921207 | SIESP CHIETI - DRIVE IN LANCIANO | Istituto Zooprofilattico Sperimentale dell'Abruzzo e Molise "G. Caporale" | Lorusso A, Marcacci M, Di Domenico M, Ancora M, Curini V, Di Lollo Valeria, Mangone I, Rinaldi A, Delli Compagni E, Scialabba S, Caporale M, Di Pasquale A, Cammà C, Puglia I, Calistri P, Savini G |
| EPI_ISL_2921208, EPI_ISL_2921209 | OSPEDALE SAN SALVATORE - MEDICINA DI LABORATORIO | Istituto Zooprofilattico Sperimentale dell'Abruzzo e Molise "G. Caporale" | Lorusso A, Marcacci M, Di Domenico M, Ancora M, Curini V, Di Lollo Valeria, Mangone I, Rinaldi A, Delli Compagni E, Scialabba S, Caporale M, Di Pasquale A, Cammà C, Puglia I, Calistri P, Savini G |
| EPI_ISL_2921210, EPI_ISL_2921211, EPI_ISL_2921212, EPI_ISL_2921213, EPI_ISL_2921214, EPI_ISL_2921215, EPI_ISL_2921216, EPI_ISL_2921217, EPI_ISL_2921218, EPI_ISL_2921219, EPI_ISL_2921220, EPI_ISL_2921221, EPI_ISL_2921222, EPI_ISL_2921223, EPI_ISL_2921224 | | | |
| see above | PO AVEZZANO - U.O.C. Laboratorio Analisi | Istituto Zooprofilattico Sperimentale dell'Abruzzo e Molise "G. Caporale" | Lorusso A, Marcacci M, Di Domenico M, Ancora M, Curini V, Di Lollo Valeria, Mangone I, Rinaldi A, Delli Compagni E, Scialabba S, Caporale M, Di Pasquale A, Cammà C, Puglia I, Calistri P, Savini G |
| EPI_ISL_2921225, EPI_ISL_2921226, EPI_ISL_2921227, EPI_ISL_2921228, EPI_ISL_2921229, EPI_ISL_2921230, EPI_ISL_2921231, EPI_ISL_2921232, EPI_ISL_2921233, EPI_ISL_2921234, EPI_ISL_2921235, EPI_ISL_2921236, EPI_ISL_2921237, EPI_ISL_2921238, EPI_ISL_2921239, EPI_ISL_2921240, EPI_ISL_2921241, EPI_ISL_2921242 | | | |
| see above | OSPEDALE CIVILE TERAMO - CENTRO TRASFUSIONALE | Istituto Zooprofilattico Sperimentale dell'Abruzzo e Molise "G. Caporale" | Lorusso A, Marcacci M, Di Domenico M, Ancora M, Curini V, Di Lollo Valeria, Mangone I, Rinaldi A, Delli Compagni E, Scialabba S, Caporale M, Di Pasquale A, Cammà C, Puglia I, Calistri P, Savini G |
| EPI_ISL_2921243, EPI_ISL_2921244, EPI_ISL_2921245, EPI_ISL_2921246, EPI_ISL_2921247, EPI_ISL_2921248, EPI_ISL_2921249, EPI_ISL_2921250, EPI_ISL_2921251, EPI_ISL_2921252, EPI_ISL_2921253, EPI_ISL_2921254, EPI_ISL_2921255, EPI_ISL_2921256, EPI_ISL_2921257 | | | |
| see above | PO AVEZZANO - U.O.C. Laboratorio Analisi | Istituto Zooprofilattico Sperimentale dell'Abruzzo e Molise "G. Caporale" | Lorusso A, Marcacci M, Di Domenico M, Ancora M, Curini V, Di Lollo Valeria, Mangone I, Rinaldi A, Delli Compagni E, Scialabba S, Caporale M, Di Pasquale A, Cammà C, Puglia I, Calistri P, Savini G |
| EPI_ISL_2921258 | SIESP DIPARTIMENTO DI PREVENZIONE CHIETI | Istituto Zooprofilattico Sperimentale dell'Abruzzo e Molise "G. Caporale" | Lorusso A, Marcacci M, Di Domenico M, Ancora M, Curini V, Di Lollo Valeria, Mangone I, Rinaldi A, Delli Compagni E, Scialabba S, Caporale M, Di Pasquale A, Cammà C, Puglia I, Calistri P, Savini G |
| EPI_ISL_2921259 | SIESP CHIETI - DRIVE IN LANCIANO | Istituto Zooprofilattico Sperimentale dell'Abruzzo e Molise "G. Caporale" | Lorusso A, Marcacci M, Di Domenico M, Ancora M, Curini V, Di Lollo Valeria, Mangone I, Rinaldi A, Delli Compagni E, Scialabba S, Caporale M, Di Pasquale A, Cammà C, Puglia I, Calistri P, Savini G |
| EPI_ISL_2921260, EPI_ISL_2921261 | Terengganu State Health Department | Institute for Medical Research, Infectious Disease Research Centre, National Institutes of Health, Ministry of Health Malaysia | Suppiah J, Kamel K, Mohd Zawawi Z, Azizan MA, Ramly N, Robert F, Anasir MI, Thayan R |
| EPI_ISL_2921262 | Pantai Hospital Kuala Lumpur | Institute for Medical Research, Infectious Disease Research Centre, National Institutes of Health, Ministry of Health Malaysia | Suppiah J, Kamel K, Mohd Zawawi Z, Azizan MA, Ramly N, Robert F, Anasir MI, Thayan R |
| EPI_ISL_2921263, EPI_ISL_2921265 | Pahang State Health Department | Institute for Medical Research, Infectious Disease Research Centre, National Institutes of Health, Ministry of Health Malaysia | Suppiah J, Kamel K, Mohd Zawawi Z, Azizan MA, Ramly N, Robert F, Anasir MI, Thayan R |
| EPI_ISL_2921611 | Labuan Hospital | Institute for Medical Research, Infectious Disease Research Centre, National Institutes of Health, Ministry of Health Malaysia | Suppiah J, Kamel K, Mohd Zawawi Z, Azizan MA, Ramly N, Robert F, Anasir MI, Thayan R |
| EPI_ISL_2921612 | Sultan Haji Ahmad Shah Hospital | Institute for Medical Research, Infectious Disease Research Centre, National Institutes of Health, Ministry of Health Malaysia | Suppiah J, Kamel K, Mohd Zawawi Z, Azizan MA, Ramly N, Robert F, Anasir MI, Thayan R |
| EPI_ISL_2921996 | Melaka Hospital | Institute for Medical Research, Infectious Disease Research Centre, National Institutes of Health, Ministry of Health Malaysia | Suppiah J, Kamel K, Mohd Zawawi Z, Azizan MA, Ramly N, Robert F, Anasir MI, Thayan R |
| EPI_ISL_2921997 | Tengku Ampuan Rahimah Hospital | Institute for Medical Research, Infectious Disease Research | Suppiah J, Kamel K, Mohd Zawawi Z, Azizan MA, Ramly N, Robert F, Anasir MI, Thayan R |

| | | | |
|---|---|---|---|
| | | Centre, National Institutes of Health, Ministry of Health Malaysia | |
| EPI_ISL_2923113 | Labo Analyses Med | National Reference Center for Viruses of Respiratory Infections, Institut Pasteur, Paris | Marion Barbet, Sylvie Behillil, Méline Bizard, Angela Brisebarre, Camille Capel, Vincent Enouf, Louise Lefrançois, Frédéric Lemoine, Christophe Malabat, Corinne Maufrais, Hub Bioinformatique Biostatistiques, Etienne Simon-Lorière, Maud Vanpeene, Sylvie Van der Werf ,GERSOHN Marc |
| EPI_ISL_2923114 | Labo Analyses Med | National Reference Center for Viruses of Respiratory Infections, Institut Pasteur, Paris | Marion Barbet, Sylvie Behillil, Méline Bizard, Angela Brisebarre, Camille Capel, Vincent Enouf, Louise Lefrançois, Frédéric Lemoine, Christophe Malabat, Corinne Maufrais, Hub Bioinformatique Biostatistiques, Etienne Simon-Lorière, Maud Vanpeene, Sylvie Van der Werf ,DES GARETS Marie-Caroline |
| EPI_ISL_2923115 | Hospital | National Reference Center for Viruses of Respiratory Infections, Institut Pasteur, Paris | Marion Barbet, Sylvie Behillil, Méline Bizard, Angela Brisebarre, Camille Capel, Vincent Enouf, Louise Lefrançois, Frédéric Lemoine, Christophe Malabat, Corinne Maufrais, Hub Bioinformatique Biostatistiques, Etienne Simon-Lorière, Maud Vanpeene, Sylvie Van der Werf ,FARFOUR Eric |
| EPI_ISL_2923116 | Hospital | National Reference Center for Viruses of Respiratory Infections, Institut Pasteur, Paris | Marion Barbet, Sylvie Behillil, Méline Bizard, Angela Brisebarre, Camille Capel, Vincent Enouf, Louise Lefrançois, Frédéric Lemoine, Christophe Malabat, Corinne Maufrais, Hub Bioinformatique Biostatistiques, Etienne Simon-Lorière, Maud Vanpeene, Sylvie Van der Werf ,GUINARD Jérome |
| EPI_ISL_2923117, EPI_ISL_2923118 | Labo Analyses Med | National Reference Center for Viruses of Respiratory Infections, Institut Pasteur, Paris | Marion Barbet, Sylvie Behillil, Méline Bizard, Angela Brisebarre, Camille Capel, Vincent Enouf, Louise Lefrançois, Frédéric Lemoine, Christophe Malabat, Corinne Maufrais, Damien Mornico, Etienne Simon-Lorière, Maud Vanpeene, Sylvie Van der Werf ,TROMBERT Sabine |
| EPI_ISL_2923119 | Labo Analyses Med | National Reference Center for Viruses of Respiratory Infections, Institut Pasteur, Paris | Marion Barbet, Sylvie Behillil, Méline Bizard, Angela Brisebarre, Camille Capel, Vincent Enouf, Louise Lefrançois, Frédéric Lemoine, Christophe Malabat, Corinne Maufrais, Hub Bioinformatique Biostatistiques, Etienne Simon-Lorière, Maud Vanpeene, Sylvie Van der Werf ,TROMBERT Sabine |
| EPI_ISL_2923120 | Hospital | National Reference Center for Viruses of Respiratory Infections, Institut Pasteur, Paris | Marion Barbet, Sylvie Behillil, Méline Bizard, Angela Brisebarre, Camille Capel, Vincent Enouf, Louise Lefrançois, Frédéric Lemoine, Christophe Malabat, Corinne Maufrais, Damien Mornico, Etienne Simon-Lorière, Maud Vanpeene, Sylvie Van der Werf ,RAMIREZ José-Manuel |
| EPI_ISL_2923121 | Labo Analyses Med | National Reference Center for Viruses of Respiratory Infections, Institut Pasteur, Paris | Marion Barbet, Sylvie Behillil, Méline Bizard, Angela Brisebarre, Camille Capel, Vincent Enouf, Louise Lefrançois, Frédéric Lemoine, Christophe Malabat, Corinne Maufrais, Damien Mornico, Etienne Simon-Lorière, Maud Vanpeene, Sylvie Van der Werf ,LABORATOIRE Palissy |
| EPI_ISL_2923122 | Labo Analyses Med | National Reference Center for Viruses of Respiratory Infections, Institut Pasteur, Paris | Marion Barbet, Sylvie Behillil, Méline Bizard, Angela Brisebarre, Camille Capel, Vincent Enouf, Louise Lefrançois, Frédéric Lemoine, Christophe Malabat, Corinne Maufrais, Victoire Baillet, Etienne Simon-Lorière, Maud Vanpeene, Sylvie Van der Werf ,LABORATOIRE Palissy |
| EPI_ISL_2923123, EPI_ISL_2923124 | Labo Analyses Med | National Reference Center for Viruses of Respiratory Infections, Institut Pasteur, Paris | Marion Barbet, Sylvie Behillil, Méline Bizard, Angela Brisebarre, Camille Capel, Vincent Enouf, Louise Lefrançois, Frédéric Lemoine, Christophe Malabat, Corinne Maufrais, Damien Mornico, Etienne Simon-Lorière, Maud Vanpeene, Sylvie Van der Werf ,LABORATOIRE Palissy |
| EPI_ISL_2923125, EPI_ISL_2923126 | Hospital | National Reference Center for Viruses of Respiratory Infections, Institut Pasteur, Paris | Marion Barbet, Sylvie Behillil, Méline Bizard, Angela Brisebarre, Camille Capel, Vincent Enouf, Louise Lefrançois, Frédéric Lemoine, Christophe Malabat, Corinne Maufrais, Hub Bioinformatique Biostatistiques, Etienne Simon-Lorière, Maud Vanpeene, Sylvie Van der Werf ,HOLSTEIN Anne |
| EPI_ISL_2923127 | Hospital | National Reference Center for Viruses of Respiratory Infections, Institut Pasteur, Paris | Marion Barbet, Sylvie Behillil, Méline Bizard, Angela Brisebarre, Camille Capel, Vincent Enouf, Louise Lefrançois, Frédéric Lemoine, Christophe Malabat, Corinne Maufrais, Victoire Baillet, Etienne Simon-Lorière, Maud Vanpeene, Sylvie Van der Werf ,HOLSTEIN Anne |
| EPI_ISL_2923128 | Hospital | National Reference Center for Viruses of Respiratory Infections, Institut Pasteur, Paris | Marion Barbet, Sylvie Behillil, Méline Bizard, Angela Brisebarre, Camille Capel, Vincent Enouf, Louise Lefrançois, Frédéric Lemoine, Christophe Malabat, Corinne Maufrais, Victoire Baillet, Etienne Simon-Lorière, Maud Vanpeene, Sylvie Van der Werf ,SERAZIN Valérie |
| EPI_ISL_2923130 | Labo Analyses Med | National Reference Center for Viruses of Respiratory Infections, Institut Pasteur, Paris | Marion Barbet, Sylvie Behillil, Méline Bizard, Angela Brisebarre, Camille Capel, Vincent Enouf, Louise Lefrançois, Frédéric Lemoine, Christophe Malabat, Corinne Maufrais, Victoire Baillet, Etienne Simon-Lorière, Maud Vanpeene, Sylvie Van der Werf ,TROMBERT Sabine |
| EPI_ISL_2923131, EPI_ISL_2923132 | Labo Analyses Med | National Reference Center for Viruses of Respiratory Infections, Institut Pasteur, Paris | Marion Barbet, Sylvie Behillil, Méline Bizard, Angela Brisebarre, Camille Capel, Vincent Enouf, Louise Lefrançois, Frédéric Lemoine, Christophe Malabat, Corinne Maufrais, Hub Bioinformatique Biostatistiques, Etienne Simon-Lorière, Maud Vanpeene, Sylvie Van der Werf ,TROMBERT Sabine |
| EPI_ISL_2923134, EPI_ISL_2923135 | Labo Analyses Med | National Reference Center for Viruses of Respiratory Infections, Institut Pasteur, Paris | Marion Barbet, Sylvie Behillil, Méline Bizard, Angela Brisebarre, Camille Capel, Vincent Enouf, Louise Lefrançois, Frédéric Lemoine, Christophe Malabat, Corinne Maufrais, Damien Mornico, Etienne Simon-Lorière, Maud Vanpeene, Sylvie Van der Werf ,TROMBERT Sabine |
| EPI_ISL_2923138 | Hospital | National Reference Center for Viruses of Respiratory Infections, Institut Pasteur, Paris | Marion Barbet, Sylvie Behillil, Méline Bizard, Angela Brisebarre, Camille Capel, Vincent Enouf, Louise Lefrançois, Frédéric Lemoine, Christophe Malabat, Corinne Maufrais, Hub Bioinformatique Biostatistiques, Etienne Simon-Lorière, Maud Vanpeene, Sylvie Van der Werf ,MARTRES Pascale |
| EPI_ISL_2923139 | Labo Analyses Med | National Reference Center for Viruses of Respiratory Infections, Institut Pasteur, Paris | Marion Barbet, Sylvie Behillil, Méline Bizard, Angela Brisebarre, Camille Capel, Vincent Enouf, Louise Lefrançois, Frédéric Lemoine, Christophe Malabat, Corinne Maufrais, Hub Bioinformatique Biostatistiques, Etienne Simon-Lorière, Maud Vanpeene, Sylvie Van der Werf ,POTIRON Grégoire |
| EPI_ISL_2923140 | Labo Analyses Med | National Reference Center for Viruses of Respiratory Infections, Institut Pasteur, Paris | Marion Barbet, Sylvie Behillil, Méline Bizard, Angela Brisebarre, Camille Capel, Vincent Enouf, Louise Lefrançois, Frédéric Lemoine, Christophe Malabat, Corinne Maufrais, Amaury Vaysse, Etienne Simon-Lorière, Maud Vanpeene, Sylvie Van der Werf ,POTIRON Grégoire |
| EPI_ISL_2923141, EPI_ISL_2923142 | Labo Analyses Med | National Reference Center for Viruses of Respiratory Infections, Institut Pasteur, Paris | Marion Barbet, Sylvie Behillil, Méline Bizard, Angela Brisebarre, Camille Capel, Vincent Enouf, Louise Lefrançois, Frédéric Lemoine, Christophe Malabat, Corinne Maufrais, Hub Bioinformatique Biostatistiques, Etienne Simon-Lorière, Maud Vanpeene, Sylvie Van der Werf ,POTIRON Grégoire |
| EPI_ISL_2923143 | Labo Analyses Med | National Reference Center for Viruses of Respiratory Infections, Institut Pasteur, Paris | Marion Barbet, Sylvie Behillil, Méline Bizard, Angela Brisebarre, Camille Capel, Vincent Enouf, Louise Lefrançois, Frédéric Lemoine, Christophe Malabat, Corinne Maufrais, Hub Bioinformatique Biostatistiques, Etienne Simon-Lorière, Maud Vanpeene, Sylvie Van der Werf ,LEONARD Pierre-Yves |
| EPI_ISL_2923144, EPI_ISL_2923145, EPI_ISL_2923146, EPI_ISL_2923147 | Labo Analyses Med | National Reference Center for Viruses of Respiratory Infections, Institut Pasteur, Paris | Marion Barbet, Sylvie Behillil, Méline Bizard, Angela Brisebarre, Camille Capel, Vincent Enouf, Louise Lefrançois, Frédéric Lemoine, Christophe Malabat, Corinne Maufrais, Hub Bioinformatique Biostatistiques, Etienne Simon-Lorière, Maud Vanpeene, Sylvie Van der Werf ,HAGUENOER Eve |
| EPI_ISL_2923148, EPI_ISL_2923149, EPI_ISL_2923150, EPI_ISL_2923151, EPI_ISL_2923152, EPI_ISL_2923155, EPI_ISL_2923156 | Labo Analyses Med | National Reference Center for Viruses of Respiratory Infections, Institut Pasteur, Paris | Marion Barbet, Sylvie Behillil, Méline Bizard, Angela Brisebarre, Camille Capel, Vincent Enouf, Louise Lefrançois, Frédéric Lemoine, Christophe Malabat, Corinne Maufrais, Hub Bioinformatique Biostatistiques, Etienne Simon-Lorière, Maud Vanpeene, Sylvie Van der Werf ,DURIVAULT Jérôme |
| EPI_ISL_2923157 | Labo Analyses Med | National Reference Center for Viruses of Respiratory Infections, Institut Pasteur, Paris | Marion Barbet, Sylvie Behillil, Méline Bizard, Angela Brisebarre, Camille Capel, Vincent Enouf, Louise Lefrançois, Frédéric Lemoine, Christophe Malabat, Corinne Maufrais,Hub Bioinformatique Biostatistiques, Etienne Simon-Lorière, Maud Vanpeene, Sylvie Van der Werf ,DURIVAULT Jérôme |
| EPI_ISL_2923158 | Labo Analyses Med | National Reference Center for Viruses of Respiratory Infections, Institut Pasteur, Paris | Marion Barbet, Sylvie Behillil, Méline Bizard, Angela Brisebarre, Camille Capel, Vincent Enouf, Louise Lefrançois, Frédéric Lemoine, Christophe Malabat, Corinne Maufrais, Hub Bioinformatique Biostatistiques, Etienne Simon-Lorière, Maud Vanpeene, Sylvie Van der Werf ,DURIVAULT Jérôme |
| EPI_ISL_2923159 | Labo Analyses Med | National Reference Center for Viruses of Respiratory Infections, Institut Pasteur, Paris | Marion Barbet, Sylvie Behillil, Méline Bizard, Angela Brisebarre, Camille Capel, Vincent Enouf, Louise Lefrançois, Frédéric Lemoine, Christophe Malabat, Corinne Maufrais,Hub Bioinformatique Biostatistiques, Etienne Simon-Lorière, Maud Vanpeene, Sylvie Van der Werf ,DURIVAULT Jérôme |
| EPI_ISL_2923160, EPI_ISL_2923161 | Labo Analyses Med | National Reference Center for Viruses of Respiratory Infections, Institut Pasteur, Paris | Marion Barbet, Sylvie Behillil, Méline Bizard, Angela Brisebarre, Camille Capel, Vincent Enouf, Louise Lefrançois, Frédéric Lemoine, Christophe Malabat, Corinne Maufrais, Amaury Vaysse, Etienne Simon-Lorière, Maud Vanpeene, Sylvie Van der Werf ,DURIVAULT Jérôme |
| EPI_ISL_2923162, EPI_ISL_2923163 | Hospital | National Reference Center for Viruses of Respiratory Infections, Institut Pasteur, Paris | Marion Barbet, Sylvie Behillil, Méline Bizard, Angela Brisebarre, Camille Capel, Vincent Enouf, Louise Lefrançois, Frédéric Lemoine, Christophe Malabat, Corinne Maufrais, Hub Bioinformatique Biostatistiques, Etienne Simon-Lorière, Maud Vanpeene, Sylvie Van der Werf ,DJAMDJIAN Laura |
| EPI_ISL_2923165, EPI_ISL_2923166 | Hospital | National Reference Center for Viruses of Respiratory Infections, Institut Pasteur, Paris | Marion Barbet, Sylvie Behillil, Méline Bizard, Angela Brisebarre, Camille Capel, Vincent Enouf, Louise Lefrançois, Frédéric Lemoine, Christophe Malabat, Corinne Maufrais, Hub Bioinformatique Biostatistiques, Etienne Simon-Lorière, Maud Vanpeene, Sylvie Van der Werf ,GUINARD Jérome |
| EPI_ISL_2923167 | Hospital | National Reference Center for Viruses of Respiratory Infections, Institut Pasteur, Paris | Marion Barbet, Sylvie Behillil, Méline Bizard, Angela Brisebarre, Camille Capel, Vincent Enouf, Louise Lefrançois, Frédéric Lemoine, Christophe Malabat, Corinne Maufrais, Amaury Vaysse, Etienne Simon-Lorière, Maud Vanpeene, Sylvie Van der Werf ,GALEAZZI GUY |
| EPI_ISL_2923168 | Labo Analyses Med | National Reference Center for Viruses of Respiratory Infections, Institut Pasteur, Paris | Marion Barbet, Sylvie Behillil, Méline Bizard, Angela Brisebarre, Camille Capel, Vincent Enouf, Louise Lefrançois, Frédéric Lemoine, Christophe Malabat, Corinne Maufrais, Hub Bioinformatique Biostatistiques, Etienne Simon-Lorière, Maud Vanpeene, Sylvie Van der Werf ,VAN AGT S. |
| EPI_ISL_2923169, EPI_ISL_2923170 | Labo Analyses Med | National Reference Center for Viruses of Respiratory Infections, Institut Pasteur, Paris | Marion Barbet, Sylvie Behillil, Méline Bizard, Angela Brisebarre, Camille Capel, Vincent Enouf, Louise Lefrançois, Frédéric Lemoine, Christophe Malabat, Corinne Maufrais, Hub Bioinformatique Biostatistiques, Etienne Simon-Lorière, Maud Vanpeene, Sylvie Van der Werf ,MIARA Philippe |
| EPI_ISL_2923171, EPI_ISL_2923172 | Labo Analyses Med | National Reference Center for Viruses of Respiratory Infections, Institut Pasteur, Paris | Marion Barbet, Sylvie Behillil, Méline Bizard, Angela Brisebarre, Camille Capel, Vincent Enouf, Louise Lefrançois, Frédéric Lemoine, Christophe Malabat, Corinne Maufrais, Hub Bioinformatique Biostatistiques, Etienne Simon-Lorière, Maud Vanpeene, Sylvie Van der Werf ,TROMBERT Sabine |
| EPI_ISL_2923173 | Labo Analyses Med | National Reference Center for Viruses of Respiratory | Marion Barbet, Sylvie Behillil, Méline Bizard, Angela Brisebarre, Camille Capel, Vincent Enouf, Louise Lefrançois, Frédéric Lemoine, Christophe Malabat, |

| | | | |
|---|---|---|---|
| | | Infections, Institut Pasteur, Paris | Corinne Maufrais, Gael Millot, Etienne Simon-Lorière, Maud Vanpeene, Sylvie Van der Werf ,TROMBERT Sabine |
| EPI_ISL_2923174, EPI_ISL_2923175 | Labo Analyses Med | National Reference Center for Viruses of Respiratory Infections, Institut Pasteur, Paris | Marion Barbet, Sylvie Behillil, Méline Bizard, Angela Brisebarre, Camille Capel, Vincent Enouf, Louise Lefrançois, Frédéric Lemoine, Christophe Malabat, Corinne Maufrais, Hub Bioinformatique Biostatistiques, Etienne Simon-Lorière, Maud Vanpeene, Sylvie Van der Werf ,TROMBERT Sabine |
| EPI_ISL_2923176 | Labo Analyses Med | National Reference Center for Viruses of Respiratory Infections, Institut Pasteur, Paris | Marion Barbet, Sylvie Behillil, Méline Bizard, Angela Brisebarre, Camille Capel, Vincent Enouf, Louise Lefrançois, Frédéric Lemoine, Christophe Malabat, Corinne Maufrais, Gael Millot, Etienne Simon-Lorière, Maud Vanpeene, Sylvie Van der Werf ,TROMBERT Sabine |
| EPI_ISL_2923177, EPI_ISL_2923178 | Labo Analyses Med | National Reference Center for Viruses of Respiratory Infections, Institut Pasteur, Paris | Marion Barbet, Sylvie Behillil, Méline Bizard, Angela Brisebarre, Camille Capel, Vincent Enouf, Louise Lefrançois, Frédéric Lemoine, Christophe Malabat, Corinne Maufrais, Hub Bioinformatique Biostatistiques, Etienne Simon-Lorière, Maud Vanpeene, Sylvie Van der Werf ,TROMBERT Sabine |
| EPI_ISL_2923179 | Labo Analyses Med | National Reference Center for Viruses of Respiratory Infections, Institut Pasteur, Paris | Marion Barbet, Sylvie Behillil, Méline Bizard, Angela Brisebarre, Camille Capel, Vincent Enouf, Louise Lefrançois, Frédéric Lemoine, Christophe Malabat, Corinne Maufrais, Amaury Vaysse, Etienne Simon-Lorière, Maud Vanpeene, Sylvie Van der Werf ,TROMBERT Sabine |
| EPI_ISL_2923180 | Labo Analyses Med | National Reference Center for Viruses of Respiratory Infections, Institut Pasteur, Paris | Marion Barbet, Sylvie Behillil, Méline Bizard, Angela Brisebarre, Camille Capel, Vincent Enouf, Louise Lefrançois, Frédéric Lemoine, Christophe Malabat, Corinne Maufrais, Hub Bioinformatique Biostatistiques, Etienne Simon-Lorière, Maud Vanpeene, Sylvie Van der Werf ,TROMBERT Sabine |
| EPI_ISL_2923181, EPI_ISL_2923182 | Labo Analyses Med | National Reference Center for Viruses of Respiratory Infections, Institut Pasteur, Paris | Marion Barbet, Sylvie Behillil, Méline Bizard, Angela Brisebarre, Camille Capel, Vincent Enouf, Louise Lefrançois, Frédéric Lemoine, Christophe Malabat, Corinne Maufrais, Amaury Vaysse, Etienne Simon-Lorière, Maud Vanpeene, Sylvie Van der Werf ,TROMBERT Sabine |
| EPI_ISL_2923183, EPI_ISL_2923184, EPI_ISL_2923185, EPI_ISL_2923186 | Labo Analyses Med | National Reference Center for Viruses of Respiratory Infections, Institut Pasteur, Paris | Marion Barbet, Sylvie Behillil, Méline Bizard, Angela Brisebarre, Camille Capel, Vincent Enouf, Louise Lefrançois, Frédéric Lemoine, Christophe Malabat, Corinne Maufrais, Hub Bioinformatique Biostatistiques, Etienne Simon-Lorière, Maud Vanpeene, Sylvie Van der Werf ,TROMBERT Sabine |
| EPI_ISL_2923187 | Labo Analyses Med | National Reference Center for Viruses of Respiratory Infections, Institut Pasteur, Paris | Marion Barbet, Sylvie Behillil, Méline Bizard, Angela Brisebarre, Camille Capel, Vincent Enouf, Louise Lefrançois, Frédéric Lemoine, Christophe Malabat, Corinne Maufrais, Amaury Vaysse, Etienne Simon-Lorière, Maud Vanpeene, Sylvie Van der Werf ,TROMBERT Sabine |
| EPI_ISL_2923188, EPI_ISL_2923189 | Labo Analyses Med | National Reference Center for Viruses of Respiratory Infections, Institut Pasteur, Paris | Marion Barbet, Sylvie Behillil, Méline Bizard, Angela Brisebarre, Camille Capel, Vincent Enouf, Louise Lefrançois, Frédéric Lemoine, Christophe Malabat, Corinne Maufrais, Hub Bioinformatique Biostatistiques, Etienne Simon-Lorière, Maud Vanpeene, Sylvie Van der Werf ,TROMBERT Sabine |
| EPI_ISL_2923190 | Labo Analyses Med | National Reference Center for Viruses of Respiratory Infections, Institut Pasteur, Paris | Marion Barbet, Sylvie Behillil, Méline Bizard, Angela Brisebarre, Camille Capel, Vincent Enouf, Louise Lefrançois, Frédéric Lemoine, Christophe Malabat, Corinne Maufrais, Amaury Vaysse, Etienne Simon-Lorière, Maud Vanpeene, Sylvie Van der Werf ,TROMBERT Sabine |
| EPI_ISL_2923191, EPI_ISL_2923192, EPI_ISL_2923193 | Labo Analyses Med | National Reference Center for Viruses of Respiratory Infections, Institut Pasteur, Paris | Marion Barbet, Sylvie Behillil, Méline Bizard, Angela Brisebarre, Camille Capel, Vincent Enouf, Louise Lefrançois, Frédéric Lemoine, Christophe Malabat, Corinne Maufrais, Hub Bioinformatique Biostatistiques, Etienne Simon-Lorière, Maud Vanpeene, Sylvie Van der Werf ,TROMBERT Sabine |
| EPI_ISL_2923194 | Labo Analyses Med | National Reference Center for Viruses of Respiratory Infections, Institut Pasteur, Paris | Marion Barbet, Sylvie Behillil, Méline Bizard, Angela Brisebarre, Camille Capel, Vincent Enouf, Louise Lefrançois, Frédéric Lemoine, Christophe Malabat, Corinne Maufrais, Hub Bioinformatique Biostatistiques, Etienne Simon-Lorière, Maud Vanpeene, Sylvie Van der Werf ,VIEILLEFOND VINCENT |
| EPI_ISL_2923195 | Labo Analyses Med | National Reference Center for Viruses of Respiratory Infections, Institut Pasteur, Paris | Marion Barbet, Sylvie Behillil, Méline Bizard, Angela Brisebarre, Camille Capel, Vincent Enouf, Louise Lefrançois, Frédéric Lemoine, Christophe Malabat, Corinne Maufrais, Amaury Vaysse, Etienne Simon-Lorière, Maud Vanpeene, Sylvie Van der Werf ,VIEILLEFOND VINCENT |
| EPI_ISL_2923196, EPI_ISL_2923198, EPI_ISL_2923199 | Labo Analyses Med | National Reference Center for Viruses of Respiratory Infections, Institut Pasteur, Paris | Marion Barbet, Sylvie Behillil, Méline Bizard, Angela Brisebarre, Camille Capel, Vincent Enouf, Louise Lefrançois, Frédéric Lemoine, Christophe Malabat, Corinne Maufrais, Hub Bioinformatique Biostatistiques, Etienne Simon-Lorière, Maud Vanpeene, Sylvie Van der Werf ,VIEILLEFOND VINCENT |
| EPI_ISL_2923200, EPI_ISL_2923201 | Labo Analyses Med | National Reference Center for Viruses of Respiratory Infections, Institut Pasteur, Paris | Marion Barbet, Sylvie Behillil, Méline Bizard, Angela Brisebarre, Camille Capel, Vincent Enouf, Louise Lefrançois, Frédéric Lemoine, Christophe Malabat, Corinne Maufrais, Amaury Vaysse, Etienne Simon-Lorière, Maud Vanpeene, Sylvie Van der Werf ,VIEILLEFOND VINCENT |
| EPI_ISL_2923202, EPI_ISL_2923203, EPI_ISL_2923204, EPI_ISL_2923205 | Labo Analyses Med | National Reference Center for Viruses of Respiratory Infections, Institut Pasteur, Paris | Marion Barbet, Sylvie Behillil, Méline Bizard, Angela Brisebarre, Camille Capel, Vincent Enouf, Louise Lefrançois, Frédéric Lemoine, Christophe Malabat, Corinne Maufrais, Hub Bioinformatique Biostatistiques, Etienne Simon-Lorière, Maud Vanpeene, Sylvie Van der Werf ,VIEILLEFOND VINCENT |
| EPI_ISL_2923206 | Labo Analyses Med | National Reference Center for Viruses of Respiratory Infections, Institut Pasteur, Paris | Marion Barbet, Sylvie Behillil, Méline Bizard, Angela Brisebarre, Camille Capel, Vincent Enouf, Louise Lefrançois, Frédéric Lemoine, Christophe Malabat, Corinne Maufrais, Pierre Lechat, Etienne Simon-Lorière, Maud Vanpeene, Sylvie Van der Werf ,VIEILLEFOND VINCENT |
| EPI_ISL_2923207, EPI_ISL_2923208 | Labo Analyses Med | National Reference Center for Viruses of Respiratory Infections, Institut Pasteur, Paris | Marion Barbet, Sylvie Behillil, Méline Bizard, Angela Brisebarre, Camille Capel, Vincent Enouf, Louise Lefrançois, Frédéric Lemoine, Christophe Malabat, Corinne Maufrais, Hub Bioinformatique Biostatistiques, Etienne Simon-Lorière, Maud Vanpeene, Sylvie Van der Werf ,VIEILLEFOND VINCENT |
| EPI_ISL_2923210 | Labo Analyses Med | National Reference Center for Viruses of Respiratory Infections, Institut Pasteur, Paris | Marion Barbet, Sylvie Behillil, Méline Bizard, Angela Brisebarre, Camille Capel, Vincent Enouf, Louise Lefrançois, Frédéric Lemoine, Christophe Malabat, Corinne Maufrais, Pierre Lechat, Etienne Simon-Lorière, Maud Vanpeene, Sylvie Van der Werf ,VIEILLEFOND VINCENT |
| EPI_ISL_2923211, EPI_ISL_2923212 | Labo Analyses Med | National Reference Center for Viruses of Respiratory Infections, Institut Pasteur, Paris | Marion Barbet, Sylvie Behillil, Méline Bizard, Angela Brisebarre, Camille Capel, Vincent Enouf, Louise Lefrançois, Frédéric Lemoine, Christophe Malabat, Corinne Maufrais, Hub Bioinformatique Biostatistiques, Etienne Simon-Lorière, Maud Vanpeene, Sylvie Van der Werf ,VIEILLEFOND VINCENT |
| EPI_ISL_2923213, EPI_ISL_2923214 | Labo Analyses Med | National Reference Center for Viruses of Respiratory Infections, Institut Pasteur, Paris | Marion Barbet, Sylvie Behillil, Méline Bizard, Angela Brisebarre, Camille Capel, Vincent Enouf, Louise Lefrançois, Frédéric Lemoine, Christophe Malabat, Corinne Maufrais, Hub Bioinformatique Biostatistiques, Etienne Simon-Lorière, Maud Vanpeene, Sylvie Van der Werf ,FOULQUIER Jean-Baptiste |
| EPI_ISL_2923216, EPI_ISL_2923217 | Labo Analyses Med | National Reference Center for Viruses of Respiratory Infections, Institut Pasteur, Paris | Marion Barbet, Sylvie Behillil, Méline Bizard, Angela Brisebarre, Camille Capel, Vincent Enouf, Louise Lefrançois, Frédéric Lemoine, Christophe Malabat, Corinne Maufrais, Hub Bioinformatique Biostatistiques, Etienne Simon-Lorière, Maud Vanpeene, Sylvie Van der Werf ,PASQUIER Emmanuel |
| EPI_ISL_2923218 | Labo Analyses Med | National Reference Center for Viruses of Respiratory Infections, Institut Pasteur, Paris | Marion Barbet, Sylvie Behillil, Méline Bizard, Angela Brisebarre, Camille Capel, Vincent Enouf, Louise Lefrançois, Frédéric Lemoine, Christophe Malabat, Corinne Maufrais, Hub Bioinformatique Biostatistiques, Etienne Simon-Lorière, Maud Vanpeene, Sylvie Van der Werf ,FELLONI Claire |
| EPI_ISL_2923219 | Labo Analyses Med | National Reference Center for Viruses of Respiratory Infections, Institut Pasteur, Paris | Marion Barbet, Sylvie Behillil, Méline Bizard, Angela Brisebarre, Camille Capel, Vincent Enouf, Louise Lefrançois, Frédéric Lemoine, Christophe Malabat, Corinne Maufrais, Gael Millot, Etienne Simon-Lorière, Maud Vanpeene, Sylvie Van der Werf ,FELLONI Claire |
| EPI_ISL_2923220 | Labo Analyses Med | National Reference Center for Viruses of Respiratory Infections, Institut Pasteur, Paris | Marion Barbet, Sylvie Behillil, Méline Bizard, Angela Brisebarre, Camille Capel, Vincent Enouf, Louise Lefrançois, Frédéric Lemoine, Christophe Malabat, Corinne Maufrais, Hub Bioinformatique Biostatistiques, Etienne Simon-Lorière, Maud Vanpeene, Sylvie Van der Werf ,FELLONI Claire |
| EPI_ISL_2923221 | Labo Analyses Med | National Reference Center for Viruses of Respiratory Infections, Institut Pasteur, Paris | Marion Barbet, Sylvie Behillil, Méline Bizard, Angela Brisebarre, Camille Capel, Vincent Enouf, Louise Lefrançois, Frédéric Lemoine, Christophe Malabat, Corinne Maufrais, Gael Millot, Etienne Simon-Lorière, Maud Vanpeene, Sylvie Van der Werf ,HOLSTEIN Anne |
| EPI_ISL_2923222, EPI_ISL_2923223, EPI_ISL_2923224, EPI_ISL_2923225 | Labo Analyses Med | National Reference Center for Viruses of Respiratory Infections, Institut Pasteur, Paris | Marion Barbet, Sylvie Behillil, Méline Bizard, Angela Brisebarre, Camille Capel, Vincent Enouf, Louise Lefrançois, Frédéric Lemoine, Christophe Malabat, Corinne Maufrais, Hub Bioinformatique Biostatistiques, Etienne Simon-Lorière, Maud Vanpeene, Sylvie Van der Werf ,HOLSTEIN Anne |
| EPI_ISL_2923226 | Labo Analyses Med | National Reference Center for Viruses of Respiratory Infections, Institut Pasteur, Paris | Marion Barbet, Sylvie Behillil, Méline Bizard, Angela Brisebarre, Camille Capel, Vincent Enouf, Louise Lefrançois, Frédéric Lemoine, Christophe Malabat, Corinne Maufrais, Gael Millot, Etienne Simon-Lorière, Maud Vanpeene, Sylvie Van der Werf ,HOLSTEIN Anne |
| EPI_ISL_2923227 | Labo Analyses Med | National Reference Center for Viruses of Respiratory Infections, Institut Pasteur, Paris | Marion Barbet, Sylvie Behillil, Méline Bizard, Angela Brisebarre, Camille Capel, Vincent Enouf, Louise Lefrançois, Frédéric Lemoine, Christophe Malabat, Corinne Maufrais, Hub Bioinformatique Biostatistiques, Etienne Simon-Lorière, Maud Vanpeene, Sylvie Van der Werf ,HOLSTEIN Anne |
| EPI_ISL_2923228, EPI_ISL_2923229, EPI_ISL_2923230 | Labo Analyses Med | National Reference Center for Viruses of Respiratory Infections, Institut Pasteur, Paris | Marion Barbet, Sylvie Behillil, Méline Bizard, Angela Brisebarre, Camille Capel, Vincent Enouf, Louise Lefrançois, Frédéric Lemoine, Christophe Malabat, Corinne Maufrais, Hub Bioinformatique Biostatistiques, Etienne Simon-Lorière, Maud Vanpeene, Sylvie Van der Werf ,GESTIN Brieuc |
| EPI_ISL_2923231 | Labo Analyses Med | National Reference Center for Viruses of Respiratory Infections, Institut Pasteur, Paris | Marion Barbet, Sylvie Behillil, Méline Bizard, Angela Brisebarre, Camille Capel, Vincent Enouf, Louise Lefrançois, Frédéric Lemoine, Christophe Malabat, Corinne Maufrais, Gael Millot, Etienne Simon-Lorière, Maud Vanpeene, Sylvie Van der Werf ,GESTIN Brieuc |
| EPI_ISL_2923232, EPI_ISL_2923233, EPI_ISL_2923234, EPI_ISL_2923235, EPI_ISL_2923236, EPI_ISL_2923237, EPI_ISL_2923238 | Labo Analyses Med | National Reference Center for Viruses of Respiratory Infections, Institut Pasteur, Paris | Marion Barbet, Sylvie Behillil, Méline Bizard, Angela Brisebarre, Camille Capel, Vincent Enouf, Louise Lefrançois, Frédéric Lemoine, Christophe Malabat, Corinne Maufrais, Hub Bioinformatique Biostatistiques, Etienne Simon-Lorière, Maud Vanpeene, Sylvie Van der Werf ,GESTIN Brieuc |
| EPI_ISL_2923239, EPI_ISL_2923241 | Labo Analyses Med | National Reference Center for Viruses of Respiratory Infections, Institut Pasteur, Paris | Marion Barbet, Sylvie Behillil, Méline Bizard, Angela Brisebarre, Camille Capel, Vincent Enouf, Louise Lefrançois, Frédéric Lemoine, Christophe Malabat, Corinne Maufrais, Hub Bioinformatique Biostatistiques, Etienne Simon-Lorière, Maud Vanpeene, Sylvie Van der Werf ,CERBALLIANCE Ile de France sud |
| EPI_ISL_2923242 | Labo Analyses Med | National Reference Center for Viruses of Respiratory Infections, Institut Pasteur, Paris | Marion Barbet, Sylvie Behillil, Méline Bizard, Angela Brisebarre, Camille Capel, Vincent Enouf, Louise Lefrançois, Frédéric Lemoine, Christophe Malabat, Corinne Maufrais, Pierre Lechat, Etienne Simon-Lorière, Maud Vanpeene, Sylvie Van der Werf ,CERBALLIANCE Ile de France sud |

| | | | |
|---|---|---|---|
| EPI_ISL_2923243 | Labo Analyses Med | National Reference Center for Viruses of Respiratory Infections, Institut Pasteur, Paris | Marion Barbet, Sylvie Behillil, Méline Bizard, Angela Brisebarre, Camille Capel, Vincent Enouf, Louise Lefrançois, Frédéric Lemoine, Christophe Malabat, Corinne Maufrais, Gael Millot, Etienne Simon-Lorière, Maud Vanpeene, Sylvie Van der Werf ,BESSON Jérôme |
| EPI_ISL_2923244 | Labo Analyses Med | National Reference Center for Viruses of Respiratory Infections, Institut Pasteur, Paris | Marion Barbet, Sylvie Behillil, Méline Bizard, Angela Brisebarre, Camille Capel, Vincent Enouf, Louise Lefrançois, Frédéric Lemoine, Christophe Malabat, Corinne Maufrais, Pierre Lechat, Etienne Simon-Lorière, Maud Vanpeene, Sylvie Van der Werf ,BESSON Jérôme |
| EPI_ISL_2923245 | Labo Analyses Med | National Reference Center for Viruses of Respiratory Infections, Institut Pasteur, Paris | Marion Barbet, Sylvie Behillil, Méline Bizard, Angela Brisebarre, Camille Capel, Vincent Enouf, Louise Lefrançois, Frédéric Lemoine, Christophe Malabat, Corinne Maufrais, Gael Millot, Etienne Simon-Lorière, Maud Vanpeene, Sylvie Van der Werf ,BESSON Jérôme |
| EPI_ISL_2923246 | Labo Analyses Med | National Reference Center for Viruses of Respiratory Infections, Institut Pasteur, Paris | Marion Barbet, Sylvie Behillil, Méline Bizard, Angela Brisebarre, Camille Capel, Vincent Enouf, Louise Lefrançois, Frédéric Lemoine, Christophe Malabat, Corinne Maufrais, Pierre Lechat, Etienne Simon-Lorière, Maud Vanpeene, Sylvie Van der Werf ,BESSON Jérôme |
| EPI_ISL_2923247 | Labo Analyses Med | National Reference Center for Viruses of Respiratory Infections, Institut Pasteur, Paris | Marion Barbet, Sylvie Behillil, Méline Bizard, Angela Brisebarre, Camille Capel, Vincent Enouf, Louise Lefrançois, Frédéric Lemoine, Christophe Malabat, Corinne Maufrais, Hub Bioinformatique Biostatistiques, Etienne Simon-Lorière, Maud Vanpeene, Sylvie Van der Werf ,BESSON Jérôme |
| EPI_ISL_2923248, EPI_ISL_2923249, EPI_ISL_2923250 | Labo Analyses Med | National Reference Center for Viruses of Respiratory Infections, Institut Pasteur, Paris | Marion Barbet, Sylvie Behillil, Méline Bizard, Angela Brisebarre, Camille Capel, Vincent Enouf, Louise Lefrançois, Frédéric Lemoine, Christophe Malabat, Corinne Maufrais, Hub Bioinformatique Biostatistiques, Etienne Simon-Lorière, Maud Vanpeene, Sylvie Van der Werf ,COCCO ARDITI |
| EPI_ISL_2923251 | Labo Analyses Med | National Reference Center for Viruses of Respiratory Infections, Institut Pasteur, Paris | Marion Barbet, Sylvie Behillil, Méline Bizard, Angela Brisebarre, Camille Capel, Vincent Enouf, Louise Lefrançois, Frédéric Lemoine, Christophe Malabat, Corinne Maufrais, Pierre Lechat, Etienne Simon-Lorière, Maud Vanpeene, Sylvie Van der Werf ,COCCO ARDITI |
| EPI_ISL_2923252, EPI_ISL_2923253, EPI_ISL_2923254, EPI_ISL_2923256, EPI_ISL_2923257 | Labo Analyses Med | National Reference Center for Viruses of Respiratory Infections, Institut Pasteur, Paris | Marion Barbet, Sylvie Behillil, Méline Bizard, Angela Brisebarre, Camille Capel, Vincent Enouf, Louise Lefrançois, Frédéric Lemoine, Christophe Malabat, Corinne Maufrais, Hub Bioinformatique Biostatistiques, Etienne Simon-Lorière, Maud Vanpeene, Sylvie Van der Werf ,COCCO ARDITI |
| EPI_ISL_2923258 | Labo Analyses Med | National Reference Center for Viruses of Respiratory Infections, Institut Pasteur, Paris | Marion Barbet, Sylvie Behillil, Méline Bizard, Angela Brisebarre, Camille Capel, Vincent Enouf, Louise Lefrançois, Frédéric Lemoine, Christophe Malabat, Corinne Maufrais, Gael Millot, Etienne Simon-Lorière, Maud Vanpeene, Sylvie Van der Werf ,COCCO ARDITI |
| EPI_ISL_2923259, EPI_ISL_2923260 | Labo Analyses Med | National Reference Center for Viruses of Respiratory Infections, Institut Pasteur, Paris | Marion Barbet, Sylvie Behillil, Méline Bizard, Angela Brisebarre, Camille Capel, Vincent Enouf, Louise Lefrançois, Frédéric Lemoine, Christophe Malabat, Corinne Maufrais, Hub Bioinformatique Biostatistiques, Etienne Simon-Lorière, Maud Vanpeene, Sylvie Van der Werf ,COCCO ARDITI |
| EPI_ISL_2923261 | Labo Analyses Med | National Reference Center for Viruses of Respiratory Infections, Institut Pasteur, Paris | Marion Barbet, Sylvie Behillil, Méline Bizard, Angela Brisebarre, Camille Capel, Vincent Enouf, Louise Lefrançois, Frédéric Lemoine, Christophe Malabat, Corinne Maufrais, Hub Bioinformatique Biostatistiques, Etienne Simon-Lorière, Maud Vanpeene, Sylvie Van der Werf ,COMES Jean-François |
| EPI_ISL_2923262, EPI_ISL_2923263 | Labo Analyses Med | National Reference Center for Viruses of Respiratory Infections, Institut Pasteur, Paris | Marion Barbet, Sylvie Behillil, Méline Bizard, Angela Brisebarre, Camille Capel, Vincent Enouf, Louise Lefrançois, Frédéric Lemoine, Christophe Malabat, Corinne Maufrais, Gael Millot, Etienne Simon-Lorière, Maud Vanpeene, Sylvie Van der Werf ,COMES Jean-François |
| EPI_ISL_2923264, EPI_ISL_2923265, EPI_ISL_2923266 | Labo Analyses Med | National Reference Center for Viruses of Respiratory Infections, Institut Pasteur, Paris | Marion Barbet, Sylvie Behillil, Méline Bizard, Angela Brisebarre, Camille Capel, Vincent Enouf, Louise Lefrançois, Frédéric Lemoine, Christophe Malabat, Corinne Maufrais, Hub Bioinformatique Biostatistiques, Etienne Simon-Lorière, Maud Vanpeene, Sylvie Van der Werf ,COMES Jean-François |
| EPI_ISL_2923267 | Hospital | National Reference Center for Viruses of Respiratory Infections, Institut Pasteur, Paris | Marion Barbet, Sylvie Behillil, Méline Bizard, Angela Brisebarre, Camille Capel, Vincent Enouf, Louise Lefrançois, Frédéric Lemoine, Christophe Malabat, Corinne Maufrais, Gael Millot, Etienne Simon-Lorière, Maud Vanpeene, Sylvie Van der Werf ,FARFOUR Eric |
| EPI_ISL_2923268 | Hospital | National Reference Center for Viruses of Respiratory Infections, Institut Pasteur, Paris | Marion Barbet, Sylvie Behillil, Méline Bizard, Angela Brisebarre, Camille Capel, Vincent Enouf, Louise Lefrançois, Frédéric Lemoine, Christophe Malabat, Corinne Maufrais, Gael Millot, Etienne Simon-Lorière, Maud Vanpeene, Sylvie Van der Werf ,GAUDY GRAFFIN Catherine |
| EPI_ISL_2923269 | Hospital | National Reference Center for Viruses of Respiratory Infections, Institut Pasteur, Paris | Marion Barbet, Sylvie Behillil, Méline Bizard, Angela Brisebarre, Camille Capel, Vincent Enouf, Louise Lefrançois, Frédéric Lemoine, Christophe Malabat, Corinne Maufrais, Hub Bioinformatique Biostatistiques, Etienne Simon-Lorière, Maud Vanpeene, Sylvie Van der Werf ,GAUDY GRAFFIN Catherine |
| EPI_ISL_2923271 | Labo Analyses Med | National Reference Center for Viruses of Respiratory Infections, Institut Pasteur, Paris | Marion Barbet, Sylvie Behillil, Méline Bizard, Angela Brisebarre, Camille Capel, Vincent Enouf, Louise Lefrançois, Frédéric Lemoine, Christophe Malabat, Corinne Maufrais, Hub Bioinformatique Biostatistiques, Etienne Simon-Lorière, Maud Vanpeene, Sylvie Van der Werf ,WEBER Valérie |
| EPI_ISL_2923272 | Labo Analyses Med | National Reference Center for Viruses of Respiratory Infections, Institut Pasteur, Paris | Marion Barbet, Sylvie Behillil, Méline Bizard, Angela Brisebarre, Camille Capel, Vincent Enouf, Louise Lefrançois, Frédéric Lemoine, Christophe Malabat, Corinne Maufrais, Gael Millot, Etienne Simon-Lorière, Maud Vanpeene, Sylvie Van der Werf ,WEBER Valérie |
| EPI_ISL_2923273, EPI_ISL_2923274 | Labo Analyses Med | National Reference Center for Viruses of Respiratory Infections, Institut Pasteur, Paris | Marion Barbet, Sylvie Behillil, Méline Bizard, Angela Brisebarre, Camille Capel, Vincent Enouf, Louise Lefrançois, Frédéric Lemoine, Christophe Malabat, Corinne Maufrais, Hub Bioinformatique Biostatistiques, Etienne Simon-Lorière, Maud Vanpeene, Sylvie Van der Werf ,TROMBERT Sabine |
| EPI_ISL_2923275 | Labo Analyses Med | National Reference Center for Viruses of Respiratory Infections, Institut Pasteur, Paris | Marion Barbet, Sylvie Behillil, Méline Bizard, Angela Brisebarre, Camille Capel, Vincent Enouf, Louise Lefrançois, Frédéric Lemoine, Christophe Malabat, Corinne Maufrais, Gael Millot, Etienne Simon-Lorière, Maud Vanpeene, Sylvie Van der Werf ,TROMBERT Sabine |
| EPI_ISL_2923276, EPI_ISL_2923277 | Labo Analyses Med | National Reference Center for Viruses of Respiratory Infections, Institut Pasteur, Paris | Marion Barbet, Sylvie Behillil, Méline Bizard, Angela Brisebarre, Camille Capel, Vincent Enouf, Louise Lefrançois, Frédéric Lemoine, Christophe Malabat, Corinne Maufrais, Hub Bioinformatique Biostatistiques, Etienne Simon-Lorière, Maud Vanpeene, Sylvie Van der Werf ,TROMBERT Sabine |
| EPI_ISL_2923278 | Labo Analyses Med | National Reference Center for Viruses of Respiratory Infections, Institut Pasteur, Paris | Marion Barbet, Sylvie Behillil, Méline Bizard, Angela Brisebarre, Camille Capel, Vincent Enouf, Louise Lefrançois, Frédéric Lemoine, Christophe Malabat, Corinne Maufrais, Hub Bioinformatique Biostatistiques, Etienne Simon-Lorière, Maud Vanpeene, Sylvie Van der Werf ,MICHEZ Karine |
| EPI_ISL_2923279, EPI_ISL_2923280 | Labo Analyses Med | National Reference Center for Viruses of Respiratory Infections, Institut Pasteur, Paris | Marion Barbet, Sylvie Behillil, Méline Bizard, Angela Brisebarre, Camille Capel, Vincent Enouf, Louise Lefrançois, Frédéric Lemoine, Christophe Malabat, Corinne Maufrais, Gael Millot, Etienne Simon-Lorière, Maud Vanpeene, Sylvie Van der Werf ,MICHEZ Karine |
| EPI_ISL_2923281 | Labo Analyses Med | National Reference Center for Viruses of Respiratory Infections, Institut Pasteur, Paris | Marion Barbet, Sylvie Behillil, Méline Bizard, Angela Brisebarre, Camille Capel, Vincent Enouf, Louise Lefrançois, Frédéric Lemoine, Christophe Malabat, Corinne Maufrais, Hub Bioinformatique Biostatistiques, Etienne Simon-Lorière, Maud Vanpeene, Sylvie Van der Werf ,MICHEZ Karine |
| EPI_ISL_2923282 | Labo Analyses Med | National Reference Center for Viruses of Respiratory Infections, Institut Pasteur, Paris | Marion Barbet, Sylvie Behillil, Méline Bizard, Angela Brisebarre, Camille Capel, Vincent Enouf, Louise Lefrançois, Frédéric Lemoine, Christophe Malabat, Corinne Maufrais, Hub Bioinformatique Biostatistiques, Etienne Simon-Lorière, Maud Vanpeene, Sylvie Van der Werf ,POTIRON Grégoire |
| EPI_ISL_2924057 | Labuan Hospital | Institute for Medical Research, Infectious Disease Research Centre, National Institutes of Health, Ministry of Health Malaysia | Suppiah J, Kamel K, Mohd Zawawi Z, Azizan MA, Ramly N, Robert F, Anasir MI, Thayan R |
| EPI_ISL_2924058 | Instituto Adolfo Lutz Central | Instituto Adolfo Lutz, Rapid Response Center, Strategic Laboratory | Claudio Tavares Sacchi, Claudia Regina Gonçalves, Erica Valessa Ramos Gomes, Karoline Rodrigues Campos, Caio Vinicius Dias Lopes, Leonardo Jose Tadeu de Araujo |
| EPI_ISL_2924059, EPI_ISL_2924060 | P.O. Cannizzaro | AOU Polclinico "G Rodolico-S.Marco" PO Rodolico and BIOMETEC UNICT CATANIA | STEFANI AND SCALIA |
| EPI_ISL_2924137 | Sultan Haji Ahmad Shah Hospital | Institute for Medical Research, Infectious Disease Research Centre, National Institutes of Health, Ministry of Health Malaysia | Suppiah J, Kamel K, Mohd Zawawi Z, Azizan MA, Ramly N, Robert F, Anasir MI, Thayan R |
| EPI_ISL_2924138 | AOU Polclinico "G Rodolico-S.Marco" PO Rodolico Catania | AOU Polclinico "G Rodolico-S.Marco" PO Rodolico and BIOMETEC UNICT CATANIA | STEFANI AND SCALIA |
| EPI_ISL_2924139 | Sultan Haji Ahmad Shah Hospital | Institute for Medical Research, Infectious Disease Research Centre, National Institutes of Health, Ministry of Health Malaysia | Suppiah J, Kamel K, Mohd Zawawi Z, Azizan MA, Ramly N, Robert F, Anasir MI, Thayan R |
| EPI_ISL_2924140 | AOU Polclinico "G Rodolico-S.Marco" PO Rodolico CATANIA | AOU Polclinico "G Rodolico-S.Marco" PO Rodolico and BIOMETEC UNICT CATANIA | STEFANI AND SCALIA |
| EPI_ISL_2924141 | P.O. Cannizzaro | AOU Polclinico "G Rodolico-S.Marco" PO Rodolico and BIOMETEC UNICT CATANIA | STEFANI AND SCALIA |
| EPI_ISL_2925475, EPI_ISL_2925477 | National Center of Infectious and Parasitic Diseases | National Center of Infectious and Parasitic Diseases | Alexiev et al |

| | | | |
|---|---|---|---|
| EPI_ISL_2925481 | AOU Polclinico "G Rodolico-S.Marco" PO Rodolico CATANIA | AOU Polclinico "G Rodolico-S.Marco" PO Rodolico and BIOMETEC UNICT CATANIA | STEFANI AND SCALIA |
| EPI_ISL_2925823 | Clinical Microbiology, Nationwide Children's Hospital | James Molecular Laboratory | Huanyu Wang, Pam Snyder, Zachary Rudy, Dan Jones, Amy Leber |
| EPI_ISL_2927083 | Cliniques universitaires Saint-Luc | UCLouvain/IREC/MBLG | Jean Ruelle, Bertrand Bearzatto, Benoit Kabamba Mukadi |
| EPI_ISL_2927972, EPI_ISL_2927973 | Imágenes Diagnósticas de Mazatlán | Microbial Genomics Laboratory | Consorcio Mexicano de Vigilancia Genómica (CoViGen-Mex). Authors (in alphabetical order): Julio Elias Alvarado-Yaah, Carlos F. Arias, Santiago Ávila-Ríos, Eduardo Becerril-Vargas, Víctor Hugo Borja-Aburto, Celia Boukadida, Cristóbal Cháidez-Quiróz, Juan Bautista Chale-Dzul, Ricardo Ciria Merce, Andreu Comas-García, Célida Duque Molina, Julissa Enciso-Ibarra, José Antonio Enciso-Moreno, Gloria Elena Espinosa-Ayala, Fernando Fontove-Herrera, Daniel Fregoso-Rueda, Victor Eduardo García-Arias, Alejandra García-Gasca, Bruno Gómez-Gil, Jean Pierre González, Irvin González-López, Concepción Grajales-Muñiz, Ricardo Grande, Rosa María Gutiérrez Rios, Jesús Hernández, Alejandra Hernández-Terán, Alfredo Herrera-Estrella, Carla Ivón Herrera-Najera, Pavel Isa, Verónica Mata-Haro, Daniel Lira Morales, Susana Lopez, Antonio Loza Román, Brenda Irasema Maldonado-Meza, Bernardo Martínez-Miguel, José Arturo Martínez-Orozco, Célida Martínez- Rodríguez, Margarita Matías-Florentino, Fidencio Mejía-Nepomuceno, María Guadalupe de Jesús Mireles-Rivera, Gloria María Molina-Salinas, Hector Montoya-Fuentes, , Mario Mújica-Sánchez, José Esteban Muñoz-Medina, José de Jesús Nuñez-Contreras, Alicia Ocaña-Mondragón, Luis Alberto Ochoa-Carrera, Hector Esteban Paz-Juárez, Marissa Perez-Garcia, Francisco Pulido, Helen Haydee Fernanda Ramírez-Plascencia, Jorge Salas-Hernández, Angel Gustavo Salas-Lais, Alejandro Sánchez-Flores, Clara Esperanza Santacruz-Tinoco, María Guadalupe Santiago-Mauricio, Selene Zárate, Nelly Sélem-Mojica, Blanca Taboada, Gloria Vazquez, Joel Armando Vázquez-Pérez. |
| EPI_ISL_2927975 | Hospital Sharp | Microbial Genomics Laboratory | Consorcio Mexicano de Vigilancia Genómica (CoViGen-Mex). Authors (in alphabetical order): Julio Elias Alvarado-Yaah, Carlos F. Arias, Santiago Ávila-Ríos, Eduardo Becerril-Vargas, Víctor Hugo Borja-Aburto, Celia Boukadida, Cristóbal Cháidez-Quiróz, Juan Bautista Chale-Dzul, Ricardo Ciria Merce, Andreu Comas-García, Célida Duque Molina, Julissa Enciso-Ibarra, José Antonio Enciso-Moreno, Gloria Elena Espinosa-Ayala, Fernando Fontove-Herrera, Daniel Fregoso-Rueda, Victor Eduardo García-Arias, Alejandra García-Gasca, Bruno Gómez-Gil, Jean Pierre González, Irvin González-López, Concepción Grajales-Muñiz, Ricardo Grande, Rosa María Gutiérrez Rios, Jesús Hernández, Alejandra Hernández-Terán, Alfredo Herrera-Estrella, Carla Ivón Herrera-Najera, Pavel Isa, Verónica Mata-Haro, Daniel Lira Morales, Susana Lopez, Antonio Loza Román, Brenda Irasema Maldonado-Meza, Bernardo Martínez-Miguel, José Arturo Martínez-Orozco, Célida Martínez- Rodríguez, Margarita Matías-Florentino, Fidencio Mejía-Nepomuceno, María Guadalupe de Jesús Mireles-Rivera, Gloria María Molina-Salinas, Hector Montoya-Fuentes, , Mario Mújica-Sánchez, José Esteban Muñoz-Medina, José de Jesús Nuñez-Contreras, Alicia Ocaña-Mondragón, Luis Alberto Ochoa-Carrera, Hector Esteban Paz-Juárez, Marissa Perez-Garcia, Francisco Pulido, Helen Haydee Fernanda Ramírez-Plascencia, Jorge Salas-Hernández, Angel Gustavo Salas-Lais, Alejandro Sánchez-Flores, Clara Esperanza Santacruz-Tinoco, María Guadalupe Santiago-Mauricio, Selene Zárate, Nelly Sélem-Mojica, Blanca Taboada, Gloria Vazquez, Joel Armando Vázquez-Pérez. |
| EPI_ISL_2927976, EPI_ISL_2927977 | Hospital Margarita Maza de Juárez | Microbial Genomics Laboratory | Consorcio Mexicano de Vigilancia Genómica (CoViGen-Mex). Authors (in alphabetical order): Julio Elias Alvarado-Yaah, Carlos F. Arias, Santiago Ávila-Ríos, Eduardo Becerril-Vargas, Víctor Hugo Borja-Aburto, Celia Boukadida, Cristóbal Cháidez-Quiróz, Juan Bautista Chale-Dzul, Ricardo Ciria Merce, Andreu Comas-García, Célida Duque Molina, Julissa Enciso-Ibarra, José Antonio Enciso-Moreno, Gloria Elena Espinosa-Ayala, Fernando Fontove-Herrera, Daniel Fregoso-Rueda, Victor Eduardo García-Arias, Alejandra García-Gasca, Bruno Gómez-Gil, Jean Pierre González, Irvin González-López, Concepción Grajales-Muñiz, Ricardo Grande, Rosa María Gutiérrez Rios, Jesús Hernández, Alejandra Hernández-Terán, Alfredo Herrera-Estrella, Carla Ivón Herrera-Najera, Pavel Isa, Verónica Mata-Haro, Daniel Lira Morales, Susana Lopez, Antonio Loza Román, Brenda Irasema Maldonado-Meza, Bernardo Martínez-Miguel, José Arturo Martínez-Orozco, Célida Martínez- Rodríguez, Margarita Matías-Florentino, Fidencio Mejía-Nepomuceno, María Guadalupe de Jesús Mireles-Rivera, Gloria María Molina-Salinas, Hector Montoya-Fuentes, , Mario Mújica-Sánchez, José Esteban Muñoz-Medina, José de Jesús Nuñez-Contreras, Alicia Ocaña-Mondragón, Luis Alberto Ochoa-Carrera, Hector Esteban Paz-Juárez, Marissa Perez-Garcia, Francisco Pulido, Helen Haydee Fernanda Ramírez-Plascencia, Jorge Salas-Hernández, Angel Gustavo Salas-Lais, Alejandro Sánchez-Flores, Clara Esperanza Santacruz-Tinoco, María Guadalupe Santiago-Mauricio, Selene Zárate, Nelly Sélem-Mojica, Blanca Taboada, Gloria Vazquez, Joel Armando Vázquez-Pérez. |
| EPI_ISL_2927979, EPI_ISL_2927980 | CIAD Hermosillo | CIAD Hermosillo | Consorcio Mexicano de Vigilancia Genómica (CoViGen-Mex). Authors (in alphabetical order): Julio Elias Alvarado-Yaah, Carlos F. Arias, Santiago Ávila-Ríos, Eduardo Becerril-Vargas, Víctor Hugo Borja-Aburto, Celia Boukadida, Cristóbal Cháidez-Quiróz, Juan Bautista Chale-Dzul, Ricardo Ciria Merce, Andreu Comas-García, Célida Duque Molina, Julissa Enciso-Ibarra, José Antonio Enciso-Moreno, Gloria Elena Espinosa-Ayala, Fernando Fontove-Herrera, Daniel Fregoso-Rueda, Victor Eduardo García-Arias, Alejandra García-Gasca, Bruno Gómez-Gil, Jean Pierre González, Irvin González-López, Concepción Grajales-Muñiz, Ricardo Grande, Rosa María Gutiérrez Rios, Jesús Hernández, Alejandra Hernández-Terán, Alfredo Herrera-Estrella, Carla Ivón Herrera-Najera, Pavel Isa, Verónica Mata-Haro, Daniel Lira Morales, Susana Lopez, Antonio Loza Román, Brenda Irasema Maldonado-Meza, Bernardo Martínez-Miguel, José Arturo Martínez-Orozco, Célida Martínez- Rodríguez, Margarita Matías-Florentino, Fidencio Mejía-Nepomuceno, María Guadalupe de Jesús Mireles-Rivera, Gloria María Molina-Salinas, Hector Montoya-Fuentes, , Mario Mújica-Sánchez, José Esteban Muñoz-Medina, José de Jesús Nuñez-Contreras, Alicia Ocaña-Mondragón, Luis Alberto Ochoa-Carrera, Hector Esteban Paz-Juárez, Marissa Perez-Garcia, Francisco Pulido, Helen Haydee Fernanda Ramírez-Plascencia, Jorge Salas-Hernández, Angel Gustavo Salas-Lais, Alejandro Sánchez-Flores, Clara Esperanza Santacruz-Tinoco, María Guadalupe Santiago-Mauricio, Selene Zárate, Nelly Sélem-Mojica, Blanca Taboada, Gloria Vazquez, Joel Armando Vázquez-Pérez. |
| EPI_ISL_2928225 | Biomédicos de Mérida | Biomédicos de Mérida | CANCHE-PECH JOSE REYES, MENDOZA-OLIVEROS TAHALI, CHIN-HAU LETICIA, SAURI-MEDINA EMILY, CAUICH-VERA KRISTIAN, CEBALLOS-LOPEZ ADRIAN, LOPEZ-NOVELO MARIA E, GARCIA-GONZALEZ IGRID |
| EPI_ISL_2928226, EPI_ISL_2928228, EPI_ISL_2928229, EPI_ISL_2928230, EPI_ISL_2928234, EPI_ISL_2928235, EPI_ISL_2928237, EPI_ISL_2928241, EPI_ISL_2928242, EPI_ISL_2928244, EPI_ISL_2928246, EPI_ISL_2928247, EPI_ISL_2928248, EPI_ISL_2928251, EPI_ISL_2928253, EPI_ISL_2928254, EPI_ISL_2928255, EPI_ISL_2928257, EPI_ISL_2928258, EPI_ISL_2928260, EPI_ISL_2928261, EPI_ISL_2928262, EPI_ISL_2928263 | | | |
| see above | Baylor Scott & White-Temple | Baylor Scott & White-Temple | Ari Rao, Linden Morales, Kimberly Walker, Marcus Volz, Shelby Johnson |
| EPI_ISL_2928264 | Baylor Scott & White-Waxahachie | Baylor Scott & White-Temple | Ari Rao, Linden Morales, Kimberly Walker, Marcus Volz, Shelby Johnson |
| EPI_ISL_2928268, EPI_ISL_2928283, EPI_ISL_2928284, EPI_ISL_2928285, EPI_ISL_2928286, EPI_ISL_2928292, EPI_ISL_2928293, EPI_ISL_2928294, EPI_ISL_2928295, EPI_ISL_2928296 | Baylor Scott & White-Temple | Baylor Scott & White-Temple | Ari Rao, Linden Morales, Kimberly Walker, Marcus Volz, Shelby Johnson |
| EPI_ISL_2928297 | Biomédicos de Mérida | Biomédicos de Mérida | CANCHE-PECH JOSE REYES, MENDOZA-OLIVEROS TAHALI, CHIN-HAU LETICIA, SAURI-MEDINA EMILY, CAUICH-VERA KRISTIAN, CEBALLOS-LOPEZ ADRIAN, LOPEZ-NOVELO MARIA E, GARCIA-GONZALEZ IGRID |
| EPI_ISL_2928507 | Laboratório de Microbiologia Molecular - Universidade FEEVALE | Laboratório de Microbiologia Molecular - Universidade FEEVALE | Alana Witt Hansen, Fágner Henrique Heldt, Fernando Rosado Spilki, Flávio Silveira, Juliana Schons Gulart, Juliane Deise Fleck, Mariana Soares da Silva, Meriane Demoliner, Matheus Nunes Weber, Paula Rodrigues de Almeida, Michele Filippi. |
| EPI_ISL_2928510 | Secretaria Municipal da Saúde- Porto Alegre | Laboratório de Microbiologia Molecular - Universidade FEEVALE | Alana Witt Hansen, Fágner Henrique Heldt, Fernando Rosado Spilki, Flávio Silveira, Juliana Schons Gulart, Juliane Deise Fleck, Mariana Soares da Silva, Meriane Demoliner, Matheus Nunes Weber, Paula Rodrigues de Almeida, Michele Filippi. |
| EPI_ISL_2931262 | Labo Analyses Med | National Reference Center for Viruses of Respiratory Infections, Institut Pasteur, Paris | Marion Barbet, Sylvie Behillil, Méline Bizard, Angela Brisebarre, Camille Capel, Vincent Enouf, Louise Lefrançois, Frédéric Lemoine, Christophe Malabat, Corinne Maufrais, Damien Mornico, Etienne Simon-Lorière, Maud Vanpeene, Sylvie Van der Werf ,TROMBERT Sabine |
| EPI_ISL_2931263 | Hospital | National Reference Center for Viruses of Respiratory Infections, Institut Pasteur, Paris | Marion Barbet, Sylvie Behillil, Méline Bizard, Angela Brisebarre, Camille Capel, Vincent Enouf, Louise Lefrançois, Frédéric Lemoine, Christophe Malabat, Corinne Maufrais, Victoire Baillet, Etienne Simon-Lorière, Maud Vanpeene, Sylvie Van der Werf ,DUCANCELLE Alexandra |
| EPI_ISL_2931264 | Hospital | National Reference Center for Viruses of Respiratory Infections, Institut Pasteur, Paris | Marion Barbet, Sylvie Behillil, Méline Bizard, Angela Brisebarre, Camille Capel, Vincent Enouf, Louise Lefrançois, Frédéric Lemoine, Christophe Malabat, Corinne Maufrais, Gael Millot, Etienne Simon-Lorière, Maud Vanpeene, Sylvie Van der Werf ,FARFOUR Eric |
| EPI_ISL_2931265 | Labo Analyses Med | National Reference Center for Viruses of Respiratory Infections, Institut Pasteur, Paris | Marion Barbet, Sylvie Behillil, Méline Bizard, Angela Brisebarre, Camille Capel, Vincent Enouf, Louise Lefrançois, Frédéric Lemoine, Christophe Malabat, Corinne Maufrais, Gael Millot, Etienne Simon-Lorière, Maud Vanpeene, Sylvie Van der Werf ,HAGUENOER Eve |
| EPI_ISL_2931266 | Labo Analyses Med | National Reference Center for Viruses of Respiratory | Marion Barbet, Sylvie Behillil, Méline Bizard, Angela Brisebarre, Camille Capel, Vincent Enouf, Louise Lefrançois, Frédéric Lemoine, Christophe Malabat, |

| | | | |
|---|---|---|---|
| | | Infections, Institut Pasteur, Paris | Corinne Maufrais, Gael Millot, Etienne Simon-Lorière, Maud Vanpeene, Sylvie Van der Werf ,TROMBERT Sabine |
| EPI_ISL_2931267 | Labo Analyses Med | National Reference Center for Viruses of Respiratory Infections, Institut Pasteur, Paris | Marion Barbet, Sylvie Behillil, Méline Bizard, Angela Brisebarre, Camille Capel, Vincent Enouf, Louise Lefrançois, Frédéric Lemoine, Christophe Malabat, Corinne Maufrais, Amaury Vaysse, Etienne Simon-Lorière, Maud Vanpeene, Sylvie Van der Werf ,TROMBERT Sabine |
| EPI_ISL_2931268 | Labo Analyses Med | National Reference Center for Viruses of Respiratory Infections, Institut Pasteur, Paris | Marion Barbet, Sylvie Behillil, Méline Bizard, Angela Brisebarre, Camille Capel, Vincent Enouf, Louise Lefrançois, Frédéric Lemoine, Christophe Malabat, Corinne Maufrais, Hub Bioinformatique Biostatistiques, Etienne Simon-Lorière, Maud Vanpeene, Sylvie Van der Werf ,TROMBERT Sabine |
| EPI_ISL_2931269 | Labo Analyses Med | National Reference Center for Viruses of Respiratory Infections, Institut Pasteur, Paris | Marion Barbet, Sylvie Behillil, Méline Bizard, Angela Brisebarre, Camille Capel, Vincent Enouf, Louise Lefrançois, Frédéric Lemoine, Christophe Malabat, Corinne Maufrais, Hub Bioinformatique Biostatistiques, Etienne Simon-Lorière, Maud Vanpeene, Sylvie Van der Werf ,VIEILLEFOND VINCENT |
| EPI_ISL_2931270 | Labo Analyses Med | National Reference Center for Viruses of Respiratory Infections, Institut Pasteur, Paris | Marion Barbet, Sylvie Behillil, Méline Bizard, Angela Brisebarre, Camille Capel, Vincent Enouf, Louise Lefrançois, Frédéric Lemoine, Christophe Malabat, Corinne Maufrais, Pierre Lechat, Etienne Simon-Lorière, Maud Vanpeene, Sylvie Van der Werf ,HOLSTEIN Anne |
| EPI_ISL_2931271 | Labo Analyses Med | National Reference Center for Viruses of Respiratory Infections, Institut Pasteur, Paris | Marion Barbet, Sylvie Behillil, Méline Bizard, Angela Brisebarre, Camille Capel, Vincent Enouf, Louise Lefrançois, Frédéric Lemoine, Christophe Malabat, Corinne Maufrais, Pierre Lechat, Etienne Simon-Lorière, Maud Vanpeene, Sylvie Van der Werf ,CERBALLIANCE Ile de France sud |
| EPI_ISL_2931360 | Home Quarantine Taskforce | Hong Kong Department of Health | Alan K.L. Tsang, Peter C.W. Yip, Ken H.L. Ng, Edman T.K. Lam, Rickjason C.W. Chan |
| EPI_ISL_2931361, EPI_ISL_2931364 | Temporary Specimen Collection Centre at the AsiaWorld-Expo | Hong Kong Department of Health | Alan K.L. Tsang, Peter C.W. Yip, Ken H.L. Ng, Edman T.K. Lam, Rickjason C.W. Chan |
| EPI_ISL_2931366 | Quarantine Centre Silka Tsuen Wan | Hong Kong Department of Health | Alan K.L. Tsang, Peter C.W. Yip, Ken H.L. Ng, Edman T.K. Lam, Rickjason C.W. Chan |
| EPI_ISL_2931367, EPI_ISL_2931368, EPI_ISL_2931369 | North Lantau Hospital | Hong Kong Department of Health | Alan K.L. Tsang, Peter C.W. Yip, Ken H.L. Ng, Edman T.K. Lam, Rickjason C.W. Chan |
| EPI_ISL_2931370, EPI_ISL_2931371, EPI_ISL_2931374, EPI_ISL_2931376, EPI_ISL_2931378, EPI_ISL_2931380, EPI_ISL_2931383, EPI_ISL_2931385 | Laboratori de Referencia de Catalunya | Laboratori de Referencia de Catalunya | Ramirez A., Bellosillo B., Padilla E., Vilas A., Hernandez JJ., Canal M. |
| EPI_ISL_2931689, EPI_ISL_2931691 | I.R.C.C.S. "S. De Bellis" - Ente Ospedaliero | Istituto Zooprofilattico Sperimentale della Puglia e della Basilicata | Parisi A., Bianco A., Capozzi L., Del Sambro L., Simone D., Difato L., Notarnicola M., Lippolis A. |
| EPI_ISL_2931692, EPI_ISL_2931694, EPI_ISL_2931695, EPI_ISL_2931699, EPI_ISL_2931700, EPI_ISL_2931701 | Ospedale Santa Caterina Novella | Istituto Zooprofilattico Sperimentale della Puglia e della Basilicata | Parisi A., Bianco A., Capozzi L., Del Sambro L., Simone D., Difato L., Bruno A. R. |
| EPI_ISL_2931702, EPI_ISL_2931703, EPI_ISL_2931704, EPI_ISL_2931705, EPI_ISL_2931706 | Ospedale Vito Fazzi | Istituto Zooprofilattico Sperimentale della Puglia e della Basilicata | Parisi A., Bianco A., Capozzi L., Del Sambro L., Simone D., Difato L., Lobreglio G. |
| EPI_ISL_2931707, EPI_ISL_2931708, EPI_ISL_2931709, EPI_ISL_2931710, EPI_ISL_2931711, EPI_ISL_2931712 | Ospedale Santissima Annunziata | Istituto Zooprofilattico Sperimentale della Puglia e della Basilicata | Parisi A., Bianco A., Capozzi L., Del Sambro L., Simone D., Difato L., Ridolfi D., Giannico A, Tampoia M. |
| EPI_ISL_2931716, EPI_ISL_2931717, EPI_ISL_2931719, EPI_ISL_2931720, EPI_ISL_2931721, EPI_ISL_2931722, EPI_ISL_2931723, EPI_ISL_2931724 | Presidio di Brindisi Di Summa - Perrino | Istituto Zooprofilattico Sperimentale della Puglia e della Basilicata | Parisi A., Bianco A., Capozzi L., Del Sambro L., Simone D., Difato L., Santoro A., Ridolfi D., Giannico A. |
| EPI_ISL_2931729 | RSUD dr Soediran Mangun Sumarso, Wonogiri | National Institute of Health Research and Development | Vivi Setiawaty, Hana Apsari Pawestri, Subangkit, Kartika Dewi Puspa, Arie Ardiansyah Nugraha, Hartanti Dian Ikawati, Krisna Pangesti, Triyani Soekarso, Nelly Puspandari |
| EPI_ISL_2931756 | RSUD dr Soediran Mangun Sumarso, Wonogiri | National Institute of Health Research and Development | Hana Apsari Pawestri, Subangkit, Kartika Dewi Puspa, Arie Ardiansyah Nugraha, Hartanti Dian Ikawati, Krisna Pangesti, Triyani Soekarso, Nelly Puspandari, Vivi Setiawaty |
| EPI_ISL_2931764 | Dinkes Musi Banyu Asin | National Institute of Health Research and Development | Hana Apsari Pawestri, Subangkit, Kartika Dewi Puspa, Arie Ardiansyah Nugraha, Hartanti Dian Ikawati, Krisna Pangesti, Triyani Soekarso, Nelly Puspandari, Vivi Setiawaty |
| EPI_ISL_2931781 | RSKIA Kota Bandung | National Institute of Health Research and Development | Subangkit, Hana Apsari Pawestri, , Kartika Dewi Puspa, Arie Ardiansyah Nugraha, Hartanti Dian Ikawati, Krisna Pangesti, Triyani Soekarso, Nelly Puspandari, Vivi Setiawaty |
| EPI_ISL_2931790 | Dinkes Musi Banyu Asin | National Institute of Health Research and Development | Subangkit, Hana Apsari Pawestri, , Kartika Dewi Puspa, Arie Ardiansyah Nugraha, Hartanti Dian Ikawati, Krisna Pangesti, Triyani Soekarso, Nelly Puspandari, Vivi Setiawaty |
| EPI_ISL_2931840, EPI_ISL_2931841, EPI_ISL_2931842, EPI_ISL_2931843, EPI_ISL_2931844, EPI_ISL_2931845, EPI_ISL_2931846, EPI_ISL_2931847, EPI_ISL_2931848, EPI_ISL_2931849, EPI_ISL_2931850, EPI_ISL_2931852, EPI_ISL_2931854, EPI_ISL_2931855, EPI_ISL_2931856, EPI_ISL_2931857, EPI_ISL_2931859, EPI_ISL_2931861, EPI_ISL_2931862, EPI_ISL_2931863, EPI_ISL_2931864, EPI_ISL_2931865, EPI_ISL_2931868, EPI_ISL_2931872 | | | |
| see above | Microbiology Department. Complexo Hospitalario Universitario de Vigo | Microbiology Department. Complexo Hospitalario Universitario de Vigo | Alfaya N, Alonso I, Alvarez M, Cabrera JJ, Carballo R, Cores O, Cortizo S, del-Campo V, Martinez L, Mediero G, Perez S, Davina C, Potel C, Regueiro B, Rey S, Vasallo FJ |
| EPI_ISL_2931920 | Sultanah Fatimah Specialist Hospital, Muar | Institute for Medical Research, Infectious Disease Research Centre, National Institutes of Health, Ministry of Health Malaysia | Suppiah J, Kamel K, Mohd Zawawi Z, Azizan MA, Ramly N, Robert F, Anasir MI, Thayan R |
| EPI_ISL_2931921 | Labuan Hospital | Institute for Medical Research, Infectious Disease Research Centre, National Institutes of Health, Ministry of Health Malaysia | Suppiah J, Kamel K, Mohd Zawawi Z, Azizan MA, Ramly N, Robert F, Anasir MI, Thayan R |
| EPI_ISL_2931922 | Tuanku Fauziah Hospital | Institute for Medical Research, Infectious Disease Research Centre, National Institutes of Health, Ministry of Health Malaysia | Suppiah J, Kamel K, Mohd Zawawi Z, Azizan MA, Ramly N, Robert F, Anasir MI, Thayan R |
| EPI_ISL_2931923 | Sultanah Fatimah Specialist Hospital, Muar | Institute for Medical Research, Infectious Disease Research Centre, National Institutes of Health, Ministry of Health Malaysia | Suppiah J, Kamel K, Mohd Zawawi Z, Azizan MA, Ramly N, Robert F, Anasir MI, Thayan R |
| EPI_ISL_2932519, EPI_ISL_2932521, EPI_ISL_2932526 | National Institute of Infectious Diseases-Prof. Dr. Matei Bals Molecular Diagnostics Laboratory | National Institute of Infectious Diseases-Prof. Dr. Matei Bals Molecular Diagnostics Laboratory | Corina Casangiu, Leontina Banica, Marius Surleac, Petre Milu, Robert Hohan, Simona Paraschiv, Dan Otelea |
| EPI_ISL_2932530, EPI_ISL_2932531 | Institute of Public Health Kraljevo | Veterinary Specialized Institute Kraljevo | Vidanovic,D., Tesovic,B., Knezevic,A., Jovanovic,T., Jankovic,M., Sekler,M., Djukic V,., Banovic Djeri,B., Petrovic,T., Volkening,J., Afonso,C. |
| EPI_ISL_2932610 | KOTA Yogya DKK Gondokusuman II | Genetics Working Group (Pokja Genetik) Faculty of Medicine, Public Health and Nursing Universitas Gadjah Mada (FK-KMK UGM); Disease Investigation Center Wates Ministry of Agriculture Indonesia; Department of Microbiology FK-KMK UGM; Laboratorium Diagnostik Yayasan Tahija World Mosquito Program (WMP) Yogyakarta Center for | Gunadi, Hendra Wibawa, Marcellus, Mohamad S. Hakim, Edwin W. Daniwijaya, Ludhang P. Rizki, Endah Supriyati, Eggi Arguni, Titik Nuryastuti, Tri Wibawa, Dwi AA Nugrahaningsih, Afiahayati, Siswanto,Kristy Iskandar,Nungki Anggorowati, Pramesti G Dewi, Dwi Indaryati,Indarto Sulistiyono, Khanza a Vujira, Alvina A Setiawan |
| EPI_ISL_2932613 | DKK Sleman/Puskesmas Depok 1 | Genetics Working Group (Pokja Genetik) Faculty of Medicine, Public Health and Nursing Universitas Gadjah Mada (FK-KMK UGM); Disease Investigation Center Wates Ministry of | Gunadi, Hendra Wibawa, Marcellus, Mohamad S. Hakim, Edwin W. Daniwijaya, Ludhang P. Rizki, Endah Supriyati, Eggi Arguni, Titik Nuryastuti, Tri Wibawa, Dwi AA Nugrahaningsih, Afiahayati, Siswanto,Kristy Iskandar,Nungki Anggorowati, Pramesti G Dewi, Dwi Indaryati,Indarto Sulistiyono, Irene Tania, Hana F Hanifin |

| | | | |
|---|---|---|---|
| | | Agriculture Indonesia; Department of Microbiology FK-KMK UGM; Laboratorium Diagnostik Yayasan Tahija World Mosquito Program (WMP) Yogyakarta Center for Tropical Medicine FK-KMK UGM; Integrated Research Center FK-KMK UGM; Department of Computer Science and Electronics FMIPA UGM; RSUP Dr. Sardjito | |
| EPI_ISL_2932618 | Jordan University of Science Technology/ Irbid filed Hospital/ MOH | Princess Haya Biotechnology Center/ Jordan University of Science Technology | Saied Jaradat, Hazem Haddad, Areej Alquran, Maha Karam, Shereen Issa,Suha Hasan, Emad Abu Yaqeen |
| EPI_ISL_2932708 | Cliniques universitaires Saint-Luc | UCLouvain/IREC/MBLG | Jean Ruelle, Bertrand Bearzatto, Benoit Kabamba Mukadi |
| EPI_ISL_2933109 | DKK Sleman/Puskesmas Depok 1 | Genetics Working Group (Pokja Genetik) Faculty of Medicine, Public Health and Nursing Universitas Gadjah Mada (FK-KMK UGM); Disease Investigation Center Wates Ministry of Agriculture Indonesia; Department of Microbiology FK-KMK UGM; Laboratorium Diagnostik Yayasan Tahija World Mosquito Program (WMP) Yogyakarta Center for Tropical Medicine FK-KMK UGM; Integrated Research Center FK-KMK UGM; Department of Computer Science and Electronics FMIPA UGM; RSUP Dr. Sardjito | Gunadi, Hendra Wibawa, Marcellus, Mohamad S. Hakim, Edwin W. Daniwijaya, Ludhang P. Rizki, Endah Supriyati, Eggi Arguni, Titik Nuryastuti, Tri Wibawa, Dwi AA Nugrahaningsih, Afiahayati, Siswanto,Kristy Iskandar,Nungki Anggorowati, Pramesti G Dewi, Dwi Indaryati,Indarto Sulistiyono, Susan Simanjaya, Dyah A Puspitarani |
| EPI_ISL_2933110 | DKK Sleman/Puskesmas Depok 1 | Genetics Working Group (Pokja Genetik) Faculty of Medicine, Public Health and Nursing Universitas Gadjah Mada (FK-KMK UGM); Disease Investigation Center Wates Ministry of Agriculture Indonesia; Department of Microbiology FK-KMK UGM; Laboratorium Diagnostik Yayasan Tahija World Mosquito Program (WMP) Yogyakarta Center for Tropical Medicine FK-KMK UGM; Integrated Research Center FK-KMK UGM; Department of Computer Science and | Gunadi, Hendra Wibawa, Marcellus, Mohamad S. Hakim, Edwin W. Daniwijaya, Ludhang P. Rizki, Endah Supriyati, Eggi Arguni, Titik Nuryastuti, Tri Wibawa, Dwi AA Nugrahaningsih, Afiahayati, Siswanto,Kristy Iskandar,Nungki Anggorowati, Pramesti G Dewi, Dwi Indaryati,Indarto Sulistiyono, Khanza a Vujira, Alvina A Setiawan |
| EPI_ISL_2934573, EPI_ISL_2934574 | SECP MHU | The Institute of Molecular Biology and Genetics of NASU | M.Tukalo et al. |
| EPI_ISL_2937751 | LESP Jalisco | Instituto de Diagnostico y Referencia Epidemiologicos (INDRE) | Claudia Wong-Arambula, Abril Rodriguez-Maldonado, Vanessa Rivero-Arredondo, Ariadna Medina-Benitez, Joaquin Quiroz-Mercado, Sergio Rangel-Guerrero, Natividad Cruz-Ortiz, Tatiana Nunez-Garcia, Gisela Barrera-Badillo, Lucia Hernandez-Rivas, Irma Lopez-Martinez, Ernesto Ramirez-Gonzalez. |
| EPI_ISL_2937752 | LESP Ciudad de Mexico | Instituto de Diagnostico y Referencia Epidemiologicos (INDRE) | Claudia Wong-Arambula, Abril Rodriguez-Maldonado, Vanessa Rivero-Arredondo, Ariadna Medina-Benitez, Joaquin Quiroz-Mercado, Sergio Rangel-Guerrero, Maribel Gonzalez-Villa, Natividad Cruz-Ortiz, Tatiana Nunez-Garcia, Gisela Barrera-Badillo, Lucia Hernandez-Rivas, Irma Lopez-Martinez, Ernesto Ramirez-Gonzalez. |
| EPI_ISL_2937753, EPI_ISL_2937754, EPI_ISL_2937755 | LESP Tabasco | Instituto de Diagnostico y Referencia Epidemiologicos (INDRE) | Claudia Wong-Arambula, Abril Rodriguez-Maldonado, Vanessa Rivero-Arredondo, Ariadna Medina-Benitez, Joaquin Quiroz-Mercado, Sergio Rangel-Guerrero, Maribel Gonzalez-Villa, Natividad Cruz-Ortiz, Tatiana Nunez-Garcia, Gisela Barrera-Badillo, Lucia Hernandez-Rivas, Irma Lopez-Martinez, Ernesto Ramirez-Gonzalez. |
| EPI_ISL_2937756 | LESP Puebla | Instituto de Diagnostico y Referencia Epidemiologicos (INDRE) | Claudia Wong-Arambula, Abril Rodriguez-Maldonado, Vanessa Rivero-Arredondo, Ariadna Medina-Benitez, Joaquin Quiroz-Mercado, Sergio Rangel-Guerrero, Maribel Gonzalez-Villa, Natividad Cruz-Ortiz, Tatiana Nunez-Garcia, Gisela Barrera-Badillo, Lucia Hernandez-Rivas, Irma Lopez-Martinez, Ernesto Ramirez-Gonzalez. |
| EPI_ISL_2937757 | LESP Jalisco | Instituto de Diagnostico y Referencia Epidemiologicos (INDRE) | Claudia Wong-Arambula, Abril Rodriguez-Maldonado, Vanessa Rivero-Arredondo, Ariadna Medina-Benitez, Joaquin Quiroz-Mercado, Sergio Rangel-Guerrero, Maribel Gonzalez-Villa, Natividad Cruz-Ortiz, Tatiana Nunez-Garcia, Gisela Barrera-Badillo, Lucia Hernandez-Rivas, Irma Lopez-Martinez, Ernesto Ramirez-Gonzalez. |
| EPI_ISL_2937758, EPI_ISL_2937759, EPI_ISL_2937760, EPI_ISL_2937761, EPI_ISL_2937762, EPI_ISL_2937763, EPI_ISL_2937764, EPI_ISL_2937765, EPI_ISL_2937766 | LESP Quintana Roo | Instituto de Diagnostico y Referencia Epidemiologicos (INDRE) | Claudia Wong-Arambula, Abril Rodriguez-Maldonado, Vanessa Rivero-Arredondo, Ariadna Medina-Benitez, Joaquin Quiroz-Mercado, Sergio Rangel-Guerrero, Maribel Gonzalez-Villa, Natividad Cruz-Ortiz, Tatiana Nunez-Garcia, Gisela Barrera-Badillo, Lucia Hernandez-Rivas, Irma Lopez-Martinez, Ernesto Ramirez-Gonzalez. |
| EPI_ISL_2937767 | LESP Sonora | Instituto de Diagnostico y Referencia Epidemiologicos (INDRE) | Claudia Wong-Arambula, Abril Rodriguez-Maldonado, Vanessa Rivero-Arredondo, Ariadna Medina-Benitez, Joaquin Quiroz-Mercado, Sergio Rangel-Guerrero, Maribel Gonzalez-Villa, Natividad Cruz-Ortiz, Tatiana Nunez-Garcia, Gisela Barrera-Badillo, Lucia Hernandez-Rivas, Irma Lopez-Martinez, Ernesto Ramirez-Gonzalez. |
| EPI_ISL_2937768 | LESP Hidalgo | Instituto de Diagnostico y Referencia Epidemiologicos (INDRE) | Claudia Wong-Arambula, Abril Rodriguez-Maldonado, Vanessa Rivero-Arredondo, Ariadna Medina-Benitez, Joaquin Quiroz-Mercado, Sergio Rangel-Guerrero, Maribel Gonzalez-Villa, Natividad Cruz-Ortiz, Tatiana Nunez-Garcia, Gisela Barrera-Badillo, Lucia Hernandez-Rivas, Irma Lopez-Martinez, Ernesto Ramirez-Gonzalez. |
| EPI_ISL_2937769, EPI_ISL_2937770 | LESP Ciudad de Mexico | Instituto de Diagnostico y Referencia Epidemiologicos (INDRE) | Claudia Wong-Arambula, Abril Rodriguez-Maldonado, Vanessa Rivero-Arredondo, Ariadna Medina-Benitez, Joaquin Quiroz-Mercado, Sergio Rangel-Guerrero, Maribel Gonzalez-Villa, Natividad Cruz-Ortiz, Tatiana Nunez-Garcia, Gisela Barrera-Badillo, Lucia Hernandez-Rivas, Irma Lopez-Martinez, Ernesto Ramirez-Gonzalez. |
| EPI_ISL_2937771 | LESP Estado de Mexico | Instituto de Diagnostico y Referencia Epidemiologicos (INDRE) | Claudia Wong-Arambula, Abril Rodriguez-Maldonado, Vanessa Rivero-Arredondo, Ariadna Medina-Benitez, Joaquin Quiroz-Mercado, Sergio Rangel-Guerrero, Maribel Gonzalez-Villa, Natividad Cruz-Ortiz, Tatiana Nunez-Garcia, Gisela Barrera-Badillo, Lucia Hernandez-Rivas, Irma Lopez-Martinez, Ernesto Ramirez-Gonzalez. |
| EPI_ISL_2937772 | LESP Guanajuato | Instituto de Diagnostico y Referencia Epidemiologicos (INDRE) | Claudia Wong-Arambula, Abril Rodriguez-Maldonado, Vanessa Rivero-Arredondo, Ariadna Medina-Benitez, Joaquin Quiroz-Mercado, Sergio Rangel-Guerrero, Maribel Gonzalez-Villa, Natividad Cruz-Ortiz, Tatiana Nunez-Garcia, Gisela Barrera-Badillo, Lucia Hernandez-Rivas, Irma Lopez-Martinez, Ernesto Ramirez-Gonzalez. |
| EPI_ISL_2937773 | LESP San Luis Potosi | Instituto de Diagnostico y Referencia Epidemiologicos (INDRE) | Claudia Wong-Arambula, Abril Rodriguez-Maldonado, Vanessa Rivero-Arredondo, Ariadna Medina-Benitez, Joaquin Quiroz-Mercado, Sergio Rangel-Guerrero, Maribel Gonzalez-Villa, Natividad Cruz-Ortiz, Tatiana Nunez-Garcia, Gisela Barrera-Badillo, Lucia Hernandez-Rivas, Irma Lopez-Martinez, Ernesto Ramirez-Gonzalez. |
| EPI_ISL_2937774 | LESP Nayarit | Instituto de Diagnostico y Referencia Epidemiologicos (INDRE) | Claudia Wong-Arambula, Abril Rodriguez-Maldonado, Vanessa Rivero-Arredondo, Ariadna Medina-Benitez, Joaquin Quiroz-Mercado, Sergio Rangel-Guerrero, Maribel Gonzalez-Villa, Natividad Cruz-Ortiz, Tatiana Nunez-Garcia, Gisela Barrera-Badillo, Lucia Hernandez-Rivas, Irma Lopez-Martinez, Ernesto Ramirez-Gonzalez. |
| EPI_ISL_2937775, EPI_ISL_2937776, EPI_ISL_2937777 | LESP Tamaulipas | Instituto de Diagnostico y Referencia Epidemiologicos (INDRE) | Claudia Wong-Arambula, Abril Rodriguez-Maldonado, Vanessa Rivero-Arredondo, Ariadna Medina-Benitez, Joaquin Quiroz-Mercado, Sergio Rangel-Guerrero, Maribel Gonzalez-Villa, Natividad Cruz-Ortiz, Tatiana Nunez-Garcia, Gisela Barrera-Badillo, Lucia Hernandez-Rivas, Irma Lopez-Martinez, Ernesto Ramirez-Gonzalez. |
| EPI_ISL_2937778, EPI_ISL_2937779, EPI_ISL_2937780, EPI_ISL_2937781, EPI_ISL_2937782, EPI_ISL_2937783 | LESP Yucatan | Instituto de Diagnostico y Referencia Epidemiologicos (INDRE) | Claudia Wong-Arambula, Abril Rodriguez-Maldonado, Vanessa Rivero-Arredondo, Ariadna Medina-Benitez, Joaquin Quiroz-Mercado, Sergio Rangel-Guerrero, Maribel Gonzalez-Villa, Natividad Cruz-Ortiz, Tatiana Nunez-Garcia, Gisela Barrera-Badillo, Lucia Hernandez-Rivas, Irma Lopez-Martinez, Ernesto Ramirez-Gonzalez. |
| EPI_ISL_2937784 | LESP Campeche | Instituto de Diagnostico y Referencia Epidemiologicos (INDRE) | Claudia Wong-Arambula, Abril Rodriguez-Maldonado, Vanessa Rivero-Arredondo, Ariadna Medina-Benitez, Joaquin Quiroz-Mercado, Sergio Rangel-Guerrero, Maribel Gonzalez-Villa, Natividad Cruz-Ortiz, Tatiana Nunez-Garcia, Gisela Barrera-Badillo, Lucia Hernandez-Rivas, Irma Lopez-Martinez, Ernesto Ramirez-Gonzalez. |

| | | | |
|---|---|---|---|
| EPI_ISL_2937785, EPI_ISL_2937786, EPI_ISL_2937787, EPI_ISL_2937788 | LESP Yucatan | Instituto de Diagnostico y Referencia Epidemiologicos (INDRE) | Claudia Wong-Arambula, Abril Rodriguez-Maldonado, Vanessa Rivero-Arredondo, Ariadna Medina-Benitez, Joaquin Quiroz-Mercado, Sergio Rangel-Guerrero, Maribel Gonzalez-Villa, Natividad Cruz-Ortiz, Tatiana Nunez-Garcia, Gisela Barrera-Badillo, Lucia Hernandez-Rivas, Irma Lopez-Martinez, Ernesto Ramirez-Gonzalez. |
| EPI_ISL_2937789, EPI_ISL_2937790, EPI_ISL_2937791, EPI_ISL_2937792 | LESP Campeche | Instituto de Diagnostico y Referencia Epidemiologicos (INDRE) | Claudia Wong-Arambula, Abril Rodriguez-Maldonado, Vanessa Rivero-Arredondo, Ariadna Medina-Benitez, Joaquin Quiroz-Mercado, Sergio Rangel-Guerrero, Maribel Gonzalez-Villa, Natividad Cruz-Ortiz, Tatiana Nunez-Garcia, Gisela Barrera-Badillo, Lucia Hernandez-Rivas, Irma Lopez-Martinez, Ernesto Ramirez-Gonzalez. |
| EPI_ISL_2937793, EPI_ISL_2937794 | LESP Veracruz | Instituto de Diagnostico y Referencia Epidemiologicos (INDRE) | Claudia Wong-Arambula, Abril Rodriguez-Maldonado, Vanessa Rivero-Arredondo, Ariadna Medina-Benitez, Joaquin Quiroz-Mercado, Sergio Rangel-Guerrero, Maribel Gonzalez-Villa, Natividad Cruz-Ortiz, Tatiana Nunez-Garcia, Gisela Barrera-Badillo, Lucia Hernandez-Rivas, Irma Lopez-Martinez, Ernesto Ramirez-Gonzalez. |
| EPI_ISL_2937795, EPI_ISL_2937796, EPI_ISL_2937797 | LESP Sonora | Instituto de Diagnostico y Referencia Epidemiologicos (INDRE) | Claudia Wong-Arambula, Abril Rodriguez-Maldonado, Vanessa Rivero-Arredondo, Ariadna Medina-Benitez, Joaquin Quiroz-Mercado, Sergio Rangel-Guerrero, Maribel Gonzalez-Villa, Natividad Cruz-Ortiz, Tatiana Nunez-Garcia, Gisela Barrera-Badillo, Lucia Hernandez-Rivas, Irma Lopez-Martinez, Ernesto Ramirez-Gonzalez. |
| EPI_ISL_2937798, EPI_ISL_2937799, EPI_ISL_2937800 | LESP Quintana Roo | Instituto de Diagnostico y Referencia Epidemiologicos (INDRE) | Claudia Wong-Arambula, Abril Rodriguez-Maldonado, Vanessa Rivero-Arredondo, Ariadna Medina-Benitez, Joaquin Quiroz-Mercado, Sergio Rangel-Guerrero, Maribel Gonzalez-Villa, Natividad Cruz-Ortiz, Tatiana Nunez-Garcia, Gisela Barrera-Badillo, Lucia Hernandez-Rivas, Irma Lopez-Martinez, Ernesto Ramirez-Gonzalez. |
| EPI_ISL_2937801 | LESP Jalisco | Instituto de Diagnostico y Referencia Epidemiologicos (INDRE) | Claudia Wong-Arambula, Abril Rodriguez-Maldonado, Vanessa Rivero-Arredondo, Ariadna Medina-Benitez, Joaquin Quiroz-Mercado, Sergio Rangel-Guerrero, Maribel Gonzalez-Villa, Natividad Cruz-Ortiz, Tatiana Nunez-Garcia, Gisela Barrera-Badillo, Lucia Hernandez-Rivas, Irma Lopez-Martinez, Ernesto Ramirez-Gonzalez. |
| EPI_ISL_2937802, EPI_ISL_2937803, EPI_ISL_2937804, EPI_ISL_2937805 | LESP Ciudad de Mexico | Instituto de Diagnostico y Referencia Epidemiologicos (INDRE) | Claudia Wong-Arambula, Abril Rodriguez-Maldonado, Vanessa Rivero-Arredondo, Ariadna Medina-Benitez, Joaquin Quiroz-Mercado, Sergio Rangel-Guerrero, Maribel Gonzalez-Villa, Natividad Cruz-Ortiz, Tatiana Nunez-Garcia, Gisela Barrera-Badillo, Lucia Hernandez-Rivas, Irma Lopez-Martinez, Ernesto Ramirez-Gonzalez. |
| EPI_ISL_2937806 | LESP Tabasco | Instituto de Diagnostico y Referencia Epidemiologicos (INDRE) | Claudia Wong-Arambula, Abril Rodriguez-Maldonado, Vanessa Rivero-Arredondo, Ariadna Medina-Benitez, Joaquin Quiroz-Mercado, Sergio Rangel-Guerrero, Maribel Gonzalez-Villa, Natividad Cruz-Ortiz, Tatiana Nunez-Garcia, Gisela Barrera-Badillo, Lucia Hernandez-Rivas, Irma Lopez-Martinez, Ernesto Ramirez-Gonzalez. |
| EPI_ISL_2937807 | LESP Chiapas | Instituto de Diagnostico y Referencia Epidemiologicos (INDRE) | Claudia Wong-Arambula, Abril Rodriguez-Maldonado, Vanessa Rivero-Arredondo, Ariadna Medina-Benitez, Joaquin Quiroz-Mercado, Sergio Rangel-Guerrero, Maribel Gonzalez-Villa, Natividad Cruz-Ortiz, Tatiana Nunez-Garcia, Gisela Barrera-Badillo, Lucia Hernandez-Rivas, Irma Lopez-Martinez, Ernesto Ramirez-Gonzalez. |
| EPI_ISL_2937808, EPI_ISL_2937809, EPI_ISL_2937810, EPI_ISL_2937811, EPI_ISL_2937812, EPI_ISL_2937813, EPI_ISL_2937814 | LESP Tabasco | Instituto de Diagnostico y Referencia Epidemiologicos (INDRE) | Claudia Wong-Arambula, Abril Rodriguez-Maldonado, Vanessa Rivero-Arredondo, Ariadna Medina-Benitez, Joaquin Quiroz-Mercado, Sergio Rangel-Guerrero, Maribel Gonzalez-Villa, Natividad Cruz-Ortiz, Tatiana Nunez-Garcia, Gisela Barrera-Badillo, Lucia Hernandez-Rivas, Irma Lopez-Martinez, Ernesto Ramirez-Gonzalez. |
| EPI_ISL_2937815 | LESP Jalisco | Instituto de Diagnostico y Referencia Epidemiologicos (INDRE) | Claudia Wong-Arambula, Abril Rodriguez-Maldonado, Vanessa Rivero-Arredondo, Ariadna Medina-Benitez, Joaquin Quiroz-Mercado, Sergio Rangel-Guerrero, Maribel Gonzalez-Villa, Natividad Cruz-Ortiz, Tatiana Nunez-Garcia, Gisela Barrera-Badillo, Lucia Hernandez-Rivas, Irma Lopez-Martinez, Ernesto Ramirez-Gonzalez. |
| EPI_ISL_2937816, EPI_ISL_2937817, EPI_ISL_2937818, EPI_ISL_2937819, EPI_ISL_2937820, EPI_ISL_2937821, EPI_ISL_2937822, EPI_ISL_2937823, EPI_ISL_2937824, EPI_ISL_2937825, EPI_ISL_2937826, EPI_ISL_2937827 | | | |
| see above | LESP Tabasco | Instituto de Diagnostico y Referencia Epidemiologicos (INDRE) | Claudia Wong-Arambula, Abril Rodriguez-Maldonado, Vanessa Rivero-Arredondo, Ariadna Medina-Benitez, Joaquin Quiroz-Mercado, Sergio Rangel-Guerrero, Maribel Gonzalez-Villa, Natividad Cruz-Ortiz, Tatiana Nunez-Garcia, Gisela Barrera-Badillo, Lucia Hernandez-Rivas, Irma Lopez-Martinez, Ernesto Ramirez-Gonzalez. |
| EPI_ISL_2937828, EPI_ISL_2937829, EPI_ISL_2937830 | LESP Tamaulipas | Instituto de Diagnostico y Referencia Epidemiologicos (INDRE) | Claudia Wong-Arambula, Abril Rodriguez-Maldonado, Vanessa Rivero-Arredondo, Ariadna Medina-Benitez, Joaquin Quiroz-Mercado, Sergio Rangel-Guerrero, Maribel Gonzalez-Villa, Natividad Cruz-Ortiz, Tatiana Nunez-Garcia, Gisela Barrera-Badillo, Lucia Hernandez-Rivas, Irma Lopez-Martinez, Ernesto Ramirez-Gonzalez. |
| EPI_ISL_2937831 | LESP Sonora | Instituto de Diagnostico y Referencia Epidemiologicos (INDRE) | Claudia Wong-Arambula, Abril Rodriguez-Maldonado, Vanessa Rivero-Arredondo, Ariadna Medina-Benitez, Joaquin Quiroz-Mercado, Sergio Rangel-Guerrero, Maribel Gonzalez-Villa, Natividad Cruz-Ortiz, Tatiana Nunez-Garcia, Gisela Barrera-Badillo, Lucia Hernandez-Rivas, Irma Lopez-Martinez, Ernesto Ramirez-Gonzalez. |
| EPI_ISL_2937832 | LESP Ciudad de Mexico | Instituto de Diagnostico y Referencia Epidemiologicos (INDRE) | Claudia Wong-Arambula, Abril Rodriguez-Maldonado, Vanessa Rivero-Arredondo, Ariadna Medina-Benitez, Joaquin Quiroz-Mercado, Sergio Rangel-Guerrero, Maribel Gonzalez-Villa, Natividad Cruz-Ortiz, Tatiana Nunez-Garcia, Gisela Barrera-Badillo, Lucia Hernandez-Rivas, Irma Lopez-Martinez, Ernesto Ramirez-Gonzalez. |
| EPI_ISL_2937836 | LESP Baja California Sur | Instituto de Diagnostico y Referencia Epidemiologicos (INDRE) | Claudia Wong-Arambula, Abril Rodriguez-Maldonado, Vanessa Rivero-Arredondo, Ariadna Medina-Benitez, Joaquin Quiroz-Mercado, Sergio Rangel-Guerrero, Maribel Gonzalez-Villa, Natividad Cruz-Ortiz, Tatiana Nunez-Garcia, Gisela Barrera-Badillo, Lucia Hernandez-Rivas, Irma Lopez-Martinez, Ernesto Ramirez-Gonzalez. |
| EPI_ISL_2937837 | LESP Zacatecas | Instituto de Diagnostico y Referencia Epidemiologicos (INDRE) | Claudia Wong-Arambula, Abril Rodriguez-Maldonado, Vanessa Rivero-Arredondo, Ariadna Medina-Benitez, Joaquin Quiroz-Mercado, Sergio Rangel-Guerrero, Maribel Gonzalez-Villa, Natividad Cruz-Ortiz, Tatiana Nunez-Garcia, Gisela Barrera-Badillo, Lucia Hernandez-Rivas, Irma Lopez-Martinez, Ernesto Ramirez-Gonzalez. |
| EPI_ISL_2937838, EPI_ISL_2937839, EPI_ISL_2937840, EPI_ISL_2937841, EPI_ISL_2937842, EPI_ISL_2937843 | LESP Ciudad de Mexico | Instituto de Diagnostico y Referencia Epidemiologicos (INDRE) | Claudia Wong-Arambula, Abril Rodriguez-Maldonado, Vanessa Rivero-Arredondo, Ariadna Medina-Benitez, Joaquin Quiroz-Mercado, Sergio Rangel-Guerrero, Maribel Gonzalez-Villa, Natividad Cruz-Ortiz, Tatiana Nunez-Garcia, Gisela Barrera-Badillo, Lucia Hernandez-Rivas, Irma Lopez-Martinez, Ernesto Ramirez-Gonzalez. |
| EPI_ISL_2937844 | LESP Nuevo Leon | Instituto de Diagnostico y Referencia Epidemiologicos (INDRE) | Claudia Wong-Arambula, Abril Rodriguez-Maldonado, Vanessa Rivero-Arredondo, Ariadna Medina-Benitez, Joaquin Quiroz-Mercado, Sergio Rangel-Guerrero, Maribel Gonzalez-Villa, Natividad Cruz-Ortiz, Tatiana Nunez-Garcia, Gisela Barrera-Badillo, Lucia Hernandez-Rivas, Irma Lopez-Martinez, Ernesto Ramirez-Gonzalez. |
| EPI_ISL_2937845 | LESP Tlaxcala | Instituto de Diagnostico y Referencia Epidemiologicos (INDRE) | Claudia Wong-Arambula, Abril Rodriguez-Maldonado, Vanessa Rivero-Arredondo, Ariadna Medina-Benitez, Joaquin Quiroz-Mercado, Sergio Rangel-Guerrero, Maribel Gonzalez-Villa, Natividad Cruz-Ortiz, Tatiana Nunez-Garcia, Gisela Barrera-Badillo, Lucia Hernandez-Rivas, Irma Lopez-Martinez, Ernesto Ramirez-Gonzalez. |
| EPI_ISL_2937846 | LESP Michoacan | Instituto de Diagnostico y Referencia Epidemiologicos (INDRE) | Claudia Wong-Arambula, Abril Rodriguez-Maldonado, Vanessa Rivero-Arredondo, Ariadna Medina-Benitez, Joaquin Quiroz-Mercado, Sergio Rangel-Guerrero, Maribel Gonzalez-Villa, Natividad Cruz-Ortiz, Tatiana Nunez-Garcia, Gisela Barrera-Badillo, Lucia Hernandez-Rivas, Irma Lopez-Martinez, Ernesto Ramirez-Gonzalez. |
| EPI_ISL_2937847, EPI_ISL_2937848 | LESP Baja California Sur | Instituto de Diagnostico y Referencia Epidemiologicos (INDRE) | Claudia Wong-Arambula, Abril Rodriguez-Maldonado, Vanessa Rivero-Arredondo, Ariadna Medina-Benitez, Joaquin Quiroz-Mercado, Sergio Rangel-Guerrero, Maribel Gonzalez-Villa, Natividad Cruz-Ortiz, Tatiana Nunez-Garcia, Gisela Barrera-Badillo, Lucia Hernandez-Rivas, Irma Lopez-Martinez, Ernesto Ramirez-Gonzalez. |
| EPI_ISL_2937849, EPI_ISL_2937850 | LESP Veracruz | Instituto de Diagnostico y Referencia Epidemiologicos (INDRE) | Claudia Wong-Arambula, Abril Rodriguez-Maldonado, Vanessa Rivero-Arredondo, Ariadna Medina-Benitez, Joaquin Quiroz-Mercado, Sergio Rangel-Guerrero, Maribel Gonzalez-Villa, Natividad Cruz-Ortiz, Tatiana Nunez-Garcia, Gisela Barrera-Badillo, Lucia Hernandez-Rivas, Irma Lopez-Martinez, Ernesto Ramirez-Gonzalez. |
| EPI_ISL_2937851, EPI_ISL_2937852, EPI_ISL_2937853, EPI_ISL_2937854, EPI_ISL_2937855 | LESP Sonora | Instituto de Diagnostico y Referencia Epidemiologicos (INDRE) | Claudia Wong-Arambula, Abril Rodriguez-Maldonado, Vanessa Rivero-Arredondo, Ariadna Medina-Benitez, Joaquin Quiroz-Mercado, Sergio Rangel-Guerrero, Maribel Gonzalez-Villa, Natividad Cruz-Ortiz, Tatiana Nunez-Garcia, Gisela Barrera-Badillo, Lucia Hernandez-Rivas, Irma Lopez-Martinez, Ernesto Ramirez-Gonzalez. |
| EPI_ISL_2937856 | LESP Quintana Roo | Instituto de Diagnostico y Referencia Epidemiologicos | Claudia Wong-Arambula, Abril Rodriguez-Maldonado, Vanessa Rivero-Arredondo, Ariadna Medina-Benitez, Joaquin Quiroz-Mercado, Sergio |

| | | | |
|---|---|---|---|
| | | (INDRE) | Rangel-Guerrero, Maribel Gonzalez-Villa, Natividad Cruz-Ortiz, Tatiana Nunez-Garcia, Gisela Barrera-Badillo, Lucia Hernandez-Rivas, Irma Lopez-Martinez, Ernesto Ramirez-Gonzalez. |
| EPI_ISL_2937857, EPI_ISL_2937858, EPI_ISL_2937859, EPI_ISL_2937860 | LESP Ciudad de Mexico | Instituto de Diagnostico y Referencia Epidemiologicos (INDRE) | Claudia Wong-Arambula, Abril Rodriguez-Maldonado, Vanessa Rivero-Arredondo, Ariadna Medina-Benitez, Joaquin Quiroz-Mercado, Sergio Rangel-Guerrero, Maribel Gonzalez-Villa, Natividad Cruz-Ortiz, Tatiana Nunez-Garcia, Gisela Barrera-Badillo, Lucia Hernandez-Rivas, Irma Lopez-Martinez, Ernesto Ramirez-Gonzalez. |
| EPI_ISL_2937861 | LESP Estado de Mexico | Instituto de Diagnostico y Referencia Epidemiologicos (INDRE) | Claudia Wong-Arambula, Abril Rodriguez-Maldonado, Vanessa Rivero-Arredondo, Ariadna Medina-Benitez, Joaquin Quiroz-Mercado, Sergio Rangel-Guerrero, Maribel Gonzalez-Villa, Natividad Cruz-Ortiz, Tatiana Nunez-Garcia, Gisela Barrera-Badillo, Lucia Hernandez-Rivas, Irma Lopez-Martinez, Ernesto Ramirez-Gonzalez. |
| EPI_ISL_2937862, EPI_ISL_2937863, EPI_ISL_2937864, EPI_ISL_2937865, EPI_ISL_2937866, EPI_ISL_2937867, EPI_ISL_2937868, EPI_ISL_2937869, EPI_ISL_2937870, EPI_ISL_2937871, EPI_ISL_2937872, EPI_ISL_2937873, EPI_ISL_2937874, EPI_ISL_2937875, EPI_ISL_2937876, EPI_ISL_2937877, EPI_ISL_2937878, EPI_ISL_2937879, EPI_ISL_2937880, EPI_ISL_2937881, EPI_ISL_2937882, EPI_ISL_2937883, EPI_ISL_2937884, EPI_ISL_2937885, EPI_ISL_2937886, EPI_ISL_2937887, EPI_ISL_2937888, EPI_ISL_2937889, EPI_ISL_2937890, EPI_ISL_2937891, EPI_ISL_2937892, EPI_ISL_2937893, EPI_ISL_2937894, EPI_ISL_2937895 | | | |
| see above | LESP Ciudad de Mexico | Instituto de Diagnostico y Referencia Epidemiologicos (INDRE) | Claudia Wong-Arambula, Abril Rodriguez-Maldonado, Vanessa Rivero-Arredondo, Ariadna Medina-Benitez, Joaquin Quiroz-Mercado, Sergio Rangel-Guerrero, Maribel Gonzalez-Villa, Natividad Cruz-Ortiz, Tatiana Nunez-Garcia, Gisela Barrera-Badillo, Lucia Hernandez-Rivas, Irma Lopez-Martinez, Ernesto Ramirez-Gonzalez. |
| EPI_ISL_2937896, EPI_ISL_2937897, EPI_ISL_2937898 | LESP Jalisco | Instituto de Diagnostico y Referencia Epidemiologicos (INDRE) | Claudia Wong-Arambula, Abril Rodriguez-Maldonado, Vanessa Rivero-Arredondo, Ariadna Medina-Benitez, Joaquin Quiroz-Mercado, Sergio Rangel-Guerrero, Maribel Gonzalez-Villa, Natividad Cruz-Ortiz, Tatiana Nunez-Garcia, Gisela Barrera-Badillo, Lucia Hernandez-Rivas, Irma Lopez-Martinez, Ernesto Ramirez-Gonzalez. |
| EPI_ISL_2937899 | LESP Tabasco | Instituto de Diagnostico y Referencia Epidemiologicos (INDRE) | Claudia Wong-Arambula, Abril Rodriguez-Maldonado, Vanessa Rivero-Arredondo, Ariadna Medina-Benitez, Joaquin Quiroz-Mercado, Sergio Rangel-Guerrero, Maribel Gonzalez-Villa, Natividad Cruz-Ortiz, Tatiana Nunez-Garcia, Gisela Barrera-Badillo, Lucia Hernandez-Rivas, Irma Lopez-Martinez, Ernesto Ramirez-Gonzalez. |
| EPI_ISL_2937901, EPI_ISL_2937902, EPI_ISL_2937905 | LESP Aguascalientes | Instituto de Diagnostico y Referencia Epidemiologicos (INDRE) | Claudia Wong-Arambula, Abril Rodriguez-Maldonado, Vanessa Rivero-Arredondo, Ariadna Medina-Benitez, Joaquin Quiroz-Mercado, Sergio Rangel-Guerrero, Maribel Gonzalez-Villa, Natividad Cruz-Ortiz, Tatiana Nunez-Garcia, Gisela Barrera-Badillo, Lucia Hernandez-Rivas, Irma Lopez-Martinez, Ernesto Ramirez-Gonzalez. |
| EPI_ISL_2937907 | LESP Coahuila | Instituto de Diagnostico y Referencia Epidemiologicos (INDRE) | Claudia Wong-Arambula, Abril Rodriguez-Maldonado, Vanessa Rivero-Arredondo, Ariadna Medina-Benitez, Joaquin Quiroz-Mercado, Sergio Rangel-Guerrero, Natividad Cruz-Ortiz, Tatiana Nunez-Garcia, Gisela Barrera-Badillo, Lucia Hernandez-Rivas, Irma Lopez-Martinez, Ernesto Ramirez-Gonzalez. |
| EPI_ISL_2937908, EPI_ISL_2937909, EPI_ISL_2937910, EPI_ISL_2937911, EPI_ISL_2937912, EPI_ISL_2937913, EPI_ISL_2937914, EPI_ISL_2937915, EPI_ISL_2937916, EPI_ISL_2937917, EPI_ISL_2937918, EPI_ISL_2937919, EPI_ISL_2937920, EPI_ISL_2937921, EPI_ISL_2937922, EPI_ISL_2937923, EPI_ISL_2937924, EPI_ISL_2937925 | | | |
| see above | LESP Ciudad de Mexico | Instituto de Diagnostico y Referencia Epidemiologicos (INDRE) | Claudia Wong-Arambula, Abril Rodriguez-Maldonado, Vanessa Rivero-Arredondo, Ariadna Medina-Benitez, Joaquin Quiroz-Mercado, Sergio Rangel-Guerrero, Maribel Gonzalez-Villa, Natividad Cruz-Ortiz, Tatiana Nunez-Garcia, Gisela Barrera-Badillo, Lucia Hernandez-Rivas, Irma Lopez-Martinez, Ernesto Ramirez-Gonzalez. |
| EPI_ISL_2937926, EPI_ISL_2937927 | LESP Queretaro | Instituto de Diagnostico y Referencia Epidemiologicos (INDRE) | Claudia Wong-Arambula, Abril Rodriguez-Maldonado, Vanessa Rivero-Arredondo, Ariadna Medina-Benitez, Joaquin Quiroz-Mercado, Sergio Rangel-Guerrero, Maribel Gonzalez-Villa, Natividad Cruz-Ortiz, Tatiana Nunez-Garcia, Gisela Barrera-Badillo, Lucia Hernandez-Rivas, Irma Lopez-Martinez, Ernesto Ramirez-Gonzalez. |
| EPI_ISL_2937928 | LESP Ciudad de Mexico | Instituto de Diagnostico y Referencia Epidemiologicos (INDRE) | Claudia Wong-Arambula, Abril Rodriguez-Maldonado, Vanessa Rivero-Arredondo, Ariadna Medina-Benitez, Joaquin Quiroz-Mercado, Sergio Rangel-Guerrero, Natividad Cruz-Ortiz, Tatiana Nunez-Garcia, Gisela Barrera-Badillo, Lucia Hernandez-Rivas, Irma Lopez-Martinez, Ernesto Ramirez-Gonzalez. |
| EPI_ISL_2937929 | LESP Tamaulipas | Instituto de Diagnostico y Referencia Epidemiologicos (INDRE) | Claudia Wong-Arambula, Abril Rodriguez-Maldonado, Vanessa Rivero-Arredondo, Ariadna Medina-Benitez, Joaquin Quiroz-Mercado, Sergio Rangel-Guerrero, Maribel Gonzalez-Villa, Natividad Cruz-Ortiz, Tatiana Nunez-Garcia, Gisela Barrera-Badillo, Lucia Hernandez-Rivas, Irma Lopez-Martinez, Ernesto Ramirez-Gonzalez. |
| EPI_ISL_2937930 | LESP Veracruz | Instituto de Diagnostico y Referencia Epidemiologicos (INDRE) | Claudia Wong-Arambula, Abril Rodriguez-Maldonado, Vanessa Rivero-Arredondo, Ariadna Medina-Benitez, Joaquin Quiroz-Mercado, Sergio Rangel-Guerrero, Maribel Gonzalez-Villa, Natividad Cruz-Ortiz, Tatiana Nunez-Garcia, Gisela Barrera-Badillo, Lucia Hernandez-Rivas, Irma Lopez-Martinez, Ernesto Ramirez-Gonzalez. |
| EPI_ISL_2938097 | Health Center Velika Kladuša | Clinical Center, University of Sarajevo; Unit for Clinical Microbiology | Irma Salimovi-Beši, Amela Dedei-Ljubovi, Edina Zahirovi, Sandra Vegar-Zubovi, Sebija Izetbegovi |
| EPI_ISL_2938098 | Health Center Novi Grad Sarajevo | Clinical Center, University of Sarajevo; Unit for Clinical Microbiology | Irma Salimovi-Beši, Amela Dedei-Ljubovi, Edina Zahirovi, Sandra Vegar-Zubovi, Sebija Izetbegovi |
| EPI_ISL_2938099, EPI_ISL_2938100, EPI_ISL_2938101 | Health Center Cazin | Clinical Center, University of Sarajevo; Unit for Clinical Microbiology | Irma Salimovi-Beši, Amela Dedei-Ljubovi, Edina Zahirovi, Sandra Vegar-Zubovi, Sebija Izetbegovi |
| EPI_ISL_2938103 | Health Center Novo Sarajevo | Clinical Center, University of Sarajevo; Unit for Clinical Microbiology | Irma Salimovi-Beši, Amela Dedei-Ljubovi, Edina Zahirovi, Sandra Vegar-Zubovi, Sebija Izetbegovi |
| EPI_ISL_2938104 | Clinical Center, University of Sarajevo; Unit for Clinical Microbiology | Clinical Center, University of Sarajevo; Unit for Clinical Microbiology | Irma Salimovi-Beši, Amela Dedei-Ljubovi, Edina Zahirovi, Sandra Vegar-Zubovi, Sebija Izetbegovi |
| EPI_ISL_2938105 | Health Center Novi Grad Sarajevo | Clinical Center, University of Sarajevo; Unit for Clinical Microbiology | Irma Salimovi-Beši, Amela Dedei-Ljubovi, Edina Zahirovi, Sandra Vegar-Zubovi, Sebija Izetbegovi |
| EPI_ISL_2938106 | Clinical Center, University of Sarajevo; Unit for Clinical Microbiology | Clinical Center, University of Sarajevo; Unit for Clinical Microbiology | Irma Salimovi-Beši, Amela Dedei-Ljubovi, Edina Zahirovi, Sandra Vegar-Zubovi, Sebija Izetbegovi |
| EPI_ISL_2938107 | Health Center Sarajevo- Center | Clinical Center, University of Sarajevo; Unit for Clinical Microbiology | Irma Salimovi-Beši, Amela Dedei-Ljubovi, Edina Zahirovi, Sandra Vegar-Zubovi, Sebija Izetbegovi |
| EPI_ISL_2938108 | Clinical Center, University of Sarajevo; Unit for Clinical Microbiology | Clinical Center, University of Sarajevo; Unit for Clinical Microbiology | Irma Salimovi-Beši, Amela Dedei-Ljubovi, Edina Zahirovi, Sandra Vegar-Zubovi, Sebija Izetbegovi |
| EPI_ISL_2938109 | Health Center Sarajevo- Center | Clinical Center, University of Sarajevo; Unit for Clinical Microbiology | Irma Salimovi-Beši, Amela Dedei-Ljubovi, Edina Zahirovi, Sandra Vegar-Zubovi, Sebija Izetbegovi |
| EPI_ISL_2938110 | Health Center Novi Grad Sarajevo | Clinical Center, University of Sarajevo; Unit for Clinical Microbiology | Irma Salimovi-Beši, Amela Dedei-Ljubovi, Edina Zahirovi, Sandra Vegar-Zubovi, Sebija Izetbegovi |
| EPI_ISL_2938111 | Health Center Velika Kladuša | Clinical Center, University of Sarajevo; Unit for Clinical Microbiology | Irma Salimovi-Beši, Amela Dedei-Ljubovi, Edina Zahirovi, Sandra Vegar-Zubovi, Sebija Izetbegovi |
| EPI_ISL_2938112 | Health Center Cazin | Clinical Center, University of Sarajevo; Unit for Clinical Microbiology | Irma Salimovi-Beši, Amela Dedei-Ljubovi, Edina Zahirovi, Sandra Vegar-Zubovi, Sebija Izetbegovi |
| EPI_ISL_2938114 | Health Center Sarajevo- Center | Clinical Center, University of Sarajevo; Unit for Clinical Microbiology | Irma Salimovi-Beši, Amela Dedei-Ljubovi, Edina Zahirovi, Sandra Vegar-Zubovi, Sebija Izetbegovi |
| EPI_ISL_2938115 | Dr. Irfan Ljubijankic Cantonal Hospital Biha, Laboratory for Microbiology | Clinical Center, University of Sarajevo; Unit for Clinical Microbiology | Irma Salimovi-Beši, Amela Dedei-Ljubovi, Edina Zahirovi, Sandra Vegar-Zubovi, Sebija Izetbegovi |
| EPI_ISL_2938116 | Health Center Vogoša | Clinical Center, University of Sarajevo; Unit for Clinical | Irma Salimovi-Beši, Amela Dedei-Ljubovi, Edina Zahirovi, Sandra Vegar-Zubovi, Sebija Izetbegovi |

| | | | |
|---|---|---|---|
| | | Microbiology | |
| EPI_ISL_2938117 | Health Center Novo Sarajevo | Clinical Center, University of Sarajevo; Unit for Clinical Microbiology | Irma Salimovi-Beši, Amela Dedei-Ljubovi, Edina Zahirovi, Sandra Vegar-Zubovi, Sebija Izetbegovi |
| EPI_ISL_2938119 | Health Center Novi Grad Sarajevo | Clinical Center, University of Sarajevo; Unit for Clinical Microbiology | Irma Salimovi-Beši, Amela Dedei-Ljubovi, Edina Zahirovi, Sandra Vegar-Zubovi, Sebija Izetbegovi |
| EPI_ISL_2938311, EPI_ISL_2938312, EPI_ISL_2938313 | St James's Hospital, Dublin 8, Ireland. | ELDA biotech | Deirdre M. Broderick, Peter R. Flanagan, Brendan Crowley |
| EPI_ISL_2940309, EPI_ISL_2940312, EPI_ISL_2940314, EPI_ISL_2940316, EPI_ISL_2940319, EPI_ISL_2940325, EPI_ISL_2940328, EPI_ISL_2940333, EPI_ISL_2940334, EPI_ISL_2940336, EPI_ISL_2940339, EPI_ISL_2940342, EPI_ISL_2940344, EPI_ISL_2940345, EPI_ISL_2940351, EPI_ISL_2940359, EPI_ISL_2940360, EPI_ISL_2940392, EPI_ISL_2940394 | | | |
| see above | NJDOH, Public Health and Environmental Laboratories | NJ_PHEL | Lindsey Bodnar, Shiv K. Verma, Ryan Pachucki, Chelsea San Filippo, Jacquelyn Deverell, Dana Woell, Allison Roder, Byeong Jeong |
| EPI_ISL_2941577 | Baylor Scott & White-Irving | Baylor Scott & White-Temple | Ari Rao, Linden Morales, Kimberly Walker, Marcus Volz, Shelby Johnson |
| EPI_ISL_2942326, EPI_ISL_2942327, EPI_ISL_2942329, EPI_ISL_2942330, EPI_ISL_2942331, EPI_ISL_2942332, EPI_ISL_2942333, EPI_ISL_2942336, EPI_ISL_2942337, EPI_ISL_2942339, EPI_ISL_2942340, EPI_ISL_2942341, EPI_ISL_2942344, EPI_ISL_2942347, EPI_ISL_2942350, EPI_ISL_2942351, EPI_ISL_2942352, EPI_ISL_2942353, EPI_ISL_2942354, EPI_ISL_2942355, EPI_ISL_2942356, EPI_ISL_2942357, EPI_ISL_2942359, EPI_ISL_2942362, EPI_ISL_2942364, EPI_ISL_2942367, EPI_ISL_2942370, EPI_ISL_2942371, EPI_ISL_2942372, EPI_ISL_2942373, EPI_ISL_2942374, EPI_ISL_2942375, EPI_ISL_2942376, EPI_ISL_2942377, EPI_ISL_2942378, EPI_ISL_2942379, EPI_ISL_2942380, EPI_ISL_2942382, EPI_ISL_2942383, EPI_ISL_2942384, EPI_ISL_2942385, EPI_ISL_2942386, EPI_ISL_2942387, EPI_ISL_2942389, EPI_ISL_2942390, EPI_ISL_2942392, EPI_ISL_2942393, EPI_ISL_2942394, EPI_ISL_2942395, EPI_ISL_2942396, EPI_ISL_2942397, EPI_ISL_2942398, EPI_ISL_2942400, EPI_ISL_2942402, EPI_ISL_2942405, EPI_ISL_2942406 | | | |
| see above | Unidad de Investigacion Medica de Yucatan (UIMY) | Unidad de Genomica Avanzada | Consorcio Mexicano de Vigilancia Genomica (CoViGen-Mex). Authors (in alphabetical order): Julio Elias Alvarado-Yaah, Carlos F. Arias, Santiago avila-Rios, Eduardo Becerril-Vargas, Victor Hugo Borja-Aburto, Celia Boukadida, Cristobal Chaidez-Quiroz, Juan Bautista Chale-Dzul, Ricardo Ciria Merce, Andreu Comas-Garcia, Celida Duque Molina, Julissa Enciso-Ibarra, Jose Antonio Enciso-Moreno, Gloria Elena Espinosa-Ayala, Fernando Fontove-Herrera, Daniel Fregoso-Rueda, Victor Eduardo Garcia-Arias, Alejandra Garcia-Gasca, Bruno Gomez-Gil, Jean Pierre Gonzalez, Irvin Gonzalez-Lopez, Concepcion Grajales-Muñiz, Ricardo Grande, Rosa Maria Gutierrez Rios, Jesus Hernandez, Alejandra Hernandez-Teran, Alfredo Herrera-Estrella, Carla Ivon Herrera-Najera, Pavel Isa, Veronica Mata-Haro, Daniel Lira Morales, Susana Lopez, Antonio Loza Roman, Brenda Irasema Maldonado-Meza, Bernardo Martinez-Miguel, Jose Arturo Martinez-Orozco, Celida Martinez- Rodriguez, Margarita Matias-Florentino, Fidencio Mejia-Nepomuceno, Maria Guadalupe de Jesus Mireles-Rivera, Gloria Maria Molina-Salinas, Hector Montoya-Fuentes, , Mario Mujica-Sanchez, Jose Esteban Muñoz-Medina, Jose de Jesus Nuñez-Contreras, Alicia Ocaña-Mondragon, Luis Alberto Ochoa-Carrera, Hector Esteban Paz-Juarez, Marissa Perez-Garcia, Francisco Pulido, Helen Haydee Fernanda Ramirez-Plascencia, Jorge Salas-Hernandez, Angel Gustavo Salas-Lais, Alejandro Sanchez-Flores, Clara Esperanza Santacruz-Tinoco, Maria Guadalupe Santiago-Mauricio, Selene Zarate, Nelly Selem-Mojica, Blanca Taboada, Gloria Vazquez, Joel Armando Vazquez-Perez. |
| EPI_ISL_2942408, EPI_ISL_2942409, EPI_ISL_2942410, EPI_ISL_2942411, EPI_ISL_2942412, EPI_ISL_2942413, EPI_ISL_2942414, EPI_ISL_2942415, EPI_ISL_2942416, EPI_ISL_2942418, EPI_ISL_2942419, EPI_ISL_2942420, EPI_ISL_2942421, EPI_ISL_2942422, EPI_ISL_2942424, EPI_ISL_2942425, EPI_ISL_2942426, EPI_ISL_2942427, EPI_ISL_2942428, EPI_ISL_2942429, EPI_ISL_2942430, EPI_ISL_2942431, EPI_ISL_2942432, EPI_ISL_2942433, EPI_ISL_2942434, EPI_ISL_2942435, EPI_ISL_2942436, EPI_ISL_2942437, EPI_ISL_2942438, EPI_ISL_2942439, EPI_ISL_2942440, EPI_ISL_2942442, EPI_ISL_2942443, EPI_ISL_2942444, EPI_ISL_2942445, EPI_ISL_2942446, EPI_ISL_2942447, EPI_ISL_2942451, EPI_ISL_2942452, EPI_ISL_2942453, EPI_ISL_2942454, EPI_ISL_2942455, EPI_ISL_2942456, EPI_ISL_2942457, EPI_ISL_2942458, EPI_ISL_2942459, EPI_ISL_2942460, EPI_ISL_2942461, EPI_ISL_2942462, EPI_ISL_2942463, EPI_ISL_2942464, EPI_ISL_2942465, EPI_ISL_2942466 | | | |
| see above | Centro de Investigacion Biomedica de Occidente (CIBO) | Unidad de Genomica Avanzada | Consorcio Mexicano de Vigilancia Genomica (CoViGen-Mex). Authors (in alphabetical order): Julio Elias Alvarado-Yaah, Carlos F. Arias, Santiago avila-Rios, Eduardo Becerril-Vargas, Victor Hugo Borja-Aburto, Celia Boukadida, Cristobal Chaidez-Quiroz, Juan Bautista Chale-Dzul, Ricardo Ciria Merce, Andreu Comas-Garcia, Celida Duque Molina, Julissa Enciso-Ibarra, Jose Antonio Enciso-Moreno, Gloria Elena Espinosa-Ayala, Fernando Fontove-Herrera, Daniel Fregoso-Rueda, Victor Eduardo Garcia-Arias, Alejandra Garcia-Gasca, Bruno Gomez-Gil, Jean Pierre Gonzalez, Irvin Gonzalez-Lopez, Concepcion Grajales-Muñiz, Ricardo Grande, Rosa Maria Gutierrez Rios, Jesus Hernandez, Alejandra Hernandez-Teran, Alfredo Herrera-Estrella, Carla Ivon Herrera-Najera, Pavel Isa, Veronica Mata-Haro, Daniel Lira Morales, Susana Lopez, Antonio Loza Roman, Brenda Irasema Maldonado-Meza, Bernardo Martinez-Miguel, Jose Arturo Martinez-Orozco, Celida Martinez- Rodriguez, Margarita Matias-Florentino, Fidencio Mejia-Nepomuceno, Maria Guadalupe de Jesus Mireles-Rivera, Gloria Maria Molina-Salinas, Hector Montoya-Fuentes, , Mario Mujica-Sanchez, Jose Esteban Muñoz-Medina, Jose de Jesus Nuñez-Contreras, Alicia Ocaña-Mondragon, Luis Alberto Ochoa-Carrera, Hector Esteban Paz-Juarez, Marissa Perez-Garcia, Francisco Pulido, Helen Haydee Fernanda Ramirez-Plascencia, Jorge Salas-Hernandez, Angel Gustavo Salas-Lais, Alejandro Sanchez-Flores, Clara Esperanza Santacruz-Tinoco, Maria Guadalupe Santiago-Mauricio, Selene Zarate, Nelly Selem-Mojica, Blanca Taboada, Gloria Vazquez, Joel Armando Vazquez-Perez. |
| EPI_ISL_2942467, EPI_ISL_2942468, EPI_ISL_2942469, EPI_ISL_2942470, EPI_ISL_2942471, EPI_ISL_2942472, EPI_ISL_2942474, EPI_ISL_2942475, EPI_ISL_2942476, EPI_ISL_2942478, EPI_ISL_2942479, EPI_ISL_2942480 | | | |
| see above | Unidad de Investigacion Biomedica de Zacatecas (UIBZ) | Unidad de Genomica Avanzada | Consorcio Mexicano de Vigilancia Genomica (CoViGen-Mex). Authors (in alphabetical order): Julio Elias Alvarado-Yaah, Carlos F. Arias, Santiago avila-Rios, Eduardo Becerril-Vargas, Victor Hugo Borja-Aburto, Celia Boukadida, Cristobal Chaidez-Quiroz, Juan Bautista Chale-Dzul, Ricardo Ciria Merce, Andreu Comas-Garcia, Celida Duque Molina, Julissa Enciso-Ibarra, Jose Antonio Enciso-Moreno, Gloria Elena Espinosa-Ayala, Fernando Fontove-Herrera, Daniel Fregoso-Rueda, Victor Eduardo Garcia-Arias, Alejandra Garcia-Gasca, Bruno Gomez-Gil, Jean Pierre Gonzalez, Irvin Gonzalez-Lopez, Concepcion Grajales-Muñiz, Ricardo Grande, Rosa Maria Gutierrez Rios, Jesus Hernandez, Alejandra Hernandez-Teran, Alfredo Herrera-Estrella, Carla Ivon Herrera-Najera, Pavel Isa, Veronica Mata-Haro, Daniel Lira Morales, Susana Lopez, Antonio Loza Roman, Brenda Irasema Maldonado-Meza, Bernardo Martinez-Miguel, Jose Arturo Martinez-Orozco, Celida Martinez- Rodriguez, Margarita Matias-Florentino, Fidencio Mejia-Nepomuceno, Maria Guadalupe de Jesus Mireles-Rivera, Gloria Maria Molina-Salinas, Hector Montoya-Fuentes, , Mario Mujica-Sanchez, Jose Esteban Muñoz-Medina, Jose de Jesus Nuñez-Contreras, Alicia Ocaña-Mondragon, Luis Alberto Ochoa-Carrera, Hector Esteban Paz-Juarez, Marissa Perez-Garcia, Francisco Pulido, Helen Haydee Fernanda Ramirez-Plascencia, Jorge Salas-Hernandez, Angel Gustavo Salas-Lais, Alejandro Sanchez-Flores, Clara Esperanza Santacruz-Tinoco, Maria Guadalupe Santiago-Mauricio, Selene Zarate, Nelly Selem-Mojica, Blanca Taboada, Gloria Vazquez, Joel Armando Vazquez-Perez. |
| EPI_ISL_2942482, EPI_ISL_2942484, EPI_ISL_2942486, EPI_ISL_2942487, EPI_ISL_2942490, EPI_ISL_2942491, EPI_ISL_2942492, EPI_ISL_2942493, EPI_ISL_2942494, EPI_ISL_2942495, EPI_ISL_2942496, EPI_ISL_2942497, EPI_ISL_2942498, EPI_ISL_2942500, EPI_ISL_2942501, EPI_ISL_2942502, EPI_ISL_2942503, EPI_ISL_2942504, EPI_ISL_2942505, EPI_ISL_2942506, EPI_ISL_2942508, EPI_ISL_2942509, EPI_ISL_2942511 | | | |
| see above | Centro de Investigacion Biomedica del Noreste (CIBIN) | Unidad de Genomica Avanzada | Consorcio Mexicano de Vigilancia Genomica (CoViGen-Mex). Authors (in alphabetical order): Julio Elias Alvarado-Yaah, Carlos F. Arias, Santiago avila-Rios, Eduardo Becerril-Vargas, Victor Hugo Borja-Aburto, Celia Boukadida, Cristobal Chaidez-Quiroz, Juan Bautista Chale-Dzul, Ricardo Ciria Merce, Andreu Comas-Garcia, Celida Duque Molina, Julissa Enciso-Ibarra, Jose Antonio Enciso-Moreno, Gloria Elena Espinosa-Ayala, Fernando Fontove-Herrera, Daniel Fregoso-Rueda, Victor Eduardo Garcia-Arias, Alejandra Garcia-Gasca, Bruno Gomez-Gil, Jean Pierre Gonzalez, Irvin Gonzalez-Lopez, Concepcion Grajales-Muñiz, Ricardo Grande, Rosa Maria Gutierrez Rios, Jesus Hernandez, Alejandra Hernandez-Teran, Alfredo Herrera-Estrella, Carla Ivon Herrera-Najera, Pavel Isa, Veronica Mata-Haro, Daniel Lira Morales, Susana Lopez, Antonio Loza Roman, Brenda Irasema Maldonado-Meza, Bernardo Martinez-Miguel, Jose Arturo Martinez-Orozco, Celida Martinez- Rodriguez, Margarita Matias-Florentino, Fidencio Mejia-Nepomuceno, Maria Guadalupe de Jesus Mireles-Rivera, Gloria Maria Molina-Salinas, Hector Montoya-Fuentes, , Mario Mujica-Sanchez, Jose Esteban Muñoz-Medina, Jose de Jesus Nuñez-Contreras, Alicia Ocaña-Mondragon, Luis Alberto Ochoa-Carrera, Hector Esteban Paz-Juarez, Marissa Perez-Garcia, Francisco Pulido, Helen Haydee Fernanda Ramirez-Plascencia, Jorge Salas-Hernandez, Angel Gustavo Salas-Lais, Alejandro Sanchez-Flores, Clara Esperanza Santacruz-Tinoco, Maria Guadalupe Santiago-Mauricio, Selene Zarate, Nelly Selem-Mojica, Blanca Taboada, Gloria Vazquez, Joel Armando Vazquez-Perez. |
| EPI_ISL_2942513, EPI_ISL_2942516, EPI_ISL_2942518, EPI_ISL_2942520, EPI_ISL_2942521, EPI_ISL_2942522, EPI_ISL_2942523, EPI_ISL_2942524, EPI_ISL_2942525, EPI_ISL_2942526, EPI_ISL_2942527, EPI_ISL_2942528, EPI_ISL_2942529, EPI_ISL_2942531, EPI_ISL_2942532, EPI_ISL_2942533, EPI_ISL_2942534, EPI_ISL_2942535, EPI_ISL_2942536, EPI_ISL_2942537, EPI_ISL_2942538, EPI_ISL_2942539, EPI_ISL_2942540, EPI_ISL_2942541, EPI_ISL_2942542, EPI_ISL_2942543, EPI_ISL_2942544, EPI_ISL_2942545, EPI_ISL_2942546, EPI_ISL_2942547, EPI_ISL_2942548, EPI_ISL_2942549, EPI_ISL_2942550, EPI_ISL_2942552, EPI_ISL_2942553, EPI_ISL_2942554, EPI_ISL_2942555, EPI_ISL_2942556, EPI_ISL_2942557, EPI_ISL_2942558, EPI_ISL_2942559, EPI_ISL_2942560, EPI_ISL_2942561, EPI_ISL_2942562, EPI_ISL_2942563, EPI_ISL_2942564, EPI_ISL_2942565, EPI_ISL_2942566, EPI_ISL_2942568, EPI_ISL_2942569, EPI_ISL_2942570, EPI_ISL_2942571, EPI_ISL_2942572, EPI_ISL_2942573, EPI_ISL_2942575, EPI_ISL_2942576, EPI_ISL_2942577, EPI_ISL_2942578, EPI_ISL_2942579, EPI_ISL_2942580, EPI_ISL_2942581, EPI_ISL_2942582, EPI_ISL_2942583, EPI_ISL_2942584, EPI_ISL_2942585, EPI_ISL_2942586, EPI_ISL_2942587, EPI_ISL_2942588, EPI_ISL_2942589, EPI_ISL_2942591, EPI_ISL_2942592, EPI_ISL_2942593, EPI_ISL_2942594, EPI_ISL_2942596, EPI_ISL_2942598, EPI_ISL_2942599, EPI_ISL_2942600, EPI_ISL_2942601, EPI_ISL_2942602, EPI_ISL_2942603, EPI_ISL_2942604, EPI_ISL_2942605, EPI_ISL_2942606, EPI_ISL_2942607, EPI_ISL_2942608, EPI_ISL_2942609, EPI_ISL_2942610, EPI_ISL_2942611, EPI_ISL_2942612, EPI_ISL_2942613, EPI_ISL_2942614, EPI_ISL_2942615, EPI_ISL_2942616, EPI_ISL_2942617, EPI_ISL_2942619, EPI_ISL_2942621, EPI_ISL_2942622, EPI_ISL_2942623, EPI_ISL_2942624 | | | |
| see above | Laboratorio Central de Epidemiologia (LCE) | Unidad de Genomica Avanzada | Consorcio Mexicano de Vigilancia Genomica (CoViGen-Mex). Authors (in alphabetical order): Julio Elias Alvarado-Yaah, Carlos F. Arias, Santiago avila-Rios, Eduardo Becerril-Vargas, Victor Hugo Borja-Aburto, Celia Boukadida, Cristobal Chaidez-Quiroz, Juan Bautista Chale-Dzul, Ricardo Ciria Merce, Andreu Comas-Garcia, Celida Duque Molina, Julissa Enciso-Ibarra, Jose Antonio Enciso-Moreno, Gloria Elena Espinosa-Ayala, Fernando |

| | | | |
|---|---|---|---|
| | | | Fontove-Herrera, Daniel Fregoso-Rueda, Victor Eduardo Garcia-Arias, Alejandra Garcia-Gasca, Bruno Gomez-Gil, Jean Pierre Gonzalez, Irvin Gonzalez-Lopez, Concepcion Grajales-Muñiz, Ricardo Grande, Rosa Maria Gutierrez Rios, Jesus Hernandez, Alejandra Hernandez-Teran, Alfredo Herrera-Estrella, Carla Ivon Herrera-Najera, Pavel Isa, Veronica Mata-Haro, Daniel Lira Morales, Susana Lopez, Antonio Loza Roman, Brenda Irasema Maldonado-Meza, Bernardo Martinez-Miguel, Jose Arturo Martinez-Orozco, Celida Martinez- Rodriguez, Margarita Matias-Florentino, Fidencio Mejia-Nepomuceno, Maria Guadalupe de Jesus Mireles-Rivera, Gloria Maria Molina-Salinas, Hector Montoya-Fuentes, , Mario Mujica-Sanchez, Jose Esteban Muñoz-Medina, Jose de Jesus Nuñez-Contreras, Alicia Ocaña-Mondragon, Luis Alberto Ochoa-Carrera, Hector Esteban Paz-Juarez, Marissa Perez-Garcia, Francisco Pulido, Helen Haydee Fernanda Ramirez-Plascencia, Jorge Salas-Hernandez, Angel Gustavo Salas-Lais, Alejandro Sanchez-Flores, Clara Esperanza Santacruz-Tinoco, Maria Guadalupe Santiago-Mauricio, Selene Zarate, Nelly Selem-Mojica, Blanca Taboada, Gloria Vazquez, Joel Armando Vazquez-Perez. |
| EPI_ISL_2942626, EPI_ISL_2942627, EPI_ISL_2942628, EPI_ISL_2942629, EPI_ISL_2942630, EPI_ISL_2942631, EPI_ISL_2942632, EPI_ISL_2942633, EPI_ISL_2942634, EPI_ISL_2942635, EPI_ISL_2942636, EPI_ISL_2942637, EPI_ISL_2942638, EPI_ISL_2942639, EPI_ISL_2942640, EPI_ISL_2942641, EPI_ISL_2942642, EPI_ISL_2942643, EPI_ISL_2942644, EPI_ISL_2942645, EPI_ISL_2942646, EPI_ISL_2942647, EPI_ISL_2942648, EPI_ISL_2942649, EPI_ISL_2942650, EPI_ISL_2942651, EPI_ISL_2942652, EPI_ISL_2942653, EPI_ISL_2942654, EPI_ISL_2942655, EPI_ISL_2942656, EPI_ISL_2942657, EPI_ISL_2942658, EPI_ISL_2942659, EPI_ISL_2942660, EPI_ISL_2942661, EPI_ISL_2942662, EPI_ISL_2942663, EPI_ISL_2942664, EPI_ISL_2942665, EPI_ISL_2942666, EPI_ISL_2942667, EPI_ISL_2942668, EPI_ISL_2942669, EPI_ISL_2942670, EPI_ISL_2942671, EPI_ISL_2942672, EPI_ISL_2942673, EPI_ISL_2942674, EPI_ISL_2942675, EPI_ISL_2942676, EPI_ISL_2942677, EPI_ISL_2942678, EPI_ISL_2942679, EPI_ISL_2942680, EPI_ISL_2942681, EPI_ISL_2942682, EPI_ISL_2942683, EPI_ISL_2942684, EPI_ISL_2942685, EPI_ISL_2942686, EPI_ISL_2942687, EPI_ISL_2942688, EPI_ISL_2942689, EPI_ISL_2942690, EPI_ISL_2942691, EPI_ISL_2942692, EPI_ISL_2942693, EPI_ISL_2942694, EPI_ISL_2942695, EPI_ISL_2942696, EPI_ISL_2942697, EPI_ISL_2942698, EPI_ISL_2942699, EPI_ISL_2942700, EPI_ISL_2942701, EPI_ISL_2942702, EPI_ISL_2942703, EPI_ISL_2942704, EPI_ISL_2942705, EPI_ISL_2942706, EPI_ISL_2942707, EPI_ISL_2942708, EPI_ISL_2942709, EPI_ISL_2942710, EPI_ISL_2942711, EPI_ISL_2942713, EPI_ISL_2942714, EPI_ISL_2942715, EPI_ISL_2942716, EPI_ISL_2942717, EPI_ISL_2942718, EPI_ISL_2942719 | | | |
| see above | Unidad de Investigacion Medica de Yucatan (UIMY) | Unidad de Genomica Avanzada | Consorcio Mexicano de Vigilancia Genomica (CoViGen-Mex). Authors (in alphabetical order): Julio Elias Alvarado-Yaah, Carlos F. Arias, Santiago avila-Rios, Eduardo Becerril-Vargas, Victor Hugo Borja-Aburto, Celia Boukadida, Cristobal Chaidez-Quiroz, Juan Bautista Chale-Dzul, Ricardo Ciria Merce, Andreu Comas-Garcia, Celida Duque Molina, Julissa Enciso-Ibarra, Jose Antonio Enciso-Moreno, Gloria Elena Espinosa-Ayala, Fernando Fontove-Herrera, Daniel Fregoso-Rueda, Victor Eduardo Garcia-Arias, Alejandra Garcia-Gasca, Bruno Gomez-Gil, Jean Pierre Gonzalez, Irvin Gonzalez-Lopez, Concepcion Grajales-Muñiz, Ricardo Grande, Rosa Maria Gutierrez Rios, Jesus Hernandez, Alejandra Hernandez-Teran, Alfredo Herrera-Estrella, Carla Ivon Herrera-Najera, Pavel Isa, Veronica Mata-Haro, Daniel Lira Morales, Susana Lopez, Antonio Loza Roman, Brenda Irasema Maldonado-Meza, Bernardo Martinez-Miguel, Jose Arturo Martinez-Orozco, Celida Martinez- Rodriguez, Margarita Matias-Florentino, Fidencio Mejia-Nepomuceno, Maria Guadalupe de Jesus Mireles-Rivera, Gloria Maria Molina-Salinas, Hector Montoya-Fuentes, , Mario Mujica-Sanchez, Jose Esteban Muñoz-Medina, Jose de Jesus Nuñez-Contreras, Alicia Ocaña-Mondragon, Luis Alberto Ochoa-Carrera, Hector Esteban Paz-Juarez, Marissa Perez-Garcia, Francisco Pulido, Helen Haydee Fernanda Ramirez-Plascencia, Jorge Salas-Hernandez, Angel Gustavo Salas-Lais, Alejandro Sanchez-Flores, Clara Esperanza Santacruz-Tinoco, Maria Guadalupe Santiago-Mauricio, Selene Zarate, Nelly Selem-Mojica, Blanca Taboada, Gloria Vazquez, Joel Armando Vazquez-Perez. |
| EPI_ISL_2942720, EPI_ISL_2942721, EPI_ISL_2942722, EPI_ISL_2942723, EPI_ISL_2942724, EPI_ISL_2942726, EPI_ISL_2942727, EPI_ISL_2942728, EPI_ISL_2942730, EPI_ISL_2942731, EPI_ISL_2942732, EPI_ISL_2942733, EPI_ISL_2942734, EPI_ISL_2942735, EPI_ISL_2942736, EPI_ISL_2942737, EPI_ISL_2942738, EPI_ISL_2942740, EPI_ISL_2942741, EPI_ISL_2942742, EPI_ISL_2942743, EPI_ISL_2942744, EPI_ISL_2942745, EPI_ISL_2942746, EPI_ISL_2942747, EPI_ISL_2942748, EPI_ISL_2942749 | | | |
| see above | Centro de Investigacion Biomedica de Occidente (CIBO) | Unidad de Genomica Avanzada | Consorcio Mexicano de Vigilancia Genomica (CoViGen-Mex). Authors (in alphabetical order): Julio Elias Alvarado-Yaah, Carlos F. Arias, Santiago avila-Rios, Eduardo Becerril-Vargas, Victor Hugo Borja-Aburto, Celia Boukadida, Cristobal Chaidez-Quiroz, Juan Bautista Chale-Dzul, Ricardo Ciria Merce, Andreu Comas-Garcia, Celida Duque Molina, Julissa Enciso-Ibarra, Jose Antonio Enciso-Moreno, Gloria Elena Espinosa-Ayala, Fernando Fontove-Herrera, Daniel Fregoso-Rueda, Victor Eduardo Garcia-Arias, Alejandra Garcia-Gasca, Bruno Gomez-Gil, Jean Pierre Gonzalez, Irvin Gonzalez-Lopez, Concepcion Grajales-Muñiz, Ricardo Grande, Rosa Maria Gutierrez Rios, Jesus Hernandez, Alejandra Hernandez-Teran, Alfredo Herrera-Estrella, Carla Ivon Herrera-Najera, Pavel Isa, Veronica Mata-Haro, Daniel Lira Morales, Susana Lopez, Antonio Loza Roman, Brenda Irasema Maldonado-Meza, Bernardo Martinez-Miguel, Jose Arturo Martinez-Orozco, Celida Martinez- Rodriguez, Margarita Matias-Florentino, Fidencio Mejia-Nepomuceno, Maria Guadalupe de Jesus Mireles-Rivera, Gloria Maria Molina-Salinas, Hector Montoya-Fuentes, , Mario Mujica-Sanchez, Jose Esteban Muñoz-Medina, Jose de Jesus Nuñez-Contreras, Alicia Ocaña-Mondragon, Luis Alberto Ochoa-Carrera, Hector Esteban Paz-Juarez, Marissa Perez-Garcia, Francisco Pulido, Helen Haydee Fernanda Ramirez-Plascencia, Jorge Salas-Hernandez, Angel Gustavo Salas-Lais, Alejandro Sanchez-Flores, Clara Esperanza Santacruz-Tinoco, Maria Guadalupe Santiago-Mauricio, Selene Zarate, Nelly Selem-Mojica, Blanca Taboada, Gloria Vazquez, Joel Armando Vazquez-Perez. |
| EPI_ISL_2942750, EPI_ISL_2942752, EPI_ISL_2942753 | Unidad de Investigacion Biomedica de Zacatecas (UIBZ) | Unidad de Genomica Avanzada | Consorcio Mexicano de Vigilancia Genomica (CoViGen-Mex). Authors (in alphabetical order): Julio Elias Alvarado-Yaah, Carlos F. Arias, Santiago avila-Rios, Eduardo Becerril-Vargas, Victor Hugo Borja-Aburto, Celia Boukadida, Cristobal Chaidez-Quiroz, Juan Bautista Chale-Dzul, Ricardo Ciria Merce, Andreu Comas-Garcia, Celida Duque Molina, Julissa Enciso-Ibarra, Jose Antonio Enciso-Moreno, Gloria Elena Espinosa-Ayala, Fernando Fontove-Herrera, Daniel Fregoso-Rueda, Victor Eduardo Garcia-Arias, Alejandra Garcia-Gasca, Bruno Gomez-Gil, Jean Pierre Gonzalez, Irvin Gonzalez-Lopez, Concepcion Grajales-Muñiz, Ricardo Grande, Rosa Maria Gutierrez Rios, Jesus Hernandez, Alejandra Hernandez-Teran, Alfredo Herrera-Estrella, Carla Ivon Herrera-Najera, Pavel Isa, Veronica Mata-Haro, Daniel Lira Morales, Susana Lopez, Antonio Loza Roman, Brenda Irasema Maldonado-Meza, Bernardo Martinez-Miguel, Jose Arturo Martinez-Orozco, Celida Martinez- Rodriguez, Margarita Matias-Florentino, Fidencio Mejia-Nepomuceno, Maria Guadalupe de Jesus Mireles-Rivera, Gloria Maria Molina-Salinas, Hector Montoya-Fuentes, , Mario Mujica-Sanchez, Jose Esteban Muñoz-Medina, Jose de Jesus Nuñez-Contreras, Alicia Ocaña-Mondragon, Luis Alberto Ochoa-Carrera, Hector Esteban Paz-Juarez, Marissa Perez-Garcia, Francisco Pulido, Helen Haydee Fernanda Ramirez-Plascencia, Jorge Salas-Hernandez, Angel Gustavo Salas-Lais, Alejandro Sanchez-Flores, Clara Esperanza Santacruz-Tinoco, Maria Guadalupe Santiago-Mauricio, Selene Zarate, Nelly Selem-Mojica, Blanca Taboada, Gloria Vazquez, Joel Armando Vazquez-Perez. |
| EPI_ISL_2942755, EPI_ISL_2942756, EPI_ISL_2942757, EPI_ISL_2942758, EPI_ISL_2942759, EPI_ISL_2942760, EPI_ISL_2942761, EPI_ISL_2942762, EPI_ISL_2942763, EPI_ISL_2942764, EPI_ISL_2942766, EPI_ISL_2942768, EPI_ISL_2942770, EPI_ISL_2942771, EPI_ISL_2942772, EPI_ISL_2942773, EPI_ISL_2942774, EPI_ISL_2942775, EPI_ISL_2942776, EPI_ISL_2942778, EPI_ISL_2942779, EPI_ISL_2942780, EPI_ISL_2942781, EPI_ISL_2942784, EPI_ISL_2942785, EPI_ISL_2942786, EPI_ISL_2942788, EPI_ISL_2942789, EPI_ISL_2942790, EPI_ISL_2942792 | | | |
| see above | Centro de Investigacion Biomedica del Noreste (CIBIN) | Unidad de Genomica Avanzada | Consorcio Mexicano de Vigilancia Genomica (CoViGen-Mex). Authors (in alphabetical order): Julio Elias Alvarado-Yaah, Carlos F. Arias, Santiago avila-Rios, Eduardo Becerril-Vargas, Victor Hugo Borja-Aburto, Celia Boukadida, Cristobal Chaidez-Quiroz, Juan Bautista Chale-Dzul, Ricardo Ciria Merce, Andreu Comas-Garcia, Celida Duque Molina, Julissa Enciso-Ibarra, Jose Antonio Enciso-Moreno, Gloria Elena Espinosa-Ayala, Fernando Fontove-Herrera, Daniel Fregoso-Rueda, Victor Eduardo Garcia-Arias, Alejandra Garcia-Gasca, Bruno Gomez-Gil, Jean Pierre Gonzalez, Irvin Gonzalez-Lopez, Concepcion Grajales-Muñiz, Ricardo Grande, Rosa Maria Gutierrez Rios, Jesus Hernandez, Alejandra Hernandez-Teran, Alfredo Herrera-Estrella, Carla Ivon Herrera-Najera, Pavel Isa, Veronica Mata-Haro, Daniel Lira Morales, Susana Lopez, Antonio Loza Roman, Brenda Irasema Maldonado-Meza, Bernardo Martinez-Miguel, Jose Arturo Martinez-Orozco, Celida Martinez- Rodriguez, Margarita Matias-Florentino, Fidencio Mejia-Nepomuceno, Maria Guadalupe de Jesus Mireles-Rivera, Gloria Maria Molina-Salinas, Hector Montoya-Fuentes, , Mario Mujica-Sanchez, Jose Esteban Muñoz-Medina, Jose de Jesus Nuñez-Contreras, Alicia Ocaña-Mondragon, Luis Alberto Ochoa-Carrera, Hector Esteban Paz-Juarez, Marissa Perez-Garcia, Francisco Pulido, Helen Haydee Fernanda Ramirez-Plascencia, Jorge Salas-Hernandez, Angel Gustavo Salas-Lais, Alejandro Sanchez-Flores, Clara Esperanza Santacruz-Tinoco, Maria Guadalupe Santiago-Mauricio, Selene Zarate, Nelly Selem-Mojica, Blanca Taboada, Gloria Vazquez, Joel Armando Vazquez-Perez. |
| EPI_ISL_2942793, EPI_ISL_2942795, EPI_ISL_2942797, EPI_ISL_2942798, EPI_ISL_2942799, EPI_ISL_2942800, EPI_ISL_2942803, EPI_ISL_2942804, EPI_ISL_2942805, EPI_ISL_2942807, EPI_ISL_2942808, EPI_ISL_2942809, EPI_ISL_2942810, EPI_ISL_2942813, EPI_ISL_2942814, EPI_ISL_2942815, EPI_ISL_2942816, EPI_ISL_2942817, EPI_ISL_2942818, EPI_ISL_2942819, EPI_ISL_2942820, EPI_ISL_2942821, EPI_ISL_2942822, EPI_ISL_2942823, EPI_ISL_2942824, EPI_ISL_2942826, EPI_ISL_2942827, EPI_ISL_2942828, EPI_ISL_2942829, EPI_ISL_2942830, EPI_ISL_2942831, EPI_ISL_2942832, EPI_ISL_2942833, EPI_ISL_2942834, EPI_ISL_2942835, EPI_ISL_2942837, EPI_ISL_2942838, EPI_ISL_2942839, EPI_ISL_2942840, EPI_ISL_2942841, EPI_ISL_2942842, EPI_ISL_2942843, EPI_ISL_2942844, EPI_ISL_2942845, EPI_ISL_2942846, EPI_ISL_2942847, EPI_ISL_2942848, EPI_ISL_2942849, EPI_ISL_2942850, EPI_ISL_2942854, EPI_ISL_2942857, EPI_ISL_2942858, EPI_ISL_2942859, EPI_ISL_2942860, EPI_ISL_2942861, EPI_ISL_2942862, EPI_ISL_2942863, EPI_ISL_2942864, EPI_ISL_2942865, EPI_ISL_2942867, EPI_ISL_2942868, EPI_ISL_2942869, EPI_ISL_2942870, EPI_ISL_2942871, EPI_ISL_2942872, EPI_ISL_2942873, EPI_ISL_2942874, EPI_ISL_2942875, EPI_ISL_2942876, EPI_ISL_2942877, EPI_ISL_2942878, EPI_ISL_2942879, EPI_ISL_2942880, EPI_ISL_2942881, EPI_ISL_2942882, EPI_ISL_2942883, EPI_ISL_2942885, EPI_ISL_2942887, EPI_ISL_2942888, EPI_ISL_2942889, EPI_ISL_2942890, EPI_ISL_2942891, EPI_ISL_2942892, EPI_ISL_2942893, EPI_ISL_2942894, EPI_ISL_2942895, EPI_ISL_2942896, EPI_ISL_2942898, EPI_ISL_2942899, EPI_ISL_2942900, EPI_ISL_2942901, EPI_ISL_2942902, EPI_ISL_2942903, EPI_ISL_2942904, EPI_ISL_2942905, EPI_ISL_2942906, EPI_ISL_2942907, EPI_ISL_2942908, EPI_ISL_2942910, EPI_ISL_2942912, EPI_ISL_2942913, EPI_ISL_2942915, EPI_ISL_2942916, EPI_ISL_2942917, EPI_ISL_2942918, EPI_ISL_2942921, EPI_ISL_2942923, EPI_ISL_2942924, EPI_ISL_2942925, EPI_ISL_2942927, EPI_ISL_2942928, EPI_ISL_2942929, EPI_ISL_2942930, EPI_ISL_2942931, EPI_ISL_2942932, EPI_ISL_2942933, EPI_ISL_2942934, EPI_ISL_2942935 | | | |
| see above | Laboratorio Central de Epidemiologia (LCE) | Unidad de Genomica Avanzada | Consorcio Mexicano de Vigilancia Genomica (CoViGen-Mex). Authors (in alphabetical order): Julio Elias Alvarado-Yaah, Carlos F. Arias, Santiago avila-Rios, Eduardo Becerril-Vargas, Victor Hugo Borja-Aburto, Celia Boukadida, Cristobal Chaidez-Quiroz, Juan Bautista Chale-Dzul, Ricardo Ciria Merce, Andreu Comas-Garcia, Celida Duque Molina, Julissa Enciso-Ibarra, Jose Antonio Enciso-Moreno, Gloria Elena Espinosa-Ayala, Fernando Fontove-Herrera, Daniel Fregoso-Rueda, Victor Eduardo Garcia-Arias, Alejandra Garcia-Gasca, Bruno Gomez-Gil, Jean Pierre Gonzalez, Irvin Gonzalez-Lopez, Concepcion Grajales-Muñiz, Ricardo Grande, Rosa Maria Gutierrez Rios, Jesus Hernandez, Alejandra Hernandez-Teran, Alfredo Herrera-Estrella, Carla Ivon Herrera-Najera, Pavel Isa, Veronica Mata-Haro, Daniel Lira Morales, Susana Lopez, Antonio Loza Roman, Brenda Irasema Maldonado-Meza, Bernardo Martinez-Miguel, Jose Arturo Martinez-Orozco, Celida Martinez- Rodriguez, Margarita Matias-Florentino, Fidencio Mejia-Nepomuceno, Maria Guadalupe de Jesus Mireles-Rivera, Gloria Maria Molina-Salinas, Hector Montoya-Fuentes, , Mario Mujica-Sanchez, Jose Esteban Muñoz-Medina, Jose de Jesus Nuñez-Contreras, Alicia Ocaña-Mondragon, Luis Alberto Ochoa-Carrera, Hector Esteban Paz-Juarez, Marissa |

| | | | |
|---|---|---|---|
| | | | Perez-Garcia, Francisco Pulido, Helen Haydee Fernanda Ramirez-Plascencia, Jorge Salas-Hernandez, Angel Gustavo Salas-Lais, Alejandro Sanchez-Flores, Clara Esperanza Santacruz-Tinoco, Maria Guadalupe Santiago-Mauricio, Selene Zarate, Nelly Selem-Mojica, Blanca Taboada, Gloria Vazquez, Joel Armando Vazquez-Perez. |
| EPI_ISL_2943074 | National Institute of Infectious Diseases-Prof. Dr. Matei Bals Molecular Diagnostics Laboratory | National Institute of Infectious Diseases-Prof. Dr. Matei Bals Molecular Diagnostics Laboratory | Corina Casangiu, Leontina Banica, Marius Surleac, Petre Milu, Robert Hohan, Simona Paraschiv, Dan Otelea |
| EPI_ISL_2943173 | RSUP Dr. Sardjito | Genetics Working Group (Pokja Genetik) Faculty of Medicine, Public Health and Nursing Universitas Gadjah Mada (FK-KMK UGM); Disease Investigation Center Wates Ministry of Agriculture Indonesia; Department of Microbiology FK-KMK UGM; Laboratorium Diagnostik Yayasan Tahija World Mosquito Program | Gunadi, Hendra Wibawa, . Marcellus, Mohamad S. Hakim, Edwin W. Daniwijaya, Ludhang P. Rizki, Endah Supriyati, Eggi Arguni, Titik Nuryastuti, Tri Wibawa, Dwi AA Nugrahaningsih, Afiahayati, Siswanto, Kristy Iskandar, Nungki Anggorowati, Ika Trisnawati, Sumardi, Eko Budiono, Bambang Sigit Riyanto, Heni Retnowulan, Munawar Gani, Satria Maulana, Nur Rahmi Ananda, Riat El Khair, Yunika Puspadewi, Osman Sianipar, Umi Solekhah Intansari, Elizabeth Henny Herningtiyas, Ira Puspitawati, Nur Imma Fatimah Harahap, Irene Tania, Hana F Hanifin |
| EPI_ISL_2943231 | CENTRO MEDICO NAVAL SEMAR LAB BIOLOGIA MOLECULAR | CENTRO MEDICO NAVAL SEMAR LAB PATOLOGIA MOLECULAR | MEIXUEIRO CALDERON CLAUDIA. MORALES RAMIREZ OMAR. BALTIERRA URIBE SHANTAL. ZUNO CORONADO TEUTLI. GARCIA ALEGRIA IRMA.ALVAREZ JIMENEZ VIOLETA DEJANIRA. LOPEZ BUSTOS ABRAHAM. CABRERA GONZALEZ MOISES. ROMERO ROMAN EDGAR. LIEVANO ESQUINCA JAZMIN. FIGUEROA ANTONIO VICTOR. DIAZ PONCE MEDRANO JUAN ALBERTO. |
| EPI_ISL_2943232 | RS Panti Rapih | Genetics Working Group (Pokja Genetik) Faculty of Medicine, Public Health and Nursing Universitas Gadjah Mada (FK-KMK UGM); Disease Investigation Center Wates Ministry of Agriculture Indonesia; Department of Microbiology FK-KMK UGM; Laboratorium Diagnostik Yayasan Tahija World Mosquito Program (WMP) Yogyakarta Center for Tropical Medicine FK-KMK UGM; Integrated Research Center FK-KMK UGM; Department of Computer Science and Electronics FMIPA UGM; RSUP Dr. Sardjito | Gunadi, Hendra Wibawa, Marcellus, Mohamad S. Hakim, Edwin W. Daniwijaya, Ludhang P. Rizki, Endah Supriyati, Eggi Arguni, Titik Nuryastuti, Tri Wibawa, Dwi AA Nugrahaningsih, Afiahayati, Siswanto,Kristy Iskandar,Nungki Anggorowati, Pramesti G Dewi, Dwi Indaryati,Indarto Sulistiyono, Cita S Amalia, Alvina A Setiawan |
| EPI_ISL_2943233 | CENTRO MEDICO NAVAL SEMAR LAB BIOLOGIA MOLECULAR | CENTRO MEDICO NAVAL SEMAR LABORATORIO DE PATOLOGIA MOLECULAR | MEIXUEIRO CALDERON CLAUDIA. MORALES RAMIREZ OMAR. BALTIERRA URIBE SHANTAL. ZUNO CORONADO TEUTLI. GARCIA ALEGRIA IRMA.ALVAREZ JIMENEZ VIOLETA DEJANIRA. LOPEZ BUSTOS ABRAHAM. CABRERA GONZALEZ MOISES. ROMERO ROMAN EDGAR. LIEVANO ESQUINCA JAZMIN. FIGUEROA ANTONIO VICTOR. DIAZ PONCE MEDRANO JUAN ALBERTO. |
| EPI_ISL_2943235 | BBTKLPP Yogyakarta | Genetics Working Group (Pokja Genetik) Faculty of Medicine, Public Health and Nursing Universitas Gadjah Mada (FK-KMK UGM); Disease Investigation Center Wates Ministry of Agriculture Indonesia; Department of Microbiology FK-KMK UGM; Laboratorium Diagnostik Yayasan Tahija World Mosquito Program (WMP) Yogyakarta Center for Tropical Medicine FK-KMK UGM; Integrated Research Center FK-KMK UGM; Department of Computer Science and Electronics FMIPA UGM; RSUP Dr. Sardjito | Gunadi, Hendra Wibawa, Marcellus, Mohamad S. Hakim, Edwin W. Daniwijaya, Ludhang P. Rizki, Endah Supriyati, Eggi Arguni, Titik Nuryastuti, Tri Wibawa, Dwi AA Nugrahaningsih, Afiahayati, Siswanto,Kristy Iskandar,Nungki Anggorowati, Pramesti G Dewi, Dwi Indaryati,Indarto Sulistiyono, Irene Tania, Hana F Hanifin |
| EPI_ISL_2943237 | DKK Sleman/Puskesmas Depok 1 | Genetics Working Group (Pokja Genetik) Faculty of Medicine, Public Health and Nursing Universitas Gadjah Mada (FK-KMK UGM); Disease Investigation Center Wates Ministry of Agriculture Indonesia; Department of Microbiology FK-KMK UGM; Laboratorium Diagnostik Yayasan Tahija World Mosquito Program (WMP) Yogyakarta Center for Tropical Medicine FK-KMK UGM; Integrated Research Center FK-KMK UGM; Department of Computer Science and Electronics FMIPA UGM; RSUP Dr. Sardjito | Gunadi, Hendra Wibawa, Marcellus, Mohamad S. Hakim, Edwin W. Daniwijaya, Ludhang P. Rizki, Endah Supriyati, Eggi Arguni, Titik Nuryastuti, Tri Wibawa, Dwi AA Nugrahaningsih, Afiahayati, Siswanto,Kristy Iskandar,Nungki Anggorowati, Pramesti G Dewi, Dwi Indaryati,Indarto Sulistiyono, Cita S Amalia, Alvina A Setiawan |
| EPI_ISL_2951626 | DKK Sleman/Puskesmas Depok 1 | Genetics Working Group (Pokja Genetik) Faculty of Medicine, Public Health and Nursing Universitas Gadjah Mada (FK-KMK UGM); Disease Investigation Center Wates Ministry of Agriculture Indonesia; Department of Microbiology FK-KMK UGM; Laboratorium Diagnostik Yayasan Tahija World Mosquito Program (WMP) Yogyakarta Center for Tropical Medicine FK-KMK UGM; Integrated Research Center FK-KMK UGM; Department of Computer Science and Electronics FMIPA UGM; RSUP Dr. | Gunadi, Hendra Wibawa, Marcellus, Mohamad S. Hakim, Edwin W. Daniwijaya, Ludhang P. Rizki, Endah Supriyati, Eggi Arguni, Titik Nuryastuti, Tri Wibawa, Dwi AA Nugrahaningsih, Afiahayati, Siswanto,Kristy Iskandar,Nungki Anggorowati, Pramesti G Dewi, Dwi Indaryati,Indarto Sulistiyono, Irene Tania, Hana F Hanifin |
| EPI_ISL_2952510 | DKK Sleman/Puskesmas Depok 1 | Genetics Working Group (Pokja Genetik) Faculty of Medicine, Public Health and Nursing Universitas Gadjah Mada (FK-KMK UGM); Disease Investigation Center Wates Ministry of Agriculture Indonesia; Department of Microbiology FK-KMK UGM; Laboratorium Diagnostik Yayasan Tahija World Mosquito Program (WMP) Yogyakarta Center for Tropical Medicine FK-KMK UGM; Integrated Research Center FK-KMK UGM; Department of Computer Science and Electronics FMIPA UGM; RSUP Dr. Sardjito | Gunadi, Hendra Wibawa, Marcellus, Mohamad S. Hakim, Edwin W. Daniwijaya, Ludhang P. Rizki, Endah Supriyati, Eggi Arguni, Titik Nuryastuti, Tri Wibawa, Dwi AA Nugrahaningsih, Afiahayati, Siswanto,Kristy Iskandar,Nungki Anggorowati, Pramesti G Dewi, Dwi Indaryati,Indarto Sulistiyono, Susan Simanjaya, Dyah A Puspitarani |
| EPI_ISL_2953393 | DKK Sleman/Puskesmas Depok 1 | Genetics Working Group (Pokja Genetik) Faculty of Medicine, Public Health and Nursing Universitas Gadjah Mada (FK-KMK UGM); Disease Investigation Center Wates Ministry of Agriculture Indonesia; Department of Microbiology FK-KMK UGM; Laboratorium Diagnostik Yayasan Tahija World Mosquito Program (WMP) Yogyakarta Center for Tropical Medicine FK-KMK UGM; Integrated Research Center FK-KMK UGM; Department of Computer Science and Electronics FMIPA UGM; RSUP Dr. Sardjito | Gunadi, Hendra Wibawa, Marcellus, Mohamad S. Hakim, Edwin W. Daniwijaya, Ludhang P. Rizki, Endah Supriyati, Eggi Arguni, Titik Nuryastuti, Tri Wibawa, Dwi AA Nugrahaningsih, Afiahayati, Siswanto,Kristy Iskandar,Nungki Anggorowati, Pramesti G Dewi, Dwi Indaryati,Indarto Sulistiyono, Khanza a Vujira, Alvina A Setiawan |
| EPI_ISL_2954064 | RSUD Panembahan Senopati, Bantul | Genetics Working Group (Pokja Genetik) Faculty of Medicine, Public Health and Nursing Universitas Gadjah Mada (FK-KMK UGM); Disease Investigation Center Wates Ministry of Agriculture Indonesia; Department of Microbiology FK-KMK UGM; Laboratorium Diagnostik Yayasan Tahija World Mosquito Program (WMP) Yogyakarta Center for Tropical Medicine FK-KMK UGM; Integrated Research Center FK-KMK UGM; Department of Computer | Gunadi, Hendra Wibawa, Marcellus, Mohamad S. Hakim, Edwin W. Daniwijaya, Ludhang P. Rizki, Endah Supriyati, Eggi Arguni, Titik Nuryastuti, Tri Wibawa, Dwi AA Nugrahaningsih, Afiahayati, Siswanto,Kristy Iskandar,Nungki Anggorowati, Pramesti G Dewi, Dwi Indaryati,Indarto Sulistiyono, Irene Tania, Hana F Hanifin |
| EPI_ISL_2955330 | Labkesda Bantul | Genetics Working Group (Pokja Genetik) Faculty of Medicine, Public Health and Nursing Universitas Gadjah Mada (FK-KMK | Gunadi, Hendra Wibawa, Marcellus, Mohamad S. Hakim, Edwin W. Daniwijaya, Ludhang P. Rizki, Endah Supriyati, Eggi Arguni, Titik Nuryastuti, Tri Wibawa, Dwi AA Nugrahaningsih, Afiahayati, Siswanto,Kristy Iskandar,Nungki Anggorowati, Pramesti G Dewi, Dwi Indaryati,Indarto Sulistiyono, Cita S |

| | | | |
|---|---|---|---|
| | | UGM); Disease Investigation Center Wates Ministry of Agriculture Indonesia; Department of Microbiology FK-KMK UGM; Laboratorium Diagnostik Yayasan Tahija World Mosquito Program (WMP) Yogyakarta Center for Tropical Medicine FK-KMK UGM; Integrated Research Center FK-KMK UGM; Department of Computer Science and Electronics FMIPA UGM; RSUP Dr. Sardjito | Amalia, Alvina A Setiawan |
| EPI_ISL_2955335 | DKK Gunungkidul/ Puskesmas Karangmojo I | Genetics Working Group (Pokja Genetik) Faculty of Medicine, Public Health and Nursing Universitas Gadjah Mada (FK-KMK UGM); Disease Investigation Center Wates Ministry of Agriculture Indonesia; Department of Microbiology FK-KMK UGM; Laboratorium Diagnostik Yayasan Tahija World Mosquito Program (WMP) Yogyakarta Center for Tropical Medicine FK-KMK UGM; Integrated Research Center FK-KMK UGM; Department of Computer Science and Electronics FMIPA UGM; RSUP Dr. Sardjito | Gunadi, Hendra Wibawa, Marcellus, Mohamad S. Hakim, Edwin W. Daniwijaya, Ludhang P. Rizki, Endah Supriyati, Eggi Arguni, Titik Nuryastuti, Tri Wibawa, Dwi AA Nugrahaningsih, Afiahayati, Siswanto,Kristy Iskandar,Nungki Anggorowati, Pramesti G Dewi, Dwi Indaryati,Indarto Sulistiyono, Irene Tania, Hana F Hanifin |
| EPI_ISL_2955437 | ZARV/NHLS, Department Medical Virology, University of Pretoria | KRISP, KZn Research Innovation and Sequencing Platform | Amy Strydom, Adriano Mendes, Micheala Davids, Sim Mayaphi and Marietjie Venter, Giandhari J, Pillay S, Naidoo Y, Ramphal U, Lessells R, Emmanuel SJ, Tegally H, Wilkinson E, de Oliveira T |
| EPI_ISL_2955597, EPI_ISL_2955598 | DKK Gunungkidul/ Puskesmas Karangmojo I | Genetics Working Group (Pokja Genetik) Faculty of Medicine, Public Health and Nursing Universitas Gadjah Mada (FK-KMK UGM); Disease Investigation Center Wates Ministry of Agriculture Indonesia; Department of Microbiology FK-KMK UGM; Laboratorium Diagnostik Yayasan Tahija World Mosquito Program (WMP) Yogyakarta Center for Tropical Medicine FK-KMK UGM; Integrated Research Center FK-KMK UGM; Department of Computer Science and Electronics FMIPA UGM; RSUP Dr. Sardjito | Gunadi, Hendra Wibawa, Marcellus, Mohamad S. Hakim, Edwin W. Daniwijaya, Ludhang P. Rizki, Endah Supriyati, Eggi Arguni, Titik Nuryastuti, Tri Wibawa, Dwi AA Nugrahaningsih, Afiahayati, Siswanto,Kristy Iskandar,Nungki Anggorowati, Pramesti G Dewi, Dwi Indaryati,Indarto Sulistiyono, Khanza a Vujira, Alvina A Setiawan |
| EPI_ISL_2955605 | COVID LAB | Cayman Islands Forensic Science Lab | Jonathan Smellie, Karla Montes, Brittany Balcewich, Allen Grolla and Tanisha Gilbert |
| EPI_ISL_2955606 | RSUP Sardjito | Genetics Working Group (Pokja Genetik) Faculty of Medicine, Public Health and Nursing Universitas Gadjah Mada (FK-KMK UGM); Disease Investigation Center Wates Ministry of Agriculture Indonesia; Department of Microbiology FK-KMK UGM; Laboratorium Diagnostik Yayasan Tahija World Mosquito Program (WMP) Yogyakarta Center for Tropical Medicine FK-KMK UGM; Integrated Research Center FK-KMK UGM; Department of Computer Science and Electronics FMIPA UGM; RSUP Dr. Sardjito | Gunadi, Hendra Wibawa, . Marcellus, Mohamad S. Hakim, Edwin W. Daniwijaya, Ludhang P. Rizki, Endah Supriyati, Eggi Arguni, Titik Nuryastuti, Tri Wibawa, Dwi AA Nugrahaningsih, Afiahayati, Siswanto, Kristy Iskandar, Nungki Anggorowati, Ika Trisnawati, Sumardi, Eko Budiono, Bambang Sigit Riyanto, Heni Retnowulan, Munawar Gani, Satria Maulana, Nur Rahmi Ananda, Riat El Khair, Yunika Puspadewi, Osman Sianipar, Umi Solekhah Intansari, Elizabeth Henny Herningtiyas, Ira Puspitawati, Nur Imma Fatimah Harahap, Khanza a Vujira, Alvina A Setiawan |
| EPI_ISL_2955608 | RSA UGM | Genetics Working Group (Pokja Genetik) Faculty of Medicine, Public Health and Nursing Universitas Gadjah Mada (FK-KMK UGM); Disease Investigation Center Wates Ministry of Agriculture Indonesia; Department of Microbiology FK-KMK UGM; Laboratorium Diagnostik Yayasan Tahija World Mosquito Program (WMP) Yogyakarta Center for Tropical Medicine FK-KMK UGM; Integrated Research Center FK-KMK UGM; Department of Computer Science and Electronics FMIPA UGM; RSUP Dr. Sardjito | Gunadi, Hendra Wibawa, Marcellus, Mohamad S. Hakim, Edwin W. Daniwijaya, Ludhang P. Rizki, Endah Supriyati, Eggi Arguni, Titik Nuryastuti, Tri Wibawa, Dwi AA Nugrahaningsih, Afiahayati, Siswanto,Kristy Iskandar,Nungki Anggorowati, Pramesti G Dewi, Dwi Indaryati,Indarto Sulistiyono, Susan Simanjaya, Dyah A Puspitarani |
| EPI_ISL_2955609 | DKK Sleman/Puskesmas Depok 1 | Genetics Working Group (Pokja Genetik) Faculty of Medicine, Public Health and Nursing Universitas Gadjah Mada (FK-KMK UGM); Disease Investigation Center Wates Ministry of Agriculture Indonesia; Department of Microbiology FK-KMK UGM; Laboratorium Diagnostik Yayasan Tahija World Mosquito Program (WMP) Yogyakarta Center for Tropical Medicine FK-KMK UGM; Integrated Research Center FK-KMK UGM; Department of Computer Science and Electronics FMIPA UGM; RSUP Dr. Sardjito | Gunadi, Hendra Wibawa, Marcellus, Mohamad S. Hakim, Edwin W. Daniwijaya, Ludhang P. Rizki, Endah Supriyati, Eggi Arguni, Titik Nuryastuti, Tri Wibawa, Dwi AA Nugrahaningsih, Afiahayati, Siswanto,Kristy Iskandar,Nungki Anggorowati, Pramesti G Dewi, Dwi Indaryati,Indarto Sulistiyono, Cita S Amalia, Alvina A Setiawan |
| EPI_ISL_2955627 | DKK Sleman/Puskesmas Ngemplak 2 | Genetics Working Group (Pokja Genetik) Faculty of Medicine, Public Health and Nursing Universitas Gadjah Mada (FK-KMK UGM); Disease Investigation Center Wates Ministry of Agriculture Indonesia; Department of Microbiology FK-KMK UGM; Laboratorium Diagnostik Yayasan Tahija World Mosquito Program (WMP) Yogyakarta Center for Tropical Medicine FK-KMK UGM; Integrated Research Center FK-KMK UGM; Department of Computer Science and Electronics FMIPA UGM; RSUP Dr. Sardjito | Gunadi, Hendra Wibawa, Marcellus, Mohamad S. Hakim, Edwin W. Daniwijaya, Ludhang P. Rizki, Endah Supriyati, Eggi Arguni, Titik Nuryastuti, Tri Wibawa, Dwi AA Nugrahaningsih, Afiahayati, Siswanto,Kristy Iskandar,Nungki Anggorowati, Pramesti G Dewi, Dwi Indaryati,Indarto Sulistiyono, Irene Tania, Hana F Hanifin |
| EPI_ISL_2955628 | DKK Sleman/Puskesmas Ngemplak 2 | Genetics Working Group (Pokja Genetik) Faculty of Medicine, Public Health and Nursing Universitas Gadjah Mada (FK-KMK UGM); Disease Investigation Center Wates Ministry of Agriculture Indonesia; Department of Microbiology FK-KMK UGM; Laboratorium Diagnostik Yayasan Tahija World Mosquito Program (WMP) Yogyakarta Center for Tropical Medicine FK-KMK UGM; Integrated Research Center FK-KMK UGM; Department of Computer Science and Electronics FMIPA UGM; RSUP Dr. Sardjito | Gunadi, Hendra Wibawa, Marcellus, Mohamad S. Hakim, Edwin W. Daniwijaya, Ludhang P. Rizki, Endah Supriyati, Eggi Arguni, Titik Nuryastuti, Tri Wibawa, Dwi AA Nugrahaningsih, Afiahayati, Siswanto,Kristy Iskandar,Nungki Anggorowati, Pramesti G Dewi, Dwi Indaryati,Indarto Sulistiyono, Susan Simanjaya, Dyah A Puspitarani |
| EPI_ISL_2958743, EPI_ISL_2958744, EPI_ISL_2958745, EPI_ISL_2958746, EPI_ISL_2958747, EPI_ISL_2958748, EPI_ISL_2958749, EPI_ISL_2958750, EPI_ISL_2958751 | labo Analyses Med | National Reference Center for Viruses of Respiratory Infections, Institut Pasteur, Paris | Marion Barbet, Sylvie Behillil, Méline Bizard, Angela Brisebarre, Camille Capel, Vincent Enouf, Louise Lefrançois, Frédéric Lemoine, Christophe Malabat, Corinne Maufrais, Hub Bioinformatique Biostatistiques, Etienne Simon-Lorière, Maud Vanpeene, Sylvie Van der Werf ,GLAUDON LOUVEAU DE LA GUIGNERAYE Marie-Hélène |
| EPI_ISL_2958752, EPI_ISL_2958753, EPI_ISL_2958754, EPI_ISL_2958755, EPI_ISL_2958756 | labo Analyses Med | National Reference Center for Viruses of Respiratory Infections, Institut Pasteur, Paris | Marion Barbet, Sylvie Behillil, Méline Bizard, Angela Brisebarre, Camille Capel, Vincent Enouf, Louise Lefrançois, Frédéric Lemoine, Christophe Malabat, Corinne Maufrais, Hub Bioinformatique Biostatistiques, Etienne Simon-Lorière, Maud Vanpeene, Sylvie Van der Werf ,LEONARD Pierre-Yves |

| | | | |
|---|---|---|---|
| EPI_ISL_2958757 | labo Analyses Med | National Reference Center for Viruses of Respiratory Infections, Institut Pasteur, Paris | Marion Barbet, Sylvie Behillil, Méline Bizard, Angela Brisebarre, Camille Capel, Vincent Enouf, Louise Lefrançois, Frédéric Lemoine, Christophe Malabat, Corinne Maufrais, Victoire Baillet, Etienne Simon-Lorière, Maud Vanpeene, Sylvie Van der Werf ,LEONARD Pierre-Yves |
| EPI_ISL_2958758, EPI_ISL_2958759, EPI_ISL_2958760 | labo Analyses Med | National Reference Center for Viruses of Respiratory Infections, Institut Pasteur, Paris | Marion Barbet, Sylvie Behillil, Méline Bizard, Angela Brisebarre, Camille Capel, Vincent Enouf, Louise Lefrançois, Frédéric Lemoine, Christophe Malabat, Corinne Maufrais, Hub Bioinformatique Biostatistiques, Etienne Simon-Lorière, Maud Vanpeene, Sylvie Van der Werf ,LEONARD Pierre-Yves |
| EPI_ISL_2958761, EPI_ISL_2958762 | Labo Analyses Med | National Reference Center for Viruses of Respiratory Infections, Institut Pasteur, Paris | Marion Barbet, Sylvie Behillil, Méline Bizard, Angela Brisebarre, Camille Capel, Vincent Enouf, Louise Lefrançois, Frédéric Lemoine, Christophe Malabat, Corinne Maufrais, Victoire Baillet, Etienne Simon-Lorière, Maud Vanpeene, Sylvie Van der Werf ,POTIRON Grégoire |
| EPI_ISL_2958763 | Labo Analyses Med | National Reference Center for Viruses of Respiratory Infections, Institut Pasteur, Paris | Marion Barbet, Sylvie Behillil, Méline Bizard, Angela Brisebarre, Camille Capel, Vincent Enouf, Louise Lefrançois, Frédéric Lemoine, Christophe Malabat, Corinne Maufrais, Hub Bioinformatique Biostatistiques, Etienne Simon-Lorière, Maud Vanpeene, Sylvie Van der Werf ,POTIRON Grégoire |
| EPI_ISL_2958764 | Labo Analyses Med | National Reference Center for Viruses of Respiratory Infections, Institut Pasteur, Paris | Marion Barbet, Sylvie Behillil, Méline Bizard, Angela Brisebarre, Camille Capel, Vincent Enouf, Louise Lefrançois, Frédéric Lemoine, Christophe Malabat, Corinne Maufrais, Hub Bioinformatique Biostatistiques, Etienne Simon-Lorière, Maud Vanpeene, Sylvie Van der Werf ,HOLSTEIN Anne |
| EPI_ISL_2958765 | Hospital | National Reference Center for Viruses of Respiratory Infections, Institut Pasteur, Paris | Marion Barbet, Sylvie Behillil, Méline Bizard, Angela Brisebarre, Camille Capel, Vincent Enouf, Louise Lefrançois, Frédéric Lemoine, Christophe Malabat, Corinne Maufrais, Victoire Baillet, Etienne Simon-Lorière, Maud Vanpeene, Sylvie Van der Werf ,FARRUGIA Cécile |
| EPI_ISL_2958766 | Hospital | National Reference Center for Viruses of Respiratory Infections, Institut Pasteur, Paris | Marion Barbet, Sylvie Behillil, Méline Bizard, Angela Brisebarre, Camille Capel, Vincent Enouf, Louise Lefrançois, Frédéric Lemoine, Christophe Malabat, Corinne Maufrais, Hub Bioinformatique Biostatistiques, Etienne Simon-Lorière, Maud Vanpeene, Sylvie Van der Werf ,FARRUGIA Cécile |
| EPI_ISL_2958767 | Hospital | National Reference Center for Viruses of Respiratory Infections, Institut Pasteur, Paris | Marion Barbet, Sylvie Behillil, Méline Bizard, Angela Brisebarre, Camille Capel, Vincent Enouf, Louise Lefrançois, Frédéric Lemoine, Christophe Malabat, Corinne Maufrais, Victoire Baillet, Etienne Simon-Lorière, Maud Vanpeene, Sylvie Van der Werf ,PILORGE Léa |
| EPI_ISL_2958768 | Hospital | National Reference Center for Viruses of Respiratory Infections, Institut Pasteur, Paris | Marion Barbet, Sylvie Behillil, Méline Bizard, Angela Brisebarre, Camille Capel, Vincent Enouf, Louise Lefrançois, Frédéric Lemoine, Christophe Malabat, Corinne Maufrais, Hub Bioinformatique Biostatistiques, Etienne Simon-Lorière, Maud Vanpeene, Sylvie Van der Werf ,PILORGE Léa |
| EPI_ISL_2958769 | Hospital | National Reference Center for Viruses of Respiratory Infections, Institut Pasteur, Paris | Marion Barbet, Sylvie Behillil, Méline Bizard, Angela Brisebarre, Camille Capel, Vincent Enouf, Louise Lefrançois, Frédéric Lemoine, Christophe Malabat, Corinne Maufrais, Hub Bioinformatique Biostatistiques, Etienne Simon-Lorière, Maud Vanpeene, Sylvie Van der Werf ,BURGARD Marianne |
| EPI_ISL_2958770, EPI_ISL_2958771, EPI_ISL_2958772 | labo Analyses Med | National Reference Center for Viruses of Respiratory Infections, Institut Pasteur, Paris | Marion Barbet, Sylvie Behillil, Méline Bizard, Angela Brisebarre, Camille Capel, Vincent Enouf, Louise Lefrançois, Frédéric Lemoine, Christophe Malabat, Corinne Maufrais, Hub Bioinformatique Biostatistiques, Etienne Simon-Lorière, Maud Vanpeene, Sylvie Van der Werf ,GUSSOND Thierry |
| EPI_ISL_2958773 | labo Analyses Med | National Reference Center for Viruses of Respiratory Infections, Institut Pasteur, Paris | Marion Barbet, Sylvie Behillil, Méline Bizard, Angela Brisebarre, Camille Capel, Vincent Enouf, Louise Lefrançois, Frédéric Lemoine, Christophe Malabat, Corinne Maufrais, Victoire Baillet, Etienne Simon-Lorière, Maud Vanpeene, Sylvie Van der Werf ,GUSSOND Thierry |
| EPI_ISL_2958774, EPI_ISL_2958775 | labo Analyses Med | National Reference Center for Viruses of Respiratory Infections, Institut Pasteur, Paris | Marion Barbet, Sylvie Behillil, Méline Bizard, Angela Brisebarre, Camille Capel, Vincent Enouf, Louise Lefrançois, Frédéric Lemoine, Christophe Malabat, Corinne Maufrais, Hub Bioinformatique Biostatistiques, Etienne Simon-Lorière, Maud Vanpeene, Sylvie Van der Werf ,GUSSOND Thierry |
| EPI_ISL_2958776 | Labo Analyses Med | National Reference Center for Viruses of Respiratory Infections, Institut Pasteur, Paris | Marion Barbet, Sylvie Behillil, Méline Bizard, Angela Brisebarre, Camille Capel, Vincent Enouf, Louise Lefrançois, Frédéric Lemoine, Christophe Malabat, Corinne Maufrais, Hub Bioinformatique Biostatistiques, Etienne Simon-Lorière, Maud Vanpeene, Sylvie Van der Werf ,ABS Gilles |
| EPI_ISL_2958777 | Labo Analyses Med | National Reference Center for Viruses of Respiratory Infections, Institut Pasteur, Paris | Marion Barbet, Sylvie Behillil, Méline Bizard, Angela Brisebarre, Camille Capel, Vincent Enouf, Louise Lefrançois, Frédéric Lemoine, Christophe Malabat, Corinne Maufrais, Emmanuelle Permal, Etienne Simon-Lorière, Maud Vanpeene, Sylvie Van der Werf ,GOUBARD Agathe |
| EPI_ISL_2958778 | Labo Analyses Med | National Reference Center for Viruses of Respiratory Infections, Institut Pasteur, Paris | Marion Barbet, Sylvie Behillil, Méline Bizard, Angela Brisebarre, Camille Capel, Vincent Enouf, Louise Lefrançois, Frédéric Lemoine, Christophe Malabat, Corinne Maufrais, Hub Bioinformatique Biostatistiques, Etienne Simon-Lorière, Maud Vanpeene, Sylvie Van der Werf ,BLANC-GALIBY M.-J. |
| EPI_ISL_2958779 | labo Analyses Med | National Reference Center for Viruses of Respiratory Infections, Institut Pasteur, Paris | Marion Barbet, Sylvie Behillil, Méline Bizard, Angela Brisebarre, Camille Capel, Vincent Enouf, Louise Lefrançois, Frédéric Lemoine, Christophe Malabat, Corinne Maufrais, Hub Bioinformatique Biostatistiques, Etienne Simon-Lorière, Maud Vanpeene, Sylvie Van der Werf ,LEONARD Pierre-Yves |
| EPI_ISL_2958781, EPI_ISL_2958782, EPI_ISL_2958783 | Labo Analyses Med | National Reference Center for Viruses of Respiratory Infections, Institut Pasteur, Paris | Marion Barbet, Sylvie Behillil, Méline Bizard, Angela Brisebarre, Camille Capel, Vincent Enouf, Louise Lefrançois, Frédéric Lemoine, Christophe Malabat, Corinne Maufrais, Hub Bioinformatique Biostatistiques, Etienne Simon-Lorière, Maud Vanpeene, Sylvie Van der Werf ,LEONARD Pierre-Yves |
| EPI_ISL_2958784, EPI_ISL_2958785 | Labo Analyses Med | National Reference Center for Viruses of Respiratory Infections, Institut Pasteur, Paris | Marion Barbet, Sylvie Behillil, Méline Bizard, Angela Brisebarre, Camille Capel, Vincent Enouf, Louise Lefrançois, Frédéric Lemoine, Christophe Malabat, Corinne Maufrais, Hub Bioinformatique Biostatistiques, Etienne Simon-Lorière, Maud Vanpeene, Sylvie Van der Werf |
| EPI_ISL_2958786 | Labo Analyses Med | National Reference Center for Viruses of Respiratory Infections, Institut Pasteur, Paris | Marion Barbet, Sylvie Behillil, Méline Bizard, Angela Brisebarre, Camille Capel, Vincent Enouf, Louise Lefrançois, Frédéric Lemoine, Christophe Malabat, Corinne Maufrais, Victoire Baillet, Etienne Simon-Lorière, Maud Vanpeene, Sylvie Van der Werf |
| EPI_ISL_2958787 | Labo Analyses Med | National Reference Center for Viruses of Respiratory Infections, Institut Pasteur, Paris | Marion Barbet, Sylvie Behillil, Méline Bizard, Angela Brisebarre, Camille Capel, Vincent Enouf, Louise Lefrançois, Frédéric Lemoine, Christophe Malabat, Corinne Maufrais, Hub Bioinformatique Biostatistiques, Etienne Simon-Lorière, Maud Vanpeene, Sylvie Van der Werf ,TROMBERT Sabine |
| EPI_ISL_2958788, EPI_ISL_2958789 | Labo Analyses Med | National Reference Center for Viruses of Respiratory Infections, Institut Pasteur, Paris | Marion Barbet, Sylvie Behillil, Méline Bizard, Angela Brisebarre, Camille Capel, Vincent Enouf, Louise Lefrançois, Frédéric Lemoine, Christophe Malabat, Corinne Maufrais, Hub Bioinformatique Biostatistiques, Etienne Simon-Lorière, Maud Vanpeene, Sylvie Van der Werf ,GLAUDON LOUVEAU DE LA GUIGNERAYE Marie-Hélène |
| EPI_ISL_2958791 | Labo Analyses Med | National Reference Center for Viruses of Respiratory Infections, Institut Pasteur, Paris | Marion Barbet, Sylvie Behillil, Méline Bizard, Angela Brisebarre, Camille Capel, Vincent Enouf, Louise Lefrançois, Frédéric Lemoine, Christophe Malabat, Corinne Maufrais, Hub Bioinformatique Biostatistiques, Etienne Simon-Lorière, Maud Vanpeene, Sylvie Van der Werf ,POTIRON Grégoire |
| EPI_ISL_2958792, EPI_ISL_2958793 | Labo Analyses Med | National Reference Center for Viruses of Respiratory Infections, Institut Pasteur, Paris | Marion Barbet, Sylvie Behillil, Méline Bizard, Angela Brisebarre, Camille Capel, Vincent Enouf, Louise Lefrançois, Frédéric Lemoine, Christophe Malabat, Corinne Maufrais, Victoire Baillet, Etienne Simon-Lorière, Maud Vanpeene, Sylvie Van der Werf ,POTIRON Grégoire |
| EPI_ISL_2958794, EPI_ISL_2958795, EPI_ISL_2958796, EPI_ISL_2958797, EPI_ISL_2958798, EPI_ISL_2958799 | Labo Analyses Med | National Reference Center for Viruses of Respiratory Infections, Institut Pasteur, Paris | Marion Barbet, Sylvie Behillil, Méline Bizard, Angela Brisebarre, Camille Capel, Vincent Enouf, Louise Lefrançois, Frédéric Lemoine, Christophe Malabat, Corinne Maufrais, Hub Bioinformatique Biostatistiques, Etienne Simon-Lorière, Maud Vanpeene, Sylvie Van der Werf ,NESTOUR François |
| EPI_ISL_2958800 | Labo Analyses Med | National Reference Center for Viruses of Respiratory Infections, Institut Pasteur, Paris | Marion Barbet, Sylvie Behillil, Méline Bizard, Angela Brisebarre, Camille Capel, Vincent Enouf, Louise Lefrançois, Frédéric Lemoine, Christophe Malabat, Corinne Maufrais, Victoire Baillet, Etienne Simon-Lorière, Maud Vanpeene, Sylvie Van der Werf ,NESTOUR François |
| EPI_ISL_2958801, EPI_ISL_2958802, EPI_ISL_2958804, EPI_ISL_2958805, EPI_ISL_2958806 | Labo Analyses Med | National Reference Center for Viruses of Respiratory Infections, Institut Pasteur, Paris | Marion Barbet, Sylvie Behillil, Méline Bizard, Angela Brisebarre, Camille Capel, Vincent Enouf, Louise Lefrançois, Frédéric Lemoine, Christophe Malabat, Corinne Maufrais, Hub Bioinformatique Biostatistiques, Etienne Simon-Lorière, Maud Vanpeene, Sylvie Van der Werf ,NESTOUR François |
| EPI_ISL_2958807 | Hospital | National Reference Center for Viruses of Respiratory Infections, Institut Pasteur, Paris | Marion Barbet, Sylvie Behillil, Méline Bizard, Angela Brisebarre, Camille Capel, Vincent Enouf, Louise Lefrançois, Frédéric Lemoine, Christophe Malabat, Corinne Maufrais, Hub Bioinformatique Biostatistiques, Etienne Simon-Lorière, Maud Vanpeene, Sylvie Van der Werf ,GUINARD Jérome |
| EPI_ISL_2958808, EPI_ISL_2958809, EPI_ISL_2958810, EPI_ISL_2958811 | Hospital | National Reference Center for Viruses of Respiratory Infections, Institut Pasteur, Paris | Marion Barbet, Sylvie Behillil, Méline Bizard, Angela Brisebarre, Camille Capel, Vincent Enouf, Louise Lefrançois, Frédéric Lemoine, Christophe Malabat, Corinne Maufrais, Hub Bioinformatique Biostatistiques, Etienne Simon-Lorière, Maud Vanpeene, Sylvie Van der Werf ,FAGOUR Laurence |
| EPI_ISL_2958812 | Hospital | National Reference Center for Viruses of Respiratory Infections, Institut Pasteur, Paris | Marion Barbet, Sylvie Behillil, Méline Bizard, Angela Brisebarre, Camille Capel, Vincent Enouf, Louise Lefrançois, Frédéric Lemoine, Christophe Malabat, Corinne Maufrais, Victoire Baillet, Etienne Simon-Lorière, Maud Vanpeene, Sylvie Van der Werf ,FAGOUR Laurence |
| EPI_ISL_2958813 | Labo Analyses Med | National Reference Center for Viruses of Respiratory Infections, Institut Pasteur, Paris | Marion Barbet, Sylvie Behillil, Méline Bizard, Angela Brisebarre, Camille Capel, Vincent Enouf, Louise Lefrançois, Frédéric Lemoine, Christophe Malabat, Corinne Maufrais, Hub Bioinformatique Biostatistiques, Etienne Simon-Lorière, Maud Vanpeene, Sylvie Van der Werf ,POTIRON Grégoire |
| EPI_ISL_2958814 | Labo Analyses Med | National Reference Center for Viruses of Respiratory Infections, Institut Pasteur, Paris | Marion Barbet, Sylvie Behillil, Méline Bizard, Angela Brisebarre, Camille Capel, Vincent Enouf, Louise Lefrançois, Frédéric Lemoine, Christophe Malabat, Corinne Maufrais, Hub Bioinformatique Biostatistiques, Etienne Simon-Lorière, Maud Vanpeene, Sylvie Van der Werf ,FAGOUR Laurence |
| EPI_ISL_2958815, EPI_ISL_2958816, EPI_ISL_2958817, EPI_ISL_2958818 | Labo Analyses Med | National Reference Center for Viruses of Respiratory Infections, Institut Pasteur, Paris | Marion Barbet, Sylvie Behillil, Méline Bizard, Angela Brisebarre, Camille Capel, Vincent Enouf, Louise Lefrançois, Frédéric Lemoine, Christophe Malabat, Corinne Maufrais, Hub Bioinformatique Biostatistiques, Etienne Simon-Lorière, Maud Vanpeene, Sylvie Van der Werf ,POTIRON Grégoire |
| EPI_ISL_2958819 | Labo Analyses Med | National Reference Center for Viruses of Respiratory Infections, Institut Pasteur, Paris | Marion Barbet, Sylvie Behillil, Méline Bizard, Angela Brisebarre, Camille Capel, Vincent Enouf, Louise Lefrançois, Frédéric Lemoine, Christophe Malabat, Corinne Maufrais, Victoire Baillet, Etienne Simon-Lorière, Maud Vanpeene, Sylvie Van der Werf ,POTIRON Grégoire |

| | | | |
|---|---|---|---|
| EPI_ISL_2958820 | Labo Analyses Med | National Reference Center for Viruses of Respiratory Infections, Institut Pasteur, Paris | Marion Barbet, Sylvie Behillil, Méline Bizard, Angela Brisebarre, Camille Capel, Vincent Enouf, Louise Lefrançois, Frédéric Lemoine, Christophe Malabat, Corinne Maufrais, Hub Bioinformatique Biostatistiques, Etienne Simon-Lorière, Maud Vanpeene, Sylvie Van der Werf ,POTIRON Grégoire |
| EPI_ISL_2958821 | Labo Analyses Med | National Reference Center for Viruses of Respiratory Infections, Institut Pasteur, Paris | Marion Barbet, Sylvie Behillil, Méline Bizard, Angela Brisebarre, Camille Capel, Vincent Enouf, Louise Lefrançois, Frédéric Lemoine, Christophe Malabat, Corinne Maufrais, Victoire Baillet, Etienne Simon-Lorière, Maud Vanpeene, Sylvie Van der Werf ,POTIRON Grégoire |
| EPI_ISL_2958822 | Labo Analyses Med | National Reference Center for Viruses of Respiratory Infections, Institut Pasteur, Paris | Marion Barbet, Sylvie Behillil, Méline Bizard, Angela Brisebarre, Camille Capel, Vincent Enouf, Louise Lefrançois, Frédéric Lemoine, Christophe Malabat, Corinne Maufrais, Hub Bioinformatique Biostatistiques, Etienne Simon-Lorière, Maud Vanpeene, Sylvie Van der Werf ,POTIRON Grégoire |
| EPI_ISL_2958988, EPI_ISL_2958991, EPI_ISL_2958994, EPI_ISL_2958995, EPI_ISL_2959000, EPI_ISL_2959001, EPI_ISL_2959011 | Microbiology Department. Complexo Hospitalario Universitario de Vigo | Microbiology Department. Complexo Hospitalario Universitario de Vigo | Alfaya N, Alonso I, Alvarez M, Cabrera JJ, Carballo R, Cores O, Cortizo S, del-Campo V, Martinez L, Mediero G, Perez S, Davina C, Potel C, Regueiro B, Rey S, Vasallo FJ |
| EPI_ISL_2964613, EPI_ISL_2964614, EPI_ISL_2964618, EPI_ISL_2964620, EPI_ISL_2964622, EPI_ISL_2964624 | Juntendo University Hospital | Juntendo University Hospital | Koji Tsuchiya, Yoshie Hosaka,Yoko Tabe |
| EPI_ISL_2964677 | RSUD DR. MOEWARDI Surakarta | Genetics Working Group (Pokja Genetik) Faculty of Medicine, Public Health and Nursing Universitas Gadjah Mada (FK-KMK UGM); Disease Investigation Center Wates Ministry of Agriculture Indonesia; Department of Microbiology FK-KMK UGM; Laboratorium Diagnostik Yayasan Tahija World Mosquito Program (WMP) Yogyakarta Center for Tropical Medicine FK-KMK UGM; Integrated Research Center FK-KMK UGM; Department of Computer Science and Electronics FMIPA UGM; RSUP Dr. Sardjito | Gunadi, Hendra Wibawa, Marcellus, Mohamad S. Hakim, Edwin W. Daniwijaya, Ludhang P. Rizki, Endah Supriyati, Eggi Arguni, Titik Nuryastuti, Tri Wibawa, Dwi AA Nugrahaningsih, Afiahayati, Siswanto,Kristy Iskandar,Nungki Anggorowati, Pramesti G Dewi, Dwi Indaryati,Indarto Sulistiyono, Khanza a Vujira, Alvina A Setiawan |
| EPI_ISL_2964681 | Puskesmas Colomadu 1 DKK Karanganyar | Genetics Working Group (Pokja Genetik) Faculty of Medicine, Public Health and Nursing Universitas Gadjah Mada (FK-KMK UGM); Disease Investigation Center Wates Ministry of Agriculture Indonesia; Department of Microbiology FK-KMK UGM; Laboratorium Diagnostik Yayasan Tahija World Mosquito Program (WMP) Yogyakarta Center for Tropical Medicine FK-KMK UGM; Integrated Research Center FK-KMK UGM; Department of Computer Science and Electronics FMIPA UGM; RSUP Dr. Sardjito | Gunadi, Hendra Wibawa, Marcellus, Mohamad S. Hakim, Edwin W. Daniwijaya, Ludhang P. Rizki, Endah Supriyati, Eggi Arguni, Titik Nuryastuti, Tri Wibawa, Dwi AA Nugrahaningsih, Afiahayati, Siswanto,Kristy Iskandar,Nungki Anggorowati, Pramesti G Dewi, Dwi Indaryati,Indarto Sulistiyono, Khanza a Vujira, Alvina A Setiawan |
| EPI_ISL_2964684 | RSUD DR. MOEWARDI | Genetics Working Group (Pokja Genetik) Faculty of Medicine, Public Health and Nursing Universitas Gadjah Mada (FK-KMK UGM); Disease Investigation Center Wates Ministry of Agriculture Indonesia; Department of Microbiology FK-KMK UGM; Laboratorium Diagnostik Yayasan Tahija World Mosquito Program (WMP) Yogyakarta Center for Tropical Medicine FK-KMK UGM; Integrated Research Center FK-KMK UGM; Department of Computer Science and Electronics FMIPA UGM; RSUP Dr. Sardjito | Gunadi, Hendra Wibawa, Marcellus, Mohamad S. Hakim, Edwin W. Daniwijaya, Ludhang P. Rizki, Endah Supriyati, Eggi Arguni, Titik Nuryastuti, Tri Wibawa, Dwi AA Nugrahaningsih, Afiahayati, Siswanto,Kristy Iskandar,Nungki Anggorowati, Pramesti G Dewi, Dwi Indaryati,Indarto Sulistiyono, Cita S Amalia, Alvina A Setiawan |
| EPI_ISL_2964685 | NILMRC | Genomic Research Laboratory, BSMMU Bangabandhu Sheikh Mujib Medical University (BSMMU) i | Laila Anjuman Banu, Md.Saydur Rahman, Sharadindu Kanti Sinha, Arifa Akram, Md.Sharfuddin Ahmed |
| EPI_ISL_2964688 | RSUD DR. MOEWARDI | Genetics Working Group (Pokja Genetik) Faculty of Medicine, Public Health and Nursing Universitas Gadjah Mada (FK-KMK UGM); Disease Investigation Center Wates Ministry of Agriculture Indonesia; Department of Microbiology FK-KMK UGM; Laboratorium Diagnostik Yayasan Tahija World Mosquito Program (WMP) Yogyakarta Center for Tropical Medicine FK-KMK UGM; Integrated Research Center FK-KMK UGM; Department of Computer Science and Electronics FMIPA UGM; RSUP Dr. Sardjito | Gunadi, Hendra Wibawa, Marcellus, Mohamad S. Hakim, Edwin W. Daniwijaya, Ludhang P. Rizki, Endah Supriyati, Eggi Arguni, Titik Nuryastuti, Tri Wibawa, Dwi AA Nugrahaningsih, Afiahayati, Siswanto,Kristy Iskandar,Nungki Anggorowati, Pramesti G Dewi, Dwi Indaryati,Indarto Sulistiyono, Irene Tania, Hana F Hanifin |
| EPI_ISL_2964692 | RSUD DR. MOEWARDI | Genetics Working Group (Pokja Genetik) Faculty of Medicine, Public Health and Nursing Universitas Gadjah Mada (FK-KMK UGM); Disease Investigation Center Wates Ministry of Agriculture Indonesia; Department of Microbiology FK-KMK UGM; Laboratorium Diagnostik Yayasan Tahija World Mosquito Program (WMP) Yogyakarta Center for Tropical Medicine FK-KMK UGM; Integrated Research Center FK-KMK UGM; Department of Computer Science and Electronics FMIPA UGM; RSUP Dr. Sardjito | Gunadi, Hendra Wibawa, Marcellus, Mohamad S. Hakim, Edwin W. Daniwijaya, Ludhang P. Rizki, Endah Supriyati, Eggi Arguni, Titik Nuryastuti, Tri Wibawa, Dwi AA Nugrahaningsih, Afiahayati, Siswanto,Kristy Iskandar,Nungki Anggorowati, Pramesti G Dewi, Dwi Indaryati,Indarto Sulistiyono, Susan Simanjaya, Dyah A Puspitarani |
| EPI_ISL_2964697 | DINKES KAB. JEPARA | Genetics Working Group (Pokja Genetik) Faculty of Medicine, Public Health and Nursing Universitas Gadjah Mada (FK-KMK UGM); Disease Investigation Center Wates Ministry of Agriculture Indonesia; Department of Microbiology FK-KMK UGM; Laboratorium Diagnostik Yayasan Tahija World Mosquito Program (WMP) Yogyakarta Center for Tropical Medicine FK-KMK UGM; Integrated Research Center FK-KMK UGM; Department of Computer Science and Electronics FMIPA UGM; RSUP Dr. Sardjito | Gunadi, Hendra Wibawa, Marcellus, Mohamad S. Hakim, Edwin W. Daniwijaya, Ludhang P. Rizki, Endah Supriyati, Eggi Arguni, Titik Nuryastuti, Tri Wibawa, Dwi AA Nugrahaningsih, Afiahayati, Siswanto,Kristy Iskandar,Nungki Anggorowati, Pramesti G Dewi, Dwi Indaryati,Indarto Sulistiyono, Cita S Amalia, Alvina A Setiawan |
| EPI_ISL_2964700 | DINKES KAB. JEPARA | Genetics Working Group (Pokja Genetik) Faculty of Medicine, Public Health and Nursing Universitas Gadjah Mada (FK-KMK UGM); Disease Investigation Center Wates Ministry of Agriculture Indonesia; Department of Microbiology FK-KMK UGM; Laboratorium Diagnostik Yayasan Tahija World Mosquito Program (WMP) Yogyakarta Center for Tropical Medicine FK-KMK UGM; Integrated Research Center FK-KMK UGM; Department of Computer Science and Electronics FMIPA UGM; RSUP Dr. Sardjito | Gunadi, Hendra Wibawa, Marcellus, Mohamad S. Hakim, Edwin W. Daniwijaya, Ludhang P. Rizki, Endah Supriyati, Eggi Arguni, Titik Nuryastuti, Tri Wibawa, Dwi AA Nugrahaningsih, Afiahayati, Siswanto,Kristy Iskandar,Nungki Anggorowati, Pramesti G Dewi, Dwi Indaryati,Indarto Sulistiyono, Khanza a Vujira, Alvina A Setiawan |
| EPI_ISL_2964704 | DINKES KAB. GROBOGAN | Genetics Working Group (Pokja Genetik) Faculty of Medicine, Public Health and Nursing Universitas Gadjah Mada (FK-KMK UGM); Disease Investigation Center Wates Ministry of | Gunadi, Hendra Wibawa, Marcellus, Mohamad S. Hakim, Edwin W. Daniwijaya, Ludhang P. Rizki, Endah Supriyati, Eggi Arguni, Titik Nuryastuti, Tri Wibawa, Dwi AA Nugrahaningsih, Afiahayati, Siswanto,Kristy Iskandar,Nungki Anggorowati, Pramesti G Dewi, Dwi Indaryati,Indarto Sulistiyono, Susan Simanjaya, Dyah A Puspitarani |

| | | | |
|---|---|---|---|
| | | Agriculture Indonesia; Department of Microbiology FK-KMK UGM; Laboratorium Diagnostik Yayasan Tahija World Mosquito Program (WMP) Yogyakarta Center for Tropical Medicine FK-KMK UGM; Integrated Research Center FK-KMK UGM; Department of Computer Science and Electronics FMIPA UGM; RSUP Dr. Sardjito | |
| EPI_ISL_2964707 | RS Dr. Oen Kandang Sapi Solo | Genetics Working Group (Pokja Genetik) Faculty of Medicine, Public Health and Nursing Universitas Gadjah Mada (FK-KMK UGM); Disease Investigation Center Wates Ministry of Agriculture Indonesia; Department of Microbiology FK-KMK UGM; Laboratorium Diagnostik Yayasan Tahija World Mosquito Program (WMP) Yogyakarta Center for Tropical Medicine FK-KMK UGM; Integrated Research Center FK-KMK UGM; Department of Computer Science and Electronics FMIPA UGM; RSUP Dr. Sardjito | Gunadi, Hendra Wibawa, Marcellus, Mohamad S. Hakim, Edwin W. Daniwijaya, Ludhang P. Rizki, Endah Supriyati, Eggi Arguni, Titik Nuryastuti, Tri Wibawa, Dwi AA Nugrahaningsih, Afiahayati, Siswanto,Kristy Iskandar,Nungki Anggorowati, Pramesti G Dewi, Dwi Indaryati,Indarto Sulistiyono, Susan Simanjaya, Dyah A Puspitarani |
| EPI_ISL_2964872 | RS Dr. Oen Kandang Sapi Solo | Genetics Working Group (Pokja Genetik) Faculty of Medicine, Public Health and Nursing Universitas Gadjah Mada (FK-KMK UGM); Disease Investigation Center Wates Ministry of Agriculture Indonesia; Department of Microbiology FK-KMK UGM; Laboratorium Diagnostik Yayasan Tahija World Mosquito Program (WMP) Yogyakarta Center for Tropical Medicine FK-KMK UGM; Integrated Research Center FK-KMK UGM; Department of Computer Science and Electronics FMIPA UGM; RSUP Dr. Sardjito | Gunadi, Hendra Wibawa, Marcellus, Mohamad S. Hakim, Edwin W. Daniwijaya, Ludhang P. Rizki, Endah Supriyati, Eggi Arguni, Titik Nuryastuti, Tri Wibawa, Dwi AA Nugrahaningsih, Afiahayati, Siswanto,Kristy Iskandar,Nungki Anggorowati, Pramesti G Dewi, Dwi Indaryati,Indarto Sulistiyono, Cita S Amalia, Alvina A Setiawan |
| EPI_ISL_2964917 | RS Dr. OEN KANDANG SAPI SOLO | Genetics Working Group (Pokja Genetik) Faculty of Medicine, Public Health and Nursing Universitas Gadjah Mada (FK-KMK UGM); Disease Investigation Center Wates Ministry of Agriculture Indonesia; Department of Microbiology FK-KMK UGM; Laboratorium Diagnostik Yayasan Tahija World Mosquito Program (WMP) Yogyakarta Center for Tropical Medicine FK-KMK UGM; Integrated Research Center FK-KMK UGM; Department of Computer Science and Electronics FMIPA UGM; RSUP Dr. Sardjito | Gunadi, Hendra Wibawa, Marcellus, Mohamad S. Hakim, Edwin W. Daniwijaya, Ludhang P. Rizki, Endah Supriyati, Eggi Arguni, Titik Nuryastuti, Tri Wibawa, Dwi AA Nugrahaningsih, Afiahayati, Siswanto,Kristy Iskandar,Nungki Anggorowati, Pramesti G Dewi, Dwi Indaryati,Indarto Sulistiyono, Irene Tania, Hana F Hanifin |
| EPI_ISL_2964918, EPI_ISL_2964919 | RS Dr. OEN KANDANG SAPI SOLO | Genetics Working Group (Pokja Genetik) Faculty of Medicine, Public Health and Nursing Universitas Gadjah Mada (FK-KMK UGM); Disease Investigation Center Wates Ministry of Agriculture Indonesia; Department of Microbiology FK-KMK UGM; Laboratorium Diagnostik Yayasan Tahija World Mosquito Program (WMP) Yogyakarta Center for Tropical Medicine FK-KMK UGM; Integrated Research Center FK-KMK UGM; Department of Computer Science and Electronics FMIPA UGM; RSUP Dr. Sardjito | Gunadi, Hendra Wibawa, Marcellus, Mohamad S. Hakim, Edwin W. Daniwijaya, Ludhang P. Rizki, Endah Supriyati, Eggi Arguni, Titik Nuryastuti, Tri Wibawa, Dwi AA Nugrahaningsih, Afiahayati, Siswanto,Kristy Iskandar,Nungki Anggorowati, Pramesti G Dewi, Dwi Indaryati,Indarto Sulistiyono, Cita S Amalia, Alvina A Setiawan |
| EPI_ISL_2964920 | RS Dr. OEN KANDANG SAPI SOLO | Genetics Working Group (Pokja Genetik) Faculty of Medicine, Public Health and Nursing Universitas Gadjah Mada (FK-KMK UGM); Disease Investigation Center Wates Ministry of Agriculture Indonesia; Department of Microbiology FK-KMK UGM; Laboratorium Diagnostik Yayasan Tahija World Mosquito Program (WMP) Yogyakarta Center for Tropical Medicine FK-KMK UGM; Integrated Research Center FK-KMK UGM; Department of Computer Science and Electronics FMIPA UGM; RSUP Dr. Sardjito | Gunadi, Hendra Wibawa, Marcellus, Mohamad S. Hakim, Edwin W. Daniwijaya, Ludhang P. Rizki, Endah Supriyati, Eggi Arguni, Titik Nuryastuti, Tri Wibawa, Dwi AA Nugrahaningsih, Afiahayati, Siswanto,Kristy Iskandar,Nungki Anggorowati, Pramesti G Dewi, Dwi Indaryati,Indarto Sulistiyono, Susan Simanjaya, Dyah A Puspitarani |
| EPI_ISL_2964921 | RS Dr. OEN KANDANG SAPI SOLO | Genetics Working Group (Pokja Genetik) Faculty of Medicine, Public Health and Nursing Universitas Gadjah Mada (FK-KMK UGM); Disease Investigation Center Wates Ministry of Agriculture Indonesia; Department of Microbiology FK-KMK UGM; Laboratorium Diagnostik Yayasan Tahija World Mosquito Program (WMP) Yogyakarta Center for Tropical Medicine FK-KMK UGM; Integrated Research Center FK-KMK UGM; Department of Computer Science and Electronics FMIPA UGM; RSUP Dr. Sardjito | Gunadi, Hendra Wibawa, Marcellus, Mohamad S. Hakim, Edwin W. Daniwijaya, Ludhang P. Rizki, Endah Supriyati, Eggi Arguni, Titik Nuryastuti, Tri Wibawa, Dwi AA Nugrahaningsih, Afiahayati, Siswanto,Kristy Iskandar,Nungki Anggorowati, Pramesti G Dewi, Dwi Indaryati,Indarto Sulistiyono, Khanza a Vujira, Alvina A Setiawan |
| EPI_ISL_2964943 | RSUD DR. MOEWARDI | Genetics Working Group (Pokja Genetik) Faculty of Medicine, Public Health and Nursing Universitas Gadjah Mada (FK-KMK UGM); Disease Investigation Center Wates Ministry of Agriculture Indonesia; Department of Microbiology FK-KMK UGM; Laboratorium Diagnostik Yayasan Tahija World Mosquito Program (WMP) Yogyakarta Center for Tropical Medicine FK-KMK UGM; Integrated Research Center FK-KMK UGM; Department of Computer Science and Electronics FMIPA UGM; RSUP Dr. Sardjito | Gunadi, Hendra Wibawa, Marcellus, Mohamad S. Hakim, Edwin W. Daniwijaya, Ludhang P. Rizki, Endah Supriyati, Eggi Arguni, Titik Nuryastuti, Tri Wibawa, Dwi AA Nugrahaningsih, Afiahayati, Siswanto,Kristy Iskandar,Nungki Anggorowati, Pramesti G Dewi, Dwi Indaryati,Indarto Sulistiyono, Susan Simanjaya, Dyah A Puspitarani |
| EPI_ISL_2964944 | RSUD DR. MOEWARDI | Genetics Working Group (Pokja Genetik) Faculty of Medicine, Public Health and Nursing Universitas Gadjah Mada (FK-KMK UGM); Disease Investigation Center Wates Ministry of Agriculture Indonesia; Department of Microbiology FK-KMK UGM; Laboratorium Diagnostik Yayasan Tahija World Mosquito Program (WMP) Yogyakarta Center for Tropical Medicine FK-KMK UGM; Integrated Research Center FK-KMK UGM; Department of Computer Science and Electronics FMIPA UGM; RSUP Dr. Sardjito | Gunadi, Hendra Wibawa, Marcellus, Mohamad S. Hakim, Edwin W. Daniwijaya, Ludhang P. Rizki, Endah Supriyati, Eggi Arguni, Titik Nuryastuti, Tri Wibawa, Dwi AA Nugrahaningsih, Afiahayati, Siswanto,Kristy Iskandar,Nungki Anggorowati, Pramesti G Dewi, Dwi Indaryati,Indarto Sulistiyono, Susan Simanjaya, I Putu Aditio A |
| EPI_ISL_2965763 | Dinkes Kabupaten Merauke | National Institute of Health Research and Development | Arie Ardiansyah Nugraha, Hana Apsari Pawestri, Subangkit, Kartika Dewi Puspa, Hartanti Dian Ikawati, Krisna Pangesti, Triyani Soekarso, Fajar Nur Sulistiyohadi, Nelly Puspandari, Vivi Setiawaty |
| EPI_ISL_2965772 | Dinkes Provinsi Kalimantan Utara | National Institute of Health Research and Development | Hana Apsari Pawestri, Subangkit, Arie Ardiansyah Nugraha, Kartika Dewi Puspa, Hartanti Dian Ikawati, Krisna Pangesti, Triyani Soekarso, Fajar Nur Sulistiyohadi, Nelly Puspandari, Vivi Setiawaty |

| | | | |
|---|---|---|---|
| EPI_ISL_2965781 | Dinkes Provinsi Kalimantan Timur | National Institute of Health Research and Development | Kartika Dewi Puspa, Arie Ardiansyah Nugraha, Hana Apsari Pawestri, Subangkit, Hartanti Dian Ikawati, Krisna Pangesti, Triyani Soekarso, Fajar Nur Sulistiyohadi, Nelly Puspandari, Vivi Setiawaty |
| EPI_ISL_2966246 | Dipartimento di Medicina di Laboratorio, Azienda sanitaria universitaria Friuli Centrale (ASU FC) | Dipartimento di Medicina di Laboratorio, Azienda sanitaria universitaria Friuli Centrale (ASU FC) | Corrado Pipan, Catia Mio, Chiara Dal Secco, Stefania Marzinotto, Francesco Curcio |
| EPI_ISL_2966248, EPI_ISL_2966249, EPI_ISL_2966250, EPI_ISL_2966251, EPI_ISL_2966252, EPI_ISL_2966253, EPI_ISL_2966254, EPI_ISL_2966255, EPI_ISL_2966256, EPI_ISL_2966257, EPI_ISL_2966258, EPI_ISL_2966259, EPI_ISL_2966260, EPI_ISL_2966262, EPI_ISL_2966263, EPI_ISL_2966265, EPI_ISL_2966266, EPI_ISL_2966267, EPI_ISL_2966268, EPI_ISL_2966269, EPI_ISL_2966270, EPI_ISL_2966271, EPI_ISL_2966272, EPI_ISL_2966273, EPI_ISL_2966274, EPI_ISL_2966275, EPI_ISL_2966276, EPI_ISL_2966277, EPI_ISL_2966279, EPI_ISL_2966280, EPI_ISL_2966281, EPI_ISL_2966282, EPI_ISL_2966283, EPI_ISL_2966284, EPI_ISL_2966286, EPI_ISL_2966287, EPI_ISL_2966288, EPI_ISL_2966290, EPI_ISL_2966291, EPI_ISL_2966292, EPI_ISL_2966293, EPI_ISL_2966294, EPI_ISL_2966296, EPI_ISL_2966297, EPI_ISL_2966298, EPI_ISL_2966300, EPI_ISL_2966301, EPI_ISL_2966304, EPI_ISL_2966305, EPI_ISL_2966306, EPI_ISL_2966307, EPI_ISL_2966308, EPI_ISL_2966309, EPI_ISL_2966312, EPI_ISL_2966313, EPI_ISL_2966314, EPI_ISL_2966315, EPI_ISL_2966317, EPI_ISL_2966318, EPI_ISL_2966321, EPI_ISL_2966322, EPI_ISL_2966323, EPI_ISL_2966325, EPI_ISL_2966326, EPI_ISL_2966327, EPI_ISL_2966331, EPI_ISL_2966332, EPI_ISL_2966338 | | | |
| see above | Cliniques universitaires Saint-Luc | UCLouvain/IREC/MBLG | Jean Ruelle, Bertrand Bearzatto, Benoit Kabamba Mukadi |
| EPI_ISL_2966354 | LESP Tabasco | Instituto de Diagnostico y Referencia Epidemiologicos (INDRE) | Claudia Wong-Arambula, Abril Rodriguez-Maldonado, Vanessa Rivero-Arredondo, Ariadna Medina-Benitez, Joaquin Quiroz-Mercado, Sergio Rangel-Guerrero, Natividad Cruz-Ortiz, Tatiana Nunez-Garcia, Gisela Barrera-Badillo, Lucia Hernandez-Rivas, Irma Lopez-Martinez, Ernesto Ramirez-Gonzalez. |
| EPI_ISL_2966355 | LESP Zacatecas | Instituto de Diagnostico y Referencia Epidemiologicos (INDRE) | Claudia Wong-Arambula, Abril Rodriguez-Maldonado, Vanessa Rivero-Arredondo, Ariadna Medina-Benitez, Joaquin Quiroz-Mercado, Sergio Rangel-Guerrero, Natividad Cruz-Ortiz, Tatiana Nunez-Garcia, Gisela Barrera-Badillo, Lucia Hernandez-Rivas, Irma Lopez-Martinez, Ernesto Ramirez-Gonzalez. |
| EPI_ISL_2966356, EPI_ISL_2966357 | LESP Puebla | Instituto de Diagnostico y Referencia Epidemiologicos (INDRE) | Claudia Wong-Arambula, Abril Rodriguez-Maldonado, Vanessa Rivero-Arredondo, Ariadna Medina-Benitez, Joaquin Quiroz-Mercado, Sergio Rangel-Guerrero, Natividad Cruz-Ortiz, Tatiana Nunez-Garcia, Gisela Barrera-Badillo, Lucia Hernandez-Rivas, Irma Lopez-Martinez, Ernesto Ramirez-Gonzalez. |
| EPI_ISL_2966358 | LESP Sonora | Instituto de Diagnostico y Referencia Epidemiologicos (INDRE) | Claudia Wong-Arambula, Abril Rodriguez-Maldonado, Vanessa Rivero-Arredondo, Ariadna Medina-Benitez, Joaquin Quiroz-Mercado, Sergio Rangel-Guerrero, Natividad Cruz-Ortiz, Tatiana Nunez-Garcia, Gisela Barrera-Badillo, Lucia Hernandez-Rivas, Irma Lopez-Martinez, Ernesto Ramirez-Gonzalez. |
| EPI_ISL_2966359 | LESP Morelos | Instituto de Diagnostico y Referencia Epidemiologicos (INDRE) | Claudia Wong-Arambula, Abril Rodriguez-Maldonado, Vanessa Rivero-Arredondo, Ariadna Medina-Benitez, Joaquin Quiroz-Mercado, Sergio Rangel-Guerrero, Natividad Cruz-Ortiz, Tatiana Nunez-Garcia, Gisela Barrera-Badillo, Lucia Hernandez-Rivas, Irma Lopez-Martinez, Ernesto Ramirez-Gonzalez. |
| EPI_ISL_2966360, EPI_ISL_2966361, EPI_ISL_2966362, EPI_ISL_2966363, EPI_ISL_2966364, EPI_ISL_2966365, EPI_ISL_2966366 | LESP Ciudad de Mexico | Instituto de Diagnostico y Referencia Epidemiologicos (INDRE) | Claudia Wong-Arambula, Abril Rodriguez-Maldonado, Vanessa Rivero-Arredondo, Ariadna Medina-Benitez, Joaquin Quiroz-Mercado, Sergio Rangel-Guerrero, Natividad Cruz-Ortiz, Tatiana Nunez-Garcia, Gisela Barrera-Badillo, Lucia Hernandez-Rivas, Irma Lopez-Martinez, Ernesto Ramirez-Gonzalez. |
| EPI_ISL_2966367 | LESP Queretaro | Instituto de Diagnostico y Referencia Epidemiologicos (INDRE) | Claudia Wong-Arambula, Abril Rodriguez-Maldonado, Vanessa Rivero-Arredondo, Ariadna Medina-Benitez, Joaquin Quiroz-Mercado, Sergio Rangel-Guerrero, Natividad Cruz-Ortiz, Tatiana Nunez-Garcia, Gisela Barrera-Badillo, Lucia Hernandez-Rivas, Irma Lopez-Martinez, Ernesto Ramirez-Gonzalez. |
| EPI_ISL_2966368 | LESP Estado de Mexico | Instituto de Diagnostico y Referencia Epidemiologicos (INDRE) | Claudia Wong-Arambula, Abril Rodriguez-Maldonado, Vanessa Rivero-Arredondo, Ariadna Medina-Benitez, Joaquin Quiroz-Mercado, Sergio Rangel-Guerrero, Natividad Cruz-Ortiz, Tatiana Nunez-Garcia, Gisela Barrera-Badillo, Lucia Hernandez-Rivas, Irma Lopez-Martinez, Ernesto Ramirez-Gonzalez. |
| EPI_ISL_2966369 | LESP Estado de Mexico | Instituto de Diagnostico y Referencia Epidemiologicos (INDRE) | Claudia Wong-Arambula, Abril Rodriguez-Maldonado, Vanessa Rivero-Arredondo, Ariadna Medina-Benitez, Joaquin Quiroz-Mercado, Sergio Rangel-Guerrero, Maribel Gonzalez-Villa, Natividad Cruz-Ortiz, Tatiana Nunez-Garcia, Gisela Barrera-Badillo, Lucia Hernandez-Rivas, Irma Lopez-Martinez, Ernesto Ramirez-Gonzalez. |
| EPI_ISL_2966370 | LESP Coahuila | Instituto de Diagnostico y Referencia Epidemiologicos (INDRE) | Claudia Wong-Arambula, Abril Rodriguez-Maldonado, Vanessa Rivero-Arredondo, Ariadna Medina-Benitez, Joaquin Quiroz-Mercado, Sergio Rangel-Guerrero, Maribel Gonzalez-Villa, Natividad Cruz-Ortiz, Tatiana Nunez-Garcia, Gisela Barrera-Badillo, Lucia Hernandez-Rivas, Irma Lopez-Martinez, Ernesto Ramirez-Gonzalez. |
| EPI_ISL_2966371, EPI_ISL_2966372, EPI_ISL_2966373, EPI_ISL_2966374 | LESP Nuevo Leon | Instituto de Diagnostico y Referencia Epidemiologicos (INDRE) | Claudia Wong-Arambula, Abril Rodriguez-Maldonado, Vanessa Rivero-Arredondo, Ariadna Medina-Benitez, Joaquin Quiroz-Mercado, Sergio Rangel-Guerrero, Maribel Gonzalez-Villa, Natividad Cruz-Ortiz, Tatiana Nunez-Garcia, Gisela Barrera-Badillo, Lucia Hernandez-Rivas, Irma Lopez-Martinez, Ernesto Ramirez-Gonzalez. |
| EPI_ISL_2966375 | LESP Tamaulipas | Instituto de Diagnostico y Referencia Epidemiologicos (INDRE) | Claudia Wong-Arambula, Abril Rodriguez-Maldonado, Vanessa Rivero-Arredondo, Ariadna Medina-Benitez, Joaquin Quiroz-Mercado, Sergio Rangel-Guerrero, Maribel Gonzalez-Villa, Natividad Cruz-Ortiz, Tatiana Nunez-Garcia, Gisela Barrera-Badillo, Lucia Hernandez-Rivas, Irma Lopez-Martinez, Ernesto Ramirez-Gonzalez. |
| EPI_ISL_2966376 | LESP Chihuahua | Instituto de Diagnostico y Referencia Epidemiologicos (INDRE) | Claudia Wong-Arambula, Abril Rodriguez-Maldonado, Vanessa Rivero-Arredondo, Ariadna Medina-Benitez, Joaquin Quiroz-Mercado, Sergio Rangel-Guerrero, Maribel Gonzalez-Villa, Natividad Cruz-Ortiz, Tatiana Nunez-Garcia, Gisela Barrera-Badillo, Lucia Hernandez-Rivas, Irma Lopez-Martinez, Ernesto Ramirez-Gonzalez. |
| EPI_ISL_2966377, EPI_ISL_2966378 | LESP Chiapas | Instituto de Diagnostico y Referencia Epidemiologicos (INDRE) | Claudia Wong-Arambula, Abril Rodriguez-Maldonado, Vanessa Rivero-Arredondo, Ariadna Medina-Benitez, Joaquin Quiroz-Mercado, Sergio Rangel-Guerrero, Maribel Gonzalez-Villa, Natividad Cruz-Ortiz, Tatiana Nunez-Garcia, Gisela Barrera-Badillo, Lucia Hernandez-Rivas, Irma Lopez-Martinez, Ernesto Ramirez-Gonzalez. |
| EPI_ISL_2966379 | LESP Veracruz | Instituto de Diagnostico y Referencia Epidemiologicos (INDRE) | Claudia Wong-Arambula, Abril Rodriguez-Maldonado, Vanessa Rivero-Arredondo, Ariadna Medina-Benitez, Joaquin Quiroz-Mercado, Sergio Rangel-Guerrero, Maribel Gonzalez-Villa, Natividad Cruz-Ortiz, Tatiana Nunez-Garcia, Gisela Barrera-Badillo, Lucia Hernandez-Rivas, Irma Lopez-Martinez, Ernesto Ramirez-Gonzalez. |
| EPI_ISL_2966380 | LESP Sonora | Instituto de Diagnostico y Referencia Epidemiologicos (INDRE) | Claudia Wong-Arambula, Abril Rodriguez-Maldonado, Vanessa Rivero-Arredondo, Ariadna Medina-Benitez, Joaquin Quiroz-Mercado, Sergio Rangel-Guerrero, Maribel Gonzalez-Villa, Natividad Cruz-Ortiz, Tatiana Nunez-Garcia, Gisela Barrera-Badillo, Lucia Hernandez-Rivas, Irma Lopez-Martinez, Ernesto Ramirez-Gonzalez. |
| EPI_ISL_2966381, EPI_ISL_2966382, EPI_ISL_2966383, EPI_ISL_2966384, EPI_ISL_2966385, EPI_ISL_2966386, EPI_ISL_2966387, EPI_ISL_2966388, EPI_ISL_2966389, EPI_ISL_2966390, EPI_ISL_2966391, EPI_ISL_2966392, EPI_ISL_2966393, EPI_ISL_2966394, EPI_ISL_2966395, EPI_ISL_2966396, EPI_ISL_2966397, EPI_ISL_2966398, EPI_ISL_2966399, EPI_ISL_2966400, EPI_ISL_2966401, EPI_ISL_2966402 | | | |
| see above | LESP Ciudad de Mexico | Instituto de Diagnostico y Referencia Epidemiologicos (INDRE) | Claudia Wong-Arambula, Abril Rodriguez-Maldonado, Vanessa Rivero-Arredondo, Ariadna Medina-Benitez, Joaquin Quiroz-Mercado, Sergio Rangel-Guerrero, Maribel Gonzalez-Villa, Natividad Cruz-Ortiz, Tatiana Nunez-Garcia, Gisela Barrera-Badillo, Lucia Hernandez-Rivas, Irma Lopez-Martinez, Ernesto Ramirez-Gonzalez. |
| EPI_ISL_2966403, EPI_ISL_2966404, EPI_ISL_2966405, EPI_ISL_2966406, EPI_ISL_2966407, EPI_ISL_2966408 | LESP Queretaro | Instituto de Diagnostico y Referencia Epidemiologicos (INDRE) | Claudia Wong-Arambula, Abril Rodriguez-Maldonado, Vanessa Rivero-Arredondo, Ariadna Medina-Benitez, Joaquin Quiroz-Mercado, Sergio Rangel-Guerrero, Maribel Gonzalez-Villa, Natividad Cruz-Ortiz, Tatiana Nunez-Garcia, Gisela Barrera-Badillo, Lucia Hernandez-Rivas, Irma Lopez-Martinez, Ernesto Ramirez-Gonzalez. |
| EPI_ISL_2966409, EPI_ISL_2966410, EPI_ISL_2966411, EPI_ISL_2966412 | LESP Tabasco | Instituto de Diagnostico y Referencia Epidemiologicos (INDRE) | Claudia Wong-Arambula, Abril Rodriguez-Maldonado, Vanessa Rivero-Arredondo, Ariadna Medina-Benitez, Joaquin Quiroz-Mercado, Sergio Rangel-Guerrero, Maribel Gonzalez-Villa, Natividad Cruz-Ortiz, Tatiana Nunez-Garcia, Gisela Barrera-Badillo, Lucia Hernandez-Rivas, Irma Lopez-Martinez, Ernesto Ramirez-Gonzalez. |
| EPI_ISL_2966413 | LESP Jalisco | Instituto de Diagnostico y Referencia Epidemiologicos (INDRE) | Claudia Wong-Arambula, Abril Rodriguez-Maldonado, Vanessa Rivero-Arredondo, Ariadna Medina-Benitez, Joaquin Quiroz-Mercado, Sergio Rangel-Guerrero, Maribel Gonzalez-Villa, Natividad Cruz-Ortiz, Tatiana Nunez-Garcia, Gisela Barrera-Badillo, Lucia Hernandez-Rivas, Irma Lopez-Martinez, Ernesto Ramirez-Gonzalez. |
| EPI_ISL_2966415 | LESP Aguascalientes | Instituto de Diagnostico y Referencia Epidemiologicos (INDRE) | Claudia Wong-Arambula, Abril Rodriguez-Maldonado, Vanessa Rivero-Arredondo, Ariadna Medina-Benitez, Joaquin Quiroz-Mercado, Sergio Rangel-Guerrero, Maribel Gonzalez-Villa, Natividad Cruz-Ortiz, Tatiana Nunez-Garcia, Gisela Barrera-Badillo, Lucia Hernandez-Rivas, Irma Lopez-Martinez, Ernesto Ramirez-Gonzalez. |

| | | | |
|---|---|---|---|
| EPI_ISL_2966416 | LESP Ciudad de Mexico | Instituto de Diagnostico y Referencia Epidemiologicos (INDRE) | Claudia Wong-Arambula, Abril Rodriguez-Maldonado, Vanessa Rivero-Arredondo, Ariadna Medina-Benitez, Joaquin Quiroz-Mercado, Sergio Rangel-Guerrero, Maribel Gonzalez-Villa, Natividad Cruz-Ortiz, Tatiana Nunez-Garcia, Gisela Barrera-Badillo, Lucia Hernandez-Rivas, Irma Lopez-Martinez, Ernesto Ramirez-Gonzalez. |
| EPI_ISL_2966417 | LESP Coahuila | Instituto de Diagnostico y Referencia Epidemiologicos (INDRE) | Claudia Wong-Arambula, Abril Rodriguez-Maldonado, Vanessa Rivero-Arredondo, Ariadna Medina-Benitez, Joaquin Quiroz-Mercado, Sergio Rangel-Guerrero, Natividad Cruz-Ortiz, Tatiana Nunez-Garcia, Gisela Barrera-Badillo, Lucia Hernandez-Rivas, Irma Lopez-Martinez, Ernesto Ramirez-Gonzalez. |
| EPI_ISL_2966512 | SECP MHU | The Institute of Molecular Biology and Genetics of NASU | M.Tukalo et al. |
| EPI_ISL_2966613, EPI_ISL_2966614, EPI_ISL_2966615, EPI_ISL_2966616, EPI_ISL_2966618, EPI_ISL_2966619 | Ochsner Health | Bioinfoexperts, LLC | Judy Crabtree, Amy Feehan, Rebecca Rose, David J. Nolan, Tessa LaFleur, Julia-Garcia-Diaz, Lucio Miele, Susanna L. Lamers |
| EPI_ISL_2967035, EPI_ISL_2967036, EPI_ISL_2967037, EPI_ISL_2967038, EPI_ISL_2967039, EPI_ISL_2967040, EPI_ISL_2967041, EPI_ISL_2967042, EPI_ISL_2967043, EPI_ISL_2967044, EPI_ISL_2967045, EPI_ISL_2967046, EPI_ISL_2967047, EPI_ISL_2967050, EPI_ISL_2967051, EPI_ISL_2967052, EPI_ISL_2967053, EPI_ISL_2967055, EPI_ISL_2967056, EPI_ISL_2967057, EPI_ISL_2967059, EPI_ISL_2967060, EPI_ISL_2967061, EPI_ISL_2967062, EPI_ISL_2967063, EPI_ISL_2967064, EPI_ISL_2967065, EPI_ISL_2967066, EPI_ISL_2967067, EPI_ISL_2967068, EPI_ISL_2967069, EPI_ISL_2967070, EPI_ISL_2967071, EPI_ISL_2967072, EPI_ISL_2967073, EPI_ISL_2967074, EPI_ISL_2967075, EPI_ISL_2967076, EPI_ISL_2967077, EPI_ISL_2967078, EPI_ISL_2967079, EPI_ISL_2967080, EPI_ISL_2967081, EPI_ISL_2967082, EPI_ISL_2967084, EPI_ISL_2967085, EPI_ISL_2967086, EPI_ISL_2967087, EPI_ISL_2967088, EPI_ISL_2967089, EPI_ISL_2967093, EPI_ISL_2967094, EPI_ISL_2967096, EPI_ISL_2967097, EPI_ISL_2967098, EPI_ISL_2967099, EPI_ISL_2967100, EPI_ISL_2967101, EPI_ISL_2967102, EPI_ISL_2967103, EPI_ISL_2967105, EPI_ISL_2967106, EPI_ISL_2967107, EPI_ISL_2967108, EPI_ISL_2967109, EPI_ISL_2967110, EPI_ISL_2967112, EPI_ISL_2967113, EPI_ISL_2967114, EPI_ISL_2967115, EPI_ISL_2967118, EPI_ISL_2967127, EPI_ISL_2967128, EPI_ISL_2967129, EPI_ISL_2967130, EPI_ISL_2967131, EPI_ISL_2967133, EPI_ISL_2967134, EPI_ISL_2967135, EPI_ISL_2967137, EPI_ISL_2967138, EPI_ISL_2967140, EPI_ISL_2967142, EPI_ISL_2967143, EPI_ISL_2967144, EPI_ISL_2967146, EPI_ISL_2967147, EPI_ISL_2967149, EPI_ISL_2967151, EPI_ISL_2967152, EPI_ISL_2967155, EPI_ISL_2967156, EPI_ISL_2967158, EPI_ISL_2967159, EPI_ISL_2967160, EPI_ISL_2967161, EPI_ISL_2967163, EPI_ISL_2967178, EPI_ISL_2967181, EPI_ISL_2967182, EPI_ISL_2967183, EPI_ISL_2967184, EPI_ISL_2967186 | | | |
| see above | National Center of Infectious and Parasitic Diseases | National Center of Infectious and Parasitic Diseases | Alexiev et al |
| EPI_ISL_2967820, EPI_ISL_2967822, EPI_ISL_2967823, EPI_ISL_2967826, EPI_ISL_2967827, EPI_ISL_2967830, EPI_ISL_2967831, EPI_ISL_2967832, EPI_ISL_2967835, EPI_ISL_2967836, EPI_ISL_2967841, EPI_ISL_2967842, EPI_ISL_2967864, EPI_ISL_2967865, EPI_ISL_2967866, EPI_ISL_2967867, EPI_ISL_2967868, EPI_ISL_2967869, EPI_ISL_2967873, EPI_ISL_2967874, EPI_ISL_2967875, EPI_ISL_2967876, EPI_ISL_2967877, EPI_ISL_2967893, EPI_ISL_2967894, EPI_ISL_2967895, EPI_ISL_2967896 | | | |
| see above | Microbiology Department. Complexo Hospitalario Universitario de Vigo | Microbiology Department. Complexo Hospitalario Universitario de Vigo | Alfaya N, Alonso I, Alvarez M, Cabrera JJ, Carballo R, Cores O, Cortizo S, del-Campo V, Martinez L, Mediero G, Perez S, Davina C, Potel C, Regueiro B, Rey S, Vasallo FJ |
| EPI_ISL_2968072, EPI_ISL_2968074, EPI_ISL_2968076, EPI_ISL_2968078, EPI_ISL_2968079, EPI_ISL_2968080, EPI_ISL_2968081, EPI_ISL_2968084, EPI_ISL_2968085, EPI_ISL_2968087, EPI_ISL_2968092 | | | |
| see above | NILMRC | Genomic Research Laboratory, Bangabandhu Sheikh Mujib Medical University | Laila Anjuman Banu, Md.Saydur Rahman, Sharadindu Kanti Sinha, Arifa Akram, Md.Sharfuddin Ahmed |
| EPI_ISL_2978347 | NJDOH, Public Health and Environmental Laboratories | NJ_PHEL | Lindsey Bodnar, Shiv K. Verma, Ryan Pachucki, Chelsea San Filippo, Jacquelyn Deverell, Dana Woell, Allison Roder, Byeong Jeong |
| EPI_ISL_2978731 | Genome Analysis Center, Yamanashi Central Hospital | Genome Analysis Center, Yamanashi Central Hospital | Yosuke Hirotsu |
| EPI_ISL_2982121, EPI_ISL_2982122, EPI_ISL_2982123, EPI_ISL_2982124, EPI_ISL_2982125, EPI_ISL_2982126, EPI_ISL_2982127, EPI_ISL_2982128, EPI_ISL_2982129, EPI_ISL_2982130, EPI_ISL_2982131, EPI_ISL_2982132, EPI_ISL_2982133, EPI_ISL_2982134, EPI_ISL_2982135 | | | |
| see above | Microbiology Department. Complexo Hospitalario Universitario de Vigo | Microbiology Department. Complexo Hospitalario Universitario de Vigo | Alfaya N, Alonso I, Alvarez M, Cabrera JJ, Carballo R, Cores O, Cortizo S, del-Campo V, Martinez L, Mediero G, Perez S, Davina C, Potel C, Regueiro B, Rey S, Vasallo FJ |
| EPI_ISL_2982136, EPI_ISL_2982137, EPI_ISL_2982138, EPI_ISL_2982139, EPI_ISL_2982140, EPI_ISL_2982141, EPI_ISL_2982142, EPI_ISL_2982143, EPI_ISL_2982144, EPI_ISL_2982145, EPI_ISL_2982146, EPI_ISL_2982147, EPI_ISL_2982148, EPI_ISL_2982149, EPI_ISL_2982150, EPI_ISL_2982151, EPI_ISL_2982152, EPI_ISL_2982153, EPI_ISL_2982154 | | | |
| see above | OSPEDALE CIVILE TERAMO - CENTRO TRASFUSIONALE | Istituto Zooprofilattico Sperimentale dell'Abruzzo e Molise "G. Caporale" | Lorusso A, Marcacci M, Di Domenico M, Ancora M, Curini V, Di Lollo Valeria, Mangone I, Rinaldi A, Delli Compagni E, Scialabba S, Caporale M, Di Pasquale A, Cammà C, Puglia I, Calistri P, Savini G |
| EPI_ISL_2982155 | OSPEDALE SAN SALVATORE - U.O.C. Malattie infettive | Istituto Zooprofilattico Sperimentale dell'Abruzzo e Molise "G. Caporale" | Lorusso A, Marcacci M, Di Domenico M, Ancora M, Curini V, Di Lollo Valeria, Mangone I, Rinaldi A, Delli Compagni E, Scialabba S, Caporale M, Di Pasquale A, Cammà C, Puglia I, Calistri P, Savini G |
| EPI_ISL_2982156 | OSPEDALE CIVILE TERAMO - CENTRO TRASFUSIONALE | Istituto Zooprofilattico Sperimentale dell'Abruzzo e Molise "G. Caporale" | Lorusso A, Marcacci M, Di Domenico M, Ancora M, Curini V, Di Lollo Valeria, Mangone I, Rinaldi A, Delli Compagni E, Scialabba S, Caporale M, Di Pasquale A, Cammà C, Puglia I, Calistri P, Savini G |
| EPI_ISL_2982157, EPI_ISL_2982158, EPI_ISL_2982159 | OSPEDALE SAN SALVATORE - MEDICINA DI LABORATORIO | Istituto Zooprofilattico Sperimentale dell'Abruzzo e Molise "G. Caporale" | Lorusso A, Marcacci M, Di Domenico M, Ancora M, Curini V, Di Lollo Valeria, Mangone I, Rinaldi A, Delli Compagni E, Scialabba S, Caporale M, Di Pasquale A, Cammà C, Puglia I, Calistri P, Savini G |
| EPI_ISL_2982160, EPI_ISL_2982161, EPI_ISL_2982162, EPI_ISL_2982163, EPI_ISL_2982164, EPI_ISL_2982165, EPI_ISL_2982166 | PO AVEZZANO - U.O.C. Laboratorio Analisi | Istituto Zooprofilattico Sperimentale dell'Abruzzo e Molise "G. Caporale" | Lorusso A, Marcacci M, Di Domenico M, Ancora M, Curini V, Di Lollo Valeria, Mangone I, Rinaldi A, Delli Compagni E, Scialabba S, Caporale M, Di Pasquale A, Cammà C, Puglia I, Calistri P, Savini G |
| EPI_ISL_2982167 | OSPEDALE CIVILE TERAMO - CENTRO TRASFUSIONALE | Istituto Zooprofilattico Sperimentale dell'Abruzzo e Molise "G. Caporale" | Lorusso A, Marcacci M, Di Domenico M, Ancora M, Curini V, Di Lollo Valeria, Mangone I, Rinaldi A, Delli Compagni E, Scialabba S, Caporale M, Di Pasquale A, Cammà C, Puglia I, Calistri P, Savini G |
| EPI_ISL_2982168, EPI_ISL_2982169, EPI_ISL_2982170, EPI_ISL_2982171, EPI_ISL_2982172, EPI_ISL_2982173, EPI_ISL_2982174, EPI_ISL_2982175, EPI_ISL_2982176 | PO AVEZZANO - U.O.C. Laboratorio Analisi | Istituto Zooprofilattico Sperimentale dell'Abruzzo e Molise "G. Caporale" | Lorusso A, Marcacci M, Di Domenico M, Ancora M, Curini V, Di Lollo Valeria, Mangone I, Rinaldi A, Delli Compagni E, Scialabba S, Caporale M, Di Pasquale A, Cammà C, Puglia I, Calistri P, Savini G |
| EPI_ISL_2982177, EPI_ISL_2982178, EPI_ISL_2982179, EPI_ISL_2982180, EPI_ISL_2982181, EPI_ISL_2982182, EPI_ISL_2982183, EPI_ISL_2982184, EPI_ISL_2982185, EPI_ISL_2982186, EPI_ISL_2982187, EPI_ISL_2982188, EPI_ISL_2982189, EPI_ISL_2982190, EPI_ISL_2982191, EPI_ISL_2982192, EPI_ISL_2982193, EPI_ISL_2982194, EPI_ISL_2982195, EPI_ISL_2982196, EPI_ISL_2982197, EPI_ISL_2982198, EPI_ISL_2982199, EPI_ISL_2982200, EPI_ISL_2982201, EPI_ISL_2982202, EPI_ISL_2982203, EPI_ISL_2982204, EPI_ISL_2982205, EPI_ISL_2982206, EPI_ISL_2982207, EPI_ISL_2982208, EPI_ISL_2982209, EPI_ISL_2982220 | | | |
| see above | OSPEDALE CIVILE TERAMO - CENTRO TRASFUSIONALE | Istituto Zooprofilattico Sperimentale dell'Abruzzo e Molise "G. Caporale" | Lorusso A, Marcacci M, Di Domenico M, Ancora M, Curini V, Di Lollo Valeria, Mangone I, Rinaldi A, Delli Compagni E, Scialabba S, Caporale M, Di Pasquale A, Cammà C, Puglia I, Calistri P, Savini G |
| EPI_ISL_2982762, EPI_ISL_2982763, EPI_ISL_2982764, EPI_ISL_2982765, EPI_ISL_2982766, EPI_ISL_2982767, EPI_ISL_2982768, EPI_ISL_2982769, EPI_ISL_2982776, EPI_ISL_2982802, EPI_ISL_2982807, EPI_ISL_2982815, EPI_ISL_2982817, EPI_ISL_2982826, EPI_ISL_2982827 | | | |
| see above | Laboratorio Central de Saude Publica do Estado do Rio Grande do Sul (LACEN-RS) | Laboratory of Respiratory Viruses and Measles, Oswaldo Cruz Institute, FIOCRUZ | Paola Resende, Luciana Appolinario, Fernando Motta, Anna Carolina Paixao, Ana Carolina Mendonca, Alice Sampaio Rocha, Taina Venas, Elisa Cavalcante Pereira, Renata Serrano Lopes, Anderson Brandao Leite, Marilda Siqueira on behalf of the Fiocruz COVID-19 Genomic Surveillance Network |
| EPI_ISL_2983187 | Laboratory of Respiratory Viruses and Measles, Oswaldo Cruz Institute, FIOCRUZ | Laboratory of Respiratory Viruses and Measles, Oswaldo Cruz Institute, FIOCRUZ | Paola Resende, Luciana Appolinario, Fernando Motta, Anna Carolina Paixao, Ana Carolina Mendonca, Ighor Leonardo Arantes Gomes, Agatha Cristinne Prudencio, Alice Sampaio Rocha, Taina Venas, Elisa Cavalcante Pereira, Renata Serrano Lopes, Marilda Siqueira on behalf of the Fiocruz COVID-19 Genomic Surveillance Network |
| EPI_ISL_2983373, EPI_ISL_2983426, EPI_ISL_2983431, EPI_ISL_2983432, EPI_ISL_2983437, EPI_ISL_2983438, EPI_ISL_2983439, EPI_ISL_2983443, EPI_ISL_2983444, EPI_ISL_2983449, EPI_ISL_2983452 | | | |
| see above | Laboratorio Central de Saude Publica do Estado do Parana (LACEN/PR) | Laboratory of Respiratory Viruses and Measles, Oswaldo Cruz Institute, FIOCRUZ | Paola Resende, Luciana Appolinario, Fernando Motta, Anna Carolina Paixao, Ana Carolina Mendonca, Alice Sampaio Rocha, Taina Venas, Elisa Cavalcante Pereira, Renata Serrano Lopes, Irina Riediger, Marilda Siqueira on behalf of the Fiocruz COVID-19 Genomic Surveillance Network |
| EPI_ISL_2983480 | CNR Institut Pasteur de la Guyane | Institut Pasteur de la Guyane | Anne Lavergne, Dominique Rousset, Antoine Enfissi, Arielle Salmier |
| EPI_ISL_2983866 | GH A.CHENEVIER-H.MONDOR | Department of Virology, Henri Mondor University Hospital, Assistance Publique Hôpitaux de Paris, Université Paris-Est Créteil, INSERM U955 | Christophe Rodriguez, Slim Fourati, Vanessa Demontant, Guillaume Gricourt, Melissa N'Debi, Alexandre Soulier, Elisabeth Trawinski, Jean-Michel Pawlotsky |
| EPI_ISL_2984224, EPI_ISL_2984226, EPI_ISL_2984227, EPI_ISL_2984228, EPI_ISL_2984231, EPI_ISL_2984232, EPI_ISL_2984233, EPI_ISL_2984234, EPI_ISL_2984236, EPI_ISL_2984237, EPI_ISL_2984238, EPI_ISL_2984239, EPI_ISL_2984240, EPI_ISL_2984242, EPI_ISL_2984243, EPI_ISL_2984244, EPI_ISL_2984245, EPI_ISL_2984246 | | | |
| see above | Jahra Hospital | Kuwait Cancer Control Center | Shakir Bahzad, Mona Alateeqi |

| | | | |
|---|---|---|---|
| EPI_ISL_2985348 | Department of Virology, Istituto Zooprofilattico Sperimentale del Lazio e della Toscana (IZSLT) | Department of General Diagnostics; Department of Virology; Istituto Zooprofilattico Sperimentale del Lazio e della Toscana (IZSLT) | Patricia Alba, Giuseppe Manna, Elena L. Diaconu, Fabiola Feltrin, Raffaela Conti, Teresa Scicluna, Virginia Carfora, Antonella Cersini, Alessia Franco, Antonio Battisti. |
| EPI_ISL_2985351, EPI_ISL_2985362, EPI_ISL_2985364, EPI_ISL_2985375, EPI_ISL_2985376 | Ospedale San Filippo Neri | Department of General Diagnostics; Department of Virology; Istituto Zooprofilattico Sperimentale del Lazio e della Toscana (IZSLT) | Patricia Alba, Giuseppe Manna, Elena L. Diaconu, Fabiola Feltrin, Raffaela Conti, Teresa Scicluna, Virginia Carfora, Antonella Cersini, Alessia Franco, Antonio Battisti. |
| EPI_ISL_2985383, EPI_ISL_2985385, EPI_ISL_2985387, EPI_ISL_2985389, EPI_ISL_2985390 | Ospedale Sandro Pertini ASL Roma2 | Department of General Diagnostics; Department of Virology; Istituto Zooprofilattico Sperimentale del Lazio e della Toscana (IZSLT) | Patricia Alba, Giuseppe Manna, Elena L. Diaconu, Fabiola Feltrin, Raffaela Conti, Teresa Scicluna, Virginia Carfora, Antonella Cersini, Alessia Franco, Antonio Battisti. |
| EPI_ISL_2985398, EPI_ISL_2985399, EPI_ISL_2985400, EPI_ISL_2985401, EPI_ISL_2985402, EPI_ISL_2985403, EPI_ISL_2985404, EPI_ISL_2985405, EPI_ISL_2985406, EPI_ISL_2985407, EPI_ISL_2985408, EPI_ISL_2985409, EPI_ISL_2985410, EPI_ISL_2985411, EPI_ISL_2985412, EPI_ISL_2985413, EPI_ISL_2985414, EPI_ISL_2985415, EPI_ISL_2985416, EPI_ISL_2985418, EPI_ISL_2985419, EPI_ISL_2985420, EPI_ISL_2985421, EPI_ISL_2985422, EPI_ISL_2985423, EPI_ISL_2985424, EPI_ISL_2985425, EPI_ISL_2985426, EPI_ISL_2985427, EPI_ISL_2985428, EPI_ISL_2985429, EPI_ISL_2985430, EPI_ISL_2985431, EPI_ISL_2985432, EPI_ISL_2985433, EPI_ISL_2985434, EPI_ISL_2985435, EPI_ISL_2985436, EPI_ISL_2985437, EPI_ISL_2985438, EPI_ISL_2985439, EPI_ISL_2985440, EPI_ISL_2985442, EPI_ISL_2985443, EPI_ISL_2985444, EPI_ISL_2985445, EPI_ISL_2985447, EPI_ISL_2985448, EPI_ISL_2985449, EPI_ISL_2985451, EPI_ISL_2985452, EPI_ISL_2985453, EPI_ISL_2985454, EPI_ISL_2985455, EPI_ISL_2985456, EPI_ISL_2985457, EPI_ISL_2985458, EPI_ISL_2985460, EPI_ISL_2985461, EPI_ISL_2985462, EPI_ISL_2985464, EPI_ISL_2985465, EPI_ISL_2985467, EPI_ISL_2985468, EPI_ISL_2985470, EPI_ISL_2985471, EPI_ISL_2985473, EPI_ISL_2985475, EPI_ISL_2985477, EPI_ISL_2985478, EPI_ISL_2985479, EPI_ISL_2985480, EPI_ISL_2985481, EPI_ISL_2985482, EPI_ISL_2985483, EPI_ISL_2985484, EPI_ISL_2985485, EPI_ISL_2985487, EPI_ISL_2985489, EPI_ISL_2985491, EPI_ISL_2985492, EPI_ISL_2985493, EPI_ISL_2985494, EPI_ISL_2985496, EPI_ISL_2985497, EPI_ISL_2985498, EPI_ISL_2985499, EPI_ISL_2985500, EPI_ISL_2985501, EPI_ISL_2985502, EPI_ISL_2985503, EPI_ISL_2985504, EPI_ISL_2985505, EPI_ISL_2985506, EPI_ISL_2985507, EPI_ISL_2985508, EPI_ISL_2985509, EPI_ISL_2985510, EPI_ISL_2985511, EPI_ISL_2985512, EPI_ISL_2985513, EPI_ISL_2985514, EPI_ISL_2985515, EPI_ISL_2985516, EPI_ISL_2985517, EPI_ISL_2985519, EPI_ISL_2985520, EPI_ISL_2985521, EPI_ISL_2985523, EPI_ISL_2985524, EPI_ISL_2985527, EPI_ISL_2985528, EPI_ISL_2985530, EPI_ISL_2985531, EPI_ISL_2985532, EPI_ISL_2985533, EPI_ISL_2985535, EPI_ISL_2985537, EPI_ISL_2985538, EPI_ISL_2985539, EPI_ISL_2985540, EPI_ISL_2985541, EPI_ISL_2985542, EPI_ISL_2985543, EPI_ISL_2985544, EPI_ISL_2985545, EPI_ISL_2985546, EPI_ISL_2985547, EPI_ISL_2985548, EPI_ISL_2985549, EPI_ISL_2985550, EPI_ISL_2985551, EPI_ISL_2985552, EPI_ISL_2985554, EPI_ISL_2985555, EPI_ISL_2985557, EPI_ISL_2985558, EPI_ISL_2985559, EPI_ISL_2985561, EPI_ISL_2985563, EPI_ISL_2985564, EPI_ISL_2985565, EPI_ISL_2985567, EPI_ISL_2985568, EPI_ISL_2985569, EPI_ISL_2985570, EPI_ISL_2985571, EPI_ISL_2985572, EPI_ISL_2985573, EPI_ISL_2985574, EPI_ISL_2985578, EPI_ISL_2985579, EPI_ISL_2985580, EPI_ISL_2985581, EPI_ISL_2985582, EPI_ISL_2985584, EPI_ISL_2985585, EPI_ISL_2985587, EPI_ISL_2985588, EPI_ISL_2985589, EPI_ISL_2985590, EPI_ISL_2985591, EPI_ISL_2985592, EPI_ISL_2985593, EPI_ISL_2985594, EPI_ISL_2985595, EPI_ISL_2985597, EPI_ISL_2985598, EPI_ISL_2985599, EPI_ISL_2985601 | | | |
| see above | Austrian Agency for Health and Food Safety (AGES) | Bergthaler laboratory, CeMM Research Center for Molecular Medicine of the Austrian Academy of Sciences | Lukas Endler, Anna Schedl, Fabian Amman, Petr Triska, Matthew Thornton, Thomas Penz, Benedikt Agerer, Maelle Le Moing, Michael Schuster, Bekir Erguner, Jan Laine, Martin Senekowitsch, Christoph Bock, Andreas Bergthaler |
| EPI_ISL_2985606 | Johannes Kepler Universität Linz | Bergthaler laboratory, CeMM Research Center for Molecular Medicine of the Austrian Academy of Sciences | Lukas Endler, Anna Schedl, Fabian Amman, Petr Triska, Matthew Thornton, Thomas Penz, Benedikt Agerer, Maelle Le Moing, Michael Schuster, Bekir Erguner, Jan Laine, Martin Senekowitsch, Christoph Bock, Andreas Bergthaler |
| EPI_ISL_2986982, EPI_ISL_2986983, EPI_ISL_2986984, EPI_ISL_2986985 | Ayass Bioscience LLC | Ayass Bioscience LLC | Mohamad Ammar Ayass, Natalya Griko, Kevin Zhu, Nazanin Taheri, Lina Abi Mosleh |
| EPI_ISL_2987599 | LAB RSUD Koja Jakut | National Institute of Health Research and Development | Hana Apsari Pawestri, Subangkit, Arie Ardiansyah Nugraha, Kartika Dewi Puspa, Hartanti Dian Ikawati, Krisna Pangesti, Triyani Soekarso, Fajar Nur Sulistiyohadi, Nelly Puspandari, Vivi Setiawaty |
| EPI_ISL_2987608 | RSU Mitra Keluarga Kalideres | National Institute of Health Research and Development | Hana Apsari Pawestri, Subangkit, Arie Ardiansyah Nugraha, Kartika Dewi Puspa, Hartanti Dian Ikawati, Krisna Pangesti, Triyani Soekarso, Fajar Nur Sulistiyohadi, Nelly Puspandari, Vivi Setiawaty |
| EPI_ISL_2987617 | PUSKESMAS BOJONGSARI | National Institute of Health Research and Development | Kartika Dewi Puspa, Arie Ardiansyah Nugraha, Hana Apsari Pawestri, Subangkit, Hartanti Dian Ikawati, Krisna Pangesti, Triyani Soekarso, Fajar Nur Sulistiyohadi, Nelly Puspandari, Vivi Setiawaty |
| EPI_ISL_2987618 | RS Mata Cicendo Bandung | National Institute of Health Research and Development | Hana Apsari Pawestri, Subangkit, Arie Ardiansyah Nugraha, Kartika Dewi Puspa, Hartanti Dian Ikawati, Krisna Pangesti, Triyani Soekarso, Fajar Nur Sulistiyohadi, Nelly Puspandari, Vivi Setiawaty |
| EPI_ISL_2987631 | RSUD KRMT Wongsonegoro | National Institute of Health Research and Development | Kartika Dewi Puspa, Arie Ardiansyah Nugraha, Hana Apsari Pawestri, Subangkit, Hartanti Dian Ikawati, Krisna Pangesti, Triyani Soekarso, Fajar Nur Sulistiyohadi, Nelly Puspandari, Vivi Setiawaty |
| EPI_ISL_2987652 | BTKL PP Kelas I Batam | National Institute of Health Research and Development | Vivi Setiawaty, Hana Apsari Pawestri, Subangkit, Kartika Dewi Puspa, Arie Ardiansyah Nugraha, Hartanti Dian Ikawati, Krisna Pangesti, Triyani Soekarso, Fajar Nur Sulistiyohadi, Nelly Puspandari |
| EPI_ISL_2987662 | Dinkes Kabupaten Merauke | National Institute of Health Research and Development | Vivi Setiawaty, Hana Apsari Pawestri, Subangkit, Kartika Dewi Puspa, Arie Ardiansyah Nugraha, Hartanti Dian Ikawati, Krisna Pangesti, Triyani Soekarso, Fajar Nur Sulistiyohadi, Nelly Puspandari |
| EPI_ISL_2987697, EPI_ISL_2987699, EPI_ISL_2987700, EPI_ISL_2987705, EPI_ISL_2987706, EPI_ISL_2987707, EPI_ISL_2987708, EPI_ISL_2987709, EPI_ISL_2987710 | Hospital Margarita Maza de Juárez | Microbial Genomics Laboratory | Consorcio Mexicano de Vigilancia Genómica (CoViGen-Mex). Authors (in alphabetical order): Julio Elias Alvarado-Yaah, Carlos F. Arias, Santiago Ávila-Ríos, Eduardo Becerril-Vargas, Víctor Hugo Borja-Aburto, Celia Boukadida, Cristóbal Cháidez-Quiróz, Juan Bautista Chale-Dzul, Ricardo Ciria Merce, Andreu Comas-García, Célida Duque Molina, Julissa Enciso-Ibarra, José Antonio Enciso-Moreno, Gloria Elena Espinosa-Ayala, Fernando Fontove-Herrera, Daniel Fregoso-Rueda, Victor Eduardo García-Arias, Alejandra García-Gasca, Bruno Gómez-Gil, Jean Pierre González, Irvin González-López, Concepción Grajales-Muñiz, Ricardo Grande, Rosa María Gutiérrez Rios, Jesús Hernández, Alejandra Hernández-Terán, Alfredo Herrera-Estrella, Carla Ivón Herrera-Najera, Pavel Isa, Verónica Mata-Haro, Daniel Lira Morales, Susana Lopez, Antonio Loza Román, Brenda Irasema Maldonado-Meza, Bernardo Martínez-Miguel, José Arturo Martínez-Orozco, Célida Martínez- Rodríguez, Margarita Matías-Florentino, Fidencio Mejía-Nepomuceno, María Guadalupe de Jesús Mireles-Rivera, Gloria María Molina-Salinas, Hector Montoya-Fuentes, , Mario Mújica-Sánchez, José Esteban Muñoz-Medina, José de Jesús Nuñez-Contreras, Alicia Ocaña-Mondragón, Luis Alberto Ochoa-Carrera, Hector Esteban Paz-Juárez, Marissa Perez-Garcia, Francisco Pulido, Helen Haydee Fernanda Ramírez-Plascencia, Jorge Salas-Hernández, Angel Gustavo Salas-Lais, Alejandro Sánchez-Flores, Clara Esperanza Santacruz-Tinoco, María Guadalupe Santiago-Mauricio, Selene Zárate, Nelly Sélem-Mojica, Blanca Taboada, Gloria Vazquez, Joel Armando Vázquez-Pérez. |
| EPI_ISL_2987722 | Hospital Sharp | Microbial Genomics Laboratory | Consorcio Mexicano de Vigilancia Genómica (CoViGen-Mex). Authors (in alphabetical order): Julio Elias Alvarado-Yaah, Carlos F. Arias, Santiago Ávila-Ríos, Eduardo Becerril-Vargas, Víctor Hugo Borja-Aburto, Celia Boukadida, Cristóbal Cháidez-Quiróz, Juan Bautista Chale-Dzul, Ricardo Ciria Merce, Andreu Comas-García, Célida Duque Molina, Julissa Enciso-Ibarra, José Antonio Enciso-Moreno, Gloria Elena Espinosa-Ayala, Fernando Fontove-Herrera, Daniel Fregoso-Rueda, Victor Eduardo García-Arias, Alejandra García-Gasca, Bruno Gómez-Gil, Jean Pierre González, Irvin González-López, Concepción Grajales-Muñiz, Ricardo Grande, Rosa María Gutiérrez Rios, Jesús Hernández, Alejandra Hernández-Terán, Alfredo Herrera-Estrella, Carla Ivón Herrera-Najera, Pavel Isa, Verónica Mata-Haro, Daniel Lira Morales, Susana Lopez, Antonio Loza Román, Brenda Irasema Maldonado-Meza, Bernardo Martínez-Miguel, José Arturo Martínez-Orozco, Célida Martínez- Rodríguez, Margarita Matías-Florentino, Fidencio Mejía-Nepomuceno, María Guadalupe de Jesús Mireles-Rivera, Gloria María Molina-Salinas, Hector Montoya-Fuentes, , Mario Mújica-Sánchez, José Esteban Muñoz-Medina, José de Jesús Nuñez-Contreras, Alicia Ocaña-Mondragón, Luis Alberto Ochoa-Carrera, Hector Esteban Paz-Juárez, Marissa Perez-Garcia, Francisco Pulido, Helen Haydee Fernanda Ramírez-Plascencia, Jorge Salas-Hernández, Angel Gustavo Salas-Lais, Alejandro Sánchez-Flores, Clara Esperanza Santacruz-Tinoco, María Guadalupe Santiago-Mauricio, Selene Zárate, Nelly Sélem-Mojica, Blanca Taboada, Gloria Vazquez, Joel Armando Vázquez-Pérez. |
